# Maleic acid adjuvates BCG to induce CD4^+^CD8^+^ T cells and sterilize tuberculosis

**DOI:** 10.64898/2026.09.20.752978

**Authors:** Chenyue Shi, Hongyu Cheng, Xiaoli Feng, Xinyu Cao, Hongjie Liu, Qi Liu, Lu Zhang, Yuanna Cheng, Cheng Peng, Jinya Zhang, Qiaojiang Yang, Kejing Dong, Yuli Gao, Xinyi Hu, Yudan Liu, Yuhan Bian, Shenzhi Li, Yifan Yang, Lin Wang, Jie Wang, Xiaochen Huang, Xia Cai, Di Qu, Minghua Li, Hua Yang, Baoxue Ge

## Abstract

Anti-tuberculosis (TB) immunity typically emphasizes CD4⁺ and CD8⁺ T cells that only provide limited protection against *Mycobacterium tuberculosis* (*Mtb*) infection. The anti-TB function of CD4^+^CD8^+^ T cells remains unexplored, mostly due to the dogma that ThPOK and Runx3 antagonize each other to block their development and their scarcity. Here, we show that immunization of BCG (Bacillus Calmette–Guérin) with Complete Freund’s Adjuvant (CFA) markedly induced CD4^+^CD8^+^ T cells through the gut microbiota *Clostridium butyricum* or its metabolite maleic acid. *Mtb*-specific CD4^+^CD8^+^ T cells exhibited a broad T-cell receptor repertoire and sterilized *Mtb* infection by inducing SLAMF1 and neutrophil responses. Mechanistically, maleic acid interacted with PAK2 to promote STAT5 phosphorylation at Ser127, thus upregulating Ets1 expression. Direct binding of Ets1 to the *Runx3* and *ThPOK* promoters orchestrated their expression to redirect the CD8^+^ T-cell differentiation towards the CD4^+^CD8^+^ lineage. Moreover, immunization of BCG plus *Clostridium butyricum* or maleic acid sterilized *Mtb* infection in mouse models. These findings establish CD4^+^CD8^+^ T cells as superior immune effectors against *Mtb*, a concept that could fundamentally redirect the current paradigm for anti-TB vaccine development.

## Introduction

Tuberculosis (TB) is a global health issue and the leading cause of death ^1^. Bacillus Calmette-Guérin (BCG; live attenuated *Mycobacterium bovis*), administered intradermally at birth, remains the only clinically licensed TB vaccine. While it confers protection against disseminated TB in infants, its efficacy against pulmonary TB declines significantly in adolescent and adult populations ^2, 3, 4^ ^5, 6^, which is responsible for the majority of mortality and TB transmission^2, 7^. Three large randomized trials indicate the ineffectiveness of BCG revaccination ^8, 9, 10^. As a live attenuated vaccine, BCG vaccination result in a disseminated infection in infants with immunodeficiency^11^. Therefore, a safer, efficacious and durable vaccine is urgently needed to prevent TB disease in these populations.

During past decade, tuberculosis vaccine research and development has produced a number of novel vaccine candidates in the clinical trial pipeline with some encouraging results^12, 13^. Currently, 16 vaccine candidates are being investigated in clinical trials, but their reported efficacies plateau at approximately 50%, a threshold insufficient to meet global targets for TB elimination ^14, 15^. For instance, the recombinant BCG vaccine VPM1002 reduced lung *Mycobacterium tuberculosis* (*Mtb*) loads by nearly 10-fold relative to naive mice by day 30 post-challenge ^16, 17^, but recent Phase III clinical trials failed to demonstrate statistically significant protection against TB in humans (per-protocol vaccine efficacy 21.4%, 95% CI –8.9% to 43.2% against all TB; not significant) ^18^. Novel approaches are actively being explored. A trivalent mRNA-lipid-nanoparticle (LNP) vaccine (comprising PPE20/Rv1387, EsxG/Rv0287, and PE18/Rv1788) augmented and even exceeded BCG protection in multiple mouse models, achieving a durable 10.7-fold reduction in lung bacterial loads when mice were challenged three months post-immunization (P= 0.0079) ^19^. Another two mRNA–LNP-based vaccine candidates also reduced the bacterial burden in *Mtb*-challenged mice ^20^. In parallel, studies in rhesus macaques showed that BCG efficacy can be greatly enhanced by switching from conventional low-dose intradermal injection to high-dose intravenous administration^21^, but the intravenous route poses substantial safety challenges. Therefore, superior protective effects greater than what current candidates have demonstrated is required for the TB vaccine strategy to eventually achieve clinical success.

Though a number of immune cells including innate immune cells and humoral or B cells have protective roles in TB^22, 23, 24, 25, 26, 27, 28, 29, 30, 31^ ^32^, central immune protective mediators are CD4⁺ and CD8⁺ T cells that mediate anti-TB immunity through the coordinated secretion of pro-inflammatory cytokines such as the IL-12-IFNγ axis and TNF^23, 33, 34, 35, 36, 37, 38^, thus serving as essential components of adaptive immunity against *Mtb* infection ^39, 40^. In a murine model, the adoptive transfer of BCG ΔureC::hly vaccine (rBCG) induces mycobacteria-specific central memory CD4^+^ T cells, thus validating their moderate protective role against pulmonary TB (<10-fold total colony forming units (CFUs))^41^. Similarly, CD8^+^ T cell responses have been shown to be critical for controlling acute and chronic *Mtb* infection in non-human primates (NHPs), and their magnitude correlates with the moderate protection afforded by BCG (also roughly <10-fold CFU reduction) ^42, 43^. CD4^+^CD8^+^ double-positive (DP) T cells have been found in intravenously (i.v.) BCG immunization models and other diseases^43, 44, 45^, but their functional role in the immune protection or pathogenesis of diseases and the mechanism underlying their lineage differentiation remain unclear, which is primarily attributable to the dogma that ThPOK and Runx3 antagonize each other to induce functionally distinct “helper” CD4^+^ or “cytotoxic” CD8^+^ T cell fates and their scarcity **_46_**.

Adjuvants enhance immune responses while also protecting antigens from degradation and facilitating their delivery to target tissues ^47^. TB vaccine adjuvants have progressed from traditional alum, which fails to induce robust Th1/CD8⁺ T cell responses against intracellular *Mtb*, toward clinically advanced liposomal and emulsion systems ^48^. Among these, AS01 (MPL+QS-21) demonstrates the highest efficacy in humans, conferring 54% protection over 3 years in patients with latent TB in a phase Ⅲ clinical trial (NCT01755598) ^14, 15^. CAF01 (DDA/TDB) safely primes Th1/Th17 responses but exhibits limited pulmonary control post-boosting, while GLA-SE (TLR4 agonist+squalene) ^49^ and IC31 (KLK/ODN1a) ^50^ enhance polyfunctional CD4⁺ T cell magnitude and durability in early-phase trials. Novel platforms such as dextran/CpG nanoparticles (GamTBVac, phase IIa) and Advax microparticles (preclinical) have shown promising immunogenicity, but AS01 remains the only adjuvant with positive phase III efficacy data, which underscores the persistent challenge of translating preclinical immunogenicity into clinical protection^51^. Complete Freund’s Adjuvant (CFA) is primarily composed of inactivated H37Ra and mineral oil ^52, 53^, and serves as a broad-spectrum immune potentiator in animal models^54, 55, 56^. However, the protective efficacy of the combined subcutaneous administration of BCG and CFA against TB remains unexplored.

In this study, we demonstrate that combined immunization with BCG and CFA markedly induced CD4^+^CD8^+^ T cells through the gut microbiota *Clostridium butyricum* or its metabolite maleic acid. *Mtb*-specific CD4^+^CD8^+^ T cells exhibited a broad TCR repertoire and conferred sterilizing protection against *Mtb* challenge by eliciting SLAMF1 and neutrophil responses. Mechanistically, interaction of maleic acid with PAK2 promoted STAT5 phosphorylation at Ser127 and subsequently upregulated the transcription factor Ets1, which directly bound to *Runx3* and *ThPOK* promoter loci, thereby upregulating their expression and driving the differentiation of CD8^+^ T-cell towards the CD4^+^CD8^+^ lineage.

## Results

### BCG plus CFA immunization provide sterilizing immunity against *Mtb*

To investigate the adjuvant effects of CFA on BCG efficacy against *Mtb* infection, the mice were subcutaneously vaccinated with BCG plus CFA and other control treatments (**Fig 1a**). 4 weeks after immunization, the mice were challenged with a nominal dose of 200 CFUs of *Mtb* H37Rv in three separate cohorts, with a pre-defined study end point of 4 weeks after challenge. The primary measure of protection was a comprehensive quantification of the *Mtb* burden (CFUs) in the lungs and spleen, combined with histopathological analysis of the lung tissue. The mean total lung CFUs for subcutaneous (SC) BCG (4.11 log10 CFU) was slightly lower than that of unvaccinated mice (5.80 log10 CFU). However, the bacterial burden in the lung tissues of mice immunized with BCG plus CFA (0.46 log10 CFU) was much lower (>100,000-fold) than unvaccinated mice (5.80 log10 CFU) or (>1000-fold) mice immunized with BCG (4.11 log10 CFU). Eight out of ten mice had no detectable *Mtb* in their lung tissues **(Fig. 1b)**. The mean of total spleen CFUs for BCG plus CFA (0.92 log10 CFU) was reduced by approximately 1000-fold relative to that in unvaccinated mice (3.66 log10 CFU) **(Fig. 1c)**. Consistent with this, lung tissues from mice immunized with BCG plus CFA showed markedly reduced immune-cell infiltration and fewer inflammatory lesions than those from mice immunized with parental BCG **(Fig. 1d–f)**. Six out of ten mice immunized with BCG plus CFA showed no detectable histopathological damage in their lung tissues **(Fig. 1d–f)**. In addition, subcutaneous immunization of BCG plus CFA resulted in drastically reduced expression of effector molecules Granzyme B, IL-12p70, IFN-γ and IL-2 in mice serum at 4 weeks after *Mtb* challenge, relative to control animals, consistent with the resolution of inflammation post-clearance **(Fig. 1g).** To evaluate the long-term protective efficacy of SC BCG plus CFA immunization, mice were challenged with approximately 200 CFUs of *Mtb* H37Rv at 3 months (12 weeks) post-vaccination, with predefined experimental endpoints assessed 4 weeks post-challenge **(Extended Data Fig. 1a)**. Notably, the mean total lung CFUs for SC BCG (5.15 log10 CFU) was slightly lower than that of unvaccinated mice (5.74 log10 CFU). However, the bacterial burden in the lung tissues of mice immunized with BCG plus CFA (0.48 log10 CFU) was markedly reduced (>100,000-fold) than unvaccinated mice (5.74 log10 CFU) or (>10,000-fold) mice immunized with parental BCG (5.15 log10 CFU). Six out of seven mice had no detectable *Mtb* in their lung tissues (P<0.0001; **Extended Data Fig. 1b**). Consistent with this, lung tissues from mice immunized with BCG plus CFA showed markedly reduced immune-cell infiltration and fewer inflammatory lesions than those from mice immunized with parental BCG at 4 weeks post-infection (wpi) **(Extended Data Fig. 1c, d)**. Two out of four mice immunized with BCG plus CFA showed no detectable histopathological damage in their lung tissues **(Extended Data Fig. 1c, d)**. To evaluate the safety of subcutaneous BCG combined with CFA vaccination, key clinical indices were monitored. The BCG plus CFA combination demonstrated a safety profile comparable to that of BCG alone **(Extended Data Fig. 2)**. These results suggest that subcutaneous BCG plus CFA immunization may confer an unprecedented degree of protection against *Mtb* infection.

**Fig. 1:**
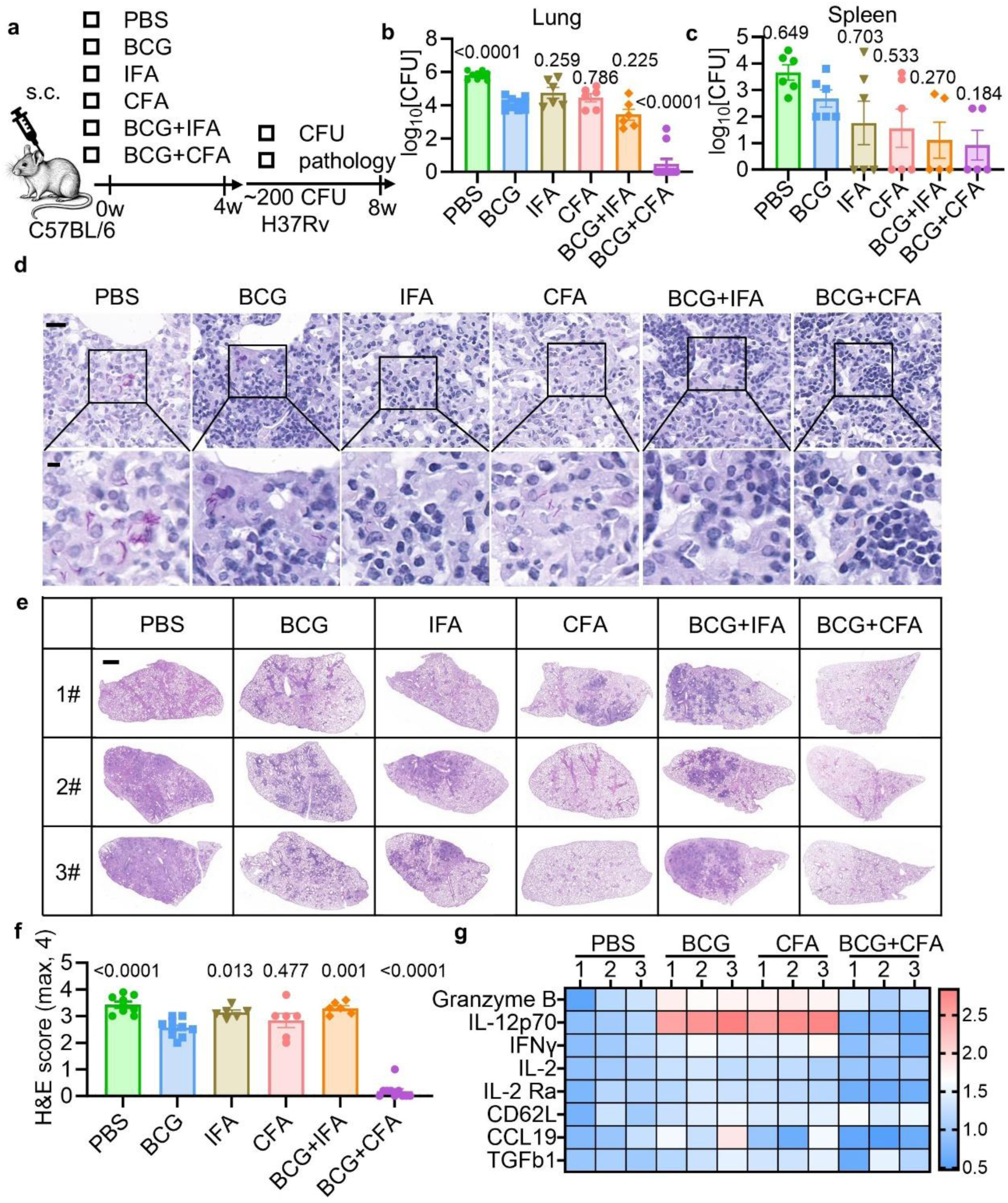
BCG plus CFA immunization provide robust protection against *Mtb*. **a-f**, Experiment design **(a)**, C57BL/6 mice from each vaccine group were aerosol-infected with ∼200 c.f.u. per mouse of *Mtb* H37Rv. After 4 weeks of infection, the following were assessed: bacterial load of lung **(b)**, bacterial load of spleen **(c)**, lung sections acid-fast staining (**d**; scale bar, 20 μm (top) and 5 μm (bottom)), lung sections H&E staining (**e**; scale bar=1mm) and histology score **(f)**. **g**, Heatmap showing serum cytokine levels in *Mtb* H37Rv-infected mice at 4 weeks post-infection across vaccine groups, quantified by cytometric bead-based immunoassays. Data represent one experiment with at least three independent biological replicates and are shown as the mean ± s.e.m.. Ordinary one-way ANOVA with Dunnett’s multiple comparison test comparing each group to the BCG group (**b, c, f**) was used for statistical analyses.

### BCG plus CFA vaccination induces *Mtb*-specific CD4^+^CD8^+^ T cells

To delineate the protective immunity conferred by subcutaneous BCG plus CFA vaccination, we performed multiparametric flow cytometry (FCM) to systematically profile innate and adaptive immune cell dynamics in the lungs, spleen and lymph nodes at 4 weeks post-vaccination. Compared to BCG-vaccinated, CFA-vaccinated, or unvaccinated controls, subcutaneous co-administration of BCG with CFA did not significantly alter the proportions of lung CD4⁺ T cells **(Extended Data Fig. 3a)**, CD8⁺ T cells **(Extended Data Fig. 3a)**, CD4^+^CD8^+^ T cells **(Extended Data Fig. 3a)**, dendritic cells (DCs) **(Extended Data Fig. 3b)**, macrophages **(Extended Data Fig. 3b)**, or monocytes **(Extended Data Fig. 3b)**. Similarly, splenic compartment analysis revealed no significant BCG plus CFA vaccination-induced changes in the frequencies of CD4⁺ T cells **(Extended Data Fig. 3c)**, CD8⁺ T cells **(Extended Data Fig. 3c)**, CD4^+^CD8^+^ T cells **(Extended Data Fig. 3c)**, macrophages **(Extended Data Fig. 3d)**, monocytes **(Extended Data Fig. 3d)**, B cells **(Extended Data Fig. 3e)**, NK cells **(Extended Data Fig. 3e)**, or NKT cells **(Extended Data Fig. 3f)**. Notably, however, BCG plus CFA vaccination induced a compartment-specific expansion of CD4⁺CD8⁺ T cells, increasing their frequency to up to 50% of total T cells exclusively within the lung-draining lymph nodes (LNs) at 4 weeks post-immunization **(Fig. 2a)**. This spatial selectivity contrasted sharply with unaltered CD4^+^CD8^+^ T cell proportions in the axillary LNs **(Extended Data Fig. 4a)**, mesenteric LNs **(Extended Data Fig. 4b)**, and Peyer’s patches **(Extended Data Fig. 4c)**. Spatially resolved immunofluorescence analysis revealed the compartmentalized enrichment of CD4⁺CD8⁺ T cells within the deep cortical regions of lung-draining LNs **(Fig. 2b)**, along with markedly higher CD44 expression in CD4^+^CD8^+^ T cells than in CD4^+^ and CD8^+^ T cells **(Fig. 2c)**, indicative of an activated effector-memory phenotype.

**Fig. 2:**
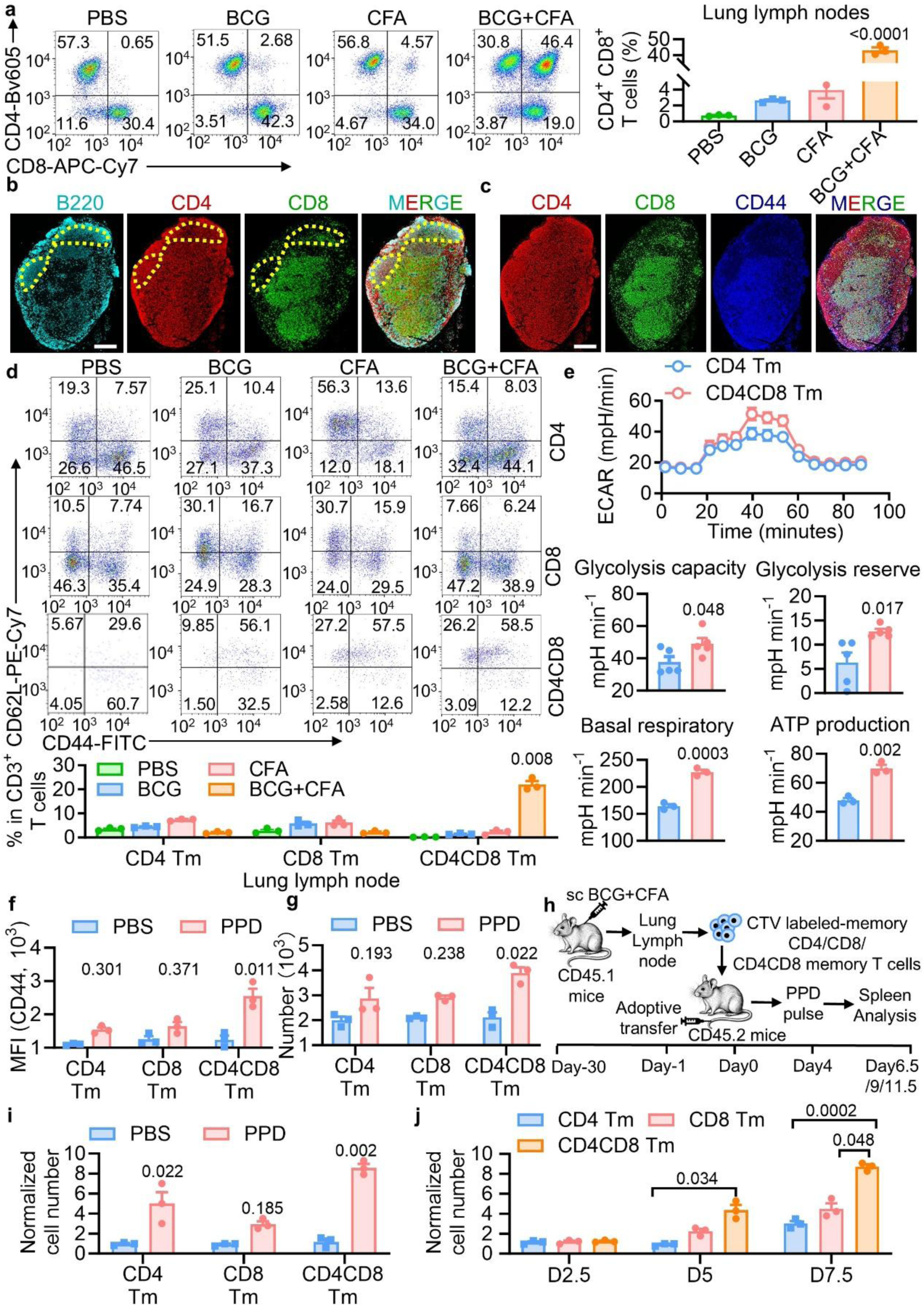
BCG plus CFA vaccination induces *Mtb*-specific CD4^+^CD8^+^ T cells. **a,** FCM analyses of CD4^+^CD8^+^ T cells (gated from CD3^+^ T cells) in the lung lymph nodes of each vaccine group, at 4 weeks post vaccination. Frequency of each population is summarized beside. **b-c**, Immunofluorescence data showing B220 (amber), CD4 (red), CD8 (green) staining. The yellow dashed box outlines the superficial cortex (B cell zone), CD4^+^CD8^+^ T cells reside in the surrounding deep cortex (T cell zone) **(b)** and CD4 (green), CD8 (red), CD44 (blue) staining (**c**) in lung lymph nodes sections of 4 weeks post BCG plus CFA vaccinated mice. (**b-c**, scale bar=1mm). **d**, FCM analyses of CD4^+^, CD8^+^, CD4^+^CD8^+^ memory T cells (CD44^+^CD62L^+^, gated from CD4^+^, CD8^+^, CD4^+^CD8^+^ T cells) in the lung lymph nodes of each vaccine group, at 4 weeks post vaccination. Frequency of each population is summarized below. **e**, ECAR, OCR were measured and analysed via Seahorse metabolic assay. **f-g**, Recall assay in vitro. Geometric mean fluorescence intensities (MFIs) of CD44 (**f**) and cell number of each group (**g**). **h-j**, Experiment design (**h**), we labeled FACS-isolated CD4^+^ memory T cell, CD8^+^ memory T cell and CD4^+^CD8^+^ memory T cells from BCG+CFA-induced CD45.1 donor mice with Cell Trace Violet (CTV). These labeled cell populations were adoptively transferred into syngeneic CD45.2 recipient mice. After in vivo PPD stimulation at day 4 post-transfer, donor-derived proliferation in spleen was quantified by flow cytometry at 2.5, 5-, and 7.5-days post-stimulation, fold changes relatively to initial cell number of each group at 7.5 days post-stimulation (**i**) and cell number of each group at 2.5, 5-, and 7.5-days post-stimulation **(j)**. Data represent one experiment with at least three independent biological replicates and are shown as the mean ± s.e.m.. Ordinary one-way ANOVA with Dunnett’s multiple comparison test comparing BCG+CFA group to the BCG group (**a**), two-way ANOVA with Dunnett‘s multiple comparisons test (**d, j**), two-tailed unpaired Student’s t-tests (**e**) and two-way ANOVA with Šídák’s multiple comparisons test (**f, g, i**) were used for statistical analyses.

Memory immune responses are mounted more rapidly than naïve counterparts, enabling effective containment of pathogen rechallenge ^57^. This accelerated response underpins vaccine efficacy, which is largely mediated by adaptive immune memory that confers durable protection ^58^. Notably, approximately 50% of the CD4^+^CD8^+^ T cells induced by BCG plus CFA co-expressed CD44 and CD62L **(Fig. 2d)**, a surface phenotype characteristic of central memory T cells (T_CM_) ^59, 60^. Memory T cells preferentially rely on mitochondrial oxidative phosphorylation (OXPHOS), metabolizing glucose, fatty acids, and amino acids to efficiently generate adenosine triphosphate (ATP)^61^. Using the Seahorse XF Analyzer, we found that the CD4^+^CD8^+^CD44^+^CD62L^+^ T cells induced by BCG plus CFA exhibited higher OXPHOS and glycolytic capacities than did conventional CD4^+^CD8^−^CD44^+^CD62L^+^ T cells (CD4^+^ T_CM_) **(Fig. 2e)**. This metabolic profile, quantified by glycolytic rate, glycolytic reserve, basal respiration and ATP production capacity, identifies CD4⁺CD8⁺CD44⁺CD62L⁺ T cells as memory cells with enhanced metabolic plasticity, suggesting their unique bioenergetic adaptations.

Memory T cells possess hallmark functional properties, including heightened self-renewal, durable persistence and robust antigen-specific recall responses, that collectively provide acute antimicrobial defense and sustained protection against recurrent infections ^62, 63^. To delineate the self-renewal capacity of CD4^+^CD8^+^ memory T cells, we developed a co-culture model combining BCG plus CFA-induced CD4^+^CD8^+^ memory T cells with purified protein derivative (PPD)-pulsed bone marrow-derived macrophages (BMDMs). Quantitative cell counting assays demonstrated significant proliferation of BCG plus CFA-primed CD4^+^CD8^+^ memory T cells following PPD stimulation, while FCM analysis revealed concomitant upregulation of CD44 expression in these antigen-experienced cells **(Fig. 2f, g)**. To further investigate the self-renewal dynamics of CD4^+^CD8^+^ memory T cells, we isolated CD4^+^, CD8^+^, and CD4^+^CD8^+^ memory T cells from BCG plus CFA-immunized CD45.1 donor mice using fluorescence-activated cell sorting (FACS) and labeled them with Cell Trace Violet (CTV). These labeled cell populations were adoptively transferred into syngeneic CD45.2 recipient mice. Following PPD pulse at day 4 post-transfer, splenic proliferation kinetics of donor-derived cells were quantified using multiparametric FCM on days 2.5, 5, and 7.5 **(Fig. 2h)**. As a result, 7.5 days post-PPD pulse, CD4^+^CD8^+^ memory T cells exhibited significantly greater proliferative potential than conventional CD4^+^ or CD8^+^ memory subsets, as evidenced by a 3.2-fold increase in fluorescence-diluted progeny, indicative of robust antigen-specific clonal expansion **(Fig. 2i, j)**. Collectively, these data indicate that BCG+CFA immunization elicits CD4^+^CD8^+^ memory T cells with superior functional competence, antigen-specific proliferative capacity and long-term persistence compared with conventional CD4^+^ or CD8^+^ memory T cell subsets.

To further characterize the antigen specificity of BCG plus CFA-induced CD4⁺CD8⁺ T cells, we performed the Pan-Peptide Meta Learning (PanPep) framework ^64^ analysis in conjunction with single-cell RNA sequencing analysis paired with TCR sequencing (scRNA-seq/TCR-seq) analysis of CD4^+^ T cells and CD8^+^ T cells isolated from lung-draining lymph nodes (LNs) of mice one month after subcutaneous immunization with BCG+CFA **(Extended Data Fig. 5a)**. Enriched cells were then subjected to scRNA-seq using the BD Rhapsody platform (SRA, SRP719462). Unsupervised clustering with the Louvain algorithm (resolution 0.7) resolved 17 distinct T cell clusters, as visualized with uniform manifold approximation and projection (UMAP) **(Extended Data Fig. 5a)**. Clusters 2, 3, and 15 co-expressed *Cd4*, *Cd8a*, and *Cd8b1* at the transcript level, confirming their identity as CD4⁺CD8⁺ T cells **(Extended Data Fig. 5b)**. Notably, BCG plus CFA-induced CD4⁺CD8⁺ T cells exhibited a remarkable breadth of antigen specificity against *Mtb*, and the TCR repertoire analysis revealed a polyclonal response without skewing toward a single immunodominant epitope, indicating a broad and unbiased antigen recognition profile of CD4⁺CD8⁺ T cells **(Extended Data Fig. 5c, d and Table S1)**.

### CD4^+^CD8^+^ memory T cells control *Mtb* infection

Given the phenotypic heterogeneity of BCG+CFA-induced CD4^+^CD8^+^ T cells **(Fig. 2d)**, we sought to test their functional protective capacity in vivo. To examine whether CD4^+^CD8^+^ memory T cells are functionally required for BCG+CFA-mediated protection, we adoptively transferred FACS-purified CD4^+^CD8^+^, CD4^+^ or CD8^+^ memory T cells (from lung-draining LNs of BCG+CFA-immunized mice) into *Rag1*^⁻/⁻^ recipients. The recipient mice were subsequently challenged with the *Mtb* H37Rv strain, and subjected to quantitative analysis of pulmonary and splenic bacterial burdens at 28 days **(Fig. 3a)**. The mean total lung CFUs in recipient mice that received CD4⁺CD8⁺ memory T cells (no detectable CFUs) were much lower (decreased by >1, 000,000-fold) than those in control *Rag1*^⁻/⁻^ mice (6.69 log10 CFU). All six mice had no detectable *Mtb* in their lung tissues **(Fig. 3b, c)**. The mean total spleen CFUs in recipient mice that received CD4⁺CD8⁺ memory T cells (2.41 log10 CFU) were much lower (decreased by >1, 000-fold) than those in control *Rag1*^⁻/⁻^ mice (5.65 log10 CFU). Two out of five mice exhibited no detectable *Mtb* in their spleen tissues **(Extended Data Fig. 6a)**. Consistent with these findings, lung tissues from recipient mice that received CD4⁺CD8⁺ memory T cells showed markedly reduced immune cell infiltration and fewer inflammatory lesions than those from control *Rag1*^⁻/⁻^ mice **(Fig. 3d, e)**. Collectively, these data suggest that BCG plus CFA-induced CD4^+^CD8^+^ memory T cell may confer sterilizing protection against *Mtb* infection *in vivo*.

**Fig. 3:**
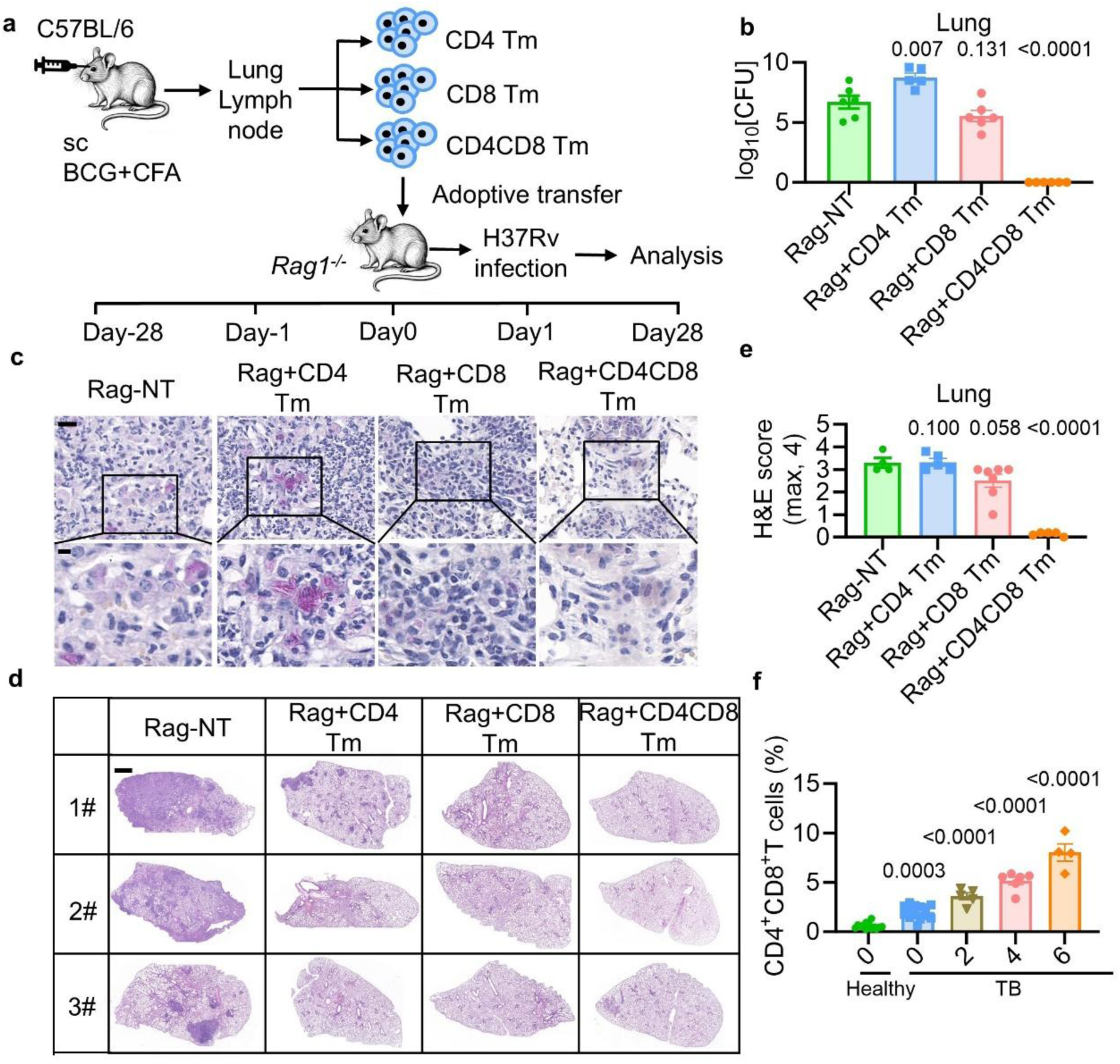
CD4^+^CD8^+^ memory T cells conferred sterilizing protection against *Mtb*. **a-e**, Experimental design **(a)**. CD4⁺ memory T cells, CD8⁺ memory T cells, and CD4⁺CD8⁺ memory T cells were sorted from lung lymph nodes of mice vaccinated with BCG plus CFA at 4 weeks post-vaccination. These cells were then transferred into *Rag1*^⁻/⁻^ mice. Recipient mice were subsequently aerosol-infected with ∼200 CFU per mouse of *Mtb* H37Rv. At 4 weeks post-infection, the following parameters were assessed: bacterial load in the lung **(b)**, acid-fast staining (**c**; scale bar, 20 μm (top) and 5 μm (bottom)), histopathology of lung sections via H&E staining (**d**; scale bar, 1 mm) and histology score **(e). f,** Proportions of CD4^+^CD8^+^ T cells in the peripheral blood of each group. Data represent one experiment with at least three independent biological replicates and are shown as the mean ± s.e.m.. Ordinary one-way ANOVA with Dunnett’s multiple comparison test comparing each group to the Rag-NT group (**b, e**) or healthy group (**f**) was used for statistical analyses.

We next determined the presence of the CD4^+^CD8^+^ T cell population in patients with clinical TB, wherein they were detected in the peripheral blood of infected individuals **(Extended Data Fig. 6b)**. Notably, the percentage of CD4⁺CD8⁺ T cells increased progressively over the course of anti-TB treatment **(Fig. 3f and Extended Data Fig. 6c).** At baseline (before treatment), patients with TB had higher percentages of Tem and Temra CD4⁺CD8⁺ T cells than healthy controls **(Extended Data Fig. 6d-h)**. After completion of standard therapy, however, CD4⁺CD8⁺ T cells predominantly exhibited Tcm and naive phenotypes **(Extended Data Fig. 6d–h)**. Collectively, these observations suggest that during active TB, CD4⁺CD8⁺ T cells are enriched in effector-memory and terminally differentiated subsets, which may contribute to the ongoing immune response; following clinical resolution, the compartment shifts towards Tcm and naive phenotypes, potentially reflecting the restoration and maintenance of immunological memory.

### CD4^+^CD8^+^ T cells induce SLAMF1 and neutrophil responses

To further investigate how CD4^+^CD8^+^ memory T cell populations confer robust protection against *Mtb* infection *in vivo,* we isolated CD4^+^CD8^+^ memory T cells from lung-draining LNs of mice subcutaneously immunized with BCG plus CFA, and adoptively transferred them into *Rag1^-/-^* mice. The recipient mice were subsequently challenged with the *Mtb* H37Rv. Quantification of pulmonary cytokines at 3 wpi revealed that the adoptive transfer recipients of CD4^+^CD8^+^ memory T cells exhibited significantly upregulated levels of signaling lymphocytic activation molecule family 1 (SLAMF1) compared with control recipients **(Extended Data Fig. 7a, b)**. Following *Mtb* infection, BCG+CFA immunization markedly increased SLAMF1 expression in CD4⁺CD8⁺ T cells, whereas no such upregulation was observed in CD4⁺ or CD8⁺ single-positive subsets relative to all control groups **(Extended Data Fig. 7c–e)**. This suggests that BCG+CFA immunization selectively induces CD4⁺CD8⁺ T cells that express SLAMF1.

As a cell surface molecule predominantly expressed on immune cells, SLAMF1 interacts with adaptor proteins SLAM-associated protein (SAP) and Ewing’s sarcoma-associated transcript 2 (EAT-2) through homophilic SLAMF1–SLAMF1 binding, and subsequently activating downstream signaling pathways to modulate diverse immune cell functions^65, 66^. To further investigate which subsets of immune cells are affected by SLAMF1-expressing CD4^+^CD8^+^ memory T cells, we sorted CD4^+^CD8^+^ memory T cells from CD45.1 mice immunized with BCG plus CFA and adoptively transferred them into CD45.2 C57BL/6 mice. Three weeks after *Mtb* infection, we performed single-cell RNA sequencing analysis of CD45^+^ cells in lungs from recipient mice (SRA, SRP719608). Notably, we observed a significant increase in the number of CD45.2⁺ neutrophils in the lungs of mice that had received CD45.1⁺ CD4⁺CD8⁺ T cells **(Extended Data Fig. 7f, g)**. The neutrophil population highly expressed genes associated with bactericidal function, including direct antimicrobial effectors (*Bri3, Nos2, Plac8, Rab37, Sod2*) and immunoregulatory molecules that modulate chemotaxis and inflammation (*Cxcl10, Fpr1, Fpr2, Il1a, Pnp*) **(Extended Data Fig. 7h)**.

To determine whether the protective effect of CD4^+^CD8^+^ memory T cells against *Mtb* infection depends on SLAMF1 or neutrophil responses, we isolated CD4^+^CD8^+^ memory T cells from BCG plus CFA-immunized *Slamf1*^⁻/⁻^ (knockout) and Wild type (WT) mice and adoptively transferred them into *Rag1*^⁻/⁻^ recipients. After *Mtb* H37Rv challenge, neutrophils were depleted using an anti-Ly6G antibody, with an isotype IgG antibody as a control. Four weeks post-infection, we assessed bacterial burden (CFU) and histopathological damage in the lungs and spleens **(Fig. 4a)** of the *Mtb* H37Rv-infected mice. In the mice treated with isotype control IgG, adoptive transfer of WT CD4^+^CD8^+^ memory T cells significantly reduced bacterial burden in the lungs and spleens **(Fig. 4b-d)** as well as pulmonary pathology **(Fig. 4e, f)** compared to those non-transferred (NT) controls. In contrast, transfer of *Slamf1*^⁻/⁻^ CD4^+^CD8^+^ memory T cells did not significantly change the bacterial burden in the lungs and spleens or lung pathology of the recipient mice compared to those control mice, suggesting an essential role of SLAMF1 for the protective anti-*Mtb* effects of CD4^+^CD8^+^ memory T cells. Notably, administration of anti-Ly6G neutralization antibody attenuated the protective effects of CD4^+^CD8^+^ memory T cells on the bacterial loads in the lung tissues of *Mtb* H37Rv-infected mice **(Fig. 4b-f)**. Consistently, the differential survival of the *Mtb* H37Rv in the mice transferred with WT or *Slamf1*⁻/⁻ CD4^+^CD8^+^ memory T cells was eliminated upon neutrophil depletion **(Fig. 4b-f)**. Collectively, these results suggest that CD4⁺CD8⁺ memory T cells may control *Mtb* infection by engaging SLAMF1 and eliciting neutrophil responses.

**Fig. 4:**
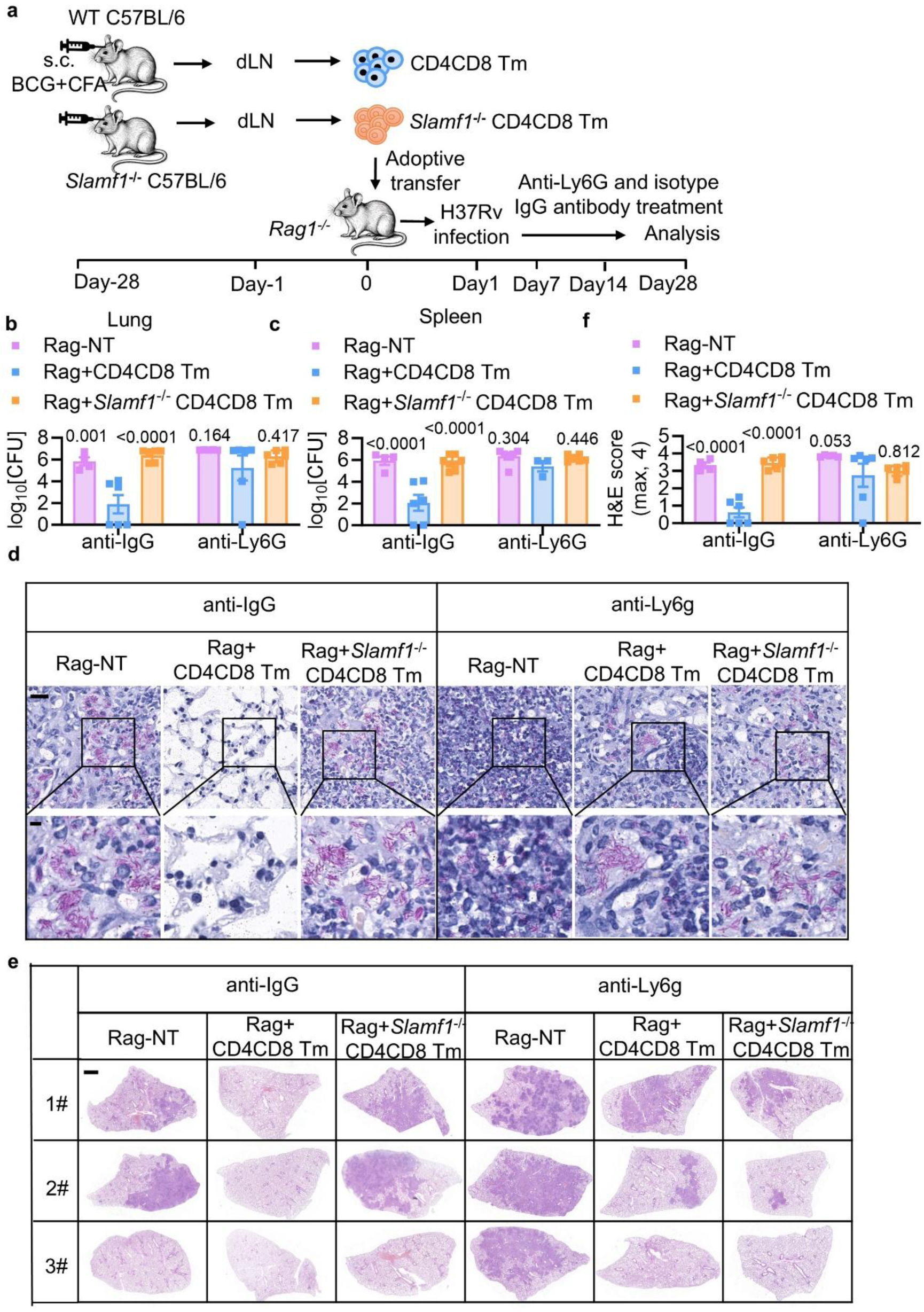
CD4^+^CD8^+^ memory T cells control *Mtb* infection by SLAMF1-dependent stimulation of neutrophils. **a-f,** Experimental design **(a)**, CD4⁺CD8⁺ memory T cells were sorted from lung lymph nodes of mice vaccinated with BCG plus CFA at 4 weeks post-vaccination. *SLAMF1*^-/-^ CD4⁺CD8⁺ memory T cells were sorted from lung lymph nodes of *SLAMF1*^-/-^ mice vaccinated with BCG plus CFA at 4 weeks post-vaccination. These cells were then transferred into *Rag1*^⁻/⁻^ mice. Recipient mice were subsequently aerosol-infected with ∼200 CFU per mouse of *Mtb* H37Rv. At 4 weeks post-infection, the following parameters were assessed: bacterial load in the lung **(b)**, bacterial load in the spleen **(c),** acid-fast staining (**d**; scale bar, 20 μm (top) and 5 μm (bottom)), histopathology of lung sections via H&E staining (**e**; scale bar, 1 mm) and histology score **(f)**. Data represent one experiment with at least three independent biological replicates and are shown as the mean ± s.e.m.. Statistical significance in **(b, c and f)** was assessed by two-way ANOVA with Dunnett’s multiple comparison test comparing each group to the Rag+CD4CD8 Tm group.

### Maleic acid adjuvates BCG to provide sterilizing protection

Given the safety concern of CFA, we next sought to identify the safer CFA-derivatives. Vaccination triggers a systemic host metabolic recalibration through a process of immunometabolism reprogramming^67, 68^, a fundamental mechanism governing immune cell identity and functional capacity ^69, 70^. To investigate whether the enhanced anti-TB immune protection conferred by BCG plus CFA immunization is mediated through metabolic reprogramming, we performed metabolomic sequencing on serum samples collected from mice 4 weeks after immunization with PBS, BCG alone, or BCG combined with CFA (MetaboLights, MTBLS15097). Comparative metabolomic analysis revealed that, compared to the PBS and BCG-only immunization groups, the BCG plus CFA group exhibited significant upregulation of the Nicotinate and nicotinamide metabolism and Butanoate metabolism pathways **(Extended Data Fig. 8a)**, which include the key metabolite Pyruvate, Fumaric acid, Malic acid, Butanoic acid, β-Muricholic acid, Orotic acid, Cafestol, Maleic Acid, α-Linolenic acid, Gamma-linolenic acid, and 16:1 (DELTA 9-Cis) PC **(Fig. 5a)**. To elucidate which specific upregulated metabolite primarily mediates the enhanced anti-TB protection observed with BCG plus CFA immunization, we established a mouse model. In this model, mice subcutaneously vaccinated with BCG received intravenous supplementation of metabolites found to be elevated in the serum following BCG plus CFA immunization. FCM analysis performed 28 days post-immunization revealed an increase in CD4^+^CD8^+^ T cells within the lung-draining LN of mice receiving BCG plus maleic acid, compared to the BCG-only group **(Fig. 5b, c)**. To directly investigate whether the immune protective effect of BCG plus CFA is mediated by the induction of maleic acid, we infected four groups of mice, PBS, BCG alone, maleic acid alone, and BCG plus maleic acid, with *Mtb* H37Rv 4 weeks after immunization. Bacterial burden and pathological damage in the lungs were assessed 4 wpi. The mean total lung CFUs for s.c. BCG plus i.v. maleic acid group (no detectable CFUs) was reduced by >100,000-fold compared with unvaccinated controls (5.95 log10 CFU) or >1,000-fold compared with mice immunized with parental BCG (3.76 log10 CFU). All four mice had no detectable *Mtb* in their lung tissues **(Fig. 5d)**. The mean total spleen CFUs for s.c. BCG plus i.v. maleic acid group (0.86 log10 CFU) was reduced by approximately 1000-fold relative to that in unvaccinated mice (4.18 log10 CFU) **(Extended Data Fig. 8b)**. Four out of six mice had no detectable *Mtb* in their spleen tissues **(Extended Data Fig. 8b)**. Consistently, lung histopathological damage in the s.c. BCG plus i.v. maleic acid group was significantly attenuated relative to that in the other groups **(Fig. 5e-g)**. Collectively, these results indicate that the immunoprotected effect of BCG plus CFA is mechanistically mediated by the induction of maleic acid, which promotes the formation of CD4^+^CD8^+^ T cells.

**Fig. 5:**
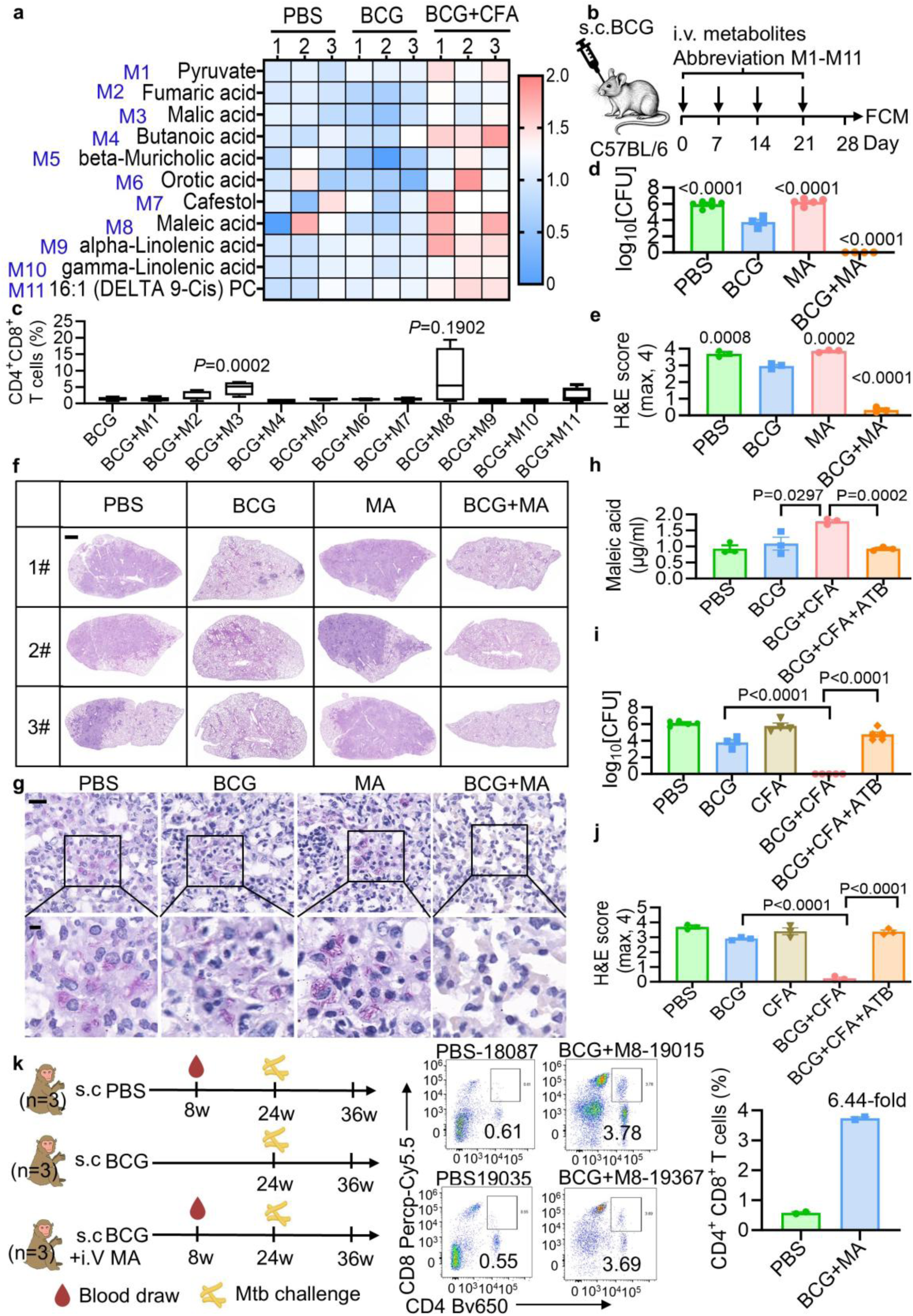
Maleic acid adjuvates BCG to provide sterilizing protection. **a**, Heatmap showing the top 11 differentially abundant metabolites identified by untargeted metabolomics. Blue text denotes metabolite abbreviations. **b-c,** Experimental design **(b)**, mice received subcutaneous vaccination and were injected intravenously with metabolites (abbreviations were defined in Fig. 5a) for 4 weeks once a week. FCM analyses of CD4^+^CD8^+^ T cells (gated from CD3^+^ cells) in the lung lymph nodes of each vaccine group, at 4 weeks post vaccination **(c)**. **d-g,** C57BL/6 mice, of each vaccine group, were aerosol-infected with ∼200 c.f.u. per mouse of *Mtb* H37Rv. After 4 weeks of infection, bacterial load of lung **(d)**, lung sections histology score **(e)**, lung sections H&E staining (**f**; scale bar=1mm), and lung sections acid-fast staining (**g**; scale bar, 20 μm (top) and 5 μm (bottom)). **h,** C57BL/6 mice were received indicated subcutaneous vaccination and treatment. Maleic acid (MA) concentration in serum across all groups was analysed at 4 weeks post vaccination. **i-j,** C57BL/6 mice, of each vaccine group, were aerosol-infected with ∼200 c.f.u. per mouse of *Mtb* H37Rv. After 4 weeks of infection, bacterial load of lung **(i)** and histology score **(j)** were analysed. **k,** FCM analyses of CD4^+^CD8^+^ T cells in the serum of each vaccine group, at 8 weeks post vaccination. Frequency of each group is summarized beside. Data represent one experiment with at least three independent biological replicates and are shown as the mean ± s.e.m.. Statistical significance in **c**, **h-j** was assessed by two-tailed unpaired Student’s t-tests. Statistical significance in **d**, **e** was assessed by ordinary one-way ANOVA with Dunnett’s multiple comparison test comparing each group to the BCG group.

Maleic acid is a specialized metabolite that cannot be synthesized by host eukaryotic cells; its production is restricted to certain microbial pathways, such as gentisate catabolism in *Pseudomonas alcaligenes*^71^. Thus, we hypothesize that BCG plus CFA immunization may elevate maleic acid levels by altering the composition or metabolic output of the gut microbiota. To investigate the causal role of the gut microbiota in the BCG plus CFA-induced immunometabolism reprogramming and anti-TB protection, we depleted the resident intestinal flora using a combination of antibiotics (ampicillin, metronidazole, neomycin, vancomycin). Mice immunized with BCG plus CFA received this regimen for 4 weeks prior to challenge with *Mtb* H37Rv **(Extended Data Fig. 8c)**. Targeted Liquid Chromatography-Tandem Mass Spectrometry (LC-MS/MS) quantification revealed that BCG plus CFA immunization significantly elevated serum levels of maleic acid, compared to BCG-only and PBS control groups. Notably, this vaccine-induced increase in circulating maleic acid was completely abrogated by antibiotic-mediated gut microbiota depletion **(Fig. 5h).** Along with the loss of maleic acid induction, the protective efficacy conferred by BCG plus CFA vaccination was entirely abolished. At 4 wpi, the mean lung *Mtb* H37Rv burden in the microbiota-depleted, BCG plus CFA-immunized group was 4.61 log10 CFU, representing a > 10, 000-fold increase relative to the non-antibiotic-treated BCG plus CFA controls (0 log10 CFU) **(Fig. 5i)**. Similarly, in the spleen, the mean *Mtb* H37Rv burden in the microbiota-depleted, BCG plus CFA-immunized group was 3.54 log10 CFU, representing a > 100-fold increase relative to the non-antibiotic-treated BCG plus CFA controls (0.67 log10 CFU) **(Extended Data Fig. 8d)**. Consistent with this loss of *Mtb* H37Rv control, lung histopathological damage was significantly exacerbated in the antibiotic-treated group compared to the immunized controls at 4 wpi **(Fig. 5j and Extended Data Fig. 8e, f)**. Collectively, these results suggest that BCG+CFA immunization may enhance anti-TB protection in a gut microbiota-dependent manner, through the elevation of immunomodulatory maleic acid.

We have demonstrated that BCG combined with maleic acid immunization confers protection against TB in a mouse model and induces immune-protective CD4^+^CD8^+^ T cells. To determine whether this observation extends to a higher animal model that more closely recapitulates human TB pathogenesis, we next evaluated the immune response in rhesus macaques. Notably, at 8 weeks post-immunization, the frequency of peripheral blood CD4⁺CD8⁺ T cells were significantly higher in macaques immunized with BCG+maleic acid than in non-vaccinated controls **(Fig. 5k)**. These findings support the translational potential of the immunization strategy and may provide a basis for further evaluation of its protective efficacy against *Mtb* challenge in macaques.

### Maleic acid is derived from *Clostridium butyricum*

To determine the gut microbial origin of maleic acid, we performed 16S rRNA sequencing to analyze the composition and dynamics of the intestinal microbiota in mice immunized for 4 weeks with PBS, BCG, CFA, or a combination of BCG and CFA (Sequence Read Archive (SRA), SRP719314). Principal coordinate analysis (PCoA) showed that the intestinal microbiota of BCG plus CFA-immunized mice formed distinct clusters, which were significantly separated from those of control animals, albeit with some overlap **(Fig. 6a)**. Furthermore, at the bacterial genus level, the gut microbiome of BCG plus CFA-immunized mice exhibited a significant increase in the abundance of *Parasutterella*, *Bacteroides*, *Rikenella*, *Phocaeicola*, and *Clostridium* in the cecum **(Fig. 6b and Extended Data Fig. 9a-e)**.

**Fig. 6:**
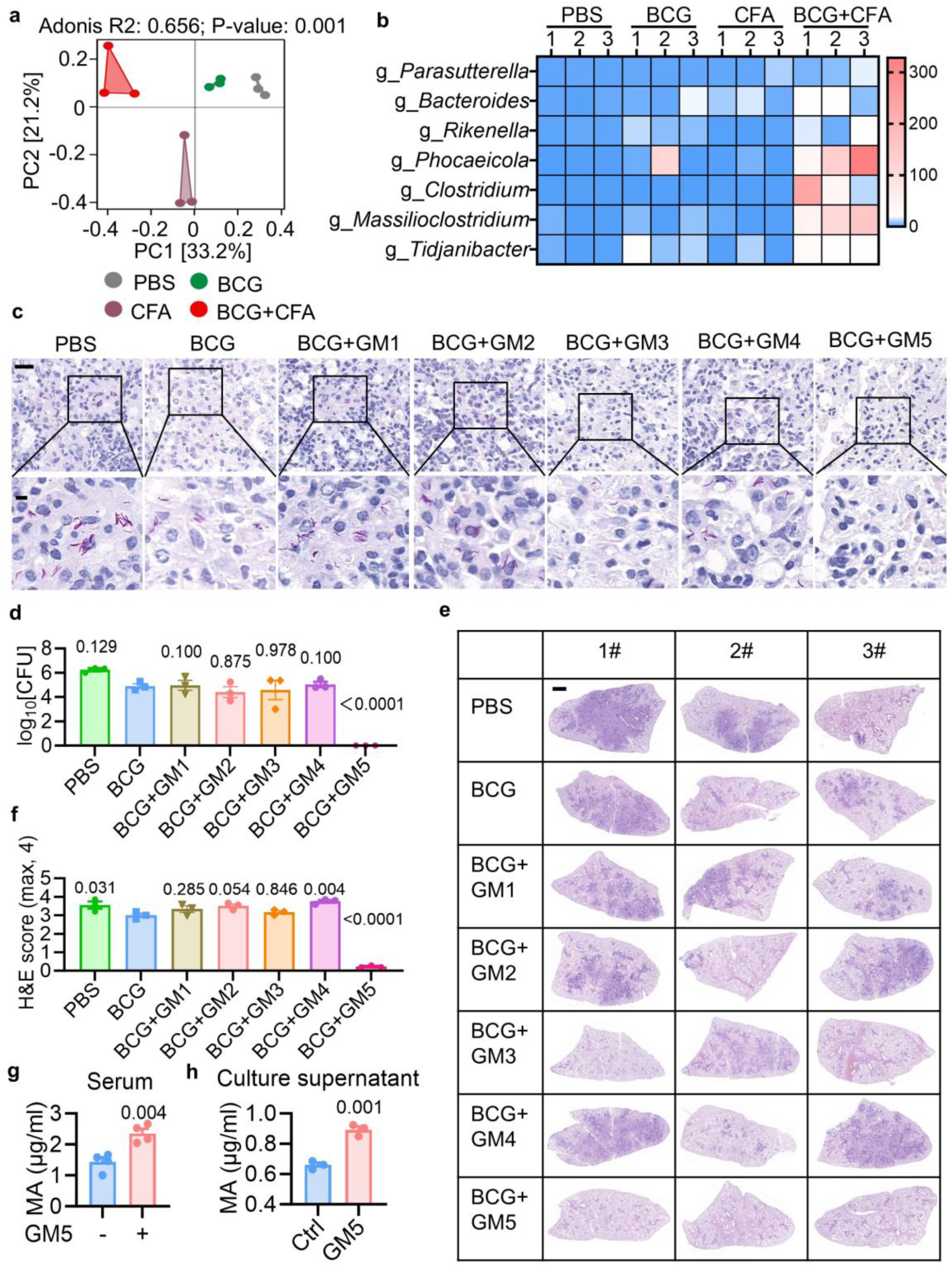
Maleic acid is derived from *Clostridium butyricum*. **a**, PCoA indicates the similarity or difference in the species composition of the gut microbiota in different treatment groups based on OTU levels. **b**, heat map of the most abundant top 7 bacteria at the genus level among four groups. Blue denotes a low relative abundance across a taxon (row); red denotes a high relative abundance. The color key indicates correspondence between the red-blue coloring and standard deviations from the mean abundance of each taxon. **c-f**, C57BL/6 mice, of each vaccine group, were aerosol-infected with ∼200 c.f.u. per mouse of *Mtb* H37Rv. After 4 weeks of infection, lung sections acid-fast staining (**c**; scale bar, 20 μm (top) and 5 μm (bottom), bacterial load of lung **(d)**, lung sections H&E staining (**e**; scale bar=1mm), and histology score **(f)**. **g**, C57BL/6 mice were received indicated subcutaneous vaccination and treatment. Maleic acid (MA) concentrations in serum were analyzed at 4 weeks post vaccination. **h**, Maleic acid (MA) concentrations in GM5 culture supernatant were analyzed. Data represent one experiment with at least three independent biological replicates and are shown as the mean ± s.e.m.. Ordinary one-way ANOVA with Dunn’s multiple comparison test comparing each group to the BCG group **(d, f)**, and two-tailed unpaired Student’s t-tests **(g, h)** were used for statistical analyses.

We next employed a fecal microbiota transplantation (FMT) approach to investigate the gut microbial origin of maleic acid. BCG-vaccinated mice were treated with a defined antibiotic cocktail for 2 weeks to deplete the native microbiota, followed by transplantation with a defined bacterial consortium comprising *Parasutterella excrementihominis*, *Bacteroides fragilis*, *Rikenella microfusus*, *Phocaeicola vulgatus*, and *Clostridium butyricum*, which was allowed to colonize for 5 weeks **(Extended Data Fig. 9f)**. These mice were subsequently infected with *Mtb* H37Rv for 4 weeks. At 4 wpi, the mean total lung CFUs for the *C. butyricum*-colonized BCG-vaccinated group (no detectable CFUs) was much lower (>10,000-fold) than that of BCG-vaccinated controls (4.91 log10 CFU). All mice had no detectable *Mtb* in their lung tissues **(Fig. 6c, d)**. Consistent with this improved bacterial control, lung histopathological damage was significantly attenuated in the *C. butyricum*-colonized BCG group compared to BCG-immunized controls at 4 wpi **(Fig. 6e, f)**. Targeted LC-MS/MS analysis revealed that colonization with *Clostridium butyricum* in BCG-vaccinated mice significantly elevated serum levels of maleic acid compared to BCG-only **(Fig. 6g)**. Furthermore, maleic acid was also detected in the *in vitro* culture supernatant of this bacterium **(Fig. 6h)**. Taken together, these findings demonstrate that colonization with *Clostridium butyricum* in BCG-vaccinated mice enhances protective immunity against *Mtb* infection by elevating serum levels of maleic acid.

### Maleic acid interacts with PAK2

To investigate mechanisms underlying the differentiation of CD4^+^CD8^+^ T cell subsets, we performed the FACS purification of CD4^+^CD8^+^ T cells along with conventional CD4^+^ or CD8^+^ T cells isolated from the lung-draining LNs of BCG plus CFA-immunized mice, and subjected them to comparative phospho-proteomic analysis (iProX, PXD081332). Kyoto Encyclopedia of Genes and Genomes (KEGG) pathway analysis revealed that CD4^+^CD8^+^ T cells exhibited significant upregulation of STAT5 signaling pathway components, compared with CD4^+^ and CD8^+^ T cell populations **(Fig. 7a, b)**. The STAT5 signaling axis is a pivotal regulator of lymphoid development, orchestrating lineage specification through phosphorylation-dependent transcriptional activation^72, 73^, yet the regulatory role of STAT5 signaling in mature T cell differentiation remains poorly understood. Notably, phospho-proteomic profiling identified enhanced phosphorylation at serine 127 (S127), a previously unreported regulatory site, of STAT5b in CD4^+^CD8^+^ T cells, compared to that in CD4^+^ or CD8^+^ T cell subsets **(Fig. 7c)**. To detect phosphorylation at Ser127 of STAT5b in CD4^+^CD8^+^ T cells, we generated a phospho-specific antibody against this site. FCM analyses revealed that, compared to CD4⁺ or CD8⁺ T cells, CD4⁺CD8⁺ T cells from BCG plus CFA-immunized mice exhibited markedly increased in STAT5b phosphorylation at Ser127 **(Fig. 7d)**.

**Fig. 7:**
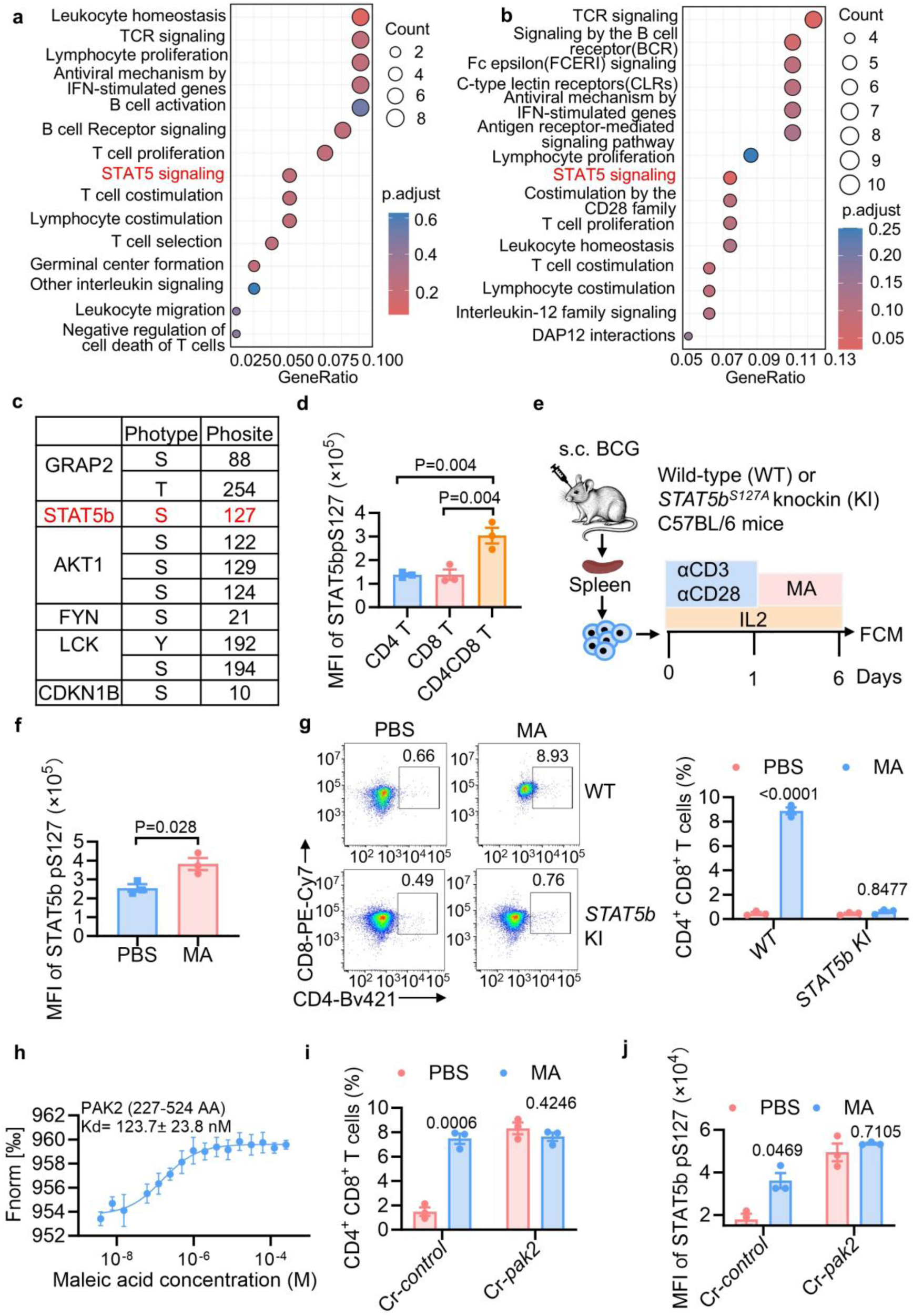
Maleic acid interacts with PAK2. **a-b**, The result of KEGG enrichment analysis by ImmPort database, CD4^+^CD8^+^ T cells V.S. CD4^+^ T cell **(a)**; CD4^+^CD8^+^ T cells V.S. CD8^+^ T cell **(b)**. **c**, The phosphorylation site of the associated protein in enrichment STAT5 signaling pathway detected by DIA-MS (data-independent acquisition mass spectrometry). **d.** Mean fluorescence intensity (MFI) of STAT5b pS127 levels in CD4^+^, CD8^+^ and CD4^+^CD8^+^ T cells (gated from CD3^+^ cells) in the lung lymph nodes were measured by FCM. **e-g**, Experiment design **(e)**, CD8^+^ T cells isolated from BCG-immuned mice spleen were stimulated with anti-CD3 antibody (1 day), anti-CD28 antibody (1 day) and IL-2 (6 days), and treated with maleic acid (MA, 1uM) and solvent control. MFI of STAT5b pS127 levels **(f)** and percentage of CD4^+^CD8^+^ T cells **(g)** of each group were measured by FCM. **h**, Binding affinity determination of recombinant PAK2 protein (expression Region: 227-524) with maleic acid assessed by microscale thermophoresis. **i-j**, CD8 T cells isolated from BCG-immuned mice spleen were stimulated with anti-CD3 antibody, anti-CD28 antibody and IL-2 for 1 day, then transfected with *Pak2* crRNA by Easy editing^TM^ kit. After 1 day, cells were treated with IL-2 and MA for 4 days. Percentage of CD4^+^CD8^+^ T cells **(i)** and MFI of STAT5b pS127 levels **(j)** of each group were measured by FCM. Data represent one experiment with at least three independent biological replicates and are shown as the mean ± s.e.m.. Ordinary one-way ANOVA with Dunn’s multiple comparison test comparing each group to the CD4^+^CD8^+^ T cells group **(d)**, two-tailed unpaired Student’s t-tests **(f)** and two-way ANOVA with Šídák’s multiple comparisons test **(g, i, j)** were used for statistical analyses.

To delineate the cellular origin of CD4⁺ CD8⁺ T cells, CD3⁺ T cells were enriched from lung-draining LNs of mice immunized with BCG plus CFA by magnetic-activated cell sorting (MACS) on day 28 post-immunization. The enriched cells were subjected to single-cell RNA sequencing (scRNA-seq) on the BD Rhapsody platform. Unsupervised clustering using the Louvain algorithm with a resolution of 0.7 resolved 17 distinct T cell clusters, as visualized by uniform manifold approximation and projection (UMAP) **(Extended Data Fig. 5a)**. Clusters 2, 3, and 15 exhibited co-expression of *Cd4*, *Cd8a*, and *Cd8b1* at the transcript level, confirming their identity as CD4⁺CD8⁺ T cells **(Extended Data Fig. 5b)**. To reconstruct the developmental trajectory of CD4⁺CD8⁺ T cells, we performed pseudotime analysis. Cells were ordered along a trajectory originating from CD8⁺ single-positive clusters and progressing toward CD4⁺CD8⁺ T cells **(Extended Data Fig. 10a)**. This analysis indicated that CD4⁺CD8⁺ T cells arise predominantly from CD8⁺ T cells, suggesting a lineage conversion event during T cell differentiation under BCG plus CFA immunization conditions.

To determine the dependence of CD4⁺CD8⁺ T-cell development on STAT5 signaling, we established an *in vitro* differentiation model. CD8⁺ T cells isolated from the spleens of BCG-vaccinated mice were treated with maleic acid. FCM analysis revealed a pronounced increase in phosphorylation at Ser127 of STAT5b **(Fig. 7e, f)**, which was paralleled by the generation of CD4^+^CD8^+^ T cells **(Fig. 7g)**. Furthermore, we generated knock-in mice harboring a STAT5b Ser127A mutation. Splenic CD8⁺ T cells were isolated from BCG-immunized STAT5b Ser127A knock-in mice and their littermate negative controls. Using an *in vitro* differentiation model followed by flow cytometry, we found that while maleic acid increased the percentage of CD4⁺CD8⁺ T cells in wild-type CD8⁺ T cells, the effect was completely abrogated in STAT5b Ser127A knock-in CD8⁺ T cells **(Fig. 7g)**. These results suggest that maleic acid may drive the generation of CD4⁺CD8⁺ T cells from CD8⁺ T cells by promoting phosphorylation at Ser127 of STAT5b.

To further delineate the pathway through which maleic acid regulates the phosphorylation of STAT5b, we performed limited proteolysis-coupled mass spectrometry (LiP-MS) and identified p21-activated kinase 2 (PAK2) ^74^ as a protein directly interacting with maleic acid **(Extended Data Fig. 10b, c and Table S2)**. This interaction was further confirmed by microscale thermophoresis (MST) assay, yielding a dissociation constant (Kd) of 123.7 ± 23.8 nM **(Fig. 7h)**. To determine whether maleic acid modulates STAT5b phosphorylation at Ser127 via PAK2, we knocked out PAK2 expression in CD8⁺ T cells using CRISPR-Cas12 with a specific CRISPR RNA (crRNA). Deletion of *Pak2* abolished the enhanced effect of maleic acid on CD4⁺CD8⁺ T cell differentiation **(Fig. 7i)**. Consistently, FCM analysis confirmed that PAK2 deletion rescued the effect of maleic acid on STAT5b phosphorylation, as it attenuated the promotion at Ser127 **(Fig. 7j)**. These results suggest that interaction of maleic acid with PAK2 may drive the STAT5b-dependent differentiation of CD4⁺CD8⁺ T cells.

### Ets1 is a key mediator of CD4⁺CD8⁺ T cell development

Current characterization of CD4^+^CD8^+^ T cell biomarkers has focused primarily on transcriptional regulators controlling lineage commitment, particularly the ThPOK-Runx3 axis ^75^ and cytokines conventionally linked to mature CD4^+^ or CD8^+^ T cell populations (such as IFN-γ, TNF, IL-10, and granzymes) ^75^. However, the specific transcription factors that govern CD4⁺CD8⁺ T cell differentiation remain unidentified. Our single-cell RNA sequencing analysis of lung-draining LNs from BCG plus CFA-immunized mice revealed that the transcription factors among the marker genes (top 110) of clusters 2, 3, and 15 (CD4⁺CD8⁺ T cells) included *Ets1*, *Sox4*, *Myb*, *Tcf7*, *Ets2* and *Tcf12* **(Extended Data Fig. 11a)**. To identify which transcription factor directly regulates the transcriptional programs related to CD4⁺CD8⁺ T cell differentiation, we performed Cleavage Under Targets and Tagmentation (CUT&Tag) to identify the genome-wide binding sites for ETS1, SOX4, MYB, TCF1, ETS2 and TCF12 (SRA, SRP719386), which was then compared with gene expression changes established by single-cell RNA sequencing. The data revealed that Ets1 bound to about 80% of the marker genes characteristic of CD4⁺CD8⁺ T cells **(Extended Data Fig. 11b)**. Ets1 is a proto-oncogene member of the ETS family of transcription factors that plays key regulatory roles in immune cell differentiation, angiogenesis, and tumorigenesis ^76^. SCENIC analysis also identified *Ets1* regulon activity was specifically enriched in clusters 2 and 3 (CD4⁺CD8⁺ T cells) **(Extended Data Fig. 11c)**. These results suggest that Ets1 may serves as a putative transcription factor for CD4^+^CD8^+^ T cells.

To further determine whether Ets1 acts as a key transcriptional regulator of CD4⁺CD8⁺ T cell differentiation, we performed FCM analysis. The data revealed that in BCG plus CFA-immunized mice, Ets1 expression in CD4⁺CD8⁺ T cells was significantly higher than that in CD8⁺ T cells but significantly lower than that in CD4⁺

T cells in vivo **(Fig. 8a)**. Notably, we also found that maleic acid promoted Ets1 expression in CD8⁺ T cells isolated from the spleens of BCG-vaccinated mice (**Fig. 8b**). We then overexpressed *Ets1* mRNA in CD8⁺ T cells isolated from the spleens of BCG-vaccinated mice. Five days later, *Ets1*-overexpressing CD8⁺ T cells showed a significantly higher frequency of CD4⁺CD8⁺ T cells compared to the control **(Fig. 8c)**. Notably, *Ets1*-overexpressing group exhibited markedly increased expression of Runx3 and ThPOK, two lineage-defining transcription factors associated with CD4⁺ or CD8⁺ T cell development **(Fig. 8d, e)**. Furthermore, we knocked out *Ets1* expression in CD8⁺ T cells using CRISPR-Cas12 with a specific crRNA. Deletion of *Ets1* abolished the enhancing effect of maleic acid on CD4⁺CD8⁺ T cell differentiation from CD8⁺ T cell **(Fig. 8f)**. Notably, maleic acid-treated group exhibited markedly increased expression of RUNX3 and ThPOK, two lineage-defining transcription factors associated with CD4⁺ or CD8⁺ T cell development **(Fig. 8d, e)**. However, FCM analysis confirmed that Ets1 deletion also abolished the enhanced effects of maleic acid on the expression of Runx3 and ThPOK **(Fig. 8g)**. These results suggest that maleic acid may promote CD4⁺CD8⁺ T cell differentiation through inducing Ets1, which may mediate the development of overriding the lineage-defining antagonism between ThPOK and Runx3. To determine how Ets1 regulate the differentiation of CD4⁺CD8⁺ T cell, CUT-Tag assay indicated that Ets1 directly bound to gene loci for *Cd4* **(Fig. 8h)**, *Cd8a* **(Fig. 8i)**, *Runx3* **(Fig. 8j)** and *Zbtb7b* (the gene encoding ThPOK) **(Fig. 8k)**, functionally linking Ets1 binding to the promotion of this differentiation pathway. These results suggest that Ets1 may directly bind to the related genes to drive CD4⁺CD8⁺ T cell differentiation, overriding antagonism between ThPOK and Runx3.

**Fig. 8:**
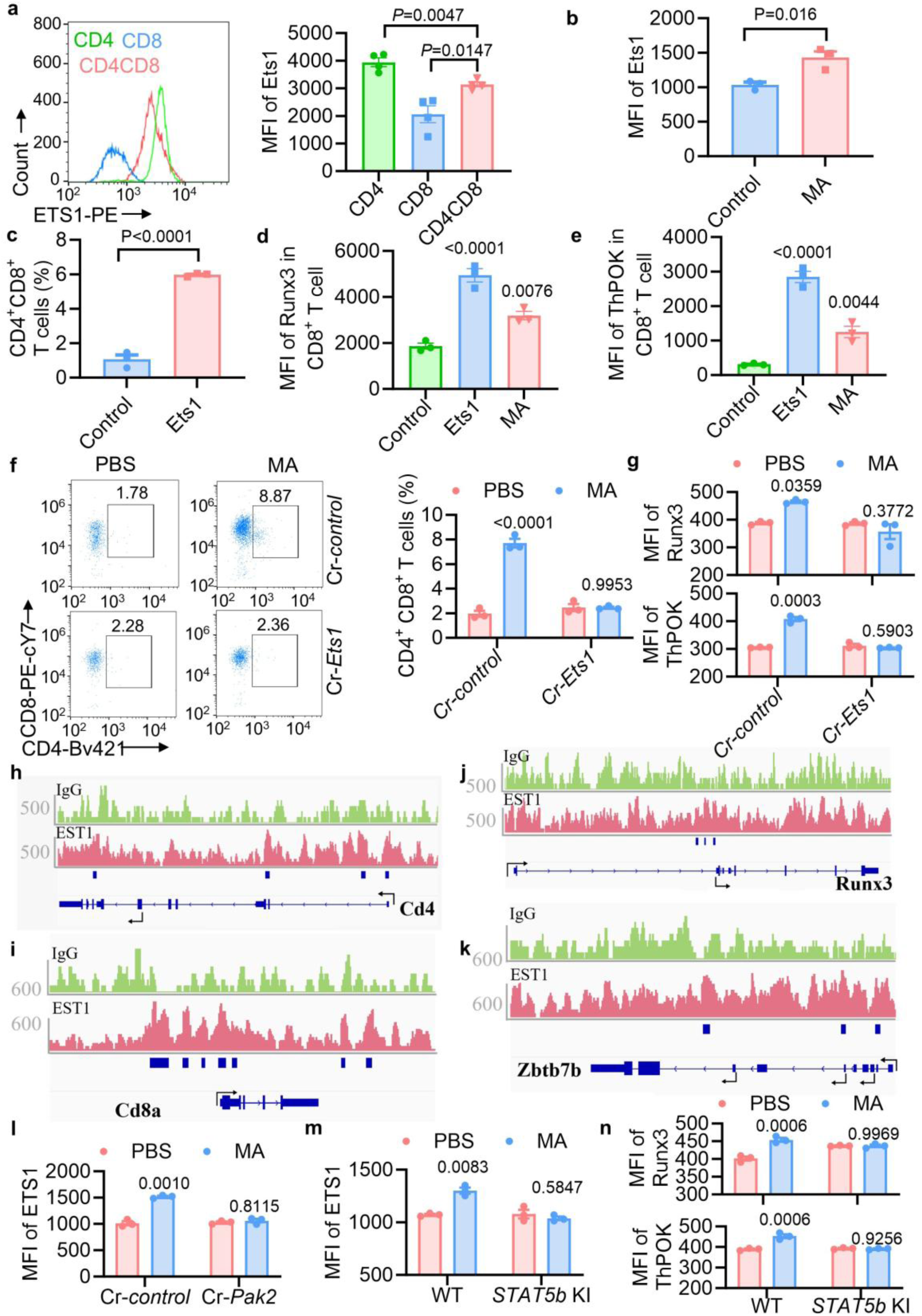
Ets1 is central for the development of CD4⁺CD8⁺ T cell. **a,** MFI of Ets1 in CD4^+^, CD8^+^ and CD4^+^CD8^+^ T cells (gated from CD3^+^ T cells) in the lung lymph nodes of 4 weeks post BCG plus CFA vaccinated mice was measured by FCM. **b**, MFI of Ets1 in maleic acid treated CD8^+^ T cell isolated from BCG-immuned mice spleen after 5 days. **c,** CD8^+^T cells isolated from BCG-immuned mice spleen were stimulated with anti-CD3 antibody, anti-CD28 antibody and IL-2 for 1 day, then transfected with *Ets1* mRNA by ProteanFect Max Mouse Immunocyte Transfection Kit. After 1 day, cells were treated with IL-2 for expansion by 4 days. Percentage of CD4^+^CD8^+^ T cells in non-transduced CD8^+^T cells and Ets1-overexpressing CD8^+^T cells was measured by FCM **(c)**. **d-e**, MFI of Thpok **(d)** and Runx3 **(e)** of each group were measured by flow FCM. **f-g,** CD8^+^T cells were transfected with Ets1 crRNA and treated with MA, percentage of CD4^+^CD8^+^ T cells **(f)** and MFI of Runx3 and Thpok **(g)** of each group were measured by FCM. **h-k**, CUT&Tag peak distribution for ETS1 at *Cd4* **(h)**, *Cd8* **(i)**, *Runx3* **(j)**, *zbtb7b* **(k)** gene loci. **l,** MFI of Ets1 was measured by FCM in MA-treated CD8^+^T cells transfected with PAK2 crRNA. **m-n**, MFI of Ets1 **(m)**, Runx3 and Thpok **(n)** were measured by FCM in MA-treated CD8^+^T cells from WT and *Stat5b*^S127A^ knockin mice. Data represent one experiment with at least three independent biological replicates and are shown as the mean ± s.e.m.. Two-tailed unpaired Student’s t-tests **(a-e)** and two-way ANOVA with Šídák’s multiple comparisons test **(f-g, l-n)** were used for statistical analyses.

### Maleic acid upregulates Ets1 expression through PAK2/STAT5 axis

Given that interaction of maleic acid with PAK2 drives the STAT5b-dependent differentiation of CD4⁺CD8⁺ T cells and maleic acid induces Ets1 to mediate the generation of CD4⁺CD8⁺ T cells, we next examined whether maleic acid induces Ets1 expression through PAK2-mediated phosphorylation of STAT5b at Ser127. Firstly, using the same in vitro differentiation model, we found that deletion of *Pak2* abolished the enhancing effect of maleic acid on Ets1 expression **(Fig. 8l)**. Furthermore, splenic CD8⁺ T cells were isolated from BCG-immunized STAT5b S127A knock-in mice and their littermate negative controls. Using an in vitro differentiation model followed by FCM analysis, we found that maleic acid promoted Ets1 expression in wild type CD8⁺ T cells; however, in STAT5b S127A knock-in CD8⁺ T cells, maleic acid failed to induce Ets1 expression **(Fig. 8m)**. Consistently, FCM analysis confirmed that the STAT5b S127A mutation also abolished the enhanced effects of maleic acid on expression of Runx3 and ThPOK **(Fig. 8n)**. Together, these results suggest that maleic acid may induce Ets1 expression through PAK2-mediated phosphorylation of STAT5b at Ser127, thus leading to the generation of CD4⁺CD8⁺ T cells.

## Discussion

Despite limited durable immunity against TB disease, BCG has been used for over 100 years to protect children^77^. However, there has been a lack of success by alternative vaccine candidates and regimens to improve protection in adults^77^. Identification and dissection of the immune factors required for the prevention of TB is urgently needed for the development of effective anti-TB vaccine or drugs. Here, our study establishes CD4^+^CD8^+^ T cells as a previously uncharacterized anti-TB immune factor, which is markedly induced by BCG plus CFA, *Clostridium butyricum* or maleic acid. *Mtb*-specific CD4^+^CD8^+^ T cells had a broad TCR repertoire and conferred sterilizing protection against *Mtb* challenge through inducing SLAMF1 and neutrophil responses, providing a paradigm-shifting strategy for enhancing TB vaccine efficacy by promoting the generation of CD4^+^CD8^+^ T cells. Furthermore, identification of Ets1 as a pivotal mediator of CD4⁺CD8⁺ T cell development subverts the dogma of the lineage-defining antagonism between ThPOK and Runx3, suggesting a degree of plasticity and variability for the differentiation of CD4^+^CD8^+^ T cells in response to disease and environmental stimuli **(Extended Data Fig. 12)**. Moreover, Co-immunization of BCG with *Clostridium butyricum* or maleic acid, which confers sterilizing protection against *Mtb* infection, introduces a potentially actionable strategy for improving vaccine performance through targeted expansion of CD4^+^CD8^+^ T cells—a paradigm not previously explored in the anti-TB vaccine development. Notably, considering the extremely low percentage of CD4^+^CD8^+^ T cells and their association with tumors or inflammatory diseases ^44, 45, 78, 79, 80^, similar strategies should be used for promoting CD4^+^CD8^+^ T cell differentiation for functional studies and potential clinical translation in other types of related diseases in response to different stresses or stimuli.

Majority of the adjuvants effective in TB animal models, including the ones in clinical development, rely on vaccine adjuvant-delivery systems. CFA, primarily composed of inactivated H37Ra and mineral oil ^52, 53^, has been widely used in animal models to potentiate vaccine-induced immune responses. Its protective efficacy has been demonstrated in diverse contexts, including the following: (1) antiviral protection against group B coxsackievirus ^54^, dengue virus ^81^, and respiratory syncytial virus infections ^82^; (2) antibacterial defense against *Helicobacter pylori* ^55^ and virulent *Brucella* species ^83^; (3) antiparasitic activity in *Neospora caninum* infections ^84^; and (4) neuroprotective effects evidenced by prolonged survival in murine prion disease models ^56^. However, the adjuvant efficacy of CFA on BCG-induced protection and immune responses against TB remains unexplored. Our findings reveal that subcutaneous BCG plus CFA immunization markedly increased antigen-responsive T cells, including CD4^+^CD8^+^ T cells, and unprecedented sterilizing protection against *Mtb* challenge. Furthermore, we found that BCG plus CFA immunization actually increases the abundance of *Clostridium butyricum*^85, 86^ and its derivative maleic acid. Unexpectedly, immunization of BCG plus *Clostridium butyricum*^85, 86^ or maleic acid also achieved sterilizing immunity against *Mtb* in 80% of vaccinated mice. Given the safety concerns of CFA, utilization of *Clostridium butyricum* or maleic acid as the novel adjuvants of BCG may represent major progress in the field of anti-TB vaccine strategies. However, the underlying mechanisms of how CFA, *Clostridium butyricum* or maleic acid stimulate BCG-induced innate immunity, potentially through DC cross presentation, to trigger the *Mtb-*specific T cell responses in lung LNs need further investigation.

Accumulating data from rodent or macaque models and human observational studies demonstrate a key role of CD4^+^ T cells in anti-mycobacterial protection ^35, 87, 88^.

CD8α^+^ lymphocytes also play a key role in the control of early *Mtb* infection in macaques ^35, 42, 43^. Indeed, our correlation analysis also observed increased number of CD4^+^ and CD8^+^ T cells as well as the production of protective cytokines following BCG plus CFA immunization. However, BCG plus CFA immunization dramatically increased the number of CD4^+^CD8^+^ T cells. The presence of CD4^+^CD8^+^ T cells in i.v. BCG immunization models and other diseases is also observed^43, 44, 45^ but their functional role in the immune protection or pathogenesis of diseases remains unclear, which is primarily attributable to their extreme scarcity. In our *Mtb* challenge model, we found that the adoptive transfer of mycobacteria-specific CD4^+^CD8^+^ T cells, not CD4^+^ or CD8^+^ T cells into *Rag1^-/-^*mice significantly reduced *Mtb* loads in both the lungs and spleen, validating their critical role in protection against *Mtb* infection. Notably, *Mtb*-specific CD4⁺CD8⁺ T cells exhibited a broad and unbiased antigen recognition profile, which may resolve the heterogeneity in tuberculosis ^89^. To our knowledge, this study is the first to define the protective role of CD4^+^CD8^+^ T cells in host immunity. However, further mechanistic studies are warranted to elucidate how CD4⁺CD8⁺ T cells exert their functions in anti-TB immunity and potentially in other disease settings.

Baena and Porcelli proposed that certain mycobacterial antigens capable of eliciting immunodominant T-cell responses may in fact function as immunological “decoys,” diverting immunity away from subdominant epitopes critical for protective immunity ^90^. This view is further supported by work from the Andersen group, which showed that although cryptic epitopes of ESAT-6 trigger only modest responses during natural infection, vaccines designed to prime CD4⁺ T cells against these subdominant epitopes confer stronger and more durable protection against tuberculosis ^91, 92, 93, 94^. Moreover, a parallel mechanism appears to operate in CD8⁺ T cell immunity. Most *Mtb*-specific CD8⁺ T cells recognize only a single antigen, TB10.4, encoded by the esxH gene ^95^. However, CD8⁺ T cells specific for the immunodominant epitope TB10.4₄-₁₁ poorly recognize *Mtb-*infected macrophages, suggesting that *Mtb* may evade CD8⁺ T cell detection by focusing the response toward this non-protective “decoy” antigen ^37, 96^. In this study, we found that BCG+CFA-induced CD4⁺CD8⁺ T cells exhibit a remarkable broad and unbiased antigen recognition profile, which may induce a broader repertoire of antigen-specific T-cell responses to overcome the pathogen’s strategic use of immunodominant decoy antigens. However, the detailed mechanism underlying the *Mtb*-specific antigen diversity of BCG plus CFA-induced CD4⁺CD8⁺ T cells remain to be explored.

CD8^+^ or CD4^+^ T cells respond to *Mtb* by producing pro-inflammatory cytokines such as IFN-γ that is critical for the control of *Mtb* clearance by stimulating macrophages^23, 33, 34^. We found that CD4^+^CD8^+^ T cells, not CD4^+^ or CD8^+^ T cell, had highly expressed SLAMF1, which is required for the CD4^+^CD8^+^ T cells to exert their anti-TB functions. SLAMF1 interacts with adaptor proteins SAP and EAT-2 through homophilic SLAMF1–SLAMF1 binding, and subsequently activating downstream signaling pathways to modulate diverse immune cell functions ^66, 97, 98^. It has been shown that SLAMF1/CD150 is highly and uniquely induced in macrophages by antigen-specific interactions with CD4^+^ T cells, and is specifically required in macrophages to restrict mycobacterial growth and limit IL-1β production for enhanced TB defense^65^. However, our single-cell RNA sequencing analysis observed a significant expansion of a subpopulation of CD45.2⁺ neutrophils in the lungs of mice that received CD45.1⁺ CD4^+^CD8^+^ memory T cells. The neutrophil subset highly expressed genes associated with bactericidal function, including direct antimicrobial effectors (*Nos2*, *Plac8*, *Rab37*, *Bri3*, *Sod2*) and immunoregulatory molecules that modulate chemotaxis and inflammation (*Fpr1*, *Fpr2*, *Cxcl10*, *Il1a*, *Pnp*). Recent studies have demonstrated that autophagy in human neutrophils during active tuberculosis is tightly regulated by SLAMF1 signaling^99^. Thus, CD4^+^CD8^+^ memory T cells may control *Mtb* infection by engaging SLAMF1 to drive neutrophil responses, highlighting a novel T cell-neutrophil crosstalk mechanism in infectious immunity. However, the detailed mechanism underlying how the CD4^+^CD8^+^ T cells drive the expansion of neutrophils and how the neutrophils control the *Mtb* infection need further investigation.

Conventionally, CD4 and CD8 expression is fixed and mutually exclusive in mature T cells ^46^, which is derived from the concept that key transcription factors ThPOK and Runx3 negatively regulate each other during T cell development in the thymus to induce functionally distinct “helper” CD4^+^ or “cytotoxic” CD8^+^ single positive (SP) T cell fates. However, peripheral expression of these transcription factors is further maintained, suggesting that their lineage defining roles remain active outside the thymus ^100^. In our study, we found that BCG plus CFA vaccination induced higher abundance of *Mtb*-specific CD4^+^CD8^+^ T cells in lung-draining lymph nodes. Ets1 a proto-oncogene member of the ETS family of transcription factors ^76^ was identified as a critical regulator for the generation of *Mtb*-specific CD4^+^CD8^+^ T cells. Interaction of maleic acid with PAK2 activates phosphorylation of STAT5b at Ser127, leading to upregulation of Ets1 expression. Ets1 directly binds to the promoter loci of the related genes, *Cd4*, *Cd8a*, *Runx3* and *Zbtb7b* (encoding ThPOK), to drive their expression, thereby overriding the antagonism between ThPOK and Runx3 and driving the generation of CD4⁺CD8⁺ T cells. Thus, our data demonstrate a degree of plasticity and variability of ThPOK and Runx3-independent lineage determination for the differentiation of CD4^+^CD8^+^ T cells, which may potentially be due to ETS1-dependent transcriptional modulation of the related genes during differentiation in response to antigen stimulation ^75, 101^. However, further analysis of the differentiation of these cells and whether ETS1-mediated development of CD4^+^CD8^+^ T cell is involved in other types of contexts is warranted.

Despite these advances, several important questions remain unanswered. First, mechanisms linking the adjuvant to gut microbiome restructuring (skin-gut axis) has not been elucidated. It is unclear whether BCG plus CFA alter the host cytokine profile to reshape gut microbiota ^102^. Second, Intriguingly, despite the role of intestinal maleic acid in priming CD4^+^CD8^+^ T cells formation, these cells are predominantly enriched within the lung-draining lymph nodes rather than the gut-associated lymphoid tissues. It remains to be elucidated whether the gut serves as the primary inductive site for CD4^+^CD8^+^ T cells, and whether the lung-draining lymph nodes possess a unique microenvironmental architecture that selectively orchestrates their local proliferation and maintenance. Third, beyond its canonical role in governing the survival, activation, and memory formation of CD8^+^ T cells ^103, 104^, we identify Ets1 as a master regulator of CD4^+^CD8^+^ T cell differentiation. Central to this expanded role is the hitherto unrecognized regulatory axis involving STAT5b phosphorylation at S127, a modification whose precise control over Ets1 activity represents a compelling frontier for further mechanistic exploration.

## Methods

### MDR-TB patient and healthy people serum samples

All the patients with TB and healthy volunteers providing blood samples were from the Shanghai Pulmonary Hospital. Patients with multidrug-resistant (MDR)-TB (18–50 years old) were diagnosed based on chest radiographs, acid-fast bacillus staining of biofluid samples, culture on Lowenstein–Jensen media, and clinical symptoms. Their Mycobacteria Growth Indicator Tube (MGIT) drug sensitivity test and results are resistant to isoniazid and rifampicin, including pre-XDR-TB and XDR-TB patients. All patients received standardized anti-MDR-TB regimens. Controls were recruited from a pool of individuals who participated in a health examination program. The control population was also subjected to a physical examination, blood testing, and chest X-rays and was screened for a medical or family history of TB. All cases and controls with a history of alcoholism, diabetes, chronic use of corticosteroids, or immunodeficiency were excluded from the study. During 0/2/4/6 months after treatment, blood samples were collected in the presence of EDTA from each donor, to analyze CD4^+^CD8^+^ T cells. All participants provided written informed consent. This study was conducted according to the Declaration of Helsinki principles. The protocol was approved by the local ethics committee of Shanghai Pulmonary Hospital and written informed consent was obtained from all participants (Ethics Number: No. L25-495).

### Experimental animals

SCID mice (Cat. NO. SM-015) were obtained from Shanghai Model Organisms Center, Inc. C57BL/6, CD45.1 mice, *Stat5b*^S127A^ mice and *Rag1^−/−^* mice were purchased from Cyagen Biosciences (Suzhou, China). SLAMF1-deficient (*Slamf1*^−/−^) mice^105^ were a gift from Shasha Chen (Anhui Medical University, Hefei, China). *Stat5b*^S127A^ mice were generated by Cyagen Company via CRISPR/Cas9 technology. Briefly, Cas9 protein, a specific gRNA (gRNA-A1: TCCAGCTGGAGAGCTGCCCTGGG) and a donor oligonucleotide containing p.S127A mutation (AGC to GCA) were co-injected into zygotes. The embryos were transferred to recipient female mice to obtain F0 mice. The genotype of point mutation mice was confirmed by PCR using a pair of primers (F1: 5’-GCATGTGGAGTTCTAGTTTCTCAA-3’, R1:5’- TAAGCTACCCAGCTCCTTCTTCAG-3’) and sequencing. F0 founder mice were crossed to C57BL/6J mice to produce heterozygous F1. All genetic models were of the C57BL/6J background. Female mice aged between 6–8 weeks were used for the study. All mice were housed in specific pathogen-free conditions at the Shanghai Pulmonary Hospital Laboratory Animal Center (temperature, 20–26 °C, relative humidity, 50–60%, light intensity in the feeding room 15–20 lx with 12 h:12 h light: dark cycle) and fed 25 kGy-irradiated feed (SLAC, P1101F), in accordance with the hospital guidelines. All animal experiments were reviewed and approved by the Animal Experiment Administration Committee of the Tongji University School of Medicine (Ethics Number: No. TJAA06522101), and conducted in accordance with the National Institutes of Health Guidelines for the Care and Use of Laboratory Animals.

### Bacterial strains

Bacterial strains used in this study are described in **Table S3**. The *Mtb* H37Rv strains and BCG strains were cultured in Middlebrook 7H9 broth (BD Biosciences, 271310) plus 10% oleic acid-albumin-dextrose-catalase (OADC, BD Biosciences, 212351) and 0.05% tyloxapol (MedChemExpress, HY-B1068).

Strains were cultured to mid-log phase (OD_600_ nm = 0.6). CFUs/mL were determined by plating serial dilutions on 7H10 agar and enumerating colonies after incubation at 37°C for 4 weeks. Bacterial aliquots were prepared in glycerol (20%) and Middlebrook 7H9 medium, stored at -80°C.

### Vaccination

BCG strain cultures were centrifuged at 12,000 ×*g* for 3 min at 4°C, washed once with ice-cold phosphate buffered saline (PBS), and resuspended in PBS to a concentration of 2 mg ml^−1^.

For BCG plus CFA immunization: Equal volumes of 2 mg ml^−1^ BCG and CFA (CFA, MedChemExpress, HY-153808) were mixed in a sonication tube, yielding a final BCG concentration of 1 mg ml^−1^ (containing 5×10^6^ CFU BCG using plate counting). For CFA immunization: Equal volumes of PBS and CFA were mixed in a sonication tube.

For BCG plus IFA immunization: Equal volumes of 2 mg ml^−1^ BCG and incomplete Freund’s adjuvant (IFA, MedChemExpress, HY-153808A) were mixed in a sonication tube, yielding a final BCG concentration of 1 mg ml^−1^ (containing 5×10^6^ CFU BCG quantified using plate counting).

For IFA immunization: Equal volumes of PBS and IFA were mixed in a sonication tube.

For BCG immunization: Equal volumes of 2 mg ml^−1^ BCG and PBS were mixed in a sonication tube, yielding a final BCG concentration of 1 mg ml^−1^ (containing 5×10^6^ CFU BCG quantified using plate counting).

The mixtures underwent two sonication cycles for primary emulsification, followed by 10 min of mechanical homogenization using an electric emulsifier (Biodragon, BDYQ1001) immediately prior to immunization until achieving water-stable consistency. The mice received 100 μL of emulsion via subcutaneous injection in the frontal scalp region.

### ABX cocktail treatment

Antibiotics (Abx) treatment was performed to normalize the gut microbiota in animal experiments. A quadruple antibiotics containing 0.2 g/L Ampicillin (MedChemExpress, HY-B0522), 0.2 g/L Metronidazole (MedChemExpress, HY-B0318), 0.2 g/L Neomycin (MedChemExpress, HY-B0470) and 0.1 g/L Vancomycin (MedChemExpress, HY-B0671) was used in drinking water for two weeks ^106, 107^.

### Microbial colonization

Parasutterella excrementihominis (5×10^8^ CFU), Bacteroides fragilis (5×10^8^ CFU), Rikenella microfusus (5×10^8^ CFU), Phocaeicola vulgatus (5×10^8^ CFU), Clostridium butyricum (5×10^8^ CFU) or the same volume of medium were administered via intragastric gavage in 200 μL of PBS once.

### Administration of metabolites to BCG-immuned mice

For metabolite treatment, after BCG vaccination, the mice were intravenously injected with Pyruvic acid, Fumaric acid, Malic acid, Butyric acid, β-Muricholic aci, Orotic acid, Cafestol, Maleic Acid, α-Linolenic acid, Gamma-linolenic acid and 16:1 (DELTA 9-Cis) PC.

### Mice infection

Female mice were divided randomly into cages and infected using an aerosol method with approximately 200 CFUs of *Mtb* H37Rv strains for 4 weeks (using an inhalation exposure system from Glas-col) at the Biosafety Level-3 Laboratory. Mice were euthanized at 4 weeks after infection. All the mice were age-, weight-, and sex-matched in each experiment.

### Neutrophil depletion

In neutrophil depletion experiments, 100 μg anti-Ly6G (clone 1A8, Bio X Cell) or isotype control (clone 2A3, Bio X Cell) was administered intravenously (i.v.) one day before infection, followed by weekly administrations post-infection.

### CFU assay

Lung and spleen tissues were homogenized in 1mL PBS. Homogenates in 10-fold serial dilutions were plated on 7H10 agar supplemented with 10% OADC enrichment medium and incubated at 37°C. Colonies were counted after 4weeks of incubation at 37°C in 5% CO_2_ and displayed on a log scale.

### Histopathological analysis

Lung tissues were fixed in 4% neutral-buffered paraformaldehyde solution for 24h and then embedded in paraffin. A series of 4-7μm-thick sections were then cut and stained with hematoxylin and eosin (H&E) or Ziehl–Neelsen (acid-fast bacilli) stain, in accordance with standard protocols. Imaging was performed using an automated whole-slide scanning device (3DHISTECH, Sysmex). Individual H&E slides were scored in a single-blinded fashion with a maximum score of four using leucocyte infiltration, hemorrhage, alveolar wall thickening, and alveolar edema as scoring indicators.

### Single cell isolation in mouse experiments

LNs were mechanically disrupted and filtered through a 70-μm cell strainer. Mouse spleens were mechanically disrupted and filtered through a 70-μm cell strainer, and then red blood cells were lysed with red cell lysis buffer. Lung tissue was digested using 0.2% collagenase, Type Ⅱ (Diamond) and 120 U/ml DNase (Sigma-Aldrich) for 30-45 min at 37°C with shaking, followed by passing through a 70-μm cell strainer and then the red blood cells were lysed.

### Non-human primates (NHPs) studies

#### Rhesus macaques

This research involved the use of Chinese-origin rhesus macaques (n=9), between 3 and 5 years of age, weighing 2.5 to 5 kg, and split approximately evenly between males and females. All experimental procedures involving the care of animals complied with ethical regulations. Prior to the experiments, all macaques underwent rigorous physical examinations to rule out potential underlying diseases. Absence of previous exposure to mycobacterial antigens was confirmed by a tuberculin skin test and chest X-rays.

#### BCG vaccination

Macaques were randomized into vaccinated (n=6) and unvaccinated (n=3) groups based on age, weight, and gender. The macaques were vaccinated under sedation. BCG was diluted in cold PBS to a target dose of 5 × 10⁶ CFUs. The solution was delivered intradermally in a volume of 0.5 ml. Maleic acid (5 mg/kg) was dissolved in saline, and the solution was delivered intravenously in a volume of 0.5 ml once a week.

#### *Mtb* challenge

Twenty-four weeks after vaccination, macaques were challenged by bronchoscope with 5000 CFUs of *Mtb* into the right lung as previously described ^108^. The challenge dose was verified by plating the bacterial suspensions from the last three dilutions on the 7H10 plates for CFU determination. Animals were euthanized at the pre-determined endpoint of the study. Animals that reached the humane endpoint prior to pre-determined endpoint were humanely euthanized.

#### Clinical assessments

Body weight, temperature, blood samples and chest X-rays were acquired prior to infection; 4, 8, and 12 weeks post infection. Blood specimens were used to detect the erythrocyte sedimentation rate (ESR), C-reactive protein (CRP), clinical hematology, biochemistry and flow cytometry. For chest X-rays, an experienced clinician who was blinded to the treatment groups evaluated the radiographs using a relative scoring system.

#### Flow cytometry

PBMCs were isolated from whole blood draws using Ficoll-Paque Plus (Cytavia) gradient separation, as previously described ^109^. Additionally, cryopreserved PBMCs were processed as a batch. After incubated with Fc Block (BD Biosciences, 553142), samples were stained with a fixable viability dye. Then, cells received a surface marker antibody stain for 30 min at 4°C. Surface marker antibody were used as follows: anti-CD4 (BV650, BD Biosciences, 563737), anti-CD8 (RB705, BD Biosciences, 757674). Cells were then washed and acquired on a a four laser BD FACSCelesta™ flow cytometer. Analysis was performed using FlowJo.

### Immunofluorescence staining

Lung LNs was fixed using 4% paraformaldehyde before embedding in paraffin. A series of 4 μm-thick sections were then cut. The prepared tissue sections (4 μm) were baked for 2 h at 65°C, tissue slices were deparaffinized in xylene, rehydrated in a series of ethanol concentrations (100,100, 95,90, and 85%), incubated with 3% hydrogen peroxide, antigen retrieved with Tris-EDTA pH 9.0, and blocked with 5% bovine serum albumin (BSA). The Opal 7-Color Manual IHC Kit (AKOYA Biosciences, USA, NEL820001KT) was then used to stain the samples using the primary antibodies from the following panel: anti-CD45R/B220 (Biolegend, 103201, 1:100), anti-CD4 (Abcam, ab288724, 1:200), anti-mouse CD8 alpha (Abcam, ab217344, 1:100), anti-CD44 (Abcam, ab232556, 1:200). Slice was then incubated with the corresponding horseradish peroxidase-conjugated goat anti-mouse or goat anti-rabbit secondary antibodies at 37°C for 10 min. Subsequently, the slide was again placed in Tris-EDTA pH 9.0, to remove redundant antibodies before the next step. Finally, the slice was incubated with DAPI solution at 37°C for 10 min in the dark. Scanning images of tissue samples were obtained using the Akoya Phenocycler Fusion (AKOYA Biosciences, USA).

### Cell sorting and adoptive transfer

Single-cell suspensions prepared from murine lung LNs of BCG+CFA-immuned WT or *slamf1*^-/-^ mice were stained with BV421-conjugated anti-CD4, FITC-conjugated anti-CD8α, PE-conjugated anti-CD44, and APC-conjugated anti-CD62L antibodies in PBS containing 0.5% (w/v) BSA. Cell sorting was performed on an FACSAriaⅢ system (BD Biosciences). The purity for all the populations was > 95%. For memory T cells adoptive transfer experiments, equal numbers (5×10^4^) of CD4^+^ (CD4^+^CD8^-^ CD44^+^CD62L^+^), CD8^+^ (CD4^-^CD8^+^CD44^+^CD62L^+^), and CD4^+^CD8^+^ (CD4^+^CD8^+^CD44^+^CD62L^+^) memory T cells were adoptively transferred i.v. to each recipient mouse infected with *Mtb* H37Rv strains on the following day^110^.

### Isolation and stimulation of CD8^+^ T cells

CD8^+^ T cells obtained from spleens of BCG-immuned C57BL/6 mice were stimulated with 5 µg ml^−1^ of plate-bound anti-CD3 (BioXcell, BE0001-1) in the presence of 1 µg ml^−1^ anti-CD28 antibodies (BioXcell, BE0001-1) and 20 ng ml^−1^ IL-2 (Peprotech, 212-12) for 24 hours.

Activated CD8^+^ T cells were treated with PBS, Maleic acid (1 uM) in complete RPMI 1640 plus FBS (10%, v/v) and penicillin–streptomycin (1%, v/v) and mouse IL-2 (20 ng ml^−1^) for 120 h.

Activated CD8^+^ T cells were transfected with Ets1 or control mRNAs with the ProteanFect Max Mouse Immunocyte Transfection Kit (Nanoportal Biotech, PT03) or transfected with PAK2, Ets1 and control crRNAs with Easy editing^TM^ kit (Tongxin Technology, Beijing, China, TX-EE12). Cells were then cultured with mouse IL-2 in complete RPMI 1640 plus FBS (10%, v/v) and penicillin–streptomycin (1%, v/v) for 120 h.

### Bone marrow-derived macrophages

Mice were euthanized and surface-sterilized in 75% ethanol. The femurs and tibias were then harvested and the bone marrow cells from all bones were flushed out. After centrifugation for 5 min at 400g, the erythrocytes were eliminated using Red Blood Cell Lysing Buffer (C3702; Beyotime). Cells were plated in Roswell Park Memorial Institute Medium 1640 (C11875500; Thermo Fisher) supplemented with 10% (v/v) fetal bovine serum (FBS), 1% (v/v) penicillin–streptomycin, and 20 ng ml^−1^ macrophage colony-stimulating factor (ABclonal, RP01216). Cultures were maintained at 37°C in 5% CO_2_ for 6 d, with medium replacement every 72 h. All cells were routinely tested for mycoplasma contamination.

### *In vitro* recall assay

CD4^+^, CD8^+^, and CD4^+^CD8^+^ memory T cells were sorted from the lymph nodes of C57BL/6 mice immunized with BCG plus CFA. For co-culture assays, approximately 1×10^5^ BMDMs/well were pulsed with 5 ng/mL Purified Protein Derivative of Tuberculin (TB-PPD, Simcere). After 1 h, CD4^+^, CD8^+^, and CD4^+^CD8^+^ memory T cells were co-cultured with BMDMs (5×10^4^/well) for 72 h. After co-culturing, CD44 levels and cell number were determined.

### CTV Labeling and Transgenic T Cell Transfer

CD4^+^, CD8^+^, and CD4^+^CD8^+^ memory T cells were sorted from the lymph nodes of CD45.1 donor mice immunized with BCG plus CFA, and were then labeled using CTV following the manufacturer’s protocol (C34557; Thermo Fisher). In brief, single-cell suspensions were incubated with CTV at a final concentration of 5 μM for 10 min before quenching with complete growth medium supplemented with 10% FBS. Labelled cells were adoptively transferred i.v. into recipient mice^110^. 4 days later, the recipient mice were pulsed with PPD. At 6.5, 9, and 11.5 d post-transfer, the mice were euthanized and spleens were harvested for flow cytometric (FCM) analysis of proliferation indices in CTV-labelled cells^111^.

### Seahorse metabolic assay

Oxygen consumption (OCR) and extracellular acidification rate (ECAR) were measured using the Seahorse XF Cell Mito Stress Test kit (Agilent), following the manufacturer’s instructions. In brief, different T cells (5 × 10^4^) obtained from mice lymph nodes or spleen were suspended in XF medium and subsequently seeded in a poly-l-lysine-coated XF96 plate. The OCR and ECAR were measured under basal conditions and in response to 1 μM oligomycin, 1.5 μM carbonyl cyanide 4- (trifluoromethoxy) phenylhydrazone (FCCP), and a mixture of rotenone and antimycin A (0.5 μM each) using an XF96 Extracellular Flux Analyzer (Seahorse Bioscience). Data were analyzed using the Seahorse Wave software (v2.6) and normalized to the exact cell number per well.

### PBMC isolation

Whole blood samples were collected in either heparin or EDTA containing blood tubes. PBMCs were subsequently isolated using Ficoll-Paque Plus (Cytavia) according to the manufacturer’s instruction. Briefly, whole blood was first centrifuged to allow the collection of plasma, and the serum-depleted blood was thereafter diluted in PBS, layered on Ficoll-Paque Plus, and centrifuged at 400 g for 30 min with the brakes off. The interphase cell layer resulting from this spin was collected, washed with PBS for flow staining.

### Generation of phospho-specific STAT5b antibodies

Rabbit polyclonal antibodies targeting STAT5b phosphorylated at Ser127 were raised against a KLH-conjugated synthetic peptide (C-RLVREANNGS(p)SPAGS; residues 118–132). To ensure mono-specificity, the crude antiserum was subjected to a sequential two-step affinity purification protocol. Initial positive selection was performed using phosphopeptide-immobilized resin to capture the total peptide-reactive IgG fraction. The resulting eluate was subsequently subjected to negative selection (depletion) by passage through a secondary column conjugated with the non-phosphorylated peptide counterpart (C-RLVREANNGSSPAGS). The flow-through, containing high-affinity Ser127 phospho-specific antibodies, was neutralized and stored at -80°C for downstream applications.

### Flow cytometry

Single cell suspensions prepared as described above were incubated with Fc Block (BD Biosciences, 553142). Then, cells received a surface marker antibody (Ab) stain for 20 min at 4°C. Surface Abs were used as follows: anti-CD3e (PE, BD Biosciences,553063; Alexa Fluor 700, Biolegend, 100216; Percp-Cy5.5, BD Biosciences, 560835), anti-CD4 (BV605, BD Biosciences, 563151; FITC, BD Biosciences, 553046; BV421, BD Biosciences, 562891; APC-Cy7, Biolegend, 100414; BV421, BD Biosciences, 562842), anti-CD8a (Percp-Cy5.5, BD Biosciences, 551162; APC-Cy7, BD Biosciences, 557654; PE-Cy7, BD Biosciences, 552877; FITC, BD Biosciences, 561966; APC-Cy7, BD Biosciences, 557760; Alexa Fluor 700, BD Biosciences, 561453), anti-CD44 (PE, BD Biosciences, 553134; FITC, BD Biosciences, 553133), anti-CD62L (PE-Cy7, Biolegend, 560516; APC, Biolegend, 104412; BV650, Biolegend, 104453; APC-Cy7, Biolegend, 104428), anti-CD69 (FITC, Invitrogen, 11-0691-85), anti-CD103 (Percp-Cy5.5, Biolegend, 121416), anti-CD127 (BV421, BD Biosciences, 566377), anti-KLRG1 (APC, Invitrogen, 17-5893-82), anti-CD45 (APC-Cy7, BD Biosciences, 557659), anti-CD11b (Alexa Fluor 700, BD Biosciences, 557960), anti-LY-6C (PE-Cy7, BD Biosciences, 560593), anti-F4/80 (APC, BD Biosciences, 566787), anti-CD86 (FITC, Biolegend, 105006), anti-CD206 (BV421, Biolegend, 141717), anti-MHCI-A/I-E (PE, Biolegend, 107608), anti-CD11c (BV605, BD Biosciences, 563057), anti-NK1.1 (APC, BD Biosciences, 550627), anti-B220 (PE-Cy7, Biolegend, 103236), anti-CD45RA (BV605, BD Biosciences, 562886), anti-CCR7(PE, BD Biosciences, 552176), anti-CD150/SLAMF1 (TC15-12F12.2; Biolegend). For dead cell exclusion, cells were stained with Zombie NIR Fixable Viability Kit (BioLegend, 423105) for 10 min at 4℃ and washed in FACS buffer. For intracellular cytokine and transcription factor staining, surface Ab-stained cells were first fixed and permeabilized using the fixation/permeabilization kit (BD Biosciences, 554714) following manufacturer’s instructions. Cells were further stained with Abs against intracellular proteins for 30 min at 4℃. Intracelluar Abs were used as follows: anti-Ets1 (PE, Santa Cruz, sc-55581PE), anti-Runx3 (Alexa 488, BD Biosciences, 565740), anti-Zbtb7b/ThPOK (Alexa 647, BD Biosciences, 565500; R718, BD Biosciences, 566942), anti-Phospho-STA5B-Ser127 (ABclonal) and anti-rabbit IgG (PE, Biolegend, 406421). Samples were collected using a Beckman Coulter cytoFLEX and analyzed with FlowJo 10 Software (Tree Star).

### Microscale thermophoresis

His-tagged PAK2 protein was labeled with a fluorescent dye (NanoTemper, MO-L018) according to the manufacturer’s instructions. Binding was performed in standard capillaries in MST buffer (25 mM HEPES pH 7.4, 50 mM NaCl, 2.5 mM MgCl₂, and 0.025% NP-40). Molecular interaction experiments were performed using a Monolith NT.115 instrument, with data collection managed via MO. Control software.

### Cytometric beads-based immunoassays

Analyte-specific antibodies are pre-coated onto magnetic beads with fluorophores at set ratios for each unique microparticle region. Coupled beads, standards and samples are pipetted into wells and the immobilized antibodies bind the analytes of interest. After washing away any unbound substances, a biotinylated antibody cocktail specific to the analytes of interest is added to each well. Following a wash to remove any unbound biotinylated antibody, streptavidin-phycoerythrin conjugate (SA-PE), which binds to the biotinylated antibody, is added to each well. Final washes remove unbound Streptavidin-PE, the beads are resuspended in buffer and read using the ABplex-100 Analyzer. A Coupled bead in the analyzer captures and holds the superparamagnetic microparticles in a monolayer. Two spectrally distinct Light Emitting Diodes (LEDs) illuminate the microparticles. One LED excites the dyes inside each microparticle to identify the region and the second LED excites the PE to measure the amount of analyte bound to the microparticle, PE serves as a fluorescent indicator, or reporter. A sample from each well is imaged with a CCD camera with a set of filters to differentiate excitation levels.

### Single-cell RNA sequencing

Lung LNs of BCG plus CFA-immunized mice were ground using a syringe rubber and 70 μm cell strainer. Flow-through containing immune cells and tissue debris was collected. The single cell solution was isolated using the CD4^+^ T Cell Isolation Kit (Cat# 130-117-043; Miltenyi Biotec) and CD8^+^ T Cell Isolation Kit (Cat# 130-116-478; Miltenyi Biotec).

CD45.2 recipient mice and control WT mice lung tissue were collected at 4 weeks post-infection. The single cell solution was isolated using the CD45^+^ Cell Isolation Kit (Cat# 130-052-301; Miltenyi Biotec).

Cells were then stained with the fluorescent dyes Calcein AM and Draq7, to accurately determine the cell concentration and viability using the BD Rhapsody™ Scanner (BD Biosciences). Cells were loaded in one BD Rhapsody micro-well cartridge. Cell capture beads were then loaded excessively to ensure that every micro-well contains one bead; the excess beads were washed away from the cartridge. After lysing the cells with lysis buffer, the cell capture beads were retrieved and washed prior to performing reverse transcription. The single cell transcriptome was generated into a cDNA library containing cell labels and unique molecular identifiers (UMI) information. All procedures were performed using the BD Rhapsody cDNA Kit (Cat. No. 633773; BD Biosciences) and BD Rhapsody Targeted mRNA & AbSeq Amplification Kit (Cat No. 633801BD; Biosciences), strictly following the manufacturers’ protocol. All the libraries were sequenced in a PE150 mode (Pair-End for 150bp read) on the NovaSeq-6000 platform (Illumina).

### Analysis of scRNA-seq data

Raw sequencing reads of the cDNA library were processed through the BD Rhapsody Whole Transcriptome Assay Analysis Pipeline (v1.11), which included filtering by reads quality, annotating reads, annotating molecules, determining putative cells, and generating single-cell expression matrix. Among all the output files, the matrix of UMI counts for each gene per cell was used for downstream analysis. Genome Reference Consortium Mouse Build 38 was used as reference for the BD pipeline. We utilized Seurat for subsequent clustering analysis and visualization. Gene expression matrices for each sample were read and converted to Seurat objects. Cells with more than 25% mitochondrial UMI, less than 500 UMI, or 200 genes were excluded from the downstream analysis. After log-normalization according to the total cellular UMI count, principal component analysis (PCA) was performed based on the top 2000 highly variable features after scaling the data with respect to the UMI counts. Clustering was then performed at a resolution of 0.6 and data visualized using either t-Distributed Stochastic Neighbor Embedding or Uniform Manifold Approximation and Projection.

### Pan-Peptide Meta Learning (PanPep) analysis

Antigen Peptide Extraction and Filtering

C57BL/6 mice were infected with *Mtb* or immunized with BCG plus CFA. Four weeks post-infection or immunization, single-cell suspensions were prepared from the lungs and subjected to sequencing analysis. MHC class I-restricted peptides were identified from the Mtb-infected model, whereas both MHC class I-and class II-restricted peptides were identified from the BCG plus CFA immunized model. Candidate peptides were filtered to retain sequences composed exclusively of the 20 standard amino acids with lengths of 25 or fewer residues. This yielded a finalized library of 389 *Mtb*-associated antigen pool for downstream analysis.

### TCR Repertoire Profiling and Preprocessing

For T cell receptor (TCR) profiling, lymph nodes were harvested from mice four weeks post-immunization with BCG plus CFA for single-cell RNA and TCR sequencing. TCR sequences from CD4^+^CD8^+^ T cell subsets were filtered for the 20 standard amino acids and independently deduplicated within each cluster. The final dataset retained unique TCRs across three distinct clusters: cluster 2 (n = 794), cluster 3 (n = 646), and cluster 15 (n = 65).

### TCR-Antigen Pairing and Prediction

The preprocessed TCR sequences from each cluster were systematically paired against the 389-antigen pool. Specific binding interactions for these pairs were computationally predicted using PanPep.

### 4D DIA phosphoproteomic analysis

#### Sample Preparation for Mass Spectrometry

CD4^+^, CD8^+^, and CD4^+^CD8^+^ memory T cells sorted from the LNs of C57BL/6 mice immunized with BCG plus CFA were loaded into PCT tubes, with each tube containing 30 μL of lysis buffer (6 M urea, 2 M thiourea), 1.5 μL of 25× phosphatase inhibitor, and 1.5 μL of 25× complete protease inhibitor. Subsequently, 5 μL of 0.2 M TCEP and 2.5 μL of 0.8 M IAA were added, followed by incubation in the dark. The samples were subjected to high-pressure cycling (45 kpsi, 30 s high pressure, 10 s ambient pressure) for 90 cycles at 30°C. For enzymatic digestion, trypsin (100 μg/vial) was dissolved in 200 μL of 1 mM HCl or 50 mM acetic acid (final concentration: 0.5 μg/μL), while rLys-C protease (50 μg) was dissolved in 100 μL of 1 mM HCl or 50 mM acetic acid (final concentration: 0.5 μg/μL). Subsequently, 75 μL of 0.1 M TEAB was added to each sample tube, followed by 10 μL (5 μg) of trypsin and 2.5 μL (1.25 μg) of rLys-C protease. The volume was adjusted with 0.1 M TEAB to obtain a final pH of 8. The samples were processed under high-pressure conditions (20 kpsi, 50 s high pressure, 10 s ambient pressure) for 120 cycles at 30°C. Enzymatic digestion was terminated by adding 15 μL of 10% trifluoroacetic acid (TFA), achieving a final TFA concentration of 1%. The pH was verified to range between 2–3. An equal volume of enrichment buffer was added to the digest, followed by an appropriate amount of enrichment material based on protein quantification. The mixture was incubated on a rotary shaker at 25 °C for 30 min. Subsequently, each tube was treated with washing buffer 1 and incubated on a rotary shaker for 30 min, followed by centrifugation to separate the supernatant. This step was repeated twice using washing buffer 2, with centrifugation (25°C, 3 min, 15,000 ×*g*), to discard the supernatant. For elution, 100 μL of elution buffer was added, and the samples were subjected to ice-bath sonication for 5 min, followed by rotary shaking for 15 min. After centrifugation (25°C, 3 min, 18,000 ×*g*), the supernatant was collected, and the elution step was repeated once. The combined eluates were dried under vacuum. Prior to mass spectrometry (MS) analysis, the samples were desalted and reconstituted in 98% water, 2% ACN, and 0.1% formic acid for liquid chromatography-tandem mass spectrometry (LC-MS/MS) acquisition.

#### Proteome sample analysis

LC-MS/MS analysis was performed using a Vanquish Neo UHPLC system coupled to an Orbitrap Astral mass spectrometer (Thermo Scientific, San Jose, USA) in the data-independent acquisition (DIA) mode. Mobile phase B consisted of 80% ACN, 20% water, and 0.1% formic acid (all reagents were MS-grade). For sample loading, peptides were injected at 800 bar onto a pre-column (5 µm, 5 mm × 300 µm i.d.) and then separated on an analytical column (1.9 µm, 120 Å, 150 mm × 75 µm i.d.) at a flow rate of 400 nL/min. An optimized 18-min LC gradient was applied, with mobile phase B increasing from 5% to 35%. The FAIMS voltage was set at -45 V. Full MS scans covered a range of 380–980 m/z with a resolution of 240,000 (at 200 m/z), a normalized AGC target of 500%, and a maximum ion injection time (max IT) of 3 ms. MS/MS scans were acquired over a range of 150-2000 m/z with a normalized AGC target of 500%, a normalized collision energy of 25%, and a max IT of 3 ms. Precursor ions were selected within 380–980 m/z using an isolation window of 2 m/z (no overlap), resulting in 299 total windows.

#### Mass spectrometry data analysis

MS data were processed using DIA-NN (version 1.8.1) for spectral library-free analysis. The search was conducted against the reviewed mouse proteome database (UP000000589_PD_mouse. fasta, downloaded on June 19, 2023). Precursor mass accuracy was set to 0.0 ppm, and identifications were filtered at both precursor and protein levels using a 1% false discovery rate threshold. The analysis considered fragment ions in the m/z range of 200–1800 and precursors within 300–1800 m/z, allowing for charge states from +1 to +4. Trypsin digestion specificity was enforced while permitting up to two missed cleavages, and only peptides with lengths between 7–30 amino acids were included.

Post-translational modifications were carefully accounted for, with variable modifications including oxidation (+15.994915 Da) of methionine residues and phosphorylation (+79.966331 Da) of serine, threonine, and tyrosine residues. Carbamidomethylation (+57.021464 Da) of cysteine was set as a static modification. The DIA-NN workflow specifically monitored phosphorylation events (UniMod:21) while enabling the detection of both oxidation (UniMod:35) and phosphorylation modifications during database searching.

### Metabolomics analysis

#### Metabolites Extraction

Samples were extracted for metabolites by using the Starlid™workstation. The 50 ul serum sample was mixed with 200 ul of extraction solution (methanol:acetonitrile=1:1 (v/v), including internal standards), shaken at 750 rpm for 5 min and settled down for 5 min, then filtered through a protein precipitation plate and the filtrate was collected.

#### LC-MS/MS Analysis

For polar metabolites, LC-MS/MS analyses were performed using an UHPLC system (Vanquish, Thermo Fisher Scientific) with a Waters ACQUITY UPLC BEH Amide (2.1 mm × 100 mm, 1.7 μm) coupled to Orbitrap Exploris 120 mass spectrometer (Orbitrap MS, Thermo). The mobile phase consisted of 25 mmol/L ammonium acetate and 25 mmol/L ammonia hydroxide in water (pH= 9.75) (A) and acetonitrile (B). The auto-sampler temperature was 4 °C, and the injection volume was 2 uL.

For non-polar metabolites, LC-MS/MS analyses were performed using an UHPLC system (Vanquish, Thermo Fisher Scientific) with a Phenomenex Kinetex C18 column (2.1 mm × 100 mm, 2.6 μm) coupled to Orbitrap Exploris 120 mass spectrometer (Thermo). The mobile phase consisted of 0.01% acetic acid in water (A) and isopropanol:acetonitrile (1:1, v/v) (B). The auto-sampler temperature was 4 °C, and the injection volume was 2 uL.

The Orbitrap Exploris 120 mass spectrometer was used to acquire MS and MS/MS data under the control of the Xcalibur software (version 4.4, Thermo). The detailed parameters were set as following: sheath gas flow rate as 50 Arb, Aux gas flow rate as 15 Arb, capillary temperature 320 °C, full MS resolution as 60000, MS/MS resolution as 15000, collision energy: SNCE 20/30/40, spray voltage as 3.8 kV (positive) or -3.4 kV (negative).

#### Data preprocessing and annotation

The raw data were converted to the mzXML format using ProteoWizard and processed with an in-house program. which was developed using R and based on XCMS, for feature detection, extraction, alignment, and integration. The R package and the BiotreeDB (V3.0) were applied in metabolite identification^112^.

Due to the small cohort size (n = 9) processed within a single stable analytical batch, pooled quality controls were omitted. Instrument stability was instead confirmed via consistent total ion chromatogram (TIC) overlays. To minimize systematic bias, raw data were filtered to remove interquartile range (IQR) outliers and features with >50% missing values. Remaining missing data were imputed using half the minimum detected peak area and data were normalized to internal standards (IS).

### Maleic acid assays

#### Sample preparation and extraction

For serum sample preparation, A aliquot of 50 μL from each serum sample was transferred into a pre-labeled 1.5 mL microcentrifuge tube after thawing on ice and vortex-mixing for 10 s to ensure homogeneity, followed by addition of 250 μL of 20% acetonitrile/methanol extraction solvent. The mixture was vortexed for 3 min and centrifuged at 12,000 rpm for 10 min at 4 °C. Subsequently, 250 μL of the supernatant was transferred to a second pre-labeled 1.5 mL microcentrifuge tube and incubated at −20 °C for 30 min, followed by a second centrifugation at 12,000 rpm for 10 min at 4 °C. Finally, 180 μL of the clarified supernatant was transferred into a fresh vial for analysis.

#### UPLC–MS/MS analysis

Chromatographic separation was carried out on a Waters ACQUITY H-Class UPLC system using an ACQUITY UPLC BEH Amide column (2.1 × 100 mm, 1.7 μm). Mobile phase A was water containing 10 mM ammonium acetate and 0.3% ammonium hydroxide, and mobile phase B was 90% acetonitrile in water (v/v). The gradient was programmed as follows: 0-1.2 min, 95% B; 8.0 min, 70% B; 9.0–11.0 min, 50% B; 11.1–15.0 min, 95% B. The flow rate was 0.4 mL/min, the column temperature was maintained at 40 °C, and the injection volume was 2 μL.

MS/MS detection was performed on a QTRAP 6500 + LC-MS/MS system (SCIEX) equipped with an electrospray ionization (ESI) source operated in positive and negative modes. Source settings were: ion spray voltage +5500 V (positive) and -4500 V (negative), source temperature 550 °C, and curtain gas 35 psi. Metabolites were quantified using scheduled MRM, with declustering potential (DP) and collision energy (CE) optimized for each transition. Data acquisition and quantification were conducted using Analyst (v1.6.3, SCIEX) and MultiQuant (v3.0.3, SCIEX), respectively.

#### Quantification and quality control

In total, 65 organic acid-related metabolites were quantified based on an in-house database (MWDB, Metware). Absolute concentrations were determined using external calibration curves with 20 levels (0.01–15,000 ng/mL for most analytes). QC samples were generated by pooling equal aliquots of all extracts and injected in triplicate across the run. Instrument performance was assessed by total ion chromatograms (TICs) overlap of QC injections. The Pearson correlation coefficients among QC samples were > 0.99, and the coefficient of variation (CV) was < 30% for > 80% of detected metabolites.

### 16S rRNA gene sequencing

Fresh stool samples were collected from animals placed in a clean empty cage without bedding for 15 min and immediately stored at −80 °C until analysis. Microbial DNA was extracted from mouse feces samples using the OMEGA Soil DNA Kit (D5625-01; Omega Bio-Tek, USA). DNA concentration and purity were measured with a NanoDrop ND-1000 spectrophotometer (Thermo Fisher Scientific), and integrity was checked by agarose gel electrophoresis. The V3–V4 region of the bacterial 16S rRNA gene was amplified using primers 338F (5′-ACTCCTACGGGAGGCAGCA-3′) and 806R (5′-GGACTACHVGGGTWTCTAAT-3′). Amplicons were sequenced on an Illumina HiSeq PE250 platform (2 × 250 bp) by a commercial provider (Personalbio). Demultiplexed, quality-filtered reads with barcodes and host-derived sequences removed were used for downstream analyses. OTU clustering was performed using UCLUST in QIIME (http://qiime.org/scripts/pick_otus.html), and taxonomic assignment was conducted against the Greengene database.

Sequence processing and statistical analyses were conducted primarily with QIIME2 (v2019.4) and R packages (v3.2.0). Alpha diversity and Bray–Curtis-based beta diversity were calculated in QIIME2 using the standard workflow. Differential taxa abundance at the ASV level was tested using metagenomeSeq and visualized with Manhattan plots.

### CUT&Tag assay

The CUT&Tag assay was performed with the Hyperactive Universal CUT&Tag Assay Kit for Illumina (Vazyme, TD903) as described by the manufacturer with some modifications. For each sample, 1×10⁵ cells were harvested, and the nuclei were extracted and incubated with ConA beads at room temperature for 10 min according to the manufacturer’s instructions. The mixed complexes of nuclei and ConA beads were incubated with Rabbit Monoclonal anti-TCF1/TCF7 antibody (CST, 2203), Rabbit Monoclonal anti-ETS1 antibody (CST, 14069), Rabbit monoclonal Anti-c-Myb (phospho S11) antibody (Abcam, ab45150), Rabbit polyclonal anti-SOX4 antibody (Diagenode, C15310129), Rabbit Polyclonal anti-TCF12 antibody (Bethyl, A300-754A), Mouse monoclonal PE anti-Ets2 antibody (SCBT, sc-365666) and mouse IgG (with IgG antibodies serving as the control) and then incubated with the secondary antibody the next day. A diluted pA-Tn5 adapter complex was subsequently added to the above system and incubated at room temperature for 1 h. After extensive washing, the system was sheared by adding 5 × TTBL, and proteinase K and DNA extract beads were then used to obtain eluted DNA. Following this procedure, CUT&Tag libraries were established after 12 cycles of PCR amplification and sequenced by Illumina Novaseq 6000 platform with paired-end 2×150 as the sequencing mode.

CUT&Tag sequencing reads were trimmed to remove adapters, short reads (length <36 bp) and low-quality reads using Trimmomatic v0.38 (non-default parameters: SLIDINGWINDOW:4:15 LEADING:10 TRAILING:10 MINLEN:36) after which FastQC (with default parameters) was used to ensure high read quality. The clean reads were aligned to the mouse genome (assembly GRCm38) using Bowtie2 v2.4.1 (non-default parameters: -I 10 -X 700 -no-discordant -no-mixed -local -very-sensitive-local)^113^. Duplicate reads were removed to generate uniquely mapped reads via Picard MarkDuplicates. All the genome peaks were identified via MACS2 v2.1.2 (non-default parameters: -f BAMPE -g hs/mm -q 0.05) with 0.05 set as the q-value cutoff ^114^. Annotation of peak sites to gene features was performed using the ChIPseeker R package^115^.

### LIP-MS

LiP-SMap approach was performed as previously described^116^. Briefly, vehicle- or maleic acid-treated CD8^+^ T cells were lysed by bead-beating in PBS at 4°C, followed by centrifugation at 16,000 × g for 10 minutes at 4°C to collect the supernatant. Limited proteolysis was carried out by adding Proteinase K (Sangon Biotech, China) at a 1:100 enzyme-to-substrate ratio and incubating for 5 minutes at 25°C. The reaction was stopped by heating the samples at 98°C for 5 minutes, followed by the addition of sodium deoxycholate (DOC) to a final concentration of 2%. Protein fragments were reduced with 10 mM dithiothreitol (DTT) for 30 minutes at 37°C and then alkylated with 40 mM iodoacetamide for 45 minutes at 25°C in the dark. The samples were digested overnight at 37°C with trypsin at a 1:50 enzyme-to-substrate ratio under agitation at 800 rpm. After digestion, the DOC was precipitated by adding a 50% trifluoroacetic acid (TFA) solution to a final concentration of 2%, and the pH of the samples was adjusted to less than 3 with formic acid. Peptide mixtures were desalted using Sep-Pak C18 cartridges and eluted with 70% acetonitrile–0.1% formic acid. The samples were dried in a vacuum centrifuge, resuspended in 0.1% formic acid, and analyzed by mass spectrometry. Peptide fragments were analyzed using Nano Acuity Ultra-High-Pressure Liquid Chromatography coupled with a Thermo Q Exactive mass spectrometer (Thermo Fisher, USA). Proteins and peptides were identified using a target-decoy approach with a reversed database, queried against the Mouse UniProt FASTA database, and quantified using MaxQuant software for label-free quantification (LFQ).

### Cytokine antibody array

The cytokine profiles of the lung supernatants of Rag-NT, Rag+CD4CD8 Tm groups were analyzed using a mouse cytokine antibody array (cat. QAM-CAA-4000; RayBiotech, ATL, USA) and tested according to the experimental protocols. A combination of factors, including ranking by fold change (< 0.67 and > 1.5) and signal intensity (> 150), were used to identify robust changes, and the meta-ranking of proteins was generated. The relative signal intensity of the indicated cytokines was presented in a heatmap. The detailed protocol of the assay using the Quantibody® mouse cytokine antibody array 4000 has been described previously^117^.

### Quantification and statistical analysis

All experimental data were analyzed using GraphPad Prism 10.1.2 software. Data distribution was assumed to be normal, but this was not formally tested. Data collection was randomized, and animals or samples were assigned to experimental groups through a randomized process. The data collection and analysis were not performed blind to the conditions of the experiments, except for the H&E slides scoring. No animals or data points were excluded from the analyses for any reason. The statistical significance of differences between two groups was determined by two-tailed unpaired Student’s t-test. One-way ANOVA with Dunnett’s multiple comparison test, two-way ANOVA with Dunnett’s multiple comparison test, two-way ANOVA with Šídák’s multiple comparisons test and Multiple unpaired t tests were used for statistical analysis when comparing more than two groups. *P* < 0.05 was considered significant. All data were expressed as mean ± s.e.m. of the averages of technical replicates from the indicated number of independent experiments.

## Acknowledgments

We thank Prof. Zhongjun Dong (Anhui Medical University, Hefei), Prof. Shasha Chen (Anhui Medical University, Hefei) for *slamf1*^−/−^ mice, Prof. Hui Hu (University of Alabama) for critical reading of the manuscript, and members of B. Ge’s laboratory (Shanghai Key Laboratory of Tuberculosis, Shanghai Pulmonary Hospital, Tongji University School of Medicine, Shanghai, China) for helpful discussions and technical assistance. This project was supported by grants from National Natural Science Foundation of China (32188101 to B.G.; 82470001, 82270006 to H.Y.), National Science and Technology Major Project (2025ZD1802400, 2025ZD1802401 to H.Y.), the National Key R&D Program of China (2021YFA1300902 to H.Y.), National Program on Key Basic Research Project (2017YFA0505900 to B.G.), The Science and Technology Innovation Action Plan (STCSM, 24YF2735400 to H.C.), Prevention and Control of Emerging and Major Infectious Diseases-National Science and Technology Major Project (2025ZD01900902 to M.L.), and The Science and Technology Department of Yunnan Province (202602AS100003 to M.L.).

## Contributions

H.C., C.S., H.Y. and B.G. designed the study. H.C. and C.S. performed most experiments and analyzed data. X.F., J.Z., Q.Y., and M.L. performed Non-human primates (NHPs) experiment. X.C., H.L., Y.C., C.P., Y.L., Y.B., S.L., Y.Y., L.W., J.W., X.H., X.C. and D.Q. assisted with manuscript preparation and performed mice infection experiments. Q.L., K.D., and Y.G. performed the Pan-Peptide Meta Learning (PanPep) analysis. L.Z. and X.H. performed the microscale thermophoresis experiment. J.W. and X.H. preserved the *Mtb* and BCG strains. All authors discussed the results and commented on the manuscript.

## Data availability

Data supporting the findings of this study are available from the following public repositories. Metabolomics datasets from mouse serum samples collected 4 weeks after immunization with PBS, BCG, or BCG+CFA are deposited in MetaboLights under accession MTBLS15097. 16S rRNA gene sequences of gut microbiota from 6- to 8-week-old female C57BL/6 mice subcutaneously immunized with PBS, BCG, CFA, or BCG+CFA are available in the Sequence Read Archive (SRA) under SRP719314. CUT&Tag data for genome-wide binding sites of ETS1, SOX4, MYB, TCF1, ETS2 and TCF12 are deposited in the SRA under SRP719386. Single-cell transcriptome and TCR sequencing data of T cells from lung-draining lymph nodes of mice immunized with BCG+CFA for 4 weeks are deposited in the SRA under accession SRP719462. Single-cell transcriptome data of lung T cells from naive C57BL/6 mice and from mice adoptively transferred with CD4⁺CD8⁺ T cells are available in the SRA under accession SRP719608. Phosphoproteomics data of CD4⁺CD8⁺, CD4⁺, and CD8⁺ T cells isolated from lung-draining lymph nodes of BCG+CFA-immunized mice are deposited in the ProteomeXchange Consortium (https://proteomecentral.proteomexchange.org) via the iProX partner repository with the dataset identifier PXD081332. The LiP-MS datasets for profiling maleic acid-interacting proteins are presented in Table S2. Additional data are available from the corresponding authors upon reasonable request.

## Code availability

All analyses were done reproducibly using publicly available R scripts.

