## Extended data figure and figure legend for "Maleic acid adjuvates BCG to induce CD4^+^CD8^+^ T cells and sterilize tuberculosis"

**Extended Data Figure legend**

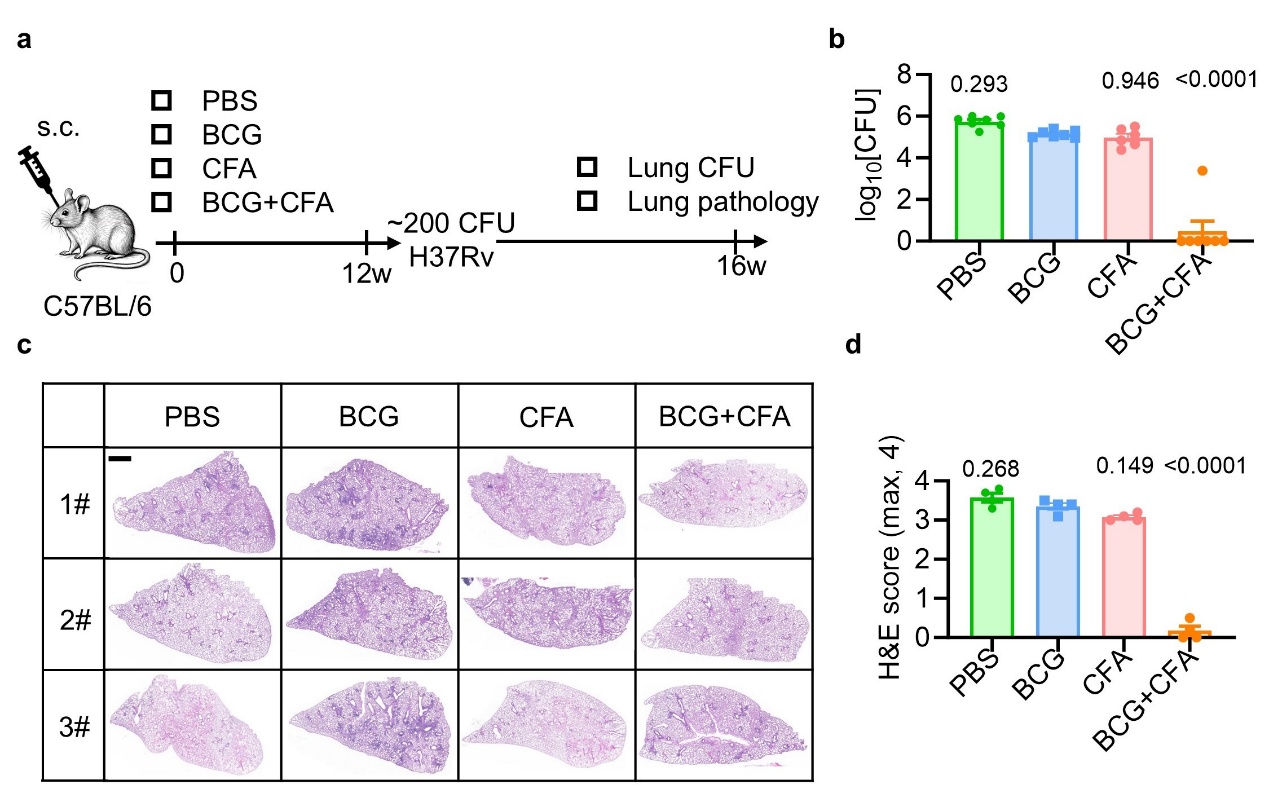

**Extended Data Fig. 1: BCG plus CFA immunization provides long-term, robust protection against *Mtb***

**a-d**, Experimental design **(a)**, C57BL/6 mice (vaccinated at 12 weeks of age per group) were aerosol-infected with ~200 CFU per mouse of *Mtb* H37Rv. At 4 weeks post-infection, the following assessments were performed: bacterial load in the lungs **(b)**, histopathology of lung sections via H&E staining (**c**; scale bar, 1 mm), and histology score **(d)**. Data in **b, d** represent one experiment with at least three independent biological replicates; mean ± SEM. Ordinary one-way ANOVA with Dunnett’s multiple comparison test comparing each group to the BCG group **(b, d)** was used for statistical analyses.

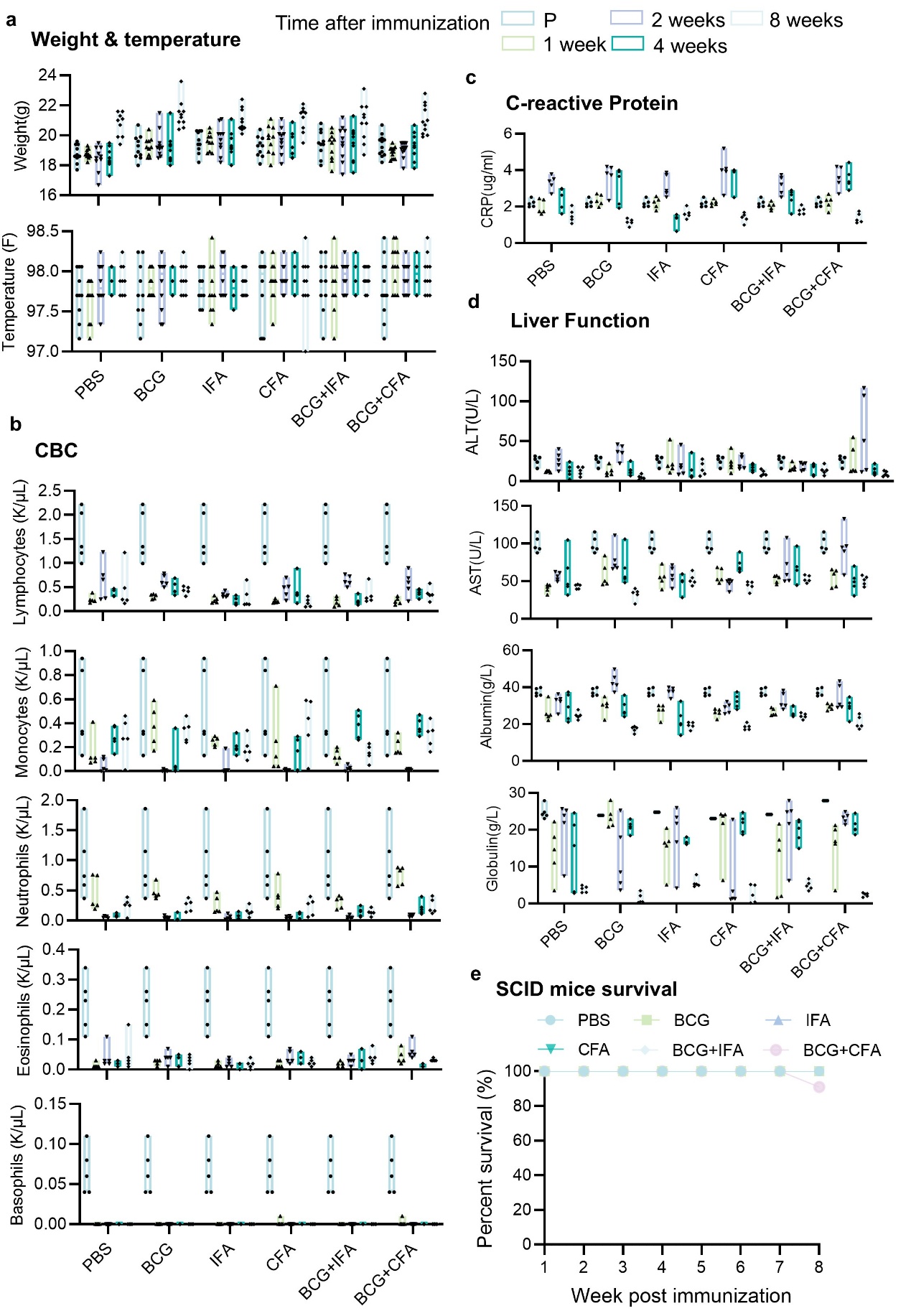

**Extended Data Fig. 2: The safety of subcutaneous BCG combined with CFA vaccination**

**a-b**, to assess safety of each vaccine group, all SCID mice were monitored for changes in several clinical parameters at various time points after immunization: Weight and temperature **(a)**, Complete blood counts (CBC) **(b)**, C-reactive protein **(c)**, liver Function **(d)**, kidney Function **(e)** and SCID mice survival **(f)**. For each parameter, pre-vaccination (P) measurements for all mice were combined and compared against distributions from every vaccine group at every time point using Dunnett’s test for multiple comparisons.

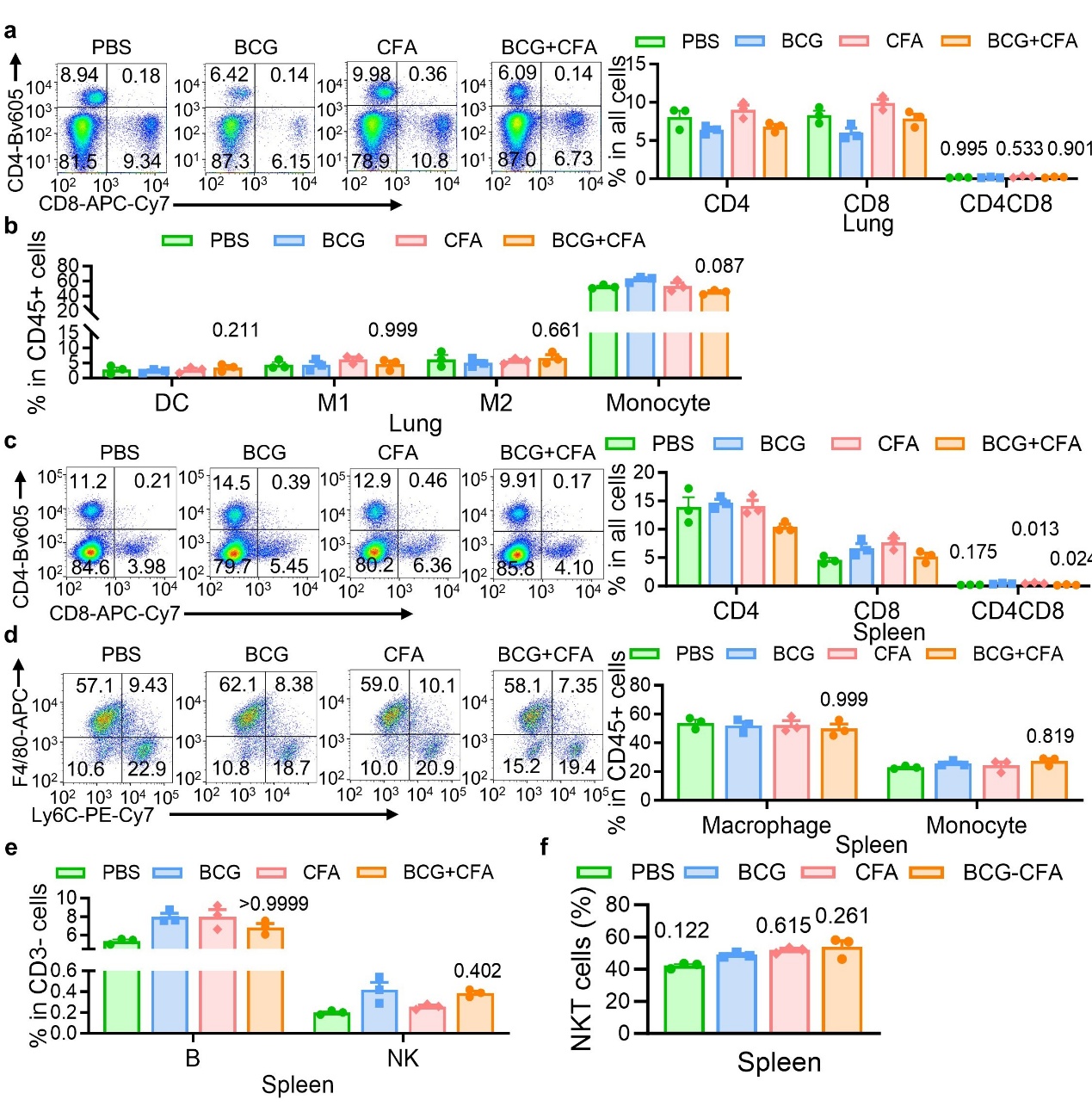

**Extended Data Fig. 3: Cellular composition and immune analysis after vaccination**

**a-f**, Proportions of cells of indicated leukocyte population in the lung and spleen of each vaccine group at 4 weeks post vaccination, as determined by flow cytometry (FCM). The proportions of CD4^+^, CD8^+^ and CD4^+^CD8^+^ T cells in lung **(a)**; the proportions of DCs, monocytes, M1 and M2 macrophages in lung **(b)**; the proportions of CD4^+^, CD8^+^ and CD4^+^CD8^+^ T cells in spleen **(c)**; the proportions of macrophage, monocytes in spleen **(d)**; the proportions of B cells and NK cells in spleen **(e)**; the proportions of NKT cells in spleen **(f)**. Data in **a-f** represent one experiment with at least three independent biological replicates; mean ± SEM. Two-way ANOVA with Tukey’s multiple comparisons test comparing the group to respective BCG group **(a-e)** and one-way ANOVA with Dunnett’s multiple comparison test comparing each group to the BCG group **(f)** were used for statistical analyses.

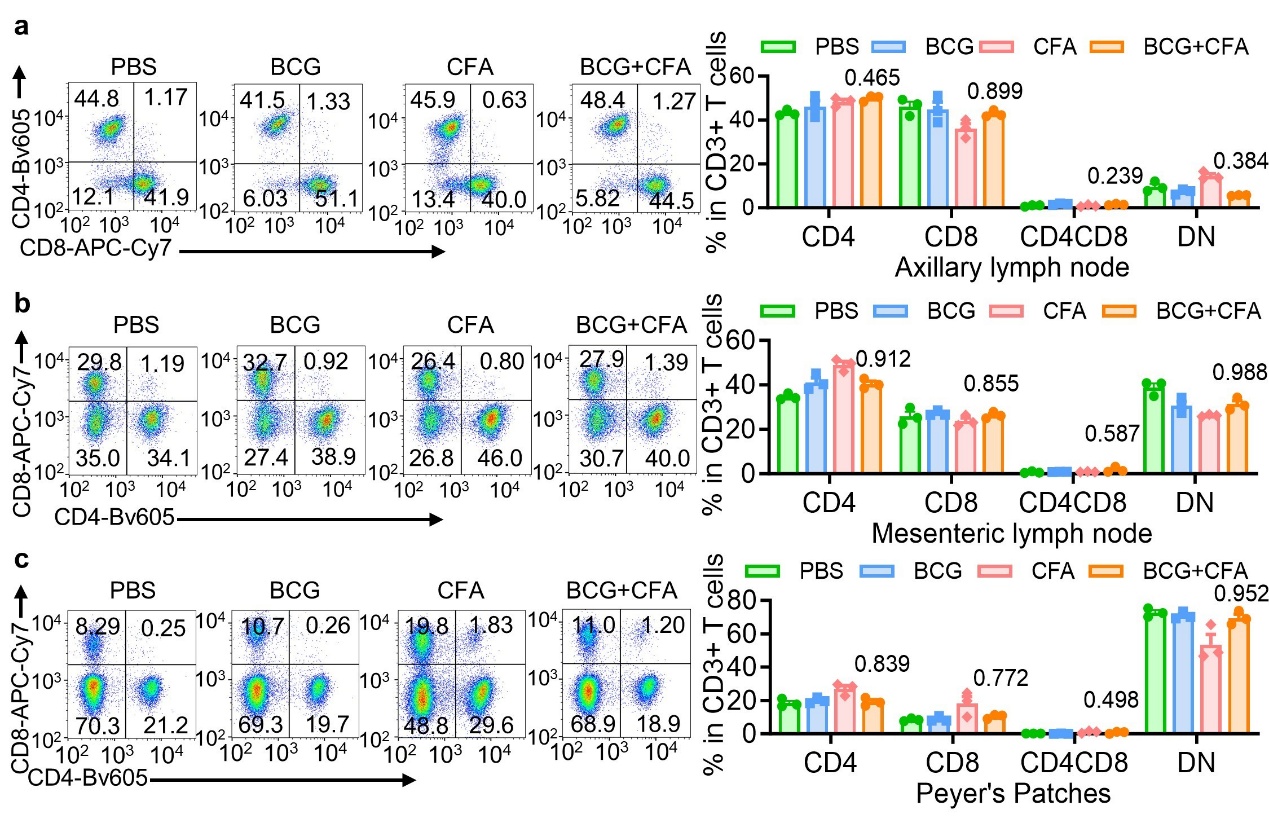

**Extended Data Fig. 4: Proportions of CD4^+^CD8^+^ T cells in lymph nodes after vaccination**

**a-c**, Proportions of CD4^+^, CD8^+^, CD4^+^CD8^+^ and CD4^-^CD8^-^ double negative (DN) T cells (gated from CD3^+^ T cells) in axillary lymph nodes **(a)**, mesenteric lymph nodes **(b)**, and Peyer’s patches **(c)** of each vaccine group at 4 weeks post vaccination, as determined by FCM. Data in **a-c** represent one experiment with at least three independent biological replicates; mean ± SEM. Two-way ANOVA with Tukey’s multiple comparisons test comparing the BCG+CFA group to the respective BCG group **(a-c)** was used for statistical analyses.

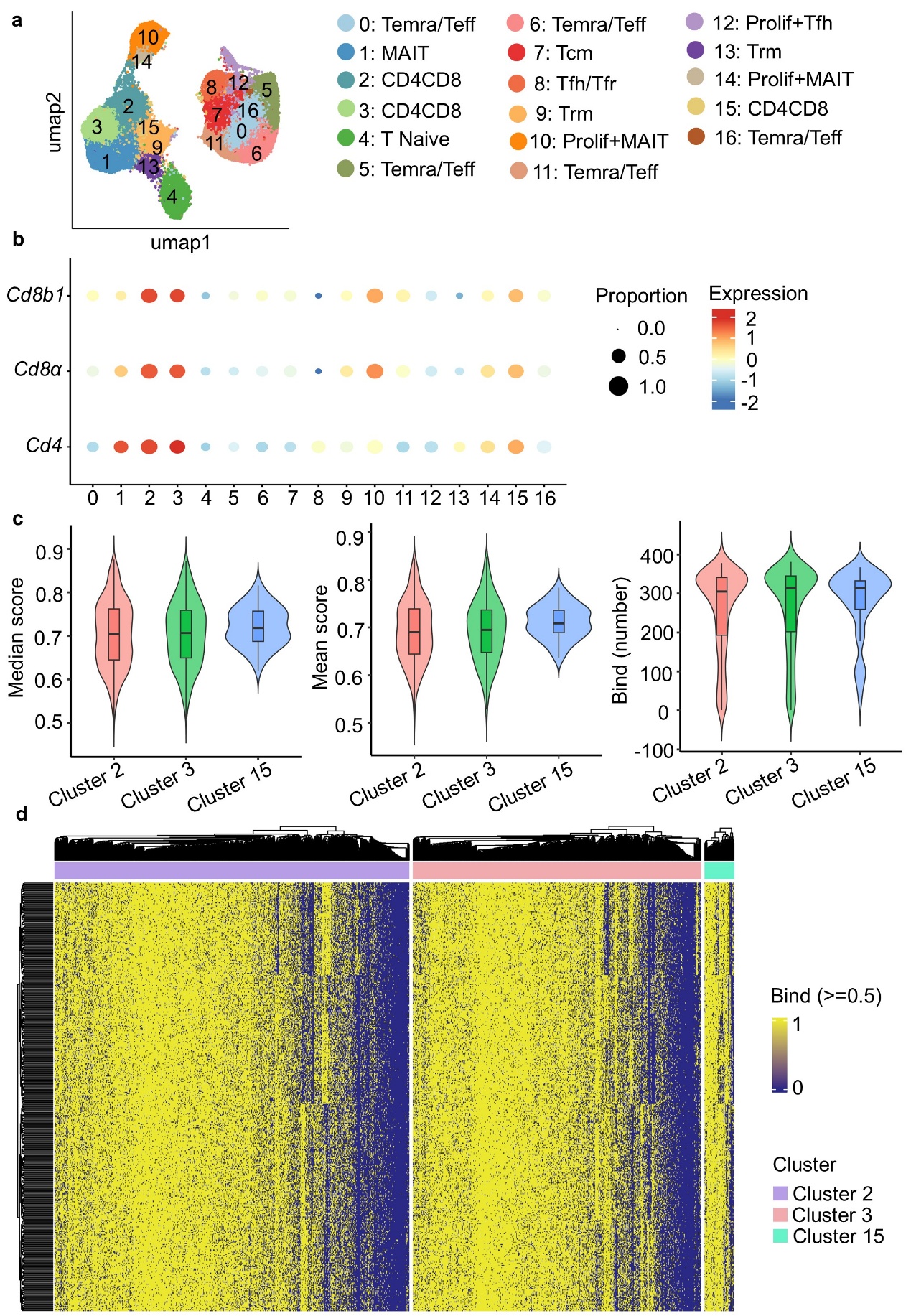

**Extended Data Fig. 5: Polyclonal CD4^+^CD8^+^ T cells recognize a broad repertoire of *Mtb* antigens**

**a,** Single-cell transcriptional analysis of lung lymph nodes cells from 4 weeks post BCG plus CFA vaccinated mice. **b,** Dot plot representing the relative average expression of *Cd4*, *Cd8a* and *Cd8b1* genes across all clusters. As indicated on the legend, dot size denotes the percentage of cells in a cluster expressing each gene. Dot color represents the relative average expression levels. **c,** Median and mean binding fractions and number of binding peptides of TCRs with binding capacity at CD4^+^CD8^+^ T cell clusters in **(a). d,** Heatmap showing the binding of TCR and Peptide at CD4^+^CD8^+^ T cells clusters in **(a)**, each row is a peptide, each column is a TCR, and the corresponding value is the binding. Yellow represents binding (score greater than 0.5), blue represents no binding.

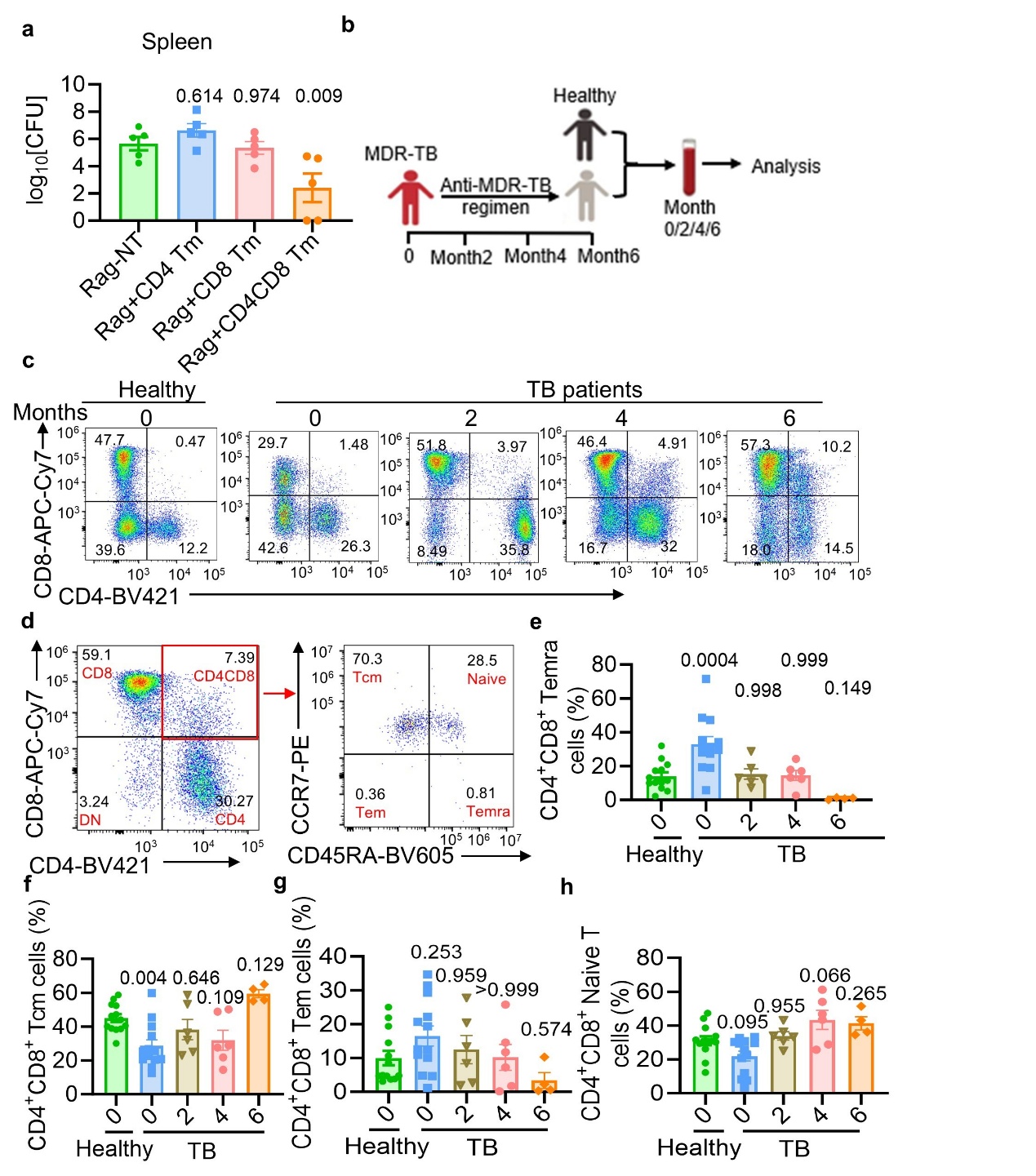

**Extended Data Fig. 6: CD4^+^CD8^+^ T cells conferred sterilizing protection against *Mtb***

**a,** Analysis of spleen bacterial load at 4 weeks post-infection from the experiment shown in **Fig. 3a**. **b-h**, Experimental design **(b),** MDR‑TB patients received standardized anti‑MDR‑TB regimens. Blood samples were collected in EDTA at 0, 2, 4, and 6 months after treatment initiation from each donor (TB patients and healthy controls) for FCM analysis. Proportions of CD4^+^CD8^+^ T cells in the peripheral blood of each group was determined by FCM analysis **(c)**. Central memory CD4^+^CD8^+^ T cells (Tcm) (CCR7^+^CD45RA^-^), naive CD4^+^CD8^+^ T cells (CCR7^+^CD45RA^+^), effector memory CD4^+^CD8^+^ T cells (Tem) (CCR7^-^CD45RA^-^), terminally differentiated CD4^+^CD8^+^ T cells (Temra) (CCR7^-^CD45RA^+^) (gated from CD4^+^CD8^+^ T cells) as determined by FCM analysis **(d)**: Proportions of CD4^+^CD8^+^ Temra cells **(e)**, CD4^+^CD8^+^ Tcm cells **(f)**, CD4^+^CD8^+^ Tem cells **(g)** and CD4^+^CD8^+^ naive T cells **(h)** in the peripheral blood of each group. Data in **a, e-h** represent one experiment with at least three independent biological replicates; mean ± SEM. Ordinary one-way ANOVA with Dunnett’s multiple comparison test comparing each group to the Rag-NT group **(a)** or healthy group (**e-h**) was used for statistical analyses.

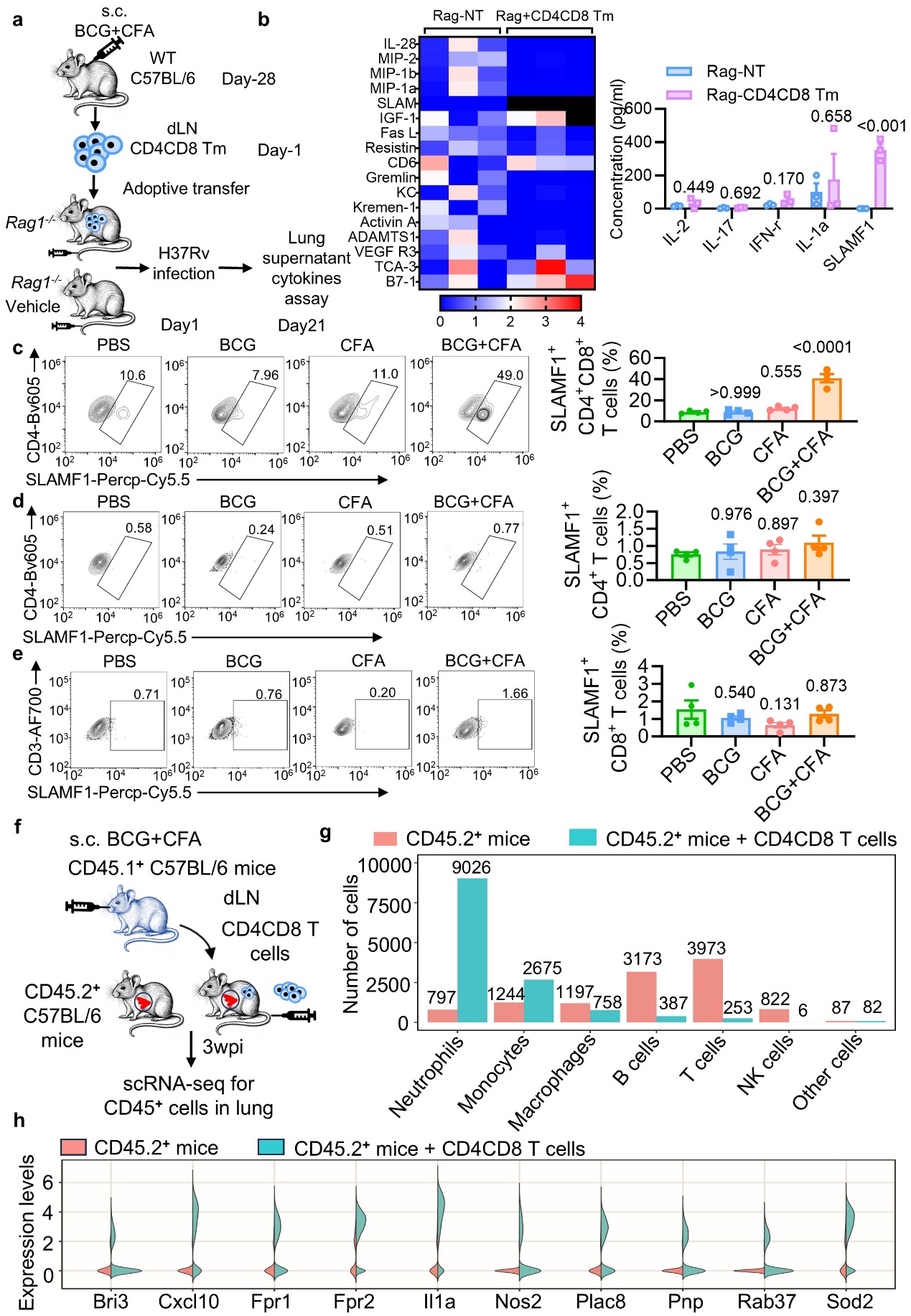

**Extended Data Fig. 7: BCG plus CFA induces** **SLAMF1-expressing CD4^+^CD8^+^ T cells.**

**a-b,** Experimental design **(a)**, heatmap showing the normalized expression of the indicated cytokines in the lung supernatants of PBS- or memory CD4^+^CD8^+^ T cells (CD4CD8 Tm) adoptive transferred *rag1*^-/-^ mice after three weeks post H37Rv infection. Results were analyzed with the Raybiotech 200 mouse cytokine array (three independent experiments), black indicates values above the maximum in the heatmap. Absolute quantification statistical of indicate cytokine concentrations is summarized beside **(b)**. **c-e,** Proportions of SLAMF1^+^ T cells in CD4^+^CD8^+^ T cells **(c)**, CD4^+^ T cells **(d)**, CD8^+^ T cells **(e)** in lung of H37Rv infected mice of each vaccine group, as determined by FCM. **f-g**, Experimental design **(f),** CD4⁺CD8⁺ T cells were sorted from lung lymph nodes of CD45.1^+^ C57BL/6 donor mice vaccinated with BCG plus CFA at 4 weeks post‑vaccination. These cells were then transferred into CD45.2^+^ C57BL/6 recipient mice. Recipient mice and wild type CD45.2^+^ C57BL/6 mice were subsequently aerosol‑infected with ~200 CFU per mouse of *Mtb* H37Rv. At 3 weeks post‑infection, lung tissue from two groups of mice was collected for single-cell sequencing. The number of different cell types in the lung tissue of the two groups of mice **(g). h,** Violin plots comparing gene expression levels in neutrophil cell types between the two groups of mice in **(f)**. The Y-axis represents log-normalised expression values, and the width of each violin indicates the density of cells at each expression level. Data represent one experiment with at least three independent biological replicates and are shown as the mean ± s.e.m.. Statistical significance in **(b)** was assessed by Multiple unpaired t tests. Statistical significance in **(c-e)** was assessed by one-way ANOVA with Dunn’s multiple comparison test comparing each group to the PBS group.

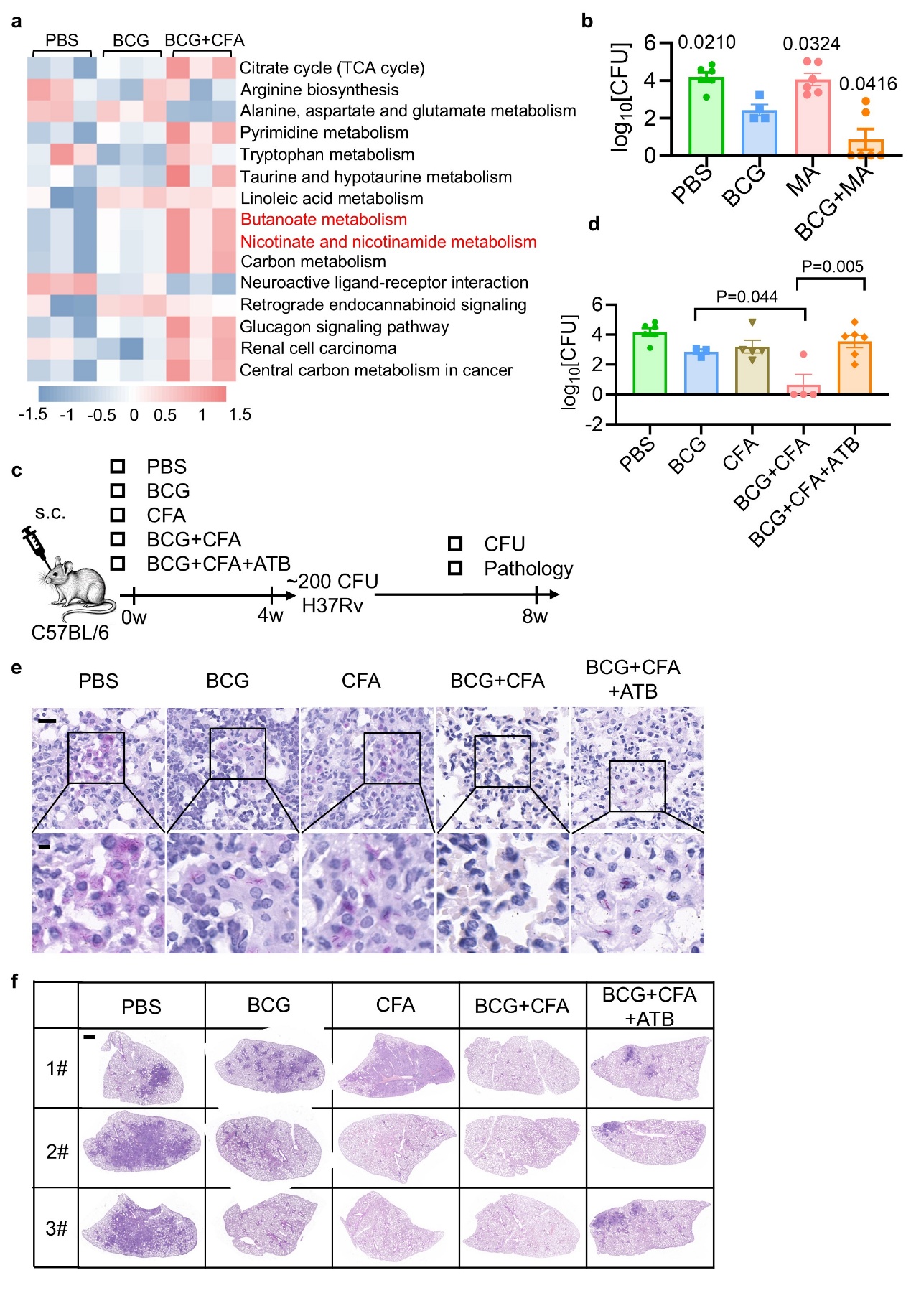

**Extended Data Fig. 8: Gut microbiota mediates vaccine protection via maleic acid**

**a**, Metabolomics sequencing was performed on serum metabolites from mice immunized for 4 weeks with PBS, BCG, or BCG+CFA. Enrichment analysis of significantly differentially secreted metabolites was then conducted. **b**, C57BL/6 mice from each vaccine group were aerosol-infected with ~200 CFU per mouse of *Mtb* H37Rv. At 4 weeks post-infection, bacterial load in the spleen was analyzed. **c-f**, Experimental design of the TB preventive vaccine model **(c)**. Mice received subcutaneous vaccination and ABX cocktail treatment, then were challenged via aerosol with ~200 CFU of *Mtb* H37Rv. At 4 weeks post‑infection, the following parameters were assessed: bacterial load in the spleen **(d)**; acid‑fast staining (**e**; scale bar, 20 μm (top) and 5 μm (bottom)); and histopathology of lung sections via H&E staining (**f**; scale bar, 1 mm). Data in **b, d,** represent one experiment with at least three independent biological replicates; mean±SEM. Ordinary one-way ANOVA with Dunnett’s multiple comparison test comparing each group to the BCG group **(b)**, and two-tailed unpaired Student’s t-tests **(d)** were used for statistical analyses.

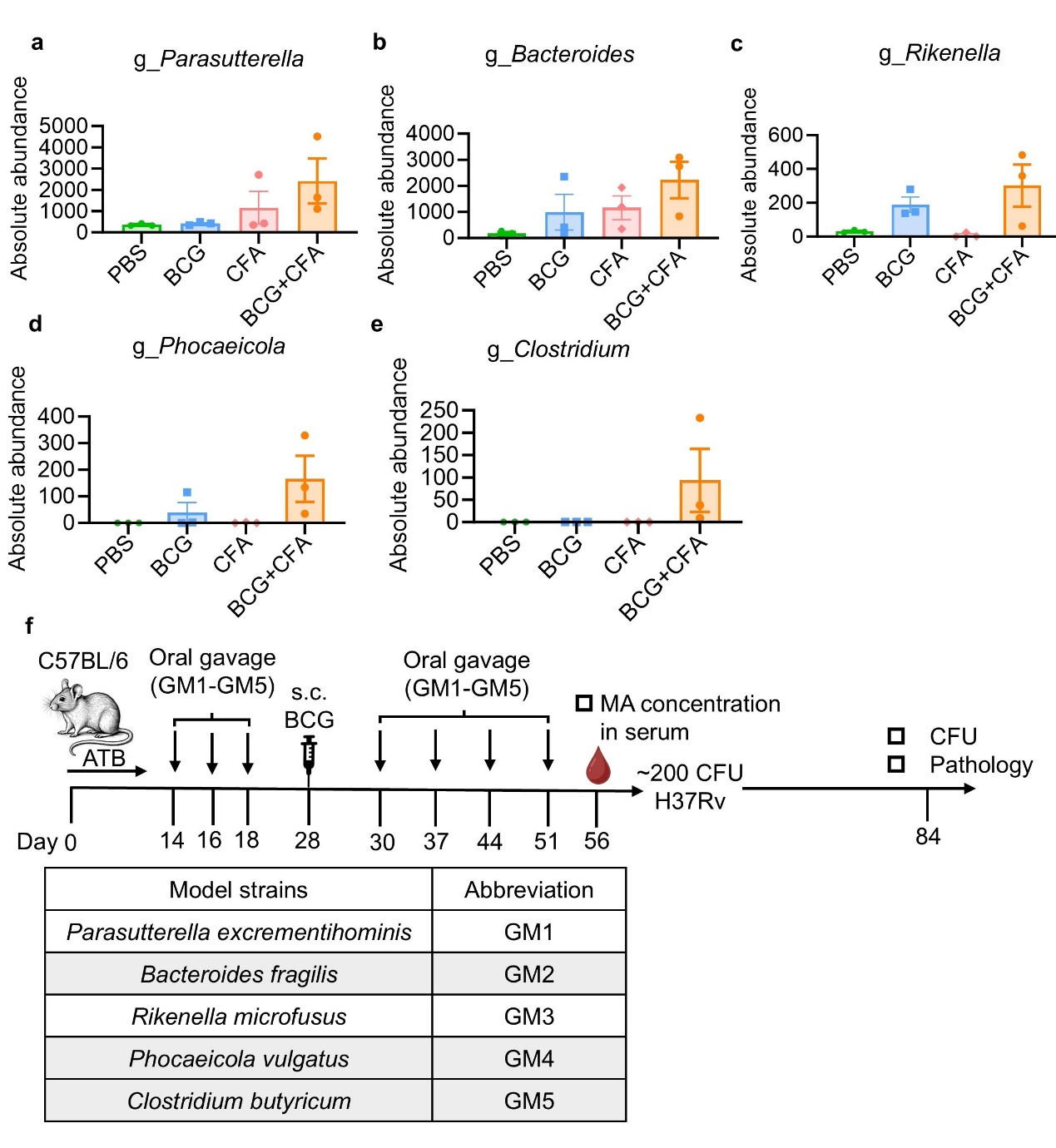

**Extended Data Fig. 9: Maleic acid is derived from *Clostridium butyricum***

**a-e**, Abundances of selected bacterial taxa. **f**, Experimental scheme of the TB preventive vaccine with bacterial transplantation model.

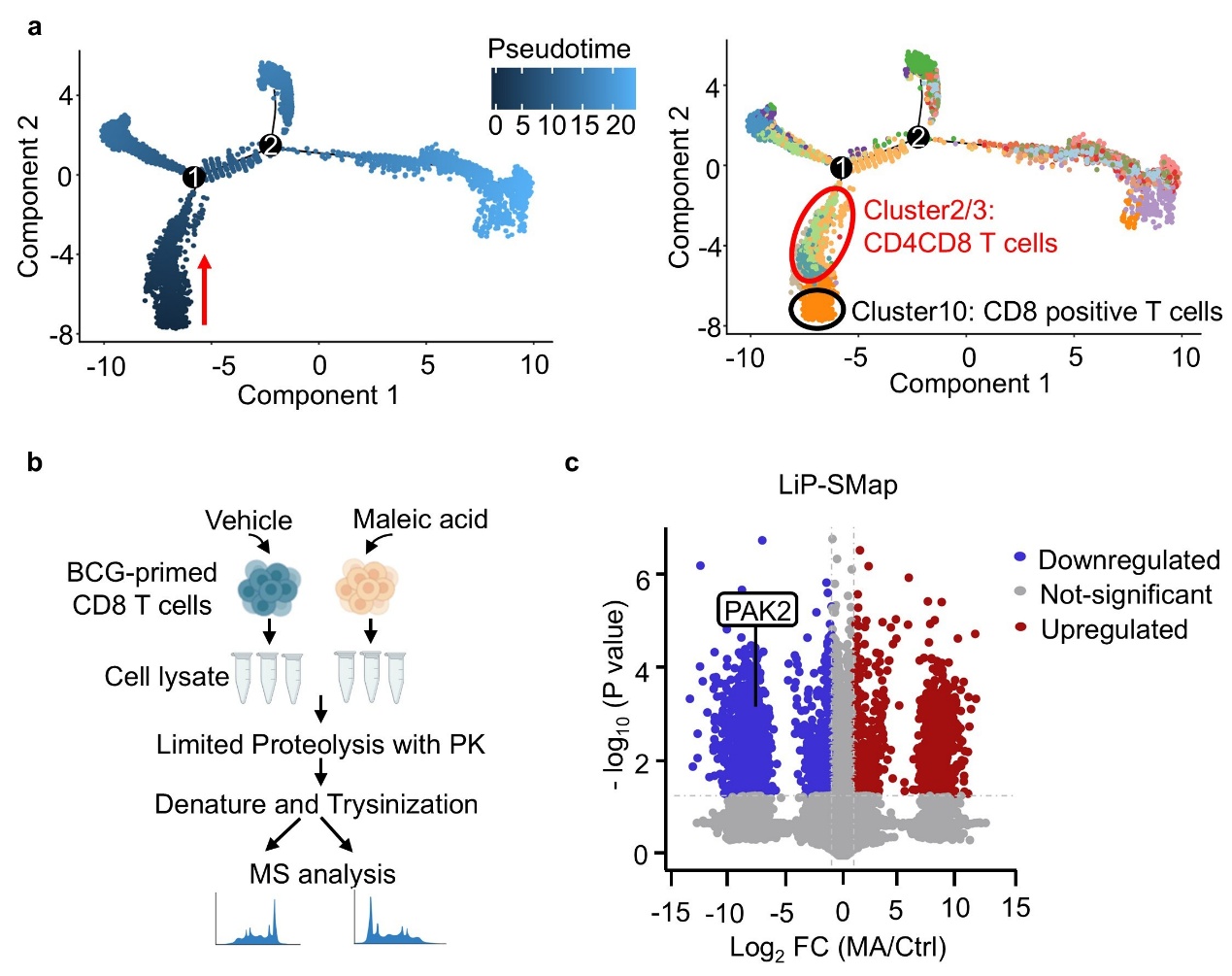

**Extended Data Fig. 10: Maleic acid interacts with PAK2**

**a,** Monocle pseudotime trajectory across all clusters in **Extended Data Fig. 5a**. Cells are labeled by pseudotime (left) and cell subtypes (right). **b.** Scheme of chemical proteomics for target identification. **c**. Volcano plots of LiP-SMap experiments with maleic acid treatment.

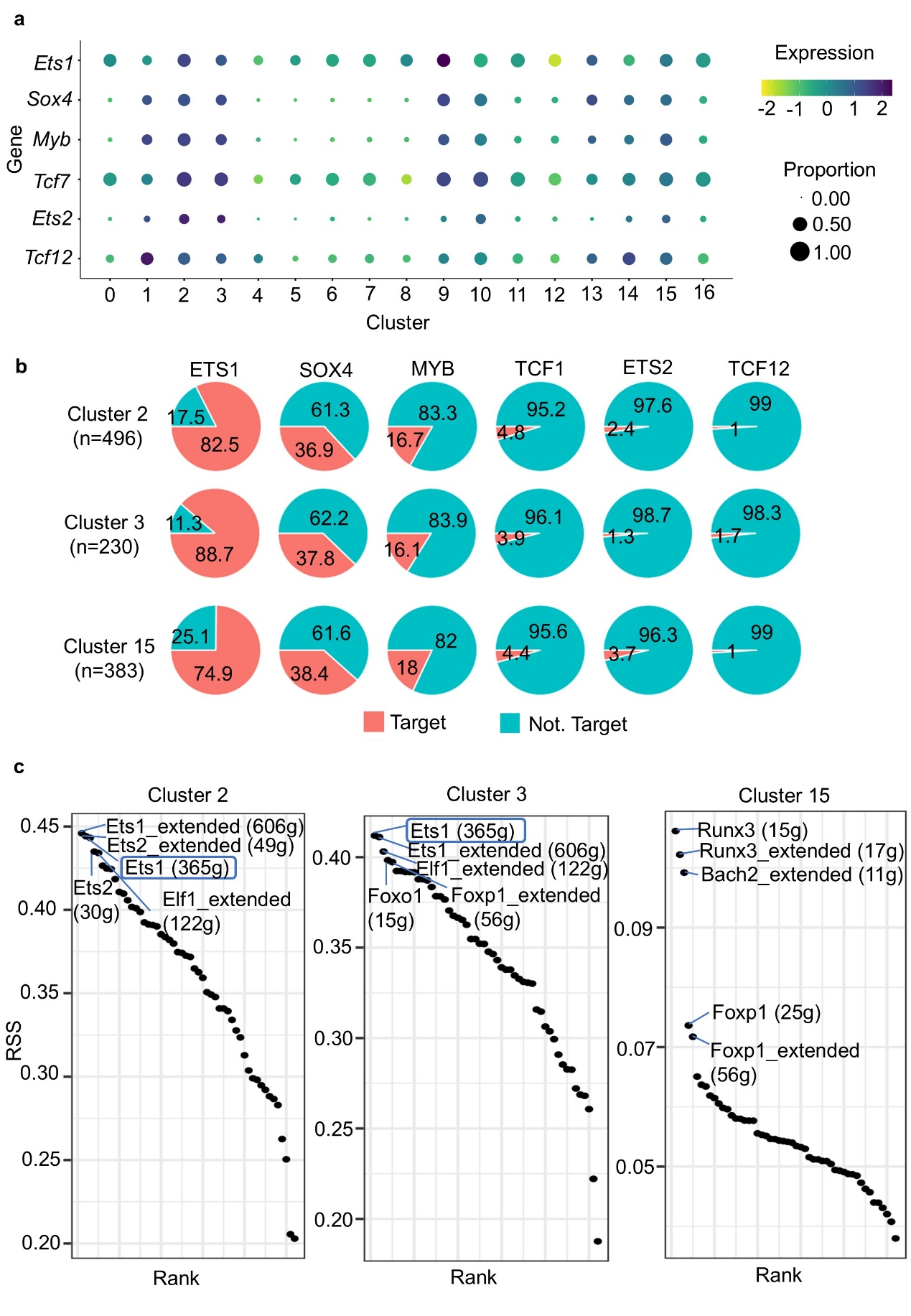

**Extended Data Fig. 11: Ets1 is central to the development of CD4⁺CD8⁺ T cell**

**a**, Heatmap showing the relative average expression of a subset of genes (y axis) across the clusters (x axis). As indicated on the legend, dot size denotes the percentage of cells in a cluster expressing each gene. Dot color represents the relative average expression levels. **b,** Conjoint analysis of CUT&Tag assay and Single cell sequencing for each TF regulating the proportion of CD4^+^CD8^+^ T cell (cluster2, 3, 15) marker gene transcription. **c**, Dot plot showing regulon specificity scores (RSS) in CD4^+^CD8^+^ T cell (cluster2, 3, 15) by Single-cell regulatory network inference and clustering (SCENIC) analysis.

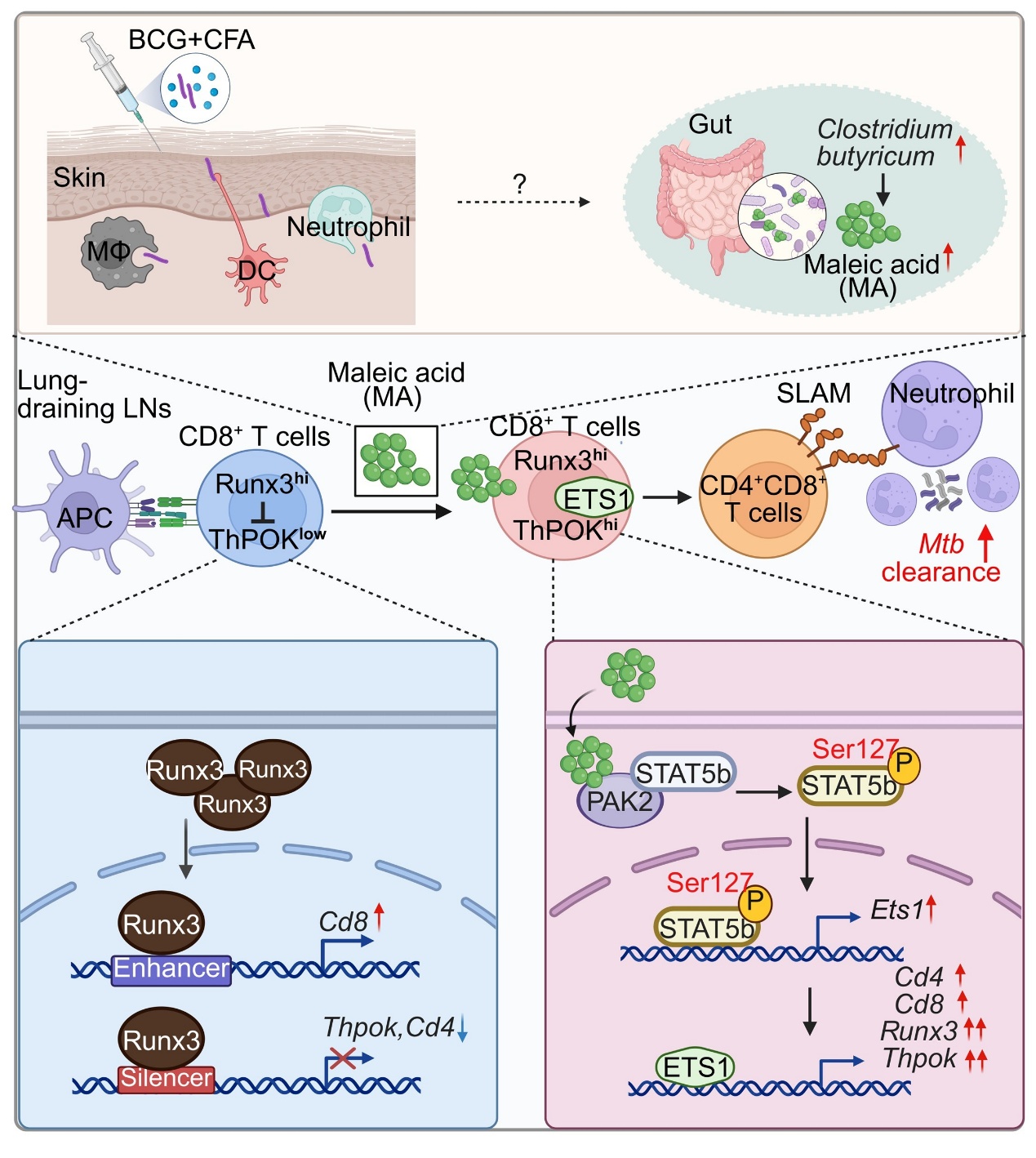

**Extended Data Fig. 12: Summary diagram**

Subcutaneous BCG+CFA immunization promotes gut *Clostridium butyricum* abundance, leading to increased production of maleic acid. Maleic acid directly interacts with PAK2, promoting STAT5 phosphorylation on Ser127 and subsequent upregulation of Ets1 transcription. Ets1 directly binds to the *Runx3* and *Zbtb7b* (encoding ThPOK) loci, upregulating both Runx3 and ThPOK to override their lineage‑defining antagonism, thereby promoting CD4⁺CD8⁺ T cells differentiation. CD4⁺CD8⁺ T cells highly express SLAMF1, which activates host neutrophils and further contributes to *Mtb* clearance.
