## Supplementary material for "Maleic acid adjuvates BCG to induce CD4^+^CD8^+^ T cells and sterilize tuberculosis": Table S2

| | | | | | | | X.PEP.Label-Free Quant | | | | | |
|---|---|---|---|---|---|---|---|---|---|---|---|---|
| PG.Genes | PEP.StrippedSequence | PEP.StartingPositions | PEP.EndingPositions | PEP.NrOfMissedCleavages | PEP.QValue | PEP.PeptideLength | MA1 | MA2 | MA3 | NT1 | NT2 | NT3 |
| Naca | NILFVITKPDVYK | (2073);(101) | (2085);(113) | 1 | 0.000431655 | 13 | 3073.5295 | 3510.6785 | 3585.9897 | 2604.9114 | 3316.8984 | 3545.8118 |
| Naca | FVITKPDVYK | (2076);(104) | (2085);(113) | 1 | 0.000907206 | 10 | 1136.9294 | 1261.9191 | 1315.3112 | 2721.1282 | 2529.067 | 2475.3745 |
| Naca | AAAEIDEEPVSK | (2027);(55) | (2038);(66) | 0 | 0 | 12 | 1057.4084 | 887.85095 | 842.4646 | 881.2449 | 1293.0388 | 1313.1454 |
| Naca | PGEATETVPATEQELPQPQ | (2);(2) | (20);(20) | 0 | 0.002972566 | 19 | 618.38763 | 598.2701 | 614.97925 | 1057.9215 | 1219.7917 | 1223.0801 |
| Naca | VQGEAVSNIQENTQTPTVQ | (2117);(145) | (2135);(163) | 0 | 0.002265552 | 19 | 624.72485 | 707.379 | 591.97723 | NaN | NaN | NaN |
| Naca | SEEEEVDETGVEVK | (2138);(166) | (2151);(179) | 0 | 0.00078661 | 14 | 612.17914 | 450.8213 | 522.9147 | NaN | NaN | NaN |
| Naca | AEIDEEPVSK | (2029);(57) | (2038);(66) | 0 | 0.005946382 | 10 | 1168.0084 | NaN | NaN | NaN | NaN | 1429.8538 |
| Mcm6 | ELRDEEQTAESIK | -314 | -326 | 1 | 0.00025015 | 13 | 6616.5796 | 4491.098 | 5187.888 | 5416.387 | 2860.8225 | 3653.9119 |
| Mcm6 | YTQPNICR | -174 | -181 | 0 | 0 | 8 | 3297.0078 | 4099.362 | 2036.609 | 3530.1484 | 4186.6025 | 2059.334 |
| Mcm6 | LSTTIQEEFYR | -75 | -85 | 0 | 0 | 11 | 2424.9912 | 2578.857 | 2087.723 | 2206.562 | 1058.621 | 1018.00543 |
| Mcm6 | HVDEFSPR | -408 | -415 | 0 | 0 | 8 | 2178.5999 | 2199.9585 | 2273.4053 | 1818.7457 | 2245.3413 | 791.0075 |
| Mcm6 | AEPGAGSQHPEVR | -7 | -19 | 0 | 0 | 13 | 1606.8264 | 1746.2842 | 2076.5908 | 86.62462 | 1950.809 | 2122.5037 |
| Mcm6 | LFPTIHGNDEVK | -354 | -365 | 0 | 0 | 12 | 1514.9552 | 1544.1232 | 1549.4459 | 1637.7114 | 1472.9436 | 1577.1193 |
| Mcm6 | FLEEFQGSDGEIK | -33 | -45 | 0 | 0 | 13 | 1505.1987 | 661.97174 | 955.27856 | 431.79077 | 1222.4543 | 1173.4211 |
| Mcm6 | EIESEIDSEEELINKK | -755 | -770 | 1 | 0 | 16 | 1023.83875 | 1202.9614 | 1246.8992 | 1182.4806 | 1838.7828 | 1698.7303 |
| Mcm6 | EEEEDESALKR | -735 | -745 | 1 | 0 | 11 | 888.6936 | 944.6739 | 1191.6674 | 929.4609 | 1517.5531 | 1474.8444 |
| Mcm6 | THPVHPELVSGTFL | -144 | -157 | 0 | 0 | 14 | 793.4778 | 657.6941 | 943.459 | 1192.333 | 986.2532 | 1040.1885 |
| Mcm6 | THPVHPELVSGTFL | -144 | -157 | 0 | 0 | 14 | 793.4778 | 657.6941 | 943.459 | 1192.333 | 986.2532 | 1040.1885 |
| Mcm6 | AAAEPGAGSQHPEVR | -5 | -19 | 0 | 0 | 15 | 582.4403 | 758.6723 | 709.04034 | 772.284 | 765.2646 | 649.5759 |
| Mcm6 | THPVHPELVSGT | -144 | -155 | 0 | 0.00025015 | 12 | 479.29526 | 154.78035 | 1356.8691 | 1837.4263 | 1289.0898 | 1376.3021 |
| Mcm6 | THPVHPELVSGT | -144 | -155 | 0 | 0.00025015 | 12 | 479.29526 | 154.78035 | 1356.8691 | 1837.4263 | 1289.0898 | 1376.3021 |
| Mcm6 | GHADSPAPVNR | -685 | -695 | 0 | 0 | 11 | 210.36893 | 1588.2932 | 1751.0997 | 959.3485 | 1423.1562 | 1446.6931 |
| Mcm6 | IVDLHSR | -552 | -558 | 0 | 0.000627943 | 7 | 1543.5677 | NaN | 793.0861 | 1455.4932 | 1153.8971 | 1195.0911 |
| Mcm6 | VYSLDDIR | -566 | -573 | 0 | 0 | 8 | 1047.9623 | 1002.1445 | NaN | 539.27606 | 913.29803 | 532.17633 |
| Mcm6 | TVSKPSLR | -707 | -714 | 1 | 0 | 8 | 850.085 | 349.43384 | NaN | 307.3386 | 615.6582 | 599.07886 |
| Mcm6 | THPVHPELVSGTF | -144 | -156 | 0 | 0 | 13 | 837.3487 | 802.41974 | 833.0319 | 574.243 | 824.56226 | NaN |
| Mcm6 | FNGSSEDASQETVSKPSLR | -696 | -714 | 1 | 0.000230707 | 19 | 468.92786 | 384.54938 | 412.18524 | NaN | 583.1233 | 700.33954 |
| Mcm6 | GSSEGSESYEEDPYLVVNPN | -797 | -816 | 0 | 0 | 20 | 327.1429 | NaN | 257.62976 | 447.3846 | 236.839 | 282.6928 |
| Mcm6 | RIVDLHSR | -551 | -558 | 1 | 0.000431655 | 8 | 244.60773 | 221.13435 | 260.97284 | NaN | 61.49812 | 299.92703 |
| Mcm6 | LIVVPDVSK | -248 | -256 | 0 | 0 | 9 | 1230.4727 | 1023.565 | 1104.2031 | 814.3698 | NaN | NaN |
| Mcm6 | STTIQEEFYR | -76 | -85 | 0 | 0.002595029 | 10 | 1108.825 | 1390.8048 | NaN | NaN | 1195.3345 | 1350.7765 |
| Mcm6 | FQDLPTR | -114 | -120 | 0 | 0.00122419 | 7 | 863.6687 | NaN | 825.66223 | NaN | 740.02625 | 402.06058 |
| Mcm6 | AAANPVSGHYDR | -501 | -512 | 0 | 0 | 12 | 647.04364 | 1513.3486 | 1275.4646 | 1099.5253 | NaN | NaN |
| Mcm6 | DEESHEFVIE | -436 | -445 | 0 | 0.003133457 | 10 | 512.0894 | 640.7568 | NaN | 558.155 | NaN | 498.2842 |
| Mcm6 | SELVNWYLK | -746 | -754 | 0 | 0 | 9 | 803.65216 | 93.115585 | 985.3406 | NaN | NaN | NaN |
| Mcm6 | RSELVNWYLK | -745 | -754 | 1 | 0.000550307 | 10 | 509.16074 | 554.4752 | 608.57495 | NaN | NaN | NaN |
| Mcm6 | IQEEFYR | -79 | -85 | 0 | 0.000826806 | 7 | 452.94095 | 217.30188 | NaN | 484.62146 | NaN | NaN |
| Mcm6 | EIESEIDSEEELINK | -755 | -769 | 0 | 0 | 15 | 378.69217 | 346.44666 | NaN | NaN | 597.00574 | NaN |
| Fh | QFDEWVKPK | -493 | -501 | 1 | 0.00025015 | 9 | 5768.093 | 6692.837 | 5744.9395 | 6927.536 | 159.35025 | 6464.138 |
| Fh | ALNPHIGYDK | -455 | -464 | 0 | 0 | 10 | 1928.5597 | 1360.0109 | 1314.1897 | 1493.4039 | 841.6289 | 1355.1265 |
| Fh | LNDHFPLVVW | -129 | -138 | 0 | 0 | 10 | 1487.4216 | 1686.5302 | 1804.5629 | 2065.54 | 2402.5808 | 2246.3225 |
| Fh | AAAEVNQEYGLDPK | -99 | -112 | 0 | 0 | 14 | 1192.7838 | 1292.4299 | 1252.7401 | 998.88 | 978.528 | 1113.5151 |
| Fh | AAEVNQEYGLDPK | -100 | -112 | 0 | 0 | 13 | 674.6423 | 683.6174 | 623.903 | 952.7212 | 1399.0093 | 1184.6255 |
| Fh | ALTGLPFVTAPNK | -296 | -308 | 0 | 0 | 13 | 650.15356 | 370.82028 | 553.29535 | 495.0139 | 611.24786 | 644.64514 |
| Fh | SGLGELILPENEPGSSIMPGK | -348 | -368 | 0 | 0 | 21 | 439.75507 | 503.37732 | 480.19922 | 574.8824 | 813.568 | 753.3591 |
| Fh | SGLGELILPENEPGSSIMPGK | -348 | -368 | 0 | 0 | 21 | 439.75507 | 503.37732 | 480.19922 | 574.8824 | 813.568 | 753.3591 |
| Fh | KVLLPGLQK | -202 | -210 | 1 | 0.000550307 | 9 | 320.9837 | NaN | 369.9041 | 501.97278 | 537.8042 | 489.38016 |
| Fh | AEQFDEWVKPK | -491 | -501 | 1 | 0 | 11 | 3635.9568 | 3587.1277 | NaN | NaN | 4679.2744 | 4543.518 |
| Fh | LHDALSAK | -211 | -218 | 0 | 0 | 8 | 759.7894 | NaN | 1283.977 | NaN | 618.7778 | 689.17676 |
| Fh | VLLPGLQK | -203 | -210 | 0 | 0.00025015 | 8 | 102.50996 | 237.36377 | 336.67072 | 67.95374 | NaN | NaN |
| Fh | FDTFGELK | -51 | -58 | 0 | 0.002466863 | 8 | 616.0298 | NaN | NaN | NaN | NaN | NaN |
| H1-3 | SETAPAAPAAPAPVEK | -2 | -17 | 0 | 0 | 16 | 1988.3462 | 2221.1433 | 2307.9648 | 2154.648 | 2162.4424 | 2077.5264 |
| H1-3 | SETAPAAPAAPAPVEK | -2 | -17 | 0 | 0 | 16 | 1988.3462 | 2221.1433 | 2307.9648 | 2154.648 | 2162.4424 | 2077.5264 |
| H1-3 | TAPAAPAAPAPVEK | -4 | -17 | 0 | 0.00025015 | 14 | 1342.118 | 1276.4672 | 1254.6362 | 1270.5892 | 828.0883 | 996.859 |
| H1-3 | SETAPAAPAAPAPVEKTPVK | -2 | -21 | 1 | 0 | 20 | 452.17773 | 446.76974 | 544.11304 | 562.2554 | 613.94196 | 599.04956 |
| H1-3 | APAAPAPVEK | -8 | -17 | 0 | 0.00111006 | 10 | 946.9937 | NaN | 901.0178 | 1073.47 | 1219.7632 | 1069.4191 |
| H1-3 | PAAPAPVEK | -9 | -17 | 0 | 0.002502116 | 9 | 775.3193 | 1112.4567 | 1063.7179 | 1242.8043 | NaN | NaN |
| Mcm3 | HDSLLHGT | -548 | -555 | 0 | 0 | 8 | 4528.81 | 3652.401 | 3556.862 | 420.06967 | 864.1986 | 426.5732 |
| Mcm3 | ELISDNQYR | -38 | -46 | 0 | 0 | 9 | 4071.7114 | 3958.8904 | 3725.203 | 1340.231 | 2011.3494 | 311.02332 |
| Mcm3 | AGTVVLDDVELR | -2 | -13 | 0 | 0 | 12 | 1746.2885 | 1811.6632 | 1797.2831 | 375.26837 | 2176.5684 | 2323.071 |
| Mcm3 | TAIHEVMEQGR | -420 | -430 | 0 | 0 | 11 | 1704.7766 | 1709.5791 | 1813.7698 | 4802.753 | 281.091 | 754.3271 |
| Mcm3 | EISDHVLR | -496 | -503 | 0 | 0 | 8 | 1502.954 | 1303.2682 | 1469.4934 | 1775.7966 | 1520.5863 | 1698.9675 |
| Mcm3 | NREEPFSSEEIQAC | -779 | -792 | 1 | 0.00025015 | 14 | 1497.9677 | 1326.7239 | 1336.1274 | 280.1308 | 1252.7239 | 1202.5098 |
| Mcm3 | ATDDPDFTQDDQQDTR | -528 | -543 | 0 | 0 | 16 | 1274.7734 | 1362.8169 | 1417.8396 | 1647.4117 | 1727.178 | 1755.3735 |
| Mcm3 | TPMENIGLQDSLLSR | -464 | -478 | 0 | 0 | 15 | 1209.6282 | 1341.4043 | 1377.4392 | 1443.5034 | 1494.9307 | 1557.259 |
| Mcm3 | TPMENIGLQDSLLSR | -464 | -478 | 0 | 0 | 15 | 1209.6282 | 1341.4043 | 1377.4392 | 1443.5034 | 1494.9307 | 1557.259 |
| Mcm3 | VVCVEGIVTK | -124 | -133 | 0 | 0 | 10 | 1012.12787 | 860.30206 | 761.0934 | 537.0203 | 576.1621 | 420.15024 |
| Mcm3 | APSIHGHDYVK | -304 | -314 | 0 | 0 | 11 | 476.28946 | 648.9341 | 787.9334 | 671.6258 | 615.5785 | 515.2868 |
| Mcm3 | APSIHGHDYVK | -304 | -314 | 0 | 0 | 11 | 476.28946 | 648.9341 | 787.9334 | 671.6258 | 615.5785 | 515.2868 |
| Mcm3 | DIQPAFSADDIAK | -271 | -283 | 0 | 0 | 13 | 722.014 | 789.00104 | 817.9205 | 583.2745 | 701.3688 | 1097.1547 |
| Mcm3 | AVTTDQETGER | -381 | -391 | 0 | 0 | 11 | 533.21063 | 616.12915 | 418.46213 | 713.8038 | 810.94226 | 687.63745 |
| Mcm3 | REISDHVLR | -495 | -503 | 1 | 0.000471409 | 9 | 372.3966 | 452.37753 | 632.3561 | 522.2843 | 153.3735 | 614.55597 |
| Mcm3 | LDQMDPEQDR | -486 | -495 | 0 | 0 | 10 | 297.69812 | 853.9281 | 791.0309 | 295.51346 | 992.88654 | 1815.8455 |
| Mcm3 | DYLDFLDDEEDQGIYQNK | -18 | -35 | 0 | 0 | 18 | 287.88748 | 314.68274 | 190.3133 | 281.36905 | 289.62695 | 219.52475 |
| Mcm3 | SLAPSIHGHDYVK | -302 | -314 | 0 | 0 | 13 | 1543.4685 | 1602.8838 | 1208.4546 | 849.159 | NaN | 1273.2598 |
| Mcm3 | APAGQLPR | -208 | -215 | 0 | 0.001868731 | 8 | 1454.6729 | 460.9605 | NaN | 868.92566 | 758.82336 | 677.6553 |

| | | | | | | | | | | | | |
|---|---|---|---|---|---|---|---|---|---|---|---|---|
| Mcm3 | PAGQLPR | -209 | -215 | 0 | 0.000230707 | 7 | 698.3629 | 624.4617 | 678.36255 | 473.2833 | 546.545 | NaN |
| Mcm3 | SVHYCPATK | -144 | -152 | 0 | 0.00078661 | 9 | 688.3074 | 731.6329 | 638.2741 | NaN | NaN | 561.5119 |
| Mcm3 | SVDVILDDDLVDK | -216 | -228 | 0 | 0 | 13 | 3088.384 | NaN | NaN | 860.68915 | NaN | 1234.0747 |
| Mcm3 | FLDDEEDQGIYQNK | -22 | -35 | 0 | 0.00025015 | 14 | 842.20654 | 555.6383 | NaN | NaN | NaN | NaN |
| Mcm3 | AAANPVYGR | -450 | -458 | 0 | 0.000947459 | 9 | 644.54425 | 124.29126 | NaN | NaN | NaN | NaN |
| Mcm3 | DDPDFTQDDQQDTR | -530 | -543 | 0 | 0.001092529 | 14 | 585.66815 | NaN | NaN | 422.1043 | NaN | NaN |
| Mcm3 | YLDFLDDEEDQGIYQNK | -19 | -35 | 0 | 0.001092529 | 17 | 498.9693 | NaN | NaN | NaN | NaN | 568.8098 |
| Mcm3 | IIKPTLTQESAAY | -577 | -589 | 1 | 0 | 13 | 347.97546 | NaN | NaN | 435.54123 | NaN | NaN |
| Cox20 | AAAPEPHETEK | -2 | -12 | 0 | 0.000230707 | 11 | 447.90753 | 451.8321 | NaN | NaN | 536.7637 | 691.8145 |
| Ddx20 | AAAPVQAVEPTPASPWTQR | -24 | -42 | 0 | 0 | 19 | 697.6963 | 528.04016 | 711.1077 | 635.44617 | 395.56772 | 944.14453 |
| Hnrnpd | IFVGGLSPDTPEEK | -184 | -197 | 0 | 0 | 14 | 5528.608 | 5523.2686 | 5831.6836 | 4379.646 | 4196.155 | 4346.685 |
| Hnrnpd | YHNVGLSK | -244 | -251 | 0 | 0 | 8 | 2525.782 | 2653.5564 | 2549.9387 | 2426.8186 | 2554.1523 | 2472.0825 |
| Hnrnpd | ITFKEEEPVKK | -228 | -238 | 2 | 0 | 11 | 1554.7422 | 569.1011 | 1372.3419 | 1012.61096 | 1086.8431 | 1302.4028 |
| Hnrnpd | ITFKEEEPVKK | -228 | -238 | 2 | 0 | 11 | 1554.7422 | 569.1011 | 1372.3419 | 1012.61096 | 1086.8431 | 1302.4028 |
| Hnrnpd | FKEEEPVKK | -230 | -238 | 2 | 0 | 9 | 2195.443 | 2264.7166 | 2240.0942 | 2404.464 | 2588.494 | 2645.6807 |
| Hnrnpd | FKEEEPVKK | -230 | -238 | 2 | 0 | 9 | 2195.443 | 2264.7166 | 2240.0942 | 2404.464 | 2588.494 | 2645.6807 |
| Hnrnpd | KLDPITGR | -129 | -136 | 1 | 0 | 8 | 1758.7872 | 1654.7067 | 1621.5856 | 1109.533 | 1373.3593 | 1760.5913 |
| Hnrnpd | FKESESVDK | -145 | -153 | 1 | 0 | 9 | 1556.1078 | 1596.8904 | 2148.2131 | 2183.623 | 2283.4158 | 2480.4592 |
| Hnrnpd | FKEEEPVK | -230 | -237 | 1 | 0 | 8 | 1343.5743 | 1088.106 | 1420.7498 | 1062.0164 | 476.60886 | 1414.4602 |
| Hnrnpd | RGFCFITF | -223 | -230 | 1 | 0.000591124 | 8 | 521.5561 | 125.02362 | 567.7648 | 560.75433 | 530.8288 | 498.30606 |
| Hnrnpd | KYHNVGLS | -243 | -250 | 1 | 0.002265552 | 8 | 1501.1967 | 2110.055 | 2612.381 | 2288.2297 | NaN | 3257.329 |
| Hnrnpd | KYHNVGLSK | -243 | -251 | 1 | 0.00025015 | 9 | 678.31616 | 804.00507 | 1179.9604 | 802.05493 | NaN | 521.1375 |
| Hnrnpd | KYHNVGLSK | -243 | -251 | 1 | 0.00025015 | 9 | 678.31616 | 804.00507 | 1179.9604 | 802.05493 | NaN | 521.1375 |
| Hnrnpd | NEEDEGHSNSSPR | -73 | -85 | 0 | 0.00078661 | 13 | 654.8725 | 581.8827 | 563.93304 | NaN | 16.645004 | 328.4793 |
| Hnrnpd | GSGSGGGGSAAGGTEGGSAEAEGAK | -43 | -67 | 0 | 0.001392869 | 25 | 365.02954 | NaN | 331.14856 | NaN | 169.10141 | 325.05725 |
| Hnrnpd | AAQGPAAAAGSGSGGGGSAAGGTEGGSAEAEGAK | -34 | -67 | 0 | 0.00078661 | 34 | 353.91855 | 449.2356 | 368.7704 | NaN | NaN | NaN |
| Hnrnpd | EYFGGFGEVE | -200 | -209 | 0 | 0.000627943 | 10 | 838.4987 | NaN | NaN | 1179.5845 | NaN | NaN |
| Hnrnpd | AAAQGPAAAAGSGSGGGGSAAGGTEGGSAEAEGA | -33 | -67 | 0 | 0.001846793 | 35 | 308.34195 | NaN | 346.51584 | NaN | NaN | NaN |
| Lcp1 | IKVPVDWNR | -433 | -441 | 1 | 0 | 9 | 12812.617 | 14510.097 | 15625.541 | 14605.65 | 12157.624 | 12693.39 |
| Lcp1 | IKVPVDWNR | -433 | -441 | 1 | 0 | 9 | 12812.617 | 14510.097 | 15625.541 | 14605.65 | 12157.624 | 12693.39 |
| Lcp1 | ALPEDLVEVNPK | -599 | -610 | 0 | 0 | 12 | 9918.392 | 9373.058 | 11214.096 | 11250.251 | 8686.648 | 9116.98 |
| Lcp1 | ALPEDLVEVNPK | -599 | -610 | 0 | 0 | 12 | 9918.392 | 9373.058 | 11214.096 | 11250.251 | 8686.648 | 9116.98 |
| Lcp1 | AACLPLPGYR | -40 | -49 | 0 | 0.00025015 | 10 | 8336.044 | 8630.511 | 9041.08 | 8166.67 | 8697.713 | 8596.365 |
| Lcp1 | HVVNIGAEDLK | -207 | -217 | 0 | 0 | 11 | 6789.803 | 7056.15 | 7186.955 | 8496.898 | 8237.332 | 7888.794 |
| Lcp1 | VPVDWNR | -435 | -441 | 0 | 0.001536401 | 7 | 6651.37 | 6534.986 | 5808.3555 | 5502.3545 | 5246.4287 | 5093.393 |
| Lcp1 | ISFDEFIK | -69 | -76 | 0 | 0 | 8 | 5100.636 | 6050.2856 | 6069.697 | 5974.822 | 7997.2646 | 8097.843 |
| Lcp1 | WANYHLENAGCTK | -273 | -285 | 0 | 0 | 13 | 3769.8958 | 4334.7505 | 4485.6597 | 5837.1406 | 6304.5757 | 6091.4175 |
| Lcp1 | WANYHLENAGCTK | -273 | -285 | 0 | 0 | 13 | 3769.8958 | 4334.7505 | 4485.6597 | 5837.1406 | 6304.5757 | 6091.4175 |
| Lcp1 | NLDDEEKLNNAK | -573 | -584 | 1 | 0.000712251 | 12 | 3367.9307 | 3648.602 | 3810.209 | 3422.9846 | 3738.4004 | 3829.8923 |
| Lcp1 | ENCNYAVDLGK | -458 | -468 | 0 | 0 | 11 | 2932.4167 | 3025.4553 | 1110.5466 | 945.25934 | 3795.2266 | 3785.72 |
| Lcp1 | ENCNYAVDLGK | -458 | -468 | 0 | 0 | 11 | 2932.4167 | 3025.4553 | 1110.5466 | 945.25934 | 3795.2266 | 3785.72 |
| Lcp1 | HVIPMNPNTDDLFNAV | -142 | -157 | 0 | 0 | 16 | 2814.456 | 2733.4868 | 2785.9504 | 2863.9158 | 2208.059 | 1692.8365 |
| Lcp1 | HVIPMNPNTDDLFNAV | -142 | -157 | 0 | 0 | 16 | 2814.456 | 2733.4868 | 2785.9504 | 2863.9158 | 2208.059 | 1692.8365 |
| Lcp1 | TENLDDEEKLNNAK | -571 | -584 | 1 | 0 | 14 | 2535.498 | 2641.7214 | 2743.4048 | 1924.9513 | 2104.277 | 2134.548 |
| Lcp1 | GSVSDEEMMELR | -4 | -15 | 0 | 0 | 12 | 1229.2506 | 1230.7745 | 1196.3318 | 1342.4073 | 1260.5912 | 1188.0565 |
| Lcp1 | GSVSDEEMMELR | -4 | -15 | 0 | 0 | 12 | 1229.2506 | 1230.7745 | 1196.3318 | 1342.4073 | 1260.5912 | 1188.0565 |
| Lcp1 | EGESLEDLMK | -254 | -263 | 0 | 0 | 10 | 2293.626 | 2311.7798 | 2156.3064 | 2432.447 | 2129.7278 | 2177.008 |
| Lcp1 | EGESLEDLMK | -254 | -263 | 0 | 0 | 10 | 2293.626 | 2311.7798 | 2156.3064 | 2432.447 | 2129.7278 | 2177.008 |
| Lcp1 | CNELNDLFK | -31 | -39 | 0 | 0.000230707 | 9 | 2251.1038 | 2365.4622 | 2513.8052 | 2024.8442 | 2135.9873 | 2160.289 |
| Lcp1 | QFVTATDVVR | -348 | -357 | 0 | 0 | 10 | 1969.2358 | 1971.2998 | 2142.4653 | 1865.9365 | 2301.7144 | 2422.0056 |
| Lcp1 | YHLLEQVAPK | -300 | -309 | 0 | 0 | 10 | 1907.3365 | 1932.7491 | 3156.824 | 3775.8918 | 707.02954 | 3124.0925 |
| Lcp1 | GLLWQVIK | -226 | -233 | 0 | 0.00025015 | 8 | 1611.3745 | 1758.8439 | 1810.3579 | 1507.7001 | 1819.834 | 2046.8867 |
| Lcp1 | AYYHLLEQVAPK | -298 | -309 | 0 | 0 | 12 | 1565.861 | 2215.0286 | 1733.9452 | 984.4263 | 2218.1963 | 1922.3085 |
| Lcp1 | FSLVGIAGQDLNEGNR | -473 | -488 | 0 | 0 | 16 | 1480.7211 | 1630.1187 | 1751.939 | 2140.6917 | 2802.4253 | 2729.076 |
| Lcp1 | WANYHLENAGCT | -273 | -284 | 0 | 0 | 12 | 1435.9159 | 1504.822 | 1385.8145 | 1442.2971 | 1217.1451 | 1231.352 |
| Lcp1 | YPALHKPENQDIDWGALEGETR | -374 | -395 | 1 | 0 | 22 | 1405.5643 | 1358.8167 | 1436.8198 | 1565.3738 | 1883.0695 | 1831.4072 |
| Lcp1 | YPALHKPENQDIDWGALEGETR | -374 | -395 | 1 | 0 | 22 | 1405.5643 | 1358.8167 | 1436.8198 | 1565.3738 | 1883.0695 | 1831.4072 |
| Lcp1 | GQDLNEGNR | -480 | -488 | 0 | 0.00025015 | 9 | 1338.578 | 1325.101 | 1476.514 | 1316.8118 | 1243.5472 | 1284.3922 |
| Lcp1 | GDEEGIPAVVIDMSGLR | -310 | -326 | 0 | 0 | 17 | 1189.8053 | 759.9156 | 1409.4225 | 731.7924 | 921.6356 | 1109.487 |
| Lcp1 | GDEEGIPAVVIDMSGLR | -310 | -326 | 0 | 0 | 17 | 1189.8053 | 759.9156 | 1409.4225 | 731.7924 | 921.6356 | 1109.487 |
| Lcp1 | EGICAIGGTSEQSSVGTQH | -98 | -116 | 0 | 0 | 19 | 1173.3113 | 1314.8335 | 1446.8171 | 1803.3563 | 2259.2986 | 2386.782 |
| Lcp1 | EITENLMATGDLDQDGK | -52 | -68 | 0 | 0 | 17 | 1103.4194 | 1238.3657 | 1153.5485 | 1683.228 | 1852.5975 | 1832.8297 |
| Lcp1 | NEALIALLR | -245 | -253 | 0 | 0 | 9 | 991.02136 | 974.95905 | 1054.2517 | 2097.7673 | 2710.5916 | 2645.7456 |
| Lcp1 | GIAGQDLNEGNR | -477 | -488 | 0 | 0 | 12 | 887.05225 | 616.4747 | 599.85864 | 829.2383 | 835.3014 | 841.5633 |
| Lcp1 | KLENCNYAVDLGK | -456 | -468 | 1 | 0 | 13 | 882.9805 | 341.0138 | 962.1755 | 393.4905 | 944.8589 | 940.9829 |
| Lcp1 | KTENLDDEEK | -570 | -579 | 1 | 0 | 10 | 836.80835 | 599.14844 | 735.9422 | 1119.8492 | 1148.7606 | 1021.9833 |
| Lcp1 | TLNILEDIGGGQK | -503 | -515 | 0 | 0 | 13 | 721.08264 | 747.04767 | 673.31415 | 1648.9268 | 1461.3983 | 1368.6703 |
| Lcp1 | AIGCHVVNIGAEDLK | -203 | -217 | 0 | 0 | 15 | 477.52377 | 463.23264 | 570.7751 | 599.61346 | 1521.2302 | 1381.9725 |
| Lcp1 | KTENLDDEEKLNNAK | -570 | -584 | 2 | 0 | 15 | 461.5991 | 584.24005 | 423.6428 | 776.3292 | 1636.3685 | 1403.8259 |
| Lcp1 | KTENLDDEEKLNNAK | -570 | -584 | 2 | 0 | 15 | 461.5991 | 584.24005 | 423.6428 | 776.3292 | 1636.3685 | 1403.8259 |
| Lcp1 | SVGTQHSYSEEEK | -111 | -123 | 0 | 0 | 13 | 426.5085 | 666.96436 | 764.8337 | 484.83075 | 707.8804 | 343.5228 |
| Lcp1 | EGICAIGGTSEQSSVGTQHSYSEEEK | -98 | -123 | 0 | 0 | 26 | 404.12088 | 438.19397 | 473.7447 | 464.28772 | 501.21982 | 496.7345 |
| Lcp1 | VNHLYSDLSDAL | -413 | -424 | 0 | 0.00051306 | 12 | 313.661 | 1351.1915 | 1308.016 | 936.42444 | 1138.185 | 1324.5627 |
| Lcp1 | VYALPEDLVEVNPK | -597 | -610 | 0 | 0 | 14 | 276.9264 | 1500.0979 | 1514.4583 | 1173.9056 | 1574.3232 | 1463.0111 |
| Lcp1 | EGICAIGGTSEQSSVGTQHSY | -98 | -118 | 0 | 0 | 21 | 199.52354 | 324.66742 | 321.10352 | 386.2365 | 356.34726 | 428.97595 |
| Lcp1 | NILEDIGGGQK | -505 | -515 | 0 | 0.000712251 | 11 | 5502.614 | 6138.429 | NaN | 4709.8486 | 432.8106 | 5763.883 |
| Lcp1 | HKPENQDIDWGALEGETR | -378 | -395 | 1 | 0 | 18 | 2232.4617 | 535.6897 | 1534.2562 | NaN | 675.9512 | 838.48474 |
| Lcp1 | WVNTTLK | -524 | -530 | 0 | 0.00111006 | 7 | 1677.1492 | 1302.1981 | 1194.6995 | 1769.4368 | NaN | 1229.3105 |

| | | | | | | | | | | | | |
|---|---|---|---|---|---|---|---|---|---|---|---|---|
| Lcp1 | VNHLYSDL | -413 | -420 | 0 | 0.002068213 | 8 | 1057.3147 | 1094.5032 | 1101.6537 | 280.39502 | NaN | 908.77716 |
| Lcp1 | LREGESLEDLMK | -252 | -263 | 1 | 0.00142999 | 12 | 673.4901 | NaN | 606.7068 | 491.8505 | 698.96136 | 593.95264 |
| Lcp1 | SAIGCHVVNIGAEDLK | -202 | -217 | 0 | 0 | 16 | 417.7642 | NaN | 686.6415 | 1402.6459 | 1109.1063 | 1209.6156 |
| Lcp1 | EGICAIGGTSEQS | -98 | -110 | 0 | 0 | 13 | 80.68804 | 286.29443 | 203.65778 | 209.64496 | NaN | 249.34885 |
| Lcp1 | TENLDDEEK | -571 | -579 | 0 | 0.00051306 | 9 | 2518.4646 | 2679.7551 | 2754.496 | NaN | NaN | 461.67752 |
| Lcp1 | WANYHLENA | -273 | -281 | 0 | 0.000712251 | 9 | 1043.8971 | NaN | 1094.4098 | 1313.9093 | 1458.9478 | NaN |
| Lcp1 | NELNDLFK | -32 | -39 | 0 | 0.002265552 | 8 | 1607.0804 | NaN | 1727.6039 | NaN | NaN | 1115.8481 |
| Lcp1 | AIGGTSEQSSVGTQHSYSEEEK | -102 | -123 | 0 | 0.000230707 | 22 | 1574.8787 | 1619.2625 | 1941.7904 | NaN | NaN | NaN |
| Lcp1 | YHLENAGCTK | -276 | -285 | 0 | 0 | 10 | 777.69037 | 171.3162 | NaN | 246.75105 | NaN | NaN |
| Lcp1 | AIQPGSINYDLLK | -558 | -570 | 0 | 0 | 13 | 712.22437 | 869.7186 | 852.7683 | NaN | NaN | NaN |
| Lcp1 | SEQSSVGTQHSYSEEEK | -107 | -123 | 0 | 0.001636916 | 17 | 352.78983 | NaN | 327.2305 | NaN | NaN | 368.6882 |
| Lcp1 | ILEDIGGGQK | -506 | -515 | 0 | 0.006677519 | 10 | 615.6699 | NaN | NaN | 505.79547 | NaN | NaN |
| Lcp1 | VTATDVVR | -350 | -357 | 0 | 0.002780285 | 8 | 507.7325 | 446.3681 | NaN | NaN | NaN | NaN |
| Lcp1 | YPALHKPENQDIDWGALE | -374 | -391 | 1 | 0.000712251 | 18 | 402.43442 | NaN | 593.3142 | NaN | NaN | NaN |
| Lcp1 | ISCNELNDLFK | -29 | -39 | 0 | 0.001092529 | 11 | 115.21358 | NaN | NaN | NaN | 841.7302 | NaN |
| Rps6 | DIPGLTDTTVPR | -120 | -131 | 0 | 0 | 12 | 8571.19 | 9691.747 | 9736.875 | 9447.473 | 10162.455 | 10150.272 |
| Rps6 | LNISFPATGCQK | -3 | -14 | 0 | 0 | 12 | 5109.378 | 133.39758 | 5627.2217 | 5776.0977 | 6535.9277 | 6119.733 |
| Rps6 | LNISFPATGCQK | -3 | -14 | 0 | 0 | 12 | 5109.378 | 133.39758 | 5627.2217 | 5776.0977 | 6535.9277 | 6119.733 |
| Rps6 | LIEVDDER | -15 | -22 | 0 | 0 | 8 | 1807.5399 | 1723.1001 | 2296.782 | 2684.8047 | 1570.4437 | 1328.0123 |
| Rps6 | KLNISFPATGCQK | -2 | -14 | 1 | 0 | 13 | 1647.1294 | 715.7768 | 1368.9878 | 1371.5626 | 1292.3767 | 1353.1501 |
| Rps6 | KLNISFPATGCQK | -2 | -14 | 1 | 0 | 13 | 1647.1294 | 715.7768 | 1368.9878 | 1371.5626 | 1292.3767 | 1353.1501 |
| Rps6 | NISFPATGCQK | -4 | -14 | 0 | 0 | 11 | 1131.614 | 1765.0146 | 2342.4663 | 2195.1636 | 1870.4924 | 1908.2972 |
| Rps6 | ISFPATGCQK | -5 | -14 | 0 | 0 | 10 | 1063.4994 | 1011.4762 | 1395.3823 | 1122.7634 | 1224.6923 | 866.05676 |
| Rps6 | PGLTDTTVPR | -122 | -131 | 0 | 0.001776473 | 10 | 1032.9225 | 2136.5017 | 2098.3489 | 760.76917 | 1253.8796 | 4214.22 |
| Rps6 | VAADALGEEWK | -36 | -46 | 0 | 0 | 11 | 525.9402 | 2363.893 | 525.81384 | 2520.0576 | 168.57199 | 2940.377 |
| Rps6 | LIEVDDERK | -15 | -23 | 1 | 0.00025015 | 9 | 6960.527 | 6954.968 | NaN | 2918.474 | 1715.7474 | 3671.938 |
| Rps6 | ADALGEEWK | -38 | -46 | 0 | 0 | 9 | 462.02866 | 785.4548 | NaN | NaN | 878.94617 | 881.523 |
| Rps6 | AADALGEEWK | -37 | -46 | 0 | 0.00025015 | 10 | 1111.6995 | NaN | NaN | NaN | NaN | 1214.3303 |
| Rps6 | NKEEAAEYAK | -202 | -211 | 1 | 0.000431655 | 10 | 858.02374 | 577.4547 | NaN | NaN | NaN | NaN |
| Hmgb1 | IKGEHPGLSIGDVAK | -113 | -127 | 1 | 0 | 15 | 6643.2925 | 6945.4023 | 7585.3926 | 6974.918 | 9154.592 | 6561.7007 |
| Hmgb1 | IKGEHPGLSIGDVAK | -113 | -127 | 1 | 0 | 15 | 6643.2925 | 6945.4023 | 7585.3926 | 6974.918 | 9154.592 | 6561.7007 |
| Hmgb1 | GEHPGLSIGDVAK | -115 | -127 | 0 | 0 | 13 | 6225.532 | 6841.548 | 7136.584 | 7124.0205 | 94.89296 | 8031.5464 |
| Hmgb1 | GEHPGLSIGDVAK | -115 | -127 | 0 | 0 | 13 | 6225.532 | 6841.548 | 7136.584 | 7124.0205 | 94.89296 | 8031.5464 |
| Hmgb1 | FKDPNAPK | -89 | -96 | 1 | 0 | 8 | 5455.738 | 5314.2446 | 5454.783 | 5441.6733 | 5897.104 | 5745.537 |
| Hmgb1 | IKGEHPGLSIGDVA | -113 | -126 | 1 | 0 | 14 | 2773.454 | 3230.0742 | 3273.721 | 6604.056 | 1734.9263 | 6454.7603 |
| Hmgb1 | TYIPPKGETK | -77 | -86 | 1 | 0 | 10 | 1691.4851 | 3056.2463 | 3100.343 | 2255.6528 | 3484.4414 | 3901.4675 |
| Hmgb1 | TAADDKQPYEK | -136 | -146 | 1 | 0 | 11 | 1433.758 | 1515.296 | 1877.3947 | 2108.6697 | 160.40715 | 1246.0656 |
| Hmgb1 | WNNTAADDKQPYEK | -133 | -146 | 1 | 0 | 14 | 1043.7426 | 1008.66455 | 1415.7284 | 955.9239 | 987.351 | 1282.8124 |
| Hmgb1 | WNNTAADDKQPYEK | -133 | -146 | 1 | 0 | 14 | 1043.7426 | 1008.66455 | 1415.7284 | 955.9239 | 987.351 | 1282.8124 |
| Hmgb1 | GKFEDMAK | -58 | -65 | 1 | 0.002265552 | 8 | 964.3016 | 1090.093 | 934.76544 | 217.7631 | 488.84402 | 334.9013 |
| Hmgb1 | KFKDPNAPK | -88 | -96 | 2 | 0.002329279 | 9 | 350.58957 | 367.84882 | 378.98785 | 259.3143 | 280.6328 | 280.37543 |
| Hmgb1 | KHPDASVNFSEF | -30 | -41 | 1 | 0 | 12 | 3034.8389 | NaN | 3672.7422 | 325.5442 | 5276.72 | 4609.924 |
| Hmgb1 | GEHPGLSIGDVA | -115 | -126 | 0 | 0.000298805 | 12 | 765.48303 | 856.4592 | 818.5205 | 1188.8342 | NaN | 1447.245 |
| Hmgb1 | KHPDASVNF | -30 | -38 | 1 | 0.000947459 | 9 | 754.0313 | NaN | 763.3495 | 700.0546 | 307.58316 | 530.91077 |
| Hmgb1 | KKEEEDDEEDEEDEEEEEEEDEDEEEDDDDE | -184 | -215 | 2 | 0.001868731 | 32 | 586.45764 | NaN | 662.88104 | 888.18353 | 926.4217 | 1034.6124 |
| Hmgb1 | AADDKQPYEK | -137 | -146 | 1 | 0 | 10 | 497.0875 | NaN | 635.8167 | 544.2208 | 853.65125 | 740.5833 |
| Hmgb1 | FCSEYRPK | -105 | -112 | 1 | 0 | 8 | 1372.2449 | 1538.0482 | NaN | NaN | 1798.0333 | 804.0537 |
| Hmgb1 | KTYIPPKGETK | -76 | -86 | 2 | 0.00830926 | 11 | 300.29288 | NaN | NaN | NaN | NaN | NaN |
| Nap1l1 | ADFEIGHFLR | -322 | -331 | 0 | 0.000431655 | 10 | 2588.0671 | 2344.629 | 335.37924 | 2652.704 | 3061.9446 | 3233.7576 |
| Nap1l1 | DFEIGHFLR | -323 | -331 | 0 | 0 | 9 | 1979.6381 | 2343.928 | 2404.6814 | 1569.3014 | 2001.6589 | 2128.8816 |
| Nap1l1 | SDMVQEHDEPILK | -182 | -194 | 0 | 0 | 13 | 1789.3828 | 1762.8641 | 3034.1519 | 3279.817 | 3345.807 | 2751.9841 |
| Nap1l1 | SDMVQEHDEPILK | -182 | -194 | 0 | 0 | 13 | 1789.3828 | 1762.8641 | 3034.1519 | 3279.817 | 3345.807 | 2751.9841 |
| Nap1l1 | WKPDEEDEVSEELKEK | -134 | -149 | 2 | 0 | 16 | 728.3172 | 850.7066 | 797.4487 | 693.27783 | 866.3907 | 228.91202 |
| Nap1l1 | NVDLLSDMVQEHDEPILK | -177 | -194 | 0 | 0 | 18 | 567.3447 | 755.4553 | 568.26483 | 821.55646 | 545.8987 | 470.61182 |
| Nap1l1 | NVDLLSDMVQEHDEPILK | -177 | -194 | 0 | 0 | 18 | 567.3447 | 755.4553 | 568.26483 | 821.55646 | 545.8987 | 470.61182 |
| Nap1l1 | TVSNDSFFNF | -291 | -300 | 0 | 0 | 10 | 506.12567 | 599.61816 | 622.44385 | 520.5047 | 880.67694 | 851.9542 |
| Nap1l1 | GEEEGDEENDPDYDPK | -365 | -380 | 0 | 0.000471409 | 16 | 393.8334 | 383.17175 | 362.4123 | 425.0769 | 316.17142 | 302.1211 |
| Nap1l1 | AADFEIGHFLR | -321 | -331 | 0 | 0 | 11 | 314.76093 | 391.15106 | 762.05817 | 178.82195 | 978.8163 | 370.63477 |
| Nap1l1 | LDGLVDTPTGYIE | -56 | -68 | 0 | 0.00025015 | 13 | 204.64903 | 172.09142 | 200.11024 | 215.36119 | 196.13394 | 212.57513 |
| Nap1l1 | EWKPDEEDEVSEELKEK | -133 | -149 | 2 | 0.000230707 | 17 | 121.301315 | 216.52602 | 99.112236 | 1233.921 | 1441.0916 | 1359.3542 |
| Nap1l1 | HFEPNDYFTNEVLTK | -217 | -231 | 0 | 0.001023236 | 15 | 637.9058 | NaN | 751.9286 | 521.42694 | 1039.113 | 1071.525 |
| Nap1l1 | EFHFEPNDYFTNEVLTK | -215 | -231 | 0 | 0 | 17 | 375.19476 | 447.4157 | NaN | 332.72147 | 433.14954 | 434.7359 |
| Nap1l1 | EWKPDEEDEVSEELK | -133 | -147 | 1 | 0.000431655 | 15 | 783.6317 | 1036.568 | 924.36285 | NaN | NaN | 1096.5249 |
| Nap1l1 | EVEEEETGEETK | -21 | -32 | 0 | 0 | 12 | 561.51337 | 696.6416 | 603.83636 | NaN | NaN | NaN |
| Nap1l1 | EDVEEVEEEETGEETK | -17 | -32 | 0 | 0.00025015 | 16 | 417.3081 | 398.95172 | 412.52228 | NaN | NaN | NaN |
| Nap1l1 | LDGLVDTPTGYIESLPK | -56 | -72 | 0 | 0 | 17 | 2676.213 | 2905.3296 | NaN | NaN | NaN | NaN |
| Nap1l1 | EQSELDQDLEDVE | -8 | -20 | 0 | 0.000471409 | 13 | 927.8045 | NaN | 915.9062 | NaN | NaN | NaN |
| Canx | AVKPDDWDEDAPSK | -299 | -312 | 1 | 0 | 14 | 4274.194 | 5535.1533 | 5069.8745 | 3534.8796 | 3081.4043 | 2919.0767 |
| Canx | VVDDWANDGWGLK | -447 | -459 | 0 | 0 | 13 | 4087.387 | 4896.3774 | 4844.805 | 1090.9397 | 5155.5537 | 5157.82 |
| Canx | DEEEEEEKLEEK | -539 | -550 | 1 | 0 | 12 | 4070.6047 | 4342.7417 | 4615.708 | 324.7903 | 309.52695 | 5064.546 |
| Canx | TAELSLDQFHDK | -172 | -183 | 0 | 0 | 12 | 3926.3862 | 4064.969 | 4432.571 | 5077.547 | 6034.5615 | 5895.2974 |
| Canx | KPEDWDERPK | -284 | -293 | 2 | 0 | 10 | 3818.2524 | 4475.216 | 4461.03 | 3595.7712 | 4472.076 | 4632.6025 |
| Canx | KPEDWDERPK | -284 | -293 | 2 | 0 | 10 | 3818.2524 | 4475.216 | 4461.03 | 3595.7712 | 4472.076 | 4632.6025 |
| Canx | DAPQPDVKDEEGKEEEK | -519 | -535 | 2 | 0 | 17 | 3481.104 | 3881.5352 | 3807.8828 | 3125.299 | 4382.553 | 4658.251 |
| Canx | DAPQPDVKDEEGKEEEK | -519 | -535 | 2 | 0 | 17 | 3481.104 | 3881.5352 | 3807.8828 | 3125.299 | 4382.553 | 4658.251 |
| Canx | AEEDEILNR | -573 | -581 | 0 | 0 | 9 | 1862.3423 | 2015.473 | 2075.5552 | 1979.421 | 2074.871 | 257.74167 |
| Canx | SDASTPPSPK | -49 | -58 | 0 | 0 | 10 | 1402.8213 | 1471.3784 | 1547.7753 | 1517.3636 | 456.10864 | 991.9678 |
| Canx | ELSLDQFHDK | -174 | -183 | 0 | 0 | 10 | 1214.6958 | 1879.8574 | 302.38306 | 2226.9128 | 978.5318 | 1173.2274 |

| | | | | | | | | | | | | |
|---|---|---|---|---|---|---|---|---|---|---|---|---|
| Canx | SAPGCGVWQR | -363 | -372 | 0 | 0 | 10 | 1182.4088 | 354.35297 | 1077.0073 | 1181.2406 | 1237.417 | 1149.5737 |
| Canx | RPDADLK | -222 | -228 | 1 | 0.006677519 | 7 | 1077.4731 | 828.1176 | 1261.0756 | 708.42145 | 715.71924 | 804.70776 |
| Canx | KDDTDDEIAK | -91 | -100 | 1 | 0.000298805 | 10 | 1022.37427 | 967.3081 | 1180.2157 | 1056.787 | 1100.4131 | 990.6015 |
| Canx | CESAPGCGVWQR | -361 | -372 | 0 | 0 | 12 | 1004.7202 | 1425.5619 | 1006.628 | 1047.2704 | 793.00793 | 811.14014 |
| Canx | RDEEEEEEKLEEK | -538 | -550 | 2 | 0 | 13 | 950.8415 | 1015.4053 | 1114.5674 | 936.4568 | 1385.7059 | 1324.9275 |
| Canx | KAADGAAEPGVVLQ | -460 | -473 | 1 | 0.00025015 | 14 | 888.421 | 821.0051 | 803.2666 | 855.82196 | 802.4216 | 792.94086 |
| Canx | KDDTDDEIAKY | -91 | -101 | 2 | 0 | 11 | 833.08105 | 839.9548 | 859.76685 | 487.52197 | 1339.0685 | 1427.3018 |
| Canx | TDAPQPDVKDEEGKEEEK | -518 | -535 | 2 | 0 | 18 | 736.0565 | 799.4781 | 803.9117 | 420.1223 | 575.212 | 621.3272 |
| Canx | LNKPFLFDTKPLIVQ | -136 | -150 | 2 | 0 | 15 | 640.17914 | 706.7208 | 698.3701 | 947.88544 | 1310.2433 | 1146.8862 |
| Canx | LNKPFLFDTKPLIVQ | -136 | -150 | 2 | 0 | 15 | 640.17914 | 706.7208 | 698.3701 | 947.88544 | 1310.2433 | 1146.8862 |
| Canx | KIPNPDFFEDLEPFK | -402 | -416 | 1 | 0 | 15 | 559.5587 | 1399.3909 | 784.4527 | 930.52936 | 1395.5688 | 1273.7346 |
| Canx | TSDIFFDNFIISGDR | -431 | -445 | 0 | 0 | 15 | 559.42664 | 454.4466 | 518.9243 | 458.0639 | 591.9125 | 474.96872 |
| Canx | IADPDAVKPDDWDEDAPSK | -294 | -312 | 1 | 0 | 19 | 508.2909 | 590.81116 | 592.19366 | 564.08417 | 661.664 | 643.7696 |
| Canx | TAELSLDQFHDKTPY | -172 | -186 | 1 | 0.00025015 | 15 | 447.37424 | 451.81442 | 519.0382 | 641.6873 | 757.1326 | 742.1856 |
| Canx | SVVNSGNLLNDMTPPVNPSR | -256 | -275 | 0 | 0 | 20 | 437.82532 | 447.42725 | 449.62988 | 353.1629 | 698.3798 | 621.20825 |
| Canx | LNKPFLFDTKPLIVQY | -136 | -151 | 2 | 0.00025015 | 16 | 381.37366 | 461.64236 | 361.9123 | 414.4322 | 509.6474 | 593.5765 |
| Canx | LNKPFLFDTKPLIVQY | -136 | -151 | 2 | 0.00025015 | 16 | 381.37366 | 461.64236 | 361.9123 | 414.4322 | 509.6474 | 593.5765 |
| Canx | QKSDAEEDGVTGSQDEEDSKPK | -551 | -572 | 2 | 0 | 22 | 266.74045 | 257.46484 | 280.74454 | 278.73004 | 298.1451 | 281.6188 |
| Canx | VDQSVVNSGNLLNDMTPPVNPSR | -253 | -275 | 0 | 0 | 23 | 258.75647 | 97.76054 | 306.20508 | 96.577065 | 266.69824 | 128.2648 |
| Canx | VVDDWANDGWGLKK | -447 | -460 | 1 | 0 | 14 | 1190.6064 | 778.2366 | 784.07446 | 695.8686 | 850.85364 | NaN |
| Canx | LNKPFLFDTK | -136 | -145 | 1 | 0 | 10 | 698.76953 | NaN | 741.53644 | 822.2525 | 863.2652 | 926.65125 |
| Canx | LNKPFLFDTKPLIV | -136 | -149 | 2 | 0.00025015 | 14 | 337.7676 | 287.55444 | 376.10458 | NaN | 311.83035 | 381.7204 |
| Canx | LNKPFLFDTKPLIV | -136 | -149 | 2 | 0.00025015 | 14 | 337.7676 | 287.55444 | 376.10458 | NaN | 311.83035 | 381.7204 |
| Canx | TYFTDKK | -229 | -235 | 1 | 0.00051306 | 7 | 1303.3362 | 2337.3079 | 1457.9513 | NaN | NaN | 319.35168 |
| Canx | SLDQFHDK | -176 | -183 | 0 | 0 | 8 | 1029.4783 | 1536.5251 | 1287.0159 | NaN | NaN | 1398.6908 |
| Canx | SDAEEDGVTGSQDEEDSKPK | -553 | -572 | 1 | 0.000627943 | 20 | 277.9681 | NaN | 198.82822 | 177.93797 | NaN | 192.30692 |
| Canx | IADPDAVKPD | -294 | -303 | 1 | 0.00025015 | 10 | 1356.2424 | 618.2985 | 758.2586 | NaN | NaN | NaN |
| Canx | DDLDDVIEEVEDSK | -33 | -46 | 0 | 0 | 14 | 652.3392 | 604.7089 | 668.43256 | NaN | NaN | NaN |
| Canx | LNKPFLF | -136 | -142 | 1 | 0.00437931 | 7 | 459.97183 | NaN | NaN | 386.7271 | 339.91443 | NaN |
| Canx | TSDIFFDNFIISGDRR | -431 | -446 | 1 | 0.00078661 | 16 | 306.66974 | NaN | NaN | 389.15836 | 744.28455 | NaN |
| Canx | GVTGSQDEEDSKPK | -559 | -572 | 1 | 0.00025015 | 14 | 290.99686 | 290.69144 | 323.05472 | NaN | NaN | NaN |
| Canx | IPNPDFFEDLEPFK | -403 | -416 | 0 | 0.00025015 | 14 | 210.47878 | NaN | 370.9319 | NaN | 358.87927 | NaN |
| Canx | NDMTPPVNPSR | -265 | -275 | 0 | 0.001610812 | 11 | 1159.0808 | NaN | NaN | 1396.2484 | NaN | NaN |
| Canx | AADGAAEPGVVLQ | -461 | -473 | 0 | 0.003966045 | 13 | 1019.0317 | NaN | NaN | NaN | NaN | NaN |
| Hspa9 | RYDDPEVQK | -127 | -135 | 1 | 0 | 9 | 7620.575 | 7741.306 | 905.41785 | 1593.5754 | 11922.3125 | 782.8962 |
| Hspa9 | AENYLGHTAK | -178 | -187 | 0 | 0 | 10 | 4153.579 | 70.81048 | 3824.6248 | 4018.62 | 1111.3889 | 4001.6904 |
| Hspa9 | VQQTVQDLFGR | -395 | -405 | 0 | 0 | 11 | 4093.1416 | 4404.9746 | 4400.7114 | 3067.0774 | 3447.8489 | 3358.4172 |
| Hspa9 | ETAENYLGHTAK | -176 | -187 | 0 | 0 | 12 | 3143.8335 | 1419.7982 | 2556.7778 | 5830.4844 | 7108.7065 | 6919.05 |
| Hspa9 | AEGIIHDTETK | -585 | -595 | 0 | 0 | 11 | 2942.3674 | 3750.3037 | 691.1325 | 3453.6343 | 4292.7573 | 4645.341 |
| Hspa9 | FEGIVTDLIK | -351 | -360 | 0 | 0 | 10 | 2244.7258 | 4478.9067 | 4333.318 | 2186.0825 | 5030.081 | 4999.0244 |
| Hspa9 | TLIGIPPAPR | -504 | -513 | 0 | 0 | 10 | 1630.8247 | 1463.3754 | 3302.7358 | 2596.2034 | 2643.028 | 2776.6873 |
| Hspa9 | AQFEGIVTDLIK | -349 | -360 | 0 | 0 | 12 | 1454.231 | 1292.5433 | 1270.1956 | 508.0766 | 1638.7717 | 1626.6099 |
| Hspa9 | QTVQDLFGR | -397 | -405 | 0 | 0 | 9 | 1267.1034 | 1385.589 | 1812.2408 | 2012.4155 | 2152.898 | 2090.392 |
| Hspa9 | STAADGQTQVEIK | -473 | -485 | 0 | 0 | 13 | 1120.1307 | 1060.0885 | 1012.3236 | 1073.1267 | 623.2453 | 1341.4009 |
| Hspa9 | ETAENYLGHTA | -176 | -186 | 0 | 0 | 11 | 1030.2557 | 171.99757 | 1108.9297 | 462.88293 | 229.66058 | 1300.5719 |
| Hspa9 | DIGEVILVGGMTR | -379 | -391 | 0 | 0 | 13 | 1001.72406 | 1047.0482 | 1046.7297 | 1142.2855 | 1274.053 | 1250.4812 |
| Hspa9 | FTLIGIPPAPR | -503 | -513 | 0 | 0 | 11 | 964.0648 | 1829.2614 | 964.55566 | 1836.4122 | 2843.5295 | 2812.6816 |
| Hspa9 | LLGQFTLIGIPPAPR | -499 | -513 | 0 | 0 | 15 | 901.82025 | 642.0805 | 757.9558 | 1313.6678 | 1579.5219 | 1488.356 |
| Hspa9 | RAQFEGIVTDLIK | -348 | -360 | 1 | 0.00051306 | 13 | 854.1515 | 1088.4216 | 981.6841 | 657.1488 | 982.4012 | 1580.0192 |
| Hspa9 | AADGQTQVEIK | -475 | -485 | 0 | 0 | 11 | 800.1562 | 1849.0771 | 656.4552 | 1149.9938 | 1917.0032 | 1285.9944 |
| Hspa9 | STNGDTFLGGEDFDQALLR | -266 | -284 | 0 | 0.000230707 | 19 | 695.468 | 615.84784 | 776.31323 | 738.9597 | 924.22675 | 894.62415 |
| Hspa9 | MEEFKDQLPADECNK | -596 | -610 | 1 | 0 | 15 | 555.89874 | 562.0511 | 610.04767 | 626.86395 | 735.1141 | 722.23456 |
| Hspa9 | MEEFKDQLPADECNK | -596 | -610 | 1 | 0 | 15 | 555.89874 | 562.0511 | 610.04767 | 626.86395 | 735.1141 | 722.23456 |
| Hspa9 | KNAVITVPAYFNDSQR | -187 | -202 | 1 | 0 | 16 | 543.2251 | 669.9218 | 966.6802 | 907.2175 | 1289.8978 | 1228.248 |
| Hspa9 | KNAVITVPAYFNDSQR | -187 | -202 | 1 | 0 | 16 | 543.2251 | 669.9218 | 966.6802 | 907.2175 | 1289.8978 | 1228.248 |
| Hspa9 | SSGGLSKDDIENMVK | -549 | -563 | 1 | 0 | 15 | 190.29086 | 1158.0297 | 1205.3643 | 694.5397 | 2070.9805 | 2254.3657 |
| Hspa9 | DQLPADECNK | -601 | -610 | 0 | 0 | 10 | 167.73758 | 1054.1749 | 1089.5343 | 1063.754 | 827.3946 | 423.55975 |
| Hspa9 | ADGQTQVEIK | -476 | -485 | 0 | 0.00025015 | 10 | 254.49265 | 1271.2982 | 1065.5114 | 945.5817 | NaN | 828.3848 |
| Hspa9 | VEAVNMAEGIIHDTETK | -579 | -595 | 0 | 0 | 17 | 693.33966 | 815.2607 | 781.39856 | NaN | NaN | 874.91516 |
| Hspa9 | QAVTNPNNTFYATK | -108 | -121 | 0 | 0 | 14 | 997.66565 | 1097.2577 | 981.14325 | NaN | NaN | NaN |
| Hspa9 | DDIENMVK | -556 | -563 | 0 | 0.007343498 | 8 | 1374.8214 | NaN | NaN | NaN | NaN | NaN |
| Hspa9 | QAVTNPNNTFYAT | -108 | -120 | 0 | 0.007256894 | 13 | 1130.0591 | NaN | NaN | NaN | NaN | NaN |
| Hspa9 | ETGVDLTK | -293 | -300 | 0 | 0.001743443 | 8 | 720.17163 | NaN | NaN | NaN | NaN | NaN |
| Hspa9 | LIGIPPAPR | -505 | -513 | 0 | 0 | 9 | 582.2416 | NaN | NaN | NaN | NaN | NaN |
| Prpf3 | ELDELKPWIEK | -7 | -17 | 1 | 0 | 11 | 498.21265 | 1194.9854 | 1154.1404 | 945.39233 | 1478.3003 | 1339.6385 |
| Prpf3 | QLSFISPPAPQPK | -159 | -171 | 0 | 0 | 13 | 577.3143 | 487.17453 | 467.58197 | 410.77002 | 1311.1976 | NaN |
| Prpf3 | AADHLKPFLDDSTLR | -47 | -61 | 1 | 0.000627943 | 15 | 554.12524 | NaN | 169.95709 | NaN | 671.6395 | 754.3426 |
| Ybx1 | RPQYSNPPVQGE | -203 | -214 | 1 | 0.000431655 | 12 | 5834.4326 | 5546.4844 | 5471.049 | 5449.393 | 5083.1064 | 5459.3286 |
| Ybx1 | NVTGPGGVPVQGSK | -122 | -135 | 0 | 0 | 14 | 3046.728 | 3019.0623 | 2908.1663 | 3118.2588 | 2880.3005 | 2848.1836 |
| Ybx1 | NVTGPGGVPVQGSK | -122 | -135 | 0 | 0 | 14 | 3046.728 | 3019.0623 | 2908.1663 | 3118.2588 | 2880.3005 | 2848.1836 |
| Ybx1 | RPQYSNPPVQGEVM | -203 | -216 | 1 | 0 | 14 | 944.83923 | 871.26624 | 367.08392 | 1035.985 | 698.88684 | 849.61017 |
| Ybx1 | SSEAETQQPPAAPAAAL | -2 | -18 | 0 | 0.000550307 | 17 | 844.1731 | 1063.3177 | 1221.1829 | 1047.8064 | 851.13525 | 885.44965 |
| Ybx1 | KAADPPAENSSAPEAEQGGAE | -302 | -322 | 1 | 0.000431655 | 21 | 676.1084 | 760.88025 | 779.4362 | 685.93005 | 784.1668 | 891.64276 |
| Ybx1 | EDGNEEDKENQGDETQGQQPPQR | -255 | -277 | 1 | 0 | 23 | 565.82 | 679.7414 | 694.5693 | 1162.0634 | 1046.7726 | 916.2265 |
| Ybx1 | KENQGDETQGQQPPQR | -262 | -277 | 1 | 0 | 16 | 560.81866 | 401.4797 | 526.92993 | 355.85657 | 529.01227 | 606.93585 |
| Ybx1 | EGADNQGAGEQGRPVR | -217 | -232 | 1 | 0 | 16 | 342.84497 | 235.68817 | 317.1819 | 467.28113 | 420.08014 | 396.23843 |
| Ybx1 | EGADNQGAGEQGRPVR | -217 | -232 | 1 | 0 | 16 | 342.84497 | 235.68817 | 317.1819 | 467.28113 | 420.08014 | 396.23843 |
| Ybx1 | AADPPAENSSAPEAEQGGAE | -303 | -322 | 0 | 0 | 20 | 418.31073 | 505.5375 | 374.23114 | 812.5059 | 697.77954 | 594.7117 |

| | | | | | | | | | | | | |
|---|---|---|---|---|---|---|---|---|---|---|---|---|
| Ybx1 | EDGNEEDKENQGDETQ | -255 | -270 | 1 | 0.000431655 | 16 | 361.8444 | 420.13123 | 367.55667 | 228.76996 | 352.25262 | 440.38727 |
| Ybx1 | EGADNQGAGEQGR | -217 | -229 | 0 | 0 | 13 | 79.83908 | 619.31934 | 817.94037 | 352.6773 | 573.4892 | 19.867975 |
| Ybx1 | RPENPKPQDGK | -289 | -299 | 2 | 0.001868731 | 11 | 357.22003 | 413.5733 | 463.7336 | 307.1235 | NaN | 345.1091 |
| Ybx1 | SAADTKPGSTGSGAGSGGPGGLTSAAPAGGDKK | -19 | -51 | 2 | 0.001092529 | 33 | 341.8999 | 360.24014 | 332.3614 | NaN | NaN | NaN |
| Ybx1 | SAADTKPGSTGSGAGSGGPGGLTSAAPAGGDKK | -19 | -51 | 2 | 0.001092529 | 33 | 341.8999 | 360.24014 | 332.3614 | NaN | NaN | NaN |
| Ybx1 | ANVTGPGGVPVQGSK | -121 | -135 | 0 | 0 | 15 | 734.67896 | 713.2115 | 844.34607 | NaN | NaN | NaN |
| Ybx1 | DTKPGSTGSGAGSGGPGGLTSAAPAGGDKK | -22 | -51 | 2 | 0.003814256 | 30 | 527.8792 | 468.50018 | 504.63126 | NaN | NaN | NaN |
| Ybx1 | NYQQNYQNSESGEK | -155 | -168 | 0 | 0.00025015 | 14 | 242.31018 | 212.4159 | 342.64 | NaN | NaN | NaN |
| Ybx1 | GAEAANVTGPGGVPVQGSK | -117 | -135 | 0 | 0.00025015 | 19 | 1350.3496 | NaN | NaN | NaN | 907.0519 | NaN |
| Ybx1 | SAADTKPGSTGSGAGSGGPGGLTSAAPAGGDK | -19 | -50 | 1 | 0.000989247 | 32 | 266.45245 | NaN | 577.9762 | NaN | NaN | NaN |
| Stxbp2 | AYDLLDIEQDTYR | -253 | -265 | 0 | 0 | 13 | 374.0158 | 536.3428 | 343.76617 | 685.6985 | 676.9846 | 757.0552 |
| Stxbp2 | AADPVSPLLHEL | -236 | -247 | 0 | 0.00051306 | 12 | 2370.805 | 2426.5352 | 2779.8289 | NaN | 2679.6348 | 2356.1372 |
| Pkm | LDIDSAPITAR | -33 | -43 | 0 | 0 | 11 | 10022.277 | 9440.758 | 9671.287 | 9814.023 | 7383.187 | 242.32803 |
| Pkm | GDYPLEAVR | -368 | -376 | 0 | 0.00025015 | 9 | 8601.167 | 8250.427 | 8112.631 | 7248.09 | 10293.802 | 10513.884 |
| Pkm | GIFPVLCK | -468 | -475 | 0 | 0.001941082 | 8 | 7132.175 | 238.7815 | 6088.464 | 5584.0244 | 3478.3384 | 4078.7515 |
| Pkm | KCDENILWLDYK | -151 | -162 | 1 | 0.000230707 | 12 | 6612.439 | 6997.639 | 7268.1074 | 10869.181 | 6494.736 | 6673.7075 |
| Pkm | DIDSAPITAR | -34 | -43 | 0 | 0.00025015 | 10 | 6412.0186 | 4544.583 | 5056.94 | 3580.3604 | 2207.9146 | 1095.7379 |
| Pkm | LNFSHGTHEYHAETIK | -74 | -89 | 0 | 0 | 16 | 898.8173 | 636.0616 | 812.20795 | 808.4943 | 191.13763 | 1163.1514 |
| Pkm | LNFSHGTHEYHAETIK | -74 | -89 | 0 | 0 | 16 | 898.8173 | 636.0616 | 812.20795 | 808.4943 | 191.13763 | 1163.1514 |
| Pkm | RGIFPVLCK | -467 | -475 | 1 | 0.002595029 | 9 | 5677.9053 | 6334.0586 | 6822.9927 | 4603.4346 | 7298.169 | 7247.113 |
| Pkm | KGVNLPGAAVDLPAVSEK | -207 | -224 | 1 | 0 | 18 | 5288.667 | 7486.3857 | 7003.737 | 7631.2104 | 10380.324 | 10147.633 |
| Pkm | KGVNLPGAAVDLPAVSEK | -207 | -224 | 1 | 0 | 18 | 5288.667 | 7486.3857 | 7003.737 | 7631.2104 | 10380.324 | 10147.633 |
| Pkm | EATESFASDPILYR | -93 | -106 | 0 | 0 | 14 | 4953.183 | 6260.4146 | 5974.78 | 6251.008 | 7373.651 | 7180.7324 |
| Pkm | GVNLPGAAVDLPAVSEK | -208 | -224 | 0 | 0 | 17 | 4759.9355 | 5219.7124 | 5335.9043 | 6112.3516 | 6494.9917 | 6528.699 |
| Pkm | GSGTAEVELKK | -126 | -136 | 1 | 0 | 11 | 4523.2847 | 5296.282 | 6065.4395 | 5845.4062 | 6500.0312 | 2650.463 |
| Pkm | GSGTAEVELK | -126 | -135 | 0 | 0 | 10 | 4245.2725 | 3760.843 | 3569.49 | 3695.0989 | 3046.9446 | 76.221924 |
| Pkm | ITLDNAYMEK | -142 | -151 | 0 | 0 | 10 | 3935.083 | 1569.7233 | 1558.6316 | 508.38074 | 1507.8031 | 505.9606 |
| Pkm | ITLDNAYMEK | -142 | -151 | 0 | 0 | 10 | 3935.083 | 1569.7233 | 1558.6316 | 508.38074 | 1507.8031 | 505.9606 |
| Pkm | PKPHSEAGTAF | -2 | -12 | 1 | 0 | 11 | 3716.617 | 4819.327 | 4959.5386 | 272.84384 | 5205.141 | 5137.188 |
| Pkm | LNFSHGTHEYHAE | -74 | -86 | 0 | 0 | 13 | 3460.1758 | 3956.3633 | 3907.1926 | 3032.8203 | 5415.7437 | 5128.0693 |
| Pkm | LNFSHGTHEYHAE | -74 | -86 | 0 | 0 | 13 | 3460.1758 | 3956.3633 | 3907.1926 | 3032.8203 | 5415.7437 | 5128.0693 |
| Pkm | RLDIDSAPITAR | -32 | -43 | 1 | 0.000230707 | 12 | 3185.2483 | 3405.246 | 3833.9053 | 3103.8953 | 3465.2476 | 3102.8467 |
| Pkm | RLDIDSAPITAR | -32 | -43 | 1 | 0.000230707 | 12 | 3185.2483 | 3405.246 | 3833.9053 | 3103.8953 | 3465.2476 | 3102.8467 |
| Pkm | HLQLFEELR | -391 | -399 | 0 | 0 | 9 | 3097.369 | 901.14557 | 3426.5994 | 4761.5986 | 5566.993 | 5303.1265 |
| Pkm | EATESFASDPILYRPV | -93 | -108 | 1 | 0 | 16 | 2883.442 | 3458.225 | 3666.5864 | 5148.602 | 6576.0645 | 6425.41 |
| Pkm | RLAPITSDPTEAAAVGAVE | -400 | -418 | 1 | 0 | 19 | 2451.6396 | 2741.5007 | 2891.6794 | 2335.9697 | 2761.0432 | 2622.9993 |
| Pkm | PKPHSEAGTAFIQ | -2 | -14 | 1 | 0 | 13 | 2220.3928 | 2371.5513 | 2425.8347 | 3444.4333 | 4043.1338 | 3722.5063 |
| Pkm | PKPHSEAGTAFIQ | -2 | -14 | 1 | 0 | 13 | 2220.3928 | 2371.5513 | 2425.8347 | 3444.4333 | 4043.1338 | 3722.5063 |
| Pkm | VDLPAVSEK | -216 | -224 | 0 | 0 | 9 | 2218.7207 | 1840.5936 | 1417.4099 | 1353.6204 | 933.86395 | 896.05115 |
| Pkm | GIFPVLCKD | -468 | -476 | 1 | 0.001190916 | 9 | 2132.6194 | 2153.3594 | 2064.3281 | 1426.5706 | 1508.1901 | 1572.521 |
| Pkm | IENHEGVR | -271 | -278 | 0 | 0 | 8 | 1788.092 | 1450.832 | 1897.7234 | 1424.8596 | 1548.0557 | 1763.0271 |
| Pkm | GADFLVTEVENGGSLGSK | -189 | -206 | 0 | 0 | 18 | 1606.0355 | 2091.4258 | 2004.7076 | 2596.797 | 3836.8352 | 3920.2012 |
| Pkm | DENILWLDYK | -153 | -162 | 0 | 0 | 10 | 1532.9906 | 1544.9618 | 1703.3434 | 2194.1897 | 1087.4592 | 2581.3887 |
| Pkm | DENILWLDYK | -153 | -162 | 0 | 0 | 10 | 1532.9906 | 1544.9618 | 1703.3434 | 2194.1897 | 1087.4592 | 2581.3887 |
| Pkm | KGDYPLEAVR | -367 | -376 | 1 | 0 | 10 | 1436.3138 | 1640.9583 | 1632.525 | 1152.6378 | 148.71188 | 1912.9093 |
| Pkm | CDENILWLDYK | -152 | -162 | 0 | 0 | 11 | 1244.5259 | 3179.2952 | 1757.7584 | 1123.9845 | 1778.8959 | 1436.7289 |
| Pkm | RLAPITSDPTEAAAV | -400 | -414 | 1 | 0.00025015 | 15 | 1224.1976 | 1035.396 | 1056.8744 | 1030.9235 | 1632.4891 | 1073.0032 |
| Pkm | KITLDNAYMEK | -141 | -151 | 1 | 0 | 11 | 1135.8948 | 1001.7122 | 1170.6041 | 1510.443 | 1704.4441 | 1721.866 |
| Pkm | GIICTIGPASR | -46 | -56 | 0 | 0 | 11 | 1000.74567 | 966.2384 | 894.84247 | 1073.3411 | 1073.0803 | 1121.9469 |
| Pkm | HLQLFEELRR | -391 | -400 | 1 | 0 | 10 | 945.7272 | 1367.0747 | 1558.01 | 1435.1017 | 1927.5309 | 2709.5474 |
| Pkm | EATESFASDPILYRPVAV | -93 | -110 | 1 | 0 | 18 | 914.929 | 947.6771 | 900.4399 | 1571.2943 | 2158.6667 | 1960.247 |
| Pkm | ADTFLEHMCR | -23 | -32 | 0 | 0 | 10 | 861.6993 | 922.144 | 829.6423 | 474.54208 | 716.484 | 203.90158 |
| Pkm | EATESFASDPILYRPVAVAL | -93 | -112 | 1 | 0 | 20 | 841.0672 | 994.0439 | 898.8998 | 1458.1879 | 2182.7437 | 2062.2808 |
| Pkm | CCSGAIIVLTK | -423 | -433 | 0 | 0 | 11 | 784.80963 | 587.81885 | 1337.8973 | 533.3322 | 1094.8982 | 1205.3044 |
| Pkm | LDNAYMEK | -144 | -151 | 0 | 0 | 8 | 743.05426 | 180.60727 | 157.11511 | 1243.1084 | 886.67706 | 994.44135 |
| Pkm | MADTFLEHMCR | -22 | -32 | 0 | 0 | 11 | 438.77863 | 681.1026 | 519.8714 | 583.61945 | 853.86334 | 398.218 |
| Pkm | MADTFLEHMCR | -22 | -32 | 0 | 0 | 11 | 438.77863 | 681.1026 | 519.8714 | 583.61945 | 853.86334 | 398.218 |
| Pkm | RLAPITSDPTEAAAVGAVEA | -400 | -419 | 1 | 0.000627943 | 20 | 650.04913 | 801.6215 | 1455.3329 | 1630.3593 | 2282.4358 | 1892.152 |
| Pkm | AADVHEVR | -248 | -255 | 0 | 0.001291103 | 8 | 569.10364 | 541.04065 | 771.4953 | 484.63614 | 750.75183 | 696.7192 |
| Pkm | EATESFASDPILYRPVAVA | -93 | -111 | 1 | 0 | 19 | 552.7474 | 582.9633 | 578.0019 | 950.17834 | 1133.7604 | 1051.371 |
| Pkm | KGVNLPGAAVDLPAVS | -207 | -222 | 1 | 0 | 16 | 543.05426 | 577.24414 | 619.3844 | 466.93134 | 468.60138 | 721.64233 |
| Pkm | LAPITSDPTEAAAVGAVEAS | -401 | -420 | 0 | 0.00078661 | 20 | 469.17252 | 443.81384 | 352.07523 | 1052.4965 | 1011.56415 | 1030.3806 |
| Pkm | LAPITSDPTEAAAVGAVEASFK | -401 | -422 | 0 | 0 | 22 | 469.09595 | 436.23135 | 464.83218 | 746.371 | 925.13025 | 837.67755 |
| Pkm | LAPITSDPTEAAAVGAVEASFK | -401 | -422 | 0 | 0 | 22 | 469.09595 | 436.23135 | 464.83218 | 746.371 | 925.13025 | 837.67755 |
| Pkm | IYVDDGLISL | -174 | -183 | 0 | 0 | 10 | 437.0959 | 429.84576 | 436.44135 | 530.952 | 676.3341 | 552.41974 |
| Pkm | SGETAKGDYPLEAVR | -362 | -376 | 1 | 0 | 15 | 429.24524 | 572.38007 | 753.02905 | 897.8811 | 808.7068 | 755.93524 |
| Pkm | LAPITSDPTEAAAV | -401 | -414 | 0 | 0 | 14 | 419.95966 | 1114.2687 | 1141.5477 | 1375.4083 | 328.91733 | 1006.20325 |
| Pkm | AWAEDVDLR | -481 | -489 | 0 | 0 | 9 | 408.41116 | 369.99274 | 4931.3438 | 6048.13 | 6946.8726 | 6482.496 |
| Pkm | IYVDDGLISLQ | -174 | -184 | 0 | 0.00025015 | 11 | 280.5709 | 245.51433 | 295.37842 | 350.0806 | 416.367 | 392.70364 |
| Pkm | IYVDDGLISLQ | -174 | -184 | 0 | 0.00025015 | 11 | 280.5709 | 245.51433 | 295.37842 | 350.0806 | 416.367 | 392.70364 |
| Pkm | RLAPITSDPTEAAAVGAVEAS | -400 | -420 | 1 | 0.000550307 | 21 | 144.02614 | 208.9059 | 277.03607 | 472.29578 | 445.74127 | 430.99448 |
| Pkm | ADVHEVR | -249 | -255 | 0 | 0.001708008 | 7 | 34.022217 | 518.54694 | 697.6145 | 578.1118 | 480.7789 | 521.3538 |
| Pkm | FGVEQDVDMVFAS | -231 | -243 | 0 | 0 | 13 | 2039.8761 | 2070.2927 | 2222.1833 | 1980.762 | NaN | 2156.277 |
| Pkm | LNFSHGTH | -74 | -81 | 0 | 0.001392869 | 8 | 1163.8575 | 1145.0886 | 557.89905 | NaN | 255.04463 | 1604.7402 |
| Pkm | RLAPITSDPTEAA | -400 | -412 | 1 | 0 | 13 | 961.5119 | 1039.0662 | 1118.6907 | NaN | 559.6943 | 519.19165 |
| Pkm | RLAPITSDPTEAAAVGA | -400 | -416 | 1 | 0.001023236 | 17 | 636.4601 | 716.9953 | NaN | 974.1039 | 1031.7404 | 877.9375 |
| Pkm | IDSAPITAR | -35 | -43 | 0 | 0.007493591 | 9 | 567.431 | 565.4363 | 819.92645 | NaN | 971.22003 | 1014.50684 |
| Pkm | GSGTAEVELKKG | -126 | -137 | 2 | 0 | 12 | 483.03964 | 2366.7004 | NaN | 2036.9409 | 2841.467 | 3061.038 |

| | | | | | | | | | | | | |
|---|---|---|---|---|---|---|---|---|---|---|---|---|
| Pkm | MQHLIAR | -377 | -383 | 0 | 0.001636916 | 7 | 5813.271 | 3959.695 | 2957.6887 | 4172.395 | NaN | NaN |
| Pkm | ASDPILYR | -99 | -106 | 0 | 0 | 8 | 957.87573 | 507.17813 | 985.2998 | 967.9938 | NaN | NaN |
| Pkm | LAPITSDPTEAAAVGAVEA | -401 | -419 | 0 | 0.001291103 | 19 | 803.2323 | 1028.4197 | 1064.2786 | NaN | 1553.1349 | NaN |
| Pkm | EVENGGSLGSK | -196 | -206 | 0 | 0.00025015 | 11 | 396.11734 | 349.2802 | 352.175 | 686.7067 | NaN | NaN |
| Pkm | RFDEILEA | -279 | -286 | 1 | 0.006677519 | 8 | 240.77142 | 3009.627 | 2761.118 | 1787.7098 | NaN | NaN |
| Pkm | FASDPILYR | -98 | -106 | 0 | 0 | 9 | 1300.1475 | 1303.1184 | 1556.1111 | NaN | NaN | NaN |
| Pkm | RLAPITSDPTEA | -400 | -411 | 1 | 0 | 12 | 994.4413 | 1754.9508 | 727.075 | NaN | NaN | NaN |
| Pkm | LNFSHGTHEY | -74 | -83 | 0 | 0 | 10 | 645.53296 | 889.49554 | 940.68 | NaN | NaN | NaN |
| Pkm | KIENHEGVR | -270 | -278 | 1 | 0.004412728 | 9 | 611.06445 | NaN | NaN | NaN | 355.867 | 350.15912 |
| Pkm | EKDIQDLK | -223 | -230 | 1 | 0.001291103 | 8 | 1331.4264 | 1366.6016 | NaN | NaN | NaN | NaN |
| Pkm | SFASDPILYR | -97 | -106 | 0 | 0 | 10 | 1190.9849 | NaN | 1206.5046 | NaN | NaN | NaN |
| Pkm | AAVGAVEASFK | -412 | -422 | 0 | 0 | 11 | 1033.5892 | NaN | 1028.3087 | NaN | NaN | NaN |
| Pkm | LAPITSDPTEAAAVGAVE | -401 | -418 | 0 | 0.000394361 | 18 | 577.6484 | NaN | NaN | NaN | NaN | 976.5507 |
| Pkm | TFLEHMCR | -25 | -32 | 0 | 0.003521747 | 8 | 423.7943 | NaN | 281.09348 | NaN | NaN | NaN |
| Pkm | VNLAMDVGK | -490 | -498 | 0 | 0.001392869 | 9 | 4010.801 | NaN | NaN | NaN | NaN | NaN |
| Pkm | LAPITSDPTEAAAVG | -401 | -415 | 0 | 0 | 15 | 1239.5586 | NaN | NaN | NaN | NaN | NaN |
| Pkm | SDGIMVAR | -287 | -294 | 0 | 0.007880195 | 8 | 865.6646 | NaN | NaN | NaN | NaN | NaN |
| Pkm | APIIAVTR | -448 | -455 | 0 | 0 | 8 | 550.4595 | NaN | NaN | NaN | NaN | NaN |
| Pdcd6ip | LVKPTPVNVPVSQK | -337 | -350 | 1 | 0 | 14 | 2120.5642 | 2265.895 | 2190.221 | 2574.8503 | 2876.3435 | 2618.3691 |
| Pdcd6ip | LVKPTPVNVPVSQK | -337 | -350 | 1 | 0 | 14 | 2120.5642 | 2265.895 | 2190.221 | 2574.8503 | 2876.3435 | 2618.3691 |
| Pdcd6ip | LLDEEEATDNDLR | -457 | -469 | 0 | 0 | 13 | 1420.0447 | 1607.7925 | 1534.7798 | 1813.947 | 1265.6027 | 1521.0073 |
| Pdcd6ip | TPSNDLYKPLR | -479 | -489 | 1 | 0 | 11 | 1411.2993 | 1563.1389 | 1672.6859 | 1478.4641 | 1888.6082 | 1802.394 |
| Pdcd6ip | SLLSNLDEIKK | -554 | -564 | 1 | 0 | 11 | 1166.0934 | 1223.5753 | 2732.904 | 2249.2214 | 1001.6285 | 980.4415 |
| Pdcd6ip | YYDQICSIEPK | -71 | -81 | 0 | 0 | 11 | 931.6179 | 1461.4229 | 512.0054 | 946.7785 | 1113.9487 | 1079.3223 |
| Pdcd6ip | FTDLFEK | -351 | -357 | 0 | 0 | 7 | 761.26227 | 694.00977 | 620.9719 | 566.1339 | 657.7957 | 391.62326 |
| Pdcd6ip | FYNELTEILVR | -676 | -686 | 0 | 0 | 11 | 634.40015 | 647.61066 | 582.85876 | 903.28253 | 1100.7842 | 877.9314 |
| Pdcd6ip | AQDGVINEEALSVTELDR | -589 | -606 | 0 | 0 | 18 | 498.45102 | 517.21326 | 517.4985 | 633.56134 | 767.4524 | 827.3614 |
| Pdcd6ip | DKHEGALETLLR | -59 | -70 | 1 | 0.000230707 | 12 | 489.60538 | 421.42133 | 381.19754 | 408.92462 | 322.21716 | 506.46872 |
| Pdcd6ip | NLPAAIEDVSGDTVPQSILTK | -400 | -420 | 0 | 0 | 21 | 455.43512 | 551.41907 | 432.74902 | 660.78033 | 784.7986 | 784.963 |
| Pdcd6ip | AADYFGDAFK | -220 | -229 | 0 | 0.00025015 | 10 | 358.37967 | 1585.0814 | 998.27057 | 1575.4031 | 1674.8066 | 786.70056 |
| Pdcd6ip | TPSNDLYKPL | -479 | -488 | 1 | 0.000471409 | 10 | 211.13803 | 565.5396 | 213.2373 | 966.5609 | 952.8519 | 905.18097 |
| Pdcd6ip | LANQAADYFGDAFK | -216 | -229 | 0 | 0 | 14 | 130.8329 | 820.92267 | 917.24664 | 544.01074 | 668.9111 | 640.4477 |
| Pdcd6ip | TSEVDLAKPLVK | -12 | -23 | 1 | 0 | 12 | 737.9394 | 742.96545 | 876.057 | NaN | 457.72382 | 510.65018 |
| Pdcd6ip | KDNDFIYHDR | -313 | -322 | 1 | 0 | 10 | 502.63644 | 507.70026 | 640.7736 | NaN | 653.5455 | 460.28296 |
| Pdcd6ip | HEGALETLLR | -61 | -70 | 0 | 0 | 10 | 1421.4976 | 2674.354 | NaN | 1231.0885 | 1161.7583 | NaN |
| Pdcd6ip | IAAEQNLDNDEGLK | -134 | -147 | 0 | 0 | 14 | 980.1011 | NaN | NaN | 585.6422 | 609.2223 | 320.49365 |
| Pdcd6ip | ELPELLQR | -439 | -446 | 0 | 0.004923919 | 8 | 1930.0925 | 1912.091 | NaN | NaN | 1242.2954 | NaN |
| Pdcd6ip | NIQVSHQEFSK | -628 | -638 | 0 | 0.000230707 | 11 | 1697.998 | 2030.4377 | 2140.747 | NaN | NaN | NaN |
| Pdcd6ip | TSEVDLAKPL | -12 | -21 | 1 | 0 | 10 | 515.03754 | NaN | NaN | NaN | 434.5623 | 638.22815 |
| Pdcd6ip | ATLVKPTPVNVPVSQK | -335 | -350 | 1 | 0.001023236 | 16 | 337.35486 | 417.5855 | NaN | NaN | NaN | 259.24194 |
| Pdcd6ip | TMQGSEVVSVLK | -542 | -553 | 0 | 0 | 12 | 3870.8538 | 4216.551 | NaN | NaN | NaN | NaN |
| Dhx9 | AAECNIVVTQPR | -437 | -448 | 0 | 0 | 12 | 4919.947 | 7222.8403 | 1890.2844 | 1707.4078 | 2325.9412 | 2206.8616 |
| Dhx9 | KVFDPVPDGVTK | -699 | -710 | 1 | 0 | 12 | 2422.242 | 2539.477 | 2771.9922 | 3087.2886 | 3186.1013 | 3282.5322 |
| Dhx9 | GISHVIVDEIHER | -506 | -518 | 0 | 0 | 13 | 2011.2579 | 2868.6504 | 2887.8464 | 2579.1304 | 2820.344 | 3426.3384 |
| Dhx9 | GISHVIVDEIHER | -506 | -518 | 0 | 0 | 13 | 2011.2579 | 2868.6504 | 2887.8464 | 2579.1304 | 2820.344 | 3426.3384 |
| Dhx9 | LAHFEPSQR | -317 | -325 | 0 | 0 | 9 | 1660.4534 | 1851.0756 | 2054.1777 | 1124.5718 | 680.71814 | 2417.3257 |
| Dhx9 | DALDANDELTPLGR | -842 | -855 | 0 | 0 | 14 | 1628.8488 | 2136.3718 | 2138.4094 | 1881.6316 | 2377.9429 | 2595.1282 |
| Dhx9 | YGDGPRPPK | -1156 | -1164 | 1 | 0.000431655 | 9 | 1618.7976 | 1328.085 | 1520.9698 | 1475.8235 | 1540.3367 | 1508.0669 |
| Dhx9 | AATCFPEPFISEGK | -887 | -900 | 0 | 0 | 14 | 1467.8696 | 1578.7642 | 1496.1508 | 1866.3701 | 1552.0029 | 1495.374 |
| Dhx9 | AKLPIEPR | -858 | -865 | 1 | 0.000394361 | 8 | 1371.4941 | 1481.816 | 1498.5186 | 1339.4639 | 1450.5103 | 1408.0933 |
| Dhx9 | VFDPVPDGVTK | -700 | -710 | 0 | 0 | 11 | 1255.946 | 112.086395 | 1287.7844 | 1144.7737 | 1139.5579 | 1312.1532 |
| Dhx9 | LICGDEYGPETK | -612 | -623 | 0 | 0 | 12 | 985.5879 | 1077.0411 | 1093.146 | 1813.1594 | 399.04434 | 1895.3807 |
| Dhx9 | GYGSPGPTWDR | -133 | -143 | 0 | 0 | 11 | 936.6076 | 899.6863 | 1000.2877 | 865.85754 | 902.8185 | 751.11554 |
| Dhx9 | LQISHEAAAC | -1092 | -1101 | 0 | 0.00025015 | 10 | 867.32996 | 1337.932 | 1049.329 | 645.4772 | 799.8558 | 524.29047 |
| Dhx9 | TTQVPQYILDDFIQNDR | -420 | -436 | 0 | 0.00025015 | 17 | 822.9343 | 1021.2517 | 683.8255 | 1013.1233 | 1212.3038 | 1080.1204 |
| Dhx9 | TPLHEIALSIK | -798 | -808 | 0 | 0.00025015 | 11 | 759.93475 | 811.0582 | 793.6051 | 793.3762 | 897.5109 | 913.5039 |
| Dhx9 | QDHNLQSVLQER | -376 | -387 | 0 | 0 | 12 | 729.1965 | 636.94135 | 753.9028 | 811.3175 | 1310.4437 | 894.843 |
| Dhx9 | QDHNLQSVLQER | -376 | -387 | 0 | 0 | 12 | 729.1965 | 636.94135 | 753.9028 | 811.3175 | 1310.4437 | 894.843 |
| Dhx9 | FCTVGVLLR | -490 | -498 | 0 | 0 | 9 | 705.3241 | 816.5123 | 753.3696 | 240.7064 | 1233.4205 | 1213.4867 |
| Dhx9 | QVPQYILDDFIQNDR | -422 | -436 | 0 | 0 | 15 | 662.04834 | 586.34625 | 535.4882 | 943.05566 | 919.5134 | 852.5002 |
| Dhx9 | TCFPEPFISEGK | -889 | -900 | 0 | 0 | 12 | 641.0186 | 1035.7516 | 373.83252 | 737.7302 | 804.43134 | 510.3178 |
| Dhx9 | AFGVYPNVCYHK | -998 | -1009 | 0 | 0 | 12 | 574.7215 | 388.10147 | 317.60587 | 355.04236 | 894.40607 | 684.5686 |
| Dhx9 | YILDDFIQNDR | -426 | -436 | 0 | 0 | 11 | 558.1546 | 525.156 | 597.04346 | 709.32544 | 882.5896 | 730.5219 |
| Dhx9 | DKEDDGGEDDDANCNLIC | -597 | -614 | 1 | 0 | 18 | 447.53717 | 396.14432 | 374.5596 | 405.8059 | 371.94354 | 411.43408 |
| Dhx9 | YTQVGPDHNR | -202 | -211 | 0 | 0 | 10 | 388.3197 | 862.33795 | 751.39496 | 776.32434 | 1037.3948 | 795.91833 |
| Dhx9 | FESILPR | -477 | -483 | 0 | 0.00025015 | 7 | 1635.378 | 1251.4128 | 1117.0656 | 710.6042 | 543.3462 | NaN |
| Dhx9 | HLENNSHFGSHR | -671 | -682 | 0 | 0 | 12 | 412.12204 | NaN | 433.51883 | 278.10263 | 464.1756 | 547.70276 |
| Dhx9 | HLENNSHFGSHR | -671 | -682 | 0 | 0 | 12 | 412.12204 | NaN | 433.51883 | 278.10263 | 464.1756 | 547.70276 |
| Dhx9 | DVVLAYPEVR | -531 | -540 | 0 | 0 | 10 | 6682.9956 | 776.6438 | 7048.6865 | 5607.197 | NaN | NaN |
| Dhx9 | DANDELTPLGR | -845 | -855 | 0 | 0 | 11 | 2299.8357 | NaN | 2896.3987 | 3220.7854 | 3724.5874 | NaN |
| Dhx9 | KLEAGIR | -499 | -505 | 1 | 0.00644558 | 7 | 1128.9585 | 1088.309 | 996.0657 | NaN | 582.1209 | NaN |
| Dhx9 | LGGIGQFLAK | -812 | -821 | 0 | 0 | 10 | 916.2308 | NaN | 1044.1014 | NaN | 323.44937 | 892.3254 |
| Dhx9 | GVYPNVCYHK | -1000 | -1009 | 0 | 0 | 10 | 1235.7081 | NaN | NaN | NaN | 802.89215 | 1243.7124 |
| Dhx9 | LNQYFQK | -187 | -193 | 0 | 0.000394361 | 7 | 411.0047 | 1556.2509 | 958.4144 | NaN | NaN | NaN |
| Dhx9 | FDRLETH | -784 | -790 | 1 | 0.002199523 | 7 | 576.8737 | NaN | NaN | 1206.1317 | NaN | NaN |
| Dhx9 | MSGEEAEIR | -932 | -940 | 0 | 0.000550307 | 9 | 401.01 | NaN | 451.9265 | NaN | NaN | NaN |
| Hspa4 | FVSEDDRNTFTLK | -640 | -652 | 1 | 0 | 13 | 4583.5444 | 3679.7563 | 4881.814 | 1299.2897 | 1987.1884 | 1708.12 |
| Hspa4 | FVSEDDRNTFTLK | -640 | -652 | 1 | 0 | 13 | 4583.5444 | 3679.7563 | 4881.814 | 1299.2897 | 1987.1884 | 1708.12 |

| | | | | | | | | | | | | |
|---|---|---|---|---|---|---|---|---|---|---|---|---|
| Hspa4 | DLPALEEKPR | -187 | -196 | 1 | 0 | 10 | 5003.8413 | 3371.0623 | 2742.4675 | 12593.133 | 4125.376 | 3820.478 |
| Hspa4 | FQESEERPK | -690 | -698 | 1 | 0 | 9 | 3539.367 | 3416.6191 | 3521.6765 | 1838.1206 | 1898.4026 | 3850.4204 |
| Hspa4 | QDLPALEEKPR | -186 | -196 | 1 | 0 | 11 | 3363.0227 | 3901.7021 | 3828.9116 | 3128.1047 | 4362.9424 | 4356.2363 |
| Hspa4 | ELTSICSPIISK | -775 | -786 | 0 | 0 | 12 | 2534.174 | 1153.5729 | 2776.786 | 2199.289 | 2094.1177 | 1987.429 |
| Hspa4 | LKETAESVLK | -125 | -134 | 1 | 0 | 10 | 1820.5037 | 1844.4958 | 1882.2081 | 2252.8364 | 2120.7712 | 2083.4941 |
| Hspa4 | WNSPAEEGLSDCEVFPK | -406 | -422 | 0 | 0 | 17 | 1275.5679 | 1513.1675 | 1596.0663 | 1593.4574 | 2096.9636 | 1942.3816 |
| Hspa4 | VEPPKEEPK | -791 | -799 | 1 | 0.00025015 | 9 | 1222.0472 | 1159.7677 | 1430.8353 | 1303.5192 | 1105.5771 | 1872.0696 |
| Hspa4 | EHGADTAVPSDGDK | -820 | -833 | 0 | 0 | 14 | 1205.6936 | 1701.3093 | 1615.4789 | 1895.0892 | 1769.1155 | 2405.1646 |
| Hspa4 | KPVVDCVVSVPSFYTD | -135 | -150 | 1 | 0.00025015 | 16 | 1132.393 | 1670.12 | 1512.3795 | 1331.9385 | 1235.8293 | 1366.94 |
| Hspa4 | AGGIETIANEYSDR | -20 | -33 | 0 | 0 | 14 | 1092.1705 | 1231.0082 | 1173.9402 | 1184.4929 | 1456.6927 | 1304.9526 |
| Hspa4 | QLPTGLTGIK | -93 | -102 | 0 | 0 | 10 | 1058.5234 | 1048.6926 | 984.4398 | 407.9783 | 106.9815 | 601.09894 |
| Hspa4 | HFCEEFGK | -243 | -250 | 0 | 0.000298805 | 8 | 1037.5906 | 1071.3729 | 1091.6597 | 340.73856 | 213.51587 | 1808.265 |
| Hspa4 | KLEDTENWLYEDGEDQPK | -652 | -669 | 1 | 0 | 18 | 420.4332 | 420.77664 | 449.5159 | 994.0129 | 591.4555 | 645.29926 |
| Hspa4 | KLEDTENWLYEDGEDQPK | -652 | -669 | 1 | 0 | 18 | 420.4332 | 420.77664 | 449.5159 | 994.0129 | 591.4555 | 645.29926 |
| Hspa4 | NKEDQYEHLDAA | -719 | -730 | 1 | 0.000230707 | 12 | 946.75146 | 975.46655 | 528.8576 | 815.13763 | 909.20044 | 925.90674 |
| Hspa4 | FVSEDDRNTFTL | -640 | -651 | 1 | 0.00025015 | 12 | 667.84436 | 611.0747 | 687.067 | 1726.6309 | 1343.0164 | 1369.26 |
| Hspa4 | NKEDQYEHLDAADVTK | -719 | -734 | 1 | 0 | 16 | 603.95685 | 5577.575 | 7318.9395 | 4574.7007 | 5453.2544 | 6457.242 |
| Hspa4 | NKEDQYEHLDAADVTK | -719 | -734 | 1 | 0 | 16 | 603.95685 | 5577.575 | 7318.9395 | 4574.7007 | 5453.2544 | 6457.242 |
| Hspa4 | KPVVDCVVSVPSFYTDAER | -135 | -153 | 1 | 0 | 19 | 570.4246 | 629.0172 | 633.3825 | 124.18141 | 317.66193 | 248.08084 |
| Hspa4 | KPVVDCVVSVPSFYTDAER | -135 | -153 | 1 | 0 | 19 | 570.4246 | 629.0172 | 633.3825 | 124.18141 | 317.66193 | 248.08084 |
| Hspa4 | AAEHGADTAVPSDGDK | -818 | -833 | 0 | 0 | 16 | 472.00403 | 480.12665 | 550.71027 | 631.4788 | 835.72 | 804.0877 |
| Hspa4 | SNLAYDIVQLPTGLTGIK | -85 | -102 | 0 | 0 | 18 | 423.40793 | 422.72955 | 312.64188 | 452.5418 | 541.5771 | 493.70123 |
| Hspa4 | HAEQNGPVDGQGDNPGSQ | -800 | -817 | 0 | 0 | 18 | 384.15884 | 403.08087 | 421.2727 | 394.73285 | 542.2714 | 536.29956 |
| Hspa4 | HAEQNGPVDGQGDNPGSQAA | -800 | -819 | 0 | 0 | 20 | 317.88867 | 299.20435 | 303.81943 | 453.48297 | 515.2379 | 549.484 |
| Hspa4 | SITDVVPYPISLR | -393 | -405 | 0 | 0 | 13 | 3383.4912 | 3621.8665 | 3835.3071 | 5259.086 | 6033.88 | NaN |
| Hspa4 | VVSVPSFYTDAER | -141 | -153 | 0 | 0 | 13 | 2839.4539 | 3256.473 | 3666.6807 | NaN | 4670.699 | 4792.0874 |
| Hspa4 | DVVPYPISLR | -396 | -405 | 0 | 0 | 10 | 1168.4237 | 1274.5127 | 1267.176 | NaN | 186.95982 | 818.65063 |
| Hspa4 | QSLTVDPVVK | -756 | -765 | 0 | 0 | 10 | 1427.9111 | NaN | NaN | 1171.6707 | 1520.751 | 1406.9825 |
| Hspa4 | CTPACVSFGPK | -34 | -44 | 0 | 0 | 11 | 1158.4983 | 1156.3129 | NaN | NaN | 1224.5479 | 1217.4459 |
| Hspa4 | FVSEDDRNTF | -640 | -649 | 1 | 0 | 10 | 807.58685 | 747.1487 | 266.5757 | 577.1369 | NaN | NaN |
| Hspa4 | LEDTENWLYEDGEDQPK | -653 | -669 | 0 | 0 | 17 | 670.7169 | 648.8888 | 620.1947 | 707.63513 | NaN | NaN |
| Hspa4 | EDQYEHLDAADVTK | -721 | -734 | 0 | 0 | 14 | 562.64874 | 918.5358 | 910.40564 | 906.5921 | NaN | NaN |
| Hspa4 | NLAYDIVQLPTGLTGIK | -86 | -102 | 0 | 0 | 17 | 361.84512 | NaN | NaN | 392.50806 | 493.9353 | 515.13635 |
| Hspa4 | IVQLPTGLTGIK | -91 | -102 | 0 | 0.001392869 | 12 | 232.46384 | 246.76797 | 83.79257 | 95.77029 | NaN | NaN |
| Hspa4 | STTLNADEAVTR | -363 | -374 | 0 | 0 | 12 | 1146.81 | 1224.5927 | NaN | 359.41614 | NaN | NaN |
| Hspa4 | NAVEEYVYEMR | -620 | -630 | 0 | 0.000431655 | 11 | 1582.7144 | 1615.8668 | NaN | NaN | NaN | NaN |
| Hspa4 | EDIYAVEIVGGATR | -333 | -346 | 0 | 0 | 14 | 894.66394 | 1075.0881 | NaN | NaN | NaN | NaN |
| Hspa4 | HAEQNGPVDGQGDNPGSQAAEHGADTAVPSDGD | -800 | -833 | 0 | 0 | 34 | 171.34784 | 1315.8455 | NaN | NaN | NaN | NaN |
| Hspa4 | ATAFDTTLGGR | -224 | -234 | 0 | 0.00025015 | 11 | 512.7867 | NaN | NaN | NaN | NaN | NaN |
| Hspa4 | ALVEVHK | -496 | -502 | 0 | 0.000712251 | 7 | 343.87784 | NaN | NaN | NaN | NaN | NaN |
| Stat1 | QEQLLLHK | -194 | -201 | 0 | 0 | 8 | 4808.323 | 4705.8184 | 5084.158 | 4790.2754 | 2836.2056 | 2773.3435 |
| Stat1 | YLYPNIDKDHAFGK | -666 | -679 | 1 | 0 | 14 | 4078.5828 | 4178.315 | 4098.2246 | 9213.656 | 4543.2573 | 5267.0063 |
| Stat1 | YLYPNIDKDHAFGK | -666 | -679 | 1 | 0 | 14 | 4078.5828 | 4178.315 | 4098.2246 | 9213.656 | 4543.2573 | 5267.0063 |
| Stat1 | FTYEPDPITK | -287 | -296 | 0 | 0 | 10 | 3607.4673 | 3780.1848 | 3888.8677 | 4204.379 | 5129.4775 | 4974.9224 |
| Stat1 | DQQPGTFLLR | -593 | -602 | 0 | 0 | 10 | 2383.6802 | 2424.2651 | 2543.3743 | 1372.7198 | 313.27936 | 1608.1016 |
| Stat1 | FHDLLSQLDDQYSR | -57 | -70 | 0 | 0 | 14 | 2214.7332 | 2273.837 | 3012.0044 | 1380.9591 | 3635.2473 | 2121.7444 |
| Stat1 | FHDLLSQLDDQYSR | -57 | -70 | 0 | 0 | 14 | 2214.7332 | 2273.837 | 3012.0044 | 1380.9591 | 3635.2473 | 2121.7444 |
| Stat1 | ELSAVTFPDIIR | -638 | -649 | 0 | 0 | 12 | 2191.6033 | 2294.3213 | 2503.67 | 2524.8164 | 3172.485 | 3224.6248 |
| Stat1 | EELQDEYDFK | -164 | -173 | 0 | 0 | 10 | 2107.6528 | 2349.2854 | 2593.2012 | 3962.9255 | 4304.942 | 4083.4087 |
| Stat1 | EELQDEYDFK | -164 | -173 | 0 | 0 | 10 | 2107.6528 | 2349.2854 | 2593.2012 | 3962.9255 | 4304.942 | 4083.4087 |
| Stat1 | TLEELQDEYDFK | -162 | -173 | 0 | 0 | 12 | 1804.407 | 2550.723 | 2437.034 | 1573.9886 | 261.34256 | 2074.2214 |
| Stat1 | EAPEPMELDDPKR | -686 | -698 | 1 | 0 | 13 | 1623.6458 | 1565.756 | 1539.6062 | 1660.386 | 150.03058 | 1903.0356 |
| Stat1 | EAPEPMELDDPKR | -686 | -698 | 1 | 0 | 13 | 1623.6458 | 1565.756 | 1539.6062 | 1660.386 | 150.03058 | 1903.0356 |
| Stat1 | LLGPNAGPDGLIPWTR | -526 | -541 | 0 | 0 | 16 | 1568.9805 | 1791.3771 | 1760.9657 | 1621.2847 | 1973.6682 | 2231.0662 |
| Stat1 | AAENIPENPLK | -655 | -665 | 0 | 0 | 11 | 1527.8549 | 1534.2825 | 1774.2339 | 340.76254 | 2073.7402 | 386.00446 |
| Stat1 | AGPDGLIPWTR | -531 | -541 | 0 | 0 | 11 | 1354.6832 | 1599.3064 | 1296.2389 | 245.30505 | 2386.0132 | 2336.6943 |
| Stat1 | YLYPNIDK | -666 | -673 | 0 | 0 | 8 | 1102.9608 | 1115.2229 | 1080.8146 | 1146.3273 | 1033.4427 | 1011.56323 |
| Stat1 | DQVMCIEQEIK | -151 | -161 | 0 | 0 | 11 | 942.1832 | 954.8709 | 895.5392 | 627.7148 | 1449.4465 | 781.8974 |
| Stat1 | NTLINDELVEWK | -229 | -240 | 0 | 0 | 12 | 807.861 | 814.6667 | 707.39996 | 1165.8367 | 3862.3882 | 879.52 |
| Stat1 | SDQKQEQLLLHK | -190 | -201 | 1 | 0 | 12 | 757.2187 | 910.07227 | 950.80994 | 672.133 | 1088.5973 | 1188.3545 |
| Stat1 | SDQKQEQLLLHK | -190 | -201 | 1 | 0 | 12 | 757.2187 | 910.07227 | 950.80994 | 672.133 | 1088.5973 | 1188.3545 |
| Stat1 | KLEELEQK | -279 | -286 | 1 | 0.003133457 | 8 | 711.87506 | 2908.537 | 2944.63 | 3804.2366 | 3861.2634 | 3177.6863 |
| Stat1 | YLYPNIDKDHAF | -666 | -677 | 1 | 0.000230707 | 12 | 2990.0928 | 668.3873 | 3158.867 | NaN | 3164.2273 | 3413.2634 |
| Stat1 | FLEQVHQL | -14 | -21 | 0 | 0.00025015 | 8 | 2903.0564 | 3149.7407 | 3257.8323 | 3430.868 | 643.6448 | NaN |
| Stat1 | LSQLDDQYSR | -61 | -70 | 0 | 0 | 10 | 1154.0581 | 1194.6375 | 1373.0237 | 2025.1464 | 1655.399 | NaN |
| Stat1 | KTGVQFTVK | -336 | -344 | 1 | 0.000431655 | 9 | 1079.0583 | NaN | 1390.5175 | 1194.1161 | 1320.7145 | 1272.8258 |
| Stat1 | KLEELEQKF | -279 | -287 | 2 | 0.000431655 | 9 | 192.97815 | 118.11856 | 533.00525 | NaN | 317.59888 | 487.59515 |
| Stat1 | INDELVEWK | -232 | -240 | 0 | 0.000550307 | 9 | 1293.189 | 663.06885 | NaN | NaN | 1556.5497 | 2301.2139 |
| Stat1 | NLQDNFQEDPVQMS | -89 | -102 | 0 | 0 | 14 | 887.037 | 916.69965 | 779.5586 | 732.49084 | NaN | NaN |
| Stat1 | LKDQQPGTFLLR | -591 | -602 | 1 | 0 | 12 | 678.38385 | 668.9421 | 708.57135 | 561.04584 | NaN | NaN |
| Stat1 | QKQEQLLLHK | -192 | -201 | 1 | 0 | 10 | 514.64514 | 494.2135 | 427.60327 | 407.17297 | NaN | NaN |
| Stat1 | FHDLLSQL | -57 | -64 | 0 | 0.005605646 | 8 | 1045.9882 | NaN | NaN | 1491.9686 | 1476.9722 | NaN |
| Stat1 | YDDSFPMEIR | -22 | -31 | 0 | 0 | 10 | 884.655 | NaN | 923.6251 | 972.2177 | NaN | NaN |
| Stat1 | SWQFSSVTK | -503 | -511 | 0 | 0 | 9 | 832.4595 | 720.0492 | 363.67767 | NaN | NaN | NaN |
| Stat1 | QDWEHAAYDVSFA | -41 | -53 | 0 | 0.00025015 | 13 | 566.7257 | 595.11847 | 697.81775 | NaN | NaN | NaN |
| Stat1 | SAVTFPDIIR | -640 | -649 | 0 | 0 | 10 | 341.82288 | NaN | 334.72852 | NaN | NaN | 86.20054 |
| Stat1 | GLNADQLSMLGEK | -513 | -525 | 0 | 0.001092529 | 13 | 1356.3563 | NaN | NaN | NaN | 3185.745 | NaN |
| Nemf | DQQQNEIIVK | -544 | -553 | 0 | 0.008736733 | 10 | 894.35986 | 1153.4021 | 1288.9344 | NaN | NaN | NaN |

| | | | | | | | | | | | | |
|---|---|---|---|---|---|---|---|---|---|---|---|---|
| Nemf | AAEPLLTLER | -160 | -169 | 0 | 0.004439092 | 10 | 287.5654 | 441.33536 | NaN | NaN | NaN | NaN |
| Plrg1 | AAEPQIITGSHDTTIR | -339 | -354 | 0 | 0 | 16 | 714.2194 | 673.339 | 622.193 | 698.63525 | 698.50305 | 952.85126 |
| Plrg1 | VISGHLGWVR | -198 | -207 | 0 | 0 | 10 | 178.5706 | 1382.3927 | 1049.3607 | 980.54553 | 1067.0164 | 832.8235 |
| Plrg1 | TGYNFQR | -442 | -448 | 0 | 0.000431655 | 7 | 1119.1605 | 1078.2529 | NaN | 1087.3267 | 639.01996 | NaN |
| Plrg1 | TGPAGSEYR | -141 | -149 | 0 | 0.003490217 | 9 | 680.90784 | 673.65314 | NaN | NaN | 718.4169 | NaN |
| Nup93 | SEPSYIGDVNPPGR | -162 | -175 | 0 | 0 | 14 | 4008.3003 | 2983.8948 | 2858.9507 | 3358.214 | 4336.0474 | 3670.8574 |
| Nup93 | LNQVCFDDDGTSSPQDR | -418 | -434 | 0 | 0.000712251 | 17 | 1353.1218 | 1479.7579 | 1455.4954 | 2216.4143 | 2729.9612 | 2805.6929 |
| Nup93 | FTSDTKPIINK | -601 | -611 | 1 | 0 | 11 | 1334.022 | 1260.6934 | 1355.4918 | 1302.7374 | 1448.4121 | 1277.0458 |
| Nup93 | LLSPVVPQISAPQSNK | -645 | -660 | 0 | 0 | 16 | 1179.2871 | 866.08167 | 819.35034 | 770.6043 | 653.07874 | 756.0007 |
| Nup93 | AAETEGISELPHVER | -17 | -31 | 0 | 0 | 15 | 1058.1335 | 1144.9978 | 1213.2753 | 1389.3367 | 1667.563 | 1689.7395 |
| Nup93 | GLDISHISQR | -68 | -77 | 0 | 0 | 10 | 682.5485 | 991.5407 | 926.6784 | 1276.9084 | 879.8981 | 873.04315 |
| Nup93 | QVCFDDDGTSSPQDR | -420 | -434 | 0 | 0 | 15 | 452.2058 | 519.6793 | 465.80096 | 718.5526 | 497.81015 | 492.64624 |
| Nup93 | AEQLAAETEGISELPHVER | -13 | -31 | 0 | 0 | 19 | 337.40695 | 418.19168 | 460.1731 | 425.16455 | 585.8546 | 462.7465 |
| Nup93 | RNFSDEIR | -735 | -742 | 1 | 0.000591124 | 8 | 690.3285 | 1204.622 | 811.09924 | NaN | 1128.4792 | 1162.8126 |
| Nup93 | LPAPSPGLQDGEVEGHPVWAL | -301 | -321 | 0 | 0.00149795 | 21 | 327.5307 | NaN | 382.81885 | 308.98898 | 427.09756 | 497.7388 |
| Nup93 | NLQEIQQAGER | -32 | -42 | 0 | 0 | 11 | 662.1979 | 855.0484 | 981.5445 | NaN | 1048.0951 | NaN |
| Nup93 | LVPLNQESVEER | -719 | -730 | 0 | 0 | 12 | 1125.505 | 1016.311 | NaN | 1078.1467 | NaN | NaN |
| Nup93 | AVYCIIGR | -389 | -396 | 0 | 0.00111006 | 8 | 1639.3488 | NaN | NaN | 1133.47 | NaN | NaN |
| Nup93 | FTQESEPSYIGDVNPPGR | -158 | -175 | 0 | 0.005698678 | 18 | 136.08707 | NaN | NaN | NaN | NaN | 218.90636 |
| Sarnp | AAETVELHK | -2 | -10 | 0 | 0 | 9 | 2614.7673 | 2710.4316 | 2677.102 | 585.0681 | 3047.4731 | 3143.5327 |
| Sarnp | DVLGDETEEEEPKPIELPVKEEEPPEK | -54 | -80 | 2 | 0.001392869 | 27 | 646.7945 | 728.2592 | 692.5519 | 706.88336 | 839.07 | 832.67804 |
| Sarnp | ANEEDVLGDETEEEEPKPIELPVKEEEPPEK | -50 | -80 | 2 | 0.000230707 | 31 | 442.89435 | 406.21448 | 380.18683 | 530.6964 | 449.51382 | 776.7287 |
| Sarnp | ANEEDVLGDETEEEEPKPIELPVKEEEPPEK | -50 | -80 | 2 | 0.000230707 | 31 | 442.89435 | 406.21448 | 380.18683 | 530.6964 | 449.51382 | 776.7287 |
| Sarnp | GLSSDTKPMVNLDK | -136 | -149 | 1 | 0 | 14 | 439.3802 | 600.2801 | 792.46686 | 488.68597 | 262.4877 | NaN |
| Snrnp200 | GSEEQLLPVEK | -456 | -466 | 0 | 0 | 11 | 6134.5557 | 6752.265 | 6934.883 | 4535.7866 | 7159.496 | 7827.1733 |
| Snrnp200 | AAEYENIPIR | -1852 | -1861 | 0 | 0 | 10 | 1525.1901 | 1356.4476 | 580.5454 | 1539.4989 | 645.7882 | 760.2353 |
| Snrnp200 | VYNSDDNVFVGAPTGSGK | -1339 | -1356 | 0 | 0 | 18 | 1401.2487 | 1675.6907 | 1319.9755 | 813.64856 | 1383.4578 | 1472.0211 |
| Snrnp200 | LDFVAPATGGHNY | -2092 | -2104 | 0 | 0 | 13 | 1329.9878 | 1179.6553 | 1071.9056 | 1474.8477 | 1454.1122 | 1467.8004 |
| Snrnp200 | PFGSEEQLLPVEK | -454 | -466 | 0 | 0 | 13 | 1225.0367 | 2211.001 | 1670.2655 | 684.9148 | 1586.6527 | 1722.8584 |
| Snrnp200 | YAQDEHLITF | -1245 | -1254 | 0 | 0.000230707 | 10 | 1179.6965 | 1350.0149 | 1264.6172 | 1421.4312 | 1426.7225 | 1384.6873 |
| Snrnp200 | WLSCETQLPVSFR | -1275 | -1287 | 0 | 0 | 13 | 1111.9113 | 1171.8298 | 1558.1307 | 2093.4143 | 1272.5599 | 1261.9391 |
| Snrnp200 | KLPEEVVK | -1134 | -1141 | 1 | 0.00025015 | 8 | 936.5081 | 1022.7306 | 1027.3181 | 817.349 | 995.2087 | 1083.6344 |
| Snrnp200 | ELTGDHQLCK | -568 | -577 | 0 | 0 | 10 | 897.3667 | 1104.72 | 179.35664 | 1166.457 | 1266.483 | 1064.8684 |
| Snrnp200 | TLVEDLFADK | -795 | -804 | 0 | 0 | 10 | 847.3713 | 1231.3738 | 682.77234 | 1970.0293 | 230.60909 | 2235.4585 |
| Snrnp200 | GVNLPAHTVIIK | -818 | -829 | 0 | 0.000431655 | 12 | 818.0923 | 963.28796 | 1061.8198 | 1027.4326 | 1222.0037 | 1198.3048 |
| Snrnp200 | VAELTGDHQLCK | -566 | -577 | 0 | 0.000230707 | 12 | 796.3279 | 945.6098 | 798.06104 | 730.6912 | 951.6723 | 841.458 |
| Snrnp200 | EEEVTGPVIAPLFPQK | -2044 | -2059 | 0 | 0 | 16 | 693.9012 | 880.7463 | 990.0352 | 774.44464 | 1126.8129 | 1100.4902 |
| Snrnp200 | NTVYNSDDNVFVGAPTGSGK | -1337 | -1356 | 0 | 0 | 20 | 566.4297 | 399.37656 | 296.58423 | 1802.815 | 375.33856 | 165.1124 |
| Snrnp200 | SPTLYGISHDDLK | -932 | -944 | 0 | 0 | 13 | 501.28085 | 676.5781 | 696.40674 | 302.26425 | 1009.6659 | 833.3854 |
| Snrnp200 | EEGWWVVIGDAK | -2061 | -2072 | 0 | 0 | 12 | 417.99738 | 343.82297 | 360.6676 | 147.45612 | 664.70435 | 956.9169 |
| Snrnp200 | LYDLNHNEIGELIR | -1153 | -1166 | 0 | 0 | 14 | 374.12027 | 636.6489 | 522.9872 | 783.21106 | 509.64758 | 423.96347 |
| Snrnp200 | LYDLNHNEIGELIR | -1153 | -1166 | 0 | 0 | 14 | 374.12027 | 636.6489 | 522.9872 | 783.21106 | 509.64758 | 423.96347 |
| Snrnp200 | VFNTVYNSDDNVFVGAPTGSGK | -1335 | -1356 | 0 | 0 | 22 | 337.47385 | 307.77255 | 146.54312 | 128.99132 | 350.80084 | 146.89578 |
| Snrnp200 | LHETEKEDLIR | -363 | -373 | 1 | 0 | 11 | 212.17076 | 674.40546 | 656.8387 | 671.6779 | 911.07874 | 850.3575 |
| Snrnp200 | LTITPDFQWDEK | -1202 | -1213 | 0 | 0.00025015 | 12 | 3410.6997 | NaN | 3715.1028 | 3942.4841 | 591.9361 | 5207.7153 |
| Snrnp200 | EKDLIPYLEK | -1594 | -1603 | 1 | 0 | 10 | 2045.7488 | 2254.5007 | 2489.2664 | NaN | 507.432 | 2377.1514 |
| Snrnp200 | GNIIISTPEK | -1422 | -1431 | 0 | 0.00025015 | 10 | 1201.3484 | 1069.4912 | 1115.5021 | 964.8667 | NaN | 499.79733 |
| Snrnp200 | NLPAHTVIIK | -820 | -829 | 0 | 0.000347792 | 10 | 513.46014 | 596.59155 | 548.05566 | 439.52283 | 428.00165 | NaN |
| Snrnp200 | ANRPLQDDEGR | -1691 | -1701 | 1 | 0.00025015 | 11 | 78.96153 | 473.37784 | 652.28937 | NaN | 186.03323 | 767.0742 |
| Snrnp200 | LTDSQIADVAR | -1999 | -2009 | 0 | 0 | 11 | 2503.4395 | 1783.6329 | 1915.2534 | NaN | NaN | 904.31256 |
| Snrnp200 | QIIVCTPEK | -584 | -592 | 0 | 0 | 9 | 1061.1011 | 349.8444 | 378.91827 | 232.144 | NaN | NaN |
| Snrnp200 | HHEDNLLR | -1862 | -1869 | 0 | 0.000947459 | 8 | 962.44336 | 530.6768 | 842.8342 | 921.04675 | NaN | NaN |
| Snrnp200 | FLYEPLPVESHLDHC | -1717 | -1731 | 0 | 0 | 15 | 558.07684 | 450.36267 | 210.23788 | NaN | NaN | 460.38843 |
| Snrnp200 | FVPVFEPLPPQYFIR | -1255 | -1269 | 0 | 0 | 15 | 165.47281 | NaN | NaN | 252.65335 | 182.80962 | 195.58078 |
| Snrnp200 | RLDLVHTA | -953 | -960 | 1 | 0.006735683 | 8 | 1256.3417 | NaN | NaN | NaN | 1270.6362 | 1329.8049 |
| Snrnp200 | EEEKLELQK | -1031 | -1039 | 1 | 0.008680443 | 9 | 3517.0923 | NaN | NaN | NaN | 2704.0393 | NaN |
| Snrnp200 | DLDQGGEALAPR | -395 | -406 | 0 | 0 | 12 | 2257.9053 | 1400.4819 | NaN | NaN | NaN | NaN |
| Snrnp200 | YVHLFPK | -1177 | -1183 | 0 | 0.00078661 | 7 | 401.52972 | 376.89713 | NaN | NaN | NaN | NaN |
| Snrnp200 | TQLPVSFR | -1280 | -1287 | 0 | 0.00025015 | 8 | 341.50665 | NaN | 459.3041 | NaN | NaN | NaN |
| Snrnp200 | NQLLKPTLSEIELFR | -1002 | -1016 | 1 | 0 | 15 | 239.22763 | 259.7318 | NaN | NaN | NaN | NaN |
| Snrnp200 | NQLLKPTLSEIELFR | -1002 | -1016 | 1 | 0 | 15 | 239.22763 | 259.7318 | NaN | NaN | NaN | NaN |
| Snrnp200 | YISSQIERPIR | -1476 | -1486 | 1 | 0.006078721 | 11 | 2273.1467 | NaN | NaN | NaN | NaN | NaN |
| Snrnp200 | HEYFLLK | -1236 | -1242 | 0 | 0.001811524 | 7 | 1271.698 | NaN | NaN | NaN | NaN | NaN |
| Snrnp200 | CQLPDGSFR | -428 | -436 | 0 | 0.006260252 | 9 | 493.47086 | NaN | NaN | NaN | NaN | NaN |
| Snrnp200 | SATLPNYEDVATFLR | -652 | -666 | 0 | 0.008129024 | 15 | 372.78726 | NaN | NaN | NaN | NaN | NaN |
| Tln1 | ENEEIGAHIK | -1908 | -1917 | 0 | 0 | 10 | 7442.7095 | 4557.076 | 8339.186 | 7620.6865 | 5402.6523 | 4895.0137 |
| Tln1 | GVEEHETLLLR | -184 | -194 | 0 | 0 | 11 | 2891.0806 | 2694.2678 | 2994.2112 | 2096.9739 | 1995.7109 | 2150.8987 |
| Tln1 | NHDEYSLVR | -123 | -131 | 0 | 0 | 9 | 2290.0513 | 2150.9607 | 831.1981 | 1753.1646 | 1585.3208 | 1623.7781 |
| Tln1 | EDQENETVVVK | -2481 | -2491 | 0 | 0 | 11 | 2216.5518 | 2270.8726 | 2657.1948 | 755.5104 | 3613.3152 | 2870.128 |
| Tln1 | AAFEDQENETVVVK | -2478 | -2491 | 0 | 0 | 14 | 1775.9418 | 1815.7135 | 1992.9174 | 2065.829 | 2085.2659 | 2059.7598 |
| Tln1 | AIVLDTASDVLDK | -1149 | -1161 | 0 | 0 | 13 | 1720.1307 | 2189.2063 | 2139.669 | 1863.0441 | 2292.4023 | 2323.5154 |
| Tln1 | ALNELLQHVK | -777 | -786 | 0 | 0 | 10 | 1564.5084 | 1637.6439 | 1689.3947 | 1834.5358 | 1834.1434 | 1839.5583 |
| Tln1 | AENEEIGAHIK | -1907 | -1917 | 0 | 0 | 11 | 1512.7876 | 1549.6458 | 1510.3732 | 1105.1538 | 250.74188 | 1716.1251 |
| Tln1 | ASNPEFSSVPAQISPEGR | -1576 | -1593 | 0 | 0 | 18 | 1051.0209 | 1378.7781 | 1566.8917 | 1600.2443 | 1999.621 | 1858.3549 |
| Tln1 | LHTDDELNWLDHGR | -165 | -178 | 0 | 0 | 14 | 307.26703 | 1052.3392 | 1056.5596 | 1092.8977 | 1464.9293 | 1353.225 |
| Tln1 | LHTDDELNWLDHGR | -165 | -178 | 0 | 0 | 14 | 307.26703 | 1052.3392 | 1056.5596 | 1092.8977 | 1464.9293 | 1353.225 |
| Tln1 | ADAEGESDLENSR | -842 | -854 | 0 | 0 | 13 | 990.257 | 937.70013 | 275.7445 | 853.6406 | 713.81036 | 798.6574 |
| Tln1 | VVAPTISSPVCQEQLVEAGR | -722 | -741 | 0 | 0 | 20 | 900.576 | 728.12067 | 604.6239 | 594.27905 | 869.49023 | 820.5767 |

| | | | | | | | | | | | | |
|---|---|---|---|---|---|---|---|---|---|---|---|---|
| Tln1 | AFEDQENETVVVK | -2479 | -2491 | 0 | 0 | 13 | 788.5818 | 833.3744 | 442.36356 | 708.7224 | 982.7127 | 341.56165 |
| Tln1 | ALDGDFTEENR | -1545 | -1555 | 0 | 0 | 11 | 781.94025 | 113.28813 | 377.1092 | 3081.3093 | 5166.4175 | 828.0913 |
| Tln1 | VGDDPAVWQLK | -2105 | -2115 | 0 | 0 | 11 | 745.2744 | 1061.6896 | 1024.3209 | 669.2356 | 794.78656 | 795.32227 |
| Tln1 | EAAFHPEVAPDVR | -2221 | -2233 | 0 | 0 | 13 | 716.583 | 1439.6345 | 1638.1339 | 1184.2737 | 1289.25 | 1341.7823 |
| Tln1 | QATLDDFETLPPLGQDAASK | -510 | -529 | 0 | 0 | 20 | 668.59064 | 519.27576 | 667.11053 | 852.09283 | 897.0404 | 793.3218 |
| Tln1 | AVAEQIPLLVQ | -958 | -968 | 0 | 0 | 11 | 630.829 | 803.8242 | 965.6944 | 640.74866 | 895.83276 | 1150.2114 |
| Tln1 | GLAGAVSELLR | -614 | -624 | 0 | 0 | 11 | 552.95544 | 567.7958 | 548.4091 | 681.262 | 701.7826 | 710.5589 |
| Tln1 | ASGHPGDPESQQR | -1172 | -1184 | 0 | 0 | 13 | 479.75064 | 103.21589 | 486.9909 | 458.894 | 84.29317 | 575.2704 |
| Tln1 | GLVEPTQFAR | -1460 | -1469 | 0 | 0 | 10 | 382.9012 | 1027.1766 | 866.2498 | 999.54126 | 313.73535 | 1197.9745 |
| Tln1 | LNEAAAGLNQAATELVQASR | -1242 | -1261 | 0 | 0 | 20 | 210.32292 | 246.54485 | 242.06847 | 440.80753 | 524.2855 | 527.4595 |
| Tln1 | FLPSELRDEH | -2532 | -2541 | 1 | 0 | 10 | 1996.4851 | 1139.3527 | 1838.5891 | NaN | 157.99168 | 1925.4825 |
| Tln1 | AQATSDLVNAIK | -830 | -841 | 0 | 0 | 12 | 1010.69104 | 883.4251 | 1051.5509 | NaN | 752.3608 | 278.5372 |
| Tln1 | IGITNHDEYSLVR | -119 | -131 | 0 | 0 | 13 | 832.4337 | 959.5097 | NaN | 646.60754 | 851.0185 | 1483.3145 |
| Tln1 | IGITNHDEYSLVR | -119 | -131 | 0 | 0 | 13 | 832.4337 | 959.5097 | NaN | 646.60754 | 851.0185 | 1483.3145 |
| Tln1 | DAEGESDLENSR | -843 | -854 | 0 | 0 | 12 | 417.08762 | NaN | 565.5986 | 945.40826 | 934.1229 | 845.3978 |
| Tln1 | LLGEIAQGNENYAGIA | -1105 | -1120 | 0 | 0 | 16 | 390.59604 | 344.9552 | 359.8751 | NaN | 301.79553 | 401.4827 |
| Tln1 | GAAAHPDSEEQQQR | -876 | -889 | 0 | 0.001092529 | 14 | 329.44067 | 358.3178 | 261.6977 | NaN | 282.78336 | 319.34662 |
| Tln1 | QAATEDGQLLR | -755 | -765 | 0 | 0 | 11 | 2985.4917 | 1663.5143 | 3459.5354 | NaN | 4686.798 | NaN |
| Tln1 | CQIQFGPHNEQK | -243 | -254 | 0 | 0.00078661 | 12 | 1188.6935 | NaN | 1773.0435 | NaN | 2830.817 | 2303.222 |
| Tln1 | LKPLPGETMEK | -1076 | -1086 | 1 | 0 | 11 | 4985.9287 | NaN | NaN | 5104.006 | NaN | 4706.4033 |
| Tln1 | ALSTDPASPNLK | -1321 | -1332 | 0 | 0 | 12 | 3303.7993 | 3478.379 | NaN | 1332.9974 | NaN | NaN |
| Tln1 | STDPASPNLK | -1323 | -1332 | 0 | 0.000298805 | 10 | 1481.3478 | 598.6972 | 1237.6254 | NaN | NaN | NaN |
| Tln1 | ERELEEAR | -2512 | -2519 | 1 | 0.003681885 | 8 | 1201.325 | NaN | NaN | 898.018 | NaN | 962.3066 |
| Tln1 | ASQSFLQPGGK | -989 | -999 | 0 | 0 | 11 | 1062.863 | 1474.3005 | 1640.3693 | NaN | NaN | NaN |
| Tln1 | SKDHFGLEGDEE | -405 | -416 | 1 | 0.000230707 | 12 | 653.97296 | 607.18945 | 255.72054 | NaN | NaN | NaN |
| Tln1 | NQAATELVQASR | -1250 | -1261 | 0 | 0 | 12 | 448.98236 | NaN | NaN | NaN | 1429.0759 | 1359.3086 |
| Tln1 | DKAPGQLECETAIAA | -1653 | -1667 | 1 | 0.000471409 | 15 | 2658.6777 | NaN | NaN | 3465.253 | NaN | NaN |
| Tln1 | TSTPEDFIR | -2169 | -2177 | 0 | 0 | 9 | 769.3429 | NaN | NaN | 334.06146 | NaN | NaN |
| Tln1 | ATSDLVNAIK | -832 | -841 | 0 | 0 | 10 | 621.3877 | NaN | NaN | NaN | 863.14905 | NaN |
| Tln1 | DGDFTEENR | -1547 | -1555 | 0 | 0.004321969 | 9 | 340.76544 | NaN | NaN | NaN | 120.29137 | NaN |
| Tln1 | AGFLDLK | -257 | -263 | 0 | 0.003745449 | 7 | 193.20702 | 438.9615 | NaN | NaN | NaN | NaN |
| Dld | SEEQLKEEGIEFK | -405 | -417 | 1 | 0 | 13 | 2201.331 | 2439.8394 | 2482.279 | 2410.7446 | 2586.8254 | 2578.8354 |
| Dld | AIGDVVAGPMLAHK | -352 | -365 | 0 | 0 | 14 | 1175.72 | 1321.0836 | 1190.8287 | 956.19147 | 1218.6488 | 1049.1609 |
| Dld | HAHPTLSEAFR | -485 | -495 | 0 | 0 | 11 | 1160.5834 | 1121.9943 | 990.4655 | 579.6333 | 579.9882 | 1020.1621 |
| Dld | CVPSVIYTHPEVAWVGK | -388 | -404 | 0 | 0 | 17 | 921.89355 | 1054.8351 | 1112.8225 | 424.72025 | 1442.7988 | 1540.8337 |
| Dld | ALTGGIAHLFK | -133 | -143 | 0 | 0 | 11 | 583.7004 | 2680.006 | 2512.5017 | 3203.9238 | 3221.4814 | 2845.9006 |
| Dld | VCHAHPTLSEAFR | -483 | -495 | 0 | 0 | 13 | 581.1261 | 738.45844 | 742.8714 | 723.57434 | 1024.795 | 941.9256 |
| Dld | AAFGKPINF | -501 | -509 | 1 | 0 | 9 | 1183.394 | 1142.3984 | 1191.8164 | 956.8314 | 766.28754 | NaN |
| Dld | ADGSTQVIDTK | -167 | -177 | 0 | 0 | 11 | 324.19318 | 710.6074 | 878.30347 | 1003.39453 | 944.79205 | NaN |
| Dld | NLGLEELGIELDPK | -321 | -334 | 0 | 0 | 14 | 381.64935 | 530.0295 | NaN | NaN | 448.8659 | NaN |
| Dld | THPEVAWVGK | -395 | -404 | 0 | 0.00051306 | 10 | 1055.6616 | NaN | 1070.2866 | NaN | NaN | NaN |
| Dld | EYGASCEDIAR | -472 | -482 | 0 | 0.000230707 | 11 | 1367.3469 | NaN | NaN | NaN | NaN | NaN |
| Srm | VLIIGGGDGGVLR | -97 | -109 | 0 | 0.000230707 | 13 | 2389.2507 | 2532.8289 | 2537.3425 | 3198.1921 | 3505.45 | 3805.711 |
| Srm | IANLPLCSHPNPR | -83 | -95 | 0 | 0 | 13 | 1957.1743 | 2147.8757 | 2431.4414 | 578.7724 | 1914.4404 | 2017.4857 |
| Srm | AAFVLPEFTR | -286 | -295 | 0 | 0 | 10 | 1098.0609 | 1392.1818 | 1296.4071 | 666.79913 | 286.99573 | 671.8408 |
| Srm | AFVLPEFTR | -287 | -295 | 0 | 0 | 9 | 1071.0386 | 1146.7897 | 1157.5858 | 1056.5662 | 1181.6543 | 1197.0205 |
| Srm | EIDEDVIEVSKK | -124 | -135 | 1 | 0.000394361 | 12 | 961.16565 | 1111.5604 | 411.00107 | 1241.7667 | 1200.1105 | 565.4126 |
| Srm | MEPGPDGPAAPGPAAIR | -1 | -17 | 0 | 0 | 17 | 667.10486 | 645.0327 | 629.2678 | 891.3246 | 1047.4163 | 702.653 |
| Srm | GECQWLHLDLIK | -207 | -218 | 0 | 0 | 12 | 490.49838 | 891.81226 | 567.03986 | 490.80685 | 685.4228 | 710.19495 |
| Srm | FVLPEFTR | -288 | -295 | 0 | 0 | 8 | 2173.8794 | 2499.1865 | NaN | 2732.159 | 3289.6128 | 3449.815 |
| Srm | YQDILVFR | -48 | -55 | 0 | 0 | 8 | 1148.4595 | 1168.6523 | 1234.083 | 648.6295 | 655.85223 | NaN |
| Srm | NLPLCSHPNPR | -85 | -95 | 0 | 0 | 11 | 15122.883 | 12761.779 | 13416.639 | NaN | 15378.249 | NaN |
| Srm | EIDEDVIEVSK | -124 | -134 | 0 | 0 | 11 | 3770.5208 | 1559.6154 | 1771.5942 | NaN | NaN | 1975.0197 |
| Srm | TALKEDGILCCQ | -195 | -206 | 1 | 0.00025015 | 12 | 937.13245 | 1426.1323 | 1382.9658 | 668.02673 | NaN | NaN |
| Srm | KVLIIGGGDGGVL | -96 | -108 | 1 | 0.000394361 | 13 | 512.4742 | 527.4634 | 580.7458 | 596.065 | NaN | NaN |
| Srm | SKNPSTNFR | -252 | -260 | 1 | 0.00078661 | 9 | 395.84225 | NaN | 818.2418 | 163.95816 | NaN | 523.85315 |
| Srm | SLQVEQLLHHR | -34 | -44 | 0 | 0 | 11 | 364.9029 | 401.69278 | 434.60406 | NaN | NaN | NaN |
| Srm | HPSVESVVQC | -114 | -123 | 0 | 0.001023236 | 10 | 1501.3231 | 1764.4642 | NaN | NaN | NaN | NaN |
| Srm | NPSTNFR | -254 | -260 | 0 | 0.00453359 | 7 | 204.17114 | NaN | NaN | NaN | NaN | 968.9717 |
| Tomm22 | AAGAGEPLSPEELLPK | -7 | -22 | 0 | 0.00025015 | 16 | 1052.917 | 1278.0009 | 1033.289 | 352.20673 | 1134.0323 | 972.4874 |
| Tomm22 | VLPVVFETEK | -96 | -105 | 0 | 0.00025015 | 10 | 876.4753 | 968.399 | 936.15497 | 1873.987 | 2185.311 | 622.884 |
| Dnajc2 | AFNSVDPTFDNSVPSK | -156 | -171 | 0 | 0.000230707 | 16 | 1171.9684 | 1201.0415 | 1239.736 | 1279.3971 | 1486.3091 | 1150.1897 |
| Dnajc2 | AAGEPIKEGDNDYFTC | -125 | -140 | 1 | 0.000826806 | 16 | 623.4408 | 545.7556 | 488.8117 | 420.56903 | 424.56104 | 390.0715 |
| Dnajc2 | FEGPCIDSTPWTTEEQK | -544 | -560 | 0 | 0 | 17 | 562.9018 | 613.97833 | 672.82 | 597.4336 | 677.1728 | 661.10785 |
| Dnajc2 | NHFSDNEADR | -364 | -373 | 0 | 0.001708008 | 10 | 380.90344 | NaN | 649.8369 | 595.583 | 756.7874 | 762.7996 |
| Dnajc2 | EFSYLDEEEKEK | -226 | -237 | 1 | 0 | 12 | 515.7758 | 757.07446 | 550.2392 | NaN | 482.7561 | NaN |
| Dnajc2 | NQDHYAVL | -86 | -93 | 0 | 0.001811524 | 8 | 997.51404 | NaN | NaN | 334.9752 | 533.603 | NaN |
| Dnajc2 | AVNLFPAGTNSR | -465 | -476 | 0 | 0 | 12 | 410.48587 | NaN | 413.7096 | NaN | NaN | NaN |
| Dnajc2 | TYPVNTPER | -568 | -576 | 0 | 0.000431655 | 9 | 491.75522 | NaN | NaN | NaN | NaN | NaN |
| Gart | DQAEQVLHDVR | -744 | -754 | 0 | 0 | 11 | 2673.6135 | 2957.3013 | 3468.2788 | 3652.1694 | 3974.3887 | 4007.736 |
| Gart | FGDPECQVILPLLK | -293 | -306 | 0 | 0 | 14 | 1476.9248 | 1512.7361 | 1088.2103 | 1303.5409 | 1532.2969 | 1302.8071 |
| Gart | NIHPSLLPSFK | -913 | -923 | 0 | 0 | 11 | 1419.5359 | 1647.5715 | 1611.7899 | 274.86505 | 1995.919 | 1849.5182 |
| Gart | SHSLLPIIR | -663 | -671 | 0 | 0.000471409 | 9 | 1182.4417 | 1551.5168 | 1183.7075 | 1325.7161 | 1398.8153 | 1310.6299 |
| Gart | AHITGGGLLENIPR | -679 | -692 | 0 | 0 | 14 | 734.77075 | 859.0183 | 1044.6434 | 1002.71436 | 1329.7339 | 1390.6467 |
| Gart | AHITGGGLLENIPR | -679 | -692 | 0 | 0 | 14 | 734.77075 | 859.0183 | 1044.6434 | 1002.71436 | 1329.7339 | 1390.6467 |
| Gart | GFKDPLLASGTDGVGTK | -483 | -499 | 1 | 0 | 17 | 614.7085 | 746.6143 | 822.89655 | 732.2044 | 912.5864 | 866.3174 |
| Gart | FFLDYFSCGK | -533 | -542 | 0 | 0 | 10 | 440.02768 | 494.2648 | 497.1579 | 548.9555 | 549.8771 | 559.854 |
| Gart | SANFPALVVK | -139 | -148 | 0 | 0 | 10 | 353.52515 | 925.7565 | 1001.799 | 860.9047 | 964.0531 | 649.0286 |

| | | | | | | | | | | | | |
|---|---|---|---|---|---|---|---|---|---|---|---|---|
| Gart | VVACPEDSPR | -766 | -775 | 0 | 0.001190916 | 10 | 1519.5203 | 1195.9537 | 981.6793 | 467.83813 | NaN | 356.74652 |
| Gart | HEIPTAQWR | -117 | -125 | 0 | 0.00025015 | 9 | 1406.844 | 1347.708 | 624.03094 | NaN | 1163.0217 | 1059.9122 |
| Gart | EHTLAWK | -14 | -20 | 0 | 0.001636916 | 7 | 1376.0111 | 1606.9504 | 273.67273 | 1620.5115 | NaN | NaN |
| Gart | AAGFKDPLLASGTDGVGTK | -481 | -499 | 1 | 0 | 19 | 1324.4847 | 1041.925 | 1222.5851 | NaN | NaN | 1143.4768 |
| Gart | AAGFKDPLLASGTDGVGTK | -481 | -499 | 1 | 0 | 19 | 1324.4847 | 1041.925 | 1222.5851 | NaN | NaN | 1143.4768 |
| Gart | NFPALVVK | -141 | -148 | 0 | 0.001868731 | 8 | 351.43326 | 594.1101 | 590.3415 | 436.13953 | NaN | NaN |
| Gart | SSPAPGGCGDQTLGDLLLTPTR | -639 | -660 | 0 | 0 | 22 | 389.02878 | NaN | NaN | 396.59692 | 370.8751 | NaN |
| Gart | TVAEMPPAQDHK | -208 | -219 | 0 | 0 | 12 | 1802.635 | NaN | 1170.2516 | NaN | NaN | NaN |
| Gart | ILSGPFVR | -898 | -905 | 0 | 0.00383302 | 8 | 1292.0461 | NaN | 1050.8287 | NaN | NaN | NaN |
| Gart | AVQEIMQEK | -170 | -178 | 0 | 0.000550307 | 9 | 119.74795 | NaN | NaN | NaN | NaN | NaN |
| Gart | LQQEGELSEEEMAR | -716 | -729 | 0 | 0.00025015 | 14 | 116.88137 | NaN | NaN | NaN | NaN | NaN |
| Nolc1 | AAGGAVSTPAPGK | -497 | -509 | 0 | 0 | 13 | 3707.6328 | 259.24515 | 4307.911 | 3862.606 | 7888.1865 | 4184.7188 |
| Nolc1 | LETPNTFPK | -608 | -616 | 0 | 0.00025015 | 9 | 2391.1033 | 2266.4924 | 2332.409 | 442.87103 | 1991.3248 | 1885.8783 |
| Nolc1 | VVPSDLYPLVLR | -9 | -20 | 0 | 0 | 12 | 1220.426 | 1228.2618 | 1359.608 | 1083.4624 | 1335.509 | 1285.5372 |
| Nolc1 | SSVPPPSVPLPK | -299 | -310 | 0 | 0 | 12 | 1070.8407 | 1108.2338 | 1204.5891 | 625.09753 | 762.1575 | 725.1161 |
| Nolc1 | SEEEEEEEETEEK | -563 | -575 | 0 | 0 | 13 | 317.9285 | 360.54462 | 353.4944 | 338.8147 | 329.27948 | 353.8947 |
| Nolc1 | ESEEEEEEEETEEK | -562 | -575 | 0 | 0 | 14 | 271.6736 | 359.2014 | 386.7745 | 305.76852 | 341.1101 | 342.28668 |
| Nolc1 | AGPYSSVPPPSVPLPK | -295 | -310 | 0 | 0 | 16 | 3188.9905 | 2583.77 | 2405.5388 | NaN | 2727.6162 | 2658.5735 |
| Nolc1 | AEAKPGTPAK | -206 | -215 | 1 | 0.001868731 | 10 | 230.35788 | NaN | 226.7042 | 127.2891 | 285.3083 | 167.00537 |
| Nolc1 | SPKPAVTPK | -394 | -402 | 1 | 0 | 9 | 498.88754 | 451.04114 | 290.81165 | NaN | NaN | NaN |
| Nolc1 | VADNSFDAK | -642 | -650 | 0 | 0.00025015 | 9 | 244.10623 | 681.0111 | NaN | NaN | NaN | NaN |
| Uqcrc1 | GYGPIEQLPDYNR | -458 | -470 | 0 | 0 | 13 | 2510.9302 | 3087.8237 | 3544.8733 | 3645.5913 | 5490.259 | 5398.2354 |
| Uqcrc1 | TDLTDYLNR | -214 | -222 | 0 | 0.00025015 | 9 | 1940.9746 | 1845.2389 | 1871.6312 | 2419.9507 | 2209.7832 | 2175.256 |
| Uqcrc1 | VYEEDAVPGLTPCR | -256 | -269 | 0 | 0 | 14 | 1919.1262 | 2215.8887 | 2254.2358 | 2161.9648 | 3494.5647 | 3231.5852 |
| Uqcrc1 | GGVEHQQLLDLAQK | -235 | -248 | 0 | 0 | 14 | 1696.0254 | 2338.1372 | 2340.6592 | 1323.9991 | 1859.2467 | 1794.3275 |
| Uqcrc1 | GGVEHQQLLDLAQK | -235 | -248 | 0 | 0 | 14 | 1696.0254 | 2338.1372 | 2340.6592 | 1323.9991 | 1859.2467 | 1794.3275 |
| Uqcrc1 | LVSHLDGTTPVCEDIGR | -399 | -415 | 0 | 0 | 17 | 1596.5717 | 1996.8156 | 1708.6451 | 2223.65 | 2229.7334 | 1869.3777 |
| Uqcrc1 | LVSHLDGTTPVCEDIGR | -399 | -415 | 0 | 0 | 17 | 1596.5717 | 1996.8156 | 1708.6451 | 2223.65 | 2229.7334 | 1869.3777 |
| Uqcrc1 | GPIEQLPDYNR | -460 | -470 | 0 | 0 | 11 | 1592.3414 | 1032.178 | 1440.7952 | 1696.246 | 1696.2869 | 1481.9575 |
| Uqcrc1 | RFTGSEIR | -269 | -276 | 1 | 0 | 8 | 1394.9426 | 1826.4833 | 2075.6187 | 1676.242 | 2135.0386 | 1876.1686 |
| Uqcrc1 | HLDGTTPVCEDIGR | -402 | -415 | 0 | 0 | 14 | 889.19135 | 1141.3651 | 982.05444 | 2300.2524 | 3120.8687 | 3150.3823 |
| Uqcrc1 | NALVSHLDGTTPVCEDIGR | -397 | -415 | 0 | 0 | 19 | 825.1864 | 596.4672 | 566.97034 | 609.80774 | 811.9362 | 767.0807 |
| Uqcrc1 | NALVSHLDGTTPVCEDIGR | -397 | -415 | 0 | 0 | 19 | 825.1864 | 596.4672 | 566.97034 | 609.80774 | 811.9362 | 767.0807 |
| Uqcrc1 | AAGGVEHQQLLDLAQK | -233 | -248 | 0 | 0 | 16 | 751.4264 | 868.58655 | 892.3786 | 1211.3779 | 1278.4098 | 1188.3862 |
| Uqcrc1 | AAGGVEHQQLLDLAQK | -233 | -248 | 0 | 0 | 16 | 751.4264 | 868.58655 | 892.3786 | 1211.3779 | 1278.4098 | 1188.3862 |
| Uqcrc1 | AGGVEHQQLLDLAQK | -234 | -248 | 0 | 0 | 15 | 539.62866 | 311.89755 | 630.4223 | 437.68726 | 367.50653 | 308.84753 |
| Uqcrc1 | AGGVEHQQLLDLAQK | -234 | -248 | 0 | 0 | 15 | 539.62866 | 311.89755 | 630.4223 | 437.68726 | 367.50653 | 308.84753 |
| Uqcrc1 | AVEGPSENVR | -200 | -209 | 0 | 0 | 10 | 4264.596 | 4283.231 | 5116.6387 | NaN | 4874.2544 | 6707.3955 |
| Uqcrc1 | LCTSATESEVTR | -379 | -390 | 0 | 0 | 12 | 3466.674 | 3526.7961 | 3722.7607 | 3173.0056 | 3073.903 | NaN |
| Uqcrc1 | EVESIGAHLN | -112 | -121 | 0 | 0.000550307 | 10 | 1908.3932 | 1918.0297 | NaN | 2261.0352 | 2989.4944 | NaN |
| Uqcrc1 | SSLEDSQIEK | -154 | -163 | 0 | 0.000947459 | 10 | 2106.7302 | 1669.7014 | NaN | NaN | 2753.735 | NaN |
| Uqcrc1 | NRPGNALEK | -103 | -111 | 1 | 0.00149795 | 9 | 958.79535 | NaN | NaN | 132.89047 | NaN | NaN |
| Uqcrc1 | EHTAYLIK | -127 | -134 | 0 | 0.000591124 | 8 | 1919.1622 | NaN | NaN | NaN | NaN | NaN |
| Ilf2 | EISTWDGVIVTPSEK | -342 | -356 | 0 | 0 | 15 | 1844.411 | 2519.2573 | 2365.96 | 2387.8176 | 3077.5237 | 3531.8442 |
| Ilf2 | WFEENASQSTVK | -210 | -221 | 0 | 0 | 12 | 1039.2703 | 762.176 | 784.1147 | 1000.5008 | 847.7379 | 1157.6091 |
| Ilf2 | SVGITDPCESGNFR | -284 | -297 | 0 | 0.000230707 | 14 | 805.9943 | 634.919 | 681.816 | 1204.7668 | 504.38055 | 512.0091 |
| Ilf2 | KLDPELHLDIK | -186 | -196 | 1 | 0 | 11 | 731.61017 | 613.0656 | 525.70447 | 1035.6871 | 489.26645 | 548.01465 |
| Ilf2 | KLDPELHLDIK | -186 | -196 | 1 | 0 | 11 | 731.61017 | 613.0656 | 525.70447 | 1035.6871 | 489.26645 | 548.01465 |
| Ilf2 | AAGLFLPGSVGITDPCESGNFR | -276 | -297 | 0 | 0.000347792 | 22 | 330.5734 | 385.5358 | 331.4148 | 414.69122 | 396.47522 | 431.7997 |
| Ilf2 | GLFLPGSVGITDPCESGNFR | -278 | -297 | 0 | 0 | 20 | 289.17645 | 411.7028 | 274.03644 | 414.3726 | 552.05994 | 464.0992 |
| Ilf2 | VKPAPDETSFSEALLK | -44 | -59 | 1 | 0 | 16 | 195.37163 | 481.8797 | 546.74066 | 512.4181 | 717.1075 | 710.3821 |
| Ilf2 | VKPAPDETSFSEA | -44 | -56 | 1 | 0.00025015 | 13 | 355.8519 | 2064.23 | 1995.3096 | NaN | 924.4887 | 1057.7832 |
| Ilf2 | TWDGVIVTPSEK | -345 | -356 | 0 | 0 | 12 | 558.5936 | NaN | 1663.7642 | 1667.2863 | NaN | 1450.4255 |
| Ilf2 | ASEISTWDGVIVTPSEK | -340 | -356 | 0 | 0 | 17 | 508.7654 | 1055.1997 | NaN | 1018.56635 | 1275.5748 | NaN |
| Kpnb1 | LLETTDRPDGHQNNLR | -510 | -525 | 1 | 0 | 16 | 2784.6218 | 2683.64 | 2987.5347 | 2102.7214 | 4145.4023 | 3784.8276 |
| Kpnb1 | LLETTDRPDGHQNNLR | -510 | -525 | 1 | 0 | 16 | 2784.6218 | 2683.64 | 2987.5347 | 2102.7214 | 4145.4023 | 3784.8276 |
| Kpnb1 | VLANPGNSQVAR | -43 | -54 | 0 | 0 | 12 | 2381.219 | 2405.4592 | 2316.2688 | 1615.4664 | 2405.5952 | 2310.941 |
| Kpnb1 | AVGLVGDLCR | -670 | -679 | 0 | 0 | 10 | 2212.9424 | 2308.5076 | 2338.0432 | 2207.3909 | 2590.686 | 493.6974 |
| Kpnb1 | TVSPDRLELEAAQK | -10 | -23 | 1 | 0 | 14 | 2041.3323 | 2913.1624 | 2871.8577 | 3596.2556 | 3889.1316 | 3996.211 |
| Kpnb1 | TVSPDRLELEAAQK | -10 | -23 | 1 | 0 | 14 | 2041.3323 | 2913.1624 | 2871.8577 | 3596.2556 | 3889.1316 | 3996.211 |
| Kpnb1 | VVCEATQCPDTR | -221 | -232 | 0 | 0 | 12 | 1518.002 | 1569.4225 | 1583.3804 | 1386.6471 | 1354.4067 | 236.5701 |
| Kpnb1 | QDIDPEQLQDK | -159 | -169 | 0 | 0 | 11 | 1494.2422 | 858.5242 | 1536.9221 | 1446.5282 | 1366.9153 | 1300.4574 |
| Kpnb1 | EAFKPFLGIGLK | -648 | -659 | 1 | 0.00025015 | 12 | 1481.0812 | 1554.1295 | 1668.5676 | 904.08356 | 1417.1168 | 1350.8096 |
| Kpnb1 | LLENLGNENVHR | -696 | -707 | 0 | 0 | 12 | 1473.9591 | 1614.629 | 1851.951 | 1106.7122 | 1211.4601 | 1122.326 |
| Kpnb1 | LLENLGNENVHR | -696 | -707 | 0 | 0 | 12 | 1473.9591 | 1614.629 | 1851.951 | 1106.7122 | 1211.4601 | 1122.326 |
| Kpnb1 | NSLTSKDPDIK | -63 | -73 | 1 | 0 | 11 | 1097.3816 | 817.39655 | 1137.6818 | 1039.7762 | 1178.2258 | 459.4277 |
| Kpnb1 | ICQDIDPEQLQDK | -157 | -169 | 0 | 0 | 13 | 1074.0869 | 969.53876 | 1084.2788 | 1331.3757 | 1231.0719 | 952.44543 |
| Kpnb1 | WLAIDANAR | -80 | -88 | 0 | 0 | 9 | 1052.1781 | 1162.9978 | 1241.9135 | 1854.9677 | 1296.9247 | 592.5676 |
| Kpnb1 | ENLGNENVHR | -698 | -707 | 0 | 0 | 10 | 882.11896 | 906.61743 | 831.6592 | 505.97858 | 654.7187 | 677.84863 |
| Kpnb1 | DENDDDDDWNPCK | -334 | -346 | 0 | 0 | 13 | 676.6752 | 699.096 | 700.59735 | 793.2554 | 837.80597 | 883.1779 |
| Kpnb1 | LQQVLQMESHIQ | -558 | -569 | 0 | 0 | 12 | 661.93396 | 609.342 | 565.20483 | 470.45483 | 339.8074 | 744.05237 |
| Kpnb1 | EAAEQGRPPEHTSK | -299 | -312 | 1 | 0 | 14 | 648.0458 | 391.91 | 736.9764 | 429.3118 | 666.8458 | 805.0255 |
| Kpnb1 | YMEAFKPFLGIGLK | -646 | -659 | 1 | 0 | 14 | 618.7487 | 829.27966 | 805.5459 | 686.03827 | 703.5557 | 642.7343 |
| Kpnb1 | YMEAFKPFLGIGLK | -646 | -659 | 1 | 0 | 14 | 618.7487 | 829.27966 | 805.5459 | 686.03827 | 703.5557 | 642.7343 |
| Kpnb1 | EHIKNPDWR | -373 | -381 | 1 | 0 | 9 | 573.2229 | 667.99396 | 931.6053 | 500.85382 | 2171.3618 | 1041.7607 |
| Kpnb1 | AAVENLPTFLVELSR | -28 | -42 | 0 | 0 | 15 | 542.53107 | 104.7545 | 415.4854 | 614.4496 | 435.7045 | 605.3376 |
| Kpnb1 | PFLGIGLK | -652 | -659 | 0 | 0.00111006 | 8 | 510.34628 | 383.26886 | 344.36758 | 389.75134 | 557.19434 | 527.83563 |
| Kpnb1 | LENLGNENVHR | -697 | -707 | 0 | 0 | 11 | 384.17487 | 1380.5527 | 1453.5438 | 1002.3139 | 1212.3152 | 1167.0582 |

| | | | | | | | | | | | |
|---|---|---|---|---|---|---|---|---|---|---|---|
| Kpnb1 | GDQENVHPDVMLVQPR | -778 | -793 | 0 | 0 | 16 | 340.5352 | 687.5299 | 306.18665 | 573.6466 | 552.244 | 724.7852 |
| Kpnb1 | SEAAEQGRPPEHTSK | -298 | -312 | 1 | 0 | 15 | 331.56335 | 403.20947 | 445.82892 | 38.899582 | 435.56952 | 411.71515 |
| Kpnb1 | IHELLTEGR | -843 | -851 | 0 | 0 | 9 | 1446.7812 | 1637.3783 | NaN | 2244.9546 | 1866.379 | 850.0253 |
| Kpnb1 | LLETTDRPDGHQN | -510 | -522 | 1 | 0 | 13 | 862.3252 | 597.39374 | 786.2204 | 740.3493 | NaN | 1040.5735 |
| Kpnb1 | GLIGDLCTAFGK | -820 | -831 | 0 | 0 | 12 | 719.992 | 1785.3049 | NaN | 1334.6886 | 1460.347 | 422.25607 |
| Kpnb1 | KDPSVVVR | -418 | -425 | 1 | 0 | 8 | 717.1653 | NaN | 1218.7444 | 794.0187 | 856.01807 | NaN |
| Kpnb1 | AAGLIGDLCTAFGK | -818 | -831 | 0 | 0 | 14 | 1782.6952 | NaN | 1879.729 | NaN | NaN | 824.08026 |
| Kpnb1 | LAAVGLVGDLCR | -668 | -679 | 0 | 0 | 12 | 884.2467 | 1051.6875 | 708.9927 | NaN | NaN | NaN |
| Kpnb1 | SDFDMVDYLNELR | -750 | -762 | 0 | 0 | 13 | 435.02097 | NaN | NaN | NaN | NaN | 491.2748 |
| Kpnb1 | SSSFELIVQK | -500 | -509 | 0 | 0.00025015 | 10 | 372.65186 | NaN | NaN | NaN | NaN | NaN |
| Bola1 | NESGGHAVPAGSETHFR | -53 | -69 | 0 | 0 | 17 | 1143.1826 | 1043.4352 | 1143.2023 | 1004.62616 | 646.2574 | 1192.5311 |
| Bola1 | NESGGHAVPAGSETHFR | -53 | -69 | 0 | 0 | 17 | 1143.1826 | 1043.4352 | 1143.2023 | 1004.62616 | 646.2574 | 1192.5311 |
| Bola1 | LEQALSPEVLELR | -40 | -52 | 0 | 0 | 13 | 1115.6146 | NaN | NaN | 1545.8907 | 1403.4224 | 1421.958 |
| Bola1 | AAGPVEAAIR | -28 | -37 | 0 | 0.00201047 | 10 | 632.19257 | 625.18256 | NaN | NaN | NaN | NaN |
| Eef1a1 | SFSDYPPLGR | -414 | -423 | 0 | 0 | 10 | 24284.922 | 23031.125 | 24161.982 | 20468.357 | 18923.229 | 915.4715 |
| Eef1a1 | RYEEIVK | -166 | -172 | 1 | 0.00504706 | 7 | 16576.402 | 14064.341 | 14924.567 | 12462.588 | 14966.555 | 14795.2705 |
| Eef1a1 | FAPVNVTTEVK | -280 | -290 | 0 | 0 | 11 | 15217.69 | 1220.2551 | 7204.9673 | 6292.8535 | 7528.963 | 7562.0493 |
| Eef1a1 | APVNVTTEVK | -281 | -290 | 0 | 0 | 10 | 11435.851 | 12207.114 | 12035.902 | 445.00766 | 13321.795 | 12924.635 |
| Eef1a1 | DCILPPTRPTDKPLR | -233 | -247 | 2 | 0.00025015 | 15 | 10667.168 | 11793.58 | 25552.73 | 15363.901 | 12699.661 | 12325.981 |
| Eef1a1 | DCILPPTRPTDKPLR | -233 | -247 | 2 | 0.00025015 | 15 | 10667.168 | 11793.58 | 25552.73 | 15363.901 | 12699.661 | 12325.981 |
| Eef1a1 | DCILPPTRPTDKPLR | -233 | -247 | 2 | 0.00025015 | 15 | 10667.168 | 11793.58 | 25552.73 | 15363.901 | 12699.661 | 12325.981 |
| Eef1a1 | YYVTIIDAPGHR | -85 | -96 | 0 | 0 | 12 | 9928.298 | 11164.631 | 11565.285 | 6979.145 | 8729.311 | 8216.773 |
| Eef1a1 | YYVTIIDAPGHR | -85 | -96 | 0 | 0 | 12 | 9928.298 | 11164.631 | 11565.285 | 6979.145 | 8729.311 | 8216.773 |
| Eef1a1 | MDSTEPPYSQK | -155 | -165 | 0 | 0 | 11 | 7305.405 | 7248.169 | 7739.9966 | 7070.357 | 8050.1455 | 7837.546 |
| Eef1a1 | VETGVLKPGMVVTF | -267 | -280 | 1 | 0 | 14 | 6775.3403 | 8489.918 | 8068.1626 | 10658.306 | 12166.838 | 12372.32 |
| Eef1a1 | VETGVLKPGMVVTF | -267 | -280 | 1 | 0 | 14 | 6775.3403 | 8489.918 | 8068.1626 | 10658.306 | 12166.838 | 12372.32 |
| Eef1a1 | DSTEPPYSQK | -156 | -165 | 0 | 0 | 10 | 6724.506 | 3116.3413 | 3475.3047 | 3318.7986 | 3553.3525 | 3880.4265 |
| Eef1a1 | DSTEPPYSQK | -156 | -165 | 0 | 0 | 10 | 6724.506 | 3116.3413 | 3475.3047 | 3318.7986 | 3553.3525 | 3880.4265 |
| Eef1a1 | DCILPPTRPTDK | -233 | -244 | 1 | 0.000230707 | 12 | 4426.2905 | 412.65002 | 4446.425 | 4162.3833 | 3667.465 | 4237.113 |
| Eef1a1 | ALDCILPPTRPTDKPLR | -231 | -247 | 2 | 0 | 17 | 3407.674 | 3946.558 | 3994.4907 | 1675.2837 | 2554.0244 | 2969.5662 |
| Eef1a1 | LEPSANMPWFK | -202 | -212 | 0 | 0 | 11 | 3017.9602 | 3038.7483 | 3138.1333 | 337.55127 | 3103.4858 | 2984.1868 |
| Eef1a1 | LEPSANMPWFK | -202 | -212 | 0 | 0 | 11 | 3017.9602 | 3038.7483 | 3138.1333 | 337.55127 | 3103.4858 | 2984.1868 |
| Eef1a1 | FSDYPPLGR | -415 | -423 | 0 | 0 | 9 | 2933.6436 | 32383.45 | 27708.195 | 35694.16 | 38704.094 | 37991.363 |
| Eef1a1 | TLLEALDCILPPTRPTDKPLR | -227 | -247 | 2 | 0 | 21 | 2637.1633 | 3673.9456 | 2958.0935 | 2586.4705 | 3749.1746 | 324.94208 |
| Eef1a1 | EALDCILPPTRPTDKPLR | -230 | -247 | 2 | 0 | 18 | 1581.8542 | 2046.5593 | 1874.5508 | 2112.1897 | 2647.3198 | 2973.3477 |
| Eef1a1 | VETGVLKPGMVVT | -267 | -279 | 1 | 0 | 13 | 1476.3646 | 1612.8632 | 1166.4226 | 4946.552 | 4824.0063 | 4364.203 |
| Eef1a1 | VETGVLKPGMVVT | -267 | -279 | 1 | 0 | 13 | 1476.3646 | 1612.8632 | 1166.4226 | 4946.552 | 4824.0063 | 4364.203 |
| Eef1a1 | LEALDCILPPTRPTDK | -229 | -244 | 1 | 0 | 16 | 1461.1299 | 1874.8594 | 1585.9968 | 2193.4202 | 2406.532 | 3342.9004 |
| Eef1a1 | ANMPWFK | -206 | -212 | 0 | 0 | 7 | 1428.5859 | 1175.4459 | 1256.7506 | 869.7855 | 755.38947 | 904.96344 |
| Eef1a1 | ANMPWFK | -206 | -212 | 0 | 0 | 7 | 1428.5859 | 1175.4459 | 1256.7506 | 869.7855 | 755.38947 | 904.96344 |
| Eef1a1 | VTIIDAPGHR | -87 | -96 | 0 | 0 | 10 | 1109.2955 | 1155.044 | 892.7338 | 1718.3568 | 1615.7296 | 1498.459 |
| Eef1a1 | IGYNPDTVAFVPIS | -181 | -194 | 0 | 0 | 14 | 1036.6923 | 950.12146 | 1053.9022 | 113.88458 | 1557.6075 | 1542.3475 |
| Eef1a1 | CVESFSDYPPLGR | -411 | -423 | 0 | 0 | 13 | 951.3638 | 1165.541 | 887.45886 | 2393.5742 | 2908.5818 | 2727.3384 |
| Eef1a1 | YAPVLDCHTAHIACK | -357 | -371 | 0 | 0 | 15 | 938.07874 | 1143.7607 | 833.6064 | 1840.1726 | 2175.7764 | 2020.3009 |
| Eef1a1 | YAPVLDCHTAHIACK | -357 | -371 | 0 | 0 | 15 | 938.07874 | 1143.7607 | 833.6064 | 1840.1726 | 2175.7764 | 2020.3009 |
| Eef1a1 | KMDSTEPPYSQK | -154 | -165 | 1 | 0 | 12 | 805.25244 | 772.09717 | 834.46454 | 770.81445 | 1373.8427 | 1250.0421 |
| Eef1a1 | KDGSASGTTLLEAL | -219 | -232 | 1 | 0.00201047 | 14 | 609.4144 | 1490.1873 | 517.5369 | 870.60144 | 2482.3374 | 388.187 |
| Eef1a1 | GYAPVLDCHTAHIACK | -356 | -371 | 0 | 0 | 16 | 571.59576 | 765.5469 | 731.62823 | 578.8759 | 1097.1748 | 798.30725 |
| Eef1a1 | PVNVTTEVK | -282 | -290 | 0 | 0 | 9 | 483.93268 | 1379.0989 | 1052.061 | 2643.6445 | 3554.5737 | 3402.758 |
| Eef1a1 | KIGYNPDTVAFVPISGW | -180 | -196 | 1 | 0 | 17 | 319.99304 | 286.4932 | 327.059 | 282.64563 | 397.24347 | 376.59412 |
| Eef1a1 | LEALDCILPPTRPTDKPLR | -229 | -247 | 2 | 0 | 19 | 141.71246 | 4409.0493 | 4434.963 | 5084.0806 | 7353.383 | 7643.312 |
| Eef1a1 | SDYPPLGR | -416 | -423 | 0 | 0.001023236 | 8 | 2014.6742 | 1881.3943 | NaN | 1358.1797 | 1376.1257 | 1186.8264 |
| Eef1a1 | TFAPVNVTTEVK | -279 | -290 | 0 | 0 | 12 | 740.76764 | 94.26724 | 871.9224 | NaN | 867.4127 | 815.3745 |
| Eef1a1 | RYEEIVKE | -166 | -173 | 2 | 0.001636916 | 8 | 419.3485 | 2054.9048 | 1997.837 | 1308.4473 | 459.36066 | NaN |
| Eef1a1 | EALDCILPPTRPTDK | -230 | -244 | 1 | 0 | 15 | 1785.9331 | 2146.6143 | NaN | NaN | 2718.0334 | 2513.0466 |
| Rplp1 | ILHDDEVTVTEDK | -15 | -27 | 0 | 0 | 13 | 2578.0027 | 2803.5842 | 2770.2192 | 2355.148 | 2604.7153 | 2530.5671 |
| Rplp1 | GVSVEPFWPGLFAK | -36 | -49 | 0 | 0 | 14 | 1744.2429 | 1674.2144 | 1703.9199 | 3232.304 | 3304.295 | 3135.3164 |
| Rplp1 | HDDEVTVTEDK | -17 | -27 | 0 | 0 | 11 | 1716.0602 | 324.43246 | 1875.9595 | 1135.2216 | 1236.1914 | 1370.3477 |
| Rplp1 | VEPFWPGLFAK | -39 | -49 | 0 | 0 | 11 | 1244.7643 | 1295.7205 | 1470.4933 | 688.1261 | 992.9238 | 961.2884 |
| Rplp1 | SALILHDDEVTVTEDK | -12 | -27 | 0 | 0.000712251 | 16 | 1026.0416 | 1220.1063 | 1229.4512 | 1044.5929 | 1254.2722 | 1177.7164 |
| Rplp1 | EDKINALIK | -25 | -33 | 1 | 0.00025015 | 9 | 1018.83044 | 1050.3674 | 1092.1829 | 1674.4458 | 1804.8809 | 1972.0637 |
| Rplp1 | NVGAGGPAPAAGAAPAGGAAPSTAAAPAEEK | -62 | -92 | 0 | 0.001055699 | 31 | 998.1743 | 1073.4924 | 1086.4962 | 604.23926 | 620.7973 | 616.83856 |
| Rplp1 | NVGAGGPAPAAGAAPAGGAAPSTAAAPAEEK | -62 | -92 | 0 | 0.001055699 | 31 | 998.1743 | 1073.4924 | 1086.4962 | 604.23926 | 620.7973 | 616.83856 |
| Rplp1 | SVEPFWPGLFAK | -38 | -49 | 0 | 0 | 12 | 833.55707 | 861.64954 | 863.00543 | 1013.36127 | 1442.1058 | 1384.9171 |
| Rplp1 | AAGVSVEPFWPGLFAK | -34 | -49 | 0 | 0 | 16 | 750.5318 | 767.00806 | 789.6917 | 634.6444 | 831.5806 | 825.16266 |
| Rplp1 | LILHDDEVTVTEDK | -14 | -27 | 0 | 0 | 14 | 459.76938 | 202.19598 | 401.9375 | 568.46893 | 587.81915 | 600.6607 |
| Rplp1 | SLICNVGAGGPAPAAGAAPAGGAAPSTAAAPAEEK | -58 | -92 | 0 | 0.00122419 | 35 | 297.0818 | 257.41095 | 242.56079 | 221.15063 | 253.05972 | 213.07315 |
| Rplp1 | KEESEESEDDMGFGL | -98 | -112 | 1 | 0.000431655 | 15 | 342.04114 | NaN | NaN | 541.61487 | 475.32257 | 411.24454 |
| Rplp1 | ALANVNIGSLIC | -50 | -61 | 0 | 0 | 12 | 667.5537 | 713.9479 | 701.0217 | NaN | NaN | NaN |
| Rplp1 | KEESEESEDDMGFGLFD | -98 | -114 | 1 | 0.001392869 | 17 | 410.61713 | 422.64435 | NaN | 357.75693 | NaN | NaN |
| Rplp1 | HDDEVTVTEDKINALIK | -17 | -33 | 1 | 0.000712251 | 17 | 267.39948 | NaN | NaN | NaN | NaN | NaN |
| Fus | HDSEQDNSDNNTIFVQ | -268 | -283 | 0 | 0 | 16 | 4583.661 | 4407.134 | 4784.635 | 4286.078 | 3642.379 | 3706.5098 |
| Fus | AAIDWFDGK | -342 | -350 | 0 | 0 | 9 | 3208.2598 | 2926.7085 | 2928.7952 | 3260.5015 | 2466.5063 | 3543.709 |
| Fus | VSFDDPPSAK | -332 | -341 | 0 | 0.000230707 | 10 | 2963.805 | 3173.929 | 3317.4343 | 3281.301 | 3166.128 | 3423.7976 |
| Fus | ATVSFDDPPSAK | -330 | -341 | 0 | 0 | 12 | 1527.037 | 1475.168 | 1344.7535 | 2099.8213 | 1148.0748 | 715.9072 |
| Fus | GEATVSFDDPPSAK | -328 | -341 | 0 | 0 | 14 | 714.75433 | 734.1781 | 865.24603 | 135.38663 | 702.2916 | 784.79175 |
| Fus | FDDPPSAK | -334 | -341 | 0 | 0 | 8 | 293.88126 | 897.03925 | 439.94638 | 503.1419 | 848.98 | 353.51102 |
| Fus | APKPDGPGGGPGGSHM | -442 | -457 | 1 | 0 | 16 | 809.23676 | NaN | 832.4084 | 299.23737 | 611.975 | 828.82104 |

| | | | | | | | | | | | | |
|---|---|---|---|---|---|---|---|---|---|---|---|---|
| Ddx21 | DAQELSQNTCIK | -522 | -533 | 0 | 0 | 12 | 5430.212 | 6063.4844 | 5199.0503 | 6439.739 | 6091.923 | 6527.9062 |
| Ddx21 | LKDGLSQPSEPK | -221 | -232 | 1 | 0 | 12 | 3920.0981 | 1850.7678 | 4358.7607 | 3113.292 | 267.3604 | 3105.1587 |
| Ddx21 | LLDSVPPTAISHFK | -654 | -667 | 0 | 0 | 14 | 3604.8374 | 3185.425 | 4726.1284 | 4434.2715 | 5993.54 | 3166.1794 |
| Ddx21 | VATEQPELEGPPDGYR | -779 | -794 | 0 | 0 | 16 | 3418.8416 | 10302.98 | 10623.482 | 8518.57 | 4210.727 | 9753.557 |
| Ddx21 | SLHGDIPQK | -538 | -546 | 0 | 0 | 9 | 3224.0588 | 3310.1174 | 3871.207 | 2558.0818 | 1929.778 | 2449.1272 |
| Ddx21 | EGAFSNFPISEETVK | -257 | -271 | 0 | 0 | 15 | 2841.8877 | 3860.8057 | 3482.8079 | 3971.937 | 4890.027 | 4801.209 |
| Ddx21 | AAVIGDVIR | -496 | -504 | 0 | 0 | 9 | 2739.3293 | 1513.4374 | 1740.4208 | 979.9301 | 760.1483 | 619.66235 |
| Ddx21 | SGIDILVGTPGR | -379 | -390 | 0 | 0 | 12 | 2367.3706 | 2646.07 | 2736.4058 | 2279.6538 | 2199.8105 | 2211.9395 |
| Ddx21 | IGVPSATEIIK | -635 | -645 | 0 | 0 | 11 | 2357.407 | 2473.9106 | 2551.9028 | 2977.8296 | 2833.9446 | 2702.1938 |
| Ddx21 | KGPSEDDVDPPK | -61 | -72 | 1 | 0 | 12 | 2048.0308 | 2112.9585 | 2176.7717 | 390.5231 | 2172.6377 | 563.57275 |
| Ddx21 | SKDFSDITK | -349 | -357 | 1 | 0 | 9 | 1920.5481 | 732.5103 | 2279.0232 | 2992.5054 | 3209.532 | 3326.8735 |
| Ddx21 | TEQPELEGPPDGYR | -781 | -794 | 0 | 0 | 14 | 1550.5559 | 1582.6354 | 2022.2673 | 1892.2433 | 1638.5114 | 1637.7452 |
| Ddx21 | GVNFLFPIQAK | -277 | -287 | 0 | 0 | 11 | 1207.1447 | 575.00726 | 1451.7329 | 985.2774 | 1292.1338 | 1111.836 |
| Ddx21 | AAITVEHLAIK | -479 | -489 | 0 | 0.000230707 | 11 | 919.81 | 982.0354 | 1035.4487 | 1577.7682 | 1170.6407 | 1035.8127 |
| Ddx21 | SFAIPLIEK | -311 | -319 | 0 | 0.00025015 | 9 | 918.3877 | 1061.5818 | 1039.2235 | 862.8096 | 1105.1562 | 1160.8192 |
| Ddx21 | ELKEQLGESIDAK | -728 | -740 | 1 | 0 | 13 | 840.09375 | 469.99542 | 995.47925 | 988.3517 | 1027.4089 | 1028.1855 |
| Ddx21 | NGLSQPSEEEADIPKPK | -149 | -165 | 1 | 0 | 17 | 791.5125 | 860.61816 | 886.0727 | 1117.002 | 1078.8208 | 942.3022 |
| Ddx21 | CFYGGTPYGGQIER | -363 | -376 | 0 | 0 | 14 | 767.5364 | 454.11124 | 667.1973 | 351.21008 | 533.302 | 765.43274 |
| Ddx21 | NGLSQPSEEEVDIPKPK | -186 | -202 | 1 | 0.00025015 | 17 | 703.42773 | 732.0764 | 733.78345 | 739.59875 | 877.6555 | 1307.2616 |
| Ddx21 | FYGGTPYGGQIER | -364 | -376 | 0 | 0 | 13 | 376.1521 | 599.88684 | 692.523 | 648.6174 | 794.03406 | 560.68744 |
| Ddx21 | TFSFAIPLIEK | -309 | -319 | 0 | 0 | 11 | 255.97565 | 268.39093 | 340.6753 | 321.59906 | 271.547 | 136.9734 |
| Ddx21 | YEQVDLIGK | -466 | -474 | 0 | 0 | 9 | 2157.7292 | NaN | 6877.6045 | 5240.1167 | 7043.006 | 6977.845 |
| Ddx21 | SNFPISEETVK | -261 | -271 | 0 | 0 | 11 | 1448.3921 | 2147.1255 | 2390.545 | NaN | 3168.0735 | 2724.586 |
| Ddx21 | KDAQELSQNTCIK | -521 | -533 | 1 | 0.00025015 | 13 | 900.3059 | NaN | 713.93146 | 683.1613 | 806.9549 | 882.88086 |
| Ddx21 | LGVCFDVR | -751 | -758 | 0 | 0 | 8 | 810.4311 | 770.80316 | NaN | 918.1256 | 381.51428 | 817.0443 |
| Ddx21 | LGSDGAEESMETLPKPSEK | -11 | -29 | 1 | 0 | 19 | 380.78607 | 480.5788 | NaN | 659.2088 | 639.49365 | 455.64737 |
| Ddx21 | TFHHVYSGK | -288 | -296 | 0 | 0.00025015 | 9 | 340.0754 | 423.17438 | 217.23184 | 227.57275 | 188.55383 | NaN |
| Ddx21 | SNSSDAPGEESSSETEK | -233 | -249 | 0 | 0 | 17 | 323.84528 | 277.08948 | 376.0011 | NaN | 337.26642 | 398.20932 |
| Ddx21 | EQLGESIDAK | -731 | -740 | 0 | 0 | 10 | 1590.6354 | 1409.1774 | 802.45123 | 1063.2206 | NaN | NaN |
| Ddx21 | TEAVTEIQEK | -759 | -768 | 0 | 0 | 10 | 72.6977 | 986.01227 | 2281.0781 | NaN | 1046.6593 | NaN |
| Ddx21 | VEHLAIK | -483 | -489 | 0 | 0.005333786 | 7 | 1815.9032 | NaN | 1704.382 | 1153.6562 | NaN | NaN |
| Ddx21 | EQPELEGPPDGYR | -782 | -794 | 0 | 0 | 13 | 244.1257 | 999.7151 | NaN | NaN | 607.491 | NaN |
| Ddx21 | STYEQVDLIGK | -464 | -474 | 0 | 0 | 11 | 16796 | NaN | 20394.434 | NaN | NaN | NaN |
| Ddx21 | DSVPPTAISHFK | -656 | -667 | 0 | 0 | 12 | 1671.1548 | NaN | 2537.0737 | NaN | NaN | NaN |
| Ddx21 | KEQLGESIDAK | -730 | -740 | 1 | 0.000550307 | 11 | 479.31412 | NaN | 541.77545 | NaN | NaN | NaN |
| Ddx21 | APQVLVLAPTR | -332 | -342 | 0 | 0.003966045 | 11 | 259.24692 | NaN | NaN | NaN | NaN | NaN |
| Cap1 | NQENVSNLVIDDTELK | -332 | -347 | 0 | 0 | 16 | 5584.803 | 7367.258 | 6577.3755 | 9081.668 | 11367.124 | 11011.124 |
| Cap1 | KEPALLELEGK | -316 | -326 | 1 | 0.000907206 | 11 | 5219.62 | 5436.1816 | 5896.5703 | 6926.267 | 7218.6387 | 7123.426 |
| Cap1 | CVNTTLQIK | -355 | -363 | 0 | 0 | 9 | 3494.443 | 3442.8132 | 688.67896 | 3712.1167 | 395.9866 | 3867.6458 |
| Cap1 | KTDGCHAYLSK | -411 | -421 | 1 | 0.00078661 | 11 | 2812.508 | 498.75955 | 813.7445 | 981.79346 | 1329.7434 | 1720.6824 |
| Cap1 | NSLDCEIVSAK | -422 | -432 | 0 | 0 | 11 | 2253.0344 | 2396.1296 | 2304.0068 | 2275.109 | 1123.9861 | 2065.2825 |
| Cap1 | EPALLELEGK | -317 | -326 | 0 | 0 | 10 | 2104.7175 | 2243.4297 | 2345.3083 | 2234.0073 | 90.24429 | 2224.8147 |
| Cap1 | LANPVAEYLK | -50 | -59 | 0 | 0.000550307 | 10 | 1613.9398 | 1746.4866 | 1684.5345 | 1422.6484 | 94.40479 | 1625.3386 |
| Cap1 | DSLLANPVAEYLK | -47 | -59 | 0 | 0.000230707 | 13 | 1536.0482 | 1512.4541 | 1515.799 | 1696.3594 | 2052.255 | 1992.6641 |
| Cap1 | HAEMVHTGLK | -71 | -80 | 0 | 0.000550307 | 10 | 1380.1343 | 747.46387 | 597.7179 | 1222.2727 | 841.79645 | 1171.9385 |
| Cap1 | HAEMVHTGLK | -71 | -80 | 0 | 0.000550307 | 10 | 1380.1343 | 747.46387 | 597.7179 | 1222.2727 | 841.79645 | 1171.9385 |
| Cap1 | KCVNTTLQIK | -354 | -363 | 1 | 0 | 10 | 1370.835 | 763.9684 | 722.5183 | 1369.2701 | 2556.0388 | 1732.1267 |
| Cap1 | AAKPGPFVK | -146 | -154 | 1 | 0.000591124 | 9 | 1273.3843 | 1259.4397 | 1353.0813 | 1255.2552 | 1081.7491 | 1132.0273 |
| Cap1 | AFDSLLANPVAEYLK | -45 | -59 | 0 | 0 | 15 | 1105.867 | 1014.0017 | 956.3199 | 1546.7814 | 102.67535 | 1589.3738 |
| Cap1 | VENQENVSNLVIDDTELK | -330 | -347 | 0 | 0 | 18 | 929.14197 | 1084.5555 | 1075.2927 | 1463.2112 | 1942.5054 | 1963.5956 |
| Cap1 | SAPKPQTSPSPK | -300 | -311 | 1 | 0 | 12 | 921.12573 | 1052.4579 | 1169.3958 | 810.72144 | 986.08936 | 858.79034 |
| Cap1 | SLDCEIVSAK | -423 | -432 | 0 | 0 | 10 | 906.1045 | 1035.0796 | 798.8603 | 910.73987 | 1028.3231 | 1145.0166 |
| Cap1 | INQGESITHALK | -260 | -271 | 0 | 0 | 12 | 833.42523 | 925.8419 | 961.9452 | 149.35115 | 681.3906 | 751.3307 |
| Cap1 | KNSLDCEIVSAK | -421 | -432 | 1 | 0 | 12 | 487.20737 | 579.97833 | 730.3251 | 1911.1165 | 219.06606 | 1518.4006 |
| Cap1 | INSITVDNCK | -366 | -375 | 0 | 0 | 10 | 1689.2627 | 1639.7617 | 1633.1699 | 696.9267 | NaN | 632.0172 |
| Cap1 | EGGDFNEFPVPEQFK | -443 | -457 | 0 | 0 | 15 | 658.04095 | 417.49783 | NaN | 136.2236 | 883.9242 | 661.02277 |
| Cap1 | LEAVSHTSDMHCGYGDSPSK | -18 | -37 | 0 | 0.000826806 | 20 | 389.14136 | 433.58234 | 412.55655 | NaN | 359.41846 | 352.5074 |
| Cap1 | VLIPTEGGDFNEFPVPEQFK | -438 | -457 | 0 | 0.002629273 | 20 | 279.39685 | NaN | 110.92605 | 285.81674 | 302.6373 | 259.5636 |
| Cap1 | NVLIPTEGGDFNEFPVPEQFK | -437 | -457 | 0 | 0 | 21 | 164.5659 | 215.80699 | 218.7143 | 269.8363 | 218.26953 | NaN |
| Cap1 | LSDLLAPISEQIQ | -100 | -112 | 0 | 0.00025015 | 13 | 717.63306 | 899.45886 | 806.6554 | NaN | 728.9817 | NaN |
| Cap1 | EFPVPEQFK | -449 | -457 | 0 | 0 | 9 | 1686.5344 | 858.71423 | NaN | 1652.726 | NaN | NaN |
| Cap1 | INSITVDNCKK | -366 | -376 | 1 | 0 | 11 | 747.9135 | 594.7592 | 708.9864 | NaN | NaN | NaN |
| Cap1 | VIDDTELK | -340 | -347 | 0 | 0.001636916 | 8 | 536.0802 | 550.66113 | NaN | 630.14923 | NaN | NaN |
| Cap1 | GKVPTISINK | -402 | -411 | 1 | 0.00670918 | 10 | 414.08484 | NaN | 411.16656 | 255.30922 | NaN | NaN |
| Cap1 | ADMQNLVER | -2 | -10 | 0 | 0 | 9 | 692.90656 | 743.5424 | NaN | NaN | NaN | NaN |
| Cap1 | DDVVGIVEIINSR | -382 | -394 | 0 | 0 | 13 | 680.7077 | NaN | 568.5634 | NaN | NaN | NaN |
| Cap1 | ENVSNLVIDDTELK | -334 | -347 | 0 | 0 | 14 | 124.05523 | NaN | NaN | NaN | NaN | 904.9875 |
| Hars1 | HGAEVIDTPVFELK | -87 | -100 | 0 | 0 | 14 | 9709.523 | 10953.384 | 10559.919 | 11322.78 | 10640.283 | 12925.26 |
| Hars1 | VAIIGEQELK | -463 | -472 | 0 | 0 | 10 | 2388.303 | 2441.6155 | 2316.3464 | 1857.0326 | 2015.6669 | 2175.5308 |
| Hars1 | REDLVEEIR | -491 | -499 | 1 | 0.001023236 | 9 | 2361.6387 | 299.81644 | 2423.2842 | 2315.8 | 2652.6428 | 2812.399 |
| Hars1 | EKVFDVIIR | -74 | -82 | 1 | 0.00025015 | 9 | 621.62573 | 626.26306 | 479.26508 | 293.69525 | 653.02856 | 703.40405 |
| Hars1 | IFSIVEQR | -389 | -396 | 0 | 0 | 8 | 590.29224 | 126.05031 | 893.2777 | 2632.2466 | 2074.8608 | 2410.173 |
| Hars1 | REDLVEEIRR | -491 | -500 | 2 | 0.000471409 | 10 | 373.35208 | 441.7661 | 456.31284 | 422.62573 | 563.50476 | 652.6194 |
| Hars1 | AALEELVR | -5 | -12 | 0 | 0 | 8 | 1793.717 | 3048.976 | 2431.382 | 4330.001 | 5612.2266 | NaN |
| Hars1 | IGDYVQQHGGVS | -267 | -278 | 0 | 0 | 12 | 813.74695 | 1109.6832 | NaN | NaN | 247.57846 | 581.2215 |
| Hars1 | GGVSLVEQLLQDPK | -275 | -288 | 0 | 0 | 14 | 476.39847 | NaN | NaN | 588.3186 | 545.45026 | 564.8507 |
| Hars1 | QAVEGLGDLK | -294 | -303 | 0 | 0.000230707 | 10 | 1870.2799 | NaN | NaN | 1302.2457 | 1290.363 | NaN |
| Hars1 | LVEQLLQDPK | -279 | -288 | 0 | 0 | 10 | 640.1258 | 655.7275 | NaN | 832.3356 | NaN | NaN |

| | | | | | | | | | | | | |
|---|---|---|---|---|---|---|---|---|---|---|---|---|
| Hars1 | TICSSVDKLDK | -233 | -243 | 1 | 0 | 11 | 2632.442 | NaN | NaN | NaN | NaN | NaN |
| Hars1 | VSWEEVK | -244 | -250 | 0 | 0.003745449 | 7 | 513.8884 | NaN | NaN | NaN | NaN | NaN |
| Krt19 | TKFETEHALR | -170 | -179 | 1 | 0.00025015 | 10 | 4437.42 | NaN | 3047.9937 | NaN | 240.14285 | 889.3252 |
| Krt19 | AALEGTLAETEAR | -321 | -333 | 0 | 0 | 13 | 1434.458 | NaN | NaN | NaN | NaN | NaN |
| Rplp2 | AAPGSAAPAAGSAPAAAEEK | -75 | -94 | 0 | 0 | 20 | 5030.027 | 5483.5776 | 5560.934 | 4571.5225 | 4899.4995 | 4966.54 |
| Rplp2 | ILDSVGIEADDDRLNK | -26 | -41 | 1 | 0.00025015 | 16 | 4749.3086 | 1364.9784 | 1370.5784 | 5308.5146 | 6705.8735 | 3184.773 |
| Rplp2 | ILDSVGIEADDDRLNK | -26 | -41 | 1 | 0.00025015 | 16 | 4749.3086 | 1364.9784 | 1370.5784 | 5308.5146 | 6705.8735 | 3184.773 |
| Rplp2 | ILDSVGIEADDDR | -26 | -38 | 0 | 0 | 13 | 3749.474 | 4644.541 | 4318.3657 | 531.0982 | 4348.1396 | 235.59688 |
| Rplp2 | ILDSVGIEADDDR | -26 | -38 | 0 | 0 | 13 | 3749.474 | 4644.541 | 4318.3657 | 531.0982 | 4348.1396 | 235.59688 |
| Rplp2 | NIEDVIAQGVGK | -50 | -61 | 0 | 0 | 12 | 3216.6438 | 3737.6868 | 3640.2441 | 3048.2593 | 974.6572 | 226.38081 |
| Rplp2 | SAAPGSAAPAAGSAPAAAEEK | -74 | -94 | 0 | 0.00149795 | 21 | 3215.3064 | 3583.6907 | 3794.7322 | 3309.2703 | 3356.9963 | 3449.027 |
| Rplp2 | LAALGGNSSPSAK | -9 | -21 | 0 | 0 | 13 | 2180.0095 | 2317.8262 | 1291.268 | 3587.481 | 2241.5337 | 2113.2712 |
| Rplp2 | VSAAPGSAAPAAGSAPAAAEEK | -73 | -94 | 0 | 0.000298805 | 22 | 1409.6332 | 1529.1132 | 1602.7811 | 1604.4985 | 1968.1039 | 1924.1201 |
| Rplp2 | KILDSVGIEADDDRLNK | -25 | -41 | 2 | 0.000230707 | 17 | 1303.3511 | 2615.7368 | 2107.708 | 1716.7281 | 2350.7522 | 2191.5212 |
| Rplp2 | KILDSVGIEADDDR | -25 | -38 | 1 | 0 | 14 | 1125.2144 | 1218.1692 | 1708.4543 | 2645.9011 | 1430.2185 | 98.96587 |
| Rplp2 | ALGGNSSPSAK | -11 | -21 | 0 | 0 | 11 | 1041.2103 | 829.2693 | 368.38156 | 1686.5984 | 869.34766 | 1052.3126 |
| Rplp2 | AALGGNSSPSAK | -10 | -21 | 0 | 0 | 12 | 1033.8113 | 1155.7699 | 239.60954 | 1115.5887 | 931.7736 | 1279.6003 |
| Rplp2 | APGSAAPAAGSAPAAAEEK | -76 | -94 | 0 | 0.00025015 | 19 | 871.9492 | 559.05334 | 688.09033 | 707.81647 | 912.2031 | 555.34076 |
| Rplp2 | LLAALGGNSSPSAK | -8 | -21 | 0 | 0 | 14 | 736.89575 | 671.9691 | 842.34515 | 896.89716 | 1217.418 | 1259.9531 |
| Rplp2 | AVSAAPGSAAPAAGSAPAAAEEK | -72 | -94 | 0 | 0.000989247 | 23 | 349.87698 | 383.80676 | 404.40167 | 375.4287 | 455.81284 | 428.449 |
| Rplp2 | SVGIEADDDRLNK | -29 | -41 | 1 | 0.00078661 | 13 | 792.0692 | NaN | 829.8022 | 834.5909 | 653.41144 | 680.4891 |
| Rplp2 | KEESEESDDDMGFGLFD | -99 | -115 | 1 | 0 | 17 | 436.8046 | 486.93994 | 542.5238 | 302.74704 | NaN | 349.81113 |
| Rplp2 | ILDSVGIEADDDRLN | -26 | -40 | 1 | 0.005363933 | 15 | 9740.073 | NaN | NaN | 9277.827 | NaN | 11069.66 |
| Rplp2 | GIEADDDRLNK | -31 | -41 | 1 | 0.002199523 | 11 | 679.0817 | 702.5873 | NaN | NaN | NaN | NaN |
| Rplp2 | KEESEESDDDMGF | -99 | -111 | 1 | 0.001392869 | 13 | 142.42776 | NaN | NaN | NaN | NaN | NaN |
| Npepl1 | ELGITPTIIR | -207 | -216 | 0 | 0.00025015 | 10 | 633.7642 | NaN | 704.89124 | 578.78986 | NaN | 654.44995 |
| Npepl1 | LDIAAPVHAGER | -463 | -474 | 0 | 0 | 12 | 1176.7686 | 262.9647 | 431.1809 | NaN | NaN | NaN |
| Npepl1 | VHLDIAAPVHAGER | -461 | -474 | 0 | 0 | 14 | 676.4231 | NaN | NaN | 838.02625 | NaN | NaN |
| Npepl1 | AALHPPALAIL | -234 | -244 | 0 | 0.000550307 | 11 | 392.41953 | 643.71765 | NaN | NaN | NaN | NaN |
| Dnaaf5 | EGSVLFPDVLVR | -813 | -824 | 0 | 0 | 12 | 868.61 | 806.0538 | 894.3048 | 867.6344 | 1185.1066 | 992.9028 |
| Dnaaf5 | AALLDPFAAVR | -171 | -181 | 0 | 0 | 11 | 223.08966 | 283.67307 | 393.91602 | NaN | 406.7909 | 389.37714 |
| Mcm5 | EVADEVTRPR | -105 | -114 | 1 | 0.000230707 | 10 | 5987.1226 | 690.718 | 6229.8594 | 693.7427 | 7451.7754 | 5857.221 |
| Mcm5 | HYNLGEYWIE | -62 | -71 | 0 | 0 | 10 | 3762.3298 | 2818.0193 | 3821.7227 | 191.39856 | 3868.0542 | 3762.9758 |
| Mcm5 | ELPDAVPHGEMPR | -231 | -243 | 0 | 0 | 13 | 2656.7222 | 3060.291 | 2661.5938 | 4273.051 | 4121.196 | 585.45416 |
| Mcm5 | SFGGDPGAEEGQAR | -14 | -27 | 0 | 0 | 14 | 1513.7101 | 1554.2888 | 1485.5465 | 1471.6824 | 1485.3442 | 113.900925 |
| Mcm5 | RLPDGLTR | -364 | -371 | 1 | 0.00078661 | 8 | 1431.3795 | 1436.1597 | 365.17462 | 1208.3074 | 1575.7842 | 1345.1619 |
| Mcm5 | SVSPQEEEFR | -313 | -323 | 0 | 0 | 11 | 1246.0016 | 1221.1093 | 1311.1808 | 1303.4496 | 1378.8197 | 765.7338 |
| Mcm5 | AGSVSPQEEEEFR | -311 | -323 | 0 | 0 | 13 | 1176.1064 | 1116.8936 | 972.2876 | 1374.2991 | 1361.8784 | 1397.7905 |
| Mcm5 | TQTQAVEGEIDLAK | -545 | -558 | 0 | 0 | 14 | 1124.0311 | 1199.388 | 1092.5723 | 490.5225 | 561.4762 | 659.4284 |
| Mcm5 | ATEADVEEALR | -632 | -642 | 0 | 0 | 11 | 1093.1332 | 912.6998 | 1029.8408 | 1570.2108 | 1123.9618 | 529.6089 |
| Mcm5 | AALPNIYELISK | -326 | -337 | 0 | 0 | 12 | 995.76715 | 1005.8191 | 902.2085 | 1499.6285 | 1586.7219 | 1710.3373 |
| Mcm5 | ASFDEELADHLHK | -78 | -90 | 0 | 0 | 13 | 950.6259 | 982.24927 | 82.41477 | 1172.1611 | 1478.362 | 75.677795 |
| Mcm5 | LAALPNIYELISK | -325 | -337 | 0 | 0 | 13 | 395.63895 | 438.68127 | 389.20596 | 432.8125 | 434.14374 | 492.20953 |
| Mcm5 | KYPEHAIR | -702 | -709 | 1 | 0 | 8 | 384.20294 | 400.01794 | 1036.6653 | 424.36765 | 500.4669 | 519.47064 |
| Mcm5 | KFGLNPSK | -273 | -280 | 1 | 0.00111006 | 8 | 2419.0361 | 2203.7964 | 2188.5889 | NaN | 1584.1842 | 1750.7521 |
| Mcm5 | QKYPEHAIR | -701 | -709 | 1 | 0.000431655 | 9 | 1262.4845 | 1551.0942 | 2487.202 | NaN | NaN | 1379.223 |
| Mcm5 | SFDEELADHLHK | -79 | -90 | 0 | 0 | 12 | 547.9229 | 371.43835 | 517.45996 | 372.99887 | NaN | NaN |
| Mcm5 | ADGGVVCIDEFDK | -437 | -449 | 0 | 0 | 13 | 490.68503 | 489.36542 | 474.87296 | NaN | NaN | 610.0924 |
| Mcm5 | QPAEHLQLLEEAAK | -91 | -104 | 0 | 0 | 14 | 2091.3784 | 2933.3262 | 3118.276 | NaN | NaN | NaN |
| Mcm5 | LQPFATEADVEEALR | -628 | -642 | 0 | 0 | 15 | 1080.86 | 1378.198 | 1424.3368 | NaN | NaN | NaN |
| Mcm5 | VSPQEEEFR | -314 | -323 | 0 | 0 | 10 | 899.89215 | 707.7223 | 800.7553 | NaN | NaN | NaN |
| Mcm5 | AGSVSPQEEEEFRR | -311 | -324 | 1 | 0 | 14 | 504.62558 | 391.84543 | 158.4002 | NaN | NaN | NaN |
| Septin1 | VNIIPVIGK | -163 | -171 | 0 | 0 | 9 | 2191.537 | 1901.3969 | 2155.0344 | 1841.556 | 4732.601 | 1859.4688 |
| Septin1 | RGVEIEEGGIK | -72 | -82 | 1 | 0.00025015 | 11 | 1226.0933 | 1003.49927 | 983.4965 | 1268.1108 | 1328.597 | 1301.7727 |
| Septin1 | VENPHHCDFLNLR | -253 | -265 | 0 | 0 | 13 | 909.5214 | 1009.2912 | 1031.384 | 2403.222 | 862.42145 | 930.85583 |
| Septin1 | VENPHHCDFLNLR | -253 | -265 | 0 | 0 | 13 | 909.5214 | 1009.2912 | 1031.384 | 2403.222 | 862.42145 | 930.85583 |
| Septin1 | EVTHDLLYEGY | -278 | -288 | 0 | 0.001190916 | 11 | 830.9549 | 666.4743 | 742.3087 | 1006.3513 | 830.0657 | 570.3618 |
| Septin1 | FIEEQFEQYLR | -109 | -119 | 0 | 0 | 11 | 582.424 | 599.27356 | 628.8622 | 657.5834 | 517.01715 | 602.294 |
| Septin1 | GLRPLDVAFLR | -147 | -157 | 1 | 0.000230707 | 11 | 327.7715 | 595.04156 | 609.6171 | 596.8907 | 707.23364 | 804.8611 |
| Septin1 | FLTNLYEDR | -46 | -54 | 0 | 0.00025015 | 9 | 261.21655 | 336.03784 | 776.6475 | 730.09485 | 1009.9016 | 1031.9476 |
| Septin1 | AALPNQLHR | -9 | -17 | 0 | 0.00025015 | 9 | 660.41077 | 584.3518 | 782.8357 | NaN | 280.0459 | NaN |
| Septin1 | GVEIEEGGIK | -73 | -82 | 0 | 0 | 10 | 1231.9526 | NaN | NaN | 1349.4421 | NaN | NaN |
| Rps25 | AALQELLSK | -86 | -94 | 0 | 0.000230707 | 9 | 2768.9175 | 2715.8308 | 2896.3455 | 111.92235 | 951.4872 | 1111.6597 |
| Rps25 | DKLNNLVLFD | -42 | -51 | 1 | 0.001291103 | 10 | 896.1467 | 1048.9585 | 1084.1934 | 733.9695 | 751.73486 | 861.32776 |
| Rps25 | DKLNNLVLFDK | -42 | -52 | 1 | 0 | 11 | 800.0793 | 834.95654 | 877.08044 | 441.83588 | 989.5993 | NaN |
| Rps25 | LNNLVLFDK | -44 | -52 | 0 | 0 | 9 | 524.03973 | NaN | NaN | 496.3287 | 414.69058 | 538.1067 |
| Rps25 | LITPAVVSER | -67 | -76 | 0 | 0 | 10 | 1388.7076 | 1077.8705 | 1042.4576 | NaN | NaN | NaN |
| Eif4g1 | KVEYTLGEESEAPGQR | -1421 | -1436 | 1 | 0 | 16 | 1319.4493 | 1218.221 | 1023.823 | 826.4631 | 813.321 | 780.6837 |
| Eif4g1 | IHNAENIQPGEQK | -585 | -597 | 0 | 0 | 13 | 1211.8408 | 1143.0392 | 1295.4143 | 775.05945 | 1182.0931 | 1166.6893 |
| Eif4g1 | DKDDDEVFEK | -858 | -867 | 1 | 0.00025015 | 10 | 1069.7722 | 1143.135 | 1235.8177 | 1456.793 | 1331.1276 | 1678.1951 |
| Eif4g1 | LGIESTLER | -1281 | -1289 | 0 | 0 | 9 | 987.01935 | 563.00256 | 756.6615 | 1243.6422 | 762.863 | 687.8094 |
| Eif4g1 | GLPLVDDGGWNTVPISK | -1047 | -1063 | 0 | 0 | 17 | 676.198 | 585.7667 | 644.1823 | 452.39365 | 730.0791 | 621.42126 |
| Eif4g1 | TLAFEELR | -1437 | -1444 | 0 | 0 | 8 | 637.5481 | 648.0128 | 743.80817 | 341.74185 | 357.9215 | 860.98126 |
| Eif4g1 | VVTLEQPANLLR | -1532 | -1543 | 0 | 0 | 12 | 518.0128 | 636.80505 | 602.39856 | 643.6721 | 737.5654 | 761.139 |
| Eif4g1 | ERPSQPEGLR | -1198 | -1207 | 1 | 0 | 10 | 3624.4812 | 4983.649 | NaN | 918.45355 | 4478.388 | 2550.7349 |
| Eif4g1 | TIDQIHKE | -1005 | -1012 | 1 | 0 | 8 | 1260.1732 | 912.2208 | NaN | 984.1467 | 1323.1421 | 1723.4978 |
| Eif4g1 | GVIDLIFEK | -800 | -808 | 0 | 0.00025015 | 9 | 1077.3826 | 466.99802 | 745.4647 | NaN | 558.1558 | 568.2813 |
| Eif4g1 | ITKPGSIDSNNQLFAPGGR | -1076 | -1094 | 1 | 0 | 19 | 592.2053 | 625.15106 | 664.15436 | NaN | 226.48544 | 666.8288 |

| | | | | | | | | | | | | |
|---|---|---|---|---|---|---|---|---|---|---|---|---|
| Eif4g1 | ITKPGSIDSNNQLFAPGGR | -1076 | -1094 | 1 | 0 | 19 | 592.2053 | 625.15106 | 664.15436 | NaN | 226.48544 | 666.8288 |
| Eif4g1 | AALSVDEVEKK | -1235 | -1245 | 1 | 0 | 11 | 660.1983 | 755.4125 | 553.467 | NaN | 490.18787 | NaN |
| Eif4g1 | AIIFETPLR | -1490 | -1498 | 0 | 0.00204437 | 9 | 589.5627 | 504.27808 | 614.32166 | NaN | NaN | 556.4763 |
| Eif4g1 | PGGELPR | -692 | -698 | 0 | 0.001055699 | 7 | 658.144 | 723.14294 | NaN | NaN | NaN | NaN |
| Eif4g1 | SDQWKPLNLEEK | -602 | -613 | 1 | 0 | 12 | 373.80292 | NaN | 1726.3839 | NaN | NaN | NaN |
| Eif4g1 | QQTLPAENTDNR | -1129 | -1140 | 0 | 0 | 12 | 132.4001 | 1323.745 | NaN | NaN | NaN | NaN |
| Eif4g1 | YLCDEQK | -1514 | -1520 | 0 | 0.005605646 | 7 | 127.19439 | NaN | NaN | 411.8233 | NaN | NaN |
| Cad | LGYPVLVR | -548 | -555 | 0 | 0 | 8 | 2191.493 | 378.2988 | 1875.5386 | 1594.6106 | 1118.7535 | 1392.4432 |
| Cad | PLAPEVSIK | -157 | -165 | 0 | 0.00025015 | 9 | 1872.1844 | 1910.2418 | 1915.7438 | 1614.8572 | 1619.862 | 1352.672 |
| Cad | GEVRPELGSR | -1658 | -1667 | 1 | 0.000230707 | 10 | 1557.4688 | 1886.308 | 2263.7744 | 3391.4397 | 3942.9036 | 2843.2358 |
| Cad | EITTTPERPR | -1829 | -1838 | 1 | 0 | 10 | 1470.8969 | 806.6891 | 1298.9785 | 1155.1185 | 1915.7957 | 975.1188 |
| Cad | GTPDGPCYPAPPVPR | -1883 | -1897 | 0 | 0 | 15 | 1393.9156 | 1208.8254 | 1103.9156 | 1119.007 | 921.9196 | 1091.7059 |
| Cad | KWPQGVVPQPPPSTPATT | -1811 | -1828 | 1 | 0 | 18 | 1194.7856 | 1140.5018 | 639.8349 | 1689.384 | 653.27454 | 1582.4967 |
| Cad | RVIPGLPDGR | -1839 | -1848 | 1 | 0.00025015 | 10 | 997.24347 | 1033.1428 | 207.73154 | 1111.9873 | 1169.4443 | 656.37305 |
| Cad | GTPVETIELTEDRR | -503 | -516 | 1 | 0 | 14 | 986.5612 | 1062.1946 | 1222.6245 | 1019.11816 | 1467.4948 | 1985.8494 |
| Cad | LSLDDLLQR | -1723 | -1731 | 0 | 0 | 9 | 973.6462 | 1061.0095 | 1096.7025 | 956.8091 | 1295.9594 | 1413.0127 |
| Cad | KEEILLIK | -1618 | -1625 | 1 | 0 | 8 | 947.7242 | 1066.8578 | 1105.2726 | 1335.0172 | 1252.7052 | 1292.009 |
| Cad | IDCWFLHR | -832 | -839 | 0 | 0 | 8 | 930.6174 | 969.55457 | 1000.997 | 1579.3278 | 1156.4342 | 946.35144 |
| Cad | IDCWFLHR | -832 | -839 | 0 | 0 | 8 | 930.6174 | 969.55457 | 1000.997 | 1579.3278 | 1156.4342 | 946.35144 |
| Cad | VDPNARPLAPEVSIK | -151 | -165 | 1 | 0 | 15 | 146.92917 | 781.99054 | 805.43 | 599.4906 | 620.69073 | 635.39935 |
| Cad | VDPNARPLAPEVSIK | -151 | -165 | 1 | 0 | 15 | 146.92917 | 781.99054 | 805.43 | 599.4906 | 620.69073 | 635.39935 |
| Cad | YWGNTHDLDFR | -921 | -931 | 0 | 0 | 11 | 906.2224 | 1707.9467 | 1660.4446 | 2510.3862 | 3561.952 | 3281.7988 |
| Cad | AALVLEDGSVLQGR | -2 | -15 | 0 | 0 | 14 | 694.6551 | 827.06177 | 719.88525 | 860.8143 | 1072.7031 | 809.62775 |
| Cad | LLDTIGISQPQWR | -1054 | -1066 | 0 | 0 | 13 | 657.47943 | 757.4435 | 786.23157 | 910.3646 | 1181.1807 | 1163.8396 |
| Cad | VYFLPITLH | -450 | -458 | 0 | 0 | 9 | 577.90204 | 471.21368 | 512.58374 | 569.1981 | 583.0614 | 629.2208 |
| Cad | LCPPELPIPGSGLPPPR | -378 | -394 | 0 | 0 | 17 | 273.29504 | 297.6386 | 832.4692 | 492.01337 | 747.5761 | 706.634 |
| Cad | GTSPEAIDSAENR | -1036 | -1048 | 0 | 0 | 13 | 5124.407 | 5750.983 | 2104.924 | NaN | 13931.266 | 232.68994 |
| Cad | THDLDFR | -925 | -931 | 0 | 0.001708008 | 7 | 2034.461 | 1364.6301 | 1028.0413 | NaN | 944.21844 | 899.44214 |
| Cad | GTADFYTEHGVK | -1347 | -1358 | 0 | 0 | 12 | 790.11993 | NaN | 1664.8695 | 1173.0417 | 1110.9259 | 1064.6443 |
| Cad | VPQFSFSR | -1263 | -1270 | 0 | 0 | 8 | 756.49194 | 769.08435 | 875.5891 | 630.97296 | NaN | 230.14514 |
| Cad | NERPDGVLLTF | -466 | -476 | 1 | 0.000431655 | 11 | 745.1442 | 841.68274 | NaN | 727.501 | 875.9035 | 890.1969 |
| Cad | YIDGQVLVPPGYGQDVR | -1794 | -1810 | 0 | 0.00025015 | 17 | 580.97986 | NaN | 817.38367 | 923.54803 | 939.2304 | 966.4021 |
| Cad | LHEWLQQR | -104 | -111 | 0 | 0 | 8 | 519.1632 | 320.41376 | 753.90393 | 753.30176 | 591.0134 | NaN |
| Cad | VVPWDHELDSQK | -202 | -213 | 0 | 0 | 12 | 835.50073 | NaN | NaN | 1302.0621 | 729.9471 | 1095.3156 |
| Cad | EVAPHHLFLNR | -1637 | -1647 | 0 | 0 | 11 | 637.77136 | 749.97864 | 986.4829 | NaN | 619.7022 | NaN |
| Cad | FYTEHGVK | -1351 | -1358 | 0 | 0 | 8 | 943.3085 | 1043.0875 | 881.84106 | NaN | NaN | NaN |
| Cad | VTQHLGIVGECNVQ | -655 | -668 | 0 | 0 | 14 | 587.6853 | 686.8453 | 698.4883 | NaN | NaN | NaN |
| Cad | SFSEATSSVQK | -1993 | -2003 | 0 | 0 | 11 | 323.82596 | 638.07465 | NaN | 1533.9755 | NaN | NaN |
| Cad | SVPLIIDIK | -1423 | -1431 | 0 | 0.001610812 | 9 | 261.05853 | 335.39246 | 387.19574 | NaN | NaN | NaN |
| Cad | ARPLAPEVSIK | -155 | -165 | 1 | 0.00383302 | 11 | 843.9617 | NaN | NaN | 623.35284 | NaN | NaN |
| Cad | GQIGPAPPLK | -1441 | -1450 | 0 | 0.00078661 | 10 | 409.7006 | NaN | NaN | 478.0234 | NaN | NaN |
| Cad | ALGQIGPAPPLK | -1439 | -1450 | 0 | 0.00025015 | 12 | 277.45004 | NaN | NaN | 785.149 | NaN | NaN |
| Cad | SELLPTVR | -1326 | -1333 | 0 | 0.001776473 | 8 | 463.64124 | NaN | NaN | NaN | NaN | NaN |
| Rab3gap1 | AANPGCFLEDFVR | -688 | -700 | 0 | 0 | 13 | 701.3455 | 811.8575 | 870.0821 | 289.32614 | NaN | 360.7032 |
| Rab3gap1 | SDEISFADFR | -62 | -71 | 0 | 0 | 10 | 423.0722 | 423.33887 | 394.7358 | NaN | 489.35202 | 414.8643 |
| Ftsj3 | TSVTDFLR | -252 | -259 | 0 | 0 | 8 | 824.66895 | 824.4009 | 1125.6002 | 1170.7899 | 682.8049 | 1082.6978 |
| Ftsj3 | LACDFLAR | -143 | -150 | 0 | 0 | 8 | 599.14575 | 537.45416 | 512.2578 | 449.11697 | 324.12668 | NaN |
| Ftsj3 | AANPVDFLSK | -260 | -269 | 0 | 0.001904399 | 10 | 1266.4325 | NaN | 500.73495 | 723.3712 | NaN | 453.71173 |
| Tpi1 | HVFGESDELIGQK | -101 | -113 | 0 | 0 | 13 | 7434.3257 | 7385.665 | 7721.565 | 8764.827 | 8698.458 | 8466.112 |
| Tpi1 | SLKPEFVDIINAK | -236 | -248 | 1 | 0 | 13 | 7407.5073 | 9124.469 | 9046.01 | 6998.6436 | 9658.942 | 721.01404 |
| Tpi1 | SLKPEFVDIINAK | -236 | -248 | 1 | 0 | 13 | 7407.5073 | 9124.469 | 9046.01 | 6998.6436 | 9658.942 | 721.01404 |
| Tpi1 | LKPEFVDIINAK | -237 | -248 | 1 | 0 | 12 | 3205.261 | 3660.3625 | 3745.3179 | 2727.6382 | 3329.1128 | 3332.632 |
| Tpi1 | LKPEFVDIINAK | -237 | -248 | 1 | 0 | 12 | 3205.261 | 3660.3625 | 3745.3179 | 2727.6382 | 3329.1128 | 3332.632 |
| Tpi1 | WVVLGHSER | -91 | -99 | 0 | 0.00025015 | 9 | 3136.6448 | 3354.8093 | 3335.7737 | 5370.736 | 5293.704 | 5303.6 |
| Tpi1 | VVLGHSER | -92 | -99 | 0 | 0.00025015 | 8 | 2732.8787 | 2724.356 | 2944.938 | 2764.0735 | 2698.9316 | 2880.466 |
| Tpi1 | ASLKPEFVDIINAK | -235 | -248 | 1 | 0 | 14 | 2459.9006 | 2771.6409 | 3100.5808 | 2857.193 | 2522.2488 | 4206.2744 |
| Tpi1 | HVFGESDELIGQ | -101 | -112 | 0 | 0 | 12 | 2414.0667 | 237.31091 | 2292.4805 | 2079.3884 | 1139.6731 | 949.7139 |
| Tpi1 | HVFGESDELIGQ | -101 | -112 | 0 | 0 | 12 | 2414.0667 | 237.31091 | 2292.4805 | 2079.3884 | 1139.6731 | 949.7139 |
| Tpi1 | VVLAYEPVWAIGTGK | -161 | -175 | 0 | 0 | 15 | 2320.0347 | 313.4677 | 2639.4287 | 2726.3486 | 112.03789 | 4097.9277 |
| Tpi1 | TATPQQAQEVHEK | -176 | -188 | 0 | 0 | 13 | 2273.8003 | 2401.6196 | 2472.8328 | 2522.7449 | 4127.0103 | 3950.1045 |
| Tpi1 | TATPQQAQEVHEK | -176 | -188 | 0 | 0 | 13 | 2273.8003 | 2401.6196 | 2472.8328 | 2522.7449 | 4127.0103 | 3950.1045 |
| Tpi1 | AYEPVWAIGTGK | -164 | -175 | 0 | 0 | 12 | 1999.861 | 1806.8832 | 365.3312 | 509.3487 | 833.718 | 477.53198 |
| Tpi1 | RHVFGESDELIGQK | -100 | -113 | 1 | 0 | 14 | 1891.3087 | 1831.8105 | 2110.4985 | 2900.988 | 2121.0352 | 2173.9028 |
| Tpi1 | RHVFGESDELIGQK | -100 | -113 | 1 | 0 | 14 | 1891.3087 | 1831.8105 | 2110.4985 | 2900.988 | 2121.0352 | 2173.9028 |
| Tpi1 | ELASQPDVDGFLVGGA | -220 | -235 | 0 | 0.000230707 | 16 | 882.0324 | 886.5919 | 995.22864 | 1228.5186 | 1590.4392 | 1582.9723 |
| Tpi1 | NVPAGTEVVCAPPTAYIDFAR | -33 | -53 | 0 | 0 | 21 | 797.5364 | 870.4239 | 818.84845 | 1440.4226 | 2205.5293 | 1916.9105 |
| Tpi1 | ELASQPDVDGFLVGGAS | -220 | -236 | 0 | 0.00078661 | 17 | 766.78784 | 876.8802 | 782.6334 | 778.85474 | 931.83673 | 811.83936 |
| Tpi1 | NAANVPAGTEVVCAPPTAYIDFAR | -30 | -53 | 0 | 0 | 24 | 498.73334 | 602.66565 | 547.70416 | 733.04456 | 989.04474 | 942.2871 |
| Tpi1 | LVGGASLKPEFVDIINAK | -231 | -248 | 1 | 0 | 18 | 466.21844 | 568.9917 | 513.7856 | 657.16406 | 821.4048 | 858.2792 |
| Tpi1 | VVLAYEPVWAIGT | -161 | -173 | 0 | 0.001291103 | 13 | 366.4871 | 546.802 | 716.5488 | 363.8354 | 633.5638 | 490.4424 |
| Tpi1 | AANVPAGTEVVCAPPTAYIDFAR | -31 | -53 | 0 | 0 | 23 | 271.58856 | 292.23413 | 268.5096 | 381.34726 | 484.71704 | 554.015 |
| Tpi1 | SNVNDGVAQSTR | -195 | -206 | 0 | 0 | 12 | 2114.339 | 1485.7411 | 1353.1742 | NaN | 1649.2805 | 1533.0078 |
| Tpi1 | FFVGGNWK | -7 | -14 | 0 | 0.00025015 | 8 | 197.03798 | 2521.3413 | 1932.5232 | NaN | 675.2225 | 1140.0533 |
| Tpi1 | APPTAYIDFAR | -43 | -53 | 0 | 0 | 11 | 8298.953 | 8363.711 | 9296.98 | NaN | NaN | 7537.501 |
| Tpi1 | RHVFGESDELIGQ | -100 | -112 | 1 | 0.000230707 | 13 | 4313.062 | NaN | 430.7498 | 4110.1675 | NaN | 5683.9473 |
| Tpi1 | RHVFGESDELIGQ | -100 | -112 | 1 | 0.000230707 | 13 | 4313.062 | NaN | 430.7498 | 4110.1675 | NaN | 5683.9473 |
| Tpi1 | VTNGAFTGEISPGMIK | -70 | -85 | 0 | 0 | 16 | 2255.9387 | NaN | NaN | 1414.4883 | 1468.1843 | 1120.5793 |
| Tpi1 | KFFVGGNWK | -6 | -14 | 1 | 0.000471409 | 9 | 1669.4526 | 1073.2485 | 966.44226 | NaN | NaN | 720.482 |

| | | | | | | | | | | | | |
|---|---|---|---|---|---|---|---|---|---|---|---|---|
| Tpi1 | EKVVFEQTK | -141 | -149 | 1 | 0 | 9 | 1184.1737 | 1072.9255 | 1169.9333 | 1190.3696 | NaN | NaN |
| Tpi1 | FVDIINAK | -241 | -248 | 0 | 0.001636916 | 8 | 689.8055 | 772.6313 | 709.6429 | NaN | NaN | NaN |
| Tpi1 | SDELIGQK | -106 | -113 | 0 | 0.002810811 | 8 | 837.40735 | 921.11926 | NaN | NaN | NaN | NaN |
| Vim | ETNLESLPLVDTHSK | -425 | -439 | 0 | 0 | 15 | 8013.239 | 9355.334 | 9628.002 | 9442.648 | 10896.674 | 10873.443 |
| Vim | ETNLESLPLVDTHSK | -425 | -439 | 0 | 0 | 15 | 8013.239 | 9355.334 | 9628.002 | 9442.648 | 10896.674 | 10873.443 |
| Vim | FANYIDK | -114 | -120 | 0 | 0.001811524 | 7 | 6788.3906 | 2410.623 | 7569.778 | 1943.0557 | 201.32855 | 1886.0779 |
| Vim | LQEEMLQR | -189 | -196 | 0 | 0 | 8 | 6450.6724 | 472.10565 | 2275.7576 | 5095.176 | 4322.097 | 2495.3372 |
| Vim | FADLSEAANR | -295 | -304 | 0 | 0 | 10 | 6245.1484 | 6649.683 | 5130.859 | 7267.1177 | 5107.996 | 308.8963 |
| Vim | KLHDEEIQEL | -236 | -245 | 1 | 0.00025015 | 10 | 5189.1265 | 5563.0947 | 5291.372 | 7076.6895 | 785.443 | 1062.483 |
| Vim | KLHDEEIQEL | -236 | -245 | 1 | 0.00025015 | 10 | 5189.1265 | 5563.0947 | 5291.372 | 7076.6895 | 785.443 | 1062.483 |
| Vim | DGQVINETSQHHDDLE | -451 | -466 | 0 | 0 | 16 | 4792.63 | 5020.6724 | 1447.9534 | 3262.4788 | 4234.019 | 4159.5474 |
| Vim | DGQVINETSQHHDDLE | -451 | -466 | 0 | 0 | 16 | 4792.63 | 5020.6724 | 1447.9534 | 3262.4788 | 4234.019 | 4159.5474 |
| Vim | KVESLQEEIAFLK | -223 | -235 | 1 | 0 | 13 | 4759.193 | 5941.004 | 6036.2656 | 7973.075 | 9119.447 | 9460.991 |
| Vim | KVESLQEEIAFLK | -223 | -235 | 1 | 0 | 13 | 4759.193 | 5941.004 | 6036.2656 | 7973.075 | 9119.447 | 9460.991 |
| Vim | KLHDEEIQELQAQ | -236 | -248 | 1 | 0.000394361 | 13 | 4158.961 | 4932.832 | 4941.469 | 3624.9714 | 4529.4263 | 4401.098 |
| Vim | KLHDEEIQELQAQ | -236 | -248 | 1 | 0.000394361 | 13 | 4158.961 | 4932.832 | 4941.469 | 3624.9714 | 4529.4263 | 4401.098 |
| Vim | ILLAELEQLK | -130 | -139 | 0 | 0 | 10 | 4055.795 | 4402.8633 | 4199.156 | 4352.3076 | 5260.859 | 316.05414 |
| Vim | LGDLYEEEMR | -146 | -155 | 0 | 0 | 10 | 3854.7908 | 1159.7506 | 4215.846 | 1458.4302 | 1252.0714 | 1075.7506 |
| Vim | LGDLYEEEMR | -146 | -155 | 0 | 0 | 10 | 3854.7908 | 1159.7506 | 4215.846 | 1458.4302 | 1252.0714 | 1075.7506 |
| Vim | QEEIAFLK | -228 | -235 | 0 | 0.00025015 | 8 | 3663.206 | 3405.3176 | 3605.5898 | 2558.936 | 2317.1235 | 2555.9976 |
| Vim | LQDEIQNMK | -365 | -373 | 0 | 0 | 9 | 2352.4858 | 1849.7899 | 2368.03 | 2185.0059 | 2547.0369 | 2833.059 |
| Vim | LQDEIQNMK | -365 | -373 | 0 | 0 | 9 | 2352.4858 | 1849.7899 | 2368.03 | 2185.0059 | 2547.0369 | 2833.059 |
| Vim | KLHDEEIQELQAQIQ | -236 | -250 | 1 | 0 | 15 | 3502.372 | 4489.0586 | 4330.3164 | 4363.504 | 5579.4673 | 5170.9717 |
| Vim | KLHDEEIQELQAQIQ | -236 | -250 | 1 | 0 | 15 | 3502.372 | 4489.0586 | 4330.3164 | 4363.504 | 5579.4673 | 5170.9717 |
| Vim | DNLAEDIMR | -176 | -184 | 0 | 0 | 9 | 3387.2517 | 1210.466 | 3636.302 | 3463.5635 | 1623.8392 | 1717.8038 |
| Vim | DNLAEDIMR | -176 | -184 | 0 | 0 | 9 | 3387.2517 | 1210.466 | 3636.302 | 3463.5635 | 1623.8392 | 1717.8038 |
| Vim | EKLQEEMLQR | -187 | -196 | 1 | 0 | 10 | 2630.6077 | 1571.566 | 1683.1079 | 1413.1383 | 1898.6497 | 1936.9099 |
| Vim | EKLQEEMLQR | -187 | -196 | 1 | 0 | 10 | 2630.6077 | 1571.566 | 1683.1079 | 1413.1383 | 1898.6497 | 1936.9099 |
| Vim | TCEVDALK | -327 | -334 | 0 | 0 | 8 | 2509.6306 | 3058.7468 | 1918.4012 | 127.368065 | 140.68893 | 471.84616 |
| Vim | NLESLPLVDTHSK | -427 | -439 | 0 | 0 | 13 | 2393.5615 | 2756.4895 | 2674.969 | 3087.934 | 3146.2834 | 3091.594 |
| Vim | RQDVDNASLAR | -207 | -217 | 1 | 0 | 11 | 2226.901 | 2276.9814 | 525.87415 | 2014.9932 | 275.07632 | 2898.6775 |
| Vim | RQVDQLTNDK | -159 | -168 | 1 | 0.000471409 | 10 | 1916.9233 | 2434.9233 | 2165.5579 | 1538.1166 | 2244.2356 | 2335.3447 |
| Vim | NYQDTIGR | -357 | -364 | 0 | 0 | 8 | 1829.4408 | 1613.8209 | 1393.014 | 1394.6406 | 940.5217 | 1399.8906 |
| Vim | ANYQDTIGR | -356 | -364 | 0 | 0 | 9 | 1599.7645 | 1416.9403 | 1460.9646 | 622.8078 | 686.178 | 922.88086 |
| Vim | SLPLVDTHSK | -430 | -439 | 0 | 0.000550307 | 10 | 1579.821 | 1429.3114 | 1202.8129 | 1218.9795 | 1403.3048 | 1419.7589 |
| Vim | FSLADAINTEFK | -86 | -97 | 0 | 0 | 12 | 1556.4688 | 1523.0598 | 189.73192 | 1642.5221 | 2061.158 | 275.12036 |
| Vim | LHDEEIQEL | -237 | -245 | 0 | 0.00025015 | 9 | 1355.6984 | 963.88513 | 1143.6128 | 1190.8413 | 1788.0061 | 1840.3032 |
| Vim | KLHDEEIQELQA | -236 | -247 | 1 | 0.00111006 | 12 | 1290.6019 | 1397.779 | 1774.9431 | 1269.0709 | 1801.2904 | 2049.2666 |
| Vim | ETNLESLPLVDTH | -425 | -437 | 0 | 0 | 13 | 1237.3195 | 1499.6556 | 1411.2294 | 377.45517 | 1894.5378 | 1855.99 |
| Vim | SLQEEIAFLK | -226 | -235 | 0 | 0 | 10 | 1097.678 | 1077.8711 | 1136.0706 | 257.64227 | 762.18933 | 796.30786 |
| Vim | SLYSSSPGGAYVTR | -51 | -64 | 0 | 0 | 14 | 1022.9934 | 885.7793 | 1120.7379 | 928.75854 | 928.3121 | 1059.4116 |
| Vim | EAANYQDTIGR | -354 | -364 | 0 | 0 | 11 | 972.16406 | 781.08997 | 2901.2454 | 4651.1025 | 183.19092 | 816.2861 |
| Vim | VESLQEEIAFLK | -224 | -235 | 0 | 0 | 12 | 928.802 | 861.84143 | 786.8909 | 919.3711 | 1013.4734 | 1033.5428 |
| Vim | KVESLQEEIAFL | -223 | -234 | 1 | 0.00025015 | 12 | 766.9153 | 444.26263 | 579.70825 | 1086.6976 | 1412.8721 | 730.547 |
| Vim | VSKPDLTAALR | -260 | -270 | 1 | 0 | 11 | 759.8596 | 747.8022 | 800.30457 | 1062.3826 | 678.49585 | 560.31537 |
| Vim | IDVDVSKPDLTAALR | -256 | -270 | 1 | 0 | 15 | 635.3728 | 569.64276 | 602.823 | 1187.7753 | 1370.2769 | 1416.3799 |
| Vim | ILLAELEQLKG | -130 | -140 | 1 | 0 | 11 | 621.1473 | 538.71594 | 662.47906 | 585.0218 | 753.1023 | 757.58496 |
| Vim | SLTCEVDALK | -325 | -334 | 0 | 0 | 10 | 593.9525 | 966.8255 | 670.52844 | 1075.4454 | 1285.8385 | 1182.0541 |
| Vim | VDVSKPDLTAALR | -258 | -270 | 1 | 0 | 13 | 582.52844 | 653.3209 | 658.33374 | 713.3019 | 799.8031 | 831.39685 |
| Vim | VDVSKPDLTAALR | -258 | -270 | 1 | 0 | 13 | 582.52844 | 653.3209 | 658.33374 | 713.3019 | 799.8031 | 831.39685 |
| Vim | LQDEIQNMKEEMAR | -365 | -378 | 1 | 0 | 14 | 331.83212 | 386.69757 | 409.80646 | 373.90814 | 569.6821 | 525.70197 |
| Vim | LHDEEIQELQAQ | -237 | -248 | 0 | 0.000471409 | 12 | 90.67808 | 1815.1633 | 1960.018 | 2564.4775 | 1499.5393 | 1580.1788 |
| Vim | KLHDEEIQELQ | -236 | -246 | 1 | 0.00051306 | 11 | 5764.7354 | 3370.9941 | NaN | 5485.4497 | 546.877 | 2289.617 |
| Vim | AANYQDTIGR | -355 | -364 | 0 | 0 | 10 | 2916.4978 | 191.51671 | 1439.7815 | NaN | 2399.5874 | 2463.9805 |
| Vim | QVDQLTNDK | -160 | -168 | 0 | 0 | 9 | 2715.7415 | 3810.505 | 1544.2034 | 2619.6729 | 1950.0771 | NaN |
| Vim | ILLAELEQLKGQ | -130 | -141 | 1 | 0 | 12 | 575.3815 | NaN | 529.70233 | 618.0955 | 621.2874 | 693.81464 |
| Vim | QQYESVAAK | -274 | -282 | 0 | 0 | 9 | 3062.4207 | 2674.8076 | 2613.8066 | 2182.5562 | NaN | NaN |
| Vim | YSSSPGGAYVTR | -53 | -64 | 0 | 0 | 12 | 1904.956 | NaN | 948.71844 | 950.3093 | 340.42572 | NaN |
| Vim | ISLPLPTFS | -411 | -419 | 0 | 0 | 9 | 394.37677 | NaN | NaN | 481.83643 | 770.63086 | 816.00653 |
| Vim | QDVDNASLAR | -208 | -217 | 0 | 0 | 10 | 3093.2898 | 335.05612 | NaN | 2377.3008 | NaN | NaN |
| Vim | LYSSSPGGAYVTR | -52 | -64 | 0 | 0 | 13 | 1011.91785 | 1436.1423 | NaN | 1302.2833 | NaN | NaN |
| Vim | LHDEEIQELQA | -237 | -247 | 0 | 0.001055699 | 11 | 1045.0364 | NaN | NaN | NaN | NaN | NaN |
| Rplp0 | SVETEYTFPLTEK | -252 | -264 | 0 | 0 | 13 | 6212.1055 | 1907.257 | 7796.104 | 7860.5127 | 9875.376 | 9470.069 |
| Rplp0 | IIQLLDDYPK | -17 | -26 | 0 | 0.000230707 | 10 | 4646.9033 | 5475.5205 | 5220.1025 | 4646.805 | 5297.6284 | 5532.5146 |
| Rplp0 | CFIVGADNVGSK | -27 | -38 | 0 | 0 | 12 | 2638.1965 | 205.74698 | 2931.0322 | 362.69003 | 2938.87 | 2798.9324 |
| Rplp0 | EVTVPAQNTGLGPEK | -120 | -134 | 0 | 0 | 15 | 2552.9937 | 2601.361 | 5558.129 | 1929.6119 | 1946.0958 | 1971.1046 |
| Rplp0 | AGAIAPCEVTVPAQNTGLGPEK | -113 | -134 | 0 | 0 | 22 | 905.6507 | 1018.57697 | 1065.3098 | 782.4769 | 1147.8203 | 1149.7083 |
| Rplp0 | AIAPCEVTVPAQNTGLGPEK | -115 | -134 | 0 | 0 | 20 | 891.4576 | 969.9798 | 876.8291 | 1107.1724 | 967.0688 | 858.1523 |
| Rplp0 | RAGAIAPCEVTVPAQNTGLGPEK | -112 | -134 | 1 | 0.000230707 | 23 | 550.5312 | 67.40383 | 684.3518 | 582.6827 | 817.4759 | 589.7953 |
| Rplp0 | GAIAPCEVTVPAQNTGLGPEK | -114 | -134 | 0 | 0.000431655 | 21 | 529.4041 | 513.05084 | 508.2533 | 510.627 | 632.41187 | 621.3841 |
| Rplp0 | GHLENNPALEK | -67 | -77 | 0 | 0 | 11 | 370.76712 | 6417.7515 | 6265.88 | 5662.741 | 6628.9307 | 6464.8013 |
| Rplp0 | IVGADNVGSK | -29 | -38 | 0 | 0.00025015 | 10 | 363.05154 | 378.45825 | 354.86362 | 609.07635 | 852.4182 | 612.18445 |
| Rplp0 | TKEDLTEIR | -91 | -99 | 1 | 0 | 9 | 290.4523 | 6894.343 | 6894.8955 | 8216.797 | 8187.597 | 8617.622 |
| Rplp0 | SVPHSIINGYK | -236 | -246 | 0 | 0 | 11 | 2268.9841 | 2076.0615 | 1857.1061 | NaN | NaN | NaN |
| Rplp0 | TTAAPAAAAAPAK | -285 | -297 | 0 | 0.00111006 | 13 | 963.0672 | 1208.7413 | NaN | NaN | NaN | NaN |
| Rplp0 | AAPAAAAAPAK | -287 | -297 | 0 | 0.006788754 | 11 | 1074.5598 | NaN | NaN | NaN | NaN | NaN |
| Ppp1r14b | AAPAPGPGSGSTGPR | -13 | -27 | 0 | 0 | 15 | 1735.3328 | 1879.5983 | 2416.6929 | 794.2088 | 2057.5027 | 2170.699 |
| Ppp1r14b | FQSPPGAAGEGPGGADDDGPVR | -30 | -51 | 0 | 0 | 22 | 620.8234 | 693.1083 | 523.21106 | 791.1699 | 832.7776 | 912.90094 |

| | | | | | | | | | | | | |
|---|---|---|---|---|---|---|---|---|---|---|---|---|
| Ppp1r14b | VYFQSPPGAAGEGPGGADDDGPVR | -28 | -51 | 0 | 0 | 24 | 346.09106 | 473.35577 | 451.63043 | 374.67896 | 391.80594 | 420.88617 |
| Mrpl12 | KLVESLPQEIK | -166 | -176 | 1 | 0 | 11 | 1169.1215 | 1033.2603 | 1416.5322 | 1021.3041 | 1410.876 | 1401.939 |
| Mrpl12 | SEALAGAPLDNAPK | -46 | -59 | 0 | 0 | 14 | 1125.3517 | 1218.7493 | 1297.4141 | 422.85068 | 1483.6797 | 1341.6213 |
| Mrpl12 | LVESLPQEIK | -167 | -176 | 0 | 0 | 10 | 568.20636 | 543.41833 | 680.6383 | 623.37415 | 977.1826 | NaN |
| Mrpl12 | LTEAKPVDK | -137 | -145 | 1 | 0.002099344 | 9 | 218.79921 | 203.28464 | NaN | 374.5824 | 377.06055 | 519.3006 |
| Mrpl12 | AAPASEAAEEEDVPK | -112 | -126 | 0 | 0.001463897 | 15 | 11361.123 | NaN | NaN | 5410.516 | NaN | 8072.7437 |
| Nptn | AAPDITGHK | -235 | -243 | 0 | 0 | 9 | 1276.7312 | NaN | NaN | 1501.0941 | 615.36084 | 1304.975 |
| Faf2 | AAPEEQDLTQEQTEK | -2 | -16 | 0 | 0.000230707 | 15 | 608.43445 | 545.8939 | 439.08453 | 550.74384 | 472.63525 | 379.297 |
| Faf2 | LHGDDHQDSDEFCR | -177 | -190 | 0 | 0 | 14 | 581.6036 | 608.3071 | 545.82385 | 362.4311 | 151.85948 | 537.4539 |
| Nhp2 | VKPHEEYQETYDK | -129 | -141 | 1 | 0 | 13 | 10020.854 | 10368.95 | 8339.692 | 996.0927 | 7106.9644 | 10227.239 |
| Nhp2 | VKPHEEYQETYDK | -129 | -141 | 1 | 0 | 13 | 10020.854 | 10368.95 | 8339.692 | 996.0927 | 7106.9644 | 10227.239 |
| Nhp2 | AAPEESEAQAEGCSEER | -6 | -22 | 0 | 0 | 17 | 238.0548 | NaN | 176.96957 | NaN | NaN | NaN |
| Me2 | SIVDNWPENHVK | -145 | -156 | 0 | 0 | 12 | 1226.6605 | 742.25885 | 706.4039 | 1235.5259 | 1238.4247 | 528.31177 |
| Me2 | TTQLTDAELAQGR | -499 | -511 | 0 | 0 | 13 | 717.23486 | 634.9508 | 672.90643 | 1291.8467 | 1029.5483 | 1135.3202 |
| Me2 | AAPESIPATFEDAVNK | -368 | -383 | 0 | 0 | 16 | 656.5702 | 1428.4977 | 615.08075 | 1194.493 | 661.4406 | 1462.5693 |
| Me2 | LKPSVIIGVA | -384 | -393 | 1 | 0.00133086 | 10 | 940.06604 | NaN | 1036.1315 | 888.0245 | 920.4366 | NaN |
| Me2 | GAGPLFTHGVIK | -394 | -405 | 0 | 0 | 12 | 934.32983 | NaN | NaN | 836.12024 | 764.2818 | 1789.6927 |
| Me2 | SIPATFEDAVNK | -372 | -383 | 0 | 0 | 12 | 666.14636 | 718.2656 | 925.2195 | NaN | NaN | 370.3656 |
| Me2 | VINKPVSEHK | -297 | -306 | 1 | 0.00111006 | 10 | 188.45757 | 193.45844 | NaN | NaN | NaN | NaN |
| Uqcrc2 | IIENLHDVAYK | -173 | -183 | 0 | 0 | 11 | 2773.7551 | 2530.9312 | 2811.7202 | 2412.1587 | 2710.4902 | 2696.623 |
| Uqcrc2 | GNLGHTPFLDEL | -442 | -453 | 0 | 0 | 12 | 1564.1162 | 1714.1077 | 1702.9503 | 2107.7827 | 2584.3845 | 2348.7705 |
| Uqcrc2 | LVGLGVSHSVLK | -220 | -231 | 0 | 0 | 12 | 1466.1802 | 1790.5105 | 1691.9148 | 2121.0476 | 1867.8331 | 2137.7 |
| Uqcrc2 | AAPGGVPLQPQDLEFTK | -26 | -42 | 0 | 0 | 17 | 1445.8412 | 1970.3088 | 1861.3798 | 2044.0115 | 2413.4727 | 2393.03 |
| Uqcrc2 | IIENLHDVAYKN | -173 | -184 | 1 | 0 | 12 | 1173.5865 | 1088.4397 | 1003.11835 | 1250.8197 | 1307.5177 | 1280.3215 |
| Uqcrc2 | GSHQPFDVSAF | -316 | -326 | 0 | 0.00025015 | 11 | 1140.4987 | 1549.3359 | 1656.5103 | 1484.0842 | 149.08176 | 1100.088 |
| Uqcrc2 | YEDSNNLGTSHLLR | -71 | -84 | 0 | 0 | 14 | 1030.7134 | 1163.0408 | 1177.8953 | 1575.0342 | 273.17133 | 1624.2266 |
| Uqcrc2 | SGNLGHTPFLDEL | -441 | -453 | 0 | 0 | 13 | 985.07764 | 695.36163 | 430.8607 | 1509.6134 | 1258.3046 | 843.16284 |
| Uqcrc2 | QHLLGAGPHIK | -290 | -300 | 0 | 0.00025015 | 11 | 842.0952 | 1235.0529 | 1188.6451 | 708.8919 | 976.71436 | 1005.0948 |
| Uqcrc2 | YEDSNNLGTSHLL | -71 | -83 | 0 | 0 | 13 | 571.31433 | 2397.6008 | 2317.808 | 561.4008 | 2542.5881 | 2628.5947 |
| Uqcrc2 | SLENYAPLSR | -51 | -60 | 0 | 0 | 10 | 1511.8429 | 1491.3754 | 206.18094 | 1327.152 | NaN | 1741.9956 |
| Uqcrc2 | ITSEELHYFVQ | -200 | -210 | 0 | 0 | 11 | 321.5138 | 404.18668 | 1513.1328 | 301.71762 | NaN | 1599.2823 |
| Uqcrc2 | TSAAPGGVPLQPQDLEFTK | -24 | -42 | 0 | 0 | 19 | 1685.4296 | 2045.1428 | 1865.999 | NaN | 2688.157 | NaN |
| Uqcrc2 | AASGNLGHTPFLDEL | -439 | -453 | 0 | 0.00025015 | 15 | 625.0497 | NaN | 1181.6991 | 1021.8405 | 1443.321 | NaN |
| Uqcrc2 | YEDSNNLGTSH | -71 | -81 | 0 | 0 | 11 | 664.13464 | NaN | NaN | 915.5096 | NaN | NaN |
| Uqcrc2 | LNVTTAPEFR | -138 | -147 | 0 | 0 | 10 | 445.49426 | NaN | NaN | NaN | NaN | NaN |
| Med25 | AAPPALLEPLQQPADVSQDPR | -198 | -218 | 0 | 0 | 21 | 106.313354 | NaN | NaN | NaN | 239.18852 | 253.40324 |
| Wars1 | FLEDDDRLEQIR | -410 | -421 | 1 | 0 | 12 | 1339.7994 | 1770.7002 | 1852.2079 | 1957.0763 | 1043.1948 | 733.50696 |
| Wars1 | SEDFVDPWTVR | -85 | -95 | 0 | 0 | 11 | 1044.6896 | 1182.4908 | 1085.3264 | 1241.0461 | 1458.1042 | 1426.2065 |
| Wars1 | AAPSFSNSFPK | -289 | -299 | 0 | 0 | 11 | 854.84814 | 891.924 | 800.84985 | 1161.947 | 1249.3625 | 833.9313 |
| Wars1 | FFLEDDDRLEQIR | -409 | -421 | 1 | 0 | 13 | 786.63257 | 685.28644 | 717.4159 | 732.04376 | 901.33154 | 962.78064 |
| Wars1 | FFLEDDDRLEQIR | -409 | -421 | 1 | 0 | 13 | 786.63257 | 685.28644 | 717.4159 | 732.04376 | 901.33154 | 962.78064 |
| Wars1 | ASEDFVDPWTVR | -84 | -95 | 0 | 0 | 12 | 667.21356 | 757.74817 | 768.50555 | 664.5018 | 957.8955 | 1055.061 |
| Wars1 | KPFYLYTGR | -158 | -166 | 1 | 0 | 9 | 511.099 | 395.2808 | 399.1519 | 898.9256 | 929.8475 | 975.84784 |
| Wars1 | FFPALQGAQTK | -343 | -353 | 0 | 0 | 11 | 832.6959 | NaN | 845.48816 | 826.9965 | 590.4715 | NaN |
| Wars1 | IATQGELVR | -20 | -28 | 0 | 0.002910424 | 9 | 613.1434 | 686.2831 | 650.0949 | NaN | 3188.6504 | NaN |
| Wars1 | IGHPKPALLH | -331 | -340 | 1 | 0 | 10 | 1013.08295 | NaN | NaN | 561.6352 | 823.1824 | NaN |
| Wars1 | FVDPWTVR | -88 | -95 | 0 | 0 | 8 | 818.9503 | 766.99756 | 897.90027 | NaN | NaN | NaN |
| Wars1 | IGHPKPALLHST | -331 | -342 | 1 | 0 | 12 | 634.8976 | NaN | 491.1661 | NaN | NaN | 618.78424 |
| Eif2a | APSTPLLTVR | -2 | -11 | 0 | 0 | 10 | 736.6643 | 759.4252 | 809.44745 | 706.1435 | 738.02606 | 748.7046 |
| Eif2a | KVNDFNLSPGTQPYK | -169 | -183 | 1 | 0 | 15 | 934.43854 | NaN | 945.88135 | NaN | 1123.903 | 1106.033 |
| Eif2a | NAAFYSPHGHIL | -318 | -329 | 0 | 0 | 12 | 1848.5651 | NaN | NaN | 1968.965 | 2028.2722 | NaN |
| Eif2a | AVPSEVPSEEPK | -430 | -441 | 0 | 0.002296977 | 12 | 1406.4497 | 1668.903 | NaN | 1293.3618 | NaN | NaN |
| Eif2a | AAPTPVPQSAPR | -505 | -516 | 0 | 0.000826806 | 12 | 606.6151 | 732.6757 | NaN | NaN | NaN | NaN |
| Tcp1 | SLHDALCVVK | -391 | -400 | 0 | 0 | 10 | 3239.4333 | 3190.4412 | 3407.4946 | 655.2388 | 3792.8694 | 3763.8674 |
| Tcp1 | SLHDALCVVK | -391 | -400 | 0 | 0 | 10 | 3239.4333 | 3190.4412 | 3407.4946 | 655.2388 | 3792.8694 | 3763.8674 |
| Tcp1 | GQAEEVVQER | -346 | -355 | 0 | 0 | 10 | 2046.1738 | 1548.284 | 1427.9628 | 1441.016 | 1415.986 | 206.47087 |
| Tcp1 | VVITDPEKLDQIR | -252 | -264 | 1 | 0 | 13 | 1788.5377 | 2310.9807 | 2048.5215 | 1857.6066 | 5076.8027 | 2701.3862 |
| Tcp1 | VVITDPEKLDQIR | -252 | -264 | 1 | 0 | 13 | 1788.5377 | 2310.9807 | 2048.5215 | 1857.6066 | 5076.8027 | 2701.3862 |
| Tcp1 | SFGPVGLDK | -35 | -43 | 0 | 0 | 9 | 1757.4935 | 1631.5826 | 1753.6821 | 1560.5636 | 1579.1642 | 1633.0254 |
| Tcp1 | LLEVEHPAAK | -64 | -73 | 0 | 0 | 10 | 1681.7924 | 1676.4514 | 1649.5428 | 143.38564 | 1325.3787 | 1376.4457 |
| Tcp1 | KLLEVEHPAAK | -63 | -73 | 1 | 0 | 11 | 1533.0293 | 1584.7482 | 1560.2529 | 1855.1332 | 2166.621 | 2258.9243 |
| Tcp1 | KLLEVEHPAAK | -63 | -73 | 1 | 0 | 11 | 1533.0293 | 1584.7482 | 1560.2529 | 1855.1332 | 2166.621 | 2258.9243 |
| Tcp1 | AAQDSTDLVAK | -456 | -466 | 0 | 0.00025015 | 11 | 1437.1189 | 1011.7981 | 1058.3201 | 1026.1342 | 1056.1255 | 1222.5156 |
| Tcp1 | WIGLDLVHGK | -485 | -494 | 0 | 0 | 10 | 1413.1616 | 890.55615 | 756.29535 | 831.5699 | 1019.9386 | 1001.9345 |
| Tcp1 | ICDDELILIK | -356 | -365 | 0 | 0 | 10 | 1236.459 | 2118.307 | 1152.5156 | 1697.5001 | 1758.7803 | 1776.848 |
| Tcp1 | HNEAQVNPER | -471 | -480 | 0 | 0 | 10 | 977.69385 | 1117.7599 | 1255.5931 | 84.27415 | 1304.8806 | 1316.2128 |
| Tcp1 | WIGLDLVHGKPR | -485 | -496 | 1 | 0 | 12 | 911.35486 | 1210.6495 | 1204.4525 | 1880.83 | 1222.8784 | 1546.4143 |
| Tcp1 | RICDDELILIK | -355 | -365 | 1 | 0 | 11 | 815.8583 | 2411.3142 | 2482.4346 | 942.3706 | 1311.9535 | 4034.2085 |
| Tcp1 | AFHNEAQVNPER | -469 | -480 | 0 | 0 | 12 | 732.0596 | 1357.4946 | 1099.5862 | 1215.1805 | 699.0586 | 731.8275 |
| Tcp1 | AFHNEAQVNPER | -469 | -480 | 0 | 0 | 12 | 732.0596 | 1357.4946 | 1099.5862 | 1215.1805 | 699.0586 | 731.8275 |
| Tcp1 | MEGPLSVFGDR | -1 | -11 | 0 | 0 | 11 | 500.6043 | 466.43198 | 178.66972 | 375.02652 | 776.9521 | 824.4541 |
| Tcp1 | RIDDLIK | -526 | -532 | 1 | 0.003003218 | 7 | 2270.8057 | NaN | 2304.177 | 3656.0486 | 3471.238 | 3254.3018 |
| Tcp1 | MLGQAEEVVQER | -344 | -355 | 0 | 0 | 12 | 630.2301 | 1663.4739 | 1020.8065 | NaN | 1040.7808 | NaN |
| Tcp1 | MLGQAEEVVQER | -344 | -355 | 0 | 0 | 12 | 630.2301 | 1663.4739 | 1020.8065 | NaN | 1040.7808 | NaN |
| Tcp1 | IACLDFSLQK | -234 | -243 | 0 | 0 | 10 | 1139.2444 | 548.7178 | 550.8715 | NaN | NaN | 525.9748 |
| Tcp1 | YINENLIINTDELGR | -131 | -145 | 0 | 0 | 15 | 569.8173 | NaN | NaN | NaN | 2092.2334 | 2144.1895 |
| Tcp1 | VLCELADLQDK | -74 | -84 | 0 | 0 | 11 | 717.32355 | 307.55112 | NaN | NaN | NaN | NaN |
| Tcp1 | QDSTDLVAK | -458 | -466 | 0 | 0.002566359 | 9 | 1868.3708 | NaN | NaN | NaN | NaN | NaN |

| | | | | | | | | | | | | |
|---|---|---|---|---|---|---|---|---|---|---|---|---|
| Rps14 | DVTPIPSDSTR | -131 | -141 | 0 | 0 | 11 | 7770.392 | 8445.687 | 6986.597 | 7774.6367 | 7837.407 | 6669.4204 |
| Rps14 | EKKEEQVISL | -8 | -17 | 2 | 0.001392869 | 10 | 4631.842 | 4905.3774 | 4624.3657 | 4929.464 | 6157.3794 | 5262.765 |
| Rps14 | RIEDVTPIPSDSTR | -128 | -141 | 1 | 0 | 14 | 3672.2964 | 3779.1072 | 3873.1865 | 3160.4075 | 3218.083 | 3395.0017 |
| Rps14 | RIEDVTPIPSDSTR | -128 | -141 | 1 | 0 | 14 | 3672.2964 | 3779.1072 | 3873.1865 | 3160.4075 | 3218.083 | 3395.0017 |
| Rps14 | ADRDESSPYAA | -64 | -74 | 1 | 0.000471409 | 11 | 1511.6385 | 1733.271 | 1436.8082 | 1924.8978 | 1370.7352 | 1500.6459 |
| Rps14 | FNDTFVHVTDLSGK | -37 | -50 | 0 | 0 | 14 | 1332.8427 | 1696.4048 | 1650.9108 | 1842.4156 | 3031.3616 | 2683.271 |
| Rps14 | LAAQDVAQR | -76 | -84 | 0 | 0 | 9 | 1230.437 | 1373.371 | 1406.5308 | 988.9925 | 213.65536 | 1082.5339 |
| Rps14 | ASFNDTFVHVTDLSGK | -35 | -50 | 0 | 0 | 16 | 840.0958 | 1183.342 | 888.7324 | 1114.4985 | 1333.8828 | 1351.3823 |
| Rps14 | IEDVTPIPSDSTR | -129 | -141 | 0 | 0 | 13 | 518.3657 | 293.01886 | 1833.7306 | 2014.2489 | 1564.1392 | 527.0442 |
| Rps14 | NDTFVHVTDLSGK | -38 | -50 | 0 | 0.000230707 | 13 | 6306.3525 | 6572.468 | 6931.709 | NaN | 5806.889 | 5532.48 |
| Rps14 | VKADRDESSPYAA | -62 | -74 | 2 | 0 | 13 | 3257.6282 | 2778.212 | 3007.7598 | NaN | 2688.7717 | 2593.9229 |
| Rps14 | EKKEEQVISLGPQ | -8 | -20 | 2 | 0.00025015 | 13 | 1144.7975 | 415.09668 | 1323.7235 | NaN | 995.2781 | 553.4424 |
| Rps14 | ADRDESSPY | -64 | -72 | 1 | 0.001392869 | 9 | 680.67114 | 637.19727 | 627.2624 | NaN | 797.1453 | 783.63586 |
| Rps14 | KEEQVISLGPQ | -10 | -20 | 1 | 0 | 11 | 3192.8264 | 2800.665 | NaN | 3599.938 | 1001.42444 | NaN |
| Rps14 | GRIEDVTPIPSDSTR | -127 | -141 | 1 | 0.001092529 | 15 | 967.896 | NaN | 1040.1476 | NaN | 2021.7225 | 2047.2799 |
| Rps14 | AAQDVAQR | -77 | -84 | 0 | 0 | 8 | 656.2411 | NaN | 513.7285 | 398.81992 | NaN | NaN |
| Prpf6 | AVIGIGIEEEDRK | -528 | -540 | 1 | 0 | 13 | 907.9894 | 1024.101 | 1030.0354 | 555.08575 | 854.8452 | 933.3559 |
| Prpf6 | ETNPHHPPAWIA | -303 | -314 | 0 | 0 | 12 | 1415.0537 | NaN | 1579.4839 | 631.929 | 1239.6104 | 1358.124 |
| Prpf6 | LTPVPDSFFAK | -179 | -189 | 0 | 0 | 11 | 399.19067 | 383.79645 | 569.64185 | NaN | 837.6687 | 738.5131 |
| Prpf6 | FELQHGTEEQQEEVR | -884 | -898 | 0 | 0 | 15 | 383.8647 | 352.4484 | 385.69162 | NaN | 838.70166 | 794.01105 |
| Prpf6 | ELCEEALR | -696 | -703 | 0 | 0.001636916 | 8 | 2073.785 | NaN | 2151.1882 | 1886.2344 | 1940.8008 | NaN |
| Prpf6 | HLQTGENHTSVDPR | -190 | -203 | 0 | 0 | 14 | 500.7969 | 608.96814 | NaN | NaN | 343.3982 | 544.73975 |
| Prpf6 | VELEEPEDAR | -411 | -420 | 0 | 0.008014596 | 10 | 4025.912 | NaN | NaN | NaN | NaN | 3497.3896 |
| Prpf6 | WLAGDVPAAR | -619 | -628 | 0 | 0 | 10 | 1409.1221 | NaN | NaN | NaN | 1439.9165 | NaN |
| Prpf6 | AAQELCEEALR | -693 | -703 | 0 | 0.00025015 | 11 | 392.82602 | NaN | 1293.5254 | NaN | NaN | NaN |
| Ran | NVPNWHR | -100 | -106 | 0 | 0.00111006 | 7 | 9758.331 | 9222.596 | 9114.656 | 5961.8003 | 9105.91 | 5149.0044 |
| Ran | SNYNFEKPFLWLAR | -153 | -166 | 1 | 0 | 14 | 1221.5044 | 1228.3113 | 1023.1466 | 517.0347 | 2274.5583 | 1858.8107 |
| Ran | SNYNFEKPFLWLAR | -153 | -166 | 1 | 0 | 14 | 1221.5044 | 1228.3113 | 1023.1466 | 517.0347 | 2274.5583 | 1858.8107 |
| Ran | LVLVGDGGTGK | -13 | -23 | 0 | 0 | 11 | 955.1627 | 673.33246 | 1026.0774 | 5990.188 | 3968.5369 | 4429.794 |
| Ran | SNYNFEKPFLW | -153 | -163 | 1 | 0.000230707 | 11 | 666.3629 | 1059.0034 | 776.72864 | 706.35565 | 1038.3046 | 1126.25 |
| Ran | SNYNFEKPFLWLA | -153 | -165 | 1 | 0 | 13 | 565.07495 | 647.7964 | 712.42926 | 1058.9529 | 1233.3545 | 1143.2711 |
| Dpp3 | SVLNTDPALDSELTSK | -218 | -233 | 0 | 0 | 16 | 3313.0955 | 4461.764 | 4587.846 | 4782.335 | 6095.6416 | 6173.6196 |
| Dpp3 | LTFLEEEDKDLYIR | -424 | -437 | 1 | 0 | 14 | 2283.7788 | 3416.6675 | 2484.8223 | 4042.6196 | 5938.84 | 5263.1753 |
| Dpp3 | DKFLTPDFTSL | -366 | -376 | 1 | 0.000230707 | 11 | 727.86145 | 543.2428 | 963.2538 | 815.4715 | 1012.6006 | 313.42456 |
| Dpp3 | VGLHELLGHGSGK | -447 | -459 | 0 | 0 | 13 | 717.2183 | 739.1422 | NaN | 596.1712 | 591.4347 | 602.0216 |
| Dpp3 | VGLHELLGHGSGK | -447 | -459 | 0 | 0 | 13 | 717.2183 | 739.1422 | NaN | 596.1712 | 591.4347 | 602.0216 |
| Dpp3 | AAQQRPEEVR | -132 | -141 | 1 | 0.00025015 | 10 | 1311.1373 | 1180.0538 | 1522.7097 | 1411.2739 | 1975.3169 | NaN |
| Dpp3 | NTDPALDSELTSK | -221 | -233 | 0 | 0 | 13 | 1065.5347 | 850.4881 | 1131.277 | NaN | 616.0142 | 660.7178 |
| Dpp3 | VTPTTGSDGRPDAR | -589 | -602 | 1 | 0 | 14 | 428.20084 | 416.22244 | NaN | 393.9332 | 557.5344 | 501.19467 |
| Dpp3 | ELPWPLAFEK | -356 | -365 | 0 | 0 | 10 | 602.35315 | 509.3469 | NaN | 709.36053 | 718.8595 | NaN |
| Dpp3 | EFYTPEAANWR | -555 | -565 | 0 | 0 | 11 | 286.46933 | 193.53693 | NaN | NaN | NaN | NaN |
| Dpp3 | EKLTFLEEEDKDLYIR | -422 | -437 | 2 | 0.000230707 | 16 | 814.50824 | NaN | NaN | NaN | NaN | NaN |
| Igbp1 | AASEDELLLPR | -2 | -12 | 0 | 0 | 11 | 1155.1211 | 1265.8793 | 1319.32 | 1417.9299 | 1707.043 | 1806.9764 |
| Igbp1 | LLEDVEVATEPTGSR | -23 | -37 | 0 | 0 | 15 | 416.94952 | 585.4747 | 615.0741 | 322.6535 | NaN | 742.0596 |
| Esd | ASEHGLVVIAPDTSPR | -71 | -86 | 0 | 0 | 16 | 11962.147 | 11609.465 | 3696.379 | 11671.7 | 8939.461 | 800.83923 |
| Esd | ASEHGLVVIAPDTSPR | -71 | -86 | 0 | 0 | 16 | 11962.147 | 11609.465 | 3696.379 | 11671.7 | 8939.461 | 800.83923 |
| Esd | AASEHGLVVIAPDTSPR | -70 | -86 | 0 | 0 | 17 | 3391.8633 | 3785.1892 | 4055.5088 | 3825.5828 | 5240.3755 | 4817.5806 |
| Esd | AASEHGLVVIAPDTSPR | -70 | -86 | 0 | 0 | 17 | 3391.8633 | 3785.1892 | 4055.5088 | 3825.5828 | 5240.3755 | 4817.5806 |
| Esd | GLTCTEQNFISK | -53 | -64 | 0 | 0 | 12 | 2452.2134 | 2644.8525 | 3479.5925 | 1750.0405 | 2277.8892 | 3286.4646 |
| Esd | KIPVVFR | -247 | -253 | 1 | 0.000627943 | 7 | 2416.8398 | 1816.5762 | 1690.4377 | 792.1763 | 636.6321 | 638.4144 |
| Esd | TFIADHIR | -267 | -274 | 0 | 0 | 8 | 1882.4199 | 1998.2302 | 1673.7576 | 2830.5823 | 344.11035 | 2583.6216 |
| Esd | FIADHIR | -268 | -274 | 0 | 0.002265552 | 7 | 1502.3129 | 1522.049 | 1413.3425 | 1578.6753 | 1643.4908 | 1712.1447 |
| Esd | VFEHSSVELK | -18 | -27 | 0 | 0 | 10 | 1018.15765 | 1223.8398 | 1264.738 | 1573.411 | 971.2276 | 1388.6189 |
| Esd | SGYQQAASEHGLVVIAPDTSPR | -65 | -86 | 0 | 0 | 22 | 231.4122 | 299.89166 | 246.42513 | 369.79974 | 461.6742 | 440.67126 |
| Esd | SGYQQAASEHGLVVIAPDTSPR | -65 | -86 | 0 | 0 | 22 | 231.4122 | 299.89166 | 246.42513 | 369.79974 | 461.6742 | 440.67126 |
| Esd | VTEELPQLINANFPVDPQR | -124 | -142 | 0 | 0 | 19 | 370.80997 | 299.52966 | 254.70908 | 509.36755 | 528.6053 | 490.9663 |
| Esd | SEHGLVVIAPDTSPR | -72 | -86 | 0 | 0 | 15 | 620.68494 | 947.8413 | 1118.488 | 830.5098 | 1273.4224 | NaN |
| Esd | SEHGLVVIAPDTSPR | -72 | -86 | 0 | 0 | 15 | 620.68494 | 947.8413 | 1118.488 | 830.5098 | 1273.4224 | NaN |
| Esd | AFSGYLGPDESK | -187 | -198 | 0 | 0 | 12 | 542.10956 | 580.3589 | 1343.625 | 1116.5367 | NaN | 519.5558 |
| Esd | AFSGYLGPDESK | -187 | -198 | 0 | 0 | 12 | 542.10956 | 580.3589 | 1343.625 | 1116.5367 | NaN | 519.5558 |
| Esd | EELPQLINANFPVDPQR | -126 | -142 | 0 | 0 | 17 | 386.05545 | 572.6515 | 336.3015 | NaN | 540.65704 | 664.96106 |
| Esd | GEDDSWDFGTGA | -92 | -103 | 0 | 0 | 12 | 255.59705 | 217.33145 | 256.6213 | NaN | 238.32256 | 222.71951 |
| Esd | TCTEQNFISK | -55 | -64 | 0 | 0 | 10 | 2003.5636 | 1972.8066 | 1837.9406 | NaN | NaN | 1316.9343 |
| Esd | FAVYLPPQAESGK | -32 | -44 | 0 | 0 | 13 | 379.74457 | 630.4219 | 611.89276 | NaN | NaN | 1313.973 |
| Thop1 | LKPLGEQER | -302 | -310 | 1 | 0.00111006 | 9 | 4395.4253 | 2829.7498 | 5031.675 | 1561.2837 | 2254.8735 | 2806.0889 |
| Thop1 | LLEEAFNCR | -239 | -247 | 0 | 0.000230707 | 9 | 670.83167 | 590.568 | 470.9331 | 1283.9181 | 560.2039 | 1555.1295 |
| Thop1 | AASPASTVNHLR | -14 | -25 | 0 | 0 | 12 | 2047.5698 | NaN | NaN | NaN | 1904.3766 | 750.7436 |
| Thop1 | FKQEGVLSPK | -627 | -636 | 1 | 0 | 10 | 744.4386 | 711.6216 | 838.0498 | NaN | NaN | NaN |
| Rpsa | QMKEEDVLK | -9 | -17 | 1 | 0 | 9 | 8115.6675 | 364.49155 | 408.9363 | 4339.157 | 8212.611 | 8151.2085 |
| Rpsa | YVDIAIPCNNK | -156 | -166 | 0 | 0 | 11 | 3700.5598 | 3173.251 | 4555.6953 | 4175.822 | 3801.715 | 3743.7283 |
| Rpsa | YVDIAIPCNNK | -156 | -166 | 0 | 0 | 11 | 3700.5598 | 3173.251 | 4555.6953 | 4175.822 | 3801.715 | 3743.7283 |
| Rpsa | LLVVTDPR | -121 | -128 | 0 | 0.001055699 | 8 | 3672.399 | 2499.7615 | 2690.8784 | 237.95654 | 1111.3699 | 1213.5468 |
| Rpsa | FYRDPEEIEK | -203 | -212 | 1 | 0 | 10 | 3149.6995 | 3301.245 | 3542.309 | 488.85245 | 2064.1777 | 1720.1489 |
| Rpsa | FYRDPEEIEK | -203 | -212 | 1 | 0 | 10 | 3149.6995 | 3301.245 | 3542.309 | 488.85245 | 2064.1777 | 1720.1489 |
| Rpsa | RDPEEIEKEEQAAAEK | -205 | -220 | 2 | 0 | 16 | 1675.1167 | 1705.6168 | 1938.3479 | 4066.549 | 4813.2227 | 4616.9326 |
| Rpsa | RDPEEIEKEEQAAAEK | -205 | -220 | 2 | 0 | 16 | 1675.1167 | 1705.6168 | 1938.3479 | 4066.549 | 4813.2227 | 4616.9326 |
| Rpsa | AIVAIENPADVSVIS | -64 | -78 | 0 | 0 | 15 | 1298.329 | 1281.6418 | 1316.2751 | 1431.2429 | 1775.59 | 1910.2727 |
| Rpsa | DPEEIEKEEQAAAEK | -206 | -220 | 1 | 0 | 15 | 1227.9951 | 1552.0381 | 1755.7372 | 1351.0249 | 1507.4305 | 1452.858 |

| | | | | | | | | | | | |
|---|---|---|---|---|---|---|---|---|---|---|---|
| Rpsa | FYRDPEEIEKEEQAAAEK | -203 | -220 | 2 | 0 | 18 | 1108.8673 | 1144.7902 | 1136.3542 | 1910.0153 | 2025.8037 | 1809.6084 |
| Rpsa | ADHQPLTEAS | -129 | -138 | 0 | 0 | 10 | 1061.838 | 903.73224 | 678.157 | 693.4206 | 527.89557 | 680.83606 |
| Rpsa | LCNTDSPLR | -147 | -155 | 0 | 0 | 9 | 937.2235 | 2009.948 | 1979.4058 | 2224.2073 | 2625.8066 | 2852.787 |
| Rpsa | FTPGTFTNQIQ | -103 | -113 | 0 | 0 | 11 | 896.31287 | 1008.324 | 165.1391 | 1025.2395 | 263.95197 | 1035.3279 |
| Rpsa | FTPGTFTNQIQ | -103 | -113 | 0 | 0 | 11 | 896.31287 | 1008.324 | 165.1391 | 1025.2395 | 263.95197 | 1035.3279 |
| Rpsa | DPEEIEKEEQAAAE | -206 | -219 | 1 | 0 | 14 | 892.3893 | 917.81665 | 936.3962 | 981.0648 | 997.2645 | 855.8412 |
| Rpsa | ADHQPLTEASYVNLPTIA | -129 | -146 | 0 | 0.00122419 | 18 | 640.1311 | 683.0504 | 678.8794 | 934.94824 | 1215.7607 | 1234.608 |
| Rpsa | TGATPIAGR | -94 | -102 | 0 | 0 | 9 | 864.72485 | 1085.2542 | 1062.1125 | 797.88055 | 647.71173 | NaN |
| Rpsa | FTPGTFTNQIQAA | -103 | -115 | 0 | 0.00078661 | 13 | 517.4062 | 498.4597 | NaN | 569.18713 | 764.0881 | 1092.2687 |
| Rpsa | EHPWEVMPDLY | -192 | -202 | 0 | 0 | 11 | 437.60214 | NaN | 85.6723 | 404.7829 | 349.60422 | 446.62494 |
| Rpsa | CNTDSPLR | -148 | -155 | 0 | 0.001092529 | 8 | 1229.464 | 706.91925 | NaN | 395.22556 | 827.2833 | NaN |
| Rpsa | KSDGIYIINLK | -42 | -52 | 1 | 0.001092529 | 11 | 490.17166 | NaN | NaN | 550.4047 | 850.1555 | 866.19214 |
| Rpsa | ADHQPLTEA | -129 | -137 | 0 | 0.002068213 | 9 | 168.86606 | NaN | 284.10492 | 992.6987 | NaN | 725.74677 |
| Rpsa | LYFYRDPEEIEKEEQAAAEK | -201 | -220 | 2 | 0.005605646 | 20 | 1293.4592 | 1375.1743 | NaN | NaN | NaN | 1311.3683 |
| Rpsa | AIVAIENPADVSVISSR | -64 | -80 | 0 | 0 | 17 | 790.0941 | NaN | NaN | NaN | 849.7001 | 836.07697 |
| Rpsa | AATGATPIAGR | -92 | -102 | 0 | 0.000431655 | 11 | 3202.2112 | 1406.9292 | NaN | NaN | NaN | NaN |
| Rpsa | YVDIAIPCNNKG | -156 | -167 | 1 | 0 | 12 | 317.15762 | NaN | NaN | 6968.5415 | NaN | NaN |
| Nosip | AATLPNTEGEQPGPSVGPVGK | -133 | -153 | 0 | 0 | 21 | 446.80286 | 348.25793 | 386.246 | 365.45712 | 426.47833 | 322.16394 |
| Vcp | LAGESESNLR | -278 | -287 | 0 | 0 | 10 | 3926.2397 | 3530.0881 | 3364.5442 | 2911.2427 | 3348.371 | 2827.5957 |
| Vcp | ELQELVQYPVEHPDK | -488 | -502 | 0 | 0 | 15 | 3767.4873 | 4631.2905 | 781.7363 | 5031.229 | 6184.0957 | 5842.671 |
| Vcp | ELQELVQYPVEHPDK | -488 | -502 | 0 | 0 | 15 | 3767.4873 | 4631.2905 | 781.7363 | 5031.229 | 6184.0957 | 5842.671 |
| Vcp | EVDIGIPDATGR | -366 | -377 | 0 | 0 | 12 | 3412.8762 | 3479.6033 | 3649.162 | 3951.8796 | 2956.8337 | 7005.276 |
| Vcp | LGDVISIQPCPDVK | -96 | -109 | 0 | 0 | 14 | 3043.25 | 3796.601 | 3510.1978 | 236.5695 | 4744.565 | 4702.891 |
| Vcp | GVLFYGPPGCGK | -513 | -524 | 0 | 0 | 12 | 2645.4583 | 2573.0452 | 2651.4277 | 2752.6274 | 2708.6902 | 2522.6675 |
| Vcp | GVLFYGPPGCGK | -513 | -524 | 0 | 0 | 12 | 2645.4583 | 2573.0452 | 2651.4277 | 2752.6274 | 2708.6902 | 2522.6675 |
| Vcp | LDQLIYIPLPDEK | -639 | -651 | 0 | 0 | 13 | 2199.558 | 2176.516 | 2118.2334 | 2171.0952 | 2589.9858 | 2663.7598 |
| Vcp | GFSGADLTEICQR | -681 | -693 | 0 | 0 | 13 | 2099.0107 | 3919.6782 | 2403.3074 | 3421.8584 | 575.9275 | 3416.3574 |
| Vcp | IDELDAIAPK | -303 | -312 | 0 | 0 | 10 | 1978.0741 | 1354.5411 | 2347.6543 | 2303.1829 | 1494.9984 | 2576.5745 |
| Vcp | FLEAYRPIR | -139 | -147 | 1 | 0.00025015 | 9 | 1772.1628 | 2048.4883 | 343.7983 | 2947.9253 | 3269.8857 | 3203.6082 |
| Vcp | EDEEESLNEVGYDDIGGCR | -192 | -210 | 0 | 0.000230707 | 19 | 1440.0648 | 1542.7762 | 1642.7142 | 1882.4409 | 1760.5111 | 1642.945 |
| Vcp | FFDELDSIAK | -575 | -584 | 0 | 0 | 10 | 1302.8644 | 5032.6 | 5500.5947 | 2050.1387 | 1970.0519 | 443.0642 |
| Vcp | IYIPLPDEK | -643 | -651 | 0 | 0 | 9 | 1232.4689 | 1236.8799 | 1252.6573 | 1418.0345 | 1617.4673 | 1594.7037 |
| Vcp | AIGVKPPR | -232 | -239 | 1 | 0 | 8 | 1159.8611 | 1350.0479 | 1233.4792 | 1059.9025 | 1493.9159 | 1584.126 |
| Vcp | VLFFDELDSIAK | -573 | -584 | 0 | 0 | 12 | 1044.6692 | 1023.5357 | 1009.809 | 239.0959 | 1483.3015 | 1494.7268 |
| Vcp | LLTLMDGLK | -328 | -336 | 0 | 0 | 9 | 873.7638 | 1015.18787 | 1014.1548 | 764.91003 | 1393.6006 | 73.66971 |
| Vcp | LLTLMDGLK | -328 | -336 | 0 | 0 | 9 | 873.7638 | 1015.18787 | 1014.1548 | 764.91003 | 1393.6006 | 73.66971 |
| Vcp | AIANECQANFISIK | -530 | -543 | 0 | 0 | 14 | 691.22687 | 538.68286 | 444.73068 | 728.051 | 989.961 | 613.07886 |
| Vcp | IHVLPIDDTVEGIT | -114 | -127 | 0 | 0 | 14 | 615.0852 | 1129.7504 | 973.5384 | 1085.6351 | 1461.6603 | 1308.5804 |
| Vcp | ATNRPNSIDPALR | -346 | -358 | 1 | 0.000230707 | 13 | 602.6209 | 429.67752 | 357.43555 | 349.34128 | 764.0546 | 768.5029 |
| Vcp | EDDPVPEIR | -724 | -732 | 0 | 0 | 9 | 555.4514 | 2161.7087 | 1885.352 | 570.5388 | 748.8043 | 3482.7454 |
| Vcp | LADDVDLEQVANETHGHVGAD | -390 | -410 | 0 | 0 | 21 | 489.8398 | 679.4131 | 849.49805 | 755.26355 | 978.0798 | 906.08325 |
| Vcp | LADDVDLEQVANETHGHVG | -390 | -408 | 0 | 0 | 19 | 339.3942 | 409.97168 | 263.4358 | 467.45895 | 473.91312 | 415.817 |
| Vcp | GGNIGDGGGAADR | -587 | -599 | 0 | 0.00025015 | 13 | 282.47528 | 414.70435 | 1190.632 | 213.7232 | 1907.2208 | 433.24628 |
| Vcp | AATNRPNSIDPALR | -345 | -358 | 1 | 0.000826806 | 14 | 5294.0347 | 1075.4487 | 310.81094 | NaN | 2976.907 | 1009.44244 |
| Vcp | EMVELPLR | -218 | -225 | 0 | 0.000591124 | 8 | 2911.5422 | 2321.0369 | 2129.3008 | NaN | 2403.7937 | 1487.6052 |
| Vcp | NRPNSIDPALR | -348 | -358 | 1 | 0 | 11 | 2048.6594 | 1860.4678 | 1896.7546 | 2490.9685 | NaN | 2410.9336 |
| Vcp | ESIESEIR | -701 | -708 | 0 | 0.00025015 | 8 | 1293.0254 | 3439.7214 | 3135.2688 | 4022.4707 | NaN | 1205.7015 |
| Vcp | FPSGNQGGAGPSQGSGGGTGGSVY | -773 | -796 | 0 | 0 | 24 | 1080.6759 | 1155.1035 | 1153.1915 | NaN | 485.10147 | 201.30414 |
| Vcp | AGESESNLR | -279 | -287 | 0 | 0 | 9 | 983.40955 | 939.01526 | NaN | 948.16943 | 650.4204 | 705.5145 |
| Vcp | IHVLPIDDTVEGITGN | -114 | -129 | 0 | 0.000907206 | 16 | 937.3064 | 926.1542 | NaN | 1098.8271 | 1244.6085 | 1313.092 |
| Vcp | MDELQLFR | -46 | -53 | 0 | 0 | 8 | 589.9886 | 550.55084 | 585.768 | NaN | 562.46655 | 538.4745 |
| Vcp | MDELQLFR | -46 | -53 | 0 | 0 | 8 | 589.9886 | 550.55084 | 585.768 | NaN | 562.46655 | 538.4745 |
| Vcp | CIVAPDTVIHCEGEPIKR | -174 | -191 | 1 | 0 | 18 | 433.09344 | 525.47375 | 482.36362 | NaN | 573.5668 | 555.60474 |
| Vcp | EVEEDDPVPEIR | -721 | -732 | 0 | 0 | 12 | 146.45724 | 3437.282 | 174.83672 | 3199.8567 | 3999.8022 | NaN |
| Vcp | WFGESEANVR | -551 | -560 | 0 | 0 | 10 | 131.69766 | NaN | 1098.2971 | 372.46347 | 145.21672 | 1522.3345 |
| Vcp | TNRPNSIDPALR | -347 | -358 | 1 | 0.001291103 | 12 | 1362.042 | 940.52673 | NaN | 577.934 | 1226.6724 | NaN |
| Vcp | RSVSDNDIR | -745 | -753 | 1 | 0.000431655 | 9 | 785.0953 | NaN | 379.8994 | NaN | 734.088 | 897.06134 |
| Vcp | VVETDPSPYCIVAPDTVIHCE | -165 | -185 | 0 | 0.001463897 | 21 | 190.77922 | 200.8013 | 199.83272 | NaN | 190.03343 | NaN |
| Vcp | IHVLPIDDTVE | -114 | -124 | 0 | 0.000347792 | 11 | 1407.8771 | 2954.7424 | 2978.2544 | NaN | NaN | NaN |
| Vcp | VVETDPSPYCIVAPDTVIH | -165 | -183 | 0 | 0.006100831 | 19 | 727.32153 | NaN | NaN | NaN | 1857.2838 | NaN |
| Vcp | DVDLEFLAK | -669 | -677 | 0 | 0.008075282 | 9 | 131.82213 | NaN | 506.10336 | NaN | NaN | NaN |
| Vcp | SGADLTEICQR | -683 | -693 | 0 | 0 | 11 | 1962.5515 | NaN | NaN | NaN | NaN | NaN |
| Vcp | VVETDPSPYCIVAPDTVIHCEGEPIKR | -165 | -191 | 1 | 0.00122419 | 27 | 385.3858 | NaN | NaN | NaN | NaN | NaN |
| Erap1 | LQWLLDQSFK | -803 | -812 | 0 | 0 | 10 | 1345.6125 | 1305.9945 | 1534.429 | 1492.1707 | 1731.0199 | 1717.0232 |
| Erap1 | AATQFEPTAAR | -167 | -177 | 0 | 0 | 11 | 1176.7231 | 1034.168 | 900.8583 | 963.3862 | 1331.2915 | 778.47986 |
| Erap1 | NTEGWDFLYSK | -767 | -777 | 0 | 0 | 11 | 1032.3112 | 1170.7605 | 1215.8612 | 1205.8035 | 1459.8514 | 1304.403 |
| Erap1 | AFPCFDEPALK | -179 | -189 | 0 | 0 | 11 | 433.68805 | 1314.8416 | 1288.7324 | 1135.855 | 1500.7267 | 1313.5747 |
| Erap1 | NPVGYPLAWK | -831 | -840 | 0 | 0.00025015 | 10 | 227.61687 | 688.5525 | 681.3733 | 560.4529 | 835.5302 | 603.49725 |
| Erap1 | QTWTDEGSVSER | -703 | -714 | 0 | 0 | 12 | 97.568016 | 283.0411 | 1166.8573 | 891.5107 | 1425.3959 | 1410.1476 |
| Erap1 | TIETIEENIR | -900 | -909 | 0 | 0.000550307 | 10 | 986.0696 | NaN | 1382.6797 | NaN | 1575.7903 | 1413.038 |
| Erap1 | EFYEDYFNIPYPLPK | -280 | -294 | 0 | 0.00078661 | 15 | 302.82208 | NaN | NaN | 497.5985 | 577.9016 | 501.48697 |
| Erap1 | TQEFPHIL | -818 | -825 | 0 | 0.00111006 | 8 | 897.42633 | 680.94055 | 718.80817 | NaN | NaN | NaN |
| Erap1 | NYQPCVQR | -728 | -735 | 0 | 0.00122419 | 8 | 525.8836 | 609.2079 | NaN | 705.57574 | NaN | NaN |
| Erap1 | FIISDFK | -238 | -244 | 0 | 0.00830926 | 7 | 559.3651 | NaN | 664.53107 | NaN | NaN | NaN |
| Dync1h1 | SIPLDEGEDEAQR | -2382 | -2394 | 0 | 0 | 13 | 2205.328 | 2529.7026 | 2400.159 | 5793.1094 | 5533.3857 | 5590.6846 |
| Dync1h1 | LLNTFLER | -4262 | -4269 | 0 | 0 | 8 | 2175.6218 | 2870.6611 | 2411.2285 | 2883.0266 | 3084.5532 | 935.0242 |
| Dync1h1 | EVLDHVLR | -2912 | -2919 | 0 | 0 | 8 | 1738.6143 | 676.9962 | 1338.0173 | 1077.9575 | 1409.9434 | 1909.6895 |
| Dync1h1 | KYTGEDFDEDLR | -2964 | -2975 | 1 | 0 | 12 | 1443.4177 | 1521.9307 | 395.91434 | 1231.6477 | 132.829 | 1416.8276 |

| | | | | | | | | | | | | |
|---|---|---|---|---|---|---|---|---|---|---|---|---|
| Dync1h1 | EWTDGLFTHVLR | -2272 | -2283 | 0 | 0 | 12 | 1421.469 | 1302.452 | 1543.0542 | 1821.0295 | 535.133 | 686.5537 |
| Dync1h1 | KYEFGESDLR | -4202 | -4211 | 1 | 0.00025015 | 10 | 1365.406 | 1307.4132 | 875.7809 | 587.5365 | 1836.338 | 810.1111 |
| Dync1h1 | DESNVLDSGFLER | -2993 | -3005 | 0 | 0 | 13 | 1339.5557 | 1552.6481 | 1352.0933 | 1710.2756 | 2034.6667 | 1744.7458 |
| Dync1h1 | AATSPALFNR | -3077 | -3086 | 0 | 0 | 10 | 1279.7444 | 2010.448 | 1940.4907 | 1056.5989 | 1406.2197 | 1379.102 |
| Dync1h1 | TVENIKDPLFR | -4399 | -4409 | 1 | 0 | 11 | 1178.7408 | 1386.4375 | 1301.2471 | 1519.6411 | 1410.8112 | 1343.8411 |
| Dync1h1 | EDLDKVEPAVIEAQ | -3278 | -3291 | 1 | 0 | 14 | 1009.794 | 2462.984 | 2824.6875 | 780.89746 | 1946.009 | 1271.5564 |
| Dync1h1 | DVESYDPVLNPVLNR | -3635 | -3649 | 0 | 0 | 15 | 875.1361 | 99.32283 | 1004.7369 | 1006.70667 | 1264.0635 | 1301.4813 |
| Dync1h1 | SHYDFGLR | -2082 | -2089 | 0 | 0 | 8 | 868.3928 | 836.8008 | 884.81647 | 872.96606 | 934.06946 | 882.7908 |
| Dync1h1 | HLLPVETQR | -1557 | -1565 | 0 | 0 | 9 | 859.3726 | 798.48236 | 946.66174 | 743.40015 | 665.931 | 705.4601 |
| Dync1h1 | LWSEETPTTPIGQAIHR | -3971 | -3987 | 0 | 0 | 17 | 801.24786 | 835.6005 | 813.693 | 792.8479 | 478.2644 | 884.3407 |
| Dync1h1 | FNYGFEYLGVQDK | -1864 | -1876 | 0 | 0 | 13 | 779.4216 | 486.42224 | 653.42847 | 699.8528 | 824.3053 | 818.29126 |
| Dync1h1 | SRQELEQHSVDTA | -1147 | -1159 | 1 | 0 | 13 | 736.9769 | 807.4224 | 904.46185 | 580.83026 | 742.12805 | 609.4743 |
| Dync1h1 | AVDDLNLHSY | -882 | -891 | 0 | 0 | 10 | 648.5217 | 649.54114 | 558.66956 | 490.78818 | 650.3402 | 668.9042 |
| Dync1h1 | NYMSNPSYNYEIVNR | -3368 | -3382 | 0 | 0 | 15 | 629.977 | 80.24136 | 625.921 | 564.72375 | 672.6728 | 617.4314 |
| Dync1h1 | VSFDTDTGLK | -312 | -321 | 0 | 0 | 10 | 605.12915 | 899.93915 | 1119.0249 | 1345.6089 | 1881.9287 | 646.70734 |
| Dync1h1 | TPVSITEHPK | -1686 | -1695 | 0 | 0 | 10 | 303.09525 | 1557.9609 | 670.76666 | 1234.3514 | 606.53644 | 978.70984 |
| Dync1h1 | TTDLLTDWEK | -1252 | -1261 | 0 | 0 | 10 | 250.18062 | 1415.4062 | 1526.5641 | 452.83023 | 1790.4255 | 1812.517 |
| Dync1h1 | DTQPHYIYSPR | -2785 | -2795 | 0 | 0 | 11 | 4258.8647 | 4681.472 | NaN | 2811.6167 | 2433.8308 | 2111.6946 |
| Dync1h1 | FVGACNPPTDPGR | -2706 | -2718 | 0 | 0 | 13 | 2963.7546 | NaN | 3230.1836 | 3014.1223 | 732.3605 | 3028.769 |
| Dync1h1 | SNLPIWVNK | -892 | -900 | 0 | 0.00025015 | 9 | 1300.1245 | 541.13684 | 719.5832 | 884.3319 | NaN | 512.41296 |
| Dync1h1 | LGGSPFGPAGTGK | -1898 | -1910 | 0 | 0 | 13 | 1001.39716 | 1090.8759 | NaN | 539.0282 | 116.38132 | 2405.6592 |
| Dync1h1 | LVQTPLTDR | -1877 | -1885 | 0 | 0 | 9 | 812.757 | 830.3144 | 1812.8148 | 420.62 | 738.1976 | NaN |
| Dync1h1 | WVSDFSER | -4476 | -4483 | 0 | 0.000591124 | 8 | 746.0205 | NaN | 772.1117 | 766.204 | 787.20026 | 837.8312 |
| Dync1h1 | TSAPITCELLNK | -1991 | -2002 | 0 | 0 | 12 | 439.66382 | 365.57068 | NaN | 861.24005 | 491.20605 | 347.6011 |
| Dync1h1 | GWDDLFNK | -1487 | -1494 | 0 | 0 | 8 | 369.65466 | 458.7597 | 350.5884 | 291.63858 | NaN | 463.8987 |
| Dync1h1 | YTGEDFDEDLR | -2965 | -2975 | 0 | 0 | 11 | 733.78156 | NaN | 624.96545 | 1206.1598 | 1132.8208 | NaN |
| Dync1h1 | LQGEFQLR | -3732 | -3739 | 0 | 0 | 8 | 702.806 | 699.548 | NaN | NaN | 627.694 | 602.8004 |
| Dync1h1 | LEGVEGVAHIIDPK | -2242 | -2255 | 0 | 0 | 14 | 431.63312 | 534.9238 | NaN | NaN | 458.53802 | 421.8849 |
| Dync1h1 | NVVHELR | -955 | -961 | 0 | 0.008890469 | 7 | 1350.2563 | 527.67096 | NaN | NaN | 1072.0732 | NaN |
| Dync1h1 | DYIPVDQEELR | -2878 | -2888 | 0 | 0 | 11 | 980.7905 | 1040.8215 | 774.65265 | NaN | NaN | NaN |
| Dync1h1 | ETPTTPIGQAIHR | -3975 | -3987 | 0 | 0 | 13 | 496.39194 | 653.0532 | 601.5392 | NaN | NaN | NaN |
| Dync1h1 | SEETPTTPIGQAIHR | -3973 | -3987 | 0 | 0 | 15 | 468.52408 | 259.5521 | NaN | 737.66736 | NaN | NaN |
| Dync1h1 | FNRYPLIIDPSGQAT | -3581 | -3595 | 1 | 0 | 15 | 1512.9591 | NaN | NaN | NaN | NaN | NaN |
| Dync1h1 | SAEEISDAIR | -3353 | -3362 | 0 | 0.000431655 | 10 | 1333.8129 | NaN | NaN | NaN | NaN | NaN |
| Dync1h1 | EQPWVSVQPR | -1346 | -1355 | 0 | 0.001392869 | 10 | 677.7311 | NaN | NaN | NaN | NaN | NaN |
| Dync1h1 | EALELTDTGLLSGSEER | -1300 | -1316 | 0 | 0.00025015 | 17 | 82.4256 | NaN | NaN | NaN | NaN | NaN |
| Khsrp | SVSLTGAPESVQK | -192 | -204 | 0 | 0 | 13 | 2862.2466 | 2862.57 | 1835.581 | 2990.9429 | 2696.1907 | 2090.2778 |
| Khsrp | DGSQNTNVDKPLR | -273 | -285 | 1 | 0.000712251 | 13 | 2359.044 | 2182.566 | 2285.9978 | 1814.9567 | 1873.8242 | 1888.6539 |
| Khsrp | ISPDSGGLPER | -181 | -191 | 0 | 0 | 11 | 2266.8564 | 2163.4624 | 2007.8342 | 1126.8245 | 2408.4402 | 621.4172 |
| Khsrp | VGGGIDVPVPR | -322 | -332 | 0 | 0 | 11 | 1986.2723 | 2114.8525 | 2194.228 | 1752.3866 | 2026.2549 | 2027.3875 |
| Khsrp | HSVGVVIGR | -333 | -341 | 0 | 0 | 9 | 1791.8632 | 1855.9952 | 1869.7772 | 1981.5708 | 1936.5444 | 1923.7343 |
| Khsrp | IGQQPQQPGAPPQQDYTK | -630 | -647 | 0 | 0 | 18 | 1761.0449 | 1822.7921 | 1969.4647 | 1847.7434 | 2404.2144 | 2212.2383 |
| Khsrp | VPDGMVGLIIGR | -152 | -163 | 0 | 0 | 12 | 309.04254 | 304.11807 | 290.6339 | 358.65717 | 411.83395 | 448.003 |
| Khsrp | VPDGMVGLIIGR | -152 | -163 | 0 | 0 | 12 | 309.04254 | 304.11807 | 290.6339 | 358.65717 | 411.83395 | 448.003 |
| Khsrp | VQISPDSGGLPER | -179 | -191 | 0 | 0 | 13 | 1353.5107 | 1263.7383 | 1312.0977 | 1222.7231 | 891.3795 | 971.1899 |
| Khsrp | AINQQTGAFVEISR | -450 | -463 | 0 | 0 | 14 | 983.22235 | 1056.6777 | 1017.06616 | 1266.8456 | 1464.5652 | 1261.6565 |
| Khsrp | GGGGPGGGPGGGGASGGPSQPPGGGGPGIR | -41 | -71 | 0 | 0 | 31 | 980.80005 | 1419.1212 | 953.8966 | 820.59814 | 2686.8403 | 2540.459 |
| Khsrp | GGGGPGGGPGGGGASGGPSQPPGGGGPGIR | -41 | -71 | 0 | 0 | 31 | 980.80005 | 1419.1212 | 953.8966 | 820.59814 | 2686.8403 | 2540.459 |
| Khsrp | KDAFADAVQR | -72 | -81 | 1 | 0 | 10 | 856.4527 | 782.73254 | 844.9059 | 233.5734 | 1121.8828 | 603.7352 |
| Khsrp | GSPQQIDHAK | -480 | -489 | 0 | 0 | 10 | 645.32587 | 544.2155 | 678.4628 | 639.27814 | 539.88403 | 630.02185 |
| Khsrp | LASQGDSIGSQLGPIHPPPR | -124 | -143 | 0 | 0.00025015 | 20 | 445.8057 | 491.57227 | 655.7986 | 487.6258 | 441.27997 | 370.31787 |
| Khsrp | IGGDAATTVNNNTPDFGFGGQK | -89 | -110 | 0 | 0 | 22 | 289.75436 | 310.0593 | 377.86682 | 539.9227 | 575.68774 | 532.23016 |
| Khsrp | MILIQDGSQNTNVDKPLR | -268 | -285 | 1 | 0.000230707 | 18 | 286.36548 | 19895.043 | 21166.898 | 24536.943 | 27666.062 | 26896.59 |
| Khsrp | GDSIGSQLGPIHPPPR | -128 | -143 | 0 | 0 | 16 | 933.6307 | 1230.3616 | 1759.4291 | NaN | 2887.5688 | 4519.0483 |
| Khsrp | FKQDDGTGPEK | -359 | -369 | 1 | 0.00025015 | 11 | 921.3507 | 1043.4473 | NaN | 845.37946 | 919.03674 | 1069.3267 |
| Khsrp | DQGGFGDRNEYGSR | -308 | -321 | 1 | 0 | 14 | 426.07187 | 378.2528 | 425.3514 | NaN | NaN | NaN |
| Khsrp | TTVNNNTPDFGFGGQK | -95 | -110 | 0 | 0.00111006 | 16 | 649.9379 | NaN | NaN | NaN | NaN | NaN |
| Khsrp | DAFADAVQR | -73 | -81 | 0 | 0.005668207 | 9 | 610.5874 | NaN | NaN | NaN | NaN | NaN |
| Khsrp | AATTVNNNTPDFGFGGQK | -93 | -110 | 0 | 0.000907206 | 18 | 388.015 | NaN | NaN | NaN | NaN | NaN |
| Sri | SGTVDPQELQK | -117 | -127 | 0 | 0 | 11 | 11432.945 | 11104.696 | 9815.941 | 7582.1016 | 8206.643 | 8338.655 |
| Sri | GQDGQIDADELQR | -44 | -56 | 0 | 0 | 13 | 1799.711 | 1871.6107 | 2080.0046 | 1391.3699 | 382.81598 | 2397.6746 |
| Sri | VAGQDGQIDADELQR | -42 | -56 | 0 | 0 | 15 | 1556.3661 | 1613.3264 | 842.1048 | 1639.3007 | 802.3698 | 1035.9915 |
| Sri | AVAGQDGQIDADELQR | -41 | -56 | 0 | 0 | 16 | 1365.0304 | 1393.0581 | 943.51 | 1098.696 | 908.5067 | 894.6505 |
| Sri | QHFISFDSDR | -107 | -116 | 0 | 0 | 10 | 982.70654 | 326.96237 | 998.9661 | 1477.7341 | 156.64111 | 1091.4949 |
| Sri | AAVAGQDGQIDADELQR | -40 | -56 | 0 | 0 | 17 | 473.92322 | 548.3209 | 529.02795 | 719.96643 | 779.331 | 638.7862 |
| Sri | GIAGGYKPFNLETCR | -62 | -76 | 1 | 0 | 15 | 757.0317 | 953.749 | 945.2795 | NaN | 1196.3733 | 991.6035 |
| Sri | RDSGQQGVVNF | -176 | -186 | 1 | 0.001023236 | 11 | 841.1972 | 91.61447 | 775.7579 | NaN | NaN | NaN |
| Sri | SGIAGGYKPFNLETCR | -61 | -76 | 1 | 0.00111006 | 16 | 1351.6603 | NaN | 1242.3358 | NaN | NaN | NaN |
| Sri | GYKPFNLETCR | -66 | -76 | 1 | 0.009362332 | 11 | 806.8545 | NaN | NaN | NaN | NaN | NaN |
| Ets1 | AAVDLKPTLTIIK | -3 | -15 | 1 | 0 | 13 | 272.7753 | 344.86295 | 337.5355 | 317.80612 | 354.67365 | 361.2438 |
| Acat1 | AVDPIDFPLAPAYAVPK | -316 | -332 | 0 | 0.000471409 | 17 | 2080.8455 | 2605.0547 | 2814.4365 | 1555.9255 | 4292.7773 | 2700.5364 |
| Acat1 | VDPIDFPLAPAYAVPK | -317 | -332 | 0 | 0 | 16 | 858.0304 | 830.12756 | 926.85956 | 907.7147 | 959.61237 | 914.91455 |
| Acat1 | QATLGAGLPISTPCTTVNK | -103 | -121 | 0 | 0 | 19 | 704.37775 | 807.31354 | 690.4671 | 880.45795 | 837.50305 | 880.40735 |
| Acat1 | EVYMGNVIQGGEGQAPTR | -85 | -102 | 0 | 0 | 18 | 554.5802 | 404.9627 | 692.2973 | 609.97284 | 475.49478 | 573.0985 |
| Acat1 | ADAAVDPIDFPLAPAYAVPK | -313 | -332 | 0 | 0 | 20 | 357.8377 | 439.99908 | 476.59244 | 615.68286 | 706.312 | 671.7619 |
| Acat1 | AAVDPIDFPLAPAYAVPK | -315 | -332 | 0 | 0 | 18 | 354.46252 | 300.38272 | 354.09106 | 263.32593 | 317.05807 | 323.4321 |
| Acat1 | VIQGGEGQAPTR | -91 | -102 | 0 | 0 | 12 | 684.0172 | 669.77576 | 606.3379 | 285.9058 | NaN | 515.2315 |
| Acat1 | NVIQGGEGQAPTR | -90 | -102 | 0 | 0 | 13 | 10383.357 | NaN | NaN | NaN | 188.36638 | 6831.055 |

| | | | | | | | | | | | | |
|---|---|---|---|---|---|---|---|---|---|---|---|---|
| Rrp12 | EGGGDEPLNFLDPK | -1110 | -1123 | 0 | 0 | 14 | 1725.247 | 1908.2938 | 1959.1501 | 613.89966 | 1108.7104 | 1032.537 |
| Rrp12 | FGSNQEDALQR | -892 | -902 | 0 | 0 | 11 | 1118.0702 | 984.8502 | 1197.0227 | 639.74805 | 702.9801 | 648.4411 |
| Rrp12 | FVQSHLDDLKK | -810 | -820 | 1 | 0.000230707 | 11 | 673.0897 | 672.89813 | 411.68927 | 1519.7203 | 343.78992 | 600.2257 |
| Rrp12 | FVQSHLDDLKK | -810 | -820 | 1 | 0.000230707 | 11 | 673.0897 | 672.89813 | 411.68927 | 1519.7203 | 343.78992 | 600.2257 |
| Rrp12 | VLDPASSDFTR | -741 | -751 | 0 | 0 | 11 | 577.3706 | 807.15405 | 878.97253 | 620.29944 | 1402.9385 | 1778.3458 |
| Rrp12 | DLGLGHLAR | -408 | -416 | 0 | 0 | 9 | 345.9849 | 722.0251 | 720.9814 | 384.62018 | 708.88855 | 666.1865 |
| Rrp12 | AAVEEEEEEEEEEPVQSK | -1052 | -1070 | 0 | 0.00025015 | 19 | 203.18024 | 401.41177 | 380.72354 | 365.33237 | 492.51678 | 464.46255 |
| Rrp12 | AVEEEEEEEEEEPVQSK | -1053 | -1070 | 0 | 0 | 18 | 226.39561 | NaN | 231.61879 | 320.14203 | 276.70428 | 408.31036 |
| Rrp12 | SWLLPVIR | -568 | -575 | 0 | 0.000471409 | 8 | 440.2805 | 449.33466 | NaN | NaN | 461.44098 | 480.8124 |
| Rrp12 | EAQLVNSFLEK | -726 | -736 | 0 | 0 | 11 | 358.6006 | 1463.346 | 1534.5508 | NaN | NaN | 1805.1873 |
| Rrp12 | GTAINERPDLR | -649 | -659 | 1 | 0.001291103 | 11 | 2869.3574 | NaN | NaN | NaN | 1871.2506 | 2811.6807 |
| Rrp12 | SSASGPPQYITK | -460 | -471 | 0 | 0.000394361 | 12 | 1826.0225 | NaN | NaN | NaN | NaN | NaN |
| Rrp12 | LFIKFTR | -1008 | -1014 | 1 | 0.00272025 | 7 | 580.0766 | NaN | NaN | NaN | NaN | NaN |
| Ewsr1 | GDATVSYEDPPTAK | -410 | -423 | 0 | 0 | 14 | 1835.4941 | 2347.2808 | 2107.5127 | 2364.4028 | 951.0558 | 2521.4785 |
| Ewsr1 | GVYGQESGGFSGPGENR | -276 | -292 | 0 | 0.000712251 | 17 | 844.1155 | 911.1438 | 986.2911 | 1329.9316 | 1460.6642 | 1489.2534 |
| Ewsr1 | GQESGGFSGPGENR | -279 | -292 | 0 | 0 | 14 | 741.5532 | 1373.25 | 1595.1833 | 1142.6287 | 932.84656 | 1241.9984 |
| Ewsr1 | APKPEGFLPPPFPPPGGD | -544 | -561 | 1 | 0 | 18 | 716.52527 | 360.87057 | 927.88434 | 746.3256 | 1237.8713 | 1118.0353 |
| Ewsr1 | GGFNKPGGPMDEGPDLD | -330 | -346 | 1 | 0.000230707 | 17 | 703.3717 | 745.82416 | 809.6821 | 823.6049 | 958.67285 | 933.26154 |
| Ewsr1 | AAVEWFDGK | -424 | -432 | 0 | 0 | 9 | 696.619 | 563.9784 | 659.271 | 805.39594 | 685.4713 | 642.1255 |
| Ewsr1 | AGDWQCPNPGCGNQNFAWR | -518 | -536 | 0 | 0 | 19 | 546.6227 | 647.3843 | 654.15894 | 1067.0804 | 1550.3562 | 1431.3618 |
| Ewsr1 | ESGGFSGPGENR | -281 | -292 | 0 | 0 | 12 | 302.25134 | 987.3898 | 875.3643 | 262.9167 | 1009.8321 | 582.40875 |
| Ewsr1 | VSYEDPPTAK | -414 | -423 | 0 | 0.00025015 | 10 | 1663.3711 | NaN | 169.91794 | 257.86267 | NaN | NaN |
| Ewsr1 | QCPNPGCGNQNFAWR | -522 | -536 | 0 | 0.000431655 | 15 | 562.744 | 503.5827 | 480.67545 | NaN | NaN | NaN |
| Ndufv2 | INDNYYEDLTPK | -187 | -198 | 0 | 0 | 12 | 1706.7701 | 1729.7985 | 1866.8629 | 840.9978 | 1203.1595 | 1009.39435 |
| Ndufv2 | AVLPVLDLAQR | -77 | -87 | 0 | 0 | 11 | 812.0943 | 924.86 | 1007.77594 | 751.21893 | 909.8225 | 933.31006 |
| Ndufv2 | AAVLPVLDLAQR | -76 | -87 | 0 | 0 | 12 | 771.0241 | 896.73956 | 825.114 | 722.2117 | 784.14886 | 878.8126 |
| Ndufv2 | DTPENNPDTPFDFTPENYK | -42 | -60 | 0 | 0 | 19 | 377.0749 | 382.28888 | 444.69467 | 418.3164 | 117.261734 | 374.6888 |
| Ndufv2 | DIEEIIDELK | -199 | -208 | 0 | 0.00025015 | 10 | 326.33664 | 214.63991 | 243.19102 | 544.1747 | 498.90173 | 486.51868 |
| Ndufv2 | DSDSILETLQR | -143 | -153 | 0 | 0 | 11 | 252.68303 | 830.2113 | 894.3862 | 321.76 | 1013.0243 | 1152.0404 |
| Hsp90ab1 | IDIIPNPQER | -73 | -82 | 0 | 0 | 10 | 11631.666 | 12371.055 | 12637.618 | 10297.037 | 10938.152 | 11157.821 |
| Hsp90ab1 | TNDWEDHLAVK | -309 | -319 | 0 | 0 | 11 | 11569.732 | 11610.275 | 12529.248 | 11813.297 | 12869.227 | 13019.361 |
| Hsp90ab1 | TNDWEDHLAVK | -309 | -319 | 0 | 0 | 11 | 11569.732 | 11610.275 | 12529.248 | 11813.297 | 12869.227 | 13019.361 |
| Hsp90ab1 | TNDWEDHLAVK | -309 | -319 | 0 | 0 | 11 | 11569.732 | 11610.275 | 12529.248 | 11813.297 | 12869.227 | 13019.361 |
| Hsp90ab1 | RAPFDLFENK | -338 | -347 | 1 | 0.000550307 | 10 | 10589.624 | 11180.985 | 11607.672 | 7680.9746 | 9181.495 | 9181.553 |
| Hsp90ab1 | NPDDITQEEYGEFYK | -292 | -306 | 0 | 0 | 15 | 9175.704 | 10415.843 | 10654.778 | 9097.097 | 11076.552 | 120.58989 |
| Hsp90ab1 | EIPPLEGDEDASR | -707 | -719 | 0 | 0 | 13 | 6522.541 | 5872.381 | 5904.081 | 3136.1758 | 1781.6016 | 1687.2987 |
| Hsp90ab1 | HLEINPDHPIVETL | -625 | -638 | 0 | 0 | 14 | 6462.8843 | 7071.9863 | 6823.6646 | 8182.93 | 7984.136 | 8438.431 |
| Hsp90ab1 | HLEINPDHPIVETL | -625 | -638 | 0 | 0 | 14 | 6462.8843 | 7071.9863 | 6823.6646 | 8182.93 | 7984.136 | 8438.431 |
| Hsp90ab1 | PEEVHHGEEEVETF | -2 | -15 | 0 | 0 | 14 | 6102.4365 | 7674.4478 | 7366.605 | 7942.3267 | 8724.09 | 8651.997 |
| Hsp90ab1 | PEEVHHGEEEVETF | -2 | -15 | 0 | 0 | 14 | 6102.4365 | 7674.4478 | 7366.605 | 7942.3267 | 8724.09 | 8651.997 |
| Hsp90ab1 | GFSLEDPQTHSNR | -667 | -679 | 0 | 0 | 13 | 5967.4517 | 6218.224 | 870.0639 | 645.28485 | 6048.8774 | 5934.744 |
| Hsp90ab1 | SLEDPQTHSNR | -669 | -679 | 0 | 0 | 11 | 5146.0615 | 5443.865 | 6268.35 | 3539.0864 | 5513.1704 | 5366.2256 |
| Hsp90ab1 | SLTNDWEDHLAVK | -307 | -319 | 0 | 0 | 13 | 4705.8384 | 5304.401 | 5311.8315 | 3384.3643 | 3808.8477 | 3801.0098 |
| Hsp90ab1 | DWEDHLAVK | -311 | -319 | 0 | 0.000394361 | 9 | 4697.401 | 392.96997 | 445.22397 | 4287.9688 | 5070.1313 | 181.35143 |
| Hsp90ab1 | RAPFDLFENKK | -338 | -348 | 2 | 0 | 11 | 4030.9304 | 4837.727 | 4931.78 | 1631.2612 | 2613.1523 | 2671.1394 |
| Hsp90ab1 | ADHGEPIGR | -169 | -177 | 0 | 0 | 9 | 3851.3928 | 3773.4617 | 620.57886 | 4077.366 | 4161.2915 | 4451.43 |
| Hsp90ab1 | ADHGEPIGR | -169 | -177 | 0 | 0 | 9 | 3851.3928 | 3773.4617 | 620.57886 | 4077.366 | 4161.2915 | 4451.43 |
| Hsp90ab1 | LGIHEDSTNR | -439 | -448 | 0 | 0 | 10 | 3744.9753 | 3796.1326 | 5176.5796 | 2740.5737 | 5301.555 | 4939.754 |
| Hsp90ab1 | LGIHEDSTNR | -439 | -448 | 0 | 0 | 10 | 3744.9753 | 3796.1326 | 5176.5796 | 2740.5737 | 5301.555 | 4939.754 |
| Hsp90ab1 | KIDIIPNPQER | -72 | -82 | 1 | 0.00025015 | 11 | 3632.8171 | 3651.8196 | 3827.8633 | 3914.1514 | 3666.5474 | 3643.581 |
| Hsp90ab1 | KIDIIPNPQER | -72 | -82 | 1 | 0.00025015 | 11 | 3632.8171 | 3651.8196 | 3827.8633 | 3914.1514 | 3666.5474 | 3643.581 |
| Hsp90ab1 | HSQFIGYPITLY | -205 | -216 | 0 | 0 | 12 | 3480.2993 | 3417.9814 | 3884.5146 | 2996.9036 | 3958.736 | 3878.1096 |
| Hsp90ab1 | HSQFIGYPITLY | -205 | -216 | 0 | 0 | 12 | 3480.2993 | 3417.9814 | 3884.5146 | 2996.9036 | 3958.736 | 3878.1096 |
| Hsp90ab1 | HLEINPDHPIVE | -625 | -636 | 0 | 0 | 12 | 3098.7317 | 3519.075 | 3517.4138 | 3205.6477 | 1342.1144 | 1583.0587 |
| Hsp90ab1 | SELAEDKENYK | -417 | -427 | 1 | 0 | 11 | 2745.885 | 2804.6492 | 2828.2837 | 2663.6174 | 4139.412 | 4001.9385 |
| Hsp90ab1 | HLEINPDHPIVETLR | -625 | -639 | 0 | 0 | 15 | 2718.2495 | 2955.8542 | 2680.013 | 3340.5654 | 5109.294 | 4252.783 |
| Hsp90ab1 | HLEINPDHPIVETLR | -625 | -639 | 0 | 0 | 15 | 2718.2495 | 2955.8542 | 2680.013 | 3340.5654 | 5109.294 | 4252.783 |
| Hsp90ab1 | LLSSGFSLEDPQTHSNR | -663 | -679 | 0 | 0 | 17 | 2552.4214 | 3060.334 | 2859.1985 | 2443.6992 | 2813.0776 | 2925.8403 |
| Hsp90ab1 | LLSSGFSLEDPQTHSNR | -663 | -679 | 0 | 0 | 17 | 2552.4214 | 3060.334 | 2859.1985 | 2443.6992 | 2813.0776 | 2925.8403 |
| Hsp90ab1 | AKFENLCK | -558 | -565 | 1 | 0.00025015 | 8 | 2495.9192 | 4445.4272 | 4496.23 | 6085.364 | 5418.2676 | 271.36932 |
| Hsp90ab1 | SGFSLEDPQTHSNR | -666 | -679 | 0 | 0 | 14 | 2285.4832 | 2405.5063 | 2543.2332 | 3418.8435 | 3545.7185 | 862.88666 |
| Hsp90ab1 | SGFSLEDPQTHSNR | -666 | -679 | 0 | 0 | 14 | 2285.4832 | 2405.5063 | 2543.2332 | 3418.8435 | 3545.7185 | 862.88666 |
| Hsp90ab1 | KHLEINPDHPIVETL | -624 | -638 | 1 | 0 | 15 | 1880.4225 | 2885.6191 | 3409.587 | 3273.8218 | 3713.3677 | 4473.5273 |
| Hsp90ab1 | KHLEINPDHPIVETL | -624 | -638 | 1 | 0 | 15 | 1880.4225 | 2885.6191 | 3409.587 | 3273.8218 | 3713.3677 | 4473.5273 |
| Hsp90ab1 | AAVPDEIPPLEGDEDASR | -702 | -719 | 0 | 0 | 18 | 1796.0046 | 2013.2316 | 2252.5154 | 3284.323 | 4368.2363 | 2687.9556 |
| Hsp90ab1 | APFDLFENK | -339 | -347 | 0 | 0 | 9 | 1689.5378 | 2728.0952 | 2683.931 | 1803.4565 | 178.2322 | 1314.9307 |
| Hsp90ab1 | QSGDEMTSLSEYVSR | -461 | -475 | 0 | 0 | 15 | 1548.9536 | 152.94019 | 712.18427 | 917.4991 | 141.38701 | 1028.5211 |
| Hsp90ab1 | KHLEINPDHPIVE | -624 | -636 | 1 | 0 | 13 | 1516.3057 | 1624.4238 | 5164.709 | 1133.1398 | 1187.8214 | 1212.1139 |
| Hsp90ab1 | KHLEINPDHPIVE | -624 | -636 | 1 | 0 | 13 | 1516.3057 | 1624.4238 | 5164.709 | 1133.1398 | 1187.8214 | 1212.1139 |
| Hsp90ab1 | SSGFSLEDPQTHSNR | -665 | -679 | 0 | 0 | 15 | 1475.4597 | 1638.6641 | 1659.9849 | 2617.7722 | 2706.9353 | 2694.7275 |
| Hsp90ab1 | SSGFSLEDPQTHSNR | -665 | -679 | 0 | 0 | 15 | 1475.4597 | 1638.6641 | 1659.9849 | 2617.7722 | 2706.9353 | 2694.7275 |
| Hsp90ab1 | LTNDWEDHLAVK | -308 | -319 | 0 | 0 | 12 | 1147.5203 | 564.09174 | 4160.8843 | 674.1767 | 6936.0547 | 1082.9215 |
| Hsp90ab1 | LTNDWEDHLAVK | -308 | -319 | 0 | 0 | 12 | 1147.5203 | 564.09174 | 4160.8843 | 674.1767 | 6936.0547 | 1082.9215 |
| Hsp90ab1 | KLGIHEDSTNR | -438 | -448 | 1 | 0 | 11 | 1088.3358 | 503.05545 | 594.1758 | 304.26926 | 699.57446 | 822.56433 |
| Hsp90ab1 | SELAEDKENYKK | -417 | -428 | 2 | 0 | 12 | 1001.19293 | 1302.7433 | 1348.4567 | 1192.7837 | 1851.0546 | 2018.4003 |
| Hsp90ab1 | SELAEDKENYKK | -417 | -428 | 2 | 0 | 12 | 1001.19293 | 1302.7433 | 1348.4567 | 1192.7837 | 1851.0546 | 2018.4003 |
| Hsp90ab1 | AEEPSAAVPDEIPPLEGDEDASR | -697 | -719 | 0 | 0 | 23 | 992.5028 | 1102.5358 | 1197.0272 | 1171.0869 | 1635.5262 | 1515.3972 |
| Hsp90ab1 | LSSGFSLEDPQTHSNR | -664 | -679 | 0 | 0 | 16 | 984.2815 | 770.51025 | 800.2434 | 1042.2959 | 732.72375 | 729.96106 |

| | | | | | | | | | | | | |
|---|---|---|---|---|---|---|---|---|---|---|---|---|
| Hsp90ab1 | GIHEDSTNR | -440 | -448 | 0 | 0.002265552 | 9 | 956.39435 | 1335.4308 | 1181.6823 | 1279.841 | 2069.3142 | 1689.8688 |
| Hsp90ab1 | KHSQFIGYPITLY | -204 | -216 | 1 | 0.00025015 | 13 | 945.5657 | 1261.4902 | 1338.5294 | 1640.3938 | 1567.9988 | 1680.6604 |
| Hsp90ab1 | KHSQFIGYPITLY | -204 | -216 | 1 | 0.00025015 | 13 | 945.5657 | 1261.4902 | 1338.5294 | 1640.3938 | 1567.9988 | 1680.6604 |
| Hsp90ab1 | KHLEINPDHPIVETLR | -624 | -639 | 1 | 0 | 16 | 924.1476 | 815.8435 | 1123.8864 | 1411.2125 | 1457.8002 | 1377.2246 |
| Hsp90ab1 | KHLEINPDHPIVETLR | -624 | -639 | 1 | 0 | 16 | 924.1476 | 815.8435 | 1123.8864 | 1411.2125 | 1457.8002 | 1377.2246 |
| Hsp90ab1 | KHLEINPDHPIVETLR | -624 | -639 | 1 | 0 | 16 | 924.1476 | 815.8435 | 1123.8864 | 1411.2125 | 1457.8002 | 1377.2246 |
| Hsp90ab1 | HSQFIGYPITLYLEK | -205 | -219 | 0 | 0 | 15 | 849.79675 | 939.311 | 876.53284 | 664.81866 | 1093.5823 | 970.10956 |
| Hsp90ab1 | IEDVGSDEEDDSGK | -250 | -263 | 0 | 0 | 14 | 836.3751 | 1050.3589 | 1524.1132 | 854.13354 | 922.2045 | 862.9717 |
| Hsp90ab1 | EEPSAAVPDEIPPLEGDEDASR | -698 | -719 | 0 | 0 | 22 | 804.4254 | 935.7319 | 974.7992 | 932.6049 | 1173.8774 | 1133.6963 |
| Hsp90ab1 | KSLTNDWEDHLAVK | -306 | -319 | 1 | 0 | 14 | 800.7486 | 1007.54 | 1089.1301 | 1278.5544 | 1930.299 | 2025.1257 |
| Hsp90ab1 | DVGSDEEDDSGKDK | -252 | -265 | 1 | 0 | 14 | 797.8357 | 1021.9234 | 1123.3522 | 467.36945 | 580.1596 | 571.1444 |
| Hsp90ab1 | TEPIDEYCVQQLK | -514 | -526 | 0 | 0 | 13 | 715.8897 | 857.3645 | 763.2718 | 1270.9688 | 1352.8859 | 1591.1255 |
| Hsp90ab1 | SCDELIPEYLNFIR | -365 | -378 | 0 | 0 | 14 | 663.6131 | 713.363 | 623.69336 | 559.5381 | 718.71936 | 696.6771 |
| Hsp90ab1 | IEDVGSDEEDDSGKDK | -250 | -265 | 1 | 0.00025015 | 16 | 662.2713 | 103.68402 | 744.8782 | 1111.6239 | 1368.534 | 1253.136 |
| Hsp90ab1 | APFDLFENKK | -339 | -348 | 1 | 0 | 10 | 651.9418 | 551.34296 | 308.98358 | 129.22858 | 684.782 | 577.9464 |
| Hsp90ab1 | FSELAEDKENYKK | -416 | -428 | 2 | 0 | 13 | 647.11554 | 816.4855 | 910.55273 | 625.3216 | 894.7237 | 1303.6432 |
| Hsp90ab1 | IEDVGSDEEDDSGKDKK | -250 | -266 | 2 | 0 | 17 | 581.447 | 524.2636 | 629.88196 | 483.97473 | 814.3312 | 619.77374 |
| Hsp90ab1 | IEDVGSDEEDDSGKDKK | -250 | -266 | 2 | 0 | 17 | 581.447 | 524.2636 | 629.88196 | 483.97473 | 814.3312 | 619.77374 |
| Hsp90ab1 | NPDDITQEEYGEFY | -292 | -305 | 0 | 0.00025015 | 14 | 541.4282 | 476.02493 | 405.23743 | 584.62555 | 365.18256 | 374.07687 |
| Hsp90ab1 | LGIHEDSTNRR | -439 | -449 | 1 | 0 | 11 | 496.28345 | 554.14825 | 576.6675 | 1951.1781 | 651.1079 | 632.05676 |
| Hsp90ab1 | PDDITQEEYGEFYK | -293 | -306 | 0 | 0.000230707 | 14 | 432.92145 | 573.64154 | 563.0857 | 540.6653 | 502.89972 | 542.8907 |
| Hsp90ab1 | ALLSSGFSLEDPQTHSNR | -662 | -679 | 0 | 0 | 18 | 369.80054 | 555.7812 | 575.33875 | 670.22327 | 915.88434 | 808.3267 |
| Hsp90ab1 | DSCDELIPEYLNFIR | -364 | -378 | 0 | 0 | 15 | 294.68106 | 280.74542 | 278.90308 | 400.53864 | 469.73755 | 431.04443 |
| Hsp90ab1 | LKEFDGK | -525 | -531 | 1 | 0.00149795 | 7 | 1544.8774 | 1552.5659 | NaN | 2174.713 | 1363.3313 | 1613.2242 |
| Hsp90ab1 | HLEINPDHPIVET | -625 | -637 | 0 | 0 | 13 | 1351.053 | 1028.2035 | 1042.9525 | NaN | 887.3096 | 687.0166 |
| Hsp90ab1 | NDWEDHLAVK | -310 | -319 | 0 | 0 | 10 | 964.73486 | 997.8313 | 543.053 | NaN | 753.2563 | 759.8163 |
| Hsp90ab1 | LVSSPCCIVTS | -584 | -594 | 0 | 0.00025015 | 11 | 429.01224 | NaN | 426.12787 | 123.68624 | 689.03467 | 626.88055 |
| Hsp90ab1 | SIYYITGESK | -482 | -491 | 0 | 0 | 10 | 1085.7374 | 613.98553 | NaN | NaN | 857.1043 | 792.05707 |
| Hsp90ab1 | LELFSELAEDKENYK | -413 | -427 | 1 | 0 | 15 | 478.4015 | NaN | 741.5974 | NaN | 947.6223 | 902.0478 |
| Hsp90ab1 | DVGSDEEDDSGKDKK | -252 | -266 | 2 | 0.001811524 | 15 | 341.85712 | 479.30225 | 4.3994355 | NaN | NaN | 471.4047 |
| Hsp90ab1 | IIPNPQER | -75 | -82 | 0 | 0.000712251 | 8 | 2210.8582 | 1420.2798 | 1006.0013 | NaN | NaN | NaN |
| Hsp90ab1 | DDITQEEYGEFYK | -294 | -306 | 0 | 0.00078661 | 13 | 616.58185 | 652.9816 | 629.69434 | NaN | NaN | NaN |
| Hsp90ab1 | TSLSEYVSR | -467 | -475 | 0 | 0 | 9 | 2931.1018 | NaN | 3224.8252 | NaN | NaN | NaN |
| Snx5 | AAVPELLEQQEEDR | -2 | -15 | 0 | 0 | 14 | 548.24634 | 572.8497 | 504.76434 | 731.0493 | 1050.5001 | 910.4201 |
| Snx5 | NVADDYIHTA | -239 | -248 | 0 | 0.001055699 | 10 | 473.05853 | 891.4851 | 559.84436 | NaN | NaN | NaN |
| Eno1 | VVGDDLTVTNPK | -315 | -326 | 0 | 0 | 12 | 16957.395 | 17101.465 | 916.6972 | 16851.41 | 1319.9733 | 1077.8574 |
| Eno1 | VVGDDLTVTNPK | -315 | -326 | 0 | 0 | 12 | 16957.395 | 17101.465 | 916.6972 | 16851.41 | 1319.9733 | 1077.8574 |
| Eno1 | YITPDQLADLYK | -270 | -281 | 0 | 0 | 12 | 7990.6196 | 8966.052 | 10049.255 | 435.13293 | 14914.887 | 14534.749 |
| Eno1 | YITPDQLADLYK | -270 | -281 | 0 | 0 | 12 | 7990.6196 | 8966.052 | 10049.255 | 435.13293 | 14914.887 | 14534.749 |
| Eno1 | SCNCLLLK | -336 | -343 | 0 | 0.000230707 | 8 | 7749.582 | 8926.803 | 175.51503 | 6893.6094 | 5992.468 | 6198.902 |
| Eno1 | DATNVGDEGGFAPNILENK | -203 | -221 | 0 | 0 | 19 | 7659.9116 | 8991.27 | 9038.387 | 10869.27 | 13226.194 | 12386.119 |
| Eno1 | SGKYDLDFK | -254 | -262 | 1 | 0 | 9 | 6097.518 | 6295.4907 | 1745.1132 | 7054.2227 | 5655.042 | 2327.1907 |
| Eno1 | KVNVVEQEK | -81 | -89 | 1 | 0.000550307 | 9 | 6041.332 | 6089.115 | 7049.1045 | 6327.9624 | 6057.031 | 6858.327 |
| Eno1 | KVNVVEQEK | -81 | -89 | 1 | 0.000550307 | 9 | 6041.332 | 6089.115 | 7049.1045 | 6327.9624 | 6057.031 | 6858.327 |
| Eno1 | GVSQAVEHINK | -61 | -71 | 0 | 0 | 11 | 5259.1055 | 5213.855 | 5170.475 | 4455.5225 | 3838.188 | 412.1186 |
| Eno1 | IGAEVYHNLK | -184 | -193 | 0 | 0 | 10 | 4029.0195 | 4268.131 | 4374.2866 | 6910.816 | 6488.4717 | 1875.9237 |
| Eno1 | IGAEVYHNLK | -184 | -193 | 0 | 0 | 10 | 4029.0195 | 4268.131 | 4374.2866 | 6910.816 | 6488.4717 | 1875.9237 |
| Eno1 | AGIQVVGDDLTVTNPK | -311 | -326 | 0 | 0 | 16 | 3812.9436 | 4603.5693 | 4455.6973 | 5177.471 | 6058.93 | 5981.696 |
| Eno1 | RIEEELGSK | -412 | -420 | 1 | 0.002810811 | 9 | 3050.5698 | 437.23163 | 231.94705 | 5124.7437 | 7325.692 | 339.2544 |
| Eno1 | VNVVEQEK | -82 | -89 | 0 | 0 | 8 | 2962.707 | 3372.7603 | 3108.0076 | 3686.1914 | 3448.3857 | 3062.9358 |
| Eno1 | ATNVGDEGGFAPNILENK | -204 | -221 | 0 | 0 | 18 | 2224.2937 | 1865.6802 | 1126.908 | 1420.525 | 924.48303 | 797.05145 |
| Eno1 | VVGDDLTVTNPKR | -315 | -327 | 1 | 0 | 13 | 1779.0841 | 1802.3605 | 1367.2577 | 2687.7825 | 4308.9927 | 2875.8223 |
| Eno1 | GNPEVILPVPAFNVINGGSHAGNK | -139 | -162 | 0 | 0 | 24 | 1095.2736 | 849.53204 | 935.9304 | 667.91614 | 722.99884 | 827.1282 |
| Eno1 | HIADLAGNPEVILPVPAFNVINGGSHAGNK | -133 | -162 | 0 | 0 | 30 | 974.89734 | 882.84204 | 743.79407 | 1232.7979 | 1177.3108 | 1061.1272 |
| Eno1 | HIADLAGNPEVILPVPA | -133 | -149 | 0 | 0.00272025 | 17 | 738.1275 | 723.1686 | 717.21967 | 739.6145 | 914.5682 | 922.1008 |
| Eno1 | GIQVVGDDLTVTNPK | -312 | -326 | 0 | 0 | 15 | 658.3191 | 860.85034 | 683.86194 | 486.03024 | 542.0261 | 855.3463 |
| Eno1 | YITPDQLADLY | -270 | -280 | 0 | 0.00025015 | 11 | 637.20844 | 617.5654 | 699.56995 | 1017.65283 | 1045.187 | 984.61804 |
| Eno1 | HIADLAGNPEVILPVPAFN | -133 | -151 | 0 | 0 | 19 | 486.91138 | 513.52734 | 446.80768 | 512.5656 | 564.09973 | 530.3129 |
| Eno1 | TIAPALVSK | -72 | -80 | 0 | 0 | 9 | 294.94308 | 298.8543 | 307.5985 | 230.69037 | 336.13885 | 386.58206 |
| Eno1 | IGAEVYHN | -184 | -191 | 0 | 0 | 8 | 220.36893 | 1278.5378 | 975.2211 | 1984.7634 | 1299.3754 | 959.4271 |
| Eno1 | HIADLAGNPEVILPVPAF | -133 | -150 | 0 | 0 | 18 | 195.90326 | 197.77063 | 167.60403 | 167.77841 | 232.03835 | 190.79192 |
| Eno1 | IGAEVYHNL | -184 | -192 | 0 | 0.00025015 | 9 | 5180.321 | NaN | 803.5498 | 4337.7227 | 6311.716 | 5338.2036 |
| Eno1 | TNVGDEGGFAPNILENK | -205 | -221 | 0 | 0 | 17 | 1150.7761 | NaN | 267.84552 | 838.5173 | 931.66296 | 2431.7256 |
| Eno1 | LMIEMDGTENK | -93 | -103 | 0 | 0 | 11 | 785.41547 | NaN | NaN | 1135.3851 | 971.27856 | 662.57825 |
| Eno1 | EALELLKT | -222 | -229 | 1 | 0.006788754 | 8 | 543.3124 | 362.71735 | 322.84024 | 706.6195 | NaN | NaN |
| Eno1 | KTIAPALVSK | -71 | -80 | 1 | 0.000471409 | 10 | 1202.0841 | 1143.5409 | 1210.0295 | NaN | NaN | NaN |
| Eno1 | KYNQILR | -406 | -412 | 1 | 0 | 7 | 148.402 | 209.66869 | 184.78029 | NaN | NaN | NaN |
| Eno1 | KVNVVEQEKID | -81 | -91 | 2 | 0.000907206 | 11 | 765.5391 | 678.8028 | NaN | NaN | NaN | NaN |
| Eno1 | IQVVGDDLTVTNPK | -313 | -326 | 0 | 0.003681885 | 14 | 3170.8972 | NaN | NaN | NaN | NaN | NaN |
| Rpl5 | RFPGYDSESK | -179 | -188 | 1 | 0 | 10 | 11390.755 | 11860.411 | 12144.98 | 13473.194 | 13843.355 | 13984.294 |
| Rpl5 | GAVDGGLSIPHSTK | -165 | -178 | 0 | 0 | 14 | 8373.253 | 8388.585 | 8188.6045 | 13287.896 | 8216.082 | 7428.406 |
| Rpl5 | GAVDGGLSIPHSTK | -165 | -178 | 0 | 0 | 14 | 8373.253 | 8388.585 | 8188.6045 | 13287.896 | 8216.082 | 7428.406 |
| Rpl5 | GAVDGGLSIPH | -165 | -175 | 0 | 0 | 11 | 6826.9683 | 7802.4585 | 6591.292 | 1007.04974 | 5355.1665 | 6827.0547 |
| Rpl5 | AYAHELPK | -78 | -85 | 0 | 0 | 8 | 4561.4644 | 4360.0063 | 3645.2515 | 6655.6357 | 4222.5757 | 3763.4668 |
| Rpl5 | IRENPVYEK | -247 | -255 | 1 | 0 | 9 | 2730.5698 | 2867.0217 | 1552.1492 | 2794.739 | 2881.7124 | 232.94106 |
| Rpl5 | KGAVDGGLSIPHSTK | -164 | -178 | 1 | 0 | 15 | 2597.608 | 2644.9688 | 2682.017 | 3130.9084 | 3859.6814 | 3509.6248 |
| Rpl5 | KGAVDGGLSIPHSTK | -164 | -178 | 1 | 0 | 15 | 2597.608 | 2644.9688 | 2682.017 | 3130.9084 | 3859.6814 | 3509.6248 |
| Rpl5 | FPGYDSESK | -180 | -188 | 0 | 0 | 9 | 1790.3127 | 232.95978 | 1878.0822 | 2056.3906 | 1967.7379 | 1923.1395 |

| | | | | | | | | | | | |
|---|---|---|---|---|---|---|---|---|---|---|---|
| Rpl5 | TCYLDAGLAR | -143 | -152 | 0 | 0 | 10 | 1552.7676 | 111.75811 | 128.9667 | 1811.035 | 1816.4042 | 1719.6761 |
| Rpl5 | AAYAHELPK | -77 | -85 | 0 | 0 | 9 | 1416.0299 | 1482.5901 | 1487.0208 | 1404.04 | 1428.547 | 1463.1034 |
| Rpl5 | YLDAGLAR | -145 | -152 | 0 | 0.002265552 | 8 | 1092.0142 | 680.3947 | 753.6793 | 760.74524 | 486.31375 | 529.79443 |
| Rpl5 | YAHELPK | -79 | -85 | 0 | 0.006915721 | 7 | 756.7821 | 2041.595 | 2115.5415 | 578.2695 | 792.0795 | 1972.6294 |
| Rpl5 | MEEDEDAYKK | -212 | -221 | 1 | 0 | 10 | 387.38925 | 329.1232 | 466.92377 | 366.41583 | 639.6129 | 535.3918 |
| Rpl5 | MEEDEDAYKK | -212 | -221 | 1 | 0 | 10 | 387.38925 | 329.1232 | 466.92377 | 366.41583 | 639.6129 | 535.3918 |
| Rpl5 | CTGLLLAR | -100 | -107 | 0 | 0 | 8 | 634.0709 | 673.823 | 439.21667 | 704.0745 | 525.0112 | 653.3907 |
| Rpl5 | ENPVYEK | -249 | -255 | 0 | 0.000431655 | 7 | 1263.8268 | 1238.887 | 968.1617 | 1418.661 | 701.1584 | NaN |
| Rpl5 | EFNAEVHR | -189 | -196 | 0 | 0 | 8 | 419.164 | 2448.8232 | 758.9751 | 3012.8594 | 2125.2842 | NaN |
| Rpl5 | YCTGLLLAR | -99 | -107 | 0 | 0 | 9 | 1465.3705 | NaN | NaN | 2901.9814 | 905.0787 | 1328.041 |
| Rpl5 | YLMEEDEDAYKK | -210 | -221 | 1 | 0 | 12 | 1273.4753 | 926.0833 | 1095.9027 | 89.97674 | NaN | NaN |
| Rpl5 | QFSQYIK | -222 | -228 | 0 | 0.004350979 | 7 | 2419.1882 | 2152.3323 | NaN | NaN | 1367.0558 | NaN |
| Rpl5 | GGLSIPHSTK | -169 | -178 | 0 | 0.00025015 | 10 | 1111.2474 | NaN | 887.06305 | 728.84045 | NaN | NaN |
| Slc25a5 | QIFLGGVDKR | -97 | -106 | 1 | 0 | 10 | 4462.692 | 149.24345 | 4878.0776 | 4015.7673 | 4338.069 | 140.13881 |
| Slc25a5 | GLTSYPFDTVR | -225 | -235 | 0 | 0 | 11 | 2141.602 | 2062.4915 | 1038.2115 | 1639.8585 | 1649.4974 | 1499.9963 |
| Slc25a5 | AAYFGIYDTAK | -189 | -199 | 0 | 0 | 11 | 2092.7805 | 2123.4792 | 2137.197 | 2031.842 | 2589.498 | 2522.3013 |
| Slc25a5 | EQGVLSFWR | -64 | -72 | 0 | 0 | 9 | 770.2027 | 812.1009 | 387.61877 | 2859.1316 | 3537.6897 | 3734.38 |
| Slc25a5 | TSYPFDTVR | -227 | -235 | 0 | 0 | 9 | 250.63637 | 1411.6062 | 1607.9045 | 1401.1356 | 1441.7075 | 1537.3789 |
| Slc25a5 | QIFLGGVDK | -97 | -105 | 0 | 0 | 9 | 236.90808 | 3165.3838 | 3458.0288 | 2181.1824 | 1254.0991 | 1653.8522 |
| Slc25a5 | KGTDIMYTG | -245 | -253 | 1 | 0.008129024 | 9 | 537.7499 | NaN | NaN | NaN | NaN | NaN |
| Rars1 | DLLEEGLKR | -464 | -472 | 1 | 0 | 9 | 13090.039 | 12042.722 | 1427.7615 | 11437.413 | 9473.524 | 9040.893 |
| Rars1 | DKVLTEEELK | -483 | -492 | 1 | 0.00025015 | 10 | 1353.4502 | 1312.8589 | 1524.3733 | 1927.2803 | 1795.1029 | 1744.5071 |
| Rars1 | AYPDLENPPLIVTPSQQPK | -91 | -109 | 0 | 0 | 19 | 827.4729 | 920.0663 | 844.4929 | 953.0462 | 1186.9564 | 1276.0979 |
| Rars1 | NCGCLEASPSLEQLR | -31 | -45 | 0 | 0 | 15 | 698.4163 | 1244.3506 | 1149.5012 | 900.86847 | 1128.3093 | 1001.0834 |
| Rars1 | VIVDFSSPNIAK | -194 | -205 | 0 | 0 | 12 | 676.7394 | 867.3754 | 813.4662 | 825.9333 | 975.5 | 1062.054 |
| Rars1 | IVFVPGCSVPLTIVK | -363 | -377 | 0 | 0 | 15 | 619.09955 | 627.4687 | 722.6699 | 436.2816 | 1229.659 | 1231.82 |
| Rars1 | ILDDLFLHTL | -588 | -597 | 0 | 0 | 10 | 584.44855 | 528.5155 | 524.6526 | 470.41168 | 34.952766 | 680.54156 |
| Rars1 | YIDKVEIAGPGFIN | -150 | -163 | 1 | 0.000394361 | 14 | 574.92957 | 815.3426 | 666.0309 | 392.41058 | 1654.1459 | 183.3984 |
| Rars1 | AAYPDLENPPLIVTPSQQPK | -90 | -109 | 0 | 0 | 20 | 530.84766 | 1038.287 | 808.8064 | 656.8148 | 903.5429 | 1323.0417 |
| Rars1 | KDFVSEQLTSL | -168 | -178 | 1 | 0.00025015 | 11 | 235.7088 | 240.05585 | 273.81473 | 144.33621 | 898.0902 | 804.8745 |
| Rars1 | SDGGYTYDTSDLAAIK | -378 | -393 | 0 | 0 | 16 | 1241.9197 | 1511.402 | 1664.7589 | 1848.8397 | 2099.0364 | NaN |
| Rars1 | NLICDVSR | -309 | -316 | 0 | 0 | 8 | 1064.382 | 1230.1152 | NaN | 1126.8163 | 1455.6741 | 975.42163 |
| Rars1 | RFDADEEFKK | -279 | -288 | 2 | 0 | 10 | 717.78314 | 585.19305 | 736.5502 | NaN | 1126.7434 | 611.1554 |
| Rars1 | YADLSHNR | -505 | -512 | 0 | 0.00025015 | 8 | 715.8711 | 1301.6753 | NaN | 973.3985 | 1147.2894 | 1253.7466 |
| Rars1 | FDADEEFK | -280 | -287 | 0 | 0 | 8 | 542.09644 | 588.08386 | NaN | 628.4726 | 609.6745 | 650.73834 |
| Rars1 | ILDDLFLH | -588 | -595 | 0 | 0 | 8 | 851.32874 | 851.9626 | 1100.1931 | NaN | NaN | NaN |
| Rars1 | AWNLICDVSR | -307 | -316 | 0 | 0 | 10 | 597.4324 | 669.4605 | 643.1593 | NaN | NaN | NaN |
| Rars1 | GVQLPVLGDK | -182 | -191 | 0 | 0 | 10 | 1127.295 | NaN | NaN | NaN | 1332.933 | NaN |
| Vps35 | LSQLEGVNVER | -227 | -237 | 0 | 0 | 11 | 2291.2126 | 1915.6173 | 1585.4253 | 4538.7856 | 3813.7056 | 3676.7053 |
| Vps35 | LLDEAIQAVK | -15 | -24 | 0 | 0 | 10 | 985.55206 | 373.39355 | 455.5237 | 375.54352 | 506.94034 | 776.8848 |
| Vps35 | AELAELPLR | -574 | -582 | 0 | 0 | 9 | 943.1993 | 845.4379 | 962.51263 | 1318.6742 | 1177.5721 | 1092.606 |
| Vps35 | GILEQVVNCR | -245 | -254 | 0 | 0 | 10 | 768.61127 | 1013.0419 | 854.17487 | 391.53937 | 1575.9713 | 839.0146 |
| Vps35 | AFSLYEDEISDSK | -610 | -622 | 0 | 0 | 13 | 541.2722 | 1156.022 | 1044.0433 | 992.4256 | 711.32526 | 948.894 |
| Vps35 | ACAELHQNVNVK | -285 | -296 | 0 | 0 | 12 | 291.37732 | 507.83868 | 634.4031 | 495.8593 | 540.6815 | 430.6351 |
| Vps35 | AELHQNVNVK | -287 | -296 | 0 | 0 | 10 | 647.95416 | 663.69366 | NaN | 1042.5778 | 961.8627 | 1107.395 |
| Vps35 | TLNPFLR | -278 | -284 | 0 | 0.003620005 | 7 | 1209.429 | 1309.3812 | 1159.3633 | 880.35986 | NaN | NaN |
| Vps35 | LALFAHR | -306 | -312 | 0 | 0.003745449 | 7 | 785.2892 | 414.1844 | NaN | 689.0515 | NaN | NaN |
| Vps35 | QYAGNIIPR | -99 | -107 | 0 | 0.000989247 | 9 | 634.8882 | NaN | NaN | NaN | NaN | NaN |
| Gars1 | GITIDFDTVNK | -659 | -669 | 0 | 0 | 11 | 3369.1265 | 1310.6425 | 3067.8425 | 723.14795 | 1059.213 | 1610.8842 |
| Gars1 | APQVDVDR | -76 | -83 | 0 | 0.000394361 | 8 | 1706.3988 | 362.2947 | 1013.4105 | 2152.8853 | 1159.8828 | 1158.3079 |
| Gars1 | EVVPSVIEPSFGLGR | -559 | -573 | 0 | 0 | 15 | 1493.0535 | 1579.8195 | 1748.4677 | 1402.4749 | 2024.4756 | 1800.9724 |
| Gars1 | EHTFHVR | -580 | -586 | 0 | 0.00025015 | 7 | 1189.1812 | 1098.2485 | 1958.8738 | 194.62929 | 622.8589 | 1170.6282 |
| Gars1 | SCYDLSCHAR | -455 | -464 | 0 | 0 | 10 | 1097.7134 | 1052.4312 | 1086.5049 | 1523.5892 | 1903.24 | 1927.4878 |
| Gars1 | GGVSGLYDFGPVGCALK | -132 | -148 | 0 | 0 | 17 | 1095.906 | 1186.9586 | 1362.0565 | 1703.3284 | 2912.2878 | 2659.288 |
| Gars1 | VIEPSFGLGR | -564 | -573 | 0 | 0 | 10 | 1073.8804 | 1312.0128 | 557.4483 | 1124.6285 | 293.3585 | 1207.2709 |
| Gars1 | EIEHFVDPTEKDHPK | -340 | -354 | 1 | 0 | 15 | 674.2215 | 799.9973 | 932.8867 | 854.1405 | 652.6494 | 904.7005 |
| Gars1 | EIEHFVDPTEKDHPK | -340 | -354 | 1 | 0 | 15 | 674.2215 | 799.9973 | 932.8867 | 854.1405 | 652.6494 | 904.7005 |
| Gars1 | SVLGYFIGR | -394 | -402 | 0 | 0 | 9 | 406.87326 | 555.27356 | 495.2244 | 591.0708 | 315.53177 | 859.6726 |
| Gars1 | TFFSFPAVVAPFK | -593 | -605 | 0 | 0 | 13 | 390.6109 | 317.2547 | 259.00278 | 397.23474 | 1039.3845 | 482.215 |
| Gars1 | FSFPAVVAPFK | -595 | -605 | 0 | 0 | 11 | 352.55493 | 381.51828 | 237.32584 | 527.4369 | 444.24744 | 229.82964 |
| Gars1 | VDDSSGSIGR | -637 | -646 | 0 | 0 | 10 | 113.73074 | 1212.0677 | 1324.516 | 366.3363 | 435.23685 | 1475.7899 |
| Gars1 | IYLYLTK | -403 | -409 | 0 | 0 | 7 | 3973.1206 | 3185.7322 | NaN | 937.46497 | 799.9215 | 1316.772 |
| Gars1 | TPHTATLR | -670 | -677 | 0 | 0.001092529 | 8 | 2741.045 | NaN | 2905.2285 | 2764.0735 | 2691.5642 | 2860.6755 |
| Gars1 | YPLFEGQETGKK | -713 | -724 | 1 | 0 | 12 | 2415.174 | NaN | 2566.8784 | 2283.5989 | 2167.7063 | 2069.8333 |
| Gars1 | NNIIQAWR | -149 | -156 | 0 | 0 | 8 | 1587.0348 | 1457.4731 | 1473.3264 | NaN | 2010.721 | 1684.9827 |
| Gars1 | HFVDPTEKDHPK | -343 | -354 | 1 | 0 | 12 | 258.07864 | 389.74072 | 232.09265 | NaN | 376.2582 | 442.8304 |
| Gars1 | HFVDPTEKDHPK | -343 | -354 | 1 | 0 | 12 | 258.07864 | 389.74072 | 232.09265 | NaN | 376.2582 | 442.8304 |
| Gars1 | TAQGIFLNFK | -290 | -299 | 0 | 0 | 10 | 607.47754 | NaN | 876.592 | 263.5554 | 71.50157 | 182.22249 |
| Gars1 | LLEFNQGK | -301 | -308 | 0 | 0.00437931 | 8 | 2089.9512 | 2113.3901 | 619.07556 | NaN | NaN | 1481.2882 |
| Gars1 | VPLVAEKPLKEPK | -468 | -480 | 2 | 0 | 13 | 1006.07654 | 457.1554 | 559.25116 | NaN | 530.83765 | NaN |
| Gars1 | EIEHFVDPTEK | -340 | -350 | 0 | 0.00025015 | 11 | 846.51654 | 987.5212 | 873.6487 | 762.24695 | NaN | NaN |
| Gars1 | VLGYFIGR | -395 | -402 | 0 | 0.00111006 | 8 | 509.36395 | 479.4107 | NaN | 353.04214 | NaN | NaN |
| Gars1 | ACDCWDAESK | -431 | -440 | 0 | 0.002068213 | 10 | 310.61542 | 281.0955 | NaN | 488.48233 | NaN | NaN |
| Gars1 | CDCWDAESK | -432 | -440 | 0 | 0.000907206 | 9 | 249.31279 | 212.93771 | NaN | NaN | NaN | 205.98175 |
| Gars1 | AEVSELPNVVR | -687 | -697 | 0 | 0.000230707 | 11 | 1304.8193 | NaN | NaN | NaN | 3278.8657 | NaN |
| Gars1 | DLANGNITWADVEAR | -698 | -712 | 0 | 0.000230707 | 15 | 608.7994 | NaN | NaN | NaN | NaN | 902.98236 |
| Gars1 | HFVDPTEK | -343 | -350 | 0 | 0 | 8 | 1457.2776 | NaN | NaN | NaN | NaN | NaN |
| Sacm1l | NFVETEQIVHYSGNR | -236 | -250 | 0 | 0 | 15 | 754.36224 | 921.03656 | 907.2673 | 1031.8512 | 507.99567 | 1175.76 |

| | | | | | | | | | | | | |
|---|---|---|---|---|---|---|---|---|---|---|---|---|
| Sacm1l | LEEQDEFEK | -424 | -432 | 0 | 0 | 9 | 3106.7493 | NaN | 1383.4987 | 3279.6914 | 1601.1931 | NaN |
| Sacm1l | ACDDGADDVLIIDR | -22 | -35 | 0 | 0 | 14 | 1143.504 | 1327.0491 | NaN | NaN | 1261.1351 | 1353.1256 |
| Sacm1l | FALPVLHGF | -185 | -193 | 0 | 0.004224523 | 9 | 152.90288 | 332.024 | NaN | NaN | NaN | NaN |
| Ttc38 | ACDLWEQILR | -123 | -132 | 0 | 0 | 10 | 1498.6224 | 876.7744 | NaN | 1048.0408 | 1187.3899 | 1207.0527 |
| Ttc38 | VLELLLPIR | -390 | -398 | 0 | 0.00025015 | 9 | 178.65677 | NaN | NaN | 222.02675 | NaN | 221.63307 |
| Ttc38 | WTPDIPLNSYVK | -168 | -179 | 0 | 0 | 12 | 152.73253 | NaN | 399.61758 | NaN | NaN | 638.24756 |
| Itgal | TFVGQEPLTSDVR | -386 | -398 | 0 | 0 | 13 | 2107.6992 | 1625.5096 | 1936.3481 | 1638.2505 | 1581.6713 | 1785.407 |
| Itgal | YQHVGQVL | -425 | -432 | 0 | 0.001291103 | 8 | 1336.5404 | 454.7926 | 1395.5643 | 1264.889 | 659.2146 | 1284.9315 |
| Itgal | TLSNSGEDAYWVR | -802 | -814 | 0 | 0 | 13 | 1278.9232 | 1138.6844 | 1260.5833 | 1206.9156 | 1056.7096 | 921.38666 |
| Itgal | LFQAPEAGGR | -433 | -442 | 0 | 0 | 10 | 1269.9126 | 1465.9927 | 1257.4841 | 1233.047 | 1179.7515 | 127.29728 |
| Itgal | IKPLTPQFQGR | -654 | -664 | 1 | 0.00025015 | 11 | 1238.5293 | 1371.1531 | 1365.2104 | 1359.1544 | 1378.6494 | 1405.5469 |
| Itgal | RPSSEAEQPCLPGVQFR | -1001 | -1017 | 1 | 0 | 17 | 1162.5117 | 1279.9005 | 1454.4359 | 1724.3905 | 1578.1578 | 1545.8198 |
| Itgal | ASEPVEEFVK | -301 | -310 | 0 | 0 | 10 | 954.6272 | 913.15314 | 913.42377 | 1095.3629 | 1204.5969 | 919.05035 |
| Itgal | LADVVVGAEGR | -594 | -604 | 0 | 0 | 11 | 952.4782 | 996.6734 | 1145.8197 | 1011.0687 | 1053.5198 | 1057.2039 |
| Itgal | TEFTFLDYVK | -201 | -210 | 0 | 0 | 10 | 720.32825 | 663.2707 | 717.4168 | 857.80365 | 1038.9102 | 956.83575 |
| Itgal | TEFTFLDYVK | -201 | -210 | 0 | 0 | 10 | 720.32825 | 663.2707 | 717.4168 | 857.80365 | 1038.9102 | 956.83575 |
| Itgal | LIGAPLFFGEQR | -477 | -488 | 0 | 0 | 12 | 621.6145 | 139.38568 | 606.1538 | 1007.18066 | 1240.2239 | 1234.7656 |
| Itgal | SFSPEEIPVHEVECSYSAR | -621 | -639 | 0 | 0 | 19 | 600.3674 | 522.4875 | 524.9874 | 573.5033 | 180.4522 | 947.8452 |
| Itgal | SFSPEEIPVHEVECSYSAR | -621 | -639 | 0 | 0 | 19 | 600.3674 | 522.4875 | 524.9874 | 573.5033 | 180.4522 | 947.8452 |
| Itgal | DWAGGFLDLR | -370 | -379 | 0 | 0 | 10 | 588.86395 | 673.4192 | 600.258 | 716.7847 | 754.3284 | 720.2399 |
| Itgal | SVQTDPLVTCHSEDLK | -985 | -1000 | 0 | 0 | 16 | 576.9907 | 587.348 | 187.70947 | 963.4006 | 1103.4453 | 1091.351 |
| Itgal | VKVDLIHEK | -1076 | -1084 | 1 | 0 | 9 | 544.0328 | 574.01215 | 514.9434 | 640.10455 | 570.37445 | 545.7599 |
| Itgal | IYAIEGTNR | -329 | -337 | 0 | 0 | 9 | 463.28336 | 2323.453 | 2198.4976 | 1602.6993 | 1260.7139 | 2573.554 |
| Itgal | EDLQGATFVGQEPLTSDVR | -380 | -398 | 0 | 0 | 19 | 387.9599 | 410.2663 | 427.28546 | 748.28925 | 929.8125 | 975.90717 |
| Itgal | LDLDFPR | -815 | -821 | 0 | 0.001776473 | 7 | 1475.7102 | 1452.8905 | 1588.2213 | 1112.8334 | NaN | 1021.8039 |
| Itgal | LKDLFTDLQR | -318 | -327 | 1 | 0 | 10 | 1003.6284 | 1142.3324 | NaN | 1293.0013 | 1636.0071 | 1553.2505 |
| Itgal | SNSGEDAYWVR | -804 | -814 | 0 | 0 | 11 | 1123.6388 | 494.43283 | 1384.5436 | NaN | 349.65182 | NaN |
| Itgal | ACDPGLSR | -107 | -114 | 0 | 0.00025015 | 8 | 907.3641 | NaN | 300.44177 | 444.22928 | NaN | 389.88055 |
| Itgal | LTDVAVGAPLEEQGAVY | -534 | -550 | 0 | 0.002068213 | 17 | 801.8171 | 781.947 | NaN | 1210.4763 | NaN | 1422.0104 |
| Itgal | FVGQEPLTSDVR | -387 | -398 | 0 | 0.001055699 | 12 | 771.0621 | NaN | NaN | 1126.7858 | 954.8205 | NaN |
| Itgal | VLVIITDGEASDK | -256 | -268 | 0 | 0.00111006 | 13 | 758.4981 | 1061.76 | NaN | NaN | 1666.6975 | NaN |
| Itgal | DMGCLEPLR | -1149 | -1157 | 0 | 0.005363933 | 9 | 977.3321 | 800.3458 | NaN | NaN | NaN | NaN |
| Itgal | AMQPILRPSIH | -746 | -756 | 1 | 0.000907206 | 11 | 355.66995 | NaN | NaN | NaN | NaN | 182.75882 |
| Itgal | EESGARPDATK | -245 | -255 | 1 | 0.002099344 | 11 | 214.68953 | NaN | NaN | NaN | NaN | 509.72635 |
| C3 | QDFFIDLR | -827 | -834 | 0 | 0 | 8 | 3366.9424 | 3551.0435 | 3575.567 | 3593.6968 | 4281.7993 | 4255.596 |
| C3 | KIWDVVEK | -614 | -621 | 1 | 0.000471409 | 8 | 2665.259 | 3132.373 | 3081.2349 | 1116.1495 | 1300.8014 | 2263.002 |
| C3 | TIYTPGSTVLYR | -138 | -149 | 0 | 0 | 12 | 2356.4482 | 2907.929 | 2923.6829 | 3575.6562 | 3431.3572 | 3403.1392 |
| C3 | ACEPGVDYVYK | -1536 | -1546 | 0 | 0 | 11 | 1466.9294 | 1410.5704 | 1522.6061 | 3720.0684 | 1611.6174 | 1751.6287 |
| C3 | NHFISDGVK | -918 | -926 | 0 | 0.000431655 | 9 | 944.4671 | 773.4661 | 917.0104 | 802.7008 | 1006.9762 | 985.6788 |
| C3 | TSQGLQTEQR | -647 | -656 | 0 | 0.000471409 | 10 | 472.62015 | 436.99335 | 659.68115 | 982.67505 | 1265.0769 | 773.2805 |
| C3 | NEQVEIR | -842 | -848 | 0 | 0.003201138 | 7 | 963.26404 | 1111.9497 | 909.4137 | NaN | NaN | 295.8487 |
| C3 | IETPDGIPVK | -168 | -177 | 0 | 0 | 10 | 798.8442 | 866.5747 | NaN | 909.56903 | NaN | NaN |
| C3 | IETPDGIPVKR | -168 | -178 | 1 | 0 | 11 | 501.76617 | 321.55664 | 907.16846 | NaN | NaN | NaN |
| Gnb2 | ACGDSTLTQITAGLDPVGR | -24 | -42 | 0 | 0 | 19 | 547.9162 | 500.68048 | 491.03333 | 687.74506 | 818.2983 | 732.7289 |
| Paics | GLNNHLVWAK | -392 | -401 | 0 | 0 | 10 | 990.6059 | 1059.3613 | 1100.8718 | 1047.9012 | 1291.8197 | 1322.869 |
| Paics | NPGVQEGYK | -112 | -120 | 0 | 0.000230707 | 9 | 800.06384 | 956.16895 | 775.5238 | 698.3548 | 716.8632 | 680.81024 |
| Paics | EIVLADVIDNDSWR | -202 | -215 | 0 | 0 | 14 | 725.0094 | 632.88715 | 680.7732 | 66.65502 | 745.4402 | 696.06024 |
| Paics | NFEWVADR | -248 | -255 | 0 | 0.000230707 | 8 | 704.54034 | 293.62125 | 664.75934 | 669.06024 | 723.6129 | 701.7634 |
| Paics | SWLPQDCTLVDMK | -179 | -191 | 0 | 0 | 13 | 610.4713 | 226.70805 | 710.3359 | 515.2381 | 1131.3264 | 725.9215 |
| Paics | ACGNFGIPCELR | -287 | -298 | 0 | 0 | 12 | 538.52026 | 610.0958 | 619.86005 | 683.4472 | 821.2258 | 781.57043 |
| Paics | DDANNDPQWSEEQLIAAK | -132 | -149 | 0 | 0 | 18 | 525.1575 | 590.8141 | 780.1024 | 373.53625 | 767.2988 | 782.39465 |
| Paics | DANNDPQWSEEQLIAAK | -133 | -149 | 0 | 0 | 17 | 504.21222 | 663.7464 | 723.64795 | 431.7387 | 524.95746 | 641.6676 |
| Paics | GSTSDLGHCEK | -273 | -283 | 0 | 0 | 11 | 377.5678 | 529.4358 | 465.91205 | 533.361 | 466.99057 | 470.5898 |
| Paics | RNPGVQEGYK | -111 | -120 | 1 | 0.000431655 | 10 | 374.94797 | 369.56766 | 436.46613 | 745.7305 | 556.1426 | 302.07178 |
| Paics | TKEVYELLDTPGR | -18 | -30 | 1 | 0 | 13 | 333.87506 | 839.73944 | 833.6223 | 741.89685 | 1039.5382 | 560.69745 |
| Paics | EVYELLDTPGR | -20 | -30 | 0 | 0.000230707 | 11 | 1947.292 | 1802.0586 | 2087.9082 | NaN | 2203.9302 | 2108.7178 |
| Paics | VIDNDSWR | -208 | -215 | 0 | 0.001291103 | 8 | 1109.3671 | 1239.4712 | 1346.9666 | 1132.5067 | 968.3033 | NaN |
| Paics | KNFEWVADR | -247 | -255 | 1 | 0 | 9 | 1108.9438 | 1603.2699 | 1577.8279 | NaN | 691.0189 | 2516.538 |
| Paics | EVTPEGLQMVK | -236 | -246 | 0 | 0 | 11 | 729.43134 | 717.14795 | NaN | 287.75018 | 494.12604 | 600.0681 |
| Paics | DVIDNDSWR | -207 | -215 | 0 | 0.00025015 | 9 | 328.1929 | 745.1444 | 692.23517 | 663.1641 | 713.98083 | NaN |
| Paics | SKDQITAGNAAR | -35 | -46 | 1 | 0.00025015 | 12 | 288.2509 | NaN | NaN | NaN | NaN | 407.19806 |
| Aco2 | FKLEAPDADELPR | -522 | -534 | 1 | 0 | 13 | 5615.296 | 6599.4526 | 6870.379 | 5621.1494 | 6364.4683 | 6249.078 |
| Aco2 | FKLEAPDADELPR | -522 | -534 | 1 | 0 | 13 | 5615.296 | 6599.4526 | 6870.379 | 5621.1494 | 6364.4683 | 6249.078 |
| Aco2 | FTITPGSEQIR | -414 | -424 | 0 | 0 | 11 | 3340.9275 | 3741.0156 | 3707.5813 | 4613.545 | 3975.5684 | 1452.4053 |
| Aco2 | KFNPETDFLTGK | -506 | -517 | 1 | 0 | 12 | 2552.9836 | 2949.123 | 3114.6804 | 3234.3474 | 464.0318 | 3661.4746 |
| Aco2 | EFGPVPDTAR | -639 | -648 | 0 | 0 | 10 | 2552.2285 | 2131.9998 | 2127.8357 | 1068.6964 | 1481.2723 | 423.64905 |
| Aco2 | SDFDPGQDTYQHPPK | -535 | -549 | 0 | 0 | 15 | 2355.4675 | 2446.28 | 2563.6104 | 2791.9302 | 2705.376 | 2667.331 |
| Aco2 | SDFDPGQDTYQHPPK | -535 | -549 | 0 | 0 | 15 | 2355.4675 | 2446.28 | 2563.6104 | 2791.9302 | 2705.376 | 2667.331 |
| Aco2 | EHAALEPR | -672 | -679 | 0 | 0 | 8 | 2306.255 | 3212.2866 | 3193.2214 | 2427.752 | 1957.1702 | 2202.9873 |
| Aco2 | LTGSLSGWTSPK | -234 | -245 | 0 | 0 | 12 | 2147.971 | 979.88684 | 1073.8499 | 1949.917 | 2139.9612 | 2130.763 |
| Aco2 | VDVSPTSQR | -556 | -564 | 0 | 0 | 9 | 2119.622 | 2299.7937 | 2408.5752 | 2129.6536 | 519.7187 | 2425.2773 |
| Aco2 | DFAPGKPLK | -731 | -739 | 1 | 0.00025015 | 9 | 2093.0613 | 2115.6836 | 2100.123 | 2037.483 | 2153.8008 | 314.722 |
| Aco2 | SQFTITPGSEQIR | -412 | -424 | 0 | 0 | 13 | 2030.6345 | 891.4737 | 2298.418 | 1292.4259 | 1687.8073 | 1665.3007 |
| Aco2 | IVYGHLDDPANQEIER | -69 | -84 | 0 | 0 | 16 | 1727.6769 | 2181.2197 | 2297.4648 | 1614.5781 | 1805.6029 | 2735.567 |
| Aco2 | IVYGHLDDPANQEIER | -69 | -84 | 0 | 0 | 16 | 1727.6769 | 2181.2197 | 2297.4648 | 1614.5781 | 1805.6029 | 2735.567 |
| Aco2 | WVVIGDENYGEGSSR | -657 | -671 | 0 | 0 | 15 | 1636.2703 | 1707.5209 | 1870.3512 | 1853.7487 | 2376.1301 | 2178.0806 |
| Aco2 | FNPETDFLTGK | -507 | -517 | 0 | 0 | 11 | 1529.5085 | 1636.8303 | 1067.655 | 953.9837 | 1079.1338 | 937.7846 |
| Aco2 | LEAPDADELPR | -524 | -534 | 0 | 0 | 11 | 1462.3457 | 462.683 | 1789.9849 | 712.45276 | 3715.9573 | 1672.6171 |

| | | | | | | | | | | | | |
|---|---|---|---|---|---|---|---|---|---|---|---|---|
| Aco2 | NDANPETHAF | -480 | -489 | 0 | 0 | 10 | 1419.733 | 1358.4451 | 1294.7583 | 1116.7737 | 332.28897 | 996.61176 |
| Aco2 | GHLDNISNNLLIGA | -608 | -621 | 0 | 0 | 14 | 1238.5109 | 151.03789 | 468.80243 | 606.92773 | 686.69147 | 1142.2598 |
| Aco2 | EGWPLDIR | -371 | -378 | 0 | 0 | 8 | 1049.4939 | 969.7062 | 1078.4113 | 512.6044 | 722.9742 | 735.2365 |
| Aco2 | LQLLEPFDKWDGK | -565 | -577 | 1 | 0 | 13 | 706.144 | 739.89056 | 934.0653 | 624.0959 | 859.90393 | 919.2152 |
| Aco2 | VAVPSTIHCDHLIEAQ | -118 | -133 | 0 | 0 | 16 | 544.04517 | 656.5495 | 596.5282 | 602.01086 | 1133.0376 | 1008.7058 |
| Aco2 | VAVPSTIHCDHLIEAQ | -118 | -133 | 0 | 0 | 16 | 544.04517 | 656.5495 | 596.5282 | 602.01086 | 1133.0376 | 1008.7058 |
| Aco2 | QGLLPLTFADPSDYNK | -702 | -717 | 0 | 0.000591124 | 16 | 569.7361 | 523.5594 | 521.27344 | 821.2048 | 849.6234 | 713.3136 |
| Aco2 | NDANPETHAFVTSPEIVTAL | -480 | -499 | 0 | 0 | 20 | 412.43774 | 559.839 | 384.6294 | 397.91122 | 528.7008 | 463.65112 |
| Aco2 | VVIGDENYGEGSSR | -658 | -671 | 0 | 0 | 14 | 1284.2125 | 1185.32 | NaN | 2545.8657 | 665.7239 | 1679.6335 |
| Aco2 | LNRPLTL | -59 | -65 | 1 | 0.000230707 | 7 | 1228.3312 | 696.52167 | 645.7195 | 644.959 | 522.1358 | NaN |
| Aco2 | NETQIEWFR | -759 | -767 | 0 | 0 | 9 | 672.2582 | 2192.3865 | NaN | 147.40527 | 938.15735 | 793.8124 |
| Aco2 | SHFEPSEYIR | -35 | -44 | 0 | 0 | 10 | 608.9993 | 827.7185 | 2018.6487 | NaN | 1018.9615 | 1012.379 |
| Aco2 | LDDPANQEIER | -74 | -84 | 0 | 0 | 11 | 554.1691 | 597.6245 | 886.162 | 806.6316 | 937.8249 | NaN |
| Aco2 | MSHFEPSEYIR | -34 | -44 | 0 | 0 | 11 | 312.01932 | NaN | 731.6321 | 3077.8274 | 531.7356 | 411.2126 |
| Aco2 | ACGPCIGQWDR | -447 | -457 | 0 | 0 | 11 | 831.4269 | 835.31964 | 888.0057 | 85.059906 | NaN | NaN |
| Aco2 | GHLDNISNNLLIGAIN | -608 | -623 | 0 | 0.000431655 | 16 | 685.2014 | 771.46106 | 796.6709 | 516.8779 | NaN | NaN |
| Aco2 | VTQEFGPVPDTAR | -636 | -648 | 0 | 0 | 13 | 933.43604 | NaN | NaN | 907.8973 | NaN | 1060.1697 |
| Aco2 | TFADPSDYNK | -708 | -717 | 0 | 0 | 10 | 878.47156 | 749.8319 | 765.48706 | NaN | NaN | NaN |
| Aco2 | KQGLLPLTF | -701 | -709 | 1 | 0.005015524 | 9 | 453.6751 | 545.48224 | 478.1005 | NaN | NaN | NaN |
| Aco2 | HFEPSEYIR | -36 | -44 | 0 | 0.000394361 | 9 | 1475.5701 | NaN | NaN | 811.33997 | NaN | NaN |
| Aco2 | IANLAEEFK | -315 | -323 | 0 | 0.002595029 | 9 | 351.84467 | 557.6132 | NaN | NaN | NaN | NaN |
| Cct3 | EIQVQHPAAK | -69 | -78 | 0 | 0 | 10 | 7572.323 | 6523.1177 | 10619.917 | 8849.79 | 11728.679 | 11065.041 |
| Cct3 | IDDIVSGHK | -519 | -527 | 0 | 0 | 9 | 3660.58 | 191.34592 | 2915.763 | 2955.137 | 2527.5605 | 2671.5688 |
| Cct3 | TGVEQWPYR | -430 | -438 | 0 | 0 | 9 | 3161.9597 | 3078.009 | 3113.3887 | 3511.4004 | 192.88771 | 213.5857 |
| Cct3 | TNDGNAILR | -60 | -68 | 0 | 0 | 9 | 2558.4634 | 1633.8224 | 1471.2913 | 1925.3892 | 695.5935 | 2524.7983 |
| Cct3 | AMTGVEQWPYR | -428 | -438 | 0 | 0 | 11 | 2098.4414 | 2323.5093 | 2442.6865 | 1076.0625 | 1316.8274 | 186.44858 |
| Cct3 | KTIADIIR | -31 | -38 | 1 | 0.00504706 | 8 | 2009.3629 | 2052.961 | 2244.5266 | 2273.0356 | 2198.6907 | 2316.1309 |
| Cct3 | IPGGIIEDSCVLR | -204 | -216 | 0 | 0 | 13 | 1657.0779 | 2015.9822 | 2000.5565 | 2158.4302 | 2495.1326 | 2689.2832 |
| Cct3 | TLIQNCGASTIR | -450 | -461 | 0 | 0 | 12 | 1490.1467 | 1419.1511 | 1748.7494 | 900.47595 | 1553.5342 | 1585.84 |
| Cct3 | KIPGGIIEDSCVLR | -203 | -216 | 1 | 0 | 14 | 1347.0444 | 1093.7026 | 1732.2853 | 2111.93 | 2236.4531 | 2197.0535 |
| Cct3 | KIPGGIIEDSCVLR | -203 | -216 | 1 | 0 | 14 | 1347.0444 | 1093.7026 | 1732.2853 | 2111.93 | 2236.4531 | 2197.0535 |
| Cct3 | IVSRPEELREDDVGTGA | -331 | -347 | 2 | 0.00078661 | 17 | 1071.0032 | 1166.9098 | 1424.7894 | 1121.7404 | 1126.7616 | 1171.4348 |
| Cct3 | IVSRPEELREDDVGTGA | -331 | -347 | 2 | 0.00078661 | 17 | 1071.0032 | 1166.9098 | 1424.7894 | 1121.7404 | 1126.7616 | 1171.4348 |
| Cct3 | IGDEYFTFITDCKDPK | -355 | -370 | 1 | 0 | 16 | 1203.8477 | 1390.2806 | 1485.709 | 675.5058 | 804.55273 | 824.05426 |
| Cct3 | REEDFTR | -260 | -266 | 1 | 0.000394361 | 7 | 1193.4327 | 317.31393 | 1204.9961 | 1367.3492 | 1290.8066 | 1934.7522 |
| Cct3 | ACNIALDAVK | -172 | -181 | 0 | 0 | 10 | 1080.9717 | 1342.878 | 1199.8689 | 678.27014 | 1146.4758 | 1144.0822 |
| Cct3 | EILSEVER | -382 | -389 | 0 | 0.00025015 | 8 | 991.7414 | 892.89636 | 879.46655 | 994.9406 | 1059.4856 | 1256.5791 |
| Cct3 | NLQDAMQVCR | -390 | -399 | 0 | 0 | 10 | 751.3501 | 682.4137 | 700.07623 | 679.8994 | 602.21277 | 581.607 |
| Cct3 | KIGDEYFTFITDCKDPK | -354 | -370 | 2 | 0.000230707 | 17 | 729.5492 | 746.6845 | 833.98724 | 1167.5809 | 1563.2285 | 1657.756 |
| Cct3 | SKEILSEVER | -380 | -389 | 1 | 0.00025015 | 10 | 707.038 | 762.03516 | 1001.7367 | 1956.7579 | 1833.2067 | 1476.555 |
| Cct3 | ELGIWEPLAVK | -492 | -502 | 0 | 0 | 11 | 696.0564 | 756.4849 | 783.24146 | 424.17633 | 1384.8715 | 1282.2941 |
| Cct3 | LRIDDIVSGHK | -517 | -527 | 1 | 0 | 11 | 460.67236 | 443.28134 | 420.74725 | 535.1908 | 436.19028 | 343.72644 |
| Cct3 | ISTPVDVNNR | -140 | -149 | 0 | 0.000394361 | 10 | 447.02148 | 2054.755 | 406.59583 | 605.69006 | 1681.5463 | 212.81621 |
| Cct3 | GIWEPLAVK | -494 | -502 | 0 | 0 | 9 | 387.72083 | 839.47 | 828.1013 | 449.5042 | 654.6387 | 694.5015 |
| Cct3 | KGESQTDIEITR | -249 | -260 | 1 | 0 | 12 | 1225.6946 | 1269.7067 | 1273.5588 | NaN | 1297.7971 | 1109.1737 |
| Cct3 | KISTPVDVNNR | -139 | -149 | 1 | 0 | 11 | 988.3386 | 1203.9795 | 1189.7128 | 1132.2915 | 1370.8608 | NaN |
| Cct3 | NVLLDPQLVPGGGASE | -400 | -415 | 0 | 0 | 16 | 345.7349 | 111.42098 | 653.7494 | NaN | 677.2335 | 379.9808 |
| Cct3 | GISDLAQHY | -295 | -303 | 0 | 0.000471409 | 9 | 215.26454 | 716.35376 | NaN | NaN | 901.80066 | 347.4381 |
| Cct3 | TVQFEENGR | -182 | -190 | 0 | 0 | 9 | 583.6991 | 540.52576 | 378.6115 | NaN | NaN | NaN |
| Cct3 | KGESQTDIEIT | -249 | -259 | 1 | 0.001868731 | 11 | 1120.943 | NaN | 1405.2832 | NaN | NaN | NaN |
| Cct3 | ACTILLR | -371 | -377 | 0 | 0.001708008 | 7 | 737.9779 | NaN | NaN | NaN | NaN | NaN |
| Cct3 | HTQESCETW | -470 | -478 | 0 | 0.00078661 | 9 | 602.94806 | NaN | NaN | NaN | NaN | NaN |
| Itgb2 | SRGDCDGVQINNPVTF | -397 | -412 | 1 | 0 | 16 | 5491.1777 | 834.2633 | 795.51154 | 580.1299 | 1180.4283 | 1308.1074 |
| Itgb2 | SNEFDYPSVGQLAHK | -297 | -311 | 0 | 0 | 15 | 3347.4724 | 4729.7627 | 4355.328 | 5040.535 | 5901.5967 | 5069.274 |
| Itgb2 | LGAILTPNDGR | -276 | -286 | 0 | 0.00025015 | 11 | 3059.4678 | 3098.923 | 3976.7773 | 3461.0127 | 2900.8594 | 4341.9 |
| Itgb2 | SIANPEFDQR | -84 | -93 | 0 | 0 | 10 | 2356.5 | 906.3214 | 2320.0776 | 2659.3982 | 2195.1047 | 754.6401 |
| Itgb2 | VFLDHSTLPDTLK | -368 | -380 | 0 | 0 | 13 | 2098.001 | 2476.4275 | 2278.8027 | 2536.4885 | 2949.9692 | 2810.4062 |
| Itgb2 | VFLDHSTLPDTLK | -368 | -380 | 0 | 0 | 13 | 2098.001 | 2476.4275 | 2278.8027 | 2536.4885 | 2949.9692 | 2810.4062 |
| Itgb2 | FDKGPFEK | -636 | -643 | 1 | 0 | 8 | 1779.5255 | 510.11993 | 1664.6677 | 1764.6919 | 1439.4221 | 1405.7769 |
| Itgb2 | LTDNSNQFQTEVGK | -211 | -224 | 0 | 0 | 14 | 1269.1956 | 1325.135 | 1137.4509 | 1296.2168 | 1385.5251 | 1388.0519 |
| Itgb2 | ACPEEIGWR | -246 | -254 | 0 | 0 | 9 | 1047.1416 | 1053.4232 | 1080.2406 | 1323.9446 | 1443.8385 | 1400.1663 |
| Itgb2 | VTLYLRPGQAA | -104 | -114 | 1 | 0 | 11 | 1032.821 | 1026.0352 | 959.129 | 1456.7439 | 1230.102 | 1251.6814 |
| Itgb2 | DDGFHFAGDGK | -265 | -275 | 0 | 0 | 11 | 811.2084 | 845.65765 | 882.6857 | 788.45233 | 785.5676 | 738.13995 |
| Itgb2 | ATDDGFHFAGDGK | -263 | -275 | 0 | 0 | 13 | 766.0735 | 768.93036 | 815.9678 | 287.73947 | 944.84143 | 908.20154 |
| Itgb2 | DCIQSGPGCSWCQK | -36 | -49 | 0 | 0 | 14 | 710.7151 | 719.5285 | 736.0687 | 1014.77997 | 1058.1478 | 1073.0022 |
| Itgb2 | SAVGELSDDSSNVVQLIK | -341 | -358 | 0 | 0 | 18 | 594.6287 | 711.06006 | 647.4685 | 789.5804 | 935.3343 | 955.73065 |
| Itgb2 | YCECDNVNCER | -534 | -544 | 0 | 0 | 11 | 565.4325 | 592.05164 | 695.83264 | 633.0463 | 707.4592 | 714.6986 |
| Itgb2 | VAACPEEIGWR | -244 | -254 | 0 | 0 | 11 | 459.18546 | 147.26056 | 1243.7766 | 1433.6504 | 1508.0284 | 259.96268 |
| Itgb2 | SDDSSNVVQLIK | -347 | -358 | 0 | 0 | 12 | 444.72406 | 339.40955 | 417.7906 | 428.97525 | 315.9412 | 571.23303 |
| Itgb2 | VFATDDGFHFAGDGK | -261 | -275 | 0 | 0 | 15 | 419.98273 | 738.787 | 348.50583 | 771.4339 | 796.2363 | 990.8866 |
| Itgb2 | NIYNIHVEDSLECVK | -685 | -699 | 0 | 0 | 15 | 391.86597 | 667.11176 | 554.95905 | 579.04175 | 873.5469 | 468.57083 |
| Itgb2 | ACQPPFAFR | -198 | -206 | 0 | 0.001392869 | 9 | 1053.7289 | 1200.5098 | 1412.1835 | 1204.3092 | NaN | 1307.1237 |
| Itgb2 | SRGDCDGVQINNPVTFQ | -397 | -413 | 1 | 0.00111006 | 17 | 611.58325 | 613.14496 | 606.81964 | 554.7018 | 754.2764 | NaN |
| Itgb2 | EKLTEIIPK | -332 | -340 | 1 | 0 | 9 | 981.22107 | NaN | 1033.4574 | 129.18636 | 1242.6348 | NaN |
| Itgb2 | SQWNNDNPLFK | -747 | -757 | 0 | 0 | 11 | 880.7286 | NaN | NaN | 881.10547 | 1320.016 | 789.4835 |
| Itgb2 | KLGGDLLQAL | -149 | -158 | 1 | 0.003299042 | 10 | 424.77014 | 539.0946 | 553.00714 | 497.5007 | NaN | NaN |
| Itgb2 | DNSNQFQTEVGK | -213 | -224 | 0 | 0 | 12 | 2257.2961 | NaN | NaN | NaN | 671.2743 | 1098.0033 |
| Itgb2 | SESNIQPIFAVTK | -313 | -325 | 0 | 0 | 13 | 1032.212 | NaN | NaN | 1492.6935 | NaN | NaN |

| | | | | | | | | | | | | |
|---|---|---|---|---|---|---|---|---|---|---|---|---|
| Itgb2 | GCPADDIMDPR | -73 | -83 | 0 | 0 | 11 | 552.9411 | NaN | NaN | NaN | NaN | NaN |
| Prkcsh | SKFEEVER | -368 | -375 | 1 | 0 | 8 | 3062.4207 | 2568.0686 | 662.55023 | 523.21875 | 1967.5686 | 545.1276 |
| Prkcsh | GTWGSWAGPDHDK | -440 | -452 | 0 | 0 | 13 | 2820.9565 | 2900.7505 | 2801.9143 | 3430.3284 | 391.20578 | 3170.211 |
| Prkcsh | SLGTWGSWAGPDHDK | -438 | -452 | 0 | 0 | 15 | 913.20526 | 948.7441 | 760.8669 | 897.1846 | 875.58325 | 877.2975 |
| Prkcsh | ACPEPPPEAPSDGDHDEL | -504 | -521 | 0 | 0 | 18 | 732.45276 | 837.61816 | 737.1516 | 1165.3326 | 1192.5513 | 1084.9128 |
| Prkcsh | KYEQGTGCWQGPNR | -457 | -470 | 1 | 0 | 14 | 702.43134 | 856.4432 | 866.26575 | 754.14075 | 1225.0106 | 875.71606 |
| Prkcsh | KYEQGTGCWQGPNR | -457 | -470 | 1 | 0 | 14 | 702.43134 | 856.4432 | 866.26575 | 754.14075 | 1225.0106 | 875.71606 |
| Prkcsh | SLEDQVETLR | -168 | -177 | 0 | 0 | 10 | 849.96533 | 882.38367 | 700.3869 | NaN | 1400.7583 | 1374.1232 |
| Prkcsh | ETVVTSTTEPSR | -481 | -492 | 0 | 0 | 12 | 1194.9868 | 1081.6597 | 592.8285 | 558.6876 | NaN | NaN |
| Prkcsh | EKESLQQLAEVTR | -123 | -135 | 1 | 0 | 13 | 437.34528 | NaN | 235.34698 | 658.00073 | 744.19775 | NaN |
| Prkcsh | ATIPFDQVNDDYCDCK | -44 | -59 | 0 | 0 | 16 | 239.69809 | 569.0646 | 350.90286 | 110.891716 | NaN | NaN |
| Prkcsh | SEVPPTDIPVPEETEPK | -282 | -298 | 0 | 0.006472924 | 17 | 983.7722 | NaN | NaN | 561.6316 | NaN | 1832.1367 |
| Prkcsh | EKESLQQLAEVT | -123 | -134 | 1 | 0.002329279 | 12 | 2456.4536 | NaN | NaN | NaN | NaN | NaN |
| Rpa2 | NQVLNLIK | -210 | -217 | 0 | 0.00025015 | 8 | 2397.0762 | 1600.2277 | 1460.0132 | 1168.6365 | 2112.5781 | 793.6728 |
| Rpa2 | ACPRPEGLNFQDLR | -218 | -231 | 1 | 0 | 14 | 841.02246 | 870.564 | 810.54 | 929.37384 | 1779.9359 | 842.9836 |
| Rpa2 | APTNIVYK | -86 | -93 | 0 | 0 | 8 | 725.9918 | 757.1088 | 728.4479 | 811.6413 | 916.2322 | 864.76886 |
| Rpa2 | VDTDDASGENAVVPPETYVK | -108 | -127 | 0 | 0 | 20 | 676.6596 | 1187.8682 | 604.8849 | 995.31006 | 823.98584 | 1079.776 |
| Rpa2 | STVDDDHFK | -257 | -265 | 0 | 0 | 9 | 469.24002 | 143.72293 | 841.39484 | 296.62265 | 932.3706 | 702.23785 |
| Rpa2 | QAVDFLCNEGHIY | -244 | -256 | 0 | 0 | 13 | 435.586 | 592.04297 | 544.27966 | 506.35776 | 748.6355 | 696.19965 |
| Rpa2 | TVVQNQVLNLIK | -206 | -217 | 0 | 0 | 12 | 261.43137 | 259.69403 | NaN | 347.93454 | 329.90573 | 351.00613 |
| Rpa2 | SPGGFGSPTPSQAEK | -23 | -37 | 0 | 0 | 15 | 4803.63 | 2265.2986 | 1633.6667 | NaN | NaN | 4035.1228 |
| Rpa2 | SQLQHMPVPSIK | -232 | -243 | 0 | 0.002099344 | 12 | 2282.6653 | 590.9878 | NaN | 563.90485 | 484.0957 | NaN |
| Rpa2 | LTDEVFR | -59 | -65 | 0 | 0.00383302 | 7 | 1483.9473 | NaN | NaN | NaN | NaN | NaN |
| Hnrnpm | QGGGGAGGSVPGIER | -388 | -402 | 0 | 0 | 15 | 2833.4785 | 2205.229 | 2300.4778 | 2247.288 | 379.23822 | 2462.6597 |
| Hnrnpm | ADILEDKDGK | -232 | -241 | 1 | 0 | 10 | 2783.5059 | 3073.3413 | 573.5895 | 2943.893 | 2885.824 | 2932.68 |
| Hnrnpm | DKFNECGHVL | -670 | -679 | 1 | 0.001055699 | 10 | 1656.0957 | 1824.3556 | 1809.0896 | 1946.4592 | 213.1174 | 1988.7625 |
| Hnrnpm | HSLSGRPL | -134 | -141 | 1 | 0.00025015 | 8 | 1471.7158 | 979.4651 | 1395.13 | 907.0254 | 1145.5443 | 1481.5212 |
| Hnrnpm | GLGPGGQPIDANHLSK | -306 | -321 | 0 | 0 | 16 | 1251.5967 | 1365.4226 | 1349.0887 | 1209.8625 | 1382.4088 | 2064.2031 |
| Hnrnpm | GLGPGGQPIDANHLSK | -306 | -321 | 0 | 0 | 16 | 1251.5967 | 1365.4226 | 1349.0887 | 1209.8625 | 1382.4088 | 2064.2031 |
| Hnrnpm | ITNIPFDVK | -74 | -82 | 0 | 0 | 9 | 1062.2908 | 984.25507 | 1153.4633 | 1363.8252 | 1514.9348 | 381.6175 |
| Hnrnpm | FNECGHVL | -672 | -679 | 0 | 0.00025015 | 8 | 706.80493 | 745.7639 | 641.7753 | 977.62 | 831.5482 | 165.80309 |
| Hnrnpm | GGAGGHAPGVAR | -638 | -649 | 0 | 0.00025015 | 12 | 630.8648 | 635.3589 | 789.31714 | 482.79404 | 601.1801 | 641.83276 |
| Hnrnpm | GGAGGHAPGVAR | -638 | -649 | 0 | 0.00025015 | 12 | 630.8648 | 635.3589 | 789.31714 | 482.79404 | 601.1801 | 641.83276 |
| Hnrnpm | VKEDPDGEHAR | -143 | -153 | 1 | 0 | 11 | 44.385902 | 444.77365 | 386.0817 | 304.6574 | 459.5199 | 401.82822 |
| Hnrnpm | VKEDPDGEHAR | -143 | -153 | 1 | 0 | 11 | 44.385902 | 444.77365 | 386.0817 | 304.6574 | 459.5199 | 401.82822 |
| Hnrnpm | NLPFDFTWK | -658 | -666 | 0 | 0 | 9 | 97.47554 | 1110.7277 | 1304.3423 | 1350.862 | 1628.104 | 1683.949 |
| Hnrnpm | NLPFDFTWK | -658 | -666 | 0 | 0 | 9 | 97.47554 | 1110.7277 | 1304.3423 | 1350.862 | 1628.104 | 1683.949 |
| Hnrnpm | DRVGSEIER | -427 | -435 | 1 | 0.001392869 | 9 | 2090.0647 | 1831.7869 | NaN | 1578.1962 | 1430.5886 | 541.0741 |
| Hnrnpm | GGGGAGGSVPGIER | -389 | -402 | 0 | 0 | 14 | 1479.2037 | 1395.5093 | NaN | 2454.0862 | 527.4055 | 2602.9087 |
| Hnrnpm | VGQTIER | -503 | -509 | 0 | 0.001392869 | 7 | 1158.8324 | 595.203 | 1266.3605 | 702.9376 | NaN | 455.17 |
| Hnrnpm | FESPEVAER | -698 | -706 | 0 | 0 | 9 | 3258.901 | 3081.1882 | NaN | 605.9622 | NaN | 1711.3403 |
| Hnrnpm | ACQIFVR | -651 | -657 | 0 | 0 | 7 | 1280.6576 | 1548.3837 | 1742.5537 | 1747.3269 | NaN | NaN |
| Hnrnpm | AFITNIPFDVK | -72 | -82 | 0 | 0 | 11 | 848.31836 | 830.7849 | 1812.6193 | 1552.7891 | NaN | NaN |
| Hnrnpm | GGAGGASFDR | -611 | -620 | 0 | 0.00051306 | 10 | 630.86035 | 1049.8013 | NaN | NaN | 1297.842 | 617.22406 |
| Rps3a | TTDGYLLR | -129 | -136 | 0 | 0 | 8 | 6209.138 | 6308.824 | 5784.793 | 5099.5225 | 4004.4414 | 4018.409 |
| Rps3a | RADGYEPPVQESV | -252 | -264 | 1 | 0 | 13 | 3420.6477 | 2635.0344 | 2437.8577 | 4369.603 | 1010.9949 | 2322.5664 |
| Rps3a | ACQSIYPLHDVFVR | -200 | -213 | 0 | 0 | 14 | 2797.942 | 3659.4456 | 3781.1406 | 3622.83 | 4418.1807 | 4434.526 |
| Rps3a | ACQSIYPLHDVFVR | -200 | -213 | 0 | 0 | 14 | 2797.942 | 3659.4456 | 3781.1406 | 3622.83 | 4418.1807 | 4434.526 |
| Rps3a | DLQNDEVAFR | -73 | -82 | 0 | 0 | 10 | 2595.522 | 383.48123 | 3277.816 | 469.71457 | 252.60207 | 539.86145 |
| Rps3a | SIYPLHDVFVR | -203 | -213 | 0 | 0 | 11 | 2408.0298 | 2698.3762 | 2892.7085 | 2325.2598 | 3648.198 | 3442.0686 |
| Rps3a | SIYPLHDVFVR | -203 | -213 | 0 | 0 | 11 | 2408.0298 | 2698.3762 | 2892.7085 | 2325.2598 | 3648.198 | 3442.0686 |
| Rps3a | IYPLHDVFVR | -204 | -213 | 0 | 0 | 10 | 1380.0002 | 1503.6251 | 1584.1267 | 1676.9829 | 1815.0383 | 2089.1633 |
| Rps3a | IYPLHDVFVR | -204 | -213 | 0 | 0 | 10 | 1380.0002 | 1503.6251 | 1584.1267 | 1676.9829 | 1815.0383 | 2089.1633 |
| Rps3a | EVSLADLQNDEVAFR | -68 | -82 | 0 | 0 | 15 | 1842.4065 | 2000.0525 | 2105.2334 | 1912.9187 | 2865.7632 | 2790.723 |
| Rps3a | CQSIYPLHDVFVR | -201 | -213 | 0 | 0 | 13 | 1658.6506 | 1725.808 | 1839.1521 | 2942.5056 | 472.707 | 3344.0706 |
| Rps3a | ADLQNDEVAFR | -72 | -82 | 0 | 0 | 11 | 1570.7389 | 1230.883 | 1141.2385 | 2212.4707 | 1643.2206 | 1617.3188 |
| Rps3a | ADGYEPPVQESV | -253 | -264 | 0 | 0 | 12 | 390.30328 | 374.01212 | 344.15704 | 363.57428 | 434.17896 | 456.22177 |
| Rps3a | ADGYEPPVQESV | -253 | -264 | 0 | 0 | 12 | 390.30328 | 374.01212 | 344.15704 | 363.57428 | 434.17896 | 456.22177 |
| Rps3a | QSIYPLHDVFVR | -202 | -213 | 0 | 0 | 12 | 1200.3134 | 1799.1321 | 1767.8813 | 1767.0765 | 1836.5801 | 580.2495 |
| Rps3a | QSIYPLHDVFVR | -202 | -213 | 0 | 0 | 12 | 1200.3134 | 1799.1321 | 1767.8813 | 1767.0765 | 1836.5801 | 580.2495 |
| Rps3a | KDWYDVK | -28 | -34 | 1 | 0 | 7 | 1044.3679 | 1185.2798 | 941.03424 | 1255.3608 | 1602.435 | 1671.719 |
| Rps3a | LADLQNDEVAFR | -71 | -82 | 0 | 0 | 12 | 388.83014 | 4055.749 | 3753.8772 | 827.2735 | 3409.5552 | 3245.3452 |
| Rps3a | YPLHDVFVR | -205 | -213 | 0 | 0.002972566 | 9 | 196.0215 | 491.14432 | 377.47186 | 1459.7378 | 442.77914 | 971.28625 |
| Rps3a | APAMFNIR | -35 | -42 | 0 | 0.000627943 | 8 | 1742.959 | NaN | 824.1563 | 1217.8773 | 1275.3938 | 1130.4452 |
| Rps3a | SLADLQNDEVAFR | -70 | -82 | 0 | 0 | 13 | 1065.5212 | 1058.7574 | 1508.1614 | 435.8597 | 1145.0519 | NaN |
| Rps3a | NCLTNFHGM | -95 | -103 | 0 | 0.000230707 | 9 | 739.9863 | 283.37604 | 675.1662 | 772.3453 | 754.1157 | NaN |
| Rps3a | LFCVGFTK | -137 | -144 | 0 | 0 | 8 | 1279.7457 | 1383.8134 | 1431.201 | 1288.314 | NaN | NaN |
| Rps3a | LITEDVQGK | -86 | -94 | 0 | 0.00051306 | 9 | 1135.7468 | 1289.905 | 1355.7922 | 874.71686 | NaN | NaN |
| Rps3a | KVVDPFSK | -20 | -27 | 1 | 0.000712251 | 8 | 404.95712 | 553.448 | 724.1492 | 731.3213 | NaN | NaN |
| Rps3a | LMELHGEGGSSGK | -228 | -240 | 0 | 0.00025015 | 13 | 1245.3954 | NaN | NaN | NaN | NaN | NaN |
| Fasn | TLEAVQDLLEQGR | -430 | -442 | 0 | 0 | 13 | 22206.79 | 20111.639 | 20353.125 | 19826.139 | 17359.053 | 18227.049 |
| Fasn | VGDPQELNGITR | -299 | -310 | 0 | 0 | 12 | 3553.2332 | 2608.5027 | 4524.559 | 424.29114 | 4139.617 | 355.2686 |
| Fasn | VLEALLPLK | -2376 | -2384 | 0 | 0 | 9 | 2553.3313 | 3148.9978 | 3452.3958 | 3665.8496 | 5258.575 | 850.99805 |
| Fasn | AIPQEKPIFL | -1400 | -1409 | 1 | 0 | 10 | 1968.6361 | 2445.5435 | 205.05554 | 2450.742 | 2948.5237 | 3904.5576 |
| Fasn | LQQHDVAQGQ | -1282 | -1291 | 0 | 0 | 10 | 1898.2307 | 2067.5454 | 2070.5261 | 254.66069 | 2382.5693 | 1229.5284 |
| Fasn | LFDHPEVPTPPESA | -968 | -981 | 0 | 0 | 14 | 1781.3567 | 1992.6978 | 1852.047 | 742.9203 | 3087.5388 | 1943.696 |
| Fasn | VSVHIIEGDHR | -2465 | -2475 | 0 | 0 | 11 | 1741.7089 | 1633.0687 | 1173.752 | 911.4767 | 921.55896 | 1065.7461 |
| Fasn | ACVDTALENLSTLK | -1221 | -1234 | 0 | 0 | 14 | 1672.3462 | 537.52435 | 492.58582 | 3200.9243 | 115.99004 | 146.91327 |

| | | | | | | | | | | | | |
|---|---|---|---|---|---|---|---|---|---|---|---|---|
| Fasn | RLTQGEVYK | -985 | -993 | 1 | 0.000869338 | 9 | 1583.8008 | 828.93884 | 1117.6182 | 1260.6573 | 845.4252 | 952.5669 |
| Fasn | RQQEQLVPTLEK | -1106 | -1117 | 1 | 0.00025015 | 12 | 1564.8358 | 1730.565 | 2140.4343 | 1563.6127 | 1971.7885 | 2035.7241 |
| Fasn | ALFPPVEFPAPR | -814 | -825 | 0 | 0 | 12 | 1498.3612 | 897.03467 | 880.7801 | 1570.8741 | 596.2055 | 1777.733 |
| Fasn | AIYIDPATHR | -1055 | -1064 | 0 | 0 | 10 | 1436.415 | 1580.3062 | 1292.607 | 1928.0444 | 762.80994 | 1901.3707 |
| Fasn | DTSFEQHVLLHTGGK | -1718 | -1732 | 0 | 0 | 15 | 1275.7231 | 512.17053 | 527.5733 | 1253.4263 | 1634.411 | 812.4208 |
| Fasn | DTSFEQHVLLHTGGK | -1718 | -1732 | 0 | 0 | 15 | 1275.7231 | 512.17053 | 527.5733 | 1253.4263 | 1634.411 | 812.4208 |
| Fasn | DTSFQWVDSLK | -1413 | -1423 | 0 | 0 | 11 | 1209.2644 | 1340.3425 | 1422.0662 | 1436.6332 | 1516.8055 | 141.07263 |
| Fasn | VVDRPLPVR | -376 | -384 | 1 | 0.003037016 | 9 | 1175.7933 | 1214.341 | 1273.1844 | 902.815 | 1056.4355 | 1062.5559 |
| Fasn | GGLGGFGLELAR | -1884 | -1895 | 0 | 0 | 12 | 1133.7167 | 1311.3737 | 1279.164 | 893.1696 | 1219.3308 | 1310.1252 |
| Fasn | LFDHPEVPTPPESASVS | -968 | -984 | 0 | 0 | 17 | 931.68024 | 921.5283 | 915.8661 | 1045.7883 | 1264.932 | 1266.9152 |
| Fasn | EAVQDLLEQGR | -432 | -442 | 0 | 0 | 11 | 877.6974 | 507.19525 | 4405.513 | 4419.644 | 958.81537 | 4517.1353 |
| Fasn | SQEEWESLFSR | -1367 | -1377 | 0 | 0.000471409 | 11 | 805.8619 | 806.0723 | 781.99005 | 1038.6382 | 1133.2811 | 1033.2317 |
| Fasn | SIDASPESPDHYLVDHCIDGR | -863 | -883 | 0 | 0 | 21 | 801.98047 | 890.86224 | 1093.3408 | 336.66168 | 1182.1082 | 1145.7804 |
| Fasn | TFDDIVHAF | -545 | -553 | 0 | 0 | 9 | 761.6452 | 797.27216 | 152.3013 | 807.7651 | 1044.5953 | 105.660095 |
| Fasn | ALHLVGLK | -1379 | -1386 | 0 | 0.000550307 | 8 | 706.18225 | 719.9179 | 768.2145 | 391.1339 | 474.68222 | 548.91254 |
| Fasn | VDTALENLSTLK | -1223 | -1234 | 0 | 0 | 12 | 605.5182 | 797.19763 | 866.32245 | 2442.0657 | 289.52222 | 1374.365 |
| Fasn | NVHVILQPNTR | -399 | -409 | 0 | 0 | 11 | 599.4809 | 521.39954 | 479.74738 | 388.52737 | 529.1032 | 408.35904 |
| Fasn | LQVVDRPLPVR | -374 | -384 | 1 | 0.000627943 | 11 | 553.80786 | 364.41592 | 679.61884 | 760.42676 | 802.2886 | 771.7093 |
| Fasn | LQVVDRPLPVR | -374 | -384 | 1 | 0.000627943 | 11 | 553.80786 | 364.41592 | 679.61884 | 760.42676 | 802.2886 | 771.7093 |
| Fasn | LDPGSPELQQVLK | -1479 | -1491 | 0 | 0 | 13 | 485.34116 | 1211.6075 | 1218.496 | 1201.8691 | 1628.3384 | 1745.9185 |
| Fasn | GINVNPNALFPPVEFPAPR | -807 | -825 | 0 | 0 | 19 | 296.6164 | 407.04996 | 425.3016 | 428.76752 | 428.7154 | 428.57794 |
| Fasn | GQEDPPQHGLPR | -1164 | -1175 | 0 | 0 | 12 | 280.88474 | 583.5149 | 584.15326 | 688.3472 | 853.4003 | 785.31573 |
| Fasn | DYGPQFQGICEATLEGEQGK | -1001 | -1020 | 0 | 0 | 20 | 188.00676 | 124.02855 | 200.61082 | 277.46738 | 281.36838 | 316.3207 |
| Fasn | VSDLLLSTDER | -534 | -544 | 0 | 0 | 11 | 1488.6866 | 1629.5637 | 1679.0457 | 403.3051 | NaN | 1600.211 |
| Fasn | RHDGLPGLAVQW | -2042 | -2053 | 1 | 0.000591124 | 12 | 876.81665 | NaN | 1235.6746 | 1896.6237 | 1285.4346 | 844.1877 |
| Fasn | DTSFEQHVL | -1718 | -1726 | 0 | 0 | 9 | 871.72974 | 943.3479 | 1077.8743 | 1010.11285 | NaN | 268.96765 |
| Fasn | DHKDNLEFFLT | -788 | -798 | 1 | 0 | 11 | 687.67834 | 812.62305 | 885.66125 | NaN | 978.55176 | 931.58673 |
| Fasn | SVEDTSFQWVDSLK | -1410 | -1423 | 0 | 0 | 14 | 637.28455 | 581.8365 | NaN | 236.24995 | 737.1435 | 753.74414 |
| Fasn | FEEANDSWR | -1793 | -1801 | 0 | 0.000230707 | 9 | 598.837 | 514.46967 | 385.41415 | 379.39645 | NaN | 508.40604 |
| Fasn | VHIIEGDHR | -2467 | -2475 | 0 | 0.00025015 | 9 | 279.10883 | 375.20297 | 543.4665 | NaN | 841.5229 | 572.3728 |
| Fasn | HTQPSSSGAQLC | -1546 | -1557 | 0 | 0.003299042 | 12 | 193.37132 | 595.2079 | 1013.08344 | 292.50583 | 133.37456 | NaN |
| Fasn | SNLSNTSHAPK | -1468 | -1478 | 0 | 0 | 11 | 76.62761 | NaN | 749.21466 | 194.55673 | 622.2014 | 159.82605 |
| Fasn | QAPLLIGSTK | -317 | -326 | 0 | 0 | 10 | 2714.959 | 2793.9595 | NaN | NaN | 2141.0205 | 2167.4207 |
| Fasn | ASPESPDHYLVDHCIDGR | -866 | -883 | 0 | 0 | 18 | 311.40607 | 426.10184 | 1093.8391 | NaN | NaN | 1344.9927 |
| Fasn | PPQHGLPR | -1168 | -1175 | 0 | 0.008828666 | 8 | 699.3809 | 615.0423 | NaN | NaN | NaN | 306.89566 |
| Fasn | NQTPELFQDVNKPK | -1975 | -1988 | 1 | 0 | 14 | 678.74835 | 928.1531 | 978.6748 | NaN | NaN | NaN |
| Fasn | VSTSNVSSLEGAR | -1934 | -1946 | 0 | 0 | 13 | 403.84064 | NaN | NaN | 984.22705 | 1615.4968 | NaN |
| Fasn | ILNAGTNTDGSK | -246 | -257 | 0 | 0 | 12 | 400.51187 | NaN | 279.35297 | 349.21115 | NaN | NaN |
| Fasn | NDTVIGGTLPQR | -2069 | -2080 | 0 | 0 | 12 | 319.69516 | 359.89432 | 657.24884 | NaN | NaN | NaN |
| Fasn | VYLWEDPNSK | -958 | -967 | 0 | 0 | 10 | 666.9818 | NaN | 555.35565 | NaN | NaN | NaN |
| Fasn | TGTVALEVR | -928 | -936 | 0 | 0 | 9 | 398.34882 | NaN | NaN | NaN | NaN | 620.7462 |
| Fasn | VTAIYIDPATHR | -1053 | -1064 | 0 | 0 | 12 | 320.22656 | NaN | NaN | NaN | NaN | NaN |
| Fasn | YNGTLNLDR | -1989 | -1997 | 0 | 0.000627943 | 9 | 144.35509 | NaN | NaN | NaN | NaN | NaN |
| Cct8 | DIALHYANK | -299 | -307 | 0 | 0 | 9 | 12279.205 | 13107.764 | 12139.868 | 13314.991 | 11326.595 | 11625.68 |
| Cct8 | KAHEILPELVCC | -138 | -149 | 1 | 0.000230707 | 12 | 757.56635 | 754.9111 | 758.9022 | 460.34338 | 580.1645 | 642.0982 |
| Cct8 | KAHEILPELVCC | -138 | -149 | 1 | 0.000230707 | 12 | 757.56635 | 754.9111 | 758.9022 | 460.34338 | 580.1645 | 642.0982 |
| Cct8 | QYGSETFLAK | -172 | -181 | 0 | 0.00025015 | 10 | 3552.5803 | 185.23055 | 502.29404 | 3127.0435 | 2668.5293 | 550.37573 |
| Cct8 | LVPGGGATEIELAK | -408 | -421 | 0 | 0 | 14 | 3380.8284 | 3931.506 | 4409.6396 | 4658.7446 | 4104.8926 | 4668.685 |
| Cct8 | RELEVQHPAAK | -74 | -84 | 1 | 0 | 11 | 2532.8142 | 2899.9402 | 3203.931 | 3077.5535 | 4191.645 | 4300.479 |
| Cct8 | RELEVQHPAAK | -74 | -84 | 1 | 0 | 11 | 2532.8142 | 2899.9402 | 3203.931 | 3077.5535 | 4191.645 | 4300.479 |
| Cct8 | AIAGTGANVIVTGGK | -282 | -296 | 0 | 0 | 15 | 1668.4913 | 1910.8767 | 2032.9442 | 1591.1779 | 1827.8202 | 1878.9773 |
| Cct8 | ELEVQHPAAK | -75 | -84 | 0 | 0.000230707 | 10 | 1541.6521 | 1671.7784 | 1747.6118 | 994.6336 | 1433.3809 | 1351.482 |
| Cct8 | RVDQIIMAK | -520 | -528 | 1 | 0.00025015 | 9 | 1453.6902 | 1286.4025 | 1625.7363 | 1083.5283 | 1357.0106 | 1125.8468 |
| Cct8 | IFPDSGNFNVDNIR | -190 | -203 | 0 | 0 | 14 | 1442.6279 | 1934.428 | 1834.4482 | 2005.0768 | 2810.2832 | 2789.296 |
| Cct8 | GETCPGLEQYAIK | -427 | -439 | 0 | 0 | 13 | 1307.6307 | 1230.4968 | 924.80664 | 1681.7002 | 1063.041 | 1210.2682 |
| Cct8 | LYSVHQEGNK | -467 | -476 | 0 | 0 | 10 | 1298.1415 | 1389.8662 | 1400.7767 | 696.349 | 2012.1068 | 2033.7913 |
| Cct8 | DVDEVSSLLR | -156 | -165 | 0 | 0 | 10 | 1221.3617 | 201.844 | 1204.822 | 1301.9185 | 1398.176 | 1378.8718 |
| Cct8 | VYSCPFDGMITETK | -241 | -254 | 0 | 0 | 14 | 1202.1886 | 858.78796 | 1022.6157 | 1245.6028 | 890.4501 | 1048.0195 |
| Cct8 | GSTDNLMDDIER | -379 | -390 | 0 | 0 | 12 | 1093.0972 | 656.12665 | 533.6266 | 1084.8986 | 962.0515 | 707.9555 |
| Cct8 | AHEILPELVCCS | -139 | -150 | 0 | 0 | 12 | 1055.0128 | 584.272 | 995.892 | 680.9662 | 1324.8969 | 1157.8093 |
| Cct8 | VSIFPDSGNFNVDNIR | -188 | -203 | 0 | 0 | 16 | 950.6767 | 937.5099 | 985.1782 | 1006.8473 | 1292.1206 | 1286.0078 |
| Cct8 | LELAEELLR | -112 | -120 | 0 | 0 | 9 | 855.4819 | 885.7822 | 956.5095 | 719.28735 | 731.8994 | 475.99493 |
| Cct8 | HFSGLEEAVYR | -21 | -31 | 0 | 0 | 11 | 501.36322 | 2972.6228 | 3423.666 | 2443.8743 | 1052.8928 | 3203.4897 |
| Cct8 | AHEILPELVCC | -139 | -149 | 0 | 0 | 11 | 368.23315 | 399.905 | 1251.814 | 1044.7239 | 677.26465 | 739.91534 |
| Cct8 | LLELAEELLR | -111 | -120 | 0 | 0.00025015 | 10 | 263.59308 | 273.53818 | 317.64825 | 334.02365 | 299.938 | 412.38968 |
| Cct8 | PAGGPKPPSGK | -529 | -539 | 1 | 0 | 11 | 567.6362 | 545.72736 | 597.6526 | NaN | 69.303665 | 447.2673 |
| Cct8 | PAGGPKPPSGK | -529 | -539 | 1 | 0 | 11 | 567.6362 | 545.72736 | 597.6526 | NaN | 69.303665 | 447.2673 |
| Cct8 | ACVSIFPDSGNFNVDNIR | -186 | -203 | 0 | 0 | 18 | 534.6946 | 574.2978 | NaN | 706.2676 | 863.3044 | 777.4016 |
| Cct8 | AHEILPELVCCSAK | -139 | -152 | 0 | 0 | 14 | 481.00647 | NaN | 537.7353 | 1136.3269 | 1121.257 | 1054.3213 |
| Cct8 | AEAFEAIPR | -442 | -450 | 0 | 0 | 9 | 801.2659 | NaN | 1214.2568 | 1196.5415 | 853.8526 | NaN |
| Cct8 | SILDTYLGK | -496 | -504 | 0 | 0.00025015 | 9 | 565.5927 | 548.39484 | NaN | NaN | 791.0305 | 901.0261 |
| Cct8 | ILDTYLGK | -497 | -504 | 0 | 0.001392869 | 8 | 253.08072 | 638.5627 | 508.35025 | NaN | NaN | 377.27188 |
| Cct8 | AVDDGVNTFK | -391 | -400 | 0 | 0 | 10 | 1461.9498 | NaN | 1641.0707 | 1833.6887 | NaN | NaN |
| Cct8 | EVGDTQVVVFK | -354 | -364 | 0 | 0 | 11 | 346.36914 | 1353.0087 | 1386.6753 | NaN | NaN | NaN |
| Atp1a1 | SPDFTNENPLETR | -228 | -240 | 0 | 0 | 13 | 2863.7986 | 3065.4167 | 3186.997 | 2741.081 | 3575.0588 | 3396.626 |
| Atp1a1 | LSLDELHR | -46 | -53 | 0 | 0.00025015 | 8 | 2270.2122 | 2226.9749 | 2297.442 | 3681.565 | 2696.982 | 2853.0159 |
| Atp1a1 | IVEIPFNSTNK | -477 | -487 | 0 | 0 | 11 | 1846.2715 | 2225.5845 | 2263.5142 | 2533.8252 | 229.79338 | 2700.2173 |
| Atp1a1 | LNIPVNQVNPR | -648 | -658 | 0 | 0 | 11 | 1782.4047 | 1755.8516 | 1914.6809 | 1412.3326 | 1590.6073 | 829.26733 |

| | | | | | | | | | | | | |
|---|---|---|---|---|---|---|---|---|---|---|---|---|
| Atp1a1 | YHTEIVFAR | -684 | -692 | 0 | 0 | 9 | 1721.4248 | 2021.9967 | 2069.7256 | 1853.7001 | 2039.2048 | 842.4623 |
| Atp1a1 | QENLPILK | -437 | -444 | 0 | 0 | 8 | 1326.979 | 1483.8136 | 1241.4447 | 1217.614 | 1009.3032 | 1203.6979 |
| Atp1a1 | AVAGDASESALLK | -446 | -458 | 0 | 0 | 13 | 1142.1646 | 1147.7495 | 198.05829 | 1071.969 | 1137.497 | 1270.3303 |
| Atp1a1 | AEDVVVGDLVEVK | -182 | -194 | 0 | 0 | 13 | 912.9396 | 2370.0945 | 822.685 | 1958.4395 | 3366.4368 | 3118.7788 |
| Atp1a1 | ANQENLPILK | -435 | -444 | 0 | 0 | 10 | 875.72626 | 450.62045 | 374.27966 | 640.51556 | 569.03766 | 780.7501 |
| Atp1a1 | EQPLDEELKDAFQN | -527 | -540 | 1 | 0 | 14 | 817.023 | 1011.46375 | 1100.0967 | 1184.6437 | 1330.6976 | 1381.219 |
| Atp1a1 | IHEADTTENQSGVSFDK | -397 | -413 | 0 | 0 | 17 | 808.36273 | 848.2587 | 883.9645 | 130.15955 | 929.6595 | 532.43414 |
| Atp1a1 | DMTSEELDDILR | -672 | -683 | 0 | 0 | 12 | 787.3196 | 625.2293 | 915.09485 | 903.10297 | 903.9317 | 617.2977 |
| Atp1a1 | NIPVNQVNPR | -649 | -658 | 0 | 0.001023236 | 10 | 955.8482 | 945.9941 | 2343.77 | NaN | NaN | NaN |
| Atp1a1 | YQLSIHK | -488 | -494 | 0 | 0.000347792 | 7 | 696.6491 | NaN | NaN | 283.61804 | 628.8306 | NaN |
| Atp1a1 | DTTENQSGVSFDK | -401 | -413 | 0 | 0 | 13 | 222.27356 | 567.86865 | 649.17804 | NaN | NaN | NaN |
| Atp1a1 | IFVYDEVR | -998 | -1005 | 0 | 0.00025015 | 8 | 386.23395 | NaN | NaN | NaN | NaN | NaN |
| Rps3 | ELNEFLTR | -20 | -27 | 0 | 0 | 8 | 16662.287 | 13424.868 | 14269.5 | 11050.086 | 8917.576 | 18583.713 |
| Rps3 | ELAEDGYSGVEVR | -28 | -40 | 0 | 0 | 13 | 6716.1436 | 6801.9595 | 6962.604 | 8075.7236 | 8447.136 | 8442.691 |
| Rps3 | KPLPDHVSIVEPK | -202 | -214 | 1 | 0 | 13 | 5402.5186 | 6017.545 | 6294.9214 | 6544.2607 | 7678.601 | 7724.344 |
| Rps3 | KPLPDHVSIVEPK | -202 | -214 | 1 | 0 | 13 | 5402.5186 | 6017.545 | 6294.9214 | 6544.2607 | 7678.601 | 7724.344 |
| Rps3 | GGKPEPPAMPQPVPTA | -228 | -243 | 1 | 0 | 16 | 4839.9146 | 3219.4673 | 4580.8984 | 5596.0347 | 3823.6714 | 5453.6743 |
| Rps3 | LNEFLTR | -21 | -27 | 0 | 0.000230707 | 7 | 4277.292 | 3519.3118 | 3636.49 | 2452.667 | 1595.8573 | 2491.4707 |
| Rps3 | IMLPWDPSGK | -188 | -197 | 0 | 0 | 10 | 2894.9197 | 3455.8389 | 3474.6477 | 3698.2107 | 4521.383 | 4443.0806 |
| Rps3 | IMLPWDPSGK | -188 | -197 | 0 | 0 | 10 | 2894.9197 | 3455.8389 | 3474.6477 | 3698.2107 | 4521.383 | 4443.0806 |
| Rps3 | DEILPTTPISEQK | -215 | -227 | 0 | 0 | 13 | 2824.8618 | 4918.4653 | 4661.1987 | 12376.306 | 5132.9146 | 4763.3154 |
| Rps3 | KGGKPEPPAMPQPVPTA | -227 | -243 | 2 | 0.00078661 | 17 | 2009.8846 | 2059.3987 | 2148.1904 | 1925.0695 | 2006.9116 | 1959.1193 |
| Rps3 | SGDPVNYYVDTAVR | -160 | -173 | 0 | 0 | 14 | 1904.4003 | 1939.8914 | 1899.4484 | 2714.1943 | 3010.2302 | 2862.849 |
| Rps3 | RFIMESGAK | -124 | -132 | 1 | 0.00025015 | 9 | 1719.1987 | 547.9626 | 1554.0286 | 1156.0962 | 519.6546 | 485.3625 |
| Rps3 | KFVADGIFK | -10 | -18 | 1 | 0 | 9 | 1419.3247 | 1326.09 | 1420.9731 | 9311.525 | 4543.1846 | 403.71207 |
| Rps3 | AEDGYSGVEVR | -30 | -40 | 0 | 0 | 11 | 1185.6588 | 478.8927 | 1045.2081 | 706.3056 | 727.25885 | 526.4194 |
| Rps3 | HVSIVEPK | -207 | -214 | 0 | 0 | 8 | 984.80914 | 1076.1154 | 1118.0576 | 687.46747 | 940.29333 | 1075.4773 |
| Rps3 | RTQNVLGEK | -54 | -62 | 1 | 0.000627943 | 9 | 622.2012 | 919.7366 | 796.94727 | 808.2713 | 781.435 | 453.31097 |
| Rps3 | FGFPEGSVELY | -77 | -87 | 0 | 0 | 11 | 424.21207 | 320.16928 | 298.54718 | 409.9883 | 532.4852 | 526.12537 |
| Rps3 | RFGFPEGSVELY | -76 | -87 | 1 | 0 | 12 | 273.5239 | 242.27863 | 460.53262 | 2323.0989 | 782.6408 | 665.1577 |
| Rps3 | AELNEFLTR | -19 | -27 | 0 | 0 | 9 | 3465.6523 | 3441.503 | 3704.4014 | NaN | 1714.6447 | 1624.0071 |
| Rps3 | RFGFPEGSVE | -76 | -85 | 1 | 0.000627943 | 10 | 1011.3178 | 1084.0929 | 993.4362 | NaN | 380.50214 | 488.64532 |
| Rps3 | TQNVLGEK | -55 | -62 | 0 | 0.00025015 | 8 | 656.24036 | 554.4293 | 502.8501 | 527.91406 | 576.683 | NaN |
| Rps3 | LAEDGYSGVEVR | -29 | -40 | 0 | 0 | 12 | 1068.315 | NaN | NaN | 998.9824 | 1178.9872 | 1127.1874 |
| Rps3 | DEILPTTPISEQ | -215 | -226 | 0 | 0.00025015 | 12 | 786.30835 | 708.45807 | 518.6429 | NaN | 159.18364 | NaN |
| Rps3 | DEILPTTPISEQKG | -215 | -228 | 1 | 0.000230707 | 14 | 591.681 | NaN | 698.1595 | 862.1011 | 631.5821 | NaN |
| Rps3 | FVADGIFK | -11 | -18 | 0 | 0 | 8 | 540.69366 | 490.2246 | 675.4949 | NaN | NaN | NaN |
| Rps3 | ACYGVLR | -118 | -124 | 0 | 0.00111006 | 7 | 3927.991 | 3318.6714 | NaN | NaN | NaN | NaN |
| Api5 | ELPQFATGENLPR | -85 | -97 | 0 | 0 | 13 | 1977.309 | 1850.4158 | 1679.5824 | 1149.926 | 1738.966 | 1814.8873 |
| Api5 | HFPELADSAINAQ | -55 | -67 | 0 | 0 | 13 | 1454.9413 | 2570.391 | 2935.9421 | 1478.8304 | 1713.8917 | 1755.2256 |
| Api5 | KLPDFLTAK | -356 | -364 | 1 | 0 | 9 | 1316.3068 | 1458.2434 | 1449.663 | 1963.5267 | 2135.7852 | 2250.9067 |
| Api5 | ADATEQVGQHK | -15 | -25 | 0 | 0 | 11 | 603.84814 | 106.54563 | 626.80237 | 535.26624 | 537.42584 | 576.51544 |
| Api5 | PTVEELYR | -2 | -9 | 0 | 0.000298805 | 8 | 536.1221 | 461.088 | 388.40103 | 974.54706 | 537.2247 | 1414.8265 |
| Api5 | DAYQVILDGVK | -26 | -36 | 0 | 0 | 11 | 386.50436 | 438.29468 | 676.87067 | 400.74515 | 695.1316 | 917.573 |
| Api5 | LVNNALLSIFK | -116 | -126 | 0 | 0 | 11 | 384.48914 | 401.7717 | 391.71423 | 402.81616 | 457.96307 | 509.8419 |
| Api5 | QADLEQAFSPSDPDCVDR | -220 | -237 | 0 | 0 | 18 | 158.38245 | 397.6267 | NaN | 276.99084 | 556.15515 | 522.1 |
| Api5 | EVEELILTESK | -170 | -180 | 0 | 0 | 11 | 4142.149 | 4449.25 | 4423.131 | 3170.612 | NaN | NaN |
| Api5 | TLPDEVLTK | -161 | -169 | 0 | 0 | 9 | 837.03796 | 908.64136 | 942.72864 | 747.2866 | NaN | NaN |
| Api5 | ILQGEDIVR | -140 | -148 | 0 | 0 | 9 | 1680.7439 | 1703.7086 | NaN | NaN | 1422.3218 | NaN |
| Api5 | QAFSPSDPDCVDR | -225 | -237 | 0 | 0 | 13 | 1083.3094 | NaN | NaN | NaN | 370.3353 | NaN |
| Api5 | ADLEQAFSPSDPDCVDR | -221 | -237 | 0 | 0 | 17 | 304.11304 | NaN | NaN | NaN | 108.386986 | NaN |
| Api5 | SQILQGEDIVR | -138 | -148 | 0 | 0.00078661 | 11 | 7481.194 | NaN | NaN | NaN | NaN | NaN |
| Api5 | TGEALKTEENK | -399 | -409 | 1 | 0.001708008 | 11 | 550.82214 | NaN | NaN | NaN | NaN | NaN |
| Api5 | HFPELADSAINA | -55 | -66 | 0 | 0.004439092 | 12 | 490.2669 | NaN | NaN | NaN | NaN | NaN |
| Api5 | KITNNINVLIK | -416 | -426 | 1 | 0.00025015 | 11 | 440.72293 | NaN | NaN | NaN | NaN | NaN |
| Snd1 | DTPDEPWAFPAR | -72 | -83 | 0 | 0 | 12 | 2496.5698 | 1464.735 | 2106.58 | 2518.5308 | 2765.096 | 2450.4387 |
| Snd1 | RDYVPPTANLDQK | -327 | -339 | 1 | 0.00111006 | 13 | 1926.5575 | 455.01746 | 2092.4453 | 2099.89 | 1836.6913 | 1886.442 |
| Snd1 | KYTIENPR | -177 | -184 | 1 | 0 | 8 | 1846.4763 | 1843.5686 | 1922.9163 | 1659.0452 | 1949.7322 | 2127.4807 |
| Snd1 | SASYKPVFVT | -679 | -688 | 1 | 0.000550307 | 10 | 1659.5392 | 1449.7681 | 1732.181 | 1405.862 | 1286.9497 | 1297.335 |
| Snd1 | IQVPQDEDAR | -801 | -810 | 0 | 0 | 10 | 1105.2747 | 325.18207 | 816.5061 | 2813.917 | 3513.5896 | 3489.1755 |
| Snd1 | SSGSSGGPAVPTVQR | -7 | -21 | 0 | 0 | 15 | 1097.6365 | 815.4463 | 994.5657 | 1433.9358 | 722.5361 | 840.8471 |
| Snd1 | FVDGEWYR | -740 | -747 | 0 | 0 | 8 | 999.3965 | 1809.5131 | 1861.7916 | 1579.201 | 1542.9636 | 1653.0872 |
| Snd1 | NDISSHPPVEGSYAPR | -716 | -731 | 0 | 0 | 16 | 976.7314 | 944.6822 | 926.21423 | 1036.3467 | 969.3042 | 899.03107 |
| Snd1 | NDISSHPPVEGSYAPR | -716 | -731 | 0 | 0 | 16 | 976.7314 | 944.6822 | 926.21423 | 1036.3467 | 969.3042 | 899.03107 |
| Snd1 | QFLPFLQR | -516 | -523 | 0 | 0 | 8 | 799.3474 | 378.68887 | 926.8211 | 474.9955 | 609.3385 | 679.15027 |
| Snd1 | SPATETVPAFSER | -426 | -438 | 0 | 0 | 13 | 718.86035 | 1344.2584 | 1947.4287 | 1246.7068 | 1780.3705 | 1637.0833 |
| Snd1 | PASPATETVPAFSER | -424 | -438 | 0 | 0 | 15 | 584.79266 | 929.6249 | 467.7249 | 518.6414 | 484.39227 | 397.19174 |
| Snd1 | LLAGIECPR | -554 | -562 | 0 | 0 | 9 | 488.66235 | 809.7764 | 478.38312 | 715.35626 | 849.24426 | 749.7223 |
| Snd1 | DIQNTQCLLNVEHLSA | -820 | -835 | 0 | 0 | 16 | 456.24194 | 480.24353 | 602.52905 | 837.38306 | 713.17804 | 679.19214 |
| Snd1 | LSECEEQAK | -149 | -157 | 0 | 0 | 9 | 240.28989 | 288.5794 | 1167.9907 | 1041.2203 | 938.8067 | 1439.6887 |
| Snd1 | GSSGGPAVPTVQR | -9 | -21 | 0 | 0 | 13 | 4191.6636 | 4364.4893 | 4453.5366 | NaN | 6935.415 | 7156.447 |
| Snd1 | LGTLPPAFSTR | -777 | -787 | 0 | 0.000394361 | 11 | 2634.128 | 2482.6606 | 3052.345 | NaN | 2806.693 | 4203.145 |
| Snd1 | QSSGSSGGPAVPTVQR | -6 | -21 | 0 | 0 | 16 | 1447.3975 | 1579.8193 | 1696.3248 | NaN | 1351.3177 | 1224.9397 |
| Snd1 | KFVDGEWYR | -739 | -747 | 1 | 0.00025015 | 9 | 1145.8068 | 1366.2948 | 1518.7906 | NaN | 1726.1976 | 1727.3555 |
| Snd1 | VHFTAER | -630 | -636 | 0 | 0.003166358 | 7 | 1075.2255 | 394.03415 | 1131.0985 | 986.5434 | NaN | 1108.4684 |
| Snd1 | LEGDNIQDK | -378 | -386 | 0 | 0.000712251 | 9 | 2326.323 | NaN | 2094.7837 | 2033.7181 | NaN | 2024.0529 |
| Snd1 | YIRPASPATETVPAFSER | -421 | -438 | 1 | 0 | 18 | 1411.7108 | 331.916 | 1241.2021 | NaN | NaN | 776.89453 |
| Snd1 | ADDADEFGYSR | -900 | -910 | 0 | 0 | 11 | 539.62585 | 591.8966 | 545.1651 | NaN | 595.898 | NaN |

| | | | | | | | | | | | | |
|---|---|---|---|---|---|---|---|---|---|---|---|---|
| Snd1 | SAYYKPLLSAE | -637 | -647 | 1 | 0 | 11 | 393.9382 | 420.644 | NaN | 333.79913 | 427.86786 | NaN |
| Snd1 | DVETGTQLEK | -700 | -709 | 0 | 0 | 10 | 851.9806 | 911.2986 | NaN | 781.20105 | NaN | NaN |
| Snd1 | HFVDSHHQKPVNAIIE | -185 | -200 | 1 | 0 | 16 | 360.3368 | 305.12442 | 471.53903 | NaN | NaN | NaN |
| Snd1 | YIDYGNR | -763 | -769 | 0 | 0.001636916 | 7 | 688.1662 | 1195.076 | NaN | NaN | NaN | NaN |
| Snd1 | RETDGSETPEPFAAE | -233 | -247 | 1 | 0.001811524 | 15 | 812.7158 | NaN | NaN | NaN | NaN | NaN |
| Snd1 | GDVGLGLVK | -849 | -857 | 0 | 0.002099344 | 9 | 565.064 | NaN | NaN | NaN | NaN | NaN |
| Mcm4 | FSNKVEAIDVEEAK | -748 | -761 | 1 | 0 | 14 | 3707.622 | 2866.6472 | 2467.9487 | 2028.5409 | 587.81323 | 1170.411 |
| Mcm4 | QRPDLGSAR | -113 | -121 | 1 | 0 | 9 | 2941.1343 | 3032.6782 | 3057.8186 | 5405.92 | 5185.853 | 5291.377 |
| Mcm4 | THIDVIHYR | -413 | -421 | 0 | 0 | 9 | 1429.6016 | 1500.2538 | 1335.6172 | 2434.8726 | 2045.7559 | 2371.7185 |
| Mcm4 | THIDVIHYR | -413 | -421 | 0 | 0 | 9 | 1429.6016 | 1500.2538 | 1335.6172 | 2434.8726 | 2045.7559 | 2371.7185 |
| Mcm4 | VEAIDVEEAKR | -752 | -762 | 1 | 0 | 11 | 1005.6512 | 974.44324 | 344.47324 | 1184.8583 | 1208.5902 | 1077.1355 |
| Mcm4 | GEPFLNVNCEHIK | -203 | -215 | 0 | 0 | 13 | 988.5563 | 1091.1631 | 1108.6516 | 1407.8418 | 1715.2114 | 1810.4071 |
| Mcm4 | NIQLPHTLLSR | -632 | -642 | 0 | 0.000627943 | 11 | 951.0165 | 944.82477 | 1079.9923 | 1000.96643 | 1101.2809 | 1107.4408 |
| Mcm4 | SDGAAAEDIVPSEQSLGQK | -130 | -148 | 0 | 0 | 19 | 942.7737 | 917.00574 | 924.43475 | 1103.9828 | 1209.7593 | 1314.073 |
| Mcm4 | APSIYEHEDIKK | -466 | -477 | 1 | 0 | 12 | 896.5406 | 1072.3895 | 1036.7649 | 828.8519 | 994.39655 | 946.3515 |
| Mcm4 | YVYNLVPR | -521 | -528 | 0 | 0 | 8 | 797.0064 | 250.71341 | 959.49866 | 859.8016 | 905.5215 | 787.72815 |
| Mcm4 | ADDDFLTVTGK | -847 | -857 | 0 | 0 | 11 | 741.006 | 1269.3351 | 1170.9083 | 1361.4343 | 386.04333 | 1591.7203 |
| Mcm4 | QLFEDIR | -821 | -827 | 0 | 0.00078661 | 7 | 594.6169 | 553.1643 | 523.63007 | 526.82336 | 343.33722 | 631.43835 |
| Mcm4 | TTIENIQLPHTLLSR | -628 | -642 | 0 | 0 | 15 | 473.68713 | 467.2005 | 479.906 | 783.5633 | 641.41516 | 751.38873 |
| Mcm4 | RLHGLDEEAEQK | -427 | -438 | 1 | 0 | 12 | 446.81384 | 343.78015 | 343.82492 | 954.2277 | 649.81146 | 595.5444 |
| Mcm4 | THIDVIHY | -413 | -420 | 0 | 0 | 8 | 1409.7294 | 1577.4835 | 1082.1586 | NaN | 1746.7151 | 268.31656 |
| Mcm4 | LHGLDEEAEQK | -428 | -438 | 0 | 0 | 11 | 1253.2133 | 1363.2085 | 1548.1611 | NaN | 2156.522 | 2147.6177 |
| Mcm4 | LDPQDEAYDR | -650 | -659 | 0 | 0 | 10 | 1249.9332 | 1066.1156 | 904.2127 | NaN | 839.62744 | NaN |
| Mcm4 | LAHHLVSL | -661 | -668 | 0 | 0.00383302 | 8 | 962.59344 | NaN | 502.46106 | NaN | 1167.0852 | 1143.4889 |
| Mcm4 | ANPIESQWNPK | -616 | -626 | 0 | 0 | 11 | 3398.1675 | NaN | NaN | 2586.904 | NaN | NaN |
| Mcm4 | DIVPSEQSLGQK | -137 | -148 | 0 | 0.001055699 | 12 | 1908.1943 | NaN | NaN | NaN | NaN | NaN |
| Mcm4 | AHNDLVDK | -373 | -380 | 0 | 0.003003218 | 8 | 417.62625 | NaN | NaN | NaN | NaN | NaN |
| Samhd1 | KEQVSQLLPEK | -576 | -586 | 1 | 0.000431655 | 11 | 3123.5981 | 3437.7078 | 3387.1746 | 3798.6052 | 3872.5896 | 1063.2377 |
| Samhd1 | RICEVEYK | -371 | -378 | 1 | 0.00025015 | 8 | 2912.013 | 2920.5894 | 3026.612 | 553.8206 | 4294.612 | 4208.3896 |
| Samhd1 | LSADDDFQNTDLR | -63 | -75 | 0 | 0 | 13 | 2881.381 | 3133.4233 | 3299.4065 | 2524.818 | 2746.1846 | 2855.7822 |
| Samhd1 | IIDTPQFQR | -167 | -175 | 0 | 0 | 9 | 2872.2612 | 3002.1843 | 565.5042 | 3095.0603 | 1245.632 | 3722.8872 |
| Samhd1 | KVLDIFR | -95 | -101 | 1 | 0 | 7 | 2736.1494 | 2435.804 | 2431.4995 | 2313.4773 | 2407.1143 | 2281.3381 |
| Samhd1 | QPELQISER | -218 | -226 | 0 | 0 | 9 | 2331.409 | 2100.8909 | 1797.8912 | 2118.7527 | 1159.3198 | 1219.0618 |
| Samhd1 | FAEQLIR | -587 | -593 | 0 | 0 | 7 | 1717.6857 | 1371.3779 | 1551.4072 | 1154.072 | 1092.1826 | 1020.289 |
| Samhd1 | FEHSLGVGYLA | -197 | -207 | 0 | 0 | 11 | 1672.505 | 1959.7203 | 1945.8108 | 206.05392 | 2597.8823 | 2798.307 |
| Samhd1 | GLVPEEDITFIK | -290 | -301 | 0 | 0 | 12 | 1601.5715 | 1793.0399 | 1581.0381 | 336.57907 | 2610.2068 | 2801.6797 |
| Samhd1 | EQIMGPPITPVK | -302 | -313 | 0 | 0 | 12 | 1029.8865 | 204.56725 | 1447.7233 | 1119.7565 | 1428.3912 | 1378.2327 |
| Samhd1 | MFEHLVNSNELK | -272 | -283 | 0 | 0 | 12 | 1000.4876 | 1067.7927 | 1113.1471 | 3096.8975 | 1407.3231 | 1290.6984 |
| Samhd1 | EQVSQLLPEK | -577 | -586 | 0 | 0 | 10 | 988.73315 | 1034.313 | 751.84937 | 11069.542 | 1076.7476 | 941.7648 |
| Samhd1 | NLSADDDFQNTDLR | -62 | -75 | 0 | 0 | 14 | 854.42786 | 898.5396 | 877.1268 | 993.59564 | 1077.8945 | 253.84541 |
| Samhd1 | NYGLVPEEDITFIK | -288 | -301 | 0 | 0 | 14 | 760.23737 | 831.126 | 1606.1661 | 548.3643 | 572.38916 | 454.87146 |
| Samhd1 | WKHEQGSIE | -263 | -271 | 1 | 0.00149795 | 9 | 661.42035 | 1104.4855 | 1316.9316 | 565.98645 | 2550.2793 | 2439.7686 |
| Samhd1 | NGIDVDKWDYFAR | -338 | -350 | 1 | 0 | 13 | 588.49603 | 809.028 | 782.98267 | 864.89276 | 1303.7207 | 1241.8301 |
| Samhd1 | IAGSFLPFLDEDR | -105 | -117 | 0 | 0 | 13 | 519.4738 | 465.10724 | 443.67072 | 475.54303 | 511.24054 | 589.5327 |
| Samhd1 | GGGYYVFPGASHNR | -183 | -196 | 0 | 0 | 14 | 248.98221 | 752.535 | 561.1228 | 475.87775 | 411.0849 | 567.8768 |
| Samhd1 | FEHSLGVGY | -197 | -205 | 0 | 0.00111006 | 9 | 1912.1133 | 1964.2953 | 1861.5184 | 1809.6631 | NaN | 1771.1595 |
| Samhd1 | NDPIHGHIEFHPLLIR | -151 | -166 | 0 | 0 | 16 | 801.6548 | 491.1402 | NaN | 425.58603 | 762.2 | 748.31134 |
| Samhd1 | DFTKPQDGDIIAPLITPL | -619 | -636 | 1 | 0.001092529 | 18 | 378.43506 | 169.04076 | 240.18536 | NaN | 370.61902 | 335.83765 |
| Samhd1 | FLPFLDEDR | -109 | -117 | 0 | 0.00025015 | 9 | 376.7587 | 263.10025 | 487.3603 | 692.4106 | 697.5619 | NaN |
| Samhd1 | TWEPEDVCSF | -76 | -85 | 0 | 0.000230707 | 10 | 205.38637 | 253.54245 | NaN | 280.00198 | 275.12363 | 247.47324 |
| Samhd1 | VINVDYGMEDKNPIDR | -544 | -559 | 1 | 0 | 16 | 4059.3923 | NaN | 5686.298 | NaN | 4107.844 | 4131.723 |
| Samhd1 | ADDDFQNTDLR | -65 | -75 | 0 | 0.00025015 | 11 | 1760.4854 | 1935.2296 | 1867.5854 | NaN | 1479.9762 | NaN |
| Samhd1 | VFPGASHNR | -188 | -196 | 0 | 0.003299042 | 9 | 1430.5165 | 1456.8813 | 1839.6456 | 1528.5127 | NaN | NaN |
| Samhd1 | DFTKPQDGDIIAPL | -619 | -632 | 1 | 0.000550307 | 14 | 1160.5862 | NaN | NaN | 3538.4592 | 4374.691 | 2721.0051 |
| Samhd1 | EHLVNSNELK | -274 | -283 | 0 | 0 | 10 | 1064.0391 | NaN | NaN | 93.1207 | 977.8998 | 1243.3599 |
| Samhd1 | HSTDPQLSEAQSILR | -475 | -489 | 0 | 0 | 15 | 714.1049 | NaN | 882.8703 | 918.53046 | NaN | 1396.972 |
| Samhd1 | SADDDFQNTDLR | -64 | -75 | 0 | 0.000431655 | 12 | 691.9247 | 645.17224 | NaN | 963.0182 | 843.04205 | NaN |
| Samhd1 | TWEPEDVCSFLENR | -76 | -89 | 0 | 0 | 14 | 669.7 | 758.6984 | NaN | NaN | 950.47754 | 924.10315 |
| Samhd1 | QVSQLLPEK | -578 | -586 | 0 | 0 | 9 | 454.156 | 943.6521 | 1609.8381 | 409.7466 | NaN | NaN |
| Samhd1 | NYGLVPEEDITFIKE | -288 | -302 | 1 | 0 | 15 | 352.43756 | 340.6013 | NaN | 258.3934 | NaN | NaN |
| Samhd1 | GIQNNFDYK | -356 | -364 | 0 | 0.003966045 | 9 | 195.56688 | NaN | NaN | NaN | NaN | NaN |
| Pds5a | VINDECYQVR | -919 | -928 | 0 | 0 | 10 | 2659.3276 | 783.2399 | 954.61365 | 995.12756 | 915.7831 | 959.3205 |
| Pds5a | YLVPHNLETEER | -471 | -482 | 0 | 0 | 12 | 1396.0172 | 1101.3757 | 1210.1676 | 494.42322 | 1353.3806 | 1067.2511 |
| Pds5a | LWSPDEEVSPEVLAK | -820 | -834 | 0 | 0 | 15 | 1088.4103 | 1198.2908 | 1291.6892 | 1604.2279 | 1918.1499 | 1842.4094 |
| Pds5a | VDESGPPAPSKPR | -1255 | -1267 | 1 | 0 | 13 | 780.0659 | 787.8904 | 707.90643 | 626.7326 | 868.4102 | 814.4229 |
| Pds5a | TLFSVINNSHNTK | -169 | -181 | 0 | 0 | 13 | 768.7083 | 874.14703 | 987.7092 | 917.4802 | 1417.155 | 1134.225 |
| Pds5a | IETDLPQIR | -708 | -716 | 0 | 0 | 9 | 387.32675 | 520.2604 | 2146.0728 | 1424.457 | 1896.6326 | 1860.9954 |
| Pds5a | GAENIQKPDEK | -1244 | -1254 | 1 | 0 | 11 | 384.25885 | 554.3764 | 659.45404 | 448.9909 | 589.68304 | 719.49133 |
| Pds5a | FNDIHVPVR | -331 | -339 | 0 | 0 | 9 | 1824.1958 | 1958.0098 | 2234.0015 | 1633.3676 | NaN | 2227.057 |
| Pds5a | EQSSEASETGVSENEENPVR | -1183 | -1202 | 0 | 0 | 20 | 207.67064 | 308.25668 | 277.0456 | NaN | 360.69287 | 258.4087 |
| Pds5a | ADDEEEGVSPDSAIR | -642 | -656 | 0 | 0 | 15 | 705.5784 | NaN | 584.36993 | NaN | 327.82303 | NaN |
| Pds5a | ELLDLHK | -517 | -523 | 0 | 0.009541249 | 7 | 617.13635 | NaN | NaN | NaN | 939.9586 | 1021.74365 |
| Ppia | FEDENFILK | -83 | -91 | 0 | 0 | 9 | 16538.654 | 17400.135 | 462.27725 | 17383.168 | 19534.633 | 19800.535 |
| Ppia | SIYGEKFEDENFILK | -77 | -91 | 1 | 0 | 15 | 12893.467 | 13147.394 | 13862.564 | 13679.065 | 16271.422 | 13232.446 |
| Ppia | SIYGEKFEDENFILK | -77 | -91 | 1 | 0 | 15 | 12893.467 | 13147.394 | 13862.564 | 13679.065 | 16271.422 | 13232.446 |
| Ppia | FDITADDEPLGR | -8 | -19 | 0 | 0 | 12 | 11324.49 | 12714.821 | 12941.119 | 300.26266 | 16015.611 | 15655.762 |
| Ppia | ADDEPLGR | -12 | -19 | 0 | 0 | 8 | 6587.5293 | 4687.948 | 3608.9019 | 3242.5945 | 1677.592 | 2117.5693 |
| Ppia | VSFELFADK | -20 | -28 | 0 | 0 | 9 | 3855.5847 | 4474.906 | 4334.4087 | 4448.9263 | 5195.8604 | 5291.7305 |

| | | | | | | | | | | | | |
|---|---|---|---|---|---|---|---|---|---|---|---|---|
| Ppia | AKTEWLDGK | -117 | -125 | 1 | 0.000907206 | 9 | 3834.1616 | 3639.3757 | 3844.2695 | 4868.585 | 5034.833 | 4908.905 |
| Ppia | VSFELFADKVPK | -20 | -31 | 1 | 0 | 12 | 2249.011 | 3028.155 | 3027.2402 | 4156.6953 | 5514.2915 | 5369.35 |
| Ppia | FELFADKVPK | -22 | -31 | 1 | 0 | 10 | 1696.7599 | 1855.1311 | 1825.6224 | 4332.1855 | 5047.778 | 4762.596 |
| Ppia | KFEDENFILK | -82 | -91 | 1 | 0 | 10 | 1566.1929 | 10095.215 | 10424.45 | 525.26636 | 7321.3423 | 6986.3433 |
| Ppia | KFEDENFILK | -82 | -91 | 1 | 0 | 10 | 1566.1929 | 10095.215 | 10424.45 | 525.26636 | 7321.3423 | 6986.3433 |
| Ppia | GEKFEDENFILK | -80 | -91 | 1 | 0 | 12 | 1506.2925 | 1857.2949 | 1963.2483 | 2243.2417 | 737.71014 | 2172.2224 |
| Ppia | KTEWLDGK | -118 | -125 | 1 | 0.00025015 | 8 | 621.4309 | 662.99194 | 2620.4624 | 2840.5269 | 3054.8662 | 628.4447 |
| Ppia | SIYGEKFEDENFIL | -77 | -90 | 1 | 0.00025015 | 14 | 538.1436 | 738.7335 | 711.4561 | 531.07324 | 652.6722 | 628.5077 |
| Ppia | VNPTVFFDIT | -2 | -11 | 0 | 0 | 10 | 205.45027 | 222.58041 | 207.30421 | 81.35664 | 272.37332 | 296.98682 |
| Ppia | CQGGDFTR | -62 | -69 | 0 | 0.000471409 | 8 | 4452.8887 | 3860.4878 | 292.21494 | NaN | 4516.019 | 5273.228 |
| Ppia | VFFDITADDEPLGR | -6 | -19 | 0 | 0 | 14 | 678.8994 | 520.75085 | 579.33826 | 453.81577 | 989.1116 | NaN |
| Ppia | EDENFILK | -84 | -91 | 0 | 0 | 8 | 846.29016 | 3474.9011 | NaN | 784.4035 | 2273.1257 | NaN |
| Ppia | FELFADK | -22 | -28 | 0 | 0.002780285 | 7 | 2270.664 | NaN | NaN | 2994.0796 | 2150.6177 | NaN |
| Ppia | TEWLDGK | -119 | -125 | 0 | 0.00111006 | 7 | 1505.4067 | 1225.8217 | NaN | NaN | 982.7716 | NaN |
| Psme2 | WIQHLIPK | -124 | -131 | 0 | 0 | 8 | 4283.363 | 4451.6816 | 4392.6665 | 4163.6333 | 3978.9072 | 3593.185 |
| Psme2 | WIQHLIPK | -124 | -131 | 0 | 0 | 8 | 4283.363 | 4451.6816 | 4392.6665 | 4163.6333 | 3978.9072 | 3593.185 |
| Psme2 | HIISSNLEK | -218 | -226 | 0 | 0.000431655 | 9 | 3595.821 | 3082.4458 | 2733.631 | 4844.228 | 3111.9631 | 2831.2842 |
| Psme2 | ALVKPEVWTLK | -105 | -115 | 1 | 0 | 11 | 2408.007 | 2803.0212 | 2946.8467 | 2075.1643 | 2639.531 | 2607.9607 |
| Psme2 | ALVKPEVWTLK | -105 | -115 | 1 | 0 | 11 | 2408.007 | 2803.0212 | 2946.8467 | 2075.1643 | 2639.531 | 2607.9607 |
| Psme2 | KQVDVFR | -15 | -21 | 1 | 0.000230707 | 7 | 2392.1975 | 2153.1382 | 2214.354 | 1698.9379 | 1556.349 | 1754.5504 |
| Psme2 | VKPEVWTLK | -107 | -115 | 1 | 0.000230707 | 9 | 1701.9429 | 995.9553 | 2066.658 | 693.6979 | 633.11285 | 513.1801 |
| Psme2 | LVKPEVWTLK | -106 | -115 | 1 | 0 | 10 | 1506.9827 | 1189.3619 | 1710.1753 | 1175.8142 | 953.7821 | 1273.43 |
| Psme2 | LVKPEVWTLK | -106 | -115 | 1 | 0 | 10 | 1506.9827 | 1189.3619 | 1710.1753 | 1175.8142 | 953.7821 | 1273.43 |
| Psme2 | CGYLPGNEK | -91 | -99 | 0 | 0 | 9 | 1303.4327 | 1269.9646 | 1234.6304 | 1553.1798 | 1242.2367 | 1251.812 |
| Psme2 | IEDGNDFGVAIQEK | -132 | -145 | 0 | 0 | 14 | 892.5887 | 1014.61487 | 1028.6042 | 1442.6764 | 1630.716 | 1586.3474 |
| Psme2 | DEAAYGALR | -195 | -203 | 0 | 0 | 9 | 600.78894 | 410.1972 | 423.3766 | 1068.313 | 482.8821 | 557.7352 |
| Psme2 | LALVKPEVWTLK | -104 | -115 | 1 | 0 | 12 | 595.4428 | 743.73114 | 776.54395 | 753.1816 | 974.8106 | 1071.7324 |
| Psme2 | LALVKPEVWTLK | -104 | -115 | 1 | 0 | 12 | 595.4428 | 743.73114 | 776.54395 | 753.1816 | 974.8106 | 1071.7324 |
| Psme2 | ADDFLCTFLPR | -28 | -38 | 0 | 0 | 11 | 1884.9313 | 1624.4171 | 1995.1968 | NaN | 2076.6133 | 1861.8644 |
| Psme2 | LYHIISSNLEK | -216 | -226 | 0 | 0 | 11 | 1621.5853 | 2694.364 | 2653.726 | 2534.0718 | 2491.2603 | NaN |
| Psme2 | IEDGNDFGVAIQ | -132 | -143 | 0 | 0.004133093 | 12 | 635.10645 | 385.3049 | 649.07043 | NaN | NaN | NaN |
| Psme2 | VHERDEAAYGALR | -191 | -203 | 1 | 0 | 13 | 124.87598 | NaN | NaN | 655.47125 | NaN | NaN |
| Psme2 | VHERDEAAYGALR | -191 | -203 | 1 | 0 | 13 | 124.87598 | NaN | NaN | 655.47125 | NaN | NaN |
| Aldoa | PHPYPALTPEQK | -2 | -13 | 0 | 0 | 12 | 10782.831 | 11841.2295 | 11183.872 | 11134.3955 | 12219.063 | 11913.5205 |
| Aldoa | PHPYPALTPEQK | -2 | -13 | 0 | 0 | 12 | 10782.831 | 11841.2295 | 11183.872 | 11134.3955 | 12219.063 | 11913.5205 |
| Aldoa | DDGRPFPQVIK | -89 | -99 | 1 | 0 | 11 | 8428.431 | 8315.553 | 8934.292 | 9027.95 | 9759.174 | 1097.5903 |
| Aldoa | DDGRPFPQVIK | -89 | -99 | 1 | 0 | 11 | 8428.431 | 8315.553 | 8934.292 | 9027.95 | 9759.174 | 1097.5903 |
| Aldoa | IGEHTPSAL | -154 | -162 | 0 | 0.00025015 | 9 | 7526.605 | 8089.9272 | 6973.834 | 6497.5405 | 5138.974 | 5116.1826 |
| Aldoa | KELSDIAHR | -14 | -22 | 1 | 0 | 9 | 7413.1426 | 3882.8264 | 4592.075 | 3558.1301 | 5676.936 | 5861.5005 |
| Aldoa | KELSDIAHR | -14 | -22 | 1 | 0 | 9 | 7413.1426 | 3882.8264 | 4592.075 | 3558.1301 | 5676.936 | 5861.5005 |
| Aldoa | CPLLKPW | -290 | -296 | 1 | 0 | 7 | 5813.8994 | 5802.2603 | 5985.8535 | 6778.962 | 7161.144 | 7053.9116 |
| Aldoa | ADDGRPFPQVIK | -88 | -99 | 1 | 0 | 12 | 5735.537 | 5366.499 | 1376.6854 | 5178.428 | 4180.2407 | 3839.3853 |
| Aldoa | ADDGRPFPQVIK | -88 | -99 | 1 | 0 | 12 | 5735.537 | 5366.499 | 1376.6854 | 5178.428 | 4180.2407 | 3839.3853 |
| Aldoa | LQSIGTENTEENRR | -44 | -57 | 1 | 0 | 14 | 4625.7134 | 5628.495 | 5574.7603 | 3287.5833 | 4276.339 | 3972.6526 |
| Aldoa | LQSIGTENTEENRR | -44 | -57 | 1 | 0 | 14 | 4625.7134 | 5628.495 | 5574.7603 | 3287.5833 | 4276.339 | 3972.6526 |
| Aldoa | QLLLTADDR | -61 | -69 | 0 | 0 | 9 | 3045.2793 | 3005.0835 | 1641.7039 | 807.0045 | 1194.2888 | 1155.9214 |
| Aldoa | QSIGTENTEENRR | -45 | -57 | 1 | 0 | 13 | 2422.3596 | 2670.8857 | 2793.6116 | 2159.97 | 2373.23 | 2743.8865 |
| Aldoa | QSIGTENTEENRR | -45 | -57 | 1 | 0 | 13 | 2422.3596 | 2670.8857 | 2793.6116 | 2159.97 | 2373.23 | 2743.8865 |
| Aldoa | LQSIGTENTEENR | -44 | -56 | 0 | 0 | 13 | 2097.552 | 2285.4038 | 2436.8855 | 1241.5762 | 3775.2778 | 3733.9097 |
| Aldoa | ADDGRPFPQ | -88 | -96 | 1 | 0.00078661 | 9 | 1671.7097 | 1621.853 | 1711.4283 | 2594.0532 | 1632.371 | 1628.4242 |
| Aldoa | SIGTENTEENRR | -46 | -57 | 1 | 0 | 12 | 1470.6288 | 644.4944 | 1711.7994 | 2607.4714 | 2989.3145 | 3262.6921 |
| Aldoa | SIGTENTEENRR | -46 | -57 | 1 | 0 | 12 | 1470.6288 | 644.4944 | 1711.7994 | 2607.4714 | 2989.3145 | 3262.6921 |
| Aldoa | SIGTENTEENR | -46 | -56 | 0 | 0 | 11 | 873.54095 | 692.2077 | 744.46716 | 1337.616 | 1038.2627 | 1083.9907 |
| Aldoa | VNPCIGGVIL | -70 | -79 | 0 | 0 | 10 | 786.6636 | 971.677 | 36.092236 | 840.88165 | 428.5912 | 993.39685 |
| Aldoa | QSIGTENTEENR | -45 | -56 | 0 | 0 | 12 | 785.4695 | 800.1517 | 747.9667 | 517.3904 | 863.50696 | 881.5032 |
| Aldoa | CPLLKPWAL | -290 | -298 | 1 | 0.00025015 | 9 | 743.3452 | 807.90955 | 828.57635 | 593.31274 | 684.86127 | 587.6692 |
| Aldoa | ADESTGSIAK | -33 | -42 | 0 | 0 | 10 | 714.7801 | 627.0075 | 526.95953 | 738.10944 | 345.17142 | 602.98334 |
| Aldoa | VNPCIGGVILFH | -70 | -81 | 0 | 0 | 12 | 711.847 | 536.47363 | 803.9031 | 726.93866 | 777.2247 | 983.3105 |
| Aldoa | GILAADESTGSIA | -29 | -41 | 0 | 0 | 13 | 607.82263 | 725.80695 | 721.70325 | 1193.5562 | 1303.4922 | 1232.5038 |
| Aldoa | KADDGRPFPQVIK | -87 | -99 | 2 | 0 | 13 | 501.2681 | 412.42062 | 738.85504 | 1155.7217 | 502.9259 | 1375.2207 |
| Aldoa | GILAADESTGSIAK | -29 | -42 | 0 | 0.000230707 | 14 | 6022.312 | 5969.0176 | 6181.5034 | 5351.975 | 1501.1588 | NaN |
| Aldoa | DGRPFPQVIK | -90 | -99 | 1 | 0.00078661 | 10 | 4976.541 | 5038.4634 | 5113.058 | NaN | 6667.3564 | 6613.865 |
| Aldoa | ELSDIAHR | -15 | -22 | 0 | 0 | 8 | 4905.3438 | 3473.1357 | NaN | 4706.358 | 1644.8885 | 2117.4453 |
| Aldoa | GVVPLAGTNGETTTQGLDGLSER | -112 | -134 | 0 | 0 | 23 | 633.4372 | 680.8338 | 653.38635 | 965.7431 | NaN | 895.7716 |
| Aldoa | VNPCIGGVILFHET | -70 | -83 | 0 | 0 | 14 | 425.91226 | 424.81943 | NaN | 509.47476 | 241.31595 | 712.53674 |
| Aldoa | LLLTADDR | -62 | -69 | 0 | 0.00025015 | 8 | 3881.6304 | 3322.405 | 3377.561 | 3079.4954 | NaN | NaN |
| Aldoa | PFPQVIK | -93 | -99 | 0 | 0.003681885 | 7 | 1209.1779 | 955.4446 | 1025.9082 | 804.6719 | NaN | NaN |
| Aldoa | GETTTQGLDGLSER | -121 | -134 | 0 | 0 | 14 | 1018.01935 | 1201.0074 | NaN | NaN | 1392.1913 | 1421.7903 |
| Aldoa | LSDIAHR | -16 | -22 | 0 | 0.008947895 | 7 | 730.7514 | 489.94702 | 713.64575 | 603.5061 | NaN | NaN |
| Aldoa | SGGQSEEEASINLNAINK | -272 | -289 | 0 | 0 | 18 | 665.78125 | NaN | NaN | 1317.0205 | 875.17316 | 1098.9335 |
| Aldoa | MVTPGHACTQK | -233 | -243 | 0 | 0 | 11 | 530.7543 | 413.0755 | 397.7225 | NaN | NaN | NaN |
| Aldoa | MVTPGHACTQK | -233 | -243 | 0 | 0 | 11 | 530.7543 | 413.0755 | 397.7225 | NaN | NaN | NaN |
| Aldoa | YTPSGQSGAAA | -343 | -353 | 0 | 0.000471409 | 11 | 1392.8397 | NaN | NaN | 1276.9708 | NaN | NaN |
| Aldoa | SKGGVVGIK | -100 | -108 | 1 | 0.005668207 | 9 | 1104.7971 | NaN | NaN | NaN | NaN | 1216.1779 |
| Aldoa | QEEYIKR | -325 | -331 | 1 | 0.004957095 | 7 | 800.3573 | NaN | NaN | NaN | 1106.2144 | NaN |
| Uhrf1 | ALSEDEPCSSSAVK | -159 | -172 | 0 | 0 | 14 | 8035.7607 | 7282.156 | 641.81104 | 7019.7886 | 5962.5605 | 173.91498 |
| Uhrf1 | VNQPLQTILNQ | -763 | -773 | 0 | 0 | 11 | 2909.0505 | 3047.7224 | 3136.9548 | 1296.6409 | 1336.1921 | 1204.5603 |
| Uhrf1 | FELDHSSPTR | -753 | -762 | 0 | 0 | 10 | 2225.7864 | 881.15344 | 433.17993 | 495.9971 | 2555.14 | 1983.6829 |

| | | | | | | | | | | | | |
|---|---|---|---|---|---|---|---|---|---|---|---|---|
| Uhrf1 | IEEVFHVEPQLQR | -32 | -44 | 0 | 0 | 13 | 850.0784 | 889.9846 | 1130.2766 | 1028.3011 | 550.67334 | 942.69226 |
| Uhrf1 | KIEEVFHVEPQLQR | -31 | -44 | 1 | 0 | 14 | 381.40833 | 375.84717 | 443.7226 | 368.59338 | 478.95474 | 402.67896 |
| Uhrf1 | KIEEVFHVEPQLQR | -31 | -44 | 1 | 0 | 14 | 381.40833 | 375.84717 | 443.7226 | 368.59338 | 478.95474 | 402.67896 |
| Uhrf1 | YDDYPEHGVDIVK | -184 | -196 | 0 | 0.00025015 | 13 | 496.80707 | 3831.9036 | 3719.159 | 4614.8125 | 3264.3691 | 3662.7903 |
| Uhrf1 | RDDTEPEPWTR | -586 | -596 | 1 | 0.00025015 | 11 | 2032.4579 | 1575.5188 | 2143.8867 | NaN | 6875.147 | 6482.023 |
| Uhrf1 | NCHSPINEK | -516 | -524 | 0 | 0.00142999 | 9 | 1275.3185 | NaN | 1311.3782 | 872.4354 | 100.40241 | NaN |
| Uhrf1 | LLNDSQLNNCR | -257 | -267 | 0 | 0 | 11 | 2083.7446 | 2229.9749 | 2275.5557 | NaN | NaN | NaN |
| Uhrf1 | STGPTLSSPR | -649 | -658 | 0 | 0 | 10 | 976.11414 | NaN | 1270.1206 | NaN | NaN | 339.37646 |
| Uhrf1 | ADDKTVWEDTDLGLYK | -117 | -132 | 1 | 0.00025015 | 16 | 846.7975 | NaN | NaN | NaN | 1186.4451 | NaN |
| Uhrf1 | SPSQPPPALR | -289 | -298 | 0 | 0.00383302 | 10 | 1492.0037 | NaN | NaN | NaN | NaN | NaN |
| Mrrf | ADDTVAELDQHLAAK | -242 | -256 | 0 | 0 | 15 | 438.24014 | 674.1613 | 664.9111 | 498.9592 | 531.7268 | 721.28705 |
| Dars1 | IHDPQLLTER | -432 | -441 | 0 | 0 | 10 | 5045.552 | 5852.318 | 2327.9373 | 8618.639 | 5691.8057 | 7178.7793 |
| Dars1 | LPLQLDDAIRPE | -152 | -163 | 1 | 0 | 12 | 3711.489 | 3895.4534 | 4081.6963 | 3392.3403 | 3250.093 | 3130.901 |
| Dars1 | FGAPPHAGGGIGLER | -461 | -475 | 0 | 0 | 15 | 2355.023 | 2793.2793 | 1692.7177 | 2298.6167 | 3196.2107 | 3678.5085 |
| Dars1 | FGAPPHAGGGIGLER | -461 | -475 | 0 | 0 | 15 | 2355.023 | 2793.2793 | 1692.7177 | 2298.6167 | 3196.2107 | 3678.5085 |
| Dars1 | VFCIGPVFR | -265 | -273 | 0 | 0 | 9 | 2268.2273 | 2867.5554 | 2647.3376 | 2303.7205 | 3071.5325 | 3102.694 |
| Dars1 | ALHHGIDLEK | -442 | -451 | 0 | 0 | 10 | 1933.7773 | 1852.8523 | 1701.3108 | 1483.5002 | 1834.4606 | 1711.666 |
| Dars1 | ALHHGIDLEK | -442 | -451 | 0 | 0 | 10 | 1933.7773 | 1852.8523 | 1701.3108 | 1483.5002 | 1834.4606 | 1711.666 |
| Dars1 | ALVAVGDHASK | -89 | -99 | 0 | 0 | 11 | 1669.4348 | 1526.5115 | 1904.7239 | 1353.3677 | 1343.2162 | 1475.3511 |
| Dars1 | FLGLHNVR | -480 | -487 | 0 | 0 | 8 | 1081.251 | 728.3834 | 793.656 | 1537.2362 | 190.88654 | 434.92603 |
| Dars1 | EIVDAAEDYAK | -16 | -26 | 0 | 0 | 11 | 1001.59247 | 395.22324 | 1951.7874 | 1825.4087 | 730.8164 | 1380.0942 |
| Dars1 | LPLQLDDAIRPEVEGE | -152 | -167 | 1 | 0 | 16 | 866.152 | 1020.75336 | 995.3191 | 1075.6979 | 1173.194 | 1140.5566 |
| Dars1 | EKYDTDFYVL | -382 | -391 | 1 | 0.000550307 | 10 | 244.19136 | 286.60892 | 161.01497 | 313.56818 | 287.18234 | 331.03027 |
| Dars1 | EKYDTDFYVL | -382 | -391 | 1 | 0.000550307 | 10 | 244.19136 | 286.60892 | 161.01497 | 313.56818 | 287.18234 | 331.03027 |
| Dars1 | IGSCTQQDVELHVQ | -127 | -140 | 0 | 0 | 14 | 629.12634 | 728.5167 | 684.82513 | 705.3204 | 714.10645 | 793.7553 |
| Dars1 | ESIIDVEGVVR | -111 | -121 | 0 | 0 | 11 | 505.57886 | 574.6186 | 763.5426 | 569.6344 | 652.26666 | 654.06445 |
| Dars1 | LPLQLDDAIRPEVEGEEDGR | -152 | -171 | 1 | 0 | 20 | 443.99048 | 310.04602 | 503.39655 | 383.55493 | 1397.2947 | 2062.1821 |
| Dars1 | SGICHLFR | -200 | -207 | 0 | 0 | 8 | 1692.7705 | 1806.155 | NaN | 1516.0864 | 1895.5219 | 1913.0327 |
| Dars1 | TQQDVELHVQK | -131 | -141 | 0 | 0 | 11 | 1053.4114 | 1128.7003 | 572.4496 | NaN | 1010.65155 | 1015.8764 |
| Dars1 | SIIDVEGVVR | -112 | -121 | 0 | 0 | 10 | 718.806 | NaN | 1095.91 | 495.32138 | 661.03046 | 313.122 |
| Dars1 | IGSCTQQDVELHVQK | -127 | -141 | 0 | 0 | 15 | 541.4661 | 531.9347 | 641.9146 | 702.3924 | 472.10693 | NaN |
| Dars1 | IGSCTQQDVELHVQK | -127 | -141 | 0 | 0 | 15 | 541.4661 | 531.9347 | 641.9146 | 702.3924 | 472.10693 | NaN |
| Dars1 | RIHDPQLLTER | -431 | -441 | 1 | 0 | 11 | 667.82104 | NaN | NaN | 1201.5513 | 1993.9246 | 747.1108 |
| Dars1 | FGAPPHAGGGIGLE | -461 | -474 | 0 | 0.00025015 | 14 | 1408.6732 | 970.73846 | NaN | NaN | 102.759995 | NaN |
| Dars1 | QCFLVLR | -75 | -81 | 0 | 0 | 7 | 943.9132 | 520.0379 | 643.9188 | NaN | NaN | NaN |
| Dars1 | GEEILSGAQR | -422 | -431 | 0 | 0.001055699 | 10 | 5373.5244 | 5340.9873 | NaN | NaN | NaN | NaN |
| Dars1 | ADDVVWVR | -56 | -63 | 0 | 0 | 8 | 1035.9359 | 1082.2855 | NaN | NaN | NaN | NaN |
| Snx27 | ADEDGEGIHPSAPHR | -2 | -16 | 0 | 0 | 15 | 945.1804 | NaN | NaN | 528.23047 | 5523.095 | 5856.014 |
| Pdcd5 | HGDPGDAAQQEAK | -21 | -33 | 0 | 0 | 13 | 556.11426 | 528.8463 | 553.36676 | 258.0066 | 163.24225 | 530.90826 |
| Pdcd5 | ADEELEALRK | -2 | -11 | 1 | 0.00025015 | 10 | 1475.9169 | 1040.5299 | 649.71326 | NaN | 1472.3369 | NaN |
| Ipo7 | VLTGVAGEDAECHAAK | -725 | -740 | 0 | 0 | 16 | 1135.6792 | 1277.8518 | 911.4724 | 1169.3928 | 1799.8583 | 1970.6726 |
| Ipo7 | ALTHGLNEEQR | -979 | -989 | 0 | 0 | 11 | 826.88074 | 693.4999 | 858.2973 | 521.1945 | 947.36755 | 898.5099 |
| Ipo7 | EDDRPELPWWK | -249 | -259 | 1 | 0 | 11 | 811.4632 | 912.38983 | 926.84375 | 334.73502 | 949.366 | 832.12445 |
| Ipo7 | THGLNEEQR | -981 | -989 | 0 | 0.000298805 | 9 | 741.41956 | 680.79865 | 725.0573 | 579.2433 | 848.8983 | 666.7505 |
| Ipo7 | ADEELWQEDPYEYIR | -363 | -377 | 0 | 0 | 15 | 712.42114 | 448.80606 | 600.9036 | 433.67215 | 447.13248 | 470.35248 |
| Ipo7 | HYFCEVK | -481 | -487 | 0 | 0 | 7 | 537.88226 | 956.5019 | 444.5063 | 601.51843 | 501.0318 | 521.04083 |
| Ipo7 | SDQSDQSVLIQK | -190 | -201 | 0 | 0 | 12 | 2287.4045 | 1274.5029 | NaN | 2530.7415 | 1426.8779 | 843.1712 |
| Ipo7 | ETENDDLTNVIQK | -552 | -564 | 0 | 0 | 13 | 1716.3998 | 2478.4578 | 592.084 | NaN | 1661.5774 | 2111.279 |
| Ipo7 | NQVIQTGPDEEGSDDK | -589 | -604 | 0 | 0.00025015 | 16 | 1053.0463 | 781.29956 | 856.2994 | 847.40265 | 932.47314 | NaN |
| Ipo7 | QHFLPVLK | -175 | -182 | 0 | 0 | 8 | 612.1612 | 1046.2362 | 1098.125 | 733.83246 | 827.34393 | NaN |
| Ipo7 | YGSPGNVSK | -275 | -283 | 0 | 0.000230707 | 9 | 567.9949 | NaN | 1311.3811 | 547.74585 | 675.0813 | 584.9915 |
| Ipo7 | MDPNTIIEALR | -1 | -11 | 0 | 0 | 11 | 309.64993 | NaN | 389.12225 | 149.1328 | 389.13016 | 378.63852 |
| Ipo7 | VLTGVAGEDAECH | -725 | -737 | 0 | 0.000230707 | 13 | 1052.7603 | NaN | 1153.3536 | 1358.1709 | NaN | NaN |
| Ipo7 | AIFQTIQNR | -964 | -972 | 0 | 0.000431655 | 9 | 856.43915 | 808.89233 | 724.0369 | NaN | NaN | NaN |
| Ipo7 | STIIDDEDNPVDEYQIFK | -946 | -963 | 0 | 0.000298805 | 18 | 258.66022 | 270.72232 | 268.77902 | NaN | NaN | NaN |
| Ipo7 | FPNNVEPVTNHFITQ | -806 | -820 | 0 | 0.001092529 | 15 | 800.3546 | NaN | 1412.1656 | NaN | NaN | NaN |
| Prdx1 | IGYPAPNFK | -8 | -16 | 0 | 0 | 9 | 11367.719 | 11072.2705 | 11687.75 | 10805.012 | 10455.783 | 10506.056 |
| Prdx1 | KADEGISFR | -120 | -128 | 1 | 0 | 9 | 7672.287 | 7753.7227 | 8236.058 | 8719.206 | 523.1648 | 9904.415 |
| Prdx1 | GWKPGSDTIKPDVNK | -176 | -190 | 2 | 0 | 15 | 6121.716 | 6679.0713 | 6863.9326 | 7350.14 | 7927.407 | 7469.5938 |
| Prdx1 | GWKPGSDTIKPDVNK | -176 | -190 | 2 | 0 | 15 | 6121.716 | 6679.0713 | 6863.9326 | 7350.14 | 7927.407 | 7469.5938 |
| Prdx1 | NIPLISDPK | -101 | -109 | 0 | 0.00025015 | 9 | 5962.077 | 5887.2104 | 6253.1143 | 5759.8604 | 3674.7764 | 4253.3096 |
| Prdx1 | TIAQDYGVLK | -111 | -120 | 0 | 0 | 10 | 616.046 | 369.16528 | 777.477 | 453.90594 | 1538.298 | 637.816 |
| Prdx1 | TIKPDVNK | -183 | -190 | 1 | 0 | 8 | 1572.631 | 1138.0605 | NaN | 1206.3693 | 1127.793 | 1254.7666 |
| Prdx1 | SDRADEFK | -60 | -67 | 1 | 0.002199523 | 8 | 819.1381 | NaN | 851.00684 | 1500.7048 | 1035.1152 | 973.14746 |
| Prdx1 | ADEGISFR | -121 | -128 | 0 | 0 | 8 | 401.43634 | 683.2472 | 7579.017 | 718.4593 | NaN | 611.4387 |
| Prdx1 | AVMPDGQFK | -19 | -27 | 0 | 0.00111006 | 9 | 2823.3435 | 2774.1206 | 2799.7834 | 4692.742 | NaN | NaN |
| Prdx1 | AVMPDGQFK | -19 | -27 | 0 | 0.00111006 | 9 | 2823.3435 | 2774.1206 | 2799.7834 | 4692.742 | NaN | NaN |
| Prdx1 | AWINTPK | -86 | -92 | 0 | 0.005605646 | 7 | 488.27567 | NaN | NaN | NaN | NaN | NaN |
| Prdx1 | ATAVMPDGQFK | -17 | -27 | 0 | 0.006195994 | 11 | 390.7303 | NaN | NaN | NaN | NaN | NaN |
| Rps5 | HAFEIIHL | -95 | -102 | 0 | 0.00025015 | 8 | 4871.327 | 495.87503 | 4204.857 | 4736.296 | 3568.3425 | 286.7416 |
| Rps5 | CLADELINAAK | -172 | -182 | 0 | 0 | 11 | 3738.1243 | 548.4911 | 398.1936 | 4586.564 | 99.98472 | 4796.366 |
| Rps5 | HAFEIIHLL | -95 | -103 | 0 | 0.00204437 | 9 | 2887.0876 | 2948.1921 | 2669.255 | 2059.7449 | 2826.338 | 2846.2104 |
| Rps5 | QCPIVER | -65 | -71 | 0 | 0.00025015 | 7 | 2529.9182 | 288.95224 | 1850.6025 | 2131.6162 | 1529.1294 | 1553.7926 |
| Rps5 | TEWEAATPAVAETPDIK | -2 | -18 | 0 | 0 | 17 | 1697.5725 | 1232.3776 | 1078.6704 | 721.5029 | 856.5632 | 886.44995 |
| Rps5 | LADELINAAK | -173 | -182 | 0 | 0 | 10 | 1462.3801 | 1384.7839 | 1449.5994 | 1758.6545 | 1139.4698 | 1496.2919 |
| Rps5 | RVNQAIWLL | -146 | -154 | 1 | 0.002199523 | 9 | 1144.8695 | 1336.8112 | 1347.3799 | 1584.9922 | 2179.555 | 2255.0913 |
| Rps5 | TIAECLADELINAAK | -168 | -182 | 0 | 0 | 15 | 884.8043 | 679.9834 | 761.71155 | 899.78033 | 1119.7917 | 1064.0736 |
| Rps5 | ADELINAAK | -174 | -182 | 0 | 0 | 9 | 1538.1772 | 749.10034 | 1197.4489 | NaN | 1019.67224 | 787.1307 |

| | | | | | | | | | | | | |
|---|---|---|---|---|---|---|---|---|---|---|---|---|
| Rps5 | VAETPDIK | -11 | -18 | 0 | 0 | 8 | 940.61176 | 506.6046 | 750.1266 | 692.8086 | 487.3999 | NaN |
| Rps5 | VNAIINSGPR | -113 | -122 | 0 | 0 | 10 | 4345.471 | 4637.674 | NaN | 4049.7632 | 3922.3809 | NaN |
| Rps5 | QAVDVSPLR | -137 | -145 | 0 | 0 | 9 | 5029.471 | 3775.4238 | 4031.4065 | NaN | NaN | NaN |
| Rps5 | VNQAIWLL | -147 | -154 | 0 | 0.000826806 | 8 | 723.00305 | NaN | NaN | 898.4589 | 1534.9755 | NaN |
| Rps5 | AQCPIVER | -64 | -71 | 0 | 0.000627943 | 8 | 1925.6484 | 2053.2458 | NaN | NaN | NaN | NaN |
| Rps5 | HAFEIIHLLT | -95 | -104 | 0 | 0.00111006 | 10 | 298.46738 | NaN | NaN | NaN | NaN | 171.25935 |
| Slc25a20 | ADEPKPISPFK | -2 | -12 | 1 | 0.001055699 | 11 | 4646.3706 | 4045.695 | 4170.6875 | 826.29614 | 2526.668 | NaN |
| Slc25a20 | LVFVGHPLDTVK | -24 | -35 | 0 | 0 | 12 | 1009.2014 | 1040.4907 | 1034.6443 | NaN | 1752.5535 | 1759.4343 |
| Slc25a20 | NWAVAIPPDVLK | -223 | -234 | 0 | 0 | 12 | 701.3151 | 916.0062 | NaN | 352.15063 | 302.89374 | 310.28506 |
| Rcc2 | NHTLVLDSQK | -338 | -347 | 0 | 0 | 10 | 36838.633 | 36295.742 | 36554.926 | 29425.957 | 30399.143 | 32037.812 |
| Rcc2 | RIEYDCELVPR | -298 | -308 | 1 | 0.00025015 | 11 | 2656.3423 | 3071.5745 | 3151.9531 | 3036.1804 | 3152.4678 | 779.6942 |
| Rcc2 | AIITEPEHTK | -81 | -90 | 0 | 0 | 10 | 2361.0696 | 506.4345 | 2081.005 | 1499.8707 | 1806.6222 | 1746.095 |
| Rcc2 | LPEYTPR | -512 | -518 | 0 | 0.004133093 | 7 | 2131.585 | 2115.0835 | 2173.3855 | 2343.7861 | 2139.1711 | 1991.8135 |
| Rcc2 | DGQILPVPNVVVR | -319 | -331 | 0 | 0 | 13 | 1922.0385 | 2189.9863 | 2193.5994 | 2350.2856 | 2801.027 | 2701.3682 |
| Rcc2 | GQNLWGPHR | -131 | -139 | 0 | 0 | 9 | 1718.7098 | 1581.9944 | 1694.4015 | 1491.795 | 2187.928 | 2143.7988 |
| Rcc2 | GAAAIITEPEHTK | -78 | -90 | 0 | 0 | 13 | 1555.4391 | 1616.9989 | 1715.374 | 1831.4979 | 1969.6609 | 2068.6846 |
| Rcc2 | NHTLALTDTGSVF | -210 | -222 | 0 | 0.00078661 | 13 | 1312.6896 | 1489.9357 | 1347.6917 | 1367.2036 | 1973.756 | 1650.4926 |
| Rcc2 | SHSLVIAR | -492 | -499 | 0 | 0 | 8 | 1307.1154 | 1189.5507 | 1103.0581 | 339.60565 | 861.9733 | 942.18005 |
| Rcc2 | GATNWDLIGR | -108 | -117 | 0 | 0 | 10 | 1122.78 | 1075.3696 | 1076.1389 | 1322.032 | 1391.3932 | 1421.1116 |
| Rcc2 | IEYDCELVPR | -299 | -308 | 0 | 0 | 10 | 994.26764 | 954.19507 | 1688.7538 | 1612.6311 | 888.3992 | 795.6089 |
| Rcc2 | NLGQNLWGPHR | -129 | -139 | 0 | 0 | 11 | 954.3205 | 2671.3757 | 3225.5398 | 1426.9376 | 1610.9916 | 1619.5363 |
| Rcc2 | SFGCPEYGQLGHNSDGK | -275 | -291 | 0 | 0 | 17 | 871.39246 | 893.9383 | 278.6423 | 659.9199 | 677.0875 | 648.11725 |
| Rcc2 | GCPEYGQLGHNSDGK | -277 | -291 | 0 | 0 | 15 | 871.13794 | 894.2879 | 892.1488 | 886.4699 | 1067.0859 | 1073.5554 |
| Rcc2 | TKDGQILPVPNVVVR | -317 | -331 | 1 | 0 | 15 | 415.96915 | 456.79932 | 460.4177 | 502.99255 | 527.1596 | 547.3265 |
| Rcc2 | TKDGQILPVPNVVVR | -317 | -331 | 1 | 0 | 15 | 415.96915 | 456.79932 | 460.4177 | 502.99255 | 527.1596 | 547.3265 |
| Rcc2 | AVQDLCGWR | -421 | -429 | 0 | 0 | 9 | 727.19446 | 1125.8048 | 1080.256 | 226.59268 | 856.8861 | 934.4567 |
| Rcc2 | DESTISWGPSPTFGELGYGDHKPK | -445 | -468 | 1 | 0 | 24 | 1269.8494 | NaN | 1433.0184 | 1043.626 | 1614.8588 | 1790.2196 |
| Rcc2 | RVAIFIEK | -309 | -316 | 1 | 0.001392869 | 8 | 575.485 | 598.6073 | NaN | 958.73505 | 847.526 | 863.2291 |
| Rcc2 | LFDFPGR | -376 | -382 | 0 | 0.002068213 | 7 | 2245.7654 | 2055.4104 | NaN | 1346.11 | 1093.875 | NaN |
| Rcc2 | TNWDLIGR | -110 | -117 | 0 | 0.000550307 | 8 | 847.1204 | NaN | 873.96277 | NaN | 689.25995 | NaN |
| Rcc2 | ADESTISWGPSPTFGELGYGDHKPK | -444 | -468 | 1 | 0 | 25 | 297.67203 | NaN | 331.35226 | NaN | NaN | 412.04718 |
| Myh9 | ELEEQLEEEESAR | -947 | -959 | 0 | 0 | 13 | 11294.85 | 11562.665 | 10169.676 | 14406.881 | 2590.4167 | 2360.2812 |
| Myh9 | NLQIDQINTDLNLER | -1756 | -1770 | 0 | 0 | 15 | 9410.589 | 10593.438 | 11025.435 | 9917.688 | 12464.419 | 12389.34 |
| Myh9 | ISELQEDLESER | -1113 | -1124 | 0 | 0 | 12 | 5000.8027 | 5510.025 | 5889.7104 | 5842.4614 | 5818.148 | 5856.1675 |
| Myh9 | EQLEEEEAKR | -1343 | -1353 | 1 | 0 | 11 | 5000.1323 | 1344.2202 | 5412.1553 | 3732.554 | 5197.677 | 5019.879 |
| Myh9 | HEDELLAK | -843 | -850 | 0 | 0.003069674 | 8 | 4607.519 | 4496.905 | 4210.514 | 840.5859 | 4787.7544 | 5007.7705 |
| Myh9 | FVSELWK | -607 | -613 | 0 | 0.00122419 | 7 | 4416.4604 | 4716.531 | 5032.718 | 3839.3147 | 3617.3398 | 3818.067 |
| Myh9 | VAAYDKLEK | -1405 | -1413 | 1 | 0.000550307 | 9 | 3903.1167 | 3256.737 | 330.8781 | 1421.1306 | 1512.0043 | 1859.762 |
| Myh9 | TTNLMEEEEK | -1005 | -1014 | 0 | 0 | 10 | 3661.8984 | 762.78094 | 2923.4639 | 804.1017 | 5502.7344 | 4894.9355 |
| Myh9 | QLEEQLDNETK | -1818 | -1828 | 0 | 0 | 11 | 3550.6594 | 3367.0535 | 275.5587 | 3333.6243 | 453.6661 | 3238.6113 |
| Myh9 | QLEEQLDNETK | -1818 | -1828 | 0 | 0 | 11 | 3550.6594 | 3367.0535 | 275.5587 | 3333.6243 | 453.6661 | 3238.6113 |
| Myh9 | YYLLSGAGEHLK | -278 | -289 | 0 | 0 | 12 | 3496.9238 | 3312.9827 | 3960.31 | 3456.7488 | 5050.302 | 4637.245 |
| Myh9 | GIPEDEQMGLLR | -330 | -341 | 0 | 0 | 12 | 3140.5042 | 3313.748 | 3564.8066 | 2959.8115 | 3340.943 | 3252.7705 |
| Myh9 | GIPEDEQMGLLR | -330 | -341 | 0 | 0 | 12 | 3140.5042 | 3313.748 | 3564.8066 | 2959.8115 | 3340.943 | 3252.7705 |
| Myh9 | MEDGVGCLETAEEAK | -1373 | -1387 | 0 | 0 | 15 | 3006.3572 | 3061.3625 | 3125.9153 | 4018.7026 | 237.30373 | 3785.0884 |
| Myh9 | VVFQEFR | -712 | -718 | 0 | 0 | 7 | 2922.7893 | 2860.095 | 2905.5142 | 1398.0875 | 1187.4432 | 1298.2837 |
| Myh9 | KLEGDSTDLSDQIAELQ | -1052 | -1068 | 1 | 0.000712251 | 17 | 2480.0928 | 2656.2488 | 2661.763 | 3056.2373 | 3893.4258 | 3719.711 |
| Myh9 | ELEEICHDLEAR | -912 | -923 | 0 | 0 | 12 | 2329.969 | 2760.3052 | 2766.5994 | 3088.1758 | 3812.29 | 3627.2498 |
| Myh9 | HSQAVEELADQLEQTK | -1194 | -1209 | 0 | 0 | 16 | 2283.2646 | 2732.3718 | 2608.6406 | 3706.2622 | 3690.3467 | 3555.5586 |
| Myh9 | HSQAVEELADQLEQTK | -1194 | -1209 | 0 | 0 | 16 | 2283.2646 | 2732.3718 | 2608.6406 | 3706.2622 | 3690.3467 | 3555.5586 |
| Myh9 | HSQAVEELADQLEQTKR | -1194 | -1210 | 1 | 0 | 17 | 2199.4507 | 2693.223 | 2520.242 | 3571.3735 | 4625.258 | 4171.3896 |
| Myh9 | HSQAVEELADQLEQTKR | -1194 | -1210 | 1 | 0 | 17 | 2199.4507 | 2693.223 | 2520.242 | 3571.3735 | 4625.258 | 4171.3896 |
| Myh9 | KLVWVPSSK | -30 | -38 | 1 | 0 | 9 | 2157.9373 | 2189.6716 | 2299.948 | 722.17334 | 797.0523 | 780.8116 |
| Myh9 | KLEEDQIIMEDQNCK | -975 | -989 | 1 | 0 | 15 | 1169.6848 | 1348.1261 | 1356.5825 | 1288.3441 | 1525.6428 | 1550.324 |
| Myh9 | KLEEDQIIMEDQNCK | -975 | -989 | 1 | 0 | 15 | 1169.6848 | 1348.1261 | 1356.5825 | 1288.3441 | 1525.6428 | 1550.324 |
| Myh9 | KLEEDQIIMEDQNCK | -975 | -989 | 1 | 0 | 15 | 1169.6848 | 1348.1261 | 1356.5825 | 1288.3441 | 1525.6428 | 1550.324 |
| Myh9 | TDLLLEPYNK | -290 | -299 | 0 | 0 | 10 | 2051.7148 | 2731.7595 | 133.27098 | 454.39722 | 3212.747 | 137.9857 |
| Myh9 | ALELDSNLYR | -746 | -755 | 0 | 0 | 10 | 2045.4294 | 2145.1313 | 2399.9934 | 152.68681 | 1342.4382 | 1409.6221 |
| Myh9 | QLGNIAFK | -348 | -355 | 0 | 0 | 8 | 2015.7975 | 1748.0104 | 1965.6503 | 808.8806 | 651.1466 | 722.85767 |
| Myh9 | HSQAVEELADQL | -1194 | -1205 | 0 | 0 | 12 | 1974.9209 | 1716.884 | 1755.405 | 1093.0342 | 1210.4031 | 1100.9669 |
| Myh9 | ESQLQDTQELLQEENR | -1307 | -1322 | 0 | 0 | 16 | 1938.3895 | 2370.6055 | 2398.884 | 1536.9009 | 1661.0566 | 1810.7616 |
| Myh9 | KQELEEICHDL | -910 | -920 | 1 | 0 | 11 | 1825.9756 | 2088.0251 | 2122.7747 | 2406.0571 | 751.5263 | 2490.2048 |
| Myh9 | AVEELADQLEQTKR | -1197 | -1210 | 1 | 0 | 14 | 1731.675 | 800.7092 | 1187.544 | 829.0665 | 1034.2578 | 918.5936 |
| Myh9 | DELADEIANSSGK | -1704 | -1716 | 0 | 0 | 13 | 1685.3522 | 1870.7555 | 2209.527 | 1131.559 | 154.22571 | 2178.796 |
| Myh9 | NFINNPLAQADWAAK | -15 | -29 | 0 | 0 | 15 | 1657.4253 | 1781.2299 | 1991.9679 | 1756.4358 | 2595.774 | 2749.006 |
| Myh9 | DVLLQVEDERR | -1846 | -1856 | 1 | 0 | 11 | 1624.3026 | 1847.6693 | 2078.1812 | 1870.8247 | 1618.0981 | 1762.616 |
| Myh9 | KMEDGVGCLETAEEAK | -1372 | -1387 | 1 | 0 | 16 | 1563.2816 | 1526.1063 | 1263.9207 | 3132.163 | 1830.2399 | 1305.8148 |
| Myh9 | QELEEICHDLEAR | -911 | -923 | 0 | 0 | 13 | 1544.1287 | 842.74146 | 820.6729 | 1891.8777 | 1437.4178 | 1309.7386 |
| Myh9 | NKHEAMITDLEER | -1023 | -1035 | 1 | 0 | 13 | 1538.0657 | 1599.9167 | 908.3345 | 1182.7587 | 2071.5618 | 1139.3933 |
| Myh9 | NKHEAMITDLEER | -1023 | -1035 | 1 | 0 | 13 | 1538.0657 | 1599.9167 | 908.3345 | 1182.7587 | 2071.5618 | 1139.3933 |
| Myh9 | NKHEAMITDLEER | -1023 | -1035 | 1 | 0 | 13 | 1538.0657 | 1599.9167 | 908.3345 | 1182.7587 | 2071.5618 | 1139.3933 |
| Myh9 | NKHEAMITDLEER | -1023 | -1035 | 1 | 0 | 13 | 1538.0657 | 1599.9167 | 908.3345 | 1182.7587 | 2071.5618 | 1139.3933 |
| Myh9 | KQELEEICHDLEAR | -910 | -923 | 1 | 0 | 14 | 1520.847 | 2189.8948 | 2235.064 | 1361.9259 | 1159.3252 | 2140.7864 |
| Myh9 | KQELEEICHDLEAR | -910 | -923 | 1 | 0 | 14 | 1520.847 | 2189.8948 | 2235.064 | 1361.9259 | 1159.3252 | 2140.7864 |
| Myh9 | THEAQIQEMR | -1182 | -1191 | 0 | 0 | 10 | 1514.1393 | 1038.2277 | 1579.721 | 1068.2769 | 1116.1575 | 1362.4807 |
| Myh9 | IAQLEEQLDNETK | -1816 | -1828 | 0 | 0 | 13 | 1427.4812 | 1437.0574 | 1541.5469 | 1131.9495 | 1307.752 | 1327.3838 |
| Myh9 | QAQQERDELADEIANSS | -1698 | -1714 | 1 | 0 | 17 | 1413.3889 | 1682.4629 | 1303.6272 | 1710.8335 | 836.33276 | 774.225 |
| Myh9 | ANLQIDQINTDLNLER | -1755 | -1770 | 0 | 0 | 16 | 1371.8497 | 1353.3907 | 1221.3339 | 1282.0582 | 1562.1528 | 1639.2451 |

| | | | | | | | | | | | | |
|---|---|---|---|---|---|---|---|---|---|---|---|---|
| Myh9 | RGDLPFVVTR | -1923 | -1932 | 1 | 0.00025015 | 10 | 1334.7826 | 1606.9092 | 1885.8678 | 1383.3663 | 1512.8579 | 1542.0466 |
| Myh9 | GHVTIPGQQDK | -306 | -316 | 0 | 0 | 11 | 1315.5173 | 2170.4192 | 2374.449 | 2725.7288 | 2990.397 | 3055.0425 |
| Myh9 | KTELEDTLDSTAAQQELR | -1145 | -1162 | 1 | 0 | 18 | 1251.257 | 1834.783 | 1698.8362 | 1998.4435 | 3074.0999 | 2984.5554 |
| Myh9 | KTELEDTLDSTAAQQELR | -1145 | -1162 | 1 | 0 | 18 | 1251.257 | 1834.783 | 1698.8362 | 1998.4435 | 3074.0999 | 2984.5554 |
| Myh9 | LKDVLLQVEDERR | -1844 | -1856 | 2 | 0 | 13 | 1221.2228 | 1516.1013 | 1325.3623 | 1117.9795 | 1380.4025 | 1183.7831 |
| Myh9 | LKDVLLQVEDERR | -1844 | -1856 | 2 | 0 | 13 | 1221.2228 | 1516.1013 | 1325.3623 | 1117.9795 | 1380.4025 | 1183.7831 |
| Myh9 | IDQINTDLNLER | -1759 | -1770 | 0 | 0 | 12 | 1202.8011 | 1958.6696 | 2188.513 | 1022.8353 | 1537.8549 | 1638.442 |
| Myh9 | QELDDLLVDLDHQR | -1420 | -1433 | 0 | 0 | 14 | 1172.8673 | 1188.666 | 1327.8341 | 652.92883 | 880.08594 | 817.08105 |
| Myh9 | QTLENERGELANEVK | -1220 | -1234 | 1 | 0 | 15 | 1145.5184 | 851.93225 | 1410.9065 | 1671.4457 | 1803.286 | 1739.5044 |
| Myh9 | QTLENERGELANEVK | -1220 | -1234 | 1 | 0 | 15 | 1145.5184 | 851.93225 | 1410.9065 | 1671.4457 | 1803.286 | 1739.5044 |
| Myh9 | ELQEDLESER | -1115 | -1124 | 0 | 0 | 10 | 1070.7744 | 220.04004 | 237.27176 | 1086.033 | 844.8814 | 965.4007 |
| Myh9 | LCAEAEELR | -895 | -903 | 0 | 0 | 9 | 1050.0604 | 1063.319 | 992.0934 | 639.5193 | 848.9426 | 817.5126 |
| Myh9 | EMEAELEDERK | -1593 | -1603 | 1 | 0 | 11 | 996.6584 | 1176.6028 | 979.0192 | 1037.5557 | 1577.3992 | 1481.8413 |
| Myh9 | LEEEEEAKR | -1345 | -1353 | 1 | 0.000550307 | 9 | 899.9638 | 882.709 | 177.83235 | 1217.2719 | 1339.1659 | 1268.029 |
| Myh9 | LQQELDDLLVDLDHQR | -1418 | -1433 | 0 | 0 | 16 | 885.6468 | 990.1077 | 743.5423 | 1145.8698 | 1431.9995 | 1413.8995 |
| Myh9 | DLEAHIDTANK | -1621 | -1631 | 0 | 0 | 11 | 871.0567 | 873.55634 | 398.96494 | 1447.467 | 869.5869 | 873.92194 |
| Myh9 | KLEMDLK | -1614 | -1620 | 1 | 0.001636916 | 7 | 859.1875 | 1640.8655 | 1815.5189 | 1514.4673 | 1682.7025 | 2048.744 |
| Myh9 | ELETQISELQEDLESER | -1108 | -1124 | 0 | 0 | 17 | 837.21747 | 786.3376 | 775.7745 | 1109.283 | 1192.2344 | 1117.0844 |
| Myh9 | SDKFVSELWK | -604 | -613 | 1 | 0 | 10 | 821.11694 | 3138.6523 | 1156.8585 | 1410.4354 | 4820.208 | 148.38986 |
| Myh9 | NIQELEEQLEEEESAR | -944 | -959 | 0 | 0 | 16 | 796.44507 | 797.1922 | 841.4497 | 1187.678 | 1444.5005 | 1424.4686 |
| Myh9 | QAQQERDELADEIANSSGK | -1698 | -1716 | 1 | 0 | 19 | 790.68164 | 514.7532 | 728.2595 | 871.91437 | 700.06464 | 676.64026 |
| Myh9 | HEAMITDLEER | -1025 | -1035 | 0 | 0 | 11 | 785.8093 | 730.8906 | 1071.7747 | 1119.0979 | 1187.899 | 1229.1257 |
| Myh9 | KLEGDSTDLSDQIAEL | -1052 | -1067 | 1 | 0.000471409 | 16 | 780.5807 | 1035.2416 | 931.47015 | 554.6784 | 564.706 | 714.8048 |
| Myh9 | QSVSNLEK | -1434 | -1441 | 0 | 0 | 8 | 780.1028 | 885.58765 | 1007.1198 | 772.77124 | 1251.8877 | 289.24234 |
| Myh9 | RKLEGDSTDLSDQIAELQ | -1051 | -1068 | 2 | 0.000230707 | 18 | 773.83813 | 648.45355 | 749.9243 | 689.5621 | 804.6812 | 716.413 |
| Myh9 | RKLEGDSTDLSDQIAELQ | -1051 | -1068 | 2 | 0.000230707 | 18 | 773.83813 | 648.45355 | 749.9243 | 689.5621 | 804.6812 | 716.413 |
| Myh9 | HQSSDKFVSELWK | -601 | -613 | 1 | 0.000431655 | 13 | 728.6339 | 1089.848 | 1099.2516 | 1290.1122 | 1292.0541 | 1196.6722 |
| Myh9 | LQDTQELLQEENR | -1310 | -1322 | 0 | 0 | 13 | 714.2506 | 593.7532 | 662.19855 | 948.584 | 720.8711 | 748.81226 |
| Myh9 | VAEFTTNLMEEEEK | -1001 | -1014 | 0 | 0 | 14 | 690.20746 | 742.6007 | 868.7288 | 831.1104 | 905.235 | 757.0742 |
| Myh9 | ALEQQVEEMK | -1529 | -1538 | 0 | 0 | 10 | 688.3995 | 1287.8518 | 1314.7155 | 700.6654 | 850.5732 | 1052.055 |
| Myh9 | ALEQQVEEMK | -1529 | -1538 | 0 | 0 | 10 | 688.3995 | 1287.8518 | 1314.7155 | 700.6654 | 850.5732 | 1052.055 |
| Myh9 | RQLEEAEEEAQR | -1877 | -1888 | 1 | 0.000712251 | 12 | 678.19745 | 1364.7386 | 1366.554 | 1021.7025 | 1607.99 | 1482.1329 |
| Myh9 | NMDPLNDNIATLL | -588 | -600 | 0 | 0.00111006 | 13 | 625.7716 | 663.80725 | 703.79724 | 713.3943 | 849.1263 | 846.3021 |
| Myh9 | QLEEELEEEQGNTELINDR | -1733 | -1751 | 0 | 0 | 19 | 616.79663 | 692.90924 | 691.98645 | 871.58606 | 1157.9463 | 1177.72 |
| Myh9 | KANLQIDQINTDLNLER | -1754 | -1770 | 1 | 0 | 17 | 525.0529 | 625.05865 | 635.3049 | 735.9236 | 666.6154 | 689.6023 |
| Myh9 | KANLQIDQINTDLNLER | -1754 | -1770 | 1 | 0 | 17 | 525.0529 | 625.05865 | 635.3049 | 735.9236 | 666.6154 | 689.6023 |
| Myh9 | QVELDSVTGLLSQSDSK | -1279 | -1295 | 0 | 0 | 17 | 479.78403 | 552.36884 | 558.86 | 1235.0896 | 1207.7723 | 1301.2781 |
| Myh9 | AETELCAEAEELR | -891 | -903 | 0 | 0 | 13 | 475.71048 | 171.63492 | 523.7886 | 374.99252 | 659.55023 | 751.23865 |
| Myh9 | ELDDLLVDLDHQR | -1421 | -1433 | 0 | 0 | 13 | 428.5078 | 485.8271 | 486.27234 | 1125.8467 | 1226.9674 | 1247.2019 |
| Myh9 | VISGVLQLGNIAFK | -342 | -355 | 0 | 0 | 14 | 394.87784 | 388.58002 | 489.4735 | 809.51544 | 671.6992 | 883.28754 |
| Myh9 | MQQNIQELEEQLEEEESAR | -941 | -959 | 0 | 0 | 19 | 382.09335 | 399.47662 | 332.57156 | 402.91003 | 340.37585 | 373.52237 |
| Myh9 | MQQNIQELEEQLEEEESAR | -941 | -959 | 0 | 0 | 19 | 382.09335 | 399.47662 | 332.57156 | 402.91003 | 340.37585 | 373.52237 |
| Myh9 | TELEDTLDSTAAQQELR | -1146 | -1162 | 0 | 0 | 17 | 381.11868 | 636.96027 | 707.1102 | 1035.0441 | 1465.6196 | 446.8349 |
| Myh9 | NAEQFKDQADK | -1857 | -1867 | 1 | 0.000230707 | 11 | 180.4981 | 429.87253 | 510.9999 | 344.16953 | 525.3323 | 611.0091 |
| Myh9 | RGDLPFVVT | -1923 | -1931 | 1 | 0.00272025 | 9 | 3498.1304 | 2490.046 | 257.143 | 1640.6738 | NaN | 1452.3475 |
| Myh9 | SSDKFVSELWK | -603 | -613 | 1 | 0 | 11 | 1823.0857 | 582.6893 | 3050.4805 | NaN | 817.6899 | 1690.1364 |
| Myh9 | CQYLQAEK | -931 | -938 | 0 | 0 | 8 | 1671.3909 | 800.7537 | 852.0548 | 1102.2606 | 722.5745 | NaN |
| Myh9 | ELEDATETADAMNR | -1899 | -1912 | 0 | 0.000230707 | 14 | 868.9409 | 801.3915 | 1020.08215 | NaN | 987.51404 | 1015.225 |
| Myh9 | ELDSNLYR | -748 | -755 | 0 | 0 | 8 | 863.9767 | 680.74164 | 585.76385 | 755.0484 | 486.44702 | NaN |
| Myh9 | DTQELLQEENR | -1312 | -1322 | 0 | 0 | 11 | 789.804 | 1004.45715 | 1130.585 | 1916.6525 | 1532.2852 | NaN |
| Myh9 | LKDVLLQVEDER | -1844 | -1855 | 1 | 0 | 12 | 783.64166 | 1016.33026 | 456.93314 | 2063.531 | NaN | 1492.8016 |
| Myh9 | SVHELEK | -1519 | -1525 | 0 | 0.001463897 | 7 | 675.888 | 865.4551 | NaN | 753.84045 | 776.30286 | 700.2065 |
| Myh9 | NFINNPLAQAD | -15 | -25 | 0 | 0.00025015 | 11 | 364.08316 | 370.75262 | NaN | 485.0168 | 627.28107 | 616.9523 |
| Myh9 | NAEQFKDQADKA | -1857 | -1868 | 2 | 0 | 12 | 334.1409 | 539.76013 | 191.14796 | NaN | 794.00256 | 910.1307 |
| Myh9 | LEEAEEEAQR | -1879 | -1888 | 0 | 0 | 10 | 303.3541 | 1862.8588 | NaN | 1275.8007 | 1547.974 | 269.1584 |
| Myh9 | IAQLEEELEEEQGNTELINDR | -1731 | -1751 | 0 | 0 | 21 | 297.2057 | 227.91045 | 318.7739 | NaN | 424.74994 | 380.34976 |
| Myh9 | AVEELADQLEQTK | -1197 | -1209 | 0 | 0 | 13 | 1531.1083 | 1541.1432 | 1463.4313 | 1507.378 | NaN | NaN |
| Myh9 | QINTDLNLER | -1761 | -1770 | 0 | 0 | 10 | 1166.395 | NaN | NaN | 1389.738 | 1049.049 | 806.62756 |
| Myh9 | ELEDTLDSTAAQQELR | -1147 | -1162 | 0 | 0 | 16 | 1109.1084 | NaN | NaN | 1272.398 | 1419.2112 | 1387.345 |
| Myh9 | TELCAEAEELR | -893 | -903 | 0 | 0 | 11 | 961.5249 | 824.1322 | 836.4314 | 1746.5176 | NaN | NaN |
| Myh9 | YEILTPNSIPK | -721 | -731 | 0 | 0 | 11 | 652.9596 | 663.01794 | 759.14453 | 1274.7338 | NaN | NaN |
| Myh9 | LEEDQIIMEDQNCK | -976 | -989 | 0 | 0 | 14 | 652.1833 | 669.52844 | NaN | 1247.856 | NaN | 1349.5077 |
| Myh9 | QEELAAAER | -1686 | -1694 | 0 | 0 | 9 | 543.8472 | NaN | 827.5305 | 927.3419 | 873.0818 | NaN |
| Myh9 | EEQLEEEESAR | -949 | -959 | 0 | 0 | 11 | 521.89276 | 532.0894 | 673.6279 | NaN | NaN | 587.98425 |
| Myh9 | SIRHEDELLAK | -840 | -850 | 1 | 0 | 11 | 268.59027 | NaN | NaN | 554.53784 | 565.38904 | 571.9388 |
| Myh9 | ALEEAMEQK | -1484 | -1492 | 0 | 0.001610812 | 9 | 257.4007 | NaN | 1992.9349 | 507.4099 | 2629.4875 | NaN |
| Myh9 | RALEEAMEQK | -1483 | -1492 | 1 | 0.00111006 | 10 | 220.05646 | 381.92047 | NaN | 310.07297 | NaN | 502.28827 |
| Myh9 | IQELEEQLEEEESAR | -945 | -959 | 0 | 0 | 15 | 178.62604 | NaN | NaN | 1164.2467 | 1331.5828 | 1276.8308 |
| Myh9 | EELAAAER | -1687 | -1694 | 0 | 0.00453359 | 8 | 2026.457 | NaN | 1958.7001 | 2381.182 | NaN | NaN |
| Myh9 | EQLEEEEEAK | -1343 | -1352 | 0 | 0 | 10 | 1523.3281 | 1409.0814 | 671.26526 | NaN | NaN | NaN |
| Myh9 | DLEGLSQR | -1393 | -1400 | 0 | 0.002329279 | 8 | 858.37885 | 836.9219 | NaN | NaN | 402.94275 | NaN |
| Myh9 | KTQLEELEDELQATEDAK | -1538 | -1555 | 1 | 0.000230707 | 18 | 684.93195 | 670.3711 | 624.12915 | NaN | NaN | NaN |
| Myh9 | KTQLEELEDELQATEDAK | -1538 | -1555 | 1 | 0.000230707 | 18 | 684.93195 | 670.3711 | 624.12915 | NaN | NaN | NaN |
| Myh9 | EQEVSILK | -1166 | -1173 | 0 | 0.00266297 | 8 | 1789.3562 | NaN | 1016.9665 | NaN | NaN | NaN |
| Myh9 | FLSNGHVTIPGQQDK | -302 | -316 | 0 | 0 | 15 | 1372.3909 | NaN | NaN | 2800.191 | NaN | NaN |
| Myh9 | NAEQFKDQ | -1857 | -1864 | 1 | 0.002810811 | 8 | 720.6557 | NaN | NaN | 816.911 | NaN | NaN |
| Myh9 | QLQDTQELLQEENR | -1309 | -1322 | 0 | 0 | 14 | 473.62122 | NaN | NaN | NaN | 822.2413 | NaN |
| Myh9 | SVTGLLSQSDSK | -1284 | -1295 | 0 | 0 | 12 | 1545.1322 | NaN | NaN | NaN | NaN | NaN |

| | | | | | | | | | | | | |
|---|---|---|---|---|---|---|---|---|---|---|---|---|
| Myh9 | LQEMESAVK | -1794 | -1802 | 0 | 0.001868731 | 9 | 1052.9409 | NaN | NaN | NaN | NaN | NaN |
| Myh9 | IMGIPEDEQMGLLR | -328 | -341 | 0 | 0.00025015 | 14 | 173.62173 | NaN | NaN | NaN | NaN | NaN |
| Slc3a2 | WLQDGVDGFQFR | -231 | -242 | 0 | 0.000298805 | 12 | 3211.3691 | 3447.44 | 3866.9534 | 2215.0413 | 2636.2769 | 2689.0354 |
| Slc3a2 | SLLHGDFHAL | -428 | -437 | 0 | 0 | 10 | 2614.5571 | 222.65363 | 2956.9978 | 1803.5098 | 3123.4468 | 3177.3828 |
| Slc3a2 | EEDTSLKLENL | -500 | -510 | 1 | 0.00025015 | 11 | 2253.7102 | 2440.453 | 2587.0762 | 2691.946 | 2887.9922 | 2827.5632 |
| Slc3a2 | GLVLGPIHK | -151 | -159 | 0 | 0.00025015 | 9 | 2196.339 | 2195.189 | 2331.367 | 1837.2532 | 1914.8591 | 1843.9128 |
| Slc3a2 | ADFIPDHLLR | -336 | -345 | 0 | 0 | 10 | 1841.0356 | 2069.5085 | 1998.7119 | 441.29288 | 1702.7943 | 1632.0917 |
| Slc3a2 | QINPTLGSQEDFK | -172 | -184 | 0 | 0 | 13 | 1483.0162 | 1406.9574 | 1443.0159 | 1456.552 | 1624.7048 | 1523.7635 |
| Slc3a2 | NLPAGISLPASAK | -475 | -487 | 0 | 0 | 13 | 1369.2585 | 1562.154 | 1626.8721 | 1940.1079 | 1792.642 | 2011.4475 |
| Slc3a2 | NQKDEINETDLK | -160 | -171 | 1 | 0 | 12 | 1283.2855 | 1562.9662 | 119.92351 | 1989.6708 | 2027.9027 | 2083.4849 |
| Slc3a2 | VAEDETEAGVK | -43 | -53 | 0 | 0 | 11 | 1270.9276 | 1222.3359 | 1300.5557 | 962.15515 | 724.6849 | 614.2375 |
| Slc3a2 | GDELGLQGALPGQPAK | -363 | -378 | 0 | 0 | 16 | 1214.6473 | 1763.1709 | 1328.0953 | 1079.0776 | 1013.60596 | 2734.9058 |
| Slc3a2 | SHLEYLSTL | -139 | -147 | 0 | 0.00025015 | 9 | 967.48706 | 755.27484 | 967.5282 | 1281.8899 | 1239.279 | 1432.3168 |
| Slc3a2 | HWDQNER | -449 | -455 | 0 | 0.00025015 | 7 | 892.491 | 928.4715 | 1019.5536 | 1005.86743 | 1066.3492 | 995.939 |
| Slc3a2 | SSSPDLFSYIR | -438 | -448 | 0 | 0 | 11 | 622.48975 | 2885.9646 | 2836.58 | 2952.505 | 181.22797 | 3708.4233 |
| Slc3a2 | LGASNLPAGISLPASAK | -471 | -487 | 0 | 0 | 17 | 577.2678 | 859.4768 | 834.0516 | 733.52563 | 905.26215 | 951.09827 |
| Slc3a2 | QINPTLGSQEDFKDLLQ | -172 | -188 | 1 | 0.00025015 | 17 | 570.1024 | 563.31854 | 566.22546 | 579.0312 | 841.9247 | 1511.6786 |
| Slc3a2 | QINPTLGSQEDFKDLLQSAK | -172 | -191 | 1 | 0 | 20 | 362.3278 | 537.63855 | 915.6926 | 755.94183 | 709.8888 | 743.6344 |
| Slc3a2 | GQNEDPGSLLTQF | -404 | -416 | 0 | 0.000627943 | 13 | 105.00957 | 684.1804 | 474.06293 | 518.3255 | 559.9972 | 507.14023 |
| Slc3a2 | DVELNELEPEK | -11 | -21 | 0 | 0 | 11 | 4251.9644 | 2932.608 | 4295.9937 | 3086.0977 | 3312.2798 | NaN |
| Slc3a2 | SHLEYLSTLK | -139 | -148 | 0 | 0.00025015 | 10 | 2267.5808 | 1106.7961 | 2319.536 | NaN | 1954.2399 | 1932.0587 |
| Slc3a2 | NEDPGSLLTQFR | -406 | -417 | 0 | 0 | 12 | 714.63666 | 800.6696 | NaN | 909.638 | 794.80194 | 980.2306 |
| Slc3a2 | WFLPAQADIVATK | -211 | -223 | 0 | 0 | 13 | 563.59784 | 2246.3506 | 3583.7988 | 5378.4688 | NaN | 5952.456 |
| Slc3a2 | QKDEINETDLK | -161 | -171 | 1 | 0 | 11 | 561.4161 | NaN | 853.6099 | 1629.8182 | 1165.783 | 953.4679 |
| Slc3a2 | GQNEDPGSLLTQFR | -404 | -417 | 0 | 0 | 14 | 2385.6719 | NaN | 1995.89 | 1730.2983 | 3348.0857 | NaN |
| Slc3a2 | LGASNLPAGISLPA | -471 | -484 | 0 | 0 | 14 | 590.2043 | 640.3126 | NaN | 455.90814 | 538.6226 | NaN |
| Slc3a2 | FLPAQADIVATK | -212 | -223 | 0 | 0 | 12 | 1055.6144 | NaN | 361.9632 | NaN | NaN | 477.4882 |
| Slc3a2 | SSPDLFSYIR | -439 | -448 | 0 | 0.00111006 | 10 | 475.12714 | 307.7646 | 276.2399 | NaN | NaN | NaN |
| Slc3a2 | VAGSPGWVR | -65 | -73 | 0 | 0.00025015 | 9 | 855.81537 | NaN | NaN | 1429.3643 | NaN | NaN |
| Vars1 | YVSPHPDAFPSLR | -5 | -17 | 0 | 0 | 13 | 811.4255 | 114.264084 | 2186.3914 | 1259.0994 | 1221.378 | 458.5041 |
| Vars1 | YVSPHPDAFPSLR | -5 | -17 | 0 | 0 | 13 | 811.4255 | 114.264084 | 2186.3914 | 1259.0994 | 1221.378 | 458.5041 |
| Vars1 | ICLQPPPSSR | -40 | -49 | 0 | 0 | 10 | 2838.3962 | 3606.639 | 663.651 | 3016.4692 | 2632.678 | 3611.7847 |
| Vars1 | VPLEVQEADEAK | -1230 | -1241 | 0 | 0 | 12 | 1851.5989 | 1950.952 | 2140.2651 | 2539.9602 | 2421.7244 | 2397.8076 |
| Vars1 | TLLPVPGYK | -498 | -506 | 0 | 0 | 9 | 1822.27 | 1792.6663 | 2120.1423 | 1674.3829 | 1188.9417 | 1442.9635 |
| Vars1 | VHDPAVPPGEDPDGR | -737 | -751 | 0 | 0 | 15 | 1479.9908 | 2006.1449 | 2096.3296 | 959.0007 | 1899.0374 | 1732.4302 |
| Vars1 | YGEAGDGPGWGGPHPR | -24 | -39 | 0 | 0 | 16 | 1455.6262 | 1485.4407 | 1412.1821 | 1373.6528 | 1539.2346 | 1563.8628 |
| Vars1 | YGEAGDGPGWGGPHPR | -24 | -39 | 0 | 0 | 16 | 1455.6262 | 1485.4407 | 1412.1821 | 1373.6528 | 1539.2346 | 1563.8628 |
| Vars1 | NSAISDIEVDKK | -481 | -492 | 1 | 0 | 12 | 1304.3656 | 1528.5538 | 1364.0336 | 1445.85 | 921.81085 | 1045.7821 |
| Vars1 | SKDVVEPLLRPQ | -665 | -676 | 2 | 0.00025015 | 12 | 1221.5009 | 1001.5421 | 1846.6787 | 1235.2013 | 1229.943 | 1193.5981 |
| Vars1 | GALINVPPPFLGLPR | -618 | -632 | 0 | 0 | 15 | 1204.3309 | 1152.8279 | 1174.6892 | 1427.161 | 1753.9583 | 1665.5919 |
| Vars1 | ALNPLEDWLR | -138 | -147 | 0 | 0 | 10 | 1163.9012 | 1269.1803 | 1344.997 | 1540.426 | 1796.424 | 163.40813 |
| Vars1 | ILPEAHQR | -699 | -706 | 0 | 0.001291103 | 8 | 1048.7803 | 907.3741 | 904.51196 | 1320.3591 | 1008.0567 | 1028.589 |
| Vars1 | ISDIEVDKK | -484 | -492 | 1 | 0.00266297 | 9 | 982.9438 | 992.2123 | 978.9249 | 1163.8622 | 1095.5446 | 1059.7977 |
| Vars1 | ITPAHDQNDYEVGQR | -591 | -605 | 0 | 0 | 15 | 980.8479 | 996.73785 | 1257.4901 | 1876.9381 | 1275.4072 | 1169.6285 |
| Vars1 | ITPAHDQNDYEVGQR | -591 | -605 | 0 | 0 | 15 | 980.8479 | 996.73785 | 1257.4901 | 1876.9381 | 1275.4072 | 1169.6285 |
| Vars1 | DFPAGIPECGTDALR | -908 | -922 | 0 | 0 | 15 | 867.17413 | 998.82104 | 498.19638 | 1624.3496 | 1795.5107 | 789.2895 |
| Vars1 | SAISDIEVDKK | -482 | -492 | 1 | 0.000431655 | 11 | 705.9648 | 893.76154 | 1532.6466 | 1809.5714 | 1124.7455 | 1686.8634 |
| Vars1 | SVTQQPGSEVIAPQK | -215 | -229 | 0 | 0 | 15 | 567.6216 | 651.646 | 721.82056 | 307.25647 | 647.5164 | 592.6423 |
| Vars1 | GATLPALGLR | -113 | -122 | 0 | 0 | 10 | 562.9009 | 130.53935 | 1307.9238 | 534.527 | 475.04346 | 548.84625 |
| Vars1 | SILYVSPHPDAFPSLR | -2 | -17 | 0 | 0 | 16 | 361.76703 | 341.52655 | 407.30432 | 685.6329 | 797.6755 | 422.58133 |
| Vars1 | LGSSLDWDR | -433 | -441 | 0 | 0 | 9 | 13734.58 | 22961.898 | 1297.7257 | NaN | 26440.213 | 706.6191 |
| Vars1 | LHEEGVIYR | -461 | -469 | 0 | 0 | 9 | 2823.088 | 291.26886 | 2473.2407 | NaN | 1952.0886 | 3053.737 |
| Vars1 | LLWPAGLGGPGGSR | -78 | -91 | 0 | 0 | 14 | 1801.9684 | NaN | 2068.964 | 1705.8978 | 1764.7915 | 1902.9167 |
| Vars1 | SKPEGHESLVDR | -973 | -984 | 1 | 0 | 12 | 1455.7498 | NaN | 1471.0875 | 837.2212 | 1755.7429 | 1478.5244 |
| Vars1 | TNAIQDSLTR | -356 | -365 | 0 | 0.000347792 | 10 | 1412.9103 | 1390.3705 | 1157.632 | 1118.462 | NaN | 1086.9989 |
| Vars1 | TPFPPPR | -50 | -56 | 0 | 0 | 7 | 936.78973 | 709.7298 | 627.7847 | 725.49054 | 402.5866 | NaN |
| Vars1 | ADFPAGIPECGTDALR | -907 | -922 | 0 | 0 | 16 | 805.02625 | 1148.2703 | 1187.3817 | NaN | 1249.8164 | 1142.0647 |
| Vars1 | HDQNDYEVGQR | -595 | -605 | 0 | 0 | 11 | 376.84628 | NaN | 407.38855 | 369.5153 | 566.1249 | 593.5374 |
| Vars1 | SLGNVIDPLDVIHGV | -865 | -879 | 0 | 0 | 15 | 331.57034 | 292.17084 | NaN | 321.25687 | 269.40594 | 279.75894 |
| Vars1 | SLGNVIDPLDVIH | -865 | -877 | 0 | 0 | 13 | 3054.3252 | 2083.1624 | 3042.8435 | NaN | NaN | NaN |
| Vars1 | DVVEPLLRPQ | -667 | -676 | 1 | 0 | 10 | 2400.8762 | 1347.325 | 1959.4086 | NaN | NaN | NaN |
| Vars1 | QLWWGHR | -723 | -729 | 0 | 0.000394361 | 7 | 1803.3816 | 1852.8636 | NaN | 1810.3145 | NaN | NaN |
| Vars1 | LQQTEAELR | -1242 | -1250 | 0 | 0 | 9 | 1438.5695 | 1404.7925 | 1635.8618 | NaN | NaN | NaN |
| Vars1 | QGFFKPEYGR | -315 | -324 | 1 | 0 | 10 | 1316.5685 | 1688.1544 | 1697.0522 | NaN | NaN | NaN |
| Vars1 | PSVSAPNPR | -325 | -333 | 0 | 0 | 9 | 587.42303 | 805.6331 | 765.35333 | NaN | NaN | NaN |
| Vars1 | GLYDQLLNSNLDPSEVEK | -883 | -900 | 0 | 0.00025015 | 18 | 500.3306 | 529.16034 | 449.622 | NaN | NaN | NaN |
| Dad1 | FLEEYLSSTPQR | -12 | -23 | 0 | 0 | 12 | 2844.181 | 3396.6514 | 3469.295 | 3075.7288 | 3560.4158 | 3924.0022 |
| Dad1 | ADFQGISPER | -83 | -92 | 0 | 0 | 10 | 1801.4462 | 1807.5604 | 1916.1316 | 1605.7285 | 1380.6111 | 1371.1954 |
| Dad1 | IQINPQNK | -75 | -82 | 0 | 0 | 8 | 677.93677 | 97.135765 | 559.50354 | NaN | NaN | 918.9722 |
| Got1 | IGADFLGR | -115 | -122 | 0 | 0.00025015 | 8 | 2199.971 | 1819.2214 | 1815.2655 | 1100.1265 | 1251.0348 | 1289.1537 |
| Got1 | AHNPTGTDPTPEQWK | -193 | -207 | 0 | 0 | 15 | 1145.2521 | 1089.2847 | 257.0019 | 997.8344 | 960.2003 | 1001.97485 |
| Got1 | AHNPTGTDPTPEQWK | -193 | -207 | 0 | 0 | 15 | 1145.2521 | 1089.2847 | 257.0019 | 997.8344 | 960.2003 | 1001.97485 |
| Got1 | LVLGDNSLAIR | -87 | -97 | 0 | 0 | 11 | 1623.7211 | 1788.4545 | 1840.115 | 2132.1863 | 2077.7556 | 1963.6196 |
| Got1 | HIYLLPSGR | -379 | -387 | 0 | 0 | 9 | 1556.5234 | 1437.0996 | 1513.5225 | 1610.4198 | 1706.7792 | 1673.1134 |
| Got1 | RTDESQPWVLPVVR | -42 | -55 | 1 | 0 | 14 | 1338.1566 | 277.01785 | 952.4731 | 6833.9644 | 600.652 | 480.05826 |
| Got1 | RTDESQPWVLPVVR | -42 | -55 | 1 | 0 | 14 | 1338.1566 | 277.01785 | 952.4731 | 6833.9644 | 600.652 | 480.05826 |
| Got1 | TDESQPWVLPVVR | -43 | -55 | 0 | 0 | 13 | 1196.5973 | 1450.7882 | 1350.0579 | 2414.4673 | 1518.6581 | 2939.0044 |
| Got1 | AQVPQAPPVLVFK | -8 | -20 | 0 | 0 | 13 | 743.84644 | 983.6816 | 892.97394 | 748.5642 | 327.3049 | 634.73834 |

| | | | | | | | | | | | | |
|---|---|---|---|---|---|---|---|---|---|---|---|---|
| Got1 | IANDNSLNHEYLPILGLA | -61 | -78 | 0 | 0.000869338 | 18 | 653.37476 | 532.4839 | 496.02267 | 624.06213 | 850.5318 | 728.49695 |
| Got1 | NTPIYVSSPTWENHNAVFSAA | -131 | -151 | 0 | 0 | 21 | 500.8649 | 549.38367 | 652.72144 | 770.139 | 945.6813 | 1034.265 |
| Got1 | NTPIYVSSPTWENHNAVF | -131 | -148 | 0 | 0 | 18 | 326.0891 | 349.63025 | 653.97186 | 465.89636 | 656.46277 | 382.92212 |
| Got1 | NTPIYVSSPTWENHNAVFS | -131 | -149 | 0 | 0.000627943 | 19 | 303.39542 | 228.31845 | 390.79004 | 279.29062 | 350.4643 | 351.19183 |
| Got1 | APPSVFAQVPQAPPVLVFK | -2 | -20 | 0 | 0.00025015 | 19 | 232.23184 | 307.9726 | 238.02599 | 278.8257 | 364.88556 | 332.0036 |
| Got1 | NFGLYNER | -260 | -267 | 0 | 0 | 8 | 1163.2155 | NaN | 1279.6345 | 1174.935 | 278.24155 | 1391.4105 |
| Got1 | IANDNSLNHEYLPILGL | -61 | -77 | 0 | 0.000230707 | 17 | 308.3393 | 318.618 | NaN | 148.036 | 415.9582 | 399.05103 |
| Got1 | TLSDPELFK | -310 | -318 | 0 | 0.002099344 | 9 | 3875.8782 | NaN | NaN | NaN | NaN | NaN |
| Got1 | ADFRDDPDPR | -23 | -32 | 1 | 0.004595494 | 10 | 1075.8157 | NaN | NaN | NaN | NaN | NaN |
| Icam1 | ADGALLPIGVVK | -438 | -449 | 0 | 0 | 12 | 1068.213 | 1216.8632 | 984.7943 | 1137.2412 | 951.50616 | 1040.9415 |
| Icam1 | TFDLPATIPK | -213 | -222 | 0 | 0 | 10 | 3577.1123 | 3960.379 | 3939.8625 | 2881.0789 | 3244.4976 | NaN |
| Icam1 | WLKDELESGPNWK | -64 | -76 | 1 | 0 | 13 | 575.1771 | 764.2271 | 632.9396 | 1176.1029 | 735.48987 | NaN |
| Icam1 | TELDLRPQGLAL | -190 | -201 | 1 | 0.00025015 | 12 | 363.33377 | NaN | NaN | 745.9588 | 555.5968 | 815.1501 |
| Icam1 | KADGALLPIGVVK | -437 | -449 | 1 | 0 | 13 | 162.07037 | 212.44556 | NaN | NaN | NaN | NaN |
| Icam1 | EGLFPASEAR | -242 | -251 | 0 | 0.001291103 | 10 | 373.54904 | NaN | NaN | NaN | NaN | NaN |
| Cdkn2aipnl | HLPDYRDPPDGGGR | -47 | -60 | 1 | 0 | 14 | 1706.3441 | 1298.324 | 1185.617 | 1002.60693 | 704.4879 | 843.2332 |
| Cdkn2aipnl | ADGIEVEDLPQFTTR | -93 | -107 | 0 | 0 | 15 | 466.2314 | 522.4845 | 515.0847 | 447.3099 | 227.28523 | 542.5273 |
| Mrpl24 | ADGIVPETWTDGPK | -162 | -175 | 0 | 0 | 14 | 1838.184 | 2165.259 | 1797.4266 | 1274.3018 | 1869.6173 | 1600.6447 |
| Rps8 | SLLEEQFQQGK | -160 | -170 | 0 | 0 | 11 | 8561.634 | 8825.213 | 9339.807 | 9148.893 | 9107.308 | 9708.791 |
| Rps8 | SLLEEQFQQGK | -160 | -170 | 0 | 0 | 11 | 8561.634 | 8825.213 | 9339.807 | 9148.893 | 9107.308 | 9708.791 |
| Rps8 | KYELGRPA | -26 | -33 | 2 | 0.00122419 | 8 | 6197.31 | 5435.938 | 5181.2383 | 2373.6855 | 5698.526 | 5669.034 |
| Rps8 | SWGSECCTR | -66 | -74 | 0 | 0 | 9 | 2817.5767 | 2764.7036 | 2796.6938 | 3819.408 | 3350.487 | 3714.191 |
| Rps8 | IVLIDSTPYR | -101 | -110 | 0 | 0 | 10 | 1841.353 | 1982.8656 | 2055.1812 | 2819.7532 | 2717.9136 | 3044.8618 |
| Rps8 | NCIVLIDSTPYR | -99 | -110 | 0 | 0 | 12 | 1585.1929 | 1847.6565 | 2214.1885 | 1660.9983 | 1903.9088 | 1870.7355 |
| Rps8 | QWYESHYALPL | -111 | -121 | 0 | 0 | 11 | 1153.2281 | 296.06332 | 1313.1571 | 350.84482 | 300.77194 | 2012.2455 |
| Rps8 | LTPEEEEILNK | -129 | -139 | 0 | 0 | 11 | 833.55145 | 3616.9607 | 3826.361 | 2478.378 | 4394.951 | 3921.5193 |
| Rps8 | LEEQFQQGK | -162 | -170 | 0 | 0 | 9 | 1749.3319 | NaN | 1670.8602 | 1918.1891 | 1665.3212 | 2009.952 |
| Rps8 | ISSLLEEQFQQGK | -158 | -170 | 0 | 0 | 13 | 1557.6849 | 1385.4939 | 1676.7244 | 2302.4585 | NaN | 557.02716 |
| Rps8 | VGNFSWGSECCTR | -62 | -74 | 0 | 0 | 13 | 742.7053 | 770.3115 | 776.00165 | 827.862 | 657.9706 | NaN |
| Rps8 | IIDVVYNASNNELVR | -78 | -92 | 0 | 0 | 15 | 457.62012 | NaN | 390.72467 | 186.92245 | 416.22827 | 446.68784 |
| Rps8 | LDVGNFSWGSECCTR | -60 | -74 | 0 | 0 | 15 | 313.0165 | 366.5858 | 331.5301 | 723.41364 | 888.25146 | NaN |
| Rps8 | CIVLIDSTPYR | -100 | -110 | 0 | 0 | 11 | 974.8396 | NaN | NaN | 1567.199 | 949.5572 | 1368.2576 |
| Rps8 | ADGYVLEGK | -185 | -193 | 0 | 0 | 9 | 299.41187 | 1727.6613 | NaN | NaN | NaN | NaN |
| Ppp1cc | NVQLQENEIR | -27 | -36 | 0 | 0 | 10 | 1010.693 | 869.3175 | 1197.5784 | 1166.0259 | 1154.6333 | 687.02795 |
| Ppp1cc | ADIDKLNIDSIIQR | -2 | -15 | 1 | 0 | 14 | 187.66505 | 481.17227 | 404.04547 | 466.2886 | 480.8079 | 461.88303 |
| Ppp1cc | LNIDSIIQR | -7 | -15 | 0 | 0.000471409 | 9 | 777.92957 | NaN | 275.27167 | NaN | 740.5312 | NaN |
| Hnrnpk | RPAEDMEEEQAFK | -22 | -34 | 1 | 0.00025015 | 13 | 10134.256 | 5585.0005 | 1186.0333 | 6059.615 | 3550.6687 | 3899.897 |
| Hnrnpk | RPAEDMEEEQAFK | -22 | -34 | 1 | 0.00025015 | 13 | 10134.256 | 5585.0005 | 1186.0333 | 6059.615 | 3550.6687 | 3899.897 |
| Hnrnpk | VVLIGGKPDR | -192 | -201 | 1 | 0.00051306 | 10 | 9630.16 | 9992.725 | 9821.428 | 10152.814 | 9811.944 | 10072.963 |
| Hnrnpk | VSVPDSSGPER | -76 | -86 | 0 | 0 | 11 | 6170.8105 | 6372.3984 | 6068.2617 | 5177.6343 | 5047.6074 | 5363.621 |
| Hnrnpk | IDEPLEGSEDR | -423 | -433 | 0 | 0 | 11 | 5793.292 | 5849.4727 | 6364.098 | 6137.522 | 6344.9307 | 6637.4214 |
| Hnrnpk | IILDLISESPIK | -208 | -219 | 0 | 0 | 12 | 3720.7983 | 3427.636 | 3345.3198 | 4049.8997 | 4581.365 | 4471.2886 |
| Hnrnpk | IILDLISESPIK | -208 | -219 | 0 | 0 | 12 | 3720.7983 | 3427.636 | 3345.3198 | 4049.8997 | 4581.365 | 4471.2886 |
| Hnrnpk | SLAGGIIGVK | -154 | -163 | 0 | 0 | 10 | 3576.3489 | 2988.6335 | 2939.4817 | 1959.8351 | 1259.7925 | 117.69823 |
| Hnrnpk | ADIETIGEILK | -92 | -102 | 0 | 0 | 11 | 3055.336 | 3548.2896 | 3279.8518 | 3054.124 | 3994.7644 | 3995.2207 |
| Hnrnpk | GSDFDCELR | -140 | -148 | 0 | 0 | 9 | 2558.713 | 2877.1306 | 2119.134 | 2644.1226 | 1725.1543 | 1415.4331 |
| Hnrnpk | NLPLPPPPPR | -306 | -316 | 0 | 0 | 11 | 2224.1804 | 2456.1975 | 2385.751 | 2128.202 | 1370.9885 | 2185.3809 |
| Hnrnpk | LFQECCPHSTDR | -180 | -191 | 0 | 0 | 12 | 1654.5944 | 1799.7887 | 1736.7362 | 1938.7571 | 1867.6406 | 1841.5938 |
| Hnrnpk | LFQECCPHSTDR | -180 | -191 | 0 | 0 | 12 | 1654.5944 | 1799.7887 | 1736.7362 | 1938.7571 | 1867.6406 | 1841.5938 |
| Hnrnpk | GSYGDLGGPIITTQ | -378 | -391 | 0 | 0 | 14 | 1092.5303 | 1086.3854 | 1086.4359 | 1030.1305 | 221.2301 | 228.04102 |
| Hnrnpk | IHQSLAGGIIGVK | -151 | -163 | 0 | 0 | 13 | 971.75494 | 1020.41644 | 1070.0236 | 1835.5808 | 1930.8074 | 1901.0199 |
| Hnrnpk | ADIETIGEILKK | -92 | -103 | 1 | 0 | 12 | 951.32904 | 1040.7798 | 1004.9706 | 1852.0508 | 2429.0342 | 2294.1858 |
| Hnrnpk | KGSDFDCELR | -139 | -148 | 1 | 0.00025015 | 10 | 887.2731 | 758.3083 | 807.11163 | 2298.4114 | 518.32587 | 2483.267 |
| Hnrnpk | AYEPQGGSGYDYSYAGGR | -360 | -377 | 0 | 0 | 18 | 762.6729 | 861.6681 | 893.21936 | 883.2165 | 1017.0699 | 1001.7648 |
| Hnrnpk | QSLAGGIIGVK | -153 | -163 | 0 | 0 | 11 | 751.9644 | 615.9463 | 733.65564 | 918.4987 | 524.34357 | 592.18243 |
| Hnrnpk | RTDYNASVSVPDSSGPER | -69 | -86 | 1 | 0 | 18 | 749.9264 | 516.2289 | 454.9098 | 584.72485 | 1300.4294 | 1287.0304 |
| Hnrnpk | IIPTLEEGLQLPSPTAT | -104 | -120 | 0 | 0 | 17 | 566.61725 | 671.2055 | 610.79846 | 882.5359 | 1058.2571 | 985.64526 |
| Hnrnpk | KIIPTLEEGLQLPSPTAT | -103 | -120 | 1 | 0 | 18 | 438.78503 | 608.6538 | 676.5735 | 749.67303 | 1077.5239 | 1009.27814 |
| Hnrnpk | AQPYDPNFYDE | -222 | -232 | 0 | 0 | 11 | 255.82109 | 192.60889 | 249.78745 | 262.46616 | 194.2345 | 218.44667 |
| Hnrnpk | AQPYDPNFYDE | -222 | -232 | 0 | 0 | 11 | 255.82109 | 192.60889 | 249.78745 | 262.46616 | 194.2345 | 218.44667 |
| Hnrnpk | IIPTLEEGLQLPSPT | -104 | -118 | 0 | 0 | 15 | 336.44638 | 352.30743 | 195.21875 | 341.15088 | 534.28076 | 530.7249 |
| Hnrnpk | GSYGDLGGPIITT | -378 | -390 | 0 | 0 | 13 | 261.59216 | 216.50607 | 327.10342 | 375.40414 | 395.56476 | 336.1275 |
| Hnrnpk | GGSGYDYSYAGGR | -365 | -377 | 0 | 0 | 13 | 223.31354 | 910.9296 | 170.63487 | 522.92584 | 1096.673 | 164.24846 |
| Hnrnpk | RPAEDMEEEQAFKR | -22 | -35 | 2 | 0 | 14 | 1628.6105 | 732.8933 | 1361.4055 | NaN | 1262.9956 | 2516.801 |
| Hnrnpk | NTDEMVELR | -38 | -46 | 0 | 0 | 9 | 1474.1451 | 1431.0979 | 1185.1508 | NaN | 1983.1139 | 2054.0393 |
| Hnrnpk | IETIGEILK | -94 | -102 | 0 | 0.001055699 | 9 | 1132.1691 | NaN | 1225.4528 | 1196.0204 | 803.18854 | 1171.8146 |
| Hnrnpk | LAGGIIGVK | -155 | -163 | 0 | 0.001291103 | 9 | 1092.8555 | 556.97974 | 576.14856 | 514.10657 | 478.68927 | NaN |
| Hnrnpk | IKIDEPLEGSEDR | -421 | -433 | 1 | 0 | 13 | 802.0886 | NaN | 704.8469 | 1265.0865 | 995.4605 | 1048.1809 |
| Hnrnpk | TDYNASVSVPDSSGPER | -70 | -86 | 0 | 0 | 17 | 775.21277 | 860.07336 | NaN | 1192.8373 | 1261.9585 | 1211.8225 |
| Hnrnpk | YLLQNSVK | -449 | -456 | 0 | 0 | 8 | 766.8187 | NaN | 802.8379 | 753.12494 | 779.5119 | 746.02924 |
| Hnrnpk | IILDLISESPIKG | -208 | -220 | 1 | 0 | 13 | 578.0264 | 712.9035 | 482.39288 | NaN | 763.2008 | 865.0836 |
| Hnrnpk | SISADIETIGEILK | -89 | -102 | 0 | 0 | 14 | 395.37787 | NaN | 402.55862 | 352.71906 | 368.88806 | 462.50864 |
| Hnrnpk | YEPQGGSGYDYSYAGGR | -361 | -377 | 0 | 0 | 17 | 541.53564 | NaN | NaN | 796.8478 | 683.6468 | 665.10443 |
| Hnrnpk | QECCPHSTDR | -182 | -191 | 0 | 0.002329279 | 10 | 279.53284 | NaN | 617.07166 | 358.99966 | 472.70685 | NaN |
| Hnrnpk | LIGGKPDR | -194 | -201 | 1 | 0.007609801 | 8 | 964.4845 | 1000.6544 | NaN | 1018.6282 | NaN | NaN |
| Glg1 | LSSDCEDQIR | -976 | -985 | 0 | 0 | 10 | 1402.7233 | 1101.3733 | 1553.3193 | 2165.8682 | 5153.3145 | 1953.9396 |
| Glg1 | FCHGILTK | -945 | -952 | 0 | 0 | 8 | 1282.109 | 992.5199 | 1108.8759 | 806.4109 | 1455.7058 | 873.10645 |
| Glg1 | ADIFVDPVLHTA | -1055 | -1066 | 0 | 0 | 12 | 294.32278 | 505.79474 | 843.1136 | NaN | 901.31665 | NaN |

| | | | | | | | | | | | | |
|---|---|---|---|---|---|---|---|---|---|---|---|---|
| Glg1 | ELSSDDFHLDR | -296 | -306 | 0 | 0 | 11 | 935.5641 | NaN | NaN | NaN | 688.24194 | 677.17456 |
| Atp6v0a2 | ADIPLPEGEASPPAPPLK | -75 | -92 | 0 | 0.000712251 | 18 | 638.91016 | NaN | 359.53644 | NaN | NaN | NaN |
| Atp6v0a2 | LSQSQEILR | -437 | -445 | 0 | 0.001392869 | 9 | 777.3809 | NaN | NaN | NaN | NaN | NaN |
| Mrps23 | AFDLFNPNFK | -84 | -93 | 0 | 0 | 10 | 1475.572 | 702.22534 | 540.3978 | NaN | 1777.8523 | 2192.552 |
| Mrps23 | ADIQDIFYQEDQIR | -58 | -71 | 0 | 0 | 14 | 332.9358 | NaN | 460.97815 | NaN | 1240.8414 | 1041.3424 |
| Mrps23 | SEGHEPQEDDDLAQR | -149 | -163 | 0 | 0.001868731 | 15 | 188.40656 | NaN | 233.13802 | NaN | 334.3584 | 337.55368 |
| Mrps23 | YTELQNLGETDEEK | -103 | -116 | 0 | 0 | 14 | 995.9609 | NaN | 909.29944 | NaN | NaN | NaN |
| Csnk2a1 | NQHLVSPEALDFLDK | -289 | -303 | 0 | 0 | 15 | 1412.5934 | 1734.0549 | 1652.8204 | 2058.8013 | 2796.1804 | 2882.3108 |
| Csnk2a1 | NQHLVSPEALDFLDK | -289 | -303 | 0 | 0 | 15 | 1412.5934 | 1734.0549 | 1652.8204 | 2058.8013 | 2796.1804 | 2882.3108 |
| Csnk2a1 | TPALVFEHVNNTDFK | -108 | -122 | 0 | 0 | 15 | 506.17834 | 1007.4339 | 620.1108 | 1228.155 | 1300.7103 | 1331.2273 |
| Csnk2a1 | ADIVKDPVSR | -98 | -107 | 1 | 0 | 10 | 964.5791 | NaN | 664.02936 | 1101.6932 | 933.96265 | 129.52596 |
| Csnk2a1 | TDVNTHRPR | -13 | -21 | 1 | 0.001392869 | 9 | 290.96423 | 236.24055 | 303.93686 | 378.63718 | 231.99918 | NaN |
| Csnk2a1 | TDVNTHRPR | -13 | -21 | 1 | 0.001392869 | 9 | 290.96423 | 236.24055 | 303.93686 | 378.63718 | 231.99918 | NaN |
| Csnk2a1 | EPFFHGHDNYDQLVR | -230 | -244 | 0 | 0 | 15 | 264.99487 | 1311.5477 | NaN | 1201.6686 | 1772.9377 | 1446.5892 |
| Csnk2a1 | DIVKDPVSR | -99 | -107 | 1 | 0 | 9 | 2043.1775 | NaN | 2081.2847 | 2447.7014 | NaN | NaN |
| Csnk2a1 | VVEWGNQDDYQLVR | -30 | -43 | 0 | 0.000230707 | 14 | 571.29846 | NaN | 725.4491 | NaN | NaN | NaN |
| Npm1 | VDNDENEHQLSLR | -33 | -45 | 0 | 0 | 13 | 12080.013 | 12168.011 | 12275.687 | 13151.829 | 13114.389 | 12404.994 |
| Npm1 | VDNDENEHQLSLR | -33 | -45 | 0 | 0 | 13 | 12080.013 | 12168.011 | 12275.687 | 13151.829 | 13114.389 | 12404.994 |
| Npm1 | DLKPSTPR | -212 | -219 | 1 | 0.000394361 | 8 | 11267.219 | 11517.437 | 10769.524 | 11366.647 | 8984.816 | 10550.482 |
| Npm1 | DENEHQLSLR | -36 | -45 | 0 | 0 | 10 | 5169.4 | 3905.3801 | 4323.681 | 6281.68 | 6730.819 | 6027.2725 |
| Npm1 | ADKDYHFK | -25 | -32 | 1 | 0.000230707 | 8 | 4085.8208 | 3112.9712 | 3754.9705 | 365.3946 | 3269.196 | 4034.895 |
| Npm1 | ADKDYHFK | -25 | -32 | 1 | 0.000230707 | 8 | 4085.8208 | 3112.9712 | 3754.9705 | 365.3946 | 3269.196 | 4034.895 |
| Npm1 | ADKDYHFK | -25 | -32 | 1 | 0.000230707 | 8 | 4085.8208 | 3112.9712 | 3754.9705 | 365.3946 | 3269.196 | 4034.895 |
| Npm1 | EITPPVVLR | -93 | -101 | 0 | 0.000550307 | 9 | 3537.9414 | 3365.311 | 3378.6545 | 2403.9524 | 2017.2365 | 2231.1267 |
| Npm1 | VDNDENEHQL | -33 | -42 | 0 | 0 | 10 | 3300.4805 | 1980.1115 | 190.24625 | 2215.6846 | 1644.6537 | 1019.0767 |
| Npm1 | NDENEHQLSLR | -35 | -45 | 0 | 0 | 11 | 3256.5518 | 3289.086 | 3480.937 | 2801.8418 | 2924.3481 | 2812.63 |
| Npm1 | NDENEHQLSLR | -35 | -45 | 0 | 0 | 11 | 3256.5518 | 3289.086 | 3480.937 | 2801.8418 | 2924.3481 | 2812.63 |
| Npm1 | CGSGPVHIS | -104 | -112 | 0 | 0.00051306 | 9 | 2925.1345 | 2944.6458 | 2852.333 | 244.60175 | 2676.518 | 2899.9595 |
| Npm1 | DQEAIQDLWQWR | -278 | -289 | 0 | 0 | 12 | 2876.1482 | 2856.8872 | 3332.38 | 3432.053 | 5193.274 | 4430.1855 |
| Npm1 | GGFEITPPVVLR | -90 | -101 | 0 | 0 | 12 | 2188.4985 | 2474.5942 | 2526.9177 | 2006.7698 | 2522.309 | 2603.1711 |
| Npm1 | EAIQDLWQWR | -280 | -289 | 0 | 0 | 10 | 2008.2612 | 1930.3553 | 2210.711 | 1885.3234 | 2313.7983 | 1654.3008 |
| Npm1 | VSLGGFEITPPVVLR | -87 | -101 | 0 | 0 | 15 | 1904.4407 | 1756.2356 | 1497.9719 | 1457.1421 | 1521.1533 | 1385.6505 |
| Npm1 | AMNYEGSPIK | -64 | -73 | 0 | 0 | 10 | 1462.6887 | 1497.7919 | 634.13416 | 923.23645 | 1776.0482 | 2175.1482 |
| Npm1 | AMNYEGSPIK | -64 | -73 | 0 | 0 | 10 | 1462.6887 | 1497.7919 | 634.13416 | 923.23645 | 1776.0482 | 2175.1482 |
| Npm1 | NYLFGCELK | -16 | -24 | 0 | 0 | 9 | 1351.7557 | 617.30884 | 1392.6094 | 3771.7034 | 3532.6577 | 3370.6223 |
| Npm1 | SAPGGGNKVPQK | -143 | -154 | 1 | 0 | 12 | 1324.261 | 1419.3591 | 1661.1754 | 1383.478 | 1794.6587 | 1801.2625 |
| Npm1 | LGGFEITPPVVLR | -89 | -101 | 0 | 0 | 13 | 1224.6016 | 1089.6809 | 1300.5875 | 1093.3809 | 1257.2587 | 1248.9779 |
| Npm1 | GFEITPPVVLR | -91 | -101 | 0 | 0 | 11 | 1105.3373 | 1155.0599 | 1157.8702 | 862.4007 | 160.67857 | 1044.5559 |
| Npm1 | DAESEDEDEEDVK | -122 | -134 | 0 | 0.000230707 | 13 | 1070.8701 | 1186.1472 | 1339.4304 | 622.0479 | 737.0666 | 715.1615 |
| Npm1 | DELHIVEAEAM | -55 | -65 | 0 | 0 | 11 | 997.7153 | 1106.7076 | 1770.3665 | 2193.0186 | 2362.5864 | 1927.8208 |
| Npm1 | TVSLGAGAKDELHIV | -46 | -60 | 1 | 0 | 15 | 874.4511 | 1011.65564 | 939.019 | 948.98914 | 1016.61566 | 1016.7194 |
| Npm1 | DNDENEHQLSLR | -34 | -45 | 0 | 0 | 12 | 690.864 | 734.4328 | 555.8232 | 1174.0076 | 409.9567 | 1427.6202 |
| Npm1 | EDAESEDEDEEDVK | -121 | -134 | 0 | 0 | 14 | 636.0902 | 789.5495 | 742.81775 | 587.9121 | 695.2742 | 669.6842 |
| Npm1 | VAVEEDAESEDEDEEDVK | -117 | -134 | 0 | 0.00111006 | 18 | 629.49585 | 615.1764 | 576.33466 | 1096.9093 | 1362.4766 | 1287.4858 |
| Npm1 | KVDNDENEHQLSLR | -32 | -45 | 1 | 0 | 14 | 578.56805 | 859.54175 | 857.28546 | 1424.6326 | 1837.8943 | 1761.8737 |
| Npm1 | KVDNDENEHQLSLR | -32 | -45 | 1 | 0 | 14 | 578.56805 | 859.54175 | 857.28546 | 1424.6326 | 1837.8943 | 1761.8737 |
| Npm1 | TVSLGAGAKDELHIVEAE | -46 | -63 | 1 | 0 | 18 | 570.9547 | 603.39465 | 527.2162 | 742.0969 | 596.86414 | 464.01654 |
| Npm1 | VEEDAESEDEDEEDVK | -119 | -134 | 0 | 0 | 16 | 562.849 | 557.8695 | 289.7308 | 1086.4617 | 1094.7281 | 1157.1409 |
| Npm1 | MTDQEAIQDLWQWR | -276 | -289 | 0 | 0 | 14 | 471.20435 | 309.05237 | 568.1937 | 704.6338 | 758.7341 | 668.50507 |
| Npm1 | VQPTVSLGGFEITPPVVLR | -83 | -101 | 0 | 0 | 19 | 385.12375 | 388.46024 | 395.62354 | 468.87198 | 548.5697 | 582.456 |
| Npm1 | EEDAESEDEDEEDVK | -120 | -134 | 0 | 0 | 15 | 360.37338 | 432.9854 | 310.4096 | 536.98645 | 434.9879 | 519.672 |
| Npm1 | GQHLVAVEEDAESEDEDEEDVK | -113 | -134 | 0 | 0.00111006 | 22 | 353.01422 | 347.72287 | 328.2422 | 232.32639 | 198.75003 | 271.9668 |
| Npm1 | AVEEDAESEDEDEEDVK | -118 | -134 | 0 | 0.000869338 | 17 | 346.78497 | 243.92557 | 230.74442 | 305.8709 | 390.92514 | 415.51425 |
| Npm1 | VDNDENEHQLSL | -33 | -44 | 0 | 0 | 12 | 340.1935 | 639.3569 | 394.801 | 554.4612 | 582.68506 | 278.84222 |
| Npm1 | SVQPTVSLGGFEITPPVVLR | -82 | -101 | 0 | 0 | 20 | 320.30884 | 287.94415 | 271.44086 | 370.4054 | 464.2194 | 446.74155 |
| Npm1 | GPSSVEDIK | -238 | -246 | 0 | 0.000394361 | 9 | 309.28662 | 4135.352 | 2611.2917 | 1894.7197 | 3445.6777 | 2565.4976 |
| Npm1 | DELHIVEAE | -55 | -63 | 0 | 0.001291103 | 9 | 1886.1917 | 2330.2668 | 946.04315 | 1703.6122 | 1348.0767 | NaN |
| Npm1 | LFGCELK | -18 | -24 | 0 | 0 | 7 | 882.09564 | 725.01794 | NaN | 907.99927 | 622.2813 | 675.77966 |
| Npm1 | VKLDEDDEDDDEDDEDDEDDDDDDFDEEETEEKVP | -156 | -192 | 2 | 0.006654467 | 37 | 935.4224 | NaN | NaN | 1028.3644 | 1246.6707 | 1152.6998 |
| Npm1 | AIQDLWQWR | -281 | -289 | 0 | 0 | 9 | 796.15265 | 676.5532 | 769.61896 | NaN | 1306.8135 | NaN |
| Npm1 | TDQEAIQDLWQWR | -277 | -289 | 0 | 0 | 13 | 366.1998 | 437.75366 | NaN | NaN | 813.42444 | 820.1189 |
| Npm1 | DELHIVEAEAMNYEGSPIK | -55 | -73 | 0 | 0.001392869 | 19 | 253.39899 | NaN | 326.03952 | 585.8886 | 659.76764 | NaN |
| Npm1 | NYEGSPIK | -66 | -73 | 0 | 0.001463897 | 8 | 1595.5067 | 1745.6382 | NaN | 928.26575 | NaN | NaN |
| Npm1 | TEEKVPVK | -185 | -192 | 1 | 0.008129024 | 8 | 1040.1884 | 811.4921 | 867.4627 | NaN | NaN | NaN |
| Npm1 | EAMNYEGSPIK | -63 | -73 | 0 | 0.005605646 | 11 | 1303.6423 | NaN | NaN | NaN | NaN | NaN |
| Rbbp4 | WLPDVTRPEGK | -49 | -59 | 1 | 0 | 11 | 4092.8904 | 4973.196 | 3957.9417 | 4867.934 | 6144.546 | 6255.054 |
| Rbbp4 | ADKEAAFDDAVEER | -2 | -15 | 1 | 0 | 14 | 2029.8384 | 807.89594 | 2208.1895 | 2746.5027 | 3092.0183 | 2937.4194 |
| Rbbp4 | EAAFDDAVEER | -5 | -15 | 0 | 0 | 11 | 1289.7878 | 1477.7566 | 1474.7432 | 1810.8472 | 1820.2812 | 1807.3256 |
| Rbbp4 | HPSKPDPSGECNPDLR | -157 | -172 | 1 | 0 | 16 | 1027.9692 | 1083.4226 | 1276.4146 | 843.958 | 1335.1605 | 1161.6338 |
| Rbbp4 | HPSKPDPSGECNPDLR | -157 | -172 | 1 | 0 | 16 | 1027.9692 | 1083.4226 | 1276.4146 | 843.958 | 1335.1605 | 1161.6338 |
| Rbbp4 | LWDISAVPK | -204 | -212 | 0 | 0 | 9 | 980.8105 | 905.0467 | 748.4584 | 916.45355 | 871.8959 | 928.79944 |
| Rbbp4 | IGEEQSPEDAEDGPPELLFIH | -350 | -370 | 0 | 0 | 21 | 866.89026 | 1023.4543 | 1053.3527 | 969.10236 | 1512.7683 | 1447.6782 |
| Rbbp4 | GEFGGFGSVSGK | -103 | -114 | 0 | 0 | 12 | 837.7729 | 904.3694 | 473.02014 | 123.704735 | 458.77576 | 1380.52 |
| Rbbp4 | LVLGTHTSDEQNHLVIA | -66 | -82 | 0 | 0 | 17 | 539.73694 | 926.397 | 474.59686 | 412.3308 | 706.82733 | 674.3924 |
| Rbbp4 | SNNTSKPSHSVDAHTA | -259 | -274 | 1 | 0.000550307 | 16 | 380.68762 | 451.80154 | 514.4676 | 854.70557 | 418.46838 | 691.4665 |
| Rbbp4 | SNNTSKPSHSVDAHTA | -259 | -274 | 1 | 0.000550307 | 16 | 380.68762 | 451.80154 | 514.4676 | 854.70557 | 418.46838 | 691.4665 |
| Rbbp4 | LHSFESHKDEIFQVQ | -310 | -324 | 1 | 0 | 15 | 495.26566 | 441.20715 | 354.91345 | 930.668 | 409.97302 | NaN |
| Rbbp4 | LHSFESHKDEIFQVQ | -310 | -324 | 1 | 0 | 15 | 495.26566 | 441.20715 | 354.91345 | 930.668 | 409.97302 | NaN |

| | | | | | | | | | | | | |
|---|---|---|---|---|---|---|---|---|---|---|---|---|
| Rbbp4 | IGEEQSPEDAEDGPPELL | -350 | -367 | 0 | 0 | 18 | 350.2446 | NaN | 321.8863 | 524.4163 | 618.5643 | 643.24286 |
| Rbbp4 | IGEEQSPEDAEDGPPELLFIHG | -350 | -371 | 0 | 0 | 22 | 309.3678 | 319.67755 | 279.0408 | 300.2439 | 348.93222 | NaN |
| Rbbp4 | AFDDAVEER | -7 | -15 | 0 | 0 | 9 | 656.02594 | 700.9549 | 625.10175 | NaN | NaN | NaN |
| Capza2 | AQYNLDQFTPVK | -55 | -66 | 0 | 0 | 12 | 931.7491 | 981.272 | 1000.8996 | 876.0417 | 711.26794 | 892.9339 |
| Capza2 | DIQDSLTVSNEVQTAK | -211 | -226 | 0 | 0 | 16 | 785.6869 | 842.5631 | 809.5518 | 856.9654 | 1036.181 | 1003.471 |
| Capza2 | ADLEEQLSDEEK | -2 | -13 | 0 | 0 | 12 | 731.7092 | 2591.2307 | 2809.1746 | 194.06796 | 2674.337 | 2447.3318 |
| Capza2 | FTVTPSTTQVVGILK | -179 | -193 | 0 | 0 | 15 | 360.52368 | 438.53738 | 435.451 | 480.54837 | 541.94336 | 565.0138 |
| Capza2 | YNLDQFTPVK | -57 | -66 | 0 | 0 | 10 | 1069.2257 | 1211.6761 | 1055.2699 | NaN | 1343.8752 | 1402.7986 |
| Capza2 | IESHQFQAK | -158 | -166 | 0 | 0.000230707 | 9 | 1019.7305 | 624.78827 | 875.3984 | NaN | NaN | 1070.88 |
| Spcs3 | YFFFDDGNGLK | -127 | -137 | 0 | 0 | 11 | 1325.7979 | 1297.2252 | 1348.9126 | 1141.8159 | 884.94165 | 968.8837 |
| Spcs3 | ADLENIFDWNVK | -72 | -83 | 0 | 0 | 12 | 443.9635 | 526.8688 | 435.9361 | 572.12476 | 566.5447 | 610.1985 |
| Spcs3 | NVEDFTGPR | -50 | -58 | 0 | 0 | 9 | 1294.9806 | 1392.9338 | NaN | 1868.5303 | 956.02747 | 1040.9298 |
| Spcs3 | ERSDLGFITF | -59 | -68 | 1 | 0.00122419 | 10 | 384.5079 | 395.95853 | 409.7558 | 490.46274 | NaN | NaN |
| Spcs3 | NNALNQVVLWDK | -97 | -108 | 0 | 0 | 12 | 1124.3112 | 1170.2336 | NaN | NaN | NaN | NaN |
| Ap1s1 | ADLLQEEDESPR | -138 | -149 | 0 | 0 | 12 | 226.3221 | 777.5969 | 768.2439 | 790.6158 | NaN | NaN |
| Efhd2 | ADLNQGIGEPQSPSR | -63 | -77 | 0 | 0 | 15 | 1099.2335 | 1605.7555 | 713.5258 | 324.41275 | 1336.04 | 1269.1305 |
| Efhd2 | VFNPYTEFK | -79 | -87 | 0 | 0 | 9 | 455.52927 | 69.767365 | 809.93146 | 435.66437 | 252.74347 | 328.39 |
| Adsl | LEHLLDPSSFTGR | -440 | -452 | 0 | 0 | 13 | 1774.6461 | 1906.5247 | 1956.6714 | 1624.9104 | 2125.1216 | 1510.4574 |
| Adsl | NAFDLLLPK | -126 | -134 | 0 | 0 | 9 | 992.85144 | 62.57235 | 1171.7523 | 789.0109 | 975.1542 | 1007.1439 |
| Adsl | FLEEEVRPL | -460 | -468 | 1 | 0.00025015 | 9 | 972.1088 | 1540.3348 | 1340.5381 | 3306.0981 | 3081.2605 | 4464.0054 |
| Adsl | YVGDNTDLIILR | -114 | -125 | 0 | 0 | 12 | 949.23224 | 963.66284 | 1038.1464 | 1219.119 | 1449.449 | 1468.1345 |
| Adsl | DRADLPTLGFTH | -148 | -159 | 1 | 0.00025015 | 12 | 809.4266 | 835.7806 | 779.3401 | 1127.3094 | 1165.5216 | 1142.3605 |
| Adsl | DRADLPTLGFT | -148 | -158 | 1 | 0.00025015 | 11 | 672.1777 | 792.3383 | 684.35443 | 411.91684 | 92.97209 | 633.4366 |
| Adsl | ADLPTLGFTHFQPA | -150 | -163 | 0 | 0.000431655 | 14 | 461.78247 | 189.21245 | 1611.703 | 1440.403 | 2073.9885 | 1783.1174 |
| Adsl | QEGGDNDLIER | -416 | -426 | 0 | 0 | 11 | 245.92523 | 4934.568 | 4855.606 | 5124.8794 | 5478.693 | 5001.3877 |
| Adsl | QNISEGLVVYPK | -356 | -367 | 0 | 0 | 12 | 632.30396 | 726.6889 | 674.189 | 742.81415 | NaN | 1012.7075 |
| Adsl | SCYVGDNTDLIILR | -112 | -125 | 0 | 0 | 14 | 399.41107 | 377.62692 | 350.96695 | 655.7071 | 490.68677 | NaN |
| Mical1 | ADLQSENQVLR | -1012 | -1022 | 0 | 0 | 11 | 1276.5984 | 303.03903 | NaN | 320.82956 | NaN | NaN |
| Mical1 | FTGLQPPPR | -181 | -189 | 0 | 0.001868731 | 9 | 739.55835 | 654.2961 | 163.92325 | NaN | NaN | NaN |
| Mdh2 | VNVPVIGGHAGK | -192 | -203 | 0 | 0 | 12 | 8880.797 | 9312.192 | 9094.548 | 9700.752 | 7926.169 | 8028.7437 |
| Mdh2 | DLSHIETR | -67 | -74 | 0 | 0 | 8 | 3909.8975 | 3954.5312 | 3766.5706 | 3939.533 | 2741.6262 | 2930.1875 |
| Mdh2 | GYLGPEQLPDCLK | -79 | -91 | 0 | 0 | 13 | 3675.7683 | 301.9211 | 4552.476 | 4666.0786 | 5766.647 | 5452.714 |
| Mdh2 | GCDVVVIPAGVPR | -92 | -104 | 0 | 0 | 13 | 2808.1562 | 3239.5527 | 3342.4624 | 456.3736 | 4061.2393 | 3941.6208 |
| Mdh2 | VDFPQDQLATLT | -216 | -227 | 0 | 0.000431655 | 12 | 2240.152 | 1781.9299 | 2017.6348 | 349.04337 | 1742.7814 | 2784.5823 |
| Mdh2 | KGEDFVK | -329 | -335 | 1 | 0 | 7 | 2159.039 | 1925.8091 | 2036.8466 | 2155.4275 | 1779.6172 | 1762.0989 |
| Mdh2 | VDFPQDQLATLTGR | -216 | -229 | 0 | 0 | 14 | 2090.0046 | 2594.258 | 141.50475 | 3208.0413 | 4492.7617 | 4582.316 |
| Mdh2 | ADLSHIETR | -66 | -74 | 0 | 0 | 9 | 1735.7839 | 657.3902 | 1943.1149 | 2729.0957 | 2994.1096 | 3139.9158 |
| Mdh2 | IQEAGTEVVK | -230 | -239 | 0 | 0 | 10 | 1252.7549 | 1278.6389 | 1079.1584 | 1065.5354 | 1037.5382 | 1601.1078 |
| Mdh2 | HTPGVAADLSHIETR | -60 | -74 | 0 | 0 | 15 | 826.80035 | 716.44183 | 661.7583 | 1281.4326 | 413.2156 | 1732.8282 |
| Mdh2 | TIIPLISQCTPK | -204 | -215 | 0 | 0 | 12 | 747.60297 | 753.319 | 810.7161 | 1069.2477 | 1041.374 | 1005.37585 |
| Mdh2 | LGPEQLPDCLK | -81 | -91 | 0 | 0 | 11 | 745.70044 | 378.87158 | 1067.6874 | 992.1843 | 542.34753 | 286.85416 |
| Mdh2 | LTLYDIAHTPGVAADL | -53 | -68 | 0 | 0.00025015 | 16 | 634.4612 | 152.82042 | 482.68 | 597.1648 | 798.7765 | 812.43695 |
| Mdh2 | GASGGIGQPLSLLLK | -31 | -45 | 0 | 0 | 15 | 573.80585 | 577.4846 | 633.4135 | 891.3196 | 729.8911 | 866.3038 |
| Mdh2 | ETECTYFSTPLLLGK | -282 | -296 | 0 | 0 | 15 | 524.4367 | 566.71075 | 571.1121 | 463.0068 | 517.4168 | 517.8829 |
| Mdh2 | DIAHTPGVAADLSHIETR | -57 | -74 | 0 | 0 | 18 | 430.83472 | 370.44803 | 504.4539 | 1101.7996 | 1337.7604 | 1278.3722 |
| Mdh2 | IIANPVNSTIPITAEVFK | -139 | -156 | 0 | 0.000230707 | 18 | 374.5037 | 353.35193 | 317.427 | 269.06494 | 308.80417 | 370.2767 |
| Mdh2 | YFSTPLLLGK | -287 | -296 | 0 | 0 | 10 | 170.05002 | 573.21234 | 561.1648 | 477.29755 | 474.69476 | 581.2462 |
| Mdh2 | GYLGPEQLPDCLKG | -79 | -92 | 1 | 0 | 14 | 2048.7268 | 2509.2822 | 2451.9116 | NaN | 2394.9944 | 2216.967 |
| Mdh2 | LTLYDIAHTPGVA | -53 | -65 | 0 | 0 | 13 | 1030.01 | 1685.7501 | 1289.904 | NaN | 2183.5024 | 1646.6063 |
| Mdh2 | EGVVECSFVQSK | -270 | -281 | 0 | 0 | 12 | 422.4405 | 313.513 | 499.21115 | 1341.4019 | 1336.4032 | NaN |
| Mdh2 | DVVVIPAGVPR | -94 | -104 | 0 | 0 | 11 | 1822.6299 | 2051.5103 | 468.56912 | 1021.80414 | NaN | NaN |
| Mdh2 | VAVLGASGGIGQPL | -27 | -40 | 0 | 0.000627943 | 14 | 1383.3438 | 1359.0928 | NaN | NaN | 1137.2028 | NaN |
| Ndufs3 | ILTDYGFEGHPFR | -186 | -198 | 0 | 0 | 13 | 688.34375 | 907.01294 | 1013.5861 | 1091.2802 | 233.02457 | 1338.5718 |
| Ndufs3 | IHPDGVIPTLTFLR | -87 | -100 | 0 | 0 | 14 | 410.14246 | 379.42392 | 434.84296 | 382.72757 | 441.9747 | 459.16644 |
| Ndufs3 | FFNHPDLR | -177 | -184 | 0 | 0.001055699 | 8 | 5140.951 | NaN | 5736.0137 | 117.887985 | 6122.4316 | 6401.6377 |
| Ndufs3 | DLNSPWEAFPAYR | -233 | -245 | 0 | 0 | 13 | 424.3355 | NaN | 474.0946 | 54.05116 | 731.99506 | 733.3985 |
| Ndufs3 | DFPLTGYVELR | -200 | -210 | 0 | 0 | 11 | 373.81244 | NaN | 345.87253 | 464.9437 | 184.02786 | 518.1158 |
| Ndufs3 | ADLTAVDVPTR | -111 | -121 | 0 | 0 | 11 | 713.8583 | 693.8395 | NaN | 485.39728 | NaN | 729.06335 |
| Ndufs3 | ILTDYGFEGHPF | -186 | -197 | 0 | 0.002068213 | 12 | 369.98587 | NaN | 423.61874 | NaN | 654.8612 | 582.7826 |
| Ndufs3 | GFEGHPFR | -191 | -198 | 0 | 0 | 8 | 674.3286 | 783.20496 | 744.6652 | NaN | NaN | NaN |
| Cct6a | KKDEPIDLF | -181 | -189 | 2 | 0.003133457 | 9 | 3385.3818 | 3701.4736 | 3964.407 | 3851.7046 | 4727.718 | 338.5013 |
| Cct6a | ETLIDVAR | -146 | -153 | 0 | 0 | 8 | 3093.4375 | 2828.253 | 2118.83 | 2068.9731 | 1216.473 | 1012.229 |
| Cct6a | ALQFLEQVK | -130 | -138 | 0 | 0 | 9 | 1898.3463 | 2191.4692 | 2193.5754 | 2269.3452 | 2267.4658 | 2445.7139 |
| Cct6a | HTLTQIK | -382 | -388 | 0 | 0.003270762 | 7 | 1842.0388 | 1696.7245 | 1663.4874 | 1044.1 | 1067.1844 | 803.91907 |
| Cct6a | TEVNSGFFYK | -242 | -251 | 0 | 0 | 10 | 1794.728 | 1981.2594 | 1836.9446 | 760.10187 | 2051.4238 | 1897.136 |
| Cct6a | SETDTSLIR | -200 | -208 | 0 | 0 | 9 | 1743.8499 | 1622.7097 | 1451.0908 | 214.20258 | 1239.589 | 1411.965 |
| Cct6a | HKSETDTSLIR | -198 | -208 | 1 | 0 | 11 | 1504.9244 | 1775.66 | 2628.6248 | 1422.46 | 1583.9341 | 1458.2538 |
| Cct6a | HKSETDTSLIR | -198 | -208 | 1 | 0 | 11 | 1504.9244 | 1775.66 | 2628.6248 | 1422.46 | 1583.9341 | 1458.2538 |
| Cct6a | GFDLQETLVK | -456 | -465 | 0 | 0 | 10 | 975.15234 | 935.66986 | 1038.3059 | 807.9194 | 315.1714 | 324.07162 |
| Cct6a | GIDPFSLDALAK | -296 | -307 | 0 | 0 | 12 | 699.17883 | 233.51428 | 738.01953 | 835.2275 | 1006.70667 | 292.70093 |
| Cct6a | AVVDSILAIR | -171 | -180 | 0 | 0.000431655 | 10 | 630.4686 | 699.6873 | 636.06824 | 447.39288 | 549.6949 | 664.2331 |
| Cct6a | TEAVVDSILAIR | -169 | -180 | 0 | 0 | 12 | 529.6713 | 645.6719 | 641.3014 | 377.53235 | 476.62195 | 474.69788 |
| Cct6a | VLAQNSGFDLQETLVK | -450 | -465 | 0 | 0 | 16 | 447.48642 | 658.04126 | 622.70496 | 477.27164 | 616.30365 | 675.52826 |
| Cct6a | IITEGFEAAK | -118 | -127 | 0 | 0.000230707 | 10 | 1990.0844 | 2247.0332 | 2219.458 | 5351.5903 | NaN | 3970.666 |
| Cct6a | NSGFDLQETLVK | -454 | -465 | 0 | 0.000230707 | 12 | 1759.3162 | NaN | 621.471 | 4584.326 | 1442.8596 | 5328.5425 |
| Cct6a | SGFDLQETLVK | -455 | -465 | 0 | 0 | 11 | 282.2569 | 440.71298 | 487.3934 | NaN | 629.95874 | 772.07 |
| Cct6a | AQNSGFDLQETLVK | -452 | -465 | 0 | 0 | 14 | 1386.854 | NaN | NaN | 873.20764 | 1921.9264 | 2149.769 |
| Cct6a | EVNSGFFYK | -243 | -251 | 0 | 0.000431655 | 9 | 249.53088 | 405.9261 | 374.39505 | NaN | NaN | NaN |

| | | | | | | | | | | | | |
|---|---|---|---|---|---|---|---|---|---|---|---|---|
| Hdgfl2 | ADNEKLEEQPGEQAPR | -567 | -582 | 1 | 0.001392869 | 16 | 346.69635 | 597.0486 | 566.1695 | 458.25885 | 489.54257 | 422.4447 |
| Phb2 | IVQAEGEAEAAK | -225 | -236 | 0 | 0 | 12 | 2754.7996 | 2492.9321 | 2435.1626 | 2842.5864 | 2045.0436 | 2086.9458 |
| Phb2 | LGLDYEER | -124 | -131 | 0 | 0 | 8 | 2078.7336 | 476.33682 | 2161.3928 | 92.63355 | 1946.8702 | 1976.1746 |
| Phb2 | ESVFTVEGGHR | -38 | -48 | 0 | 0 | 11 | 1808.2765 | 2015.2251 | 709.8027 | 1214.8765 | 4125.052 | 3615.1333 |
| Phb2 | VLPSIVNEVLK | -132 | -142 | 0 | 0 | 11 | 1419.4681 | 1486.8507 | 1472.1775 | 1501.0879 | 1711.2205 | 1768.8821 |
| Phb2 | DNLVLNLQDESFTR | -276 | -289 | 0 | 0 | 14 | 582.1442 | 514.78345 | 709.0214 | 889.6454 | 499.2277 | 679.2355 |
| Phb2 | ADNLVLNLQDESFTR | -275 | -289 | 0 | 0 | 15 | 416.77252 | 452.09656 | 401.63806 | 644.6347 | 824.32367 | 790.9418 |
| Phb2 | NLVLNLQDESFTR | -277 | -289 | 0 | 0 | 13 | 1955.4907 | 1593.9269 | NaN | 1733.8235 | 1440.4208 | 1808.6133 |
| Phb2 | NLQDESFTR | -281 | -289 | 0 | 0 | 9 | 867.87177 | 842.1267 | 850.56866 | 810.9855 | 655.2282 | NaN |
| Phb2 | FNASQLITQR | -148 | -157 | 0 | 0 | 10 | 127.00005 | NaN | NaN | 1035.5891 | 680.13135 | 952.9954 |
| Phb2 | SKNPGYIK | -243 | -250 | 1 | 0.001392869 | 8 | 1222.5205 | NaN | NaN | 1016.71545 | NaN | 1401.5905 |
| Prpf8 | SHLVPVYDVEPLEK | -903 | -916 | 0 | 0 | 14 | 3014.1094 | 3995.802 | 3871.112 | 4710.998 | 5353.012 | 5018.41 |
| Prpf8 | FSPIPFPPLSYK | -822 | -833 | 0 | 0 | 12 | 2245.1248 | 2265.2673 | 2633.3152 | 145.35106 | 3457.47 | 3022.2004 |
| Prpf8 | DGVWNLQNEVTK | -1211 | -1222 | 0 | 0 | 12 | 2209.4885 | 3006.7583 | 2667.0676 | 3313.62 | 3426.0398 | 371.56702 |
| Prpf8 | LVVDSHVQYR | -556 | -565 | 0 | 0 | 10 | 1591.9463 | 1861.5789 | 2204.9595 | 1545.9313 | 1384.483 | 1286.5431 |
| Prpf8 | IDLTLLNR | -988 | -995 | 0 | 0 | 8 | 1300.0372 | 1322.0382 | 1144.2937 | 1760.1973 | 2095.521 | 2217.2534 |
| Prpf8 | VHWLESR | -814 | -820 | 0 | 0.000591124 | 7 | 1280.759 | 1002.42883 | 834.1468 | 776.2965 | 875.8832 | 750.45874 |
| Prpf8 | YLTEHPDPNNENIVGY | -1113 | -1128 | 0 | 0 | 16 | 1262.3853 | 494.36508 | 486.2378 | 1211.5925 | 1215.4558 | 1130.47 |
| Prpf8 | IFIFNPR | -1807 | -1813 | 0 | 0.000347792 | 7 | 1032.9799 | 882.9986 | 892.3389 | 796.7219 | 604.5581 | 658.6212 |
| Prpf8 | GYLPSHYER | -2250 | -2258 | 0 | 0 | 9 | 1014.0864 | 1241.081 | 1269.8387 | 1443.3625 | 2010.5168 | 2243.3765 |
| Prpf8 | WYHTPNVVFIK | -331 | -341 | 0 | 0 | 11 | 736.4873 | 743.49603 | 817.251 | 815.3672 | 1122.7507 | 1114.2012 |
| Prpf8 | WYHTPNVVFIK | -331 | -341 | 0 | 0 | 11 | 736.4873 | 743.49603 | 817.251 | 815.3672 | 1122.7507 | 1114.2012 |
| Prpf8 | EHCPAGQPVK | -433 | -442 | 0 | 0 | 10 | 786.66876 | 698.4533 | 921.8384 | 495.2042 | 911.0102 | 524.0591 |
| Prpf8 | YIQPWESEFIDSQR | -1371 | -1384 | 0 | 0 | 14 | 732.4125 | 879.38513 | 929.2034 | 476.02515 | 697.24316 | 814.9808 |
| Prpf8 | GYLYGVSPPDNPQVK | -2126 | -2140 | 0 | 0 | 15 | 713.3241 | 799.0557 | 1031.2704 | 994.88696 | 755.15845 | 978.2307 |
| Prpf8 | WGDYDSHDIER | -1668 | -1678 | 0 | 0 | 11 | 700.2401 | 790.1189 | 751.06213 | 824.5306 | 853.5603 | 863.57166 |
| Prpf8 | TVHLPSQLPQHEYLK | -2156 | -2170 | 0 | 0 | 15 | 606.26086 | 1297.0848 | 1239.9237 | 929.24164 | 889.99066 | 1507.2609 |
| Prpf8 | TVHLPSQLPQHEYLK | -2156 | -2170 | 0 | 0 | 15 | 606.26086 | 1297.0848 | 1239.9237 | 929.24164 | 889.99066 | 1507.2609 |
| Prpf8 | LVPVYDVEPLEK | -905 | -916 | 0 | 0 | 12 | 523.02954 | 507.5695 | 451.96896 | 494.16074 | 521.32025 | 612.2343 |
| Prpf8 | LTLEDLEDSWDR | -1403 | -1414 | 0 | 0 | 12 | 363.9445 | 366.67865 | 544.9163 | 388.0634 | 310.73056 | 510.30075 |
| Prpf8 | DINLQDEDWNEFNDINK | -285 | -301 | 0 | 0 | 17 | 362.32178 | 299.42236 | 347.48083 | 433.71936 | 459.14523 | 420.2424 |
| Prpf8 | QAYDNPHEALSR | -869 | -880 | 0 | 0 | 12 | 2114.0876 | 1906.9989 | 850.30945 | 1826.7677 | 2648.26 | NaN |
| Prpf8 | FICISDLR | -2114 | -2121 | 0 | 0.000230707 | 8 | 884.9588 | NaN | 823.9977 | 266.55182 | 816.9845 | 826.897 |
| Prpf8 | VVDSHVQYR | -557 | -565 | 0 | 0 | 9 | 601.60016 | 3110.3718 | NaN | 299.5677 | 5844.2124 | 3014.66 |
| Prpf8 | WLGNLLAR | -651 | -658 | 0 | 0 | 8 | 589.55096 | NaN | 606.29816 | 538.3377 | 887.154 | 1052.3308 |
| Prpf8 | HLDYNFNLKPVK | -517 | -528 | 1 | 0 | 12 | 547.16895 | 241.35335 | 561.5649 | NaN | 842.1999 | 842.24634 |
| Prpf8 | ALDIPLVK | -421 | -428 | 0 | 0 | 8 | 474.20032 | 392.4695 | 401.71024 | 358.04382 | 452.51828 | NaN |
| Prpf8 | SGLNQIPNR | -1524 | -1532 | 0 | 0 | 9 | 1529.3284 | 1335.2568 | 1381.2487 | 2154.1855 | NaN | NaN |
| Prpf8 | LYGVSPPDNPQVK | -2128 | -2140 | 0 | 0 | 13 | 1386.5968 | 1767.6853 | 2207.2856 | NaN | NaN | 2334.13 |
| Prpf8 | FYDHQPLR | -207 | -214 | 0 | 0 | 8 | 1232.5332 | 1161.0353 | 1167.3171 | NaN | 993.29175 | NaN |
| Prpf8 | TILQHLSEAWR | -708 | -718 | 0 | 0 | 11 | 565.45154 | NaN | NaN | 1136.0593 | 1066.2219 | 2025.9199 |
| Prpf8 | LNLLIHR | -504 | -510 | 0 | 0 | 7 | 441.22852 | 455.39377 | 463.74585 | 299.2444 | NaN | NaN |
| Prpf8 | TSYEEFTHK | -1202 | -1210 | 0 | 0 | 9 | 155.71808 | 3689.3694 | NaN | 5011.4585 | 9502.315 | NaN |
| Prpf8 | WWSPTINR | -1537 | -1544 | 0 | 0 | 8 | 4056.2483 | 4078.9966 | 4365.931 | NaN | NaN | NaN |
| Prpf8 | ADNPSWDGEK | -2201 | -2210 | 0 | 0.000907206 | 10 | 694.2538 | NaN | NaN | NaN | 4433.122 | 246.83255 |
| Prpf8 | VILKPDK | -1952 | -1958 | 1 | 0.007987481 | 7 | 672.0244 | 653.4634 | NaN | NaN | 638.8547 | NaN |
| Prpf8 | LTPSGYEWGR | -2230 | -2239 | 0 | 0 | 10 | 476.3802 | 1073.4808 | 999.7131 | NaN | NaN | NaN |
| Prpf8 | FGDLILK | -1902 | -1908 | 0 | 0.002434616 | 7 | 394.75754 | 227.0362 | 297.0575 | NaN | NaN | NaN |
| Prpf8 | KQNPFWWTHQR | -1449 | -1459 | 1 | 0 | 11 | 5124.881 | 4213.465 | NaN | NaN | NaN | NaN |
| Prpf8 | AYDNPHEALSR | -870 | -880 | 0 | 0 | 11 | 825.7942 | NaN | NaN | NaN | 938.9818 | NaN |
| Prpf8 | QNPFWWTHQ | -1450 | -1458 | 0 | 0.003069674 | 9 | 755.96625 | NaN | NaN | NaN | 922.086 | NaN |
| Prpf8 | TVHWLESR | -813 | -820 | 0 | 0.003003218 | 8 | 726.3592 | NaN | NaN | 211.91493 | NaN | NaN |
| Prpf8 | LYDDWLK | -1919 | -1925 | 0 | 0.002329279 | 7 | 543.13995 | 559.916 | NaN | NaN | NaN | NaN |
| Glod4 | KLEWPLSK | -102 | -109 | 1 | 0.00025015 | 8 | 3726.1113 | 2447.4385 | 3360.7515 | 1606.9011 | 938.03815 | 1258.2638 |
| Glod4 | VAEGIFETEAPGGYK | -110 | -124 | 0 | 0 | 15 | 918.62134 | 937.01605 | 939.1668 | 1098.6599 | 1108.265 | 1106.0762 |
| Glod4 | ALLGYADNQCK | -173 | -183 | 0 | 0 | 11 | 733.13306 | 582.8738 | 709.0178 | 686.6511 | 566.40283 | 790.8288 |
| Glod4 | ESHSILTPLVSLDTPGK | -221 | -237 | 0 | 0 | 17 | 671.49615 | 747.60187 | 600.87524 | 940.6347 | 1192.7983 | 1114.5979 |
| Glod4 | ESHSILTPLVSLDTPGK | -221 | -237 | 0 | 0 | 17 | 671.49615 | 747.60187 | 600.87524 | 940.6347 | 1192.7983 | 1114.5979 |
| Glod4 | TYNYGIGDYK | -72 | -81 | 0 | 0 | 10 | 400.61533 | 1095.8585 | 555.7264 | 791.3968 | 773.552 | 643.5265 |
| Glod4 | ADPDGHEICFVGDEAFR | -246 | -262 | 0 | 0 | 17 | 340.77832 | 341.26212 | 464.752 | NaN | 1331.86 | 1173.8403 |
| Glod4 | FQTVHFFR | -17 | -24 | 0 | 0 | 8 | 2472.444 | 209.22339 | NaN | NaN | NaN | NaN |
| Uba2 | KLSDFGIR | -503 | -510 | 1 | 0.000550307 | 8 | 4013.3257 | 4265.0137 | 4192.335 | 4334.481 | 4471.6797 | 4314.3286 |
| Uba2 | NTPSEPIHCIVWAK | -177 | -190 | 0 | 0 | 14 | 1475.9885 | 1757.5984 | 1727.383 | 1354.2548 | 1441.5652 | 1304.4341 |
| Uba2 | VVGAGGIGCELLK | -22 | -34 | 0 | 0 | 13 | 1302.1868 | 1583.8466 | 1640.3135 | 2301.5833 | 2691.424 | 2663.4678 |
| Uba2 | SQGEANADQQNEPQLGLK | -289 | -306 | 0 | 0 | 18 | 1135.8678 | 1046.2305 | 930.19855 | 1847.7511 | 1384.0533 | 2161.8267 |
| Uba2 | ISSEEGETEANNPK | -489 | -502 | 0 | 0 | 14 | 583.7891 | 602.56256 | 628.76624 | 612.3665 | 755.7237 | 863.2852 |
| Uba2 | KLEENEAASTK | -611 | -621 | 1 | 0.000627943 | 11 | 580.4921 | 624.94995 | 722.8366 | 666.1201 | 734.79175 | 612.4845 |
| Uba2 | GLIVLEGLK | -394 | -402 | 0 | 0.00025015 | 9 | 4912.5923 | 5632.634 | NaN | 4435.675 | 5153.3853 | 5386.7876 |
| Uba2 | KPPVPLDWAEVQ | -277 | -288 | 1 | 0.00025015 | 12 | 723.43286 | 601.3678 | 801.13477 | 740.5368 | NaN | 1009.2115 |
| Uba2 | ELAEAVSGGR | -10 | -19 | 0 | 0 | 10 | 499.1145 | 2192.7356 | 606.94824 | 2500.7747 | NaN | 1864.3577 |
| Uba2 | DVEFEVVGDSPEK | -539 | -551 | 0 | 0 | 13 | 1036.4202 | 1387.5338 | 1368.0859 | 1458.0555 | NaN | NaN |
| Uba2 | ADPEAAWEPTEAEAR | -211 | -225 | 0 | 0 | 15 | 632.84406 | 599.6317 | 651.1925 | NaN | NaN | 836.7938 |
| Uba2 | ASKPEVTVR | -443 | -451 | 1 | 0.002068213 | 9 | 568.45917 | NaN | 447.6145 | NaN | 600.61615 | 301.54926 |
| Uba2 | SEEGETEANNPK | -491 | -502 | 0 | 0 | 12 | 424.71588 | 297.39398 | 343.543 | 294.07477 | NaN | NaN |
| Uba2 | FAMVAPDVQIEDGK | -471 | -484 | 0 | 0 | 14 | 209.98808 | NaN | 322.17456 | NaN | 404.362 | 533.581 |
| Uba2 | VLVVGAGGIGCELLK | -20 | -34 | 0 | 0 | 15 | 2310.1782 | 2332.0364 | 2541.8628 | NaN | NaN | NaN |
| Uba2 | QLGLKDQQVLDVK | -302 | -314 | 1 | 0.003102268 | 13 | 893.38586 | NaN | NaN | 795.86145 | 978.3857 | NaN |
| Uba2 | WEPTEAEAR | -217 | -225 | 0 | 0 | 9 | 500.96783 | NaN | NaN | NaN | NaN | 1234.2604 |

| | | | | | | | | | | | | |
|---|---|---|---|---|---|---|---|---|---|---|---|---|
| Stk26 | NNPPTLIGDFTK | -234 | -245 | 0 | 0 | 12 | 1124.53 | 1292.0194 | 608.4457 | 499.4646 | 1362.9124 | 1282.1415 |
| Stk26 | ESNPHPEWSFTT | -317 | -328 | 0 | 0.003069674 | 12 | 487.31973 | NaN | NaN | NaN | 706.7008 | NaN |
| Stk24 | NNPPTLEGNYSKPLK | -234 | -248 | 1 | 0 | 15 | 4866.3867 | 6409.896 | 7093.569 | 13334.453 | 11432.78 | 11745.331 |
| Stk24 | NVLLSEHGEVK | -149 | -159 | 0 | 0 | 11 | 1155.3956 | 1128.1769 | 4571.7705 | 1236.1263 | 1189.0491 | 1286.5305 |
| Stk24 | ADPEELFTKL | -18 | -27 | 1 | 0 | 10 | 238.39499 | 460.54724 | 543.1927 | 472.3179 | 574.66437 | 579.3563 |
| Pabpn1 | ESEELEPGELLPEPEPEEEPPRPR | -51 | -74 | 1 | 0.000230707 | 24 | 977.6726 | 958.8573 | 1012.87463 | 928.79016 | 1022.64087 | 1088.0355 |
| Pabpn1 | LALDESLFR | -226 | -234 | 0 | 0 | 9 | 853.0941 | 1059.0474 | 1104.4724 | 1621.1167 | 129.83769 | 1871.9144 |
| Pabpn1 | ADPGDGAIEDPELEAIK | -103 | -119 | 0 | 0 | 17 | 702.0877 | 881.0369 | 905.34424 | 1003.4237 | 1201.5417 | 1210.7996 |
| Pabpn1 | HLVPGAGGEAGEGDPGGAGDYGNGL | -26 | -50 | 0 | 0 | 25 | 578.64624 | 486.72003 | 484.6858 | 352.7015 | 335.59558 | 283.39072 |
| Pabpn1 | HLVPGAGGEAGEGDPGGAGDY | -26 | -46 | 0 | 0 | 21 | 359.3309 | 560.9405 | 383.591 | 498.35226 | 452.2425 | 518.8385 |
| Pabpn1 | TSLALDESLFR | -224 | -234 | 0 | 0 | 11 | 2217.711 | 2619.5164 | 2351.1023 | NaN | 1741.3099 | NaN |
| Pabpn1 | LKELQNEVEK | -132 | -141 | 1 | 0.000431655 | 10 | 735.70776 | 616.7074 | 921.54785 | NaN | NaN | NaN |
| Arhgdia | VAVSADPNVPNVIVTR | -59 | -74 | 0 | 0 | 16 | 6961.787 | 7264.616 | 6942.9634 | 5940.8096 | 6825.178 | 465.04865 |
| Arhgdia | AENEEDEHSVNYKPPAQK | -16 | -33 | 1 | 0 | 18 | 6520.754 | 7004.494 | 7335.566 | 5181.0645 | 8044.324 | 7927.306 |
| Arhgdia | AENEEDEHSVNYKPPAQK | -16 | -33 | 1 | 0 | 18 | 6520.754 | 7004.494 | 7335.566 | 5181.0645 | 8044.324 | 7927.306 |
| Arhgdia | VSADPNVPNVIVTR | -61 | -74 | 0 | 0 | 14 | 5329.1733 | 568.6817 | 6044.446 | 6635.2666 | 7098.989 | 7053.3936 |
| Arhgdia | SIQEIQELDKDDESLRK | -34 | -50 | 2 | 0 | 17 | 4475.7217 | 5713.4336 | 5448.433 | 5284.657 | 7808.1343 | 7614.2124 |
| Arhgdia | SIQEIQELDKDDESLRK | -34 | -50 | 2 | 0 | 17 | 4475.7217 | 5713.4336 | 5448.433 | 5284.657 | 7808.1343 | 7614.2124 |
| Arhgdia | ADPNVPNVIVTR | -63 | -74 | 0 | 0 | 12 | 2584.1436 | 2677.6968 | 201.91245 | 3289.688 | 3222.6504 | 3042.523 |
| Arhgdia | SIQEIQELDKDDESLR | -34 | -49 | 1 | 0 | 16 | 2532.2422 | 3350.424 | 3328.2124 | 3322.1294 | 4794.157 | 4594.3643 |
| Arhgdia | SIQEIQELDKDDESLR | -34 | -49 | 1 | 0 | 16 | 2532.2422 | 3350.424 | 3328.2124 | 3322.1294 | 4794.157 | 4594.3643 |
| Arhgdia | FTDDDKTDHL | -181 | -190 | 1 | 0 | 10 | 2485.5469 | 2650.5073 | 2963.3171 | 1057.2095 | 1562.8346 | 1658.7035 |
| Arhgdia | ELDKDDESLR | -40 | -49 | 1 | 0 | 10 | 2223.6726 | 2324.1775 | 2428.0398 | 2844.2104 | 3443.8774 | 335.70197 |
| Arhgdia | PNVPNVIVTR | -65 | -74 | 0 | 0 | 10 | 1354.1089 | 1413.8918 | 1404.5098 | 1568.7281 | 1458.0347 | 1194.7388 |
| Arhgdia | FTDDDKTDHLSW | -181 | -192 | 1 | 0.000230707 | 12 | 1091.1486 | 1099.0927 | 1530.9215 | 959.0007 | 476.15118 | 748.25366 |
| Arhgdia | IAAENEEDEHSVNYKPPAQK | -14 | -33 | 1 | 0 | 20 | 1002.97406 | 995.231 | 906.2069 | 1217.5166 | 1359.7676 | 1437.5967 |
| Arhgdia | AQIAAENEEDEHSVNYKPPAQK | -12 | -33 | 1 | 0 | 22 | 872.7228 | 776.95764 | 771.3024 | 932.92975 | 1030.1295 | 1092.6367 |
| Arhgdia | AEEYEFLTPMEEAPK | -153 | -167 | 0 | 0 | 15 | 839.7112 | 977.6 | 1246.3169 | 1443.6818 | 1671.4899 | 1686.1864 |
| Arhgdia | AEEYEFLTPMEEAPK | -153 | -167 | 0 | 0 | 15 | 839.7112 | 977.6 | 1246.3169 | 1443.6818 | 1671.4899 | 1686.1864 |
| Arhgdia | STAPGPLELDLTGDLESFK | -80 | -98 | 0 | 0 | 19 | 604.7223 | 786.9752 | 740.53265 | 380.65988 | 952.3388 | 388.64746 |
| Arhgdia | YIQHTYR | -128 | -134 | 0 | 0.001055699 | 7 | 602.93445 | 1018.47705 | 1715.0623 | 804.1128 | 1144.517 | 1127.102 |
| Arhgdia | STAPGPLELDLTGDLESFKK | -80 | -99 | 1 | 0 | 20 | 360.63025 | 406.42896 | 366.96893 | 415.5611 | 502.83234 | 475.9908 |
| Arhgdia | EYEFLTPMEEAPK | -155 | -167 | 0 | 0 | 13 | 1398.0901 | 1162.1228 | 1106.7201 | NaN | 408.1268 | 216.34094 |
| Arhgdia | FTDDDKTDHLSWEW | -181 | -194 | 1 | 0.000712251 | 14 | 390.7527 | 400.4411 | 477.83957 | NaN | 424.39297 | 488.0688 |
| Arhgdia | SADPNVPNVIVTR | -62 | -74 | 0 | 0 | 13 | 3260.7021 | NaN | NaN | 4214.8237 | 4729.0776 | NaN |
| Arhgdia | AEQEPTAEQLAQIA | -2 | -15 | 0 | 0.002168954 | 14 | 936.95294 | NaN | NaN | 517.8242 | NaN | 403.95007 |
| Arhgdia | ELDKDDESLRK | -40 | -50 | 2 | 0 | 11 | 834.70197 | 806.9291 | NaN | NaN | 646.0822 | NaN |
| Arhgdia | ELDKDDESLRK | -40 | -50 | 2 | 0 | 11 | 834.70197 | 806.9291 | NaN | NaN | 646.0822 | NaN |
| Rps6ka3 | NSIQFTDGYEVK | -414 | -425 | 0 | 0 | 12 | 3054.8682 | 2212.7368 | 2475.454 | 1427.6597 | 1604.1896 | 1685.4674 |
| Rps6ka3 | NQSPVLEPVGR | -713 | -723 | 0 | 0 | 11 | 555.3246 | 1074.2935 | 1458.6818 | 1101.4685 | 1973.1663 | 1332.1313 |
| Rps6ka3 | LGAGPDGVEEIK | -311 | -322 | 0 | 0 | 12 | 2388.383 | 1987.7272 | 469.65796 | 2220.4143 | 1547.2181 | NaN |
| Rps6ka3 | PLAQLADPWQK | -2 | -12 | 0 | 0 | 11 | 769.5166 | 747.42944 | 729.25885 | 1112.2637 | NaN | NaN |
| Rps6ka3 | SIQFTDGYEVK | -415 | -425 | 0 | 0 | 11 | 380.96307 | NaN | NaN | 982.2898 | 1050.8622 | NaN |
| Copb2 | QYPLVTPNEER | -823 | -833 | 0 | 0 | 11 | 3784.6218 | 1681.3964 | 1798.6965 | 1157.9644 | 720.0761 | 1290.4679 |
| Copb2 | GSNNVALGYDEGSIIVK | -282 | -298 | 0 | 0 | 17 | 3726.1345 | 4272.261 | 4855.18 | 6246.099 | 5778.4014 | 8211.692 |
| Copb2 | ETHADLWPAK | -813 | -822 | 0 | 0 | 10 | 2172.1528 | 3179.5247 | 2837.21 | 1230.9507 | 1610.2699 | 1774.648 |
| Copb2 | AWAHDSSEYAIR | -395 | -406 | 0 | 0 | 12 | 1912.8882 | 2134.6711 | 666.25397 | 1498.3254 | 1469.7308 | 1870.5553 |
| Copb2 | VWNHETQTLVK | -41 | -51 | 0 | 0 | 11 | 853.4197 | 953.1649 | 966.3623 | 1132.7987 | 993.95264 | 972.3543 |
| Copb2 | NHETQTLVK | -43 | -51 | 0 | 0 | 9 | 757.59955 | 1138.9945 | 1305.8126 | 875.7641 | 1308.1769 | 773.0269 |
| Copb2 | ADPTEYENLFPGLK | -789 | -802 | 0 | 0 | 14 | 732.5561 | 773.49835 | 817.29785 | 731.56946 | 1053.3031 | 1013.5207 |
| Copb2 | ALTVSTDPEHR | -634 | -644 | 0 | 0 | 11 | 695.64777 | 575.01465 | 655.1406 | 778.6068 | 870.7778 | 907.52637 |
| Copb2 | TCVQTLEGHAQNVSCA | -220 | -235 | 0 | 0.001092529 | 16 | 684.3946 | 732.2802 | 291.2931 | 712.2756 | 706.8105 | 772.7621 |
| Copb2 | TEESFFILK | -481 | -489 | 0 | 0.00025015 | 9 | 676.6552 | 705.8895 | 787.5254 | 365.93695 | 365.2898 | 630.64954 |
| Copb2 | AFYDWENTELIR | -448 | -459 | 0 | 0 | 12 | 625.84265 | 782.1019 | 739.1798 | 682.0066 | 794.61975 | 858.9122 |
| Copb2 | YDWENTELIR | -450 | -459 | 0 | 0 | 10 | 378.46643 | NaN | 841.1863 | 407.79333 | 903.1158 | 821.1952 |
| Copb2 | TFEVCDLPVR | -52 | -61 | 0 | 0 | 10 | 1545.5464 | NaN | 1723.477 | 226.70914 | 1762.9237 | NaN |
| Copb2 | LADPTEYENLFPGLK | -788 | -802 | 0 | 0.00025015 | 15 | 182.9892 | 572.8994 | 524.6627 | NaN | NaN | NaN |
| Copb2 | IWDYQNK | -213 | -219 | 0 | 0.006472924 | 7 | 557.3312 | NaN | NaN | NaN | NaN | NaN |
| Zc3h18 | TAPGPEPAKPGDLR | -649 | -662 | 1 | 0.000230707 | 14 | 359.9872 | 684.06665 | 1103.9512 | 1108.0278 | 1115.8577 | 577.06195 |
| Zc3h18 | ADPYYDYEIER | -364 | -374 | 0 | 0.001776473 | 11 | 851.1454 | NaN | NaN | NaN | NaN | NaN |
| Coro1a | LDRLEETVQAK | -451 | -461 | 1 | 0 | 11 | 8918.207 | 8937.38 | 9114.071 | 10292.188 | 10816.864 | 10670.336 |
| Coro1a | LDRLEETVQAK | -451 | -461 | 1 | 0 | 11 | 8918.207 | 8937.38 | 9114.071 | 10292.188 | 10816.864 | 10670.336 |
| Coro1a | ATPEPSGTPSSDTVSR | -417 | -432 | 0 | 0 | 16 | 7843.8955 | 8115.3413 | 8718.488 | 8362.338 | 7865.6416 | 8631.163 |
| Coro1a | EPVITLEGHTK | -122 | -132 | 0 | 0 | 11 | 7499.319 | 8408.909 | 922.74896 | 9375.079 | 8606.875 | 8431.715 |
| Coro1a | HVFGQPAK | -13 | -20 | 0 | 0 | 8 | 5706.001 | 5676.1074 | 5964.0127 | 5192.732 | 4770.656 | 4685.532 |
| Coro1a | QTTWDSGFCAVNPK | -32 | -45 | 0 | 0 | 14 | 5561.0684 | 5826.9077 | 5858.143 | 377.78952 | 6534.0063 | 6357.845 |
| Coro1a | QCYEDVR | -23 | -29 | 0 | 0.001291103 | 7 | 4267.234 | 4006.1135 | 207.88348 | 4220.668 | 4189.8057 | 4343.261 |
| Coro1a | LDRLEETVQA | -451 | -460 | 1 | 0.00111006 | 10 | 2691.3906 | 3000.897 | 3024.8594 | 4974.9673 | 3159.3586 | 3111.5105 |
| Coro1a | EIPDGGLVLPLR | -110 | -121 | 0 | 0 | 12 | 2615.2395 | 3109.1533 | 2987.5083 | 2835.7642 | 3532.07 | 3293.0635 |
| Coro1a | HLEEPLSLQEL | -254 | -264 | 0 | 0.00025015 | 11 | 2607.611 | 3138.8154 | 833.5406 | 2028.2051 | 2654.4265 | 2623.7607 |
| Coro1a | ADQCYEDVR | -21 | -29 | 0 | 0.00025015 | 9 | 2513.2185 | 2151.3442 | 1944.7727 | 2101.6418 | 1300.569 | 1530.6299 |
| Coro1a | NVPLVCGHTAPVLDIA | -73 | -88 | 0 | 0 | 16 | 2399.3945 | 3048.8596 | 2918.2527 | 2299.0488 | 3504.9797 | 3663.6597 |
| Coro1a | KSDLFQEDLYPPTAGPDPALTAE | -355 | -377 | 1 | 0.00025015 | 23 | 1570.4249 | 1606.7185 | 1608.1257 | 1842.5106 | 2528.6724 | 2269.9395 |
| Coro1a | VSQTTWDSGFCAVNPK | -30 | -45 | 0 | 0 | 16 | 1390.5416 | 1685.9811 | 1708.8462 | 699.3096 | 781.0227 | 825.47565 |
| Coro1a | TTWDSGFCAVNPK | -33 | -45 | 0 | 0 | 13 | 1351.9277 | 1541.2195 | 1496.7534 | 1677.3074 | 1592.6976 | 1497.0833 |
| Coro1a | WDSGFCAVNPK | -35 | -45 | 0 | 0 | 11 | 1182.9897 | 1410.6923 | 1410.9532 | 1711.3965 | 2137.9573 | 2112.5881 |
| Coro1a | PVHAVFVSEGK | -223 | -233 | 0 | 0 | 11 | 972.7537 | 1139.877 | 1059.7065 | 1742.7429 | 1744.3074 | 1557.8295 |
| Coro1a | VWEIPDGGLVLPLR | -108 | -121 | 0 | 0 | 14 | 883.0414 | 164.0265 | 794.2291 | 992.3187 | 1412.2574 | 1260.3981 |

| | | | | | | | | | | | | |
|---|---|---|---|---|---|---|---|---|---|---|---|---|
| Coro1a | DRPHEGTRPVHAV | -215 | -227 | 2 | 0.000230707 | 13 | 192.75656 | 519.16504 | 1214.3545 | 509.30185 | 369.1738 | 1824.5701 |
| Coro1a | DRPHEGTRPVHAV | -215 | -227 | 2 | 0.000230707 | 13 | 192.75656 | 519.16504 | 1214.3545 | 509.30185 | 369.1738 | 1824.5701 |
| Coro1a | DAGPLLISLK | -384 | -393 | 0 | 0 | 10 | 863.45874 | 1021.1534 | 1012.9571 | 1673.8162 | 1940.5266 | 1956.4254 |
| Coro1a | KDGYVPPK | -393 | -400 | 1 | 0.002566359 | 8 | 788.9694 | 1595.7623 | 1818.1646 | 2212.7976 | 1603.6431 | 1671.1981 |
| Coro1a | FDPDTNIVYLCGK | -275 | -287 | 0 | 0 | 13 | 653.2048 | 898.38806 | 1150.0458 | 2658.769 | 2519.8953 | 3402.0317 |
| Coro1a | KSDLFQEDLYPPTAGPD | -355 | -371 | 1 | 0.00051306 | 17 | 558.86304 | 567.2631 | 567.8702 | 539.138 | 836.27106 | 682.71985 |
| Coro1a | KSDLFQEDLYPPTAGPDPALT | -355 | -375 | 1 | 0.000431655 | 21 | 533.7594 | 611.1636 | 621.91406 | 916.5084 | 1046.1239 | 1016.01166 |
| Coro1a | KADQCYEDVR | -20 | -29 | 1 | 0 | 10 | 494.3499 | 2902.8708 | 3319.7886 | 199.05664 | 4941.9297 | 4863.843 |
| Coro1a | FLVLPLGK | -59 | -66 | 0 | 0.001023236 | 8 | 380.97186 | 444.95343 | 412.32123 | 276.71497 | 321.87485 | 345.78632 |
| Coro1a | KSDLFQEDLYPPTAGPDPALTA | -355 | -376 | 1 | 0.000230707 | 22 | 371.66132 | 368.47015 | 279.18146 | 509.88834 | 625.4761 | 585.5924 |
| Coro1a | QVALWDTK | -246 | -253 | 0 | 0 | 8 | 3584.6233 | 2062.3726 | 2304.1685 | 1452.2544 | 965.1323 | NaN |
| Coro1a | HLEEPLSLQELDTS | -254 | -267 | 0 | 0.00025015 | 14 | 1616.1323 | NaN | 260.91418 | 1890.1493 | 551.513 | 1774.3911 |
| Coro1a | TWDSGFCAVNPK | -34 | -45 | 0 | 0 | 12 | 1277.2047 | 1214.8933 | 1274.0012 | 969.9907 | 265.17578 | NaN |
| Coro1a | HLEEPLSLQELDTSS | -254 | -268 | 0 | 0 | 15 | 1242.4674 | 1335.227 | 1455.3201 | NaN | 1970.5507 | 2051.1816 |
| Coro1a | LKDGYVPPK | -392 | -400 | 1 | 0.00025015 | 9 | 999.88464 | NaN | 1127.9664 | 1543.7085 | 1976.3976 | 931.2661 |
| Coro1a | PVHAVFVSE | -223 | -231 | 0 | 0 | 9 | 915.2477 | 646.48846 | 787.2015 | 900.34656 | 1098.2848 | NaN |
| Coro1a | HLEEPLSLQ | -254 | -262 | 0 | 0.00025015 | 9 | 1415.7145 | 1380.1329 | 2511.7964 | 1013.28 | NaN | NaN |
| Coro1a | LEETVQAK | -454 | -461 | 0 | 0.001055699 | 8 | 2965.2515 | 2584.0776 | 3048.9587 | NaN | NaN | NaN |
| Coro1a | LDRLEETVQ | -451 | -459 | 1 | 0.007791272 | 9 | 529.8998 | 597.6861 | 1007.28186 | NaN | NaN | NaN |
| Coro1a | DQCYEDVR | -22 | -29 | 0 | 0.001291103 | 8 | 517.33636 | NaN | NaN | 446.12497 | NaN | NaN |
| Calm1;Calm2;( | FDKDGDGTITTK | (20);(20);(20) | (31);(31);(31) | 1 | 0 | 12 | 8877.513 | 9576.708 | 9490.284 | 8193.916 | 10714.622 | 10639.315 |
| Calm1;Calm2;( | FDKDGDGTITTK | (20);(20);(20) | (31);(31);(31) | 1 | 0 | 12 | 8877.513 | 9576.708 | 9490.284 | 8193.916 | 10714.622 | 10639.315 |
| Calm1;Calm2;( | FDKDGDGTITTK | (20);(20);(20) | (31);(31);(31) | 1 | 0 | 12 | 8877.513 | 9576.708 | 9490.284 | 8193.916 | 10714.622 | 10639.315 |
| Calm1;Calm2;( | FSLFDKDGDGTITTK | (17);(17);(17) | (31);(31);(31) | 1 | 0 | 15 | 6867.6606 | 8757.116 | 8594.789 | 5145.631 | 2892.524 | 6479.5107 |
| Calm1;Calm2;( | FSLFDKDGDGTITTK | (17);(17);(17) | (31);(31);(31) | 1 | 0 | 15 | 6867.6606 | 8757.116 | 8594.789 | 5145.631 | 2892.524 | 6479.5107 |
| Calm1;Calm2;( | LTDEEVDEMIR | (117);(117);(117) | (127);(127);(127) | 0 | 0 | 11 | 1656.2587 | 1765.278 | 1666.1414 | 1444.9825 | 698.6751 | 479.12732 |
| Calm1;Calm2;( | LTDEEVDEMIR | (117);(117);(117) | (127);(127);(127) | 0 | 0 | 11 | 1656.2587 | 1765.278 | 1666.1414 | 1444.9825 | 698.6751 | 479.12732 |
| Calm1;Calm2;( | SLFDKDGDGTITTK | (18);(18);(18) | (31);(31);(31) | 1 | 0 | 14 | 1261.3112 | 880.6054 | 1384.4502 | 1832.5472 | 1302.488 | 1237.2014 |
| Calm1;Calm2;( | SLFDKDGDGTITTK | (18);(18);(18) | (31);(31);(31) | 1 | 0 | 14 | 1261.3112 | 880.6054 | 1384.4502 | 1832.5472 | 1302.488 | 1237.2014 |
| Calm1;Calm2;( | VFDKDGNGYIS | (92);(92);(92) | (102);(102);(102) | 1 | 0.000471409 | 11 | 1055.052 | 2724.3157 | 2519.359 | 2425.76 | 1738.3356 | 1517.1133 |
| Calm1;Calm2;( | VFDKDGNGYISAA | (92);(92);(92) | (104);(104);(104) | 1 | 0.000431655 | 13 | 896.6256 | 918.74634 | 485.22522 | 706.31396 | 1026.5371 | 912.14764 |
| Calm1;Calm2;( | EAFSLFDKDGDGTITTK | (15);(15);(15) | (31);(31);(31) | 1 | 0 | 17 | 861.09467 | 687.61694 | 1102.2057 | 1627.1628 | 2183.2554 | 2176.3384 |
| Calm1;Calm2;( | EAFSLFDKDGDGTITTK | (15);(15);(15) | (31);(31);(31) | 1 | 0 | 17 | 861.09467 | 687.61694 | 1102.2057 | 1627.1628 | 2183.2554 | 2176.3384 |
| Nup133 | ECEVPNSLTPADVFFR | -675 | -690 | 0 | 0 | 16 | 412.7854 | 357.93988 | 440.09305 | 478.07153 | 412.11975 | 418.71588 |
| Nup133 | EYLPEVTDLLR | -1116 | -1126 | 0 | 0 | 11 | 368.3799 | 585.2323 | 634.7351 | 647.5102 | 128.37277 | 880.26196 |
| Nup133 | ADQNFSDFLFR | -892 | -902 | 0 | 0 | 11 | 265.76303 | 515.46124 | 215.44528 | 141.32706 | 342.06897 | 277.8153 |
| Nup133 | DDWSGSDGKDDPIEVSK | -1080 | -1096 | 1 | 0.000431655 | 17 | 1151.5524 | 1395.0122 | NaN | 636.00446 | 805.23175 | 676.37787 |
| Nup133 | EDTYSDTCVDLGGEK | -192 | -206 | 0 | 0.000591124 | 15 | 644.48315 | 531.9055 | NaN | NaN | 360.06976 | 496.5774 |
| Nup133 | FADQNFSDFLFR | -891 | -902 | 0 | 0 | 12 | 401.28778 | NaN | NaN | 426.6669 | 520.91345 | 490.48428 |
| Nup133 | TLPEQLLTER | -1006 | -1015 | 0 | 0.000431655 | 10 | 652.4007 | 593.5226 | 711.19507 | NaN | NaN | NaN |
| Cdk6 | NHPYFQDLER | -296 | -305 | 0 | 0 | 10 | 3411.8271 | 687.8457 | 3785.0547 | 3887.5483 | 3882.3374 | 640.6208 |
| Cdk6 | YKDNLNSHLPSNQ | -306 | -318 | 1 | 0 | 13 | 1942.9307 | 2159.4639 | 1995.9014 | 1529.6945 | 2391.4355 | 1097.117 |
| Cdk6 | DIIGLPGEEDWPR | -233 | -245 | 0 | 0 | 13 | 1489.1193 | 1503.9082 | 1559.0029 | 1700.1449 | 1928.2842 | 1878.7291 |
| Cdk6 | GALNHPYFQDLER | -293 | -305 | 0 | 0 | 13 | 1219.6003 | 16490.13 | 1543.5391 | 1185.0741 | 1477.2466 | 1474.7383 |
| Cdk6 | HLETFEHPNVVR | -67 | -78 | 0 | 0 | 12 | 928.9296 | 888.51953 | 1160.3705 | 958.5542 | 922.35474 | 1520.1177 |
| Cdk6 | HLETFEHPNVVR | -67 | -78 | 0 | 0 | 12 | 928.9296 | 888.51953 | 1160.3705 | 958.5542 | 922.35474 | 1520.1177 |
| Cdk6 | LDKVPEPGVPTETIK | -109 | -123 | 1 | 0 | 15 | 768.1421 | 759.3419 | 821.4123 | 760.9045 | 861.2293 | 855.3777 |
| Cdk6 | ADQQYECVAEIGEGAYGK | -9 | -26 | 0 | 0 | 18 | 495.7621 | 595.33057 | 595.92816 | 188.09286 | 744.44336 | 714.4629 |
| Cdk6 | DNLNSHLPSNQSTSELNTA | -308 | -326 | 0 | 0 | 19 | 310.96674 | 380.1548 | 343.0428 | 478.46387 | 331.8906 | 389.52167 |
| Cdk6 | GLDFLHSHR | -132 | -140 | 0 | 0.000550307 | 9 | 133.31766 | NaN | NaN | NaN | NaN | NaN |
| Rab1a | ADSLGIPFLETSAK | -143 | -156 | 0 | 0 | 14 | 142.77603 | 604.976 | 426.77826 | 539.4547 | 343.7383 | 366.01135 |
| Rab1a | KVVDYTTAK | -132 | -140 | 1 | 0 | 9 | 486.76935 | 500.03998 | NaN | NaN | 572.1559 | 617.18146 |
| Rab1a | DVTDQESFNNVK | -92 | -103 | 0 | 0 | 12 | 676.3858 | NaN | 612.07654 | NaN | NaN | NaN |
| Rab1b | DVTDQESYANVK | -89 | -100 | 0 | 0 | 12 | 4304.8657 | 2422.918 | 2125.9512 | 2771.828 | 1193.9595 | 1191.9664 |
| Rab1b | ADSLGVPFLETSAK | -140 | -153 | 0 | 0 | 14 | 627.2101 | 713.23145 | 988.10376 | 1058.6914 | 857.0049 | 868.0575 |
| Rab1b | MGPGAASGGERPNLK | -173 | -187 | 1 | 0 | 15 | 318.1939 | NaN | 550.86926 | NaN | 569.5782 | NaN |
| Rab1b | VYDVTDQESYANVK | -87 | -100 | 0 | 0.00025015 | 14 | 662.9527 | NaN | NaN | NaN | 432.67618 | NaN |
| Plcg1 | FSDVLHTIK | -390 | -398 | 0 | 0 | 9 | 473.9078 | 1040.2075 | 1160.3114 | 1092.3241 | 912.122 | 1041.0142 |
| Plcg1 | EHLDENSPLGDLLR | -860 | -873 | 0 | 0 | 14 | 981.25757 | 558.8002 | 673.1781 | NaN | 536.8344 | 695.1087 |
| Plcg1 | ISNPEFAFLR | -1149 | -1158 | 0 | 0 | 10 | 283.2251 | 243.28918 | NaN | NaN | NaN | 388.88983 |
| Plcg1 | ADSQEELQDWVK | -913 | -924 | 0 | 0 | 12 | 1118.3656 | NaN | NaN | NaN | 1275.4414 | NaN |
| Cct2 | AIEHADFAGVER | -311 | -322 | 0 | 0 | 12 | 3466.3083 | 3659.7385 | 3994.337 | 4185.6206 | 4476.22 | 4419.4937 |
| Cct2 | GSGNLEAIHVIK | -192 | -203 | 0 | 0 | 12 | 2713.6692 | 3161.102 | 3502.5962 | 3148.4695 | 4286.951 | 4376.025 |
| Cct2 | KNIGVDNPAAK | -72 | -82 | 1 | 0.00025015 | 11 | 2095.4854 | 2146.5208 | 365.75906 | 2403.805 | 2875.3757 | 3038.3945 |
| Cct2 | ADSYLDEGFLLDKK | -210 | -223 | 1 | 0 | 14 | 1637.0287 | 1147.7634 | 1539.9513 | 2091.6265 | 3408.4875 | 4325.7065 |
| Cct2 | LDEGFLLDK | -214 | -222 | 0 | 0 | 9 | 1620.1571 | 1783.5958 | 1519.4286 | 1700.0431 | 1631.3767 | 1613.4282 |
| Cct2 | DHPELVK | -336 | -342 | 0 | 0.00025015 | 7 | 1606.5399 | 1270.6283 | 1108.8596 | 1559.3824 | 1214.667 | 1236.7568 |
| Cct2 | LDEGFLLDKK | -214 | -223 | 1 | 0 | 10 | 1444.7681 | 2972.064 | 3308.4558 | 4218.6494 | 7655.054 | 6221.3027 |
| Cct2 | ADSYLDEGFLLDK | -210 | -222 | 0 | 0 | 13 | 1165.1251 | 1250.6305 | 1307.9376 | 1388.1764 | 1550.2225 | 1512.0457 |
| Cct2 | GATQQILDEAER | -377 | -388 | 0 | 0 | 12 | 1163.6202 | 1274.6051 | 1365.9583 | 839.59186 | 918.9421 | 849.91174 |
| Cct2 | STFDHPELVK | -333 | -342 | 0 | 0 | 10 | 1051.0845 | 1159.6034 | 1547.6487 | 1293.402 | 207.05144 | 1191.3859 |
| Cct2 | GNLEAIHVIK | -194 | -203 | 0 | 0 | 10 | 1012.26154 | 873.7499 | 1113.3785 | 1540.0696 | 1456.2847 | 1444.6559 |
| Cct2 | SAVDHGSDEAR | -144 | -154 | 0 | 0 | 11 | 819.24146 | 636.87384 | 424.89258 | 834.9464 | 561.77234 | 796.5665 |
| Cct2 | QLIYNYPEQLFGAA | -294 | -307 | 0 | 0 | 14 | 737.1289 | 685.159 | 750.6504 | 983.55365 | 1057.4695 | 1083.0233 |
| Cct2 | GGSLADSYLDEGFLLDKK | -206 | -223 | 1 | 0 | 18 | 594.20496 | 741.25024 | 723.62 | 447.33365 | 1210.8438 | 1284.3938 |
| Cct2 | SFIGAIAIGDLVK | -28 | -40 | 0 | 0 | 13 | 576.5757 | 560.5243 | 505.35602 | 911.346 | 952.32684 | 920.1037 |
| Cct2 | SLHDALCVLAQ | -389 | -399 | 0 | 0 | 11 | 290.77606 | 2619.7856 | 1029.4961 | 2511.0752 | 3253.9983 | 1266.4855 |
| Cct2 | GVMAIEHADFAGVER | -308 | -322 | 0 | 0 | 15 | 650.2661 | 701.633 | 826.5432 | NaN | 1019.34326 | 503.59528 |

| | | | | | | | | | | | | |
|---|---|---|---|---|---|---|---|---|---|---|---|---|
| Cct2 | GGSLADSYLDEGFLLDK | -206 | -222 | 0 | 0 | 17 | 399.90158 | NaN | 338.9116 | 644.49207 | 1166.728 | 904.4693 |
| Cct2 | LIHFSGVAL | -359 | -367 | 0 | 0.007546675 | 9 | 372.42804 | NaN | 520.874 | 330.14923 | 969.7623 | 1102.3446 |
| Cct2 | LIEEVMIGEDK | -348 | -358 | 0 | 0 | 11 | 1421.3839 | 816.1037 | NaN | 1242.8439 | NaN | 729.57526 |
| Cct2 | GEACTIVLR | -368 | -376 | 0 | 0 | 9 | 1396.4908 | NaN | NaN | 1338.4636 | 1491.1663 | 1591.6649 |
| Cct2 | TQQILDEAER | -379 | -388 | 0 | 0 | 10 | 2470.1482 | NaN | NaN | 2953.1616 | NaN | 1269.8457 |
| Cct2 | IAIGDLVK | -33 | -40 | 0 | 0.006820063 | 8 | 772.5218 | 279.0265 | NaN | NaN | NaN | NaN |
| Cct2 | GYDSADLVAQLR | -455 | -466 | 0 | 0 | 12 | 477.68457 | NaN | NaN | NaN | NaN | NaN |
| Pfkp | DLLFQPVAELK | -725 | -735 | 0 | 0 | 11 | 3305.843 | 3712.2988 | 9825.772 | 9554.943 | 4925.829 | 4954.9673 |
| Pfkp | ALNTITDTCDR | -554 | -564 | 0 | 0 | 11 | 1160.9128 | 1134.3741 | 548.1758 | 1270.4329 | 1045.8188 | 1300.8214 |
| Pfkp | SNVVHLTEK | -616 | -624 | 0 | 0 | 9 | 1144.6444 | 3092.9004 | 3428.2417 | 2685.651 | 1186.3696 | 1181.3359 |
| Pfkp | QGLVDGGSNIVEAK | -64 | -77 | 0 | 0 | 14 | 1075.1674 | 1611.3519 | 1155.0367 | 1711.1589 | 1290.0132 | 1287.6581 |
| Pfkp | EATPETPACVVSLR | -334 | -347 | 0 | 0 | 14 | 848.9263 | 917.05566 | 1099.5272 | 1067.2878 | 1080.1228 | 1354.6326 |
| Pfkp | KEWSGLLEELAR | -138 | -149 | 1 | 0.00025015 | 12 | 801.80206 | 744.3359 | 651.01917 | 835.1453 | 1169.1385 | 1147.4702 |
| Pfkp | VGIADGHK | -434 | -441 | 0 | 0 | 8 | 772.1273 | 555.5462 | 635.7566 | 752.32043 | 605.83527 | 378.72485 |
| Pfkp | EWSGLLEELAR | -139 | -149 | 0 | 0 | 11 | 755.3294 | 729.89874 | 819.5404 | 1134.1979 | 1201.277 | 1222.4523 |
| Pfkp | QGGAPSPFDR | -673 | -682 | 0 | 0 | 10 | 704.1113 | 529.33484 | 1051.2039 | 134.4887 | 423.97864 | 817.7458 |
| Pfkp | EGYQGLVDGGSNIVEAK | -61 | -77 | 0 | 0 | 17 | 693.9361 | 786.35455 | 784.4938 | 91.82442 | 94.990814 | 1175.2034 |
| Pfkp | NGDIDNDTVQKY | -150 | -161 | 1 | 0 | 12 | 656.62256 | 619.6714 | 641.7468 | 911.0583 | 848.9484 | 887.4312 |
| Pfkp | YLEHLSGDGK | -15 | -24 | 0 | 0 | 10 | 618.86316 | 549.7089 | 3540.812 | 3313.4204 | 571.75024 | 375.09787 |
| Pfkp | LLEATPETPACVVSLR | -332 | -347 | 0 | 0 | 16 | 618.18005 | 695.2274 | 720.18225 | 840.9599 | 1188.706 | 1009.58057 |
| Pfkp | ADTALNTITDTCDR | -551 | -564 | 0 | 0 | 14 | 425.3887 | 677.1925 | 912.2273 | 795.16315 | 784.2919 | 851.6492 |
| Pfkp | EIGWADVGGWTGQ | -459 | -471 | 0 | 0.000627943 | 13 | 311.17325 | 458.45233 | 333.18597 | 201.91306 | 573.57587 | 156.62202 |
| Pfkp | TALNTITDTCDR | -553 | -564 | 0 | 0 | 12 | 1719.0153 | 1948.095 | 1788.1965 | NaN | 1428.9683 | 1429.703 |
| Pfkp | VDGGSNIVEAK | -67 | -77 | 0 | 0 | 11 | 1459.111 | 1266.6837 | 1009.2839 | 1017.70825 | NaN | 720.835 |
| Pfkp | FIYEGYQGLVDGGSNIVEAK | -58 | -77 | 0 | 0 | 20 | 497.1485 | 275.9861 | NaN | 947.8618 | 935.9833 | 1007.0762 |
| Pfkp | QVGGTIIGSAR | -86 | -96 | 0 | 0 | 11 | 1613.8252 | 1445.1758 | NaN | 959.69464 | 973.1531 | NaN |
| Pfkp | SDLDSSSSSAYPK | -2 | -14 | 0 | 0 | 13 | 1212.2084 | 526.24023 | 858.42505 | 663.63 | NaN | NaN |
| Pfkp | SLQHHEEL | -777 | -784 | 0 | 0.007701635 | 8 | 1159.8771 | 1271.3223 | 706.29297 | 405.13083 | NaN | NaN |
| Pfkp | KEWSGLLEEL | -138 | -147 | 1 | 0.00201047 | 10 | 1060.4756 | 934.3957 | 1105.0778 | NaN | NaN | 375.1441 |
| Pfkp | VTDFEHR | -737 | -743 | 0 | 0.00078661 | 7 | 2567.8057 | NaN | 2633.0508 | NaN | NaN | NaN |
| Pfkp | RFEGNLNTY | -385 | -393 | 1 | 0.00025015 | 9 | 306.56625 | 578.22485 | NaN | NaN | NaN | NaN |
| Nucb1 | RFEEELAAR | -348 | -356 | 1 | 0.00025015 | 9 | 1429.6394 | 1465.6223 | 1771.4512 | 1422.536 | 77.70018 | 1823.9048 |
| Nucb1 | KVPEQPPELPQLDSQHL | -443 | -459 | 1 | 0 | 17 | 674.3721 | 906.93414 | 918.37756 | 716.41595 | 932.0106 | 740.19727 |
| Nucb1 | TKLDELK | -97 | -103 | 1 | 0.00025015 | 7 | 636.40753 | 634.1041 | 561.0603 | 551.06396 | 716.16425 | 666.98303 |
| Nucb1 | LSQETEALGR | -367 | -376 | 0 | 0 | 10 | 608.596 | 599.01276 | 676.77203 | 772.7552 | 1224.3757 | 588.70905 |
| Nucb1 | ADTDDAPVPAPAGDQK | -420 | -435 | 0 | 0 | 16 | 1724.1838 | 1655.4434 | 1949.2534 | 2142.8564 | NaN | 1921.686 |
| Nucb1 | EVWEELDGLDPNR | -229 | -241 | 0 | 0 | 13 | 652.04407 | 962.3816 | 722.2924 | 622.8366 | NaN | 1305.68 |
| Tars1 | GISQGLADNTVVAK | -107 | -120 | 0 | 0 | 14 | 2219.2007 | 386.05005 | 560.3336 | 821.9423 | 2341.8745 | 709.8031 |
| Tars1 | LNLSTRPEK | -495 | -503 | 1 | 0 | 9 | 2215.1196 | 2431.2134 | 2060.4502 | 1569.6896 | 2236.8389 | 2256.04 |
| Tars1 | AELNPWPEYINTR | -44 | -56 | 0 | 0 | 13 | 1052.4164 | 1368.4353 | 1381.9137 | 1143.2838 | 1335.7366 | 1315.7285 |
| Tars1 | ADTDLDPGCTLNK | -647 | -659 | 0 | 0 | 13 | 987.3866 | 1378.4115 | 300.47998 | 489.59366 | 909.9163 | 1313.839 |
| Tars1 | LKAEHDSILAEK | -63 | -74 | 1 | 0 | 12 | 860.76984 | 887.12744 | 850.12915 | 859.293 | 1188.5627 | 946.2239 |
| Tars1 | QLENSLNEFGEK | -517 | -528 | 0 | 0 | 12 | 678.2858 | 316.7266 | 928.08014 | 1303.9884 | 797.2134 | 1238.2924 |
| Tars1 | FNLTYVSHDGDDK | -570 | -582 | 0 | 0 | 13 | 534.97284 | 640.39246 | 607.12366 | 159.44794 | 672.0513 | 693.0751 |
| Tars1 | DQELYFFHE | -328 | -336 | 0 | 0 | 9 | 332.90427 | 372.90558 | 427.1177 | 397.13654 | 400.87106 | 435.80695 |
| Tars1 | CGPLIDLCR | -253 | -261 | 0 | 0 | 9 | 134.87578 | 479.3427 | 401.71884 | 456.6822 | 731.5556 | 542.90344 |
| Tars1 | WELNPGDGAFYGPK | -529 | -542 | 0 | 0 | 14 | 1482.648 | 1656.8137 | NaN | 1905.2207 | 1382.7523 | 859.32513 |
| Tars1 | RTVEETVR | -698 | -705 | 1 | 0.009569685 | 8 | 1134.7716 | 1141.8794 | NaN | 895.61194 | 1254.0837 | 724.2979 |
| Tars1 | QLENSLNEFGEKW | -517 | -529 | 1 | 0.000471409 | 13 | 984.06903 | 1200.6686 | NaN | 1055.4047 | 1192.2445 | 958.1159 |
| Tars1 | WPFWLSPR | -611 | -618 | 0 | 0.00025015 | 8 | 865.9996 | 885.2016 | 1184.8578 | 1177.0902 | 1289.5563 | NaN |
| Tars1 | LSPGSCFFLPK | -337 | -347 | 0 | 0 | 11 | 3074.906 | NaN | NaN | 1606.5206 | 1561.4489 | 1919.3275 |
| Tars1 | GLADNTVVAK | -111 | -120 | 0 | 0.00025015 | 10 | 658.0748 | 755.8053 | 928.6914 | NaN | NaN | NaN |
| Tars1 | LKAEHDSILAE | -63 | -73 | 1 | 0.005333786 | 11 | 2745.2722 | 3084.754 | NaN | NaN | NaN | NaN |
| Psmc4 | KDEQEHEFYK | -409 | -418 | 1 | 0 | 10 | 1692.1873 | 1589.1116 | 1605.6846 | 917.75635 | 1611.3057 | 1067.9961 |
| Psmc4 | KDEQEHEFYK | -409 | -418 | 1 | 0 | 10 | 1692.1873 | 1589.1116 | 1605.6846 | 917.75635 | 1611.3057 | 1067.9961 |
| Psmc4 | EVQEEYIKDEQK | -55 | -66 | 1 | 0 | 12 | 1667.9983 | 2699.156 | 1057.3782 | 590.1314 | 1399.6676 | 1107.7463 |
| Psmc4 | ADTLDPALLRPGR | -314 | -326 | 1 | 0 | 13 | 1146.6948 | 1125.5095 | 1185.6842 | 1039.796 | 1008.514 | 1055.4861 |
| Psmc4 | EEYIKDEQK | -58 | -66 | 1 | 0 | 9 | 923.5638 | 909.2461 | 779.88116 | 1857.4216 | 661.6911 | 2478.3806 |
| Psmc4 | SFLGPEPEDLEDLYSR | -28 | -43 | 0 | 0 | 16 | 614.8462 | 546.5606 | 753.1049 | 573.0958 | 913.614 | 885.7809 |
| Psmc4 | LGPEPEDLEDLYSR | -30 | -43 | 0 | 0 | 14 | 420.3394 | 644.9149 | 753.548 | 823.8421 | 338.41083 | 987.9507 |
| Psmc4 | EAVELPLTHFELYK | -179 | -192 | 0 | 0 | 14 | 363.12354 | 347.3129 | 547.77356 | 633.8679 | 757.822 | 650.8977 |
| Psmc4 | IDEIDAIATK | -264 | -273 | 0 | 0 | 10 | 2029.3868 | NaN | 619.43066 | 1843.5184 | 1448.888 | 1480.144 |
| Psmc4 | NQMDGFDQNVNVK | -294 | -306 | 0 | 0.00122419 | 13 | 432.5824 | 430.94727 | NaN | NaN | NaN | NaN |
| Ldha | SLNPELGTDADKEQWK | -213 | -228 | 1 | 0 | 16 | 13471.673 | 14247.535 | 14416.967 | 14733.534 | 15459.109 | 15153.059 |
| Ldha | SLNPELGTDADKEQWK | -213 | -228 | 1 | 0 | 16 | 13471.673 | 14247.535 | 14416.967 | 14733.534 | 15459.109 | 15153.059 |
| Ldha | SADTLWGIQK | -319 | -328 | 0 | 0 | 10 | 13205.74 | 12993.592 | 193.49377 | 13227.4 | 14082.036 | 14501.379 |
| Ldha | SADTLWGIQK | -319 | -328 | 0 | 0 | 10 | 13205.74 | 12993.592 | 193.49377 | 13227.4 | 14082.036 | 14501.379 |
| Ldha | KFIIPNIVK | -118 | -126 | 1 | 0.000431655 | 9 | 6544.407 | 6792.719 | 6581.2983 | 5436.586 | 5719.2456 | 5351.134 |
| Ldha | DLQHGSLFLK | -64 | -73 | 0 | 0 | 10 | 6050.4062 | 5793.1504 | 6079.8145 | 571.95953 | 5557.0557 | 4913.3975 |
| Ldha | FIIPNIVK | -119 | -126 | 0 | 0.000591124 | 8 | 5562.9966 | 6490.99 | 6509.076 | 5054.1914 | 6061.0044 | 6135.229 |
| Ldha | VTLTPEEEAR | -306 | -315 | 0 | 0 | 10 | 4171.7236 | 3891.2322 | 3038.7483 | 1836.7954 | 317.48404 | 1582.2639 |
| Ldha | KISGFPK | -149 | -155 | 1 | 0.000755556 | 7 | 3981.971 | 3427.5347 | 2799.439 | 1710.8693 | 1614.9722 | 2146.6726 |
| Ldha | QVVDSAYEVIK | -233 | -243 | 0 | 0 | 11 | 3786.477 | 4177.7944 | 4461.089 | 5021.658 | 4666.146 | 4360.2153 |
| Ldha | QVVDSAYEVIK | -233 | -243 | 0 | 0 | 11 | 3786.477 | 4177.7944 | 4461.089 | 5021.658 | 4666.146 | 4360.2153 |
| Ldha | SLNPELGTDADK | -213 | -224 | 0 | 0 | 12 | 3185.6953 | 1761.3604 | 1701.0154 | 4447.5103 | 947.6298 | 1622.12 |
| Ldha | SLNPELGTDADK | -213 | -224 | 0 | 0 | 12 | 3185.6953 | 1761.3604 | 1701.0154 | 4447.5103 | 947.6298 | 1622.12 |
| Ldha | LLKEEQAPQNK | -12 | -22 | 1 | 0 | 11 | 2687.0557 | 2548.8125 | 2767.0776 | 2733.7307 | 3989.878 | 3815.5176 |
| Ldha | LLKEEQAPQNK | -12 | -22 | 1 | 0 | 11 | 2687.0557 | 2548.8125 | 2767.0776 | 2733.7307 | 3989.878 | 3815.5176 |

| | | | | | | | | | | | | |
|---|---|---|---|---|---|---|---|---|---|---|---|---|
| Ldha | KSADTLWGIQK | -318 | -328 | 1 | 0 | 11 | 1934.2047 | 2197.7656 | 2270.2803 | 2645.3496 | 2823.8027 | 2822.1763 |
| Ldha | DQLIVNLLK | -6 | -14 | 0 | 0 | 9 | 1644.7816 | 1844.4323 | 1786.7413 | 2230.74 | 3151.1172 | 3274.4265 |
| Ldha | VHPISTMIK | -270 | -278 | 0 | 0 | 9 | 1461.8646 | 1331.5813 | 1356.1455 | 2030.5496 | 2188.784 | 2054.3987 |
| Ldha | VHPISTMIK | -270 | -278 | 0 | 0 | 9 | 1461.8646 | 1331.5813 | 1356.1455 | 2030.5496 | 2188.784 | 2054.3987 |
| Ldha | LKEEQAPQNK | -13 | -22 | 1 | 0 | 10 | 1326.8687 | 1282.1884 | 1569.1326 | 1457.3345 | 1681.2463 | 1665.4019 |
| Ldha | ADTLWGIQK | -320 | -328 | 0 | 0 | 9 | 1119.3473 | 826.20337 | 1099.7932 | 542.82745 | 712.64 | 1280.0686 |
| Ldha | LTPEEEAR | -308 | -315 | 0 | 0.001392869 | 8 | 976.95184 | 6352.6753 | 5839.899 | 352.20346 | 2302.5063 | 4421.6357 |
| Ldha | SVADLAESIMK | -255 | -265 | 0 | 0 | 11 | 426.89267 | 3560.2517 | 572.12756 | 3941.519 | 3967.5134 | 3940.054 |
| Ldha | SVADLAESIMK | -255 | -265 | 0 | 0 | 11 | 426.89267 | 3560.2517 | 572.12756 | 3941.519 | 3967.5134 | 3940.054 |
| Ldha | AIGLSVADLAESIMK | -251 | -265 | 0 | 0 | 15 | 894.23645 | 1028.1925 | 404.0311 | 256.78793 | 1526.3893 | 1317.5831 |
| Ldha | LNPELGTDADKEQWK | -214 | -228 | 1 | 0 | 15 | 812.18115 | 808.4988 | 793.57306 | 611.15485 | 700.3949 | 733.09265 |
| Ldha | DLADELALVDVMEDK | -43 | -57 | 0 | 0 | 15 | 699.7101 | 935.19965 | 859.30066 | 654.8999 | 889.84204 | 729.87524 |
| Ldha | DLADELALVDVMEDK | -43 | -57 | 0 | 0 | 15 | 699.7101 | 935.19965 | 859.30066 | 654.8999 | 889.84204 | 729.87524 |
| Ldha | PELGTDADKEQWK | -216 | -228 | 1 | 0 | 13 | 662.11755 | 751.56213 | 768.5916 | 684.179 | 651.30273 | 409.84515 |
| Ldha | PELGTDADKEQWK | -216 | -228 | 1 | 0 | 13 | 662.11755 | 751.56213 | 768.5916 | 684.179 | 651.30273 | 409.84515 |
| Ldha | SLNPELGTDADKE | -213 | -225 | 1 | 0 | 13 | 581.37994 | 401.7984 | 1313.39 | 1023.3464 | 1039.4591 | 946.21405 |
| Ldha | ILGQNGISDVVK | -294 | -305 | 0 | 0 | 12 | 443.4388 | 553.1055 | 684.8513 | 636.93964 | 890.8459 | 965.8084 |
| Ldha | HGSLFLK | -67 | -73 | 0 | 0.00025015 | 7 | 435.48767 | 623.96 | 571.6293 | 1025.1549 | 596.58 | 517.4953 |
| Ldha | ALVDVMEDK | -49 | -57 | 0 | 0 | 9 | 1988.9202 | 2310.8616 | NaN | 2198.5269 | 1960.0287 | 489.1986 |
| Ldha | ALVDVMEDK | -49 | -57 | 0 | 0 | 9 | 1988.9202 | 2310.8616 | NaN | 2198.5269 | 1960.0287 | 489.1986 |
| Ldha | QHGSLFLK | -66 | -73 | 0 | 0 | 8 | 2003.8025 | NaN | 1823.1263 | 1533.102 | 1152.0619 | 1160.874 |
| Ldha | RVHPISTMIK | -269 | -278 | 1 | 0 | 10 | 361.08136 | NaN | 571.1889 | 662.44183 | 432.1757 | 469.5753 |
| Ldha | GQNGISDVVK | -296 | -305 | 0 | 0.00051306 | 10 | 1773.3296 | 996.94794 | NaN | 397.43774 | NaN | 1585.8629 |
| Ldha | SLNPELGTDADKEQ | -213 | -226 | 1 | 0.00025015 | 14 | 789.4248 | 869.5462 | 761.74194 | NaN | NaN | NaN |
| Safb2 | ADTVNQILHAF | -144 | -154 | 0 | 0 | 11 | 217.65991 | NaN | 729.80365 | 395.62833 | 890.07477 | NaN |
| Safb2 | LNEGQGLPPPPR | -872 | -883 | 0 | 0.003166358 | 12 | 243.26648 | NaN | NaN | 226.64438 | NaN | NaN |
| Sun2 | SALLDDHLHSEPY | -92 | -104 | 0 | 0.000627943 | 13 | 1498.0221 | 1985.745 | 2014.7583 | 1763.4308 | 2673.7544 | 2856.7458 |
| Sun2 | EGIVGVTEEQVHR | -532 | -544 | 0 | 0 | 13 | 1193.6074 | 1389.1089 | 1631.3025 | 1356.565 | 1697.6907 | 1957.2677 |
| Sun2 | ADVESQFPDWIR | -465 | -476 | 0 | 0 | 12 | 953.9528 | NaN | 1653.194 | 995.2718 | 1299.5648 | 1434.9885 |
| Sun2 | VILQPDVHPGNCWAF | -604 | -618 | 0 | 0.000431655 | 15 | 823.3517 | 991.59875 | 1012.46106 | 806.45953 | NaN | 1041.5819 |
| Sun2 | VWTLVTFPGR | -177 | -186 | 0 | 0 | 10 | 299.53632 | 265.9629 | NaN | 431.6098 | 462.32513 | 447.92578 |
| Sun2 | QNSVADEVGLLPQK | -446 | -459 | 0 | 0.000471409 | 14 | 3688.2473 | 6181.9614 | 3251.2854 | NaN | NaN | NaN |
| Sun2 | AEHHQDSEDLFK | -378 | -389 | 0 | 0.000591124 | 12 | 1227.2788 | NaN | NaN | 1093.4014 | 1055.3702 | NaN |
| Sun2 | HLSPAPQLGPSSDSHTS | -53 | -69 | 0 | 0.00149795 | 17 | 5083.8145 | NaN | 451.009 | NaN | NaN | NaN |
| Pcna | GIPEQEYSCVIK | -127 | -138 | 0 | 0 | 12 | 3409.215 | 2811.7588 | 527.67017 | 2895.4846 | 2460.2854 | 2440.5107 |
| Pcna | LGIPEQEYSCVIK | -126 | -138 | 0 | 0 | 13 | 2278.587 | 3021.4844 | 2968.5156 | 1352.1389 | 1699.1235 | 1631.6582 |
| Pcna | SEGFDTYR | -54 | -61 | 0 | 0 | 8 | 1977.3619 | 2022.0269 | 1844.7178 | 1926.8234 | 1375.487 | 1516.6617 |
| Pcna | YLNFFTK | -211 | -217 | 0 | 0.001392869 | 7 | 1901.7649 | 1758.8439 | 1862.9922 | 1019.68445 | 984.3309 | 953.301 |
| Pcna | RYLNFFTK | -210 | -217 | 1 | 0.00078661 | 8 | 1801.8796 | 2935.208 | 3008.403 | 1134.0422 | 658.2722 | 2629.5618 |
| Pcna | PEQEYSCVIK | -129 | -138 | 0 | 0.00025015 | 10 | 1714.0327 | 1611.4847 | 1502.8479 | 687.7227 | 2011.5635 | 574.4686 |
| Pcna | DLSHIGDAVVIS | -150 | -161 | 0 | 0 | 12 | 1692.3866 | 1948.3285 | 1957.7758 | 1514.7 | 1427.8923 | 1799.4738 |
| Pcna | CAGNEDIITLR | -81 | -91 | 0 | 0 | 11 | 1596.4375 | 1523.5398 | 1539.351 | 976.27594 | 1177.324 | 1189.1411 |
| Pcna | SADVPLVVEYK | -230 | -240 | 0 | 0 | 11 | 1585.0596 | 1545.5386 | 1599.6177 | 1141.6913 | 557.63153 | 555.16156 |
| Pcna | ALVFEAPNQEK | -100 | -110 | 0 | 0 | 11 | 1413.9612 | 1861.7509 | 484.35242 | 3229.2117 | 550.3008 | 1951.5328 |
| Pcna | AGNEDIITLR | -82 | -91 | 0 | 0 | 10 | 1086.5348 | 1129.4296 | 1089.613 | 844.1753 | 911.7309 | 1027.9755 |
| Pcna | LSQTSNVDKEEEAVTIE | -182 | -198 | 1 | 0 | 17 | 912.8132 | 1051.4279 | 1079.4556 | 577.6846 | 719.1901 | 714.1227 |
| Pcna | DLSHIGDAVVISC | -150 | -162 | 0 | 0 | 13 | 464.10767 | 855.42053 | 799.71106 | 835.5421 | 953.9405 | 1030.967 |
| Pcna | DLDVEQLGIPEQEYSCVIK | -120 | -138 | 0 | 0.00025015 | 19 | 463.14233 | 329.15417 | 359.3985 | 382.34708 | 434.9077 | 450.49213 |
| Pcna | LIQGSILK | -6 | -13 | 0 | 0 | 8 | 2107.3643 | 1118.2913 | 989.4188 | 946.0349 | NaN | 754.29205 |
| Pcna | FSASGELGNGNIK | -169 | -181 | 0 | 0 | 13 | 1425.3062 | 909.94867 | 1164.4557 | 1497.8691 | NaN | 1101.2151 |
| Pcna | ADVPLVVEYK | -231 | -240 | 0 | 0 | 10 | 1292.8888 | NaN | 1448.1298 | NaN | 1391.1609 | NaN |
| Pcna | VFEAPNQEK | -102 | -110 | 0 | 0.00025015 | 9 | 7758.451 | NaN | NaN | 4969.0645 | NaN | NaN |
| Sptan1 | SSEEIESAFR | -2405 | -2414 | 0 | 0 | 10 | 2796.731 | 2709.448 | 3051.0024 | 374.5495 | 2787.8474 | 612.2592 |
| Sptan1 | KHEAFETDFTVHK | -1911 | -1923 | 1 | 0 | 13 | 2365.5886 | 1002.96436 | 4218.108 | 1118.1221 | 686.0764 | 903.12933 |
| Sptan1 | ISEGHFASETIR | -110 | -121 | 0 | 0 | 12 | 2198.358 | 930.6941 | 1929.7931 | 2481.3525 | 1993.0945 | 2241.5896 |
| Sptan1 | ELPTAFDYVEFTR | -2455 | -2467 | 0 | 0 | 13 | 2194.555 | 1929.2275 | 2751.1062 | 2299.2195 | 3485.1177 | 2129.4282 |
| Sptan1 | SDENYKDPTNLQGK | -68 | -81 | 1 | 0 | 14 | 2039.8405 | 1475.2676 | 2280.1387 | 4340.644 | 1716.7611 | 795.5696 |
| Sptan1 | GEIDAHEDSFK | -408 | -418 | 0 | 0 | 11 | 1460.6946 | 1545.5132 | 1322.069 | 1238.1058 | 576.73987 | 1024.4269 |
| Sptan1 | FEEFQTDLAAHEER | -191 | -204 | 0 | 0.00025015 | 14 | 1215.8265 | 2274.53 | 2254.6116 | 1660.7515 | 1758.231 | 1904.198 |
| Sptan1 | LGDSHDLQR | -1335 | -1343 | 0 | 0 | 9 | 891.01196 | 442.8854 | 949.49603 | 976.17004 | 928.4822 | 1569.6432 |
| Sptan1 | LGDSHDLQR | -1335 | -1343 | 0 | 0 | 9 | 891.01196 | 442.8854 | 949.49603 | 976.17004 | 928.4822 | 1569.6432 |
| Sptan1 | LIQEQHPEEELIK | -215 | -227 | 0 | 0 | 13 | 878.081 | 1147.4929 | 991.5568 | 4009.3933 | 1060.3651 | 5254.595 |
| Sptan1 | SDESYKDPTNIQSK | -1575 | -1588 | 1 | 0 | 14 | 832.3127 | 1808.3815 | 1601.9713 | 548.9658 | 1577.2404 | 1371.2886 |
| Sptan1 | EELYQNLTR | -2427 | -2435 | 0 | 0 | 9 | 806.3713 | 718.4329 | 750.23566 | 539.4013 | 575.1538 | 575.69135 |
| Sptan1 | ITALDEFATK | -523 | -532 | 0 | 0 | 10 | 616.0527 | 674.7694 | 686.612 | 1240.3163 | 1812.5464 | 817.5183 |
| Sptan1 | NQALNTDNYGHDLASVQ | -1253 | -1269 | 0 | 0 | 17 | 607.42236 | 454.55725 | 486.71915 | 527.4702 | 515.40564 | 509.5246 |
| Sptan1 | EKLDILDQER | -1418 | -1427 | 1 | 0 | 10 | 602.7074 | 530.5213 | 727.8682 | 477.69247 | 1818.088 | 1763.74 |
| Sptan1 | KFEEFQTDLAAHEER | -190 | -204 | 1 | 0 | 15 | 549.39465 | 977.22003 | 621.0533 | 477.73676 | 753.1441 | 687.32324 |
| Sptan1 | SQLLGSAHEVQR | -1226 | -1237 | 0 | 0 | 12 | 538.011 | 531.77136 | 677.6664 | 756.5448 | 1164.1448 | 1212.096 |
| Sptan1 | TATDEAYKDPSNLQGK | -595 | -610 | 1 | 0 | 16 | 394.9414 | 447.0214 | 508.43542 | 452.8686 | 545.3731 | 482.25287 |
| Sptan1 | RFLADFR | -365 | -371 | 1 | 0 | 7 | 2066.9878 | 1538.4143 | 880.2311 | 1219.392 | NaN | 768.44354 |
| Sptan1 | SHPESAEDLKEK | -1302 | -1313 | 1 | 0 | 12 | 450.03735 | 369.46188 | NaN | 439.08105 | 457.1713 | 474.2979 |
| Sptan1 | LIDVNHYAK | -638 | -646 | 0 | 0 | 9 | 816.5185 | 377.38916 | NaN | 452.6688 | 1134.3575 | NaN |
| Sptan1 | QADFNQLAELDR | -2144 | -2155 | 0 | 0 | 12 | 844.7693 | NaN | NaN | 1494.262 | NaN | 1340.4153 |
| Sptan1 | EANELQQWITEK | -1099 | -1110 | 0 | 0 | 12 | 673.2657 | 938.9978 | 946.6262 | NaN | NaN | NaN |
| Sptan1 | RQDLEDSLQAQ | -883 | -893 | 1 | 0.001023236 | 11 | 2043.515 | NaN | NaN | 339.86127 | NaN | NaN |
| Sptan1 | ETAEDIQER | -10 | -18 | 0 | 0.000907206 | 9 | 380.70535 | NaN | NaN | 668.4663 | NaN | NaN |
| Gbp2 | LQTDQSLTEAAK | -474 | -485 | 0 | 0 | 12 | 1476.2422 | 2731.1052 | 2227.1096 | 2255.7578 | 2236.4287 | 2271.0193 |

| | | | | | | | | | | | | |
|---|---|---|---|---|---|---|---|---|---|---|---|---|
| Gbp2 | FQTELGNLLVAK | -378 | -389 | 0 | 0 | 12 | 1233.3945 | 1486.5936 | 1382.3479 | 171.11526 | 2637.852 | 2534.5713 |
| Gbp2 | QLLAEQENIIAAK | -539 | -551 | 0 | 0 | 13 | 953.1159 | 996.98627 | 1204.7979 | 935.2627 | 1087.9625 | 1055.3839 |
| Gbp2 | AITHYEEQMNQK | -329 | -340 | 0 | 0 | 12 | 715.901 | 691.68134 | 798.0626 | 137.38135 | 825.0487 | 838.6208 |
| Gbp2 | AITHYEEQMNQK | -329 | -340 | 0 | 0 | 12 | 715.901 | 691.68134 | 798.0626 | 137.38135 | 825.0487 | 838.6208 |
| Gbp2 | LETLREEELCGEFVEQ | -248 | -263 | 1 | 0.001392869 | 16 | 578.3241 | 629.5126 | 525.03723 | 526.1628 | 468.68518 | 439.11026 |
| Gbp2 | SSPDQSGVDDSANFVGFFPT | -156 | -175 | 0 | 0 | 20 | 537.9334 | 426.80215 | 495.96655 | 483.9107 | 687.03143 | 576.06006 |
| Gbp2 | ADVVETLLQTDQSLTEAAK | -467 | -485 | 0 | 0 | 19 | 534.17706 | 581.4449 | 533.08075 | 341.50497 | 436.81604 | 378.4169 |
| Gbp2 | SYQEHVK | -521 | -527 | 0 | 0.002265552 | 7 | 485.62994 | 450.02136 | 416.09253 | 412.95786 | 386.25317 | 387.17947 |
| Gbp2 | QLLAEQENIIAA | -539 | -550 | 0 | 0 | 12 | 312.46204 | 387.36792 | 201.1151 | 553.1314 | 194.16919 | 329.923 |
| Gbp2 | LLQTDQSLTEAAK | -473 | -485 | 0 | 0 | 13 | 2497.1396 | 2839.5356 | 1831.8792 | NaN | NaN | NaN |
| Erh | SHTILLVQPTK | -2 | -12 | 0 | 0 | 11 | 1648.2559 | 1689.7286 | 1610.3992 | 2434.3875 | 3106.1401 | 2548.6562 |
| Erh | TYADYESVNECMEGVCK | -18 | -34 | 0 | 0 | 17 | 411.63333 | 527.7587 | 500.21927 | 463.64233 | 626.00964 | 541.25604 |
| Erh | TYADYESVNECMEGVCK | -18 | -34 | 0 | 0 | 17 | 411.63333 | 527.7587 | 500.21927 | 463.64233 | 626.00964 | 541.25604 |
| Erh | ADYESVNECMEGVCK | -20 | -34 | 0 | 0 | 15 | 396.60333 | 387.93207 | 402.60132 | 822.1075 | 821.7761 | 822.80597 |
| Fnbp1 | HTQWGIDILEK | -17 | -27 | 0 | 0 | 11 | 1864.3535 | 1974.2375 | 2034.9209 | 1960.697 | 1606.7467 | 1634.4915 |
| Fnbp1 | SVIEEDKGDGWTR | -577 | -589 | 1 | 0 | 13 | 998.36365 | 1062.8475 | 1053.4976 | 866.292 | 915.42224 | 864.6492 |
| Fnbp1 | LTEVTQNIEK | -457 | -466 | 0 | 0.00025015 | 10 | 770.3737 | 726.89667 | 858.2926 | 843.5916 | 774.0292 | 835.18787 |
| Fnbp1 | QVIPIIGK | -240 | -247 | 0 | 0.002068213 | 8 | 490.65878 | 567.55145 | 509.953 | 478.8667 | NaN | NaN |
| Fnbp1 | NPQMGDPASLDQK | -444 | -456 | 0 | 0 | 13 | 365.74954 | 242.48218 | 423.52875 | 463.12753 | NaN | NaN |
| Fnbp1 | FNQEQWEYYHT | -197 | -207 | 0 | 0.00025015 | 11 | 857.08966 | 261.66016 | 704.3867 | NaN | NaN | NaN |
| Fnbp1 | ADYSLILQR | -188 | -196 | 0 | 0 | 9 | 679.6921 | NaN | NaN | NaN | NaN | NaN |
| Gcn1 | SWCQEELPVAVR | -937 | -948 | 0 | 0 | 12 | 5847.154 | 6917.8193 | 4515.594 | 14708.4795 | 2754.2456 | 1442.8414 |
| Gcn1 | YLLDNCAPLLR | -241 | -251 | 0 | 0 | 11 | 733.7318 | 582.55225 | 481.2125 | 1769.4054 | 1119.9718 | 840.97235 |
| Gcn1 | AIAPVLLDALTDPSR | -1576 | -1590 | 0 | 0 | 15 | 459.64337 | 460.20447 | 491.11813 | 480.1313 | 632.60406 | 570.99603 |
| Gcn1 | GVTSILPVLR | -2260 | -2269 | 0 | 0 | 10 | 434.9751 | 307.7825 | 486.92316 | 395.72403 | 564.2945 | 512.8504 |
| Gcn1 | LYRPPPVLDAL | -1209 | -1219 | 1 | 0 | 11 | 392.9653 | 295.54807 | 556.15076 | 590.1732 | 652.8545 | 525.43274 |
| Gcn1 | VDIAPHVR | -1746 | -1753 | 0 | 0 | 8 | 260.61453 | 386.72876 | 340.34924 | 266.62643 | 486.9345 | 497.65326 |
| Gcn1 | ADYSSHLR | -2184 | -2191 | 0 | 0 | 8 | 1505.3771 | 932.9838 | 852.2516 | 408.2818 | 1028.6865 | NaN |
| Gcn1 | DLAPYLPSVTPGLK | -1650 | -1663 | 0 | 0 | 14 | 670.19586 | 695.5431 | 680.2526 | 598.4442 | 981.0127 | NaN |
| Gcn1 | KLTDWFK | -424 | -430 | 1 | 0.000431655 | 7 | 584.71185 | NaN | 704.8559 | 918.31976 | 658.0175 | 609.75336 |
| Gcn1 | GEHVPGFCLPK | -2248 | -2258 | 0 | 0 | 11 | 632.85986 | NaN | NaN | 381.38397 | 468.10007 | 355.59146 |
| Gcn1 | LLKPECALDK | -927 | -936 | 1 | 0 | 10 | 630.7612 | NaN | 444.36176 | 1029.3354 | 1131.619 | NaN |
| Gcn1 | FQFFVPDALNDR | -1257 | -1268 | 0 | 0 | 12 | 270.37003 | 367.75705 | 376.921 | 234.98958 | NaN | NaN |
| Gcn1 | VQNPALCLVTR | -750 | -760 | 0 | 0.000591124 | 11 | 726.2835 | 1133.0427 | 1172.9738 | NaN | NaN | NaN |
| Gcn1 | FDKEDEIR | -1135 | -1142 | 1 | 0 | 8 | 621.0326 | 751.1754 | 902.6278 | NaN | NaN | NaN |
| Gcn1 | DLILPTIQK | -260 | -268 | 0 | 0.002068213 | 9 | 523.4303 | 285.55603 | NaN | NaN | NaN | NaN |
| Gcn1 | DAPNDASYDAVR | -1305 | -1316 | 0 | 0.001636916 | 12 | 814.8749 | NaN | NaN | NaN | NaN | NaN |
| Tkt | TSRPENAIIY | -472 | -481 | 1 | 0.001868731 | 10 | 6206.1943 | 6258.716 | 6353.7847 | 5043.5884 | 332.48187 | 3833.5767 |
| Tkt | ISSDLDGHPVPK | -103 | -114 | 0 | 0 | 12 | 5346.278 | 1550.4496 | 5598.6274 | 5769.725 | 6208.0615 | 5880.5703 |
| Tkt | ISSDLDGHPVPK | -103 | -114 | 0 | 0 | 12 | 5346.278 | 1550.4496 | 5598.6274 | 5769.725 | 6208.0615 | 5880.5703 |
| Tkt | VFYPSDGVATEK | -445 | -456 | 0 | 0 | 12 | 205.27846 | 433.55746 | 473.3489 | 500.56433 | 516.85846 | 754.23004 |
| Tkt | VFYPSDGVATEK | -445 | -456 | 0 | 0 | 12 | 205.27846 | 433.55746 | 473.3489 | 500.56433 | 516.85846 | 754.23004 |
| Tkt | VLDPFTIKPLDR | -531 | -542 | 1 | 0 | 12 | 4271.413 | 5351.092 | 5410.3096 | 5806.9536 | 7193.244 | 6959.176 |
| Tkt | VLDPFTIKPLDR | -531 | -542 | 1 | 0 | 12 | 4271.413 | 5351.092 | 5410.3096 | 5806.9536 | 7193.244 | 6959.176 |
| Tkt | NSTFSELFK | -344 | -352 | 0 | 0.00025015 | 9 | 4049.1946 | 3900.0754 | 4264.3857 | 4868.942 | 5434.8394 | 5247.5757 |
| Tkt | SNNEDFQVGQAK | -482 | -493 | 0 | 0 | 12 | 3829.635 | 3805.49 | 3685.7668 | 1402.4058 | 3570.801 | 3521.787 |
| Tkt | SNNEDFQVGQAK | -482 | -493 | 0 | 0 | 12 | 3829.635 | 3805.49 | 3685.7668 | 1402.4058 | 3570.801 | 3521.787 |
| Tkt | STVFYPSDGVATEK | -443 | -456 | 0 | 0 | 14 | 3454.2446 | 3638.2708 | 3781.3018 | 4798.3916 | 5144.4185 | 5919.1494 |
| Tkt | EAWHGKPLPK | -255 | -264 | 1 | 0.001055699 | 10 | 3047.2246 | 2961.9253 | 2840.5293 | 3240.4104 | 3763.1572 | 3466.2039 |
| Tkt | EAWHGKPLPK | -255 | -264 | 1 | 0.001055699 | 10 | 3047.2246 | 2961.9253 | 2840.5293 | 3240.4104 | 3763.1572 | 3466.2039 |
| Tkt | LGQSDPAPLQHQVDIY | -187 | -202 | 0 | 0 | 16 | 2959.3599 | 4166.8823 | 3988.927 | 3423.5515 | 4194.704 | 4147.571 |
| Tkt | LGQSDPAPLQHQ | -187 | -198 | 0 | 0 | 12 | 2745.4976 | 172.2124 | 3773.9482 | 2005.2938 | 2242.917 | 1900.3416 |
| Tkt | AVVGEPGVTVTR | -575 | -586 | 0 | 0.000431655 | 12 | 2457.7625 | 359.73822 | 2968.1533 | 452.73392 | 2239.9368 | 285.4159 |
| Tkt | LGQSDPAPLQHQVDIYQK | -187 | -204 | 0 | 0 | 18 | 2157.0486 | 3713.261 | 1969.282 | 1150.8159 | 1535.2104 | 1554.9674 |
| Tkt | LGQSDPAPLQHQVDIYQK | -187 | -204 | 0 | 0 | 18 | 2157.0486 | 3713.261 | 1969.282 | 1150.8159 | 1535.2104 | 1554.9674 |
| Tkt | SDLDGHPVPK | -105 | -114 | 0 | 0 | 10 | 2135.8237 | 2087.5044 | 2274.9631 | 2352.529 | 2629.4595 | 1550.4414 |
| Tkt | KISSDLDGHPVPK | -102 | -114 | 1 | 0 | 13 | 2052.7004 | 2388.4727 | 2668.0837 | 2333.956 | 3098.5664 | 3225.0542 |
| Tkt | KISSDLDGHPVPK | -102 | -114 | 1 | 0 | 13 | 2052.7004 | 2388.4727 | 2668.0837 | 2333.956 | 3098.5664 | 3225.0542 |
| Tkt | VLDPFTIKPL | -531 | -540 | 1 | 0.002068213 | 10 | 1813.0573 | 2020.9493 | 2068.4487 | 2052.7302 | 2434.0461 | 2606.5356 |
| Tkt | IIALDGDTK | -335 | -343 | 0 | 0 | 9 | 1752.7786 | 1366.3348 | 1017.9017 | 1475.9784 | 735.97437 | 1059.4924 |
| Tkt | GWHTIIVDGHSVEELCK | -210 | -226 | 0 | 0 | 17 | 1556.2427 | 909.1743 | 1210.9656 | 2176.4646 | 2455.1113 | 1799.6116 |
| Tkt | GWHTIIVDGHSVEELCK | -210 | -226 | 0 | 0 | 17 | 1556.2427 | 909.1743 | 1210.9656 | 2176.4646 | 2455.1113 | 1799.6116 |
| Tkt | VVGEPGVTVTR | -576 | -586 | 0 | 0.00025015 | 11 | 1359.2458 | 1436.2908 | 1247.3423 | 1257.2472 | 1131.9767 | 1181.4818 |
| Tkt | SVPMSTVFYPSDGVATEK | -439 | -456 | 0 | 0 | 18 | 1080.7214 | 1264.27 | 1113.0819 | 1253.2295 | 1316.0225 | 1426.2432 |
| Tkt | SVPMSTVFYPSDGVATEK | -439 | -456 | 0 | 0 | 18 | 1080.7214 | 1264.27 | 1113.0819 | 1253.2295 | 1316.0225 | 1426.2432 |
| Tkt | HTIIVDGHSVEELCK | -212 | -226 | 0 | 0 | 15 | 1009.2541 | 1310.2026 | 1243.3181 | 1982.4158 | 2516.1475 | 2527.3474 |
| Tkt | HTIIVDGHSVEELCK | -212 | -226 | 0 | 0 | 15 | 1009.2541 | 1310.2026 | 1243.3181 | 1982.4158 | 2516.1475 | 2527.3474 |
| Tkt | ATPPQEDAPSVDIANIR | -286 | -302 | 0 | 0 | 17 | 800.6166 | 1089.5779 | 1155.3109 | 1383.9688 | 1874.4225 | 1689.9646 |
| Tkt | SKDDQVTVIGAGVTL | -498 | -512 | 1 | 0 | 15 | 711.38074 | 858.085 | 739.68304 | 845.1112 | 1052.129 | 784.97504 |
| Tkt | MEGYHKPDQQK | -1 | -11 | 1 | 0 | 11 | 638.3719 | 333.3166 | 832.63336 | 554.6917 | 819.00244 | 783.458 |
| Tkt | MEGYHKPDQQK | -1 | -11 | 1 | 0 | 11 | 638.3719 | 333.3166 | 832.63336 | 554.6917 | 819.00244 | 783.458 |
| Tkt | MEGYHKPDQQK | -1 | -11 | 1 | 0 | 11 | 638.3719 | 333.3166 | 832.63336 | 554.6917 | 819.00244 | 783.458 |
| Tkt | ILATPPQEDAPSVDIA | -284 | -299 | 0 | 0.00025015 | 16 | 634.3673 | 782.6903 | 871.276 | 1034.7548 | 729.97656 | 911.8214 |
| Tkt | FLPEAELLNLR | -91 | -101 | 0 | 0 | 11 | 603.678 | 483.53876 | 563.64954 | 169.18587 | 660.029 | 611.3304 |
| Tkt | QAFTDVATGSLGQGL | -115 | -129 | 0 | 0 | 15 | 462.3429 | 2943.5312 | 2937.4268 | 2769.2378 | 3471.125 | 2993.9036 |
| Tkt | AEAGFLPEAELLNLR | -87 | -101 | 0 | 0 | 15 | 433.44965 | 422.06415 | 317.01117 | 622.1285 | 665.3571 | 656.20264 |
| Tkt | QAFTDVATGSLGQGLGAAC | -115 | -133 | 0 | 0 | 19 | 337.54196 | 403.46423 | 264.00983 | 1019.4921 | 1070.744 | 952.35315 |
| Tkt | KLILDSAR | -543 | -550 | 1 | 0 | 8 | 10574.611 | 11415.934 | 13402.182 | 17012.318 | 18215.63 | NaN |

| | | | | | | | | | | | | |
|---|---|---|---|---|---|---|---|---|---|---|---|---|
| Tkt | NEDFQVGQAK | -484 | -493 | 0 | 0 | 10 | 1950.4044 | 2421.8733 | 405.8348 | 2129.9094 | NaN | 1345.2142 |
| Tkt | GIDKDAIVQAVK | -606 | -617 | 1 | 0 | 12 | 528.41364 | 632.87213 | 583.39594 | 574.2938 | 587.7075 | NaN |
| Tkt | SAAVVGEPGVTVTR | -573 | -586 | 0 | 0 | 14 | 509.16724 | 419.32306 | 707.4409 | 755.49243 | 718.4586 | NaN |
| Tkt | SKDDQVTVIGAGVTLH | -498 | -513 | 1 | 0 | 16 | 488.38333 | 695.4805 | NaN | 821.70526 | 940.80756 | 734.5092 |
| Tkt | AGFLPEAELLNLR | -89 | -101 | 0 | 0 | 13 | 420.93668 | 654.5521 | NaN | 221.92676 | 316.9514 | 282.20898 |
| Tkt | LDNLVAIFDINR | -175 | -186 | 0 | 0 | 12 | 269.3064 | 210.84698 | NaN | 385.09326 | 405.7149 | 390.86945 |
| Tkt | SKDDQVTVIGA | -498 | -508 | 1 | 0.002068213 | 11 | 2916.6353 | NaN | 4367.1304 | 2890.1875 | 3323.0962 | NaN |
| Tkt | SKDDQVTVIGAGVT | -498 | -511 | 1 | 0.001291103 | 14 | 2335.25 | NaN | 2391.824 | 1914.4523 | 2162.102 | NaN |
| Tkt | HQPTAIIAK | -233 | -241 | 0 | 0.00025015 | 9 | 1569.5974 | 1447.1978 | NaN | 1786.931 | 1631.0526 | NaN |
| Tkt | DKEAWHGKPLPK | -253 | -264 | 2 | 0.000230707 | 12 | 3348.7236 | 794.0504 | 861.9435 | NaN | NaN | NaN |
| Tkt | MPTPPSYK | -303 | -310 | 0 | 0.00025015 | 8 | 466.34726 | NaN | NaN | NaN | 629.4067 | NaN |
| Tkt | MPTPPSYK | -303 | -310 | 0 | 0.00025015 | 8 | 466.34726 | NaN | NaN | NaN | 629.4067 | NaN |
| Zbp1 | ALGDSSPQTTEPVLR | -379 | -393 | 0 | 0 | 15 | 562.0788 | 402.94223 | 535.89453 | 1139.196 | 1228.6447 | 1453.9913 |
| Zbp1 | AEAPVDLSTGDNLEQK | -2 | -17 | 0 | 0.00122419 | 16 | 492.9003 | 591.35376 | 497.3517 | 498.64404 | 858.8247 | 692.57556 |
| Zbp1 | GDGAPAIPENSSAQPSLDER | -74 | -93 | 0 | 0.00111006 | 20 | 341.53113 | 626.6987 | 595.96686 | 319.62338 | 738.54675 | 624.76086 |
| Zbp1 | EHEVQDIESSQDTGLSK | -394 | -410 | 0 | 0.000394361 | 17 | 559.7776 | 671.3173 | NaN | 602.2975 | 637.49414 | 658.12756 |
| Zbp1 | SGDGAPAIPENSSAQPSLDER | -73 | -93 | 0 | 0.000826806 | 21 | 420.59982 | 536.9574 | 427.1627 | 573.0989 | 556.64655 | NaN |
| Zbp1 | TSHPLPLAWDASAQ | -213 | -226 | 0 | 0.007493591 | 14 | 867.3426 | NaN | NaN | NaN | 5671.7397 | NaN |
| Vdac2 | YQLDPTASISAK | -237 | -248 | 0 | 0 | 12 | 1199.1803 | 1396.039 | 1596.2325 | 1861.2167 | 2375.2124 | 2207.8875 |
| Vdac2 | LTFDTTFSPNTGK | -109 | -121 | 0 | 0 | 13 | 894.26416 | 909.50903 | 552.2866 | 659.0571 | 1463.61 | 1333.3846 |
| Vdac2 | WCEYGLTFTEK | -76 | -86 | 0 | 0 | 11 | 666.99133 | 722.45355 | 688.123 | 255.91733 | 950.72754 | 989.5114 |
| Vdac2 | SCSGVEFSTSGSSNTDTGK | -47 | -65 | 0 | 0 | 19 | 135.17459 | 643.0273 | 679.6537 | 865.3144 | 870.39984 | 883.49316 |
| Vdac2 | AECCVPVCPR | -2 | -11 | 0 | 0 | 10 | 803.6572 | 696.83167 | NaN | 824.83716 | NaN | NaN |
| Vdac2 | GFGFGLVK | -33 | -40 | 0 | 0.00713295 | 8 | 526.7038 | NaN | NaN | NaN | NaN | NaN |
| Ptma | VAEDDEDDDVDTK | -91 | -103 | 0 | 0 | 13 | 7144.6997 | 8321.947 | 8359.123 | 6159.997 | 7768.763 | 1533.2742 |
| Ptma | SDAAVDTSSEITTK | -2 | -15 | 0 | 0 | 14 | 7040.7666 | 6054.5957 | 6782.543 | 5822.9185 | 5647.587 | 5892.552 |
| Ptma | EVVEEAENGR | -22 | -31 | 0 | 0 | 10 | 5404.8696 | 4186.4756 | 2488.8284 | 3488.0183 | 5682.136 | 7817.666 |
| Ptma | RVAEDDEDDDVDTK | -90 | -103 | 1 | 0 | 14 | 4865.1665 | 4876.1626 | 5193.2617 | 7837.0967 | 8376.053 | 11063.509 |
| Ptma | RVAEDDEDDDVDTK | -90 | -103 | 1 | 0 | 14 | 4865.1665 | 4876.1626 | 5193.2617 | 7837.0967 | 8376.053 | 11063.509 |
| Ptma | RVAEDDEDDDVDTKK | -90 | -104 | 2 | 0.000230707 | 15 | 3772.8652 | 4772.9995 | 4871.081 | 3785.083 | 4423.253 | 4773.3438 |
| Ptma | RVAEDDEDDDVDTKK | -90 | -104 | 2 | 0.000230707 | 15 | 3772.8652 | 4772.9995 | 4871.081 | 3785.083 | 4423.253 | 4773.3438 |
| Ptma | RVAEDDEDDDVD | -90 | -101 | 1 | 0.000230707 | 12 | 3755.4763 | 4005.9155 | 3780.1353 | 3411.666 | 3658.2107 | 3961.4297 |
| Ptma | RVAEDDEDDDVD | -90 | -101 | 1 | 0.000230707 | 12 | 3755.4763 | 4005.9155 | 3780.1353 | 3411.666 | 3658.2107 | 3961.4297 |
| Ptma | VAEDDEDDDVDTKK | -91 | -104 | 1 | 0 | 14 | 2053.4973 | 2223.8203 | 2332.106 | 1010.38824 | 1403.4564 | 1320.7844 |
| Ptma | VAEDDEDDDVDTKK | -91 | -104 | 1 | 0 | 14 | 2053.4973 | 2223.8203 | 2332.106 | 1010.38824 | 1403.4564 | 1320.7844 |
| Ptma | KEVVEEAENGR | -21 | -31 | 1 | 0 | 11 | 1967.5642 | 1419.5338 | 2077.928 | 2398.4316 | 3026.6326 | 2805.831 |
| Ptma | KEVVEEAENGR | -21 | -31 | 1 | 0 | 11 | 1967.5642 | 1419.5338 | 2077.928 | 2398.4316 | 3026.6326 | 2805.831 |
| Ptma | EDDEDDDVDTKK | -93 | -104 | 1 | 0 | 12 | 555.2396 | 482.88757 | 470.0243 | 432.75488 | 475.33987 | 496.51514 |
| Ptma | SDAAVDTSSEITTKDLK | -2 | -18 | 1 | 0 | 17 | 475.6247 | 538.82275 | 252.54259 | 797.76825 | 325.6018 | 572.59503 |
| Ptma | AEDDEDDDVDTKK | -92 | -104 | 1 | 0.000431655 | 13 | 337.5585 | 381.2079 | 349.23767 | 319.75943 | 408.13007 | 368.92764 |
| Ptma | AEDDEDDDVDTK | -92 | -103 | 0 | 0 | 12 | 220.93121 | NaN | NaN | 292.02252 | 359.18414 | 279.72955 |
| Tbca | RVVLDSVK | -98 | -105 | 1 | 0.00025015 | 8 | 1658.041 | 1659.1807 | 1751.5734 | 2352.1697 | 2835.105 | 2693.3735 |
| Tbca | ESEKDLEEAEEYK | -83 | -95 | 1 | 0 | 13 | 614.36633 | 710.2149 | 643.2469 | 543.5287 | 493.00806 | 544.18866 |
| Tbca | KAEDGENYAIK | -41 | -51 | 1 | 0 | 11 | 1405.7703 | 1571.7427 | 1560.4907 | 1676.0848 | NaN | 1977.4777 |
| Tbca | AEDGENYAIK | -42 | -51 | 0 | 0 | 10 | 517.8559 | 287.46216 | NaN | NaN | NaN | NaN |
| Anxa2 | GSVKPYTNFDAER | -25 | -37 | 1 | 0 | 13 | 4594.2803 | 4830.1797 | 4750.354 | 7996.207 | 11996.12 | 11758.534 |
| Anxa2 | TPAQYDASELK | -105 | -115 | 0 | 0 | 11 | 2663.3513 | 2717.9026 | 3198.6296 | 4112.335 | 3683.5752 | 3567.4868 |
| Anxa2 | TNQELQEINR | -136 | -145 | 0 | 0 | 10 | 2646.6665 | 2666.6726 | 66.39502 | 2788.406 | 110.75934 | 1984.3713 |
| Anxa2 | STVHEILCK | -2 | -10 | 0 | 0 | 9 | 2237.1938 | 2616.2578 | 2607.5767 | 2718.4663 | 3446.2961 | 3158.4893 |
| Anxa2 | AEDGSVIDYELIDQDAR | -180 | -196 | 0 | 0 | 17 | 1331.9658 | 1447.0779 | 1517.0647 | 1421.6539 | 1561.0038 | 1485.4109 |
| Anxa2 | DIISDTSGDFR | -158 | -168 | 0 | 0 | 11 | 1288.5908 | 107.13669 | 1339.9775 | 1137.0043 | 983.24585 | 1131.8191 |
| Anxa2 | RAEDGSVIDYELIDQDAR | -179 | -196 | 1 | 0 | 18 | 1207.672 | 1471.6919 | 1505.9646 | 1517.5833 | 2126.3552 | 2077.4058 |
| Anxa2 | RAEDGSVIDYELIDQDAR | -179 | -196 | 1 | 0 | 18 | 1207.672 | 1471.6919 | 1505.9646 | 1517.5833 | 2126.3552 | 2077.4058 |
| Anxa2 | TDLEKDIISDTSGDFR | -153 | -168 | 1 | 0 | 16 | 919.611 | 988.39825 | 1457.9786 | 1146.0275 | 1182.6484 | 1327.0529 |
| Anxa2 | SYSPYDMLESIK | -234 | -245 | 0 | 0 | 12 | 336.12006 | 3088.0469 | 2505.7053 | 435.2142 | 2539.7607 | 508.44318 |
| Anxa2 | SYSPYDMLESIK | -234 | -245 | 0 | 0 | 12 | 336.12006 | 3088.0469 | 2505.7053 | 435.2142 | 2539.7607 | 508.44318 |
| Anxa2 | DALNIETAVK | -38 | -47 | 0 | 0 | 10 | 652.5753 | 1108.3136 | 1182.3131 | 972.95844 | 694.33844 | 445.4571 |
| Anxa2 | GLGTDEDSLIEIICSR | -120 | -135 | 0 | 0 | 16 | 578.2612 | 679.09265 | 188.7282 | 525.5637 | 564.2016 | 507.93604 |
| Anxa2 | GTDEDSLIEIICSR | -122 | -135 | 0 | 0 | 14 | 456.42523 | 471.68378 | 507.9791 | 725.0092 | 440.66672 | 522.1794 |
| Anxa2 | RTNQELQEINR | -135 | -145 | 1 | 0.001636916 | 11 | 956.19025 | 443.13867 | NaN | 883.1591 | 938.0165 | 260.7575 |
| Anxa2 | CIQNKPLYFADR | -262 | -273 | 1 | 0 | 12 | 544.0094 | NaN | 453.5813 | 495.11276 | 862.54456 | 767.0268 |
| Anxa2 | LSLEGDHSTPPSAY | -11 | -24 | 0 | 0 | 14 | 1917.1205 | 2242.5208 | 2176.73 | NaN | NaN | 1317.1324 |
| Hdlbp | VFHVPLEER | -79 | -87 | 0 | 0 | 9 | 2614.846 | 1946.7864 | 1294.8502 | 3158.3274 | 269.29077 | 3641.9326 |
| Hdlbp | VFHVPLEER | -79 | -87 | 0 | 0 | 9 | 2614.846 | 1946.7864 | 1294.8502 | 3158.3274 | 269.29077 | 3641.9326 |
| Hdlbp | AEDKDQDLITIIGK | -766 | -779 | 1 | 0 | 14 | 908.31384 | 799.487 | 926.8005 | 636.83215 | 797.85065 | 640.24634 |
| Hdlbp | TLNSEEENDPPTYK | -28 | -41 | 0 | 0 | 14 | 590.6415 | 492.34192 | 584.6776 | 682.2786 | 476.0569 | 483.8011 |
| Hdlbp | GNSLQEILER | -315 | -324 | 0 | 0 | 10 | 211.77701 | 239.04353 | 278.7503 | 551.99207 | 546.1132 | 648.38055 |
| Hdlbp | SVSAPSWLHR | -368 | -377 | 0 | 0.003133457 | 10 | 6718.202 | 6615.7764 | NaN | NaN | NaN | NaN |
| Rps2 | SPYQEFTDHLVK | -264 | -275 | 0 | 0 | 12 | 5957.492 | 7271.4336 | 6989.797 | 7680.245 | 9129.237 | 290.87042 |
| Rps2 | SPYQEFTDHLVK | -264 | -275 | 0 | 0 | 12 | 5957.492 | 7271.4336 | 6989.797 | 7680.245 | 9129.237 | 290.87042 |
| Rps2 | DKEWIPVTK | -57 | -65 | 1 | 0 | 9 | 5370.406 | 7219.7915 | 5231.3027 | 4104.4966 | 6029.837 | 4285.235 |
| Rps2 | SLKDEVLK | -101 | -108 | 1 | 0.000431655 | 8 | 5028.9087 | 5642.6187 | 5978.531 | 6111.838 | 5781.4404 | 5534.7725 |
| Rps2 | TYSYLTPDLWK | -247 | -257 | 0 | 0 | 11 | 3348.6562 | 3813.7104 | 3895.7246 | 1999.9817 | 1982.0599 | 1769.7341 |
| Rps2 | TYSYLTPDLWK | -247 | -257 | 0 | 0 | 11 | 3348.6562 | 3813.7104 | 3895.7246 | 1999.9817 | 1982.0599 | 1769.7341 |
| Rps2 | GIDDCYTSAR | -218 | -227 | 0 | 0 | 10 | 2997.4421 | 2750.5278 | 150.9617 | 2860.2908 | 2167.1292 | 2389.2095 |
| Rps2 | GCTATLGNFAK | -228 | -238 | 0 | 0 | 11 | 2104.2847 | 1289.7595 | 1810.585 | 1556.9241 | 1315.3718 | 1232.2856 |
| Rps2 | VAIGDYNGHVGLGVK | -128 | -142 | 0 | 0 | 15 | 1839.3749 | 2577.9292 | 2598.7534 | 2496.4697 | 3337.6914 | 3332.925 |
| Rps2 | VAIGDYNGHVGLGVK | -128 | -142 | 0 | 0 | 15 | 1839.3749 | 2577.9292 | 2598.7534 | 2496.4697 | 3337.6914 | 3332.925 |

| | | | | | | | | | | | | |
|---|---|---|---|---|---|---|---|---|---|---|---|---|
| Rps2 | IGDYNGHVGLGVK | -130 | -142 | 0 | 0 | 13 | 819.0602 | 638.5025 | 755.40076 | 218.01257 | 333.12915 | 1054.2772 |
| Rps2 | IGKPHTVPCK | -174 | -183 | 1 | 0 | 10 | 802.57776 | 944.631 | 128.98666 | 278.71625 | 403.9025 | 560.6931 |
| Rps2 | IGKPHTVPCK | -174 | -183 | 1 | 0 | 10 | 802.57776 | 944.631 | 128.98666 | 278.71625 | 403.9025 | 560.6931 |
| Rps2 | MAGIDDCYTSAR | -216 | -227 | 0 | 0 | 12 | 798.8561 | 1027.2404 | 1029.2821 | 1077.1238 | 911.60175 | 924.353 |
| Rps2 | KLSIVPVR | -159 | -166 | 1 | 0 | 8 | 512.3481 | 597.2399 | 630.89075 | 562.7665 | 747.64 | 763.7918 |
| Rps2 | KAEDKEWIPVTK | -54 | -65 | 2 | 0 | 12 | 479.63406 | 525.8885 | 513.44275 | 310.21683 | 1116.5475 | 1115.1272 |
| Rps2 | SYLTPDLWK | -249 | -257 | 0 | 0 | 9 | 430.11832 | 1448.6495 | 386.17523 | 1396.0272 | 1715.9647 | 1738.386 |
| Rps2 | FLGASLKDEVLK | -97 | -108 | 1 | 0 | 12 | 334.55273 | 515.4344 | 513.32965 | 872.1801 | 1129.5721 | 1163.9794 |
| Rps2 | EDKEWIPVTK | -56 | -65 | 1 | 0.00025015 | 10 | 2143.0713 | NaN | 2266.687 | 4358.1895 | 4497.0093 | 4569.683 |
| Rps2 | AGIDDCYTSAR | -217 | -227 | 0 | 0.00025015 | 11 | 1451.3135 | 214.74998 | 1582.8745 | 1801.7589 | 1723.8207 | NaN |
| Rps2 | SPYQEFTDHLV | -264 | -274 | 0 | 0 | 11 | 493.98224 | 504.32205 | 624.77295 | NaN | 422.50266 | 290.1477 |
| Rps2 | AFVAIGDYNGHVGLGVK | -126 | -142 | 0 | 0 | 17 | 1884.7733 | 3436.478 | 2363.3418 | NaN | 3299.0837 | NaN |
| Rps2 | TATLGNFAK | -230 | -238 | 0 | 0 | 9 | 383.0838 | 4275.089 | 225.24698 | 229.6148 | NaN | NaN |
| Rps2 | AEDKEWIPVTK | -55 | -65 | 1 | 0.00025015 | 11 | 3903.775 | NaN | 591.6141 | NaN | NaN | 186.24525 |
| Rps2 | RTQAPAVATT | -284 | -293 | 1 | 0.002880184 | 10 | 1985.9178 | 1926.8744 | NaN | 2874.721 | NaN | NaN |
| Rps2 | GTGIVSAPVPK | -201 | -211 | 0 | 0 | 11 | 1047.3411 | 1992.2383 | 1903.9667 | NaN | NaN | NaN |
| Rps2 | EWIPVTK | -59 | -65 | 0 | 0.000591124 | 7 | 1079.4908 | 904.431 | NaN | NaN | NaN | NaN |
| Trmt2a | AEDLVPGLVSR | -479 | -489 | 0 | 0.002068213 | 11 | 1624.8456 | NaN | 1249.3623 | NaN | 856.8843 | NaN |
| Trmt2a | DTVHIPEATK | -267 | -276 | 0 | 0.001463897 | 10 | 1247.497 | NaN | NaN | 1963.5958 | NaN | NaN |
| Ccar1 | LLLPTPTIK | -1000 | -1008 | 0 | 0 | 9 | 954.6639 | 44.97121 | 972.2064 | 1165.613 | 702.06726 | 1289.7167 |
| Ccar1 | LEDNKEHSFEVS | -754 | -765 | 1 | 0 | 12 | 510.93613 | 675.7987 | 721.39514 | 691.4804 | 806.8824 | 865.42395 |
| Ccar1 | TILNLENSNK | -1064 | -1073 | 0 | 0 | 10 | 938.09393 | 429.893 | NaN | 939.6377 | 698.9175 | 832.3495 |
| Ccar1 | TLPNQNQSQTQPLLK | -204 | -218 | 0 | 0.000471409 | 15 | 711.9368 | 477.09064 | 496.03455 | 472.8391 | NaN | 631.0036 |
| Ccar1 | IQTLPNQNQSQTQPLLK | -202 | -218 | 0 | 0.001392869 | 17 | 400.4722 | 351.96143 | 434.78207 | 429.6613 | 355.92224 | NaN |
| Ccar1 | AEDPQDLR | -465 | -472 | 0 | 0.000431655 | 8 | 837.6465 | NaN | NaN | NaN | NaN | NaN |
| Arfgap1 | EWSLESSPAQNWTPPQPK | -123 | -140 | 0 | 0 | 18 | 350.84457 | 368.20886 | 453.78113 | 498.0655 | 480.86127 | 524.2776 |
| Arfgap1 | AEDSSDRPLEGHSY | -298 | -311 | 1 | 0 | 14 | 333.4502 | 699.90125 | 501.0693 | NaN | 288.813 | 304.78085 |
| Arfgap1 | NENVLKPAQEK | -248 | -258 | 1 | 0 | 11 | 1770.4451 | 680.71295 | NaN | NaN | 1592.1857 | 910.5175 |
| Hsph1 | AGGIETIANEFSDR | -20 | -33 | 0 | 0 | 14 | 2081.3308 | 2423.5623 | 2303.0442 | 2161.2295 | 3189.0935 | 3044.1306 |
| Hsph1 | YSDPQGVPYPEAK | -446 | -458 | 0 | 0 | 13 | 1702.6042 | 1595.5486 | 972.2374 | 2331.5005 | 1041.8636 | 1005.28174 |
| Hsph1 | FICEQEHEK | -657 | -665 | 0 | 0 | 9 | 1441.4949 | 848.9395 | 1397.6167 | 1355.2908 | 1394.066 | 1406.2981 |
| Hsph1 | AFNDPFIQK | -74 | -82 | 0 | 0 | 9 | 1300.2136 | 1530.21 | 1597.4249 | 123.88996 | 1968.3485 | 2119.7678 |
| Hsph1 | GTAFDPFLGGK | -224 | -234 | 0 | 0.000230707 | 11 | 1236.8875 | 1243.3887 | 1309.3289 | 359.78903 | 1518.2175 | 187.57095 |
| Hsph1 | ELNNVCEPVVTQPKPK | -792 | -807 | 1 | 0.000550307 | 16 | 348.49432 | 389.5138 | 413.39172 | 459.31192 | 509.85187 | 499.112 |
| Hsph1 | ELNNVCEPVVTQPKPK | -792 | -807 | 1 | 0.000550307 | 16 | 348.49432 | 389.5138 | 413.39172 | 459.31192 | 509.85187 | 499.112 |
| Hsph1 | AFDPFLGGK | -226 | -234 | 0 | 0 | 9 | 812.5297 | 794.8767 | 893.0259 | 390.4312 | 615.1859 | 695.0701 |
| Hsph1 | FQEAEERPK | -707 | -715 | 1 | 0 | 9 | 788.21643 | 705.3076 | 855.26514 | 592.1318 | 322.24655 | 775.0809 |
| Hsph1 | ELNNVCEPVVTQPK | -792 | -805 | 0 | 0 | 14 | 670.64606 | 566.0468 | 650.5133 | 756.5974 | 392.4186 | 735.20966 |
| Hsph1 | TETEDWLYEEGEDQAK | -671 | -686 | 0 | 0 | 16 | 629.9141 | 627.21606 | 775.32635 | 827.05505 | 982.80536 | 929.1273 |
| Hsph1 | LKETAENNLK | -125 | -134 | 1 | 0 | 10 | 517.02374 | 522.8241 | 534.82135 | 512.3051 | 622.5953 | 674.8495 |
| Hsph1 | DKLCGPYEK | -648 | -656 | 1 | 0 | 9 | 1773.7551 | 2009.8405 | 616.89825 | 2169.7817 | 2254.414 | NaN |
| Hsph1 | DLPNAEEKPR | -187 | -196 | 1 | 0 | 10 | 1620.7701 | 1713.2489 | 1731.6178 | NaN | 1887.3057 | 1920.7963 |
| Hsph1 | VWNHDSEETEGVHEVFSR | -405 | -422 | 0 | 0 | 18 | 1164.6727 | 1327.3245 | 1523.0537 | NaN | 2769.1104 | 2865.6714 |
| Hsph1 | IEVPLHSL | -317 | -324 | 0 | 0 | 8 | 788.8956 | 668.9088 | 774.9847 | 560.84656 | NaN | 419.06305 |
| Hsph1 | LTETEDWLYEEGEDQAK | -670 | -686 | 0 | 0 | 17 | 649.0901 | NaN | 504.27902 | 269.06195 | 309.64444 | 284.37314 |
| Hsph1 | AFYSDPQGVPYPEAK | -444 | -458 | 0 | 0 | 15 | 645.6647 | 1111.4996 | 968.52985 | 986.1826 | NaN | 918.4744 |
| Hsph1 | GQQTSQSPPSPELTSEESK | -549 | -567 | 0 | 0.000431655 | 19 | 546.49805 | 491.35876 | NaN | 440.8797 | 560.6816 | 479.60095 |
| Hsph1 | VLEELGQR | -716 | -723 | 0 | 0.000431655 | 8 | 9867.155 | 10399.428 | 9701.357 | 10598.19 | NaN | NaN |
| Hsph1 | KQDLPNAEEKPR | -185 | -196 | 2 | 0 | 12 | 461.42892 | 447.09784 | NaN | 456.80652 | NaN | 572.153 |
| Hsph1 | AEDVSAIEIVGGATR | -332 | -346 | 0 | 0 | 15 | 1110.5999 | 1088.0353 | 1149.9812 | NaN | NaN | NaN |
| Hsph1 | SQFEELCAELLQK | -304 | -316 | 0 | 0 | 13 | 1086.2705 | NaN | 1402.0878 | 1258.2512 | NaN | NaN |
| Cdc37 | EGEEAGPGDPLLEAVPK | -354 | -370 | 0 | 0 | 17 | 2089.4722 | 2676.0325 | 2575.5925 | 2575.8452 | 3411.0376 | 3290.0332 |
| Cdc37 | CIDSGLWVPNSK | -337 | -348 | 0 | 0 | 12 | 976.71313 | 849.3925 | 916.0486 | 999.41473 | 1052.402 | 1083.7769 |
| Cdc37 | ELEVAESDGQVELER | -70 | -84 | 0 | 0 | 15 | 834.7185 | 1030.8467 | 1135.8235 | 764.2967 | 720.554 | 690.5797 |
| Cdc37 | AEEDSEEAREQ | -137 | -147 | 1 | 0 | 11 | 387.56097 | 317.3498 | 497.0063 | 65.04573 | 371.6517 | 478.44086 |
| Cdc37 | ESLPEELQK | -300 | -308 | 0 | 0.000230707 | 9 | 1216.6 | 616.9561 | 445.8181 | 1109.7344 | 855.5615 | NaN |
| Cdc37 | LGPGGLDPVEVYESLPEELQK | -288 | -308 | 0 | 0.00025015 | 21 | 330.79938 | 362.13928 | NaN | 425.04352 | 362.907 | 323.46854 |
| Dbnl | NGPALQEAYVR | -8 | -18 | 0 | 0 | 11 | 5591.4087 | 4566.841 | 4464.499 | 2917.3967 | 1556.6665 | 2514.8516 |
| Dbnl | WTGEGVNDVR | -84 | -93 | 0 | 0.00025015 | 10 | 1934.5145 | 2025.821 | 2070.8994 | 2305.7932 | 2319.1846 | 2273.3025 |
| Dbnl | QEWESAGQQAPHPR | -253 | -266 | 0 | 0 | 14 | 1543.042 | 2093.6646 | 2881.679 | 2779.54 | 2419.8418 | 1932.6855 |
| Dbnl | TGEPEQEAVSR | -240 | -250 | 0 | 0 | 11 | 1528.3584 | 1404.0538 | 1530.3745 | 1680.8616 | 719.65497 | 1970.2384 |
| Dbnl | QLTQPETSYGR | -297 | -307 | 0 | 0 | 11 | 1035.186 | 1152.7635 | 1090.4998 | 989.5495 | 888.0658 | 1029.6237 |
| Dbnl | VKDPNSGLPK | -69 | -78 | 1 | 0.00025015 | 10 | 728.2826 | 631.5732 | 692.1491 | 642.5225 | 707.08405 | 755.20886 |
| Dbnl | ESTSFQDVGPQAPVGSVY | -145 | -162 | 0 | 0.000230707 | 18 | 694.3062 | 362.93564 | 400.13242 | 461.4144 | 840.77356 | 972.59436 |
| Dbnl | SPTDWALFTYEGNSNDIR | -24 | -41 | 0 | 0 | 18 | 559.0462 | 593.8085 | 550.93506 | 769.6316 | 1171.7085 | 1021.7929 |
| Dbnl | GTGEGGLEELVEELNSGK | -44 | -61 | 0 | 0 | 18 | 514.9949 | 511.15955 | 441.83798 | 584.1264 | 603.7941 | 592.15326 |
| Dbnl | AEEDVEPECIMEK | -119 | -131 | 0 | 0 | 13 | 615.82446 | 601.0764 | NaN | 118.70218 | NaN | 603.4914 |
| Dbnl | GANYSFHK | -137 | -144 | 0 | 0.005112649 | 8 | 4504.16 | NaN | NaN | NaN | 3591.1206 | 450.75278 |
| Dbnl | GIEVIDEGWWR | -406 | -416 | 0 | 0 | 11 | 615.7847 | 323.2074 | NaN | NaN | 493.0069 | NaN |
| Rabep2 | AEELIQEIQR | -160 | -169 | 0 | 0 | 10 | 12336.544 | NaN | NaN | NaN | NaN | NaN |
| Clic1 | NSNPALNDNLEK | -120 | -131 | 0 | 0 | 12 | 5035.1533 | 5043.9463 | 4848.5234 | 2989.6892 | 2433.5886 | 39.680405 |
| Clic1 | ALNPESNTSGLDIFAK | -98 | -113 | 0 | 0 | 16 | 1514.7609 | 2346.005 | 2331.9956 | 3092.015 | 3936.401 | 3926.3542 |
| Clic1 | LHIVQVVCK | -184 | -192 | 0 | 0 | 9 | 1237.251 | 1357.602 | 1467.2076 | 1025.214 | 1840.0164 | 1744.7882 |
| Clic1 | LNPESNTSGLDIFAK | -99 | -113 | 0 | 0 | 15 | 1034.4845 | 1093.4943 | 1066.4087 | 1672.1974 | 1561.0219 | 1704.4142 |
| Clic1 | LAALNPESNTSGLDIFAK | -96 | -113 | 0 | 0 | 18 | 906.06024 | 1026.8215 | 1115.3728 | 658.267 | 874.5217 | 873.3776 |
| Clic1 | FLDGNELTLADCNLLPK | -167 | -183 | 0 | 0 | 17 | 837.8051 | 765.05206 | 676.2167 | 1063.4336 | 260.88297 | 854.0705 |
| Clic1 | KVLDNYLTSPLPEEVDETSAEDEGISQR | -138 | -165 | 1 | 0 | 28 | 485.6884 | 500.18567 | 643.44214 | 679.42975 | 883.7605 | 768.12726 |
| Clic1 | LAALNPESNTSGLDIF | -96 | -111 | 0 | 0.00025015 | 16 | 305.9842 | 290.79077 | 273.89465 | 333.18478 | 465.6972 | 429.48053 |

| | | | | | | | | | | | | |
|---|---|---|---|---|---|---|---|---|---|---|---|---|
| Clic1 | GFTIPEAFR | -196 | -204 | 0 | 0 | 9 | 274.98572 | 993.08057 | 912.47986 | 336.21906 | 465.09564 | 363.6132 |
| Clic1 | SAEDEGISQR | -156 | -165 | 0 | 0 | 10 | 1640.3212 | NaN | 880.6043 | 1681.4681 | 844.80316 | 1888.7067 |
| Clic1 | AEEQPQVELFVK | -2 | -13 | 0 | 0 | 12 | 622.5536 | 665.8017 | NaN | 381.43832 | 783.26855 | 692.89636 |
| Clic1 | NPESNTSGLDIFAK | -100 | -113 | 0 | 0.00025015 | 14 | 590.53784 | NaN | 1059.4135 | NaN | 1215.2526 | 1098.26 |
| Ppp4r2 | AEETETAAPPPDKDR | -273 | -287 | 1 | 0 | 15 | 723.38324 | NaN | 925.7305 | 1602.8849 | 2323.0417 | 2310.4573 |
| Ipo5 | NDSCYQNDDSVLK | -226 | -238 | 0 | 0 | 13 | 3881.1086 | 2994.1047 | 3685.973 | 2838.1812 | 3108.5115 | 3133.4927 |
| Ipo5 | ANEHNVALFK | -203 | -212 | 0 | 0 | 10 | 2826.1611 | 3159.3328 | 2888.9866 | 464.88568 | 1913.6445 | 1421.6481 |
| Ipo5 | AVNDSCYQNDDSVLK | -224 | -238 | 0 | 0 | 15 | 2076.1982 | 2306.5974 | 2365.5884 | 2332.7896 | 1387.0889 | 2350.5352 |
| Ipo5 | EHNVALFK | -205 | -212 | 0 | 0 | 8 | 1766.9861 | 1392.3193 | 2061.513 | 977.7625 | 1506.4008 | 1102.5222 |
| Ipo5 | NLIDEDGNNQWPEGLK | -119 | -134 | 0 | 0 | 16 | 1420.7985 | 1481.437 | 1693.4773 | 2125.783 | 3349.6123 | 3692.2278 |
| Ipo5 | YVCDNSPEVR | -913 | -922 | 0 | 0 | 10 | 1401.1729 | 1069.236 | 959.25134 | 1174.8092 | 112.43658 | 1129.9884 |
| Ipo5 | ICDIAAELAR | -109 | -118 | 0 | 0 | 10 | 1327.9856 | 1558.9418 | 1348.82 | 1500.7228 | 1591.5795 | 1293.9897 |
| Ipo5 | GNLLSPDNVVR | -16 | -26 | 0 | 0 | 11 | 1091.8184 | 1415.24 | 1736.8577 | 1403.5613 | 1632.4695 | 2416.2314 |
| Ipo5 | GNQQQHYLDVIK | -163 | -174 | 0 | 0 | 12 | 1068.435 | 1588.7091 | 1842.5898 | 1320.2286 | 1806.489 | 1851.1997 |
| Ipo5 | GNQQQHYLDVIK | -163 | -174 | 0 | 0 | 12 | 1068.435 | 1588.7091 | 1842.5898 | 1320.2286 | 1806.489 | 1851.1997 |
| Ipo5 | YLRPHLEATLQ | -250 | -260 | 1 | 0.00078661 | 11 | 707.43994 | 1025.5118 | 1190.6102 | 1826.3267 | 2537.1592 | 2774.0098 |
| Ipo5 | ENVNATENCISAVGK | -964 | -978 | 0 | 0 | 15 | 688.73755 | 1113.5544 | 647.3981 | 843.08325 | 835.5706 | 950.897 |
| Ipo5 | HIVENAVQK | -540 | -548 | 0 | 0 | 9 | 682.8986 | 737.73096 | 935.2865 | 787.61005 | 814.15216 | 937.4669 |
| Ipo5 | LQYVCDNSPEVR | -911 | -922 | 0 | 0 | 12 | 640.3826 | 525.8542 | 649.9602 | 541.7134 | 896.8966 | 800.32263 |
| Ipo5 | LGNLLSPDNVVR | -15 | -26 | 0 | 0 | 12 | 606.8764 | 693.7156 | 732.2007 | 456.54608 | 572.2946 | 722.83325 |
| Ipo5 | VSDILHSIFSSYK | -842 | -854 | 0 | 0 | 13 | 499.3335 | 501.5201 | 699.1686 | 626.39294 | 1543.6875 | 1403.7902 |
| Ipo5 | FYFHDGVR | -713 | -720 | 0 | 0 | 8 | 390.08472 | 3364.4316 | 3391.856 | 801.4913 | 677.8827 | 4138.53 |
| Ipo5 | HFADLLPGFLQ | -213 | -223 | 0 | 0.000907206 | 11 | 382.69714 | 151.18027 | 354.7843 | 331.2697 | 385.06723 | 320.36575 |
| Ipo5 | SLLIPYLDNLVK | -476 | -487 | 0 | 0.00025015 | 12 | 259.45425 | 203.95668 | 180.59302 | 345.32327 | 339.3142 | 306.21143 |
| Ipo5 | ELEDDDFDSNAVAGESALDR | -326 | -345 | 0 | 0 | 20 | 238.65869 | 246.58003 | 255.31534 | 284.36566 | 320.81345 | 302.71536 |
| Ipo5 | AEETYENIPGR | -29 | -39 | 0 | 0 | 11 | 206.91893 | 282.0481 | 342.22293 | 1548.4956 | 237.4198 | 1241.9243 |
| Ipo5 | INFTEDCPK | -467 | -475 | 0 | 0 | 9 | 1195.8883 | 3112.12 | 2656.6707 | NaN | 1678.7607 | 1775.6727 |
| Ipo5 | QQHYLDVIK | -166 | -174 | 0 | 0.000431655 | 9 | 982.5484 | 675.2098 | 877.557 | 617.30225 | 673.53064 | NaN |
| Ipo5 | SAFDEVYPALPSDVQTAIK | -72 | -90 | 0 | 0.00025015 | 19 | 210.14064 | 288.28098 | NaN | 328.05023 | 337.88135 | 285.36554 |
| Ipo5 | TKENVNATENCISAVGK | -962 | -978 | 1 | 0.00078661 | 17 | 1114.0056 | 1200.7251 | NaN | 1113.8773 | 1496.4852 | NaN |
| Ipo5 | IFDDIVEHCSPASFK | -887 | -901 | 0 | 0 | 15 | 772.47754 | 950.14056 | 441.18677 | NaN | 757.1717 | NaN |
| Ipo5 | LCGDTNLNNMQR | -265 | -276 | 0 | 0 | 12 | 1183.2039 | NaN | NaN | 1223.2233 | 506.48492 | NaN |
| Ipo5 | LCGDTNLNNMQR | -265 | -276 | 0 | 0 | 12 | 1183.2039 | NaN | NaN | 1223.2233 | 506.48492 | NaN |
| Ipo5 | CTDALPLLVR | -944 | -953 | 0 | 0 | 10 | 1020.84814 | 294.9281 | NaN | 377.90738 | NaN | NaN |
| Ipo5 | FKPDCVNVE | -982 | -990 | 1 | 0.00078661 | 9 | 780.9314 | 2445.701 | 906.5745 | NaN | NaN | NaN |
| Ipo5 | CIFDDIVEHCSPASFK | -886 | -901 | 0 | 0.002329279 | 16 | 350.97937 | 355.0136 | NaN | NaN | NaN | 353.52527 |
| Ipo5 | YLRPHLEAT | -250 | -258 | 1 | 0 | 9 | 10007.498 | 7754.3364 | NaN | NaN | NaN | NaN |
| Ipo5 | SMPLLLECAR | -726 | -735 | 0 | 0.006677519 | 10 | 3016.465 | NaN | NaN | NaN | NaN | NaN |
| Ipo5 | TASIKPEVALL | -633 | -643 | 1 | 0.005301614 | 11 | 1270.3123 | NaN | NaN | NaN | NaN | NaN |
| Exosc8 | FIADVVDNSQVIK | -108 | -120 | 0 | 0 | 13 | 125.60307 | 612.7394 | 609.87506 | 618.73694 | 608.41504 | 761.1412 |
| Exosc8 | AEFAAPPVDAPDR | -64 | -76 | 0 | 0 | 13 | 1165.7781 | 1925.0531 | 2213.349 | 1649.2289 | NaN | 1647.5 |
| Exosc8 | TVEPLEYYR | -7 | -15 | 0 | 0.000431655 | 9 | 1053.4822 | 882.41455 | 1071.4229 | NaN | NaN | NaN |
| Pa2g4 | LVKPGNQNTQVTEA | -156 | -169 | 1 | 0.001463897 | 14 | 7311.2627 | 7684.068 | 7515.8613 | 3358.7483 | 7757.8125 | 7507.5625 |
| Pa2g4 | SGPFEPDLYK | -335 | -344 | 0 | 0 | 10 | 5631.0513 | 5879.27 | 5702.575 | 313.3167 | 6180.1816 | 349.36353 |
| Pa2g4 | TIIQNPTDQQK | -200 | -210 | 0 | 0 | 11 | 5156.5034 | 5188.1333 | 6178.068 | 5980.588 | 5545.821 | 5739.3223 |
| Pa2g4 | TIIQNPTDQQK | -200 | -210 | 0 | 0 | 11 | 5156.5034 | 5188.1333 | 6178.068 | 5980.588 | 5545.821 | 5739.3223 |
| Pa2g4 | QQEQTIAEDLVVTK | -7 | -20 | 0 | 0 | 14 | 5076.16 | 5495.876 | 5719.5645 | 3334.9275 | 3188.59 | 6041.346 |
| Pa2g4 | LVKPGNQNTQVT | -156 | -167 | 1 | 0.001055699 | 12 | 3992.6167 | 3692.1394 | 3850.3572 | 3152.9878 | 3650.2007 | 3572.6843 |
| Pa2g4 | LVKPGNQNTQVTEAW | -156 | -170 | 1 | 0 | 15 | 2347.0063 | 3041.5168 | 3187.0027 | 4150.7734 | 4820.1953 | 4688.3027 |
| Pa2g4 | SDQDYILK | -94 | -101 | 0 | 0 | 8 | 2286.8335 | 2317.1472 | 2375.6335 | 3723.476 | 2464.297 | 2566.695 |
| Pa2g4 | NCVCHFSPLK | -84 | -93 | 0 | 0 | 10 | 2027.5729 | 2224.2737 | 2207.8584 | 2619.0425 | 2347.486 | 627.55566 |
| Pa2g4 | AFFSEVER | -264 | -271 | 0 | 0 | 8 | 1635.9327 | 762.7202 | 1697.2632 | 1903.9213 | 1704.6909 | 1895.5315 |
| Pa2g4 | ITSGPFEPDLYK | -333 | -344 | 0 | 0 | 12 | 1130.8636 | 1223.0723 | 1363.3052 | 1036.9191 | 752.35876 | 863.2702 |
| Pa2g4 | NCTPIEGMLSHQLK | -178 | -191 | 0 | 0 | 14 | 1103.058 | 1218.0496 | 1641.4425 | 1759.4071 | 1382.4669 | 1462.6349 |
| Pa2g4 | NCTPIEGMLSHQLK | -178 | -191 | 0 | 0 | 14 | 1103.058 | 1218.0496 | 1641.4425 | 1759.4071 | 1382.4669 | 1462.6349 |
| Pa2g4 | AEFEVHEVY | -216 | -224 | 0 | 0 | 9 | 1069.0018 | 1357.1576 | 1500.5928 | 2376.0034 | 1835.1816 | 1391.909 |
| Pa2g4 | TIAEDLVVTK | -11 | -20 | 0 | 0 | 10 | 885.9427 | 136.4771 | 976.415 | 867.76196 | 882.2138 | 722.9729 |
| Pa2g4 | HELLQPFNVL | -299 | -308 | 0 | 0 | 10 | 761.03656 | 822.402 | 882.97266 | 143.1252 | 1036.9948 | 1048.7008 |
| Pa2g4 | LVKPGNQNTQVTEAWNK | -156 | -172 | 1 | 0 | 17 | 751.74695 | 746.49774 | 868.77625 | 935.0233 | 965.8199 | 966.469 |
| Pa2g4 | LVKPGNQNTQVTEAWNK | -156 | -172 | 1 | 0 | 17 | 751.74695 | 746.49774 | 868.77625 | 935.0233 | 965.8199 | 966.469 |
| Pa2g4 | QHVIDGEK | -192 | -199 | 0 | 0.00078661 | 8 | 688.9651 | 536.84 | 734.0342 | 348.53143 | 548.38116 | 700.5276 |
| Pa2g4 | SGEDEQQEQTIAEDLVVTK | -2 | -20 | 0 | 0 | 19 | 317.05942 | 510.48547 | 544.16364 | 690.8116 | 888.896 | 860.82263 |
| Pa2g4 | VCHFSPLK | -86 | -93 | 0 | 0 | 8 | 697.77734 | 671.5161 | NaN | 620.54785 | 322.308 | 341.6108 |
| Pa2g4 | AVDVLVSSGEGK | -225 | -236 | 0 | 0 | 12 | 412.24814 | NaN | 415.23013 | 651.17896 | 1093.9688 | 561.4575 |
| Pa2g4 | LVKPGNQNTQVTE | -156 | -168 | 1 | 0.00416551 | 13 | 1511.4087 | NaN | 752.5479 | NaN | NaN | 878.60114 |
| Pa2g4 | CVCHFSPLK | -85 | -93 | 0 | 0 | 9 | 830.7099 | 2523.4229 | NaN | 495.8676 | NaN | NaN |
| Pa2g4 | AEFEVHEVYAV | -216 | -226 | 0 | 0.001941082 | 11 | 1002.2186 | NaN | 957.50934 | NaN | NaN | NaN |
| Phf6 | AEFGDFDIK | -258 | -266 | 0 | 0 | 9 | 575.88873 | 568.08704 | 580.71735 | 525.9603 | 517.2182 | 624.60455 |
| Twf2 | AEFLYDEVHPK | -305 | -315 | 0 | 0 | 11 | 1312.0068 | 1949.5033 | 1202.573 | 1322.4287 | 931.5834 | 914.2447 |
| Twf2 | WSPDNSPVR | -88 | -96 | 0 | 0 | 9 | 689.1261 | 1223.8862 | 397.56693 | 1349.6371 | 1147.6127 | 1004.8568 |
| Twf2 | EFGGGHIKDELFGTVK | -111 | -126 | 1 | 0 | 16 | 419.80643 | 676.70276 | 907.271 | 819.44836 | 519.7033 | 645.40186 |
| Twf2 | EFGGGHIKDELFGTVK | -111 | -126 | 1 | 0 | 16 | 419.80643 | 676.70276 | 907.271 | 819.44836 | 519.7033 | 645.40186 |
| Twf2 | AHQTGIHATEELK | -2 | -14 | 0 | 0 | 13 | 92.76079 | 651.2116 | 681.77594 | 1273.1101 | 924.66693 | 907.1577 |
| Twf2 | LDAQEPCYLLFR | -61 | -72 | 0 | 0 | 12 | 1013.7401 | NaN | 1282.2786 | 1193.9152 | 1251.6779 | 1588.4347 |
| Twf2 | EELKEFFAK | -11 | -19 | 1 | 0 | 9 | 659.9027 | 1861.5151 | 1841.1974 | NaN | 1756.0581 | 721.6066 |
| Twf2 | HTHEGDALESVVF | -243 | -255 | 0 | 0.000431655 | 13 | 488.73938 | 817.07874 | NaN | NaN | 1541.317 | 784.0052 |
| Twf2 | HTHEGDALESVVFIY | -243 | -257 | 0 | 0.003620005 | 15 | 280.80057 | NaN | NaN | 285.50958 | NaN | 329.99033 |
| Dhx15 | TDIKPEWLVK | -751 | -760 | 1 | 0 | 10 | 8829.738 | 8618.766 | 9913.74 | 9922.272 | 11231.124 | 7098.3125 |

| | | | | | | | | | | | | |
|---|---|---|---|---|---|---|---|---|---|---|---|---|
| Dhx15 | DRFNLPR | -669 | -675 | 1 | 0 | 7 | 1933.5989 | 1152.2979 | 1187.3854 | 3507.6975 | 4821.733 | 4154.4 |
| Dhx15 | YGVIILDEAHER | -254 | -265 | 0 | 0 | 12 | 1867.7849 | 2180.8564 | 2275.0757 | 2653.5854 | 3696.1829 | 3534.472 |
| Dhx15 | YGVIILDEAHER | -254 | -265 | 0 | 0 | 12 | 1867.7849 | 2180.8564 | 2275.0757 | 2653.5854 | 3696.1829 | 3534.472 |
| Dhx15 | TNLPHTPR | -119 | -126 | 0 | 0.000230707 | 8 | 1752.9435 | 1801.587 | 1734.3394 | 1807.8779 | 1997.9432 | 1987.3032 |
| Dhx15 | EVDDLGPEVGDIK | -372 | -384 | 0 | 0 | 13 | 1542.3379 | 1831.2084 | 1804.8412 | 2534.729 | 2165.3386 | 1162.7626 |
| Dhx15 | AEFPLDPQLAK | -563 | -573 | 0 | 0 | 11 | 1525.0675 | 1444.1233 | 1687.8806 | 842.9872 | 1559.8431 | 2135.1804 |
| Dhx15 | TCTDIKPEWLVK | -749 | -760 | 1 | 0 | 12 | 1382.0854 | 1536.6448 | 1700.8247 | 1731.8423 | 1882.8314 | 1869.2501 |
| Dhx15 | LQLPVWEYK | -135 | -143 | 0 | 0 | 9 | 1348.4644 | 1530.554 | 1093.6749 | 998.4824 | 1230.853 | 937.77826 |
| Dhx15 | IYFDNCPLLTIPGR | -302 | -315 | 0 | 0 | 14 | 1090.6964 | 540.5339 | 696.7012 | 583.964 | 579.962 | 557.38513 |
| Dhx15 | IFEPPPPK | -400 | -407 | 0 | 0.000347792 | 8 | 883.7085 | 760.9623 | 769.0459 | 392.8548 | 736.923 | 603.073 |
| Dhx15 | SNLGSVVLQLK | -505 | -515 | 0 | 0 | 11 | 741.4185 | 707.58374 | 737.6826 | 653.5291 | 856.5779 | 878.9279 |
| Dhx15 | LDLGEDYPSGK | -7 | -17 | 0 | 0 | 11 | 692.7524 | 675.32385 | 1103.2157 | 1025.7572 | 1188.6111 | 1131.3837 |
| Dhx15 | FAHIDGDHLTLL | -617 | -628 | 0 | 0 | 12 | 632.8842 | 1243.5624 | 680.9459 | 1686.7919 | 895.2881 | 730.793 |
| Dhx15 | FAHIDGDHLTLL | -617 | -628 | 0 | 0 | 12 | 632.8842 | 1243.5624 | 680.9459 | 1686.7919 | 895.2881 | 730.793 |
| Dhx15 | SFVLVGETGSGK | -155 | -166 | 0 | 0 | 12 | 1547.6746 | 442.4755 | 1444.2595 | 1771.8354 | NaN | 1332.2682 |
| Dhx15 | GQEEIDEACKR | -358 | -368 | 1 | 0.000550307 | 11 | 969.67584 | NaN | 1197.4563 | 1059.9265 | 1511.4148 | 1459.1781 |
| Dhx15 | HQSFVLVGETGSGK | -153 | -166 | 0 | 0 | 14 | 1371.4382 | 1392.1294 | 1173.1017 | 1143.9745 | NaN | NaN |
| Dhx15 | LGEDYPSGK | -9 | -17 | 0 | 0 | 9 | 1054.1425 | 245.52174 | 1168.506 | NaN | NaN | 1469.6832 |
| Dhx15 | VFVIDPGFAK | -434 | -443 | 0 | 0 | 10 | 318.5228 | 481.0582 | 413.54324 | NaN | NaN | NaN |
| Dhx15 | DNQVVQLHPSTVL | -715 | -727 | 0 | 0.00025015 | 13 | 761.05023 | NaN | NaN | NaN | 667.4404 | NaN |
| Dhx15 | MNDPLLER | -246 | -253 | 0 | 0 | 8 | 531.28674 | NaN | NaN | NaN | NaN | 1262.5955 |
| Dhx15 | DNQVVQLHPS | -715 | -724 | 0 | 0.000431655 | 10 | 609.3981 | NaN | NaN | NaN | NaN | NaN |
| Fip1l1 | AEFTSPPSLFK | -231 | -241 | 0 | 0 | 11 | 945.45526 | 729.78 | 313.27704 | 815.03015 | 1224.0885 | 221.01352 |
| Fip1l1 | RLPGAIDVIGQ | -285 | -295 | 1 | 0.000550307 | 11 | 722.5697 | 841.48773 | 802.15924 | 763.65686 | NaN | 895.30615 |
| Fip1l1 | GTAPVNLNIK | -113 | -122 | 0 | 0.00078661 | 10 | 429.8422 | 389.14368 | 368.78894 | NaN | 416.9093 | 392.35904 |
| Fip1l1 | QWDYYAR | -435 | -441 | 0 | 0.000431655 | 7 | 764.90686 | 512.22565 | NaN | NaN | NaN | NaN |
| Atp5pd | SWNETFHAR | -33 | -41 | 0 | 0 | 9 | 2841.8613 | 2932.2466 | 278.53296 | 2855.1282 | 3160.5315 | 3040.9924 |
| Atp5pd | VAKPGLVDDFEK | -61 | -72 | 1 | 0 | 12 | 2339.4644 | 2545.6228 | 2697.8298 | 3416.9307 | 3103.2305 | 3181.6667 |
| Atp5pd | VAKPGLVDDFEK | -61 | -72 | 1 | 0 | 12 | 2339.4644 | 2545.6228 | 2697.8298 | 3416.9307 | 3103.2305 | 3181.6667 |
| Atp5pd | KYPYWPHQPIENL | -149 | -161 | 1 | 0 | 13 | 1098.6418 | 1075.7677 | 1020.7106 | 975.7269 | 1248.6663 | 958.2081 |
| Atp5pd | KYPYWPHQPIENL | -149 | -161 | 1 | 0 | 13 | 1098.6418 | 1075.7677 | 1020.7106 | 975.7269 | 1248.6663 | 958.2081 |
| Atp5pd | ANVAKPGLVDDFEK | -59 | -72 | 1 | 0 | 14 | 1679.5549 | 1612.7788 | 1758.1703 | 2630.165 | 1087.1901 | 1146.4252 |
| Atp5pd | SEKPPAIDWAYYR | -46 | -58 | 1 | 0 | 13 | 1591.8608 | 2028.6218 | 1958.8278 | 1954.5973 | 2192.193 | 2238.4 |
| Atp5pd | KSWNETFHAR | -32 | -41 | 1 | 0.00025015 | 10 | 1406.6909 | 1377.8828 | 1274.5093 | 848.199 | 1340.7792 | 1500.7974 |
| Atp5pd | SLSEKPPAIDWAYYR | -44 | -58 | 1 | 0 | 15 | 1329.8976 | 1519.8413 | 1681.0571 | 1785.8918 | 776.0103 | 2446.6719 |
| Atp5pd | SLSEKPPAIDWAYYR | -44 | -58 | 1 | 0 | 15 | 1329.8976 | 1519.8413 | 1681.0571 | 1785.8918 | 776.0103 | 2446.6719 |
| Atp5pd | NVAKPGLVDDFEK | -60 | -72 | 1 | 0 | 13 | 1305.8079 | 1295.0775 | 1665.8265 | 1504.6044 | 1251.9431 | 1737.7557 |
| Atp5pd | YPYWPHQPIENL | -150 | -161 | 0 | 0 | 12 | 1024.0938 | 1033.4999 | 1208.6836 | 1205.4734 | 1296.8224 | 1235.6117 |
| Atp5pd | VDQEEKEDVK | -90 | -99 | 1 | 0 | 10 | 881.579 | 961.62976 | 1045.3225 | 830.89545 | 985.03613 | 729.60815 |
| Atp5pd | TIDDLNEIFPETK | -132 | -144 | 0 | 0 | 13 | 273.30807 | 1942.3354 | 1571.4147 | 466.3955 | 191.78358 | 246.9679 |
| Atp5pd | KIPVPEDK | -78 | -85 | 1 | 0 | 8 | 1105.3405 | NaN | 1032.1522 | 1243.1737 | 813.1149 | 1014.1752 |
| Atp5pd | SCAEFVSGSQLR | -100 | -111 | 0 | 0 | 12 | 792.09985 | 715.31573 | 798.62366 | 779.7183 | NaN | 719.56104 |
| Atp5pd | VAKPGLVDDFEKK | -61 | -73 | 2 | 0 | 13 | 537.37256 | NaN | 676.48865 | 398.63177 | 655.7835 | 749.63214 |
| Atp5pd | VAKPGLVDDFEKK | -61 | -73 | 2 | 0 | 13 | 537.37256 | NaN | 676.48865 | 398.63177 | 655.7835 | 749.63214 |
| Atp5pd | LASLSEKPPAIDWAYYR | -42 | -58 | 1 | 0 | 17 | 3651.6775 | NaN | NaN | 4626.285 | 4225.3496 | 4997.753 |
| Atp5pd | EKPPAIDWAYYR | -47 | -58 | 1 | 0 | 12 | 891.2818 | 440.35803 | 960.11835 | NaN | NaN | 606.9781 |
| Atp5pd | YTALVDQEEKEDVK | -86 | -99 | 1 | 0.000826806 | 14 | 244.1785 | 748.5408 | 580.5722 | NaN | NaN | NaN |
| Atp5pd | AEFVSGSQLR | -102 | -111 | 0 | 0.00122419 | 10 | 1011.18396 | NaN | 267.33398 | NaN | NaN | NaN |
| Srsf4 | SKPNVPAESR | -445 | -454 | 1 | 0 | 10 | 479.44855 | 508.62143 | 584.79895 | 773.93176 | 445.5484 | 577.2547 |
| Srsf4 | AEGESEAPNPEPR | -424 | -436 | 0 | 0 | 13 | 464.78534 | 667.5916 | 675.3809 | 706.03375 | 732.6147 | 771.53894 |
| Srsf4 | DKYGPPTR | -94 | -101 | 1 | 0.000591124 | 8 | 782.7864 | NaN | NaN | NaN | 597.4888 | 455.77322 |
| Usp15 | AEGGAADLDTQR | -2 | -13 | 0 | 0.00025015 | 12 | 427.6471 | NaN | NaN | NaN | NaN | NaN |
| Tnpo1 | SECLNNIGDSSPLIR | -101 | -115 | 0 | 0 | 15 | 3416.2305 | 4433.0986 | 4598.3457 | 1846.2156 | 2659.229 | 2810.0044 |
| Tnpo1 | FSDQFPLPLK | -880 | -889 | 0 | 0 | 10 | 1735.5958 | 1719.321 | 528.5101 | 1820.198 | 1722.6624 | 1831.429 |
| Tnpo1 | GELQNWPDLLPK | -129 | -140 | 0 | 0 | 12 | 867.5047 | 757.0064 | 596.24677 | 578.1464 | 915.2162 | 934.21655 |
| Tnpo1 | AEGLGGNIEQLVAR | -660 | -673 | 0 | 0 | 14 | 812.09174 | 870.724 | 962.47046 | 861.7602 | 1035.8297 | 1018.8045 |
| Tnpo1 | NQVGDENWR | -870 | -878 | 0 | 0.00025015 | 9 | 749.2026 | 882.1703 | 583.45825 | 864.2988 | 622.8901 | 656.80786 |
| Tnpo1 | DELLPHILPLLK | -400 | -411 | 0 | 0 | 12 | 288.65656 | 359.8892 | 372.32962 | 340.17252 | 404.47455 | 386.13358 |
| Tnpo1 | ELLFHHEWVVK | -412 | -422 | 0 | 0 | 11 | 666.0054 | 1045.4315 | NaN | 822.0499 | 1511.0742 | 1195.3931 |
| Tnpo1 | ELLFHHEWVVK | -412 | -422 | 0 | 0 | 11 | 666.0054 | 1045.4315 | NaN | 822.0499 | 1511.0742 | 1195.3931 |
| Tnpo1 | DSDVLDRPLNIMIPK | -172 | -186 | 1 | 0.00025015 | 15 | 502.08542 | NaN | NaN | NaN | 227.75331 | 426.32898 |
| Ddb1 | YYLQIHPQELR | -517 | -527 | 0 | 0 | 11 | 3058.487 | 3285.017 | 3032.2876 | 1644.8009 | 1989.1497 | 1612.0485 |
| Ddb1 | LEELHVIDVK | -159 | -168 | 0 | 0 | 10 | 1702.9468 | 1986.7533 | 1944.2854 | 774.09656 | 2484.779 | 2322.9102 |
| Ddb1 | IEHSFWR | -1068 | -1074 | 0 | 0.00111006 | 7 | 1575.8531 | 1494.2927 | 1476.1774 | 1794.4019 | 1927.2098 | 1807.0906 |
| Ddb1 | LYDGLFK | -135 | -141 | 0 | 0 | 7 | 1080.8259 | 1153.9648 | 1085.8501 | 897.0526 | 1058.1017 | 849.93695 |
| Ddb1 | YLAIAPPIIK | -245 | -254 | 0 | 0 | 10 | 881.38434 | 991.4411 | 1058.5247 | 968.3977 | 1219.7415 | 1226.6938 |
| Ddb1 | IGRPSETGIIGIIDPECR | -112 | -129 | 1 | 0.00025015 | 18 | 769.87476 | 669.95337 | 737.1079 | 834.2638 | 941.01263 | 1011.636 |
| Ddb1 | LVFSNVNLK | -666 | -674 | 0 | 0 | 9 | 767.49524 | 849.17303 | 867.646 | 1302.996 | 1327.7999 | 1488.9502 |
| Ddb1 | LPSFELLHK | -571 | -579 | 0 | 0 | 9 | 706.3253 | 823.1465 | 805.02386 | 870.82086 | 981.2643 | 932.0672 |
| Ddb1 | IEVQDSSGGTTALRPSASTQ | -740 | -759 | 1 | 0.00025015 | 20 | 538.8918 | 686.06177 | 665.5526 | 62.833363 | 436.51953 | 399.41125 |
| Ddb1 | KLPSFELLHK | -570 | -579 | 1 | 0 | 10 | 341.99875 | 388.39194 | 71.549164 | 613.0737 | 353.53162 | 354.18655 |
| Ddb1 | AEGLRPVK | -46 | -53 | 1 | 0.00025015 | 8 | 630.97876 | 668.644 | 756.48505 | NaN | 681.3116 | 630.4241 |
| Ddb1 | LGKDPNTYFIV | -821 | -831 | 1 | 0.000591124 | 11 | 837.2174 | 164.8432 | 1295.2594 | NaN | 1154.6918 | NaN |
| Ddb1 | GDLIESFLDISRPK | -1091 | -1104 | 1 | 0 | 14 | 342.79083 | NaN | NaN | 315.42932 | 508.05725 | 411.09595 |
| Ddb1 | SCNSSQVVVAVGR | -502 | -514 | 0 | 0 | 13 | 10956.422 | 10972.437 | NaN | 221.60156 | NaN | NaN |
| Ddb1 | TVPLYESPR | -714 | -722 | 0 | 0.00659934 | 9 | 1282.0599 | 1737.1874 | NaN | 909.0009 | NaN | NaN |
| Ddb1 | NIETGLLSDR | -617 | -626 | 0 | 0 | 10 | 984.9291 | NaN | NaN | 1048.2257 | 1206.8151 | NaN |
| Ddb1 | VTLGTQPTVLR | -629 | -639 | 0 | 0.000230707 | 11 | 396.96234 | 503.0205 | NaN | NaN | NaN | 80.723404 |

| | | | | | | | | | | | | |
|---|---|---|---|---|---|---|---|---|---|---|---|---|
| Ddb1 | PSETGIIGIIDPECR | -115 | -129 | 0 | 0 | 15 | 511.6616 | NaN | NaN | NaN | 852.3054 | NaN |
| Flna | VEPGLGADNSVVR | -1020 | -1032 | 0 | 0 | 13 | 3481.3809 | 2996.9297 | 4842.267 | 4390.3154 | 3871.5166 | 4500.041 |
| Flna | TPCEEILVK | -2599 | -2607 | 0 | 0 | 9 | 2991.1394 | 3258.8982 | 2340.949 | 1591.7565 | 3214.443 | 2655.0012 |
| Flna | EAPQDFHPDR | -665 | -674 | 0 | 0 | 10 | 2925.655 | 1494.6501 | 2746.225 | 3233.0806 | 2069.751 | 1687.7905 |
| Flna | TGVELGKPTHF | -885 | -895 | 1 | 0 | 11 | 2883.8906 | 2454.529 | 747.97266 | 2334.343 | 2956.5398 | 432.06598 |
| Flna | NVTYGGHQVPGSPFK | -1425 | -1439 | 0 | 0 | 15 | 2123.2087 | 2242.755 | 716.36694 | 1901.544 | 1960.4633 | 3471.85 |
| Flna | VATVPQHATSGPGPADVSK | -2541 | -2559 | 0 | 0 | 19 | 414.92575 | 364.96036 | 474.51154 | 425.651 | 481.89328 | 578.59766 |
| Flna | VATVPQHATSGPGPADVSK | -2541 | -2559 | 0 | 0 | 19 | 414.92575 | 364.96036 | 474.51154 | 425.651 | 481.89328 | 578.59766 |
| Flna | DKGEYTLVVK | -2622 | -2631 | 1 | 0 | 10 | 2001.5281 | 246.70258 | 2097.2917 | 696.12695 | 2055.544 | 1956.6052 |
| Flna | VTYGGDHIPK | -949 | -958 | 0 | 0 | 10 | 1798.1023 | 1612.687 | 1596.6133 | 1416.099 | 675.67816 | 599.0044 |
| Flna | VPVHDVTDASK | -1440 | -1450 | 0 | 0 | 11 | 1790.2375 | 1625.6537 | 1681.1107 | 469.73233 | 1438.9728 | 317.37207 |
| Flna | VLFADQATPTSPIR | -850 | -863 | 0 | 0 | 14 | 1668.1001 | 1618.0789 | 1728.8611 | 1295.1982 | 2287.9412 | 1857.632 |
| Flna | YVPEVTGTHK | -346 | -355 | 0 | 0.000431655 | 10 | 1649.0339 | 1486.337 | 1552.6031 | 1948.9897 | 1896.3125 | 1581.4991 |
| Flna | FGGEHVPNSPFQVT | -1726 | -1739 | 0 | 0 | 14 | 1582.0477 | 1797.4443 | 1915.032 | 1599.303 | 2128.5015 | 1488.867 |
| Flna | YNDQHIPGSPFTAR | -1938 | -1951 | 0 | 0 | 14 | 1575.1022 | 1586.0192 | 1655.7515 | 1402.4764 | 1180.9064 | 1364.0021 |
| Flna | YNDQHIPGSPFTAR | -1938 | -1951 | 0 | 0 | 14 | 1575.1022 | 1586.0192 | 1655.7515 | 1402.4764 | 1180.9064 | 1364.0021 |
| Flna | VYVEDPAGHQEEAK | -318 | -331 | 0 | 0 | 14 | 1499.063 | 1455.2968 | 1727.4213 | 1427.129 | 1709.2798 | 1648.9127 |
| Flna | VYVEDPAGHQEEAK | -318 | -331 | 0 | 0 | 14 | 1499.063 | 1455.2968 | 1727.4213 | 1427.129 | 1709.2798 | 1648.9127 |
| Flna | NIGSHCDLSLK | -2155 | -2165 | 0 | 0 | 11 | 1465.9944 | 1672.8165 | 1634.0193 | 1576.8766 | 1597.0068 | 1712.1486 |
| Flna | VDVGKDQEFTVK | -983 | -994 | 1 | 0 | 12 | 1444.2185 | 1289.8422 | 1585.2004 | 2576.135 | 2491.276 | 2445.4963 |
| Flna | ALAGDQPTVQTPLR | -1740 | -1753 | 0 | 0 | 14 | 1360.611 | 1445.661 | 1223.1089 | 1442.4529 | 1322.734 | 1359.5507 |
| Flna | ETGEHLVHVK | -2015 | -2024 | 0 | 0 | 10 | 1354.3618 | 1377.4518 | 1325.3546 | 1290.793 | 1459.1204 | 1507.2361 |
| Flna | SPFSVGVSPSLDLSK | -959 | -973 | 0 | 0 | 15 | 1301.6896 | 1500.9731 | 1729.6077 | 2026.2495 | 2353.6663 | 2819.062 |
| Flna | TGVELGKPTHFTVN | -885 | -898 | 1 | 0.000230707 | 14 | 1255.7158 | 1406.612 | 1454.266 | 1065.7362 | 1395.743 | 1298.0219 |
| Flna | TGVELGKPTHFTVN | -885 | -898 | 1 | 0.000230707 | 14 | 1255.7158 | 1406.612 | 1454.266 | 1065.7362 | 1395.743 | 1298.0219 |
| Flna | AFGPGLQGGNAGSPAR | -1072 | -1087 | 0 | 0 | 16 | 1200.7272 | 1205.4792 | 1334.0691 | 2425.048 | 2306.8757 | 2386.651 |
| Flna | VGEPGHGGDPGLVSAY | -2420 | -2435 | 0 | 0 | 16 | 1152.12 | 1000.40894 | 1041.7643 | 1331.5093 | 1913.7722 | 1704.9956 |
| Flna | FADQHVPGSPFSVK | -2120 | -2133 | 0 | 0 | 14 | 1130.731 | 933.54663 | 1287.8651 | 1429.924 | 1041.1838 | 1851.6222 |
| Flna | FADQHVPGSPFSVK | -2120 | -2133 | 0 | 0 | 14 | 1130.731 | 933.54663 | 1287.8651 | 1429.924 | 1041.1838 | 1851.6222 |
| Flna | AWGPGLEGGIVGK | -581 | -593 | 0 | 0 | 13 | 1120.1111 | 1263.7866 | 1128.4889 | 1519.4865 | 569.6453 | 2224.5132 |
| Flna | WGDEHIPGSPYR | -2632 | -2643 | 0 | 0 | 12 | 1075.6504 | 1243.7555 | 1153.1367 | 1344.5271 | 1103.8127 | 1230.7516 |
| Flna | WGDEHIPGSPYR | -2632 | -2643 | 0 | 0 | 12 | 1075.6504 | 1243.7555 | 1153.1367 | 1344.5271 | 1103.8127 | 1230.7516 |
| Flna | GGVSIPNSPFR | -750 | -760 | 0 | 0 | 11 | 1061.7065 | 1128.9498 | 1298.0812 | 1253.6765 | 1491.9335 | 1288.5072 |
| Flna | SLNVTYGGHQVPGSPFK | -1423 | -1439 | 0 | 0 | 17 | 1060.7013 | 1227.6539 | 1243.9661 | 1108.9883 | 1598.7882 | 1444.5204 |
| Flna | YGGDEIPFSPYR | -1622 | -1633 | 0 | 0 | 12 | 1006.926 | 1142.405 | 1065.5658 | 646.3844 | 817.4907 | 485.9871 |
| Flna | ANLPQSFQVDTSK | -1465 | -1477 | 0 | 0 | 13 | 978.63745 | 552.2818 | 255.21555 | 336.095 | 274.84692 | 475.58224 |
| Flna | GAGTGGLGLAVEGPSEAK | -1382 | -1399 | 0 | 0 | 18 | 878.84283 | 821.54364 | 840.14215 | 851.2958 | 930.0016 | 857.0436 |
| Flna | SPFEVYVDK | -368 | -376 | 0 | 0 | 9 | 863.5005 | 475.15884 | 971.85254 | 1211.0645 | 371.74365 | 1300.107 |
| Flna | THEAEIVEGENHTY | -2185 | -2198 | 0 | 0 | 14 | 859.6189 | 873.8917 | 816.18054 | 567.6046 | 586.5683 | 589.17834 |
| Flna | THEAEIVEGENHTY | -2185 | -2198 | 0 | 0 | 14 | 859.6189 | 873.8917 | 816.18054 | 567.6046 | 586.5683 | 589.17834 |
| Flna | VDINTEDLEDGTCR | -2090 | -2103 | 0 | 0 | 14 | 849.32495 | 932.1413 | 904.29626 | 906.0004 | 1040.7552 | 986.4114 |
| Flna | VLPTHDASK | -1539 | -1547 | 0 | 0.001092529 | 9 | 797.6768 | 833.1641 | 766.83813 | 662.77057 | 852.10535 | 890.1561 |
| Flna | DVDIIDHHDNTYTVK | -922 | -936 | 0 | 0 | 15 | 794.7228 | 942.46686 | 920.2079 | 773.99866 | 1394.6478 | 1117.0255 |
| Flna | DVDIIDHHDNTYTVK | -922 | -936 | 0 | 0 | 15 | 794.7228 | 942.46686 | 920.2079 | 773.99866 | 1394.6478 | 1117.0255 |
| Flna | GFSVEGPSQAK | -610 | -620 | 0 | 0 | 11 | 774.01636 | 875.9126 | 787.827 | 887.7916 | 753.71155 | 691.81433 |
| Flna | FNEEHIPDSPFVVPVASPSGDAR | -2311 | -2333 | 0 | 0 | 23 | 767.9669 | 665.8208 | 743.87634 | 721.27325 | 806.321 | 887.11835 |
| Flna | DCTEAGQGDVSIGIK | -795 | -809 | 0 | 0 | 15 | 742.6604 | 753.5731 | 807.7816 | 757.1855 | 1014.8476 | 715.3285 |
| Flna | AEISFEDRK | -2281 | -2289 | 1 | 0 | 9 | 713.8 | 699.631 | 841.0255 | 584.78345 | 829.13885 | 241.69174 |
| Flna | KTHIQDNHDGTYTVA | -1593 | -1607 | 1 | 0.000591124 | 15 | 667.70953 | 864.08875 | 843.7322 | 1665.4447 | 1038.8289 | 895.12177 |
| Flna | KTHIQDNHDGTYTVA | -1593 | -1607 | 1 | 0.000591124 | 15 | 667.70953 | 864.08875 | 843.7322 | 1665.4447 | 1038.8289 | 895.12177 |
| Flna | YGGQPVPNFPSKL | -1235 | -1247 | 1 | 0.00025015 | 13 | 651.9558 | 960.4455 | 987.6211 | 1980.5011 | 3143.2927 | 2768.0022 |
| Flna | VKVEPSHDASK | -864 | -874 | 1 | 0 | 11 | 558.61566 | 723.6878 | 470.09937 | 559.4366 | 704.1982 | 422.13593 |
| Flna | THEAEIVEGENHTYCIR | -2185 | -2201 | 0 | 0 | 17 | 311.32513 | 317.85217 | 323.13077 | 328.39578 | 411.67914 | 476.13605 |
| Flna | THEAEIVEGENHTYCIR | -2185 | -2201 | 0 | 0 | 17 | 311.32513 | 317.85217 | 323.13077 | 328.39578 | 411.67914 | 476.13605 |
| Flna | VTAQGPGLEPSGNIANK | -384 | -400 | 0 | 0 | 17 | 534.8141 | 556.26825 | 499.30093 | 533.47064 | 169.2234 | 744.0401 |
| Flna | ITDPEGKPK | -1584 | -1592 | 1 | 0 | 9 | 528.45917 | 567.8657 | 599.0611 | 458.78323 | 475.92328 | 496.88812 |
| Flna | WGGVSIPNSPFR | -749 | -760 | 0 | 0 | 12 | 520.6234 | 704.084 | 823.0972 | 623.5669 | 751.42847 | 750.3644 |
| Flna | TGVAVNKPAEFTVDAK | -685 | -700 | 1 | 0.00025015 | 16 | 454.63278 | 613.8933 | 537.0081 | 697.7149 | 721.74884 | 714.6212 |
| Flna | DAGEGLLAVQITDPEGKPK | -1574 | -1592 | 1 | 0 | 19 | 429.9935 | 456.62488 | 457.96683 | 769.1727 | 700.93866 | 991.9021 |
| Flna | VEDPAGHQEEAK | -320 | -331 | 0 | 0 | 12 | 379.88388 | 363.52014 | 424.64484 | 165.28745 | 546.1318 | 510.91385 |
| Flna | EADIDFDIIR | -819 | -828 | 0 | 0 | 10 | 334.82367 | 419.0564 | 406.73828 | 572.45325 | 418.24237 | 200.35663 |
| Flna | YDGVPVPGSPFPLEAVAPTKPSK | -1047 | -1069 | 1 | 0 | 23 | 135.42276 | 771.4032 | 627.61273 | 759.1915 | 785.9778 | 689.9981 |
| Flna | YDGVPVPGSPFPLEAVAPTKPSK | -1047 | -1069 | 1 | 0 | 23 | 135.42276 | 771.4032 | 627.61273 | 759.1915 | 785.9778 | 689.9981 |
| Flna | GLVEPVDVVDNADGTGTQTVNYVPSR | -1492 | -1515 | 0 | 0 | 24 | 265.01636 | 291.0045 | 258.15692 | 299.93878 | 272.7765 | 297.9637 |
| Flna | VANPSGNLTDTYVQDCGDGTYK | -1297 | -1318 | 0 | 0.00025015 | 22 | 203.3726 | 151.90982 | 251.52338 | 266.52182 | 272.9948 | 233.29141 |
| Flna | YGGQPVPNFPSK | -1235 | -1246 | 0 | 0 | 12 | 4326.2427 | 3548.0005 | NaN | 1222.698 | 1815.0936 | 1863.853 |
| Flna | YVVQEPGDYEVSVK | -2297 | -2310 | 0 | 0 | 14 | 2688.4548 | 3420.7732 | 1048.893 | 527.2904 | NaN | 541.34705 |
| Flna | GTVEPQLEAR | -428 | -437 | 0 | 0 | 10 | 2467.9697 | 8842.367 | 6237.5503 | 11717.943 | NaN | 5304.351 |
| Flna | YSPSEAGLHEMDIR | -1832 | -1845 | 0 | 0 | 14 | 1224.9573 | NaN | 1545.6732 | 1522.202 | 1708.7625 | 1678.4205 |
| Flna | AEGPGLNR | -877 | -884 | 0 | 0 | 8 | 848.5636 | 678.211 | 545.52875 | 590.4218 | 795.6899 | NaN |
| Flna | ITWGGQNIGR | -554 | -563 | 0 | 0 | 10 | 697.0292 | 837.2443 | 679.4847 | NaN | 1157.3223 | 683.6333 |
| Flna | YGGPYHIGGSPF | -2501 | -2512 | 0 | 0.001055699 | 12 | 623.0346 | NaN | 712.5657 | 467.39426 | 558.82166 | 581.11743 |
| Flna | FAGQHIAK | -360 | -367 | 0 | 0 | 8 | 505.24875 | 514.52124 | 527.48224 | NaN | 497.59583 | 634.24646 |
| Flna | SHCDLSLK | -2158 | -2165 | 0 | 0.00025015 | 8 | 409.71472 | NaN | 663.253 | 523.26605 | 457.396 | 347.20905 |
| Flna | EATTEFSVDAR | -1273 | -1283 | 0 | 0 | 11 | 239.50436 | 350.80234 | 610.1789 | 1302.4073 | 171.08528 | NaN |
| Flna | GPGLQGGNAGSPAR | -1074 | -1087 | 0 | 0 | 14 | 2022.787 | NaN | NaN | 2073.0522 | 779.5983 | 2933.999 |
| Flna | VSGQGLHEGHTFEPA | -2052 | -2066 | 0 | 0.00078661 | 15 | 557.32764 | 492.7049 | 481.1846 | 431.84048 | NaN | NaN |
| Flna | TGVAVNKPAEFTVD | -685 | -698 | 1 | 0 | 14 | 2304.0225 | 1839.2587 | 1981.6326 | NaN | NaN | NaN |

| | | | | | | | | | | | | |
|---|---|---|---|---|---|---|---|---|---|---|---|---|
| Flna | AHVAPCFDASK | -1152 | -1162 | 0 | 0 | 11 | 2105.885 | 2054.4326 | 2357.009 | NaN | NaN | NaN |
| Flna | FADQHVPGSPF | -2120 | -2130 | 0 | 0 | 11 | 939.1329 | NaN | NaN | NaN | 1533.2517 | 1655.6389 |
| Flna | AGDQPTVQTPLR | -1742 | -1753 | 0 | 0 | 12 | 710.75574 | 1524.2917 | NaN | 1527.6737 | NaN | NaN |
| Flna | WDEEEDEEAKK | -155 | -165 | 1 | 0.000230707 | 11 | 696.7036 | 697.3001 | NaN | 814.6034 | NaN | NaN |
| Flna | ALTQTGGPHVK | -1284 | -1294 | 0 | 0 | 11 | 567.60004 | 496.69257 | 518.2394 | NaN | NaN | NaN |
| Flna | VVIQDPTGQK | -418 | -427 | 0 | 0.007670622 | 10 | 1369.8655 | 1179.9899 | NaN | NaN | NaN | NaN |
| Flna | VGEPGHGGDPGLVS | -2420 | -2433 | 0 | 0.000712251 | 14 | 1354.3862 | NaN | NaN | NaN | NaN | 1986.7886 |
| Flna | HIGGSPFK | -2506 | -2513 | 0 | 0 | 8 | 1129.7241 | 1405.2771 | NaN | NaN | NaN | NaN |
| Flna | THIPGSPFK | (1143)(2409) | (1151)(2417) | 0 | 0 | 9 | 686.82794 | NaN | NaN | NaN | 1089.3315 | NaN |
| Flna | YIPVQQGPVGVN | -937 | -948 | 0 | 0.003490217 | 12 | 676.97015 | NaN | NaN | NaN | 117.18193 | NaN |
| Flna | THIQDNHDGTYTVA | -1594 | -1607 | 0 | 0.002168954 | 14 | 1821.2573 | NaN | NaN | NaN | NaN | NaN |
| Flna | IVSPSGAAVPCK | -1008 | -1019 | 0 | 0.004224523 | 12 | 1447.4099 | NaN | NaN | NaN | NaN | NaN |
| Flna | LVGVHGPR | -2591 | -2598 | 0 | 0.002595029 | 8 | 1254.6626 | NaN | NaN | NaN | NaN | NaN |
| Flna | LLVGVHGPR | -2590 | -2598 | 0 | 0.000471409 | 9 | 690.84607 | NaN | NaN | NaN | NaN | NaN |
| Flna | LKPGAPLRPK | -269 | -278 | 2 | 0.008014596 | 10 | 376.91068 | NaN | NaN | NaN | NaN | NaN |
| Rexo2 | AEGPNLIIK | -76 | -84 | 0 | 0.000230707 | 9 | 2414.028 | 2225.2131 | 2352.8687 | 2048.3037 | 1812.445 | 1781.2473 |
| Rexo2 | RALDDISESIK | -195 | -205 | 1 | 0.00025015 | 11 | 861.45984 | 847.383 | 973.3086 | 1310.9774 | 1363.3048 | 1379.2014 |
| Rexo2 | RWYPEDYEFAPK | -178 | -189 | 1 | 0.00025015 | 12 | 411.85825 | 550.89453 | 841.5383 | 489.3587 | 328.66562 | 629.51086 |
| Rexo2 | ALDDISESIK | -196 | -205 | 0 | 0 | 10 | 1217.4519 | 1445.5438 | 746.6853 | 1235.5586 | NaN | 1649.0374 |
| Kif5b | AEGSTYVPYR | -269 | -278 | 0 | 0 | 10 | 857.7358 | 827.303 | NaN | 1343.7511 | NaN | NaN |
| Lmnb1 | ASPPTDLIWK | -508 | -517 | 0 | 0 | 10 | 5317.052 | 5516.916 | 5755.591 | 3634.8489 | 3847.8535 | 3798.909 |
| Lmnb1 | IGDTSVSYK | -476 | -484 | 0 | 0 | 9 | 3685.0322 | 3957.6028 | 3531.607 | 1300.7592 | 1927.4437 | 2762.4563 |
| Lmnb1 | NQNSWGTGEDVK | -518 | -529 | 0 | 0 | 12 | 3451.0547 | 3369.0042 | 4014.383 | 2918.3784 | 3876.931 | 3190.5427 |
| Lmnb1 | NQNSWGTGEDVK | -518 | -529 | 0 | 0 | 12 | 3451.0547 | 3369.0042 | 4014.383 | 2918.3784 | 3876.931 | 3190.5427 |
| Lmnb1 | LREYEAALNSK | -136 | -146 | 1 | 0 | 11 | 3125.0825 | 2491.1218 | 1971.1177 | 2592.3696 | 1722.0024 | 2363.6362 |
| Lmnb1 | LYKEELEQTYHAK | -260 | -272 | 1 | 0 | 13 | 2564.149 | 2932.6152 | 3249.2249 | 2562.4932 | 4057.953 | 3367.7473 |
| Lmnb1 | LYKEELEQTYHAK | -260 | -272 | 1 | 0 | 13 | 2564.149 | 2932.6152 | 3249.2249 | 2562.4932 | 4057.953 | 3367.7473 |
| Lmnb1 | QLADETLLK | -184 | -192 | 0 | 0.000230707 | 9 | 2549.8003 | 2385.787 | 2308.1965 | 2338.8687 | 2991.1829 | 2705.5273 |
| Lmnb1 | SLSSQLSNLQK | -303 | -313 | 0 | 0.00025015 | 11 | 2365.5671 | 2123.7651 | 2161.9033 | 2940.5908 | 1776.9688 | 2387.5776 |
| Lmnb1 | SLTEDLEFR | -201 | -209 | 0 | 0 | 9 | 1983.3171 | 2475.7744 | 2794.197 | 2583.3357 | 5289.869 | 1219.7195 |
| Lmnb1 | HDQLLLNYAK | -115 | -124 | 0 | 0 | 10 | 1904.7115 | 1782.6974 | 1755.3839 | 1948.3164 | 1707.868 | 1151.2764 |
| Lmnb1 | LSDYEQLLDVK | -358 | -368 | 0 | 0 | 11 | 1864.8534 | 1611.1552 | 2207.0837 | 4020.173 | 3341.631 | 2881.0378 |
| Lmnb1 | IQELEDMLAK | -322 | -331 | 0 | 0 | 10 | 1821.0034 | 1355.8091 | 2771.3713 | 2597.3757 | 2593.9717 | 2766.9602 |
| Lmnb1 | IQELEDMLAK | -322 | -331 | 0 | 0 | 10 | 1821.0034 | 1355.8091 | 2771.3713 | 2597.3757 | 2593.9717 | 2766.9602 |
| Lmnb1 | KQLADETLLK | -183 | -192 | 1 | 0 | 10 | 1529.1927 | 1678.329 | 1664.6263 | 1617.9084 | 1576.7749 | 1582.5549 |
| Lmnb1 | FKAEHDQLLL | -111 | -120 | 1 | 0.000471409 | 10 | 1513.5869 | 1023.6962 | 978.13116 | 1051.7776 | 1424.3726 | 2411.2378 |
| Lmnb1 | ASAPATPLSPTR | -16 | -27 | 0 | 0 | 12 | 1479.3087 | 1600.9795 | 1657.4882 | 499.69012 | 1256.9061 | 1204.7177 |
| Lmnb1 | FKAEHDQLLLNYAK | -111 | -124 | 1 | 0 | 14 | 1391.8992 | 1259.2198 | 1347.9592 | 835.69476 | 1760.2349 | 1363.7646 |
| Lmnb1 | FKAEHDQLLLNYAK | -111 | -124 | 1 | 0 | 14 | 1391.8992 | 1259.2198 | 1347.9592 | 835.69476 | 1760.2349 | 1363.7646 |
| Lmnb1 | AEHDQLLLNYAK | -113 | -124 | 0 | 0 | 12 | 1244.0913 | 1594.1954 | 1781.2518 | 1458.7975 | 1990.1069 | 2525.6328 |
| Lmnb1 | AEHDQLLLNYAK | -113 | -124 | 0 | 0 | 12 | 1244.0913 | 1594.1954 | 1781.2518 | 1458.7975 | 1990.1069 | 2525.6328 |
| Lmnb1 | GNVCIEEIDVDGK | -441 | -453 | 0 | 0 | 13 | 1114.0043 | 1225.7646 | 1217.7981 | 2103.6072 | 980.72565 | 1593.1648 |
| Lmnb1 | NSQGEEVAQR | -534 | -543 | 0 | 0 | 10 | 1079.9794 | 814.1026 | 1467.4576 | 784.8181 | 1451.2761 | 1081.8159 |
| Lmnb1 | PPTDLIWK | -510 | -517 | 0 | 0.00025015 | 8 | 1017.1871 | 1075.9128 | 1190.4766 | 2680.7695 | 3495.055 | 3235.7573 |
| Lmnb1 | SAPATPLSPTR | -17 | -27 | 0 | 0 | 11 | 987.7652 | 1143.7949 | 1485.5916 | 1238.4302 | 1258.8276 | 1295.647 |
| Lmnb1 | KEELEQTYHAK | -262 | -272 | 1 | 0.000471409 | 11 | 914.1386 | 1000.8318 | 1181.0728 | 1265.9044 | 1905.9207 | 1732.4197 |
| Lmnb1 | KEELEQTYHAK | -262 | -272 | 1 | 0.000471409 | 11 | 914.1386 | 1000.8318 | 1181.0728 | 1265.9044 | 1905.9207 | 1732.4197 |
| Lmnb1 | KALYETELADAR | -80 | -91 | 1 | 0 | 12 | 679.81616 | 911.8014 | 878.9007 | 227.61658 | 1102.6818 | 1367.1726 |
| Lmnb1 | ALYETELADAR | -81 | -91 | 0 | 0 | 11 | 668.302 | 3033.4219 | 3138.1248 | 1970.4247 | 1798.1082 | 1202.719 |
| Lmnb1 | QNSWGTGEDVK | -519 | -529 | 0 | 0 | 11 | 548.7199 | 552.3798 | 573.48096 | 1244.5496 | 453.1855 | 969.19434 |
| Lmnb1 | MYEEEINETR | -212 | -221 | 0 | 0 | 10 | 547.5984 | 515.8423 | 624.5069 | 116.88798 | 839.03784 | 837.70526 |
| Lmnb1 | SATGNVCIEEIDVDGK | -438 | -453 | 0 | 0 | 16 | 514.5605 | 574.5313 | 680.9165 | 471.83902 | 834.37665 | 830.413 |
| Lmnb1 | MQQQLSDYEQLLDVK | -354 | -368 | 0 | 0 | 15 | 482.52237 | 614.22205 | 598.3404 | 930.71375 | 855.64594 | 859.6222 |
| Lmnb1 | MQQQLSDYEQLLDVK | -354 | -368 | 0 | 0 | 15 | 482.52237 | 614.22205 | 598.3404 | 930.71375 | 855.64594 | 859.6222 |
| Lmnb1 | DQMQQQLSDYEQLLDVK | -352 | -368 | 0 | 0 | 17 | 414.12726 | 298.29388 | 261.1817 | 295.2326 | 322.7942 | 373.19318 |
| Lmnb1 | KESDLSGAQIK | -125 | -135 | 1 | 0 | 11 | 387.67694 | 272.93155 | 1634.332 | 2078.702 | 2326.4238 | 425.1523 |
| Lmnb1 | IESLSSQLSNLQK | -301 | -313 | 0 | 0 | 13 | 384.8137 | 233.51062 | 505.09924 | 572.1677 | 609.576 | 633.2357 |
| Lmnb1 | TIPEEEEEEEEPIGVAVEEER | -550 | -571 | 0 | 0 | 22 | 374.36172 | 459.21777 | 411.31497 | 358.07755 | 370.37885 | 316.5858 |
| Lmnb1 | EELEQTYHAK | -263 | -272 | 0 | 0.00025015 | 10 | 2210.402 | 3232.0752 | 2396.7637 | NaN | 2440.2808 | 7955.481 |
| Lmnb1 | FKAEHDQLLLNY | -111 | -122 | 1 | 0 | 12 | 1839.1581 | 1984.0779 | 2000.997 | NaN | 1027.5131 | 1278.7623 |
| Lmnb1 | LYKEELEQTYH | -260 | -270 | 1 | 0 | 11 | 1231.4995 | 1038.67 | 1500.9773 | 1699.9669 | NaN | 1132.6941 |
| Lmnb1 | QLSDYEQLLDVK | -357 | -368 | 0 | 0 | 12 | 973.4082 | 656.8862 | NaN | 1518.4121 | 1796.3142 | 99.41834 |
| Lmnb1 | KIGDTSVSYK | -475 | -484 | 1 | 0.00025015 | 10 | 687.46405 | 1397.6809 | 1740.6676 | 1335.9069 | NaN | 1435.4382 |
| Lmnb1 | LSSEMNTSTVNSAR | -278 | -291 | 0 | 0 | 14 | 541.7833 | 638.91693 | 235.42566 | 419.29376 | 825.0445 | NaN |
| Lmnb1 | NAGVTASPPTDLIWK | -503 | -517 | 0 | 0 | 15 | 529.0983 | NaN | 740.78656 | 1376.3912 | 1437.5339 | 1405.174 |
| Lmnb1 | ESDLSGAQIK | -126 | -135 | 0 | 0.000627943 | 10 | 2441.724 | 2176.1345 | 1779.9036 | 1959.4731 | NaN | NaN |
| Lmnb1 | AEHDQLLLNY | -113 | -122 | 0 | 0.001811524 | 10 | 2379.6782 | 1546.9763 | 4133.698 | NaN | NaN | NaN |
| Lmnb1 | KTTIPEEEEEEEEPIGVAVEEER | -548 | -571 | 1 | 0.001868731 | 24 | 617.44147 | 551.7911 | 557.9418 | NaN | NaN | NaN |
| Lmnb1 | NMYEEEINETR | -211 | -221 | 0 | 0 | 11 | 1824.3091 | 2104.095 | NaN | NaN | NaN | NaN |
| Lmnb1 | YETELADAR | -83 | -91 | 0 | 0 | 9 | 519.8111 | NaN | NaN | NaN | NaN | NaN |
| Tmbim6 | KINFDALLK | -7 | -15 | 1 | 0.000471409 | 9 | 1195.3492 | 1453.3704 | 2784.0789 | 1016.34656 | 1431.1179 | 1307.1658 |
| Tmbim6 | FSHITPSTQQHLK | -16 | -28 | 0 | 0 | 13 | 457.17365 | 635.96844 | 571.5341 | 906.2853 | 961.4368 | 997.0606 |
| Tmbim6 | AEHGDKDYIWHC | -196 | -207 | 1 | 0 | 12 | 500.08 | 468.80963 | 554.2461 | NaN | 373.2981 | 324.1613 |
| Tgtp2;Tgtp1 | HISSVTGGPVAAVT | (355);(355) | (368);(368) | 0 | 0 | 14 | 4185.6685 | 4420.73 | 5427.5684 | 5198.779 | 5719.2524 | 5778.779 |
| Tgtp2;Tgtp1 | HISSVTGGPVAAVT | (355);(355) | (368);(368) | 0 | 0 | 14 | 4185.6685 | 4420.73 | 5427.5684 | 5198.779 | 5719.2524 | 5778.779 |
| Tgtp2;Tgtp1 | TKIDSDLDNEQK | (170);(170) | (181);(181) | 1 | 0 | 12 | 1973.4716 | 2097.318 | 2286.1108 | 1934.8909 | 2414.182 | 2459.4668 |
| Tgtp2;Tgtp1 | LLQDLPAHK | (234);(234) | (242);(242) | 0 | 0 | 9 | 1873.784 | 1933.9135 | 1877.6235 | 127.546364 | 2156.6223 | 1768.2646 |
| Tgtp2;Tgtp1 | FKENDAQLAK | (147);(147) | (156);(156) | 1 | 0 | 10 | 1662.6124 | 1652.0748 | 1784.56 | 1503.566 | 577.5379 | 1733.5621 |

| | | | | | | | | | | | | |
|---|---|---|---|---|---|---|---|---|---|---|---|---|
| Tgtp2;Tgtp1 | DIESAPLHIA | (51);(51) | (60);(60) | 0 | 0.00025015 | 10 | 1493.3367 | 1446.5953 | 180.69574 | 1521.2107 | 1313.6976 | 1249.9609 |
| Tgtp2;Tgtp1 | TPYPHPK | (99);(99) | (105);(105) | 0 | 0.00025015 | 7 | 1329.1335 | 1113.7095 | 623.1735 | 1360.2715 | 1423.5306 | 1329.1467 |
| Tgtp2;Tgtp1 | SFNKEEVLK | (186);(186) | (194);(194) | 1 | 0 | 9 | 1054.8606 | 1354.383 | 1009.07935 | 1580.2998 | 1489.879 | 1568.4564 |
| Tgtp2;Tgtp1 | KVGPYISEPPEYWEA | (401);(401) | (415);(415) | 1 | 0.000230707 | 15 | 509.15344 | 741.5713 | 706.7888 | 193.64687 | 747.85846 | 788.68695 |
| Tgtp2;Tgtp1 | IDSDLDNEQK | (172);(172) | (181);(181) | 0 | 0 | 10 | 1535.4816 | 1184.2437 | NaN | 851.469 | 884.23865 | 1408.7229 |
| Tgtp2;Tgtp1 | AEHNDESLEDK | (338);(338) | (348);(348) | 0 | 0 | 11 | 369.596 | 436.5622 | 373.49103 | 332.24063 | NaN | 377.8558 |
| Tgtp2;Tgtp1 | FPHLFAE | (333);(333) | (339);(339) | 0 | 0.005459848 | 7 | 1477.8783 | 1559.8718 | 1624.1389 | 68.642334 | NaN | NaN |
| Tgtp2;Tgtp1 | ENLDETFNLYR | (293);(293) | (303);(303) | 0 | 0.00025015 | 11 | 1094.1256 | NaN | NaN | NaN | NaN | NaN |
| Uchl5 | NQDDWITAVRPVIEK | -192 | -206 | 1 | 0.00025015 | 15 | 2580.8408 | 3603.651 | 2855.0142 | 4549.8325 | 6042.922 | 6870.939 |
| Uchl5 | LYELDGLR | -175 | -182 | 0 | 0 | 8 | 1697.3345 | 1680.1427 | 1766.7773 | 1256.5417 | 1381.5638 | 1366.8942 |
| Uchl5 | TPAKEEDAFHF | -155 | -165 | 1 | 0 | 11 | 859.71985 | 897.02527 | 1103.1047 | 1155.4766 | 1093.7429 | 1420.5237 |
| Uchl5 | AEHQQLIPLVEK | -304 | -315 | 0 | 0 | 12 | 848.68994 | 854.1708 | 813.3214 | 1284.7815 | 994.88367 | 1084.5503 |
| Uchl5 | WQPGEEPAGSVVQDSR | -58 | -73 | 0 | 0 | 16 | 678.5734 | 687.66675 | 662.1929 | 699.2211 | 956.4815 | 940.8335 |
| Uchl5 | TPAKEEDAFHFV | -155 | -166 | 1 | 0.001023236 | 12 | 1518.0098 | 932.00476 | 1061.1793 | NaN | 3274.959 | NaN |
| Uchl5 | LETIFFAK | -74 | -81 | 0 | 0 | 8 | 758.15826 | 873.49286 | NaN | 990.0665 | 740.32654 | NaN |
| Uchl5 | GAQVEEIWSLEPESFEK | -29 | -45 | 0 | 0.00025015 | 17 | 277.70215 | 357.33258 | NaN | NaN | 546.3019 | 584.74817 |
| Uchl5 | QVEEIWSLEPESFEK | -31 | -45 | 0 | 0.000627943 | 15 | 306.55356 | NaN | NaN | NaN | 1268.0431 | 1031.4971 |
| Uchl5 | ALSNSDVIR | -129 | -137 | 0 | 0.008129024 | 9 | 1225.4227 | NaN | 1098.6572 | NaN | NaN | NaN |
| Stat5b | AWDSIDLDNPQENIK | -42 | -56 | 0 | 0 | 15 | 820.92896 | 906.4729 | 941.0787 | 1065.2887 | NaN | 1200.3677 |
| Mthfd1l | LSGGSPGVPVDK | -299 | -310 | 0 | 0 | 12 | 1899.7255 | 144.92073 | 1631.394 | 1474.9104 | 1828.557 | 1764.4579 |
| Mthfd1l | DFTLPISDVR | -925 | -934 | 0 | 0.00025015 | 10 | 1876.4291 | 2387.549 | 2392.1074 | 2391.3027 | 3593.7534 | 2980.9236 |
| Mthfd1l | VLVVGAHGPLEAALQ | -222 | -236 | 0 | 0 | 15 | 1629.1188 | 1748.601 | 1618.1554 | 1834.2395 | 1956.9055 | 1988.3561 |
| Mthfd1l | VLNALKPEK | -168 | -176 | 1 | 0 | 9 | 1584.6881 | 1359.076 | 1514.7314 | 1498.0323 | 1151.747 | 1142.1118 |
| Mthfd1l | TDPSTLTEEEVR | -547 | -558 | 0 | 0 | 12 | 1472.5763 | 1024.1125 | 917.4819 | 2046.3964 | 1618.1555 | 235.39796 |
| Mthfd1l | QPSQGPTFGVK | -453 | -463 | 0 | 0 | 11 | 1367.108 | 1502.8824 | 1763.2035 | 2014.5173 | 2465.9614 | 2332.0623 |
| Mthfd1l | FHDFLSGK | -291 | -298 | 0 | 0 | 8 | 1293.4708 | 833.7909 | 856.9842 | 1610.2653 | 1271.9622 | 1318.3813 |
| Mthfd1l | DVDGVTDINLGK | -177 | -188 | 0 | 0 | 12 | 963.1029 | 1249.2363 | 1247.3888 | 1173.555 | 781.49176 | 912.0803 |
| Mthfd1l | FYDIDLDTETEQVK | -961 | -974 | 0 | 0 | 14 | 811.2038 | 960.42395 | 1038.8795 | 2002.2145 | 2494.9712 | 2400.0408 |
| Mthfd1l | SSSGGGDPEGLR | -36 | -47 | 0 | 0 | 12 | 783.7346 | 819.1663 | 849.2914 | 913.0333 | 890.0512 | 895.0175 |
| Mthfd1l | SCPWATPQLPDK | -248 | -259 | 0 | 0 | 12 | 697.57465 | 1083.0653 | 627.98004 | 1006.21454 | 835.9213 | 826.1561 |
| Mthfd1l | ILHESTQTDK | -506 | -515 | 0 | 0 | 10 | 628.167 | 760.7183 | 678.14966 | 742.7203 | 835.31006 | 806.9961 |
| Mthfd1l | LQPLSPVPSDIEISR | -352 | -366 | 0 | 0 | 15 | 558.42413 | 637.97595 | 618.53 | 583.3096 | 640.4982 | 658.3335 |
| Mthfd1l | AEIDLVCELAK | -807 | -817 | 0 | 0 | 11 | 477.4485 | 485.607 | 528.818 | 414.63364 | 276.10126 | 872.93976 |
| Mthfd1l | THLSLSHEPDKK | -909 | -920 | 1 | 0 | 12 | 419.16647 | 913.49976 | 548.84735 | 98.89749 | 1088.5099 | 794.31903 |
| Mthfd1l | NIDPATITWQR | -564 | -574 | 0 | 0 | 11 | 341.69333 | 1102.8401 | 1118.3112 | 758.0497 | 287.19885 | 1161.2126 |
| Mthfd1l | NPAFKPVLVVIQ | -92 | -103 | 1 | 0.000550307 | 12 | 106.21119 | 1167.3171 | 1135.281 | 1317.2561 | 1692.7573 | 1562.834 |
| Mthfd1l | NCFHDFLSGK | -289 | -298 | 0 | 0.000230707 | 10 | 1931.5192 | 1948.1108 | 2053.6902 | 1073.199 | 3925.782 | NaN |
| Mthfd1l | DIELSPEAQSK | -880 | -890 | 0 | 0 | 11 | 1576.7855 | 390.98654 | 1053.7131 | 1812.3853 | 790.90106 | NaN |
| Mthfd1l | KITIGQGSTEK | -585 | -595 | 1 | 0 | 11 | 1241.81 | 1078.9897 | 922.8088 | 2264.0063 | 983.4273 | NaN |
| Mthfd1l | GDAPECFVSPLAK | -192 | -204 | 0 | 0 | 13 | 1240.6737 | 1121.7616 | 1269.8785 | 1309.9354 | NaN | 1543.7474 |
| Mthfd1l | LNIDPATITWQR | -563 | -574 | 0 | 0 | 12 | 657.4482 | 1018.5365 | 831.7831 | 523.8592 | NaN | 486.81616 |
| Mthfd1l | YHWSAGGK | -828 | -835 | 0 | 0.000230707 | 8 | 541.78705 | 3530.0261 | 646.8994 | NaN | 2849.421 | 2786.8403 |
| Mthfd1l | IAHGNSSVLADK | -684 | -695 | 0 | 0 | 12 | 1933.6343 | NaN | NaN | NaN | 3139.308 | 2922.4316 |
| Anxa6 | TNYDIEHVIK | -570 | -579 | 0 | 0 | 10 | 9374.085 | 9345.659 | 9711.284 | 6322.5386 | 5263.1035 | 5117.8213 |
| Anxa6 | SLHQAIEGDTSGDFMK | -648 | -663 | 0 | 0 | 16 | 7139.4277 | 7201.8765 | 7295.973 | 2938.5369 | 2540.0732 | 2611.4458 |
| Anxa6 | SLHQAIEGDTSGDFMK | -648 | -663 | 0 | 0 | 16 | 7139.4277 | 7201.8765 | 7295.973 | 2938.5369 | 2540.0732 | 2611.4458 |
| Anxa6 | SLEDALSSDTSGHFR | -484 | -498 | 0 | 0 | 15 | 6884.4087 | 8671.938 | 8438.135 | 8511.235 | 10575.701 | 10304.305 |
| Anxa6 | SLEDALSSDTSGHFR | -484 | -498 | 0 | 0 | 15 | 6884.4087 | 8671.938 | 8438.135 | 8511.235 | 10575.701 | 10304.305 |
| Anxa6 | DLESDIIGDTSGHFQK | -141 | -156 | 0 | 0 | 16 | 4194.5317 | 5121.0767 | 4622.761 | 6112.7876 | 7289.431 | 7192.3696 |
| Anxa6 | TTGKPIEASIR | -221 | -231 | 1 | 0 | 11 | 4032.6538 | 4021.3267 | 3918.255 | 3400.3843 | 3880.8843 | 3830.808 |
| Anxa6 | GFGSDKESILELITSR | -35 | -50 | 1 | 0 | 16 | 468.4568 | 607.66974 | 439.65125 | 788.81067 | 765.5914 | 708.6454 |
| Anxa6 | GFGSDKESILELITSR | -35 | -50 | 1 | 0 | 16 | 468.4568 | 607.66974 | 439.65125 | 788.81067 | 765.5914 | 708.6454 |
| Anxa6 | NKPLFFADK | -599 | -607 | 1 | 0 | 9 | 3536.416 | 560.6267 | 4114.387 | 4522.5938 | 4422.6426 | 4053.8035 |
| Anxa6 | NYDIEHVIK | -571 | -579 | 0 | 0 | 9 | 3513.191 | 3577.5715 | 4049.705 | 5274.863 | 236.41415 | 5226.8506 |
| Anxa6 | EATIIDIVTHR | -383 | -393 | 0 | 0.000431655 | 11 | 2211.5474 | 2067.2366 | 2050.9666 | 1407.6558 | 1466.4218 | 1473.4492 |
| Anxa6 | GIGTDEATIIDIVTHR | -378 | -393 | 0 | 0 | 16 | 2204.4727 | 2106.284 | 1730.0034 | 2492.7925 | 326.88867 | 2801.3584 |
| Anxa6 | GIGTDEATIIDIVTHR | -378 | -393 | 0 | 0 | 16 | 2204.4727 | 2106.284 | 1730.0034 | 2492.7925 | 326.88867 | 2801.3584 |
| Anxa6 | GSVHDFPEFDANQDAEAL | -12 | -29 | 0 | 0 | 18 | 2192.7852 | 2562.6792 | 2747.3215 | 1024.6179 | 1247.9321 | 1247.4678 |
| Anxa6 | SELDMLDIR | -282 | -290 | 0 | 0 | 9 | 2092.8674 | 2027.0645 | 2119.445 | 224.56563 | 2481.1326 | 2336.964 |
| Anxa6 | SELDMLDIR | -282 | -290 | 0 | 0 | 9 | 2092.8674 | 2027.0645 | 2119.445 | 224.56563 | 2481.1326 | 2336.964 |
| Anxa6 | DVQDLYEAGELK | -180 | -191 | 0 | 0 | 12 | 2064.5884 | 2354.931 | 2040.8 | 298.3224 | 258.0342 | 1612.7014 |
| Anxa6 | DVQDLYEAGELK | -180 | -191 | 0 | 0 | 12 | 2064.5884 | 2354.931 | 2040.8 | 298.3224 | 258.0342 | 1612.7014 |
| Anxa6 | NDFNPDADAK | -361 | -370 | 0 | 0 | 10 | 1938.6201 | 1268.0566 | 912.19745 | 1478.6104 | 436.1264 | 709.6969 |
| Anxa6 | QEICQNYK | -56 | -63 | 0 | 0 | 8 | 1621.6244 | 325.6209 | 1537.3221 | 873.86993 | 1570.843 | 1553.8396 |
| Anxa6 | KTNYDIEHVIK | -569 | -579 | 1 | 0 | 11 | 1376.9121 | 1582.5873 | 1553.0105 | 1777.0763 | 1733.205 | 1603.7869 |
| Anxa6 | KTNYDIEHVIK | -569 | -579 | 1 | 0 | 11 | 1376.9121 | 1582.5873 | 1553.0105 | 1777.0763 | 1733.205 | 1603.7869 |
| Anxa6 | AINEAYKEDYHK | -472 | -483 | 1 | 0 | 12 | 1331.4893 | 1612.9536 | 1782.3317 | 1549.5537 | 2521.9966 | 2410.532 |
| Anxa6 | AINEAYKEDYHK | -472 | -483 | 1 | 0 | 12 | 1331.4893 | 1612.9536 | 1782.3317 | 1549.5537 | 2521.9966 | 2410.532 |
| Anxa6 | SEIDLLNIR | -630 | -638 | 0 | 0 | 9 | 1287.7867 | 1991.3787 | 1554.6478 | 3735.1838 | 4391.866 | 4402.634 |
| Anxa6 | PEYFAER | -253 | -259 | 0 | 0.00111006 | 7 | 1216.053 | 2088.6194 | 2242.9224 | 2265.4695 | 2899.7961 | 2970.6416 |
| Anxa6 | GKPIEASIR | -223 | -231 | 1 | 0.00133086 | 9 | 1200.2855 | 572.3737 | 991.6195 | 1413.4504 | 859.2867 | 1604.7764 |
| Anxa6 | LVFDEYLK | -213 | -220 | 0 | 0 | 8 | 1155.4375 | 145.4054 | 1119.5007 | 450.9679 | 1124.6514 | 1147.028 |
| Anxa6 | ESILELITSR | -41 | -50 | 0 | 0 | 10 | 869.89026 | 69.64205 | 859.0793 | 940.4092 | 221.43513 | 1300.6683 |
| Anxa6 | TNEQMHQL | -123 | -130 | 0 | 0 | 8 | 775.63904 | 626.06305 | 902.63855 | 335.963 | 307.37048 | 943.5154 |
| Anxa6 | GELSGDFEKL | -232 | -241 | 1 | 0 | 10 | 739.9825 | 711.2684 | 740.2843 | 1174.3492 | 1223.7507 | 1233.9442 |
| Anxa6 | EIADTPSGDKTSLETR | -531 | -546 | 1 | 0.001291103 | 16 | 661.2511 | 550.79034 | 597.94226 | 514.6758 | 560.4082 | 953.0506 |
| Anxa6 | EIKDAISGVGTDEK | -100 | -113 | 1 | 0 | 14 | 615.92444 | 678.8336 | 676.7238 | 911.97534 | 849.7439 | 845.96686 |
| Anxa6 | AEILEIADTPSGDK | -527 | -540 | 0 | 0 | 14 | 603.73956 | 1329.6694 | 1430.3881 | 1415.2224 | 989.9516 | 1244.1199 |

| | | | | | | | | | | | | |
|---|---|---|---|---|---|---|---|---|---|---|---|---|
| Anxa6 | GSDKESILELITSR | -37 | -50 | 1 | 0.00078661 | 14 | 599.2856 | 660.0168 | 664.7438 | 1069.1492 | 1397.7666 | 1428.8916 |
| Anxa6 | GIGTDEATIIDIVTH | -378 | -392 | 0 | 0 | 15 | 544.20056 | 477.95294 | 422.14462 | 791.4001 | 898.08545 | 846.51294 |
| Anxa6 | EEGGENRDQAQEDAQVA | -510 | -526 | 1 | 0.00025015 | 17 | 532.98114 | 593.8479 | 821.0179 | 695.62646 | 799.8309 | 99.02241 |
| Anxa6 | AINEAYKE | -472 | -479 | 1 | 0.000591124 | 8 | 519.5421 | 900.00104 | 800.8797 | 830.97687 | 796.31195 | 766.3124 |
| Anxa6 | DLIEDLKYELTGK | -69 | -81 | 1 | 0 | 13 | 511.65564 | 534.7276 | 604.4435 | 679.5592 | 833.03 | 789.4546 |
| Anxa6 | EDALSSDTSGHFR | -486 | -498 | 0 | 0 | 13 | 342.19446 | 762.8781 | 698.03 | 652.4933 | 818.14667 | 647.19824 |
| Anxa6 | STPEYFAER | -251 | -259 | 0 | 0.000394361 | 9 | 289.47174 | 1091.0605 | 436.0479 | 1065.7786 | 713.7672 | 391.01025 |
| Anxa6 | IKNDTSGEYK | -305 | -314 | 1 | 0 | 10 | 146.70036 | 150.6912 | 759.4871 | 185.95796 | 134.6405 | 667.4684 |
| Anxa6 | RVFQEFIK | -561 | -568 | 1 | 0.009388464 | 8 | 1574.5962 | NaN | 1906.4335 | 404.55756 | 2094.5056 | 2508.8413 |
| Anxa6 | ATIIDIVTHR | -384 | -393 | 0 | 0.000431655 | 10 | 1359.075 | 1436.2759 | 1502.1814 | NaN | 691.99036 | 717.08704 |
| Anxa6 | DLIEDLKYELT | -69 | -79 | 1 | 0.00025015 | 11 | 1388.153 | NaN | NaN | 2307.2378 | 2723.4368 | 2579.7039 |
| Anxa6 | GELSGDFEK | -232 | -240 | 0 | 0 | 9 | 1064.6263 | 1023.7853 | 930.35046 | 1051.091 | NaN | NaN |
| Anxa6 | PLAYCDAK | -92 | -99 | 0 | 0 | 8 | 320.8735 | NaN | 1407.8505 | 545.2037 | 568.5959 | NaN |
| Anxa6 | IIDIVTHR | -386 | -393 | 0 | 0.000907206 | 8 | 286.96326 | 902.6404 | 643.723 | NaN | 228.62624 | NaN |
| Anxa6 | NKPLFFAD | -599 | -606 | 1 | 0.009060424 | 8 | 2367.638 | 1173.3372 | 1668.2594 | NaN | NaN | NaN |
| Anxa6 | DAISGVGTDEK | -103 | -113 | 0 | 0.000431655 | 11 | 605.86926 | 288.92294 | NaN | 1591.8967 | NaN | NaN |
| Anxa6 | YKDAYER | -134 | -140 | 1 | 0.000627943 | 7 | 559.9704 | NaN | 454.44724 | NaN | NaN | 631.0699 |
| Anxa6 | EIADTPSGDK | -531 | -540 | 0 | 0 | 10 | 727.0527 | NaN | NaN | 587.8474 | NaN | NaN |
| Anxa6 | GFGSDKESILELIT | -35 | -48 | 1 | 0.000431655 | 14 | 634.16565 | NaN | NaN | 651.1329 | NaN | NaN |
| Ganab | IILTAQPFR | -149 | -157 | 0 | 0 | 9 | 4743.869 | 5598.8096 | 5157.3057 | 3874.4976 | 177.35004 | 5355.33 |
| Ganab | TGDNTAEWDHLK | -616 | -627 | 0 | 0 | 12 | 2532.3176 | 1184.3168 | 1968.857 | 623.4328 | 1744.9927 | 1446.7836 |
| Ganab | VPDVLVADPPTAR | -115 | -127 | 0 | 0 | 13 | 2053.8865 | 2225.286 | 1632.3209 | 3528.9714 | 2761.5283 | 2880.8416 |
| Ganab | NPEPELLVR | -654 | -662 | 0 | 0 | 9 | 1652.2365 | 2161.3108 | 1744.1368 | 1374.1716 | 1789.4918 | 1731.2098 |
| Ganab | AEKDEPGAWEETFK | -218 | -231 | 1 | 0 | 14 | 1561.9221 | 530.99054 | 1826.412 | 1869.8376 | 2073.9302 | 2083.486 |
| Ganab | GHLETPIWIER | -881 | -891 | 0 | 0 | 11 | 1210.167 | 1187.8123 | 1374.6978 | 1521.1003 | 1784.4467 | 1797.6261 |
| Ganab | MLDYLQGSGETPQTDIR | -338 | -354 | 0 | 0 | 17 | 1025.2123 | 1074.4777 | 874.3933 | 1144.8501 | 928.9151 | 1307.2021 |
| Ganab | YFTWDPTR | -438 | -445 | 0 | 0 | 8 | 989.7973 | 267.4991 | 1038.6477 | 891.0822 | 1343.3042 | 1336.8795 |
| Ganab | RLDLLEDR | -157 | -164 | 1 | 0.00111006 | 8 | 981.4082 | 855.07043 | 849.7711 | 1952.6927 | 2199.4517 | 2133.386 |
| Ganab | LSFQHDPETSVLIL | -915 | -928 | 0 | 0.00025015 | 14 | 804.49396 | 808.565 | 671.09204 | 881.61664 | 1249.8118 | 909.9655 |
| Ganab | YRVPDVLVADPPTAR | -113 | -127 | 1 | 0 | 15 | 782.2274 | 825.61865 | 971.6627 | 822.5071 | 902.1506 | 823.66547 |
| Ganab | WTGDNTAEWDHLK | -615 | -627 | 0 | 0 | 13 | 767.1109 | 890.5535 | 928.48157 | 999.457 | 1302.9407 | 532.8972 |
| Ganab | HHGPQTLYLPVTLS | -785 | -798 | 0 | 0 | 14 | 692.5032 | 532.36786 | 371.74625 | 706.371 | 756.4421 | 865.9772 |
| Ganab | HHGPQTLYLPVTLS | -785 | -798 | 0 | 0 | 14 | 692.5032 | 532.36786 | 371.74625 | 706.371 | 756.4421 | 865.9772 |
| Ganab | LGPSVFDVFR | -368 | -377 | 0 | 0 | 10 | 616.4621 | 568.54346 | 644.11316 | 622.62537 | 806.49774 | 794.1071 |
| Ganab | SFCGADVGGFFK | -642 | -653 | 0 | 0 | 12 | 539.36237 | 473.63336 | 477.41623 | 522.0388 | 692.01495 | 605.7318 |
| Ganab | GHLETPIWIE | -881 | -890 | 0 | 0.00111006 | 10 | 514.15674 | 1334.713 | 547.8335 | 1014.6894 | 1033.8225 | 504.7446 |
| Ganab | TCDESSFCK | -40 | -48 | 0 | 0 | 9 | 444.69547 | 447.48578 | 420.3634 | 536.6455 | 568.461 | 513.21277 |
| Ganab | DYLQGSGETPQTDIR | -340 | -354 | 0 | 0 | 15 | 165.13757 | 827.35004 | 821.0324 | 1452.1422 | 1139.7699 | 1010.4381 |
| Ganab | SDWSIHLR | -937 | -944 | 0 | 0 | 8 | 132.17284 | 678.17535 | 4661.739 | 5522.966 | 5180.831 | 4111.4214 |
| Ganab | SGGIERPFVLS | -591 | -601 | 1 | 0 | 11 | 25622.18 | 28004.824 | 1069.4233 | NaN | 28923.703 | 28362.227 |
| Ganab | SIRPGLSPY | -52 | -60 | 1 | 0.002068213 | 9 | 1449.4421 | 1287.4755 | 1371.2334 | 1379.3708 | NaN | 1069.0854 |
| Ganab | LSFQHDPETSVLILR | -915 | -929 | 0 | 0 | 15 | 742.61145 | 908.7932 | 926.92346 | NaN | 1430.5283 | 1411.8558 |
| Ganab | LSFQHDPETSVLILR | -915 | -929 | 0 | 0 | 15 | 742.61145 | 908.7932 | 926.92346 | NaN | 1430.5283 | 1411.8558 |
| Ganab | SYPDFTNPR | -508 | -516 | 0 | 0.000826806 | 9 | 998.9685 | 1085.7703 | NaN | 661.90875 | 856.7716 | NaN |
| Ganab | TVHLIHEVTK | -75 | -84 | 0 | 0 | 10 | 593.09235 | NaN | 732.29504 | NaN | 146.45245 | 393.97504 |
| Ganab | TRHEFLLR | -856 | -863 | 1 | 0.00111006 | 8 | 445.93372 | 497.48877 | 292.22852 | 284.2588 | NaN | NaN |
| Ganab | IVDPHIK | -466 | -472 | 0 | 0.004321969 | 7 | 3199.9766 | 3166.4783 | NaN | NaN | 2610.5034 | NaN |
| Ganab | LSFQHDPETSVL | -915 | -926 | 0 | 0.000627943 | 12 | 892.62744 | NaN | 1007.99365 | 1004.6304 | NaN | NaN |
| Ganab | ATPGDGDKPEETQEK | -203 | -217 | 1 | 0 | 15 | 461.17737 | NaN | NaN | 309.7969 | 625.14594 | NaN |
| Ganab | VTEGGEPYR | -270 | -278 | 0 | 0.000230707 | 9 | 147.24251 | NaN | NaN | 362.37048 | NaN | 495.05075 |
| Ganab | SASYPDFTNPR | -506 | -516 | 0 | 0 | 11 | 483.3073 | NaN | NaN | 679.87396 | NaN | NaN |
| Eef2 | EDLYLKPIQR | -440 | -449 | 1 | 0 | 10 | 7107.766 | 7264.703 | 7768.905 | 7414.3027 | 158.96507 | 7382.0596 |
| Eef2 | KEDLYLKPIQR | -439 | -449 | 2 | 0 | 11 | 6829.3115 | 7435.556 | 7746.122 | 6395.1494 | 7550.343 | 7809.916 |
| Eef2 | KEDLYLKPIQR | -439 | -449 | 2 | 0 | 11 | 6829.3115 | 7435.556 | 7746.122 | 6395.1494 | 7550.343 | 7809.916 |
| Eef2 | STLTDSLVCK | -33 | -42 | 0 | 0.000431655 | 10 | 5819.232 | 5589.6484 | 5836.671 | 6096.3706 | 4916.747 | 5081.604 |
| Eef2 | QILPGDPFDNSSR | -818 | -830 | 0 | 0 | 13 | 5180.9595 | 4713.832 | 4929.234 | 3879.9817 | 3203.1611 | 1498.7224 |
| Eef2 | KYEWDVAEAR | -638 | -647 | 1 | 0.00025015 | 10 | 4766.0527 | 5328.5537 | 5483.375 | 4604.3423 | 867.53015 | 1037.2262 |
| Eef2 | ILDPIFK | -293 | -299 | 0 | 0.000471409 | 7 | 4584.1206 | 3958.8743 | 4404.5234 | 1107.2598 | 883.64197 | 692.34875 |
| Eef2 | KFSVSPVVR | -498 | -506 | 1 | 0 | 9 | 4536.148 | 4849.1206 | 4482.0454 | 4129.1562 | 4258.652 | 3940.3848 |
| Eef2 | DLEEDHACIPIKK | -560 | -572 | 1 | 0 | 13 | 4436.8223 | 4836.616 | 4744.281 | 5111.632 | 5826.033 | 5415.249 |
| Eef2 | DLEEDHACIPIKK | -560 | -572 | 1 | 0 | 13 | 4436.8223 | 4836.616 | 4744.281 | 5111.632 | 5826.033 | 5415.249 |
| Eef2 | FDVHDVTLHAD | -702 | -712 | 0 | 0.00025015 | 11 | 4257.3184 | 5170.8643 | 5184.744 | 798.0894 | 3443.7114 | 3044.1511 |
| Eef2 | FDVHDVTLHAD | -702 | -712 | 0 | 0.00025015 | 11 | 4257.3184 | 5170.8643 | 5184.744 | 798.0894 | 3443.7114 | 3044.1511 |
| Eef2 | VGVDQFLVK | -473 | -481 | 0 | 0 | 9 | 4105.41 | 5040.2095 | 5133.132 | 5219.246 | 5131.13 | 5234.2114 |
| Eef2 | ARPFPDGLAEDIDKGEVS | -606 | -623 | 2 | 0 | 18 | 3808.2095 | 4297.201 | 4503.5684 | 4470.142 | 5995.849 | 5945.8584 |
| Eef2 | ARPFPDGLAEDIDKGEVS | -606 | -623 | 2 | 0 | 18 | 3808.2095 | 4297.201 | 4503.5684 | 4470.142 | 5995.849 | 5945.8584 |
| Eef2 | VNFTVDQIR | -2 | -10 | 0 | 0 | 9 | 3407.0586 | 3578.6572 | 3590.744 | 834.2896 | 1414.6788 | 1392.7518 |
| Eef2 | YEWDVAEAR | -639 | -647 | 0 | 0 | 9 | 3376.5857 | 2824.4116 | 2748.6882 | 187.4836 | 2148.2078 | 2221.7356 |
| Eef2 | SENDLNFIK | -82 | -90 | 0 | 0 | 9 | 3272.451 | 2873.8845 | 3246.8557 | 1043.9623 | 2534.6816 | 2219.6309 |
| Eef2 | GPNYTPGK | -431 | -438 | 0 | 0 | 8 | 2772.2666 | 2413.7996 | 2799.2908 | 2367.319 | 938.1199 | 2930.36 |
| Eef2 | GLVGVDQFLVK | -471 | -481 | 0 | 0 | 11 | 2671.0063 | 3175.241 | 2979.5195 | 2545.406 | 3138.1396 | 3106.7314 |
| Eef2 | ARPFPDGLAEDIDK | -606 | -619 | 1 | 0.00025015 | 14 | 2477.1758 | 2819.1038 | 2643.3801 | 2423.5247 | 214.33635 | 2921.2878 |
| Eef2 | YEGPPDDEAAMGIK | -373 | -386 | 0 | 0 | 14 | 2249.2695 | 2457.3335 | 2222.6748 | 2254.8167 | 2714.3533 | 1511.6532 |
| Eef2 | ARPFPDGLAED | -606 | -616 | 1 | 0.000627943 | 11 | 2244.284 | 385.08783 | 1698.7025 | 1523.3945 | 916.2202 | 407.37292 |
| Eef2 | FYELSENDLNFIK | -78 | -90 | 0 | 0 | 13 | 2034.396 | 2128.5542 | 2172.7764 | 2288.528 | 3422.6157 | 3272.2334 |
| Eef2 | VFDAIMNFR | -300 | -308 | 0 | 0 | 9 | 1914.4169 | 1980.0808 | 1060.8777 | 2119.4429 | 2301.9058 | 2516.722 |
| Eef2 | VFDAIMNFR | -300 | -308 | 0 | 0 | 9 | 1914.4169 | 1980.0808 | 1060.8777 | 2119.4429 | 2301.9058 | 2516.722 |
| Eef2 | GPDGTGPNILTDITK | -653 | -667 | 0 | 0 | 15 | 1890.9376 | 1511.7412 | 2032.0542 | 1467.5741 | 2612.298 | 2549.3264 |

| | | | | | | | | | | | | |
|---|---|---|---|---|---|---|---|---|---|---|---|---|
| Eef2 | SVIAHVDHGK | -23 | -32 | 0 | 0 | 10 | 1662.8209 | 1538.0409 | 1544.9866 | 2362.7288 | 2138.6912 | 2484.622 |
| Eef2 | SVIAHVDHGK | -23 | -32 | 0 | 0 | 10 | 1662.8209 | 1538.0409 | 1544.9866 | 2362.7288 | 2138.6912 | 2484.622 |
| Eef2 | AEKYEWDVAEAR | -636 | -647 | 1 | 0 | 12 | 1658.6633 | 1642.3362 | 1850.392 | 2226.0654 | 2436.4192 | 2551.575 |
| Eef2 | KIWCFGPDGTGPN | -648 | -660 | 1 | 0 | 13 | 1563.6711 | 560.3771 | 1709.5975 | 2006.9996 | 555.62006 | 2058.5547 |
| Eef2 | IVGLVGVDQFLVK | -469 | -481 | 0 | 0 | 13 | 1530.4548 | 1554.858 | 1570.3188 | 2401.0994 | 2562.0896 | 2808.075 |
| Eef2 | DLEEDHACIPIK | -560 | -571 | 0 | 0 | 12 | 1429.8738 | 3329.335 | 3361.3303 | 4499.9067 | 4037.5752 | 3759.5686 |
| Eef2 | LILDPIFK | -292 | -299 | 0 | 0.00025015 | 8 | 1416.3154 | 1589.0635 | 1679.527 | 1441.9175 | 2013.6383 | 1944.2289 |
| Eef2 | CELLYEGPPDDEAAMGIK | -369 | -386 | 0 | 0 | 18 | 1412.2926 | 2020.5867 | 1710.497 | 148.18707 | 2558.5017 | 2305.6887 |
| Eef2 | TFCQLILDPIFK | -288 | -299 | 0 | 0 | 12 | 1396.4255 | 1350.792 | 1239.3242 | 2450.5867 | 2672.5137 | 2596.0107 |
| Eef2 | LKDLEEDHACIPIKK | -558 | -572 | 2 | 0 | 15 | 1318.3308 | 1614.5576 | 1841.64 | 1954.3446 | 2864.9944 | 2275.5315 |
| Eef2 | LKDLEEDHACIPIKK | -558 | -572 | 2 | 0 | 15 | 1318.3308 | 1614.5576 | 1841.64 | 1954.3446 | 2864.9944 | 2275.5315 |
| Eef2 | ARPFPDGLAEDIDKGEVSAR | -606 | -625 | 2 | 0.000230707 | 20 | 1240.9303 | 10226.626 | 10140.84 | 11531.882 | 11779.682 | 15481.523 |
| Eef2 | EKYEWDVAEAR | -637 | -647 | 1 | 0 | 11 | 1193.8276 | 1242.4192 | 1249.9846 | 754.31 | 1123.9689 | 483.54733 |
| Eef2 | CFGPDGTGPNILTDITK | -651 | -667 | 0 | 0 | 17 | 1161.258 | 1113.483 | 1105.4707 | 1293.5063 | 1425.5768 | 1256.7427 |
| Eef2 | YELSENDLNFIK | -79 | -90 | 0 | 0 | 12 | 1160.7268 | 1354.8995 | 1376.6267 | 1466.7068 | 2035.8401 | 2271.6025 |
| Eef2 | EGIPALDNFLDK | -846 | -857 | 0 | 0 | 12 | 1022.4459 | 1005.63165 | 1191.0204 | 1281.5135 | 1025.7753 | 2029.5068 |
| Eef2 | LKDLEEDHACIPIK | -558 | -571 | 1 | 0 | 14 | 1000.68256 | 1168.4114 | 1161.9364 | 1142.438 | 1324.0177 | 1331.4635 |
| Eef2 | KSDPVVSYR | -572 | -580 | 1 | 0 | 9 | 978.2316 | 1744.188 | 348.63364 | 1134.6415 | 1330.6814 | 411.14258 |
| Eef2 | EGALCEENMR | -689 | -698 | 0 | 0 | 10 | 940.08496 | 1031.2568 | 1022.37354 | 319.1644 | 960.8088 | 879.3662 |
| Eef2 | DIKLDSEDKDK | -320 | -330 | 2 | 0 | 11 | 917.9962 | 716.3218 | 786.9201 | 941.0958 | 849.63855 | 853.5166 |
| Eef2 | EEDHACIPIKK | -562 | -572 | 1 | 0 | 11 | 914.58356 | 697.1211 | 1003.3165 | 477.70364 | 1544.4152 | 446.61905 |
| Eef2 | LLYEGPPDDEAAMGIK | -371 | -386 | 0 | 0 | 16 | 914.53937 | 1867.4786 | 1806.4813 | 1846.3496 | 1852.5571 | 2007.6599 |
| Eef2 | LLYEGPPDDEAAMGIK | -371 | -386 | 0 | 0 | 16 | 914.53937 | 1867.4786 | 1806.4813 | 1846.3496 | 1852.5571 | 2007.6599 |
| Eef2 | EDLYLKPIQ | -440 | -448 | 1 | 0 | 9 | 886.5524 | 496.95425 | 1417.1707 | 562.0024 | 582.7861 | 748.8231 |
| Eef2 | CQLILDPIFK | -290 | -299 | 0 | 0 | 10 | 859.7908 | 877.2195 | 933.4851 | 582.24243 | 560.6317 | 1655.4374 |
| Eef2 | YVEPIEDVPCGNIV | -457 | -470 | 0 | 0 | 14 | 839.9835 | 867.58105 | 1048.8352 | 797.7264 | 2450.3184 | 972.04016 |
| Eef2 | DLYLKPIQR | -441 | -449 | 1 | 0 | 9 | 826.7653 | 291.90796 | 931.76514 | 882.08234 | 947.0848 | 882.2744 |
| Eef2 | KNPADLPK | -512 | -519 | 1 | 0.000230707 | 8 | 739.7315 | 861.7694 | 736.3832 | 403.15134 | 506.76575 | 508.29733 |
| Eef2 | LDNFLDKL | -851 | -858 | 1 | 0.00025015 | 8 | 700.084 | 502.6674 | 816.20856 | 1551.0688 | 1918.5812 | 1960.5702 |
| Eef2 | LEPEELYQTFQR | -169 | -180 | 0 | 0 | 12 | 592.1492 | 706.42377 | 750.4426 | 835.0077 | 918.17804 | 1062.1034 |
| Eef2 | IDKGEVSAR | -617 | -625 | 1 | 0.000394361 | 9 | 508.2494 | 524.8297 | 525.326 | 420.65204 | 568.70667 | 609.0688 |
| Eef2 | YVEPIEDVPCGNIVGL | -457 | -472 | 0 | 0 | 16 | 495.09055 | 424.06595 | 417.67365 | 532.9392 | 731.87213 | 661.9132 |
| Eef2 | AYLPVNESFGF | -786 | -796 | 0 | 0 | 11 | 307.35822 | 387.80576 | 311.3922 | 403.22104 | 448.56268 | 458.81488 |
| Eef2 | IWCFGPDGTGPNILTDITK | -649 | -667 | 0 | 0 | 19 | 292.13794 | 355.36307 | 269.69568 | 352.86188 | 485.4732 | 367.5513 |
| Eef2 | EESQVAGTPMFVVK | -772 | -785 | 0 | 0 | 14 | 274.72305 | 1650.9746 | 1773.0336 | 2446.7966 | 2577.942 | 2320.505 |
| Eef2 | FDVHDVTLH | -702 | -710 | 0 | 0.00051306 | 9 | 210.83614 | 1867.674 | 1867.0767 | 1205.5575 | 1574.3785 | 1593.0017 |
| Eef2 | VFSGVVSTGLK | -416 | -426 | 0 | 0 | 11 | 6814.9976 | NaN | 6584.4316 | 5689.119 | 5575.475 | 5414.5835 |
| Eef2 | FSVSPVVR | -499 | -506 | 0 | 0 | 8 | 2015.3308 | 1744.8632 | 1954.7689 | 288.9505 | 649.86127 | NaN |
| Eef2 | AKNPADLPK | -511 | -519 | 1 | 0.00025015 | 9 | 1295.8545 | 1118.4419 | 1130.8093 | 367.16473 | 725.12244 | NaN |
| Eef2 | IHLPSPVTAQK | -356 | -366 | 0 | 0 | 11 | 1142.6547 | 1231.1647 | 1216.4602 | 607.3521 | 495.8125 | NaN |
| Eef2 | QILPGDPFDNSSRPSQVVAETR | -818 | -839 | 1 | 0 | 22 | 825.8458 | 965.893 | 1042.7864 | 1508.5632 | NaN | 1325.7311 |
| Eef2 | YVEPIEDVPCGN | -457 | -468 | 0 | 0 | 12 | 789.19745 | NaN | 626.1719 | 176.47163 | 168.84511 | 206.98108 |
| Eef2 | FGPDGTGPNILTDITK | -652 | -667 | 0 | 0 | 16 | 635.9258 | NaN | 1174.7427 | 521.45935 | 435.06354 | 550.35913 |
| Eef2 | VTDGALVVVDCVSGV | -121 | -135 | 0 | 0 | 15 | 500.0116 | 869.1638 | 801.37994 | NaN | 529.792 | 764.39197 |
| Eef2 | EEDHACIPIK | -562 | -571 | 0 | 0 | 10 | 275.29187 | 2418.2495 | 2689.6614 | NaN | 2898.3027 | 639.26874 |
| Eef2 | PSQVVAETR | -831 | -839 | 0 | 0.000907206 | 9 | 1185.6013 | NaN | NaN | 1369.1002 | 1581.2966 | 1574.4232 |
| Eef2 | RWLPAGDALLQ | -342 | -352 | 1 | 0 | 11 | 1161.6057 | 960.3819 | 1115.5706 | NaN | NaN | 1391.2758 |
| Eef2 | KEDLYLKPIQ | -439 | -448 | 2 | 0.000755556 | 10 | 926.523 | NaN | NaN | 917.183 | 1008.9519 | 1153.4583 |
| Eef2 | VTDGALVVVDCVSGVC | -121 | -136 | 0 | 0.001811524 | 16 | 791.0181 | 872.6726 | 726.1486 | NaN | NaN | 935.23804 |
| Eef2 | DIDKGEVSAR | -616 | -625 | 1 | 0 | 10 | 701.33405 | 256.57837 | 303.7097 | NaN | 606.43005 | NaN |
| Eef2 | YFDPANGK | -265 | -272 | 0 | 0.00051306 | 8 | 406.4821 | 424.13 | 420.5178 | 529.5891 | NaN | NaN |
| Eef2 | ARPFPDGLAE | -606 | -615 | 1 | 0.00122419 | 10 | 2812.3035 | 373.90204 | 527.85834 | NaN | NaN | NaN |
| Eef2 | ARPFPDGLAEDID | -606 | -618 | 1 | 0.000431655 | 13 | 1723.1039 | NaN | NaN | 1534.9966 | NaN | 1686.6699 |
| Eef2 | STLTDSLVCKA | -33 | -43 | 1 | 0.000431655 | 11 | 1249.5271 | NaN | 299.1178 | 288.52005 | NaN | NaN |
| Eef2 | KIWCFGPDGTGPNIL | -648 | -662 | 1 | 0.001463897 | 15 | 1251.3849 | NaN | NaN | NaN | NaN | 2121.4866 |
| Eef2 | GVQYLNEIKDSVVA | -668 | -681 | 1 | 0.000431655 | 14 | 1032.846 | NaN | 1435.8281 | NaN | NaN | NaN |
| Eef2 | VLTAQPR | -733 | -739 | 0 | 0.000907206 | 7 | 976.575 | NaN | NaN | 1107.7426 | NaN | NaN |
| Eef2 | EWDVAEAR | -640 | -647 | 0 | 0.000712251 | 8 | 866.29083 | NaN | NaN | NaN | NaN | NaN |
| Got2 | KQWLQEVK | -338 | -345 | 1 | 0.00025015 | 8 | 5618.0884 | 7472.6763 | 6482.774 | 5516.39 | 7841.1685 | 6809.965 |
| Got2 | HFIEQGINVC | -263 | -272 | 0 | 0 | 10 | 3092.4436 | 237.98323 | 210.71709 | 1282.4246 | 383.03217 | 994.412 |
| Got2 | AHNPTGVDPRPEQWK | -213 | -227 | 1 | 0 | 15 | 2727.824 | 3055.1392 | 3340.6077 | 2709.5884 | 3456.7559 | 3355.7004 |
| Got2 | AHNPTGVDPRPEQWK | -213 | -227 | 1 | 0 | 15 | 2727.824 | 3055.1392 | 3340.6077 | 2709.5884 | 3456.7559 | 3355.7004 |
| Got2 | PSWGNHTPIFR | -160 | -170 | 0 | 0 | 11 | 2579.8174 | 2551.1455 | 2694.9365 | 3725.56 | 2719.661 | 2918.2905 |
| Got2 | PSWGNHTPIFR | -160 | -170 | 0 | 0 | 11 | 2579.8174 | 2551.1455 | 2694.9365 | 3725.56 | 2719.661 | 2918.2905 |
| Got2 | GDGDKDAWAVR | -252 | -262 | 1 | 0 | 11 | 2229.7305 | 303.31042 | 196.82637 | 1262.8448 | 1798.78 | 1725.1879 |
| Got2 | ATILTSPDLR | -328 | -337 | 0 | 0 | 10 | 2129.155 | 1790.9879 | 2306.5579 | 1559.7853 | 2069.454 | 1527.533 |
| Got2 | ASGDGDKDAWAVR | -250 | -262 | 1 | 0 | 13 | 2018.2186 | 1971.6785 | 2152.492 | 690.83673 | 1098.9534 | 581.52704 |
| Got2 | ASGDGDKDAWAVR | -250 | -262 | 1 | 0 | 13 | 2018.2186 | 1971.6785 | 2152.492 | 690.83673 | 1098.9534 | 581.52704 |
| Got2 | AELALGENNEVLK | -110 | -122 | 0 | 0 | 13 | 1627.5731 | 1890.0752 | 1964.3672 | 2914.4844 | 3130.3062 | 3217.4207 |
| Got2 | CFTGLKPEQVER | -382 | -393 | 1 | 0 | 12 | 1551.0452 | 1601.1348 | 1620.2151 | 1182.7437 | 1357.9816 | 505.06674 |
| Got2 | QWLQEVK | -339 | -345 | 0 | 0.000712251 | 7 | 1503.5461 | 1420.3154 | 1405.0955 | 1614.2971 | 1556.063 | 1279.5543 |
| Got2 | LALGENNEVLK | -112 | -122 | 0 | 0 | 11 | 1470.6829 | 1326.9731 | 1435.9773 | 1324.7988 | 859.66656 | 858.8304 |
| Got2 | FTGLKPEQVER | -383 | -393 | 1 | 0.000394361 | 11 | 1081.7252 | 919.22833 | 1120.0587 | 2245.0183 | 1188.4707 | 1170.6982 |
| Got2 | FTGLKPEQVER | -383 | -393 | 1 | 0.000394361 | 11 | 1081.7252 | 919.22833 | 1120.0587 | 2245.0183 | 1188.4707 | 1170.6982 |
| Got2 | DDNGKPYVLPSVR | -69 | -81 | 1 | 0 | 13 | 1303.7406 | 1424.9846 | 1617.6803 | 960.3074 | 2038.0356 | 1911.9526 |
| Got2 | IPEQSVLLLH | -201 | -210 | 0 | 0 | 10 | 1241.527 | 1362.031 | 1367.6317 | 1800.6326 | 2022.0762 | 1951.1377 |
| Got2 | NLDKEYLPIGGLA | -91 | -103 | 1 | 0 | 13 | 1112.2028 | 1216.6956 | 131.37454 | 1035.3906 | 1191.8115 | 987.59344 |
| Got2 | GLKPEQVER | -385 | -393 | 1 | 0.000347792 | 9 | 1051.5809 | 538.4274 | 1098.0968 | 639.45 | 947.56146 | 621.45435 |

| | | | | | | | | | | | |
|---|---|---|---|---|---|---|---|---|---|---|---|
| Got2 | NLDKEYLPIGGL | -91 | -102 | 1 | 0.00025015 | 12 | 951.49677 | 1006.1687 | 1027.1112 | 725.1398 | 395.98154 | 1047.2828 |
| Got2 | GFDFSGALEDISK | -188 | -200 | 0 | 0 | 13 | 938.619 | 950.5513 | 962.1882 | 1643.7627 | 1874.7952 | 1779.457 |
| Got2 | GFASGDGDKDAWAVR | -248 | -262 | 1 | 0 | 15 | 808.41595 | 2374.5178 | 2250.9307 | 2055.4082 | 1634.0618 | 1847.8762 |
| Got2 | TCGFDFSGALEDISK | -186 | -200 | 0 | 0 | 15 | 732.34576 | 857.413 | 832.5002 | 532.91626 | 800.04987 | 845.3981 |
| Got2 | RDDNGKPYVLPSVR | -68 | -81 | 2 | 0 | 14 | 634.73145 | 1067.7872 | 310.41623 | 882.0797 | 291.68802 | 155.39496 |
| Got2 | NLDKEYLPIGGLAEFCK | -91 | -107 | 1 | 0 | 17 | 410.79047 | 453.3446 | 356.2017 | 377.8615 | 409.39658 | 491.40015 |
| Got2 | NLDKEYLPIGGLAEFC | -91 | -106 | 1 | 0.000471409 | 16 | 361.55103 | 365.52515 | 415.0454 | 500.43427 | 607.553 | 795.9052 |
| Got2 | TGLKPEQVER | -384 | -393 | 1 | 0 | 10 | 2328.807 | 1838.313 | 2284.1812 | NaN | 2024.8855 | 2429.0767 |
| Got2 | SAELALGENNEVLK | -109 | -122 | 0 | 0 | 14 | 1004.3333 | 1112.0618 | 1052.8003 | 924.87823 | NaN | NaN |
| Got2 | RDVFLPK | -153 | -159 | 1 | 0 | 7 | 414.87274 | 534.49506 | NaN | 836.9481 | 263.63232 | NaN |
| Nudc | ELTDEEAERL | -106 | -115 | 1 | 0 | 10 | 1628.6423 | 1752.2479 | 137.60593 | 2055.1116 | 365.62976 | 285.6065 |
| Nudc | GQPPVVDGELYNEVK | -214 | -228 | 0 | 0 | 15 | 1596.495 | 1671.2279 | 1700.215 | 1092.8424 | 1146.3827 | 1025.2725 |
| Nudc | DAEDQEAQLK | -124 | -133 | 0 | 0 | 10 | 968.8715 | 771.5899 | 1506.7706 | 714.36707 | 1383.0454 | 794.56964 |
| Nudc | KDAEDQEAQLK | -123 | -133 | 1 | 0 | 11 | 725.5757 | 963.2301 | 876.5939 | 639.826 | 850.26434 | 936.39246 |
| Nudc | LSDLDSETR | -277 | -285 | 0 | 0.00025015 | 9 | 691.5061 | 1593.2133 | 1661.4845 | 1357.5817 | 967.39667 | 812.7323 |
| Nudc | VEESSWLIEDGK | -229 | -240 | 0 | 0 | 12 | 545.4363 | 323.94696 | 1665.8519 | 1314.5264 | 487.28702 | 1427.6295 |
| Nudc | LQLEIDQK | -115 | -122 | 0 | 0.00025015 | 8 | 540.8762 | 832.0737 | 965.89923 | 666.29663 | 559.38995 | 561.6844 |
| Nudc | KAETPGPQIK | -96 | -105 | 1 | 0 | 10 | 331.27478 | 197.52512 | 443.07977 | 209.9975 | 387.5458 | 226.18579 |
| Nudc | DAEDEEDEEDEKDK | -144 | -157 | 1 | 0.000627943 | 14 | 271.83615 | 299.6191 | 325.85468 | 232.19504 | 375.63004 | 377.784 |
| Nudc | ELTDEEAER | -106 | -114 | 0 | 0.000471409 | 9 | 828.9565 | 798.94086 | 743.6978 | 670.1707 | NaN | 862.0979 |
| Nudc | LKPNLGNGADLPNYR | -160 | -174 | 1 | 0 | 15 | 730.7117 | NaN | 874.06146 | 579.00446 | 770.2719 | NaN |
| Nudc | LKPNLGNGADLPNYR | -160 | -174 | 1 | 0 | 15 | 730.7117 | NaN | 874.06146 | 579.00446 | 770.2719 | NaN |
| Nudc | ESSWLIEDGK | -231 | -240 | 0 | 0 | 10 | 657.69305 | 560.4071 | 608.1453 | 548.58813 | NaN | NaN |
| Nudc | SMGLPTSDEQK | -299 | -309 | 0 | 0.001055699 | 11 | 4713.3184 | NaN | 3943.9136 | NaN | NaN | NaN |
| Nudc | GKDVVVDIQR | -196 | -205 | 1 | 0.00025015 | 10 | 558.9228 | 650.3457 | NaN | NaN | NaN | NaN |
| Nudc | AELDLAVPFR | -180 | -189 | 0 | 0.002810811 | 10 | 271.5163 | NaN | NaN | NaN | NaN | NaN |
| Iars2 | AELEYNPEHVSR | -256 | -267 | 0 | 0 | 12 | 889.31354 | 913.2904 | 971.47266 | 868.51465 | 841.9966 | NaN |
| Grap2 | DSSNPSWWTGR | -294 | -304 | 0 | 0.000230707 | 11 | 918.35474 | 1100.2573 | 1161.2482 | 817.6049 | 473.9125 | 1028.9221 |
| Grap2 | VRHEDDVQHFK | -96 | -106 | 1 | 0 | 11 | 737.679 | 605.28723 | 608.8771 | 565.7713 | 101.82012 | 692.71796 |
| Grap2 | VRHEDDVQHFK | -96 | -106 | 1 | 0 | 11 | 737.679 | 605.28723 | 608.8771 | 565.7713 | 101.82012 | 692.71796 |
| Grap2 | HEDDVQHFK | -98 | -106 | 0 | 0 | 9 | 384.80322 | 451.23642 | 459.01855 | 644.6512 | 597.31165 | 705.57196 |
| Grap2 | HEDDVQHFK | -98 | -106 | 0 | 0 | 9 | 384.80322 | 451.23642 | 459.01855 | 644.6512 | 597.31165 | 705.57196 |
| Grap2 | SQGGPHPSGTVGEEIRPSVNR | -163 | -183 | 1 | 0.000230707 | 21 | 702.62463 | 763.084 | 874.687 | 788.23376 | 886.0758 | 826.9044 |
| Grap2 | GEDELSFR | -13 | -20 | 0 | 0 | 8 | 684.9188 | 781.2523 | 710.1106 | 293.6458 | 926.363 | 819.4374 |
| Grap2 | SQGGPHPSGTVGEEIRPS | -163 | -180 | 1 | 0.000230707 | 18 | 647.0342 | 747.6096 | 671.0416 | 102.73286 | 430.65396 | 564.49036 |
| Grap2 | DIGFFIIR | -76 | -83 | 0 | 0.00025015 | 8 | 384.80753 | 367.73547 | 94.89143 | 319.9598 | 412.72153 | 375.65222 |
| Grap2 | YLQHFHQDR | -218 | -226 | 0 | 0 | 9 | 352.2365 | 1491.579 | 338.40112 | NaN | 1503.9478 | 487.89087 |
| Grap2 | KFPSLNK | -121 | -127 | 1 | 0.00111006 | 7 | 286.748 | 290.82535 | 300.53485 | 615.95013 | 164.9314 | NaN |
| Grap2 | AELGSQEGYVPK | -37 | -48 | 0 | 0 | 12 | 3124.6104 | NaN | 1206.9008 | 2582.5479 | 728.2064 | NaN |
| Grap2 | SNQEEWLK | -29 | -36 | 0 | 0.00025015 | 8 | 640.4844 | 725.33685 | NaN | NaN | NaN | 642.9465 |
| Grap2 | RHTDPVQLQ | -252 | -260 | 1 | 0.002780285 | 9 | 901.2102 | NaN | NaN | NaN | NaN | NaN |
| Eif2s2 | FPEEDEILEKD | -133 | -143 | 1 | 0 | 11 | 2928.246 | 3640.94 | 3579.8105 | 339.13147 | 4143.535 | 4020.8286 |
| Eif2s2 | RSPDTILQK | -283 | -291 | 1 | 0.00025015 | 9 | 2214.9636 | 2431.2134 | 2394.5217 | 1532.2637 | 2310.5952 | 2283.497 |
| Eif2s2 | AELGTSGSIDGNNQLVIK | -240 | -257 | 0 | 0 | 18 | 1960.9691 | 3830.0518 | 4705.8574 | 5184.2207 | 6314.261 | 5959.3525 |
| Eif2s2 | DYTYEELLNR | -171 | -180 | 0 | 0 | 10 | 1613.0295 | 388.61267 | 556.2173 | 573.4056 | 2246.7852 | 1998.2065 |
| Eif2s2 | TSFVNFTDICK | -215 | -225 | 0 | 0 | 11 | 1404.9308 | 1338.1074 | 1453.1548 | 1530.9664 | 1456.8367 | 1425.8696 |
| Eif2s2 | SIDGNNQLVIK | -247 | -257 | 0 | 0 | 11 | 860.00757 | 810.0457 | 957.8639 | 639.5148 | 652.4271 | 2072.7268 |
| Eif2s2 | DASDDLDDLNFFNQK | -65 | -79 | 0 | 0 | 15 | 842.5364 | 976.9036 | 948.7723 | 193.57063 | 795.61646 | 752.60266 |
| Eif2s2 | EVEPEPTEEKDVDADEEDSR | -43 | -62 | 1 | 0.00025015 | 20 | 610.3971 | 695.8289 | 748.9324 | 634.6947 | 1149.5663 | 1089.265 |
| Eif2s2 | DIDEAEEAIKDVK | -90 | -102 | 1 | 0 | 13 | 1537.2562 | 562.4269 | 674.6003 | NaN | 1466.1659 | 1595.7316 |
| Eif2s2 | NVKFPEEDEILEKD | -130 | -143 | 2 | 0.00051306 | 14 | 1567.9941 | NaN | 1484.1803 | NaN | 1029.9121 | 1132.9717 |
| Eif2s2 | SPDTILQK | -284 | -291 | 0 | 0 | 8 | 1122.8705 | 302.47107 | NaN | 562.6312 | 649.4909 | NaN |
| Eif2s2 | SDAQEPAEPEDDLDIMLGNK | -105 | -124 | 0 | 0 | 20 | 302.45193 | NaN | NaN | 321.08643 | 423.11517 | 349.06207 |
| Eif2s2 | IFDIDEAEEAIK | -88 | -99 | 0 | 0 | 12 | 2863.5925 | 1460.8301 | 1752.0525 | NaN | NaN | NaN |
| Eif2s2 | TGFQAVTGK | -314 | -322 | 0 | 0 | 9 | 1061.4012 | 1081.601 | 284.15375 | NaN | NaN | NaN |
| Eif2s2 | GSIDGNNQLVIK | -246 | -257 | 0 | 0 | 12 | 930.2272 | 977.34814 | NaN | NaN | 1437.6727 | NaN |
| Eif2s2 | IFDIDEAEEAIKDVK | -88 | -102 | 1 | 0 | 15 | 752.7701 | 356.58282 | 705.584 | NaN | NaN | NaN |
| Eif2s2 | KIFDIDEAEEAIK | -87 | -99 | 1 | 0.00025015 | 13 | 735.8773 | 745.60345 | 649.38824 | NaN | NaN | NaN |
| Eif2s2 | FPEEDEILEKDEAL | -133 | -146 | 1 | 0.00383302 | 14 | 595.1273 | NaN | 811.5161 | NaN | NaN | NaN |
| Eif2s2 | DIDEAEEAIK | -90 | -99 | 0 | 0.000431655 | 10 | 525.81146 | NaN | NaN | 341.5882 | NaN | NaN |
| Umps | GLQEVGLPLHR | -353 | -363 | 0 | 0 | 11 | 1002.4174 | 1024.3738 | 1056.4196 | 1113.0807 | 1075.6472 | 1261.6996 |
| Umps | RGSDVIIVGR | -442 | -451 | 1 | 0.00051306 | 10 | 876.4414 | 777.7406 | 1062.3887 | 1777.7269 | 1869.6779 | 1637.2562 |
| Umps | QEVGLPLHR | -355 | -363 | 0 | 0 | 9 | 743.0661 | 823.933 | 628.8451 | 1409.423 | 820.9885 | 724.4452 |
| Umps | VTDAIVLLDR | -146 | -155 | 0 | 0 | 10 | 560.6152 | 608.3801 | 331.5556 | 573.4057 | 444.7242 | 436.20636 |
| Umps | AELPGTHPLASK | -231 | -242 | 0 | 0 | 12 | 548.3335 | 1005.56146 | 1115.7345 | 934.30096 | 1125.365 | 1030.6764 |
| Umps | SGLSSPVYIDLR | -30 | -41 | 0 | 0 | 12 | 149.07028 | 661.1619 | 636.3828 | 560.04767 | 147.91911 | 493.27737 |
| Umps | FADIGNTVK | -315 | -323 | 0 | 0 | 9 | 653.67035 | NaN | 589.7413 | 628.86707 | 711.08405 | 648.8048 |
| Umps | RLVEGEINPGQTC | -107 | -119 | 1 | 0.00025015 | 13 | 1358.7594 | NaN | NaN | 279.72568 | 3447.1372 | NaN |
| Tmed3 | AELTFELPDNAK | -31 | -42 | 0 | 0 | 12 | 704.27454 | 1363.1521 | NaN | 2328.352 | 887.4919 | 879.68835 |
| Tmed3 | QCFHEEVEQGVK | -43 | -54 | 0 | 0.00078661 | 12 | 361.1896 | NaN | NaN | 2821.6628 | NaN | NaN |
| Nasp | SLQEQYLEAHDR | -556 | -567 | 0 | 0 | 12 | 4457.857 | 4294.404 | 4181.7373 | 3819.7432 | 3680.5266 | 3936.1506 |
| Nasp | LSLQEQYLEAHDR | -555 | -567 | 0 | 0 | 13 | 1985.6132 | 2602.3833 | 2848.9165 | 2501.7747 | 1715.6814 | 1636.8932 |
| Nasp | LSLQEQYLEAHDR | -555 | -567 | 0 | 0 | 13 | 1985.6132 | 2602.3833 | 2848.9165 | 2501.7747 | 1715.6814 | 1636.8932 |
| Nasp | NCVTDISHLVR | -693 | -703 | 0 | 0 | 11 | 1047.7493 | 1078.0867 | 1271.0137 | 972.95355 | 1758.006 | 1257.1185 |
| Nasp | KIEDAPAPSTSADK | -20 | -33 | 1 | 0 | 14 | 692.86224 | 701.42444 | 766.47015 | 622.5104 | 1056.4177 | 998.295 |
| Nasp | KYGETANECGEAF | -76 | -88 | 1 | 0.000230707 | 13 | 616.1783 | 658.90814 | 696.5527 | 696.9178 | 622.89325 | 645.4822 |
| Nasp | KLLGLGQK | -45 | -52 | 1 | 0.001392869 | 8 | 578.07623 | 501.33167 | 508.81573 | 673.8731 | 593.1867 | 579.41974 |
| Nasp | SLQENEEEEIGNLELA | -489 | -504 | 0 | 0 | 16 | 412.8765 | 461.75397 | 449.65674 | 608.8632 | 859.9758 | 769.2064 |

| | | | | | | | | | | | |
|---|---|---|---|---|---|---|---|---|---|---|---|
| Nasp | AELVSADKIEDAPAPSTSADK | -13 | -33 | 1 | 0.001291103 | 21 | 97.65843 | 365.69687 | 231.28232 | 243.41406 | 228.6814 | 322.9181 |
| Nasp | SGNVAELALK | -648 | -657 | 0 | 0 | 10 | 641.49554 | 568.6011 | 547.5244 | 470.84814 | NaN | NaN |
| Nasp | EAEGSFTEYEK | -613 | -623 | 0 | 0 | 11 | 1531.14 | 2493.2163 | 2224.7478 | NaN | NaN | NaN |
| Nasp | QEQYLEAHDR | -558 | -567 | 0 | 0 | 10 | 585.0221 | NaN | NaN | 375.02844 | 566.48346 | NaN |
| Nasp | GYNSQYDEAVAQFGK | -581 | -595 | 0 | 0.000471409 | 15 | 140.0609 | NaN | 141.09306 | NaN | NaN | NaN |
| Nasp | LLAETHYQL | -568 | -576 | 0 | 0.006568856 | 9 | 864.45105 | NaN | NaN | NaN | NaN | NaN |
| Eloc | LISSDGHEFIVK | -21 | -32 | 0 | 0.000230707 | 12 | 2962.2888 | 3147.0486 | 3416.9556 | 2881.325 | 3409.2576 | 3499.4065 |
| Eloc | LISSDGHEFIVK | -21 | -32 | 0 | 0.000230707 | 12 | 2962.2888 | 3147.0486 | 3416.9556 | 2881.325 | 3409.2576 | 3499.4065 |
| Eloc | EIPSHVLSK | -64 | -72 | 0 | 0.00025015 | 9 | 1887.3854 | 1518.078 | 1807.9641 | 1675.183 | 1318.5242 | 1667.9276 |
| Eloc | EHALTSGTIK | -34 | -43 | 0 | 0 | 10 | 1681.0116 | 1986.042 | 1830.0535 | 2410.9192 | 2194.044 | 3047.353 |
| Eloc | LSGPGQFAENETNEVNFR | -46 | -63 | 0 | 0 | 18 | 948.4871 | 971.9708 | 1135.6831 | 1418.6166 | 1918.5544 | 1800.8613 |
| Eloc | SDGHEFIVK | -24 | -32 | 0 | 0 | 9 | 804.66455 | 341.7533 | 707.8136 | 1361.8718 | 1276.8566 | 961.03796 |
| Eloc | TYGGCEGPDAMYVK | -7 | -20 | 0 | 0 | 14 | 385.11774 | 379.24197 | NaN | 440.51477 | 406.44263 | 361.4016 |
| Eloc | AENETNEVNFR | -53 | -63 | 0 | 0 | 11 | 820.9081 | 888.61316 | 721.43054 | NaN | NaN | 1030.5917 |
| Gvin1 | WINQSEYGEEQVK | -623 | -635 | 0 | 0 | 13 | 3897.3887 | 5334.1084 | 5858.388 | 5675.1343 | 8114.9243 | 4920.445 |
| Gvin1 | YIQKPEEFFR | -2152 | -2161 | 1 | 0 | 10 | 2043.4028 | 2107.3623 | 2172.907 | 1210.6606 | 870.3417 | 1419.3878 |
| Gvin1 | SLVGPEHGTEIK | -218 | -229 | 0 | 0 | 12 | 1912.8784 | 1432.9877 | 1918.7496 | 1975.261 | 1813.2897 | 1918.2961 |
| Gvin1 | HIDEIVPDIEK | -2272 | -2282 | 0 | 0 | 11 | 1672.8645 | 1272.2483 | 1665.0516 | 1042.3744 | 1196.1893 | 1341.4933 |
| Gvin1 | GFIIDEDQR | -1241 | -1249 | 0 | 0 | 9 | 1220.5826 | 642.07983 | 335.70364 | 1033.3588 | 889.70526 | 538.8319 |
| Gvin1 | TIEELLEQTTDHR | -323 | -335 | 0 | 0.000298805 | 13 | 1112.4877 | 1538.6685 | 1453.1179 | 1306.7034 | 1756.0145 | 1789.9364 |
| Gvin1 | TLGIEHLLR | -1412 | -1420 | 0 | 0 | 9 | 1070.9996 | 1051.7877 | 1043.0958 | 1009.3004 | 1075.2683 | 1123.709 |
| Gvin1 | DYSQNYFHEILR | -2001 | -2012 | 0 | 0 | 12 | 960.40076 | 1979.4493 | 1741.3866 | 716.82434 | 1030.8145 | 1143.8667 |
| Gvin1 | YNEWTWELR | -1757 | -1765 | 0 | 0 | 9 | 664.77747 | 677.1677 | 706.33093 | 654.59955 | 731.77325 | 719.0271 |
| Gvin1 | LGLDNYYPK | -872 | -880 | 0 | 0 | 9 | 654.2247 | 1308.741 | 1482.678 | 736.27997 | 461.95987 | 736.4474 |
| Gvin1 | SITPDSSTQPYWK | -2373 | -2385 | 0 | 0 | 13 | 571.63715 | 915.1113 | 909.86816 | 1043.9275 | 1194.7976 | 1090.4573 |
| Gvin1 | TLAHSPNLSDGDLVR | -169 | -183 | 0 | 0 | 15 | 514.06964 | 737.26013 | 893.0708 | 1181.3462 | 641.67957 | 169.77641 |
| Gvin1 | TLAHSPNLSDGDLVR | -169 | -183 | 0 | 0 | 15 | 514.06964 | 737.26013 | 893.0708 | 1181.3462 | 641.67957 | 169.77641 |
| Gvin1 | FGSHANQGPLQLGGIY | -352 | -367 | 0 | 0 | 16 | 379.60446 | 348.5672 | 356.46475 | 840.87463 | 928.36957 | 1175.0737 |
| Gvin1 | FKNEAPETVEEAER | -658 | -671 | 1 | 0 | 14 | 815.59143 | 1026.4258 | NaN | 539.8507 | 738.3963 | 670.4793 |
| Gvin1 | DFHLIYK | -886 | -892 | 0 | 0 | 7 | 812.9172 | 738.5139 | NaN | 704.6174 | 604.96765 | 592.88635 |
| Gvin1 | AENFFHR | -345 | -351 | 0 | 0.00025015 | 7 | 729.2488 | 616.7689 | 654.6666 | 961.5086 | 836.2598 | NaN |
| Gvin1 | TTVEITDISTEEKR | -761 | -774 | 1 | 0 | 14 | 455.89716 | 495.78177 | 618.42523 | NaN | 574.16693 | 630.3432 |
| Gvin1 | SNLLEEPLSR | -1789 | -1798 | 0 | 0.000230707 | 10 | 451.00705 | NaN | 2195.1704 | 1783.3123 | 1239.3483 | 1218.1299 |
| Gvin1 | SEPCEGDYTFTK | -2020 | -2031 | 0 | 0 | 12 | 159.2337 | 1066.3082 | 1041.9991 | 1115.675 | NaN | 1067.023 |
| Gvin1 | TANDPVNYLER | -2058 | -2068 | 0 | 0.00111006 | 11 | 323.6358 | 362.8742 | 1058.1307 | NaN | 581.40173 | NaN |
| Gvin1 | SQNWDHELATL | -1561 | -1571 | 0 | 0.004224523 | 11 | 1641.7809 | 595.2283 | 1191.7084 | NaN | NaN | NaN |
| Gvin1 | YFHEILR | -2006 | -2012 | 0 | 0.003681885 | 7 | 1133.133 | 1185.9862 | 1353.9006 | NaN | NaN | NaN |
| Arpc4 | HNKPEVEVR | -33 | -41 | 1 | 0.00025015 | 9 | 2346.0852 | 2637.2954 | 2467.4805 | 2308.0198 | 2482.8088 | 2648.9402 |
| Arpc4 | HNKPEVEVR | -33 | -41 | 1 | 0.00025015 | 9 | 2346.0852 | 2637.2954 | 2467.4805 | 2308.0198 | 2482.8088 | 2648.9402 |
| Arpc4 | ELLLQPVTISR | -45 | -55 | 0 | 0.00025015 | 11 | 1910.6636 | 2291.5261 | 2062.3796 | 2019.1677 | 1384.0895 | 1449.029 |
| Arpc4 | AENFFILR | -98 | -105 | 0 | 0 | 8 | 1471.8132 | 1909.7168 | 1817.7134 | 1820.7446 | 1495.1135 | 2246.685 |
| Arpc4 | TATLRPYLSAVR | -2 | -13 | 1 | 0 | 12 | 1248.813 | 1182.8607 | 1285.1385 | 660.1005 | 864.49677 | 830.3536 |
| Arpc4 | IVAEEFLK | -159 | -166 | 0 | 0 | 8 | 1006.54706 | 1066.454 | 190.40527 | 1513.4683 | 272.90234 | 1230.8823 |
| Arpc4 | RKPVEGYDISFL | -106 | -117 | 2 | 0.002199523 | 12 | 984.11743 | NaN | 599.83105 | 857.5519 | 767.4312 | 777.1183 |
| Arpc4 | TATLRPYLS | -2 | -10 | 1 | 0 | 9 | 817.51544 | NaN | 900.8646 | 783.7024 | 544.7085 | 798.95605 |
| Arpc4 | RKPVEGY | -106 | -112 | 2 | 0.008222027 | 7 | 808.0011 | 515.2991 | NaN | 665.4391 | 984.3476 | 961.45044 |
| Smarcc1 | EKPIDLQNFGLR | -590 | -601 | 1 | 0 | 12 | 1247.9845 | 1188.7064 | 1358.4966 | 1278.8086 | 1381.1223 | 1423.3328 |
| Smarcc1 | DVDAEIEDAPIPEKPWK | -247 | -263 | 1 | 0 | 17 | 482.7063 | 520.2967 | 518.4896 | 684.9482 | 787.40894 | 734.24335 |
| Smarcc1 | KVEHEISEGNVA | -855 | -866 | 1 | 0 | 12 | 1305.238 | 1151.5836 | NaN | 1393.6532 | 485.47247 | 812.63605 |
| Smarcc1 | HVTNPAFTK | -94 | -102 | 0 | 0 | 9 | 1143.271 | 1008.5815 | NaN | NaN | 985.4397 | 311.5275 |
| Smarcc1 | VENESDEGDKIQDR | -796 | -809 | 1 | 0 | 14 | 822.40753 | NaN | NaN | NaN | 835.31995 | 809.1306 |
| Smarcc1 | EQDSDVSEDVKPEEK | -818 | -832 | 1 | 0 | 15 | 658.7602 | 712.11975 | 848.61523 | NaN | NaN | NaN |
| Smarcc1 | AENKVENESDEGDKIQDR | -792 | -809 | 2 | 0.000394361 | 18 | 451.79187 | 570.23 | 544.3898 | NaN | NaN | NaN |
| Smarcc1 | RFDLQNPSR | -135 | -143 | 1 | 0.001868731 | 9 | 850.72107 | NaN | 1384.907 | NaN | NaN | NaN |
| Hexim1 | AENLQLLTENELHR | -331 | -344 | 0 | 0 | 14 | 6414.5815 | 6995.7744 | 7168.367 | NaN | NaN | 7483.101 |
| Kpna3 | SGILPILVK | -113 | -121 | 0 | 0 | 9 | 879.0056 | 985.0829 | 1047.2819 | 852.256 | 1035.585 | 1051.5546 |
| Kpna3 | NVPQEESLEDSDVDADFK | -50 | -67 | 0 | 0 | 18 | 350.52295 | 358.85486 | 401.56757 | 548.2639 | 205.16434 | 728.8905 |
| Kpna3 | AENPGLENHR | -2 | -11 | 0 | 0 | 10 | 1439.8419 | 1408.0519 | 1380.8774 | NaN | 801.7642 | 1347.593 |
| Kpna3 | DSQVVQVVLDGLK | -424 | -436 | 0 | 0 | 13 | 1191.6117 | 526.1631 | 1327.2954 | 800.95355 | NaN | 699.79193 |
| Kpna3 | SHFPNLLSHPK | -329 | -339 | 0 | 0 | 11 | 582.1958 | NaN | NaN | 141.78181 | 540.761 | NaN |
| Nap1l4 | SELVQEYDEPILK | -174 | -186 | 0 | 0 | 13 | 3351.7473 | 4231.2563 | 3899.2908 | 4877.965 | 5284.8115 | 5324.333 |
| Nap1l4 | AENSLSDGGPADSVEAAK | -2 | -19 | 0 | 0 | 18 | 2071.659 | 1652.028 | 1370.2157 | 1990.7637 | 2432.4182 | 2336.057 |
| Nap1l4 | LDNVSHTPSSYIE | -45 | -57 | 0 | 0.00025015 | 13 | 1467.5416 | 1426.7001 | 1332.4802 | 1057.78 | 942.666 | 198.76834 |
| Nap1l4 | SDFEIGHFFR | -315 | -324 | 0 | 0 | 10 | 1128.1084 | 1131.7314 | 1204.7267 | 1002.02014 | 1132.0422 | 1194.6434 |
| Nap1l4 | EKEAATVEELNPK | -145 | -157 | 1 | 0 | 13 | 993.0342 | 878.8576 | 1408.9967 | 2439.6711 | 2391.2266 | 2364.4084 |
| Nap1l4 | EFHFEPNDYFTNPVLTK | -207 | -223 | 0 | 0 | 17 | 475.4845 | 559.516 | 507.4116 | 602.53394 | 794.1019 | 715.9208 |
| Nap1l4 | EFHFEPNDYFTNPVLTK | -207 | -223 | 0 | 0 | 17 | 475.4845 | 559.516 | 507.4116 | 602.53394 | 794.1019 | 715.9208 |
| Nap1l4 | SFEGPEIVDCDGCTIDWK | -238 | -255 | 0 | 0.00025015 | 18 | 431.73523 | 460.75262 | 504.0904 | 609.69257 | 707.5885 | 693.01355 |
| Nap1l4 | LDNVSHTPSSYIETLPK | -45 | -61 | 0 | 0 | 17 | 634.04047 | 1077.469 | 1407.0061 | NaN | 1304.2651 | 2028.0682 |
| Nap1l4 | LDNVSHTPSSYIETLPK | -45 | -61 | 0 | 0 | 17 | 634.04047 | 1077.469 | 1407.0061 | NaN | 1304.2651 | 2028.0682 |
| Nap1l4 | HFEPNDYFTNPVLTK | -209 | -223 | 0 | 0 | 15 | 480.05673 | 569.45667 | 873.30945 | 505.90317 | NaN | 490.97293 |
| Nap1l4 | EIGHFFR | -318 | -324 | 0 | 0.000947459 | 7 | 597.5044 | NaN | NaN | NaN | NaN | NaN |
| Rrbp1 | LKGELESSDQVR | -1383 | -1394 | 1 | 0 | 12 | 914.6255 | 865.6837 | 871.29047 | 698.6703 | 1014.9669 | 995.2898 |
| Rrbp1 | HLEDIVEK | -1375 | -1382 | 0 | 0 | 8 | 526.92975 | 739.30286 | 679.1896 | 697.4862 | 665.20074 | 688.1964 |
| Rrbp1 | TEATLEAEQTR | -1492 | -1502 | 0 | 0.00025015 | 11 | 761.5737 | 745.80664 | 937.30975 | 773.90295 | NaN | 653.0602 |
| Rrbp1 | AENSQLTER | -1151 | -1159 | 0 | 0.001392869 | 9 | 685.81696 | 418.92984 | 367.28116 | NaN | NaN | 407.90305 |
| Rrbp1 | EAEETQNSLQAECDQYR | -1315 | -1331 | 0 | 0 | 17 | 186.34822 | 252.00143 | 172.08145 | 154.09192 | NaN | NaN |
| Rrbp1 | LLATEQEDAAVAK | -911 | -923 | 0 | 0.00078661 | 13 | 1721.3552 | NaN | 1812.1965 | NaN | NaN | 2277.8535 |

| | | | | | | | | | | | | |
|---|---|---|---|---|---|---|---|---|---|---|---|---|
| Rrbp1 | EHTSHLEAELEK | -1395 | -1406 | 0 | 0 | 12 | 736.4863 | NaN | 1204.2589 | 685.04395 | NaN | NaN |
| Rrbp1 | VFSEGEAQR | -864 | -872 | 0 | 0 | 9 | 428.83408 | NaN | NaN | NaN | NaN | NaN |
| Uba1 | DNPGVVTCLDEAR | -227 | -239 | 0 | 0 | 13 | 10074.817 | 8385.8545 | 8485.032 | 12791.684 | 10901.371 | 10119.409 |
| Uba1 | QFLFRPWDVTK | -516 | -526 | 1 | 0 | 11 | 356.64624 | 5074.911 | 432.8378 | 377.99973 | 555.59656 | 235.3313 |
| Uba1 | QFLFRPWDVTK | -516 | -526 | 1 | 0 | 11 | 356.64624 | 5074.911 | 432.8378 | 377.99973 | 555.59656 | 235.3313 |
| Uba1 | SLVLQRPQTW | -694 | -703 | 1 | 0 | 10 | 4596.3813 | 5008.648 | 5082.269 | 4957.0815 | 5260.991 | 5131.7803 |
| Uba1 | LAGTQPLEVLEAVQR | -679 | -693 | 0 | 0 | 15 | 3739.4692 | 4142.538 | 4172.578 | 3038.0234 | 4257.7524 | 4659.1807 |
| Uba1 | SVQVPEFTPK | -793 | -802 | 0 | 0.00025015 | 10 | 2871.3948 | 1772.2428 | 3808.8755 | 4423.3286 | 3489.8503 | 3530.8525 |
| Uba1 | IHVSDQELQSA | -807 | -817 | 0 | 0.00025015 | 11 | 2413.8335 | 2475.5212 | 2185.087 | 325.86398 | 1086.988 | 629.24054 |
| Uba1 | ATLPSPDKLPGFK | -831 | -843 | 1 | 0 | 13 | 2215.4062 | 2424.669 | 2432.4333 | 2745.106 | 3187.0317 | 3186.3196 |
| Uba1 | ATLPSPDKLPGFK | -831 | -843 | 1 | 0 | 13 | 2215.4062 | 2424.669 | 2432.4333 | 2745.106 | 3187.0317 | 3186.3196 |
| Uba1 | TNSPLEAQLR | -165 | -174 | 0 | 0 | 10 | 2004.3143 | 2162.993 | 2109.214 | 1286.5206 | 1841.236 | 2209.014 |
| Uba1 | VVQGHQQLDSY | -912 | -922 | 0 | 0 | 11 | 1874.8601 | 1536.5472 | 1514.0565 | 3356.4365 | 1243.8263 | 1140.1176 |
| Uba1 | SLDEDLIR | -377 | -384 | 0 | 0 | 8 | 1760.8328 | 1600.8972 | 1498.7216 | 1256.6866 | 1284.3569 | 909.7361 |
| Uba1 | NFPNAIEHTLQ | -636 | -646 | 0 | 0 | 11 | 1628.1118 | 1837.0306 | 327.3846 | 2815.4307 | 2853.3362 | 2718.541 |
| Uba1 | PFWSGPK | -740 | -746 | 0 | 0.00111006 | 7 | 1603.2551 | 1811.4307 | 1952.1931 | 1184.3617 | 1012.2504 | 1481.5522 |
| Uba1 | TYGLTGSQDR | -775 | -784 | 0 | 0 | 10 | 1581.2388 | 1594.0234 | 1691.2267 | 1849.6945 | 1434.7812 | 425.3632 |
| Uba1 | VVQGHQQLDSYK | -912 | -923 | 0 | 0 | 12 | 1505.4482 | 1672.8872 | 1776.804 | 1323.8494 | 1857.9064 | 1860.3628 |
| Uba1 | VVQGHQQLDSYK | -912 | -923 | 0 | 0 | 12 | 1505.4482 | 1672.8872 | 1776.804 | 1323.8494 | 1857.9064 | 1860.3628 |
| Uba1 | DTSNFSDYIR | -279 | -288 | 0 | 0 | 10 | 1450.2245 | 95.71309 | 1874.7914 | 599.43195 | 1041.7725 | 959.4664 |
| Uba1 | HGFETGDFVSF | -240 | -250 | 0 | 0 | 11 | 1345.3817 | 1410.394 | 1497.4772 | 1464.4032 | 1670.083 | 1747.5605 |
| Uba1 | HGFETGDFVSF | -240 | -250 | 0 | 0 | 11 | 1345.3817 | 1410.394 | 1497.4772 | 1464.4032 | 1670.083 | 1747.5605 |
| Uba1 | PAQLHIGFQ | -326 | -334 | 0 | 0 | 9 | 1311.7852 | 1129.5413 | 1118.0142 | 1550.7615 | 850.41095 | 998.2439 |
| Uba1 | QNSLDEDLIR | -375 | -384 | 0 | 0 | 10 | 1220.4072 | 604.9548 | 1074.3347 | 900.10376 | 94.23031 | 1551.969 |
| Uba1 | NGSEADIDESLYSR | -44 | -57 | 0 | 0 | 14 | 1164.6975 | 762.8046 | 614.80426 | 468.82578 | 770.5748 | 609.7126 |
| Uba1 | NASVDDSRLEELK | -818 | -830 | 1 | 0 | 13 | 1083.2421 | 1172.3575 | 1249.2584 | 1269.8424 | 1381.8131 | 1339.7885 |
| Uba1 | ECLPEDKEALTEEK | -430 | -443 | 1 | 0 | 14 | 1079.9974 | 1236.4872 | 1206.4193 | 1510.1687 | 1719.0823 | 1704.6946 |
| Uba1 | VFGSDFQEK | -457 | -465 | 0 | 0 | 9 | 1056.6249 | 1679.6204 | 1469.2328 | 1279.645 | 1549.188 | 1579.7169 |
| Uba1 | SICDTSNFSDYIR | -276 | -288 | 0 | 0 | 13 | 1027.9484 | 1068.6151 | 1175.6318 | 1462.3105 | 1216.2643 | 1189.754 |
| Uba1 | NFPNAIEHTLQWAR | -636 | -649 | 0 | 0 | 14 | 987.4433 | 1306.0175 | 977.7893 | 1144.5372 | 1853.5391 | 1808.1299 |
| Uba1 | KATLPSPDKLPGFK | -830 | -843 | 2 | 0 | 14 | 982.5692 | 1181.3641 | 1148.9983 | 1123.1571 | 851.87946 | 1181.7435 |
| Uba1 | KATLPSPDKLPGFK | -830 | -843 | 2 | 0 | 14 | 982.5692 | 1181.3641 | 1148.9983 | 1123.1571 | 851.87946 | 1181.7435 |
| Uba1 | CNDESGEDVEVPYVR | -1040 | -1054 | 0 | 0 | 15 | 777.17957 | 875.80176 | 844.3452 | 889.7997 | 1066.9763 | 982.50385 |
| Uba1 | PAQLHIGF | -326 | -333 | 0 | 0.001636916 | 8 | 753.84595 | 723.501 | 777.6018 | 618.7888 | 627.34314 | 649.0233 |
| Uba1 | QGHQQLDSYK | -914 | -923 | 0 | 0.000230707 | 10 | 630.0938 | 762.7211 | 716.97015 | 462.3057 | 706.87384 | 982.49915 |
| Uba1 | VGEFCHSR | -175 | -182 | 0 | 0.000230707 | 8 | 595.9205 | 618.94775 | 2280.2627 | 1827.1306 | 2396.827 | 1994.6886 |
| Uba1 | VGEFCHSR | -175 | -182 | 0 | 0.000230707 | 8 | 595.9205 | 618.94775 | 2280.2627 | 1827.1306 | 2396.827 | 1994.6886 |
| Uba1 | IYDDDFFQNLDGVAN | -559 | -573 | 0 | 0.00025015 | 15 | 555.8739 | 521.739 | 501.18982 | 760.65686 | 870.30554 | 813.17194 |
| Uba1 | ELCCNDESGEDVEVPYVR | -1037 | -1054 | 0 | 0 | 18 | 397.53647 | 468.42242 | 546.1394 | 445.58923 | 574.6703 | 515.1831 |
| Uba1 | ALPFFGFSEPLAAPR | -930 | -944 | 0 | 0 | 15 | 273.5453 | 256.43365 | 265.67365 | 420.50803 | 477.91705 | 572.3953 |
| Uba1 | QSVQVPEFTPK | -792 | -802 | 0 | 0 | 11 | 10117.558 | 4592.849 | NaN | 6140.4536 | 7482.3413 | 1677.014 |
| Uba1 | QPAENVNQYLTDSK | -658 | -671 | 0 | 0 | 14 | 1532.4556 | NaN | 1482.9572 | 1326.8518 | 1206.654 | 1199.9401 |
| Uba1 | FGFSEPLAAPR | -934 | -944 | 0 | 0 | 11 | 1134.324 | 1907.5597 | 1180.3776 | NaN | 2077.2717 | 2067.2527 |
| Uba1 | NYDISPADR | -872 | -880 | 0 | 0 | 9 | 736.8185 | 809.5249 | NaN | 977.496 | 678.38336 | 1206.9847 |
| Uba1 | YSRPAQLHIGFQ | -323 | -334 | 1 | 0.000471409 | 12 | 630.31305 | 949.9335 | 2359.156 | NaN | 1875.8481 | 997.5408 |
| Uba1 | QLLHNFPPDQLTSS | -724 | -737 | 0 | 0.000471409 | 14 | 577.50214 | 1164.5342 | 1127.2559 | NaN | 1083.8107 | 1187.2295 |
| Uba1 | GTQPLEVLEAVQR | -681 | -693 | 0 | 0 | 13 | 1891.5933 | 1740.241 | 1296.5151 | 2604.1208 | NaN | NaN |
| Uba1 | TEHKLEIT | -981 | -988 | 1 | 0.000627943 | 8 | 1273.4507 | 380.09903 | 1324.8616 | 318.73767 | NaN | NaN |
| Uba1 | NFPNAIEHTL | -636 | -645 | 0 | 0 | 10 | 1198.7638 | 1254.6401 | 1202.9395 | NaN | NaN | 860.1693 |
| Uba1 | AENYDISPADR | -870 | -880 | 0 | 0 | 11 | 1754.2551 | 1187.1528 | 456.47208 | NaN | NaN | NaN |
| Uba1 | RCPHPLTFDVN | -747 | -757 | 1 | 0 | 11 | 769.7476 | NaN | 836.642 | 1011.5279 | NaN | NaN |
| Uba1 | AVTLHDQGTTQ | -98 | -108 | 0 | 0.002910424 | 11 | 496.41415 | NaN | 919.28955 | 1403.7363 | NaN | NaN |
| Mybbp1a | SPAESCDVLGDIQTCIK | -607 | -623 | 0 | 0 | 17 | 3667.1782 | 3981.842 | 3975.1877 | 3892.1104 | 5158.903 | 2214.5996 |
| Mybbp1a | QHFSFPLDDR | -497 | -506 | 0 | 0 | 10 | 2210.5476 | 2514.9963 | 2654.4119 | 804.6295 | 2267.9922 | 1786.6206 |
| Mybbp1a | SVFGHICPHLTPR | -670 | -682 | 0 | 0 | 13 | 2108.074 | 2403.289 | 2822.7527 | 1832.2505 | 3037.416 | 2847.6445 |
| Mybbp1a | SVFGHICPHLTPR | -670 | -682 | 0 | 0 | 13 | 2108.074 | 2403.289 | 2822.7527 | 1832.2505 | 3037.416 | 2847.6445 |
| Mybbp1a | LPNVALDLLR | -262 | -271 | 0 | 0 | 10 | 2046.561 | 2179.847 | 2210.8694 | 1947.9644 | 2343.1133 | 2439.6775 |
| Mybbp1a | PQIPETK | -490 | -496 | 0 | 0 | 7 | 2032.0859 | 1808.7882 | 1627.2555 | 1011.2932 | 658.0194 | 676.911 |
| Mybbp1a | SPAPSNPTLSPSTPAK | -1244 | -1259 | 0 | 0.000826806 | 16 | 1873.1245 | 2041.8538 | 2061.7544 | 1388.8547 | 1815.4155 | 2020.0007 |
| Mybbp1a | LVQLADMLLNHNR | -543 | -555 | 0 | 0 | 13 | 1685.278 | 1453.8427 | 1789.8217 | 1610.4944 | 1108.0819 | 1175.5657 |
| Mybbp1a | VGTFLEGCQDDPKR | -354 | -367 | 1 | 0 | 14 | 1554.7892 | 1009.4659 | 1408.1217 | 2781.5303 | 1186.8589 | 1530.4509 |
| Mybbp1a | KATPQIPETK | -487 | -496 | 1 | 0 | 10 | 1441.1262 | 1514.0796 | 1590.7368 | 1678.8779 | 1678.1422 | 1809.5463 |
| Mybbp1a | VISKPQNLFK | -336 | -345 | 1 | 0.00025015 | 10 | 1140.0491 | 1159.3276 | 1239.0116 | 1004.7296 | 1013.86224 | 70.3507 |
| Mybbp1a | AEPATPAEAAQSDR | -10 | -23 | 0 | 0 | 14 | 1002.29846 | 1582.8679 | 1583.6119 | 1167.0034 | 1565.0323 | 1600.7917 |
| Mybbp1a | SQHPNHLQGQPIK | -165 | -177 | 0 | 0 | 13 | 984.53906 | 1518.4847 | 2438.9448 | 1358.3364 | 1161.5869 | 1135.2456 |
| Mybbp1a | SQHPNHLQGQPIK | -165 | -177 | 0 | 0 | 13 | 984.53906 | 1518.4847 | 2438.9448 | 1358.3364 | 1161.5869 | 1135.2456 |
| Mybbp1a | KSPAPSNPTLSPSTPAK | -1243 | -1259 | 1 | 0 | 17 | 939.56055 | 1012.67426 | 1095.107 | 1047.5195 | 1338.3419 | 1298.6818 |
| Mybbp1a | KSPAPSNPTLSPSTPAK | -1243 | -1259 | 1 | 0 | 17 | 939.56055 | 1012.67426 | 1095.107 | 1047.5195 | 1338.3419 | 1298.6818 |
| Mybbp1a | KVLEEGLLK | -285 | -293 | 1 | 0.000431655 | 9 | 905.60767 | 908.2059 | 889.7622 | 1110.7959 | 925.4423 | 922.5898 |
| Mybbp1a | QEQDLLHK | -867 | -874 | 0 | 0 | 8 | 865.92896 | 2355.5989 | 2140.1765 | 2821.0496 | 920.16693 | 1114.9651 |
| Mybbp1a | SASLESLLTK | -974 | -983 | 0 | 0 | 10 | 820.4098 | 791.37714 | 915.63385 | 882.641 | 1164.3771 | 1312.1506 |
| Mybbp1a | HSLLEHSR | -24 | -31 | 0 | 0 | 8 | 818.5405 | 866.9888 | 134.97392 | 653.51044 | 682.4972 | 735.2579 |
| Mybbp1a | RYCHEVGPC | -888 | -896 | 1 | 0.001023236 | 9 | 754.99615 | 813.8397 | 841.85724 | 562.4877 | 773.5865 | 822.77234 |
| Mybbp1a | GATPVSPIEPESK | -1275 | -1287 | 0 | 0 | 13 | 711.53424 | 541.5024 | 798.0583 | 2165.2832 | 692.547 | 683.3782 |
| Mybbp1a | FCLFHAF | -476 | -482 | 0 | 0.000627943 | 7 | 575.50366 | 696.1692 | 664.41626 | 601.94666 | 958.251 | 867.64264 |
| Mybbp1a | LDQNLASLFK | -788 | -797 | 0 | 0.000230707 | 10 | 446.47693 | 1171.2141 | 1134.0837 | 884.2608 | 985.6019 | 1048.6351 |
| Mybbp1a | GLPVMPTFWR | -382 | -391 | 0 | 0 | 10 | 426.47644 | 287.39633 | 504.24585 | 494.48636 | 653.4808 | 667.5708 |
| Mybbp1a | VGTFLEGCQDDPK | -354 | -366 | 0 | 0 | 13 | 1111.344 | 1307.5623 | 1277.0905 | NaN | 1522.1797 | 1406.0948 |

| | | | | | | | | | | | | |
|---|---|---|---|---|---|---|---|---|---|---|---|---|
| Mybbp1a | DQNLASLFK | -789 | -797 | 0 | 0.000431655 | 9 | 837.0921 | 927.3354 | 829.18866 | 1071.7439 | NaN | 947.1414 |
| Mybbp1a | SVFGHICPH | -670 | -678 | 0 | 0 | 9 | 625.2476 | 2019.1566 | 2248.0933 | NaN | 1221.0958 | 3640.1978 |
| Mybbp1a | ILSQHPNHLQGQPIK | -163 | -177 | 0 | 0.000230707 | 15 | 394.7042 | 353.04495 | NaN | 47.27999 | 587.8999 | 340.1378 |
| Mybbp1a | ILSQHPNHLQGQPIK | -163 | -177 | 0 | 0.000230707 | 15 | 394.7042 | 353.04495 | NaN | 47.27999 | 587.8999 | 340.1378 |
| Mybbp1a | ELLNTLLR | -1084 | -1091 | 0 | 0.001636916 | 8 | 78.16429 | 1064.1006 | 1045.3716 | 60.155254 | 940.98065 | NaN |
| Mybbp1a | DIPSDTQSPVSTK | -1157 | -1169 | 0 | 0 | 13 | 2561.5747 | NaN | NaN | 3311.6929 | 1999.6492 | 1817.0599 |
| Mybbp1a | VDHLHLEK | -458 | -465 | 0 | 0 | 8 | 849.72186 | NaN | 957.82764 | 1436.9352 | 1136.3109 | NaN |
| Mybbp1a | EHKLPNVALDLLR | -259 | -271 | 1 | 0 | 13 | 612.9258 | NaN | NaN | 639.0474 | 652.501 | 739.0221 |
| Mybbp1a | EDIPLCDILDQIQEK | -102 | -116 | 0 | 0.00051306 | 15 | 320.5861 | 296.32666 | NaN | NaN | 232.98819 | 201.37767 |
| Mybbp1a | VLEEGLLK | -286 | -293 | 0 | 0.00025015 | 8 | 2620.795 | 1539.498 | 1565.0067 | NaN | NaN | NaN |
| Mybbp1a | FWDIAKPDQETR | -37 | -48 | 1 | 0 | 12 | 1026.0212 | 1371.7826 | 1409.2302 | NaN | NaN | NaN |
| Mybbp1a | HFSFPLDDR | -498 | -506 | 0 | 0.00025015 | 9 | 858.62 | NaN | 828.9212 | 735.60516 | NaN | NaN |
| Mybbp1a | YCHEVGPCAEAL | -889 | -900 | 0 | 0.000230707 | 12 | 521.85034 | NaN | NaN | 643.1394 | 653.08936 | NaN |
| Mybbp1a | ALDQNLASLFK | -787 | -797 | 0 | 0.000591124 | 11 | 314.51022 | 331.2061 | 356.6797 | NaN | NaN | NaN |
| Mybbp1a | SEDNIPSLVNILK | -238 | -250 | 0 | 0.000431655 | 13 | 143.92065 | NaN | 872.3671 | 548.93994 | NaN | NaN |
| Mybbp1a | WDIAKPDQETR | -38 | -48 | 1 | 0.00149795 | 11 | 1026.356 | NaN | NaN | 621.11993 | NaN | NaN |
| Mybbp1a | ICPHLTPR | -675 | -682 | 0 | 0 | 8 | 446.10394 | NaN | NaN | NaN | NaN | 1259.271 |
| Fam107b | YIEDDNPELIRPQK | -6 | -19 | 1 | 0 | 14 | 3483.6309 | 3904.569 | 3830.2493 | 4384.5225 | 5096.397 | 5001.714 |
| Fam107b | LQEEQENAPEFVK | -101 | -113 | 0 | 0 | 13 | 2037.1782 | 2233.9253 | 2367.3262 | 2321.388 | 2409.69 | 2511.8765 |
| Fam107b | AEPDYIEDDNPELIRPQK | -2 | -19 | 1 | 0 | 18 | 1145.4889 | 1419.3477 | 1416.1482 | 1416.8977 | 2075.779 | 2092.0188 |
| Fam107b | KSDLEIELLK | -77 | -86 | 1 | 0 | 10 | 853.4531 | 541.76025 | 1001.6992 | 754.93353 | 845.5479 | 845.87744 |
| Fam107b | GLAPQNKPELQK | -44 | -55 | 1 | 0 | 12 | 774.0723 | 937.0976 | 893.60547 | 402.4818 | 531.441 | 755.7184 |
| Fam107b | QKLQEEQENAPEFVK | -99 | -113 | 1 | 0 | 15 | 490.0156 | 627.3556 | 757.10284 | 732.3755 | 493.75595 | 505.05817 |
| Fam107b | KLQEEQENAPEFVK | -100 | -113 | 1 | 0.000471409 | 14 | 420.0465 | 639.1665 | NaN | 854.766 | 957.98346 | 418.43488 |
| Actr3 | NTEPQFIIPSCIAIK | -24 | -38 | 0 | 0 | 15 | 6444.1074 | 6573.2246 | 7343.6597 | 464.2228 | 8847.766 | 8445.691 |
| Actr3 | YSYVCPDLVK | -231 | -240 | 0 | 0 | 10 | 2498.9963 | 2653.9736 | 2731.5994 | 3402.111 | 3425.2285 | 3394.9524 |
| Actr3 | YSYVCPDLVK | -231 | -240 | 0 | 0 | 10 | 2498.9963 | 2653.9736 | 2731.5994 | 3402.111 | 3425.2285 | 3394.9524 |
| Actr3 | LPACVVDCGTGYTK | -5 | -18 | 0 | 0 | 14 | 2292.4946 | 2631.5806 | 2877.8115 | 2853.4526 | 3629.8857 | 3730.409 |
| Actr3 | FMEQVIFK | -92 | -99 | 0 | 0 | 8 | 1438.8584 | 1578.3373 | 1118.6366 | 1466.7705 | 1766.4346 | 804.61896 |
| Actr3 | FMEQVIFK | -92 | -99 | 0 | 0 | 8 | 1438.8584 | 1578.3373 | 1118.6366 | 1466.7705 | 1766.4346 | 804.61896 |
| Actr3 | DYEEIGPSICR | -399 | -409 | 0 | 0 | 11 | 973.34955 | 1052.4767 | 1043.5352 | 629.2435 | 864.921 | 775.8076 |
| Actr3 | DREVGIPPEQSLETA | -210 | -224 | 1 | 0.000471409 | 15 | 860.5331 | 837.2633 | 808.40137 | 1382.776 | 931.71924 | 1317.0747 |
| Actr3 | KDYEEIGPSICR | -398 | -409 | 1 | 0 | 12 | 860.0302 | 835.29114 | 849.42224 | 895.5971 | 1020.901 | 964.1784 |
| Actr3 | EFSIDVGYER | -266 | -275 | 0 | 0 | 10 | 784.44366 | 808.1159 | 743.4778 | 795.377 | 854.89545 | 358.60544 |
| Actr3 | TLTGTVIDSGDGVTH | -162 | -176 | 0 | 0 | 15 | 604.40625 | 403.09238 | 805.47076 | 704.40295 | 1336.5117 | 1756.1876 |
| Actr3 | TLTGTVIDSGDGVTHVIPVAEGY | -162 | -184 | 0 | 0.00025015 | 23 | 254.03017 | 364.42337 | 249.27591 | 421.48865 | 464.92377 | 422.79694 |
| Actr3 | LPACVVDCGTGY | -5 | -16 | 0 | 0.00025015 | 12 | 163.42383 | 205.64627 | 202.32297 | 163.57286 | 215.70787 | 198.92094 |
| Actr3 | LKPKPIDVQ | -358 | -366 | 2 | 0.002265552 | 9 | 1282.5343 | 1123.722 | 1046.9872 | 1245.6606 | 608.1345 | NaN |
| Actr3 | AIEKPTYATK | -66 | -75 | 1 | 0 | 10 | 2565.913 | 3250.4895 | 3347.3303 | NaN | NaN | NaN |
| Actr3 | LLTEPPLNTPENR | -111 | -123 | 0 | 0 | 13 | 2450.3027 | 2639.7236 | 2758.4185 | NaN | NaN | NaN |
| Actr3 | AEPEDHYFLL | -103 | -112 | 0 | 0.002401448 | 10 | 1947.0582 | 2135.712 | 269.43997 | NaN | NaN | NaN |
| Actr3 | EAIEKPTYATK | -65 | -75 | 1 | 0.00025015 | 11 | 799.6599 | 790.02655 | 751.3559 | NaN | NaN | NaN |
| Actr3 | QYTGVNAISK | -255 | -264 | 0 | 0 | 10 | 1972.5576 | 504.57318 | NaN | NaN | NaN | NaN |
| Actr3 | TLTGTVIDSGDGVTHVIPVAE | -162 | -182 | 0 | 0.001023236 | 21 | 172.75555 | 237.92213 | NaN | NaN | NaN | NaN |
| Actr3 | NCPIDVR | -306 | -312 | 0 | 0.002199523 | 7 | 816.95844 | NaN | NaN | NaN | NaN | NaN |
| Hyou1 | YGDLGFLGPEDLR | -497 | -509 | 0 | 0 | 13 | 2835.7578 | 3144.3423 | 3030.7363 | 3944.481 | 4203.4126 | 1584.3334 |
| Hyou1 | SRFPEHELIVDPQR | -118 | -131 | 1 | 0 | 14 | 2122.5583 | 1962.5728 | 1328.2345 | 2024.6511 | 3009.7036 | 3122.2668 |
| Hyou1 | LPATEKPVLLSK | -878 | -889 | 1 | 0 | 12 | 1594.0724 | 1689.5432 | 1785.4077 | 6520.425 | 2496.412 | 2907.9294 |
| Hyou1 | VEDSPEEESTLTK | -564 | -576 | 0 | 0 | 13 | 981.36633 | 971.8657 | 940.26245 | 441.38895 | 938.8444 | 951.274 |
| Hyou1 | FPEHELIVDPQR | -120 | -131 | 0 | 0 | 12 | 978.8528 | 136.9554 | 553.03455 | 663.23944 | 927.1447 | 599.2124 |
| Hyou1 | SLAEDFAEQPIK | -159 | -170 | 0 | 0 | 12 | 813.64124 | 933.078 | 836.0075 | 623.2409 | 678.3599 | 311.5897 |
| Hyou1 | LYQPEYQEVSTEEQR | -754 | -768 | 0 | 0 | 15 | 779.4297 | 859.548 | 1151.9463 | 919.91 | 888.4778 | 865.63605 |
| Hyou1 | DLPDLPEDELAR | -707 | -718 | 0 | 0 | 12 | 705.19965 | 528.5735 | 789.8597 | 757.54486 | 666.2458 | 818.3899 |
| Hyou1 | AVLDLPDLPEDELAR | -704 | -718 | 0 | 0 | 15 | 686.0898 | 703.4226 | 589.7645 | 596.0059 | 846.7373 | 622.88055 |
| Hyou1 | LPATEKPVLL | -878 | -887 | 1 | 0 | 10 | 599.20917 | 481.08557 | 613.9525 | 472.4764 | 642.1497 | 614.8529 |
| Hyou1 | GFLGPEDLR | -501 | -509 | 0 | 0 | 9 | 565.923 | 689.3595 | 401.97495 | 358.23224 | 903.6063 | 259.00397 |
| Hyou1 | DAVITVPAFFNQAER | -171 | -185 | 0 | 0 | 15 | 377.30035 | 428.92087 | 392.399 | 483.1479 | 604.5493 | 709.9949 |
| Hyou1 | LAVLDLPDLPEDELAR | -703 | -718 | 0 | 0 | 16 | 276.58176 | 348.2018 | 309.88605 | 345.2305 | 343.85892 | 352.58554 |
| Hyou1 | ESGDKPEAQKPNEK | -656 | -669 | 2 | 0.000591124 | 14 | 553.4319 | 558.6315 | 731.17395 | NaN | 335.04623 | 385.02374 |
| Hyou1 | ESGDKPEAQKPNEK | -656 | -669 | 2 | 0.000591124 | 14 | 553.4319 | 558.6315 | 731.17395 | NaN | 335.04623 | 385.02374 |
| Hyou1 | LVEDSPEEESTLTK | -563 | -576 | 0 | 0 | 14 | 623.2997 | 1246.8297 | 714.1212 | NaN | NaN | 623.0176 |
| Hyou1 | SLDQIEQVILVGGATR | -373 | -388 | 0 | 0.00025015 | 16 | 352.98224 | 282.7165 | 427.794 | NaN | NaN | NaN |
| Hyou1 | AEPPAEETSQPPPSEPK | -626 | -642 | 0 | 0.001023236 | 17 | 268.89673 | 257.27954 | 480.75485 | NaN | NaN | NaN |
| Hyou1 | EVEEEPGLR | -452 | -460 | 0 | 0 | 9 | 588.2774 | 648.09454 | NaN | NaN | NaN | NaN |
| Hyou1 | AEPPLNASAGDQEEK | -926 | -940 | 0 | 0.00897527 | 15 | 1608.3511 | NaN | NaN | NaN | NaN | NaN |
| Tap1 | VEFQDVSFAYPNQPK | -479 | -493 | 0 | 0 | 15 | 6880.8066 | 6912.9385 | 6637.089 | 4906.025 | 5690.7866 | 6106.002 |
| Tap1 | ITDWILQDK | -187 | -195 | 0 | 0 | 9 | 1965.6652 | 1399.6875 | 1615.5007 | 1996.7444 | 1634.9128 | 2103.4175 |
| Tap1 | VAAVGQEPLLFGR | -557 | -569 | 0 | 0 | 13 | 1465.448 | 1757.0328 | 1562.9983 | 1720.4805 | 1829.9182 | 1412.55 |
| Tap1 | GYDTEVGETGNQLSGGQR | -607 | -624 | 0 | 0 | 18 | 1247.8496 | 1529.0624 | 1475.4415 | 1667.2207 | 1707.1235 | 1559.1665 |
| Tap1 | LGSLWAPSGNR | -133 | -143 | 0 | 0 | 11 | 1118.6332 | 1353.8578 | 1300.9407 | 1238.2218 | 2868.3445 | 2925.74 |
| Tap1 | AVGQEPLLFGR | -559 | -569 | 0 | 0 | 11 | 983.86 | 1056.5552 | 1095.4613 | 1918.0387 | 625.7367 | 819.71027 |
| Tap1 | SLLVPEVPLLR | -29 | -39 | 0 | 0 | 11 | 975.3957 | 985.5625 | 995.23627 | 876.57556 | 1094.9517 | 1131.6709 |
| Tap1 | MLGFLGPK | -152 | -159 | 0 | 0.00051306 | 8 | 689.53876 | 629.5475 | 539.7648 | 528.7709 | 356.05463 | 527.161 |
| Tap1 | GLVEFQDVSFAYPNQPK | -477 | -493 | 0 | 0 | 17 | 651.35785 | 623.7693 | 775.66455 | 884.6497 | 1060.2614 | 1167.2075 |
| Tap1 | TPCSPLSGSLAPSNMK | -461 | -476 | 0 | 0 | 16 | 616.9134 | 596.3814 | 523.278 | 623.7093 | 648.8096 | 712.94727 |
| Tap1 | DDATSALDAGNQLR | -643 | -656 | 0 | 0 | 14 | 563.7733 | 749.0692 | 1243.2561 | 879.0481 | 721.253 | 1131.0239 |
| Tap1 | RLLYESPK | -659 | -666 | 1 | 0 | 8 | 457.74426 | 365.48798 | 487.24683 | 729.3078 | 539.79425 | 866.5627 |
| Tap1 | SLALPGLALFR | -80 | -90 | 0 | 0 | 11 | 330.9512 | 332.64084 | 266.1582 | 300.2084 | 367.32797 | 412.74622 |

| | | | | | | | | | | | | |
|---|---|---|---|---|---|---|---|---|---|---|---|---|
| Tap1 | ALPGLALFR | -82 | -90 | 0 | 0.00266297 | 9 | 443.1664 | NaN | 568.2792 | 400.1933 | 456.0206 | 412.52585 |
| Tap1 | FQDVSFAYPNQPK | -481 | -493 | 0 | 0 | 13 | 1582.1584 | NaN | 1276.0974 | 1061.8721 | NaN | 1333.4014 |
| Tap1 | AEQAHHILFLR | -682 | -692 | 0 | 0.00025015 | 11 | 514.7036 | NaN | NaN | NaN | NaN | 408.92282 |
| Tap1 | EVGETGNQLSGGQR | -611 | -624 | 0 | 0.000230707 | 14 | 939.90765 | NaN | NaN | NaN | NaN | NaN |
| Tap1 | LDAGNQLR | -649 | -656 | 0 | 0.004199202 | 8 | 732.90076 | NaN | NaN | NaN | NaN | NaN |
| Csk | AEQDLPFCK | -24 | -32 | 0 | 0 | 9 | 1154.9708 | 1057.2258 | 1123.7672 | 1250.9542 | 1225.9016 | 981.37256 |
| Csk | LLYPPETGLFLVR | -95 | -107 | 0 | 0 | 13 | 622.56506 | 819.29706 | 925.9201 | 717.11633 | 945.8162 | 744.0321 |
| Csk | IPLKDVVPR | -390 | -398 | 1 | 0.000230707 | 9 | 383.82782 | 179.13036 | 207.00612 | 733.99603 | 504.2839 | 292.94644 |
| Csk | LYPPETGLFLVR | -96 | -107 | 0 | 0 | 12 | 665.15 | NaN | 622.29535 | 1164.2562 | 1426.258 | 1132.2339 |
| Csk | GSLVDYLR | -272 | -279 | 0 | 0.002296977 | 8 | 738.3784 | 720.14923 | 763.07245 | 576.5527 | NaN | NaN |
| Csk | EQLEHIK | -438 | -444 | 0 | 0.00635324 | 7 | 2722.494 | NaN | NaN | 2483.688 | 538.1826 | NaN |
| Csk | NVLVSEDNVAK | -319 | -329 | 0 | 0.000230707 | 11 | 2489.0898 | 2700.0137 | 2397.2534 | NaN | NaN | NaN |
| Csk | TTDADGLCTR | -157 | -166 | 0 | 0.000550307 | 10 | 628.3304 | NaN | NaN | NaN | NaN | NaN |
| Luc7l | AEQLGAEGNVDESQK | -140 | -154 | 0 | 0 | 15 | 1002.0352 | 886.4833 | 701.28656 | NaN | 1282.0201 | 909.12494 |
| Luc7l | LHLGFIQ | -214 | -220 | 0 | 0.004350979 | 7 | 386.50867 | NaN | 429.17712 | NaN | NaN | NaN |
| Ubqln1 | AESAESGGPPGAQDSAADGGPAEPK | -2 | -26 | 0 | 0.000431655 | 25 | 180.0219 | 248.84389 | 307.2029 | 250.85825 | 208.50693 | 475.47256 |
| Ubqln1 | EKEEFAVPENSSVQ | -37 | -50 | 1 | 0.000230707 | 14 | 19853.01 | 21963.846 | 22701.146 | 1410.0647 | 1776.0006 | NaN |
| Ubqln1 | ESGGPPGAQDSAADGGPAEPK | -6 | -26 | 0 | 0 | 21 | 495.889 | 460.86813 | NaN | 671.3778 | 449.3456 | NaN |
| Ostf1 | QAESIDNPLHEAAK | -69 | -82 | 0 | 0 | 14 | 1358.7501 | 1783.6504 | 1786.2089 | 2048.7048 | 2287.0205 | 2374.943 |
| Ostf1 | FTQPNVELNQQNK | -126 | -138 | 0 | 0 | 13 | 1245.0776 | 1392.0293 | 1436.9635 | 1488.3339 | 1961.8895 | 1642.2638 |
| Ostf1 | AESIDNPLHEAAK | -70 | -82 | 0 | 0 | 13 | 1093.4829 | 1001.67163 | 1075.686 | 975.45764 | 999.73474 | 964.0451 |
| Ostf1 | VAEQAESIDNPLHEAAK | -66 | -82 | 0 | 0 | 17 | 458.1942 | 481.0393 | 477.4145 | 337.36487 | 546.7351 | 267.49078 |
| Smc1a | EEDTQFNYHR | -179 | -188 | 0 | 0 | 10 | 3062.7095 | 3070.2761 | 3134.7788 | 3945.5798 | 4109.379 | 3999.2126 |
| Smc1a | SGELAQEYDKR | -161 | -171 | 1 | 0 | 11 | 1565.0695 | 837.6234 | 1566.5039 | 858.25073 | 1536.0758 | 1851.2053 |
| Smc1a | DYLEVKPTDEK | -574 | -584 | 1 | 0 | 11 | 1197.7809 | 2538.314 | 2433.5754 | 1014.6698 | 1547.6831 | 1491.2958 |
| Smc1a | SKLESELANFGPR | -735 | -747 | 1 | 0 | 13 | 1016.55853 | 1152.9741 | 929.13354 | 1012.8003 | 1155.9333 | 1126.6486 |
| Smc1a | VVQLHEYSEELEK | -111 | -123 | 0 | 0 | 13 | 505.04785 | 647.0821 | 773.2225 | 1164.9122 | 1130.7838 | 651.3135 |
| Smc1a | AESLIGVYPEQGDCVISK | -1197 | -1214 | 0 | 0 | 18 | 332.06427 | 406.66888 | 463.4461 | 662.37585 | 120.76796 | 156.05428 |
| Smc1a | DKFQETSDEFEAA | -1036 | -1048 | 1 | 0.001291103 | 13 | 7822.3706 | 2754.7131 | NaN | 9643.811 | 4778.8716 | 1913.1156 |
| Smc1a | DLIHGAPVGK | -63 | -72 | 0 | 0 | 10 | 956.1778 | 746.8298 | 744.5647 | NaN | 364.05844 | 641.72266 |
| Smc1a | LESELANFGPR | -737 | -747 | 0 | 0 | 11 | 1485.6158 | 548.54315 | 3225.9387 | NaN | NaN | 693.1367 |
| Smc1a | LKDEVVR | -212 | -218 | 1 | 0 | 7 | 8203.315 | 7829.627 | NaN | NaN | 7359.5483 | NaN |
| Smc1a | LNEQQSVLQR | -1010 | -1019 | 0 | 0 | 10 | 2866.2305 | 2956.2776 | NaN | 3451.5537 | NaN | NaN |
| Smc1a | EDTQFNYHR | -180 | -188 | 0 | 0 | 9 | 883.85535 | NaN | 816.06226 | NaN | 1897.4138 | NaN |
| Smc1a | LYHNEVEIEK | -228 | -237 | 0 | 0.00051306 | 10 | 554.09015 | NaN | 2286.6282 | NaN | NaN | NaN |
| Smc1a | RDLIHGAPVGK | -62 | -72 | 1 | 0 | 11 | 444.61447 | 476.5404 | NaN | NaN | NaN | NaN |
| Dis3 | IIFSEHLPLSK | -226 | -236 | 0 | 0 | 11 | 1289.8114 | 1490.1377 | 1338.334 | 2395.2515 | 1364.2549 | 1196.4268 |
| Dis3 | LAYDDEIPSLR | -885 | -895 | 0 | 0 | 11 | 1208.6136 | NaN | 1381.6598 | 1715.292 | 1756.7899 | 1316.3622 |
| Dis3 | SVDPPGCTDIDDALHCR | -477 | -493 | 0 | 0 | 17 | 371.3319 | 2495.1272 | 2646.5767 | NaN | 652.77515 | 619.32556 |
| Dis3 | AESPDFPYLNTLLR | -728 | -741 | 0 | 0 | 14 | 318.5285 | 243.03323 | NaN | 295.44992 | 359.5516 | 347.0821 |
| Dis3 | LHQIDVLEHPAIR | -74 | -86 | 0 | 0 | 13 | 462.91156 | NaN | NaN | 515.687 | 466.65073 | 319.6048 |
| Dis3 | YADIIVHR | -782 | -789 | 0 | 0.00025015 | 8 | 2041.7683 | NaN | 1702.0347 | 1209.5859 | NaN | NaN |
| Dis3 | NAIVVLIPK | -858 | -866 | 0 | 0.002099344 | 9 | 954.747 | 1056.6271 | NaN | NaN | NaN | NaN |
| Dis3 | SETHDPIDLQTK | -643 | -654 | 0 | 0.00111006 | 12 | 998.2596 | NaN | NaN | NaN | NaN | NaN |
| Cds2 | TAPLPTSVDDTPEVLNR | -40 | -56 | 0 | 0 | 17 | 451.54733 | 799.2717 | 842.55664 | 562.5975 | 700.7611 | 600.9746 |
| Cds2 | AETAPLPTSVDDTPEVLNR | -38 | -56 | 0 | 0 | 19 | 349.9497 | 216.49487 | 465.9854 | 189.80872 | 234.10193 | 207.82832 |
| Cds2 | TVDCEPSDLFR | -300 | -310 | 0 | 0 | 11 | 595.2302 | NaN | NaN | 670.82367 | 213.36737 | NaN |
| Cds2 | SVDDTPEVLNR | -46 | -56 | 0 | 0 | 11 | 1442.1796 | NaN | NaN | NaN | NaN | NaN |
| F5 | DIHSGLIGPL | -1699 | -1708 | 0 | 0.00025015 | 10 | 1368.4043 | 1598.4525 | 1588.0895 | 1035.8365 | 1206.1534 | 1273.0736 |
| F5 | IYNLPGLR | -1771 | -1778 | 0 | 0 | 8 | 1190.1351 | 1298.2944 | 1285.6624 | 1207.7969 | 392.37714 | 1182.0768 |
| F5 | GVWPLLPGSFK | -1819 | -1829 | 0 | 0 | 11 | 738.6697 | 948.9721 | 939.0084 | 426.1588 | 423.87854 | 294.31348 |
| F5 | SGPENPGSACR | -1676 | -1686 | 0 | 0 | 11 | 302.8989 | 269.5065 | 259.89914 | 940.911 | 159.43822 | 624.41205 |
| F5 | AEVDDVIQVR | -1610 | -1619 | 0 | 0 | 10 | 998.1655 | 1073.9246 | 949.64185 | 1307.551 | 1103.0865 | NaN |
| F5 | DIHVVHFHGQ | -1797 | -1806 | 0 | 0.000627943 | 10 | 732.46014 | 1262.5073 | NaN | 508.72034 | 461.3554 | 639.6437 |
| F5 | NFFNPPIISR | -2149 | -2158 | 0 | 0 | 10 | 606.23425 | 1106.5826 | 1147.4768 | NaN | NaN | NaN |
| Bzw1 | GFEDEVKK | -127 | -134 | 1 | 0 | 8 | 6537.772 | 5080.077 | 4374.0312 | 4114.238 | 494.97076 | 5276.1763 |
| Bzw1 | AEVLSEEPILK | -369 | -379 | 0 | 0 | 11 | 1189.8241 | 1736.1436 | 2470.6343 | 3367.5703 | 4396.4697 | 4396.017 |
| Bzw1 | TETGTDLEAVAK | -41 | -52 | 0 | 0 | 12 | 1097.0547 | 701.9813 | 1437.8689 | 1166.0476 | 851.8824 | 1344.6533 |
| Bzw1 | IQEYCYDNIHFMK | -345 | -357 | 0 | 0.00025015 | 13 | 1081.4843 | 1104.9111 | 964.0908 | 1617.7051 | 1327.2262 | 561.5867 |
| Bzw1 | TTQGQSELTLLLK | -332 | -344 | 0 | 0 | 13 | 871.5597 | 1578.7632 | 752.71436 | 790.0773 | 936.37726 | 1141.3867 |
| Bzw1 | GQSELTLLLK | -335 | -344 | 0 | 0 | 10 | 536.35895 | 598.7357 | 651.26154 | 573.1527 | 574.3977 | 560.35223 |
| Bzw1 | ILNSLYNENLVK | -167 | -178 | 0 | 0 | 12 | 404.90805 | 481.91675 | 92.43902 | NaN | 630.9893 | 599.0829 |
| Bzw1 | NSLYNENLVK | -169 | -178 | 0 | 0 | 10 | 1096.4274 | 948.0477 | NaN | NaN | NaN | NaN |
| Rpn1 | VLALEPELESR | -71 | -81 | 0 | 0 | 11 | 3000.4758 | 2197.9824 | 1928.6473 | 3762.129 | 1147.8846 | 3848.6157 |
| Rpn1 | NLVEQHIQDIVVH | -416 | -428 | 0 | 0 | 13 | 2457.3035 | 2466.7864 | 2385.7751 | 2022.2617 | 1901.5631 | 1545.898 |
| Rpn1 | LPVALDPGSK | -118 | -127 | 0 | 0 | 10 | 2417.9604 | 2480.331 | 2573.714 | 2316.2507 | 2767.857 | 2731.016 |
| Rpn1 | DIPAYSQDTFK | -206 | -216 | 0 | 0 | 11 | 2010.5535 | 1920.1204 | 2064.0469 | 1872.054 | 1596.0226 | 1664.8248 |
| Rpn1 | RHFDETVNR | -495 | -503 | 1 | 0.000230707 | 9 | 1983.3346 | 1681.3668 | 1214.0999 | 527.0308 | 1248.8873 | 1602.6313 |
| Rpn1 | RHFDETVNR | -495 | -503 | 1 | 0.000230707 | 9 | 1983.3346 | 1681.3668 | 1214.0999 | 527.0308 | 1248.8873 | 1602.6313 |
| Rpn1 | TVDLSSHLAK | -40 | -49 | 0 | 0 | 10 | 1902.9503 | 1868.5417 | 1822.3474 | 2627.7285 | 2462.8867 | 2593.145 |
| Rpn1 | SEDVLDYGPFK | -195 | -205 | 0 | 0 | 11 | 1568.7013 | 1819.038 | 1760.2937 | 1854.629 | 2087.0085 | 2208.2446 |
| Rpn1 | SEDVLDYGPFK | -195 | -205 | 0 | 0 | 11 | 1568.7013 | 1819.038 | 1760.2937 | 1854.629 | 2087.0085 | 2208.2446 |
| Rpn1 | AEVVLVHPGGGSTSR | -52 | -66 | 0 | 0 | 15 | 1479.5406 | 1637.4749 | 1690.4191 | 2387.3962 | 2757.3953 | 2675.049 |
| Rpn1 | FPLFGGWK | -321 | -328 | 0 | 0.00025015 | 8 | 1388.6354 | 1659.3961 | 1873.2242 | 1216.4965 | 1803.0084 | 1861.2194 |
| Rpn1 | VTAEVVLVHPGGGSTSR | -50 | -66 | 0 | 0 | 17 | 1213.4113 | 1405.3232 | 1472.1578 | 1062.5846 | 1464.4243 | 1437.0975 |
| Rpn1 | VHYENNSPFLTIT | -217 | -229 | 0 | 0 | 13 | 1172.8402 | 1479.6122 | 1491.1708 | 807.9072 | 1213.4419 | 1153.032 |
| Rpn1 | LKTEGSDLCDR | -538 | -548 | 1 | 0 | 11 | 1076.7997 | 1359.9907 | 1450.1139 | 1470.8157 | 1472.3685 | 1528.4944 |
| Rpn1 | TEGSDLCDR | -540 | -548 | 0 | 0 | 9 | 1020.63983 | 1175.1797 | 1383.8029 | 1349.2191 | 180.47437 | 2070.9397 |

| | | | | | | | | | | | | |
|---|---|---|---|---|---|---|---|---|---|---|---|---|
| Rpn1 | SSEAPPLVNEDVKR | -26 | -39 | 1 | 0 | 14 | 912.99304 | 1031.3691 | 1086.4984 | 1158.1868 | 1269.9584 | 1144.0746 |
| Rpn1 | VLHPYPTQITQSEK | -139 | -152 | 0 | 0.000230707 | 14 | 750.65466 | 799.6191 | 833.67737 | 732.10333 | 876.967 | 919.6852 |
| Rpn1 | IKGEDEEDNNLEVR | -89 | -102 | 1 | 0 | 14 | 656.0315 | 573.9253 | 762.192 | 1074.8794 | 874.2261 | 892.6947 |
| Rpn1 | IKGEDEEDNNLEVR | -89 | -102 | 1 | 0 | 14 | 656.0315 | 573.9253 | 762.192 | 1074.8794 | 874.2261 | 892.6947 |
| Rpn1 | LAHLGVQIK | -82 | -90 | 0 | 0 | 9 | 602.8445 | 656.3725 | 694.0768 | 545.6813 | 698.2682 | 510.2316 |
| Rpn1 | NLVEQHIQDIVVHY | -416 | -429 | 0 | 0 | 14 | 584.62646 | 612.76117 | 865.9722 | 746.0597 | 942.79065 | 1142.9026 |
| Rpn1 | KLPVALDPGSK | -117 | -127 | 1 | 0 | 11 | 423.5737 | 484.85684 | 477.11987 | 596.6243 | 553.7013 | 561.30707 |
| Rpn1 | FVDHVFDEQVIDSL | -356 | -369 | 0 | 0 | 14 | 412.8648 | 436.65967 | 470.23602 | 877.1648 | 343.91867 | 969.44226 |
| Rpn1 | GEDEEDNNLEVR | -91 | -102 | 0 | 0.00025015 | 12 | 167.14432 | 91.67562 | 1177.6099 | 186.6593 | 226.7543 | 42.44239 |
| Rpn1 | SSEAPPLVNEDVK | -26 | -38 | 0 | 0 | 13 | 1662.0547 | 870.1717 | 1137.0447 | 1992.216 | NaN | 2241.7021 |
| Rpn1 | YLDTFGRPVIVAYK | -401 | -414 | 1 | 0 | 14 | 786.67303 | 1112.8578 | 986.2855 | NaN | NaN | 720.1816 |
| Rpn1 | VVLVHPGGGSTSR | -54 | -66 | 0 | 0.00025015 | 13 | 978.4063 | NaN | 6813.5063 | 3124.2585 | NaN | NaN |
| Cope | SQHLGKPPEVTNR | -260 | -272 | 1 | 0 | 13 | 780.54065 | 822.2905 | 303.33817 | 346.81677 | 962.08514 | 848.6882 |
| Cope | FHDQNPDAALR | -128 | -138 | 0 | 0 | 11 | 521.69946 | 483.82056 | 1514.1974 | 1212.4761 | 1481.4473 | 1322.634 |
| Cope | SYQQCINEAQR | -30 | -40 | 0 | 0 | 11 | 495.0619 | 542.3669 | 496.00528 | 271.656 | 329.08682 | 633.2378 |
| Cope | APPVPGAVSGGSGEVDELFDVK | -2 | -23 | 0 | 0 | 22 | 407.21292 | 511.53894 | 492.54617 | 583.39484 | 580.3342 | 597.43536 |
| Cope | KYGVVLDEIKPS | -65 | -76 | 2 | 0 | 12 | 1652.3164 | 2334.9048 | 1973.7765 | NaN | 2969.8894 | 3888.9224 |
| Cope | AEYLASENQR | -88 | -97 | 0 | 0.00025015 | 10 | 793.21423 | NaN | NaN | NaN | NaN | NaN |
| Eif3e | NLYSDDIPHALR | -60 | -71 | 0 | 0 | 12 | 2705.213 | 4148.183 | 3729.1143 | 770.90234 | 2030.5221 | 2140.6655 |
| Eif3e | QEYLDTLYR | -125 | -133 | 0 | 0 | 9 | 1923.7662 | 2012.1222 | 1890.1139 | 463.2263 | 613.052 | 424.64944 |
| Eif3e | AEYLYFFR | -148 | -155 | 0 | 0 | 8 | 1660.0013 | 1411.4437 | 2024.7471 | 819.2183 | 1122.3506 | 1630.3634 |
| Eif3e | YSDDIPHALR | -62 | -71 | 0 | 0 | 10 | 1531.0233 | 3257.0789 | 3731.8418 | 555.8497 | 1622.5565 | 816.3942 |
| Eif3e | YVNFDFDGAQK | -300 | -310 | 0 | 0 | 11 | 496.5968 | 598.13727 | 411.1239 | 700.3197 | 603.61 | 662.8241 |
| Eif3e | AVSPYQQVIEK | -397 | -407 | 0 | 0 | 11 | 409.4073 | 836.8808 | 769.298 | 956.8494 | 782.78235 | 1063.3755 |
| Eif3e | LEDFIENAR | -329 | -337 | 0 | 0 | 9 | 350.86484 | 503.02075 | 830.3584 | 551.7727 | 815.48584 | 496.25214 |
| Eif3e | CLEDFIENAR | -328 | -337 | 0 | 0 | 10 | 313.36734 | 709.53394 | 674.85895 | 724.9823 | 702.17377 | 642.015 |
| Eif3e | IAHFLDR | -10 | -16 | 0 | 0.001023236 | 7 | 1652.3916 | 1904.6355 | NaN | 1284.2086 | 1291.2881 | 1213.3544 |
| Eif3e | QLQAETEPIVK | -83 | -93 | 0 | 0 | 11 | 532.75665 | 150.21388 | 489.25055 | NaN | 908.00757 | 602.91156 |
| Eif3e | FDFDGAQK | -303 | -310 | 0 | 0.00383302 | 8 | 644.1468 | 536.4631 | 534.91766 | NaN | NaN | 433.70096 |
| Eif3e | NWDAAMEDLTR | -181 | -191 | 0 | 0 | 11 | 497.4102 | NaN | NaN | 584.18445 | 473.67273 | 387.85278 |
| Eif3e | NLYSDDIPHAL | -60 | -70 | 0 | 0 | 11 | 492.08075 | 526.9443 | 609.81885 | NaN | NaN | NaN |
| Eif3e | EYLYFFR | -149 | -155 | 0 | 0.000230707 | 7 | 450.65955 | 357.4407 | 391.736 | NaN | NaN | NaN |
| Eif3e | NAVSPYQQVIEK | -396 | -407 | 0 | 0 | 12 | 341.47995 | NaN | NaN | 911.4019 | 1282.2223 | NaN |
| Mgat2 | AEYPDSFGHYR | -208 | -218 | 0 | 0 | 11 | 566.14514 | 203.90529 | 653.57135 | NaN | 447.0174 | 518.4117 |
| Mgat2 | SDALGPPLLDAEPVR | -34 | -48 | 0 | 0.001055699 | 15 | 3815.9917 | NaN | NaN | NaN | 3925.8428 | NaN |
| Ddx27 | AFALPVLER | -237 | -245 | 0 | 0 | 9 | 458.01764 | 569.91766 | 350.54935 | 874.68036 | 328.7315 | 317.23203 |
| Ddx27 | ILPQDVILK | -556 | -564 | 0 | 0.001868731 | 9 | 5980.2495 | NaN | 3091.7505 | 2887.4846 | 901.45337 | 888.8532 |
| Ddx27 | LIDHLHNCPSFHLS | -316 | -329 | 0 | 0 | 14 | 259.22617 | 1874.3646 | 1341.8359 | NaN | NaN | 2042.37 |
| Ddx27 | VLVLVPTR | -258 | -265 | 0 | 0.000826806 | 8 | 547.04767 | 373.1326 | 432.77124 | NaN | NaN | NaN |
| Ndufb9 | EAEEEFWQNQHPQPY | -66 | -80 | 0 | 0 | 15 | 468.6533 | 634.33124 | 534.2269 | 545.3368 | 620.483 | 554.58606 |
| Ndufb9 | AFCAPPAYLTHQ | -2 | -13 | 0 | 0.00025015 | 12 | 888.3344 | 884.8662 | 903.25977 | NaN | 1145.8824 | 1068.9486 |
| Ndufb9 | IFPDSPGGTSFER | -81 | -93 | 0 | 0 | 13 | 924.90594 | NaN | NaN | 1328.6904 | 1676.1467 | NaN |
| Ywhag | KNVTELNEPLSNEER | -28 | -42 | 1 | 0 | 15 | 2226.4404 | 2598.2285 | 2538.4873 | 2924.31 | 2993.832 | 2984.6128 |
| Ywhag | KNVTELNEPLSNEER | -28 | -42 | 1 | 0 | 15 | 2226.4404 | 2598.2285 | 2538.4873 | 2924.31 | 2993.832 | 2984.6128 |
| Ywhag | AYSEAHEISK | -153 | -162 | 0 | 0 | 10 | 860.3517 | 326.3607 | 417.2126 | 436.20557 | 1519.1206 | 458.35577 |
| Ywhag | NAPEQACHLAK | -188 | -198 | 0 | 0 | 11 | 502.44543 | 1183.8391 | 1278.4989 | 203.1147 | 2418.596 | 2505.6475 |
| Ywhag | NVTELNEPLSNEER | -29 | -42 | 0 | 0 | 14 | 490.99966 | 1320.759 | 1206.1273 | 671.29456 | 517.4817 | 432.50842 |
| Ywhag | NCSETQYESK | -111 | -120 | 0 | 0 | 10 | 397.12 | 413.4594 | 446.77792 | 144.09535 | 366.40756 | 394.8406 |
| Ywhag | YLAEVATGEK | -133 | -142 | 0 | 0 | 10 | 8485.692 | 8546.113 | NaN | 6967.4062 | 325.8728 | 6416.3755 |
| Ywhag | LDNYLIK | -104 | -110 | 0 | 0.00930008 | 7 | 786.1925 | 910.5389 | NaN | NaN | NaN | NaN |
| Ywhae | HLIPAANTGESK | -107 | -118 | 0 | 0 | 12 | 5466.215 | 5353.7646 | 5423.2344 | 3995.8098 | 4783.1455 | 5071.0796 |
| Ywhae | YLAEFATGNDR | -131 | -141 | 0 | 0 | 11 | 2170.671 | 1950.0385 | 2234.4124 | 2356.059 | 2074.5696 | 2030.9436 |
| Ywhae | ELPPTHPIR | -162 | -170 | 0 | 0.001055699 | 9 | 1705.1364 | 1772.3286 | 1674.6461 | 2197.8958 | 1721.9532 | 2386.1572 |
| Ywhae | YYEILNSPDR | -181 | -190 | 0 | 0 | 10 | 1543.421 | 1454.6755 | 1468.8413 | 3213.5227 | 3226.6707 | 3082.0544 |
| Ywhae | LICCDILDVLDK | -95 | -106 | 0 | 0 | 12 | 1489.6018 | 1412.0558 | 1197.2462 | 1414.7849 | 1539.4805 | 281.3163 |
| Ywhae | EALQDVEDENQ | -245 | -255 | 0 | 0 | 11 | 253.12708 | 235.09746 | 273.47098 | 295.2743 | 298.24512 | 294.5555 |
| Ywhae | EALQDVEDENQ | -245 | -255 | 0 | 0 | 11 | 253.12708 | 235.09746 | 273.47098 | 295.2743 | 298.24512 | 294.5555 |
| Ywhae | MDDREDLVYQAK | -1 | -12 | 1 | 0 | 12 | 98.90339 | 2866.1357 | 2801.5288 | 2377.7983 | 2507.2812 | 2652.6228 |
| Ywhae | MDDREDLVYQAK | -1 | -12 | 1 | 0 | 12 | 98.90339 | 2866.1357 | 2801.5288 | 2377.7983 | 2507.2812 | 2652.6228 |
| Ywhae | PPTHPIR | -164 | -170 | 0 | 0.001092529 | 7 | 518.86743 | 504.53534 | 408.2487 | 400.80707 | 726.2163 | 378.71265 |
| Ywhae | CDILDVLDK | -98 | -106 | 0 | 0 | 9 | 507.12628 | 598.73145 | 567.48987 | 523.3434 | 611.8858 | 644.1187 |
| Ywhae | FDDAIAELDTLSEESYK | -199 | -215 | 0 | 0 | 17 | 501.3081 | 443.70535 | 482.68988 | 372.32678 | 413.8399 | 419.98093 |
| Ywhae | AFDDAIAELDTLSEESYK | -198 | -215 | 0 | 0 | 18 | 498.49606 | 520.0936 | 462.52902 | 559.41797 | 176.8631 | 693.6089 |
| Ywhae | VAGMDVELTVEER | -30 | -42 | 0 | 0.000230707 | 13 | 305.71286 | 1094.5087 | 956.81696 | 1130.5457 | 279.26056 | 1330.1222 |
| Ywhae | SDIAMTELPPTHPIR | -156 | -170 | 0 | 0.001190916 | 15 | 638.00336 | 599.6497 | NaN | 736.30774 | 769.78015 | 498.81946 |
| Ywhae | DMQGDGEEQNKEALQDVEDENQ | -234 | -255 | 1 | 0.000471409 | 22 | 126.21239 | NaN | 178.19617 | 174.34244 | 157.38036 | 159.99576 |
| Ywhae | DILDVLDK | -99 | -106 | 0 | 0.001868731 | 8 | 576.4345 | NaN | NaN | 519.5949 | 648.8218 | NaN |
| Ywhaq | EKVESELR | -84 | -91 | 1 | 0.000907206 | 8 | 2220.5396 | 2456.9001 | 2516.51 | 231.60393 | 2276.6602 | 2546.878 |
| Ywhaq | KAVTEQGAELSNEER | -27 | -41 | 1 | 0 | 15 | 1799.2881 | 1381.6721 | 1581.4418 | 1608.9562 | 2039.3436 | 1945.3828 |
| Ywhaq | KAVTEQGAELSNEER | -27 | -41 | 1 | 0 | 15 | 1799.2881 | 1381.6721 | 1581.4418 | 1608.9562 | 2039.3436 | 1945.3828 |
| Ywhaq | AVTEQGAELSNEER | -28 | -41 | 0 | 0 | 14 | 1616.1488 | 2095.9785 | 2066.9753 | 1580.3665 | 1681.6818 | 1590.6211 |
| Ywhaq | YLAEVACGDDRK | -128 | -139 | 1 | 0.000230707 | 12 | 1477.5659 | 2054.6692 | 2981.806 | 311.17776 | 2473.75 | 2576.964 |
| Ywhaq | KQTIENSQGAYQE | -139 | -151 | 1 | 0.00025015 | 13 | 870.21136 | 844.3739 | 830.658 | 1013.4092 | 1099.337 | 1118.1743 |
| Ywhaq | SICTTVLELLDK | -92 | -103 | 0 | 0 | 12 | 679.19055 | 431.9055 | 608.5201 | 919.02435 | 926.16254 | 816.04425 |
| Ywhaq | KQTIENSQGAYQEAF | -139 | -153 | 1 | 0 | 15 | 532.9481 | 644.88477 | 687.0548 | 729.966 | 787.78314 | 808.6514 |
| Ywhaq | FDEAIAELDTLNEDSYK | -196 | -212 | 0 | 0 | 17 | 469.45972 | 489.34814 | 435.0377 | 482.4914 | 491.03452 | 387.75986 |
| Ywhaq | AFDEAIAELDTLNEDSYK | -195 | -212 | 0 | 0 | 18 | 213.09192 | 183.3943 | 342.05203 | 256.1575 | 477.4226 | 254.76836 |
| Ywhaq | QTIENSQGAYQEAFDISK | -140 | -157 | 0 | 0 | 18 | 1145.4911 | NaN | 1338.4607 | 1181.2094 | 133.96895 | 1631.7244 |

| | | | | | | | | | | | | |
|---|---|---|---|---|---|---|---|---|---|---|---|---|
| Ywhab | YLSEVASGENK | -130 | -140 | 0 | 0 | 11 | 3163.9878 | 176.35307 | 875.37103 | 2536.4785 | 1198.6256 | 2427.1162 |
| Ywhab | KAVTEQGHELSNEER | -29 | -43 | 1 | 0 | 15 | 2062.4294 | 966.83856 | 3114.5513 | 916.8196 | 883.78687 | 2530.4363 |
| Ywhab | AVTEQGHELSNEER | -30 | -43 | 0 | 0 | 14 | 1776.4153 | 1995.9424 | 2060.5593 | 1649.1862 | 2352.7744 | 754.8529 |
| Ywhab | AVTEQGHELSNEER | -30 | -43 | 0 | 0 | 14 | 1776.4153 | 1995.9424 | 2060.5593 | 1649.1862 | 2352.7744 | 754.8529 |
| Ywhab | LDTLNEESYK | -205 | -214 | 0 | 0 | 10 | 1520.4692 | 1751.9144 | 1584.5704 | 1762.0739 | 658.9935 | 1678.7114 |
| Ywhab | AFDEAIAELDTLNEESYK | -197 | -214 | 0 | 0 | 18 | 535.861 | 477.3666 | NaN | 571.2682 | 737.8084 | 597.9215 |
| Ywhab | IAELDTLNEESYK | -202 | -214 | 0 | 0 | 13 | 562.8007 | 599.9099 | 1154.7623 | NaN | NaN | 403.83838 |
| Ywhab | DICNDVLELLDK | -94 | -105 | 0 | 0 | 12 | 239.442 | NaN | NaN | 317.37265 | 158.31021 | 462.73767 |
| Ywhab | TAFDEAIAELDTLNEESYK | -196 | -214 | 0 | 0 | 19 | 182.87717 | 224.11127 | NaN | NaN | 251.50859 | 254.20181 |
| Ywhab | AIAELDTLNEESYK | -201 | -214 | 0 | 0 | 14 | 1679.7179 | 1942.8041 | 1947.0546 | NaN | NaN | NaN |
| Ywhab | FDEAIAELDTLNEESYK | -198 | -214 | 0 | 0 | 17 | 628.60236 | 682.2719 | 664.58044 | NaN | NaN | NaN |
| Ywhaz | FLIPNASQPESK | -104 | -115 | 0 | 0.00025015 | 12 | 13362.724 | 13190.772 | 13559.06 | 11708.9375 | 11692.365 | 11490.586 |
| Ywhaz | FLIPNASQPESK | -104 | -115 | 0 | 0.00025015 | 12 | 13362.724 | 13190.772 | 13559.06 | 11708.9375 | 11692.365 | 11490.586 |
| Ywhaz | SVTEQGAELSNEER | -28 | -41 | 0 | 0 | 14 | 6872.1177 | 6882.1787 | 1159.0016 | 7904.0176 | 8261.018 | 798.36865 |
| Ywhaz | YLAEVAAGDDKK | -128 | -139 | 1 | 0 | 12 | 6010.174 | 6329.414 | 6227.738 | 5088.49 | 5800.3823 | 6060.238 |
| Ywhaz | YLAEVAAGDDKK | -128 | -139 | 1 | 0 | 12 | 6010.174 | 6329.414 | 6227.738 | 5088.49 | 5800.3823 | 6060.238 |
| Ywhaz | EKIETELR | -84 | -91 | 1 | 0.000471409 | 8 | 5162.1694 | 880.8927 | 4771.3184 | 4950.7803 | 4287.157 | 162.30725 |
| Ywhaz | YLAEVAAGDDK | -128 | -138 | 0 | 0 | 11 | 1791.8132 | 1838.4994 | 162.42227 | 147.76845 | 1392.0181 | 1635.8398 |
| Ywhaz | KSVTEQGAELSNEER | -27 | -41 | 1 | 0 | 15 | 1703.5559 | 2442.2383 | 1957.4429 | 2729.402 | 3662.0928 | 3386.8665 |
| Ywhaz | MDKNELVQK | -1 | -9 | 1 | 0.000230707 | 9 | 1522.3038 | 4431.287 | 1643.6492 | 4367.729 | 4417.6523 | 1967.6368 |
| Ywhaz | MDKNELVQK | -1 | -9 | 1 | 0.000230707 | 9 | 1522.3038 | 4431.287 | 1643.6492 | 4367.729 | 4417.6523 | 1967.6368 |
| Ywhaz | YLAEVAAGDDKKG | -128 | -140 | 2 | 0 | 13 | 1424.3395 | 909.4977 | 1198.3983 | 1313.3645 | 1248.5066 | 1024.7183 |
| Ywhaz | DICNDVLSLLEK | -92 | -103 | 0 | 0 | 12 | 457.77637 | 382.03207 | 409.3124 | 616.57697 | 815.5468 | 849.0662 |
| Ywhaz | AFDEAIAELDTLSEESYK | -195 | -212 | 0 | 0 | 18 | 371.5021 | 466.60034 | 415.20447 | 574.78125 | 778.01105 | 537.01605 |
| Ywhaz | WTSDTQGDEAEAGEGGEN | -228 | -245 | 0 | 0 | 18 | 290.0254 | 281.82086 | 247.51793 | 299.94482 | 260.31345 | 245.52872 |
| Ywhaz | GIVDQSQQAYQE | -140 | -151 | 0 | 0 | 12 | 187.8411 | 171.4755 | 202.47183 | 304.53027 | 255.79578 | 248.34985 |
| Ywhaz | GIVDQSQQAYQEAFEISK | -140 | -157 | 0 | 0 | 18 | 469.1541 | NaN | 563.407 | 1090.452 | 1253.4833 | 1117.7004 |
| Ywhaz | FDEAIAELDTLSEESYK | -196 | -212 | 0 | 0 | 17 | 427.6408 | 527.57733 | 473.59827 | 299.7604 | 233.31683 | NaN |
| Ywhaz | VVSSIEQK | -61 | -68 | 0 | 0.001776473 | 8 | 1606.2424 | NaN | 1272.2474 | NaN | NaN | NaN |
| Ywhaz | TAFDEAIAELDTLSEESYK | -194 | -212 | 0 | 0.001291103 | 19 | 68.597725 | NaN | NaN | NaN | NaN | NaN |
| Cct5 | IADGYEQAAR | -133 | -142 | 0 | 0 | 10 | 8759.865 | 5912.4834 | 7184.273 | 5745.4067 | 5079.0625 | 4877.6216 |
| Cct5 | TCPFEPPKPK | -252 | -261 | 1 | 0 | 10 | 6927.7686 | 4850.231 | 4037.1296 | 1678.3453 | 3383.8032 | 4261.506 |
| Cct5 | TCPFEPPKPK | -252 | -261 | 1 | 0 | 10 | 6927.7686 | 4850.231 | 4037.1296 | 1678.3453 | 3383.8032 | 4261.506 |
| Cct5 | ESNPALGIDCLHK | -484 | -496 | 0 | 0 | 13 | 2602.9531 | 2722.904 | 2851.79 | 3532.4038 | 3284.5703 | 3235.3562 |
| Cct5 | DVDFELIKVE | -203 | -212 | 1 | 0.00025015 | 10 | 1517.1437 | 1180.2759 | 1574.0835 | 1043.2837 | 194.36594 | 1398.1763 |
| Cct5 | HVIETLIGK | -505 | -513 | 0 | 0.000471409 | 9 | 1194.0834 | 1728.8411 | 1711.3928 | 1115.2045 | 2010.6519 | 2163.7344 |
| Cct5 | LEEAEQLLDR | -117 | -126 | 0 | 0 | 10 | 1191.2859 | 827.2576 | 368.54346 | 2342.0889 | 1697.5789 | 1079.488 |
| Cct5 | IAIQHLDK | -143 | -150 | 0 | 0.000627943 | 8 | 1053.8618 | 2577.8303 | 2346.5562 | 2861.8462 | 1408.809 | 1735.0931 |
| Cct5 | VLVDINNPEPLIQ | -155 | -167 | 0 | 0.00025015 | 13 | 895.84534 | 837.3197 | 821.1352 | 1018.3175 | 1165.0719 | 1722.9893 |
| Cct5 | ILTCPFEPPKPK | -250 | -261 | 1 | 0 | 12 | 500.48315 | 939.0464 | 1396.3558 | 667.5179 | 95.74367 | 1207.3412 |
| Cct5 | ILTCPFEPPKPK | -250 | -261 | 1 | 0 | 12 | 500.48315 | 939.0464 | 1396.3558 | 667.5179 | 95.74367 | 1207.3412 |
| Cct5 | GALLEEAEQLLDR | -114 | -126 | 0 | 0 | 13 | 548.1722 | 366.65158 | 538.00574 | 396.86496 | 750.0654 | 691.748 |
| Cct5 | AGALLEEAEQLLDR | -113 | -126 | 0 | 0 | 14 | 390.69012 | 321.31598 | 411.97192 | 258.38892 | 86.46074 | 324.08893 |
| Cct5 | AFDEYGRPFLIIK | -8 | -20 | 1 | 0 | 13 | 795.6607 | 949.0742 | 826.01733 | NaN | 781.9548 | 805.69196 |
| Cct5 | VLVDINNPEPLIQTA | -155 | -169 | 0 | 0.001055699 | 15 | 555.5531 | NaN | 408.60315 | 783.48346 | 754.4436 | 867.9309 |
| Cct5 | SQDDEIGDGTTGVVVLA | -97 | -113 | 0 | 0 | 17 | 55.059 | 536.3179 | NaN | 911.86365 | 361.93427 | 443.04474 |
| Cct5 | ESNPALGIDCLHKG | -484 | -497 | 1 | 0 | 14 | 1679.6843 | 1214.6611 | NaN | NaN | 1401.1633 | 937.4028 |
| Cct5 | AVLTVADMER | -192 | -201 | 0 | 0.000298805 | 10 | 770.1817 | NaN | NaN | NaN | 1105.3723 | NaN |
| Cct5 | VVQEISFGTTK | -358 | -368 | 0 | 0 | 11 | 730.9852 | 1385.3595 | NaN | NaN | NaN | NaN |
| Uqcr10 | GADAIYEHINEGK | -39 | -51 | 0 | 0 | 13 | 15990.321 | 16329.21 | 16490.201 | 1894.5214 | 13470.85 | 13883.3545 |
| Uqcr10 | AFDQGADAIYEHINEGK | -35 | -51 | 0 | 0 | 17 | 7215.3403 | 9088.691 | 10150.767 | 14957.612 | 9300.487 | 7419.0303 |
| Uqcr10 | LYSLLFR | -10 | -16 | 0 | 0 | 7 | 580.8698 | 641.7976 | 731.3631 | 625.456 | 596.31104 | 705.0113 |
| Eif3i | LFDSTTLEHQK | -214 | -224 | 0 | 0 | 11 | 2539.8386 | 2562.5984 | 2715.0046 | 2674.736 | 2768.3271 | 2541.3928 |
| Eif3i | SYSSGGEDGYVR | -299 | -310 | 0 | 0 | 12 | 1484.8738 | 1279.7959 | 1278.1877 | 1553.4398 | 77.047264 | 1201.411 |
| Eif3i | VSFFDLR | -122 | -128 | 0 | 0.000591124 | 7 | 640.8158 | 615.30896 | 629.49945 | 115.356804 | 230.7098 | 710.85455 |
| Eif3i | GHESGELNQYSAK | -164 | -176 | 0 | 0 | 13 | 611.85205 | 758.68463 | 734.78015 | 901.2251 | 1123.2186 | 1113.7289 |
| Eif3i | QINDIQLSR | -190 | -198 | 0 | 0.00025015 | 9 | 601.6484 | 1729.3109 | 1628.214 | 1739.7034 | 300.8877 | 1495.0673 |
| Eif3i | CVDADWDTK | -57 | -65 | 0 | 0 | 9 | 1059.1588 | 741.30194 | 799.45544 | 930.4245 | 797.9575 | NaN |
| Eif3i | HVLTGSADNSCR | -66 | -77 | 0 | 0 | 12 | 605.0241 | 588.7153 | 835.3601 | 822.94806 | NaN | 1072.0002 |
| Eif3i | GHFGPINSVAFHPDGK | -283 | -298 | 0 | 0 | 16 | 536.92065 | 2203.0198 | NaN | 1125.3441 | 3270.8877 | 3594.4702 |
| Eif3i | GHFGPINSV | -283 | -291 | 0 | 0.000826806 | 9 | 1223.0853 | 881.0782 | 957.7683 | NaN | NaN | NaN |
| Eif3i | FEEEFGR | -274 | -280 | 0 | 0.000431655 | 7 | 1204.6127 | 695.6242 | 390.17892 | NaN | NaN | NaN |
| Eif3i | AFEEEFGR | -273 | -280 | 0 | 0 | 8 | 734.00964 | 548.00714 | NaN | 724.94037 | NaN | NaN |
| Rnpep | YLAHLVGDQDQFDK | -423 | -436 | 0 | 0 | 14 | 2005.3971 | 3818.1104 | 3602.42 | 268.74353 | 3728.2512 | 3497.8926 |
| Rnpep | YLAHLVGDQDQFDK | -423 | -436 | 0 | 0 | 14 | 2005.3971 | 3818.1104 | 3602.42 | 268.74353 | 3728.2512 | 3497.8926 |
| Rnpep | IEPGVDPDDTYNETPYEK | -399 | -416 | 0 | 0 | 18 | 78.59336 | 530.5261 | 655.11426 | 570.37573 | 762.3681 | 727.99084 |
| Rnpep | GVDSIPGFEFDR | -470 | -481 | 0 | 0 | 12 | 461.78568 | 281.58307 | 538.3313 | 925.5798 | NaN | 1244.6552 |
| Rnpep | AFEILHL | -33 | -39 | 0 | 0 | 7 | 411.17023 | NaN | 418.9659 | 673.0389 | 81.18733 | NaN |
| Rnpep | WGQIILK | -575 | -581 | 0 | 0.001291103 | 7 | 912.3404 | NaN | NaN | 740.28723 | NaN | NaN |
| Stmn1 | SHEAEVLK | -63 | -70 | 0 | 0.000230707 | 8 | 2760.6077 | 2567.1208 | 2666.749 | 1576.614 | 3647.8496 | 2298.192 |
| Stmn1 | SKESVPDFPLSPPK | -28 | -41 | 1 | 0 | 14 | 2490.2256 | 2032.7548 | 554.36316 | 2409.4158 | 1120.1576 | 917.19525 |
| Stmn1 | AFELILSPR | -19 | -27 | 0 | 0 | 9 | 1389.3656 | 1287.5375 | 1209.8556 | 1360.7264 | 1401.1588 | 1558.0504 |
| Stmn1 | ESKDPADETEAD | -138 | -149 | 1 | 0 | 12 | 172.00269 | 191.65897 | 1038.5408 | 797.27057 | 200.82567 | 165.18893 |
| Stmn1 | ESVPDFPLSPPK | -30 | -41 | 0 | 0 | 12 | 2950.4854 | 3359.8171 | 3228.584 | NaN | 3619.7725 | 3773.5508 |
| Stmn1 | DKHVEEVR | -127 | -134 | 1 | 0.004350979 | 8 | 678.2673 | NaN | NaN | NaN | NaN | NaN |
| Ccdc134 | AFFGDVVLR | -113 | -121 | 0 | 0.006100831 | 9 | 593.47266 | NaN | 707.2308 | NaN | 655.8922 | NaN |
| Hmox1 | FTFPNIDSPTK | -167 | -177 | 0 | 0 | 11 | 538.1168 | 383.14722 | 630.55914 | 554.09576 | 439.4568 | 495.8407 |
| Hmox1 | TALEEEIER | -59 | -67 | 0 | 0 | 9 | 449.50504 | 764.14465 | 1065.1233 | 1156.1854 | 1041.0801 | 986.54407 |

| | | | | | | | | | | | | |
|---|---|---|---|---|---|---|---|---|---|---|---|---|
| Hmox1 | YLGDLSGGQVLK | -137 | -148 | 0 | 0 | 12 | 6977.2183 | 7283.6 | 1841.8875 | NaN | 5775.726 | 6385.2803 |
| Hmox1 | THPELLVAH | -124 | -132 | 0 | 0.00025015 | 9 | 105.685455 | NaN | 1547.7876 | NaN | 2178.5962 | NaN |
| Hmox1 | AFFTFPNIDSPTK | -165 | -177 | 0 | 0.000471409 | 13 | 616.89545 | NaN | NaN | NaN | NaN | 513.74634 |
| Erp44 | AFGDLSKPER | -194 | -203 | 1 | 0 | 10 | 1670.1273 | 1864.723 | 2132.807 | 1908.6798 | 508.40387 | 2221.4272 |
| Erp44 | NIIGYFEQK | -162 | -170 | 0 | 0 | 9 | 1111.0426 | 1065.8712 | 1352.079 | 1139.1636 | 1295.5403 | 1230.3318 |
| Erp44 | FYADWCR | -53 | -59 | 0 | 0.000712251 | 7 | 840.8527 | 924.52734 | 1021.46924 | 813.2468 | 947.28796 | 1002.9248 |
| Erp44 | YNGDNVIYKPPGR | -204 | -216 | 1 | 0 | 13 | 661.4766 | 830.0158 | 922.0872 | 1295.402 | 1049.8916 | 889.7014 |
| Erp44 | VDCDQHSDIAQR | -90 | -101 | 0 | 0 | 12 | 120.53837 | 560.999 | 579.0946 | 425.63507 | 801.20447 | 693.1757 |
| Erp44 | DDTESLEIFQNEVAR | -271 | -285 | 0 | 0 | 15 | 449.46472 | 331.94476 | NaN | 359.54413 | 638.09296 | 798.673 |
| Erp44 | RVDCDQHSDIAQR | -89 | -101 | 1 | 0 | 13 | 335.43167 | 664.9315 | NaN | 365.53796 | 288.73093 | 536.00183 |
| Erp44 | QFVFDLHSGK | -346 | -355 | 0 | 0 | 10 | 1068.2979 | 1337.2051 | 1182.8744 | NaN | NaN | 1419.7891 |
| Leprotl1 | AFGLPVVFAR | -81 | -90 | 0 | 0 | 10 | 287.74084 | 251.3625 | 311.53473 | 369.2281 | 348.51245 | 416.19138 |
| Cul4b | AFGSTIVINPEK | -557 | -568 | 0 | 0 | 12 | 1505.2982 | 1550.1128 | 1730.2294 | 664.40735 | 710.23425 | 2033.8591 |
| Cul4b | LPENYTDETWQK | -265 | -276 | 0 | 0 | 12 | 950.95795 | NaN | 807.4484 | 1094.3293 | 738.93524 | 410.17184 |
| Cul4b | EVPEYLHHVNK | -466 | -476 | 0 | 0 | 11 | 497.43982 | 554.79834 | 622.442 | NaN | 697.7182 | 673.7127 |
| Cul4b | HATGIEDGELR | -826 | -836 | 0 | 0 | 11 | 6424.7363 | 4878.5513 | 4930.7173 | NaN | NaN | 5725.0073 |
| Cul4b | GLNSLLDENR | -517 | -526 | 0 | 0 | 10 | 4164.19 | 3513.0527 | 6937.147 | NaN | 306.82642 | NaN |
| Cul4b | FPVKPADLK | -936 | -944 | 1 | 0.003360691 | 9 | 674.2567 | 696.7286 | NaN | NaN | NaN | NaN |
| Nrm | AFGTGVEFVR | -18 | -27 | 0 | 0 | 10 | 529.78577 | NaN | 573.03754 | NaN | 521.9133 | 456.6347 |
| Psmd6 | PLENLEEEGLPK | -2 | -13 | 0 | 0 | 12 | 2696.552 | 3340.3374 | 3000.6409 | 4397.157 | 4695.066 | 5920.5576 |
| Psmd6 | RLDEELEDAEK | -83 | -93 | 1 | 0 | 11 | 2143.0266 | 2285.9243 | 2289.5186 | 2080.295 | 1821.2828 | 1829.5677 |
| Psmd6 | VNEIVETNRPDSK | -350 | -362 | 1 | 0 | 13 | 1463.1967 | 1326.2067 | 1304.2188 | 1299.4122 | 1786.1647 | 1862.6068 |
| Psmd6 | KDWLFAPHYR | -284 | -293 | 1 | 0 | 10 | 936.8269 | 751.59485 | 658.2488 | 108.92361 | 692.27203 | 593.4646 |
| Psmd6 | KDWLFAPHYR | -284 | -293 | 1 | 0 | 10 | 936.8269 | 751.59485 | 658.2488 | 108.92361 | 692.27203 | 593.4646 |
| Psmd6 | EVLHSLPAVR | -248 | -257 | 0 | 0 | 10 | 772.4475 | 1414.7842 | 1425.9369 | 1071.4943 | 1242.2166 | 1582.8458 |
| Psmd6 | IGLFYMDNDLITR | -147 | -159 | 0 | 0 | 13 | 585.6904 | 525.9812 | 203.5571 | 947.40063 | 943.405 | 909.5846 |
| Psmd6 | KVNEIVETNRPDSK | -349 | -362 | 2 | 0.000431655 | 14 | 500.7704 | 578.3741 | 546.73376 | 481.06268 | 565.32135 | 559.647 |
| Psmd6 | IDKVNEIVETNRPDSK | -347 | -362 | 2 | 0 | 16 | 468.92743 | 505.28406 | 501.67828 | 549.91223 | 896.0146 | 728.4824 |
| Psmd6 | IDKVNEIVETNRPDSK | -347 | -362 | 2 | 0 | 16 | 468.92743 | 505.28406 | 501.67828 | 549.91223 | 896.0146 | 728.4824 |
| Psmd6 | IALERPDLR | -229 | -237 | 1 | 0.002265552 | 9 | 3738.463 | 967.57764 | 4110.236 | NaN | 4114.8784 | 4895.868 |
| Psmd6 | IEEGGDWDRR | -168 | -177 | 1 | 0.00025015 | 10 | 1931.0559 | NaN | 1681.3259 | 1308.7087 | 1416.8373 | 2124.7224 |
| Psmd6 | NWQYQETIK | -363 | -371 | 0 | 0 | 9 | 1837.3235 | 1247.5289 | NaN | 1140.8002 | 388.079 | 534.9308 |
| Psmd6 | GVGVDFIDQELSR | -324 | -336 | 0 | 0 | 13 | 655.62683 | NaN | 727.2444 | 996.7744 | 247.65356 | 1025.7678 |
| Psmd6 | SQLLESYR | -305 | -312 | 0 | 0 | 8 | 966.57214 | 954.1472 | NaN | NaN | 747.0232 | NaN |
| Psmd6 | FLLSLPEHR | -24 | -32 | 0 | 0.000550307 | 9 | 662.6016 | NaN | NaN | 541.07794 | NaN | NaN |
| Psmd6 | AFGVGVDFIDQELSR | -322 | -336 | 0 | 0.00383302 | 15 | 336.66513 | NaN | NaN | NaN | NaN | NaN |
| Srsf3 | AFGYYGPLR | -29 | -37 | 0 | 0 | 9 | 329.99957 | 3761.16 | 3844.9067 | 2830.8813 | 2769.2493 | 3007.5269 |
| Srsf3 | DSCPLDCK | -4 | -11 | 0 | 0 | 8 | 810.48883 | 712.93994 | 635.30927 | 616.5764 | NaN | 426.85132 |
| Srsf3 | SCPLDCK | -5 | -11 | 0 | 0 | 7 | 791.1649 | NaN | 708.7215 | 710.2343 | 726.69775 | 773.82465 |
| Srsf3 | VYVGNLGNNGNK | -12 | -23 | 0 | 0 | 12 | 1110.33 | NaN | NaN | 901.8397 | 1313.2368 | 949.1609 |
| Rpl29 | ASAPAQAPK | -137 | -145 | 0 | 0.00025015 | 9 | 612.97546 | 717.015 | 651.9439 | 538.3144 | 771.708 | 676.9772 |
| Rpl29 | AFIAHPK | -95 | -101 | 0 | 0.00025015 | 7 | 1078.498 | 1007.9153 | 891.73224 | 941.44196 | 799.9744 | NaN |
| Rpl29 | LAFIAHPK | -94 | -101 | 0 | 0 | 8 | 960.95905 | 910.318 | 869.9076 | NaN | NaN | NaN |
| Actn1 | ANVIGPWIQTK | -639 | -649 | 0 | 0 | 11 | 14566.992 | 16551.82 | 16476.676 | 18585.002 | 23638.078 | 23921.803 |
| Actn1 | VIGPWIQTK | -641 | -649 | 0 | 0 | 9 | 3224.2031 | 4283.421 | 4682.7837 | 4209.2314 | 5172.1445 | 4697.6353 |
| Actn1 | DHSGTLGPEEFK | -761 | -772 | 0 | 0 | 12 | 3043.6306 | 1077.8334 | 2984.6445 | 3182.8386 | 2960.2983 | 2793.842 |
| Actn1 | DHSGTLGPEEFK | -761 | -772 | 0 | 0 | 12 | 3043.6306 | 1077.8334 | 2984.6445 | 3182.8386 | 2960.2983 | 2793.842 |
| Actn1 | ATLPDADKER | -556 | -565 | 1 | 0.00111006 | 10 | 2570.6018 | 2478.959 | 2694.912 | 1853.9092 | 2637.499 | 2349.1453 |
| Actn1 | HRPELIDYGK | -186 | -195 | 1 | 0 | 10 | 1825.548 | 1948.4429 | 2053.588 | 1735.011 | 1947.7607 | 1959.5881 |
| Actn1 | HRPELIDYGK | -186 | -195 | 1 | 0 | 10 | 1825.548 | 1948.4429 | 2053.588 | 1735.011 | 1947.7607 | 1959.5881 |
| Actn1 | IDQLECDHQLIQ | -685 | -696 | 0 | 0.001023236 | 12 | 1786.184 | 1442.7269 | 1599.5416 | 1579.5446 | 1289.404 | 1317.1124 |
| Actn1 | NYITEDELR | -841 | -849 | 0 | 0 | 9 | 1241.5708 | 1238.9684 | 972.15607 | 1269.8364 | 1886.8082 | 995.23596 |
| Actn1 | QKDYETATLSEIK | -419 | -431 | 1 | 0 | 13 | 1206.8037 | 837.5937 | 587.45636 | 260.14755 | 835.4404 | 677.0545 |
| Actn1 | GYDIGNDPQGEAEFAR | -779 | -794 | 0 | 0 | 16 | 851.5496 | 1191.7263 | 1190.766 | 1008.1279 | 1329.0779 | 1266.8414 |
| Actn1 | GISQEQMNEFR | -742 | -752 | 0 | 0 | 11 | 795.4306 | 210.15211 | 125.84042 | 911.83527 | 140.48032 | 725.30554 |
| Actn1 | DAEDIVGTARPDEK | -223 | -236 | 1 | 0 | 14 | 635.0563 | 228.8858 | 1835.2891 | 1133.0386 | 652.72 | 1189.969 |
| Actn1 | LAILGIHNEVSK | -566 | -577 | 0 | 0 | 12 | 527.8499 | 562.1892 | 550.14795 | 491.66055 | 528.14355 | 517.5704 |
| Actn1 | HTNYNMEHIR | -705 | -714 | 0 | 0 | 10 | 517.2685 | 677.074 | 563.7552 | 1482.6731 | 623.4128 | 634.4115 |
| Actn1 | ELNELDYYDSPSVNAR | -460 | -475 | 0 | 0 | 16 | 477.3946 | 513.6722 | 535.469 | 1253.6002 | 1260.6104 | 1107.7135 |
| Actn1 | MLDAEDIVGTARPDEK | -221 | -236 | 1 | 0 | 16 | 476.2986 | 1447.9663 | 1297.1506 | 887.41235 | 978.7892 | 464.37695 |
| Actn1 | RDQALTEEHAR | -614 | -624 | 1 | 0 | 11 | 471.64862 | 572.9786 | 578.69806 | 597.97375 | 855.49304 | 904.9793 |
| Actn1 | LISLGYDIGNDPQGEAEFAR | -775 | -794 | 0 | 0 | 20 | 406.76147 | 387.60474 | 374.23065 | 405.18372 | 480.07803 | 471.86487 |
| Actn1 | QELNELDYYDSPSVNAR | -459 | -475 | 0 | 0.004791731 | 17 | 362.49963 | 386.28555 | 442.55212 | 477.23352 | 706.4054 | 688.012 |
| Actn1 | MHGTLEDQLSHLR | -660 | -672 | 0 | 0 | 13 | 71.579605 | 960.7177 | 883.24884 | 852.3557 | 978.9176 | 1143.4158 |
| Actn1 | HRPELIDY | -186 | -193 | 1 | 0.001392869 | 8 | 688.0859 | 2172.1882 | 4789.1895 | 1242.6251 | NaN | 4718.552 |
| Actn1 | DQALTEEHAR | -615 | -624 | 0 | 0 | 10 | 306.552 | 3719.7744 | NaN | 1169.4486 | 3187.938 | 3704.8328 |
| Actn1 | IDQLECDHQLIQE | -685 | -697 | 0 | 0.001636916 | 13 | 1314.9381 | 3495.6511 | 3574.2788 | NaN | 372.8907 | NaN |
| Actn1 | AVNQENEQLMEDYEK | -267 | -281 | 0 | 0.00025015 | 15 | 1155.8057 | 1220.4124 | 1120.4071 | NaN | NaN | 602.2792 |
| Actn1 | SIVNYKPK | -677 | -684 | 1 | 0.002068213 | 8 | 593.3305 | 681.9248 | 688.3556 | NaN | NaN | NaN |
| Actn1 | KDDPLTNLNTAFDVAER | -198 | -214 | 1 | 0.002068213 | 17 | 1361.9061 | NaN | NaN | NaN | NaN | 2384.0325 |
| Actn1 | KDDPLTNLNTAF | -198 | -209 | 1 | 0.004628201 | 12 | 672.4187 | NaN | NaN | NaN | NaN | NaN |
| Lck | ITFPGLHDLVR | -197 | -207 | 0 | 0.000550307 | 11 | 1426.106 | 502.95062 | 422.87015 | 471.32962 | 2240.924 | 482.88083 |
| Lck | ERPEDRPTFDYLR | -479 | -491 | 2 | 0 | 13 | 1219.2645 | 1452.7618 | 1552.4735 | 1452.9604 | 1484.9323 | 1580.1974 |
| Lck | ERPEDRPTFDYLR | -479 | -491 | 2 | 0 | 13 | 1219.2645 | 1452.7618 | 1552.4735 | 1452.9604 | 1484.9323 | 1580.1974 |
| Lck | TTGQEGFIPFNFVAK | -104 | -118 | 0 | 0 | 15 | 741.8027 | 119.08011 | 801.522 | 915.1499 | 992.94525 | 1023.6624 |
| Lck | QLLAPGNTHGSFLIR | -140 | -154 | 0 | 0 | 15 | 497.34412 | 355.5114 | 631.01917 | 581.8146 | 760.97644 | 895.23486 |
| Lck | QLLAPGNTHGSFLIR | -140 | -154 | 0 | 0 | 15 | 497.34412 | 355.5114 | 631.01917 | 581.8146 | 760.97644 | 895.23486 |
| Lck | HYTNASDGLCTK | -208 | -219 | 0 | 0 | 12 | 669.7166 | 589.7102 | 676.2467 | 890.85315 | 1005.2673 | 847.54596 |

| | | | | | | | | | | | | |
|---|---|---|---|---|---|---|---|---|---|---|---|---|
| Lck | AFIEEQNYIHR | -353 | -363 | 0 | 0 | 11 | 584.742 | 772.3162 | 788.738 | 1036.8208 | 1049.7074 | 1023.62146 |
| Lck | GILLTEIVTHGR | -427 | -438 | 0 | 0.00025015 | 12 | 542.8396 | 805.36847 | 815.72766 | 1286.9672 | 1719.9197 | 1557.8655 |
| Lck | HSYEPSHDGDLGFEK | -70 | -84 | 0 | 0 | 15 | 535.26855 | 848.84705 | 816.2012 | 714.7384 | 999.66547 | 1016.66364 |
| Lck | HSYEPSHDGDLGFEK | -70 | -84 | 0 | 0 | 15 | 535.26855 | 848.84705 | 816.2012 | 714.7384 | 999.66547 | 1016.66364 |
| Lck | ANSLEPEPWFFK | -119 | -130 | 0 | 0 | 12 | 526.10254 | 479.64838 | 540.3284 | 769.1442 | 1178.5974 | 794.0088 |
| Lck | ILEQSGEWWK | -90 | -99 | 0 | 0 | 10 | 363.94852 | 1098.4628 | 542.3958 | 238.90863 | 472.90585 | 326.08575 |
| Lck | SLEPEPWFFK | -121 | -130 | 0 | 0.000347792 | 10 | 328.25595 | 1005.03644 | 437.96014 | 1066.2103 | 562.9801 | 419.31546 |
| Lck | SLTTGQEGFIPFNFVAK | -102 | -118 | 0 | 0 | 17 | 282.44113 | 164.63239 | 354.17358 | 368.0345 | 452.15234 | 442.22342 |
| Lck | LHSYEPSHDGDLGFEK | -69 | -84 | 0 | 0.00025015 | 16 | 1508.9193 | 1534.5568 | NaN | 1886.8492 | 609.87634 | 1511.9553 |
| Lck | DFDQNQGEVVK | -169 | -179 | 0 | 0 | 11 | 1300.5457 | 1339.3375 | 195.62967 | NaN | 206.2589 | 1317.4391 |
| Lck | AQSLTTGQEGFIPFNFVAK | -100 | -118 | 0 | 0 | 19 | 377.27634 | NaN | 341.87268 | 556.31604 | 650.8607 | 621.7052 |
| Lck | NLDNGGFYISPR | -185 | -196 | 0 | 0 | 12 | 896.7428 | 794.5056 | 1047.599 | NaN | NaN | NaN |
| Lck | ITFPGLHDL | -197 | -205 | 0 | 0.002752526 | 9 | 512.2082 | 365.89484 | 276.82642 | NaN | NaN | NaN |
| Snrnp70 | DPIPYLPPLEK | -17 | -27 | 0 | 0 | 11 | 2463.3823 | 2540.5837 | 2781.8206 | 2896.7568 | 3333.6792 | 3505.617 |
| Snrnp70 | YDERPGPSPLPHR | -219 | -231 | 1 | 0 | 13 | 2026.1465 | 2161.94 | 2212.6365 | 1260.8203 | 1648.9386 | 1890.0812 |
| Snrnp70 | YDERPGPSPLPHR | -219 | -231 | 1 | 0 | 13 | 2026.1465 | 2161.94 | 2212.6365 | 1260.8203 | 1648.9386 | 1890.0812 |
| Snrnp70 | RQQEVETELK | -78 | -87 | 1 | 0.00025015 | 10 | 1371.4679 | 1205.2444 | 1647.2607 | 1378.4039 | 1788.5868 | 1886.888 |
| Snrnp70 | DERPGPSPLPHR | -220 | -231 | 1 | 0 | 12 | 1353.7246 | 879.78595 | 752.20264 | 1089.1998 | 748.55023 | 823.698 |
| Snrnp70 | HHNQPYCGIAPY | -33 | -44 | 0 | 0 | 12 | 1003.6672 | 901.51764 | 989.5632 | 1739.5975 | 913.3688 | 5049.0215 |
| Snrnp70 | IEYEHER | -149 | -155 | 0 | 0.00204437 | 7 | 794.72424 | 964.98224 | 335.72836 | 346.02924 | 888.1143 | 727.61163 |
| Snrnp70 | EAGDGAPDDGPPGELGPEGPDGPEEK | -333 | -358 | 0 | 0 | 26 | 482.39505 | 479.25693 | 533.47107 | 648.9747 | 728.3238 | 608.47424 |
| Snrnp70 | AFIEYEHER | -147 | -155 | 0 | 0 | 9 | 1241.8718 | 1366.3462 | NaN | 1827.397 | 1956.6919 | 1888.578 |
| Snrnp70 | GGGGSGQDNGLEGLGSDGR | -404 | -422 | 0 | 0 | 19 | 829.9957 | 2093.4055 | 526.5316 | NaN | NaN | 618.4932 |
| Snrnp70 | EFEVYGPIK | -122 | -130 | 0 | 0.000826806 | 9 | 676.55176 | NaN | 548.0794 | 686.9555 | 454.55234 | NaN |
| Syncrip | AGPIWDLR | -185 | -192 | 0 | 0 | 8 | 7149.896 | 7183.928 | 7653.5166 | 6642.2705 | 6356.4087 | 6643.536 |
| Syncrip | VTEGLTDVILYHQPDDK | -266 | -282 | 0 | 0.00025015 | 17 | 1979.6064 | 289.17856 | 1909.6859 | 136.82298 | 3332.0469 | 3693.8638 |
| Syncrip | EQILEEFSK | -257 | -265 | 0 | 0 | 9 | 1511.1473 | 926.3746 | 1314.6503 | 2123.009 | 1957.7764 | 308.26593 |
| Syncrip | GLVAHSDLDER | -50 | -60 | 0 | 0 | 11 | 1471.0292 | 3554.557 | 3385.3113 | 2391.6492 | 2010.3815 | 1141.5682 |
| Syncrip | TKEQILEEFSK | -255 | -265 | 1 | 0 | 11 | 1191.9712 | 1164.209 | 1230.2802 | 1749.8646 | 2257.4905 | 105.12161 |
| Syncrip | TVTEEILEK | -348 | -356 | 0 | 0 | 9 | 916.5567 | 440.3275 | 941.4456 | 875.1751 | 906.00916 | 971.38324 |
| Syncrip | LEYEDHK | -291 | -297 | 0 | 0.00078661 | 7 | 884.86316 | 906.03674 | 1000.25964 | 1137.2328 | 1166.2885 | 1104.1702 |
| Syncrip | DSDLSHVQNK | -82 | -91 | 0 | 0.000471409 | 10 | 873.8557 | 1749.2697 | 1097.767 | 1124.7163 | 401.81824 | 240.99638 |
| Syncrip | VAHSDLDER | -52 | -60 | 0 | 0 | 9 | 806.96564 | 238.18097 | 988.2178 | 268.61823 | 401.9927 | 1199.6747 |
| Syncrip | KYGGPPPDSVYSGQQPSVGTE | -143 | -163 | 1 | 0 | 21 | 693.9499 | 709.7405 | 625.5482 | 1161.956 | 943.00336 | 855.46875 |
| Syncrip | AFIHFDER | -374 | -381 | 0 | 0 | 8 | 131.7172 | 1570.3173 | 1563.7314 | 1209.9324 | 1283.3417 | 1606.9004 |
| Syncrip | TGYTLDVTTGQR | -131 | -142 | 0 | 0 | 12 | 1377.0714 | NaN | 1063.182 | 832.8898 | 1250.091 | 1056.7242 |
| Syncrip | NLANTVTEEILEK | -344 | -356 | 0 | 0 | 13 | 8771.025 | NaN | 7857.967 | NaN | 1792.0857 | 1800.0253 |
| Syncrip | LYNNHEIR | -222 | -229 | 0 | 0.00025015 | 8 | 2351.187 | 2612.793 | NaN | 1349.2407 | NaN | 1814.1136 |
| Syncrip | TKEQILEEF | -255 | -263 | 1 | 0.002880184 | 9 | 1345.3606 | NaN | NaN | 1215.0317 | 1483.0588 | 1527.0969 |
| Syncrip | ANTVTEEILEK | -346 | -356 | 0 | 0 | 11 | 1005.2104 | 1113.4067 | NaN | 2217.275 | NaN | 503.58026 |
| Syncrip | NFQTLLDAGLPQK | -26 | -38 | 0 | 0 | 13 | 479.7888 | 460.35883 | NaN | 611.8231 | NaN | 348.05835 |
| Syncrip | LFVGSIPK | -245 | -252 | 0 | 0.002780285 | 8 | 790.73474 | 685.4028 | NaN | NaN | 258.17126 | NaN |
| Syncrip | LDAGLPQK | -31 | -38 | 0 | 0.002168954 | 8 | 648.64075 | 634.6127 | NaN | NaN | NaN | NaN |
| Syncrip | EFNEDGALAVLQ | -67 | -78 | 0 | 0.000431655 | 12 | 216.26479 | NaN | 178.8761 | NaN | NaN | NaN |
| Dnm2 | NFRPDDPTR | -316 | -324 | 1 | 0 | 9 | 2301.272 | 1151.7793 | 2286.2322 | 2083.365 | 2246.6597 | 154.48708 |
| Dnm2 | VPVGDQPPDIEYQIK | -143 | -157 | 0 | 0 | 15 | 1156.0057 | 1213.1287 | 1370.8284 | 1121.7877 | 1226.385 | 1292.3105 |
| Dnm2 | IEGSGDQVDTLELSGGAR | -344 | -361 | 0 | 0 | 18 | 450.36743 | 271.87436 | 723.18 | 538.67334 | 625.8473 | 442.15628 |
| Dnm2 | AFIHHELLAY | -689 | -698 | 0 | 0.002434616 | 10 | 459.10413 | 1006.0812 | NaN | NaN | NaN | 433.91135 |
| Dnm2 | NLVDSYVAIINK | -658 | -669 | 0 | 0.001868731 | 12 | 227.42857 | NaN | NaN | NaN | NaN | NaN |
| Ide | SEKPPHYLESR | -852 | -862 | 1 | 0 | 11 | 1058.614 | 1101.9062 | 341.59607 | 737.2551 | 1211.7494 | 1192.0198 |
| Ide | SEKPPHYLESR | -852 | -862 | 1 | 0 | 11 | 1058.614 | 1101.9062 | 341.59607 | 737.2551 | 1211.7494 | 1192.0198 |
| Ide | LRPENVR | -471 | -477 | 1 | 0.00025015 | 7 | 1043.3778 | 1043.7307 | 1092.4934 | 525.33887 | 1050.1357 | 511.787 |
| Ide | DELKEALDDVTLPR | -698 | -711 | 1 | 0 | 14 | 670.82837 | 368.07666 | 617.84705 | 367.76215 | 674.1229 | 809.8851 |
| Ide | EVNAVDSEHEK | -182 | -192 | 0 | 0 | 11 | 627.72296 | 831.167 | 830.98083 | 591.27313 | 678.0469 | 740.8832 |
| Ide | ENEYSQFLSEHAGSSN | -124 | -139 | 0 | 0.00025015 | 16 | 337.38718 | 500.45663 | 410.6782 | 507.37265 | 516.9424 | 627.13574 |
| Ide | AFIPQLLSR | -714 | -722 | 0 | 0 | 9 | 994.48486 | 1111.4957 | 1150.0117 | 1347.9934 | NaN | 1587.6189 |
| Ide | VLLISDPTTDK | -75 | -85 | 0 | 0 | 11 | 1491.7316 | 1626.6844 | 1674.3922 | 1513.1953 | NaN | NaN |
| Ide | NVPLPEFPEHPFQEE | -282 | -296 | 0 | 0 | 15 | 412.44568 | 500.7348 | 481.80923 | NaN | NaN | 847.20996 |
| Ide | HTKPLLPSQLVR | -754 | -765 | 1 | 0.00025015 | 12 | 332.45145 | NaN | 321.2073 | NaN | 243.16321 | 442.33658 |
| Ide | ALLHGNITK | -727 | -735 | 0 | 0.000712251 | 9 | 1383.2004 | NaN | NaN | 611.2804 | 1118.5807 | NaN |
| Ide | TKDDIIR | -932 | -938 | 1 | 0.000230707 | 7 | 736.4414 | NaN | NaN | 2292.1062 | 1737.5193 | NaN |
| Ide | DATPYPALIK | -543 | -552 | 0 | 0.00149795 | 10 | 729.1559 | 256.74146 | NaN | 1864.3123 | NaN | NaN |
| Ogt | IHQDGIHIL | -628 | -636 | 0 | 0.000550307 | 9 | 335.70715 | 350.71967 | 1562.1294 | 1206.7051 | NaN | 919.09625 |
| Ogt | IIFSPVAPK | -900 | -908 | 0 | 0 | 9 | 354.77298 | 612.293 | 489.92358 | NaN | NaN | 614.61414 |
| Ogt | ISSPLFNTK | -1002 | -1010 | 0 | 0.000755556 | 9 | 916.6592 | 2260.6907 | NaN | NaN | NaN | NaN |
| Ogt | AFLDSLPDVK | -743 | -752 | 0 | 0.00555405 | 10 | 859.9361 | NaN | NaN | 933.223 | NaN | NaN |
| Ogt | ALEVFPEFAAAH | -353 | -364 | 0 | 0 | 12 | 343.29285 | 242.0701 | NaN | NaN | NaN | NaN |
| Gbp6 | KLVELIGENK | -380 | -389 | 1 | 0.00051306 | 10 | 561.99445 | 562.2015 | 853.1985 | 662.61 | 922.9053 | 657.0855 |
| Gbp6 | AFLEEGFK | -554 | -561 | 0 | 0 | 8 | 1804.3721 | 1504.5656 | NaN | 1898.7477 | 1709.0345 | NaN |
| Gbp6 | DSITEDQLDPK | -248 | -258 | 0 | 0 | 11 | 4888.1875 | 5173.293 | NaN | 6005.316 | NaN | NaN |
| Gbp6 | LVELIGENK | -381 | -389 | 0 | 0.00266297 | 9 | 724.4663 | NaN | NaN | NaN | NaN | 1026.2382 |
| Ppp5c | LRDYETVVK | -116 | -124 | 1 | 0 | 9 | 1071.6046 | 1091.1246 | 1241.6713 | 1287.0737 | 1472.8364 | 1350.1322 |
| Ppp5c | LLYPDHFHLLR | -291 | -301 | 0 | 0 | 11 | 625.1636 | 829.43677 | 505.94366 | 524.51154 | 756.1296 | 724.3511 |
| Ppp5c | AFLEENQLDYIIR | -413 | -425 | 0 | 0 | 13 | 479.3167 | 537.3497 | 568.8742 | 122.4086 | 357.10193 | 568.2638 |
| Ppp5c | AIELNPGNAIYYGNR | -54 | -68 | 0 | 0 | 15 | 1108.4727 | NaN | 474.069 | 1407.6946 | 1425.2322 | 1592.358 |
| Ppp5c | SEDGVTLDDIR | -357 | -367 | 0 | 0 | 11 | 170.35567 | 290.5349 | 497.95007 | 622.79315 | NaN | 396.3995 |
| Ppp5c | GGLFSEDGVTLDDIR | -353 | -367 | 0 | 0 | 15 | 1596.8387 | NaN | 1955.458 | 1929.2659 | NaN | 2584.489 |
| Ppp5c | AKDYENAIK | -41 | -49 | 1 | 0 | 9 | 2541.3467 | 2485.7588 | NaN | 3042.035 | NaN | NaN |

| | | | | | | | | | | | | |
|---|---|---|---|---|---|---|---|---|---|---|---|---|
| Rragd;Rragc | AFLFDVVSK | (309);(258) | (317);(266) | 0 | 0 | 9 | 665.8151 | NaN | 626.2806 | NaN | 525.78 | 461.18616 |
| Tubb5 | YHGDSDLQLDR | -36 | -46 | 0 | 0 | 11 | 1378.2463 | 1431.0096 | 1698.718 | 1598.9203 | 3263.363 | 3087.1228 |
| Tubb5 | ISVYYNEATGGK | -47 | -58 | 0 | 0 | 12 | 1031.9282 | 979.74524 | 929.71436 | 1091.7885 | 1151.7217 | 1272.1885 |
| Tubb5 | GTYHGDSDLQLDR | -34 | -46 | 0 | 0 | 13 | 1023.1299 | 1422.9425 | 1031.7885 | 1771.794 | 3749.7964 | 1953.6946 |
| Tubb5 | GDSDLQLDR | -38 | -46 | 0 | 0 | 9 | 841.7837 | 685.391 | 955.6895 | 1205.0906 | 991.6276 | 848.4757 |
| Tubb5 | EVISDEHGIDPTGTYHGDSDLQLDR | -22 | -46 | 0 | 0 | 25 | 649.7114 | 707.4128 | 747.0836 | 585.4234 | 679.0063 | 635.39435 |
| Tubb5 | EHGIDPTGTYHGDSDLQLDR | -27 | -46 | 0 | 0 | 20 | 540.5215 | 653.25653 | 479.26913 | 590.56226 | 633.0909 | 852.9442 |
| Tubb5 | TAEEEEDFGEEAEEEA | -429 | -444 | 0 | 0 | 16 | 344.40668 | 417.15637 | 431.2744 | 859.86926 | 819.08875 | 708.9969 |
| Rpl34 | SACGVCPGR | -44 | -52 | 0 | 0.00025015 | 9 | 183.32089 | 1210.7051 | 1338.5156 | 1624.3622 | 1362.3583 | 1562.8187 |
| Rpl34 | AFLIEEQK | -94 | -101 | 0 | 0 | 8 | 1396.83 | 1193.8511 | 975.9871 | 567.74097 | NaN | NaN |
| Nfkb1 | RLEPVVSDAIYDSK | -228 | -241 | 1 | 0 | 14 | 1251.8037 | 1211.326 | 1344.9364 | 1329.6178 | 1401.5873 | 1282.6384 |
| Nfkb1 | HCEDGVCTVTAGPK | -115 | -128 | 0 | 0 | 14 | 1099.0087 | 1116.7502 | 1126.8455 | 1161.1603 | 1010.9237 | 1214.2672 |
| Nfkb1 | FTAFLPDSTGSFTR | -214 | -227 | 0 | 0 | 14 | 940.1888 | 642.81213 | 1030.8026 | 354.27347 | 717.42426 | 858.6465 |
| Nfkb1 | AFLPDSTGSFTR | -216 | -227 | 0 | 0 | 12 | 556.5894 | 619.7244 | 1136.321 | 358.63885 | 898.9144 | 994.1095 |
| Nfkb1 | FLPDSTGSFTR | -217 | -227 | 0 | 0 | 11 | 413.45288 | 909.1122 | 903.1201 | 373.85287 | 921.07336 | 1215.3416 |
| Nfkb1 | LYYPEIK | -346 | -352 | 0 | 0.003360691 | 7 | 2069.659 | 1907.029 | 1361.699 | NaN | 725.49 | 878.0638 |
| Nfkb1 | KSDLETSEPKPF | -334 | -345 | 2 | 0.000230707 | 12 | 903.1299 | NaN | 948.6633 | 870.86176 | 516.0275 | 1007.78766 |
| Nfkb1 | YKDVNITKPA | -316 | -325 | 2 | 0.00025015 | 10 | 700.38184 | NaN | NaN | 721.8974 | 737.0261 | 672.14417 |
| Nfkb1 | YVCEGPSHGGLPGAS | -57 | -71 | 0 | 0.000230707 | 15 | 389.77423 | NaN | NaN | NaN | 657.8025 | 847.5226 |
| Nfkb1 | VGADLSLLDR | -600 | -609 | 0 | 0 | 10 | 1766.0696 | NaN | NaN | NaN | 1983.1621 | NaN |
| Nfkb1 | ICNYVGPAK | -84 | -92 | 0 | 0.002168954 | 9 | 1342.071 | NaN | NaN | NaN | NaN | NaN |
| Cd2ap | LGLFPSNFVK | -154 | -163 | 0 | 0 | 10 | 799.6863 | 817.28845 | 830.7712 | 591.4404 | 802.64575 | 615.74866 |
| Cd2ap | AFLTPLELK | -560 | -568 | 0 | 0 | 9 | 400.19696 | NaN | 214.84833 | 243.21658 | 242.63084 | NaN |
| Cd2ap | TQNVEVTKPDVDGK | -254 | -267 | 1 | 0.00025015 | 14 | 1672.8424 | NaN | 2092.6323 | NaN | 2243.0002 | NaN |
| Cd2ap | ISELDKDFPKPK | -328 | -339 | 2 | 0.003998192 | 12 | 98.22389 | NaN | NaN | NaN | NaN | NaN |
| Hnrnpu | DLPEHAVLK | -603 | -611 | 0 | 0.00025015 | 9 | 9201.731 | 9993.532 | 9951.493 | 548.52844 | 605.89355 | 10608.226 |
| Hnrnpu | DIDIHEVR | -329 | -336 | 0 | 0 | 8 | 6509.118 | 730.1267 | 7122.192 | 8742.903 | 9463.302 | 9199.433 |
| Hnrnpu | EVLADRPLFPHVL | -413 | -425 | 1 | 0 | 13 | 5213.268 | 5914.618 | 5843.1655 | 6220.013 | 8093.24 | 7992.4595 |
| Hnrnpu | EVLADRPLFPHVL | -413 | -425 | 1 | 0 | 13 | 5213.268 | 5914.618 | 5843.1655 | 6220.013 | 8093.24 | 7992.4595 |
| Hnrnpu | EKPYFPIPEDCTFIQ | -439 | -453 | 1 | 0 | 15 | 3536.5935 | 3294.9707 | 3108.2007 | 4422.33 | 5721.334 | 4866.818 |
| Hnrnpu | GYFEYIEENKYSR | -232 | -244 | 1 | 0 | 13 | 3275.1165 | 382.85358 | 4009.8723 | 557.7942 | 5103.343 | 4657.982 |
| Hnrnpu | ANFETDEVELSYAK | -386 | -399 | 0 | 0 | 14 | 3019.0217 | 3484.8313 | 3625.979 | 2690.677 | 3397.0225 | 3210.9304 |
| Hnrnpu | AVEFNFGQK | -430 | -438 | 0 | 0.000431655 | 9 | 2972.5686 | 2581.6907 | 3114.5188 | 2422.2463 | 627.39246 | 336.61194 |
| Hnrnpu | GKDLPEHAVLK | -601 | -611 | 1 | 0 | 11 | 2891.6116 | 81.56971 | 2850.11 | 2150.4492 | 2544.8628 | 2025.9219 |
| Hnrnpu | GKDLPEHAVLK | -601 | -611 | 1 | 0 | 11 | 2891.6116 | 81.56971 | 2850.11 | 2150.4492 | 2544.8628 | 2025.9219 |
| Hnrnpu | HYHQGYY | -794 | -800 | 0 | 0.001092529 | 7 | 2313.2974 | 2340.314 | 1989.1781 | 2430.5764 | 2516.356 | 2298.438 |
| Hnrnpu | LLEQYKEESKK | -641 | -651 | 2 | 0 | 11 | 2079.1992 | 566.3772 | 732.68256 | 574.62726 | 627.67255 | 703.0473 |
| Hnrnpu | LLEQYKEESKK | -641 | -651 | 2 | 0 | 11 | 2079.1992 | 566.3772 | 732.68256 | 574.62726 | 627.67255 | 703.0473 |
| Hnrnpu | EVLADRPLFPH | -413 | -423 | 1 | 0 | 11 | 1928.437 | 587.77936 | 2494.2026 | 2252.2427 | 2943.6943 | 2912.8389 |
| Hnrnpu | TCNCETEDYGEKF | -364 | -376 | 1 | 0 | 13 | 1566.9573 | 1648.6669 | 1656.8732 | 2421.2861 | 1896.1237 | 2008.6974 |
| Hnrnpu | LNTLLQR | -528 | -534 | 0 | 0.001291103 | 7 | 1368.3922 | 861.2453 | 787.2486 | 451.1665 | 514.7237 | 498.46246 |
| Hnrnpu | FETDEVELSYAK | -388 | -399 | 0 | 0 | 12 | 1281.6075 | 1407.957 | 1372.4465 | 366.35733 | 1016.3409 | 223.38705 |
| Hnrnpu | LLEQYKEESK | -641 | -650 | 1 | 0 | 10 | 1245.0906 | 1321.2434 | 1162.836 | 1239.4404 | 1133.2474 | 1250.6558 |
| Hnrnpu | GYFEYIEENK | -232 | -241 | 0 | 0 | 10 | 1055.3394 | 1121.7859 | 1187.4911 | 1025.0881 | 277.6497 | 940.4167 |
| Hnrnpu | EVLADRPLFPHVLCHN | -413 | -428 | 1 | 0.00111006 | 16 | 994.36255 | 1190.9425 | 1149.0275 | 1027.2186 | 759.9004 | 1711.8558 |
| Hnrnpu | EVLADRPLFPHVLCHN | -413 | -428 | 1 | 0.00111006 | 16 | 994.36255 | 1190.9425 | 1149.0275 | 1027.2186 | 759.9004 | 1711.8558 |
| Hnrnpu | PLFPHVL | -419 | -425 | 0 | 0.001868731 | 7 | 934.02954 | 156.32846 | 991.85376 | 968.87164 | 1241.4758 | 1236.401 |
| Hnrnpu | PEHAVLK | -605 | -611 | 0 | 0.000431655 | 7 | 686.8783 | 463.44073 | 596.90704 | 671.20044 | 1117.2712 | 1188.4712 |
| Hnrnpu | NCAVEFNFGQK | -428 | -438 | 0 | 0 | 11 | 676.3057 | 665.75476 | 585.32007 | 901.8142 | 224.43953 | 995.5106 |
| Hnrnpu | DEITYVELQK | -626 | -635 | 0 | 0 | 10 | 665.16815 | 908.7627 | 7941.7744 | 465.0141 | 5145.367 | 5607.043 |
| Hnrnpu | DEITYVELQKEEAQK | -626 | -640 | 1 | 0 | 15 | 544.9913 | 610.72595 | 581.48065 | 436.94534 | 657.9533 | 685.6105 |
| Hnrnpu | EKPYFPIPEDCTF | -439 | -451 | 1 | 0.00025015 | 13 | 472.69455 | 374.71432 | 409.90625 | 147.67372 | 408.79712 | 300.80157 |
| Hnrnpu | FANFETDEVELSYAK | -385 | -399 | 0 | 0 | 15 | 417.94592 | 2839.0986 | 3035.4556 | 2055.2659 | 3992.874 | 3753.767 |
| Hnrnpu | HAAENPGKY | -493 | -501 | 1 | 0.001055699 | 9 | 407.7072 | 770.68097 | 837.4877 | 543.71844 | 866.69073 | 1051.3265 |
| Hnrnpu | SFAFLWAGGR | -291 | -300 | 0 | 0 | 10 | 396.0829 | 446.364 | 462.76337 | 393.48663 | 588.89923 | 560.6471 |
| Hnrnpu | TCNCETEDYGEK | -364 | -375 | 0 | 0 | 12 | 299.13397 | 275.79996 | 307.0077 | 335.3367 | 290.22333 | 405.12387 |
| Hnrnpu | KAVVVCPK | -578 | -585 | 1 | 0 | 8 | 298.4143 | 683.73883 | 252.41658 | 1019.09503 | 241.04256 | 189.25356 |
| Hnrnpu | LQAALDNEAGGRPA | -38 | -51 | 1 | 0 | 14 | 3144.814 | 3374.589 | 2112.7183 | NaN | 539.78406 | 2122.733 |
| Hnrnpu | AFLWAGGR | -293 | -300 | 0 | 0.001776473 | 8 | 2699.4 | 2615.6338 | 2135.758 | 1347.6995 | NaN | 1097.7336 |
| Hnrnpu | GGIGYPYPR | -722 | -730 | 0 | 0 | 9 | 1624.5197 | NaN | 1418.2548 | 2636.5564 | 1764.9296 | 1884.0525 |
| Hnrnpu | GEEEFSYGYSLK | -349 | -360 | 0 | 0 | 12 | 1065.0934 | NaN | 1343.0767 | 109.87535 | 1644.5613 | 1407.9327 |
| Hnrnpu | LLEQYKEE | -641 | -648 | 1 | 0.003521747 | 8 | 683.5035 | 1468.9185 | 1369.2324 | 899.80853 | NaN | 1034.2526 |
| Hnrnpu | NGQDLGVAFK | -400 | -409 | 0 | 0 | 10 | 3768.1416 | NaN | 3227.2043 | 2667.388 | NaN | 2035.6199 |
| Hnrnpu | SSGPTSLFAVT | -182 | -192 | 0 | 0.00111006 | 11 | 1880.7228 | 2647.8845 | 2555.906 | 1314.1068 | NaN | NaN |
| Hnrnpu | FEYIEENK | -234 | -241 | 0 | 0 | 8 | 1212.7576 | 1219.0414 | 873.7127 | NaN | NaN | 1038.7041 |
| Hnrnpu | VSELKEELK | -13 | -21 | 1 | 0 | 9 | 858.9071 | 676.9253 | 710.1417 | NaN | NaN | 605.49506 |
| Hnrnpu | LQAALDNEAGGR | -38 | -49 | 0 | 0 | 12 | 1672.5723 | 1295.5084 | 1294.7107 | NaN | NaN | NaN |
| Hnrnpu | FDEITYVELQK | -625 | -635 | 0 | 0.000947459 | 11 | 995.6964 | 1260.4568 | 1212.2444 | NaN | NaN | NaN |
| Hnrnpu | VSELKEELKK | -13 | -22 | 2 | 0 | 10 | 633.3634 | 671.52014 | 622.2572 | NaN | NaN | NaN |
| Hnrnpu | VSELKEELKK | -13 | -22 | 2 | 0 | 10 | 633.3634 | 671.52014 | 622.2572 | NaN | NaN | NaN |
| Hnrnpu | GQKPWSQHYHQGYY | -787 | -800 | 1 | 0.001463897 | 14 | 714.1648 | NaN | 456.2221 | NaN | NaN | NaN |
| Hnrnpu | ETDEVELSYAK | -389 | -399 | 0 | 0.003037016 | 11 | 1165.151 | NaN | NaN | NaN | NaN | NaN |
| Rpl3 | DDASKPVHLT | -35 | -44 | 1 | 0.00025015 | 10 | 5437.7524 | 5456.333 | 5676.567 | 1517.8265 | 6155.5986 | 6041.3613 |
| Rpl3 | KYCQVIR | -155 | -161 | 1 | 0 | 7 | 2817.528 | 2713.2827 | 2634.079 | 6412.6235 | 3024.2134 | 1983.6389 |
| Rpl3 | SINPLGGFVHY | -313 | -323 | 0 | 0.00025015 | 11 | 2226.615 | 2540.31 | 2700.0652 | 2556.2466 | 3219.0798 | 3260.9958 |
| Rpl3 | ERLEQQVPVNQVFGQ | -199 | -213 | 1 | 0.00051306 | 15 | 1807.0292 | 2430.1716 | 2323.6064 | 1394.9465 | 1776.8441 | 1905.6375 |
| Rpl3 | LEQQVPVNQVFGQ | -201 | -213 | 0 | 0 | 13 | 1478.767 | 1251.7582 | 881.49493 | 1277.7437 | 1670.2603 | 1623.21 |
| Rpl3 | ERLEQQVPVNQ | -199 | -209 | 1 | 0.00025015 | 11 | 1258.1345 | 1382.3556 | 545.61096 | 2189.6565 | 1716.4143 | 1740.7048 |

| | | | | | | | | | | | | |
|---|---|---|---|---|---|---|---|---|---|---|---|---|
| Rpl3 | EKLDWAR | -192 | -198 | 1 | 0.000627943 | 7 | 1246.9767 | 1030.2135 | 1178.1936 | 1759.9447 | 967.34247 | 1078.1001 |
| Rpl3 | TVFAEHISDECK | -104 | -115 | 0 | 0 | 12 | 1115.4568 | 1231.623 | 1230.3398 | 728.0733 | 324.2527 | 627.4447 |
| Rpl3 | NASTDYDLSDK | -302 | -312 | 0 | 0 | 11 | 1043.7455 | 970.6101 | 841.1216 | 148.12833 | 336.95645 | 1139.9475 |
| Rpl3 | SINPLGGFVHYG | -313 | -324 | 0 | 0.00025015 | 12 | 938.21985 | 1130.4282 | 1114.6519 | 1084.153 | 1225.8027 | 464.76965 |
| Rpl3 | GEVTNDFIMLK | -324 | -334 | 0 | 0 | 11 | 857.6294 | 2809.3748 | 2783.816 | 2804.418 | 2561.3938 | 170.83362 |
| Rpl3 | ASTDYDLSDK | -303 | -312 | 0 | 0 | 10 | 726.8683 | 1119.9431 | 926.20654 | 769.761 | 979.38135 | 961.1327 |
| Rpl3 | ERLEQQVPVNQVF | -199 | -211 | 1 | 0 | 13 | 635.5263 | 1349.2203 | 1271.4769 | 946.0554 | 1099.1416 | 1200.45 |
| Rpl3 | NNASTDYDLSDK | -301 | -312 | 0 | 0 | 12 | 114.924255 | 1665.6077 | 1588.9435 | 1395.7756 | 126.88541 | 382.01163 |
| Rpl3 | DEMIDVIGVTK | -214 | -224 | 0 | 0 | 11 | 1042.165 | 1101.9921 | 587.12537 | 1196.3613 | NaN | 1343.8313 |
| Rpl3 | TVFAEHISDECKR | -104 | -116 | 1 | 0 | 13 | 1904.9083 | 1781.489 | 1732.4762 | NaN | NaN | 2040.4396 |
| Rpl3 | TVFAEHISDECKR | -104 | -116 | 1 | 0 | 13 | 1904.9083 | 1781.489 | 1732.4762 | NaN | NaN | 2040.4396 |
| Rpl3 | HGSLGFLPR | -11 | -19 | 0 | 0 | 9 | 1478.7518 | 1674.907 | 1733.4646 | NaN | 485.56653 | NaN |
| Rpl3 | FAEHISDECK | -106 | -115 | 0 | 0 | 10 | 901.98303 | NaN | NaN | 115.869675 | 125.28399 | 1204.7434 |
| Rpl3 | LEQQVPVNQVFGQDE | -201 | -215 | 0 | 0.00025015 | 15 | 1291.333 | 841.0424 | 118.83038 | NaN | NaN | NaN |
| Rpl3 | IGQGYLIK | -287 | -294 | 0 | 0 | 8 | 670.22186 | 537.9367 | 535.7706 | NaN | NaN | NaN |
| Rpl3 | ERLEQQVPVNQVFGQDE | -199 | -215 | 1 | 0.000230707 | 17 | 667.1401 | 509.23737 | 633.5582 | NaN | NaN | NaN |
| Rpl3 | SINPLGGFVHYGEVTND | -313 | -329 | 0 | 0 | 17 | 515.19165 | 485.84985 | 505.3877 | NaN | NaN | NaN |
| Rpl3 | AFMGPLKK | -387 | -394 | 1 | 0.00635324 | 8 | 668.8389 | NaN | 401.82437 | NaN | NaN | NaN |
| Rpl3 | KAFMGPLK | -386 | -393 | 1 | 0 | 8 | 194.39487 | NaN | NaN | NaN | 476.40524 | NaN |
| Hbb-b1 | AFNDGLNHLDSLK | -71 | -83 | 0 | 0 | 13 | 1900.7812 | 3043.3586 | 2490.9487 | 556.45667 | 3476.404 | 3916.0876 |
| Hbb-b1 | NDGLNHLDSLK | -73 | -83 | 0 | 0 | 11 | 724.46686 | 745.25366 | 4514.4424 | 4438.6616 | 1155.9496 | 4740.2163 |
| Rpa1 | FDSEGQALDGVSISDHR | -424 | -440 | 0 | 0 | 17 | 1875.2827 | 1315.7848 | 1511.5298 | 1808.5161 | 2097.1245 | 1426.9016 |
| Rpa1 | DVKPVDFR | -602 | -609 | 1 | 0 | 8 | 1619.7227 | 1669.7357 | 1690.7048 | 1744.4069 | 1554.9781 | 1501.0225 |
| Rpa1 | FFPLIEVNK | -254 | -262 | 0 | 0 | 9 | 1209.5597 | 1219.2278 | 1200.7671 | 836.0547 | 1498.3057 | 1717.7045 |
| Rpa1 | LWGEDADKFDGSR | -369 | -381 | 1 | 0 | 13 | 1016.28534 | 648.66125 | 838.9728 | 659.41455 | 588.3024 | 1225.8656 |
| Rpa1 | LWGEDADKFDGSR | -369 | -381 | 1 | 0 | 13 | 1016.28534 | 648.66125 | 838.9728 | 659.41455 | 588.3024 | 1225.8656 |
| Rpa1 | TVIVNPDIPEAYK | -407 | -419 | 0 | 0 | 13 | 889.02655 | 816.7451 | 680.9341 | 1112.2885 | 1256.063 | 1170.3572 |
| Rpa1 | FDFTGIGDLESK | -309 | -320 | 0 | 0 | 12 | 884.20905 | 833.4119 | 880.5017 | 988.6511 | 1312.4012 | 1260.1655 |
| Rpa1 | TLWGEDADKFDGSR | -368 | -381 | 1 | 0 | 14 | 642.65546 | 773.61865 | 810.1205 | 222.52585 | 989.0055 | 1099.7101 |
| Rpa1 | TLWGEDADKFDGSR | -368 | -381 | 1 | 0 | 14 | 642.65546 | 773.61865 | 810.1205 | 222.52585 | 989.0055 | 1099.7101 |
| Rpa1 | SLELVDESGEIR | -232 | -243 | 0 | 0 | 12 | 516.51843 | 524.74414 | 955.8321 | 1204.7708 | 953.7497 | 414.93018 |
| Rpa1 | AKDALVDIIGIC | -321 | -332 | 1 | 0 | 12 | 515.14435 | 453.37857 | 428.8817 | 277.15662 | 783.99786 | 322.83752 |
| Rpa1 | SSSTVIVNPDIPEAYK | -404 | -419 | 0 | 0 | 16 | 507.95865 | 633.625 | 668.5917 | 592.2745 | 705.3489 | 769.77057 |
| Rpa1 | VIDQQNGLYR | -499 | -508 | 0 | 0 | 10 | 385.16486 | 429.51822 | 426.91962 | 876.55853 | 630.2726 | 532.4994 |
| Rpa1 | EKNEQAFEEVFQNA | -561 | -574 | 1 | 0.00025015 | 14 | 350.3047 | 525.80164 | 492.64282 | 603.01733 | 736.546 | 707.05365 |
| Rpa1 | VKLETYNDESR | -585 | -595 | 1 | 0 | 11 | 3777.7615 | 1969.8949 | 1657.1981 | NaN | 745.818 | 648.95526 |
| Rpa1 | AFNEQVDK | -246 | -253 | 0 | 0 | 8 | 3703.5728 | 2182.7395 | 1592.8993 | 1905.3688 | NaN | 850.9029 |
| Rpa1 | NEQAFEEVFQNA | -563 | -574 | 0 | 0.000431655 | 12 | 858.422 | NaN | 1101.5283 | 938.6235 | 895.6787 | 1195.451 |
| Rpa1 | DALVDIIGICK | -323 | -333 | 0 | 0 | 11 | 691.16956 | NaN | 748.58624 | 763.3438 | 976.58826 | 158.30966 |
| Rpa1 | ELVDESGEIR | -234 | -243 | 0 | 0 | 10 | 561.6842 | 660.52014 | 491.1598 | 664.1199 | NaN | NaN |
| Rpa1 | SGGAGGGNTNWK | -441 | -452 | 0 | 0.00644558 | 12 | 836.8655 | NaN | 252.8609 | 218.03432 | NaN | NaN |
| Rpa1 | VVPIASLTPYQSK | -193 | -205 | 0 | 0 | 13 | 371.30045 | 444.38385 | NaN | 451.60037 | NaN | NaN |
| Rpa1 | SENLGQGDKADYF | -459 | -471 | 1 | 0.000591124 | 13 | 614.4201 | NaN | NaN | NaN | 756.59314 | NaN |
| Rpa1 | GWFDSEGQALDGVSISDHR | -422 | -440 | 0 | 0.006624942 | 19 | 265.68835 | NaN | NaN | NaN | NaN | NaN |
| Actr2 | HLWDYTFGPEK | -87 | -97 | 0 | 0 | 11 | 4940.7046 | 5765.2695 | 5752.439 | 6041.5166 | 7710.3687 | 7487.0103 |
| Actr2 | NFPEHIFPALVGR | -26 | -38 | 0 | 0 | 13 | 1894.9115 | 2190.7417 | 2168.7039 | 1358.9114 | 2896.0955 | 2335.363 |
| Actr2 | NFPEHIFPALVGR | -26 | -38 | 0 | 0 | 13 | 1894.9115 | 2190.7417 | 2168.7039 | 1358.9114 | 2896.0955 | 2335.363 |
| Actr2 | FEAPEALFQPH | -259 | -269 | 0 | 0 | 11 | 1406.205 | 1698.7253 | 1639.9966 | 2631.1323 | 2357.1235 | 2398.3853 |
| Actr2 | FEAPEALFQPH | -259 | -269 | 0 | 0 | 11 | 1406.205 | 1698.7253 | 1639.9966 | 2631.1323 | 2357.1235 | 2398.3853 |
| Actr2 | SNFPEHIFPALVGR | -25 | -38 | 0 | 0 | 14 | 1254.3417 | 1477.59 | 1660.1713 | 878.8754 | 1630.4806 | 1268.5475 |
| Actr2 | ILLTEPPMNPTK | -107 | -118 | 0 | 0 | 12 | 1224.5082 | 1128.292 | 735.88116 | 602.19104 | 698.7324 | 745.1683 |
| Actr2 | AFNHSADFETVR | -203 | -214 | 0 | 0 | 12 | 453.57156 | 560.5588 | 453.17957 | 477.6406 | 628.31134 | 549.93304 |
| Actr2 | AFNHSADFETVR | -203 | -214 | 0 | 0 | 12 | 453.57156 | 560.5588 | 453.17957 | 477.6406 | 628.31134 | 549.93304 |
| Actr2 | VVVCDNGTGFVK | -8 | -19 | 0 | 0 | 12 | 372.0087 | 877.771 | 813.2016 | 322.57315 | 2423.372 | 2412.325 |
| Actr2 | LCYVGYNIEQEQK | -220 | -232 | 0 | 0 | 13 | 5747.095 | 7358.4 | 7810.043 | 8660.947 | 9409.322 | NaN |
| Actr2 | HLWDYTFGPEKL | -87 | -98 | 1 | 0 | 12 | 1071.2372 | NaN | 1298.671 | 470.04633 | 1324.9518 | 591.86707 |
| Actr2 | DKDNFWMTR | -367 | -375 | 1 | 0 | 9 | 857.4195 | 871.08514 | 968.19415 | 823.2285 | NaN | 220.4333 |
| Actr2 | YVGYNIEQEQK | -222 | -232 | 0 | 0 | 11 | 857.0868 | NaN | 1197.2611 | 1158.8597 | 1101.8768 | 980.35535 |
| Actr2 | NHSADFETVR | -205 | -214 | 0 | 0.00025015 | 10 | 807.69495 | 1328.2009 | 1252.0919 | NaN | 1713.4791 | 1127.129 |
| Actr2 | VESYTLPDGR | -241 | -250 | 0 | 0.00025015 | 10 | 1040.6133 | NaN | NaN | 915.82227 | 1042.9152 | 1158.3903 |
| Actr2 | FEAPEALFQPHLINVE | -259 | -274 | 0 | 0.000298805 | 16 | 420.90985 | 460.35638 | NaN | NaN | NaN | 605.5028 |
| Actr2 | VGDEASELR | -57 | -65 | 0 | 0.007987481 | 9 | 2607.0261 | 2555.2375 | NaN | NaN | NaN | NaN |
| Actr2 | MYPGLPSR | -309 | -316 | 0 | 0.005015524 | 8 | 1064.5015 | NaN | NaN | 587.5709 | NaN | NaN |
| Krtcap2 | AFNNLENLVFGK | -54 | -65 | 0 | 0 | 12 | 535.5766 | 450.9092 | 426.80212 | 519.8163 | 456.01874 | 672.0389 |
| Lancl1 | AFPNPYADYNK | -5 | -15 | 0 | 0.001868731 | 11 | 636.562 | NaN | 526.7267 | 764.5536 | 672.7529 | NaN |
| Slc9a1 | AFPVLDIDYPHVR | -92 | -104 | 0 | 0.00111006 | 13 | 491.31198 | NaN | NaN | 626.30334 | NaN | NaN |
| Gzma | AFPYPCYDEYTR | -99 | -110 | 0 | 0 | 12 | 2010.0994 | 2489.5942 | 2738.456 | 2872.1814 | 3677.8792 | 3558.472 |
| Gzma | HYNFHPVIGLN | -183 | -193 | 0 | 0.000230707 | 11 | 1854.6006 | 2099.857 | 2198.3467 | 611.6161 | 2832.0425 | 2583.8953 |
| Gzma | IIGGDTVVPH | -29 | -38 | 0 | 0 | 10 | 1617.1953 | 1494.3177 | 1377.8875 | 1498.6177 | 461.61993 | 1396.3605 |
| Gzma | IIGGDTVVPHSR | -29 | -40 | 0 | 0 | 12 | 1540.7804 | 1612.2059 | 1664.4006 | 1540.2058 | 1681.8219 | 1672.9341 |
| Gzma | KAFPYPCYDEYTR | -98 | -110 | 1 | 0 | 13 | 812.8323 | 1017.30524 | 988.6809 | 954.8248 | 1346.6049 | 1267.4004 |
| Gzma | KAFPYPCYDEYTR | -98 | -110 | 1 | 0 | 13 | 812.8323 | 1017.30524 | 988.6809 | 954.8248 | 1346.6049 | 1267.4004 |
| Gzma | DSCNGDSGSPLLCDGILR | -205 | -222 | 0 | 0 | 18 | 765.9927 | 763.68036 | 1021.18994 | 1041.052 | 1315.7247 | 1103.8878 |
| Gzma | SINKEPEQQILTVK | -84 | -97 | 1 | 0 | 14 | 521.444 | 607.0453 | 618.23267 | 1295.0504 | 1425.4061 | 1489.0469 |
| Gzma | HYNFHPVIGL | -183 | -192 | 0 | 0 | 10 | 215.41425 | 245.4297 | 1937.2592 | 1471.969 | 196.91618 | 1479.3436 |
| Gzma | EGDLQLVR | -111 | -118 | 0 | 0 | 8 | 1813.1184 | 1706.8059 | 1597.2448 | 811.2357 | NaN | 728.6995 |
| Gzma | LCDGILR | -216 | -222 | 0 | 0.001868731 | 7 | 959.0569 | NaN | NaN | NaN | NaN | NaN |
| Uggt1 | IVPEWQDYDQEIK | -1509 | -1521 | 0 | 0 | 13 | 1490.0442 | 1825.6041 | 1907.062 | 406.20715 | 2828.0544 | 2748.745 |

| | | | | | | | | | | | |
|---|---|---|---|---|---|---|---|---|---|---|---|
| Uggt1 | NIQPSETDYAVDIR | -441 | -454 | 0 | 0 | 14 | 1329.4703 | 526.13934 | 2701.361 | 492.9553 | 501.5778 | 479.8539 |
| Uggt1 | SILGIDSAYDQNR | -604 | -616 | 0 | 0 | 13 | 1225.6339 | 2386.4592 | 2359.7046 | 742.4775 | 2886.9275 | 2930.6245 |
| Uggt1 | AFQQIAVDEPPPEGCK | -133 | -148 | 0 | 0.000550307 | 16 | 1205.1501 | 796.7193 | 889.3285 | 1529.0372 | 1489.4613 | 1007.6396 |
| Uggt1 | ELYPALEGQLK | -292 | -302 | 0 | 0 | 11 | 946.6176 | 898.9688 | 876.84216 | 884.09863 | 798.6099 | 870.67725 |
| Uggt1 | WIVGDFDSPSGR | -764 | -775 | 0 | 0 | 12 | 837.51483 | 873.3834 | 894.9314 | 626.58484 | 827.7456 | 996.46045 |
| Uggt1 | FLFVDADQIVR | -1354 | -1364 | 0 | 0 | 11 | 800.0652 | 977.1891 | 849.4901 | 1359.9121 | 1513.2134 | 1591.604 |
| Uggt1 | KDISQNFPTK | -347 | -356 | 1 | 0 | 10 | 1101.7794 | NaN | 3645.3506 | 3061.547 | 181.81729 | 2356.2368 |
| Uggt1 | GTIGLQPGDSALFIN | -382 | -396 | 0 | 0.003166358 | 15 | 656.8105 | 635.23016 | NaN | 527.4476 | 674.31415 | 857.76404 |
| Uggt1 | ETQEGSQKHEEL | -1540 | -1551 | 1 | 0 | 12 | 348.33102 | 305.38745 | 852.95917 | 804.18304 | 576.5977 | NaN |
| Uggt1 | LNIQPSETDYAVDIR | -440 | -454 | 0 | 0.000230707 | 15 | 560.5888 | 330.21185 | 696.9628 | NaN | NaN | 268.95752 |
| Uggt1 | KNYLSPTFK | -1292 | -1300 | 1 | 0 | 9 | 1307.5388 | NaN | 858.7551 | NaN | 751.61426 | NaN |
| Uggt1 | QLLYDAIK | -776 | -783 | 0 | 0 | 8 | 1589.1367 | NaN | NaN | 1198.2603 | NaN | NaN |
| Uggt1 | GYTPFCDSR | -1381 | -1389 | 0 | 0 | 9 | 342.84906 | NaN | 509.23633 | NaN | NaN | NaN |
| Uggt1 | VEHVVSVL | -586 | -593 | 0 | 0.002910424 | 8 | 1121.7874 | NaN | NaN | NaN | NaN | NaN |
| Uggt1 | ASNNFYVDDFAR | -711 | -722 | 0 | 0.000431655 | 12 | 541.5054 | NaN | NaN | NaN | NaN | NaN |
| Pes1 | SHDHPTDVDYR | -208 | -218 | 0 | 0 | 11 | 2377.9424 | 2214.0955 | 2765.1033 | 1995.4747 | 1785.1996 | 1853.479 |
| Pes1 | SHDHPTDVDYR | -208 | -218 | 0 | 0 | 11 | 2377.9424 | 2214.0955 | 2765.1033 | 1995.4747 | 1785.1996 | 1853.479 |
| Pes1 | WVFDCVNAR | -396 | -404 | 0 | 0 | 9 | 1199.3318 | 1199.7861 | 1265.0212 | 1237.1846 | 1056.4836 | 1144.4142 |
| Pes1 | LHEPIVNK | -74 | -81 | 0 | 0 | 8 | 1080.1407 | 1261.3984 | 1625.1809 | 1523.9713 | 1657.3027 | 1179.8353 |
| Pes1 | AFSHDHPTDVDYR | -206 | -218 | 0 | 0 | 13 | 675.83746 | 654.42676 | 664.4054 | 717.42896 | 718.10834 | 684.4846 |
| Pes1 | LLGFVNFR | -230 | -237 | 0 | 0 | 8 | 479.8435 | 468.52972 | 383.2451 | 409.90405 | 423.2777 | 477.10025 |
| Pes1 | IVDRPGQQTPIIGR | -376 | -389 | 1 | 0.000471409 | 14 | 791.85 | 883.0209 | 1009.483 | NaN | 928.32965 | 949.74164 |
| Pes1 | KVFLSIK | -180 | -186 | 1 | 0 | 7 | 491.962 | 353.34665 | 326.97778 | 400.94614 | NaN | NaN |
| Pes1 | LAQEEESEAK | -520 | -529 | 0 | 0.00025015 | 10 | 915.42017 | NaN | 924.6592 | 140.57597 | NaN | NaN |
| Pes1 | SEWNAVER | -99 | -106 | 0 | 0 | 8 | 698.4998 | 484.36783 | 1765.3599 | NaN | NaN | NaN |
| Pes1 | SFGGDVSWDK | -348 | -357 | 0 | 0 | 10 | 650.2888 | NaN | NaN | NaN | NaN | 4281.4517 |
| Sec61a1 | CVILPEIQKPER | -13 | -24 | 1 | 0 | 12 | 987.60254 | 1119.0093 | 1071.1979 | 423.0822 | 1332.3821 | 1445.5311 |
| Sec61a1 | GTWSDTSSGGPAR | -322 | -334 | 0 | 0 | 13 | 952.6862 | 513.4381 | 950.65564 | 774.51105 | 614.6803 | 618.077 |
| Sec61a1 | VILPEIQKPER | -14 | -24 | 1 | 0.000394361 | 11 | 581.0578 | 627.0834 | 595.1105 | 682.0373 | 507.51352 | 500.35327 |
| Sec61a1 | AFSPTTVNTGR | -195 | -205 | 0 | 0 | 11 | 325.16858 | 445.18005 | 646.90234 | 682.4622 | 549.64935 | NaN |
| Sfpq | FGQGGAGPVGGQGPR | -659 | -673 | 0 | 0 | 15 | 2646.511 | 2562.331 | 2640.5278 | 2603.775 | 2401.104 | 2241.9746 |
| Sfpq | HHQGPPPGGPGPR | -247 | -259 | 0 | 0 | 13 | 1538.3234 | 1461.8193 | 1084.6705 | 987.34265 | 1449.9224 | 1597.8086 |
| Sfpq | HHQGPPPGGPGPR | -247 | -259 | 0 | 0 | 13 | 1538.3234 | 1461.8193 | 1084.6705 | 987.34265 | 1449.9224 | 1597.8086 |
| Sfpq | LFVGNLPADITEDEFK | -291 | -306 | 0 | 0 | 16 | 1123.1791 | 1037.5886 | 798.8499 | 1461.9597 | 1468.6123 | 1377.6602 |
| Sfpq | NLPADITEDEFKR | -295 | -307 | 1 | 0 | 13 | 891.8641 | 472.7203 | 765.9793 | 616.3254 | 1595.4724 | 1463.2706 |
| Sfpq | TTTPRPVIVEPLEQLDDEDGLPEK | -431 | -454 | 1 | 0.001055699 | 24 | 764.5307 | 986.6355 | 787.7693 | 1248.2614 | 1284.6426 | 981.07635 |
| Sfpq | LFVGNLPADITEDEFKR | -291 | -307 | 1 | 0 | 17 | 759.5238 | 713.1099 | 846.5894 | 702.6884 | 860.7525 | 852.61273 |
| Sfpq | LFVGNLPADITEDEFKR | -291 | -307 | 1 | 0 | 17 | 759.5238 | 713.1099 | 846.5894 | 702.6884 | 860.7525 | 852.61273 |
| Sfpq | VGNLPADITEDEFKR | -293 | -307 | 1 | 0 | 15 | 719.6981 | 923.2002 | 1243.0016 | 1314.6365 | 1495.6818 | 1258.0216 |
| Sfpq | VGNLPADITEDEFKR | -293 | -307 | 1 | 0 | 15 | 719.6981 | 923.2002 | 1243.0016 | 1314.6365 | 1495.6818 | 1258.0216 |
| Sfpq | PGPGPGGPK | -192 | -200 | 0 | 0.002068213 | 9 | 706.1837 | 572.3808 | 738.0178 | 392.1186 | 155.94624 | 873.26996 |
| Sfpq | GREEYEGPNK | -686 | -695 | 1 | 0.00025015 | 10 | 611.18195 | 749.433 | 544.00635 | 711.2396 | 627.8851 | 624.4861 |
| Sfpq | PVIVEPLEQLDDEDGLPEK | -436 | -454 | 0 | 0 | 19 | 577.87085 | 559.13617 | 619.1404 | 786.1854 | 937.8119 | 946.48157 |
| Sfpq | TTPRPVIVEPLEQLDDEDGLPEK | -432 | -454 | 1 | 0 | 23 | 547.7995 | 644.16925 | 681.19446 | 708.62726 | 779.75555 | 750.5556 |
| Sfpq | FAQHGTFEYEYSQR | -472 | -485 | 0 | 0 | 14 | 544.8726 | 3179.1108 | 1732.2325 | 4417.653 | 3285.2126 | 1868.9302 |
| Sfpq | FAQHGTFEYEYSQR | -472 | -485 | 0 | 0 | 14 | 544.8726 | 3179.1108 | 1732.2325 | 4417.653 | 3285.2126 | 1868.9302 |
| Sfpq | ELLEEAFSQFGPIER | -377 | -391 | 0 | 0 | 15 | 380.62933 | 360.21063 | 304.2635 | 409.76202 | 492.23718 | 438.07007 |
| Sfpq | GTFEYEYSQR | -476 | -485 | 0 | 0.00025015 | 10 | 2680.2305 | 2179.616 | NaN | 311.31995 | NaN | 1116.2467 |
| Sfpq | AFSQFGPIER | -382 | -391 | 0 | 0 | 10 | 1099.8823 | 961.17975 | 1124.9708 | 1150.7351 | NaN | NaN |
| Sfpq | EQLDDEDGLPEK | -443 | -454 | 0 | 0 | 12 | 827.31323 | 1288.5458 | 465.90277 | 945.4856 | NaN | NaN |
| Sfpq | FSQFGPIER | -383 | -391 | 0 | 0.00025015 | 9 | 1405.6155 | NaN | 403.0325 | NaN | NaN | NaN |
| Sfpq | FGQGGAGPVGGQGPRG | -659 | -674 | 1 | 0.002972566 | 16 | 948.9817 | NaN | NaN | NaN | NaN | NaN |
| Sfpq | LEEAFSQFGPIER | -379 | -391 | 0 | 0 | 13 | 309.74805 | NaN | NaN | NaN | NaN | NaN |
| Pspc1 | LFVGNLPTDITEEDFKR | -83 | -99 | 1 | 0 | 17 | 3231.802 | 405.084 | 3960.463 | 3717.1418 | 4958.2046 | 5122.714 |
| Pspc1 | LFVGNLPTDITEEDFKR | -83 | -99 | 1 | 0 | 17 | 3231.802 | 405.084 | 3960.463 | 3717.1418 | 4958.2046 | 5122.714 |
| Pspc1 | SFLKPGEK | -68 | -75 | 1 | 0 | 8 | 881.4719 | 1145.7917 | 1121.1582 | 838.1636 | 691.13806 | 1150.1753 |
| Pspc1 | LFVGNLPTDITEEDFK | -83 | -98 | 0 | 0 | 16 | 453.74432 | 353.50275 | 752.2452 | 546.01855 | 903.0943 | 792.7273 |
| Pspc1 | AFSQFGPVEK | -174 | -183 | 0 | 0 | 10 | 694.4228 | NaN | 917.2354 | NaN | NaN | NaN |
| Pspc1 | EQFDDEDGLPEK | -235 | -246 | 0 | 0.000550307 | 12 | 554.9846 | NaN | NaN | NaN | NaN | NaN |
| Srsf7 | RPFDPNDR | -127 | -134 | 1 | 0.00025015 | 8 | 5904.26 | 5673.253 | 5606.296 | 4270.015 | 6392.4277 | 6012.2354 |
| Srsf7 | VYVGNLGTGAGKG | -42 | -54 | 1 | 0.000471409 | 13 | 6893.522 | 7187.1177 | 4044.7725 | 7717.001 | NaN | 515.82556 |
| Srsf7 | FDRPPAR | -120 | -126 | 1 | 0.008129024 | 7 | 2772.2126 | 2767.034 | 2881.5933 | 832.2379 | 2435.427 | NaN |
| Srsf7 | VYVGNLGTGAGK | -42 | -53 | 0 | 0 | 12 | 1969.9348 | 2281.9937 | 1069.9426 | 4521.723 | 2606.3477 | NaN |
| Srsf7 | AFSYYGPLR | -59 | -67 | 0 | 0.00111006 | 9 | 1212.022 | 1103.758 | 1139.0715 | NaN | NaN | NaN |
| Prpf31 | VGYELKDEIER | -316 | -326 | 1 | 0 | 11 | 1423.2649 | 1532.1406 | 1604.5903 | 1817.1704 | 1861.7693 | 1913.5106 |
| Prpf31 | VDSFHESTEGK | -305 | -315 | 0 | 0 | 11 | 1145.0166 | 944.63983 | 912.25555 | 276.2852 | 731.1534 | 512.5834 |
| Prpf31 | AFTPLQGLEIVNPQAAEK | -453 | -470 | 0 | 0 | 18 | 373.2003 | 514.7823 | 574.37976 | 218.24918 | 522.2686 | 528.3784 |
| Prpf31 | SVAFTPLQGLEIVNPQAAEK | -451 | -470 | 0 | 0 | 20 | 181.79181 | 219.92844 | 261.07727 | 364.0358 | 488.7371 | 371.70746 |
| Prpf31 | EVMGPVEAAPEYR | -79 | -91 | 0 | 0.00025015 | 13 | 1043.5076 | 1454.0944 | 1765.8876 | 1351.7351 | 1310.9995 | NaN |
| Prpf31 | DKWQEPPPVK | -329 | -338 | 1 | 0.003461185 | 10 | 514.69275 | NaN | 779.10974 | NaN | NaN | NaN |
| Prpf31 | GPVEAAPEYR | -82 | -91 | 0 | 0.00450079 | 10 | 1934.4708 | NaN | NaN | NaN | NaN | NaN |
| Prpf31 | GQQLSDEELER | -176 | -186 | 0 | 0.002595029 | 11 | 1275.149 | NaN | NaN | NaN | NaN | NaN |
| Rpl7a | TNYNDRYDEIR | -224 | -234 | 1 | 0 | 11 | 9324.351 | 9185.093 | 7742.8623 | 9258.524 | 9932.519 | 6838.8315 |
| Rpl7a | TNYNDRYDEIR | -224 | -234 | 1 | 0 | 11 | 9324.351 | 9185.093 | 7742.8623 | 9258.524 | 9932.519 | 6838.8315 |
| Rpl7a | HWGGNVLGPK | -236 | -245 | 0 | 0 | 10 | 6185.4565 | 7935.819 | 7738.9365 | 3303.7935 | 5116.282 | 4703.5967 |
| Rpl7a | FTQVNSEDKGALAK | -204 | -217 | 1 | 0 | 14 | 2448.938 | 703.8015 | 655.3808 | 451.98758 | 852.19464 | 470.764 |
| Rpl7a | YNDRYDEIR | -226 | -234 | 1 | 0.00025015 | 9 | 2021.3958 | 2133.9592 | 1968.8496 | 2454.9556 | 345.65308 | 2546.5195 |
| Rpl7a | VNSEDKGALAK | -207 | -217 | 1 | 0 | 11 | 1899.6761 | 1472.2146 | 1479.3497 | 70.58102 | 2424.748 | 3051.3955 |

| | | | | | | | | | | | | |
|---|---|---|---|---|---|---|---|---|---|---|---|---|
| Rpl7a | KVAPAPAVVK | -11 | -20 | 1 | 0.00051306 | 10 | 1524.998 | 1489.2656 | 1462.2463 | 1576.2767 | 1126.5897 | 1202.2864 |
| Rpl7a | KAQLVVIAH | -151 | -159 | 1 | 0.00142999 | 9 | 1470.235 | 1636.86 | 1481.6238 | 1733.1669 | 1869.4105 | 1659.5066 |
| Rpl7a | VFLPALCR | -168 | -175 | 0 | 0 | 8 | 1352.5399 | 1100.8036 | 1162.5326 | 1257.1998 | 1451.8533 | 1169.7123 |
| Rpl7a | AFTQVNSEDKGALAK | -203 | -217 | 1 | 0 | 15 | 1173.3164 | 1204.9697 | 1455.2014 | 1681.7412 | 1541.6638 | 1520.8903 |
| Rpl7a | TQVNSEDKGALAK | -205 | -217 | 1 | 0 | 13 | 583.81354 | 1172.7861 | 958.70325 | 452.15958 | 887.6762 | 1156.8625 |
| Rpl7a | RHWGGNVL | -235 | -242 | 1 | 0.001291103 | 8 | 412.3074 | 844.8839 | 1001.3893 | 1626.8251 | 994.1737 | 1193.342 |
| Rpl7a | QVNSEDKGALAK | -206 | -217 | 1 | 0 | 12 | 293.64758 | 339.80847 | 130.22064 | 117.60373 | 397.20502 | 351.99985 |
| Rpl7a | KVVNPLFEK | -26 | -34 | 1 | 0 | 9 | 3564.1187 | 3515.2827 | 3701.2349 | NaN | 532.04767 | 573.10583 |
| Rpl7a | QFTQALDR | -82 | -89 | 0 | 0 | 8 | 1379.8705 | 1391.7827 | NaN | 314.52277 | 1212.8964 | 1807.9811 |
| Rpl7a | LGHLVHR | -190 | -196 | 0 | 0 | 7 | 1347.3027 | 1301.2513 | 1503.2649 | 370.4171 | NaN | 174.88998 |
| Rpl7a | NYNDRYDEIR | -225 | -234 | 1 | 0.000826806 | 10 | 1011.7565 | NaN | 1033.7549 | 1258.8607 | 1340.04 | 1361.8801 |
| Rpl7a | AQLVVIAH | -152 | -159 | 0 | 0.003069674 | 8 | 936.31177 | NaN | NaN | NaN | NaN | NaN |
| Tfrc | LNSIEFADTIK | -137 | -147 | 0 | 0 | 11 | 1509.5507 | 1648.2532 | 1743.0056 | 1710.0846 | 2089.1003 | 259.1042 |
| Tfrc | YHFLSPYVSPR | -686 | -696 | 0 | 0 | 11 | 1479.53 | 1334.7367 | 1434.49 | 1585.4146 | 1786.4453 | 1557.4257 |
| Tfrc | FSNLFGGEPLSYTR | -9 | -22 | 0 | 0 | 14 | 1431.154 | 1385.186 | 1294.3198 | 848.99585 | 5104.349 | 1064.3809 |
| Tfrc | YIENQFHEFK | -170 | -179 | 0 | 0 | 10 | 1097.3855 | 197.91605 | 957.13763 | 1774.5491 | 1511.4087 | 1438.8923 |
| Tfrc | WNIDSSCK | -359 | -366 | 0 | 0 | 8 | 1005.2818 | 1086.169 | 931.0114 | 1402.2529 | 1316.5955 | 1305.9712 |
| Tfrc | LFGGEPLSYTR | -12 | -22 | 0 | 0 | 11 | 963.6549 | 2580.5195 | 1272.336 | 5472.0806 | 6659.5005 | 1203.1007 |
| Tfrc | LVHANFGTK | -234 | -242 | 0 | 0.000230707 | 9 | 955.69257 | 1038.6781 | 903.5367 | 821.3114 | 1077.3593 | 1230.2378 |
| Tfrc | LTTDFHNAEK | -659 | -668 | 0 | 0 | 10 | 613.72473 | 1166.2006 | 1357.3682 | 1267.1436 | 375.47333 | 608.77057 |
| Tfrc | FCEDADYPYLGTR | -560 | -572 | 0 | 0 | 13 | 579.8404 | 609.7595 | 646.01556 | 508.65604 | 512.72534 | 536.20917 |
| Tfrc | LYWADLK | -124 | -130 | 0 | 0 | 7 | 553.6864 | 663.9542 | 708.226 | 505.33804 | 419.35352 | 381.45264 |
| Tfrc | CFCEDADYPYLGTR | -559 | -572 | 0 | 0 | 14 | 330.6381 | 537.88605 | 598.4885 | 493.8274 | 480.27356 | 515.4207 |
| Tfrc | GLPNIPVQTISR | -330 | -341 | 0 | 0 | 12 | 1527.8401 | 1379.6733 | 1798.6676 | 1706.5154 | 408.60495 | NaN |
| Tfrc | QLSQNTYTPR | -148 | -157 | 0 | 0 | 10 | 1247.5177 | NaN | NaN | 1996.2229 | 2627.6438 | 2170.0908 |
| Tfrc | SQNTYTPR | -150 | -157 | 0 | 0.004595494 | 8 | 452.43054 | NaN | NaN | 956.6793 | 815.20154 | 893.3994 |
| Tfrc | VEYHFLSPYVSPR | -684 | -696 | 0 | 0 | 13 | 797.06573 | 973.012 | 211.74467 | NaN | NaN | NaN |
| Tfrc | AGEITFAEK | -261 | -269 | 0 | 0 | 9 | 493.05798 | NaN | 609.602 | 635.32947 | NaN | NaN |
| Tfrc | EAGSQKDESLAY | -158 | -169 | 1 | 0.007905138 | 12 | 193.09421 | 213.29013 | 733.95337 | NaN | NaN | NaN |
| Tfrc | AFTYINLDK | -481 | -489 | 0 | 0 | 9 | 1164.2026 | NaN | NaN | NaN | NaN | NaN |
| Acadl | LPANALLGEENK | -268 | -279 | 0 | 0 | 12 | 1754.2732 | 1764.7113 | 2040.2471 | 1741.297 | 2111.3958 | 2092.531 |
| Acadl | IFSSEHDIFR | -52 | -61 | 0 | 0 | 10 | 1276.8529 | 3034.8777 | 3184.8647 | 1747.21 | 1674.3787 | 1707.0868 |
| Acadl | DTAELFFEDVR | -257 | -267 | 0 | 0 | 11 | 941.62085 | 994.6568 | 1093.2146 | 1243.3134 | 1321.6229 | 1264.6777 |
| Acadl | FFQEEVIPHHTEWEK | -67 | -81 | 0 | 0 | 15 | 842.8756 | 602.5611 | 513.6153 | 848.8995 | 606.7565 | 825.9211 |
| Acadl | FFQEEVIPHHTEWEK | -67 | -81 | 0 | 0 | 15 | 842.8756 | 602.5611 | 513.6153 | 848.8995 | 606.7565 | 825.9211 |
| Acadl | AFVDSCLQLHETK | -346 | -358 | 0 | 0 | 13 | 652.20465 | 769.2714 | 880.6122 | 1093.1385 | 445.45792 | 1497.17 |
| Acadl | SSEHDIFR | -54 | -61 | 0 | 0 | 8 | 621.1314 | 1155.2885 | 1274.3529 | 1253.8861 | 1521.0292 | 1554.0945 |
| Acadl | VQPIYGGTNEIMK | -407 | -419 | 0 | 0 | 13 | 344.6197 | 483.93842 | 457.1162 | 564.32 | 695.1297 | 608.3292 |
| Acadl | SCLQLHETK | -350 | -358 | 0 | 0 | 9 | 1285.2424 | 2030.1412 | 1540.1438 | NaN | 1022.3796 | 1122.6505 |
| Acadl | TEPGAGSDLQGVR | -173 | -185 | 0 | 0 | 13 | 3405.4282 | 919.4051 | 4221.925 | 4600.4536 | NaN | NaN |
| Acadl | EVIPHHTEWEK | -71 | -81 | 0 | 0.002199523 | 11 | 2202.0042 | NaN | NaN | NaN | NaN | NaN |
| Acadl | QELPQER | -286 | -292 | 0 | 0 | 7 | 980.12067 | NaN | NaN | NaN | NaN | NaN |
| Acadl | MTEPGAGSDLQGVR | -172 | -185 | 0 | 0.00025015 | 14 | 674.67096 | NaN | NaN | NaN | NaN | NaN |
| C1qbp | FLTDEIKEEK | -82 | -91 | 1 | 0 | 10 | 3965.1018 | 301.45325 | 4298.5156 | 286.87387 | 549.50867 | 4868.6943 |
| C1qbp | AFVEFLTDEIKEEK | -78 | -91 | 1 | 0 | 14 | 690.0077 | 733.10956 | 827.7925 | 1170.0635 | 1082.6099 | 1190.7734 |
| C1qbp | ATGDSEWR | -209 | -216 | 0 | 0 | 8 | 500.7134 | 381.88538 | 302.15222 | 666.60425 | 611.3263 | 623.26483 |
| C1qbp | NSIPPTFDGEEEPSQGQK | -134 | -151 | 0 | 0 | 18 | 900.803 | 1193.6809 | 1191.1901 | NaN | 694.76624 | 726.01416 |
| C1qbp | AFVEFLTDEIK | -78 | -88 | 0 | 0 | 11 | 441.9049 | 585.782 | NaN | 722.18823 | 717.5419 | 736.56854 |
| C1qbp | FLTDEIK | -82 | -88 | 0 | 0.001392869 | 7 | 1095.9818 | NaN | 569.6255 | 651.1878 | NaN | 708.1482 |
| C1qbp | NINNSIPPTFDGEEEPSQGQK | -131 | -151 | 0 | 0 | 21 | 298.80148 | 397.22403 | 354.98663 | NaN | 108.02775 | NaN |
| Ddx49 | AFVLPILQK | -55 | -63 | 0 | 0 | 9 | 1716.9486 | 2018.389 | 2111.563 | 476.77942 | 2176.5032 | 2169.7722 |
| Ddx49 | TAAFVLPILQK | -53 | -63 | 0 | 0 | 11 | 303.6403 | NaN | NaN | NaN | 248.86205 | 160.72037 |
| Ddx49 | VVINHNTPGLPK | -319 | -330 | 0 | 0.001463897 | 12 | 359.2831 | NaN | NaN | 544.5573 | NaN | NaN |
| Upf2 | AFVPAILFK | -475 | -483 | 0 | 0 | 9 | 302.12646 | 165.87265 | 407.72797 | 405.61127 | 434.65646 | 501.26685 |
| Pcnp | SAEDEAADLPTKPTK | -53 | -67 | 1 | 0 | 15 | 2002.4695 | 1915.4453 | 1960.4125 | 2058.0112 | 2373.604 | 2502.392 |
| Pcnp | AGAAGGPEEEAEKPVK | -17 | -32 | 1 | 0 | 16 | 791.2895 | 1111.7139 | 1276.5146 | 666.209 | 774.895 | 892.9983 |
| Pcnp | AGAAGGPEEEAEKPVK | -17 | -32 | 1 | 0 | 16 | 791.2895 | 1111.7139 | 1276.5146 | 666.209 | 774.895 | 892.9983 |
| Pcnp | AGEEKPEKPQR | -6 | -16 | 2 | 0.00111006 | 11 | 202.77165 | 303.3014 | 351.0068 | NaN | 217.55087 | 246.07565 |
| Pcnp | AGEEKPEKPQR | -6 | -16 | 2 | 0.00111006 | 11 | 202.77165 | 303.3014 | 351.0068 | NaN | 217.55087 | 246.07565 |
| Pcnp | SHLGNVHDQDN | -168 | -178 | 0 | 0 | 11 | 414.77786 | 355.84488 | NaN | 481.24442 | 350.49594 | NaN |
| Cyb5b | ATESFEDVGHSPDAR | -70 | -84 | 0 | 0 | 15 | 1935.147 | 1652.877 | 2292.5886 | 1281.5604 | 1697.0305 | 1648.5906 |
| Cyb5b | YIGDVHPSDLKPK | -91 | -103 | 1 | 0 | 13 | 1295.6716 | 633.5775 | 2008.996 | 1884.9127 | 4940.2964 | 3383.6892 |
| Cyb5b | YIGDVHPSDLKPK | -91 | -103 | 1 | 0 | 13 | 1295.6716 | 633.5775 | 2008.996 | 1884.9127 | 4940.2964 | 3383.6892 |
| Cyb5b | HFWADSK | -138 | -144 | 0 | 0.000550307 | 7 | 1107.5424 | 1265.7654 | 1203.3347 | 1210.0221 | 1153.7084 | 1146.1245 |
| Cyb5b | GADATESFEDVGHSPDAR | -67 | -84 | 0 | 0 | 18 | 990.59204 | 921.8776 | 892.63477 | 764.0174 | 494.80188 | 87.17507 |
| Cyb5b | VEGSEPSVTYYR | -13 | -24 | 0 | 0 | 12 | 825.9387 | 4199.7637 | 3962.962 | 1273.4557 | 3052.8184 | 3085.2776 |
| Cyb5b | AGADATESFEDVGHSPDAR | -66 | -84 | 0 | 0 | 19 | 611.89276 | 608.9793 | 618.25775 | 813.8892 | 709.62366 | 718.0788 |
| Cyb5b | AGADATESFEDVGHSPDAR | -66 | -84 | 0 | 0 | 19 | 611.89276 | 608.9793 | 618.25775 | 813.8892 | 709.62366 | 718.0788 |
| Cyb5b | YYIGDVHPSDLKPK | -90 | -103 | 1 | 0 | 14 | 7780.0273 | 7754.4067 | NaN | 8082.9365 | 10009.655 | 10589.267 |
| Cyb5b | FLSEHPGGEEVLLEQA | -51 | -66 | 0 | 0.00078661 | 16 | 703.8724 | 819.20703 | 934.0277 | 160.62256 | NaN | 985.7766 |
| Cyb5b | FLSEHPGGEEVLLEQAGAD | -51 | -69 | 0 | 0.001776473 | 19 | 1379.509 | 1374.3225 | NaN | NaN | NaN | NaN |
| Cyb5b | SFEDVGHSPDAR | -73 | -84 | 0 | 0 | 12 | 618.3537 | 1048.1887 | NaN | NaN | NaN | NaN |
| Atp5mk | AGAESDGQFQFT | -2 | -13 | 0 | 0.00025015 | 12 | 300.01498 | NaN | 293.66687 | 400.3061 | 367.43155 | 411.6958 |
| Rbm42 | AGAGPAPGLPVAGGPVVPGPGVGIPGK | -6 | -32 | 0 | 0.00025015 | 27 | 553.4098 | 668.7396 | 658.4693 | 829.11896 | 746.23706 | 975.97754 |
| Rbm42 | ALRPPLEEPAAPR | -221 | -233 | 1 | 0 | 13 | 1028.0001 | NaN | NaN | 3974.7766 | 3477.3816 | NaN |
| Noc3l | DLSSSEPVHIK | -113 | -123 | 0 | 0 | 11 | 2588.033 | 2706.5532 | 2710.1077 | 1853.9258 | 1839.2112 | 2062.6453 |
| Noc3l | ELIHLLPIK | -148 | -156 | 0 | 0 | 9 | 793.1299 | 784.62146 | 753.3222 | 610.7217 | 633.4774 | 661.7967 |
| Noc3l | AGAPSEGSEALKPELSR | -712 | -728 | 1 | 0 | 17 | 566.35706 | 587.0488 | 797.43616 | 448.61697 | 815.1675 | 941.70593 |

| | | | | | | | | | | | | |
|---|---|---|---|---|---|---|---|---|---|---|---|---|
| Noc3l | YCNEAAPETPLDFAK | -778 | -792 | 0 | 0 | 15 | 390.34933 | 436.83746 | 526.9233 | 573.2944 | 851.45496 | 961.6394 |
| Noc3l | SPLLPAVLEGLAK | -511 | -523 | 0 | 0 | 13 | 284.26776 | 309.84814 | NaN | 328.20026 | 386.44403 | 415.11478 |
| Noc3l | HYHPIVR | -698 | -704 | 0 | 0.00078661 | 7 | 160.49751 | NaN | NaN | NaN | NaN | NaN |
| Ube2k | VYAGAPVSSPEYTK | -151 | -164 | 0 | 0 | 14 | 1530.8949 | 1147.6857 | 2023.5616 | 1422.1932 | 1395.7964 | 1397.9205 |
| Ube2k | IPETYPFNPPK | -62 | -72 | 0 | 0 | 11 | 1367.048 | 1299.7883 | 1691.8921 | 1341.4788 | 1189.7106 | 1427.8163 |
| Ube2k | AHVYAGAPVSSPEYTK | -149 | -164 | 0 | 0.000230707 | 16 | 1044.0156 | 1811.6508 | 1959.3068 | NaN | 1706.815 | 1492.7068 |
| Ube2k | AGAPVSSPEYTK | -153 | -164 | 0 | 0 | 12 | 3130.3745 | 2716.3755 | 3571.1404 | NaN | NaN | 5779.3115 |
| Ube2k | IAGPPDTPYEGGR | -43 | -55 | 0 | 0 | 13 | 694.938 | 657.29675 | 980.5248 | 1378.7305 | NaN | NaN |
| Ube2k | VDLVDENFTELR | -29 | -40 | 0 | 0 | 12 | 609.2346 | NaN | NaN | 1594.8705 | 707.1633 | 794.59784 |
| Ube2k | GEIAGPPDTPYEGGR | -41 | -55 | 0 | 0 | 15 | 438.37143 | 719.1044 | 452.67883 | NaN | NaN | NaN |
| Taldo1 | LSSTWEGIQAGK | -143 | -154 | 0 | 0 | 12 | 4813.169 | 1714.4712 | 5454.032 | 6799.8545 | 6379.9634 | 3518.576 |
| Taldo1 | LSSTWEGIQAGK | -143 | -154 | 0 | 0 | 12 | 4813.169 | 1714.4712 | 5454.032 | 6799.8545 | 6379.9634 | 3518.576 |
| Taldo1 | AGCDFLTISPK | -248 | -258 | 0 | 0 | 11 | 3448.1504 | 749.5658 | 3977.7637 | 4678.5435 | 4888.299 | 5014.574 |
| Taldo1 | SYEPQEDPGVK | -205 | -215 | 0 | 0 | 11 | 2365.75 | 2028.1587 | 1849.8798 | 1609.3898 | 1501.8834 | 1803.79 |
| Taldo1 | SYEPQEDPGVK | -205 | -215 | 0 | 0 | 11 | 2365.75 | 2028.1587 | 1849.8798 | 1609.3898 | 1501.8834 | 1803.79 |
| Taldo1 | ELEEQHGIHCN | -155 | -165 | 0 | 0 | 11 | 1548.195 | 236.48512 | 1744.2334 | 1661.7028 | 2102.6702 | 2008.7163 |
| Taldo1 | KFAADAIK | -314 | -321 | 1 | 0 | 8 | 1411.4215 | 2022.4542 | 2498.9834 | 1624.2703 | 2389.014 | 2285.863 |
| Taldo1 | MESALDQLK | -11 | -19 | 0 | 0 | 9 | 832.53766 | 1183.2152 | 1213.7947 | 1720.1162 | 776.778 | 1694.3523 |
| Taldo1 | ALAGCDFLTISPK | -246 | -258 | 0 | 0 | 13 | 803.8501 | 1210.8917 | 1005.2498 | 1139.7557 | 1195.4412 | 1129.7542 |
| Taldo1 | WLHNEDQMAVEK | -296 | -307 | 0 | 0 | 12 | 605.38324 | 503.6396 | 573.4712 | 457.2211 | 472.28552 | 472.20447 |
| Taldo1 | DWHVANTDKK | -195 | -204 | 1 | 0.000230707 | 10 | 564.3109 | 454.4485 | 529.3746 | 514.19446 | 524.9252 | 468.40558 |
| Taldo1 | LGGPQEEQIK | -72 | -81 | 0 | 0.00111006 | 10 | 261.843 | 863.9941 | 4168.142 | 774.54974 | 3915.3186 | 1781.0386 |
| Taldo1 | RLIELYK | -124 | -130 | 1 | 0.004857192 | 7 | 1027.1791 | 1061.8724 | 980.9315 | NaN | 1153.8501 | 1096.3888 |
| Taldo1 | AGVTLISPFVGR | -181 | -192 | 0 | 0.001291103 | 12 | 576.49396 | 600.45856 | NaN | 733.0621 | 211.94826 | 498.4069 |
| Taldo1 | LISPFVGR | -185 | -192 | 0 | 0 | 8 | 1965.5499 | NaN | 773.33154 | 1033.8885 | 447.79578 | NaN |
| Taldo1 | ILDWHVANTDKK | -193 | -204 | 1 | 0 | 12 | 1702.4904 | NaN | NaN | 2297.595 | 2255.804 | 2354.821 |
| Taldo1 | ILDWHVANTDKK | -193 | -204 | 1 | 0 | 12 | 1702.4904 | NaN | NaN | 2297.595 | 2255.804 | 2354.821 |
| Txnl1 | TEDDIKEDGIVPLR | -221 | -234 | 1 | 0 | 14 | 895.0136 | 1075.6869 | 1074.8528 | 1231.4696 | 1020.97943 | 1140.5818 |
| Txnl1 | QHLENDPGSNEDADIPK | -105 | -121 | 0 | 0 | 17 | 837.17584 | 874.2398 | 865.26324 | 919.9006 | 1023.4694 | 967.9746 |
| Txnl1 | IDQYQGADAVGLEEK | -88 | -102 | 0 | 0 | 15 | 774.77783 | 844.4821 | 796.4584 | 185.1313 | 1246.9579 | 853.82104 |
| Txnl1 | VGVKPVGSDPDFQPELSGAGSR | -2 | -23 | 1 | 0 | 22 | 721.4054 | 813.5239 | 863.49347 | 1853.4048 | 1012.3716 | 812.4974 |
| Txnl1 | NISATPTFLFFR | -72 | -83 | 0 | 0 | 12 | 587.7036 | 581.47363 | 612.1074 | 693.7897 | 832.3681 | 804.651 |
| Txnl1 | ELTEDDIKEDGIVPLR | -219 | -234 | 1 | 0 | 16 | 490.93237 | 333.25476 | 506.8279 | 571.74774 | 467.2358 | 494.44177 |
| Txnl1 | IFINLPR | -196 | -202 | 0 | 0.00025015 | 7 | 1950.3 | 1535.3141 | 1255.0457 | NaN | NaN | NaN |
| Txnl1 | AGCECLNESDEHGFDNCLR | -133 | -151 | 0 | 0.000907206 | 19 | 635.075 | 1208.6556 | NaN | NaN | 402.0809 | NaN |
| Txnl1 | ATPTFLFFR | -75 | -83 | 0 | 0.001776473 | 9 | 492.6035 | 484.09692 | 147.36092 | NaN | NaN | NaN |
| Rpf2 | THLASDDLYK | -228 | -237 | 0 | 0.000471409 | 10 | 1679.3074 | 1620.1719 | 1501.6912 | 574.24084 | 850.6649 | 932.5941 |
| Rpf2 | AGDDFDVTEDFRR | -146 | -158 | 1 | 0.000431655 | 13 | 1316.752 | 114.6932 | 1179.0726 | 2455.8076 | 2794.6099 | 3337.2986 |
| Rpf2 | NLLIDFFR | -161 | -168 | 0 | 0.000947459 | 8 | 335.25513 | 260.00772 | 180.09647 | 315.0857 | 376.4064 | 128.63548 |
| Rpf2 | AGDDFDVTEDFR | -146 | -157 | 0 | 0 | 12 | 446.48337 | 576.0898 | 519.1752 | NaN | NaN | NaN |
| Hadh | LKNELFQR | -126 | -133 | 1 | 0 | 8 | 1044.2549 | 1079.2333 | 1061.914 | 1014.65173 | 1064.5784 | 1071.095 |
| Hadh | AGDEFVEK | -88 | -95 | 0 | 0.000471409 | 8 | 1278.5364 | 870.6514 | 786.4848 | 1918.106 | NaN | NaN |
| Hadh | LLVPYLIEAVR | -222 | -232 | 0 | 0 | 11 | 593.3607 | NaN | NaN | 882.46 | 1192.2064 | 959.0011 |
| Ermp1 | AGDNILAVLK | -365 | -374 | 0 | 0 | 10 | 956.98303 | 744.7142 | 921.8651 | NaN | 1044.2125 | 1237.4177 |
| Ermp1 | ILIDSIQR | -357 | -364 | 0 | 0.000591124 | 8 | 399.071 | 528.70746 | NaN | NaN | 221.24582 | NaN |
| Ermp1 | TSGLQGEFDAR | -94 | -104 | 0 | 0 | 11 | 1573.4891 | 1751.1613 | NaN | NaN | NaN | NaN |
| Hspa1l | NQVAMNPQNTVFDAK | -59 | -73 | 0 | 0.00149795 | 15 | 2665.036 | 588.4641 | NaN | NaN | NaN | NaN |
| Fermt3 | DEAPGDPTQQLNLK | -388 | -401 | 0 | 0 | 14 | 1642.4834 | 2261.9897 | 2360.289 | 2073.9114 | 2655.4668 | 2095.68 |
| Fermt3 | YYSFFDLDPK | -253 | -262 | 0 | 0 | 10 | 1447.7382 | 1344.5942 | 1404.4124 | 149.48514 | 1166.176 | 1516.0419 |
| Fermt3 | VGEEDPEAQSVTLR | -22 | -35 | 0 | 0 | 14 | 1396.9645 | 1500.0684 | 1335.8374 | 1805.5435 | 1780.5126 | 1725.0004 |
| Fermt3 | LFFGPQHR | -91 | -98 | 0 | 0 | 8 | 1340.1952 | 1422.1488 | 1066.73 | 1573.5598 | 1158.6447 | 1035.9961 |
| Fermt3 | IDLAVGDVVK | -579 | -588 | 0 | 0.000431655 | 10 | 1250.403 | 1290.7883 | 1258.2834 | 1233.1984 | 1399.5042 | 1425.0823 |
| Fermt3 | KQDWSDHAIWWEQK | -56 | -69 | 1 | 0 | 14 | 425.98773 | 411.61346 | 377.5235 | 1806.2642 | 411.06137 | 415.88092 |
| Fermt3 | KQDWSDHAIWWEQK | -56 | -69 | 1 | 0 | 14 | 425.98773 | 411.61346 | 377.5235 | 1806.2642 | 411.06137 | 415.88092 |
| Fermt3 | AGDVLWLR | -243 | -250 | 0 | 0 | 8 | 995.97986 | 1059.9407 | 956.2699 | 1237.823 | 1137.0308 | 1204.25 |
| Fermt3 | LAFLSLQR | -470 | -477 | 0 | 0 | 8 | 492.4796 | 537.87396 | 507.3768 | 426.0275 | 464.9012 | 512.34143 |
| Fermt3 | QWNVNWDIR | -597 | -605 | 0 | 0 | 9 | 142.74414 | 1289.6997 | 1367.0938 | 800.10925 | 918.40393 | 275.87897 |
| Fermt3 | SGDYIDSSWELR | -8 | -19 | 0 | 0 | 12 | 138.63846 | 1102.7898 | 1067.2633 | 1314.743 | 1397.4753 | 183.88377 |
| Fermt3 | VFVGEEDPEAQSVTLR | -20 | -35 | 0 | 0 | 16 | 3419.2947 | 3224.544 | 3368.9429 | 1824.0946 | 4939.549 | NaN |
| Fermt3 | EKEPEEEVHDL | -154 | -164 | 1 | 0 | 11 | 1716.8026 | 1610.1615 | 2785.8606 | 2170.847 | 1931.7396 | NaN |
| Fermt3 | GCEVVPDVNVSGQK | -402 | -415 | 0 | 0 | 14 | 1293.4308 | 1317.5117 | 1268.5542 | NaN | NaN | 1363.4766 |
| Fermt3 | LTQLYEQAR | -268 | -276 | 0 | 0 | 9 | 2623.5835 | 559.08276 | 2917.1846 | NaN | NaN | NaN |
| Fermt3 | EVVPDVNVSGQK | -404 | -415 | 0 | 0 | 12 | 2021.1992 | NaN | 944.9721 | 2153.8716 | NaN | NaN |
| Fermt3 | SQDEAPGDPTQQLNLK | -386 | -401 | 0 | 0 | 16 | 935.75433 | NaN | NaN | 1381.9584 | 855.38245 | NaN |
| Fermt3 | QYWVVFK | -371 | -377 | 0 | 0.005176406 | 7 | 654.37476 | 660.7537 | 755.3105 | NaN | NaN | NaN |
| Fermt3 | EKEPEEEVHDLTK | -154 | -166 | 1 | 0.00025015 | 13 | 295.41855 | NaN | NaN | 417.31235 | NaN | 1020.88605 |
| Fermt3 | ASGDYIDSSWELR | -7 | -19 | 0 | 0 | 13 | 377.12216 | NaN | NaN | 755.73285 | NaN | NaN |
| Fermt3 | VTGESHIGGVL | -36 | -46 | 0 | 0.001055699 | 11 | 1026.8271 | NaN | NaN | NaN | NaN | NaN |
| Ybx3 | NVTGPDGVPVEGSR | -148 | -161 | 0 | 0 | 14 | 846.8076 | 2046.2078 | 2052.5808 | 3138.5032 | 2225.59 | 2411.3372 |
| Ybx3 | SEAGEATTGGTTLPQAAADAPAAAPPDPAPK | -2 | -32 | 0 | 0.002534621 | 31 | 331.98965 | 369.85623 | 404.77893 | 491.7836 | 411.5677 | 1003.4201 |
| Ybx3 | LAGSPGGDAAPGPAPASSAPAGGEDAEK | -49 | -76 | 0 | 0.00051306 | 28 | 234.36229 | 250.70815 | 263.4539 | 208.44066 | 222.19305 | 205.49971 |
| Ybx3 | AGEAPTENPAPATEQSSAE | -343 | -361 | 0 | 0 | 19 | 393.47647 | 622.1012 | NaN | 779.3763 | 411.20157 | 624.33704 |
| Ybx3 | GSPGGDAAPGPAPASSAPAGGEDAEK | -51 | -76 | 0 | 0.00149795 | 26 | 365.43637 | 168.25719 | 457.11664 | 93.48715 | 212.07988 | NaN |
| Ybx3 | ANVTGPDGVPVEGSR | -147 | -161 | 0 | 0.00051306 | 15 | 781.54565 | NaN | NaN | NaN | NaN | NaN |
| Xpo1 | NVDILKDPETVK | -675 | -686 | 1 | 0.000431655 | 12 | 1963.3535 | 1926.2335 | 1966.7646 | 2117.685 | 2143.3962 | 2094.4128 |
| Xpo1 | FSLNQDIPAFK | -1002 | -1012 | 0 | 0 | 11 | 1510.1707 | 1013.0629 | 1520.3164 | 1185.7078 | 1262.3723 | 1031.3668 |
| Xpo1 | SAFPHLQDAQVK | -984 | -995 | 0 | 0 | 12 | 862.0337 | 1125.5839 | 1058.5712 | 1318.7308 | 571.11957 | 1542.7561 |
| Xpo1 | AGEDTSDLFLEER | -1026 | -1038 | 0 | 0 | 13 | 590.4213 | 602.8342 | 539.748 | 495.10312 | 699.849 | 853.9039 |

| | | | | | | | | | | | | |
|---|---|---|---|---|---|---|---|---|---|---|---|---|
| Xpo1 | SVPGILNPHEIPEEMCD | -1055 | -1071 | 0 | 0 | 17 | 419.4713 | 511.09033 | 463.77982 | 539.1101 | 605.6113 | 569.35236 |
| Xpo1 | HLDYVDTEIIMTK | -466 | -478 | 0 | 0 | 13 | 950.13043 | 859.50256 | NaN | 1059.378 | 660.2076 | 925.3486 |
| Xpo1 | FLNVPMFR | -254 | -261 | 0 | 0.00025015 | 8 | 556.69824 | NaN | 636.24677 | 693.98926 | 684.6053 | 1088.0111 |
| Xpo1 | VSEVEETEIFK | -357 | -367 | 0 | 0 | 11 | 3961.5798 | 4446.59 | NaN | NaN | 1369.6466 | 3934.5576 |
| Xpo1 | GLFSLNQDIPAFK | -1000 | -1012 | 0 | 0 | 13 | 240.49557 | 232.62158 | NaN | NaN | 687.3963 | 610.33984 |
| Xpo1 | EFAGEDTSDLFLEER | -1024 | -1038 | 0 | 0 | 15 | 3036.7039 | NaN | 2789.0679 | 6511.0044 | NaN | NaN |
| Xpo1 | EHPDAWTR | -55 | -62 | 0 | 0.005789881 | 8 | 1587.9492 | NaN | NaN | 1479.5891 | NaN | NaN |
| Xpo1 | ISTPLNPGNPVNNQ | -958 | -971 | 0 | 0.000550307 | 14 | 951.68567 | NaN | NaN | NaN | NaN | NaN |
| Xpo1 | LYHGEGAQQR | -35 | -44 | 0 | 0.005789881 | 10 | 536.2506 | NaN | NaN | NaN | NaN | NaN |
| Fdx2 | AGEEAADSPELPR | -44 | -56 | 0 | 0 | 13 | 1095.437 | 1220.988 | NaN | NaN | NaN | 480.28787 |
| Rps21 | MNVAEVDR | -34 | -41 | 0 | 0 | 8 | 801.4109 | 331.1087 | 481.5255 | 736.45013 | 930.5415 | 839.0558 |
| Rps21 | MQNDAGEFVDLYVPR | -1 | -15 | 0 | 0 | 15 | 548.05493 | 585.43164 | 532.30664 | 823.275 | 810.5671 | 835.1542 |
| Rps21 | DAGEFVDLYVPR | -4 | -15 | 0 | 0 | 12 | 508.47476 | 578.7421 | 653.6878 | 971.4025 | 1098.9518 | 1042.2252 |
| Rps21 | MGESDDSILR | -62 | -71 | 0 | 0 | 10 | 398.83917 | 824.8371 | 1458.2042 | 1668.6685 | 1862.6316 | 411.43945 |
| Rps21 | AGEFVDLYVPR | -5 | -15 | 0 | 0 | 11 | 525.471 | NaN | 5677.6978 | 752.67926 | 5373.0454 | 5132.544 |
| Rps21 | GESDDSILR | -63 | -71 | 0 | 0 | 9 | 1977.6614 | 1831.6633 | 1957.679 | 2211.3066 | NaN | NaN |
| Rps21 | VDLYVPR | -9 | -15 | 0 | 0.001868731 | 7 | 998.99133 | 58.81647 | 501.6189 | 457.02234 | NaN | NaN |
| Rps18 | IPDWFLNR | -79 | -86 | 0 | 0.000471409 | 8 | 6250.4907 | 7534.805 | 7427.864 | 3129.3284 | 4110.626 | 4111.572 |
| Rps18 | LREDLER | -107 | -113 | 1 | 0.004439092 | 7 | 5746.271 | 1607.7599 | 6434.1694 | 6401.1724 | 8196.121 | 8113.272 |
| Rps18 | RAGELTEDEVER | -55 | -66 | 1 | 0 | 12 | 3677.3298 | 3695.5674 | 3949.7314 | 4649.4907 | 4796.767 | 5131.4727 |
| Rps18 | YAHVVLR | -40 | -46 | 0 | 0.00142999 | 7 | 2719.1077 | 1419.9208 | 1489.8163 | 2165.2908 | 2272.457 | 2389.0667 |
| Rps18 | VLNTNIDGR | -15 | -23 | 0 | 0 | 9 | 1451.4019 | 1723.9617 | 2558.6936 | 1699.087 | 2065.8284 | 1957.3804 |
| Rps18 | KADIDLTK | -47 | -54 | 1 | 0 | 8 | 1036.9243 | 1104.8716 | 1259.3893 | 1016.8531 | 856.2521 | 930.01917 |
| Rps18 | AGELTEDEVER | -56 | -66 | 0 | 0.000627943 | 11 | 654.5067 | 5016.0312 | 4976.529 | 5315.3677 | 5744.2583 | 5170.477 |
| Rps18 | KIPDWFLNR | -78 | -86 | 1 | 0.000627943 | 9 | 8793.522 | 9296.807 | 287.6466 | 11168.86 | 11682.646 | NaN |
| Ssrp1 | LFLLPHK | -242 | -248 | 0 | 0 | 7 | 2610.9731 | 2326.608 | 2299.6787 | 1702.2777 | 1832.77 | 1854.2405 |
| Ssrp1 | NIQAGELTEGIWR | -42 | -54 | 0 | 0 | 13 | 2127.004 | 2762.9521 | 2426.5508 | 2501.9373 | 2818.5002 | 2841.222 |
| Ssrp1 | ASSGLLYPLER | -347 | -357 | 0 | 0 | 11 | 2073.61 | 1961.4872 | 2189.707 | 1215.5637 | 813.7909 | 563.9067 |
| Ssrp1 | VDNIQAGELTEGIWR | -40 | -54 | 0 | 0 | 15 | 1504.3345 | 2398.5327 | 2252.6787 | 195.3382 | 2829.9663 | 2504.3225 |
| Ssrp1 | VHKPPVHIR | -362 | -370 | 1 | 0.00025015 | 9 | 996.838 | 1473.1456 | 1348.5787 | 929.5884 | 384.03073 | 1203.516 |
| Ssrp1 | VHKPPVHIR | -362 | -370 | 1 | 0.00025015 | 9 | 996.838 | 1473.1456 | 1348.5787 | 929.5884 | 384.03073 | 1203.516 |
| Ssrp1 | IYPTFLHLHGK | -218 | -228 | 0 | 0 | 11 | 956.7462 | 497.50934 | 132.80685 | 1250.8235 | 975.3002 | 931.40063 |
| Ssrp1 | KITVPGNFQGHSGAQ | -325 | -339 | 1 | 0 | 15 | 820.52094 | 731.1775 | 911.2767 | 720.04956 | 1161.2715 | 757.0709 |
| Ssrp1 | RELQCLTPR | -203 | -211 | 1 | 0.00122419 | 9 | 798.8454 | 970.99335 | 935.993 | 1119.7314 | 584.22296 | 1125.8142 |
| Ssrp1 | FYVPPTQEDGVDPVEAF | -165 | -181 | 0 | 0 | 17 | 730.40155 | 785.1424 | 799.621 | 697.6311 | 924.14685 | 880.2871 |
| Ssrp1 | IGDQPVFEIPLSNVSQCTTGK | -123 | -143 | 0 | 0 | 21 | 422.31696 | 405.03717 | 436.20963 | 460.9801 | 584.1573 | 641.0608 |
| Ssrp1 | EFNDIFQEVK | -6 | -15 | 0 | 0 | 10 | 376.9909 | 1728.8398 | 1799.885 | 2013.1178 | 2032.4543 | 2076.1472 |
| Ssrp1 | FLHLHGK | -222 | -228 | 0 | 0 | 7 | 325.3329 | 222.34317 | 185.39793 | 1135.2258 | 591.60114 | 798.321 |
| Ssrp1 | AGELTEGIWR | -45 | -54 | 0 | 0.000298805 | 10 | 1304.915 | 1258.3892 | 1352.3754 | NaN | 1458.6965 | 1264.031 |
| Ssrp1 | SGLLYPLER | -349 | -357 | 0 | 0 | 9 | 1066.1587 | 1026.3842 | NaN | 1675.4459 | 2233.8555 | 2378.5938 |
| Ssrp1 | ITVPGNFQGHSGAQ | -326 | -339 | 0 | 0 | 14 | 940.956 | 963.49756 | 1066.7665 | 2286.0566 | 2215.254 | NaN |
| Ssrp1 | SDHPGISITDL | -567 | -577 | 0 | 0.00272025 | 11 | 1135.7302 | 919.8608 | NaN | NaN | 948.4006 | 1362.7589 |
| Ssrp1 | DYADSDEDQHDAYLER | -440 | -455 | 0 | 0.000826806 | 16 | 1081.1846 | 1255.0292 | NaN | 1178.8032 | 1548.7891 | NaN |
| Ssrp1 | IKSDHPGISIT | -565 | -575 | 1 | 0.001868731 | 11 | 767.1903 | 783.96484 | 674.44183 | 1207.9713 | NaN | NaN |
| Ssrp1 | GDAICIFR | -196 | -203 | 0 | 0.00111006 | 8 | 410.36017 | 640.9588 | 560.5127 | NaN | NaN | NaN |
| Ssrp1 | SDHPGISITDLSK | -567 | -579 | 0 | 0 | 13 | 2091.8337 | NaN | 2587.974 | NaN | NaN | NaN |
| Ssrp1 | IPYTTVLR | -234 | -241 | 0 | 0.001291103 | 8 | 868.2261 | NaN | NaN | NaN | 1078.0741 | NaN |
| Ssrp1 | LFDFVNAK | -414 | -421 | 0 | 0.00078661 | 8 | 520.3203 | 634.3485 | NaN | NaN | NaN | NaN |
| Ssrp1 | IYPTFLHLH | -218 | -226 | 0 | 0.009002904 | 9 | 471.59766 | 434.7859 | NaN | NaN | NaN | NaN |
| Ecpas | VASYLIKPEVQVR | -480 | -492 | 1 | 0 | 13 | 599.5459 | 263.7149 | 511.36917 | 154.43288 | 358.5246 | 338.82993 |
| Ecpas | VIGDNPWTPEQLEQCK | -224 | -239 | 0 | 0 | 16 | 274.92877 | 410.2534 | 357.39218 | 558.6102 | 991.61237 | 1104.6501 |
| Ecpas | EGAVLKPELK | -311 | -320 | 1 | 0 | 10 | 1156.6577 | 912.26624 | NaN | 1505.5242 | 1963.2268 | 1598.6886 |
| Ecpas | VWQENVPGSFGGIR | -1504 | -1517 | 0 | 0 | 14 | 808.4327 | NaN | 804.2053 | 540.5579 | 393.09445 | 918.49866 |
| Ecpas | HIISPNPHVR | -945 | -954 | 0 | 0 | 10 | 622.2358 | 555.2849 | 554.0519 | NaN | 423.71875 | 633.14685 |
| Ecpas | TTQCPQDLTPYSGK | -1382 | -1395 | 0 | 0 | 14 | 492.17938 | 688.91925 | 588.53485 | NaN | 1010.70667 | NaN |
| Ecpas | QSLIDWNNPAIINK | -283 | -296 | 0 | 0 | 14 | 249.95305 | NaN | 462.34372 | NaN | NaN | 343.8802 |
| Ecpas | AGEQLAPFLPQLVPR | -1107 | -1121 | 0 | 0 | 15 | 441.2992 | NaN | 354.0217 | NaN | NaN | NaN |
| Numa1 | EGQQILQQPLPER | -48 | -60 | 0 | 0 | 13 | 3986.4683 | 4550.1914 | 4779.472 | 2632.0955 | 868.31323 | 2400.2993 |
| Numa1 | LQEQLQDLEELQK | -1613 | -1625 | 0 | 0 | 13 | 3345.981 | 3524.8455 | 3760.0007 | 520.4664 | 1460.4534 | 692.36316 |
| Numa1 | ALVSELLPAK | -1351 | -1360 | 0 | 0 | 10 | 2187.4736 | 2327.2104 | 2283.3901 | 1677.4825 | 1696.5264 | 1578.5957 |
| Numa1 | IDHLALLNEK | -256 | -265 | 0 | 0 | 10 | 2103.5693 | 2520.5273 | 2235.1785 | 423.79886 | 3424.9167 | 3327.596 |
| Numa1 | FREELEQSK | -1377 | -1385 | 1 | 0.000947459 | 9 | 1247.0978 | 1302.024 | 722.6927 | 1138.9347 | 1593.2822 | 1276.8265 |
| Numa1 | ALETLQGQLEEK | -1159 | -1170 | 0 | 0 | 12 | 987.4301 | 1140.8826 | 1213.2474 | 952.3882 | 316.33682 | 602.15094 |
| Numa1 | LKEQQLEEAAK | -556 | -566 | 1 | 0 | 11 | 882.9888 | 960.3668 | 895.47736 | 794.23584 | 870.6136 | 868.00494 |
| Numa1 | DNAQTSVTQAQQEK | -626 | -639 | 0 | 0 | 14 | 858.5977 | 939.06335 | 1061.3419 | 874.4107 | 1080.6389 | 1219.9662 |
| Numa1 | AGEQQETASLELLK | -913 | -926 | 0 | 0 | 14 | 805.7339 | 893.62006 | 1044.4265 | 887.7623 | 329.78354 | 727.1783 |
| Numa1 | TQPDGTSVPGEPASPISQR | -1726 | -1744 | 0 | 0 | 19 | 745.4196 | 691.89026 | 555.7766 | 815.1353 | 548.74176 | 976.76044 |
| Numa1 | LATDNTQLQTR | -423 | -433 | 0 | 0 | 11 | 742.664 | 891.41736 | 888.62787 | 1848.3412 | 1527.2982 | 1014.26495 |
| Numa1 | QAASSQEPSELEELR | -266 | -280 | 0 | 0.00025015 | 15 | 708.4884 | 1064.1218 | 668.1138 | 1053.2217 | 1008.0655 | 1102.625 |
| Numa1 | ELQQTLEQLK | -1069 | -1078 | 0 | 0 | 10 | 685.9613 | 121.99422 | 1004.96356 | 1016.9026 | 455.4685 | 1015.0139 |
| Numa1 | LSQLEDQATR | -385 | -394 | 0 | 0 | 10 | 725.09796 | 408.19574 | 965.96466 | NaN | 1016.8495 | 1062.0754 |
| Numa1 | LDFVCSFLQK | -61 | -70 | 0 | 0 | 10 | 688.8594 | 478.39545 | 680.8693 | NaN | 420.86508 | 699.2928 |
| Numa1 | IINTIHDTK | -39 | -47 | 0 | 0 | 9 | 1604.4187 | NaN | NaN | 782.6394 | 1426.4799 | 1417.816 |
| Numa1 | TDAETHLAEMR | -1481 | -1491 | 0 | 0.000989247 | 11 | 711.82556 | 867.11383 | 602.346 | NaN | NaN | NaN |
| Numa1 | LQESPAPEKGEVL | -395 | -407 | 1 | 0.000627943 | 13 | 765.6363 | NaN | 738.6222 | NaN | NaN | NaN |
| Numa1 | KLELEEPDSANSS | -1804 | -1816 | 1 | 0.008736733 | 13 | 202.01393 | NaN | 18519.895 | NaN | NaN | NaN |
| Numa1 | YVQELAAVR | -1472 | -1480 | 0 | 0.00025015 | 9 | 472.18878 | NaN | NaN | NaN | NaN | NaN |
| Bax | AGETPELTLEQPPQDASTK | -39 | -57 | 0 | 0 | 19 | 1289.0139 | 1456.5951 | 1288.2357 | 998.8349 | 711.70105 | 1237.895 |

| | | | | | | | | | | | | |
|---|---|---|---|---|---|---|---|---|---|---|---|---|
| Bax | GETPELTLEQPPQDASTK | -40 | -57 | 0 | 0 | 18 | 870.8924 | 1038.2737 | 988.54443 | 1023.2256 | 1235.9215 | 1215.5868 |
| Bax | IGDELDSNMELQR | -66 | -78 | 0 | 0 | 13 | 403.85046 | 953.9612 | 993.3356 | 1001.274 | 1001.6524 | 945.906 |
| Nono | DKGFGFIR | -110 | -117 | 1 | 0 | 8 | 1911.063 | 1771.3275 | 278.54593 | 1771.138 | 1242.169 | 1239.2178 |
| Nono | DQLDDEEGLPEK | -230 | -241 | 0 | 0 | 12 | 964.4226 | 906.8212 | 834.81165 | 339.69086 | 472.56647 | 580.2127 |
| Nono | FAQPGSFEYEYAMR | -259 | -272 | 0 | 0 | 14 | 930.37335 | 1141.2751 | 980.08545 | 150.10652 | 1400.6813 | 1542.4102 |
| Nono | FAQPGSFEYEYAMR | -259 | -272 | 0 | 0 | 14 | 930.37335 | 1141.2751 | 980.08545 | 150.10652 | 1400.6813 | 1542.4102 |
| Nono | LFVGNLPPDITEEEMR | -78 | -93 | 0 | 0 | 16 | 748.5656 | 1276.0691 | 1154.5654 | 776.2748 | 1667.0321 | 822.40924 |
| Nono | NEGLTIDLK | -54 | -62 | 0 | 0 | 9 | 608.5327 | 632.0145 | 927.1908 | 767.20575 | 1250.7732 | 1243.7385 |
| Nono | AGEVFIHK | -102 | -109 | 0 | 0.005758933 | 8 | 1021.41547 | NaN | NaN | 1151.3496 | 1800.7456 | 1687.708 |
| Actr1b | SEEHPVLLTEAPLNPSK | -103 | -119 | 0 | 0.00078661 | 17 | 307.54486 | NaN | NaN | NaN | NaN | 546.28033 |
| Actbl2 | GIHETTFNSIM | -274 | -284 | 0 | 0 | 11 | 774.3502 | 928.9285 | 1200.8625 | 1223.846 | 90.93809 | 1388.1487 |
| Actbl2 | GIHETTFNSIM | -274 | -284 | 0 | 0 | 11 | 774.3502 | 928.9285 | 1200.8625 | 1223.846 | 90.93809 | 1388.1487 |
| Actbl2 | GIHETTFNSI | -274 | -283 | 0 | 0.001291103 | 10 | 1645.1642 | NaN | NaN | 872.2113 | 3313.6028 | 405.2729 |
| Actbl2 | AGFGGDDAPR | -20 | -29 | 0 | 0.009897819 | 10 | 314.12756 | NaN | 3865.9385 | 3951.8296 | NaN | 178.86049 |
| Ly6e | NNINCLWPVSCQEK | -37 | -50 | 0 | 0 | 14 | 1393.2827 | 1774.343 | 1614.4877 | 2305.165 | 2767.532 | 113.11929 |
| Ly6e | AGFGNVNLGYTLNK | -61 | -74 | 0 | 0 | 14 | 948.99255 | 1069.0424 | 904.587 | 1283.5298 | 1621.0387 | 1543.7999 |
| Ly6e | GNVNLGYTLNK | -64 | -74 | 0 | 0 | 11 | 764.90186 | 969.83264 | 843.6648 | 1053.2959 | NaN | NaN |
| Bud23 | RPEHSGPPELFYDQ | -7 | -20 | 1 | 0.000431655 | 14 | 547.02704 | NaN | 829.5285 | 772.1731 | 672.6104 | 1064.9583 |
| Bud23 | AGFTGGVVVDFPNSAK | -181 | -196 | 0 | 0.007670622 | 16 | 240.73961 | NaN | NaN | NaN | NaN | NaN |
| Nat10 | KDLPPLLLK | -689 | -697 | 1 | 0 | 9 | 842.8232 | 884.7869 | 915.6918 | 337.67413 | 442.52762 | 895.58875 |
| Nat10 | AGFVPVYLR | -726 | -734 | 0 | 0 | 9 | 525.49493 | 472.44913 | 469.82175 | 567.6512 | 505.90967 | 254.25656 |
| Nat10 | TVSEQFQDPDFGGLSGGR | -608 | -625 | 0 | 0 | 18 | 354.0148 | 576.57043 | 574.63257 | 539.5632 | 883.5248 | 809.31573 |
| Nat10 | QTIQYIHPADAVK | -366 | -378 | 0 | 0 | 13 | 1232.2137 | 311.45184 | 1408.833 | 613.4745 | NaN | 605.6322 |
| Nat10 | YCYYNETHK | -104 | -112 | 0 | 0 | 9 | 357.25363 | 398.14716 | NaN | 208.93317 | 406.53784 | 462.6412 |
| Nat10 | SLFVVVGDR | -24 | -32 | 0 | 0 | 9 | 1533.2128 | 681.3593 | 1614.0304 | 1215.2133 | NaN | NaN |
| Nat10 | VCLEGEISR | -579 | -587 | 0 | 0.00025015 | 9 | 622.22876 | NaN | 473.28723 | 538.731 | 649.8863 | NaN |
| Nat10 | IVSGCPLPEACELYYVNR | -495 | -512 | 0 | 0.000431655 | 18 | 275.96573 | NaN | 379.26605 | NaN | 451.19904 | 262.52448 |
| Nat10 | DFEALTPNLLAR | -125 | -136 | 0 | 0 | 12 | 684.69775 | 547.8183 | 1396.5187 | NaN | NaN | NaN |
| Nat10 | LFNDVQEK | -905 | -912 | 0 | 0 | 8 | 3877.1606 | 3907.3613 | NaN | NaN | NaN | NaN |
| Nat10 | GDDEEWNEVLSK | -967 | -978 | 0 | 0 | 12 | 514.8437 | NaN | NaN | NaN | NaN | NaN |
| Cyb5a | TYIIGELHPDDR | -78 | -89 | 0 | 0 | 12 | 3921.4084 | 4499.306 | 4446.5786 | 4796.7383 | 5171.813 | 5013.6206 |
| Cyb5a | FLEEHPGGEEVLR | -40 | -52 | 0 | 0 | 13 | 2853.5588 | 3127.7087 | 6029.282 | 2837.2874 | 2092.1199 | 2133.6045 |
| Cyb5a | FLEEHPGGEEVLR | -40 | -52 | 0 | 0 | 13 | 2853.5588 | 3127.7087 | 6029.282 | 2837.2874 | 2092.1199 | 2133.6045 |
| Cyb5a | YYTLEEIQK | -11 | -19 | 0 | 0 | 9 | 1477.1716 | 1370.8735 | 438.0423 | 1407.972 | 1559.7367 | 1310.3077 |
| Cyb5a | IAKPSDTLITT | -92 | -102 | 1 | 0.00051306 | 11 | 1367.0332 | 1314.2709 | 1411.4493 | 1523.3518 | 1379.137 | 1442.3306 |
| Cyb5a | AGGDATENFEDVGHSTDAR | -55 | -73 | 0 | 0 | 19 | 812.69354 | 804.09607 | 821.5701 | 1144.2205 | 1432.5972 | 1364.9117 |
| Cyb5a | AGGDATENFEDVGHSTDAR | -55 | -73 | 0 | 0 | 19 | 812.69354 | 804.09607 | 821.5701 | 1144.2205 | 1432.5972 | 1364.9117 |
| Cyb5a | IAKPSDTLIT | -92 | -101 | 1 | 0.001055699 | 10 | 3126.4224 | NaN | 3576.8894 | 105.984184 | NaN | 5221.477 |
| Cyb5a | EQAGGDATENFEDVGHSTDAR | -53 | -73 | 0 | 0.001023236 | 21 | 295.43033 | 324.50494 | 310.29718 | NaN | 432.20654 | NaN |
| Cyb5a | EQAGGDATENFEDVGHST | -53 | -70 | 0 | 0.000230707 | 18 | 420.49976 | 419.0969 | 449.9372 | NaN | NaN | NaN |
| Cyb5a | IIGELHPDDR | -80 | -89 | 0 | 0 | 10 | 2085.5686 | NaN | 1597.8701 | NaN | NaN | NaN |
| Ppp2r1a | LAGGDWFTSR | -135 | -144 | 0 | 0 | 10 | 5094.7007 | 4166.4556 | 4330.4893 | 5878.203 | 5819.599 | 5075.4966 |
| Ppp2r1a | VLSEVCGQDITTK | -507 | -519 | 0 | 0 | 13 | 2679.9373 | 1986.9698 | 2289.985 | 3302.0947 | 2709.2056 | 1392.5186 |
| Ppp2r1a | ELVSDANQHVK | -332 | -342 | 0 | 0 | 11 | 2460.762 | 2659.0742 | 2715.3037 | 2340.3862 | 3288.015 | 3005.7144 |
| Ppp2r1a | SGDPNYLHR | -490 | -498 | 0 | 0 | 9 | 2138.9253 | 1944.7125 | 2410.87 | 1831.5284 | 1596.642 | 2596.9998 |
| Ppp2r1a | SEVKPILEK | -558 | -566 | 1 | 0 | 9 | 1524.6105 | 1006.72656 | 1538.5862 | 2065.6555 | 2289.414 | 2049.1863 |
| Ppp2r1a | AISHEHSPSDLEAH | -114 | -127 | 0 | 0 | 14 | 1350.6122 | 1518.7924 | 1590.5117 | 780.5948 | 1304.7742 | 1292.4117 |
| Ppp2r1a | AISHEHSPSDLEAH | -114 | -127 | 0 | 0 | 14 | 1350.6122 | 1518.7924 | 1590.5117 | 780.5948 | 1304.7742 | 1292.4117 |
| Ppp2r1a | IGPILDNSTLQSEVKPILEK | -547 | -566 | 1 | 0 | 20 | 1119.2678 | 1169.7227 | 983.9719 | 1581.5822 | 1479.9105 | 1403.1108 |
| Ppp2r1a | IGPILDNSTLQSEVKPILEK | -547 | -566 | 1 | 0 | 20 | 1119.2678 | 1169.7227 | 983.9719 | 1581.5822 | 1479.9105 | 1403.1108 |
| Ppp2r1a | AGGDWFTSR | -136 | -144 | 0 | 0 | 9 | 848.1007 | 820.76917 | 192.63164 | 984.5454 | 995.62665 | 1061.3259 |
| Ppp2r1a | EFCENLSADCR | -308 | -318 | 0 | 0 | 11 | 527.2735 | 521.8022 | 472.90546 | 671.48627 | 111.23556 | 470.51077 |
| Ppp2r1a | DNTIEHLLPL | -359 | -368 | 0 | 0.00111006 | 10 | 491.99866 | 613.18964 | 596.51013 | 527.57275 | 212.83029 | 189.64636 |
| Ppp2r1a | VADKFTELQK | -263 | -272 | 1 | 0 | 10 | 955.93 | 898.54553 | 1922.5178 | NaN | 2680.9373 | 1391.9767 |
| Ppp2r1a | AMSGDPNYLHR | -488 | -498 | 0 | 0 | 11 | 2409.9963 | NaN | NaN | 3171.1355 | 3412.7993 | 3602.0098 |
| Ppp2r1a | FSNLASDEQDSVR | -209 | -221 | 0 | 0 | 13 | 2252.8206 | NaN | 2411.8381 | 1583.3337 | NaN | 1561.1965 |
| Ppp2r1a | KFTELQK | -266 | -272 | 1 | 0 | 7 | 560.5906 | 651.0052 | NaN | NaN | NaN | NaN |
| Eif2s3x | LTPLSHEVISR | -28 | -38 | 0 | 0 | 11 | 2425.755 | 2474.9277 | 2452.0046 | 2909.0576 | 2737.7363 | 2761.332 |
| Eif2s3x | GSSTPDEFPTDIPGTK | -106 | -121 | 0 | 0 | 16 | 2076.417 | 215.06544 | 1836.3633 | 1094.9299 | 1087.5151 | 1209.8138 |
| Eif2s3x | AGGEGGVTLGQPHLSR | -2 | -17 | 0 | 0 | 16 | 1784.0879 | 2305.9143 | 1954.8328 | 4918.695 | 5126.899 | 5084.2437 |
| Eif2s3x | AGGEGGVTLGQPHLSR | -2 | -17 | 0 | 0 | 16 | 1784.0879 | 2305.9143 | 1954.8328 | 4918.695 | 5126.899 | 5084.2437 |
| Eif2s3x | GVTIKPTVDDD | -462 | -472 | 1 | 0.000550307 | 11 | 968.8653 | 1170.9167 | 1285.4559 | 1202.8829 | 650.4197 | 579.86633 |
| Eif2s3x | PLSHEVISR | -30 | -38 | 0 | 0 | 9 | 866.7859 | 946.3916 | 937.84485 | 960.86926 | 1349.8975 | 1297.8519 |
| Eif2s3x | PLSHEVISR | -30 | -38 | 0 | 0 | 9 | 866.7859 | 946.3916 | 937.84485 | 960.86926 | 1349.8975 | 1297.8519 |
| Eif2s3x | RGVTIKPTVDDD | -461 | -472 | 2 | 0.000230707 | 12 | 639.55646 | 637.5513 | 661.5724 | 875.68536 | 762.5183 | 730.7081 |
| Eif2s3x | SCGSSTPDEFPTDIPGTK | -104 | -121 | 0 | 0 | 18 | 475.87833 | 505.19788 | 468.57596 | NaN | NaN | NaN |
| Ndufb2 | AGGGVHIQPR | -34 | -43 | 0 | 0 | 10 | 748.4693 | 874.2922 | 787.9612 | 804.8713 | 934.494 | 783.8456 |
| Ndufb2 | FWHDSDAVLGHF | -73 | -84 | 0 | 0 | 12 | 694.84357 | 531.818 | NaN | NaN | 336.28714 | 297.1271 |
| H2az1;H2az2 | GDEELDSLIK | (93);(93) | (102);(102) | 0 | 0 | 10 | 1292.9841 | 2129.8943 | 1362.7793 | 2199.9863 | 2321.5864 | 2208.5552 |
| H2az1;H2az2 | GDEELDSLIK | (93);(93) | (102);(102) | 0 | 0 | 10 | 1292.9841 | 2129.8943 | 1362.7793 | 2199.9863 | 2321.5864 | 2208.5552 |
| H2az1;H2az2 | IAGGGVIPHIHK | (105);(105) | (116);(116) | 0 | 0 | 12 | 2650.173 | 3439.5945 | 3942.6877 | 3613.1016 | 5804.076 | 6032.571 |
| H2az1;H2az2 | IAGGGVIPHIHK | (105);(105) | (116);(116) | 0 | 0 | 12 | 2650.173 | 3439.5945 | 3942.6877 | 3613.1016 | 5804.076 | 6032.571 |
| H2az1;H2az2 | GGGVIPHIHK | (107);(107) | (116);(116) | 0 | 0 | 10 | 2458.8088 | 2721.684 | 2520.0845 | 1126.4233 | 2994.5625 | 3041.7593 |
| H2az1;H2az2 | AIRGDEELDSLIK | (90);(90) | (102);(102) | 1 | 0 | 13 | 2279.9543 | 3089.186 | 3146.8164 | 3461.098 | 4066.8208 | 4115.151 |
| H2az1;H2az2 | AIRGDEELDSLIK | (90);(90) | (102);(102) | 1 | 0 | 13 | 2279.9543 | 3089.186 | 3146.8164 | 3461.098 | 4066.8208 | 4115.151 |
| H2az1;H2az2 | IRGDEELDSLIK | (91);(91) | (102);(102) | 1 | 0.000431655 | 12 | 1422.782 | 1952.2826 | 236.93398 | 1527.1556 | 2193.7458 | 2169.3457 |
| H2az1;H2az2 | ATIAGGGVIPHIHK | (103);(103) | (116);(116) | 0 | 0 | 14 | 986.1395 | 1088.8695 | 965.5465 | 433.14352 | 776.94244 | 870.6349 |
| H2az1;H2az2 | AGGGVIPHIHK | (106);(106) | (116);(116) | 0 | 0 | 11 | 930.2856 | 966.52954 | NaN | 1764.792 | 802.9208 | 1528.6615 |

| | | | | | | | | | | | | |
|---|---|---|---|---|---|---|---|---|---|---|---|---|
| H2az1;H2az2 | ATIAGGGVIPHIH | (103);(103) | (115);(115) | 0 | 0 | 13 | 386.65823 | 915.10315 | 999.17706 | NaN | 115.23209 | 1094.2449 |
| H2az1;H2az2 | RGDEELDSLIK | (92);(92) | (102);(102) | 1 | 0.000431655 | 11 | 223.35925 | 1230.081 | 1416.1343 | NaN | 1407.7903 | NaN |
| H2az1;H2az2 | TAEVLELAGNASK | (63);(63) | (75);(75) | 0 | 0 | 13 | 1127.4225 | NaN | NaN | 1769.5732 | NaN | 1916.3577 |
| Ttll12 | LPLAISPVAR | -285 | -294 | 0 | 0 | 10 | 526.57874 | 572.6329 | 505.9583 | 113.07456 | 572.05896 | 593.2568 |
| Ttll12 | AGGPEGPPWLPR | -373 | -384 | 0 | 0 | 12 | 834.04126 | 1045.5088 | NaN | 4666.6245 | 4870.0537 | 2032.5709 |
| Ttll12 | EVNFNPDCER | -600 | -609 | 0 | 0 | 10 | 457.6708 | 273.78253 | NaN | 418.5372 | NaN | NaN |
| Ttll12 | DLDTGEEVTR | -226 | -235 | 0 | 0.005015524 | 10 | 1039.483 | NaN | NaN | NaN | NaN | NaN |
| Cybc1 | SFLELHR | -159 | -165 | 0 | 0 | 7 | 1978.5923 | 1900.1667 | 1974.1392 | 1744.7899 | 1772.7273 | 1720.8788 |
| Cybc1 | LKDIQDVNVEEEK | -104 | -116 | 1 | 0 | 13 | 611.5203 | 672.0806 | 559.7638 | 874.32263 | 782.48083 | 801.5418 |
| Cybc1 | RSDVEAIAK | -147 | -155 | 1 | 0.002752526 | 9 | 581.5177 | 957.8443 | 1318.6243 | 1336.0842 | 1689.6553 | 1351.2449 |
| Cybc1 | DIQDVNVEEEK | -106 | -116 | 0 | 0 | 11 | 2588.3108 | 3582.0127 | NaN | 402.03418 | NaN | NaN |
| Cybc1 | AGHDQVVVL | -95 | -103 | 0 | 0.009002904 | 9 | 895.3224 | NaN | 857.2177 | NaN | NaN | NaN |
| Nup155 | SPFLEPHLVR | -1057 | -1066 | 0 | 0 | 10 | 1791.1967 | 1767.854 | 2112.9348 | 909.8842 | 761.2704 | 550.813 |
| Nup155 | NVGGDGEEIER | -527 | -537 | 0 | 0 | 11 | 1060.8193 | 204.77858 | 1017.1721 | 1301.5984 | 848.9574 | 272.25232 |
| Nup155 | LRPVDQLR | -514 | -521 | 1 | 0.00025015 | 8 | 678.16437 | 646.9754 | 687.0422 | 222.08102 | 813.8426 | 716.2643 |
| Nup155 | DHIPITDSPVVVQ | -479 | -491 | 0 | 0 | 13 | 590.20087 | 916.9159 | 1046.0013 | 671.6488 | 516.6119 | 869.84564 |
| Nup155 | RVPLPPELVEQF | -86 | -97 | 1 | 0 | 12 | 437.4397 | 319.20023 | 405.16223 | 272.50418 | 804.7377 | 657.5798 |
| Nup155 | NSQFSGGPLGNPNTTAR | -724 | -740 | 0 | 0 | 17 | 424.07062 | 472.68735 | 500.03705 | 556.82776 | 615.55176 | 474.06088 |
| Nup155 | NEIGVPLPR | -1290 | -1298 | 0 | 0 | 9 | 5858.653 | 5701.2183 | 5914.4165 | 5457.278 | NaN | NaN |
| Nup155 | SVFKPIVQIA | -337 | -346 | 1 | 0.00111006 | 10 | 313.34082 | 332.12692 | 356.5564 | NaN | NaN | 232.26848 |
| Nup155 | SRDPFWNR | -1309 | -1316 | 1 | 0.001636916 | 8 | 1422.3855 | NaN | NaN | NaN | 1518.3955 | 1377.6323 |
| Nup155 | AGIFQPHVR | -154 | -162 | 0 | 0.001904399 | 9 | 1877.1504 | NaN | NaN | NaN | NaN | 808.6098 |
| Nup155 | ALSAIDELKVDK | -460 | -471 | 1 | 0.002099344 | 12 | 687.3354 | NaN | NaN | NaN | NaN | 911.67957 |
| Nup155 | LLEVYDQLFK | -1299 | -1308 | 0 | 0 | 10 | 465.68143 | NaN | NaN | NaN | NaN | 490.154 |
| Nup155 | LYGEFADPFK | -1188 | -1197 | 0 | 0.000431655 | 10 | 267.81308 | NaN | 225.0092 | NaN | NaN | NaN |
| Ahcy | WLNENAVEKV | -310 | -319 | 1 | 0.000230707 | 10 | 8495.592 | 778.6228 | 11142.475 | 1576.0804 | 1558.693 | 2006.2084 |
| Ahcy | SKFDNLYGCR | -187 | -196 | 1 | 0 | 10 | 5840.236 | 6498.12 | 6392.296 | 5947.7803 | 6441.8896 | 6741.146 |
| Ahcy | YPQLLSGIR | -143 | -151 | 0 | 0.00149795 | 9 | 4326.0386 | 4780.106 | 4777.3296 | 4413.596 | 5925.442 | 5948.6865 |
| Ahcy | GISEETTTGVHNLYK | -152 | -166 | 0 | 0 | 15 | 3398.1482 | 3863.7063 | 3668.1846 | 4795.3975 | 5162.145 | 4861.585 |
| Ahcy | VPAINVNDSVTK | -175 | -186 | 0 | 0 | 12 | 2853.2263 | 3073.287 | 3087.5332 | 3416.4043 | 3764.5596 | 3708.3442 |
| Ahcy | SDKLPYK | -2 | -8 | 1 | 0.001055699 | 7 | 2764.1782 | 2960.3638 | 2915.5469 | 1966.397 | 2007.279 | 1986.8903 |
| Ahcy | AGIPVFAWK | -95 | -103 | 0 | 0 | 9 | 2242.5247 | 2736.5635 | 2775.6108 | 1167.8088 | 1584.5183 | 1685.4271 |
| Ahcy | FDNLYGCR | -189 | -196 | 0 | 0 | 8 | 1592.2407 | 1714.6068 | 1757.2424 | 744.03375 | 1431.8186 | 1525.3672 |
| Ahcy | VNIKPQVDR | -319 | -327 | 1 | 0.00025015 | 9 | 1543.0195 | 1453.7504 | 1487.2749 | 1177.5399 | 1170.1273 | 1141.5231 |
| Ahcy | KFDNLYGCR | -188 | -196 | 1 | 0.000591124 | 9 | 1145.9518 | 1288.1864 | 1206.5282 | 1747.357 | 1690.3896 | 1488.7563 |
| Ahcy | DDGGDLTNLIHTK | -130 | -142 | 0 | 0 | 13 | 883.6395 | 1003.60394 | 917.8567 | 955.60754 | 392.85226 | 674.66 |
| Ahcy | TTGCVDIILGR | -275 | -285 | 0 | 0 | 11 | 2160.7078 | 2139.1135 | 2258.9124 | NaN | 2255.3618 | 2550.783 |
| Ahcy | LWTHPDKYPVGVHFLPK | -372 | -388 | 1 | 0 | 17 | 749.1437 | 777.07666 | NaN | 640.8534 | 1277.2452 | 1056.075 |
| Ahcy | HFEQMKDDAIVC | -286 | -297 | 1 | 0 | 12 | 507.65817 | NaN | 616.59216 | 700.58563 | 613.6056 | 686.7944 |
| Ahcy | IELWTHPDK | -370 | -378 | 0 | 0 | 9 | 466.86908 | 410.79153 | 539.45496 | 499.19397 | NaN | 1885.5562 |
| Ahcy | THPDKYPVGVHFLPK | -374 | -388 | 1 | 0 | 15 | 538.5137 | NaN | 623.56824 | 172.07529 | NaN | 604.9169 |
| Ahcy | THPDKYPVGVHFLPK | -374 | -388 | 1 | 0 | 15 | 538.5137 | NaN | 623.56824 | 172.07529 | NaN | 604.9169 |
| Ahcy | TTTGVHNLYK | -157 | -166 | 0 | 0.00025015 | 10 | 2674.5093 | 2769.0825 | 3165.0286 | NaN | NaN | NaN |
| Ahcy | GISEETTTGVHN | -152 | -163 | 0 | 0.00025015 | 12 | 1083.0468 | NaN | 959.78217 | 1236.7302 | NaN | NaN |
| Ahcy | GCVDIILGR | -277 | -285 | 0 | 0 | 9 | 264.20773 | 3040.6313 | NaN | 3032.2622 | NaN | NaN |
| Ahcy | LWTHPDK | -372 | -378 | 0 | 0.003620005 | 7 | 880.6203 | 637.1781 | NaN | NaN | NaN | NaN |
| Ahcy | YSASKPLK | -39 | -46 | 1 | 0.004224523 | 8 | 306.7494 | NaN | 641.24426 | NaN | NaN | NaN |
| Ahcy | YPVGVHFLPK | -379 | -388 | 0 | 0.00025015 | 10 | 2280.0273 | NaN | NaN | NaN | NaN | NaN |
| Gsn | QTQVSVLPEGGETPLFK | -372 | -388 | 0 | 0 | 17 | 2912.274 | 3373.685 | 3344.7527 | 967.1697 | 1130.8864 | 1152.2754 |
| Gsn | LDEELGGTPVQSR | -511 | -523 | 0 | 0.000431655 | 13 | 1745.9575 | 1863.538 | 2078.7878 | 1822.8354 | 2178.5247 | 2147.1653 |
| Gsn | AQLDEELGGTPVQSR | -509 | -523 | 0 | 0 | 15 | 1288.2051 | 1847.9813 | 1780.9852 | 1289.7463 | 2306.0093 | 1880.3777 |
| Gsn | EPGLQIWR | -63 | -70 | 0 | 0 | 8 | 816.11066 | 1534.8162 | 1455.6938 | 1326.7534 | 1471.4436 | 1756.3793 |
| Gsn | LVADENPFAQGALR | -312 | -325 | 0 | 0 | 14 | 792.17896 | 956.238 | 739.80884 | 1034.8901 | 3989.5469 | 4543.42 |
| Gsn | SVSLVADENPFAQGALR | -309 | -325 | 0 | 0 | 17 | 515.02765 | 814.6174 | 673.1601 | 567.1468 | 938.33167 | 835.456 |
| Gsn | VLGPKPALPEGTEDTAK | -271 | -287 | 1 | 0.001743443 | 17 | 444.99777 | 526.6779 | 600.5177 | 497.4306 | 770.75696 | 822.466 |
| Gsn | VLGPKPALPEGTEDTAK | -271 | -287 | 1 | 0.001743443 | 17 | 444.99777 | 526.6779 | 600.5177 | 497.4306 | 770.75696 | 822.466 |
| Gsn | QLDEELGGTPVQSR | -510 | -523 | 0 | 0 | 14 | 1003.7435 | 342.0395 | 572.71924 | 346.20496 | 193.85999 | NaN |
| Gsn | RYIETDPANR | -727 | -736 | 1 | 0 | 10 | 756.7804 | NaN | 932.37134 | 880.5571 | 1214.938 | 922.66455 |
| Gsn | TAQLDEELGGTPVQSR | -508 | -523 | 0 | 0.00111006 | 16 | 815.8597 | NaN | NaN | 1177.931 | 786.6273 | 751.7316 |
| Gsn | DSQEEEKTEALTSAK | -712 | -726 | 1 | 0 | 15 | 3227.6177 | 2546.079 | NaN | NaN | NaN | 3867.5037 |
| Gsn | SHIANVER | -410 | -417 | 0 | 0 | 8 | 726.25366 | 710.922 | NaN | NaN | 404.09097 | NaN |
| Gsn | AGKEPGLQIW | -60 | -69 | 1 | 0.001636916 | 10 | 448.52466 | 481.67093 | NaN | NaN | 707.48895 | NaN |
| Gsn | PALPEGTEDTAK | -276 | -287 | 0 | 0.002099344 | 12 | 2071.882 | NaN | NaN | NaN | 4680.7617 | NaN |
| Gsn | YIETDPANR | -728 | -736 | 0 | 0.00025015 | 9 | 953.4977 | NaN | 605.5434 | NaN | NaN | NaN |
| Gsn | NSNDAFVLK | -587 | -595 | 0 | 0.004255025 | 9 | 526.6151 | NaN | NaN | NaN | NaN | NaN |
| Rpl10a | RDTLYEAVR | -7 | -15 | 1 | 0.00078661 | 9 | 1353.6892 | 395.6814 | 1263.2617 | 1839.7651 | 1376.2212 | 2502.036 |
| Rpl10a | MTDDELVYNIH | -174 | -184 | 0 | 0 | 11 | 361.89725 | 1025.6354 | 1126.7421 | 1126.938 | 1168.6006 | 1051.7758 |
| Rpl10a | MTDDELVYNIH | -174 | -184 | 0 | 0 | 11 | 361.89725 | 1025.6354 | 1126.7421 | 1126.938 | 1168.6006 | 1051.7758 |
| Rpl10a | AGKFPSLL | -131 | -138 | 1 | 0.00111006 | 8 | 551.9725 | NaN | NaN | 352.13742 | 380.44415 | 407.67783 |
| Rpl10a | AVDIPHMDIEAL | -79 | -90 | 0 | 0.000627943 | 12 | 2031.6708 | 2252.883 | NaN | NaN | NaN | 1726.8195 |
| Rpl10a | FPSLLTHN | -134 | -141 | 0 | 0.00272025 | 8 | 1763.8762 | NaN | 1599.7582 | NaN | NaN | NaN |
| Rpl10a | AVDIPHMDIE | -79 | -88 | 0 | 0.007493591 | 10 | 676.50195 | NaN | 706.39594 | NaN | NaN | NaN |
| Nifk | PSLALNPQEDSQFEK | -10 | -24 | 0 | 0 | 15 | 4526.858 | 1929.3401 | 3199.1812 | 6435.517 | 5885.747 | 6003.9263 |
| Nifk | RDSEGNQVLPDQK | -203 | -215 | 1 | 0 | 13 | 710.51434 | 600.83844 | 662.34424 | 548.0076 | 892.85767 | 900.1912 |
| Nifk | GIDYSFPSLVLPK | -182 | -194 | 0 | 0 | 13 | 623.6372 | 364.41302 | NaN | 333.5412 | 902.7095 | 807.4573 |
| Nifk | AGLAGPAKPSLALNPQEDSQFEK | -2 | -24 | 1 | 0.001055699 | 23 | 418.91125 | 396.83325 | NaN | 490.7434 | 704.1498 | 679.23645 |
| Nifk | AGLAGPAKPSLALNPQEDSQFEK | -2 | -24 | 1 | 0.001055699 | 23 | 418.91125 | 396.83325 | NaN | 490.7434 | 704.1498 | 679.23645 |
| Nifk | DSEGNQVLPDQK | -204 | -215 | 0 | 0.001636916 | 12 | 1828.3707 | NaN | NaN | NaN | NaN | NaN |

| | | | | | | | | | | | | |
|---|---|---|---|---|---|---|---|---|---|---|---|---|
| Flnb | IGNLQTDLSDGLR | -37 | -49 | 0 | 0.000550307 | 13 | 560.56213 | 463.8549 | 606.96704 | NaN | NaN | NaN |
| Flnb | EVGEHLVSIK | -1970 | -1979 | 0 | 0.00078661 | 10 | 288.6076 | NaN | NaN | 438.57385 | NaN | NaN |
| Flnb | AGLAPLEVR | -1450 | -1458 | 0 | 0.002068213 | 9 | 522.1421 | NaN | NaN | NaN | NaN | NaN |
| Psmb3 | GVIVHVIEK | -184 | -192 | 0 | 0 | 9 | 1641.1389 | 1670.0828 | 1652.5668 | 1511.5851 | 1430.9519 | 1383.6215 |
| Psmb3 | YTEPVIAGLDPK | -104 | -115 | 0 | 0 | 12 | 1239.9722 | 3244.491 | 905.5171 | 4527.385 | 7148.0293 | 2753.6582 |
| Psmb3 | AGLATDVQTVAQR | -54 | -66 | 0 | 0 | 13 | 596.0043 | 1486.2516 | 1764.0134 | 1394.9928 | 1649.1819 | 1975.3121 |
| Psmb3 | GLATDVQTVAQR | -55 | -66 | 0 | 0 | 12 | 1469.7444 | 501.6389 | 1598.165 | 1218.4551 | NaN | 1239.3916 |
| Psmb3 | FGPYYTEPVIA | -100 | -110 | 0 | 0 | 11 | 182.03325 | 176.58496 | 170.94244 | 280.28915 | 308.77438 | NaN |
| Psmb3 | FGPYYTEPVIAGLDPK | -100 | -115 | 0 | 0 | 16 | 438.7869 | NaN | 1151.9362 | NaN | 653.57043 | 1102.9323 |
| Sdha | LEYRPVIDK | -639 | -647 | 1 | 0 | 9 | 2968.2424 | 2695.4324 | 1883.5486 | 2232.8076 | 693.91534 | 696.0894 |
| Sdha | WHFYDTVK | -121 | -128 | 0 | 0 | 8 | 1622.9951 | 1737.5808 | 1861.332 | 2373.043 | 2278.1401 | 2124.5444 |
| Sdha | LHHLPPEQLATR | -368 | -379 | 0 | 0 | 12 | 1349.422 | 891.7901 | 1841.4248 | 2900.4595 | 397.53494 | 906.5978 |
| Sdha | EYRPVIDK | -640 | -647 | 1 | 0.000627943 | 8 | 1329.7183 | 553.5258 | 1296.4808 | 931.9562 | 877.75464 | 1234.1871 |
| Sdha | ISQLYGDLK | -539 | -547 | 0 | 0 | 9 | 899.79877 | 1119.389 | 1049.5769 | 758.57294 | 1364.3945 | 1225.0096 |
| Sdha | GSDWLGDQDAIHY | -129 | -141 | 0 | 0 | 13 | 881.28467 | 747.90204 | 676.7427 | 686.3263 | 572.6836 | 689.6094 |
| Sdha | TLNEADCATVPPAIR | -648 | -662 | 0 | 0 | 15 | 792.0087 | 149.0379 | 858.83154 | 1397.8958 | 1026.2207 | 1435.3357 |
| Sdha | GIYGAGCLITEGCR | -299 | -312 | 0 | 0 | 14 | 789.0175 | 1061.1254 | 1058.3293 | 1086.9685 | 789.29736 | 697.6493 |
| Sdha | AGLPCQDLEFVQFHPT | -283 | -298 | 0 | 0 | 16 | 412.2559 | 454.4305 | 432.60785 | 928.3348 | 603.4475 | 550.0016 |
| Sdha | GCGPEKDHVYLQ | -356 | -367 | 1 | 0 | 12 | 1281.988 | 1075.2236 | 1372.9778 | NaN | 5516.1196 | 5276.368 |
| Sdha | CIEDGSIHR | -238 | -246 | 0 | 0 | 9 | 1252.5469 | NaN | 268.57495 | 1533.984 | 1795.0048 | 718.5042 |
| Sdha | ANAGEESVMNLDK | -486 | -498 | 0 | 0 | 13 | 979.8689 | 924.0218 | 935.0317 | 966.1467 | 989.41315 | NaN |
| Sdha | EPIPVLPTVHYN | -398 | -409 | 0 | 0.002068213 | 12 | 1057.6719 | 490.8387 | 555.4027 | 1101.008 | NaN | NaN |
| Sdha | VTLEYRPVIDK | -637 | -647 | 1 | 0 | 11 | 1053.731 | 1241.4633 | NaN | 1553.5363 | 1731.3042 | NaN |
| Sdha | KPFGEHWR | -616 | -623 | 1 | 0 | 8 | 605.81537 | 1353.6534 | 215.3353 | NaN | 1411.0529 | NaN |
| Sdha | KHTLSYVDIK | -624 | -633 | 1 | 0.00644558 | 10 | 519.42865 | NaN | NaN | NaN | NaN | NaN |
| Dnajc11 | AGLPGFYDPCVGEEK | -509 | -523 | 0 | 0 | 15 | 573.9461 | 230.64311 | 659.9766 | 515.09143 | 504.12534 | 497.0952 |
| Cstf2 | GTGAPVIESPYGESISPEDAPESISK | -105 | -130 | 0 | 0.000230707 | 26 | 228.10826 | 249.94092 | NaN | 320.7015 | 348.14285 | 336.23343 |
| Cstf2 | IVDPEIALK | -181 | -189 | 0 | 0 | 9 | 1927.8513 | 1492.0017 | 1410.2074 | 629.24225 | NaN | NaN |
| Cstf2 | AGLPVRDPAVDR | -2 | -13 | 1 | 0.002099344 | 12 | 1106.6951 | NaN | 1568.4426 | NaN | NaN | 1237.8594 |
| Pdhb | VVSPWNSEDAK | -174 | -184 | 0 | 0 | 11 | 6722.0234 | 86.14706 | 7751.7944 | 9762.87 | 539.93274 | 811.57794 |
| Pdhb | GLQPVPIVFR | -136 | -145 | 0 | 0 | 10 | 1470.0493 | 1494.5895 | 1371.656 | 1123.5509 | 1387.733 | 1442.2506 |
| Pdhb | EGPAFNFLDAPAVR | -311 | -324 | 0 | 0 | 14 | 1169.646 | 1279.0221 | 1383.3077 | 1999.6763 | 2426.6445 | 2265.2114 |
| Pdhb | EGIECEVINLR | -259 | -269 | 0 | 0 | 11 | 1015.3756 | 1054.2185 | 1220.5729 | 1200.1904 | 1385.464 | 2971.422 |
| Pdhb | WYGHCPGLK | -165 | -173 | 0 | 0 | 9 | 929.49396 | 1232.9092 | 1662.5045 | 1511.9441 | 1180.7628 | 1949.3848 |
| Pdhb | SKDFLIPIGK | -218 | -227 | 1 | 0 | 10 | 648.5846 | 1993.5413 | 2078.96 | 1730.4545 | 3486.0164 | 1672.0013 |
| Pdhb | AGLQPVPIVFR | -135 | -145 | 0 | 0 | 11 | 572.835 | 644.71515 | 643.52106 | 632.0631 | 716.7345 | 727.0073 |
| Pdhb | SKEGIECEVINLR | -257 | -269 | 1 | 0 | 13 | 315.0186 | 857.0045 | 500.8555 | 408.10504 | 345.75162 | 713.17285 |
| Pdhb | YGHCPGLK | -166 | -173 | 0 | 0.006624942 | 8 | 649.47626 | NaN | NaN | NaN | 337.10547 | NaN |
| Ddx54 | DQEFYVPYRPK | -669 | -679 | 1 | 0 | 11 | 1480.7825 | 3731.499 | 2966.6648 | 2145.867 | 3961.1074 | 2748.51 |
| Ddx54 | AGLTEPVLIR | -290 | -299 | 0 | 0 | 10 | 610.04254 | 497.56732 | 557.1496 | 549.92725 | 712.1055 | 477.08548 |
| Runx3 | VDVLADHAGELVR | -57 | -69 | 0 | 0 | 13 | 462.78436 | 741.3813 | 722.6531 | 573.62537 | 678.59937 | 733.122 |
| Runx3 | AQTPIQGSSDLNPFSDPR | -227 | -244 | 0 | 0 | 18 | 387.1131 | 614.6066 | 586.68195 | 439.30215 | 656.3193 | 350.93964 |
| Top1 | GPVFAPPYEPLPESVK | -226 | -241 | 0 | 0 | 16 | 4190.493 | 4623.2666 | 4744.904 | 3670.9773 | 4444.2812 | 3856.6978 |
| Top1 | ELTAPDENVPAK | -606 | -617 | 0 | 0 | 12 | 1239.9619 | 1011.672 | 352.65802 | 1512.8165 | 941.4687 | 1266.3527 |
| Top1 | SGDHLHNDSQIEAD | -2 | -15 | 0 | 0.00272025 | 14 | 929.4656 | 940.72424 | 951.96875 | 947.5985 | 1338.9746 | 892.7226 |
| Top1 | IKDEPEDDGYFAPPK | -118 | -132 | 1 | 0 | 15 | 744.6242 | 780.9282 | 836.7555 | 804.54156 | 812.9898 | 842.68 |
| Top1 | LRDEDDADYKPK | -141 | -152 | 2 | 0 | 12 | 355.4359 | 1104.2747 | 1082.2902 | 1314.7726 | 1175.5121 | 1470.3153 |
| Top1 | LRDEDDADYKPK | -141 | -152 | 2 | 0 | 12 | 355.4359 | 1104.2747 | 1082.2902 | 1314.7726 | 1175.5121 | 1470.3153 |
| Top1 | AGNEKEEGETADTVGCC | -491 | -507 | 1 | 0 | 17 | 375.28107 | 363.4468 | 353.54117 | 371.06735 | 351.17505 | 327.00986 |
| Top1 | VPSPPPGHK | -394 | -402 | 0 | 0.002972566 | 9 | 328.55188 | NaN | NaN | 395.51697 | 364.07892 | 2510.595 |
| Top1 | KWGVPIEK | -737 | -744 | 1 | 0.002199523 | 8 | 288.49347 | 833.06213 | 147.16182 | NaN | NaN | 653.45135 |
| Top1 | KQPEDDLFDR | -560 | -569 | 1 | 0.000230707 | 10 | 5361.844 | NaN | NaN | NaN | 6310.885 | 6564.431 |
| Top1 | WTENIQGSIK | -418 | -427 | 0 | 0 | 10 | 3682.2214 | 3743.615 | 1365.3113 | NaN | NaN | NaN |
| Top1 | WWEEERYPEGIK | -207 | -218 | 1 | 0.001463897 | 12 | 502.3104 | NaN | NaN | NaN | NaN | 8394.009 |
| Rpl7 | KVPAVPETL | -32 | -40 | 1 | 0.002910424 | 9 | 2655.6294 | 1750.2036 | 1869.5477 | 1503.67 | 884.10596 | 936.40436 |
| Rpl7 | VEGGDAGNREDQINR | -250 | -264 | 1 | 0 | 15 | 2135.9517 | 381.56833 | 2932.8088 | 1605.3569 | 2070.4644 | 2095.7136 |
| Rpl7 | VEGGDAGNREDQINR | -250 | -264 | 1 | 0 | 15 | 2135.9517 | 381.56833 | 2932.8088 | 1605.3569 | 2070.4644 | 2095.7136 |
| Rpl7 | KVLQLLR | -129 | -135 | 1 | 0.000550307 | 7 | 2095.4912 | 2060.6897 | 1970.7826 | 525.08124 | 532.08636 | 490.4287 |
| Rpl7 | IAWGYPNLK | -162 | -170 | 0 | 0.000230707 | 9 | 1769.2391 | 1863.7788 | 1928.2601 | 1695.8214 | 1911.7551 | 1871.4946 |
| Rpl7 | TTHFVEGGDAGNR | -246 | -258 | 0 | 0 | 13 | 1312.7374 | 1349.5 | 1308.0778 | 453.84918 | 773.7857 | 788.7986 |
| Rpl7 | TTHFVEGGDAGNR | -246 | -258 | 0 | 0 | 13 | 1312.7374 | 1349.5 | 1308.0778 | 453.84918 | 773.7857 | 788.7986 |
| Rpl7 | HFVEGGDAGNREDQINR | -248 | -264 | 1 | 0 | 17 | 1410.4534 | 1438.9178 | 1436.648 | 2845.5642 | 3625.0957 | 3535.6401 |
| Rpl7 | HFVEGGDAGNR | -248 | -258 | 0 | 0 | 11 | 959.36505 | 1084.6559 | 881.71045 | 351.28217 | 376.05417 | 1118.4419 |
| Rpl7 | LTDNSLIAR | -191 | -199 | 0 | 0.00025015 | 9 | 950.6378 | 958.28107 | 1103.9213 | 1551.7421 | 1570.0475 | 1425.3987 |
| Rpl7 | TTHFVEGGDAGNREDQINR | -246 | -264 | 1 | 0 | 19 | 798.90356 | 831.7776 | 893.08716 | 588.8207 | 601.2136 | 558.9983 |
| Rpl7 | TTHFVEGGDAGNREDQINR | -246 | -264 | 1 | 0 | 19 | 798.90356 | 831.7776 | 893.08716 | 588.8207 | 601.2136 | 558.9983 |
| Rpl7 | EDLIHEIYTVGK | -210 | -221 | 0 | 0 | 12 | 650.4284 | 1272.2445 | 728.0855 | 1353.0112 | 886.0535 | 789.39233 |
| Rpl7 | GGDAGNREDQINR | -252 | -264 | 1 | 0 | 13 | 473.30988 | 304.90033 | 392.9841 | 352.1357 | 402.4133 | 395.12082 |
| Rpl7 | GGDAGNREDQINR | -252 | -264 | 1 | 0 | 13 | 473.30988 | 304.90033 | 392.9841 | 352.1357 | 402.4133 | 395.12082 |
| Rpl7 | KKVPAVPETL | -31 | -40 | 2 | 0.000591124 | 10 | 407.73645 | 506.97104 | 491.2475 | 583.32196 | 342.45093 | 490.91357 |
| Rpl7 | GNFYVPAEPK | -101 | -110 | 0 | 0 | 10 | 3783.0234 | 3851.9285 | 3294.0608 | 4254.737 | 2461.3735 | NaN |
| Rpl7 | MEDLIHEIYTVGK | -209 | -221 | 0 | 0 | 13 | 3256.0842 | 3237.384 | NaN | 4267.0225 | 5122.2295 | 4568.708 |
| Rpl7 | AGNFYVPAEPK | -100 | -110 | 0 | 0 | 11 | 3172.8716 | 1463.6492 | 1466.9945 | NaN | 1765.1138 | 941.86566 |
| Rpl7 | IALTDNSLIAR | -189 | -199 | 0 | 0 | 11 | 1586.9327 | 1805.683 | 1860.1112 | 793.5521 | 1204.3635 | NaN |
| Rpl7 | TTHFVEGGDAGNREDQIN | -246 | -263 | 1 | 0.00025015 | 18 | 1377.4584 | 1419.7161 | 1430.7118 | 947.39813 | 563.16547 | NaN |
| Rpl7 | TTHFVEGGDAGNREDQIN | -246 | -263 | 1 | 0.00025015 | 18 | 1377.4584 | 1419.7161 | 1430.7118 | 947.39813 | 563.16547 | NaN |
| Rpl7 | KAGNFYVPAEPK | -99 | -110 | 1 | 0 | 12 | 1462.5421 | 1072.0222 | 1387.0809 | NaN | NaN | 711.3494 |
| Rpl7 | DLIHEIYTVGK | -211 | -221 | 0 | 0.000826806 | 11 | 1250.4967 | 1055.5437 | 1165.2922 | NaN | NaN | NaN |

| | | | | | | | | | | | | |
|---|---|---|---|---|---|---|---|---|---|---|---|---|
| Rpl7 | KVPAVPETLK | -32 | -41 | 1 | 0 | 10 | 899.07526 | 920.1383 | 191.16257 | NaN | NaN | NaN |
| Rpl7 | VPAVPETLK | -33 | -41 | 0 | 0.00025015 | 9 | 224.13977 | 585.3483 | 771.53845 | NaN | NaN | NaN |
| Rpl7 | KVATVPGTLK | -10 | -19 | 1 | 0.001291103 | 10 | 276.8311 | NaN | NaN | NaN | NaN | NaN |
| Ndufb3 | AGNITFPSVILK | -64 | -75 | 0 | 0 | 12 | 585.91504 | 511.0389 | NaN | 673.44696 | 561.01117 | 569.8233 |
| Ndufb3 | NITFPSVILK | -66 | -75 | 0 | 0 | 10 | 409.99948 | NaN | 606.6997 | NaN | NaN | NaN |
| Rpl22 | NFEQFLQER | -38 | -46 | 0 | 0 | 9 | 6357.757 | 6886.102 | 6613.465 | 6542.9307 | 7065.1914 | 7103.9375 |
| Rpl22 | AGNLGGGVVTIER | -53 | -65 | 0 | 0 | 13 | 799.982 | 1161.3701 | 1447.8711 | 293.86874 | 621.3337 | 549.26855 |
| Rpl22 | ANFEQFLQER | -37 | -46 | 0 | 0 | 10 | 1167.9961 | 1375.0154 | 1600.3136 | NaN | NaN | NaN |
| Eno3 | TLGPALLEK | -72 | -80 | 0 | 0.00025015 | 9 | 835.74097 | 758.30945 | 899.0235 | 635.32715 | NaN | NaN |
| Eno3 | AGNPDLVLPVPAFNVINGGSHAGNK | -138 | -162 | 0 | 0.00122419 | 25 | 648.63367 | NaN | 498.23035 | NaN | NaN | NaN |
| Srrt | HKDEEWFR | -191 | -198 | 1 | 0 | 8 | 745.6527 | 856.58405 | 735.88007 | 321.07632 | 501.58344 | 662.7091 |
| Srrt | HKDEEWFR | -191 | -198 | 1 | 0 | 8 | 745.6527 | 856.58405 | 735.88007 | 321.07632 | 501.58344 | 662.7091 |
| Srrt | EEKEEAEEALKEK | -380 | -392 | 2 | 0 | 13 | 2690.2349 | 1999.4584 | 1564.81 | 2030.2174 | 3399.0366 | 2753.4978 |
| Srrt | KDPEQEVEKF | -690 | -699 | 2 | 0.00025015 | 10 | 2304.888 | 1834.1313 | 3339.3306 | 816.7026 | 557.17615 | 3048.299 |
| Srrt | SLDDSVDETEAVKR | -158 | -171 | 1 | 0 | 14 | 1120.9666 | 983.69147 | 1007.4294 | 1010.3889 | 1986.7914 | 2111.3596 |
| Srrt | KDPEQEVEK | -690 | -698 | 1 | 0.000298805 | 9 | 907.03766 | 924.158 | 546.3685 | 813.8069 | 980.8639 | 1101.3649 |
| Srrt | ESLSEEEAQK | -677 | -686 | 0 | 0 | 10 | 800.7486 | 836.10657 | 934.81323 | 602.3142 | 519.2042 | 531.5753 |
| Srrt | RPALPEIKPAQPPGPA | -758 | -773 | 2 | 0.00078661 | 16 | 1249.5498 | 1663.3727 | 43.183384 | NaN | 1751.4283 | 1957.3176 |
| Srrt | RPALPEIKPAQPPGPA | -758 | -773 | 2 | 0.00078661 | 16 | 1249.5498 | 1663.3727 | 43.183384 | NaN | 1751.4283 | 1957.3176 |
| Srrt | DDSVDETEAVKR | -160 | -171 | 1 | 0 | 12 | 578.5857 | 500.56302 | 590.0827 | 519.4724 | NaN | NaN |
| Srrt | CGIIHVR | -639 | -645 | 0 | 0 | 7 | 973.80005 | 620.45715 | NaN | NaN | 641.81305 | NaN |
| Srrt | EEKEEAEEALK | -380 | -390 | 1 | 0 | 11 | 1186.4398 | NaN | NaN | NaN | 1624.4996 | NaN |
| Srrt | AGPALGEGER | -291 | -300 | 0 | 0.00122419 | 10 | 948.9045 | NaN | NaN | NaN | NaN | NaN |
| Arpc5 | GFESPSDNSSAVLL | -113 | -126 | 0 | 0 | 14 | 856.9639 | 1419.2871 | 950.043 | 992.2795 | 2843.5886 | 2490.917 |
| Arpc5 | AGPDEGEVDSCLR | -35 | -47 | 0 | 0 | 13 | 820.9635 | 758.5281 | 788.4688 | 1778.3083 | 2001.7675 | 1879.567 |
| Arpc5 | ALAAGGVGSIVR | -132 | -143 | 0 | 0 | 12 | 1291.0896 | 969.71844 | NaN | 1061.183 | NaN | 894.5015 |
| Arpc5 | EDGGDGQAGPDEGEVDSCLR | -28 | -47 | 0 | 0.000471409 | 20 | 496.0638 | 483.8619 | 469.65735 | NaN | NaN | NaN |
| Sart3 | GYDYNCHVELIR | -110 | -121 | 0 | 0 | 12 | 854.31104 | 647.56036 | 452.93558 | 257.8354 | 570.312 | 580.3463 |
| Sart3 | AGPEAEGEEDEAKPAGVQR | -18 | -36 | 1 | 0.00025015 | 19 | 337.02042 | 322.43478 | NaN | 644.27545 | 780.6262 | 533.5331 |
| Sart3 | LRPLFEVC | -722 | -729 | 1 | 0.00659934 | 8 | 970.45624 | 1173.3794 | 1094.0472 | NaN | NaN | 1251.6953 |
| Sart3 | EHVYELFER | -168 | -176 | 0 | 0 | 9 | 1153.0149 | NaN | NaN | 509.40717 | NaN | 1079.8662 |
| Sart3 | YKPYEEALLQ | -296 | -305 | 1 | 0.00122419 | 10 | 618.50836 | NaN | NaN | NaN | 670.49475 | 1107.8015 |
| Sart3 | EYLQQEVEER | -455 | -464 | 0 | 0.00025015 | 10 | 2376.8794 | 1881.5563 | NaN | NaN | NaN | NaN |
| Ddx6 | NLVCTDLFTR | -387 | -396 | 0 | 0 | 10 | 5353.1406 | 5601.659 | 360.36044 | 492.278 | 378.75476 | 981.4895 |
| Ddx6 | EDIILTLPK | -261 | -269 | 0 | 0 | 9 | 3630.834 | 3749.395 | 3927.1304 | 3135.8367 | 3203.8953 | 3384.1643 |
| Ddx6 | LITYDDRFNLK | -437 | -447 | 1 | 0.000230707 | 11 | 3510.343 | 1975.2205 | 2590.9312 | 2621.9758 | 2851.325 | 3398.4607 |
| Ddx6 | AGPGGGGTQPQPQLNQLK | -29 | -46 | 0 | 0 | 18 | 3099.8887 | 3522.9858 | 3649.082 | 3209.5176 | 4146.95 | 3671.3345 |
| Ddx6 | EYHSEPAEDEKP | -472 | -483 | 1 | 0.00025015 | 12 | 1832.2739 | 1804.7158 | 2137.9443 | 400.61417 | 1302.6376 | 1383.5756 |
| Ddx6 | SGAYLIPLLER | -147 | -157 | 0 | 0 | 11 | 1489.0209 | 1701.1785 | 1961.2504 | 1267.028 | 1516.1202 | 1307.0393 |
| Ddx6 | GNEFEDYCLK | -95 | -104 | 0 | 0 | 10 | 632.2243 | 630.34406 | 659.37317 | 646.04016 | 460.54507 | 488.80832 |
| Ddx6 | SIEEQLGTEIKPIPSNIDK | -448 | -466 | 1 | 0 | 19 | 607.83923 | 682.2897 | 661.7073 | 769.6605 | 1145.9706 | 1174.8372 |
| Ddx6 | SIEEQLGTEIKPIPSNIDK | -448 | -466 | 1 | 0 | 19 | 607.83923 | 682.2897 | 661.7073 | 769.6605 | 1145.9706 | 1174.8372 |
| Ddx6 | AYLIPLLER | -149 | -157 | 0 | 0 | 9 | 386.1212 | 335.21783 | 408.44342 | 378.20532 | 468.027 | 522.21387 |
| Ddx6 | ASAGPGGGGTQPQPQLNQLK | -27 | -46 | 0 | 0.00111006 | 20 | 646.83813 | 694.1589 | 749.6198 | NaN | NaN | NaN |
| Ddx6 | SAGPGGGGTQPQPQLNQLK | -28 | -46 | 0 | 0.000347792 | 19 | 442.58316 | 375.70612 | NaN | NaN | NaN | NaN |
| Smg9 | AGPGSSPLFSLLPGYR | -446 | -461 | 0 | 0 | 16 | 275.58948 | NaN | 452.69336 | 120.238235 | 491.2041 | 420.8432 |
| Sod1 | HVGDLGNVTAGK | -81 | -92 | 0 | 0 | 12 | 6851.909 | 7216.109 | 7359.953 | 8222.982 | 9009.005 | 8961.871 |
| Sod1 | VISLSGEHSIIGR | -104 | -116 | 0 | 0 | 13 | 6333.9854 | 6855.9463 | 7292.0225 | 8963.349 | 9835.37 | 9574.889 |
| Sod1 | VISLSGEHSIIGR | -104 | -116 | 0 | 0 | 13 | 6333.9854 | 6855.9463 | 7292.0225 | 8963.349 | 9835.37 | 9574.889 |
| Sod1 | GDGPVQGTIHFEQK | -11 | -24 | 0 | 0 | 14 | 3165.129 | 3560.5269 | 3617.871 | 228.65495 | 532.8238 | 6705.033 |
| Sod1 | SGEHSIIGR | -108 | -116 | 0 | 0 | 9 | 2825.3848 | 2735.0874 | 2808.7612 | 3034.414 | 2930.4863 | 3040.9934 |
| Sod1 | DGVANVSIEDR | -93 | -103 | 0 | 0 | 11 | 2626.5938 | 2293.1448 | 1571.6337 | 2813.525 | 453.81638 | 885.42224 |
| Sod1 | ASGEPVVLSGQIT | -25 | -37 | 0 | 0.00111006 | 13 | 352.20438 | 379.45834 | 365.40366 | 316.85086 | 316.10056 | 292.02118 |
| Sod1 | AGPHFNPHSK | -61 | -70 | 0 | 0.00025015 | 10 | 616.0063 | 454.04895 | 264.409 | 578.35516 | 195.67577 | NaN |
| Pnp | DHINLPGFCGQNPLR | -134 | -148 | 0 | 0 | 15 | 2255.0767 | 1906.3907 | 2756.2898 | 1901.696 | 2390.5842 | 3126.82 |
| Pnp | DHINLPGFCGQNPLR | -134 | -148 | 0 | 0 | 15 | 2255.0767 | 1906.3907 | 2756.2898 | 1901.696 | 2390.5842 | 3126.82 |
| Pnp | AGPNFETVAESR | -196 | -207 | 0 | 0 | 12 | 1624.0413 | NaN | 1796.4945 | 2110.8103 | 1622.5339 | 808.052 |
| Pnp | VVMDYENLEK | -245 | -254 | 0 | 0 | 10 | 883.28705 | NaN | NaN | 1042.2285 | 927.5761 | NaN |
| Pcbp2 | IANPVEGSTDR | -320 | -330 | 0 | 0 | 11 | 4245.2397 | 4686.849 | 4596.714 | 637.28595 | 3457.9656 | 3939.419 |
| Pcbp2 | GIPQSIIECVK | -150 | -160 | 0 | 0 | 11 | 2899.6077 | 3908.4814 | 3228.567 | 2000.9401 | 3985.895 | 3818.2314 |
| Pcbp2 | GVTIPYRPKPSSSPVIF | -173 | -189 | 2 | 0.000431655 | 17 | 600.81006 | 769.09625 | 665.1344 | 704.1506 | 809.5365 | 779.51764 |
| Pcbp2 | GVTIPYRPKPSSSPVIF | -173 | -189 | 2 | 0.000431655 | 17 | 600.81006 | 769.09625 | 665.1344 | 704.1506 | 809.5365 | 779.51764 |
| Pcbp2 | IITLAGPTNAIFK | -58 | -70 | 0 | 0 | 13 | 119.68282 | 1539.601 | 1611.9413 | 1773.4083 | 2272.742 | 2271.041 |
| Pcbp2 | YAIPQPDLTK | -232 | -241 | 0 | 0 | 10 | 115.20023 | 2507.291 | 2615.7466 | 949.82056 | 1817.8667 | 2210.3005 |
| Pcbp2 | GVTIPYRPK | -173 | -181 | 1 | 0 | 9 | 1130.1519 | NaN | 1281.705 | 997.2631 | 982.01135 | 1081.2543 |
| Pcbp2 | PKPSSSPVIF | -180 | -189 | 1 | 0.001904399 | 10 | 1113.0354 | 564.4232 | 591.267 | 887.5465 | 1196.518 | NaN |
| Pcbp2 | GQYAIPQPDLTK | -230 | -241 | 0 | 0.00078661 | 12 | 13362.724 | 13200.076 | NaN | NaN | 11705.694 | 11556.804 |
| Pcbp2 | AGPTNAIFK | -62 | -70 | 0 | 0.00111006 | 9 | 1941.7899 | 1515.2985 | NaN | NaN | NaN | NaN |
| Bhmt | AGPWTPEAAVEHPEAVR | -41 | -57 | 0 | 0 | 17 | 786.2965 | 1045.2239 | 867.2068 | 810.4692 | 1092.9614 | 1167.9009 |
| Pgd | AGQAVDDFIEK | -77 | -87 | 0 | 0 | 11 | 1784.7571 | 1253.2671 | 2075.7087 | 1546.4257 | 1198.176 | 1417.5449 |
| Pgd | ALSFYDGYR | -426 | -434 | 0 | 0 | 9 | 1271.3844 | 1477.041 | 1456.8623 | 1373.318 | 1517.6255 | 1520.6577 |
| Pgd | VDNCQDSWR | -399 | -407 | 0 | 0 | 9 | 920.17255 | 979.5505 | 927.9314 | 1144.7063 | 1027.6632 | 1018.1748 |
| Pgd | GQAVDDFIEK | -78 | -87 | 0 | 0 | 10 | 624.30743 | 714.9665 | 639.10236 | 1508.1505 | 1122.3113 | 1243.837 |
| Pgd | AVDNCQDSWR | -398 | -407 | 0 | 0 | 10 | 583.0441 | 819.5575 | 74.61555 | 229.00494 | 685.1444 | 822.63995 |
| Pgd | VGTGEPCCDWVGDEGAGHFVK | -164 | -184 | 0 | 0 | 21 | 533.73444 | 655.5208 | 647.9702 | 571.3059 | 830.7501 | 713.343 |
| Pgd | VGTGEPCCDWVGDEGAGHFVK | -164 | -184 | 0 | 0 | 21 | 533.73444 | 655.5208 | 647.9702 | 571.3059 | 830.7501 | 713.343 |
| Pgd | LLDDFFK | -390 | -396 | 0 | 0.00111006 | 7 | 479.52066 | 428.42004 | 496.20734 | 290.57867 | 284.8538 | 290.1234 |
| Pgd | EYGMPVTLIGEAVFAR | -273 | -288 | 0 | 0 | 16 | 477.95792 | 459.7383 | 519.26794 | 358.83005 | 836.0136 | 754.35535 |

| | | | | | | | | | | | | |
|---|---|---|---|---|---|---|---|---|---|---|---|---|
| Pgd | SAVDNCQDSWR | -397 | -407 | 0 | 0 | 11 | 364.1626 | 724.8316 | 679.8407 | 677.9472 | 557.80615 | 696.5193 |
| Pgd | NLLLDDFFK | -388 | -396 | 0 | 0 | 9 | 246.27318 | 451.33096 | 373.2025 | 71.454094 | 484.74838 | 318.8661 |
| Pgd | IKDAFER | -376 | -382 | 1 | 0.000591124 | 7 | 1005.29346 | NaN | 175.84764 | 940.4683 | 1017.4713 | 492.95972 |
| Pgd | LSFYDGYR | -427 | -434 | 0 | 0.000755556 | 8 | 815.6121 | 410.07626 | 743.87866 | 906.7947 | 705.8517 | NaN |
| Pgd | KSFLEDIR | -317 | -324 | 1 | 0 | 8 | 467.89194 | 757.46075 | 916.0022 | NaN | 678.6957 | 689.48627 |
| Pgd | SFLEDIRK | -318 | -325 | 1 | 0.00025015 | 8 | 1618.4005 | 1543.3364 | 1378.4315 | 1497.6812 | NaN | NaN |
| Pgd | WVGDEGAGHFVK | -173 | -184 | 0 | 0 | 12 | 1566.0077 | 1641.7083 | 1671.6194 | NaN | NaN | NaN |
| Pgd | VGTGEPCCDWVGDEGAGHF | -164 | -182 | 0 | 0.000230707 | 19 | 269.53986 | NaN | 333.57993 | NaN | 582.6374 | NaN |
| Pgd | CLSSLKEER | -289 | -297 | 1 | 0 | 9 | 518.7944 | NaN | NaN | NaN | NaN | NaN |
| Tubb4b | INVYYNEATGGK | -47 | -58 | 0 | 0 | 12 | 74.53076 | 1057.9631 | 1028.7478 | NaN | NaN | 1108.5247 |
| Arid1a | EFDSGLLHWR | -1834 | -1843 | 0 | 0 | 10 | 847.7046 | 971.51715 | 891.3119 | NaN | 813.3736 | 755.81683 |
| Arid1a | AGQESEGPAVGPPQPLGK | -52 | -69 | 0 | 0 | 18 | 487.11758 | 400.52008 | 640.14185 | NaN | 630.38293 | 809.61017 |
| Arid1a | GYQGYPGGDYGGGPQDGGAGK | -308 | -328 | 0 | 0.00025015 | 21 | 222.03308 | NaN | 61.947952 | 452.48346 | 417.71423 | 152.0532 |
| Acaca | EEFLIPIYHQ | -2171 | -2180 | 0 | 0 | 10 | 1210.056 | 1091.043 | 1424.8805 | 950.7384 | 183.90051 | 997.98175 |
| Acaca | AGQVWFPDSAFK | -2013 | -2024 | 0 | 0.00025015 | 12 | 295.29724 | 520.23816 | 456.47015 | 524.61304 | 633.14355 | 569.77686 |
| Acaca | VNNADDFPNLFR | -323 | -334 | 0 | 0 | 12 | 692.73975 | 473.78854 | 746.533 | 285.39523 | 1010.7544 | NaN |
| Acaca | LPELLLK | -217 | -223 | 0 | 0.005605646 | 7 | 187.3242 | NaN | NaN | 122.67595 | NaN | 94.80355 |
| Acaca | EEFLIPIYHQV | -2171 | -2181 | 0 | 0.003681885 | 11 | 438.22107 | 481.2942 | NaN | NaN | NaN | NaN |
| Acaca | TVELSIPADPANLDSEAK | -1991 | -2008 | 0 | 0.001291103 | 18 | 385.73828 | NaN | NaN | NaN | 694.08826 | NaN |
| Tmpo | YSDNDEDSKIELK | -182 | -194 | 1 | 0 | 13 | 925.94696 | 1046.3567 | 1264.4172 | 1210.6663 | 1467.3182 | 1763.5806 |
| Tmpo | GAAGRPLELSDFR | -368 | -380 | 1 | 0 | 13 | 915.67004 | 1064.827 | 1293.4553 | 928.23486 | 1042.7443 | 1002.0428 |
| Tmpo | GAVISESTPIAETIK | -271 | -285 | 0 | 0 | 15 | 897.1296 | 723.745 | 1285.6489 | 1381.9891 | 1594.5804 | 1352.8082 |
| Tmpo | HASSILPITEF | -303 | -313 | 0 | 0.00025015 | 11 | 854.0846 | 928.4094 | 1730.3646 | 1353.2837 | 749.45154 | 1424.3348 |
| Tmpo | HASSILPITEF | -303 | -313 | 0 | 0.00025015 | 11 | 854.0846 | 928.4094 | 1730.3646 | 1353.2837 | 749.45154 | 1424.3348 |
| Tmpo | IDGAVISESTPIAETIK | -269 | -285 | 0 | 0 | 17 | 837.13055 | 548.113 | 716.39056 | 840.208 | 1030.6942 | 1163.3433 |
| Tmpo | GAAGRPLELSDF | -368 | -379 | 1 | 0.001291103 | 12 | 963.07495 | 1343.4053 | 1233.7024 | NaN | 1726.5059 | NaN |
| Tmpo | HASSILPITEFSDITR | -303 | -318 | 0 | 0 | 16 | 274.1792 | NaN | 421.36765 | 326.98212 | NaN | 351.43576 |
| Tmpo | ESTPIAETIK | -276 | -285 | 0 | 0.000230707 | 10 | 1414.814 | 1377.2767 | 965.7782 | NaN | NaN | NaN |
| Tmpo | AGRPLELSDFR | -370 | -380 | 1 | 0 | 11 | 333.194 | NaN | NaN | 237.06735 | NaN | NaN |
| Farsb | SSTFPELPYR | -296 | -305 | 0 | 0 | 10 | 4264.605 | 3514.4924 | 3508.2573 | 1375.623 | 344.97733 | 1464.2615 |
| Farsb | LGVLHPDVITK | -561 | -571 | 0 | 0 | 11 | 1461.2443 | 1570.7502 | 1597.6191 | 1500.3438 | 2010.178 | 1971.5175 |
| Farsb | TTLLPGLLK | -452 | -460 | 0 | 0 | 9 | 771.15845 | 829.7251 | 784.07874 | 670.19934 | 795.71533 | 793.56976 |
| Farsb | TPGFEIIHGL | -505 | -514 | 0 | 0 | 10 | 649.68024 | 807.1271 | 789.2047 | 676.72064 | 642.8352 | 248.568 |
| Farsb | TPGFEIIHGLLDR | -505 | -517 | 0 | 0 | 13 | 604.25586 | 637.8415 | 699.6388 | 3598.2678 | 5132.4233 | 4961.6777 |
| Farsb | DRYDSFIEL | -134 | -142 | 1 | 0.006850902 | 9 | 464.94974 | 436.97473 | 619.13025 | NaN | NaN | NaN |
| Farsb | AGSAFFPGR | -539 | -547 | 0 | 0.000627943 | 9 | 2160.8835 | NaN | NaN | NaN | NaN | NaN |
| Clic4 | NSRPEANEALER | -131 | -142 | 1 | 0 | 12 | 2369.9001 | 989.23206 | 2436.2158 | 2717.412 | 2926.3557 | 2774.1428 |
| Clic4 | NSRPEANEALER | -131 | -142 | 1 | 0 | 12 | 2369.9001 | 989.23206 | 2436.2158 | 2717.412 | 2926.3557 | 2774.1428 |
| Clic4 | DEFTNTCPSDKEVEIA | -228 | -243 | 1 | 0.00025015 | 16 | 1869.6794 | 1153.8402 | 2031.5123 | 1845.7981 | 234.74387 | 1026.8699 |
| Clic4 | AGSDGESIGNCPFSQR | -25 | -40 | 0 | 0 | 16 | 804.5513 | 899.6866 | 859.22595 | 926.6181 | 930.8832 | 961.70886 |
| Clic4 | EEDKEPLIELF | -12 | -22 | 1 | 0.00111006 | 11 | 942.04926 | 977.2592 | 1215.1729 | NaN | NaN | 1278.1112 |
| Ythdf2 | AGSGSTPSEPHPVLEK | -376 | -391 | 0 | 0 | 16 | 616.2898 | 543.4651 | 582.755 | 568.8256 | 501.20837 | 585.90845 |
| Ythdf2 | LGSTEVASSVPK | -194 | -205 | 0 | 0.000431655 | 12 | 648.5687 | 719.4317 | 150.4516 | 757.7814 | NaN | NaN |
| Ythdf2 | DGLNDDDFEPYLSPQAR | -27 | -43 | 0 | 0 | 17 | 502.70862 | NaN | NaN | 474.0253 | 697.8272 | NaN |
| Arhgap45 | SQASPHDISNVLK | -819 | -831 | 0 | 0 | 13 | 4010.8877 | 2916.7046 | 4494.7817 | 3602.1372 | 4016.7798 | 3811.5708 |
| Arhgap45 | AGSPNPQSSSGELPR | -21 | -35 | 0 | 0 | 15 | 3748.0361 | 3334.7456 | 3731.6138 | 3698.782 | 3882.1216 | 4278.7905 |
| Arhgap45 | LSSLVDYPHQAR | -947 | -958 | 0 | 0 | 12 | 1219.1823 | 1629.626 | 2218.839 | 1810.3625 | 949.33624 | 983.40125 |
| Arhgap45 | SLVDYPHQAR | -949 | -958 | 0 | 0 | 10 | 1169.9812 | 1946.2756 | 2118.454 | 1213.401 | 2075.4595 | 2054.5352 |
| Arhgap45 | KDWTEAPGLEPPATSL | -36 | -51 | 1 | 0.00111006 | 16 | 985.7488 | 1187.9672 | 1215.5818 | 858.1105 | 1093.7736 | 1088.1437 |
| Arhgap45 | QLPEPLISFR | -836 | -845 | 0 | 0 | 10 | 752.19836 | 932.10925 | 1075.8104 | 851.3981 | 918.1795 | 1702.5814 |
| Arhgap45 | ALSTPDGVPFIVK | -764 | -776 | 0 | 0 | 13 | 651.94147 | 774.56555 | 672.5228 | 708.0181 | 804.7514 | 817.02405 |
| Arhgap45 | LYDPGQQYASHVR | -532 | -544 | 0 | 0 | 13 | 646.10516 | 186.03014 | 1034.2972 | 582.0029 | 893.18585 | 1147.546 |
| Arhgap45 | KGSFNPGDASGPEAAGSPPEEGGTS | -575 | -599 | 1 | 0.00078661 | 25 | 146.86357 | 265.81207 | 275.1728 | 266.37466 | 154.95625 | 155.56847 |
| Arhgap45 | ECVLQDDLLEAR | -134 | -145 | 0 | 0 | 12 | 791.7522 | NaN | 727.4374 | 562.94116 | 396.4991 | 1366.0236 |
| Arhgap45 | RPLAHECLGEAL | -146 | -157 | 1 | 0 | 12 | 1522.9817 | NaN | NaN | 856.1586 | 2613.2878 | 992.1341 |
| Eef1d | SLEVENQNLR | -86 | -95 | 0 | 0 | 10 | 3758.454 | 4256.245 | 4296.188 | 4761.4272 | 4386.8125 | 4164.7754 |
| Eef1d | ATNFLAHEK | -2 | -10 | 0 | 0.00149795 | 9 | 3494.7603 | 1324.7751 | 1397.8912 | 2874.945 | 3584.4714 | 3418.3086 |
| Eef1d | SLAGSSGPGASSGPGGDHSELIVR | -60 | -83 | 0 | 0 | 24 | 1684.2275 | 1712.5189 | 1737.2362 | 1581.4518 | 1992.2219 | 1953.0929 |
| Eef1d | SLAGSSGPGASSGPGGDHSELIVR | -60 | -83 | 0 | 0 | 24 | 1684.2275 | 1712.5189 | 1737.2362 | 1581.4518 | 1992.2219 | 1953.0929 |
| Eef1d | SGPGASSGPGGDHSELIVR | -65 | -83 | 0 | 0 | 19 | 1581.827 | 1759.9613 | 1845.3041 | 1790.8617 | 2134.7852 | 2014.3773 |
| Eef1d | SGPGASSGPGGDHSELIVR | -65 | -83 | 0 | 0 | 19 | 1581.827 | 1759.9613 | 1845.3041 | 1790.8617 | 2134.7852 | 2014.3773 |
| Eef1d | GVVQDLQQAISK | -96 | -107 | 0 | 0.00025015 | 12 | 1459.0082 | 1586.7688 | 1506.8025 | 1937.2269 | 1395.9326 | 1560.7896 |
| Eef1d | SGPGGDHSELIVR | -71 | -83 | 0 | 0 | 13 | 1166.8523 | 1092.2504 | 1390.2406 | 1441.1145 | 1431.0796 | 1690.9086 |
| Eef1d | FKYDDAER | -16 | -23 | 1 | 0.000431655 | 8 | 943.8363 | 944.3547 | 914.8727 | 706.69324 | 615.5675 | 854.97705 |
| Eef1d | AGSSGPGASSGPGGDHSELIVR | -62 | -83 | 0 | 0 | 22 | 740.812 | 884.2419 | 926.67126 | 810.94073 | 918.5584 | 891.2251 |
| Eef1d | AGSSGPGASSGPGGDHSELIVR | -62 | -83 | 0 | 0 | 22 | 740.812 | 884.2419 | 926.67126 | 810.94073 | 918.5584 | 891.2251 |
| Eef1d | FYEQMNGPVTSGSR | -25 | -38 | 0 | 0 | 14 | 720.5683 | 372.73065 | 793.99493 | 917.2295 | 891.9907 | 904.3496 |
| Eef1d | SSGPGASSGPGGDHSELIVR | -64 | -83 | 0 | 0 | 20 | 567.4645 | 646.02795 | 459.4841 | 1231.7667 | 178.51425 | 874.4093 |
| Eef1d | SSGPGASSGPGGDHSELIVR | -64 | -83 | 0 | 0 | 20 | 567.4645 | 646.02795 | 459.4841 | 1231.7667 | 178.51425 | 874.4093 |
| Eef1d | GATPAEDDEDKDIDLF | -145 | -160 | 1 | 0.00025015 | 16 | 499.2314 | 708.88446 | 586.78973 | 232.55843 | 1321.2589 | 1312.2837 |
| Eef1d | LVPVGYGIR | -233 | -241 | 0 | 0.002534621 | 9 | 3433.0652 | 1976.4541 | 2145.93 | NaN | 2607.5342 | NaN |
| Eef1d | ATAPQTQHVSPMR | -124 | -136 | 0 | 0 | 13 | 463.95993 | 505.0639 | 504.49 | NaN | NaN | 417.09564 |
| Eef1d | ITSLEVENQNLR | -84 | -95 | 0 | 0 | 12 | 2553.719 | 440.03522 | 523.5954 | NaN | NaN | NaN |
| Pld3 | LFVVPTDESQAR | -395 | -406 | 0 | 0 | 12 | 2495.537 | 2743.8333 | 2429.4514 | 2102.8977 | 2493.0725 | 2942.2688 |
| Pld3 | SQLEAVFLR | -456 | -464 | 0 | 0 | 9 | 697.9453 | NaN | 755.00275 | 1030.5524 | 1027.8511 | NaN |
| Pld3 | DWESPYSHDLDTSAN | -465 | -479 | 0 | 0.000230707 | 15 | 422.90723 | 411.69705 | 411.58084 | NaN | NaN | 330.62726 |
| Pld3 | ALLNVVDSAR | -308 | -317 | 0 | 0 | 10 | 1409.291 | NaN | NaN | 769.0733 | NaN | 1523.5563 |
| Pld3 | AGSSIPSTWPR | -255 | -265 | 0 | 0.00025015 | 11 | 721.2912 | NaN | 892.9993 | NaN | NaN | 554.1437 |

| | | | | | | | | | | | | |
|---|---|---|---|---|---|---|---|---|---|---|---|---|
| Nop2 | VLLDAPCSGTGVISK | -442 | -456 | 0 | 0 | 15 | 2073.445 | 2042.354 | 1625.0935 | 1914.0908 | 1946.1956 | 1834.2981 |
| Nop2 | IQDIVGVLR | -228 | -236 | 0 | 0 | 9 | 933.0861 | 1057.1045 | 1035.5687 | 750.87695 | 719.4462 | 716.51935 |
| Nop2 | TIISHYDGR | -422 | -430 | 0 | 0 | 9 | 814.37286 | 6225.4873 | 3940.4272 | 921.81433 | 6870.312 | 4587.631 |
| Nop2 | KDDKAEGDLQIN | -186 | -197 | 2 | 0.00025015 | 12 | 610.37024 | 432.03107 | 748.29663 | 1096.482 | 1072.584 | 951.8508 |
| Nop2 | EVPRPITLR | -290 | -298 | 1 | 0.000394361 | 9 | 1312.5189 | 1297.972 | 1313.0305 | 956.6838 | 1131.84 | NaN |
| Nop2 | SVVGNLHR | -409 | -416 | 0 | 0 | 8 | 969.0034 | 1302.6641 | 1209.4545 | 1207.7554 | 971.15704 | NaN |
| Nop2 | SYGDFLLSK | -264 | -272 | 0 | 0.00025015 | 9 | 1384.1163 | NaN | 1500.0157 | NaN | 973.719 | 1372.678 |
| Nop2 | FLPAAGDENSK | -32 | -42 | 0 | 0.00051306 | 11 | 961.23395 | 1773.5804 | NaN | 3428.3 | NaN | 2143.2913 |
| Nop2 | AGSVDVPKPNK | -57 | -67 | 1 | 0 | 11 | 754.9521 | 948.4038 | 770.5011 | NaN | NaN | 1314.234 |
| Nop2 | GVTNTIISHYDGR | -418 | -430 | 0 | 0 | 13 | 503.49545 | NaN | 404.43472 | NaN | 658.44385 | 699.59344 |
| Nop2 | NTIISHYDGR | -421 | -430 | 0 | 0 | 10 | 442.1434 | NaN | NaN | 1316.8761 | 818.46375 | 418.77457 |
| Fmnl1 | SKPLDQSVEDLSK | -178 | -190 | 1 | 0 | 13 | 763.5314 | 821.1096 | 807.1026 | 736.30835 | 780.1797 | 870.14545 |
| Fmnl1 | HSDLHFLDK | -893 | -901 | 0 | 0 | 9 | 662.5311 | 863.5803 | 432.74722 | 265.0781 | 833.91016 | 889.97864 |
| Fmnl1 | AGSVSLDSVLGDVR | -902 | -915 | 0 | 0 | 14 | 648.0537 | 687.16016 | 794.2556 | NaN | 947.31616 | 1034.3604 |
| Fmnl1 | TAQEAYESVVEYFGENPK | -957 | -974 | 0 | 0.00078661 | 18 | 305.43118 | NaN | 255.10406 | 287.7078 | 354.27197 | 347.87888 |
| Fmnl1 | SLDSVLGDVR | -906 | -915 | 0 | 0 | 10 | 749.8593 | 198.66478 | 456.87695 | 897.47406 | NaN | NaN |
| Fmnl1 | FSESTPMGTSR | -446 | -456 | 0 | 0 | 11 | 538.60614 | 564.7697 | 564.9515 | NaN | NaN | NaN |
| Fmnl1 | FLPTDYER | -750 | -757 | 0 | 0.003620005 | 8 | 2539.127 | 2590.4888 | NaN | NaN | NaN | NaN |
| Fmnl1 | TNHIGWVQ | -130 | -137 | 0 | 0.007987481 | 8 | 1295.4011 | 1433.2521 | NaN | NaN | NaN | NaN |
| Fmnl1 | EFLNEENR | -138 | -145 | 0 | 0.000627943 | 8 | 808.5002 | NaN | 672.3408 | NaN | NaN | NaN |
| Fmnl1 | SKPLDQSVEDL | -178 | -188 | 1 | 0.001743443 | 11 | 1875.5878 | NaN | NaN | NaN | NaN | NaN |
| Lman2 | DITDGNSEHLK | -47 | -57 | 0 | 0 | 11 | 3192.6763 | 1435.962 | 3710.9324 | 3132.4348 | 3784.5962 | 3804.003 |
| Lman2 | AIFLDTYPNDETTER | -160 | -174 | 0 | 0 | 15 | 2021.2084 | 2351.2493 | 2388.49 | 2321.5112 | 2654.1248 | 2674.493 |
| Lman2 | PGPVFGSK | -146 | -153 | 0 | 0.001055699 | 8 | 1308.1515 | 1832.1589 | 1902.6296 | 1353.4004 | 985.77783 | 1469.9844 |
| Lman2 | DHDTFLAVR | -212 | -220 | 0 | 0 | 9 | 1300.2123 | 1105.6578 | 991.22626 | 1130.2574 | 1143.8345 | 401.121 |
| Lman2 | TDLEDKNEWK | -230 | -239 | 1 | 0 | 10 | 1271.3296 | 1329.6587 | 486.12067 | 142.54315 | 1493.7769 | 1493.9838 |
| Lman2 | SAGTGDLSDNHDIISIK | -258 | -274 | 0 | 0 | 17 | 1024.597 | 1105.4907 | 1047.5183 | 1128.1831 | 1081.5782 | 1052.7532 |
| Lman2 | SAGTGDLSDNHDIISIK | -258 | -274 | 0 | 0 | 17 | 1024.597 | 1105.4907 | 1047.5183 | 1128.1831 | 1081.5782 | 1052.7532 |
| Lman2 | DNVDDPTGNFR | -306 | -316 | 0 | 0 | 11 | 896.8213 | 856.6438 | 925.5809 | 970.2151 | 451.64148 | 1154.8558 |
| Lman2 | NHQPCFLK | -105 | -112 | 0 | 0.000550307 | 8 | 816.9424 | 1217.1919 | 1243.0465 | 570.2997 | 1375.4615 | 2134.924 |
| Lman2 | TPEEESIDWTK | -283 | -293 | 0 | 0 | 11 | 704.1042 | 725.012 | 448.26526 | 962.35284 | 732.4022 | 770.6025 |
| Lman2 | AGTGDLSDNHDIISIK | -259 | -274 | 0 | 0 | 16 | 3737.0093 | 4044.2239 | NaN | 3702.4304 | 4290.4863 | 4098.7847 |
| Lman2 | DITDGNSEHLKR | -47 | -58 | 1 | 0 | 12 | 918.0602 | NaN | 617.63385 | 644.9074 | 1374.6842 | 1644.2686 |
| Lman2 | RTPEEESIDWTK | -282 | -293 | 1 | 0.00142999 | 12 | 1218.4845 | 909.34973 | NaN | 283.44962 | NaN | 1642.1735 |
| Lman2 | LVPGPVFGSK | -144 | -153 | 0 | 0.000947459 | 10 | 3017.0852 | 2470.4148 | 873.06 | NaN | NaN | NaN |
| Lman2 | EHSLIKPYQGVGSS | -59 | -72 | 1 | 0 | 14 | 2159.729 | NaN | 1989.2771 | NaN | NaN | 2400.691 |
| Lman2 | IEPGVNFLK | -294 | -302 | 0 | 0.008045078 | 9 | 1349.0154 | NaN | NaN | NaN | NaN | NaN |
| Baz1b | DHSNLPDDDYCPR | -944 | -956 | 0 | 0.001291103 | 13 | 913.28455 | 524.74097 | 519.89154 | 664.6979 | 679.3556 | 908.04224 |
| Baz1b | IISNVPADSLIR | -232 | -243 | 0 | 0 | 12 | 793.18555 | 860.6768 | 881.63367 | 1062.2682 | 1026.5668 | 996.68536 |
| Baz1b | FSDFLLDPYK | -288 | -297 | 0 | 0 | 10 | 563.6932 | 611.106 | 628.01117 | 649.35944 | 719.48505 | 687.249 |
| Baz1b | LSEIPLTLH | -679 | -687 | 0 | 0 | 9 | 535.80475 | 600.74725 | 663.35187 | 845.482 | 674.095 | 605.391 |
| Baz1b | YAVGEECDFEVGK | -114 | -126 | 0 | 0 | 13 | 504.19626 | 1061.5448 | 796.99744 | 83.23066 | 636.1324 | 757.03796 |
| Baz1b | LYEVPDGEWQCPACQPPTAR | -1218 | -1237 | 0 | 0 | 20 | 320.60876 | 328.8162 | 324.80963 | 349.19305 | 351.95557 | 359.89603 |
| Baz1b | AGTGENAPWVVEDELVK | -264 | -280 | 0 | 0 | 17 | 646.47614 | 730.1906 | 745.7068 | 918.22925 | NaN | 1277.3229 |
| Baz1b | KEEFPNWYEK | -77 | -86 | 1 | 0 | 10 | 494.29172 | 1004.917 | 1115.7449 | NaN | 1641.8857 | 1381.612 |
| Baz1b | SCDGNQELLNFLR | -1044 | -1056 | 0 | 0.000230707 | 13 | 219.88823 | 461.76117 | 337.2819 | 219.58305 | 752.55457 | NaN |
| Baz1b | LKEEFPNWYEK | -76 | -86 | 1 | 0.00025015 | 11 | 1734.7852 | 1968.3452 | NaN | NaN | 1391 | 2366.3853 |
| Baz1b | GGFLYLNR | -647 | -654 | 0 | 0 | 8 | 609.785 | 614.02136 | 614.2101 | NaN | NaN | 202.9273 |
| Baz1b | ARLPEELR | -513 | -520 | 1 | 0 | 8 | 346.83444 | 877.7311 | NaN | 746.3363 | 1180.8647 | NaN |
| Baz1b | HLPGHPYVR | -1439 | -1447 | 0 | 0 | 9 | 733.5769 | NaN | NaN | NaN | 554.94037 | 440.3916 |
| Baz1b | EVPDGEWQCPACQPPTAR | -1220 | -1237 | 0 | 0.001392869 | 18 | 663.1645 | 386.91998 | NaN | NaN | 323.69446 | NaN |
| Baz1b | KPNLGLK | -1037 | -1043 | 1 | 0.001846793 | 7 | 151.60165 | NaN | NaN | NaN | NaN | NaN |
| Phgdh | AGTGVDNVDLEAATR | -76 | -90 | 0 | 0 | 15 | 4358.318 | 4487.651 | 4916.735 | 4421.7847 | 5490.5386 | 5414.421 |
| Phgdh | GGIVDEGALLR | -237 | -247 | 0 | 0 | 11 | 3741.0535 | 4059.9148 | 4135.8896 | 3812.214 | 4202.4097 | 4161.8115 |
| Phgdh | ILQDGGLQVVEK | -22 | -33 | 0 | 0 | 12 | 2930.673 | 3187.0093 | 3334.8474 | 3536.348 | 3074.9192 | 2868.6492 |
| Phgdh | VLISDSLDPCCR | -9 | -20 | 0 | 0 | 12 | 2906.7185 | 9728.831 | 7506.414 | 169.18669 | 3168.3477 | 8631.094 |
| Phgdh | KVLISDSLDPCCR | -8 | -20 | 1 | 0 | 13 | 2827.78 | 2968.0884 | 3716.9065 | 4734.3027 | 2825.407 | 4108.887 |
| Phgdh | ALVDHENVISCPHLGASTK | -271 | -289 | 0 | 0 | 19 | 2709.7786 | 3237.5085 | 3162.4233 | 2929.8171 | 3554.51 | 3280.2542 |
| Phgdh | ALVDHENVISCPHLGASTK | -271 | -289 | 0 | 0 | 19 | 2709.7786 | 3237.5085 | 3162.4233 | 2929.8171 | 3554.51 | 3280.2542 |
| Phgdh | ALVDHENVISCPHLGASTK | -271 | -289 | 0 | 0 | 19 | 2709.7786 | 3237.5085 | 3162.4233 | 2929.8171 | 3554.51 | 3280.2542 |
| Phgdh | ALDVFTEEPPRDR | -258 | -270 | 1 | 0 | 13 | 2435.102 | 2348.8525 | 2167.8704 | 2098.1165 | 2305.1763 | 2111.306 |
| Phgdh | LDVFTEEPPRDR | -259 | -270 | 1 | 0 | 12 | 2366.6226 | 2254.1333 | 2365.6462 | 2201.792 | 2378.3623 | 941.5669 |
| Phgdh | DVFTEEPPRDR | -260 | -270 | 1 | 0.00025015 | 11 | 2268.8953 | 2442.8794 | 2219.5598 | 2294.1248 | 2160.8381 | 2143.0122 |
| Phgdh | KILQDGGLQVVEK | -21 | -33 | 1 | 0.00025015 | 13 | 1487.793 | 1631.3477 | 1745.99 | 1963.5161 | 2046.5299 | 1999.3944 |
| Phgdh | VTADVINAAEK | -59 | -69 | 0 | 0 | 11 | 1201.3699 | 2004.6268 | 848.5176 | 1030.2979 | 658.9208 | 756.62476 |
| Phgdh | LSPAVIVGLLR | -370 | -380 | 0 | 0 | 11 | 1058.7563 | 1123.9291 | 1033.2521 | 735.0879 | 913.72034 | 921.77686 |
| Phgdh | ALVDHENVISCPHLGAS | -271 | -287 | 0 | 0 | 17 | 873.3209 | 1091.6609 | 972.6013 | 881.1092 | 1005.8131 | 1001.20874 |
| Phgdh | ALVDHENVISCPHLGAS | -271 | -287 | 0 | 0 | 17 | 873.3209 | 1091.6609 | 972.6013 | 881.1092 | 1005.8131 | 1001.20874 |
| Phgdh | SPAVIVGLLR | -371 | -380 | 0 | 0 | 10 | 466.3645 | 473.18866 | 495.60486 | 516.9301 | 571.8601 | 626.9065 |
| Phgdh | DCEGLIVR | -47 | -54 | 0 | 0 | 8 | 406.79816 | 687.93787 | 460.51614 | 763.60925 | 548.585 | 445.81067 |
| Phgdh | NAGTCLSPAVIVGL | -365 | -378 | 0 | 0.000230707 | 14 | 308.6308 | 366.12097 | 378.9819 | 482.35852 | 579.55035 | 404.8345 |
| Phgdh | VDHENVISCPHLGASTK | -273 | -289 | 0 | 0 | 17 | 253.53836 | 423.82376 | 372.51547 | 365.29993 | 476.9824 | 538.5011 |
| Phgdh | NAGTCLSPAVIVGLLR | -365 | -380 | 0 | 0 | 16 | 566.2013 | 509.45825 | NaN | 594.8816 | 617.5967 | 588.8481 |
| Phgdh | GTCLSPAVIVGLLR | -367 | -380 | 0 | 0 | 14 | 81.000656 | 261.40628 | NaN | 486.4311 | 288.8002 | 418.48953 |
| Phgdh | ELQDCEGLIVR | -44 | -54 | 0 | 0 | 11 | 821.4192 | 928.6069 | 993.40607 | NaN | NaN | 877.0797 |
| Phgdh | ALDVFTEEPPR | -258 | -268 | 0 | 0 | 11 | 774.48804 | 814.57605 | NaN | NaN | 1972.1216 | 1653.7152 |
| Phgdh | HENVISCPHLGASTK | -275 | -289 | 0 | 0 | 15 | 631.20264 | 529.5539 | 433.10886 | NaN | NaN | 1296.0105 |
| Phgdh | DVFTEEPPR | -260 | -268 | 0 | 0 | 9 | 1860.2319 | NaN | 1982.3094 | NaN | NaN | NaN |

| | | | | | | | | | | | | |
|---|---|---|---|---|---|---|---|---|---|---|---|---|
| Phgdh | EKLQVVGR | -68 | -75 | 1 | 0.004658491 | 8 | 857.9634 | NaN | NaN | 558.92694 | NaN | NaN |
| Phgdh | HAWAGSPK | -344 | -351 | 0 | 0.002068213 | 8 | 747.7966 | NaN | NaN | NaN | NaN | 652.2554 |
| Phgdh | QADVNLVNAK | -385 | -394 | 0 | 0.00025015 | 10 | 1013.56726 | NaN | NaN | NaN | NaN | NaN |
| Phgdh | ALVDHENVISCPHLGA | -271 | -286 | 0 | 0 | 16 | 781.84485 | NaN | NaN | NaN | NaN | NaN |
| Actn4 | HRPELIEYDK | -206 | -215 | 1 | 0.00025015 | 10 | 1482.9186 | 1705.2415 | 1307.6162 | 1534.5813 | 1858.768 | 1817.8015 |
| Actn4 | HRPELIEYDK | -206 | -215 | 1 | 0.00025015 | 10 | 1482.9186 | 1705.2415 | 1307.6162 | 1534.5813 | 1858.768 | 1817.8015 |
| Actn4 | LSGSNPYTTVTPQIINSK | -606 | -623 | 0 | 0 | 18 | 2382.6023 | 1002.26874 | 900.1465 | 755.9327 | 900.8667 | 869.15326 |
| Actn4 | MLDAEDIVNTARPDEK | -241 | -256 | 1 | 0 | 16 | 1905.0428 | 2141.7876 | 2175.4082 | 3140.115 | 205.25075 | 1379.8032 |
| Actn4 | MLDAEDIVNTARPDEK | -241 | -256 | 1 | 0 | 16 | 1905.0428 | 2141.7876 | 2175.4082 | 3140.115 | 205.25075 | 1379.8032 |
| Actn4 | DAEDIVNTARPDEK | -243 | -256 | 1 | 0 | 14 | 1534.0942 | 1475.5449 | 1504.2538 | 1754.9712 | 1799.968 | 1846.866 |
| Actn4 | LISLGYDVENDR | -795 | -806 | 0 | 0 | 12 | 1443.8878 | 1399.1769 | 1608.0563 | 189.17868 | 1472.171 | 1330.6241 |
| Actn4 | LVSIGAEEIVDGNAK | -127 | -141 | 0 | 0 | 15 | 1354.6539 | 1096.155 | 673.3991 | 1727.4106 | 1331.7538 | 1667.8755 |
| Actn4 | GTQIENIDEDFRDGLK | -69 | -84 | 1 | 0 | 16 | 1301.8501 | 968.6402 | 2052.1685 | 1362.7113 | 937.2199 | 1509.1139 |
| Actn4 | GTQIENIDEDFR | -69 | -80 | 0 | 0 | 12 | 1254.045 | 1045.1984 | 154.98582 | 1464.6394 | 1539.159 | 1383.893 |
| Actn4 | NFITAEELR | -861 | -869 | 0 | 0 | 9 | 1104.0206 | 1102.5245 | 1131.9717 | 307.16467 | 679.5852 | 1127.8322 |
| Actn4 | ISAHDQFK | -568 | -575 | 0 | 0 | 8 | 959.7535 | 740.787 | 932.8739 | 1335.0538 | 2525.974 | 2395.5828 |
| Actn4 | RDHALLEEQSK | -634 | -644 | 1 | 0 | 11 | 892.15137 | 928.1169 | 405.9387 | 882.00494 | 1637.7327 | 1430.2625 |
| Actn4 | FNHFDKDHGGALGPEEFK | -775 | -792 | 1 | 0 | 18 | 755.75305 | 874.94464 | 1021.27747 | 783.0305 | 995.3711 | 109.28101 |
| Actn4 | FNHFDKDHGGALGPEEFK | -775 | -792 | 1 | 0 | 18 | 755.75305 | 874.94464 | 1021.27747 | 783.0305 | 995.3711 | 109.28101 |
| Actn4 | SIGAEEIVDGNAK | -129 | -141 | 0 | 0 | 13 | 699.8339 | 876.67346 | 996.93176 | 1093.541 | 1348.2004 | 1331.9346 |
| Actn4 | QELNELDYYDSHNVNTR | -479 | -495 | 0 | 0 | 17 | 687.0057 | 684.68 | 661.8477 | 676.6646 | 839.69324 | 849.2162 |
| Actn4 | QELNELDYYDSHNVNTR | -479 | -495 | 0 | 0 | 17 | 687.0057 | 684.68 | 661.8477 | 676.6646 | 839.69324 | 849.2162 |
| Actn4 | ELNELDYYDSHNVNTR | -480 | -495 | 0 | 0 | 16 | 681.7322 | 858.67834 | 743.44324 | 690.8092 | 848.66223 | 888.19617 |
| Actn4 | QLETIDQLHLEYAK | -523 | -536 | 0 | 0 | 14 | 557.48724 | 710.7409 | 615.07495 | 748.2971 | 894.0115 | 974.22284 |
| Actn4 | ICDQWDNLGSLTHSR | -499 | -513 | 0 | 0 | 15 | 466.0257 | 537.5513 | 521.20306 | 806.1993 | 995.4725 | 990.44147 |
| Actn4 | TIEEIEGLISAHDQFK | -560 | -575 | 0 | 0.00025015 | 16 | 400.48767 | 445.13727 | 332.70493 | 499.66092 | 665.93726 | 611.7411 |
| Actn4 | GISQEQMQEFR | -762 | -772 | 0 | 0 | 11 | 317.96863 | 1948.334 | 2140.5862 | 267.7314 | 6325.851 | 7140.405 |
| Actn4 | GYDVENDR | -799 | -806 | 0 | 0 | 8 | 4922.372 | 5112.6426 | 5308.367 | 3716.688 | NaN | 5189.874 |
| Actn4 | KDDPVTNLNNAFEVA | -218 | -232 | 1 | 0 | 15 | 1623.0519 | 1769.3655 | NaN | 1837.5248 | 1537.7185 | 1982.0975 |
| Actn4 | AGTQIENIDEDFR | -68 | -80 | 0 | 0 | 13 | 676.3667 | 819.02814 | NaN | 365.0473 | 714.6059 | 628.3666 |
| Actn4 | ETTDTDTADQVIASFK | -839 | -854 | 0 | 0 | 16 | 651.3772 | 388.2727 | 788.77905 | NaN | 895.608 | 764.39874 |
| Actn4 | ICDQWDNLGSLTH | -499 | -511 | 0 | 0 | 13 | 482.10022 | 423.49646 | 557.5005 | NaN | 445.3936 | 577.6294 |
| Actn4 | RDHALLEEQ | -634 | -642 | 1 | 0.000431655 | 9 | 1357.8627 | 1364.2523 | 1167.064 | 2487.421 | NaN | NaN |
| Actn4 | EAILAIHK | -586 | -593 | 0 | 0 | 8 | 1225.2417 | NaN | NaN | 1212.7323 | 1104.2979 | 1182.9802 |
| Actn4 | DYYDSHNVNTR | -485 | -495 | 0 | 0 | 11 | 1170.9849 | 1084.4884 | 1054.4086 | 1142.6885 | NaN | NaN |
| Actn4 | IMSVVDPNHSGLVTFQ | -815 | -830 | 0 | 0.00122419 | 16 | 480.88596 | 583.6106 | 470.56418 | NaN | NaN | 842.94 |
| Actn4 | SIVDYKPSLDLLEQQHQLIQ | -697 | -716 | 1 | 0.002810811 | 20 | 170.23528 | NaN | NaN | 258.80902 | 253.83229 | 310.22934 |
| Actn4 | NIDEDFRDGLK | -74 | -84 | 1 | 0 | 11 | 1732.117 | 900.83307 | 923.448 | NaN | NaN | NaN |
| Actn4 | STLPDADRER | -576 | -585 | 1 | 0.006537474 | 10 | 6790.991 | NaN | NaN | NaN | NaN | 273.56256 |
| Actn4 | GSNPYTTVTPQIINSK | -608 | -623 | 0 | 0.001092529 | 16 | 737.3797 | NaN | NaN | NaN | NaN | 1316.6956 |
| Fubp1 | IGGNEGIDVPIPR | -268 | -280 | 0 | 0.000230707 | 13 | 3520.075 | 3834.626 | 4253.767 | 4155.4463 | 5191.7246 | 4874.9883 |
| Fubp1 | RPLEDGDQPDAK | -61 | -72 | 1 | 0 | 12 | 1923.9108 | 1993.5074 | 1172.0431 | 885.02203 | 2459.5984 | 1033.5002 |
| Fubp1 | FKPDDGTTPDR | -307 | -317 | 1 | 0.00051306 | 11 | 1430.4205 | 1774.4219 | 1893.6433 | 1542.8954 | 2121.34 | 1950.6592 |
| Fubp1 | IAQITGPPDR | -318 | -327 | 0 | 0 | 10 | 1386.6921 | 525.33295 | 1457.284 | 1177.8207 | 1134.4008 | 1085.3457 |
| Fubp1 | IAPDSGGLPER | -132 | -142 | 0 | 0.000230707 | 11 | 1271.7229 | 1317.4414 | 1533.8665 | 2526.6697 | 1249.2515 | 1321.7609 |
| Fubp1 | RLLDQIVEK | -158 | -166 | 1 | 0.00025015 | 9 | 1179.9412 | 1114.4017 | 1298.316 | 1467.0408 | 1490.5372 | 1537.1508 |
| Fubp1 | IQIAPDSGGLPER | -130 | -142 | 0 | 0 | 13 | 889.42346 | 1651.6052 | 1637.8073 | 486.83502 | 2264.3403 | 2328.3408 |
| Fubp1 | SVQAGNPGGPGPGGR | -341 | -355 | 0 | 0 | 15 | 517.40546 | 528.3987 | 607.29944 | 591.9238 | 822.0986 | 834.79443 |
| Fubp1 | GTPQQIDYAR | -427 | -436 | 0 | 0.000431655 | 10 | 9514.49 | 4896.3105 | 11203.056 | 5377.1846 | NaN | NaN |
| Fubp1 | DGPQNTGADKPLR | -220 | -232 | 1 | 0.000627943 | 13 | 1979.5457 | 1806.1818 | 2272.613 | NaN | NaN | 849.2259 |
| Fubp1 | AGTSLNSNDYGYGGQK | -45 | -60 | 0 | 0 | 16 | 751.52 | NaN | NaN | 681.026 | 464.4764 | 293.761 |
| Fubp1 | EIITDLLR | -333 | -340 | 0 | 0 | 8 | 293.85077 | NaN | 401.31747 | 229.70573 | 295.55835 | NaN |
| Fubp1 | IQFKPDDGTTPDR | -305 | -317 | 1 | 0 | 13 | 4558.203 | 576.1956 | 4095.4387 | NaN | NaN | NaN |
| Fubp1 | IQFKPDDGTTPDR | -305 | -317 | 1 | 0 | 13 | 4558.203 | 576.1956 | 4095.4387 | NaN | NaN | NaN |
| Fubp1 | TSLNSNDYGYGGQK | -47 | -60 | 0 | 0 | 14 | 3534.606 | NaN | NaN | 699.34174 | 3386.959 | NaN |
| Fubp1 | SCMLTGTPESVQSAK | -143 | -157 | 0 | 0 | 15 | 921.81616 | NaN | 771.1438 | 783.09174 | NaN | NaN |
| Fubp1 | EFNFIVPTGK | -374 | -383 | 0 | 0 | 10 | 613.95966 | NaN | NaN | NaN | NaN | NaN |
| Pgk1 | VADKIQLINNMLDK | -217 | -230 | 1 | 0 | 14 | 13735.784 | 14212.8955 | 13630.232 | 5982.432 | 590.06946 | 8382.982 |
| Pgk1 | VADKIQLINNMLDK | -217 | -230 | 1 | 0 | 14 | 13735.784 | 14212.8955 | 13630.232 | 5982.432 | 590.06946 | 8382.982 |
| Pgk1 | MVGVNLPQK | -176 | -184 | 0 | 0 | 9 | 6396.448 | 6161.909 | 6927.1016 | 6260.259 | 6245.033 | 7799.7124 |
| Pgk1 | MVGVNLPQK | -176 | -184 | 0 | 0 | 9 | 6396.448 | 6161.909 | 6927.1016 | 6260.259 | 6245.033 | 7799.7124 |
| Pgk1 | YSLEPVAAELK | -76 | -86 | 0 | 0 | 11 | 5877.934 | 6808.9175 | 6945.8657 | 7124.433 | 7778.0996 | 7903.324 |
| Pgk1 | ALESPERPFLAIL | -200 | -212 | 1 | 0 | 13 | 2619.9348 | 2770.1897 | 2864.8486 | 3646.532 | 4037.0894 | 3926.6995 |
| Pgk1 | ANPAAGTVILLENLR | -109 | -123 | 0 | 0 | 15 | 2190.7483 | 2651.0195 | 2492.7834 | 2745.8286 | 3328.5747 | 3253.5825 |
| Pgk1 | IQLINNMLDK | -221 | -230 | 0 | 0 | 10 | 2058.0513 | 2184.6484 | 2344.5771 | 3397.1296 | 3366.3945 | 3358.901 |
| Pgk1 | IQLINNMLDK | -221 | -230 | 0 | 0 | 10 | 2058.0513 | 2184.6484 | 2344.5771 | 3397.1296 | 3366.3945 | 3358.901 |
| Pgk1 | EWEAFAR | -344 | -350 | 0 | 0 | 7 | 1751.0305 | 1419.9012 | 1505.7982 | 1232.8607 | 281.4838 | 966.1035 |
| Pgk1 | SLMDEVVK | -354 | -361 | 0 | 0 | 8 | 1636.1903 | 758.14966 | 1064.841 | 547.8867 | 578.1653 | 634.665 |
| Pgk1 | ITLPVDFVTADKFDENAK | -280 | -297 | 1 | 0 | 18 | 1572.1824 | 1101.017 | 1606.4509 | 2381.184 | 2985.965 | 2897.0269 |
| Pgk1 | ITLPVDFVTADKFDENAK | -280 | -297 | 1 | 0 | 18 | 1572.1824 | 1101.017 | 1606.4509 | 2381.184 | 2985.965 | 2897.0269 |
| Pgk1 | ITLPVDFVTADK | -280 | -291 | 0 | 0 | 12 | 1415.0638 | 1191.3773 | 1534.1332 | 1877.8925 | 1308.9054 | 2064.2979 |
| Pgk1 | SLEPVAAELK | -77 | -86 | 0 | 0 | 10 | 1331.2782 | 1290.1522 | 1386.4355 | 1109.6592 | 1144.8104 | 1194.8467 |
| Pgk1 | KYAEAVGR | -323 | -330 | 1 | 0.003133457 | 8 | 1245.2583 | 1288.1007 | 1331.447 | 614.00116 | 1701.3647 | 1695.2786 |
| Pgk1 | YAEAVGR | -324 | -330 | 0 | 0.00078661 | 7 | 1199.2463 | 954.5349 | 1513.9475 | 1368.3191 | 1188.3387 | 811.0183 |
| Pgk1 | SHLGRPDGVPMPDK | -62 | -75 | 1 | 0.00025015 | 14 | 887.2694 | 745.8015 | 792.5867 | 843.1508 | 975.63885 | 1017.12897 |
| Pgk1 | SHLGRPDGVPMPDK | -62 | -75 | 1 | 0.00025015 | 14 | 887.2694 | 745.8015 | 792.5867 | 843.1508 | 975.63885 | 1017.12897 |
| Pgk1 | SHLGRPDGVPMPDK | -62 | -75 | 1 | 0.00025015 | 14 | 887.2694 | 745.8015 | 792.5867 | 843.1508 | 975.63885 | 1017.12897 |
| Pgk1 | SGIPAGWMGLDCGTESSK | -305 | -322 | 0 | 0 | 18 | 1110.3889 | 1158.6389 | 1182.8057 | 1718.7101 | 1842.2827 | 1815.5728 |

| | | | | | | | | | | | | |
|---|---|---|---|---|---|---|---|---|---|---|---|---|
| Pgk1 | SGIPAGWMGLDCGTESSK | -305 | -322 | 0 | 0 | 18 | 1110.3889 | 1158.6389 | 1182.8057 | 1718.7101 | 1842.2827 | 1815.5728 |
| Pgk1 | GCITIIGGGDTATCC | -366 | -380 | 0 | 0 | 15 | 1110.0935 | 1086.7189 | 1098.0457 | 1085.5944 | 715.21893 | 848.96765 |
| Pgk1 | ITLPVDFVTADKF | -280 | -292 | 1 | 0 | 13 | 1094.5947 | 909.4089 | 916.77325 | 1391.1587 | 1585.9526 | 1574.8873 |
| Pgk1 | ESPERPFLAILGGAK | -202 | -216 | 1 | 0 | 15 | 654.0869 | 748.3783 | 798.07916 | 1940.851 | 3015.5645 | 2666.2576 |
| Pgk1 | ESPERPFLAILGGAK | -202 | -216 | 1 | 0 | 15 | 654.0869 | 748.3783 | 798.07916 | 1940.851 | 3015.5645 | 2666.2576 |
| Pgk1 | QIVWNGPVGVF | -333 | -343 | 0 | 0 | 11 | 632.0373 | 631.1043 | 796.00696 | 603.9949 | 947.6663 | 1199.4473 |
| Pgk1 | QIVWNGPVGVF | -333 | -343 | 0 | 0 | 11 | 632.0373 | 631.1043 | 796.00696 | 603.9949 | 947.6663 | 1199.4473 |
| Pgk1 | VASGIPAGWMGLDCGTESSK | -303 | -322 | 0 | 0.00025015 | 20 | 605.2207 | 874.13824 | 628.2429 | 588.75726 | 853.34344 | 893.6667 |
| Pgk1 | WNTEDKVSHVSTGGGA | -383 | -398 | 1 | 0 | 16 | 496.52435 | 558.4344 | 556.7264 | 493.11682 | 605.8245 | 1636.8439 |
| Pgk1 | WNTEDKVSHVSTGGGA | -383 | -398 | 1 | 0 | 16 | 496.52435 | 558.4344 | 556.7264 | 493.11682 | 605.8245 | 1636.8439 |
| Pgk1 | AGTVILLENLR | -113 | -123 | 0 | 0.00025015 | 11 | 451.46234 | 511.885 | 486.17636 | 410.67856 | 496.76886 | 471.60178 |
| Pgk1 | ITLPVDFVTAD | -280 | -290 | 0 | 0.00142999 | 11 | 384.80527 | 430.75894 | 482.08142 | 365.8003 | 519.3217 | 530.5344 |
| Pgk1 | ITLPVDFVTAD | -280 | -290 | 0 | 0.00142999 | 11 | 384.80527 | 430.75894 | 482.08142 | 365.8003 | 519.3217 | 530.5344 |
| Pgk1 | WNTEDKVSHVS | -383 | -393 | 1 | 0.00111006 | 11 | 308.4125 | 1145.4905 | 1304.5999 | 1312.3545 | 1474.7462 | 1605.888 |
| Pgk1 | GCITIIGGGDTATCCAK | -366 | -382 | 0 | 0 | 17 | 594.22174 | NaN | 787.47754 | 1488.701 | 1420.4021 | 1385.976 |
| Pgk1 | LYDEEGAK | -257 | -264 | 0 | 0.001392869 | 8 | 424.762 | 313.89102 | 4167.7607 | 3806.917 | 3368.9402 | NaN |
| Pgk1 | KELNYFAK | -192 | -199 | 1 | 0 | 8 | 1238.7418 | 1049.8428 | 1073.7089 | 1088.2584 | NaN | NaN |
| Pgk1 | CANPAAGTVILLENLR | -108 | -123 | 0 | 0 | 16 | 1073.2668 | NaN | 1089.4277 | NaN | 1085.6074 | 1085.4802 |
| Pgk1 | VILLENLR | -116 | -123 | 0 | 0 | 8 | 602.01294 | 554.7956 | NaN | 461.7543 | 639.80725 | NaN |
| Pgk1 | VLPGVDALSNV | -407 | -417 | 0 | 0 | 11 | 550.8906 | NaN | 568.9173 | 1499.073 | 3551.401 | NaN |
| Prkcb | VLALPGKPPFLTQ | -393 | -405 | 1 | 0 | 13 | 838.1348 | 767.6244 | 782.72406 | 822.663 | 808.9287 | 693.23846 |
| Prkcb | KGTDELYAVK | -362 | -371 | 1 | 0 | 10 | 699.10297 | 1463.6548 | 1727.39 | 1637.6066 | 1092.1166 | 1966.2784 |
| Prkcb | AGVDGWFK | -269 | -276 | 0 | 0.00122419 | 8 | 927.58435 | 187.09172 | 698.2651 | 704.6996 | NaN | 501.24057 |
| Prkcb | KENIWDGVTTK | -489 | -499 | 1 | 0.000431655 | 11 | 1250.4268 | NaN | NaN | NaN | NaN | NaN |
| Prkcb | EIQPPYKPK | -612 | -620 | 1 | 0.009242859 | 9 | 325.8598 | NaN | NaN | NaN | NaN | NaN |
| Fto | AGVGPSCDDEVDLK | -178 | -191 | 0 | 0 | 14 | 1593.9054 | 2504.2004 | 2920.1074 | 1452.8185 | 375.6243 | 1318.2216 |
| Fto | LFTVPWPVK | -113 | -121 | 0 | 0 | 9 | 382.15085 | 491.36072 | 475.58896 | 311.39478 | 365.23163 | 403.9895 |
| Fto | STGTLDYILER | -324 | -334 | 0 | 0.003003218 | 11 | 502.01978 | 573.41235 | NaN | NaN | 606.57715 | NaN |
| Macroh2a1 | SIAFPSIGSGR | -305 | -315 | 0 | 0 | 11 | 6969.7363 | 7122.7686 | 7487.0234 | 6267.3745 | 6534.294 | 6616.851 |
| Macroh2a1 | SGGVLPNIHPELLAK | -102 | -116 | 0 | 0 | 15 | 3168.741 | 740.51825 | 3184.8157 | 3790.8982 | 3961.551 | 3849.7854 |
| Macroh2a1 | AISAGHGLPAK | -258 | -268 | 0 | 0 | 11 | 2554.9258 | 2673.5864 | 2717.1692 | 2295.9922 | 2497.6072 | 2592.2136 |
| Macroh2a1 | ISAGHGLPAK | -259 | -268 | 0 | 0 | 10 | 2519.5945 | 2316.9456 | 2387.2595 | 2188.9788 | 2107.806 | 2093.603 |
| Macroh2a1 | KAGVIFPVGR | -18 | -27 | 1 | 0 | 10 | 2314.427 | 2287.8992 | 2499.8218 | 2235.0166 | 2620.5745 | 2588.2366 |
| Macroh2a1 | NCLALADDRK | -293 | -302 | 1 | 0.00025015 | 10 | 2092.3005 | 2580.4973 | 2797.0044 | 583.8743 | 3186.8542 | 1664.7277 |
| Macroh2a1 | QTAAQLILK | -321 | -329 | 0 | 0.000826806 | 9 | 1734.7249 | 1663.5807 | 1805.6804 | 1486.3185 | 1353.9972 | 1385.3483 |
| Macroh2a1 | SPVWGADKCEELLEK | -275 | -289 | 1 | 0 | 15 | 1440.7086 | 1269.391 | 1374.909 | 1200.2456 | 1330.7814 | 1258.5759 |
| Macroh2a1 | SAGHGLPAK | -260 | -268 | 0 | 0.001370415 | 9 | 1407.999 | 1776.0706 | 1304.9164 | 11713.338 | 1967.1257 | 2070.6814 |
| Macroh2a1 | CNSPVWGADKCEELLEK | -273 | -289 | 1 | 0 | 17 | 503.78268 | 551.3534 | 527.4742 | 613.001 | 243.98062 | 354.2528 |
| Macroh2a1 | IAFPSIGSGR | -306 | -315 | 0 | 0.00025015 | 10 | 2248.1213 | 1888.7279 | 328.98447 | 1465.2192 | 697.01636 | NaN |
| Macroh2a1 | NSPVWGADKCEELLEK | -274 | -289 | 1 | 0 | 16 | 557.2714 | 787.3749 | 752.8778 | 671.0392 | NaN | 856.3663 |
| Macroh2a1 | FVIHCNSPVWGADK | -269 | -282 | 0 | 0 | 14 | 502.05692 | 557.76227 | NaN | 845.8235 | 432.18893 | 1042.0494 |
| Macroh2a1 | NCLALADDR | -293 | -301 | 0 | 0 | 9 | 451.33157 | 155.99762 | NaN | 82.80482 | 1018.7648 | 1063.8728 |
| Macroh2a1 | CNSPVWGADK | -273 | -282 | 0 | 0 | 10 | 518.8782 | 569.68225 | 1057.2688 | 655.2166 | NaN | NaN |
| Macroh2a1 | AIITPPPAK | -126 | -134 | 0 | 0.007374243 | 9 | 949.861 | 895.1821 | NaN | 833.58405 | NaN | NaN |
| Slc25a12 | FGLYLPK | -653 | -659 | 0 | 0 | 7 | 984.0918 | 906.4604 | 1072.3944 | 774.369 | 638.3323 | 801.5359 |
| Slc25a12 | LVTYELLQR | -595 | -603 | 0 | 0 | 9 | 670.214 | 699.06946 | 688.0007 | 866.3521 | 1105.4755 | 950.7307 |
| Slc25a12 | AGVPAASLVTPADVIK | -528 | -543 | 0 | 0 | 16 | 598.0581 | 1285.3026 | 745.86664 | 741.1855 | 1013.2233 | 964.44666 |
| Slc25a12 | LTLADIER | -284 | -291 | 0 | 0 | 8 | 568.5153 | 597.24994 | 735.30237 | 520.9403 | 867.5674 | 464.70953 |
| Slc25a12 | YLGLYNDPNSNPK | -43 | -55 | 0 | 0 | 13 | 436.8959 | 343.803 | 1635.9001 | 1879.9271 | 1701.039 | 382.80847 |
| Slc25a12 | IVQLLAGVADQTK | -56 | -68 | 0 | 0 | 13 | 401.92612 | 519.1715 | 466.0095 | 551.84705 | 567.86914 | 585.5922 |
| Metap1 | AGVTTEEIDHAVHLA | -159 | -173 | 0 | 0.000471409 | 15 | 680.7656 | 749.1891 | 424.08908 | 408.57507 | 392.2285 | 496.5034 |
| Metap1 | LFHTAPNVPHYA | -299 | -310 | 0 | 0.000298805 | 12 | 414.23422 | 219.23062 | 437.61392 | NaN | NaN | NaN |
| Metap1 | LLVTDTGCEILTR | -361 | -373 | 0 | 0.002434616 | 13 | 3103.6035 | NaN | NaN | NaN | NaN | NaN |
| Pfn1 | EGVHGGLINK | -117 | -126 | 0 | 0 | 10 | 11843.672 | 10965.406 | 10971.226 | 11075.377 | 9349.776 | 10280.6045 |
| Pfn1 | DSLLQDGEFTMDLR | -76 | -89 | 0 | 0 | 14 | 3632.8127 | 4654.676 | 4558.7593 | 1769.891 | 9913.92 | 9708.3125 |
| Pfn1 | SLMADGTCQDAAIVGYK | -10 | -26 | 0 | 0 | 17 | 3129.2383 | 4545.5596 | 2984.2595 | 4921.509 | 4621.8433 | 4065.7273 |
| Pfn1 | SLMADGTCQDAAIVGYK | -10 | -26 | 0 | 0 | 17 | 3129.2383 | 4545.5596 | 2984.2595 | 4921.509 | 4621.8433 | 4065.7273 |
| Pfn1 | KDSPSVWAAVPGK | -26 | -38 | 1 | 0 | 13 | 2793.3904 | 2881.1055 | 3088.6067 | 3764.6533 | 4275.164 | 4218.3003 |
| Pfn1 | TFVSITPAEVGVLVGK | -39 | -54 | 0 | 0 | 16 | 1424.9027 | 1428.2433 | 1368.057 | 2225.2383 | 2305.655 | 2315.6519 |
| Pfn1 | EGVHGGLINKK | -117 | -127 | 1 | 0 | 11 | 1290.4288 | 1451.8593 | 1617.977 | 1307.3673 | 1727.6183 | 1989.0824 |
| Pfn1 | EGVHGGLINKK | -117 | -127 | 1 | 0 | 11 | 1290.4288 | 1451.8593 | 1617.977 | 1307.3673 | 1727.6183 | 1989.0824 |
| Pfn1 | TFVSITPAEVGVL | -39 | -51 | 0 | 0 | 13 | 1049.4048 | 1189.3232 | 1131.8143 | 1384.0243 | 1674.8097 | 1857.4841 |
| Pfn1 | TFVSITPAEVGVL | -39 | -51 | 0 | 0 | 13 | 1049.4048 | 1189.3232 | 1131.8143 | 1384.0243 | 1674.8097 | 1857.4841 |
| Pfn1 | GKEGVHGGLINK | -115 | -126 | 1 | 0 | 12 | 919.95325 | 598.14777 | 665.3557 | 655.391 | 1451.8456 | 1326.7542 |
| Pfn1 | GKEGVHGGLINK | -115 | -126 | 1 | 0 | 12 | 919.95325 | 598.14777 | 665.3557 | 655.391 | 1451.8456 | 1326.7542 |
| Pfn1 | MADGTCQDAAIVGYK | -12 | -26 | 0 | 0 | 15 | 914.2442 | 744.4202 | 637.63354 | 809.4943 | 833.16736 | 960.9433 |
| Pfn1 | VSITPAEVGVLVGK | -41 | -54 | 0 | 0 | 14 | 865.42175 | 1066.3779 | 1023.76666 | 1897.3774 | 2467.1182 | 2218.289 |
| Pfn1 | ITPAEVGVLVGK | -43 | -54 | 0 | 0.000471409 | 12 | 857.19037 | 921.68317 | 848.2683 | 746.1763 | 904.5901 | 816.1691 |
| Pfn1 | SLLQDGEFTMDLR | -77 | -89 | 0 | 0 | 13 | 744.96747 | 1130.6414 | 860.70886 | 1071.6298 | 1197.5605 | 800.0301 |
| Pfn1 | GTCQDAAIVGYK | -15 | -26 | 0 | 0 | 12 | 618.4143 | 1020.12177 | 1081.6343 | 763.9486 | 903.485 | 453.0239 |
| Pfn1 | DSPSVWAAVPGK | -27 | -38 | 0 | 0 | 12 | 195.63812 | 5825.583 | 4941.749 | 7634.5513 | 5952.1914 | 6442.0767 |
| Pfn1 | KEGVHGGLINK | -116 | -126 | 1 | 0 | 11 | 1292.8241 | 1447.1167 | 1616.0188 | NaN | 1718.9674 | 1989.0737 |
| Pfn1 | AGWNAYID | -2 | -9 | 0 | 0.000550307 | 8 | 1129.5455 | NaN | NaN | 1277.1493 | 1968.4186 | 1532.7607 |
| Pfn1 | CQDAAIVGYK | -17 | -26 | 0 | 0 | 10 | 901.3883 | 324.2058 | 844.51917 | NaN | 329.4062 | NaN |
| Pfn1 | SSFFVNGLTLGGQK | -57 | -70 | 0 | 0.002199523 | 14 | 866.814 | 2302.0972 | 2241.7312 | NaN | NaN | NaN |
| Pfn1 | EGVHGGLIN | -117 | -125 | 0 | 0.009272078 | 9 | 2137.991 | 1827.0448 | NaN | NaN | NaN | NaN |
| Selenoi | WSPSDILEIHPR | -279 | -290 | 0 | 0 | 12 | 738.1952 | 890.41833 | 926.4984 | 2056.0393 | 2697.2974 | 415.24445 |
| Selenoi | AGYEYVSPEQLSGFDK | -2 | -17 | 0 | 0 | 16 | 297.75058 | 324.33307 | 402.58063 | 473.41922 | 564.5977 | 449.2729 |

| | | | | | | | | | | | | |
|---|---|---|---|---|---|---|---|---|---|---|---|---|
| Prmt5 | LSPWIHPDSK | -86 | -95 | 0 | 0 | 10 | 6029.2817 | 4102.419 | 6591.679 | 3643.2375 | 2808.7656 | 2854.08 |
| Prmt5 | LHNFHQL | -506 | -512 | 0 | 0.001941082 | 7 | 1450.9209 | 2516.01 | 2736.6685 | 1120.2335 | 5060.326 | 4607.7236 |
| Prmt5 | FLLPLNQEDNTNLAR | -122 | -136 | 0 | 0 | 15 | 1292.7428 | 2437.6296 | 717.2147 | 2782.338 | 4224.521 | 3632.8953 |
| Prmt5 | VPLVAPEDLR | -155 | -164 | 0 | 0 | 10 | 1201.8895 | 1179.1425 | 1226.8168 | 1165.9608 | 1308.4889 | 1284.6309 |
| Prmt5 | TYEVFEKDPIK | -323 | -333 | 1 | 0 | 11 | 681.3291 | 1799.7178 | 2233.6624 | 1090.3734 | 579.54675 | 887.4823 |
| Prmt5 | EIGADLPSNHVIDR | -207 | -220 | 0 | 0.00025015 | 14 | 491.23102 | 2891.373 | 3208.4202 | 2878.362 | 2969.163 | 2967.1963 |
| Prmt5 | AGYFETVLYR | -552 | -561 | 0 | 0 | 10 | 188.80556 | 175.50845 | 420.23264 | 498.1523 | 540.19574 | 367.4331 |
| Prmt5 | VPEEEKETNVQVL | -349 | -361 | 1 | 0 | 13 | 1729.8416 | 1548.7615 | 227.8137 | NaN | 1603.7288 | 1335.6226 |
| Prmt5 | QPITVHEGQNICVR | -588 | -601 | 0 | 0.00025015 | 14 | 709.72687 | NaN | 824.85114 | 144.74033 | 1647.2318 | 1565.5923 |
| Prmt5 | GLPAFLLPLNQEDNTNLAR | -118 | -136 | 0 | 0.002068213 | 19 | 307.9965 | NaN | 315.2217 | 143.14609 | 301.96335 | 326.41913 |
| Sgpl1 | AGYPLEKPFDFR | -329 | -340 | 1 | 0 | 12 | 1751.9595 | 1235.1936 | 1014.54333 | 1951.6581 | 1037.2205 | 2544.579 |
| Sgpl1 | TPEIVAPESAHAA | -232 | -244 | 0 | 0 | 13 | 2830.1785 | 89.67704 | NaN | NaN | 864.7596 | NaN |
| Rack1 | FSPNSSNPIIVSC | -156 | -168 | 0 | 0 | 13 | 5770.7705 | 5576.81 | 5687.252 | 4082.3918 | 231.20468 | 3160.9878 |
| Rack1 | TNHIGHTGYLN | -186 | -196 | 0 | 0.000591124 | 11 | 3937.498 | 1442.13 | 4148.7974 | 3607.4126 | 3306.6797 | 260.12527 |
| Rack1 | DETNYGIPQR | -48 | -57 | 0 | 0 | 10 | 3428.053 | 492.68634 | 334.5586 | 419.83026 | 2156.8193 | 2045.5836 |
| Rack1 | TRDETNYGIPQR | -46 | -57 | 1 | 0.00025015 | 12 | 2389.0925 | 2360.1548 | 2310.1545 | 2451.4927 | 2671.4094 | 2585.3108 |
| Rack1 | DESHSEWVSCVR | -144 | -155 | 0 | 0 | 12 | 2032.2024 | 2319.8875 | 2203.078 | 379.44934 | 3395.2522 | 3447.917 |
| Rack1 | TLFAGYTDNLVR | -297 | -308 | 0 | 0 | 12 | 1967.3746 | 2343.1648 | 2538.5186 | 3019.656 | 3271.6804 | 3416.2473 |
| Rack1 | TVTVSPDGSLCASGGK | -197 | -212 | 0 | 0 | 16 | 1578.9705 | 1691.2522 | 1708.3738 | 875.5791 | 3835.4863 | 3773.3765 |
| Rack1 | VQDESHSEWVSCVR | -142 | -155 | 0 | 0 | 14 | 1480.9183 | 1236.6418 | 1061.769 | 645.318 | 716.96796 | 757.4452 |
| Rack1 | IIVDELKQEVIST | -265 | -277 | 1 | 0.00025015 | 13 | 1256.6887 | 1508.5643 | 1515.4315 | 1979.2258 | 2057.0457 | 2244.5557 |
| Rack1 | IIVDELKQEVISTS | -265 | -278 | 1 | 0 | 14 | 1033.891 | 1199.9883 | 1148.1433 | 1574.2865 | 1711.6632 | 1727.5575 |
| Rack1 | SGSWDGTLR | -80 | -88 | 0 | 0 | 9 | 1029.739 | 695.11426 | 612.44025 | 1106.2765 | 173.32994 | 635.8265 |
| Rack1 | HLYTLDGGDIINAL | -226 | -239 | 0 | 0 | 14 | 960.6599 | 1017.16895 | 931.18616 | 732.45776 | 935.51184 | 909.3009 |
| Rack1 | SDGQFALSGSWDGTLR | -73 | -88 | 0 | 0 | 16 | 738.32654 | 986.41 | 1014.6227 | 1102.2408 | 921.49146 | 979.6088 |
| Rack1 | YTVQDESHSEWVSCVR | -140 | -155 | 0 | 0 | 16 | 713.9619 | 839.6679 | 944.75476 | 799.7349 | 921.4705 | 873.68713 |
| Rack1 | YTVQDESHSEWVSCVR | -140 | -155 | 0 | 0 | 16 | 713.9619 | 839.6679 | 944.75476 | 799.7349 | 921.4705 | 873.68713 |
| Rack1 | FAGYTDNLVR | -299 | -308 | 0 | 0 | 10 | 649.771 | 2830.9456 | 3041.7627 | 2672.8948 | 2893.729 | 2879.073 |
| Rack1 | SADGQTLFAGYTDNLVR | -292 | -308 | 0 | 0 | 17 | 545.21423 | 556.7949 | 423.14706 | 841.0549 | 993.315 | 932.57587 |
| Rack1 | GHSHFVSDVVIS | -61 | -72 | 0 | 0 | 12 | 281.9873 | 2508.2036 | 2319.1216 | 2701.0635 | 744.946 | 824.23303 |
| Rack1 | GHSHFVSDVVIS | -61 | -72 | 0 | 0 | 12 | 281.9873 | 2508.2036 | 2319.1216 | 2701.0635 | 744.946 | 824.23303 |
| Rack1 | LWDLTTGTTTR | -89 | -99 | 0 | 0 | 11 | 199.68037 | 3643.1433 | 229.17993 | 2740.0337 | 2467.4817 | 2946.1948 |
| Rack1 | TVSPDGSLCASGGK | -199 | -212 | 0 | 0 | 14 | 1824.2522 | 2119.5378 | 2325.9531 | 847.9551 | NaN | 2043.1747 |
| Rack1 | IIVDELKQE | -265 | -273 | 1 | 0 | 9 | 1660.0721 | 1680.6066 | 1954.0386 | 1195.6283 | NaN | 1128.4379 |
| Rack1 | LWNTLGVCK | -131 | -139 | 0 | 0 | 9 | 1128.0201 | 1193.2356 | NaN | 1181.6938 | NaN | 1816.1694 |
| Rack1 | YTVQDESHSEWVS | -140 | -152 | 0 | 0.00025015 | 13 | 2921.284 | 3335.4312 | 3718.0076 | NaN | NaN | NaN |
| Rack1 | IWDLEGK | -258 | -264 | 0 | 0.000826806 | 7 | 2731.323 | NaN | 2421.3494 | NaN | 1184.5518 | NaN |
| Rack1 | VWQVTIGTR | -309 | -317 | 0 | 0 | 9 | 1065.9064 | 1133.4818 | 1118.5771 | NaN | NaN | NaN |
| Rack1 | AGYTDNLVR | -300 | -308 | 0 | 0.001291103 | 9 | 816.9323 | NaN | NaN | 1020.6835 | NaN | NaN |
| Rack1 | IWDLEGKI | -258 | -265 | 1 | 0.005821388 | 8 | 984.4984 | NaN | NaN | NaN | NaN | NaN |
| Tmed2 | AHAEECFFER | -27 | -36 | 0 | 0 | 10 | 1839.7327 | 2079.988 | 2071.3662 | 1814.1272 | 1935.7957 | 60.398983 |
| Tmed2 | TIDIGEAPK | -109 | -117 | 0 | 0.001811524 | 9 | 729.8755 | 696.0255 | 640.9785 | 862.5035 | NaN | NaN |
| Tmed2 | HEQEYMEVR | -146 | -154 | 0 | 0 | 9 | 425.61285 | NaN | NaN | 675.7539 | NaN | NaN |
| Preb | NLEAVQTDFSNEPLQK | -143 | -158 | 0 | 0 | 16 | 572.37274 | 588.4461 | 674.7024 | 402.91956 | 734.0469 | 486.93936 |
| Preb | AHEGEIGDLTLGPDGK | -194 | -209 | 0 | 0 | 16 | 577.7393 | 927.6086 | 593.6446 | NaN | 1183.6627 | 1126.9131 |
| Cebpz | AHELLCDKPEEEK | -369 | -381 | 1 | 0 | 13 | 424.69427 | 1959.0906 | 974.34515 | 1545.9818 | NaN | 3147.047 |
| Cebpz | ETEPESSAEAEKPK | -665 | -678 | 1 | 0 | 14 | 1403.4513 | 1721.8914 | NaN | NaN | 2439.3843 | 1918.172 |
| Cebpz | SHEESELANK | -451 | -460 | 0 | 0.008736733 | 10 | 606.5264 | NaN | NaN | 447.79974 | NaN | NaN |
| Rpl13a | LAHEVGWK | -141 | -148 | 0 | 0 | 8 | 2013.3032 | 1129.4125 | 1684.543 | 1385.06 | 1095.0696 | 2057.8384 |
| Rpl13a | AHEVGWK | -142 | -148 | 0 | 0 | 7 | 1159.5381 | 862.76263 | 1055.089 | 1600.4723 | 1225.4999 | 1306.7133 |
| Rpl13a | VLDGIPPPYDK | -104 | -114 | 0 | 0 | 11 | 1085.8248 | 1468.525 | 696.9913 | 1707.75 | NaN | 894.02466 |
| Prmt1 | SPESPYTHWK | -304 | -313 | 0 | 0 | 10 | 5163.321 | 1181.1831 | 5001.3965 | 4408.6045 | 3496.3943 | 3545.064 |
| Prmt1 | TGFSTSPESPYTHWK | -299 | -313 | 0 | 0 | 15 | 1845.4358 | 2741.1516 | 2789.8228 | 2000.1107 | 2396.51 | 2251.2656 |
| Prmt1 | TGFSTSPESPYTHWK | -299 | -313 | 0 | 0 | 15 | 1845.4358 | 2741.1516 | 2789.8228 | 2000.1107 | 2396.51 | 2251.2656 |
| Prmt1 | KLDHVVTIIK | -134 | -143 | 1 | 0 | 10 | 1570.7782 | 1777.2754 | 1753.695 | 1654.499 | 2572.7034 | 2334.4321 |
| Prmt1 | LNTVLHAR | -174 | -181 | 0 | 0 | 8 | 1196.0704 | 715.7985 | 1071.9934 | 1245.5287 | 917.5527 | 1091.4554 |
| Prmt1 | LDHVVTIIK | -135 | -143 | 0 | 0 | 9 | 1166.4911 | 1202.775 | 1255.579 | 1061.0316 | 1219.6815 | 1127.5264 |
| Prmt1 | WLAPDGLIFPDR | -184 | -195 | 0 | 0 | 12 | 945.28564 | 928.51434 | 973.516 | 1342.4132 | 1600.7258 | 1635.8135 |
| Prmt1 | TGEEIFGTIGMRPN | -327 | -340 | 1 | 0 | 14 | 867.46466 | 1064.519 | 1074.6284 | 601.20337 | 896.22736 | 1272.5221 |
| Prmt1 | TGFSTSPESPYTHW | -299 | -312 | 0 | 0.00025015 | 14 | 739.45374 | 814.7322 | 692.6194 | 1290.8406 | 1488.1744 | 1734.7501 |
| Prmt1 | DVAIKEPLVDVVDPK | -229 | -243 | 1 | 0 | 15 | 720.5331 | 1056.5205 | 955.08563 | 1378.1553 | 1595.6782 | 1748.0726 |
| Prmt1 | LCELSCSTDYR | -359 | -369 | 0 | 0 | 11 | 703.002 | 466.37292 | 455.72034 | 1075.0767 | 1131.6039 | 1113.7369 |
| Prmt1 | DKWLAPDGLIFPDR | -182 | -195 | 1 | 0 | 14 | 518.1464 | 595.0591 | 687.53107 | 751.043 | 724.94543 | 874.063 |
| Prmt1 | DKWLAPDGLIFPDR | -182 | -195 | 1 | 0 | 14 | 518.1464 | 595.0591 | 687.53107 | 751.043 | 724.94543 | 874.063 |
| Prmt1 | RNDYVHAL | -277 | -284 | 1 | 0.001536401 | 8 | 273.59384 | 1053.7295 | 1343.5337 | 3654.4473 | 2805.5112 | 1994.9056 |
| Prmt1 | DKWLAPDGLIFPD | -182 | -194 | 1 | 0.00025015 | 13 | 401.55505 | 327.0044 | 348.31085 | 376.47748 | 360.1922 | NaN |
| Prmt1 | TGEEIFGTIGMRPNAK | -327 | -342 | 1 | 0 | 16 | 1774.0227 | NaN | NaN | 2374.2302 | 2715.694 | 2916.4692 |
| Prmt1 | GQLCELSCSTDYR | -357 | -369 | 0 | 0.000826806 | 13 | 903.22473 | 443.05017 | NaN | 1541.5825 | 627.8863 | NaN |
| Prmt1 | EPLVDVVDPK | -234 | -243 | 0 | 0 | 10 | 1810.2954 | NaN | 1802.8616 | 1644.9261 | NaN | NaN |
| Pafah1b3 | AHFLDADPGFVHSD | -170 | -183 | 0 | 0.007343498 | 14 | 1815.6221 | 1901.0908 | NaN | NaN | NaN | NaN |
| Atp5f1b | IPVGPETLGR | -134 | -143 | 0 | 0.000347792 | 10 | 9966.312 | 9471.214 | 10073.355 | 10410.286 | 11677.947 | 11228.062 |
| Atp5f1b | NVIGEPIDER | -146 | -155 | 0 | 0.00025015 | 10 | 7009.6855 | 6803.0913 | 5985.248 | 7021.668 | 6672.2983 | 624.8227 |
| Atp5f1b | KVLDSGAPIK | -124 | -133 | 1 | 0.00078661 | 10 | 4842.6177 | 4639.629 | 4964.186 | 3086.6428 | 3198.4824 | 3949.8323 |
| Atp5f1b | VGPIEEAVAK | -510 | -519 | 0 | 0 | 10 | 4583.0303 | 5110.03 | 4991.1694 | 6971.837 | 6859.742 | 6831.158 |
| Atp5f1b | TIAMDGTEGLVR | -110 | -121 | 0 | 0 | 12 | 3070.416 | 3116.6274 | 3157.296 | 5006.1494 | 1192.5042 | 4445.37 |
| Atp5f1b | TIAMDGTEGLVR | -110 | -121 | 0 | 0 | 12 | 3070.416 | 3116.6274 | 3157.296 | 5006.1494 | 1192.5042 | 4445.37 |
| Atp5f1b | EVFTGHMGK | -472 | -480 | 0 | 0 | 9 | 1257.0652 | 1375.5146 | 1470.4866 | 2657.0283 | 1767.3467 | 3417.0276 |
| Atp5f1b | EVFTGHMGK | -472 | -480 | 0 | 0 | 9 | 1257.0652 | 1375.5146 | 1470.4866 | 2657.0283 | 1767.3467 | 3417.0276 |

| | | | | | | | | | | | | |
|---|---|---|---|---|---|---|---|---|---|---|---|---|
| Atp5f1b | QHLGESTVR | -101 | -109 | 0 | 0 | 9 | 2848.0593 | 2911.508 | 3068.3071 | 2873.2173 | 96.25676 | 1896.8917 |
| Atp5f1b | FTQAGSEVSALLGR | -311 | -324 | 0 | 0 | 14 | 2460.6963 | 2892.259 | 456.18436 | 2773.6704 | 2681.6562 | 2890.3103 |
| Atp5f1b | AIAELGIYPAVDPLDSTSR | -388 | -406 | 0 | 0 | 19 | 2144.1902 | 2198.7246 | 2012.1528 | 2357.3816 | 2789.2297 | 2928.0498 |
| Atp5f1b | IESGVINLK | -251 | -259 | 0 | 0 | 9 | 2044.8893 | 1867.3501 | 623.89197 | 1176.3236 | 1176.4203 | 1347.773 |
| Atp5f1b | IGLFGGAGVGK | -202 | -212 | 0 | 0 | 11 | 1758.5212 | 2174.4578 | 2155.8975 | 1847.9523 | 1684.0231 | 1849.34 |
| Atp5f1b | IMDPNIVGNEHYDVAR | -407 | -422 | 0 | 0 | 16 | 1652.4249 | 1871.6733 | 1946.1489 | 2253.783 | 3429.0774 | 3364.9084 |
| Atp5f1b | IMDPNIVGNEHYDVAR | -407 | -422 | 0 | 0 | 16 | 1652.4249 | 1871.6733 | 1946.1489 | 2253.783 | 3429.0774 | 3364.9084 |
| Atp5f1b | IMDPNIVGNEHYDVAR | -407 | -422 | 0 | 0 | 16 | 1652.4249 | 1871.6733 | 1946.1489 | 2253.783 | 3429.0774 | 3364.9084 |
| Atp5f1b | IMDPNIVGNEHYDVAR | -407 | -422 | 0 | 0 | 16 | 1652.4249 | 1871.6733 | 1946.1489 | 2253.783 | 3429.0774 | 3364.9084 |
| Atp5f1b | GNEHYDVAR | -414 | -422 | 0 | 0.001536401 | 9 | 1628.4312 | 1500.39 | 1790.8503 | 652.4111 | 1881.2563 | 1970.207 |
| Atp5f1b | VVDLLAPYAK | -189 | -198 | 0 | 0.000230707 | 10 | 1533.9749 | 456.30548 | 1634.486 | 207.73027 | 1336.0491 | 2009.9482 |
| Atp5f1b | IVGNEHYDVAR | -412 | -422 | 0 | 0 | 11 | 1532.4502 | 646.93726 | 1564.0122 | 2049.6511 | 1666.3354 | 1663.7632 |
| Atp5f1b | IVGNEHYDVAR | -412 | -422 | 0 | 0 | 11 | 1532.4502 | 646.93726 | 1564.0122 | 2049.6511 | 1666.3354 | 1663.7632 |
| Atp5f1b | QAGSEVSALLGR | -313 | -324 | 0 | 0 | 12 | 1129.4934 | 2688.4692 | 1647.5325 | 2665.755 | 386.4182 | 2670.4604 |
| Atp5f1b | EVAQHLGESTVR | -98 | -109 | 0 | 0 | 12 | 1106.9326 | 1469.8926 | 1923.7905 | 1406.2104 | 1706.3468 | 2572.5059 |
| Atp5f1b | VLDSGAPIKIPVGPETLGR | -125 | -143 | 1 | 0 | 19 | 826.49274 | 858.2871 | 906.24915 | 1044.5126 | 1649.0886 | 1506.4729 |
| Atp5f1b | VLDSGAPIKIPVGPETLGR | -125 | -143 | 1 | 0 | 19 | 826.49274 | 858.2871 | 906.24915 | 1044.5126 | 1649.0886 | 1506.4729 |
| Atp5f1b | AHGGYSVFAGVGER | -226 | -239 | 0 | 0 | 14 | 727.3926 | 782.558 | 728.6268 | 2026.6074 | 550.1758 | 1835.285 |
| Atp5f1b | IAELGIYPAVDPLDSTSR | -389 | -406 | 0 | 0 | 18 | 715.0532 | 701.0349 | 741.80774 | 1671.3821 | 2274.4722 | 2079.7305 |
| Atp5f1b | DPNIVGNEHYDVAR | -409 | -422 | 0 | 0 | 14 | 634.7884 | 2269.8604 | 1019.0705 | 6240.3135 | 490.51984 | 719.1563 |
| Atp5f1b | KVLDSGAPIKIPVGPETLGR | -124 | -143 | 2 | 0.00025015 | 20 | 539.3882 | 463.88083 | 540.5354 | 566.78534 | 803.79584 | 760.2097 |
| Atp5f1b | QFAPIHAEAPEFIEM | -162 | -176 | 0 | 0.001023236 | 15 | 469.53583 | 524.68005 | 416.03397 | 297.98425 | 388.0899 | 405.07553 |
| Atp5f1b | VLDSGAPIKIPVGPETL | -125 | -141 | 1 | 0 | 17 | 210.70831 | 281.98624 | 399.39594 | 282.68237 | 236.97458 | 290.77658 |
| Atp5f1b | MVGPIEEAVAK | -509 | -519 | 0 | 0.001708008 | 11 | 931.45856 | 294.42523 | 610.8649 | NaN | 862.1043 | 665.59705 |
| Atp5f1b | IPSAVGYQPTLATD | -325 | -338 | 0 | 0.000471409 | 14 | 534.7181 | 979.43243 | NaN | 1083.4551 | 851.1823 | 901.8368 |
| Atp5f1b | EGNDLYHEMIESGVIN | -242 | -257 | 0 | 0.00025015 | 16 | 526.4912 | NaN | 540.4333 | 738.2769 | 750.495 | 667.56415 |
| Atp5f1b | LFIDNIFR | -303 | -310 | 0 | 0 | 8 | 420.37747 | 494.07208 | NaN | 212.75543 | 379.52155 | 159.24936 |
| Atp5f1b | ELGIYPAVDPLDSTSR | -391 | -406 | 0 | 0 | 16 | 398.56677 | NaN | 792.5432 | 1035.478 | 1181.5481 | 1282.9536 |
| Atp5f1b | GFQQILAGEYDHLPEQAF | -490 | -507 | 0 | 0 | 18 | 358.54868 | 548.751 | 441.86472 | NaN | 623.4504 | 505.16693 |
| Atp5f1b | VLDSGAPIK | -125 | -133 | 0 | 0.002296977 | 9 | 1990.7834 | 1922.7035 | 1825.6066 | 1589.6299 | NaN | NaN |
| Atp5f1b | LGIYPAVDPLDSTSR | -392 | -406 | 0 | 0 | 15 | 1343.0157 | 1453.916 | 1455.6083 | NaN | 1761.5032 | NaN |
| Atp5f1b | YMVGPIEEAVAK | -508 | -519 | 0 | 0 | 12 | 2061.2886 | NaN | NaN | NaN | 1138.3037 | 1046.3835 |
| Atp5f1b | FIDNIFR | -304 | -310 | 0 | 0.002972566 | 7 | 719.923 | 859.37616 | NaN | NaN | NaN | 863.19507 |
| Atp5f1b | EHYDVAR | -416 | -422 | 0 | 0 | 7 | 679.9087 | NaN | 862.5143 | NaN | 852.7374 | NaN |
| Atp5f1b | VDPLDSTSR | -398 | -406 | 0 | 0.002265552 | 9 | 480.53064 | 298.39655 | NaN | 573.40234 | NaN | NaN |
| Atp5f1b | IMNVIGEPIDER | -144 | -155 | 0 | 0 | 12 | 330.1861 | 1397.1594 | 1631.8088 | NaN | NaN | NaN |
| Atp5f1b | FTQAGSEVSAL | -311 | -321 | 0 | 0.003490217 | 11 | 1394.0334 | NaN | NaN | NaN | NaN | NaN |
| Tgm2 | SVEVSDPVPAGDLVK | -634 | -648 | 0 | 0 | 15 | 3297.6096 | 2983.9 | 3719.2258 | 1698.9984 | 5044.1025 | 5926.1235 |
| Tgm2 | YSGCLTESNLIK | -550 | -561 | 0 | 0 | 12 | 1992.8633 | 1363.9701 | 1278.7489 | 3446.5952 | 841.15497 | 3438.2734 |
| Tgm2 | VDLFPTDIGLHK | -651 | -662 | 0 | 0 | 12 | 1872.4763 | 1771.511 | 2082.6907 | 2363.5195 | 4261.1445 | 572.5576 |
| Tgm2 | SLTFGAVTGPDPSEEAGTK | -56 | -74 | 0 | 0 | 19 | 1683.5242 | 2108.5718 | 1996.4547 | 1654.0164 | 1399.7036 | 1145.488 |
| Tgm2 | SEGTYCCGPVSVR | -365 | -377 | 0 | 0 | 13 | 1085.5524 | 136.76733 | 886.9751 | 1163.2828 | 1147.5273 | 1126.5249 |
| Tgm2 | KYPEGSPEER | -444 | -453 | 1 | 0 | 10 | 982.9258 | 972.62683 | 330.19608 | 831.09796 | 1317.9625 | 1284.3849 |
| Tgm2 | DDREDITHTYK | -434 | -444 | 1 | 0 | 11 | 594.1057 | 754.11145 | 845.03595 | 794.4222 | 600.3292 | 860.4891 |
| Tgm2 | DDREDITHTYK | -434 | -444 | 1 | 0 | 11 | 594.1057 | 754.11145 | 845.03595 | 794.4222 | 600.3292 | 860.4891 |
| Tgm2 | YPEGSPEER | -445 | -453 | 0 | 0 | 9 | 751.8516 | 145.58113 | 841.39484 | 932.2417 | 526.59814 | 950.6273 |
| Tgm2 | TFGAVTGPDPSEEAGTK | -58 | -74 | 0 | 0 | 17 | 675.9133 | 755.1522 | 766.3473 | 645.79407 | 1056.7231 | 733.1267 |
| Tgm2 | GAVTGPDPSEEAGTK | -60 | -74 | 0 | 0 | 15 | 591.4687 | 664.3709 | 690.91156 | 351.41376 | 1026.4042 | 1034.8292 |
| Tgm2 | VNADVVDWIR | -397 | -406 | 0 | 0 | 10 | 569.0664 | 712.8257 | 776.9918 | 732.24243 | 798.22076 | 817.84125 |
| Tgm2 | DLYLENPEIK | -580 | -589 | 0 | 0 | 10 | 408.35626 | 851.59625 | 1110.7565 | 847.3164 | 778.32135 | 448.69247 |
| Tgm2 | AHIGNDTSETR | -493 | -503 | 0 | 0 | 11 | 82.01685 | 2033.1625 | 661.4595 | 1575.112 | 1921.7196 | 1669.4193 |
| Tgm2 | NEFGELESNK | -318 | -327 | 0 | 0 | 10 | 743.068 | 651.9757 | 675.7836 | 503.2537 | NaN | NaN |
| Tgm2 | DDREDITHTY | -434 | -443 | 1 | 0.00122419 | 10 | 86.60374 | NaN | 737.06934 | NaN | 999.9325 | 900.35986 |
| Tgm2 | VNCNDDQGVLLGR | -228 | -240 | 0 | 0 | 13 | 306.76416 | NaN | 605.2237 | NaN | NaN | 932.9566 |
| Cnpy2 | TDLCDHALHR | -168 | -177 | 0 | 0 | 10 | 1165.6389 | 1015.2217 | 1062.1726 | 1278.0499 | 230.85309 | 154.7302 |
| Cnpy2 | EYGEQIDPSTHR | -91 | -102 | 0 | 0 | 12 | 1107.8397 | 1030.4166 | 937.68365 | 1242.2667 | 675.79834 | 762.98724 |
| Cnpy2 | INPDGSQSVVEVPYAR | -58 | -73 | 0 | 0 | 16 | 1053.0509 | 1154.6699 | 869.0056 | 1335.0916 | 1421.7275 | 1550.9717 |
| Cnpy2 | AHLTELLEEVCDR | -76 | -88 | 0 | 0 | 13 | 922.6076 | 1125.9458 | 723.91534 | 4411.97 | 781.5706 | 1117.8838 |
| Cnpy2 | RTDLCDHALHR | -167 | -177 | 1 | 0 | 11 | 352.64966 | 669.16595 | 979.5305 | 510.21582 | 561.3763 | 878.30804 |
| Cnpy2 | SQDLHCGACR | -23 | -32 | 0 | 0.000627943 | 10 | 595.27576 | NaN | 617.7446 | NaN | 192.24783 | 813.18286 |
| Nampt | YDGHLPIEVK | -108 | -117 | 0 | 0 | 10 | 6421.9893 | 32738.92 | 21263.05 | 15195.944 | 36799.504 | 33188.016 |
| Nampt | EHFQDDVFNER | -89 | -99 | 0 | 0 | 11 | 2216.0737 | 2001.1371 | 2026.533 | 2425.7544 | 1835.4069 | 1975.886 |
| Nampt | NAQLNIEQDVAPH | -479 | -491 | 0 | 0 | 13 | 1475.6609 | 1576.1483 | 1618.3604 | 1983.1196 | 2058.8672 | 1939.0607 |
| Nampt | LHDFGYR | -190 | -196 | 0 | 0.001092529 | 7 | 1343.3265 | 1439.0033 | 1289.7125 | 1674.2733 | 1427.8988 | 1505.5127 |
| Nampt | KFPVTENSK | -331 | -339 | 1 | 0 | 9 | 979.9066 | 999.3681 | 1432.2479 | 420.0885 | 1036.1304 | 406.11096 |
| Nampt | DAFEHIVTQF | -260 | -269 | 0 | 0.00051306 | 10 | 762.5823 | 808.4803 | 830.24084 | 619.6612 | 1014.989 | 501.69678 |
| Nampt | STEAPLIIRPDSGNPLDTVLK | -303 | -323 | 1 | 0 | 21 | 665.5708 | 703.853 | 646.811 | 814.8945 | 790.51044 | 839.4382 |
| Nampt | DPVPGYSVPAAEHSTIT | -235 | -251 | 0 | 0.002566359 | 17 | 827.72375 | 814.6877 | 886.4182 | NaN | 875.59094 | 904.56177 |
| Nampt | AHLVNFK | -210 | -216 | 0 | 0.00111006 | 7 | 610.13727 | 573.4981 | 429.72244 | NaN | 463.08374 | 432.28873 |
| Nampt | PEGSVIPR | -120 | -127 | 0 | 0 | 8 | 537.25024 | 590.22125 | 1204.7971 | NaN | NaN | 1069.3529 |
| Nampt | IWGEDLR | -290 | -296 | 0 | 0.00204437 | 7 | 641.678 | 737.5935 | NaN | 732.75995 | NaN | NaN |
| Cand1 | KHTVDDGLDIR | -1072 | -1082 | 1 | 0 | 11 | 4669.7764 | 1294.2203 | 4922.9077 | 1079.6906 | 3761.731 | 3634.5164 |
| Cand1 | YHISNLLEK | -6 | -14 | 0 | 0.00051306 | 9 | 3262.3982 | 2622.2256 | 5303.497 | 2755.2217 | 3728.9075 | 3376.0286 |
| Cand1 | DHPQPIDPLLK | -995 | -1005 | 0 | 0 | 11 | 1789.437 | 1726.7356 | 1883.6617 | 2600.8696 | 2462.783 | 2441.112 |
| Cand1 | VIRPLDQPSSF | -549 | -559 | 1 | 0.000230707 | 11 | 1702.9581 | 1951.6788 | 1758.9232 | 1925.24 | 1823.8098 | 2242.514 |
| Cand1 | HTVDDGLDIR | -1073 | -1082 | 0 | 0 | 10 | 1179.7653 | 1171.8678 | 829.1765 | 600.88257 | 835.03265 | 512.1968 |
| Cand1 | LTLIDPETLLPR | -958 | -969 | 0 | 0 | 12 | 813.0387 | 934.4021 | 866.7655 | 1050.0115 | 805.828 | 695.8176 |
| Cand1 | LKNEITR | -619 | -625 | 1 | 0.000550307 | 7 | 741.127 | 528.3035 | 728.4542 | 603.58435 | 591.531 | 657.82654 |

| | | | | | | | | | | | | |
|---|---|---|---|---|---|---|---|---|---|---|---|---|
| Cand1 | NHVEDGLKDHYDIK | -1107 | -1120 | 1 | 0 | 14 | 735.7302 | 673.441 | 626.22546 | 598.92535 | 723.5236 | 753.8742 |
| Cand1 | NHVEDGLKDHYDIK | -1107 | -1120 | 1 | 0 | 14 | 735.7302 | 673.441 | 626.22546 | 598.92535 | 723.5236 | 753.8742 |
| Cand1 | TISDHPQPIDPLLK | -992 | -1005 | 0 | 0.000230707 | 14 | 573.50385 | 648.6788 | 474.55438 | 634.44885 | 822.96173 | 770.5415 |
| Cand1 | IDLRPVLGEGVPILAS | -642 | -657 | 1 | 0.002168954 | 16 | 573.047 | 577.95154 | 628.4849 | 559.0561 | 759.0201 | 533.59814 |
| Cand1 | FTISDHPQPIDPLLK | -991 | -1005 | 0 | 0 | 15 | 497.48563 | 582.36176 | 629.83234 | 506.60648 | 591.05743 | 653.5557 |
| Cand1 | FTISDHPQPIDPLLK | -991 | -1005 | 0 | 0 | 15 | 497.48563 | 582.36176 | 629.83234 | 506.60648 | 591.05743 | 653.5557 |
| Cand1 | ALTLIAGSPLK | -631 | -641 | 0 | 0 | 11 | 469.3276 | 456.06683 | 389.17105 | 767.9633 | 483.979 | 539.4148 |
| Cand1 | VIRPLDQPSSFDATPY | -549 | -564 | 1 | 0.00025015 | 16 | 399.29807 | 476.9836 | 512.3313 | 1061.1278 | 866.65234 | 608.14435 |
| Cand1 | LSTLCPSAVLQR | -1130 | -1141 | 0 | 0 | 12 | 269.68274 | 731.83655 | 447.26346 | 812.45715 | 909.0845 | 426.97073 |
| Cand1 | NCIGDFLK | -1006 | -1013 | 0 | 0 | 8 | 895.0205 | 923.54346 | NaN | 988.49744 | 1006.6173 | 992.03394 |
| Cand1 | TCLLPQLTSPR | -178 | -188 | 0 | 0 | 11 | 769.11145 | 747.1309 | 704.163 | 631.2123 | 748.39136 | NaN |
| Cand1 | AHNKPSLIR | -1034 | -1042 | 1 | 0.000627943 | 9 | 567.92676 | 586.64136 | 634.79456 | 351.4958 | NaN | 493.88492 |
| Cand1 | IDLRPVLGEGVPILA | -642 | -656 | 1 | 0.00111006 | 15 | 162.61917 | 188.3511 | NaN | 187.8247 | 78.68499 | 201.91034 |
| Cand1 | RQYLLLHSLK | -905 | -914 | 1 | 0 | 10 | 215.10757 | NaN | 205.51436 | NaN | NaN | 378.8055 |
| Cand1 | NELIGLVR | -736 | -743 | 0 | 0 | 8 | 4206.635 | NaN | 3318.571 | NaN | NaN | NaN |
| Cand1 | IGEYLEK | -249 | -255 | 0 | 0.004983786 | 7 | 3132.5227 | 2778.3748 | NaN | NaN | NaN | NaN |
| Cand1 | LLTIPEAEK | -1183 | -1191 | 0 | 0.00025015 | 9 | 1087.8168 | NaN | NaN | 898.28375 | NaN | NaN |
| Cand1 | CLGPLVSK | -71 | -78 | 0 | 0.002068213 | 8 | 668.9461 | NaN | NaN | NaN | NaN | NaN |
| Polr2a | IFHINPR | -993 | -999 | 0 | 0.00025015 | 7 | 949.1268 | 861.7304 | 628.7324 | 425.69095 | 325.75674 | 305.88068 |
| Polr2a | AHNNELEPTPGNTLR | -720 | -734 | 0 | 0 | 15 | 609.16205 | 639.66394 | 679.07544 | 1314.6736 | 1192.1403 | 847.5864 |
| Polr2a | TVITPDPNLSIDQVGVPR | -365 | -382 | 0 | 0 | 18 | 366.8874 | 383.53476 | 580.31226 | 554.0517 | 708.57764 | 540.3839 |
| Polr2a | DWLLGEIESK | -1066 | -1075 | 0 | 0.000230707 | 10 | 434.38657 | 397.4777 | 519.7394 | NaN | 664.81665 | 374.0402 |
| Polr2a | RIPFGFK | -797 | -803 | 1 | 0.000627943 | 7 | 772.5085 | 657.7819 | 599.2709 | NaN | 423.6175 | NaN |
| Polr2a | YSPTSPTYSPTTPK | -1874 | -1887 | 0 | 0 | 14 | 669.1523 | 813.22833 | 867.2745 | NaN | 941.34766 | NaN |
| Polr2a | FGVEQPEGDEDLTK | -164 | -177 | 0 | 0 | 14 | 650.9752 | 713.78455 | 838.552 | NaN | 974.81683 | NaN |
| Polr2a | INAGFGDDLNCIFNDDNAEK | -1235 | -1254 | 0 | 0.00078661 | 20 | 196.447 | 218.00482 | NaN | 218.48543 | 282.7945 | NaN |
| Polr2a | YSPTSPTYSPTSPVYTPTSPK | -1888 | -1908 | 0 | 0.000627943 | 21 | 331.80865 | NaN | 346.85333 | NaN | NaN | NaN |
| Snrpg | AHPPELK | -4 | -10 | 0 | 0 | 7 | 3357.021 | 689.3108 | 1158.7064 | 849.5585 | 1064.2615 | 962.98425 |
| Snrpg | HVQGILR | -26 | -32 | 0 | 0.000947459 | 7 | 3163.2666 | 3201.4307 | 3235.7642 | 2749.5488 | 2674.3865 | 2443.7695 |
| Snrpg | TSGQQNNIGMVVIR | -50 | -63 | 0 | 0 | 14 | 951.6162 | 969.00824 | 1000.9118 | 1462.3932 | 1414.355 | 1450.3752 |
| Snrpg | ATSGQQNNIGMVVIR | -49 | -63 | 0 | 0 | 15 | 800.63007 | 694.6502 | 594.7557 | 933.02106 | 773.25464 | 673.0428 |
| Snrpg | ATSGQQNNIGMVVIR | -49 | -63 | 0 | 0 | 15 | 800.63007 | 694.6502 | 594.7557 | 933.02106 | 773.25464 | 673.0428 |
| Snrpg | NIGMVVIR | -56 | -63 | 0 | 0.007224385 | 8 | 291.42395 | NaN | NaN | NaN | NaN | NaN |
| Nmi | GIPDELSEEQTR | -164 | -175 | 0 | 0 | 12 | 853.66956 | 612.2776 | 866.0844 | 844.1301 | 709.9419 | 610.893 |
| Nmi | AHPVPLNTGVR | -138 | -148 | 0 | 0 | 11 | 588.48334 | 575.35333 | 655.37286 | 872.0951 | 656.46375 | 690.224 |
| Nmi | DKLELSFCK | -176 | -184 | 1 | 0 | 9 | 123.65719 | 575.99896 | NaN | NaN | 937.3721 | 327.12595 |
| Nmi | KLEAELQSDAR | -47 | -57 | 1 | 0 | 11 | 4942.662 | NaN | NaN | NaN | NaN | NaN |
| Vwa5a | YVQELPLETDGALR | -129 | -142 | 0 | 0 | 14 | 1550.4998 | 1589.916 | 1647.7073 | 1626.3342 | 1863.9517 | 1974.6478 |
| Vwa5a | YLLPAILNPR | -143 | -152 | 0 | 0 | 10 | 690.6195 | 438.37686 | 615.66943 | 473.90027 | 402.0888 | 628.89825 |
| Vwa5a | AHSEYEEALSQ | -83 | -93 | 0 | 0.000907206 | 11 | 3072.2046 | NaN | NaN | NaN | NaN | NaN |
| B2m | AHTEFTPTETDTYACR | -86 | -101 | 0 | 0 | 16 | 2995.6924 | 3618.739 | 3597.5715 | 4877.492 | 5681.121 | 5623.082 |
| B2m | AHTEFTPTETDTYACR | -86 | -101 | 0 | 0 | 16 | 2995.6924 | 3618.739 | 3597.5715 | 4877.492 | 5681.121 | 5623.082 |
| B2m | FTPTETDTYACR | -90 | -101 | 0 | 0 | 12 | 1166.0779 | 1368.2806 | 1123.2234 | 1392.7732 | 602.02075 | 1558.7448 |
| B2m | EFTPTETDTYACR | -89 | -101 | 0 | 0 | 13 | 403.31488 | 570.901 | 504.0258 | 551.4495 | 560.27673 | 576.3594 |
| B2m | HTEFTPTETDTYACR | -87 | -101 | 0 | 0.00025015 | 15 | 359.64453 | 397.5243 | NaN | 300.112 | 239.10416 | 613.33936 |
| B2m | ETDTYACR | -94 | -101 | 0 | 0.00719662 | 8 | 396.37216 | NaN | NaN | NaN | NaN | NaN |
| Dock8 | KIPELYKPIFR | -2065 | -2075 | 2 | 0.000230707 | 11 | 466.2876 | 727.19714 | 708.2339 | 781.54333 | 680.97394 | 837.73254 |
| Dock8 | KIPELYKPIFR | -2065 | -2075 | 2 | 0.000230707 | 11 | 466.2876 | 727.19714 | 708.2339 | 781.54333 | 680.97394 | 837.73254 |
| Dock8 | AHTPSIDPSSQAR | -319 | -331 | 0 | 0 | 13 | 1197.161 | 1128.2734 | 1239.9552 | 1252.1917 | 1093.5248 | 1119.6885 |
| Dock8 | LVIPILEAHR | -1755 | -1764 | 0 | 0 | 10 | 939.4893 | 925.9668 | 958.12476 | 911.72156 | 886.46747 | 898.6807 |
| Dock8 | STGGLYETVNEVYK | -1741 | -1754 | 0 | 0 | 14 | 907.6453 | 915.073 | 1115.3619 | 959.16864 | 1226.3538 | 1097.9331 |
| Dock8 | ITESTLEHELK | -765 | -775 | 0 | 0 | 11 | 740.2844 | 800.43536 | 823.7535 | 1200.8966 | 837.0281 | 356.3991 |
| Dock8 | SWLPILLNER | -670 | -679 | 0 | 0 | 10 | 517.1033 | 646.7086 | 679.4182 | 446.12115 | 589.40283 | 583.0123 |
| Dock8 | SLPEEGVELDPHVR | -106 | -119 | 0 | 0 | 14 | 6595.3535 | 5726.3467 | NaN | 7858.7314 | 5032.67 | 5870.6504 |
| Dock8 | TLQHSLPEEGVELDPHVR | -102 | -119 | 0 | 0 | 18 | 685.69855 | 689.8672 | 844.44055 | NaN | 192.40674 | 814.7927 |
| Dock8 | QHSLPEEGVELDPHVR | -104 | -119 | 0 | 0.00025015 | 16 | 211.62119 | 355.47858 | 316.30377 | NaN | NaN | 675.2141 |
| Dock8 | AKLEEALLR | -1363 | -1371 | 1 | 0.00201047 | 9 | 2173.8772 | NaN | NaN | NaN | 805.8878 | 774.90326 |
| Dock8 | LQTGSYCLPVALEK | -680 | -693 | 0 | 0 | 14 | 486.72742 | NaN | NaN | NaN | 531.2879 | 546.00476 |
| Dock8 | EWLIVNR | -128 | -134 | 0 | 0.008222027 | 7 | 343.10022 | 305.84784 | NaN | 995.1406 | NaN | NaN |
| Irf4 | EGIDKPDPPTWK | -83 | -94 | 1 | 0 | 12 | 1829.064 | 1820.0475 | 994.0272 | 1959.1068 | 1075.9275 | 961.13916 |
| Irf4 | ELPEHVTTPDYHR | -429 | -441 | 0 | 0 | 13 | 1094.6052 | 1131.5944 | 928.9425 | 1198.0432 | 586.0161 | 427.36493 |
| Irf4 | FYACAPPESQAPGIPIEPSIR | -220 | -240 | 0 | 0 | 21 | 851.1559 | 983.9424 | 888.9401 | 1360.5033 | 1101.4673 | NaN |
| Irf4 | APPESQAPGIPIEPSIR | -224 | -240 | 0 | 0.000431655 | 17 | 798.7748 | 830.95105 | 736.0966 | NaN | 869.91284 | 825.58075 |
| Irf4 | SNDFEELVER | -104 | -113 | 0 | 0 | 10 | 389.25598 | 767.4053 | NaN | 1057.0901 | 822.4501 | NaN |
| Irf4 | SQLDISDPYK | -114 | -123 | 0 | 0 | 10 | 920.426 | NaN | NaN | 1317.0245 | NaN | NaN |
| Irf4 | AHVEPLLAR | -403 | -411 | 0 | 0 | 9 | 376.05307 | 1081.8755 | NaN | NaN | NaN | NaN |
| Htt | AHVLDDVTPGPAIK | -1133 | -1146 | 0 | 0 | 14 | 2447.4492 | NaN | NaN | NaN | 4394.5728 | 919.0209 |
| Mdh1 | EALKDDSWLK | -211 | -220 | 1 | 0 | 10 | 5756.336 | 6175.687 | 797.81305 | 6140.467 | 6358.558 | 6463.4097 |
| Mdh1 | AIADHIR | -249 | -255 | 0 | 0.002466863 | 7 | 3943.8652 | 4028.0388 | 4071.1943 | 3749.6423 | 3254.3367 | 3322.8223 |
| Mdh1 | LQDVIATDKEEIAFK | -65 | -79 | 1 | 0 | 15 | 2907.1782 | 1703.2714 | 848.9192 | 1399.5391 | 1760.1567 | 1653.8435 |
| Mdh1 | LQDVIATDKEEIAFK | -65 | -79 | 1 | 0 | 15 | 2907.1782 | 1703.2714 | 848.9192 | 1399.5391 | 1760.1567 | 1653.8435 |
| Mdh1 | STQYPDVNHAK | -189 | -199 | 0 | 0 | 11 | 1954.5468 | 1389.3588 | 1931.8038 | 2000.9016 | 2126.3315 | 2138.4023 |
| Mdh1 | GNHSSTQYPDVNHAK | -185 | -199 | 0 | 0 | 15 | 1738.6521 | 1789.8663 | 2013.6013 | 1857.2352 | 1946.3481 | 2079.9639 |
| Mdh1 | GNHSSTQYPDVNHAK | -185 | -199 | 0 | 0 | 15 | 1738.6521 | 1789.8663 | 2013.6013 | 1857.2352 | 1946.3481 | 2079.9639 |
| Mdh1 | KDDSWLK | -214 | -220 | 1 | 0.000431655 | 7 | 1309.7683 | 1267.6002 | 1194.1427 | 1281.6669 | 1284.0167 | 1248.3457 |
| Mdh1 | DVIATDKEEIAFK | -67 | -79 | 1 | 0.000431655 | 13 | 1184.8937 | 1258.7833 | 1284.1882 | 1056.3348 | 1159.9788 | 1124.9951 |
| Mdh1 | ELTEEKETAFEF | -319 | -330 | 1 | 0.000431655 | 12 | 768.7484 | 826.46704 | 943.4906 | 981.728 | 1054.7958 | 948.3983 |
| Mdh1 | FVEGLPINDFSR | -299 | -310 | 0 | 0 | 12 | 615.1917 | 594.3999 | 6302.074 | 1318.0988 | 831.268 | 758.44965 |

| | | | | | | | | | | | | |
|---|---|---|---|---|---|---|---|---|---|---|---|---|
| Mdh1 | GVPDDLLYSFPVVIK | -279 | -293 | 0 | 0 | 15 | 610.40356 | 530.469 | 374.51544 | 690.317 | 724.636 | 648.53455 |
| Mdh1 | HSSTQYPDVNHAK | -187 | -199 | 0 | 0 | 13 | 585.0995 | 540.04517 | 462.439 | 671.51807 | 677.4541 | 807.2848 |
| Mdh1 | NVIIWGNHSSTQYPDVNHAK | -180 | -199 | 0 | 0 | 20 | 533.1395 | 552.9704 | 534.3783 | 1238.3251 | 1586.8531 | 1693.8046 |
| Mdh1 | KSAPSIPK | -142 | -149 | 1 | 0.002068213 | 8 | 890.1831 | 851.91473 | 967.2525 | NaN | 942.2213 | 921.13293 |
| Mdh1 | KAIADHIR | -248 | -255 | 1 | 0.000394361 | 8 | 707.86084 | 603.00916 | NaN | 791.26855 | 833.24805 | 799.33813 |
| Mdh1 | ELTEEKETAFEFLSSA | -319 | -334 | 1 | 0 | 16 | 366.34354 | 327.83472 | NaN | 612.865 | 740.66187 | 492.3956 |
| Mdh1 | ELTEEKETAFEFL | -319 | -331 | 1 | 0 | 13 | 3098.8486 | NaN | 3634.8582 | 2571.7017 | 3360.4062 | NaN |
| Mdh1 | TDKEEIAFK | -71 | -79 | 1 | 0.001708008 | 9 | 3878.1956 | 2651.8787 | NaN | NaN | NaN | 2175.065 |
| Mdh1 | LGVTADDVK | -171 | -179 | 0 | 0.000550307 | 9 | 1420.361 | 1089.7688 | NaN | NaN | NaN | NaN |
| Mdh1 | VIVVGNPANTNCLTASK | -126 | -142 | 0 | 0 | 17 | 713.3868 | 943.7763 | NaN | NaN | NaN | NaN |
| Mdh1 | NVIIWGNHS | -180 | -188 | 0 | 0.002534621 | 9 | 688.3483 | 817.6258 | NaN | NaN | NaN | NaN |
| Mdh1 | VIVVGNPANTN | -126 | -136 | 0 | 0.002265552 | 11 | 108.87618 | NaN | NaN | NaN | NaN | NaN |
| Psmb1 | NMQNVEHVPLTLDR | -184 | -197 | 0 | 0 | 14 | 3132.4475 | 3643.288 | 3974.3662 | 1548.9213 | 3600.257 | 3434.6848 |
| Psmb1 | NMQNVEHVPLTLDR | -184 | -197 | 0 | 0 | 14 | 3132.4475 | 3643.288 | 3974.3662 | 1548.9213 | 3600.257 | 3434.6848 |
| Psmb1 | QNVEHVPLTLDR | -186 | -197 | 0 | 0 | 12 | 341.41165 | 618.0134 | 444.81738 | 396.31207 | 750.08386 | 756.29004 |
| Psmb1 | QNVEHVPLTLDR | -186 | -197 | 0 | 0 | 12 | 341.41165 | 618.0134 | 444.81738 | 396.31207 | 750.08386 | 756.29004 |
| Psmb1 | SFDPVGSYQR | -150 | -159 | 0 | 0 | 10 | 2864.3206 | 2778.2595 | 2945.589 | 3450.2705 | 658.40906 | 679.6655 |
| Psmb1 | EGIREETVPLR | -228 | -238 | 1 | 0 | 11 | 2834.615 | 2847.8455 | 2871.76 | 353.12067 | 4756.711 | 3896.2063 |
| Psmb1 | EGIREETVPLR | -228 | -238 | 1 | 0 | 11 | 2834.615 | 2847.8455 | 2871.76 | 353.12067 | 4756.711 | 3896.2063 |
| Psmb1 | SGFHGDCLTLTK | -82 | -93 | 0 | 0 | 12 | 2363.0396 | 2342.2273 | 3089.2942 | 5540.29 | 967.2776 | 5391.5747 |
| Psmb1 | LSEGFSIHTR | -56 | -65 | 0 | 0 | 10 | 1499.3477 | 1448.8485 | 1694.6271 | 2262.1357 | 904.7195 | 821.53406 |
| Psmb1 | AIAGEDFSIVASDTR | -41 | -55 | 0 | 0 | 15 | 1291.062 | 980.44824 | 3618.3645 | 291.69357 | 1080.4747 | 992.21643 |
| Psmb1 | YVYNIIGGLDEEGK | -132 | -145 | 0 | 0 | 14 | 874.9769 | 890.69196 | 1091.5111 | 1246.0896 | 1613.9741 | 2846.6226 |
| Psmb1 | LQPLLDNQVGFK | -172 | -183 | 0 | 0 | 12 | 546.00397 | 3463.3477 | 3504.5771 | 3977.4673 | 4288.6597 | 4209.3984 |
| Psmb1 | EGIREETVPLRK | -228 | -239 | 2 | 0 | 12 | 459.30994 | 524.34265 | 567.0197 | 639.1028 | 864.122 | 890.89166 |
| Psmb1 | VYSFDPVGSYQR | -148 | -159 | 0 | 0 | 12 | 729.3046 | 618.52936 | 866.0631 | 425.4393 | 14866.332 | NaN |
| Psmb1 | NIIGGLDEEGK | -135 | -145 | 0 | 0 | 11 | 486.86255 | 2161.0542 | 2194.7292 | NaN | 1770.4182 | 1866.8773 |
| Psmb1 | AMLQPLLDNQVGFK | -170 | -183 | 0 | 0.000230707 | 14 | 294.8423 | 596.95856 | NaN | 437.2758 | 613.78485 | 444.89243 |
| Psmb1 | IIGGLDEEGK | -136 | -145 | 0 | 0 | 10 | 1217.7681 | NaN | NaN | 254.51027 | NaN | NaN |
| Psmb1 | EETVPLRK | -232 | -239 | 1 | 0.002068213 | 8 | 1029.3185 | NaN | NaN | 2096.7207 | NaN | NaN |
| Mthfd1 | THLSLSHNPEQK | -867 | -878 | 0 | 0 | 12 | 3060.8477 | 4200.2744 | 347.11038 | 671.4961 | 4910.01 | 110.15574 |
| Mthfd1 | GVPTGFVLPIR | -879 | -889 | 0 | 0 | 11 | 1173.4895 | 1318.6141 | 1361.2253 | 1108.184 | 1524.8464 | 1451.7776 |
| Mthfd1 | QPSQGPTFGIK | -411 | -421 | 0 | 0 | 11 | 1014.4661 | 956.5139 | 482.09296 | 386.31946 | 409.0095 | 866.66223 |
| Mthfd1 | KGEPISCEDLGVSGAL | -596 | -611 | 1 | 0.000591124 | 16 | 890.54596 | 946.0451 | 921.9501 | 994.50696 | 1160.3768 | 1105.9429 |
| Mthfd1 | KITIGQSPTEK | -543 | -553 | 1 | 0 | 11 | 846.83826 | 576.15015 | 709.47723 | 575.1443 | 614.0966 | 569.18555 |
| Mthfd1 | LDIDPETITWQR | -521 | -532 | 0 | 0 | 12 | 761.38806 | 815.93994 | 851.2591 | 504.429 | 1511.3788 | 1761.6221 |
| Mthfd1 | AIAPEKDVDGLTSVSAGK | -117 | -134 | 1 | 0.000471409 | 18 | 644.73425 | 751.122 | 725.56274 | 730.0503 | 786.98615 | 854.4691 |
| Mthfd1 | GDLNDCFIPCTPK | -138 | -150 | 0 | 0 | 13 | 372.95874 | 616.731 | 676.0866 | 868.0368 | 706.8373 | 215.93022 |
| Mthfd1 | KTDTDAELDLVSR | -760 | -772 | 1 | 0 | 13 | 114.107925 | 1605.9686 | 1050.8805 | 1468.4824 | 1290.1187 | 1323.9908 |
| Mthfd1 | KVVGDVAYDEAK | -251 | -262 | 1 | 0 | 12 | 2259.2336 | NaN | 2406.1685 | 2788.3718 | 462.67212 | 3333.471 |
| Mthfd1 | TPVPSDIAISR | -314 | -324 | 0 | 0 | 11 | 1217.8602 | 476.51968 | 1931.8763 | 872.91895 | NaN | 264.63644 |
| Mthfd1 | IFHELTQTDK | -464 | -473 | 0 | 0 | 10 | 759.1168 | 327.51743 | 848.63666 | 1381.1781 | 1640.4528 | NaN |
| Mthfd1 | IFHELTQTDK | -464 | -473 | 0 | 0 | 10 | 759.1168 | 327.51743 | 848.63666 | 1381.1781 | 1640.4528 | NaN |
| Mthfd1 | EHGAFDAVK | -776 | -784 | 0 | 0 | 9 | 981.6122 | NaN | 785.9359 | 1698.2129 | 664.1474 | 652.15625 |
| Mthfd1 | KTDPTTLTDDEINR | -504 | -517 | 1 | 0.00025015 | 14 | 586.7801 | 629.455 | 626.6754 | NaN | 530.8075 | 506.62994 |
| Mthfd1 | VVGDVAYDEAK | -252 | -262 | 0 | 0 | 11 | 488.0437 | 705.5284 | NaN | 252.706 | 813.2743 | 780.4005 |
| Mthfd1 | TDPTTLTDDEINR | -505 | -517 | 0 | 0 | 13 | 1714.1812 | NaN | NaN | 1186.3048 | 882.1329 | 1369.4526 |
| Mthfd1 | RIYGADDIELLPEAQNK | -832 | -848 | 1 | 0.00025015 | 17 | 657.9976 | NaN | 684.49457 | NaN | 823.8901 | 757.9662 |
| Mthfd1 | RIYGADDIELLPEAQNK | -832 | -848 | 1 | 0.00025015 | 17 | 657.9976 | NaN | 684.49457 | NaN | 823.8901 | 757.9662 |
| Mthfd1 | DIDPETITWQR | -522 | -532 | 0 | 0 | 11 | 1057.728 | NaN | NaN | 1210.3358 | NaN | NaN |
| Mthfd1 | IYGADDIELLPEAQNK | -833 | -848 | 0 | 0.002296977 | 16 | 17285.672 | NaN | NaN | NaN | NaN | NaN |
| Ier3ip1 | AIAVLHEER | -18 | -26 | 0 | 0 | 9 | 858.98505 | 2053.386 | 1963.5894 | 2597.0608 | 3391.6738 | 4216.63 |
| Ier3ip1 | GIGGFGEEPGIK | -38 | -49 | 0 | 0.00025015 | 12 | 21077.547 | 20727.287 | 21463.922 | NaN | 21492.607 | NaN |
| Qdpr | AIAVLPVTLDTPMNR | -174 | -188 | 0 | 0 | 15 | 364.5172 | 436.28503 | 466.1101 | 372.21155 | 485.74393 | 562.54016 |
| Qdpr | TISSHLATK | -113 | -121 | 0 | 0.000431655 | 9 | 3317.6082 | NaN | NaN | 2980.1296 | NaN | 2577.6462 |
| Gabpa | HITTISDETSEQVTR | -151 | -165 | 0 | 0.001392869 | 15 | 571.8305 | 709.7577 | 1271.5686 | 826.9688 | 703.6684 | 772.7364 |
| Gabpa | LNILEIVK | -108 | -115 | 0 | 0 | 8 | 460.6112 | 555.54565 | 632.84186 | NaN | 1058.697 | 1147.6904 |
| Gabpa | AIDINEPIGNLK | -41 | -52 | 0 | 0 | 12 | 466.37152 | 227.14232 | 592.73376 | NaN | NaN | NaN |
| St13 | LQKPNAAIR | -160 | -168 | 1 | 0.000431655 | 9 | 958.9233 | 1007.4552 | 1036.8297 | 1724.6115 | 2155.461 | 1909.2941 |
| St13 | LQKPNAAIR | -160 | -168 | 1 | 0.000431655 | 9 | 958.9233 | 1007.4552 | 1036.8297 | 1724.6115 | 2155.461 | 1909.2941 |
| St13 | AIEINPDSAQPY | -173 | -184 | 0 | 0 | 12 | 822.48145 | 695.978 | 910.10706 | 576.21643 | 184.46472 | 216.47203 |
| St13 | AIDLFTDAIK | -132 | -141 | 0 | 0 | 10 | 741.3109 | 800.9926 | 756.8406 | 861.2828 | 1187.1981 | 1128.41 |
| Abhd14b | AIDLPGLGR | -66 | -74 | 0 | 0.000755556 | 9 | 281.0526 | NaN | NaN | NaN | NaN | NaN |
| Larp7 | AIEFLNNPPEEAPR | -176 | -189 | 0 | 0 | 14 | 678.18854 | 831.2832 | 341.80292 | 1246.1453 | 2038.8003 | 2096.2998 |
| Larp7 | IVSGEPLPGR | -447 | -456 | 0 | 0 | 10 | 1194.6361 | 1204.766 | 1140.7823 | 298.04102 | 2011.3141 | NaN |
| Larp7 | LNNPPEEAPR | -180 | -189 | 0 | 0.000712251 | 10 | 507.06552 | 1466.038 | NaN | NaN | NaN | NaN |
| Set | LNEQASEEILKVE | -57 | -69 | 1 | 0 | 13 | 5832.601 | 6469.707 | 6572.386 | 5453.6694 | 6268.4507 | 6387.492 |
| Set | FYFDENPYFENK | -138 | -149 | 0 | 0.00025015 | 12 | 3669.3877 | 3783.7544 | 4138.966 | 4629.9844 | 5577.27 | 5670.625 |
| Set | LGEEDEEALHYLTR | -108 | -121 | 0 | 0 | 14 | 3476.6184 | 4522.69 | 4672.171 | 1892.7435 | 2739.5806 | 2839.6948 |
| Set | LGEEDEEALHYLTR | -108 | -121 | 0 | 0 | 14 | 3476.6184 | 4522.69 | 4672.171 | 1892.7435 | 2739.5806 | 2839.6948 |
| Set | QHEEPESFFTW | -194 | -204 | 0 | 0.001291103 | 11 | 3405.6362 | 3121.0889 | 3655.6882 | 3615.407 | 42.016853 | 4237.0986 |
| Set | LNEQASEEILK | -57 | -67 | 0 | 0 | 11 | 2863.3337 | 2849.02 | 2868.1477 | 2644.1338 | 2535.4358 | 2472.1714 |
| Set | LNEQASEEILK | -57 | -67 | 0 | 0 | 11 | 2863.3337 | 2849.02 | 2868.1477 | 2644.1338 | 2535.4358 | 2472.1714 |
| Set | RVEVTEFEDIK | -121 | -131 | 1 | 0 | 11 | 2468.9941 | 2144.6797 | 2787.6296 | 1505.5919 | 1592.0396 | 1487.995 |
| Set | EFHLNESGDPSSK | -154 | -166 | 0 | 0 | 13 | 2000.2151 | 3159.0073 | 3242.855 | 10397.803 | 4571.517 | 734.9502 |
| Set | EFHLNESGDPSSK | -154 | -166 | 0 | 0 | 13 | 2000.2151 | 3159.0073 | 3242.855 | 10397.803 | 4571.517 | 734.9502 |
| Set | EFHLNESGDPSSK | -154 | -166 | 0 | 0 | 13 | 2000.2151 | 3159.0073 | 3242.855 | 10397.803 | 4571.517 | 734.9502 |
| Set | VEVTEFEDIK | -122 | -131 | 0 | 0 | 10 | 1968.9677 | 1956.1395 | 1914.7869 | 467.56104 | 1986.9238 | 1828.6781 |

| | | | | | | | | | | | | |
|---|---|---|---|---|---|---|---|---|---|---|---|---|
| Set | FTDHSDAGADELGEVIK | -205 | -221 | 0 | 0 | 17 | 1367.4137 | 1597.7021 | 1514.6342 | 1500.626 | 1644.7804 | 1679.1033 |
| Set | EQQEAIEHIDEVQNEIDR | -39 | -56 | 0 | 0 | 18 | 1193.3538 | 1447.2247 | 1564.6566 | 1811.1666 | 2688.2612 | 2413.8618 |
| Set | EQQEAIEHIDEVQNEIDR | -39 | -56 | 0 | 0 | 18 | 1193.3538 | 1447.2247 | 1564.6566 | 1811.1666 | 2688.2612 | 2413.8618 |
| Set | LRQPFFQK | -75 | -82 | 1 | 0.00078661 | 8 | 1168.6658 | 1322.1694 | 1396.695 | 708.9443 | 785.3578 | 940.3579 |
| Set | ALLGEEDEEALHYLTR | -106 | -121 | 0 | 0 | 16 | 876.51184 | 1081.3713 | 878.64746 | 1065.4514 | 1409.5441 | 1218.7314 |
| Set | RQHEEPESFFTW | -193 | -204 | 1 | 0.001868731 | 12 | 854.81476 | 1316.1129 | 1418.7622 | 1072.6384 | 82.29048 | 2072.9736 |
| Set | RQHEEPESFFTW | -193 | -204 | 1 | 0.001868731 | 12 | 854.81476 | 1316.1129 | 1418.7622 | 1072.6384 | 82.29048 | 2072.9736 |
| Set | IDFYFDENPYFENK | -136 | -149 | 0 | 0 | 14 | 674.7158 | 588.01874 | 539.8344 | 750.05817 | 799.8175 | 741.57526 |
| Set | EQQEAIEHID | -39 | -48 | 0 | 0.00025015 | 10 | 1456.4011 | 1117.6023 | 1068.968 | 736.88434 | NaN | NaN |
| Set | AIEHIDEVQNEIDR | -43 | -56 | 0 | 0 | 14 | 1212.4099 | 1195.6367 | 1366.0414 | NaN | 1659.9613 | NaN |
| Set | LRQPFFQ | -75 | -81 | 1 | 0.003681885 | 7 | 993.485 | 473.0112 | 791.84015 | NaN | 403.75397 | NaN |
| Set | QSAILPQPK | -6 | -14 | 0 | 0 | 9 | 1495.2849 | 1398.5449 | 1375.5819 | NaN | NaN | NaN |
| Set | RQSAILPQPK | -5 | -14 | 1 | 0.000712251 | 10 | 1058.8032 | 1043.0422 | 1113.7864 | NaN | NaN | NaN |
| Set | GEKEQQEAIEHID | -36 | -48 | 1 | 0 | 13 | 964.7941 | 1051.5503 | 994.52 | NaN | NaN | NaN |
| Set | EVQNEIDR | -49 | -56 | 0 | 0.000431655 | 8 | 1403.4967 | NaN | 724.57916 | NaN | NaN | NaN |
| Oxsr1 | LLSGGSVLDIIK | -94 | -105 | 0 | 0 | 12 | 1639.1327 | 469.91177 | 506.04462 | 522.3819 | 621.3788 | 639.2089 |
| Oxsr1 | IPISLVLR | -432 | -439 | 0 | 0.000230707 | 8 | 440.81894 | 535.7535 | 530.3939 | NaN | 336.2193 | NaN |
| Oxsr1 | IVEEPQSNR | -489 | -497 | 0 | 0.006820063 | 9 | 2614.717 | NaN | NaN | 2596.511 | NaN | 3416.9956 |
| Fxr1 | AIELDEDTGTFR | -252 | -263 | 0 | 0 | 12 | 2649.8582 | 597.10693 | 2686.2695 | 1626.275 | 518.15753 | 788.7826 |
| Fxr1 | DVIEEHGPSEK | -624 | -634 | 0 | 0 | 11 | 992.0966 | 872.3912 | 877.1759 | 935.65076 | 865.186 | 589.5006 |
| Fxr1 | HEEFVVR | -220 | -226 | 0 | 0 | 7 | 944.6481 | 1370.5363 | NaN | 1135.965 | 2050.4915 | 1122.1305 |
| Fxr1 | LEFVEDFIQVPR | -279 | -290 | 0 | 0 | 12 | 596.18494 | NaN | NaN | 455.31924 | 472.24875 | 399.38928 |
| Hnrnpab | IFVGGLNPEATEEK | -161 | -174 | 0 | 0 | 14 | 5804.3735 | 6505.5312 | 6703.698 | 5400.973 | 5970.217 | 5973.205 |
| Hnrnpab | FKEEDPVKK | -207 | -215 | 2 | 0 | 9 | 3977.1335 | 4204.509 | 4885.9697 | 4201.437 | 5105.749 | 5316.9214 |
| Hnrnpab | AIELPIDPK | -187 | -195 | 0 | 0 | 9 | 3530.9526 | 3649.2222 | 3744.5376 | 3456.3625 | 2985.0554 | 3059.561 |
| Hnrnpab | ITFKEEDPVKK | -205 | -215 | 2 | 0 | 11 | 3345.5037 | 2661.405 | 229.84802 | 1594.4109 | 2939.524 | 3094.0305 |
| Hnrnpab | DLKDYFTK | -90 | -97 | 1 | 0.000627943 | 8 | 2414.734 | 2509.7654 | 136.17697 | 2633.8423 | 2802.756 | 2860.524 |
| Hnrnpab | ITFKEEDPVK | -205 | -214 | 1 | 0 | 10 | 1371.0682 | 1031.9438 | 2534.558 | 2961.8394 | 2280.023 | 3574.9348 |
| Hnrnpab | FKEEDPVK | -207 | -214 | 1 | 0.000431655 | 8 | 1323.0273 | 1477.9164 | 1611.0278 | 1285.0514 | 1681.3541 | 1550.4108 |
| Hnrnpab | FKDSSSVEK | -123 | -131 | 1 | 0.001055699 | 9 | 1010.2966 | 217.15776 | 942.00415 | 936.53 | 392.41815 | 1169.4398 |
| Hnrnpab | FGEIEAIELPIDPK | -182 | -195 | 0 | 0 | 14 | 524.4239 | 369.85147 | 412.50803 | 619.6186 | 961.70404 | 809.622 |
| Hnrnpab | GEIEAIELPIDPK | -183 | -195 | 0 | 0 | 13 | 416.77652 | 359.68182 | 415.5163 | 434.16418 | 376.59076 | 691.1656 |
| Hnrnpab | EYFGQFGEIE | -177 | -186 | 0 | 0.00025015 | 10 | 398.24075 | 365.93256 | 366.89694 | 404.25763 | 480.54584 | 406.22498 |
| Hnrnpab | VGGLNPEATEEK | -163 | -174 | 0 | 0 | 12 | 2434.8909 | NaN | 2365.337 | 230.25363 | 1760.656 | 1902.9568 |
| Hnrnpab | GLNPEATEEK | -165 | -174 | 0 | 0.000431655 | 10 | 2085.822 | 1930.8501 | NaN | 1962.3513 | NaN | NaN |
| Hnrnpab | IFVGGLNPEATEE | -161 | -173 | 0 | 0 | 13 | 812.0025 | 218.58073 | 805.35596 | NaN | NaN | NaN |
| Rmdn3 | AIELQPEDPR | -356 | -365 | 0 | 0.001536401 | 10 | 449.8973 | 626.89636 | 273.71747 | NaN | NaN | NaN |
| Otulin | AIELYDDKEK | -240 | -249 | 1 | 0 | 10 | 645.9357 | 553.7025 | 581.9195 | NaN | 1112.7797 | 1460.2583 |
| Otulin | ELLIHER | -67 | -73 | 0 | 0.000907206 | 7 | 715.76086 | NaN | NaN | 749.77356 | NaN | NaN |
| Reep5 | HESQVDSVVK | -144 | -153 | 0 | 0 | 10 | 1236.0481 | 1005.58746 | 1140.5825 | 1121.1573 | 1221.1381 | 1242.8923 |
| Reep5 | AIESPNKDDDTQWLTY | -70 | -85 | 1 | 0.00025015 | 16 | 551.67395 | 520.0444 | 504.04773 | 1057.1707 | 1093.6156 | 1298.7988 |
| Jak1 | NVLVESEHQVK | -1007 | -1017 | 0 | 0 | 11 | 1371.6359 | 977.24774 | 1601.2946 | 2318.663 | 1797.0901 | 2446.287 |
| Jak1 | AIETDKEYYTVK | -1026 | -1037 | 1 | 0 | 12 | 566.6677 | 633.9304 | 623.6455 | 974.6368 | 762.7979 | 621.88934 |
| Jak1 | IPWIAPECVEDSK | -755 | -767 | 0 | 0 | 13 | 97.20625 | 491.15857 | NaN | 539.43054 | 885.5607 | 839.4154 |
| Jak1 | QPTTEVDPTHFEK | -859 | -871 | 0 | 0 | 13 | 1390.2842 | NaN | NaN | 1323.0724 | 1429.1335 | 1348.5555 |
| Jak1 | DIIQGEHLGR | -581 | -590 | 0 | 0 | 10 | 1224.3589 | NaN | NaN | NaN | 1602.7972 | NaN |
| Jak1 | YDPEGDNTGEQVAVK | -893 | -907 | 0 | 0 | 15 | 540.91406 | NaN | 776.99786 | NaN | NaN | NaN |
| Ppm1g | SSQEVVDFIQSK | -445 | -456 | 0 | 0 | 12 | 939.5497 | 830.1464 | 960.8808 | 657.37805 | 385.3842 | 1052.5192 |
| Ppm1g | ALQDAFLAIDAK | -93 | -104 | 0 | 0 | 12 | 430.54755 | 580.58325 | 586.3391 | 464.32935 | 470.8686 | 525.0135 |
| Ppm1g | AIGDHFYK | -396 | -403 | 0 | 0 | 8 | 1068.649 | 948.97626 | NaN | 677.6095 | 1503.7734 | 1605.7605 |
| Ppm1g | KLEEALSTEGAEDTGNSDK | -518 | -536 | 1 | 0.00025015 | 19 | 370.96005 | 1358.952 | NaN | 928.1491 | 837.30426 | 826.357 |
| Ppm1g | SGIGTGDEPGPQGLNGEAGPEDPSR | -173 | -197 | 0 | 0.000550307 | 25 | 300.53195 | 445.4183 | 339.40552 | 333.493 | 405.9616 | NaN |
| Ppm1g | SYDHKPEDEVELAR | -360 | -373 | 1 | 0.00025015 | 14 | 2422.8044 | NaN | 1080.141 | 2062.9868 | NaN | 1952.9805 |
| Ppm1g | YLPDIIK | -74 | -80 | 0 | 0.004658491 | 7 | 241.61679 | 670.2439 | 459.52856 | 691.86237 | NaN | NaN |
| Atp6v0c | AIGIVGDAGVR | -109 | -119 | 0 | 0 | 11 | 2781.038 | 2532.9407 | 2633.57 | 1356.183 | 1318.2198 | 1237.6335 |
| Atp6v0c | GFAIGIVGDAGVR | -107 | -119 | 0 | 0.000550307 | 13 | 803.9493 | 900.54407 | NaN | NaN | NaN | NaN |
| Kpna2 | EKQPPIDNIIR | -107 | -117 | 1 | 0 | 11 | 3624.9858 | 3665.1626 | 3821.2031 | 3943.9985 | 3724.2573 | 3642.7017 |
| Kpna2 | NVSSFPDDATSPLQENR | -52 | -68 | 0 | 0 | 17 | 1727.6792 | 1528.7213 | 1418.2888 | 1342.4186 | 1229.9274 | 836.93713 |
| Kpna2 | AVFPSLLTNPK | -338 | -348 | 0 | 0.00025015 | 11 | 1669.9725 | 1654.295 | 1641.9625 | 219.4318 | 1609.1388 | 1423.3065 |
| Kpna2 | AIGNIVTGTDEQTQK | -316 | -330 | 0 | 0 | 15 | 1390.7163 | 1461.315 | 1402.9044 | 1348.7361 | 1301.5769 | 1301.1462 |
| Kpna2 | GATELPIVTPALR | -303 | -315 | 0 | 0 | 13 | 1182.7395 | 1244.0063 | 1156.0076 | 1232.2821 | 1299.3691 | 1314.0038 |
| Kpna2 | WSVEDIVK | -76 | -83 | 0 | 0.000550307 | 8 | 1141.4515 | 1336.4462 | 1307.0474 | 1314.9152 | 1377.2661 | 1356.7969 |
| Kpna2 | HGAIDPLLAL | -203 | -212 | 0 | 0.00025015 | 10 | 788.6327 | 999.7241 | 1019.76385 | 595.237 | 750.1615 | 833.0161 |
| Kpna2 | QDQIQQVVNHGLVPF | -367 | -381 | 0 | 0 | 15 | 611.9867 | 643.9711 | 534.0813 | 704.94995 | 842.0544 | 902.71326 |
| Kpna2 | LLGATELPIVTPALR | -301 | -315 | 0 | 0 | 15 | 433.38763 | 451.66144 | 402.76746 | 396.51364 | 389.75153 | 460.62347 |
| Kpna2 | NKNPAPPLDAVEQILPT | -239 | -255 | 1 | 0.00025015 | 17 | 362.54556 | 252.66011 | 438.59186 | 658.0318 | 663.69604 | 714.72205 |
| Kpna2 | KGVVPQLVK | -292 | -300 | 1 | 0.000826806 | 9 | 247.7912 | 568.7355 | 634.2911 | 579.5794 | 550.3483 | NaN |
| Kpna2 | NKNPAPPLDAVEQ | -239 | -251 | 1 | 0.00122419 | 13 | 2442.7488 | NaN | 702.2262 | NaN | 2118.3362 | NaN |
| Kpna2 | QPPIDNIIR | -109 | -117 | 0 | 0 | 9 | 882.50757 | 731.9375 | NaN | 845.2115 | NaN | NaN |
| Kpna2 | GNIAGDGSAFR | -187 | -197 | 0 | 0 | 11 | 663.54175 | NaN | 1323.8802 | 476.6782 | NaN | NaN |
| Kpna2 | FVSFLGK | -124 | -130 | 0 | 0.001463897 | 7 | 537.15717 | NaN | 447.6907 | NaN | NaN | NaN |
| Kpna2 | LLHHNDPEVLAD | -259 | -270 | 0 | 0.000230707 | 12 | 975.78766 | NaN | NaN | NaN | NaN | NaN |
| Kpna2 | STNENANLPAAR | -2 | -13 | 0 | 0.003003218 | 12 | 204.61755 | NaN | NaN | NaN | NaN | NaN |
| Sf3b1 | VLPPPAGYVPIR | -414 | -425 | 0 | 0 | 12 | 7491.7764 | 7466.426 | 5159.003 | 2832.861 | 5873.227 | 2307.2656 |
| Sf3b1 | DQAETPGHTPSLR | -219 | -231 | 0 | 0 | 13 | 2084.2578 | 1364.5907 | 1467.2578 | 741.4299 | 913.32263 | 631.9943 |
| Sf3b1 | THEDIEAQIR | -7 | -16 | 0 | 0 | 10 | 1917.6377 | 2303.7322 | 3495.1433 | 1902.3599 | 2379.838 | 2613.9192 |
| Sf3b1 | LDDLVRPY | -563 | -570 | 1 | 0 | 8 | 1335.4839 | 1336.0798 | 1410.8341 | 324.4624 | 897.3876 | 998.3276 |
| Sf3b1 | IVDDLKDEAEQYR | -853 | -865 | 1 | 0 | 13 | 1301.8868 | 1733.7952 | 1800.2742 | 1563.7206 | 2006.1926 | 1929.099 |

| | | | | | | | | | | | | |
|---|---|---|---|---|---|---|---|---|---|---|---|---|
| Sf3b1 | IYNDDKNTYIR | -1287 | -1297 | 1 | 0 | 11 | 997.5217 | 966.39197 | 1079.1743 | 791.34375 | 853.6674 | 841.8422 |
| Sf3b1 | TEILPPFFK | -808 | -816 | 0 | 0 | 9 | 832.89453 | 1215.8622 | 1378.0671 | 2034.172 | 1586.8126 | 1378.1726 |
| Sf3b1 | IWDPTPSHTPAGAATPGR | -253 | -270 | 0 | 0 | 18 | 752.13605 | 880.1936 | 719.5111 | 787.50616 | 848.40906 | 904.12415 |
| Sf3b1 | IWDPTPSHTPAGAATPGR | -253 | -270 | 0 | 0 | 18 | 752.13605 | 880.1936 | 719.5111 | 787.50616 | 848.40906 | 904.12415 |
| Sf3b1 | LIDEDYYAR | -582 | -590 | 0 | 0 | 9 | 751.84796 | 712.2838 | 529.20276 | 896.47705 | 420.82147 | 556.67377 |
| Sf3b1 | RGGDSIGETPTPGASK | -318 | -333 | 1 | 0 | 16 | 686.34216 | 945.84045 | 703.49023 | 1243.4647 | 917.76263 | 802.74475 |
| Sf3b1 | IADREDEYKK | -112 | -121 | 2 | 0.00111006 | 10 | 684.6736 | 430.5093 | 512.2048 | 684.196 | 606.3163 | 447.124 |
| Sf3b1 | KTHEDIEAQIR | -6 | -16 | 1 | 0 | 11 | 594.1929 | 622.61566 | 817.8518 | 340.9473 | 932.6912 | 664.20105 |
| Sf3b1 | KTHEDIEAQIR | -6 | -16 | 1 | 0 | 11 | 594.1929 | 622.61566 | 817.8518 | 340.9473 | 932.6912 | 664.20105 |
| Sf3b1 | LSSWDQAETPGHTPSLR | -215 | -231 | 0 | 0 | 17 | 367.39465 | 399.95792 | 355.32144 | 447.21298 | 416.70007 | 527.5432 |
| Sf3b1 | SFDSVLKPLWK | -723 | -733 | 1 | 0 | 11 | 344.5375 | 1749.3998 | 1705.7397 | 231.0185 | 404.06128 | 247.14996 |
| Sf3b1 | SLVEIIEHGLVDEQQK | -685 | -700 | 0 | 0 | 16 | 336.17786 | 354.057 | 479.33807 | 561.36707 | 396.13922 | 441.14893 |
| Sf3b1 | GIPSLLPFLK | -640 | -649 | 0 | 0 | 10 | 329.78235 | 320.22742 | 297.35004 | 253.71986 | 367.3654 | 392.2296 |
| Sf3b1 | GDTPGHATPGHGGAT | -271 | -285 | 0 | 0 | 15 | 272.47946 | 292.6575 | 284.52808 | 363.83414 | 363.84326 | 331.41492 |
| Sf3b1 | GDTPGHATPGHGGAT | -271 | -285 | 0 | 0 | 15 | 272.47946 | 292.6575 | 284.52808 | 363.83414 | 363.84326 | 331.41492 |
| Sf3b1 | ATELEDDDDDYSSTSLLGQK | -60 | -80 | 0 | 0 | 21 | 287.24872 | 319.79535 | 271.0607 | 405.08026 | 476.66458 | 418.76846 |
| Sf3b1 | LVDVDESTLSPEEQK | -479 | -493 | 0 | 0 | 15 | 1987.4775 | 2360.0293 | 1936.0264 | 2596.9932 | 3045.2817 | NaN |
| Sf3b1 | LKPDDIQYFDK | -467 | -477 | 1 | 0 | 11 | 1571.6814 | 248.6505 | 387.1221 | NaN | 161.98068 | 1042.0985 |
| Sf3b1 | EFQSPDEEMKK | -776 | -786 | 1 | 0 | 11 | 782.615 | NaN | 857.3416 | 675.2833 | 967.38965 | 824.23016 |
| Sf3b1 | NDDKNTYIR | -1289 | -1297 | 1 | 0 | 9 | 655.1426 | 623.13934 | 574.7908 | 641.9206 | NaN | 611.64496 |
| Sf3b1 | SWDQAETPGHTPSLR | -217 | -231 | 0 | 0 | 15 | 1344.0104 | NaN | 966.6778 | 1493.0703 | 1710.0905 | NaN |
| Sf3b1 | GCAILPHLR | -676 | -684 | 0 | 0 | 9 | 691.8582 | 1419.7755 | 472.66547 | 927.6643 | NaN | NaN |
| Sf3b1 | KPGYHAPVAL | -81 | -90 | 1 | 0.00111006 | 10 | 679.70935 | 587.9521 | 1018.3741 | NaN | 2149.909 | NaN |
| Sf3b1 | SFLFEYIGEMGK | -1152 | -1163 | 0 | 0.000431655 | 12 | 398.30927 | NaN | 367.1486 | 427.742 | 349.21622 | NaN |
| Sf3b1 | VTPLLEDALMDR | -1169 | -1180 | 0 | 0.000471409 | 12 | 396.13403 | NaN | NaN | 727.0002 | 522.3955 | 538.35645 |
| Sf3b1 | DTPGHGSGWAETPR | -302 | -315 | 0 | 0 | 14 | 209.10303 | NaN | NaN | 689.239 | 1095.2738 | 1006.38464 |
| Sf3b1 | STGYYDQEIYGGSDSR | -35 | -50 | 0 | 0 | 16 | 1617.9812 | NaN | NaN | 1842.4598 | 1329.5304 | NaN |
| Sf3b1 | VQENCIDLVGR | -1031 | -1041 | 0 | 0 | 11 | 753.09155 | NaN | NaN | 926.4834 | NaN | 885.6552 |
| Sf3b1 | KLTATPTPLGGMT | -430 | -442 | 1 | 0.001392869 | 13 | 720.2536 | 965.66626 | 857.8127 | NaN | NaN | NaN |
| Sf3b1 | SSWDQAETPGHTPSLR | -216 | -231 | 0 | 0 | 16 | 718.8199 | 787.30206 | NaN | NaN | 581.0392 | NaN |
| Sf3b1 | AIGPHDVLATL | -1087 | -1097 | 0 | 0.003133457 | 11 | 563.82806 | NaN | NaN | 566.125 | NaN | 656.7948 |
| Sf3b1 | KAALDEAQGVGLD | -22 | -34 | 1 | 0.00025015 | 13 | 469.6366 | 1667.168 | 1671.5895 | NaN | NaN | NaN |
| Sf3b1 | ILPPFFK | -810 | -816 | 0 | 0.005946382 | 7 | 426.86908 | 472.33112 | 207.35538 | NaN | NaN | NaN |
| Sf3b1 | LLVDVDESTLSPEEQK | -478 | -493 | 0 | 0 | 16 | 419.85068 | 487.58725 | 473.95178 | NaN | NaN | NaN |
| Sf3b1 | WDQTADQTPGATPK | -200 | -213 | 0 | 0 | 14 | 1586.9912 | 1642.8242 | NaN | NaN | NaN | NaN |
| Sf3b1 | VGAAEIISR | -844 | -852 | 0 | 0 | 9 | 1568.2654 | NaN | 1335.9895 | NaN | NaN | NaN |
| Psma5 | SSLQEVYHK | -179 | -187 | 0 | 0 | 9 | 2756.0154 | 1254.329 | 1315.7067 | 2445.4492 | 747.4813 | 2527.9543 |
| Psma5 | TKEELEEVIK | -230 | -239 | 1 | 0 | 10 | 2731.792 | 3191.1208 | 2955.8586 | 3429.1006 | 3183.7246 | 3069.9194 |
| Psma5 | SLQEVYHK | -180 | -187 | 0 | 0 | 8 | 1827.7076 | 2351.24 | 247.78477 | 1698.672 | 2395.639 | 2361.1284 |
| Psma5 | GPQLFHMDPSGTFVQC | -150 | -165 | 0 | 0 | 16 | 1606.5205 | 2198.8157 | 1702.9088 | 2097.4749 | 1913.1748 | 2514.705 |
| Psma5 | GPQLFHMDPSGTFVQC | -150 | -165 | 0 | 0 | 16 | 1606.5205 | 2198.8157 | 1702.9088 | 2097.4749 | 1913.1748 | 2514.705 |
| Psma5 | TSEGVCLAVEK | -42 | -52 | 0 | 0 | 11 | 1606.3602 | 902.4674 | 833.39136 | 912.5872 | 461.21222 | 1342.9004 |
| Psma5 | TKEELEEVIKDI | -230 | -241 | 2 | 0.00025015 | 12 | 1518.2164 | 1663.0477 | 1652.6876 | 2547.6084 | 3151.6284 | 3173.2786 |
| Psma5 | SEGAQSSLQEVYHK | -174 | -187 | 0 | 0 | 14 | 1095.3901 | 284.757 | 1280.0109 | 977.323 | 1182.8369 | 1210.4628 |
| Psma5 | ASEGAQSSLQEVYHK | -173 | -187 | 0 | 0 | 15 | 1024.7808 | 1097.9254 | 558.28534 | 879.9076 | 932.5411 | 940.6174 |
| Psma5 | VETQNHWFTYN | -94 | -104 | 0 | 0 | 11 | 939.9014 | 1171.4432 | 1127.3203 | 882.0949 | 877.7335 | 773.86707 |
| Psma5 | AIGSASEGAQSSLQEVYHK | -169 | -187 | 0 | 0 | 19 | 802.303 | 894.4653 | 956.3139 | 1412.2216 | 1913.6826 | 1873.9462 |
| Psma5 | AIGSASEGAQSSLQEVYHK | -169 | -187 | 0 | 0 | 19 | 802.303 | 894.4653 | 956.3139 | 1412.2216 | 1913.6826 | 1873.9462 |
| Psma5 | ITSPLMEPSSIEK | -54 | -66 | 0 | 0 | 13 | 610.105 | 776.75854 | 793.75366 | 590.785 | 1135.3002 | 1059.6829 |
| Psma5 | QFGEEDADPGAMSR | -122 | -135 | 0 | 0.000230707 | 14 | 458.6295 | 537.92786 | 446.47815 | 552.02576 | 554.3505 | 575.15686 |
| Psma5 | GVNTFSPEGR | -11 | -20 | 0 | 0 | 10 | 1359.6522 | 1416.9343 | 1033.4443 | 506.19943 | NaN | 878.06714 |
| Psma5 | EGAQSSLQEVYHK | -175 | -187 | 0 | 0 | 13 | 2625.989 | 2517.043 | 1091.1008 | 2607.974 | NaN | NaN |
| Psma5 | VETQNHWFTYNET | -94 | -106 | 0 | 0 | 13 | 2088.5356 | 1954.0604 | 1838.0002 | NaN | NaN | NaN |
| Psma5 | EELEEVIKDI | -232 | -241 | 1 | 0.00025015 | 10 | 838.27783 | 832.2303 | 842.2802 | NaN | NaN | NaN |
| Psma5 | VETQNHWFTY | -94 | -103 | 0 | 0.00111006 | 10 | 839.5484 | NaN | NaN | 843.97046 | NaN | NaN |
| Psma5 | ALQFGEEDADPGAMSR | -120 | -135 | 0 | 0.00111006 | 16 | 434.60944 | NaN | NaN | 555.7852 | NaN | NaN |
| Nfs1 | AIGTDEDLAHSSIR | -396 | -409 | 0 | 0 | 14 | 773.5089 | 792.1216 | 837.8462 | 980.7306 | 801.3197 | 617.10815 |
| Impdh2 | HGFCGIPITDTGR | -137 | -149 | 0 | 0 | 13 | 2823.853 | 3145.5674 | 3439.9727 | 3103.2664 | 3672.2637 | 3459.11 |
| Impdh2 | DIDFLKEEEHDR | -162 | -173 | 1 | 0 | 12 | 2456.688 | 3021.7432 | 3191.9856 | 1642.6549 | 1875.6415 | 3356.4958 |
| Impdh2 | DIDFLKEEEHDR | -162 | -173 | 1 | 0 | 12 | 2456.688 | 3021.7432 | 3191.9856 | 1642.6549 | 1875.6415 | 3356.4958 |
| Impdh2 | GIQHSCQDIGAK | -463 | -474 | 0 | 0 | 12 | 1249.264 | 1324.104 | 1336.3561 | 1042.6299 | 1592.7318 | 1494.1696 |
| Impdh2 | GAAIGTHEDDKYR | -247 | -259 | 1 | 0 | 13 | 1080.3776 | 969.38934 | 1114.7144 | 377.71124 | 2056.6958 | 809.7072 |
| Impdh2 | AIGTHEDDKYR | -249 | -259 | 1 | 0 | 11 | 1075.1301 | 779.52625 | 1098.7739 | 545.14856 | 1154.5867 | 1067.471 |
| Impdh2 | AIGTHEDDKYR | -249 | -259 | 1 | 0 | 11 | 1075.1301 | 779.52625 | 1098.7739 | 545.14856 | 1154.5867 | 1067.471 |
| Impdh2 | EQGFITDPVVLSPK | -111 | -124 | 0 | 0 | 14 | 831.2154 | 1040.3083 | 998.41956 | 1790.3444 | 2317.1577 | 2586.5623 |
| Impdh2 | FLEEIMTK | -174 | -181 | 0 | 0 | 8 | 779.1453 | 797.1263 | 824.284 | 866.8727 | 622.1088 | 466.89417 |
| Impdh2 | FLEEIMTK | -174 | -181 | 0 | 0 | 8 | 779.1453 | 797.1263 | 824.284 | 866.8727 | 622.1088 | 466.89417 |
| Impdh2 | REDLVVAPAGVTLK | -182 | -195 | 1 | 0.00025015 | 14 | 774.19727 | 889.41394 | 852.07153 | 824.41174 | 978.82544 | 948.595 |
| Impdh2 | REDLVVAPAGVTLK | -182 | -195 | 1 | 0.00025015 | 14 | 774.19727 | 889.41394 | 852.07153 | 824.41174 | 978.82544 | 948.595 |
| Impdh2 | EKYPSLQVIGGNVVTAA | -292 | -308 | 1 | 0.00025015 | 17 | 706.60913 | 873.60034 | 829.9402 | 672.4251 | 852.13367 | 843.1382 |
| Impdh2 | KYEQGFITDPVVLSPK | -109 | -124 | 1 | 0 | 16 | 638.4402 | 736.989 | 735.3509 | 597.03845 | 627.4886 | 612.67126 |
| Impdh2 | KYEQGFITDPVVLSPK | -109 | -124 | 1 | 0 | 16 | 638.4402 | 736.989 | 735.3509 | 597.03845 | 627.4886 | 612.67126 |
| Impdh2 | YEQGFITDPVVLSPK | -110 | -124 | 0 | 0 | 15 | 630.82275 | 813.2093 | 738.4115 | 619.19037 | 764.47455 | 1191.1554 |
| Impdh2 | VRDVFEAK | -127 | -134 | 1 | 0.00025015 | 8 | 618.0075 | 1047.1873 | 427.77988 | 1164.4938 | 1218.2252 | 1085.9664 |
| Impdh2 | EKYPSLQVIGGNVVTA | -292 | -307 | 1 | 0.00025015 | 16 | 445.2067 | 509.43332 | 583.0645 | 444.47333 | 762.4789 | 760.98157 |
| Impdh2 | NLIDAGVDALR | -312 | -322 | 0 | 0 | 11 | 1675.8982 | 2044.3724 | 2093.418 | 1387.0106 | 1978.1202 | NaN |
| Impdh2 | EKYPSLQVIGGN | -292 | -303 | 1 | 0.00201047 | 12 | 1156.7135 | 602.194 | 1200.59 | 492.9126 | 493.2293 | NaN |
| Sf3b3 | LNENLPESIFGAPK | -843 | -856 | 0 | 0 | 14 | 3267.9407 | 3854.3506 | 3951.6455 | 4837.794 | 6444.3564 | 6359.17 |

| | | | | | | | | | | | | |
|---|---|---|---|---|---|---|---|---|---|---|---|---|
| Sf3b3 | NFGDQPDIR | -260 | -268 | 0 | 0 | 9 | 3078.031 | 3060.6348 | 2858.7092 | 2108.0647 | 3121.4275 | 3025.147 |
| Sf3b3 | ELLRPDPNTGK | -39 | -49 | 1 | 0 | 11 | 2731.612 | 2651.2004 | 2904.939 | 3510.5935 | 4269.3696 | 4514.818 |
| Sf3b3 | EEGDTFFFQPR | -380 | -390 | 0 | 0 | 11 | 2199.3127 | 2421.8994 | 2405.978 | 2429.4978 | 2074.0444 | 1778.7611 |
| Sf3b3 | TVLDPVTGDLSDTR | -677 | -690 | 0 | 0 | 14 | 1754.9062 | 2345.874 | 2288.1707 | 1654.2998 | 2220.193 | 2382.9612 |
| Sf3b3 | LPPNTNDEVDEDPTGNK | -1058 | -1074 | 0 | 0.000230707 | 17 | 1429.0043 | 1872.31 | 1903.1683 | 1921.2036 | 127.62826 | 2010.8389 |
| Sf3b3 | SEHPPLCGR | -1150 | -1158 | 0 | 0 | 9 | 1340.7571 | 408.46136 | 1342.7092 | 724.8791 | 2551.2053 | 1465.3357 |
| Sf3b3 | TPVEEVPAAIAPFQGR | -943 | -958 | 0 | 0 | 16 | 1168.2705 | 1803.2003 | 1700.5359 | 1944.9318 | 2543.58 | 2332.1707 |
| Sf3b3 | AVSELPGNPNAVWTVR | -448 | -463 | 0 | 0.000471409 | 16 | 932.59863 | 1155.6625 | 1162.2083 | 1289.4066 | 1397.8514 | 1542.363 |
| Sf3b3 | LTISSPLEAHK | -152 | -162 | 0 | 0 | 11 | 870.436 | 1338.93 | 1362.4954 | 923.51575 | 1148.4851 | 1089.8081 |
| Sf3b3 | SLLDYDTVAGADK | -1037 | -1049 | 0 | 0 | 13 | 854.00574 | 1026.8651 | 1085.3715 | 1005.91394 | 522.4104 | 943.2527 |
| Sf3b3 | NVIDGDLCEQFNSMEPNK | -1172 | -1189 | 0 | 0 | 18 | 795.78064 | 959.2544 | 753.2874 | 1178.0917 | 1526.281 | 1384.589 |
| Sf3b3 | YHVGETVLSLQK | -1095 | -1106 | 0 | 0 | 12 | 682.31134 | 780.3854 | 568.6095 | 1428.8145 | 1351.7844 | 1312.8619 |
| Sf3b3 | ELPGNPNAVWTVR | -451 | -463 | 0 | 0 | 13 | 519.307 | 872.079 | 973.58093 | 1064.4875 | 1185.3362 | 816.5678 |
| Sf3b3 | FLAVGLVDNTVR | -604 | -615 | 0 | 0 | 12 | 294.5097 | 329.2821 | 527.30035 | 639.4183 | 764.3308 | 785.0303 |
| Sf3b3 | VSELPGNPNAVWTVR | -449 | -463 | 0 | 0 | 15 | 725.1498 | NaN | 687.8272 | 1349.6495 | 1333.512 | 1207.427 |
| Sf3b3 | DVQESFIWVR | -1004 | -1013 | 0 | 0 | 10 | 516.19824 | 632.3875 | NaN | 836.4322 | 891.4231 | 826.431 |
| Sf3b3 | IISLDPSDCLQPL | -616 | -628 | 0 | 0.001868731 | 13 | 468.48126 | 254.87613 | NaN | 734.02167 | 655.4621 | 330.21283 |
| Sf3b3 | AIHGNFSGTK | -17 | -26 | 0 | 0.00078661 | 10 | 299.34055 | NaN | 1032.0265 | 509.29373 | 930.3171 | 1006.98914 |
| Sf3b3 | DYIVVGSDSGR | -76 | -86 | 0 | 0 | 11 | 3301.3352 | 3354.3027 | 137.90263 | 3587.6282 | NaN | NaN |
| Sf3b3 | AVGLVDNTVR | -606 | -615 | 0 | 0.003681885 | 10 | 875.6218 | 818.78735 | NaN | NaN | 794.16705 | 974.55005 |
| Sf3b3 | FLNENLPESIFGAPK | -842 | -856 | 0 | 0.00025015 | 15 | 672.6711 | 906.53503 | NaN | 797.76575 | 1046.2992 | NaN |
| Sf3b3 | TISSPLEAHK | -153 | -162 | 0 | 0 | 10 | 639.1382 | 730.45386 | 717.81885 | 523.68066 | NaN | NaN |
| Sf3b3 | IVPGQFLAVDPK | -115 | -126 | 0 | 0 | 12 | 636.69244 | 275.92258 | 329.43307 | 620.257 | NaN | NaN |
| Sf3b3 | MDPSGQLNEYTER | -568 | -580 | 0 | 0 | 13 | 312.36456 | NaN | NaN | 268.63074 | 430.8556 | 316.62302 |
| Sf3b3 | KQQEIVVSR | -26 | -34 | 1 | 0 | 9 | 1804.6677 | NaN | NaN | NaN | 1344.5747 | 1584.9939 |
| Sf3b3 | LAQTEQGDIFK | -302 | -312 | 0 | 0.000230707 | 11 | 501.26276 | NaN | NaN | NaN | 890.87524 | 749.81616 |
| Sf3b3 | QDELGER | -651 | -657 | 0 | 0.006100831 | 7 | 492.61896 | NaN | NaN | NaN | NaN | NaN |
| Psmb4 | GAYLAQPLLR | -187 | -196 | 0 | 0 | 10 | 1539.7816 | 1604.3799 | 1746.6471 | 1884.7534 | 1349.2253 | 1576.6135 |
| Psmb4 | QPVLSQTEAR | -202 | -211 | 0 | 0 | 10 | 1289.457 | 729.87366 | 387.1268 | 1727.3129 | 636.4285 | 716.13385 |
| Psmb4 | VIDEELLGDGHSYSPR | -116 | -131 | 0 | 0 | 16 | 1078.4341 | 1262.3032 | 1280.4379 | 1515.0879 | 1594.0542 | 1306.849 |
| Psmb4 | SGDYADFQYLK | -99 | -109 | 0 | 0 | 11 | 783.2238 | 843.82574 | 860.5484 | 1112.2335 | 1034.3202 | 970.2427 |
| Psmb4 | MVIDEELLGDGHSYSPR | -115 | -131 | 0 | 0 | 17 | 740.61035 | 996.4304 | 990.4397 | 1169.2131 | 725.2472 | 909.53204 |
| Psmb4 | MVIDEELLGDGHSYSPR | -115 | -131 | 0 | 0 | 17 | 740.61035 | 996.4304 | 990.4397 | 1169.2131 | 725.2472 | 909.53204 |
| Psmb4 | GASGDYADFQYLK | -97 | -109 | 0 | 0 | 13 | 754.78485 | 940.48425 | 761.163 | 426.76593 | NaN | 861.2865 |
| Psmb4 | AIHSWLTR | -132 | -139 | 0 | 0.000394361 | 8 | 9538.527 | NaN | NaN | NaN | 6747.518 | 6926.603 |
| Psmb4 | YLAQPLLR | -189 | -196 | 0 | 0.000471409 | 8 | 465.79083 | NaN | NaN | NaN | NaN | NaN |
| Emp3 | AIHTEEILAK | -119 | -128 | 0 | 0.000394361 | 10 | 3197.0405 | 2645.0679 | 2358.9128 | 3242.8943 | 3163.2651 | 2482.2039 |
| Pip4k2b | AIIDILTPYDAK | -366 | -377 | 0 | 0.00078661 | 12 | 2389.9336 | NaN | 2856.1365 | NaN | NaN | NaN |
| Rpl30 | AIIDPGDSDIIR | -95 | -106 | 0 | 0 | 12 | 8548.345 | 9883.124 | 9736.875 | 9371.486 | 10042.146 | 10121.963 |
| Rpl30 | AIIDPGDSDIIR | -95 | -106 | 0 | 0 | 12 | 8548.345 | 9883.124 | 9736.875 | 9371.486 | 10042.146 | 10121.963 |
| Rpl30 | ANNCPALR | -49 | -56 | 0 | 0.000550307 | 8 | 3550.5916 | 3514.2441 | 3484.4438 | 3859.4944 | 4207.338 | 378.88934 |
| Rpl30 | TLAIIDPGDSDIIR | -93 | -106 | 0 | 0 | 14 | 2338.2314 | 2718.9453 | 2774.0198 | 2935.3416 | 3630.4436 | 3911.1262 |
| Rpl30 | TGVHHYSGNNIELGTAC | -69 | -85 | 0 | 0 | 17 | 993.5297 | 1083.1664 | 1005.07733 | 1705.5216 | 1858.2844 | 1831.865 |
| Rpl30 | TGVHHYSGNNIELGTAC | -69 | -85 | 0 | 0 | 17 | 993.5297 | 1083.1664 | 1005.07733 | 1705.5216 | 1858.2844 | 1831.865 |
| Rpl30 | TGVHHYSGNNIELGTA | -69 | -84 | 0 | 0 | 16 | 481.07248 | 529.9448 | 623.97186 | 690.51294 | 487.88596 | 618.6428 |
| Rpl30 | KSLESINSR | -9 | -17 | 1 | 0.00025015 | 9 | 888.4377 | 1039.4751 | NaN | 528.16693 | NaN | 665.9764 |
| Rpl30 | PGDSDIIR | -99 | -106 | 0 | 0.00111006 | 8 | 10248.128 | NaN | NaN | NaN | 792.44763 | 850.39966 |
| Rpl30 | TGVHHYSGNNIELGT | -69 | -83 | 0 | 0 | 15 | 1773.5436 | 1520.0864 | NaN | NaN | NaN | NaN |
| Washc2 | AIIFDDDEGDLFK | -474 | -486 | 0 | 0.000471409 | 13 | 972.1954 | NaN | 1201.8961 | NaN | 979.92523 | 866.304 |
| Rps7 | KEIEVGGGR | -49 | -57 | 1 | 0.001092529 | 9 | 4484.661 | 4387.153 | 5078.42 | 9855.99 | 7188.4795 | 7144.211 |
| Rps7 | VETFSGVYK | -170 | -178 | 0 | 0 | 9 | 1570.6805 | 1241.7262 | 681.1904 | 670.4128 | 1750.6697 | 1209.2709 |
| Rps7 | LELEMNSDLK | -28 | -37 | 0 | 0.00025015 | 10 | 1558.8055 | 1974.2666 | 384.54663 | 2377.8499 | 3792.0752 | 387.35074 |
| Rps7 | IVKPNGEKPDEFESGISQ | -8 | -25 | 2 | 0.001092529 | 18 | 1506.971 | 1445.1396 | 1648.8555 | 1335.581 | 1663.7449 | 1643.5748 |
| Rps7 | IVKPNGEKPDEFESGISQ | -8 | -25 | 2 | 0.001092529 | 18 | 1506.971 | 1445.1396 | 1648.8555 | 1335.581 | 1663.7449 | 1643.5748 |
| Rps7 | AILEDLVFPSEIVGK | -128 | -142 | 0 | 0 | 15 | 365.42645 | 298.3887 | 391.70853 | 304.48065 | 325.69977 | 356.92776 |
| Rps7 | AIIIFVPVPQLK | -59 | -70 | 0 | 0 | 12 | 328.77808 | 306.75696 | 213.53 | 444.6111 | 346.01544 | 298.977 |
| Rps7 | VHDAILEDLVFPSEIVGK | -125 | -142 | 0 | 0.000230707 | 18 | 275.74774 | 334.23303 | 516.182 | 390.54468 | 542.7981 | 602.5405 |
| Rps7 | IVKPNGEKPDEFESGIS | -8 | -24 | 2 | 0.00025015 | 17 | 413.61874 | 640.43396 | 592.8542 | NaN | 884.1402 | 363.49173 |
| Rps7 | HDAILEDLVFPSEIVGK | -126 | -142 | 0 | 0.00025015 | 17 | 211.39824 | 350.50296 | NaN | 381.5455 | 532.28827 | 361.023 |
| Rps7 | KAIIIFVPVPQLK | -58 | -70 | 1 | 0 | 13 | 526.53876 | 492.58978 | 463.69968 | NaN | NaN | 549.7873 |
| Rps7 | KAIIIFVPVPQ | -58 | -68 | 1 | 0.00025015 | 11 | 444.85638 | NaN | NaN | 621.28564 | 530.07947 | 619.46356 |
| Rps7 | LVFPSEIVGK | -133 | -142 | 0 | 0.00025015 | 10 | 505.94385 | 562.7576 | 507.48782 | NaN | NaN | NaN |
| Rps7 | VFPSEIVGK | -134 | -142 | 0 | 0.00025015 | 9 | 4286.049 | 3447.4895 | NaN | NaN | NaN | NaN |
| Cstf3 | AIIPDPVVAPSIVPVLK | -556 | -572 | 0 | 0 | 17 | 338.34134 | 342.06586 | 279.08447 | 141.65898 | 375.10266 | 367.31735 |
| Cstf3 | DYNKYEEGINIH | -189 | -200 | 1 | 0.000947459 | 12 | 626.4634 | NaN | NaN | NaN | 764.52454 | NaN |
| Msto1 | AIIPFPLVR | -358 | -366 | 0 | 0 | 9 | 410.9209 | 531.3776 | NaN | 427.23877 | 501.50757 | 482.79205 |
| Srp68 | KVTEELLTDNR | -111 | -121 | 1 | 0 | 11 | 1774.4167 | 1878.451 | 2438.5598 | 1835.1942 | 2201.9185 | 1421.0564 |
| Srp68 | SPRPQDLIR | -417 | -425 | 1 | 0.002068213 | 9 | 3534.1091 | 3383.3147 | NaN | 1938.0547 | 2318.9678 | 2258.7295 |
| Srp68 | AILDANDSHQTDTSSQVK | -530 | -547 | 0 | 0 | 18 | 963.5555 | 890.65045 | 823.52277 | NaN | 874.40497 | 723.787 |
| Srp68 | FEHQEWK | -198 | -204 | 0 | 0 | 7 | 1199.9706 | 1547.3628 | 1522.866 | NaN | 557.0889 | NaN |
| Srp68 | QIPGGGSGGGGSGSGGGGGGSGGGR | -6 | -30 | 0 | 0.003490217 | 25 | 437.8737 | NaN | NaN | NaN | NaN | NaN |
| Zwint | AILEPVGLQEEAELPAK | -27 | -43 | 0 | 0 | 17 | 487.69666 | 246.04416 | 231.83792 | 411.158 | 256.51407 | 321.3765 |
| Fbl | VIFADVAQPDQTR | -238 | -250 | 0 | 0 | 13 | 6818.1553 | 889.4069 | 7616.699 | 4563.854 | 5384.0483 | 4830.706 |
| Fbl | GGVDQIHIKPGAK | -156 | -168 | 1 | 0 | 13 | 3264.79 | 3416.5227 | 3288.9265 | 2860.5483 | 2801.884 | 2846.3254 |
| Fbl | GGVDQIHIKPGAK | -156 | -168 | 1 | 0 | 13 | 3264.79 | 3416.5227 | 3288.9265 | 2860.5483 | 2801.884 | 2846.3254 |
| Fbl | NLVPGESVYGEK | -116 | -127 | 0 | 0 | 12 | 2799.8804 | 2424.3423 | 2247.1052 | 1044.7845 | 1824.8956 | 2846.9412 |
| Fbl | NLVPGESVYGEK | -116 | -127 | 0 | 0 | 12 | 2799.8804 | 2424.3423 | 2247.1052 | 1044.7845 | 1824.8956 | 2846.9412 |
| Fbl | KNLVPGESVYGEK | -115 | -127 | 1 | 0 | 13 | 1779.4486 | 1794.013 | 1787.3562 | 2471.336 | 2827.5151 | 2724.955 |

| | | | | | | | | | | | | |
|---|---|---|---|---|---|---|---|---|---|---|---|---|
| Fbl | VDVIFADVAQPDQTR | -236 | -250 | 0 | 0 | 15 | 1224.0806 | 1575.0084 | 1463.3108 | 2622.0476 | 3253.9746 | 3080.0493 |
| Fbl | ILGGVDQIHIKPGAK | -154 | -168 | 1 | 0 | 15 | 1087.9081 | 1333.2427 | 1499.8269 | 1634.6327 | 1806.7592 | 1879.0167 |
| Fbl | ILGGVDQIHIKPGAK | -154 | -168 | 1 | 0 | 15 | 1087.9081 | 1333.2427 | 1499.8269 | 1634.6327 | 1806.7592 | 1879.0167 |
| Fbl | MVDVIFADVAQPDQTR | -235 | -250 | 0 | 0 | 16 | 752.13165 | 846.27435 | 818.5272 | 967.58374 | 835.3248 | 942.3851 |
| Fbl | MVDVIFADVAQPDQTR | -235 | -250 | 0 | 0 | 16 | 752.13165 | 846.27435 | 818.5272 | 967.58374 | 835.3248 | 942.3851 |
| Fbl | VSISEGDDKIEYR | -129 | -141 | 1 | 0 | 13 | 728.09357 | 2720.1165 | 2705.276 | 2278.7766 | 2625.11 | 2553.3748 |
| Fbl | VSISEGDDKIEYR | -129 | -141 | 1 | 0 | 13 | 728.09357 | 2720.1165 | 2705.276 | 2278.7766 | 2625.11 | 2553.3748 |
| Fbl | VAQPDQTR | -243 | -250 | 0 | 0 | 8 | 716.0968 | 698.1028 | 749.74 | 1042.4852 | 807.4051 | 1216.2017 |
| Fbl | PGESVYGEK | -119 | -127 | 0 | 0 | 9 | 646.68585 | 971.90045 | 980.4339 | 895.6606 | 1428.2954 | 1598.9618 |
| Fbl | MKPQEQLTLEPYER | -297 | -310 | 1 | 0 | 14 | 517.5509 | 541.8594 | 486.7409 | 413.78494 | 677.21655 | 514.5715 |
| Fbl | HEGVFICR | -99 | -106 | 0 | 0 | 8 | 501.78717 | 175.5541 | 5986.9087 | 3740.274 | 7284.268 | 6274.3833 |
| Fbl | AILGGVDQIHIKPGAK | -153 | -168 | 1 | 0 | 16 | 279.99664 | 368.8485 | 361.19415 | 445.70062 | 475.14035 | 463.9112 |
| Fbl | EGDDKIEYR | -133 | -141 | 1 | 0 | 9 | 238.47876 | 1422.063 | 358.13608 | 2113.585 | 1654.7642 | 1629.884 |
| Fbl | NMKPQEQLTLEPYER | -296 | -310 | 1 | 0.001092529 | 15 | 2250.1433 | 432.74506 | 456.88098 | 471.47913 | 552.4842 | NaN |
| Fbl | NGGHFVISIK | -262 | -271 | 0 | 0.003490217 | 10 | 17850.49 | 291.4638 | 15453.972 | NaN | NaN | NaN |
| Fbl | LVPGESVYGEK | -117 | -127 | 0 | 0.00025015 | 11 | 3661.0452 | 1771.9901 | NaN | NaN | NaN | NaN |
| Fbl | QQENMKPQEQLTLEPYER | -293 | -310 | 1 | 0.001291103 | 18 | 235.25703 | NaN | NaN | NaN | NaN | NaN |
| Copz1 | AILILDNDGDR | -15 | -25 | 0 | 0 | 11 | 1165.1644 | 1249.3312 | 1174.5574 | 1189.9963 | 1521.6558 | 1385.4515 |
| Copz1 | YYDDTYPSVK | -30 | -39 | 0 | 0 | 10 | 1003.1032 | 562.34247 | 1025.6392 | 1003.8483 | 1026.8765 | 220.3124 |
| Sec23b | ERPDLPPVQYEPVLC | -47 | -61 | 1 | 0 | 15 | 1089.0641 | 1029.7913 | 988.47705 | 888.7738 | 828.81244 | 1175.099 |
| Sec23b | VVDTCLEEDDLQALK | -134 | -148 | 0 | 0 | 15 | 945.0716 | 682.88556 | 520.0348 | 347.77386 | 765.2253 | 619.0192 |
| Sec23b | LFTGGPPTQGPGMVVGDELK | -290 | -309 | 0 | 0 | 20 | 643.9491 | 503.5142 | 659.29 | 1046.2053 | 724.11255 | 1127.1893 |
| Sec23b | SIAVGLLEGTFPNTGAR | -271 | -287 | 0 | 0 | 17 | 480.51633 | 301.92267 | 327.93408 | 357.363 | 402.84824 | 390.48596 |
| Sec23b | LEGTFPNTGAR | -277 | -287 | 0 | 0 | 11 | 988.35974 | 1191.7751 | 813.1818 | 951.23395 | NaN | 763.55975 |
| Sec23b | FLQPIHK | -231 | -237 | 0 | 0.001392869 | 7 | 824.5624 | 473.38647 | 612.7753 | 574.27765 | NaN | NaN |
| Sec23b | AILNPLCQVDYR | -68 | -79 | 0 | 0 | 12 | 643.7767 | NaN | NaN | 759.13196 | 857.73846 | 2385.063 |
| Lta4h | APLPLGHIK | -511 | -519 | 0 | 0 | 9 | 4130.707 | 4008.821 | 3960.2773 | 4779.4136 | 4843.1143 | 5249.3076 |
| Lta4h | DGEAPDPEDPSR | -176 | -187 | 0 | 0 | 12 | 2297.2446 | 980.1767 | 1239.638 | 541.29535 | 781.6187 | 693.88855 |
| Lta4h | SSALQWLTPEQTSGK | -113 | -127 | 0 | 0 | 15 | 1785.4009 | 876.59283 | 862.6755 | 1940.0479 | 1336.6111 | 2212.5977 |
| Lta4h | SFLYSHFK | -427 | -434 | 0 | 0 | 8 | 1483.7512 | 953.9745 | 1232.6818 | 1639.2244 | 1798.37 | 1739.5841 |
| Lta4h | SHDQAVHTY | -581 | -589 | 0 | 0.00078661 | 9 | 1469.967 | 1487.7999 | 1701.5974 | 1239.3595 | 1966.7601 | 1948.9083 |
| Lta4h | DLSSHQLNEFLAQ | -494 | -506 | 0 | 0 | 13 | 1386.286 | 937.2194 | 997.4996 | 1874.8157 | 1949.344 | 1338.0359 |
| Lta4h | SLSNVIAHEISHS | -289 | -301 | 0 | 0.00025015 | 13 | 1332.9508 | 1385.0665 | 1186.8091 | 1045.8873 | 1464.862 | 1355.2452 |
| Lta4h | GGWGELQNTIK | -345 | -355 | 0 | 0 | 11 | 1266.7996 | 112.41832 | 890.4949 | 924.6025 | 797.05084 | 1055.4054 |
| Lta4h | AILPCQDTPSVK | -143 | -154 | 0 | 0 | 12 | 1147.378 | 974.47815 | 929.8853 | 2154.9036 | 1204.8323 | 1996.0182 |
| Lta4h | WTGNLVTNK | -302 | -310 | 0 | 0 | 9 | 992.27734 | 439.15634 | 427.55786 | 225.60217 | 376.65494 | 511.28787 |
| Lta4h | FTRPLFK | -567 | -573 | 1 | 0.001743443 | 7 | 873.5913 | 837.6527 | 729.44965 | 696.48346 | 617.9556 | 591.89026 |
| Lta4h | DVDPDVAYSSIPYEK | -372 | -386 | 0 | 0 | 15 | 778.7601 | 1003.99493 | NaN | 841.6619 | 673.87225 | 547.9842 |
| Lta4h | NAINNSEIR | -528 | -536 | 0 | 0 | 9 | 383.03125 | 209.7676 | 677.60376 | 1004.9969 | NaN | 550.45605 |
| Lta4h | TWDHFWLN | -311 | -318 | 0 | 0.001463897 | 8 | 372.2595 | 383.8293 | NaN | NaN | 348.9717 | 490.6403 |
| Lta4h | TQHLHLR | -19 | -25 | 0 | 0.003681885 | 7 | 551.22406 | 432.43634 | 570.22485 | NaN | NaN | NaN |
| Pds5b | LVPPVVDDDEEEER | -1317 | -1330 | 0 | 0 | 14 | 1579.6427 | 1781.7758 | 1825.1082 | 305.38574 | 1778.4863 | 1778.1837 |
| Pds5b | IEEDFPHIR | -699 | -707 | 0 | 0 | 9 | 1227.311 | 603.16583 | 256.51965 | 859.31323 | 1339.003 | 1262.3511 |
| Pds5b | FNDIHVPIR | -322 | -330 | 0 | 0 | 9 | 722.8767 | 697.5897 | 817.55286 | 606.55725 | 1014.52466 | 759.3661 |
| Pds5b | AINDECYQVR | -910 | -919 | 0 | 0 | 10 | 616.6765 | 659.53467 | 606.9866 | 757.16895 | 338.20462 | 498.11987 |
| Pds5b | SIDGTADDEDEGVPTDQAIR | -628 | -647 | 0 | 0 | 20 | 392.9354 | 438.0219 | 435.89832 | 442.7321 | 468.0216 | 439.2557 |
| Pds5b | EDPDLSELEKPR | -1202 | -1213 | 1 | 0 | 12 | 1896.8722 | NaN | 2459.581 | 2236.5 | 1732.6968 | 1821.4417 |
| Pds5b | LLAPDQFAAPLK | -775 | -786 | 0 | 0.00025015 | 12 | 608.795 | 614.84875 | 677.4622 | NaN | 805.48676 | 651.74725 |
| Pds5b | SVLPQLEFK | -272 | -280 | 0 | 0.003521747 | 9 | 483.78708 | 589.3124 | 547.37836 | NaN | 306.6969 | 398.1815 |
| Pds5b | IFEPLHK | -747 | -753 | 0 | 0.004628201 | 7 | 2638.2385 | NaN | NaN | NaN | NaN | 1270.0039 |
| Iqgap1 | VAADTFTALK | -232 | -241 | 0 | 0 | 10 | 2010.9326 | 820.17017 | 1248.8776 | 826.22516 | 1064.9269 | 835.2086 |
| Iqgap1 | FPDAGEDELLK | -1175 | -1185 | 0 | 0 | 11 | 1812.9075 | 1859.6959 | 2056.8438 | 2364.3682 | 2198.9277 | 2887.9673 |
| Iqgap1 | AINEAVDSGDVGR | -629 | -641 | 0 | 0 | 13 | 1663.93 | 1603.6962 | 1619.6006 | 1699.0004 | 1436.7072 | 1623.0806 |
| Iqgap1 | DHINDIIK | -834 | -841 | 0 | 0.000869338 | 8 | 1238.7228 | 1274.885 | 319.14743 | 1191.5923 | 1219.9358 | 1186.309 |
| Iqgap1 | VDHVNLVVHEEHER | -525 | -538 | 0 | 0 | 14 | 1110.3717 | 3518.5283 | 3643.793 | 1002.72266 | 1128.2505 | 2564.7979 |
| Iqgap1 | VDHVNLVVHEEHER | -525 | -538 | 0 | 0 | 14 | 1110.3717 | 3518.5283 | 3643.793 | 1002.72266 | 1128.2505 | 2564.7979 |
| Iqgap1 | KLGNFFSPK | -80 | -88 | 1 | 0.00025015 | 9 | 964.86774 | 812.36584 | 415.5305 | 918.596 | 1200.4397 | 1346.8066 |
| Iqgap1 | YQELINDIAK | -1478 | -1487 | 0 | 0 | 10 | 924.0494 | 815.8374 | 234.80423 | 1149.4296 | 1220.0511 | 1119.7506 |
| Iqgap1 | VDQIQEIVTGNPTVIK | -1038 | -1053 | 0 | 0 | 16 | 818.2052 | 961.26227 | 898.7583 | 1134.9803 | 1682.0355 | 1386.935 |
| Iqgap1 | KLPYDVTPEQALSHEEVK | -1111 | -1128 | 1 | 0 | 18 | 773.2195 | 875.27985 | 848.4695 | 279.979 | 195.54585 | 1145.4971 |
| Iqgap1 | LGLAPQIQDLYGK | -162 | -174 | 0 | 0 | 13 | 631.10144 | 773.93994 | 719.4452 | 772.5116 | 927.1265 | 954.1593 |
| Iqgap1 | ETPATNEQEAEHQR | -1409 | -1422 | 0 | 0 | 14 | 490.2694 | 477.13474 | 728.37286 | 407.4513 | 486.4379 | 582.18463 |
| Iqgap1 | RLIVDVIR | -1390 | -1397 | 1 | 0.000230707 | 8 | 444.25226 | 520.5908 | 544.9238 | 394.59485 | 574.3242 | 559.8536 |
| Iqgap1 | SKVDQIQEIVTGNPTVIK | -1036 | -1053 | 1 | 0 | 18 | 394.34708 | 567.58185 | 560.2854 | 567.7696 | 926.54974 | 864.14825 |
| Iqgap1 | GVIPECGETYQSDLAEAK | -655 | -672 | 0 | 0 | 18 | 278.14465 | 403.50348 | 221.47395 | 449.2301 | 567.14594 | 558.8503 |
| Iqgap1 | LVNLEEGLAPTYQDVLYQAK | -247 | -266 | 0 | 0.00025015 | 20 | 234.845 | 291.1056 | 161.69135 | 307.25165 | 338.91275 | 237.87659 |
| Iqgap1 | IFYPETTDIYDR | -131 | -142 | 0 | 0 | 12 | 3676.181 | 4328.299 | 4172.6763 | NaN | 4596.909 | 1493.9508 |
| Iqgap1 | INEALDEGDAQK | -545 | -556 | 0 | 0 | 12 | 1684.1055 | 931.58984 | 766.10803 | 819.61743 | 846.01794 | NaN |
| Iqgap1 | REEYLLLR | -1017 | -1024 | 1 | 0.00025015 | 8 | 1034.0988 | 706.1247 | 705.08264 | NaN | 486.04337 | 466.61346 |
| Iqgap1 | GGYHYYHNL | -692 | -700 | 0 | 0.001055699 | 9 | 777.94135 | 692.1804 | 689.9017 | NaN | 568.5087 | 539.5505 |
| Iqgap1 | LPYDVTPEQALSHEEVK | -1112 | -1128 | 0 | 0.000471409 | 17 | 419.06088 | NaN | 490.5298 | 356.8262 | 555.271 | 800.67993 |
| Iqgap1 | QSGQTDPLQK | -359 | -368 | 0 | 0 | 10 | 3734.4485 | NaN | NaN | 1468.4379 | 790.9929 | 1515.1859 |
| Iqgap1 | HTDNVIQWLN | -112 | -121 | 0 | 0.000230707 | 10 | 3509.597 | NaN | 319.32745 | 759.52026 | NaN | 1053.5645 |
| Iqgap1 | SNQQLENDLNLMDIK | -902 | -916 | 0 | 0 | 15 | 2779.8943 | NaN | 3056.918 | NaN | 2949.9387 | 2928.2598 |
| Iqgap1 | EALDEGDAQK | -547 | -556 | 0 | 0 | 10 | 1703.134 | 1442.7195 | 565.12787 | 1023.24414 | NaN | NaN |
| Iqgap1 | QHYQDTLIR | -577 | -585 | 0 | 0.000826806 | 9 | 1347.104 | 1510.9758 | 1385.4543 | 1744.4608 | NaN | NaN |
| Iqgap1 | LGNFFSPK | -81 | -88 | 0 | 0 | 8 | 932.15356 | 863.67505 | 364.415 | 854.3502 | NaN | NaN |
| Iqgap1 | SAGGQLTTDQR | -1212 | -1222 | 0 | 0 | 11 | 898.7666 | NaN | 1078.9003 | NaN | 1192.4851 | 492.70734 |

| | | | | | | | | | | | | |
|---|---|---|---|---|---|---|---|---|---|---|---|---|
| Iqgap1 | GLTNIEEENCQR | -485 | -496 | 0 | 0 | 12 | 769.4986 | 726.60425 | NaN | 1466.089 | NaN | 829.26135 |
| Iqgap1 | QLQSDLQQK | -349 | -357 | 0 | 0 | 9 | 407.925 | 472.07956 | 809.46655 | 866.2023 | NaN | NaN |
| Iqgap1 | RDHINDIIK | -833 | -841 | 1 | 0.00025015 | 9 | 210.8736 | NaN | NaN | 1456.0596 | 742.9092 | 787.5384 |
| Iqgap1 | ITLQDVVSHSK | -925 | -935 | 0 | 0 | 11 | 2036.7874 | 664.436 | 2532.9211 | NaN | NaN | NaN |
| Iqgap1 | EIVTGNPTVIK | -1043 | -1053 | 0 | 0 | 11 | 556.1222 | 779.3404 | 571.35284 | NaN | NaN | NaN |
| Iqgap1 | ALESGDMTTVWK | -466 | -477 | 0 | 0 | 12 | 1299.5713 | NaN | NaN | NaN | NaN | NaN |
| Srrm2 | TKPDSSIYPLVDSK | -1103 | -1116 | 1 | 0.000230707 | 14 | 3472.9683 | 2833.7822 | 2621.1812 | 3358.623 | 3020.4812 | 3074.4 |
| Srrm2 | TPQAPTPANLVVGPR | -2281 | -2295 | 0 | 0 | 15 | 1097.1628 | 1435.5487 | 1555.1067 | 1073.5256 | 1469.4138 | 1466.6024 |
| Srrm2 | RPNPDILDHER | -52 | -62 | 1 | 0 | 11 | 913.08264 | 1289.3414 | 1199.9395 | 931.8752 | 1109.7825 | 1190.9983 |
| Srrm2 | SRLEPSELK | -1122 | -1130 | 1 | 0.000431655 | 9 | 732.88257 | 790.7396 | 793.7911 | 922.5406 | 897.16876 | 920.8979 |
| Srrm2 | EGRPQEPTPAK | -2546 | -2556 | 1 | 0.000591124 | 11 | 501.61554 | 581.4588 | 619.1567 | 582.082 | 679.7457 | 650.585 |
| Srrm2 | TAVAPSAVNLADPR | -2227 | -2240 | 0 | 0 | 14 | 1512.6635 | 1536.4963 | 1470.7623 | NaN | 1626.1904 | 1775.8206 |
| Srrm2 | ENSFESSLEFK | -1282 | -1292 | 0 | 0 | 11 | 386.74307 | 408.9016 | 437.97122 | NaN | 305.0755 | 352.935 |
| Srrm2 | NSGPVSEVNTGFSPEVK | -1293 | -1309 | 0 | 0 | 17 | 1923.163 | NaN | 2160.7776 | 1986.7131 | NaN | 1518.4662 |
| Srrm2 | TPAGLAPTNLSSSR | -2310 | -2323 | 0 | 0 | 14 | 1145.6204 | 1121.5245 | 1379.0903 | NaN | NaN | NaN |
| Srrm2 | GYEEQQIQEK | -81 | -90 | 0 | 0.000230707 | 10 | 945.07733 | 404.09912 | NaN | NaN | 961.1787 | NaN |
| Srrm2 | KPIDSLR | -2634 | -2640 | 1 | 0.000394361 | 7 | 899.30927 | NaN | NaN | 544.98834 | NaN | NaN |
| Srrm2 | SLTGSGTPPTAANYPSSSR | -2262 | -2280 | 0 | 0.002595029 | 19 | 288.12482 | NaN | NaN | NaN | NaN | 257.21927 |
| Srrm2 | AIPASVNLADSR | -2201 | -2212 | 0 | 0.000550307 | 12 | 1457.2299 | NaN | NaN | NaN | NaN | NaN |
| Rsl1d1 | ETPEASAPGTSGK | -398 | -410 | 0 | 0 | 13 | 3940.3013 | 1035.4949 | 247.38226 | 1082.7925 | 3538.159 | 3269.0198 |
| Rsl1d1 | VTSQDENAVSLR | -267 | -278 | 0 | 0 | 12 | 2635.635 | 1957.9093 | 1815.7577 | 1920.1417 | 2953.0793 | 327.32935 |
| Rsl1d1 | LVPIGETPDKENVK | -354 | -367 | 1 | 0 | 14 | 1794.1737 | 2314.081 | 2521.444 | 3392.273 | 3797.9634 | 3817.0608 |
| Rsl1d1 | HLPSHIGR | -163 | -170 | 0 | 0.001023236 | 8 | 743.5534 | 784.7439 | 688.8922 | 211.79573 | 446.68024 | 506.87854 |
| Rsl1d1 | HLPSHIGR | -163 | -170 | 0 | 0.001023236 | 8 | 743.5534 | 784.7439 | 688.8922 | 211.79573 | 446.68024 | 506.87854 |
| Rsl1d1 | TKDEFDSPEQTEGFYK | -99 | -114 | 1 | 0 | 16 | 1402.0657 | 1711.3973 | 1674.3379 | 2047.9564 | 2397.874 | 2476.3154 |
| Rsl1d1 | TKDEFDSPEQTEGFYK | -99 | -114 | 1 | 0 | 16 | 1402.0657 | 1711.3973 | 1674.3379 | 2047.9564 | 2397.874 | 2476.3154 |
| Rsl1d1 | EFDSPEQTEGFYK | -102 | -114 | 0 | 0 | 13 | 946.9988 | 1077.2604 | 986.10046 | 1020.77826 | 1140.7297 | 1127.6516 |
| Rsl1d1 | HGVNTISQIIPF | -120 | -131 | 0 | 0 | 12 | 428.38562 | 337.80997 | 363.02606 | 891.8171 | 419.69214 | 366.30652 |
| Rsl1d1 | TETPEASAPGTSGK | -397 | -410 | 0 | 0 | 14 | 122.87634 | 823.26953 | 570.0778 | 813.0018 | 950.43726 | 639.184 |
| Rsl1d1 | ALLATETPEASAPGTSGK | -393 | -410 | 0 | 0 | 18 | 3062.923 | 3626.5757 | 3585.1443 | NaN | NaN | NaN |
| Rsl1d1 | AIPQLVPIGETPDKENVK | -350 | -367 | 1 | 0.00025015 | 18 | 1519.5233 | 1691.3651 | 1608.3036 | NaN | NaN | NaN |
| Rsl1d1 | VPLPHSILSE | -82 | -91 | 0 | 0.005789881 | 10 | 1226.0714 | NaN | NaN | NaN | NaN | NaN |
| Psmd3 | ELDTVTLEDIK | -62 | -72 | 0 | 0 | 11 | 3777.9658 | 4632.3745 | 4467.1 | 239.16272 | 4446.1685 | 255.01732 |
| Psmd3 | SVFPEQANNNEWAR | -270 | -283 | 0 | 0 | 14 | 1392.7294 | 1421.4142 | 1404.8784 | 1746.1339 | 1760.5328 | 1646.6426 |
| Psmd3 | EREQQDLEFAK | -509 | -519 | 1 | 0 | 11 | 1998.1648 | NaN | 333.4643 | 3020.2683 | 3262.681 | 3268.6707 |
| Psmd3 | EQQDLEFAK | -511 | -519 | 0 | 0 | 9 | 2694.1245 | 2753.2317 | 1117.1791 | NaN | NaN | 981.15326 |
| Psmd3 | LQLDSPEDAEFIVAK | -422 | -436 | 0 | 0 | 15 | 1067.1285 | 1173.1168 | 1258.6346 | NaN | 1452.6359 | NaN |
| Psmd3 | EPQLAFHQR | -467 | -475 | 0 | 0 | 9 | 2327.116 | 1817.2631 | 2254.079 | NaN | NaN | NaN |
| Psmd3 | KAPQHTAVGFK | -311 | -321 | 1 | 0 | 11 | 2012.9868 | 2250.7273 | 686.0877 | NaN | NaN | NaN |
| Psmd3 | AIQLEYSEAR | -293 | -302 | 0 | 0.000471409 | 10 | 980.88513 | NaN | NaN | NaN | 2667.1409 | 2503.2322 |
| Psmd3 | VYEFLDKL | -213 | -220 | 1 | 0.002265552 | 8 | 258.4978 | NaN | 314.30347 | NaN | NaN | 277.1717 |
| H2ac20;Hist2h | AIRNDEELNKLLGK | (87);(87) | (100);(100) | 2 | 0.000230707 | 14 | 2310.589 | 831.55225 | 560.7197 | 836.28174 | 252.74211 | 1032.4524 |
| H2ac20;Hist2h | NDEELNKLLGKVT | (90);(90) | (102);(102) | 2 | 0.000907206 | 13 | 1024.7635 | 1161.4697 | 857.7764 | 721.8049 | 1136.4207 | 921.9452 |
| Mta1 | VWEAHNPLVDK | -198 | -208 | 0 | 0 | 11 | 724.0668 | 703.3895 | 744.2872 | 531.86066 | 1065.1763 | 753.7106 |
| Mta1 | QLESLPATHIR | -113 | -123 | 0 | 0 | 11 | 443.93448 | 516.56165 | 497.4106 | 554.4015 | 557.2711 | 538.6515 |
| Mta1 | YQADITDLLK | -173 | -182 | 0 | 0 | 10 | 1494.4183 | 1748.7394 | 1590.0394 | NaN | 1971.8174 | 2042.2031 |
| Mta1 | AISALVPQGGPVLCR | -268 | -282 | 0 | 0.000431655 | 15 | 528.0272 | NaN | NaN | 364.36383 | NaN | NaN |
| Zpr1 | AISGLEQDQPTR | -153 | -164 | 0 | 0 | 12 | 545.0167 | 800.03064 | 712.9225 | 1215.9291 | 855.2422 | 777.2457 |
| Zpr1 | NPFTLGDSSNPDQSEK | -368 | -383 | 0 | 0 | 16 | 1211.9009 | NaN | NaN | 879.1046 | 877.96075 | 905.4364 |
| Nop58 | MDGLIPGVEPR | -123 | -133 | 0 | 0 | 11 | 2639.631 | 2685.307 | 2528.5522 | 1166.7902 | 1419.5691 | 1644.8467 |
| Nop58 | MDGLIPGVEPR | -123 | -133 | 0 | 0 | 11 | 2639.631 | 2685.307 | 2528.5522 | 1166.7902 | 1419.5691 | 1644.8467 |
| Nop58 | IEEVDKEDEITEK | -445 | -457 | 1 | 0 | 13 | 1911.7722 | 1780.2548 | 2655.1982 | 5144.8286 | 5655.46 | 5587.5503 |
| Nop58 | IISDNLTYCK | -197 | -206 | 0 | 0 | 10 | 1551.815 | 1515.6838 | 1588.8646 | 1823.6107 | 767.6965 | 616.1005 |
| Nop58 | TYDPSGDSTLPTCSK | -427 | -441 | 0 | 0 | 15 | 1543.7719 | 1793.0078 | 1741.4241 | 164.66643 | 1966.5178 | 1937.6377 |
| Nop58 | KIEEVDKEDEITEK | -444 | -457 | 2 | 0.00025015 | 14 | 1457.83 | 1530.2576 | 1708.096 | 2330.7417 | 3005.34 | 3081.5527 |
| Nop58 | KIEEVDKEDEITEK | -444 | -457 | 2 | 0.00025015 | 14 | 1457.83 | 1530.2576 | 1708.096 | 2330.7417 | 3005.34 | 3081.5527 |
| Nop58 | WKEFETPEK | -29 | -37 | 1 | 0 | 9 | 1085.3202 | 249.35312 | 1211.1766 | 1581.0471 | 1483.6619 | 433.92654 |
| Nop58 | AISLLDDLDK | -165 | -174 | 0 | 0 | 10 | 1058.065 | 1249.3782 | 1247.9879 | 1202.2738 | 1347.732 | 1302.4225 |
| Nop58 | LQEVDSLWK | -22 | -30 | 0 | 0 | 9 | 1027.5063 | 1070.8448 | 1193.6671 | 945.7337 | 1266.7251 | 1168.2507 |
| Nop58 | LNLSCIHSPVVNEL | -102 | -115 | 0 | 0.000230707 | 14 | 1016.4493 | 1086.5209 | 1069.7738 | 1245.3777 | 1557.8302 | 1579.6067 |
| Nop58 | LYEYLQNR | -271 | -278 | 0 | 0 | 8 | 546.1126 | 1482.9833 | 1959.1775 | 2178.3853 | 2100.898 | 1988.1621 |
| Nop58 | EAEEEQVVEEEPTVK | -478 | -492 | 0 | 0 | 15 | 421.219 | 537.15326 | 466.58844 | 512.9509 | 548.2324 | 562.23083 |
| Nop58 | EVDKEDEITEK | -447 | -457 | 1 | 0 | 11 | 210.43176 | 854.49286 | 893.877 | 453.26263 | 408.23724 | 845.0459 |
| Nop58 | HIKEEPLSEEEPCTST | -502 | -517 | 1 | 0.000550307 | 16 | 10170.531 | 12591.658 | 10551.057 | NaN | 12561.454 | 10603.95 |
| Nop58 | TQLYEYLQNR | -269 | -278 | 0 | 0 | 10 | 3370.1562 | 2691.533 | 2304.5803 | 3634.479 | NaN | 3658.0784 |
| Nop58 | EWYGWHFPELGK | -185 | -196 | 0 | 0 | 12 | 485.20798 | NaN | 686.5954 | 475.70624 | 1017.1665 | 943.87476 |
| Nop58 | KLQEVDSLWK | -21 | -30 | 1 | 0.000230707 | 10 | 768.32153 | 717.0567 | 817.19086 | NaN | NaN | NaN |
| Nop58 | KIEEVDKEDEIT | -444 | -455 | 2 | 0.005112649 | 12 | 795.89044 | NaN | NaN | NaN | NaN | NaN |
| Cdv3 | KTPQGPPEIYSDTQFPSLQ | -204 | -222 | 1 | 0 | 19 | 516.9801 | 532.0374 | 674.25635 | 674.6611 | 759.43036 | 844.00745 |
| Cdv3 | EDPGDNWEEGGGGSGAEK | -142 | -159 | 0 | 0 | 18 | 512.2655 | 475.21918 | 491.27347 | 496.66162 | 467.06738 | 467.33487 |
| Cdv3 | AITKDENEWK | -102 | -111 | 1 | 0 | 10 | 1862.8607 | 385.72995 | 1322.5051 | NaN | 2353.449 | 2131.1929 |
| Cdv3 | PGDGGSLGSGAR | -46 | -57 | 0 | 0 | 12 | 836.46466 | 359.0213 | 930.6286 | NaN | 1045.2977 | 418.2824 |
| Cdv3 | KTPQGPPEIY | -204 | -213 | 1 | 0.00025015 | 10 | 1254.6335 | 1288.9135 | NaN | NaN | NaN | NaN |
| Cdv3 | KTPQGPPEIYSDT | -204 | -216 | 1 | 0.003521747 | 13 | 534.67395 | NaN | NaN | NaN | NaN | NaN |
| Psma7 | ILNPEEIEK | -219 | -227 | 0 | 0 | 9 | 3480.0562 | 3486.8877 | 3863.3904 | 2596.3962 | 2909.4878 | 2767.5662 |
| Psma7 | NYTDDAIETDDLTIK | -175 | -189 | 0 | 0 | 15 | 2631.6768 | 3187.765 | 3333.0806 | 3489.6025 | 137.36185 | 4064.3818 |
| Psma7 | IVGFDFDGTPR | -133 | -143 | 0 | 0 | 11 | 226.75552 | 2269.4475 | 2196.5784 | 2274.1768 | 1752.1222 | 247.56245 |
| Psma7 | YTDDAIETDDLTIK | -176 | -189 | 0 | 0 | 14 | 2100.702 | 2466.0764 | NaN | 2487.528 | 2896.486 | NaN |

| | | | | | | | | | | | | |
|---|---|---|---|---|---|---|---|---|---|---|---|---|
| Ppa1 | VDDPDAANYK | -161 | -170 | 0 | 0 | 10 | 1975.5153 | 1334.8562 | 1881.732 | 331.65912 | 2052.3945 | 1453.2192 |
| Ppa1 | TWEDPGHSDK | -100 | -109 | 0 | 0 | 10 | 1964.1938 | 2149.568 | 2026.6598 | 1506.2478 | 2175.8699 | 3439.2915 |
| Ppa1 | NKDFAVDIIK | -212 | -221 | 1 | 0 | 10 | 1759.6636 | 1892.4878 | 2023.0029 | 2045.385 | 2256.9844 | 2257.3342 |
| Ppa1 | GAIPQTWEDPGHSDK | -95 | -109 | 0 | 0 | 15 | 1288.8301 | 1417.4021 | 1291.263 | 1567.4497 | 1365.5872 | 1343.0652 |
| Ppa1 | YKVPDGKPENEFAFNA | -193 | -208 | 2 | 0.00025015 | 16 | 1171.8016 | 1356.557 | 1440.1125 | 1497.3368 | 1833.103 | 2020.4309 |
| Ppa1 | YKVPDGKPENEFAFNA | -193 | -208 | 2 | 0.00025015 | 16 | 1171.8016 | 1356.557 | 1440.1125 | 1497.3368 | 1833.103 | 2020.4309 |
| Ppa1 | NTTVSESPFK | -244 | -253 | 0 | 0 | 10 | 1088.8121 | 1621.2797 | 1563.3657 | 1153.1232 | 569.7538 | 1157.3427 |
| Ppa1 | HTGCCGDNDPIDVCEIGSK | -110 | -128 | 0 | 0 | 19 | 914.01794 | 783.3592 | 908.97125 | 953.8282 | 1126.5875 | 1136.4692 |
| Ppa1 | HTGCCGDNDPIDVCEIGSK | -110 | -128 | 0 | 0 | 19 | 914.01794 | 783.3592 | 908.97125 | 953.8282 | 1126.5875 | 1136.4692 |
| Ppa1 | HTGCCGDNDPIDVC | -110 | -123 | 0 | 0 | 14 | 719.9373 | 743.2665 | 662.5869 | 535.13586 | 411.612 | 365.04245 |
| Ppa1 | VIAINVDDPDAANYK | -156 | -170 | 0 | 0 | 15 | 710.22034 | 860.09955 | 825.72314 | 653.663 | 596.77386 | 621.31476 |
| Ppa1 | GQYISPFHDVPIYADK | -26 | -41 | 0 | 0 | 16 | 672.37634 | 988.37836 | 856.91516 | 1959.0608 | 2313.7332 | 2327.951 |
| Ppa1 | NYGAIPQTWEDPGHSDK | -93 | -109 | 0 | 0 | 17 | 663.9974 | 686.28375 | 718.1302 | 679.8489 | 773.3859 | 806.1891 |
| Ppa1 | AIVDALPPPCESACSLPTDVDK | -261 | -282 | 0 | 0 | 22 | 419.83054 | 465.02344 | 406.66815 | 602.92535 | 839.9656 | 774.5988 |
| Ppa1 | ALPPPCESACSLPTDVDK | -265 | -282 | 0 | 0.000627943 | 18 | 559.4645 | NaN | 155.61812 | 935.482 | 898.82086 | 974.40155 |
| Ppa1 | HDVPIYADK | -33 | -41 | 0 | 0 | 9 | 440.00644 | 1307.0536 | 1833.4368 | NaN | 786.2693 | 477.36295 |
| Ppa1 | IDEGETDWK | -147 | -155 | 0 | 0 | 9 | 1456.5918 | 1450.1204 | NaN | 1737.4915 | NaN | NaN |
| Ppa1 | AMIDEGETDWK | -145 | -155 | 0 | 0 | 11 | 1053.1473 | NaN | 970.968 | NaN | NaN | NaN |
| Ppa1 | DFAVDIIK | -214 | -221 | 0 | 0 | 8 | 473.87488 | NaN | 427.9788 | NaN | NaN | NaN |
| Ddx39b | NCPHIVVGTPGR | -164 | -175 | 0 | 0 | 12 | 2979.8892 | 3200.8665 | 3149.2168 | 3359.3005 | 3722.4285 | 3522.6345 |
| Ddx39b | NCPHIVVGTPGR | -164 | -175 | 0 | 0 | 12 | 2979.8892 | 3200.8665 | 3149.2168 | 3359.3005 | 3722.4285 | 3522.6345 |
| Ddx39b | HFILDECDK | -192 | -200 | 0 | 0 | 9 | 2208.8599 | 2452.3289 | 2488.2588 | 1717.0231 | 2122.3696 | 2072.8975 |
| Ddx39b | LDYEDDEVETAAGADGTEAPAK | -11 | -32 | 0 | 0.00025015 | 22 | 126.22969 | 175.30551 | 339.79602 | NaN | 241.9325 | NaN |
| Bin2 | ASPDASSNPEPAETR | -442 | -456 | 0 | 0 | 15 | 1519.734 | 1781.2856 | 1887.8021 | 723.3989 | 1621.0482 | 1704.2837 |
| Bin2 | DWDGHEDLK | -92 | -100 | 0 | 0 | 9 | 1266.5492 | 255.28123 | 1173.0028 | 434.61304 | 446.77188 | 1109.8082 |
| Bin2 | EGAGSSGPEEPR | -459 | -470 | 0 | 0 | 12 | 1119.9176 | 465.5197 | 794.4267 | 395.4919 | 664.0088 | 543.58344 |
| Bin2 | PSPGDVVLR | -379 | -387 | 0 | 0.00025015 | 9 | 615.857 | 1208.4949 | 1016.3662 | 1191.6223 | 307.03537 | 238.63309 |
| Bin2 | EKEGAGSSGPEEPR | -457 | -470 | 1 | 0.00025015 | 14 | 546.9379 | 676.5803 | 287.03766 | 511.1926 | 611.3703 | 625.55133 |
| Bin2 | AIVGNNDLLWEDYEEK | -101 | -116 | 0 | 0.000230707 | 16 | 532.22144 | 670.7629 | 625.04425 | 342.64374 | 477.69855 | 684.3513 |
| Bin2 | ASGSGSCNAPGSPEGSSQLCSPR | -419 | -441 | 0 | 0.00025015 | 23 | 306.99643 | 428.81818 | 367.8579 | 427.82272 | 411.56122 | 539.4578 |
| Bin2 | NNDLLWEDYEEK | -105 | -116 | 0 | 0 | 12 | 118.53342 | 420.71646 | 591.8876 | 889.43604 | 1085.2054 | 1228.1582 |
| Bin2 | DTPSPGDVVLR | -377 | -387 | 0 | 0 | 11 | 1247.5684 | 508.50574 | 548.1616 | NaN | 444.50586 | 538.469 |
| Hnrnpul1 | HLPSTEPDPHVVR | -272 | -284 | 0 | 0 | 13 | 588.57904 | 764.19385 | 784.9878 | 687.8263 | 1017.29553 | 721.9019 |
| Hnrnpul1 | HLPSTEPDPHVVR | -272 | -284 | 0 | 0 | 13 | 588.57904 | 764.19385 | 784.9878 | 687.8263 | 1017.29553 | 721.9019 |
| Hnrnpul1 | QNQFYETPVIK | -107 | -117 | 0 | 0 | 11 | 1370.3109 | 1390.4532 | 1309.2333 | 1340.2009 | 1797.9138 | 309.04562 |
| Hnrnpul1 | AIVICPTDEDLKDR | -529 | -542 | 1 | 0 | 14 | 1059.009 | 937.92004 | 917.41614 | 859.68933 | 1046.6263 | 970.6162 |
| Hnrnpul1 | AIVICPTDEDLKDR | -529 | -542 | 1 | 0 | 14 | 1059.009 | 937.92004 | 917.41614 | 859.68933 | 1046.6263 | 970.6162 |
| Hnrnpul1 | STQLGEEPFSYGYGGTGK | -293 | -310 | 0 | 0 | 18 | 589.5314 | 1196.8619 | 689.01587 | 7726.332 | 1639.8896 | 1271.8094 |
| Hnrnpul1 | RTDEEGKDVPDHAVLE | -546 | -561 | 2 | 0.002068213 | 16 | 555.29785 | 379.864 | 550.63525 | NaN | 277.7623 | 342.48456 |
| Hnrnpul1 | ALYPHVLVK | -368 | -376 | 0 | 0.000471409 | 9 | 1157.1226 | NaN | NaN | NaN | 1657.5588 | 1453.561 |
| Hnrnpul1 | TDEEGKDVPDHAVLE | -547 | -561 | 1 | 0.001392869 | 15 | 577.1888 | NaN | 767.6924 | NaN | 127.45592 | NaN |
| Hnrnpul1 | HLPLSER | -403 | -409 | 0 | 0.006417541 | 7 | 1167.8254 | 1053.1149 | NaN | NaN | NaN | NaN |
| Hnrnpul1 | AVEFNFGQR | -379 | -387 | 0 | 0.00025015 | 9 | 1023.89124 | NaN | NaN | NaN | 963.78503 | NaN |
| Psmc1 | SKENVLYK | -422 | -429 | 1 | 0 | 8 | 1010.0505 | 1030.8711 | 1010.8502 | 899.50134 | 1070.0355 | 1276.3544 |
| Psmc1 | AKDDLSGADIK | -386 | -396 | 1 | 0.000550307 | 11 | 988.32056 | 947.4665 | 956.3499 | 1229.0588 | 1434.0022 | 1208.4856 |
| Psmc1 | IGGLDNQIQEIK | -187 | -198 | 0 | 0 | 12 | 937.2138 | 1034.7018 | 1131.9207 | 859.5692 | 1406.258 | 1222.9476 |
| Psmc1 | IETLDPALIRPGR | -334 | -346 | 1 | 0 | 13 | 905.8286 | 1027.5104 | 1037.1104 | 819.365 | 813.91125 | 752.2945 |
| Psmc1 | IETLDPALIRPGR | -334 | -346 | 1 | 0 | 13 | 905.8286 | 1027.5104 | 1037.1104 | 819.365 | 813.91125 | 752.2945 |
| Psmc1 | KIEFPLPDEK | -350 | -359 | 1 | 0 | 10 | 832.71136 | 1352.6714 | 1367.7136 | 1531.4329 | 857.5826 | 1333.1321 |
| Psmc1 | APQETYADIGGLDNQIQEIK | -179 | -198 | 0 | 0 | 20 | 662.50116 | 719.6025 | 734.16376 | 921.15137 | 399.75827 | 781.5133 |
| Psmc1 | SKVDDLR | -97 | -103 | 1 | 0.00025015 | 7 | 288.99228 | 269.91425 | 291.61084 | 237.47476 | 397.01144 | 260.09216 |
| Psmc1 | KQEGTPEGLYL | -430 | -440 | 1 | 0.002534621 | 11 | 842.9692 | 3494.4922 | NaN | 323.4037 | 1306.4315 | NaN |
| Psmc1 | KYEPPVPTR | -24 | -32 | 1 | 0.00025015 | 9 | 1718.0746 | 1356.5986 | 1169.5404 | NaN | NaN | NaN |
| Psmc1 | VVGSELIQK | -250 | -258 | 0 | 0 | 9 | 1104.5594 | NaN | 595.7333 | 538.43445 | NaN | NaN |
| Psmc1 | LPLVTPHTQ | -49 | -57 | 0 | 0.00025015 | 9 | 977.3367 | NaN | 834.1275 | NaN | NaN | NaN |
| Eef1g | GLLDTHLK | -140 | -147 | 0 | 0 | 8 | 8163.6196 | 2715.7356 | 4272.779 | 3352.607 | 3203.011 | 3933.9285 |
| Eef1g | KLDPGSEETQTLVR | -401 | -414 | 1 | 0 | 14 | 5656.8804 | 6159.5737 | 5800.216 | 6941.193 | 7221.651 | 6817.789 |
| Eef1g | KLDPGSEETQTLVR | -401 | -414 | 1 | 0 | 14 | 5656.8804 | 6159.5737 | 5800.216 | 6941.193 | 7221.651 | 6817.789 |
| Eef1g | STFVLDEFKR | -286 | -295 | 1 | 0 | 10 | 5079.5576 | 4064.506 | 4341.901 | 5820.1167 | 6009.367 | 396.74356 |
| Eef1g | AKDPFAHLPK | -276 | -285 | 1 | 0 | 10 | 4873.984 | 4760.9263 | 4478.669 | 7759.1396 | 5349.3145 | 4516.8345 |
| Eef1g | AKDPFAHLPK | -276 | -285 | 1 | 0 | 10 | 4873.984 | 4760.9263 | 4478.669 | 7759.1396 | 5349.3145 | 4516.8345 |
| Eef1g | YTYPENWR | -7 | -14 | 0 | 0 | 8 | 4193.334 | 639.8841 | 4847.284 | 4542.2305 | 4183.8 | 4306.482 |
| Eef1g | SAPPHFHFGQTNR | -33 | -45 | 0 | 0 | 13 | 3641.7979 | 3908.4658 | 3658.4016 | 3672.2178 | 4324.188 | 4174.2593 |
| Eef1g | SAPPHFHFGQTNR | -33 | -45 | 0 | 0 | 13 | 3641.7979 | 3908.4658 | 3658.4016 | 3672.2178 | 4324.188 | 4174.2593 |
| Eef1g | FWEHFDK | -310 | -316 | 0 | 0 | 7 | 3275.2593 | 3461.8706 | 3371.1545 | 3605.9495 | 4168.667 | 4206.5854 |
| Eef1g | QVLEPSFR | -174 | -181 | 0 | 0 | 8 | 3185.9946 | 2943.6726 | 2954.314 | 2788.201 | 1864.6664 | 1906.2756 |
| Eef1g | LDPGSEETQTLVR | -402 | -414 | 0 | 0 | 13 | 3161.3462 | 3315.8303 | 3478.5417 | 3286.9917 | 3444.59 | 3564.3904 |
| Eef1g | ILGLLDTHLK | -138 | -147 | 0 | 0 | 10 | 3077.2673 | 3965.919 | 3877.0088 | 1772.5779 | 3533.4473 | 3775.007 |
| Eef1g | YVSNEELR | -78 | -85 | 0 | 0 | 8 | 2577.0632 | 2254.736 | 1835.8834 | 1484.4662 | 1171.6752 | 1667.9285 |
| Eef1g | KYSNEDTLSVALPY | -296 | -309 | 1 | 0.001055699 | 14 | 1799.0258 | 2358.3877 | 2314.808 | 2411.2407 | 3128.8147 | 3177.5586 |
| Eef1g | FVLDEFKR | -288 | -295 | 1 | 0.00025015 | 8 | 1469.3167 | 1474.8595 | 332.6264 | 327.34323 | 2984.9536 | 314.27838 |
| Eef1g | DPFAHLPK | -278 | -285 | 0 | 0 | 8 | 1412.0184 | 1543.1812 | 1123.7866 | 1738.1667 | 1811.991 | 1640.4241 |
| Eef1g | QAFPNTNR | -182 | -189 | 0 | 0 | 8 | 1336.5347 | 1070.1517 | 566.8612 | 1260.5784 | 2437.295 | 1370.2036 |
| Eef1g | EYFSWEGTFQHVGK | -415 | -428 | 0 | 0 | 14 | 1207.6958 | 1391.5912 | 1504.9886 | 2979.0864 | 3626.7366 | 3566.265 |
| Eef1g | LYTYPENWR | -6 | -14 | 0 | 0 | 9 | 1009.0885 | 1063.5906 | 542.18713 | 1486.5518 | 926.01587 | 647.1083 |
| Eef1g | VPAFEGDDGFCVF | -58 | -70 | 0 | 0 | 13 | 979.073 | 1116.566 | 1147.7917 | 1551.8354 | 1780.5616 | 1571.0828 |
| Eef1g | FVLDEFK | -288 | -294 | 0 | 0.000431655 | 7 | 699.31366 | 877.0255 | 743.02637 | 756.8088 | 783.90607 | 794.9934 |
| Eef1g | PFAHLPK | -279 | -285 | 0 | 0.003299042 | 7 | 550.0273 | 2836.8567 | 865.61456 | 2979.6077 | 207.78961 | 1013.0728 |

| | | | | | | | | | | | | |
|---|---|---|---|---|---|---|---|---|---|---|---|---|
| Eef1g | SPDWQVDYESYTWR | -387 | -400 | 0 | 0 | 14 | 544.7113 | 530.03516 | 572.8423 | 596.8954 | 724.4752 | 579.1957 |
| Eef1g | KAAAPAPEEEMDECEQAL | -253 | -270 | 1 | 0 | 18 | 326.93997 | 690.7512 | 772.5567 | 814.0033 | 879.6253 | 589.99945 |
| Eef1g | RTPEFLR | -45 | -51 | 1 | 0.005242018 | 7 | 158.04776 | 4647.782 | 4804.08 | 3590.2073 | 4667.421 | 4781.6157 |
| Eef1g | KLDPGSEETQTL | -401 | -412 | 1 | 0.000627943 | 12 | 3019.1362 | 2332.5757 | 3012.7627 | NaN | 221.33328 | 3487.8066 |
| Eef1g | KFAESQPK | -220 | -227 | 1 | 0.00078661 | 8 | 2396.9014 | 1973.9473 | 2514.6238 | 2243.2998 | 2339.9036 | NaN |
| Eef1g | STFVLDEFK | -286 | -294 | 0 | 0 | 9 | 1792.3137 | 1873.2009 | 1983.7921 | NaN | 1582.1749 | 1767.2312 |
| Eef1g | TLLWLYK | -167 | -173 | 0 | 0.002466863 | 7 | 1610.484 | 1940.5566 | 1888.4775 | NaN | 1800.0654 | 1770.692 |
| Eef1g | KAAAPAPEEEMDECEQA | -253 | -269 | 1 | 0.000712251 | 17 | 314.02792 | 232.47975 | NaN | 387.5125 | 412.78476 | 483.40967 |
| Eef1g | SISGVWVFR | -370 | -378 | 0 | 0 | 9 | 579.44476 | 426.3991 | 601.465 | 428.6636 | NaN | NaN |
| Eef1g | INQPQFR | -195 | -201 | 0 | 0.004031417 | 7 | 270.26822 | 1447.8679 | 1275.5101 | NaN | 735.0646 | NaN |
| Eef1g | KAAAPAPEEEMDECEQ | -253 | -268 | 1 | 0.00272025 | 16 | 218.8766 | 320.2387 | 276.2641 | NaN | NaN | NaN |
| Eef1g | VLSAPPHFHFGQTNR | -31 | -45 | 0 | 0 | 15 | 3553.1885 | 1301.1647 | NaN | NaN | NaN | NaN |
| Ezr | SQEQLAAELAEYTAK | -413 | -427 | 0 | 0 | 15 | 2532.5955 | 1616.3083 | 2808.452 | 2591.511 | 2862.4373 | 2967.922 |
| Ezr | RKEDEVEEWQHR | -437 | -448 | 2 | 0 | 12 | 1207.5524 | 1541.7109 | 1651.7777 | 902.1191 | 130.01924 | 1736.8228 |
| Ezr | RKEDEVEEWQHR | -437 | -448 | 2 | 0 | 12 | 1207.5524 | 1541.7109 | 1651.7777 | 902.1191 | 130.01924 | 1736.8228 |
| Ezr | RKEDEVEEWQHR | -437 | -448 | 2 | 0 | 12 | 1207.5524 | 1541.7109 | 1651.7777 | 902.1191 | 130.01924 | 1736.8228 |
| Ezr | AKEAQDDLVK | -449 | -458 | 1 | 0 | 10 | 1020.81226 | 585.5542 | 1213.6904 | 1120.1366 | 1531.7861 | 1470.1405 |
| Ezr | KEDEVEEWQHR | -438 | -448 | 1 | 0 | 11 | 818.9893 | 836.90546 | 808.1492 | 1101.0209 | 679.1357 | 908.2567 |
| Ezr | KEDEVEEWQHR | -438 | -448 | 1 | 0 | 11 | 818.9893 | 836.90546 | 808.1492 | 1101.0209 | 679.1357 | 908.2567 |
| Ezr | EQLAAELAEYTAK | -415 | -427 | 0 | 0 | 13 | 353.9791 | 557.08307 | 512.5495 | 820.58374 | 767.37256 | 753.9638 |
| Ezr | EDEVEEWQHR | -439 | -448 | 0 | 0 | 10 | 300.3957 | 1065.5594 | 371.25278 | 821.0129 | 929.5232 | 977.60956 |
| Ezr | KENPVQFK | -72 | -79 | 1 | 0 | 8 | 2040.5233 | 1987.8651 | 1895.8937 | 1798.2225 | NaN | 1430.4777 |
| Ezr | ELSSEGILDDR | -502 | -512 | 0 | 0 | 11 | 1105.5493 | 752.8917 | 922.64294 | NaN | 862.141 | 851.3982 |
| Ezr | THNDIIHNENMR | -548 | -559 | 0 | 0 | 12 | 827.83765 | 483.0938 | 79.49807 | NaN | 746.80164 | 764.18414 |
| Ezr | AQEEAERLEADR | -382 | -393 | 1 | 0 | 12 | 735.2002 | NaN | 740.12305 | 651.30634 | NaN | NaN |
| Prkacb | DLKPENLLIDHQ | -167 | -178 | 1 | 0 | 12 | 685.25586 | 790.8445 | 779.8955 | 930.7489 | 758.44385 | 1332.3381 |
| Prkacb | QAVEFPFLVR | -97 | -106 | 0 | 0 | 10 | 363.90036 | 592.70355 | 484.51288 | 748.246 | 945.02216 | 851.724 |
| Prkacb | AKEDFLR | -23 | -29 | 1 | 0 | 7 | 1034.512 | 669.4697 | 617.52435 | 1109.3645 | 482.8667 | NaN |
| Prkacb | AVEFPFLVR | -98 | -106 | 0 | 0 | 9 | 430.8647 | 494.4802 | 481.72968 | NaN | 546.36993 | NaN |
| Prkacb | VEFPFLVR | -99 | -106 | 0 | 0 | 8 | 1256.1763 | NaN | NaN | 62.709282 | 247.62065 | NaN |
| Gng2 | EDPLLTPVPASENPFR | -47 | -62 | 0 | 0 | 16 | 172.90155 | 1002.2263 | 1053.1024 | 970.198 | 1270.3594 | 1155.4967 |
| Gng2 | AKEDPLLTPVPASENPFR | -45 | -62 | 1 | 0 | 18 | 331.69296 | 352.4427 | NaN | 437.9915 | 446.8087 | 559.0702 |
| Rrp7a | EEEGVPDEEGWVK | -193 | -205 | 0 | 0 | 13 | 608.404 | 873.78864 | 642.4172 | 996.0866 | 741.8958 | 612.1224 |
| Rrp7a | TVELQEKPDLAESPTEPK | -87 | -104 | 1 | 0.000550307 | 18 | 886.6766 | 994.0246 | 1004.7273 | NaN | 866.5641 | NaN |
| Rrp7a | AKEEEGVPDEEGWVK | -191 | -205 | 1 | 0 | 15 | 327.63443 | 623.34796 | 503.04465 | NaN | NaN | NaN |
| Cotl1 | WIGEDVSGLQR | -81 | -91 | 0 | 0 | 11 | 4500.3706 | 4579.695 | 4687.562 | 666.26746 | 5383.849 | 4861.646 |
| Cotl1 | KELEEDFIR | -118 | -126 | 1 | 0 | 9 | 3973.182 | 4159.091 | 4235.016 | 4749.811 | 5100.61 | 4991.888 |
| Cotl1 | ELEEDFIR | -119 | -126 | 0 | 0 | 8 | 3318.404 | 3466.7786 | 3493.4653 | 593.0229 | 3174.5488 | 3310.8354 |
| Cotl1 | YDGATIVPGDQGADYQHFIQ | -31 | -50 | 0 | 0 | 20 | 1291.1844 | 1439.903 | 1588.4574 | 1779.6677 | 2104.8022 | 1950.1687 |
| Cotl1 | YDGATIVPGDQGADYQHF | -31 | -48 | 0 | 0 | 18 | 956.8507 | 1103.4093 | 1114.0509 | 1078.4275 | 1174.2483 | 1219.1567 |
| Cotl1 | YDGATIVPGDQGADYQHFIQQ | -31 | -51 | 0 | 0 | 21 | 729.3492 | 747.21204 | 789.3905 | 946.0593 | 1207.9875 | 1117.7833 |
| Cotl1 | AKEFVISDR | -109 | -117 | 1 | 0.00078661 | 9 | 1592.2689 | NaN | NaN | NaN | NaN | NaN |
| Hspa5 | VMEHFIK | -263 | -269 | 0 | 0 | 7 | 7661.929 | 8466.047 | 7081.0615 | 1465.3801 | 1308.8187 | 4678.656 |
| Hspa5 | VMEHFIK | -263 | -269 | 0 | 0 | 7 | 7661.929 | 8466.047 | 7081.0615 | 1465.3801 | 1308.8187 | 4678.656 |
| Hspa5 | VYEGERPLTK | -466 | -475 | 1 | 0.000550307 | 10 | 7303.622 | 7714.4336 | 7647.392 | 6194.359 | 7310.873 | 7274.483 |
| Hspa5 | FEELNMDLFR | -328 | -337 | 0 | 0 | 10 | 6999.7188 | 6244.9565 | 6727.4253 | 2383.0466 | 1300.2749 | 1356.447 |
| Hspa5 | FEELNMDLFR | -328 | -337 | 0 | 0 | 10 | 6999.7188 | 6244.9565 | 6727.4253 | 2383.0466 | 1300.2749 | 1356.447 |
| Hspa5 | NQLTSNPENTVFDAK | -83 | -97 | 0 | 0 | 15 | 6116.401 | 6041.8345 | 6330.3267 | 6065.491 | 5126.0747 | 4961.591 |
| Hspa5 | DIDEIVLVGGSTR | -356 | -368 | 0 | 0 | 13 | 4759.43 | 540.91437 | 4636.9883 | 5657.456 | 4986.652 | 5083.9614 |
| Hspa5 | KSDIDEIVLVGGSTR | -354 | -368 | 1 | 0 | 15 | 4157.846 | 4892.1475 | 4821.5493 | 3395.0146 | 3329.6143 | 3483.2708 |
| Hspa5 | KSDIDEIVLVGGSTR | -354 | -368 | 1 | 0 | 15 | 4157.846 | 4892.1475 | 4821.5493 | 3395.0146 | 3329.6143 | 3483.2708 |
| Hspa5 | ITITNDQNR | -525 | -533 | 0 | 0 | 9 | 4118.3306 | 3979.562 | 2049.5073 | 2974.833 | 1639.3801 | 1636.8495 |
| Hspa5 | LYGSGGPPPTGEEDTSEKDEL | -635 | -655 | 1 | 0 | 21 | 3997.144 | 5097.996 | 5047.4434 | 2331.2708 | 3357.239 | 3397.689 |
| Hspa5 | VVTVPAYFNDAQR | -170 | -182 | 0 | 0 | 13 | 3363.4937 | 4112.874 | 4049.4333 | 5309.848 | 5774.4175 | 5720.212 |
| Hspa5 | ELEEIVQPIISK | -623 | -634 | 0 | 0 | 12 | 3229.22 | 3949.5312 | 3850.6655 | 3770.6719 | 4305.34 | 4496.9272 |
| Hspa5 | ELEEIVQPIISK | -623 | -634 | 0 | 0 | 12 | 3229.22 | 3949.5312 | 3850.6655 | 3770.6719 | 4305.34 | 4496.9272 |
| Hspa5 | TWNDPSVQQDIK | -103 | -114 | 0 | 0 | 12 | 2964.4893 | 3248.07 | 3320.8706 | 1838.0062 | 1998.6392 | 2070.5908 |
| Hspa5 | SFFEGEDFSETLTR | -312 | -325 | 0 | 0 | 14 | 2953.145 | 3356.6174 | 3471.453 | 3366.5193 | 4682.942 | 4591.3486 |
| Hspa5 | YVAFTPEGER | -66 | -75 | 0 | 0 | 10 | 2695.8953 | 2514.589 | 2468.8577 | 2775.5193 | 2022.3353 | 2077.4539 |
| Hspa5 | ITPSYVAFTPEGER | -62 | -75 | 0 | 0 | 14 | 2637.4417 | 2606.2244 | 2646.2666 | 2693.3347 | 2706 | 2739.8064 |
| Hspa5 | VDIGGGQTK | -131 | -139 | 0 | 0.00025015 | 9 | 1803.0292 | 1427.9752 | 1328.1552 | 7006.5654 | 1256.4562 | 1627.8071 |
| Hspa5 | ATNGDTHLGGEDFDQR | -247 | -262 | 0 | 0 | 16 | 1655.2184 | 3840.5483 | 470.31717 | 488.2286 | 529.6325 | 5156.1865 |
| Hspa5 | STASDNQPTVTIK | -453 | -465 | 0 | 0 | 13 | 1625.3497 | 1547.6078 | 1597.7983 | 2272.491 | 1814.2177 | 1782.5886 |
| Hspa5 | IEWLESHQDADIEDFK | -603 | -618 | 0 | 0 | 16 | 1625.0066 | 1708.8018 | 1954.0817 | 1737.3815 | 217.28679 | 2386.0056 |
| Hspa5 | IEWLESHQDADIEDFK | -603 | -618 | 0 | 0 | 16 | 1625.0066 | 1708.8018 | 1954.0817 | 1737.3815 | 217.28679 | 2386.0056 |
| Hspa5 | VLEDSDLKK | -346 | -354 | 1 | 0 | 9 | 1549.0726 | 1589.2294 | 1606.5261 | 1452.8727 | 1468.8241 | 1587.5168 |
| Hspa5 | KIEWLESHQDADIEDFK | -602 | -618 | 1 | 0 | 17 | 1465.3455 | 1429.4335 | 1880.99 | 1920.9169 | 1321.9895 | 460.26282 |
| Hspa5 | SHQDADIEDFK | -608 | -618 | 0 | 0 | 11 | 1460.7018 | 1404.0299 | 1349.0931 | 1439.7667 | 1494.8148 | 1478.9303 |
| Hspa5 | SHQDADIEDFK | -608 | -618 | 0 | 0 | 11 | 1460.7018 | 1404.0299 | 1349.0931 | 1439.7667 | 1494.8148 | 1478.9303 |
| Hspa5 | TKPYIQVDIGGGQTK | -125 | -139 | 1 | 0 | 15 | 1346.6195 | 1682.3837 | 1703.0957 | 638.2015 | 821.0472 | 742.73663 |
| Hspa5 | TKPYIQVDIGGGQTK | -125 | -139 | 1 | 0 | 15 | 1346.6195 | 1682.3837 | 1703.0957 | 638.2015 | 821.0472 | 742.73663 |
| Hspa5 | AKFEELNMDLFR | -326 | -337 | 1 | 0 | 12 | 1252.2622 | 1554.3054 | 1625.0867 | 1735.01 | 2151.6777 | 702.7682 |
| Hspa5 | AKFEELNMDLFR | -326 | -337 | 1 | 0 | 12 | 1252.2622 | 1554.3054 | 1625.0867 | 1735.01 | 2151.6777 | 702.7682 |
| Hspa5 | KKELEEIVQPIISK | -621 | -634 | 2 | 0.00025015 | 14 | 1057.393 | 602.73883 | 539.46405 | 409.55087 | 1156.2181 | 709.6028 |
| Hspa5 | DNHLLGTFDLT | -476 | -486 | 0 | 0 | 11 | 1012.52313 | 1050.7035 | 1129.3634 | 949.60236 | 1069.491 | 1128.3317 |
| Hspa5 | GGPPPTGEEDTSEKDEL | -639 | -655 | 1 | 0 | 17 | 1012.19904 | 1115.8956 | 1175.3788 | 2605.147 | 2766.753 | 614.6547 |
| Hspa5 | PSVQQDIK | -107 | -114 | 0 | 0.000230707 | 8 | 846.44806 | 1025.8438 | 1177.3794 | 1092.5164 | 1514.6077 | 1499.1532 |
| Hspa5 | QDADIEDFK | -610 | -618 | 0 | 0 | 9 | 738.57574 | 889.2217 | 384.943 | 669.6644 | 559.24133 | 465.6823 |

| | | | | | | | | | | | | |
|---|---|---|---|---|---|---|---|---|---|---|---|---|
| Hspa5 | SDIDEIVLVGGSTR | -355 | -368 | 0 | 0 | 14 | 409.13937 | 1327.0665 | 295.3191 | 1287.8671 | 1149.5878 | 1059.113 |
| Hspa5 | YEGERPLTK | -467 | -475 | 1 | 0 | 9 | 380.82474 | 594.8691 | 754.9429 | 441.7841 | 560.84937 | 773.5696 |
| Hspa5 | GDTHLGGEDFDQR | -250 | -262 | 0 | 0 | 13 | 365.64948 | 433.7859 | 2403.6653 | 533.05646 | 2667.9006 | 197.86172 |
| Hspa5 | LTPEEIER | -534 | -541 | 0 | 0.000712251 | 8 | 3736.6782 | 3563.9128 | 3610.331 | 212.45204 | 2516.2324 | NaN |
| Hspa5 | ASDNQPTVTIK | -455 | -465 | 0 | 0 | 11 | 1816.0789 | NaN | 1891.0037 | 1959.2208 | 2092.8225 | 1532.3126 |
| Hspa5 | NGDTHLGGEDFDQR | -249 | -262 | 0 | 0 | 14 | 1022.0494 | 410.40353 | 1021.6168 | 488.35907 | 1079.1819 | NaN |
| Hspa5 | IESFFEGEDFSETLTR | -310 | -325 | 0 | 0 | 16 | 587.617 | 534.49634 | 439.39423 | NaN | 644.54877 | 653.43286 |
| Hspa5 | NDPSVQQDIK | -105 | -114 | 0 | 0 | 10 | 3505.281 | 2561.8423 | 2398.112 | 2773.3704 | NaN | NaN |
| Hspa5 | AVVTVPAYFNDAQR | -169 | -182 | 0 | 0 | 14 | 1303.6821 | 1709.3727 | NaN | 1762.4626 | NaN | 2299.2683 |
| Hspa5 | EFFNGKEPSR | -378 | -387 | 1 | 0.001868731 | 10 | 1257.9293 | 1143.8799 | NaN | 1805.4647 | NaN | 705.78503 |
| Hspa5 | VTHAVVTVPAYFNDAQR | -166 | -182 | 0 | 0 | 17 | 554.40643 | 202.51358 | 539.6668 | NaN | 1132.6163 | NaN |
| Hspa5 | SDNQPTVTIK | -456 | -465 | 0 | 0.00025015 | 10 | 4650.0786 | 4637.821 | 383.95435 | NaN | NaN | NaN |
| Hspa5 | WLESHQDADIEDFK | -605 | -618 | 0 | 0.00025015 | 14 | 466.98828 | 368.92072 | 440.21524 | NaN | NaN | NaN |
| Hspa5 | WLESHQDADIEDFK | -605 | -618 | 0 | 0.00025015 | 14 | 466.98828 | 368.92072 | 440.21524 | NaN | NaN | NaN |
| Hspa5 | VTVPAYFNDAQR | -171 | -182 | 0 | 0.001291103 | 12 | 717.6659 | 1023.2478 | NaN | NaN | NaN | NaN |
| Hspa5 | YFNDAQR | -176 | -182 | 0 | 0.007523243 | 7 | 515.9484 | 529.8254 | NaN | NaN | NaN | NaN |
| Grpel1 | EEISNNNPHLK | -128 | -138 | 0 | 0 | 11 | 1363.6967 | 1381.9902 | 1427.2432 | 1125.3997 | 1520.6586 | 1454.501 |
| Grpel1 | FDPYEHEALFHTPVEGK | -170 | -186 | 0 | 0 | 17 | 657.1337 | 678.74884 | 586.80804 | 699.84375 | 605.9532 | 879.648 |
| Grpel1 | FDPYEHEALFHTPVEGK | -170 | -186 | 0 | 0 | 17 | 657.1337 | 678.74884 | 586.80804 | 699.84375 | 605.9532 | 879.648 |
| Grpel1 | DLLEVADILEK | -110 | -120 | 0 | 0 | 11 | 446.65628 | 431.95953 | 523.89685 | 509.96674 | 534.5081 | 554.7659 |
| Grpel1 | ALADTENLR | -81 | -89 | 0 | 0 | 9 | 1188.6218 | 1093.9939 | 1128.1829 | 856.22595 | 1045.0543 | NaN |
| Grpel1 | AKLEEQLR | -65 | -72 | 1 | 0.000230707 | 8 | 606.2881 | 763.53015 | NaN | 885.2188 | 896.0468 | 878.3661 |
| Grpel1 | FDPYEHEAL | -170 | -178 | 0 | 0.005605646 | 9 | 2063.5085 | NaN | NaN | NaN | 5174.4873 | 1369.2279 |
| Mapre1 | KLTVEDLEK | -204 | -212 | 1 | 0 | 9 | 1690.4447 | 1800.5857 | 2221.587 | 2419.9844 | 2103.607 | 2609.6792 |
| Mapre1 | STSVTSDNLSR | -7 | -17 | 0 | 0 | 11 | 1153.8811 | 1275.3317 | 960.28394 | 917.5982 | 1926.5201 | 873.78296 |
| Mapre1 | LEHEYIQNFK | -67 | -76 | 0 | 0 | 10 | 1117.369 | 1270.5161 | 1221.766 | 1701.212 | 971.83075 | 950.4761 |
| Mapre1 | LEHEYIQNFK | -67 | -76 | 0 | 0 | 10 | 1117.369 | 1270.5161 | 1221.766 | 1701.212 | 971.83075 | 950.4761 |
| Mapre1 | NIELICQENEGENDPVLQR | -223 | -241 | 0 | 0 | 19 | 894.37103 | 1037.5334 | 967.7831 | 828.2939 | 975.1742 | 986.9374 |
| Mapre1 | ESLQLNLTK | -26 | -34 | 0 | 0 | 9 | 829.5199 | 795.0585 | 856.76807 | 804.4918 | 232.33545 | 717.682 |
| Mapre1 | QENEGENDPVLQR | -229 | -241 | 0 | 0 | 13 | 776.81024 | 793.5751 | 866.04474 | 861.0429 | 725.17737 | 942.4104 |
| Mapre1 | AKLEHEYIQNFK | -65 | -76 | 1 | 0 | 12 | 1047.755 | NaN | 592.5 | 1031.3329 | 1070.5082 | 1042.9558 |
| Mapre1 | LVAPALSKPK | -141 | -150 | 1 | 0 | 10 | 334.80835 | NaN | NaN | NaN | NaN | NaN |
| Plec | RPELEDSTLR | -659 | -668 | 1 | 0.002068213 | 10 | 4088.7085 | 4461.1353 | 5329.9146 | 4505.337 | 1002.68475 | 5077.425 |
| Plec | SGFLLDPVR | -2880 | -2888 | 0 | 0 | 9 | 3494.573 | 2490.046 | 395.97842 | 677.2786 | 1563.7202 | 1327.8796 |
| Plec | YLTGGLIEPDTPGR | -4536 | -4549 | 0 | 0 | 14 | 1637.348 | 2256.591 | 424.593 | 1679.4154 | 2269.8037 | 2311.6375 |
| Plec | LAAEQELIR | -1854 | -1862 | 0 | 0 | 9 | 1478.3801 | 564.08594 | 1904.4408 | 674.19794 | 591.12305 | 815.0693 |
| Plec | RQELEAELAK | -1895 | -1904 | 1 | 0 | 10 | 1038.9591 | 986.6728 | 907.62134 | 903.8299 | 1295.1923 | 1264.4996 |
| Plec | GLVGPELHDR | -3895 | -3904 | 0 | 0 | 10 | 961.06085 | 15717.742 | 1060.6592 | 10441.1875 | 1890.4158 | 20213.723 |
| Plec | AKLEQLFQDEVAK | -2659 | -2671 | 1 | 0 | 13 | 762.7191 | 844.9649 | 1033.8046 | 801.53845 | 904.23505 | 675.72815 |
| Plec | VLALPEPSPAAPTLR | -1195 | -1209 | 0 | 0 | 15 | 602.53815 | 573.9124 | 510.5646 | 413.01923 | 464.17526 | 451.0936 |
| Plec | RAPVPASELLDAK | -3453 | -3465 | 1 | 0 | 13 | 492.7486 | 854.85864 | 1066.5496 | 1189.9907 | 1242.5636 | 1122.8146 |
| Plec | LPVDVAYQR | -3627 | -3635 | 0 | 0 | 9 | 385.71732 | 849.5989 | 868.539 | 707.91785 | 501.9206 | 786.1495 |
| Plec | VSLDEALQR | -4550 | -4558 | 0 | 0 | 9 | 384.1552 | 736.0888 | 823.3965 | 416.60547 | 699.30084 | 759.76263 |
| Plec | SQVEEELFSVR | -2369 | -2379 | 0 | 0 | 11 | 326.09277 | 590.6198 | 802.34064 | 737.51245 | 769.9333 | 760.5081 |
| Plec | QLLEEELAR | -1875 | -1883 | 0 | 0.000591124 | 9 | 1114.2843 | 1177.2202 | 1200.7588 | 1047.0453 | 1204.0542 | NaN |
| Plec | DTHDQLSEPSEVR | -3976 | -3988 | 0 | 0 | 13 | 839.6215 | 835.9955 | 1195.7404 | NaN | 218.67113 | 621.1859 |
| Plec | RLEEQAALHK | -2024 | -2033 | 1 | 0 | 10 | 794.5847 | 814.0457 | 787.18 | 406.98828 | NaN | 641.1059 |
| Plec | VPLDVAYAR | -3296 | -3304 | 0 | 0.000230707 | 9 | 768.0369 | NaN | 978.57007 | 714.5413 | 703.36066 | 961.12415 |
| Plec | VPGAQDGVR | -413 | -421 | 0 | 0 | 9 | 751.7889 | 515.0081 | 548.73785 | 706.7903 | NaN | 421.28168 |
| Plec | KQEELQHLEQQR | -2714 | -2725 | 1 | 0 | 12 | 723.7858 | 1044.469 | 719.88074 | NaN | 882.54785 | 892.76196 |
| Plec | SKEQAELEAAR | -2106 | -2116 | 1 | 0 | 11 | 567.47943 | 912.96375 | 424.23203 | 619.6871 | NaN | 517.2599 |
| Plec | LLDPEDVDVPQPDEK | -383 | -397 | 0 | 0 | 15 | 564.6758 | 658.8799 | 694.36316 | NaN | 707.4183 | 167.65814 |
| Plec | LGFHLPLEVA | -3958 | -3967 | 0 | 0.000471409 | 10 | 544.4157 | 567.33435 | 657.2025 | NaN | 763.43384 | 871.1645 |
| Plec | LPLDKDPAR | -1154 | -1162 | 1 | 0 | 9 | 1908.5992 | NaN | 1478.8508 | 2691.6155 | 1876.1069 | NaN |
| Plec | QLAEEDLAQQR | -2440 | -2450 | 0 | 0 | 11 | 1901.8683 | 588.532 | 2364.853 | 1972.5413 | NaN | NaN |
| Plec | FPVTEAVNK | -4474 | -4482 | 0 | 0.00025015 | 9 | 814.2018 | NaN | 759.594 | NaN | 650.7084 | 576.1275 |
| Plec | ALVPAAELLDSGVISHE | -3123 | -3139 | 0 | 0 | 17 | 435.43066 | NaN | 308.34622 | 402.34348 | NaN | 521.76337 |
| Plec | LATGGIVDPR | -3948 | -3957 | 0 | 0.002434616 | 10 | 1535.1714 | 1809.2556 | 1760.8264 | NaN | NaN | NaN |
| Plec | LVEETQGFQR | -2502 | -2511 | 0 | 0.000230707 | 10 | 675.71985 | 778.80896 | NaN | 479.78174 | NaN | NaN |
| Plec | IVITVVEEHER | -3102 | -3112 | 0 | 0 | 11 | 180.40176 | NaN | NaN | NaN | 1934.7391 | 1940.3307 |
| Plec | GGELVYTDTEAR | -3035 | -3046 | 0 | 0.007987481 | 12 | 4935.6104 | NaN | NaN | 2604.346 | NaN | NaN |
| Plec | ELIPAEEALR | -3933 | -3942 | 0 | 0.001846793 | 10 | 2528.9932 | NaN | NaN | NaN | NaN | NaN |
| Plec | LQLEETDHQK | -2338 | -2347 | 0 | 0.00294445 | 10 | 610.53815 | NaN | NaN | NaN | NaN | NaN |
| Ncl | ETLEEVFEK | -501 | -509 | 0 | 0.000230707 | 9 | 5326.3906 | 5584.573 | 5863.998 | 4143.7983 | 4320.4966 | 4498.051 |
| Ncl | ETLEEVFEK | -501 | -509 | 0 | 0.000230707 | 9 | 5326.3906 | 5584.573 | 5863.998 | 4143.7983 | 4320.4966 | 4498.051 |
| Ncl | KVIPTPGK | -102 | -109 | 1 | 0 | 8 | 7473.922 | 7794.4077 | 7413.639 | 7661.154 | 7597.2397 | 7987.8687 |
| Ncl | KETLEEVFEK | -500 | -509 | 1 | 0 | 10 | 5585.8105 | 5583.198 | 601.24915 | 7632.412 | 8519.537 | 7817.477 |
| Ncl | KETLEEVFEK | -500 | -509 | 1 | 0 | 10 | 5585.8105 | 5583.198 | 601.24915 | 7632.412 | 8519.537 | 7817.477 |
| Ncl | SFNITEDELK | -403 | -412 | 0 | 0 | 10 | 4264.9077 | 4787.616 | 4487.9673 | 3827.968 | 2549.4958 | 3313.0127 |
| Ncl | VTLDWAKPK | -637 | -645 | 1 | 0 | 9 | 3844.1477 | 3969.2456 | 4281.0176 | 6789.8345 | 7450.24 | 7819.5083 |
| Ncl | GLSEDTTEETLK | -575 | -586 | 0 | 0 | 12 | 3835.0366 | 3903.492 | 3569.6777 | 3232.8396 | 193.0942 | 170.40578 |
| Ncl | GLSEDTTEETLK | -575 | -586 | 0 | 0 | 12 | 3835.0366 | 3903.492 | 3569.6777 | 3232.8396 | 193.0942 | 170.40578 |
| Ncl | GFGFVDFNSEEDAK | -608 | -621 | 0 | 0 | 14 | 3557.1367 | 4053.1553 | 4066.2524 | 2417.945 | 2924.8105 | 3119.247 |
| Ncl | GLSEDTTEETLKESFEGSVR | -575 | -594 | 1 | 0 | 20 | 476.46155 | 522.0269 | 479.7056 | 522.44495 | 696.80927 | 796.68207 |
| Ncl | GLSEDTTEETLKESFEGSVR | -575 | -594 | 1 | 0 | 20 | 476.46155 | 522.0269 | 479.7056 | 522.44495 | 696.80927 | 796.68207 |
| Ncl | NDLAVVDVR | -336 | -344 | 0 | 0 | 9 | 3090.6655 | 2477.999 | 2986.754 | 2854.731 | 1546.7761 | 2491.181 |
| Ncl | SEDTTEETLK | -577 | -586 | 0 | 0 | 10 | 2975.147 | 3007.1287 | 3492.2378 | 3313.2825 | 3506.5913 | 3994.2283 |
| Ncl | SEDTTEETLK | -577 | -586 | 0 | 0 | 10 | 2975.147 | 3007.1287 | 3492.2378 | 3313.2825 | 3506.5913 | 3994.2283 |

| | | | | | | | | | | | | |
|---|---|---|---|---|---|---|---|---|---|---|---|---|
| Ncl | NLFIGNLNPNK | -310 | -320 | 0 | 0 | 11 | 2922.9958 | 3647.7163 | 3050 | 2896.3608 | 4300.3003 | 3771.523 |
| Ncl | KFGYVDFESAEDLEK | -350 | -364 | 1 | 0 | 15 | 2886.4844 | 3336.333 | 3530.115 | 4435.803 | 5017.2036 | 4853.006 |
| Ncl | KFGYVDFESAEDLEK | -350 | -364 | 1 | 0 | 15 | 2886.4844 | 3336.333 | 3530.115 | 4435.803 | 5017.2036 | 4853.006 |
| Ncl | EVFEDAMEIR | -413 | -422 | 0 | 0 | 10 | 2801.8433 | 2431.959 | 2388.2788 | 3042.1167 | 1080.2225 | 499.56125 |
| Ncl | EVFEDAMEIR | -413 | -422 | 0 | 0 | 10 | 2801.8433 | 2431.959 | 2388.2788 | 3042.1167 | 1080.2225 | 499.56125 |
| Ncl | TKETLEEVFEK | -499 | -509 | 1 | 0 | 11 | 2319.1692 | 1331.7703 | 1530.3135 | 2092.6294 | 1495.0359 | 1719.9813 |
| Ncl | KFAISELFAK | -326 | -335 | 1 | 0.001055699 | 10 | 2005.5839 | 2189.2188 | 2386.2212 | 800.1529 | 2082.7227 | 2492.1133 |
| Ncl | FAISELFAK | -327 | -335 | 0 | 0 | 9 | 1972.3776 | 2014.7035 | 2240.3655 | 2051.5063 | 3032.5544 | 3152.0073 |
| Ncl | SEDTTEETLKESFEGSVR | -577 | -594 | 1 | 0 | 18 | 1741.1135 | 2112.1648 | 2150.0854 | 2388.7505 | 3754.9695 | 3294.161 |
| Ncl | SEDTTEETLKESFEGSVR | -577 | -594 | 1 | 0 | 18 | 1741.1135 | 2112.1648 | 2150.0854 | 2388.7505 | 3754.9695 | 3294.161 |
| Ncl | FIGNLNPNK | -312 | -320 | 0 | 0.00025015 | 9 | 1611.867 | 1178.0764 | 1172.6039 | 1094.5297 | 1395.369 | 1140.5726 |
| Ncl | KVTLDWAKPK | -636 | -645 | 2 | 0.00051306 | 10 | 1305.6057 | 1189.423 | 1415.1665 | 1034.5558 | 1415.2927 | 1151.1235 |
| Ncl | FGYVDFESAEDLEK | -351 | -364 | 0 | 0 | 14 | 1297.8601 | 1570.3793 | 1546.7555 | 2689.213 | 3158.6104 | 3191.1572 |
| Ncl | FKSEADAEK | -438 | -446 | 1 | 0.000431655 | 9 | 1248.7986 | 181.6687 | 1329.7197 | 1125.875 | 615.94495 | 1325.7358 |
| Ncl | IEFASFEDAK | -529 | -538 | 0 | 0 | 10 | 1193.0568 | 1441.0892 | 1244.7919 | 1814.0707 | 1965.127 | 2014.1958 |
| Ncl | KVEGSEPTTPFNLFIGNLNPNK | -299 | -320 | 1 | 0 | 22 | 1090.8252 | 1082.1495 | 1090.4906 | 1584.1152 | 2198.414 | 2253.3657 |
| Ncl | KVEGSEPTTPFNLFIGNLNPNK | -299 | -320 | 1 | 0 | 22 | 1090.8252 | 1082.1495 | 1090.4906 | 1584.1152 | 2198.414 | 2253.3657 |
| Ncl | FASFEDAK | -531 | -538 | 0 | 0 | 8 | 1074.3798 | 1102.7797 | 850.8998 | 451.6224 | 830.8696 | 305.67746 |
| Ncl | VDFNSEEDAK | -612 | -621 | 0 | 0 | 10 | 921.35236 | 1046.6431 | 4519.547 | 3856.6882 | 168.45674 | 5115.885 |
| Ncl | GFVDFNSEEDAK | -610 | -621 | 0 | 0 | 12 | 736.3944 | 2874.6433 | 3116.5688 | 733.3799 | 596.7194 | 4124.947 |
| Ncl | GFVDFNSEEDAK | -610 | -621 | 0 | 0 | 12 | 736.3944 | 2874.6433 | 3116.5688 | 733.3799 | 596.7194 | 4124.947 |
| Ncl | GYVDFESAEDLEK | -352 | -364 | 0 | 0 | 13 | 632.30994 | 667.0827 | 717.03534 | 351.4884 | 936.8786 | 890.01965 |
| Ncl | IKVPQNPHGK | -513 | -522 | 1 | 0 | 10 | 632.02386 | 600.8681 | 750.6868 | 799.7942 | 1001.59314 | 957.4909 |
| Ncl | IKVPQNPHGK | -513 | -522 | 1 | 0 | 10 | 632.02386 | 600.8681 | 750.6868 | 799.7942 | 1001.59314 | 957.4909 |
| Ncl | VEGSEPTTPFNLFIGNLNPNK | -300 | -320 | 0 | 0 | 21 | 528.0949 | 558.6294 | 588.7473 | 456.2457 | 641.58154 | 624.2273 |
| Ncl | QGAEIDGR | -452 | -459 | 0 | 0.000230707 | 8 | 506.25714 | 2870.3228 | 3099.165 | 2692.2383 | 3115.9214 | 759.8439 |
| Ncl | VPQNPHGKPK | -515 | -524 | 1 | 0.00025015 | 10 | 469.7 | 292.36118 | 506.89017 | 428.0908 | 437.6279 | 396.41602 |
| Ncl | ETLEEVFEKA | -501 | -510 | 1 | 0.001636916 | 10 | 414.59064 | 2808.2114 | 790.2666 | 155.59593 | 935.4439 | 229.70195 |
| Ncl | ETLEEVFEKA | -501 | -510 | 1 | 0.001636916 | 10 | 414.59064 | 2808.2114 | 790.2666 | 155.59593 | 935.4439 | 229.70195 |
| Ncl | GSEPTTPFNLFIGNLNPNK | -302 | -320 | 0 | 0.000230707 | 19 | 297.72012 | 231.75635 | 274.76346 | 294.31573 | 336.62387 | 387.6529 |
| Ncl | GIAYIEFK | -432 | -439 | 0 | 0 | 8 | 857.0827 | 659.7569 | 628.44073 | 830.2903 | 844.6466 | NaN |
| Ncl | KAAVPTPAK | -71 | -79 | 1 | 0.00025015 | 9 | 844.6144 | 388.63895 | 909.407 | NaN | 654.7637 | 646.11707 |
| Ncl | AKNDLAVVDVR | -334 | -344 | 1 | 0 | 11 | 679.10205 | 690.2268 | NaN | 1126.405 | 1369.5085 | 1205.3043 |
| Ncl | QKVEGSEPTTPFNLFIGN | -298 | -315 | 1 | 0.003681885 | 18 | 476.3983 | 725.9809 | NaN | 803.2525 | 592.8418 | 617.6149 |
| Ncl | QKVEGSEPTTPFNLFIG | -298 | -314 | 1 | 0.007343498 | 17 | 354.24756 | NaN | 410.49847 | 293.10785 | 392.7718 | 369.93298 |
| Ncl | SVSLYYTGEK | -460 | -469 | 0 | 0.000431655 | 10 | 2922.3445 | NaN | NaN | 862.6168 | 1375.9453 | 1215.4047 |
| Ncl | NLSFNITEDELK | -401 | -412 | 0 | 0.00025015 | 12 | 634.8068 | NaN | 768.2937 | 400.2309 | NaN | 289.00247 |
| Ncl | FESAEDLEK | -356 | -364 | 0 | 0 | 9 | 284.8864 | 2015.6095 | 1780.8359 | NaN | NaN | 1218.703 |
| Ncl | EDELKEVFEDAMEIR | -408 | -422 | 1 | 0.00025015 | 15 | 260.54132 | NaN | NaN | 471.87402 | 539.478 | 479.86295 |
| Ncl | EDELKEVFEDAMEIR | -408 | -422 | 1 | 0.00025015 | 15 | 260.54132 | NaN | NaN | 471.87402 | 539.478 | 479.86295 |
| Ncl | IKVPQNPHGKPK | -513 | -524 | 2 | 0 | 12 | 259.4631 | NaN | NaN | 375.56158 | 372.7443 | 372.06903 |
| Ncl | LELQGSNSR | -556 | -564 | 0 | 0.00025015 | 9 | 2278.6838 | 1432.7368 | NaN | 1140.937 | NaN | NaN |
| Ncl | KEDSDEDEDEEDEDDSDEDEDDEEEDEFEPPIVK | -142 | -175 | 1 | 0.007288197 | 34 | 992.8274 | 938.3765 | NaN | 880.63074 | NaN | NaN |
| Ncl | VPQNPHGKPKG | -515 | -525 | 2 | 0.005605646 | 11 | 689.31726 | NaN | 688.506 | NaN | NaN | 706.75116 |
| Ncl | SEDEDDSSGEEEVVIPQK | -34 | -51 | 0 | 0 | 18 | 660.5773 | 612.506 | 609.2468 | NaN | NaN | NaN |
| Ncl | QKVEGSEPTTPFNLFIGNLNPNK | -298 | -320 | 1 | 0 | 23 | 589.71716 | 479.71725 | 503.70013 | NaN | NaN | NaN |
| Ncl | EDDEDDDEEEEEEEPVK | -260 | -276 | 0 | 0.001023236 | 17 | 344.44427 | 349.65268 | 348.1527 | NaN | NaN | NaN |
| Ncl | DDEDDDEEEEEEEPVK | -261 | -276 | 0 | 0 | 16 | 316.95285 | 387.21027 | 364.2696 | NaN | NaN | NaN |
| Ncl | EDDSSGEEEVVIPQKK | -37 | -52 | 1 | 0 | 16 | 292.91495 | 678.82983 | 661.1749 | NaN | NaN | NaN |
| Ncl | SEDEDDSSGEEEVVIPQKK | -34 | -52 | 1 | 0.002329279 | 19 | 282.33545 | 229.79118 | 363.03177 | NaN | NaN | NaN |
| Ncl | FNSEEDAK | -614 | -621 | 0 | 0 | 8 | 254.07248 | NaN | 425.05502 | NaN | 279.06046 | NaN |
| Ncl | QKVEGSEPTTPFNLF | -298 | -312 | 1 | 0.001636916 | 15 | 352.3189 | NaN | 552.551 | NaN | NaN | NaN |
| Ncl | QKVEGSEPTTPF | -298 | -309 | 1 | 0.00272025 | 12 | 374.76276 | NaN | NaN | NaN | NaN | NaN |
| Cs | SVLDPDEGIR | -83 | -92 | 0 | 0 | 10 | 2694.4114 | 2289.6091 | 379.82837 | 2234.2139 | 1949.5637 | 2118.8828 |
| Cs | IVPNILLEQGK | -383 | -393 | 0 | 0 | 11 | 1537.7052 | 1600.318 | 1532.2015 | 1741.0713 | 1861.1638 | 1799.1531 |
| Cs | ALNSESNFAR | -174 | -183 | 0 | 0 | 10 | 1508.0186 | 453.18777 | 513.0353 | 1485.0713 | 855.78644 | 473.44995 |
| Cs | VTGQMPTEEQVSWLSR | -124 | -139 | 0 | 0 | 16 | 1243.5199 | 1304.3379 | 1290.5276 | 1530.8027 | 1363.4738 | 1229.4866 |
| Cs | VTGQMPTEEQVSWLSR | -124 | -139 | 0 | 0 | 16 | 1243.5199 | 1304.3379 | 1290.5276 | 1530.8027 | 1363.4738 | 1229.4866 |
| Cs | EGSSIGAIDSR | -223 | -233 | 0 | 0 | 11 | 1156.328 | 1263.6676 | 1003.2626 | 1739.2527 | 781.6243 | 549.65137 |
| Cs | VLDPDEGIR | -84 | -92 | 0 | 0 | 9 | 1135.5168 | 1131.8469 | 931.91095 | 1094.4584 | 624.9963 | 877.27454 |
| Cs | VYETSVLDPDEGIR | -79 | -92 | 0 | 0 | 14 | 805.6232 | 914.902 | 1366.4774 | 1274.3745 | 1589.4696 | 1539.6034 |
| Cs | VVPGYGHAVLR | -341 | -351 | 0 | 0.00025015 | 11 | 795.8627 | 830.94885 | 908.072 | 321.83487 | 629.4951 | 221.08875 |
| Cs | NPWPNVDAHSGVLL | -396 | -409 | 0 | 0 | 14 | 678.45044 | 716.45935 | 89.344666 | 145.208 | 282.32462 | 773.39966 |
| Cs | AKNPWPNVDAHSGVLL | -394 | -409 | 1 | 0 | 16 | 437.5899 | 577.5171 | 526.9413 | 422.75644 | 616.74054 | 709.09375 |
| Cs | ALGFPLERPK | -441 | -450 | 1 | 0 | 10 | 427.00916 | 3023.9807 | 3073.5474 | 4068.2524 | 4666.822 | 4918.687 |
| Cs | GQMPTEEQVSWLSR | -126 | -139 | 0 | 0 | 14 | 316.30014 | 560.7654 | 608.3908 | 639.7719 | 735.6185 | 514.64374 |
| Cs | AKNPWPNVDAHSGVLLQ | -394 | -410 | 1 | 0.000907206 | 17 | 925.8776 | NaN | 1340.9772 | 906.49084 | 249.74396 | 224.45486 |
| Cs | AKNPWPNVDAHSGVLLQ | -394 | -410 | 1 | 0.000907206 | 17 | 925.8776 | NaN | 1340.9772 | 906.49084 | 249.74396 | 224.45486 |
| Cs | ETSVLDPDEGIR | -81 | -92 | 0 | 0 | 12 | 1392.1906 | 1419.6442 | NaN | 2105.8171 | 1375.6414 | 1447.0856 |
| Cs | NPWPNVDAHSGVLLQ | -396 | -410 | 0 | 0.000550307 | 15 | 855.33655 | NaN | 1045.0679 | 1046.8955 | 1157.427 | 1226.7797 |
| Cs | ALGFPLER | -441 | -448 | 0 | 0 | 8 | 117.576675 | 1053.5311 | 1223.7891 | NaN | 305.06824 | 626.292 |
| Cs | NPWPNVDAH | -396 | -404 | 0 | 0 | 9 | 770.10486 | NaN | NaN | 241.75156 | 772.57776 | NaN |
| Cs | IYEDCMDLIAK | -198 | -208 | 0 | 0.009569685 | 11 | 625.9933 | NaN | 1178.7726 | NaN | NaN | NaN |
| Cs | TVLFGVSR | -421 | -428 | 0 | 0.003166358 | 8 | 504.77002 | NaN | NaN | NaN | NaN | NaN |
| Eif3b | DAEEDAEAGPEPEVR | -47 | -61 | 0 | 0 | 15 | 1659.4235 | 1868.302 | 1732.0781 | 1823.3705 | 1896.8358 | 1968.9989 |
| Eif3b | LEYASPAHAVDAVK | -224 | -237 | 0 | 0 | 14 | 1546.3057 | 2187.4666 | 2741.2437 | 3770.487 | 3397.0908 | 4049.2349 |
| Eif3b | FAVLHGEAPR | -566 | -575 | 0 | 0 | 10 | 1411.4879 | 1108.6228 | 3108.1987 | 3100.6172 | 1372.8168 | 2831.1768 |
| Eif3b | NLFNVVDCK | -497 | -505 | 0 | 0 | 9 | 1383.6765 | 1491.8098 | 1467.5254 | 1424.0665 | 1670.9457 | 1540.6562 |

| | | | | | | | | | | | | |
|---|---|---|---|---|---|---|---|---|---|---|---|---|
| Eif3b | IINDYYPEEDGK | -206 | -217 | 0 | 0 | 12 | 1017.2582 | 412.66745 | 1111.8086 | 1330.6199 | 906.96747 | 937.3976 |
| Eif3b | WNDVKDPVSIEER | -309 | -321 | 1 | 0 | 13 | 895.4213 | 1157.4551 | 1034.8073 | 813.3984 | 983.9496 | 943.26624 |
| Eif3b | WNDVKDPVSIEER | -309 | -321 | 1 | 0 | 13 | 895.4213 | 1157.4551 | 1034.8073 | 813.3984 | 983.9496 | 943.26624 |
| Eif3b | SEEETATSPAASPTPQSAER | -68 | -87 | 0 | 0 | 20 | 875.0541 | 753.64746 | 1043.5807 | 449.9154 | 1159.8718 | 1246.9054 |
| Eif3b | DVEWDPTGR | -645 | -653 | 0 | 0 | 9 | 842.4583 | 758.5131 | 797.7489 | 664.58325 | 80.965866 | 621.3205 |
| Eif3b | SHQGVQLIDFSPCER | -361 | -375 | 0 | 0 | 15 | 611.68774 | 718.89374 | 778.6344 | 647.34924 | 775.00885 | 743.82166 |
| Eif3b | SHQGVQLIDFSPCER | -361 | -375 | 0 | 0 | 15 | 611.68774 | 718.89374 | 778.6344 | 647.34924 | 775.00885 | 743.82166 |
| Eif3b | GHPSAGAEEEGGSDGSAA | -108 | -125 | 0 | 0 | 18 | 429.5195 | 350.99667 | 354.90686 | 406.8341 | 382.72092 | 452.42377 |
| Eif3b | SSAHWPIFK | -411 | -419 | 0 | 0.00025015 | 9 | 3752.538 | NaN | 3922.336 | 2455.0776 | 2668.9832 | 2572.6975 |
| Eif3b | WVPEDKDIPAR | -471 | -481 | 1 | 0 | 11 | 1948.9526 | NaN | 1832.9064 | 3121.9824 | 526.79156 | 2642.3 |
| Eif3b | SVSWWSHK | -658 | -665 | 0 | 0 | 8 | 1696.3429 | NaN | 1895.8104 | 2371.8057 | 2706.0186 | 2575.1023 |
| Eif3b | YWLEEAECR | -284 | -292 | 0 | 0 | 9 | 659.6414 | 819.7299 | 856.9863 | 229.71753 | NaN | 911.63806 |
| Eif3b | ASPAHAVDAVK | -227 | -237 | 0 | 0 | 11 | 4951.7466 | 1756.9972 | 3272.3994 | NaN | NaN | NaN |
| Eif3b | AVLHGEAPR | -567 | -575 | 0 | 0.000989247 | 9 | 984.65643 | NaN | NaN | 1011.48724 | 849.68396 | NaN |
| Eif3b | PRPPTLL | -695 | -701 | 1 | 0.001392869 | 7 | 934.68536 | NaN | 783.8556 | 592.3219 | NaN | NaN |
| Eif3b | TSIFWNDVK | -305 | -313 | 0 | 0.00025015 | 9 | 485.65784 | 357.43466 | 283.7917 | NaN | NaN | NaN |
| Eif3b | AKPAAQSEEETATSPAASPTPQSAER | -62 | -87 | 1 | 0 | 26 | 382.70096 | 432.5327 | 421.69714 | NaN | NaN | NaN |
| Eif3b | MQDAENVAVPEAAEER | -1 | -16 | 0 | 0.002329279 | 16 | 259.58972 | 421.74957 | NaN | NaN | 193.04442 | NaN |
| Eif3b | QQPASESPPTDEAAGSGGSEVGQTE | -22 | -46 | 0 | 0.002168954 | 25 | 192.61356 | 238.19914 | 257.11444 | NaN | NaN | NaN |
| Eif3b | DVSEEELLGDVLK | -151 | -163 | 0 | 0 | 13 | 1072.3262 | NaN | 1032.7958 | NaN | NaN | NaN |
| Eif3b | NVPQVGPDR | -180 | -188 | 0 | 0.001291103 | 9 | 1019.24603 | NaN | NaN | 849.41223 | NaN | NaN |
| Dlst | TPAFAESVTEGDVR | -76 | -89 | 0 | 0.000431655 | 14 | 2398.606 | 2248.4478 | 401.83325 | 2635.657 | 2461.9988 | 2476.3193 |
| Dlst | VEGGTPLFTLR | -135 | -145 | 0 | 0 | 11 | 1172.513 | 1142.1672 | 247.60336 | 828.14185 | 1055.433 | 1138.7509 |
| Dlst | AKPAETPAPAHK | -154 | -165 | 1 | 0.00025015 | 12 | 240.46863 | 227.37044 | 356.0361 | 226.90584 | NaN | 195.36934 |
| Dlst | PPLAEAGAAK | -209 | -218 | 0 | 0.002329279 | 10 | 1226.7662 | NaN | NaN | 931.9378 | NaN | NaN |
| Dlst | GLVVPVIR | -327 | -334 | 0 | 0.003299042 | 8 | 216.26828 | NaN | NaN | NaN | NaN | NaN |
| Rrp9 | LWQCGEGFR | -402 | -410 | 0 | 0 | 9 | 2273.0042 | 457.82278 | 2150.9336 | 854.49786 | 685.99963 | 730.9172 |
| Rrp9 | AKPAPGPGAAPVAGK | -11 | -25 | 1 | 0.00078661 | 15 | 175.327 | 1582.2559 | 1501.5334 | 2790.1582 | 2571.0278 | 3163.3323 |
| Rrp9 | EIQAPAPTDIR | -130 | -140 | 0 | 0 | 11 | 2634.8762 | 2687.5427 | 2837.7607 | 3214.0898 | 1618.1927 | NaN |
| Rrp9 | LKEDVLEQR | -112 | -120 | 1 | 0.00025015 | 9 | 1076.9917 | 1148.264 | 1121.1842 | 1159.8606 | NaN | 1267.3167 |
| Rrp9 | NSVCIIPLR | -457 | -465 | 0 | 0.00025015 | 9 | 304.94836 | 371.96912 | 309.24063 | NaN | 601.8464 | NaN |
| Rrp9 | EAHGLHGEPGLEQPFWVS | -363 | -380 | 0 | 0.00383302 | 18 | 4265.254 | NaN | NaN | NaN | 667.10114 | 2084.6448 |
| Hmgn2 | AKPAPPKPEPK | -30 | -40 | 2 | 0 | 11 | 3234.0757 | 3525.8313 | 3620.3245 | 3593.9326 | 4047.288 | 3719.3171 |
| Hmgn2 | AKPAPPKPEPK | -30 | -40 | 2 | 0 | 11 | 3234.0757 | 3525.8313 | 3620.3245 | 3593.9326 | 4047.288 | 3719.3171 |
| Hmgn2 | PAPPKPEPK | -32 | -40 | 1 | 0.00111006 | 9 | 381.47147 | 377.70743 | 486.45148 | 433.30005 | 421.12415 | 490.2154 |
| Hmgn2 | PAPPKPEPK | -32 | -40 | 1 | 0.00111006 | 9 | 381.47147 | 377.70743 | 486.45148 | 433.30005 | 421.12415 | 490.2154 |
| Hmgn2 | SAKPAPPKPEPK | -29 | -40 | 2 | 0 | 12 | 450.66895 | 519.175 | 569.69714 | 510.70496 | 640.04614 | 193.97308 |
| Fabp5 | FDETTADGR | -73 | -81 | 0 | 0 | 9 | 1167.8768 | 1171.9084 | 1139.7882 | 1098.626 | 1064.0023 | 1113.9332 |
| Fabp5 | AKPDCIITCDGNNITVK | -39 | -55 | 1 | 0 | 17 | 424.1302 | 580.17664 | 535.5732 | 639.25195 | 945.31946 | 372.77908 |
| Pygb | QAVDQISSGFFSPK | -741 | -754 | 0 | 0 | 14 | 2892.8115 | 1328.1039 | 3417.9177 | 610.9551 | 2731.8264 | 3132.114 |
| Pygb | VEHTPDGVLWLDTQ | -207 | -220 | 0 | 0 | 14 | 902.21765 | 1024.7688 | 1111.3695 | 573.77264 | 1308.5582 | 1481.4526 |
| Pygb | LLSLVDDEAFIR | -522 | -533 | 0 | 0 | 12 | 713.9406 | 681.31256 | 647.0952 | 183.06473 | 512.7052 | 120.05167 |
| Pygb | SIGDVVNHDPVVGDR | -626 | -640 | 0 | 0 | 15 | 423.78836 | 557.1401 | 465.58606 | 696.07715 | 683.5696 | 662.4623 |
| Pygb | LVTSIGDVVNHDPVVGDR | -623 | -640 | 0 | 0 | 18 | 892.8929 | 864.5928 | 104.09268 | NaN | 674.4384 | 198.8244 |
| Pygb | LVTSIGDVVNHDPVVGDR | -623 | -640 | 0 | 0 | 18 | 892.8929 | 864.5928 | 104.09268 | NaN | 674.4384 | 198.8244 |
| Pygb | AKPLTDSER | -2 | -10 | 1 | 0.000471409 | 9 | 2029.7352 | 1550.4385 | NaN | 2861.7107 | 3029.4412 | NaN |
| Serbp1 | FEKPLEEK | -138 | -145 | 1 | 0.001868731 | 8 | 7313.358 | 6662.172 | 6814.4287 | 233.36633 | 5828.5405 | 6272.488 |
| Serbp1 | NKENEVEEVKEEGPK | -271 | -285 | 2 | 0 | 15 | 857.0099 | 225.7566 | 1145.8257 | 1625.1467 | 2472.256 | 2295.8618 |
| Serbp1 | NKENEVEEVKEEGPK | -271 | -285 | 2 | 0 | 15 | 857.0099 | 225.7566 | 1145.8257 | 1625.1467 | 2472.256 | 2295.8618 |
| Serbp1 | RFEKPLEEK | -137 | -145 | 2 | 0.00078661 | 9 | 4569.2686 | 4294.7466 | 4044.1726 | 5719.346 | 5418.233 | 5093.349 |
| Serbp1 | RFEKPLEEK | -137 | -145 | 2 | 0.00078661 | 9 | 4569.2686 | 4294.7466 | 4044.1726 | 5719.346 | 5418.233 | 5093.349 |
| Serbp1 | ENEVEEVKEEGPK | -273 | -285 | 1 | 0 | 13 | 2964.0576 | 3273.6165 | 3338.0232 | 3196.8762 | 3666.1125 | 3727.4155 |
| Serbp1 | SVDRPIIER | -152 | -160 | 1 | 0.00078661 | 9 | 2749.6633 | 2778.9773 | 2708.8313 | 2482.4414 | 2916.8193 | 2887.7202 |
| Serbp1 | TENKENEVEEVKEEGPK | -269 | -285 | 2 | 0 | 17 | 2575.9329 | 2640.2249 | 2939.5117 | 2658.3794 | 3422.3186 | 3372.5767 |
| Serbp1 | GGSGSHNWGTVK | -217 | -228 | 0 | 0 | 12 | 2501.0305 | 2479.0767 | 2418.637 | 1877.0074 | 1208.4475 | 1924.1133 |
| Serbp1 | KPNEGADGQWK | -309 | -319 | 1 | 0 | 11 | 2072.5283 | 2215.6013 | 2627.1062 | 1783.1163 | 433.81018 | 2292.5232 |
| Serbp1 | KNPLPPSVGVADK | -79 | -91 | 1 | 0 | 13 | 1519.4249 | 1496.8319 | 1626.4241 | 2681.387 | 1478.8752 | 2861.343 |
| Serbp1 | DSVMDHHFR | -336 | -344 | 0 | 0 | 9 | 1447.6824 | 133.78517 | 1598.5498 | 726.6998 | 1091.5736 | 1108.5629 |
| Serbp1 | DSVMDHHFR | -336 | -344 | 0 | 0 | 9 | 1447.6824 | 133.78517 | 1598.5498 | 726.6998 | 1091.5736 | 1108.5629 |
| Serbp1 | PGHLQEGFGCVVT | -2 | -14 | 0 | 0.00025015 | 13 | 1394.4536 | 1140.812 | 1099.3801 | 1638.997 | 1700.1996 | 1546.463 |
| Serbp1 | AKVEFNIR | -301 | -308 | 1 | 0 | 8 | 1231.0798 | 1444.3174 | 1505.9893 | 1654.8289 | 1895.0967 | 1800.2103 |
| Serbp1 | PGHLQEGFGC | -2 | -11 | 0 | 0.001868731 | 10 | 1092.9475 | 5566.9307 | 5130.9775 | 6498.818 | 1249.8425 | 5766.5815 |
| Serbp1 | EAGGGGVGGPGAK | -40 | -52 | 0 | 0 | 13 | 1045.3184 | 1100.2449 | 1360.0706 | 1020.1673 | 803.8632 | 1199.2563 |
| Serbp1 | NVTEETPEGEEHPVADTENKENEVEEVKEEGPK | -253 | -285 | 2 | 0.00025015 | 33 | 822.8583 | 979.18317 | 1120.201 | 1776.1693 | 2274.462 | 2098.9744 |
| Serbp1 | PGHLQEGFGCVVTNR | -2 | -16 | 0 | 0 | 15 | 764.88916 | 744.2113 | 831.7763 | 806.7806 | 909.90234 | 554.21576 |
| Serbp1 | PGHLQEGFGCVVTNR | -2 | -16 | 0 | 0 | 15 | 764.88916 | 744.2113 | 831.7763 | 806.7806 | 909.90234 | 554.21576 |
| Serbp1 | PGHLQEGFGCVV | -2 | -13 | 0 | 0 | 12 | 687.6292 | 1579.8163 | 1834.004 | 802.7872 | 1401.5659 | 236.58415 |
| Serbp1 | KNPLPPSVGVA | -79 | -89 | 1 | 0.000471409 | 11 | 630.73535 | 368.02042 | 752.5129 | 1052.9489 | 773.66235 | 840.34204 |
| Serbp1 | SSASAPDVDDPEAFPALA | -390 | -407 | 0 | 0 | 18 | 569.729 | 688.8028 | 770.7971 | 594.3954 | 962.80115 | 848.74316 |
| Serbp1 | KNPLPPSVGVADKK | -79 | -92 | 2 | 0.000230707 | 14 | 479.97562 | 366.23273 | 425.08032 | 679.1118 | 1058.9795 | 293.27368 |
| Serbp1 | KSSASAPDVDDPEAFPALA | -389 | -407 | 1 | 0 | 19 | 358.64117 | 252.65614 | 460.64084 | 550.06323 | 624.1854 | 736.2512 |
| Serbp1 | RPDQQLQGDGK | -112 | -122 | 1 | 0.000230707 | 11 | 267.22678 | 302.67896 | 1565.0598 | 601.88275 | 788.75055 | 782.46954 |
| Serbp1 | FDQLFDDESDPFEVLK | -17 | -32 | 0 | 0 | 16 | 68.32267 | 621.2079 | 668.5451 | 504.91608 | 620.66565 | 594.35785 |
| Serbp1 | KPANDITSQ | -345 | -353 | 1 | 0.002168954 | 9 | 1310.0857 | 247.87735 | NaN | 1081.7588 | 975.0955 | 940.76855 |
| Serbp1 | SSFSHYSGLK | -202 | -211 | 0 | 0 | 10 | 1162.1274 | 299.34326 | 1025.1481 | NaN | 770.3814 | 337.7177 |
| Serbp1 | EETQPPVALK | -93 | -102 | 0 | 0 | 10 | 1084.7947 | 995.6367 | 1107.6367 | 761.8724 | NaN | 449.56656 |
| Serbp1 | ENKENEVEEVKEEGPK | -270 | -285 | 2 | 0.000298805 | 16 | 600.80975 | NaN | 604.2653 | 832.749 | 1039.068 | 1102.1859 |
| Serbp1 | KNPLPPSVGVAD | -79 | -90 | 1 | 0 | 12 | 489.31955 | 557.4525 | 602.3845 | NaN | 1085.6692 | 1380.316 |

| | | | | | | | | | | | | |
|---|---|---|---|---|---|---|---|---|---|---|---|---|
| Serbp1 | FEKPLEEKGEGGEF | -138 | -151 | 2 | 0.00051306 | 14 | 314.44714 | 1123.4762 | NaN | 1087.3007 | 1324.7894 | 1268.1498 |
| Serbp1 | PGHLQEGFGCVVTN | -2 | -15 | 0 | 0.000230707 | 14 | 1485.686 | NaN | NaN | 1964.6671 | 2352.239 | 438.21036 |
| Serbp1 | RPDQQLQ | -112 | -118 | 1 | 0.003393785 | 7 | 897.9475 | NaN | NaN | 1135.3844 | 316.52548 | 600.5198 |
| Serbp1 | KPANDITSQLEIN | -345 | -357 | 1 | 0.002199523 | 13 | 441.9621 | 2563.9448 | 2763.3586 | 2707.1133 | NaN | NaN |
| Serbp1 | EETQPPVALKK | -93 | -103 | 1 | 0.002296977 | 11 | 554.3679 | NaN | 448.86774 | 433.03195 | NaN | NaN |
| Serbp1 | DQLFDDESDPFEVLK | -18 | -32 | 0 | 0 | 15 | 469.44373 | 471.9449 | 477.92508 | NaN | NaN | NaN |
| Serbp1 | EINFGDLGR | -355 | -363 | 0 | 0.008284062 | 9 | 628.47076 | NaN | 627.77704 | NaN | NaN | NaN |
| Serbp1 | DDESDPFEVLK | -22 | -32 | 0 | 0.001291103 | 11 | 528.9713 | 435.25412 | NaN | NaN | NaN | NaN |
| Serbp1 | KEETQPPVALK | -92 | -102 | 1 | 0 | 11 | 450.66974 | NaN | 1172.3341 | NaN | NaN | NaN |
| Luc7l2 | RVHELNEEIGK | -125 | -135 | 1 | 0.000230707 | 11 | 455.41315 | 718.9519 | 760.08093 | 732.46344 | 1445.7767 | 932.2646 |
| Luc7l2 | RVHELNEEIGK | -125 | -135 | 1 | 0.000230707 | 11 | 455.41315 | 718.9519 | 760.08093 | 732.46344 | 1445.7767 | 932.2646 |
| Luc7l2 | QLGAEGNVEESQK | -142 | -154 | 0 | 0 | 13 | 1059.701 | 1134.5896 | 995.64044 | 868.6559 | 910.8584 | 1008.50366 |
| Luc7l2 | AKVEQLGAEGNVEESQK | -138 | -154 | 1 | 0.00201047 | 17 | 962.3969 | 1292.875 | 1536.0211 | 1817.5208 | 1046.9083 | 1612.8297 |
| Luc7l2 | EKLEELKR | -223 | -230 | 2 | 0.000230707 | 8 | 683.29974 | 636.91077 | 769.3148 | 982.6488 | 938.6884 | 350.9406 |
| Luc7l2 | SHLLNCCPHDVLSGTR | -38 | -53 | 0 | 0 | 16 | 257.2259 | 351.9453 | 432.1277 | 577.7848 | 511.3686 | 463.04468 |
| Luc7l2 | SHLLNCCPHDVLSGTR | -38 | -53 | 0 | 0 | 16 | 257.2259 | 351.9453 | 432.1277 | 577.7848 | 511.3686 | 463.04468 |
| Luc7l2 | VEQLGAEGNVEESQK | -140 | -154 | 0 | 0 | 15 | 800.0103 | 835.30145 | NaN | 771.4601 | 1014.95715 | 1038.7255 |
| Luc7l2 | LHLGFIEIR | -214 | -222 | 0 | 0 | 9 | 1818.8901 | NaN | NaN | 1020.9402 | 923.76404 | 940.9087 |
| Luc7l2 | VHDLALR | -62 | -68 | 0 | 0.002199523 | 7 | 914.1083 | 422.04657 | 485.09418 | NaN | NaN | NaN |
| Luc7l2 | NCCPHDVLSGTR | -42 | -53 | 0 | 0.00025015 | 12 | 414.5067 | 499.0504 | NaN | NaN | NaN | NaN |
| Dbi | TVGDVNTDRPGLLDLK | -36 | -51 | 1 | 0 | 16 | 1706.9424 | 1838.3496 | 2043.3905 | 1833.4222 | 2626.0994 | 2555.465 |
| Dbi | TVGDVNTDRPGLLDLK | -36 | -51 | 1 | 0 | 16 | 1706.9424 | 1838.3496 | 2043.3905 | 1833.4222 | 2626.0994 | 2555.465 |
| Dbi | AKWDSWNK | -54 | -61 | 1 | 0 | 8 | 995.22754 | 212.76486 | 1216.2543 | 701.29675 | 516.68427 | 228.3432 |
| Dbi | QATVGDVNTDRPGLLDLK | -34 | -51 | 1 | 0.004350979 | 18 | 530.1308 | NaN | NaN | NaN | NaN | NaN |
| H1-5 | ATGPPVSELITK | -35 | -46 | 0 | 0 | 12 | 8891.999 | 9175.896 | 9287.001 | 6536.578 | 7435.254 | 7342.6104 |
| H1-5 | KATGPPVSELITK | -34 | -46 | 1 | 0 | 13 | 2798.3164 | 2794.261 | 2988.4788 | 1694.7269 | 1775.4658 | 1768.8024 |
| H1-5 | SETAPAETAAPAPVEK | -2 | -17 | 0 | 0.000230707 | 16 | 1640.742 | 1631.6256 | 1575.7113 | 1404.8723 | 1177.1454 | 1264.9933 |
| H1-5 | SETAPAETAAPAPVEK | -2 | -17 | 0 | 0.000230707 | 16 | 1640.742 | 1631.6256 | 1575.7113 | 1404.8723 | 1177.1454 | 1264.9933 |
| H1-5 | ALAAGGYDVEK | -65 | -75 | 0 | 0 | 11 | 1442.7284 | 1412.8638 | 1163.1788 | 853.52594 | 855.4435 | 550.161 |
| H1-5 | TAPAETAAPAPVEK | -4 | -17 | 0 | 0.00025015 | 14 | 783.70807 | 733.7846 | 771.546 | 974.21185 | 775.9842 | 598.5215 |
| H1-5 | KATGPPVSELIT | -34 | -45 | 1 | 0.000627943 | 12 | 8647.846 | 8168.184 | 8568.033 | 6575.89 | 7534.466 | NaN |
| H1-5 | GGVSLPALK | -55 | -63 | 0 | 0.001392869 | 9 | 1274.3783 | 843.8144 | NaN | 600.26447 | 659.4271 | 503.5659 |
| H1-5 | KATGPPVSEL | -34 | -43 | 1 | 0.001092529 | 10 | 514.2404 | 1780.2925 | 1785.921 | 1258.8156 | NaN | 265.84573 |
| Pigs | FVQPFLNR | -254 | -261 | 0 | 0 | 8 | 774.7046 | 617.53394 | 643.25836 | 768.0249 | 772.2555 | 108.1585 |
| Pigs | ALADHLPEDK | -203 | -212 | 0 | 0.003360691 | 10 | 1771.3651 | 1965.9362 | NaN | NaN | 1322.2205 | NaN |
| H2-K1 | HKWEQAGEAER | -166 | -176 | 1 | 0 | 11 | 531.8082 | 530.9344 | 830.1181 | 756.71173 | 610.3161 | 671.39636 |
| H2-K1 | HKWEQAGEAER | -166 | -176 | 1 | 0 | 11 | 531.8082 | 530.9344 | 830.1181 | 756.71173 | 610.3161 | 671.39636 |
| H2-K1 | VISGCEVGSDGR | -118 | -129 | 0 | 0 | 12 | 1582.7747 | 1697.7102 | 230.43007 | 1814.3838 | 1734.8861 | 1823.1372 |
| H2-K1 | WEQAGEAER | -168 | -176 | 0 | 0 | 9 | 770.4696 | 889.68286 | 805.6757 | 948.3815 | 845.3157 | 965.6519 |
| H2-K1 | YMEVGYVDDTEFVR | -43 | -56 | 0 | 0 | 14 | 692.5242 | 883.00116 | 929.96216 | 852.1374 | 1015.74426 | 981.47845 |
| H2-K1 | EVGYVDDTEFVR | -45 | -56 | 0 | 0 | 12 | 585.0502 | 535.9424 | 556.3182 | 729.9001 | 275.82617 | 717.54834 |
| H2-K1 | GGSHTIQVISGCEVGSDGR | -111 | -129 | 0 | 0 | 19 | 464.00708 | 507.93646 | 430.26254 | 925.0415 | 1049.408 | 1044.567 |
| H2-K1 | HKWEQAGEAERL | -166 | -177 | 2 | 0 | 12 | 441.44586 | 752.05206 | 840.0666 | 442.58145 | 894.96576 | 887.1707 |
| Nop56 | IVNDNATYCR | -203 | -212 | 0 | 0 | 10 | 8357.401 | 563.57227 | 286.33368 | 7550.175 | 8908.403 | 567.32666 |
| Nop56 | SEGVVHEDLR | -68 | -77 | 0 | 0 | 10 | 4471.912 | 4079.7727 | 4954.9775 | 1845.1912 | 1808.19 | 1853.0024 |
| Nop56 | SEPEEAASPTTPK | -536 | -548 | 0 | 0 | 13 | 3596.145 | 3451.6309 | 3618.3054 | 1411.2904 | 1762.8145 | 3742.5515 |
| Nop56 | SEVPTSVFGEK | -386 | -396 | 0 | 0 | 11 | 2961.9077 | 1736.6671 | 2200.1802 | 2130.28 | 1218.4965 | 2652.345 |
| Nop56 | AVSEGVVHEDLR | -66 | -77 | 0 | 0 | 12 | 1708.6759 | 1525.4213 | 999.8375 | 709.4242 | 880.9569 | 1485.1345 |
| Nop56 | EWYGYHFPELVK | -191 | -202 | 0 | 0 | 12 | 1603.7858 | 2119.7026 | 1528.1107 | 1614.2377 | 339.38306 | 2126.725 |
| Nop56 | LSFYETGEIPR | -405 | -415 | 0 | 0 | 11 | 1436.2356 | 1574.8191 | 1352.493 | 1583.0737 | 1173.6676 | 1577.7755 |
| Nop56 | ELNEEKLEK | -222 | -230 | 1 | 0 | 9 | 1280.0222 | 1339.7548 | 1558.0011 | 1637.0162 | 1857.588 | 1767.0759 |
| Nop56 | NANAVSEGVVHEDLR | -63 | -77 | 0 | 0 | 15 | 1102.7891 | 975.18085 | 2324.8047 | 607.08673 | 1101.3756 | 1454.8652 |
| Nop56 | ANAVSEGVVHEDLR | -64 | -77 | 0 | 0 | 14 | 1033.1049 | 1242.5211 | 1220.0487 | 869.8824 | 952.0985 | 990.2261 |
| Nop56 | IDCFSEVPTSVFGEK | -382 | -396 | 0 | 0 | 15 | 864.5195 | 953.90643 | 1070.6134 | 1139.4757 | 1576.9036 | 1489.5653 |
| Nop56 | SENSSTPEECEEVNEK | -462 | -477 | 0 | 0 | 16 | 596.99536 | 729.72656 | 814.883 | 560.3181 | 803.72565 | 547.9402 |
| Nop56 | NSSTPEECEEVNEK | -464 | -477 | 0 | 0 | 14 | 510.3995 | 548.3474 | 699.85614 | 73.1766 | 641.092 | 726.2643 |
| Nop56 | QTGGVIAEILR | -113 | -123 | 0 | 0.00025015 | 11 | 382.90417 | 981.71405 | 383.37982 | 877.0284 | 979.57733 | 1014.1193 |
| Nop56 | TGGVIAEILR | -114 | -123 | 0 | 0.000230707 | 10 | 227.21637 | 502.13345 | 392.40033 | 425.6438 | 430.8465 | 425.46338 |
| Nop56 | ASSENSSTPEECEEVNEK | -460 | -477 | 0 | 0 | 18 | 117.31363 | 187.05702 | 195.35054 | 213.91986 | 196.4262 | 198.04562 |
| Nop56 | LHFHNLVK | -127 | -134 | 0 | 0 | 8 | 4624.1455 | 4562.161 | 7484.0205 | 1755.1676 | 5370.7935 | NaN |
| Nop56 | GVVHEDLR | -70 | -77 | 0 | 0.00025015 | 8 | 1894.2198 | 2036.6691 | 1796.4315 | 775.16254 | NaN | 962.29517 |
| Nop56 | EEVASEPEEAASPTTPK | -532 | -548 | 0 | 0 | 17 | 1444.2991 | 1419.9924 | NaN | 1269.8588 | 505.62863 | 1471.2036 |
| Nop56 | KFSEEPEVAAN | -552 | -562 | 1 | 0.000230707 | 11 | 832.7477 | 483.7957 | NaN | 2647.1355 | NaN | 1324.7869 |
| Nop56 | ENANAVSEGVVHEDLR | -62 | -77 | 0 | 0 | 16 | 745.8312 | NaN | 667.5526 | NaN | 679.1725 | 671.9154 |
| Nop56 | ALASSENSSTPEECEEVNEK | -458 | -477 | 0 | 0.00025015 | 20 | 433.28577 | NaN | 457.5578 | 291.67596 | 334.2058 | NaN |
| Nop56 | LLLETYLPSK | -78 | -87 | 0 | 0 | 10 | 944.93176 | 1267.6127 | 1153.3148 | NaN | NaN | NaN |
| Nop56 | ALIGEAVGAR | -298 | -307 | 0 | 0.001023236 | 10 | 731.506 | 540.3109 | 668.38104 | NaN | NaN | NaN |
| Nop56 | KVLLGVGDPK | -90 | -99 | 1 | 0.00133086 | 10 | 636.36096 | 712.19244 | 662.65344 | NaN | NaN | NaN |
| Nop56 | SISLLDQLDK | -171 | -180 | 0 | 0.000394361 | 10 | 475.28506 | 647.55347 | 648.85254 | NaN | NaN | NaN |
| Nop56 | VLLGVGDPK | -91 | -99 | 0 | 0.00025015 | 9 | 894.8674 | 594.37964 | NaN | NaN | NaN | NaN |
| Nop56 | LEEITMDGAK | -231 | -240 | 0 | 0.00122419 | 10 | 546.6554 | NaN | NaN | 574.00305 | NaN | NaN |
| Atp2a2 | WGSNELPAEEGK | -36 | -47 | 0 | 0 | 12 | 3024.543 | 3061.2446 | 3436.3604 | 3535.7988 | 3929.0757 | 3779.287 |
| Atp2a2 | EFDELSPSAQR | -656 | -666 | 0 | 0 | 11 | 1163.814 | 1385.8358 | 926.84174 | 414.24252 | 1153.6923 | 1080.7002 |
| Atp2a2 | EWGSGSDTLR | -550 | -559 | 0 | 0 | 10 | 1139.4398 | 1036.3096 | 1132.5686 | 975.8682 | 704.6592 | 971.9748 |
| Atp2a2 | TVEEVLGHF | -8 | -16 | 0 | 0.000550307 | 9 | 970.67114 | NaN | 1261.91 | 1211.5348 | 1486.057 | 1460.7678 |
| Atp2a2 | ALATHDNPLKR | -562 | -572 | 1 | 0.00025015 | 11 | 478.08932 | NaN | 464.67502 | NaN | 472.60892 | 544.2029 |
| Atp2a2 | IGIFGQDEDVTSK | -638 | -650 | 0 | 0 | 13 | 2200.2444 | NaN | NaN | 3532.5308 | NaN | 3740.155 |
| Atp2a2 | GSTYAPIGEVQK | -386 | -397 | 0 | 0 | 12 | 1234.2092 | NaN | 1046.1011 | NaN | NaN | 2965.309 |

| | | | | | | | | | | | | |
|---|---|---|---|---|---|---|---|---|---|---|---|---|
| Ubr4 | WFDFPFTR | -2382 | -2389 | 0 | 0.00025015 | 8 | 241.53954 | 262.50958 | 289.58844 | 295.1804 | 338.80127 | 447.04858 |
| Ubr4 | LNLPYFLR | -261 | -268 | 0 | 0.00111006 | 8 | 1400.975 | NaN | NaN | 824.48193 | 198.6679 | 1303.183 |
| Ubr4 | SNPSVLQGLLR | -4605 | -4615 | 0 | 0 | 11 | 1753.3413 | 1921.2623 | 1924.532 | NaN | NaN | NaN |
| Ubr4 | SKEDDYESDAATIVQK | -365 | -380 | 1 | 0.002099344 | 16 | 582.93756 | NaN | NaN | 995.03577 | 796.4838 | NaN |
| Ubr4 | QLFPINIK | -2143 | -2150 | 0 | 0 | 8 | 317.66092 | NaN | NaN | 205.46085 | 491.87265 | NaN |
| Ubr4 | GHWANPDLASSL | -3954 | -3965 | 0 | 0.002199523 | 12 | 1004.2211 | NaN | NaN | NaN | NaN | NaN |
| Ubr4 | ALATNPALR | -3886 | -3894 | 0 | 0.009897819 | 9 | 700.9626 | NaN | NaN | NaN | NaN | NaN |
| Arpc5l | GFEKPTENSSAVLL | -115 | -128 | 1 | 0 | 14 | 428.3218 | 665.16656 | 551.3156 | 882.55725 | 968.36206 | 1182.4894 |
| Arpc5l | ALAVGGLGSIIR | -134 | -145 | 0 | 0 | 12 | 801.732 | 817.16925 | 906.81824 | NaN | 836.8039 | 834.83234 |
| Vkorc1l1 | ALCDLGPWVK | -48 | -57 | 0 | 0 | 10 | 295.37985 | 168.68283 | 289.82394 | NaN | 664.3602 | 139.85419 |
| Rnh1 | LLCEGLQDPQCR | -127 | -138 | 0 | 0 | 12 | 2950.3071 | 2830.6865 | 2709.199 | 1372.4386 | 931.9932 | 929.55023 |
| Rnh1 | ALCPGLLLPSCK | -241 | -252 | 0 | 0 | 12 | 1011.78503 | 928.2654 | 989.08234 | 1362.2726 | 952.2841 | 993.5461 |
| Rnh1 | ELVLSNNDLHEPGVR | -169 | -183 | 0 | 0.00025015 | 15 | 751.6875 | 876.2811 | 938.1399 | 772.2869 | 907.70435 | 964.859 |
| Rnh1 | ALSQPDTVLR | -359 | -368 | 0 | 0 | 10 | 741.32275 | 833.6472 | 1007.94855 | 775.48083 | 596.5009 | 545.39026 |
| Rnh1 | SLASNELKDEGAR | -285 | -297 | 1 | 0 | 13 | 1309.8622 | 1448.0669 | 1522.2112 | 1711.0286 | 1514.2036 | NaN |
| Rnh1 | LDDCGLTEVR | -30 | -39 | 0 | 0 | 10 | 1261.5005 | 1397.1622 | 502.83734 | NaN | 1163.1476 | 1303.8181 |
| Rnh1 | SNPLGDEGVQELCK | -345 | -358 | 0 | 0 | 14 | 998.9403 | NaN | 845.974 | NaN | 1170.4038 | 923.2404 |
| Rnh1 | SCEPLASVLR | -153 | -162 | 0 | 0.000550307 | 10 | 662.2929 | 462.2705 | 913.18414 | NaN | NaN | NaN |
| Rnh1 | ALEEERPSLR | -444 | -453 | 1 | 0.000550307 | 10 | 1011.75903 | 1037.1797 | NaN | NaN | NaN | NaN |
| Praf2 | ALDDFVLGSAR | -11 | -21 | 0 | 0 | 11 | 520.8535 | NaN | NaN | 630.2462 | NaN | NaN |
| Praf2 | LAAPDPGDPQR | -22 | -32 | 0 | 0.000431655 | 11 | 720.6351 | NaN | NaN | NaN | NaN | NaN |
| Pml | SAIGQLDHAR | -267 | -276 | 0 | 0 | 10 | 2391.6333 | 2197.3706 | 3371.7998 | 2615.7195 | 2082.3384 | 2299.7097 |
| Pml | NPALTDIYCR | -200 | -209 | 0 | 0 | 10 | 1063.3981 | 909.13654 | 846.97687 | 983.71844 | 549.3523 | 622.78033 |
| Pml | LAVLPLIR | -717 | -724 | 0 | 0.002099344 | 8 | 716.4809 | 849.1063 | 820.5582 | 571.6037 | 683.05365 | 700.80554 |
| Pml | STHPVTTSTPEDLEQPK | -416 | -432 | 0 | 0.000230707 | 17 | 439.3498 | 641.95795 | 549.15985 | 286.19196 | 474.08646 | 558.94025 |
| Pml | CSGCLEAPGLQCPICK | -82 | -97 | 0 | 0.000907206 | 16 | 1791.8604 | 698.5108 | NaN | NaN | 691.9256 | NaN |
| Pml | ELLEAVNDR | -300 | -308 | 0 | 0 | 9 | 1348.0322 | 1251.8142 | NaN | 1433.3596 | NaN | NaN |
| Pml | LCLQDFISCITQR | -385 | -397 | 0 | 0 | 13 | 343.64932 | NaN | NaN | NaN | 527.4483 | 573.09766 |
| Pml | SCLEAVLQR | -321 | -329 | 0 | 0 | 9 | 1372.5825 | 1056.7526 | NaN | NaN | NaN | NaN |
| Pml | ALDESLAEPHLEDR | -605 | -618 | 0 | 0 | 14 | 753.7602 | NaN | 1900.1895 | NaN | NaN | NaN |
| Pml | DNSVSSFLDSTR | -176 | -187 | 0 | 0.00025015 | 12 | 586.589 | NaN | NaN | NaN | NaN | NaN |
| Prpf19 | VAHPIRPKPPSAT | -57 | -69 | 2 | 0.000431655 | 13 | 5395.0557 | 3975.2678 | 2042.3474 | 897.12946 | 5031.581 | 4500.139 |
| Prpf19 | VAHPIRPKPPSAT | -57 | -69 | 2 | 0.000431655 | 13 | 5395.0557 | 3975.2678 | 2042.3474 | 897.12946 | 5031.581 | 4500.139 |
| Prpf19 | TVPEELVKPEELSK | -193 | -206 | 1 | 0 | 14 | 2824.526 | 2942.778 | 3339.6602 | 3647.575 | 3702.4658 | 3342.0632 |
| Prpf19 | TVPEELVKPEEL | -193 | -204 | 1 | 0 | 12 | 1022.65967 | 1093.1229 | 1079.6478 | 1318.1245 | 1364.9869 | 1319.8856 |
| Prpf19 | QWTEILHF | -464 | -471 | 0 | 0.00111006 | 8 | 664.2811 | 573.66724 | 592.0314 | 616.5542 | 769.1349 | 771.03516 |
| Prpf19 | SISNEVPEHPCVSPVSNHVYER | -6 | -27 | 0 | 0.001023236 | 22 | 661.0076 | 736.5964 | 665.73975 | 2408.9128 | 902.4715 | 865.947 |
| Prpf19 | LYQHDAACR | -107 | -115 | 0 | 0 | 9 | 657.16437 | 498.98474 | 607.36206 | 598.66785 | 404.17676 | 411.4468 |
| Prpf19 | TNVANFPGHSGPITSIAF | -383 | -400 | 0 | 0 | 18 | 441.74695 | 713.42346 | 611.4309 | 709.4517 | 690.25885 | 674.84564 |
| Prpf19 | QLDNNFEVK | -431 | -439 | 0 | 0 | 9 | 1883.0875 | 133.13684 | 2260.4087 | NaN | 1918.4714 | 1954.5498 |
| Prpf19 | KVTSVVFHPS | -266 | -275 | 1 | 0 | 10 | 1038.4955 | 193.37787 | 308.0916 | 658.91144 | 457.98633 | NaN |
| Prpf19 | ALDLCPSDTNK | -226 | -236 | 0 | 0 | 11 | 529.61597 | 1789.0886 | 2138.8376 | NaN | 1331.7526 | 1373.6774 |
| Prpf19 | QVASHVGLH | -209 | -217 | 0 | 0.000627943 | 9 | 1006.4328 | 917.17773 | NaN | NaN | NaN | NaN |
| Prpf19 | TLQLDNNFEVK | -429 | -439 | 0 | 0 | 11 | 915.0429 | 719.5869 | NaN | NaN | NaN | NaN |
| Stip1 | HEANNLQLK | -101 | -109 | 0 | 0.000347792 | 9 | 5418.2056 | 3981.428 | 5212.6836 | 4402.0957 | 4073.2925 | 4511.6714 |
| Stip1 | DPQALSEHLK | -514 | -523 | 0 | 0 | 10 | 4794.4873 | 4484.969 | 4459.969 | 5744.7666 | 5408.001 | 4830.457 |
| Stip1 | TVDLKPDWGK | -64 | -73 | 1 | 0.00025015 | 10 | 4299.347 | 4704.8447 | 4985.7866 | 4805.943 | 4821.808 | 306.09485 |
| Stip1 | LAYINPDLALEEK | -352 | -364 | 0 | 0 | 13 | 3444.7405 | 4334.5244 | 4219.424 | 5377.299 | 6485.8164 | 6500.6284 |
| Stip1 | YKDAIHFYNK | -316 | -325 | 1 | 0 | 10 | 243.49234 | 668.46344 | 773.323 | 1402.4664 | 1017.30005 | 2628.1687 |
| Stip1 | YKDAIHFYNK | -316 | -325 | 1 | 0 | 10 | 243.49234 | 668.46344 | 773.323 | 1402.4664 | 1017.30005 | 2628.1687 |
| Stip1 | SLLSDPTYR | -145 | -153 | 0 | 0 | 9 | 2926.5793 | 2367.2947 | 330.73215 | 2499.1704 | 1556.4862 | 1442.5247 |
| Stip1 | LDPQNHVLY | -33 | -41 | 0 | 0 | 9 | 2443.8496 | 1026.7588 | 1342.4446 | 842.78534 | 2004.5262 | 2134.804 |
| Stip1 | LDPQNHVLYSN | -33 | -43 | 0 | 0 | 11 | 2290.9192 | 2148.5813 | 2642.9478 | 2767.5928 | 2599.0798 | 2401.5444 |
| Stip1 | ELIEQLQNKPSDLGTK | -154 | -169 | 1 | 0 | 16 | 1855.9132 | 2241.0874 | 2232.173 | 2983.8425 | 3912.9229 | 3795.3708 |
| Stip1 | ELIEQLQNKPSDLGTK | -154 | -169 | 1 | 0 | 16 | 1855.9132 | 2241.0874 | 2232.173 | 2983.8425 | 3912.9229 | 3795.3708 |
| Stip1 | LDPQNHVLYSNR | -33 | -44 | 0 | 0 | 12 | 1820.3381 | 2125.9385 | 1547.589 | 2333.789 | 2296.7773 | 2612.6902 |
| Stip1 | LDPQNHVLYSNR | -33 | -44 | 0 | 0 | 12 | 1820.3381 | 2125.9385 | 1547.589 | 2333.789 | 2296.7773 | 2612.6902 |
| Stip1 | TYEEGLKHEANNL | -94 | -106 | 1 | 0 | 13 | 1704.0356 | 1917.2585 | 1932.3136 | 2827.1707 | 2946.762 | 2112.151 |
| Stip1 | ALDLDSSCK | -454 | -462 | 0 | 0 | 9 | 1466.6968 | 1241.4686 | 1010.59357 | 1577.4185 | 997.78534 | 1011.3248 |
| Stip1 | NLPNLYQK | -129 | -136 | 0 | 0.00025015 | 8 | 1449.4421 | 1382.4076 | 1377.1064 | 1397.571 | 1005.7982 | 1084.9266 |
| Stip1 | YKDAIHFY | -316 | -323 | 1 | 0.00025015 | 8 | 1031.5455 | 934.10516 | 112.24546 | 803.35333 | 1318.1498 | 1241.5588 |
| Stip1 | DAIHFYNK | -318 | -325 | 0 | 0 | 8 | 1014.95654 | 476.26117 | 869.99976 | 731.91754 | 1140.2133 | 1116.0566 |
| Stip1 | ALSAGNIDDALQCY | -14 | -27 | 0 | 0 | 14 | 997.11426 | 739.0237 | 212.4125 | 723.01996 | 957.8181 | 921.58997 |
| Stip1 | LMDVGLIAIR | -534 | -543 | 0 | 0 | 10 | 982.74164 | 1041.3468 | 828.2677 | 1531.2397 | 1814.343 | 1879.6405 |
| Stip1 | LMDVGLIAIR | -534 | -543 | 0 | 0 | 10 | 982.74164 | 1041.3468 | 828.2677 | 1531.2397 | 1814.343 | 1879.6405 |
| Stip1 | DCEECIQLEPTFIK | -416 | -429 | 0 | 0 | 14 | 947.6811 | 1070.6787 | 1012.52167 | 944.75854 | 1097.288 | 1065.646 |
| Stip1 | FMNPFNLPNLYQK | -124 | -136 | 0 | 0 | 13 | 678.46515 | 803.8417 | 574.17694 | 820.42474 | 835.24963 | 1013.81177 |
| Stip1 | TYEEGLKHEANNLQLK | -94 | -109 | 1 | 0 | 16 | 609.7533 | 801.32196 | 694.9815 | 731.3296 | 1049.5466 | 1007.4974 |
| Stip1 | ALSAGNIDDALQCYSEAIK | -14 | -32 | 0 | 0 | 19 | 414.73062 | 298.51328 | 274.43262 | 342.6046 | 194.869 | 332.50583 |
| Stip1 | PQNHVLYSNR | -35 | -44 | 0 | 0.000591124 | 10 | 357.24515 | 6567.1973 | 5683.313 | 366.32452 | 456.6632 | 8362.354 |
| Stip1 | HDSPEDVKR | -479 | -487 | 1 | 0.00025015 | 9 | 275.91235 | 240.36853 | 338.13095 | 311.41928 | 298.83997 | 319.74515 |
| Stip1 | EGLQNMEAR | -110 | -118 | 0 | 0 | 9 | 256.06464 | 1051.9917 | 1501.076 | 2646.4768 | 146.47437 | 2243.6436 |
| Stip1 | TPDVLKK | -332 | -338 | 1 | 0 | 7 | 491.58002 | 630.328 | 793.02686 | 546.9904 | NaN | 460.09375 |
| Stip1 | QNKPSDLGTK | -160 | -169 | 1 | 0.001291103 | 10 | 854.46906 | 705.24945 | 832.2264 | 733.5436 | NaN | NaN |
| Stip1 | TYEEGLKHEAN | -94 | -104 | 1 | 0 | 11 | 1477.8396 | NaN | NaN | NaN | 1512.9021 | 1273.9034 |
| Stip1 | KEPKPEPMEEDLPENK | -207 | -222 | 2 | 0.001868731 | 16 | 524.3539 | NaN | NaN | NaN | 584.61206 | 474.29477 |
| Stip1 | YINPDLALEEK | -354 | -364 | 0 | 0 | 11 | 115.3582 | 806.8912 | 800.60333 | NaN | NaN | NaN |
| Stip1 | QAAVHFEK | -265 | -272 | 0 | 0.007791272 | 8 | 1847.3301 | NaN | NaN | 2125.3374 | NaN | NaN |

| | | | | | | | | | | | | |
|---|---|---|---|---|---|---|---|---|---|---|---|---|
| Nt5dc1 | ALDLEDGTFIK | -67 | -77 | 0 | 0 | 11 | 3232.9905 | NaN | 3499.8474 | NaN | 826.9249 | 411.40347 |
| Hk1 | HIDLVEGDEGR | -300 | -310 | 0 | 0 | 11 | 1883.9321 | 2399.481 | 155.07889 | 1855.1736 | 1794.3771 | 1893.6903 |
| Hk1 | GLNSTCDDSILVK | -864 | -876 | 0 | 0 | 13 | 998.29755 | 1005.6647 | 1255.2797 | 919.87195 | 1208.593 | 911.6043 |
| Hk1 | DVAAIETDKEGVQNAK | -394 | -409 | 1 | 0.001023236 | 16 | 692.8312 | 696.56757 | 836.0292 | 811.3234 | 1063.4937 | 1007.61383 |
| Hk1 | ISDFLDYMGIK | -638 | -648 | 0 | 0 | 11 | 651.0284 | 433.72437 | NaN | 593.5504 | 638.876 | 791.9864 |
| Hk1 | GDDGSLEDIR | -321 | -330 | 0 | 0 | 10 | 993.53754 | 880.3635 | 886.7115 | NaN | 1561.009 | NaN |
| Hk1 | TEWGAFGDDGSLEDIR | -315 | -330 | 0 | 0 | 16 | 392.81268 | 751.2337 | 563.10266 | NaN | NaN | NaN |
| Hk1 | ESLLFEGR | -372 | -379 | 0 | 0.000627943 | 8 | 797.6022 | NaN | NaN | NaN | NaN | NaN |
| Rnf213 | SILLDCATPDAVVR | -3215 | -3228 | 0 | 0 | 14 | 403.38422 | 463.41608 | 455.09912 | 372.98044 | 973.5488 | 1210.416 |
| Rnf213 | ETVPVLWHF | -4706 | -4714 | 0 | 0.000471409 | 9 | 549.2614 | 603.6706 | 622.31805 | NaN | 619.355 | 612.1777 |
| Rnf213 | ALDNDQHLPDK | -1461 | -1471 | 0 | 0.00025015 | 11 | 613.1503 | NaN | 729.44836 | NaN | 867.34753 | NaN |
| Rnf213 | SLSPFDDVVDQTPVIR | -4066 | -4081 | 0 | 0 | 16 | 974.14886 | NaN | 1063.2242 | NaN | NaN | NaN |
| Rnf213 | LRDETVPLK | -1975 | -1983 | 1 | 0.001291103 | 9 | 638.8487 | NaN | NaN | NaN | 1905.4045 | NaN |
| Rnf213 | IIDNITEEGR | -1047 | -1056 | 0 | 0.001023236 | 10 | 553.6877 | 568.75397 | NaN | NaN | NaN | NaN |
| Rnf213 | LLLPDGHGYPEALR | -1528 | -1541 | 0 | 0.004628201 | 14 | 388.38025 | NaN | NaN | NaN | NaN | NaN |
| Cmc1 | ALDPAEQHLR | -2 | -11 | 0 | 0 | 10 | 292.82208 | 2224.6729 | 2486.8809 | NaN | 1446.6747 | 2816.812 |
| Ccar2 | FAEFQYLQPGPPR | -405 | -417 | 0 | 0 | 13 | 764.7669 | 809.97705 | 804.27606 | 637.5133 | 493.13397 | 1312.4221 |
| Ccar2 | SPAPPLLHVA | -124 | -133 | 0 | 0.00025015 | 10 | 391.38873 | 443.44437 | 247.38736 | 437.98123 | 781.42737 | 339.02847 |
| Ccar2 | EAAPDTGAEPSPEDSDPTYSSK | -288 | -309 | 0 | 0 | 22 | 257.58707 | 363.5968 | 368.8632 | 331.98978 | 423.186 | 376.87424 |
| Ccar2 | SSPGLEEFYR | -314 | -323 | 0 | 0 | 10 | 775.4904 | NaN | 95.96539 | 976.33606 | 109.742096 | 1049.1187 |
| Ccar2 | TVDSPTCDFLELQR | -232 | -245 | 0 | 0 | 14 | 1220.7682 | NaN | 1188.5691 | NaN | 1462.5575 | 1259.0635 |
| Ccar2 | ALDPDLLLLR | -646 | -655 | 0 | 0 | 10 | 540.8251 | 400.05408 | 690.43427 | NaN | NaN | NaN |
| Ccar2 | TLSNQPLLK | -115 | -123 | 0 | 0.00025015 | 9 | 680.4963 | NaN | 353.9307 | NaN | NaN | NaN |
| Ccar2 | VSKDEVQNEGTAA | -596 | -608 | 1 | 0.002697088 | 13 | 497.96625 | NaN | NaN | NaN | NaN | NaN |
| Nup210 | ELYLEDSPLELK | -127 | -138 | 0 | 0 | 12 | 1486.5728 | 1610.6001 | 1307.4587 | 1803.6903 | 2089.7935 | 2092.3064 |
| Nup210 | STPQQIEVFPPFR | -1070 | -1082 | 0 | 0 | 13 | 752.5547 | 716.0005 | 865.83496 | 642.04224 | 1526.101 | 1765.6598 |
| Nup210 | SIEPLGSSEQQCSQK | -65 | -79 | 0 | 0 | 15 | 720.40515 | 689.8728 | 461.09836 | 1914.4355 | 2280.0984 | 2022.1793 |
| Nup210 | VHDLCLAFPAPAK | -958 | -970 | 0 | 0 | 13 | 656.2 | 1617.8192 | 1998.2825 | 1356.605 | 1106.484 | 2028.7528 |
| Nup210 | VHDLCLAFPAPAK | -958 | -970 | 0 | 0 | 13 | 656.2 | 1617.8192 | 1998.2825 | 1356.605 | 1106.484 | 2028.7528 |
| Nup210 | ALDPTAGQLHGL | -1259 | -1270 | 0 | 0 | 12 | 645.09106 | 1249.5083 | 1006.99365 | 1142.9486 | 750.58765 | 1291.8411 |
| Nup210 | SSNLLGECSPAQR | -1585 | -1597 | 0 | 0 | 13 | 438.5813 | 3412.9133 | 12959.016 | 3051.6333 | 411.71487 | 2149.8433 |
| Nup210 | VFDWTIVK | -156 | -163 | 0 | 0.00201047 | 8 | 1901.8032 | 2159.6848 | 2104.7878 | 2489.3 | NaN | 3140.7583 |
| Nup210 | KVTVSVGDR | -1576 | -1584 | 1 | 0.00025015 | 9 | 1174.5554 | 980.8556 | NaN | 1613.7751 | 2222.9277 | 2183.5764 |
| Nup210 | WSSTRPEVA | -56 | -64 | 1 | 0.00025015 | 9 | 1078.1824 | 967.7234 | 612.8164 | 629.9462 | 720.2024 | NaN |
| Nup210 | VVDKVEIGK | -985 | -993 | 1 | 0 | 9 | 755.4259 | 570.10944 | 520.18726 | 486.3664 | NaN | 539.4913 |
| Nup210 | VSSTPQQIEVFPPFR | -1068 | -1082 | 0 | 0 | 15 | 597.9163 | NaN | 853.94586 | 694.1201 | 872.9254 | 187.82094 |
| Nup210 | LSLLGPPASR | -692 | -701 | 0 | 0.00025015 | 10 | 1159.5612 | 1204.8496 | NaN | NaN | NaN | 1523.7527 |
| Nup210 | NQGVFQPLPGR | -587 | -597 | 0 | 0.001776473 | 11 | 1086.3914 | NaN | 941.8461 | NaN | NaN | NaN |
| Nup210 | PWVLEPSK | -670 | -677 | 0 | 0.002595029 | 8 | 873.2105 | 921.8397 | NaN | NaN | NaN | NaN |
| Nup210 | VSPVSYLR | -1371 | -1378 | 0 | 0.006100831 | 8 | 1893.5961 | NaN | NaN | NaN | NaN | NaN |
| Nup210 | VYPSDNIR | -380 | -387 | 0 | 0.001636916 | 8 | 1325.3846 | NaN | NaN | NaN | NaN | NaN |
| Nup210 | APIVHTDEK | -1329 | -1337 | 0 | 0 | 9 | 1033.9476 | NaN | NaN | NaN | NaN | NaN |
| Nup210 | SGDTGLSVIR | -506 | -515 | 0 | 0.006820063 | 10 | 467.86652 | NaN | NaN | NaN | NaN | NaN |
| Rps17 | DQEIIEVDPDTK | -92 | -103 | 0 | 0 | 12 | 3026.6853 | 2205.2083 | 2497.966 | 1760.9734 | 1174.8054 | 1745.5702 |
| Rps17 | DQEIIEVDPDTK | -92 | -103 | 0 | 0 | 12 | 3026.6853 | 2205.2083 | 2497.966 | 1760.9734 | 1174.8054 | 1745.5702 |
| Rps17 | RDNYVPEVSALD | -81 | -92 | 1 | 0 | 12 | 1816.1357 | 1756.4128 | 958.2652 | 698.3058 | 773.81714 | 1001.2763 |
| Rps17 | SNLQVTQPTVGMNFK | -115 | -129 | 0 | 0 | 15 | 1063.1747 | 1125.1389 | 1089.3909 | 1323.9775 | 1323.124 | 1459.8037 |
| Rps17 | SNLQVTQPTVGMNFK | -115 | -129 | 0 | 0 | 15 | 1063.1747 | 1125.1389 | 1089.3909 | 1323.9775 | 1323.124 | 1459.8037 |
| Rps17 | VCEEIAIIPSK | -34 | -44 | 0 | 0 | 11 | 1283.8817 | 1686.1823 | 822.0355 | 1314.8623 | 576.2412 | 2067.6426 |
| Rps17 | RDNYVPEVSAL | -81 | -91 | 1 | 0.00078661 | 11 | 1107.7397 | 1189.0464 | 1193.4608 | 1330.149 | 959.3564 | 373.2381 |
| Rps17 | ALDQEIIEVDPDTK | -90 | -103 | 0 | 0 | 14 | 940.08514 | 409.51138 | 765.48267 | 747.096 | 805.7244 | 905.1914 |
| Rps17 | DQEIIEVDPDTKEMLK | -92 | -107 | 1 | 0 | 16 | 561.841 | 697.4159 | 635.56067 | 486.5161 | 531.4102 | 578.8733 |
| Rps17 | QVTQPTVGMNFK | -118 | -129 | 0 | 0.000230707 | 12 | 3356.6938 | 3739.249 | 3134.4363 | NaN | 4290.416 | 2815.6223 |
| Rps17 | QVTQPTVGMNFK | -118 | -129 | 0 | 0.000230707 | 12 | 3356.6938 | 3739.249 | 3134.4363 | NaN | 4290.416 | 2815.6223 |
| Rps17 | LGNDFHTN | -24 | -31 | 0 | 0.002595029 | 8 | 1008.56573 | 990.4795 | 938.43896 | NaN | 777.66626 | NaN |
| Rps17 | VCEEIAIIPSKK | -34 | -45 | 1 | 0 | 12 | 531.98236 | 556.207 | 593.99036 | 339.871 | NaN | NaN |
| Rps17 | LGNDFHTNK | -24 | -32 | 0 | 0 | 9 | 3553.3723 | 3792.9773 | NaN | NaN | 3734.824 | NaN |
| Myg1 | EHLYHLE | -273 | -279 | 0 | 0.00025015 | 7 | 2836.3755 | 3966.29 | 4128.2217 | 4160.64 | 4412.7334 | 4084.1777 |
| Myg1 | LPLPEPWR | -316 | -323 | 0 | 0 | 8 | 1316.7068 | 1356.9028 | 1304.264 | 1210.4641 | 1293.4393 | 1311.3105 |
| Myg1 | LLPEYANAEIVR | -66 | -77 | 0 | 0 | 12 | 1052.7727 | 948.37225 | 1069.7834 | 1302.4753 | 1446.7769 | 1312.9984 |
| Myg1 | LNPTWNQPNQDTEAGFR | -201 | -217 | 0 | 0 | 17 | 167.30113 | 1323.7512 | 1649.3505 | 1471.8954 | 2290.7783 | 2053.7458 |
| Myg1 | VIYTDQAGQWR | -291 | -301 | 0 | 0 | 11 | 787.4779 | 711.4862 | NaN | 808.8249 | 697.59155 | 558.6993 |
| Myg1 | ALDQVSGIPGCIFVH | -329 | -343 | 0 | 0 | 15 | 299.2168 | NaN | 376.10828 | 340.69873 | 463.97687 | 407.06363 |
| Myg1 | GTSEEDSVVDTIYDK | -148 | -162 | 0 | 0 | 15 | 262.92865 | 472.13403 | 335.9965 | 507.00943 | 510.9997 | NaN |
| Gspt1 | PFVPNVHAA | -75 | -83 | 0 | 0 | 9 | 3096.2834 | 2241.903 | 2372.0154 | 2353.4504 | 1862.737 | 122.375786 |
| Gspt1 | EFVPSFLR | -84 | -91 | 0 | 0 | 8 | 1035.0244 | 1140.4844 | 1245.0537 | 139.90181 | 1191.2094 | 1230.961 |
| Gspt1 | SDDVETDSVAPGENLK | -485 | -500 | 0 | 0.00025015 | 16 | 702.952 | 1111.6646 | 1054.0338 | 1681.6014 | 1455.8944 | 799.8384 |
| Gspt1 | SVDGPIRLPIVDK | -434 | -446 | 1 | 0 | 13 | 629.0009 | 671.3268 | 637.7151 | 750.7199 | 823.87585 | 775.16656 |
| Gspt1 | GILSDDVETDSVAPGENLK | -482 | -500 | 0 | 0 | 19 | 458.25638 | 612.1919 | 542.7708 | 510.41415 | 449.0532 | 543.6744 |
| Gspt1 | DDPTVNWSNER | -360 | -370 | 0 | 0 | 11 | 937.56775 | 1020.8932 | 467.29373 | 1347.1798 | NaN | NaN |
| Gspt1 | LDNLPNFNR | -425 | -433 | 0 | 0.000471409 | 9 | 936.62244 | NaN | NaN | 999.8483 | NaN | NaN |
| Nip7 | HQADIGEYVR | -165 | -174 | 0 | 0.00025015 | 10 | 1964.1218 | NaN | 1828.2919 | 232.17067 | NaN | NaN |
| Nip7 | ALDYLAPYAK | -85 | -94 | 0 | 0.000471409 | 10 | 256.573 | NaN | NaN | NaN | NaN | NaN |
| Hprt1 | DLNHVCVISETGK | -201 | -213 | 0 | 0 | 13 | 3943.0637 | 1529.4058 | 1572.0447 | 2782.3708 | 2864.8809 | 2946.6963 |
| Hprt1 | VIGGDDLSTLTGK | -116 | -128 | 0 | 0 | 13 | 2405.7178 | 2756.1934 | 2633.5916 | 2563.9707 | 478.41223 | 539.1259 |
| Hprt1 | SVGYRPDFVGF | -171 | -181 | 1 | 0.001092529 | 11 | 1887.8483 | 2147.2546 | 2290.8489 | 911.37195 | 1769.5032 | 1736.0486 |
| Hprt1 | ALDYNEYFR | -192 | -200 | 0 | 0 | 9 | 1854.3627 | 2172.823 | 2120.3472 | 2624.258 | 3010.56 | 2837.3618 |
| Hprt1 | SYCNDQSTGDIK | -104 | -115 | 0 | 0 | 12 | 1123.1381 | 89.920876 | 466.7754 | 1243.2395 | 1727.6353 | 1823.4443 |

| | | | | | | | | | | | | |
|---|---|---|---|---|---|---|---|---|---|---|---|---|
| Hprt1 | CIPNHYAEDLEK | -23 | -34 | 0 | 0 | 12 | 684.4983 | 653.7525 | 663.0682 | 991.06934 | 2844.2915 | 708.78406 |
| Hprt1 | NVLIVEDIIDTGK | -129 | -141 | 0 | 0 | 13 | 622.6189 | 670.65985 | 703.4543 | 581.4029 | 832.134 | 889.1245 |
| Hprt1 | HYAEDLEK | -27 | -34 | 0 | 0.001636916 | 8 | 316.4191 | 307.32382 | 611.1268 | 1101.1268 | 496.26855 | 640.77563 |
| Hprt1 | SVGYRPDFVGFEIPDK | -171 | -186 | 1 | 0 | 16 | 269.0108 | 457.27878 | 494.11288 | 705.2452 | 742.9423 | 884.63306 |
| Hprt1 | FFADLLDYIK | -74 | -83 | 0 | 0 | 10 | 181.0311 | 198.53465 | 174.37439 | 226.43808 | 201.12054 | 236.29604 |
| Cndp2 | LGGSVELVDIGK | -55 | -66 | 0 | 0 | 12 | 1839.7197 | 284.34293 | 2001.1936 | 2247.0435 | 2617.4463 | 2670.1846 |
| Cndp2 | AVFQYIDENQDR | -6 | -17 | 0 | 0 | 12 | 1817.9667 | 1661.0411 | 1645.205 | 1379.8328 | 1525.2406 | 1589.1735 |
| Cndp2 | GSTDDKGPVAGW | -129 | -140 | 1 | 0 | 12 | 1456.2457 | 1443.6365 | 1439.0184 | 1314.3115 | 1390.179 | 1267.0773 |
| Cndp2 | VFGVEPDLTR | -404 | -413 | 0 | 0 | 10 | 1432.3848 | 1866.1785 | 2052.7593 | 961.98267 | 1564.3531 | 2381.0537 |
| Cndp2 | ISDNYWLGK | -193 | -201 | 0 | 0 | 9 | 1272.918 | 2107.469 | 2065.6191 | 1660.796 | 2816.8892 | 1963.5192 |
| Cndp2 | DFNHPHYQAGR | -388 | -398 | 0 | 0 | 11 | 1227.5972 | 1309.8524 | 1376.035 | 721.0467 | 1094.3494 | 1037.3618 |
| Cndp2 | DFNHPHYQAGR | -388 | -398 | 0 | 0 | 11 | 1227.5972 | 1309.8524 | 1376.035 | 721.0467 | 1094.3494 | 1037.3618 |
| Cndp2 | VSDFNHPHYQAGR | -386 | -398 | 0 | 0 | 13 | 1148.1775 | 789.7966 | 1102.4476 | 606.92163 | 1450.328 | 1472.2349 |
| Cndp2 | VSDFNHPHYQAGR | -386 | -398 | 0 | 0 | 13 | 1148.1775 | 789.7966 | 1102.4476 | 606.92163 | 1450.328 | 1472.2349 |
| Cndp2 | NVMLLPVGSADDGAHSQ | -431 | -447 | 0 | 0 | 17 | 1024.0417 | 1041.4055 | 900.7834 | 1110.8198 | 1133.2863 | 1161.2567 |
| Cndp2 | ALEDGWDSEPFTLVER | -106 | -121 | 0 | 0 | 16 | 959.41675 | 790.8153 | 814.2615 | 870.7532 | 1119.5104 | 1022.59375 |
| Cndp2 | QKLPDGSEIPLPPILLGK | -67 | -84 | 1 | 0 | 18 | 894.76416 | 823.6763 | 933.72424 | 954.28076 | 1002.84656 | 971.4132 |
| Cndp2 | QKLPDGSEIPLPPILLGK | -67 | -84 | 1 | 0 | 18 | 894.76416 | 823.6763 | 933.72424 | 954.28076 | 1002.84656 | 971.4132 |
| Cndp2 | TVFGVEPDLTR | -403 | -413 | 0 | 0 | 11 | 573.6194 | 717.1129 | 863.2958 | 1171.7217 | 1764.7316 | 1760.58 |
| Cndp2 | ILIPGINDAVAPVTDEEHAL | -257 | -276 | 0 | 0 | 20 | 528.5392 | 443.98062 | 387.5067 | 498.9669 | 690.41797 | 609.3023 |
| Cndp2 | NVMLLPVGSADDGAHSQNEK | -431 | -450 | 0 | 0 | 20 | 385.61298 | 419.7413 | 416.0052 | 490.53036 | 647.7627 | 651.79974 |
| Cndp2 | NVMLLPVGSADDGAHSQNEK | -431 | -450 | 0 | 0 | 20 | 385.61298 | 419.7413 | 416.0052 | 490.53036 | 647.7627 | 651.79974 |
| Cndp2 | SVSAWPEKR | -30 | -38 | 1 | 0.00025015 | 9 | 305.04218 | 946.4762 | 670.96564 | 664.16583 | 1072.7306 | 1043.2605 |
| Cndp2 | LPDGSEIPLPPILLGK | -69 | -84 | 0 | 0.000230707 | 16 | 273.13275 | 246.88269 | 326.5794 | 288.85083 | 353.48306 | 383.91876 |
| Cndp2 | NKPCITYGLR | -202 | -211 | 1 | 0.00025015 | 10 | 168.12595 | 1917.5675 | 2004.9512 | 5113.8564 | 1495.6775 | 2144.0017 |
| Cndp2 | TGQEIPVNLR | -150 | -159 | 0 | 0.000230707 | 10 | 3847.3342 | 2856.6804 | NaN | 4829.5293 | 2974.507 | 4435.703 |
| Cndp2 | LNYIEGTK | -454 | -461 | 0 | 0 | 8 | 1163.015 | 1184.2446 | 1185.8809 | NaN | 957.11444 | 1045.8602 |
| Cndp2 | YIDENQDR | -10 | -17 | 0 | 0 | 8 | 524.3486 | 563.517 | 687.9862 | NaN | 577.13245 | 610.725 |
| Krt10 | LENEIQTYR | -440 | -448 | 0 | 0 | 9 | 632.2431 | 1226.5471 | 1356.7233 | 1191.0706 | 2408.2505 | NaN |
| Krt10 | ALEESNYELEGK | -164 | -175 | 0 | 0 | 12 | 331.5782 | 287.70688 | 410.7483 | NaN | 556.2023 | 709.3734 |
| Krt10 | ISALEEQLQQIR | -409 | -420 | 0 | 0 | 12 | 291.18896 | NaN | NaN | NaN | NaN | 437.86404 |
| Cltc | GQFSTDELVAEVEKR | -838 | -852 | 1 | 0 | 15 | 5884.2275 | 5495.3247 | 6605.0137 | 6734.203 | 7252.349 | 6738.6025 |
| Cltc | QLPLVKPY | -1444 | -1451 | 1 | 0.00204437 | 8 | 3489.1565 | 241.16676 | 165.41177 | 3237.776 | 2727.3933 | 2519.1924 |
| Cltc | EDKLECSEELGDLVK | -454 | -468 | 1 | 0 | 15 | 3371.1711 | 1220.333 | 1192.3993 | 1228.2493 | 1263.8427 | 1282.9255 |
| Cltc | FQEHLQLQ | -9 | -16 | 0 | 0.003490217 | 8 | 2609.385 | 5553.574 | 5613.733 | 5454.7744 | 5356.0293 | 1811.9221 |
| Cltc | LTDQLPLIIVCDR | -768 | -780 | 0 | 0 | 13 | 2544.4792 | 2532.0874 | 2695.56 | 3136.9644 | 4000.8625 | 4162.033 |
| Cltc | HELIEFR | -1502 | -1508 | 0 | 0.000471409 | 7 | 2357.676 | 2440.46 | 2616.503 | 2462.686 | 2762.3025 | 2743.3103 |
| Cltc | IVLDNSVFSEHR | -1011 | -1022 | 0 | 0 | 12 | 2221.5479 | 2603.633 | 2662.9963 | 2711.9048 | 3769.8032 | 3585.0847 |
| Cltc | IVLDNSVFSEHR | -1011 | -1022 | 0 | 0 | 12 | 2221.5479 | 2603.633 | 2662.9963 | 2711.9048 | 3769.8032 | 3585.0847 |
| Cltc | NNRPSEGPLQTR | -572 | -583 | 1 | 0 | 12 | 2112.666 | 2413.4165 | 2549.3967 | 1303.1486 | 2322.204 | 2586.3638 |
| Cltc | NNRPSEGPLQTR | -572 | -583 | 1 | 0 | 12 | 2112.666 | 2413.4165 | 2549.3967 | 1303.1486 | 2322.204 | 2586.3638 |
| Cltc | VLQNPDLALR | -341 | -350 | 0 | 0.00025015 | 10 | 2064.485 | 1928.7368 | 1990.6715 | 1751.9939 | 2071.5503 | 1991.2166 |
| Cltc | ALSETQDPEEVSVTVK | -978 | -993 | 0 | 0.000230707 | 16 | 2022.4828 | 1885.7391 | 1447.9673 | 2775.2444 | 3445.1118 | 3188.9663 |
| Cltc | YNNVSNFGR | -1218 | -1226 | 0 | 0 | 9 | 1951.2639 | 3469.3052 | 3531.2417 | 422.70172 | 667.20325 | 336.27203 |
| Cltc | NNLAGAEELFAR | -355 | -366 | 0 | 0 | 12 | 1701.7439 | 2069.343 | 1923.3147 | 347.34573 | 2790.2632 | 543.4898 |
| Cltc | RPLIDQVVQT | -968 | -977 | 1 | 0.001846793 | 10 | 1518.9021 | 1675.1735 | 1572.3206 | 1536.8722 | 1703.7402 | 1760.492 |
| Cltc | LHIIEVGTPPTGNQPFPK | -228 | -245 | 0 | 0 | 18 | 1502.9576 | 1735.4836 | 1772.2906 | 1860.3467 | 2525.3423 | 2476.926 |
| Cltc | LHIIEVGTPPTGNQPFPK | -228 | -245 | 0 | 0 | 18 | 1502.9576 | 1735.4836 | 1772.2906 | 1860.3467 | 2525.3423 | 2476.926 |
| Cltc | ALEHFTDLYDIK | -626 | -637 | 0 | 0 | 12 | 1487.7128 | 1903.0189 | 1997.1982 | 1993.4882 | 2492.6887 | 2330.597 |
| Cltc | VLIEHIGNLDR | -1084 | -1094 | 0 | 0 | 11 | 1441.0995 | 1293.4834 | 1533.1094 | 366.1754 | 1692.3226 | 1539.613 |
| Cltc | IEHIGNLDR | -1086 | -1094 | 0 | 0.001190916 | 9 | 1352.5277 | 1385.8892 | 1406.5326 | 1247.5664 | 1071.858 | 1161.5918 |
| Cltc | SVEDSLECLR | -659 | -668 | 0 | 0 | 10 | 1314.4348 | 1447.4017 | 1488.8882 | 344.76865 | 122.96205 | 1551.1635 |
| Cltc | LDNYDAPDIANIA | -1047 | -1059 | 0 | 0.00025015 | 13 | 1207.5188 | 1349.9448 | 1353.5106 | 1619.6954 | 811.462 | 2138.637 |
| Cltc | EHFTDLYDIK | -628 | -637 | 0 | 0.000230707 | 10 | 1025.1038 | 1376.4304 | 474.48386 | 1668.1029 | 1734.8812 | 458.7986 |
| Cltc | RPLIDQVVQTA | -968 | -978 | 1 | 0.00025015 | 11 | 1004.38855 | 1262.9844 | 1107.1539 | 1190.262 | 1101.98 | 1330.8655 |
| Cltc | IDSNNNPER | -884 | -892 | 0 | 0 | 9 | 947.6266 | 793.4749 | 987.3951 | 886.01465 | 1111.8188 | 918.15216 |
| Cltc | IHEGCEEPATHNAL | -866 | -879 | 0 | 0 | 14 | 902.93304 | 850.4749 | 832.5653 | 574.238 | 451.66177 | 389.52908 |
| Cltc | ITNVLQNPDLALR | -338 | -350 | 0 | 0 | 13 | 880.79736 | 957.7521 | 855.28546 | 935.54877 | 1317.9072 | 1217.8851 |
| Cltc | KVSQPIEGHAA | -189 | -199 | 1 | 0 | 11 | 870.4029 | 572.47864 | 730.2609 | 735.1903 | 698.0717 | 832.738 |
| Cltc | AVQVLIEHIGNLDR | -1081 | -1094 | 0 | 0 | 14 | 760.338 | 927.5706 | 875.108 | 1025.4971 | 1143.6013 | 1176.92 |
| Cltc | LLLPWLEAR | -857 | -865 | 0 | 0 | 9 | 756.0168 | 918.9005 | 962.9589 | 833.098 | 1183.6791 | 1298.158 |
| Cltc | ESNCYDPER | -750 | -758 | 0 | 0 | 9 | 750.0792 | 802.51086 | 736.35114 | 66.03331 | 744.7815 | 805.6918 |
| Cltc | GLLDVDCSEDVIK | -818 | -830 | 0 | 0 | 13 | 717.12506 | 687.82275 | 836.81866 | 1160.003 | 1072.9231 | 1267.2711 |
| Cltc | LAELEEFINGPNNAHIQ | -1183 | -1199 | 0 | 0 | 17 | 591.612 | 600.93494 | 698.28265 | 760.9209 | 861.2536 | 786.0252 |
| Cltc | ISGETIFVTAPHEATAG | -298 | -314 | 0 | 0 | 17 | 591.34076 | 625.191 | 832.4243 | 274.47516 | 704.1427 | 722.6566 |
| Cltc | TAFLLDALK | -563 | -571 | 0 | 0 | 9 | 524.37665 | 256.24347 | 612.74756 | 649.1092 | 629.312 | 672.4348 |
| Cltc | RALEHFTDLYDIK | -625 | -637 | 1 | 0 | 13 | 272.4924 | 519.16736 | 464.91992 | 506.65973 | 678.4534 | 555.27216 |
| Cltc | NHPTDAWK | -1380 | -1387 | 0 | 0 | 8 | 198.52158 | 247.18404 | 231.41933 | 1506.898 | 1342.5603 | 1968.4453 |
| Cltc | HDVVFLITK | -270 | -278 | 0 | 0 | 9 | 4751.2407 | 2007.9215 | 1622.6852 | NaN | 4210.2354 | 4105.5884 |
| Cltc | LDQGQLNK | -422 | -429 | 0 | 0.00051306 | 8 | 1909.683 | 2229.3489 | 2148.3845 | 2171.6946 | NaN | 2841.31 |
| Cltc | RKDPELWGSVL | -950 | -960 | 2 | 0.002168954 | 11 | 1880.5913 | 861.2282 | 996.80176 | 4050.0415 | NaN | 1184.0199 |
| Cltc | LECSEELGDLVK | -457 | -468 | 0 | 0 | 12 | 1062.0565 | 222.52686 | 925.345 | 1615.6162 | 1054.3042 | NaN |
| Cltc | VANVELYYK | -1398 | -1406 | 0 | 0 | 9 | 816.7597 | 719.01465 | 1002.3025 | 1225.7225 | 603.1076 | NaN |
| Cltc | GILLDQGQLNK | -419 | -429 | 0 | 0 | 11 | 676.4006 | 627.53754 | 448.49646 | NaN | 466.96426 | 462.62094 |
| Cltc | KDPELWGSVL | -951 | -960 | 1 | 0.00111006 | 10 | 581.80975 | 629.1482 | 618.1227 | NaN | 601.6614 | 516.5806 |
| Cltc | NVLQNPDLALR | -340 | -350 | 0 | 0 | 11 | 505.70737 | 707.00323 | 471.77396 | NaN | 1195.4797 | 795.7419 |
| Cltc | HSSLAGCQIINYR | -145 | -157 | 0 | 0 | 13 | 318.4777 | 521.60986 | NaN | 124.84186 | 773.29956 | 777.21136 |
| Cltc | VSQPIEGHAA | -190 | -199 | 0 | 0.003620005 | 10 | 165.75322 | 4493.025 | 282.3846 | 6162.352 | 5452.0967 | NaN |

| | | | | | | | | | | | | |
|---|---|---|---|---|---|---|---|---|---|---|---|---|
| Cltc | NFSQDPDVHFK | -720 | -730 | 0 | 0 | 11 | 2070.4756 | 407.48975 | 2091.36 | 414.02835 | NaN | NaN |
| Cltc | WFLQEEK | -1556 | -1562 | 0 | 0 | 7 | 1333.3593 | 1626.5524 | 1318.2821 | NaN | 1014.3281 | NaN |
| Cltc | EGCEEPATHNALAK | -868 | -881 | 0 | 0.000230707 | 14 | 1260.3164 | NaN | NaN | 663.8158 | 1296.0322 | 1302.8257 |
| Cltc | RKDPELWGS | -950 | -958 | 2 | 0.002534621 | 9 | 1165.004 | 1191.7827 | 1328.1451 | NaN | NaN | 1606.2202 |
| Cltc | ISGETIFVTAPHEATA | -298 | -313 | 0 | 0 | 16 | 845.26514 | 730.90967 | 1379.2334 | NaN | 1298.7654 | NaN |
| Cltc | YHEQLSTQ | -689 | -696 | 0 | 0.00025015 | 8 | 378.45117 | 712.0564 | 780.5184 | 567.3148 | NaN | NaN |
| Cltc | IYIDSNNNPER | -882 | -892 | 0 | 0 | 11 | 202.78607 | NaN | NaN | 1391.9258 | 1226.8738 | 1455.2449 |
| Cltc | TSIDAYDNFDNISLAQR | -1482 | -1498 | 0 | 0 | 17 | 3221.2168 | NaN | 3481.2637 | NaN | 451.1639 | NaN |
| Cltc | QNPDLALR | -343 | -350 | 0 | 0.001392869 | 8 | 2151.773 | NaN | NaN | 2701.9712 | NaN | 2088.3926 |
| Cltc | ENPYYDSR | -896 | -903 | 0 | 0 | 8 | 709.4202 | 661.5298 | 692.57367 | NaN | NaN | NaN |
| Cltc | SIDAYDNFDNISLAQR | -1483 | -1498 | 0 | 0 | 16 | 496.34778 | 853.2865 | 758.1483 | NaN | NaN | NaN |
| Cltc | IHEGCEEPATHNALAK | -866 | -881 | 0 | 0.00025015 | 16 | 330.87964 | 279.3134 | 277.8781 | NaN | NaN | NaN |
| Cltc | FAQGNYSEAAK | -372 | -382 | 0 | 0 | 11 | 257.59998 | NaN | NaN | 880.73895 | NaN | NaN |
| Sgta | ALELDPDNDTYK | -186 | -197 | 0 | 0 | 12 | 892.27136 | 856.72986 | 936.95245 | 718.0675 | 716.0552 | NaN |
| Rdx | EIHKPGYLANDR | -140 | -151 | 1 | 0 | 12 | 809.985 | 941.9897 | 866.9191 | 931.0543 | 1045.5454 | 899.73206 |
| Rdx | KTQNDVLHAE | -544 | -553 | 1 | 0.001392869 | 10 | 779.91425 | 812.6266 | 758.04486 | 836.7881 | 968.8259 | 1119.8633 |
| Rdx | NQEQLAAELAEFTAK | -413 | -427 | 0 | 0 | 15 | 544.23724 | 534.6184 | 750.6487 | 751.9117 | 910.0586 | 969.03046 |
| Rdx | KALELEQER | -371 | -379 | 1 | 0 | 9 | 474.5939 | 760.20856 | 1043.5853 | 521.5396 | 364.49295 | 848.22095 |
| Rdx | IQNWHEEHR | -172 | -180 | 0 | 0 | 9 | 430.52664 | 563.8528 | 505.0623 | 854.15753 | 459.0515 | 1184.643 |
| Pcmt1 | ELLFDQLHEGAK | -70 | -81 | 0 | 0 | 12 | 2461.8804 | 2760.7842 | 2812.2317 | 2770.3318 | 2382.4465 | 2262.343 |
| Pcmt1 | ALELLFDQLHEGAK | -68 | -81 | 0 | 0 | 14 | 1294.4624 | 1575.436 | 1523.3029 | 657.51385 | 818.0927 | 888.2863 |
| Pcmt1 | ALIDQLKPGGR | -168 | -178 | 1 | 0 | 11 | 948.72314 | 1077.916 | 976.22046 | 1054.0303 | 1368.9951 | 1327.0366 |
| Pcmt1 | SGGASHSELIHNL | -5 | -17 | 0 | 0 | 13 | 623.9968 | NaN | NaN | 469.30093 | 561.4749 | NaN |
| Pcmt1 | ELVDDSITNVK | -114 | -124 | 0 | 0 | 11 | 1447.6508 | NaN | 991.94324 | NaN | NaN | NaN |
| Itih3 | FAHNVVTTR | -58 | -66 | 0 | 0.00025015 | 9 | 2766.0083 | 2952.929 | 2975.9224 | 1762.237 | 2611.7373 | 2745.8684 |
| Itih3 | YHFVTPLTS | -592 | -600 | 0 | 0.001055699 | 9 | 2063.2847 | 2046.1454 | 438.44946 | 724.57 | 596.1053 | 1998.0374 |
| Itih3 | ALENHGLAR | -440 | -448 | 0 | 0 | 9 | 1050.4875 | 1090.7332 | 1023.4748 | 971.3219 | 1179.5333 | 1232.0956 |
| Itih3 | RIYEDSDANLQLQ | -449 | -461 | 1 | 0.000298805 | 13 | 1023.751 | 905.1029 | 1198.8408 | NaN | 1347.1963 | 862.2572 |
| Itih3 | FVHFFAPQGLPVVPK | -269 | -283 | 0 | 0.00122419 | 15 | 582.2366 | NaN | 652.4821 | 1384.9419 | NaN | NaN |
| Htatsf1 | ALEPEEGNPQTEAQENGPER | -481 | -500 | 0 | 0 | 20 | 378.02045 | 331.3608 | 392.6724 | NaN | 472.82407 | 451.35333 |
| Sugt1 | AYCHILLGK | -52 | -60 | 0 | 0 | 9 | 3557.22 | 2137.5273 | 330.60056 | 1301.8696 | 4034.1594 | 328.49506 |
| Sugt1 | IYSDGSDEVKR | -287 | -297 | 1 | 0 | 11 | 2878.5027 | 2582.436 | 2370.5037 | 1645.5557 | 825.0153 | 793.5312 |
| Sugt1 | VKIPAGEDYSLK | -181 | -192 | 1 | 0 | 12 | 2365.8845 | 869.8354 | 2344.7966 | 1528.77 | 1373.132 | 2841.7195 |
| Sugt1 | LEGQGDEPTPK | -228 | -238 | 0 | 0 | 11 | 728.7948 | 885.45605 | 744.41394 | 745.6508 | 590.3641 | 147.02686 |
| Sugt1 | LLHPIIPEQSTFK | -195 | -207 | 0 | 0 | 13 | 707.3886 | 893.194 | 445.5121 | 747.9713 | 980.0911 | 941.72723 |
| Sugt1 | LVGEIKEEEK | -261 | -270 | 1 | 0 | 10 | 684.02814 | 928.39374 | 904.1771 | 1545.8231 | 2355.1663 | 1556.9285 |
| Sugt1 | SLELNPNNCTALLR | -71 | -84 | 0 | 0 | 14 | 681.131 | 568.4035 | 826.1069 | 2680.7527 | 3078.6855 | 2240.064 |
| Sugt1 | ALEQNPDDAQYYCQR | -37 | -51 | 0 | 0 | 15 | 651.9521 | 589.81177 | 297.5146 | 384.68445 | 471.90924 | 253.0385 |
| Sugt1 | LDSTDTNFDTWIK | -108 | -120 | 0 | 0 | 13 | 651.2128 | 701.58154 | 329.03534 | 651.5986 | 259.50174 | 709.97144 |
| Sugt1 | SDALIDGDPQAALEELTK | -19 | -36 | 0 | 0 | 18 | 533.5686 | 589.801 | 522.0478 | 549.64197 | 778.8509 | 685.0901 |
| Sugt1 | LVGEIKEEEKNEK | -261 | -273 | 2 | 0 | 13 | 473.97327 | 785.1229 | 731.2832 | 626.95776 | 1084.8566 | 757.73444 |
| Sugt1 | LVGEIKEEEKNEK | -261 | -273 | 2 | 0 | 13 | 473.97327 | 785.1229 | 731.2832 | 626.95776 | 1084.8566 | 757.73444 |
| Sugt1 | LLHPIIPEQSTF | -195 | -206 | 0 | 0 | 12 | 467.01965 | 431.7067 | 603.181 | 976.95105 | 907.3957 | 940.88367 |
| Sugt1 | EKLEGQGDEPTPK | -226 | -238 | 1 | 0 | 13 | 393.28085 | 163.1684 | 513.88916 | 412.58902 | 449.41095 | 409.27686 |
| Sugt1 | YRDGIADVK | -61 | -69 | 1 | 0.00025015 | 9 | 2305.1702 | 1550.788 | 2073.4636 | 1900.3257 | 1449.0404 | NaN |
| Sugt1 | IPAGEDYSLK | -183 | -192 | 0 | 0.00111006 | 10 | 1416.3079 | 1453.469 | NaN | NaN | 707.8047 | 668.35144 |
| Sugt1 | LEGDAALNK | -274 | -282 | 0 | 0 | 9 | 1129.6031 | NaN | NaN | 1035.5151 | NaN | NaN |
| Sugt1 | STNWSDVGK | -312 | -320 | 0 | 0.002697088 | 9 | 623.2673 | NaN | NaN | NaN | NaN | NaN |
| Nans | EIGQNHQGDIDVAK | -24 | -37 | 0 | 0 | 14 | 1978.6919 | 1028.7834 | 992.7879 | 1070.696 | 1151.7219 | 1208.2944 |
| Nans | EIGQNHQGDIDVAK | -24 | -37 | 0 | 0 | 14 | 1978.6919 | 1028.7834 | 992.7879 | 1070.696 | 1151.7219 | 1208.2944 |
| Nans | VGSGDTNNFPYLEK | -132 | -145 | 0 | 0 | 14 | 1563.763 | 1964.7998 | 1893.7233 | 1951.6733 | 2047.945 | 2305.1406 |
| Nans | ALERPYTSK | -66 | -74 | 1 | 0 | 9 | 1454.9567 | 759.66876 | 834.5095 | 626.4868 | 697.50055 | 662.23206 |
| Nans | HLEFSHDQYK | -87 | -96 | 0 | 0 | 10 | 1249.585 | 1223.9158 | 1436.543 | 1332.309 | 1792.3712 | 1682.3861 |
| Nans | SAYPLQPEDANLR | -186 | -198 | 0 | 0 | 13 | 924.12506 | 1007.76917 | 657.6351 | 1059.0588 | 859.11554 | 808.2474 |
| Nans | GSDHSASLEPGELAELVR | -247 | -264 | 0 | 0 | 18 | 571.61017 | 668.5608 | 685.0316 | 761.4708 | 1046.3766 | 1093.648 |
| Nans | GSDHSASLEPGELAELVR | -247 | -264 | 0 | 0 | 18 | 571.61017 | 668.5608 | 685.0316 | 761.4708 | 1046.3766 | 1093.648 |
| Nans | CTSAYPLQPEDANLR | -184 | -198 | 0 | 0 | 15 | 427.58694 | 696.4813 | 375.75824 | 550.637 | 565.6564 | 468.3623 |
| Nans | PLNPNFCFLQ | -174 | -183 | 0 | 0 | 10 | 657.0971 | 627.4469 | NaN | 953.4611 | 1045.8434 | 506.75882 |
| Nans | PLELELCPGR | -2 | -11 | 0 | 0 | 10 | 595.70636 | 310.3399 | 1374.2856 | 1205.1306 | NaN | 1644.8467 |
| Nans | QVYQIVKPLNPNFCFLQ | -167 | -183 | 1 | 0.002099344 | 17 | 207.43301 | NaN | 176.35147 | 192.36948 | 273.4875 | 227.6055 |
| Nans | AYPLQPEDANLR | -187 | -198 | 0 | 0.000550307 | 12 | 521.1446 | NaN | 487.23807 | 1242.5917 | NaN | NaN |
| Nans | KSELEFK | -55 | -61 | 1 | 0 | 7 | 629.7676 | NaN | NaN | NaN | NaN | 610.5253 |
| Nans | VKIPAGTTL | -299 | -307 | 1 | 0 | 9 | 525.05396 | NaN | NaN | NaN | NaN | NaN |
| Kif11 | EEDPLAGIIPR | -127 | -137 | 0 | 0.000471409 | 11 | 808.8396 | 844.8007 | 338.26166 | 305.4404 | 763.06476 | 763.3479 |
| Kif11 | ALESLVSIPENVSAR | -584 | -598 | 0 | 0 | 15 | 437.38153 | 450.41864 | 701.16113 | 474.91364 | 597.24475 | 568.6486 |
| Kif11 | NEELFDLLSPSSDVSER | -164 | -180 | 0 | 0 | 17 | 299.9751 | NaN | NaN | NaN | 198.09323 | 334.9886 |
| Kif11 | EQLLDQLQK | -944 | -952 | 0 | 0 | 9 | 798.41754 | NaN | 827.923 | NaN | NaN | NaN |
| Oat | KTEQGPPSSEYIFER | -32 | -46 | 1 | 0 | 15 | 1702.8884 | 1985.4706 | 2306.7976 | 627.0459 | 152.29373 | 2905.099 |
| Oat | KTEQGPPSSEYIFER | -32 | -46 | 1 | 0 | 15 | 1702.8884 | 1985.4706 | 2306.7976 | 627.0459 | 152.29373 | 2905.099 |
| Oat | NYHPLPVALER | -54 | -64 | 0 | 0 | 11 | 2031.7168 | 2068.7153 | 2173.5862 | 1611.0177 | 2336.2517 | 2020.2372 |
| Oat | NYHPLPVALER | -54 | -64 | 0 | 0 | 11 | 2031.7168 | 2068.7153 | 2173.5862 | 1611.0177 | 2336.2517 | 2020.2372 |
| Oat | TEQGPPSSEYIFER | -33 | -46 | 0 | 0 | 14 | 1113.8079 | 1011.8118 | 811.4783 | 1253.3735 | 1423.7626 | 1574.5063 |
| Oat | NVLGEYEEYITK | -118 | -129 | 0 | 0 | 12 | 940.29193 | 916.1533 | 415.8051 | 549.16077 | 649.165 | 956.03815 |
| Oat | YNNVLGEYEEYITK | -116 | -129 | 0 | 0 | 14 | 825.6565 | 972.85754 | 853.3094 | 494.935 | 612.39996 | 751.02795 |
| Oat | EVLEEENLAENADK | -338 | -351 | 0 | 0.000471409 | 14 | 1401.9764 | 1620.1577 | 1828.9781 | NaN | 1315.8679 | 1264.0044 |
| Oat | LAPPLVIK | -414 | -421 | 0 | 0.00201047 | 8 | 737.724 | 629.3582 | 617.347 | 608.0314 | NaN | 397.67075 |
| Oat | IVFADGNFWGR | -170 | -180 | 0 | 0 | 11 | 518.80493 | 332.09647 | 595.23395 | 216.20618 | NaN | NaN |
| Oat | GLLAKPTHGDIIR | -401 | -413 | 1 | 0.000550307 | 13 | 216.83452 | NaN | 804.7001 | NaN | 345.6432 | 557.887 |

| | | | | | | | | | | | | |
|---|---|---|---|---|---|---|---|---|---|---|---|---|
| Oat | GLLNAIVIR | -375 | -383 | 0 | 0 | 9 | 6097.268 | 4731.9854 | 5449.234 | NaN | NaN | NaN |
| Oat | IADEIQTGLAR | -261 | -271 | 0 | 0 | 11 | 1184.114 | NaN | NaN | NaN | 1486.0823 | NaN |
| Oat | ALEVLEEENLAENADK | -336 | -351 | 0 | 0 | 16 | 988.66675 | NaN | 1049.9419 | NaN | NaN | NaN |
| Psmd2 | FGGSGSQVDSAR | -358 | -369 | 0 | 0 | 12 | 12394.5 | 1327.3153 | 10693.285 | 9759.841 | 12510.051 | 573.773 |
| Psmd2 | YVPEPENSALLR | -239 | -250 | 0 | 0 | 12 | 2581.6738 | 1826.129 | 1814.163 | 1605.0894 | 1392.7024 | 1650.4896 |
| Psmd2 | TFGHLLR | -674 | -680 | 0 | 0 | 7 | 1910.5432 | 1610.4452 | 1759.6567 | 1726.8568 | 1049.9487 | 1536.9138 |
| Psmd2 | YLYSSEDYIK | -432 | -441 | 0 | 0 | 10 | 1780.9495 | 948.0477 | 734.96173 | 1655.4761 | 1586.1871 | 1471.0874 |
| Psmd2 | DEELRPLPVSVR | -834 | -845 | 1 | 0 | 12 | 1725.2786 | 1136.215 | 1182.2073 | 485.12207 | 1287.2507 | 1373.7975 |
| Psmd2 | DEELRPLPVSVR | -834 | -845 | 1 | 0 | 12 | 1725.2786 | 1136.215 | 1182.2073 | 485.12207 | 1287.2507 | 1373.7975 |
| Psmd2 | WLPLGLGLNHLGK | -557 | -569 | 0 | 0 | 13 | 1038.7771 | 1079.1313 | 1131.1968 | 149.6159 | 544.6385 | 635.4992 |
| Psmd2 | ALEVVSEPFR | -579 | -588 | 0 | 0 | 10 | 850.9727 | 764.8126 | 932.4877 | 610.48083 | 619.36096 | 763.56464 |
| Psmd2 | VGQAVDVVGQAGKPK | -846 | -860 | 1 | 0 | 15 | 751.81683 | 658.9501 | 765.297 | 677.07306 | 799.04816 | 739.41895 |
| Psmd2 | DKTPVQSQQPSATTPSGADEK | -7 | -27 | 1 | 0 | 21 | 502.62152 | 546.6712 | 568.0756 | 506.555 | 599.54346 | 618.8837 |
| Psmd2 | DKTPVQSQQPSATTPSGADEK | -7 | -27 | 1 | 0 | 21 | 502.62152 | 546.6712 | 568.0756 | 506.555 | 599.54346 | 618.8837 |
| Psmd2 | RYGEPTLR | -680 | -687 | 1 | 0.000230707 | 8 | 426.5247 | 1130.61 | 1156.1582 | 7355.6006 | 1471.3044 | 1495.8749 |
| Psmd2 | LVGSQEELASWGHEYVR | -144 | -160 | 0 | 0 | 17 | 248.08356 | 449.24362 | 521.5417 | 544.21106 | 829.0078 | 675.6536 |
| Psmd2 | VNYVPEPENSALLR | -237 | -250 | 0 | 0 | 14 | 2157.437 | 2544.767 | NaN | 2528.7905 | 3168.4028 | 3095.7212 |
| Psmd2 | DKEQELSEEDK | -40 | -50 | 1 | 0 | 11 | 1130.0093 | 1307.6416 | 1233.0984 | NaN | 1152.97 | 1231.4023 |
| Psmd2 | FDEELRPLPVSVR | -833 | -845 | 1 | 0 | 13 | 578.5382 | 738.33234 | NaN | 857.26904 | 1164.7935 | 1228.6742 |
| Psmd2 | TPVQSQQPSATTPSGADEK | -9 | -27 | 0 | 0.000712251 | 19 | 382.81125 | 473.22034 | NaN | 418.53033 | 586.9225 | 550.0727 |
| Psmd2 | EWQELDDAEK | -169 | -178 | 0 | 0 | 10 | 87.072266 | 76.10632 | 1789.009 | 1182.7788 | 651.3389 | NaN |
| Psmd2 | LWDVDGGLTQIDK | -419 | -431 | 0 | 0 | 13 | 2636.9287 | 2117.801 | NaN | NaN | 1834.1077 | 2673.156 |
| Psmd2 | LNILDTLSK | -704 | -712 | 0 | 0 | 9 | 327.72488 | 763.8398 | NaN | NaN | 625.3548 | 281.35867 |
| Psmd2 | FLRPHYGK | -98 | -105 | 1 | 0.002752526 | 8 | 901.5298 | 308.9548 | 542.1552 | NaN | NaN | NaN |
| Psmd2 | EPLLTLVK | -182 | -189 | 0 | 0 | 8 | 529.07074 | 521.52216 | 512.7394 | NaN | NaN | NaN |
| Psmd2 | LEVVSEPFR | -580 | -588 | 0 | 0.000712251 | 9 | 794.18115 | 1437.2517 | NaN | NaN | NaN | NaN |
| Psmd2 | CVNYVPEPENSALLR | -236 | -250 | 0 | 0.00025015 | 15 | 998.1243 | NaN | NaN | NaN | NaN | NaN |
| Grb2 | FDFDPQEDGELGFR | -165 | -178 | 0 | 0 | 14 | 1571.3947 | 89.12467 | 1752.8867 | 2079.9526 | 132.45447 | 2622.5652 |
| Grb2 | FNSLNELVDYHR | -125 | -136 | 0 | 0 | 12 | 652.8419 | 1172.0645 | 962.4532 | 655.1062 | 1015.98444 | 1284.904 |
| Grb2 | NEECDQNWYK | -29 | -38 | 0 | 0 | 10 | 648.5555 | 612.0377 | 598.64905 | 58.831646 | 598.0377 | 605.0217 |
| Grb2 | NSLNELVDYHR | -126 | -136 | 0 | 0 | 11 | 503.25372 | 627.5646 | 1064.9216 | 522.66534 | 1064.1776 | 1163.8491 |
| Grb2 | LFDFDPQEDGELGFR | -164 | -178 | 0 | 0 | 15 | 294.72266 | 230.50253 | 368.30228 | 403.3539 | 369.91837 | 344.55673 |
| Grb2 | NQQIFLR | -143 | -149 | 0 | 0 | 7 | 1731.3339 | 1420.4744 | 1651.0776 | 1492.344 | 1892.9221 | NaN |
| Grb2 | SLNELVDYHR | -127 | -136 | 0 | 0 | 10 | 938.2191 | NaN | 980.5439 | 1424.6041 | 1499.1815 | 563.95905 |
| Grb2 | FGNDVQHFK | -101 | -109 | 0 | 0 | 9 | 435.65683 | 1833.2571 | 696.1157 | NaN | 1237.8547 | 622.91724 |
| Grb2 | ALFDFDPQEDGELGFR | -163 | -178 | 0 | 0 | 16 | 220.67674 | 356.1355 | 145.84161 | NaN | 333.1177 | 333.827 |
| Grb2 | FGNDVQHF | -101 | -108 | 0 | 0 | 8 | 1881.516 | 1559.513 | 1170.0767 | NaN | 2123.3186 | NaN |
| Atic | SDICDVPTAK | -322 | -331 | 0 | 0 | 10 | 5418.475 | 280.8298 | 3405.9788 | 3354.8738 | 3416.7058 | 3151.993 |
| Atic | EVSDGIVAPGYEEEALK | -336 | -352 | 0 | 0.00025015 | 17 | 2212.0334 | 2582.197 | 2925.7102 | 2837.1528 | 3731.0808 | 3262.6816 |
| Atic | HVSPAGAAVGVPL | -267 | -279 | 0 | 0.00078661 | 13 | 2033.3 | 2231.985 | 729.1583 | 3013.2476 | 2827.1375 | 3230.1787 |
| Atic | SDAFFPFR | -538 | -545 | 0 | 0 | 8 | 1768.0938 | 2003.9241 | 1952.3896 | 1486.6986 | 1609.6157 | 1726.2101 |
| Atic | IVAPSGSTADK | -559 | -569 | 0 | 0.00025015 | 11 | 1724.0543 | 1884.0334 | 1824.5544 | 1603.776 | 1414.9573 | 1690.5896 |
| Atic | LDFNLVR | -91 | -97 | 0 | 0 | 7 | 1545.0278 | 1404.8044 | 1365.3506 | 920.4968 | 599.37213 | 632.426 |
| Atic | TLHPAVHAGIL | -67 | -77 | 0 | 0 | 11 | 1376.6677 | 1409.8254 | 1630.9938 | 2426.8782 | 1717.8588 | 741.9552 |
| Atic | FEEVPELLTEAEKK | -512 | -525 | 1 | 0 | 14 | 1319.5594 | 1279.1666 | 1395.0405 | 1502.045 | 1544.0532 | 1357.3923 |
| Atic | YGMNPHQTPAQLY | -208 | -220 | 0 | 0 | 13 | 1079.3125 | 1007.67303 | 1075.9923 | 563.6201 | 1036.2251 | 969.22296 |
| Atic | YGMNPHQTPAQLY | -208 | -220 | 0 | 0 | 13 | 1079.3125 | 1007.67303 | 1075.9923 | 563.6201 | 1036.2251 | 969.22296 |
| Atic | HVSPAGAAVGVPLSEDEAR | -267 | -285 | 0 | 0 | 19 | 1018.5383 | 1080.4868 | 1052.5293 | 1330.1074 | 1584.3774 | 1557.276 |
| Atic | NAIDQYVTGTIGEGEDLVK | -489 | -507 | 0 | 0 | 19 | 789.55585 | 1104.491 | 973.9153 | 724.89667 | 1158.2297 | 128.03491 |
| Atic | QYDEAISDYFR | -184 | -194 | 0 | 0 | 11 | 538.91656 | 533.92053 | 568.7982 | 379.74603 | 686.95575 | 685.53186 |
| Atic | YDEAISDYFR | -185 | -194 | 0 | 0 | 10 | 388.48425 | 353.93515 | 251.3905 | 590.04987 | 618.2473 | 552.9398 |
| Atic | TAQYDEAISDYFR | -182 | -194 | 0 | 0.000550307 | 13 | 346.14795 | 295.3493 | 287.65717 | 620.43823 | 831.0585 | 686.7357 |
| Atic | EALFEEVPELLTEAEKK | -509 | -525 | 1 | 0 | 17 | 277.78323 | 204.51851 | 335.9584 | 465.27927 | 209.2157 | 476.18927 |
| Atic | EALFEEVPELLTEAEK | -509 | -524 | 0 | 0 | 16 | 170.79617 | 270.0935 | 167.72852 | 316.2732 | 360.06287 | 435.40936 |
| Atic | NIPEDAADMAR | -80 | -90 | 0 | 0.002099344 | 11 | 1446.013 | 525.222 | 1373.2983 | 1776.5994 | 1683.7938 | NaN |
| Atic | SYKPDENEVR | -370 | -379 | 1 | 0 | 10 | 433.88376 | 411.94763 | 768.99176 | NaN | 4061.0864 | 3130.2888 |
| Atic | TIGEGEDLVK | -498 | -507 | 0 | 0.000826806 | 10 | 2176.1233 | NaN | 2114.4128 | 2038.8418 | 1877.3257 | NaN |
| Atic | DKTGLVEFAR | -13 | -22 | 1 | 0.000431655 | 10 | 1862.5847 | NaN | NaN | 2241.9143 | 2370.8862 | 2170 |
| Atic | TGLVEFAR | -15 | -22 | 0 | 0 | 8 | 1405.1184 | 619.61145 | 549.806 | 677.881 | NaN | NaN |
| Atic | SGVAYIVAPSGSTADK | -554 | -569 | 0 | 0 | 16 | 885.90326 | NaN | NaN | 1045.5096 | 1366.185 | 1150.4702 |
| Atic | NLYPFVK | -102 | -108 | 0 | 0.002502116 | 7 | 721.5129 | NaN | 636.6642 | 494.28098 | NaN | 135.91771 |
| Atic | ALFEEVPELLTEAEK | -510 | -524 | 0 | 0.00025015 | 15 | 488.15213 | 407.29254 | 534.2182 | NaN | 786.86395 | NaN |
| Atic | EVPELLTEAEK | -514 | -524 | 0 | 0 | 11 | 804.9735 | 1348.3082 | 582.2833 | NaN | NaN | NaN |
| Atic | LFEEVPELLTEAEK | -511 | -524 | 0 | 0.000394361 | 14 | 387.0526 | 345.2365 | 352.81415 | NaN | NaN | NaN |
| Atic | YIVAPSGSTADK | -558 | -569 | 0 | 0.000431655 | 12 | 1160.5599 | NaN | NaN | NaN | NaN | NaN |
| Abcf2 | EFGIDLDTR | -422 | -430 | 0 | 0 | 9 | 792.3025 | 820.4567 | 804.9854 | 1124.0664 | 1137.1932 | 835.7777 |
| Abcf2 | ETTGDGEVDLLTK | -51 | -63 | 0 | 0 | 13 | 2168.8562 | 1450.3767 | 2050.88 | 1998.0061 | 1800.7675 | NaN |
| Abcf2 | EVPIPEHIDIY | -140 | -150 | 0 | 0 | 11 | 670.6951 | NaN | 771.5851 | 1214.0891 | 875.33685 | 355.71246 |
| Abcf2 | ALFIRPF | -247 | -253 | 1 | 0.004791731 | 7 | 393.21823 | 456.85132 | NaN | 400.91873 | 403.94263 | NaN |
| Cdk2 | ALTGIPLPLIK | -95 | -105 | 0 | 0 | 11 | 251.42137 | 298.8365 | 287.49866 | 305.00467 | 337.21527 | 284.60248 |
| Cdk2 | EFLHQDLK | -81 | -88 | 0 | 0.003681885 | 8 | 1290.2576 | 2560.7104 | NaN | 1865.196 | 2000.7958 | 2437.4434 |
| Cdk2 | LDTETEGVPSTAIR | -37 | -50 | 0 | 0 | 14 | 1215.8431 | 829.29785 | 830.6439 | NaN | 1053.884 | 845.1446 |
| Cdk2 | TGIPLPLIK | -97 | -105 | 0 | 0.000230707 | 9 | 245.94846 | 1713.9313 | NaN | 1232.502 | 2055.9216 | 2102.2522 |
| Cdk2 | APEILLGCK | -170 | -178 | 0 | 0.001536401 | 9 | 936.83954 | NaN | NaN | NaN | NaN | 1367.124 |
| Ftl1 | LLEFQNDR | -66 | -73 | 0 | 0.00025015 | 8 | 1654.5537 | 1455.922 | 1628.1583 | 56.53649 | 1387.3186 | 1367.65 |
| Ftl1 | ALFQDVQKPSQDEWGK | -77 | -92 | 1 | 0 | 16 | 879.2434 | 1072.6648 | 949.5756 | 797.9657 | 977.67633 | 997.4747 |
| Ftl1 | ALFQDVQKPSQDEWGK | -77 | -92 | 1 | 0 | 16 | 879.2434 | 1072.6648 | 949.5756 | 797.9657 | 977.67633 | 997.4747 |
| Ftl1 | NLNQALLDLHAL | -106 | -117 | 0 | 0 | 12 | 563.325 | 536.3632 | 580.0902 | 466.8963 | 496.72235 | 496.6235 |

| | | | | | | | | | | | | |
|---|---|---|---|---|---|---|---|---|---|---|---|---|
| Ftl1 | APQGSLGEYLFER | -165 | -177 | 0 | 0 | 13 | 1155.0166 | 1379.8802 | NaN | 1562.2922 | 1770.4976 | 1786.4977 |
| Ftl1 | GAPQGSLGEYLFER | -164 | -177 | 0 | 0 | 14 | 353.46838 | 7777.7026 | 7341.8086 | NaN | 14994.053 | 2072.4895 |
| Ftl1 | VAGPQPAQTGAPQGSLGEYLFER | -155 | -177 | 0 | 0.000431655 | 23 | 183.99898 | NaN | NaN | 262.3502 | 306.74234 | 338.78705 |
| Tbcd | ALGALPGFLLR | -780 | -790 | 0 | 0 | 11 | 335.69412 | 353.68558 | 355.59592 | 348.9889 | 140.9434 | 425.87183 |
| Dcps | VNEDSGDTHGEDAVVIL | -67 | -83 | 0 | 0 | 17 | 1920.6685 | 3062.1472 | 2184.969 | 1352.22 | 805.7768 | 1312.2673 |
| Dcps | RDLTPEHLPLLR | -230 | -241 | 1 | 0 | 12 | 1605.1118 | 1833.1344 | 1771.5403 | 2006.8064 | 2485.3489 | 2468.6418 |
| Dcps | DLTPEHLPLLR | -231 | -241 | 0 | 0 | 11 | 1316.5 | 1532.9751 | 1533.7817 | 1508.7832 | 1891.1542 | 1940.921 |
| Dcps | DLTPEHLPLLR | -231 | -241 | 0 | 0 | 11 | 1316.5 | 1532.9751 | 1533.7817 | 1508.7832 | 1891.1542 | 1940.921 |
| Dcps | ALGFEAPGSGVER | -281 | -293 | 0 | 0 | 13 | 1067.3505 | 1150.6239 | 1126.2034 | 1475.1519 | 1351.8423 | 728.98755 |
| Dcps | AQVIENLECDPK | -298 | -309 | 0 | 0 | 12 | 824.2528 | 1120.3981 | 988.99176 | 1186.1431 | 1407.2255 | 1645.8328 |
| Dcps | EQEAEEAETPSTEEK | -13 | -27 | 0 | 0 | 15 | 736.7927 | 782.37317 | 795.1638 | 808.0917 | 1018.87366 | 998.3502 |
| Dcps | TDDPLLQLLQK | -322 | -332 | 0 | 0.00025015 | 11 | 731.62213 | 791.2903 | 729.809 | 888.06384 | 1460.1873 | 1474.95 |
| Dcps | WVYNILDK | -174 | -181 | 0 | 0 | 8 | 908.5598 | NaN | 1014.1479 | 665.8597 | 549.374 | 646.7285 |
| Dcps | TPFQVEHVAQLL | -86 | -97 | 0 | 0 | 12 | 724.9752 | 615.67456 | NaN | 2178.2776 | 2944.9858 | NaN |
| Dcps | VNEDSGDTHGEDAVVILEK | -67 | -85 | 0 | 0 | 19 | 335.74954 | NaN | 396.76898 | NaN | 403.85596 | 411.50488 |
| Dcps | IVFENPDPSDGFVLIPDLK | -188 | -206 | 0 | 0 | 19 | 246.57121 | NaN | NaN | 55.400658 | 337.44824 | 453.46155 |
| Dcps | GFEAPGSGVER | -283 | -293 | 0 | 0 | 11 | 978.7517 | 568.1417 | NaN | NaN | NaN | NaN |
| Dcps | IIFLHGK | -60 | -66 | 0 | 0.007987481 | 7 | 827.6337 | NaN | NaN | NaN | NaN | NaN |
| Rtn3 | TQIDHYVGIAR | -931 | -941 | 0 | 0 | 11 | 806.8752 | 644.6249 | 628.5704 | 779.8387 | 716.6945 | 788.15137 |
| Rtn3 | TQIDHYVGIAR | -931 | -941 | 0 | 0 | 11 | 806.8752 | 644.6249 | 628.5704 | 779.8387 | 716.6945 | 788.15137 |
| Rtn3 | GGGGGSPGACPALGAK | -26 | -41 | 0 | 0 | 16 | 1868.6079 | 1620.4797 | 1983.7467 | 504.79004 | 1550.916 | 1647.8997 |
| Rtn3 | ALGGGGGSPGACPALGAK | -24 | -41 | 0 | 0 | 18 | 1079.278 | 1219.5023 | 1622.0806 | 1669.0569 | 1788.6978 | 1747.4548 |
| Rtn3 | SSGAEPSALGGGGGSPGACPALGAK | -17 | -41 | 0 | 0.000394361 | 25 | 274.9105 | 324.33902 | 334.002 | NaN | 2341.9648 | NaN |
| Rtn3 | SEEGHPFK | -838 | -845 | 0 | 0.001743443 | 8 | 712.79565 | NaN | NaN | NaN | NaN | NaN |
| Msn | EDAVLEYLK | -185 | -193 | 0 | 0 | 9 | 3465.2593 | 3273.775 | 3158.6863 | 199.5472 | 2473.0217 | 189.34418 |
| Msn | IQVWHEEHR | -172 | -180 | 0 | 0 | 9 | 1740.145 | 1623.1082 | 214.95703 | 1295.488 | 1491.8098 | 1509.7839 |
| Msn | IQVWHEEHR | -172 | -180 | 0 | 0 | 9 | 1740.145 | 1623.1082 | 214.95703 | 1295.488 | 1491.8098 | 1509.7839 |
| Msn | KAQQELEEQTR | -360 | -370 | 1 | 0 | 11 | 1324.7229 | 1512.6667 | 1695.6282 | 842.885 | 1627.6703 | 1704.4471 |
| Msn | KAQQELEEQTR | -360 | -370 | 1 | 0 | 11 | 1324.7229 | 1512.6667 | 1695.6282 | 842.885 | 1627.6703 | 1704.4471 |
| Msn | GLNIYEQNDR | -224 | -233 | 0 | 0 | 10 | 1090.6425 | 981.4456 | 1081.7317 | 1291.8778 | 757.4637 | 1119.835 |
| Msn | AQQELEEQTR | -361 | -370 | 0 | 0 | 10 | 879.06836 | 1022.5496 | 4654.8584 | 3579.6882 | 4177.045 | 4712.5576 |
| Msn | ESEAVEWQQK | -439 | -448 | 0 | 0 | 10 | 477.0613 | 494.33978 | 447.4931 | 407.6252 | 467.75623 | 404.32394 |
| Msn | FYPEDVSEELIQDITQR | -84 | -100 | 0 | 0 | 17 | 183.70789 | 145.54947 | 54.640705 | 203.4591 | 338.35077 | 350.62622 |
| Msn | KESPLLFK | -72 | -79 | 1 | 0.00025015 | 8 | 1900.3108 | 1968.4626 | 2140.5923 | NaN | 1552.9421 | 1546.2308 |
| Msn | KTANDMIHAE | -538 | -547 | 1 | 0 | 10 | 1179.0573 | 920.2068 | 1046.7395 | 467.83813 | NaN | 936.8437 |
| Msn | GLQYQDTK | -46 | -53 | 0 | 0.001092529 | 8 | 1102.5696 | 847.2739 | 1149.6033 | 894.3233 | 1228.0894 | NaN |
| Msn | RALELEQER | -371 | -379 | 1 | 0.000591124 | 9 | 814.6456 | 956.54333 | 1257.9047 | NaN | 971.6756 | 1230.9277 |
| Msn | VWHEEHR | -174 | -180 | 0 | 0.002972566 | 7 | 465.3333 | 415.7531 | 399.66223 | 223.70448 | NaN | 336.06555 |
| Msn | ALTSELANAR | -524 | -533 | 0 | 0 | 10 | 1762.0034 | 1826.0391 | 524.35126 | 815.93414 | NaN | NaN |
| Msn | DVSEELIQDITQR | -88 | -100 | 0 | 0 | 13 | 1006.6437 | NaN | 1300.1733 | 304.82315 | 1109.1473 | NaN |
| Msn | KTQEQLASEMAELT | -412 | -425 | 1 | 0.00025015 | 14 | 566.09033 | NaN | 669.8354 | NaN | 894.58124 | 1067.767 |
| Msn | ALLENEKK | -321 | -328 | 1 | 0.002199523 | 8 | 1166.4985 | NaN | 1242.7804 | 461.67346 | NaN | NaN |
| Msn | AQMVQEDLEK | -449 | -458 | 0 | 0.000230707 | 10 | 70.74431 | 445.68732 | 474.15594 | NaN | NaN | NaN |
| Msn | ALGLNIYEQNDR | -222 | -233 | 0 | 0.009272078 | 12 | 821.514 | NaN | NaN | NaN | NaN | NaN |
| Lrrc47 | VVDLEGSLK | -553 | -561 | 0 | 0.00025015 | 9 | 4392.6816 | 579.42334 | 3994.1423 | 3489.5186 | 3174.718 | 3143.5352 |
| Lrrc47 | ELLLTGPGLEER | -25 | -36 | 0 | 0 | 12 | 1029.6215 | 1234.1362 | 1317.4806 | 1183.1561 | 975.753 | 1121.6018 |
| Lrrc47 | NALGPGLSPELGPLPALR | -87 | -104 | 0 | 0 | 18 | 581.69745 | 495.46182 | 565.29877 | 558.05994 | 435.408 | 615.1637 |
| Lrrc47 | ALGPGLSPELGPLPALR | -88 | -104 | 0 | 0 | 17 | 512.8218 | 502.56848 | 484.62198 | 551.74097 | 633.00775 | 620.7638 |
| Lrrc47 | HVSENPTPLTVR | -312 | -323 | 0 | 0 | 12 | 424.02386 | 747.3658 | 770.32446 | 522.6188 | 860.2992 | 857.2018 |
| Lrrc47 | AVSEAWPELELAER | -7 | -20 | 0 | 0 | 14 | 1358.5131 | 1234.9901 | 1648.6117 | 607.04956 | NaN | 2967.9758 |
| Myl6 | EAFQLFDR | -14 | -21 | 0 | 0 | 8 | 5822.825 | 5788.39 | 5842.309 | 3894.27 | 2821.523 | 3596.3335 |
| Myl6 | CDFTEDQTAEFK | -2 | -13 | 0 | 0 | 12 | 2476.9368 | 2982.7285 | 2810.1096 | 2826.9353 | 2703.3323 | 2883.5488 |
| Myl6 | NKDQGTYEDYVEGLR | -80 | -94 | 1 | 0 | 15 | 1996.6865 | 2302.489 | 2293.5674 | 730.4728 | 1389.6019 | 883.9422 |
| Myl6 | NKDQGTYEDYVEGLR | -80 | -94 | 1 | 0 | 15 | 1996.6865 | 2302.489 | 2293.5674 | 730.4728 | 1389.6019 | 883.9422 |
| Myl6 | HVLVTLGEK | -111 | -119 | 0 | 0.00025015 | 9 | 710.0177 | 609.17377 | 779.3862 | 462.49658 | 511.79245 | 784.47565 |
| Myl6 | DQGTYEDYVEGLR | -82 | -94 | 0 | 0 | 13 | 470.1781 | 2427.3254 | 2680.3208 | 1522.6959 | 1042.4733 | 983.4104 |
| Myl6 | NKDQGTYEDYVEGL | -80 | -93 | 1 | 0 | 14 | 147.28632 | 1829.6934 | 1885.4011 | 2727.8682 | 3050.8374 | 3358.927 |
| Myl6 | DQGTYEDYVEGL | -82 | -93 | 0 | 0 | 12 | 464.21408 | NaN | 371.96335 | 572.7608 | 671.2631 | 640.2729 |
| Myl6 | NKDQGTYEDYVE | -80 | -91 | 1 | 0.001392869 | 12 | 580.86084 | NaN | NaN | NaN | 550.62573 | 596.97687 |
| Apex1 | EAAGEGPVLYEDPPDQK | -35 | -51 | 0 | 0 | 17 | 2014.2996 | 1776.2932 | 1965.1368 | 2063.3477 | 2287.3591 | 2351.7656 |
| Apex1 | SHSLLPALCDSK | -287 | -298 | 0 | 0 | 12 | 1964.8091 | 2107.4265 | 813.8034 | 1970.3655 | 2387.0317 | 142.30917 |
| Apex1 | VSYGIGEEEHDQEGR | -141 | -155 | 0 | 0 | 15 | 1735.5933 | 1762.3945 | 1979.0596 | 1904.3899 | 2145.4539 | 2069.7275 |
| Apex1 | SWNVDGLR | -65 | -72 | 0 | 0 | 8 | 972.91144 | 5475.8726 | 8968.521 | 9842.941 | 8382.645 | 14054.057 |
| Apex1 | LPAELQELPGLTHQY | -103 | -117 | 0 | 0 | 15 | 943.6626 | 971.5311 | 982.0003 | 48.104694 | 289.6631 | 1044.6783 |
| Apex1 | YGIGEEEHDQEGR | -143 | -155 | 0 | 0 | 13 | 634.6577 | 639.38116 | 587.4059 | 508.15274 | 627.68243 | 707.72925 |
| Apex1 | QAVPLADSFR | -244 | -253 | 0 | 0 | 10 | 608.7549 | 2376.4446 | 3305.1746 | 3360.4282 | 3189.7297 | 411.53296 |
| Apex1 | LPAELQELPGLTHQ | -103 | -116 | 0 | 0 | 14 | 574.05493 | 995.5525 | 772.39215 | 935.0372 | 1459.4604 | 1043.084 |
| Apex1 | ALGSDHCPITLY | -303 | -314 | 0 | 0.00025015 | 12 | 523.6981 | 1797.6736 | 1933.2728 | 1958.7125 | 1895.2761 | 1704.569 |
| Apex1 | EKEAAGEGPVLYEDPPDQK | -33 | -51 | 1 | 0.000471409 | 19 | 428.69452 | 428.74832 | 399.9676 | 570.2447 | 645.4839 | 587.1205 |
| Apex1 | KGLDWVK | -78 | -84 | 1 | 0.000627943 | 7 | 932.04236 | 866.10944 | 1508.905 | 1532.0436 | NaN | 1024.8723 |
| Apex1 | AVPLADSFR | -245 | -253 | 0 | 0.007791272 | 9 | 676.7791 | NaN | NaN | NaN | NaN | NaN |
| Apex1 | HEEIDLR | -214 | -220 | 0 | 0.003003218 | 7 | 343.25842 | NaN | NaN | NaN | NaN | NaN |
| Cers2 | ALHDASDYLLESAK | -237 | -250 | 0 | 0 | 14 | 5295.2285 | 6281.755 | 6328.564 | 3887.6255 | 4662.82 | 4459.6504 |
| Pycr2 | ILDEIGADVQER | -78 | -89 | 0 | 0 | 12 | 1673.2441 | 1474.8776 | 1634.2618 | 2341.7002 | 2243.2148 | 1935.59 |
| Pycr2 | LDSEDHPGQLK | -218 | -228 | 0 | 0 | 11 | 1496.6825 | 1075.0242 | 850.78467 | 606.28687 | 1530.0046 | 1529.5784 |
| Pycr2 | ALHFLESGGFR | -241 | -251 | 0 | 0 | 11 | 4803.8843 | NaN | 3584.4937 | 2846.6558 | 1695.2437 | 545.1714 |
| Cct4 | TIGTKPVAHID | -339 | -349 | 1 | 0.00078661 | 11 | 5487.414 | 4885.1323 | 4947.5146 | 5575.7197 | 5325.678 | 5339.519 |
| Cct4 | GIHPTIISESFQK | -127 | -139 | 0 | 0 | 13 | 3044.2258 | 3324.522 | 3474.0718 | 3334.6096 | 4539.939 | 4667.383 |

| | | | | | | | | | | | | |
|---|---|---|---|---|---|---|---|---|---|---|---|---|
| Cct4 | GGGAPEIELALR | -424 | -435 | 0 | 0 | 12 | 2900.2197 | 2221.367 | 2014.3833 | 2162.699 | 2801.781 | 2477.511 |
| Cct4 | DVEREDIEFICK | -327 | -338 | 1 | 0.000230707 | 12 | 2470.5376 | 249.87082 | 3090.6492 | 3437.3281 | 3560.5295 | 311.21588 |
| Cct4 | IDDVVNTR | -532 | -539 | 0 | 0 | 8 | 2430.3792 | 1883.5406 | 1267.14 | 1834.5065 | 1616.2012 | 773.6895 |
| Cct4 | EDIEFICK | -331 | -338 | 0 | 0 | 8 | 2337.7192 | 1135.3622 | 2849.6628 | 1158.8253 | 1104.3916 | 1366.2019 |
| Cct4 | TGCNVLLIQK | -293 | -302 | 0 | 0 | 10 | 1213.6941 | 719.4685 | 1743.7891 | 1301.7378 | 1773.319 | 1577.8815 |
| Cct4 | FCLSAPK | -251 | -257 | 0 | 0 | 7 | 905.6046 | 820.78516 | 666.5789 | 866.1829 | 817.2915 | 799.6302 |
| Cct4 | SGAPTAGPGSR | -9 | -19 | 0 | 0 | 11 | 819.4656 | 832.72437 | 806.17474 | 561.3357 | 723.35187 | 258.80518 |
| Cct4 | DALSDLALHFLNK | -307 | -319 | 0 | 0 | 13 | 639.6463 | 569.5947 | 499.3062 | 715.3484 | 815.31116 | 775.08234 |
| Cct4 | DALSDLALHF | -307 | -316 | 0 | 0.00025015 | 10 | 350.2021 | 85.4776 | 299.37152 | 333.64734 | 104.68275 | 540.42474 |
| Cct4 | ALTLATETVR | -518 | -527 | 0 | 0.000230707 | 10 | 3690.1658 | NaN | 2793.332 | 3478.3425 | 2028.487 | 1124.2695 |
| Cct4 | QVLHPAAR | -82 | -89 | 0 | 0 | 8 | 708.5561 | NaN | 879.0932 | 704.4287 | 507.1477 | 632.3717 |
| Cct4 | VIDPATATSVDLR | -194 | -206 | 0 | 0 | 13 | 528.94476 | 529.6248 | 1440.4403 | NaN | NaN | 1710.7037 |
| Cct4 | ALHFLNK | -313 | -319 | 0 | 0 | 7 | 2167.9592 | 1486.7662 | 1461.5697 | NaN | NaN | NaN |
| Cct4 | LVIEEAER | -396 | -403 | 0 | 0 | 8 | 1694.5851 | NaN | 1366.3997 | NaN | 1376.635 | NaN |
| Cct4 | GSLLDSCTK | -114 | -122 | 0 | 0.00025015 | 9 | 2209.6318 | 1942.7513 | NaN | NaN | NaN | NaN |
| Cct4 | KTGCNVLLIQK | -292 | -302 | 1 | 0.00025015 | 11 | 425.64014 | NaN | NaN | NaN | 377.84106 | NaN |
| Ctsa | APDQDEIDCLPGLAK | -24 | -38 | 0 | 0 | 15 | 736.81683 | 732.6755 | 932.6792 | 809.7787 | 978.34314 | 1145.9885 |
| Ctsa | ALHIPESLPR | -343 | -352 | 0 | 0 | 10 | 185.53297 | 592.2811 | 605.89056 | 430.2572 | NaN | NaN |
| Ppa2 | TWEDPHLR | -143 | -150 | 0 | 0 | 8 | 703.93085 | 769.66675 | 940.5803 | 711.1828 | 699.61255 | 786.1231 |
| Ppa2 | ALIDQSETDWK | -188 | -198 | 0 | 0.00025015 | 11 | 1483.3602 | NaN | NaN | NaN | NaN | NaN |
| Ppa2 | FHDIDDVKK | -212 | -220 | 1 | 0.001868731 | 9 | 484.4304 | NaN | NaN | NaN | NaN | NaN |
| Ptbp1 | KLPSDVTEGEVISL | -64 | -77 | 1 | 0.000298805 | 14 | 3404.689 | 4379.6665 | 3961.5854 | 4373.603 | 5418.8853 | 5389.7383 |
| Ptbp1 | YYTSVAPVLR | -112 | -121 | 0 | 0 | 10 | 3387.0637 | 2276.2363 | 3694.0103 | 1800.1921 | 918.27136 | 999.2798 |
| Ptbp1 | HQSVQLPR | -435 | -442 | 0 | 0 | 8 | 2861.1257 | 2382.735 | 2118.4102 | 2098.0598 | 1459.8474 | 1518.2163 |
| Ptbp1 | NIPPSVSEDDLK | -484 | -495 | 0 | 0 | 12 | 1038.4755 | 5756.099 | 1593.8395 | 1363.873 | 1482.5696 | 1373.938 |
| Ptbp1 | YGSSPLHR | -454 | -461 | 0 | 0 | 8 | 993.77625 | 1120.6555 | 979.11865 | 1168.7767 | 1119.9067 | 824.2206 |
| Ptbp1 | LQYADPVSAQHAK | -225 | -237 | 0 | 0 | 13 | 969.5692 | 350.7419 | 382.32983 | 1244.4487 | 438.02097 | 1014.9365 |
| Ptbp1 | DYGSSPLHR | -453 | -461 | 0 | 0 | 9 | 302.0235 | 3957.6802 | 4360.685 | 2611.4885 | 2420.8145 | 4144.3403 |
| Ptbp1 | MAGQSPVLR | -176 | -184 | 0 | 0 | 9 | 1251.6584 | 1056.7158 | 1095.9625 | 953.08527 | NaN | 1317.0387 |
| Ptbp1 | RGSDELFSTC | -14 | -23 | 1 | 0.00025015 | 10 | 460.1338 | 686.953 | 662.0966 | NaN | 804.58234 | 636.5741 |
| Ptbp1 | YTSVAPVLR | -113 | -121 | 0 | 0.00025015 | 9 | 1538.9412 | NaN | 940.3943 | NaN | NaN | NaN |
| Top2a | DSTTIEISELPIR | -916 | -928 | 0 | 0 | 13 | 10086.737 | 9107.824 | 8250.963 | 3387.6118 | 1100.4486 | 8003.764 |
| Top2a | ALIENPTFDSQTK | -365 | -377 | 0 | 0 | 13 | 1971.2861 | 1043.7717 | 2242.7693 | 1632.6422 | 1512.7397 | 1537.8627 |
| Top2a | TTLPFKPVK | -1274 | -1282 | 1 | 0 | 9 | 1919.124 | 1961.5642 | 2089.9233 | 2197.327 | 2500.7656 | 2322.5554 |
| Top2a | KYDTVLDILR | -1010 | -1019 | 1 | 0 | 10 | 1896.6091 | 2132.822 | 2226.159 | 1755.213 | 2601.6816 | 314.78024 |
| Top2a | SIPSMVDGLKPGQR | -713 | -726 | 1 | 0 | 14 | 1403.9827 | 1821.6376 | 1684.7281 | 1471.9636 | 2151.1604 | 2086.5242 |
| Top2a | SIPSMVDGLKPGQR | -713 | -726 | 1 | 0 | 14 | 1403.9827 | 1821.6376 | 1684.7281 | 1471.9636 | 2151.1604 | 2086.5242 |
| Top2a | HVDYVADQIVSK | -324 | -335 | 0 | 0 | 12 | 1303.8707 | 1301.0892 | 1340.7921 | 2097.2712 | 2171.8638 | 2276.894 |
| Top2a | DKYGVFPLR | -478 | -486 | 1 | 0 | 9 | 1195.3496 | 1237.6364 | 1137.8097 | 637.48926 | 562.8287 | 202.7379 |
| Top2a | NVEGTPAEDGAEPGSLR | -1241 | -1257 | 0 | 0.00078661 | 17 | 1129.2383 | 1133.2231 | 1108.58 | 656.4932 | 851.5495 | 357.98166 |
| Top2a | LDDANDAGSR | -440 | -449 | 0 | 0 | 10 | 958.6176 | 900.1709 | 1185.2511 | 705.0712 | 799.85535 | 222.47876 |
| Top2a | KLLGLPEDYLYGQ | -675 | -687 | 1 | 0 | 13 | 841.4597 | 896.9112 | 835.9112 | 739.4612 | 932.40436 | 632.1351 |
| Top2a | VLFDHVGCLK | -1000 | -1009 | 0 | 0 | 10 | 797.6911 | 664.9007 | 667.26697 | 674.0266 | 786.9522 | 741.3733 |
| Top2a | VIHEQVNPR | -287 | -295 | 0 | 0 | 9 | 559.49133 | 505.22824 | 659.87177 | 541.8942 | 614.3347 | 557.0349 |
| Top2a | VIHEQVNPR | -287 | -295 | 0 | 0 | 9 | 559.49133 | 505.22824 | 659.87177 | 541.8942 | 614.3347 | 557.0349 |
| Top2a | NSTECTLILTEGDSAK | -450 | -465 | 0 | 0 | 16 | 590.81165 | 681.9951 | 303.5703 | 638.44855 | 576.80975 | 769.6417 |
| Top2a | SGEDYTCITFQPDLSK | -209 | -224 | 0 | 0 | 16 | 486.42548 | 484.1115 | 492.22308 | 659.865 | 910.8884 | 654.18176 |
| Top2a | GEEQDFPVDLEDTIAPR | -1489 | -1505 | 0 | 0 | 17 | 437.97098 | 661.97314 | 505.0625 | 944.9234 | 1102.2379 | 853.0354 |
| Top2a | LDMPLWYLTK | -1126 | -1135 | 0 | 0 | 10 | 305.49307 | 252.42065 | 188.00336 | 719.89746 | 338.45578 | 314.83643 |
| Top2a | STSYLTYNDFINK | -688 | -700 | 0 | 0 | 13 | 266.96725 | 746.4388 | 716.4142 | 1026.5181 | 2514.6245 | 4270.2285 |
| Top2a | EKQDEQVGLPGK | -1182 | -1193 | 1 | 0 | 12 | 10679.843 | 11783.479 | NaN | 13567.766 | 13994.902 | 14272.41 |
| Top2a | DFPVDLEDTIAPR | -1493 | -1505 | 0 | 0 | 13 | 918.6499 | 865.4503 | 911.20483 | NaN | 1012.3254 | 1150.6018 |
| Top2a | KVIHEQVNPR | -286 | -295 | 1 | 0.00025015 | 10 | 302.33017 | NaN | 396.26263 | 265.21173 | 323.7506 | 394.70532 |
| Top2a | KVIHEQVNPR | -286 | -295 | 1 | 0.00025015 | 10 | 302.33017 | NaN | 396.26263 | 265.21173 | 323.7506 | 394.70532 |
| Top2a | RGYDSDPVK | -1079 | -1087 | 1 | 0.00272025 | 9 | 690.27795 | NaN | 1224.3699 | 1026.4592 | 1207.2311 | 1311.0498 |
| Top2a | YDTVLDILR | -1011 | -1019 | 0 | 0 | 9 | 274.24207 | 883.6491 | 951.40704 | NaN | 1116.3497 | 882.65045 |
| Top2a | EKQDEQVGLPGKA | -1182 | -1194 | 2 | 0 | 13 | 8877.871 | NaN | NaN | 10894.1455 | 6755.2124 | 7534.24 |
| Top2a | TPSLITDYR | -949 | -957 | 0 | 0 | 9 | 1617.7101 | 1646.3364 | 2057.2458 | 1446.0455 | NaN | NaN |
| Top2a | NYEDEDSLK | -520 | -528 | 0 | 0 | 9 | 476.425 | 438.1583 | 526.1532 | 507.06528 | NaN | NaN |
| Top2a | SLPEFEEWK | -590 | -598 | 0 | 0.000431655 | 9 | 441.38492 | 468.3118 | NaN | 504.3811 | NaN | 557.4939 |
| Top2a | DTVLDILR | -1012 | -1019 | 0 | 0.00111006 | 8 | 321.7545 | 362.99854 | NaN | 433.06998 | NaN | 222.53452 |
| Top2a | IVIENKPK | -1062 | -1069 | 1 | 0.003745449 | 8 | 1237.4813 | 465.15063 | 523.74915 | NaN | NaN | NaN |
| Top2a | SENVEGTPAEDGAEPGSLR | -1239 | -1257 | 0 | 0.00111006 | 19 | 775.5123 | NaN | 574.53735 | NaN | NaN | NaN |
| Top2a | DKVDETGNSL | -276 | -285 | 1 | 0.002697088 | 10 | 320.79398 | NaN | NaN | NaN | NaN | NaN |
| Bak1 | ALIGDDINR | -77 | -85 | 0 | 0.00111006 | 9 | 1707.2664 | 695.41077 | 1956.0614 | 2265.993 | 2971.9429 | 3091.727 |
| Rps12 | TALIHDGLAR | -24 | -33 | 0 | 0 | 10 | 5222.654 | 734.6808 | 5569.378 | 3031.6267 | 3495.1782 | 3522.1233 |
| Rps12 | LIHDGLAR | -26 | -33 | 0 | 0 | 8 | 3185.0107 | 3166.4778 | 3120.0698 | 3009.1226 | 2560.7014 | 3359.8438 |
| Rps12 | CAEHQINLIK | -69 | -78 | 0 | 0 | 10 | 3025.507 | 3198.8823 | 3165.9907 | 545.3611 | 337.35098 | 3685.9211 |
| Rps12 | ALIHDGLAR | -25 | -33 | 0 | 0 | 9 | 2792.299 | 443.87122 | 2877.8096 | 3795.7905 | 4386.9307 | 3953.7869 |
| Rps12 | DVNTALQEVLK | -13 | -23 | 0 | 0 | 11 | 1489.7664 | 1801.3894 | 1748.0955 | 972.7769 | 1237.9515 | 1962.1649 |
| Rps12 | VLASNCDEPMYVK | -51 | -63 | 0 | 0 | 13 | 1351.9712 | 1157.2661 | 1935.0178 | 2095.524 | 2082.4587 | 2401.9316 |
| Rps12 | DVIEEYFK | -122 | -129 | 0 | 0 | 8 | 1162.0958 | 1039.3386 | 665.3594 | 1654.1007 | 1524.1241 | 1869.0216 |
| Rps12 | HQINLIK | -72 | -78 | 0 | 0.001636916 | 7 | 867.76917 | 888.453 | 765.78577 | 594.5206 | 607.3295 | 637.7808 |
| Rps12 | ASNCDEPMYVK | -53 | -63 | 0 | 0 | 11 | 783.8448 | 864.69904 | 801.68243 | 541.5326 | 856.1812 | 858.65967 |
| Rps12 | GGVMDVNTALQEVLK | -9 | -23 | 0 | 0 | 15 | 688.1605 | 870.64905 | 757.4189 | 1196.5432 | 893.3326 | 1455.0112 |
| Rps12 | KLGEWVGLC | -84 | -92 | 1 | 0.001291103 | 9 | 631.8779 | 707.5492 | 747.96893 | 954.873 | 898.51825 | 770.4298 |
| Rps12 | LGEWVGLCK | -85 | -93 | 0 | 0 | 9 | 436.2166 | 3966.7104 | 4132.357 | 330.01135 | 4496.949 | 4566.865 |
| Rps12 | KLGEWVGLCK | -84 | -93 | 1 | 0 | 10 | 1176.608 | 479.08575 | 1300.1439 | NaN | 1180.1603 | 1105.5548 |

| | | | | | | | | | | | |
|---|---|---|---|---|---|---|---|---|---|---|---|
| Rps12 | EHQINLIK | -71 | -78 | 0 | 0.000907206 | 8 | 893.4753 | 771.45905 | NaN | 949.42126 | NaN | NaN |
| Rps12 | TALQEVLK | -16 | -23 | 0 | 0.002199523 | 8 | 221.96184 | NaN | 346.2613 | 1236.6941 | NaN | NaN |
| Lypla1 | WFDIVGLSPDSQEDESGIK | -72 | -90 | 0 | 0 | 19 | 256.2834 | 232.55379 | 291.32642 | 322.46637 | 391.8228 | 350.1579 |
| Lypla1 | INPANVTFK | -193 | -201 | 0 | 0.00025015 | 9 | 2130.4832 | NaN | 1940.4486 | NaN | 482.3733 | NaN |
| Lypla1 | ALINPANVTFK | -191 | -201 | 0 | 0 | 11 | 1079.0142 | 1625.1488 | NaN | NaN | NaN | 2174.393 |
| Lypla1 | SCWLPLR | -143 | -149 | 0 | 0.004439092 | 7 | 917.7438 | 219.3748 | 838.9343 | NaN | NaN | NaN |
| Lypla1 | SPDSQEDESGIK | -79 | -90 | 0 | 0.002434616 | 12 | 827.9627 | NaN | NaN | NaN | 748.80316 | 1056.883 |
| Rrm1 | SIPEIPDDLK | -668 | -677 | 0 | 0.00025015 | 10 | 1311.5941 | 1380.1838 | 1410.1497 | 1052.868 | 1737.3668 | 1444.9459 |
| Rrm1 | VTPEHTYDFEK | -455 | -465 | 0 | 0 | 11 | 868.6356 | 1300.5399 | 1609.5543 | 930.02313 | 729.569 | 976.58374 |
| Rrm1 | EYGPYETYEGSPVSK | -549 | -563 | 0 | 0.00025015 | 15 | 838.13354 | 902.2308 | 949.9516 | 670.1165 | 1234.7162 | 2661.3103 |
| Rrm1 | TRPAANPIQFT | -741 | -751 | 1 | 0.000230707 | 11 | 779.1295 | 719.141 | 993.62976 | 1416.1675 | 998.2904 | 948.8634 |
| Rrm1 | TRPAANPIQF | -741 | -750 | 1 | 0.00025015 | 10 | 715.99426 | 582.7783 | 864.3788 | 652.73254 | 696.8692 | 892.5814 |
| Rrm1 | LNSAIIYDR | -131 | -139 | 0 | 0 | 9 | 690.1066 | 175.11858 | 748.2131 | 772.34784 | 839.2875 | 819.2912 |
| Rrm1 | ECPGLDEVWGEEFEK | -355 | -369 | 0 | 0 | 15 | 646.573 | 614.2137 | 834.3909 | 1082.8439 | 1088.0759 | 1495.7491 |
| Rrm1 | YVTPEHTYDFEK | -454 | -465 | 0 | 0 | 12 | 644.14166 | 682.66797 | 768.04175 | 738.88055 | 685.017 | 608.16 |
| Rrm1 | VLSGEFQIVNPHLLK | -629 | -643 | 0 | 0 | 15 | 470.88474 | 500.71246 | 471.92883 | 591.76843 | 723.9273 | 784.1095 |
| Rrm1 | NVAPTDLWDWKPLK | -572 | -585 | 1 | 0 | 14 | 366.0878 | 402.34247 | 460.84903 | 594.38336 | 863.08563 | 718.76886 |
| Rrm1 | ALKEEEEKER | -763 | -772 | 2 | 0.000431655 | 10 | 328.79684 | 371.78748 | 368.35522 | 303.0755 | 340.35175 | 385.87405 |
| Rrm1 | WFTHASPTLF | -197 | -206 | 0 | 0.000471409 | 10 | 517.677 | NaN | 553.6933 | 389.83432 | 316.4155 | 448.51367 |
| Rrm1 | TVWEISQK | -682 | -689 | 0 | 0 | 8 | 429.95007 | 580.5591 | 548.3782 | 986.4279 | NaN | 731.77026 |
| Rrm1 | KDDSIEGIYDTLK | -224 | -236 | 1 | 0 | 13 | 2049.8667 | NaN | 2138.9375 | 2108.1968 | 2546.2644 | NaN |
| Rrm1 | HRPIGIGVQ | -498 | -506 | 1 | 0.002780285 | 9 | 1227.2692 | 1465.0708 | NaN | 1442.3286 | 1690.7986 | NaN |
| Rrm1 | IVNPHLLK | -636 | -643 | 0 | 0 | 8 | 525.8238 | 457.1151 | 568.9788 | NaN | 373.59018 | NaN |
| Rrm1 | TRPAANPIQFTL | -741 | -752 | 1 | 0.003998192 | 12 | 797.80225 | 876.1931 | 840.1117 | NaN | NaN | NaN |
| Rrm1 | EFLDLKK | -310 | -316 | 1 | 0.002296977 | 7 | 1758.6108 | NaN | NaN | 2236.8965 | NaN | NaN |
| Rrm1 | KSNQQNLGTIK | -414 | -424 | 1 | 0.00111006 | 11 | 473.22778 | NaN | 617.0538 | NaN | NaN | NaN |
| Acsl5 | GAAPISTPVLTFFR | -419 | -432 | 0 | 0 | 14 | 584.9496 | 496.1756 | 515.2427 | 559.01984 | 646.01874 | 603.13727 |
| Acsl5 | TQEVLDKDGWLHTG | -511 | -524 | 1 | 0.000230707 | 14 | 928.68616 | NaN | 1086.7361 | 979.2158 | 1238.737 | 1201.9489 |
| Acsl5 | ALKPTVFPTVPR | -342 | -353 | 1 | 0 | 12 | 396.8623 | NaN | NaN | 426.65497 | 532.5222 | NaN |
| Acsl5 | EAILEDLQK | -617 | -625 | 0 | 0 | 9 | 1793.8073 | NaN | 1901.5737 | NaN | NaN | NaN |
| Acsl5 | SLHDAENIGK | -232 | -241 | 0 | 0.00078661 | 10 | 205.58005 | NaN | 1120.9 | NaN | NaN | NaN |
| Gltp | SICDGERDENHPNLIR | -110 | -125 | 1 | 0 | 16 | 562.7886 | 2659.5066 | 737.0526 | 2624.3044 | 994.74457 | 793.518 |
| Gltp | GQNVTEEECLEK | -168 | -179 | 0 | 0 | 12 | 562.6226 | 587.0263 | 543.02264 | 387.24268 | 609.4935 | 715.1741 |
| Gltp | ALLAEHLLKPLPADR | -2 | -16 | 1 | 0 | 15 | 376.01373 | 432.56836 | 405.46027 | 481.605 | 667.6349 | 637.4315 |
| Gltp | ALLAEHLLKPLPADR | -2 | -16 | 1 | 0 | 15 | 376.01373 | 432.56836 | 405.46027 | 481.605 | 667.6349 | 637.4315 |
| Gltp | SKGQNVTEEECLEK | -166 | -179 | 1 | 0 | 14 | 1055.366 | 1035.8623 | NaN | 1076.5522 | 1291.1593 | 1317.7914 |
| Gltp | HLLKPLPADR | -7 | -16 | 1 | 0.00025015 | 10 | 447.7123 | NaN | NaN | NaN | NaN | NaN |
| Eif3d | TLNEWDSR | -395 | -402 | 0 | 0 | 8 | 1281.3794 | 1257.3025 | 1259.0712 | 1297.4779 | 1229.9528 | 192.1951 |
| Eif3d | ALLAGSEYLK | -439 | -448 | 0 | 0 | 10 | 1074.9131 | 846.8831 | 683.62866 | 887.42175 | 116.90869 | 470.04672 |
| Eif3d | ETANEPPQDEGNSFNSPR | -293 | -310 | 0 | 0 | 18 | 913.54065 | 826.0111 | 882.8083 | 941.67236 | 1207.3174 | 1122.427 |
| Eif3d | LGDDIDLIVR | -365 | -374 | 0 | 0 | 10 | 450.27756 | 494.9545 | 500.68 | 445.7963 | 829.0525 | 488.24332 |
| Eif3d | HCNGVDWR | -403 | -410 | 0 | 0.002780285 | 8 | 656.0529 | NaN | NaN | NaN | NaN | NaN |
| Eif3d | DETSFQLVDTAR | -76 | -87 | 0 | 0 | 12 | 652.79254 | NaN | NaN | NaN | NaN | NaN |
| Psmb8 | SGCAADCQYWER | -118 | -129 | 0 | 0 | 12 | 2190.335 | 2059.3606 | 2001.3073 | 2265.7559 | 1785.4237 | 1304.14 |
| Psmb8 | QDLSPEEAYDLGR | -217 | -229 | 0 | 0.000298805 | 13 | 1212.3506 | 1142.9008 | 1172.5454 | 1011.92773 | 603.4231 | 1075.0482 |
| Psmb8 | SSDVSDLLYK | -261 | -270 | 0 | 0 | 10 | 1110.3105 | 2132.08 | 2198.2742 | 3037.4583 | 2597.6929 | 2522.021 |
| Psmb8 | ALLDLCGAAR | -2 | -11 | 0 | 0 | 10 | 255.47643 | 337.0622 | 391.3167 | 300.21408 | 418.48846 | 406.04388 |
| Psmb8 | VESSDVSDLLYK | -259 | -270 | 0 | 0 | 12 | 1206.8643 | NaN | 1155.6077 | NaN | NaN | NaN |
| Psmb8 | FQHGVIVAV | -80 | -88 | 0 | 0.00111006 | 9 | 552.65717 | 769.9401 | NaN | NaN | NaN | NaN |
| Psmb8 | FQHGVIV | -80 | -86 | 0 | 0.008105996 | 7 | 483.59174 | 513.4452 | NaN | NaN | NaN | NaN |
| Eif3h | HYQEEGQGTEVVQGVLL | -52 | -68 | 0 | 0 | 17 | 1650.2411 | 1936.9156 | 1962.9615 | 2452.9832 | 3031.886 | 3066.9905 |
| Eif3h | ALLDSQFSYQH | -132 | -142 | 0 | 0 | 11 | 1478.8192 | 1458.3363 | 1523.2523 | 1173.8042 | 1518.4509 | 447.30695 |
| Eif3h | LFKPHQAPAR | -304 | -313 | 1 | 0 | 10 | 864.81323 | 900.4802 | 992.75336 | 251.54265 | 685.982 | 743.1384 |
| Eif3h | LFKPHQAPAR | -304 | -313 | 1 | 0 | 10 | 864.81323 | 900.4802 | 992.75336 | 251.54265 | 685.982 | 743.1384 |
| Eif3h | SRGEPPLPEEDLSK | -290 | -303 | 1 | 0.000230707 | 14 | 569.27075 | 1359.173 | 1399.44 | 902.29425 | 1803.6799 | 1831.6943 |
| Eif3h | HVNIDHLHVG | -109 | -118 | 0 | 0.00111006 | 10 | 2315.6965 | 2180.3362 | 1830.7861 | 374.64752 | NaN | 1747.1375 |
| Eif3h | HVNIDHLHVG | -109 | -118 | 0 | 0.00111006 | 10 | 2315.6965 | 2180.3362 | 1830.7861 | 374.64752 | NaN | 1747.1375 |
| Eif3h | SAVADKHELL | -222 | -231 | 1 | 0.001392869 | 10 | 1970.1088 | 2002.4922 | 1785.0304 | 2057.5422 | NaN | 1236.3466 |
| Eif3h | SVVLIYDPIK | -147 | -156 | 0 | 0 | 10 | 787.5524 | 458.32574 | 332.319 | NaN | 448.55984 | 407.76178 |
| Eif3h | YYGSFVTR | -124 | -131 | 0 | 0 | 8 | 1825.6411 | 1735.3124 | NaN | NaN | 1638.0348 | 1699.1617 |
| Eif3h | HYQEEGQGTEVVQ | -52 | -64 | 0 | 0.00201047 | 13 | 1015.2806 | NaN | NaN | NaN | NaN | NaN |
| Hsp90aa1 | TNDWEEHLAVK | -318 | -328 | 0 | 0 | 11 | 8528.253 | 974.1533 | 9495.821 | 544.5571 | 123.32853 | 667.9098 |
| Hsp90aa1 | TNDWEEHLAVK | -318 | -328 | 0 | 0 | 11 | 8528.253 | 974.1533 | 9495.821 | 544.5571 | 123.32853 | 667.9098 |
| Hsp90aa1 | TNDWEEHLAVK | -318 | -328 | 0 | 0 | 11 | 8528.253 | 974.1533 | 9495.821 | 544.5571 | 123.32853 | 667.9098 |
| Hsp90aa1 | LGIHEDSQNR | -448 | -457 | 0 | 0 | 10 | 7283.1963 | 2621.2065 | 7573.9556 | 5605.6523 | 6802.031 | 7093.613 |
| Hsp90aa1 | LGIHEDSQNR | -448 | -457 | 0 | 0 | 10 | 7283.1963 | 2621.2065 | 7573.9556 | 5605.6523 | 6802.031 | 7093.613 |
| Hsp90aa1 | HLEINPDHSIIETLR | -634 | -648 | 0 | 0 | 15 | 6813.474 | 10086.6 | 6239.1045 | 1450.89 | 6643.449 | 5962.3213 |
| Hsp90aa1 | HLEINPDHSIIETLR | -634 | -648 | 0 | 0 | 15 | 6813.474 | 10086.6 | 6239.1045 | 1450.89 | 6643.449 | 5962.3213 |
| Hsp90aa1 | RAPFDLFENR | -347 | -356 | 1 | 0.001291103 | 10 | 6387.1206 | 5339.3604 | 8871.66 | 3552.6436 | 6462.3203 | 5640.68 |
| Hsp90aa1 | NPDDITNEEYGEFYK | -301 | -315 | 0 | 0 | 15 | 5200.7163 | 5562.531 | 5433.7812 | 397.71057 | 4002.1104 | 4244.2515 |
| Hsp90aa1 | ELISNSSDALDK | -47 | -58 | 0 | 0 | 12 | 4915.986 | 4700.3545 | 4590.7793 | 5272.476 | 3923.5417 | 344.81766 |
| Hsp90aa1 | ELISNSSDALDK | -47 | -58 | 0 | 0 | 12 | 4915.986 | 4700.3545 | 4590.7793 | 5272.476 | 3923.5417 | 344.81766 |
| Hsp90aa1 | SLEDPQTHANR | -678 | -688 | 0 | 0 | 11 | 4625.797 | 4496.2593 | 5188.826 | 2639.6184 | 4236.6416 | 4310.218 |
| Hsp90aa1 | HLEINPDHSIIETL | -634 | -647 | 0 | 0 | 14 | 2612.7058 | 3212.534 | 3094.3994 | 4196.0596 | 4825.044 | 4982.7856 |
| Hsp90aa1 | HLEINPDHSIIETL | -634 | -647 | 0 | 0 | 14 | 2612.7058 | 3212.534 | 3094.3994 | 4196.0596 | 4825.044 | 4982.7856 |
| Hsp90aa1 | GFSLEDPQTHANR | -676 | -688 | 0 | 0 | 13 | 2441.7874 | 2644.3066 | 2472.6504 | 1970.3798 | 797.71326 | 1962.2172 |
| Hsp90aa1 | SLTNDWEEHLAVK | -316 | -328 | 0 | 0 | 13 | 2007.5554 | 2733.851 | 2802.5757 | 1674.5431 | 2256.3994 | 2343.3538 |
| Hsp90aa1 | DWEEHLAVK | -320 | -328 | 0 | 0 | 9 | 1628.6582 | 1751.7042 | 1873.9285 | 1538.7814 | 1875.702 | 1907.2775 |

| | | | | | | | | | | | |
|---|---|---|---|---|---|---|---|---|---|---|---|
| Hsp90aa1 | SGFSLEDPQTHANR | -675 | -688 | 0 | 0 | 14 | 1449.3601 | 1723.9093 | 2023.9065 | 765.87787 | 2641.0293 | 2512.1482 |
| Hsp90aa1 | SGFSLEDPQTHANR | -675 | -688 | 0 | 0 | 14 | 1449.3601 | 1723.9093 | 2023.9065 | 765.87787 | 2641.0293 | 2512.1482 |
| Hsp90aa1 | SSGFSLEDPQTHANR | -674 | -688 | 0 | 0 | 15 | 1424.4543 | 1561.4128 | 1506.4338 | 1526.7799 | 1552.0703 | 1495.722 |
| Hsp90aa1 | SSGFSLEDPQTHANR | -674 | -688 | 0 | 0 | 15 | 1424.4543 | 1561.4128 | 1506.4338 | 1526.7799 | 1552.0703 | 1495.722 |
| Hsp90aa1 | FTELAEDKENYK | -425 | -436 | 1 | 0 | 12 | 1166.9482 | 1129.433 | 1124.746 | 1951.4553 | 1713.316 | 2453.49 |
| Hsp90aa1 | LLSSGFSLEDPQTHANR | -672 | -688 | 0 | 0 | 17 | 1128.4796 | 1206.2711 | 1156.7208 | 1405.9403 | 575.90125 | 479.10266 |
| Hsp90aa1 | HIYFITGETK | -491 | -500 | 0 | 0 | 10 | 915.1733 | 938.9799 | 778.0119 | 1061.6708 | 1157.0769 | 1177.885 |
| Hsp90aa1 | KHLEINPDHSIIE | -633 | -645 | 1 | 0 | 13 | 903.28937 | 994.71234 | 920.8919 | 685.98517 | 538.1901 | 583.0271 |
| Hsp90aa1 | KHLEINPDHSIIE | -633 | -645 | 1 | 0 | 13 | 903.28937 | 994.71234 | 920.8919 | 685.98517 | 538.1901 | 583.0271 |
| Hsp90aa1 | ASGDEMVSLK | -470 | -479 | 0 | 0 | 10 | 700.6166 | 920.3392 | 883.5896 | 585.8058 | 886.41425 | 996.11536 |
| Hsp90aa1 | ASGDEMVSLK | -470 | -479 | 0 | 0 | 10 | 700.6166 | 920.3392 | 883.5896 | 585.8058 | 886.41425 | 996.11536 |
| Hsp90aa1 | LTNDWEEHLAVK | -317 | -328 | 0 | 0 | 12 | 848.5536 | 661.0109 | 777.2103 | 1130.8931 | 1023.166 | 490.75412 |
| Hsp90aa1 | KHLEINPDHSIIETL | -633 | -647 | 1 | 0 | 15 | 824.73755 | 1217.372 | 1323.2095 | 1324.7183 | 1992.2325 | 2326.576 |
| Hsp90aa1 | KHLEINPDHSIIETL | -633 | -647 | 1 | 0 | 15 | 824.73755 | 1217.372 | 1323.2095 | 1324.7183 | 1992.2325 | 2326.576 |
| Hsp90aa1 | HSQFIGYPITLF | -210 | -221 | 0 | 0 | 12 | 779.09485 | 774.4 | 855.3648 | 903.59753 | 952.2953 | 291.02765 |
| Hsp90aa1 | KHLEINPDHSIIETLR | -633 | -648 | 1 | 0 | 16 | 648.6613 | 247.85152 | 480.81525 | 550.388 | 1568.1277 | 1012.28094 |
| Hsp90aa1 | FTELAEDKENYKK | -425 | -437 | 2 | 0 | 13 | 501.69507 | 754.9908 | 474.34174 | 575.04504 | 755.0415 | 504.30197 |
| Hsp90aa1 | FTELAEDKENYKK | -425 | -437 | 2 | 0 | 13 | 501.69507 | 754.9908 | 474.34174 | 575.04504 | 755.0415 | 504.30197 |
| Hsp90aa1 | KSLTNDWEEHLAVK | -315 | -328 | 1 | 0 | 14 | 437.17355 | 86.34669 | 687.86804 | 102.13473 | 966.2064 | 912.2827 |
| Hsp90aa1 | EEMPPLEGDDDTSR | -715 | -728 | 0 | 0 | 14 | 396.30002 | 1261.7017 | 1173.8666 | 1587.5432 | 1540.8839 | 1561.0111 |
| Hsp90aa1 | ESDDKPEIEDVGSDEEEEKK | -251 | -271 | 2 | 0.000947459 | 21 | 369.81244 | 389.2753 | 361.72635 | 454.84122 | 412.3615 | 448.2453 |
| Hsp90aa1 | ALLSSGFSLEDPQTHANR | -671 | -688 | 0 | 0 | 18 | 314.91852 | 383.88788 | 476.38208 | 438.12927 | 751.7172 | 677.4015 |
| Hsp90aa1 | DNCEELIPEYLNFIR | -373 | -387 | 0 | 0 | 15 | 240.81256 | 183.7377 | 217.05457 | 79.369286 | 337.5416 | 329.60693 |
| Hsp90aa1 | ISNSSDALDK | -49 | -58 | 0 | 0 | 10 | 1937.4968 | 1063.3403 | NaN | 839.3451 | 803.7574 | 1952.0889 |
| Hsp90aa1 | LKEFEGK | -534 | -540 | 1 | 0.001776473 | 7 | 1649.4473 | 1700.4574 | 1619.1785 | 762.29315 | NaN | 998.9402 |
| Hsp90aa1 | LGIHEDSQNRK | -448 | -458 | 1 | 0 | 11 | 893.0986 | 119.731064 | 1043.6207 | NaN | 708.80414 | 740.9856 |
| Hsp90aa1 | LVTSPCCIVT | -593 | -602 | 0 | 0 | 10 | 532.571 | 595.31494 | 576.4658 | 630.3835 | 619.2263 | NaN |
| Hsp90aa1 | DDITNEEYGEFYK | -303 | -315 | 0 | 0 | 13 | 380.67145 | 397.47333 | 368.37558 | 224.3229 | 264.3925 | NaN |
| Hsp90aa1 | DKEVSDDEAEEK | -227 | -238 | 1 | 0.000550307 | 12 | 1035.8921 | 891.802 | NaN | 831.6933 | 1121.3031 | NaN |
| Hsp90aa1 | EMPPLEGDDDTSR | -716 | -728 | 0 | 0 | 13 | 833.11346 | 894.6589 | 903.54346 | 555.7996 | NaN | NaN |
| Hsp90aa1 | KVTVITK | -147 | -153 | 1 | 0.009416377 | 7 | 310.78098 | 240.999 | 311.41348 | 226.30046 | NaN | NaN |
| Hsp90aa1 | HSQFIGYPITLFVEK | -210 | -224 | 0 | 0 | 15 | 659.07495 | 733.2506 | 717.5197 | NaN | NaN | NaN |
| Hsp90aa1 | ESDDKPEIE | -251 | -259 | 1 | 0.007343498 | 9 | 1004.1256 | 855.585 | NaN | NaN | NaN | NaN |
| Hsp90aa1 | HLEINPDHSIIE | -634 | -645 | 0 | 0 | 12 | 3467.219 | NaN | NaN | NaN | NaN | NaN |
| Hsp90aa1 | ALLFVPR | -340 | -346 | 0 | 0.002534621 | 7 | 1183.7025 | NaN | NaN | NaN | NaN | NaN |
| Hsp90aa1 | LSSGFSLEDPQTHANR | -673 | -688 | 0 | 0.003620005 | 16 | 307.42667 | NaN | NaN | NaN | NaN | NaN |
| Xpot | DGPVGFADFVYK | -861 | -872 | 0 | 0 | 12 | 306.95334 | 475.6748 | 432.25494 | 360.6788 | 470.51584 | 564.973 |
| Xpot | ALLGLNPNADSDFR | -5 | -18 | 0 | 0 | 14 | 1395.0533 | 282.38672 | 1772.0438 | NaN | 389.02658 | 2529.348 |
| Xpot | LVELWGGK | -853 | -860 | 0 | 0.00025015 | 8 | 582.156 | NaN | 613.25604 | 517.32996 | 615.9122 | 529.8672 |
| Xpot | IVNSEYPAENK | -604 | -614 | 0 | 0.002068213 | 11 | 1442.2755 | 1233.3923 | NaN | 987.6445 | NaN | NaN |
| Xpot | QMNPYIEEILNR | -559 | -570 | 0 | 0.002099344 | 12 | 418.71768 | NaN | NaN | NaN | NaN | 1207.2798 |
| Gimap4 | SHELGNQDQGIPQLR | -18 | -32 | 0 | 0 | 15 | 382.81857 | 813.70575 | 822.4816 | 1354.8566 | 1205.6925 | 1210.6663 |
| Gimap4 | SHELGNQDQGIPQLR | -18 | -32 | 0 | 0 | 15 | 382.81857 | 813.70575 | 822.4816 | 1354.8566 | 1205.6925 | 1210.6663 |
| Gimap4 | SSHELGNQDQGIPQLR | -17 | -32 | 0 | 0 | 16 | 2501.0002 | 3188.155 | 3526.0164 | 2467.3752 | 3290.4036 | 3494.1475 |
| Gimap4 | SSHELGNQDQGIPQLR | -17 | -32 | 0 | 0 | 16 | 2501.0002 | 3188.155 | 3526.0164 | 2467.3752 | 3290.4036 | 3494.1475 |
| Gimap4 | KDDLEDTDIHEY | -156 | -167 | 1 | 0.000672736 | 12 | 2140.5027 | 2333.4253 | 2390.2615 | 1293.1088 | 1393.7549 | 1387.733 |
| Gimap4 | FFQEVMHEFQNR | -174 | -185 | 0 | 0 | 12 | 2082.7693 | 1669.3589 | 2656.8665 | 2284.5212 | 1042.8132 | 1334.1499 |
| Gimap4 | FFQEVMHEFQNR | -174 | -185 | 0 | 0 | 12 | 2082.7693 | 1669.3589 | 2656.8665 | 2284.5212 | 1042.8132 | 1334.1499 |
| Gimap4 | SSTGNSILGEK | -44 | -54 | 0 | 0 | 11 | 1843.6274 | 819.0268 | 1413.8414 | 1176.04 | 335.42487 | 413.67883 |
| Gimap4 | KDDLEDTDIHEYLEK | -156 | -170 | 1 | 0.00025015 | 15 | 1528.5984 | 1445.7195 | 1825.3878 | 2539.154 | 3132.1858 | 2777.9773 |
| Gimap4 | KDDLEDTDIHEYLEK | -156 | -170 | 1 | 0.00025015 | 15 | 1528.5984 | 1445.7195 | 1825.3878 | 2539.154 | 3132.1858 | 2777.9773 |
| Gimap4 | MYESAECVIQK | -225 | -235 | 0 | 0 | 11 | 1021.129 | 522.6604 | 775.3026 | 1313.9912 | 1174.4565 | 1236.8787 |
| Gimap4 | VVVDTPGIFDTEVPDADTQR | -82 | -101 | 0 | 0 | 20 | 752.0148 | 829.1642 | 831.30145 | 894.48083 | 1046.9609 | 1095.0574 |
| Gimap4 | ELVVVDTPGIFDTEVPDADTQR | -80 | -101 | 0 | 0 | 22 | 403.7193 | 390.53223 | 240.64622 | 389.67157 | 419.23547 | 377.6456 |
| Gimap4 | FFQEVMHEF | -174 | -182 | 0 | 0.00025015 | 9 | 383.90405 | 339.3155 | 390.6183 | 380.41113 | 357.68707 | 420.4143 |
| Gimap4 | ALLLVVPLGR | -116 | -125 | 0 | 0 | 10 | 369.54416 | 386.84625 | 415.65918 | 534.00116 | 635.3213 | 621.3183 |
| Gimap4 | LLVVPLGR | -118 | -125 | 0 | 0.00025015 | 8 | 4181.27 | 2291.2307 | 2113.837 | 1230.757 | 537.67834 | NaN |
| Gimap4 | YCLFNNR | -186 | -192 | 0 | 0.002780285 | 7 | 1550.8951 | 1689.8916 | 1815.7421 | 1703.922 | 1231.3857 | NaN |
| Gimap4 | CGGAGFIPESSR | -5 | -16 | 0 | 0 | 12 | 1374.6647 | NaN | 1385.7142 | 1174.3795 | 1492.089 | 1523.3624 |
| Gimap4 | DDLEDTDIHEYLEK | -157 | -170 | 0 | 0 | 14 | 634.7945 | NaN | 705.85596 | 424.45047 | 908.28845 | 1624.9497 |
| Gimap4 | DTEVPDADTQR | -91 | -101 | 0 | 0 | 11 | 1291.0598 | 1125.876 | 1098.5732 | NaN | NaN | 747.8391 |
| Gimap4 | MEVQCGGAGFIPESSR | -1 | -16 | 0 | 0.000431655 | 16 | 577.6186 | NaN | 676.90314 | 320.386 | NaN | NaN |
| Gimap4 | NQDQGIPQLR | -23 | -32 | 0 | 0 | 10 | 524.519 | 680.48206 | NaN | 143.02902 | NaN | NaN |
| Gimap4 | ILDMFGK | -137 | -143 | 0 | 0.009955823 | 7 | 335.54324 | NaN | 512.80444 | NaN | NaN | NaN |
| Hnrnpul2 | VVVVVPNEEDWK | -557 | -568 | 0 | 0 | 12 | 1044.7506 | 1191.1694 | 1198.7644 | 1247.7616 | 1262.5787 | 1334.5126 |
| Hnrnpul2 | NFILDQCNVYNSGQR | -530 | -544 | 0 | 0 | 15 | 965.5494 | 1236.1794 | 1111.1302 | 840.938 | 855.8511 | 915.6208 |
| Hnrnpul2 | SGDETPGSEAPGDK | -159 | -172 | 0 | 0 | 14 | 679.41693 | 686.7248 | 886.8795 | 722.94025 | 882.5838 | 1017.1713 |
| Hnrnpul2 | QLGEDEFSYGFDGR | -324 | -337 | 0 | 0 | 14 | 650.326 | 682.9224 | 677.7257 | 656.7471 | 799.35944 | 888.5649 |
| Hnrnpul2 | VVVVVPNEEDWKR | -557 | -569 | 1 | 0 | 13 | 497.44836 | 556.7112 | 691.546 | 802.3344 | 937.3327 | 790.36066 |
| Hnrnpul2 | SCSQLGEDEFSYGFDGR | -321 | -337 | 0 | 0.000230707 | 17 | 404.09225 | 442.85785 | 425.21048 | 573.41895 | 496.21555 | 603.5559 |
| Hnrnpul2 | AVEEQGDDQDSEK | -173 | -185 | 0 | 0 | 13 | 381.16003 | 461.46683 | 481.79196 | 376.24677 | 471.81543 | 432.55588 |
| Hnrnpul2 | DWQNYYYHHQ | -719 | -728 | 0 | 0 | 10 | 71.81769 | 761.67523 | 95.11015 | 591.8113 | 532.3425 | 600.67303 |
| Hnrnpul2 | EKEEAQPIVTK | -612 | -622 | 1 | 0 | 11 | 1445.6515 | 919.71423 | 1432.775 | NaN | 1290.7107 | 1310.3021 |
| Hnrnpul2 | EGCTEVSLLR | -304 | -313 | 0 | 0 | 10 | 886.6858 | 1960.3083 | 1048.5553 | 1837.5256 | NaN | 652.75995 |
| Hnrnpul2 | ALLPHVLCK | -396 | -404 | 0 | 0 | 9 | 575.2577 | 705.13684 | 868.1332 | NaN | 831.04944 | 1182.1688 |
| Hnrnpul2 | YGGQPLFSEK | -262 | -271 | 0 | 0 | 10 | 685.02167 | 797.2026 | 1148.5386 | 888.4653 | NaN | NaN |
| Hnrnpul2 | KVVVVVPNEEDWK | -556 | -568 | 1 | 0 | 13 | 750.5153 | 643.7713 | NaN | 629.4323 | NaN | NaN |
| Gpi | KINYTENR | -89 | -96 | 1 | 0.000591124 | 8 | 9301.395 | 7694.831 | 181.1849 | 10536.491 | 10222.788 | 3628.3528 |

| | | | | | | | | | | | | |
|---|---|---|---|---|---|---|---|---|---|---|---|---|
| Gpi | HWMDQHFLK | -301 | -309 | 0 | 0 | 9 | 5456.667 | 521.19293 | 848.4741 | 2794.0166 | 6300.82 | 6417.4717 |
| Gpi | ELFEADPER | -28 | -36 | 0 | 0 | 9 | 4725.6455 | 201.63997 | 4865.453 | 5002.5654 | 4508.853 | 313.83347 |
| Gpi | VFEGNRPTNSIVFTK | -467 | -481 | 1 | 0 | 15 | 4623.7686 | 5223.935 | 5845.442 | 4114.549 | 4755.2046 | 4729.166 |
| Gpi | VFEGNRPTNSIVFTK | -467 | -481 | 1 | 0 | 15 | 4623.7686 | 5223.935 | 5845.442 | 4114.549 | 4755.2046 | 4729.166 |
| Gpi | DFLIPVQTQHPIR | -405 | -417 | 0 | 0 | 13 | 4461.0967 | 5116.8467 | 5068.7524 | 2862.9878 | 3464.997 | 3175.8 |
| Gpi | DFLIPVQTQHPIR | -405 | -417 | 0 | 0 | 13 | 4461.0967 | 5116.8467 | 5068.7524 | 2862.9878 | 3464.997 | 3175.8 |
| Gpi | NIDGTHIAK | -186 | -194 | 0 | 0 | 9 | 3655.8162 | 1828.9382 | 4534.3086 | 6381.165 | 1602.8529 | 4140.765 |
| Gpi | FVSNIDGTHIAK | -183 | -194 | 0 | 0 | 12 | 3571.1982 | 1091.8307 | 3763.3127 | 4224.373 | 4811.157 | 4676.459 |
| Gpi | INYTENR | -90 | -96 | 0 | 0.001636916 | 7 | 2347.8706 | 2257.4282 | 2159.431 | 2042.3135 | 1911.7549 | 2098.62 |
| Gpi | SFDQWGVELGK | -509 | -519 | 0 | 0 | 11 | 1928.0648 | 732.50964 | 2418.6914 | 2147.6099 | 2621.3022 | 2724.5505 |
| Gpi | SFDQWGVELGK | -509 | -519 | 0 | 0 | 11 | 1928.0648 | 732.50964 | 2418.6914 | 2147.6099 | 2621.3022 | 2724.5505 |
| Gpi | GKSPEDLEK | -453 | -461 | 1 | 0.00051306 | 9 | 1798.1313 | 1974.9523 | 2039.9174 | 1859.2322 | 1920.9451 | 2020.9633 |
| Gpi | ETITNAETAK | -217 | -226 | 0 | 0 | 10 | 1528.3595 | 1294.292 | 341.35083 | 1626.0732 | 847.04846 | 1766.4471 |
| Gpi | VFEGNRPTNSIVF | -467 | -479 | 1 | 0 | 13 | 1387.0568 | 1776.1205 | 1767.5874 | 2196.2637 | 2534.5952 | 2563.7676 |
| Gpi | SHDSSTNGLISFIK | -537 | -550 | 0 | 0.00025015 | 14 | 1226.7257 | 1621.297 | 1602.8015 | 1531.7253 | 2307.8071 | 2159.3342 |
| Gpi | DQWGVELGK | -511 | -519 | 0 | 0 | 9 | 1068.5282 | 874.5418 | 553.83453 | 794.0536 | 760.3266 | 687.6097 |
| Gpi | EWFLEAAK | -227 | -234 | 0 | 0.00111006 | 8 | 1027.1459 | 1004.56384 | 1117.3245 | 1371.4949 | 234.88326 | 1468.0349 |
| Gpi | ALLPYDQYMHR | -337 | -347 | 0 | 0 | 11 | 508.67822 | 1084.7805 | 705.0783 | 1377.9745 | 1744.7041 | 1556.2816 |
| Gpi | ALLPYDQYMHR | -337 | -347 | 0 | 0 | 11 | 508.67822 | 1084.7805 | 705.0783 | 1377.9745 | 1744.7041 | 1556.2816 |
| Gpi | VFEGNRPTNSIVFT | -467 | -480 | 1 | 0.00025015 | 14 | 802.16064 | 878.4836 | 744.33435 | 1306.6619 | 1542.1687 | 1399.6091 |
| Gpi | SNIDGTHIAK | -185 | -194 | 0 | 0.001868731 | 10 | 772.39905 | 1322.4518 | 1052.9298 | 1286.5288 | 1364.6427 | 1507.2485 |
| Gpi | TFTTQETITNAETAK | -212 | -226 | 0 | 0 | 15 | 480.1087 | 620.924 | 522.78925 | 1011.1 | 1094.3265 | 1201.0642 |
| Gpi | VDHQTGPIVW | -371 | -380 | 0 | 0.001392869 | 10 | 380.95496 | 577.2176 | 846.4436 | 3329.7407 | 491.81573 | 954.429 |
| Gpi | KIEPELEGSSAVT | -524 | -536 | 1 | 0.000431655 | 13 | 4583.65 | 3124.2717 | 3722.0718 | 3624.4155 | 2394.9792 | NaN |
| Gpi | GKLPEEAR | -439 | -446 | 1 | 0.001092529 | 8 | 3077.1472 | 2956.0615 | 3263.8396 | 3355.625 | 418.55466 | NaN |
| Gpi | KIEPELEGSSA | -524 | -534 | 1 | 0.001092529 | 11 | 1282.052 | 1700.2062 | 1608.9998 | 647.1112 | NaN | 1334.2941 |
| Gpi | VDHQTGPIVWGEPGTNGQHAF | -371 | -391 | 0 | 0.000431655 | 21 | 9148.078 | NaN | 11458.642 | NaN | 286.80304 | 7866.8276 |
| Gpi | WDWVGGR | -267 | -273 | 0 | 0.001868731 | 7 | 1050.9661 | 1000.9412 | 940.295 | NaN | NaN | 669.26215 |
| Gpi | VDHQTGPIVWGEPGTN | -371 | -386 | 0 | 0 | 16 | 583.25757 | 613.86816 | 647.2475 | NaN | 565.5356 | NaN |
| Gpi | SNTPIKVDGK | -107 | -116 | 1 | 0.00025015 | 10 | 2625.2847 | 1300.2609 | NaN | 2311.3088 | NaN | NaN |
| Gpi | YQLIHQGTK | -392 | -400 | 0 | 0.00025015 | 9 | 1044.8651 | 1091.0796 | 1126.6937 | NaN | NaN | NaN |
| Gpi | KIEPELEGS | -524 | -532 | 1 | 0.002329279 | 9 | 765.876 | NaN | NaN | NaN | NaN | NaN |
| Shmt2 | SFLLKDPETSQR | -470 | -481 | 1 | 0 | 12 | 3708.8237 | 3705.62 | 3830.3933 | 3718.3503 | 4216.9824 | 4208.6914 |
| Shmt2 | LLKDPETSQR | -472 | -481 | 1 | 0 | 10 | 3486.621 | 3815.2395 | 3659.19 | 3355.889 | 4866.2573 | 4852.1216 |
| Shmt2 | TALLQPHDR | -152 | -160 | 0 | 0.00025015 | 9 | 2871.5647 | 2977.397 | 3123.4402 | 2797.6812 | 2357.5728 | 3332.1255 |
| Shmt2 | VSGGTDTHLVLVDLRPK | -373 | -389 | 1 | 0 | 17 | 2322.0864 | 2548.1025 | 1061.3629 | 3667.7 | 3710.9966 | 3441.776 |
| Shmt2 | VSGGTDTHLVLVDLRPK | -373 | -389 | 1 | 0 | 17 | 2322.0864 | 2548.1025 | 1061.3629 | 3667.7 | 3710.9966 | 3441.776 |
| Shmt2 | GGTDTHLVLVDLRPK | -375 | -389 | 1 | 0 | 15 | 1922.5532 | 2436.5178 | 2470.897 | 1784.3787 | 921.434 | 2565.3916 |
| Shmt2 | GGTDTHLVLVDLRPK | -375 | -389 | 1 | 0 | 15 | 1922.5532 | 2436.5178 | 2470.897 | 1784.3787 | 921.434 | 2565.3916 |
| Shmt2 | ELIASENFCSR | -72 | -82 | 0 | 0 | 11 | 1661.9564 | 1644.5162 | 1756.341 | 158.36887 | 2878.6353 | 161.00269 |
| Shmt2 | GFPMPGFDER | -495 | -504 | 0 | 0 | 10 | 1487.4377 | 1575.5867 | 1706.1534 | 2264.0688 | 2584.7788 | 2632.2295 |
| Shmt2 | GFPMPGFDER | -495 | -504 | 0 | 0 | 10 | 1487.4377 | 1575.5867 | 1706.1534 | 2264.0688 | 2584.7788 | 2632.2295 |
| Shmt2 | LNPQTGLIDYDQLALTAR | -197 | -214 | 0 | 0 | 18 | 410.35834 | 394.40408 | 398.56146 | 429.7046 | 452.8589 | 462.21796 |
| Shmt2 | LGAPALTSR | -426 | -434 | 0 | 0.00025015 | 9 | 361.04834 | 978.03564 | 677.2336 | 222.04953 | 568.87195 | 620.3438 |
| Shmt2 | INFAVFPSLQGGPHNHAIAA | -315 | -334 | 0 | 0 | 20 | 355.61835 | 311.02502 | 391.0986 | 465.925 | 570.5537 | 663.1204 |
| Shmt2 | YSEGYPGK | -96 | -103 | 0 | 0.000550307 | 8 | 348.79022 | 1185.7357 | 1225.0731 | 1319.3809 | 1224.1124 | 1268.9061 |
| Shmt2 | KVIPSPFK | -262 | -269 | 1 | 0 | 8 | 4141.7856 | 4463.3438 | NaN | 4342.4546 | 4014.008 | 3854.2412 |
| Shmt2 | ALLQPHDR | -153 | -160 | 0 | 0.000298805 | 8 | 845.5819 | 800.083 | 384.94415 | 688.95636 | NaN | 1774.4044 |
| Shmt2 | IMGLDLPDGGHLTHGY | -161 | -176 | 0 | 0.00025015 | 16 | 627.1593 | 553.1973 | NaN | 282.5462 | 469.80923 | 822.84344 |
| Shmt2 | LNPQTGLIDYDQLAL | -197 | -211 | 0 | 0.001536401 | 15 | 591.0722 | 409.08478 | NaN | 446.8899 | 691.54443 | 556.31384 |
| Shmt2 | EIPYTFEDR | -306 | -314 | 0 | 0.000627943 | 9 | 478.88965 | NaN | 1489.6742 | 2155.1494 | 1333.9816 | 1647.4436 |
| Shmt2 | LNPQTGLIDYDQLALTA | -197 | -213 | 0 | 0.00025015 | 17 | 343.89322 | NaN | 412.59845 | 487.79852 | 613.8808 | 586.4738 |
| Shmt2 | LNPQTGLIDYDQL | -197 | -209 | 0 | 0 | 13 | 530.0055 | NaN | NaN | 584.8824 | 1026.3091 | 672.48175 |
| Shmt2 | GYSLVSGGTDTHLVLVDLRPK | -369 | -389 | 1 | 0 | 21 | 119.01141 | 1078.0782 | NaN | NaN | 1728.6843 | 621.12366 |
| Shmt2 | QQVEQFAR | -487 | -494 | 0 | 0.00025015 | 8 | 4035.664 | 1613.9662 | NaN | NaN | NaN | NaN |
| Shmt2 | LLQPHDR | -154 | -160 | 0 | 0.001463897 | 7 | 2150.2349 | NaN | NaN | 1633.5018 | NaN | NaN |
| Shmt2 | SGLIFYR | -287 | -293 | 0 | 0.00122419 | 7 | 365.69614 | NaN | NaN | NaN | NaN | NaN |
| Shmt2 | HLVLVDLRPK | -380 | -389 | 1 | 0.00078661 | 10 | 228.60333 | NaN | NaN | NaN | NaN | NaN |
| Ube2n | DKWSPALQIR | -93 | -102 | 1 | 0 | 10 | 2768.6025 | 2805.6448 | 3062.9824 | 2977.9768 | 3385.2253 | 3078.183 |
| Ube2n | WSPALQIR | -95 | -102 | 0 | 0 | 8 | 2093.6392 | 1650.2074 | 1946.3676 | 1537.576 | 1592.7594 | 1552.629 |
| Ube2n | KTNEAQAIETAR | -130 | -141 | 1 | 0 | 12 | 1613.4823 | 2732.4824 | 1370.567 | 749.33185 | 3547.2056 | 3256.502 |
| Ube2n | LLAEPVPGIK | -15 | -24 | 0 | 0 | 10 | 1452.161 | 1454.5037 | 1645.5383 | 1172.7749 | 1367.5469 | 1272.3259 |
| Ube2n | LFLPEEYPMAAPK | -56 | -68 | 0 | 0 | 13 | 564.254 | 962.53656 | 701.47656 | 142.29018 | 843.2633 | 796.6836 |
| Ube2n | IAGPQDSPFEGGTFK | -39 | -53 | 0 | 0 | 15 | 187.7216 | 2041.6077 | 2101.7622 | 1973.6642 | 1986.8716 | 1709.4338 |
| Ube2n | HVVIAGPQDSPFEGGTFK | -36 | -53 | 0 | 0.001392869 | 18 | 607.39667 | NaN | 631.38965 | 764.5378 | 929.1399 | 975.1851 |
| Ube2n | LELFLPEEYPMAAPK | -54 | -68 | 0 | 0 | 15 | 590.0551 | NaN | 553.1745 | 308.4426 | 381.14038 | 444.37207 |
| Ube2n | TNEAQAIETAR | -131 | -141 | 0 | 0.00025015 | 11 | 1454.8225 | 526.9127 | NaN | 1303.8396 | 690.2097 | NaN |
| Ube2n | LSAPNPDDPLANDVAEQWK | -112 | -130 | 0 | 0 | 19 | 495.37906 | 543.9894 | 577.78015 | NaN | 395.84863 | NaN |
| Ube2n | ALLSAPNPDDPLANDVAEQWK | -110 | -130 | 0 | 0.000907206 | 21 | 215.94691 | NaN | 338.75174 | 252.19118 | 426.6223 | NaN |
| Lrpap1 | ALNEDTQDELGDPR | -140 | -153 | 0 | 0.000471409 | 14 | 2180.2256 | 2307.7222 | 2425.48 | 3289.1765 | 3380.2063 | 169.1643 |
| Lrpap1 | ELESFREELK | -276 | -285 | 1 | 0.000627943 | 10 | 473.66473 | 343.2913 | 787.97943 | 1135.6226 | 1351.5116 | 1643.8711 |
| Lrpap1 | HVESIGDPEHISR | -310 | -322 | 0 | 0 | 13 | 327.64987 | NaN | 740.17004 | 605.6561 | 701.8543 | 657.64514 |
| Anxa7 | GFGTDEQAIVDVVSNR | -175 | -190 | 0 | 0 | 16 | 1290.51 | 1341.1886 | 1383.6774 | 1207.4618 | 1840.0514 | 817.4777 |
| Anxa7 | EFSGYVESGLK | -373 | -383 | 0 | 0 | 11 | 695.96875 | 363.00476 | 837.49506 | 1233.9531 | 1138.0903 | 644.91675 |
| Anxa7 | SDTSGHFER | -288 | -296 | 0 | 0 | 9 | 580.687 | 617.544 | 643.16943 | 229.30042 | 593.4221 | 265.01074 |
| Anxa7 | IASDTSGDYR | -445 | -454 | 0 | 0.002910424 | 10 | 280.42578 | 4970.225 | NaN | 3502.9612 | NaN | 382.85983 |
| Anxa7 | GAGTDDSTLVR | -406 | -416 | 0 | 0 | 11 | 2167.689 | 1109.2449 | 628.00006 | NaN | NaN | NaN |
| Anxa7 | SEIDLVQIK | -422 | -430 | 0 | 0 | 9 | 132.06825 | NaN | 3593.0532 | 3369.9697 | NaN | NaN |

| | | | | | | | | | | | | |
|---|---|---|---|---|---|---|---|---|---|---|---|---|
| Anxa7 | ALNRPAFFAER | -389 | -399 | 1 | 0.00025015 | 11 | 290.0349 | 290.00598 | NaN | NaN | NaN | NaN |
| Anxa7 | CYQLEFGR | -273 | -280 | 0 | 0.000712251 | 8 | 82.81357 | NaN | 656.8288 | NaN | NaN | NaN |
| Gtpbp1 | VGDNDFLEVR | -153 | -162 | 0 | 0 | 10 | 1513.3518 | 1671.4357 | 1684.9917 | 1316.8201 | 1490.424 | 1434.3639 |
| Gtpbp1 | VITFIDLAGHEK | -247 | -258 | 0 | 0 | 12 | 704.6484 | 329.50702 | 330.4546 | 235.76418 | 839.58875 | 957.2819 |
| Gtpbp1 | IDLAGHEK | -251 | -258 | 0 | 0.001023236 | 8 | 1113.6218 | 1249.2599 | 1226.2865 | NaN | 1232.0511 | 1316.6566 |
| Gtpbp1 | VSPTSEQYDSLLR | -68 | -80 | 0 | 0 | 13 | 809.6001 | 768.8557 | 541.73474 | 912.20734 | NaN | 577.01245 |
| Gtpbp1 | ALNVPVFVVVTK | -298 | -309 | 0 | 0 | 12 | 596.31525 | 272.63403 | 1036.7831 | 210.0088 | 751.9567 | NaN |
| Gtpbp1 | VLHHPTTISPR | -498 | -508 | 0 | 0.00025015 | 11 | 706.79803 | NaN | NaN | NaN | NaN | NaN |
| Gtf2i | ALPAEDEESSESR | -8 | -20 | 0 | 0 | 13 | 2013.5391 | 1641.2075 | 1399.4272 | 1389.146 | 863.3203 | 1021.70557 |
| Gtf2i | SPSWYGIPR | -515 | -523 | 0 | 0 | 9 | 1190.1919 | 730.0988 | 1354.4949 | 635.196 | 910.9446 | 1478.314 |
| Gtf2i | EDLQLDKPASGVK | -444 | -456 | 1 | 0.000431655 | 13 | 657.745 | 537.3429 | 1025.2808 | 1331.7821 | 1662.3757 | 1651.8696 |
| Gtf2i | SPTWFGIPR | -620 | -628 | 0 | 0.00025015 | 9 | 626.1677 | 643.79895 | 748.69147 | 680.3301 | 586.50397 | 746.12585 |
| Gtf2i | EVPVTEEIKETDGSSQIK | -974 | -991 | 1 | 0 | 18 | 1104.048 | 1126.7249 | NaN | 481.40564 | 1006.64276 | 1133.5446 |
| Gtf2i | ILDSAEFIK | -929 | -937 | 0 | 0.000907206 | 9 | 866.4265 | NaN | NaN | NaN | 767.9132 | 793.2165 |
| Gtf2i | GLPEGIPFR | (401)(611) | (409)(619) | 0 | 0.007320451 | 9 | 5525.913 | NaN | 4133.738 | NaN | NaN | NaN |
| Gtf2i | PENIPFR | -508 | -514 | 0 | 0.009745776 | 7 | 608.40765 | NaN | NaN | NaN | 209.78429 | NaN |
| Chd1l | ALPAEHGPEPLR | -11 | -22 | 0 | 0.008498216 | 12 | 821.94806 | NaN | NaN | NaN | NaN | 2015.3395 |
| Wipf1 | SAPILDKPK | -55 | -63 | 1 | 0.002566359 | 9 | 3449.3596 | 3459.5774 | 2687.3503 | 5561.2354 | 2619.8384 | 4198.5464 |
| Wipf1 | ALPATPQLPSR | -384 | -394 | 0 | 0 | 11 | 3022.7864 | 2692.1218 | 2749.3303 | 2765.8699 | 2034.9985 | 2281.337 |
| Wipf1 | SGPLPPPPPINR | -366 | -377 | 0 | 0 | 12 | 1567.1766 | 1764.0728 | 1338.8461 | 1845.6294 | 1512.9492 | 1412.0619 |
| Wipf1 | GAPPLPPIPR | -484 | -493 | 0 | 0 | 10 | 230.91512 | 1074.0536 | 1246.7084 | 1072.64 | 702.53253 | 873.98315 |
| Wipf1 | FPAPSPGHR | -151 | -159 | 0 | 0.002752526 | 9 | 719.7292 | NaN | NaN | NaN | NaN | NaN |
| Mphosph10 | ALPDDEAEDTSPLAVK | -336 | -351 | 0 | 0 | 16 | 1087.1019 | 1054.6405 | 2242.6472 | 4515.92 | 4379.2256 | 3234.3657 |
| Mphosph10 | TEEEDNPEHVEIQK | -486 | -499 | 0 | 0 | 14 | 704.7825 | 655.5802 | 586.7622 | 161.35135 | 552.1207 | 438.0343 |
| Il16 | GSLHGDKPLTIN | -1245 | -1256 | 1 | 0 | 12 | 1372.2424 | 1076.3345 | 1626.9768 | 1117.4739 | 2092.1921 | 2149.5898 |
| Il16 | ALPDGPVTIVIR | -1296 | -1307 | 0 | 0 | 12 | 559.5644 | 458.9616 | 461.2915 | 566.45355 | 531.3712 | 566.87805 |
| Il16 | SEGLTEPGETEDR | -1051 | -1063 | 0 | 0 | 13 | 489.685 | 625.1551 | 517.90906 | 630.7907 | 308.87512 | 603.5201 |
| Il16 | GSLHGDKPLTINR | -1245 | -1257 | 1 | 0 | 13 | 508.2224 | 734.7408 | 521.04755 | 496.00177 | 468.13806 | NaN |
| Il16 | GSLHGDKPLTINR | -1245 | -1257 | 1 | 0 | 13 | 508.2224 | 734.7408 | 521.04755 | 496.00177 | 468.13806 | NaN |
| Il16 | EYSEGLTEPGETEDR | -1049 | -1063 | 0 | 0 | 15 | 516.5833 | 495.90958 | 443.72025 | NaN | NaN | 412.38474 |
| Il16 | SEDTQGPGLK | -942 | -951 | 0 | 0 | 10 | 1084.2047 | 929.59564 | NaN | NaN | NaN | 1756.5667 |
| Il16 | TSLGEEDGTQGCPEGGLSK | -739 | -757 | 0 | 0.000550307 | 19 | 912.7823 | NaN | NaN | 567.33905 | NaN | 560.7646 |
| Il16 | ENFGSSQLPDR | -841 | -851 | 0 | 0.000627943 | 11 | 1689.4386 | NaN | NaN | NaN | 730.83264 | NaN |
| Il16 | DPGLPESPPPGQR | -911 | -923 | 0 | 0.002068213 | 13 | 702.5425 | NaN | NaN | NaN | NaN | NaN |
| Il16 | GATHNDALAIL | -1165 | -1175 | 0 | 0.000230707 | 11 | 680.62195 | NaN | NaN | NaN | NaN | NaN |
| Pgam1 | YDADLSPAGHEEAK | -26 | -39 | 0 | 0 | 14 | 6412.5117 | 6596.085 | 6711.359 | 7666.467 | 7966.6855 | 5634.317 |
| Pgam1 | HGESAWNLENR | -11 | -21 | 0 | 0 | 11 | 4621.76 | 266.3534 | 131.30453 | 6202.1694 | 7089.5933 | 6899.901 |
| Pgam1 | YADLTEDQLPSCESLK | -142 | -157 | 0 | 0 | 16 | 2258.0815 | 2986.1775 | 2933.958 | 3956.2864 | 4976.8623 | 4627.089 |
| Pgam1 | MWLPVVR | -77 | -83 | 0 | 0 | 7 | 2175.0278 | 1997.273 | 2522.9312 | 2420.428 | 1390.7986 | 1385.6964 |
| Pgam1 | MWLPVVR | -77 | -83 | 0 | 0 | 7 | 2175.0278 | 1997.273 | 2522.9312 | 2420.428 | 1390.7986 | 1385.6964 |
| Pgam1 | FSGWYDADLSPAGHEEAK | -22 | -39 | 0 | 0 | 18 | 1988.8467 | 2426.259 | 2373.3652 | 2908.511 | 3834.0503 | 3751.9478 |
| Pgam1 | NLPTGIPIVYELDK | -209 | -222 | 0 | 0 | 14 | 1756.0353 | 1744.7001 | 1551.5452 | 1194.2429 | 1509.2428 | 1499.2754 |
| Pgam1 | WNEEIVPQIK | -167 | -176 | 0 | 0 | 10 | 1074.0277 | 5494.9536 | 5486.9814 | 5770.97 | 6858.381 | 7128.2036 |
| Pgam1 | WNEEIVPQIK | -167 | -176 | 0 | 0 | 10 | 1074.0277 | 5494.9536 | 5486.9814 | 5770.97 | 6858.381 | 7128.2036 |
| Pgam1 | YDADLSPAGHEEAKR | -26 | -40 | 1 | 0 | 15 | 861.0687 | 769.8884 | 1954.415 | 1464.5242 | 2782.323 | 1249.467 |
| Pgam1 | YDADLSPAGHEEAKR | -26 | -40 | 1 | 0 | 15 | 861.0687 | 769.8884 | 1954.415 | 1464.5242 | 2782.323 | 1249.467 |
| Pgam1 | LNLPTGIPIVYELDK | -208 | -222 | 0 | 0 | 15 | 855.0288 | 786.9334 | 759.02716 | 702.6153 | 772.2376 | 675.86566 |
| Pgam1 | LSPAGHEEAK | -30 | -39 | 0 | 0 | 10 | 631.97656 | 662.43945 | 732.8332 | 595.13385 | 569.35156 | 538.73376 |
| Pgam1 | DAIDQMWLPVVR | -72 | -83 | 0 | 0 | 12 | 579.9518 | 682.8085 | 632.2969 | 1059.6748 | 804.17694 | 789.7525 |
| Pgam1 | DAIDQMWLPVVR | -72 | -83 | 0 | 0 | 12 | 579.9518 | 682.8085 | 632.2969 | 1059.6748 | 804.17694 | 789.7525 |
| Pgam1 | QFLGDEETVR | -231 | -240 | 0 | 0 | 10 | 449.36954 | 385.48837 | 159.9651 | 6739.904 | 4794.9053 | 5087.179 |
| Pgam1 | SYDVPPPPMEPDHPFYSN | -118 | -135 | 0 | 0 | 18 | 435.88525 | 456.86282 | 538.95593 | 673.03033 | 548.5002 | 574.7627 |
| Pgam1 | YADLTEDQLPSCESLKDTIA | -142 | -161 | 1 | 0 | 20 | 299.11398 | 288.55408 | 243.92671 | 251.71863 | 275.26056 | 284.08466 |
| Pgam1 | FSGWYDADLSPAGHEEAKR | -22 | -40 | 1 | 0 | 19 | 218.65417 | 684.1591 | 591.6665 | 846.0142 | 144.6947 | 925.82574 |
| Pgam1 | QFLGDEETVRK | -231 | -241 | 1 | 0 | 11 | 2767.5464 | NaN | 2876.0771 | 2021.4644 | 3619.167 | 3615.369 |
| Pgam1 | YADLTEDQLPSCESLKDTIAR | -142 | -162 | 1 | 0 | 21 | 1089.5659 | NaN | 1025.8826 | 1371.091 | 1213.4451 | 512.3496 |
| Pgam1 | YADLTEDQLPSCESLKDTIAR | -142 | -162 | 1 | 0 | 21 | 1089.5659 | NaN | 1025.8826 | 1371.091 | 1213.4451 | 512.3496 |
| Pgam1 | ALPFWNEEIVPQIK | -163 | -176 | 0 | 0 | 14 | 900.2107 | 788.68896 | 778.93866 | NaN | 840.48596 | 812.3694 |
| Pgam1 | RYADLTEDQLPSCE | -141 | -154 | 1 | 0.00025015 | 14 | 816.40784 | 170.54002 | 168.9026 | NaN | 904.56744 | 649.14795 |
| Pgam1 | TEDQLPSCESLK | -146 | -157 | 0 | 0 | 12 | 537.7246 | 1076.8024 | 1140.1118 | 1095.1104 | NaN | NaN |
| Pgam1 | ELNLPTGIPIVYELDK | -207 | -222 | 0 | 0.000230707 | 16 | 295.00015 | 275.963 | 262.9657 | NaN | 225.67581 | NaN |
| Pgam1 | EDQLPSCESLK | -147 | -157 | 0 | 0 | 11 | 2764.6384 | 518.41205 | 3085.6072 | NaN | NaN | NaN |
| Pgam1 | LSPAGHEEAKR | -30 | -40 | 1 | 0.000230707 | 11 | 599.912 | 455.85245 | 483.98505 | NaN | NaN | NaN |
| Pgam1 | FSGWYDADLSPAGHEE | -22 | -37 | 0 | 0.000230707 | 16 | 570.6256 | 679.14606 | 710.15533 | NaN | NaN | NaN |
| Pgam1 | YADLTEDQLPSCE | -142 | -154 | 0 | 0.00025015 | 13 | 508.47797 | 606.4421 | 573.06024 | NaN | NaN | NaN |
| Gstp1 | TIVYFPVR | -5 | -12 | 0 | 0.000298805 | 8 | 318.69086 | 488.44635 | 474.8027 | 114.464386 | 414.9482 | 342.24835 |
| Gstp1 | PPYTIVYFPVR | -2 | -12 | 0 | 0 | 11 | 623.3084 | 459.54633 | 641.0547 | NaN | 577.0021 | 883.46564 |
| Rel | ALPPIVSNPIYDNR | -166 | -179 | 0 | 0.005363933 | 14 | 462.27393 | NaN | NaN | NaN | NaN | NaN |
| Utp14a | NKQPVTDPLLTPVEK | -195 | -209 | 1 | 0 | 15 | 651.2611 | 514.29926 | 511.59418 | 452.42175 | NaN | 566.8432 |
| Utp14a | ALPTTVEELEDEGAR | -572 | -586 | 0 | 0 | 15 | 560.2777 | 812.8172 | 1891.3171 | NaN | 1052.5935 | 1011.58154 |
| Utp14a | VVTKPGHIIKPITAE | -718 | -732 | 2 | 0.00555405 | 15 | 296.63724 | 419.95294 | NaN | NaN | NaN | 323.71402 |
| Bcas2 | ALPYFDQGYEAPGVR | -15 | -29 | 0 | 0 | 15 | 870.2596 | 772.47186 | 170.96875 | 768.9972 | 473.27402 | 487.31927 |
| Cse1l | TGNIPALVR | -698 | -706 | 0 | 0 | 9 | 1289.0703 | 946.4623 | 1085.797 | 406.88797 | 850.5427 | 694.16583 |
| Cse1l | ALQEIFDGIQPK | -805 | -816 | 0 | 0 | 12 | 1194.4445 | 561.01117 | 1303.3936 | 2300.332 | 949.5153 | 1482.212 |
| Cse1l | TLDPDPAIR | -18 | -26 | 0 | 0 | 9 | 826.6255 | 852.5497 | 884.7113 | 890.90063 | 959.9889 | 666.7342 |
| Cse1l | ANDHQGFYLLN | -742 | -752 | 0 | 0 | 11 | 702.6109 | 781.8464 | 790.6024 | 407.30588 | 821.19073 | 324.7081 |
| Cse1l | NLFEDQNTLTSICEK | -332 | -346 | 0 | 0 | 15 | 670.67676 | 858.5938 | 863.6557 | 1309.2498 | 1661.8567 | 1482.2358 |
| Cse1l | QLSDAISIIGR | -110 | -120 | 0 | 0 | 11 | 587.717 | 767.7206 | 691.2618 | 727.74133 | 788.5819 | 747.042 |

| | | | | | | | | | | | | |
|---|---|---|---|---|---|---|---|---|---|---|---|---|
| Cse1l | LSDAISIIGR | -111 | -120 | 0 | 0 | 10 | 485.52704 | NaN | 130.42845 | 323.3421 | 413.35663 | 417.1024 |
| Cse1l | HILPDLK | -464 | -470 | 0 | 0.003133457 | 7 | 2076.2979 | 2065.44 | 1794.5361 | NaN | NaN | 2808.4048 |
| Cse1l | IVEDEPNKICEADR | -76 | -89 | 1 | 0 | 14 | 1486.8115 | NaN | 3687.4717 | NaN | 3933.5002 | 3830.8594 |
| Cse1l | IIIPEIQK | -825 | -832 | 0 | 0.001291103 | 8 | 1446.7112 | 1502.5306 | NaN | 1019.80597 | 777.7492 | NaN |
| Cse1l | NPSKPHFNHY | -608 | -617 | 1 | 0 | 10 | 1042.8229 | 1755.315 | 438.1809 | NaN | NaN | 162.81851 |
| Cse1l | NPSKPHFNHY | -608 | -617 | 1 | 0 | 10 | 1042.8229 | 1755.315 | 438.1809 | NaN | NaN | 162.81851 |
| Cse1l | FQSGDFHVIN | -138 | -147 | 0 | 0 | 10 | 768.51447 | NaN | 966.90643 | 428.0115 | NaN | 3332.2393 |
| Cse1l | TDDEEEAGLLELLK | -255 | -268 | 0 | 0 | 14 | 611.5006 | 506.98877 | NaN | 593.1758 | NaN | 543.6897 |
| Cse1l | ANDHQGFYLL | -742 | -751 | 0 | 0.001743443 | 10 | 1584.8824 | NaN | NaN | 998.38336 | NaN | 1137.9705 |
| Cse1l | LSSPEQIQK | -101 | -109 | 0 | 0 | 9 | 634.90936 | NaN | NaN | 882.9782 | NaN | 669.93964 |
| Cse1l | STHANDASALR | -196 | -206 | 0 | 0 | 11 | 568.3271 | 324.36578 | 372.8116 | NaN | NaN | NaN |
| Cse1l | KTLDPDPAIR | -17 | -26 | 1 | 0 | 10 | 1280.182 | NaN | NaN | NaN | 483.0708 | NaN |
| Cse1l | DLEGSDIDTR | -373 | -382 | 0 | 0 | 10 | 546.2648 | NaN | NaN | 4760.9766 | NaN | NaN |
| Igtp | FSPSFHDFPELR | -243 | -254 | 0 | 0 | 12 | 14603.165 | 11853.227 | 9375.656 | 705.2771 | 8934.924 | 7486.663 |
| Igtp | FSPSFHDFPELR | -243 | -254 | 0 | 0 | 12 | 14603.165 | 11853.227 | 9375.656 | 705.2771 | 8934.924 | 7486.663 |
| Igtp | LPQNIWK | -7 | -13 | 0 | 0 | 7 | 5239.0566 | 5577.2974 | 4964.13 | 4226.1924 | 4399.418 | 4728.5684 |
| Igtp | FIGHEEEDSAPTGVVR | -106 | -121 | 0 | 0 | 16 | 3968.044 | 4450.6025 | 4631.2974 | 4929.7803 | 5422.615 | 5683.238 |
| Igtp | FIGHEEEDSAPTGVVR | -106 | -121 | 0 | 0 | 16 | 3968.044 | 4450.6025 | 4631.2974 | 4929.7803 | 5422.615 | 5683.238 |
| Igtp | FHDFPELR | -247 | -254 | 0 | 0 | 8 | 2182.9033 | 2370.438 | 2363.1318 | 2231.413 | 1778.7367 | 2337.4543 |
| Igtp | ALQFQDLIK | -345 | -353 | 0 | 0.000230707 | 9 | 2166.2615 | 2501.9885 | 2626.0054 | 2278.1787 | 2756.552 | 2875.634 |
| Igtp | VSSEDDTANLER | -302 | -313 | 0 | 0.00025015 | 12 | 2026.3435 | 2078.7197 | 2152.851 | 2396.5298 | 2289.9087 | 2354.5645 |
| Igtp | QNSSSPEVIEDIGK | -46 | -59 | 0 | 0 | 14 | 1995.365 | 131.5755 | 2189.5903 | 2816.791 | 2377.3052 | 3101.385 |
| Igtp | YRDPLEIISQ | -266 | -275 | 1 | 0.001743443 | 10 | 1054.4363 | 2695.452 | 2792.1292 | 2466.3848 | 2548.162 | 2598.7444 |
| Igtp | VIGIVKDEIQSK | -69 | -80 | 1 | 0 | 12 | 968.14685 | 1081.2211 | 1109.0051 | 1325.9844 | 1285.9058 | 1439.0652 |
| Igtp | CFSPSFHDFPELR | -242 | -254 | 0 | 0 | 13 | 957.7495 | 537.8191 | 580.3137 | 725.4988 | 834.66125 | 859.31384 |
| Igtp | LFGVDDGSLQQVAR | -321 | -334 | 0 | 0 | 14 | 190.02768 | 1956.1038 | 1853.57 | 2190.9468 | 2724.2834 | 2562.4832 |
| Igtp | EAAVSSEDDTANLER | -299 | -313 | 0 | 0 | 15 | 1127.6302 | NaN | 1005.71985 | 1311.4641 | 711.0024 | 644.9635 |
| Igtp | NSSSPEVIEDIGK | -47 | -59 | 0 | 0.00025015 | 13 | 3205.902 | NaN | NaN | 3202.0107 | NaN | NaN |
| Igtp | NSSSPEVIEDIGK | -47 | -59 | 0 | 0.00025015 | 13 | 3205.902 | NaN | NaN | 3202.0107 | NaN | NaN |
| Ppp2r1b | FGTEWAQNTIVPK | -485 | -497 | 0 | 0 | 13 | 2560.8345 | 3110.3396 | 2910.8787 | 6192.766 | 4862.021 | 5618.056 |
| Ppp2r1b | ELVSDTNQHVK | -344 | -354 | 0 | 0 | 11 | 923.204 | 1057.3003 | 950.2228 | 1185.7255 | 1042.845 | 1211.4342 |
| Ppp2r1b | ALQGEVKPVLQK | -567 | -578 | 1 | 0 | 12 | 337.67554 | 370.68875 | 318.08478 | 342.69998 | 304.32712 | NaN |
| Ppp2r1b | IGPILDTNALQGEVKPVLQK | -559 | -578 | 1 | 0.001776473 | 20 | 279.22894 | NaN | NaN | 283.6585 | 331.0353 | 348.93073 |
| Ppp2r1b | ASGDWFTSR | -148 | -156 | 0 | 0 | 9 | 94.38352 | NaN | NaN | 484.22528 | 543.0966 | 519.9666 |
| Ppp2r1b | LASGDWFTSR | -147 | -156 | 0 | 0 | 10 | 408.1833 | NaN | NaN | NaN | NaN | 353.81662 |
| Copg1 | TLEPSEKPFDLK | -566 | -577 | 1 | 0 | 12 | 3606.263 | 2916.282 | 3144.7412 | 2544.6426 | 2100.784 | 1692.8164 |
| Copg1 | GGHDILVR | -836 | -843 | 0 | 0 | 8 | 1286.9722 | 741.3792 | 1010.73254 | 901.81934 | 812.4729 | 840.00714 |
| Copg1 | NLDLENLVTDSNR | -326 | -338 | 0 | 0.000431655 | 13 | 1250.4988 | 634.08746 | 1040.5762 | 192.47542 | 1480.1644 | 1397.3446 |
| Copg1 | VVLEHEEVR | -480 | -488 | 0 | 0 | 9 | 1173.6455 | 1157.8695 | 1122.7472 | 938.91986 | 975.16736 | 1216.4948 |
| Copg1 | DEESGGGSNPLQHLEK | -9 | -24 | 0 | 0 | 16 | 1001.14014 | 684.3893 | 1148.9988 | 851.336 | 934.4133 | 1201.4166 |
| Copg1 | LFQSNDPTLR | -76 | -85 | 0 | 0 | 10 | 906.0597 | 880.4936 | 965.5412 | 846.5744 | 797.9314 | 791.9435 |
| Copg1 | KDEESGGGSNPLQHLEK | -8 | -24 | 1 | 0 | 17 | 675.813 | 782.29474 | 783.4624 | 610.3756 | 668.75714 | 678.053 |
| Copg1 | KDEESGGGSNPLQHLEK | -8 | -24 | 1 | 0 | 17 | 675.813 | 782.29474 | 783.4624 | 610.3756 | 668.75714 | 678.053 |
| Copg1 | ALQQYTLEPSEKPFDLK | -561 | -577 | 1 | 0 | 17 | 453.3407 | 529.0276 | 469.1958 | 502.89166 | 202.50357 | 563.5258 |
| Copg1 | LAAVPEFQGLGPLFK | -617 | -631 | 0 | 0 | 15 | 188.14445 | 417.6833 | 405.44656 | 644.5944 | 846.69293 | 528.87427 |
| Copg1 | GLLYHVR | -198 | -204 | 0 | 0.002697088 | 7 | 513.5832 | 447.588 | NaN | 558.06824 | 813.56744 | 903.3168 |
| Copg1 | RVVLEHEEVR | -479 | -488 | 1 | 0 | 10 | 612.20325 | NaN | NaN | 527.3893 | 1527.3121 | 1211.8508 |
| Abhd11 | ALQRPDVVER | -143 | -152 | 1 | 0.00078661 | 10 | 761.837 | NaN | 790.86884 | NaN | 942.6604 | 932.2244 |
| Abhd11 | LHGLFGSK | -64 | -71 | 0 | 0.000869338 | 8 | 1024.7183 | NaN | NaN | NaN | NaN | NaN |
| Tnpo2 | GLLFHPEWVVK | -401 | -411 | 0 | 0 | 11 | 267.03842 | 1297.648 | 1251.7756 | NaN | NaN | 1354.3276 |
| Tnpo2 | EELLPHLLPLLK | -389 | -400 | 0 | 0 | 12 | 447.43872 | 488.68875 | NaN | NaN | NaN | 547.3242 |
| Psmc6 | HGEIDYEAIVK | -323 | -333 | 0 | 0 | 11 | 8439.395 | 2762.553 | 10889.422 | 1604.5186 | 11271.796 | 3014.8396 |
| Psmc6 | NRPDTLDPALLRPGR | -280 | -294 | 2 | 0 | 15 | 2189.1304 | 2299.021 | 1862.3593 | 1894.5243 | 1626.155 | 303.9395 |
| Psmc6 | NRPDTLDPALLRPGR | -280 | -294 | 2 | 0 | 15 | 2189.1304 | 2299.021 | 1862.3593 | 1894.5243 | 1626.155 | 303.9395 |
| Psmc6 | GCLLYGPPGTGK | -169 | -180 | 0 | 0 | 12 | 1502.1821 | 1659.1129 | 2076.704 | 718.232 | 933.66156 | 2206.4707 |
| Psmc6 | KIHIDLPNEQAR | -298 | -309 | 1 | 0 | 12 | 1160.8962 | 1304.8582 | 1348.1947 | 1086.431 | 1481.4392 | 1511.2244 |
| Psmc6 | KIHIDLPNEQAR | -298 | -309 | 1 | 0 | 12 | 1160.8962 | 1304.8582 | 1348.1947 | 1086.431 | 1481.4392 | 1511.2244 |
| Psmc6 | ATNRPDTLDPALLRPGR | -278 | -294 | 2 | 0 | 17 | 987.36487 | 1101.4093 | 80.76836 | 574.3933 | 843.09784 | 890.35504 |
| Psmc6 | EVIELPLTNPELFQR | -147 | -161 | 0 | 0 | 15 | 529.181 | 648.89935 | 619.9294 | 605.9 | 778.11816 | 651.07697 |
| Psmc6 | KIHAGPITK | -314 | -322 | 1 | 0.00142999 | 9 | 352.7298 | 445.01147 | 413.16876 | 510.7075 | 360.42416 | 406.22577 |
| Psmc6 | QSVGQIVGEVLK | -51 | -62 | 0 | 0 | 12 | 309.54193 | 303.7056 | 343.52438 | 563.2284 | 560.8554 | 566.4293 |
| Psmc6 | TNRPDTLDPALLRPGR | -279 | -294 | 2 | 0 | 16 | 708.99384 | 815.1259 | 483.06888 | NaN | 938.31036 | 139.66342 |
| Psmc6 | IHAGPITK | -315 | -322 | 0 | 0.00025015 | 8 | 430.66385 | 266.52875 | 210.05327 | NaN | 337.47873 | 418.75058 |
| Psmc6 | ALQSVGQIVGEVLK | -49 | -62 | 0 | 0.000230707 | 14 | 392.55383 | 269.3825 | 286.90308 | NaN | NaN | NaN |
| Psmc6 | LSDGFNGADLR | -334 | -344 | 0 | 0 | 11 | 292.74466 | 693.6307 | 607.7685 | NaN | NaN | NaN |
| Ap2a1 | DFLTPPLLSVR | -833 | -843 | 0 | 0 | 11 | 549.1834 | 658.3407 | 605.6579 | 593.92755 | 715.51373 | 670.50055 |
| Ap2a1 | ALQVGCLLR | -936 | -944 | 0 | 0.002168954 | 9 | 432.30276 | NaN | NaN | NaN | 570.7696 | NaN |
| Shmt1 | YDQLEENASLFHPK | -177 | -190 | 0 | 0 | 14 | 10253.334 | 11791.633 | 11600.567 | 10422.813 | 11827.447 | 11159.654 |
| Shmt1 | VAHFIHR | -415 | -421 | 0 | 0 | 7 | 5770.235 | 5410.5186 | 776.33325 | 1801.4813 | 5588.5327 | 5783.787 |
| Shmt1 | ALSDALTELGYK | -331 | -342 | 0 | 0 | 12 | 826.90137 | 683.226 | 849.38995 | 760.08795 | 990.2035 | 822.35223 |
| Shmt1 | YSEGYPGQR | -67 | -75 | 0 | 0 | 9 | 682.85876 | 345.8794 | 396.7199 | 532.797 | 111.02977 | 326.12268 |
| Shmt1 | IVTGGSDNHLIL | -343 | -354 | 0 | 0 | 12 | 978.0626 | 964.71045 | 1029.8219 | NaN | NaN | 909.68604 |
| Shmt1 | TALVEPHGR | -123 | -131 | 0 | 0.005821388 | 9 | 506.4394 | NaN | NaN | NaN | NaN | 4054.1858 |
| Mtch1 | ALSHPLLYVK | -94 | -103 | 0 | 0.000907206 | 10 | 261.5563 | NaN | NaN | NaN | NaN | NaN |
| Unc45a | ALSLGATPQDQAILHR | -47 | -62 | 0 | 0.000550307 | 16 | 743.54535 | NaN | NaN | NaN | 904.23553 | NaN |
| Itga2b | FGSAIAPLGDLNR | -386 | -398 | 0 | 0 | 13 | 687.6359 | 1227.5858 | 719.66833 | 710.95886 | 1035.8154 | 1522.4752 |
| Itga2b | GAVEILDSYYQPLHR | -295 | -309 | 0 | 0 | 15 | 677.5664 | 622.95337 | 878.54297 | 825.8047 | 241.20883 | 336.1758 |
| Itga2b | FTYDNSNPVFFDGYR | -251 | -265 | 0 | 0 | 15 | 378.16183 | 448.56256 | 466.40256 | 392.38647 | 553.30945 | 561.78424 |

| | | | | | | | | | | | | |
|---|---|---|---|---|---|---|---|---|---|---|---|---|
| Itga2b | SLPYSVPVVSLPSGQAR | -960 | -976 | 0 | 0 | 17 | 13457.934 | 14240.593 | 14382.118 | 16161.135 | NaN | 16805.752 |
| Itga2b | IILDCGEDDLCVPQLR | -628 | -643 | 0 | 0 | 16 | 490.7811 | 564.95215 | 349.6464 | NaN | 588.7367 | NaN |
| Itga2b | ALSNIEGFER | -692 | -701 | 0 | 0.00025015 | 10 | 907.94135 | 1010.1456 | 3145.0725 | NaN | NaN | NaN |
| Itga2b | PSQVLDSPFPTGSGF | -433 | -447 | 0 | 0.000755556 | 15 | 447.50464 | NaN | NaN | NaN | NaN | 713.29724 |
| Ubqln4 | ALSNLESVPGGYNALR | -263 | -278 | 0 | 0 | 16 | 897.34436 | 819.88434 | 855.2268 | NaN | 1418.8135 | NaN |
| Ubqln4 | DGDTLSQHGIK | -63 | -73 | 0 | 0 | 11 | 1736.4852 | NaN | NaN | NaN | NaN | NaN |
| Lactb | IALEFDKDR | -541 | -549 | 1 | 0.000394361 | 9 | 738.83264 | 690.117 | 959.2416 | 235.93388 | NaN | 853.47406 |
| Lactb | ALSPGSPHTPAPPAAR | -85 | -100 | 0 | 0.009388464 | 16 | 1240.8827 | NaN | NaN | NaN | NaN | NaN |
| Ogfr | SQVGPEDPQSQVGPEQAASK | -581 | -600 | 0 | 0 | 20 | 383.54718 | 562.041 | 524.7796 | 697.7113 | 497.0501 | 585.50366 |
| Ogfr | DNYDLLEENHSYIQ | -109 | -122 | 0 | 0 | 14 | 303.86725 | 437.40314 | 543.7829 | NaN | 615.4154 | 896.7723 |
| Ogfr | ALSPTPSECPESQK | -450 | -463 | 0 | 0.00025015 | 14 | 1399.7942 | NaN | NaN | NaN | 1509.1849 | 1476.2448 |
| Ogfr | SQVEPEDPK | (554)(563)(572) | (562)(571)(580) | 0 | 0 | 9 | 1298.2565 | NaN | NaN | 900.9927 | NaN | NaN |
| Rbm10 | TYIPALEQSADGHK | -614 | -627 | 0 | 0 | 14 | 5750.6914 | 5857.562 | 6347.928 | 5944.9595 | 5152.3696 | 5423.1562 |
| Rbm10 | INEDWLCNK | -213 | -221 | 0 | 0 | 9 | 379.25815 | 836.5688 | 894.0478 | 266.3691 | 612.1937 | 493.1799 |
| Rbm10 | ALSQGSEPSSENANDTIILR | -286 | -305 | 0 | 0.00111006 | 20 | 3365.2795 | 3715.1711 | 4296.9834 | NaN | NaN | NaN |
| Itpa | ALSTGDPSQPVLLFR | -119 | -133 | 0 | 0 | 15 | 890.5828 | 1090.2439 | 1072.2406 | 940.521 | 90.65023 | 1217.1906 |
| Itpa | STGDPSQPVLLFR | -121 | -133 | 0 | 0 | 13 | 1123.3871 | 1063.2738 | 1086.4379 | NaN | 1391.3472 | 1392.7366 |
| Itpa | NALGGLPGPYIK | -78 | -89 | 0 | 0 | 12 | 691.1447 | 658.8458 | NaN | 717.77356 | 547.79034 | 791.26135 |
| Itpa | LKPEGLHQL | -95 | -103 | 1 | 0.00201047 | 9 | 1356.5725 | 1418.2129 | NaN | NaN | NaN | NaN |
| Itpa | GDPSQPVLLFR | -123 | -133 | 0 | 0 | 11 | 281.17218 | NaN | 482.3805 | NaN | NaN | NaN |
| Psmb10 | IHFIAPK | -73 | -79 | 0 | 0.000627943 | 7 | 906.22205 | 1030.6746 | 968.72455 | 865.8781 | 852.7996 | 790.7571 |
| Psmb10 | ALSTPTEPVQR | -227 | -237 | 0 | 0 | 11 | 883.0029 | 1003.3644 | 983.03516 | 1328.685 | 1569.9379 | 1470.6204 |
| Psmb10 | FAPGTTPVLTR | -243 | -253 | 0 | 0 | 11 | 396.4483 | 1993.7711 | 2113.8523 | 2015.8223 | 2481.7593 | 2440.42 |
| Psmb10 | YQGHVGASL | -129 | -137 | 0 | 0 | 9 | 11637.598 | 11122.751 | 9981.604 | 177.45502 | NaN | NaN |
| Tomm70 | NVDLSTFYQNR | -152 | -162 | 0 | 0 | 11 | 813.8661 | 875.36084 | 271.6374 | 889.6404 | 1029.5608 | 944.16376 |
| Tomm70 | SDEDKDKEGEAL | -282 | -293 | 2 | 0.00025015 | 12 | 630.56726 | 661.47076 | 655.82324 | 946.08655 | 761.58673 | 729.60944 |
| Tomm70 | ALTDQQQFGK | -488 | -497 | 0 | 0 | 10 | 8440.892 | NaN | NaN | NaN | NaN | NaN |
| Tomm70 | YGLKPPTL | -604 | -611 | 1 | 0.003998192 | 8 | 1701.691 | NaN | NaN | NaN | NaN | NaN |
| Elmo1 | ALTTKPSSLDQFK | -491 | -503 | 1 | 0 | 13 | 2388.303 | 1023.326 | 1868.2244 | 714.10736 | 1343.9816 | 1520.6206 |
| Elmo1 | TTKPSSLDQFK | -493 | -503 | 1 | 0 | 11 | 1705.2368 | 2031.8496 | 2254.8198 | 1613.1512 | 1617.0082 | 3142.1423 |
| Elmo1 | KPLSAIIK | -28 | -35 | 1 | 0 | 8 | 937.22516 | 928.9506 | 789.4282 | 1073.216 | 884.39154 | 791.2802 |
| Elmo1 | LLDLENIQIPDAPPPIPK | -698 | -715 | 0 | 0 | 18 | 432.33862 | 412.05545 | 498.5469 | 511.46762 | 700.534 | 553.6938 |
| Elmo1 | VAIEWPGAYPK | -10 | -20 | 0 | 0 | 11 | 617.98755 | 504.2781 | 668.0563 | 1264.6556 | NaN | 1443.3993 |
| Elmo1 | TSPAQNAQQLHER | -79 | -91 | 0 | 0 | 13 | 573.7877 | NaN | 535.7079 | 463.58224 | 711.687 | 696.2621 |
| Elmo1 | VLNSHDLYQK | -209 | -218 | 0 | 0 | 10 | 2641.5908 | 2160.5588 | 3786.7993 | NaN | NaN | 3189.4453 |
| Elmo1 | LQNLSYTEILK | -506 | -516 | 0 | 0 | 11 | 571.5079 | 451.59055 | NaN | NaN | 518.9601 | 1085.4163 |
| Elmo1 | LTTSPAQNAQQLHER | -77 | -91 | 0 | 0 | 15 | 510.82938 | NaN | 1460.4138 | NaN | 351.22632 | NaN |
| Rpl14 | LVAIVDVIDQNR | -24 | -35 | 0 | 0 | 12 | 2992.508 | 3279.1692 | 157.01976 | 2240.6636 | 2793.2505 | 2850.481 |
| Rpl14 | YISFGPHAGK | -14 | -23 | 0 | 0 | 10 | 313.37662 | 6376.03 | 6252.5117 | 7108.0557 | 6359.487 | 5940.256 |
| Rpl14 | ALVDGPCTR | -36 | -44 | 0 | 0 | 9 | 2777.264 | 2383.0469 | 1956.2651 | 1857.54 | NaN | 1642.4652 |
| Rpl14 | VAYISFGPHAGK | -12 | -23 | 0 | 0 | 12 | 1741.1499 | 1762.3857 | 1918.1271 | 696.07526 | NaN | 1164.652 |
| Rpl14 | VAYISFGPHAGK | -12 | -23 | 0 | 0 | 12 | 1741.1499 | 1762.3857 | 1918.1271 | 696.07526 | NaN | 1164.652 |
| Rpl14 | IVDVIDQNR | -27 | -35 | 0 | 0 | 9 | 1556.3921 | 1104.0929 | NaN | 1793.4761 | 1704.2133 | 1882.4286 |
| Map4k1 | ALVDPDIFNKDPR | -2 | -14 | 1 | 0 | 13 | 334.2776 | 344.3317 | 752.1421 | 814.2163 | 896.1584 | 926.33374 |
| Map4k1 | LVTPEGAPAPGLR | -737 | -749 | 0 | 0 | 13 | 1013.6322 | 305.84772 | 626.5104 | 576.8171 | NaN | NaN |
| Upf1 | ALVEGPLNNLR | -903 | -913 | 0 | 0 | 11 | 2755.1287 | 2237.2285 | 2606.2078 | 2031.425 | 2582.9888 | 2590.393 |
| Upf1 | LLGHEVEDVVIK | -451 | -462 | 0 | 0 | 12 | 1866.2722 | 3595.852 | 3794.5645 | 3793.2131 | 4744.594 | 4639.8477 |
| Upf1 | APSNIAVDQLTEK | -527 | -539 | 0 | 0 | 13 | 1286.7043 | 866.32513 | 1391.9479 | 1134.3883 | 507.92786 | 1294.9285 |
| Upf1 | IPSEQEQLR | -240 | -248 | 0 | 0 | 9 | 407.45074 | 2866.878 | 2977.0593 | 2660.6467 | 3047.674 | 3275.4497 |
| Upf1 | NFEEDEEDTYYTK | -99 | -111 | 0 | 0 | 13 | 307.32794 | 599.292 | 582.0353 | 536.0604 | 512.1396 | 591.7227 |
| Upf1 | DINWDSSQWQPLIQDR | -216 | -231 | 0 | 0 | 16 | 288.1438 | 288.62805 | 273.31766 | 312.98566 | 503.8802 | 486.50974 |
| Upf1 | YGVIIVGNPK | -874 | -883 | 0 | 0 | 10 | 730.85974 | 761.6937 | 835.0057 | 647.5709 | NaN | 595.01794 |
| Upf1 | IAFFTLPK | -337 | -344 | 0 | 0.00078661 | 8 | 558.62445 | 705.44806 | 598.6728 | NaN | NaN | NaN |
| Upf1 | QVDFVWK | -416 | -422 | 0 | 0.004439092 | 7 | 545.2395 | 543.2274 | NaN | NaN | NaN | NaN |
| Upf1 | DLEKPGVDEEPQHVLLR | -274 | -290 | 1 | 0.00025015 | 17 | 521.1678 | NaN | 369.67328 | NaN | NaN | NaN |
| Upf1 | TVLQRPLSLIQ | -487 | -497 | 1 | 0.00078661 | 11 | 474.02737 | NaN | NaN | NaN | 452.204 | NaN |
| Upf1 | ENPSATLEDLEKPGVDEEPQHVLLR | -266 | -290 | 1 | 0.00272025 | 25 | 405.6815 | 348.76178 | NaN | NaN | NaN | NaN |
| Upf1 | WDLGLNK | -328 | -334 | 0 | 0.005668207 | 7 | 154.20259 | NaN | NaN | NaN | NaN | NaN |
| Phb1 | ATAGDGLIELR | -229 | -239 | 0 | 0 | 11 | 3288.5432 | 3209.4026 | 3269.1003 | 3245.6345 | 3544.0054 | 3410.718 |
| Phb1 | LYNVDAGHR | -27 | -35 | 0 | 0 | 9 | 2558.2122 | 2686.497 | 2697.1372 | 2433.977 | 2362.932 | 2385.129 |
| Phb1 | TSIGEDYDER | -108 | -117 | 0 | 0 | 10 | 1955.5579 | 168.85045 | 1694.5618 | 1761.1112 | 1370.4447 | 506.563 |
| Phb1 | GVQDIVVGEGTHF | -44 | -56 | 0 | 0 | 13 | 1497.4957 | 1780.2438 | 1565.1041 | 1693.2324 | 1788.1245 | 1631.4126 |
| Phb1 | VLPSITTEILK | -118 | -128 | 0 | 0 | 11 | 1446.9181 | 1827.383 | 1701.3462 | 1823.6487 | 2529.809 | 2558.793 |
| Phb1 | DLQNVNITLR | -84 | -93 | 0 | 0 | 10 | 1435.6239 | 1719.3228 | 1806.2235 | 1528.3461 | 1705.9382 | 1771.6931 |
| Phb1 | SLATAGDGLIELR | -227 | -239 | 0 | 0 | 13 | 1223.751 | 1512.0393 | 1743.2026 | 1709.0034 | 1927.3673 | 1862.3617 |
| Phb1 | SALYNVDAGHR | -25 | -35 | 0 | 0 | 11 | 1184.9984 | 1519.2864 | 2172.653 | 1538.5664 | 1475.5461 | 1353.1521 |
| Phb1 | IYTSIGEDYDER | -106 | -117 | 0 | 0 | 12 | 1078.931 | 1499.912 | 1246.2089 | 1597.5795 | 1376.52 | 1242.9326 |
| Phb1 | FDAGELITQR | -134 | -143 | 0 | 0.00025015 | 10 | 445.51398 | 3281.388 | 3500.0278 | 259.88788 | 3399.2656 | 3342.823 |
| Phb1 | LIPWVQKPIIFDCR | -57 | -70 | 1 | 0 | 14 | 171.7713 | 287.58304 | 445.34515 | 242.51797 | 343.62955 | 341.89557 |
| Phb1 | NITYLPAGQSVLL | -256 | -268 | 0 | 0 | 13 | 380.34177 | 497.70627 | NaN | 453.0673 | 497.71826 | 460.86853 |
| Phb1 | PVASQLPR | -98 | -105 | 0 | 0.000591124 | 8 | 994.631 | 1048.3138 | NaN | 972.65405 | 648.0017 | NaN |
| Phb1 | QVAQQEAER | -187 | -195 | 0 | 0 | 9 | 604.25574 | NaN | NaN | 496.6897 | 873.8965 | 796.9579 |
| Phb1 | ALYNVDAGHR | -26 | -35 | 0 | 0.00051306 | 10 | 1262.9858 | 612.338 | NaN | 595.9878 | NaN | NaN |
| Phb1 | KLEAAEDIAYQL | -240 | -251 | 1 | 0 | 12 | 891.10645 | 814.22485 | 710.5124 | NaN | NaN | NaN |
| Phb1 | EFTEAVEAK | -178 | -186 | 0 | 0.000627943 | 9 | 368.16165 | 331.86475 | NaN | NaN | NaN | NaN |
| Phb1 | QVSDDLTER | -149 | -157 | 0 | 0.004983786 | 9 | 1385.6392 | NaN | NaN | NaN | NaN | NaN |
| Slc25a3 | SCGLTHTAVVPLDLVK | -70 | -85 | 0 | 0 | 16 | 2556.389 | 2900.6243 | 2964.7607 | 2941.2756 | 3562.6104 | 3684.2002 |
| Slc25a3 | GEENTYLWR | -148 | -156 | 0 | 0 | 9 | 2115.6475 | 132.15404 | 2139.4124 | 923.9781 | 900.6568 | 1234.6388 |

| | | | | | | | | | | | | |
|---|---|---|---|---|---|---|---|---|---|---|---|---|
| Slc25a3 | FGFYEVFK | -133 | -140 | 0 | 0 | 8 | 1863.9128 | 1798.4254 | 1911.0488 | 1249.286 | 1547.8127 | 1507.4733 |
| Slc25a3 | QWFIYDSVK | -324 | -332 | 0 | 0 | 9 | 1625.0322 | 1995.6522 | 1042.074 | 1002.54395 | 284.7445 | 1965.8972 |
| Slc25a3 | FVVPKPR | -243 | -249 | 1 | 0 | 7 | 1622.0712 | 1166.0486 | 1552.4993 | 1132.5518 | 1088.5723 | 1128.0181 |
| Slc25a3 | LPRPPPPEMPESLKK | -337 | -351 | 2 | 0.001023236 | 15 | 894.3675 | 894.2854 | 824.27155 | 819.6644 | 743.2586 | 770.6775 |
| Slc25a3 | TAVVPLDLVK | -76 | -85 | 0 | 0 | 10 | 715.00415 | 716.0208 | 625.61914 | 856.23834 | 947.34766 | 1045.6956 |
| Slc25a3 | LPRPPPPEMPESLK | -337 | -350 | 1 | 0 | 14 | 702.15735 | 1003.72 | 271.0044 | 549.17847 | 482.0347 | 854.4066 |
| Slc25a3 | LPRPPPPEMPESLK | -337 | -350 | 1 | 0 | 14 | 702.15735 | 1003.72 | 271.0044 | 549.17847 | 482.0347 | 854.4066 |
| Slc25a3 | YSNILGEENTYLWR | -143 | -156 | 0 | 0 | 14 | 670.5181 | 754.1315 | 822.7334 | 1500.6927 | 2076.7861 | 1959.9211 |
| Slc25a3 | VVPLDLVK | -78 | -85 | 0 | 0.00025015 | 8 | 1706.899 | 1185.0383 | 1028.9379 | 654.08026 | 413.77457 | NaN |
| Slc25a3 | AVVPLDLVK | -77 | -85 | 0 | 0 | 9 | 1547.5356 | 1061.0892 | 1118.6962 | 1255.1322 | 1515.2263 | NaN |
| Slc25a3 | IQTQPGYANTLR | -185 | -196 | 0 | 0 | 12 | 887.4863 | 532.52466 | 1059.7552 | 891.57623 | 1880.9661 | NaN |
| Slc25a3 | GWAPTLIGY | -117 | -125 | 0 | 0.00025015 | 9 | 1538.8325 | NaN | NaN | 2163.722 | 3691.806 | 3699.6816 |
| Slc25a3 | GSTASQVLQR | -291 | -300 | 0 | 0 | 10 | 1103.2664 | 1053.0327 | 1074.0135 | NaN | NaN | 757.58606 |
| Slc25a3 | KFACFER | -229 | -235 | 1 | 0 | 7 | 686.7794 | NaN | 700.2784 | NaN | 1375.229 | 965.70087 |
| Slc25a3 | ALYSNILGEENTYLWR | -141 | -156 | 0 | 0 | 16 | 388.4166 | NaN | NaN | 717.36523 | 542.58246 | 832.514 |
| Slc25a3 | NILGEENTYLWR | -145 | -156 | 0 | 0.003003218 | 12 | 2351.9983 | NaN | 2679.445 | 2406.3062 | NaN | NaN |
| Slc25a3 | ILGEENTYLWR | -146 | -156 | 0 | 0 | 11 | 1209.1544 | 1054.2462 | NaN | 491.81635 | NaN | NaN |
| Slc25a3 | WFIYDSVK | -325 | -332 | 0 | 0.000550307 | 8 | 858.54834 | 565.08154 | NaN | NaN | NaN | NaN |
| Slc25a3 | AVEEYSCEFGSMK | -46 | -58 | 0 | 0 | 13 | 320.96622 | NaN | NaN | 881.2852 | NaN | NaN |
| Slc25a3 | THTAVVPLDLVK | -74 | -85 | 0 | 0.000431655 | 12 | 587.1873 | NaN | NaN | NaN | NaN | NaN |
| Pdia4 | EVSQPDWTPPPEVTLS | -159 | -174 | 0 | 0.000471409 | 16 | 11604.827 | 11611.553 | 12012.158 | 24977.852 | 289.02167 | 26519.469 |
| Pdia4 | FIDEHATK | -623 | -630 | 0 | 0 | 8 | 3256.4358 | 3183.6716 | 2899.305 | 3996.4006 | 3935.319 | 4349.438 |
| Pdia4 | FHHTFSPEIAK | -329 | -339 | 0 | 0 | 11 | 1295.6036 | 1718.9645 | 5664.8105 | 1903.2715 | 1274.4746 | 1505.8215 |
| Pdia4 | FHHTFSPEIAK | -329 | -339 | 0 | 0 | 11 | 1295.6036 | 1718.9645 | 5664.8105 | 1903.2715 | 1274.4746 | 1505.8215 |
| Pdia4 | IADEEDYATEVK | -445 | -456 | 0 | 0 | 12 | 1008.453 | 1041.4264 | 1072.4489 | 410.67685 | 296.03497 | 1872.0027 |
| Pdia4 | AMEPEEFDSDTLR | -480 | -492 | 0 | 0 | 13 | 751.40546 | 911.6307 | 932.6765 | 1129.2228 | 597.77734 | 1095.7924 |
| Pdia4 | KDNDPPIAVAK | -102 | -112 | 1 | 0.00025015 | 11 | 640.788 | 575.53235 | 435.00647 | 657.6677 | 673.65173 | 660.91956 |
| Pdia4 | FQGDGDPAYLQYQDAANNLR | -305 | -324 | 0 | 0.000394361 | 20 | 289.0216 | 452.46625 | 353.1851 | 484.4096 | 602.8442 | 594.10864 |
| Pdia4 | GQAVDYDGSR | -139 | -148 | 0 | 0 | 10 | 81.86082 | 1117.0706 | 1262.2238 | 1142.8348 | 985.4292 | 1104.8932 |
| Pdia4 | DLEHLSK | -616 | -622 | 0 | 0.004224523 | 7 | 2148.5286 | 2211.0667 | 2033.5181 | 2366.233 | NaN | 1779.7607 |
| Pdia4 | KLAPEYEK | -204 | -211 | 1 | 0 | 8 | 999.9713 | 975.6734 | NaN | 822.8504 | 1109.4685 | 1237.8096 |
| Pdia4 | LFQGDGDPAYLQYQDAANNLR | -304 | -324 | 0 | 0.000627943 | 21 | 200.44144 | 233.88638 | NaN | 336.01846 | 339.48618 | 360.42157 |
| Pdia4 | KGQAVDYDGSR | -138 | -148 | 1 | 0 | 11 | 857.6651 | 744.0453 | 816.5546 | 475.78122 | NaN | NaN |
| Pdia4 | SPPIPLAK | -220 | -227 | 0 | 0.000826806 | 8 | 797.19696 | 591.7329 | 607.571 | 374.27094 | NaN | NaN |
| Pdia4 | GLSESGEDVNAAILDESGK | -459 | -477 | 0 | 0 | 19 | 600.8622 | 74.32995 | NaN | 1068.4724 | 946.3088 | NaN |
| Pdia4 | FHHTFSPEIA | -329 | -338 | 0 | 0.00111006 | 10 | 313.43268 | 679.1085 | NaN | NaN | 690.38153 | 988.09827 |
| Pdia4 | VDATEQTDLAK | -228 | -238 | 0 | 0 | 11 | 4765.589 | NaN | NaN | NaN | 5736.519 | 6232.6543 |
| Pdia4 | QFAPEYEK | -89 | -96 | 0 | 0 | 8 | 3838.5054 | 4118.9077 | NaN | 2392.1245 | NaN | NaN |
| Pdia4 | ATEQTDLAK | -230 | -238 | 0 | 0.002780285 | 9 | 975.3295 | NaN | NaN | NaN | NaN | NaN |
| Pabpc1 | EFTNVYIK | -189 | -196 | 0 | 0.00051306 | 8 | 4398.834 | 595.9696 | 6801.338 | 5321.3784 | 5500.339 | 6706.91 |
| Pabpc1 | AMHPSLAGK | -572 | -580 | 0 | 0 | 9 | 1420.3511 | 1512.6934 | 1712.8022 | 1767.5875 | 1767.5161 | 1990.0195 |
| Pabpc1 | AMHPSLAGK | -572 | -580 | 0 | 0 | 9 | 1420.3511 | 1512.6934 | 1712.8022 | 1767.5875 | 1767.5161 | 1990.0195 |
| Pabpc1 | FSPAGPILSIR | -31 | -41 | 0 | 0 | 11 | 3620.849 | 4168.5635 | 4226.5435 | 3939.0332 | 4339.9697 | 342.81976 |
| Pabpc1 | SKVDEAVAVLQ | -605 | -615 | 1 | 0.000347792 | 11 | 3413.396 | 3463.0073 | 3493.3992 | 3833.3381 | 3816.9 | 3500.416 |
| Pabpc1 | NLDDGIDDER | -300 | -309 | 0 | 0 | 10 | 2254.1882 | 1939.2405 | 1748.9591 | 2928.6792 | 2353.2124 | 2416.2222 |
| Pabpc1 | KEFSPFGTIT | -312 | -321 | 1 | 0 | 10 | 2154.5823 | 2238.478 | 2316.4702 | 288.46277 | 1588.7761 | 2010.5305 |
| Pabpc1 | IVATKPLYVAL | -357 | -367 | 1 | 0.002566359 | 11 | 1923.1875 | 2358.9978 | 2195.6016 | 1989.3993 | 1757.159 | 2004.9166 |
| Pabpc1 | VNFQQPADAER | -57 | -67 | 0 | 0 | 11 | 1624.054 | 1714.0717 | 1803.0598 | 2160.2427 | 1764.9708 | 749.58057 |
| Pabpc1 | VHFETQEAAER | -143 | -153 | 0 | 0 | 11 | 1361.341 | 1367.1476 | 1470.9583 | 1295.4655 | 1939.515 | 1941.8356 |
| Pabpc1 | NPQQHLNAQPQVT | -519 | -531 | 0 | 0.001055699 | 13 | 1343.9576 | 1419.7867 | 1555.8274 | 1281.9766 | 1309.1565 | 1264.3958 |
| Pabpc1 | IVATKPLYVA | -357 | -366 | 1 | 0.00025015 | 10 | 1313.6736 | 1242.7405 | 1394.7955 | 1340.7554 | 1254.3337 | 1212.745 |
| Pabpc1 | FSSPEEATK | -340 | -348 | 0 | 0 | 9 | 1249.5767 | 1581.7062 | 936.5764 | 1059.4308 | 958.84686 | 1023.78564 |
| Pabpc1 | SPAGPILSIR | -32 | -41 | 0 | 0 | 10 | 1197.8995 | 1168.7095 | 608.6407 | 1507.5088 | 1402.5167 | 1378.5267 |
| Pabpc1 | FETQEAAER | -145 | -153 | 0 | 0 | 9 | 982.5516 | 1016.4413 | 1258.866 | 1265.4629 | 798.6656 | 1120.0289 |
| Pabpc1 | NPQQHLNAQPQ | -519 | -529 | 0 | 0.00111006 | 11 | 820.35046 | 915.2084 | 1027.1975 | 815.8789 | 335.3945 | 994.263 |
| Pabpc1 | AVPNPVINPYQPAPPSGY | -391 | -408 | 0 | 0.00025015 | 18 | 759.69885 | 670.75494 | 869.3605 | 917.50385 | 1261.364 | 956.87506 |
| Pabpc1 | EKFSPAGPILSIR | -29 | -41 | 1 | 0 | 13 | 288.46765 | 411.28378 | 470.64392 | 443.96802 | 441.67737 | 513.6638 |
| Pabpc1 | EKFSPAGPILSIR | -29 | -41 | 1 | 0 | 13 | 288.46765 | 411.28378 | 470.64392 | 443.96802 | 441.67737 | 513.6638 |
| Pabpc1 | VCFSSPEEATK | -338 | -348 | 0 | 0 | 11 | 636.45715 | 1211.5188 | 1345.7354 | 1725.7028 | 1616.3416 | 1565.628 |
| Pabpc1 | NFGEDMDDER | -197 | -206 | 0 | 0 | 10 | 634.4935 | 704.817 | 584.17096 | 545.4271 | 475.2321 | 452.1747 |
| Pabpc1 | FQQPADAER | -59 | -67 | 0 | 0 | 9 | 168.11835 | 1854.7695 | 1871.2545 | 2211.1968 | 1410.1394 | 1551.3608 |
| Pabpc1 | FGPALSVK | -214 | -221 | 0 | 0.002168954 | 8 | 1669.9146 | 1360.8663 | NaN | 879.18884 | 836.94666 | 1083.9977 |
| Pabpc1 | AVPNPVINPYQPAPPSGYFMA | -391 | -411 | 0 | 0 | 21 | 160.59174 | NaN | 310.1645 | 352.219 | 373.41074 | 383.90363 |
| Pabpc1 | SKVDEAVAVL | -605 | -614 | 1 | 0.002199523 | 10 | 2504.1704 | 665.5986 | 1728.3595 | NaN | NaN | NaN |
| Pabpc1 | YQGVNLYVK | -291 | -299 | 0 | 0 | 9 | 1586.6669 | 1313.099 | 1139.4027 | NaN | NaN | NaN |
| Satb1 | EYKDEELLK | -719 | -727 | 1 | 0.00025015 | 9 | 1966.1987 | 1811.5223 | 1609.3833 | 231.19493 | 2138.0234 | 236.1578 |
| Satb1 | LLNQQYAVNR | -326 | -335 | 0 | 0 | 10 | 1591.7325 | 1790.6235 | 2018.5259 | 1380.347 | 2384.2456 | 2063.7605 |
| Satb1 | WKEDPSPENR | -534 | -543 | 1 | 0 | 10 | 791.98615 | 463.96182 | 1356.6676 | 1878.5884 | 1654.8632 | 1480.4498 |
| Satb1 | AMQNFLQLPEAER | -428 | -440 | 0 | 0 | 13 | 597.03394 | 493.73 | 826.665 | 1376.553 | 1501.6075 | 1306.7317 |
| Satb1 | TPLPNLHPGLVSTPISPQ | -298 | -315 | 0 | 0 | 18 | 504.00473 | 684.57587 | 642.1547 | 769.9118 | 1137.804 | 1043.8087 |
| Satb1 | TPLPNLHPGLVSTPISPQLVNQ | -298 | -319 | 0 | 0.00122419 | 22 | 368.6402 | 272.98886 | 276.61682 | 355.30377 | 349.72275 | 339.55203 |
| Satb1 | QNFLQLPEAER | -430 | -440 | 0 | 0 | 11 | 357.9257 | 1425.1785 | 343.53278 | 1768.0408 | 1518.6669 | 792.1238 |
| Satb1 | MDHLNEATQGK | -1 | -11 | 0 | 0 | 11 | 306.0582 | 3823.6963 | 2319.3643 | 373.0917 | 3425.7563 | 1177.8799 |
| Satb1 | TPLPNLHPGLVSTPISPQLVNQQ | -298 | -320 | 0 | 0.000431655 | 23 | 261.7801 | 307.26538 | 256.4211 | 323.64032 | 360.07614 | 373.35678 |
| Satb1 | YLNHPPPVSR | -346 | -355 | 0 | 0 | 10 | 5059.061 | 4932.3643 | 623.8308 | NaN | 3751.1257 | 3711.0203 |
| Satb1 | FLSLPQPER | -555 | -563 | 0 | 0 | 9 | 2270.8694 | 2634.2288 | NaN | 2639.2903 | 1635.3392 | 1574.3254 |
| Satb1 | GPAPLLSTPPSRPPQVK | -459 | -475 | 1 | 0.00025015 | 17 | 200.95528 | 1609.4266 | 1547.6355 | NaN | 2055.2615 | 1970.9906 |
| Satb1 | GPAPLLSTPPSRPPQVK | -459 | -475 | 1 | 0.00025015 | 17 | 200.95528 | 1609.4266 | 1547.6355 | NaN | 2055.2615 | 1970.9906 |

| | | | | | | | | | | | | |
|---|---|---|---|---|---|---|---|---|---|---|---|---|
| Satb1 | EHYENAIEYDCK | -81 | -92 | 0 | 0 | 12 | 355.80322 | 365.15012 | 342.11 | 294.30963 | 223.26709 | NaN |
| Satb1 | FLQLPEAER | -432 | -440 | 0 | 0 | 9 | 1015.4204 | 918.6125 | 1044.137 | 5125.7812 | NaN | NaN |
| Satb1 | NANTLFSVK | -737 | -745 | 0 | 0 | 9 | 897.6971 | NaN | NaN | 1217.0715 | 772.45734 | 644.77576 |
| Satb1 | EYKDEELLKDLEESVQDK | -719 | -736 | 2 | 0.00025015 | 18 | 677.1307 | NaN | NaN | 504.18713 | 770.9296 | 613.64905 |
| Satb1 | EYKDEELLKDLEESVQDK | -719 | -736 | 2 | 0.00025015 | 18 | 677.1307 | NaN | NaN | 504.18713 | 770.9296 | 613.64905 |
| Satb1 | IQLHSCPK | -168 | -175 | 0 | 0 | 8 | 656.46954 | 685.327 | 770.7607 | NaN | NaN | NaN |
| Satb1 | LKDNSGLEVD | -707 | -716 | 1 | 0.003966045 | 10 | 1419.2461 | NaN | NaN | NaN | NaN | NaN |
| Psmd4 | LANDCEVLTTLTPDTGR | -54 | -70 | 0 | 0 | 17 | 1745.7471 | 2123.7642 | 2238.0615 | 2341.1218 | 3066.2334 | 2916.23 |
| Psmd4 | EAGIATPGTEDSDDALLK | -245 | -262 | 0 | 0.00025015 | 18 | 890.2969 | 129.38484 | 184.1435 | 219.40338 | 547.5345 | 955.7323 |
| Psmd4 | SVLENLPGVDPNNAAIR | -336 | -352 | 0 | 0.000471409 | 17 | 627.6484 | 681.56476 | 910.654 | 519.2673 | 699.5602 | 736.0534 |
| Psmd4 | VGSPVEDNEKDLVK | -113 | -126 | 1 | 0 | 14 | 592.3174 | 935.6489 | 1004.5348 | 873.0836 | 866.77454 | 888.9617 |
| Psmd4 | IINFGEEEVNTEK | -140 | -152 | 0 | 0 | 13 | 1070.1771 | 1514.2163 | NaN | NaN | 1332.6079 | 678.28516 |
| Psmd4 | NDCEVLTTLTPDTGR | -56 | -70 | 0 | 0.000591124 | 15 | 1508.2573 | NaN | 1451.1433 | NaN | NaN | 1649.6924 |
| Psmd4 | ANDCEVLTTLTPDTGR | -55 | -70 | 0 | 0.000550307 | 16 | 652.231 | NaN | 421.17715 | NaN | 637.77844 | NaN |
| Psmd4 | SAAEAGIATPGTEDSDDALLK | -242 | -262 | 0 | 0.000230707 | 21 | 522.7227 | 522.6161 | 586.1381 | NaN | NaN | NaN |
| Psmd4 | ENLPGVDPNNAAIR | -339 | -352 | 0 | 0.002199523 | 14 | 776.4501 | NaN | NaN | NaN | 1316.366 | NaN |
| Trir | ANDGSFLELFK | -56 | -66 | 0 | 0 | 11 | 199.09052 | 842.3784 | 819.42334 | 838.4856 | 245.48537 | 173.23311 |
| Trir | QKTEDEVLTSK | -133 | -143 | 1 | 0 | 11 | 618.57446 | 610.63574 | NaN | 505.20233 | 4685.429 | 4476.3286 |
| Trir | QEEPPPGPQRPDPPASAA | -77 | -94 | 1 | 0.002566359 | 18 | 1770.7533 | NaN | NaN | 1775.8312 | 2216.3303 | 2195.7588 |
| Abcf1 | SFNLPYQDAR | -706 | -715 | 0 | 0 | 10 | 1622.6075 | 1606.2473 | 280.50467 | 1240.7517 | 1427.6742 | 282.7332 |
| Abcf1 | LQGQLEQGDDTAAEK | -389 | -403 | 0 | 0 | 15 | 1026.6462 | 1032.7017 | 1182.4791 | 447.46375 | 657.46747 | 599.6383 |
| Abcf1 | NQDEESQEPPELLK | -582 | -595 | 0 | 0 | 14 | 909.2724 | 861.47424 | 538.8153 | 1227.9844 | 863.4928 | 807.891 |
| Abcf1 | GVSQIDGFDDYKR | -808 | -821 | 1 | 0 | 14 | 636.30524 | 584.78955 | 655.774 | 619.8411 | 755.8632 | 471.8648 |
| Abcf1 | IQDDSEEEEEEENR | -134 | -148 | 0 | 0.000230707 | 15 | 271.93747 | 274.77835 | 257.94983 | 357.26306 | 324.7707 | 326.81024 |
| Abcf1 | GYEGQKPLFK | -625 | -634 | 1 | 0 | 10 | 4580.3486 | NaN | 4651.6504 | 3438.9087 | 3961.3796 | 4012.4224 |
| Abcf1 | RLQGQLEQGDDTAAEK | -388 | -403 | 1 | 0.004224523 | 16 | 6584.525 | 6942.825 | 5917.199 | NaN | NaN | 3739.1567 |
| Abcf1 | GAVIVVSHDAR | -782 | -792 | 0 | 0 | 11 | 1341.8728 | 1666.9707 | 1660.8799 | 1549.4086 | NaN | NaN |
| Abcf1 | IDGDFDDYKR | -812 | -821 | 1 | 0.000230707 | 10 | 579.42676 | NaN | NaN | 423.60275 | 1076.0049 | 1297.2399 |
| Abcf1 | SVPASDEEDEVPAPIPR | -103 | -119 | 0 | 0 | 17 | 911.787 | NaN | 992.0967 | NaN | 1090.0049 | NaN |
| Abcf1 | GFDPEMQNRPTQK | -431 | -443 | 1 | 0.009122264 | 13 | 875.9283 | NaN | NaN | 548.9731 | 487.70233 | NaN |
| Abcf1 | ANDPYANLSK | -238 | -247 | 0 | 0 | 10 | 1884.6724 | NaN | 833.6225 | NaN | NaN | NaN |
| Abcf1 | LNNYLQGWR | -482 | -490 | 0 | 0.00122419 | 9 | 643.1913 | NaN | 199.78145 | NaN | NaN | NaN |
| Pdia3 | SEPIPESNEGPVK | -367 | -379 | 0 | 0 | 13 | 5076.3154 | 5067.7246 | 4684.5664 | 4209.954 | 4124.298 | 4274.68 |
| Pdia3 | SEPIPESNEGPVK | -367 | -379 | 0 | 0 | 13 | 5076.3154 | 5067.7246 | 4684.5664 | 4209.954 | 4124.298 | 4274.68 |
| Pdia3 | SDVLELTDENFESR | -25 | -38 | 0 | 0 | 14 | 4811.119 | 5871.2627 | 6012.952 | 10264.759 | 6977.0967 | 6899.2437 |
| Pdia3 | TADGIVSHLK | -120 | -129 | 0 | 0 | 10 | 4629.4893 | 4703.8457 | 4520.41 | 4520.2944 | 4252.8696 | 4237.1953 |
| Pdia3 | STTGEVPVVAIR | -318 | -329 | 0 | 0 | 12 | 3379.8186 | 3387.8542 | 3473.2825 | 1983.3365 | 1498.6725 | 1785.3518 |
| Pdia3 | RDGEEAGAYDGPR | -107 | -119 | 1 | 0 | 13 | 3002.9863 | 2978.3293 | 3172.8772 | 3165.0142 | 936.9154 | 1515.192 |
| Pdia3 | RDGEEAGAYDGPR | -107 | -119 | 1 | 0 | 13 | 3002.9863 | 2978.3293 | 3172.8772 | 3165.0142 | 936.9154 | 1515.192 |
| Pdia3 | FLDAGHKL | -290 | -297 | 1 | 0.000230707 | 8 | 2720.3726 | 2615.4937 | 2303.8518 | 2392.235 | 2186.0627 | 2310.0369 |
| Pdia3 | LAPEYEAAATR | -63 | -73 | 0 | 0 | 11 | 2315.7307 | 2418.0474 | 817.4793 | 540.8469 | 811.03705 | 973.1017 |
| Pdia3 | YGVSGYPTLK | -95 | -104 | 0 | 0 | 10 | 2252.441 | 2661.6636 | 2848.4287 | 2082.6274 | 2531.8672 | 2618.4006 |
| Pdia3 | FVMQEEFSR | -336 | -344 | 0 | 0 | 9 | 655.76135 | 72.63696 | 704.73773 | 195.63168 | 86.3069 | 379.45868 |
| Pdia3 | FVMQEEFSR | -336 | -344 | 0 | 0 | 9 | 655.76135 | 72.63696 | 704.73773 | 195.63168 | 86.3069 | 379.45868 |
| Pdia3 | EYDDNGEGITIFRPL | -195 | -209 | 1 | 0 | 15 | 2008.895 | 2393.8813 | 2714.9402 | 1869.9824 | 101.67154 | 2843.5752 |
| Pdia3 | DASVVGFFR | -153 | -161 | 0 | 0 | 9 | 1970.2766 | 2240.482 | 2178.0557 | 353.65997 | 2843.4094 | 297.2572 |
| Pdia3 | GLESTTGEVPVVAIR | -315 | -329 | 0 | 0 | 15 | 1678.6221 | 1060.5856 | 1456.4581 | 1980.8036 | 1073.9446 | 1142.4717 |
| Pdia3 | DGEEAGAYDGPR | -108 | -119 | 0 | 0 | 12 | 1661.2931 | 1408.9196 | 1072.2163 | 1626.2665 | 793.57733 | 1059.6444 |
| Pdia3 | TFSHELSDFGLE | -306 | -317 | 0 | 0 | 12 | 1353.0142 | 1314.2119 | 1380.9388 | 155.95279 | 171.19556 | 1045.0636 |
| Pdia3 | EATNPPIIQEEKPK | -483 | -496 | 1 | 0 | 14 | 1214.3518 | 1318.3177 | 1459.4526 | 1137.4668 | 1206.7003 | 1154.5756 |
| Pdia3 | FISDKDASVV | -148 | -157 | 1 | 0.00025015 | 10 | 991.01263 | 3860.9226 | 4396.123 | 951.9588 | 3789.115 | 4162.5527 |
| Pdia3 | FISDKDASVVGF | -148 | -159 | 1 | 0 | 12 | 932.0864 | 985.6924 | 907.21136 | 1147.7429 | 1297.4255 | 1168.7765 |
| Pdia3 | TFSHELSDF | -306 | -314 | 0 | 0 | 9 | 899.0924 | 973.4653 | 1024.945 | 778.52966 | 756.033 | 777.1645 |
| Pdia3 | FAHTNIESL | -184 | -192 | 0 | 0.002434616 | 9 | 889.7736 | 471.75342 | 779.4088 | 1879.7983 | 1085.6632 | 582.3384 |
| Pdia3 | DATANDVPSPYEVK | -435 | -448 | 0 | 0 | 14 | 869.4182 | 1192.217 | 447.54398 | 954.09485 | 522.9002 | 1150.6998 |
| Pdia3 | FAHTNIESLVK | -184 | -194 | 0 | 0.00025015 | 11 | 734.87427 | 724.3729 | 3225.1155 | 2303.6084 | 3456.7163 | 3404.576 |
| Pdia3 | FAHTNIESLVK | -184 | -194 | 0 | 0.00025015 | 11 | 734.87427 | 724.3729 | 3225.1155 | 2303.6084 | 3456.7163 | 3404.576 |
| Pdia3 | SVPLRTEEEFK | -136 | -146 | 1 | 0 | 11 | 682.1299 | 671.42975 | 686.90607 | 707.5347 | 832.9028 | 979.7368 |
| Pdia3 | YDVDYEK | -265 | -271 | 0 | 0 | 7 | 636.0799 | 613.4906 | 625.65027 | 602.86487 | 566.9174 | 541.0969 |
| Pdia3 | LVEFFAPWCGHCK | -49 | -61 | 0 | 0 | 13 | 589.61053 | 910.6175 | 969.97455 | 1166.2029 | 1097.9584 | 1253.5089 |
| Pdia3 | FISDKDASVVGFFR | -148 | -161 | 1 | 0 | 14 | 584.01135 | 509.04028 | 407.76666 | 477.88055 | 594.9037 | 529.0777 |
| Pdia3 | DLFSDGHSEFLK | -162 | -173 | 0 | 0 | 12 | 497.0438 | 6093.019 | 6006.2744 | 1119.376 | 888.3793 | 7909.7134 |
| Pdia3 | DLFSDGHSEFLK | -162 | -173 | 0 | 0 | 12 | 497.0438 | 6093.019 | 6006.2744 | 1119.376 | 888.3793 | 7909.7134 |
| Pdia3 | DLFSDGHSEFL | -162 | -172 | 0 | 0 | 11 | 495.2009 | 499.78232 | 551.183 | 483.01703 | 587.8251 | 567.5715 |
| Pdia3 | EQFLQEYFDGNLK | -350 | -362 | 0 | 0 | 13 | 473.813 | 497.45337 | 589.36676 | 670.36127 | 434.39832 | 877.4598 |
| Pdia3 | VDCTANTNTCNK | -83 | -94 | 0 | 0 | 12 | 409.03757 | 428.01257 | 522.2091 | 376.02338 | 460.49127 | 451.9633 |
| Pdia3 | GFPTIYFSPANK | -449 | -460 | 0 | 0 | 12 | 395.6117 | 689.0247 | 413.29123 | 831.2376 | 790.05865 | 441.34418 |
| Pdia3 | SDFGLESTTGEVPVVAIR | -312 | -329 | 0 | 0 | 18 | 324.11108 | 336.81744 | 473.60715 | 634.0909 | 439.74954 | 324.9488 |
| Pdia3 | LSDFGLESTTGEVPVVAIR | -311 | -329 | 0 | 0 | 19 | 239.46275 | 280.69434 | 371.72696 | 346.61267 | 595.6674 | 526.8037 |
| Pdia3 | LSKDPNIVIAK | -423 | -433 | 1 | 0.000431655 | 11 | 3033.1882 | 4040.2432 | 3756.4775 | NaN | 5240.898 | 5539.018 |
| Pdia3 | EDNKDLIQGK | -249 | -258 | 1 | 0 | 10 | 2460.7515 | NaN | 2644.4893 | 2013.7687 | 2213.5505 | 2184.0989 |
| Pdia3 | MTEDNKDLIQGK | -247 | -258 | 1 | 0 | 12 | 1339.1749 | 1505.9489 | NaN | 1090.956 | 1452.0126 | 1564.1029 |
| Pdia3 | TEEEFKK | -141 | -147 | 1 | 0 | 7 | 1063.8053 | 828.65564 | 957.61914 | 983.17847 | NaN | 924.93054 |
| Pdia3 | SVPLRTEEEFKK | -136 | -147 | 2 | 0.002466863 | 12 | 772.8771 | 670.9064 | 712.31964 | NaN | 545.63727 | 748.7573 |
| Pdia3 | HLANKFEDK | -210 | -218 | 1 | 0.001092529 | 9 | 559.6676 | 372.46747 | 638.3676 | NaN | 4354.0894 | 742.25616 |
| Pdia3 | TTGEVPVVAIR | -319 | -329 | 0 | 0 | 11 | 518.77856 | 450.42123 | 482.54248 | 232.14133 | NaN | 463.35187 |
| Pdia3 | TFSHELSDFGLES | -306 | -318 | 0 | 0.000230707 | 13 | 940.64984 | 1016.2532 | 856.2556 | NaN | 430.90204 | NaN |
| Pdia3 | EYFDGNLKR | -355 | -363 | 1 | 0 | 9 | 590.71313 | 1797.6298 | 2072.872 | 2166.9678 | NaN | NaN |

| | | | | | | | | | | | | |
|---|---|---|---|---|---|---|---|---|---|---|---|---|
| Pdia3 | PSPYEVK | -442 | -448 | 0 | 0.001392869 | 7 | 419.87976 | NaN | NaN | 789.7648 | 982.4853 | 933.1304 |
| Pdia3 | ANDVPSPYEVK | -438 | -448 | 0 | 0 | 11 | 398.77966 | 600.07263 | NaN | 417.7189 | NaN | 876.4048 |
| Pdia3 | DPNIVIAK | -426 | -433 | 0 | 0.001023236 | 8 | 202.6211 | NaN | NaN | 343.36115 | 226.35925 | 285.6415 |
| Pdia3 | FLDAGHK | -290 | -296 | 0 | 0.003620005 | 7 | 1968.1642 | 936.13617 | NaN | NaN | NaN | 636.50726 |
| Pdia3 | EYFDGNLK | -355 | -362 | 0 | 0.00025015 | 8 | 557.15875 | 315.14743 | 450.5808 | NaN | NaN | NaN |
| Pdia3 | KFLDAGHK | -289 | -296 | 1 | 0.00025015 | 8 | 1533.8584 | NaN | NaN | NaN | NaN | 3120.3887 |
| Pdia3 | ATANDVPSPYEVK | -436 | -448 | 0 | 0.00025015 | 13 | 1202.8844 | NaN | 1124.9607 | NaN | NaN | NaN |
| Pdia3 | DVPSPYEVK | -440 | -448 | 0 | 0.004224523 | 9 | 992.0171 | 736.3591 | NaN | NaN | NaN | NaN |
| Pdia3 | EYDDNGEGITIFR | -195 | -207 | 0 | 0 | 13 | 313.53537 | NaN | NaN | NaN | NaN | NaN |
| Pdap1 | KGVEGLIDIENPNR | -75 | -88 | 1 | 0 | 14 | 3501.7937 | 1291.0719 | 3145.414 | 2504.4758 | 3151.1157 | 2773.9304 |
| Pdap1 | KGVEGLIDIENPNR | -75 | -88 | 1 | 0 | 14 | 3501.7937 | 1291.0719 | 3145.414 | 2504.4758 | 3151.1157 | 2773.9304 |
| Pdap1 | ANEEDEQEEGGDGASGDPK | -34 | -52 | 0 | 0 | 19 | 292.01288 | 339.03308 | 452.87134 | 209.5248 | 223.80577 | 371.0849 |
| Pdap1 | KVTQLDLDGPK | -95 | -105 | 1 | 0 | 11 | 607.4155 | 650.81915 | 711.7806 | NaN | NaN | 777.0761 |
| Pdap1 | SLDSDESEDEDDDYQQK | -57 | -73 | 0 | 0 | 17 | 427.64758 | 475.24164 | 454.3752 | 393.9411 | NaN | NaN |
| Pdap1 | GVEGLIDIENPNR | -76 | -88 | 0 | 0 | 13 | 1955.0815 | NaN | NaN | NaN | 2903.0596 | NaN |
| Pdap1 | QYTSPEEIDAQLQAEK | -16 | -31 | 0 | 0.00830926 | 16 | 121.00668 | NaN | NaN | NaN | NaN | NaN |
| Prdx3 | NWTPESPTIKPSPTASK | -233 | -249 | 1 | 0 | 17 | 2457.6243 | 2771.6604 | 2996.2593 | 2793.1228 | 3223.4675 | 3277.2107 |
| Prdx3 | NWTPESPTIKPSPTASK | -233 | -249 | 1 | 0 | 17 | 2457.6243 | 2771.6604 | 2996.2593 | 2793.1228 | 3223.4675 | 3277.2107 |
| Prdx3 | HLSVNDLPVGR | -198 | -208 | 0 | 0 | 11 | 2230.9792 | 2303.2405 | 2413.8315 | 1884.1726 | 1412.3485 | 1948.0623 |
| Prdx3 | ANEFHDVNCEVVAV | -120 | -133 | 0 | 0 | 14 | 1193.5463 | 1501.1047 | 1460.4674 | 1570.6351 | 1711.2815 | 1606.9681 |
| Prdx3 | GLFIIDPNGVVK | -186 | -197 | 0 | 0 | 12 | 1087.9779 | 1190.4117 | 1167.6786 | 1126.718 | 1273.6106 | 734.91254 |
| Prdx3 | LESAGIALR | -177 | -185 | 0 | 0 | 9 | 854.1412 | 789.4071 | 645.0136 | 565.6672 | 460.63116 | 540.9007 |
| Prdx3 | SVNDLPVGR | -200 | -208 | 0 | 0.000471409 | 9 | 1378.8816 | 1220.2688 | NaN | NaN | NaN | 987.37494 |
| Ptbp3 | TDNLPNQAR | -109 | -117 | 0 | 0 | 9 | 5704.633 | 6031.09 | 6100.7144 | 5113.1953 | 4875.084 | 5494.7163 |
| Ptbp3 | DFSNSPLHR | -421 | -429 | 0 | 0 | 9 | 2856.615 | 2766.17 | 1835.5056 | 1831.8381 | 2128.2922 | 1732.2577 |
| Ptbp3 | HQAVQLPR | -403 | -410 | 0 | 0 | 8 | 1299.4757 | 1150.3398 | 1037.0216 | 983.178 | 837.0134 | 864.3599 |
| Ptbp3 | ANEGTLLPGQSPVLR | -140 | -154 | 0 | 0 | 15 | 1134.9926 | 1260.8438 | 1363.2927 | 1302.015 | 1394.3593 | 1379.9922 |
| Ptbp3 | EGTLLPGQSPVLR | -142 | -154 | 0 | 0.002845514 | 13 | 888.1383 | NaN | NaN | NaN | NaN | NaN |
| Ptbp3 | NLFTEAGCSVK | -464 | -474 | 0 | 0 | 11 | 182.66045 | NaN | NaN | NaN | NaN | NaN |
| Psat1 | IIGNTENLVR | -52 | -61 | 0 | 0 | 10 | 4182.0283 | 3539.4248 | 3859.5833 | 3561.081 | 3015.1436 | 3111.7441 |
| Psat1 | FLSRPVDVSK | -181 | -190 | 1 | 0.001904399 | 10 | 2388.603 | 2376.5784 | 2278.254 | 270.66174 | 2249.7666 | 2372.1558 |
| Psat1 | NETVHGVEFDFVPDVK | -154 | -169 | 0 | 0 | 16 | 1966.3594 | 666.2547 | 1355.7474 | 451.38138 | 520.68585 | 1425.8334 |
| Psat1 | VVNFGPGPAK | -7 | -16 | 0 | 0 | 10 | 1258.0786 | 746.42065 | 688.93396 | 604.4151 | 952.1352 | 773.21436 |
| Psat1 | AVPLNLIGLK | -85 | -94 | 0 | 0 | 10 | 1242.8367 | 1410.2117 | 1284.9575 | 597.6608 | 628.469 | 733.1098 |
| Psat1 | ANETVHGVEFDFVPDVK | -153 | -169 | 0 | 0 | 17 | 896.1047 | 993.04 | 1061.6823 | 1007.261 | 135.59229 | 1094.9207 |
| Psat1 | ANETVHGVEFDFVPDVK | -153 | -169 | 0 | 0 | 17 | 896.1047 | 993.04 | 1061.6823 | 1007.261 | 135.59229 | 1094.9207 |
| Psat1 | GTVNIVHPK | -119 | -127 | 0 | 0 | 9 | 885.2366 | 658.10175 | 675.13293 | 828.11755 | 1133.6421 | 857.91077 |
| Psat1 | KQLLDYR | -27 | -33 | 1 | 0 | 7 | 830.29224 | 742.3228 | 861.1954 | 1082.8374 | 1098.1293 | 1019.0823 |
| Psat1 | QVVNFGPGPAK | -6 | -16 | 0 | 0 | 11 | 823.0161 | 758.4895 | 917.7147 | 928.09106 | 802.5677 | 602.9149 |
| Psat1 | ETVHGVEFDFVPDVK | -155 | -169 | 0 | 0 | 15 | 700.2827 | 654.2597 | 1075.4333 | 993.3171 | 943.4452 | 853.82684 |
| Psat1 | RECPSVLDYK | -222 | -231 | 1 | 0.00025015 | 10 | 641.83734 | 913.75116 | 1070.9104 | 774.6616 | 1178.312 | 1078.7175 |
| Psat1 | LPHSVLLEIQK | -17 | -27 | 0 | 0 | 11 | 548.9295 | 625.8822 | 592.7907 | 586.81836 | 746.48254 | 719.9183 |
| Psat1 | LPHSVLLEIQK | -17 | -27 | 0 | 0 | 11 | 548.9295 | 625.8822 | 592.7907 | 586.81836 | 746.48254 | 719.9183 |
| Psat1 | ECPSVLDYK | -223 | -231 | 0 | 0 | 9 | 320.30713 | 2445.6672 | 2201.67 | 2964.225 | 2581.2046 | 2736.0137 |
| Psat1 | NFLEMHQL | -363 | -370 | 0 | 0 | 8 | 2142.3347 | NaN | 2993.8872 | 2449.5798 | 2615.0742 | 2364.983 |
| Psat1 | EIIDNSQGFYVCPVER | -280 | -295 | 0 | 0 | 16 | 696.5985 | 762.5801 | 705.1018 | 599.8828 | 409.15262 | NaN |
| Psat1 | GFYVCPVER | -287 | -295 | 0 | 0.001463897 | 9 | 849.50745 | 788.67236 | 872.66943 | NaN | NaN | NaN |
| Psat1 | YVCPVER | -289 | -295 | 0 | 0.000627943 | 7 | 710.3819 | NaN | NaN | 741.32227 | NaN | 692.67615 |
| Psat1 | FSAVPLNLIGLK | -83 | -94 | 0 | 0.001291103 | 12 | 446.30682 | NaN | 435.6823 | NaN | NaN | NaN |
| Lactb2 | ANIIYPGHGPVIHNA | -192 | -206 | 0 | 0.00078661 | 15 | 484.62677 | NaN | NaN | NaN | 895.465 | NaN |
| Sae1 | TFANLGEHEFVEEK | -169 | -182 | 0 | 0 | 14 | 2463.0815 | 160.4642 | 3714.673 | 1528.8304 | 2905.271 | 2646.6453 |
| Sae1 | GISPDLLPDDFVR | -285 | -297 | 0 | 0 | 13 | 968.307 | 873.1198 | 1019.22205 | 990.10046 | 1143.1997 | 1132.3218 |
| Sae1 | EEAGGGGGGGISEEEAAQYDR | -5 | -25 | 0 | 0.001392869 | 21 | 530.33386 | 597.973 | 626.57153 | 640.819 | 723.43805 | 745.55444 |
| Sae1 | KVSQGVEDGPEAK | -187 | -199 | 1 | 0.00025015 | 13 | 465.50403 | 307.20435 | 586.12085 | 218.2811 | 694.83215 | 486.42398 |
| Sae1 | VEKEEAGGGGGGGISEEEAAQYDR | -2 | -25 | 1 | 0 | 24 | 386.2228 | 464.11667 | 819.47925 | 595.80084 | 625.1504 | 554.13873 |
| Sae1 | KPESFFTK | -122 | -129 | 1 | 0 | 8 | 373.46912 | 1715.5051 | 1707.9307 | 2063.281 | 1610.939 | 1878.5425 |
| Sae1 | ANLGEHEFVEEK | -171 | -182 | 0 | 0 | 12 | 336.10712 | 523.9104 | 7378.586 | 5721.0396 | 1015.3095 | 11849.332 |
| Sae1 | VDQICHR | -146 | -152 | 0 | 0 | 7 | 396.90643 | 492.69855 | 510.63992 | 575.94507 | 525.0598 | NaN |
| Sae1 | DPPHNNFFFFD | -326 | -336 | 0 | 0.001868731 | 11 | 344.20944 | NaN | 369.7566 | NaN | NaN | NaN |
| Ttn | ANLIVEEEDLR | -14514 | -14524 | 0 | 0.00025015 | 11 | 2830.2444 | 662.33044 | 145.36151 | 897.5779 | 3636.3936 | 3514.198 |
| Septin2 | GVVEVENPEHNDFLK | -261 | -275 | 0 | 0 | 15 | 1425.6722 | 887.44116 | 2127.2097 | 1056.2676 | 1502.7859 | 1480.5432 |
| Septin2 | SLFLTDLYPER | -56 | -66 | 0 | 0 | 11 | 1156.9081 | 677.70667 | 500.2128 | 926.8731 | 781.5577 | 499.95325 |
| Septin2 | YLHDESGLNR | -129 | -138 | 0 | 0 | 10 | 889.62744 | 822.917 | 738.2668 | 722.33575 | 806.9116 | 1261.4152 |
| Septin2 | TIISYIDEQFER | -117 | -128 | 0 | 0 | 12 | 742.2175 | 1394.9517 | 1397.4385 | 1201.8992 | 1470.2697 | 297.9077 |
| Septin2 | VVDTPGYGDAINCR | -99 | -112 | 0 | 0 | 14 | 731.035 | 583.0977 | 739.4148 | 869.7683 | 929.72034 | 778.7307 |
| Septin2 | LTVVDTPGYGDAINCR | -97 | -112 | 0 | 0 | 16 | 690.42773 | 781.3674 | 745.2632 | 727.69135 | 981.2587 | 643.87463 |
| Septin2 | ILDEIEEHSIK | -199 | -209 | 0 | 0 | 11 | 2940.4429 | 3256.7993 | NaN | 3064.027 | NaN | NaN |
| Septin2 | ANLPNQVHR | -21 | -29 | 0 | 0 | 9 | 679.49036 | 267.92017 | 685.4999 | NaN | NaN | NaN |
| Septin7 | LYSPEYPGPSHR | -74 | -85 | 0 | 0 | 12 | 1981.0029 | 1673.3593 | 1640.6473 | 1242.0796 | 1074.0796 | 1061.4186 |
| Septin7 | DTLTPEECQQFK | -196 | -207 | 0 | 0 | 12 | 1814.8339 | 4957.806 | 730.4095 | 2823.254 | 725.74817 | 715.5288 |
| Septin7 | TLTPEECQQFK | -197 | -207 | 0 | 0 | 11 | 1000.642 | 953.41034 | 964.73035 | 804.76904 | 907.0486 | 1304.3193 |
| Septin7 | FLTDLYSPEYPGPSHR | -70 | -85 | 0 | 0 | 16 | 776.41095 | 960.2842 | 1011.0056 | 1072.5736 | 1403.637 | 1492.3098 |
| Septin7 | FLTDLYSPEYPGPSHR | -70 | -85 | 0 | 0 | 16 | 776.41095 | 960.2842 | 1011.0056 | 1072.5736 | 1403.637 | 1492.3098 |
| Septin7 | DVTNNVHYENYR | -298 | -309 | 0 | 0 | 12 | 666.3805 | 713.28613 | 648.4035 | 894.44977 | 892.4813 | 915.0945 |
| Septin7 | KADTLTPEECQQFK | -194 | -207 | 1 | 0.000550307 | 14 | 581.72015 | 727.2779 | 672.2613 | 721.21136 | 748.59204 | 764.8866 |
| Septin7 | FEDYLNAESR | -137 | -146 | 0 | 0 | 10 | 553.15283 | 693.12695 | 595.793 | 745.0674 | 753.1191 | 800.63855 |
| Septin7 | SLFLTDLYSPEYPGPSHR | -68 | -85 | 0 | 0 | 18 | 295.8061 | 308.70346 | 345.30188 | 608.3017 | 575.3719 | 478.2948 |
| Septin7 | DRLPLAVVGSN | -241 | -251 | 1 | 0.001023236 | 11 | 937.3416 | 981.4228 | 934.769 | 893.1434 | NaN | 989.6323 |

| | | | | | | | | | | | | |
|---|---|---|---|---|---|---|---|---|---|---|---|---|
| Septin7 | DLYSPEYPGPSHR | -73 | -85 | 0 | 0 | 13 | 601.1134 | 1054.6825 | 531.996 | NaN | 1114.5277 | 727.5657 |
| Septin7 | ANLPNQVYR | -33 | -41 | 0 | 0 | 9 | 283.4705 | 470.7144 | 681.598 | 1141.7256 | 828.4871 | NaN |
| Septin7 | EFPETDDEEENKLVK | -223 | -237 | 1 | 0 | 15 | 187.24385 | 961.4854 | NaN | 1194.2134 | 1142.3042 | 984.8766 |
| Septin7 | YSPEYPGPSHR | -75 | -85 | 0 | 0 | 11 | 17517.713 | 17928.559 | 17869.607 | NaN | NaN | 21741.814 |
| Septin7 | LKDSEAELQR | -371 | -380 | 1 | 0.000627943 | 10 | 545.6123 | NaN | 809.5602 | 890.93445 | NaN | NaN |
| Septin7 | NLPNQVYR | -34 | -41 | 0 | 0.00025015 | 8 | 381.6366 | 636.13727 | 572.96185 | NaN | NaN | NaN |
| Cd247 | REEYDVLEK | -80 | -88 | 1 | 0.000907206 | 9 | 1564.577 | 1260.4458 | 333.7037 | 1795.6847 | 1750.2057 | 1761.4154 |
| Cd247 | ANLQDPNQLYNELNLGR | -63 | -79 | 0 | 0.000347792 | 17 | 106.84416 | 1114.8026 | 928.726 | 927.8007 | 1132.6475 | 1189.7751 |
| Nrdc | NFDLNPDLHLPAENK | -658 | -672 | 0 | 0 | 15 | 1733.1603 | 1675.5615 | 1858.2646 | 317.9475 | 2619.7559 | 2469.1008 |
| Nrdc | LNFAPLER | -892 | -899 | 0 | 0 | 8 | 1128.6151 | 1092.8345 | 1171.3922 | 1160.7235 | 1195.2805 | 189.5714 |
| Nrdc | LVAGEHGLIIR | -763 | -773 | 0 | 0 | 11 | 624.9894 | 692.8411 | 741.88715 | 745.7171 | 741.8272 | 724.7794 |
| Nrdc | FHLISPLIQK | -719 | -728 | 0 | 0 | 10 | 463.68634 | 383.3121 | 473.0119 | 523.24927 | 419.1593 | 439.62408 |
| Nrdc | SNFDLNPDLHLPAENK | -657 | -672 | 0 | 0 | 16 | 555.58954 | 1063.6038 | 951.5 | 345.35577 | NaN | NaN |
| Nrdc | FHLISPLIQ | -719 | -727 | 0 | 0.001392869 | 9 | 190.67946 | 710.8541 | NaN | NaN | 966.77313 | 291.58374 |
| Nrdc | HTVFQYLK | -534 | -541 | 0 | 0.001055699 | 8 | 1055.9048 | 841.56537 | NaN | NaN | NaN | 946.98376 |
| Nrdc | ANLVLLSGANEGR | -618 | -630 | 0 | 0.005270854 | 13 | 3031.9587 | NaN | NaN | NaN | NaN | NaN |
| Nrdc | LAHEIEALK | -1064 | -1072 | 0 | 0.001904399 | 9 | 1367.0146 | NaN | NaN | NaN | NaN | NaN |
| Ncbp1 | ANNYNEAVYLVR | -118 | -129 | 0 | 0 | 12 | 1294.8007 | 1378.3022 | 1554.9381 | 1728.4272 | 206.71211 | 557.0542 |
| Ncbp1 | NYNEAVYLVR | -120 | -129 | 0 | 0 | 10 | 707.9658 | 624.6377 | 737.64453 | 608.589 | 670.5153 | 482.6699 |
| Ncbp1 | KDVPNPNQVDDDDEGFR | -522 | -538 | 1 | 0.00078661 | 17 | 692.1304 | 723.7292 | 706.7334 | 609.10406 | 391.91263 | 1048.5992 |
| Ncbp1 | CETDGTSILTPWYK | -728 | -741 | 0 | 0 | 14 | 623.9711 | 1062.1852 | NaN | 1817.6274 | 565.62 | 631.8297 |
| Ncbp1 | LSSLPWVGK | -174 | -182 | 0 | 0.001392869 | 9 | 11748.608 | 9966.905 | 12342.102 | NaN | NaN | 7242.972 |
| Ncbp1 | FLSDLVNCHVIAAPS | -130 | -144 | 0 | 0.000712251 | 15 | 444.55237 | NaN | 483.52783 | NaN | 792.21655 | 849.9502 |
| Ncbp1 | IPLNYHIV | -345 | -352 | 0 | 0.008736733 | 8 | 1436.7383 | 1590.7657 | NaN | NaN | 1820.9795 | NaN |
| Ncbp1 | TDGTSILTPWYK | -730 | -741 | 0 | 0 | 12 | 980.0132 | 923.00146 | 884.7771 | NaN | NaN | NaN |
| Ncbp1 | YGDESSNSLPGHSVAL | -487 | -502 | 0 | 0 | 16 | 5932.9126 | NaN | 9869.97 | NaN | NaN | NaN |
| Aimp2 | SVLQQTGGSSGAAPTNVQR | -283 | -301 | 0 | 0.000755556 | 19 | 821.6408 | 698.1255 | 730.7049 | 696.081 | 785.7477 | 713.8055 |
| Aimp2 | FLFSLFGQK | -216 | -224 | 0 | 0 | 9 | 150.69826 | 126.80346 | 238.64557 | 340.03113 | 343.9321 | 348.23047 |
| Aimp2 | ANPASPPLSLLVLHR | -126 | -140 | 0 | 0 | 15 | 64.91749 | 840.88104 | 684.50543 | 717.34326 | 953.798 | 1027.1953 |
| Aimp2 | TTSPATDAGHVQ | -34 | -45 | 0 | 0.00025015 | 12 | 1205.618 | 2513.1565 | NaN | 1023.87537 | 785.6009 | NaN |
| Aimp2 | TMCPIEGEGNIAR | -203 | -215 | 0 | 0 | 13 | 1462.2281 | 1542.0294 | 1314.1927 | NaN | NaN | NaN |
| Tra2b | GYDDRDYYSR | -229 | -238 | 1 | 0 | 10 | 1082.7578 | 1127.2155 | 1178.9797 | 1183.8556 | 2243.5747 | 1451.2368 |
| Tra2b | ANPDPNCCLGVF | -112 | -123 | 0 | 0 | 12 | 824.7283 | 941.4049 | 888.11426 | 900.3752 | 120.08486 | 137.96101 |
| Tra2b | VYFENVDDAK | -164 | -173 | 0 | 0 | 10 | 1838.3512 | 213.7179 | 1676.1993 | 432.53552 | NaN | 1376.0886 |
| Rps23 | KGHAVGDIPGVR | -108 | -119 | 1 | 0 | 12 | 2312.829 | 2032.2485 | 1963.8018 | 2868.7817 | NaN | NaN |
| Rps23 | GHAVGDIPGVR | -109 | -119 | 0 | 0 | 11 | 983.1383 | 221.57956 | NaN | 246.53874 | NaN | 816.36694 |
| Rps23 | ANPFGGASHAK | -38 | -48 | 0 | 0 | 11 | 1994.5658 | 2136.7156 | 2490.5 | NaN | NaN | NaN |
| Rps23 | KITAFVPNDGCLN | -80 | -92 | 1 | 0 | 13 | 1184.4547 | 1058.8427 | 1077.4854 | NaN | NaN | NaN |
| Cbl | EGFYLFPDGR | -332 | -341 | 0 | 0 | 10 | 894.4651 | 1009.20105 | 1014.62415 | 796.0634 | 994.11646 | 376.66855 |
| Cbl | LFQPWSSLLR | -245 | -254 | 0 | 0 | 10 | 2366.1772 | 2726.8164 | NaN | 2253.6877 | 2441.787 | 2370.8855 |
| Cbl | ANPVATAPSPQLSSEIER | -853 | -870 | 0 | 0 | 18 | 999.2607 | NaN | 995.8855 | NaN | 881.8136 | 1072.9718 |
| mt-Nd3 | ANPYECGFDPTSSAR | -34 | -48 | 0 | 0.00025015 | 15 | 364.00317 | 1219.9626 | NaN | NaN | NaN | NaN |
| Nup107 | SGSSLPGHLLR | -539 | -549 | 0 | 0 | 11 | 1810.4714 | 1828.8132 | 1899.3456 | 2203.8164 | 2230.0854 | 2169.1978 |
| Nup107 | VLEENQEHYHIV | -505 | -516 | 0 | 0 | 12 | 837.9704 | 838.7208 | 1608.0245 | 731.65967 | 878.12683 | 1979.6206 |
| Nup107 | VLEENQEHYHIVQK | -505 | -518 | 0 | 0.000989247 | 14 | 381.60922 | 155.26932 | 276.08142 | 457.6067 | 774.514 | 785.6859 |
| Nup107 | ANQEDNFGTATPR | -36 | -48 | 0 | 0 | 13 | 466.6286 | 515.6147 | 537.79767 | NaN | NaN | NaN |
| Smarca5 | LDGQTPHDER | -528 | -537 | 0 | 0 | 10 | 1000.1495 | 970.2463 | 775.2077 | 864.26587 | 1055.1685 | 1124.3246 |
| Smarca5 | ANRFEYLLK | -88 | -96 | 1 | 0 | 9 | 715.5916 | 992.9613 | 873.8742 | 1033.3365 | 1378.6115 | 1374.5033 |
| Smarca5 | GFDKENVYDELR | -962 | -973 | 1 | 0 | 12 | 711.8583 | 812.1442 | 780.1036 | 837.7418 | 982.54675 | 1007.5365 |
| Smarca5 | EIQEPDPTYEEK | -71 | -82 | 0 | 0 | 12 | 111.3988 | 2109.5505 | 1982.7631 | 129.55495 | 2121.7837 | 2110.9229 |
| Smarca5 | RTEQEEDEELLTE | -144 | -156 | 1 | 0.000947459 | 13 | 2776.4402 | NaN | 968.82666 | 5953.2407 | 6685.661 | 6077.482 |
| Smarca5 | FEDSPSYVK | -167 | -175 | 0 | 0 | 9 | 1152.6843 | 1287.4681 | 1413.8053 | NaN | 1213.8708 | 1145.6675 |
| Smarca5 | RFITDNTVEER | -605 | -615 | 1 | 0 | 11 | 1080.019 | 1376.4562 | 1204.8754 | NaN | 1552.8336 | 1528.5658 |
| Smarca5 | EAEPLNDEELEEK | -821 | -833 | 0 | 0 | 13 | 619.1208 | 465.11755 | 525.6001 | NaN | 696.543 | 780.7224 |
| Smarca5 | LGFDKENVYDELR | -961 | -973 | 1 | 0 | 13 | 485.58423 | 556.09924 | 642.9522 | NaN | 721.762 | 1280.5089 |
| Smarca5 | RTEQEEDEELLTES | -144 | -157 | 1 | 0.00025015 | 14 | 845.91815 | 871.30023 | 1089.1405 | 1020.35504 | NaN | NaN |
| Smarca5 | TEQEEDEELLTESSK | -145 | -159 | 0 | 0 | 15 | 818.2674 | NaN | 895.18353 | 390.19055 | 1005.84094 | NaN |
| Smarca5 | IDEAEPLNDEELEEK | -819 | -833 | 0 | 0.001610812 | 15 | 690.9387 | 778.6979 | 823.2523 | NaN | NaN | 654.49585 |
| Smarca5 | QKEIQEPDPTYEEK | -69 | -82 | 1 | 0.002068213 | 14 | 505.64612 | 510.70282 | 703.01013 | NaN | 627.2424 | NaN |
| Smarca5 | LVDQNLNK | -636 | -643 | 0 | 0 | 8 | 400.06403 | 595.0791 | 853.315 | NaN | NaN | NaN |
| Serpinc1 | TSDQIHFF | -148 | -155 | 0 | 0.000431655 | 8 | 2041.6101 | 1955.6434 | 2045.0886 | 433.85907 | 1333.9052 | 1322.1698 |
| Serpinc1 | TSDQIHFFFAK | -148 | -158 | 0 | 0 | 11 | 1755.704 | 1788.3009 | 1724.5012 | 2068.7383 | 1784.8723 | 296.7375 |
| Serpinc1 | FYQHLADSK | -95 | -103 | 0 | 0.000298805 | 9 | 826.5858 | 801.27783 | 856.5653 | 789.6681 | 851.7616 | 822.6032 |
| Serpinc1 | ANRPFLVLIR | -437 | -446 | 1 | 0 | 10 | 761.3003 | 938.57684 | 990.28485 | 462.97824 | 803.74176 | 784.5607 |
| Serpinc1 | TSDQIHFFFA | -148 | -157 | 0 | 0 | 10 | 601.77466 | 574.5731 | 693.25635 | 820.2918 | 885.09467 | 834.74725 |
| Serpinc1 | VLVNTIYFK | -247 | -255 | 0 | 0 | 9 | 523.0843 | 590.6111 | 599.5063 | 780.8926 | 968.44855 | 949.3573 |
| Serpinc1 | KLQPLDFK | -202 | -209 | 1 | 0.000431655 | 8 | 863.9313 | NaN | 217.64157 | 974.77936 | 1107.6954 | 1002.84375 |
| Serpinc1 | GDDITMVLILPK | -309 | -320 | 0 | 0 | 12 | 831.4609 | 817.93756 | 585.27893 | NaN | 634.207 | 636.82385 |
| Serpinc1 | KFDTISEK | -140 | -147 | 1 | 0.002068213 | 8 | 1004.92456 | NaN | NaN | 1204.9302 | 1083.589 | 1751.5082 |
| Acly | LTKPVVCW | -726 | -733 | 1 | 0.000471409 | 8 | 2980.5076 | 2318.5166 | 2474.798 | 4017.2153 | 1883.0299 | 1725.6548 |
| Acly | YQDTPGVK | -694 | -701 | 0 | 0.00025015 | 8 | 2250.378 | 2290.3977 | 2422.9956 | 1449.5564 | 2213.7046 | 2339.9941 |
| Acly | EILIPVFK | -537 | -544 | 0 | 0.00025015 | 8 | 1416.2861 | 1563.0343 | 1690.358 | 1444.6141 | 2045.8894 | 2004.2722 |
| Acly | NFLIEPFVPHSQ | -104 | -115 | 0 | 0 | 12 | 996.2656 | 836.03235 | 589.033 | 1019.2449 | 1639.5679 | 1355.662 |
| Acly | VTPDTDWAHLL | -34 | -44 | 0 | 0 | 11 | 867.0927 | 939.3659 | 784.2524 | 605.15936 | 839.07294 | 903.7348 |
| Acly | TVGGIKPGCFK | -615 | -625 | 1 | 0 | 11 | 864.90295 | 1203.5812 | 1308.0436 | 765.74115 | 1101.7002 | 1323.4862 |
| Acly | WGDIEFPPPFGR | -233 | -244 | 0 | 0 | 12 | 762.7142 | 829.17554 | 703.50854 | 817.13885 | 1169.526 | 776.4993 |
| Acly | NFLIEPFVPH | -104 | -113 | 0 | 0.00025015 | 10 | 695.0154 | 788.07855 | 877.1945 | 415.51917 | 498.09357 | 547.16016 |
| Acly | HHEGGVDVGDVDAK | -135 | -148 | 0 | 0 | 14 | 678.1876 | 774.2385 | 812.26416 | 711.1442 | 803.2094 | 883.833 |

| | | | | | | | | | | | | |
|---|---|---|---|---|---|---|---|---|---|---|---|---|
| Acly | HHEGGVDVGDVDAK | -135 | -148 | 0 | 0 | 14 | 678.1876 | 774.2385 | 812.26416 | 711.1442 | 803.2094 | 883.833 |
| Acly | TTGIPIHVF | -397 | -405 | 0 | 0 | 9 | 597.1229 | 612.38654 | 705.2427 | 560.426 | 726.39856 | 683.05035 |
| Acly | HLLVHAPEDKK | -168 | -178 | 1 | 0 | 11 | 491.3507 | 576.7308 | 678.63165 | 298.858 | 403.03363 | 672.6152 |
| Acly | DGVYILDLAAK | -210 | -220 | 0 | 0 | 11 | 429.03415 | 719.0091 | 343.29584 | 285.58603 | 284.28464 | 186.81265 |
| Acly | ELANYGEYSGAPSEQQTYDYAK | -300 | -321 | 0 | 0 | 22 | 381.14215 | 419.55585 | 284.47244 | 545.0768 | 237.00705 | 246.14961 |
| Acly | ANYGEYSGAPSEQQTYDYAK | -302 | -321 | 0 | 0 | 20 | 303.91226 | 327.26016 | 387.38785 | 487.0492 | 419.95023 | 434.39062 |
| Acly | LEINPLVVTK | -200 | -209 | 0 | 0 | 10 | 280.46622 | 384.84714 | 958.66644 | 295.2791 | 1117.6929 | 289.3456 |
| Acly | LNTEDIKR | -160 | -167 | 1 | 0 | 8 | 130.2867 | 1413.8367 | 1402.5381 | 1280.907 | 1732.7982 | 1349.6355 |
| Acly | HLLVHAPEDK | -168 | -177 | 0 | 0.000431655 | 10 | 919.7862 | NaN | 555.3318 | 1217.1064 | 1338.9314 | 1030.2999 |
| Acly | KLLVGVDEK | -151 | -159 | 1 | 0 | 9 | 611.3513 | NaN | 636.0681 | 914.7619 | 885.1297 | 882.67377 |
| Acly | TSGLLTIGDR | -915 | -924 | 0 | 0 | 10 | 1635.063 | 1747.1884 | 1912.0728 | 1436.7856 | NaN | NaN |
| Acly | SGAPSEQQTYDYAK | -308 | -321 | 0 | 0 | 14 | 1847.3987 | 2333.275 | NaN | NaN | NaN | 2377.6816 |
| Acly | QHFPATPLLDYAL | -986 | -998 | 0 | 0.00025015 | 13 | 962.5382 | 506.2752 | 1029.5065 | NaN | NaN | NaN |
| Ada | SSFLPEEEKK | -332 | -341 | 1 | 0 | 10 | 1661.6404 | 1771.467 | 2241.3699 | 1344.9717 | 1215.5607 | 1356.8318 |
| Ada | TVHAGEVGSPEVVR | -212 | -225 | 0 | 0 | 14 | 1599.1099 | 1100.5996 | 1434.4507 | 1299.0319 | 1289.6399 | 1044.0741 |
| Ada | HTIEDEALYNR | -241 | -251 | 0 | 0 | 11 | 2085.9402 | 2016.1569 | 2030.9038 | 2039.2219 | NaN | 1865.4531 |
| Ada | ANYSLNTDDPLIFK | -288 | -301 | 0 | 0 | 14 | 977.2057 | 1013.3346 | 808.26184 | NaN | 859.7794 | 1178.8617 |
| Ada | GIALPADTVEELR | -35 | -47 | 0 | 0 | 13 | 965.5373 | 1267.7166 | 909.3672 | 820.8209 | NaN | NaN |
| Mars1 | SFICTLPAGHR | -786 | -796 | 0 | 0 | 11 | 1484.433 | 1743.7812 | 1645.9683 | 1727.4661 | 192.36095 | 130.16765 |
| Mars1 | QHLFLDLPK | -450 | -458 | 0 | 0 | 9 | 1219.9784 | 1065.3481 | 1091.1246 | 844.20013 | 1486.4718 | 1577.9525 |
| Mars1 | DTGIPADIWR | -611 | -620 | 0 | 0 | 10 | 1187.1401 | 1471.7473 | 658.6188 | 633.6152 | 943.0738 | 1214.5599 |
| Mars1 | VLTHIDHSLSR | -120 | -130 | 0 | 0 | 11 | 805.38403 | 912.7494 | 959.594 | 677.61896 | 710.9887 | 901.2377 |
| Mars1 | VLTHIDHSLSR | -120 | -130 | 0 | 0 | 11 | 805.38403 | 912.7494 | 959.594 | 677.61896 | 710.9887 | 901.2377 |
| Mars1 | LQQHPVLPVPGER | -254 | -266 | 0 | 0 | 13 | 893.4382 | 941.25397 | 956.0083 | 675.1769 | 1563.585 | 1419.2194 |
| Mars1 | WGTPVPLEGFEDK | -503 | -515 | 0 | 0 | 13 | 736.78394 | 578.6328 | 1034.4524 | 229.28363 | 820.7385 | 625.08655 |
| Mars1 | FVEGVCPFCGYEEAR | -402 | -416 | 0 | 0 | 15 | 516.1251 | 903.3626 | 849.8285 | 751.4965 | 648.1771 | 999.7475 |
| Mars1 | KGEDILGPLR | -109 | -118 | 1 | 0 | 10 | 1549.4006 | 2099.0806 | 2137.674 | NaN | 1655.4308 | 1688.691 |
| Mars1 | ANYTDQWEK | -532 | -540 | 0 | 0.000230707 | 9 | 1102.8092 | 1232.3274 | 251.62424 | 924.8852 | NaN | 505.62506 |
| Mars1 | SVLGAEDNYTLVK | -569 | -581 | 0 | 0 | 13 | 861.03345 | 358.90173 | 887.17664 | 5414.146 | 4552.65 | NaN |
| Mars1 | NNSELLNNLGNFINR | -644 | -658 | 0 | 0 | 15 | 580.5128 | 732.119 | 936.85236 | 700.58923 | NaN | 1389.9637 |
| Mars1 | YHAIHADIYR | -338 | -347 | 0 | 0 | 10 | 470.00372 | 625.6007 | 719.12506 | NaN | 566.4454 | 608.0989 |
| Mars1 | TVPGSDWTPNAR | -470 | -481 | 0 | 0 | 12 | 467.47937 | 1332.3999 | NaN | 1951.5588 | 507.34314 | 1503.1113 |
| Mars1 | ITQDIFQR | -367 | -374 | 0 | 0 | 8 | 330.85226 | 1079.138 | NaN | 376.8459 | 854.2602 | 902.8723 |
| Mars1 | IGTVSPLFQK | -797 | -806 | 0 | 0.00111006 | 10 | 5581.803 | 6063.8125 | NaN | 4558.107 | 7249.602 | NaN |
| Mars1 | QHPVLPVPGER | -256 | -266 | 0 | 0.00025015 | 11 | 10685.6 | NaN | NaN | 13663.082 | 15266.897 | NaN |
| Mars1 | ICTLPAGHR | -788 | -796 | 0 | 0.007523243 | 9 | 982.4337 | NaN | NaN | NaN | NaN | NaN |
| Mars1 | DNVPFHGL | -557 | -564 | 0 | 0.005428329 | 8 | 672.1873 | NaN | NaN | NaN | NaN | NaN |
| H1-2 | APAAAPPAEK | -8 | -17 | 0 | 0.00078661 | 10 | 2607.965 | 2478.164 | 1953.9277 | 2037.8807 | 2021.8429 | 1289.4752 |
| H1-2 | SEAAPAAPAAAPPAEK | -2 | -17 | 0 | 0 | 16 | 299.77472 | 1349.4967 | 1298.7444 | 990.12256 | 802.58826 | 982.5035 |
| Mrpl41 | FQETVAPAIEK | -90 | -100 | 0 | 0.00111006 | 11 | 3039.2869 | NaN | 3055.6638 | NaN | NaN | NaN |
| Mrpl41 | APAGIDTPLTAK | -76 | -87 | 0 | 0.001904399 | 12 | 1334.0557 | NaN | NaN | NaN | NaN | NaN |
| Gtf2b | VTCPNHPDAILVEDYR | -13 | -28 | 0 | 0 | 16 | 1431.124 | 1622.1061 | 1454.717 | 647.91895 | 790.2171 | 782.1095 |
| Gtf2b | APDLFPSDFK | -296 | -305 | 0 | 0 | 10 | 679.0674 | NaN | 1183.6675 | 474.01517 | 1449.0404 | 1340.1836 |
| Rpl23a | TLIRPDGEK | -126 | -134 | 1 | 0 | 9 | 4815.9927 | 5332.389 | 5363.8433 | 4550.8213 | 5587.9307 | 5114.4453 |
| Rpl23a | LAPDYDALDVANK | -140 | -152 | 0 | 0 | 13 | 3240.7798 | 3569.5466 | 3323.7666 | 3973.6748 | 1563.2257 | 1314.7888 |
| Rpl23a | APDYDALDVANK | -141 | -152 | 0 | 0 | 12 | 1063.2012 | 1317.308 | 1344.9572 | 760.4409 | 781.2226 | 795.3253 |
| Rpl23a | KEAPAPPKAE | -7 | -16 | 2 | 0 | 10 | 1046.7451 | 1129.7944 | 221.98546 | 692.5729 | 1205.3116 | 1037.6372 |
| Rpl23a | TLIRPDGEKK | -126 | -135 | 2 | 0 | 10 | 570.3555 | 833.2137 | 941.73285 | 711.6124 | 616.69824 | 821.4435 |
| Rpl23a | TLIRPDGEKK | -126 | -135 | 2 | 0 | 10 | 570.3555 | 833.2137 | 941.73285 | 711.6124 | 616.69824 | 821.4435 |
| Rpl23a | LDHYAIIK | -71 | -78 | 0 | 0 | 8 | 2026.0815 | 2117.0522 | 1684.6187 | 1451.2847 | 1865.648 | NaN |
| Rpl23a | VNTLIRPDGEK | -124 | -134 | 1 | 0 | 11 | 1809.7812 | 1308.2587 | 1055.5731 | 1514.8032 | 1424.9425 | NaN |
| Rpl23a | KEAPAPPK | -7 | -14 | 1 | 0.000591124 | 8 | 362.51456 | NaN | 398.06055 | 284.22272 | 303.22443 | 441.84326 |
| Rpl23a | NKLDHYA | -69 | -75 | 1 | 0.007791272 | 7 | 835.2265 | 831.60754 | 683.3563 | NaN | NaN | NaN |
| Rpl23a | KIEDNNTLVFIV | -89 | -100 | 1 | 0.002168954 | 12 | 571.7274 | NaN | 426.72092 | NaN | 464.95795 | NaN |
| Anp32e | EITNLEDYR | -124 | -132 | 0 | 0 | 9 | 1395.8234 | 1253.4473 | 949.0174 | 1681.5394 | 1214.9368 | 1166.6655 |
| Anp32e | CPNLTYLNLSGNK | -87 | -99 | 0 | 0 | 13 | 1392.8375 | 1538.1987 | 1156.7966 | 1603.343 | 1838.3245 | 2127.2078 |
| Anp32e | GEIEGLNDTFK | -32 | -42 | 0 | 0 | 11 | 1241.226 | 763.5364 | 1269.1119 | 1250.7325 | 770.4198 | 1100.4954 |
| Anp32e | GEEEEEEEEGLRGEK | -231 | -246 | 1 | 0.00025015 | 16 | 1158.1875 | 1279.4885 | 1470.5592 | 1007.45886 | 147.55084 | 1293.2385 |
| Anp32e | NRAPEEVTEL | -13 | -22 | 1 | 0.000591124 | 10 | 842.7999 | 682.123 | 889.3245 | 454.6922 | 956.45245 | 1145.596 |
| Anp32e | EGEEEEEEEEGLRGEK | -230 | -246 | 1 | 0 | 17 | 788.30774 | 870.99036 | 945.4542 | 440.5369 | 693.3371 | 535.01416 |
| Anp32e | APEEVTELVLDNC | -15 | -27 | 0 | 0 | 13 | 589.6792 | 640.5661 | 728.319 | 819.62335 | 948.53 | 1103.8683 |
| Anp32e | NRAPEEVTELVLDNC | -13 | -27 | 1 | 0.00051306 | 15 | 337.19904 | 4033.1123 | 4045.6235 | 4133.973 | 5950.6094 | 5624.445 |
| Anp32e | DLSTVEALQNLK | -102 | -113 | 0 | 0 | 12 | 545.9758 | NaN | 875.19 | 951.7573 | 914.3248 | 1023.61163 |
| Anp32e | EEEEEEEEGLRGEK | -233 | -246 | 1 | 0 | 14 | 374.5084 | 363.06332 | 606.7464 | NaN | NaN | NaN |
| Anp32e | LTYLNLSGNK | -90 | -99 | 0 | 0 | 10 | 599.9921 | NaN | NaN | 541.2251 | NaN | NaN |
| Anp32e | SLDLFNCEITNLEDYR | -117 | -132 | 0 | 0.003900536 | 16 | 314.56677 | NaN | NaN | NaN | NaN | 113.96233 |
| Anp32e | NRAPEEVTELVLDNCL | -13 | -28 | 1 | 0.004439092 | 16 | 1122.3296 | NaN | NaN | NaN | NaN | NaN |
| Cdk7 | APELLFGAR | -180 | -188 | 0 | 0.005363933 | 9 | 799.1469 | NaN | NaN | NaN | 605.5754 | 622.4702 |
| G3bp2 | TEELKPQVEEK | -206 | -216 | 1 | 0 | 11 | 2026.624 | 2195.7476 | 2499.8066 | 675.15753 | 2537.3018 | 2629.7566 |
| G3bp2 | HLEELEEK | -217 | -224 | 0 | 0 | 8 | 1118.6725 | 697.5419 | 1141.8314 | 330.85916 | 617.9857 | 1268.4519 |
| G3bp2 | NKAPEYLHR | -24 | -32 | 1 | 0.000230707 | 9 | 500.56445 | 491.9463 | 452.15613 | 542.1744 | 490.89783 | 130.667 |
| G3bp2 | APEYLHR | -26 | -32 | 0 | 0.00025015 | 7 | 1319.756 | 1550.1162 | 1328.6642 | 1048.0636 | 1161.1711 | NaN |
| G3bp2 | VDAKPEVQ | -278 | -285 | 1 | 0 | 8 | 1295.3779 | NaN | 1257.3113 | 275.27048 | NaN | 1022.4015 |
| G3bp2 | VLAPEGSVPNK | -113 | -123 | 0 | 0.001392869 | 11 | 262.1337 | 321.80545 | NaN | 490.81 | NaN | NaN |
| Psmd12 | LAGVINFQR | -409 | -417 | 0 | 0 | 9 | 2416.488 | 2840.118 | 2649.5852 | 1510.4114 | 1907.1367 | 2069.0242 |
| Psmd12 | YVILAPFDNEQSDLVHR | -273 | -289 | 0 | 0 | 17 | 1913.4741 | 2583.46 | 2022.6866 | 1675.8417 | 1442.5609 | 2002.3262 |
| Psmd12 | DPNNLLNDWSQK | -420 | -431 | 0 | 0 | 12 | 1593.8492 | 1685.6499 | 1521.302 | 1223.7915 | 2484.1755 | 680.1086 |
| Psmd12 | FFQEENTENLK | -213 | -223 | 0 | 0.00025015 | 11 | 1419.317 | 1609.4249 | 1496.1216 | 2043.2914 | 851.08215 | 1859.0259 |

| | | | | | | | | | | | | |
|---|---|---|---|---|---|---|---|---|---|---|---|---|
| Psmd12 | APFDNEQSDLVHR | -277 | -289 | 0 | 0 | 13 | 1327.4448 | 805.33545 | 1015.97455 | 896.8284 | 76.765686 | 1163.8376 |
| Psmd12 | APFDNEQSDLVHR | -277 | -289 | 0 | 0 | 13 | 1327.4448 | 805.33545 | 1015.97455 | 896.8284 | 76.765686 | 1163.8376 |
| Psmd12 | VILAPFDNEQSDLVHR | -274 | -289 | 0 | 0 | 16 | 1152.8489 | 1413.6764 | 1440.1833 | 889.3718 | 1174.7765 | 1194.2208 |
| Psmd12 | VILAPFDNEQSDLVHR | -274 | -289 | 0 | 0 | 16 | 1152.8489 | 1413.6764 | 1440.1833 | 889.3718 | 1174.7765 | 1194.2208 |
| Psmd12 | LDQHEGSYLSICK | -233 | -245 | 0 | 0 | 13 | 733.41846 | 744.295 | 735.54083 | 717.57776 | 679.3812 | 865.18176 |
| Psmd12 | VVEHNIR | -358 | -364 | 0 | 0.001023236 | 7 | 584.4884 | 178.13904 | 638.8603 | 412.68762 | 114.21286 | NaN |
| Psmd12 | KGSSETPATDVFSST | -330 | -344 | 1 | 0.001023236 | 15 | 518.48425 | 492.69092 | 561.996 | NaN | NaN | NaN |
| Psmd12 | LAGVINFQRPK | -409 | -419 | 1 | 0.00025015 | 11 | 427.07678 | NaN | NaN | 498.2073 | 515.0448 | NaN |
| Tmem201 | APFQVILLR | -211 | -219 | 0 | 0 | 9 | 450.34045 | 592.2428 | NaN | NaN | 516.0781 | 525.1896 |
| Tmem43 | LEDPHVDIIR | -211 | -220 | 0 | 0 | 10 | 1908.3037 | 1664.846 | 1557.6597 | 1132.4331 | 1529.8207 | 1266.1448 |
| Tmem43 | YSYNTEWR | -145 | -152 | 0 | 0 | 8 | 563.7867 | 543.8362 | 506.6033 | 629.40607 | 761.5445 | 494.838 |
| Tmem43 | QWVETEESSEYTEDGQVK | -123 | -140 | 0 | 0 | 18 | 311.32864 | 328.40637 | 231.94106 | 401.41742 | 471.29047 | 412.46445 |
| Tmem43 | SLVVSPDSIHSVAPENEGR | -69 | -87 | 0 | 0 | 19 | 425.73093 | 535.8808 | 481.45926 | NaN | 692.3413 | 584.00903 |
| Tmem43 | SEPQPGFLER | -19 | -28 | 0 | 0.004439092 | 10 | 1094.4434 | NaN | NaN | 430.1322 | NaN | 1023.6478 |
| Tmem43 | APFVQIGR | -182 | -189 | 0 | 0 | 8 | 3033.663 | 2876.3105 | NaN | NaN | NaN | NaN |
| Tmem43 | RGDFFYHSENPK | -221 | -232 | 1 | 0.00025015 | 12 | 375.84235 | NaN | NaN | 538.49414 | NaN | NaN |
| Tmem43 | VDWFPVFR | -334 | -341 | 0 | 0.000907206 | 8 | 368.30768 | NaN | NaN | 333.2337 | NaN | NaN |
| Tmem43 | QRGDQLIPY | -267 | -275 | 1 | 0.009388464 | 9 | 1186.3248 | NaN | NaN | NaN | NaN | NaN |
| Hspd1 | QSKPVTTPEEIAQVA | -158 | -172 | 1 | 0 | 15 | 4418.0713 | 4766.9287 | 4847.156 | 5161.6987 | 4681.9126 | 5081.95 |
| Hspd1 | KNAGVEGSLIVEK | -481 | -493 | 1 | 0.00025015 | 13 | 3782.151 | 4100.035 | 3909.286 | 4817.823 | 5261.5938 | 4989.434 |
| Hspd1 | VGLQVVAVK | -293 | -301 | 0 | 0 | 9 | 2784.926 | 2803.167 | 2991.098 | 1239.1868 | 786.10077 | 795.35144 |
| Hspd1 | APGFGDNR | -302 | -309 | 0 | 0.00025015 | 8 | 2619.3284 | 2110.1533 | 2171.8064 | 2038.2688 | 1402.839 | 1719.6703 |
| Hspd1 | VGGTSDVEVNEK | -406 | -417 | 0 | 0 | 12 | 2540.0376 | 2697.047 | 2744.8264 | 2184.6177 | 406.17734 | 74.92585 |
| Hspd1 | VGGTSDVEVNEK | -406 | -417 | 0 | 0 | 12 | 2540.0376 | 2697.047 | 2744.8264 | 2184.6177 | 406.17734 | 74.92585 |
| Hspd1 | KVGGTSDVEVNEK | -405 | -417 | 1 | 0 | 13 | 2276.4607 | 2317.6443 | 2495.6587 | 2152.224 | 3276.4158 | 3253.5688 |
| Hspd1 | KVGGTSDVEVNEK | -405 | -417 | 1 | 0 | 13 | 2276.4607 | 2317.6443 | 2495.6587 | 2152.224 | 3276.4158 | 3253.5688 |
| Hspd1 | TLNDELEIIEGMK | -206 | -218 | 0 | 0 | 13 | 2084.0283 | 2173.232 | 2202.5408 | 1590.552 | 1507.7046 | 434.3192 |
| Hspd1 | CIPALDSLKPANEDQK | -447 | -462 | 1 | 0 | 16 | 1526.9427 | 1613.393 | 1543.3145 | 1789.3086 | 1999.5398 | 1816.2345 |
| Hspd1 | CIPALDSLKPANEDQK | -447 | -462 | 1 | 0 | 16 | 1526.9427 | 1613.393 | 1543.3145 | 1789.3086 | 1999.5398 | 1816.2345 |
| Hspd1 | NLEDVQAHDLGK | -333 | -344 | 0 | 0.000347792 | 12 | 1359.9069 | 1508.77 | 1531.3673 | 1815.6973 | 600.8005 | 2178.6145 |
| Hspd1 | EAVVTEIPKEEK | -543 | -554 | 1 | 0 | 12 | 1356.0912 | 1412.4736 | 1502.1123 | 1963.776 | 1154.7332 | 1174.7133 |
| Hspd1 | KAPGFGDNR | -301 | -309 | 1 | 0 | 9 | 1289.0884 | 1319.9222 | 416.9972 | 1971.0352 | 1879.3108 | 1881.248 |
| Hspd1 | GEEGLNLNLEDVQAHDLGK | -326 | -344 | 0 | 0 | 19 | 1269.6836 | 1591.0573 | 1665.3473 | 1365.4656 | 927.0886 | 1874.219 |
| Hspd1 | GEEGLNLNLEDVQAHDLGK | -326 | -344 | 0 | 0 | 19 | 1269.6836 | 1591.0573 | 1665.3473 | 1365.4656 | 927.0886 | 1874.219 |
| Hspd1 | AVDAVIAELK | -147 | -156 | 0 | 0 | 10 | 1251.9956 | 1235.8397 | 1258.6161 | 1269.1237 | 1362.1061 | 1371.061 |
| Hspd1 | TTAEAVVTEIPKEEK | -540 | -554 | 1 | 0 | 15 | 931.3742 | 1039.7458 | 1026.9062 | 441.41797 | 578.4871 | 606.1606 |
| Hspd1 | SIVPALEIANAHR | -256 | -268 | 0 | 0 | 13 | 639.13226 | 197.39265 | 884.1802 | 945.6433 | 1100.0322 | 1073.582 |
| Hspd1 | LVQDVANNTNEEAGDGTTT | -97 | -115 | 0 | 0.000550307 | 19 | 628.5902 | 687.28735 | 757.4558 | 641.74567 | 718.6325 | 734.51746 |
| Hspd1 | LDITTSEYEK | -378 | -387 | 0 | 0 | 10 | 442.82254 | 837.4054 | 955.10516 | 1325.811 | 1158.4312 | 1195.6687 |
| Hspd1 | LVQDVANNTNEEAGDGTTTAT | -97 | -117 | 0 | 0.000627943 | 21 | 442.00525 | 583.40314 | 513.4395 | 493.6397 | 589.70483 | 531.5566 |
| Hspd1 | LVQDVANNTNEEAGDGTTTATVL | -97 | -119 | 0 | 0.00025015 | 23 | 419.94412 | 498.74615 | 463.4257 | 184.83058 | 629.9858 | 531.45264 |
| Hspd1 | KQSKPVTTPEEIAQVA | -157 | -172 | 2 | 0.00025015 | 16 | 2166.094 | 2371.9895 | NaN | 1814.3654 | 1763.5977 | 1622.1222 |
| Hspd1 | KPLVIIAE | -269 | -276 | 1 | 0.002099344 | 8 | 1836.731 | 1258.7029 | 1219.4974 | 403.80963 | NaN | 349.83063 |
| Hspd1 | QSKPVTTPEEIAQ | -158 | -170 | 1 | 0 | 13 | 1310.8799 | 1418.1794 | 1372.5621 | NaN | 2137.2583 | 148.92702 |
| Hspd1 | IGIEIIK | -463 | -469 | 0 | 0.006100831 | 7 | 4230.331 | 3120.5042 | NaN | NaN | 1779.2928 | 1707.1775 |
| Hspd1 | LSDGVAVLK | -397 | -405 | 0 | 0 | 9 | 2166.4045 | 122.26462 | 1669.255 | 2185.2822 | NaN | NaN |
| Hspd1 | KDDAMLLK | -352 | -359 | 1 | 0.00272025 | 8 | 665.7132 | NaN | NaN | 738.61365 | 649.5333 | 589.4887 |
| Hspd1 | AVDAVIAELKK | -147 | -157 | 1 | 0 | 11 | 517.8928 | NaN | NaN | 1023.9595 | 1025.3468 | 1069.3978 |
| Hspd1 | VTDALNATR | -421 | -429 | 0 | 0.000431655 | 9 | 3221.6155 | NaN | 6945.272 | 1025.1705 | NaN | NaN |
| Hspd1 | TTAEAVVTEIPK | -540 | -551 | 0 | 0 | 12 | 624.61334 | 5524.0396 | 7110.9683 | NaN | NaN | NaN |
| Hspd1 | AVIAELKK | -150 | -157 | 1 | 0.007493591 | 8 | 183.18253 | NaN | 159.99937 | NaN | NaN | 135.45338 |
| Hspd1 | GLNLNLEDVQAHDLGK | -329 | -344 | 0 | 0.000431655 | 16 | 485.17923 | NaN | 612.4031 | NaN | NaN | NaN |
| Hspd1 | TVIIEQSWGSPK | -61 | -72 | 0 | 0 | 12 | 1169.9529 | NaN | NaN | NaN | NaN | NaN |
| Hspd1 | DRVTDALNATR | -419 | -429 | 1 | 0.003102268 | 11 | 1024.0305 | NaN | NaN | NaN | NaN | NaN |
| Mki67 | EITQISDHSEK | -2866 | -2876 | 0 | 0 | 11 | 1342.6948 | 1901.4059 | 1114.1904 | 847.95374 | 642.53326 | 1748.5007 |
| Mki67 | HGDIITIIDR | -84 | -93 | 0 | 0 | 10 | 965.9729 | 1134.4298 | 1124.4038 | 1370.2074 | 1416.0386 | 1393.9495 |
| Mki67 | SQETELQNPDDGAK | -3061 | -3074 | 0 | 0 | 14 | 827.7444 | 773.4723 | 849.522 | 594.91705 | 1084.7208 | 784.2132 |
| Mki67 | VDQQVQDNENAPQR | -854 | -867 | 0 | 0 | 14 | 355.30292 | 386.0791 | 392.20032 | 405.17087 | 400.04327 | 401.14612 |
| Mki67 | DGVQPLEVLGDSK | -2853 | -2865 | 0 | 0 | 13 | 2466.7783 | 2606.7478 | 2737.807 | 3789.2517 | 3260.7947 | NaN |
| Mki67 | NVHNITDLEK | -786 | -795 | 0 | 0.00025015 | 10 | 1493.1078 | 316.91788 | 1445.5681 | NaN | 1784.4597 | 1422.9314 |
| Mki67 | APGTPAPVQEENDSTAF | (1156)(1986)(2224) | (1172)(2002)(2240) | 0 | 0.00025015 | 17 | 896.137 | 1030.2386 | 1219.8293 | 1033.9033 | 1175.5181 | NaN |
| Mki67 | ISGEIVDLPR | -2463 | -2472 | 0 | 0 | 10 | 685.46893 | 1103.6436 | 793.24286 | NaN | 926.6991 | 1235.7762 |
| Mki67 | SSPTQSLSNSNEK | -212 | -224 | 0 | 0 | 13 | 551.53705 | 712.81134 | 621.8021 | NaN | 959.5834 | 625.77637 |
| Mki67 | VHTTQEQEDNAIK | -1627 | -1639 | 0 | 0.00025015 | 13 | 534.138 | 528.97925 | 446.44324 | 232.97696 | NaN | 249.57874 |
| Mki67 | FISPVPTNIIK | -1098 | -1108 | 0 | 0.00025015 | 11 | 465.46896 | NaN | 544.35205 | 470.12155 | 401.27576 | 415.10126 |
| Mki67 | IQLPVVSK | -39 | -46 | 0 | 0.00078661 | 8 | 207.02875 | 672.8197 | 832.3558 | 540.154 | 216.2837 | NaN |
| Mki67 | VVHIPIGPEDDTENK | -1746 | -1760 | 0 | 0 | 15 | 64.07771 | 930.6442 | 1121.8225 | NaN | NaN | NaN |
| Mki67 | KTPVSETEPLK | -796 | -806 | 1 | 0.004439092 | 11 | 877.35297 | NaN | 997.21436 | NaN | NaN | NaN |
| Mki67 | SPQPGFVR | -1704 | -1711 | 0 | 0.000230707 | 8 | 2252.3372 | NaN | NaN | NaN | NaN | NaN |
| Mki67 | SPQLGPVR | -1587 | -1594 | 0 | 0.007670622 | 8 | 923.5595 | NaN | NaN | NaN | NaN | NaN |
| Mki67 | NSGDIPEPITTEILGEK | -732 | -748 | 0 | 0.001291103 | 17 | 876.46716 | NaN | NaN | NaN | NaN | NaN |
| Mki67 | RHSVNLDEGESAQ | -335 | -347 | 1 | 0.002972566 | 13 | 402.17798 | NaN | NaN | NaN | NaN | NaN |
| Rpl18 | TNRPPLSL | -56 | -63 | 1 | 0.001743443 | 8 | 6857.5605 | 7033.5356 | 7087.061 | 7297.141 | 6304.1406 | 6799.558 |
| Rpl18 | TAVVVGTVTDDVR | -79 | -91 | 0 | 0 | 13 | 2425.6155 | 2823.597 | 2781.2788 | 1943.9319 | 1886.3518 | 1740.8157 |
| Rpl18 | TFDQLALESPK | -122 | -132 | 0 | 0 | 11 | 2204.3083 | 2290.2922 | 2428.957 | 4281.2334 | 3992.965 | 3878.2917 |
| Rpl18 | TNRPPLSLS | -56 | -64 | 1 | 0.00025015 | 9 | 2010.725 | 2005.5658 | 2063.7854 | 2159.0603 | 835.1543 | 1939.479 |
| Rpl18 | VVVGTVTDDVR | -81 | -91 | 0 | 0 | 11 | 1148.325 | 1134.4961 | 1188.7728 | 1561.396 | 1606.8838 | 1618.7765 |
| Rpl18 | RTNSTFNQVVLK | -38 | -49 | 1 | 0.00025015 | 12 | 621.2492 | 643.7442 | 636.2954 | 624.85516 | 581.8729 | 568.85815 |

| | | | | | | | | | | | | |
|---|---|---|---|---|---|---|---|---|---|---|---|---|
| Rpl18 | NSTFNQVVLK | -40 | -49 | 0 | 0 | 10 | 579.9104 | 1264.7372 | 797.34155 | 1085.8129 | 1792.9116 | 1802.5488 |
| Rpl18 | APGTPHSHTKPY | -155 | -166 | 1 | 0 | 12 | 514.8733 | 509.9031 | 548.51245 | 725.96063 | 139.49426 | 609.2526 |
| Rpl18 | STFNQVVLK | -41 | -49 | 0 | 0 | 9 | 7843.9053 | NaN | 8114.2246 | 9653.924 | 6842.153 | 8018.876 |
| Rpl18 | TNSTFNQVVLK | -39 | -49 | 0 | 0 | 11 | 4406.0845 | 4692.8643 | 5051.567 | 1504.0743 | 1692.5525 | NaN |
| Rpl18 | TNSTFNQVVLK | -39 | -49 | 0 | 0 | 11 | 4406.0845 | 4692.8643 | 5051.567 | 1504.0743 | 1692.5525 | NaN |
| Rpl18 | SQDIYLR | -20 | -26 | 0 | 0 | 7 | 2515.0815 | 2427.7065 | 1918.3588 | 641.8485 | 327.87393 | NaN |
| Rpl18 | APGTPHSHTKPYVR | -155 | -168 | 1 | 0.00025015 | 14 | 2335.508 | 1431.1079 | 2053.266 | NaN | NaN | NaN |
| Rpl18 | APGTPHSHTKPYVR | -155 | -168 | 1 | 0.00025015 | 14 | 2335.508 | 1431.1079 | 2053.266 | NaN | NaN | NaN |
| Rpl18 | TNRPPLSLSR | -56 | -65 | 1 | 0.00051306 | 10 | 996.84863 | 1025.7191 | 121.256454 | NaN | NaN | NaN |
| Rpl18 | QLALESPK | -125 | -132 | 0 | 0 | 8 | 284.24847 | 179.90921 | NaN | NaN | NaN | NaN |
| Eif4a1 | EELTLEGIR | -239 | -247 | 0 | 0 | 9 | 5720.77 | 5797.178 | 5636.9844 | 6703.591 | 6739.1196 | 5903.3564 |
| Eif4a1 | KEELTLEGIR | -238 | -247 | 1 | 0 | 10 | 4522.3174 | 4590.469 | 4845.609 | 4841.7495 | 4850.508 | 4671.386 |
| Eif4a1 | QQIELDLK | -92 | -99 | 0 | 0.00111006 | 8 | 2784.1042 | 2702.6084 | 2929.9204 | 3441.583 | 2623.2976 | 3904.4946 |
| Eif4a1 | SATMPSDVLEVTK | -213 | -225 | 0 | 0 | 13 | 2022.1904 | 1976.242 | 2013.7871 | 1887.3447 | 2599.6523 | 2147.1997 |
| Eif4a1 | SATMPSDVLEVTK | -213 | -225 | 0 | 0 | 13 | 2022.1904 | 1976.242 | 2013.7871 | 1887.3447 | 2599.6523 | 2147.1997 |
| Eif4a1 | EAPHIIVGTPGR | -150 | -161 | 0 | 0 | 12 | 1428.8075 | 1548.6633 | 1641.9564 | 1592.2601 | 1457.1484 | 1532.7993 |
| Eif4a1 | ATQALVLAPTR | -100 | -110 | 0 | 0 | 11 | 1385.7231 | 1195.1354 | 1370.1597 | 1386.1515 | 991.9032 | 1138.7609 |
| Eif4a1 | QMEAPHIIVGTPGR | -148 | -161 | 0 | 0 | 14 | 1117.8989 | 968.2677 | 1047.4728 | 1378.8909 | 1372.5417 | 1382.568 |
| Eif4a1 | QMEAPHIIVGTPGR | -148 | -161 | 0 | 0 | 14 | 1117.8989 | 968.2677 | 1047.4728 | 1378.8909 | 1372.5417 | 1382.568 |
| Eif4a1 | QMEAPHIIVGTPGR | -148 | -161 | 0 | 0 | 14 | 1117.8989 | 968.2677 | 1047.4728 | 1378.8909 | 1372.5417 | 1382.568 |
| Eif4a1 | GFKDQIYDIF | -191 | -200 | 1 | 0.000431655 | 10 | 1103.4124 | 1328.7788 | 1071.9327 | 1076.9181 | 1123.8956 | 1189.4297 |
| Eif4a1 | LQMEAPHIIVGTPGR | -147 | -161 | 0 | 0 | 15 | 1032.9647 | 1161.5322 | 1285.1143 | 1118.0997 | 1356.303 | 1342.8134 |
| Eif4a1 | LQMEAPHIIVGTPGR | -147 | -161 | 0 | 0 | 15 | 1032.9647 | 1161.5322 | 1285.1143 | 1118.0997 | 1356.303 | 1342.8134 |
| Eif4a1 | GFKDQIYDIFQK | -191 | -202 | 1 | 0.000230707 | 12 | 826.46124 | 1000.4497 | 1391.4576 | 967.6855 | 2537.5537 | 2991.0627 |
| Eif4a1 | KLQMEAPHIIVGTPGR | -146 | -161 | 1 | 0 | 16 | 737.1225 | 587.32526 | 623.4835 | 1019.2633 | 698.3851 | 693.72406 |
| Eif4a1 | KLQMEAPHIIVGTPGR | -146 | -161 | 1 | 0 | 16 | 737.1225 | 587.32526 | 623.4835 | 1019.2633 | 698.3851 | 693.72406 |
| Eif4a1 | SRDNGPDGMEPEGVIESN | -9 | -26 | 1 | 0.000431655 | 18 | 600.53094 | 576.5068 | 511.93472 | 620.6018 | 585.3185 | 947.83545 |
| Eif4a1 | APHIIVGTPGR | -151 | -161 | 0 | 0 | 11 | 254.28018 | 131.2819 | 549.2192 | 264.62314 | 267.70474 | NaN |
| Eif4a1 | SRDNGPDGMEPEGVIE | -9 | -24 | 1 | 0.001291103 | 16 | 724.3406 | NaN | 661.7853 | 711.3577 | 644.2173 | NaN |
| Eif4a1 | SIEEMPLNVADLI | -394 | -406 | 0 | 0 | 13 | 394.9154 | NaN | NaN | 479.40707 | 387.77435 | 487.71008 |
| Eif4a1 | MEAPHIIVGTPGR | -149 | -161 | 0 | 0 | 13 | 229.47098 | NaN | NaN | 792.0657 | 954.9984 | NaN |
| Eif4a1 | ELTLEGIR | -240 | -247 | 0 | 0.000394361 | 8 | 410.12912 | NaN | 352.65366 | NaN | NaN | NaN |
| Idh3a | KAPIQWEER | -58 | -66 | 1 | 0 | 9 | 4727.931 | 4998.9536 | 1609.8171 | 3516.2334 | 4009.5728 | 5300.135 |
| Idh3a | APIQWEER | -59 | -66 | 0 | 0 | 8 | 1688.3855 | 1543.9564 | 627.3404 | 2157.2222 | 1879.4612 | 2475.226 |
| Idh3a | TPYTDVNIVTIR | -135 | -146 | 0 | 0 | 12 | 1169.6605 | 1420.0066 | 1358.9996 | 1403.2972 | 1616.9108 | 1479.4287 |
| Idh3a | CSDFTEEICR | -351 | -360 | 0 | 0 | 10 | 806.6004 | 524.89703 | 681.05896 | 765.1343 | 742.52997 | 706.6702 |
| Idh3a | ENTEGEYSGIEHVIVDGVVQ | -147 | -166 | 0 | 0.00078661 | 20 | 497.82556 | 417.853 | 355.44638 | 529.96204 | 277.00345 | 517.04956 |
| Idh3a | IAEFAFEYAR | -179 | -188 | 0 | 0 | 10 | 236.57011 | 199.7446 | 4030.85 | 4667.2153 | 395.7735 | 4154.0845 |
| Idh3a | LITEEASKR | -170 | -178 | 1 | 0.00025015 | 9 | 396.75168 | 354.91235 | 348.63977 | NaN | 491.91614 | 418.46127 |
| Idh3a | MSDGLFLQK | -206 | -214 | 0 | 0 | 9 | 1300.0889 | 802.2913 | 1517.9088 | NaN | NaN | NaN |
| Idh3a | NVTAIQGPGGK | -67 | -77 | 0 | 0 | 11 | 1008.56335 | 894.869 | NaN | NaN | NaN | 1001.03687 |
| Idh3a | HMGLFDHAAK | -317 | -326 | 0 | 0.00111006 | 10 | 239.89365 | 297.87097 | 472.43542 | NaN | NaN | NaN |
| Idh3a | HMGLFDHAAK | -317 | -326 | 0 | 0.00111006 | 10 | 239.89365 | 297.87097 | 472.43542 | NaN | NaN | NaN |
| Rpl4 | APIRPDIVNF | -30 | -39 | 1 | 0.000550307 | 10 | 7041.0933 | 8162.65 | 7886.2334 | 6928.0547 | 7844.486 | 8089.0474 |
| Rpl4 | NVTLPAVFK | -21 | -29 | 0 | 0 | 9 | 4038.4268 | 3973.0356 | 9322.813 | 6571.41 | 6967.279 | 8917.007 |
| Rpl4 | LAPGGHVGR | -240 | -248 | 0 | 0.00025015 | 9 | 2477.5872 | 3241.267 | 2956.2412 | 2657.1562 | 2265.1196 | 2267.1091 |
| Rpl4 | KLDELYGTWR | -259 | -268 | 1 | 0 | 10 | 1636.1945 | 1967.111 | 497.80203 | 1491.0817 | 1457.2682 | 1261.489 |
| Rpl4 | IEEVPELPLVVEDK | -144 | -157 | 0 | 0 | 14 | 1365.0935 | 1392.9727 | 1404.4113 | 1881.0256 | 2697.5952 | 2576.4548 |
| Rpl4 | KLAPGGHVGR | -239 | -248 | 1 | 0.00025015 | 10 | 812.04803 | 763.93085 | 1065.853 | 906.34503 | 590.8913 | 764.5457 |
| Rpl4 | KLAPGGHVGR | -239 | -248 | 1 | 0.00025015 | 10 | 812.04803 | 763.93085 | 1065.853 | 906.34503 | 590.8913 | 764.5457 |
| Rpl4 | IEEVPELPLVVEDKVEGYK | -144 | -162 | 1 | 0 | 19 | 443.7306 | 451.4018 | 443.11102 | 584.71185 | 753.5152 | 726.9082 |
| Rpl4 | IEEVPELPLVVEDKVEGYK | -144 | -162 | 1 | 0 | 19 | 443.7306 | 451.4018 | 443.11102 | 584.71185 | 753.5152 | 726.9082 |
| Rpl4 | SNYNLPMHK | -275 | -283 | 0 | 0 | 9 | 521.85834 | 457.36545 | 573.57263 | 1522.8623 | 1596.516 | 1818.0874 |
| Rpl4 | IEEVPELPLVVEDKVEGY | -144 | -161 | 1 | 0.001092529 | 18 | 411.79327 | 368.6914 | 400.38763 | 718.8504 | 777.50775 | 683.94806 |
| Rpl4 | PGGHVGR | -242 | -248 | 0 | 0.00025015 | 7 | 87.17946 | 324.73877 | 553.49176 | 365.25906 | 306.71735 | 583.4114 |
| Rpl4 | IWTESAFR | -251 | -258 | 0 | 0.001392869 | 8 | 3371.213 | 1386.263 | 1398.4258 | 1482.7084 | 1178.0403 | NaN |
| Rpl4 | DELYGTWR | -261 | -268 | 0 | 0 | 8 | 1224.0642 | 944.69135 | 1336.3069 | NaN | 1852.6471 | 901.70966 |
| Rpl4 | GPCIIYNEDNGIIK | -206 | -219 | 0 | 0 | 14 | 962.1726 | 1113.7515 | 1235.0797 | NaN | 1063.1571 | 1355.7523 |
| Rpl4 | KLDELYGTW | -259 | -267 | 1 | 0.000550307 | 9 | 443.33154 | 1456.9558 | 1634.8583 | NaN | NaN | 475.7178 |
| Rpl4 | IEEVPELPLVVED | -144 | -156 | 0 | 0 | 13 | 2750.1062 | 1082.6119 | 1800.6693 | NaN | NaN | NaN |
| Rpl4 | KSNYNLPMHK | -274 | -283 | 1 | 0.001463897 | 10 | 590.0867 | NaN | 310.92532 | NaN | NaN | 727.73114 |
| Rpl4 | IEEVPELPLVVEDKVE | -144 | -159 | 1 | 0 | 16 | 527.3463 | 387.62296 | 387.9378 | NaN | NaN | NaN |
| Rpl4 | YNLPMHK | -277 | -283 | 0 | 0.00025015 | 7 | 877.89874 | NaN | NaN | NaN | NaN | NaN |
| Srrm1 | SAQEEPAAAPEPR | -898 | -910 | 0 | 0 | 13 | 2554.857 | 2588.9478 | 138.20972 | 2749.3604 | 168.37036 | 1806.52 |
| Srrm1 | TTTSAQEEPAAAPEPR | -895 | -910 | 0 | 0 | 16 | 1293.2395 | 8576.661 | 9850.435 | 8236.323 | 8154.1343 | 1624.6711 |
| Srrm1 | KETESEAEDDNLDDLER | -911 | -927 | 1 | 0 | 17 | 801.5947 | 856.3045 | 944.8233 | 985.87384 | 934.07324 | 955.58887 |
| Srrm1 | VNLEVIKPWITK | -43 | -54 | 1 | 0 | 12 | 530.29016 | 665.162 | 712.21924 | 623.7571 | 809.35785 | 836.0623 |
| Srrm1 | KVNLEVIKPWITK | -42 | -54 | 2 | 0.000230707 | 13 | 496.2641 | 414.64966 | 472.18567 | 663.34674 | 486.35425 | 547.6606 |
| Srrm1 | KVNLEVIKPWITK | -42 | -54 | 2 | 0.000230707 | 13 | 496.2641 | 414.64966 | 472.18567 | 663.34674 | 486.35425 | 547.6606 |
| Srrm1 | NLEVIKPWITK | -44 | -54 | 1 | 0 | 11 | 246.19946 | 339.43106 | 537.47345 | 431.25955 | 813.22174 | 741.78345 |
| Srrm1 | APKPEPVPEPK | -247 | -257 | 1 | 0 | 11 | 901.1757 | 1159.3215 | NaN | 479.58926 | 1227.9539 | 1282.9485 |
| Srrm1 | ENIAGIPSAFLELK | -114 | -127 | 0 | 0 | 14 | 276.43652 | 318.893 | NaN | 339.1339 | 464.02005 | 398.4866 |
| Btf3 | TNQGTVIHFNNPK | -108 | -120 | 0 | 0 | 13 | 4235.4653 | 4407.4346 | 4260.9272 | 4607.176 | 364.56638 | 4944.4917 |
| Btf3 | NQLGADSLTSLR | -150 | -161 | 0 | 0 | 12 | 2331.1228 | 1705.888 | 1646.3948 | 1692.2721 | 1569.2448 | 1400.2402 |
| Btf3 | KLQFSLK | -85 | -91 | 1 | 0.00025015 | 7 | 1113.143 | 1243.5914 | 1158.1138 | 322.8589 | 187.13371 | 236.37141 |
| Btf3 | GKAPLATGEDDDDEVPDLVENFDEASK | -174 | -200 | 1 | 0.001291103 | 27 | 548.8468 | 510.04532 | 444.47476 | NaN | 149.92085 | NaN |
| Btf3 | GKAPLATGEDDDDEVPDLVENFDEASKNEAN | -174 | -204 | 2 | 0.00078661 | 31 | 350.45874 | 396.97897 | 346.09714 | NaN | NaN | NaN |
| Btf3 | APLATGEDDDDEVPDLVENFDEASK | -176 | -200 | 0 | 0.002910424 | 25 | 276.22855 | NaN | 238.12914 | NaN | 220.88782 | NaN |

| | | | | | | | | | | | | |
|---|---|---|---|---|---|---|---|---|---|---|---|---|
| Psme1 | TENLLGSYFPK | -25 | -35 | 0 | 0.00025015 | 11 | 5669.379 | 6525.215 | 6405.9985 | 685.5557 | 6510.0522 | 6739.787 |
| Psme1 | TENLLGSYFPK | -25 | -35 | 0 | 0.00025015 | 11 | 5669.379 | 6525.215 | 6405.9985 | 685.5557 | 6510.0522 | 6739.787 |
| Psme1 | KQPHVGDYR | -190 | -198 | 1 | 0.000907206 | 9 | 5304.5054 | 5100.7437 | 5448.454 | 5127.59 | 3168.077 | 6693.7256 |
| Psme1 | APLDIPVPDPVK | -59 | -70 | 0 | 0 | 12 | 4600.852 | 5625.822 | 5410.5107 | 5787.377 | 6980.3003 | 7239.795 |
| Psme1 | APLDIPVPDPVK | -59 | -70 | 0 | 0 | 12 | 4600.852 | 5625.822 | 5410.5107 | 5787.377 | 6980.3003 | 7239.795 |
| Psme1 | VHELDEAEYQEIR | -201 | -213 | 0 | 0 | 13 | 4415.1206 | 5048.098 | 4702.1084 | 6062.374 | 6675.143 | 6384.2646 |
| Psme1 | VHELDEAEYQEIR | -201 | -213 | 0 | 0 | 13 | 4415.1206 | 5048.098 | 4702.1084 | 6062.374 | 6675.143 | 6384.2646 |
| Psme1 | WLQLQIPR | -134 | -141 | 0 | 0 | 8 | 3806.9727 | 4595.9937 | 4491.92 | 3552.4102 | 4355.7036 | 4575.1763 |
| Psme1 | LQLQIPR | -135 | -141 | 0 | 0.001023236 | 7 | 2208.1614 | 1903.691 | 1818.3483 | 1337.5878 | 963.99243 | 1020.45123 |
| Psme1 | KISELDAFLK | -36 | -45 | 1 | 0.000471409 | 10 | 2172.2324 | 2591.0889 | 2851.521 | 2168.3489 | 2950.5457 | 2871.5042 |
| Psme1 | ISELDAFLK | -37 | -45 | 0 | 0 | 9 | 1892.4098 | 2309.622 | 2302.3472 | 2431.7827 | 2734.7505 | 2938.854 |
| Psme1 | EPALNEANLSNLK | -46 | -58 | 0 | 0 | 13 | 1542.3997 | 1438.3917 | 1256.7948 | 2110.367 | 2202.1355 | 2671.9753 |
| Psme1 | APLDIPVPDPVKEK | -59 | -72 | 1 | 0 | 14 | 988.3114 | 1392.7916 | 1211.8387 | 1411.852 | 1777.0238 | 1891.8877 |
| Psme1 | LKEPALNEANLSNLK | -44 | -58 | 1 | 0 | 15 | 984.58203 | 1174.0671 | 1182.2659 | 1020.128 | 1526.3987 | 1445.6917 |
| Psme1 | LKEPALNEANLSNLK | -44 | -58 | 1 | 0 | 15 | 984.58203 | 1174.0671 | 1182.2659 | 1020.128 | 1526.3987 | 1445.6917 |
| Psme1 | IEDGNNFGVAVQEK | -142 | -155 | 0 | 0 | 14 | 871.14557 | 837.29224 | 1021.7678 | 1463.4022 | 1473.7136 | 1501.5779 |
| Psme1 | KAPLDIPVPDPVK | -58 | -70 | 1 | 0.000230707 | 13 | 775.7638 | 936.4541 | 914.9718 | 1070.9427 | 1215.2319 | 1154.473 |
| Psme1 | LVHELDEAEYQEIR | -200 | -213 | 0 | 0 | 14 | 703.41034 | 873.67084 | 707.1322 | 814.6284 | 771.8621 | 835.8181 |
| Psme1 | ELDEAEYQEIR | -203 | -213 | 0 | 0 | 11 | 608.1046 | 1078.1375 | 714.4669 | 1226.5725 | 718.2543 | 352.0053 |
| Psme1 | TWLQLQIPR | -133 | -141 | 0 | 0.000394361 | 9 | 207.3383 | 853.1973 | 927.7839 | 831.584 | 1003.3053 | 1017.8636 |
| Psme1 | LYDIILK | -226 | -232 | 0 | 0.00025015 | 7 | 1775.7361 | NaN | 1948.5234 | 1170.7799 | 975.7263 | 996.735 |
| Psme1 | QPHVGDYR | -191 | -198 | 0 | 0.000826806 | 8 | 905.36395 | 2491.5542 | 2749.3582 | 2044.0846 | NaN | 741.25116 |
| Psme1 | ENLLGSYFPK | -26 | -35 | 0 | 0 | 10 | 702.3088 | 433.98914 | 570.43396 | 1635.0193 | 368.71844 | NaN |
| Psme1 | QLVHELDEAEYQEIR | -199 | -213 | 0 | 0.00025015 | 15 | 607.05597 | 729.9011 | 792.7653 | 60.898666 | 429.9783 | NaN |
| Psme1 | QLVHELDEAEYQEIR | -199 | -213 | 0 | 0.00025015 | 15 | 607.05597 | 729.9011 | 792.7653 | 60.898666 | 429.9783 | NaN |
| Psme1 | KGDEDDKGPPCGPVNCNEK | -91 | -109 | 2 | 0.00025015 | 19 | 265.71994 | 783.7236 | NaN | 692.1413 | 296.2772 | 821.5481 |
| Psme1 | ISELDAFLKEPALNEANLS | -37 | -55 | 1 | 0.001392869 | 19 | 243.32552 | 205.75705 | 275.22604 | 189.82562 | 403.21353 | NaN |
| Psme1 | DEAEYQEIR | -205 | -213 | 0 | 0 | 9 | 989.46924 | 377.37738 | 721.90717 | NaN | NaN | NaN |
| Psme1 | DVTEQLNLVTT | -123 | -133 | 0 | 0.001392869 | 11 | 967.8389 | 250.39265 | 791.7916 | NaN | NaN | NaN |
| Psme1 | NLLGSYFPK | -27 | -35 | 0 | 0.002880184 | 9 | 179.77637 | 472.2474 | 503.6463 | NaN | NaN | NaN |
| Nckap1l | APLHLHENPDLAK | -501 | -513 | 0 | 0 | 13 | 713.66833 | 721.61633 | 791.527 | 792.8986 | 788.2258 | 799.17914 |
| Nckap1l | APLHLHENPDLAK | -501 | -513 | 0 | 0 | 13 | 713.66833 | 721.61633 | 791.527 | 792.8986 | 788.2258 | 799.17914 |
| Nckap1l | NSTQHLGPVHR | -69 | -79 | 0 | 0 | 11 | 576.7078 | 849.749 | 282.34198 | 530.2475 | 899.56824 | 733.30554 |
| Nckap1l | EILRPSELLAGVK | -733 | -745 | 1 | 0 | 13 | 442.75626 | 563.9157 | 559.695 | 590.8304 | 818.2184 | 814.4944 |
| Nckap1l | LLPQLTGAENVLK | -894 | -906 | 0 | 0 | 13 | 312.64554 | 358.58475 | 417.85086 | 310.95572 | 439.74646 | 477.31168 |
| Nckap1l | DEVTWLVR | -378 | -385 | 0 | 0 | 8 | 94.315155 | 416.31195 | 836.80493 | 691.19836 | 778.2664 | 942.6414 |
| Nckap1l | SLAQFLGADASR | -753 | -764 | 0 | 0 | 12 | 490.4401 | 388.55823 | NaN | 215.37323 | 601.1338 | 338.82953 |
| Nckap1l | VTEDLFSSLK | -304 | -313 | 0 | 0.000471409 | 10 | 406.40277 | NaN | 357.74164 | NaN | 544.0514 | 627.8281 |
| Nckap1l | ELETVLNDEPGLLGPK | -350 | -365 | 0 | 0 | 16 | 361.96274 | NaN | 408.1786 | NaN | 1084.0449 | 977.1861 |
| Nckap1l | EYLSSHLEAR | -707 | -716 | 0 | 0 | 10 | 1196.1324 | 1467.6698 | 1445.026 | NaN | NaN | NaN |
| Nckap1l | SLHFLFVR | -207 | -214 | 0 | 0.000869338 | 8 | 732.22394 | 599.0796 | NaN | NaN | 389.03723 | NaN |
| Nckap1l | SKPPFLLEK | -41 | -49 | 1 | 0.00078661 | 9 | 524.4155 | NaN | 737.7046 | 813.9177 | NaN | NaN |
| Nckap1l | EDVLQVHK | -296 | -303 | 0 | 0 | 8 | 1238.1212 | 1263.9263 | NaN | NaN | NaN | NaN |
| Nckap1l | FLLEEIR | -409 | -415 | 0 | 0.002910424 | 7 | 471.94525 | NaN | NaN | NaN | 223.71228 | NaN |
| Iars1 | SGSIVVEGHELHEEDIR | -930 | -946 | 0 | 0 | 17 | 1147.345 | 1260.8186 | 1083.7086 | 1621.1383 | 1592.1976 | 1528.2013 |
| Iars1 | SGSIVVEGHELHEEDIR | -930 | -946 | 0 | 0 | 17 | 1147.345 | 1260.8186 | 1083.7086 | 1621.1383 | 1592.1976 | 1528.2013 |
| Iars1 | VQQVPENISFPAEEEK | -2 | -17 | 0 | 0 | 16 | 827.58936 | 822.95087 | 877.0681 | 1012.0723 | 278.9062 | 966.0042 |
| Iars1 | WIDFDNDYK | -141 | -149 | 0 | 0.00025015 | 9 | 788.4249 | 816.9067 | 832.086 | 803.5854 | 113.99798 | 1029.696 |
| Iars1 | VREELIDKK | -809 | -817 | 2 | 0.00025015 | 9 | 626.46747 | 649.65234 | 813.952 | 325.89536 | 426.40613 | 708.84814 |
| Iars1 | GHELHEEDIR | -937 | -946 | 0 | 0 | 10 | 431.36502 | 652.3591 | 1337.6913 | 1657.8253 | 2631.0803 | 594.85895 |
| Iars1 | GHELHEEDIR | -937 | -946 | 0 | 0 | 10 | 431.36502 | 652.3591 | 1337.6913 | 1657.8253 | 2631.0803 | 594.85895 |
| Iars1 | DVLLPWYNAYR | -651 | -661 | 0 | 0 | 11 | 474.54752 | 400.854 | 575.66223 | 486.3374 | 617.37225 | 299.1685 |
| Iars1 | APLKPYPVPTSDNILIQ | -1039 | -1055 | 1 | 0.00078661 | 17 | 433.2391 | 486.3241 | 593.97125 | 635.1018 | 754.4585 | 740.2485 |
| Iars1 | QAPYFGADDHR | -324 | -334 | 0 | 0 | 11 | 1863.8959 | 826.78357 | 617.7383 | 3506.1777 | 977.2993 | NaN |
| Iars1 | LLIDPASLR | -786 | -794 | 0 | 0.001743443 | 9 | 1624.3658 | 1922.982 | 1822.0178 | 1803.6993 | 1846.5448 | NaN |
| Iars1 | SIDHLTIPSR | -502 | -511 | 0 | 0 | 10 | 733.83636 | 753.8748 | 1322.177 | NaN | 825.88544 | 607.8836 |
| Iars1 | RLSNEELER | -918 | -926 | 1 | 0.00051306 | 9 | 1087.3599 | NaN | 1103.8368 | 1077.3964 | 1274.5938 | NaN |
| Iars1 | VHYPFESK | -538 | -545 | 0 | 0.00025015 | 8 | 800.0057 | NaN | 367.84033 | 1318.7605 | 1499.3387 | NaN |
| Iars1 | QDPEALEDIR | -855 | -864 | 0 | 0.00025015 | 10 | 4325.8447 | 1176.8741 | NaN | NaN | 2611.777 | NaN |
| Iars1 | FKEEGVR | -640 | -646 | 1 | 0.002068213 | 7 | 2193.711 | NaN | NaN | 3340.0713 | 2736.2917 | NaN |
| Iars1 | NNDLCYWVPEFVR | -429 | -441 | 0 | 0 | 13 | 297.07306 | NaN | NaN | 273.99796 | NaN | 343.6898 |
| Iars1 | ESIDHLTIPSR | -501 | -511 | 0 | 0 | 11 | 914.45496 | NaN | NaN | NaN | 1056.6036 | NaN |
| Iars1 | LSNEELER | -919 | -926 | 0 | 0.008890469 | 8 | 2112.6604 | NaN | NaN | NaN | NaN | NaN |
| Chd4 | SAVDLTPIVVEDKEEK | -1651 | -1666 | 1 | 0 | 16 | 923.18604 | 980.69653 | 1010.03284 | 1127.2235 | 1409.914 | 1370.0603 |
| Chd4 | EESVDPDYWEK | -1299 | -1309 | 0 | 0 | 11 | 832.50446 | 701.55963 | 751.76764 | 874.9738 | 958.03186 | 958.47046 |
| Chd4 | EVAPPPPPVEVPIR | -238 | -251 | 0 | 0 | 14 | 753.6468 | 802.50037 | 682.15155 | 741.0275 | 677.89026 | 639.65216 |
| Chd4 | EEEMGEEEEVER | -1282 | -1293 | 0 | 0 | 12 | 743.1553 | 733.51575 | 900.3914 | 334.06787 | 529.8087 | 528.182 |
| Chd4 | GTTEVEKVEEK | -1640 | -1650 | 1 | 0 | 11 | 706.5555 | 976.3301 | 828.45306 | 447.4856 | 240.35059 | 621.5592 |
| Chd4 | SLKEEESTEGEK | -1563 | -1574 | 1 | 0 | 12 | 459.6291 | 505.16284 | 491.26825 | 348.841 | 443.37833 | 430.56708 |
| Chd4 | EGEDSSVIHYDDK | -1233 | -1245 | 0 | 0 | 13 | 220.61375 | 929.6904 | 1136.3455 | 1074.5012 | 1148.1694 | 1115.6395 |
| Chd4 | ERPPETPTVDPTVK | -691 | -704 | 1 | 0.000230707 | 14 | 908.46826 | 1202.1498 | NaN | 1824.0038 | 2037.5825 | 1805.0038 |
| Chd4 | ENSLKEEESTEGEK | -1561 | -1574 | 1 | 0 | 14 | 1031.1409 | 1195.5249 | NaN | NaN | 1703.5677 | 1459.8529 |
| Chd4 | FHNLEGFLE | -917 | -925 | 0 | 0.000230707 | 9 | 831.6989 | 832.2852 | 862.9409 | NaN | NaN | 1104.0143 |
| Chd4 | ATDGGGDNKEGEDSSVIHYDDK | -1224 | -1245 | 1 | 0 | 22 | 569.7929 | NaN | NaN | 692.0363 | 643.86487 | NaN |
| Chd4 | TYEIWHR | -1728 | -1734 | 0 | 0.00025015 | 7 | 296.73563 | NaN | NaN | 488.74072 | 229.46115 | NaN |
| Chd4 | EFSTNNPFK | -204 | -212 | 0 | 0.00078661 | 9 | 799.662 | 466.36154 | NaN | NaN | NaN | NaN |
| Wdr12 | APLYDLAAHEDK | -377 | -388 | 0 | 0 | 12 | 3533.0317 | 3787.3809 | 3550.6072 | 4409.92 | 4355.4287 | 1196.7192 |
| Wdr12 | GHAGSVDAIAVDSSGAK | -187 | -203 | 0 | 0 | 17 | 581.5465 | 536.7669 | 487.71182 | 754.6566 | 663.5803 | 753.31195 |

| | | | | | | | | | | | | |
|---|---|---|---|---|---|---|---|---|---|---|---|---|
| Wdr12 | SYSPTTSHVGA | -413 | -423 | 0 | 0.003681885 | 11 | 5395.014 | 6382.27 | 6047.2476 | NaN | NaN | NaN |
| Rpl18a | IFAPNHVVAK | -32 | -41 | 0 | 0 | 10 | 3919.2917 | 3612.6763 | 3465.1072 | 3332.5142 | 3216.679 | 3141.644 |
| Rpl18a | IFAPNHVVAK | -32 | -41 | 0 | 0 | 10 | 3919.2917 | 3612.6763 | 3465.1072 | 3332.5142 | 3216.679 | 3141.644 |
| Rpl18a | CHTPPLYR | -22 | -29 | 0 | 0 | 8 | 2170.4482 | 294.169 | 2620.9253 | 1996.6758 | 382.80118 | 1268.6744 |
| Rpl18a | APNHVVAK | -34 | -41 | 0 | 0.004350979 | 8 | 438.3043 | 505.68433 | NaN | 658.9416 | 624.7979 | 743.1928 |
| Rpl18a | FAPNHVVAK | -33 | -41 | 0 | 0.008105996 | 9 | 621.70935 | 745.7396 | 724.80664 | NaN | 664.05884 | NaN |
| Rpl18a | IKFPLPH | -150 | -156 | 1 | 0.003037016 | 7 | 399.1793 | NaN | NaN | 425.1642 | NaN | NaN |
| Ass1 | WHSPECEFVR | -326 | -335 | 0 | 0 | 10 | 3957.1455 | 2749.3896 | 2349.117 | 3207.9663 | 1812.3975 | 1324.4728 |
| Ass1 | WHSPECEFVR | -326 | -335 | 0 | 0 | 10 | 3957.1455 | 2749.3896 | 2349.117 | 3207.9663 | 1812.3975 | 1324.4728 |
| Ass1 | KVFIEDVSK | -58 | -66 | 1 | 0 | 9 | 4867.124 | NaN | 5351.0527 | 2941.574 | 3400.5122 | 3462.1558 |
| Ass1 | APNSPDVLEIEFK | -216 | -228 | 0 | 0 | 13 | 1380.2789 | NaN | 2184.95 | 970.0886 | 892.5389 | 1596.4935 |
| Ak2 | NLETPSCK | -86 | -93 | 0 | 0 | 8 | 1092.7551 | 854.1007 | 926.91406 | 1126.1144 | 837.856 | 996.0295 |
| Ak2 | NGFLLDGFPR | -94 | -103 | 0 | 0 | 10 | 1082.3412 | 835.3038 | 925.5506 | 1128.2335 | 1430.2615 | 1514.3224 |
| Ak2 | AVLLGPPGAGK | -18 | -28 | 0 | 0 | 11 | 1283.0378 | 815.04224 | 875.0314 | 671.9405 | NaN | 428.1109 |
| Ak2 | HTQTTPLVEYYR | -191 | -202 | 0 | 0 | 12 | 1740.3741 | NaN | NaN | 1132.3767 | 1822.2249 | 1677.6525 |
| Ak2 | APNVLASEPEIPK | -2 | -14 | 0 | 0 | 13 | 1171.9105 | 1303.3447 | NaN | NaN | NaN | 1291.5172 |
| Ak2 | SIQDSLLIR | -129 | -137 | 0 | 0 | 9 | 453.43533 | NaN | NaN | 381.38095 | NaN | NaN |
| Ak2 | TTPLVEYYR | -194 | -202 | 0 | 0.001055699 | 9 | 1052.4569 | NaN | NaN | NaN | NaN | NaN |
| Dync1i2 | EFLSFFDHSTR | -195 | -205 | 0 | 0.000431655 | 11 | 741.17035 | 771.2317 | 680.5626 | 63.2314 | 776.13794 | 724.1366 |
| Dync1i2 | SVSTPSEAGSQDSGDGAVGSR | -86 | -106 | 0 | 0 | 21 | 561.18225 | 497.16318 | 570.6943 | 433.17856 | 576.5834 | 681.17725 |
| Dync1i2 | DVGEQIAVPR | -572 | -581 | 0 | 0 | 10 | 2544.5886 | 2496.1692 | 3064.3257 | 2013.1877 | 2173.746 | NaN |
| Dync1i2 | RADAEEEAATR | -599 | -609 | 1 | 0.000230707 | 11 | 930.6896 | NaN | 966.87964 | 1278.1941 | 1216.324 | 1155.0117 |
| Dync1i2 | APPHELTEEEK | -177 | -187 | 0 | 0 | 11 | 812.6675 | 1084.6793 | 1052.6185 | 706.6881 | NaN | NaN |
| Gnl2 | SWENAEDFLEK | -414 | -424 | 0 | 0 | 11 | 728.5796 | 888.2319 | 1418.7975 | 986.7345 | NaN | 869.7713 |
| Gnl2 | APQDHIGAVL | -389 | -398 | 0 | 0.000431655 | 10 | 6328.5083 | NaN | NaN | NaN | 437.8178 | NaN |
| Capn1 | VLSEEEIDDNFK | -536 | -547 | 0 | 0 | 12 | 1272.4423 | 1541.3447 | 399.42987 | 1697.6337 | 1986.0798 | 2103.4272 |
| Capn1 | ELDDQIQANLPDEK | -522 | -535 | 0 | 0 | 14 | 1077.3523 | 1213.4623 | 838.01575 | 734.04114 | 1119.201 | 429.07474 |
| Capn1 | LEEVDDADDYDNR | -401 | -413 | 0 | 0.000230707 | 13 | 372.05487 | 359.9341 | 407.46674 | 554.92566 | 588.51666 | 590.8347 |
| Capn1 | SLTLNETILHR | -123 | -133 | 0 | 0 | 11 | 5849.8115 | NaN | 6297.8037 | 2104.5952 | 2027.121 | 2527.1155 |
| Capn1 | ELAGQPVHLK | -451 | -460 | 0 | 0 | 10 | 2241.3276 | 2042.8922 | NaN | 903.9114 | 1003.3164 | 1393.1208 |
| Capn1 | KAPSDLYQIILK | -229 | -240 | 1 | 0.00078661 | 12 | 787.08826 | NaN | 939.37726 | 1042.7919 | 1588.0822 | 1557.3522 |
| Capn1 | APSDLYQIILK | -230 | -240 | 0 | 0 | 11 | 721.72644 | 744.8991 | 676.68805 | 811.6625 | 881.1071 | NaN |
| Capn1 | LPPGEYIVVPSTFEPNK | -488 | -504 | 0 | 0 | 17 | 4677.0386 | NaN | NaN | 6126.858 | 9076.037 | 793.9682 |
| Capn1 | TFEPNKEGDFLLR | -499 | -511 | 1 | 0 | 13 | 1065.714 | 1104.6211 | 1070.6642 | NaN | 1846.6776 | NaN |
| Capn1 | DKELGLGR | -23 | -30 | 1 | 0.005270854 | 8 | 1822.6211 | NaN | 1781.9547 | NaN | NaN | NaN |
| Capn1 | ELQTILNR | -564 | -571 | 0 | 0.002566359 | 8 | 957.181 | NaN | NaN | NaN | NaN | NaN |
| Capn1 | TLNETILHR | -125 | -133 | 0 | 0 | 9 | 692.89966 | NaN | NaN | NaN | NaN | NaN |
| Aifm1 | DGEQHEDLNEVAK | -593 | -605 | 0 | 0 | 13 | 880.04156 | 746.5324 | 1058.4055 | 1532.2529 | 981.1769 | 1075.929 |
| Aifm1 | TGGLEIDSDFGGFR | -408 | -421 | 0 | 0 | 14 | 627.8569 | 430.83258 | 439.31775 | 397.87088 | 804.4289 | 494.5443 |
| Aifm1 | ELWFSDDPNVTK | -177 | -188 | 0 | 0 | 12 | 615.53094 | 717.7099 | 696.62506 | 857.1789 | 992.4745 | 791.7392 |
| Aifm1 | APSHVPFLLIGGGTAA | -127 | -142 | 0 | 0 | 16 | 470.63962 | 183.8393 | 451.30832 | 435.54916 | NaN | 443.93536 |
| Aifm1 | KVETDHIVTAV | -387 | -397 | 1 | 0.003201138 | 11 | 2115.4197 | NaN | NaN | 573.10767 | 3073.6338 | NaN |
| Aifm1 | GLEPNVELAK | -398 | -407 | 0 | 0.003966045 | 10 | 833.59174 | 605.8377 | 449.70923 | NaN | NaN | NaN |
| Aifm1 | APSHVPFLLIGGGT | -127 | -140 | 0 | 0.003681885 | 14 | 631.7348 | 669.48376 | NaN | 534.5729 | NaN | NaN |
| Aifm1 | ATEGGSVPQIR | -116 | -126 | 0 | 0.003069674 | 11 | 529.5499 | NaN | NaN | NaN | NaN | NaN |
| Apbb1ip | KNPQNFYLDNK | -271 | -281 | 1 | 0 | 11 | 751.90643 | 730.6017 | 786.2279 | 622.71515 | 908.039 | 913.478 |
| Apbb1ip | APTDHCFVLK | -379 | -388 | 0 | 0 | 10 | 6707.838 | NaN | 8930.238 | 8665.192 | 6589.6187 | 8094.284 |
| Hint1 | VAQPGGDTIFGK | -10 | -21 | 0 | 0 | 12 | 3745.682 | 3802.5273 | 2655.9258 | 7761.101 | 4042.5117 | 3966.0686 |
| Hint1 | HDISPQAPTHFLVIPK | -42 | -57 | 0 | 0 | 16 | 1860.8252 | 1963.5048 | 1810.4928 | 2059.3513 | 1884.1628 | 1868.6672 |
| Hint1 | HDISPQAPTHFLVIPK | -42 | -57 | 0 | 0 | 16 | 1860.8252 | 1963.5048 | 1810.4928 | 2059.3513 | 1884.1628 | 1868.6672 |
| Hint1 | APTHFLVIPK | -48 | -57 | 0 | 0.000347792 | 10 | 476.35437 | 494.28186 | 503.58783 | 336.6654 | NaN | 405.37677 |
| Hint1 | MVVNEGADGGQSVYHIH | -96 | -112 | 0 | 0.00416551 | 17 | 987.2671 | 1086.5677 | NaN | NaN | 1860.4597 | NaN |
| Rrm2 | DIQHWEALKPDER | -113 | -125 | 1 | 0 | 13 | 1117.7605 | 1251.1046 | 1279.7838 | 1502.9021 | 1613.5393 | 1709.2719 |
| Rrm2 | DIQHWEALKPDER | -113 | -125 | 1 | 0 | 13 | 1117.7605 | 1251.1046 | 1279.7838 | 1502.9021 | 1613.5393 | 1709.2719 |
| Rrm2 | APTNPSVEDEPLLR | -62 | -75 | 0 | 0 | 14 | 1198.0419 | 1251.8914 | 1273.0225 | 1961.4836 | 2599.1475 | 1532.0516 |
| Rrm2 | LIDTYIKDPK | -180 | -189 | 1 | 0.000230707 | 10 | 1196.866 | 1251.5043 | 1414.313 | 1193.0912 | 1409.8883 | 1385.6093 |
| Rrm2 | IEQEFLTEALPVK | -300 | -312 | 0 | 0 | 13 | 766.3832 | 937.3256 | 860.75836 | 1107.596 | 1532.1715 | 1543.1733 |
| Rrm2 | IFQDSAELESK | -51 | -61 | 0 | 0 | 11 | 436.1515 | 5157.97 | 5425.587 | 4466.8525 | 4924.339 | 920.8706 |
| Rrm2 | FSQEVQVTEAR | -150 | -160 | 0 | 0 | 11 | 2044.0851 | 2350.5962 | NaN | 2260.4902 | 3077.6824 | 2329.845 |
| Rrm2 | HFISHVL | -126 | -132 | 0 | 0.00025015 | 7 | 1546.7135 | 1877.1841 | 1914.2311 | NaN | 922.66565 | 1309.2336 |
| Rrm2 | SKAPTNPSVEDEPLLR | -60 | -75 | 1 | 0 | 16 | 660.2039 | NaN | 443.70663 | NaN | 1390.8911 | 1504.3597 |
| Rrm2 | DQQQLQLSPLK | -13 | -23 | 0 | 0.000591124 | 11 | 593.1925 | 504.60602 | 468.72437 | NaN | NaN | NaN |
| Rrm2 | EALKPDER | -118 | -125 | 1 | 0 | 8 | 567.59393 | NaN | 531.79663 | NaN | NaN | 432.4872 |
| Pfas | YVPLSPGQHR | -852 | -861 | 0 | 0.002972566 | 10 | 5859.7563 | 6496.8833 | 5745.849 | 10795.16 | 7179.3677 | 6875.908 |
| Pfas | GDNTSDLDFGAVQR | -473 | -486 | 0 | 0 | 14 | 1137.2589 | 1045.7343 | 1141.7665 | 274.27594 | 902.3953 | 1120.3903 |
| Pfas | SVPCKPGGPVPR | -1052 | -1063 | 1 | 0 | 12 | 854.06305 | 812.72833 | 885.0724 | 789.4155 | 528.5206 | 583.6618 |
| Pfas | SQLGEHPPDLDLPENLVR | -872 | -889 | 0 | 0 | 18 | 528.2349 | 548.71814 | 407.72522 | 582.672 | 1183.9906 | 874.59973 |
| Pfas | SLGLQLPDGQR | -407 | -417 | 0 | 0 | 11 | 1336.4341 | 3034.1519 | 2225.2915 | 1624.5205 | NaN | 1968.5537 |
| Pfas | AVVVEEPVGELR | -996 | -1007 | 0 | 0 | 12 | 492.75983 | 346.11902 | 708.7043 | 1073.8365 | NaN | 1241.38 |
| Pfas | LNFSTPASTNIVSVC | -94 | -108 | 0 | 0.00272025 | 15 | 1927.0294 | NaN | NaN | NaN | 1862.2728 | 1834.326 |
| Pfas | APVLHFY | -2 | -8 | 0 | 0.003620005 | 7 | 815.097 | NaN | NaN | 418.19476 | 422.30435 | NaN |
| Yars1 | NLQEVLGEEK | -17 | -26 | 0 | 0 | 10 | 2829.2966 | 3059.948 | 3300.8289 | 3078.8184 | 3206.2256 | 3368.1548 |
| Yars1 | DFAAEVVHPGDLK | -298 | -310 | 0 | 0 | 13 | 2013.4117 | 2388.1816 | 2313.5352 | 1408.328 | 1772.9916 | 1684.1217 |
| Yars1 | KIDVGEAEPR | -391 | -400 | 1 | 0 | 10 | 1477.574 | 1403.2144 | 1575.1658 | 1562.6626 | 371.33072 | 1688.9177 |
| Yars1 | VDAQFGGVDQR | -179 | -189 | 0 | 0 | 11 | 1446.7667 | 1683.0643 | 1046.7162 | 401.75583 | 1928.7454 | 431.77264 |
| Yars1 | APWELLELR | -85 | -93 | 0 | 0 | 9 | 580.63745 | 556.6589 | 620.3652 | 681.0786 | 767.2946 | 751.2301 |
| Yars1 | QVEPLDPPAGSAPGER | -451 | -466 | 0 | 0.00025015 | 16 | 566.2856 | 529.41565 | 1335.7051 | 823.7565 | 149.44633 | 613.92664 |
| Yars1 | GQPDEELKPK | -475 | -484 | 1 | 0 | 10 | 292.13712 | 1271.0918 | 377.23123 | 1203.9468 | 1646.3473 | 1454.6975 |

| | | | | | | | | | | | |
|---|---|---|---|---|---|---|---|---|---|---|---|
| Yars1 | HPDADSLYVEK | -381 | -391 | 0 | 0.00025015 | 11 | 3874.3247 | NaN | 4265.747 | 5384.6167 | 6769.4043 | 1188.9893 |
| Yars1 | LYPGLQALDEEYLK | -165 | -178 | 0 | 0 | 14 | 297.14154 | NaN | 459.214 | 438.36963 | 569.339 | 762.09827 |
| Yars1 | NSEPEEVIPSR | -357 | -367 | 0 | 0 | 11 | 1386.0363 | 1423.9332 | 1008.59607 | NaN | NaN | NaN |
| H2-D1 | AVSRPGLEEPR | -35 | -45 | 1 | 0 | 11 | 3767.7961 | 3201.1814 | 3625.4775 | 3136.7107 | 3688.0442 | 3437.613 |
| H2-D1 | SRPGLEEPR | -37 | -45 | 1 | 0.00025015 | 9 | 1502.6576 | 2170.639 | 4160.932 | 2020.5209 | 1049.4895 | 1974.9647 |
| H2-D1 | SGCDLGSDWR | -123 | -132 | 0 | 0.000627943 | 10 | 1354.8286 | 1305.9227 | 1379.3464 | 1693.9178 | 1532.4344 | 1711.9674 |
| H2-D1 | WEQSGAAEHYK | -171 | -181 | 0 | 0 | 11 | 632.818 | NaN | 898.9536 | 428.53064 | 569.5776 | 1066.3586 |
| H2-D1 | PGLEEPR | -39 | -45 | 0 | 0.003360691 | 7 | 1292.3259 | NaN | 1039.0972 | NaN | 1038.0994 | NaN |
| H2-D1 | VSRPGLEEPR | -36 | -45 | 1 | 0.000431655 | 10 | 1396.703 | NaN | NaN | 433.78305 | NaN | NaN |
| Zdhhc20 | APWTLWR | -2 | -8 | 0 | 0.001392869 | 7 | 314.99017 | NaN | 390.43317 | 355.24045 | 585.958 | NaN |
| Plac8 | AQAPTVIVTQPGFVR | -2 | -16 | 0 | 0 | 15 | 2861.1216 | 3225.6147 | 3344.0388 | 4947.6313 | 6348.646 | 5035.8896 |
| Hadhb | DQLLLGPTYATPK | -350 | -362 | 0 | 0 | 13 | 1426.2034 | 1094.0037 | 1173.6411 | 1280.331 | 1595.1414 | 1574.036 |
| Hadhb | AQDEGHLSDIVPF | -256 | -268 | 0 | 0.00025015 | 13 | 659.21027 | 155.15091 | 719.1987 | 788.5623 | 821.9847 | 819.82404 |
| Hadhb | KAQDEGHLSDIVPF | -255 | -268 | 1 | 0 | 14 | 475.9756 | 630.02905 | 526.4441 | 429.3576 | 525.03424 | 163.39946 |
| Hadhb | KAQDEGHLSDIVPF | -255 | -268 | 1 | 0 | 14 | 475.9756 | 630.02905 | 526.4441 | 429.3576 | 525.03424 | 163.39946 |
| Hadhb | LKPAFIKPY | -293 | -301 | 2 | 0.004350979 | 9 | 580.14874 | 570.1723 | 632.1883 | NaN | NaN | NaN |
| Sec31a | AQDGSSPLSLQDLIEK | -707 | -722 | 0 | 0 | 16 | 1359.6333 | 859.4524 | 1262.8876 | 475.1159 | 682.9029 | 1188.8352 |
| Sec31a | VSGDIDGLITR | -564 | -574 | 0 | 0 | 11 | 1009.12555 | 1222.4398 | 1314.4186 | 824.7566 | 916.74835 | 1079.9762 |
| Sec31a | RLEFLYDK | -1159 | -1166 | 1 | 0.002910424 | 8 | 953.79584 | 1266.8881 | 1290.2982 | 269.56213 | 1382.0289 | 1340.7466 |
| Sec31a | KPIPEEHLILK | -1120 | -1130 | 1 | 0 | 11 | 935.9717 | 1105.111 | 1047.5116 | 820.42706 | 992.7272 | 1032.9032 |
| Sec31a | TTFEDLIQR | -1131 | -1139 | 0 | 0 | 9 | 641.8579 | 811.9367 | 261.34985 | NaN | NaN | NaN |
| Sec31a | ESVAVPLQQGAEQQR | -408 | -422 | 0 | 0.00025015 | 15 | 4852.104 | NaN | NaN | 3982.7236 | NaN | NaN |
| Sec31a | VNFEEDSR | -478 | -485 | 0 | 0.006788754 | 8 | 1440.2423 | NaN | NaN | NaN | NaN | NaN |
| Nudcd2 | AQDIQCGLQSR | -40 | -50 | 0 | 0 | 11 | 902.8327 | 2622.633 | 893.3109 | 534.4305 | 2185.0925 | 2412.1567 |
| Nudcd2 | IADEGTWTLEDR | -71 | -82 | 0 | 0 | 12 | 1164.1571 | 1926.956 | 1925.6874 | NaN | 2429.259 | 4433.642 |
| Nudcd2 | GGPDFSNLEK | -148 | -157 | 0 | 0 | 10 | 595.97485 | 1095.8517 | NaN | 492.1112 | 917.144 | 909.27313 |
| Esyt1 | AQDLPLKK | -477 | -484 | 1 | 0.00025015 | 8 | 16632.717 | 16534.416 | 15933.185 | 13183.517 | 14358.215 | 18196.432 |
| Esyt1 | GITSRPEPPSAAIL | -457 | -470 | 1 | 0 | 14 | 2843.85 | 3753.5098 | 3693.104 | 2270.461 | 2776.2747 | 3132.2012 |
| Esyt1 | RQLLDDEER | -96 | -104 | 1 | 0.00025015 | 9 | 1689.7369 | 1525.8121 | 1566.0388 | 1630.1267 | 1857.8795 | 1805.1741 |
| Esyt1 | FFLQDPR | -521 | -527 | 0 | 0 | 7 | 1574.6205 | 1633.9742 | 242.25456 | 784.7813 | 783.83545 | 740.00354 |
| Esyt1 | VELGEKPLR | -174 | -182 | 1 | 0.000230707 | 9 | 1542.1431 | 1575.0157 | 818.21313 | 1927.8079 | 1606.4664 | 1567.4807 |
| Esyt1 | LLAETVAPAVR | -150 | -160 | 0 | 0 | 11 | 1408.7837 | 1443.0447 | 1542.9232 | 1671.1254 | 1816.4054 | 1667.8662 |
| Esyt1 | RVELGEKPLR | -173 | -182 | 2 | 0.00025015 | 10 | 1300.8827 | 1247.5146 | 1304.493 | 1275.9629 | 1416.764 | 1461.05 |
| Esyt1 | LTHGDSPSEAPVGPLGQVK | -946 | -964 | 0 | 0 | 19 | 1259.0215 | 1230.0967 | 1373.3142 | 1068.6057 | 1269.2128 | 1418.4275 |
| Esyt1 | LTHGDSPSEAPVGPLGQVK | -946 | -964 | 0 | 0 | 19 | 1259.0215 | 1230.0967 | 1373.3142 | 1068.6057 | 1269.2128 | 1418.4275 |
| Esyt1 | VFDKDPDKDDFLGR | -392 | -405 | 2 | 0 | 14 | 1170.6995 | 1247.0701 | 1419.7776 | 1140.1849 | 1069.0017 | 1518.9484 |
| Esyt1 | VFDKDPDKDDFLGR | -392 | -405 | 2 | 0 | 14 | 1170.6995 | 1247.0701 | 1419.7776 | 1140.1849 | 1069.0017 | 1518.9484 |
| Esyt1 | GGPTHPASPVLEVR | -930 | -943 | 0 | 0 | 14 | 1118.0914 | 1169.03 | 1271.32 | 736.1453 | 941.6237 | 1197.5764 |
| Esyt1 | KGTKPPSPYATIT | -801 | -813 | 2 | 0.000712251 | 13 | 986.1269 | 1121.5453 | 1119.4718 | 823.11646 | 751.87415 | 994.02606 |
| Esyt1 | VIDEELNPHWGETY | -363 | -376 | 0 | 0 | 14 | 962.9547 | 1220.9795 | 1245.9452 | 1006.58685 | 1224.4054 | 1116.9375 |
| Esyt1 | ILYLDYSEIR | -585 | -594 | 0 | 0 | 10 | 932.8268 | 983.31854 | 1046.8988 | 988.20874 | 633.30096 | 916.8095 |
| Esyt1 | ELPAWVSFPDVEK | -116 | -128 | 0 | 0 | 13 | 916.19324 | 877.64087 | 914.73566 | 1067.856 | 1281.4944 | 1297.7549 |
| Esyt1 | HTTPNSHFGTENVLR | -623 | -637 | 0 | 0.00025015 | 15 | 814.2961 | 962.8246 | 1314.0579 | 2139.8413 | 1372.3412 | 1648.1682 |
| Esyt1 | HTTPNSHFGTENVLR | -623 | -637 | 0 | 0.00025015 | 15 | 814.2961 | 962.8246 | 1314.0579 | 2139.8413 | 1372.3412 | 1648.1682 |
| Esyt1 | VFDKDLDKDDFLGR | -707 | -720 | 2 | 0 | 14 | 793.113 | 936.2146 | 949.2087 | 917.1738 | 213.017 | 199.94492 |
| Esyt1 | DNWYPLQGGQGQVHLR | -420 | -435 | 0 | 0 | 16 | 660.7848 | 627.08594 | 778.7038 | 576.8003 | 831.42163 | 764.14484 |
| Esyt1 | EWDLPLDGTLR | -1030 | -1040 | 0 | 0 | 11 | 647.59656 | 836.73944 | 833.6276 | 815.6968 | 867.5208 | 785.1146 |
| Esyt1 | YSTNSPVWEEAFR | -508 | -520 | 0 | 0 | 13 | 555.7449 | 693.38275 | 602.69653 | 1304.281 | 791.2668 | 761.65497 |
| Esyt1 | DLDKDDFLGR | -711 | -720 | 1 | 0 | 10 | 544.1715 | 1224.291 | 1282.1287 | 1177.8977 | 589.1159 | 698.5734 |
| Esyt1 | LEWLSLLPDAEK | -436 | -447 | 0 | 0 | 12 | 534.83435 | 478.24536 | 538.0004 | 538.366 | 730.7911 | 659.08984 |
| Esyt1 | FEWDLPLDGTLR | -1029 | -1040 | 0 | 0 | 12 | 375.77682 | 470.23465 | 372.51733 | 364.72348 | 407.34747 | 378.6888 |
| Esyt1 | LTHGDSPSEAPVGPL | -946 | -960 | 0 | 0 | 15 | 250.0685 | 1394.2771 | 1654.8849 | 2453.249 | 2144.3784 | 2213.5679 |
| Esyt1 | LISIIHSCR | -976 | -984 | 0 | 0 | 9 | 1075.6973 | 1504.6189 | 1316.8995 | NaN | 543.28326 | 606.90985 |
| Esyt1 | DKDLDKDDFLGR | -709 | -720 | 2 | 0 | 12 | 580.2809 | 2865.0515 | NaN | 833.8695 | 1453.4634 | 1481.919 |
| Esyt1 | DKDLDKDDFLGR | -709 | -720 | 2 | 0 | 12 | 580.2809 | 2865.0515 | NaN | 833.8695 | 1453.4634 | 1481.919 |
| Esyt1 | TLNPEFNER | -1020 | -1028 | 0 | 0 | 9 | 1800.0706 | 2034.3209 | 2100.3047 | 557.5421 | NaN | NaN |
| Esyt1 | GANPHLQTF | -161 | -169 | 0 | 0.002629273 | 9 | 1205.7318 | 1128.295 | 1903.1564 | NaN | 2769.4958 | NaN |
| Esyt1 | YLDYSEIR | -587 | -594 | 0 | 0 | 8 | 708.74817 | NaN | NaN | 810.8281 | 862.6922 | 883.1704 |
| Esyt1 | STNSPVWEEAFR | -509 | -520 | 0 | 0 | 12 | 706.1718 | NaN | 643.3269 | 583.3153 | NaN | NaN |
| Esyt1 | TNSPVWEEAFR | -510 | -520 | 0 | 0.000431655 | 11 | 1060.691 | NaN | NaN | NaN | 1170.6973 | NaN |
| Nsun2 | DGQWFADWHEVPQGR | -370 | -384 | 0 | 0 | 15 | 2524.6465 | 2579.118 | 2838.1384 | 1326.6727 | 1706.5758 | 1617.479 |
| Nsun2 | FYALDPSFPR | -535 | -544 | 0 | 0 | 10 | 2094.4453 | 2635.3726 | 2454.0984 | 2196.9956 | 2212.942 | 2184.6829 |
| Nsun2 | LTQENPFFR | -630 | -638 | 0 | 0 | 9 | 1244.8417 | 1106.3649 | 986.03253 | 1247.924 | 1327.7249 | 1218.5209 |
| Nsun2 | AQDPCVPDSVPGCDAAAAEPSR | -736 | -757 | 0 | 0 | 22 | 1217.789 | 1419.3524 | 1437.1672 | 1276.8346 | 762.77026 | 1237.7103 |
| Nsun2 | GQAGWEGGYPEIVK | -30 | -43 | 0 | 0 | 14 | 1101.8595 | 1264.0736 | 1264.4366 | 1624.4598 | 1036.359 | 1340.2924 |
| Nsun2 | NNSGEEFDCAFR | -590 | -601 | 0 | 0 | 12 | 1068.7509 | 1082.9474 | 1160.2017 | 1050.6088 | 1326.7537 | 1320.7397 |
| Nsun2 | TLLTQENPFFR | -628 | -638 | 0 | 0 | 11 | 890.19244 | 794.3702 | 1088.1084 | 776.9038 | 1424.9402 | 1433.06 |
| Nsun2 | VVNHDASSIPR | -238 | -248 | 0 | 0 | 11 | 846.9771 | 940.8641 | 835.5575 | 576.7667 | 836.33795 | 820.10004 |
| Nsun2 | VVNHDASSIPR | -238 | -248 | 0 | 0 | 11 | 846.9771 | 940.8641 | 835.5575 | 576.7667 | 836.33795 | 820.10004 |
| Nsun2 | REPLPATLR | -71 | -79 | 1 | 0.001023236 | 9 | 777.68024 | 879.17993 | 974.4382 | 396.7084 | 871.7842 | 856.11304 |
| Nsun2 | FQQPPQPEGEEDASDGGR | -10 | -27 | 0 | 0.00025015 | 18 | 735.1225 | 780.4516 | 678.82 | 854.84454 | 258.25058 | 800.9692 |
| Nsun2 | ENAASPEQPGDEDAK | -719 | -733 | 0 | 0.00025015 | 15 | 666.935 | NaN | 846.04047 | 356.4049 | 1268.9655 | 1219.9904 |
| Nsun2 | LFEHYYQELK | -47 | -56 | 0 | 0 | 10 | 3784.0364 | 214.4373 | 1707.1577 | NaN | NaN | 894.9052 |
| Nsun2 | ELEDLEVDGQK | -101 | -111 | 0 | 0 | 11 | 1681.0686 | NaN | 1125.1343 | NaN | 1789.551 | 1744.9954 |
| Nsun2 | ILPHHQNTGGFFVA | -412 | -425 | 0 | 0 | 14 | 1235.7045 | 1292.5099 | 2133.4092 | 1979.3411 | NaN | NaN |
| Nsun2 | GAEQLAEGGR | -306 | -315 | 0 | 0 | 10 | 928.4059 | 1297.2866 | 990.73737 | 694.7988 | NaN | NaN |
| Nsun2 | HDASSIPR | -241 | -248 | 0 | 0.000230707 | 8 | 805.9439 | 827.40326 | NaN | NaN | 965.45795 | 649.0149 |
| Nsun2 | ILPHHQNTGGFF | -412 | -423 | 0 | 0 | 12 | 760.3355 | 609.8997 | 920.49176 | 177.32106 | NaN | NaN |

| | | | | | | | | | | | | |
|---|---|---|---|---|---|---|---|---|---|---|---|---|
| Nsun2 | SEGALELADVSAELPGLK | -339 | -356 | 0 | 0 | 18 | 369.02365 | NaN | NaN | 435.66443 | 465.79626 | 479.4081 |
| Nsun2 | VSSHVAATEGNPSDQSELESQ | -454 | -474 | 0 | 0 | 21 | 252.76802 | 346.8287 | 315.90192 | NaN | NaN | NaN |
| Nsun2 | RFQQPPQPEGEEDASDGGR | -9 | -27 | 1 | 0.00051306 | 19 | 594.52106 | NaN | 1084.0562 | NaN | NaN | NaN |
| Nsun2 | LAQEGIYTLYPFINSR | -602 | -617 | 0 | 0 | 16 | 192.22092 | NaN | NaN | NaN | 231.10776 | NaN |
| Nsun2 | KEILFYDR | -257 | -264 | 1 | 0.002265552 | 8 | 1253.4617 | NaN | NaN | NaN | NaN | NaN |
| Zbtb8os | KYEYLDHTADVQ | -30 | -41 | 1 | 0 | 12 | 591.9558 | 1030.11 | 1937.9908 | 1353.5242 | 932.0817 | 2115.0615 |
| Zbtb8os | SADEYFIPR | -101 | -109 | 0 | 0 | 9 | 91.529915 | 842.9248 | 737.4652 | 989.10693 | 1061.539 | 857.7923 |
| Zbtb8os | HFLDEWLYK | -91 | -99 | 0 | 0.00078661 | 9 | 3821.311 | NaN | NaN | NaN | 7173.8965 | NaN |
| Zbtb8os | FSADEYFIPR | -100 | -109 | 0 | 0 | 10 | 573.08 | NaN | 338.37125 | NaN | NaN | NaN |
| Zbtb8os | AQEEEDVRDYNLTEEQK | -2 | -18 | 1 | 0.00272025 | 17 | 340.31796 | NaN | NaN | NaN | NaN | 336.05573 |
| Ppid | EYDQALADLKK | -322 | -332 | 1 | 0 | 11 | 2665.9026 | 575.05237 | 1616.6432 | 740.6035 | 1891.2872 | 1868.6158 |
| Ppid | FEDENFHYK | -103 | -111 | 0 | 0 | 9 | 1486.5868 | 267.25766 | 1613.5547 | 1414.7637 | 483.81622 | 1554.1367 |
| Ppid | DGSGDSHPDFPEDADIDLK | -196 | -214 | 0 | 0.000627943 | 19 | 1386.7604 | 1677.2434 | 1634.2007 | 1526.3417 | 1818.7673 | 1721.122 |
| Ppid | EGDDWGIFPK | -186 | -195 | 0 | 0 | 10 | 1166.6096 | 1344.1431 | 1179.2745 | 1184.4929 | 1238.5144 | 1422.9845 |
| Ppid | AQEIAPGDK | -333 | -341 | 0 | 0 | 9 | 911.08844 | 721.3451 | 790.5291 | 676.61615 | 733.73285 | 893.23395 |
| Ppid | LCVIAECGELK | -175 | -185 | 0 | 0 | 11 | 1243.5969 | 1109.4307 | 1153.6526 | NaN | 752.36053 | 659.0868 |
| Ppid | KFEDENFHYK | -102 | -111 | 1 | 0 | 10 | 1056.0039 | NaN | 842.88153 | 290.85416 | 926.92975 | 232.79651 |
| Ppid | KFEDENFHYK | -102 | -111 | 1 | 0 | 10 | 1056.0039 | NaN | 842.88153 | 290.85416 | 926.92975 | 232.79651 |
| Ppid | PTPHLDGK | -138 | -145 | 0 | 0 | 8 | 509.50314 | 1384.9199 | 1455.2993 | 753.8683 | 530.53015 | NaN |
| Ppid | ILLISEDLK | -219 | -227 | 0 | 0.000230707 | 9 | 1359.2449 | 1345.7904 | 1224.4244 | NaN | NaN | NaN |
| Ppid | FDVDIGGER | -20 | -28 | 0 | 0.00025015 | 9 | 1210.9546 | NaN | 747.949 | NaN | NaN | 857.67255 |
| Ppid | ENVEVNGEKPAK | -163 | -174 | 1 | 0.00025015 | 12 | 769.8057 | NaN | NaN | NaN | 867.42523 | 942.2308 |
| Echs1 | AQFGQPEILLGTIPGAGGTQR | -158 | -178 | 0 | 0 | 21 | 479.4289 | 559.7833 | 640.5728 | 623.8066 | 980.9522 | 844.59485 |
| Echs1 | FLSHWDHITR | -116 | -125 | 0 | 0 | 10 | 831.4517 | NaN | 19898.924 | 609.48535 | 828.4708 | 659.65405 |
| Echs1 | TFQDCYSSK | -107 | -115 | 0 | 0.007987481 | 9 | 461.32822 | NaN | NaN | NaN | NaN | NaN |
| Atp5f1a | GIRPAINVGL | -403 | -412 | 1 | 0.001150701 | 10 | 11727.058 | 12196.599 | 11677.407 | 9881.253 | 8245.135 | 9581.154 |
| Atp5f1a | AVDSLVPIGR | -195 | -204 | 0 | 0 | 10 | 6606.205 | 6255.9023 | 6530.469 | 6431.775 | 5031.1313 | 5326.887 |
| Atp5f1a | HALIIYDDLSK | -306 | -316 | 0 | 0 | 11 | 4945 | 5788.5645 | 5830.3145 | 5919.748 | 7848.5483 | 7785.3887 |
| Atp5f1a | HALIIYDDLSK | -306 | -316 | 0 | 0 | 11 | 4945 | 5788.5645 | 5830.3145 | 5919.748 | 7848.5483 | 7785.3887 |
| Atp5f1a | HQSLLGNIR | -519 | -527 | 0 | 0 | 9 | 4289.83 | 4602.4517 | 5008.3135 | 3965.9795 | 4249.363 | 4396.3496 |
| Atp5f1a | TGAIVDVPVGEELLGR | -134 | -149 | 0 | 0 | 16 | 4016.4595 | 4516.69 | 4284.4487 | 4466.6377 | 4932.908 | 5397.402 |
| Atp5f1a | RELIIGDR | -207 | -214 | 1 | 0.000826806 | 8 | 3688.5437 | 3791.1177 | 4062.7212 | 4500.583 | 5761.946 | 5521.0073 |
| Atp5f1a | EAYPGDVFYLH | -335 | -345 | 0 | 0 | 11 | 3407.2886 | 4016.9053 | 3970.4583 | 2901.4695 | 2207.7786 | 2425.7478 |
| Atp5f1a | EAYPGDVFYLH | -335 | -345 | 0 | 0 | 11 | 3407.2886 | 4016.9053 | 3970.4583 | 2901.4695 | 2207.7786 | 2425.7478 |
| Atp5f1a | GYLDKLEPSK | -494 | -503 | 1 | 0 | 10 | 2677.4387 | 2945.1848 | 2982.0166 | 2697.2239 | 750.24884 | 3079.7532 |
| Atp5f1a | IAIDTIINQK | -221 | -230 | 0 | 0 | 10 | 2168.3545 | 2260.6028 | 2144.3022 | 2754.062 | 2897.162 | 2489.5122 |
| Atp5f1a | TSVDLEETGR | -64 | -73 | 0 | 0 | 10 | 1750.1775 | 1776.0587 | 3263.7756 | 2246.1914 | 1669.8375 | 1893.9989 |
| Atp5f1a | LELAQYR | -435 | -441 | 0 | 0 | 7 | 1515.8778 | 1274.6279 | 897.7101 | 708.70056 | 589.56146 | 759.09186 |
| Atp5f1a | HVISQHQSLLGNIR | -514 | -527 | 0 | 0 | 14 | 1215.7352 | 1059.326 | 685.1529 | 1336.0831 | 1759.9973 | 1499.4081 |
| Atp5f1a | HVISQHQSLLGNIR | -514 | -527 | 0 | 0 | 14 | 1215.7352 | 1059.326 | 685.1529 | 1336.0831 | 1759.9973 | 1499.4081 |
| Atp5f1a | VREPMQTGIK | -185 | -194 | 1 | 0 | 10 | 1120.2573 | 943.5452 | 621.1098 | 1094.2605 | 1142.8435 | 1145.8888 |
| Atp5f1a | VREPMQTGIK | -185 | -194 | 1 | 0 | 10 | 1120.2573 | 943.5452 | 621.1098 | 1094.2605 | 1142.8435 | 1145.8888 |
| Atp5f1a | ILGADTSVDLEETGR | -59 | -73 | 0 | 0 | 15 | 1103.847 | 1306.0421 | 1215.8668 | 858.35583 | 908.0191 | 954.1727 |
| Atp5f1a | GAIVDVPVGEELLGR | -135 | -149 | 0 | 0 | 15 | 1013.8756 | 789.7379 | 1579.5402 | 359.90268 | 1856.7999 | 1635.957 |
| Atp5f1a | FENAFLSHVISQHQ | -507 | -520 | 0 | 0.000431655 | 14 | 988.7194 | 1031.1995 | 1246.2745 | 2337.4285 | 1618.5392 | 2630.0386 |
| Atp5f1a | LDKLEPSK | -496 | -503 | 1 | 0.00025015 | 8 | 695.2058 | 704.43726 | 673.7962 | 1074.0773 | 1157.134 | 1280.9637 |
| Atp5f1a | VISQHQSLLGNIR | -515 | -527 | 0 | 0 | 13 | 586.1233 | 506.8795 | 485.95715 | 503.02844 | 460.6726 | 117.03196 |
| Atp5f1a | VISQHQSLLGNIR | -515 | -527 | 0 | 0 | 13 | 586.1233 | 506.8795 | 485.95715 | 503.02844 | 460.6726 | 117.03196 |
| Atp5f1a | FENAFLSHVISQ | -507 | -518 | 0 | 0.00025015 | 12 | 558.6602 | 705.68524 | 1193.6991 | 906.3718 | 817.9491 | 262.16 |
| Atp5f1a | AQFGSDLDAATQQLLSR | -447 | -463 | 0 | 0 | 17 | 349.4988 | 317.86896 | 333.87717 | 701.35785 | 846.53815 | 733.5809 |
| Atp5f1a | FENAFLSH | -507 | -514 | 0 | 0.000591124 | 8 | 210.40076 | 645.36426 | 711.6419 | 479.94128 | 749.59357 | 539.2541 |
| Atp5f1a | LIIYDDLSK | -308 | -316 | 0 | 0 | 9 | 2131.11 | 2458.6152 | NaN | 2576.1323 | 2639.8503 | 3013.681 |
| Atp5f1a | LKEIVTNF | -540 | -547 | 1 | 0.00025015 | 8 | 1600.7085 | 1887.3118 | NaN | 1200.508 | 1047.4513 | 1097.751 |
| Atp5f1a | QSLLGNIR | -520 | -527 | 0 | 0.000230707 | 8 | 1122.2324 | 1530.0936 | 1219.584 | 952.29315 | 1404.2069 | NaN |
| Atp5f1a | SMGEYFR | -295 | -301 | 0 | 0 | 7 | 804.1544 | 778.22406 | 782.4959 | 729.7363 | 629.6236 | NaN |
| Atp5f1a | RFNDGTDEK | -231 | -239 | 1 | 0.000431655 | 9 | 737.83954 | NaN | 854.7071 | 388.5795 | 818.73334 | 893.0275 |
| Atp5f1a | QHQSLLGNIR | -518 | -527 | 0 | 0 | 10 | 678.6421 | 648.3081 | 862.44653 | 642.76776 | 1178.4528 | NaN |
| Atp5f1a | SLNLEPDNVGVVVFGNDK | -106 | -123 | 0 | 0 | 18 | 323.99326 | 331.62784 | 535.8567 | NaN | 393.8062 | 372.81232 |
| Atp5f1a | SIAIDTIINQK | -220 | -230 | 0 | 0 | 11 | 296.60095 | 225.8269 | NaN | 460.17017 | 473.26654 | 530.5863 |
| Atp5f1a | VVDALGNAIDGK | -150 | -161 | 0 | 0 | 12 | 1268.3083 | 1204.6146 | 1117.1616 | NaN | 1211.9329 | NaN |
| Atp5f1a | TSIAIDTIINQK | -219 | -230 | 0 | 0 | 12 | 543.9119 | 757.9851 | 247.21094 | 462.72498 | NaN | NaN |
| Atp5f1a | VLSIGDGIAR | -74 | -83 | 0 | 0.00025015 | 10 | 3564.1943 | 3278.4683 | 3566.059 | NaN | NaN | NaN |
| Atp5f1a | INVGLSVSR | -408 | -416 | 0 | 0 | 9 | 941.7451 | 975.21716 | 1066.8629 | NaN | NaN | NaN |
| Atp5f1a | FNDGTDEKK | -232 | -240 | 1 | 0.008129024 | 9 | 803.7513 | 838.7361 | NaN | NaN | NaN | 763.71356 |
| Atp5f1a | VDSLVPIGR | -196 | -204 | 0 | 0.00025015 | 9 | 1368.4637 | NaN | NaN | 946.7233 | NaN | NaN |
| Atp5f1a | IDTIINQK | -223 | -230 | 0 | 0.00111006 | 8 | 879.1261 | NaN | NaN | 556.23315 | NaN | NaN |
| Atp5f1a | RTGAIVDVPVGEELLGR | -133 | -149 | 1 | 0 | 17 | 835.7006 | NaN | NaN | 670.4429 | NaN | NaN |
| Bin1 | TFQSIAGLEENFHK | -241 | -254 | 0 | 0 | 14 | 922.46045 | 1011.1467 | 1035.0626 | 71.12849 | 904.3723 | 1153.7673 |
| Bin1 | AQHDYTATDTDELQLK | -522 | -537 | 0 | 0 | 16 | 498.60904 | 472.19342 | 375.3858 | 536.7985 | 559.1869 | 338.24802 |
| Bin1 | GVFPENFTER | -577 | -586 | 0 | 0 | 10 | 63.716904 | 727.94867 | 585.2886 | 708.1348 | 793.08826 | 1040.395 |
| Rab11b | SALDSTNVEEAFK | -154 | -166 | 0 | 0 | 13 | 993.1502 | 835.33264 | 866.51526 | 841.28815 | 444.7189 | 563.9712 |
| Rab11b | NILTEIYR | -167 | -174 | 0 | 0 | 8 | 732.5425 | 642.14 | 1003.84644 | NaN | 834.2344 | 866.64264 |
| Trap1 | AQLLQPTLEINPR | -637 | -649 | 0 | 0 | 13 | 998.0724 | 809.1856 | 965.33496 | 1044.1394 | 1201.4718 | 1219.2068 |
| Trap1 | DISEFQHEEFYR | -307 | -318 | 0 | 0 | 12 | 963.2792 | 1119.2262 | 1283.1887 | 484.61838 | 1538.6971 | 1501.7432 |
| Trap1 | ELISNASDALEK | -117 | -128 | 0 | 0 | 12 | 897.11316 | 667.7632 | 968.1953 | 1362.3416 | 419.10635 | 1112.5894 |
| Trap1 | KLLDIVAR | -98 | -105 | 1 | 0 | 8 | 605.7021 | 589.8407 | 649.724 | 717.3276 | 908.7412 | 786.02136 |
| Trap1 | KFEDTSPADER | -565 | -575 | 1 | 0 | 11 | 1925.5477 | 2114.5283 | NaN | NaN | 2396.936 | 2541.3486 |
| Trap1 | TDAPLNIR | -335 | -342 | 0 | 0.00111006 | 8 | 869.2764 | NaN | NaN | 746.0957 | NaN | NaN |

| | | | | | | | | | | | | |
|---|---|---|---|---|---|---|---|---|---|---|---|---|
| Trap1 | TAEQDIKEDIAK | -457 | -468 | 1 | 0 | 12 | 2246.6526 | NaN | NaN | NaN | NaN | NaN |
| Trap1 | HLAEHSPYYEA | -508 | -518 | 0 | 0.00111006 | 11 | 1572.3268 | NaN | NaN | NaN | NaN | NaN |
| Pola2 | AQLPVTPDVFIVPSELR | -529 | -545 | 0 | 0 | 17 | 312.84814 | 368.8174 | 349.42548 | 405.1828 | 375.41318 | 362.75604 |
| Pola2 | VVGDPEPLTGSYK | -195 | -207 | 0 | 0 | 13 | 1154.5114 | 940.6677 | NaN | 740.617 | NaN | NaN |
| Sfxn3 | NLLLSGEQLEASR | -35 | -47 | 0 | 0 | 13 | 1372.23 | 1401.7079 | 474.82895 | 791.0059 | 2028.8477 | 1487.5979 |
| Sfxn3 | ELQVGIPVTDEAGQR | -199 | -213 | 0 | 0 | 15 | 1058.8973 | 1163.4272 | 1156.4856 | 1277.3529 | 1274.9556 | 1297.061 |
| Sfxn3 | AQNPSIDVVYYNK | -307 | -319 | 0 | 0 | 13 | 1637.1011 | NaN | 1840.2554 | 1997.5496 | 2072.0398 | 1863.4346 |
| Sfxn3 | GDLPLNINIQEPR | -2 | -14 | 0 | 0.00025015 | 13 | 843.0082 | NaN | 759.1689 | 731.41125 | 984.3201 | 331.65613 |
| Adi1 | LDADKYENDPELEK | -41 | -54 | 1 | 0 | 14 | 542.86414 | 491.14026 | 443.10065 | 444.79352 | 506.96606 | 584.5844 |
| Adi1 | DADKYENDPELEK | -42 | -54 | 1 | 0.00025015 | 13 | 1539.5782 | 1850.8617 | 1058.4692 | 1885.9923 | NaN | 1921.0453 |
| Adi1 | AQPDRPVSLEQLR | -20 | -32 | 1 | 0 | 13 | 200.23734 | 426.4917 | 272.75922 | 442.4429 | NaN | NaN |
| Sp110 | REVPCTPANK | -236 | -245 | 1 | 0.00025015 | 10 | 665.63684 | 1348.2858 | 521.0292 | 840.7002 | 1379.8549 | 1704.0074 |
| Sp110 | FSPTLPVTCGK | -358 | -368 | 0 | 0 | 11 | 642.6726 | 455.9551 | 632.0632 | 634.3142 | 161.892 | 573.4259 |
| Sp110 | ATAQPIIEILDEQPSPSPR | -161 | -179 | 0 | 0 | 19 | 592.78687 | 884.9249 | 629.0582 | 585.7374 | 820.1177 | 775.4523 |
| Sp110 | EYPSLVAIFR | -92 | -101 | 0 | 0 | 10 | 552.974 | 412.93054 | 277.95087 | 874.2812 | 1087.6603 | 1035.0803 |
| Sp110 | AQPIIEILDEQPSPSPR | -163 | -179 | 0 | 0 | 17 | 523.5128 | 683.43427 | 685.1785 | 971.207 | 1046.2882 | 1046.6105 |
| Sp110 | TKNDAVDFSPTLPVTC | -351 | -366 | 1 | 0 | 16 | 516.3109 | 784.58105 | 667.9557 | 642.5201 | 759.07544 | 740.2418 |
| Sp110 | CIQNEAGDWLTVK | -386 | -398 | 0 | 0 | 13 | 746.56213 | 1489.9834 | 1285.0157 | 1063.39 | NaN | 466.97693 |
| Sp110 | AVPLLGCIQEGK | -180 | -191 | 0 | 0 | 12 | 531.3228 | NaN | NaN | 994.68823 | 1449.5896 | NaN |
| Sp110 | NDAVDFSPTLPVTCGK | -353 | -368 | 0 | 0 | 16 | 1072.0579 | 1349.5468 | NaN | NaN | NaN | NaN |
| Sp110 | VASPGHGVQEK | -275 | -285 | 0 | 0.000627943 | 11 | 436.68365 | NaN | NaN | NaN | NaN | NaN |
| Kdm1a | AQPITPGPSIPGAPQPIPR | -778 | -796 | 0 | 0 | 19 | 374.47226 | 350.52948 | 410.4735 | 690.5263 | 493.54443 | 490.8393 |
| Kdm1a | FVPPLPEWK | -640 | -648 | 0 | 0 | 9 | 1722.6077 | 1790.5278 | NaN | 1650.4728 | NaN | 2220.5642 |
| Rbm14 | KPGAGDTAFPGTGGFSA | -167 | -183 | 1 | 0.000591124 | 17 | 683.5505 | 874.67505 | 1352.9961 | 768.47003 | 799.32996 | 658.0417 |
| Rbm14 | KPGAGDTAFPGTGGFSAT | -167 | -184 | 1 | 0.000712251 | 18 | 520.84314 | 1151.6534 | 1570.6283 | 2788.7515 | 297.8 | 1482.74 |
| Rbm14 | QPTPPFFGR | -204 | -212 | 0 | 0 | 9 | 939.8036 | 1380.4078 | NaN | 2325.7085 | 2575.1245 | 2272.897 |
| Rbm14 | ASYDDPYKK | -586 | -594 | 1 | 0.00025015 | 9 | 756.9878 | 839.23865 | 464.89587 | NaN | 713.5808 | NaN |
| Rbm14 | LSESQLSFR | -617 | -625 | 0 | 0 | 9 | 813.2444 | NaN | 771.8539 | NaN | NaN | NaN |
| Rbm14 | AQPSVSLGAPYR | -275 | -286 | 0 | 0.000230707 | 12 | 1807.8 | NaN | NaN | NaN | NaN | NaN |
| Fkbp8 | AQQGEYSEAIPILR | -307 | -320 | 0 | 0 | 14 | 631.0187 | 810.78674 | 908.0597 | 1111.3585 | 1131.7422 | 1223.0704 |
| Fkbp8 | SCSQVLEHQPDNIK | -284 | -297 | 0 | 0 | 14 | 566.12805 | 648.13275 | 708.6218 | 603.4575 | 705.9914 | 608.01996 |
| Fkbp8 | LKLEPSNK | -323 | -330 | 1 | 0.002068213 | 8 | 370.9983 | NaN | 261.8906 | NaN | 823.8178 | 505.8185 |
| Fkbp8 | ASWAEPSEPAALR | -2 | -14 | 0 | 0 | 13 | 1115.27 | 1160.831 | 1113.8859 | NaN | NaN | NaN |
| Stat5a | AVDGYVKPQIK | -690 | -700 | 1 | 0 | 11 | 1124.9995 | 1416.7032 | 1376.9326 | 1844.5878 | 1632.7827 | 1599.1044 |
| Stat5a | NLWNLKPF | -639 | -646 | 1 | 0.001636916 | 8 | 722.7089 | 176.36047 | 927.1042 | 588.54956 | 905.278 | 1039.1647 |
| Stat5a | AQQLQGDALR | -7 | -16 | 0 | 0.00122419 | 10 | 798.9124 | 768.93835 | 772.40454 | 909.2437 | NaN | NaN |
| Stat5a | YYTPVLAK | -682 | -689 | 0 | 0 | 8 | 782.39844 | NaN | NaN | 353.24612 | NaN | NaN |
| Atp5f1d | AQSELSGAADEAAR | -137 | -150 | 0 | 0 | 14 | 6671.5713 | 6508.6904 | 6752.1113 | 6905.516 | 1710.5688 | 7940.0786 |
| Atp5f1d | IEANEALVK | -157 | -165 | 0 | 0 | 9 | 1685.3639 | 1830.5099 | 1856.1093 | 1594.9524 | 1509.5371 | 1352.8695 |
| Atp5f1d | PGLVVVHTEDGTTTK | -82 | -96 | 0 | 0 | 15 | 1832.7734 | 1903.3918 | NaN | 1562.8494 | 1818.4617 | 1641.6827 |
| Atp5f1d | ASPTQVFFDSANVK | -43 | -56 | 0 | 0 | 14 | 685.78595 | 2034.4966 | 3164.772 | NaN | 3167.896 | 852.3052 |
| Atp5f1d | ASHVPTLQVLR | -71 | -81 | 0 | 0 | 11 | 338.17606 | 749.8134 | 693.13477 | 635.7374 | 729.96 | NaN |
| Atp5f1d | QVFFDSANVK | -47 | -56 | 0 | 0.000230707 | 10 | 428.9076 | 896.1508 | NaN | NaN | 916.72363 | 862.5496 |
| Atp5f1d | QVDVPTLTGAF | -57 | -67 | 0 | 0 | 11 | 337.13434 | NaN | NaN | 460.16306 | 558.0101 | 574.9501 |
| Atp5f1d | DMLDLGAAR | -123 | -131 | 0 | 0.00272025 | 9 | 1697.6405 | NaN | NaN | 1435.5844 | 1520.1829 | NaN |
| Rap1b | VKDTDDVPMILVGN | -103 | -116 | 1 | 0 | 14 | 859.8952 | 1059.7871 | NaN | 1080.1869 | 907.7678 | 830.1448 |
| Rap1b | QWNNCAFLESSAK | -137 | -149 | 0 | 0 | 13 | 802.373 | NaN | NaN | 1229.01 | 1106.7457 | 1092.1813 |
| Rpl12 | SVGCNVDGR | -138 | -146 | 0 | 0 | 9 | 2476.7478 | 2748.759 | 3456.1638 | 2423.7627 | 3244.0212 | 2473.5186 |
| Rpl12 | HPHDIIDDINSGAVECPAS | -147 | -165 | 0 | 0 | 19 | 1703.4288 | 2084.6987 | 2072.1025 | 3264.641 | 3227.1257 | 3250.271 |
| Rpl12 | HPHDIIDDINSGAVECPAS | -147 | -165 | 0 | 0 | 19 | 1703.4288 | 2084.6987 | 2072.1025 | 3264.641 | 3227.1257 | 3250.271 |
| Rpl12 | KVGDDIAK | -41 | -48 | 1 | 0.000431655 | 8 | 1533.014 | 1246.0869 | 1709.0802 | 1949.6171 | 1970.1165 | 332.34354 |
| Rpl12 | AQSVGCNVDGR | -136 | -146 | 0 | 0 | 11 | 801.0763 | 1366.2886 | 1008.456 | 315.32562 | 1374.4786 | 1314.0881 |
| Rpl12 | QSVGCNVDGR | -137 | -146 | 0 | 0 | 10 | 1932.5781 | 548.312 | 1544.8694 | NaN | 93.150116 | 921.62427 |
| Rpl12 | IGPLGLSPK | -32 | -40 | 0 | 0.00078661 | 9 | 1751.2717 | 1412.6733 | 1372.4329 | NaN | NaN | NaN |
| Grwd1 | AQTGAPCLSFDIVR | -60 | -73 | 0 | 0 | 14 | 1579.9327 | 1185.3367 | 1256.1376 | 2237.9497 | 3320.994 | 3451.507 |
| Grwd1 | NVDQRPFVGHTR | -251 | -262 | 1 | 0 | 12 | 551.5513 | 681.7611 | 186.2723 | 428.50714 | 585.98267 | 258.85995 |
| Grwd1 | NVHLWTPTEGGSW | -238 | -250 | 0 | 0 | 13 | 630.9173 | 699.81445 | 942.17676 | NaN | 584.89465 | 964.154 |
| Grwd1 | EPFLLSGGDDGALK | -320 | -333 | 0 | 0.00025015 | 14 | 1470.9325 | NaN | NaN | 1549.1523 | NaN | NaN |
| Grwd1 | TCDPGTEGPSQVYLPGR | -21 | -37 | 0 | 0 | 17 | 1371.7607 | NaN | NaN | NaN | NaN | NaN |
| Stat3 | VVENLQDDFDFNYK | -164 | -177 | 0 | 0 | 14 | 1167.2844 | 1198.5736 | 942.9502 | 1541.1427 | 2006.1445 | 1701.4858 |
| Stat3 | TLTDEELADWK | -234 | -244 | 0 | 0 | 11 | 1077.3306 | 1171.7717 | 1148.7313 | 150.3138 | 1148.0372 | 953.9585 |
| Stat3 | LLGEIDQQYSR | -60 | -70 | 0 | 0 | 11 | 768.9234 | 1541.5697 | 1535.0548 | 2365.6255 | 2847.907 | 2832.4783 |
| Stat3 | AQWNQLQQLDTR | -2 | -13 | 0 | 0 | 12 | 2797.3481 | 3324.3293 | 3732.9385 | 3483.5945 | NaN | NaN |
| Stat3 | FLQESNVLYQHNLR | -71 | -84 | 0 | 0 | 14 | 619.07404 | NaN | NaN | 721.5336 | 780.0021 | 721.2836 |
| Stat3 | LYPDIPKEEAFGK | -673 | -685 | 1 | 0 | 13 | 1015.1289 | NaN | NaN | NaN | 1133.7748 | 1426.4463 |
| Stat3 | IDLETHSLPVVVIS | -452 | -465 | 0 | 0.001055699 | 14 | 644.2522 | NaN | NaN | NaN | 963.77435 | 987.30695 |
| Stat3 | YLEQLHQL | -14 | -21 | 0 | 0.00025015 | 8 | 536.0304 | NaN | NaN | 396.33752 | 616.55273 | NaN |
| Stat3 | SLAESQLQTR | -269 | -278 | 0 | 0 | 10 | 480.83377 | NaN | NaN | 966.30273 | NaN | NaN |
| Por | SYENQKPPFDAK | -268 | -279 | 1 | 0 | 12 | 755.54156 | 839.0618 | 844.60767 | 903.52747 | 869.5186 | 867.07825 |
| Por | AQYASEPSEQEHLHK | -390 | -404 | 0 | 0 | 15 | 1232.9747 | NaN | 1639.1504 | NaN | NaN | NaN |
| Por | AQYASEPSEQEHLHK | -390 | -404 | 0 | 0 | 15 | 1232.9747 | NaN | 1639.1504 | NaN | NaN | NaN |
| Por | SDEDYLYREE | -569 | -578 | 1 | 0.007224385 | 10 | 628.0827 | NaN | NaN | NaN | NaN | NaN |
| Rpl35a | DETEFYLGK | -37 | -45 | 0 | 0 | 9 | 3002.525 | 2953.6665 | 3132.3323 | 2740.1655 | 893.6291 | 3191.6733 |
| Rpl35a | NNTVTPGGKPNK | -55 | -66 | 1 | 0 | 12 | 436.26947 | 459.0042 | 485.93652 | 329.9092 | 447.1486 | 498.63965 |
| Rpl35a | ARDETEFYLGK | -35 | -45 | 1 | 0 | 11 | 1023.9832 | NaN | 1250.1047 | 1305.5975 | 650.6894 | 1429.0728 |
| Rpl35a | EHTALLK | -23 | -29 | 0 | 0.000431655 | 7 | 1379.1246 | 1394.3032 | 1359.5721 | NaN | NaN | NaN |
| Hspa8 | SQIHDIVLVGGSTR | -329 | -342 | 0 | 0 | 14 | 12902.929 | 13864.277 | 14678.162 | 11376.966 | 13433.8955 | 13668.895 |
| Hspa8 | SQIHDIVLVGGSTR | -329 | -342 | 0 | 0 | 14 | 12902.929 | 13864.277 | 14678.162 | 11376.966 | 13433.8955 | 13668.895 |

| | | | | | | | | | | | | |
|---|---|---|---|---|---|---|---|---|---|---|---|---|
| Hspa8 | VCNPIITK | -602 | -609 | 0 | 0 | 8 | 9695.131 | 9766.602 | 9717.391 | 10475.49 | 9536.845 | 9344.972 |
| Hspa8 | VVTVPAYFNDSQR | -143 | -155 | 0 | 0 | 13 | 9103.452 | 10570.26 | 10757.242 | 8331.582 | 8159.263 | 7659.0786 |
| Hspa8 | NQVAMNPTNTVFDAK | -57 | -71 | 0 | 0 | 15 | 6209.9146 | 6558.65 | 6917.741 | 7042.768 | 7509.0986 | 7602.86 |
| Hspa8 | NQVAMNPTNTVFDAK | -57 | -71 | 0 | 0 | 15 | 6209.9146 | 6558.65 | 6917.741 | 7042.768 | 7509.0986 | 7602.86 |
| Hspa8 | NQTAEKEEFEHQQK | -584 | -597 | 1 | 0 | 14 | 5282.699 | 5481.822 | 5836.527 | 5994.9907 | 6614.435 | 6430.709 |
| Hspa8 | NQTAEKEEFEHQQK | -584 | -597 | 1 | 0 | 14 | 5282.699 | 5481.822 | 5836.527 | 5994.9907 | 6614.435 | 6430.709 |
| Hspa8 | VAMNPTNTVFDAK | -59 | -71 | 0 | 0 | 13 | 5109.4395 | 10052.085 | 9642.07 | 5250.6084 | 13256.355 | 10697.785 |
| Hspa8 | VAMNPTNTVFDAK | -59 | -71 | 0 | 0 | 13 | 5109.4395 | 10052.085 | 9642.07 | 5250.6084 | 13256.355 | 10697.785 |
| Hspa8 | FELTGIPPAPR | -459 | -469 | 0 | 0 | 11 | 4353.4917 | 4463.5557 | 5192.787 | 4456.047 | 4936.985 | 5043.014 |
| Hspa8 | KNQVAMNPTNTVFDAK | -56 | -71 | 1 | 0 | 16 | 4174.8125 | 4537.4424 | 4258.228 | 4395.3135 | 4956.633 | 5046.906 |
| Hspa8 | KNQVAMNPTNTVFDAK | -56 | -71 | 1 | 0 | 16 | 4174.8125 | 4537.4424 | 4258.228 | 4395.3135 | 4956.633 | 5046.906 |
| Hspa8 | HWPFMVVN | -89 | -96 | 0 | 0.00078661 | 8 | 116.858894 | 1644.3436 | 1789.9209 | 939.94464 | 1125.5442 | 1213.6923 |
| Hspa8 | HWPFMVVN | -89 | -96 | 0 | 0.00078661 | 8 | 116.858894 | 1644.3436 | 1789.9209 | 939.94464 | 1125.5442 | 1213.6923 |
| Hspa8 | TVTNAVVTVPAYFNDSQR | -138 | -155 | 0 | 0 | 18 | 3530.4673 | 4249.2295 | 3643.3665 | 6922.074 | 8977.169 | 7500.0605 |
| Hspa8 | TVTNAVVTVPAYFNDSQR | -138 | -155 | 0 | 0 | 18 | 3530.4673 | 4249.2295 | 3643.3665 | 6922.074 | 8977.169 | 7500.0605 |
| Hspa8 | SQIHDIVLVGG | -329 | -339 | 0 | 0.00051306 | 11 | 3428.0278 | 251.01865 | 3944.123 | 4131.563 | 3779.4062 | 3908.9705 |
| Hspa8 | LDKNQTAEKEEFEHQQK | -581 | -597 | 2 | 0 | 17 | 3347.3977 | 3658.0369 | 3537.6155 | 3377.4934 | 5305.1816 | 5070.2607 |
| Hspa8 | LDKNQTAEKEEFEHQQK | -581 | -597 | 2 | 0 | 17 | 3347.3977 | 3658.0369 | 3537.6155 | 3377.4934 | 5305.1816 | 5070.2607 |
| Hspa8 | SQIHDIVLVGGST | -329 | -341 | 0 | 0 | 13 | 2945.7 | 2781.779 | 3144.2065 | 4018.0642 | 2752.853 | 2247.1294 |
| Hspa8 | MNPTNTVFDAK | -61 | -71 | 0 | 0 | 11 | 2690.4233 | 822.7649 | 3181.6047 | 2226.33 | 2768.3523 | 2563.2595 |
| Hspa8 | MNPTNTVFDAK | -61 | -71 | 0 | 0 | 11 | 2690.4233 | 822.7649 | 3181.6047 | 2226.33 | 2768.3523 | 2563.2595 |
| Hspa8 | HWPFMVVNDAGRPK | -89 | -102 | 1 | 0 | 14 | 1218.3405 | 1226.919 | 966.59827 | 957.7147 | 1102.7919 | 1112.7008 |
| Hspa8 | CNEIISWLDK | -574 | -583 | 0 | 0 | 10 | 1216.7544 | 1204.1647 | 1263.192 | 1269.5618 | 1494.2542 | 1207.9254 |
| Hspa8 | LSKEDIER | -510 | -517 | 1 | 0.001023236 | 8 | 1205.8982 | 1286.7625 | 1447.9518 | 410.0819 | 1744.547 | 1610.5361 |
| Hspa8 | PTNTVFDAK | -63 | -71 | 0 | 0 | 9 | 1199.9282 | 1235.487 | 1018.6632 | 1042.0999 | 1263.6243 | 1003.39795 |
| Hspa8 | NQTAEKEEFEHQ | -584 | -595 | 1 | 0 | 12 | 1113.1841 | 1203.2224 | 1384.9888 | 865.3667 | 1220.5342 | 1168.1311 |
| Hspa8 | ELTGIPPAPR | -460 | -469 | 0 | 0 | 10 | 840.639 | 1804.8849 | 1707.0958 | 1147.2483 | 1387.8359 | 1086.7654 |
| Hspa8 | SQIHDIVLVGGS | -329 | -340 | 0 | 0 | 12 | 687.92175 | 668.4201 | 882.15454 | 741.27124 | 819.90125 | 785.6214 |
| Hspa8 | EIAEAYLGK | -129 | -137 | 0 | 0.00025015 | 9 | 585.56934 | 468.66086 | 725.6964 | 1062.0575 | 703.8783 | 660.0026 |
| Hspa8 | NEIISWLDK | -575 | -583 | 0 | 0 | 9 | 473.76083 | 478.82196 | 537.6889 | 736.0751 | 916.44763 | 943.9392 |
| Hspa8 | AVVTVPAYFNDSQR | -142 | -155 | 0 | 0 | 14 | 426.50592 | 7567.25 | 7072.5415 | 9109.107 | 10394.78 | 10576.364 |
| Hspa8 | QTAEKEEFEHQQK | -585 | -597 | 1 | 0 | 13 | 179.8954 | 448.85437 | 425.8634 | 563.41724 | 214.74072 | 197.5429 |
| Hspa8 | RGTLDPVEK | -311 | -319 | 1 | 0.002068213 | 9 | 883.08453 | 1412.5272 | 1230.504 | NaN | 1288.2197 | 431.31207 |
| Hspa8 | QVAMNPTNTVFDAK | -58 | -71 | 0 | 0 | 14 | 844.381 | NaN | 693.5863 | 1388.752 | 1370.3928 | 1204.3457 |
| Hspa8 | TNAVVTVPAYFNDSQR | -140 | -155 | 0 | 0.001846793 | 16 | 571.33514 | 718.32416 | NaN | 835.1956 | 1164.4224 | 1046.2866 |
| Hspa8 | VQVEYKGETK | -103 | -112 | 1 | 0 | 10 | 1051.4498 | 363.6497 | 846.1931 | NaN | 1942.2499 | NaN |
| Hspa8 | EEFEHQQK | -590 | -597 | 0 | 0 | 8 | 941.9793 | 908.1322 | NaN | NaN | 202.31055 | 429.38528 |
| Hspa8 | SQIHDIVLV | -329 | -337 | 0 | 0.002780285 | 9 | 735.59607 | 689.8071 | NaN | 835.7423 | 667.8217 | NaN |
| Hspa8 | MVVNDAGRPK | -93 | -102 | 1 | 0 | 10 | 674.2286 | NaN | NaN | 271.70996 | 320.71973 | 984.496 |
| Hspa8 | RFDDAVVQ | -77 | -84 | 1 | 0.00450079 | 8 | 581.34717 | 1318.492 | 1127.8112 | 711.2721 | NaN | NaN |
| Hspa8 | SQIHDIVLVG | -329 | -338 | 0 | 0.001463897 | 10 | 2191.4573 | NaN | NaN | 2638.5435 | 1176.48 | NaN |
| Hspa8 | GTLDPVEK | -312 | -319 | 0 | 0.002466863 | 8 | 853.39105 | 864.9037 | NaN | 454.36557 | NaN | NaN |
| Hspa8 | DKSQIHDIVLVGGSTR | -327 | -342 | 1 | 0 | 16 | 334.54294 | 425.82858 | 548.8624 | NaN | NaN | NaN |
| Ndufa8 | ARPEPNPVIEGDLKPAK | -146 | -162 | 2 | 0.000230707 | 17 | 1433.473 | 1763.8312 | 1921.9619 | 1760.827 | 2337.4285 | 2376.932 |
| Ndufa8 | TDRPLPENPYHSR | -133 | -145 | 1 | 0 | 13 | 1398.4755 | 1391.7689 | 1458.7312 | 1278.503 | 1644.9801 | 1797.9169 |
| Ndufa8 | LGWVRPDLGQLSK | -115 | -127 | 1 | 0 | 13 | 692.1828 | 909.49255 | 1111.2838 | 1220.1072 | 677.6639 | 1231.9135 |
| Ndufa8 | FDQCVLDKL | -107 | -115 | 1 | 0 | 9 | 408.19684 | 489.67078 | 338.86145 | NaN | 903.9687 | 435.12466 |
| Ndufa8 | LGWVRPDLGQLS | -115 | -126 | 1 | 0.000712251 | 12 | 821.29724 | NaN | 223.85765 | NaN | 968.39185 | 889.9536 |
| Kbtbd11 | VLNLPEKPDR | -618 | -627 | 1 | 0 | 10 | 2242.7708 | 2302.6785 | 2333.982 | NaN | 2538.7073 | 2403.2083 |
| Kbtbd11 | ARPSDQVYCYNPVTDSW | -382 | -398 | 1 | 0.00025015 | 17 | 286.16855 | NaN | 278.60782 | 47.690132 | 386.12387 | 367.8531 |
| Kbtbd11 | SDHYLEVLR | -266 | -274 | 0 | 0.00025015 | 9 | 1511.7299 | NaN | 1838.2594 | NaN | NaN | 2963.0493 |
| Kbtbd11 | LLLADAYSGR | -192 | -201 | 0 | 0.001636916 | 10 | 679.1723 | NaN | NaN | NaN | NaN | NaN |
| Kif2a | FSLIDLAGNER | -452 | -462 | 0 | 0 | 11 | 369.20444 | 231.7926 | 416.7292 | NaN | NaN | 230.20613 |
| Kif2a | ARPSQLPEQSSSAQ | -117 | -130 | 1 | 0 | 14 | 754.1406 | 600.83844 | 695.9399 | NaN | NaN | NaN |
| Oas1a | ARPVILDPADPTGNVA | -296 | -311 | 1 | 0 | 16 | 2178.1023 | 2346.9197 | 2315.2288 | NaN | 3276.106 | NaN |
| Oas1a | YYDFQHQEVSK | -277 | -287 | 0 | 0.007880195 | 11 | 586.19073 | NaN | NaN | NaN | NaN | NaN |
| Ints3 | ASCLQELFK | -618 | -626 | 0 | 0.005270854 | 9 | 922.9009 | NaN | NaN | NaN | NaN | NaN |
| Ints3 | LIVDHHGTAQ | -214 | -223 | 0 | 0.007546675 | 10 | 552.4828 | NaN | NaN | NaN | NaN | NaN |
| Gapdh | LISWYDNEYGYSNR | -308 | -321 | 0 | 0 | 14 | 7044.304 | 8241.971 | 8172.819 | 2568.3652 | 3226.1677 | 3216.0334 |
| Gapdh | NIIPASTGAAK | -203 | -213 | 0 | 0 | 11 | 5847.5825 | 5430.708 | 5076.6226 | 3059.4907 | 1503.9034 | 1780.0518 |
| Gapdh | QERDPTNIK | -76 | -84 | 1 | 0.001055699 | 9 | 4840.8413 | 4839.301 | 5338.991 | 5151.496 | 5834.574 | 6186.0244 |
| Gapdh | QERDPTNIK | -76 | -84 | 1 | 0.001055699 | 9 | 4840.8413 | 4839.301 | 5338.991 | 5151.496 | 5834.574 | 6186.0244 |
| Gapdh | GAAQNIIPASTGAAK | -199 | -213 | 0 | 0 | 15 | 3304.3347 | 3176.7808 | 2680.4067 | 1737.7283 | 1201.713 | 1334.8663 |
| Gapdh | GILGYTEDQVVSC | -270 | -282 | 0 | 0 | 13 | 209.82368 | 241.50435 | 187.02977 | 244.77747 | 197.92043 | 211.94344 |
| Gapdh | GILGYTEDQVVSC | -270 | -282 | 0 | 0 | 13 | 209.82368 | 241.50435 | 187.02977 | 244.77747 | 197.92043 | 211.94344 |
| Gapdh | DAGAGIALNDNFVK | -294 | -307 | 0 | 0 | 14 | 2764.116 | 2835.3623 | 2894.1794 | 4105.2183 | 5082.8164 | 4620.827 |
| Gapdh | MFQYDSTHGK | -44 | -53 | 0 | 0 | 10 | 2590.264 | 2860.9014 | 3017.9187 | 3052.084 | 4567.6265 | 4898.0034 |
| Gapdh | MFQYDSTHGK | -44 | -53 | 0 | 0 | 10 | 2590.264 | 2860.9014 | 3017.9187 | 3052.084 | 4567.6265 | 4898.0034 |
| Gapdh | MFQYDSTHGK | -44 | -53 | 0 | 0 | 10 | 2590.264 | 2860.9014 | 3017.9187 | 3052.084 | 4567.6265 | 4898.0034 |
| Gapdh | LVINGKPITIF | -65 | -75 | 1 | 0 | 11 | 2177.182 | 722.75775 | 336.42636 | 1200.4053 | 1398.843 | 76.56158 |
| Gapdh | FQYDSTHGK | -45 | -53 | 0 | 0 | 9 | 1499.3063 | 11205.556 | 10593.182 | 7248.6553 | 5273.97 | 1775.1111 |
| Gapdh | RVIISAPSADAPMFVM | -116 | -131 | 1 | 0 | 16 | 1025.5002 | 1085.7603 | 1177.8379 | 1385.5016 | 1664.6438 | 1673.3304 |
| Gapdh | RVIISAPSADAPMFVM | -116 | -131 | 1 | 0 | 16 | 1025.5002 | 1085.7603 | 1177.8379 | 1385.5016 | 1664.6438 | 1673.3304 |
| Gapdh | RVIISAPSADAPMFVM | -116 | -131 | 1 | 0 | 16 | 1025.5002 | 1085.7603 | 1177.8379 | 1385.5016 | 1664.6438 | 1673.3304 |
| Gapdh | SHSSTFDAGAGIALNDNFVK | -288 | -307 | 0 | 0 | 20 | 829.3287 | 956.6332 | 876.76544 | 1104.0857 | 1473.7627 | 1549.955 |
| Gapdh | SHSSTFDAGAGIALNDNFVK | -288 | -307 | 0 | 0 | 20 | 829.3287 | 956.6332 | 876.76544 | 1104.0857 | 1473.7627 | 1549.955 |
| Gapdh | VIHDNFGIVEGL | -161 | -172 | 0 | 0 | 12 | 479.0798 | 7739.0938 | 7628.261 | 10204.409 | 12019.178 | 574.30334 |
| Gapdh | VIHDNFGIVEGL | -161 | -172 | 0 | 0 | 12 | 479.0798 | 7739.0938 | 7628.261 | 10204.409 | 12019.178 | 574.30334 |

| | | | | | | | | | | | |
|---|---|---|---|---|---|---|---|---|---|---|---|
| Gapdh | GILGYTEDQVVSCDFNSN | -270 | -287 | 0 | 0 | 18 | 453.288 | 504.6942 | 388.5022 | 1112.1229 | 1276.4028 | 1177.0447 |
| Gapdh | VIHDNFGIVE | -161 | -170 | 0 | 0.000989247 | 10 | 203.97641 | 16565.29 | 16810.91 | 11196.723 | 10840.856 | 11814.185 |
| Gapdh | LISWYDNEYGYSN | -308 | -320 | 0 | 0.00025015 | 13 | 889.00836 | 804.5154 | 913.786 | NaN | 453.85538 | 463.46484 |
| Gapdh | VYMFQYDSTHGK | -42 | -53 | 0 | 0 | 12 | 641.3059 | NaN | 1060.8894 | 1235.4377 | 1406.2751 | 1586.0817 |
| Gapdh | ISWYDNEYGYSNR | -309 | -321 | 0 | 0 | 13 | 1140.0674 | NaN | NaN | 905.72626 | 994.82025 | 846.1256 |
| Gapdh | RVIISAPSADAPM | -116 | -128 | 1 | 0 | 13 | 706.42395 | 486.08887 | 312.56503 | 672.8174 | NaN | NaN |
| Gapdh | TFDAGAGIALNDNFVK | -292 | -307 | 0 | 0 | 16 | 432.69986 | NaN | NaN | 211.75958 | 616.7952 | 422.82166 |
| Gapdh | RVIISAPSAD | -116 | -125 | 1 | 0.001463897 | 10 | 1125.8042 | 962.1752 | 823.2662 | NaN | NaN | NaN |
| Gapdh | RVIISAPSADAPMFV | -116 | -130 | 1 | 0.00025015 | 15 | 575.9407 | 556.77563 | 661.3541 | NaN | NaN | NaN |
| Gapdh | SWYDNEYGYSNR | -310 | -321 | 0 | 0.000431655 | 12 | 625.2752 | NaN | NaN | NaN | NaN | NaN |
| Gapdh | GILGYTEDQVVS | -270 | -281 | 0 | 0.000869338 | 12 | 166.75096 | NaN | NaN | NaN | NaN | NaN |
| NaN | ASDDLGEPDVFATAPFR | -114 | -130 | 0 | 0 | 17 | 811.3742 | 844.8457 | NaN | 935.8937 | 1495.0884 | NaN |
| NaN | LSQSNEGPLLAGR | -194 | -206 | 0 | 0 | 13 | 402.31757 | 1271.6912 | 398.8885 | NaN | NaN | NaN |
| NaN | GLPLYPDPCR | -5 | -14 | 0 | 0.001055699 | 10 | 1362.728 | NaN | NaN | 549.472 | NaN | NaN |
| Nfkb2 | GDFSPTDVHK | -274 | -283 | 0 | 0.00025015 | 10 | 1249.0514 | NaN | NaN | 870.27875 | NaN | NaN |
| Nfkb2 | ASDGSFSLPLKPVISQPIHDSK | -200 | -221 | 1 | 0 | 22 | 939.79694 | NaN | NaN | 708.66833 | NaN | NaN |
| Capza1 | EASDPQPEDVDGGLK | -104 | -118 | 0 | 0 | 15 | 2649.641 | 2748.702 | 2614.4287 | 2651.5227 | 2586.813 | 312.4598 |
| Capza1 | ASDPQPEDVDGGLK | -105 | -118 | 0 | 0 | 14 | 1548.736 | 1612.3518 | 1509.2026 | 1312.9415 | 517.699 | 1045.0674 |
| Capza1 | FITHAPPGEFNEVFNDVR | -20 | -37 | 0 | 0 | 18 | 1001.324 | 838.4196 | 1071.9093 | 1461.8608 | 1469.4167 | 1457.4978 |
| Capza1 | FITHAPPGEFNEVFNDVR | -20 | -37 | 0 | 0 | 18 | 1001.324 | 838.4196 | 1071.9093 | 1461.8608 | 1469.4167 | 1457.4978 |
| Capza1 | KEASDPQPEDVDGGLK | -103 | -118 | 1 | 0 | 16 | 969.1842 | 1530.233 | 1738.2905 | 2206.8235 | 841.88513 | 1940.7178 |
| Capza1 | ESCDSALR | -122 | -129 | 0 | 0.000230707 | 8 | 625.12305 | NaN | NaN | 464.26733 | 119.91762 | NaN |
| Capza1 | YNMDQFTPVK | -57 | -66 | 0 | 0 | 10 | 361.35208 | NaN | NaN | NaN | NaN | NaN |
| Hsp90b1 | EFEPLLNWMK | -614 | -623 | 0 | 0 | 10 | 658.43317 | 666.3192 | 694.07874 | 579.9175 | 467.42236 | 452.4672 |
| Hsp90b1 | EFEPLLNWMK | -614 | -623 | 0 | 0 | 10 | 658.43317 | 666.3192 | 694.07874 | 579.9175 | 467.42236 | 452.4672 |
| Hsp90b1 | YNDTFWK | -480 | -486 | 0 | 0.00111006 | 7 | 3272.6519 | 3300.5742 | 3480.9214 | 283.47485 | 3953.8955 | 3713.6284 |
| Hsp90b1 | EAESSPFVER | -548 | -557 | 0 | 0 | 10 | 2731.5747 | 2689.555 | 204.50833 | 3136.1423 | 3046.4373 | 2897.385 |
| Hsp90b1 | KEAESSPFVER | -547 | -557 | 1 | 0 | 11 | 2461.4492 | 351.78357 | 2808.8596 | 2760.4194 | 3458.591 | 2949.6865 |
| Hsp90b1 | SGYLLPDTK | -725 | -733 | 0 | 0 | 9 | 2268.6958 | 1907.8347 | 1977.2477 | 1464.764 | 1522.6233 | 1678.1178 |
| Hsp90b1 | LGVIEDHSNR | -494 | -503 | 0 | 0 | 10 | 2265.3318 | 4177.194 | 2261.6177 | 1692.6587 | 4310.1343 | 1570.4933 |
| Hsp90b1 | LGVIEDHSNR | -494 | -503 | 0 | 0 | 10 | 2265.3318 | 4177.194 | 2261.6177 | 1692.6587 | 4310.1343 | 1570.4933 |
| Hsp90b1 | IKEDEDDKTVMDL | -701 | -713 | 2 | 0 | 13 | 2009.8226 | 2149.3362 | 2194.5251 | 2761.2656 | 2252.96 | 1891.5702 |
| Hsp90b1 | IKEDEDDKTVMDL | -701 | -713 | 2 | 0 | 13 | 2009.8226 | 2149.3362 | 2194.5251 | 2761.2656 | 2252.96 | 1891.5702 |
| Hsp90b1 | ESDSNEFSVIADPR | -224 | -237 | 0 | 0 | 14 | 1984.5631 | 2142.103 | 2176.222 | 2839.8728 | 2856.468 | 2992.6926 |
| Hsp90b1 | GVVDSDDLPLNVSR | -435 | -448 | 0 | 0 | 14 | 1932.6985 | 2462.9668 | 2244.104 | 2968.014 | 3365.092 | 3421.0063 |
| Hsp90b1 | SILFVPTSAPR | -385 | -395 | 0 | 0 | 11 | 1691.9281 | 1633.9801 | 1505.6831 | 1661.2336 | 1475.6364 | 1488.1539 |
| Hsp90b1 | FTAEGEVTFK | -375 | -384 | 0 | 0 | 10 | 1674.7268 | 1178.3207 | 1380.3903 | 1415.0703 | 577.88403 | 1455.8772 |
| Hsp90b1 | TDDEVVQR | -44 | -51 | 0 | 0 | 8 | 1640.5422 | 916.6794 | 911.36566 | 2043.1187 | 395.79385 | 1741.191 |
| Hsp90b1 | LTDENALAGNEELTVK | -120 | -135 | 0 | 0 | 16 | 1572.1865 | 1604.1791 | 1999.9385 | 2000.9531 | 1817.2081 | 1729.7748 |
| Hsp90b1 | SGTSEFLNK | -169 | -177 | 0 | 0 | 9 | 1462.0375 | 1180.1112 | 1040.114 | 101.20745 | 926.5064 | 895.7545 |
| Hsp90b1 | GLFDEYGSK | -396 | -404 | 0 | 0.000591124 | 9 | 1268.8589 | 1227.1295 | 1189.5846 | 1052.8875 | 824.41595 | 1003.019 |
| Hsp90b1 | EEASDYLELDTIK | -253 | -265 | 0 | 0 | 13 | 1166.6613 | 1391.4397 | 1294.668 | 1346.7512 | 1553.5063 | 1558.1514 |
| Hsp90b1 | KTFEINPR | -683 | -690 | 1 | 0.001092529 | 8 | 910.569 | 988.7551 | 1072.2137 | 1041.0486 | 1189.7823 | 1242.1027 |
| Hsp90b1 | EGVKFDESEK | -594 | -603 | 1 | 0 | 10 | 850.02 | 1000.8332 | 810.2673 | 945.1756 | 1120.1696 | 770.04803 |
| Hsp90b1 | TETVEEPLEEDEAAKEE | -286 | -302 | 1 | 0.001023236 | 17 | 847.52106 | 946.1745 | 910.15247 | 1288.555 | 1553.2396 | 1471.6063 |
| Hsp90b1 | TDENALAGNEELTVK | -121 | -135 | 0 | 0 | 15 | 823.0127 | 1333.1844 | 1219.3455 | 2352.5886 | 1607.5734 | 1582.1449 |
| Hsp90b1 | EASDYLELDTIK | -254 | -265 | 0 | 0 | 12 | 793.0723 | 972.70667 | 710.002 | 1009.18555 | 1467.1058 | 1809.4512 |
| Hsp90b1 | SSHHSTDITSLDQYVER | -514 | -530 | 0 | 0 | 17 | 708.2637 | 786.83923 | 817.8255 | 3427.2153 | 960.77844 | 1002.42444 |
| Hsp90b1 | SSHHSTDITSLDQYVER | -514 | -530 | 0 | 0 | 17 | 708.2637 | 786.83923 | 817.8255 | 3427.2153 | 960.77844 | 1002.42444 |
| Hsp90b1 | HHSTDITSLDQYVER | -516 | -530 | 0 | 0 | 15 | 708.1049 | 1113.0822 | 887.3563 | 910.8344 | 1365.6478 | 1222.5563 |
| Hsp90b1 | HHSTDITSLDQYVER | -516 | -530 | 0 | 0 | 15 | 708.1049 | 1113.0822 | 887.3563 | 910.8344 | 1365.6478 | 1222.5563 |
| Hsp90b1 | TETVEEPLEEDEAAKE | -286 | -301 | 1 | 0.00025015 | 16 | 698.11584 | 791.73157 | 1471.8489 | 1866.6101 | 1527.5789 | 154.64172 |
| Hsp90b1 | ASDYLELDTIK | -255 | -265 | 0 | 0 | 11 | 360.8574 | 316.02203 | 368.46112 | 265.36374 | 244.40579 | 250.73138 |
| Hsp90b1 | ASDYLELDTIK | -255 | -265 | 0 | 0 | 11 | 360.8574 | 316.02203 | 368.46112 | 265.36374 | 244.40579 | 250.73138 |
| Hsp90b1 | KEEASDYLELDTIK | -252 | -265 | 1 | 0 | 14 | 535.59326 | 626.2178 | 362.88455 | 798.553 | 1020.5467 | 953.9286 |
| Hsp90b1 | KSGTSEFLNK | -168 | -177 | 1 | 0 | 10 | 275.4856 | 405.75287 | 1959.5583 | 1750.0493 | 500.8655 | 2044.8898 |
| Hsp90b1 | TDDEVVQREEEAIQLDGLNAS | -44 | -64 | 1 | 0.000550307 | 21 | 272.04794 | 312.5855 | 274.76883 | 301.22247 | 313.22714 | 401.0332 |
| Hsp90b1 | DENALAGNEELTVK | -122 | -135 | 0 | 0 | 14 | 2241.6226 | 1475.3228 | 2189.5903 | 2816.791 | 2377.3052 | NaN |
| Hsp90b1 | TFEINPR | -684 | -690 | 0 | 0.00025015 | 7 | 1890.1287 | 1399.1245 | 1214.435 | NaN | 736.73425 | 649.68756 |
| Hsp90b1 | LISLTDENALAGNEELTVK | -117 | -135 | 0 | 0.00025015 | 19 | 1191.958 | NaN | 1321.8739 | 1243.7891 | 1630.8612 | 1631.9661 |
| Hsp90b1 | VDSDDLPLNVSR | -437 | -448 | 0 | 0 | 12 | 1044.6052 | 783.12354 | NaN | 938.4175 | 1032.9622 | 1171.8665 |
| Hsp90b1 | TDDEVVQREEEAIQL | -44 | -58 | 1 | 0.000230707 | 15 | 309.33585 | NaN | 780.9648 | 633.2543 | 729.4189 | 641.6142 |
| Hsp90b1 | TDDEVVQREEEAIQLD | -44 | -59 | 1 | 0.005522803 | 16 | 663.4244 | 656.10754 | 724.19006 | NaN | NaN | 565.8191 |
| Hsp90b1 | SQYGWSGNMER | -650 | -660 | 0 | 0.001055699 | 11 | 466.71582 | 415.13086 | 776.5354 | NaN | NaN | 661.716 |
| Hsp90b1 | SDDLPLNVSR | -439 | -448 | 0 | 0 | 10 | 121.56587 | 4813.1265 | 5364.095 | 4096.6294 | NaN | NaN |
| Hsp90b1 | DSNEFSVIADPR | -226 | -237 | 0 | 0 | 12 | 6246.191 | 6422.785 | 7082.9854 | NaN | NaN | NaN |
| Hsp90b1 | TDDEVVQREEEAIQLDGL | -44 | -61 | 1 | 0.00149795 | 18 | 303.4064 | NaN | NaN | 372.41888 | NaN | 285.87027 |
| Hsp90b1 | EVEEDEYK | -349 | -356 | 0 | 0.001055699 | 8 | 795.37854 | NaN | NaN | 550.3771 | NaN | NaN |
| Hsp90b1 | EEEAIQLDGLNAS | -52 | -64 | 0 | 0 | 13 | 963.01086 | NaN | NaN | NaN | NaN | NaN |
| Cops6 | YTTGGPPDPSDIHVHK | -122 | -137 | 0 | 0 | 16 | 14458.128 | 15420.765 | 14892.494 | 14706.555 | 14691.992 | 13234.309 |
| Cops6 | YTTGGPPDPSDIHVHK | -122 | -137 | 0 | 0 | 16 | 14458.128 | 15420.765 | 14892.494 | 14706.555 | 14691.992 | 13234.309 |
| Cops6 | ASEAGEVPFNHEILR | -241 | -255 | 0 | 0 | 15 | 674.9403 | 415.95078 | 753.5575 | 417.28244 | 451.58115 | 603.1948 |
| Cops6 | VCEIIESPLFLK | -139 | -150 | 0 | 0 | 12 | 741.1252 | 847.19806 | 712.21063 | NaN | 774.98 | 499.26428 |
| Cops6 | HTDLPVSVF | -157 | -165 | 0 | 0 | 9 | 443.83917 | 530.75977 | 278.39322 | 370.50745 | NaN | NaN |
| Cops6 | IIIDKEYYYTK | -95 | -105 | 1 | 0 | 11 | 366.2251 | NaN | NaN | 522.43677 | 478.75214 | 489.13995 |
| Cd2bp2 | GVGDEDGEDEISVPK | -10 | -24 | 0 | 0 | 15 | 1811.887 | 1645.6089 | 3009.2788 | 2595.2302 | 3602.8076 | 3528.6409 |
| Cd2bp2 | EVAEGELETPTPTQR | -255 | -269 | 0 | 0 | 15 | 1123.7462 | 1088.2845 | 1253.878 | 1198.3734 | 1457.4692 | 1227.8375 |
| Cd2bp2 | ASEDVEGQEAATLPSEGGVR | -66 | -85 | 0 | 0 | 20 | 434.63123 | 417.55566 | NaN | 340.8078 | NaN | 584.74054 |

| | | | | | | | | | | | | |
|---|---|---|---|---|---|---|---|---|---|---|---|---|
| Cd2bp2 | KLDPPGGQFYNSK | -322 | -334 | 1 | 0.000431655 | 13 | 1172.8469 | NaN | NaN | 1112.2555 | 988.8374 | NaN |
| Cd2bp2 | DSWLDNIDWVR | -116 | -126 | 0 | 0.001023236 | 11 | 326.6188 | NaN | NaN | 111.7837 | NaN | NaN |
| Cd2bp2 | LVDPVAAAGGPGSR | -27 | -40 | 0 | 0.00025015 | 14 | 546.6394 | NaN | NaN | NaN | NaN | NaN |
| Metap2 | DVGHVPIR | -388 | -395 | 0 | 0.001743443 | 8 | 826.4088 | 693.1101 | 657.05286 | 917.4001 | 540.92596 | 644.37897 |
| Metap2 | FEHTILLRPTCK | -458 | -469 | 1 | 0 | 12 | 809.2151 | 750.96747 | 670.4601 | 841.54156 | 1248.2916 | 399.42773 |
| Metap2 | TYQVKPIR | -319 | -326 | 1 | 0 | 8 | 526.8038 | 479.81323 | 493.14526 | NaN | 485.47018 | 461.40384 |
| Metap2 | IDFGTHISGR | -250 | -259 | 0 | 0.000230707 | 10 | 1687.5464 | 1225.7303 | 1654.553 | NaN | NaN | 1289.5953 |
| Metap2 | ASEEIWNDFR | -161 | -170 | 0 | 0.000471409 | 10 | 653.4381 | NaN | NaN | NaN | NaN | NaN |
| Hddc2 | ASEYEDLENKPGR | -146 | -158 | 1 | 0 | 13 | 729.81964 | 646.088 | 702.98584 | 1085.4164 | 838.7494 | 703.32947 |
| Hddc2 | EYEDLENKPGR | -148 | -158 | 1 | 0 | 11 | 424.7363 | 1297.1399 | 1093.6643 | 1292.4216 | 1324.7207 | 1342.2833 |
| Rad23b | ASFNNPDR | -213 | -220 | 0 | 0 | 8 | 3583.6565 | 2465.2385 | 2174.6235 | 2333.7986 | 1610.3798 | 1452.5223 |
| Rad23b | KIDIDPEETVK | -14 | -24 | 1 | 0 | 11 | 992.1054 | 1259.3157 | 997.9831 | 1509.7631 | 865.2362 | 1417.4407 |
| Rad23b | ATTTTTSGGHPLEFLR | -265 | -280 | 0 | 0 | 16 | 668.2316 | 736.0502 | 736.9752 | 893.12146 | 1051.2314 | 907.2183 |
| Rad23b | QIIQQNPSLLPALLQ | -290 | -304 | 0 | 0.00025015 | 15 | 288.49704 | 424.09747 | NaN | 158.88844 | 326.8257 | 211.45154 |
| Rad23b | IDIDPEETVK | -15 | -24 | 0 | 0.001055699 | 10 | 1993.2545 | NaN | 2278.3276 | NaN | NaN | NaN |
| Rad23b | TTSGGHPLEFLR | -269 | -280 | 0 | 0 | 12 | 164.68774 | NaN | NaN | 2573.0657 | NaN | NaN |
| Rad23b | ILSDDTALK | -52 | -60 | 0 | 0.001610812 | 9 | 1739.9353 | NaN | NaN | NaN | NaN | NaN |
| Tnpo3 | IVCTPGQGLGDLR | -310 | -322 | 0 | 0 | 13 | 4317.7305 | 1389.1349 | 3422.8398 | 1389.5449 | 1288.2047 | 1475.5397 |
| Tnpo3 | ASFWLGELQR | -30 | -39 | 0 | 0 | 10 | 544.751 | 667.5426 | 756.6229 | 576.905 | 691.0278 | 393.45486 |
| Tnpo3 | DHEEDFELRK | -810 | -819 | 1 | 0 | 10 | 300.82608 | 258.08737 | 374.20853 | NaN | NaN | 349.1676 |
| Cox5b | ASGGGVPTDEEQATGLER | -31 | -48 | 0 | 0 | 18 | 1911.0392 | 2055.1392 | 2977.3958 | 1006.65955 | 4066.4934 | 773.09467 |
| Cox5b | EDPNLVPSISNK | -74 | -85 | 0 | 0 | 12 | 182.73956 | 13148.224 | 14596.173 | 2278.859 | 10871.393 | 1117.1328 |
| Cox5b | EDPNLVPSISNK | -74 | -85 | 0 | 0 | 12 | 182.73956 | 13148.224 | 14596.173 | 2278.859 | 10871.393 | 1117.1328 |
| Cox5b | SGTKEDPNLVPSISNK | -70 | -85 | 1 | 0 | 16 | 662.6439 | 709.24084 | 669.9797 | 661.1708 | 672.502 | 721.18115 |
| Exosc10 | ASGGLPQFGDEYDFYR | -49 | -64 | 0 | 0.00025015 | 16 | 374.75082 | 366.51505 | 454.42072 | 427.6126 | 647.4487 | 572.89343 |
| Exosc10 | NESLTDPAIVK | -353 | -363 | 0 | 0 | 11 | 1651.5186 | 3370.4775 | 1147.1023 | 4846.8687 | NaN | NaN |
| Exosc10 | FVKPIFTDE | -481 | -489 | 1 | 0.001392869 | 9 | 754.50525 | 767.4394 | 703.4796 | NaN | NaN | 504.57147 |
| Sptbn1 | SAASGIPYHSEVPVSLK | -2241 | -2257 | 0 | 0 | 17 | 2348.662 | 2705.0815 | 2559.446 | 3769.0486 | 3751.7568 | 3390.0027 |
| Sptbn1 | LVDTGDKFR | -1907 | -1915 | 1 | 0 | 9 | 902.0833 | 799.9782 | 773.3548 | 584.41095 | 769.6148 | 968.7984 |
| Sptbn1 | LLDPEDISVDHPDEK | -250 | -264 | 0 | 0 | 15 | 710.93164 | 1654.6799 | 1806.2775 | 1571.6696 | 1130.8763 | 1674.8418 |
| Sptbn1 | ASGIPYHSEVPVSLK | -2243 | -2257 | 0 | 0 | 15 | 672.9074 | 425.44858 | 503.89456 | 606.6862 | 855.00275 | 995.2046 |
| Sptbn1 | HLLGVEDLLQK | -559 | -569 | 0 | 0 | 11 | 493.00122 | 488.26923 | 508.45416 | 899.2077 | 542.1092 | 522.40515 |
| Sptbn1 | DVEDEILWVGER | -1493 | -1504 | 0 | 0 | 12 | 476.26498 | 357.0687 | 307.558 | 685.5118 | 868.3714 | 698.6718 |
| Sptbn1 | DTGWNELHK | -1148 | -1156 | 0 | 0 | 9 | 208.94118 | NaN | 671.0855 | 570.82684 | 631.7821 | 157.54324 |
| Sptbn1 | NEIDNYEEDYQK | -1109 | -1120 | 0 | 0 | 12 | 773.6496 | 935.71765 | 1017.8235 | NaN | 232.37117 | NaN |
| Sptbn1 | EVDDLEQWIAER | -1706 | -1717 | 0 | 0 | 12 | 591.67224 | 455.2833 | NaN | 422.84265 | 500.7803 | NaN |
| Sptbn1 | LEDLEVIQHR | -881 | -890 | 0 | 0 | 10 | 516.7575 | NaN | 470.70248 | NaN | 754.1054 | NaN |
| Sptbn1 | SALPAQSAATLPAR | -2177 | -2190 | 0 | 0.00025015 | 14 | 1711.8651 | NaN | 1820.7418 | NaN | NaN | NaN |
| Sptbn1 | IVSSNDVGHDEYSTQ | -766 | -780 | 0 | 0.004439092 | 15 | 492.2001 | NaN | 1033.8096 | NaN | NaN | NaN |
| Sptbn1 | KHEAIETDIAAY | -463 | -474 | 1 | 0.00713295 | 12 | 476.97946 | NaN | NaN | NaN | NaN | NaN |
| Ruvbl1 | LLTPANLLAK | -409 | -418 | 0 | 0 | 10 | 1786.77 | 2002.7495 | 2221.9365 | 1757.2462 | 1644.2769 | 1913.3228 |
| Ruvbl1 | GTEDITSPHGIPLDLLDR | -340 | -357 | 0 | 0 | 18 | 1159.0488 | 1407.3925 | 1177.4944 | 1312.8523 | 1527.156 | 1409.1343 |
| Ruvbl1 | GTEDITSPHGIPLDLLDR | -340 | -357 | 0 | 0 | 18 | 1159.0488 | 1407.3925 | 1177.4944 | 1312.8523 | 1527.156 | 1409.1343 |
| Ruvbl1 | TISHVIIGL | -153 | -161 | 0 | 0.007343498 | 9 | 1085.2266 | 1211.7616 | 1243.1901 | 1355.6183 | 1288.6198 | 1289.7684 |
| Ruvbl1 | EHVEEISEL | -428 | -436 | 0 | 0 | 9 | 728.2303 | 577.22327 | 332.52444 | 720.6302 | 928.98193 | 729.0309 |
| Ruvbl1 | TISHVIIGLK | -153 | -162 | 0 | 0 | 10 | 702.50867 | 752.04126 | 676.4666 | 374.52673 | 568.5201 | 434.9176 |
| Ruvbl1 | GINISEEALNHLGEIGTK | -383 | -400 | 0 | 0 | 18 | 627.0133 | 764.74713 | 756.9887 | NaN | 588.24445 | 200.59387 |
| Ruvbl1 | EDITSPHGIPLDLLDR | -342 | -357 | 0 | 0 | 16 | 620.62946 | 691.7644 | 809.6969 | NaN | 686.09973 | 598.26025 |
| Ruvbl1 | EDITSPHGIPLDLLDR | -342 | -357 | 0 | 0 | 16 | 620.62946 | 691.7644 | 809.6969 | NaN | 686.09973 | 598.26025 |
| Ruvbl1 | ASGLVGQENAR | -36 | -46 | 0 | 0 | 11 | 1339.673 | 2465.308 | 3057.3447 | NaN | NaN | 2712.7688 |
| Ruvbl1 | EACGVIVELIK | -47 | -57 | 0 | 0 | 11 | 1114.8767 | NaN | 1478.6022 | 1549.3854 | 1735.5892 | NaN |
| Ruvbl1 | AVLLAGPPGTGK | -65 | -76 | 0 | 0 | 12 | 493.03656 | 468.42566 | 738.9741 | NaN | NaN | NaN |
| Xrn2 | NSLGGDVLFVGK | -677 | -688 | 0 | 0 | 12 | 2047.6309 | 842.0383 | 1118.0145 | 1271.5471 | 838.37823 | 1248.2201 |
| Xrn2 | SLPFPFDVER | -321 | -330 | 0 | 0 | 10 | 1486.0542 | 1577.1465 | 1623.5317 | 97.5877 | 1958.9568 | 1938.0493 |
| Xrn2 | GVALLPFVDER | -646 | -656 | 0 | 0 | 11 | 379.66205 | 451.1558 | 390.7516 | 487.3308 | 540.2747 | 572.05524 |
| Xrn2 | KPATVLKPGDWEK | -776 | -788 | 2 | 0.000230707 | 13 | 158.58665 | 648.078 | 635.086 | 542.04083 | NaN | 760.69086 |
| Xrn2 | KPATVLKPGDWEK | -776 | -788 | 2 | 0.000230707 | 13 | 158.58665 | 648.078 | 635.086 | 542.04083 | NaN | 760.69086 |
| Xrn2 | ATHEPNFTIIR | -242 | -252 | 0 | 0 | 11 | 2476.6926 | 2631.5293 | 2930.7556 | 3682.2444 | 3437.2375 | NaN |
| Xrn2 | LEEVYPDLTPEENR | -662 | -675 | 0 | 0 | 14 | 606.0994 | NaN | 1781.9778 | 675.5446 | 1530.3044 | 1736.3136 |
| Xrn2 | VALLPFVDER | -647 | -656 | 0 | 0 | 10 | 745.6526 | 803.8275 | 764.4833 | NaN | NaN | NaN |
| Xrn2 | ASGNFLPPTWR | -605 | -615 | 0 | 0.001055699 | 11 | 663.9446 | NaN | NaN | 1024.3185 | NaN | 1156.103 |
| Xrn2 | LNNDPGWK | -182 | -189 | 0 | 0 | 8 | 7329.277 | 6883.346 | NaN | NaN | NaN | NaN |
| Xrn2 | LFNIVRPR | -88 | -95 | 1 | 0 | 8 | 3455.808 | 3128.5166 | NaN | NaN | NaN | NaN |
| Rpl8 | DIIHDPGR | -47 | -54 | 0 | 0 | 8 | 7553.5615 | 7314.5166 | 7383.8843 | 8179.981 | 8191.324 | 1343.1403 |
| Rpl8 | IDKPILK | -175 | -181 | 1 | 0.008129024 | 7 | 3275.2754 | 2966.7876 | 2482.7007 | 3244.0728 | 2371.8477 | 2604.7588 |
| Rpl8 | VKLPSGSK | -148 | -155 | 1 | 0.000471409 | 8 | 3132.489 | 3335.2568 | 3638.7034 | 2472.449 | 2422.02 | 2922.3926 |
| Rpl8 | YATVISHNPETK | -133 | -144 | 0 | 0 | 12 | 2026.8638 | 1471.553 | 2225.7966 | 2610.9417 | 2833.9026 | 2202.1157 |
| Rpl8 | VISHNPETK | -136 | -144 | 0 | 0.00025015 | 9 | 1502.3451 | 1416.1925 | 1671.4568 | 98.53461 | 1770.932 | 1910.3857 |
| Rpl8 | TVISHNPETK | -135 | -144 | 0 | 0.00025015 | 10 | 1127.7964 | 1116.9568 | 1298.476 | 738.00995 | 1100.911 | 927.5598 |
| Rpl8 | ASGNYATVISHNPETK | -129 | -144 | 0 | 0 | 16 | 886.4943 | 713.18744 | 849.34143 | 832.5776 | 2079.591 | 1144.0576 |
| Rpl8 | CLEEKPGDR | -115 | -123 | 1 | 0 | 9 | 352.2661 | 1481.3069 | 1612.3448 | 626.8916 | 132.68979 | 2438.313 |
| Rpl8 | GNYATVISHNPETK | -131 | -144 | 0 | 0 | 14 | 10697.462 | 9999.147 | 9937.706 | NaN | 13574.745 | 13280.37 |
| Rpl8 | LEEKPGDR | -116 | -123 | 1 | 0.003620005 | 8 | 744.4843 | NaN | 885.38495 | 660.5933 | 722.10046 | 744.76544 |
| Rpl8 | KAQLNIGNVLPVGTMPEGTIV | -93 | -113 | 1 | 0.000431655 | 21 | 490.06738 | NaN | NaN | 890.77374 | 131.33646 | 1488.4926 |
| Rpl8 | GVAMNPVEHPF | -201 | -211 | 0 | 0 | 11 | 923.29346 | NaN | NaN | NaN | NaN | NaN |
| Gpd2 | ASGPFTDSVR | -289 | -298 | 0 | 0.002068213 | 10 | 2099.405 | NaN | NaN | NaN | NaN | NaN |
| Gpd2 | ELNWSELR | -580 | -587 | 0 | 0.003360691 | 8 | 714.8452 | NaN | NaN | NaN | NaN | NaN |
| Clip1 | ASGPPSSETQEEFVDDFR | -42 | -59 | 0 | 0 | 18 | 631.0466 | 466.30637 | 645.589 | 390.8051 | 902.5362 | 935.4464 |

| | | | | | | | | | | | | |
|---|---|---|---|---|---|---|---|---|---|---|---|---|
| Clip1 | EKTETLASLEDTK | -1054 | -1066 | 1 | 0.00025015 | 13 | 1117.4846 | NaN | NaN | NaN | 1182.1919 | 1114.4996 |
| Cfl1 | WAPENAPLK | -104 | -112 | 0 | 0 | 9 | 10963.211 | 9613.517 | 10724.388 | 11337.914 | 8615.491 | 9024.37 |
| Cfl1 | KNIILEEGK | -45 | -53 | 1 | 0 | 9 | 4410.369 | 4584.866 | 5312.794 | 1378.6125 | 5259.5356 | 5113.7754 |
| Cfl1 | VGQTVDDPYTTFVK | -60 | -73 | 0 | 0 | 14 | 2520.5527 | 3125.0078 | 674.5633 | 4622.712 | 5160.6826 | 5038.7324 |
| Cfl1 | NIILEEGK | -46 | -53 | 0 | 0.00149795 | 8 | 2459.572 | 1852.1313 | 954.194 | 1306.4424 | 673.2694 | 565.73395 |
| Cfl1 | MLPDKDCR | -74 | -81 | 1 | 0.00111006 | 8 | 2057.9062 | 1988.2672 | 1930.3638 | 2483.4153 | 2747.882 | 2735.8462 |
| Cfl1 | MLPDKDCR | -74 | -81 | 1 | 0.00111006 | 8 | 2057.9062 | 1988.2672 | 1930.3638 | 2483.4153 | 2747.882 | 2735.8462 |
| Cfl1 | ASGVAVSDGVIK | -2 | -13 | 0 | 0 | 12 | 1985.6802 | 118.98779 | 2798.7505 | 1337.8601 | 2927.8896 | 312.451 |
| Cfl1 | AVISLEGKPL | -157 | -166 | 1 | 0 | 10 | 1299.1996 | 1208.02 | 970.6758 | 1462.4677 | 991.96545 | 1020.3202 |
| Cfl1 | VISLEGKPL | -158 | -166 | 1 | 0.00122419 | 9 | 1285.8868 | 1049.1503 | 933.8422 | 641.0785 | 563.2777 | 440.1797 |
| Cfl1 | EILVGDVGQTVDDPYTTFVK | -54 | -73 | 0 | 0 | 20 | 1153.8865 | 1239.1167 | 1076.44 | 2016.6348 | 2531.2083 | 2153.231 |
| Cfl1 | PENAPLK | -106 | -112 | 0 | 0 | 7 | 1043.6511 | 2070.2283 | 2015.6918 | 2194.6824 | 3525.3086 | 3268.852 |
| Cfl1 | LGGSAVISLEGKPL | -153 | -166 | 1 | 0 | 14 | 828.75006 | 986.1535 | 850.589 | 1254.4706 | 1143.5973 | 1172.4989 |
| Cfl1 | NIILEEGKEILVGD | -46 | -59 | 1 | 0 | 14 | 827.9548 | 906.90564 | 822.3768 | 199.7331 | 174.42535 | 1476.5251 |
| Cfl1 | DVGQTVDDPYTTFVK | -59 | -73 | 0 | 0 | 15 | 648.27 | 854.10504 | 969.1238 | 1003.522 | 1426.1912 | 1415.8156 |
| Cfl1 | GGSAVISLEGKPL | -154 | -166 | 1 | 0 | 13 | 501.17178 | 1167.8257 | 2330.8633 | 3119.5598 | 1267.5262 | 686.9337 |
| Cfl1 | EILVGDVGQTVDDPYTTF | -54 | -71 | 0 | 0 | 18 | 440.56714 | 564.41986 | 481.253 | 551.7848 | 635.86536 | 552.1439 |
| Cfl1 | CYEEVKDR | -139 | -146 | 1 | 0 | 8 | 186.19025 | 1487.3969 | 230.86229 | 405.1824 | 2388.7734 | 2505.9963 |
| Cfl1 | AVSDGVIK | -6 | -13 | 0 | 0.00713295 | 8 | 668.32153 | 719.44275 | 885.0229 | 979.1213 | NaN | 1007.1095 |
| Cfl1 | IFWAPENAPLK | -102 | -112 | 0 | 0 | 11 | 576.787 | 457.50714 | 1351.9448 | NaN | 647.5244 | 540.5076 |
| Cfl1 | AVLFCLSEDKK | -35 | -45 | 1 | 0 | 11 | 439.42227 | NaN | 811.93445 | 2092.6755 | 1639.8656 | 3672.7969 |
| Cfl1 | GDVGQTVDDPYTTFVK | -58 | -73 | 0 | 0.001023236 | 16 | 844.533 | NaN | NaN | 1213.3607 | 753.3215 | 1357.5763 |
| Cfl1 | VDDPYTTFVK | -64 | -73 | 0 | 0 | 10 | 552.2884 | NaN | 1339.0133 | NaN | 137.90617 | 565.0284 |
| Cfl1 | NIILEEGKEILVG | -46 | -58 | 1 | 0.00025015 | 13 | 378.3676 | NaN | NaN | 530.65424 | 589.1677 | 486.20795 |
| Cfl1 | CTLAEKL | -147 | -153 | 1 | 0.00272025 | 7 | 984.0353 | NaN | NaN | NaN | 766.0273 | NaN |
| Sfxn1 | NILLTNEQLENAR | -36 | -48 | 0 | 0 | 13 | 2366.1797 | 1851.1376 | 2348.05 | 2631.124 | 487.80945 | 481.61868 |
| Sfxn1 | ASHFFTVTDPR | -25 | -35 | 0 | 0 | 11 | 1450.8889 | 1202.5779 | 1498.5774 | NaN | 1422.9359 | 855.495 |
| Sfxn1 | GLTENELWR | -62 | -70 | 0 | 0.000550307 | 9 | 1099.8062 | 1342.5228 | 2872.0674 | NaN | 1194.7673 | 1165.1919 |
| Sfxn1 | VGIPVTDENGTR | -203 | -214 | 0 | 0 | 12 | 319.0201 | 700.56934 | NaN | 531.83673 | 302.15283 | 655.928 |
| Sfxn1 | QGIVPAGLTENELWR | -56 | -70 | 0 | 0 | 15 | 437.54324 | NaN | NaN | 576.368 | 838.06006 | 696.90765 |
| Sfxn1 | CCALFPQK | -283 | -290 | 0 | 0.000826806 | 8 | 536.97125 | NaN | NaN | NaN | NaN | NaN |
| Akr1b1 | LWCTFHDK | -79 | -86 | 0 | 0 | 8 | 4488.7207 | 4419.284 | 1570.7102 | 5859.002 | 7379.163 | 7420.296 |
| Akr1b1 | MPTLGLGTWK | -13 | -22 | 0 | 0 | 10 | 4207.584 | 1700.3267 | 3278.4084 | 2857.6724 | 1145.7322 | 2208.7886 |
| Akr1b1 | MPTLGLGTWK | -13 | -22 | 0 | 0 | 10 | 4207.584 | 1700.3267 | 3278.4084 | 2857.6724 | 1145.7322 | 2208.7886 |
| Akr1b1 | QIECHPYLTQEK | -184 | -195 | 0 | 0 | 12 | 2803.7358 | 3887.9595 | 3408.7476 | 4474.036 | 2647.4668 | 2516.9436 |
| Akr1b1 | LIEYCHSK | -196 | -203 | 0 | 0.00025015 | 8 | 1922.4545 | 1133.6635 | 1599.7944 | 1063.2548 | 881.71924 | 447.50378 |
| Akr1b1 | HKDYPFHAEV | -307 | -316 | 1 | 0 | 10 | 1396.9734 | 1552.4355 | 1439.616 | 3625.616 | 664.6103 | 2169.2808 |
| Akr1b1 | HKDYPFHAEV | -307 | -316 | 1 | 0 | 10 | 1396.9734 | 1552.4355 | 1439.616 | 3625.616 | 664.6103 | 2169.2808 |
| Akr1b1 | ASHLELNNGTK | -2 | -12 | 0 | 0 | 11 | 1303.4899 | 442.9766 | 923.64105 | 1353.9341 | 900.4019 | 999.5942 |
| Akr1b1 | NFNPLQIER | -161 | -169 | 0 | 0.000230707 | 9 | 1252.5165 | 1624.901 | 483.82425 | 673.7776 | 1359.3282 | 1368.2612 |
| Akr1b1 | RQDLFIVSK | -70 | -78 | 1 | 0.000550307 | 9 | 1191.7885 | 1189.8148 | 1382.1995 | 194.72473 | 933.65753 | 888.7845 |
| Akr1b1 | VAIDLGYR | -34 | -41 | 0 | 0 | 8 | 1139.3658 | 790.62866 | 825.7863 | 675.7119 | 263.51575 | 533.1943 |
| Akr1b1 | AYSPLGSPDRPWAKPEDPSLLEDPR | -209 | -233 | 2 | 0 | 25 | 635.1023 | 519.0342 | 611.2783 | 575.4821 | 693.7005 | 733.15454 |
| Akr1b1 | SPLGSPDRPWAKPEDPSLLEDPR | -211 | -233 | 2 | 0 | 23 | 395.55154 | 426.35294 | 465.78287 | 370.9729 | 400.98892 | 412.23425 |
| Akr1b1 | LKEQVVK | -63 | -69 | 1 | 0.001055699 | 7 | 343.6484 | 281.6591 | 440.2825 | 435.39713 | 407.8525 | 371.89954 |
| Akr1b1 | SPPGQVTEAVK | -23 | -33 | 0 | 0 | 11 | 2368.1309 | NaN | 1863.6625 | 1524.9612 | 1900.1594 | 1897.9363 |
| Akr1b1 | HIDCAQVYQNEK | -42 | -53 | 0 | 0 | 12 | 524.8214 | 307.4814 | NaN | 1160.1536 | 629.3667 | 617.7862 |
| Akr1b1 | LVDEGLVK | -148 | -155 | 0 | 0 | 8 | 612.47107 | 452.84128 | 321.05725 | 487.55533 | NaN | NaN |
| Akr1b1 | GSPDRPWAKPEDPSLLEDPR | -214 | -233 | 2 | 0.001868731 | 20 | 588.2656 | 583.914 | 570.8463 | NaN | NaN | NaN |
| Arpc2 | VFKDDDDVVIGK | -168 | -179 | 1 | 0.000230707 | 12 | 3645.966 | 3539.292 | 3915.9255 | 669.17694 | 4110.501 | 4225.222 |
| Arpc2 | RDYLHYHIK | -248 | -256 | 1 | 0.001636916 | 9 | 3390.8672 | 4432.3613 | 1517.6976 | 2146.8738 | 3732.158 | 4548.1587 |
| Arpc2 | RDYLHYHIK | -248 | -256 | 1 | 0.001636916 | 9 | 3390.8672 | 4432.3613 | 1517.6976 | 2146.8738 | 3732.158 | 4548.1587 |
| Arpc2 | TVFKDDDDVVIGK | -167 | -179 | 1 | 0 | 13 | 2580.3862 | 2391.5657 | 2556.023 | 1364.2584 | 1707.5619 | 1433.9736 |
| Arpc2 | STVFKDDDDVVIGK | -166 | -179 | 1 | 0 | 14 | 2364.4673 | 2393.5178 | 2735.701 | 2513.1152 | 2761.9924 | 2538.519 |
| Arpc2 | DNTINLIHTFR | -238 | -248 | 0 | 0 | 11 | 2224.2249 | 2330.8608 | 2409.2136 | 2226.1116 | 644.72235 | 2551.56 |
| Arpc2 | ELQAHGADELLK | -66 | -77 | 0 | 0 | 12 | 2035.2407 | 2188.1638 | 2376.558 | 3153.4167 | 3585.3853 | 3770.6318 |
| Arpc2 | DNTINLIHTF | -238 | -247 | 0 | 0 | 10 | 1876.0585 | 2209.4397 | 2203.2878 | 2661.95 | 375.79077 | 544.92224 |
| Arpc2 | ASHTAPQVLFSHR | -191 | -203 | 0 | 0 | 13 | 1702.542 | 1455.7806 | 1442.9312 | 2012.9731 | 2234.2422 | 2271.9033 |
| Arpc2 | ASHTAPQVLFSHR | -191 | -203 | 0 | 0 | 13 | 1702.542 | 1455.7806 | 1442.9312 | 2012.9731 | 2234.2422 | 2271.9033 |
| Arpc2 | YFQFQEEGK | -128 | -136 | 0 | 0 | 9 | 1658.0157 | 345.3696 | 2185.0967 | 1785.7081 | 1459.5592 | 2018.7029 |
| Arpc2 | DYLHYHIK | -249 | -256 | 0 | 0 | 8 | 1642.7318 | 1135.6658 | 1634.1921 | 1741.6241 | 1608.1862 | 2024.1279 |
| Arpc2 | YDLENLPASK | -97 | -106 | 0 | 0 | 10 | 1291.0728 | 1354.9369 | 1277.59 | 1377.3748 | 1602.2153 | 134.62659 |
| Arpc2 | LLYDLENLPASK | -95 | -106 | 0 | 0 | 12 | 1048.918 | 1309.1798 | 1296.862 | 1285.6608 | 858.6329 | 1565.1044 |
| Arpc2 | IIEETLALK | -10 | -18 | 0 | 0 | 9 | 963.93005 | 867.3664 | 897.51416 | 190.87029 | 64.170654 | 844.7961 |
| Arpc2 | FADFDGVLYHISNPNGDK | -35 | -52 | 0 | 0 | 18 | 664.93933 | 604.1712 | 588.11285 | 449.21634 | 553.25073 | 619.7241 |
| Arpc2 | LYDLENLPASK | -96 | -106 | 0 | 0 | 11 | 515.2223 | 1070.215 | 1861.334 | 1779.456 | 1096.57 | 710.1347 |
| Arpc2 | EKYFQFQEEGK | -126 | -136 | 1 | 0 | 11 | 211.99654 | 273.14514 | 219.53645 | 1225.5669 | 1244.3945 | 1220.0449 |
| Arpc2 | YFQFQEEGKEGENR | -128 | -141 | 1 | 0 | 14 | 1275.6957 | 347.3861 | NaN | 77.570595 | 410.23962 | 406.39752 |
| Arpc2 | DTDAAVGDNIGYIT | -211 | -224 | 0 | 0 | 14 | 800.17255 | 515.0877 | 976.3221 | 821.5446 | 888.398 | NaN |
| Arpc2 | DTDAAVGDNIGYITF | -211 | -225 | 0 | 0.00025015 | 15 | 330.40036 | 313.34665 | NaN | 368.64902 | 410.06226 | 392.156 |
| Arpc2 | FENAAAGNKPEAVEVT | -19 | -34 | 1 | 0 | 16 | 3428.1047 | 3600.6992 | 3688.4995 | 4541.695 | NaN | NaN |
| Arpc2 | RASHTAPQVL | -190 | -199 | 1 | 0.000471409 | 10 | 1031.2974 | 893.95795 | 1159.3618 | NaN | NaN | NaN |
| Arpc2 | ASHTAPQVLF | -191 | -200 | 0 | 0.002595029 | 10 | 2814.7917 | NaN | NaN | NaN | NaN | 1870.0754 |
| Arpc2 | GADELLKR | -71 | -78 | 1 | 0.001868731 | 8 | 1169.2651 | NaN | NaN | NaN | NaN | 931.48944 |
| Sf3a1 | EDPTPSKPVVGIIYPPPEVR | -31 | -50 | 1 | 0 | 20 | 2737.3884 | 1405.7135 | 1284.0604 | 2656.819 | 2845.8965 | 2813.498 |
| Sf3a1 | EDPTPSKPVVGIIYPPPEVR | -31 | -50 | 1 | 0 | 20 | 2737.3884 | 1405.7135 | 1284.0604 | 2656.819 | 2845.8965 | 2813.498 |
| Sf3a1 | IGEEEIQKPEEK | -486 | -497 | 1 | 0 | 12 | 1434.8494 | 1550.6934 | 1705.7664 | 1263.8197 | 1367.7415 | 1631.1196 |
| Sf3a1 | TLPLTDQVSVIK | -728 | -739 | 0 | 0 | 12 | 1038.9519 | 1081.115 | 1147.45 | 1132.7783 | 1004.60406 | 1256.3562 |

| | | | | | | | | | | | | |
|---|---|---|---|---|---|---|---|---|---|---|---|---|
| Sf3a1 | RTDIFGVEETAIGK | -471 | -484 | 1 | 0 | 14 | 927.25385 | 1187.8824 | 1211.9305 | 1090.4102 | 1353.159 | 1273.1611 |
| Sf3a1 | LQYEGIFIK | -755 | -763 | 0 | 0.00025015 | 9 | 914.6265 | 1044.1768 | 1011.7784 | 1281.8236 | 1466.1448 | 1414.9325 |
| Sf3a1 | FNFLNPNDPYHAY | -81 | -93 | 0 | 0 | 13 | 896.8153 | 871.3439 | 971.25757 | 986.2258 | 1191.5443 | 1238.0834 |
| Sf3a1 | ASKPLPPAPAPDEYLVSPITGEK | -395 | -417 | 1 | 0 | 23 | 877.9478 | 948.642 | 1020.50757 | 856.70703 | 975.10254 | 978.0956 |
| Sf3a1 | EVLDQVCYR | -238 | -246 | 0 | 0 | 9 | 847.8389 | 1283.5647 | 1622.4255 | 809.3383 | 163.66478 | 1854.7415 |
| Sf3a1 | NYQFDFLRPQ | -192 | -201 | 1 | 0.00025015 | 10 | 765.2008 | 619.1614 | 761.015 | 834.8422 | 1641.5088 | 1206.101 |
| Sf3a1 | KIGEEEIQKPEEK | -485 | -497 | 2 | 0 | 13 | 591.5746 | 669.63654 | 684.70154 | 536.5249 | 956.6655 | 793.931 |
| Sf3a1 | TDIFGVEETAIGK | -472 | -484 | 0 | 0 | 13 | 582.55927 | 783.5655 | 678.61694 | 885.32874 | 571.3982 | 1118.8013 |
| Sf3a1 | QSDDEVYAPGLDIESSLK | -448 | -465 | 0 | 0 | 18 | 499.37128 | 459.45105 | 450.33704 | 469.5857 | 506.52893 | 1020.46606 |
| Sf3a1 | VFTLPLTDQVSVIK | -726 | -739 | 0 | 0 | 14 | 325.66635 | 274.1032 | 316.37872 | 267.7797 | 402.5998 | 385.3137 |
| Sf3a1 | NGPEFEAR | -63 | -70 | 0 | 0.00025015 | 8 | 506.07037 | 682.56085 | 482.97452 | NaN | 521.211 | 484.79272 |
| Sf3a1 | NYQFDFLRPQHS | -192 | -203 | 1 | 0.00025015 | 12 | 852.5864 | 657.5946 | 563.9523 | NaN | NaN | 767.00635 |
| Sf3a1 | QPIEEEASSK | -21 | -30 | 0 | 0 | 10 | 1279.4697 | 1243.1202 | 610.729 | NaN | NaN | NaN |
| Sf3a1 | KIGEEEIQK | -485 | -493 | 1 | 0.006788754 | 9 | 1270.7172 | NaN | 1535.273 | NaN | NaN | NaN |
| Sf3a1 | HSLFNYFTK | -202 | -210 | 0 | 0 | 9 | 997.4053 | NaN | NaN | 480.9225 | NaN | NaN |
| Sf3a1 | TEDSLMPEEEFLR | -685 | -697 | 0 | 0.000550307 | 13 | 881.8848 | NaN | NaN | NaN | NaN | 793.14856 |
| Sf3a1 | EQIEAIHK | -522 | -529 | 0 | 0.008919338 | 8 | 351.89133 | 5263.066 | NaN | NaN | NaN | NaN |
| Sf3a1 | NITLQEQIEAIHK | -517 | -529 | 0 | 0 | 13 | 323.3662 | NaN | NaN | NaN | 427.36047 | NaN |
| Rpn2 | ASLDRPFTDLESA | -39 | -51 | 1 | 0.00078661 | 13 | 6804.0635 | 6860.869 | 6972.936 | 4939.6777 | 4629.598 | 3369.3572 |
| Rpn2 | LSSGYYDFSVR | -341 | -351 | 0 | 0 | 11 | 3336.4705 | 327.95938 | 182.50096 | 352.47736 | 3629.9514 | 312.48016 |
| Rpn2 | LTPTHYLTK | -23 | -31 | 0 | 0.001055699 | 9 | 2234.5762 | 2028.2812 | 1868.7382 | 78.950645 | 1588.471 | 1705.4598 |
| Rpn2 | YIANTVELR | -358 | -366 | 0 | 0 | 9 | 2100.6304 | 3589.2163 | 3944.1567 | 5214.3223 | 4219.775 | 3944.0308 |
| Rpn2 | TGQEVVFVAEPDNK | -443 | -456 | 0 | 0 | 14 | 2010.8252 | 1789.6938 | 1784.0242 | 1495.632 | 1367.9152 | 1033.7544 |
| Rpn2 | LGKEETVLATVQ | -152 | -163 | 1 | 0 | 12 | 1067.4524 | 1168.0706 | 1019.29974 | 1041.7096 | 940.6647 | 1052.1953 |
| Rpn2 | TVDKDQSIAPK | -382 | -392 | 1 | 0 | 11 | 991.65717 | 1095.6606 | 1151.6766 | 1079.1357 | 1467.4374 | 1450.4146 |
| Rpn2 | YHVPVVVVPEGSTSDTQEQAILR | -267 | -289 | 0 | 0 | 23 | 713.4959 | 790.2365 | 817.83167 | 925.2641 | 1129.882 | 1098.9912 |
| Rpn2 | NVDLSTVDKDQSIAPK | -377 | -392 | 1 | 0 | 16 | 621.3386 | 479.48102 | 622.64307 | 601.72485 | 681.22253 | 582.8046 |
| Rpn2 | VDLSTVDKDQSIAPK | -378 | -392 | 1 | 0 | 15 | 518.8316 | 553.051 | 526.3623 | 504.40353 | 780.7763 | 632.6416 |
| Rpn2 | QEIQHLFR | -524 | -531 | 0 | 0 | 8 | 461.65717 | 743.58594 | 617.6009 | 1021.3521 | 626.3374 | 548.37787 |
| Rpn2 | VDVNTGAELTPHQTFVR | -421 | -437 | 0 | 0 | 17 | 420.1376 | 511.4075 | 461.43958 | 502.46777 | 681.39764 | 537.32947 |
| Rpn2 | LVDVNTGAELTPHQTFVR | -420 | -437 | 0 | 0 | 18 | 339.59045 | 435.22437 | 501.63025 | 417.73486 | 483.11322 | 440.24744 |
| Rpn2 | FELDTSER | -461 | -468 | 0 | 0 | 8 | 1010.61237 | 988.8012 | 113.75357 | NaN | 812.0896 | 568.0499 |
| Rpn2 | SSLGVQVPDVK | -59 | -69 | 0 | 0 | 11 | 3816.6406 | 4131.509 | NaN | NaN | 3722.9688 | 3714.1821 |
| Rpn2 | LIIGDATLK | -483 | -491 | 0 | 0.000591124 | 9 | 1675.508 | 1343.0403 | 979.7023 | 1273.4503 | NaN | NaN |
| Rpn2 | DTQEQAILR | -281 | -289 | 0 | 0.00025015 | 9 | 1099.6844 | NaN | 1197.3611 | 828.8616 | 689.5677 | NaN |
| Rpn2 | KNFESLSEAF | -244 | -253 | 1 | 0.002566359 | 10 | 693.27466 | 149.22223 | NaN | 342.46014 | NaN | 1494.7438 |
| Rpn2 | KTGQEVVFVAEPDNK | -442 | -456 | 1 | 0.000471409 | 15 | 246.40353 | NaN | NaN | 366.5089 | 1191.6146 | 1557.6104 |
| Rpn2 | YHVPVVVVPEGSTSDTQEQ | -267 | -285 | 0 | 0 | 19 | 668.6647 | 1098.2416 | 1027.4524 | NaN | NaN | NaN |
| Adgre5 | LLGNTPLNLEQR | -411 | -422 | 0 | 0 | 12 | 1157.5905 | 1362.2474 | 1319.1487 | 1423.2638 | 1870.393 | 1614.2916 |
| Adgre5 | DVTTVHHGQTW | -369 | -379 | 0 | 0 | 11 | 602.64343 | 1662.8578 | 179.6314 | 378.57516 | 723.16046 | 179.52225 |
| Adgre5 | SCVCKPGFSSEK | -46 | -57 | 1 | 0.00025015 | 12 | 569.1178 | 404.1566 | 727.62195 | 491.2779 | 560.2297 | 481.6019 |
| Adgre5 | ASLEDFYGSPIPSVSLK | -424 | -440 | 0 | 0 | 17 | 508.09796 | 244.4353 | NaN | 133.48672 | 566.3931 | 546.2814 |
| Psmd1 | IFLDDPQAVSDILEK | -221 | -235 | 0 | 0 | 15 | 3804.7234 | 3995.4202 | 4597.748 | 1906.6116 | 94.70204 | 1761.1118 |
| Psmd1 | LSESYNPHVR | -640 | -649 | 0 | 0 | 10 | 980.4622 | 405.1381 | 1637.2955 | 697.0015 | 642.5115 | 1004.9011 |
| Psmd1 | ASLGVIHK | -406 | -413 | 0 | 0 | 8 | 964.16724 | 856.96796 | 732.0298 | 181.78613 | 598.759 | 275.45264 |
| Psmd1 | NVNDNSEYVETIIAK | -89 | -103 | 0 | 0 | 15 | 959.9279 | 539.78534 | 642.1417 | 474.4641 | 473.74658 | 420.84726 |
| Psmd1 | PAPLEVPK | -814 | -821 | 0 | 0.000298805 | 8 | 860.01666 | 1103.1458 | 1130.8157 | 1182.5557 | 1253.8074 | 1469.494 |
| Psmd1 | TVGTPIASVPGSTNTGTVPGSEK | -270 | -292 | 0 | 0 | 23 | 631.95135 | 705.59845 | 691.34656 | 646.4859 | 771.3866 | 780.63544 |
| Psmd1 | KKEPEPNFQLLDNPAR | -868 | -883 | 2 | 0 | 16 | 605.1715 | 795.6821 | 715.20667 | 789.44464 | 455.12442 | 857.3234 |
| Psmd1 | HLQFLIR | -338 | -344 | 0 | 0.000431655 | 7 | 478.4593 | 419.6306 | 245.82114 | 431.856 | 349.1084 | 361.242 |
| Psmd1 | QDVYDLLK | -491 | -498 | 0 | 0 | 8 | 452.67166 | 844.31976 | 914.0437 | 454.5869 | 827.5575 | 815.07715 |
| Psmd1 | TPEQCPSVVSLL | -629 | -640 | 0 | 0 | 12 | 370.868 | 566.6098 | 559.6833 | 592.8772 | 657.92773 | 695.0001 |
| Psmd1 | DNSEYVETIIAK | -92 | -103 | 0 | 0 | 12 | 1439.4769 | 1538.7771 | 1598.6157 | 1284.1569 | NaN | 1199.9485 |
| Psmd1 | RLDVFEK | -159 | -165 | 1 | 0.003490217 | 7 | 1342.4373 | 1474.8092 | 1489.8142 | NaN | 1588.7057 | 1591.5012 |
| Psmd1 | GIISLLDEEEPQLK | -7 | -20 | 0 | 0 | 14 | 188.34587 | 435.53714 | 452.25854 | NaN | 575.70715 | 549.0851 |
| Psmd1 | KDTSEDVEELVEPVAAHGPK | -915 | -934 | 1 | 0 | 20 | 1283.5857 | 1911.5688 | 1389.8828 | NaN | NaN | 858.86957 |
| Psmd1 | NASNDIVR | -467 | -474 | 0 | 0.001463897 | 8 | 235.03012 | 1229.2755 | 1135.0637 | 1042.2573 | NaN | NaN |
| Psmd1 | VSDVNDDVR | -607 | -615 | 0 | 0 | 9 | 1225.2888 | NaN | NaN | NaN | 883.7929 | NaN |
| Psmd1 | LDEEEPQLK | -12 | -20 | 0 | 0 | 9 | 916.4412 | NaN | NaN | 6706.1025 | NaN | NaN |
| Psmd1 | AYPAPLEVPK | -812 | -821 | 0 | 0.00025015 | 10 | 1491.7341 | NaN | NaN | NaN | NaN | NaN |
| Psmd1 | DTSEDVEELVEPVAAHGPK | -916 | -934 | 0 | 0.00122419 | 19 | 1448.5663 | NaN | NaN | NaN | NaN | NaN |
| Lmnb2 | QGLETEVAELR | -149 | -159 | 0 | 0 | 11 | 857.37354 | 2698.841 | 2511.401 | 566.6849 | 1193.8767 | NaN |
| Lmnb2 | SQTNWGPGESFR | -526 | -537 | 0 | 0 | 12 | 794.1595 | 328.3395 | 711.71045 | 423.63483 | NaN | 713.1855 |
| Lmnb2 | NSSDKDQSLGNWR | -466 | -478 | 1 | 0 | 13 | 769.8302 | 786.93964 | 226.94807 | 381.31467 | 396.9367 | NaN |
| Lmnb2 | LVSADGEEVAVK | -540 | -551 | 0 | 0 | 12 | 2056.882 | 2183.5557 | NaN | 2879.209 | 1998.6853 | NaN |
| Lmnb2 | DLESLFHR | -129 | -136 | 0 | 0.000471409 | 8 | 779.5311 | NaN | NaN | 744.7747 | 903.5787 | 932.7162 |
| Lmnb2 | ISEKEEVTTR | -56 | -65 | 1 | 0 | 10 | 499.44843 | 508.23175 | 631.2815 | 612.27673 | NaN | NaN |
| Lmnb2 | ASLPPHAGPATPLSPTR | -2 | -18 | 0 | 0 | 17 | 1356.444 | NaN | 1727.3586 | 1881.9764 | NaN | NaN |
| Lmnb2 | TLYESELADAR | -72 | -82 | 0 | 0 | 11 | 324.7998 | NaN | NaN | 511.0932 | NaN | NaN |
| Rad21 | TQEEPYSDIIATPGPR | -616 | -631 | 0 | 0 | 16 | 843.9252 | 1079.3625 | 1145.0266 | 972.3661 | 1195.2308 | 1139.037 |
| Rad21 | QLIPELELLPEK | -518 | -529 | 0 | 0 | 12 | 729.32623 | 424.4948 | 405.14206 | 758.04425 | 496.034 | 457.04758 |
| Rad21 | EEQQPQQQQPQPQR | -428 | -441 | 0 | 0 | 14 | 362.87216 | 276.03036 | 715.6862 | 282.89468 | 330.66406 | 348.7925 |
| Rad21 | ASNLLLEPEQSTSNLNEK | -188 | -205 | 0 | 0 | 18 | 286.8272 | 586.3567 | 530.1387 | 299.0258 | 420.90518 | 464.27716 |
| Rad21 | CLTPLVPEDLR | -392 | -402 | 0 | 0 | 11 | 854.0017 | 934.3218 | NaN | 122.47843 | 723.06354 | 827.94867 |
| Rad21 | DVIDEPIIEEPSR | -442 | -454 | 0 | 0 | 13 | 749.2275 | 351.8526 | NaN | 684.28156 | 1116.5421 | 877.93115 |
| Rad21 | SASNLLLEPEQSTSNLNEK | -187 | -205 | 0 | 0.00078661 | 19 | 681.10486 | 734.2649 | 719.3483 | NaN | 721.77136 | 815.52686 |
| Rad21 | YLLADCNEAFIK | -73 | -84 | 0 | 0 | 12 | 762.8187 | 779.953 | 324.68683 | NaN | NaN | 393.93842 |
| Rad21 | TGAESISLLELCR | -578 | -590 | 0 | 0 | 13 | 801.56055 | NaN | NaN | 1259.8488 | 1688.831 | NaN |

| | | | | | | | | | | | | |
|---|---|---|---|---|---|---|---|---|---|---|---|---|
| Rad21 | SNLLLEPEQSTSNLNEK | -189 | -205 | 0 | 0.00025015 | 17 | 357.3329 | NaN | NaN | NaN | NaN | NaN |
| Atad3a | QQQLLNEENLR | -146 | -156 | 0 | 0 | 11 | 830.8909 | 950.6835 | 844.26215 | 311.04492 | 1535.5503 | 824.3398 |
| Atad3a | LVSRPQDALEGVIL | -306 | -319 | 1 | 0.001392869 | 14 | 496.95435 | 461.62735 | 487.51443 | 485.33832 | 704.1076 | 780.73724 |
| Atad3a | TAGTLLGEGFR | -227 | -237 | 0 | 0 | 11 | 1661.2374 | 2673.9763 | 1809.7328 | 1259.1816 | 1280.4581 | NaN |
| Atad3a | SNQPEQFDWAINDR | -452 | -465 | 0 | 0.000394361 | 14 | 664.0729 | NaN | 1235.4213 | NaN | 417.70782 | NaN |
| Atad3a | ASNQPEQFDWAINDR | -451 | -465 | 0 | 0 | 15 | 194.36938 | 430.3392 | 475.63107 | NaN | NaN | NaN |
| Atad3a | LGKPSLVR | -278 | -285 | 1 | 0 | 8 | 289.54575 | 270.72186 | NaN | NaN | NaN | NaN |
| Cnot3 | ASPAPSHNSGTPAPYAQ | -357 | -373 | 0 | 0 | 17 | 1561.7941 | 1294.0205 | 1790.2384 | 1297.5531 | 1523.2682 | 1896.2065 |
| Tsc22d4 | ASPEQLAQLPSSGLPR | -361 | -376 | 0 | 0.000550307 | 16 | 449.17932 | 416.3326 | 477.40067 | NaN | 376.62375 | 428.70468 |
| Hnrnpa0 | GTLTDCVVVVNPQTK | -31 | -45 | 0 | 0 | 15 | 2596.5378 | 2423.059 | 3248.7852 | 2090.4958 | 2204.608 | 2098.0276 |
| Hnrnpa0 | SPHAVDGNTVELK | -68 | -80 | 0 | 0 | 13 | 1517.4559 | 1503.237 | 1767.472 | 1406.5026 | 1677.9941 | 1540.599 |
| Hnrnpa0 | SPHAVDGNTVELK | -68 | -80 | 0 | 0 | 13 | 1517.4559 | 1503.237 | 1767.472 | 1406.5026 | 1677.9941 | 1540.599 |
| Hnrnpa0 | ASPHAVDGNTVELK | -67 | -80 | 0 | 0 | 14 | 1404.3273 | 812.0562 | 998.521 | 834.5796 | 1019.8888 | 879.27875 |
| Hnrnpa0 | ASPHAVDGNTVELK | -67 | -80 | 0 | 0 | 14 | 1404.3273 | 812.0562 | 998.521 | 834.5796 | 1019.8888 | 879.27875 |
| Hnrnpa0 | GDVAEGDLIEHF | -107 | -118 | 0 | 0 | 12 | 1314.3309 | 1363.3535 | 1441.5085 | 1537.0273 | 1556.0334 | 1474.2424 |
| Hnrnpa0 | GDVAEGDLIEHF | -107 | -118 | 0 | 0 | 12 | 1314.3309 | 1363.3535 | 1441.5085 | 1537.0273 | 1556.0334 | 1474.2424 |
| Hnrnpa0 | FHPIQGH | -160 | -166 | 0 | 0.00142999 | 7 | 1261.3236 | 1869.348 | 1065.014 | 1752.928 | 1342.1719 | 1491.7295 |
| Hnrnpa0 | GDVAEGDLIEHFS | -107 | -119 | 0 | 0 | 13 | 1079.0527 | 1391.2418 | 1210.2399 | 790.1031 | 288.65317 | 616.22076 |
| Hnrnpa0 | EDIHAGGGGAR | -177 | -187 | 0 | 0 | 11 | 946.8074 | 697.8421 | 1027.6006 | 728.78723 | 706.8179 | 782.16 |
| Hnrnpa0 | GDVAEGDLIEHFSQF | -107 | -121 | 0 | 0 | 15 | 503.30344 | 491.5766 | 489.54132 | 550.2708 | 736.2074 | 701.6097 |
| Hnrnpa0 | FHPIQGHR | -160 | -167 | 0 | 0.001092529 | 8 | 436.12415 | 1207.262 | 463.8243 | 583.48975 | 436.88437 | 956.4451 |
| Hnrnpa0 | FHPIQGHR | -160 | -167 | 0 | 0.001092529 | 8 | 436.12415 | 1207.262 | 463.8243 | 583.48975 | 436.88437 | 956.4451 |
| Hnrnpa0 | NVQTSESGLR | -15 | -24 | 0 | 0 | 10 | 1396.8317 | 1373.7941 | 1338.8665 | 1178.3698 | NaN | 362.2034 |
| Hnrnpa0 | VVVVNPQTK | -37 | -45 | 0 | 0.000431655 | 9 | 679.8191 | 2618.2488 | 3086.809 | 3006.6226 | NaN | NaN |
| Hnrnpa0 | LFVGGLKG | -100 | -107 | 1 | 0.00078661 | 8 | 420.2594 | NaN | NaN | NaN | NaN | NaN |
| Hnrnpl | SSSGLLEWDSK | -539 | -549 | 0 | 0 | 11 | 4749.508 | 4854.1816 | 5128.75 | 5056.2856 | 5422.699 | 5433.8843 |
| Hnrnpl | SSSGLLEWDSK | -539 | -549 | 0 | 0 | 11 | 4749.508 | 4854.1816 | 5128.75 | 5056.2856 | 5422.699 | 5433.8843 |
| Hnrnpl | IQHPSNVLHFF | -493 | -503 | 0 | 0.00025015 | 11 | 4054.1826 | 4837.5117 | 4939.0215 | 4748.189 | 5944.777 | 5921.218 |
| Hnrnpl | IQHPSNVLHFF | -493 | -503 | 0 | 0.00025015 | 11 | 4054.1826 | 4837.5117 | 4939.0215 | 4748.189 | 5944.777 | 5921.218 |
| Hnrnpl | IQHPSNVLHFF | -493 | -503 | 0 | 0.00025015 | 11 | 4054.1826 | 4837.5117 | 4939.0215 | 4748.189 | 5944.777 | 5921.218 |
| Hnrnpl | TPASPVVHIR | -95 | -104 | 0 | 0 | 10 | 3382.4006 | 3865.3953 | 3873.02 | 4325.164 | 3379.7764 | 3595.231 |
| Hnrnpl | TPASPVVHIR | -95 | -104 | 0 | 0 | 10 | 3382.4006 | 3865.3953 | 3873.02 | 4325.164 | 3379.7764 | 3595.231 |
| Hnrnpl | VEFDSVQSAQR | -233 | -243 | 0 | 0 | 11 | 2814.0132 | 402.74344 | 3188.7732 | 3855.5098 | 4047.8691 | 4012.9114 |
| Hnrnpl | ISRPGDSDDSR | -176 | -186 | 1 | 0 | 11 | 2519.8477 | 3279.4534 | 3252.3135 | 3038.9136 | 2080.3167 | 3344.1091 |
| Hnrnpl | ISRPGDSDDSR | -176 | -186 | 1 | 0 | 11 | 2519.8477 | 3279.4534 | 3252.3135 | 3038.9136 | 2080.3167 | 3344.1091 |
| Hnrnpl | TDVLYTICNPCGPVQR | -205 | -220 | 0 | 0 | 16 | 2017.199 | 2709.8186 | 2551.0273 | 2888.5366 | 3639.988 | 3583.313 |
| Hnrnpl | GYGPPPPHYEGR | -329 | -340 | 0 | 0 | 12 | 2003.9396 | 7708.2344 | 4364.0093 | 4009.1619 | 4212.9844 | 879.0076 |
| Hnrnpl | GYGPPPPHYEGR | -329 | -340 | 0 | 0 | 12 | 2003.9396 | 7708.2344 | 4364.0093 | 4009.1619 | 4212.9844 | 879.0076 |
| Hnrnpl | QPPLLGDHPAEYGGPHGGYHSH | -303 | -324 | 0 | 0 | 22 | 1457.5675 | 1621.5382 | 1599.7235 | 1767.3625 | 2386.2812 | 3648.3872 |
| Hnrnpl | QPPLLGDHPAEYGGPHGGYHSH | -303 | -324 | 0 | 0 | 22 | 1457.5675 | 1621.5382 | 1599.7235 | 1767.3625 | 2386.2812 | 3648.3872 |
| Hnrnpl | QPPLLGDHPAEYGGPHGGYHSH | -303 | -324 | 0 | 0 | 22 | 1457.5675 | 1621.5382 | 1599.7235 | 1767.3625 | 2386.2812 | 3648.3872 |
| Hnrnpl | QEFGPISYVVVMPK | -120 | -133 | 0 | 0 | 14 | 1405.8569 | 1434.0016 | 931.72437 | 532.2214 | 2345.0972 | 2241.5112 |
| Hnrnpl | IEYAKPTR | -262 | -269 | 1 | 0 | 8 | 1322.9635 | 1164.9012 | 1335.9569 | 902.35785 | 1031.986 | 807.50073 |
| Hnrnpl | YHDEGYGPPPPHYEGR | -325 | -340 | 0 | 0 | 16 | 1267.824 | 985.5787 | 961.38416 | 1892.583 | 1967.6523 | 2029.2687 |
| Hnrnpl | YHDEGYGPPPPHYEGR | -325 | -340 | 0 | 0 | 16 | 1267.824 | 985.5787 | 961.38416 | 1892.583 | 1967.6523 | 2029.2687 |
| Hnrnpl | TPASPVVHIRG | -95 | -105 | 1 | 0 | 11 | 1185.3418 | 951.1507 | 1101.0491 | 1199.0001 | 1218.2683 | 1205.5331 |
| Hnrnpl | KNDQDTWDYTNPNLSGQGDPGSNPNKR | -274 | -300 | 2 | 0 | 27 | 612.21277 | 771.11774 | 677.495 | 724.05676 | 196.36905 | 817.8894 |
| Hnrnpl | KNDQDTWDYTNPNLSGQGDPGSNPNKR | -274 | -300 | 2 | 0 | 27 | 612.21277 | 771.11774 | 677.495 | 724.05676 | 196.36905 | 817.8894 |
| Hnrnpl | NDQDTWDYTNPNLSGQ | -275 | -290 | 0 | 0 | 16 | 807.5603 | 873.92957 | 924.6196 | 652.6708 | 664.9693 | 645.4945 |
| Hnrnpl | NAGDQHGGGGGGGSGAAGGGGGENYDDPHKTPA | -65 | -104 | 1 | 0.00025015 | 40 | 755.4153 | 1018.6482 | 933.43353 | 964.2736 | 1144.2386 | 1132.0555 |
| Hnrnpl | NAGDQHGGGGGGGSGAAGGGGGENYDDPHKTPA | -65 | -104 | 1 | 0.00025015 | 40 | 755.4153 | 1018.6482 | 933.43353 | 964.2736 | 1144.2386 | 1132.0555 |
| Hnrnpl | NPNGPYPYTLK | -566 | -576 | 0 | 0 | 11 | 642.4079 | 1652.0642 | 1545.9512 | 1487.864 | 235.17346 | 1914.2135 |
| Hnrnpl | NPNGPYPYTLK | -566 | -576 | 0 | 0 | 11 | 642.4079 | 1652.0642 | 1545.9512 | 1487.864 | 235.17346 | 1914.2135 |
| Hnrnpl | ICNPCGPVQR | -211 | -220 | 0 | 0 | 10 | 564.63324 | 1981.402 | 2082.7551 | 507.1711 | 1711.6469 | 575.89087 |
| Hnrnpl | ASPVVHIR | -97 | -104 | 0 | 0.002434616 | 8 | 544.6474 | 556.2742 | 610.71423 | 669.7097 | 502.5411 | 576.5087 |
| Hnrnpl | QPPLLGDHPAEYGGPH | -303 | -318 | 0 | 0.000298805 | 16 | 480.6728 | 764.13245 | 898.8424 | 1225.177 | 1178.5609 | 1159.8263 |
| Hnrnpl | QPPLLGDHPAEYGGPH | -303 | -318 | 0 | 0.000298805 | 16 | 480.6728 | 764.13245 | 898.8424 | 1225.177 | 1178.5609 | 1159.8263 |
| Hnrnpl | NAGDQHGGGGGGGSGAAGGGGGENYDDPHK | -65 | -94 | 0 | 0 | 30 | 470.7266 | 488.92508 | 506.77414 | 391.8733 | 474.20486 | 524.60767 |
| Hnrnpl | NAGDQHGGGGGGGSGAAGGGGGENYDDPHK | -65 | -94 | 0 | 0 | 30 | 470.7266 | 488.92508 | 506.77414 | 391.8733 | 474.20486 | 524.60767 |
| Hnrnpl | GGGGGENYDDPHK | -82 | -94 | 0 | 0.000230707 | 13 | 346.30276 | 48.59616 | 438.64432 | 454.72797 | 485.98157 | 306.15332 |
| Hnrnpl | QPAIMPGQSYGLEDGSCSYK | -453 | -472 | 0 | 0.001023236 | 20 | 330.80194 | 316.90424 | 274.4535 | 679.6325 | 751.41516 | 671.602 |
| Hnrnpl | NPCGPVQR | -213 | -220 | 0 | 0 | 8 | 1173.516 | 2488.4312 | 1446.85 | 1736.0077 | 258.36356 | NaN |
| Hnrnpl | NDQDTWDYTNPNLSGQGDPGSNPNKR | -275 | -300 | 1 | 0 | 26 | 336.79242 | 373.104 | 384.92062 | NaN | 360.4469 | 304.91348 |
| Hnrnpl | YYGGGNEGGR | -47 | -56 | 0 | 0.00025015 | 10 | 1128.3251 | 119.8956 | 273.30432 | 700.24347 | NaN | NaN |
| Hnrnpl | SPVVHIR | -98 | -104 | 0 | 0.002401448 | 7 | 423.08298 | 680.0959 | 1532.5072 | NaN | NaN | 478.40817 |
| Hnrnpl | PASPVVHIR | -96 | -104 | 0 | 0 | 9 | 374.08463 | 341.2217 | 179.86292 | 3826.9363 | NaN | NaN |
| Hnrnpl | SYGLEDGSCSYK | -461 | -472 | 0 | 0 | 12 | 347.348 | NaN | NaN | 946.81824 | 799.6509 | 749.3242 |
| Hnrnpl | NDQDTWDYTNPNLSGQGDPGSNPNK | -275 | -299 | 0 | 0.000394361 | 25 | 209.19267 | 242.9072 | NaN | 233.84143 | 226.36153 | NaN |
| Hnrnpl | NNNFMFGQK | -437 | -445 | 0 | 0.002972566 | 9 | 578.2316 | 734.1338 | NaN | NaN | NaN | NaN |
| Hnrnpl | KISRPGDSDDSR | -175 | -186 | 2 | 0.003814256 | 12 | 455.9376 | NaN | NaN | NaN | 301.5643 | NaN |
| Hnrnpl | GLEDGSCSYK | -463 | -472 | 0 | 0 | 10 | 720.23566 | NaN | NaN | NaN | NaN | NaN |
| Gba1 | ASPWTSPTWLK | -195 | -205 | 0 | 0 | 11 | 3677.5635 | 3748.4583 | 3721.7068 | 2185.599 | 3308.8599 | 2945.096 |
| Gba1 | WAEVVLSDPEAAK | -309 | -321 | 0 | 0 | 13 | 850.7478 | 862.0342 | 583.65625 | 1163.7991 | 1259.6559 | 1178.8195 |
| Gba1 | NFVDSPIIVDIPK | -414 | -426 | 0 | 0.00025015 | 13 | 669.2614 | NaN | 433.50934 | NaN | 793.30066 | NaN |
| Zc3hav1 | ASQEFLEDGDPDGLFSR | -323 | -339 | 0 | 0 | 17 | 1251.3025 | 702.5656 | 911.1054 | 1177.0859 | 542.5434 | 816.6644 |
| Zc3hav1 | NFEWILHGNR | -892 | -901 | 0 | 0 | 10 | 484.1839 | 891.44446 | 809.8277 | 951.99023 | 587.6659 | 1108.5063 |
| Zc3hav1 | EICLDHLYR | -602 | -610 | 0 | 0 | 9 | 520.61456 | 1072.8097 | NaN | 199.42004 | 338.06018 | 1360.9672 |
| Zc3hav1 | NHFHLPYR | -620 | -627 | 0 | 0 | 8 | 500.23398 | 773.0245 | 663.416 | 450.07465 | NaN | NaN |

| | | | | | | | | | | | | |
|---|---|---|---|---|---|---|---|---|---|---|---|---|
| Zc3hav1 | YCQRPCDSLHLC | -77 | -88 | 1 | 0.000550307 | 12 | 101.18216 | NaN | NaN | NaN | 756.26715 | 775.0686 |
| Zc3hav1 | QICGQPQPCER | -160 | -170 | 0 | 0.00025015 | 11 | 515.8772 | NaN | NaN | NaN | NaN | NaN |
| Zc3hav1 | QEFLEDGDPDGLFSR | -325 | -339 | 0 | 0.000431655 | 15 | 461.459 | NaN | NaN | NaN | NaN | NaN |
| Smarcb1 | ASQPEVLVPIR | -180 | -190 | 0 | 0 | 11 | 714.67206 | 649.5825 | 641.38184 | NaN | NaN | NaN |
| Smarcb1 | LNIHVGNIS | -266 | -274 | 0 | 0.002502116 | 9 | 298.67892 | NaN | NaN | NaN | NaN | NaN |
| Psmd8 | VAEFHTELER | -208 | -217 | 0 | 0 | 10 | 2428.0532 | 297.8523 | 2556.9368 | 2918.266 | 3112.2673 | 2734.5476 |
| Psmd8 | ELNFLPTTGTK | -128 | -138 | 0 | 0 | 11 | 775.8179 | 857.72296 | 796.9802 | 1364.1782 | 1139.8196 | 1673.4691 |
| Psmd8 | YEQLKDEWNR | -98 | -107 | 1 | 0 | 10 | 1827.4238 | NaN | 633.9945 | 1716.591 | 1601.7596 | 1561.3745 |
| Psmd8 | ASQQQKPEDSTIPSTELAK | -322 | -340 | 1 | 0.00025015 | 19 | 367.04916 | 594.37195 | 450.60883 | 333.33255 | NaN | 397.59537 |
| Psmd8 | QQKPEDSTIPSTELAK | -325 | -340 | 1 | 0.000627943 | 16 | 1033.3156 | 847.9862 | 1133.9005 | NaN | NaN | 950.5522 |
| Psmd8 | EQLKDEWNR | -99 | -107 | 1 | 0.00051306 | 9 | 1814.1549 | NaN | 2340.5767 | NaN | NaN | 1709.561 |
| Ddost | SSLNPILFR | -175 | -183 | 0 | 0 | 9 | 3164.6096 | 635.0459 | 3557.391 | 3086.5361 | 3173.979 | 3793.9421 |
| Ddost | TADDPSLSLIK | -63 | -73 | 0 | 0 | 11 | 3037.183 | 2604.26 | 2808.5964 | 1060.7183 | 2952.2761 | 2799.3796 |
| Ddost | VAASSDIGDPLR | -113 | -124 | 0 | 0 | 12 | 2053.93 | 1325.4044 | 1980.2402 | 1742.5026 | 663.5311 | 653.5695 |
| Ddost | VLLDNLNVR | -34 | -42 | 0 | 0 | 9 | 1899.9224 | 1648.9528 | 446.65152 | 1103.3718 | 1786.1356 | 2455.0515 |
| Ddost | TLVLLDNLNVR | -32 | -42 | 0 | 0 | 11 | 1697.723 | 1725.3046 | 1892.7822 | 894.10614 | 1478.4736 | 1329.7697 |
| Ddost | SLNPILFR | -176 | -183 | 0 | 0 | 8 | 1125.3489 | 1476.4993 | 1378.213 | 977.4716 | 916.3513 | 1449.7179 |
| Ddost | SYSFFPDKPITQYPHAVGR | -206 | -224 | 1 | 0 | 19 | 1026.3448 | 1194.8502 | 1324.5968 | 961.0938 | 1212.6686 | 1652.307 |
| Ddost | STSYSFFPDKPITQYPHAVGR | -204 | -224 | 1 | 0 | 21 | 604.9826 | 514.8632 | 865.50964 | 1046.9614 | 900.3189 | 1312.2106 |
| Ddost | LIVADTENLLK | -157 | -167 | 0 | 0 | 11 | 602.1465 | 716.09344 | 597.28375 | 861.0251 | 979.0106 | 1313.4401 |
| Ddost | SFFPDKPITQYPHAVGR | -208 | -224 | 1 | 0 | 17 | 462.17096 | 339.96243 | 597.7944 | 426.57938 | 556.07007 | 629.0951 |
| Ddost | TAVIDHHNYDVSDLGQHTLIVA | -139 | -160 | 0 | 0 | 22 | 432.06702 | 390.2973 | 401.21432 | 321.81342 | 331.8128 | 358.42825 |
| Ddost | DTHSLFFR | -43 | -50 | 0 | 0.00025015 | 8 | 1361.6365 | NaN | 858.0735 | 1410.9838 | 1791.4058 | 1565.4353 |
| Ddost | ELGSECGIEFDEEK | -125 | -138 | 0 | 0 | 14 | 618.25336 | 749.0836 | 690.40765 | NaN | NaN | NaN |
| Ddost | ASSDIGDPLR | -115 | -124 | 0 | 0 | 10 | 586.84894 | NaN | 662.7104 | 511.86783 | NaN | NaN |
| Arpc1b | GAGLDSLHK | -318 | -326 | 0 | 0 | 9 | 16930.574 | 19131.307 | 21881.385 | 19188.51 | 28879.447 | 25395.992 |
| Arpc1b | GIDWAPESNR | -57 | -66 | 0 | 0 | 10 | 6432.293 | 6731.803 | 7043.6494 | 600.05597 | 461.84125 | 2940.8367 |
| Arpc1b | HDSTVCLVDADKK | -222 | -234 | 1 | 0 | 13 | 3150.815 | 1283.1377 | 2678.9768 | 2702.3657 | 1510.7527 | 2832.275 |
| Arpc1b | HDSTVCLVDADKK | -222 | -234 | 1 | 0 | 13 | 3150.815 | 1283.1377 | 2678.9768 | 2702.3657 | 1510.7527 | 2832.275 |
| Arpc1b | EVEERPAPTPWGSK | -175 | -188 | 1 | 0 | 14 | 2483.2798 | 2746.4153 | 2588.3828 | 2738.5674 | 2807.351 | 2744.038 |
| Arpc1b | EVEERPAPTPWGSK | -175 | -188 | 1 | 0 | 14 | 2483.2798 | 2746.4153 | 2588.3828 | 2738.5674 | 2807.351 | 2744.038 |
| Arpc1b | LVEPISCHAWNK | -7 | -18 | 0 | 0 | 12 | 1778.6869 | 1225.8304 | 1256.8822 | 3276.18 | 1512.2289 | 1066.3967 |
| Arpc1b | TWKPTLVILR | -85 | -94 | 1 | 0.000550307 | 10 | 821.7982 | 951.7024 | 951.203 | 347.98184 | 329.31934 | 345.9545 |
| Arpc1b | TQIAICPNNHEVHIY | -21 | -35 | 0 | 0 | 15 | 720.619 | 835.24585 | 826.9295 | 913.6786 | 964.8191 | 902.76544 |
| Arpc1b | TQIAICPNNHEVHIY | -21 | -35 | 0 | 0 | 15 | 720.619 | 835.24585 | 826.9295 | 913.6786 | 964.8191 | 902.76544 |
| Arpc1b | ASSEGGAATGAGLDSLHK | -309 | -326 | 0 | 0 | 18 | 602.91943 | 621.18823 | 655.0657 | 376.7753 | 552.6957 | 499.43793 |
| Arpc1b | YFEQENDWWVCK | -124 | -135 | 0 | 0 | 12 | 539.5807 | 561.0295 | 740.4282 | 1510.6683 | 1458.48 | 1449.4128 |
| Arpc1b | TWKPTLVIL | -85 | -93 | 1 | 0.003966045 | 9 | 899.1369 | 1011.599 | 1001.3462 | 242.15448 | NaN | 892.7973 |
| Arpc1b | RWAPNENK | -103 | -110 | 1 | 0.00025015 | 8 | 894.61084 | NaN | 918.15594 | 485.64157 | 1076.9521 | 1444.8358 |
| Arpc1b | HDSTVCLVDADK | -222 | -233 | 0 | 0.00025015 | 12 | 1261.2225 | 731.0658 | NaN | 720.54816 | 765.9757 | NaN |
| Arpc1b | WAPNENK | -104 | -110 | 0 | 0.002136192 | 7 | 1001.26715 | 888.5909 | NaN | 1617.6359 | NaN | 451.85822 |
| Eif4g2 | ENPLLPEEEEQR | -189 | -200 | 0 | 0 | 12 | 1221.2767 | 706.4684 | 787.18536 | 855.7996 | 1010.6309 | 918.45215 |
| Eif4g2 | LAEDAPNFDGPAAEGQPGQK | -139 | -158 | 0 | 0 | 20 | 1057.9774 | 862.33307 | 847.9037 | 509.97943 | 1127.9312 | 1018.1932 |
| Eif4g2 | LDLIHESILHK | -220 | -230 | 0 | 0 | 11 | 703.6527 | 636.24133 | 578.3459 | 869.1122 | 1244.9761 | 818.30475 |
| Eif4g2 | TQTPPLGQTPQLGLK | -505 | -519 | 0 | 0 | 15 | 620.43146 | 668.1786 | 794.2712 | 230.73041 | 813.0343 | 855.7254 |
| Eif4g2 | ASSLISLLK | -599 | -607 | 0 | 0.006654467 | 9 | 413.594 | 440.64847 | NaN | NaN | NaN | NaN |
| Tpr | FHNELNAHIK | -349 | -358 | 0 | 0 | 10 | 3250.5134 | 4315.913 | 4196.4575 | 1080.5966 | 4820.974 | 5467.151 |
| Tpr | LSQEVEYLTEDVKR | -208 | -221 | 1 | 0.000230707 | 14 | 2348.1238 | 3890.3506 | 2711.0962 | 2844.5852 | 2893.4563 | 4708.0923 |
| Tpr | QVLERPELNK | -81 | -90 | 1 | 0.00025015 | 10 | 2280.394 | 2697.7456 | 2183.5674 | 2083.207 | 2538.8438 | 2031.4911 |
| Tpr | IQQLNEEVGR | -1442 | -1451 | 0 | 0 | 10 | 2141.9436 | 238.35722 | 2285.3708 | 1941.1794 | 2332.5515 | 1954.8135 |
| Tpr | LSQEVEYLTEDVK | -208 | -220 | 0 | 0 | 13 | 2053.1382 | 1863.2847 | 1811.6576 | 1897.5215 | 1157.6149 | 1321.504 |
| Tpr | SLEGQVENLQK | -1566 | -1576 | 0 | 0 | 11 | 1931.4532 | 2217.5352 | 2220.304 | 1282.4047 | 4488.381 | 4558.07 |
| Tpr | TVPSTPTLVVPHR | -2201 | -2213 | 0 | 0 | 13 | 1488.4764 | 1558.2671 | 1788.0911 | 1848.4001 | 2074.7695 | 2072.7488 |
| Tpr | GQPGDRDDVDDLK | -1000 | -1012 | 1 | 0.00078661 | 13 | 1117.285 | 1069.0585 | 1277.655 | 1333.8295 | 1472.207 | 1416.7078 |
| Tpr | LQEQVTDLR | -774 | -782 | 0 | 0.00025015 | 9 | 1060.3131 | 1045.3782 | 948.61896 | 1113.6404 | 1150.013 | 922.24634 |
| Tpr | LENSLAELEQLR | -657 | -668 | 0 | 0 | 12 | 936.57837 | 906.63745 | 834.92786 | 1606.8436 | 1179.22 | 1154.1212 |
| Tpr | HVEDLLTK | -327 | -334 | 0 | 0 | 8 | 920.28845 | 1256.3861 | 856.4914 | 1159.1917 | 1262.4567 | 1325.0673 |
| Tpr | NIEELQQQNQR | -616 | -626 | 0 | 0 | 11 | 874.8734 | 952.7732 | 910.9977 | 1048.9668 | 1214.3683 | 1166.4044 |
| Tpr | ELQELQDSLNVER | -1308 | -1320 | 0 | 0 | 13 | 725.27594 | 900.19244 | 478.17743 | 904.1384 | 890.6551 | 895.41956 |
| Tpr | TLSTVQNEVQEALQR | -1113 | -1127 | 0 | 0.00025015 | 15 | 690.1338 | 810.6523 | 826.8279 | 758.42267 | 1048.9758 | 1092.5272 |
| Tpr | DAVQAPLNVSLNEEGK | -1250 | -1265 | 0 | 0 | 16 | 613.9113 | 659.0269 | 624.9322 | 489.63364 | 620.7983 | 572.7911 |
| Tpr | TIQDHLLQVEESK | -397 | -409 | 0 | 0 | 13 | 519.1878 | 506.3747 | 602.83685 | 714.54504 | 1427.8442 | 992.4392 |
| Tpr | TIQDHLLQVEESK | -397 | -409 | 0 | 0 | 13 | 519.1878 | 506.3747 | 602.83685 | 714.54504 | 1427.8442 | 992.4392 |
| Tpr | IKPTPVVSTPSK | -1739 | -1750 | 1 | 0.000431655 | 12 | 513.5559 | 578.5271 | 494.25833 | 550.3128 | 593.51495 | 554.2807 |
| Tpr | TVSTPAPEPVIDSTEAIEAK | -724 | -743 | 0 | 0 | 20 | 317.37625 | 415.09665 | 479.08737 | 476.5503 | 560.92236 | 417.78278 |
| Tpr | VETQDQLLLEK | -477 | -487 | 0 | 0 | 11 | 302.93118 | 1290.663 | 786.8627 | 1476.1493 | 1256.7139 | 386.3886 |
| Tpr | ELENANDLLSATK | -426 | -438 | 0 | 0 | 13 | 274.78012 | 1569.8586 | 1573.5792 | 1704.2168 | 1753.328 | 1766.407 |
| Tpr | NGALDQQKDELDVR | -1655 | -1668 | 1 | 0 | 14 | 1463.1274 | 147.48259 | 1617.8048 | 1448.6702 | 148.22163 | NaN |
| Tpr | SEQQYFEIEK | -124 | -133 | 0 | 0 | 10 | 1067.021 | 1068.6162 | 1215.072 | 247.46028 | 773.37775 | NaN |
| Tpr | KLENEVEQR | -928 | -936 | 1 | 0.00025015 | 9 | 811.8922 | NaN | 553.7551 | 1528.3046 | 1895.9681 | 603.8075 |
| Tpr | LEHEISHLK | -918 | -926 | 0 | 0 | 9 | 477.96286 | 627.7375 | 666.3219 | 716.9196 | 936.4079 | NaN |
| Tpr | VLKDEVSK | -1211 | -1218 | 1 | 0.001055699 | 8 | 411.65677 | 493.75473 | 444.2611 | NaN | 810.4296 | 653.3299 |
| Tpr | LLHDQIEK | -1232 | -1239 | 0 | 0 | 8 | 158.65918 | 283.9202 | 294.89484 | NaN | 433.8307 | 259.6859 |
| Tpr | ASTALSNEQQAR | -1128 | -1139 | 0 | 0 | 12 | 634.63727 | 639.84955 | NaN | 483.6322 | NaN | 557.13495 |
| Tpr | TSTSNVEQYR | -1031 | -1040 | 0 | 0 | 10 | 519.8121 | 625.4625 | 577.02747 | 647.8033 | NaN | NaN |
| Tpr | SQEQILEILR | -1266 | -1275 | 0 | 0 | 10 | 378.85632 | 1609.8511 | 351.4334 | NaN | 791.5848 | NaN |
| Tpr | EKGNEILELK | -288 | -297 | 1 | 0.00025015 | 10 | 333.85144 | NaN | NaN | 836.4961 | 586.33124 | 345.0625 |
| Tpr | NLDVQLLDTK | -942 | -951 | 0 | 0 | 10 | 286.50232 | 548.7046 | 234.22304 | 534.65765 | NaN | NaN |

| | | | | | | | | | | | | |
|---|---|---|---|---|---|---|---|---|---|---|---|---|
| Tpr | NEVQEALQR | -1119 | -1127 | 0 | 0.00078661 | 9 | 1664.955 | 1024.9785 | 719.582 | NaN | NaN | NaN |
| Rap2b | ASVDELFAEIVR | -151 | -162 | 0 | 0 | 12 | 265.9878 | NaN | NaN | 499.74625 | 604.79144 | 185.20087 |
| Cacybp | KPELDNEKPAAVVAPL | -53 | -68 | 2 | 0.000471409 | 16 | 1277.9155 | 1024.6045 | 1103.6965 | 922.2956 | 1188.8887 | 1237.8041 |
| Cacybp | KPELDNEKPAAVVAPL | -53 | -68 | 2 | 0.000471409 | 16 | 1277.9155 | 1024.6045 | 1103.6965 | 922.2956 | 1188.8887 | 1237.8041 |
| Cacybp | LRDTLTSEK | -27 | -35 | 1 | 0.000591124 | 9 | 842.638 | 691.3 | 722.13074 | 767.0543 | 920.2203 | 700.71265 |
| Cacybp | ASVLEELQK | -2 | -10 | 0 | 0 | 9 | 140.30724 | 1307.2234 | 86.754425 | 1314.1838 | 1616.526 | 1829.2845 |
| Cacybp | KPELDNEKPAAVVAPLTT | -53 | -70 | 2 | 0.001463897 | 18 | 323.7375 | 412.1862 | 433.31326 | NaN | NaN | 495.56985 |
| Cacybp | NYGWDQSDK | -78 | -86 | 0 | 0 | 9 | 608.7323 | NaN | NaN | 867.9028 | 468.75174 | NaN |
| Cacybp | SFDLLVK | -113 | -119 | 0 | 0.00025015 | 7 | 501.74997 | NaN | 386.44913 | NaN | NaN | NaN |
| Mif | LHISPDR | -88 | -94 | 0 | 0.000298805 | 7 | 12839.984 | 11177.187 | 10070.333 | 11973.244 | 8741.951 | 8425.75 |
| Mif | ASVPEGFLSELTQQ | -13 | -26 | 0 | 0 | 14 | 4673.4043 | 4818.8057 | 4434.9907 | 6443.6787 | 7124.963 | 6794.8057 |
| Mif | SGTNDPCALCSLHSIGK | -51 | -67 | 0 | 0 | 17 | 4237.4526 | 3878.8047 | 4162.8354 | 5742.4736 | 4554.91 | 4401.6743 |
| Mif | PAQYIAVH | -34 | -41 | 0 | 0.000712251 | 8 | 3781.0742 | 3795.8584 | 2213.475 | 3636.679 | 3016.0908 | 2368.043 |
| Mif | LLCGLLSDR | -79 | -87 | 0 | 0 | 9 | 2148.0696 | 2394.6326 | 2059.3958 | 2202.4526 | 1581.4474 | 1322.9696 |
| Mif | CSLHSIGK | -60 | -67 | 0 | 0 | 8 | 2081.217 | 2193.014 | 2229.7222 | 2527.9827 | 300.37634 | 299.43457 |
| Mif | ASVPEGFLSEL | -13 | -23 | 0 | 0 | 11 | 681.3996 | 684.04614 | 727.3943 | 526.29785 | 718.8246 | 703.68567 |
| Mif | LCGLLSDR | -80 | -87 | 0 | 0 | 8 | 597.4442 | 824.28723 | 594.98315 | 820.52606 | 883.0492 | 1175.2695 |
| Mif | SLHSIGK | -61 | -67 | 0 | 0.006850902 | 7 | 1312.9519 | 879.00256 | 1260.8163 | 1210.3276 | NaN | 1247.711 |
| Mif | ASVPEGFLSELTQ | -13 | -25 | 0 | 0 | 13 | 991.0202 | NaN | 829.3874 | 882.98096 | 481.62723 | 926.16705 |
| Mif | NVGWNGSTFA | -106 | -115 | 0 | 0.00025015 | 10 | 2874.9778 | NaN | NaN | 3716.8408 | NaN | NaN |
| Incenp | ASWGLQDSPGSTDSPWQER | -238 | -256 | 0 | 0.00025015 | 19 | 2120.952 | 2259.2097 | 2419.441 | 3023.3706 | 1867.1162 | 3375.2698 |
| Incenp | GTTAPGPICLLDLCDQK | -2 | -18 | 0 | 0 | 17 | 304.45917 | 254.96729 | 280.1378 | 423.62323 | 415.55035 | 361.92438 |
| Eprs1 | IYIPDGHTK | -700 | -708 | 0 | 0 | 9 | 20252.053 | 11632.544 | 1410.8235 | 18324.816 | 11466.148 | 14510.093 |
| Eprs1 | NPDVGLKPVWYSPK | -536 | -549 | 1 | 0 | 14 | 1666.6135 | 2210.1843 | 2149.3647 | 2827.6135 | 3098.724 | 2707.7236 |
| Eprs1 | ETVAFTDVNSILR | -44 | -56 | 0 | 0 | 13 | 1250.2482 | 405.3405 | 876.2786 | 1958.8126 | 1886.5857 | 1790.8145 |
| Eprs1 | KGDIIQLQR | -666 | -674 | 1 | 0 | 9 | 946.28357 | 961.064 | 955.8688 | 1568.7435 | 1756.7753 | 1473.1807 |
| Eprs1 | VEEGKENLLR | -32 | -41 | 1 | 0 | 10 | 887.32886 | 914.36237 | 977.0241 | 826.76654 | 1032.2133 | 955.4666 |
| Eprs1 | VYEELLAIPVVR | -1196 | -1207 | 0 | 0 | 12 | 855.7002 | 917.31934 | 1081.2279 | 918.6725 | 922.3824 | 1026.7699 |
| Eprs1 | DYVPGQPPASQNSHSNPVSNAQPAGAEKPEAK | -871 | -902 | 1 | 0.00111006 | 32 | 589.599 | 631.4006 | 644.7876 | 537.5929 | 521.70654 | 567.6964 |
| Eprs1 | DYVPGQPPASQNSHSNPVSNAQPAGAEKPEAK | -871 | -902 | 1 | 0.00111006 | 32 | 589.599 | 631.4006 | 644.7876 | 537.5929 | 521.70654 | 567.6964 |
| Eprs1 | DQVDSAVQELLQLK | -926 | -939 | 0 | 0 | 14 | 427.22766 | 260.29083 | 319.54868 | 387.77924 | 392.24033 | 418.15933 |
| Eprs1 | EKNHIEDFAPEVA | -1090 | -1102 | 1 | 0 | 13 | 2612.1785 | NaN | 2747.8254 | 1363.7374 | 1213.7732 | 4173.447 |
| Eprs1 | EFLWQEGHSA | -1166 | -1175 | 0 | 0.000826806 | 10 | 795.2463 | 945.1995 | 877.34467 | NaN | 1530.8756 | 983.90826 |
| Eprs1 | FVNEGLVDGWDDPR | -441 | -454 | 0 | 0 | 14 | 390.3595 | NaN | 429.57352 | 1453.2483 | 1811.2996 | 700.87445 |
| Eprs1 | VACQGEVVR | -908 | -916 | 0 | 0 | 9 | 1990.52 | NaN | 735.81525 | 951.89874 | 474.94662 | NaN |
| Eprs1 | FNHWELK | -1355 | -1361 | 0 | 0 | 7 | 1501.0068 | NaN | NaN | 1868.2119 | 1912.0195 | 1848.7395 |
| Eprs1 | TGQEYKPGNPSAAAVQ | -795 | -810 | 1 | 0 | 16 | 832.6129 | 879.6934 | 1193.0795 | NaN | NaN | 1096.5355 |
| Eprs1 | GLVDGWDDPR | -445 | -454 | 0 | 0 | 10 | 761.6498 | 939.30457 | 972.9796 | NaN | 682.5598 | NaN |
| Eprs1 | TDVNSILR | -49 | -56 | 0 | 0 | 8 | 552.42053 | 348.85327 | 347.47473 | 335.36102 | NaN | NaN |
| Eprs1 | EHLITKPVLGK | -627 | -637 | 1 | 0 | 11 | 399.4207 | NaN | NaN | 398.15274 | 488.1706 | 425.14267 |
| Eprs1 | GSAAWQEHLK | -130 | -139 | 0 | 0 | 10 | 1268.5529 | NaN | 1077.7515 | NaN | 816.34515 | NaN |
| Eprs1 | NHIEDFAPEVA | -1092 | -1102 | 0 | 0 | 11 | 941.8497 | 761.63684 | 717.86707 | NaN | NaN | NaN |
| Eprs1 | LNQWCNVVR | -1144 | -1152 | 0 | 0 | 9 | 727.2936 | 793.782 | NaN | NaN | NaN | NaN |
| Eprs1 | ATAEDSSVLYSR | -745 | -756 | 0 | 0.00272025 | 12 | 687.1524 | NaN | NaN | NaN | NaN | NaN |
| Stk10 | FYDVELENLER | -607 | -617 | 0 | 0 | 11 | 1323.8969 | 1205.428 | 1087.6812 | 957.1355 | 959.0999 | 893.97064 |
| Stk10 | ATCDHPYIVK | -86 | -95 | 0 | 0 | 10 | 1031.1676 | 1553.5616 | 1345.9161 | 222.17273 | 1357.5002 | 1691.5135 |
| Stk10 | KSDPPTLLTPSK | -251 | -262 | 1 | 0.001023236 | 12 | 309.79086 | 507.93527 | 553.0813 | 739.15784 | 994.1736 | 794.24976 |
| Stk10 | RFEQEINAK | -596 | -604 | 1 | 0.007493591 | 9 | 1221.1904 | 1531.4401 | NaN | 1716.6683 | NaN | NaN |
| Stk10 | NDNDLKVPVPLR | -399 | -410 | 1 | 0.001370415 | 12 | 1898.6144 | NaN | NaN | 5520.111 | NaN | NaN |
| Nsmce1 | ATDFAENELDLFR | -98 | -110 | 0 | 0.000431655 | 13 | 326.57394 | NaN | NaN | 125.44949 | 261.54678 | 396.35043 |
| Hp1bp3 | NGWLEQISGK | -394 | -403 | 0 | 0 | 10 | 3346.2544 | 3905.511 | 3556.081 | 628.1129 | 5179.836 | 642.63336 |
| Hp1bp3 | KYPSLGLER | -192 | -200 | 1 | 0 | 9 | 1084.9388 | 1842.1993 | 678.2647 | 2051.6611 | 2346.2607 | 2409.6301 |
| Hp1bp3 | LAEGEEEKPEPDGSSEESIS | -57 | -76 | 1 | 0.001150701 | 20 | 455.63187 | 628.63214 | 452.82608 | 642.40216 | 632.49677 | 600.34033 |
| Hp1bp3 | TVEEQENETPPATSSEAEQPK | -77 | -97 | 0 | 0.000230707 | 21 | 305.1563 | 331.9771 | 316.39163 | 248.85757 | 312.5562 | 352.92352 |
| Hp1bp3 | ATDMSQGELIHPK | -2 | -14 | 0 | 0 | 13 | 256.45215 | 699.1532 | 301.87634 | 367.9602 | 733.45685 | 812.66797 |
| Hp1bp3 | SFPYYPSPGVLFPK | -412 | -425 | 0 | 0 | 14 | 135.92398 | 510.07858 | 1198.0245 | 1270.2782 | 1776.0876 | 1628.8569 |
| Hp1bp3 | SGEKPLLGGSL | -331 | -341 | 1 | 0 | 11 | 1413.8018 | 1248.1791 | 1257.1836 | NaN | 1517.842 | 1595.0282 |
| Hp1bp3 | QLSFPYYPSPGVLFPK | -410 | -425 | 0 | 0.000431655 | 16 | 268.82153 | 476.83508 | NaN | 618.36414 | 588.38324 | 579.3634 |
| Hp1bp3 | KGSALDPEPQVK | -247 | -258 | 1 | 0 | 12 | 251.57806 | 1220.4755 | 313.1811 | NaN | 1480.3252 | 1298.8174 |
| Hp1bp3 | LAEGEEEKPEPDGSSEE | -57 | -73 | 1 | 0.002068213 | 17 | 305.28726 | 277.22418 | NaN | NaN | NaN | 120.46687 |
| Galnt1 | NFFYIISPGVTK | -379 | -390 | 0 | 0 | 12 | 477.54373 | 349.9679 | 584.7606 | NaN | 734.8601 | 427.42126 |
| Galnt1 | ATEEDSQVPSIR | -527 | -538 | 0 | 0 | 12 | 1350.2234 | 1581.1885 | 1346.1665 | 1525.8768 | NaN | NaN |
| Hypk | ATEGDVELELETETSGPERPPEKPR | -2 | -26 | 2 | 0 | 25 | 469.08258 | 484.80783 | 486.76086 | 514.9613 | 537.3596 | 625.2583 |
| Hypk | KHDSGAADLER | -27 | -37 | 1 | 0.00025015 | 11 | 294.92798 | NaN | 429.50708 | 375.2143 | 261.1847 | NaN |
| Hnrnph1 | GITLPVDFQGR | -140 | -150 | 0 | 0.000431655 | 11 | 1555.8237 | 2097.6272 | 1442.0096 | 1773.4333 | 1601.1064 | 1799.2589 |
| Hnrnph1 | GLPWSCSADEVQR | -17 | -29 | 0 | 0 | 13 | 487.94882 | 1694.2035 | 1620.9076 | 1961.3574 | 2198.592 | 337.30762 |
| Hnrnph1 | NVEMDWVLK | -90 | -98 | 0 | 0 | 9 | 1050.7611 | NaN | NaN | 1190.3545 | NaN | NaN |
| Hnrnph1 | SNNVEMDWVLK | -88 | -98 | 0 | 0.00025015 | 11 | 829.00006 | NaN | NaN | NaN | NaN | NaN |
| Dnajc7 | ATEPELLEDEDAKR | -13 | -26 | 1 | 0 | 14 | 1336.9579 | 1150.7961 | 997.0989 | 1566.1862 | 1232.7933 | 1290.8274 |
| Ipo4 | YVRPDDVSLAR | -190 | -200 | 1 | 0 | 11 | 678.556 | 871.5933 | 850.416 | 844.6018 | 862.52356 | 940.03613 |
| Ipo4 | ELLLPDTER | -12 | -20 | 0 | 0 | 9 | 288.4791 | 530.1105 | 622.1708 | 852.72644 | 675.0077 | 894.6824 |
| Ipo4 | AVLSDGAGDHIR | -377 | -388 | 0 | 0 | 12 | 3513.0415 | NaN | NaN | 3941.6895 | 4075.3574 | NaN |
| Ipo4 | ENFVENLGPK | -463 | -472 | 0 | 0.000591124 | 10 | 957.24915 | 1018.52997 | 451.81964 | NaN | NaN | NaN |
| Ipo4 | GLDDPSQIVR | -402 | -411 | 0 | 0.00272025 | 10 | 1932.2096 | NaN | NaN | 1729.615 | NaN | NaN |
| Ipo4 | ATEQLQTILR | -24 | -33 | 0 | 0 | 10 | 729.4952 | NaN | NaN | 648.3752 | NaN | NaN |
| Rrp1 | ATGGFTPDELLK | -43 | -54 | 0 | 0 | 12 | 905.7821 | 8762.61 | 8032.2817 | 8434.749 | 9080.8545 | 9151.92 |
| Rrp1 | TTEILNPDSQAPSGVK | -151 | -166 | 0 | 0 | 16 | 828.0395 | 798.4187 | 858.05176 | 685.82886 | 631.79236 | 607.18274 |
| Nomo1 | LDFTVEHDSLR | -301 | -311 | 0 | 0 | 11 | 2617.4346 | 2711.5635 | 411.12918 | 2745.5476 | 2619.5571 | 2355.3071 |

| | | | | | | | | | | | | |
|---|---|---|---|---|---|---|---|---|---|---|---|---|
| Nomo1 | ITFDVAPSR | -292 | -300 | 0 | 0 | 9 | 783.5975 | 892.7724 | 916.0468 | 778.08026 | 805.2714 | 998.8493 |
| Nomo1 | HHVLGTIITDK | -658 | -668 | 0 | 0 | 11 | 472.5808 | NaN | 513.23706 | 1220.3251 | 816.5816 | 569.57904 |
| Nomo1 | ATHPTWALK | -177 | -185 | 0 | 0.00025015 | 9 | 1150.31 | NaN | 582.59766 | 763.4064 | NaN | 770.7539 |
| Nomo1 | YDKGDFILK | -75 | -83 | 1 | 0.002629273 | 9 | 2684.5432 | NaN | NaN | 1669.1168 | NaN | NaN |
| Rab7a | VFDVTAPNTFK | -87 | -97 | 0 | 0 | 11 | 1502.0619 | 1561.9583 | 1662.1997 | 1692.8392 | 1465.5947 | 1523.6272 |
| Rab7a | ATIGADFLTK | -39 | -48 | 0 | 0 | 10 | 770.14966 | 954.16656 | 934.70135 | 701.1609 | 1239.7062 | 1176.2103 |
| Rab7a | EAINVEQAFQTIAR | -158 | -171 | 0 | 0 | 14 | 696.1867 | 766.2959 | 881.0265 | 727.3414 | 1204.2834 | 1159.1624 |
| Rab7a | NNIPYFETSAK | -147 | -157 | 0 | 0 | 11 | 540.9215 | 589.5314 | 329.4072 | 756.94165 | NaN | NaN |
| Rab7a | VIILGDSGVGK | -11 | -21 | 0 | 0 | 11 | 633.854 | NaN | NaN | NaN | NaN | NaN |
| Myo1g | ELYERPPHLY | -76 | -85 | 1 | 0 | 10 | 1403.342 | 1637.8326 | 1399.9851 | 1831.7245 | 1795.8965 | 1805.1423 |
| Myo1g | IIGFSPDEVESIHR | -267 | -280 | 0 | 0 | 14 | 89.892525 | 935.1835 | 961.4459 | 792.3116 | 185.87386 | 1169.3093 |
| Myo1g | IIGFSPDEVESIHR | -267 | -280 | 0 | 0 | 14 | 89.892525 | 935.1835 | 961.4459 | 792.3116 | 185.87386 | 1169.3093 |
| Myo1g | TIVELVEQPHR | -458 | -468 | 0 | 0 | 11 | 1058.681 | 1243.3225 | 1289.666 | 1318.9504 | 1511.3997 | 1410.5665 |
| Myo1g | VWPTPPAVLQPFQDTCR | -779 | -795 | 0 | 0.000230707 | 17 | 913.14923 | 942.8781 | 1028.0448 | 1036.7307 | 1356.9192 | 1346.4003 |
| Myo1g | LFEWVVNK | -369 | -376 | 0 | 0.000471409 | 8 | 703.1122 | 736.28503 | 922.5578 | 446.0399 | 632.4276 | 874.4263 |
| Myo1g | GSEDQELQGLHLER | -216 | -229 | 0 | 0 | 14 | 521.671 | 624.8884 | 698.3216 | 706.52026 | 754.937 | 907.2517 |
| Myo1g | VNPYQELPLYGPEAIAK | -55 | -71 | 0 | 0 | 17 | 507.46573 | 536.84985 | 358.82126 | 888.3214 | 958.15967 | 652.6282 |
| Myo1g | VSVNPYQELPLYGPEAIAK | -53 | -71 | 0 | 0 | 19 | 492.47894 | 370.39368 | 317.37265 | 770.4984 | 966.523 | 526.22595 |
| Myo1g | LISVEPRPEQPEPDFQSS | -996 | -1013 | 1 | 0.000230707 | 18 | 466.90396 | 528.7695 | 519.58795 | 260.90152 | 592.2246 | 470.3877 |
| Myo1g | STFTLLWPSH | -1015 | -1024 | 0 | 0 | 10 | 442.34418 | 431.4826 | 118.35832 | 637.62823 | 298.5607 | 581.39526 |
| Myo1g | ATIVELVEQPHR | -457 | -468 | 0 | 0 | 12 | 326.73285 | 852.4504 | 457.38614 | 398.86887 | 177.30934 | 630.84 |
| Myo1g | DSLFQDFK | -543 | -550 | 0 | 0.003490217 | 8 | 764.1023 | 826.5217 | 812.07275 | 688.35803 | NaN | 663.88275 |
| Myo1g | GILAVLDEACSTAGPITDR | -469 | -487 | 0 | 0 | 19 | 333.02173 | NaN | 249.98286 | 338.83353 | 314.3386 | 417.15527 |
| Myo1g | GDPVGGHIH | -180 | -188 | 0 | 0.001392869 | 9 | 3326.7173 | NaN | 4114.77 | NaN | 2710.6187 | NaN |
| Myo1g | LLYNSVDPTLR | -552 | -562 | 0 | 0.001811524 | 11 | 908.72845 | 950.48584 | NaN | NaN | NaN | 1034.0498 |
| Myo1g | QDWGCQR | -833 | -839 | 0 | 0.006677519 | 7 | 812.9009 | NaN | NaN | NaN | 1151.6646 | 1039.1969 |
| Myo1g | VSDCIPLSQR | -982 | -991 | 0 | 0.000550307 | 10 | 1750.0176 | NaN | 1651.7148 | NaN | NaN | NaN |
| Myo1g | DYLSSDTDNPTASHL | -844 | -858 | 0 | 0.001636916 | 15 | 1709.0684 | NaN | NaN | NaN | NaN | NaN |
| Myo1g | HYAGDVTYSVE | -525 | -535 | 0 | 0.007937545 | 11 | 1164.8579 | NaN | NaN | NaN | NaN | NaN |
| Cth | LVWIETPTNPTLK | -152 | -164 | 0 | 0 | 13 | 178.83643 | 393.5805 | 469.45325 | 365.1714 | 566.4811 | 557.9071 |
| Cth | ATLGINDTLIR | -364 | -374 | 0 | 0 | 11 | 835.3262 | NaN | 867.3805 | 983.48944 | NaN | 409.82916 |
| Gnpnat1 | ATLIIEHK | -101 | -108 | 0 | 0 | 8 | 663.3135 | NaN | 1215.7466 | 979.1275 | 1087.8107 | 796.65314 |
| Mtch2 | ATLITHPFHVITLR | -134 | -147 | 0 | 0 | 14 | 343.8493 | 335.92618 | 324.87036 | 287.72668 | 469.1515 | 366.04996 |
| Mtch2 | LCSGVLGTVVHGK | -78 | -90 | 0 | 0 | 13 | 921.89197 | 652.4583 | 1090.1168 | 351.62915 | NaN | NaN |
| Mtch2 | GLFTGLTPR | -69 | -77 | 0 | 0 | 9 | 814.5279 | 191.49025 | 320.8954 | 843.99426 | NaN | NaN |
| Mtch2 | VLQYYQESEKPEELGSVT | -91 | -108 | 1 | 0.001941082 | 18 | 237.83308 | NaN | 215.73709 | NaN | 323.31497 | 397.54706 |
| Mtch2 | ITHPFHVITLR | -137 | -147 | 0 | 0 | 11 | 347.07 | NaN | NaN | NaN | 604.5509 | NaN |
| Mtch2 | VLIQVGYEPLPPTIGR | -26 | -41 | 0 | 0.001636916 | 16 | 318.2501 | NaN | NaN | NaN | NaN | NaN |
| Ppp2r5a | DTTLTEPVIR | -303 | -312 | 0 | 0 | 10 | 2001.903 | 1952.14 | NaN | 1625.1246 | 2196.872 | 2221.7192 |
| Ppp2r5a | ATLNELVEYVSTNR | -99 | -112 | 0 | 0 | 14 | 753.8189 | 741.1208 | NaN | 855.2461 | 910.52094 | 668.6939 |
| Ppp2r5a | FLESPDFQPSIAK | -169 | -181 | 0 | 0 | 13 | 3706.4097 | 4867.1665 | 4845.4688 | NaN | NaN | NaN |
| Ndufb4 | VSHIEDPALIR | -57 | -67 | 0 | 0 | 11 | 1606.83 | 1781.8557 | 1604.5734 | 1851.5734 | 1807.1581 | 1700.9822 |
| Ndufb4 | ATLPSTLDPAEYDVSPETR | -12 | -30 | 0 | 0 | 19 | 798.31384 | 875.86975 | 955.6151 | 766.24713 | 1282.1776 | 1197.9412 |
| Gnb1 | LSQITNNIDPVGR | -30 | -42 | 0 | 0.000550307 | 13 | 775.0952 | 745.206 | NaN | NaN | NaN | 842.4155 |
| Gnb1 | GYDDFNCNVWDALK | -288 | -301 | 0 | 0 | 14 | 176.02472 | 272.38257 | NaN | 206.32239 | NaN | NaN |
| Gnb1 | ATLSQITNNIDPVGR | -28 | -42 | 0 | 0.00025015 | 15 | 584.6901 | NaN | NaN | 696.384 | NaN | NaN |
| Vav1 | LNPGDIVELTK | -619 | -629 | 0 | 0 | 11 | 4549.004 | 5905.646 | 6511.3223 | 6010.7354 | 7719.8267 | 7018.4414 |
| Vav1 | DRSELSLK | -797 | -804 | 1 | 0 | 8 | 1650.196 | 1804.3573 | 2014.7212 | 1102.4358 | 1503.5688 | 1500.7334 |
| Vav1 | NTATNEVGWFPCNR | -641 | -654 | 0 | 0 | 14 | 1237.4651 | 1260.5739 | 1178.6051 | 2053.484 | 1491.0044 | 406.26743 |
| Vav1 | ATNEVGWFPCNR | -643 | -654 | 0 | 0 | 12 | 467.1607 | 1173.9618 | 818.6035 | 1051.9196 | 1360.6 | 1627.4774 |
| Vav1 | RNELGLPK | -589 | -596 | 1 | 0.001636916 | 8 | 1408.7333 | 1427.488 | 1786.1418 | NaN | NaN | NaN |
| Vav1 | NLDQSLANYGR | -392 | -402 | 0 | 0 | 11 | 811.44867 | 642.7532 | 212.25516 | NaN | NaN | NaN |
| Vav1 | DLFDVQDFGK | -98 | -107 | 0 | 0 | 10 | 493.82214 | NaN | NaN | 490.94797 | NaN | 642.23004 |
| Glrx | NNTSAIQDYLQQLTGAR | -52 | -68 | 0 | 0 | 17 | 1280.0939 | 1228.6211 | 1255.6832 | 2493.3428 | 3037.8599 | 2748.6055 |
| Glrx | KTQEILSQLPFK | -28 | -39 | 1 | 0 | 12 | 1042.0526 | 1356.495 | 1104.0513 | 1033.2439 | 1417.6816 | 1529.6707 |
| Glrx | ATNNTSAIQDYLQQLTGAR | -50 | -68 | 0 | 0 | 19 | 693.7089 | 826.34106 | 876.13367 | 764.5793 | 932.82825 | 1000.6096 |
| Pnkp | FELPAFDPR | -335 | -343 | 0 | 0 | 9 | 806.11774 | 763.61206 | 417.11176 | 704.7769 | 1072.2689 | 398.6972 |
| Pnkp | NQVELIADPESR | -49 | -60 | 0 | 0 | 12 | 459.28564 | 1196.4349 | 1048.3369 | 1461.0643 | NaN | 1243.1729 |
| Pnkp | GKLPAEVFK | -224 | -232 | 1 | 0 | 9 | 395.51413 | 379.11078 | 487.77866 | 367.54077 | 491.64816 | NaN |
| Pnkp | ILYPEIPK | -193 | -200 | 0 | 0 | 8 | 611.964 | 293.4912 | NaN | NaN | 236.74416 | 173.91475 |
| Pnkp | VFPTSPSDWR | -183 | -192 | 0 | 0 | 10 | 423.29492 | 711.54285 | 597.39453 | NaN | NaN | 1089.9141 |
| Pnkp | LQEHLDPALQR | -504 | -514 | 0 | 0 | 11 | 3458.3418 | 3506.7192 | 3392.5173 | NaN | NaN | NaN |
| Pnkp | ATPEEFFLK | -321 | -329 | 0 | 0.000550307 | 9 | 1261.527 | 1447.2118 | 1629.8992 | NaN | NaN | NaN |
| Hsd17b10 | LVAGEMGQNEPDQGGQR | -131 | -147 | 0 | 0 | 17 | 661.08716 | 176.08174 | 456.24533 | 488.41003 | 731.89166 | 813.91693 |
| Hsd17b10 | LGDPAEYAHLVQ | -227 | -238 | 0 | 0 | 12 | 1259.4387 | 1381.6187 | 1367.1329 | 1042.8975 | NaN | NaN |
| Hsd17b10 | IHTLEDFQR | -108 | -116 | 0 | 0.000826806 | 9 | 714.92175 | NaN | NaN | NaN | 1014.05756 | NaN |
| Hsd17b10 | ATPLLTTLPEK | -202 | -212 | 0 | 0.000591124 | 11 | 609.4044 | NaN | 271.04166 | NaN | NaN | NaN |
| Tcof1 | VLTELLEQER | -1162 | -1171 | 0 | 0 | 10 | 5332.418 | 5881.1797 | 6504.616 | 4796.795 | 7354.134 | 7005.5464 |
| Tcof1 | ATPRPDSNSLASSAPATK | -1065 | -1082 | 1 | 0.001708008 | 18 | 498.19022 | 503.88837 | 495.75006 | 471.701 | 607.60504 | 629.0471 |
| Tcof1 | QKAEDDETLQAK | -63 | -74 | 1 | 0 | 12 | 490.1952 | 885.64856 | 899.1635 | 576.3668 | 1218.2119 | 658.4041 |
| Tcof1 | SDVDSSSESESEGAAPSTPR | -603 | -622 | 0 | 0 | 20 | 381.67682 | 424.45145 | 374.91824 | 578.7796 | 483.86395 | 642.65045 |
| Tcof1 | LGNVAPTPAKPAR | -242 | -254 | 1 | 0.00025015 | 13 | 781.8764 | 779.78735 | 787.48047 | NaN | NaN | NaN |
| Tcof1 | ATPRPTPVNSA | -103 | -113 | 1 | 0.002068213 | 11 | 1965.6631 | NaN | NaN | NaN | NaN | 1886.68 |
| Tcof1 | STSSSPAPTQTLPNSITQR | -1124 | -1142 | 0 | 0.00111006 | 19 | 318.05124 | 228.56708 | NaN | NaN | NaN | NaN |
| Ap1b1 | LVEDQGFLDTLK | -155 | -166 | 0 | 0 | 12 | 1080.9324 | 781.7892 | 800.87335 | 1449.5532 | 1751.0165 | 1658.2216 |
| Ap1b1 | ENLDSLDEPEAR | -434 | -445 | 0 | 0 | 12 | 1269.8062 | 1858.6599 | 1483.4038 | NaN | 3567.789 | 2809.8074 |
| Ap1b1 | TLCENLDSLDEPEAR | -431 | -445 | 0 | 0.00025015 | 15 | 1682.9393 | 2171.037 | NaN | NaN | 4861.969 | NaN |
| Ap1b1 | DIANENEAQFQIR | -849 | -861 | 0 | 0 | 13 | 1055.0646 | 1072.3748 | NaN | NaN | 781.7305 | NaN |

| | | | | | | | | | | | | |
|---|---|---|---|---|---|---|---|---|---|---|---|---|
| Ap1b1 | KPTETQELVQQ | -495 | -505 | 1 | 0.000989247 | 11 | 1041.8694 | NaN | NaN | NaN | NaN | 1693.4291 |
| Vasp | VKQELLEEVR | -342 | -351 | 1 | 0 | 10 | 1653.0806 | 1853.1047 | 423.81897 | 1817.3905 | 2142.7627 | 2037.3074 |
| Vasp | VKQELLEEVRK | -342 | -352 | 2 | 0 | 11 | 1070.6969 | 1333.2471 | 1415.2341 | 1110.7335 | 1138.2343 | 1563.8235 |
| Vasp | VKQELLEEVRK | -342 | -352 | 2 | 0 | 11 | 1070.6969 | 1333.2471 | 1415.2341 | 1110.7335 | 1138.2343 | 1563.8235 |
| Vasp | EAHPSTPCSSDDSDLER | -325 | -341 | 0 | 0 | 17 | 717.13086 | 825.86865 | 835.8465 | 918.09753 | 1170.7515 | 1097.1581 |
| Vasp | EAHPSTPCSSDDSDLER | -325 | -341 | 0 | 0 | 17 | 717.13086 | 825.86865 | 835.8465 | 918.09753 | 1170.7515 | 1097.1581 |
| Vasp | NGPSPEELEQQK | -131 | -142 | 0 | 0 | 12 | 5593.4883 | 5698.6714 | 5639.98 | 1165.5494 | 1439.2404 | NaN |
| Vasp | QATPIFHQWR | -74 | -83 | 0 | 0 | 10 | 889.35565 | 693.0124 | 421.1106 | 1076.6398 | 842.32025 | NaN |
| Vasp | YNQATPIFHQW | -72 | -82 | 0 | 0 | 11 | 495.7101 | 349.0562 | 541.73193 | 677.67163 | 619.8962 | NaN |
| Vasp | QELLEEVR | -344 | -351 | 0 | 0.000471409 | 8 | 603.9053 | NaN | 418.1255 | 662.2445 | NaN | 377.35367 |
| Vasp | YNQATPIFHQWR | -72 | -83 | 0 | 0 | 12 | 1027.4568 | 1079.0475 | 1071.8367 | NaN | NaN | NaN |
| Vasp | LPAQSEPVR | -294 | -302 | 0 | 0.000431655 | 9 | 811.36646 | NaN | NaN | 699.94385 | NaN | NaN |
| Vasp | ATQVGEKPPKDESASEE | -273 | -289 | 2 | 0.005668207 | 17 | 719.0976 | NaN | NaN | NaN | NaN | NaN |
| Kpna6 | ATSGGTPEQIR | -405 | -415 | 0 | 0 | 11 | 1504.1179 | NaN | 1418.7463 | NaN | NaN | NaN |
| Kpna6 | VSPCLPVLSR | -250 | -259 | 0 | 0.002972566 | 10 | 387.52933 | NaN | 578.6798 | NaN | NaN | NaN |
| Cmas | TDHDEIENVAK | -90 | -100 | 0 | 0 | 11 | 1018.2871 | 956.0454 | 982.92487 | 1058.6775 | 933.0546 | 6959.909 |
| Cmas | ATSPCLHPTDLQK | -142 | -154 | 0 | 0.000627943 | 13 | 876.3485 | 833.78644 | 829.8475 | 938.08374 | 914.06055 | 227.07 |
| Cmas | EVAYLGNEVSDEECLKR | -365 | -381 | 1 | 0.00133086 | 17 | 610.4124 | 655.91364 | 713.1434 | 738.10065 | 316.93658 | 828.97363 |
| Cmas | VGLSAVPADACSGAQK | -382 | -397 | 0 | 0 | 16 | 438.34915 | NaN | 571.05853 | 649.8501 | 746.3143 | NaN |
| Cmas | LATVDEWR | -349 | -356 | 0 | 0.000550307 | 8 | 350.25366 | NaN | NaN | 496.42548 | NaN | 684.0976 |
| Cmas | DHDEIENVAK | -91 | -100 | 0 | 0.00272025 | 10 | 900.41156 | NaN | NaN | NaN | NaN | NaN |
| Lsm5 | TNPSQLLPLELVDK | -7 | -20 | 0 | 0 | 14 | 1370.1782 | 1448.9384 | 1623.9633 | 1796.056 | 2497.9333 | 2622.7866 |
| Lsm5 | ATTNPSQLLPLELVDK | -5 | -20 | 0 | 0.000431655 | 16 | 438.06848 | 470.34256 | 426.7571 | 71.10231 | 483.45923 | 472.17746 |
| Lsm5 | VLEDVTEFEITPEGR | -50 | -64 | 0 | 0 | 15 | 382.31848 | 361.24805 | 483.13 | 364.127 | 426.87817 | 456.99973 |
| Lsm5 | SDKEIVGTLL | -32 | -41 | 1 | 0.00025015 | 10 | 1318.2894 | 1304.987 | NaN | 1122.9083 | 1099.4827 | 1248.4692 |
| Lsm5 | TTNPSQLLPLELVDK | -6 | -20 | 0 | 0 | 15 | 662.7228 | 644.078 | NaN | 685.9961 | 740.5856 | 815.69336 |
| Kdelr1;Kdelr2 | ATYDGNHDTFR | (84);(84) | (94);(94) | 0 | 0.000627943 | 11 | 1915.3593 | 2172.5083 | 2115.0535 | 1823.3127 | 1723.7421 | 3289.4692 |
| Psma2 | RLTPTEVR | -220 | -227 | 1 | 0 | 8 | 1555.0135 | 1579.9231 | 173.51624 | 1503.1348 | 1820.5399 | 1750.8071 |
| Psma2 | VAGGAPSVGIK | -29 | -39 | 0 | 0 | 11 | 1196.815 | 1155.1749 | 210.41818 | 284.22437 | 4136.754 | 1051.572 |
| Psma2 | SLTTFSPSGK | -9 | -18 | 0 | 0 | 10 | 1028.1958 | 778.2185 | 763.5102 | 1130.3947 | 1176.0059 | 1041.1035 |
| Psma2 | AVAGGAPSVGIK | -28 | -39 | 0 | 0.00025015 | 12 | 878.97437 | 697.1524 | 691.03015 | 1618.7518 | 1647.7285 | 1579.3021 |
| Psma2 | YNEDLELEDAIHT | -178 | -190 | 0 | 0 | 13 | 801.12256 | 435.03064 | 660.1763 | 676.8354 | 727.4306 | NaN |
| Psma2 | RYNEDLELEDAIHT | -177 | -190 | 1 | 0.001636916 | 14 | 1519.7808 | NaN | NaN | 730.18414 | 2513.1436 | 1695.49 |
| Psma2 | YNEDLELEDAIHTA | -178 | -191 | 0 | 0 | 14 | 569.51 | 813.09686 | 794.6225 | NaN | NaN | NaN |
| Psma2 | RYNEDLELEDAIHTA | -177 | -191 | 1 | 0.000947459 | 15 | 398.94757 | NaN | 327.06964 | 143.89204 | NaN | NaN |
| Psma2 | SGMGPDYR | -77 | -84 | 0 | 0.008129024 | 8 | 526.76135 | NaN | NaN | NaN | NaN | 611.8846 |
| Psmd14 | AVAVVVDPIQSVK | -140 | -152 | 0 | 0 | 13 | 985.92255 | 1124.0258 | 1125.0726 | 1166.6395 | 1184.1036 | 172.13098 |
| Psmd14 | QTTSNLGHLNKPSIQ | -176 | -190 | 1 | 0 | 15 | 606.2953 | 641.5267 | 776.3654 | 849.57715 | 811.92755 | 763.2155 |
| Psmd14 | RHLEEHVDVL | -282 | -291 | 1 | 0.000712251 | 10 | 1342.5731 | 1454.866 | NaN | NaN | 1428.4052 | 1347.7142 |
| Psmd14 | VVDPIQSVK | -144 | -152 | 0 | 0.000394361 | 9 | 603.9538 | 651.0529 | 565.1187 | 617.4379 | NaN | NaN |
| Psmd14 | SVEAVDPVFQAK | -83 | -94 | 0 | 0.00272025 | 12 | 1089.6987 | 1049.8126 | NaN | NaN | 333.16098 | NaN |
| Ndufa5 | AVCDTPHER | -15 | -23 | 0 | 0.000471409 | 9 | 111.6148 | 552.4261 | NaN | 563.4819 | 718.3469 | 630.1576 |
| Ddx19a | DGNPDNETYLHR | -414 | -425 | 0 | 0 | 12 | 2413.034 | 2280.284 | 1680.1898 | 3177.109 | 2751.058 | 3126.965 |
| Ddx19a | DGNPDNETYLHR | -414 | -425 | 0 | 0 | 12 | 2413.034 | 2280.284 | 1680.1898 | 3177.109 | 2751.058 | 3126.965 |
| Ddx19a | SATFEDSVWK | -273 | -282 | 0 | 0 | 10 | 1846.5244 | NaN | 2419.0024 | 1696.0092 | 793.08997 | 1712.8489 |
| Ddx19a | AVDEQEAAVK | -9 | -18 | 0 | 0 | 10 | 2450.4734 | 3118.0645 | NaN | 2385.6687 | NaN | NaN |
| Ddx19a | IVIGTPGTVLDWCSK | -212 | -226 | 0 | 0.00025015 | 15 | 725.9284 | 803.9885 | NaN | NaN | NaN | 1062.7845 |
| Thbs1 | LCNNPTPQFGGK | -518 | -529 | 0 | 0 | 12 | 1315.2711 | 3134.5635 | 3498.4517 | 3396.2395 | 3142.0908 | 3352.2517 |
| Thbs1 | FTGSQPFGR | -624 | -632 | 0 | 0 | 9 | 866.93567 | 1163.5939 | 1114.8203 | 1086.3057 | 1111.0465 | 1189.62 |
| Thbs1 | NALWHTGNTPGQVR | -1078 | -1091 | 0 | 0 | 14 | 828.0969 | 1100.5957 | 443.19476 | 903.26843 | 1182.0492 | 837.79895 |
| Thbs1 | NTDPGYNCLPCPPR | -610 | -623 | 0 | 0 | 14 | 700.7304 | 920.4827 | 849.87885 | 795.4922 | 855.52625 | 821.678 |
| Thbs1 | DCVGDVTENQVCNK | -530 | -543 | 0 | 0 | 14 | 509.36642 | 544.4418 | 462.69244 | 629.52466 | 483.77734 | 515.9712 |
| Thbs1 | TCHIQECDKR | -422 | -431 | 1 | 0.00025015 | 10 | 448.222 | 445.9753 | 653.7043 | 433.1138 | 705.9785 | 74.14635 |
| Thbs1 | TCHIQECDKR | -422 | -431 | 1 | 0.00025015 | 10 | 448.222 | 445.9753 | 653.7043 | 433.1138 | 705.9785 | 74.14635 |
| Thbs1 | FVFGTTPEDILR | -217 | -228 | 0 | 0 | 12 | 348.97153 | 1249.0714 | 1293.2952 | 1196.4457 | 1707.0175 | 1608.6287 |
| Thbs1 | DHDDDNDGIPDDR | -895 | -907 | 0 | 0 | 13 | 687.54974 | 916.916 | 870.2202 | 960.17456 | NaN | NaN |
| Thbs1 | QYDYDRDDVGDR | -762 | -773 | 1 | 0.001776473 | 12 | 676.2697 | 737.14856 | 738.4292 | 681.4234 | NaN | NaN |
| Thbs1 | AVDIDGDGILNER | -798 | -810 | 0 | 0 | 13 | 1057.0436 | 959.6792 | 1182.3522 | NaN | NaN | NaN |
| Rpl6 | AVPQLQGYLR | -271 | -280 | 0 | 0.000230707 | 10 | 2398.0708 | 2777.7039 | 2695.537 | 2754.0757 | 2536.3389 | 2552.5288 |
| Rpl6 | AVDLQILPK | -260 | -268 | 0 | 0 | 9 | 2082.1228 | 2119.5972 | 2169.7842 | 2261.1904 | 1864.7192 | 1575.971 |
| Rpl6 | SSITPGTVLIILTGR | -150 | -164 | 0 | 0 | 15 | 205.48836 | 274.34488 | 343.78192 | 308.54382 | 239.531 | 221.59645 |
| Rpl6 | RNPVLVR | -53 | -59 | 1 | 0.008498216 | 7 | 26647.469 | 24858.805 | NaN | 23018.273 | NaN | NaN |
| Rpl6 | KVDISDVK | -208 | -215 | 1 | 0.001636916 | 8 | 1048.0645 | NaN | 1190.9813 | 934.0343 | NaN | NaN |
| Rpl6 | HQEGEIFDTEK | -235 | -245 | 0 | 0 | 11 | 1035.373 | 1078.5104 | 1055.091 | NaN | NaN | NaN |
| Rpl6 | YYPTEDVPR | -123 | -131 | 0 | 0 | 9 | 700.0982 | 594.71954 | 342.54462 | NaN | NaN | NaN |
| Rpl6 | QLDSGLLLVTGPLVINR | -175 | -191 | 0 | 0.005522803 | 17 | 315.80347 | 308.33804 | 339.2874 | NaN | NaN | NaN |
| Rpl6 | EKYEITEQR | -246 | -254 | 1 | 0.001055699 | 9 | 3603.2837 | NaN | 4078.5476 | NaN | NaN | NaN |
| Rpl6 | VTGPLVINR | -183 | -191 | 0 | 0 | 9 | 1557.8042 | NaN | NaN | NaN | NaN | NaN |
| Nherf1 | EELLRPQEK | -108 | -116 | 1 | 0 | 9 | 1596.0334 | 1883.698 | 1800.2063 | 1208.0432 | 1055.3375 | 1002.723 |
| Nherf1 | LLVVDPETDER | -88 | -98 | 0 | 0 | 11 | 2447.4214 | 2321.3928 | 1423.7004 | 1553.9003 | 1528.2518 | NaN |
| Nherf1 | VVDKETDEFFK | -225 | -235 | 1 | 0 | 11 | 5915.9443 | NaN | NaN | 6251.7285 | 332.64078 | 7540.7754 |
| Nherf1 | QHGDVVSAIK | -206 | -215 | 0 | 0 | 10 | 1199.0764 | 1204.4933 | 1153.804 | 773.9994 | NaN | NaN |
| Nherf1 | SEQAEPPAAADTHEAGDQNEAEK | -117 | -139 | 0 | 0 | 23 | 409.61063 | 452.62866 | 538.5454 | 508.33295 | NaN | NaN |
| Nherf1 | ETHQQVVSR | -70 | -78 | 0 | 0.00025015 | 9 | 716.53015 | NaN | NaN | NaN | 1128.6768 | 998.3956 |
| Nherf1 | AVDPDSPAEASGLR | -176 | -189 | 0 | 0 | 14 | 654.1494 | 604.18176 | 592.1617 | NaN | NaN | NaN |
| Nherf1 | LLVVDKETDEFFK | -223 | -235 | 1 | 0 | 13 | 334.45816 | 409.93527 | 459.87732 | NaN | NaN | NaN |
| Nherf1 | EALVEPASESPR | -266 | -277 | 0 | 0 | 12 | 1360.789 | NaN | 767.31573 | NaN | NaN | NaN |
| Nherf1 | LVEPGSPAEK | -41 | -50 | 0 | 0 | 10 | 462.69348 | 798.9595 | NaN | NaN | NaN | NaN |

| | | | | | | | | | | | | |
|---|---|---|---|---|---|---|---|---|---|---|---|---|
| Lgals3bp | QTPQHPSFLFK | -454 | -464 | 0 | 0 | 11 | 1237.709 | 1250.3528 | 2044.2089 | 1281.9019 | 1871.9186 | 1963.4264 |
| Lgals3bp | EFHTVPVEVLAK | -382 | -393 | 0 | 0 | 12 | 1067.6984 | 947.7119 | 1014.42346 | 1496.3329 | 959.6907 | 912.53644 |
| Lgals3bp | AVDQWSTETIASHEDIER | -324 | -341 | 0 | 0 | 18 | 342.6426 | 300.9343 | 276.27155 | 253.81958 | 385.7089 | 343.7451 |
| Lgals3bp | SYSNPTIGYENR | -502 | -513 | 0 | 0 | 12 | 952.0569 | NaN | NaN | 2351.8406 | 2297.1318 | 767.4288 |
| Lgals3bp | GATELQDYCGR | -235 | -245 | 0 | 0 | 11 | 636.89795 | NaN | NaN | 773.14856 | 564.58655 | 133.7028 |
| Lgals3bp | TTSSYSNPTIGYENR | -499 | -513 | 0 | 0 | 15 | 446.9365 | 517.04626 | NaN | 470.16824 | NaN | 431.33087 |
| Ctla4 | AVDTGLYLCK | -121 | -130 | 0 | 0 | 10 | 495.95526 | NaN | 716.9278 | 548.12396 | NaN | NaN |
| Tufm | KGDECELLGHNK | -286 | -297 | 1 | 0 | 12 | 1127.0806 | 1158.6976 | 1167.7502 | 995.25903 | 1401.6346 | 1293.1803 |
| Tufm | KGDECELLGHNK | -286 | -297 | 1 | 0 | 12 | 1127.0806 | 1158.6976 | 1167.7502 | 995.25903 | 1401.6346 | 1293.1803 |
| Tufm | QIGVEHVVVY | -170 | -179 | 0 | 0.00025015 | 10 | 1281.5358 | 1050.789 | 992.5386 | 421.45184 | 988.8639 | 1000.44446 |
| Tufm | LLDAVDTYIPVPTR | -239 | -252 | 0 | 0 | 14 | 1043.8215 | 1174.6907 | 1042.479 | 1555.717 | 2532.2717 | 2261.9612 |
| Tufm | KYEEIDNAPEER | -91 | -102 | 1 | 0 | 12 | 705.121 | 694.056 | 678.3158 | 573.47485 | 716.9642 | 821.73816 |
| Tufm | VGTIGHVDHGK | -60 | -70 | 0 | 0 | 11 | 523.1923 | 651.01013 | 661.9817 | 947.89355 | 305.96454 | 3179.538 |
| Tufm | VGTIGHVDHGK | -60 | -70 | 0 | 0 | 11 | 523.1923 | 651.01013 | 661.9817 | 947.89355 | 305.96454 | 3179.538 |
| Tufm | YEEIDNAPEER | -92 | -102 | 0 | 0 | 11 | 439.01544 | 466.83237 | 548.6384 | 199.94609 | 622.3818 | 317.36792 |
| Tufm | DLDKPFLLPVESVY | -253 | -266 | 1 | 0 | 14 | 364.06854 | 251.64468 | 332.8862 | 333.87704 | 327.81006 | 343.05325 |
| Tufm | DKPHVNVGTIGHVD | -54 | -67 | 1 | 0 | 14 | 2274.6343 | 2284.4817 | 1893.905 | NaN | NaN | NaN |
| Tufm | DKPHVNVGTIGHVD | -54 | -67 | 1 | 0 | 14 | 2274.6343 | 2284.4817 | 1893.905 | NaN | NaN | NaN |
| Tufm | AVDTYIPVPTR | -242 | -252 | 0 | 0 | 11 | 624.32526 | 1439.6864 | 1673.4357 | NaN | NaN | NaN |
| Slc1a5 | KLGPEGELLIR | -259 | -269 | 1 | 0 | 11 | 782.9133 | 880.6985 | 836.4221 | 698.02313 | 716.7701 | 622.48663 |
| Slc1a5 | NEVSLNPLPLATEEGNPLLK | -515 | -534 | 0 | 0.00025015 | 20 | 416.8833 | 441.3425 | 341.17212 | 357.26724 | 501.03848 | 514.4856 |
| Slc1a5 | QYQGPTGDSSATFEK | -535 | -549 | 0 | 0 | 15 | 349.9537 | 468.29007 | 681.1851 | 328.90952 | 493.85275 | 506.67313 |
| Slc1a5 | AVDWLVDR | -470 | -477 | 0 | 0.000627943 | 8 | 1825.4763 | 1899.5997 | 1877.3335 | NaN | 1662.9696 | NaN |
| Slc1a5 | QYQGPTGDSSATFEKE | -535 | -550 | 1 | 0.00025015 | 16 | 742.8043 | 674.7386 | 784.7844 | 749.47046 | NaN | NaN |
| Cpsf3 | SAIPAEESDQLLIRPL | -2 | -17 | 1 | 0 | 16 | 382.8194 | NaN | 335.25058 | 479.03088 | 665.38464 | 676.87616 |
| Cpsf3 | AVECEEGSEDDESLR | -652 | -666 | 0 | 0 | 15 | 376.57162 | NaN | 373.87817 | 315.96286 | 303.3042 | 326.88107 |
| Cpsf3 | LYEALTPVH | -676 | -684 | 0 | 0 | 9 | 68.04115 | 938.16644 | 781.43506 | NaN | NaN | NaN |
| Cpsf3 | LLYTGDFSR | -174 | -182 | 0 | 0 | 9 | 2232.8765 | NaN | NaN | 1623.0684 | NaN | NaN |
| Psmb9 | AVEFDGGVVVGSDSR | -25 | -39 | 0 | 0 | 15 | 2379.1301 | 2360.164 | 2515.8413 | 2917.619 | 2742.3384 | 2575.9712 |
| Psmb9 | VILGDELPK | -207 | -215 | 0 | 0 | 9 | 2125.266 | 2205.8015 | 2546.843 | 2407.9163 | 2567.4043 | 2380.6218 |
| Psmb9 | EFDGGVVVGSDSR | -27 | -39 | 0 | 0 | 13 | 1107.5652 | 654.281 | 1277.1755 | NaN | NaN | NaN |
| Grk2 | AVELPDSFSPELR | -410 | -422 | 0 | 0 | 13 | 2840.3394 | 2919.6313 | 3424.3408 | 1483.8368 | 1836.1414 | 2540.7378 |
| Grk2 | ILLPEPSIR | -32 | -40 | 0 | 0 | 9 | 2785.1184 | 445.85852 | 2203.6885 | 361.0398 | 752.0396 | 811.8644 |
| Grk2 | NFPLTISER | -508 | -516 | 0 | 0 | 9 | 872.6201 | 829.44904 | 1429.6902 | 457.99164 | 1207.5049 | 1139.261 |
| Grk2 | LLDSDQELYR | -498 | -507 | 0 | 0 | 10 | 839.76373 | 3056.8474 | 3102.596 | 1973.8665 | 822.09393 | 1667.3823 |
| Grk2 | LGYLLFR | -63 | -69 | 0 | 0.00223472 | 7 | 400.83212 | 321.03323 | 434.10376 | 119.55506 | 231.88576 | 199.43805 |
| Psma3 | SNFGYNIPLK | -101 | -110 | 0 | 0 | 10 | 3977.39 | 4185.805 | 4379.6416 | 3535.576 | 3809.699 | 4044.7659 |
| Psma3 | IIYIVHDEVK | -197 | -206 | 0 | 0.00025015 | 10 | 3751.4836 | 3610.3215 | 3597.9758 | 3966.7136 | 3526.953 | 3375.5754 |
| Psma3 | IIYIVHDEVK | -197 | -206 | 0 | 0.00025015 | 10 | 3751.4836 | 3610.3215 | 3597.9758 | 3966.7136 | 3526.953 | 3375.5754 |
| Psma3 | AVENSSTAIGIR | -30 | -41 | 0 | 0 | 12 | 2613.196 | 2715.259 | 2428.3625 | 3008.8003 | 2716.909 | 2819.8562 |
| Psma3 | LYEEGSNKR | -58 | -66 | 1 | 0 | 9 | 802.1182 | 711.81714 | 830.92834 | 840.19666 | 909.04047 | 821.1176 |
| Psma3 | YIVHDEVK | -199 | -206 | 0 | 0.00437931 | 8 | 2457.4382 | 2345.5076 | 2374.8274 | NaN | 2168.271 | 2317.8132 |
| Psma3 | ELSWVGELTK | -213 | -222 | 0 | 0 | 10 | 1039.44 | 371.105 | 976.89087 | 178.67 | 558.1814 | NaN |
| Psma3 | GRHEIVPK | -223 | -230 | 1 | 0.007105559 | 8 | 458.72827 | 344.3036 | 375.0998 | 366.14343 | NaN | 320.30157 |
| Psma3 | RSNFGYNIPLK | -100 | -110 | 1 | 0.000431655 | 11 | 2536.4448 | NaN | NaN | 3487.5554 | NaN | NaN |
| Psmd5 | LEAPLEELR | -15 | -23 | 0 | 0 | 9 | 3644.4658 | 1489.0217 | 1403.6682 | 6591.2104 | 2068.278 | 1108.9589 |
| Psmd5 | FESNLLDDLK | -163 | -172 | 0 | 0 | 10 | 1486.5374 | 1649.9813 | 210.4878 | 521.40094 | 413.59088 | 251.67831 |
| Psmd5 | GISNQPFPELHCA | -401 | -413 | 0 | 0 | 13 | 390.53525 | 436.88763 | 418.68036 | 531.7996 | 661.78796 | 731.58124 |
| Psmd5 | TAIADQPWAQR | -419 | -429 | 0 | 0 | 11 | 1880.2483 | NaN | 2256.4043 | 2147.5469 | 1697.4248 | 1845.7302 |
| Psmd5 | AVEPIHLAR | -74 | -82 | 0 | 0 | 9 | 950.405 | 1192.6251 | 1026.312 | NaN | 1340.1805 | 1610.9631 |
| Psmd5 | IADQPWAQR | -421 | -429 | 0 | 0 | 9 | 623.8138 | 289.81042 | 351.4699 | 574.15314 | 705.171 | NaN |
| Psmd5 | LRPLFSLL | -46 | -53 | 1 | 0.006164361 | 8 | 241.34189 | 289.31738 | 320.16696 | NaN | 291.65533 | 278.93335 |
| Sh3bgrl3 | LVDISQDNALR | -37 | -47 | 0 | 0 | 11 | 5821.511 | 5784.489 | 6033.403 | 1482.3121 | 6513.5986 | 6300.6353 |
| Sh3bgrl3 | AVEQDTLQEFLK | -80 | -91 | 0 | 0 | 12 | 3107.959 | 3908.4355 | 4029.224 | 4471.8745 | 4660.2617 | 5145.393 |
| Sh3bgrl3 | IQYQLVDISQDNALR | -33 | -47 | 0 | 0.00025015 | 15 | 1334.017 | 1509.5565 | 1536.4729 | 1609.021 | 2003.3822 | 1859.0847 |
| Sh3bgrl3 | LVDISQDNALRDEMR | -37 | -51 | 1 | 0.006100831 | 15 | 555.6355 | 319.51566 | NaN | 798.77264 | NaN | NaN |
| Hirip3 | AVESTDEDHQTDLDAK | -131 | -146 | 0 | 0.000550307 | 16 | 323.5709 | 384.04105 | 446.2838 | NaN | 298.6539 | 322.11005 |
| Hirip3 | NAWNPSGEGTSPGETYR | -554 | -570 | 0 | 0 | 17 | 556.5374 | 665.4179 | 933.8873 | NaN | NaN | NaN |
| Aars1 | SLGDAFPELK | -356 | -365 | 0 | 0 | 10 | 2467.043 | 0 | 4481.7915 | 2545.9702 | 2855.4043 | 3821.7646 |
| Aars1 | SVEFCGGTHLR | -719 | -729 | 0 | 0 | 11 | 2431.376 | 3470.8672 | 4525.194 | 4732.0786 | 5110.101 | 5153.062 |
| Aars1 | ESDGVLKPLPK | -225 | -235 | 1 | 0 | 11 | 2013.5227 | 2605.129 | 2258.409 | 2283.5056 | 2468.837 | 2324.3926 |
| Aars1 | VDDSSEDKTEFTVK | -551 | -564 | 1 | 0 | 14 | 1463.4174 | 1641.43 | 1795.7522 | 1251.6467 | 1601.7144 | 1568.1936 |
| Aars1 | HNDLDDVGK | -82 | -90 | 0 | 0 | 9 | 1456.549 | 1485.6676 | 1632.3358 | 1353.1078 | 1564.7834 | 1706.431 |
| Aars1 | AVFDETYPDPVR | -684 | -695 | 0 | 0 | 12 | 1308.3586 | 587.1447 | 1282.5931 | 968.8197 | 918.6412 | 1046.7859 |
| Aars1 | GGQIYDEGYLVK | -539 | -550 | 0 | 0 | 12 | 1267.2556 | 1192.7391 | 1008.2625 | 1390.4778 | 1550.373 | 1169.1937 |
| Aars1 | YFGGDEAAGLEPDLECR | -137 | -153 | 0 | 0 | 17 | 980.1963 | 1129.0988 | 1218.2747 | 43.120792 | 1545.0916 | 1479.1635 |
| Aars1 | QIWQNLGLDEAR | -154 | -165 | 0 | 0 | 12 | 973.14294 | 1403.481 | 1274.2772 | 335.63086 | 303.96512 | 1221.6102 |
| Aars1 | VGAEDADGIDMAYR | -283 | -296 | 0 | 0 | 14 | 778.08344 | 715.4912 | 559.1717 | 655.9413 | 687.2292 | 504.05923 |
| Aars1 | LADGGRPDNTGR | -309 | -320 | 1 | 0 | 12 | 450.22934 | 453.45837 | 476.93976 | 506.4843 | 541.3698 | 461.2592 |
| Aars1 | DAAHLVNQDDPNVLEIW | -199 | -215 | 0 | 0 | 17 | 272.26944 | 284.30908 | 236.59683 | 295.15454 | 422.9351 | 343.4672 |
| Aars1 | EFGIPVER | -125 | -132 | 0 | 0.00025015 | 8 | 1469.609 | 1130.0837 | 1143.3499 | 1257.2981 | NaN | 820.75793 |
| Aars1 | DIINEEEVQFLK | -373 | -384 | 0 | 0 | 12 | 1037.7284 | 86.2594 | 947.8751 | 777.38385 | 421.73212 | NaN |
| Aars1 | AVIPQWQK | -805 | -812 | 0 | 0.000550307 | 8 | 1021.6825 | 740.24756 | NaN | 673.3386 | 588.62506 | 267.1669 |
| Aars1 | QGGQIYDEGYLVK | -538 | -550 | 0 | 0 | 13 | 932.2968 | 960.3862 | 939.8954 | NaN | NaN | 1268.8353 |
| Aars1 | TQEFGIPVER | -123 | -132 | 0 | 0.00025015 | 10 | 2323.1375 | 2423.2683 | 2533.1753 | NaN | NaN | NaN |
| Aars1 | GSWSFGDYFK | -102 | -111 | 0 | 0.00025015 | 10 | 595.1478 | NaN | NaN | NaN | NaN | NaN |
| Ilkap | SCLEDDKIQTR | -343 | -353 | 1 | 0.00025015 | 11 | 1154.8708 | 846.23706 | 659.643 | NaN | 805.9863 | 1559.8148 |
| Ilkap | AVFDGHGGIR | -149 | -158 | 0 | 0.00025015 | 10 | 2182.9683 | NaN | NaN | NaN | NaN | NaN |

| | | | | | | | | | | | | |
|---|---|---|---|---|---|---|---|---|---|---|---|---|
| Tuba4a | EIIDPVLDR | -113 | -121 | 0 | 0 | 9 | 9037.518 | 5366.3247 | 6260.1284 | 1716.3676 | 6730.208 | 6519.115 |
| Tuba4a | FVDLEPTVIDEIR | -67 | -79 | 0 | 0 | 13 | 806.33887 | 766.7656 | 680.7412 | 936.1033 | 1089.3474 | 1076.6648 |
| Tuba4a | KLSDQCTGLQGF | -124 | -135 | 1 | 0 | 12 | 806.04803 | 1196.2799 | 1084.0398 | 1319.7887 | 1536.9275 | 1005.9828 |
| Tuba4a | TIGGGDDSFTTF | -41 | -52 | 0 | 0.001055699 | 12 | 399.51697 | 496.82492 | NaN | 513.77026 | 682.6307 | 666.8635 |
| Tuba4a | AVFVDLEPTVIDEIR | -65 | -79 | 0 | 0 | 15 | 513.524 | NaN | 425.69952 | NaN | 664.08014 | 445.62622 |
| Tuba4a | LSDQCTGLQGFLVFH | -125 | -139 | 0 | 0.000431655 | 15 | 1716.215 | NaN | NaN | NaN | 1490.5754 | 1712.5989 |
| Tuba4a | FCETGAGK | -53 | -60 | 0 | 0.00051306 | 8 | 812.65924 | NaN | NaN | 570.0783 | NaN | 766.88837 |
| Tuba4a | VDLEPTVIDEIR | -68 | -79 | 0 | 0.00025015 | 12 | 780.02985 | NaN | NaN | NaN | 1165.3115 | 650.4175 |
| Tuba4a | LEPTVIDEIR | -70 | -79 | 0 | 0 | 10 | 2141.599 | NaN | 4785.389 | NaN | NaN | NaN |
| Psmf1 | SGIITPIHEQWEK | -136 | -148 | 0 | 0 | 13 | 1200.8107 | 1473.0815 | 1487.5873 | 1702.6664 | 2178.2012 | 2237.4856 |
| Psmf1 | VLIDPSSGLPNR | -220 | -231 | 0 | 0 | 12 | 766.7272 | 526.157 | 1245.8309 | 763.618 | 789.98236 | 784.1646 |
| Psmf1 | AVGGDDLDPFGCQR | -192 | -205 | 0 | 0 | 14 | 1043.7272 | NaN | NaN | NaN | 1206.2302 | 1142.0922 |
| Wrnip1 | AVGQETLLR | -240 | -248 | 0 | 0.00025015 | 9 | 668.46387 | 826.0966 | NaN | 1017.56104 | NaN | NaN |
| Bop1 | WLEVSEEEHQR | -501 | -511 | 0 | 0 | 11 | 955.03204 | 1171.3988 | 923.4916 | 1260.2152 | 1132.6458 | 1183.2816 |
| Bop1 | SYNPPPEYLPTEEER | -299 | -313 | 0 | 0 | 15 | 715.7002 | 777.959 | 824.2221 | 804.0355 | 1088.6895 | 938.8627 |
| Bop1 | YDEFPHVGYDLDGK | -130 | -143 | 0 | 0 | 14 | 694.4528 | 724.8479 | 645.9855 | 813.08704 | 873.38794 | 966.17664 |
| Bop1 | NDLLQNPLLVPVK | -682 | -694 | 0 | 0 | 13 | 365.80252 | 160.03947 | 362.80914 | 378.86838 | 368.41437 | 382.4953 |
| Bop1 | RPHDPTPSFYDL | -258 | -269 | 1 | 0.00201047 | 12 | 1104.9353 | 1546.7091 | 1529.8577 | 1378.5065 | 860.202 | NaN |
| Bop1 | SVSPGGQWLASGSDDGTLK | -406 | -424 | 0 | 0 | 19 | 806.47845 | 901.4799 | 940.63464 | NaN | 1538.724 | 978.4234 |
| Bop1 | SFIPSLVEK | -231 | -239 | 0 | 0 | 9 | 640.05646 | 726.0075 | 791.725 | 799.41626 | NaN | 856.8946 |
| Bop1 | LTDEQVALVHR | -182 | -192 | 0 | 0 | 11 | 1923.9972 | 1401.5247 | 2219.0112 | 2290.4753 | NaN | NaN |
| Bop1 | AVHPAGDNIICGSYDSK | -613 | -629 | 0 | 0.00111006 | 17 | 720.57245 | NaN | NaN | NaN | NaN | 599.624 |
| Bop1 | WAQEDPNAVLGR | -270 | -281 | 0 | 0.001708008 | 12 | 613.62646 | 562.5265 | NaN | NaN | NaN | NaN |
| Bop1 | CLDLYLCPR | -353 | -361 | 0 | 0.008045078 | 9 | 5989.7036 | NaN | NaN | NaN | NaN | NaN |
| Bop1 | TVHVGGVVR | -436 | -444 | 0 | 0.004031417 | 9 | 1171.2739 | NaN | NaN | NaN | NaN | NaN |
| Wdr1 | FTIGDHSR | -183 | -190 | 0 | 0.000627943 | 8 | 3314.3164 | 3281.2354 | 3180.1172 | 3324.3022 | 3377.2217 | 3434.149 |
| Wdr1 | DIAWTEDSKR | -107 | -116 | 1 | 0 | 10 | 2599.0889 | 693.27484 | 2784.5566 | 3188.3323 | 2596.1816 | 2681.5505 |
| Wdr1 | YWDSETGENDSFSGK | -348 | -362 | 0 | 0.00025015 | 15 | 2466.8477 | 2459.2993 | 2484.2632 | 1842.6259 | 1563.8396 | 1616.3977 |
| Wdr1 | YEYQPFAGK | -96 | -104 | 0 | 0 | 9 | 1348.2847 | 1337.6163 | 1305.5814 | 1300.6992 | 1176.3119 | 221.32587 |
| Wdr1 | IGQIVLLK | -426 | -433 | 0 | 0.00025015 | 8 | 1113.9017 | 840.7936 | 912.987 | 698.25146 | 446.4184 | 506.54865 |
| Wdr1 | LWDTGSSVGEITGHNK | -132 | -147 | 0 | 0 | 16 | 877.01166 | 1057.6057 | 1112.4138 | 1723.3201 | 103.6761 | 2167.885 |
| Wdr1 | VWTLSDPETK | -558 | -567 | 0 | 0.000431655 | 10 | 755.7137 | 1047.3354 | 1036.9343 | 1506.5498 | 957.94556 | 1163.1478 |
| Wdr1 | GPVTDVAYSHDGAFL | -490 | -504 | 0 | 0 | 15 | 651.4946 | 731.8975 | 761.3865 | 724.5575 | 612.6252 | 887.14886 |
| Wdr1 | SIDNPGYEPEVVAVHPGGDTVAVGGTDGNVR | -440 | -470 | 0 | 0 | 31 | 595.14575 | 591.846 | 637.4741 | 879.81464 | 1036.4244 | 979.607 |
| Wdr1 | SYIYSGSHDGHIN | -335 | -347 | 0 | 0 | 13 | 593.8642 | 685.5449 | 404.67303 | 928.91406 | 495.06586 | 531.77856 |
| Wdr1 | GPVTDVAYSHDGAFLAVC | -490 | -507 | 0 | 0 | 18 | 379.6804 | 435.24152 | 415.00177 | 895.1147 | 715.5246 | 670.8593 |
| Wdr1 | SVADGYSENNVFYGHHAK | -517 | -534 | 0 | 0 | 18 | 307.3124 | 366.84018 | 209.60004 | 387.2342 | 516.7281 | 515.1705 |
| Wdr1 | AVHPGGDTVAVGGTDGNVR | -452 | -470 | 0 | 0.000431655 | 19 | 779.80383 | 716.6215 | 750.9385 | NaN | 543.1092 | 671.63885 |
| Wdr1 | SENNVFYGHHAK | -523 | -534 | 0 | 0 | 12 | 537.35815 | 439.55783 | 345.27255 | NaN | 885.5998 | 754.3863 |
| Wdr1 | TEHAHQVVVAK | -55 | -65 | 0 | 0 | 11 | 442.22998 | 396.38196 | 440.24017 | 282.22287 | 286.75485 | NaN |
| Wdr1 | KVFASLPQVER | -7 | -17 | 1 | 0 | 11 | 344.7401 | NaN | 635.4211 | 401.8941 | 268.1393 | 2240.122 |
| Wdr1 | GDHFLYTN | -29 | -36 | 0 | 0.001092529 | 8 | 1843.4476 | 1510.0754 | 1744.9633 | NaN | 1155.7555 | NaN |
| Wdr1 | IKDIAWTE | -105 | -112 | 1 | 0.00713295 | 8 | 1591.7415 | 1143.0332 | 1654.145 | 1443.188 | NaN | NaN |
| Wdr1 | IKDIAWTEDSK | -105 | -115 | 1 | 0.00051306 | 11 | 768.1197 | NaN | NaN | 1025.741 | 159.521 | 947.2311 |
| Wdr1 | VFASLPQVER | -8 | -17 | 0 | 0 | 10 | 12665.455 | 148.83481 | 15822.208 | NaN | NaN | NaN |
| Wdr1 | VHPGGDTVAVGGTDGNVR | -453 | -470 | 0 | 0.00025015 | 18 | 1021.537 | NaN | 841.42377 | NaN | NaN | 357.75113 |
| Wdr1 | KYEYQPFAGK | -95 | -104 | 1 | 0.000230707 | 10 | 139.8561 | 1994.3053 | 1519.3015 | NaN | NaN | NaN |
| Wdr1 | VCALGESK | -224 | -231 | 0 | 0.00644558 | 8 | 2053.7493 | NaN | NaN | NaN | NaN | NaN |
| Wdr1 | FEGPPFK | -174 | -180 | 0 | 0.002068213 | 7 | 711.87366 | NaN | NaN | NaN | NaN | NaN |
| Wdr1 | YLDKNNPSKPLR | -303 | -314 | 2 | 0.001978766 | 12 | 261.97986 | NaN | NaN | NaN | NaN | NaN |
| Acsl4 | AVIDIPGADTLDK | -69 | -81 | 0 | 0 | 13 | 1262.5968 | 2211.4578 | 1892.1617 | 2331.5178 | 2200.2068 | 2550.4863 |
| Acsl4 | GYDAPLCNLILFK | -414 | -426 | 0 | 0 | 13 | 625.2087 | NaN | 616.6712 | 1052.339 | 746.5556 | NaN |
| Cops4 | NESTNEQLQIHYK | -176 | -188 | 0 | 0 | 13 | 2421.8447 | 1242.9747 | 2342.1746 | 1842.198 | 1606.425 | 1218.4598 |
| Cops4 | QNESTNEQLQIHYK | -175 | -188 | 0 | 0 | 14 | 1166.9585 | 1376.5729 | 1374.2872 | 1204.0216 | 1621.2543 | 1426.2059 |
| Cops4 | AVIEHNLLSASK | -303 | -314 | 0 | 0.000712251 | 12 | 563.6277 | 628.1997 | NaN | 898.2812 | NaN | 653.02106 |
| Cops4 | TIVHESER | -215 | -222 | 0 | 0.000591124 | 8 | 299.15198 | NaN | NaN | 421.68634 | 584.96344 | NaN |
| Prps1 | NCTIVSPDAGGAK | -164 | -176 | 0 | 0 | 13 | 1016.54803 | 681.99426 | 479.726 | 958.21497 | 433.62692 | 477.86865 |
| Prps1 | FSNQETCVEIGESVR | -35 | -49 | 0 | 0 | 15 | 778.1762 | 890.16895 | 867.79694 | 944.3503 | 1055.5308 | 239.02374 |
| Prps1 | FDIPVDNLYAEPAVLK | -138 | -153 | 0 | 0 | 16 | 621.12585 | 468.5194 | 464.71777 | 469.51465 | 462.652 | 539.4843 |
| Prps1 | NVDFALIHK | -186 | -194 | 0 | 0 | 9 | 503.35864 | 1409.4257 | 1418.2 | 1297.3589 | 1596.2568 | 1752.2037 |
| Prps1 | SNQETCVEIGESVR | -36 | -49 | 0 | 0 | 14 | 599.1239 | NaN | 632.8255 | 650.71545 | 665.0045 | 1126.2913 |
| Rac1 | HHCPNTPIILVGT | -103 | -115 | 0 | 0 | 13 | 6457.948 | 917.66785 | 6477.4204 | 2914.259 | 5026.881 | 4500.73 |
| Rac1 | HHCPNTPIILVGTK | -103 | -116 | 0 | 0 | 14 | 1164.533 | 1029.0502 | 1281.2775 | 896.1173 | 864.5759 | 934.64417 |
| Rac1 | HHCPNTPIILVGTK | -103 | -116 | 0 | 0 | 14 | 1164.533 | 1029.0502 | 1281.2775 | 896.1173 | 864.5759 | 934.64417 |
| Rac1 | KLTPITYPQGL | -133 | -143 | 1 | 0.00025015 | 11 | 1060.2782 | 1404.799 | 1665.1604 | 924.22894 | 1331.4174 | 2006.5093 |
| Rac2 | HHCPSTPIILVGT | -103 | -115 | 0 | 0 | 13 | 4351.8384 | 3746.1719 | 3030.554 | 4700.747 | 3671.8105 | 2779.1414 |
| Rac2 | AVLCPQPTR | -175 | -183 | 0 | 0.00025015 | 9 | 3868.57 | 3013.838 | 2420.0532 | 1810.3141 | 1596.1302 | 1833.9597 |
| Rac2 | HHCPSTPIILVGTK | -103 | -116 | 0 | 0 | 14 | 2851.217 | 3532.373 | 3542.1257 | 1741.2638 | 2747.6956 | 2667.5208 |
| Rac2 | HHCPSTPIILVGTK | -103 | -116 | 0 | 0 | 14 | 2851.217 | 3532.373 | 3542.1257 | 1741.2638 | 2747.6956 | 2667.5208 |
| Rac2 | HHCPSTPIIL | -103 | -112 | 0 | 0.001055699 | 10 | 2269.3506 | 2930.6511 | 2752.8389 | 1579.1166 | 2490.8188 | 344.71735 |
| Rac2 | VDSKPVNLGLWDTAGQEDYDR | -46 | -66 | 1 | 0 | 21 | 1089.8784 | 1128.4415 | 1209.044 | 1399.6354 | 1309.1278 | 1311.3418 |
| Rac2 | VDSKPVNLGLWDTAGQEDYDR | -46 | -66 | 1 | 0 | 21 | 1089.8784 | 1128.4415 | 1209.044 | 1399.6354 | 1309.1278 | 1311.3418 |
| Rac2 | KLAPITYPQGLAL | -133 | -145 | 1 | 0.000627943 | 13 | 626.72736 | 690.8888 | 635.4386 | 696.8393 | 927.28534 | 822.06793 |
| Rac2 | LAPITYPQGLAL | -134 | -145 | 0 | 0 | 12 | 532.83984 | 550.2641 | 621.8383 | 588.9419 | 702.7062 | 659.3579 |
| Rac2 | LAPITYPQGLALAK | -134 | -147 | 0 | 0 | 14 | 498.4091 | 351.37238 | 376.9583 | 889.4478 | 456.04883 | 460.85638 |
| Ivd | FLQENLAPK | -57 | -65 | 0 | 0 | 9 | 719.05194 | 892.65186 | 548.0184 | 606.68726 | 621.4605 | 697.44293 |
| Ivd | AVLDHTIPYLHVR | -298 | -310 | 0 | 0 | 13 | 658.79663 | 473.6212 | 435.2444 | 693.8872 | 564.80945 | 632.67224 |
| Ivd | HSILPVDDDINGLNEEQK | -31 | -48 | 0 | 0.00051306 | 18 | 536.92365 | NaN | NaN | NaN | 1043.1497 | NaN |

| | | | | | | | | | | | | |
|---|---|---|---|---|---|---|---|---|---|---|---|---|
| Cmpk2 | AVLDLVDQCPK | -235 | -245 | 0 | 0 | 11 | 1115.5155 | 1225.294 | 759.0832 | 1342.9736 | 1250.1432 | 993.85315 |
| Cmpk2 | LAELLGPPGR | -62 | -71 | 0 | 0 | 10 | 656.22064 | 540.2811 | 588.2169 | 644.11017 | 631.8307 | 692.6338 |
| Cmpk2 | ESTNFPVIVDR | -321 | -331 | 0 | 0 | 11 | 580.7132 | 86.72901 | 829.04346 | 1173.346 | 1144.0378 | 1096.3619 |
| Cmpk2 | LWELQGDR | -176 | -183 | 0 | 0 | 8 | 1064.548 | 676.1468 | 1493.912 | NaN | 1342.4238 | 777.1261 |
| Cmpk2 | AVLEECTSFIPEAR | -221 | -234 | 0 | 0 | 14 | 622.34595 | NaN | 1007.4257 | 665.71954 | 951.1224 | 888.3523 |
| Cmpk2 | HFVLGDATDHR | -48 | -58 | 0 | 0 | 11 | 421.52448 | 598.2476 | NaN | NaN | 849.5807 | NaN |
| Cmpk2 | IFDDEPTIIR | -294 | -303 | 0 | 0.001776473 | 10 | 5373.115 | NaN | 493.48566 | NaN | NaN | NaN |
| Gemin5 | VGPGAGASPGAPPFR | -41 | -55 | 0 | 0 | 15 | 1376.4377 | 1248.2268 | 1517.7495 | 1038.5544 | 957.46106 | 1792.3561 |
| Gemin5 | AVLESNPESPITITEPYR | -616 | -633 | 0 | 0 | 18 | 360.12198 | 396.93814 | 481.8289 | 432.96198 | 590.5744 | 443.78534 |
| Gemin5 | FHLGLFTDR | -890 | -898 | 0 | 0 | 9 | 1008.22784 | 992.8195 | 1031.1649 | NaN | 703.9058 | 675.2995 |
| Gemin5 | LRPEDPVLK | -1014 | -1022 | 1 | 0.000591124 | 9 | 618.8958 | 658.4714 | 586.5458 | 977.554 | 1160.8479 | NaN |
| Calr | LFPSGLDQK | -112 | -120 | 0 | 0 | 9 | 19475.28 | 19610.225 | 20438.559 | 18358.143 | 15902.163 | 698.41455 |
| Calr | FGPDICGPGTK | -132 | -142 | 0 | 0 | 11 | 8444.157 | 8571.632 | 8981.065 | 388.87738 | 9378.794 | 9341.722 |
| Calr | HEQNIDCGGGYVK | -99 | -111 | 0 | 0 | 13 | 5929.877 | 6192.355 | 6605.843 | 6502.2856 | 7460.454 | 7449.5933 |
| Calr | IVRPDNTYEVK | -175 | -185 | 1 | 0 | 11 | 5412.711 | 5027.6646 | 4926.131 | 5174.5303 | 5543.6924 | 5235.4795 |
| Calr | IVRPDNTYEVK | -175 | -185 | 1 | 0 | 11 | 5412.711 | 5027.6646 | 4926.131 | 5174.5303 | 5543.6924 | 5235.4795 |
| Calr | EFGNETWGVTK | -341 | -351 | 0 | 0 | 11 | 4150.195 | 3503.2524 | 3666.5635 | 2146.622 | 2779.637 | 1877.5742 |
| Calr | EFGNETWGVTK | -341 | -351 | 0 | 0 | 11 | 4150.195 | 3503.2524 | 3666.5635 | 2146.622 | 2779.637 | 1877.5742 |
| Calr | IKDPDAAKPEDWDER | -208 | -222 | 2 | 0 | 15 | 3592.8318 | 4348.2236 | 4515.414 | 4287.6743 | 5105.2847 | 5050.4614 |
| Calr | IKDPDAAKPEDWDER | -208 | -222 | 2 | 0 | 15 | 3592.8318 | 4348.2236 | 4515.414 | 4287.6743 | 5105.2847 | 5050.4614 |
| Calr | TLIVRPDNTYEVK | -173 | -185 | 1 | 0.00025015 | 13 | 3225.5376 | 3669.1658 | 3903.9697 | 3122.5442 | 3840.3457 | 3817.7317 |
| Calr | SLEDDWDFLPPKK | -195 | -207 | 1 | 0 | 13 | 2213.699 | 2552.9248 | 3189.8853 | 3691.483 | 4877.8325 | 4378.2515 |
| Calr | GLDLWQVK | -315 | -322 | 0 | 0.000431655 | 8 | 2126.9226 | 2206.4119 | 2332.056 | 216.44647 | 2129.9568 | 2432.341 |
| Calr | CKDDEFTHLY | -163 | -172 | 1 | 0 | 10 | 1986.4847 | 2185.5908 | 2314.4324 | 2352.56 | 2711.3704 | 2477.6582 |
| Calr | IDDPTDSKPEDWDKPEHIPDPDAK | -225 | -248 | 2 | 0 | 24 | 1819.655 | 2167.8572 | 2195.1501 | 1902.2278 | 3728.71 | 2779.377 |
| Calr | IDDPTDSKPEDWDKPEHIPDPDAK | -225 | -248 | 2 | 0 | 24 | 1819.655 | 2167.8572 | 2195.1501 | 1902.2278 | 3728.71 | 2779.377 |
| Calr | YTLIVRPDNTYEVK | -172 | -185 | 1 | 0.00025015 | 14 | 1446.2386 | 1641.3502 | 1706.9828 | 1632.803 | 2184.154 | 2288.7837 |
| Calr | EQFLDGDAWTNR | -25 | -36 | 0 | 0 | 12 | 1413.47 | 1877.5399 | 1761.566 | 1242.115 | 1470.5759 | 1601.6222 |
| Calr | NSQVESGSLEDDWDFLPPK | -188 | -206 | 0 | 0 | 19 | 1392.8162 | 1418.5465 | 1644.5696 | 1495.7588 | 1884.9385 | 1806.4032 |
| Calr | FYGDLEKDK | -56 | -64 | 1 | 0 | 9 | 1065.54 | 1477.0278 | 322.69296 | 691.02594 | 2087.8057 | 358.79868 |
| Calr | DPDAAKPEDWDER | -210 | -222 | 1 | 0 | 13 | 1023.1903 | 1125.3622 | 1129.248 | 372.20004 | 564.4622 | 1213.0787 |
| Calr | NSQVESGSLEDDWDFLPPKK | -188 | -207 | 1 | 0 | 20 | 900.563 | 999.16785 | 1150.5033 | 1013.7633 | 1246.4562 | 1203.629 |
| Calr | VKHEQNIDCGGGYVK | -97 | -111 | 1 | 0 | 15 | 897.91864 | 775.83057 | 1111.876 | 470.8991 | 1097.2623 | 1118.3475 |
| Calr | KVHVIFNYK | -143 | -151 | 1 | 0 | 9 | 895.49866 | 953.60345 | 945.3367 | 3068.3364 | 1046.356 | 1006.3871 |
| Calr | KVHVIFNYK | -143 | -151 | 1 | 0 | 9 | 895.49866 | 953.60345 | 945.3367 | 3068.3364 | 1046.356 | 1006.3871 |
| Calr | SLEDDWDFLPPK | -195 | -206 | 0 | 0 | 12 | 884.2821 | 717.75555 | 909.49646 | 1117.6537 | 1151.2721 | 1752.4167 |
| Calr | AVLGLDLWQVK | -312 | -322 | 0 | 0 | 11 | 873.16156 | 429.05927 | 930.74945 | 1012.0981 | 1061.7673 | 548.14 |
| Calr | NDEAYAEEFGNETWGVTK | -334 | -351 | 0 | 0 | 18 | 662.7461 | 575.5222 | 654.23444 | 669.7558 | 805.47186 | 685.55206 |
| Calr | KPEDWDEEMDGEWEPPVIQNPEYK | -249 | -272 | 1 | 0 | 24 | 612.0752 | 524.28033 | 562.0807 | 611.2851 | 596.77423 | 285.12122 |
| Calr | DKGLQTSQDAR | -63 | -73 | 1 | 0.00025015 | 11 | 505.0946 | 579.15674 | 584.547 | 660.205 | 622.8656 | 622.00604 |
| Calr | IDNSQVESGSLEDDWDFLPPK | -186 | -206 | 0 | 0 | 21 | 411.97156 | 647.19934 | 767.0173 | 723.24756 | 1164.5605 | 1061.5486 |
| Calr | GTWIHPEIDNPEYSPDANIY | -287 | -306 | 0 | 0.00025015 | 20 | 334.87433 | 410.94843 | 406.78806 | 302.93176 | 296.09982 | 356.9343 |
| Calr | EQFLDGDAWTN | -25 | -35 | 0 | 0 | 11 | 300.2481 | 319.8645 | 273.91113 | 220.98123 | 200.89168 | 200.05782 |
| Calr | CKDDEFTHL | -163 | -171 | 1 | 0.001291103 | 9 | 138.92139 | 1088.7649 | 1018.773 | 1268.0717 | 1509.8198 | 1257.669 |
| Calr | CKDDEFTHLYTL | -163 | -174 | 1 | 0 | 12 | 756.0112 | 963.3098 | 938.4008 | NaN | 1650.513 | 1635.1855 |
| Calr | VHVIFNYK | -144 | -151 | 0 | 0 | 8 | 685.41223 | 704.3916 | 677.9276 | 452.16553 | 555.6183 | NaN |
| Calr | GLQTSQDAR | -65 | -73 | 0 | 0 | 9 | 670.27844 | NaN | 468.54178 | 424.6637 | 720.95685 | 575.8554 |
| Calr | GLQTSQDAR | -65 | -73 | 0 | 0 | 9 | 670.27844 | NaN | 468.54178 | 424.6637 | 720.95685 | 575.8554 |
| Calr | FEPFSNK | -81 | -87 | 0 | 0.00644558 | 7 | 2050.5742 | 1394.0459 | 1255.0321 | 1255.7612 | NaN | NaN |
| Calr | DPAIYFK | -18 | -24 | 0 | 0 | 7 | 1081.9386 | 451.448 | 767.1771 | 612.35913 | NaN | NaN |
| Calr | HEQNIDCGGGY | -99 | -109 | 0 | 0 | 11 | 1057.9401 | 900.06006 | 897.11755 | NaN | NaN | 197.03635 |
| Calr | FLDGDAWTNR | -27 | -36 | 0 | 0 | 10 | 184.52507 | 854.15546 | 737.4975 | NaN | 403.5401 | NaN |
| Calr | EDEEESPGQAKDEL | -403 | -416 | 1 | 0 | 14 | 786.6182 | 721.809 | 673.6566 | NaN | NaN | NaN |
| Calr | EKEEDEEESPGQAKDEL | -400 | -416 | 2 | 0 | 17 | 503.81357 | 518.95435 | 470.2881 | NaN | NaN | NaN |
| Calr | DEKEEDEEESPGQAKDEL | -399 | -416 | 2 | 0.000431655 | 18 | 418.02988 | 435.39392 | 473.74524 | NaN | NaN | NaN |
| Calr | DEEESPGQAKDEL | -404 | -416 | 1 | 0 | 13 | 448.07892 | NaN | 908.6869 | NaN | NaN | NaN |
| Cyb5r3 | LIDKEVISPDTR | -47 | -58 | 1 | 0 | 12 | 1346.347 | 1753.768 | 1699.3428 | 1510.2134 | 1744.6715 | 1478.7277 |
| Cyb5r3 | AVLKDPNDHTVCYL | -193 | -206 | 1 | 0 | 14 | 661.29803 | 804.56 | 852.188 | 1007.98883 | 1070.4012 | 978.72015 |
| Cyb5r3 | IGDTIEFR | -136 | -143 | 0 | 0.00025015 | 8 | 536.1504 | 627.0638 | 626.9665 | 1203.0038 | 467.4883 | 979.3866 |
| Cyb5r3 | DHLPTPGEEPLIL | -260 | -272 | 0 | 0 | 13 | 1195.0533 | 1216.9535 | 1308.6147 | 278.4221 | NaN | 1609.5016 |
| Cyb5r3 | DILLRPELEELR | -215 | -226 | 1 | 0 | 12 | 729.3335 | 690.57196 | 1405.0978 | 976.74475 | 1491.7682 | NaN |
| Cyb5r3 | FALPSPQHIL | -62 | -71 | 0 | 0 | 10 | 330.22165 | 930.4326 | NaN | 798.73083 | 1014.9791 | 982.70435 |
| Cyb5r3 | DILLRPELEEL | -215 | -225 | 1 | 0.00078661 | 11 | 256.97324 | 401.7865 | 545.6912 | NaN | 213.8893 | 285.41327 |
| Cyb5r3 | IDGNLVIR | -85 | -92 | 0 | 0 | 8 | 796.2339 | 691.38403 | 652.57355 | 420.76904 | NaN | NaN |
| Cyb5r3 | KIGDTIEFR | -135 | -143 | 1 | 0.00025015 | 9 | 9404.357 | 414.7605 | NaN | NaN | NaN | NaN |
| Rhog | HHCPDVPILLVGTK | -103 | -116 | 0 | 0 | 14 | 1312.5026 | 1583.0281 | 2170.7573 | 327.15427 | 2688.6123 | 2964.2227 |
| Rhog | HHCPDVPILLVGTK | -103 | -116 | 0 | 0 | 14 | 1312.5026 | 1583.0281 | 2170.7573 | 327.15427 | 2688.6123 | 2964.2227 |
| Rhog | LWDTAGQEEYDR | -55 | -66 | 0 | 0 | 12 | 1774.6241 | 1024.5244 | 2405.0771 | 2626.1309 | 1792.4991 | 2371.4246 |
| Rhog | LKEQGQAPITPQQGQAL | -129 | -145 | 1 | 0 | 17 | 601.3398 | 603.683 | 534.65375 | 295.47473 | 608.25946 | 630.07635 |
| Rhog | TVNLNLWDTAGQEEYDR | -50 | -66 | 0 | 0 | 17 | 554.90106 | 570.64343 | 599.39606 | 445.8301 | 960.0951 | 971.28485 |
| Rhog | SIASPPSYENVR | -83 | -94 | 0 | 0.000230707 | 12 | 1854.1278 | 1552.9899 | NaN | 2822.0613 | 1918.1287 | 2404.011 |
| Rhog | AVLNPTPIK | -175 | -183 | 0 | 0.000230707 | 9 | 440.86957 | 218.71881 | 273.70624 | NaN | NaN | 438.34302 |
| Rhog | SPPSYENVR | -86 | -94 | 0 | 0.000591124 | 9 | 391.07114 | 500.26392 | 587.2082 | NaN | NaN | 644.1598 |
| Noc4l | HNLEYPDFYQK | -319 | -329 | 0 | 0 | 11 | 820.0997 | 895.2184 | 910.81775 | 815.4089 | 953.99677 | 1041.8995 |
| Noc4l | AVLQSEEPEEIEEGVR | -36 | -51 | 0 | 0 | 16 | 765.9475 | 792.3169 | 731.69476 | 899.6996 | 745.6666 | 771.82245 |
| Noc4l | LYGLLDPSIFHVK | -330 | -342 | 0 | 0 | 13 | 411.6239 | 422.89853 | 392.1406 | 788.0748 | 961.4113 | 997.2464 |
| Noc4l | FVQLEGAKPL | -125 | -134 | 1 | 0.000712251 | 10 | 1824.5483 | NaN | NaN | 2136.0127 | 2528.453 | NaN |
| Noc4l | HKLPLSLYK | -262 | -270 | 1 | 0.003393785 | 9 | 495.26605 | NaN | 1245.7002 | NaN | NaN | NaN |

| | | | | | | | | | | | | |
|---|---|---|---|---|---|---|---|---|---|---|---|---|
| Cpne1 | SVSCEHLIDK | -11 | -20 | 0 | 0 | 10 | 2806.9436 | 3528.5317 | 3144.8665 | 3852.2405 | 3366.6216 | 3655.777 |
| Cpne1 | NFAPIINHVAR | -389 | -399 | 0 | 0 | 11 | 1693.2134 | 2141.752 | 687.5362 | 943.8754 | 1126.1982 | 1225.159 |
| Cpne1 | AVPAEFECVHPEK | -235 | -247 | 0 | 0 | 13 | 1152.4333 | 1063.4249 | 1260.9226 | 1243.6339 | 125.55403 | 1327.3778 |
| Cpne1 | SDPFLEFFR | -157 | -165 | 0 | 0 | 9 | 593.1158 | 646.52014 | 759.468 | 737.2019 | 954.44385 | 962.4978 |
| Cpne1 | HFCGGDLSTPIQVR | -197 | -210 | 0 | 0 | 14 | 1152.9622 | 1099.933 | 1007.4499 | 1462.6897 | 1246.7084 | NaN |
| Cpne1 | GPTNFAPIINHVAR | -386 | -399 | 0 | 0.000431655 | 14 | 675.0185 | 522.30505 | NaN | NaN | NaN | 650.7528 |
| Cpne1 | DIVQFVPYR | -478 | -486 | 0 | 0 | 9 | 472.05023 | NaN | 1180.228 | NaN | NaN | 240.07863 |
| Add1 | SPGTPAGEGSGSPPK | -355 | -369 | 0 | 0 | 15 | 2760.968 | 2719.5105 | 2830.001 | 2630.5273 | 4063.0884 | 3966.274 |
| Add1 | VDENNPEYLR | -28 | -37 | 0 | 0 | 10 | 1411.8477 | 1409.8566 | 389.394 | 1377.2128 | 1617.2095 | 1519.111 |
| Add1 | TTGPNPFNTLTDR | -558 | -570 | 0 | 0.000230707 | 13 | 784.484 | 912.8805 | 945.36865 | 869.1842 | 1082.3301 | 1080.662 |
| Add1 | VNLQGDIVDR | -200 | -209 | 0 | 0 | 10 | 658.37354 | 475.75067 | 490.26733 | 223.95602 | NaN | 392.8554 |
| Add1 | LEEDLPQEPTSR | -618 | -629 | 0 | 0 | 12 | 1888.0286 | 1239.6022 | 1625.977 | NaN | NaN | 1057.1913 |
| Add1 | VNSEQEHFLIVPF | -173 | -185 | 0 | 0.00025015 | 13 | 520.70544 | NaN | 667.027 | 524.13586 | NaN | 557.4939 |
| Add1 | TLASAGGPDNLVLLDPGK | -331 | -348 | 0 | 0 | 18 | 368.90668 | 361.64127 | 413.19232 | NaN | NaN | 590.78784 |
| Add1 | GSEENLDETR | -585 | -594 | 0 | 0.000230707 | 10 | 1398.1997 | 1382.3115 | NaN | 681.0263 | NaN | NaN |
| Add1 | AVPNLFVPLNTNPK | -484 | -497 | 0 | 0 | 14 | 1264.2673 | NaN | NaN | NaN | NaN | NaN |
| Aip | NIAEGKDPLEGQR | -107 | -119 | 1 | 0 | 13 | 909.62396 | 851.1459 | 1261.1284 | 1451.3252 | 1017.1326 | 1061.0564 |
| Aip | AVPVIHQEGNR | -178 | -188 | 0 | 0 | 11 | 674.95984 | 812.04895 | 607.0056 | 655.3024 | 360.6468 | 323.2132 |
| Aip | HSDNEGSVIDDSR | -42 | -54 | 0 | 0 | 13 | 350.97247 | 950.42804 | 280.47977 | 906.60016 | 506.1882 | 1168.6396 |
| Aip | ELDPALAPVVSR | -293 | -304 | 0 | 0 | 12 | 932.78937 | 913.2727 | 994.7832 | 608.9862 | NaN | 622.5834 |
| Aip | NEGSVIDDSR | -45 | -54 | 0 | 0.009631128 | 10 | 8203.075 | NaN | NaN | NaN | NaN | NaN |
| G3bp1 | GLLSNNNQALR | -96 | -106 | 0 | 0 | 11 | 5520.9688 | 6665.142 | 6790.6436 | 6175.62 | 5995.2295 | 5789.7437 |
| G3bp1 | EAGEPGDVEPR | -319 | -329 | 0 | 0 | 11 | 1699.342 | 2383.4277 | 488.04227 | 3890.44 | 2164.6228 | 1468.3547 |
| G3bp1 | NLPHEVDKSELK | -344 | -355 | 1 | 0 | 12 | 1613.691 | 560.6035 | 803.07935 | 824.96063 | 479.80582 | 503.7409 |
| G3bp1 | NLPHEVDKSELK | -344 | -355 | 1 | 0 | 12 | 1613.691 | 560.6035 | 803.07935 | 824.96063 | 479.80582 | 503.7409 |
| G3bp1 | HPDSHQLFIGNLPHEVDK | -334 | -351 | 0 | 0 | 18 | 1536.5083 | 1205.4338 | 1230.6354 | 4925.4727 | 1611.928 | 1462.762 |
| G3bp1 | NLPPSGAVPVTGTPPHVVK | -256 | -274 | 0 | 0.00025015 | 19 | 1530.0947 | 1841.6332 | 1977.2395 | 1513.2683 | 2064.7375 | 1867.9034 |
| G3bp1 | FYVHNDIFR | -124 | -132 | 0 | 0 | 9 | 1436.177 | 1732.7734 | 1779.5428 | 1397.1805 | 1760.3617 | 316.44287 |
| G3bp1 | FYVHNDIFR | -124 | -132 | 0 | 0 | 9 | 1436.177 | 1732.7734 | 1779.5428 | 1397.1805 | 1760.3617 | 316.44287 |
| G3bp1 | STSPAPADVAPAQEDLR | -229 | -245 | 0 | 0.001291103 | 17 | 773.9047 | 904.16046 | 838.8121 | 897.08124 | 700.76917 | 580.76874 |
| G3bp1 | NQAPDMLHR | -24 | -32 | 0 | 0 | 9 | 669.3555 | 674.9008 | 429.27484 | 654.5554 | 228.97293 | 802.0878 |
| G3bp1 | NQAPDMLHR | -24 | -32 | 0 | 0 | 9 | 669.3555 | 674.9008 | 429.27484 | 654.5554 | 228.97293 | 802.0878 |
| G3bp1 | VTEPQEESEEEVEEPEER | -142 | -159 | 0 | 0 | 18 | 502.44867 | 529.21533 | 521.85815 | 689.1741 | 688.8047 | 721.77435 |
| G3bp1 | DFFQNFGNVVELR | -356 | -368 | 0 | 0 | 13 | 337.31702 | 302.29077 | 358.55106 | 417.1092 | 439.9554 | 404.53915 |
| G3bp1 | FIGNLPHEVDK | -341 | -351 | 0 | 0 | 11 | 780.27014 | 771.87866 | NaN | 708.0857 | 222.61674 | 248.37062 |
| G3bp1 | LPNFGFVVFDDSEPVQK | -375 | -391 | 0 | 0 | 17 | 466.10028 | 482.31058 | NaN | 667.8626 | 840.9152 | 745.42914 |
| G3bp1 | LNQAPDMLHR | -23 | -32 | 0 | 0 | 10 | 2763.4807 | NaN | 2926.726 | NaN | NaN | 3644.5208 |
| G3bp1 | GEPGDVEPR | -321 | -329 | 0 | 0.000947459 | 9 | 513.8563 | 729.4139 | NaN | 897.99384 | NaN | NaN |
| G3bp1 | YVHNDIFR | -125 | -132 | 0 | 0 | 8 | 781.4625 | NaN | NaN | 1086.407 | NaN | NaN |
| G3bp1 | VLAPEGSVANK | -113 | -123 | 0 | 0.00025015 | 11 | 664.28345 | NaN | NaN | 2162.5889 | NaN | NaN |
| G3bp1 | AVPVTGTPPHVVK | -262 | -274 | 0 | 0.002068213 | 13 | 260.30164 | 268.07974 | NaN | NaN | NaN | NaN |
| Cnot1 | AVQELVHPVVDR | -1399 | -1410 | 0 | 0 | 12 | 10306.936 | 10523.64 | 10510.043 | 10788.229 | 9206.929 | 9404.512 |
| Cnot1 | EVTYELDHPGFQIR | -356 | -369 | 0 | 0 | 14 | 2116.0361 | 2928.298 | 2966.6611 | 1527.357 | 2482.3872 | 1841.2571 |
| Cnot1 | KDFALDSEESR | -1425 | -1435 | 1 | 0 | 11 | 867.438 | 771.9479 | 773.1561 | 862.143 | 348.43893 | 475.52844 |
| Cnot1 | QLFSFPIK | -471 | -478 | 0 | 0 | 8 | 442.23477 | 441.2232 | 642.8871 | 473.34866 | 484.8831 | 527.77844 |
| Cnot1 | SPVTFLSDLR | -2187 | -2196 | 0 | 0 | 10 | 1371.6862 | 1703.3722 | NaN | NaN | NaN | NaN |
| Cnot1 | LELISHR | -2038 | -2044 | 0 | 0.00025015 | 7 | 672.7583 | 564.2259 | NaN | NaN | NaN | NaN |
| Nol6 | AVQGAHPVLR | -691 | -700 | 0 | 0 | 10 | 527.963 | 646.69257 | 483.9769 | 452.23962 | 519.4538 | 609.18994 |
| Nol6 | TEPPAEGLLQPVNLSR | -51 | -66 | 0 | 0 | 16 | 352.20737 | 730.0877 | 583.35144 | NaN | 409.79034 | 512.2458 |
| Nol6 | QQQYPAYSGVAR | -851 | -862 | 0 | 0 | 12 | 554.72437 | 486.20178 | 643.55664 | NaN | NaN | NaN |
| Eif4b | LGNLPYDVTEDSIK | -100 | -113 | 0 | 0 | 14 | 1703.209 | 2195.001 | 2203.1306 | 3443.9834 | 2286.3335 | 3010.4783 |
| Eif4b | WHSNDDDVYR | -55 | -64 | 0 | 0 | 10 | 1676.6719 | 311.1973 | 1026.3595 | 1764.151 | 1453.8728 | 1781.8579 |
| Eif4b | DSDKTDTDWR | -191 | -200 | 1 | 0 | 10 | 1245.6099 | 1201.0508 | 1303.772 | 1086.9686 | 1228.013 | 1381.604 |
| Eif4b | AVQPPEEGPSR | -514 | -524 | 0 | 0.000230707 | 11 | 1057.9241 | 1508.4753 | 528.0167 | 1897.5939 | 495.02594 | 922.09235 |
| Eif4b | SSDTEQPSPTSGGGK | -497 | -511 | 0 | 0 | 15 | 861.8259 | 852.54517 | 915.49646 | 283.5156 | 632.2037 | 491.4484 |
| Eif4b | SIFGGAKPVDTAAR | -359 | -372 | 1 | 0 | 14 | 584.2536 | 639.69336 | 1105.4302 | 640.3694 | 792.8879 | 784.1113 |
| Eif4b | TTWHSNDDDVYR | -53 | -64 | 0 | 0.00025015 | 12 | 365.893 | 384.6813 | 414.41547 | 168.04745 | 439.73676 | 384.56747 |
| Eif4b | VDVADQAQDKDRDDR | -168 | -182 | 2 | 0 | 15 | 311.47446 | 370.31122 | 565.6511 | 498.45917 | 868.77527 | 640.2798 |
| Eif4b | SLNEESLGNR | -155 | -164 | 0 | 0 | 10 | 2084.6533 | 2437.111 | 2494.0056 | 2019.8722 | 2035.2783 | NaN |
| Eif4b | VDVADQAQDKDR | -168 | -179 | 1 | 0 | 12 | 652.62213 | NaN | 654.35547 | 2157.9146 | 1796.9473 | 193.46094 |
| Eif4b | VAAVQPPEEGPSR | -512 | -524 | 0 | 0 | 13 | 2843.747 | 2998.8596 | 2990.9976 | 605.1896 | NaN | NaN |
| Eif4b | GGAKPVDTAAR | -362 | -372 | 1 | 0 | 11 | 352.50522 | NaN | 364.4497 | 52.740807 | 570.5994 | NaN |
| Eif4b | DGNKVDVVGATQGQAG | -526 | -541 | 1 | 0.001811524 | 16 | 789.97864 | NaN | NaN | 1102.9434 | 907.5072 | NaN |
| Ago2 | SIEEQQKPLTDSQR | -243 | -256 | 1 | 0.00025015 | 14 | 626.3831 | NaN | 683.68353 | 630.23157 | 995.0824 | 894.70245 |
| Ago2 | VLQPPSILYGGR | -413 | -424 | 0 | 0.00078661 | 12 | 2085.135 | NaN | NaN | 474.56445 | 759.86597 | NaN |
| H2ac21 | AVRNDEELNK | -87 | -96 | 1 | 0.000431655 | 10 | 3765.231 | 4021.9133 | 4636.9917 | 3082.6 | 4563.125 | 2430.8662 |
| H2ac21 | AVRNDEELNK | -87 | -96 | 1 | 0.000431655 | 10 | 3765.231 | 4021.9133 | 4636.9917 | 3082.6 | 4563.125 | 2430.8662 |
| Vtn | AVRPGYPK | -186 | -193 | 1 | 0.00025015 | 8 | 977.4993 | 950.72144 | 1941.3236 | 2529.0464 | 1875.237 | 1811.8259 |
| Eif2s1 | VVTDTDETELAR | -277 | -288 | 0 | 0 | 12 | 2991.8022 | 3089.2107 | 2828.797 | 2847.7979 | 1059.4264 | 1022.1984 |
| Eif2s1 | EYTKDEQLESLFQR | -120 | -133 | 1 | 0 | 14 | 2170.5166 | 2265.5767 | 2439.9053 | 646.8207 | 3057.4329 | 1741.5641 |
| Eif2s1 | EYTKDEQLESLFQR | -120 | -133 | 1 | 0 | 14 | 2170.5166 | 2265.5767 | 2439.9053 | 646.8207 | 3057.4329 | 1741.5641 |
| Eif2s1 | DEQLESLFQR | -124 | -133 | 0 | 0 | 10 | 1349.5944 | 1714.5315 | 668.56934 | 1761.108 | 1827.7152 | 1709.0942 |
| Eif2s1 | AVSDPSILDSLDLNEDER | -156 | -173 | 0 | 0 | 18 | 626.04443 | 659.5011 | NaN | 511.95483 | 608.1281 | 682.0082 |
| Eif2s1 | VSDPSILDSLDLNEDER | -157 | -173 | 0 | 0.00025015 | 17 | 346.61136 | 511.10333 | NaN | 394.41544 | 445.96176 | 445.0105 |
| Eif2s1 | INLIAPPR | -227 | -234 | 0 | 0 | 8 | 543.1786 | 170.78621 | 151.65573 | NaN | NaN | NaN |
| Rpap3 | NLDPDVFNQIIK | -580 | -591 | 0 | 0 | 12 | 378.1427 | 907.7945 | 1037.5051 | 909.2822 | 1275.6278 | 1097.3661 |
| Rpap3 | AVSDTSAPQAQVGVK | -496 | -510 | 0 | 0.001610812 | 15 | 2547.7488 | NaN | NaN | NaN | NaN | NaN |
| Ilf3 | VPTWGPLR | -240 | -247 | 0 | 0.000627943 | 8 | 5928.844 | 1316.2639 | 5642.537 | 3909.141 | 4704.3584 | 5980.1875 |

| | | | | | | | | | | | | |
|---|---|---|---|---|---|---|---|---|---|---|---|---|
| Ilf3 | SVEVDGSNFEASGPSK | -435 | -450 | 0 | 0.00025015 | 16 | 588.8128 | 628.4844 | 601.39276 | 587.0406 | 651.89435 | 949.8736 |
| Ilf3 | QLKPGLQYK | -411 | -419 | 1 | 0 | 9 | 524.4625 | 493.69724 | 464.57434 | 417.12408 | 379.82617 | 448.88983 |
| Ilf3 | EATDAIGHLDR | -298 | -308 | 0 | 0 | 11 | 1105.6906 | NaN | 1149.8865 | 1512.0602 | 1617.0571 | 1045.9818 |
| Ilf3 | SNPSSVFPSDATTEQGPILTK | -503 | -523 | 0 | 0 | 21 | 561.9624 | 647.4553 | 483.6272 | NaN | 728.39905 | 591.54694 |
| Ilf3 | EKPTTALLDK | -118 | -127 | 1 | 0.001392869 | 10 | 1300.5825 | 1207.9436 | 1394.9092 | NaN | NaN | 1604.4688 |
| Ilf3 | SVNDPPDVLDR | -190 | -200 | 0 | 0 | 11 | 1267.713 | NaN | 657.29315 | 1300.7863 | 903.865 | NaN |
| Ilf3 | HSSVYPTQEELEAVQN | -18 | -33 | 0 | 0.000230707 | 16 | 749.0054 | 1142.6724 | 978.22925 | NaN | NaN | NaN |
| Ilf3 | SVFPSDATTEQGPILTK | -507 | -523 | 0 | 0.000431655 | 17 | 6045.5684 | NaN | 7262.6357 | NaN | NaN | NaN |
| Ilf3 | AVSDWIDEQEK | -44 | -54 | 0 | 0 | 11 | 1053.1128 | NaN | 815.6041 | NaN | NaN | NaN |
| Eef1b | PVGYGIK | -179 | -185 | 0 | 0 | 7 | 1805.5736 | 1791.2775 | 1803.7382 | 1157.362 | 810.1193 | 838.7093 |
| Eef1b | TPAGLQVLNDYLADK | -8 | -22 | 0 | 0 | 15 | 1609.8584 | 1923.6086 | 1644.1793 | 2232.6396 | 4333.384 | 2794.0396 |
| Eef1b | SVEDTTGSGAADAK | -83 | -96 | 0 | 0 | 14 | 402.07343 | 643.7632 | 473.1718 | 321.23563 | 316.4279 | 578.28174 |
| Eef1b | LVPVGYGIK | -177 | -185 | 0 | 0.00025015 | 9 | 3273.5056 | 1875.723 | 1980.5986 | 1107.3871 | 430.036 | NaN |
| Eef1b | EAVSGPPPADLCHALR | -39 | -54 | 0 | 0 | 16 | 1846.7885 | 856.9081 | NaN | 2159.3767 | 2549.183 | 2401.7803 |
| Eef1b | QVLNDYLADK | -13 | -22 | 0 | 0 | 10 | 4239.1416 | 5155.2183 | 12223.744 | 7307.624 | NaN | NaN |
| Eef1b | RWYNHIK | -54 | -60 | 1 | 0.00272025 | 7 | 1360.8721 | 1419.9204 | 1750.2175 | NaN | NaN | 405.56378 |
| Eef1b | SYIEGYVPSQADVA | -23 | -36 | 0 | 0.00025015 | 14 | 963.7352 | 753.8851 | 1378.6299 | NaN | NaN | 154.56615 |
| Eef1b | LAQYESK | -123 | -129 | 0 | 0 | 7 | 1232.0773 | 389.4914 | NaN | 388.4229 | NaN | NaN |
| Eef1b | AVSGPPPADLCHALR | -40 | -54 | 0 | 0.001392869 | 15 | 796.91 | NaN | NaN | NaN | NaN | NaN |
| Ppp1r12a | LYEQILAENEK | -934 | -944 | 0 | 0 | 11 | 1150.652 | 1308.5942 | 1265.7574 | 1699.8595 | 1950.0176 | 1839.2441 |
| Ppp1r12a | SYLTPVRDEESESQ | -666 | -679 | 1 | 0.000431655 | 14 | 566.77325 | NaN | NaN | 473.58032 | 388.79755 | 550.49835 |
| Ppp1r12a | AVTAPTLSSNQGTPTSPVK | -394 | -412 | 0 | 0.00025015 | 19 | 1199.8094 | NaN | 1185.2441 | NaN | NaN | 1392.0568 |
| Ppp1r12a | DYDGWTPLHAA | -229 | -239 | 0 | 0 | 11 | 600.1654 | NaN | NaN | NaN | NaN | NaN |
| Coro1b | NVGTAEELYR | -161 | -170 | 0 | 0 | 10 | 1843.514 | 1012.013 | 1664.8905 | 1731.4722 | 1683.0824 | 1629.4883 |
| Coro1b | TAVTDVPSGNLAGAGEAGK | -429 | -447 | 0 | 0.000947459 | 19 | 737.41113 | 818.64716 | 890.9802 | 948.9613 | 860.1427 | 891.2767 |
| Coro1b | AVTDVPSGNLAGAGEAGK | -430 | -447 | 0 | 0 | 18 | 2229.1765 | 1712.0171 | 2082.4775 | 2469.3416 | NaN | NaN |
| Coro1b | HVFGQPVK | -14 | -21 | 0 | 0.00111006 | 8 | 1091.6322 | 1153.1208 | 1092.2291 | 986.17847 | NaN | NaN |
| Coro1b | DANPILISLR | -384 | -393 | 0 | 0 | 10 | 414.1589 | 490.96454 | 403.58783 | NaN | 422.29456 | NaN |
| Coro1b | REAYVPSK | -393 | -400 | 1 | 0 | 8 | 516.97595 | NaN | NaN | 799.4764 | 591.6851 | NaN |
| Itgb1 | VCGSLIGWR | -260 | -268 | 0 | 0 | 9 | 2692.7634 | 488.65405 | 2954.116 | 2655.5369 | 3050.6804 | 2866.2664 |
| Itgb1 | AVTEEFQPVYK | -336 | -346 | 0 | 0 | 11 | 1163.1835 | 490.06354 | 998.26807 | 500.75314 | 1836.4591 | 1914.4391 |
| Itgb1 | STDAGFHFAGDGK | -277 | -289 | 0 | 0 | 13 | 1294.7031 | 1370.846 | NaN | 1227.1714 | 1222.9269 | 1421.7546 |
| Itgb1 | WDTGENPIYK | -775 | -784 | 0 | 0 | 10 | 1010.4718 | 839.0111 | 671.3985 | 731.2045 | NaN | 710.6369 |
| Itgb1 | LPDGVTINYK | -389 | -398 | 0 | 0 | 10 | 642.8136 | NaN | NaN | 800.5476 | 418.34775 | NaN |
| Ywhah | KAVTELNEPLSNEDR | -28 | -42 | 1 | 0 | 15 | 2387.6562 | 2670.9094 | 2646.2766 | 2837.037 | 3001.2085 | 2917.6384 |
| Ywhah | KAVTELNEPLSNEDR | -28 | -42 | 1 | 0 | 15 | 2387.6562 | 2670.9094 | 2646.2766 | 2837.037 | 3001.2085 | 2917.6384 |
| Ywhah | AVTELNEPLSNEDR | -29 | -42 | 0 | 0 | 14 | 1518.4088 | 2338.6875 | 2661.1453 | 2111.5642 | 2126.8137 | 2227.2834 |
| Ywhah | NCNDFQYESK | -111 | -120 | 0 | 0 | 10 | 1453.3767 | 1561.4465 | 1509.9867 | 1731.1608 | 1739.8013 | 1798.2501 |
| Ywhah | NCNDFQYESK | -111 | -120 | 0 | 0 | 10 | 1453.3767 | 1561.4465 | 1509.9867 | 1731.1608 | 1739.8013 | 1798.2501 |
| Ywhah | GDREQLLQR | -2 | -10 | 1 | 0.001023236 | 9 | 1451.3834 | 1630.6499 | 648.8861 | 4139.8677 | 2148.2578 | 3582.695 |
| Ywhah | QAFDDAIAELDTLNEDSYK | -199 | -217 | 0 | 0 | 19 | 231.14508 | NaN | 210.3523 | 208.72533 | 237.41063 | 270.97614 |
| Ywhah | YLAEVASGEK | -133 | -142 | 0 | 0 | 10 | 1621.3461 | NaN | 1396.4966 | 1861.7391 | NaN | 2014.674 |
| Snrnp40 | LWNVYGDCDNYATLK | -93 | -107 | 0 | 0 | 15 | 1564.95 | 1663.0891 | 1672.86 | 412.79657 | 839.27814 | 1097.0791 |
| Snrnp40 | TFNDTSDQIISGGIDNDIK | -201 | -219 | 0 | 0.000550307 | 19 | 1554.0065 | 1517.5891 | 1970.385 | 1087.9932 | 2239.9282 | 2140.436 |
| Snrnp40 | NDTSDQIISGGIDNDIK | -203 | -219 | 0 | 0 | 17 | 1161.1995 | 1437.3727 | 342.931 | 246.28902 | 237.78545 | 239.76031 |
| Snrnp40 | KGPELPLVPVK | -8 | -18 | 1 | 0 | 11 | 939.8124 | 944.1112 | 967.4963 | 1021.25574 | 956.2667 | 978.0447 |
| Snrnp40 | NVYGDCDNYATLK | -95 | -107 | 0 | 0 | 13 | 797.05927 | 825.3235 | 498.6533 | 796.2176 | 616.6405 | 692.3796 |
| Snrnp40 | AVTFNDTSDQIISGGIDNDIK | -199 | -219 | 0 | 0 | 21 | 411.76514 | 445.80508 | 407.01453 | 417.88016 | 340.42065 | 411.0598 |
| Snrnp40 | TVAVWDSETGER | -133 | -144 | 0 | 0 | 12 | 1233.2727 | 894.99536 | 891.6085 | NaN | 928.79285 | 864.0144 |
| Snrnp40 | WNVYGDCDNYATLK | -94 | -107 | 0 | 0 | 14 | 415.03705 | 247.62262 | 392.00354 | 559.2494 | NaN | NaN |
| H4c1 | VFLENVIR | -61 | -68 | 0 | 0 | 8 | 34110.605 | 38645.35 | 35984.48 | 2135.604 | 37735.133 | 37729.47 |
| H4c1 | GITKPAIR | -29 | -36 | 1 | 0.000907206 | 8 | 27144.887 | 25385.86 | 23605.725 | 23314.146 | 206.4992 | 22575.682 |
| H4c1 | DAVTYTEHAK | -69 | -78 | 0 | 0 | 10 | 21963.416 | 21174.316 | 2525.9854 | 1565.0765 | 25821.516 | 25907.021 |
| H4c1 | DAVTYTEHAK | -69 | -78 | 0 | 0 | 10 | 21963.416 | 21174.316 | 2525.9854 | 1565.0765 | 25821.516 | 25907.021 |
| H4c1 | DNIQGITKPAIR | -25 | -36 | 1 | 0 | 12 | 15280.29 | 13441.382 | 13266.451 | 22625.719 | 12853.665 | 13116.802 |
| H4c1 | DNIQGITKPAIR | -25 | -36 | 1 | 0 | 12 | 15280.29 | 13441.382 | 13266.451 | 22625.719 | 12853.665 | 13116.802 |
| H4c1 | ISGLIYEETR | -47 | -56 | 0 | 0 | 10 | 14792.583 | 14001.941 | 14156.804 | 11002.261 | 10742.69 | 10353.153 |
| H4c1 | NIQGITKPAIR | -26 | -36 | 1 | 0 | 11 | 8984.145 | 9680.548 | 1603.4664 | 13716.101 | 14617.388 | 13659.658 |
| H4c1 | VFLENVIRD | -61 | -69 | 1 | 0 | 9 | 5532.162 | 6054.6636 | 6019.7686 | 3938.615 | 4417.261 | 4535.484 |
| H4c1 | TVTAMDVVYALK | -81 | -92 | 0 | 0 | 12 | 4253.833 | 3726.289 | 4449.667 | 3812.872 | 1475.4908 | 4309.8525 |
| H4c1 | GLIYEETR | -49 | -56 | 0 | 0.000471409 | 8 | 3797.809 | 3125.1782 | 2638.4705 | 4021.8857 | 2726.812 | 3336.2192 |
| H4c1 | IYEETRGVLK | -51 | -60 | 1 | 0.000431655 | 10 | 3397.249 | 2886.9277 | 2843.1711 | 3449.42 | 3613.811 | 3534.8577 |
| H4c1 | FLENVIR | -62 | -68 | 0 | 0.003966045 | 7 | 3240.9487 | 3903.4592 | 4161.5806 | 3774.3223 | 4673.296 | 4637.4272 |
| H4c1 | RDNIQGITK | -24 | -32 | 1 | 0.000230707 | 9 | 3025.2673 | 3009.8413 | 3029.0862 | 8733.654 | 12539.867 | 12261.116 |
| H4c1 | DNIQGITKPA | -25 | -34 | 1 | 0.00051306 | 10 | 2828.251 | 3321.2625 | 2995.7893 | 3037.117 | 3079.989 | 1888.8398 |
| H4c1 | RDNIQGITKPAIR | -24 | -36 | 2 | 0 | 13 | 2761.7043 | 2688.3286 | 2684.8926 | 934.7179 | 15270.154 | 12842.86 |
| H4c1 | RDNIQGITKPAIR | -24 | -36 | 2 | 0 | 13 | 2761.7043 | 2688.3286 | 2684.8926 | 934.7179 | 15270.154 | 12842.86 |
| H4c1 | IQGITKPAIR | -27 | -36 | 1 | 0 | 10 | 2242.5361 | 300.13428 | 2455.5408 | 2205.2957 | 2232.596 | 2140.8298 |
| H4c1 | VTYTEHAK | -71 | -78 | 0 | 0.000471409 | 8 | 1875.7208 | 1561.2957 | 1981.2595 | 1578.7233 | 192.0843 | 1564.9224 |
| H4c1 | AVTYTEHAK | -70 | -78 | 0 | 0 | 9 | 1696.0704 | 1649.7068 | 492.95547 | 1085.5645 | 1129.6824 | 1203.0325 |
| H4c1 | KVFLENVIR | -60 | -68 | 1 | 0 | 9 | 1106.389 | 878.6885 | 934.0965 | 1637.3517 | 1715.257 | 1646.4805 |
| H4c1 | TVTAMDVVYAL | -81 | -91 | 0 | 0.001811524 | 11 | 512.54285 | 566.771 | 645.36005 | 1510.1892 | 2144.5012 | 1696.4395 |
| H4c1 | KTVTAMDVVYAL | -80 | -91 | 1 | 0.001023236 | 12 | 470.60516 | 1342.0477 | 1156.1694 | 1912.1946 | 2305.2166 | 2269.5576 |
| H4c1 | LENVIRDAVTYTEHAK | -63 | -78 | 1 | 0 | 16 | 400.919 | 888.77124 | 896.76135 | 1904.342 | 3395.4473 | 3238.9565 |
| H4c1 | DNIQGITK | -25 | -32 | 0 | 0.000550307 | 8 | 1769.5171 | 1154.5377 | 1026.9749 | 898.8426 | NaN | 906.999 |
| H4c1 | RISGLIYEETR | -46 | -56 | 1 | 0 | 11 | 698.3571 | 817.6084 | NaN | 4909.5806 | 4215.943 | 4493.1 |
| H4c1 | DNIQGITKPAIRR | -25 | -37 | 2 | 0 | 13 | 455.06412 | 2673.8857 | 2697.099 | NaN | 15376.624 | 12842.861 |
| H4c1 | QGITKPAIR | -28 | -36 | 1 | 0.003360691 | 9 | 410.41412 | NaN | 792.0648 | 230.75749 | 648.5619 | 658.18884 |

| | | | | | | | | | | | | |
|---|---|---|---|---|---|---|---|---|---|---|---|---|
| H4c1 | ISGLIYEETRG | -47 | -57 | 1 | 0.003360691 | 11 | 3101.9043 | 4397.9146 | 5335.7905 | NaN | NaN | NaN |
| Adk | AVVDKDFLDK | -37 | -46 | 1 | 0 | 10 | 2559.6187 | 2617.7502 | 2769.5078 | 2384.6428 | 2578.954 | 1647.763 |
| Adk | TGCTFPEKPDFH | -350 | -361 | 1 | 0 | 12 | 2143.3223 | 308.41193 | 2542.787 | 2620.0664 | 2765.2478 | 2014.7753 |
| Adk | YSLKPNDQILAEDK | -47 | -60 | 1 | 0 | 14 | 1698.3986 | 1510.4381 | 1649.4458 | 1546.7792 | 1811.4519 | 1699.9232 |
| Adk | QWLIQEPHK | -90 | -98 | 0 | 0 | 9 | 790.4868 | 871.16895 | 907.40125 | 766.6753 | 634.44366 | 775.39856 |
| Adk | ELFDELVK | -63 | -70 | 0 | 0.000471409 | 8 | 280.17886 | 1023.6563 | 998.958 | 803.78424 | 771.61523 | 775.6144 |
| Adk | RTGCTFPEKPDFH | -349 | -361 | 2 | 0 | 13 | 630.8194 | 214.26196 | 625.16156 | NaN | 183.55173 | 785.7183 |
| Asns | VPFLDHR | -418 | -424 | 0 | 0 | 7 | 1557.8251 | 1608.7324 | 1708.7532 | 1618.4329 | 1600.3875 | 1559.9966 |
| Asns | LFPGFDLETVK | -222 | -232 | 0 | 0 | 11 | 1369.1453 | 1326.6698 | 1485.2322 | 1356.6771 | 1878.5284 | 1825.274 |
| Asns | AVVDPLFGMQPIR | -51 | -63 | 0 | 0 | 13 | 813.45856 | 884.13354 | 862.27637 | 114.41438 | 1098.7922 | 1145.7739 |
| Asns | TDEPLHAIYDSVEK | -208 | -221 | 0 | 0 | 14 | 803.95935 | 1009.43933 | 1045.5491 | 846.4177 | 1112.9423 | 1077.817 |
| Asns | HSTTPFLK | -169 | -176 | 0 | 0 | 8 | 787.4014 | 522.11237 | 1032.339 | 1122.6191 | 783.2063 | 854.96924 |
| Asns | ELYLFDVLR | -396 | -404 | 0 | 0 | 9 | 349.40292 | 162.76749 | 220.75212 | 276.5744 | 490.59854 | 480.005 |
| Asns | KYPYLWLC | -66 | -73 | 1 | 0.001776473 | 8 | 759.4199 | 116.94617 | 563.2923 | 773.2671 | 929.31433 | NaN |
| Asns | VANYIGSEHHEVLFN | -302 | -316 | 0 | 0 | 15 | 619.2699 | 349.39496 | 706.608 | NaN | 897.15967 | 743.0097 |
| Asns | EAQVQYPLQTF | -276 | -286 | 0 | 0.00025015 | 11 | 330.8716 | 335.54208 | 528.9767 | NaN | 509.5157 | 262.68628 |
| Asns | ETFEDCNLLPK | -450 | -460 | 0 | 0 | 11 | 1232.0651 | 1178.7545 | NaN | NaN | 1441.6981 | 1181.3136 |
| IntS13 | TYPVGEDSLVSDRPK | -209 | -223 | 1 | 0 | 15 | 895.6336 | 1459.2266 | 1083.8376 | 1522.1837 | 1726.7717 | 1741.2965 |
| IntS13 | AVVPLAGVIVK | -473 | -483 | 0 | 0 | 11 | 275.6993 | NaN | NaN | 267.52286 | 284.70996 | 259.76886 |
| IntS13 | SILEDPPSISEGCGGR | -394 | -409 | 0 | 0.002068213 | 16 | 783.8909 | NaN | 884.29236 | NaN | NaN | 406.62848 |
| Kdm3b | AVVSGDPGPALR | -33 | -44 | 0 | 0.005758933 | 12 | 1695.5809 | NaN | NaN | NaN | NaN | NaN |
| Dnajc9 | ELGLEEGVDNLK | -206 | -217 | 0 | 0 | 12 | 1685.2408 | 2536.6826 | 2355.3464 | 3234.7644 | 301.6617 | 242.30713 |
| Dnajc9 | KELGLEEGVDNLK | -205 | -217 | 1 | 0 | 13 | 1202.0963 | 1061.3865 | 918.24 | 650.40564 | 653.2862 | 624.5624 |
| Dnajc9 | VHPDRVEEDQKEDATR | -42 | -57 | 2 | 0 | 16 | 512.9304 | 3229.1877 | 506.0922 | 2500.8262 | 856.9157 | 767.94904 |
| Dnajc9 | AVYDEQGTVDEDSAGLNQDR | -76 | -95 | 0 | 0 | 20 | 210.01129 | 363.39505 | 306.7969 | 407.95465 | 354.3324 | 362.08322 |
| Dnajc9 | GTVDEDSAGLNQDR | -82 | -95 | 0 | 0 | 14 | 2847.639 | 433.26132 | 522.4331 | 3907.7378 | NaN | 1314.7113 |
| Dnajc9 | ISLEDIQAFEK | -108 | -118 | 0 | 0 | 11 | 519.0894 | 740.34375 | NaN | 842.2257 | 386.34378 | 607.08466 |
| Dnajc9 | VEEDQKEDATR | -47 | -57 | 1 | 0 | 11 | 305.68604 | NaN | 352.18243 | 143.5081 | 354.91415 | 425.4154 |
| Dnajc9 | QAYLDFK | -132 | -138 | 0 | 0.002752526 | 7 | 502.6776 | NaN | 621.0325 | NaN | NaN | NaN |
| Lbr | YKDGTELELK | -41 | -50 | 1 | 0 | 10 | 3887.779 | 879.46075 | 3431.427 | 949.31335 | 1332.527 | 1083.2324 |
| Lbr | LTPLVLKPF | -127 | -135 | 1 | 0 | 9 | 3373.6858 | 3861.6013 | 3973.8596 | 2787.512 | 3897.8862 | 4126.875 |
| Lbr | VAEGTPLVDGR | -291 | -301 | 0 | 0 | 11 | 2653.5894 | 337.79123 | 2572.0005 | 2030.9753 | 1722.5144 | 1647.2168 |
| Lbr | EIESEEQNLVTK | -189 | -200 | 0 | 0 | 12 | 2466.976 | 2776.66 | 2776.5493 | 2980.6228 | 3523.2349 | 3521.6953 |
| Lbr | SSGNAVYDFFIGR | -371 | -383 | 0 | 0 | 13 | 1629.1312 | 253.81682 | 1944.0232 | 1746.9806 | 2285.505 | 2478.6702 |
| Lbr | LCGYVIFR | -506 | -513 | 0 | 0 | 8 | 1618.2698 | 1286.5276 | 1456.3434 | 877.6316 | 972.30383 | 422.81992 |
| Lbr | KFVEGEVVR | -5 | -13 | 1 | 0.00025015 | 9 | 1307.464 | 1293.2299 | 1369.0804 | 419.8961 | 1024.8717 | 1064.8445 |
| Lbr | LVSGWWGFVR | -546 | -555 | 0 | 0 | 10 | 933.21356 | 666.93744 | 1112.5374 | 1148.7965 | 1476.4719 | 1652.8888 |
| Lbr | ESDIKPLK | -51 | -58 | 1 | 0 | 8 | 786.6241 | 692.474 | 781.3433 | 1025.365 | 1017.1532 | 1151.4355 |
| Lbr | LTPLVLKPFG | -127 | -136 | 1 | 0.000591124 | 10 | 753.083 | 642.1993 | 540.95386 | 516.0241 | 656.6257 | 543.02057 |
| Lbr | GNAVYDFFIGR | -373 | -383 | 0 | 0 | 11 | 631.9057 | 533.52496 | 653.18384 | 621.09216 | 564.6532 | 716.4319 |
| Lbr | LTPLVLKPFGN | -127 | -137 | 1 | 0 | 11 | 574.8954 | 592.60364 | 632.6337 | 594.3233 | 618.8723 | 575.1604 |
| Lbr | AVYDFFIGR | -375 | -383 | 0 | 0 | 9 | 503.78152 | 385.61932 | 532.08984 | 387.0371 | 473.32904 | 489.70425 |
| Lbr | YGLAWEK | -606 | -612 | 0 | 0.001904399 | 7 | 819.8939 | 924.4499 | 925.0469 | NaN | NaN | 763.06635 |
| Lbr | FVEGEVVR | -6 | -13 | 0 | 0 | 8 | 724.8325 | 664.67487 | 842.91187 | 608.9338 | NaN | NaN |
| Lbr | EELSPASSGNAVYDFFIGR | -365 | -383 | 0 | 0.00025015 | 19 | 109.55881 | 208.9919 | NaN | NaN | 315.37 | 291.60233 |
| Lbr | LTPLVLKPFGNSV | -127 | -139 | 1 | 0 | 13 | 1107.5365 | 736.7091 | 1005.0128 | NaN | NaN | NaN |
| Lbr | QVTTPQR | -209 | -215 | 0 | 0.003333688 | 7 | 912.50214 | NaN | NaN | NaN | NaN | NaN |
| Lbr | SVSASHEGDVK | -101 | -111 | 0 | 0.000755556 | 11 | 793.3143 | NaN | NaN | NaN | NaN | NaN |
| Arl6ip5 | DDFFPGSDR | -12 | -20 | 0 | 0 | 9 | 1868.4626 | 1131.036 | 1344.1459 | 923.16956 | 504.84766 | 453.94583 |
| Arl6ip5 | AWDDFFPGSDR | -10 | -20 | 0 | 0 | 11 | 731.9988 | 828.2489 | 822.3766 | 768.9005 | 889.7866 | 292.37723 |
| Arl6ip5 | MDVNLAPLR | -1 | -9 | 0 | 0 | 9 | 509.9306 | 598.9957 | 455.59583 | 544.60583 | NaN | 704.37787 |
| Arl6ip5 | DALEQQEDNINK | -167 | -178 | 0 | 0.000431655 | 12 | 2499.6729 | 2553.8545 | 2672.7373 | 2260.9268 | NaN | NaN |
| Sars1 | LLIDEAIQK | -108 | -116 | 0 | 0 | 9 | 3096.4187 | 4285.317 | 4582.348 | 4208.14 | 5054.3564 | 4472.052 |
| Sars1 | FKDPGLVDQLVK | -27 | -38 | 1 | 0 | 12 | 2098.1213 | 2977.578 | 2865.345 | 4422.8926 | 3707.6377 | 3527.9534 |
| Sars1 | FKDPGLVDQLVK | -27 | -38 | 1 | 0 | 12 | 2098.1213 | 2977.578 | 2865.345 | 4422.8926 | 3707.6377 | 3527.9534 |
| Sars1 | AWFPGSGAFR | -381 | -390 | 0 | 0 | 10 | 1502.9036 | 1388.1294 | 1258.3019 | 1071.5746 | 1055.3268 | 611.11707 |
| Sars1 | VLDLDLFR | -2 | -9 | 0 | 0 | 8 | 1443.1757 | 1734.4896 | 1610.6436 | 1535.5714 | 1947.0382 | 1974.5935 |
| Sars1 | NIVSGSLNHAASK | -363 | -375 | 0 | 0 | 13 | 1374.6097 | 1353.6311 | 1601.3574 | 1412.1147 | 1646.7104 | 1214.7589 |
| Sars1 | VKPAPIDQEPSKK | -471 | -483 | 2 | 0 | 13 | 1247.5242 | 468.25323 | 1500.7922 | 503.44104 | 1604.3636 | 1619.391 |
| Sars1 | VKPAPIDQEPSKK | -471 | -483 | 2 | 0 | 13 | 1247.5242 | 468.25323 | 1500.7922 | 503.44104 | 1604.3636 | 1619.391 |
| Sars1 | DEWLRPEDLPIK | -282 | -293 | 1 | 0.00025015 | 12 | 1234.0996 | 1210.3099 | 1274.0281 | 1999.4272 | 2111.669 | 310.15024 |
| Sars1 | DEWLRPEDLPIK | -282 | -293 | 1 | 0.00025015 | 12 | 1234.0996 | 1210.3099 | 1274.0281 | 1999.4272 | 2111.669 | 310.15024 |
| Sars1 | SQFDEELYK | -241 | -249 | 0 | 0 | 9 | 1196.4493 | 1599.592 | 1343.7572 | 297.6905 | 1382.5167 | 255.61682 |
| Sars1 | LHPSVPISNDEDADNKVER | -139 | -157 | 1 | 0 | 19 | 1056.232 | 1263.0411 | 1553.6434 | 1803.202 | 2615.6624 | 2801.0544 |
| Sars1 | LHPSVPISNDEDADNKVER | -139 | -157 | 1 | 0 | 19 | 1056.232 | 1263.0411 | 1553.6434 | 1803.202 | 2615.6624 | 2801.0544 |
| Sars1 | EIGNLLHPSVPISNDEDADNKVER | -134 | -157 | 1 | 0 | 24 | 901.17456 | 1299.2288 | 2382.88 | 526.6404 | 1186.1422 | 1329.4948 |
| Sars1 | HRDEWLRPEDLPIK | -280 | -293 | 2 | 0.001055699 | 14 | 791.50366 | 1689.512 | 1937.0374 | 1802.5591 | 2456.3633 | 2530.1328 |
| Sars1 | HRDEWLRPEDLPIK | -280 | -293 | 2 | 0.001055699 | 14 | 791.50366 | 1689.512 | 1937.0374 | 1802.5591 | 2456.3633 | 2530.1328 |
| Sars1 | QFDEELYK | -242 | -249 | 0 | 0 | 8 | 762.6861 | 1080.4668 | 1074.2777 | 1180.4039 | 837.4411 | 901.1284 |
| Sars1 | IWGDCTVR | -158 | -165 | 0 | 0.000230707 | 8 | 2587.4849 | NaN | 2806.1006 | 2770.6055 | 2879.287 | 523.4331 |
| Sars1 | IEQFVYSSPHDNK | -324 | -336 | 0 | 0 | 13 | 1602.7544 | 1534.7933 | 1328.9745 | 568.42114 | 1347.3398 | NaN |
| Sars1 | TICAILENYQAEK | -436 | -448 | 0 | 0 | 13 | 1022.87 | NaN | 999.523 | 1404.2637 | 995.13605 | 1110.7062 |
| Sars1 | DLEAWFPGSGAFR | -378 | -390 | 0 | 0 | 13 | 446.61688 | 683.8491 | 510.98187 | NaN | NaN | 639.5184 |
| Sars1 | GYTPIYTPF | -219 | -227 | 0 | 0.00025015 | 9 | 666.27686 | NaN | NaN | 864.628 | 1297.5149 | NaN |
| Sars1 | VDKGGDPALIR | -10 | -20 | 1 | 0.001868731 | 11 | 1219.1567 | NaN | NaN | 1244.482 | NaN | NaN |
| Cyrib | TSTPYSPTQHLER | -112 | -124 | 0 | 0 | 13 | 2797.0027 | 2728.3066 | 2764.9355 | 1776.2917 | 1956.3806 | 1908.584 |
| Cyrib | KFYEFSQR | -92 | -99 | 1 | 0.00051306 | 8 | 2638.5369 | 2723.5488 | 2045.3893 | 1653.5023 | 1695.1176 | 1591.5992 |
| Cyrib | NVPAEGENEVNNELANR | -170 | -186 | 0 | 0 | 17 | 1904.9951 | 2382.5723 | 2556.2866 | 2512.3115 | 3351.5422 | 3152.3125 |

| | | | | | | | | | | | | |
|---|---|---|---|---|---|---|---|---|---|---|---|---|
| Cyrib | AWGAVVPLVGK | -79 | -89 | 0 | 0 | 11 | 1430.0273 | 1670.6796 | 1736.5181 | 930.71655 | 1130.3867 | 1055.8754 |
| Cyrib | DAEGILEDLQSYR | -45 | -57 | 0 | 0 | 13 | 1211.8348 | 1127.2473 | 1106.8628 | 1365.2714 | 1859.987 | 1727.0594 |
| Cyrib | INNVPAEGENEVNNELANR | -168 | -186 | 0 | 0 | 19 | 333.58545 | 327.5894 | 412.9976 | 324.74667 | 443.44418 | 421.84857 |
| Cyrib | EAIQHPADEK | -65 | -74 | 0 | 0.00025015 | 10 | 680.8469 | 651.51953 | NaN | 804.12744 | 358.7694 | 874.3935 |
| Cyrib | STPYSPTQHLER | -113 | -124 | 0 | 0 | 12 | 618.8046 | NaN | 733.5719 | 460.69113 | 669.0396 | NaN |
| Cyrib | FENAQPTESEK | -23 | -33 | 0 | 0 | 11 | 1441.7505 | 1547.739 | 1713.925 | NaN | NaN | NaN |
| Tspo | QKPSWHPPR | -38 | -46 | 1 | 0 | 9 | 1102.4603 | 380.35037 | 1000.25507 | 844.5863 | 366.6695 | 331.16037 |
| Tspo | AWPPIFFGAR | -94 | -103 | 0 | 0 | 10 | 276.98358 | 3376.0156 | 3708.093 | 2258.5513 | 3031.732 | 3252.0032 |
| Tspo | LQKPSWHPPR | -37 | -46 | 1 | 0 | 10 | 234.40553 | 1704.8557 | 2189.7942 | 107.76955 | 216.52946 | 364.35666 |
| Vdac1 | LTFDSSFSPNTGK | -110 | -122 | 0 | 0 | 13 | 1511.5312 | 1442.5906 | 1294.9031 | 2158.835 | 1346.8678 | 1903.5903 |
| Vdac1 | WTEYGLTFTEK | -77 | -87 | 0 | 0 | 11 | 856.3832 | 760.4355 | 1024.2523 | 91.74821 | 1508.2767 | 398.0084 |
| Vdac1 | TDEFQLHTN | -188 | -196 | 0 | 0 | 9 | 4169.135 | 1424.5002 | 4504.7266 | 580.0421 | NaN | 1176.5194 |
| Vdac1 | GYTQTLKPGIK | -259 | -269 | 1 | 0.00201047 | 11 | 506.99048 | 1021.39905 | 983.086 | NaN | 1166.1182 | 569.388 |
| Vdac1 | YQVDPDACFSAK | -238 | -249 | 0 | 0 | 12 | 491.86072 | 761.3035 | 804.8751 | NaN | 962.9344 | 937.3807 |
| Vdac1 | YQVDPDACFSA | -238 | -248 | 0 | 0.000230707 | 11 | 75.415375 | 109.65052 | NaN | 168.16827 | 230.7373 | 185.08446 |
| Vdac1 | GYGFGLIK | -34 | -41 | 0 | 0 | 8 | 1694.8556 | 823.0514 | 1114.0431 | 680.1948 | NaN | NaN |
| Vdac1 | TLKPGIK | -263 | -269 | 1 | 0.00111006 | 7 | 1109.4573 | NaN | 1285.157 | 1255.2305 | NaN | 950.1108 |
| Vdac1 | AWTAGNSNTR | -222 | -231 | 0 | 0.000431655 | 10 | 2807.845 | 252.15411 | NaN | NaN | NaN | NaN |
| Vdac1 | VTQSNFAVGYK | -177 | -187 | 0 | 0.000230707 | 11 | 2170.6816 | NaN | NaN | NaN | NaN | NaN |
| Dock11 | AWVPLLK | -758 | -764 | 0 | 0 | 7 | 1825.7646 | 1142.3732 | 1013.82574 | 435.10184 | 311.6652 | 273.60764 |
| Dock11 | IELPIHLHQK | -716 | -725 | 0 | 0 | 10 | 1059.7836 | 1093.6458 | 1182.3492 | NaN | 1595.005 | 1120.608 |
| Dock11 | IELPIHLHQK | -716 | -725 | 0 | 0 | 10 | 1059.7836 | 1093.6458 | 1182.3492 | NaN | 1595.005 | 1120.608 |
| Dock11 | FPSAFFK | -1467 | -1473 | 0 | 0.001868731 | 7 | 374.34143 | 298.60034 | 610.4882 | NaN | 162.95192 | 634.03107 |
| Dock11 | KYEDFSGDFR | -118 | -127 | 1 | 0 | 10 | 630.51984 | NaN | NaN | 1444.0493 | 1317.2446 | 1229.0393 |
| Dock11 | SFFEEEDGKEYIYK | -1772 | -1785 | 1 | 0.00025015 | 14 | 599.7157 | NaN | NaN | NaN | 940.02216 | NaN |
| Hccs | YVIDYYDGGEVNK | -228 | -240 | 0 | 0 | 13 | 1173.4568 | 983.69037 | 743.6156 | 946.4665 | 356.7403 | 1003.4486 |
| Hccs | HDWIINR | -215 | -221 | 0 | 0 | 7 | 1150.9216 | 1424.108 | 1216.2152 | 1271.2391 | 1253.2135 | 1361.9034 |
| Hccs | WKDDDISQK | -141 | -149 | 1 | 0 | 9 | 738.63495 | 521.40704 | 552.7011 | 353.00705 | 618.9937 | 697.5414 |
| Hccs | AYDYVECPVTGAR | -64 | -76 | 0 | 0 | 13 | 1097.5851 | 873.646 | NaN | 943.6206 | NaN | NaN |
| Dnaja1 | DHAVFTR | -247 | -253 | 0 | 0 | 7 | 539.5536 | 1578.005 | 1707.9938 | 1311.5789 | 971.65533 | 1487.7019 |
| Dnaja1 | YHPDKNPNEGEK | -33 | -44 | 1 | 0 | 12 | 338.6488 | 339.95255 | 379.37387 | 305.61603 | 340.61838 | 319.48572 |
| Dnaja1 | YHPDKNPNEGEK | -33 | -44 | 1 | 0 | 12 | 338.6488 | 339.95255 | 379.37387 | 305.61603 | 340.61838 | 319.48572 |
| Dnaja1 | AYEDDEHHPR | -380 | -389 | 0 | 0.00025015 | 10 | 320.48163 | 224.1699 | 387.37347 | 166.85237 | 436.87665 | 401.86685 |
| Dnaja1 | LVDFDPNQER | -363 | -372 | 0 | 0 | 10 | 1547.4629 | 1735.3204 | 188.42155 | NaN | 164.13547 | NaN |
| Glo1 | AYEDKNDIPK | -74 | -83 | 1 | 0 | 10 | 1798.7048 | 1657.744 | 1441.9834 | 3157.9426 | 4460.8413 | 2793.4175 |
| Glo1 | VLGLTLLQK | -52 | -60 | 0 | 0 | 9 | 341.0828 | 364.12198 | 372.8856 | 371.1909 | 508.64944 | NaN |
| Tmed10 | LKPLEVELR | -144 | -152 | 1 | 0 | 9 | 2745.449 | 2961.8538 | 3055.9717 | 3118.3564 | 3204.816 | 3291.6301 |
| Tmed10 | AYEITDQSGGAGGLR | -57 | -71 | 0 | 0 | 15 | 1857.155 | 1134.2341 | 1997.4083 | 2689.42 | 516.9723 | 1053.937 |
| Tmed10 | ISFHLPVNSR | -32 | -41 | 0 | 0 | 10 | 1798.9458 | 2099.8604 | 894.7219 | 2182.673 | 2097.4111 | 2225.6943 |
| Tmed10 | ISFHLPVNSR | -32 | -41 | 0 | 0 | 10 | 1798.9458 | 2099.8604 | 894.7219 | 2182.673 | 2097.4111 | 2225.6943 |
| Tmed10 | GISFHLPVNSR | -31 | -41 | 0 | 0 | 11 | 847.65314 | 961.85614 | 942.6065 | 818.10956 | 3075.3315 | 3355.4673 |
| Tmed10 | GAYEITDQSGGAGGLR | -56 | -71 | 0 | 0 | 16 | 659.35364 | 2985.8455 | 3233.6562 | 3483.6897 | 233.25461 | 3889.4456 |
| Tmed10 | KLKPLEVELR | -143 | -152 | 2 | 0 | 10 | 635.41144 | 605.7974 | 536.92957 | 656.712 | 527.7529 | 480.228 |
| Tmed10 | RLEDLSESIVNDF | -153 | -165 | 1 | 0.00025015 | 13 | 578.29706 | 499.5492 | 621.10944 | 503.11987 | 329.72806 | 253.93718 |
| Tmed10 | ITDSAGHIL | -76 | -84 | 0 | 0.000431655 | 9 | 1936.0487 | 1515.9526 | 1527.6882 | 1473.4375 | NaN | NaN |
| Spcs2 | IDDKPVKID | -41 | -49 | 2 | 0.001636916 | 9 | 7572.4907 | 5881.013 | 4979.9043 | 4430.8047 | 3952.1724 | 4792.1562 |
| Spcs2 | RFDDKYTLK | -165 | -173 | 2 | 0.00122419 | 9 | 2185.5298 | 1952.7993 | 1682.331 | 1246.7474 | 2198.2146 | 1801.6395 |
| Spcs2 | RFDDKYTLK | -165 | -173 | 2 | 0.00122419 | 9 | 2185.5298 | 1952.7993 | 1682.331 | 1246.7474 | 2198.2146 | 1801.6395 |
| Spcs2 | FDDKYTLK | -166 | -173 | 1 | 0.00025015 | 8 | 1926.769 | 404.04645 | 1870.8975 | 1666.6868 | 1431.6326 | 1245.2465 |
| Spcs2 | YVENFGLIDGR | -74 | -84 | 0 | 0 | 11 | 479.23224 | 1291.6212 | 512.38885 | 514.03723 | 1427.7733 | 756.5559 |
| Spcs2 | LHDSLATER | -215 | -223 | 0 | 0 | 9 | 2427.112 | NaN | 2626.061 | 2029.8965 | 2472.8862 | 2640.6064 |
| Spcs2 | IDKWDGSAVK | -48 | -57 | 1 | 0.000550307 | 10 | 1360.8832 | NaN | NaN | 1003.013 | 2045.0159 | NaN |
| Spcs2 | AYEPEISR | -207 | -214 | 0 | 0.004595494 | 8 | 634.45624 | 1479.2634 | NaN | NaN | NaN | NaN |
| Urb1 | AYEQNNLSLISFR | -1547 | -1559 | 0 | 0.00111006 | 13 | 709.1234 | NaN | NaN | NaN | NaN | NaN |
| Urb1 | EVDPGDWQQFVK | -1421 | -1432 | 0 | 0 | 12 | 188.69519 | NaN | NaN | NaN | NaN | NaN |
| Kars1 | ILDDICVAK | -420 | -428 | 0 | 0 | 9 | 3656.1396 | 1102.906 | 3560.9783 | 3312.7407 | 1381.4725 | 1325.4344 |
| Kars1 | EVDFTPPFR | -382 | -390 | 0 | 0 | 9 | 2117.1897 | 2122.135 | 2237.0496 | 1922.5751 | 1932.3887 | 2053.675 |
| Kars1 | LPETSLFETEETR | -406 | -418 | 0 | 0 | 13 | 2046.3379 | 2056.5413 | 2190.033 | 1714.8545 | 338.21286 | 2636.958 |
| Kars1 | EICNAYTELNDPVR | -492 | -505 | 0 | 0 | 14 | 1780.9807 | 2481.3672 | 1300.4104 | 2761.3486 | 2234.0532 | 2435.5903 |
| Kars1 | SEEEFVHIN | -163 | -171 | 0 | 0 | 9 | 1166.3676 | 1161.0751 | 1231.3746 | 1303.233 | 212.02306 | 1467.8774 |
| Kars1 | LPETSLFETEETRK | -406 | -419 | 1 | 0 | 14 | 1078.2175 | 1363.7596 | 1340.7675 | 1561.6871 | 1813.4232 | 1897.5591 |
| Kars1 | KILDDICVAK | -419 | -428 | 1 | 0 | 10 | 1004.13446 | 1094.7992 | 1097.0497 | 860.81635 | 1275.4174 | 1059.8533 |
| Kars1 | NEGIDLTHNPEFTTC | -322 | -336 | 0 | 0.00025015 | 15 | 940.75543 | 1084.1544 | 1066.8884 | 1137.0402 | 1318.5322 | 1301.0065 |
| Kars1 | SHLQPGDHLTDVTLK | -111 | -125 | 0 | 0 | 15 | 910.9427 | 1118.1116 | 1423.4895 | 1395.9519 | 1296.3191 | 1436.0583 |
| Kars1 | SHLQPGDHLTDVTLK | -111 | -125 | 0 | 0 | 15 | 910.9427 | 1118.1116 | 1423.4895 | 1395.9519 | 1296.3191 | 1436.0583 |
| Kars1 | EYSHLQPGDHLTDVTLK | -109 | -125 | 0 | 0 | 17 | 624.0822 | 670.12695 | 766.3624 | 632.47437 | 719.7632 | 692.92706 |
| Kars1 | EYSHLQPGDHLTDVTLK | -109 | -125 | 0 | 0 | 17 | 624.0822 | 670.12695 | 766.3624 | 632.47437 | 719.7632 | 692.92706 |
| Kars1 | KVTGEDPYPHK | -86 | -96 | 1 | 0 | 11 | 431.7314 | 469.4934 | 559.5447 | 897.0333 | 242.64218 | 582.9712 |
| Kars1 | KGELSIIPQEITLL | -191 | -204 | 1 | 0.000230707 | 14 | 261.33032 | 258.43082 | 269.95938 | 365.9247 | 391.1998 | 356.3167 |
| Kars1 | IAPELYHK | -296 | -303 | 0 | 0.001846793 | 8 | 3831.5906 | 3687.254 | 3468.2024 | 5232.2627 | 4059.3362 | NaN |
| Kars1 | VTGEDPYPHK | -87 | -96 | 0 | 0 | 10 | 1709.3783 | 1747.2402 | 1668.8606 | 934.16315 | NaN | 887.5967 |
| Kars1 | AYEVDFTPPFR | -380 | -390 | 0 | 0 | 11 | 635.99274 | 800.30853 | NaN | 675.55585 | 701.0287 | 419.44983 |
| Kars1 | KGELSIIPQ | -191 | -199 | 1 | 0.00437931 | 9 | 758.0416 | 1054.7546 | 961.68164 | 969.9827 | NaN | NaN |
| Kars1 | ITYHPDGPEGQAY | -369 | -381 | 0 | 0.00025015 | 13 | 2006.196 | 989.3147 | 1078.3464 | NaN | NaN | NaN |
| Kars1 | EVLLFPAMKPEDK | -564 | -576 | 1 | 0 | 13 | 652.1006 | 678.3878 | 710.9307 | NaN | NaN | NaN |
| Kars1 | TLDPNQYYK | -68 | -76 | 0 | 0 | 9 | 387.43793 | 1894.8772 | 1885.4124 | NaN | NaN | NaN |
| Kars1 | FIQEYSHLQPGDHLTDVTLK | -106 | -125 | 0 | 0.00149795 | 20 | 606.0608 | NaN | NaN | 437.38336 | NaN | NaN |
| Kars1 | YLDLILNDFVR | -229 | -239 | 0 | 0.001610812 | 11 | 327.8104 | NaN | NaN | NaN | NaN | NaN |

| | | | | | | | | | | | | |
|---|---|---|---|---|---|---|---|---|---|---|---|---|
| Cars1 | FPHHDNELAQSE | -454 | -465 | 0 | 0.002810811 | 12 | 1188.8197 | 1064.5212 | 503.34756 | 932.6978 | 379.3436 | 333.7873 |
| Cars1 | DDILPELGVR | -702 | -711 | 0 | 0 | 10 | 1100.2344 | 1228.3975 | 1262.3903 | 364.8082 | 1903.5653 | 1996.7843 |
| Cars1 | FEDHEGLPTVVK | -712 | -723 | 0 | 0 | 12 | 927.7464 | 883.04956 | 968.31305 | 364.7455 | 796.0212 | 728.1071 |
| Cars1 | LSETTDPDKR | -223 | -232 | 1 | 0 | 10 | 497.29877 | 782.9954 | 678.5674 | 790.2157 | 1013.95233 | 1032.7959 |
| Cars1 | TAVHEALCDNIDTR | -582 | -595 | 0 | 0 | 14 | 414.6598 | 451.7132 | 553.5551 | 548.7741 | 424.22012 | 513.7689 |
| Cars1 | PEAVGDQK | -371 | -378 | 0 | 0.00201047 | 8 | 5015.9614 | NaN | 377.4045 | 457.25436 | 3648.9807 | 1269.5892 |
| Cars1 | WSPPAGTEPCR | -101 | -111 | 0 | 0.000230707 | 11 | 1180.0436 | 319.72668 | 291.34406 | NaN | 1062.6958 | 1102.5055 |
| Cars1 | NVLPPDVLTR | -314 | -323 | 0 | 0 | 10 | 876.8758 | 796.02655 | NaN | 617.2816 | 1504.4773 | 1972.5518 |
| Cars1 | EQKPPATQLLK | -200 | -210 | 1 | 0 | 11 | 1145.8184 | 1169.5419 | 1231.9302 | NaN | NaN | 1712.6342 |
| Cars1 | FDENGLPTHDTEGK | -784 | -797 | 0 | 0 | 14 | 1496.4872 | 367.47092 | 1582.177 | NaN | NaN | NaN |
| Cars1 | SSLSGEEVDSK | -255 | -265 | 0 | 0.00025015 | 11 | 1334.733 | 1040.5624 | 1484.0508 | NaN | NaN | NaN |
| Cars1 | AYFENDCWVR | -466 | -475 | 0 | 0 | 10 | 782.4455 | NaN | 583.90405 | 730.4251 | NaN | NaN |
| Cars1 | DVFIPQDGKK | -123 | -132 | 1 | 0.008680443 | 10 | 890.3576 | 743.17444 | NaN | NaN | NaN | NaN |
| Eif1a | TSDIILIGLR | -73 | -82 | 0 | 0 | 10 | 801.4101 | 132.35292 | 861.2552 | 611.8973 | 669.4361 | 668.0055 |
| Hspa14 | AYGIGQDHPTGK | -179 | -190 | 0 | 0 | 12 | 1081.052 | 832.3612 | 1505.9062 | 1063.9136 | 713.8637 | 634.0007 |
| Hspa14 | FTVLFPSGTPLPAR | -412 | -425 | 0 | 0 | 14 | 753.8299 | 561.61084 | 593.9466 | 533.00116 | 634.4783 | 580.96893 |
| Hspa14 | LRYEIDTGEETK | -98 | -109 | 1 | 0.000431655 | 12 | 563.46735 | NaN | 775.18243 | 1014.1123 | 884.6687 | 421.82364 |
| Hspa14 | FELLCSPLFNK | -300 | -310 | 0 | 0 | 11 | 476.58255 | 1477.8656 | 592.12506 | 1272.3438 | NaN | 1159.3447 |
| Hspa14 | TVLFPSGTPLPAR | -413 | -425 | 0 | 0 | 13 | 1445.2407 | 1642.6433 | 1612.997 | NaN | NaN | NaN |
| Bclaf1 | AYIVERPSTAK | -463 | -473 | 1 | 0 | 11 | 1427.2408 | 1712.5614 | 1723.936 | 1248.8955 | 1614.7471 | 1944.4094 |
| Bclaf1 | SSFYPEGDQETAK | -318 | -330 | 0 | 0 | 13 | 586.4426 | 254.9513 | 746.4634 | 571.0698 | 495.34726 | 713.65924 |
| Bclaf1 | FHDSEGDDTEETEDYR | -392 | -407 | 0 | 0 | 16 | 420.47125 | 50.223362 | 427.26974 | 414.824 | 520.7349 | 498.59375 |
| Bclaf1 | KAEGEPQEESPLK | -168 | -180 | 1 | 0.000550307 | 13 | 869.0684 | NaN | 716.7561 | 837.2262 | 945.60626 | 803.45386 |
| Bclaf1 | GFKEEIQK | -672 | -679 | 1 | 0.001055699 | 8 | 500.61453 | 776.79663 | 1052.5066 | NaN | 753.6677 | 943.0869 |
| Bclaf1 | DTFEHDPSESIDEFNK | -189 | -204 | 0 | 0.00025015 | 16 | 1118.1534 | 1059.0089 | 1773.1669 | NaN | 803.89014 | NaN |
| Bclaf1 | ESDGFREEK | -447 | -455 | 1 | 0.001023236 | 9 | 1108.2343 | NaN | NaN | 1533.6783 | 281.25476 | 1555.9473 |
| Snu13 | AYPLADAHLTK | -10 | -20 | 0 | 0 | 11 | 5652.198 | 6421.803 | 5867.464 | 6290.313 | 891.27576 | 5727.6436 |
| Snu13 | SIQQSIER | -118 | -125 | 0 | 0.000431655 | 8 | 2563.0337 | 5727.7935 | 1985.6444 | 1766.1517 | 3176.372 | 885.31647 |
| Snu13 | KLLDLVQQS | -21 | -29 | 1 | 0.00025015 | 9 | 627.3987 | 3637.651 | NaN | NaN | 697.8752 | 760.1726 |
| Snu13 | NVPYVFVR | -77 | -84 | 0 | 0.00025015 | 8 | 2167.8018 | 1799.4868 | 2021.223 | NaN | NaN | NaN |
| Snu13 | EIILHLPLLCEDK | -64 | -76 | 0 | 0 | 13 | 386.2959 | 229.97888 | NaN | NaN | NaN | NaN |
| Rnps1 | VEFENPDEAEK | -208 | -218 | 0 | 0 | 11 | 522.2695 | 708.98676 | 1733.9045 | 1935.6815 | 961.20935 | 197.15936 |
| Rnps1 | HMDGGQIDGQEITAT | -222 | -236 | 0 | 0.00025015 | 15 | 372.30673 | 772.9346 | 699.82837 | 426.86188 | 418.28253 | 343.09256 |
| Rnps1 | AYVEFENPDEAEK | -206 | -218 | 0 | 0 | 13 | 224.47763 | 1564.034 | 2003.6182 | 1646.4296 | 146.43677 | 1304.4227 |
| Rnps1 | FENPDEAEK | -210 | -218 | 0 | 0 | 9 | 1415.9175 | 172.69737 | 940.9525 | NaN | NaN | 1248.7898 |
| Ptprc | TLVSWPEPVSKPESASKPHGY | -392 | -412 | 2 | 0 | 21 | 5089.853 | 5751.2974 | 5683.559 | 2974.131 | 5072.0254 | 3838.1038 |
| Ptprc | FSLHDCTPK | -280 | -288 | 0 | 0 | 9 | 5075.6274 | 5369.2197 | 1507.5128 | 2314.6387 | 5428.231 | 5184.41 |
| Ptprc | HNGVYEGDSVIR | -546 | -557 | 0 | 0 | 12 | 3363.8508 | 1344.8933 | 706.4831 | 2685.4973 | 596.0674 | 1824.1844 |
| Ptprc | KRDPPSDPSPLEAEYQ | -923 | -938 | 2 | 0.000230707 | 16 | 3168.574 | 1499.8605 | 1530.8506 | 2266.8665 | 2600.5422 | 2603.6836 |
| Ptprc | KRDPPSDPSPLEAEYQ | -923 | -938 | 2 | 0.000230707 | 16 | 3168.574 | 1499.8605 | 1530.8506 | 2266.8665 | 2600.5422 | 2603.6836 |
| Ptprc | DETVDDFWR | -719 | -727 | 0 | 0 | 9 | 2728.0947 | 3046.8179 | 3067.4033 | 2949.405 | 3773.6167 | 3809.1167 |
| Ptprc | TDLGIPETPKPSCGDPAAR | -372 | -390 | 1 | 0 | 19 | 2420.7788 | 2363.9167 | 2394.9983 | 2255.7747 | 1986.5127 | 1767.1859 |
| Ptprc | TDLGIPETPKPSCGDPAAR | -372 | -390 | 1 | 0 | 19 | 2420.7788 | 2363.9167 | 2394.9983 | 2255.7747 | 1986.5127 | 1767.1859 |
| Ptprc | QELVERDDEK | -605 | -614 | 1 | 0 | 10 | 2269.4653 | 2392.5317 | 2858.3308 | 2865.2483 | 1925.7721 | 3445.9453 |
| Ptprc | KRDPPSDPSPLEAE | -923 | -936 | 2 | 0.000471409 | 14 | 2260.7637 | 601.6417 | 1041.1466 | 853.21686 | 568.7885 | 771.2765 |
| Ptprc | FHNGVYEGDSVIR | -545 | -557 | 0 | 0 | 13 | 1967.5935 | 2069.0159 | 1988.2782 | 1792.839 | 2401.2378 | 2213.4104 |
| Ptprc | SWPDHGVPEDPHLLLK | -807 | -822 | 0 | 0 | 16 | 1533.2352 | 1830.1575 | 801.0657 | 603.83295 | 2283.6365 | 2162.594 |
| Ptprc | SWPDHGVPEDPHLLLK | -807 | -822 | 0 | 0 | 16 | 1533.2352 | 1830.1575 | 801.0657 | 603.83295 | 2283.6365 | 2162.594 |
| Ptprc | QDANCVRPDGPLNK | -1244 | -1257 | 1 | 0 | 14 | 1484.7864 | 1524.8439 | 1403.7211 | 1608.1598 | 1636.6482 | 1563.4635 |
| Ptprc | QDANCVRPDGPLNK | -1244 | -1257 | 1 | 0 | 14 | 1484.7864 | 1524.8439 | 1403.7211 | 1608.1598 | 1636.6482 | 1563.4635 |
| Ptprc | CQLDNLR | -339 | -345 | 0 | 0.000712251 | 7 | 1387.6962 | 1350.0153 | 1159.0458 | 1498.5406 | 1195.9052 | 1138.0573 |
| Ptprc | SSNLDEQQELVER | -598 | -610 | 0 | 0 | 13 | 1328.9241 | 1433.909 | 1440.9727 | 1786.4843 | 1601.2338 | 1838.8021 |
| Ptprc | SYIDGFKEPR | -700 | -709 | 1 | 0.000230707 | 10 | 1187.5098 | 1512.4443 | 1685.213 | 685.579 | 1027.9459 | 1236.5509 |
| Ptprc | KRDPPSDPSPL | -923 | -933 | 2 | 0.001055699 | 11 | 1170.3438 | 896.80975 | 944.7529 | 908.07385 | 1229.6622 | 1091.3995 |
| Ptprc | SSNLDEQQELVERDDEK | -598 | -614 | 1 | 0 | 17 | 1100.2198 | 807.65356 | 687.3151 | 667.6698 | 149.64337 | 662.97534 |
| Ptprc | SSNLDEQQELVERDDEK | -598 | -614 | 1 | 0 | 17 | 1100.2198 | 807.65356 | 687.3151 | 667.6698 | 149.64337 | 662.97534 |
| Ptprc | DGSQQTGLFCAL | -1159 | -1170 | 0 | 0 | 12 | 864.65375 | 861.71796 | 918.21594 | 932.68024 | 1143.9175 | 1221.2863 |
| Ptprc | DPPSDPSPLEAEYQR | -925 | -939 | 0 | 0 | 15 | 693.4299 | 1353.9353 | 1167.8121 | 1416.3185 | 569.54645 | 1504.9761 |
| Ptprc | KTLVSWPEPVSKPE | -391 | -404 | 2 | 0.00025015 | 14 | 605.8332 | 635.5134 | 600.3756 | 489.32382 | 560.5462 | 615.49634 |
| Ptprc | TWKGEELPAEPK | -1113 | -1124 | 1 | 0 | 12 | 551.6638 | 2339.2654 | 1939.7603 | 2171.368 | 2222.124 | 2298.125 |
| Ptprc | FTSWPDHGVPEDPHLLLK | -805 | -822 | 0 | 0 | 18 | 551.0043 | 568.94836 | 641.5241 | 622.53436 | 735.1917 | 689.18774 |
| Ptprc | TNCQFYVDNLYYST | -524 | -537 | 0 | 0.000712251 | 14 | 350.43182 | 361.61945 | 392.77454 | 565.09106 | 656.932 | 777.6909 |
| Ptprc | CGNEDCENVLNNLEECSQIK | -235 | -254 | 0 | 0 | 20 | 300.05728 | 311.67923 | 337.86063 | 409.52917 | 432.41736 | 381.38666 |
| Ptprc | RNTFIPER | -331 | -338 | 1 | 0 | 8 | 273.57153 | 2856.9124 | 2585.2505 | 1809.4813 | 2311.839 | 2267.5405 |
| Ptprc | TLVSWPEPVSKPESASKPHGYVL | -392 | -414 | 2 | 0 | 23 | 163.861 | 741.1417 | 808.41614 | 571.98663 | 883.9915 | 918.38074 |
| Ptprc | YVDILPYDYNR | -672 | -682 | 0 | 0 | 11 | 897.31964 | NaN | 965.63367 | 913.15405 | 815.33875 | 694.80066 |
| Ptprc | NDSCAPATTIDLYVPPGTDK | -260 | -279 | 0 | 0 | 20 | 810.5611 | 805.02594 | 805.9589 | 778.4174 | NaN | 914.3837 |
| Ptprc | KGEELPAEPK | -1115 | -1124 | 1 | 0.000907206 | 10 | 633.35004 | NaN | 1408.4331 | 1335.9069 | 1350.1545 | 1342.9034 |
| Ptprc | DFNRVPLK | -970 | -977 | 1 | 0.002068213 | 8 | 529.41064 | 1252.9396 | 1157.4677 | NaN | 1398.103 | 1287.5923 |
| Ptprc | KTLVSWPEPVSKPESA | -391 | -406 | 2 | 0 | 16 | 453.8487 | 622.08496 | 619.63074 | 938.17395 | 697.0044 | NaN |
| Ptprc | TQHIGNQEENK | -948 | -958 | 0 | 0.00025015 | 11 | 309.43063 | 387.50558 | 213.38428 | 267.51312 | NaN | 302.63058 |
| Ptprc | TLVSWPEPVSKPESASK | -392 | -408 | 1 | 0 | 17 | 289.7473 | NaN | 506.8096 | 697.5644 | 819.9176 | 800.9304 |
| Ptprc | LGIPETPKPSCGDPAAR | -374 | -390 | 1 | 0.000230707 | 17 | 107.77222 | 3846.3267 | NaN | 395.46716 | 558.2046 | 570.8429 |
| Ptprc | TLVSWPEPVSKPE | -392 | -404 | 1 | 0 | 13 | 706.3749 | 1715.408 | 841.4691 | NaN | NaN | 622.4224 |
| Ptprc | CAEYWPSMEEGTR | -751 | -763 | 0 | 0.000230707 | 13 | 338.4105 | 255.37047 | NaN | 474.6825 | NaN | 357.11005 |
| Ptprc | EVESLKPYK | -434 | -442 | 1 | 0.002199523 | 9 | 1919.7245 | NaN | NaN | 1742.4961 | NaN | 2136.1655 |
| Ptprc | NNLEECSQIK | -245 | -254 | 0 | 0.000298805 | 10 | 1133.6119 | 1206.7642 | NaN | NaN | NaN | 743.04755 |
| Ptprc | CPDYIIQK | -778 | -785 | 0 | 0 | 8 | 1075.1028 | 1110.8916 | NaN | 1489.5394 | NaN | NaN |

| | | | | | | | | | | | | |
|---|---|---|---|---|---|---|---|---|---|---|---|---|
| Ptprc | CVRPDGPLNK | -1248 | -1257 | 1 | 0.000230707 | 10 | 1186.5408 | NaN | NaN | NaN | NaN | 2405.7837 |
| Glud1 | SEAAADREDDPNFFK | -54 | -68 | 1 | 0 | 15 | 3228.6445 | 3472.497 | 3579.7698 | 3892.4045 | 3988.319 | 4009.2754 |
| Glud1 | SEAAADREDDPNFFK | -54 | -68 | 1 | 0 | 15 | 3228.6445 | 3472.497 | 3579.7698 | 3892.4045 | 3988.319 | 4009.2754 |
| Glud1 | NLNHVSYGR | -445 | -453 | 0 | 0 | 9 | 2630.5364 | 3021.6323 | 813.2411 | 3341.7761 | 2780.8662 | 6650.3125 |
| Glud1 | HGGTIPVVPTAEFQDR | -481 | -496 | 0 | 0 | 16 | 2184.5771 | 3163.436 | 3096.3472 | 2669.7283 | 3765.872 | 3585.762 |
| Glud1 | HGGTIPVVPTAEFQDR | -481 | -496 | 0 | 0 | 16 | 2184.5771 | 3163.436 | 3096.3472 | 2669.7283 | 3765.872 | 3585.762 |
| Glud1 | YNLGLDLR | -528 | -535 | 0 | 0 | 8 | 2174.718 | 1965.0636 | 1983.7031 | 1986.7528 | 1840.6122 | 2075.9333 |
| Glud1 | EADCDILIPAASEK | -373 | -386 | 0 | 0 | 14 | 1881.4741 | 2162.2131 | 2073.311 | 1901.6825 | 1741.2089 | 2023.559 |
| Glud1 | NYTDNELEK | -192 | -200 | 0 | 0 | 9 | 1444.533 | 310.3556 | 1034.9122 | 1469.4675 | 737.2725 | 1077.0671 |
| Glud1 | DDGSWEVIEGYR | -125 | -136 | 0 | 0 | 12 | 1156.1692 | 1173.8577 | 509.39337 | 1161.2054 | 1142.9418 | 1233.8347 |
| Glud1 | SDGSIWNPDGIDPK | -333 | -346 | 0 | 0 | 14 | 1106.4711 | 1107.4567 | 1127.1072 | 1332.4065 | 1309.2528 | 1201.9772 |
| Glud1 | GFIGPGIDVPAPDMSTGER | -213 | -231 | 0 | 0 | 19 | 785.39014 | 921.38165 | 901.6222 | 779.61383 | 1049.5502 | 1031.5239 |
| Glud1 | GVFHGIENFINEA | -275 | -287 | 0 | 0 | 13 | 744.04944 | 738.8413 | 96.22832 | 593.0263 | 912.4384 | 952.7337 |
| Glud1 | CVGVGESDGSIWNPDGIDPK | -327 | -346 | 0 | 0 | 20 | 722.5598 | 776.72076 | 797.93604 | 737.1185 | 1132.3778 | 1017.1501 |
| Glud1 | GANGPTTPEADKIFLER | -404 | -420 | 1 | 0.00025015 | 17 | 595.7211 | 389.33606 | 701.69336 | 582.58307 | 578.9271 | 468.89777 |
| Glud1 | KGFIGPGIDVPAPDMSTGER | -212 | -231 | 1 | 0 | 20 | 562.0577 | 650.34106 | 675.1585 | 553.6904 | 532.40247 | 908.0329 |
| Glud1 | KGFIGPGIDVPAPDMSTGER | -212 | -231 | 1 | 0 | 20 | 562.0577 | 650.34106 | 675.1585 | 553.6904 | 532.40247 | 908.0329 |
| Glud1 | VVDVPFGGAK | -174 | -183 | 0 | 0 | 10 | 1248.6545 | 513.61835 | 847.92017 | 571.94257 | NaN | NaN |
| Glud1 | PISQGGIHGR | -259 | -268 | 0 | 0.00122419 | 10 | 484.30615 | 434.42368 | 317.91763 | NaN | NaN | NaN |
| Glud1 | CAVVDVPFGGAK | -172 | -183 | 0 | 0.00201047 | 12 | 1463.0756 | NaN | NaN | NaN | NaN | NaN |
| Alb | ECCHGDLLECADDR | -268 | -281 | 0 | 0 | 14 | 25791.047 | 27577.234 | 26898.34 | 31397.03 | 31042.236 | 30583.928 |
| Alb | ECCHGDLLECADDR | -268 | -281 | 0 | 0 | 14 | 25791.047 | 27577.234 | 26898.34 | 31397.03 | 31042.236 | 30583.928 |
| Alb | RHPYFYAPELL | -169 | -179 | 1 | 0.00025015 | 11 | 25742.941 | 30579.545 | 310.59164 | 17740.865 | 18349.725 | 19551.414 |
| Alb | RHPYFYAPELL | -169 | -179 | 1 | 0.00025015 | 11 | 25742.941 | 30579.545 | 310.59164 | 17740.865 | 18349.725 | 19551.414 |
| Alb | ECCHGDLL | -268 | -275 | 0 | 0.00644558 | 8 | 13688.337 | 12655.484 | 11723.238 | 5867.499 | 5143.079 | 5198.329 |
| Alb | ECCHGDLLECA | -268 | -278 | 0 | 0.00025015 | 11 | 5516.766 | 5921.349 | 6209.1577 | 4263.528 | 3957.008 | 3965.2253 |
| Alb | ECCHGDLLECA | -268 | -278 | 0 | 0.00025015 | 11 | 5516.766 | 5921.349 | 6209.1577 | 4263.528 | 3957.008 | 3965.2253 |
| Alb | HPYFYAPELL | -170 | -179 | 0 | 0 | 10 | 5110.555 | 5015.2617 | 5138.9053 | 1082.134 | 5268.0156 | 4607.9463 |
| Alb | HPYFYAPELL | -170 | -179 | 0 | 0 | 10 | 5110.555 | 5015.2617 | 5138.9053 | 1082.134 | 5268.0156 | 4607.9463 |
| Alb | LGEYGFQNAILVR | -422 | -434 | 0 | 0 | 13 | 3763.5505 | 4025.2517 | 4463.4014 | 5107.474 | 6967.2217 | 6566.2114 |
| Alb | RHPYFYAPE | -169 | -177 | 1 | 0.000471409 | 9 | 3059.2322 | 3069.64 | 3182.3193 | 2898.9695 | 2912.7341 | 2520.8674 |
| Alb | RHPYFYAPELLY | -169 | -180 | 1 | 0 | 12 | 2846.4727 | 2845.8635 | 2552.7805 | 4360.339 | 4273.3525 | 2626.9285 |
| Alb | GFQNAILVR | -426 | -434 | 0 | 0.000230707 | 9 | 2459.9692 | 2638.4607 | 2637.9907 | 1551.4436 | 2086.7708 | 2019.1914 |
| Alb | ECCHGDLLEC | -268 | -277 | 0 | 0 | 10 | 1901.5691 | 2075.685 | 2048.8281 | 2179.2407 | 2313.4846 | 2122.6116 |
| Alb | CCHGDLLECADDR | -269 | -281 | 0 | 0 | 13 | 1600.1284 | 1699.307 | 1691.6693 | 2053.9521 | 2308.8342 | 2283.5852 |
| Alb | CCHGDLLECADDR | -269 | -281 | 0 | 0 | 13 | 1600.1284 | 1699.307 | 1691.6693 | 2053.9521 | 2308.8342 | 2283.5852 |
| Alb | HPYFYAPELLY | -170 | -180 | 0 | 0 | 11 | 1561.6721 | 1453.1826 | 1594.0624 | 1205.7043 | 1612.8263 | 1581.2191 |
| Alb | LLECADDR | -274 | -281 | 0 | 0 | 8 | 1013.08136 | 701.06445 | 804.7712 | 666.5543 | 1239.9598 | 1260.939 |
| Alb | CHGDLLECADDR | -270 | -281 | 0 | 0 | 12 | 763.0573 | 858.28015 | 803.9675 | 853.34753 | 870.1622 | 814.5868 |
| Alb | RHPYFYAPELLYYA | -169 | -182 | 1 | 0 | 14 | 626.63544 | 533.56067 | 732.702 | 848.0713 | 833.9126 | 890.25995 |
| Alb | RHPYFYAPELLYY | -169 | -181 | 1 | 0 | 13 | 528.63885 | 780.99945 | 859.9074 | 799.3425 | 946.87823 | 540.48804 |
| Alb | HGDLLECADDR | -271 | -281 | 0 | 0 | 11 | 1848.3206 | 1812.3788 | 1776.2722 | 1794.5775 | 1900.6077 | NaN |
| Alb | HPYFYAPELLYY | -170 | -181 | 0 | 0.001055699 | 12 | 635.8114 | 650.3901 | 594.2301 | 669.8844 | 764.87305 | NaN |
| Alb | GEYGFQNAILVR | -423 | -434 | 0 | 0 | 12 | 275.83936 | NaN | 273.09647 | 1380.248 | 1726.1423 | 1621.263 |
| Alb | HPYFYAPELLYYA | -170 | -182 | 0 | 0.001708008 | 13 | 507.22495 | 660.42957 | 627.8813 | 649.48474 | NaN | NaN |
| Alb | ECCHGDLLECAD | -268 | -279 | 0 | 0 | 12 | 401.77344 | 383.80453 | 409.3012 | NaN | 420.42825 | NaN |
| Alb | NECFLQHK | -123 | -130 | 0 | 0 | 8 | 1188.6361 | 1304.1788 | 1247.017 | NaN | NaN | NaN |
| Alb | QEPERNECF | -118 | -126 | 1 | 0.000550307 | 9 | 928.24817 | 357.02255 | 317.49362 | NaN | NaN | NaN |
| Alb | TPVSEHVTK | -491 | -499 | 0 | 0.00025015 | 9 | 915.69543 | NaN | NaN | 1837.153 | 1582.383 | NaN |
| Alb | LCAIPNLR | -98 | -105 | 0 | 0.00051306 | 8 | 864.39825 | 1222.0865 | NaN | 1287.1023 | NaN | NaN |
| Alb | SLHTLFGDK | -89 | -97 | 0 | 0.001743443 | 9 | 733.05176 | 610.97327 | NaN | 239.27022 | NaN | NaN |
| Alb | RPCFSALTVDETYVPK | -509 | -524 | 1 | 0.00025015 | 16 | 6599.775 | NaN | NaN | 7564.718 | NaN | NaN |
| Alb | KYEATLEKCCAE | -376 | -387 | 2 | 0.003745449 | 12 | 3169.3733 | 2822.9836 | NaN | NaN | NaN | NaN |
| Alb | ECCHGDLLE | -268 | -276 | 0 | 0.005884925 | 9 | 1013.4227 | 968.67633 | NaN | NaN | NaN | NaN |
| Rps27a | RECPSDECGAGVF | -119 | -131 | 1 | 0.000627943 | 13 | 1136.8715 | 1196.2179 | 1276.4658 | 1506.1721 | 1617.1766 | 1602.6808 |
| Rps27a | CCLTYCFNKPEDK | -144 | -156 | 1 | 0 | 13 | 633.2929 | 522.05865 | 645.9967 | 1142.2782 | 1195.8201 | 1178.5898 |
| Rps27a | CFNKPEDK | -149 | -156 | 1 | 0 | 8 | 516.3035 | 540.4187 | 598.8517 | 887.39813 | 885.7783 | 521.9573 |
| Rps27a | LTYCFNKPEDK | -146 | -156 | 1 | 0 | 11 | 505.8589 | 816.5242 | 937.3779 | 640.8762 | 497.75848 | 714.0302 |
| Rps27a | FNKPEDK | -150 | -156 | 1 | 0.00122419 | 7 | 445.8879 | 409.292 | 585.48914 | 517.2604 | 581.4513 | 397.1015 |
| Rps27a | ECPSDECGAGVF | -120 | -131 | 0 | 0 | 12 | 275.06857 | 330.34555 | 312.0443 | 440.70618 | 481.64175 | 425.51666 |
| Rps27a | YCFNKPEDK | -148 | -156 | 1 | 0 | 9 | 808.36896 | 875.64136 | 866.3128 | NaN | 688.77563 | 694.6563 |
| Rps27a | YCFNKPEDK | -148 | -156 | 1 | 0 | 9 | 808.36896 | 875.64136 | 866.3128 | NaN | 688.77563 | 694.6563 |
| Crip1 | GKDWHRPCLK | -21 | -30 | 2 | 0 | 10 | 2654.5774 | 2976.3232 | 3612.6672 | 1569.3706 | 2882.3674 | 3525.8928 |
| Crip1 | GKDWHRPCLK | -21 | -30 | 2 | 0 | 10 | 2654.5774 | 2976.3232 | 3612.6672 | 1569.3706 | 2882.3674 | 3525.8928 |
| Crip1 | DWHRPCLK | -23 | -30 | 1 | 0.000230707 | 8 | 328.48944 | 2899.2173 | 2848.5862 | 2635.521 | 2955.1882 | 3089.996 |
| Crip1 | DWHRPCLK | -23 | -30 | 1 | 0.000230707 | 8 | 328.48944 | 2899.2173 | 2848.5862 | 2635.521 | 2955.1882 | 3089.996 |
| Crip1 | CDKEVYFAE | -7 | -15 | 1 | 0.001463897 | 9 | 819.3805 | NaN | 718.55994 | 860.64996 | 690.47363 | 522.2679 |
| Kcnh1 | CDLHVIK | -667 | -673 | 0 | 0 | 7 | 1490.6973 | NaN | NaN | NaN | 727.0304 | 607.2628 |
| Psma6 | YGYEIPVDMLCK | -105 | -116 | 0 | 0 | 12 | 3026.222 | 2950.0288 | 3073.8083 | 1081.7212 | 1442.818 | 1028.5255 |
| Psma6 | YGYEIPVDMLCK | -105 | -116 | 0 | 0 | 12 | 3026.222 | 2950.0288 | 3073.8083 | 1081.7212 | 1442.818 | 1028.5255 |
| Psma6 | IGIDEEQGPQVYK | -141 | -153 | 0 | 0 | 13 | 2689.2207 | 2381.6162 | 2752.391 | 2585.7622 | 2614.622 | 2579.8696 |
| Psma6 | LLDSSTVTHLFK | -60 | -71 | 0 | 0 | 12 | 1340.8698 | 1605.4417 | 1570.8828 | 1479.787 | 1720.2084 | 1746.0994 |
| Psma6 | TVTHLFK | -65 | -71 | 0 | 0.000230707 | 7 | 950.80054 | 953.2014 | 973.6275 | 662.8833 | 635.6118 | 729.3443 |
| Psma6 | LLDSSTVTHL | -60 | -69 | 0 | 0 | 10 | 628.8964 | 408.04358 | 434.51218 | 912.6121 | 841.42 | 830.4586 |
| Psma6 | LLDSSTVTHLF | -60 | -70 | 0 | 0 | 11 | 475.1672 | 936.1429 | 965.50854 | 1226.149 | 1486.9482 | 121.88317 |
| Psma6 | KFDWTFEQ | -186 | -193 | 1 | 0.000591124 | 8 | 326.62787 | 324.0679 | 951.217 | 888.07446 | NaN | NaN |
| Psma6 | CDPAGYYCGFK | -154 | -164 | 0 | 0 | 11 | 388.0039 | 313.30615 | 321.06842 | NaN | NaN | NaN |
| Elovl1 | CDPIDFSNSPEALR | -93 | -106 | 0 | 0 | 14 | 786.7384 | 1049.262 | 1117.7388 | 701.1501 | 803.90485 | 1057.6968 |

| | | | | | | | | | | | | |
|---|---|---|---|---|---|---|---|---|---|---|---|---|
| Hk2 | GCEGEDVVTLLK | -627 | -638 | 0 | 0 | 12 | 2188.4458 | 2542.6035 | 2544.9504 | 1292.7444 | 621.7906 | 2054.2925 |
| Hk2 | DVSDIEDDKDGIQK | -338 | -351 | 1 | 0 | 14 | 1798.7914 | 254.66956 | 1979.0607 | 1830.8358 | 2072.4727 | 1990.7661 |
| Hk2 | LFTELNQNQVQK | -10 | -21 | 0 | 0 | 12 | 1179.8762 | 1442.9609 | 1552.0594 | 879.77423 | 1102.1125 | 1154.097 |
| Hk2 | LGLSPLQEDCVATHR | -359 | -373 | 0 | 0 | 15 | 1010.76715 | 965.2638 | 1086.5043 | 905.245 | 944.9967 | 949.1692 |
| Hk2 | LGLSPLQEDCVATHR | -359 | -373 | 0 | 0 | 15 | 1010.76715 | 965.2638 | 1086.5043 | 905.245 | 944.9967 | 949.1692 |
| Hk2 | SFPCQQNSLDQSILLK | -603 | -618 | 0 | 0 | 16 | 711.94794 | 718.3675 | 628.1278 | 559.49304 | 971.06555 | 711.8123 |
| Hk2 | HLGLESTCDDSIIVK | -806 | -820 | 0 | 0.00025015 | 15 | 535.9616 | 629.1668 | 614.3343 | 572.16876 | 725.1008 | 800.1597 |
| Hk2 | GDDGTLNDIR | -265 | -274 | 0 | 0 | 10 | 793.18005 | 2205.5579 | 2175.6685 | 2282.1523 | NaN | 427.60107 |
| Hk2 | LSHEQLLEVK | -479 | -488 | 0 | 0.00025015 | 10 | 4543.0366 | 4780.1997 | 4491.121 | NaN | 6128.174 | NaN |
| Hk2 | NVELVDGEEGR | -692 | -702 | 0 | 0.00078661 | 11 | 1055.4557 | 1426.1959 | 888.88824 | NaN | NaN | NaN |
| Hk2 | CDVSFLESEDGSGK | -886 | -899 | 0 | 0 | 14 | 189.10678 | NaN | 291.60474 | NaN | NaN | 655.29846 |
| Lsp1 | SQPTLPISTIDER | -209 | -221 | 0 | 0 | 13 | 3431.6907 | 4039.3574 | 4096.8096 | 3967.48 | 4853.7983 | 4810.5034 |
| Lsp1 | EQPGQQTLISLK | -65 | -76 | 0 | 0 | 12 | 1067.905 | 1070.6068 | 234.29828 | 1154.5157 | 460.26636 | 1065.6583 |
| Lsp1 | WREEDEEEAAR | -28 | -38 | 1 | 0 | 11 | 1035.4777 | 1132.9603 | 1276.7192 | 1163.9238 | 1662.4318 | 1609.2604 |
| Lsp1 | WREEDEEEAAR | -28 | -38 | 1 | 0 | 11 | 1035.4777 | 1132.9603 | 1276.7192 | 1163.9238 | 1662.4318 | 1609.2604 |
| Lsp1 | TPSCQDIVAGDMSK | -271 | -284 | 0 | 0 | 14 | 841.0235 | 733.2669 | 787.34265 | 815.92236 | 547.5184 | 666.86646 |
| Lsp1 | SSELDEDEGFGDWSQKPEPR | -77 | -96 | 1 | 0 | 20 | 746.6106 | 648.5692 | 823.05084 | 910.1064 | 726.3482 | 709.44946 |
| Lsp1 | KSQPTLPISTIDER | -208 | -221 | 1 | 0.00025015 | 14 | 688.6683 | 665.8143 | 696.9876 | 824.1903 | 725.12006 | 701.7237 |
| Lsp1 | QLQDQDKDKEDDGGHSL | -48 | -64 | 2 | 0 | 17 | 356.7917 | 1045.8916 | 1107.1919 | 788.78253 | 1153.0431 | 1039.5345 |
| Lsp1 | QLQDQDKDKEDDGGHSL | -48 | -64 | 2 | 0 | 17 | 356.7917 | 1045.8916 | 1107.1919 | 788.78253 | 1153.0431 | 1039.5345 |
| Lsp1 | GTEPSQSERPEEK | -108 | -120 | 1 | 0.00025015 | 13 | 328.59808 | 398.43884 | 407.4673 | 585.7045 | 428.23062 | 350.73254 |
| Lsp1 | VHLEESNLSYR | -131 | -141 | 0 | 0 | 11 | 254.21144 | 1293.592 | 1053.9929 | 759.8349 | 710.7288 | 773.88776 |
| Lsp1 | LQQYTQATESSGR | -222 | -234 | 0 | 0 | 13 | 817.2565 | NaN | 1025.0093 | 685.29974 | 229.36826 | 823.06793 |
| Lsp1 | EPDPEDAVGGSGEAEEHLIR | -142 | -161 | 0 | 0 | 20 | 734.05756 | NaN | 908.37146 | 668.54803 | 133.39514 | 881.89636 |
| Lsp1 | CEEQEELHAE | -10 | -19 | 0 | 0.00111006 | 10 | 392.05673 | 348.9358 | 524.4961 | NaN | NaN | 100.06917 |
| Rhoa | QEPVKPEEGR | -136 | -145 | 1 | 0 | 10 | 1524.7498 | 1243.2145 | 1154.3245 | 347.48672 | 1471.0483 | 1338.5125 |
| Rhoa | QEPVKPEEGR | -136 | -145 | 1 | 0 | 10 | 1524.7498 | 1243.2145 | 1154.3245 | 347.48672 | 1471.0483 | 1338.5125 |
| Rhoa | MKQEPVKPEEGR | -134 | -145 | 2 | 0 | 12 | 424.2843 | 483.9399 | NaN | 629.0545 | 626.83307 | 679.7991 |
| Dock2 | VSDNLRPF | -1590 | -1597 | 1 | 0.00025015 | 8 | 3662.8135 | 3139.4343 | 2941.1772 | 1412.6908 | 2019.3569 | 1874.3848 |
| Dock2 | TVQPVLDEHPR | -1409 | -1419 | 0 | 0.00025015 | 11 | 2139.6904 | 2461.8423 | 2587.0925 | 1696.3389 | 2279.9194 | 2081.1497 |
| Dock2 | DILLPVITK | -855 | -863 | 0 | 0 | 9 | 1723.8405 | 1796.1924 | 1952.0636 | 1353.4452 | 1920.9694 | 1972.9438 |
| Dock2 | GVPGLDEANTSPR | -1768 | -1780 | 0 | 0 | 13 | 1546 | 1069.04 | 1726.584 | 2020.1938 | 1483.1646 | 1814.2255 |
| Dock2 | VSDNLRPFHDR | -1590 | -1600 | 1 | 0.000627943 | 11 | 1474.0851 | 1396.9769 | 1565.9496 | 1095.8522 | 1154.0411 | 1453.3135 |
| Dock2 | NKPVPDQIINF | -1422 | -1432 | 1 | 0 | 11 | 1394.4817 | 1181.5894 | 1479.804 | 1575.2534 | 1535.2319 | 1549.581 |
| Dock2 | NKPVPDQIINFYK | -1422 | -1434 | 1 | 0 | 13 | 895.0495 | 1127.0481 | 866.1372 | 465.3924 | 1168.0533 | 1224.0629 |
| Dock2 | LLSQFPNAEK | -1377 | -1386 | 0 | 0 | 10 | 807.76935 | 513.52136 | 828.64557 | 681.4858 | 599.7164 | 597.20184 |
| Dock2 | NIENIIPAEIPLAQ | -76 | -89 | 0 | 0 | 14 | 743.1331 | 880.19855 | 896.73035 | 874.78595 | 1318.762 | 1179.5415 |
| Dock2 | QTVISENYLVR | -250 | -260 | 0 | 0 | 11 | 695.6727 | 835.9409 | 156.21515 | 902.6279 | 1181.982 | 1389.1864 |
| Dock2 | KDYPHLVDR | -395 | -403 | 1 | 0 | 9 | 455.763 | 959.9515 | 110.68963 | 484.50763 | 944.4976 | 612.67773 |
| Dock2 | DEDGNILDPDKTSVISL | -167 | -183 | 1 | 0.00025015 | 17 | 419.9722 | 439.4888 | 369.03552 | 379.66833 | 684.5993 | 209.51585 |
| Dock2 | MLVGDIIQIR | -385 | -394 | 0 | 0 | 10 | 365.4042 | 100.35943 | 475.9013 | 619.24646 | 729.02 | 590.24286 |
| Dock2 | SAENDFLHSLLGK | -354 | -366 | 0 | 0.000471409 | 13 | 365.20804 | 325.03757 | 643.5553 | 545.9065 | 753.27185 | 660.09186 |
| Dock2 | DYPHLVDR | -396 | -403 | 0 | 0.00025015 | 8 | 3255.8137 | 3326.3306 | 240.05685 | 2974.9282 | 3080.988 | NaN |
| Dock2 | HDVETVFDAK | -804 | -813 | 0 | 0 | 10 | 2057.0103 | 1479.4209 | 1750.8097 | 1959.437 | 1653.0608 | NaN |
| Dock2 | IQETCGDWYR | -37 | -46 | 0 | 0 | 10 | 1241.1772 | 616.71576 | 516.0706 | 708.1461 | NaN | 429.74402 |
| Dock2 | VEEEPISPGSTLPEVK | -1677 | -1692 | 0 | 0 | 16 | 1061.1611 | 1438.4916 | 1120.6458 | 1416.3132 | 1686.6774 | NaN |
| Dock2 | VVFTDLGNK | -278 | -286 | 0 | 0 | 9 | 765.5597 | 700.98584 | 399.9708 | 300.1808 | 783.4915 | NaN |
| Dock2 | LWEWGSIWK | -95 | -103 | 0 | 0.00051306 | 9 | 229.844 | NaN | 346.033 | 236.45331 | 184.57242 | 114.8583 |
| Dock2 | GEQCEPILR | -727 | -735 | 0 | 0 | 9 | 1424.4172 | 1102.1147 | NaN | NaN | 441.97775 | 2410.9512 |
| Dock2 | QIPFLGAGIK | -1575 | -1584 | 0 | 0 | 10 | 668.46783 | 693.6747 | 794.2351 | NaN | 1176.4275 | NaN |
| Dock2 | TLQIGDVVR | -28 | -36 | 0 | 0 | 9 | 492.86102 | 890.8043 | 969.9741 | 868.71625 | NaN | NaN |
| Dock2 | TLIPEAELR | -1095 | -1103 | 0 | 0.003360691 | 9 | 977.0309 | NaN | 929.8855 | 685.81976 | NaN | NaN |
| Dock2 | CFTVQPVLDEHPR | -1407 | -1419 | 0 | 0.001055699 | 13 | 584.3769 | 803.4145 | 745.0922 | NaN | NaN | NaN |
| Dock2 | EDGTTLHDGYHEL | -537 | -549 | 0 | 0.00111006 | 13 | 91.596466 | 466.98016 | 242.07341 | NaN | NaN | NaN |
| Dock2 | DLHLDCENYTEAA | -1228 | -1240 | 0 | 0.009631128 | 13 | 447.4492 | NaN | 491.23383 | NaN | NaN | NaN |
| Dock2 | VLPNAICVGAGDK | -459 | -471 | 0 | 0 | 13 | 283.38434 | NaN | 926.70703 | NaN | NaN | NaN |
| Dock2 | YIPSVLHDVETVFDAK | -798 | -813 | 0 | 0.005789881 | 16 | 682.4341 | NaN | NaN | NaN | NaN | NaN |
| Dock2 | DEDGNILDPDK | -167 | -177 | 0 | 0.005301614 | 11 | 428.18802 | NaN | NaN | NaN | NaN | NaN |
| Gbp5 | DTEGLGDVEKDDK | -97 | -109 | 1 | 0 | 13 | 515.0457 | 518.31836 | 535.08386 | 502.21613 | 702.8347 | 590.92883 |
| Gbp5 | IKAEQEAQLR | -556 | -565 | 1 | 0 | 10 | 1380.4583 | 903.2399 | 496.50067 | NaN | 1316.3302 | 1818.2642 |
| Gbp5 | HSFKDEGEFFQK | -370 | -381 | 1 | 0 | 12 | 653.41614 | NaN | 688.95624 | 723.0924 | 323.11615 | NaN |
| Gbp5 | CFVFDAPALGSK | -234 | -245 | 0 | 0.00025015 | 12 | 607.2962 | 580.419 | 793.3255 | 588.1655 | NaN | NaN |
| Gbp5 | TILQTDQVLTDK | -473 | -484 | 0 | 0 | 12 | 542.8668 | 633.84644 | NaN | 548.45874 | NaN | NaN |
| Gbp5 | VAQEQQWILK | -537 | -546 | 0 | 0.002265552 | 10 | 350.7739 | NaN | 455.69464 | NaN | NaN | NaN |
| Cnbp | SLPDICYR | -49 | -56 | 0 | 0.00111006 | 8 | 3318.2341 | 3015.3696 | 2954.2354 | 3032.5276 | 1952.1277 | 1897.2257 |
| Cnbp | TSEVNCYR | -154 | -161 | 0 | 0 | 8 | 1300.6212 | 791.88074 | 854.1262 | 76.72888 | 1366.2046 | 368.00125 |
| Cnbp | CYSCGEFGHIQ | -120 | -130 | 0 | 0 | 11 | 660.9064 | 623.99976 | 670.1063 | 705.31146 | 602.8725 | 687.44183 |
| Cnbp | CYSCGEFGHIQK | -120 | -131 | 0 | 0 | 12 | 562.28375 | 1786.7529 | 2283.2908 | 1707.0475 | 1725.9783 | 1679.0273 |
| Cnbp | EQCCYNCGKPGHLAR | -96 | -110 | 1 | 0 | 15 | 161.89133 | 472.0722 | 483.4122 | 307.96262 | 793.5949 | 810.27277 |
| Cnbp | EQCCYNCGKPGHLAR | -96 | -110 | 1 | 0 | 15 | 161.89133 | 472.0722 | 483.4122 | 307.96262 | 793.5949 | 810.27277 |
| Cnbp | EQCCYNCGKPGHLAR | -96 | -110 | 1 | 0 | 15 | 161.89133 | 472.0722 | 483.4122 | 307.96262 | 793.5949 | 810.27277 |
| Cnbp | DCDLQEDEACYNCGR | -66 | -80 | 0 | 0 | 15 | 508.99802 | 546.3095 | 502.93207 | 763.12726 | 844.8604 | 837.0238 |
| Cnbp | CGETGHVAIN | -141 | -150 | 0 | 0.000431655 | 10 | 509.91476 | 565.08154 | NaN | NaN | 634.0607 | 825.7984 |
| Cnbp | GEFGHIQK | -124 | -131 | 0 | 0 | 8 | 2207.7986 | 2271.5554 | NaN | NaN | NaN | NaN |
| Cnbp | EQCCYNCGKPGHLA | -96 | -109 | 1 | 0.000712251 | 14 | 194.45695 | NaN | 195.16632 | NaN | NaN | NaN |
| Aldh9a1 | FTGSVPTGVK | -230 | -239 | 0 | 0 | 10 | 3498.5227 | 1482.5851 | 3378.5552 | 1716.8916 | 2001.0377 | 2544.1763 |
| Aldh9a1 | TFGTEAEVLER | -401 | -411 | 0 | 0 | 11 | 2924.2456 | 2137.8826 | 1726.5209 | 1965.013 | 1994.8188 | 1702.9324 |
| Aldh9a1 | GVKPITLELGGK | -247 | -258 | 1 | 0.00025015 | 12 | 1008.02893 | 1039.984 | 1131.665 | 1358.7279 | 1115.0952 | 1523.4612 |

| | | | | | | | | | | | | |
|---|---|---|---|---|---|---|---|---|---|---|---|---|
| Aldh9a1 | STGTFVVSQPLNYR | -2 | -15 | 0 | 0 | 14 | 969.9243 | 1144.5605 | 656.97925 | 683.6841 | 1655.046 | 1719.5374 |
| Aldh9a1 | NYNVSPVELPFGGYK | -447 | -461 | 0 | 0 | 15 | 610.1698 | 662.46643 | 687.2808 | 606.4495 | 845.16455 | 826.60156 |
| Aldh9a1 | INAPHLER | -331 | -338 | 0 | 0 | 8 | 717.9648 | NaN | 525.69745 | 1079.9495 | 1379.9257 | 697.03723 |
| Aldh9a1 | VVSQPLNYR | -7 | -15 | 0 | 0.00149795 | 9 | 1525.0269 | 671.913 | NaN | NaN | NaN | NaN |
| Aldh9a1 | EVNLAVENAK | -50 | -59 | 0 | 0 | 10 | 338.60568 | 496.58923 | NaN | NaN | NaN | NaN |
| Aldh9a1 | CGGEVYVPEDPK | -355 | -366 | 0 | 0.001708008 | 12 | 710.99396 | NaN | NaN | NaN | NaN | NaN |
| Rps15a | WQNNLLPSR | -89 | -97 | 0 | 0 | 9 | 9430.669 | 4877.185 | 10406.899 | 10147.459 | 12094.23 | 11951.712 |
| Rps15a | HGYIGEFEIIDDHR | -44 | -57 | 0 | 0 | 14 | 6240.4634 | 8062.9814 | 8427.679 | 7484.2783 | 10103.707 | 9459.726 |
| Rps15a | HGYIGEFEIIDDHR | -44 | -57 | 0 | 0 | 14 | 6240.4634 | 8062.9814 | 8427.679 | 7484.2783 | 10103.707 | 9459.726 |
| Rps15a | EIIDDHR | -51 | -57 | 0 | 0.000394361 | 7 | 674.4936 | 1309.3549 | 1605.2917 | 1356.5438 | 1221.9281 | 1415.0128 |
| Rps15a | EFEIIDDHR | -49 | -57 | 0 | 0 | 9 | 587.7333 | 1829.1594 | 2004.0499 | 321.61108 | 698.8034 | 1234.5702 |
| Rps15a | IGEFEIIDDHR | -47 | -57 | 0 | 0 | 11 | 388.77655 | 653.9208 | NaN | 724.1466 | 1412.6953 | 1241.6073 |
| Rps15a | IGEFEIIDDHR | -47 | -57 | 0 | 0 | 11 | 388.77655 | 653.9208 | NaN | 724.1466 | 1412.6953 | 1241.6073 |
| Rps15a | QVLIRPC | -24 | -30 | 1 | 0.000627943 | 7 | 537.5462 | 189.74823 | NaN | 87.10643 | NaN | NaN |
| Rps15a | CGVISPR | -72 | -78 | 0 | 0 | 7 | 386.51706 | NaN | NaN | NaN | NaN | NaN |
| Ampd2 | HLEEIVHVE | -387 | -395 | 0 | 0.000826806 | 9 | 1290.5027 | 501.95755 | 1259.4205 | 109.649254 | 394.90903 | 733.2892 |
| Ampd2 | YNPIGESVLR | -440 | -449 | 0 | 0 | 10 | 716.5148 | 1600.417 | 1361.4124 | 703.57336 | 1450.9707 | 1330.0303 |
| Ampd2 | LEPDILLR | -118 | -125 | 0 | 0 | 8 | 708.7736 | 150.66147 | 202.88414 | 673.1313 | 714.0607 | NaN |
| Ampd2 | STDDPLQFHFTK | -708 | -719 | 0 | 0 | 12 | 479.84717 | 592.4796 | 583.17017 | NaN | 584.94476 | NaN |
| Ampd2 | CGVPFTDLLDAAK | -176 | -188 | 0 | 0.00078661 | 13 | 390.90936 | NaN | NaN | 640.64624 | 836.44257 | NaN |
| Eif4a3 | CHACIGGTNVGEDIR | -137 | -151 | 0 | 0 | 15 | 3800.476 | 4021.248 | 3859.2588 | 4153.9365 | 4501.685 | 4546.4067 |
| Eif4a3 | DYGQHVVAGTPGR | -154 | -166 | 0 | 0 | 13 | 2433.3345 | 2590.8147 | 2572.4636 | 1880.5093 | 2814.2368 | 2872.3286 |
| Eif4a3 | IGGTNVGEDIR | -141 | -151 | 0 | 0 | 11 | 2403.5723 | 1203.0637 | 1578.3721 | 1889.4126 | 1378.6456 | 1332.7903 |
| Eif4a3 | LIINYDLPNNR | -348 | -358 | 0 | 0 | 11 | 1440.482 | 1345.9099 | 1753.6406 | 2076.495 | 2295.6738 | 2175.5051 |
| Eif4a3 | RDELTLEGIK | -243 | -252 | 1 | 0.00025015 | 10 | 1208.4208 | 1377.8207 | 1347.3668 | 878.2451 | 1534.0665 | 1389.138 |
| Eif4a3 | VLDEADEMLNK | -185 | -195 | 0 | 0 | 11 | 582.2204 | 550.3827 | 554.42206 | 844.98206 | 626.86523 | 655.3441 |
| Eif4a3 | ETQALILAPTR | -106 | -116 | 0 | 0 | 11 | 4556.7954 | 5223.014 | 4807.8867 | NaN | 726.2364 | 801.2573 |
| Eif4a3 | YLPPATQVVLIS | -207 | -218 | 0 | 0.000431655 | 12 | 908.30945 | 946.5639 | NaN | 1351.0225 | 179.21597 | 1463.7891 |
| Eif4a3 | ISTDVWAR | -332 | -339 | 0 | 0.00078661 | 8 | 788.9945 | 564.31055 | 632.80695 | 913.3067 | 365.11472 | NaN |
| Eif4a3 | DELTLEGIK | -244 | -252 | 0 | 0 | 9 | 373.5377 | 130.63623 | NaN | 1009.56445 | 1116.3987 | NaN |
| Eif4a3 | EQIYDVYR | -199 | -206 | 0 | 0.001868731 | 8 | 776.84705 | NaN | NaN | NaN | NaN | NaN |
| Tnfrsf18 | CICVTPEYHCGDPQCK | -60 | -75 | 0 | 0 | 16 | 393.88806 | 369.7401 | 400.41656 | 443.7138 | 481.55707 | 520.35065 |
| Tnfrsf18 | HYPCQPGQR | -79 | -87 | 0 | 0.00025015 | 9 | 928.8725 | NaN | NaN | 927.8079 | 1053.7081 | 1132.5055 |
| Arhgap15 | EHSTVPWFVK | -296 | -305 | 0 | 0 | 10 | 812.7725 | 855.209 | 966.3746 | 582.69867 | 811.8198 | 270.6905 |
| Arhgap15 | CIEAVEK | -307 | -313 | 0 | 0.004628201 | 7 | 659.79004 | NaN | NaN | NaN | 2966.3684 | NaN |
| Bsg | KPDQTLDEDDPGAAPLK | -354 | -370 | 1 | 0.003900536 | 17 | 1755.7073 | 1990.3145 | 2182.1929 | 2325.6543 | 2862.1443 | 2653.3828 |
| Bsg | QEDTLPDLHTK | -179 | -189 | 0 | 0 | 11 | 1578.6686 | 1666.699 | 1641.4678 | 1813.4814 | 1690.2712 | 1759.8397 |
| Bsg | VLQEDTLPDLHTK | -177 | -189 | 0 | 0 | 13 | 1385.1316 | 1573.7131 | 1626.9303 | 387.1927 | 1203.0431 | 1245.4487 |
| Bsg | SEINVEGPPR | -213 | -222 | 0 | 0 | 10 | 1315.0658 | 643.2903 | 1044.5914 | 590.3421 | 445.57462 | 179.4068 |
| Bsg | SGEYSCIFLPEPVGR | -198 | -212 | 0 | 0 | 15 | 927.1776 | 1104.7277 | 1104.0217 | 1222.6868 | 1796.5743 | 1724.3547 |
| Bsg | RKPDQTLDEDDPGAAPLK | -353 | -370 | 2 | 0.00025015 | 18 | 176.25108 | 470.63373 | 767.4914 | 419.13034 | 569.1868 | 560.3288 |
| Bsg | CIFLPEPVGR | -203 | -212 | 0 | 0 | 10 | 1797.9535 | 2110.467 | 835.52075 | NaN | NaN | NaN |
| Bsg | GVDIVGHR | -163 | -170 | 0 | 0.000431655 | 8 | 794.4749 | 195.55011 | NaN | NaN | NaN | NaN |
| Bsg | YVVVSTPEK | -278 | -286 | 0 | 0.000431655 | 9 | 1312.0194 | NaN | NaN | NaN | NaN | NaN |
| Bsg | YIVDADDR | -190 | -197 | 0 | 0.00383302 | 8 | 556.39185 | NaN | NaN | NaN | NaN | NaN |
| Timm13 | CIGKPGGSLDNSEQK | -50 | -64 | 1 | 0 | 15 | 627.29645 | 745.3278 | 780.1093 | NaN | 898.15375 | 884.48615 |
| Timm13 | PGGSLDNSEQK | -54 | -64 | 0 | 0.007546675 | 11 | 724.39166 | NaN | NaN | NaN | NaN | NaN |
| Snrpb;Snrpn | CILQDGR | (19);(19) | (25);(25) | 0 | 0 | 7 | 1614.22 | 1613.5522 | 1110.6946 | 1431.0214 | 804.068 | 190.25203 |
| Snrpb;Snrpn | IFIGTFK | (26);(26) | (32);(32) | 0 | 0.004595494 | 7 | 1406.7316 | NaN | 984.3447 | 490.7279 | 479.56458 | 454.71146 |
| Gzmb | DQQPEAICGGFLIR | -42 | -55 | 0 | 0 | 14 | 8579.982 | 10321.55 | 10388.621 | 10023.178 | 13997.229 | 13591.86 |
| Gzmb | CIPHPDYNPK | -94 | -103 | 0 | 0.000230707 | 10 | 6717.0747 | 6464.4404 | 6405.1206 | 7753.541 | 7840.4946 | 7682.248 |
| Gzmb | CIPHPDYNPK | -94 | -103 | 0 | 0.000230707 | 10 | 6717.0747 | 6464.4404 | 6405.1206 | 7753.541 | 7840.4946 | 7682.248 |
| Gzmb | NVNVKPGDVCYVA | -133 | -145 | 1 | 0 | 13 | 6372.158 | 6615.8745 | 7012.641 | 493.47916 | 5500.391 | 380.82758 |
| Gzmb | NVNVKPGDVCYVA | -133 | -145 | 1 | 0 | 13 | 6372.158 | 6615.8745 | 7012.641 | 493.47916 | 5500.391 | 380.82758 |
| Gzmb | DRECESYFK | -170 | -178 | 1 | 0.001055699 | 9 | 3300.9646 | 3599.6465 | 3536.5625 | 2921.424 | 227.52515 | 3409.379 |
| Gzmb | RNVNVKPGDVCYVA | -132 | -145 | 2 | 0 | 14 | 3105.015 | 3743.3079 | 3978.7585 | 2629.3564 | 3135.0828 | 3260.149 |
| Gzmb | RNVNVKPGDVCYVA | -132 | -145 | 2 | 0 | 14 | 3105.015 | 3743.3079 | 3978.7585 | 2629.3564 | 3135.0828 | 3260.149 |
| Gzmb | TQQVIPMVK | -85 | -93 | 0 | 0.000230707 | 9 | 2781.6448 | 2882.9521 | 3086.2288 | 3129.1296 | 3248.2837 | 3445.5461 |
| Gzmb | DQQPEAICGGF | -42 | -52 | 0 | 0.00051306 | 11 | 2142.5234 | 2755.6812 | 117.018585 | 1899.1058 | 2183.4983 | 2524.9429 |
| Gzmb | RNVNVKPGDVCYVAG | -132 | -146 | 2 | 0.00025015 | 15 | 1484.4121 | 1534.6626 | 1173.7163 | 1645.5614 | 1802.9801 | 1044.9563 |
| Gzmb | NVNVKPGDVCYVAG | -133 | -146 | 1 | 0 | 14 | 1229.5422 | 367.0917 | 1122.4674 | 1563.8324 | 659.8801 | 657.9798 |
| Gzmb | RNVNVKPGDVC | -132 | -142 | 2 | 0.000471409 | 11 | 1210.4551 | 1234.1095 | 1293.9252 | 646.02 | 835.63837 | 842.36523 |
| Gzmb | SIKDQQPEAICGGFLIR | -39 | -55 | 1 | 0 | 17 | 1156.5144 | 1222.5997 | 1376.0256 | 1388.2512 | 1738.4781 | 1725.543 |
| Gzmb | SIKDQQPEAICGGFLIR | -39 | -55 | 1 | 0 | 17 | 1156.5144 | 1222.5997 | 1376.0256 | 1388.2512 | 1738.4781 | 1725.543 |
| Gzmb | DQQPEAICGGFL | -42 | -53 | 0 | 0 | 12 | 782.92755 | 796.0123 | 804.88336 | 814.4911 | 798.552 | 805.92114 |
| Gzmb | VSSFLSWIK | -233 | -241 | 0 | 0 | 9 | 779.68463 | 900.2396 | 959.7765 | 75.29475 | 725.1789 | 838.8748 |
| Gzmb | GEIIGGHEVKPHSR | -19 | -32 | 1 | 0 | 14 | 725.1378 | 1412.441 | 940.61584 | 904.22504 | 1138.2012 | 1635.7065 |
| Gzmb | GEIIGGHEVKPHSR | -19 | -32 | 1 | 0 | 14 | 725.1378 | 1412.441 | 940.61584 | 904.22504 | 1138.2012 | 1635.7065 |
| Gzmb | YSNTLQEVELTVQ | -156 | -168 | 0 | 0.000230707 | 13 | 706.5911 | 799.5332 | 608.26215 | 783.81805 | 823.5167 | 1591.5156 |
| Gzmb | RNVNVKPGDVCYVAGW | -132 | -147 | 2 | 0 | 16 | 626.3971 | 358.94437 | 822.6275 | 941.84155 | 987.7046 | 716.4474 |
| Gzmb | RNVNVKPGDVCYVAGW | -132 | -147 | 2 | 0 | 16 | 626.3971 | 358.94437 | 822.6275 | 941.84155 | 987.7046 | 716.4474 |
| Gzmb | TNQICAGDPK | -184 | -193 | 0 | 0 | 10 | 564.4449 | 563.285 | 720.3633 | 3955.973 | 3430.4004 | 3800.7336 |
| Gzmb | TNQICAGDPK | -184 | -193 | 0 | 0 | 10 | 564.4449 | 563.285 | 720.3633 | 3955.973 | 3430.4004 | 3800.7336 |
| Gzmb | NVNVKPGDVCYVAGW | -133 | -147 | 1 | 0.00025015 | 15 | 1102.6772 | 1374.1294 | NaN | 2920.346 | 3753.5781 | 702.0709 |
| Gzmb | NQICAGDPK | -185 | -193 | 0 | 0.00025015 | 9 | 1041.5449 | 377.72174 | 1137.981 | 1675.6534 | NaN | 1316.7174 |
| Gzmb | FRGDSGGPLVCK | -199 | -210 | 1 | 0 | 12 | 850.3177 | 890.7846 | 503.86237 | NaN | 1420.3672 | 1054.3304 |
| Gzmb | YSNTLQEVELTVQK | -156 | -169 | 0 | 0 | 14 | 713.11676 | 483.14017 | 855.3283 | 593.5002 | NaN | NaN |
| Gzmb | ECESYFK | -172 | -178 | 0 | 0.002068213 | 7 | 655.16125 | 523.45306 | NaN | 513.2169 | 341.04153 | NaN |

| | | | | | | | | | | | | |
|---|---|---|---|---|---|---|---|---|---|---|---|---|
| Gzmb | NVNVKPGDVC | -133 | -142 | 1 | 0 | 10 | 1339.1982 | 3606.813 | 1877.592 | NaN | NaN | NaN |
| Gzmb | YSNTLQEVELT | -156 | -166 | 0 | 0.000230707 | 11 | 204.74995 | 240.65924 | NaN | NaN | 277.75076 | NaN |
| Gzmb | RNVNVKPGDVCY | -132 | -143 | 2 | 0.001776473 | 12 | 1218.1906 | NaN | NaN | NaN | 1831.8269 | NaN |
| Ube2i | KDHPFGF | -18 | -24 | 1 | 0.002466863 | 7 | 3293.9426 | 154.72745 | 2486.642 | 2121.5117 | 2031.3047 | 1954.7158 |
| Ube2i | EEDKDWRPAITIK | -98 | -110 | 2 | 0 | 13 | 1724.4081 | 1829.9257 | 1765.5785 | 1352.8379 | 1403.4711 | 1505.7646 |
| Ube2i | CKFEPPLFHPN | -75 | -85 | 1 | 0 | 11 | 1628.278 | 1584.5675 | 1752.0752 | 1536.1366 | 2472.538 | 2312.9287 |
| Ube2i | SILEEDKDWR | -95 | -104 | 1 | 0 | 10 | 1471.3431 | 479.13266 | 959.1182 | 2729.5698 | 1935.2671 | 2756.9614 |
| Ube2i | KGTPWEGGLF | -49 | -58 | 1 | 0.001055699 | 10 | 1270.2462 | 1386.3618 | 1451.5148 | 1546.8156 | 2105.5684 | 2135.028 |
| Ube2i | DHPFGFVAVPTK | -19 | -30 | 0 | 0 | 12 | 783.07135 | 209.99261 | 789.74756 | 967.45166 | 920.363 | 1202.3474 |
| Ube2i | FKDDYPSSPPK | -64 | -74 | 1 | 0 | 11 | 694.5175 | 2347.0725 | 2569.2966 | 2459.7366 | 2913.4353 | 2799.7664 |
| Ube2i | NWECAIPGK | -40 | -48 | 0 | 0 | 9 | 625.0819 | 1485.5679 | 296.25723 | 1613.0063 | 1476.712 | 1637.8033 |
| Ube2i | DHPFGFVAVPT | -19 | -29 | 0 | 0.00025015 | 11 | 538.6181 | 2096.9866 | 1704.138 | 2589.4514 | 326.76715 | 2793.313 |
| Ube2i | NWECAIPGKK | -40 | -49 | 1 | 0 | 10 | 333.32086 | 657.84644 | 1107.2793 | 1194.4792 | 1168.3228 | 1165.6903 |
| Ube2i | FEPPLFHPN | -77 | -85 | 0 | 0.00078661 | 9 | 263.67487 | 422.39325 | 1296.5496 | 1281.6925 | 1149.4467 | 1070.1208 |
| Ube2i | GTPWEGGLFK | -50 | -59 | 0 | 0 | 10 | 2213.274 | 2612.159 | 2670.1836 | 1339.8082 | NaN | 1764.39 |
| Ube2i | KGTPWEGGLFK | -49 | -59 | 1 | 0 | 11 | 1127.6442 | 1320.9424 | 1393.1698 | NaN | NaN | NaN |
| Ube2i | GTPWEGGLF | -50 | -58 | 0 | 0.000431655 | 9 | 987.329 | NaN | NaN | 1837.7375 | 2451.293 | NaN |
| Ube2i | KDHPFGFVAVPTK | -18 | -30 | 1 | 0 | 13 | 917.9347 | 749.5846 | 848.9823 | NaN | NaN | NaN |
| Tpt1 | DLISHDELFSDIYK | -6 | -19 | 0 | 0 | 14 | 4366.7065 | 4644.5737 | 4514.0527 | 3557.981 | 4615.666 | 4689.9976 |
| Tpt1 | DLISHDELFSDIYK | -6 | -19 | 0 | 0 | 14 | 4366.7065 | 4644.5737 | 4514.0527 | 3557.981 | 4615.666 | 4689.9976 |
| Tpt1 | ISHDELFSDIYK | -8 | -19 | 0 | 0 | 12 | 3775.1035 | 5912.5293 | 5473.9917 | 8692.976 | 4228.8184 | 3616.2708 |
| Tpt1 | FKDGLEMEK | -163 | -171 | 1 | 0 | 9 | 2606.7585 | 2325.1233 | 2686.148 | 2699.4614 | 2985.4841 | 2923.8748 |
| Tpt1 | GKLEEQKPER | -101 | -110 | 2 | 0 | 10 | 1825.4672 | 1944.0059 | 2306.777 | 813.09534 | 986.4657 | 1592.8998 |
| Tpt1 | GKLEEQKPER | -101 | -110 | 2 | 0 | 10 | 1825.4672 | 1944.0059 | 2306.777 | 813.09534 | 986.4657 | 1592.8998 |
| Tpt1 | RDLISHDELFSDIYK | -5 | -19 | 1 | 0 | 15 | 1417.0017 | 1325.0621 | 1433.8794 | 2652.2234 | 4012.2793 | 4372.233 |
| Tpt1 | RDLISHDELFSDIYK | -5 | -19 | 1 | 0 | 15 | 1417.0017 | 1325.0621 | 1433.8794 | 2652.2234 | 4012.2793 | 4372.233 |
| Tpt1 | HHLQETSFTK | -76 | -85 | 0 | 0 | 10 | 1014.2458 | 1047.4291 | 1711.3551 | 1337.2545 | 1172.5785 | 729.7252 |
| Tpt1 | HHLQETSFTK | -76 | -85 | 0 | 0 | 10 | 1014.2458 | 1047.4291 | 1711.3551 | 1337.2545 | 1172.5785 | 729.7252 |
| Tpt1 | HLQETSFTK | -77 | -85 | 0 | 0 | 9 | 873.96405 | 903.90753 | 390.17984 | 819.6304 | 652.3074 | 1052.8713 |
| Tpt1 | TEGAIDDSLIGGN | -39 | -51 | 0 | 0 | 13 | 817.73315 | 276.12674 | 331.69785 | 491.00613 | 499.8471 | 440.43463 |
| Tpt1 | LEEQKPER | -103 | -110 | 1 | 0.001023236 | 8 | 268.853 | 342.12552 | 3316.5068 | 94.67852 | 4029.061 | 4181.5747 |
| Tpt1 | TEGAIDDSLIGGNAS | -39 | -53 | 0 | 0.000230707 | 15 | 66.03177 | 1343.4691 | 1266.2947 | 159.28142 | 157.40436 | 992.78925 |
| Tpt1 | TEGAIDDSLIGGNA | -39 | -52 | 0 | 0.00078661 | 14 | 5214.812 | 6814.415 | 11191.964 | 8800.111 | NaN | 15429.414 |
| Tpt1 | CLEVEGK | -28 | -34 | 0 | 0 | 7 | 1447.3578 | 1345.1544 | NaN | 843.1702 | NaN | NaN |
| Tpt1 | EIADGLCLEVEGK | -22 | -34 | 0 | 0 | 13 | 373.1253 | 334.8177 | NaN | NaN | NaN | NaN |
| Lgals1 | DSNNLCLHFNPR | -38 | -49 | 0 | 0 | 12 | 4910.2383 | 5296.1274 | 5586.2007 | 524.60706 | 7040.822 | 6861.1406 |
| Lgals1 | DSNNLCLHFNPR | -38 | -49 | 0 | 0 | 12 | 4910.2383 | 5296.1274 | 5586.2007 | 524.60706 | 7040.822 | 6861.1406 |
| Lgals1 | LPDGHEFK | -101 | -108 | 0 | 0 | 8 | 4169.1104 | 4784.7583 | 4471.241 | 3497.2947 | 4226.648 | 241.80193 |
| Lgals1 | FDQADLTIK | -92 | -100 | 0 | 0 | 9 | 3953.6177 | 3931.216 | 3039.6028 | 2738.9724 | 981.4452 | 2713.6191 |
| Lgals1 | NLKPGECLK | -11 | -19 | 1 | 0 | 9 | 3538.4785 | 3646.0342 | 3658.4016 | 4627.169 | 6087.9766 | 2616.6033 |
| Lgals1 | SNLNLKPGECLK | -8 | -19 | 1 | 0 | 12 | 2985.797 | 3164.5032 | 3011.817 | 613.05176 | 4136.501 | 3835.392 |
| Lgals1 | NLNLKPGECLK | -9 | -19 | 1 | 0 | 11 | 2894.7505 | 1880.9323 | 3138.3396 | 2943.0225 | 3048.6882 | 2855.472 |
| Lgals1 | FNAHGDANTIVCNTK | -50 | -64 | 0 | 0 | 15 | 1070.7534 | 1146.456 | 1212.3198 | 897.2789 | 1013.9766 | 1070.8323 |
| Lgals1 | FNAHGDANTIVCNTK | -50 | -64 | 0 | 0 | 15 | 1070.7534 | 1146.456 | 1212.3198 | 897.2789 | 1013.9766 | 1070.8323 |
| Lgals1 | EDGTWGTEHR | -65 | -74 | 0 | 0 | 10 | 850.884 | 2476.6282 | 2545.2043 | 2062.7578 | 2342.976 | 2343.8276 |
| Lgals1 | VASNLNLKPGECLK | -6 | -19 | 1 | 0 | 14 | 850.6954 | 1043.6881 | 725.49506 | 1171.4712 | 1182.0493 | 1317.623 |
| Lgals1 | VASNLNLKPGECLK | -6 | -19 | 1 | 0 | 14 | 850.6954 | 1043.6881 | 725.49506 | 1171.4712 | 1182.0493 | 1317.623 |
| Lgals1 | CLHFNPR | -43 | -49 | 0 | 0 | 7 | 822.20294 | 1428.9993 | 2451.3738 | 573.28204 | 1330.1943 | 2077.8774 |
| Lgals1 | PDGHEFK | -102 | -108 | 0 | 0 | 7 | 675.4282 | 735.73157 | 843.6581 | 936.58545 | 1378.229 | 1257.3544 |
| Lgals1 | TKEDGTWGTEHR | -63 | -74 | 1 | 0 | 12 | 462.56985 | 611.2773 | 612.2268 | 1045.7616 | 659.7593 | 816.58044 |
| Lgals1 | TKEDGTWGTEHR | -63 | -74 | 1 | 0 | 12 | 462.56985 | 611.2773 | 612.2268 | 1045.7616 | 659.7593 | 816.58044 |
| Lgals1 | SFVLNLGK | -30 | -37 | 0 | 0.002595029 | 8 | 1697.5792 | 1537.0944 | 1778.9164 | 834.3699 | NaN | 599.93506 |
| Lgals1 | ITFDQADLTIK | -90 | -100 | 0 | 0 | 11 | 1566.1906 | 208.00072 | 211.19998 | NaN | 1263.3319 | 1138.282 |
| Lgals1 | VRGEVASDAK | -20 | -29 | 1 | 0.000907206 | 10 | 1205.9214 | 723.6868 | 695.47925 | 675.05475 | 940.045 | NaN |
| Lgals1 | FNAHGDANTIVC | -50 | -61 | 0 | 0 | 12 | 1064.8948 | 972.7826 | NaN | 1411.9198 | 964.63586 | 992.64954 |
| Lgals1 | EPAFPFQPGSIT | -75 | -86 | 0 | 0 | 12 | 935.0795 | NaN | 1125.0691 | 963.39734 | 671.4209 | 1029.9183 |
| Lgals1 | GDANTIVCNTK | -54 | -64 | 0 | 0 | 11 | 1207.7283 | 380.25525 | 839.4719 | 1464.2725 | NaN | NaN |
| Lgals1 | FNAHGDANTIVCN | -50 | -62 | 0 | 0.000347792 | 13 | 888.73004 | 890.6358 | 883.6834 | 1535.3553 | NaN | NaN |
| Sbds | SLKDVEEGDEKFE | -238 | -250 | 2 | 0 | 13 | 1375.9784 | 1356.5597 | 1224.2085 | 1388.5428 | 1438.9674 | 1476.3969 |
| Sbds | CLIDPGCFR | -210 | -218 | 0 | 0 | 9 | 124.23658 | 355.71414 | 341.29352 | NaN | NaN | NaN |
| Sbds | HTQLEQMFR | -101 | -109 | 0 | 0.009388464 | 9 | 3345.5789 | NaN | NaN | NaN | NaN | NaN |
| Rpl11 | EHIDLGIK | -111 | -118 | 0 | 0.000230707 | 8 | 6465.8286 | 640.10034 | 7003.3916 | 6681.7812 | 6683.738 | 7124.644 |
| Rpl11 | KYDGIILPGK | -169 | -178 | 1 | 0.001291103 | 10 | 6070.451 | 6483.576 | 6565.7056 | 7719.7456 | 8360.698 | 8487.334 |
| Rpl11 | YDGIILPGK | -170 | -178 | 0 | 0 | 9 | 4414.7827 | 4420.043 | 4525.26 | 1386.8131 | 3068.5212 | 3331.563 |
| Rpl11 | VLEQLTGQTPVFSK | -39 | -52 | 0 | 0 | 14 | 1972.0717 | 2898.991 | 2703.7947 | 3731.7593 | 4499.299 | 4400.2886 |
| Rpl11 | KNNFSDTGNFGFGIQ | -96 | -110 | 1 | 0.00025015 | 15 | 1734.5042 | 2130.5193 | 2199.3298 | 2420.5925 | 2503.295 | 3180.688 |
| Rpl11 | KAEEILEK | -78 | -85 | 1 | 0.000907206 | 8 | 1434.0404 | 1902.5425 | 1888.8575 | 3349.8042 | 3642.9001 | 4161.203 |
| Rpl11 | NNFSDTGNFGFGIQ | -97 | -110 | 0 | 0.00025015 | 14 | 684.39526 | 662.5705 | 726.4725 | 604.2921 | 675.7607 | 640.21765 |
| Rpl11 | KVLEQLTGQTPVFSK | -38 | -52 | 1 | 0 | 15 | 671.9042 | 870.2511 | 806.66583 | 1685.0585 | 2219.025 | 2387.662 |
| Rpl11 | VREYELR | -89 | -95 | 1 | 0.00025015 | 7 | 526.3552 | 720.26685 | 506.77313 | 917.6942 | 693.73047 | 634.83405 |
| Rpl11 | EQLTGQTPVFSK | -41 | -52 | 0 | 0 | 12 | 422.37445 | 6257.437 | 6595.914 | 5484.221 | 4131.3916 | 4345.5703 |
| Rpl11 | EQLTGQTPVFSK | -41 | -52 | 0 | 0 | 12 | 422.37445 | 6257.437 | 6595.914 | 5484.221 | 4131.3916 | 4345.5703 |
| Rpl11 | LTGQTPVFSK | -43 | -52 | 0 | 0 | 10 | 3615.874 | 3722.7815 | 4017.8645 | NaN | NaN | 508.44345 |
| Rpl11 | CLNICVGESGDR | -21 | -32 | 0 | 0 | 12 | 550.8271 | 615.8662 | NaN | 614.7148 | 528.0804 | NaN |
| Rpl11 | PGFSIADK | -137 | -144 | 0 | 0.00111006 | 8 | 740.5854 | 796.3648 | NaN | NaN | NaN | NaN |
| Pebp1 | SDYVGSGPPSGTGLHR | -104 | -119 | 0 | 0 | 16 | 5052.784 | 5281.921 | 5467.6465 | 5250.607 | 5724.9775 | 5690.12 |
| Pebp1 | SDYVGSGPPSGTGLHR | -104 | -119 | 0 | 0 | 16 | 5052.784 | 5281.921 | 5467.6465 | 5250.607 | 5724.9775 | 5690.12 |
| Pebp1 | YQAEWDDYVPK | -169 | -179 | 0 | 0 | 11 | 1727.1084 | 1543.1239 | 1792.4034 | 1781.7388 | 173.61746 | 1624.1354 |

| | | | | | | | | | | | | |
|---|---|---|---|---|---|---|---|---|---|---|---|---|
| Pebp1 | CLQEVDEPPQHALR | -13 | -26 | 0 | 0 | 14 | 1033.303 | 1232.5597 | 1132.3472 | 1145.8075 | 1207.0791 | 512.6508 |
| Pebp1 | EQEQPLSCDEPILSNK | -126 | -141 | 0 | 0 | 16 | 952.11084 | 1278.6206 | 1132.456 | 1053.9672 | 1383.042 | 1317.1765 |
| Pebp1 | LYTLVLTDPDAPSR | -63 | -76 | 0 | 0 | 14 | 935.1833 | 947.0735 | 963.8452 | 913.74316 | 1010.8939 | 787.6824 |
| Pebp1 | PSSISWDGLDPGK | -50 | -62 | 0 | 0 | 13 | 848.9437 | 852.7693 | 746.2807 | 654.4277 | 394.43155 | 378.093 |
| Pebp1 | FREWHHFLVVN | -81 | -91 | 1 | 0 | 11 | 688.7434 | 842.7123 | 744.6613 | 586.8032 | 1243.4573 | 982.965 |
| Pebp1 | VLTDPDAPSR | -67 | -76 | 0 | 0 | 10 | 451.1919 | 1672.2 | 1409.9141 | 1526.4612 | 1208.2325 | 1388.1707 |
| Pebp1 | LVYEQEQPLSCDEPILSNK | -123 | -141 | 0 | 0 | 19 | 306.7424 | 422.8149 | 333.9222 | 504.6484 | 728.6787 | 723.6306 |
| Pebp1 | VYEQEQPLSCDEPILSNK | -124 | -141 | 0 | 0 | 18 | 118.23562 | 320.62134 | 291.14832 | 244.5463 | 532.3132 | 558.87036 |
| Pebp1 | NRPSSISWDGLDPGK | -48 | -62 | 1 | 0 | 15 | 1098.513 | 166.76126 | 1585.9269 | 1312.9548 | NaN | 809.0584 |
| Pebp1 | NRPSSISWDGLDPGK | -48 | -62 | 1 | 0 | 15 | 1098.513 | 166.76126 | 1585.9269 | 1312.9548 | NaN | 809.0584 |
| Pebp1 | VDYAGVTVDELGK | -27 | -39 | 0 | 0 | 13 | 863.2846 | NaN | 1024.3344 | 1166.375 | 1084.5436 | 1071.8828 |
| Pebp1 | VLSDYVGSGPPSGTGLHR | -102 | -119 | 0 | 0 | 18 | 637.50555 | 581.02454 | 455.89203 | NaN | 936.3927 | 826.97546 |
| Pebp1 | KYNLGAPVAGT | -157 | -167 | 1 | 0.001636916 | 11 | 473.65472 | 340.30963 | 465.84967 | NaN | NaN | 457.46048 |
| Pebp1 | YVGSGPPSGTGLHR | -106 | -119 | 0 | 0 | 14 | 8572.06 | 8714.085 | 9495.115 | NaN | NaN | NaN |
| Pebp1 | YEQEQPLSCDEPILSNK | -125 | -141 | 0 | 0.000471409 | 17 | 255.12967 | 281.24338 | 273.44666 | NaN | NaN | NaN |
| Npm3 | DHDNQEIAVPVANLR | -88 | -102 | 0 | 0 | 15 | 2358.1045 | 2765.9182 | 2505.2468 | 2286.701 | 1926.8276 | 2544.2627 |
| Npm3 | DHDNQEIAVPVANLR | -88 | -102 | 0 | 0 | 15 | 2358.1045 | 2765.9182 | 2505.2468 | 2286.701 | 1926.8276 | 2544.2627 |
| Npm3 | FFGCELSGHTR | -38 | -48 | 0 | 0 | 11 | 1346.009 | 1470.7878 | 1690.781 | 1847.2919 | 2204.3877 | 2164.097 |
| Npm3 | DHDNQEIAVPVANL | -88 | -101 | 0 | 0 | 14 | 970.3482 | 986.95886 | 982.24725 | 997.8576 | 747.16736 | 1379.1964 |
| Npm3 | CLTEGATDECNVVEVVAR | -70 | -87 | 0 | 0 | 18 | 693.20935 | 587.6077 | 638.4759 | 878.88794 | 821.91925 | 757.2959 |
| Npm3 | TEGATDECNVVEVVAR | -72 | -87 | 0 | 0 | 16 | 1115.9529 | 999.8786 | 483.08786 | NaN | 944.0278 | 1006.81464 |
| Npm3 | VEEEDDTEHVLAL | -54 | -66 | 0 | 0 | 13 | 489.64066 | 1145.2799 | 649.9566 | 693.9269 | 923.3964 | NaN |
| Npm3 | DFQLQPPVTFR | -113 | -123 | 0 | 0 | 11 | 619.3976 | 487.76184 | 885.65753 | NaN | NaN | NaN |
| Npm3 | VEEEDDTEHVLALN | -54 | -67 | 0 | 0.001463897 | 14 | 345.3991 | NaN | 368.47986 | NaN | NaN | NaN |
| Npm3 | DHDNQEIAVPVAN | -88 | -100 | 0 | 0.00111006 | 13 | 885.63226 | NaN | NaN | NaN | NaN | NaN |
| Nutf2 | TWEGQQFQGK | -40 | -49 | 0 | 0 | 10 | 6567.8027 | 6454.9155 | 6060.825 | 6250.3247 | 6307.523 | 6121.621 |
| Nutf2 | IQHSITAQ | -64 | -71 | 0 | 0.002752526 | 8 | 1580.0011 | 1598.3271 | 1508.1316 | 1903.4291 | 1386.2993 | 1316.8767 |
| Nutf2 | HYYQLFDNDR | -17 | -26 | 0 | 0 | 10 | 951.1295 | 1074.5802 | 1152.1946 | 812.7574 | 901.2629 | 916.7063 |
| Nutf2 | IQHYYQLFDNDR | -15 | -26 | 0 | 0 | 12 | 643.8022 | 1354.2166 | 1328.3423 | 645.4405 | 1731.6545 | 1619.3066 |
| Nutf2 | CLTWEGQQFQGK | -38 | -49 | 0 | 0 | 12 | 565.551 | 465.52902 | 868.62 | 854.2449 | 1056.2512 | 1087.0621 |
| Nutf2 | GDKPIWEQIGSS | -2 | -13 | 1 | 0.000471409 | 12 | 593.61884 | 228.3018 | 597.0389 | NaN | 505.45807 | NaN |
| Nutf2 | GDKPIWEQIGSSF | -2 | -14 | 1 | 0.00025015 | 13 | 354.03006 | 320.33716 | 321.65128 | NaN | 346.26495 | NaN |
| Nutf2 | LSSLPFQK | -56 | -63 | 0 | 0.003745449 | 8 | 475.08728 | 489.87088 | 684.98083 | NaN | NaN | NaN |
| Txn | FHSLCDK | -42 | -48 | 0 | 0 | 7 | 4234.1284 | 3716.7485 | 1207.3212 | 4538.9565 | 5094.747 | 5181.9043 |
| Txn | EAFQEALAAAGDK | -9 | -21 | 0 | 0 | 13 | 3564.818 | 549.9219 | 169.36688 | 7224.171 | 7372.059 | 7248.132 |
| Txn | EVDVDDCQDVAADCEVK | -56 | -72 | 0 | 0 | 17 | 1737.7977 | 2200.9727 | 1987.9176 | 2136.9714 | 2566.3352 | 2585.8933 |
| Txn | SATWCGPCK | -28 | -36 | 0 | 0 | 9 | 1113.039 | 1053.7303 | 1074.4542 | 1008.9543 | 864.6446 | 1006.1113 |
| Txn | CMPTFQFYK | -73 | -81 | 0 | 0 | 9 | 732.7979 | 660.7514 | 764.7524 | 785.62537 | 714.0461 | 735.9149 |
| Txn | CMPTFQFYK | -73 | -81 | 0 | 0 | 9 | 732.7979 | 660.7514 | 764.7524 | 785.62537 | 714.0461 | 735.9149 |
| Txn | VGEFSGANK | -86 | -94 | 0 | 0 | 9 | 362.12875 | 5941.6323 | 5154.5977 | 6161.419 | 396.4845 | 6591.308 |
| Txn | VDVDDCQDVAADCEVK | -57 | -72 | 0 | 0.00078661 | 16 | 351.8261 | 433.60895 | 418.3593 | 494.35437 | 661.21075 | 729.395 |
| Txn | DVDDCQDVAADCEVK | -58 | -72 | 0 | 0.001776473 | 15 | 216.54459 | NaN | NaN | NaN | NaN | NaN |
| Copa | LLELGPKPEVAQQ | -1128 | -1140 | 1 | 0 | 13 | 2421.647 | 1494.5315 | 1584.8702 | 2172.4368 | 2475.573 | 1994.8771 |
| Copa | IDKFDEHDGPVR | -44 | -55 | 1 | 0 | 12 | 1460.7766 | 1529.142 | 1470.6083 | 1072.3406 | 1528.0226 | 1490.6903 |
| Copa | LLHDQVGVIQ | -942 | -951 | 0 | 0.000627943 | 10 | 1374.343 | 1506.4208 | 1566.2356 | 299.6696 | 964.6388 | 997.01776 |
| Copa | TLIDKFDEHDGPVR | -42 | -55 | 1 | 0 | 14 | 1273.3291 | 746.24146 | 806.85175 | 625.5241 | 566.5202 | 625.7865 |
| Copa | GHLDYIR | -91 | -97 | 0 | 0 | 7 | 717.1492 | 874.6958 | 785.4494 | 498.88913 | 376.82483 | 397.9898 |
| Copa | TALNLFFK | -1107 | -1114 | 0 | 0 | 8 | 609.06903 | 684.42554 | 701.6773 | 682.1431 | 768.9656 | 908.4208 |
| Copa | NLSPGAVESDVR | -171 | -182 | 0 | 0 | 12 | 4308.857 | 2143.2043 | 2773.9976 | NaN | 2342.3562 | 2949.9204 |
| Copa | LYLGDVSER | -737 | -745 | 0 | 0 | 9 | 955.3546 | NaN | 423.4112 | 799.6614 | 752.2527 | 783.63556 |
| Copa | CNIHENIR | -522 | -529 | 0 | 0.00078661 | 8 | 1556.0718 | 1597.9595 | 1424.1392 | 1634.9105 | NaN | NaN |
| Copa | GDHGIIR | -556 | -562 | 0 | 0.008498216 | 7 | 1227.839 | NaN | NaN | NaN | NaN | NaN |
| Ahsg | CNLLAEK | -219 | -225 | 0 | 0 | 7 | 10252.455 | 8272.36 | 7263.1763 | 9256.486 | 6490.164 | 6191.2305 |
| Ahsg | PDSAEDVR | -136 | -143 | 0 | 0.000907206 | 8 | 370.09442 | 383.85574 | 681.93915 | 753.032 | 818.10034 | 707.5827 |
| Snrpf | GVEEEEEDGEMRE | -74 | -86 | 1 | 0 | 13 | 1008.0793 | 1161.2122 | 1440.927 | 1233.3672 | 1599.8691 | 1585.2056 |
| Snrpf | LSGHLGEVLIR | -55 | -65 | 0 | 0 | 11 | 875.8178 | 1049.5704 | 89.577896 | 984.3355 | 880.3062 | 925.9504 |
| Snrpf | SLPLNPKPF | -2 | -10 | 1 | 0.00294445 | 9 | 804.4353 | 1051.9706 | 989.89606 | 1232.1509 | 1320.0156 | 889.6009 |
| Snrpf | GHLGEVLIR | -57 | -65 | 0 | 0 | 9 | 638.15204 | 727.598 | 803.6912 | 950.8882 | 567.30225 | 627.3005 |
| Snrpf | EYIDGALSGHLGEVLIR | -49 | -65 | 0 | 0 | 17 | 564.51886 | 459.88077 | 579.13995 | 733.676 | 700.95685 | 764.28674 |
| Snrpf | SLPLNPKPFLN | -2 | -12 | 1 | 0 | 11 | 160.11394 | 532.63416 | 220.78944 | 657.398 | 943.42755 | 847.0835 |
| Snrpf | CNNVLYIR | -66 | -73 | 0 | 0 | 8 | 1475.6436 | 333.4667 | 2319.977 | 433.4854 | 782.8408 | NaN |
| Snrpf | GALSGHLGEVLIR | -53 | -65 | 0 | 0 | 13 | 564.76794 | NaN | NaN | NaN | NaN | NaN |
| Vdac3 | LTLDTIFVPNTGK | -97 | -109 | 0 | 0 | 13 | 852.2566 | 1034.1359 | 987.63727 | 1043.3153 | 1209.671 | 1123.442 |
| Vdac3 | DTIFVPNTGK | -100 | -109 | 0 | 0 | 10 | 443.20786 | 1052.4886 | 1179.1276 | 668.20264 | 1019.0768 | 972.0512 |
| Vdac3 | WNTDNTLGTEISWENK | -75 | -90 | 0 | 0 | 16 | 538.6459 | 422.38092 | 340.10016 | NaN | 504.9546 | 1230.5283 |
| Vdac3 | KWNTDNTLGTEISWENK | -74 | -90 | 1 | 0.000431655 | 17 | 314.28394 | 303.45474 | 371.85773 | NaN | 599.521 | 433.6062 |
| Vdac3 | LSQNNFALGYK | -164 | -174 | 0 | 0 | 11 | 208.04301 | 16070.473 | 24428.947 | NaN | 19097.322 | 18012.324 |
| Vdac3 | CNTPTYCDLGK | -2 | -12 | 0 | 0 | 11 | 1316.7267 | 417.65848 | 1195.222 | 1142.1914 | NaN | NaN |
| Hectd1 | EVELPLTNFR | -1836 | -1845 | 0 | 0 | 10 | 1005.9226 | 1224.3986 | 1181.1177 | 916.9807 | 537.41846 | 1283.0647 |
| Hectd1 | HEDHQVSDGALR | -197 | -208 | 0 | 0.000550307 | 12 | 2736.6875 | NaN | 3358.1135 | NaN | 2780.9216 | 2623.9185 |
| Hectd1 | CPFLIPFETR | -2076 | -2085 | 0 | 0.000230707 | 10 | 1112.5232 | NaN | 1162.4264 | NaN | 1499.0569 | NaN |
| Hectd1 | TFLPALCK | -52 | -59 | 0 | 0.000550307 | 8 | 828.91187 | 599.52356 | NaN | NaN | NaN | NaN |
| Rps11 | DYLHYIR | -91 | -97 | 0 | 0.000394361 | 7 | 5224.809 | 5573.9004 | 5640.13 | 2541.8088 | 303.44366 | 2754.141 |
| Rps11 | RDYLHYIR | -90 | -97 | 1 | 0.00025015 | 8 | 4268.4014 | 4305.747 | 4386.456 | 4094.7102 | 2513.0771 | 2410.0178 |
| Rps11 | RDYLHYIR | -90 | -97 | 1 | 0.00025015 | 8 | 4268.4014 | 4305.747 | 4386.456 | 4094.7102 | 2513.0771 | 2410.0178 |
| Rps11 | SVHLSPCFR | -110 | -118 | 0 | 0 | 9 | 3111.8838 | 3046.4253 | 3443.2725 | 1435.7852 | 1422.3807 | 3602.0403 |
| Rps11 | SVHLSPCFR | -110 | -118 | 0 | 0 | 9 | 3111.8838 | 3046.4253 | 3443.2725 | 1435.7852 | 1422.3807 | 3602.0403 |
| Rps11 | CPFTGNVSIR | -60 | -69 | 0 | 0 | 10 | 2243.0867 | 2195.025 | 2354.4985 | 677.5076 | 1560.7169 | 1553.809 |

| | | | | | | | | | | | | |
|---|---|---|---|---|---|---|---|---|---|---|---|---|
| Rps11 | TVGECRPLSK | -127 | -136 | 1 | 0.000471409 | 10 | 1278.9026 | 1358.4747 | 1273.4547 | 1568.6932 | 2184.72 | 1950.991 |
| Rps11 | DVQIGDIVTVGECRPLSK | -119 | -136 | 1 | 0 | 18 | 679.37415 | 726.52264 | 616.72266 | 726.11255 | 851.734 | 736.7244 |
| Rps11 | EAIEGTYIDKK | -49 | -59 | 1 | 0.00025015 | 11 | 1972.9485 | 2040.5674 | 1989.5933 | 2046.9592 | NaN | NaN |
| Rps11 | KNIGLGFK | -38 | -45 | 1 | 0.001392869 | 8 | 450.49957 | 451.79633 | 444.97424 | NaN | 312.2291 | NaN |
| Rps11 | NMSVHLSPCFR | -108 | -118 | 0 | 0 | 11 | 208.48401 | NaN | NaN | NaN | NaN | NaN |
| Foxp4 | CPLIPGWK | -676 | -683 | 0 | 0.000591124 | 8 | 669.8856 | 624.8339 | NaN | NaN | 1309.4136 | 1032.9177 |
| Matr3 | SICDLPVHSNK | -294 | -304 | 0 | 0 | 11 | 1290.7363 | 1324.6587 | 1385.958 | 1110.7706 | 1053.3008 | 1102.9316 |
| Matr3 | GAPPSSNIEDFHGLLPK | -271 | -287 | 0 | 0 | 17 | 1074.1863 | 958.08246 | 1407.704 | 1412.0695 | 1876.6201 | 1876.7444 |
| Matr3 | GAPPSSNIEDFHGLLPK | -271 | -287 | 0 | 0 | 17 | 1074.1863 | 958.08246 | 1407.704 | 1412.0695 | 1876.6201 | 1876.7444 |
| Matr3 | NLPHSGYSDSAVLK | -502 | -515 | 0 | 0 | 14 | 970.45264 | 1060.4476 | 1140.5143 | 1456.425 | 2597.6523 | 2523.822 |
| Matr3 | SNLPHSGYSDSAVLK | -501 | -515 | 0 | 0 | 15 | 935.10944 | 675.29553 | 660.5192 | 748.178 | 858.35284 | 870.99115 |
| Matr3 | SNLPHSGYSDSAVLK | -501 | -515 | 0 | 0 | 15 | 935.10944 | 675.29553 | 660.5192 | 748.178 | 858.35284 | 870.99115 |
| Matr3 | IGPYQPNVPVGIDYVIPK | -780 | -797 | 0 | 0 | 18 | 817.07587 | 796.5838 | 699.88983 | 977.79694 | 1155.7574 | 1016.8341 |
| Matr3 | GVISNHLILNK | -423 | -433 | 0 | 0 | 11 | 565.3875 | 549.0371 | 493.60898 | 1158.9001 | 778.2223 | 669.3539 |
| Matr3 | DFHGLLPK | -280 | -287 | 0 | 0 | 8 | 588.3064 | 443.30127 | 587.0451 | NaN | 312.51022 | 514.2551 |
| Matr3 | LANLGDVSSDGK | -680 | -691 | 0 | 0 | 12 | 1320.6003 | 2148.0894 | NaN | 1463.5039 | NaN | 1849.9076 |
| Matr3 | NDENKEDYTIPDEYR | -765 | -779 | 1 | 0 | 15 | 412.5443 | NaN | NaN | 640.7368 | 758.694 | 754.03235 |
| Matr3 | GPGPLQER | -256 | -263 | 0 | 0.005333786 | 8 | 332.15436 | NaN | 260.41452 | 501.97504 | NaN | NaN |
| Matr3 | NHLILNK | -427 | -433 | 0 | 0 | 7 | 458.91006 | NaN | NaN | 641.19293 | NaN | NaN |
| Matr3 | CRDDSFFGET | -230 | -239 | 1 | 0.002595029 | 10 | 252.52106 | NaN | NaN | NaN | NaN | NaN |
| Tagln2 | NFSDNQLQEGK | -161 | -171 | 0 | 0 | 11 | 4595.5225 | 3312.849 | 2948.0554 | 4945.8496 | 1955.852 | 2329.3135 |
| Tagln2 | NFSDNQLQEGK | -161 | -171 | 0 | 0 | 11 | 4595.5225 | 3312.849 | 2948.0554 | 4945.8496 | 1955.852 | 2329.3135 |
| Tagln2 | DDGLFSGDPNWFPK | -140 | -153 | 0 | 0 | 14 | 3703.9788 | 3704.4836 | 4077.6938 | 3907.711 | 5163.5186 | 4937.994 |
| Tagln2 | SLYPEGQAPVK | -68 | -78 | 0 | 0.00025015 | 11 | 3451.2605 | 3023.3413 | 2534.9058 | 3087.627 | 1661.8652 | 774.14386 |
| Tagln2 | LINSLYPEGQAPVK | -65 | -78 | 0 | 0 | 14 | 2636.1257 | 3007.9514 | 2904.2317 | 1559.9684 | 1951.1129 | 2000.0216 |
| Tagln2 | SLYPEGQAPVKK | -68 | -79 | 1 | 0 | 12 | 2578.1658 | 2519.378 | 2567.4438 | 3355.1562 | 3631.3958 | 3487.2808 |
| Tagln2 | EDVGQPQPGR | -40 | -49 | 0 | 0 | 10 | 2332.9473 | 2520.1936 | 2485.8257 | 1865.2804 | 848.6329 | 1886.8296 |
| Tagln2 | RDDGLFSGDPNWFPK | -139 | -153 | 1 | 0 | 15 | 588.18134 | 626.7336 | 737.33154 | 595.30615 | 857.18286 | 803.35236 |
| Tagln2 | RDDGLFSGDPNWFPK | -139 | -153 | 1 | 0 | 15 | 588.18134 | 626.7336 | 737.33154 | 595.30615 | 857.18286 | 803.35236 |
| Tagln2 | NVIGLQMGTNR | -172 | -182 | 0 | 0.00025015 | 11 | 1034.7968 | 1448.3247 | 1355.0668 | 3242.7932 | 2796.091 | 276.6607 |
| Tagln2 | QYDADLEQILIQ | -21 | -32 | 0 | 0 | 12 | 865.03906 | 1024.0796 | 842.60376 | 1044.814 | 1220.9464 | 1160.8143 |
| Tagln2 | LINSLYPEGQAPVKK | -65 | -79 | 1 | 0 | 15 | 658.67725 | 677.15674 | 815.1891 | 913.2449 | 575.39264 | 590.2518 |
| Tagln2 | CREDVGQPQPGR | -38 | -49 | 1 | 0 | 12 | 580.9683 | 107.20488 | 618.2234 | 676.43555 | 860.725 | 933.70294 |
| Tagln2 | CREDVGQPQPGR | -38 | -49 | 1 | 0 | 12 | 580.9683 | 107.20488 | 618.2234 | 676.43555 | 860.725 | 933.70294 |
| Tagln2 | TLMNLGGLAVAR | -128 | -139 | 0 | 0 | 12 | 257.27338 | 265.90256 | NaN | 864.3494 | 1337.4014 | 1075.0287 |
| Tagln2 | GLFSGDPNWFPK | -142 | -153 | 0 | 0 | 12 | 146.01483 | 864.63586 | 1056.4545 | NaN | NaN | NaN |
| Gfus | SVGEEDEVSIK | -243 | -253 | 0 | 0 | 11 | 5533.2417 | 6107.951 | 5839.4893 | NaN | 5091.336 | 431.77264 |
| Gfus | VQPTHVIHLA | -61 | -70 | 0 | 0 | 10 | 477.66696 | 407.68982 | 530.0716 | NaN | 485.2387 | 655.22784 |
| Gfus | VTGGSGLVGR | -12 | -21 | 0 | 0.005522803 | 10 | 707.5883 | NaN | NaN | 770.3087 | NaN | NaN |
| Gfus | CTWFTDNYEQAR | -309 | -320 | 0 | 0.00111006 | 12 | 306.5593 | NaN | NaN | NaN | NaN | NaN |
| Ddx17 | VLEEANQAINPK | -457 | -468 | 0 | 0 | 12 | 4620.781 | 4946.277 | 4695.456 | 5352.608 | 5627.926 | 5660.7627 |
| Ddx17 | DYPNSSEDYVHR | -415 | -426 | 0 | 0 | 12 | 905.4592 | 993.8568 | 1005.7111 | 700.1072 | 758.1132 | 330.94794 |
| Ddx17 | YDYPNSSEDYVHR | -414 | -426 | 0 | 0 | 13 | 617.4686 | 660.371 | 895.21814 | 679.0658 | 906.1237 | 716.3256 |
| Ddx17 | NFYVEHPEVAR | -54 | -64 | 0 | 0 | 11 | 1709.681 | 1693.832 | 1762.0044 | NaN | 1504.832 | 1675.1035 |
| Ddx17 | WDLSELPK | -43 | -50 | 0 | 0.000431655 | 8 | 538.4101 | NaN | 441.54565 | 549.8539 | 480.25305 | 505.4674 |
| Ddx17 | HQPYLER | -156 | -162 | 0 | 0.000471409 | 7 | 5917.6973 | NaN | 4198.8774 | 2965.5972 | 1133.717 | NaN |
| Ddx17 | QLAEDFLR | -286 | -293 | 0 | 0 | 8 | 477.5268 | 385.00333 | 346.06406 | NaN | NaN | NaN |
| Ddx17 | LIDFLESGK | -226 | -234 | 0 | 0 | 9 | 893.5077 | NaN | NaN | NaN | NaN | 1039.2034 |
| Ddx17 | CTYLVLDEADR | -240 | -250 | 0 | 0.00201047 | 11 | 715.4038 | NaN | NaN | NaN | NaN | NaN |
| Ptges3 | KGESGQSWPR | -79 | -88 | 1 | 0.000230707 | 10 | 6528.4243 | 4965.784 | 5930.638 | 6537.6978 | 5317.561 | 7350.5986 |
| Ptges3 | FCVEDSKDVNVNFEK | -19 | -33 | 1 | 0 | 15 | 1737.3574 | 1267.3597 | 1852.5184 | 1189.8691 | 2040.2972 | 905.64417 |
| Ptges3 | HLNEIDLFHC | -49 | -58 | 0 | 0 | 10 | 992.5674 | 1137.4614 | 1125.0049 | 566.903 | 116.193306 | 1289.375 |
| Ptges3 | HCIDPNDSK | -57 | -65 | 0 | 0.001392869 | 9 | 930.91876 | 903.26404 | 929.0961 | 924.08386 | 975.1803 | 908.82446 |
| Ptges3 | HLNEIDLFHCIDPNDSK | -49 | -65 | 0 | 0 | 17 | 589.3067 | 766.2322 | 808.50366 | 1249.0236 | 1293.411 | 1107.7013 |
| Ptges3 | HLNEIDLFHCIDPNDSK | -49 | -65 | 0 | 0 | 17 | 589.3067 | 766.2322 | 808.50366 | 1249.0236 | 1293.411 | 1107.7013 |
| Ptges3 | CVEDSKDVNVNFEK | -20 | -33 | 1 | 0.000230707 | 14 | 447.8006 | 475.04837 | 482.16046 | 637.36475 | 602.72736 | 517.6108 |
| Ptges3 | VDFNNWK | -101 | -107 | 0 | 0 | 7 | 1239.6475 | 1219.6178 | NaN | 1217.7852 | NaN | 1249.8636 |
| Ptges3 | SVDFNNWK | -100 | -107 | 0 | 0.000471409 | 8 | 1448.9424 | 450.86475 | NaN | NaN | NaN | NaN |
| Ptges3 | SCLGGSDNFK | -39 | -48 | 0 | 0.00025015 | 10 | 1120.795 | NaN | NaN | 1032.6187 | NaN | NaN |
| Strap | HSAVSLEPIK | -209 | -218 | 0 | 0 | 10 | 4060.5889 | 4106.7134 | 4084.1748 | 2702.342 | 2388.362 | 1950.0437 |
| Strap | IYDLNKPEAEPK | -126 | -137 | 1 | 0 | 12 | 2234.1128 | 1829.2584 | 1397.1587 | 1908.3314 | 2805.834 | 3226.83 |
| Strap | SDSIYSSTPEVK | -338 | -349 | 0 | 0 | 12 | 1375.5283 | 1365.5208 | 1458.8164 | 694.48175 | 838.80963 | 872.27527 |
| Strap | DLNKPEAEPK | -128 | -137 | 1 | 0 | 10 | 1221.9246 | 1251.4211 | 1234.9491 | 1347.6227 | 1435.4031 | 1541.5281 |
| Strap | CVLPEEDSGELAKPK | -305 | -319 | 1 | 0 | 15 | 964.6577 | 849.8817 | 1056.4176 | 745.5264 | 1401.9563 | 842.9959 |
| Strap | GHFGPIHCVR | -263 | -272 | 0 | 0 | 10 | 802.06964 | 867.8679 | 1192.8193 | 853.93823 | 936.25 | 804.97754 |
| Strap | TVDFTQDSNYLLTGGQDK | -105 | -122 | 0 | 0 | 18 | 716.5984 | 813.81506 | 783.89954 | 896.5322 | 1137.2721 | 1129.5543 |
| Strap | VLPEEDSGELAKPK | -306 | -319 | 1 | 0 | 14 | 665.4203 | 635.282 | 858.6792 | 925.57196 | 1001.3506 | 994.2522 |
| Strap | DYNSGEELESYK | -251 | -262 | 0 | 0 | 12 | 400.83838 | 618.31604 | 670.8672 | 663.1244 | 675.5417 | 782.48505 |
| Strap | SENSDSIYSSTPEVK | -335 | -349 | 0 | 0 | 15 | 917.7183 | 394.64743 | NaN | 1256.2057 | 875.0049 | 757.4863 |
| Strap | DFTQDSNYLLTGGQDK | -107 | -122 | 0 | 0 | 16 | 662.042 | 816.2983 | 772.84503 | 769.5171 | NaN | 1163.5408 |
| Xab2 | CVTDPAYEDVNNCHER | -86 | -101 | 0 | 0 | 16 | 191.28574 | 248.84 | 266.75876 | 181.88188 | 215.0922 | 264.564 |
| Xab2 | HENYDEALK | -446 | -454 | 0 | 0.000431655 | 9 | 887.13916 | NaN | NaN | 400.51898 | 100.623405 | 1048.6299 |
| Xab2 | SHPLPETAVR | -158 | -167 | 0 | 0 | 10 | 1204.9774 | 919.42865 | 918.5185 | NaN | NaN | NaN |
| Slc7a5 | RGDGADPEGEGVTL | -34 | -47 | 1 | 0.001055699 | 14 | 2281.8865 | 2283.524 | 2506.963 | 452.86365 | 63.24021 | 392.3593 |
| Slc7a5 | RGDGADPEGEGVTLQ | -34 | -48 | 1 | 0 | 15 | 1050.543 | 403.6483 | 1167.0945 | 229.1934 | 343.40524 | 1225.4382 |
| Slc7a5 | CYAELGTTISK | -103 | -113 | 0 | 0 | 11 | 741.4933 | 759.80194 | 880.2794 | NaN | 448.5701 | 367.43597 |
| Slc7a5 | RGDGADPEGEGVTLQR | -34 | -49 | 1 | 0 | 16 | 609.5232 | NaN | 588.2838 | 756.8646 | 647.6197 | 620.0479 |
| Slc7a5 | KPELERPIK | -427 | -435 | 2 | 0.00051306 | 9 | 485.14664 | 388.2287 | NaN | 691.97534 | 394.87686 | 353.15198 |

| | | | | | | | | | | | | |
|---|---|---|---|---|---|---|---|---|---|---|---|---|
| Slc7a5 | GDGADPEGEGVTLQR | -35 | -49 | 0 | 0.000394361 | 15 | 777.83124 | 896.0882 | NaN | NaN | NaN | 733.77026 |
| Tm9sf2 | FCNPGFPIGCYITDK | -172 | -186 | 0 | 0 | 15 | 817.6731 | 842.7633 | 880.4279 | 1257.3687 | 1709.3817 | 1471.3134 |
| Tm9sf2 | NAIEHPVR | -495 | -502 | 0 | 0.00025015 | 8 | 642.1253 | 643.0701 | 592.41516 | 553.0343 | 681.1095 | 587.32117 |
| Tm9sf2 | RPSENLGQVL | -90 | -99 | 1 | 0.000550307 | 10 | 595.8093 | 708.4491 | 354.92703 | NaN | 975.051 | NaN |
| Tm9sf2 | DACVISSEFHER | -191 | -202 | 0 | 0 | 12 | 9244.048 | NaN | NaN | 8605.324 | NaN | 9038.874 |
| Pstpip1 | TTPSAPAASTETLTPTPER | -321 | -339 | 0 | 0 | 19 | 661.58264 | 1163.6774 | 391.093 | 740.56537 | 875.29126 | 842.9785 |
| Pstpip1 | EPPAPVPYQNYYDR | -277 | -290 | 0 | 0.00025015 | 14 | 601.8823 | 606.05365 | 535.8658 | 652.5198 | 984.06213 | 555.00323 |
| Pstpip1 | DADDAEQAFER | -150 | -160 | 0 | 0 | 11 | 457.96255 | 463.51776 | 659.08527 | 507.1835 | NaN | NaN |
| Pstpip1 | GFVPGSYLEK | -405 | -414 | 0 | 0 | 10 | 321.50427 | 857.9947 | 904.3445 | NaN | NaN | NaN |
| Srsf5 | GEVTFADAHRPK | -132 | -143 | 1 | 0 | 12 | 1746.8927 | 2263.1675 | 2439.1445 | 2252.3933 | 1125.0209 | 2478.7483 |
| Srsf5 | GEVTFADAHRPK | -132 | -143 | 1 | 0 | 12 | 1746.8927 | 2263.1675 | 2439.1445 | 2252.3933 | 1125.0209 | 2478.7483 |
| Srsf5 | DADDAVYELDGK | -49 | -60 | 0 | 0 | 12 | 1289.0615 | 1361.3601 | 1764.9531 | 1499.3054 | 1352.5293 | 1410.666 |
| Srsf5 | QAGEVTFADAHRPK | -130 | -143 | 1 | 0 | 14 | 1055.84 | 503.47885 | 759.5963 | 1239.1381 | 755.5179 | 718.93396 |
| Srsf5 | QAGEVTFADAHRPK | -130 | -143 | 1 | 0 | 14 | 1055.84 | 503.47885 | 759.5963 | 1239.1381 | 755.5179 | 718.93396 |
| Srsf5 | VSWQDLK | -119 | -125 | 0 | 0.003490217 | 7 | 342.9955 | 1041.6135 | 199.27684 | 993.99304 | 839.1263 | 849.54675 |
| Glyr1 | FFGTEDHAWIK | -46 | -56 | 0 | 0 | 11 | 796.28564 | 1471.3411 | 1656.4044 | 242.22174 | 697.7286 | 2485.6804 |
| Glyr1 | DADPHFHHF | -211 | -219 | 0 | 0.00025015 | 9 | 2048.5303 | 1897.5925 | NaN | 1414.0162 | 1145.032 | 1222.5334 |
| Glyr1 | DLTIPESSTVK | -179 | -189 | 0 | 0.003681885 | 11 | 804.6103 | NaN | NaN | NaN | NaN | NaN |
| Srsf10 | DAEDALHNLDR | -63 | -73 | 0 | 0 | 11 | 3010.1633 | 3144.2124 | 3045.095 | 1487.0044 | 442.8612 | 560.53015 |
| Srsf10 | RYDDYDR | -109 | -115 | 1 | 0.007987481 | 7 | 516.8583 | 529.05084 | NaN | NaN | 480.53006 | 385.8714 |
| Srsf2 | VGDVYIPR | -40 | -47 | 0 | 0.00025015 | 8 | 5275.7383 | 4500.599 | 4216.271 | 3575.5671 | 1851.7135 | 1943.705 |
| Srsf2 | RTSPDTLR | -24 | -31 | 1 | 0.000627943 | 8 | 1932.1335 | 1123.2655 | 1600.8271 | 1254.9237 | 3042.21 | 1282.8492 |
| Srsf2 | YGRPPDSHHSR | -92 | -102 | 1 | 0.00051306 | 11 | 646.5209 | 874.6627 | 906.65814 | 267.27426 | 594.45044 | 665.6982 |
| Srsf2 | DAEDAMDAMDGAVLDGR | -67 | -83 | 0 | 0 | 17 | 105.36265 | 160.55147 | NaN | 145.72047 | 152.70291 | 121.216515 |
| Srsf2 | KVDNLTYR | -17 | -24 | 1 | 0 | 8 | 6986.352 | NaN | NaN | 3001.5552 | 3347.1855 | NaN |
| Srsf2 | YGGGGYGR | -110 | -117 | 0 | 0.005242018 | 8 | 769.85815 | NaN | 662.2263 | NaN | NaN | NaN |
| Srsf1 | SHEGETAYIR | -182 | -191 | 0 | 0 | 10 | 4252.233 | 3509.1487 | 3468.3972 | 182.61255 | 3776.8474 | 3990.7986 |
| Srsf1 | TKDIEDVFYK | -29 | -38 | 1 | 0.00025015 | 10 | 3705.8677 | 3929.9675 | 4046.4229 | 4416.1216 | 5130.146 | 4495.506 |
| Srsf1 | DGTGVVEFVR | -155 | -164 | 0 | 0 | 10 | 2966.082 | 3068.6958 | 3133.4204 | 3969.853 | 4206.2134 | 4113.591 |
| Srsf1 | VGNLPPDIR | -20 | -28 | 0 | 0 | 9 | 2404.774 | 1919.3334 | 2162.2249 | 1150.5504 | 1064.6914 | 2536.502 |
| Srsf1 | IYVGNLPPDIR | -18 | -28 | 0 | 0 | 11 | 2030.7184 | 2250.756 | 2101.4714 | 2664.6655 | 4247.424 | 2307.9941 |
| Srsf1 | DAEDAVYGR | -66 | -74 | 0 | 0 | 9 | 1390.6296 | 1106.6943 | 802.7946 | 1263.7104 | 508.86237 | 571.1618 |
| Srsf1 | GGPPFAFVEFEDPR | -52 | -65 | 0 | 0 | 14 | 853.6183 | 801.1094 | 905.53424 | 406.58316 | 1486.0791 | 1697.3474 |
| Srsf1 | DGYDYDGYR | -75 | -83 | 0 | 0 | 9 | 765.8869 | 746.5015 | 744.436 | 1208.1986 | 1252.1061 | 1293.4766 |
| Srsf1 | DIEDVFYK | -31 | -38 | 0 | 0.000431655 | 8 | 373.38504 | 685.55475 | NaN | 621.77484 | 529.61414 | 337.15573 |
| Srsf1 | RGGPPFAFVE | -51 | -60 | 1 | 0.002810811 | 10 | 3024.7078 | 2941.6497 | 1901.1077 | NaN | NaN | 1150.5038 |
| Srsf1 | DGYDYDGYRL | -75 | -84 | 1 | 0 | 10 | 455.99234 | 503.525 | 422.3168 | NaN | NaN | NaN |
| Srsf1 | DAEDAVYGRD | -66 | -75 | 1 | 0.00025015 | 10 | 699.3841 | NaN | NaN | NaN | NaN | NaN |
| Ppp1r7 | DAEDVDLTHYR | -78 | -88 | 0 | 0 | 11 | 901.46326 | 3121.4272 | 5774.1567 | 5884.5615 | 6337.271 | 6648.8853 |
| Ppp1r7 | ELDLYDNQIK | -125 | -134 | 0 | 0 | 10 | 292.48474 | 948.4762 | 982.53815 | NaN | NaN | 1217.9362 |
| Ppp1r7 | GAEDPEEEHEL | -56 | -66 | 0 | 0.000230707 | 11 | 438.50925 | 450.98767 | NaN | NaN | NaN | NaN |
| Dck | DVNEDFKDKHESLVEK | -237 | -252 | 2 | 0.000431655 | 16 | 115.96564 | 857.8471 | 848.5601 | 269.5669 | 953.23535 | 796.1673 |
| Dck | QASEDWEVVPEPVAR | -43 | -57 | 0 | 0 | 15 | 901.3983 | NaN | 941.0299 | NaN | 1162.333 | 1251.7402 |
| Dck | SEDWEVVPEPVAR | -45 | -57 | 0 | 0 | 13 | 559.18945 | NaN | 742.4109 | 1973.5181 | 803.94055 | NaN |
| Dck | DAEKPVLFFER | -118 | -128 | 1 | 0 | 11 | 1272.1732 | 1626.3156 | 255.69698 | NaN | NaN | NaN |
| Dck | STQEEFEELTTSQK | -63 | -76 | 0 | 0 | 14 | 612.12445 | NaN | NaN | NaN | NaN | NaN |
| Dck | FCPSPSTSSEGTR | -8 | -20 | 0 | 0 | 13 | 477.29507 | NaN | NaN | NaN | NaN | NaN |
| Dck | HESWLLHR | -212 | -219 | 0 | 0.006788754 | 8 | 222.90175 | NaN | NaN | NaN | NaN | NaN |
| Akr1a1 | HWPYAFER | -113 | -120 | 0 | 0 | 8 | 1515.3611 | 1568.9491 | 1465.2494 | 1249.9996 | 1635.0327 | 1713.9338 |
| Akr1a1 | DAGHPLYPFNDPY | -313 | -325 | 0 | 0 | 13 | 699.784 | 902.9336 | 1005.94824 | 1004.1046 | 1013.2916 | 972.9743 |
| Akr1a1 | HPDEPVLLEEPVVL | -222 | -235 | 0 | 0 | 14 | 335.42148 | 445.40735 | 493.22382 | 513.2004 | 614.7843 | 539.7433 |
| Akr1a1 | SVYGNETEIGEALK | -48 | -61 | 0 | 0 | 14 | 5801.1943 | 5579.7256 | 5802.727 | NaN | 6909.342 | 6337.9414 |
| Akr1a1 | SVLLHTGQK | -5 | -13 | 0 | 0 | 9 | 6151.7227 | NaN | 5313.218 | NaN | 4233.497 | 4520.6123 |
| Akr1a1 | HHPEDVEPALR | -86 | -96 | 0 | 0 | 11 | 3768.498 | 3847.1606 | NaN | 4303.764 | NaN | 4042.4785 |
| Akr1a1 | HHPEDVEPALR | -86 | -96 | 0 | 0 | 11 | 3768.498 | 3847.1606 | NaN | 4303.764 | NaN | 4042.4785 |
| Eif3c | VWDLFPEADKVR | -763 | -774 | 1 | 0 | 12 | 9500.053 | 9128.803 | 7665.886 | 18281.443 | 12728.1045 | 13768.338 |
| Eif3c | VWDLFPEADKVR | -763 | -774 | 1 | 0 | 12 | 9500.053 | 9128.803 | 7665.886 | 18281.443 | 12728.1045 | 13768.338 |
| Eif3c | VWDLFPEADK | -763 | -772 | 0 | 0 | 10 | 1199.5133 | 386.9054 | 1364.1904 | 1557.7257 | 1686.4749 | 1697.557 |
| Eif3c | KIQEESLR | -780 | -787 | 1 | 0.001264841 | 8 | 969.03906 | 878.90125 | 1064.1095 | 1246.8884 | 1426.9117 | 1383.6472 |
| Eif3c | DAHNALLDIQ | -626 | -635 | 0 | 0 | 10 | 937.1096 | 1130.2943 | 1060.1665 | 863.0075 | 1403.9857 | 1006.28204 |
| Eif3c | LGSLVENNER | -861 | -870 | 0 | 0 | 10 | 815.2981 | 1513.7075 | 1426.6097 | 754.11444 | 1251.447 | 1367.0657 |
| Eif3c | NILEESENLHNFDQPLR | -421 | -437 | 0 | 0 | 17 | 460.59344 | 441.31918 | 488.51404 | 690.8689 | 825.3343 | 801.23895 |
| Eif3c | FEELTNLIR | -57 | -65 | 0 | 0 | 9 | 318.29538 | 1193.2053 | 1296.1395 | 311.9795 | 1429.4701 | 1470.9326 |
| Eif3c | GTEITHAVVIK | -319 | -329 | 0 | 0 | 11 | 2424.0176 | NaN | 853.70123 | 707.07886 | 1112.0005 | 673.08276 |
| Eif3c | SENLHNFDQPLR | -426 | -437 | 0 | 0 | 12 | 5239.7085 | 4304.1226 | 4559.6704 | 3738.2908 | NaN | NaN |
| Eif3c | SENLHNFDQPLR | -426 | -437 | 0 | 0 | 12 | 5239.7085 | 4304.1226 | 4559.6704 | 3738.2908 | NaN | NaN |
| Eif3c | KLNEILQVR | -330 | -338 | 1 | 0.00078661 | 9 | 398.86237 | NaN | NaN | 514.5701 | 557.17456 | 491.98935 |
| Eif3c | ENLHNFDQPLR | -427 | -437 | 0 | 0.00111006 | 11 | 998.3204 | NaN | 447.95715 | NaN | NaN | 933.3704 |
| Eif3c | RLDEEEEDNEGGEW | -286 | -299 | 1 | 0.00025015 | 14 | 367.58264 | 422.53528 | 395.44666 | NaN | NaN | NaN |
| Eif3c | QVPFHLHIN | -668 | -676 | 0 | 0.00025015 | 9 | 501.399 | NaN | 540.9378 | NaN | NaN | NaN |
| Eif3c | EEDNEGGEWER | -291 | -301 | 0 | 0.000230707 | 11 | 157.51569 | NaN | NaN | NaN | NaN | NaN |
| Nudt9 | DAHWSEDHAADSR | -336 | -348 | 0 | 0 | 13 | 373.7612 | 409.83218 | 409.92438 | 582.8845 | 114.51415 | 275.02603 |
| Nudt9 | FSQEHLVIYK | -257 | -266 | 0 | 0 | 10 | 1019.7279 | 327.12927 | 1021.6615 | NaN | 1150.2609 | 1247.6508 |
| Cat | LCENIAGHLK | -459 | -468 | 0 | 0 | 10 | 1690.9781 | 1929.3923 | 1926.4678 | 863.0075 | 2717.095 | 2586.2869 |
| Cat | DAILFPSFIHSQ | -157 | -168 | 0 | 0.00025015 | 12 | 649.2356 | 305.7286 | 690.6449 | 388.68323 | 288.5232 | 680.45953 |
| Cat | NFTDVHPDYGAR | -481 | -492 | 0 | 0 | 12 | 581.6412 | 756.11414 | 816.20734 | 553.3291 | 314.18042 | 548.11017 |
| Cat | EAETFPFNPFDLTK | -288 | -301 | 0 | 0 | 14 | 473.6632 | 452.0727 | 446.69138 | 525.6494 | 171.4582 | 634.3762 |
| Cat | YLQIPVNCPYR | -370 | -380 | 0 | 0 | 11 | 914.93005 | 976.3806 | 445.75403 | 880.54065 | 533.66156 | NaN |

| | | | | | | | | | | | | |
|---|---|---|---|---|---|---|---|---|---|---|---|---|
| Cat | FNSANEDNVTQVR | -432 | -444 | 0 | 0 | 13 | 1010.36316 | 1190.4171 | NaN | 1171.581 | NaN | NaN |
| Cat | NTPIFFIR | -149 | -156 | 0 | 0.001636916 | 8 | 488.00162 | 493.97302 | NaN | 301.67902 | NaN | NaN |
| Ppp1r10 | TTLPERPLTEVK | -168 | -179 | 1 | 0.000230707 | 12 | 1300.5352 | 1729.8755 | 2174.1833 | 588.2929 | 2009.7604 | 2681.4653 |
| Ppp1r10 | DAKPVDSPGDPNQLTR | -376 | -391 | 1 | 0.001392869 | 16 | 394.0761 | 420.86398 | 934.1242 | NaN | NaN | 868.6388 |
| Psmg1 | DALCCPLLEQPNIVHDL | -200 | -216 | 0 | 0 | 17 | 235.8616 | 246.78589 | 222.67883 | 312.01102 | 424.20282 | 487.39142 |
| Psmg1 | EDQQFQWLEK | -144 | -153 | 0 | 0 | 10 | 1231.9777 | 1331.1349 | NaN | NaN | 1265.0873 | 1455.2122 |
| Psmg1 | NSGVWEEVGCAK | -90 | -101 | 0 | 0 | 12 | 544.17236 | NaN | NaN | 387.68057 | 1045.4387 | NaN |
| Psmg1 | LWNEWCR | -102 | -108 | 0 | 0.00051306 | 7 | 362.2441 | NaN | 932.33673 | NaN | NaN | 1096.9503 |
| Psmg1 | IAEDQQFQWLEK | -142 | -153 | 0 | 0.00025015 | 12 | 811.6304 | NaN | 502.09613 | NaN | NaN | NaN |
| Psmg1 | VFGFQPR | -154 | -160 | 0 | 0.003620005 | 7 | 220.05522 | NaN | NaN | NaN | NaN | NaN |
| P4hb | YKPESDELTAE | -329 | -339 | 1 | 0 | 11 | 4202.2734 | 4577.4507 | 4513.6553 | 4658.9165 | 4664.1826 | 640.3233 |
| P4hb | YKPESDELTAE | -329 | -339 | 1 | 0 | 11 | 4202.2734 | 4577.4507 | 4513.6553 | 4658.9165 | 4664.1826 | 640.3233 |
| P4hb | LEEEDNVLVLK | -22 | -32 | 0 | 0 | 11 | 3563.302 | 287.17285 | 1189.6609 | 4412.027 | 1058.394 | 1171.6605 |
| P4hb | EADDIVNWLK | -123 | -132 | 0 | 0.00025015 | 10 | 3279.2493 | 3675.1423 | 3859.4019 | 3757.5764 | 4619.297 | 4616.1577 |
| P4hb | LGETYKDHENIIIAK | -412 | -426 | 1 | 0 | 15 | 3196.561 | 3687.213 | 3589.1028 | 2891.293 | 3495.4211 | 3963.0776 |
| P4hb | LGETYKDHENIIIAK | -412 | -426 | 1 | 0 | 15 | 3196.561 | 3687.213 | 3589.1028 | 2891.293 | 3495.4211 | 3963.0776 |
| P4hb | NNFEGEITK | -216 | -224 | 0 | 0 | 9 | 2548.684 | 2723.3843 | 2737.3074 | 1827.6996 | 2021.2915 | 2180.0981 |
| P4hb | YKPESDELTAEK | -329 | -340 | 1 | 0 | 12 | 2220.8633 | 2450.117 | 2603.3281 | 2925.8682 | 3869.166 | 3888.381 |
| P4hb | MDSTANEVEAVK | -427 | -438 | 0 | 0 | 12 | 2091.4932 | 1459.983 | 2439.8494 | 2293.661 | 721.62994 | 1300.9393 |
| P4hb | DHENIIIAK | -418 | -426 | 0 | 0.001092529 | 9 | 1986.3589 | 2082.489 | 2178.301 | 529.4405 | 2203.3884 | 2182.8728 |
| P4hb | TVIDYNGER | -455 | -463 | 0 | 0 | 9 | 1979.5984 | 200.11519 | 1577.7618 | 1442.8015 | 763.2811 | 1103.0358 |
| P4hb | FIFIDSDHTDNQR | -290 | -302 | 0 | 0 | 13 | 1938.7668 | 2216.115 | 2248.1191 | 2992.4146 | 3436.9756 | 3422.3835 |
| P4hb | FIFIDSDHTDNQR | -290 | -302 | 0 | 0 | 13 | 1938.7668 | 2216.115 | 2248.1191 | 2992.4146 | 3436.9756 | 3422.3835 |
| P4hb | NFEEALAAHK | -35 | -44 | 0 | 0 | 10 | 1731.4968 | 1279.5864 | 1381.4779 | 2455.3987 | 2563.0293 | 2445.1394 |
| P4hb | SQEVPEDWDKQPVK | -359 | -372 | 1 | 0 | 14 | 1625.1202 | 1626.36 | 1470.7253 | 1559.5054 | 1517.7689 | 1536.8655 |
| P4hb | SQEVPEDWDKQPVK | -359 | -372 | 1 | 0 | 14 | 1625.1202 | 1626.36 | 1470.7253 | 1559.5054 | 1517.7689 | 1536.8655 |
| P4hb | NFEEVAFDEKK | -378 | -388 | 1 | 0 | 11 | 1423.8348 | 694.5378 | 1587.9236 | 1266.6046 | 1577.6792 | 1586.8046 |
| P4hb | NFEEVAFDEK | -378 | -387 | 0 | 0 | 10 | 1282.344 | 1326.6405 | 1385.6956 | 1272.754 | 4255.2266 | 1221.4418 |
| P4hb | ILEFFGLK | -303 | -310 | 0 | 0.000431655 | 8 | 1169.1647 | 1419.3656 | 1512.7671 | 1418.9268 | 126.72655 | 2091.7695 |
| P4hb | DALEEEDNVLVLK | -20 | -32 | 0 | 0 | 13 | 1119.3237 | 2727.5173 | 2497.4146 | 1487.8323 | 2792.8184 | 2729.2173 |
| P4hb | KSNFEEALAAHK | -33 | -44 | 1 | 0 | 12 | 1078.9314 | 1059.0881 | 1167.6809 | 1152.5854 | 998.5719 | 771.892 |
| P4hb | THILLFLPK | -257 | -265 | 0 | 0.00025015 | 9 | 1020.8569 | 863.53674 | 882.6472 | 1783.5784 | 2400.9524 | 2167.7961 |
| P4hb | LKAEGSEIR | -72 | -80 | 1 | 0 | 9 | 853.5978 | 862.2811 | 979.7365 | 1107.5123 | 1180.5204 | 1196.1204 |
| P4hb | HNQLPLVIEFTEQTAPK | -233 | -249 | 0 | 0 | 17 | 812.9457 | 759.6798 | 789.0011 | 1272.3928 | 1174.7322 | 1135.7224 |
| P4hb | VGANFEEVAFDEK | -375 | -387 | 0 | 0 | 13 | 796.2293 | 897.36597 | 955.97003 | 1295.7659 | 1404.3896 | 1528.7925 |
| P4hb | HNQLPLVIEF | -233 | -242 | 0 | 0 | 10 | 772.1212 | 662.471 | 716.0425 | 687.9554 | 673.19684 | 725.1752 |
| P4hb | EVPEDWDKQPVK | -361 | -372 | 1 | 0 | 12 | 724.65985 | 1279.1984 | 2880.4075 | 2861.8306 | 371.13907 | 3395.8494 |
| P4hb | YQLDKDGVVLFK | -198 | -209 | 1 | 0 | 12 | 702.9196 | 1054.079 | 1158.5516 | 806.2338 | 1058.773 | 1152.6741 |
| P4hb | YQLDKDGVVLF | -198 | -208 | 1 | 0 | 11 | 682.37634 | 1299.9597 | 1494.064 | 1386.8394 | 1406.7521 | 1472.4733 |
| P4hb | VGANFEEVAFDEKK | -375 | -388 | 1 | 0 | 14 | 595.5837 | 585.02924 | 286.67776 | 415.30243 | 934.0488 | 1021.3317 |
| P4hb | KEECPAVR | -311 | -318 | 1 | 0 | 8 | 571.0593 | 512.954 | 532.4424 | 733.6181 | 629.14575 | 559.89606 |
| P4hb | LGETYKDHENIIIA | -412 | -425 | 1 | 0 | 14 | 570.34033 | 543.7233 | 572.00836 | 744.8389 | 678.99176 | 476.98917 |
| P4hb | FLESGGQDGAGDDEDLDLEEA | -471 | -491 | 0 | 0.000431655 | 21 | 310.60303 | 305.31314 | 263.00858 | 368.49115 | 406.95734 | 428.3659 |
| P4hb | VHSFPTLK | -439 | -446 | 0 | 0.000712251 | 8 | 1695.7692 | 1756.1406 | 1500.5378 | NaN | 993.3157 | 824.9524 |
| P4hb | QLAPIWDK | -404 | -411 | 0 | 0.000230707 | 8 | 1437.4956 | NaN | 734.06793 | 268.77698 | 448.50162 | 859.17194 |
| P4hb | DSTANEVEAVK | -428 | -438 | 0 | 0 | 11 | 1013.7662 | 262.7023 | 993.3197 | 890.012 | NaN | 944.07135 |
| P4hb | KDHENIIIAK | -417 | -426 | 1 | 0 | 10 | 897.7997 | 677.70654 | 601.15625 | 528.09326 | 340.1662 | NaN |
| P4hb | KDHENIIIAK | -417 | -426 | 1 | 0 | 10 | 897.7997 | 677.70654 | 601.15625 | 528.09326 | 340.1662 | NaN |
| P4hb | KVHSFPTLK | -438 | -446 | 1 | 0 | 9 | 633.1129 | 678.91925 | 822.53784 | 1198.0253 | NaN | 621.8814 |
| P4hb | FKDVESDSAK | -163 | -172 | 1 | 0 | 10 | 469.8822 | 665.5292 | NaN | 1130.3422 | 1267.9768 | 1185.5736 |
| P4hb | ITEFCHR | -341 | -347 | 0 | 0.000627943 | 7 | 1947.5818 | 1866.3069 | NaN | 899.9413 | NaN | 1954.898 |
| P4hb | HNQLPLVIEFTEQ | -233 | -245 | 0 | 0 | 13 | 556.00525 | 576.782 | 535.7306 | NaN | NaN | 719.6889 |
| P4hb | SQEVPEDWDK | -359 | -368 | 0 | 0 | 10 | 2422.7766 | 2689.1707 | 2666.73 | NaN | NaN | NaN |
| P4hb | SNFEEALAAHK | -34 | -44 | 0 | 0 | 11 | 1051.0758 | 1162.9037 | 1379.9902 | NaN | NaN | NaN |
| P4hb | EECPAVR | -312 | -318 | 0 | 0.00078661 | 7 | 980.27106 | 1114.2001 | NaN | 1211.0308 | NaN | NaN |
| P4hb | EVTVIGFFK | -156 | -164 | 0 | 0.00025015 | 9 | 748.5282 | 646.0826 | 772.5314 | NaN | NaN | NaN |
| P4hb | FLESGGQDGAGDDEDLDLEE | -471 | -490 | 0 | 0.000230707 | 20 | 357.32684 | 331.07858 | 357.83118 | NaN | NaN | NaN |
| P4hb | ILLFLPK | -259 | -265 | 0 | 0.009002904 | 7 | 348.8257 | NaN | NaN | NaN | NaN | 623.91254 |
| P4hb | SLVDSSEVTVIGFFK | -150 | -164 | 0 | 0 | 15 | 308.9448 | NaN | 242.97275 | NaN | NaN | NaN |
| Smyd5 | WFTPEGFR | -235 | -242 | 0 | 0 | 8 | 449.8876 | 1243.4132 | 1329.2909 | 1814.5308 | 921.0651 | 1266.1096 |
| Smyd5 | LTGKPSQILPHPELC | -87 | -101 | 1 | 0.003003218 | 15 | 3493.0432 | 542.6934 | NaN | NaN | 873.05383 | NaN |
| Smyd5 | DALELTPQDR | -268 | -277 | 0 | 0 | 10 | 1279.5637 | 825.5974 | NaN | 1675.6584 | NaN | NaN |
| Smyd5 | TANQEQAIVHK | -196 | -206 | 0 | 0.00078661 | 11 | 350.5463 | NaN | NaN | NaN | NaN | NaN |
| Nudt5 | GAEQHLTVDAK | -185 | -195 | 0 | 0 | 11 | 1620.7701 | 1713.2489 | 1869.695 | 1771.1111 | 2013.2322 | 2089.311 |
| Nudt5 | SKPFEVPFLK | -208 | -217 | 1 | 0.00111006 | 10 | 1210.7817 | 1012.99274 | 127.678604 | 873.8472 | 1418.2454 | 1313.5204 |
| Nudt5 | TLHHECVILVK | -70 | -80 | 0 | 0 | 11 | 816.6789 | 714.508 | 430.02356 | 849.83966 | 994.0896 | 680.3326 |
| Nudt5 | DALGAEQHLTVDAK | -182 | -195 | 0 | 0.000431655 | 14 | 733.52966 | 266.7716 | 1624.509 | 1894.0333 | 1168.0234 | 1343.0764 |
| Nudt5 | HANSKPFEVPFLK | -205 | -217 | 1 | 0.000989247 | 13 | 651.81573 | 911.2755 | 864.6113 | 1405.7548 | 749.6934 | 748.1666 |
| Nudt5 | HLVTSEELISEGK | -14 | -26 | 0 | 0 | 13 | 1920.8037 | 2226.7913 | NaN | 2434.614 | 2440.7864 | 2386.528 |
| Nudt5 | RLEEETGYK | -110 | -119 | 1 | 0.00025015 | 10 | 1069.5312 | 1102.448 | NaN | NaN | 1176.5872 | 285.6799 |
| Nudt5 | LDALGAEQHLTVDAK | -181 | -195 | 0 | 0 | 15 | 537.5346 | 508.50305 | 574.1769 | NaN | NaN | NaN |
| Nudt5 | SVIPVLQR | -62 | -69 | 0 | 0.004439092 | 8 | 466.34885 | NaN | NaN | NaN | NaN | NaN |
| Mia2 | LSPLLPGGGGR | -1142 | -1152 | 0 | 0 | 11 | 8094.932 | 7926.8955 | 7799.088 | NaN | 8447.209 | 8428.767 |
| Mia2 | DALNENSQLQESQK | -807 | -820 | 0 | 0 | 14 | 250.73563 | 208.49063 | 250.77953 | NaN | NaN | 246.65742 |
| Ikzf3 | SHTGERPFQCN | -138 | -148 | 1 | 0.00025015 | 11 | 988.57007 | 1018.8474 | 1205.5712 | 758.32263 | 1450.3069 | 1170.2969 |
| Ikzf3 | SVYPIALTR | -339 | -347 | 0 | 0 | 9 | 746.58374 | 587.7414 | NaN | 548.0991 | 479.94977 | 873.99634 |
| Ctbp1 | VSTLQDLLFHSD | -220 | -231 | 0 | 0.00051306 | 12 | 554.75934 | 346.9629 | 356.7953 | NaN | NaN | 524.1356 |
| Golph3 | DAPTGDVLLDEALK | -125 | -138 | 0 | 0 | 14 | 3554.8215 | NaN | 2700.2651 | 5773.5986 | 2273.1416 | 2261.2583 |

| | | | | | | | | | | | |
|---|---|---|---|---|---|---|---|---|---|---|---|
| Mier1 | GVEAIPEGSHIK | -239 | -250 | 0 | 0 | 12 | 501.52524 | 550.80914 | 278.62534 | 601.11084 | 549.51514 | 556.13165 |
| Mier1 | DAQEIIRPR | -142 | -150 | 1 | 0.000947459 | 9 | 2232.3394 | 2736.253 | 3482.1553 | NaN | 3611.1067 | 3345.6917 |
| Arhgdib | LNYKPPPQK | -21 | -29 | 1 | 0 | 9 | 8053.409 | 7780.409 | 7590.0933 | 5867.162 | 6826.439 | 6250.8403 |
| Arhgdib | ELQEMDKDDESLTK | -33 | -46 | 1 | 0 | 14 | 4800.8496 | 3543.403 | 3695.24 | 3117.2825 | 5471.559 | 3906.072 |
| Arhgdib | ELQEMDKDDESLTK | -33 | -46 | 1 | 0 | 14 | 4800.8496 | 3543.403 | 3695.24 | 3117.2825 | 5471.559 | 3906.072 |
| Arhgdib | DAQPQLEEADDDLDSK | -5 | -20 | 0 | 0 | 16 | 4334.7793 | 5076.1978 | 4974.3364 | 4326.2153 | 3100.2424 | 3082.2546 |
| Arhgdib | SFFTDDDKQDHLTW | -175 | -188 | 1 | 0.000431655 | 14 | 2747.2373 | 3236.7847 | 3615.3777 | 2350.461 | 2823.933 | 2856.829 |
| Arhgdib | SFFTDDDKQDHLTW | -175 | -188 | 1 | 0.000431655 | 14 | 2747.2373 | 3236.7847 | 3615.3777 | 2350.461 | 2823.933 | 2856.829 |
| Arhgdib | LTGDLEALKK | -86 | -95 | 1 | 0.00025015 | 10 | 1993.114 | 1889.065 | 1199.2012 | 1013.04584 | 1715.252 | 1323.4473 |
| Arhgdib | LEEADDDLDSK | -10 | -20 | 0 | 0 | 11 | 1741.2407 | 1709.3074 | 1794.4861 | 427.11194 | 470.48486 | 323.70544 |
| Arhgdib | LEEADDDLDSK | -10 | -20 | 0 | 0 | 11 | 1741.2407 | 1709.3074 | 1794.4861 | 427.11194 | 470.48486 | 323.70544 |
| Arhgdib | DLTGDLEALKK | -85 | -95 | 1 | 0 | 11 | 1581.7924 | 874.12946 | 271.31287 | 2185.7742 | 2200.7566 | 2018.6781 |
| Arhgdib | SYGPRPEEYEFLTPVEEAPK | -144 | -163 | 1 | 0 | 20 | 1470.3862 | 1527.5251 | 1781.162 | 1357.1252 | 1580.6296 | 1765.897 |
| Arhgdib | SYGPRPEEYEFLTPVEEAPK | -144 | -163 | 1 | 0 | 20 | 1470.3862 | 1527.5251 | 1781.162 | 1357.1252 | 1580.6296 | 1765.897 |
| Arhgdib | VGSYGPRPEEYEFLTPVEEAPK | -142 | -163 | 1 | 0 | 22 | 1186.7542 | 1257.6355 | 1460.6022 | 1628.4816 | 1861.1548 | 2019.3641 |
| Arhgdib | TLLGDVPVVADPTVPNVTVTR | -50 | -70 | 0 | 0.00122419 | 21 | 1184.1254 | 1154.9679 | 956.51013 | 844.71515 | 922.3928 | 896.89996 |
| Arhgdib | KDTFVLK | -95 | -101 | 1 | 0 | 7 | 584.866 | 564.3975 | 514.1947 | 666.57587 | 629.2195 | 465.60394 |
| Arhgdib | DVPVVADPTVPNVTVTR | -54 | -70 | 0 | 0 | 17 | 415.20804 | 337.2218 | 485.345 | 373.81216 | 624.62463 | 587.6902 |
| Arhgdib | LLGDVPVVADPTVPNVTVTR | -51 | -70 | 0 | 0.00025015 | 20 | 322.3362 | 250.1738 | 389.26324 | 407.98474 | 320.2025 | 349.94357 |
| Arhgdib | GSYGPRPEEYEFLTPVEEAPK | -143 | -163 | 1 | 0 | 21 | 190.0473 | 1090.8247 | 893.1233 | 1480.9523 | 979.8678 | 1735.277 |
| Arhgdib | EFLTPVEEAPK | -153 | -163 | 0 | 0 | 11 | 3554.7654 | 3838.3303 | 4088.9258 | 4014.3652 | 534.9149 | NaN |
| Arhgdib | KTLLGDVPVVADPTVPNVTVT | -49 | -69 | 1 | 0.00025015 | 21 | 305.214 | 313.8504 | NaN | 331.34494 | 427.16852 | 322.37128 |
| Arhgdib | KKDTFVLK | -94 | -101 | 2 | 0.00025015 | 8 | 166.36868 | 135.34592 | 561.39056 | NaN | 182.99443 | 438.48657 |
| Arhgdib | SFFTDDDKQDHL | -175 | -186 | 1 | 0 | 12 | 808.36053 | 941.4128 | 900.2003 | 654.1035 | NaN | NaN |
| Arhgdib | EYEFLTPVEEAPK | -151 | -163 | 0 | 0 | 13 | 1003.2066 | 1132.5038 | 1083.9331 | NaN | NaN | NaN |
| Arhgdib | PVEEAPK | -157 | -163 | 0 | 0.001868731 | 7 | 744.37305 | 354.0518 | 317.80594 | NaN | NaN | NaN |
| Arhgdib | KYVQHTYR | -123 | -130 | 1 | 0 | 8 | 157.32329 | 238.46927 | 124.721504 | NaN | NaN | NaN |
| Arhgdib | KYVQHTYR | -123 | -130 | 1 | 0 | 8 | 157.32329 | 238.46927 | 124.721504 | NaN | NaN | NaN |
| Gfpt1 | WATHGEPNPVNSHPQR | -96 | -111 | 0 | 0 | 16 | 1833.249 | 809.6174 | 177.66505 | 662.5796 | 652.73004 | 682.53033 |
| Gfpt1 | GYDFESETDTETIAK | -140 | -154 | 0 | 0 | 15 | 493.53094 | 560.3497 | 511.84882 | 761.43353 | 614.86633 | 588.81226 |
| Gfpt1 | DASAVIEHTNR | -278 | -288 | 0 | 0 | 11 | 2422.0593 | 2450.646 | NaN | 253.6775 | 2348.8818 | 458.7452 |
| Gfpt1 | VIFLEDDDVAAVVDGR | -289 | -304 | 0 | 0 | 16 | 108.124405 | NaN | NaN | 274.9826 | NaN | 340.36884 |
| Cdkn2d | DASGLTPLELAR | -136 | -147 | 0 | 0 | 12 | 940.693 | 1140.1318 | 1192.7162 | 801.7064 | 499.89615 | NaN |
| Isoc2a | DASHPQFK | -171 | -178 | 0 | 0 | 8 | 1135.3 | 1122.2053 | 1172.5193 | 1238.1824 | 1359.4066 | 1224.6023 |
| Isoc2a | NTALDLLHR | -117 | -125 | 0 | 0 | 9 | 703.3178 | 1339.5607 | 1333.41 | 1347.5189 | 1380.9785 | 1394.9725 |
| Sp100 | DHEFITGK | -52 | -59 | 0 | 0.000907206 | 8 | 2811.6377 | 1056.4381 | 1381.2775 | 2306.8276 | 2861.541 | 2651.7556 |
| Sp100 | DATPESCSLLPQNEER | -207 | -222 | 0 | 0 | 16 | 993.67175 | 1079.507 | 1159.7017 | 1348.1477 | NaN | 250.11484 |
| Sp100 | RDHEFITGK | -51 | -59 | 1 | 0.001636916 | 9 | 357.48428 | 528.7513 | NaN | 942.616 | 331.90045 | NaN |
| Sp100 | TTVPSQPAPAYSR | -288 | -300 | 0 | 0.00025015 | 13 | 1995.0504 | 2101.758 | 1051.9109 | NaN | NaN | NaN |
| Sp100 | TTTVPSQPAPAYSR | -287 | -300 | 0 | 0.00051306 | 14 | 1017.5404 | 1081.6285 | 1230.2667 | NaN | NaN | NaN |
| Sp100 | FVQLVDVK | -236 | -243 | 0 | 0.00383302 | 8 | 472.8838 | 319.0333 | 329.7577 | NaN | NaN | NaN |
| Sp100 | GGDTSDTESSIIIR | -310 | -323 | 0 | 0 | 14 | 2412.0007 | NaN | NaN | 2342.1543 | NaN | NaN |
| Wdr77 | KDTPPPLVPPAAR | -3 | -15 | 1 | 0.00025015 | 13 | 223.94167 | 189.78871 | 185.55481 | 378.52438 | 220.34962 | 204.37209 |
| Wdr77 | ILLWDTR | -192 | -198 | 0 | 0.001743443 | 7 | 500.3487 | 632.35095 | 543.7874 | NaN | 330.0148 | NaN |
| Wdr77 | DATWSPLNHSLLT | -302 | -314 | 0 | 0.003490217 | 13 | 1790.2903 | NaN | NaN | NaN | NaN | NaN |
| Eif4e | DAVTHIGR | -174 | -181 | 0 | 0 | 8 | 2923.2388 | NaN | 3134.0571 | 1849.5267 | 1911.6367 | 1930.4457 |
| Ero1a | GAVDESLSEETQK | -137 | -149 | 0 | 0 | 13 | 608.57855 | 982.75446 | 1013.41956 | 600.9489 | 1445.4458 | 1829.5071 |
| Ero1a | DCAVKPCHSDEVPDGIK | -98 | -114 | 1 | 0.00025015 | 17 | 275.37717 | 431.5464 | 472.77353 | 541.0651 | 530.42426 | 504.69318 |
| Ero1a | DIQSPDAEYVDLLLNPER | -169 | -186 | 0 | 0 | 18 | 142.61436 | 189.75465 | 180.3439 | 293.44308 | 225.65276 | 237.7976 |
| Ero1a | SFPLHFDEN | -349 | -357 | 0 | 0.00025015 | 9 | 982.3798 | NaN | 1095.5298 | 262.72736 | 836.0589 | 807.2815 |
| Ero1a | FDGILTEGEGPR | -284 | -295 | 0 | 0 | 12 | 1025.3167 | 121.62596 | 1031.5648 | NaN | 1414.9915 | NaN |
| Ero1a | LLESDYFR | -68 | -75 | 0 | 0 | 8 | 955.7383 | NaN | 913.5445 | NaN | 1018.01874 | 906.5264 |
| Ero1a | IEECEQAER | -127 | -135 | 0 | 0 | 9 | 489.33563 | 282.1311 | 454.39874 | NaN | NaN | 1168.5148 |
| Ero1a | VLPFFERPDFQ | -314 | -324 | 1 | 0 | 11 | 442.44778 | 378.27734 | 513.5695 | NaN | 721.84656 | NaN |
| Ero1a | WNDINQCGR | -88 | -96 | 0 | 0 | 9 | 381.08167 | NaN | NaN | 798.671 | 719.4076 | 688.1876 |
| Ero1a | LQDTWLEK | -264 | -271 | 0 | 0 | 8 | 717.0845 | NaN | NaN | 675.4358 | 783.038 | NaN |
| Ero1a | LGAVDESLSEETQK | -136 | -149 | 0 | 0 | 14 | 225.7319 | 402.46573 | 839.7259 | NaN | NaN | NaN |
| Ero1a | HSDEVPDGIK | -105 | -114 | 0 | 0 | 10 | 206.54372 | NaN | NaN | 3790.8464 | 3212.187 | NaN |
| Apobec3 | NVQDPETQQNLCR | -144 | -156 | 0 | 0 | 13 | 952.26556 | 848.50806 | 922.23425 | 1216.3828 | 1195.2507 | 1477.8306 |
| Apobec3 | LYNVQDPETQQNLCR | -142 | -156 | 0 | 0 | 15 | 728.01025 | 844.9653 | 812.82697 | 769.7519 | 746.4733 | 751.4719 |
| Apobec3 | SPCFECAEQIVR | -114 | -125 | 0 | 0 | 12 | 726.8851 | 1176.72 | 1005.2085 | 915.2842 | 1007.0437 | 893.3577 |
| Apobec3 | DRPDLILHIY | -340 | -349 | 1 | 0 | 10 | 317.3482 | 537.2155 | 608.66425 | 385.65802 | 269.3954 | 652.532 |
| Apobec3 | RPFWPWK | -395 | -401 | 1 | 0.001023236 | 7 | 260.82465 | 425.69003 | 253.48242 | 1253.6091 | 401.91068 | 544.679 |
| Apobec3 | DCDSPVSLHHGVFK | -63 | -76 | 0 | 0 | 14 | 955.04626 | 1049.8549 | 977.87384 | NaN | 500.59076 | 1026.3293 |
| Apobec3 | SWSPCFECAEQIVR | -112 | -125 | 0 | 0 | 14 | 139.7337 | 192.65381 | 287.6278 | NaN | NaN | NaN |
| Serinc3 | DCDVLVGFK | -88 | -96 | 0 | 0 | 9 | 912.81256 | 1849.7643 | 1919.8716 | 1476.6198 | 844.16797 | 1284.3832 |
| Serinc3 | IPGFCEGGFQIK | -68 | -79 | 0 | 0 | 12 | 633.21155 | 705.67236 | 547.8126 | NaN | 1146.019 | 643.5323 |
| Banf1 | KDEDLFR | -54 | -60 | 1 | 0 | 7 | 5582.75 | 5780.2754 | 5749.6675 | 5451.04 | 5711.13 | 5571.78 |
| Banf1 | DFVAEPMGEKPVGSLA | -9 | -24 | 1 | 0 | 16 | 5417.787 | 4863.68 | 4760.0693 | 1861.6301 | 4785.1084 | 3654.0125 |
| Banf1 | DCFGCLR | -76 | -82 | 0 | 0.001023236 | 7 | 2034.1582 | 1254.6306 | 1875.8967 | 2731.392 | 1730.1775 | 1936.8016 |
| Banf1 | LREWCDAFL | -81 | -89 | 1 | 0.00051306 | 9 | 1971.5486 | 2368.0544 | 2421.066 | 2184.836 | 377.45355 | 2636.9993 |
| Banf1 | KKDEDLFR | -53 | -60 | 2 | 0.000230707 | 8 | 1351.5625 | 1164.2727 | 297.96155 | 2525.9302 | 1249.0784 | 1358.9761 |
| Banf1 | REWCDAFL | -82 | -89 | 1 | 0.000431655 | 8 | 798.49585 | 809.4888 | 518.51044 | 516.6729 | 533.93036 | 615.03986 |
| Banf1 | GEKPVGSLAGIGDVLSK | -16 | -32 | 1 | 0 | 17 | 713.66644 | 762.62396 | 741.88293 | 1695.5361 | 2151.6746 | 2073.671 |
| Banf1 | EWCDAFL | -83 | -89 | 0 | 0.003620005 | 7 | 1885.6808 | NaN | NaN | 3245.0256 | 4403.754 | 4477.9688 |
| Sting1 | LILPGLQAR | -169 | -177 | 0 | 0.000230707 | 9 | 1699.2765 | 1766.9066 | 1835.2128 | 1265.7181 | 1264.9563 | 1366.3357 |
| Sting1 | PYSNLHPAIPR | -2 | -12 | 0 | 0 | 11 | 1164.1168 | 1186.2366 | 1227.5021 | 818.5485 | 886.3883 | 903.1945 |
| Sting1 | PYSNLHPAIPR | -2 | -12 | 0 | 0 | 11 | 1164.1168 | 1186.2366 | 1227.5021 | 818.5485 | 886.3883 | 903.1945 |

| | | | | | | | | | | | | |
|---|---|---|---|---|---|---|---|---|---|---|---|---|
| Sting1 | SHELGLLLK | -53 | -61 | 0 | 0.000431655 | 9 | 1121.9215 | 819.8092 | 952.50165 | 860.96075 | 1330.2054 | 1210.3596 |
| Sting1 | EEILEDVPESR | -295 | -305 | 0 | 0 | 11 | 6363.951 | NaN | 7344.676 | 7965.161 | 328.61783 | 130.84486 |
| Sting1 | TLEEILEDVPESR | -293 | -305 | 0 | 0 | 13 | 1068.0664 | 1059.5605 | NaN | 1036.4397 | 1358.535 | 1398.5197 |
| Sting1 | NLCCLAEELCHVQ | -62 | -74 | 0 | 0 | 13 | 967.66675 | 652.85657 | NaN | NaN | 401.08124 | 2064.8674 |
| Sting1 | DCGVPDNLSVVDPNIR | -204 | -219 | 0 | 0.00025015 | 16 | 768.656 | NaN | NaN | 730.08923 | 554.6708 | NaN |
| Dis3l2 | DCIFTIDPSTAR | -375 | -386 | 0 | 0.001392869 | 12 | 154.26779 | NaN | NaN | NaN | NaN | NaN |
| Creld2 | DCIQCEVGWAR | -211 | -221 | 0 | 0 | 11 | 919.1051 | 728.45935 | 740.7422 | 600.7082 | 661.6331 | 923.0748 |
| Creld2 | ESGQCTDIDECSLEEK | -282 | -297 | 0 | 0 | 16 | 426.5865 | 600.0735 | 498.53986 | 282.24857 | 673.7068 | 591.28986 |
| Smc3 | DLQDELAGNSEQR | -311 | -323 | 0 | 0 | 13 | 1098.8877 | 162.42 | 1013.5829 | 172.56393 | 776.65027 | 524.5348 |
| Smc3 | FVLSDEFSHLRPEQR | -47 | -61 | 1 | 0 | 15 | 760.6656 | 761.3166 | 922.276 | 693.5449 | 957.21765 | 992.9795 |
| Smc3 | SEDLDNSIDKTEAGIK | -880 | -895 | 1 | 0 | 16 | 634.16376 | 725.4219 | 668.2664 | 743.51746 | 868.0165 | 880.2386 |
| Smc3 | DCITLEGDQVSHR | -649 | -661 | 0 | 0 | 13 | 555.60895 | 619.0789 | 600.29004 | 734.67255 | 759.1752 | 887.0963 |
| Smc3 | TTSELEAINKR | -862 | -872 | 1 | 0 | 11 | 3405.8213 | NaN | 3343.3613 | 2211.583 | 3663.384 | 3453.1086 |
| Smc3 | SVPSVDQFTGVGIR | -1086 | -1099 | 0 | 0 | 14 | 879.0343 | 814.67694 | 86.725296 | NaN | 993.6465 | 992.97516 |
| Smc3 | HNVIVGR | -27 | -33 | 0 | 0.004595494 | 7 | 661.2782 | NaN | 670.14716 | 645.86835 | 181.1529 | 657.86145 |
| Smc3 | EEEKEELAQYQK | -204 | -215 | 1 | 0 | 12 | 416.05157 | 264.28705 | 554.0419 | NaN | 442.7083 | 440.51035 |
| Smc3 | LDQDLNEVK | -437 | -445 | 0 | 0 | 9 | 9193.422 | 10195.5 | 8078.9414 | NaN | NaN | 2774.4219 |
| Smc3 | TATTSELEAINKR | -860 | -872 | 1 | 0 | 13 | 3379.8494 | NaN | NaN | 775.2531 | 1963.729 | 1046.7007 |
| Smc3 | VETYLNENLR | -828 | -837 | 0 | 0 | 10 | 3309.3347 | 4492.121 | 4189.0903 | 5655.5503 | NaN | NaN |
| Smc3 | DLEDTEANKEK | -419 | -429 | 1 | 0.000471409 | 11 | 1421.0022 | NaN | 1698.874 | 945.8949 | 1195.8713 | NaN |
| Smc3 | LFYHIVDSDEVSTK | -558 | -571 | 0 | 0 | 14 | 1152.989 | 1411.816 | 1178.7559 | NaN | NaN | 794.8214 |
| Smc3 | VDALNDEIR | -797 | -805 | 0 | 0 | 9 | 5356.369 | 5516.852 | 5951.448 | NaN | NaN | NaN |
| Smc3 | DLDNSIDKTEAGIK | -882 | -895 | 1 | 0.00149795 | 14 | 539.9356 | 574.26294 | 811.7519 | NaN | NaN | NaN |
| Smc3 | IKLEGIITR | -819 | -827 | 1 | 0.001392869 | 9 | 221.06514 | 335.60193 | 380.41467 | NaN | NaN | NaN |
| Smc3 | LALLHEGTGPR | -62 | -72 | 0 | 0 | 11 | 749.78204 | 919.39496 | NaN | NaN | NaN | NaN |
| Itm2b | DCKDPGDVVPVGQR | -37 | -50 | 1 | 0 | 14 | 1187.0697 | 1232.3026 | 1380.9843 | 2085.6118 | 1739.8315 | 1526.9641 |
| Itm2b | DPGDVVPVGQR | -40 | -50 | 0 | 0 | 11 | 1358.419 | 1255.0005 | 1267.771 | 1043.9211 | 2175.2896 | NaN |
| Itm2b | DVILNEPSADAPAAR | -97 | -111 | 0 | 0.001023236 | 15 | 1328.8447 | 1349.1768 | 1248.0028 | NaN | NaN | NaN |
| Cpsf7 | DCLHGIEAK | -398 | -406 | 0 | 0.000471409 | 9 | 1180.897 | 415.3407 | NaN | NaN | 1924.8123 | NaN |
| Gnl1 | DCPGLIFPSLLPR | -411 | -423 | 0 | 0.001636916 | 13 | 296.1445 | NaN | NaN | 308.95746 | NaN | 356.44434 |
| Hpcal1 | DCPTGHLTVDEFK | -37 | -49 | 0 | 0 | 13 | 898.97754 | 1124.7739 | 551.6197 | 1166.093 | 1159.1531 | 1002.9231 |
| Hpcal1 | EHTEFTDHELQEWYK | -18 | -32 | 0 | 0 | 15 | 545.7137 | 620.099 | 646.0965 | 650.97485 | 1197.0359 | 717.3611 |
| Hpcal1 | EHTEFTDHELQEWYK | -18 | -32 | 0 | 0 | 15 | 545.7137 | 620.099 | 646.0965 | 650.97485 | 1197.0359 | 717.3611 |
| Hpcal1 | LRPEVLQDLR | -8 | -17 | 1 | 0.00025015 | 10 | 824.3336 | 989.89667 | 964.1889 | 506.155 | 822.6495 | NaN |
| Hpcal1 | LRPEVLQDL | -8 | -16 | 1 | 0.002068213 | 9 | 466.2509 | NaN | 415.84473 | 591.87415 | NaN | 202.12537 |
| Trim28 | LSPPYSSPQEFAQDVGR | -751 | -767 | 0 | 0 | 17 | 1797.4952 | 2180.6506 | 2181.0496 | 2622.4697 | 3355.3281 | 3280.225 |
| Trim28 | IVAERPGTN | -409 | -417 | 1 | 0 | 9 | 1792.1816 | 1512.555 | 1724.6237 | 1310.1439 | 1280.9734 | 1347.0957 |
| Trim28 | LDLDLTSDSQPPVFK | -493 | -507 | 0 | 0 | 15 | 1487.3953 | 1445.3325 | 1389.4607 | 1237.4453 | 1579.3545 | 2262.3198 |
| Trim28 | SAPVSGPGILDDSATICR | -612 | -629 | 0 | 0 | 18 | 1302.963 | 1761.4346 | 1344.839 | 738.0696 | 1478.7942 | 1491.6415 |
| Trim28 | GSDDPYSSAEPHVSGMK | -453 | -469 | 0 | 0 | 17 | 1276.5734 | 1039.3573 | 1120.8229 | 1192.7024 | 1333.8777 | 1399.0579 |
| Trim28 | DIVENYFMR | -129 | -137 | 0 | 0 | 9 | 500.88467 | 594.9269 | 550.5482 | 494.5208 | 615.842 | 625.51514 |
| Trim28 | FCESCDTLTCR | -221 | -231 | 0 | 0 | 11 | 328.26166 | 364.84436 | 365.4886 | 686.2064 | 737.20154 | 727.9044 |
| Trim28 | DHQYQFLEDAVR | -240 | -251 | 0 | 0 | 12 | 2329.8208 | 2825.4153 | NaN | 3094.934 | 5123.768 | 6364.514 |
| Trim28 | ELLEHCGVCR | -61 | -70 | 0 | 0 | 10 | 1005.0003 | 374.5174 | NaN | 1134.6909 | 812.66797 | 1221.0066 |
| Trim28 | VFPGSTTEDYNLIVIER | -508 | -524 | 0 | 0 | 17 | 753.37994 | NaN | 766.02386 | 791.47675 | 943.1508 | 938.4082 |
| Trim28 | EEETEAAIGAPPAAPEGPETKPVLMPLTEGPGAEGPR | -555 | -591 | 1 | 0.000826806 | 37 | 354.9022 | 370.77747 | 337.0899 | 343.3351 | NaN | 219.67992 |
| Trim28 | TVYCNVHK | -207 | -214 | 0 | 0.000230707 | 8 | 3232.4934 | 480.67154 | 3419.9177 | 3576.1792 | NaN | NaN |
| Trim28 | EQPGGTLDLTLIR | -732 | -744 | 0 | 0 | 13 | 1133.6935 | NaN | 1313.5891 | NaN | 948.18176 | 346.47485 |
| Trim28 | SAASGSPGSGEGSAGGEK | -18 | -35 | 0 | 0 | 18 | 660.73517 | 654.9502 | 715.2462 | NaN | NaN | 829.5343 |
| Trim28 | EEDGSLSLDGADSTGVVAK | -677 | -695 | 0 | 0 | 19 | 638.3547 | NaN | 465.58783 | 632.0268 | NaN | NaN |
| Trim28 | DCQLNAHK | -232 | -239 | 0 | 0.005668207 | 8 | 461.35907 | NaN | NaN | NaN | NaN | NaN |
| Magoh;Magol | SVMEELKR | (55);(55) | (62);(62) | 1 | 0.00383302 | 8 | 5998.77 | 8357.523 | 7116.117 | 5238.3833 | 2758.6926 | 4683.9966 |
| Magoh;Magol | SVMEELKR | (55);(55) | (62);(62) | 1 | 0.00383302 | 8 | 5998.77 | 8357.523 | 7116.117 | 5238.3833 | 2758.6926 | 4683.9966 |
| Magoh;Magol | SLIGLHFK | (135);(135) | (142);(142) | 0 | 0.00025015 | 8 | 1187.4452 | 1361.6573 | 1348.0825 | 1264.1135 | 1164.3663 | 1157.5929 |
| Magoh;Magol | IIDDSEITKE | (63);(63) | (72);(72) | 1 | 0.00383302 | 10 | 675.9679 | 974.19495 | 815.8525 | 1067.3672 | 850.7093 | 168.16382 |
| Magoh;Magol | IIDDSEITKEDDALWPPPDR | (63);(63) | (82);(82) | 1 | 0.00025015 | 20 | 578.65924 | 501.13248 | 682.9673 | 321.20172 | 327.69424 | 825.59186 |
| Magoh;Magol | IGSLIDVNQSK | (104);(104) | (114);(114) | 0 | 0.000298805 | 11 | 1005.2983 | 647.9034 | 604.117 | NaN | 1100.4581 | 663.905 |
| Magoh;Magol | DDALWPPPDR | (73);(73) | (82);(82) | 0 | 0.001636916 | 10 | 1960.74 | 1268.3088 | 1018.8689 | NaN | NaN | 1128.1173 |
| Hnrnpc | LKGDDLQAIK | -189 | -198 | 1 | 0.00025015 | 10 | 4729.0283 | 4728.5205 | 4678.8833 | 4555.4834 | 4860.7197 | 4830.9043 |
| Hnrnpc | KSDVEAIFSK | -30 | -39 | 1 | 0.000907206 | 10 | 2086.716 | 2162.3982 | 2211.1687 | 1347.8711 | 1245.144 | 1326.5197 |
| Hnrnpc | QKVDSLLESLEK | -206 | -217 | 1 | 0 | 12 | 2073.0933 | 2829.6956 | 2513.5188 | 3794.7637 | 4265.991 | 4328.3735 |
| Hnrnpc | VDSLLESLEK | -208 | -217 | 0 | 0 | 10 | 1702.5497 | 1773.1925 | 202.89642 | 1926.0076 | 2102.4631 | 2198.4275 |
| Hnrnpc | SFDLDYDFQR | -121 | -130 | 0 | 0.00025015 | 10 | 1195.9253 | 1227.2676 | 1323.9229 | 1533.3385 | 1755.9886 | 1679.6409 |
| Hnrnpc | QADLSFSSPVEMK | -225 | -237 | 0 | 0 | 13 | 1163.2777 | 289.0979 | 367.6305 | 1210.7284 | 1123.7135 | 966.62683 |
| Hnrnpc | DDDDNEDRGDDQLELK | -276 | -291 | 1 | 0.001868731 | 16 | 967.38745 | 985.8645 | 974.6492 | 1102.4581 | 1315.4233 | 1278.8856 |
| Hnrnpc | VLDINLAAEPK | -79 | -89 | 0 | 0 | 11 | 904.5171 | 948.907 | 956.90186 | 1577.8788 | 1188.2765 | 1118.5507 |
| Hnrnpc | LKGDDLQAIKK | -189 | -199 | 2 | 0 | 11 | 817.0559 | 882.826 | 801.52954 | 617.7076 | 643.37976 | 832.10583 |
| Hnrnpc | LKGDDLQAIKK | -189 | -199 | 2 | 0 | 11 | 817.0559 | 882.826 | 801.52954 | 617.7076 | 643.37976 | 832.10583 |
| Hnrnpc | SEEEQSSASVK | -241 | -251 | 0 | 0 | 11 | 661.1509 | 749.1886 | 820.94385 | 796.46106 | 929.8033 | 685.9335 |
| Hnrnpc | NEKSEEEQSSASVK | -238 | -251 | 1 | 0 | 14 | 269.8296 | 262.1796 | 261.60248 | 207.80132 | 249.30228 | 262.75366 |
| Hnrnpc | DDEKEPEEGEDDRD | -292 | -305 | 2 | 0 | 14 | 234.88268 | 224.7529 | 261.63287 | 331.37698 | 340.4565 | 312.56284 |
| Hnrnpc | RGDDQLELK | -283 | -291 | 1 | 0.00078661 | 9 | 2339.479 | 2481.4358 | NaN | 3550.5996 | 1947.6283 | 1817.4056 |
| Hnrnpc | SDVEAIFSK | -31 | -39 | 0 | 0 | 9 | 1320.5277 | NaN | 1577.0178 | NaN | NaN | NaN |
| Psma1 | LDGLEERPQR | -233 | -242 | 1 | 0.000230707 | 10 | 3389.0166 | 3134.0664 | 3466.7246 | 3431.346 | 2326.307 | 2516.286 |
| Psma1 | ETLPAEQDLTTK | -197 | -208 | 0 | 0 | 12 | 3137.518 | 2403.9856 | 2567.1853 | 2058.0933 | 1197.2798 | 1293.9526 |
| Psma1 | FVFDRPLPVSR | -97 | -107 | 1 | 0 | 11 | 2933.615 | 3360.5967 | 3332.6445 | 2463.5374 | 3453.6604 | 3488.673 |
| Psma1 | FVFDRPLPVSR | -97 | -107 | 1 | 0 | 11 | 2933.615 | 3360.5967 | 3332.6445 | 2463.5374 | 3453.6604 | 3488.673 |
| Psma1 | IYDDDDVSPFLDGLEERPQR | -223 | -242 | 1 | 0 | 20 | 680.55396 | 650.0789 | 688.8892 | 948.4514 | 1003.91125 | 623.05817 |

| | | | | | | | | | | | | |
|---|---|---|---|---|---|---|---|---|---|---|---|---|
| Psma1 | IYDDDDVSPFLDGLEERPQR | -223 | -242 | 1 | 0 | 20 | 680.55396 | 650.0789 | 688.8892 | 948.4514 | 1003.91125 | 623.05817 |
| Psma1 | DDDDVSPFLDGLEERPQR | -225 | -242 | 1 | 0 | 18 | 929.6586 | 962.9475 | 1110.5638 | 971.0099 | 1166.6586 | 1130.1967 |
| Psma1 | DDDDVSPFLDGLEERPQR | -225 | -242 | 1 | 0 | 18 | 929.6586 | 962.9475 | 1110.5638 | 971.0099 | 1166.6586 | 1130.1967 |
| Psma1 | YDNDVTVWSPQGR | -6 | -18 | 0 | 0 | 13 | 563.714 | 546.8505 | 713.7256 | 900.76544 | 704.1798 | 881.31696 |
| Psma1 | QYDNDVTVWSPQGR | -5 | -18 | 0 | 0 | 14 | 320.2868 | 807.927 | 203.3985 | 1558.7299 | 802.5927 | 792.31006 |
| Psma1 | RETLPAEQDLTTK | -196 | -208 | 1 | 0.000471409 | 13 | 2493.9165 | NaN | 2432.0867 | 4825.382 | 5172.891 | 4989.44 |
| Psma1 | NQYDNDVTVWSPQGR | -4 | -18 | 0 | 0 | 15 | 1486.8353 | 1549.1582 | NaN | 1618.9333 | 1879.6213 | 2014.9489 |
| Psma1 | NVSIGIVGK | -209 | -217 | 0 | 0.001092529 | 9 | 314.8464 | 452.54456 | NaN | NaN | NaN | NaN |
| Psma1 | IHQIEYAMEA | -19 | -28 | 0 | 0.002595029 | 10 | 593.9703 | NaN | NaN | NaN | NaN | NaN |
| Usp14 | GGTLKDDDWGNIK | -50 | -62 | 1 | 0 | 13 | 4720.247 | 3628.874 | 4149.7847 | 1660.6531 | 3093.6128 | 4141.6904 |
| Usp14 | FDDDKVSIVTPEDILR | -449 | -464 | 1 | 0 | 16 | 509.69907 | 520.25366 | 680.094 | 652.89526 | 835.5459 | 780.1827 |
| Usp14 | VSIVTPEDILR | -454 | -464 | 0 | 0.00025015 | 11 | 3128.3533 | 2647.9421 | NaN | 3167.176 | 2120.8247 | 3195.5637 |
| Usp14 | FKDLEDK | -374 | -380 | 1 | 0.000712251 | 7 | 1628.512 | 1474.7382 | 1538.5652 | 972.9102 | 1346.2567 | NaN |
| Usp14 | DDDKVSIVTPEDILR | -450 | -464 | 1 | 0 | 15 | 719.31354 | 757.31573 | 765.6552 | NaN | 1062.9886 | 1082.5032 |
| Usp14 | LSGGGDWHIA | -465 | -474 | 0 | 0.000431655 | 10 | 1005.3356 | 85.52613 | 1119.9143 | 1123.886 | NaN | NaN |
| Usp14 | ETDSSSAPAVTPSK | -225 | -238 | 0 | 0 | 14 | 689.18066 | NaN | 172.23921 | 590.68634 | 813.96716 | NaN |
| Usp14 | YVLLYGPR | -475 | -482 | 0 | 0.006100831 | 8 | 757.5758 | NaN | NaN | NaN | NaN | NaN |
| Tubg1;Tubg2 | IHEDIFDIIDR | (114);(114) | (124);(124) | 0 | 0 | 11 | 3048.9043 | 3221.73 | 3592.9314 | 1803.5315 | 992.222 | 2011.2664 |
| Tubg1;Tubg2 | DDEHYIPR | (56);(56) | (63);(63) | 0 | 0.002434616 | 8 | 558.42236 | 532.7159 | NaN | NaN | NaN | NaN |
| Rpl26 | HFNAPSHIR | -18 | -26 | 0 | 0 | 9 | 6802.353 | 7609.3433 | 7250.6504 | 2986.7166 | 3945.2324 | 4534.3066 |
| Rpl26 | HFNAPSHIR | -18 | -26 | 0 | 0 | 9 | 6802.353 | 7609.3433 | 7250.6504 | 2986.7166 | 3945.2324 | 4534.3066 |
| Rpl26 | TVHVGIHPSK | -94 | -103 | 0 | 0 | 10 | 5147.2827 | 5283.6978 | 5539.989 | 5196.5615 | 5398.5864 | 6014.564 |
| Rpl26 | TVHVGIHPSK | -94 | -103 | 0 | 0 | 10 | 5147.2827 | 5283.6978 | 5539.989 | 5196.5615 | 5398.5864 | 6014.564 |
| Rpl26 | KDDEVQVVR | -51 | -59 | 1 | 0.00025015 | 9 | 3084.8328 | 3780.146 | 3673.289 | 3777.0508 | 5546.54 | 5235.8633 |
| Rpl26 | YKEETIEK | -135 | -142 | 1 | 0.00025015 | 8 | 9713.109 | 9257.464 | 10287.619 | 222.81792 | NaN | 12306.867 |
| Rpl26 | VHVGIHPSK | -95 | -103 | 0 | 0 | 9 | 751.90076 | 636.6138 | NaN | 758.28723 | 660.0161 | 682.2211 |
| Rpl26 | DDEVQVVR | -52 | -59 | 0 | 0.007224385 | 8 | 1361.066 | NaN | NaN | NaN | NaN | NaN |
| Btf3l4 | SQLGADSLTSLR | -103 | -114 | 0 | 0 | 12 | 186.01796 | 190.07455 | 735.3974 | 390.59766 | 449.437 | 481.884 |
| Btf3l4 | DDGTVIHFNNPK | -62 | -73 | 0 | 0 | 12 | 210.52487 | 653.94794 | 2194.439 | 3000.212 | 745.4101 | NaN |
| Hnrnpa2b1 | FDDHDPVDK | -160 | -168 | 0 | 0 | 9 | 9028.706 | 9181.005 | 9874.395 | 8912.32 | 10106.232 | 10222.71 |
| Hnrnpa2b1 | SYNDFGNYNQQPSNYGPMK | -299 | -317 | 0 | 0 | 19 | 8720.86 | 8229.343 | 7682.967 | 10786.052 | 252.07849 | 7974.5894 |
| Hnrnpa2b1 | SFETTEESLR | -29 | -38 | 0 | 0 | 10 | 8592.51 | 9526.117 | 9263.391 | 8126.732 | 8988.932 | 8529.249 |
| Hnrnpa2b1 | IDTIEIITDR | -138 | -147 | 0 | 0 | 10 | 8184.1235 | 6908.9775 | 8888.823 | 11051.019 | 11391.243 | 10873.882 |
| Hnrnpa2b1 | TIEIITDR | -140 | -147 | 0 | 0.00025015 | 8 | 4315.8413 | 3805.344 | 3986.535 | 3398.0679 | 3118.37 | 3133.9958 |
| Hnrnpa2b1 | NYYEQWGK | -39 | -46 | 0 | 0 | 8 | 2679.923 | 2799.3179 | 2901.5515 | 569.88745 | 3873.729 | 359.78802 |
| Hnrnpa2b1 | NYYEQWGK | -39 | -46 | 0 | 0 | 8 | 2679.923 | 2799.3179 | 2901.5515 | 569.88745 | 3873.729 | 359.78802 |
| Hnrnpa2b1 | GGGNYGPGGSGGSGGYGGR | -332 | -350 | 0 | 0 | 19 | 3508.4644 | 4166.1113 | 3478.9812 | 3118.026 | 591.4869 | 3754.9658 |
| Hnrnpa2b1 | EESGKPGAHVT | -100 | -110 | 1 | 0 | 11 | 1971.3634 | 1761.4663 | 2244.483 | 2070.209 | 2057.1858 | 2282.896 |
| Hnrnpa2b1 | GGGPGGGNFGGSPGYGGGR | -248 | -266 | 0 | 0 | 19 | 1610.9116 | 1588.1627 | 1229.6147 | 1262.6879 | 1025.4856 | 964.7081 |
| Hnrnpa2b1 | GKIDTIEIITDR | -136 | -147 | 1 | 0 | 12 | 1532.2828 | 1793.4229 | 1920.7688 | 1537.4933 | 1787.7234 | 1901.6326 |
| Hnrnpa2b1 | GGNFGFGDSR | -204 | -213 | 0 | 0 | 10 | 1518.4098 | 1675.0239 | 1476.4641 | 805.71094 | 244.29596 | 869.53827 |
| Hnrnpa2b1 | GLSFETTEESLR | -27 | -38 | 0 | 0 | 12 | 1352.5443 | 1083.4752 | 1363.0487 | 1685.538 | 1104.619 | 1607.8752 |
| Hnrnpa2b1 | DYFEEYGKID | -130 | -139 | 1 | 0 | 10 | 1216.5604 | 1296.2743 | 1325.0122 | 1521.9016 | 1390.3208 | 1461.9109 |
| Hnrnpa2b1 | DYFEEYGKID | -130 | -139 | 1 | 0 | 10 | 1216.5604 | 1296.2743 | 1325.0122 | 1521.9016 | 1390.3208 | 1461.9109 |
| Hnrnpa2b1 | GGGGNFGPGPGSNF | -214 | -227 | 0 | 0 | 14 | 1169.1113 | 955.32495 | 109.66415 | 1126.782 | 903.6577 | 951.6668 |
| Hnrnpa2b1 | DDHDPVDK | -161 | -168 | 0 | 0.00078661 | 8 | 859.915 | 719.57635 | 349.46277 | 799.04584 | 926.1848 | 778.941 |
| Hnrnpa2b1 | VTFDDHDPVDK | -158 | -168 | 0 | 0 | 11 | 790.9752 | 5789.66 | 5935.258 | 7733.6235 | 8131.4243 | 8045.7876 |
| Hnrnpa2b1 | VTFDDHDPVDK | -158 | -168 | 0 | 0 | 11 | 790.9752 | 5789.66 | 5935.258 | 7733.6235 | 8131.4243 | 8045.7876 |
| Hnrnpa2b1 | VTFDDHDPVDK | -158 | -168 | 0 | 0 | 11 | 790.9752 | 5789.66 | 5935.258 | 7733.6235 | 8131.4243 | 8045.7876 |
| Hnrnpa2b1 | DYFEEYGK | -130 | -137 | 0 | 0 | 8 | 778.70416 | 695.428 | 720.6352 | 748.5707 | 595.6055 | 627.7142 |
| Hnrnpa2b1 | GGPYGGGNYGPGGSGGSGGYGGR | -328 | -350 | 0 | 0 | 23 | 744.7146 | 846.65735 | 791.52014 | 848.7047 | 859.08545 | 892.83014 |
| Hnrnpa2b1 | GGGGNFGPGPGSNFRG | -214 | -229 | 1 | 0.000755556 | 16 | 723.1837 | 4210.0874 | 510.53094 | 3869.8774 | 3946.12 | 3975.481 |
| Hnrnpa2b1 | YGPGGSGGSGGYGGR | -336 | -350 | 0 | 0 | 15 | 593.3053 | 985.57336 | 662.9575 | 858.60614 | 1122.4095 | 1061.826 |
| Hnrnpa2b1 | EESGKPGAHVTVK | -100 | -112 | 1 | 0 | 13 | 580.7454 | 552.7358 | 728.75836 | 511.47726 | 596.2703 | 627.86884 |
| Hnrnpa2b1 | NMGGPYGGGNYGPGGSGGSGGYGGR | -326 | -350 | 0 | 0 | 25 | 426.37442 | 534.2352 | 492.53378 | 528.6438 | 574.7197 | 547.148 |
| Hnrnpa2b1 | YNDFGNYNQQPSNYGPMK | -300 | -317 | 0 | 0 | 18 | 272.39093 | 362.82617 | 470.88757 | 306.8559 | 506.92865 | 154.3056 |
| Hnrnpa2b1 | GFGDGYNGYGGGPGGGNFGGSPGYGGGR | -239 | -266 | 0 | 0 | 28 | 173.88263 | 219.17169 | 228.42659 | 201.64629 | 236.54224 | 264.0853 |
| Hnrnpa2b1 | GGGGNFGPGPGSNFR | -214 | -228 | 0 | 0 | 15 | 140.06654 | 7990.483 | 7926.6216 | 246.517 | 8360.217 | 8255.667 |
| Hnrnpa2b1 | GGYGGGGPGYGNQGGGY | -267 | -283 | 0 | 0 | 17 | 1027.857 | 867.5238 | 1239.103 | 585.40234 | NaN | NaN |
| Hnrnpa2b1 | TLETVPLER | -4 | -12 | 0 | 0 | 9 | 558.1577 | 501.43237 | 709.9843 | NaN | NaN | 717.42883 |
| Hnrnpa2b1 | QQPSNYGPMK | -308 | -317 | 0 | 0.00830926 | 10 | 8216.691 | NaN | NaN | NaN | NaN | NaN |
| Hnrnpa2b1 | GGYGGGGPGYGNQGGGYG | -267 | -284 | 0 | 0.00272025 | 18 | 412.35352 | NaN | NaN | NaN | NaN | NaN |
| Hnrnpa2b1 | YGGGNYGPGGSGGSGGYGGR | -331 | -350 | 0 | 0.00025015 | 20 | 351.30133 | NaN | NaN | NaN | NaN | NaN |
| Ramac | DDKEYQEYLK | -22 | -31 | 1 | 0 | 10 | 1124.705 | 1618.3201 | 1323.5591 | NaN | 1635.4773 | 1744.2036 |
| Suclg2 | INFDDNAEFR | -261 | -270 | 0 | 0 | 10 | 1557.7732 | 1506.95 | 1552.8868 | 1721.3411 | 1891.6486 | 1567.9525 |
| Suclg2 | GLPITSAVDLEDAAK | -410 | -424 | 0 | 0 | 15 | 1322.0443 | 1426.8987 | 1315.5703 | 1888.6508 | 1714.5238 | 1754.5558 |
| Suclg2 | PANFLDLGGGVK | -328 | -339 | 0 | 0 | 12 | 1095.5127 | 599.5126 | 1420.3854 | 460.69702 | 1423.3038 | 1396.8947 |
| Suclg2 | DDKSENEPIENEAAR | -278 | -292 | 1 | 0.00025015 | 15 | 750.44336 | 596.78186 | 1239.4763 | 557.85016 | 833.1521 | 980.9784 |
| Suclg2 | LYHLFLK | -229 | -235 | 0 | 0 | 7 | 214.31984 | 2306.413 | 2564.9268 | NaN | 581.6826 | 724.4193 |
| Suclg2 | EQIDIFEGIK | -192 | -201 | 0 | 0 | 10 | 810.8108 | 755.5694 | 742.69794 | 264.8897 | NaN | NaN |
| Tsr1 | QIDAPVDPFPLNPR | -301 | -314 | 0 | 0 | 14 | 1734.8247 | 1800.0144 | 785.4017 | 891.3649 | 1092.8254 | 368.32877 |
| Tsr1 | DGPPHQVLVVPL | -79 | -90 | 0 | 0.00078661 | 12 | 1033.3412 | 1100.0365 | 1134.7761 | 969.8363 | 1353.9438 | 1473.7827 |
| Tsr1 | HVSDVPVSVVEYFR | -560 | -573 | 0 | 0 | 14 | 646.7953 | 696.49493 | 841.97235 | 887.039 | 1135.9374 | 1156.3297 |
| Tsr1 | DDLYDEKVDAEDEER | -455 | -469 | 1 | 0.002199523 | 15 | 784.19336 | NaN | 1041.1436 | NaN | 1026.4781 | 1020.81335 |
| Tsr1 | ISLPEAFK | -94 | -101 | 0 | 0.004983786 | 8 | 248.60832 | 331.0487 | NaN | NaN | NaN | NaN |
| Sub1 | ISDIDDAVR | -117 | -125 | 0 | 0 | 9 | 2204.552 | 2498.7734 | 2533.7317 | 2070.713 | 1816.3395 | 1346.6807 |
| Sub1 | GISLNMEQWSQLK | -102 | -114 | 0 | 0 | 13 | 1966.553 | 823.13446 | 2077.9692 | 1078.0134 | 1216.9801 | 1289.6702 |
| Sub1 | QAVPEKPVK | -30 | -38 | 1 | 0 | 9 | 1852.3271 | 1941.4725 | 2158.5613 | 2192.7612 | 2184.129 | 2090.111 |

| | | | | | | | | | | | | |
|---|---|---|---|---|---|---|---|---|---|---|---|---|
| Sub1 | QAVPEKPVK | -30 | -38 | 1 | 0 | 9 | 1852.3271 | 1941.4725 | 2158.5613 | 2192.7612 | 2184.129 | 2090.111 |
| Sub1 | DDNMFQIGK | -60 | -68 | 0 | 0 | 9 | 1671.1843 | 1597.4154 | 1603.6605 | 598.48065 | 554.22675 | 1776.5189 |
| Sub1 | DDNMFQIGK | -60 | -68 | 0 | 0 | 9 | 1671.1843 | 1597.4154 | 1603.6605 | 598.48065 | 554.22675 | 1776.5189 |
| Sub1 | NMEQWSQLK | -106 | -114 | 0 | 0 | 9 | 556.5664 | 540.3225 | 279.48358 | 614.95123 | 1389.3479 | 463.47443 |
| Sub1 | NMEQWSQLK | -106 | -114 | 0 | 0 | 9 | 556.5664 | 540.3225 | 279.48358 | 614.95123 | 1389.3479 | 463.47443 |
| Sub1 | MEQWSQLK | -107 | -114 | 0 | 0 | 8 | 969.1141 | 858.9346 | 858.6352 | 1121.8804 | 884.3628 | 918.3781 |
| Sub1 | EYWMDSEGEMKPGR | -87 | -100 | 1 | 0 | 14 | 561.7808 | 517.1804 | 575.76996 | 700.89514 | 808.8537 | 814.2425 |
| Sub1 | EYWMDSEGEMKPGR | -87 | -100 | 1 | 0 | 14 | 561.7808 | 517.1804 | 575.76996 | 700.89514 | 808.8537 | 814.2425 |
| Sub1 | EYWMDSEGEMKPGR | -87 | -100 | 1 | 0 | 14 | 561.7808 | 517.1804 | 575.76996 | 700.89514 | 808.8537 | 814.2425 |
| Sub1 | EQISDIDDAVR | -115 | -125 | 0 | 0 | 11 | 569.08234 | 6702.687 | 7110.0166 | 6528.5864 | 6624.486 | 6791.52 |
| Sub1 | KQAVPEKPVK | -29 | -38 | 2 | 0.00025015 | 10 | 319.53317 | 293.9525 | 247.14339 | 240.21843 | 287.96722 | 327.51337 |
| Sub1 | QAVPEKPVKK | -30 | -39 | 2 | 0 | 10 | 317.0743 | 321.88373 | 325.3432 | 369.76282 | 357.20786 | 331.90356 |
| Sub1 | LKEQISDIDDAVR | -113 | -125 | 1 | 0 | 13 | 1495.6858 | 1851.9426 | NaN | 686.3569 | 2012.3585 | 2206.37 |
| Calu | HLVYESDQNK | -272 | -281 | 0 | 0 | 10 | 2502.2793 | 591.03174 | 2667.1821 | 2013.8976 | 2509.5095 | 2866.4082 |
| Calu | WIHEDVER | -104 | -111 | 0 | 0 | 8 | 1148.7694 | 2131.2014 | 2641.9329 | 340.92438 | 3578.4163 | 2631.6545 |
| Calu | SHDGNADEPEWVK | -219 | -231 | 0 | 0 | 13 | 1002.4989 | 934.44415 | 966.284 | 1154.8546 | 1192.2443 | 1157.236 |
| Calu | LHPEEYDYMK | -180 | -189 | 0 | 0 | 10 | 800.3983 | 812.4548 | 752.38947 | 862.56384 | 767.4139 | 776.2689 |
| Calu | DWILPSDYDHAEAEAR | -256 | -271 | 0 | 0 | 16 | 723.84076 | 237.99718 | 608.38763 | 215.42758 | 1063.0474 | 93.70065 |
| Calu | DWILPSDYDHAEAEAR | -256 | -271 | 0 | 0 | 16 | 723.84076 | 237.99718 | 608.38763 | 215.42758 | 1063.0474 | 93.70065 |
| Calu | LDDPDPDDGFNYK | -138 | -150 | 0 | 0 | 13 | 709.47577 | 780.32764 | 880.193 | 962.8154 | 1052.7758 | 1048.3049 |
| Calu | VHHEPQLSDK | -28 | -37 | 0 | 0.000627943 | 10 | 581.4631 | 586.2921 | 477.85126 | 797.3208 | 781.1488 | NaN |
| Calu | DYDHAEAEAR | -262 | -271 | 0 | 0 | 10 | 101.12011 | 121.84369 | 466.83743 | 391.6293 | NaN | 425.13095 |
| Calu | DYDHDAFLGAEEAK | -46 | -59 | 0 | 0.00025015 | 14 | 321.1619 | 416.04266 | 585.84735 | NaN | 460.4601 | NaN |
| Calu | GHDLNEDGLVSWEEYK | -115 | -130 | 0 | 0 | 16 | 316.76987 | NaN | 450.4815 | 395.5067 | NaN | 571.7992 |
| Calu | SFDQLTPEESK | -60 | -70 | 0 | 0 | 11 | 1290.736 | 566.83673 | 1573.5862 | NaN | NaN | NaN |
| Calu | DDPDPDDGFNYK | -139 | -150 | 0 | 0 | 12 | 634.49164 | 541.72797 | NaN | NaN | NaN | NaN |
| Skp1 | NDFTEEEEAQVR | -143 | -154 | 0 | 0 | 12 | 1813.2583 | 2099.821 | 2000.1744 | 2037.0101 | 1428.4014 | 1357.786 |
| Skp1 | RTDDIPVWDQEFLK | -81 | -94 | 1 | 0 | 14 | 1175.0813 | 1353.7312 | 1411.2983 | 788.41797 | 1078.9008 | 1104.95 |
| Skp1 | RTDDIPVWDQEFLK | -81 | -94 | 1 | 0 | 14 | 1175.0813 | 1353.7312 | 1411.2983 | 788.41797 | 1078.9008 | 1104.95 |
| Skp1 | TDDIPVWDQEFLK | -82 | -94 | 0 | 0 | 13 | 655.40643 | 658.93005 | 666.0849 | 606.3628 | 475.3719 | 811.2178 |
| Skp1 | KENQWCEEK | -155 | -163 | 1 | 0.000230707 | 9 | 600.7109 | NaN | 826.10803 | 781.25287 | 756.48035 | 807.8063 |
| Skp1 | DDPPPPEDDENKEK | -67 | -80 | 1 | 0.00025015 | 14 | 366.43387 | 393.72766 | 488.37878 | NaN | NaN | 465.1361 |
| Skp1 | RTDDIPVWDQ | -81 | -90 | 1 | 0.007493591 | 10 | 378.14014 | NaN | NaN | NaN | NaN | NaN |
| Ptprcap | LSHASGGYYHPA | -57 | -68 | 0 | 0.00025015 | 12 | 10818.299 | 9433.161 | 10638.101 | 864.11786 | 1401.4895 | 1567.5248 |
| Ptprcap | WDDSAGGAGGQGLR | -180 | -193 | 0 | 0.000230707 | 14 | 1555.7126 | 374.66153 | 971.9618 | 1106.118 | 1910.1265 | 1765.4412 |
| Ptprcap | GGPEEAAAKEEEQR | -119 | -132 | 1 | 0 | 14 | 1194.8335 | 1258.0568 | 1148.3883 | 1018.8998 | 1332.9078 | 1233.0526 |
| Ptprcap | SGSAAWDDSAGGAGGQGLR | -175 | -193 | 0 | 0 | 19 | 1192.5424 | 1442.872 | 1350.5784 | 1236.8948 | 1166.3345 | 1425.4915 |
| Ptprcap | GGYYHPAR | -62 | -69 | 0 | 0 | 8 | 1180.3995 | 1160.466 | 1266.602 | 646.9696 | 1269.3079 | 565.283 |
| Ptprcap | LLWASPAGR | -81 | -89 | 0 | 0.00078661 | 9 | 1342.4058 | 988.1963 | 1088.3363 | 776.6526 | NaN | NaN |
| Ptprcap | DDSAGGAGGQGLR | -181 | -193 | 0 | 0 | 13 | 463.14215 | NaN | NaN | 601.8989 | NaN | 1155.256 |
| Septin11 | FESDPATHNEPGVR | -66 | -79 | 0 | 0 | 14 | 4404.886 | 4171.109 | 3891.9226 | 4184.3916 | 3863.4785 | 3808.453 |
| Septin11 | SLFNYHDTR | -138 | -146 | 0 | 0 | 9 | 980.64185 | 1045.5488 | 1008.2663 | 1220.5194 | 1344.0732 | 1277.3129 |
| Septin11 | KFESDPATHNEPGVR | -65 | -79 | 1 | 0 | 15 | 810.08356 | 810.50995 | 800.9845 | 853.3942 | 931.5767 | 944.3238 |
| Septin11 | SLSGHVGFDSLPDQLVNK | -17 | -34 | 0 | 0 | 18 | 298.42688 | 515.3955 | 459.1062 | 468.8995 | 667.34186 | 545.13214 |
| Septin11 | SLSGHVGFDSLPDQLVNK | -17 | -34 | 0 | 0 | 18 | 298.42688 | 515.3955 | 459.1062 | 468.8995 | 667.34186 | 545.13214 |
| Septin11 | DDSYKPIVEYIDAQ | -111 | -124 | 1 | 0 | 14 | 264.00113 | 231.63487 | 673.1863 | NaN | 776.00244 | 884.5642 |
| Bub3 | LNQPPEDGISSVK | -9 | -21 | 0 | 0 | 13 | 2761.5132 | 2982.3218 | 1449.8903 | 2665.25 | 3225.0127 | 1556.1309 |
| Bub3 | IWDPFNK | -260 | -266 | 0 | 0.001023236 | 7 | 2213.4868 | 2310.3938 | 2472.7234 | 2291.2188 | 2305.862 | 2534.5461 |
| Bub3 | FYDPTHAWSGGLDHQLK | -64 | -80 | 0 | 0 | 17 | 433.2239 | 510.0892 | 491.4101 | 650.73724 | 150.53238 | 670.5712 |
| Bub3 | FYDPTHAWSGGLDHQLK | -64 | -80 | 0 | 0 | 17 | 433.2239 | 510.0892 | 491.4101 | 650.73724 | 150.53238 | 670.5712 |
| Bub3 | YQHTGAVLDCA | -53 | -63 | 0 | 0 | 11 | 400.73398 | 212.09387 | 441.59982 | 553.3325 | 557.14453 | 656.14294 |
| Bub3 | QVTDAETKPK | -315 | -324 | 1 | 0 | 10 | 319.90106 | 403.5268 | 381.6204 | 292.77344 | 385.96365 | 394.27832 |
| Bub3 | VTGSWDQTVK | -112 | -121 | 0 | 0 | 10 | 1973.2571 | 1909.4543 | 2046.6481 | 2924.9812 | 2677.0007 | NaN |
| Bub3 | DDTEHPEDGIFIR | -302 | -314 | 0 | 0 | 13 | 1398.7385 | 591.98364 | NaN | 3351.3743 | 4633.676 | 1146.8491 |
| Bub3 | DDTEHPEDGIFIR | -302 | -314 | 0 | 0 | 13 | 1398.7385 | 591.98364 | NaN | 3351.3743 | 4633.676 | 1146.8491 |
| Bub3 | NTDQENLVGTHDAPIR | -85 | -100 | 0 | 0 | 16 | 527.81836 | 874.3419 | 608.0251 | NaN | 606.05865 | 472.4707 |
| Bub3 | YLDPSPEVQK | -207 | -216 | 0 | 0.000431655 | 10 | 291.815 | NaN | 585.40247 | 1292.3057 | NaN | 554.9608 |
| Bub3 | VLVWDLR | -159 | -165 | 0 | 0.001636916 | 7 | 806.063 | 577.0634 | 836.8756 | NaN | NaN | NaN |
| Bub3 | FSPNTSQFLLVS | -22 | -33 | 0 | 0.001392869 | 12 | 493.65555 | NaN | 504.83063 | NaN | 174.3631 | NaN |
| Cd6 | HFTPGQGPIHR | -214 | -224 | 0 | 0 | 11 | 540.0294 | 334.9561 | 474.00616 | NaN | 477.14017 | 556.1418 |
| Cd6 | DDTWDLQDAHVVCK | -186 | -199 | 0 | 0.000627943 | 14 | 1021.83344 | 796.58374 | 584.13983 | NaN | NaN | NaN |
| Cd6 | NYHPVPITIAK | -449 | -459 | 0 | 0 | 11 | 585.46545 | 614.39954 | 629.3671 | NaN | NaN | NaN |
| Fdps | TAPQGHVDLGR | -177 | -187 | 0 | 0 | 11 | 2239.6914 | 2403.7458 | 2623.4268 | 3378.0464 | 3439.9934 | 2503.8982 |
| Fdps | ELGHPEIGDAIAR | -32 | -44 | 0 | 0 | 13 | 1751.1803 | 1782.6313 | 1838.9656 | 1543.9902 | 1963.3003 | 1779.8972 |
| Fdps | ELGHPEIGDAIAR | -32 | -44 | 0 | 0 | 13 | 1751.1803 | 1782.6313 | 1838.9656 | 1543.9902 | 1963.3003 | 1779.8972 |
| Fdps | QILEENYGQKDPEK | -284 | -297 | 1 | 0 | 14 | 604.2906 | 621.0646 | 651.6882 | 871.85345 | 762.64417 | 674.4904 |
| Fdps | QVQDDYLDLFGDPSVTGK | -240 | -257 | 0 | 0 | 18 | 520.47455 | 489.9319 | 638.011 | 670.3082 | 807.90936 | 844.86554 |
| Fdps | DDYLDLFGDPSVTGK | -243 | -257 | 0 | 0 | 15 | 389.58575 | 492.1627 | 435.19305 | 139.96184 | 201.72372 | 268.3294 |
| Fdps | VGTDIQDNK | -258 | -266 | 0 | 0 | 9 | 967.21063 | NaN | 845.7452 | 656.57434 | 838.82056 | 1666.2722 |
| Fdps | LKEVLEYNAL | -45 | -54 | 1 | 0.000627943 | 10 | 498.49588 | 229.71283 | 643.17535 | 379.3382 | NaN | NaN |
| Fdps | YEEDSYNR | -317 | -324 | 0 | 0.000989247 | 8 | 801.5344 | NaN | 721.00323 | 454.5734 | NaN | NaN |
| Fdps | QNFIQHF | -15 | -21 | 0 | 0.004857192 | 7 | 734.5787 | NaN | NaN | 447.75162 | NaN | 934.0039 |
| Psmc3 | LKPGDLVGVNK | -148 | -158 | 1 | 0 | 11 | 3325.577 | 3390.2246 | 3570.1038 | 3742.8193 | 7818.039 | 3415.6477 |
| Psmc3 | DVDPNDQEEDGANIDLDSQR | -101 | -120 | 0 | 0 | 20 | 2435.5122 | 1543.076 | 2203.3403 | 309.7768 | 1692.0632 | 1133.4644 |
| Psmc3 | VDILDPALLR | -338 | -347 | 0 | 0 | 10 | 1520.5856 | 1529.8599 | 557.378 | 427.09534 | 377.11237 | 2461.0227 |
| Psmc3 | EVDERPTEQYSDIGGLDK | -179 | -196 | 1 | 0 | 18 | 1121.1249 | 1351.8759 | 1404.3289 | 1246.5907 | 1686.8481 | 1056.9501 |
| Psmc3 | EVDERPTEQYSDIGGLDK | -179 | -196 | 1 | 0 | 18 | 1121.1249 | 1351.8759 | 1404.3289 | 1246.5907 | 1686.8481 | 1056.9501 |
| Psmc3 | VWDEAEQDGIGEEVLK | -23 | -38 | 0 | 0 | 16 | 811.30725 | 892.1066 | 567.40204 | 1125.1866 | 1403.8557 | 1106.5038 |

| | | | | | | | | | | | | |
|---|---|---|---|---|---|---|---|---|---|---|---|---|
| Psmc3 | NRVDILDPALLR | -336 | -347 | 1 | 0 | 12 | 564.863 | 415.1592 | 499.67163 | 346.68164 | 569.78906 | 562.7759 |
| Psmc3 | DEAEQDGIGEEVLK | -25 | -38 | 0 | 0 | 14 | 352.55084 | 766.2132 | 727.4508 | 906.44617 | 611.1567 | 405.69952 |
| Psmc3 | ETLPTEYDSR | -165 | -174 | 0 | 0 | 10 | 1521.8223 | NaN | 885.3337 | 847.2798 | 1269.3367 | 851.5305 |
| Psmc3 | EKFENLGIQPPK | -213 | -224 | 1 | 0 | 12 | 770.4848 | 782.5222 | 969.07324 | NaN | 713.3173 | 834.64014 |
| Psmc3 | RVDILDPALLR | -337 | -347 | 1 | 0 | 11 | 539.13135 | 541.2051 | 616.8494 | NaN | 654.0535 | 653.94543 |
| Psmc3 | QTYFLPVIGLVDAEK | -133 | -147 | 0 | 0 | 15 | 134.96768 | 350.27386 | NaN | 194.63934 | 290.2076 | 369.002 |
| Psmc3 | MNVSPDVNYEELAR | -376 | -389 | 0 | 0 | 14 | 1914.1096 | NaN | NaN | NaN | 2681.498 | 2811.2742 |
| Nsfl1c | SYQDPSNAQFLESIR | -200 | -214 | 0 | 0 | 15 | 2791.7488 | 3876.2166 | 3840.3044 | 3430.6938 | 3819.2954 | 3624.5122 |
| Nsfl1c | DLIHDQDEEEEEEGQR | -77 | -93 | 0 | 0.00025015 | 17 | 631.68195 | 743.1876 | 726.16376 | 1035.5133 | 782.04175 | 719.208 |
| Nsfl1c | RGEVPAELR | -215 | -223 | 1 | 0.000907206 | 9 | 618.75696 | 1326.2532 | 1467.7319 | 2164.4197 | 1748.697 | 1645.8523 |
| Nsfl1c | ELADENQTLK | -347 | -356 | 0 | 0 | 10 | 1016.7505 | 826.07104 | 1348.2031 | 755.0132 | NaN | 661.0902 |
| Nsfl1c | SPGETSKPRPF | -140 | -150 | 2 | 0.001636916 | 11 | 863.54193 | 387.4548 | NaN | 699.2477 | 538.6031 | 750.3791 |
| Nsfl1c | SPNELVDDLFK | -114 | -124 | 0 | 0 | 11 | 659.44403 | 524.68787 | 444.9602 | NaN | 1368.9366 | 841.2052 |
| Nsfl1c | LGSTAPQVLNTSSPAQQAENEAK | -260 | -282 | 0 | 0 | 23 | 298.41687 | 399.8034 | 390.71378 | NaN | 516.925 | 462.71692 |
| Nsfl1c | DEDFVKPK | -240 | -247 | 1 | 0 | 8 | 3270.3257 | NaN | NaN | 1596.8341 | 2646.8386 | 1466.6326 |
| Nsfl1c | EAEPTTNIQIR | -291 | -301 | 0 | 0 | 11 | 1810.891 | 2339.6528 | 1382.6311 | 2880.4673 | NaN | NaN |
| Nsfl1c | EHGAVAVER | -128 | -136 | 0 | 0.00111006 | 9 | 737.43335 | 735.9021 | NaN | NaN | NaN | NaN |
| Rbm8a | NIHLNLDR | -101 | -108 | 0 | 0.000230707 | 8 | 2810.3892 | 3057.4995 | 2790.6206 | 3271.0408 | 2927.4424 | 2770.497 |
| Rbm8a | TLVEYETYK | -117 | -125 | 0 | 0 | 9 | 1674.7352 | 1678.6036 | 2589.587 | 1250.7898 | 613.5882 | 1645.0227 |
| Rbm8a | FAEYGEIK | -93 | -100 | 0 | 0 | 8 | 1424.104 | 570.77966 | 1120.5079 | 1234.5863 | 1216.6049 | 1351.5328 |
| Rbm8a | GVHEEATEEDIHDK | -79 | -92 | 0 | 0 | 14 | 1143.303 | 1164.2084 | 1288.9382 | 961.8996 | 1716.6962 | 1593.2792 |
| Rbm8a | GVHEEATEEDIHDK | -79 | -92 | 0 | 0 | 14 | 1143.303 | 1164.2084 | 1288.9382 | 961.8996 | 1716.6962 | 1593.2792 |
| Rbm8a | FVTGVHEEATEEDIHDK | -76 | -92 | 0 | 0.000471409 | 17 | 801.4153 | 817.6808 | 928.0409 | 1050.6047 | 1315.2544 | 449.25858 |
| Rbm8a | FVTGVHEEATEEDIHDK | -76 | -92 | 0 | 0.000471409 | 17 | 801.4153 | 817.6808 | 928.0409 | 1050.6047 | 1315.2544 | 449.25858 |
| Rbm8a | EDYDSVEQDGDEPGPQR | -52 | -68 | 0 | 0 | 17 | 292.63116 | 522.6921 | 424.2187 | 697.1846 | 892.9806 | 651.9071 |
| Rbm8a | GGEDFAMDEDGDESIHK | -11 | -27 | 0 | 0.00111006 | 17 | 249.28949 | 376.14612 | 347.74402 | 560.91003 | 612.00806 | 405.13336 |
| Rbm8a | GYTLVEYETYK | -115 | -125 | 0 | 0 | 11 | 439.7616 | 2141.8364 | 2255.7046 | NaN | 2092.9177 | 1925.82 |
| Rbm8a | GYTLVEYETYK | -115 | -125 | 0 | 0 | 11 | 439.7616 | 2141.8364 | 2255.7046 | NaN | 2092.9177 | 1925.82 |
| Rbm8a | YDSVEQDGDEPGPQR | -54 | -68 | 0 | 0 | 15 | 322.13892 | 430.01703 | 693.24 | NaN | NaN | NaN |
| Rbm8a | GFGSEEGSR | -39 | -47 | 0 | 0.007343498 | 9 | 440.9639 | 533.76855 | NaN | NaN | NaN | NaN |
| Rbm8a | DEDGDESIHK | -18 | -27 | 0 | 0.005522803 | 10 | 379.89438 | 588.06 | NaN | NaN | NaN | NaN |
| Ddx39a | NCPHVVVGTPGR | -163 | -174 | 0 | 0.00025015 | 12 | 6174.7583 | 4452.687 | 1212.4504 | 288.51346 | 3811.6482 | 3625.149 |
| Ddx39a | HFVLDECDK | -191 | -199 | 0 | 0 | 9 | 1285.8168 | 1334.8483 | 354.4756 | 236.00632 | 849.6573 | 838.9585 |
| Ddx39a | DYDEDEEPQAPQESTPAPPK | -12 | -31 | 0 | 0.00025015 | 20 | 287.761 | 287.2664 | 279.82275 | 325.58698 | 284.4475 | 373.28265 |
| Ddx39a | VENELLDYDEDEEPQAPQESTPAPPK | -6 | -31 | 0 | 0 | 26 | 403.90942 | NaN | 477.57178 | 781.3863 | 565.1684 | 598.9607 |
| Ddx39a | DEDVLKK | -156 | -162 | 1 | 0.002595029 | 7 | 1759.0687 | NaN | 1909.5787 | NaN | 1847.8525 | 1993.2561 |
| Ddx39a | EDEEPQAPQESTPAPPK | -15 | -31 | 0 | 0 | 17 | 679.0454 | 432.8891 | 283.20053 | NaN | NaN | 285.27487 |
| Ddx39a | LDYDEDEEPQAPQESTPAPPK | -11 | -31 | 0 | 0.000431655 | 21 | 321.89438 | 295.3958 | 333.83368 | NaN | NaN | NaN |
| Mtrex | DLRPVDNR | -747 | -754 | 1 | 0.000230707 | 8 | 2979.965 | 3238.884 | 2968.569 | 2673.4377 | 3177.0781 | 3767.3945 |
| Mtrex | DEDYIPLKPR | -117 | -126 | 1 | 0 | 10 | 1236.6213 | 434.7353 | 1520.8982 | 1424.1292 | 1596.8643 | 600.8012 |
| Mtrex | YCLPFLQPGR | -647 | -656 | 0 | 0 | 10 | 277.18613 | 522.0804 | 419.47122 | 601.89813 | 561.1206 | 453.44406 |
| Mtrex | EVALPADEDYIPLKPR | -111 | -126 | 1 | 0 | 16 | 957.43536 | NaN | 1093.2183 | 784.9725 | 745.9713 | 739.9317 |
| Mtrex | KLPQVEHVLPLLK | -435 | -447 | 1 | 0.00025015 | 13 | 3184.5942 | 2269.3503 | NaN | NaN | 600.3589 | 580.1011 |
| Mtrex | SNVKPNSGELDPLYVVE | -679 | -695 | 1 | 0.000230707 | 17 | 694.65784 | 940.715 | 1149.0886 | NaN | 1466.985 | NaN |
| Mtrex | QFAEWICHLHK | -289 | -299 | 0 | 0 | 11 | 349.2834 | 567.9792 | 435.04974 | NaN | NaN | 368.2077 |
| Prkcq | EGPVEIGLPCSTK | -313 | -325 | 0 | 0 | 13 | 5003.4365 | NaN | NaN | 2829.2405 | 1181.8033 | 981.66223 |
| Prkcq | DEEELFHSIR | -583 | -592 | 0 | 0 | 10 | 1091.3103 | NaN | 1141.6595 | NaN | NaN | NaN |
| Ppil2 | DLLTDEPFSR | -143 | -152 | 0 | 0 | 10 | 413.51804 | 434.2336 | 471.32446 | 458.1433 | 581.0061 | 691.0254 |
| Ppil2 | DEFRPNLSHTGR | -352 | -363 | 1 | 0 | 12 | 430.2644 | NaN | 607.9826 | NaN | 620.9883 | 671.606 |
| Thy1 | GIPEHTYR | -69 | -76 | 0 | 0.001636916 | 8 | 6974.9233 | 6907.329 | 6509.913 | 6474.737 | 5452.777 | 6463.911 |
| Thy1 | DNSIQHEFSLTR | -46 | -57 | 0 | 0 | 12 | 5367.519 | 5446.5386 | 5655.274 | 6284.85 | 1659.5543 | 7207.8677 |
| Thy1 | DNSIQHEFSLTR | -46 | -57 | 0 | 0 | 12 | 5367.519 | 5446.5386 | 5655.274 | 6284.85 | 1659.5543 | 7207.8677 |
| Thy1 | VTLSNQPYIK | -79 | -88 | 0 | 0 | 10 | 3155.6265 | 3019.2725 | 3250.2021 | 2604.2266 | 1940.0404 | 2175.0547 |
| Thy1 | DNSIQHEF | -46 | -53 | 0 | 0.001636916 | 8 | 2860.748 | 2638.0898 | 2770.3274 | 1907.9902 | 392.60583 | 1934.8849 |
| Thy1 | TLGIPEHTYR | -67 | -76 | 0 | 0 | 10 | 2443.7075 | 2707.1716 | 2342.1309 | 2676.782 | 2491.8425 | 2379.795 |
| Thy1 | TACLVNQNLR | -26 | -35 | 0 | 0 | 10 | 2357.7917 | 2234.6523 | 3146.8723 | 2873.342 | 2803.3901 | 4564.6313 |
| Thy1 | DNSIQHEFSLT | -46 | -56 | 0 | 0 | 11 | 1596.5487 | 1619.3182 | 1740.349 | 2712.7585 | 2687.7927 | 2869.0242 |
| Thy1 | DNSIQHEFSLT | -46 | -56 | 0 | 0 | 11 | 1596.5487 | 1619.3182 | 1740.349 | 2712.7585 | 2687.7927 | 2869.0242 |
| Thy1 | DEGDYFCEL | -99 | -107 | 0 | 0 | 9 | 1565.4489 | 1941.5653 | 2132.6318 | 1487.1547 | 2338.2656 | 2248.0266 |
| Thy1 | LGIPEHTYR | -68 | -76 | 0 | 0 | 9 | 1544.7462 | 1939.7896 | 1767.1736 | 2089.336 | 1856.3286 | 1838.3258 |
| Thy1 | SGTLGIPEHTYR | -65 | -76 | 0 | 0 | 12 | 1328.4662 | 1338.2448 | 859.6869 | 1217.7692 | 1234.5814 | 1042.7428 |
| Thy1 | HVLSGTLGIPEHTYR | -62 | -76 | 0 | 0 | 15 | 1286.5966 | 2842.3806 | 3081.6245 | 5237.4976 | 5094.827 | 6445.7993 |
| Thy1 | HVLSGTLGIPEHTYR | -62 | -76 | 0 | 0 | 15 | 1286.5966 | 2842.3806 | 3081.6245 | 5237.4976 | 5094.827 | 6445.7993 |
| Thy1 | LSNQPYIK | -81 | -88 | 0 | 0 | 8 | 1164.7947 | 1208.0706 | 1203.133 | 1285.3628 | 1181.8899 | 1314.9498 |
| Thy1 | GTLGIPEHTYR | -66 | -76 | 0 | 0 | 11 | 340.49554 | 662.21246 | 753.78156 | 683.18933 | 1374.8179 | 663.20026 |
| Thy1 | DNSIQHEFSL | -46 | -55 | 0 | 0.002068213 | 10 | 501.70944 | 131.99962 | 568.8953 | 475.49155 | 502.9662 | NaN |
| Thy1 | KHVLSGTL | -61 | -68 | 1 | 0.00635324 | 8 | 1874.339 | 1751.7241 | NaN | NaN | 237.89185 | 310.1645 |
| Thy1 | LVNQNLR | -29 | -35 | 0 | 0.000550307 | 7 | 1596.7366 | 1403.7775 | NaN | 319.35022 | NaN | NaN |
| Mtco2 | MIPTNDLKPGELR | -122 | -134 | 1 | 0 | 13 | 1810.1665 | 2849.1685 | 2993.4636 | 3008.5083 | 3232.883 | 3148.3208 |
| Mtco2 | MIPTNDLKPGELR | -122 | -134 | 1 | 0 | 13 | 1810.1665 | 2849.1685 | 2993.4636 | 3008.5083 | 3232.883 | 3148.3208 |
| Mtco2 | VVLPMELPIR | -142 | -151 | 0 | 0 | 10 | 951.8831 | 1061.0363 | 1044.7777 | 1322.1404 | 1481.4923 | 1433.4694 |
| Mtco2 | VVLPMELPIR | -142 | -151 | 0 | 0 | 10 | 951.8831 | 1061.0363 | 1044.7777 | 1322.1404 | 1481.4923 | 1433.4694 |
| Mtco2 | DEINNPVLTVK | -88 | -98 | 0 | 0 | 11 | 222.5618 | 691.797 | 125.84045 | 240.08765 | NaN | NaN |
| Mcts1 | DEKENVSNCIQLK | -6 | -18 | 1 | 0 | 13 | 2766.7268 | 2671.8103 | 2490.4927 | 1655.9559 | 1999.3623 | 1630.4349 |
| Dnajc3 | GKLDEAEDDFKK | -119 | -130 | 2 | 0 | 12 | 514.76306 | 529.36633 | 495.97498 | 525.20215 | 511.32062 | 533.94763 |
| Dnajc3 | DEKPVEAIR | -319 | -327 | 1 | 0 | 9 | 729.8196 | 760.59845 | 783.5391 | NaN | 1095.6406 | 1085.9735 |
| Cwc22 | NILHESEIDKR | -319 | -329 | 1 | 0 | 11 | 720.2682 | 164.72185 | 883.32874 | 753.05865 | 798.6646 | 860.50653 |
| Cwc22 | SSLDFEECAHK | -464 | -474 | 0 | 0 | 11 | 525.70996 | 559.7111 | 719.70844 | 1023.7973 | 882.6821 | 954.8099 |

| | | | | | | | | | | | | |
|---|---|---|---|---|---|---|---|---|---|---|---|---|
| Cwc22 | DELDPLLTR | -118 | -126 | 0 | 0.001055699 | 9 | 1389.7618 | 882.2427 | 964.9114 | 726.12164 | NaN | NaN |
| Cwc22 | ELLQENIVR | -182 | -190 | 0 | 0.00025015 | 9 | 347.41986 | NaN | 325.96973 | 960.7814 | NaN | 1019.54724 |
| Cwc22 | FEGLLPR | -611 | -617 | 0 | 0.00142999 | 7 | 3982.8684 | NaN | NaN | NaN | NaN | NaN |
| Parp1 | GGAAVDPDSGLEHSAHVL | -528 | -545 | 0 | 0 | 18 | 3249.8865 | 3460.5603 | 3456.1877 | 4325.542 | 4295.863 | 4233.6284 |
| Parp1 | DELGFRPEYSAS | -168 | -179 | 1 | 0 | 12 | 1958.6714 | 3104.387 | 1926.1282 | 1382.5765 | 4410.9004 | 312.83707 |
| Parp1 | GGSDDSSKDPIDVNYEK | -779 | -795 | 1 | 0 | 17 | 1452.2479 | 1462.4586 | 712.42706 | 2102.3772 | 1629.7468 | 1482.1042 |
| Parp1 | KPPLLNNADSVQAK | -747 | -760 | 1 | 0 | 14 | 1175.1146 | 1180.6603 | 1128.7148 | 1228.6532 | 1368.564 | 1269.0671 |
| Parp1 | VSQGSSESQILDLSNR | -719 | -734 | 0 | 0 | 16 | 847.7469 | 873.4276 | 788.0247 | 847.1334 | 717.9158 | 757.7269 |
| Parp1 | SLLSAEDKEALKK | -185 | -197 | 2 | 0 | 13 | 512.7485 | 632.0139 | 232.97664 | 862.9903 | 655.97266 | 650.35815 |
| Parp1 | SLLSAEDKEALKK | -185 | -197 | 2 | 0 | 13 | 512.7485 | 632.0139 | 232.97664 | 862.9903 | 655.97266 | 650.35815 |
| Parp1 | ECSGQLVFK | -297 | -305 | 0 | 0 | 9 | 622.94995 | 575.70404 | 450.7715 | 376.15564 | 5927.718 | 6177.455 |
| Parp1 | TLGLVDIVK | -555 | -563 | 0 | 0 | 9 | 523.4938 | 484.73795 | 492.63306 | 574.3005 | 463.3425 | 503.62054 |
| Parp1 | VPHWYHFS | -48 | -55 | 0 | 0 | 8 | 200.6894 | 916.62744 | 298.74457 | 761.7738 | 419.79016 | 990.12067 |
| Parp1 | LLEDDKESR | -573 | -581 | 1 | 0.00025015 | 9 | 1804.4034 | 2296.7258 | 3073.7234 | 796.4102 | NaN | 1746.4893 |
| Parp1 | GSSESQILDLSNR | -722 | -734 | 0 | 0 | 13 | 1751.7473 | 332.8277 | 2269.1987 | 1907.7715 | 1948.6007 | NaN |
| Parp1 | FYTLIPHDFGMK | -735 | -746 | 0 | 0 | 12 | 1160.0627 | NaN | NaN | 1032.7446 | 676.457 | 611.9678 |
| Parp1 | FYTLIPHDF | -735 | -743 | 0 | 0.001092529 | 9 | 428.68622 | NaN | 707.78674 | NaN | 843.2923 | 1083.7035 |
| Parp1 | EIDYGQDEEAVK | -641 | -652 | 0 | 0 | 12 | 961.9548 | NaN | NaN | 179.91077 | 1033.6512 | NaN |
| Parp1 | QVPSGESAILDR | -271 | -282 | 0 | 0.00111006 | 12 | 2999.767 | NaN | 3049.2354 | NaN | NaN | NaN |
| Eftud2 | DTPGHVNFSDEVTAGLR | -203 | -219 | 0 | 0 | 17 | 2214.576 | 2798.088 | 2345.95 | 1819.8632 | 745.4856 | 1903.9506 |
| Eftud2 | EGPLCDELIR | -775 | -784 | 0 | 0 | 10 | 1522.3347 | 1590.9604 | 1643.81 | 1612.8784 | 2315.0222 | 1798.3711 |
| Eftud2 | KYDWDLLAAR | -720 | -729 | 1 | 0 | 10 | 1349.7084 | 1358.9536 | 1246.0192 | 1414.1697 | 1416.3586 | 1202.2974 |
| Eftud2 | FSDEVTAGLR | -210 | -219 | 0 | 0 | 10 | 1162.1913 | 1109.4052 | 1254.3596 | 1168.1793 | 1341.5817 | 1460.0491 |
| Eftud2 | TLDELGIHLTK | -394 | -404 | 0 | 0 | 11 | 1036.9932 | 824.4612 | 902.30725 | 996.6111 | 1099.3906 | 1124.0189 |
| Eftud2 | QDAPIPGSPLYTIK | -858 | -871 | 0 | 0 | 14 | 847.8373 | 783.6195 | 788.9399 | 734.0703 | 2471.1729 | 2347.3215 |
| Eftud2 | FVDCLIEQTHPEIR | -144 | -157 | 0 | 0 | 14 | 779.12897 | 964.19934 | 909.0165 | 1140.085 | 1132.8519 | 1286.8372 |
| Eftud2 | IAVEPVNPSELPK | -589 | -601 | 0 | 0 | 13 | 637.31134 | 763.5792 | 704.5744 | 556.2914 | 769.5673 | 704.7797 |
| Eftud2 | SIVIRPLEPQPAPHLAR | -914 | -930 | 1 | 0 | 17 | 575.9538 | 661.3194 | 724.49835 | 611.20605 | 811.0247 | 780.7538 |
| Eftud2 | EEDTQPLTEPIIKPVK | -82 | -97 | 1 | 0 | 16 | 571.0135 | 851.1248 | 775.77673 | 592.2876 | 796.15375 | 698.2271 |
| Eftud2 | VEESGEHVIL | -620 | -629 | 0 | 0 | 10 | 550.7287 | 559.76373 | 647.3462 | 748.2676 | 659.1388 | 752.66943 |
| Eftud2 | GPDATGPNILVDDTLPSEVDK | -735 | -755 | 0 | 0.000230707 | 21 | 527.5803 | 630.6367 | 674.3908 | 639.3241 | 782.29517 | 739.2213 |
| Eftud2 | KFFGEFTGF | -421 | -429 | 1 | 0.00025015 | 9 | 492.66473 | 431.74075 | 463.50397 | 363.7456 | 257.4515 | 469.7832 |
| Eftud2 | VPAGNWVLIEGVDQPIVK | -543 | -560 | 0 | 0 | 18 | 434.35184 | 287.60983 | 532.54584 | 463.2865 | 400.6966 | 482.1487 |
| Eftud2 | TCFVDCLIEQTHPEIR | -142 | -157 | 0 | 0 | 16 | 304.33618 | 376.0332 | 305.8625 | 410.39038 | 445.3627 | 424.8948 |
| Eftud2 | VVGDVDTSLPR | -383 | -393 | 0 | 0.00025015 | 11 | 5816.7554 | NaN | 6078.725 | 3784.5083 | 209.57906 | 6130.4995 |
| Eftud2 | GNEEAQIFRPL | -569 | -579 | 1 | 0 | 11 | 1153.9159 | 1054.3206 | 1078.6722 | 316.77234 | 604.6377 | NaN |
| Eftud2 | DELGIHLTK | -396 | -404 | 0 | 0 | 9 | 507.48642 | 552.6693 | 652.12885 | 1099.1354 | 415.09988 | NaN |
| Eftud2 | GENYTLEDEEDSQICTVGR | -510 | -528 | 0 | 0 | 19 | 278.96124 | 327.02997 | NaN | 298.702 | 418.35562 | 450.28717 |
| Eftud2 | HHWQIVPGDPLDK | -901 | -913 | 0 | 0 | 13 | 2192.3313 | 863.00916 | NaN | 2492.2183 | 1133.0292 | NaN |
| Eftud2 | YHIEVNR | -536 | -542 | 0 | 0 | 7 | 1697.0637 | 1382.7762 | 1351.3815 | NaN | NaN | 1379.5187 |
| Eftud2 | STPVTVVLPDTK | -182 | -193 | 0 | 0 | 12 | 1490.989 | 1275.2509 | 1457.6919 | 1022.5145 | NaN | NaN |
| Eftud2 | QIVPGDPLDK | -904 | -913 | 0 | 0.000230707 | 10 | 792.3634 | 818.63434 | 834.4952 | 829.84436 | NaN | NaN |
| Eftud2 | VDDTLPSEVDK | -745 | -755 | 0 | 0 | 11 | 625.68835 | 890.03754 | 1079.2856 | NaN | NaN | 914.291 |
| Eftud2 | TIVQEEDTQPLTEPIIKPVK | -78 | -97 | 1 | 0.00051306 | 20 | 414.3949 | 395.90213 | 401.32965 | NaN | 185.86147 | NaN |
| Eftud2 | YDWDLLAAR | -721 | -729 | 0 | 0 | 9 | 2228.5298 | 1274.7643 | 2328.6726 | NaN | NaN | NaN |
| Eftud2 | GNEEAQIFRPLK | -569 | -580 | 1 | 0 | 12 | 864.9336 | 1545.4396 | 1549.1038 | NaN | NaN | NaN |
| Eftud2 | VEPVNPSELPK | -591 | -601 | 0 | 0.002068213 | 11 | 1915.4072 | NaN | 2271.0344 | NaN | NaN | NaN |
| Eftud2 | SIVIRPLEPQPAPH | -914 | -927 | 1 | 0.000431655 | 14 | 800.3524 | NaN | 843.4736 | NaN | NaN | NaN |
| Eftud2 | RYDQDLCYTD | -159 | -168 | 1 | 0.004472887 | 10 | 419.19788 | NaN | NaN | 322.11078 | NaN | NaN |
| Eif5b | LAHCEELR | -846 | -853 | 0 | 0 | 8 | 1613.8645 | 823.7397 | 1385.8112 | 1176.5228 | 1711.7778 | 1071.4312 |
| Eif5b | NVPLEAINEQTK | -668 | -679 | 0 | 0 | 12 | 1099.2864 | 1246.7358 | 1104.5161 | 1416.8519 | 1371.707 | 670.33167 |
| Eif5b | TSEVPYAGINIGPVHK | -996 | -1011 | 0 | 0 | 16 | 913.53723 | 1085.0756 | 835.0409 | 681.1124 | 1435.561 | 638.13434 |
| Eif5b | SFDENDSEELEDKDSK | -108 | -123 | 1 | 0 | 16 | 469.6094 | 448.14664 | 580.0256 | 478.38837 | 542.39246 | 544.50415 |
| Eif5b | IEPIPGESPK | -1157 | -1166 | 0 | 0.002595029 | 10 | 1295.2772 | 1466.8547 | 1293.3075 | 1500.7109 | NaN | NaN |
| Eif5b | LFYENKDPR | -803 | -811 | 1 | 0 | 9 | 1067.4907 | 1120.9315 | 1387.9175 | 898.6967 | NaN | NaN |
| Eif5b | DFDENDILR | -52 | -60 | 0 | 0 | 9 | 636.51416 | NaN | NaN | 646.2067 | 761.3501 | 670.7576 |
| Eif5b | DELIHELK | -957 | -964 | 0 | 0 | 8 | 556.02026 | 2292.1113 | NaN | 2055.939 | 1680.0842 | NaN |
| Eif5b | HTHVQDGEAGGITQ | -649 | -662 | 0 | 0 | 14 | 495.91888 | 466.15643 | 447.4005 | NaN | 192.79248 | NaN |
| Chp1 | DELLQVLR | -133 | -140 | 0 | 0.00272025 | 8 | 1006.9144 | NaN | NaN | 780.9348 | 1069.3802 | NaN |
| Rps19 | ELAPYDENWFYTR | -44 | -56 | 0 | 0 | 13 | 4307.8667 | 3082.9585 | 4822.312 | 4743.27 | 7238.8394 | 4971.5654 |
| Rps19 | LKVPEWVDTVK | -28 | -38 | 1 | 0 | 11 | 3991.597 | 5192.6167 | 5143.943 | 4357.5425 | 6382.23 | 6488.93 |
| Rps19 | LKVPEWVDTVK | -28 | -38 | 1 | 0 | 11 | 3991.597 | 5192.6167 | 5143.943 | 4357.5425 | 6382.23 | 6488.93 |
| Rps19 | DVNQQEFVR | -8 | -16 | 0 | 0 | 9 | 3240.2104 | 559.4788 | 3545.5945 | 4229.8687 | 3604.4404 | 3970.1528 |
| Rps19 | VKDVNQQEFVR | -6 | -16 | 1 | 0 | 11 | 675.1788 | 747.1993 | 938.1985 | 910.9322 | 887.3093 | 825.4303 |
| Rps19 | KVPEWVDTVK | -29 | -38 | 1 | 0.000230707 | 10 | 1276.9106 | 1488.9738 | NaN | 2063.6785 | 2447.7246 | 2332.6433 |
| Rps19 | LKVPEWVD | -28 | -35 | 1 | 0 | 8 | 693.24695 | 590.1075 | 1650.2952 | 875.1522 | NaN | NaN |
| Rps19 | DENWFYTR | -49 | -56 | 0 | 0.000826806 | 8 | 456.52994 | 529.84766 | 518.4636 | 608.0293 | NaN | NaN |
| Rps19 | RVLQALEGLK | -102 | -111 | 1 | 0.000627943 | 10 | 1428.0494 | NaN | 427.12622 | NaN | NaN | NaN |
| Anxa11 | STITPAVPPAQFGNR | -175 | -189 | 0 | 0 | 15 | 1652.2568 | 1546.1945 | 1436.3064 | 2448.1724 | 1930.7029 | 1835.792 |
| Anxa11 | NTPAFFAER | -429 | -437 | 0 | 0 | 9 | 1220.9982 | 560.7381 | 198.5284 | 346.21838 | 1742.6322 | 623.685 |
| Anxa11 | SDTSGHFQR | -326 | -334 | 0 | 0.00025015 | 9 | 916.2604 | 782.822 | 1075.6896 | 722.2488 | 903.00336 | 939.8351 |
| Anxa11 | SLYHDITGDTSGDYR | -478 | -492 | 0 | 0 | 15 | 771.5685 | 857.08276 | 806.2224 | 624.28015 | 928.5062 | 886.9554 |
| Anxa11 | YHDITGDTSGDYR | -480 | -492 | 0 | 0 | 13 | 514.3269 | 511.3828 | 667.5219 | 647.6396 | 118.507965 | 760.6023 |
| Anxa11 | TPVLFDVYEIK | -270 | -280 | 0 | 0 | 11 | 157.0505 | 860.3581 | 738.2515 | 1028.9056 | 1307.3484 | 1195.774 |
| Anxa11 | DESTNVDMSLVQR | -345 | -357 | 0 | 0.00025015 | 13 | 497.66098 | 411.69623 | 543.58105 | 745.0792 | NaN | 601.848 |
| Anxa11 | KSELSGNFEK | -253 | -262 | 1 | 0 | 10 | 310.1836 | NaN | 2801.1802 | NaN | 2031.4397 | 2097.1533 |
| Capzb | RLPPQQIEK | -15 | -23 | 1 | 0.001023236 | 9 | 1538.9237 | 2025.9805 | 1280.1932 | 1566.4143 | 1929.7136 | 1538.8574 |
| Capzb | DYLLCDYNR | -58 | -66 | 0 | 0 | 9 | 1482.7849 | 1792.6438 | 1438.2031 | 295.11914 | 1198.0051 | 1715.9858 |

| | | | | | | | | | | | |
|---|---|---|---|---|---|---|---|---|---|---|---|
| Capzb | YDPPLEDGAMPSAR | -79 | -92 | 0 | 0.00025015 | 14 | 1405.2661 | 961.5786 | 944.0522 | 1173.9849 | 934.69574 | 861.838 |
| Capzb | YDPPLEDGAMPSAR | -79 | -92 | 0 | 0.00025015 | 14 | 1405.2661 | 961.5786 | 944.0522 | 1173.9849 | 934.69574 | 861.838 |
| Capzb | NAFDQYR | -102 | -108 | 0 | 0 | 7 | 1347.0642 | 1502.0962 | 1335.3226 | 1120.2709 | 1243.3154 | 1098.5194 |
| Capzb | LNEIYFGK | -228 | -235 | 0 | 0.000826806 | 8 | 928.4628 | 469.38498 | 978.52246 | 1195.754 | 970.3265 | 818.4473 |
| Capzb | WDSIHVVEVQEK | -148 | -159 | 0 | 0 | 12 | 855.4576 | 577.84485 | 1408.7722 | 913.1355 | 1057.4573 | 1068.2444 |
| Capzb | GCWDSIHVVEVQEK | -146 | -159 | 0 | 0 | 14 | 785.7505 | 897.84265 | 773.6519 | 834.13135 | 916.3643 | 834.3112 |
| Capzb | SDQQLDCALDLMR | -2 | -14 | 0 | 0 | 13 | 343.90247 | 340.38123 | 442.34882 | 435.02615 | 523.9913 | 460.53815 |
| Capzb | STLNEIYFGK | -226 | -235 | 0 | 0.00025015 | 10 | 303.3205 | 734.3447 | 4754.5146 | 4121.8174 | 4652.173 | 587.4275 |
| Capzb | EDLLSSVDQPLK | -37 | -48 | 0 | 0 | 12 | 494.66098 | 1015.5947 | 1220.3788 | 927.8762 | 1031.26 | NaN |
| Capzb | DETVSDCSPHIA | -200 | -211 | 0 | 0.00025015 | 12 | 752.44275 | 617.562 | 720.75 | 310.13983 | NaN | NaN |
| Capzb | QMEKDETVSDCSPHIA | -196 | -211 | 1 | 0.007546675 | 16 | 817.4075 | NaN | 336.1503 | 1032.2891 | NaN | NaN |
| Capzb | GCWDSIHVV | -146 | -154 | 0 | 0 | 9 | 597.32635 | 639.69086 | NaN | NaN | NaN | NaN |
| Capzb | GCWDSIHVVEVQ | -146 | -157 | 0 | 0.00025015 | 12 | 713.4453 | NaN | NaN | NaN | NaN | NaN |
| Supt16h | GDKVDILYNNIK | -694 | -705 | 1 | 0 | 12 | 3936.3198 | 3724.4473 | 3939.207 | 5757.06 | 3838.5461 | 818.0758 |
| Supt16h | SINLGFSDLTNK | -378 | -389 | 0 | 0 | 12 | 1796.932 | 1950.6204 | 1718.1005 | 1176.0284 | 2089.3635 | 2095.1482 |
| Supt16h | NEGNIFPNPEATFVK | -582 | -596 | 0 | 0 | 15 | 1582.0908 | 1801.3688 | 2030.8112 | 2018.674 | 2422.4863 | 2652.807 |
| Supt16h | DEVELIHFER | -838 | -847 | 0 | 0 | 10 | 1264.9427 | 1229.568 | 1322.5715 | 1239.2493 | 1271.3712 | 1149.8058 |
| Supt16h | LDEVELIHFER | -837 | -847 | 0 | 0.000394361 | 11 | 1168.6976 | 1419.4751 | 1268.6426 | 1004.6589 | 1325.2711 | 1217.0756 |
| Supt16h | YEEEEEQSR | -1006 | -1014 | 0 | 0 | 9 | 1056.5155 | 555.4875 | 688.7723 | 140.86586 | 763.95776 | 799.10583 |
| Supt16h | TKEELEFEVPFR | -791 | -802 | 1 | 0 | 12 | 707.1572 | 757.27954 | 766.0373 | 992.31696 | 1220.5774 | 1208.209 |
| Supt16h | TKEELEFEVPFR | -791 | -802 | 1 | 0 | 12 | 707.1572 | 757.27954 | 766.0373 | 992.31696 | 1220.5774 | 1208.209 |
| Supt16h | LKNEDDEEEEEKDEAEDLLGR | -432 | -453 | 2 | 0 | 22 | 536.0968 | 507.8505 | 696.60315 | 861.3458 | 1166.2668 | 1201.6102 |
| Supt16h | INFYCPGSALGR | -570 | -581 | 0 | 0 | 12 | 386.7594 | 446.1883 | 396.04025 | 664.69696 | 526.6219 | 401.2683 |
| Supt16h | LGFSDLTNK | -381 | -389 | 0 | 0 | 9 | 165.405 | 935.12805 | 1462.155 | 627.11597 | 868.87 | 899.3155 |
| Supt16h | VEGDYTYLR | -561 | -569 | 0 | 0 | 9 | 101.095795 | 373.52515 | 2194.316 | 2352.8743 | 1563.4396 | 2180.1665 |
| Supt16h | SVEGDYTYLR | -560 | -569 | 0 | 0 | 10 | 1408.0708 | 543.3691 | NaN | 1337.4835 | 1007.35986 | 483.57004 |
| Supt16h | LKDLYIRPN | -662 | -670 | 2 | 0 | 9 | 1057.4525 | 1045.4003 | 1172.5673 | 1274.9084 | 1090.4698 | NaN |
| Supt16h | QLQEELLK | -307 | -314 | 0 | 0.001392869 | 8 | 513.9626 | 307.48145 | 427.81647 | 1510.6997 | 280.6933 | NaN |
| Supt16h | VMEIVDADEK | -202 | -211 | 0 | 0 | 10 | 517.25555 | 504.60742 | 954.9011 | NaN | NaN | NaN |
| Supt16h | KYETVIMPVFGIA | -534 | -546 | 1 | 0.004595494 | 13 | 127.538536 | 421.58408 | NaN | NaN | NaN | 941.98956 |
| Scarb2 | KLDDFVETGDIR | -391 | -402 | 1 | 0.00025015 | 12 | 1582.3606 | 1361.7587 | 1292.0419 | NaN | 2595.6956 | 2970.4062 |
| Scarb2 | FSSFENVEGLPAFR | -281 | -294 | 0 | 0 | 14 | 524.144 | 574.7542 | NaN | 607.7603 | 652.77997 | 713.9624 |
| Scarb2 | SFENVEGLPAFR | -283 | -294 | 0 | 0 | 12 | 3364.1174 | NaN | 382.73404 | NaN | 685.2825 | 630.2708 |
| Scarb2 | DEVLYLFPSDLCR | -263 | -275 | 0 | 0.00051306 | 13 | 283.19226 | NaN | NaN | NaN | NaN | 334.98553 |
| Ncbp2 | DEYREDYDAGR | -136 | -146 | 1 | 0 | 11 | 420.86612 | 494.58194 | NaN | 821.80237 | 892.8494 | 934.37463 |
| Surf4 | TFLEDGIR | -36 | -43 | 0 | 0.000471409 | 8 | 1251.0671 | 26681.43 | 26406.662 | 27116.885 | 26004.695 | 27453.611 |
| Surf4 | KQYLPHVAR | -22 | -30 | 1 | 0 | 9 | 565.2252 | 581.5364 | NaN | NaN | 471.84143 | 530.21466 |
| Surf4 | MWFQWSEQR | -44 | -52 | 0 | 0.000627943 | 9 | 350.2866 | 395.68286 | NaN | NaN | 572.77094 | 542.888 |
| Surf4 | DFADQFLR | -12 | -19 | 0 | 0 | 8 | 551.02496 | 490.01996 | 533.1841 | NaN | NaN | NaN |
| Lrrfip1 | EVEERPDKDFAEK | -51 | -63 | 2 | 0 | 13 | 812.44904 | 897.74133 | 1078.509 | 1054.3322 | 1026.6869 | 977.6523 |
| Lrrfip1 | GPQCDTSQIGSEEGHVT | -664 | -680 | 0 | 0 | 17 | 593.6573 | 566.27893 | 752.1113 | 631.972 | 712.713 | 627.4103 |
| Lrrfip1 | DFAEKGSRNMP | -59 | -69 | 2 | 0.004983786 | 11 | 373.91803 | 1137.9015 | 1017.8901 | 1167.6501 | 862.6139 | 520.6535 |
| Lrrfip1 | EIDCLSPEAQR | -11 | -21 | 0 | 0 | 11 | 2012.1423 | 1689.6115 | 1822.9458 | 2043.2762 | NaN | 433.77313 |
| Lrrfip1 | ETLQNSEQEQPKPNTGK | -260 | -276 | 1 | 0 | 17 | 276.54205 | NaN | 355.7686 | 315.46408 | 449.3758 | 344.39526 |
| Lrrfip1 | ETLQNSEQEQPKPNTGK | -260 | -276 | 1 | 0 | 17 | 276.54205 | NaN | 355.7686 | 315.46408 | 449.3758 | 344.39526 |
| Lrpprc | GSLLPELSLAER | -130 | -141 | 0 | 0 | 12 | 1661.3162 | 1365.4755 | 1816.7856 | 1006.25806 | 1119.8103 | 841.28564 |
| Lrpprc | DFAETHIK | -1090 | -1097 | 0 | 0.000712251 | 8 | 1412.1173 | 1869.4614 | 1815.0376 | 2253.2708 | 809.8406 | 692.2623 |
| Lrpprc | FSPTDFLAK | -178 | -186 | 0 | 0.00078661 | 9 | 1055.2906 | 1090.1204 | 402.61426 | 1068.7212 | 1279.9144 | 1190.9232 |
| Lrpprc | IHDVLCK | -857 | -863 | 0 | 0 | 7 | 1023.9455 | 1267.2545 | 1140.5482 | 771.4551 | 982.43304 | 890.1534 |
| Lrpprc | LELIPELR | -1306 | -1313 | 0 | 0.001392869 | 8 | 755.40295 | 894.452 | 845.72736 | 978.12 | 1295.4186 | 1272.5485 |
| Lrpprc | EEGFPIRPH | -431 | -439 | 1 | 0.00025015 | 9 | 744.3668 | 340.2097 | 976.7039 | 1648.8076 | 999.7148 | 623.56146 |
| Lrpprc | KLDDLFLK | -1348 | -1355 | 1 | 0 | 8 | 541.6007 | 459.44165 | 725.47284 | 528.8802 | 603.94794 | 474.6795 |
| Lrpprc | KIIETPGIR | -908 | -916 | 1 | 0.00025015 | 9 | 248.17409 | 250.87727 | 1215.9902 | 178.75903 | 708.80853 | 1422.7816 |
| Lrpprc | DLQEEPAPVR | -67 | -76 | 0 | 0 | 10 | 1431.5973 | 1111.0875 | 861.5052 | NaN | NaN | NaN |
| Lrpprc | GDVESIEAIQR | -1152 | -1162 | 0 | 0 | 11 | 264.15833 | 1497.1132 | 1784.1808 | NaN | NaN | NaN |
| Lrpprc | LQDAINILK | -763 | -771 | 0 | 0 | 9 | 973.6752 | NaN | NaN | NaN | NaN | NaN |
| Lrpprc | TLLELIPELR | -1304 | -1313 | 0 | 0 | 10 | 372.57462 | NaN | NaN | NaN | NaN | NaN |
| Pdlim1 | TSASGEEANSRPVVQPHPSGSLIIDK | -165 | -190 | 1 | 0.001868731 | 26 | 6689.7563 | 5905.358 | 5800.914 | 6276.2896 | 5541.0054 | 5376.9907 |
| Pdlim1 | GDPNKPSGFR | -227 | -236 | 1 | 0 | 10 | 1223.3479 | 297.33282 | 522.8898 | 731.1303 | 209.52527 | 1393.068 |
| Pdlim1 | GHFFVEDQIYCEK | -295 | -307 | 0 | 0 | 13 | 876.97406 | 1026.975 | 986.17303 | 1386.6783 | 1557.1721 | 1371.8236 |
| Pdlim1 | QIVLQGPGPWGFR | -5 | -17 | 0 | 0 | 13 | 849.3039 | 742.36206 | 684.97485 | 388.12186 | 326.8937 | 527.3749 |
| Pdlim1 | IWSPLVTEEGKR | -88 | -99 | 1 | 0 | 12 | 772.89594 | 930.6783 | 1001.228 | 1742.0876 | 1093.5287 | 991.8905 |
| Pdlim1 | IWSPLVTEEGKR | -88 | -99 | 1 | 0 | 12 | 772.89594 | 930.6783 | 1001.228 | 1742.0876 | 1093.5287 | 991.8905 |
| Pdlim1 | TSASGEEANSRPVVQPHPSGSLIIDKDSEVYK | -165 | -196 | 2 | 0.005698678 | 32 | 670.8514 | 797.61707 | NaN | 766.12054 | 849.1215 | 770.188 |
| Pdlim1 | DFEQPLAISR | -23 | -32 | 0 | 0.000230707 | 10 | 2992.516 | 1488.9867 | 1842.0199 | 1546.8391 | NaN | NaN |
| Pdlim1 | EILESDGKGDPNKPSGFR | -219 | -236 | 2 | 0.000394361 | 18 | 447.68695 | 165.46434 | 300.64746 | NaN | 760.9599 | NaN |
| Pdlim1 | TDCGINLK | -285 | -292 | 0 | 0.000907206 | 8 | 499.49298 | NaN | NaN | NaN | NaN | NaN |
| Pdlim2 | DFHTPIIVTK | -22 | -31 | 0 | 0 | 10 | 2339.5278 | 2636.7095 | 2551.9456 | NaN | NaN | NaN |
| Fkbp5 | YGFGEAGKPK | -223 | -232 | 1 | 0 | 10 | 3314.513 | 3201.2705 | 3204.3838 | 3160.55 | 2330.8513 | 2894.7288 |
| Fkbp5 | EEQCILYLGPR | -212 | -222 | 0 | 0 | 11 | 871.4282 | 165.21028 | 482.91736 | 634.86945 | 1341.0901 | 1212.3922 |
| Fkbp5 | DFKGEDLFEDSGVIR | -136 | -150 | 1 | 0 | 15 | 833.0062 | 727.45355 | 801.30426 | 1225.8105 | 1296.0226 | 1262.8384 |
| Fkbp5 | GEDLFEDSGVIR | -139 | -150 | 0 | 0 | 12 | 511.36893 | 841.4591 | 902.1988 | 604.7999 | 1128.4911 | 1011.2401 |
| Fkbp5 | VVGEGEDHDIPIGIDK | -189 | -204 | 0 | 0 | 16 | 424.27228 | 658.7922 | 517.7441 | 425.16968 | 728.6572 | 524.6436 |
| Fkbp5 | KGEGYSNPNEGATVK | -155 | -169 | 1 | 0.000230707 | 15 | 395.1879 | 385.02066 | 470.35583 | 508.05542 | 525.68024 | 520.64636 |
| Fkbp5 | VLAVNPQNR | -377 | -385 | 0 | 0 | 9 | 940.5763 | 854.01697 | 982.45886 | NaN | NaN | NaN |
| Psip1 | DFKPGDLIFAK | -4 | -14 | 1 | 0 | 11 | 2431.6104 | 2777.8943 | 3059.8254 | 2954.2588 | 3394.7454 | 3337.898 |
| Psip1 | VDEVPDGAVKPPTNK | -25 | -39 | 1 | 0 | 15 | 1065.0519 | 1163.9314 | 1263.2412 | 1755.3638 | 2039.9381 | 1969.6732 |
| Psip1 | KGFNEGLWEIDNNPK | -75 | -89 | 1 | 0.00025015 | 15 | 1022.8147 | 3467.2617 | 94.76667 | NaN | 3287.8103 | 5029.4453 |

| | | | | | | | | | | | | |
|---|---|---|---|---|---|---|---|---|---|---|---|---|
| Psip1 | GFNEGLWEIDNNPK | -76 | -89 | 0 | 0 | 14 | 554.7486 | 658.0246 | NaN | 653.6993 | 688.30396 | 655.89307 |
| Psip1 | LPIFFFGTHET | -40 | -50 | 0 | 0 | 11 | 546.4077 | NaN | 256.70032 | 718.91315 | 246.36206 | 205.96547 |
| Psip1 | LPIFFFGTH | -40 | -48 | 0 | 0.001291103 | 9 | 461.52655 | 106.32453 | 498.04022 | 377.661 | NaN | 509.49527 |
| Psip1 | QSNASSDVEVEEK | -101 | -113 | 0 | 0 | 13 | 401.37073 | 995.08905 | 826.8083 | 1038.5459 | NaN | 753.1985 |
| Psip1 | EVPDGAVKPPTNK | -27 | -39 | 1 | 0 | 13 | 1095.9714 | 1055.345 | 1221.985 | NaN | NaN | NaN |
| Psip1 | ETNVSKEDTDQEEK | -114 | -127 | 1 | 0.001291103 | 14 | 506.6801 | NaN | 531.9806 | NaN | NaN | NaN |
| Psip1 | GLWEIDNNPK | -80 | -89 | 0 | 0.001868731 | 10 | 1848.1788 | NaN | NaN | NaN | NaN | NaN |
| Tbcb | DQEDALLGSYPVDDGCR | -68 | -84 | 0 | 0 | 17 | 1427.7902 | 1347.5686 | 1545.6295 | 971.93036 | 1748.7002 | 237.1293 |
| Tbcb | LDQEDALLGSYPVDDGCR | -67 | -84 | 0 | 0 | 18 | 378.95193 | 471.69025 | 546.2022 | NaN | 859.02985 | 382.0045 |
| Tbcb | DFKPGYWVGVR | -186 | -196 | 1 | 0.00025015 | 11 | 727.59314 | 1625.9093 | NaN | NaN | 2134.8538 | NaN |
| Tbcb | IHVIDHSGVR | -85 | -94 | 0 | 0 | 10 | 514.1309 | 451.9963 | 463.3468 | NaN | NaN | NaN |
| Tbcb | IHVIDHSGVR | -85 | -94 | 0 | 0 | 10 | 514.1309 | 451.9963 | 463.3468 | NaN | NaN | NaN |
| Tbcb | LGPYNEELR | -130 | -138 | 0 | 0.00133086 | 9 | 1501.588 | NaN | 1554.4381 | NaN | NaN | NaN |
| Ebna1bp2 | LDFLEGDQKPVER | -210 | -222 | 1 | 0 | 13 | 2013.9584 | 1961.8479 | 1478.2152 | 1509.2529 | 1699.3359 | 1682.2205 |
| Ebna1bp2 | DFLEGDQKPVER | -211 | -222 | 1 | 0 | 12 | 936.5416 | 951.88293 | 993.0989 | 1204.4292 | 1151.9297 | 1138.4043 |
| Ebna1bp2 | GLLKPGLNVVL | -30 | -40 | 1 | 0.001291103 | 11 | 331.69974 | 368.05048 | 425.71432 | 497.45227 | 438.79425 | 390.725 |
| Ebna1bp2 | KGVNPEDDFQR | -94 | -104 | 1 | 0 | 11 | 4467.044 | NaN | 1187.5435 | NaN | 630.2779 | 1603.4727 |
| Ebna1bp2 | KLDFLEGDQKPVER | -209 | -222 | 2 | 0.002068213 | 14 | 543.24005 | 598.83295 | 593.3876 | NaN | 807.9518 | NaN |
| Ebna1bp2 | KAVNDVSGLK | -45 | -54 | 1 | 0 | 10 | 951.5104 | NaN | NaN | 554.2551 | NaN | 650.64734 |
| Apmap | LLEYDTVTK | -238 | -246 | 0 | 0 | 9 | 2753.1604 | 2996.0027 | 3162.6934 | 3680.8174 | 2809.028 | 3191.7092 |
| Apmap | DFLSDKPFIK | -332 | -341 | 1 | 0 | 10 | 4509.9727 | 4878.354 | 5220.28 | NaN | 435.7622 | 7485.578 |
| Apmap | SLHDPDGQVVTY | -375 | -386 | 0 | 0.001868731 | 12 | 1531.4951 | NaN | NaN | 785.7665 | 3421.3137 | NaN |
| Pi4ka | DFLVIPSPVLVK | -512 | -523 | 0 | 0 | 12 | 297.97412 | 319.55783 | 372.93143 | 342.37073 | 381.6649 | 420.78006 |
| Rpl9 | KFLDGIYVSE | -174 | -183 | 1 | 0.001636916 | 10 | 1513.2107 | 3064.9036 | 2611.702 | 4265.868 | 2334.5723 | 1784.224 |
| Rpl9 | ENGSLVEIR | -107 | -115 | 0 | 0 | 9 | 1074.0698 | 2168.5596 | 1191.6676 | 2418.3416 | 2003.1525 | 2150.7979 |
| Rpl9 | KFLDGIYVSEK | -174 | -184 | 1 | 0 | 11 | 1051.4146 | 4514.011 | 4294.6816 | 5656.3755 | 6867.4775 | 7294.222 |
| Rpl9 | SVYAHFPINVVIQ | -94 | -106 | 0 | 0 | 13 | 775.67896 | 759.4251 | 772.89966 | 897.5317 | 969.63617 | 1045.2943 |
| Rpl9 | FLDGIYVSEK | -175 | -184 | 0 | 0 | 10 | 536.79144 | 238.78624 | 3044.938 | 607.1615 | 642.98413 | 535.3891 |
| Rpl9 | RDFNHINVE | -36 | -44 | 1 | 0.000712251 | 9 | 236.86316 | 2821.1436 | 2656.079 | 2315.2427 | 3274.611 | 292.86703 |
| Rpl9 | RDFNHINVEL | -36 | -45 | 1 | 0.00111006 | 10 | 1600.6885 | 4026.0256 | NaN | 1037.7817 | 1794.8605 | 1535.2458 |
| Rpl9 | TICSHVQ | -72 | -78 | 0 | 0.00416551 | 7 | 1329.7401 | 959.98944 | NaN | 1059.714 | 3141.2595 | 3369.7422 |
| Rpl9 | KFLDGIYVSEKG | -174 | -185 | 2 | 0.000947459 | 12 | 1144.2601 | 1225.7487 | 1316.6572 | NaN | 1098.1163 | 2408.962 |
| Rpl9 | VDKWWGNR | -57 | -64 | 1 | 0.00025015 | 8 | 963.1668 | 1347.1129 | 1194.7776 | NaN | 1044.3629 | NaN |
| Rpl9 | TVDIPENVEITLK | -9 | -21 | 0 | 0 | 13 | 793.7406 | NaN | 781.2895 | 905.2124 | NaN | 940.8439 |
| Rpl9 | DFNHINVEL | -37 | -45 | 0 | 0.006915721 | 9 | 1211.4994 | 1260.6973 | NaN | 1250.049 | NaN | NaN |
| Itih2 | IQPSGGTNINEALLR | -380 | -394 | 0 | 0 | 15 | 3287.3408 | 3941.5127 | 3880.2114 | 4123.1416 | 4668.819 | 4421.8237 |
| Itih2 | VQFELHYQEVK | -177 | -187 | 0 | 0 | 11 | 2668.9136 | 2710.74 | 2794.2705 | 1140.3645 | 1843.8693 | 1774.8489 |
| Itih2 | VSDGDPTVGELK | -419 | -430 | 0 | 0 | 12 | 2281.1912 | 2364.3567 | 2322.485 | 3193.941 | 409.4452 | 449.49182 |
| Itih2 | HADPDFTK | -574 | -581 | 0 | 0 | 8 | 1006.3462 | 1021.09186 | 1022.3664 | 1219.7843 | 963.34186 | 691.31085 |
| Itih2 | LTINQLLAER | -587 | -596 | 0 | 0 | 10 | 260.5274 | 279.14957 | 355.12744 | 489.3962 | 549.43243 | 568.28937 |
| Itih2 | FKPTVAQQR | -251 | -259 | 1 | 0.000431655 | 9 | 5147.2827 | 5401.648 | 5827.9717 | NaN | 5398.5864 | 6038.9897 |
| Itih2 | KTILDDLR | -334 | -341 | 1 | 0.002810811 | 8 | 2040.1991 | 2208.3347 | NaN | 2366.1208 | 3125.858 | 2964.732 |
| Itih2 | TILDDLR | -335 | -341 | 0 | 0.008947895 | 7 | 1309.3367 | 818.0683 | 692.663 | 203.51433 | NaN | NaN |
| Itih2 | DFNHNVR | -350 | -356 | 0 | 0 | 7 | 407.56607 | 1158.6885 | 1392.3125 | NaN | 1065.0929 | NaN |
| Itih2 | FYNQVSTPLLR | -489 | -499 | 0 | 0 | 11 | 1005.958 | 744.741 | 145.57037 | NaN | NaN | NaN |
| Itih2 | GGTNINEALLR | -384 | -394 | 0 | 0 | 11 | 256.82672 | NaN | 558.35223 | NaN | NaN | NaN |
| Gmip | DFPEEVPFVITR | -558 | -569 | 0 | 0.00025015 | 12 | 1293.6672 | NaN | NaN | NaN | NaN | NaN |
| Gm12250 | SPHLLSSEPNESVADK | -345 | -360 | 0 | 0.001868731 | 16 | 2311.0828 | 2358.1914 | 1757.5917 | 1234.6165 | 1486.4243 | 1413.5365 |
| Gm12250 | NYFLDTVTEDAK | -390 | -401 | 0 | 0 | 12 | 1421.4142 | 1231.975 | 1273.4397 | 1462.9153 | 1589.1343 | 1641.3851 |
| Gm12250 | NFKPEEYLK | -133 | -141 | 1 | 0 | 9 | 1333.8771 | 1622.159 | 1534.0837 | 1760.9347 | 1618.2941 | 1706.437 |
| Gm12250 | IDSDLWNEK | -184 | -192 | 0 | 0 | 9 | 1006.47565 | 933.16364 | 1089.1796 | 820.7179 | 76.96328 | 385.80182 |
| Gm12250 | FFNDDIEEFEK | -299 | -309 | 0 | 0 | 11 | 933.645 | 890.32 | 824.2401 | 1061.478 | 751.61444 | 945.21765 |
| Gm12250 | TKIDSDLWNEK | -182 | -192 | 1 | 0.000230707 | 11 | 900.72473 | 837.08527 | 835.597 | 1937.3477 | 1388.385 | 779.30255 |
| Gm12250 | ETIDSIQSCIQEGDIQK | -36 | -52 | 0 | 0 | 17 | 589.1915 | 552.02545 | 708.5232 | 770.75543 | 891.92664 | 914.5573 |
| Gm12250 | GIGHEESESAESGAVE | -90 | -105 | 0 | 0.00078661 | 16 | 578.9197 | 442.44153 | 521.7868 | 353.58954 | 1364.3118 | 950.6987 |
| Gm12250 | SFFNDDIEEFEK | -298 | -309 | 0 | 0 | 12 | 296.1182 | 310.3503 | 327.10614 | 97.849174 | 445.36237 | 1916.3911 |
| Gm12250 | FKPEEYLK | -134 | -141 | 1 | 0 | 8 | 978.4658 | NaN | 1112.0779 | 1108.0138 | 1280.3704 | 1341.0114 |
| Gm12250 | DFPSLESTLLEELPAHK | -238 | -254 | 0 | 0 | 17 | 254.27171 | NaN | NaN | 372.63342 | 337.86926 | 437.8059 |
| Dync1li1 | SGQKPVLSDVHAE | -486 | -498 | 1 | 0 | 13 | 1376.0278 | 1475.4474 | 1771.6807 | 1241.1425 | 1658.3392 | 1527.0615 |
| Dync1li1 | KPASVSPTTPTSPTEGEAS | -505 | -523 | 1 | 0.000230707 | 19 | 1135.8525 | 1887.9303 | 1666.368 | 1623.275 | 1191.7502 | 1067.0159 |
| Dync1li1 | KVEDNFEDIITKPPVR | -344 | -359 | 2 | 0 | 16 | 631.8978 | 995.2213 | 848.12646 | 1038.8179 | 1193.6168 | 1975.5282 |
| Dync1li1 | DFQEYVEPGEDFPASPQR | -193 | -210 | 0 | 0 | 18 | 503.36023 | 693.5748 | NaN | 571.881 | 724.87274 | 773.32153 |
| Tacc3 | SKYEDLNTK | -450 | -458 | 1 | 0 | 9 | 3069.209 | 704.0228 | 962.272 | NaN | 6932.2466 | 7184.4717 |
| Tacc3 | DFQFLPPELTGR | -21 | -32 | 0 | 0 | 12 | 664.22675 | 851.65906 | 693.6787 | 921.77875 | NaN | 1302.5474 |
| Tacc3 | GLLPAEPIVDVLK | -416 | -428 | 0 | 0 | 13 | 267.96695 | 310.8181 | 234.8302 | 249.28825 | NaN | 232.47345 |
| Tacc3 | LANEEIAQVH | -570 | -579 | 0 | 0.000230707 | 10 | 585.92444 | NaN | NaN | 661.29236 | 615.51715 | 285.19272 |
| Eif4h | EALTYDGALLGDR | -97 | -109 | 0 | 0 | 13 | 1921.281 | 2180.4373 | 940.6831 | 2887.9788 | 3232.7937 | 189.42508 |
| Eif4h | DFREPTEEER | -196 | -205 | 1 | 0 | 10 | 1880.5338 | 1877.874 | 1963.063 | 2322.9143 | 2473.8096 | 2388.8748 |
| Eif4h | DFREPTEEER | -196 | -205 | 1 | 0 | 10 | 1880.5338 | 1877.874 | 1963.063 | 2322.9143 | 2473.8096 | 2388.8748 |
| Eif4h | VEFDEVDSLK | -87 | -96 | 0 | 0 | 10 | 730.538 | 779.69434 | 875.56213 | 816.3494 | 976.54626 | 919.21063 |
| Eif4h | NQVANPNSAIFGGAR | -223 | -237 | 0 | 0 | 15 | 633.8244 | NaN | 1327.9286 | 2050.3965 | 2734.831 | 1637.178 |
| Eif4h | ELPTEPPYTAY | -35 | -45 | 0 | 0.000431655 | 11 | 152.88376 | 194.83607 | 131.37688 | 215.34462 | 230.31117 | NaN |
| Eif4h | FDEVDSLK | -89 | -96 | 0 | 0.000230707 | 8 | 2433.052 | 2693.0615 | 2510.5396 | NaN | NaN | NaN |
| Mfap1a;Mfap1 | DFSAPTLEDHFNK | (359);(359) | (371);(371) | 0 | 0 | 13 | 1030.4574 | 1313.2598 | 1251.585 | 1143.9135 | 1249.9897 | 2194.737 |
| Mfap1a;Mfap1 | EAEPEEQEEDSSSDPR | (70);(70) | (85);(85) | 0 | 0 | 16 | 281.45496 | 112.18605 | 146.013 | NaN | 205.60388 | 244.47806 |
| Mfap1a;Mfap1 | ISEDVEER | (93);(93) | (100);(100) | 0 | 0 | 8 | 887.3639 | NaN | NaN | 931.765 | 1710.8192 | NaN |
| Acaa2 | LEDTLWAGLTDQHVK | -144 | -158 | 0 | 0 | 15 | 458.04443 | 497.23474 | 2252.9502 | 611.11426 | 862.9093 | 889.59204 |
| Acaa2 | KALDLDPSK | -332 | -340 | 1 | 0 | 9 | 387.6693 | 594.16797 | 662.7548 | 859.06 | 471.1284 | 684.5835 |

| | | | | | | | | | | | | |
|---|---|---|---|---|---|---|---|---|---|---|---|---|
| Acaa2 | SATDLTEFAAR | -28 | -38 | 0 | 0 | 11 | 376.1776 | 2064.0466 | 2287.5957 | 1914.6343 | 2060.4297 | 2006.8815 |
| Acaa2 | DFSATDLTEFAAR | -26 | -38 | 0 | 0 | 13 | 375.04596 | 424.46613 | 379.3296 | 472.41605 | 218.7766 | 120.89969 |
| Acaa2 | TTLEQLQK | -227 | -234 | 0 | 0.002566359 | 8 | 498.74313 | 671.9958 | NaN | 199.0163 | NaN | 633.1924 |
| Prdx6 | LPFPIIDDK | -98 | -106 | 0 | 0 | 9 | 4251.701 | 581.1459 | 4759.1157 | 532.80444 | 707.7388 | 881.44995 |
| Prdx6 | LSILYPATTGR | -145 | -155 | 0 | 0.00025015 | 11 | 3289.4902 | 2741.8828 | 2800.744 | 607.00366 | 3103.0918 | 3031.5647 |
| Prdx6 | SIDSVEDHLAWSK | -72 | -84 | 0 | 0 | 13 | 2575.7903 | 2934.4207 | 4387.3022 | 5561.841 | 9403.559 | 9190.634 |
| Prdx6 | SIDSVEDHLAWSK | -72 | -84 | 0 | 0 | 13 | 2575.7903 | 2934.4207 | 4387.3022 | 5561.841 | 9403.559 | 9190.634 |
| Prdx6 | GTKPVATPVDWK | -171 | -182 | 1 | 0 | 12 | 2194.3862 | 1700.6584 | 1771.493 | 1962.5789 | 1911.3878 | 499.934 |
| Prdx6 | VVFIFGPDKK | -133 | -142 | 1 | 0 | 10 | 1600.4991 | 2161.583 | 2098.5027 | 1915.7296 | 2435.928 | 2388.2012 |
| Prdx6 | FHDFLGDSW | -25 | -33 | 0 | 0.002697088 | 9 | 1236.1887 | 1132.2767 | 1325.7977 | 757.8849 | 1128.0791 | 1139.6235 |
| Prdx6 | KLAPEFAK | -56 | -63 | 1 | 0 | 8 | 1134.4126 | 1441.9241 | 1209.8528 | 1108.4976 | 1345.8506 | 513.2087 |
| Prdx6 | QLTGTKPVATPVDWK | -168 | -182 | 1 | 0 | 15 | 1046.0513 | 1265.9408 | 1343.5978 | 1002.3617 | 1131.6149 | 1184.4443 |
| Prdx6 | QLTGTKPVATPVDWK | -168 | -182 | 1 | 0 | 15 | 1046.0513 | 1265.9408 | 1343.5978 | 1002.3617 | 1131.6149 | 1184.4443 |
| Prdx6 | VVFIFGPDK | -133 | -141 | 0 | 0 | 9 | 569.19073 | 597.83887 | 706.3666 | 399.52924 | 709.14264 | 689.26666 |
| Prdx6 | VVDSLQLTGTKPVATPVDWK | -163 | -182 | 1 | 0 | 20 | 164.26935 | 176.04337 | 200.12764 | 404.79144 | 478.97867 | 478.38806 |
| Prdx6 | PGGLLLGDEAPNFEAN | -2 | -17 | 0 | 0 | 16 | 88.69655 | 1018.4016 | 1091.981 | 918.6205 | 1556.624 | 1531.402 |
| Prdx6 | LPFPIIDDKG | -98 | -107 | 1 | 0.00025015 | 10 | 1672.068 | 1832.8158 | 1883.0801 | 1694.5822 | NaN | 2175.2974 |
| Prdx6 | SVEDHLAWSK | -75 | -84 | 0 | 0 | 10 | 1479.663 | 1562.2777 | 1651.2485 | NaN | 1404.4517 | 1440.3544 |
| Prdx6 | DFTPVCTTELGR | -42 | -53 | 0 | 0 | 12 | 576.0339 | NaN | 1363.9943 | 1379.8865 | NaN | 1304.8041 |
| Prdx6 | VVPTLSEEEAK | -189 | -199 | 0 | 0 | 11 | 1841.9625 | 1925.83 | NaN | NaN | NaN | NaN |
| Cdc123 | DFVDLSTGEDAHK | -309 | -321 | 0 | 0 | 13 | 920.38025 | NaN | NaN | 1370.9629 | 1218.411 | 1172.8047 |
| Cdc123 | SVILPLPQNVK | -25 | -35 | 0 | 0 | 11 | 575.9201 | 242.18185 | 598.1834 | NaN | 466.83417 | NaN |
| Cdc123 | TWEELTSENNLR | -260 | -271 | 0 | 0 | 12 | 3004.0957 | NaN | NaN | NaN | 2628.9268 | 2730.096 |
| Cdc123 | WCELIPGAEFR | -169 | -179 | 0 | 0 | 11 | 695.96655 | 824.5117 | 970.45984 | NaN | NaN | NaN |
| Anapc1 | DFVKPEFLLLR | -1397 | -1407 | 1 | 0 | 11 | 311.5934 | 291.77853 | 303.51755 | NaN | 328.43915 | NaN |
| Psmd13 | VEHQAQDILT | -367 | -376 | 0 | 0 | 10 | 1321.2549 | 885.4349 | 668.76636 | 983.2001 | 682.2825 | 709.9608 |
| Psmd13 | YYQTIGNHASYYK | -162 | -174 | 0 | 0 | 13 | 1149.0668 | 1058.0494 | 1090.8536 | 1260.3289 | 133.90779 | 1188.835 |
| Psmd13 | GSIDEVDKR | -322 | -330 | 1 | 0.00025015 | 9 | 1092.1072 | 1085.1821 | 803.80255 | 1237.2498 | 959.939 | 759.18134 |
| Psmd13 | LYENFISEFEHR | -59 | -70 | 0 | 0 | 12 | 795.9151 | 1020.36676 | 153.34659 | 936.05347 | 1031.064 | 703.89764 |
| Psmd13 | DFVQDPCFAQGDGLIK | -43 | -58 | 0 | 0.001023236 | 16 | 637.3734 | 741.40924 | 793.36505 | 692.79114 | 360.64935 | 907.67004 |
| Psmd13 | HRLEELYTK | -23 | -31 | 1 | 0.001023236 | 9 | 619.19086 | 478.4935 | 432.43384 | 633.2426 | 805.7476 | 813.75665 |
| Psmd13 | LELWCTDVK | -353 | -361 | 0 | 0 | 9 | 199.66408 | 440.78543 | 322.6305 | 398.62653 | 173.69164 | 564.9389 |
| Psmd13 | ENFISEFEHR | -61 | -70 | 0 | 0 | 10 | 1605.433 | 1348.8793 | 1804.665 | 1397.274 | NaN | 1431.0764 |
| Psmd13 | FLGCVDIK | -179 | -186 | 0 | 0.00111006 | 8 | 260.0079 | 4394.1943 | 4869.1606 | NaN | NaN | NaN |
| Txnrd1 | DFVTPTPLGTR | -156 | -166 | 0 | 0 | 11 | 2370.6213 | 2502.345 | 2603.063 | 2603.8584 | 389.41693 | 2757.2368 |
| Txnrd1 | QQLDSTIGIHPVCAEIF | -577 | -593 | 0 | 0.009416377 | 17 | 1003.07947 | 861.22675 | 1126.4022 | 1269.387 | 1539.8525 | 1484.3739 |
| Txnrd1 | SDDLFSLPYCPGK | -294 | -306 | 0 | 0 | 13 | 939.397 | 961.567 | 896.0225 | 1009.52313 | 1256.406 | 1221.2522 |
| Txnrd1 | TIGLETVGVK | -413 | -422 | 0 | 0 | 10 | 921.5691 | 862.2594 | 1045.5094 | 867.48224 | 606.42334 | 679.7224 |
| Txnrd1 | IEQIEAGTPGR | -370 | -380 | 0 | 0 | 11 | 815.7302 | 783.19635 | 793.7793 | 667.1302 | 819.7196 | 676.5136 |
| Txnrd1 | LELTPVAIQAGR | -454 | -465 | 0 | 0 | 12 | 589.38934 | 687.2448 | 659.5865 | 854.82385 | 930.2133 | 867.0792 |
| Txnrd1 | IPVTDEEQTNVPYIY | -430 | -444 | 0 | 0.002810811 | 15 | 478.16376 | 414.81805 | 639.24854 | 746.0627 | 902.3004 | 765.7507 |
| Txnrd1 | FGEENIEVYHSF | -508 | -519 | 0 | 0 | 12 | 453.29053 | 375.56473 | 359.65286 | 526.3426 | 467.9146 | 335.2203 |
| Txnrd1 | VLVLDFVTPTPLGTR | -152 | -166 | 0 | 0 | 15 | 320.8286 | 284.0541 | 231.96826 | 185.18921 | NaN | 220.8385 |
| Mtx1 | QTPAAPETEEEPYR | -254 | -267 | 0 | 0 | 14 | 329.01846 | 252.31898 | 256.26685 | 250.87157 | 519.99426 | 259.41623 |
| Mtx1 | LLPVLIHTF | -104 | -112 | 0 | 0.002099344 | 9 | 580.15857 | NaN | 623.1723 | 873.2281 | 196.73726 | 110.91097 |
| Mtx1 | DGDEVPLPR | -245 | -253 | 0 | 0 | 9 | 269.81635 | NaN | NaN | NaN | NaN | NaN |
| Arhgef1 | SGLELEPEEPPGWR | -385 | -398 | 0 | 0 | 14 | 5741.409 | 6095.2197 | 6141.2983 | 137.0174 | 5823.056 | 5305.414 |
| Arhgef1 | TRPDLISEDVQR | -123 | -134 | 1 | 0.00025015 | 12 | 2162.0762 | 2314.4163 | 2377.4104 | 2179.6167 | 2376.5164 | 2299.368 |
| Arhgef1 | TVGTPGQDNPGVSLHPL | -311 | -327 | 0 | 0 | 17 | 1334.9475 | 1626.376 | 1522.0465 | 1573.7521 | 1857.5583 | 1828.4286 |
| Arhgef1 | SIGQNTEESTER | -573 | -584 | 0 | 0 | 12 | 982.0668 | 978.742 | 1061.9987 | 904.59973 | 1112.1348 | 949.9507 |
| Arhgef1 | EILHHVNQAVR | -596 | -606 | 0 | 0 | 11 | 916.2202 | 926.6736 | 886.45435 | 975.13385 | 1235.3396 | 1126.0979 |
| Arhgef1 | RLDLTHLR | -620 | -627 | 1 | 0 | 8 | 766.2421 | 440.47986 | 481.64563 | 346.54346 | 994.1517 | 904.7041 |
| Arhgef1 | RLDLTHLR | -620 | -627 | 1 | 0 | 8 | 766.2421 | 440.47986 | 481.64563 | 346.54346 | 994.1517 | 904.7041 |
| Arhgef1 | FDGAEGSWFQK | -499 | -509 | 0 | 0 | 11 | 763.35736 | 318.38098 | 627.20776 | 716.6017 | 729.06665 | 442.09927 |
| Arhgef1 | ELVPPDTLLSLPK | -399 | -411 | 0 | 0 | 13 | 698.4639 | 760.1741 | 688.76807 | 802.1238 | 1206.9991 | 1280.5 |
| Arhgef1 | QILLSTEEDSGAGPPR | -835 | -850 | 0 | 0 | 16 | 690.027 | 998.9118 | 906.045 | 821.2671 | 927.2396 | 921.055 |
| Arhgef1 | LLSHLEETQHTISTDEEK | -195 | -212 | 0 | 0 | 18 | 263.36642 | 440.85132 | 606.27673 | 598.0164 | 206.12593 | 296.12973 |
| Arhgef1 | LLSHLEETQHTISTDEEK | -195 | -212 | 0 | 0 | 18 | 263.36642 | 440.85132 | 606.27673 | 598.0164 | 206.12593 | 296.12973 |
| Arhgef1 | VEADAEKPGPADR | -287 | -299 | 1 | 0 | 13 | 651.63995 | 712.13763 | 763.5372 | 549.9809 | 813.37726 | 727.1823 |
| Arhgef1 | KAFLDFYHSF | -92 | -101 | 1 | 0 | 10 | 411.21036 | 516.1312 | 361.90564 | 495.01636 | 496.5692 | 620.0349 |
| Arhgef1 | SHLEETQHTISTDEEK | -197 | -212 | 0 | 0 | 16 | 204.54747 | 267.76837 | 218.06679 | 311.9698 | 312.21124 | 289.4171 |
| Arhgef1 | LVHEGPLTWR | -647 | -656 | 0 | 0 | 10 | 647.60614 | 701.75836 | 736.33386 | 1083.5372 | NaN | 619.96234 |
| Arhgef1 | LVHEGPLTWR | -647 | -656 | 0 | 0 | 10 | 647.60614 | 701.75836 | 736.33386 | 1083.5372 | NaN | 619.96234 |
| Arhgef1 | GGTLSPNLAAPER | -901 | -913 | 0 | 0 | 13 | 642.02496 | 485.7839 | 554.6542 | 285.97302 | NaN | 929.5948 |
| Arhgef1 | QLEELEEEFCR | -882 | -892 | 0 | 0 | 11 | 2920.283 | 259.19354 | 3514.3342 | 3896.0142 | NaN | NaN |
| Arhgef1 | LVHEGPLTW | -647 | -655 | 0 | 0.008129024 | 9 | 1168.5212 | 955.6494 | NaN | NaN | 1439.6133 | 1443.7242 |
| Arhgef1 | LRPLLSQL | -893 | -900 | 1 | 0.001055699 | 8 | 1104.1627 | 1057.4994 | NaN | 605.248 | NaN | NaN |
| Arhgef1 | LDLTHLR | -621 | -627 | 0 | 0.000869338 | 7 | 1043.929 | NaN | NaN | NaN | 720.5688 | 866.98944 |
| Arhgef1 | KNWCNLITET | -748 | -757 | 1 | 0.00111006 | 10 | 674.88684 | 947.26385 | NaN | NaN | NaN | NaN |
| Arhgef1 | EETQHTISTDEEK | -200 | -212 | 0 | 0.000627943 | 13 | 650.0587 | NaN | 450.434 | NaN | NaN | NaN |
| Arhgef1 | LLSHLEETQHTISTD | -195 | -209 | 0 | 0.004472887 | 15 | 608.87805 | NaN | NaN | NaN | NaN | 622.57324 |
| Arhgef1 | DGDGVPGGR | -851 | -859 | 0 | 0.007493591 | 9 | 824.7874 | NaN | NaN | NaN | NaN | NaN |
| Erp29 | QGQDGLLSVK | -185 | -194 | 0 | 0 | 10 | 1030.521 | 1102.484 | 1020.75726 | 1305.1462 | 589.09894 | 1482.3129 |
| Erp29 | ILDQGEDFPASEMAR | -211 | -225 | 0 | 0 | 15 | 905.0215 | 562.8108 | 925.49524 | 1253.3953 | 1026.2103 | 777.2314 |
| Erp29 | WASQYLK | -200 | -206 | 0 | 0.006164361 | 7 | 382.1526 | 1405.8351 | 1493.9393 | 1598.7913 | 2009.5461 | 1923.0026 |
| Erp29 | DGDLENPVLYNGAVK | -125 | -139 | 0 | 0.000394361 | 15 | 793.5855 | 742.96735 | NaN | 752.0178 | NaN | NaN |
| Erp29 | ESYPVFYLFR | -115 | -124 | 0 | 0.002265552 | 10 | 406.8761 | NaN | NaN | 329.5843 | NaN | NaN |
| Blmh | VYEVVVDKK | -419 | -427 | 1 | 0 | 9 | 1681.3094 | 1467.1693 | 1402.3506 | 1584.8489 | 1527.7961 | 1078.155 |

| | | | | | | | | | | | | |
|---|---|---|---|---|---|---|---|---|---|---|---|---|
| Blmh | TLYNNQPIDFLK | -298 | -309 | 0 | 0 | 12 | 923.6146 | 1041.62 | 1036.7582 | 982.3598 | 1271.6792 | 1085.1298 |
| Blmh | DGEAVWFGCDVGK | -318 | -330 | 0 | 0 | 13 | 444.51526 | 429.72653 | 409.94785 | 598.5984 | 595.2174 | 715.8075 |
| Blmh | KTLYNNQPIDFLK | -297 | -309 | 1 | 0 | 13 | 412.9359 | 577.579 | 537.21014 | 512.035 | 494.38553 | 563.6417 |
| Blmh | TAVSEKDNQEGTFVK | -377 | -391 | 1 | 0.000230707 | 15 | 338.88864 | 298.08453 | 351.387 | 371.64874 | 393.35733 | 588.3151 |
| Blmh | VENSWGEDHGHK | -394 | -405 | 0 | 0 | 12 | 140.77151 | 455.80508 | 389.94293 | 347.3515 | 379.49982 | 360.29416 |
| Blmh | SEKDNQEGTFVK | -380 | -391 | 1 | 0 | 12 | 1119.8022 | 898.30304 | 865.4759 | 1033.311 | NaN | NaN |
| Blmh | FVNDPRPQHK | -270 | -279 | 1 | 0.00025015 | 10 | 647.8037 | 662.97253 | 788.1633 | NaN | 791.72894 | NaN |
| Blmh | LNSDPQFVLAQ | -18 | -28 | 0 | 0.001023236 | 11 | 2938.5835 | NaN | NaN | NaN | NaN | NaN |
| Slc44a2 | DGECPAVITPSKPLAQR | -155 | -171 | 1 | 0.00025015 | 17 | 1377.1898 | 695.9681 | 765.2897 | 689.8092 | 913.71643 | 931.7244 |
| Slc44a2 | VVDDTACPLLR | -395 | -405 | 0 | 0 | 11 | 1461.1716 | 1074.4335 | NaN | NaN | 819.85785 | 355.04297 |
| Slc44a2 | NADKPFLFYFN | -81 | -91 | 1 | 0.000591124 | 11 | 620.9729 | NaN | 602.04944 | 427.05646 | 533.2051 | NaN |
| Slc44a2 | QFCVPGFQNNK | -137 | -147 | 0 | 0.001708008 | 11 | 2077.0251 | NaN | NaN | 2227.9688 | 1824.4932 | NaN |
| Adss2 | FIEDELQIPVK | -431 | -441 | 0 | 0 | 11 | 819.83167 | 967.6666 | 876.2969 | 927.4938 | 1031.0372 | 777.55347 |
| Adss2 | DLVSDFDGFSER | -183 | -194 | 0 | 0 | 12 | 784.53186 | 786.6571 | 937.3754 | 846.40375 | 976.1458 | 892.85547 |
| Adss2 | KGIGPVYSSK | -164 | -173 | 1 | 0 | 10 | 768.62964 | 334.848 | 299.03336 | 177.00687 | 513.62036 | 494.01443 |
| Adss2 | TLPGWNTDISNAR | -404 | -416 | 0 | 0 | 13 | 733.08075 | 2008.8943 | 1328.7285 | 1027.9261 | 1508.3873 | 1449.1058 |
| Adss2 | DGETIPHFPANQEVLNK | -381 | -397 | 0 | 0 | 17 | 562.9368 | 558.53174 | 592.262 | 493.22818 | 543.5693 | 649.97705 |
| Adss2 | VVDLLAQDADIVCR | -46 | -59 | 0 | 0 | 14 | 3451.04 | 4277.5405 | 3152.3167 | NaN | 1163.3639 | 810.5845 |
| Adss2 | LDGETIPHFPANQEVLNK | -380 | -397 | 0 | 0 | 18 | 811.596 | 710.8279 | 728.98126 | 307.45718 | 891.20374 | NaN |
| Myl12b | GNFNYIEFTR | -152 | -161 | 0 | 0 | 10 | 3669.218 | 4072.1113 | 4024.5466 | 335.14966 | 3322.497 | 3054.109 |
| Myl12b | GDRFTDEEVDELYR | -131 | -144 | 1 | 0.00051306 | 14 | 1086.779 | 1496.7681 | 1338.3827 | 683.39795 | 1853.5492 | 1878.6201 |
| Myl12b | FTDEEVDELYR | -134 | -144 | 0 | 0 | 11 | 179.62097 | 3032.7444 | 2807.0366 | 3117.0442 | 3117.5579 | 3048.4868 |
| Macf1 | SSEIDEDEIRDEK | -4936 | -4948 | 1 | 0 | 13 | 1643.1658 | 620.76697 | 963.92664 | 1363.0878 | 515.87805 | 104.85078 |
| Macf1 | VDSLIPWIR | -323 | -331 | 0 | 0 | 9 | 1390.1226 | 980.6767 | 65.24879 | 1365.6818 | 849.93286 | 586.1533 |
| Macf1 | LQCQLQENEEFQK | -4793 | -4805 | 0 | 0.000550307 | 13 | 480.50708 | 271.398 | NaN | 793.2035 | 929.737 | 321.21103 |
| Hdac1 | YHSDDYIK | -67 | -74 | 0 | 0 | 8 | 1872.9678 | 1991.521 | 1969.6003 | 1472.1711 | 1758.7231 | 2031.1915 |
| Hdac1 | DGIDDESYEAIFKPVMSK | -230 | -247 | 1 | 0 | 18 | 447.19562 | 458.21048 | 485.3409 | 571.5742 | 626.9683 | 603.7152 |
| Hdac1 | DGIDDESYEAIFKPVMSK | -230 | -247 | 1 | 0 | 18 | 447.19562 | 458.21048 | 485.3409 | 571.5742 | 626.9683 | 603.7152 |
| Hdac1 | LGCFNLTIK | -271 | -279 | 0 | 0.00051306 | 9 | 354.6872 | 351.90292 | 405.9547 | NaN | NaN | 510.88843 |
| Psmb2 | YYTPTISR | -146 | -153 | 0 | 0 | 8 | 1845.1755 | 649.74335 | 1622.1799 | 1211.6454 | 1754.5535 | 384.96164 |
| Psmb2 | KCLEELQK | -162 | -169 | 1 | 0 | 8 | 1273.267 | 1154.6667 | 1280.1014 | 358.58102 | 1226.46 | 1329.115 |
| Psmb2 | VIDKDGIHNLENIAFPK | -182 | -198 | 1 | 0 | 17 | 1191.2489 | 1245.7411 | 1345.3541 | 1623.0703 | 2247.874 | 2294.3381 |
| Psmb2 | VIDKDGIHNLENIAFPK | -182 | -198 | 1 | 0 | 17 | 1191.2489 | 1245.7411 | 1345.3541 | 1623.0703 | 2247.874 | 2294.3381 |
| Psmb2 | DGIHNLENIAFPK | -186 | -198 | 0 | 0 | 13 | 814.8574 | 1043.2793 | 680.923 | 937.8162 | 1154.6147 | 1239.7571 |
| Psmb2 | FILNLPTFSVR | -171 | -181 | 0 | 0 | 11 | 642.2682 | 639.2899 | 524.1772 | 984.16925 | 1055.2919 | 871.03937 |
| Psmb2 | NLPTFSVR | -174 | -181 | 0 | 0.00635324 | 8 | 1473.9928 | 882.4614 | 1377.506 | NaN | NaN | 586.85077 |
| Psmb7 | LDFLRPF | -226 | -232 | 1 | 0.00051306 | 7 | 3526.2478 | 3460.7095 | 4292.1797 | 2446.7249 | 3782.2095 | 4217.856 |
| Psmb7 | IHFISPNIY | -77 | -85 | 0 | 0.000230707 | 9 | 1342.9857 | 1707.689 | 1653.4677 | 1536.4734 | 1351.022 | 1446.2761 |
| Psmb7 | SKLDFLRPF | -224 | -232 | 2 | 0 | 9 | 1169.8417 | 1387.621 | 1403.8011 | 1570.5098 | 661.438 | 1126.9437 |
| Psmb7 | TTIAGVVYK | -44 | -52 | 0 | 0 | 9 | 1435.123 | 1012.3489 | 1239.2322 | 1558.3706 | 2018.7924 | NaN |
| Psmb7 | DGIVLGADTR | -53 | -62 | 0 | 0.000230707 | 10 | 587.0989 | NaN | NaN | 1802.6746 | NaN | NaN |
| Usp5 | QAEEEKVPLPELVR | -511 | -524 | 1 | 0 | 14 | 1573.9894 | 2128.5483 | 98.252914 | 1367.8525 | 2063.607 | 2352.8271 |
| Usp5 | DGLGGLPDIVR | -134 | -144 | 0 | 0 | 11 | 1110.0127 | 1316.4194 | 1323.3014 | 1224.5892 | 1390.2488 | 1341.8855 |
| Usp5 | VHKDECAFSF | -25 | -34 | 1 | 0.002199523 | 10 | 879.8228 | 632.0743 | 595.8097 | 1224.5043 | 1033.9941 | 1140.4187 |
| Usp5 | LNKEELLEYEEK | -497 | -508 | 1 | 0 | 12 | 682.00726 | 803.5618 | 781.6447 | 1056.0378 | 1126.0013 | 1102.7216 |
| Usp5 | IVILPDYLEIAR | -122 | -133 | 0 | 0 | 12 | 480.14777 | 413.72934 | 507.98682 | 302.3533 | 379.1625 | 422.57608 |
| Usp5 | GVPLKPLFGPGYTGIR | -314 | -329 | 1 | 0 | 16 | 455.60983 | 450.94897 | 463.97516 | 362.83063 | 488.88617 | 456.57376 |
| Usp5 | GVPLKPLFGPGYTGIR | -314 | -329 | 1 | 0 | 16 | 455.60983 | 450.94897 | 463.97516 | 362.83063 | 488.88617 | 456.57376 |
| Usp5 | YFDGSGGNNHAVEHYR | -223 | -238 | 0 | 0 | 16 | 363.635 | 458.6485 | 969.93665 | 548.5112 | 639.1884 | 620.9217 |
| Usp5 | YFDGSGGNNHAVEHYR | -223 | -238 | 0 | 0 | 16 | 363.635 | 458.6485 | 969.93665 | 548.5112 | 639.1884 | 620.9217 |
| Usp5 | SSENPNEVFR | -453 | -462 | 0 | 0 | 10 | 1162.8632 | 1235.6506 | 1074.9845 | 1462.5258 | 1417.8733 | NaN |
| Usp5 | IFQNAPTDPTQDFSTQ | -361 | -376 | 0 | 0 | 16 | 658.26025 | 814.4372 | 851.3893 | NaN | 785.82104 | 994.7036 |
| Usp5 | FGPGYTGIR | -321 | -329 | 0 | 0.00025015 | 9 | 650.7765 | 513.59 | 807.25806 | 480.78986 | 856.12805 | NaN |
| Usp5 | FSIPDFQR | -345 | -352 | 0 | 0.00266297 | 8 | 799.65625 | NaN | 4250.8276 | NaN | 3897.752 | NaN |
| Fuca1 | DGLIVPIFQER | -330 | -340 | 0 | 0 | 11 | 363.9826 | 394.8738 | NaN | 347.7653 | 329.94055 | 339.54114 |
| Gusb | DGLWHFR | -42 | -48 | 0 | 0.000471409 | 7 | 401.35706 | 1444.759 | 916.0794 | NaN | NaN | NaN |
| Ccdc9 | DGPAPTHELSHR | -302 | -313 | 0 | 0 | 12 | 664.6147 | NaN | NaN | NaN | NaN | 1132.047 |
| Hadha | THINYGVK | -39 | -46 | 0 | 0.000298805 | 8 | 1874.6418 | 2154.0571 | 2019.1537 | 2931.9094 | 1936.2097 | 2016.0709 |
| Hadha | HYFSPVDK | -498 | -505 | 0 | 0.000230707 | 8 | 1495.5465 | 1247.9927 | 1189.8109 | 1162.3812 | 1254.9176 | 1220.2228 |
| Hadha | LPAKPEVSSDEDVQYR | -661 | -676 | 1 | 0.00025015 | 16 | 1234.0999 | 1241.9838 | 1039.9846 | 985.0761 | 1006.5133 | 1021.97205 |
| Hadha | LPAKPEVSSDEDVQYR | -661 | -676 | 1 | 0.00025015 | 16 | 1234.0999 | 1241.9838 | 1039.9846 | 985.0761 | 1006.5133 | 1021.97205 |
| Hadha | FGGGSVELLK | -611 | -620 | 0 | 0 | 10 | 1072.8428 | 129.10324 | 1205.6243 | 1372.8013 | 1247.6245 | 1282.0492 |
| Hadha | EVESVTPEHCIFAS | -461 | -474 | 0 | 0 | 14 | 855.34766 | 728.34656 | 726.1901 | 586.14026 | 717.9084 | 677.0721 |
| Hadha | GLVDQLVEPLGPGIK | -216 | -230 | 0 | 0 | 15 | 461.2497 | 391.24127 | 301.33728 | 468.40216 | 538.2928 | 527.9233 |
| Hadha | VAQHVAEDLGK | -595 | -605 | 0 | 0 | 11 | 590.9758 | 632.13403 | 848.64343 | NaN | 643.381 | 737.7433 |
| Hadha | SSCTTPQEATR | -108 | -118 | 0 | 0 | 11 | 572.5197 | NaN | 636.0567 | 683.43585 | 666.7987 | 149.91208 |
| Hadha | EVESVTPEHCIF | -461 | -472 | 0 | 0 | 12 | 123.28159 | 3948.4004 | 3421.519 | 2733.1553 | 2997.4443 | NaN |
| Hadha | DGPGFYTTR | -541 | -549 | 0 | 0.00122419 | 9 | 464.53824 | 548.3467 | 699.4224 | 1133.3046 | NaN | NaN |
| Hadha | QLVEPLGPGIK | -220 | -230 | 0 | 0 | 11 | 273.55948 | 155.9134 | 139.62608 | NaN | NaN | 307.3913 |
| Hadha | GLGFPPCLGGPFR | -707 | -719 | 0 | 0 | 13 | 361.46375 | NaN | 542.9472 | NaN | 238.70874 | NaN |
| Hadha | QHVAEDLGK | -597 | -605 | 0 | 0.000471409 | 9 | 604.8391 | NaN | 905.26575 | NaN | NaN | NaN |
| Slc39a7 | EKPSTEEEKEVGGL | -280 | -293 | 2 | 0 | 14 | 8228.771 | 614.8366 | 652.8195 | 608.10315 | 692.26636 | 744.42737 |
| Slc39a7 | DGPVKPQSPEEEK | -305 | -317 | 1 | 0 | 13 | 826.4463 | 675.0606 | 764.4617 | NaN | 335.5766 | 345.70624 |
| Slc39a7 | FLIPVESNSPR | -162 | -172 | 0 | 0 | 11 | 1879.2064 | NaN | NaN | NaN | NaN | NaN |
| Ubap2l | SANDSTVHSPFTK | -428 | -440 | 0 | 0 | 13 | 7561.3105 | 2233.8955 | 7668.2314 | 3487.348 | 3092.4387 | 3284.335 |
| Ubap2l | DGSLASNPYSGDLTK | -870 | -884 | 0 | 0 | 15 | 6348.409 | 5027.9663 | 5394.3477 | 6940.876 | 213.15938 | 5850.3096 |
| Ubap2l | GGSTTGSQFLEQFK | -374 | -387 | 0 | 0 | 14 | 1068.7981 | 391.73965 | 1208.6511 | 1106.2424 | 1715.8071 | 1177.8097 |
| Ubap2l | LDFIGVEGSNYPR | -236 | -248 | 0 | 0 | 13 | 839.1431 | 1024.1572 | NaN | 1661.4309 | 1940.2461 | 1794.3956 |

| | | | | | | | | | | | | |
|---|---|---|---|---|---|---|---|---|---|---|---|---|
| Ubap2l | SIPFPTPTTPLTGR | -856 | -869 | 0 | 0 | 14 | 527.839 | NaN | 361.5641 | 508.0864 | 586.6056 | 378.99734 |
| Ubap2l | GNWEQPQNQNQTQHK | -12 | -26 | 0 | 0 | 15 | 405.02777 | 424.53506 | 446.41754 | NaN | 303.9632 | 288.9021 |
| Ubap2l | GSQFLEQFK | -379 | -387 | 0 | 0 | 9 | 307.67773 | 295.47165 | 295.59076 | NaN | NaN | 371.24084 |
| Ubap2l | NDSTVHSPFTK | -430 | -440 | 0 | 0 | 11 | 350.0679 | NaN | NaN | 839.7917 | NaN | NaN |
| Ptpn6 | VDDQVTHIR | -45 | -53 | 0 | 0 | 9 | 4556.1553 | 2823.824 | 2960.2844 | 3341.1462 | 3669.1604 | 5683.0454 |
| Ptpn6 | NQLLGPDENSK | -309 | -319 | 0 | 0 | 11 | 2931.318 | 3269.7034 | 2689.5608 | 3092.2249 | 2431.1997 | 2480.5574 |
| Ptpn6 | YPLNCSDPTSER | -98 | -109 | 0 | 0 | 12 | 1839.4142 | 1830.1841 | 1715.0476 | 1568.7072 | 1449.7855 | 1439.0884 |
| Ptpn6 | GQESEYGNITYPPAVR | -531 | -546 | 0 | 0 | 16 | 1826.8702 | 1906.9374 | 2736.7058 | 2952.3274 | 3844.9858 | 4680.7524 |
| Ptpn6 | DGTIIHLK | -90 | -97 | 0 | 0 | 8 | 1290.152 | 1239.2036 | 1234.1934 | 979.90515 | 1208.7228 | 926.71173 |
| Ptpn6 | QESLPHAGPIIVH | -440 | -452 | 0 | 0 | 13 | 1055.413 | 951.0917 | 961.41876 | 1405.4489 | 1168.9182 | 1082.8394 |
| Ptpn6 | NSGDFYDLYGGEK | -56 | -68 | 0 | 0 | 13 | 911.5072 | 1102.2081 | 1048.9404 | 1017.44116 | 1221.3708 | 1215.3386 |
| Ptpn6 | TDLVEHFK | -188 | -195 | 0 | 0 | 8 | 696.9829 | 639.4546 | 756.00653 | 662.76294 | 593.935 | 685.1459 |
| Ptpn6 | FWEEFESLQK | -244 | -253 | 0 | 0 | 10 | 642.57996 | 494.31302 | 263.92938 | 990.674 | 973.92236 | 847.8436 |
| Ptpn6 | QESLPHAGPIIVHC | -440 | -453 | 0 | 0.000230707 | 14 | 542.9087 | 944.46747 | 1318.4042 | 856.3923 | 1120.1458 | 866.59375 |
| Ptpn6 | SFLDQINQR | -431 | -439 | 0 | 0 | 9 | 5765.0273 | NaN | 4410.7515 | 5220.1943 | 2266.605 | 1024.6078 |
| Ptpn6 | GLDCDIDIQK | -477 | -486 | 0 | 0 | 10 | 523.5069 | 318.53766 | 157.91133 | 549.08673 | 988.3311 | NaN |
| Ptpn6 | GFWEEFESLQK | -243 | -253 | 0 | 0 | 11 | 318.2538 | 465.81683 | 363.73248 | NaN | 504.36954 | 413.424 |
| Ptpn6 | NQGDFSLSVR | -35 | -44 | 0 | 0 | 10 | 749.61755 | 1570.2166 | NaN | NaN | 953.4974 | 2543.107 |
| Ptpn6 | IQNSGDFYDLYGGEK | -54 | -68 | 0 | 0 | 15 | 444.86847 | 539.47906 | 495.05957 | NaN | NaN | NaN |
| Ptpn6 | EYGNITYPPAVR | -535 | -546 | 0 | 0.001846793 | 12 | 1576.702 | NaN | 718.9182 | NaN | NaN | NaN |
| Ptpn6 | KNQGDFSLSVR | -34 | -44 | 1 | 0.006100831 | 11 | 2378.4753 | NaN | NaN | NaN | NaN | NaN |
| Rbpj | DGYIHYGQTVK | -265 | -275 | 0 | 0 | 11 | 1090.1606 | NaN | NaN | 1351.6097 | NaN | 1675.8391 |
| Rbpj | QPVQVPVTLVR | -446 | -456 | 0 | 0 | 11 | 362.942 | NaN | NaN | 457.12448 | NaN | 702.98444 |
| Pgls | IVAPISDSPKPPPQR | -171 | -185 | 1 | 0 | 15 | 1640.3373 | 2118.4675 | 2023.8755 | 1940.1151 | 2144.6082 | 2224.8525 |
| Pgls | IVAPISDSPKPPPQR | -171 | -185 | 1 | 0 | 15 | 1640.3373 | 2118.4675 | 2023.8755 | 1940.1151 | 2144.6082 | 2224.8525 |
| Pgls | SLFPDHPLLQER | -157 | -168 | 0 | 0 | 12 | 1229.5159 | 1469.0823 | 1268.4293 | 1620.4713 | 1813.0822 | 1693.6917 |
| Pgls | TINPALPVEDAAEDYAR | -112 | -128 | 0 | 0.000230707 | 17 | 908.8827 | 1370.153 | 1087.4575 | 1259.7468 | 1466.0134 | 1487.9086 |
| Pgls | WFLDEAAAR | -237 | -245 | 0 | 0 | 9 | 813.379 | 1024.5564 | 977.0626 | 1046.5115 | 982.11456 | 960.10583 |
| Pgls | DLPAAAAPAGPASFAR | -57 | -72 | 0 | 0 | 16 | 682.90955 | 733.6456 | 941.58984 | 886.88525 | 937.8529 | 929.364 |
| Pgls | LVPFDHAESTYGLYR | -82 | -96 | 0 | 0 | 15 | 342.17294 | 526.0225 | 536.5467 | 485.53604 | 1109.3392 | 1039.1753 |
| Pgls | DHAESTYGLYR | -86 | -96 | 0 | 0 | 11 | 4406.6367 | 497.6246 | NaN | 6156.889 | 6963.0645 | 6799.308 |
| Pgls | WTLGFCDER | -73 | -81 | 0 | 0 | 9 | 1598.2743 | 1705.9602 | 1672.3491 | NaN | 2035.1025 | 2044.7692 |
| Pop4 | DHDVQELGSQR | -16 | -26 | 0 | 0 | 11 | 687.32404 | NaN | 606.1436 | NaN | 842.32697 | NaN |
| Cd69 | YNCPGLYEK | -66 | -74 | 0 | 0 | 9 | 4294.076 | 4070.6897 | 4504.6187 | 2803.236 | 434.66278 | 4812.4893 |
| Cd69 | YSGELEHWIGLK | -135 | -146 | 0 | 0 | 12 | 1078.6362 | 1099.585 | 187.86423 | 1115.3801 | 1399.07 | 1759.7301 |
| Cd69 | DHGTSIHFEK | -22 | -31 | 0 | 0 | 10 | 570.54144 | 647.0861 | 673.07227 | NaN | NaN | 524.9893 |
| Cd69 | DHGTSIHFEK | -22 | -31 | 0 | 0 | 10 | 570.54144 | 647.0861 | 673.07227 | NaN | NaN | 524.9893 |
| Ube3a | DHIIDDALVR | -525 | -534 | 0 | 0 | 10 | 435.4984 | 423.92892 | 732.442 | 1982.9069 | 591.9068 | 846.628 |
| Ube3a | VDPLETELGVK | -420 | -430 | 0 | 0 | 11 | 307.31714 | 315.1074 | 622.4897 | 693.7939 | 697.1118 | 619.9427 |
| Ube3a | NLGLYYDNR | -485 | -493 | 0 | 0 | 9 | 1163.0436 | 972.0392 | 1177.4941 | NaN | 1324.3032 | NaN |
| Aqr | LGHGEEELETEK | -879 | -890 | 0 | 0 | 12 | 1631.8002 | 1489.0186 | 1506.7513 | 1161.773 | 1063.4976 | 1309.5735 |
| Aqr | DHIKDEWEGLR | -547 | -557 | 1 | 0 | 11 | 1204.4427 | 625.9702 | 774.9352 | 1206.7217 | 1464.9491 | 1210.9764 |
| Aqr | DHIKDEWEGLR | -547 | -557 | 1 | 0 | 11 | 1204.4427 | 625.9702 | 774.9352 | 1206.7217 | 1464.9491 | 1210.9764 |
| Aqr | RIELLEEVKR | -905 | -914 | 2 | 0.00025015 | 10 | 615.0529 | 816.5588 | 863.96893 | 444.15125 | 603.1827 | 698.62213 |
| Aqr | TLHDYLLR | -461 | -468 | 0 | 0.00025015 | 8 | 600.9981 | 639.5115 | 766.4853 | NaN | NaN | NaN |
| Aqr | ENVPAWETFK | -104 | -113 | 0 | 0.00437931 | 10 | 229.30875 | NaN | 3064.7585 | 3629.3857 | NaN | NaN |
| Mrpl22 | DHNVEFR | -125 | -131 | 0 | 0.00383302 | 7 | 1113.2374 | 1390.1979 | 1151.1042 | NaN | NaN | NaN |
| Sema4d | LICSKPDSGLVF | -279 | -290 | 1 | 0.000712251 | 12 | 3171.604 | 2318.2263 | 2894.6582 | 916.3014 | 2809.5776 | 1341.3073 |
| Sema4d | SEDKDTLYVGAR | -56 | -67 | 1 | 0 | 12 | 1201.8076 | 763.0091 | 1726.1571 | 619.9926 | 2176.3823 | 1926.377 |
| Sema4d | HGSCEDCVLAR | -506 | -516 | 0 | 0 | 11 | 981.6665 | 959.54004 | 137.37067 | 1075.1038 | 1191.2883 | 1115.7623 |
| Sema4d | EVHVIEETQLFR | -457 | -468 | 0 | 0 | 12 | 610.61206 | 846.47687 | 295.4539 | 483.79688 | 610.9932 | 953.94965 |
| Sema4d | NFLGSEPIISR | -180 | -190 | 0 | 0 | 11 | 512.17456 | 544.4483 | 629.363 | 668.7087 | 672.32434 | 2015.7759 |
| Sema4d | ESFNQHFFK | -559 | -567 | 0 | 0 | 9 | 192.53554 | 1002.2647 | 2617.0527 | 2149.768 | 1161.8728 | 1437.4558 |
| Sema4d | TGYETEQDTITSK | -812 | -824 | 0 | 0 | 13 | 684.9792 | 1237.9773 | 1264.7388 | NaN | 2143.426 | 1675.9819 |
| Sema4d | SYNFLGSEPIISR | -178 | -190 | 0 | 0 | 13 | 650.0565 | 599.6386 | 510.41284 | NaN | 969.2497 | 708.5439 |
| Sema4d | DHPLMDDSVTPIDNRPK | -396 | -412 | 1 | 0.000550307 | 17 | 955.2229 | NaN | 903.0633 | 639.3615 | 770.511 | NaN |
| Sema4d | LTWEHGEVGLVQ | -30 | -41 | 0 | 0 | 12 | 222.05339 | NaN | 684.34186 | 1034.4525 | 1336.593 | NaN |
| Sema4d | YNGPVPTPR | -358 | -366 | 0 | 0.002068213 | 9 | 714.06445 | 725.7101 | 1263.0836 | NaN | NaN | NaN |
| Sema4d | QTECLNYIR | -105 | -113 | 0 | 0 | 9 | 833.69824 | NaN | NaN | 732.8733 | NaN | NaN |
| Trmt5 | DHQLPFK | -206 | -212 | 0 | 0.000230707 | 7 | 883.22034 | 862.13025 | 766.87305 | 929.6696 | NaN | 782.6866 |
| Erg28 | YLEAEPVSR | -127 | -135 | 0 | 0.000298805 | 9 | 1125.2922 | 1195.3491 | 1097.4562 | 1256.859 | 993.1323 | 948.847 |
| Erg28 | DHTFLYEK | -29 | -36 | 0 | 0 | 8 | 397.14304 | 553.18835 | 551.09125 | 565.7765 | 637.05084 | 504.26608 |
| Supt6h | HETEQFPAEPLELAK | -557 | -571 | 0 | 0.000431655 | 15 | 923.5099 | 1139.5532 | 1048.4321 | 1157.1075 | 1204.6274 | 1320.8009 |
| Supt6h | DHYQDPVPGITPS | -1513 | -1525 | 0 | 0 | 13 | 660.556 | 730.44464 | 650.14233 | NaN | 533.257 | 481.52048 |
| Supt6h | NVTGIAHR | -1170 | -1177 | 0 | 0.002329279 | 8 | 822.3958 | 538.7709 | 806.21936 | NaN | 572.8219 | NaN |
| Supt6h | GLTPEQFGENLR | -540 | -551 | 0 | 0 | 12 | 247.11511 | NaN | NaN | 1349.5176 | 969.1891 | NaN |
| Supt6h | EYVEPELHIN | -382 | -391 | 0 | 0.00830926 | 10 | 361.8217 | 596.1421 | NaN | NaN | NaN | NaN |
| Dynll1 | DIAAHIK | -37 | -43 | 0 | 0.005270854 | 7 | 299.2494 | NaN | NaN | NaN | NaN | NaN |
| Thumpd3 | LPWSDPLK | -126 | -133 | 0 | 0.00078661 | 8 | 1398.5156 | 650.8809 | 1132.1455 | 1071.3759 | 1294.6168 | 1268.0264 |
| Thumpd3 | DICNLPLR | -391 | -398 | 0 | 0.001846793 | 8 | 1075.4204 | NaN | NaN | NaN | NaN | NaN |
| Puf60 | IYYELGEDTIR | -141 | -151 | 0 | 0 | 11 | 1712.4589 | 1956.7924 | 825.5222 | 409.80527 | 2058.4924 | 2177.0896 |
| Puf60 | SVHQDLSDDDIK | -237 | -248 | 0 | 0 | 12 | 1317.2534 | 1363.5387 | 1464.0323 | 888.2035 | 1331.4716 | 1433.7804 |
| Puf60 | NLFDLGGQYLR | -293 | -303 | 0 | 0 | 11 | 1313.6982 | 670.98956 | 1154.1411 | 1455.0906 | 1852.9938 | 808.20435 |
| Puf60 | LGLPPLTPEQQEALQK | -59 | -74 | 0 | 0 | 16 | 1111.039 | 1164.1584 | 1067.0542 | 913.3346 | 1181.3901 | 1196.5054 |
| Puf60 | VHQDLSDDDIK | -238 | -248 | 0 | 0 | 11 | 875.7908 | 921.27075 | 502.1447 | 794.0032 | 946.26886 | 921.8043 |
| Puf60 | YELGEDTIR | -143 | -151 | 0 | 0 | 9 | 667.424 | 721.62494 | 645.4035 | 702.94934 | 659.4125 | 802.68835 |
| Puf60 | DIDDDLEGEVTEECGK | -479 | -494 | 0 | 0 | 16 | 495.04797 | 567.30176 | 579.4106 | 473.48532 | 532.44116 | 500.83957 |
| Puf60 | VGRPSNIGQAQPIIDQLAEEA | -207 | -227 | 1 | 0.000230707 | 21 | 313.33527 | 342.24136 | 117.69804 | 227.0696 | 216.70944 | 286.9317 |

| | | | | | | | | | | | |
|---|---|---|---|---|---|---|---|---|---|---|---|
| Puf60 | LFDLGGQYLR | -294 | -303 | 0 | 0.00078661 | 10 | 1266.1163 | 1301.9419 | 1300.2875 | NaN | 1630.0393 | 1568.2092 |
| Puf60 | QGEEEDAEIIVK | -508 | -519 | 0 | 0 | 12 | 918.29266 | 1120.6196 | 846.3715 | NaN | 3021.4055 | 1053.404 |
| Puf60 | GYGFIEYEK | -273 | -281 | 0 | 0.00025015 | 9 | 1056.9473 | NaN | 824.7106 | NaN | NaN | 301.56873 |
| Clta | DIDESSPGTEWER | -188 | -200 | 0 | 0 | 13 | 1182.1847 | 1106.227 | 1057.0735 | 850.2257 | 273.23547 | 258.05286 |
| Clta | ELEEWYAR | -141 | -148 | 0 | 0 | 8 | 896.6271 | 925.7249 | 1127.3589 | 193.63802 | NaN | 352.821 |
| Sms | LYCPVEFSK | -335 | -343 | 0 | 0 | 9 | 2628.451 | 343.6429 | 349.1342 | 599.14716 | 2919.3945 | 2189.9126 |
| Sms | TCGDVLDNLR | -236 | -245 | 0 | 0 | 10 | 513.18176 | 557.4731 | 413.1175 | 550.29694 | 939.98364 | 274.70337 |
| Sms | SGDVNLAESDLAYTR | -165 | -179 | 0 | 0 | 15 | 506.6347 | 1277.7467 | 625.49756 | 811.3212 | 930.12274 | 898.0693 |
| Sms | DIDEVVYDEDSPYQNIK | -135 | -151 | 0 | 0 | 17 | 426.36078 | 372.75958 | 391.93375 | 290.38373 | NaN | 291.0396 |
| Mrps6 | ECDGIVPVPLEEK | -104 | -116 | 0 | 0 | 13 | 1055.153 | 1035.4152 | 1051.5547 | 927.16833 | 866.8233 | 681.8251 |
| Mrps6 | DIDVVRPNIVK | -85 | -95 | 1 | 0 | 11 | 698.39856 | 920.63794 | 888.72107 | 759.37274 | 865.6837 | 438.09402 |
| Wdr3 | SFDLIHSPQGELK | -366 | -378 | 0 | 0 | 13 | 2069.6533 | 2413.6523 | 1873.0231 | 1299.7073 | 1365.4219 | 1835.0298 |
| Wdr3 | EDQPAVPGETQGDNYFTGK | -729 | -747 | 0 | 0.00025015 | 19 | 522.4529 | 552.05133 | 471.90314 | 663.24426 | 717.7544 | 564.12946 |
| Wdr3 | WWDLDNQHCFK | -175 | -185 | 0 | 0 | 11 | 457.58844 | NaN | 588.31256 | 583.39044 | 702.7823 | 702.0047 |
| Wdr3 | DIDYLQEIDDPEEPEPK | -219 | -235 | 0 | 0 | 17 | 255.3263 | 413.76904 | 343.90048 | 598.54694 | 423.25348 | NaN |
| Wdr3 | GLDFGDCHR | -618 | -626 | 0 | 0 | 9 | 403.89954 | NaN | NaN | 517.27637 | 570.97235 | 474.00345 |
| Zap70 | DIELGCGNFGSVR | -340 | -352 | 0 | 0 | 13 | 4623.154 | 5120.7334 | 6478.9614 | 187.2455 | 348.88068 | 2296.7686 |
| Zap70 | PDPAAHLPFFYGS | -2 | -14 | 0 | 0.001392869 | 13 | 1269.0518 | 1087.3392 | 1133.1385 | 1109.7375 | 1736.6622 | 1584.0413 |
| Zap70 | FHHFPIER | -56 | -63 | 0 | 0 | 8 | 679.74774 | 652.38837 | 325.08682 | 468.574 | 1186.487 | 332.0974 |
| Zap70 | FHHFPIER | -56 | -63 | 0 | 0 | 8 | 679.74774 | 652.38837 | 325.08682 | 468.574 | 1186.487 | 332.0974 |
| Zap70 | VAAPTLPAHPSTFTQPQR | -263 | -280 | 0 | 0 | 18 | 1004.0177 | 987.7046 | 660.55505 | 585.9822 | 1019.7536 | 672.54987 |
| Zap70 | VAAPTLPAHPSTFTQPQR | -263 | -280 | 0 | 0 | 18 | 1004.0177 | 987.7046 | 660.55505 | 585.9822 | 1019.7536 | 672.54987 |
| Zap70 | VDTLNSDGYTPEPAR | -282 | -296 | 0 | 0 | 15 | 927.1829 | 1025.1306 | 940.68854 | 286.78842 | 502.1112 | 923.44073 |
| Zap70 | SLVHDVR | -49 | -55 | 0 | 0 | 7 | 801.87024 | 885.6193 | 969.5038 | 651.6221 | 834.06213 | 555.0398 |
| Zap70 | YSQDPDGLPCNLR | -87 | -99 | 0 | 0 | 13 | 313.53668 | 429.60822 | 420.3785 | 467.00034 | 260.4324 | 603.2409 |
| Zap70 | HQLDNPYIVR | -390 | -399 | 0 | 0 | 10 | 264.58276 | 2607.4807 | 2157.798 | 3423.209 | 3068.8467 | 3050.4802 |
| Zap70 | ESPYSDPEELKDK | -315 | -327 | 1 | 0 | 13 | 951.639 | 896.45154 | NaN | 538.4332 | NaN | 1433.8264 |
| Zap70 | ESPYSDPEELK | -315 | -325 | 0 | 0 | 11 | 796.3735 | 809.7811 | NaN | 292.28217 | NaN | NaN |
| Exosc7 | LGHTDILVGVK | -53 | -63 | 0 | 0 | 11 | 1058.1018 | 741.6396 | 811.86633 | 1024.5647 | 800.21606 | 805.4769 |
| Exosc7 | DIELSDDPYDCIR | -179 | -191 | 0 | 0 | 13 | 636.83057 | 604.997 | 563.4987 | 596.4491 | NaN | NaN |
| Lonp1 | NVIDTIPEPLR | -629 | -639 | 0 | 0 | 11 | 2003.1342 | 2090.6038 | 2060.4312 | 686.5844 | 1544.2876 | 1477.6152 |
| Lonp1 | TANVIDTIPEPLR | -627 | -639 | 0 | 0 | 13 | 1417.5846 | 852.8335 | 1046.6696 | 965.4054 | 1172.3346 | 954.4475 |
| Lonp1 | DIIALNPLYR | -291 | -300 | 0 | 0 | 10 | 965.92914 | 1059.9105 | 1124.2389 | 1081.7177 | 1319.6697 | 1273.4718 |
| Lonp1 | ELGLEKDDKDAIEEK | -395 | -409 | 2 | 0.000712251 | 15 | 863.94714 | 901.5027 | 1289.654 | 1798.9321 | 1370.2239 | 1615.7417 |
| Lonp1 | QSDENLDLAR | -462 | -471 | 0 | 0 | 10 | 826.85126 | 409.2166 | 355.64154 | 344.92316 | 577.8435 | 1132.803 |
| Lonp1 | DIFPIAFPR | -930 | -938 | 0 | 0 | 9 | 667.61005 | 693.0463 | 814.86285 | 629.6827 | 935.43665 | 943.94006 |
| Lonp1 | NVAHEDFQVTEEVK | -266 | -279 | 0 | 0 | 14 | 575.6382 | 767.03296 | 823.8259 | 725.7026 | 772.0915 | 741.8449 |
| Lonp1 | EVEDELGPKPQLE | -234 | -246 | 1 | 0 | 13 | 277.21957 | 1271.7091 | 253.25159 | 1496.3766 | 1567.2513 | 1403.8008 |
| Lonp1 | SHIHLHVPEGATPK | -827 | -840 | 0 | 0 | 14 | 1162.3756 | 1234.3846 | 1393.1411 | NaN | 1433.0261 | 1354.8292 |
| Lonp1 | SHIHLHVPEGATPK | -827 | -840 | 0 | 0 | 14 | 1162.3756 | 1234.3846 | 1393.1411 | NaN | 1433.0261 | 1354.8292 |
| Lonp1 | TENPLVLIDEVDK | -571 | -583 | 0 | 0 | 13 | 1246.6998 | NaN | NaN | 1169.799 | 1486.968 | 1589.4907 |
| Lonp1 | ILCFHGPPGVGK | -507 | -518 | 0 | 0.00025015 | 12 | 1033.3956 | 1051.6434 | NaN | NaN | NaN | NaN |
| Edf1 | INEKPQVIADYESGR | -99 | -113 | 1 | 0 | 15 | 1478.4507 | 966.0201 | 1356.0225 | 459.94147 | 658.4696 | 751.0433 |
| Edf1 | DIGKPIEK | -136 | -143 | 1 | 0 | 8 | 1917.9524 | NaN | 233.63177 | 2220.3845 | NaN | 4727.927 |
| Edf1 | INEKPQVIAD | -99 | -108 | 1 | 0.00111006 | 10 | 637.9424 | 802.04504 | NaN | NaN | NaN | NaN |
| Edf1 | EKPQVIADYESGR | -101 | -113 | 1 | 0.001708008 | 13 | 1769.1216 | NaN | NaN | NaN | NaN | NaN |
| Son | LSEPKPIFFNL | -2163 | -2173 | 1 | 0.001023236 | 11 | 2126.404 | 1895.8213 | 2342.296 | 872.7729 | 970.8511 | 807.6978 |
| Son | ESSEDTEVPLPNK | -1726 | -1738 | 0 | 0 | 13 | 1186.3969 | 1409.1709 | 1381.5708 | 1000.87305 | 886.6156 | 290.96643 |
| Son | SIDEINEADLVRPLLPK | -1749 | -1765 | 1 | 0 | 17 | 597.8054 | 668.03723 | 591.7702 | 752.7976 | 880.3291 | 843.78204 |
| Son | EVQESHVLETLKPATK | -192 | -207 | 1 | 0 | 16 | 594.1772 | 566.4037 | 622.8483 | 606.3098 | 791.1012 | 807.6222 |
| Son | DIHLDLPSTNFVCK | -1690 | -1703 | 0 | 0 | 14 | 768.4777 | 785.35876 | 655.30035 | 758.22864 | NaN | NaN |
| Son | TTLVELPIAK | -285 | -294 | 0 | 0.001055699 | 10 | 404.04684 | NaN | 378.70734 | NaN | 175.91884 | 238.48915 |
| Usp47 | DIHQDLDWNPK | -1288 | -1298 | 0 | 0 | 11 | 569.5576 | 273.0844 | 945.43243 | 463.62378 | 147.85017 | 1138.0979 |
| Usp47 | SLLDANFEPGKK | -111 | -122 | 1 | 0.00025015 | 12 | 310.41284 | 309.63382 | NaN | 579.6058 | 306.0793 | NaN |
| Usp47 | STETSDFENIESPLNER | -900 | -916 | 0 | 0 | 17 | 425.92813 | 557.5803 | 503.20905 | NaN | NaN | NaN |
| Usp47 | LSEISGIPLEDIEFAK | -1260 | -1275 | 0 | 0.000431655 | 16 | 457.9789 | NaN | NaN | NaN | NaN | NaN |
| Rps6ka1 | DILADVNHPFVVK | -114 | -126 | 0 | 0 | 13 | 417.08557 | 408.78046 | 408.4465 | 559.0812 | 281.80322 | 646.1721 |
| Rps6ka1 | LGSGPDGAEEIKR | -294 | -306 | 1 | 0 | 13 | 1070.6221 | NaN | NaN | 2101.6958 | NaN | 2388.1873 |
| Rps6ka1 | LGSGPDGAEEIKR | -294 | -306 | 1 | 0 | 13 | 1070.6221 | NaN | NaN | 2101.6958 | NaN | 2388.1873 |
| Glrx3 | GELIGGLDIIK | -209 | -219 | 0 | 0 | 11 | 1539.5225 | 1459.5343 | 2086.169 | 2183.5056 | 2801.0535 | 677.67535 |
| Glrx3 | LDGAHAPELTK | -102 | -112 | 0 | 0 | 11 | 1461.7545 | 1585.364 | 454.36978 | 2170.8352 | 2480.0708 | 440.76004 |
| Glrx3 | GDLVGGLDIVK | -311 | -321 | 0 | 0 | 11 | 1031.4215 | 1001.93195 | 1263.0657 | 1967.2478 | 1298.308 | 1239.3347 |
| Glrx3 | EASEELDTICPK | -222 | -233 | 0 | 0 | 12 | 1002.56354 | 832.6202 | 850.73456 | 1048.0052 | 743.2977 | 881.56024 |
| Glrx3 | ELKDNGELLPILK | -322 | -334 | 1 | 0 | 13 | 985.1977 | 1053.4949 | 1019.17975 | 347.82907 | 506.97284 | 483.6616 |
| Glrx3 | EHPHVSFVK | -61 | -69 | 0 | 0 | 9 | 867.99744 | 770.70825 | 733.06964 | 782.87585 | 1159.2327 | 1123.9608 |
| Glrx3 | DNGELLPILK | -325 | -334 | 0 | 0 | 10 | 600.0891 | 698.2078 | 733.5185 | 535.0993 | 726.92017 | 688.3898 |
| Glrx3 | SVPTFLFFK | -86 | -94 | 0 | 0.00111006 | 9 | 307.83505 | 651.50134 | 110.294464 | 304.44946 | 711.92285 | 911.92804 |
| Glrx3 | SFDIFSDEEVR | -180 | -190 | 0 | 0 | 11 | 3102.8643 | 1257.0933 | NaN | 1590.5583 | 2062.426 | NaN |
| Glrx3 | TFSNWPTYPQLYVR | -297 | -310 | 0 | 0 | 14 | 2351.944 | NaN | 2812.8247 | 2881.8564 | NaN | NaN |
| Glrx3 | ELEASEELDTICPK | -220 | -233 | 0 | 0 | 14 | 1828.9639 | NaN | 3030.3557 | NaN | NaN | 3354.578 |
| Glrx3 | DILEDEEVR | -284 | -292 | 0 | 0 | 9 | 765.12787 | NaN | 744.3771 | 681.27014 | NaN | NaN |
| Glrx3 | EHLKEDLSLR | -129 | -138 | 1 | 0.003299042 | 10 | 2201.8691 | NaN | NaN | NaN | 2560.015 | NaN |
| Glrx3 | LEAEAVPEVSEK | -70 | -81 | 0 | 0.002068213 | 12 | 1095.0651 | NaN | NaN | NaN | NaN | 280.28192 |
| Pwp1 | DILGDRPSLIHSR | -413 | -425 | 1 | 0 | 13 | 142.64455 | 939.3562 | 965.9039 | 879.30975 | 1029.7498 | 1016.4629 |
| Pwp1 | ESLEDGDPQDDR | -73 | -84 | 0 | 0 | 12 | 863.6587 | NaN | NaN | 721.8543 | NaN | 484.6928 |
| Pitpnc1 | DILLIGHR | -210 | -217 | 0 | 0 | 8 | 320.52054 | NaN | NaN | 595.3905 | NaN | NaN |
| Rbm25 | LVPLDYGEDDKNATK | -707 | -721 | 1 | 0 | 15 | 763.312 | 873.8141 | 1061.3704 | 948.96796 | 1125.0464 | 932.6297 |
| Rbm25 | LVPLDYGEDDKNATK | -707 | -721 | 1 | 0 | 15 | 763.312 | 873.8141 | 1061.3704 | 948.96796 | 1125.0464 | 932.6297 |

| | | | | | | | | | | | | |
|---|---|---|---|---|---|---|---|---|---|---|---|---|
| Rbm25 | LLAEGHPDPDAELQR | -545 | -559 | 0 | 0 | 15 | 676.96545 | 613.92834 | 687.8274 | 381.48322 | 712.69025 | 478.55316 |
| Rbm25 | SSELNAPSQESDSHPR | -218 | -233 | 0 | 0 | 16 | 465.1158 | 485.46902 | 439.4959 | 443.3471 | 558.8861 | 500.72305 |
| Rbm25 | EKEELEEIR | -534 | -542 | 1 | 0.000431655 | 9 | 277.1203 | 1699.0586 | 285.65796 | 1301.581 | 1719.502 | 1894.5684 |
| Rbm25 | DREEDEEDAYER | -427 | -438 | 1 | 0.000230707 | 12 | 175.85753 | 1024.1741 | 1018.8807 | 1175.9962 | 1465.2698 | 1514.8665 |
| Rbm25 | LEDYDDDRDDPK | -495 | -506 | 1 | 0.00025015 | 12 | 80.51918 | 931.32806 | 733.2187 | 760.1379 | 997.88837 | 1005.6582 |
| Rbm25 | DINAIEMEEDKR | -262 | -273 | 1 | 0 | 12 | 1918.3484 | NaN | 1812.0715 | 142.17407 | 2235.521 | 2021.9739 |
| Rbm25 | TVTNDDEEALDEETK | -186 | -200 | 0 | 0 | 15 | 649.4531 | 831.65173 | 735.69666 | 706.5735 | 893.3164 | NaN |
| Rbm25 | GFCEYKEPESTLR | -130 | -142 | 1 | 0.000471409 | 13 | 509.95605 | 128.21848 | NaN | 706.0393 | 721.3735 | 578.67053 |
| Rbm25 | LKENDENCGPTTT | -76 | -88 | 1 | 0 | 13 | 432.85556 | 513.11035 | NaN | 587.54474 | 502.7452 | 460.9233 |
| Rbm25 | EDEDSDDVPR | -694 | -703 | 0 | 0 | 10 | 330.71213 | NaN | 338.37088 | 436.32364 | 466.1921 | 338.20465 |
| Rbm25 | EFLEDYDDDRDDPK | -493 | -506 | 1 | 0.00201047 | 14 | 304.531 | 311.08344 | NaN | 365.04248 | 427.49728 | 449.39716 |
| Rbm25 | IRPWINK | -766 | -772 | 1 | 0.00025015 | 7 | 538.0608 | NaN | 614.87897 | NaN | 290.15048 | 497.73572 |
| Rbm25 | EYSSELNAPSQESDSHPR | -216 | -233 | 0 | 0 | 18 | 303.31085 | 158.96869 | 205.69025 | NaN | 413.54816 | NaN |
| Rbm25 | LLIYETEAK | -824 | -832 | 0 | 0 | 9 | 1258.1592 | 684.752 | 693.38513 | NaN | NaN | NaN |
| Rbm25 | KLPVDSVFNKF | -683 | -693 | 2 | 0.001190916 | 11 | 2218.219 | NaN | NaN | NaN | NaN | NaN |
| Hcfc1 | DLWYLETEKPPPPAR | -355 | -369 | 1 | 0 | 15 | 1364.6775 | 1443.8396 | 1486.1973 | 1378.0635 | 2177.1873 | 1710.9414 |
| Hcfc1 | GVKPDLPPPPSK | -1802 | -1813 | 1 | 0 | 12 | 977.70575 | 1109.8392 | 921.809 | 398.6849 | 1078.475 | 837.1236 |
| Hcfc1 | TATSPTPNPVPSVPANPPK | -408 | -426 | 0 | 0.001023236 | 19 | 451.2969 | 722.85675 | 546.55444 | 484.63867 | 327.08936 | 373.84247 |
| Hcfc1 | DIPITYGVLPPPR | -188 | -200 | 0 | 0 | 13 | 1456.3126 | 1740.6506 | 351.55374 | NaN | 1928.59 | 1316.512 |
| Hcfc1 | ENQWFDVGVIK | -1819 | -1829 | 0 | 0 | 11 | 597.7385 | 4474.324 | 4449.484 | NaN | 5590.844 | 5393.8374 |
| Hcfc1 | TSPTPNPVPSVPANPPK | -410 | -426 | 0 | 0.000712251 | 17 | 684.07007 | NaN | NaN | NaN | 747.4621 | 811.68933 |
| Hcfc1 | KQELQPGTAYK | -1863 | -1873 | 1 | 0 | 11 | 198.73943 | 630.386 | 604.5509 | NaN | NaN | NaN |
| Ctc1 | DIPNVLEVPR | -862 | -871 | 0 | 0.00025015 | 10 | 921.497 | NaN | NaN | NaN | NaN | NaN |
| Parn | DIPYLNLEGPDLQPK | -422 | -436 | 0 | 0 | 15 | 1121.6621 | 1322.7273 | NaN | NaN | 123.308014 | 1798.1216 |
| Parn | VFCSGIPYLNQEEER | -126 | -140 | 0 | 0.00149795 | 15 | 128.98991 | 352.5132 | 436.14612 | NaN | NaN | NaN |
| Psmd11 | DIQENDEEAVQVK | -34 | -46 | 0 | 0 | 13 | 2866.3315 | 2678.71 | 2855.891 | 3427.922 | 2652.0518 | 3746.4382 |
| Psmd11 | FYEAFEGYDSIDSPK | -232 | -246 | 0 | 0.00025015 | 15 | 1896.9019 | 4094.096 | 2136.3408 | 3542.2424 | 1209.1134 | 820.1709 |
| Psmd11 | LYDNLLEQNLIR | -326 | -337 | 0 | 0 | 12 | 1310.3206 | 1418.5708 | 1574.2821 | 1117.0479 | 2105.8398 | 2026.2761 |
| Psmd11 | EASIDILHSIVK | -21 | -32 | 0 | 0 | 12 | 1297.9131 | 1479.0939 | 1472.7332 | 1545.3374 | 1704.9673 | 1632.5914 |
| Psmd11 | VQIEHISSLIK | -345 | -355 | 0 | 0 | 11 | 1098.9269 | 1291.8364 | 1240.2296 | 1718.6047 | 2239.8098 | 2190.6152 |
| Psmd11 | YQEALHLGSQ | -143 | -152 | 0 | 0.000431655 | 10 | 1334.3727 | 943.7423 | 363.3524 | NaN | NaN | NaN |
| Psmd11 | DNLLEQNLIR | -328 | -337 | 0 | 0 | 10 | 700.99744 | 524.3406 | NaN | 775.325 | NaN | NaN |
| Dut | IAQLICER | -127 | -134 | 0 | 0 | 8 | 1282.6664 | 1841.2589 | 1333.882 | 1908.927 | 1517.3944 | 745.18097 |
| Dut | DIQIAVPSGCYGR | -67 | -79 | 0 | 0 | 13 | 551.54 | 1110.6624 | 1134.6008 | 351.86447 | 1242.4329 | 1159.1508 |
| Dut | FIDVGAGVIDEDYR | -91 | -104 | 0 | 0 | 14 | 536.2439 | 630.14996 | 328.0375 | 659.01715 | 362.35382 | 688.3443 |
| Dut | LSEHATAPTR | -29 | -38 | 0 | 0 | 10 | 4160.4067 | 4519.964 | 4536.527 | 765.10223 | NaN | 528.7287 |
| Dut | ISYPDLEEVQTLDDTER | -135 | -151 | 0 | 0 | 17 | 694.97986 | NaN | 808.1767 | 1199.823 | 1379.2129 | 1370.9058 |
| Dut | TDIQIAVPSGCYGR | -66 | -79 | 0 | 0 | 14 | 484.9616 | 591.6187 | 550.52814 | 259.71335 | 515.1758 | NaN |
| Dut | SEHATAPTR | -30 | -38 | 0 | 0.00078661 | 9 | 448.29858 | 612.59174 | 808.753 | 302.0349 | 672.77814 | NaN |
| Dut | ISYPDLEEVQTL | -135 | -146 | 0 | 0.00201047 | 12 | 368.07095 | NaN | NaN | 642.88586 | NaN | NaN |
| Gnl3 | LLGDFQQSCGK | -238 | -248 | 0 | 0 | 11 | 429.3577 | 1836.935 | 1915.3381 | 1617.5919 | 1413.2158 | 419.66373 |
| Gnl3 | KLEVSPGDEQSNVETR | -91 | -106 | 1 | 0 | 16 | 272.32962 | 747.56006 | 744.33636 | 939.372 | 911.9203 | 1057.5083 |
| Gnl3 | DIQVGVIGFPNVGK | -249 | -262 | 0 | 0 | 14 | 87.073204 | 501.15314 | 382.45923 | 373.99973 | 576.7032 | 590.69763 |
| Gnl3 | DSLHFFTK | -355 | -362 | 0 | 0 | 8 | 1681.9635 | 1502.7218 | NaN | 1737.8408 | 1633.7406 | 1472.1982 |
| Gnl3 | KDPGVPNSAPFK | -46 | -57 | 1 | 0 | 12 | 296.94342 | 505.32928 | 615.1972 | 555.1351 | NaN | NaN |
| Gnl3 | ENLENWLNYLNK | -184 | -195 | 0 | 0 | 12 | 611.4529 | NaN | 720.725 | 542.196 | NaN | NaN |
| Gnl3 | KLEVSPGDEQSNVE | -91 | -104 | 1 | 0.00025015 | 14 | 854.8084 | 1254.9042 | NaN | NaN | NaN | NaN |
| Atp6v1g1 | DIRPEIHENYR | -105 | -115 | 1 | 0 | 11 | 972.2527 | 1226.3029 | 201.28642 | 105.45289 | 1646.5732 | 127.82979 |
| Ehd1 | EHQISSGDFPSLR | -345 | -357 | 0 | 0 | 13 | 1517.5894 | 1655.2041 | 1758.3403 | 1983.8317 | 2098.15 | 2171.602 |
| Ehd1 | EHQISSGDFPSLR | -345 | -357 | 0 | 0 | 13 | 1517.5894 | 1655.2041 | 1758.3403 | 1983.8317 | 2098.15 | 2171.602 |
| Ehd1 | GEGAGEGIDDVEWVVGK | -427 | -443 | 0 | 0 | 17 | 801.6208 | 838.5065 | NaN | 973.23474 | 997.1213 | 508.6254 |
| Ehd1 | DKPTYDEIFYTL | -444 | -455 | 1 | 0.000627943 | 12 | 602.27966 | 587.2122 | NaN | NaN | NaN | 365.43698 |
| Ehd1 | IINTPEVVR | -243 | -251 | 0 | 0 | 9 | 2272.5232 | NaN | NaN | 1061.956 | NaN | NaN |
| Ube2m | KDINELNLPK | -36 | -45 | 1 | 0 | 10 | 2076.8086 | 1937.1163 | 2430.7068 | 1693.5723 | 892.59 | 754.59467 |
| Ube2m | LVICPDEGFYK | -62 | -72 | 0 | 0 | 11 | 1161.1464 | 1348.8596 | 1288.7704 | 1663.0513 | 1912.5476 | 741.85223 |
| Ube2m | QYLFLEPNPEDPLNK | -133 | -147 | 0 | 0.001055699 | 15 | 433.3509 | 473.0125 | 819.5934 | 586.08875 | 950.0747 | 960.1842 |
| Ube2m | DISFSDPDDLLNFK | -48 | -61 | 0 | 0 | 14 | 9653.693 | 10613.151 | NaN | 13956.089 | 14488.237 | 14742.886 |
| Ube2m | EDWKPVLTIN | -117 | -126 | 1 | 0.00204437 | 10 | 175.30127 | 667.86096 | 732.4869 | 264.87546 | NaN | 858.8224 |
| Ube2m | TCDISFSDPDDLLNFK | -46 | -61 | 0 | 0.001868731 | 16 | 182.33737 | 196.91115 | 236.005 | NaN | 239.90393 | NaN |
| Aprt | DISPLLKDPDSFR | -28 | -40 | 1 | 0 | 13 | 763.2487 | NaN | 724.8285 | 949.43677 | 1671.3068 | 1010.39026 |
| Aprt | IDYIAGLDSR | -58 | -67 | 0 | 0.000394361 | 10 | 1610.9865 | NaN | NaN | 2314.3027 | 582.4063 | NaN |
| Aprt | DLLHQLR | -141 | -147 | 0 | 0.001846793 | 7 | 1403.6534 | 1398.3989 | 505.32693 | NaN | NaN | NaN |
| Aprt | LKDPDSFR | -33 | -40 | 1 | 0.001055699 | 8 | 1618.8562 | NaN | NaN | NaN | NaN | 1008.08484 |
| Anp32b | DISTLEPLKR | -102 | -111 | 1 | 0 | 10 | 5414.205 | 5671.692 | 6174.891 | 6129.379 | 7848.7466 | 7510.0293 |
| Anp32b | SLDLFGCEVTNR | -117 | -128 | 0 | 0 | 12 | 1244.6338 | 1508.0675 | 317.18512 | 562.00977 | 1255.9448 | 1430.688 |
| Anp32b | DLFGCEVTNR | -119 | -128 | 0 | 0 | 10 | 735.3031 | 881.92883 | 831.2345 | 664.40027 | 822.1546 | 823.5316 |
| Anp32b | LAEELPSLTHLNLSGNNLK | -83 | -101 | 0 | 0 | 19 | 652.82587 | 714.37616 | 1530.5377 | 1131.4757 | 1420.0405 | 1429.0175 |
| Anp32b | LAEELPSLTHLNLSGNNLK | -83 | -101 | 0 | 0 | 19 | 652.82587 | 714.37616 | 1530.5377 | 1131.4757 | 1420.0405 | 1429.0175 |
| Anp32b | LAEELPSLTHLNLS | -83 | -96 | 0 | 0 | 14 | 596.6787 | 629.6941 | 657.71545 | 654.1504 | 819.6217 | 736.9919 |
| Anp32b | LAEELPSLTHL | -83 | -93 | 0 | 0 | 11 | 322.30066 | 215.77228 | 2247.4958 | 1872.7141 | 2348.501 | 2457.6936 |
| Anp32b | LAEELPSLTHLN | -83 | -94 | 0 | 0 | 12 | 1751.6245 | 326.92578 | 445.98767 | NaN | 3492.962 | 5877.6665 |
| Anp32b | LAEELPSLTHLNLSGN | -83 | -98 | 0 | 0 | 16 | 1155.0631 | 747.5256 | 839.40375 | 637.56226 | NaN | NaN |
| Anp32b | LLPQLSYLDGYDR | -138 | -150 | 0 | 0 | 13 | 730.79456 | 799.9905 | 767.4818 | NaN | NaN | NaN |
| Anp32b | RIHLELR | -6 | -12 | 1 | 0.00266297 | 7 | 799.1981 | NaN | NaN | NaN | NaN | NaN |
| Rab5a | DITNEESFAR | -101 | -110 | 0 | 0 | 10 | 5050.958 | 4571.311 | 4847.2456 | 233.8833 | NaN | NaN |
| Rab5a | GVDLTEPAQPAR | -198 | -209 | 0 | 0.000627943 | 12 | 5052.656 | NaN | NaN | NaN | NaN | NaN |
| Rab5c | IVVYDITNTDTFAR | -98 | -111 | 0 | 0 | 14 | 2233.965 | 2865.791 | 2752.5415 | 3970.5286 | 4195.9146 | 211.37482 |
| Rab5c | DITNTDTFAR | -102 | -111 | 0 | 0 | 10 | 980.4201 | 872.10114 | 804.5806 | 1067.2356 | 1764.6852 | 1782.8788 |

| | | | | | | | | | | | | |
|---|---|---|---|---|---|---|---|---|---|---|---|---|
| Rab5c | GVDLQESNPASR | -199 | -210 | 0 | 0 | 12 | 2617.5369 | NaN | 2985.4155 | 3528.5574 | NaN | 4477.2363 |
| Cpne3 | SDPYLEFHK | -159 | -167 | 0 | 0 | 9 | 734.2032 | NaN | 828.50366 | 620.90137 | 804.83813 | 350.0235 |
| Cpne3 | LIITDGVITDLDETR | -424 | -438 | 0 | 0 | 15 | 448.02448 | 442.40298 | NaN | 581.40424 | 867.1521 | 795.8888 |
| Cpne3 | SDPYLEFHKQ | -159 | -168 | 1 | 0 | 10 | 1211.1931 | 1818.718 | 620.43054 | NaN | NaN | 575.3197 |
| Cpne3 | EFLDGDGGSLR | -465 | -475 | 0 | 0 | 11 | 773.53217 | 743.0653 | NaN | 873.05927 | NaN | NaN |
| Rbx1 | DIVVDNCAICR | -36 | -46 | 0 | 0.00025015 | 11 | 1998.7979 | NaN | NaN | NaN | NaN | NaN |
| Abi1 | DKDDELSFK | -433 | -441 | 1 | 0.00025015 | 9 | 1249.4047 | 1526.8293 | 1240.324 | 1268.4814 | 1241.8918 | 1175.1804 |
| Irf3 | VYEFVTPGAR | -106 | -115 | 0 | 0 | 10 | 1189.1957 | 699.8689 | 783.2367 | 559.0326 | 1721.6494 | 937.396 |
| Irf3 | DKDGAVFDLRPF | -307 | -318 | 2 | 0.000431655 | 12 | 600.2879 | 638.3959 | 690.8292 | NaN | 492.14365 | 641.708 |
| Irf3 | TLFCPGGLR | -213 | -221 | 0 | 0.000712251 | 9 | 1073.8159 | NaN | NaN | NaN | 2615.4355 | 3611.3013 |
| Cast | STYQDKPSTPAEK | -459 | -471 | 1 | 0 | 13 | 612.778 | 615.35004 | 604.6942 | 496.50143 | 736.95795 | 666.2273 |
| Cast | EQKPFTPASPVQSTPSKPSDK | -211 | -231 | 2 | 0 | 21 | 451.6221 | 446.49655 | 435.71805 | 387.09998 | 189.36961 | 399.53366 |
| Cast | DKDGKPLLPEPEETSK | -426 | -441 | 2 | 0.00111006 | 16 | 273.5899 | NaN | 565.34454 | 543.213 | 636.6553 | 442.06992 |
| Ash2l | DKDIIPFIDK | -189 | -198 | 1 | 0 | 10 | 545.521 | 487.48334 | 373.57004 | 563.3168 | 675.21954 | 384.85614 |
| Ash2l | FGLLDQDLSNIGPAYDNQK | -249 | -267 | 0 | 0 | 19 | 294.0247 | 401.5346 | 454.79694 | 597.5893 | 619.13385 | 446.12314 |
| Ankrd44 | DKDLNTPLHLAISK | -903 | -916 | 1 | 0 | 14 | 343.2503 | 333.7436 | 379.09302 | 392.06354 | 386.25345 | 366.15076 |
| Ankrd44 | DKNWQTPLHVA | -104 | -114 | 1 | 0 | 11 | 7018.2075 | 2350.2852 | 7045.0015 | NaN | 2822.7056 | 2297.8445 |
| Ankrd44 | IQDESLINAK | -929 | -938 | 0 | 0.00025015 | 10 | 1523.6824 | 1321.4126 | NaN | NaN | 1686.415 | 1569.8265 |
| Ankrd44 | IFSGDPEEIR | -16 | -25 | 0 | 0.00025015 | 10 | 677.37585 | NaN | NaN | 862.5146 | 851.82086 | 704.8102 |
| Naa15 | RLPLNFLSGEK | -295 | -305 | 1 | 0 | 11 | 1131.6604 | 1584.5619 | 1684.1145 | 787.7802 | 1487.756 | 985.4517 |
| Naa15 | TQQTSPDKVDYEYSELL | -177 | -193 | 1 | 0.000230707 | 17 | 671.85284 | 767.5289 | 799.1792 | 1011.67725 | 1185.6112 | 1157.0286 |
| Naa15 | VETPLEEAIKF | -650 | -660 | 1 | 0 | 11 | 466.96646 | 816.7243 | 419.91327 | 450.1101 | 795.75806 | 557.48224 |
| Naa15 | LHDNPLTDENKEHEADTA | -568 | -585 | 1 | 0.00111006 | 18 | 418.6958 | 459.44046 | 445.4446 | 506.62735 | 670.0046 | 688.6274 |
| Naa15 | LFNPNDDGKEEPPTTL | -358 | -373 | 1 | 0 | 16 | 1142.6074 | 240.98268 | 1111.0787 | 1298.1385 | NaN | NaN |
| Naa15 | QHPFYFK | -550 | -556 | 0 | 0 | 7 | 768.97253 | 721.78973 | 993.913 | 972.68787 | NaN | NaN |
| Naa15 | FKECLDR | -306 | -312 | 1 | 0 | 7 | 490.48462 | 528.0644 | 790.2755 | 1703.6255 | NaN | NaN |
| Naa15 | YYLAQHYDK | -378 | -386 | 0 | 0 | 9 | 444.1174 | NaN | NaN | 1916.4795 | 1829.645 | 209.56235 |
| Naa15 | WDKDNLQILR | -109 | -118 | 1 | 0 | 10 | 304.52676 | 451.59406 | NaN | 1465.7795 | NaN | NaN |
| Rnaseh2b | VFLLPEHLK | -16 | -24 | 0 | 0 | 9 | 194.58989 | 390.74057 | 368.7102 | 442.8571 | 518.7678 | 468.62137 |
| Rnaseh2b | DKEEDYVR | -200 | -207 | 1 | 0.00051306 | 8 | 712.68036 | 809.2626 | NaN | 1306.6154 | NaN | NaN |
| Epb41l2 | SSHETLNVVEEK | -665 | -676 | 0 | 0 | 12 | 790.34436 | 648.961 | NaN | 193.97147 | NaN | 646.17694 |
| Epb41l2 | DKEETIQDTQEEK | -165 | -177 | 1 | 0.000431655 | 13 | 266.25397 | NaN | 593.09296 | NaN | 346.65793 | 504.4751 |
| Ap1m1 | NVEIHIPVPNDADSPK | -309 | -324 | 0 | 0 | 16 | 1026.8545 | 1241.5948 | 1060.9596 | 1700.738 | 1670.3994 | 2011.524 |
| Ap1m1 | WVPENSEIVWSVK | -334 | -346 | 0 | 0 | 13 | 556.9013 | 841.54486 | 837.5271 | 772.56854 | 928.3784 | 1745.7524 |
| Ap1m1 | DKEGKPPISVK | -369 | -379 | 2 | 0.00025015 | 11 | 559.3578 | 1045.6704 | 805.50555 | NaN | NaN | NaN |
| Ap1m1 | DKEGKPPISVK | -369 | -379 | 2 | 0.00025015 | 11 | 559.3578 | 1045.6704 | 805.50555 | NaN | NaN | NaN |
| Gbp7 | GVFSVPGGHR | -424 | -433 | 0 | 0 | 10 | 1206.767 | 973.6281 | 999.5527 | NaN | 952.27136 | 907.8144 |
| Gbp7 | DKELLVHVE | -242 | -250 | 1 | 0.00025015 | 9 | 1511.6555 | NaN | NaN | NaN | 2143.0576 | 2029.2345 |
| Fkbp4 | DKFSFDLGK | -75 | -83 | 1 | 0 | 9 | 1936.3783 | 528.8261 | 2566.8677 | 2260.4673 | 2722.5532 | 628.53613 |
| Fkbp4 | VALEGYHK | -172 | -179 | 0 | 0.00025015 | 8 | 1826.3477 | 1753.6025 | 1623.508 | 1210.4524 | 1043.1395 | 1359.401 |
| Fkbp4 | VLQLYPSNK | -379 | -387 | 0 | 0 | 9 | 1639.1053 | 1685.2858 | 1499.1067 | 1365.402 | 1196.5526 | 853.6893 |
| Fkbp4 | LAVNDFDLAR | -364 | -373 | 0 | 0 | 10 | 1415.134 | 2751.0007 | 2620.7136 | 8279.213 | 8547.678 | 9132.672 |
| Fkbp4 | GEDLTEEEDGGIIR | -139 | -152 | 0 | 0 | 14 | 1364.4495 | 1682.6249 | 1689.237 | 1557.3857 | 1755.8463 | 1755.1178 |
| Fkbp4 | DLPCGLEEAIQR | -199 | -210 | 0 | 0 | 12 | 1196.9008 | 944.6721 | 282.30957 | 903.5102 | 1730.0219 | 2114.2515 |
| Fkbp4 | FDSSLDRK | -67 | -74 | 1 | 0 | 8 | 1125.6814 | 2320.046 | 2206.1477 | 1156.071 | 192.3391 | 1904.7716 |
| Fkbp4 | EFKGEDLTEEEDGGIIR | -136 | -152 | 1 | 0 | 17 | 1038.686 | 1360.0184 | 1306.9944 | 1370.3599 | 1917.7319 | 1821.5889 |
| Fkbp4 | EFKGEDLTEEEDGGIIR | -136 | -152 | 1 | 0 | 17 | 1038.686 | 1360.0184 | 1306.9944 | 1370.3599 | 1917.7319 | 1821.5889 |
| Fkbp4 | SAPLPLEGVDISPK | -15 | -28 | 0 | 0 | 14 | 821.99445 | 859.01904 | 868.0215 | 1240.4805 | 1417.464 | 1179.4028 |
| Fkbp4 | YTGWLLDGTK | -57 | -66 | 0 | 0 | 10 | 732.96423 | 317.42175 | 1033.9172 | 1388.6051 | 1654.0718 | 1083.4912 |
| Fkbp4 | VNDFDLAR | -366 | -373 | 0 | 0.000230707 | 8 | 266.0866 | 6413.827 | 6687.9565 | 5965.76 | 5405.0513 | 5576.7114 |
| Fkbp4 | EKLEQSNIVK | -265 | -274 | 1 | 0 | 10 | 1678.776 | 1730.7384 | 1674.0426 | NaN | 1995.6768 | 2084.7683 |
| Fkbp4 | FQIPPHAELR | -235 | -244 | 0 | 0 | 10 | 1073.9946 | 1836.5928 | 2111.7434 | NaN | 1067.3411 | 1346.3622 |
| Fkbp4 | ENGAQSAPLPLEGVDISPK | -10 | -28 | 0 | 0.00025015 | 19 | 693.19495 | NaN | 650.7638 | 1475.3507 | 1062.5857 | 1059.5714 |
| Fkbp4 | EGTGTETPMIGDR | -40 | -52 | 0 | 0 | 13 | 408.86768 | 553.49274 | NaN | 662.17773 | 748.7276 | 441.85422 |
| Fkbp4 | GWLLDGTK | -59 | -66 | 0 | 0.001291103 | 8 | 909.10516 | 142.92409 | 793.92236 | NaN | NaN | NaN |
| Fkbp4 | SLDLPCGLEEAIQR | -197 | -210 | 0 | 0 | 14 | 638.1033 | NaN | NaN | 468.31622 | 804.19617 | NaN |
| Fkbp4 | LKPSYAFGSVGK | -221 | -232 | 1 | 0.000550307 | 12 | 1266.3021 | NaN | NaN | NaN | NaN | NaN |
| Eif2b1 | DKGETLQGLR | -37 | -46 | 1 | 0 | 10 | 2110.3545 | NaN | NaN | 1870.7938 | 1787.6644 | 1689.4194 |
| Eif2b1 | LFPLNQEDVPDKFK | -238 | -251 | 1 | 0 | 14 | 1513.394 | NaN | NaN | NaN | 3138.5571 | 3069.5947 |
| Alad | DVPDDVQPIASLPGVAR | -39 | -55 | 0 | 0 | 17 | 830.17413 | 749.13226 | 571.8572 | 689.00525 | 1317.8317 | NaN |
| Alad | DSEDSPTIEAVR | -95 | -106 | 0 | 0.00078661 | 12 | 536.7849 | NaN | 537.5967 | NaN | NaN | 503.27844 |
| Alad | DKHPELPLAVY | -266 | -276 | 1 | 0.00051306 | 11 | 637.1659 | NaN | NaN | NaN | NaN | NaN |
| Alad | YFAPQLLK | -318 | -325 | 0 | 0.000627943 | 8 | 391.21768 | NaN | NaN | NaN | NaN | NaN |
| Mrpl38 | DKIDIGLPPPR | -86 | -96 | 1 | 0 | 11 | 1050.0641 | 855.5732 | 725.79205 | 717.67596 | 689.6761 | 804.761 |
| Mrpl38 | ETCPYLPPFPAR | -250 | -261 | 0 | 0.00383302 | 12 | 1986.5181 | NaN | NaN | 1527.6002 | NaN | NaN |
| Usp19 | DKINDLVEFPVR | -1161 | -1172 | 1 | 0 | 12 | 135.4874 | 376.17645 | 400.98297 | 326.99866 | 500.54715 | 575.50305 |
| Uso1 | LREEIEELK | -735 | -743 | 1 | 0.001743443 | 9 | 1286.7084 | 965.9778 | 954.2903 | 1002.44586 | 1271.1516 | 1031.209 |
| Uso1 | SVPVEGESEHVSAAK | -828 | -842 | 0 | 0 | 15 | 525.69824 | 658.9794 | 690.19696 | 456.72516 | 485.16516 | 429.05762 |
| Uso1 | TLEQHDNIVTHYK | -644 | -656 | 0 | 0 | 13 | 3583.8896 | 484.17352 | 3371.1567 | NaN | 3554.7122 | 4040.7603 |
| Uso1 | TLEQHDNIVTHYK | -644 | -656 | 0 | 0 | 13 | 3583.8896 | 484.17352 | 3371.1567 | NaN | 3554.7122 | 4040.7603 |
| Uso1 | TLEQHDNIVTHY | -644 | -655 | 0 | 0.00122419 | 12 | 2164.9143 | 2133.9111 | NaN | 1701.8623 | 297.90527 | 1687.9932 |
| Uso1 | SAGPQHTEAETIQK | -12 | -25 | 0 | 0.00025015 | 14 | 312.03687 | 247.98756 | 346.46872 | NaN | 340.11423 | 167.06412 |
| Uso1 | DKLDLEVTDSK | -893 | -903 | 1 | 0.001636916 | 11 | 1570.5409 | 2071.7808 | NaN | NaN | 841.38275 | 1092.4207 |
| Usp4 | DKLDTVVEFPVR | -835 | -846 | 1 | 0 | 12 | 888.9169 | 1065.3042 | 1090.3914 | 1377.4609 | 1254.8081 | 818.6039 |
| Usp4 | VLVIEPQNEDGTWPR | -212 | -226 | 0 | 0 | 15 | 429.977 | 591.17035 | 599.64307 | 633.4891 | 729.64105 | 790.2249 |
| Usp4 | KVVEHGLFVK | -120 | -129 | 1 | 0.00383302 | 10 | 262.96857 | NaN | NaN | NaN | NaN | NaN |
| Tmed9 | DKLSELQLR | -159 | -167 | 1 | 0 | 9 | 1583.0962 | 1286.1998 | 1200.2346 | 3915.0151 | 3822.1958 | 3955.4392 |
| Tmed9 | LYFHIGETEKK | -38 | -48 | 1 | 0 | 11 | 457.55438 | 513.33734 | 758.26794 | 777.017 | 427.4142 | 530.98413 |

| | | | | | | | | | | | | |
|---|---|---|---|---|---|---|---|---|---|---|---|---|
| Tmed9 | FVEVKDPEDK | -86 | -95 | 1 | 0 | 10 | 2242.573 | 1889.1908 | 1171.0073 | 979.4042 | 2637.7993 | NaN |
| Tmed9 | QLVEQVEQIQK | -170 | -180 | 0 | 0 | 11 | 833.5358 | NaN | 1220.2382 | 1717.6713 | NaN | NaN |
| Rps4x | GIPHLVTHDAR | -135 | -145 | 0 | 0 | 11 | 1391.8683 | 1214.0458 | 99.954414 | 820.3433 | 793.32416 | 651.3109 |
| Rps4x | GIPHLVTHDAR | -135 | -145 | 0 | 0 | 11 | 1391.8683 | 1214.0458 | 99.954414 | 820.3433 | 793.32416 | 651.3109 |
| Rps4x | YPDPLIK | -149 | -155 | 0 | 0.001941082 | 7 | 10005.271 | 9040.119 | 9657.776 | 8341.184 | 8403.741 | 8781.459 |
| Rps4x | GNKPWISLPR | -231 | -240 | 1 | 0.000550307 | 10 | 7461.3667 | 7448.7715 | 7855.3804 | 7373.391 | 8127.4136 | 7749.5547 |
| Rps4x | HPGSFDVVHVK | -201 | -211 | 0 | 0 | 11 | 3908.6934 | 4046.3103 | 3900.0698 | 5391.427 | 4953.2183 | 4601.4253 |
| Rps4x | HPGSFDVVHVK | -201 | -211 | 0 | 0 | 11 | 3908.6934 | 4046.3103 | 3900.0698 | 5391.427 | 4953.2183 | 4601.4253 |
| Rps4x | GIPHLVTH | -135 | -142 | 0 | 0.00025015 | 8 | 3640.6638 | 3241.293 | 3120.589 | 3741.6433 | 2806.9119 | 2686.4788 |
| Rps4x | RITPEEAK | -113 | -120 | 1 | 0.000394361 | 8 | 2840.53 | 294.65375 | 1675.9818 | 352.16257 | 2978.0667 | 3387.712 |
| Rps4x | ERHPGSFDVVHVK | -199 | -211 | 1 | 0 | 13 | 1902.8239 | 2233.2278 | 2384.77 | 2413.4019 | 2905.3135 | 222.1145 |
| Rps4x | ERHPGSFDVVHVK | -199 | -211 | 1 | 0 | 13 | 1902.8239 | 2233.2278 | 2384.77 | 2413.4019 | 2905.3135 | 222.1145 |
| Rps4x | ERHPGSFDVVHVK | -199 | -211 | 1 | 0 | 13 | 1902.8239 | 2233.2278 | 2384.77 | 2413.4019 | 2905.3135 | 222.1145 |
| Rps4x | ERHPGSFDVV | -199 | -208 | 1 | 0.001092529 | 10 | 1209.8486 | 991.731 | 1318.4491 | 421.83203 | 1006.7611 | 2354.9053 |
| Rps4x | KGNKPWISLPR | -230 | -240 | 2 | 0.00025015 | 11 | 670.271 | 823.7781 | 899.5984 | 593.3836 | 783.8084 | 887.764 |
| Rps4x | GFMDVISIDK | -85 | -94 | 0 | 0 | 10 | 537.9964 | 120.84435 | 539.32117 | 372.1752 | 450.42917 | 457.16223 |
| Rps4x | GFMDVISIDK | -85 | -94 | 0 | 0 | 10 | 537.9964 | 120.84435 | 539.32117 | 372.1752 | 450.42917 | 457.16223 |
| Rps4x | GSFDVVHVK | -203 | -211 | 0 | 0.000627943 | 9 | 1593.2524 | 1628.797 | NaN | 1535.1865 | 1408.1538 | 1315.8696 |
| Rps4x | RLIYDTK | -100 | -106 | 1 | 0 | 7 | 980.3149 | 1064.0682 | 1114.3558 | 736.95825 | NaN | 1025.3131 |
| Rps4x | PHLVTHDAR | -137 | -145 | 0 | 0.00111006 | 9 | 608.7525 | 509.78467 | NaN | 5329.6055 | 4313.3735 | 1879.8517 |
| Rps4x | PHLVTHDAR | -137 | -145 | 0 | 0.00111006 | 9 | 608.7525 | 509.78467 | NaN | 5329.6055 | 4313.3735 | 1879.8517 |
| Rps4x | LSNIFVIGK | -222 | -230 | 0 | 0.000230707 | 9 | 1440.0583 | 853.6681 | 654.1965 | 1156.7229 | NaN | NaN |
| Rps4x | GIPHLVTHDA | -135 | -144 | 0 | 0.000550307 | 10 | 2777.7124 | NaN | 3124.4268 | NaN | 5672.801 | NaN |
| Rps4x | ERHPGSF | -199 | -205 | 1 | 0.003745449 | 7 | 1473.3336 | NaN | NaN | NaN | 782.9826 | 1666.624 |
| Rps4x | KLTGVFAPR | -22 | -30 | 1 | 0.001610812 | 9 | 1320.966 | 1410.4977 | 1374.1155 | NaN | NaN | NaN |
| Rps4x | DKLTGVFAPR | -21 | -30 | 1 | 0 | 10 | 1274.4148 | 1230.9849 | 1311.074 | NaN | NaN | NaN |
| Rps4x | YALTGDEVKK | -54 | -63 | 1 | 0 | 10 | 1193.213 | 1145.3235 | 1275.7812 | NaN | NaN | NaN |
| Rps4x | SNIFVIGK | -223 | -230 | 0 | 0 | 8 | 817.16174 | 485.79446 | 640.6688 | NaN | NaN | NaN |
| Rps4x | ITPEEAK | -114 | -120 | 0 | 0.007791272 | 7 | 631.4966 | 1013.2241 | NaN | NaN | NaN | NaN |
| Rps4x | TDITYPAGFMDVISIDK | -78 | -94 | 0 | 0 | 17 | 234.64189 | NaN | NaN | NaN | 462.4114 | NaN |
| Ranbp2 | VTDEENASSGADAPSASDTTAK | -2028 | -2049 | 0 | 0 | 22 | 861.3513 | 772.3275 | 696.9382 | 708.0562 | 808.7599 | 573.1592 |
| Ranbp2 | SGEEDEEVLFK | -2754 | -2764 | 0 | 0 | 11 | 739.7925 | 846.49475 | 1015.84485 | 965.3408 | 638.75476 | 958.89154 |
| Ranbp2 | SGFEGLFPK | -1608 | -1616 | 0 | 0 | 9 | 549.49347 | 672.8412 | 471.03012 | 711.00226 | 705.3871 | 484.9521 |
| Ranbp2 | GTNTSTAPNHTLR | -1303 | -1315 | 0 | 0 | 13 | 440.8315 | 372.54144 | 385.3426 | 367.8125 | 384.19046 | 375.77036 |
| Ranbp2 | KFEECQQNIIK | -2865 | -2875 | 1 | 0 | 11 | 2543.4924 | 2771.1504 | 2846.885 | NaN | 3355.5608 | 3094.0952 |
| Ranbp2 | SQSNESAGSQVGTEFK | -1662 | -1677 | 0 | 0 | 16 | 1705.9352 | 1676.603 | 1310.5448 | NaN | 1675.654 | 1744.7214 |
| Ranbp2 | LPVPLESVK | -735 | -743 | 0 | 0.000907206 | 9 | 910.36426 | 931.114 | 900.79755 | NaN | 414.6393 | 500.05817 |
| Ranbp2 | HLENDPSFQAH | -1752 | -1762 | 0 | 0 | 11 | 815.82654 | 1629.7828 | 456.39624 | NaN | 532.6532 | 638.4653 |
| Ranbp2 | QLFHHLPQETSR | -459 | -470 | 0 | 0 | 12 | 494.15402 | 986.24585 | NaN | 2043.07 | 605.94385 | 5111.8403 |
| Ranbp2 | LLLDIPLQTPHK | -1982 | -1993 | 0 | 0 | 12 | 407.38345 | 429.75885 | NaN | 401.3394 | 492.81024 | 466.108 |
| Ranbp2 | HGLQPALLVH | -568 | -577 | 0 | 0 | 10 | 328.6924 | 340.83356 | 440.92087 | NaN | 906.2448 | 326.30157 |
| Ranbp2 | DYADELPKPEQLAIR | -1264 | -1278 | 1 | 0 | 15 | 839.2111 | 877.49176 | 949.1543 | NaN | 1190.8025 | NaN |
| Ranbp2 | GDKFDDENFDLK | -2969 | -2980 | 1 | 0.00025015 | 12 | 590.609 | 607.37445 | 442.57077 | NaN | 360.45398 | NaN |
| Ranbp2 | TGEEDEEEFFCNR | -1180 | -1192 | 0 | 0 | 13 | 1739.3186 | 1884.4421 | 2009.0743 | NaN | NaN | NaN |
| Ranbp2 | HIGETEVTNPVGCK | -2614 | -2627 | 0 | 0.00437931 | 14 | 485.7061 | 453.32745 | NaN | 750.8298 | NaN | NaN |
| Ranbp2 | DKNFQWANTGAAVF | -2701 | -2714 | 1 | 0.001868731 | 14 | 476.54794 | NaN | NaN | NaN | 593.5055 | 385.17786 |
| Ranbp2 | TGLENAQDR | -1394 | -1402 | 0 | 0.005176406 | 9 | 3150.6401 | NaN | NaN | NaN | NaN | NaN |
| Ppib | IEVEKPFAIA | -205 | -214 | 1 | 0.00025015 | 10 | 2684.147 | 2933.8127 | 2970.5706 | 2954.8232 | 3496.7756 | 3425.1472 |
| Ppib | IEVEKPFAIAK | -205 | -215 | 1 | 0 | 11 | 1770.712 | 2109.8757 | 2247.8535 | 2134.723 | 2454.8418 | 2359.1292 |
| Ppib | VVFGLFGK | -60 | -67 | 0 | 0.001941082 | 8 | 1194.7395 | 1346.8641 | 1182.2776 | 1481.8483 | 1152.2139 | 1230.0908 |
| Ppib | DKPLKDVIIVDSGK | -191 | -204 | 2 | 0 | 14 | 771.2847 | 703.8707 | 869.67096 | 968.69543 | 833.3168 | 822.76465 |
| Ppib | VLEGMDVVR | -172 | -180 | 0 | 0.00025015 | 9 | 982.9316 | NaN | 927.24585 | 879.6187 | 735.69867 | 845.2063 |
| Ppib | RFPDENFK | -122 | -129 | 1 | 0.00025015 | 8 | 1390.5781 | 1251.4088 | NaN | 942.4337 | NaN | NaN |
| Ppib | QIGDESVGR | -51 | -59 | 0 | 0 | 9 | 7419.306 | NaN | NaN | NaN | NaN | 6900.9053 |
| Pin4 | DKPVFTDPPVK | -107 | -117 | 1 | 0 | 11 | 495.58945 | 469.2924 | NaN | NaN | NaN | NaN |
| Hmgb2 | IEHPGLSIGDTAK | -115 | -127 | 0 | 0 | 13 | 11515.967 | 12051.599 | 12165.064 | 11799.52 | 12312.467 | 12338.242 |
| Hmgb2 | IEHPGLSIGDTAK | -115 | -127 | 0 | 0 | 13 | 11515.967 | 12051.599 | 12165.064 | 11799.52 | 12312.467 | 12338.242 |
| Hmgb2 | IEHPGLSIGDTAK | -115 | -127 | 0 | 0 | 13 | 11515.967 | 12051.599 | 12165.064 | 11799.52 | 12312.467 | 12338.242 |
| Hmgb2 | SKFEDLAK | -58 | -65 | 1 | 0 | 8 | 7822.876 | 7303.9087 | 7639.48 | 1121.3844 | 9030.636 | 9374.911 |
| Hmgb2 | KHPDSSVNFA | -30 | -39 | 1 | 0 | 10 | 5119.5522 | 5101.228 | 5157.5557 | 3443.924 | 4341.5933 | 4657.3535 |
| Hmgb2 | IKIEHPGLSIGDTAK | -113 | -127 | 1 | 0 | 15 | 4683.9927 | 5567.8843 | 5525.5894 | 5748.9814 | 6698.6177 | 6564.291 |
| Hmgb2 | IKIEHPGLSIGDTAK | -113 | -127 | 1 | 0 | 15 | 4683.9927 | 5567.8843 | 5525.5894 | 5748.9814 | 6698.6177 | 6564.291 |
| Hmgb2 | KHPDSSVNFAEF | -30 | -41 | 1 | 0.00149795 | 12 | 161.10722 | 194.02202 | 154.09358 | 167.62262 | 209.89586 | 209.87766 |
| Hmgb2 | KHPDSSVNFAEF | -30 | -41 | 1 | 0.00149795 | 12 | 161.10722 | 194.02202 | 154.09358 | 167.62262 | 209.89586 | 209.87766 |
| Hmgb2 | FLFCSENRPK | -103 | -112 | 1 | 0 | 10 | 1055.476 | 1634.7073 | 1967.4167 | 1050.8102 | 2235.3862 | 1992.6664 |
| Hmgb2 | KKNEPEDEEEEEEEEEEDDEEEEEDEE | -183 | -210 | 2 | 0.00133086 | 28 | 1045.2225 | 1013.96515 | 911.46234 | 2361.4573 | 1051.1144 | 913.16064 |
| Hmgb2 | IKIEHPGLSIGDT | -113 | -125 | 1 | 0 | 13 | 825.06854 | 950.11804 | 915.6503 | 689.3938 | 1161.4329 | 1252.4929 |
| Hmgb2 | DKQPYEQK | -140 | -147 | 1 | 0.00025015 | 8 | 615.23395 | 143.7301 | 497.48584 | 105.14086 | 495.22858 | 571.8319 |
| Hmgb2 | KKHPDSSVNFA | -29 | -39 | 2 | 0.00078661 | 11 | 3228.0525 | 758.8188 | 592.8574 | 5876.262 | 2180.9626 | NaN |
| Hmgb2 | NYVPPKGDK | -77 | -85 | 1 | 0 | 9 | 633.37866 | 2624.721 | NaN | NaN | 1192.7123 | 2778.8958 |
| Hmgb2 | KHPDSSVNFAEFSK | -30 | -43 | 1 | 0 | 14 | 878.30676 | 331.40588 | 385.83972 | NaN | NaN | NaN |
| Hmgb2 | KKNEPEDEEEE | -183 | -194 | 2 | 0.002434616 | 12 | 553.14355 | 567.2008 | 561.67224 | NaN | NaN | NaN |
| Hmgb2 | KKNEPEDEEEEEE | -183 | -196 | 2 | 0.00111006 | 14 | 156.11545 | NaN | 814.9821 | NaN | NaN | NaN |
| Hmgb2 | KNEPEDEEEEEEEE | -184 | -198 | 1 | 0.005301614 | 15 | 374.26157 | NaN | NaN | NaN | NaN | NaN |
| U2af1 | NPQNSSQSADGLR | -54 | -66 | 0 | 0 | 13 | 1023.329 | 993.0634 | 1063.8452 | 508.671 | 810.0339 | 660.6442 |
| U2af1 | WFNGQPIHAE | -134 | -143 | 0 | 0 | 10 | 116.2331 | 674.1738 | 760.92377 | 838.1902 | 146.55635 | 807.24023 |
| Ruvbl2 | FVQCPDGELQK | -224 | -234 | 0 | 0 | 11 | 3626.5908 | 3816.4514 | 3409.0022 | 3092.7495 | 1475.0044 | 1379.8835 |
| Ruvbl2 | VSLHEIDVINSR | -242 | -253 | 0 | 0 | 12 | 381.83914 | 491.86688 | 416.62723 | 373.19717 | 345.48987 | 438.10898 |

| | | | | | | | | | | | | |
|---|---|---|---|---|---|---|---|---|---|---|---|---|
| Ruvbl2 | VSLHEIDVINSR | -242 | -253 | 0 | 0 | 12 | 381.83914 | 491.86688 | 416.62723 | 373.19717 | 345.48987 | 438.10898 |
| Ruvbl2 | GTEVQVDDIKR | -418 | -428 | 1 | 0 | 11 | 2760.1375 | 793.7345 | 800.4761 | 831.75195 | 2672.1191 | 853.97034 |
| Ruvbl2 | LFSGDTGEIK | -260 | -269 | 0 | 0 | 10 | 1461.0004 | 1292.3732 | 1319.0056 | 1709.4163 | 1691.2239 | 1481.4965 |
| Ruvbl2 | SLHEIDVINSR | -243 | -253 | 0 | 0 | 11 | 1404.081 | 1588.5781 | 1574.8861 | 479.99597 | 1466.6494 | 1294.0438 |
| Ruvbl2 | SLHEIDVINSR | -243 | -253 | 0 | 0 | 11 | 1404.081 | 1588.5781 | 1574.8861 | 479.99597 | 1466.6494 | 1294.0438 |
| Ruvbl2 | SYQSPHGIPIDLLDR | -339 | -353 | 0 | 0 | 15 | 993.3092 | 1031.3102 | 1232.2068 | 1164.9731 | 1223.0145 | 1726.0315 |
| Ruvbl2 | SYQSPHGIPIDLLDR | -339 | -353 | 0 | 0 | 15 | 993.3092 | 1031.3102 | 1232.2068 | 1164.9731 | 1223.0145 | 1726.0315 |
| Ruvbl2 | DKVQAGDVITIDK | -185 | -197 | 1 | 0 | 13 | 863.7907 | 601.88293 | 1017.0676 | 1006.0146 | 1930.7802 | 1894.302 |
| Ruvbl2 | LLIVSTSPYSEK | -354 | -365 | 0 | 0 | 12 | 584.5517 | 755.58246 | 440.82562 | 440.0843 | 670.0135 | 587.81604 |
| Ruvbl2 | EEGKAEIIPGVL | -285 | -296 | 1 | 0 | 12 | 201.57356 | 165.35822 | 174.23418 | 827.96277 | 370.1073 | 333.4128 |
| Ruvbl2 | VQAGDVITIDK | -187 | -197 | 0 | 0 | 11 | 1019.78326 | NaN | NaN | 1327.4979 | 1954.2217 | 2010.885 |
| Ruvbl2 | GTSYQSPHGIPIDLLDR | -337 | -353 | 0 | 0.00025015 | 17 | 211.01831 | 208.32173 | 314.37585 | NaN | 340.509 | NaN |
| Ruvbl2 | VYSLFLDESR | -429 | -438 | 0 | 0 | 10 | 611.3194 | 1240.2451 | 1068.5498 | NaN | NaN | NaN |
| Pdxk | DKVVPVADIITPN | -138 | -150 | 1 | 0.000471409 | 13 | 1664.5433 | NaN | NaN | 1249.0161 | 1323.1995 | NaN |
| Cherp | LALEQQQLICK | -60 | -70 | 0 | 0 | 11 | 1003.8856 | 710.35986 | 1379.9414 | 682.3091 | 805.5916 | 810.997 |
| Cherp | LEDHEYKPLDPK | -670 | -681 | 1 | 0 | 12 | 675.216 | 476.1064 | 526.48126 | 538.48236 | 398.28107 | 221.46729 |
| Cherp | LEDHEYKPLDPK | -670 | -681 | 1 | 0 | 12 | 675.216 | 476.1064 | 526.48126 | 538.48236 | 398.28107 | 221.46729 |
| Cherp | ITADGAHFELR | -201 | -211 | 0 | 0 | 11 | 490.0323 | 1423.4552 | 1478.5109 | 1513.2252 | 1324.6588 | 1341.8447 |
| Cherp | EFDNLLQPIIDTCTK | -156 | -170 | 0 | 0.00025015 | 15 | 296.43152 | 253.00116 | 149.94267 | 393.60388 | 394.08493 | 381.33218 |
| Cherp | DKWDQYK | -887 | -893 | 1 | 0.00025015 | 7 | 1150.7355 | 1336.0009 | 1043.6797 | NaN | NaN | 1491.7002 |
| Ccdc25 | DKYENEDLIK | -22 | -31 | 1 | 0 | 10 | 487.08408 | 1013.9633 | 427.61545 | 768.8636 | 1088.243 | NaN |
| Gatad2b | SLDPADERDDVLAK | -17 | -30 | 1 | 0 | 14 | 1558.5317 | 1026.4398 | 1203.0728 | 1503.3817 | 1457.341 | 1439.0043 |
| Gatad2b | DLANLEVPHELPTK | -53 | -66 | 0 | 0 | 14 | 1047.8634 | 1350.0356 | 1321.8284 | 713.1972 | 770.58453 | 775.16113 |
| Ctps2 | LGLPINDCSSNLLFK | -270 | -284 | 0 | 0 | 15 | 941.8422 | 1039.5502 | NaN | 1297.3562 | 1620.8019 | 1882.7096 |
| Ctps2 | VPLLLEEQGVVK | -253 | -264 | 0 | 0 | 12 | 184.41574 | 296.82553 | 233.99597 | NaN | 307.2532 | 349.3864 |
| Ctps2 | DLCFVGEDVDGK | -496 | -507 | 0 | 0 | 12 | 847.00916 | 764.95557 | 758.31506 | NaN | NaN | NaN |
| Tmod3 | DRDDYVPYTGEK | -82 | -93 | 1 | 0 | 12 | 1219.8118 | 1365.2753 | 1371.769 | 991.77454 | 1019.3978 | 658.11743 |
| Tmod3 | ILPVFDEPPNPTNVEESLK | -173 | -191 | 0 | 0 | 19 | 494.6038 | 672.9783 | NaN | 772.8936 | 960.39685 | 921.4251 |
| Tmod3 | ILPVFDEPPNPTNVEESLKR | -173 | -192 | 1 | 0 | 20 | 439.08118 | 378.45334 | NaN | 206.1361 | 524.4647 | 386.95242 |
| Tmod3 | DLGDYKDLDEDELLGK | -8 | -23 | 1 | 0 | 16 | 385.58185 | 749.11523 | 480.80432 | 459.10687 | NaN | 527.84576 |
| Tmod3 | DLDEDELLGKL | -14 | -24 | 1 | 0 | 11 | 604.42914 | NaN | NaN | 633.3429 | 678.5091 | 727.8085 |
| Tmod3 | DLDEDELLGK | -14 | -23 | 0 | 0.003102268 | 10 | 1033.8815 | 392.2975 | NaN | NaN | NaN | NaN |
| Wdr26 | DLDGNLLDSWEGVR | -454 | -467 | 0 | 0 | 14 | 2402.8357 | 2250.555 | NaN | NaN | 3037.6938 | NaN |
| Bag6 | LINLVGESLR | -323 | -332 | 0 | 0 | 10 | 2209.1655 | 1841.1765 | 1771.5941 | 1598.6667 | 1185.236 | 1556.4313 |
| Bag6 | DLEAPEVQESYR | -1110 | -1121 | 0 | 0 | 12 | 1716.2743 | 1860.9115 | 1238.7731 | 1956.9385 | 2471.7722 | 2317.7961 |
| Rbm39 | HGGVIHIY | -457 | -464 | 0 | 0.001092529 | 8 | 1187.7278 | 1316.9596 | 1378.386 | 1375.2029 | 1072.0735 | 329.0646 |
| Rbm39 | SVPLAIGLTGQR | -206 | -217 | 0 | 0 | 12 | 1132.2697 | 1119.6766 | 1100.9927 | 1102.8677 | 907.40393 | 873.566 |
| Rbm39 | SSFLDSDELER | -336 | -346 | 0 | 0 | 11 | 765.04333 | 766.3282 | 780.02313 | 779.1314 | 725.36255 | 661.91125 |
| Rbm39 | HFNITEDMLR | -258 | -267 | 0 | 0 | 10 | 525.894 | 343.32974 | NaN | 1345.4406 | 634.2475 | 333.72916 |
| Rbm39 | DLEEFFSTVGK | -168 | -178 | 0 | 0 | 11 | 295.57816 | 415.84378 | 393.32724 | NaN | NaN | 266.3255 |
| Rbm39 | FLDSDELER | -338 | -346 | 0 | 0.005789881 | 9 | 554.8432 | 105.55746 | NaN | NaN | NaN | NaN |
| Ogdh | ICEEAFTR | -565 | -572 | 0 | 0 | 8 | 2295.5867 | 2685.8804 | 2640.3025 | 3101.0737 | 3487.2312 | 3464.2092 |
| Ogdh | NQGYYDYVKPR | -962 | -972 | 1 | 0 | 11 | 1154.7675 | 1772.3706 | 2464.1116 | 3044.4326 | 5693.4326 | 5393.4014 |
| Ogdh | EAQKYPNAELAWCQ | -944 | -957 | 1 | 0.000550307 | 14 | 979.43555 | 1218.2935 | 862.6473 | 1423.1057 | 405.45102 | 1354.647 |
| Ogdh | VVNAPIFHVN | -467 | -476 | 0 | 0 | 10 | 933.6245 | 1077.9976 | 951.39996 | 847.4264 | 238.93295 | 1225.7957 |
| Ogdh | RKPLIVFTPK | -851 | -860 | 2 | 0.00025015 | 10 | 412.59808 | 397.48383 | 353.99182 | 337.80444 | 312.28864 | 233.15149 |
| Ogdh | LVEDHLAVQ | -123 | -131 | 0 | 0 | 9 | 913.9996 | 1060.1907 | 1121.6294 | 956.5102 | NaN | 1130.6405 |
| Ogdh | THLTELQR | -1001 | -1008 | 0 | 0.000230707 | 8 | 478.9303 | 597.28894 | NaN | 510.89297 | 443.03052 | 602.8525 |
| Ogdh | SWDIFFR | -75 | -81 | 0 | 0.00078661 | 7 | 365.28754 | 191.26723 | 384.98593 | NaN | 383.10416 | 465.32748 |
| Ogdh | SLVEAQPNVDK | -112 | -122 | 0 | 0 | 11 | 1908.6482 | 2042.0195 | 2106.2454 | 3772.928 | NaN | NaN |
| Ogdh | FIGGQEPALPLR | -193 | -204 | 0 | 0 | 12 | 1017.109 | 1089.4482 | 1116.8451 | 947.90125 | NaN | NaN |
| Ogdh | DLEQCQWIR | -227 | -235 | 0 | 0 | 9 | 928.2308 | 811.16693 | 1117.7429 | NaN | 773.0581 | NaN |
| Ogdh | QIGFTTDPR | -445 | -453 | 0 | 0.00122419 | 9 | 1093.1741 | 1257.7507 | 796.05255 | NaN | NaN | NaN |
| Ogdh | FEEFLQR | -264 | -270 | 0 | 0.00051306 | 7 | 612.24524 | NaN | 648.7875 | 587.46185 | NaN | NaN |
| Ogdh | KPLIVFTPK | -852 | -860 | 1 | 0 | 9 | 536.4885 | 513.91284 | 573.6258 | NaN | NaN | NaN |
| Ogdh | GVVNQPEYEEEISK | -548 | -561 | 0 | 0 | 14 | 1469.9984 | NaN | 1289.348 | NaN | NaN | NaN |
| Ogdh | VNNQIGFTTDPR | -442 | -453 | 0 | 0 | 12 | 830.7947 | NaN | 1087.8407 | NaN | NaN | NaN |
| Ogdh | GRDPAAAPATGNK | -987 | -999 | 1 | 0.00437931 | 13 | 237.78922 | NaN | NaN | NaN | NaN | NaN |
| Cdc40 | SGEIVQEYDR | -407 | -416 | 0 | 0 | 10 | 1636.4445 | 179.83969 | 2058.5664 | 2556.872 | 2480.9065 | 234.23395 |
| Cdc40 | GYALDPSLDNHQVSAK | -136 | -151 | 0 | 0.00025015 | 16 | 834.34863 | 754.9176 | 809.6481 | 932.88055 | 798.6592 | 884.08453 |
| Cdc40 | TIVFVDENRR | -423 | -432 | 1 | 0.000394361 | 10 | 291.76205 | 166.18153 | 377.0901 | 402.74048 | 394.79843 | 486.1936 |
| Cdc40 | TIVFVDENR | -423 | -431 | 0 | 0.002265552 | 9 | 1231.587 | 388.73767 | 1400.0112 | 2407.993 | 3079.4033 | NaN |
| Cdc40 | FAPEFGPENPFR | -89 | -100 | 0 | 0 | 12 | 458.4969 | 906.7494 | NaN | 797.20685 | 1136.0256 | 785.26495 |
| Cdc40 | EDLETGVHLDPAVK | -64 | -77 | 0 | 0 | 14 | 368.0702 | 351.67023 | 431.36334 | NaN | 531.33594 | 650.7749 |
| Cdc40 | DLETGVHLDPAVK | -65 | -77 | 0 | 0 | 13 | 671.0351 | NaN | 585.1012 | 453.2549 | NaN | 680.0978 |
| Ckb | DLFDPIIEER | -87 | -96 | 0 | 0 | 10 | 503.87048 | NaN | 448.833 | 382.01544 | 766.9389 | 797.3473 |
| Hdgf | GLWEIENNPTVK | -85 | -96 | 0 | 0 | 12 | 4878.0977 | 5869.4424 | 5861.262 | 3251.506 | 2285.7183 | 2684.5933 |
| Hdgf | ESGDHEEEDKEIAAL | -171 | -185 | 1 | 0.000230707 | 15 | 1482.5583 | 1717.4961 | 1766.0817 | 2851.2917 | 3046.652 | 2907.258 |
| Hdgf | ESGDHEEEDKEIAAL | -171 | -185 | 1 | 0.000230707 | 15 | 1482.5583 | 1717.4961 | 1766.0817 | 2851.2917 | 3046.652 | 2907.258 |
| Hdgf | EGERPLPVEVEK | -186 | -197 | 1 | 0.000755556 | 12 | 1066.5317 | 1213.4003 | 1264.9841 | 1603.6974 | 1761.8541 | 1834.2766 |
| Hdgf | GSSDEEGKLVIDEPAK | -131 | -146 | 1 | 0 | 16 | 698.81555 | 633.1846 | 667.05096 | 991.5744 | 1274.7986 | 728.8282 |
| Hdgf | SCAAEPEVEPEAHEGDGDKK | -107 | -126 | 1 | 0.00025015 | 20 | 573.28357 | 549.7685 | 551.30396 | 601.2645 | 661.3171 | 604.2598 |
| Hdgf | SCAAEPEVEPEAHEGDGDKK | -107 | -126 | 1 | 0.00025015 | 20 | 573.28357 | 549.7685 | 551.30396 | 601.2645 | 661.3171 | 604.2598 |
| Hdgf | SCAAEPEVEPEAHEGDGDK | -107 | -125 | 0 | 0.000431655 | 19 | 468.1766 | 526.28796 | 512.864 | 341.00046 | 94.01007 | 91.90733 |
| Hdgf | RAGDVLEDSPK | -157 | -167 | 1 | 0 | 11 | 404.65735 | 1277.4263 | 996.11975 | 1224.3273 | 641.33093 | 493.7787 |
| Hdgf | GFSEGLWEIENNPTVK | -81 | -96 | 0 | 0 | 16 | 149.03276 | 456.68463 | 528.52734 | 892.9625 | 1121.8102 | 984.47064 |
| Hdgf | DLFPYEESK | -62 | -70 | 0 | 0 | 9 | 1540.4384 | NaN | 1495.1755 | 493.80786 | 1538.6523 | 1691.6309 |
| Hdgf | KGFSEGLWEIENNPTVK | -80 | -96 | 1 | 0 | 17 | 433.90393 | 534.1791 | NaN | 969.5343 | 1355.2404 | 1236.192 |

| | | | | | | | | | | | | |
|---|---|---|---|---|---|---|---|---|---|---|---|---|
| Hdgf | ESGDHEEEDKEIAALE | -171 | -186 | 1 | 0.001536401 | 16 | 360.30286 | 1252.8606 | 1245.6846 | NaN | NaN | NaN |
| Ccr7 | DLGCLSQER | -343 | -351 | 0 | 0.00025015 | 9 | 1115.7693 | 1137.2346 | 1346.6793 | NaN | 1494.9418 | 1812.0417 |
| Ttc1 | DLGNLVLRPF | -253 | -262 | 1 | 0 | 10 | 321.63034 | 405.38568 | 419.2299 | 517.99744 | 410.95697 | 92.759346 |
| Ttc1 | TDKLDEALEDYK | -202 | -213 | 1 | 0.00025015 | 12 | 655.00366 | 369.76648 | 478.51535 | NaN | NaN | 359.44495 |
| Ctcf | KEPDAEPAEGEEEEAQAA | -699 | -716 | 1 | 0.00025015 | 18 | 558.21204 | 600.8453 | 638.1868 | 861.8777 | 916.1658 | 825.9148 |
| Ctcf | EGGQEEDACHLPQNQTD | -30 | -46 | 0 | 0 | 17 | 408.89215 | 479.01706 | 383.23538 | 494.2264 | 351.87192 | 369.35852 |
| Ctcf | FHCPHCDTVIA | -437 | -447 | 0 | 0.00025015 | 11 | 655.0788 | 586.36707 | NaN | 763.13104 | 785.82043 | 888.94086 |
| Ctcf | DLGVHLR | -451 | -457 | 0 | 0.004439092 | 7 | 629.9039 | 663.6294 | 754.5303 | NaN | NaN | 516.419 |
| Ctcf | HTLPLPEGFQVVK | -156 | -168 | 0 | 0 | 13 | 481.67557 | 499.05722 | 582.23724 | NaN | 418.2489 | NaN |
| Sf3a3 | DLHDCYLK | -141 | -148 | 0 | 0.002296977 | 8 | 2754.9956 | 2739.016 | 2938.1282 | 1835.8542 | 1901.3297 | 1835.037 |
| Sf3a3 | DLYDDKDGLR | -59 | -68 | 1 | 0.00025015 | 10 | 2576.7637 | 2679.509 | 3242.486 | 3040.7188 | 3234.0486 | 3001.9958 |
| Sf3a3 | VKPLQDQNELFGK | -200 | -212 | 1 | 0 | 13 | 532.77185 | 497.2303 | 545.33813 | 441.1892 | 710.0941 | 877.66815 |
| Sf3a3 | VKPLQDQNELFGK | -200 | -212 | 1 | 0 | 13 | 532.77185 | 497.2303 | 545.33813 | 441.1892 | 710.0941 | 877.66815 |
| Sf3a3 | YLDLHDCYLK | -139 | -148 | 0 | 0 | 10 | 1105.426 | 1309.621 | 1284.7452 | 1028.5779 | 900.03 | 1562.2897 |
| Sf3a3 | DLYDDKDGLRK | -59 | -69 | 2 | 0 | 11 | 966.59906 | 526.25256 | 718.0102 | 663.8723 | 635.1451 | 733.74976 |
| Sf3a3 | NLPLGWDGKPIPY | -382 | -394 | 1 | 0 | 13 | 752.94965 | 637.77515 | 258.92044 | 1127.9974 | 1492.7136 | 1315.0237 |
| Sf3a3 | VEFTDEEGYGR | -128 | -138 | 0 | 0 | 11 | 747.0903 | 351.39465 | 827.32275 | 883.45233 | 1026.6743 | 1025.6742 |
| Sf3a3 | QLTHENVQR | -339 | -347 | 0 | 0 | 9 | 452.41818 | 252.16754 | 499.7068 | 365.8435 | 431.6198 | 467.98758 |
| Sf3a3 | WQPDTEEEYEDSSGNVVNK | -471 | -489 | 0 | 0 | 19 | 413.2411 | 576.8779 | 633.15436 | 497.30362 | 565.231 | 592.24097 |
| Sf3a3 | KEELNAISGPNEFAEFY | -69 | -85 | 1 | 0 | 17 | 309.89038 | 311.76825 | 531.80096 | 618.17175 | 709.2419 | 701.315 |
| Sf3a3 | WQPDTEEEYEDSSGNVVNKK | -471 | -490 | 1 | 0 | 20 | 253.2922 | 673.54224 | 612.8506 | 632.1946 | 773.6264 | 728.2063 |
| Sf3a3 | EYVEILGEQR | -329 | -338 | 0 | 0 | 10 | 833.43616 | NaN | NaN | 936.18866 | 5783.8936 | NaN |
| Sf3a3 | SIFDQLFDIPK | -165 | -175 | 0 | 0.00025015 | 11 | 183.22897 | NaN | NaN | NaN | 354.95804 | 244.47517 |
| Sf3a3 | SVEFEELLK | -107 | -115 | 0 | 0.000471409 | 9 | 2109.787 | NaN | NaN | NaN | NaN | NaN |
| Drg1 | SDATADDLIDVVEGNR | -223 | -238 | 0 | 0 | 16 | 1312.972 | 1323.4775 | 1307.9531 | 1599.0475 | 847.4857 | 1453.0635 |
| Drg1 | WNFDDLLEK | -276 | -284 | 0 | 0 | 9 | 670.8174 | 583.89233 | 493.1636 | 585.02704 | 681.6553 | 468.99878 |
| Drg1 | LLDLPGIIEGAK | -115 | -126 | 0 | 0 | 12 | 426.0125 | 476.27676 | 407.94608 | 464.54047 | 436.6069 | 581.383 |
| Drg1 | IGFVGFPSVGK | -67 | -77 | 0 | 0 | 11 | 368.4223 | 1021.8893 | 883.0235 | 766.7001 | 994.1905 | 1141.155 |
| Drg1 | GGGGGGPGEGFDVAK | -47 | -61 | 0 | 0 | 15 | 2033.7872 | 2225.2554 | NaN | 2654.3496 | 2286.8618 | 2221.4236 |
| Drg1 | DLIDVVEGNR | -229 | -238 | 0 | 0 | 10 | 829.05444 | 474.27853 | 507.21738 | NaN | NaN | 641.54083 |
| Cdk4 | DLIGLPPEDDWPR | -228 | -240 | 0 | 0.002595029 | 13 | 192.20058 | NaN | NaN | 486.85886 | NaN | 587.56354 |
| Lars1 | SGPADSFNDR | -788 | -797 | 0 | 0 | 10 | 5449.035 | 4825.674 | 4254.9497 | 3717.6042 | 967.1732 | 3629.4426 |
| Lars1 | SWPVAGPVDESLIR | -886 | -899 | 0 | 0 | 14 | 3152.064 | 3357.8538 | 457.36096 | 3541.6238 | 3891.654 | 4181.3857 |
| Lars1 | NSLELEHIEVK | -1031 | -1041 | 0 | 0 | 11 | 1522.0352 | 126.04763 | 775.6989 | 1834.962 | 1353.313 | 1178.633 |
| Lars1 | ELGSLPELK | -977 | -985 | 0 | 0 | 9 | 1142.7874 | 839.6326 | 725.065 | 711.6967 | 814.20874 | 896.2918 |
| Lars1 | QGDSCESIIR | -1091 | -1100 | 0 | 0 | 10 | 1117.5701 | 608.7011 | 805.56396 | 706.69244 | 663.4135 | 899.3619 |
| Lars1 | FDDPLLGPR | -1120 | -1128 | 0 | 0 | 9 | 1087.9457 | 773.1698 | 1066.085 | 1021.1744 | 1139.7253 | 1113.8427 |
| Lars1 | QTGEGVGPQEYTLVK | -256 | -270 | 0 | 0 | 15 | 1049.2637 | 804.2203 | 608.45026 | 998.09827 | 972.5027 | 832.74146 |
| Lars1 | SLELEHIEVK | -1032 | -1041 | 0 | 0 | 10 | 363.395 | 1468.6827 | 1614.8762 | 1724.6357 | 1903.4092 | 1818.9803 |
| Lars1 | NYPVWQHITLT | -945 | -955 | 0 | 0 | 11 | 281.66772 | 1591.6173 | 749.04376 | 1382.1836 | 536.8177 | 1600.7985 |
| Lars1 | LDWLQEHACSR | -567 | -577 | 0 | 0 | 11 | 1892.9137 | 1686.6345 | 1719.385 | NaN | NaN | 1191.9275 |
| Lars1 | DLIPNHLSYY | -678 | -687 | 0 | 0.00025015 | 10 | 547.4665 | 960.6572 | NaN | NaN | 1182.5442 | NaN |
| Tpp2 | GTLIEAFPVLGGK | -716 | -728 | 0 | 0 | 13 | 1167.616 | 1185.894 | 1221.8796 | 1422.3135 | 1720.7207 | 1576.3716 |
| Tpp2 | TDPRPDAATIK | -1113 | -1123 | 1 | 0.00204437 | 11 | 1004.97174 | 953.3357 | 894.0763 | 1109.0536 | 1100.0948 | 1007.27216 |
| Tpp2 | TEEPFPFHGLLPK | -6 | -18 | 0 | 0 | 13 | 720.03644 | 877.8211 | 892.9846 | 890.94324 | 1166.4595 | 1128.7749 |
| Tpp2 | WTDLFDTK | -1186 | -1193 | 0 | 0 | 8 | 719.38654 | 4703.344 | 1013.4121 | 3335.937 | 2375.4844 | 2928.2278 |
| Tpp2 | DSTDIYNELK | -1050 | -1059 | 0 | 0 | 10 | 502.62393 | 763.18243 | 892.2075 | 1062.5592 | 739.52423 | 755.5647 |
| Tpp2 | DPVQVAAPSDHGVGIEPVFPENTENSEK | -537 | -564 | 0 | 0.00025015 | 28 | 250.53601 | 504.31268 | 547.9792 | 318.2161 | 312.94202 | 415.24716 |
| Tpp2 | HEQISDLDR | -894 | -902 | 0 | 0 | 9 | 1400.457 | 1322.2744 | 733.3747 | 1267.2821 | 944.8474 | NaN |
| Tpp2 | DLKEEFTEALR | -1029 | -1039 | 1 | 0.00025015 | 11 | 996.99994 | 913.67834 | 807.71643 | 1027.0856 | 1173.8368 | NaN |
| Tpp2 | VWHDGETWR | -199 | -207 | 0 | 0 | 9 | 875.48596 | NaN | 756.5376 | 1655.5654 | 897.2247 | 841.8857 |
| Tpp2 | IWDPIHR | -137 | -143 | 0 | 0.000550307 | 7 | 1569.6082 | NaN | 1526.0569 | 1349.7557 | 1658.3944 | NaN |
| Tpp2 | EKLPANQYTWSSR | -400 | -412 | 1 | 0 | 13 | 651.2998 | NaN | NaN | 3849.0332 | 1417.5713 | 538.7911 |
| Tpp2 | FTDVHFKPGQIR | -649 | -660 | 1 | 0.001392869 | 12 | 547.5778 | 325.73102 | NaN | NaN | 290.51035 | 520.6485 |
| Tpp2 | LWIIFDQNK | -856 | -864 | 0 | 0.000712251 | 9 | 504.5861 | 492.68393 | 554.523 | NaN | 445.61032 | NaN |
| Tpp2 | SPNAGPLFR | -620 | -628 | 0 | 0.00025015 | 9 | 507.14328 | 207.14398 | 690.15857 | NaN | NaN | NaN |
| Tpp2 | LPANQYTWSSR | -402 | -412 | 0 | 0.00204437 | 11 | 1648.4685 | NaN | NaN | NaN | NaN | NaN |
| Ripk1 | IDLLNHLIR | -644 | -652 | 0 | 0.00025015 | 9 | 648.0798 | 690.52246 | 675.3314 | NaN | 483.74536 | 533.67255 |
| Ripk1 | SQIDEIDHDYER | -595 | -606 | 0 | 0 | 12 | 633.7237 | 517.41046 | 433.90717 | NaN | NaN | 389.33224 |
| Ripk1 | DLKPENILVDR | -138 | -148 | 1 | 0 | 11 | 600.0307 | 284.97382 | 533.43976 | 643.2742 | NaN | NaN |
| Ripk1 | KTQIDVPLSLK | -105 | -115 | 1 | 0.000627943 | 11 | 432.28992 | 438.43018 | NaN | NaN | 540.76526 | 401.87402 |
| Prkaca | DLKPENLLIDQQ | -167 | -178 | 1 | 0 | 12 | 613.3421 | 749.9663 | 677.1054 | 644.29944 | 604.2223 | 832.63354 |
| Prkaca | KWETPSQNTAQLDQFDR | -30 | -46 | 1 | 0 | 17 | 489.716 | NaN | 938.70886 | NaN | 421.84167 | NaN |
| Cdk1 | SEEEGVPSTAIR | -39 | -50 | 0 | 0 | 12 | 1010.32684 | 927.193 | 825.4535 | 863.01276 | 1159.7532 | 635.6922 |
| Cdk1 | NLDENGLDLLSK | -255 | -266 | 0 | 0 | 12 | 929.8639 | 968.5127 | 1019.18896 | 1064.5901 | 865.6054 | 153.59886 |
| Cdk1 | LESEEEGVPSTAIR | -37 | -50 | 0 | 0 | 14 | 663.7942 | 652.3075 | 652.3211 | 763.10004 | 1135.6565 | 1228.0311 |
| Cdk1 | KPLFHGDSEIDQLFR | -201 | -215 | 1 | 0.000431655 | 15 | 541.3309 | 727.7144 | 586.9123 | 87.593956 | 772.66895 | 903.4054 |
| Cdk1 | LKHPYFDDLDNQIK | -282 | -295 | 1 | 0.000431655 | 14 | 1602.3402 | 1715.1196 | NaN | 1771.0826 | 2333.9993 | 2281.098 |
| Cdk1 | HPYFDDLDNQIK | -284 | -295 | 0 | 0.00051306 | 12 | 791.2257 | NaN | 827.8271 | 951.01685 | NaN | 971.9355 |
| Cdk1 | DLKPQNLLIDDK | -128 | -139 | 1 | 0 | 12 | 1057.3563 | 1205.6974 | NaN | NaN | NaN | 869.1208 |
| Ripk3 | NILLDPELHAK | -148 | -158 | 0 | 0 | 11 | 1253.7056 | 854.85944 | 885.9623 | 532.727 | 563.3365 | 508.5701 |
| Ripk3 | LHLEEPSGPVPGK | -347 | -359 | 0 | 0 | 13 | 734.1952 | 764.9332 | 827.6548 | 1041.6176 | 533.53326 | 938.11053 |
| Ripk3 | DLKPSNILLDPELHAK | -143 | -158 | 1 | 0 | 16 | 484.93677 | 574.8885 | 561.6542 | 388.46146 | 317.46014 | 588.55365 |
| Ripk3 | QAQDTSVGPATPAR | -364 | -377 | 0 | 0 | 14 | 835.7545 | NaN | 921.11554 | 1130.2904 | 1215.0372 | 878.1844 |
| Ripk3 | GTTPGPVFTETPGPHPQR | -397 | -414 | 0 | 0 | 18 | 579.7038 | NaN | 731.31946 | 835.3666 | 796.14734 | 794.3589 |
| Ripk3 | KISWEVK | -57 | -63 | 1 | 0.001291103 | 7 | 608.1661 | NaN | 493.08823 | NaN | NaN | NaN |
| Mapk10;Mapk | DLKPSNIVVK | (189);(151);(151) | (198);(160);(160) | 1 | 0.00078661 | 10 | 616.6685 | NaN | 671.3827 | NaN | NaN | 1112.7579 |
| Rps27l | LHPSLEEEK | -8 | -16 | 0 | 0 | 9 | 2625.5647 | 2508.3723 | 2882.2927 | 4134.32 | 1847.15 | 3874.039 |

| | | | | | | | | | | | | |
|---|---|---|---|---|---|---|---|---|---|---|---|---|
| Rps27l | DLLHPSLEEEK | -6 | -16 | 0 | 0 | 11 | 12941.915 | 11077.221 | NaN | NaN | 9901.777 | 7827.9375 |
| Rps27l | DLLHPSLEEE | -6 | -15 | 0 | 0.008436628 | 10 | 1157.6306 | NaN | NaN | NaN | NaN | NaN |
| Rps27 | LLHPSPEEEK | -7 | -16 | 0 | 0.00025015 | 10 | 1515.7642 | 1511.9059 | 1509.3049 | 1208.7079 | 192.0332 | 1290.7396 |
| Rps27 | DLLHPSPEEEK | -6 | -16 | 0 | 0 | 11 | 1270.4469 | 1220.2949 | 1238.0374 | 1004.43005 | 1053.7759 | 980.68 |
| Rps27 | LHPSPEEEK | -8 | -16 | 0 | 0.000431655 | 9 | 1232.2847 | 1289.0323 | 2096.347 | 2148.3726 | 2367.7769 | 1679.923 |
| Rps27 | LHPSPEEEKR | -8 | -17 | 1 | 0 | 10 | 372.7803 | 552.5057 | 571.9511 | 993.303 | 943.7185 | 765.4038 |
| Rps27 | LHPSPEEEKR | -8 | -17 | 1 | 0 | 10 | 372.7803 | 552.5057 | 571.9511 | 993.303 | 943.7185 | 765.4038 |
| Sart1 | LGLKPLEVN | -130 | -138 | 1 | 0.00025015 | 9 | 366.79892 | 469.81595 | 479.75967 | 520.6397 | 467.1473 | 548.9147 |
| Sart1 | TVEHAIDSFR | -258 | -267 | 0 | 0 | 10 | 360.37756 | 175.58675 | 1773.063 | 1852.3214 | 1939.5853 | 1397.6869 |
| Sart1 | DLLPLGDQTQDGDFGSR | -422 | -438 | 0 | 0.000591124 | 17 | 641.0238 | 939.69995 | 845.1084 | NaN | NaN | NaN |
| Sart1 | YDEELEGERPHSF | -337 | -349 | 1 | 0.00025015 | 13 | 569.576 | 567.29297 | NaN | 1184.8945 | NaN | NaN |
| Trrap | DLNVVEHK | -44 | -51 | 0 | 0.001392869 | 8 | 412.32584 | 467.73224 | NaN | NaN | NaN | NaN |
| Trrap | ENWHEEVLR | -2036 | -2044 | 0 | 0 | 9 | 637.02234 | NaN | NaN | NaN | NaN | NaN |
| Pdcd4 | VVLPLDETAFEK | -155 | -166 | 0 | 0 | 12 | 693.8067 | 808.6837 | 807.1534 | 793.67395 | 1007.3476 | 568.2034 |
| Pdcd4 | IYNEIPDINLDVPHSY | -408 | -423 | 0 | 0 | 16 | 392.37796 | 73.7461 | 634.76294 | 707.12665 | 763.5699 | 624.74054 |
| Pdcd4 | GGGQQPVNHLVK | -318 | -329 | 0 | 0 | 12 | 242.48148 | 1309.7227 | 2410.7402 | 2871.716 | 1881.3687 | 3181.034 |
| Pdcd4 | VLESTGESAFK | -371 | -381 | 0 | 0.00025015 | 11 | 1197.7394 | 340.89648 | NaN | 1159.5784 | NaN | 736.6246 |
| Pdcd4 | DLPELALDTPR | -246 | -256 | 0 | 0 | 11 | 1120.9532 | 2445.8171 | NaN | 2367.72 | NaN | NaN |
| Smndc1 | LNLKPVEEGR | -120 | -129 | 1 | 0.000471409 | 10 | 2992.5754 | 3234.7524 | 3193.2336 | 2711.005 | 3829.1443 | 3396.4375 |
| Smndc1 | SFASTQPTHSWK | -63 | -74 | 0 | 0 | 12 | 2294.571 | 1917.9302 | 1813.3761 | 3533.4404 | 3437.3413 | 1145.0234 |
| Smndc1 | DLQEVIELTK | -38 | -47 | 0 | 0 | 10 | 443.16693 | 534.60046 | 491.8407 | 323.24768 | 645.3179 | 719.7036 |
| Smndc1 | SDSFASTQPTHSWK | -61 | -74 | 0 | 0 | 14 | 1365.5344 | NaN | 884.55646 | 909.9007 | 920.34424 | 718.4399 |
| Copb1 | KPITDDDVDR | -603 | -612 | 1 | 0 | 10 | 1616.3439 | 1619.19 | 1612.7241 | 2246.8499 | 1963.6726 | 1962.5056 |
| Copb1 | DLQHPNEFIR | -108 | -117 | 0 | 0 | 10 | 1518.0815 | 1539.4647 | 1783.21 | 1869.3016 | 425.68167 | 2110.8982 |
| Copb1 | EAELLEPLMPAIR | -129 | -141 | 0 | 0 | 13 | 1477.809 | 1414.8447 | 1057.0662 | 3912.1357 | 1688.0449 | 1773.2065 |
| Copb1 | KEEDRPPLR | -535 | -543 | 2 | 0.000230707 | 9 | 1012.3208 | 916.18054 | 604.10297 | 552.10205 | 544.6578 | 519.44305 |
| Copb1 | FVLPLQDHTIK | -65 | -75 | 0 | 0 | 11 | 1000.34827 | 1332.9404 | 1328.9906 | 1488.5876 | 1781.5349 | 1783.0995 |
| Copb1 | TLAPHDFANIK | -776 | -786 | 0 | 0 | 11 | 977.40894 | 1160.2317 | 1222.3666 | 734.3157 | 1296.6392 | 1364.9657 |
| Copb1 | SLGEIPIVESEIK | -482 | -494 | 0 | 0 | 13 | 580.01355 | 351.4182 | 250.97186 | 169.6647 | 609.98584 | 467.45245 |
| Copb1 | EAGELKPEEEITVGPVQK | -496 | -513 | 1 | 0 | 18 | 573.70013 | 625.59106 | 1047.3181 | 658.4883 | 806.0456 | 711.5526 |
| Copb1 | EAGELKPEEEITVGPVQK | -496 | -513 | 1 | 0 | 18 | 573.70013 | 625.59106 | 1047.3181 | 658.4883 | 806.0456 | 711.5526 |
| Copb1 | SLGEIPIVESEIKK | -482 | -495 | 1 | 0 | 14 | 456.1126 | 1152.6025 | 599.99646 | 1268.8556 | 2169.2778 | 3051.9927 |
| Copb1 | GNIVYDVSGAASDR | -801 | -814 | 0 | 0 | 14 | 858.00995 | 1006.8237 | 1112.6438 | NaN | 936.5008 | 1320.6456 |
| Copb1 | NVSEHEDTDKYR | -369 | -380 | 1 | 0 | 12 | 654.95544 | 659.5764 | 697.4805 | 616.599 | 747.47 | NaN |
| Copb1 | LVEKPSPLTLAPHDFANIK | -768 | -786 | 1 | 0 | 19 | 722.29553 | NaN | NaN | 2027.5566 | 1496.2208 | 1183.9844 |
| Copb1 | TNNVSEHEDTDKYR | -367 | -380 | 1 | 0.001463897 | 14 | 288.6381 | NaN | 526.9719 | NaN | NaN | NaN |
| G6pdx | IFGPIWNR | -220 | -227 | 0 | 0.000230707 | 8 | 3462.533 | 1938.1951 | 2112.111 | 1370.4758 | 2745.8713 | 2856.1306 |
| G6pdx | FNPEESELDLTYGNR | -413 | -427 | 0 | 0 | 15 | 1277.1702 | 1604.4127 | NaN | 1386.0557 | 1546.335 | 1583.5625 |
| G6pdx | EKPATTGSDDVR | -274 | -285 | 1 | 0.000627943 | 12 | 480.1873 | 395.50516 | NaN | 168.9137 | 435.27747 | 378.268 |
| G6pdx | DVAGDIFHQ | -375 | -383 | 0 | 0 | 9 | 1262.7539 | NaN | NaN | 520.8734 | NaN | 1169.3995 |
| G6pdx | DVAGDIFHQQ | -375 | -384 | 0 | 0 | 10 | 798.16724 | 523.5385 | 972.9796 | NaN | NaN | NaN |
| G6pdx | LEEFFAR | -98 | -104 | 0 | 0.001291103 | 7 | 632.4618 | 761.0443 | NaN | 559.71484 | NaN | NaN |
| G6pdx | VQPNEAVYTK | -394 | -403 | 0 | 0.00025015 | 10 | 70.55259 | 1103.2173 | NaN | 468.43533 | NaN | NaN |
| Gtpbp4 | LPTIDPNTR | -161 | -169 | 0 | 0 | 9 | 3470.5588 | 3648.2454 | 185.3034 | 76.19911 | 140.116 | 3991.949 |
| Gtpbp4 | KNLVDNVAK | -103 | -111 | 1 | 0 | 9 | 947.62537 | 878.26385 | 555.63226 | 578.01184 | 495.42828 | 585.2338 |
| Gtpbp4 | KLEELEKEEELR | -449 | -460 | 2 | 0.000230707 | 12 | 877.27655 | 833.0032 | 677.2903 | 559.3289 | 629.03296 | 431.07483 |
| Gtpbp4 | KLEELEKEEELR | -449 | -460 | 2 | 0.000230707 | 12 | 877.27655 | 833.0032 | 677.2903 | 559.3289 | 629.03296 | 431.07483 |
| Gtpbp4 | ERPPFIPEGVIA | -372 | -383 | 1 | 0 | 12 | 763.5221 | 901.0203 | 899.69556 | 922.32166 | 1114.332 | 1001.4657 |
| Gtpbp4 | EELEKEEELR | -451 | -460 | 1 | 0 | 10 | 299.03577 | 444.07758 | 1826.0287 | 2170.0422 | 3144.5623 | 2548.6565 |
| Gtpbp4 | IAELSEEDQK | -296 | -305 | 0 | 0 | 10 | 1430.7268 | 1427.5526 | 1463.354 | 456.52136 | 93.17555 | NaN |
| Gtpbp4 | NILYDKDHYK | -84 | -93 | 1 | 0 | 10 | 797.8554 | 873.4264 | 1049.4697 | 1579.3582 | NaN | 929.04285 |
| Gtpbp4 | NILYDKDHYK | -84 | -93 | 1 | 0 | 10 | 797.8554 | 873.4264 | 1049.4697 | 1579.3582 | NaN | 929.04285 |
| Gtpbp4 | DLSEQCGHGLK | -256 | -266 | 0 | 0 | 11 | 533.70294 | 432.6468 | 3442.4658 | 173.30641 | 7072.001 | NaN |
| Gtpbp4 | LHLAVPNK | -360 | -367 | 0 | 0 | 8 | 860.06665 | NaN | 802.9718 | NaN | 807.719 | NaN |
| Gtpbp4 | TPTVIHK | -30 | -36 | 0 | 0.002910424 | 7 | 337.17352 | 348.39944 | NaN | NaN | NaN | NaN |
| Anp32a | YLDGYDRDNK | -144 | -153 | 1 | 0 | 10 | 3815.5928 | 3523.7874 | 4638.5806 | 6196.591 | 7705.214 | 1220.5532 |
| Anp32a | DLSTIEPLKK | -102 | -111 | 1 | 0 | 10 | 1185.8965 | 1123.1685 | 1248.3933 | 1675.206 | 1694.4663 | 1650.768 |
| Anp32a | KREPDDEGEEDD | -236 | -247 | 2 | 0.001055699 | 12 | 1143.2097 | 1147.9929 | 1236.297 | 893.26556 | 896.43335 | 30.885038 |
| Anp32a | IKDLSTIEPLKK | -100 | -111 | 2 | 0 | 12 | 633.84064 | 590.99805 | 634.99384 | 845.36127 | 825.6073 | 938.19525 |
| Anp32a | STIEPLKK | -104 | -111 | 1 | 0 | 8 | 1048.7312 | 958.25616 | 1016.17487 | NaN | 842.35266 | NaN |
| Irgm1 | VKYPPVFLVS | -223 | -232 | 1 | 0.00025015 | 10 | 1913.193 | 1995.701 | 167.75165 | 1518.6656 | 1729.333 | 1818.4806 |
| Irgm1 | SEQFSSNHVK | -164 | -173 | 0 | 0 | 10 | 1659.7582 | 1784.2466 | 445.65918 | 2061.2817 | 398.61777 | 2527.9495 |
| Irgm1 | SLDPLLYDFPK | -233 | -243 | 0 | 0 | 11 | 1188.2158 | 1089.0427 | 1179.0908 | 1170.881 | 1509.9172 | 1468.9817 |
| Irgm1 | VIGHDEDASAPTGVVR | -98 | -113 | 0 | 0 | 16 | 1083.7528 | 1184.3958 | 1164.3965 | 1154.9894 | 1326.1193 | 1278.9685 |
| Irgm1 | VIGHDEDASAPTGVVR | -98 | -113 | 0 | 0 | 16 | 1083.7528 | 1184.3958 | 1164.3965 | 1154.9894 | 1326.1193 | 1278.9685 |
| Irgm1 | DLSTSVLSEVR | -195 | -205 | 0 | 0 | 11 | 981.3794 | 571.0318 | 1079.7344 | 744.53186 | 839.47107 | 328.91364 |
| Smarce1 | FLESTDSFNNELK | -259 | -271 | 0 | 0 | 13 | 729.09937 | 701.56714 | 769.66925 | 287.71008 | 602.7797 | 512.5514 |
| Smarce1 | DLTDEEKQEYLNEY | -106 | -119 | 1 | 0.006472924 | 14 | 206.41434 | NaN | NaN | NaN | NaN | NaN |
| Rbm4 | DLTEQYNEQYGAVR | -185 | -198 | 0 | 0 | 14 | 837.81055 | 870.18994 | 938.55646 | NaN | NaN | 1032.4999 |
| Pgm2 | LDQETAQWLR | -23 | -32 | 0 | 0 | 10 | 1170.9788 | 1040.3978 | 1279.187 | 1264.6792 | 74.918015 | 1170.2144 |
| Pgm2 | DLTTGYDDSQPDKK | -528 | -541 | 1 | 0 | 14 | 1069.0208 | 1011.2289 | 1103.3999 | 145.51033 | 1161.8899 | 1269.2125 |
| Pgm2 | IVLANDPDADRL | -325 | -336 | 1 | 0 | 12 | 718.7494 | 639.3393 | 775.0543 | 1080.4117 | 1332.534 | 1072.1694 |
| Pgm2 | WDNGAQIISPHDR | -186 | -198 | 0 | 0 | 13 | 534.44147 | 836.96216 | 916.9031 | 1328.7173 | NaN | 569.23425 |
| Pgm2 | WDQNPLTSESVK | -33 | -44 | 0 | 0 | 12 | 184.46349 | 1020.37054 | 1242.9946 | 165.1765 | NaN | 1275.0938 |
| Pgm2 | DPDPEFPTVK | -291 | -300 | 0 | 0 | 10 | 368.2313 | 556.8057 | NaN | NaN | 957.9861 | 736.0078 |
| Pgm2 | EHFFQPQK | -605 | -612 | 0 | 0.00111006 | 8 | 462.58368 | NaN | NaN | NaN | NaN | NaN |
| Anxa5 | GAGTDDHTLIR | -259 | -269 | 0 | 0 | 11 | 1391.9296 | 1404.7673 | 1367.3492 | 1287.2655 | 1424.929 | 1453.3877 |
| Anxa5 | VLTEIIASR | -107 | -115 | 0 | 0 | 9 | 1307.6447 | 907.7422 | 1136.7479 | 794.278 | 767.1193 | 653.5016 |

| | | | | | | | | | | | | |
|---|---|---|---|---|---|---|---|---|---|---|---|---|
| Anxa5 | RTPEELSAIK | -115 | -124 | 1 | 0.00025015 | 10 | 1263.4609 | 1122.739 | 945.17883 | 980.06506 | 1827.0615 | 1786.69 |
| Anxa5 | GTVTDFPGFDGR | -5 | -16 | 0 | 0 | 12 | 389.38177 | 1011.30237 | 421.37006 | 968.85895 | 474.3526 | 1113.2065 |
| Anxa5 | DLVDDLKSELTGK | -62 | -74 | 1 | 0 | 13 | 176.06549 | 240.91078 | 362.34238 | NaN | 479.71454 | 520.07605 |
| Anxa5 | EDDVVGDTSGYYQR | -136 | -149 | 0 | 0 | 14 | 106.93305 | 530.17664 | 333.771 | NaN | NaN | NaN |
| Eif3a | QYVPQLQNNTILR | -426 | -438 | 0 | 0 | 13 | 2946.9226 | 2501.1255 | 994.9064 | 3376.307 | 2188.9316 | 2132.548 |
| Eif3a | LLLTPWVK | -144 | -151 | 0 | 0.00025015 | 8 | 2243.1072 | 2478.8188 | 2549.9827 | 2062.2153 | 2757.9597 | 2807.8171 |
| Eif3a | ILQEHEQIK | -624 | -632 | 0 | 0.000712251 | 9 | 2162.2483 | 2305.5273 | 2404.1233 | 1908.2819 | 2477.9314 | 2336.4253 |
| Eif3a | KIDYFER | -694 | -700 | 1 | 0.00078661 | 7 | 2079.9807 | 2216.967 | 2221.66 | 2128.3088 | 2367.1687 | 1242.9414 |
| Eif3a | PGPWRPF | -1127 | -1133 | 1 | 0.00416551 | 7 | 2066.1497 | 1728.3043 | 1807.8649 | 1541.0538 | 1952.0265 | 2030.325 |
| Eif3a | RLEEIPLIK | -703 | -711 | 1 | 0.002595029 | 9 | 1843.9504 | 2018.4971 | 2043.9169 | 1711.9581 | 2126.2244 | 2119.9846 |
| Eif3a | RGTDDDRPSWR | -980 | -990 | 2 | 0.000298805 | 11 | 1565.1758 | 1780.3235 | 276.7943 | 401.31863 | 1599.806 | 1554.4215 |
| Eif3a | KLCDNLR | -196 | -202 | 1 | 0.007819401 | 7 | 1555.8616 | 1051.2631 | 1831.0636 | 802.06366 | 2335.577 | 1962.6779 |
| Eif3a | RLGDDPLSR | -877 | -885 | 1 | 0.00025015 | 9 | 1508.1099 | 2553.1558 | 3603.869 | 3031.376 | 661.6538 | 3318.9243 |
| Eif3a | PAYFQRPENALK | -2 | -13 | 1 | 0 | 12 | 1379.3806 | 1526.1755 | 1832.1058 | 2000.4849 | 1228.7462 | 1482.8485 |
| Eif3a | PAYFQRPENALK | -2 | -13 | 1 | 0 | 12 | 1379.3806 | 1526.1755 | 1832.1058 | 2000.4849 | 1228.7462 | 1482.8485 |
| Eif3a | GTDDDRPSWR | -981 | -990 | 1 | 0 | 10 | 1212.0519 | 1198.493 | 1372.087 | 1285.8959 | 1537.305 | 1655.5477 |
| Eif3a | ESSLRPDDDRIPR | -947 | -959 | 2 | 0.001055699 | 13 | 1079.7993 | 231.14342 | 1189.8756 | 1461.8177 | 1731.11 | 1493.4492 |
| Eif3a | ESSLRPDDDRIPR | -947 | -959 | 2 | 0.001055699 | 13 | 1079.7993 | 231.14342 | 1189.8756 | 1461.8177 | 1731.11 | 1493.4492 |
| Eif3a | TLSIPITPER | -326 | -335 | 0 | 0 | 10 | 981.35657 | 990.39526 | 1064.4911 | 233.24164 | 2036.29 | 2326.1228 |
| Eif3a | IGLINDMVR | -368 | -376 | 0 | 0 | 9 | 850.2736 | 619.12317 | 748.34625 | 587.1324 | 1066.8225 | 3176.1968 |
| Eif3a | IGLINDMVR | -368 | -376 | 0 | 0 | 9 | 850.2736 | 619.12317 | 748.34625 | 587.1324 | 1066.8225 | 3176.1968 |
| Eif3a | TKNETDEDGWTTVR | -1330 | -1343 | 1 | 0 | 14 | 773.6116 | 777.7496 | 763.5595 | 574.9918 | 784.6457 | 648.01917 |
| Eif3a | DMDLWEQQEEER | -721 | -732 | 0 | 0 | 12 | 471.28522 | 337.7314 | 349.10455 | 462.05304 | 98.64388 | 330.1759 |
| Eif3a | KGPEADSEWR | -901 | -910 | 1 | 0 | 10 | 357.81293 | 1456.646 | 1515.5629 | 1444.326 | 1179.9609 | 1873.3678 |
| Eif3a | DNQDREENDKDLER | -1173 | -1186 | 2 | 0 | 14 | 143.37039 | 184.03433 | 412.99048 | 407.95596 | 523.8166 | 373.38342 |
| Eif3a | SFGSDLNYATR | -492 | -502 | 0 | 0 | 11 | 112.05237 | 2241.9492 | 2319.875 | 1540.9327 | 1869.9906 | 1608.253 |
| Eif3a | TLSFGSDLNYATR | -490 | -502 | 0 | 0 | 13 | 1623.7415 | 1968.4609 | 514.7715 | 2094.9646 | NaN | 2556.9758 |
| Eif3a | QYVVPEVK | -381 | -388 | 0 | 0 | 8 | 1801.3472 | 1804.1273 | NaN | 1778.6135 | NaN | 1955.2411 |
| Eif3a | YVPQLQNNTILR | -427 | -438 | 0 | 0 | 12 | 1155.7174 | 996.8395 | NaN | 1175.5574 | NaN | 1171.9095 |
| Eif3a | HCDLQVR | -477 | -483 | 0 | 0 | 7 | 681.35596 | NaN | NaN | 786.954 | 1606.6638 | 1793.7986 |
| Eif3a | EQPEKEPELQ | -416 | -425 | 1 | 0.001463897 | 10 | 830.81854 | NaN | 1190.4014 | 2092.6143 | NaN | NaN |
| Eif3a | IYQSIEFSR | -446 | -454 | 0 | 0.00025015 | 9 | 322.49222 | 376.58215 | NaN | 2101.3555 | NaN | NaN |
| Eif3a | GADDDRGPWR | -1102 | -1111 | 1 | 0.003360691 | 10 | 1457.8542 | 683.4794 | NaN | NaN | NaN | NaN |
| Eif3a | LGDDPLSR | -878 | -885 | 0 | 0.001055699 | 8 | 1028.5845 | NaN | NaN | NaN | 217.76266 | NaN |
| Eif3a | DMDGIIVEK | -343 | -351 | 0 | 0.004224523 | 9 | 1015.9053 | NaN | NaN | 1061.8567 | NaN | NaN |
| Eif3a | NLTQEEMQR | -309 | -317 | 0 | 0 | 9 | 848.5091 | 950.1386 | NaN | NaN | NaN | NaN |
| Eif3a | HHNQSTAINLNNPESQ | -211 | -226 | 0 | 0.00025015 | 16 | 687.0747 | NaN | 923.5225 | NaN | NaN | NaN |
| Eif3a | QSIEFSR | -448 | -454 | 0 | 0.001023236 | 7 | 406.91205 | NaN | NaN | NaN | NaN | NaN |
| Map4 | NIILPTEPDESSTK | -314 | -327 | 0 | 0 | 14 | 3757.39 | 4572.6074 | 4530.7974 | 3871.097 | 4430.1226 | 4246.6445 |
| Map4 | DMSPSAETEAPLAK | -515 | -528 | 0 | 0.00025015 | 14 | 1049.2255 | 969.26935 | 840.38873 | 1055.5988 | 1932.218 | 737.01385 |
| Map4 | EFIPLLDGDEK | -46 | -56 | 0 | 0 | 11 | 663.9655 | 748.23737 | 726.22003 | 1075.9209 | 757.8006 | 937.92725 |
| Map4 | ETPGSQPSEPCSGVSR | -626 | -641 | 0 | 0 | 16 | 587.8368 | 686.3749 | 534.52466 | 589.17883 | 945.524 | 1012.21594 |
| Map4 | VGSLDNVGHLPAGGAVK | -1044 | -1060 | 0 | 0 | 17 | 541.7368 | 615.88055 | 584.7125 | 645.9016 | 580.7775 | 518.09717 |
| Map4 | VGSLDNVGHLPAGGAVK | -1044 | -1060 | 0 | 0 | 17 | 541.7368 | 615.88055 | 584.7125 | 645.9016 | 580.7775 | 518.09717 |
| Map4 | STLPIDAPSPLENLEQK | -609 | -625 | 0 | 0 | 17 | 432.01727 | 553.95984 | 530.6822 | 665.18085 | 747.3152 | 755.62555 |
| Map4 | SLDNVGHLPAGGAVK | -1046 | -1060 | 0 | 0 | 15 | 218.3655 | 286.30334 | 458.54 | 595.86536 | 634.8963 | 649.716 |
| Map4 | TSPSKPSSAPALKPGPK | -759 | -775 | 2 | 0.001291103 | 17 | 439.85117 | 490.20877 | 479.25464 | NaN | NaN | NaN |
| Map4 | TEFIPLLDGDEK | -45 | -56 | 0 | 0 | 12 | 415.24722 | 595.6863 | 98.2571 | NaN | NaN | NaN |
| Map4 | DMSPLPESEVTLGK | -473 | -486 | 0 | 0.000230707 | 14 | 1553.9021 | NaN | NaN | NaN | NaN | NaN |
| Cdc5l | GGLNTPLHESDFSGVTPQR | -381 | -399 | 0 | 0 | 19 | 591.56683 | 647.7792 | 978.32623 | 1005.8964 | 522.7813 | 1107.3882 |
| Cdc5l | GGLNTPLHESDFSGVTPQR | -381 | -399 | 0 | 0 | 19 | 591.56683 | 647.7792 | 978.32623 | 1005.8964 | 522.7813 | 1107.3882 |
| Cdc5l | WYEWLDPSIK | -50 | -59 | 0 | 0 | 10 | 547.178 | 621.33606 | 841.3273 | 793.891 | 776.68585 | 801.44116 |
| Cdc5l | EIDDTYIEDAADVDAR | -506 | -521 | 0 | 0 | 16 | 447.2129 | 518.48 | 579.8829 | 569.99036 | 553.1982 | 602.6063 |
| Cdc5l | SGTTPKPVTNATPGR | -427 | -441 | 1 | 0.00025015 | 15 | 732.874 | NaN | 754.8084 | 821.30914 | 836.58594 | 1027.3169 |
| Cdc5l | NDFEIVLPENAEK | -488 | -500 | 0 | 0 | 13 | 679.0016 | 622.50195 | 659.67395 | 632.6116 | 225.55887 | NaN |
| Cdc5l | DNEEETTDDPRK | -111 | -122 | 1 | 0 | 12 | 287.4821 | NaN | NaN | 322.46597 | 274.9144 | 245.866 |
| Cdc5l | NVEPPLTDLQK | -560 | -570 | 0 | 0 | 11 | 3048.7373 | NaN | NaN | 2428.9736 | NaN | 3322.4946 |
| Cdc5l | LGLLGLPAPK | -478 | -487 | 0 | 0 | 10 | 578.89404 | 737.79504 | 688.4582 | NaN | NaN | NaN |
| Cdc5l | DNEEETTDDPR | -111 | -121 | 0 | 0 | 11 | 218.65633 | NaN | NaN | NaN | 343.33536 | 271.60406 |
| Ube2q1 | GAAPGAGGGPGGGPGPGPCLR | -22 | -42 | 0 | 0 | 21 | 457.05383 | NaN | NaN | 426.33237 | 441.36963 | NaN |
| Dpysl2 | NQTNCPLYVTK | -244 | -254 | 0 | 0 | 11 | 1652.3722 | 1539.3954 | 1637.0852 | 1420.5354 | 340.65503 | 1392.3204 |
| Dpysl2 | KPFPDFVYK | -472 | -480 | 1 | 0 | 9 | 1536.174 | 1573.8828 | 459.86432 | 1551.1394 | 1199.1628 | 1327.4694 |
| Dpysl2 | ILDLGITGPEGHVL | -212 | -225 | 0 | 0 | 14 | 679.24194 | 583.91846 | 597.5374 | 582.46497 | 675.00696 | 644.1029 |
| Dpysl2 | GLYDGPVCEVSVTPK | -497 | -511 | 0 | 0 | 15 | 629.3714 | 781.49347 | 714.5788 | 842.60144 | 1142.1025 | 111.672035 |
| Dpysl2 | VGSDADLVIWDPDSVK | -403 | -418 | 0 | 0 | 16 | 400.91058 | 297.58405 | 437.46112 | 398.3934 | 661.7812 | 545.83124 |
| Dpysl2 | THNSALEYNIF | -424 | -434 | 0 | 0 | 11 | 111.29824 | 868.21387 | 926.1615 | 803.6942 | 614.1648 | 648.1631 |
| Dpysl2 | SRPEEVEAEAVNR | -226 | -238 | 1 | 0 | 13 | 3553.257 | 7800.187 | NaN | NaN | NaN | 8982.981 |
| Dpysl2 | DNFTLIPEGTNGTEER | -346 | -361 | 0 | 0 | 16 | 426.28662 | NaN | NaN | 273.02637 | NaN | NaN |
| Dpysl2 | VHAENGDIIAEEQQR | -197 | -211 | 0 | 0.000907206 | 15 | 411.85114 | NaN | NaN | NaN | NaN | 509.49744 |
| Rp1 | DNGEELNK | -980 | -987 | 0 | 0.000826806 | 8 | 789.1105 | 423.79324 | 1043.6003 | 572.6162 | 639.248 | 818.10254 |
| Lrrc59 | VTGLHQQPLCTSVN | -268 | -281 | 0 | 0 | 14 | 1825.0593 | 1029.5452 | 590.0633 | 1012.5638 | 404.70773 | 513.2396 |
| Lrrc59 | LQQLPADFGR | -74 | -83 | 0 | 0.000591124 | 10 | 598.5427 | 2289.5723 | 2407.8179 | NaN | 2285.3965 | 2473.0059 |
| Lrrc59 | DNPLDPVLAK | -117 | -126 | 0 | 0.002595029 | 10 | 1616.2135 | 2006.9397 | NaN | NaN | 1585.9458 | 437.88766 |
| Lrrc59 | VNTIYDNAVQGLR | -280 | -292 | 0 | 0 | 13 | 4341.463 | NaN | NaN | 2520.1592 | NaN | NaN |
| Lrrc59 | SVNTIYDNAVQGLR | -279 | -292 | 0 | 0 | 14 | 469.93756 | NaN | 656.798 | NaN | NaN | NaN |
| Ddx1 | GEDSVPDTVHHVVVPVNPK | -433 | -451 | 0 | 0 | 19 | 1232.3615 | 1301.8322 | 1393.5774 | 498.6576 | 1498.6931 | 1529.2117 |
| Ddx1 | GEDSVPDTVHHVVVPVNPK | -433 | -451 | 0 | 0 | 19 | 1232.3615 | 1301.8322 | 1393.5774 | 498.6576 | 1498.6931 | 1529.2117 |
| Ddx1 | HFPTWVDLK | -424 | -432 | 0 | 0 | 9 | 740.8813 | 753.60095 | 861.90356 | 1098.205 | 1067.9802 | 954.0706 |

| | | | | | | | | | | | | |
|---|---|---|---|---|---|---|---|---|---|---|---|---|
| Ddx1 | LDDLVSTGK | -350 | -358 | 0 | 0 | 9 | 692.51373 | 803.3055 | 1491.3629 | 1051.3328 | 905.1331 | 1099.8757 |
| Ddx1 | GIDIHGVPYVIN | -578 | -589 | 0 | 0.000471409 | 12 | 1121.3815 | 2837.3953 | NaN | 1017.7492 | 1664.0514 | 1182.5013 |
| Ddx1 | FNFGEEEFK | -240 | -248 | 0 | 0 | 9 | 392.7853 | 707.06335 | NaN | 675.6249 | 700.29803 | NaN |
| Ddx1 | FGFGFGGTGK | -163 | -172 | 0 | 0.00122419 | 10 | 474.11655 | NaN | 407.34103 | 451.37808 | NaN | NaN |
| Ddx1 | DNTRPGANSPEMWSE | -473 | -487 | 1 | 0.001941082 | 15 | 541.2383 | NaN | NaN | NaN | NaN | NaN |
| Ctsd | NYELHPDK | -344 | -351 | 0 | 0.00025015 | 8 | 4850.65 | 5200.032 | 4793.7793 | 4234.83 | 5919.638 | 5595.8984 |
| Ctsd | VSSLPTVYLK | -330 | -339 | 0 | 0 | 10 | 2942.8452 | 2443.033 | 3457.8037 | 2428.2886 | 3685.7463 | 2349.3826 |
| Ctsd | LVGPVEEVK | -299 | -307 | 0 | 0 | 9 | 1984.4619 | 1821.5594 | 2122.589 | 1102.326 | 880.83185 | 613.3665 |
| Ctsd | LWVPSIHCK | -103 | -111 | 0 | 0 | 9 | 495.65387 | 349.53784 | 410.41028 | 358.46945 | 240.14955 | 295.7074 |
| Ctsd | LWVPSIHCK | -103 | -111 | 0 | 0 | 9 | 495.65387 | 349.53784 | 410.41028 | 358.46945 | 240.14955 | 295.7074 |
| Ctsd | NYELHPDKYILK | -344 | -355 | 1 | 0 | 12 | 764.59033 | 1041.2006 | 521.50775 | 814.701 | 785.1779 | 717.5291 |
| Ctsd | NYELHPDKYILK | -344 | -355 | 1 | 0 | 12 | 764.59033 | 1041.2006 | 521.50775 | 814.701 | 785.1779 | 717.5291 |
| Ctsd | DPEGQPGGELMLGGTDSK | -234 | -251 | 0 | 0 | 18 | 498.89026 | 628.89496 | 392.04407 | 290.989 | 722.6207 | 981.5102 |
| Ctsd | GGCEAIVDTGTSLLVGPVEEVK | -286 | -307 | 0 | 0 | 22 | 363.87183 | NaN | NaN | 530.76324 | 915.70795 | 801.4812 |
| Ctsd | GGCEAIVDTGTSL | -286 | -298 | 0 | 0.001291103 | 13 | 209.17865 | NaN | 197.85846 | 279.6891 | NaN | 313.52756 |
| Emg1 | DPGEVRPDITHQ | -79 | -90 | 1 | 0 | 12 | 2349.2131 | 2311.8152 | 2637.7383 | 2387.2952 | 907.40137 | 2238.119 |
| Ltv1 | DPLAADETAPQR | -27 | -38 | 0 | 0.002810811 | 12 | 474.52295 | NaN | NaN | NaN | NaN | NaN |
| Uqcrfs1 | GAVPAASEPPVLDVK | -32 | -46 | 0 | 0 | 15 | 1035.5581 | 500.1245 | 1122.4644 | 1375.7034 | 1308.0251 | 1154.2097 |
| Uqcrfs1 | DPQHDLDR | -197 | -204 | 0 | 0 | 8 | 879.5056 | 881.3955 | 876.3989 | 1003.6442 | 1428.5897 | 1170.8853 |
| Uqcrfs1 | GVAGALRPLLQ | -21 | -31 | 1 | 0 | 11 | 522.4058 | 549.0035 | 524.3051 | 578.20844 | 500.32837 | 475.47144 |
| Uqcrfs1 | VPDFSDYR | -85 | -92 | 0 | 0.000230707 | 8 | 2467.741 | 2356.7085 | 2711.3901 | 3324.038 | 3223.6663 | NaN |
| Uqcrfs1 | TVGLNVPASVR | -67 | -77 | 0 | 0 | 11 | 1023.80804 | 917.35596 | 986.3368 | 199.57356 | NaN | 704.5494 |
| Uqcrfs1 | QLRDPQHDLDR | -194 | -204 | 1 | 0.00025015 | 11 | 680.34314 | 513.5878 | 679.0893 | NaN | 623.1299 | 699.67236 |
| Uqcrfs1 | KGPAPLNLEVPAYEFT | -251 | -266 | 1 | 0.001743443 | 16 | 1255.5289 | 1390.0367 | 1513.3275 | NaN | NaN | NaN |
| Uqcrfs1 | RDPQHDLDR | -196 | -204 | 1 | 0.001636916 | 9 | 543.0116 | NaN | 416.3379 | NaN | NaN | 784.3685 |
| Uqcrfs1 | KGPAPLNLEVPAYEF | -251 | -265 | 1 | 0.00025015 | 15 | 412.99103 | NaN | 480.68774 | NaN | NaN | NaN |
| Pitrm1 | TVPYADPDHASLK | -871 | -883 | 0 | 0 | 13 | 2469.0686 | 2477.407 | 3555.8735 | 2516.3547 | 2751.1704 | 2651.8213 |
| Pitrm1 | LIAGPNSPFYK | -353 | -363 | 0 | 0 | 11 | 1381.9033 | 1224.5824 | 1388.3359 | 168.404 | 1577.3467 | 154.56694 |
| Pitrm1 | GNFQLEGHLK | -269 | -278 | 0 | 0 | 10 | 945.29596 | 947.74585 | 914.5529 | 907.8242 | 976.97925 | 960.7142 |
| Pitrm1 | ELDFWQEGWR | -174 | -183 | 0 | 0 | 10 | 527.559 | 592.2536 | 606.41846 | 496.3989 | 540.0663 | 560.9563 |
| Pitrm1 | HQDASCLPALK | -551 | -561 | 0 | 0 | 11 | 2521.0198 | NaN | 2092.165 | 132.26726 | 1729.9631 | 1386.3513 |
| Pitrm1 | TLTPSGDLQETFSGMDQVK | -729 | -747 | 0 | 0 | 19 | 765.7725 | NaN | NaN | 1240.1664 | 1459.528 | 1146.7001 |
| Pitrm1 | DPQTPLIFK | -191 | -199 | 0 | 0.002629273 | 9 | 2186.8733 | NaN | 496.182 | NaN | 1824.1135 | NaN |
| Pitrm1 | STVDSPVAPSDK | -960 | -971 | 0 | 0 | 12 | 2142.7708 | NaN | NaN | NaN | NaN | NaN |
| Acin1 | VSDESVLPLAQK | -496 | -507 | 0 | 0 | 12 | 4695.053 | 5386.394 | 5540.9443 | 5574.0234 | 7358.2017 | 7382.8496 |
| Acin1 | WLPLTESQIVQK | -1225 | -1236 | 0 | 0 | 12 | 2541.5938 | 3732.9875 | 3584.4077 | 2547.5347 | 2356.0945 | 2435.4888 |
| Acin1 | VVDEKPLNIR | -322 | -331 | 1 | 0 | 10 | 1408.8851 | 1430.1512 | 1385.8551 | 1417.3501 | 1457.4985 | 1669.6304 |
| Acin1 | YAEQDELDYHR | -1085 | -1095 | 0 | 0 | 11 | 1355.8593 | 833.2926 | 967.97876 | 822.11285 | 922.4298 | 262.25623 |
| Acin1 | EPLASPHPAQLLR | -464 | -476 | 0 | 0 | 13 | 1268.7274 | 1022.5025 | 1135.6239 | 737.4667 | 967.40027 | 835.379 |
| Acin1 | SAPLPLTVEEFAPAK | -526 | -540 | 0 | 0 | 15 | 745.6724 | 865.9501 | 747.3337 | 940.45447 | 320.8845 | 433.51828 |
| Acin1 | ETQIVSLPQEENEVK | -364 | -378 | 0 | 0 | 15 | 592.4903 | 553.3142 | 557.521 | 416.72464 | 699.89246 | 551.967 |
| Acin1 | PLHPPPPPPVQPPPHPR | -1115 | -1131 | 0 | 0 | 17 | 340.72678 | 336.78766 | 1103.2667 | 1069.5454 | 574.35376 | 690.3826 |
| Acin1 | DPSCGQEAAAPSGPQLQVLEPK | -673 | -694 | 0 | 0 | 22 | 238.4144 | 303.43903 | 376.57953 | 306.43414 | 436.7663 | 440.25308 |
| Acin1 | GLLVDRPSETK | -1096 | -1106 | 1 | 0.000550307 | 11 | 2057.3843 | NaN | 2229.5679 | 1639.0475 | 2043.9457 | 1939.979 |
| Acin1 | VALPPPVEHEVK | -957 | -968 | 0 | 0.00111006 | 12 | 173.07817 | 3360.1748 | NaN | 275.71848 | 1699.7 | 2097.9988 |
| Acin1 | QFEDEEGEER | -305 | -314 | 0 | 0 | 10 | 279.40637 | NaN | 367.23337 | 360.1408 | 373.53607 | NaN |
| Khdrbs1 | DPSGAHPSVR | -22 | -31 | 0 | 0 | 10 | 3718.1675 | 3440.7542 | 567.44574 | 527.9349 | 3109.1738 | 3585.439 |
| Khdrbs1 | KDDEENYLDLF | -139 | -149 | 1 | 0.00025015 | 11 | 2307.1372 | 2147.835 | 2343.2556 | 2645.2852 | 3047.163 | 777.97876 |
| Ndufb8 | GGDPTKEPEPVVHYDI | -171 | -186 | 1 | 0 | 16 | 599.6273 | 661.8025 | 572.3999 | 776.82715 | 1193.4038 | 1110.9861 |
| Ndufb8 | DPWYQWDHSELR | -87 | -98 | 0 | 0 | 12 | 570.24927 | NaN | 587.6775 | 473.07785 | 646.1985 | 751.14087 |
| Cd27 | EEEGSAIPIQEDYR | -229 | -242 | 0 | 0 | 14 | 2003.063 | 1830.6016 | 2067.0916 | 2056.687 | 2322.2861 | 2125.5144 |
| Cd27 | DQECTECDPPLNPALTR | -114 | -130 | 0 | 0 | 17 | 397.33832 | 310.3863 | 365.37076 | 114.40953 | 467.82904 | 429.26663 |
| Cops3 | DQEITVNPQFVQK | -394 | -406 | 0 | 0 | 13 | 2519.2783 | 2492.3691 | 2487.67 | 2936.6382 | 2485.6655 | 2849.0051 |
| Cops3 | VQLSGPQEAEK | -326 | -336 | 0 | 0 | 11 | 793.36273 | 734.27606 | 824.5773 | 911.655 | 839.213 | 1051.9968 |
| Cops3 | STNNPSELR | -268 | -276 | 0 | 0 | 9 | 3313.0151 | 2949.724 | NaN | 1725.7616 | 3914.0369 | 2823.4128 |
| Ikzf1 | SNHSAQDAVDNLLLL | -357 | -371 | 0 | 0.000230707 | 15 | 450.05597 | 446.88318 | 582.1396 | 129.84444 | 1002.56665 | 1088.268 |
| Ikzf1 | DQGSSALSGVGGIR | -98 | -111 | 0 | 0 | 14 | 965.4579 | NaN | NaN | NaN | NaN | NaN |
| Farsa | DQHDTFFLR | -272 | -280 | 0 | 0 | 9 | 1125.7308 | 1083.7727 | 1218.7214 | 1027.2722 | 1337.4269 | 1337.5675 |
| Farsa | VVDSIEDEVQK | -128 | -138 | 0 | 0 | 11 | 966.2053 | 777.5157 | 991.79553 | 1002.38727 | 572.4513 | 386.33426 |
| Farsa | GVEHQAVVGAVK | -31 | -42 | 0 | 0.001463897 | 12 | 770.55676 | 844.0886 | 876.16565 | 660.67285 | 742.49646 | 770.3142 |
| Farsa | DQHDTFFLRDPAEAL | -272 | -286 | 1 | 0.00051306 | 15 | 342.2348 | 684.27313 | 677.47595 | 733.2221 | 723.9004 | 717.7439 |
| Farsa | DRPFKPY | -204 | -210 | 2 | 0.000550307 | 7 | 4438.13 | 4440.8687 | 4213.795 | 3188.6245 | 4466.9937 | NaN |
| Farsa | RLEVADGGLDSAELATQL | -13 | -30 | 1 | 0.002466863 | 18 | 4330.4565 | 5392.225 | 5046.199 | 4350.027 | NaN | 6068.449 |
| Farsa | VVDSIEDEVQKR | -128 | -139 | 1 | 0 | 12 | 784.88434 | 992.9255 | 796.8578 | NaN | 1172.8235 | 1077.952 |
| Farsa | FQPQQHPAR | -263 | -271 | 0 | 0.002168954 | 9 | 782.7845 | 927.6482 | 983.06555 | NaN | 130.40176 | 832.9378 |
| Farsa | GVLPDSGHLHPLLK | -216 | -229 | 0 | 0 | 14 | 914.69055 | 764.31445 | 1521.5638 | NaN | NaN | 489.946 |
| Farsa | RVVDSIEDEVQKR | -127 | -139 | 2 | 0.00025015 | 13 | 1818.2384 | NaN | NaN | NaN | 1209.2788 | NaN |
| Farsa | RVVDSIEDEVQK | -127 | -138 | 1 | 0.000471409 | 12 | 772.1113 | NaN | NaN | 600.3574 | NaN | NaN |
| Cd48 | KDPLGPYK | -45 | -52 | 1 | 0.000627943 | 8 | 509.3801 | 879.2053 | 391.22897 | 649.775 | 890.71075 | 913.9351 |
| Cd48 | ITLEVFDPVPK | -120 | -130 | 0 | 0 | 11 | 441.2112 | 739.1224 | 451.6549 | 512.5884 | 733.7714 | 722.8173 |
| Cd48 | TEASTDSCHLR | -138 | -148 | 0 | 0 | 11 | 428.8487 | 578.63824 | 810.56476 | 433.97137 | 248.64076 | 1107.283 |
| Cd48 | IHKDPLGPYK | -43 | -52 | 1 | 0.000431655 | 10 | 649.9608 | 754.62744 | 1000.63257 | 1192.9453 | NaN | 1017.8455 |
| Cd48 | WYESSGPFPK | -162 | -171 | 0 | 0 | 10 | 542.30774 | 684.3467 | 661.5307 | NaN | 848.87354 | 575.9811 |
| Cd48 | DQHVDYTWYESSGPFPK | -155 | -171 | 0 | 0 | 17 | 512.30695 | NaN | 549.7541 | 586.94934 | 729.35767 | 709.03815 |
| Txndc9 | DQIGEDELELLK | -37 | -48 | 0 | 0 | 12 | 1474.5856 | 968.54315 | 668.51984 | 2423.9192 | NaN | NaN |
| Dctn1 | IKGEELSEANVR | -961 | -972 | 1 | 0 | 12 | 3411.8855 | 4037.0566 | 4490.6475 | 6684.0503 | 3552.4285 | 3517.9114 |
| Dctn1 | VFEDGADTTSPETPDSSASK | -96 | -115 | 0 | 0.000298805 | 20 | 551.3727 | 576.43677 | 590.2836 | NaN | 487.1962 | 535.6448 |
| Dctn1 | LDETQTLLR | -998 | -1006 | 0 | 0 | 9 | 1344.5103 | NaN | 1578.2457 | NaN | NaN | NaN |

| | | | | | | | | | | | | |
|---|---|---|---|---|---|---|---|---|---|---|---|---|
| Dctn1 | EQIYGSPSSSPYECLR | -875 | -890 | 0 | 0.000712251 | 16 | 394.7401 | NaN | 373.42892 | NaN | NaN | NaN |
| Dctn1 | DQLDETVNVEPLTK | -709 | -722 | 0 | 0.006624942 | 14 | 1682.4774 | NaN | NaN | NaN | NaN | NaN |
| Cluh | FNPDIFSPGVR | -719 | -729 | 0 | 0 | 11 | 1247.7686 | 1361.3756 | 709.3247 | 1107.5264 | 1236.0222 | 158.6278 |
| Cluh | HYILDLLR | -561 | -568 | 0 | 0 | 8 | 387.5774 | 185.33026 | 684.4948 | 494.76132 | 1048.6367 | 736.4549 |
| Cluh | LGYEEHIPGQTR | -344 | -355 | 0 | 0 | 12 | 241.89908 | 489.49298 | 536.81775 | 424.55994 | 464.1532 | 518.9594 |
| Cluh | IGIGELITR | -813 | -821 | 0 | 0 | 9 | 684.12524 | 133.46902 | 813.96326 | 827.2533 | 1077.569 | NaN |
| Cluh | DWKPLQCLK | -208 | -216 | 1 | 0.00025015 | 9 | 1590.473 | 961.2495 | 828.43134 | 2011.437 | NaN | NaN |
| Cluh | DWNEELQTTR | -356 | -365 | 0 | 0 | 10 | 893.28864 | NaN | NaN | NaN | NaN | NaN |
| Cluh | DQLDHIYK | -805 | -812 | 0 | 0.007670622 | 8 | 749.7696 | NaN | NaN | NaN | NaN | NaN |
| Slbp | DRKPEDSEEGELQ | -39 | -51 | 2 | 0.000471409 | 13 | 593.78577 | 799.12805 | 923.4908 | 589.02057 | 888.4203 | 784.4216 |
| Nme2 | NIIHGSDSVESAEK | -115 | -128 | 0 | 0 | 14 | 8174.555 | 7957.357 | 7483.6455 | 9902.234 | 9571.254 | 9740.7705 |
| Nme2 | NIIHGSDSVESAEK | -115 | -128 | 0 | 0 | 14 | 8174.555 | 7957.357 | 7483.6455 | 9902.234 | 9571.254 | 9740.7705 |
| Nme2 | DRPFFPGLVK | -57 | -66 | 1 | 0.000471409 | 10 | 8010.394 | 9052.688 | 9262.92 | 9062.814 | 11145.695 | 10891.487 |
| Nme2 | FKPEELIDYK | -134 | -143 | 1 | 0 | 10 | 6782.074 | 7708.455 | 7882.16 | 6848.292 | 8343.491 | 8222.722 |
| Nme2 | FKPEELIDYK | -134 | -143 | 1 | 0 | 10 | 6782.074 | 7708.455 | 7882.16 | 6848.292 | 8343.491 | 8222.722 |
| Nme2 | QHYIDLK | -50 | -56 | 0 | 0.001055699 | 7 | 2781.5894 | 2169.9966 | 1583.0099 | 2466.4937 | 2258.5886 | 2133.444 |
| Nme2 | KDRPFFPGLVK | -56 | -66 | 2 | 0.000471409 | 11 | 1416.9545 | 1869.6741 | 2058.152 | 998.01306 | 1539.006 | 1175.1765 |
| Nme2 | KDRPFFPGLVK | -56 | -66 | 2 | 0.000471409 | 11 | 1416.9545 | 1869.6741 | 2058.152 | 998.01306 | 1539.006 | 1175.1765 |
| Nme2 | SCAHDWVYE | -144 | -152 | 0 | 0.000471409 | 9 | 1098.5494 | 1168.4949 | 1142.162 | 1066.6588 | 808.45184 | 897.03375 |
| Nme2 | WFKPEELIDYK | -133 | -143 | 1 | 0 | 11 | 777.0354 | 1203.5098 | 1186.3479 | 1185.336 | 1557.5425 | 1458.8739 |
| Nme2 | IHGSDSVESAEK | -117 | -128 | 0 | 0.00111006 | 12 | 724.7697 | 908.93335 | 994.38293 | 1245.3735 | 1786.8723 | 1514.5892 |
| Nme2 | NIIHGSDSVESA | -115 | -126 | 0 | 0.00111006 | 12 | 2197.2988 | 2074.0168 | NaN | 1646.7755 | NaN | 762.97644 |
| Nme1 | NIIHGSDSVK | -115 | -124 | 0 | 0 | 10 | 12322.126 | 11807.158 | 11553.338 | 5190.4307 | 4200.9795 | 4228.578 |
| Nme1 | DRPFFTGLVK | -57 | -66 | 1 | 0.000394361 | 10 | 2713.149 | 2901.6384 | 68.194595 | 3307.2937 | 3698.827 | 3910.8926 |
| Nme1 | DRPFFTGL | -57 | -64 | 1 | 0.000627943 | 8 | 1946.371 | 2183.7292 | 2256.806 | 2191.2798 | 2385.819 | 2637.9797 |
| Nme1 | FQPEELVEYK | -134 | -143 | 0 | 0 | 10 | 699.3008 | 4721.369 | 4878.4575 | 4771.2256 | 599.27496 | 4671.4917 |
| Nme1 | FQPEELVEYK | -134 | -143 | 0 | 0 | 10 | 699.3008 | 4721.369 | 4878.4575 | 4771.2256 | 599.27496 | 4671.4917 |
| Nme1 | KFLQASEDLLK | -39 | -49 | 1 | 0 | 11 | 416.44522 | 382.52484 | 558.30273 | 954.1494 | 1838.0771 | 1772.5599 |
| Nme1 | WFQPEELVEYK | -133 | -143 | 0 | 0.00025015 | 11 | 258.9502 | 1088.2451 | NaN | NaN | 1614.3646 | 787.5063 |
| Wnk1 | DRPVSQPSLVGSK | -161 | -173 | 1 | 0.003133457 | 13 | 2039.3171 | NaN | NaN | 2261.3816 | NaN | NaN |
| Wnk1 | SQQQDDIEELETK | -198 | -210 | 0 | 0 | 13 | 1833.3516 | 2278.9316 | NaN | NaN | NaN | NaN |
| Mrpl4 | DSAGLLDPVLR | -39 | -49 | 0 | 0 | 11 | 1576.5377 | 1983.1832 | 1830.5756 | 1433.2163 | 1198.4692 | 1569.6072 |
| Rabggta | LPENVLLR | -203 | -210 | 0 | 0 | 8 | 1417.742 | 348.4245 | 988.3937 | 720.85944 | 194.33173 | 628.2917 |
| Rabggta | DSATDEQLFR | -344 | -353 | 0 | 0.00025015 | 10 | 747.38715 | NaN | NaN | 776.85333 | NaN | NaN |
| Prss2 | TLDNDIMLIK | -103 | -112 | 0 | 0 | 10 | 5869.485 | 16530.959 | 6166.01 | 10518.723 | 3891.063 | 11606.939 |
| Prss2 | TLDNDIMLIK | -103 | -112 | 0 | 0 | 10 | 5869.485 | 16530.959 | 6166.01 | 10518.723 | 3891.063 | 11606.939 |
| Prss2 | TLDNDIMLIK | -103 | -112 | 0 | 0 | 10 | 5869.485 | 16530.959 | 6166.01 | 10518.723 | 3891.063 | 11606.939 |
| Prss2 | DSCQGDSGGPVVCNGELQ | -194 | -211 | 0 | 0 | 18 | 597.1964 | 669.6832 | 1406.2091 | 877.7347 | 689.7841 | 692.5773 |
| Try5;Try4 | MICVGFLEGGK | (183);(183) | (193);(193) | 0 | 0 | 11 | 9530.254 | 223.56464 | 6502.835 | 6080.604 | 13727.675 | 13933.94 |
| Try5;Try4 | MICVGFLEGGK | (183);(183) | (193);(193) | 0 | 0 | 11 | 9530.254 | 223.56464 | 6502.835 | 6080.604 | 13727.675 | 13933.94 |
| Try5;Try4 | LGEHNINVLEGNEQFVNSAK | (73);(73) | (92);(92) | 0 | 0.00025015 | 20 | 2638.8591 | 3750.0713 | 3848.5847 | 20256.324 | 5342.8022 | 5101.6206 |
| Try5;Try4 | LGEHNINVLEGNEQFVNSAK | (73);(73) | (92);(92) | 0 | 0.00025015 | 20 | 2638.8591 | 3750.0713 | 3848.5847 | 20256.324 | 5342.8022 | 5101.6206 |
| Try5;Try4 | ICVGFLEGGK | (184);(184) | (193);(193) | 0 | 0 | 10 | 1805.7667 | 1854.746 | 1989.7556 | 1867.6245 | 2176.4094 | 2274.1367 |
| Try5;Try4 | DSCQGDSGGPVVCNGQLQ | (194);(194) | (211);(211) | 0 | 0.00025015 | 18 | 440.03604 | 458.67047 | 1739.6742 | 557.3165 | 946.8915 | 978.50415 |
| Try5;Try4 | NNMICVGFLEGGK | (181);(181) | (193);(193) | 0 | 0 | 13 | 259.16187 | 348.39395 | 450.71207 | 1518.2925 | 1398.2175 | 1202.7169 |
| Try5;Try4 | NMICVGFLEGGK | (182);(182) | (193);(193) | 0 | 0.002296977 | 12 | 156.258 | 1663.5775 | 1718.5071 | NaN | NaN | NaN |
| Nars1 | TINEPILLCR | -441 | -450 | 0 | 0 | 10 | 4197.186 | 4679.3364 | 5053.1797 | 3753.3423 | 4829.6465 | 5176.2275 |
| Nars1 | SWDSEEILEGYK | -490 | -501 | 0 | 0 | 12 | 2382.5298 | 2334.1733 | 1981.6038 | 152.95732 | 1109.9518 | 1401.5354 |
| Nars1 | LEDLVCDVVDR | -365 | -375 | 0 | 0 | 11 | 1387.371 | 1509.9868 | 1285.3804 | 1110.402 | 1071.3851 | 1310.8073 |
| Nars1 | EPFPTIYVDSQK | -50 | -61 | 0 | 0 | 12 | 1161.6427 | 722.86444 | 1821.097 | 791.8915 | 1783.1943 | 1010.69336 |
| Nars1 | YGTCPHGGYGL | -519 | -529 | 0 | 0.000230707 | 11 | 884.4854 | 899.471 | 461.65247 | 1028.0499 | 283.52493 | 902.0438 |
| Nars1 | EFGDDIPEAPER | -425 | -436 | 0 | 0 | 12 | 875.37225 | 777.00977 | 747.9325 | 1032.7013 | 1049.9694 | 1561.0892 |
| Nars1 | SPVASIVYELNPNFKPPK | -379 | -396 | 1 | 0 | 18 | 822.753 | 816.42334 | 651.849 | 962.30414 | 976.62244 | 918.782 |
| Nars1 | YGTCPHGGYGLGLER | -519 | -533 | 0 | 0 | 15 | 544.7217 | 876.2028 | 835.1187 | 986.6543 | 825.6222 | 870.8419 |
| Nars1 | FLSWILNR | -534 | -541 | 0 | 0 | 8 | 519.0725 | 645.1344 | 715.17633 | 461.26968 | 802.12335 | 759.0233 |
| Nars1 | NDPSLPEPACVK | -116 | -127 | 0 | 0 | 12 | 11556.889 | 11368.144 | 12417.434 | NaN | 11737.641 | 1191.3558 |
| Nars1 | RFPVEIK | -450 | -456 | 1 | 0.004439092 | 7 | 2010.8743 | NaN | 2130.9512 | 2391.838 | 2842.8271 | 2863.0505 |
| Nars1 | GDDIPEAPER | -427 | -436 | 0 | 0 | 10 | 1041.056 | 1339.2999 | 1271.1055 | 342.89578 | NaN | 517.8943 |
| Nars1 | DSEEILEGYK | -492 | -501 | 0 | 0 | 10 | 265.08667 | 793.8361 | 1023.33496 | 1003.1383 | 975.2688 | NaN |
| Nars1 | DSEEILEGYKR | -492 | -502 | 1 | 0 | 11 | 1164.5707 | 1233.536 | 1215.4955 | NaN | NaN | 1035.7075 |
| Nars1 | DVCLYPR | -546 | -552 | 0 | 0.002329279 | 7 | 981.2764 | NaN | NaN | 598.2782 | NaN | 660.7242 |
| Nars1 | EGIDPAPYYWYTDQR | -503 | -517 | 0 | 0 | 15 | 576.34015 | 698.4069 | 633.22253 | NaN | NaN | NaN |
| Nars1 | LMTDTINEPILLCR | -437 | -450 | 0 | 0.001868731 | 14 | 247.32976 | 415.00983 | NaN | NaN | NaN | NaN |
| Cops5 | DSFALPVEGTETR | -94 | -106 | 0 | 0 | 13 | 2635.315 | 2811.9036 | 2509.9465 | 3665.423 | 3745.993 | 3518.7131 |
| Cops5 | LEQSEAQLGR | -273 | -282 | 0 | 0 | 10 | 1579.7472 | 1267.5077 | NaN | 1677.8683 | NaN | NaN |
| Cops5 | SEKLEQSEAQLGR | -270 | -282 | 1 | 0.002199523 | 13 | 1081.6753 | 1044.2986 | NaN | NaN | NaN | NaN |
| Ndufb7 | DSFPNFLACK | -72 | -81 | 0 | 0 | 10 | 440.1905 | 345.52902 | 636.3187 | 754.7468 | 723.6322 | 89.00415 |
| Mat2a | EQQSPDIAQGVHLDR | -111 | -125 | 0 | 0 | 15 | 6435.328 | 5686.9814 | 7233.552 | 8510.228 | 6662.8164 | 7532.4478 |
| Mat2a | FVIGGPQGDAGLTGR | -250 | -264 | 0 | 0 | 15 | 1936.3447 | 2123.2092 | 2329.3447 | 2285.475 | 2708.738 | 2546.8335 |
| Mat2a | DSFPWEVPK | -383 | -391 | 0 | 0 | 9 | 1040.6864 | 1043.7966 | 1013.06305 | 451.02582 | 406.1836 | 1130.306 |
| Mat2a | NGTLPWLRPDSK | -170 | -181 | 1 | 0 | 12 | 1007.8335 | 1394.3129 | 1153.4304 | 316.44357 | 1082.3228 | 1082.69 |
| Mat2a | NGTLPWLRPDSK | -170 | -181 | 1 | 0 | 12 | 1007.8335 | 1394.3129 | 1153.4304 | 316.44357 | 1082.3228 | 1082.69 |
| Mat2a | YLDEDTIYHLQPSGR | -235 | -249 | 0 | 0 | 15 | 564.4366 | 607.6138 | 1411.3424 | 1559.04 | 1344.2964 | 1368.0669 |
| Mat2a | YLDEDTIYHLQPSGR | -235 | -249 | 0 | 0 | 15 | 564.4366 | 607.6138 | 1411.3424 | 1559.04 | 1344.2964 | 1368.0669 |
| Mat2a | ICDQISDAVLDAHLQ | -33 | -47 | 0 | 0 | 15 | 1082.2136 | 1270.4855 | 1318.9221 | NaN | 576.85443 | 1813.3844 |
| Mat2a | ICDQISDAVLDAHLQQDPDAK | -33 | -53 | 0 | 0 | 21 | 336.90094 | 278.72632 | NaN | NaN | 509.87122 | 440.00452 |
| Sash3 | VHTDFTPSPYDHDSLK | -180 | -195 | 0 | 0 | 16 | 2154.0083 | 2287.5117 | 1966.7354 | 2516.4834 | 2336.8174 | 2214.393 |
| Sash3 | TDFTPSPYDHDSLK | -182 | -195 | 0 | 0 | 14 | 758.3147 | 740.82983 | 806.97766 | 1224.6735 | 1247.8997 | 1214.4595 |

| | | | | | | | | | | | | |
|---|---|---|---|---|---|---|---|---|---|---|---|---|
| Sash3 | SKPSSPVVSEK | -34 | -44 | 1 | 0.000431655 | 11 | 656.7556 | 854.4386 | 981.019 | NaN | 1152.4769 | 1240.0892 |
| Sash3 | DSGCFEGSESGRDEAELA | -348 | -365 | 1 | 0.000869338 | 18 | 380.74844 | 373.07227 | 386.22986 | NaN | 325.8589 | 342.9277 |
| Sash3 | EESPAPQYTGPFCGR | -163 | -177 | 0 | 0 | 15 | 943.9125 | 977.74 | 961.92126 | 766.8103 | NaN | NaN |
| Sash3 | FTPSPYDHDSLK | -184 | -195 | 0 | 0 | 12 | 1583.6128 | 1500.0264 | 1733.3114 | NaN | NaN | NaN |
| Sash3 | TLHELLER | -256 | -263 | 0 | 0.004224523 | 8 | 433.6034 | NaN | NaN | NaN | NaN | NaN |
| Ranbp1 | FASENDLPEWK | -58 | -68 | 0 | 0 | 11 | 5975.9736 | 6140.3325 | 6134.78 | 675.86584 | 6081.2925 | 182.18973 |
| Ranbp1 | FLNAENAQK | -142 | -150 | 0 | 0.00025015 | 9 | 2019.6521 | 3156.8384 | 3399.8284 | 391.81268 | 463.36414 | 3524.2854 |
| Ranbp1 | EEDEEELFK | -42 | -50 | 0 | 0 | 9 | 1683.0602 | 1566.7614 | 1545.9924 | 1505.6757 | 1055.8059 | 1291.3547 |
| Ranbp1 | NTHADFADECPKPELLAIR | -123 | -141 | 1 | 0 | 19 | 1401.5164 | 1640.3232 | 1719.4194 | 1792.7136 | 2297.512 | 1135.3552 |
| Ranbp1 | NTHADFADECPKPELLAIR | -123 | -141 | 1 | 0 | 19 | 1401.5164 | 1640.3232 | 1719.4194 | 1792.7136 | 2297.512 | 1135.3552 |
| Ranbp1 | RFASENDLPEWK | -57 | -68 | 1 | 0 | 12 | 1316.3104 | 919.1395 | 1436.55 | 1665.9775 | 2133.7246 | 2304.906 |
| Ranbp1 | RFASENDLPEWK | -57 | -68 | 1 | 0 | 12 | 1316.3104 | 919.1395 | 1436.55 | 1665.9775 | 2133.7246 | 2304.906 |
| Ranbp1 | SENDLPEWK | -60 | -68 | 0 | 0 | 9 | 1100.7903 | 1121.7922 | 1254.0112 | 1333.7645 | 1306.357 | 1277.6862 |
| Ranbp1 | TKFEECR | -153 | -159 | 1 | 0.001190916 | 7 | 1004.3057 | 294.2595 | 1210.4165 | 422.68436 | 1061.9362 | 1147.1068 |
| Ranbp1 | TLEEDEEELFK | -40 | -50 | 0 | 0 | 11 | 907.3477 | 829.0041 | 748.677 | 810.9998 | 639.82605 | 424.8011 |
| Ranbp1 | DSHEDHDTSTENADESNHDPQFEPIVSLPEQEIK | -6 | -39 | 0 | 0 | 34 | 589.7664 | 556.20905 | 553.5968 | 566.1506 | 201.34494 | 586.8592 |
| Ranbp1 | PQFEPIVSLPEQEIK | -25 | -39 | 0 | 0 | 15 | 501.57455 | 662.56696 | 598.5774 | 857.49994 | 507.0735 | 980.45984 |
| Rps13 | GLAPDLPEDLYHLIK | -79 | -93 | 0 | 0 | 15 | 2037.8201 | 2076.9836 | 1569.6201 | 2509.61 | 3132.8867 | 2663.3608 |
| Rps13 | GLAPDLPEDLYHLIK | -79 | -93 | 0 | 0 | 15 | 2037.8201 | 2076.9836 | 1569.6201 | 2509.61 | 3132.8867 | 2663.3608 |
| Rps13 | KGLTPSQIGVIL | -43 | -54 | 1 | 0.00025015 | 12 | 1243.3273 | 1401.995 | 1395.784 | 1064.0089 | 1134.2303 | 1169.6327 |
| Rps13 | GLAPDLPEDLYHL | -79 | -91 | 0 | 0 | 13 | 839.7919 | 810.2445 | 762.82983 | 1008.35693 | 1117.4276 | 1136.9036 |
| Rps13 | RVLPPNWK | -133 | -140 | 1 | 0 | 8 | 510.16104 | 1437.8558 | 1926.1599 | 1734.3191 | 1172.6616 | 2447.7722 |
| Rps13 | GLTPSQIGVIL | -44 | -54 | 0 | 0.001291103 | 11 | 453.8346 | 595.3378 | 598.72534 | 532.949 | 933.34216 | 855.8209 |
| Rps13 | DSHGVAQVR | -56 | -64 | 0 | 0.00025015 | 9 | 882.7122 | 1042.8552 | 1150.0503 | 704.2782 | NaN | 1056.0103 |
| Rps13 | GLTPSQIGVILR | -44 | -55 | 0 | 0 | 12 | 613.3469 | 661.92096 | 701.0938 | NaN | NaN | NaN |
| Zmat2 | DSINFLDHINGK | -91 | -102 | 0 | 0.000947459 | 12 | 276.28 | NaN | NaN | NaN | NaN | NaN |
| Purb | GGGGGGGGPGGFQPAPR | -11 | -28 | 0 | 0 | 18 | 896.15295 | 921.5945 | 879.8474 | 402.5435 | 1080.4424 | 1000.69824 |
| Purb | FFFDVGCNK | -244 | -252 | 0 | 0 | 9 | 658.36597 | 694.3336 | 729.86896 | 616.7767 | 675.94635 | 653.0321 |
| Purb | GGGGGGGGPGGEQETQELASK | -29 | -49 | 0 | 0 | 21 | 653.61096 | 1100.4668 | 893.8424 | 853.6742 | 797.59406 | 862.63666 |
| Purb | DSLGDFIEHY | -97 | -106 | 0 | 0 | 10 | 625.41254 | 696.9771 | 685.4715 | 515.8869 | 806.0503 | 805.90735 |
| Purb | QSGQTIALPAQGLIEFR | -179 | -195 | 0 | 0 | 17 | 376.2508 | 342.0302 | NaN | 400.75406 | 520.64526 | 556.5278 |
| Purb | RGGGSGGGDESEGEEVDED | -306 | -324 | 1 | 0.001708008 | 19 | 303.38837 | 381.47238 | 400.96594 | 326.80527 | NaN | 391.8247 |
| Msh2 | NKGEYEEAQDAIVK | -566 | -579 | 1 | 0.00025015 | 14 | 960.95764 | 1036.5151 | 1012.2335 | 1175.9591 | 1168.2051 | 1137.1803 |
| Msh2 | DSLIIIDELGR | -742 | -752 | 0 | 0 | 11 | 631.45496 | 734.5126 | 760.94855 | 126.987656 | 920.47815 | 738.76215 |
| Msh2 | SFVNEIISR | -921 | -929 | 0 | 0 | 9 | 570.54083 | 1028.3687 | 596.8031 | 1118.8972 | 1160.6941 | 1211.243 |
| Msh2 | HGEDALLAAR | -46 | -55 | 0 | 0 | 10 | 908.9866 | NaN | NaN | 635.8051 | NaN | 531.1496 |
| Msh2 | QSLQEDLLR | -374 | -382 | 0 | 0 | 9 | 1182.6937 | NaN | NaN | 444.80386 | NaN | NaN |
| Fxyd5 | ENNPFYYDDTTLR | -131 | -143 | 0 | 0 | 13 | 825.0417 | 1125.3981 | 1273.989 | 781.90454 | 576.24365 | 742.24805 |
| Fxyd5 | DSNENNPFYYDDTTLR | -128 | -143 | 0 | 0 | 16 | 368.63293 | 376.64233 | 427.76443 | NaN | NaN | NaN |
| Ddx42 | ELEPGDGPIAVIVCPTR | -320 | -336 | 0 | 0 | 17 | 617.6371 | 1028.2113 | 660.4983 | 767.9207 | 1638.236 | 1499.4839 |
| Ddx42 | NLEGANQHVSK | -616 | -626 | 0 | 0 | 11 | 500.5643 | 473.57755 | 474.0091 | 416.1907 | 496.5207 | 475.14798 |
| Ddx42 | DSNFAGDLVR | -606 | -615 | 0 | 0 | 10 | 969.86975 | 859.94434 | NaN | 1021.2779 | NaN | 1561.4452 |
| Ddx42 | GIRDDIEEEDDQEAYFR | -146 | -162 | 1 | 0.004923919 | 17 | 510.8132 | NaN | NaN | NaN | 651.9914 | NaN |
| Nup98 | EYGFLPNPVAVK | -1175 | -1186 | 0 | 0 | 12 | 1040.8296 | 557.7053 | 1141.9928 | 1205.9364 | 1493.1261 | 403.13373 |
| Nup98 | VDDNQKPPVGEGLNR | -803 | -817 | 1 | 0 | 15 | 642.0673 | 767.1383 | 854.04 | 1075.2188 | 850.3258 | 806.23706 |
| Nup98 | VDDNQKPPVGEGLNR | -803 | -817 | 1 | 0 | 15 | 642.0673 | 767.1383 | 854.04 | 1075.2188 | 850.3258 | 806.23706 |
| Nup98 | RQPGLVPLEK | -1112 | -1121 | 1 | 0.000230707 | 10 | 637.40173 | 768.94244 | 790.5837 | 660.94305 | 613.55396 | 599.33264 |
| Nup98 | DSPGDFLELPIVK | -1254 | -1266 | 0 | 0 | 13 | 441.75674 | 457.8446 | 510.28366 | 605.89325 | 700.9984 | 573.71356 |
| Nup98 | FGHIPSK | -1012 | -1018 | 0 | 0 | 7 | 3144.1233 | 2957.7087 | 3108.6277 | 1165.0995 | NaN | 406.04718 |
| Nup98 | LADINYEGR | -843 | -851 | 0 | 0 | 9 | 975.4802 | 727.4295 | NaN | 2927.1858 | 4064.3813 | 3933.6448 |
| Nup98 | EYRPETGSWVF | -865 | -875 | 1 | 0.001392869 | 11 | 593.6519 | 523.1248 | 462.7196 | NaN | 942.5374 | 995.853 |
| Nup98 | PIPQTPESVGNK | -666 | -677 | 0 | 0 | 12 | 2195.0898 | NaN | 2716.2058 | 2102.1917 | NaN | 1886.2937 |
| Nup98 | SITADPLDYR | -1513 | -1522 | 0 | 0.000230707 | 10 | 236.59435 | 886.33093 | 1169.0901 | NaN | NaN | 2154.5808 |
| Nup98 | GFLEDLAPPER | -1668 | -1678 | 0 | 0 | 11 | 709.47217 | NaN | NaN | 675.25195 | 335.6775 | NaN |
| Nup98 | LEDYQANR | -205 | -212 | 0 | 0.000907206 | 8 | 617.1011 | 1036.2579 | 795.19183 | NaN | NaN | NaN |
| Nup98 | SFIHDER | -1388 | -1394 | 0 | 0.00025015 | 7 | 1115.5503 | NaN | NaN | NaN | NaN | NaN |
| Nup98 | HCQLSETPESWAK | -1588 | -1600 | 0 | 0.009122264 | 13 | 505.89438 | NaN | NaN | NaN | NaN | NaN |
| Gls | VNDLGTEYVHR | -302 | -312 | 0 | 0 | 11 | 1053.8901 | 1197.8285 | 1412.8859 | 1355.217 | 1557.3789 | 1609.0768 |
| Gls | SSSPSEILQELGK | -73 | -85 | 0 | 0 | 13 | 915.0849 | 891.419 | 977.7276 | 1419.3843 | 1768.2174 | 1644.2036 |
| Gls | DSPGETDAFGNSEGK | -112 | -126 | 0 | 0 | 15 | 746.1189 | 691.79675 | 652.3529 | 744.34094 | 664.52094 | 708.1448 |
| Gls | SHIDELYESAK | -234 | -244 | 0 | 0 | 11 | 482.84683 | 323.483 | 586.50946 | 1181.9813 | 641.904 | 754.11816 |
| Ncapd2 | DSPSVPEPEGSQSNDELVK | -588 | -606 | 0 | 0.00122419 | 19 | 450.12146 | 490.31085 | 432.24124 | 404.02396 | 348.5298 | 386.22894 |
| Ncapd2 | GFAHPDPLWIPFK | -846 | -858 | 0 | 0 | 13 | 1277.9866 | 1342.5657 | 1355.3839 | NaN | 1788.4042 | 1835.1215 |
| Ncapd2 | SIEPGLKEDTLEFLK | -72 | -86 | 1 | 0 | 15 | 9116.321 | 9696.328 | NaN | NaN | 4680.6704 | 5666.3765 |
| Ncapd2 | LLENPTISHQK | -205 | -215 | 0 | 0 | 11 | 471.07162 | 346.21317 | 481.10864 | NaN | NaN | NaN |
| Ncapd2 | ITEAIGIISK | -624 | -633 | 0 | 0.000431655 | 10 | 411.58765 | 392.43808 | 407.74625 | NaN | NaN | NaN |
| Sdcbp | DSSGHVGFIFK | -205 | -215 | 0 | 0 | 11 | 977.3276 | NaN | 1188.1677 | 1084.3352 | 3608.125 | 3376.2751 |
| Sdcbp | DSSGHVGFIF | -205 | -214 | 0 | 0.004350979 | 10 | 396.52234 | 546.2135 | NaN | NaN | 868.7419 | 723.11096 |
| Sdcbp | SPASLVGLR | -146 | -154 | 0 | 0 | 9 | 841.3104 | NaN | 498.6017 | NaN | NaN | NaN |
| Smchd1 | GVQPEHIQYLK | -326 | -336 | 0 | 0 | 11 | 8533.3545 | 6641.5635 | 7734.569 | 3176.828 | 3018.5718 | 3221.4358 |
| Smchd1 | LCDQWDNPALVPNVK | -1286 | -1300 | 0 | 0 | 15 | 4968.935 | 5441.6514 | 3995.071 | 4003.9265 | 4265.1426 | 4468.1196 |
| Smchd1 | VVLNESNADTYAGR | -794 | -807 | 0 | 0 | 14 | 2166.135 | 2665.034 | 2587.6997 | 693.9238 | 2475.6157 | 2463.8452 |
| Smchd1 | LTILPDPEKPIR | -1360 | -1371 | 1 | 0 | 12 | 747.1046 | 896.58716 | 899.5942 | 1382.6405 | 1351.3838 | 1393.6428 |
| Smchd1 | LTILPDPEKPIR | -1360 | -1371 | 1 | 0 | 12 | 747.1046 | 896.58716 | 899.5942 | 1382.6405 | 1351.3838 | 1393.6428 |
| Smchd1 | GIITRPDLPTK | -581 | -591 | 1 | 0 | 11 | 640.8918 | 930.4743 | 988.284 | 751.8532 | 811.7777 | 864.96204 |
| Smchd1 | GLLPDVQVPTSVK | -1108 | -1120 | 0 | 0 | 13 | 392.1822 | 444.93494 | 393.29416 | 411.93646 | 577.3899 | 526.09296 |
| Smchd1 | LKDEDDDDDCFISEK | -449 | -463 | 1 | 0 | 15 | 364.97104 | 356.1151 | 401.67096 | 384.99725 | 455.82623 | 473.60742 |
| Smchd1 | FSFGLLDSPFR | -826 | -836 | 0 | 0 | 11 | 352.64868 | 128.69511 | 404.88025 | 393.8937 | 185.11946 | 392.9234 |

| | | | | | | | | | | | | |
|---|---|---|---|---|---|---|---|---|---|---|---|---|
| Smchd1 | SFGLLDSPFR | -827 | -836 | 0 | 0 | 10 | 342.36716 | 451.47565 | 503.89468 | 304.86597 | 515.2433 | 496.39886 |
| Smchd1 | DSTEYFIIFEPR | -1586 | -1597 | 0 | 0 | 12 | 320.11926 | 305.3158 | 290.78763 | 298.326 | 324.4189 | 369.0292 |
| Smchd1 | NYLHLWTR | -337 | -344 | 0 | 0 | 8 | 2556.1792 | 516.76086 | NaN | 253.93694 | 3583.9146 | 637.64465 |
| Smchd1 | SITGDGLDSSNLK | -1196 | -1208 | 0 | 0 | 13 | 941.3392 | 687.5351 | 1280.0646 | NaN | 515.94305 | 526.7937 |
| Smchd1 | HQDEVTWVAGDVIR | -1055 | -1068 | 0 | 0 | 14 | 475.29547 | 805.7625 | 696.2006 | NaN | 1373.667 | 1280.0121 |
| Smchd1 | EINITPSLAEK | -1081 | -1091 | 0 | 0.001392869 | 11 | 1786.2122 | 1197.5011 | NaN | 1337.0684 | NaN | 708.88 |
| Smchd1 | SAFTVRPLPDDPK | -1138 | -1150 | 1 | 0 | 13 | 1320.4224 | 1058.7798 | 1037.1862 | NaN | NaN | 748.35376 |
| Smchd1 | TLPLCYGSIVR | -620 | -630 | 0 | 0 | 11 | 182.64813 | 1112.0542 | 1149.9589 | NaN | NaN | NaN |
| Smchd1 | LLFDETQGKPAVA | -169 | -181 | 1 | 0.001055699 | 13 | 2183.0645 | 2444.1675 | NaN | NaN | NaN | NaN |
| Smchd1 | TSVGDFDWCAPPK | -484 | -496 | 0 | 0.000230707 | 13 | 693.3579 | NaN | NaN | 790.36475 | NaN | NaN |
| Smchd1 | GFNEEDTDTPLFIGK | -1544 | -1558 | 0 | 0 | 15 | 658.88416 | NaN | 825.1604 | NaN | NaN | NaN |
| Smchd1 | TLEFPFVK | -1561 | -1568 | 0 | 0.000230707 | 8 | 314.09042 | 377.91333 | NaN | NaN | NaN | NaN |
| Smchd1 | EFSDFLR | -1241 | -1247 | 0 | 0.00272025 | 7 | 980.06 | NaN | NaN | NaN | NaN | NaN |
| Smchd1 | VLAENSPGR | -1577 | -1585 | 0 | 0.003299042 | 9 | 934.34064 | NaN | NaN | NaN | NaN | NaN |
| Smchd1 | LLPSNQQHK | -1311 | -1319 | 0 | 0.002502116 | 9 | 324.44684 | NaN | NaN | NaN | NaN | NaN |
| Hnrnpa3 | IFVGGIKEDTEEYNLR | -128 | -143 | 1 | 0 | 16 | 10901.793 | 11845.832 | 11417.129 | 9006.77 | 15392.79 | 9064.033 |
| Hnrnpa3 | VTFDDHDTVDK | -172 | -182 | 0 | 0 | 11 | 8854.859 | 8917.49 | 8890.168 | 10768.099 | 11152.848 | 11288.888 |
| Hnrnpa3 | VTFDDHDTVDK | -172 | -182 | 0 | 0 | 11 | 8854.859 | 8917.49 | 8890.168 | 10768.099 | 11152.848 | 11288.888 |
| Hnrnpa3 | EDSVKPGAHLTVK | -114 | -126 | 1 | 0 | 13 | 6334.6777 | 6697.507 | 6477.2573 | 4420.565 | 5386.2085 | 5513.9253 |
| Hnrnpa3 | EDSVKPGAHLTVK | -114 | -126 | 1 | 0 | 13 | 6334.6777 | 6697.507 | 6477.2573 | 4420.565 | 5386.2085 | 5513.9253 |
| Hnrnpa3 | EDSVKPGAHLTVK | -114 | -126 | 1 | 0 | 13 | 6334.6777 | 6697.507 | 6477.2573 | 4420.565 | 5386.2085 | 5513.9253 |
| Hnrnpa3 | FDDHDTVDK | -174 | -182 | 0 | 0 | 9 | 4929.4204 | 4960.722 | 5180.9775 | 3650.7793 | 5211.35 | 5219.221 |
| Hnrnpa3 | FDDHDTVDK | -174 | -182 | 0 | 0 | 9 | 4929.4204 | 4960.722 | 5180.9775 | 3650.7793 | 5211.35 | 5219.221 |
| Hnrnpa3 | FDDHDTVDK | -174 | -182 | 0 | 0 | 9 | 4929.4204 | 4960.722 | 5180.9775 | 3650.7793 | 5211.35 | 5219.221 |
| Hnrnpa3 | EDSVKPGAHLT | -114 | -124 | 1 | 0.000230707 | 11 | 4517.536 | 4879.18 | 4539.4287 | 4458.704 | 2154.934 | 5385.421 |
| Hnrnpa3 | GIKEDTEEYNLR | -132 | -143 | 1 | 0 | 12 | 3712.9595 | 4107.1475 | 4288.7227 | 5600.687 | 5754.8716 | 5697.788 |
| Hnrnpa3 | GIKEDTEEYNLR | -132 | -143 | 1 | 0 | 12 | 3712.9595 | 4107.1475 | 4288.7227 | 5600.687 | 5754.8716 | 5697.788 |
| Hnrnpa3 | EDSVKPGAHL | -114 | -123 | 1 | 0.00025015 | 10 | 3500.1506 | 3627 | 2123.7886 | 2972.506 | 3424.6816 | 4146.4575 |
| Hnrnpa3 | SVKPGAHLTVK | -116 | -126 | 1 | 0 | 11 | 98.78998 | 1314.0479 | 181.10065 | 327.42645 | 404.10123 | 662.47186 |
| Hnrnpa3 | SVKPGAHLTVK | -116 | -126 | 1 | 0 | 11 | 98.78998 | 1314.0479 | 181.10065 | 327.42645 | 404.10123 | 662.47186 |
| Hnrnpa3 | GGIKEDTEEYNLR | -131 | -143 | 1 | 0 | 13 | 2139.836 | 2174.4214 | 807.432 | 183.28452 | 224.1545 | 953.11316 |
| Hnrnpa3 | SSGSPYGGGYGSGGGSGGYGSR | -356 | -377 | 0 | 0 | 22 | 1947.8505 | 2014.5979 | 2059.4878 | 1088.5776 | 1170.7114 | 1220.1682 |
| Hnrnpa3 | MEVKPPPGRPQPD | -1 | -13 | 2 | 0.00025015 | 13 | 1600.4142 | 1627.4183 | 114.94924 | 1783.7303 | 1833.9279 | 1784.2997 |
| Hnrnpa3 | SFETTDDSLR | -43 | -52 | 0 | 0 | 10 | 483.52115 | 11607.752 | 11638.826 | 11305.541 | 11872.781 | 9899.6455 |
| Hnrnpa3 | SFETTDDSLR | -43 | -52 | 0 | 0 | 10 | 483.52115 | 11607.752 | 11638.826 | 11305.541 | 11872.781 | 9899.6455 |
| Hnrnpa3 | VGGIKEDTEEYNLR | -130 | -143 | 1 | 0 | 14 | 1276.9834 | 1271.4694 | 1378.3307 | 1378.9386 | 1460.8002 | 1327.0336 |
| Hnrnpa3 | GGGYGSGGGSGGYGSR | -362 | -377 | 0 | 0 | 16 | 1234.3862 | 1285.6987 | 1310.9055 | 889.7453 | 122.21893 | 1305.9324 |
| Hnrnpa3 | SFETTDDSLREHFEK | -43 | -57 | 1 | 0 | 15 | 1049.1967 | 1289.6567 | 1334.8958 | 1130.3468 | 1895.3861 | 1840.7203 |
| Hnrnpa3 | SFETTDDSLREHFEK | -43 | -57 | 1 | 0 | 15 | 1049.1967 | 1289.6567 | 1334.8958 | 1130.3468 | 1895.3861 | 1840.7203 |
| Hnrnpa3 | GSPYGGGYGSGGGSGGYGSR | -358 | -377 | 0 | 0 | 20 | 1011.71924 | 137.07161 | 1063.2777 | 1154.503 | 1317.7372 | 1306.6978 |
| Hnrnpa3 | GLSFETTDDSLR | -41 | -52 | 0 | 0 | 12 | 773.5909 | 885.4712 | 1011.7989 | 1149.5781 | 1321.4016 | 1265.8729 |
| Hnrnpa3 | RGEEGHDPKEPEQL | -21 | -34 | 2 | 0.00025015 | 14 | 770.8077 | 895.02136 | 1087.8748 | 1045.0319 | 1429.0902 | 1238.1664 |
| Hnrnpa3 | RGEEGHDPKEPEQL | -21 | -34 | 2 | 0.00025015 | 14 | 770.8077 | 895.02136 | 1087.8748 | 1045.0319 | 1429.0902 | 1238.1664 |
| Hnrnpa3 | SGSPYGGGYGSGGGSGGYGSR | -357 | -377 | 0 | 0 | 21 | 733.5554 | 571.7761 | 313.34274 | 716.63824 | 658.7934 | 664.478 |
| Hnrnpa3 | EDTEEYNLR | -135 | -143 | 0 | 0 | 9 | 172.53062 | 6150.4956 | 4561.777 | 166.79515 | 196.96744 | 3209.8523 |
| Hnrnpa3 | RGEEGHDPKEPEQLR | -21 | -35 | 2 | 0 | 15 | 1008.8073 | 1198.0055 | 1185.4208 | NaN | 685.24774 | 870.55597 |
| Hnrnpa3 | RGEEGHDPKEPEQLR | -21 | -35 | 2 | 0 | 15 | 1008.8073 | 1198.0055 | 1185.4208 | NaN | 685.24774 | 870.55597 |
| Hnrnpa3 | GGLSFETTDDSLR | -40 | -52 | 0 | 0 | 13 | 563.9483 | 683.58966 | NaN | 770.15875 | 1136.5453 | 1085.6654 |
| Hnrnpa3 | IFVGGIKEDTEEY | -128 | -140 | 1 | 0 | 13 | 601.7445 | 563.10504 | 605.9536 | NaN | NaN | NaN |
| Hnrnpa3 | GEEGHDPKEPEQLR | -22 | -35 | 1 | 0 | 14 | 481.09116 | 865.05914 | 715.5561 | NaN | NaN | NaN |
| Hnrnpa3 | DSVKPGAHLTVK | -115 | -126 | 1 | 0.000230707 | 12 | 431.4418 | 461.8613 | 468.22467 | NaN | NaN | NaN |
| Hnrnpa3 | YHTINGHNCEVK | -188 | -199 | 0 | 0.00025015 | 12 | 214.89786 | 482.38458 | NaN | NaN | NaN | NaN |
| Get3 | VFDTAPTGHTLR | -164 | -175 | 0 | 0 | 12 | 1889.3641 | 1933.5547 | 1868.6023 | 1888.355 | 1254.9492 | 1096.2411 |
| Get3 | LLNFPTIVER | -176 | -185 | 0 | 0 | 10 | 694.67926 | 820.4281 | 770.35394 | NaN | 784.25446 | 756.64606 |
| Get3 | DTAPTGHTLR | -166 | -175 | 0 | 0 | 10 | 1883.3611 | 2111.0254 | 2386.8132 | NaN | NaN | 2529.0806 |
| Get3 | LPLLPHEVR | -318 | -326 | 0 | 0 | 9 | 722.3644 | NaN | NaN | 819.2904 | 872.4523 | 813.1046 |
| Get3 | SVSEQFKDPEQTTFIC | -231 | -246 | 1 | 0.000550307 | 16 | 669.3767 | NaN | 774.3651 | 817.8642 | NaN | NaN |
| Timm23 | DTDEFILPTGANK | -56 | -68 | 0 | 0 | 13 | 3173.685 | NaN | 3380.2693 | NaN | NaN | NaN |
| Arpc3 | VFDPQSDKPSK | -148 | -158 | 1 | 0 | 11 | 3682.3242 | 3820.3198 | 3902.1948 | 3247.8354 | 3842.1084 | 3953.8508 |
| Arpc3 | VFDPQSDKPSK | -148 | -158 | 1 | 0 | 11 | 3682.3242 | 3820.3198 | 3902.1948 | 3247.8354 | 3842.1084 | 3953.8508 |
| Arpc3 | EKVFDPQSDKPSK | -146 | -158 | 2 | 0 | 13 | 2023.8259 | 1764.9844 | 3094.1812 | 2498.0325 | 4918.735 | 3093.0508 |
| Arpc3 | NFPIPGEPGFPLNAIYAK | -102 | -119 | 0 | 0 | 18 | 666.50635 | 696.56445 | 549.5539 | 892.5276 | 1066.613 | 868.3972 |
| Arpc3 | DTDIVDEAIYYFK | -38 | -50 | 0 | 0 | 13 | 298.43735 | 335.63437 | 471.64548 | 433.35574 | 370.34705 | 413.90656 |
| Arpc3 | FKGPAPR | -28 | -34 | 1 | 0.001392869 | 7 | 1050.9125 | NaN | 1138.2299 | 759.5299 | 709.9862 | 956.09705 |
| Arpc3 | WWTCFVK | -159 | -165 | 0 | 0.001023236 | 7 | 825.40924 | 897.48376 | NaN | 720.49115 | 866.48706 | 831.356 |
| Mtdh | TVEVPEDEVVR | -131 | -141 | 0 | 0 | 11 | 2951.8584 | 2167.1062 | 1915.3993 | 1886.1326 | 2132.7908 | 1384.667 |
| Mtdh | DTEDLEKDTR | -458 | -467 | 1 | 0 | 10 | 997.3345 | 897.52966 | NaN | NaN | 1588.246 | 699.85364 |
| Mtdh | SWQDELAQQAEEGSAR | -5 | -20 | 0 | 0.00111006 | 16 | 419.50177 | 514.11145 | 483.24643 | NaN | NaN | NaN |
| Hnrnpa1 | VTFDDHDSVDK | -151 | -161 | 0 | 0 | 11 | 5497.458 | 5743.8247 | 5894.3276 | 599.38965 | 8113.1763 | 8281.858 |
| Hnrnpa1 | VTFDDHDSVDK | -151 | -161 | 0 | 0 | 11 | 5497.458 | 5743.8247 | 5894.3276 | 599.38965 | 8113.1763 | 8281.858 |
| Hnrnpa1 | VTFDDHDSVDK | -151 | -161 | 0 | 0 | 11 | 5497.458 | 5743.8247 | 5894.3276 | 599.38965 | 8113.1763 | 8281.858 |
| Hnrnpa1 | EDSQRPGAHLT | -93 | -103 | 1 | 0 | 11 | 4924.116 | 5010.382 | 5335.6294 | 586.33203 | 1048.2662 | 4933.288 |
| Hnrnpa1 | EDSQRPGAHLTVK | -93 | -105 | 1 | 0 | 13 | 3664.9243 | 2925.676 | 4214.0273 | 1822.778 | 3331.5588 | 3428.0894 |
| Hnrnpa1 | EDSQRPGAHLTVK | -93 | -105 | 1 | 0 | 13 | 3664.9243 | 2925.676 | 4214.0273 | 1822.778 | 3331.5588 | 3428.0894 |
| Hnrnpa1 | SESPKEPEQLR | -4 | -14 | 1 | 0 | 11 | 3596.325 | 3655.906 | 3652.219 | 3014.8174 | 3487.8564 | 3395.0205 |
| Hnrnpa1 | SESPKEPEQLR | -4 | -14 | 1 | 0 | 11 | 3596.325 | 3655.906 | 3652.219 | 3014.8174 | 3487.8564 | 3395.0205 |
| Hnrnpa1 | EDSQRPGAHL | -93 | -102 | 1 | 0 | 10 | 2072.015 | 2492.353 | 2529.6243 | 750.2485 | 272.85135 | 3603.228 |
| Hnrnpa1 | SSGPYGGGGQYFAK | -285 | -298 | 0 | 0 | 14 | 1535.3472 | 1705.5142 | 475.9971 | 5513.2656 | 6628.3335 | 6514.0713 |

| | | | | | | | | | | | | |
|---|---|---|---|---|---|---|---|---|---|---|---|---|
| Hnrnpa1 | SFETTDESLR | -22 | -31 | 0 | 0 | 10 | 160.82945 | 7812.1675 | 8057.885 | 504.49472 | 6735.4746 | 7413.7725 |
| Hnrnpa1 | SFETTDESLR | -22 | -31 | 0 | 0 | 10 | 160.82945 | 7812.1675 | 8057.885 | 504.49472 | 6735.4746 | 7413.7725 |
| Hnrnpa1 | SHFEQWGTLTDC | -32 | -43 | 0 | 0 | 12 | 1056.1919 | 1230.3862 | 1147.1256 | 879.1326 | 697.90247 | 55.920616 |
| Hnrnpa1 | SSNFGPMK | -270 | -277 | 0 | 0 | 8 | 850.7096 | 633.7472 | 1162.9182 | 139.44418 | 965.11884 | 985.2127 |
| Hnrnpa1 | SSGPYGGGGQYFAKPR | -285 | -300 | 1 | 0 | 16 | 664.97955 | 696.00726 | 700.3779 | 412.64224 | 427.2471 | 419.11575 |
| Hnrnpa1 | FDDHDSVDK | -153 | -161 | 0 | 0 | 9 | 530.4635 | 3342.9004 | 633.1754 | 610.285 | 546.4382 | 3580.1162 |
| Hnrnpa1 | FDDHDSVDK | -153 | -161 | 0 | 0 | 9 | 530.4635 | 3342.9004 | 633.1754 | 610.285 | 546.4382 | 3580.1162 |
| Hnrnpa1 | SHFEQWGTL | -32 | -40 | 0 | 0.00025015 | 9 | 2734.4902 | 132.76662 | 3139.7407 | 3262.838 | NaN | 2610.0437 |
| Hnrnpa1 | SHFEQWGTLT | -32 | -41 | 0 | 0 | 10 | 744.58997 | 850.0773 | 713.07495 | 938.5604 | NaN | 945.8767 |
| Hnrnpa1 | GGLSFETTDESLR | -19 | -31 | 0 | 0 | 13 | 5438.8096 | NaN | NaN | 186.59718 | 1089.8842 | 948.52295 |
| Hnrnpa1 | DYFEQYGK | -123 | -130 | 0 | 0 | 8 | 596.91943 | 510.88116 | 689.11005 | 526.7167 | NaN | NaN |
| Hnrnpa1 | SHFEQWGTLTD | -32 | -42 | 0 | 0 | 11 | 864.3431 | NaN | NaN | NaN | NaN | NaN |
| RTRAF | LLHIEELR | -206 | -213 | 0 | 0.00025015 | 8 | 12911.261 | 828.6476 | 10780.318 | 9198.94 | 5206.053 | 9624.553 |
| RTRAF | NAEPLINLDVNNPDFK | -116 | -131 | 0 | 0 | 16 | 600.03625 | 1203.9672 | 987.0059 | 638.48236 | 880.1472 | 733.5746 |
| RTRAF | NCKDETEFR | -18 | -26 | 1 | 0 | 9 | 186.21198 | NaN | 153.20375 | NaN | 541.55273 | NaN |
| RTRAF | DTGDAVLNEAAQILR | -191 | -205 | 0 | 0.002566359 | 15 | 391.9284 | NaN | NaN | NaN | NaN | NaN |
| Hmgn5 | DTGEEVEDGKIEEEGLNEKPGTAK | -143 | -166 | 2 | 0.000431655 | 24 | 458.1301 | 420.53625 | 475.19458 | 466.63782 | 123.34199 | 507.7494 |
| Hmgn5 | INEDTEEDGGEKK | -101 | -113 | 1 | 0 | 13 | 297.32147 | 370.1825 | 211.51457 | 232.22745 | 290.1153 | 340.25937 |
| Gna13 | DTILHDNLK | -364 | -372 | 0 | 0 | 9 | 1292.476 | 547.56085 | 991.1563 | 1452.0344 | 1387.0114 | 295.20926 |
| Gna13 | LGVPDYIPSQQDILLAR | -184 | -200 | 0 | 0.000230707 | 17 | 231.64857 | 288.40375 | NaN | 391.70227 | 432.62683 | 392.7971 |
| Gna13 | VFLQYLPAIR | -141 | -150 | 0 | 0 | 10 | 262.09805 | 277.9001 | NaN | NaN | 292.40985 | 244.78233 |
| Gna13 | LHIPWGDNK | -106 | -114 | 0 | 0.000627943 | 9 | 1538.7903 | 1565.6313 | NaN | NaN | NaN | NaN |
| Gna13 | YFLDNLDK | -176 | -183 | 0 | 0.00025015 | 8 | 538.1563 | NaN | NaN | NaN | NaN | NaN |
| Xpnpep1 | GHLLDSFAR | -465 | -473 | 0 | 0 | 9 | 585.6587 | 698.22986 | 855.574 | 1967.314 | 600.8717 | 1041.3701 |
| Xpnpep1 | VGVDPLIIPTDYWK | -117 | -130 | 0 | 0 | 14 | 383.50674 | 301.06342 | 376.19846 | 126.743546 | 709.2914 | 437.65295 |
| Xpnpep1 | DTPTQEDWLVSVLPEGSR | -99 | -116 | 0 | 0 | 18 | 350.7623 | 326.3017 | 123.23175 | 329.8945 | 358.45956 | 350.3263 |
| Xpnpep1 | GSDVEHNPVFFSY | -213 | -225 | 0 | 0 | 13 | 50.210575 | 408.89105 | 327.84875 | 605.86176 | 718.1914 | 596.7613 |
| Xpnpep1 | NVHEGPCGISYK | -496 | -507 | 0 | 0 | 12 | 905.9862 | NaN | 842.6594 | NaN | 738.4783 | NaN |
| Xpnpep1 | YAPVPETNR | -396 | -404 | 0 | 0.002752526 | 9 | 935.33014 | NaN | NaN | NaN | NaN | NaN |
| Ppp3r1 | TIINADKDGDGR | -136 | -147 | 1 | 0 | 12 | 940.14606 | 1065.8403 | 1081.6678 | 602.89417 | 1033.2311 | 1141.9133 |
| Ppp3r1 | DTQLQQIVDK | -126 | -135 | 0 | 0 | 10 | 1484.4629 | 822.7768 | 756.27875 | 1237.8444 | 921.3496 | NaN |
| Ppp3r1 | SHFDADEIKR | -13 | -22 | 1 | 0 | 10 | 1399.739 | 1398.5609 | 890.04535 | NaN | 315.57642 | NaN |
| Inpp5d | LLLDSDFLK | -195 | -203 | 0 | 0 | 9 | 1311.9629 | 1434.3971 | 1322.9075 | NaN | 2135.732 | 2276.7114 |
| Inpp5d | SLSETLFQR | -162 | -170 | 0 | 0 | 9 | 304.28375 | NaN | 242.18791 | 551.5955 | 371.31628 | 365.28165 |
| Inpp5d | DTSGLPEEHLK | -175 | -185 | 0 | 0 | 11 | 5488.0107 | 542.1134 | 3352.068 | NaN | NaN | NaN |
| Inpp5d | SLLHEGSESTNR | -284 | -295 | 0 | 0.00025015 | 12 | 1614.3088 | 1495.7286 | 1479.2782 | NaN | NaN | NaN |
| Inpp5d | ILPNEDDKFTVQ | -58 | -69 | 1 | 0 | 12 | 1138.9708 | NaN | 1343.556 | NaN | NaN | 1212.8796 |
| Inpp5d | TGSSNLPHLK | -204 | -213 | 0 | 0 | 10 | 628.14343 | 1080.1877 | NaN | NaN | NaN | NaN |
| Inpp5d | SQEGENEEGSEGELVVR | -766 | -782 | 0 | 0.00204437 | 17 | 480.15863 | NaN | NaN | NaN | NaN | NaN |
| Tial1 | DTSNHFHVF | -109 | -117 | 0 | 0.00111006 | 9 | 248.205 | 963.76294 | 1025.2561 | 997.0711 | 1071.0299 | 237.32361 |
| Dkc1 | LHNAIEGGTQLSR | -159 | -171 | 0 | 0 | 13 | 2901.1602 | 3525.9246 | 5604.287 | 3135.5186 | 4787.699 | 4589.494 |
| Dkc1 | IMLPGLLR | -315 | -322 | 0 | 0.00025015 | 8 | 1777.7563 | 1109.5112 | 1921.6809 | 181.33559 | 1815.9822 | 1889.9415 |
| Dkc1 | HGKPTDNTPATWK | -401 | -413 | 1 | 0 | 13 | 1529.4451 | 1732.1221 | 1945.9264 | 393.2569 | 2034.4729 | 2185.518 |
| Dkc1 | HGKPTDNTPATWK | -401 | -413 | 1 | 0 | 13 | 1529.4451 | 1732.1221 | 1945.9264 | 393.2569 | 2034.4729 | 2185.518 |
| Dkc1 | HAEEFLIKPESK | -32 | -43 | 1 | 0 | 12 | 1450.4188 | 1810.33 | 1930.8505 | 2514.3145 | 1679.9072 | 1829.6123 |
| Dkc1 | HAEEFLIKPESK | -32 | -43 | 1 | 0 | 12 | 1450.4188 | 1810.33 | 1930.8505 | 2514.3145 | 1679.9072 | 1829.6123 |
| Dkc1 | TVLESGGETGDGDNDTTK | -477 | -494 | 0 | 0 | 18 | 1075.5085 | 2146.2551 | 3050.363 | 943.5804 | 996.49536 | 1024.9354 |
| Dkc1 | KPLQEDDVAEIQ | -20 | -31 | 1 | 0.001610812 | 12 | 953.9512 | 1215.4791 | 1145.2614 | 1476.1753 | 1600.4176 | 1410.9708 |
| Dkc1 | WVSCEAGTYIR | -217 | -227 | 0 | 0 | 11 | 861.7461 | 2107.5994 | 1749.7638 | 1296.083 | 1224.0393 | 1612.7009 |
| Dkc1 | WLYDNHKDESYLR | -271 | -283 | 1 | 0 | 13 | 822.37933 | 831.2814 | 895.46094 | 749.0968 | 1263.6196 | 1229.7969 |
| Dkc1 | WLYDNHKDESYLR | -271 | -283 | 1 | 0 | 13 | 822.37933 | 831.2814 | 895.46094 | 749.0968 | 1263.6196 | 1229.7969 |
| Dkc1 | DTSQWPLLLK | -48 | -57 | 0 | 0 | 10 | 745.0809 | 864.43726 | 854.874 | 824.4365 | 926.3978 | 987.00824 |
| Dkc1 | QLDTSQWPLLLK | -46 | -57 | 0 | 0 | 12 | 403.88425 | 711.77655 | 619.7514 | 770.12695 | 1070.611 | 1059.5957 |
| Dkc1 | HYTPLPCGSNPLK | -68 | -80 | 0 | 0 | 13 | 188.97972 | 2353.4468 | 1988.9469 | 2367.7358 | 2824.1877 | 2309.9526 |
| Dkc1 | HYTPLPCGSNPLKR | -68 | -81 | 1 | 0 | 14 | 974.2793 | 1291.377 | 1575.2896 | NaN | 2272.9338 | 721.58954 |
| Dkc1 | TSQWPLLLK | -49 | -57 | 0 | 0 | 9 | 705.2624 | 550.773 | 587.86633 | NaN | 626.01154 | 598.21924 |
| Dkc1 | LHNAIEGGTQL | -159 | -169 | 0 | 0 | 11 | 610.5084 | 1411.3535 | 471.73486 | 612.9637 | 1589.2129 | NaN |
| Dkc1 | RVVYPLEK | -284 | -291 | 1 | 0.001708008 | 8 | 573.60095 | NaN | 817.42786 | 891.19934 | 1180.1318 | 1289.5082 |
| Dkc1 | TVLESGGETGDGDNDTTKK | -477 | -495 | 1 | 0.001392869 | 19 | 603.08984 | NaN | 591.938 | NaN | 764.73376 | 845.9154 |
| Dkc1 | KWGLGPK | -379 | -385 | 1 | 0.002068213 | 7 | 710.8415 | 701.88806 | NaN | NaN | NaN | NaN |
| Dkc1 | EFLIKPESK | -35 | -43 | 1 | 0 | 9 | 574.1222 | 644.7759 | NaN | NaN | NaN | NaN |
| Atp6v1f | LLGGIGELNK | -21 | -30 | 0 | 0 | 10 | 853.5564 | 790.4648 | 760.9768 | 697.6441 | 580.425 | 744.91516 |
| Atp6v1f | DTTINEIEDTFR | -42 | -53 | 0 | 0 | 12 | 263.3325 | 576.2128 | 912.0337 | 718.9966 | 835.3683 | NaN |
| Eif6 | NSLPDSVQIR | -86 | -95 | 0 | 0 | 10 | 2509.0586 | 475.75067 | 490.26733 | 561.77747 | 502.96317 | 406.94653 |
| Eif6 | HGLLVPNNTTDQELQHIR | -68 | -85 | 0 | 0 | 18 | 1896.289 | 2351.1206 | 2191.5261 | 3141.924 | 3007.8704 | 117.24656 |
| Eif6 | SNQGGLVHPK | -155 | -164 | 0 | 0 | 10 | 1445.327 | 847.0004 | 326.13672 | 1092.3209 | 1128.5339 | 1803.048 |
| Eif6 | ETEEILADVLK | -124 | -134 | 0 | 0 | 11 | 1092.0686 | 1147.9164 | 941.0425 | 1352.3054 | 1357.8944 | 1048.864 |
| Eif6 | GLDTTSTELSVVESVFK | -207 | -223 | 0 | 0 | 17 | 342.348 | 202.97095 | 310.99078 | 293.105 | 395.17023 | 357.22012 |
| Eif6 | VFSNQGGLVHPK | -153 | -164 | 0 | 0 | 12 | 1514.8939 | 1131.6943 | NaN | 285.16907 | 1270.8038 | 1879.2922 |
| Eif6 | VHPDLDR | -117 | -123 | 0 | 0.002845514 | 7 | 2973.5752 | 3166.6128 | 4031.3828 | NaN | NaN | 3565.1233 |
| Eif6 | QVPLVAGTVNR | -178 | -188 | 0 | 0.001291103 | 11 | 1165.1669 | NaN | 870.7012 | 749.2243 | NaN | NaN |
| Eif6 | DTTSTELSVVESVFK | -209 | -223 | 0 | 0 | 15 | 182.1041 | 297.94058 | 381.20602 | NaN | NaN | NaN |
| Mcm2 | HLPLVEELR | -287 | -295 | 0 | 0 | 9 | 5408.4062 | 6265.3857 | 5653.516 | 1337.1238 | 2272.7922 | 2317.0366 |
| Mcm2 | SAFYDSDLFK | -878 | -887 | 0 | 0 | 10 | 2570.582 | 2870.3215 | 2784.7646 | 2087.2734 | 2432.1855 | 2348.4211 |
| Mcm2 | NVDLTEPIISR | -646 | -656 | 0 | 0 | 11 | 1950.7471 | 2946.5232 | 2933.8706 | 2731.8518 | 4362.4883 | 4742.7085 |
| Mcm2 | VAVGELTDEDVK | -451 | -462 | 0 | 0 | 12 | 1594.3851 | 1679.9749 | 1621.2911 | 477.63553 | 1609.5663 | 1648.4342 |
| Mcm2 | IAPSIYGHEDIKR | -481 | -493 | 1 | 0 | 13 | 1160.4723 | 107.46346 | 259.41278 | 966.98596 | 1002.73267 | 1039.8313 |
| Mcm2 | THVDSHGHNVFK | -205 | -216 | 0 | 0 | 12 | 1103.0933 | 895.37604 | 379.03806 | 1096.832 | 1115.854 | 1013.2105 |

| | | | | | | | | | | | | |
|---|---|---|---|---|---|---|---|---|---|---|---|---|
| Mcm2 | THVDSHGHNVFK | -205 | -216 | 0 | 0 | 12 | 1103.0933 | 895.37604 | 379.03806 | 1096.832 | 1115.854 | 1013.2105 |
| Mcm2 | GVCLIDEFDK | -582 | -591 | 0 | 0 | 10 | 1013.8545 | 667.673 | 1066.9863 | 766.51404 | 2039.4233 | 2699.3792 |
| Mcm2 | ISHLPLVEELR | -285 | -295 | 0 | 0 | 11 | 1004.02454 | 1103.2738 | 1046.7871 | 973.7197 | 1013.03485 | 1004.69525 |
| Mcm2 | DTVDPVQDEMLAR | -665 | -677 | 0 | 0 | 13 | 941.7152 | 791.84644 | 850.3927 | 192.13217 | 1037.3286 | 611.80817 |
| Mcm2 | ESLVVNYEDLAAR | -228 | -240 | 0 | 0 | 13 | 775.99976 | 855.8437 | 124.55343 | 957.52765 | 228.37177 | 251.4259 |
| Mcm2 | SENVDLTEPIISR | -644 | -656 | 0 | 0 | 13 | 720.9166 | 793.36847 | 800.90625 | 2072.1316 | 972.98846 | 676.6476 |
| Mcm2 | ITNHIHVR | -277 | -284 | 0 | 0 | 8 | 615.2906 | 535.1865 | 525.3364 | 635.2093 | 694.0466 | 538.5585 |
| Mcm2 | ITNHIHVR | -277 | -284 | 0 | 0 | 8 | 615.2906 | 535.1865 | 525.3364 | 635.2093 | 694.0466 | 538.5585 |
| Mcm2 | GLALDDEDVEELTASQR | -94 | -110 | 0 | 0.00078661 | 17 | 441.28738 | 364.20853 | 529.22296 | 444.85086 | 560.65735 | 625.04034 |
| Mcm2 | GVEPLPQEVLK | -711 | -721 | 0 | 0 | 11 | 219.44243 | 958.0156 | 964.363 | 695.19104 | 742.5529 | 784.4205 |
| Mcm2 | ISDPLTSSPGR | -20 | -30 | 0 | 0 | 11 | 262.85825 | 1051.6064 | 1833.1012 | NaN | 1168.7065 | 432.03714 |
| Mcm2 | DYRPIPELDVYEAE | -80 | -93 | 1 | 0 | 14 | 212.40874 | 851.7627 | NaN | 528.1366 | 218.93416 | 643.6326 |
| Mcm2 | ENVDLTEPIISR | -645 | -656 | 0 | 0 | 12 | 3978.9465 | 4187.817 | 4268.728 | 4655.824 | NaN | NaN |
| Mcm2 | SIENLEDLK | -170 | -178 | 0 | 0.00025015 | 9 | 1460.0029 | 1556.8168 | 2956.505 | NaN | NaN | 1357.2052 |
| Mcm2 | SMATGSIPITVR | -754 | -765 | 0 | 0 | 12 | 216.51923 | 522.29425 | 4565.44 | 1503.4316 | NaN | NaN |
| Mcm2 | ISHLPLVEEL | -285 | -294 | 0 | 0 | 10 | 1208.2056 | 1323.173 | 1159.609 | NaN | NaN | NaN |
| Stim1 | EDLNYHDPTVK | -94 | -104 | 0 | 0 | 11 | 1238.5894 | 1176.5842 | 1719.8779 | 843.145 | 1679.5807 | 182.78294 |
| Stim1 | YAEEELEQVR | -316 | -325 | 0 | 0 | 10 | 1030.0559 | 1030.4318 | 3180.5054 | 1147.3579 | 4099.8613 | 3931.0718 |
| Stim1 | DTVLFGPPLLTR | -196 | -207 | 0 | 0 | 12 | 990.9963 | 1613.5388 | 1215.697 | 1717.2021 | 1875.2688 | 1696.4785 |
| Stim1 | LISVEDLWK | -114 | -122 | 0 | 0 | 9 | 605.3601 | 491.48724 | 562.61816 | 507.94272 | 594.3603 | 586.6491 |
| Stim1 | LIEGVHPGSLVEK | -559 | -571 | 0 | 0 | 13 | 335.282 | 876.1411 | 867.38916 | 721.6926 | 740.74445 | 821.95624 |
| Stim1 | IDKPLCHSEDEK | -51 | -62 | 1 | 0 | 12 | 2761.035 | 433.50513 | 512.4898 | NaN | 3777.1143 | 426.2619 |
| Stim1 | LRDEINLAK | -286 | -294 | 1 | 0.00437931 | 9 | 449.42694 | NaN | 317.91656 | NaN | NaN | 1383.1842 |
| Stim1 | LPDSPALAK | -572 | -580 | 0 | 0.001636916 | 9 | 9162.138 | 11925.961 | NaN | NaN | NaN | NaN |
| Ggps1 | DTYTCPTEEEYK | -134 | -145 | 0 | 0.001392869 | 12 | 305.8967 | 483.77325 | NaN | NaN | NaN | NaN |
| Mcmbp | IIQHLVPASFR | -409 | -419 | 0 | 0 | 11 | 1272.0117 | 1489.6552 | 1634.2382 | 897.4952 | 1865.673 | 1772.2202 |
| Mcmbp | DVAECGPQQELDLSSPR | -104 | -120 | 0 | 0 | 17 | 646.4749 | 761.4181 | 811.32544 | 803.4205 | 1072.3276 | 891.8353 |
| Mcmbp | LSGCPQNSTFTEHLYR | -393 | -408 | 0 | 0 | 16 | 588.9401 | 400.53735 | 462.45386 | 398.2369 | 457.03754 | 399.79486 |
| Mcmbp | NLSSPFDLNFPLPGEK | -220 | -235 | 0 | 0 | 16 | 443.60117 | 410.39835 | 442.32895 | 520.16547 | 595.407 | 528.02386 |
| Mcmbp | QTFYCVPVPGESSWVK | -127 | -142 | 0 | 0 | 16 | 290.84564 | 441.27057 | 360.02783 | 470.9387 | 486.65497 | 475.0279 |
| Mcmbp | FLDYNLSDDITK | -560 | -571 | 0 | 0 | 12 | 1449.6393 | 1560.9697 | 1604.1736 | 783.9768 | NaN | NaN |
| Mcmbp | VHSPPASLVPR | -296 | -306 | 0 | 0 | 11 | 294.75232 | 268.1546 | NaN | 564.6338 | 401.85684 | NaN |
| Mcmbp | VYEDWDCFK | -243 | -251 | 0 | 0 | 9 | 417.0324 | NaN | NaN | 401.45215 | 427.44672 | NaN |
| Ddx5 | GHNCPKPVLN | -86 | -95 | 1 | 0 | 10 | 8135.801 | 474.6848 | 553.9617 | 4715.4883 | 8212.611 | 8464.358 |
| Ddx5 | NFYQEHPDLAR | -57 | -67 | 0 | 0.000230707 | 11 | 5264.838 | 5292.3247 | 4715.0493 | 838.34827 | 3840.0679 | 3948.016 |
| Ddx5 | HQPFLER | -158 | -164 | 0 | 0 | 7 | 4850.0596 | 1066.6448 | 5161.0713 | 3920.9836 | 4340.7554 | 4376.868 |
| Ddx5 | QLAEDFLK | -288 | -295 | 0 | 0 | 8 | 4535.606 | 4855.082 | 5166.5684 | 850.7935 | 5048.748 | 5216.523 |
| Ddx5 | FYQEHPDLAR | -58 | -67 | 0 | 0 | 10 | 4181.675 | 4490.9775 | 4484.9404 | 6185.8203 | 558.0184 | 7455.875 |
| Ddx5 | LIDFLECGK | -228 | -236 | 0 | 0 | 9 | 3353.11 | 3309.024 | 3359.937 | 3060.576 | 4061.8909 | 3700.5144 |
| Ddx5 | DYPNSSEDYIHR | -417 | -428 | 0 | 0 | 12 | 2936.9602 | 744.4107 | 3360.0374 | 2239.873 | 2397.5017 | 446.16788 |
| Ddx5 | TTYLVLDEADR | -242 | -252 | 0 | 0 | 11 | 2371.3572 | 2922.1934 | 2914.4983 | 2862.5146 | 2351.2366 | 2369.343 |
| Ddx5 | TTYLVLDEADR | -242 | -252 | 0 | 0 | 11 | 2371.3572 | 2922.1934 | 2914.4983 | 2862.5146 | 2351.2366 | 2369.343 |
| Ddx5 | YDYPNSSEDYIHR | -416 | -428 | 0 | 0 | 13 | 2341.9438 | 3002.1987 | 2845.5203 | 4056.2424 | 4415.9517 | 4414.761 |
| Ddx5 | YDYPNSSEDYIHR | -416 | -428 | 0 | 0 | 13 | 2341.9438 | 3002.1987 | 2845.5203 | 4056.2424 | 4415.9517 | 4414.761 |
| Ddx5 | LLQLVEDR | -471 | -478 | 0 | 0.00025015 | 8 | 2313.7185 | 2447.6667 | 2467.3984 | 2010.2495 | 2068.0696 | 2046.9667 |
| Ddx5 | WNLDELPK | -46 | -53 | 0 | 0 | 8 | 1929.4462 | 1828.8079 | 737.96405 | 559.8487 | 1808.0155 | 1908.7532 |
| Ddx5 | HNILQIVDVCHDVEKDEK | -311 | -328 | 1 | 0 | 18 | 1852.6808 | 1660.363 | 1084.7689 | 2878.2913 | 1681.5652 | 1417.6111 |
| Ddx5 | HNILQIVDVCHDVEKDEK | -311 | -328 | 1 | 0 | 18 | 1852.6808 | 1660.363 | 1084.7689 | 2878.2913 | 1681.5652 | 1417.6111 |
| Ddx5 | KWNLDELPK | -45 | -53 | 1 | 0 | 9 | 1659.7391 | 1902.4172 | 1889.2137 | 1009.2325 | 1903.321 | 1778.1637 |
| Ddx5 | QVSDLISVLR | -452 | -461 | 0 | 0 | 10 | 1396.1664 | 1645.8011 | 1477.1711 | 1545.8872 | 1404.804 | 1352.8673 |
| Ddx5 | FVINYDYPNSSEDYIHR | -412 | -428 | 0 | 0 | 17 | 395.0232 | 348.20047 | 437.0985 | 477.99945 | 331.021 | 451.64212 |
| Ddx5 | FVINYDYPNSSEDYIHR | -412 | -428 | 0 | 0 | 17 | 395.0232 | 348.20047 | 437.0985 | 477.99945 | 331.021 | 451.64212 |
| Ddx5 | TLSYLLPAIVHIN | -145 | -157 | 0 | 0.00051306 | 13 | 723.4422 | 665.5627 | 686.99927 | 513.38196 | 818.201 | 64.18293 |
| Ddx5 | QIVDVCHDVEKDEK | -315 | -328 | 1 | 0 | 14 | 659.68866 | 752.2821 | 759.9449 | 704.9049 | 832.9283 | 831.76825 |
| Ddx5 | QIVDVCHDVEKDEK | -315 | -328 | 1 | 0 | 14 | 659.68866 | 752.2821 | 759.9449 | 704.9049 | 832.9283 | 831.76825 |
| Ddx5 | QIVDVCHDVEK | -315 | -325 | 0 | 0.000431655 | 11 | 599.32666 | 123.10718 | 743.46545 | 205.83386 | 720.5439 | 780.3953 |
| Ddx5 | DVCHDVEKDEK | -318 | -328 | 1 | 0 | 11 | 433.58273 | 524.1079 | 552.3288 | 394.22736 | 485.0816 | 496.30014 |
| Ddx5 | DVCHDVEKDEK | -318 | -328 | 1 | 0 | 11 | 433.58273 | 524.1079 | 552.3288 | 394.22736 | 485.0816 | 496.30014 |
| Ddx5 | INHQPFLER | -156 | -164 | 0 | 0.000826806 | 9 | 1093.9032 | 1234.9958 | NaN | 1045.9252 | 1045.6615 | 1057.8477 |
| Ddx5 | REANQAINPK | -461 | -470 | 1 | 0.001776473 | 10 | 824.6111 | 786.6826 | 858.2997 | NaN | 1222.937 | 1134.5458 |
| Ddx5 | DWVLNEFK | -381 | -388 | 0 | 0.00025015 | 8 | 531.3635 | 470.42126 | NaN | 118.852066 | 975.49646 | 1060.0195 |
| Ddx5 | NFPANVMDVIAR | -100 | -111 | 0 | 0 | 12 | 2064.3027 | 1093.2207 | 2267.4883 | 2278.2659 | NaN | NaN |
| Ddx5 | QNFTEPTAIQAQ | -112 | -123 | 0 | 0.000471409 | 12 | 143.47334 | 1496.535 | NaN | NaN | 230.35837 | 271.45566 |
| Ddx5 | QNFTEPTAIQAQ | -112 | -123 | 0 | 0.000471409 | 12 | 143.47334 | 1496.535 | NaN | NaN | 230.35837 | 271.45566 |
| Ddx5 | ELAQQVQQVAAEYCR | -178 | -192 | 0 | 0 | 15 | 762.68823 | NaN | NaN | 1273.685 | NaN | 598.719 |
| Ddx5 | TLSYLLPAIVH | -145 | -155 | 0 | 0.00025015 | 11 | 437.2802 | 1250.3302 | NaN | NaN | NaN | NaN |
| Snx6 | DVDDFFEHER | -205 | -214 | 0 | 0 | 10 | 773.9836 | 846.389 | 820.0912 | 803.5115 | 1050.4958 | 863.05023 |
| Snx6 | EKLEDFFK | -182 | -189 | 1 | 0 | 8 | 768.7034 | 713.3294 | 827.43555 | 657.7636 | 668.5099 | 663.6502 |
| Snx6 | LEYNQDLSVR | -167 | -176 | 0 | 0 | 10 | 807.9857 | 1270.5338 | 1011.86304 | NaN | 1685.9469 | 1642.464 |
| Snx6 | VAAHPILR | -151 | -158 | 0 | 0.002296977 | 8 | 337.89005 | NaN | NaN | NaN | NaN | NaN |
| Iah1 | DVEEAKPELSLL | -233 | -244 | 1 | 0.000550307 | 12 | 1032.5198 | NaN | 1312.3752 | 1379.2522 | NaN | 1556.8838 |
| Smu1 | DVEEEKFPTQLSR | -195 | -207 | 1 | 0 | 13 | 518.03546 | 788.02155 | 641.3732 | 447.55597 | 663.2202 | NaN |
| Carhsp1 | DVEGEYVPVEGDEVTYK | -95 | -111 | 0 | 0 | 17 | 4777.0693 | 5712.585 | 5740.618 | 5707.1167 | 5121.906 | 4151.671 |
| Carhsp1 | HETWSGHVISN | -138 | -148 | 0 | 0 | 11 | 796.0434 | 578.2635 | 569.19464 | 198.5462 | NaN | 675.11346 |
| Pdia6 | GSTAPVGGGSFPTITPR | -393 | -409 | 0 | 0 | 17 | 4809.2905 | 4896.76 | 5251.517 | 3348.0874 | 3213.9915 | 3350.3616 |
| Pdia6 | QKGESPVDYDGGR | -244 | -256 | 1 | 0 | 13 | 1657.7834 | 1731.1523 | 1756.2322 | 1499.7084 | 1966.8486 | 1978.9152 |
| Pdia6 | TAPVGGGSFPTITPR | -395 | -409 | 0 | 0 | 15 | 906.89484 | 1100.5509 | 863.7947 | 3310.3572 | 942.98517 | 893.7985 |
| Pdia6 | SSSDDVIELTPSNFNR | -22 | -37 | 0 | 0 | 16 | 343.59753 | 438.07742 | 813.7074 | 974.54193 | 1048.7324 | 881.6261 |

| | | | | | | | | | | | | |
|---|---|---|---|---|---|---|---|---|---|---|---|---|
| Pdia6 | TCEEHQLCVV | -290 | -299 | 0 | 0.000230707 | 10 | 133.24998 | 1106.907 | 1183.2572 | 1285.6025 | 1348.4429 | 1369.7327 |
| Pdia6 | SDDVIELTPSNFNR | -24 | -37 | 0 | 0 | 14 | 460.82492 | 424.7287 | 621.6461 | 513.4451 | 817.47876 | NaN |
| Pdia6 | DVELDDLEKDEL | -429 | -440 | 1 | 0 | 12 | 371.75092 | 1194.5531 | NaN | 1320.8207 | 632.9726 | 287.037 |
| Pdia6 | HQSLGGQYGVQ | -86 | -96 | 0 | 0 | 11 | 1351.6272 | 1502.8611 | 892.5962 | 1131.1921 | NaN | NaN |
| Pdia6 | FYAPWCGHCQR | -50 | -60 | 0 | 0 | 11 | 1033.0833 | 975.98944 | 1085.2753 | 799.3459 | NaN | NaN |
| Pdia6 | KTCEEHQLCVVA | -289 | -300 | 1 | 0.00122419 | 12 | 1657.8796 | 2389.7883 | 3018.1353 | NaN | NaN | NaN |
| Pdia6 | RLTPEWK | -60 | -66 | 1 | 0.003360691 | 7 | 461.92517 | NaN | NaN | 937.74756 | 604.77423 | NaN |
| Pdia6 | NKPEDYQGGR | -109 | -118 | 1 | 0.000347792 | 10 | 186.42969 | NaN | 843.3205 | 902.8499 | NaN | NaN |
| Pdia6 | VLPHILDTGAAGR | -301 | -313 | 0 | 0.002534621 | 13 | 873.85406 | NaN | NaN | NaN | NaN | NaN |
| Pdia6 | NSYLEVLLK | -314 | -322 | 0 | 0.009122264 | 9 | 491.38345 | NaN | NaN | NaN | NaN | NaN |
| Rpl27 | NIDDGTSDRPYSHAL | -28 | -42 | 1 | 0 | 15 | 6547.851 | 7259.4746 | 7171.88 | 6681.101 | 6924.126 | 6948.4375 |
| Rpl27 | NIDDGTSDRPYSHAL | -28 | -42 | 1 | 0 | 15 | 6547.851 | 7259.4746 | 7171.88 | 6681.101 | 6924.126 | 6948.4375 |
| Rpl27 | DVFRDPALK | -99 | -107 | 1 | 0 | 9 | 4778.763 | 5312.26 | 5365.322 | 4518.402 | 5392.2725 | 158.35548 |
| Rpl27 | NYNHLMPTR | -76 | -84 | 0 | 0 | 9 | 3001.9988 | 2672.6772 | 2950.9465 | 1606.6466 | 2226.2512 | 2508.4734 |
| Rpl27 | NYNHLMPTR | -76 | -84 | 0 | 0 | 9 | 3001.9988 | 2672.6772 | 2950.9465 | 1606.6466 | 2226.2512 | 2508.4734 |
| Rpl27 | NIDDGTSDRPYSH | -28 | -40 | 1 | 0 | 13 | 2944.6355 | 62.777946 | 3269.9973 | 2300.9983 | 2923.5962 | 2970.3638 |
| Rpl27 | YSVDIPLDK | -85 | -93 | 0 | 0 | 9 | 2878.3103 | 3167.566 | 3093.5244 | 3007.445 | 3205.0117 | 3003.7156 |
| Rpl27 | NIDDGTSDRPYSHA | -28 | -41 | 1 | 0.00025015 | 14 | 2083.2454 | 2397.5454 | 2412.769 | 3266.273 | 3615.0608 | 3899.5415 |
| Rpl27 | KDVFRDPALK | -98 | -107 | 2 | 0.00025015 | 10 | 952.4001 | 1000.8463 | 1202.6876 | 1931.7407 | 1690.1783 | 1751.7117 |
| Rpl27 | VFRDPALK | -100 | -107 | 1 | 0 | 8 | 943.9783 | 991.85236 | 866.7966 | 927.6298 | 715.5746 | 780.7938 |
| Rpl27 | DVFRDPALKR | -99 | -108 | 2 | 0 | 10 | 821.4425 | 783.83435 | 740.5433 | 499.96036 | 543.20337 | 393.17935 |
| Rpl27 | RYSVDIPLDK | -84 | -93 | 1 | 0 | 10 | 327.4656 | 2662.729 | 650.2484 | 3289.2554 | 3927.0698 | 3722.4956 |
| Rpl27 | NIDDGTSDRPYSHALVA | -28 | -44 | 1 | 0.00025015 | 17 | 337.89246 | 564.40784 | NaN | 812.552 | 1215.9302 | 350.27164 |
| Rpl27 | SVDIPLDK | -86 | -93 | 0 | 0.004065182 | 8 | 1067.5785 | 812.82416 | NaN | NaN | NaN | NaN |
| Rpl27 | NIDDGTSDRPY | -28 | -38 | 1 | 0.00025015 | 11 | 417.57108 | 116.69608 | NaN | NaN | NaN | NaN |
| Evl | VKEEIIDAIR | -396 | -405 | 1 | 0 | 10 | 1996.9354 | 2067.2896 | 2240.4226 | 2157.5488 | 2493.547 | 2283.3442 |
| Evl | DVGLDALDLDR | -370 | -380 | 0 | 0 | 11 | 973.7187 | 1017.62024 | 990.83484 | 811.5783 | 1069.1014 | 929.4535 |
| Evl | KEDESQTEDPSTSPSPGTR | -290 | -308 | 1 | 0 | 19 | 489.43137 | 559.4773 | 533.7044 | 527.34576 | 646.2749 | 643.1245 |
| Evl | KEDESQTEDPSTSPSPGTR | -290 | -308 | 1 | 0 | 19 | 489.43137 | 559.4773 | 533.7044 | 527.34576 | 646.2749 | 643.1245 |
| Evl | MKQEILEEVVR | -381 | -391 | 1 | 0.00025015 | 11 | 1164.2312 | NaN | 900.13763 | 1634.5648 | 1618.2921 | 2480.7886 |
| Evl | SNSVEKPVSSL | -327 | -337 | 1 | 0.001392869 | 11 | 854.2588 | 1060.7295 | NaN | 1291.8433 | 1904.3506 | 1660.1461 |
| Evl | WVPIKPGQQ | -23 | -31 | 1 | 0.001055699 | 9 | 1125.4237 | NaN | NaN | 790.3917 | 703.1736 | 738.8747 |
| Evl | VQRPEDASGGSSPSGTSK | -233 | -250 | 1 | 0.001055699 | 18 | 1162.7407 | NaN | 1796.4807 | NaN | NaN | NaN |
| Evl | QVQNGPSPEEMDIQR | -124 | -138 | 0 | 0.005668207 | 15 | 1081.4852 | NaN | NaN | NaN | 500.10223 | NaN |
| Nucks1 | QFQESDDADEDYGR | -15 | -28 | 0 | 0 | 14 | 1207.5233 | 755.68634 | 518.67126 | 1365.1012 | 1002.17236 | 1133.1593 |
| Nucks1 | SQFQESDDADEDYGR | -14 | -28 | 0 | 0 | 15 | 620.75775 | 643.9413 | 660.1977 | 489.02124 | 542.06885 | 525.76465 |
| Nucks1 | VGSEEEPEEDDEAPFQEK | -111 | -128 | 0 | 0 | 18 | 467.4666 | 451.497 | 586.36646 | 547.606 | 429.28693 | 579.2939 |
| Nucks1 | DVGSEEEPEEDDEAPFQEK | -110 | -128 | 0 | 0 | 19 | 326.0665 | 354.27448 | 388.822 | 269.9408 | 293.1763 | 462.40543 |
| Nucks1 | NSQEDSEDSEEKDVK | -53 | -67 | 1 | 0.00078661 | 15 | 227.08305 | 256.9342 | 277.95236 | 306.96506 | 431.7561 | 427.64935 |
| Sptlc1 | DVHLDLEER | -143 | -151 | 0 | 0.00272025 | 9 | 599.2395 | NaN | 1142.2961 | NaN | 1483.3086 | 1347.8617 |
| Actr1a | DVIANQPVVIDNGSGVIK | -5 | -22 | 0 | 0.001023236 | 18 | 1744.8147 | 1198.9316 | 1834.0287 | 2092.6797 | 1215.0303 | 1943.099 |
| Actr1a | YYLPDGSTIEIGPSR | -241 | -255 | 0 | 0 | 15 | 451.2916 | 521.63245 | 457.56595 | 581.07355 | 569.62537 | 817.0545 |
| Actr1a | SINPQKDETLETEK | -225 | -238 | 1 | 0.000230707 | 14 | 1827.8267 | NaN | NaN | 2411.6934 | 5359.123 | 5062.278 |
| Pold1 | IFEPILGEGR | -967 | -976 | 0 | 0 | 10 | 2597.828 | 2687.8699 | 752.1818 | 2742.0276 | 2839.7454 | 2958.7031 |
| Pold1 | DVISDLLCNR | -864 | -873 | 0 | 0 | 10 | 391.94562 | 539.2605 | 1025.2505 | 326.35678 | 1138.1368 | 1160.6283 |
| Pold1 | RFGPPGPEAW | -1096 | -1105 | 1 | 0 | 10 | 306.65826 | 641.05365 | NaN | 264.5857 | 389.8907 | 544.3664 |
| Pold1 | GLKPDEFIK | -629 | -637 | 1 | 0.001392869 | 9 | 2062.7815 | 2309.456 | NaN | NaN | 1209.3721 | 1136.9639 |
| Qars1 | SHPQDPIDTK | -96 | -105 | 0 | 0 | 10 | 2954.095 | 780.3677 | 2981.4688 | 2781.3188 | 3440.1567 | 1007.3636 |
| Qars1 | SVDPDSHQGQIVFNR | -750 | -764 | 0 | 0 | 15 | 1152.7054 | 1520.3671 | 1533.4608 | 1446.8091 | 1433.67 | 108.34758 |
| Qars1 | SVDPDSHQGQIVFNR | -750 | -764 | 0 | 0 | 15 | 1152.7054 | 1520.3671 | 1533.4608 | 1446.8091 | 1433.67 | 108.34758 |
| Qars1 | VYCPVQWEYGR | -476 | -486 | 0 | 0 | 11 | 936.1168 | 924.3753 | 963.1333 | 880.2267 | 1062.8229 | 1063.9843 |
| Qars1 | FDDTNPEKEEAK | -302 | -313 | 1 | 0 | 12 | 584.6185 | 714.3324 | 700.969 | 424.7896 | 563.4069 | 615.88416 |
| Qars1 | LYECLFQHK | -691 | -699 | 0 | 0 | 9 | 1812.1143 | 1250.0967 | 1394.8052 | 686.39594 | NaN | 834.0604 |
| Qars1 | SDFKEESEPGYK | -617 | -628 | 1 | 0 | 12 | 915.14496 | NaN | 873.7694 | 1410.5187 | 938.4083 | NaN |
| Qars1 | LASGQPVGLR | -630 | -639 | 0 | 0.00025015 | 10 | 311.26407 | 764.4442 | 586.0748 | 709.4089 | NaN | NaN |
| Qars1 | VLHLLGPK | -173 | -180 | 0 | 0.000550307 | 8 | 122.95673 | 79.43149 | 81.81044 | 567.77167 | NaN | NaN |
| Qars1 | DVLNDAAPR | -559 | -567 | 0 | 0.002099344 | 9 | 2050.8435 | 700.09985 | 2288.4995 | NaN | NaN | NaN |
| Qars1 | HTGYVIELQ | -640 | -648 | 0 | 0.001708008 | 9 | 717.12836 | 391.44638 | 370.17667 | NaN | NaN | NaN |
| QNG1 | DVLVDHEGVR | -20 | -29 | 0 | 0.00025015 | 10 | 2808.2695 | 2766.4575 | 2707.259 | NaN | 1453.5122 | 1249.512 |
| QNG1 | SLHELNPR | -49 | -56 | 0 | 0.000755556 | 8 | 2140.024 | 2238.723 | NaN | NaN | NaN | NaN |
| Park7 | VAGLAGKDPVQCSR | -35 | -48 | 1 | 0 | 14 | 4041.8413 | 3584.7034 | 3247.2053 | 3991.0361 | 3893.0156 | 3787.9346 |
| Park7 | LAHEVGFGCK | -113 | -122 | 0 | 0 | 10 | 2059.0222 | 1927.4818 | 2031.8085 | 759.3457 | 2007.696 | 2019.6125 |
| Park7 | LAHEVGFGCK | -113 | -122 | 0 | 0 | 10 | 2059.0222 | 1927.4818 | 2031.8085 | 759.3457 | 2007.696 | 2019.6125 |
| Park7 | TQGPYDVVVLPGGNLGAQ | -63 | -80 | 0 | 0 | 18 | 1380.4677 | 1548.5212 | 1516.8241 | 2084.109 | 2372.042 | 2083.6118 |
| Park7 | HYSYSESR | -138 | -145 | 0 | 0.00111006 | 8 | 1145.1831 | 1272.2843 | 137.25865 | 674.87445 | 748.33075 | 723.7523 |
| Park7 | VTTHPLAK | -123 | -130 | 0 | 0 | 8 | 716.2844 | 748.0987 | 798.3954 | 982.5513 | 786.6931 | 794.84607 |
| Park7 | DVMICPDTSLEDAK | -49 | -62 | 0 | 0 | 14 | 491.174 | 477.4187 | 1232.7117 | 1050.6587 | 1748.1653 | 1282.341 |
| Park7 | DVMICPDTSLEDAK | -49 | -62 | 0 | 0 | 14 | 491.174 | 477.4187 | 1232.7117 | 1050.6587 | 1748.1653 | 1282.341 |
| Park7 | GLAGKDPVQCSR | -37 | -48 | 1 | 0 | 12 | 2572.5454 | NaN | NaN | 1250.0608 | 1801.3478 | 1886.1829 |
| Park7 | LAIVEALVGK | -166 | -175 | 0 | 0.000712251 | 10 | 440.40787 | 458.05112 | 199.52545 | 492.54095 | NaN | NaN |
| Ciao1 | DVNCVAWNPK | -305 | -314 | 0 | 0.00383302 | 10 | 1206.083 | NaN | NaN | NaN | NaN | NaN |
| Mt-Cyb | DVNYGWLIR | -72 | -80 | 0 | 0 | 9 | 1027.0912 | 879.26324 | 720.22595 | 1105.7922 | 1433.5819 | 359.44067 |
| Mt-Cyb | LFLHVGR | -94 | -100 | 0 | 0.001868731 | 7 | 669.7034 | 622.4179 | 629.3184 | NaN | 488.5433 | 465.74008 |
| Qtrt1 | HLGLRPGPELIR | -76 | -87 | 1 | 0.00025015 | 12 | 282.39536 | 269.94852 | 275.0847 | NaN | 277.10574 | 264.45053 |
| Qtrt1 | SPYDGEETLLSPER | -129 | -142 | 0 | 0 | 14 | 1115.1185 | 1239.1052 | 1311.1288 | 672.73755 | NaN | NaN |
| Qtrt1 | DVPGFAIGGLSGGESK | -221 | -236 | 0 | 0.000712251 | 16 | 988.5005 | NaN | 1105.9121 | NaN | NaN | 1340.7306 |
| Epb41 | DVPIVHTETK | -748 | -757 | 0 | 0.001868731 | 10 | 4647.6973 | 4825.11 | NaN | NaN | 4346.274 | NaN |
| Letm1 | DVQDYSEDLQEIK | -574 | -586 | 0 | 0 | 13 | 1984.6292 | 2273.9333 | 2673.3794 | 394.30493 | 390.69257 | NaN |

| | | | | | | | | | | | | |
|---|---|---|---|---|---|---|---|---|---|---|---|---|
| Letm1 | QEHLEELKR | -477 | -485 | 1 | 0 | 9 | 1373.8802 | 574.97845 | NaN | 1112.7408 | 1526.7913 | 1687.2715 |
| Letm1 | SALDDNKDGNINIDDLVK | -672 | -689 | 1 | 0 | 18 | 547.7113 | 630.50696 | NaN | 625.4645 | 642.00446 | 504.78552 |
| Letm1 | KLEEGGPVYSPPAQVVVR | -132 | -149 | 1 | 0.001291103 | 18 | 402.54657 | NaN | 471.0187 | NaN | NaN | 551.49615 |
| Letm1 | KLEEGGPVYSPPAQVVVR | -132 | -149 | 1 | 0.001291103 | 18 | 402.54657 | NaN | 471.0187 | NaN | NaN | 551.49615 |
| Letm1 | STLQTLPEIVAK | -434 | -445 | 0 | 0 | 12 | 444.33563 | 464.68854 | 631.59125 | NaN | NaN | NaN |
| Dnaja2 | YGEQGLR | -69 | -75 | 0 | 0.000471409 | 7 | 1729.8853 | 1524.743 | 1790.2115 | 1251.8341 | 411.56802 | 1222.2545 |
| Dnaja2 | EHEVFQR | -256 | -262 | 0 | 0 | 7 | 1518.6288 | 4393.1294 | 1593.6526 | 444.19565 | 2845.8127 | 2605.9236 |
| Dnaja2 | DVQFPENNWINPDK | -334 | -347 | 0 | 0 | 14 | 1092.7539 | 1121.1512 | 1343.1544 | 1394.6888 | 1450.9236 | 1544.4868 |
| Dnaja2 | EYHPDKNPNAGDK | -34 | -46 | 1 | 0.00025015 | 13 | 516.62 | 589.13525 | 465.94385 | 369.7787 | 421.17914 | 494.11572 |
| Dnaja2 | EYHPDKNPNAGDK | -34 | -46 | 1 | 0.00025015 | 13 | 516.62 | 589.13525 | 465.94385 | 369.7787 | 421.17914 | 494.11572 |
| Dnaja2 | EGSGGGGGMDDIFSHIF | -76 | -92 | 0 | 0.00025015 | 17 | 349.43134 | 393.79214 | 430.94235 | 431.67673 | 483.47662 | 504.44443 |
| Dnaja2 | GVPPGASENELKK | -14 | -26 | 1 | 0 | 13 | 1348.1403 | 1286.1383 | 1345.3218 | NaN | 1008.5105 | 1366.0779 |
| Dnaja2 | KVSLEDLYNGK | -121 | -131 | 1 | 0 | 11 | 559.99695 | NaN | 926.8265 | 851.9117 | 1021.2663 | 1074.1963 |
| Dnaja2 | VIEPGCVR | -303 | -310 | 0 | 0.00365438 | 8 | 245.49707 | 1148.0574 | NaN | NaN | NaN | NaN |
| Dctpp1 | DWEQFHQPR | -46 | -54 | 0 | 0 | 9 | 750.7289 | 754.2029 | 815.41016 | 1184.3922 | 914.64984 | 946.6805 |
| Arf3 | DWLANQLK | -171 | -178 | 0 | 0.004065182 | 8 | 564.93756 | NaN | NaN | NaN | NaN | NaN |
| Gdi2 | LELLEPIEQK | -370 | -379 | 0 | 0 | 10 | 9277.258 | 10470.879 | 11113.564 | 8929.445 | 10907.931 | 6647.938 |
| Gdi2 | FETTCDDIKDIYK | -410 | -422 | 1 | 0 | 13 | 5681.1353 | 268.0264 | 5384.8184 | 484.99634 | 2322.707 | 2159.1533 |
| Gdi2 | MLLFTEVTR | -90 | -98 | 0 | 0 | 9 | 4535.604 | 2925.4053 | 4930.5464 | 3422.5935 | 3359.3108 | 3307.3018 |
| Gdi2 | VANFDEKDPR | -147 | -156 | 1 | 0.00025015 | 10 | 4267.1006 | 1165.629 | 3462.2783 | 700.0578 | 8221.956 | 8938.9795 |
| Gdi2 | DYLDQPCCETINR | -196 | -208 | 0 | 0 | 13 | 3814.2756 | 383.20572 | 4426.874 | 4823.1074 | 5154.047 | 5106.2534 |
| Gdi2 | NFDEKDPR | -149 | -156 | 1 | 0 | 8 | 3273.9116 | 1704.1752 | 2068.4368 | 1631.3059 | 2352.7173 | 3268.0542 |
| Gdi2 | FVSISDLFVPK | -380 | -390 | 0 | 0 | 11 | 1595.5417 | 1611.5923 | 1602.8643 | 1789.64 | 2485.427 | 2601.2192 |
| Gdi2 | THFETTCDDIKDIYK | -408 | -422 | 1 | 0 | 15 | 1441.0197 | 1934.1687 | 576.4226 | 588.09644 | 634.0898 | 577.82043 |
| Gdi2 | THFETTCDDIKDIYK | -408 | -422 | 1 | 0 | 15 | 1441.0197 | 1934.1687 | 576.4226 | 588.09644 | 634.0898 | 577.82043 |
| Gdi2 | SITPLEDLYK | -45 | -54 | 0 | 0 | 10 | 1394.3 | 1513.2886 | 1601.4749 | 1761.284 | 1845.6932 | 1689.8473 |
| Gdi2 | TDDYLDQPCCETINR | -194 | -208 | 0 | 0 | 15 | 835.9366 | 858.68854 | 815.3788 | 1059.0394 | 959.68646 | 66.592514 |
| Gdi2 | EIRPALELLEPIEQK | -365 | -379 | 1 | 0 | 15 | 757.2517 | 829.93994 | 820.9984 | 1438.6461 | 1965.9375 | 2249.2417 |
| Gdi2 | EIRPALELLEPIEQK | -365 | -379 | 1 | 0 | 15 | 757.2517 | 829.93994 | 820.9984 | 1438.6461 | 1965.9375 | 2249.2417 |
| Gdi2 | THFETTCDDIK | -408 | -418 | 0 | 0 | 11 | 538.50665 | 615.4808 | 682.0034 | 884.9622 | 965.69293 | 615.6206 |
| Gdi2 | PALELLEPIEQK | -368 | -379 | 0 | 0 | 12 | 494.66763 | 627.5802 | 607.64105 | 938.74884 | 1045.4478 | 1010.8932 |
| Gdi2 | SASITPLEDLYK | -43 | -54 | 0 | 0 | 12 | 2535.4143 | 1050.7356 | 2693.6758 | 4700.696 | NaN | 4582.412 |
| Dazap1 | GIPHNCGETELR | -119 | -130 | 0 | 0 | 12 | 8659.304 | 9957.334 | 8857.242 | 9429.292 | 5875.595 | 7670.208 |
| Dazap1 | GIPHNCGETELR | -119 | -130 | 0 | 0 | 12 | 8659.304 | 9957.334 | 8857.242 | 9429.292 | 5875.595 | 7670.208 |
| Dazap1 | PHNCGETELR | -121 | -130 | 0 | 0.000627943 | 10 | 2489.349 | 2787.7004 | 613.88904 | 1733.8909 | 5753.7715 | 7113.907 |
| Dazap1 | FKDPNCVGTVL | -58 | -68 | 1 | 0 | 11 | 1512.9661 | 1633.2234 | 991.4176 | 1453.7147 | 544.13416 | 1724.9027 |
| Dazap1 | LFVGGLDWSTTQETLR | -12 | -27 | 0 | 0 | 16 | 1135.3418 | 303.69244 | 276.24457 | 466.5214 | 673.74744 | 652.40515 |
| Dazap1 | IFVGGIPHNCGETELR | -115 | -130 | 0 | 0 | 16 | 497.32843 | 609.90125 | 523.72345 | NaN | 383.99222 | 569.3371 |
| Dazap1 | DWSTTQETLR | -18 | -27 | 0 | 0.000230707 | 10 | 947.3213 | NaN | NaN | 1287.8878 | NaN | NaN |
| Ube2s | QVTIEGPEGTPYAGGLFR | -44 | -61 | 0 | 0.000712251 | 18 | 937.54395 | 975.8645 | 685.83075 | 1200.2413 | 1318.1714 | 947.7088 |
| Ube2s | DWTAELGIR | -102 | -110 | 0 | 0.00111006 | 9 | 397.8031 | NaN | NaN | NaN | NaN | NaN |
| Rab35 | DYDHLFK | -4 | -10 | 0 | 0.00111006 | 7 | 1204.2646 | 1324.4832 | 1173.7192 | 1226.6831 | 408.1349 | 686.39417 |
| Rab35 | KVVETEDAYK | -128 | -137 | 1 | 0 | 10 | 521.803 | 437.66635 | 617.8401 | NaN | 841.8673 | 807.89014 |
| Rab35 | GTHGVIVVY | -80 | -88 | 0 | 0.002910424 | 9 | 565.0508 | 528.70496 | 379.3697 | NaN | NaN | NaN |
| Rras2 | MNVDQAFHELVR | -162 | -173 | 0 | 0 | 12 | 1441.8481 | 1390.3535 | 5156.3438 | 1059.7035 | 804.17523 | 1689.5172 |
| Rras2 | FVTDYDPTIEDSYTK | -39 | -53 | 0 | 0 | 15 | 606.6798 | 731.35443 | 780.05835 | 745.96075 | 1080.8827 | 1046.6327 |
| Kri1 | DYEDIIDDLPCR | -513 | -524 | 0 | 0 | 12 | 1536.7181 | 1673.8485 | 172.52188 | 79.33954 | 3320.683 | 3314.0317 |
| Morf4l1 | DYEVAPPEYHR | -349 | -359 | 0 | 0 | 11 | 3811.7747 | 6045.777 | 4760.1553 | 1816.4413 | 2016.654 | 1572.9402 |
| Morf4l1 | QLFYLPAK | -219 | -226 | 0 | 0 | 8 | 245.101 | 296.13263 | 222.12988 | 812.0383 | NaN | NaN |
| Npepps | KPYPDDENLVEVK | -223 | -235 | 1 | 0 | 13 | 1062.3181 | 1740.4209 | 817.3297 | 775.10736 | 847.0644 | 917.36316 |
| Npepps | DYFNVPYPLPK | -295 | -305 | 0 | 0 | 11 | 308.04916 | 237.28964 | 379.46585 | 734.62085 | 1107.4962 | 498.3874 |
| Npepps | FESHPAPSAER | -873 | -883 | 0 | 0 | 11 | 1734.2316 | 616.42725 | 649.9945 | NaN | 1487.4463 | 325.75266 |
| Npepps | IDFVGELNDK | -147 | -156 | 0 | 0 | 10 | 1173.0344 | 715.52496 | NaN | 1211.1738 | 1198.8866 | 1507.3447 |
| Npepps | NVKPDQWVK | -573 | -581 | 1 | 0.00025015 | 9 | 1514.217 | 1268.7843 | NaN | NaN | NaN | NaN |
| Npepps | VVGEYDFVETR | -250 | -260 | 0 | 0 | 11 | 1314.6921 | NaN | NaN | NaN | NaN | NaN |
| Etfa | LNVAPVSDIIEIK | -127 | -139 | 0 | 0 | 13 | 704.73724 | 845.46106 | 877.3574 | 971.98535 | 857.0744 | 1175.8912 |
| Etfa | TIVAINKDPEAPIFQVA | -295 | -311 | 1 | 0.00025015 | 17 | 156.44836 | 458.20374 | 342.27988 | 762.4107 | 491.31027 | 474.62317 |
| Etfa | SDRPELTGAK | -207 | -216 | 1 | 0 | 10 | 1223.2312 | 1436.1923 | 1117.2737 | NaN | 1600.7886 | 684.38617 |
| Etfa | DYGIVADLFK | -312 | -321 | 0 | 0 | 10 | 1021.9738 | 717.1421 | 983.23157 | 1363.8613 | 1501.9688 | NaN |
| Etfa | NVAPVSDIIEIK | -128 | -139 | 0 | 0 | 12 | 521.2365 | NaN | NaN | 787.444 | 745.09845 | 808.64984 |
| Etfa | GLLPEELTPLILETQK | -86 | -101 | 0 | 0 | 16 | 167.30789 | NaN | 276.40506 | NaN | 69.93485 | 210.52176 |
| F2 | NLPIVERPVCK | -524 | -534 | 1 | 0 | 11 | 3328.2378 | 6623.595 | 1635.3668 | 4754.8506 | 2254.2512 | 2339.727 |
| F2 | KSPQELLCGA | -381 | -390 | 1 | 0.001743443 | 10 | 1100.2971 | 282.9626 | 155.43617 | 985.4697 | 1113.2988 | 1164.9248 |
| F2 | QVVNLPIVERPVCK | -521 | -534 | 1 | 0 | 14 | 894.2597 | 1073.0469 | 1189.0269 | 1015.51416 | 1270.7567 | 1151.8649 |
| F2 | GIVSWGEGCDR | -583 | -593 | 0 | 0 | 11 | 429.02505 | 357.48712 | 729.84 | 525.7572 | 188.38736 | 287.6814 |
| F2 | DYIHPVCLPDK | -473 | -483 | 0 | 0 | 11 | 1019.8635 | 612.15796 | 508.11722 | NaN | 610.23047 | 515.89844 |
| F2 | QVVNLPIVER | -521 | -530 | 0 | 0.000230707 | 10 | 405.9277 | 739.49646 | 796.29675 | NaN | 1058.8715 | NaN |
| F2 | SDYIHPVCLPDK | -472 | -483 | 0 | 0 | 12 | 1312.2157 | NaN | 1977.36 | NaN | NaN | NaN |
| Pura | FFFDVGSNK | -230 | -238 | 0 | 0 | 9 | 1001.2552 | 1234.1418 | 1107.3147 | 975.2533 | 1162.2642 | 1114.752 |
| Pura | DYLGDFIEHY | -111 | -120 | 0 | 0.000431655 | 10 | 405.24768 | 373.90433 | 465.02426 | 511.7982 | 497.11237 | 448.30383 |
| Rps10 | DYLHLPPEIVPA | -81 | -92 | 0 | 0 | 12 | 5496.0557 | 4571.74 | 6002.55 | 6513.871 | 6069.3613 | 3556.2854 |
| Rps10 | TNEGIQYLR | -72 | -80 | 0 | 0 | 9 | 3796.7058 | 3137.9358 | 3998.115 | 4144.802 | 3102.1653 | 3615.1216 |
| Rps10 | GPEGERPAR | -108 | -116 | 1 | 0.002566359 | 9 | 846.93616 | 769.5698 | 997.932 | 599.17456 | 738.89026 | 886.1669 |
| Rps10 | LTNEGIQYLR | -71 | -80 | 0 | 0 | 10 | 769.3798 | 766.18005 | 630.48254 | 658.7651 | 587.0305 | 657.4853 |
| Rps10 | HPELADK | -32 | -38 | 0 | 0.000431655 | 7 | 516.3986 | 896.9095 | 935.5293 | 893.83563 | 699.9099 | 859.39 |
| Rps10 | IAIYELLFK | -9 | -17 | 0 | 0.001636916 | 9 | 162.23508 | NaN | 304.0308 | NaN | NaN | 296.23422 |
| Emd | QPGTSLVDADTFHHQ | -120 | -134 | 0 | 0 | 15 | 2723.8145 | 2974.308 | 2943.873 | 5368.367 | 4519.3994 | 4746.004 |
| Emd | DYNDDYYEESYLTTK | -90 | -104 | 0 | 0 | 15 | 522.9407 | 518.67883 | 586.66846 | 619.72266 | 728.24603 | 675.5345 |

| | | | | | | | | | | | | |
|---|---|---|---|---|---|---|---|---|---|---|---|---|
| Emd | KEDALLYQSK | -80 | -89 | 1 | 0 | 10 | 1590.0632 | 1786.9166 | 1656.3815 | 1213.2854 | NaN | 977.7064 |
| Cox4i1 | DYPLPDVAHVT | -43 | -53 | 0 | 0 | 11 | 3693.1108 | 3233.0632 | 3284.8992 | 515.94147 | 280.3489 | 3802.1562 |
| Cox4i1 | WDYDKNEWK | -160 | -168 | 1 | 0 | 9 | 2608.0715 | 2735.205 | 2826.2722 | 3139.2039 | 3418.0574 | 3377.4995 |
| Cox4i1 | WDYDKNEWK | -160 | -168 | 1 | 0 | 9 | 2608.0715 | 2735.205 | 2826.2722 | 3139.2039 | 3418.0574 | 3377.4995 |
| Cox4i1 | SYVYGPIPHTF | -123 | -133 | 0 | 0.00025015 | 11 | 1671.3075 | 1788.8209 | 1775.5398 | 1501.306 | 736.7429 | 1687.0017 |
| Cox4i1 | SYVYGPIPHTF | -123 | -133 | 0 | 0.00025015 | 11 | 1671.3075 | 1788.8209 | 1775.5398 | 1501.306 | 736.7429 | 1687.0017 |
| Cox4i1 | KWDYDKNEWK | -159 | -168 | 2 | 0.00025015 | 10 | 908.95435 | 1031.9603 | 913.9263 | 985.04346 | 1517.447 | 828.55493 |
| Cox4i1 | SEDYAFPTYADR | -30 | -41 | 0 | 0 | 12 | 623.02155 | 784.7661 | 776.11035 | 211.92706 | 225.65607 | 903.91864 |
| Cox4i1 | SYVYGPIPHTFDR | -123 | -135 | 0 | 0 | 13 | 352.5808 | 1590.9884 | 773.29034 | 685.7911 | 827.926 | 643.84894 |
| Cox4i1 | SYVYGPIPH | -123 | -131 | 0 | 0.00149795 | 9 | 840.78046 | NaN | NaN | 693.1095 | NaN | NaN |
| Etfdh | VDHTVGWPLDR | -290 | -300 | 0 | 0 | 11 | 1232.4639 | 551.68243 | 1352.3539 | 1401.2834 | 1350.7408 | 1070.2677 |
| Etfdh | VVGLDYQNPYLSPFR | -323 | -337 | 0 | 0.00051306 | 15 | 481.48364 | 487.6729 | NaN | 472.27322 | NaN | 639.4138 |
| Etfdh | DYQNPYLSPFR | -327 | -337 | 0 | 0 | 11 | 1347.0065 | 495.63727 | 585.601 | NaN | NaN | NaN |
| Etfdh | EVLYHEDGSVK | -201 | -211 | 0 | 0 | 11 | 1316.4492 | 1280.724 | NaN | NaN | 1863.6611 | NaN |
| Rbm3 | SFGPISEVVVVK | -28 | -39 | 0 | 0 | 12 | 4310.586 | 4965.216 | 4525.175 | 5349.0903 | 6527.336 | 6692.323 |
| Rbm3 | FTNPEHASDAMR | -54 | -65 | 0 | 0 | 12 | 3329.3452 | 3641.596 | 3980.8853 | 3484.8918 | 4486.686 | 4761.0005 |
| Rbm3 | FTNPEHASDAMR | -54 | -65 | 0 | 0 | 12 | 3329.3452 | 3641.596 | 3980.8853 | 3484.8918 | 4486.686 | 4761.0005 |
| Rbm3 | FTNPEHASDAMR | -54 | -65 | 0 | 0 | 12 | 3329.3452 | 3641.596 | 3980.8853 | 3484.8918 | 4486.686 | 4761.0005 |
| Rbm3 | YDSRPGGY | -115 | -122 | 1 | 0.00025015 | 8 | 1770.2534 | 1496.2892 | 1782.9133 | 1911.9417 | 1534.9828 | 1793.2905 |
| Rbm3 | YDSRPGGYGY | -115 | -124 | 1 | 0.001708008 | 10 | 1499.0085 | 1653.6669 | 1542.6716 | 1568.1589 | 1479.7233 | 1511.0065 |
| Rbm3 | YDSRPGGYGYGY | -115 | -126 | 1 | 0 | 12 | 784.2166 | 863.08655 | 777.9772 | 117.10905 | 869.6435 | 863.3344 |
| Rbm3 | TNPEHASDAMR | -55 | -65 | 0 | 0 | 11 | 503.2163 | 979.546 | 1092.7844 | 890.1264 | 1010.7575 | 1051.0942 |
| Rbm3 | TNPEHASDAMR | -55 | -65 | 0 | 0 | 11 | 503.2163 | 979.546 | 1092.7844 | 890.1264 | 1010.7575 | 1051.0942 |
| Rbm3 | ITFTNPEHASDAMR | -52 | -65 | 0 | 0 | 14 | 301.50342 | 341.25082 | 601.9715 | 556.8341 | 884.01276 | 556.96906 |
| Rbm3 | DYSGSQGGYDR | -131 | -141 | 0 | 0 | 11 | 560.6013 | NaN | NaN | 248.59138 | 169.57375 | 112.18106 |
| Hcls1 | VSEEHDILK | -61 | -69 | 0 | 0.000230707 | 9 | 6047.319 | 5997.26 | 4201.807 | 1672.9061 | 630.0722 | 3733.741 |
| Hcls1 | SAVGFNEMEAPTTAYK | -208 | -223 | 0 | 0 | 16 | 3600.1433 | 3032.2576 | 2850.4558 | 1281.4615 | 1230.5305 | 1433.8901 |
| Hcls1 | EYPVPSLPTR | -322 | -331 | 0 | 0 | 10 | 1207.7063 | 1225.4647 | 1788.3746 | 1133.6277 | 1520.3077 | 1253.3116 |
| Hcls1 | SAVGHEYVADVEK | -97 | -109 | 0 | 0.00025015 | 13 | 1046.1038 | 1200.0404 | 1237.8447 | 1133.0706 | 1266.8572 | 1257.2544 |
| Hcls1 | TEHINIHQLR | -49 | -58 | 0 | 0 | 10 | 453.511 | 506.7025 | 591.49054 | 473.9059 | 518.591 | 615.7607 |
| Hcls1 | EVQQPSMPVEEPAAPAQLPK | -277 | -296 | 0 | 0.000431655 | 20 | 378.66553 | 395.99274 | 451.2922 | 88.312744 | 537.9946 | 480.4075 |
| Hcls1 | VETQGDDWDTDPDFVNDISEK | -14 | -34 | 0 | 0 | 21 | 358.40744 | 364.2113 | 393.07007 | 472.88974 | 627.31274 | 583.59564 |
| Hcls1 | DYSHGFGGR | -152 | -160 | 0 | 0 | 9 | 159.81085 | 1631.5249 | 1451.4319 | 1747.3848 | 1750.3583 | 1827.2458 |
| Hcls1 | GHFGLFPANYVK | -473 | -484 | 0 | 0 | 12 | 1429.3116 | 1359.1526 | 1471.7877 | 1009.1509 | 1186.1754 | NaN |
| Hcls1 | TQGDDWDTDPDFVNDISEK | -16 | -34 | 0 | 0.00025015 | 19 | 739.6427 | 925.4827 | 1004.04626 | NaN | NaN | 506.3434 |
| Agap2 | DYSSSLPSSPNVGHR | -601 | -615 | 0 | 0 | 15 | 1250.9059 | NaN | NaN | NaN | NaN | NaN |
| Gnas | TTPEDATPEPGEDPR | (319);(1058) | (333);(1072) | 0 | 0 | 15 | 758.3106 | 650.0212 | 400.97034 | 845.60815 | 946.8788 | 568.65356 |
| Gnas | YTTPEDATPEPGEDPR | (318);(1057) | (333);(1072) | 0 | 0.000230707 | 16 | 447.53238 | 492.403 | 454.36612 | 305.5369 | 305.16803 | 265.72513 |
| Gnas | SNLVPPVELANPENQFR | (111);(850) | (127);(866) | 0 | 0 | 17 | 270.122 | 5058.9814 | 5705.339 | 3834.6338 | 1047.9924 | 349.84192 |
| Gnas | QADYVPSDQDLLR | (187);(926) | (199);(938) | 0 | 0 | 13 | 1060.6656 | 1029.1392 | NaN | 273.93817 | NaN | 2313.8374 |
| Gnas | DYVPSDQDLLR | (189);(928) | (199);(938) | 0 | 0 | 11 | 682.2587 | NaN | NaN | 303.8021 | NaN | NaN |
| Pfkl | DYVSGELEHVTR | -760 | -771 | 0 | 0 | 12 | 943.9725 | 3050.8245 | 2839.8962 | 3082.506 | 3585.1548 | 3919.2563 |
| Pfkl | FDEAIQLR | -367 | -374 | 0 | 0 | 8 | 908.1308 | 753.6893 | 1013.9181 | 1160.5944 | 1102.3119 | 907.12 |
| Pfkl | TGISEGHTVYIVHDGFEGLAK | -424 | -444 | 0 | 0 | 21 | 828.757 | 709.16907 | 821.23016 | 1233.6027 | 1426.8188 | 1277.9235 |
| Pfkl | EQWWLNLR | -738 | -745 | 0 | 0 | 8 | 725.5022 | 960.0562 | 1028.9379 | 807.27246 | 1081.5566 | 943.0741 |
| Pfkl | GHLQQGGAPTPFDR | -659 | -672 | 0 | 0 | 14 | 456.2596 | 194.91849 | 285.87766 | 370.6898 | 713.61035 | 1188.8284 |
| Pfkl | RLGFDTR | -286 | -292 | 1 | 0.000550307 | 7 | 776.9404 | NaN | NaN | 667.28107 | NaN | 666.2797 |
| Pfkl | EAYEGVLQLVEAR | -499 | -511 | 0 | 0 | 13 | 666.95795 | 372.82483 | 644.28796 | NaN | NaN | NaN |
| Pfkl | EGVLQLVEAR | -502 | -511 | 0 | 0.001392869 | 10 | 479.99802 | 462.51636 | NaN | NaN | NaN | NaN |
| Pfkl | VLGHLQQGGAPTPFDR | -657 | -672 | 0 | 0.00644558 | 16 | 1082.4515 | NaN | NaN | NaN | NaN | NaN |
| Crnkl1 | SFEDEFGTVSDKER | -599 | -612 | 1 | 0 | 14 | 789.7528 | 520.87744 | 609.8709 | 1045.0358 | 1435.8787 | 1369.7606 |
| Crnkl1 | FLEFGPENCTSWIK | -471 | -484 | 0 | 0 | 14 | 457.21747 | 517.80786 | 542.76025 | 389.30014 | 401.07013 | 439.1599 |
| Crnkl1 | EAAEQDPDKDIDESESSSF | -672 | -690 | 1 | 0.000627943 | 19 | 289.91043 | 369.95093 | 304.54987 | 362.65396 | 320.5517 | 383.54306 |
| Crnkl1 | ELELLPPPPQQK | -41 | -52 | 0 | 0 | 12 | 612.2965 | NaN | NaN | NaN | NaN | 605.0514 |
| Ly6a | QEAAVIVDSQTR | -56 | -67 | 0 | 0 | 12 | 1934.8732 | 2075.4548 | 2005.1404 | 1762.3721 | 1221.2882 | 1072.9113 |
| Ly6a | EAAVIVDSQTR | -57 | -67 | 0 | 0.00025015 | 11 | 1840.6736 | 2055.866 | 2157.8433 | 1911.1343 | 1466.0236 | NaN |
| Ly6a | NNLCLPICPPNIE | -71 | -83 | 0 | 0 | 13 | 503.37845 | NaN | NaN | NaN | 555.8259 | NaN |
| Slk | ILNEKPTTDGPEK | -380 | -392 | 1 | 0 | 13 | 803.60297 | 512.0682 | 664.193 | 1354.5326 | 1751.0394 | 474.84283 |
| Slk | EADFQAVDNEVGLTK | -506 | -520 | 0 | 0 | 15 | 1651.4524 | NaN | 1702.9384 | NaN | NaN | NaN |
| Psmd7 | EAEEVGVEHLLR | -166 | -177 | 0 | 0 | 12 | 1678.1683 | 1766.8269 | 94.56727 | 1881.7495 | 2088.8425 | 1934.6852 |
| Psmd7 | VVVHPLVLL | -9 | -17 | 0 | 0.002595029 | 9 | 568.5508 | 684.1011 | 688.1146 | 499.77496 | 597.1294 | 577.70605 |
| Psmd7 | VEEVHDDGTPTSK | -141 | -153 | 0 | 0 | 13 | 480.79974 | 505.95828 | NaN | 698.3585 | 833.6275 | 763.38116 |
| Pafah1b1 | SVQDISFDHSGK | -152 | -163 | 0 | 0 | 12 | 8096.303 | 7942.398 | 1145.3337 | 4397.6255 | 5179.55 | 1215.2493 |
| Pafah1b1 | LVGHDNWVR | -334 | -342 | 0 | 0.000550307 | 9 | 803.2749 | 2404.1025 | 466.4871 | 2595.2979 | 2591.8528 | 2984.9429 |
| Pafah1b1 | EWIPRPPEKY | -93 | -102 | 2 | 0.000471409 | 10 | 799.0089 | 2624.203 | 1115.9629 | 3222.0774 | 1176.6101 | 879.86005 |
| Pafah1b1 | EAELDMNEELDKK | -34 | -46 | 1 | 0.00025015 | 13 | 745.80316 | 549.566 | 589.6999 | 608.9742 | 599.559 | 721.8609 |
| Pafah1b1 | GHTDSVQDISFDHSGK | -148 | -163 | 0 | 0 | 16 | 657.43365 | 1120.8448 | 991.2445 | 877.04944 | 1187.7776 | 1042.3862 |
| Pafah1b1 | GHTDSVQDISFDHSGK | -148 | -163 | 0 | 0 | 16 | 657.43365 | 1120.8448 | 991.2445 | 877.04944 | 1187.7776 | 1042.3862 |
| Pafah1b1 | TAPYVVTGSVDQTVK | -391 | -405 | 0 | 0 | 15 | 358.27118 | 729.60504 | 711.92847 | 903.4865 | 997.1162 | 941.2792 |
| Pafah1b1 | LWDFQGFECIR | -176 | -186 | 0 | 0 | 11 | 205.12566 | 428.61185 | 394.8206 | 240.79657 | 519.8162 | 724.33 |
| Pafah1b1 | TLVGHDNWVR | -333 | -342 | 0 | 0 | 10 | 1805.8303 | 2001.7168 | 2174.9797 | 1648.7792 | NaN | 2499.4124 |
| Pafah1b1 | EWIPRPPEK | -93 | -101 | 1 | 0.000907206 | 9 | 931.78357 | 767.70166 | NaN | 1520.8047 | 1627.1418 | 1268.8616 |
| Pafah1b1 | EEFTSGGPLGQK | -77 | -88 | 0 | 0 | 12 | 467.7583 | 907.2183 | 465.67166 | 506.45667 | 264.64087 | NaN |
| Pafah1b1 | FTSGGPLGQK | -79 | -88 | 0 | 0 | 10 | 5985.9146 | 3626.915 | NaN | NaN | NaN | NaN |
| Pafah1b1 | VSASEDATIK | -124 | -133 | 0 | 0.000431655 | 10 | 1927.4633 | NaN | NaN | 603.9718 | NaN | NaN |
| Sell | EAENWGAGEPNNKK | -110 | -123 | 1 | 0 | 14 | 594.60144 | 638.172 | 424.38956 | 673.1339 | 910.3059 | 929.8225 |
| Sell | SPYYYWIGIR | -83 | -92 | 0 | 0 | 10 | 526.6105 | 451.59808 | 398.06696 | 660.9284 | 814.2623 | 626.0187 |
| Sell | EAENWGAGEPNNK | -110 | -122 | 0 | 0 | 13 | 481.3767 | 600.6314 | 579.72974 | NaN | 733.7323 | 829.92206 |

| | | | | | | | | | | | | |
|---|---|---|---|---|---|---|---|---|---|---|---|---|
| Inpp5b | EAFFFHDTPK | -389 | -398 | 0 | 0 | 10 | 495.21826 | NaN | 796.37866 | 596.3015 | 678.8384 | 391.0153 |
| Hibadh | EAGEQVASSPAEVAEK | -79 | -94 | 0 | 0 | 16 | 2601.2168 | 3214.2295 | 1401.419 | NaN | NaN | 2569.509 |
| Hopx | EAGLTEEQTQK | -39 | -49 | 0 | 0 | 11 | 1700.0576 | 1894.9586 | 1668.0082 | 976.73596 | 1738.3411 | 1878.8097 |
| Bysl | VRDDIAEYK | -257 | -265 | 1 | 0 | 9 | 1832.6111 | 582.4892 | 1279.7595 | 434.6731 | 1577.9938 | 992.95935 |
| Bysl | LQPHPQLSPEIR | -406 | -417 | 0 | 0 | 12 | 1446.067 | 1330.1492 | 1325.0514 | 1733.4834 | 1963.6644 | 1467.7896 |
| Bysl | QLPVLWHQ | -372 | -379 | 0 | 0.000627943 | 8 | 892.44275 | 890.6985 | 1056.6416 | 839.3896 | 983.09973 | 980.56696 |
| Bysl | EAIIVGSIITK | -303 | -313 | 0 | 0 | 11 | 465.4 | NaN | 486.58057 | 458.41782 | NaN | 509.11655 |
| Bysl | GILIPLCESGTCTLR | -288 | -302 | 0 | 0.00051306 | 15 | 486.8192 | NaN | NaN | 629.2437 | NaN | 542.189 |
| Chmp4b | EALENANTNTEVLK | -94 | -107 | 0 | 0 | 14 | 2931.1106 | 1871.8839 | 1538.1559 | 2604.1145 | 1389.4933 | 1476.0028 |
| Chmp4b | IDGTLSTIEFQR | -82 | -93 | 0 | 0.000627943 | 12 | 869.1478 | NaN | NaN | NaN | NaN | NaN |
| Zw10 | EALPDLPSPDADHK | -431 | -444 | 0 | 0 | 14 | 826.5364 | 1081.328 | 919.558 | NaN | NaN | 822.09564 |
| Zw10 | FVDLVPGFR | -574 | -582 | 0 | 0 | 9 | 486.00604 | NaN | 485.9665 | NaN | NaN | NaN |
| Alg5 | EALPSIWDSPTK | -54 | -65 | 0 | 0 | 12 | 1613.0172 | 1635.4225 | 1763.4099 | 1791.1588 | 2000.4628 | 1991.2902 |
| Alg5 | VIVVDDGSEDQTSK | -105 | -118 | 0 | 0.000550307 | 14 | 1498.1178 | NaN | 1691.8867 | NaN | 2548.1382 | NaN |
| Zc3h11a | TVLPTVPESQEEEVK | -100 | -114 | 0 | 0 | 15 | 1000.26196 | 401.0967 | 897.0031 | 721.8109 | 811.41534 | 1082.666 |
| Zc3h11a | LKAEEPSGADDSPSGTK | -356 | -372 | 1 | 0.00025015 | 17 | 600.9501 | 503.09808 | 485.3861 | 266.35626 | 532.32697 | 720.5236 |
| Zc3h11a | EANETSDETISDPTK | -516 | -530 | 0 | 0 | 15 | 523.74243 | 297.4284 | 214.02386 | 478.8497 | 482.5443 | 266.22412 |
| Zc3h11a | TLEEILLER | -336 | -344 | 0 | 0.00025015 | 9 | 92.67595 | NaN | NaN | 918.6837 | 1105.278 | NaN |
| Zc3h11a | STGKPPLSVEDDFEK | -743 | -757 | 1 | 0.005946382 | 15 | 1522.2582 | NaN | NaN | NaN | NaN | NaN |
| Aurkaip1 | EAPENWQTPK | -185 | -194 | 0 | 0.00025015 | 10 | 313.5639 | NaN | NaN | NaN | NaN | NaN |
| Eif3f | VSADNTVGR | -302 | -310 | 0 | 0.00025015 | 9 | 786.7322 | 173.47974 | 1051.7079 | 871.1095 | 1135.5737 | 1021.8476 |
| Eif3f | KVSPNELILGW | -166 | -176 | 1 | 0 | 11 | 686.6777 | 623.09595 | 646.73584 | 811.2245 | 1051.5331 | 1056.0176 |
| Eif3f | EAPNPIHLTVDTGLQHGR | -197 | -214 | 0 | 0 | 18 | 578.52924 | 922.62445 | 827.33057 | 887.3608 | 1136.7792 | 202.55154 |
| Eif3f | VDTGLQHGR | -206 | -214 | 0 | 0 | 9 | 1273.5902 | 1182.5162 | 1287.2346 | 999.63184 | 1389.8591 | NaN |
| Eif3f | QYAEDVLSGK | -292 | -301 | 0 | 0 | 10 | 1008.2325 | 1149.2831 | 794.4391 | 1133.2448 | NaN | NaN |
| Eif3f | YAYYDTER | -243 | -250 | 0 | 0 | 8 | 930.17816 | 829.9928 | 718.71643 | 665.5036 | NaN | NaN |
| Eif3f | EAPNPIHLT | -197 | -205 | 0 | 0.007819401 | 9 | 1228.9526 | 314.18195 | NaN | 1231.6064 | NaN | NaN |
| Eif3f | TCFSPNR | -259 | -265 | 0 | 0.001868731 | 7 | 775.44104 | 562.5499 | 454.0927 | NaN | NaN | NaN |
| Rps15 | TFNQVEIKPE | -101 | -110 | 1 | 0.00025015 | 10 | 2664.9504 | 475.36078 | 2626.172 | 2649.0342 | 1414.8507 | 297.04517 |
| Rps15 | EAPPMEKPEVVK | -66 | -77 | 1 | 0 | 12 | 2383.1682 | 2776.6147 | 2927.4614 | 2512.2856 | 3170.198 | 3373.4082 |
| Rps15 | EAPPMEKPEVVK | -66 | -77 | 1 | 0 | 12 | 2383.1682 | 2776.6147 | 2927.4614 | 2512.2856 | 3170.198 | 3373.4082 |
| Rps15 | HGRPGIGAT | -128 | -136 | 1 | 0.001023236 | 9 | 2248.1704 | 844.8574 | 2927.2922 | 3073.6455 | 2655.794 | 1564.9143 |
| Rps15 | TFNQVEIKPEMIGH | -101 | -114 | 1 | 0 | 14 | 1763.0978 | 2110.3909 | 1997.7404 | 2294.8608 | 2003.4498 | 2082.6033 |
| Rps15 | TFNQVEIKPEMIGH | -101 | -114 | 1 | 0 | 14 | 1763.0978 | 2110.3909 | 1997.7404 | 2294.8608 | 2003.4498 | 2082.6033 |
| Rps15 | SITYKPVK | -120 | -127 | 1 | 0.000230707 | 8 | 928.1455 | 1417.6632 | 605.1593 | 412.20987 | NaN | 582.6984 |
| Rps15 | KEAPPMEKPEVVK | -65 | -77 | 2 | 0 | 13 | 985.8526 | 1009.3432 | 555.15344 | NaN | 1002.8767 | NaN |
| Sh3kbp1 | SYLPQNDDELELK | -108 | -120 | 0 | 0 | 13 | 1115.0182 | 1376.8268 | 1096.0402 | 970.6962 | 965.27185 | 1033.5493 |
| Sh3kbp1 | SIEVENDFLPVEK | -274 | -286 | 0 | 0 | 13 | 822.4088 | 1003.8278 | 886.72076 | 741.49084 | 835.7588 | 705.2081 |
| Sh3kbp1 | KPPPPSAPVVK | -383 | -393 | 1 | 0.001055699 | 11 | 698.46155 | 925.4123 | 807.8775 | 629.3141 | 1011.2772 | 1023.7297 |
| Sh3kbp1 | RPERPVGPL | -465 | -473 | 2 | 0.001904399 | 9 | 418.0866 | 698.52466 | 760.0006 | 1225.8163 | 399.31497 | 697.5898 |
| Sh3kbp1 | SGESDELGISQDEQLSK | -156 | -172 | 0 | 0 | 17 | 1817.253 | 2086.0486 | NaN | 194.97678 | 2424.1423 | 2490.7715 |
| Sh3kbp1 | GVFPDNFVK | -360 | -368 | 0 | 0.00025015 | 9 | 998.65234 | 903.92914 | 990.91455 | 400.25998 | 895.04584 | NaN |
| Sh3kbp1 | FSYLPQNDDELELK | -107 | -120 | 0 | 0 | 14 | 720.91473 | 402.6884 | NaN | 807.5764 | 655.9201 | 732.8913 |
| Sh3kbp1 | EAQNDDELTIK | -323 | -333 | 0 | 0 | 11 | 695.1299 | 1377.0471 | NaN | 1869.8291 | 2628.7627 | 2677.4292 |
| Sh3kbp1 | LDLQKPSVPAIPPK | -432 | -445 | 1 | 0 | 14 | 527.89215 | 357.93597 | 401.8899 | 380.5331 | 352.39536 | NaN |
| Sh3kbp1 | ELSGESDELGISQDEQLSK | -154 | -172 | 0 | 0.00025015 | 19 | 744.8063 | NaN | NaN | NaN | NaN | NaN |
| Dnmt1 | NLDAPEPYR | -988 | -996 | 0 | 0 | 9 | 3094.3987 | 2745.519 | 2641.4468 | 2630.4568 | 2248.5718 | 2189.8167 |
| Dnmt1 | HIPLFPGSDWR | -1430 | -1440 | 0 | 0 | 11 | 1741.1929 | 2193.436 | 2061.5447 | 2871.8584 | 3248.4214 | 3404.162 |
| Dnmt1 | LSYCDYYRPR | -1255 | -1264 | 1 | 0.00025015 | 10 | 1480.8739 | 1666.347 | 1811.0593 | 1479.5728 | 1699.3713 | 1710.1273 |
| Dnmt1 | GHLCPVDTGLIEK | -423 | -435 | 0 | 0 | 13 | 1439.5381 | 1402.4364 | 1319.6597 | 1668.5396 | 1534.2816 | 1625.5472 |
| Dnmt1 | VLEQIEEVDGR | -914 | -924 | 0 | 0 | 11 | 1108.9303 | 1982.8523 | 1107.7592 | 1015.4203 | 1567.2739 | 969.17346 |
| Dnmt1 | LWYNQEYAR | -871 | -879 | 0 | 0 | 9 | 929.4124 | 902.7484 | 903.1158 | 450.5613 | 1099.6289 | 896.6801 |
| Dnmt1 | EGGTDPETTLPGAEDGK | -849 | -865 | 0 | 0 | 17 | 924.0011 | 827.3777 | 566.2325 | 1466.2638 | 642.9904 | 727.8072 |
| Dnmt1 | SYCDYYRPR | -1256 | -1264 | 1 | 0 | 9 | 705.05023 | 696.12585 | 257.2358 | 559.14813 | 856.7673 | 860.17584 |
| Dnmt1 | HYQPILR | -1407 | -1413 | 0 | 0 | 7 | 692.3601 | 388.8321 | 941.95056 | 1288.7811 | 1141.6243 | 1244.7084 |
| Dnmt1 | RFTEDSLLR | -550 | -558 | 1 | 0.000550307 | 9 | 683.6007 | 1742.77 | 2027.9247 | 1617.8472 | 2175.9465 | 2201.0488 |
| Dnmt1 | LPLFPEPLHVFAPR | -1327 | -1340 | 0 | 0 | 14 | 605.4939 | 658.19086 | 660.92303 | 598.2878 | 701.54395 | 697.2791 |
| Dnmt1 | LNNPGTTVFTEDCNVLLK | -1182 | -1199 | 0 | 0 | 18 | 571.8265 | 549.428 | 572.1759 | 672.6928 | 762.74347 | 825.13214 |
| Dnmt1 | EARPETHLDEDEDGKK | -261 | -276 | 2 | 0 | 16 | 401.08002 | 443.3589 | 450.93768 | 247.3325 | 387.01743 | 370.05414 |
| Dnmt1 | NFEDPPNHAR | -1098 | -1107 | 0 | 0 | 10 | 238.2852 | 1812.8247 | 1333.5388 | 583.1752 | 1405.7534 | 1683.2283 |
| Dnmt1 | WLSGFDGGEK | -471 | -480 | 0 | 0 | 10 | 188.06041 | 915.60455 | 1124.1422 | 1367.4498 | 1184.3894 | 1197.5377 |
| Dnmt1 | KLQYTFHDVK | -1457 | -1466 | 1 | 0 | 10 | 1734.4696 | 803.40625 | NaN | 1155.7007 | 1020.2761 | 1208.4629 |
| Dnmt1 | GSLPDHVR | -18 | -25 | 0 | 0.002068213 | 8 | 677.4456 | 697.6133 | 825.4051 | 920.76166 | NaN | 853.5977 |
| Dnmt1 | FFLLENVR | -1265 | -1272 | 0 | 0 | 8 | 663.81116 | 737.32385 | 741.26373 | NaN | 514.3202 | 701.84973 |
| Dnmt1 | THLDEDEDGKK | -266 | -276 | 1 | 0.00025015 | 11 | 287.24768 | NaN | 242.72993 | 317.57724 | 228.47575 | 32.062782 |
| Dnmt1 | FSDVQGR | -1056 | -1062 | 0 | 0.006677519 | 7 | 1345.633 | 1255.0828 | NaN | 605.60065 | 1382.6143 | NaN |
| Dnmt1 | GSHYQPILR | -1405 | -1413 | 0 | 0 | 9 | 1062.4425 | NaN | 462.04852 | NaN | 410.38312 | 482.99335 |
| Dnmt1 | GGTDPETTLPGAEDGK | -850 | -865 | 0 | 0.000627943 | 16 | 2619.914 | 615.50696 | 2720.9126 | NaN | NaN | NaN |
| Dnmt1 | VFTEDCNVLLK | -1189 | -1199 | 0 | 0 | 11 | 1616.3784 | 1694.1293 | 1850.6323 | NaN | NaN | NaN |
| Dnmt1 | LVMAGEVTNSLGQR | -1200 | -1213 | 0 | 0.000394361 | 14 | 925.01904 | 718.6047 | NaN | NaN | 840.84045 | NaN |
| Dnmt1 | HQVSEPKEPEAA | -1125 | -1136 | 1 | 0.00149795 | 12 | 839.5729 | 910.54987 | 882.0486 | NaN | NaN | NaN |
| Dnmt1 | KDPVNETLYPEHYR | -965 | -978 | 1 | 0.000230707 | 14 | 476.95874 | 505.42374 | 473.3689 | NaN | NaN | NaN |
| Dnmt1 | KDPVNETLYPE | -965 | -975 | 1 | 0.003966045 | 11 | 85.819336 | 2782.8362 | NaN | 3387.6553 | NaN | NaN |
| Dnmt1 | EQIEEVDGR | -916 | -924 | 0 | 0.001392869 | 9 | 5608.5063 | NaN | 5415.1626 | NaN | NaN | NaN |
| Dnmt1 | GSNLDAPEPYR | -986 | -996 | 0 | 0.002534621 | 11 | 1870.9799 | NaN | 1380.0596 | NaN | NaN | NaN |
| Dnmt1 | QVGNAVPPPLAK | -1577 | -1588 | 0 | 0.005333786 | 12 | 1103.5427 | 1223.8737 | NaN | NaN | NaN | NaN |
| Dnmt1 | QVAPETPEDRDEDEREEK | -303 | -320 | 2 | 0.000627943 | 18 | 346.40295 | NaN | 370.88196 | NaN | NaN | NaN |
| Dnmt1 | QNNPDAVYEDLINK | -523 | -536 | 0 | 0 | 14 | 237.30908 | NaN | 284.6444 | NaN | NaN | NaN |

| | | | | | | | | | | | | |
|---|---|---|---|---|---|---|---|---|---|---|---|---|
| Rnmt | GDVSEDEPSLGR | -97 | -108 | 0 | 0 | 12 | 2211.0525 | 1929.2451 | NaN | 2595.9502 | 2843.307 | 2306.4265 |
| Rnmt | EASETESFGNEIYTVK | -318 | -333 | 0 | 0 | 16 | 570.4199 | NaN | 637.87427 | 758.7318 | 164.68091 | 351.82642 |
| Rnmt | FIGTTPNSFELIR | -303 | -315 | 0 | 0 | 13 | 775.28784 | NaN | NaN | 532.66327 | 836.05994 | 1227.2441 |
| Rnmt | LNPGGYFIGTTPNSFELIR | -297 | -315 | 0 | 0.00025015 | 19 | 165.33885 | NaN | NaN | NaN | 202.09698 | 352.4517 |
| Rnmt | KLQPQDDEVPQK | -115 | -126 | 1 | 0.00025015 | 12 | 501.60547 | NaN | NaN | NaN | 298.08472 | NaN |
| Rnmt | SILIGEILEK | -170 | -179 | 0 | 0.00204437 | 10 | 205.39734 | NaN | NaN | NaN | NaN | NaN |
| Tomm6 | EASEVPDNVGDWLR | -16 | -29 | 0 | 0 | 14 | 776.99347 | 1069.4122 | 1031.5043 | 899.44 | 1115.1234 | 1127.4493 |
| Tomm6 | SEVPDNVGDWLR | -18 | -29 | 0 | 0 | 12 | 236.214 | 478.8979 | 643.6348 | 676.5746 | 920.77997 | 1212.7068 |
| Mrps12 | EAVCFIPGEGHTL | -90 | -102 | 0 | 0 | 13 | 880.05975 | NaN | 547.87244 | NaN | 836.449 | 555.8467 |
| Tsn | TKFPAEQYYR | -77 | -86 | 1 | 0 | 10 | 4665.0566 | 4597.2134 | 5333.5215 | 5109.9834 | 5429.356 | 5771.092 |
| Tsn | KVVQSLEQTAR | -26 | -36 | 1 | 0 | 11 | 3367.4087 | 4451.3975 | 4986.956 | 1070.8263 | 6592.54 | 5920.617 |
| Tsn | KVEEVVYDL | -204 | -212 | 1 | 0.00025015 | 9 | 2597.5474 | 2565.6265 | 2182.9329 | 1625.843 | 1008.5655 | 1907.4819 |
| Tsn | TFINELDSGFR | -172 | -182 | 0 | 0 | 11 | 1253.8733 | 1501.5355 | 1618.8938 | 615.9245 | 1727.2175 | 1754.7087 |
| Tsn | EAVTEILGIEPDREK | -117 | -131 | 1 | 0 | 15 | 473.79895 | 675.4088 | 439.2837 | 669.87006 | 325.51834 | 452.07034 |
| Tsn | FPAEQYYR | -79 | -86 | 0 | 0 | 8 | 2710.0051 | 2210.3635 | 2340.934 | NaN | 1288.176 | 932.54364 |
| Tsn | EHFSTVK | -63 | -69 | 0 | 0.000394361 | 7 | 1123.6504 | 880.9702 | NaN | 1127.122 | 826.6447 | 1645.7074 |
| Tsn | LETETLVTR | -108 | -116 | 0 | 0.00025015 | 9 | 1758.4545 | 2169.7605 | 1721.251 | NaN | NaN | NaN |
| Tsn | SVTAGDYSR | -158 | -166 | 0 | 0.000431655 | 9 | 612.2692 | NaN | 323.43494 | 841.8241 | NaN | NaN |
| Tsn | GFLAAEQDIR | -12 | -21 | 0 | 0 | 10 | 1562.6068 | NaN | NaN | NaN | NaN | NaN |
| Scly | EAWGNPSSSYVSGR | -44 | -57 | 0 | 0 | 14 | 775.86115 | 6081.4077 | 2869.5554 | NaN | NaN | 664.7819 |
| Stt3b | EAYFWLR | -583 | -589 | 0 | 0.00025015 | 7 | 496.58795 | 635.42474 | NaN | 549.6542 | 662.6584 | 473.06436 |
| Thyn1 | EAYPDHTQFEK | -133 | -143 | 0 | 0 | 11 | 1228.1797 | 1021.2989 | 563.01715 | 1151.4476 | 1230.7976 | 1290.6344 |
| Thyn1 | FIPLEELK | -173 | -180 | 0 | 0.00383302 | 8 | 6466.493 | NaN | 6676.705 | 4858.3374 | NaN | 3323.8728 |
| Thyn1 | SNPHYDPSSKEDDPK | -144 | -158 | 1 | 0.005946382 | 15 | 273.64157 | NaN | NaN | NaN | NaN | NaN |
| Stt3a | FLAEEGFYK | -59 | -67 | 0 | 0 | 9 | 2252.771 | 1496.5311 | 2489.1162 | 3841.531 | 5357.88 | 2830.6865 |
| Stt3a | FYSLLDPSYAK | -330 | -340 | 0 | 0 | 11 | 1896.3812 | 1855.6841 | 2059.7253 | 2411.1382 | 2671.4573 | 2666.918 |
| Stt3a | FHNWFDDR | -68 | -75 | 0 | 0.00122419 | 8 | 1454.6902 | 1402.7378 | 1451.4337 | 1198.6251 | 1189.2762 | 1173.6029 |
| Stt3a | IIFDDFR | -500 | -506 | 0 | 0.001846793 | 7 | 1246.3817 | 1293.7258 | 1340.6304 | 1237.3513 | 1116.6167 | 1117.7136 |
| Stt3a | NLDISRPDK | -432 | -440 | 1 | 0.000471409 | 9 | 755.8002 | 1192.4498 | 508.54407 | 864.17365 | 1173.172 | 583.2379 |
| Stt3a | EAYYWLR | -507 | -513 | 0 | 0.001392869 | 7 | 685.9437 | 727.65826 | 783.3331 | 777.08624 | 724.2996 | 668.16235 |
| Stt3a | LNPQQFEVLFR | -290 | -300 | 0 | 0 | 11 | 661.58936 | 564.3535 | 659.9593 | 1539.3115 | 978.34174 | 919.0003 |
| Stt3a | ENDYYTPTGEFR | -614 | -625 | 0 | 0 | 12 | 645.69214 | 879.85706 | 526.2307 | 1065.433 | 1092.0203 | 1218.4098 |
| Stt3a | EFDPYFNYR | -47 | -55 | 0 | 0 | 9 | 480.41525 | 467.2195 | 428.8054 | 296.01874 | 296.67096 | 331.16898 |
| Stt3a | GGLTGYSSDDINK | -583 | -595 | 0 | 0 | 13 | 1010.22 | NaN | 569.31 | 2331.7314 | 814.5389 | NaN |
| Stt3a | NLDISRPDKK | -432 | -441 | 2 | 0.000431655 | 10 | 1190.5388 | NaN | NaN | 226.93684 | 1820.6221 | NaN |
| Stt3a | IAVVPGYISR | -151 | -160 | 0 | 0 | 10 | 782.9816 | NaN | 837.54785 | NaN | NaN | NaN |
| Rpl23 | ECADLWPR | -124 | -131 | 0 | 0 | 8 | 4949.0947 | 4415.715 | 3847.9294 | 4357.5215 | 3648.1172 | 3748.0984 |
| Rpl23 | FEDNAGVIVNNK | -98 | -109 | 0 | 0.00025015 | 12 | 1342.6367 | 2306.6611 | 2075.5808 | 2119.3164 | 1918.1287 | 2290.7126 |
| Rpl23 | LGLPVGAVINCADNTGAK | -18 | -35 | 0 | 0 | 18 | 439.45264 | 499.83047 | 481.7269 | 1190.2222 | 1586.519 | 1490.9674 |
| Rpl23 | ISLGLPVGAVIN | -16 | -27 | 0 | 0.000230707 | 12 | 375.045 | 411.88715 | 360.67383 | 256.88028 | 341.91528 | 310.5755 |
| Rpl23 | EDNAGVIVNNK | -99 | -109 | 0 | 0 | 11 | 1053.6509 | 964.91956 | 895.3918 | 876.9828 | NaN | 929.18475 |
| Rpl23 | LYFEDNAGVIVNNK | -96 | -109 | 0 | 0 | 14 | 766.09985 | 1296.9263 | 742.8161 | NaN | 609.7041 | 1615.688 |
| Rpl23 | ISLGLPVGAVINC | -16 | -28 | 0 | 0.00025015 | 13 | 338.92365 | 451.24896 | 292.4568 | NaN | NaN | NaN |
| Rpl23 | VHPAVVIR | -76 | -83 | 0 | 0 | 8 | 620.6268 | NaN | NaN | NaN | NaN | NaN |
| Slc16a1 | FLDLSLFTHR | -245 | -254 | 0 | 0 | 10 | 3270.3845 | 3842.9634 | 3480.0232 | 3327.7004 | 652.212 | 7955.2563 |
| Slc16a1 | ECCPVLLGPPLLGR | -391 | -404 | 0 | 0 | 14 | 962.54663 | 938.3508 | 1128.7473 | 850.4502 | 1408.3639 | 1423.0364 |
| Slc16a1 | IVECCPVLLGPPLLGR | -389 | -404 | 0 | 0 | 16 | 647.4628 | 660.44507 | 614.05273 | 624.05725 | 644.0699 | 664.09344 |
| Slc16a1 | QSPQQHSSGDPTEEESPV | -476 | -493 | 0 | 0.000591124 | 18 | 634.8715 | 701.6869 | 679.27234 | 740.34686 | 815.67737 | 821.53314 |
| Slc16a1 | VTIVECCPVLLGPPLLGR | -387 | -404 | 0 | 0 | 18 | 272.10385 | 352.66327 | 266.57828 | 236.84093 | NaN | NaN |
| Ap2b1 | ILDCLSNYNPK | -222 | -232 | 0 | 0 | 11 | 4027.8591 | 4128.8677 | 4354.0103 | 461.55072 | 3465.9792 | 4052.4417 |
| Ap2b1 | SEISESHPNSNLLDLNPQNINK | -185 | -206 | 0 | 0 | 22 | 765.9712 | 790.8956 | 634.29114 | 811.5293 | 868.4878 | 606.15283 |
| Ap2b1 | ENLDSLDEPDAR | -434 | -445 | 0 | 0 | 12 | 1110.4349 | 3723.2405 | 611.736 | NaN | 2620.5332 | 2992.6865 |
| Ap2b1 | ECHLNADTVSSK | -856 | -867 | 0 | 0 | 12 | 1961.6638 | NaN | 2061.2195 | 1695.0208 | NaN | 1773.5417 |
| Ap2b1 | KPSETQELVQQ | -495 | -505 | 1 | 0.00111006 | 11 | 4648.628 | NaN | NaN | NaN | 840.2668 | NaN |
| Ap2b1 | SNLLDLNPQNINK | -194 | -206 | 0 | 0 | 13 | 595.78906 | NaN | NaN | NaN | NaN | NaN |
| Adh5 | MTDGGVDYSFECIGNVK | -257 | -273 | 0 | 0 | 17 | 3022.8938 | 3409.4177 | 3646.693 | 349.49475 | 2765.2244 | 2768.159 |
| Adh5 | MTDGGVDYSFECIGNVK | -257 | -273 | 0 | 0 | 17 | 3022.8938 | 3409.4177 | 3646.693 | 349.49475 | 2765.2244 | 2768.159 |
| Adh5 | KIDPSAPLDK | -159 | -168 | 1 | 0.001291103 | 10 | 785.3199 | 623.88495 | 478.9397 | 775.4095 | 830.8504 | 763.0067 |
| Adh5 | GHEGAGIVESVGEGVTK | -66 | -82 | 0 | 0 | 17 | 661.36945 | 727.29175 | 575.39325 | 1630.2358 | 1113.0989 | 1055.8086 |
| Adh5 | GASECISPQDFSK | -236 | -248 | 0 | 0 | 13 | 675.0382 | 613.2509 | 692.52576 | NaN | 504.96243 | 643.95825 |
| Adh5 | ECISPQDFSK | -239 | -248 | 0 | 0 | 10 | 402.31354 | NaN | 666.7421 | 1292.5892 | 695.66473 | 869.1635 |
| Nop9 | NQELIQDPFGHHVAR | -588 | -602 | 0 | 0 | 15 | 764.3236 | 1166.7195 | 1156.9701 | 530.11163 | 1249.6056 | 637.17474 |
| Nop9 | NSSADGSPLLLFLR | -311 | -324 | 0 | 0 | 14 | 229.23723 | 222.95198 | 185.97073 | 308.84943 | 266.2887 | 333.4057 |
| Nop9 | LYFSNPGLVLR | -494 | -504 | 0 | 0 | 11 | 330.52313 | NaN | NaN | 275.7224 | 242.01852 | 246.29353 |
| Nop9 | ECKPTDFEVPK | -241 | -251 | 1 | 0 | 11 | 1155.0013 | NaN | NaN | 803.3728 | 256.5507 | NaN |
| Mapkapk2 | REVELHWR | -89 | -96 | 1 | 0.00025015 | 8 | 410.7393 | 485.62878 | 303.69003 | NaN | 380.77176 | NaN |
| Mapkapk2 | ECLDGGELFSR | -125 | -135 | 0 | 0.000550307 | 11 | 589.24225 | 722.3201 | NaN | NaN | NaN | 340.88327 |
| Ifit3 | VCNQVEHLNSEEK | -38 | -50 | 0 | 0.00025015 | 13 | 787.2621 | 1185.051 | 506.90485 | 824.1276 | 993.6907 | 982.56757 |
| Ifit3 | ESPDVEEEDLYER | -352 | -364 | 0 | 0.00025015 | 13 | 601.5752 | 395.0674 | 1155.8693 | NaN | NaN | NaN |
| Ifit3 | ECLLQFPR | -382 | -389 | 0 | 0.006624942 | 8 | 838.9091 | NaN | 822.46436 | NaN | NaN | NaN |
| Mob4 | ILEPPEGQDEGVWK | -57 | -70 | 0 | 0 | 14 | 1262.8458 | 1526.3461 | 1487.0718 | 1693.2388 | 1799.419 | 1874.4849 |
| Mob4 | ECPAIDYTR | -118 | -126 | 0 | 0 | 9 | 740.79535 | 953.70667 | 860.0759 | 953.9916 | 894.85333 | 920.9985 |
| Cdca2 | ECVLNNSENDVFR | -15 | -27 | 0 | 0 | 13 | 376.74658 | NaN | NaN | NaN | NaN | NaN |
| Suz12 | NLIAPIFLH | -106 | -114 | 0 | 0 | 9 | 572.30304 | 626.92914 | 638.11 | 531.309 | 574.9171 | 603.9615 |
| Suz12 | NLIAPIFLHR | -106 | -115 | 0 | 0 | 10 | 538.6372 | 696.0441 | 672.9854 | 521.90857 | 718.3212 | 591.52484 |
| Suz12 | HGGGGGGGSGPSAGSGGGGFGGSAA | -6 | -30 | 0 | 0.001023236 | 25 | 376.84018 | 586.8712 | 66.62886 | 491.38358 | 443.4651 | 412.17593 |
| Suz12 | ECYDGSYAGNPQDIHR | -495 | -510 | 0 | 0 | 16 | 822.40533 | 1053.2212 | 945.922 | 1201.188 | 700.4516 | NaN |
| Suz12 | SQGPTLQFTLR | -352 | -362 | 0 | 0.00111006 | 11 | 709.8528 | NaN | NaN | NaN | NaN | NaN |

| | | | | | | | | | | | | |
|---|---|---|---|---|---|---|---|---|---|---|---|---|
| Ddx47 | LIDHLENTK | -153 | -161 | 0 | 0 | 9 | 1919.6083 | 1305.0377 | 1708.7032 | 1177.0336 | 1297.639 | 1053.5063 |
| Ddx47 | LQQYYLFIPSK | -238 | -248 | 0 | 0 | 11 | 899.15283 | 1063.103 | 1134.1635 | 760.201 | 997.36414 | 1039.2823 |
| Ddx47 | VTQYDVELFQR | -367 | -377 | 0 | 0 | 11 | 558.52234 | 602.4402 | 9066.788 | 4715.2686 | 3613.1404 | 537.65466 |
| Ddx47 | EDAGDDDDKEGAIGVR | -426 | -441 | 1 | 0 | 16 | 486.1118 | 549.01404 | 474.87164 | 500.40656 | 493.27344 | 502.96033 |
| Nop14 | FGEYNSNISPEEK | -88 | -100 | 0 | 0 | 13 | 441.54773 | 338.56226 | NaN | 309.63354 | 117.85329 | NaN |
| Nop14 | HTSADDLNDGFILD | -309 | -322 | 0 | 0.002296977 | 14 | 863.8701 | 981.552 | NaN | NaN | NaN | NaN |
| Nop14 | EDALELTEK | -210 | -218 | 0 | 0.00272025 | 9 | 564.2789 | NaN | NaN | NaN | NaN | NaN |
| Lsg1 | EDEDPYRPPTSEELLTAY | -482 | -499 | 1 | 0.00025015 | 18 | 294.79993 | 383.26843 | NaN | 343.03717 | 704.01245 | 615.23987 |
| Ndufab1 | SDAPPLTLDGIKDR | -69 | -82 | 1 | 0 | 14 | 3351.8035 | 3380.5046 | 3254.6675 | 3615.6335 | 3987.8442 | 4241.895 |
| Ndufab1 | EIVDYIADKK | -143 | -152 | 1 | 0.000230707 | 10 | 3199.6528 | 3273.2974 | 2636.4448 | 7562.5186 | 9443.202 | 9534.029 |
| Ndufab1 | SDAPPLTLDGIK | -69 | -80 | 0 | 0 | 12 | 3140.7212 | 3126.6887 | 3352.2869 | 3903.7615 | 3473.766 | 3971.846 |
| Ndufab1 | EDEFGFEIPDIDAEK | -123 | -137 | 0 | 0 | 15 | 720.6192 | 690.38257 | 527.535 | 520.728 | 931.24634 | 771.7517 |
| Ndufab1 | LYDKIDPEK | -89 | -97 | 1 | 0.00025015 | 9 | 1201.4419 | NaN | 1355.13 | NaN | NaN | NaN |
| Gmfg | TTDDLTETWLK | -125 | -135 | 0 | 0 | 11 | 1378.1241 | 1454.8938 | 599.64703 | 863.9985 | 599.84985 | 1812.9027 |
| Gmfg | EDELQNISPEELK | -46 | -58 | 0 | 0 | 13 | 1084.6815 | 1034.6262 | NaN | 783.8139 | 1059.8276 | NaN |
| Ola1 | IGIVGLPNVGK | -25 | -35 | 0 | 0 | 11 | 962.69995 | 1123.2391 | 1096.9342 | 908.77515 | 886.93304 | 984.2307 |
| Ola1 | EDFKDEGSENAVK | -351 | -363 | 1 | 0 | 13 | 756.9785 | 1488.2322 | 2039.0018 | 2099.2478 | 2922.2512 | 4313.6387 |
| Ola1 | KGGDGIKPPPIIGR | -5 | -18 | 2 | 0.00025015 | 14 | 102.6024 | 1103.4857 | 1219.1631 | 756.23224 | 1135.4716 | 1091.192 |
| Ola1 | YEDFKDEGSENAVK | -350 | -363 | 1 | 0.002099344 | 14 | 3262.1619 | 3516.515 | 3412.6816 | 4337.441 | 5149.6436 | NaN |
| Ola1 | IHTDFEK | -334 | -340 | 0 | 0 | 7 | 1086.4513 | 1566.3173 | 1299.1179 | 1733.2347 | NaN | 1239.4805 |
| Ola1 | KYEDFKDEGSENAVK | -349 | -363 | 2 | 0.00025015 | 15 | 1065.9122 | NaN | 813.20734 | 527.4838 | 376.6578 | 1385.7279 |
| Ola1 | FYHDWNDKEIEVL | -202 | -214 | 1 | 0.00025015 | 13 | 507.2778 | NaN | NaN | 709.09326 | 349.3154 | 511.6855 |
| Ola1 | VPVPDER | -64 | -70 | 0 | 0.00025015 | 7 | 975.1113 | 855.2655 | NaN | NaN | 698.51184 | NaN |
| Ola1 | LQELSAEER | -271 | -279 | 0 | 0 | 9 | 196.8197 | NaN | 709.45654 | NaN | NaN | NaN |
| Ctsb | EQWSNCPTIGQIR | -88 | -100 | 0 | 0 | 13 | 3192.0151 | 3137.9248 | 3265.3743 | 3326.492 | 2855.788 | 2231.184 |
| Ctsb | IDLPETFDAR | -78 | -87 | 0 | 0 | 10 | 808.56555 | 265.11194 | 799.8846 | 349.10785 | 744.7529 | 947.6361 |
| Ctsb | SCEAGYSPSYKEDK | -210 | -223 | 1 | 0 | 14 | 745.204 | 661.25946 | 713.60156 | NaN | 877.849 | 854.21436 |
| Ctsb | LPETFDAR | -80 | -87 | 0 | 0.001392869 | 8 | 2132.8643 | NaN | NaN | NaN | NaN | 3705.2563 |
| Ctsb | EDIDLPETFDAR | -76 | -87 | 0 | 0.00078661 | 12 | 1183.7322 | NaN | NaN | NaN | 1296.3285 | NaN |
| Ctsb | SCEAGYSPSYK | -210 | -220 | 0 | 0.000431655 | 11 | 671.7786 | NaN | NaN | NaN | NaN | NaN |
| Stk17b | EDKENIPEDGSLVSK | -338 | -352 | 1 | 0 | 15 | 336.5767 | 558.1741 | 439.4588 | 906.67487 | 478.04556 | 389.90332 |
| Tpm3 | VIIEGDLER | -171 | -179 | 0 | 0.00051306 | 9 | 2124.3535 | 2139.0042 | 2096.2737 | 440.24136 | 2553.607 | 2351.3137 |
| Tpm3 | KLVIIEGDLER | -169 | -179 | 1 | 0.000230707 | 11 | 1801.8048 | 2128.299 | 2086.8516 | 1356.7793 | 692.40936 | 1719.229 |
| Tpm3 | HIAEEADR | -154 | -161 | 0 | 0.00025015 | 8 | 1519.4421 | 1158.8494 | 1338.4786 | 1043.9092 | 968.7135 | 1046.1316 |
| Tpm3 | LVIIEGDLER | -170 | -179 | 0 | 0.000230707 | 10 | 920.20935 | 1157.1074 | NaN | NaN | NaN | NaN |
| Tpm3 | TIDDLEDELYAQK | -253 | -265 | 0 | 0 | 13 | 677.86646 | 595.04535 | NaN | NaN | NaN | NaN |
| Ssb | EDLHFLFSNHGEIK | -246 | -259 | 0 | 0 | 14 | 3166.0427 | 6967.5312 | 8364.425 | 4735.8794 | 11588.731 | 10081.494 |
| Ssb | KFVEIPGQK | -166 | -174 | 1 | 0 | 9 | 2436.3389 | 2470.696 | 2731.5479 | 2660.065 | 3001.7134 | 2687.6672 |
| Ssb | FSGDLDDQTCR | -235 | -245 | 0 | 0 | 11 | 2115.892 | 2023.5092 | 2326.7231 | 2411.2817 | 71.26615 | 66.60305 |
| Ssb | FVEIPGQK | -167 | -174 | 0 | 0 | 8 | 1645.622 | 1442.0122 | 1558.4629 | 1740.7471 | 1781.4489 | 1738.9707 |
| Ssb | SPSRPLPEVTDEYK | -92 | -105 | 1 | 0.000230707 | 14 | 1585.3724 | 1749.1731 | 1577.1036 | 1872.6918 | 2004.0577 | 2034.0073 |
| Ssb | SPSRPLPEVTDEYK | -92 | -105 | 1 | 0.000230707 | 14 | 1585.3724 | 1749.1731 | 1577.1036 | 1872.6918 | 2004.0577 | 2034.0073 |
| Ssb | GFPTDATLDDIK | -117 | -128 | 0 | 0 | 12 | 1099.764 | 1039.0203 | 427.72864 | 1010.4032 | 1020.692 | 269.00525 |
| Ssb | KFSGDLDDQTCR | -234 | -245 | 1 | 0 | 12 | 694.9614 | 936.50305 | 896.58545 | 912.77563 | 1135.0388 | 214.73532 |
| Ssb | KFSGDLDDQTCR | -234 | -245 | 1 | 0 | 12 | 694.9614 | 936.50305 | 896.58545 | 912.77563 | 1135.0388 | 214.73532 |
| Ssb | LDEGWVPLETMIK | -42 | -54 | 0 | 0 | 13 | 647.1106 | 660.5033 | 724.851 | 694.5696 | 996.9374 | 805.8303 |
| Ssb | LDEGWVPLETMIK | -42 | -54 | 0 | 0 | 13 | 647.1106 | 660.5033 | 724.851 | 694.5696 | 996.9374 | 805.8303 |
| Ssb | HQIEYYFGDFNLPR | -19 | -32 | 0 | 0 | 14 | 514.7338 | 512.9177 | 682.33905 | 1047.1478 | 871.69714 | 715.3781 |
| Ssb | ITDDQQESLNK | -317 | -327 | 0 | 0 | 11 | 346.8649 | 737.65326 | 177.37863 | 2309.2742 | 1051.4896 | 1955.7375 |
| Ssb | GNRPGYAGAPK | -357 | -367 | 1 | 0 | 11 | 208.40366 | 770.624 | 837.0516 | 747.69495 | 661.4987 | 748.0614 |
| Ssb | RSPSRPLPEVTDEYK | -91 | -105 | 2 | 0.000627943 | 15 | 741.3974 | NaN | 756.9966 | 1125.2767 | 1146.5673 | 429.32584 |
| Ssb | YKDTNLLILFK | -175 | -185 | 1 | 0 | 11 | 725.9687 | 724.99615 | 816.0611 | NaN | 1568.2883 | 1697.7489 |
| Ssb | SPSRPLPEVTDEYKNDVK | -92 | -109 | 2 | 0 | 18 | 702.1034 | 753.4744 | NaN | 1317.0787 | 1080.4596 | 904.85266 |
| Ssb | EWLDDKGQIL | -129 | -138 | 1 | 0.002466863 | 10 | 249.43568 | NaN | 367.2433 | 238.383 | 1094.4368 | 1043.322 |
| Ssb | KITDDQQESLNK | -316 | -327 | 1 | 0 | 12 | 972.07324 | 1121.9517 | NaN | 1032.9022 | NaN | 1473.6731 |
| Ssb | FSNHGEIK | -252 | -259 | 0 | 0.002780285 | 8 | 429.07755 | 391.568 | 437.20477 | NaN | 799.2157 | NaN |
| Ssb | FGDFNLPR | -25 | -32 | 0 | 0 | 8 | 932.1991 | 859.86835 | 911.54834 | NaN | NaN | NaN |
| Ssb | SGDLDDQTCR | -236 | -245 | 0 | 0 | 10 | 411.86346 | NaN | NaN | 388.7962 | NaN | NaN |
| Ssb | YKDTNLLILF | -175 | -184 | 1 | 0.000471409 | 10 | 342.0726 | NaN | NaN | NaN | 276.44113 | NaN |
| Eif5a | EDLRLPEGDLGK | -110 | -121 | 1 | 0.00025015 | 12 | 9082.025 | 9619.6875 | 9790.762 | 11365.267 | 12737.929 | 12207.456 |
| Eif5a | EDLRLPEGDLGK | -110 | -121 | 1 | 0.00025015 | 12 | 9082.025 | 9619.6875 | 9790.762 | 11365.267 | 12737.929 | 12207.456 |
| Eif5a | SAMTEEAAVAIK | -139 | -150 | 0 | 0 | 12 | 1249.8613 | 512.1216 | 1108.7334 | 441.2926 | 1312.9275 | 1198.4352 |
| Eif5a | SAMTEEAAVAIK | -139 | -150 | 0 | 0 | 12 | 1249.8613 | 512.1216 | 1108.7334 | 441.2926 | 1312.9275 | 1198.4352 |
| Eif5a | RNDFQLIGIQ | -86 | -95 | 1 | 0.000230707 | 10 | 1494.326 | 421.365 | 1727.2095 | 1493.1118 | 1569.7833 | 1485.124 |
| Eif5a | EDLRLPEGD | -110 | -118 | 1 | 0.00078661 | 9 | 976.36316 | 1033.7595 | 240.65973 | 974.62134 | 1098.5072 | 911.3363 |
| Eif5a | YDCGEEILIT | -127 | -136 | 0 | 0.00111006 | 10 | 807.1518 | 1003.1628 | 810.874 | 1115.5981 | 1373.2532 | 1230.1599 |
| Eif5a | RNDFQLIGIQDGYLS | -86 | -100 | 1 | 0.001291103 | 15 | 327.88757 | 303.85007 | 288.36465 | 467.47577 | 601.7692 | 233.61905 |
| Eif5a | RNDFQLIGIQD | -86 | -96 | 1 | 0.00025015 | 11 | 186.98407 | 1147.2086 | 1354.7297 | NaN | NaN | 1061.5898 |
| Eif5a | LPEGDLGK | -114 | -121 | 0 | 0.00659934 | 8 | 2739.6643 | NaN | 2078.9836 | 1735.1842 | NaN | NaN |
| Eif5a | LLQDSGEVR | -101 | -109 | 0 | 0.00025015 | 9 | 1673.401 | NaN | 872.76996 | NaN | NaN | 766.06696 |
| Eif5a | LRLPEGDLGK | -112 | -121 | 1 | 0.001392869 | 10 | 789.1566 | 710.7238 | 139.37976 | NaN | NaN | NaN |
| Eif5a | TEEAAVAIK | -142 | -150 | 0 | 0 | 9 | 1802.5864 | NaN | NaN | NaN | NaN | NaN |
| Sem1 | EDNWDDDNVEDDFSNQLR | -40 | -57 | 0 | 0 | 18 | 597.4452 | 564.0488 | 676.5711 | 659.5326 | 136.64195 | 82.01507 |
| Prex1 | EDPWNLPSSIR | -1271 | -1281 | 0 | 0.000230707 | 11 | 17118.145 | 16400.836 | 16691.262 | 7884.531 | 7151.0684 | 7397.973 |
| Prex1 | EFKDESQYFR | -580 | -589 | 1 | 0.004224523 | 10 | 183.58348 | NaN | NaN | NaN | NaN | NaN |
| Dock10 | EDQLEYQEELR | -2121 | -2131 | 0 | 0 | 11 | 1713.3114 | 231.72527 | 1507.3721 | 1149.5406 | 915.7704 | 258.99268 |
| Dock10 | NIPDPLTPSESIR | -1119 | -1131 | 0 | 0 | 13 | 884.5691 | 108.34318 | 715.30914 | 969.6177 | 830.2527 | 922.4494 |
| Dock10 | YSTVPEEAEHR | -93 | -103 | 0 | 0 | 11 | 573.20465 | 421.72156 | 556.6956 | 739.1366 | 474.55167 | 323.54578 |

| | | | | | | | | | | | | |
|---|---|---|---|---|---|---|---|---|---|---|---|---|
| Dock10 | IELPTQLHGK | -748 | -757 | 0 | 0 | 10 | 1538.5892 | 1171.2596 | 841.27893 | 1227.2344 | 1223.6938 | NaN |
| Dock10 | FHVDLNHPAVR | -442 | -452 | 0 | 0 | 11 | 358.3446 | 314.49747 | 245.63095 | NaN | 336.84995 | 365.70212 |
| Dock10 | ILQISPEGPLQGR | -288 | -300 | 0 | 0 | 13 | 1440.6172 | 1546.3243 | 1693.7081 | NaN | NaN | 1225.4116 |
| Dock10 | TLYSTVPEEAEHR | -91 | -103 | 0 | 0 | 13 | 891.68787 | 123.10676 | 1398.1825 | NaN | NaN | NaN |
| Dock10 | VTYVTPFFEEK | -1949 | -1959 | 0 | 0.005668207 | 11 | 698.44305 | NaN | NaN | NaN | NaN | 1328.6761 |
| Dock10 | SQTLPIIR | -1438 | -1445 | 0 | 0.002296977 | 8 | 300.78207 | NaN | NaN | NaN | NaN | NaN |
| Dock10 | YVTPFFEEK | -1951 | -1959 | 0 | 0.006537474 | 9 | 197.11484 | NaN | NaN | NaN | NaN | NaN |
| Jpt1 | SLGFDEPAEQPVR | -31 | -43 | 0 | 0 | 13 | 4940.84 | 5181.513 | 5641.087 | 5732.085 | 6490.614 | 1778.8257 |
| Jpt1 | IFGTPEENPPSWAK | -51 | -64 | 0 | 0 | 14 | 933.78296 | 642.1231 | 988.64136 | 925.1443 | 995.5237 | 936.9525 |
| Jpt1 | SLGFDEPAEQPVRK | -31 | -44 | 1 | 0 | 14 | 818.94214 | 1405.9688 | 1359.9203 | 1277.7551 | 1276.7946 | 1002.8925 |
| Jpt1 | MASNIFGTPEENPPSWAK | -47 | -64 | 0 | 0 | 18 | 207.2205 | 443.40637 | 438.1249 | 455.01672 | 731.07825 | 509.05273 |
| Jpt1 | VLRPPGGGSNF | -20 | -30 | 1 | 0.000230707 | 11 | 1868.5289 | 1333.4418 | 1388.2443 | NaN | 2295.3086 | 2302.4973 |
| Jpt1 | SNSSEASSGDFLDLK | -85 | -99 | 0 | 0 | 15 | 1211.6012 | 1377.7297 | 1327.4052 | 2078.0713 | NaN | 171.01694 |
| Jpt1 | GEGDMHENVDTDFQAN | -100 | -115 | 0 | 0 | 16 | 394.6228 | 412.8215 | 360.77017 | 451.12778 | NaN | 387.3822 |
| Jpt1 | LAQMEEKPVPAAPVPSPVAPAPVPSR | -116 | -141 | 1 | 0.001392869 | 26 | 545.5716 | NaN | 550.4666 | NaN | 420.2567 | 635.6362 |
| Jpt1 | EDSESPGTQR | -75 | -84 | 0 | 0 | 10 | 374.4355 | 685.6168 | NaN | NaN | NaN | 432.3601 |
| Septin6 | STVGFGDQINK | -101 | -111 | 0 | 0 | 11 | 1666.2826 | 1199.4385 | 2146.9421 | 828.7594 | 1145.4387 | 1347.9719 |
| Septin6 | SNTYDLQESNVGLK | -83 | -96 | 0 | 0 | 14 | 986.44086 | 3714.4116 | 1064.2578 | 932.8055 | 1085.2115 | 1140.752 |
| Septin6 | VENEAHCDFVK | -263 | -273 | 0 | 0 | 11 | 976.5745 | 1136.4916 | 1305.7037 | 1027.854 | 1224.9274 | 1177.9526 |
| Septin6 | TVPLAGHVGFDSLPDQLVNK | -16 | -35 | 0 | 0 | 20 | 906.21484 | 916.8583 | 916.8549 | 3871.6182 | 1219.5087 | 1290.285 |
| Septin6 | TVPLAGHVGFDSLPDQLVNK | -16 | -35 | 0 | 0 | 20 | 906.21484 | 916.8583 | 916.8549 | 3871.6182 | 1219.5087 | 1290.285 |
| Septin6 | FEGEPATHTQPGVQLQ | -67 | -82 | 0 | 0 | 16 | 499.39954 | 778.7795 | 791.3409 | 657.12427 | 564.6031 | 734.13257 |
| Septin6 | EDSYKPIVEFIDAQ | -112 | -125 | 1 | 0 | 14 | 484.7551 | 374.96707 | 436.60767 | 288.11847 | 629.0803 | 516.9159 |
| Septin6 | TYDLQESNVGLK | -85 | -96 | 0 | 0 | 12 | 951.32416 | 736.6876 | 745.32837 | 304.412 | NaN | NaN |
| Ly9 | NQVYDLVTHQDIAHAL | -558 | -573 | 0 | 0.000550307 | 16 | 601.406 | 572.4766 | 686.16675 | 709.333 | 763.42377 | 907.53546 |
| Ly9 | LQKPQIIVE | -154 | -162 | 1 | 0.002434616 | 9 | 214.34756 | NaN | 199.25053 | 116.267265 | NaN | 270.90955 |
| Ly9 | KTAQTPQQDAESPETPTYENFT | -633 | -654 | 1 | 0 | 22 | 168.43288 | 209.39041 | NaN | 135.158 | NaN | NaN |
| Ly9 | EDTHLNTYDGSHTL | -194 | -207 | 0 | 0.001392869 | 14 | 288.26343 | NaN | NaN | NaN | NaN | NaN |
| Ppme1 | SNLFLSCPIPK | -306 | -316 | 0 | 0 | 11 | 1518.021 | 1646.4047 | 2412.159 | 1274.0052 | 1365.6215 | 816.4196 |
| Ppme1 | EDVEVENETGKDTFR | -54 | -68 | 1 | 0 | 15 | 535.3176 | 496.82678 | 485.98935 | 507.61688 | 495.23883 | 494.45514 |
| Ppme1 | KDHPYTWR | -281 | -288 | 1 | 0 | 8 | 3783.7686 | 3306.2812 | NaN | NaN | NaN | 1304.1704 |
| Szrd1 | EEAADSGEIDR | -11 | -21 | 0 | 0.003393785 | 11 | 807.6262 | 888.76306 | NaN | NaN | NaN | NaN |
| Mob1b;Mob1 | ELAPLQELIEK | (200);(200) | (210);(210) | 0 | 0 | 11 | 2258.2427 | 2458.906 | 2570.4478 | 2011.9563 | 2526.777 | 2569.159 |
| Mob1b;Mob1 | WVQDQLDDETLFPSK | (121);(121) | (135);(135) | 0 | 0 | 15 | 529.8148 | 630.92487 | 556.23816 | 828.25134 | 895.2303 | 838.5738 |
| Mob1b;Mob1 | EEAHLNTSFK | (175);(175) | (184);(184) | 0 | 0 | 10 | 1920.1431 | 1313.4136 | 1134.7366 | NaN | 723.60864 | NaN |
| Mob1b;Mob1 | VQDQLDDETLFPSK | (122);(122) | (135);(135) | 0 | 0 | 14 | 368.1119 | 548.29706 | 568.8202 | NaN | NaN | NaN |
| Mob1b;Mob1 | IGVPFPK | (136);(136) | (142);(142) | 0 | 0.004983786 | 7 | 579.8222 | 318.39035 | NaN | NaN | NaN | NaN |
| Mob1b;Mob1 | KNIPEGSHQYELL | (17);(17) | (29);(29) | 1 | 0.00111006 | 13 | 162.00343 | NaN | NaN | NaN | NaN | 248.19995 |
| Usp7 | QKDDPENDNSELPTAK | -770 | -785 | 1 | 0 | 16 | 364.86728 | 462.4818 | 474.23447 | 489.55228 | 573.3075 | 336.58908 |
| Usp7 | FDDDVVSR | -481 | -488 | 0 | 0 | 8 | 1308.2183 | 1372.3188 | 1429.7313 | NaN | 1040.3032 | 1052.9082 |
| Usp7 | HQYINEDEYEVN | -1049 | -1060 | 0 | 0.00025015 | 12 | 762.5167 | 831.7616 | 416.93417 | 250.46793 | NaN | 1135.999 |
| Usp7 | HQYINEDEYEVNLK | -1049 | -1062 | 0 | 0.003360691 | 14 | 717.3977 | 692.435 | 690.6394 | NaN | 858.4592 | 1140.0317 |
| Usp7 | YDAGEHGLQEAEK | -380 | -392 | 0 | 0.000826806 | 13 | 7931.627 | 8567.187 | NaN | NaN | NaN | 7564.6323 |
| Usp7 | EEAIEHNYGGHDDDLSVR | -492 | -509 | 0 | 0.001776473 | 18 | 437.80347 | NaN | NaN | NaN | NaN | NaN |
| Tcerg1 | TLESTWEKPQELK | -454 | -466 | 1 | 0 | 13 | 1958.6411 | 2068.3406 | 2135.381 | 1979.3761 | 2414.5593 | 2445.6707 |
| Tcerg1 | ESAWTKPDGVK | -156 | -166 | 1 | 0 | 11 | 1774.5151 | 1691.9071 | 927.69586 | 1034.3916 | 1744.3535 | 1592.2883 |
| Tcerg1 | VFFYNPTTR | -546 | -554 | 0 | 0.000591124 | 9 | 1262.8 | 1184.6213 | 525.66833 | 654.91455 | 1466.1652 | 1320.0564 |
| Tcerg1 | YLVLDCVPEER | -1059 | -1069 | 0 | 0 | 11 | 1038.5338 | 629.59955 | 590.60474 | 2688.9905 | 2862.7205 | 2922.006 |
| Tcerg1 | EEAIQNFK | -899 | -906 | 0 | 0 | 8 | 1623.8167 | NaN | 1233.1993 | NaN | NaN | NaN |
| Nol12 | EEALEEADELER | -77 | -88 | 0 | 0.00111006 | 12 | 707.31134 | NaN | NaN | 714.9961 | NaN | NaN |
| Atp1b3 | EECSGVTDANFGYSK | -152 | -166 | 0 | 0 | 15 | 2705.6208 | 3208.8242 | 3453.0237 | 3570.949 | 3929.3513 | 4223.0996 |
| Atp1b3 | LFIYNPSSGEFLGR | -18 | -31 | 0 | 0 | 14 | 1246.6433 | 1206.7483 | 1330.3431 | 466.3003 | 1705.7544 | 1638.2849 |
| Atp1b3 | LVEDLESFLKPY | -106 | -117 | 1 | 0.00025015 | 12 | 465.80954 | 498.26474 | 541.2236 | 510.9859 | 162.09491 | 1152.5718 |
| Atp1b3 | IIDLIPDGYPQIS | -178 | -190 | 0 | 0 | 13 | 454.0924 | 212.95905 | 253.70767 | 398.1909 | 351.78427 | 355.96613 |
| Atp1b3 | IIDLIPDGYPQISCLPK | -178 | -194 | 0 | 0.00272025 | 17 | 348.41434 | 537.0683 | 202.03116 | 557.9373 | 703.8887 | 832.8886 |
| Atp1b3 | IYNPSSGEFLGR | -20 | -31 | 0 | 0 | 12 | 2833.997 | NaN | 1199.5173 | 2677.9707 | 2779.8416 | 2803.8896 |
| Atp1b3 | FDSGLNKK | -236 | -243 | 1 | 0 | 8 | 970.42676 | NaN | NaN | 688.0581 | NaN | NaN |
| Pak2 | SVIDPIPAPVGDSNVDSGAK | -197 | -216 | 0 | 0 | 20 | 3470.41 | 3947.7297 | 3765.4727 | 3866.5044 | 4037.4604 | 4109.4565 |
| Pak2 | SDNGELEDKPPAPPVR | -2 | -17 | 1 | 0.00111006 | 16 | 1641.0654 | 1693.5858 | 1667.6757 | 2225.3564 | 1942.2968 | 2000.2643 |
| Pak2 | SVIDPIPAPVGDSNVD | -197 | -212 | 0 | 0 | 16 | 1085.4875 | 1295.9554 | 1220.1571 | 271.97614 | 91.58397 | 1633.6617 |
| Pak2 | HSLKPLPSVPEEK | -35 | -47 | 1 | 0 | 13 | 976.8612 | 1121.3893 | 1109.0151 | 1125.4092 | 1216.2661 | 1203.201 |
| Pak2 | HSLKPLPSVPEEK | -35 | -47 | 1 | 0 | 13 | 976.8612 | 1121.3893 | 1109.0151 | 1125.4092 | 1216.2661 | 1203.201 |
| Pak2 | ERPEISPPSDFEHT | -70 | -83 | 1 | 0.000230707 | 14 | 916.2809 | 1015.38116 | 1146.6656 | 988.84186 | 767.3963 | 833.34454 |
| Pak2 | EEDDDDEDAAPPVIAPR | -170 | -186 | 0 | 0 | 17 | 268.31216 | 202.37798 | 330.06747 | NaN | 510.04773 | 403.63513 |
| Pak2 | YLSFTPPEK | -139 | -147 | 0 | 0 | 9 | 2366.526 | 145.0946 | NaN | NaN | NaN | NaN |
| Uqcrb | YEEDKFYLEPYLK | -84 | -96 | 1 | 0 | 13 | 2579.294 | 3082.5312 | 3249.2678 | 1455.0358 | 2736.7551 | 4279.049 |
| Uqcrb | EEDKFYLEPYLK | -85 | -96 | 1 | 0 | 12 | 1900.0135 | 1107.4547 | 1627.251 | 2263.3208 | 2491.4116 | 446.35608 |
| Uqcrb | YEEDKFYLEPY | -84 | -94 | 1 | 0 | 11 | 728.61316 | 646.66766 | 670.6911 | 706.8776 | 746.98047 | 870.93616 |
| Uqcrb | RDDTLHETEDVK | -34 | -45 | 1 | 0 | 12 | 1007.1804 | NaN | 1174.0986 | 1014.5116 | 1160.1482 | 1225.6462 |
| Slc2a3 | EEDQATEILQR | -218 | -228 | 0 | 0 | 11 | 1174.9578 | 981.56885 | 316.24203 | 765.02136 | 649.9452 | 742.58417 |
| Slc2a3 | KKEEDQATEILQR | -216 | -228 | 2 | 0 | 13 | 925.2561 | 1096.3562 | 1181.2756 | 1060.7823 | 1376.7792 | 1827.0399 |
| Slc2a3 | KKEEDQATEILQR | -216 | -228 | 2 | 0 | 13 | 925.2561 | 1096.3562 | 1181.2756 | 1060.7823 | 1376.7792 | 1827.0399 |
| Slc2a3 | KEEDQATEILQR | -217 | -228 | 1 | 0 | 12 | 583.7112 | 589.4688 | 730.2141 | 865.7711 | 745.87665 | NaN |
| Slc2a3 | GVINAPETILK | -29 | -39 | 0 | 0.00078661 | 11 | 494.1214 | 912.74756 | 1067.6118 | 643.45386 | NaN | 1712.8042 |
| Spag7 | HGEEWDPQKAEEK | -123 | -135 | 1 | 0.000627943 | 13 | 866.6571 | 135.38185 | 1052.449 | 850.2938 | 1062.3093 | 1104.1469 |
| Spag7 | EFAPSDEELDSYR | -110 | -122 | 0 | 0 | 13 | 604.42267 | 702.308 | 738.6601 | 614.73254 | 598.6551 | 790.248 |
| Spag7 | EVSDFIQDSGQVK | -55 | -67 | 0 | 0 | 13 | 846.2013 | NaN | NaN | 1264.7689 | NaN | 1279.7397 |
| Spag7 | SSMEKPPSLGDQESR | -10 | -24 | 1 | 0.004472887 | 15 | 542.1085 | 402.66483 | NaN | NaN | NaN | NaN |

| | | | | | | | | | | | | |
|---|---|---|---|---|---|---|---|---|---|---|---|---|
| Spag7 | EEEAAQQGPAVVSPASDYK | -146 | -164 | 0 | 0.00111006 | 19 | 519.2333 | NaN | NaN | NaN | NaN | NaN |
| Sec13 | EEEDGQWKEEQK | -196 | -207 | 1 | 0.00025015 | 12 | 427.38715 | 259.9624 | 598.3454 | 304.1438 | 529.681 | 506.8226 |
| Sec13 | FNDVVWHVS | -261 | -269 | 0 | 0 | 9 | 1419.6172 | 974.8327 | 1027.7097 | 1318.1031 | 1489.8032 | NaN |
| Sec13 | LEAHSDWVR | -208 | -216 | 0 | 0 | 9 | 493.82538 | 146.28943 | 644.24713 | NaN | 740.61536 | 378.1167 |
| Sec13 | ESVDGQWVCISDVNK | -291 | -305 | 0 | 0.000230707 | 15 | 338.1385 | NaN | 512.628 | 835.4742 | 826.90283 | 841.5476 |
| Sec13 | TYTGEGQWEVK | -133 | -143 | 0 | 0 | 11 | 1293.5872 | NaN | 132.02701 | NaN | 1435.9581 | 1128.8157 |
| Sec13 | TYTGEGQWEVK | -133 | -143 | 0 | 0 | 11 | 1293.5872 | NaN | 132.02701 | NaN | 1435.9581 | 1128.8157 |
| Sec13 | FNDVVWHV | -261 | -268 | 0 | 0 | 8 | 661.6738 | 679.07996 | 162.70355 | 601.0397 | NaN | NaN |
| Sec13 | KINNAHTIGCN | -144 | -154 | 1 | 0.000627943 | 11 | 869.4611 | NaN | NaN | 287.02213 | NaN | 821.41705 |
| Sec13 | WVCISDVNK | -297 | -305 | 0 | 0.002697088 | 9 | 893.06354 | NaN | NaN | NaN | NaN | 723.30927 |
| Mbd3 | GKPDLNTALPVR | -91 | -102 | 1 | 0 | 12 | 269.9677 | 678.09875 | 722.47144 | 881.7531 | 1040.7899 | 946.4613 |
| Mbd3 | EEEEEEEEEEPEPERV | -269 | -285 | 1 | 0.00025015 | 17 | 232.34328 | 410.7386 | 518.22626 | 410.96768 | 287.50204 | 281.4818 |
| Mbd3 | KQEELVQQVR | -227 | -236 | 1 | 0 | 10 | 894.21826 | 996.37494 | NaN | 1406.3156 | 1554.94 | 2192.355 |
| Uqcrh | ERLELCDNR | -46 | -54 | 1 | 0 | 9 | 2867.5474 | 2753.0918 | 2696.799 | 2994.1865 | 3058.589 | 2753.353 |
| Uqcrh | GDPKEEEEEELVDPLTTVR | -14 | -32 | 1 | 0 | 19 | 543.50995 | 645.21844 | 687.99115 | 698.7418 | 2282.7495 | 921.86346 |
| Uqcrh | GDPKEEEEEELVDPLTTVR | -14 | -32 | 1 | 0 | 19 | 543.50995 | 645.21844 | 687.99115 | 698.7418 | 2282.7495 | 921.86346 |
| Uqcrh | SQTEEDCTEELFDFLHAR | -59 | -76 | 0 | 0 | 18 | 528.506 | 585.7519 | 523.7264 | 594.10535 | 680.2523 | 571.9908 |
| Uqcrh | SQTEEDCTEELFDFLHAR | -59 | -76 | 0 | 0 | 18 | 528.506 | 585.7519 | 523.7264 | 594.10535 | 680.2523 | 571.9908 |
| Uqcrh | EEEEEELVDPLTTVR | -18 | -32 | 0 | 0.000627943 | 15 | 8344.0625 | 9818.643 | 9864.779 | NaN | 11244.164 | 12428.943 |
| Uqcrh | EEEELVDPLTTVR | -20 | -32 | 0 | 0 | 13 | 1559.3752 | NaN | 1270.1704 | NaN | NaN | NaN |
| Snrpd2 | SLLNKPK | -2 | -8 | 1 | 0.001392869 | 7 | 2875.0803 | 2833.2793 | 2565.9321 | 2213.831 | 1893.2219 | 1783.5662 |
| Snrpd2 | RNPLIAGK | -111 | -118 | 1 | 0.00111006 | 8 | 2337.8496 | 1765.6821 | 2378.429 | 1743.042 | 1145.2559 | 1614.5823 |
| Snrpd2 | REEEEFNTGPLSVL | -19 | -32 | 1 | 0.00025015 | 14 | 2235.68 | 2351.5637 | 2860.4202 | 2401.8281 | 3241.551 | 727.83673 |
| Snrpd2 | HCNMVLENVK | -62 | -71 | 0 | 0 | 10 | 1184.595 | 1217.7744 | 1217.4202 | 1605.6138 | 683.07623 | 1464.9222 |
| Snrpd2 | REEEEFNTGPL | -19 | -29 | 1 | 0 | 11 | 1099.0461 | 71.21139 | 1033.6531 | 982.3195 | 981.65283 | 365.39383 |
| Snrpd2 | EEEEFNTGPLSVL | -20 | -32 | 0 | 0 | 13 | 973.47095 | 920.115 | 1122.4531 | 1498.0087 | 1664.6798 | 1652.0444 |
| Snrpd2 | SEMTPEELQK | -9 | -18 | 0 | 0 | 10 | 1447.5029 | NaN | 2310.8875 | 1438.427 | 1278.2126 | 2373.7969 |
| Slc25a11 | LTGADGTPPGFLLK | -109 | -122 | 0 | 0 | 14 | 1783.2537 | 1959.2429 | 1839.6226 | 2529.2922 | 2416.252 | 2347.7336 |
| Slc25a11 | EEGVPTLWR | -174 | -182 | 0 | 0 | 9 | 1057.8757 | 1109.2769 | 1029.2775 | 1088.6655 | 1188.6613 | 1266.034 |
| Slc25a11 | GAFVGTPAEVALIR | -133 | -146 | 0 | 0 | 14 | 963.22577 | 967.1951 | 1005.9168 | 1017.7056 | 1201.5912 | 1160.778 |
| Slc25a11 | VFVQPLDLVK | -37 | -46 | 0 | 0 | 10 | 747.32367 | 893.30194 | 912.1391 | 992.2988 | 1161.6704 | 1038.3036 |
| Slc25a11 | YEGFFSLWK | -272 | -280 | 0 | 0 | 9 | 382.13232 | 122.85728 | 131.0675 | 482.66037 | 456.26727 | 490.36243 |
| Slc25a11 | VQPLDLVK | -39 | -46 | 0 | 0.000627943 | 8 | 703.25195 | 554.5058 | 550.2167 | NaN | 485.48572 | 477.84735 |
| Slc25a11 | FVGTPAEVALIR | -135 | -146 | 0 | 0 | 12 | 661.10986 | 635.46405 | 719.4789 | 181.53772 | NaN | 747.80457 |
| Slc25a11 | LGPHTVLTF | -289 | -297 | 0 | 0.002534621 | 9 | 1167.4628 | NaN | NaN | 1450.7577 | 1542.4037 | 1514.8618 |
| U2af2 | SIEIPRPVDGVEVPGCGK | -414 | -431 | 1 | 0 | 18 | 1558.8672 | 1937.2268 | 1821.5594 | 1842.189 | 2457.6492 | 2311.1338 |
| U2af2 | SIEIPRPVDGVEVPGCGK | -414 | -431 | 1 | 0 | 18 | 1558.8672 | 1937.2268 | 1821.5594 | 1842.189 | 2457.6492 | 2311.1338 |
| U2af2 | EEHGGLIR | -71 | -78 | 0 | 0.00025015 | 8 | 1451.3391 | 1409.8032 | 2854.9812 | 3064.7058 | 3420.6938 | 3158.7993 |
| U2af2 | VEFTSVFDCQK | -434 | -444 | 0 | 0 | 11 | 1055.2134 | 1143.7812 | 1145.5068 | 1312.9094 | 1421.0062 | 1484.272 |
| U2af2 | LFIGGLPNYLNDDQVK | -261 | -276 | 0 | 0 | 16 | 1046.1931 | 957.9492 | 751.3262 | 1344.8464 | 1401.01 | 1262.3002 |
| U2af2 | YCDPDSYHR | -463 | -471 | 0 | 0 | 9 | 985.2837 | 1042.7501 | 1041.4677 | 773.3461 | 1167.4773 | 1239.7703 |
| U2af2 | NFAFLEFR | -196 | -203 | 0 | 0 | 8 | 775.47424 | 752.8007 | 82.12077 | 1025.3359 | 1434.7576 | 1524.6271 |
| U2af2 | GLPNYLNDDQVK | -265 | -276 | 0 | 0 | 12 | 671.3049 | 856.91547 | 835.3911 | 861.13165 | 727.0032 | 592.4041 |
| U2af2 | KYCDPDSYHR | -462 | -471 | 1 | 0 | 10 | 553.02325 | 612.7944 | 624.34375 | 175.60966 | 784.9843 | 557.2437 |
| U2af2 | LGGLTQAPGNPVLAV | -175 | -189 | 0 | 0 | 15 | 452.16135 | 481.43344 | 572.3114 | 913.58093 | 1114.0555 | 701.8864 |
| U2af2 | ELLTSFGPLK | -277 | -286 | 0 | 0 | 10 | 285.73468 | 1866.4261 | 2172.6582 | 2048.1975 | 2595.7908 | 2798.09 |
| U2af2 | LGGLTQAPGNPVLAVQ | -175 | -190 | 0 | 0.00025015 | 16 | 133.31511 | 908.49554 | 721.92334 | 1092.1227 | 1308.0988 | 1252.3584 |
| U2af2 | ENPSVYVPGVVSTVVPDSAHK | -240 | -260 | 0 | 0.00025015 | 21 | 367.4802 | 268.37906 | NaN | 515.81616 | 574.0164 | 643.7941 |
| U2af2 | FIGGLPNYLNDDQVK | -262 | -276 | 0 | 0.00025015 | 15 | 521.3364 | 1332.3951 | 592.42975 | 1080.5055 | NaN | NaN |
| Stx6 | WTELLQGPSAATR | -28 | -40 | 0 | 0 | 13 | 1400.5878 | NaN | 691.94476 | 1107.9264 | 963.5572 | NaN |
| Stx6 | EEIDWTTNELR | -41 | -51 | 0 | 0.003133457 | 11 | 404.74554 | NaN | 3801.4966 | NaN | NaN | NaN |
| Acadm | EEIIPVAPEYDK | -58 | -69 | 0 | 0 | 12 | 477.98663 | 621.89276 | 762.1522 | 724.78186 | 724.1297 | 672.687 |
| Acadm | IVEADTPGIHIGK | -223 | -235 | 0 | 0 | 13 | 1976.8964 | NaN | 2505.0518 | 1398.4049 | 1476.4553 | 1759.335 |
| Acadm | SGEYPFPLIK | -70 | -79 | 0 | 0 | 10 | 1165.3484 | 1276.4612 | 1343.5592 | 1408.6653 | NaN | 1265.2378 |
| Pfdn6 | EELALLDGSNVVFK | -45 | -58 | 0 | 0 | 14 | 874.9951 | 834.80756 | 1008.0617 | 1065.7831 | 1498.6497 | 1301.9863 |
| Pfdn6 | ETLAQLQQEFQR | -106 | -117 | 0 | 0.00025015 | 12 | 5001.4067 | NaN | NaN | 5150.72 | 7184.068 | 4613.5713 |
| Pfdn6 | RLDYITAEIK | -79 | -88 | 1 | 0.000298805 | 10 | 566.1244 | NaN | NaN | 575.5944 | 758.1378 | 913.94275 |
| Pfdn6 | LLGPVLVK | -59 | -66 | 0 | 0 | 8 | 887.9077 | 537.60315 | 547.6091 | NaN | NaN | NaN |
| Pfdn6 | LQQEFQR | -111 | -117 | 0 | 0.002296977 | 7 | 3366.8506 | NaN | NaN | NaN | NaN | NaN |
| Utp6 | IPELEQLER | -13 | -21 | 0 | 0 | 9 | 17121.31 | 17602.934 | 15510.541 | 16125.688 | 17258.986 | 17042.576 |
| Utp6 | HTDDPLTWDYVAR | -267 | -279 | 0 | 0 | 13 | 1516.5267 | 1809.11 | 1879.571 | 1970.9679 | 2406.9617 | 1870.8771 |
| Utp6 | ELEIESQPGEEQPVSK | -281 | -296 | 0 | 0 | 16 | 233.51988 | NaN | 90.7062 | 144.50204 | 471.87857 | 473.76172 |
| Utp6 | EELNHPFGKPENCGQIY | -555 | -571 | 1 | 0.000431655 | 17 | 260.89923 | NaN | NaN | 419.84787 | 366.68503 | NaN |
| Cactin | EEQEDSFHLR | -303 | -312 | 0 | 0 | 10 | 1183.8032 | NaN | NaN | NaN | NaN | 1164.6738 |
| Cct7 | VEEGLHPQIIIR | -112 | -123 | 0 | 0 | 12 | 3807.9995 | 4459.7114 | 4372.3115 | 3493.81 | 4158.794 | 4144.162 |
| Cct7 | VEEGLHPQIIIR | -112 | -123 | 0 | 0 | 12 | 3807.9995 | 4459.7114 | 4372.3115 | 3493.81 | 4158.794 | 4144.162 |
| Cct7 | SLHDAIMIVR | -388 | -397 | 0 | 0 | 10 | 3803.466 | 5153.5356 | 11629.316 | 5692.4966 | 3972.6987 | 1620.3105 |
| Cct7 | SLHDAIMIVR | -388 | -397 | 0 | 0 | 10 | 3803.466 | 5153.5356 | 11629.316 | 5692.4966 | 3972.6987 | 1620.3105 |
| Cct7 | SLHDAIMIVR | -388 | -397 | 0 | 0 | 10 | 3803.466 | 5153.5356 | 11629.316 | 5692.4966 | 3972.6987 | 1620.3105 |
| Cct7 | YNFFTGCPK | -358 | -366 | 0 | 0.00025015 | 9 | 3188.9453 | 3211.8147 | 3206.9568 | 3512.6511 | 3538.8667 | 3606.625 |
| Cct7 | VFEETQIGGER | -347 | -357 | 0 | 0 | 11 | 1882.587 | 2267.5022 | 911.4435 | 1974.955 | 2432.8982 | 2602.6855 |
| Cct7 | LLDVVHPAAK | -68 | -77 | 0 | 0 | 10 | 1150.9884 | 1321.299 | 1460.8408 | 852.4313 | 1190.0148 | 1151.0847 |
| Cct7 | LDELLQLK | -186 | -193 | 0 | 0 | 8 | 944.1191 | 399.14514 | 1114.5884 | 1280.1019 | 1373.2374 | 1372.8306 |
| Cct7 | KLLDVVHPAAK | -67 | -77 | 1 | 0 | 11 | 864.0743 | 953.3882 | 1080.1926 | 1017.52814 | 1288.5609 | 1282.6602 |
| Cct7 | KLLDVVHPAAK | -67 | -77 | 1 | 0 | 11 | 864.0743 | 953.3882 | 1080.1926 | 1017.52814 | 1288.5609 | 1282.6602 |
| Cct7 | QLCDNAGFDATNILNK | -448 | -463 | 0 | 0 | 16 | 846.65625 | 959.9185 | 956.83344 | 989.0296 | 1128.8539 | 1056.7207 |
| Cct7 | KVQGGALEESQLVAGVA | -199 | -215 | 1 | 0.00025015 | 17 | 531.96075 | 749.53345 | 862.37866 | 1157.609 | 1434.079 | 1611.8143 |
| Cct7 | GRVPEEDLK | -312 | -320 | 1 | 0 | 9 | 3033.2764 | 3243.506 | 3528.7646 | NaN | 2815.7036 | 2918.1099 |

| | | | | | | | | | | | | |
|---|---|---|---|---|---|---|---|---|---|---|---|---|
| Cct7 | QVFEETQIGGER | -346 | -357 | 0 | 0 | 12 | 880.7553 | 671.547 | NaN | 882.5176 | 296.9731 | 769.80347 |
| Cct7 | GRVPEEDLKR | -312 | -321 | 2 | 0 | 10 | 664.981 | 2313.2664 | 538.00037 | 335.8789 | 865.1166 | NaN |
| Cct7 | VPEEDLKR | -314 | -321 | 1 | 0 | 8 | 260.87885 | 1120.3203 | NaN | 1129.8513 | 778.7525 | 832.249 |
| Cct7 | TCTIILR | -369 | -375 | 0 | 0.00025015 | 7 | 10483.987 | 8091.5103 | 8394.604 | 2289.4033 | NaN | NaN |
| Cct7 | RVPEEDLK | -313 | -320 | 1 | 0.001392869 | 8 | 1889.3708 | NaN | NaN | 144.66837 | 3180.7976 | 2961.9482 |
| Cct7 | EETQIGGER | -349 | -357 | 0 | 0 | 9 | 789.1795 | 1224.471 | 1101.4457 | NaN | NaN | NaN |
| Cct7 | GLHPQIIIR | -115 | -123 | 0 | 0.00122419 | 9 | 462.02597 | 508.05283 | 518.19604 | NaN | NaN | NaN |
| Atp6v1a | LPANHPLLTGQ | -221 | -231 | 0 | 0 | 11 | 2793.757 | 2863.5593 | 2987.1838 | 2331.7622 | 1742.5879 | 780.7512 |
| Atp6v1a | VGHSELVGEIIR | -45 | -56 | 0 | 0 | 12 | 1765.3048 | 1997.708 | 1852.7205 | 2523.3938 | 2753.3696 | 2452.252 |
| Atp6v1a | IRDEDKESTFGYVH | -9 | -22 | 2 | 0.00111006 | 14 | 591.96436 | 538.6617 | 321.03275 | 536.5821 | 627.62463 | 580.3116 |
| Atp6v1a | IRDEDKESTFGYVH | -9 | -22 | 2 | 0.00111006 | 14 | 591.96436 | 538.6617 | 321.03275 | 536.5821 | 627.62463 | 580.3116 |
| Atp6v1a | VYEETSGVSVGDPVLR | -66 | -81 | 0 | 0 | 16 | 863.3118 | 1032.6405 | 1077.7642 | 983.5646 | 1054.415 | 955.2148 |
| Atp6v1a | TQSIYIPR | -113 | -120 | 0 | 0 | 8 | 819.7211 | 595.3301 | 845.6592 | 1084.748 | 373.05087 | 911.2812 |
| Atp6v1a | EETSGVSVGDPVLR | -68 | -81 | 0 | 0 | 14 | 804.17084 | 872.56494 | 843.0371 | 1185.7457 | 906.98926 | NaN |
| Atp6v1a | HFTEFVPLR | -468 | -476 | 0 | 0 | 9 | 674.2431 | NaN | 1705.6307 | 1175.3936 | 553.1774 | NaN |
| Atp6v1a | WEFIPSK | -133 | -139 | 0 | 0.003166358 | 7 | 446.98166 | 438.08542 | NaN | 450.53143 | NaN | NaN |
| Mrto4 | QNLIEELR | -24 | -31 | 0 | 0.000471409 | 8 | 1204.7069 | 533.2179 | 586.9989 | 1163.3821 | 524.35406 | 553.927 |
| Mrto4 | TKEEVNEWFTK | -108 | -118 | 1 | 0 | 11 | 932.34906 | 1018.85345 | 1154.3749 | 1316.1886 | 1111.022 | 1132.8219 |
| Mrto4 | SPSDEYKDNLHQVSK | -80 | -94 | 1 | 0 | 15 | 601.5561 | 1119.9227 | 890.35364 | 909.03455 | 1379.9056 | 1268.1105 |
| Mrto4 | SPSDEYKDNLHQVSK | -80 | -94 | 1 | 0 | 15 | 601.5561 | 1119.9227 | 890.35364 | 909.03455 | 1379.9056 | 1268.1105 |
| Mrto4 | KEGDVLTPEQAR | -177 | -188 | 1 | 0.00025015 | 12 | 573.0686 | 709.8821 | 679.24426 | 911.2126 | 1107.675 | 1148.2222 |
| Mrto4 | VSLDPGPLK | -135 | -143 | 0 | 0 | 9 | 903.24756 | 398.13867 | 314.88086 | 1459.5848 | NaN | 224.96432 |
| Mrto4 | EGDVLTPEQAR | -178 | -188 | 0 | 0 | 11 | 1711.5084 | 1425.0791 | 1720.4064 | 952.6813 | NaN | NaN |
| Mrto4 | SPSDEYKDNLHQV | -80 | -92 | 1 | 0.00025015 | 13 | 546.2308 | NaN | NaN | 1100.7147 | NaN | 859.2249 |
| Mrto4 | WDAQSGR | -208 | -214 | 0 | 0.003867596 | 7 | 498.17123 | 547.00464 | NaN | NaN | NaN | NaN |
| Mrto4 | EEVNEWFTK | -110 | -118 | 0 | 0 | 9 | 694.30865 | NaN | NaN | NaN | NaN | NaN |
| Wdr75 | EEVQWGVFVPR | -626 | -636 | 0 | 0 | 11 | 938.719 | 740.2335 | 954.3139 | 957.77155 | 1102.9209 | 407.12354 |
| Wdr75 | LWDYVDGILIK | -83 | -93 | 0 | 0 | 11 | 564.5774 | 463.64548 | NaN | 812.6668 | 450.27405 | 494.13623 |
| Wdr75 | LGSSIEHISVSPAGDLFCT | -288 | -306 | 0 | 0.000712251 | 19 | 310.531 | 493.82547 | NaN | 525.0693 | 532.3263 | 569.34735 |
| Wdr75 | SSAGSDLFVFKPSEPR | -600 | -615 | 1 | 0.001023236 | 16 | 287.22342 | 669.45135 | 560.65375 | 801.20557 | NaN | NaN |
| Gatad2a | IPGSIIPPPLVR | -210 | -221 | 0 | 0 | 12 | 470.1975 | 598.58746 | 614.17725 | 651.9812 | 654.2709 | 725.7443 |
| Gatad2a | TLEPDLTEDDVENKK | -14 | -28 | 1 | 0 | 15 | 902.3021 | NaN | 956.13477 | 786.38135 | NaN | NaN |
| Gatad2a | EEVVQNLLETQAGR | -386 | -399 | 0 | 0 | 14 | 417.8907 | NaN | NaN | NaN | 1506.7721 | 1485.6749 |
| Rigi | KYEQWIVGVHK | -509 | -519 | 1 | 0 | 11 | 69.79959 | 1237.6284 | 1366.6442 | 789.8483 | 1386.179 | 1436.0166 |
| Rigi | NTIICAPTGCGK | -260 | -271 | 0 | 0 | 12 | 1206.1785 | NaN | 1731.0007 | 2341.5144 | 1682.917 | 2236.229 |
| Rigi | EEYHLKPETK | -621 | -630 | 1 | 0 | 10 | 1497.771 | NaN | NaN | 855.8377 | 1078.8027 | NaN |
| Rigi | SFVVEDIVSGVQNR | -892 | -905 | 0 | 0 | 14 | 479.25067 | NaN | 299.19128 | NaN | NaN | NaN |
| Rigi | NLHSPLKPR | -237 | -245 | 1 | 0.002199523 | 9 | 458.5934 | 387.8126 | NaN | NaN | NaN | NaN |
| Mien1 | EEYPGIEIESR | -48 | -58 | 0 | 0 | 11 | 957.98315 | 478.20148 | 1360.7877 | 1664.2555 | 1943.5392 | 251.57852 |
| Tmem109 | ETSADILTQIGR | -48 | -59 | 0 | 0 | 12 | 1100.9027 | 1097.1307 | 1173.7904 | 1573.7341 | 1548.5942 | 1540.3973 |
| Tmem109 | KETSADILTQIGR | -47 | -59 | 1 | 0 | 13 | 852.3008 | 332.4964 | NaN | 1269.7788 | 804.9139 | 756.7465 |
| Tmem109 | EFASPGQQK | -38 | -46 | 0 | 0.002296977 | 9 | 993.8003 | 684.79895 | NaN | 773.9432 | NaN | 703.44995 |
| Cep170b | EFFDGDTPRK | -407 | -416 | 1 | 0.00025015 | 10 | 2533.5808 | 2505.7954 | 2452.1138 | 2283.5156 | 2937.867 | 2888.0276 |
| Slc25a4 | QIFLGGVDR | -97 | -105 | 0 | 0.00025015 | 9 | 2194.7334 | 2180.1992 | 2222.447 | 1452.716 | 2216.155 | 1605.3926 |
| Slc25a4 | GLVSYPFDTVR | -225 | -235 | 0 | 0 | 11 | 1516.6088 | 1823.0059 | 2115.645 | 1862.3826 | 1212.1475 | 1314.8511 |
| Slc25a4 | VSYPFDTVR | -227 | -235 | 0 | 0 | 9 | 782.20807 | 1368.1959 | 1142.454 | 1196.9042 | 615.4508 | 1020.6568 |
| Slc25a4 | GDQALSFLK | -2 | -10 | 0 | 0.001392869 | 9 | 556.4062 | 903.7736 | 462.55777 | 667.3075 | 1129.5175 | 809.0778 |
| Slc25a4 | EQGFLSFWR | -64 | -72 | 0 | 0 | 9 | 313.6272 | 368.15195 | 407.42227 | 99.28295 | 1402.3561 | 1125.1588 |
| Slc25a4 | EFNGLGDCLTK | -153 | -163 | 0 | 0 | 11 | 756.2439 | 694.2864 | 657.6155 | 846.764 | NaN | 771.1148 |
| Slc25a4 | LVSYPFDTVR | -226 | -235 | 0 | 0 | 10 | 170.77757 | 830.1249 | 865.0453 | NaN | 459.46774 | NaN |
| Atxn2l | LSLTPTDVK | -502 | -510 | 0 | 0 | 9 | 1253.3038 | 1216.2014 | 1065.9054 | 1434.674 | 276.3889 | 181.94586 |
| Atxn2l | LQPSSSPETGLDPFPSR | -557 | -573 | 0 | 0 | 17 | 1206.8875 | 1548.8346 | 1386.3428 | 528.00836 | 1425.6503 | 1343.1 |
| Atxn2l | FELAVDAVHR | -154 | -163 | 0 | 0 | 10 | 1028.635 | 952.7157 | 854.6907 | 1582.3229 | 1754.3828 | 1325.3722 |
| Atxn2l | GDEKEEGPVTEQVK | -645 | -658 | 1 | 0 | 14 | 793.0826 | 654.4641 | 676.4764 | 610.65826 | 650.2399 | 743.8944 |
| Atxn2l | EFNPTKPLLSVN | -668 | -679 | 1 | 0.00025015 | 12 | 672.1719 | 1317.9946 | 1233.7882 | 1649.9888 | 1781.5514 | 1993.7612 |
| Atxn2l | VFKPSDVLLVHFR | -182 | -194 | 1 | 0.000230707 | 13 | 527.78436 | 529.2776 | 239.58307 | 558.24133 | 650.07434 | 695.23566 |
| Atxn2l | GKVPGLQNEQK | -531 | -541 | 1 | 0 | 11 | 702.2204 | NaN | NaN | 798.88354 | 1357.9696 | 1458.3369 |
| Atxn2l | GPHHLDNSSPGPGSEAR | -381 | -397 | 0 | 0.001055699 | 17 | 292.36365 | 520.1008 | NaN | NaN | 277.9646 | 571.4273 |
| Atxn2l | SASIPVTSSVVDPGAGSISPASPK | -478 | -501 | 0 | 0 | 24 | 362.49954 | 428.36917 | NaN | 297.84814 | NaN | NaN |
| Wbp11 | SAPPSLIQRPK | -535 | -545 | 1 | 0 | 11 | 572.17694 | 580.3367 | 555.6739 | 620.28284 | 540.29944 | 317.39078 |
| Wbp11 | KPPGPPPGPPPPQVL | -198 | -212 | 1 | 0.000591124 | 15 | 33139.38 | NaN | 34643.543 | 33652.027 | 37807.785 | 30034.215 |
| Wbp11 | EKENPDIYK | -97 | -105 | 1 | 0 | 9 | 10005.709 | 114.03744 | 1199.3812 | NaN | NaN | 10831.35 |
| Wbp11 | EFNPVQQPQLNEK | -64 | -76 | 0 | 0 | 13 | 592.14417 | 594.1171 | 737.8893 | NaN | NaN | NaN |
| Tbc1d17 | EFPFPDVLR | -503 | -511 | 0 | 0.00025015 | 9 | 404.14023 | 814.4547 | 534.5937 | 543.768 | NaN | 774.8546 |
| S100a10 | FAGDKDHLTK | -19 | -28 | 1 | 0 | 10 | 2095.6887 | 1998.357 | 2143.4648 | 888.76996 | 936.3262 | 1661.1519 |
| S100a10 | FAGDKDHLTK | -19 | -28 | 1 | 0 | 10 | 2095.6887 | 1998.357 | 2143.4648 | 888.76996 | 936.3262 | 1661.1519 |
| S100a10 | EFPGFLENQKDPLAVDK | -38 | -54 | 1 | 0 | 17 | 1335.0886 | 1363.223 | 1453.2665 | 1544.7803 | 2067.7957 | 2124.9639 |
| S100a10 | EFPGFLENQKDPLAVDK | -38 | -54 | 1 | 0 | 17 | 1335.0886 | 1363.223 | 1453.2665 | 1544.7803 | 2067.7957 | 2124.9639 |
| S100a10 | EFPGFLENQK | -38 | -47 | 0 | 0 | 10 | 1794.0128 | 2065.3782 | 2169.2092 | NaN | 2062.1453 | 2193.237 |
| Gfm1 | YQPCSPSTQEELINK | -720 | -734 | 0 | 0 | 15 | 998.73804 | 893.40063 | 942.95355 | 1286.6766 | 1391.7336 | 1533.8456 |
| Gfm1 | EFSDETFGSNVPK | -568 | -580 | 0 | 0 | 13 | 994.2367 | 846.67554 | 884.7543 | 802.8679 | 599.7715 | 819.7363 |
| Gfm1 | FVQIPIGLEGDFK | -201 | -213 | 0 | 0 | 13 | 692.5967 | 900.2902 | 708.8893 | 622.30493 | 962.5178 | 833.678 |
| Gfm1 | GIIDLIEER | -214 | -222 | 0 | 0 | 9 | 574.5548 | 261.9998 | 540.12506 | 489.88248 | 339.5554 | 268.49072 |
| Gfm1 | VDSNEISFIR | -619 | -628 | 0 | 0 | 10 | 247.0724 | 2614.5522 | 2320.8818 | 3754.7988 | 3616.749 | NaN |
| Gfm1 | VIGVLEPLPPEDYTKL | -552 | -567 | 1 | 0 | 16 | 163.458 | 329.51566 | NaN | 362.98676 | 176.75781 | 467.87985 |
| Gfm1 | LEFSDETFGSNVPK | -567 | -580 | 0 | 0.001023236 | 14 | 1119.4711 | NaN | 774.9077 | NaN | NaN | 824.09326 |
| Riox2 | EFWEQKPLLIQ | -57 | -67 | 1 | 0 | 11 | 404.01608 | 1220.6357 | 1258.7782 | 1312.044 | 1587.547 | 1421.0707 |
| Riox2 | EFWEQKPLLIQR | -57 | -68 | 1 | 0 | 12 | 345.59366 | 476.48987 | 503.80032 | 420.83636 | 398.8902 | 475.19565 |

| | | | | | | | | | | | | |
|---|---|---|---|---|---|---|---|---|---|---|---|---|
| Riox2 | LYSPTVPLAHEY | -198 | -209 | 0 | 0 | 12 | 2260.8433 | 2129.3738 | 1798.9617 | NaN | 1366.115 | 2030.869 |
| Riox2 | YKDELWR | -143 | -149 | 1 | 0 | 7 | 829.2002 | 886.5122 | NaN | 993.4809 | 1221.2133 | 1020.24817 |
| Riox2 | VQPTGDENEEASVPCK | -5 | -20 | 0 | 0.000394361 | 16 | 479.361 | 348.8861 | 390.81595 | NaN | 541.6799 | 593.99854 |
| Riox2 | FPLSHVDALK | -419 | -428 | 0 | 0.001291103 | 10 | 19466.314 | NaN | NaN | NaN | 18580.535 | NaN |
| Mcm7 | IAPEIYGHEDVKK | -340 | -352 | 1 | 0 | 13 | 5747.723 | 6198.4365 | 5967.8853 | 5235.294 | 6763.257 | 6825.2554 |
| Mcm7 | SIAPEIYGHEDVKK | -339 | -352 | 1 | 0 | 14 | 4241.6377 | 4169.6997 | 3814.4907 | 4613.6816 | 4203.6084 | 4172.481 |
| Mcm7 | SIAPEIYGHEDVKK | -339 | -352 | 1 | 0 | 14 | 4241.6377 | 4169.6997 | 3814.4907 | 4613.6816 | 4203.6084 | 4172.481 |
| Mcm7 | IAQPGDHVSVT | -264 | -274 | 0 | 0 | 11 | 3113.4312 | 3073.2004 | 3005.7563 | 3294.2173 | 3066.2056 | 3050.4194 |
| Mcm7 | EHSDQVPVGNIPR | -239 | -251 | 0 | 0 | 13 | 1964.5479 | 2114.8545 | 2105.0613 | 2187.4712 | 2514.4028 | 2814.5679 |
| Mcm7 | EFYYENELGK | -19 | -28 | 0 | 0 | 10 | 1467.9763 | 1464.2482 | 1599.1799 | 1636.3129 | 1554.2865 | 1361.9548 |
| Mcm7 | SIAPEIYGHEDVK | -339 | -351 | 0 | 0 | 13 | 1203.3243 | 1342.1753 | 1377.3496 | 646.9425 | 1085.1947 | 1499.4149 |
| Mcm7 | NPQNQYPSELMR | -121 | -132 | 0 | 0 | 12 | 1126.7163 | 522.9318 | 486.25015 | 584.6924 | 516.32654 | 474.13004 |
| Mcm7 | NPQNQYPSELMR | -121 | -132 | 0 | 0 | 12 | 1126.7163 | 522.9318 | 486.25015 | 584.6924 | 516.32654 | 474.13004 |
| Mcm7 | GIFLPVLR | -275 | -282 | 0 | 0.001776473 | 8 | 774.17365 | 1047.2134 | 981.8622 | 382.91367 | 1077.1804 | 1149.7684 |
| Mcm7 | QIAEEDFYEK | -326 | -335 | 0 | 0 | 10 | 754.4769 | 2218.1624 | 1975.9935 | 971.75415 | 599.8103 | 549.8379 |
| Mcm7 | QIAEEDFYEK | -326 | -335 | 0 | 0 | 10 | 754.4769 | 2218.1624 | 1975.9935 | 971.75415 | 599.8103 | 549.8379 |
| Mcm7 | GDVVQELLPEYK | -78 | -89 | 0 | 0 | 12 | 608.7856 | 535.2383 | 491.10895 | 1323.2521 | 1251.4194 | 988.8275 |
| Mcm7 | VLADQGVCCIDEFDK | -435 | -449 | 0 | 0 | 15 | 382.72607 | 407.0795 | 393.78143 | 1965.6466 | 2584.918 | 2340.356 |
| Mcm7 | TQRPADVIFAT | -654 | -664 | 1 | 0.00111006 | 11 | 1215.7903 | 1137.7611 | 1150.557 | 1170.0525 | 300.6754 | NaN |
| Mcm7 | LFGDVVQELLPEYK | -76 | -89 | 0 | 0 | 14 | 432.39774 | 447.89786 | 401.32452 | 340.83356 | 495.29257 | NaN |
| Mcm7 | IQEHSDQVPVGNIPR | -237 | -251 | 0 | 0 | 15 | 514.33453 | NaN | 543.4522 | 490.50778 | NaN | 1054.3306 |
| Mcm7 | QPTVPESLADYITAA | -570 | -584 | 0 | 0.00111006 | 15 | 337.7808 | 490.19897 | NaN | 467.51505 | 464.02808 | NaN |
| Zyx | VNPFRPGDSEPPVAA | -36 | -50 | 1 | 0 | 15 | 1318.3304 | 1456.8425 | 511.01123 | 592.8095 | 771.6557 | 554.23267 |
| Zyx | SPGGPGPLTLK | -336 | -346 | 0 | 0 | 11 | 757.3102 | 766.68115 | 1049.7391 | 626.2124 | 687.667 | 633.95154 |
| Zyx | EGAPYCEGCYTDTLEK | -420 | -435 | 0 | 0 | 16 | 149.55977 | 422.71408 | NaN | NaN | 399.52213 | 273.30554 |
| Lig1 | TIQDTLEEQNEDKTK | -132 | -146 | 1 | 0 | 15 | 1614.3 | 1464.7146 | 1709.7279 | 1081.1187 | 1253.6558 | 1393.2178 |
| Lig1 | KQTFCEVPDLDR | -502 | -513 | 1 | 0 | 12 | 1321.4182 | 1473.1525 | 1375.3293 | 722.2766 | 1260.4542 | 1191.4572 |
| Lig1 | EGDQLIVPSEPTK | -175 | -187 | 0 | 0 | 13 | 1246.1017 | 1685.8062 | 1671.1335 | 1844.923 | 452.90607 | 2791.2258 |
| Lig1 | NQEDNTGKYPDIISR | -588 | -602 | 1 | 0 | 15 | 1170.8156 | 1484.5238 | 1560.8889 | 1910.3456 | 2793.9797 | 2717.7654 |
| Lig1 | IIPVLLEHGLER | -514 | -525 | 0 | 0 | 12 | 853.1424 | 1081.842 | 1070.8444 | 1239.8774 | 1893.1307 | 1998.0938 |
| Lig1 | TFCEVPDLDR | -504 | -513 | 0 | 0 | 10 | 725.33655 | 743.29517 | 585.4171 | 782.59564 | 1018.0002 | 776.94037 |
| Lig1 | SCEGEDEDEAPGTPK | -65 | -79 | 0 | 0 | 15 | 631.6844 | 667.2306 | 742.96857 | 593.5002 | 746.7964 | 775.5791 |
| Lig1 | NNYHPIEDACWK | -269 | -280 | 0 | 0 | 12 | 564.75183 | 569.03046 | 593.77905 | 520.228 | 614.40515 | 544.09656 |
| Lig1 | LKPQEEEQSKPPAR | -209 | -222 | 2 | 0 | 14 | 454.8046 | 399.833 | 539.42535 | 543.2788 | 160.27742 | 583.0057 |
| Lig1 | FEEVDFTCEYK | -556 | -566 | 0 | 0.00025015 | 11 | 527.54895 | 311.1015 | 187.69139 | 1004.62964 | NaN | 434.96838 |
| Lig1 | RFEEVDFTCEYK | -555 | -566 | 1 | 0.000394361 | 12 | 485.0515 | 510.85294 | 565.9899 | 634.3169 | NaN | 693.4538 |
| Lig1 | EKIEEVSAR | -295 | -303 | 1 | 0.004099215 | 9 | 5691.2197 | NaN | NaN | 3749.7898 | 4762.439 | 4930.9087 |
| Lig1 | TIQDTLEEQNEDK | -132 | -144 | 0 | 0 | 13 | 170.38148 | 463.82303 | NaN | NaN | NaN | 1610.9335 |
| Treml1 | VSLLVLPPVPGPR | -119 | -131 | 0 | 0 | 13 | 706.21484 | 5167.96 | 5467.8813 | 6953.1523 | 7159.477 | 7836.3184 |
| Treml1 | EGEEAEDEKETYR | -132 | -144 | 1 | 0 | 13 | 446.6538 | 105.02727 | 255.37 | 333.71683 | 449.423 | 495.2483 |
| Huwe1 | HIIEDPCTLR | -1826 | -1835 | 0 | 0.000230707 | 10 | 959.2452 | 997.84106 | 867.5311 | NaN | 1163.6979 | 658.0641 |
| Huwe1 | EGEEGQDAGGLLR | -4054 | -4066 | 0 | 0 | 13 | 1174.8081 | NaN | 1355.8293 | NaN | NaN | NaN |
| Huwe1 | FNQLDLPAYESFEK | -4345 | -4358 | 0 | 0.001776473 | 14 | 799.4679 | NaN | NaN | NaN | NaN | NaN |
| Polr1a | SFLGLLPGPSLTDK | -380 | -393 | 0 | 0 | 14 | 359.2637 | 362.546 | 286.29486 | 363.76456 | 365.82568 | 445.74652 |
| Polr1a | TIDKENDSSVAIDR | -366 | -379 | 1 | 0.00025015 | 14 | 1750.7971 | 1878.3188 | 1890.3604 | NaN | NaN | 2350.1409 |
| Polr1a | EGELLCGVLDK | -748 | -758 | 0 | 0.003299042 | 11 | 437.036 | NaN | NaN | NaN | NaN | NaN |
| Ndufb10 | VDQEIMNIIQER | -98 | -109 | 0 | 0 | 12 | 4745.864 | 4812.477 | 1537.3484 | 3430.1948 | 3633.2517 | 3625.5242 |
| Ndufb10 | VDQEIMNIIQER | -98 | -109 | 0 | 0 | 12 | 4745.864 | 4812.477 | 1537.3484 | 3430.1948 | 3633.2517 | 3625.5242 |
| Ndufb10 | PDSWDKDVYPEPPSR | -2 | -16 | 1 | 0.00025015 | 15 | 1323.5316 | 1390.5819 | 1425.2922 | 796.5797 | 751.84015 | 549.3385 |
| Ndufb10 | EGENYQQNCAK | -117 | -127 | 0 | 0 | 11 | 170.82877 | 386.16644 | 193.48743 | 263.4258 | 371.3529 | 206.02103 |
| Ndufb10 | RVPDITECK | -70 | -78 | 1 | 0.001291103 | 9 | 625.2684 | 396.62637 | 871.286 | 1413.1572 | NaN | 628.01843 |
| Eif3m | EGERPSLR | -95 | -102 | 1 | 0.003003218 | 8 | 1074.6506 | 631.6389 | 1306.6249 | 1180.2941 | 1294.2164 | 1257.0201 |
| Eif3m | LFDHLLTLKPVK | -219 | -230 | 1 | 0.000230707 | 12 | 812.42267 | 981.3758 | 1023.9814 | 851.73425 | 1154.1727 | 1110.5775 |
| Eif3m | LFDHLLTLKPVK | -219 | -230 | 1 | 0.000230707 | 12 | 812.42267 | 981.3758 | 1023.9814 | 851.73425 | 1154.1727 | 1110.5775 |
| Eif3m | KWISDWK | -149 | -155 | 1 | 0.000431655 | 7 | 573.78766 | 843.45715 | 1075.2239 | 959.0029 | 509.35077 | 912.5077 |
| Eif3m | LLYEALVDCK | -167 | -176 | 0 | 0 | 10 | 167.0962 | 473.07562 | 749.8665 | 841.9675 | 878.56195 | 829.19995 |
| Eif3m | GAELSEENSEGGLHVD | -26 | -41 | 0 | 0.00025015 | 16 | 1035.9883 | 1172.9788 | NaN | NaN | NaN | 1968.1812 |
| Cdk11b | TLDEILQEK | -12 | -20 | 0 | 0 | 9 | 1328.3649 | 1428.603 | 1482.4537 | 1292.9711 | 769.77045 | NaN |
| Cdk11b | HLHDNWILHR | -541 | -550 | 0 | 0 | 10 | 584.983 | 681.1811 | 970.5229 | 489.15488 | NaN | 716.3096 |
| Cdk11b | EGFPITSLR | -465 | -473 | 0 | 0 | 9 | 240.71347 | 207.09377 | 225.73251 | 154.19356 | NaN | 261.44855 |
| Cdk11b | GELLTQKPLFPGK | -618 | -630 | 1 | 0 | 13 | 1372.5592 | 1709.594 | 1616.8292 | NaN | NaN | NaN |
| Atl3 | EHQHEEIQNVR | -240 | -250 | 0 | 0 | 11 | 748.80695 | 337.43854 | 403.2295 | 5906.6724 | 585.40173 | 437.28198 |
| Atl3 | IYQGEDLPHPK | -331 | -341 | 0 | 0 | 11 | 334.4179 | 1574.1195 | 1584.7783 | 1478.4004 | 1618.1116 | 1653.6746 |
| Atl3 | EGGHSDWLGDPEEPLTGF | -89 | -106 | 0 | 0 | 18 | 277.13226 | 281.26196 | 271.2247 | NaN | 263.16418 | 313.7067 |
| Atl3 | LAMDEIFQKPFQTL | -195 | -208 | 1 | 0.00025015 | 14 | 570.61914 | NaN | NaN | NaN | 676.46014 | NaN |
| Mogs | LGPLLDVLADSR | -706 | -717 | 0 | 0 | 12 | 5990.2925 | 5615.1836 | 5475.9727 | 5386.9805 | 4634.207 | 4942.9062 |
| Mogs | SISGHTSELGDFR | -228 | -240 | 0 | 0 | 13 | 830.2125 | 886.076 | 1142.522 | 750.54364 | 898.2932 | 2018.1737 |
| Mogs | SISGHTSELGDFR | -228 | -240 | 0 | 0 | 13 | 830.2125 | 886.076 | 1142.522 | 750.54364 | 898.2932 | 2018.1737 |
| Mogs | EGHDPDDLAFLR | -521 | -532 | 0 | 0 | 12 | 754.65674 | 1005.0516 | 991.059 | 427.43246 | 1090.6514 | 923.26074 |
| Mogs | GWEFHDGR | -146 | -153 | 0 | 0 | 8 | 693.6815 | 703.2513 | 661.1788 | 672.37274 | 743.0881 | 727.68256 |
| Mogs | LTLLPPTSPGDTVPK | -241 | -255 | 0 | 0 | 15 | 637.38416 | 781.8309 | 900.64514 | 1016.3936 | 1339.635 | 1375.6576 |
| Mogs | LWEQYSDQDGR | -802 | -812 | 0 | 0 | 11 | 1165.5404 | 1273.7749 | NaN | 1428.5148 | 1196.489 | 1104.0457 |
| Mogs | NTEHDPPYWR | -739 | -748 | 0 | 0 | 10 | 955.40393 | 1047.8491 | 924.76996 | NaN | 313.03043 | 1103.0002 |
| Mogs | RPGGQHGGDWSWR | -174 | -186 | 1 | 0 | 13 | 735.0485 | 154.47823 | 587.92474 | NaN | 600.6277 | 621.8229 |
| Mogs | RWDPHLTR | -458 | -465 | 1 | 0.00025015 | 8 | 1050.2042 | 1003.6825 | 992.43896 | NaN | 1097.4095 | NaN |
| Mogs | WDPHLTR | -459 | -465 | 0 | 0 | 7 | 170.4736 | 1786.149 | 2149.227 | NaN | 1607.5328 | NaN |
| Mogs | IHDGALR | -160 | -166 | 0 | 0.003521747 | 7 | 1049.2306 | NaN | NaN | 477.09213 | 811.1857 | NaN |
| Mogs | GFLWEQYSDQDGR | -800 | -812 | 0 | 0.00272025 | 13 | 582.65063 | 206.44968 | 202.2898 | NaN | NaN | NaN |

| | | | | | | | | | | | | |
|---|---|---|---|---|---|---|---|---|---|---|---|---|
| Mogs | TLPSGLDDYPR | -574 | -584 | 0 | 0 | 11 | 2232.5205 | 2169.6758 | NaN | NaN | NaN | NaN |
| Mogs | FVFESGSAATGGNQASGR | -331 | -348 | 0 | 0.000471409 | 18 | 614.90356 | 580.87616 | NaN | NaN | NaN | NaN |
| Mogs | HTCEQGDGVGPY | -134 | -145 | 0 | 0.006788754 | 12 | 1061.257 | NaN | NaN | NaN | NaN | NaN |
| Ensa | EGILPEKAEEAK | -25 | -36 | 1 | 0 | 12 | 563.6936 | NaN | 546.88965 | NaN | 490.75385 | 569.3934 |
| Usp10 | IHKPVSLQPR | -400 | -409 | 1 | 0.000471409 | 10 | 296.455 | 370.1777 | 316.82706 | 234.27042 | 259.29092 | NaN |
| Usp10 | EGLVPVSEDPVAIK | -376 | -389 | 0 | 0.000230707 | 14 | 604.1169 | 503.30457 | NaN | 239.16904 | NaN | 629.9151 |
| Usp10 | NIDYPVDLEISR | -697 | -708 | 0 | 0.00025015 | 12 | 419.72992 | NaN | NaN | NaN | NaN | NaN |
| Snw1 | FLPAPTQLSQDQLEAEER | -6 | -23 | 0 | 0 | 18 | 316.92023 | 390.19342 | 340.61078 | 341.88092 | 417.72516 | 381.9707 |
| Snw1 | FVPDKEFSGSDR | -472 | -483 | 1 | 0 | 12 | 1823.0466 | NaN | NaN | 663.6161 | 656.9477 | 338.6554 |
| Snw1 | EGPVQFEEDPFGLDK | -489 | -503 | 0 | 0.00078661 | 15 | 1047.5212 | NaN | NaN | NaN | NaN | 1593.4716 |
| Snw1 | IKYDAIAR | -96 | -103 | 1 | 0.00133086 | 8 | 66.1689 | NaN | NaN | NaN | NaN | NaN |
| Ddx41 | EGPYGLIICPSR | -256 | -267 | 0 | 0.000826806 | 12 | 853.53467 | NaN | NaN | NaN | 636.94824 | NaN |
| Cnot2 | EGSGNPTPLINPLAGR | -240 | -255 | 0 | 0 | 16 | 753.8442 | 519.29987 | 511.79068 | 854.5202 | 565.2758 | 629.60516 |
| mt-Co3 | EGTYQGHHTPIVQK | -64 | -77 | 0 | 0 | 14 | 1186.6582 | 984.7691 | 1087.187 | 1126.372 | 1258.3674 | 1140.7354 |
| mt-Co3 | EGTYQGHHTPIVQK | -64 | -77 | 0 | 0 | 14 | 1186.6582 | 984.7691 | 1087.187 | 1126.372 | 1258.3674 | 1140.7354 |
| mt-Co3 | HHFGFEAA | -231 | -238 | 0 | 0.001636916 | 8 | 1167.7692 | 999.98926 | 736.8895 | NaN | 754.7342 | 782.77673 |
| Spout1 | EGVVVDRPTK | -192 | -201 | 1 | 0 | 10 | 291.65103 | 273.84805 | 1503.4318 | 3310.2556 | 4760.4673 | NaN |
| Gimap5 | ELEEQEGSWIAK | -258 | -269 | 0 | 0 | 12 | 686.9797 | NaN | NaN | NaN | 219.6413 | 1233.1704 |
| Pym1 | EGYVPQEEVPVYENK | -34 | -48 | 0 | 0.000826806 | 15 | 3455.8538 | 2026.7672 | 3860.577 | 4060.687 | 4412.46 | 5278.677 |
| Pym1 | IQAGEVSQPSR | -170 | -180 | 0 | 0 | 11 | 5198.952 | 3444.1187 | 7743.348 | NaN | NaN | NaN |
| Pym1 | TQRPDGTWR | -19 | -27 | 1 | 0.00025015 | 9 | 2011.7439 | NaN | NaN | NaN | NaN | NaN |
| Smc2 | LSLVDYKPELQK | -607 | -618 | 1 | 0 | 12 | 1491.9722 | 1744.7026 | 1694.0483 | 1170.5348 | 1933.077 | 2120.1455 |
| Smc2 | TVTLGGDVFDPHGTLSGGAR | -649 | -668 | 0 | 0 | 20 | 1437.5637 | 1658.5826 | 1336.9325 | 943.9237 | 782.2946 | 756.24066 |
| Smc2 | SLVDYKPELQK | -608 | -618 | 1 | 0 | 11 | 1216.2712 | 1435.1504 | 1238.3987 | 2170.5442 | 1716.6221 | 1657.0542 |
| Smc2 | EHASNEQQLDAVNEAIK | -832 | -848 | 0 | 0.00025015 | 17 | 688.2532 | 660.08453 | 709.2195 | 730.90375 | 737.6297 | 759.71484 |
| Smc2 | ENLTELSGGQR | -1080 | -1090 | 0 | 0 | 11 | 306.73965 | 572.35187 | 381.45023 | 899.83 | 1675.7627 | NaN |
| Smc2 | HNLQNNESQLK | -899 | -909 | 0 | 0 | 11 | 63.207363 | 514.0286 | 597.0111 | NaN | 896.5778 | 625.31494 |
| Smc2 | TEEGDLLQTK | -723 | -732 | 0 | 0 | 10 | 2444.8323 | 2449.3777 | NaN | 1559.4385 | NaN | NaN |
| Smc2 | TIEESEETLK | -751 | -760 | 0 | 0.00025015 | 10 | 887.0061 | NaN | NaN | 764.41705 | NaN | NaN |
| Smc2 | SIILEGFK | -5 | -12 | 0 | 0.00437931 | 8 | 155.00691 | NaN | NaN | 301.8502 | NaN | NaN |
| Smc2 | ITDGLHGLQEASNK | -355 | -368 | 0 | 0.00025015 | 14 | 659.0409 | NaN | NaN | NaN | NaN | NaN |
| Smc2 | TILEEEITPTIQK | -197 | -209 | 0 | 0 | 13 | 594.73364 | NaN | NaN | NaN | NaN | NaN |
| Atp11b | EHDLFFK | -483 | -489 | 0 | 0.00025015 | 7 | 265.58658 | 490.14032 | 491.47156 | 383.16846 | 464.67065 | 541.4021 |
| Atp11b | WDEPWYNQK | -317 | -325 | 0 | 0 | 9 | 625.7911 | NaN | NaN | NaN | NaN | 131.65283 |
| Parp9 | EHDVLSELQTSTR | -566 | -578 | 0 | 0.000471409 | 13 | 1024.898 | NaN | NaN | 1592.7898 | 1452.7635 | 2085.0378 |
| Parp9 | NSQTAIELLK | -207 | -216 | 0 | 0.000907206 | 10 | 376.5455 | NaN | 678.4089 | NaN | NaN | NaN |
| Drg2 | TCIPGVIEYK | -98 | -107 | 0 | 0 | 10 | 1575.355 | 1665.4288 | 516.19434 | 2867.682 | 1338.3904 | 601.7041 |
| Drg2 | EHEDVIQIVK | -354 | -363 | 0 | 0 | 10 | 654.45764 | 723.3618 | 1883.7485 | 629.9763 | 849.0145 | 761.6061 |
| Drg2 | LIGFPSVGK | -67 | -75 | 0 | 0 | 9 | 164.55994 | 508.49158 | 522.4538 | 401.53787 | 408.98734 | 660.53503 |
| Drg2 | VALIGFPSVGK | -65 | -75 | 0 | 0 | 11 | 1010.4445 | 749.68787 | 934.32996 | NaN | NaN | NaN |
| Mtap | EHEEAVSVDGVLK | -226 | -238 | 0 | 0 | 13 | 1240.4077 | 1667.3602 | 1249.8567 | 2450.654 | 711.23157 | 1014.0017 |
| Mtap | GLDDPEILEGR | -19 | -29 | 0 | 0 | 11 | 995.05676 | 1157.4155 | 1069.3716 | 990.7603 | 1115.7063 | 1079.9652 |
| Mtap | YVDTPFGKPSDALIL | -33 | -47 | 1 | 0 | 15 | 557.2256 | 740.82996 | 641.9401 | 142.39473 | 975.2751 | 1031.9801 |
| Mtap | IGIIGGTGLDDPEILEGR | -12 | -29 | 0 | 0 | 18 | 461.08054 | 317.1545 | 507.82922 | 525.5492 | 562.7896 | 604.91205 |
| Polr2e | EHELVPEHVVMTK | -141 | -153 | 0 | 0.00025015 | 13 | 853.51825 | 419.2736 | 473.34143 | NaN | NaN | NaN |
| Slirp | EHFAQFGHVR | -37 | -46 | 0 | 0 | 10 | 1260.5581 | NaN | 322.76874 | 574.657 | 994.3677 | 7712.2144 |
| Slirp | SSTGRPIAF | -14 | -22 | 1 | 0.003521747 | 9 | 1825.3606 | 1418.116 | NaN | NaN | NaN | NaN |
| Lag3 | SPNLDPNFLR | -48 | -57 | 0 | 0 | 10 | 610.8784 | 514.3687 | 424.66638 | 1301.1497 | 682.53467 | 1502.0144 |
| Lag3 | EHGIQPFPAQR | -479 | -489 | 0 | 0.000627943 | 11 | 663.4867 | NaN | NaN | NaN | NaN | 640.6167 |
| Parp3 | SRPPPINSPDVLQAK | -267 | -281 | 1 | 0.000471409 | 15 | 765.7616 | 640.96497 | 481.03772 | 427.8499 | 385.01035 | 457.8821 |
| Parp3 | SPPSGFDSVIAR | -461 | -472 | 0 | 0 | 12 | 672.3136 | 839.1486 | 880.8197 | 1075.9573 | 1444.8779 | 1414.4456 |
| Parp3 | EHHITIDDPSLK | -449 | -460 | 0 | 0 | 12 | 560.90765 | 1197.4048 | NaN | 481.34897 | 729.9232 | 838.1844 |
| Cops2 | EHIEELLR | -345 | -352 | 0 | 0 | 8 | 1631.3433 | 2149.6904 | 2308.6616 | 1099.097 | 1678.5001 | 1817.1593 |
| Cops2 | SINSILDYISTSK | -111 | -123 | 0 | 0 | 13 | 624.24603 | 587.50104 | 601.6282 | 1073.7637 | 1331.8325 | 1258.336 |
| Cops2 | VLELEGEKGEWGFK | -57 | -70 | 1 | 0 | 14 | 554.2102 | 345.26254 | 889.4681 | 563.7914 | 627.8567 | 228.88496 |
| Cops2 | IHIPFISK | -369 | -376 | 0 | 0.00025015 | 8 | 989.0172 | NaN | 1318.7308 | 1304.5208 | 1500.1627 | 1448.4026 |
| Chordc1 | HDWHQTGGEVTIS | -232 | -244 | 0 | 0 | 13 | 1273.3943 | 1275.3876 | 1493.6526 | 1237.1512 | 1445.9463 | 1527.2992 |
| Chordc1 | FQEHIIQAPK | -92 | -101 | 0 | 0 | 10 | 1209.5378 | 1464.9283 | 1251.1704 | 1220.1654 | 1215.8206 | 1395.9146 |
| Chordc1 | FQEHIIQAPKPVEAIK | -92 | -107 | 1 | 0 | 16 | 1093.8229 | 1330.2904 | 1316.9972 | 1035.0839 | 1559.4807 | 1538.9979 |
| Chordc1 | FQEHIIQAPKPVEAIK | -92 | -107 | 1 | 0 | 16 | 1093.8229 | 1330.2904 | 1316.9972 | 1035.0839 | 1559.4807 | 1538.9979 |
| Chordc1 | FDPEANSDDACTYHPGVPVFHDALK | -14 | -38 | 0 | 0 | 25 | 976.35986 | 605.3211 | 1211.9823 | 631.6713 | 553.4481 | 679.04865 |
| Chordc1 | FDPEANSDDACTYHPGVPVFHDALK | -14 | -38 | 0 | 0 | 25 | 976.35986 | 605.3211 | 1211.9823 | 631.6713 | 553.4481 | 679.04865 |
| Chordc1 | LSSGSEEDKKEEDSDEIK | -135 | -152 | 2 | 0 | 18 | 909.1046 | 951.1998 | 1051.2467 | 509.4795 | 831.83856 | 831.68335 |
| Chordc1 | HNSEKPPEPVKPEVK | -64 | -78 | 2 | 0 | 15 | 695.10675 | 740.1529 | 579.1462 | 414.79742 | 759.0864 | 670.19763 |
| Chordc1 | HNSEKPPEPVKPEVK | -64 | -78 | 2 | 0 | 15 | 695.10675 | 740.1529 | 579.1462 | 414.79742 | 759.0864 | 670.19763 |
| Chordc1 | EHIIQAPKPVEAIK | -94 | -107 | 1 | 0.00025015 | 14 | 431.90417 | 463.93903 | 858.16296 | 534.1718 | 559.2317 | 1436.3687 |
| Chordc1 | NVHIVFEGEK | -265 | -274 | 0 | 0 | 10 | 293.4146 | 635.1737 | 295.0394 | 1131.8951 | 1725.1919 | 948.66925 |
| Chordc1 | QWASLELPTTK | -311 | -321 | 0 | 0 | 11 | 166.09044 | 321.12888 | 618.72034 | 748.54065 | 960.63635 | 988.0275 |
| Chordc1 | SGSEEDKKEEDSDEIK | -137 | -152 | 2 | 0.000431655 | 16 | 339.9216 | NaN | 336.73642 | NaN | 502.83398 | 528.8438 |
| Chordc1 | ELSELKPK | -84 | -91 | 1 | 0.004255025 | 8 | 871.6561 | NaN | NaN | NaN | NaN | NaN |
| Clint1 | EHIYDLR | -111 | -117 | 0 | 0 | 7 | 1233.0396 | 1888.3484 | 985.33795 | 1175.1434 | 699.75 | 933.5505 |
| Clint1 | SLENYHFVDEHGK | -118 | -130 | 0 | 0 | 13 | 448.6349 | 532.11633 | 170.95001 | 413.11993 | 547.38135 | 65.22691 |
| Clint1 | SLENYHFVDEHGK | -118 | -130 | 0 | 0 | 13 | 448.6349 | 532.11633 | 170.95001 | 413.11993 | 547.38135 | 65.22691 |
| Clint1 | GEFKDEEETVTTK | -256 | -268 | 1 | 0.00025015 | 13 | 1276.7484 | 1324.3027 | 1325.4888 | NaN | 1935.0166 | 1869.1359 |
| Dhx33 | EHKDDQLTLTPIGR | -473 | -486 | 1 | 0 | 14 | 353.53214 | 504.78992 | 312.4127 | NaN | 475.25583 | 479.79486 |
| Dhx33 | SVDSVLYNPPAR | -521 | -532 | 0 | 0.00025015 | 12 | 940.257 | NaN | 533.0573 | NaN | 1085.3092 | NaN |
| Acads | NSLYLGPILK | -120 | -129 | 0 | 0.00025015 | 10 | 352.86826 | 576.1298 | 432.0263 | 340.20013 | 278.91553 | 534.8672 |
| Acads | ITEIYEGTSEIQR | -387 | -399 | 0 | 0 | 13 | 1520.9689 | 140.5287 | 1633.8011 | 1546.1627 | 1575.3364 | NaN |

| | | | | | | | | | | | | |
|---|---|---|---|---|---|---|---|---|---|---|---|---|
| Acads | EHLFPTAQVK | -63 | -72 | 0 | 0 | 10 | 1023.69696 | 1051.4583 | NaN | 1101.7526 | 1163.8787 | 940.41144 |
| Acads | LYLGPILK | -122 | -129 | 0 | 0 | 8 | 372.22882 | 408.99213 | 354.89853 | NaN | NaN | NaN |
| Acads | EIYEGTSEIQR | -389 | -399 | 0 | 0 | 11 | 784.91785 | NaN | NaN | 727.1001 | NaN | NaN |
| Acads | LQNIQFK | -307 | -313 | 0 | 0.002401448 | 7 | 235.1749 | NaN | 365.68585 | NaN | NaN | NaN |
| Eef1e1 | EHLLGSTAEEK | -58 | -68 | 0 | 0 | 11 | 3024.1677 | 3170.4587 | 4045.7605 | 3415.5923 | 5264.3203 | 4719.2183 |
| Eef1e1 | QHLSSIVFIK | -157 | -166 | 0 | 0 | 10 | 643.09894 | 587.0074 | 344.5356 | 971.94507 | 527.1919 | 546.30835 |
| Pum3 | GGLKPLIPTLEK | -619 | -630 | 1 | 0 | 12 | 1395.6307 | 1425.8452 | 1340.0035 | 1283.2604 | 1259.7311 | 1361.6708 |
| Pum3 | EHPAGHLVLK | -549 | -558 | 0 | 0 | 10 | 999.1252 | 915.52496 | 899.06726 | NaN | 1297.429 | 1344.2623 |
| Pum3 | SDKTNYDIVVR | -126 | -136 | 1 | 0 | 11 | 972.10364 | NaN | 1102.3231 | 1001.33606 | 1189.501 | 532.4342 |
| Pum3 | TNYDIVVR | -129 | -136 | 0 | 0 | 8 | 2044.0671 | 2199.1108 | NaN | 143.69968 | 2675.0918 | NaN |
| Smap | WEAADLGNEER | -27 | -37 | 0 | 0 | 11 | 716.553 | 741.64667 | 1028.3573 | 733.5653 | 1628.6846 | 1458.771 |
| Smap | EHPDDTEDPK | -153 | -162 | 0 | 0 | 10 | 497.23242 | 387.89258 | 470.20328 | 144.97675 | 31.400711 | 541.5778 |
| Smap | SASPDDDLGSSNWEAADLGNEER | -15 | -37 | 0 | 0 | 23 | 217.78346 | 221.27502 | 236.9017 | 281.29108 | 328.07306 | 293.98343 |
| Smap | NWEAADLGNEER | -26 | -37 | 0 | 0.00025015 | 12 | 867.5016 | NaN | NaN | NaN | 728.3613 | NaN |
| Psph | VIREEGIDELAK | -25 | -36 | 1 | 0.000298805 | 12 | 2133.079 | 2057.4436 | 1630.9417 | 1666.7296 | 1238.278 | 1319.3301 |
| Psph | SIVEHVAAK | -114 | -122 | 0 | 0.00025015 | 9 | 1760.359 | 1830.4658 | 2215.9922 | 1424.6027 | 1735.8472 | 2022.3341 |
| Psph | LAEHPPHLTPGIR | -80 | -92 | 0 | 0 | 13 | 1419.4873 | 1932.1111 | 1720.4222 | 2355.2593 | 3086.4714 | 2968.6023 |
| Psph | VSHSELR | -2 | -8 | 0 | 0.001055699 | 7 | 1266.6387 | 808.9054 | 1293.6133 | 1907.9398 | 691.09546 | 1477.8208 |
| Psph | LALIQPSR | -66 | -73 | 0 | 0 | 8 | 936.89246 | 903.1477 | 837.3573 | 648.6682 | 696.6952 | 483.4746 |
| Psph | EHPPHLTPGIR | -82 | -92 | 0 | 0 | 11 | 663.988 | 346.29565 | 371.24774 | 1000.59454 | 658.3178 | 675.00824 |
| Psph | LLAEHPPHLTPGIR | -79 | -92 | 0 | 0.000826806 | 14 | 751.5804 | 611.52844 | NaN | 57.056015 | NaN | NaN |
| Cmah | EHPYEEIHSR | -509 | -518 | 0 | 0 | 10 | 277.2331 | 435.8305 | NaN | NaN | 570.3567 | 405.54025 |
| Hmox2 | ETLEDGLPVHDGK | -246 | -258 | 0 | 0 | 13 | 1194.814 | 1177.0251 | 1075.7262 | 1246.0294 | 1499.77 | 1420.8276 |
| Hmox2 | YFFGENWEEQVK | -114 | -125 | 0 | 0 | 12 | 933.2272 | 953.40985 | 960.77136 | 1764.0912 | 1823.7748 | 1941.6403 |
| Hmox2 | NKDHPAFAPL | -87 | -96 | 1 | 0.00078661 | 10 | 444.11145 | 566.88715 | 719.36096 | 706.7543 | 779.09143 | 713.7158 |
| Hmox2 | YFPTELHR | -97 | -104 | 0 | 0 | 8 | 1032.3906 | NaN | 342.37167 | 1132.8615 | 1174.8384 | 1320.8065 |
| Hmox2 | EHVDNAQQFK | -189 | -198 | 0 | 0 | 10 | 789.22284 | 782.3173 | NaN | 377.0575 | NaN | 868.4786 |
| Hmox2 | LEDGLPVHDGK | -248 | -258 | 0 | 0 | 11 | 600.4989 | 3308.022 | 2465.244 | NaN | NaN | NaN |
| Hmox2 | MADLSELLK | -29 | -37 | 0 | 0 | 9 | 568.8272 | NaN | NaN | NaN | 940.78827 | 1177.944 |
| Pdcd11 | ELSVRPSELK | -1128 | -1137 | 1 | 0.00025015 | 10 | 1322.5615 | 705.6689 | 1357.6343 | 1581.7445 | 1778.8425 | 936.9081 |
| Pdcd11 | LHQDLVNR | -910 | -917 | 0 | 0 | 8 | 798.2565 | 987.76117 | 859.547 | 899.8076 | 979.7388 | 893.6448 |
| Pdcd11 | YREGEDEVGEPK | -1496 | -1507 | 1 | 0 | 12 | 647.3467 | 591.08765 | 862.2912 | 954.5331 | 969.981 | 1205.5197 |
| Pdcd11 | VAQEEPLEDLLR | -127 | -138 | 0 | 0 | 12 | 139.53181 | 604.76465 | 905.4596 | 627.6611 | 581.11945 | 1043.5256 |
| Pdcd11 | FVTTPSEHFVEGQ | -765 | -777 | 0 | 0 | 13 | 2006.0494 | 2096.863 | 782.1026 | 1651.545 | NaN | 1327.5254 |
| Pdcd11 | EHVDVIVK | -1750 | -1757 | 0 | 0.000591124 | 8 | 1435.4114 | 501.12527 | NaN | 1301.5114 | 1293.7831 | 290.89355 |
| Pdcd11 | VQFPSGLSGLSPK | -746 | -758 | 0 | 0 | 13 | 711.65045 | 814.2046 | 820.9192 | 1014.98035 | 1139.5262 | NaN |
| Pdcd11 | FYNDVQGLVPK | -561 | -571 | 0 | 0 | 11 | 614.33276 | 540.9388 | NaN | 287.7978 | NaN | 285.3813 |
| Pdcd11 | SIIAAPFLR | -438 | -446 | 0 | 0 | 9 | 173.06038 | 373.26987 | NaN | NaN | NaN | NaN |
| Ppp2r5e | GKPIELTPLPLLK | -40 | -52 | 1 | 0 | 13 | 676.2545 | 78.26968 | 323.5492 | 880.3771 | 525.4141 | 308.9406 |
| Ppp2r5e | EHWNPAIVAL | -397 | -406 | 0 | 0.008284062 | 10 | 411.70526 | NaN | NaN | NaN | NaN | NaN |
| Ndufb1 | ELRPNEEVTWK | -47 | -57 | 1 | 0 | 11 | 593.44183 | 1696.1527 | 2024.3138 | 844.28094 | 2402.0166 | 2151.9136 |
| Ndufb1 | EHWVHILVPA | -9 | -18 | 0 | 0.002434616 | 10 | 717.37964 | 1007.1765 | 1318.5854 | NaN | 1290.6171 | 1349.7439 |
| Ndufb1 | TLFQLLR | -2 | -8 | 0 | 0.00111006 | 7 | 147.81871 | 914.9122 | NaN | 696.5993 | 955.49023 | 935.794 |
| Ndufb1 | RELRPNEEVTWK | -46 | -57 | 2 | 0.00025015 | 12 | 635.53503 | 761.0737 | NaN | NaN | 911.60834 | 1124.1617 |
| Capn2 | EIDANEEDIDDGFRR | -522 | -536 | 1 | 0 | 15 | 6388.693 | 7730.6743 | 6929.6807 | 6264.4023 | 5720.2993 | NaN |
| Capn2 | YLNQDYETLR | -27 | -36 | 0 | 0 | 10 | 880.0978 | NaN | NaN | 240.74176 | NaN | NaN |
| Capn2 | TFEPHKDGDFCIR | -488 | -500 | 1 | 0.000907206 | 13 | 2781.3506 | NaN | NaN | NaN | NaN | NaN |
| Etfb | VSVISVEEPPQR | -222 | -233 | 0 | 0 | 12 | 1609.5082 | 1612.4906 | 1766.3126 | 2101.164 | 2034.0073 | 2068.309 |
| Etfb | SCGPSQCQETIR | -65 | -76 | 0 | 0 | 12 | 1354.1864 | 1564.2786 | 1512.1218 | 979.7897 | 998.2976 | 1119.1239 |
| Etfb | EIDGGLETLR | -165 | -174 | 0 | 0 | 10 | 937.8861 | 1137.336 | 882.0554 | 413.7856 | 508.54883 | 860.00854 |
| Etfb | EKVDLLFLGK | -115 | -124 | 1 | 0.001023236 | 10 | 757.7461 | 740.83673 | 778.73047 | 852.8949 | 973.43054 | 1017.6708 |
| Etfb | VDLLFLGK | -117 | -124 | 0 | 0.00025015 | 8 | 610.80115 | 740.2293 | 853.74805 | 611.0289 | 929.5889 | 1033.6462 |
| Etfb | GIHVEIPGAQAE | -86 | -97 | 0 | 0.00025015 | 12 | 1028.3829 | 1026.7747 | 1094.3872 | NaN | 841.34955 | 870.6212 |
| Etfb | SLGPLQVAR | -98 | -106 | 0 | 0.007288197 | 9 | 823.5107 | 662.271 | 592.97125 | NaN | NaN | NaN |
| Etfb | VTLEGDKVK | -153 | -161 | 1 | 0.001392869 | 9 | 309.51675 | NaN | NaN | NaN | NaN | NaN |
| Dynlrb1 | EIDPQNDLTFLR | -59 | -70 | 0 | 0 | 12 | 3018.0408 | 2212.2903 | 5416.0977 | 327.94305 | 2481.8918 | 534.476 |
| Rangap1 | VINLNDNTFTEK | -240 | -251 | 0 | 0 | 12 | 3021.7705 | 7000.293 | 8256.676 | 288.99835 | 1001.475 | 12579.473 |
| Rangap1 | EIEEFDGLEALR | -44 | -55 | 0 | 0 | 12 | 1171.9216 | 1151.4198 | 1109.9066 | 603.30383 | 577.65814 | 554.1431 |
| Rangap1 | IVQQTDTSDPEK | -459 | -470 | 0 | 0 | 12 | 675.19336 | 679.4314 | 709.22943 | 678.7202 | 730.90106 | 835.98474 |
| Rangap1 | QVEVINFGDCLVR | -265 | -277 | 0 | 0 | 13 | 363.8832 | 365.8004 | NaN | 361.43472 | 373.55923 | 413.07495 |
| Rangap1 | LEGNTVGVEAAR | -56 | -67 | 0 | 0 | 12 | 1952.6243 | 1003.01483 | NaN | NaN | NaN | NaN |
| Rangap1 | NRLENDGATAL | -190 | -200 | 1 | 0.002972566 | 11 | 1131.6016 | NaN | NaN | 829.8637 | NaN | NaN |
| Nedd8 | EIEIDIEPTDKVER | -12 | -25 | 1 | 0.00025015 | 14 | 2233.2778 | 2269.6243 | 2247.155 | 2345.3767 | 2581.458 | 2592.7483 |
| Nedd8 | HLVLALR | -68 | -74 | 0 | 0 | 7 | 961.26776 | 1056.2368 | 1029.1333 | 530.1974 | 507.59024 | 565.0235 |
| Impa1 | LCSIPIHGIR | -182 | -191 | 0 | 0 | 10 | 2929.5886 | 3100.9856 | 3511.9656 | 3961.82 | 4092.865 | 3989.5444 |
| Impa1 | LCSIPIHGIR | -182 | -191 | 0 | 0 | 10 | 2929.5886 | 3100.9856 | 3511.9656 | 3961.82 | 4092.865 | 3989.5444 |
| Impa1 | SLLVTELGSSR | -157 | -167 | 0 | 0 | 11 | 1560.1992 | 1684.5497 | 1672.8566 | 690.383 | 2240.555 | 2122.5232 |
| Impa1 | EIEIIPLQR | -265 | -273 | 0 | 0.000591124 | 9 | 1445.0798 | 1408.1902 | 1389.9031 | 1307.7766 | 1399.3168 | 1375.4907 |
| Impa1 | LQVSQQEDITK | -146 | -156 | 0 | 0 | 11 | 1090.6721 | 1217.7396 | 189.85944 | 1369.1564 | 1384.5037 | 1544.2107 |
| Impa1 | SSPADLVTVTDQK | -37 | -49 | 0 | 0 | 13 | 1004.9908 | 1124.3191 | 1150.5045 | 1038.7328 | 1223.4923 | 1170.8838 |
| Apool | YVEEQPGNLQR | -53 | -63 | 0 | 0 | 11 | 851.6913 | NaN | NaN | 916.795 | 458.23294 | 849.5054 |
| Apool | EIHASLPDLK | -206 | -215 | 0 | 0 | 10 | 1873.4552 | NaN | NaN | NaN | NaN | 2344.5847 |
| Brk1 | EIHQDWANR | -11 | -19 | 0 | 0 | 9 | 1106.2032 | 701.22107 | 852.8209 | NaN | 854.6569 | 1044.7181 |
| Brk1 | RIEYIEAR | -60 | -67 | 1 | 0.000550307 | 8 | 1070.239 | NaN | 867.0736 | 931.58655 | 990.86145 | 1281.4229 |
| Ppp4c | EILVEESNVQR | -34 | -44 | 0 | 0 | 11 | 3900.2563 | 4171.4424 | 4189.6284 | 5209.1963 | 5266.542 | 5175.4194 |
| Ppp4c | LWSDPEDTTGWGVSPR | -196 | -211 | 0 | 0 | 16 | 455.63025 | 459.90714 | 537.82416 | 387.93088 | 505.71536 | NaN |
| Dnajc5 | EINNAHAILTDATK | -59 | -72 | 0 | 0 | 14 | 482.97226 | 1128.233 | NaN | 819.78784 | 2119.131 | NaN |
| Rbm17 | KSDSNPLTEILK | -294 | -305 | 1 | 0.00272025 | 12 | 1566.2894 | 1561.3903 | 1574.698 | 1603.4995 | 3118.2754 | 3083.4487 |

| | | | | | | | | | | | | |
|---|---|---|---|---|---|---|---|---|---|---|---|---|
| Rbm17 | QSTVLAPVIDLK | -47 | -58 | 0 | 0 | 12 | 1439.9302 | 2193.9258 | 2257.0051 | 496.52777 | 1610.0752 | 1065.1796 |
| Rbm17 | RPDPDSDEDEDYER | -150 | -163 | 1 | 0.001868731 | 14 | 835.09814 | 928.9721 | 895.14844 | 625.13605 | 811.02155 | 881.7557 |
| Rbm17 | EIPGAPDDEAVR | -347 | -358 | 0 | 0 | 12 | 2646.347 | 3239.5647 | 134.14227 | NaN | 1225.3862 | 1292.8817 |
| Rbm17 | IIVGDATEKGEAQ | -277 | -289 | 1 | 0.00025015 | 13 | 317.33627 | 944.47284 | 678.739 | NaN | 776.5325 | 704.5747 |
| Rbm17 | IAPPTSLVEK | -175 | -184 | 0 | 0.003003218 | 10 | 879.8882 | 458.29404 | 850.10895 | NaN | NaN | NaN |
| Naa25 | EIQYPFLGPVPTR | -748 | -760 | 0 | 0 | 13 | 977.4276 | 734.8917 | 960.73987 | 824.5067 | 829.4258 | 825.88776 |
| Naa25 | LISGLPSLTHPVEPK | -696 | -710 | 0 | 0 | 15 | 621.3181 | 793.4428 | 783.37616 | NaN | 1065.4934 | 1061.1218 |
| Naa25 | YQEALDVIR | -209 | -217 | 0 | 0 | 9 | 2233.8447 | NaN | 2884.8252 | NaN | NaN | 3126.7363 |
| Naa25 | NLRPEEDDVPWEDLR | -639 | -653 | 1 | 0 | 15 | 784.98425 | NaN | NaN | NaN | 917.996 | 410.13614 |
| Il2rg | GLTESLQPDYSER | -316 | -328 | 0 | 0 | 13 | 1320.5193 | 1236.8273 | 1369.0371 | 962.3738 | 831.3379 | 1085.716 |
| Il2rg | NLQNLVIPR | -147 | -155 | 0 | 0 | 9 | 577.10144 | 609.03406 | 722.34937 | 758.25903 | 764.5936 | 893.21674 |
| Il2rg | LQDPQKPQR | -132 | -140 | 1 | 0.00025015 | 9 | 596.2588 | NaN | 291.7711 | 739.3702 | 616.38635 | 599.1265 |
| Il2rg | FSLPSVDELKR | -209 | -219 | 1 | 0 | 11 | 399.05658 | 842.6306 | 929.70233 | 1280.7074 | 1285.3472 | NaN |
| Il2rg | EITSGCQIQK | -110 | -119 | 0 | 0.003166358 | 10 | 1614.5815 | 3712.5127 | NaN | 997.4779 | NaN | 4177.5996 |
| Il2rg | SWTELIVNHEPR | -197 | -208 | 0 | 0 | 12 | 1117.9761 | 1663.9946 | 689.21814 | NaN | NaN | 2700.0117 |
| Il2rg | LNLQNLVIPR | -146 | -155 | 0 | 0.000230707 | 10 | 414.7333 | 330.32364 | 414.2719 | NaN | NaN | 515.5775 |
| Il2rg | VVQLQDPQKPQR | -129 | -140 | 1 | 0 | 12 | 871.5805 | 2560.7236 | 2812.557 | NaN | NaN | NaN |
| Hnrnpf | GITLPVDPEGK | -140 | -150 | 0 | 0 | 11 | 3320.274 | 493.72073 | 3292.9424 | 3830.576 | 3410.8152 | 3669.8271 |
| Hnrnpf | TEMDWVLK | -91 | -98 | 0 | 0 | 8 | 2951.4785 | 2190.307 | 3736.736 | 1642.8984 | 670.6501 | 2196.932 |
| Hnrnpf | HSGPNSADSANDGFVR | -99 | -114 | 0 | 0 | 16 | 2750.0332 | 3018.3088 | 210.74347 | 1862.0723 | 2764.6475 | 3113.1365 |
| Hnrnpf | RYIGIVK | -218 | -224 | 1 | 0.004439092 | 7 | 2425.7742 | 1976.5231 | 1972.881 | 1384.5985 | 1130.5104 | 1164.8209 |
| Hnrnpf | EIVPNGITLPVDPEGK | -135 | -150 | 0 | 0 | 16 | 1247.9382 | 1770.8239 | 1691.8802 | 1331.7908 | 1679.4105 | 1765.355 |
| Hnrnpf | VTGEADVEFATHEEAVAA | -327 | -344 | 0 | 0 | 18 | 932.2852 | 1461.0485 | 1428.523 | 1375.2184 | 1352.1226 | 1386.01 |
| Hnrnpf | SGPNSADSANDGFVR | -100 | -114 | 0 | 0 | 15 | 578.0554 | 787.26843 | 2041.3884 | 1355.3162 | 1486.6456 | 1214.2592 |
| Hnrnpf | VELESEDDVK | -59 | -68 | 0 | 0 | 10 | 443.82767 | 2076.6047 | 1804.2056 | 2092.963 | 1461.9073 | 1337.6484 |
| Hnrnpf | SADSANDGFVR | -104 | -114 | 0 | 0 | 11 | 3138.2292 | 2636.616 | 277.78387 | 4789.932 | NaN | 1730.1345 |
| Hnrnpf | SVQRPGPYDRPGTAR | -203 | -217 | 2 | 0.00025015 | 15 | 551.7448 | 536.2059 | 575.12805 | NaN | 456.5153 | 404.36716 |
| Hnrnpf | FSGLEIVPNGITLPVDPEGK | -131 | -150 | 0 | 0 | 20 | 261.33533 | 277.72006 | 291.27267 | NaN | 319.9436 | 326.72028 |
| Srsf9 | SHEGETSYIR | -173 | -182 | 0 | 0 | 10 | 908.4035 | 951.4072 | 953.74426 | 354.29694 | 718.97406 | 887.1156 |
| Srsf9 | EKDLEDLFYK | -28 | -37 | 1 | 0.00025015 | 10 | 897.41986 | 901.8359 | 328.59955 | 175.91888 | 1319.5131 | 437.1975 |
| Srsf9 | VGNLPSDVR | -19 | -27 | 0 | 0.000230707 | 9 | 1260.2556 | NaN | NaN | 1027.1821 | NaN | NaN |
| Pwp2 | LLDVLSGHEGPVSGL | -493 | -507 | 0 | 0 | 15 | 1302.9296 | 1337.0613 | 162.22443 | 1252.5619 | 498.99713 | 349.6947 |
| Pwp2 | HFKPEIR | -698 | -704 | 1 | 0.00111006 | 7 | 631.9285 | 634.4927 | 496.1725 | 24.4058 | NaN | 1086.843 |
| Pwp2 | EKEEEEEDEEEGDRETTIR | -240 | -258 | 2 | 0 | 19 | 567.8801 | 606.7032 | 627.6436 | NaN | 799.7126 | 781.07587 |
| Ccdc86 | EKEELPVIPK | -294 | -303 | 1 | 0.00025015 | 10 | 529.88416 | 418.89584 | 493.4475 | NaN | 577.35645 | 575.2205 |
| Txlna | EQGVESPGAQPASSPR | -510 | -525 | 0 | 0 | 16 | 488.85632 | 460.19724 | 466.19235 | 475.08142 | 566.4329 | 555.39716 |
| Txlna | YAELLEEHR | -196 | -204 | 0 | 0 | 9 | 1492.3273 | 1155.651 | NaN | NaN | 1222.9491 | 1145.4868 |
| Txlna | EKEFLLK | -358 | -364 | 1 | 0.000627943 | 7 | 625.57837 | NaN | NaN | 552.4138 | NaN | NaN |
| Hsd17b12 | EKFNVETR | -96 | -103 | 1 | 0.000230707 | 8 | 2014.9224 | 1817.4636 | 1644.416 | 2368.931 | 1487.6459 | 1879.7014 |
| Hsd17b12 | IQKPTLDKPSAETF | -242 | -255 | 2 | 0.00025015 | 14 | 1279.2507 | 1168.7148 | 1159.9106 | 1302.5728 | 1209.1781 | 1049.2875 |
| Hsd17b12 | IQKPTLDKPSAE | -242 | -253 | 2 | 0.000550307 | 12 | 787.9468 | 982.9722 | 1525.6539 | 1013.15405 | 803.4214 | 990.252 |
| Sf3b2 | VGEPVALSEEER | -135 | -146 | 0 | 0 | 12 | 2271.6318 | 2406.5269 | 401.41064 | 2691.3335 | 460.1352 | 2227.982 |
| Sf3b2 | LAEIGAPIQGSR | -36 | -47 | 0 | 0 | 12 | 2259.999 | 2039.4666 | 2207.363 | 1436.2078 | 1971.4409 | 1765.959 |
| Sf3b2 | KPGDLSDELR | -588 | -597 | 1 | 0 | 10 | 1903.7051 | 2031.8019 | 1845.8489 | 3670.44 | 3351.632 | 3087.1458 |
| Sf3b2 | KGPAPELQGVEVA | -798 | -810 | 1 | 0 | 13 | 1363.0818 | 1445.0417 | 1501.0153 | 1516.4663 | 1579.1221 | 1495.1848 |
| Sf3b2 | EKKPGDLSDELR | -586 | -597 | 2 | 0 | 12 | 711.734 | 582.0329 | 716.565 | 609.6106 | 490.6203 | 411.8778 |
| Sf3b2 | GIEKPPFELPDFIK | -499 | -512 | 1 | 0 | 14 | 582.39905 | 577.42065 | 744.74805 | 908.5688 | 970.64435 | 1075.5349 |
| Sf3b2 | IPGLNSPIPESCSF | -633 | -646 | 0 | 0 | 14 | 516.7314 | 690.87524 | 589.5075 | 99.54518 | 623.6974 | 1057.4462 |
| Sf3b2 | LHDAFFK | -554 | -560 | 0 | 0.000394361 | 7 | 623.4167 | 681.6189 | 950.7187 | 491.9013 | 526.53784 | NaN |
| Sf3b2 | GPPPPPGDENR | -231 | -241 | 0 | 0 | 11 | 530.4595 | 298.7982 | 842.5039 | NaN | 705.27594 | NaN |
| Mrps27 | YVDSHQWEAR | -34 | -43 | 0 | 0 | 10 | 575.2771 | 1203.8806 | 949.79315 | 1045.8201 | 966.8074 | 1493.7513 |
| Mrps27 | EKLPACEAEDLATYEQK | -363 | -379 | 1 | 0 | 17 | 341.90802 | NaN | 348.34528 | NaN | 545.74695 | 547.6491 |
| Mrps27 | EVLDVLDGVLK | -278 | -288 | 0 | 0 | 11 | 674.46643 | NaN | NaN | NaN | NaN | 976.0589 |
| Mrps27 | TELTWEEER | -194 | -202 | 0 | 0.002780285 | 9 | 778.9915 | NaN | NaN | NaN | NaN | NaN |
| Eif3g | EKLPGELEPVQAA | -194 | -206 | 1 | 0 | 13 | 3162.1624 | 4028.2786 | 880.6715 | 4153.816 | 4902.638 | 5123.762 |
| Eif3g | ETDLQELFRPF | -252 | -262 | 1 | 0 | 11 | 931.6391 | 1057.5433 | 1250.4272 | 1341.8619 | 313.2208 | 1503.0981 |
| Eif3g | PTGDFDSKPSWAD | -2 | -14 | 1 | 0 | 13 | 581.64825 | 549.6688 | 521.3274 | 689.34985 | 543.16187 | 600.62177 |
| Eif3g | ETDLQELFR | -252 | -260 | 0 | 0 | 9 | 450.51965 | 551.44525 | 713.7725 | NaN | NaN | NaN |
| Tmsb4x | TETQEKNPLPSK | -27 | -38 | 1 | 0 | 12 | 3019.608 | 3178.014 | 3135.8174 | 2393.2034 | 3226.3503 | 3092.2166 |
| Tmsb4x | TETQEKNPLPSK | -27 | -38 | 1 | 0 | 12 | 3019.608 | 3178.014 | 3135.8174 | 2393.2034 | 3226.3503 | 3092.2166 |
| Tmsb4x | KTETQEKNPLPSK | -26 | -38 | 2 | 0 | 13 | 2239.2092 | 2322.6755 | 2588.8176 | 3575.7627 | 3538.3818 | 3867.6433 |
| Tmsb4x | KTETQEKNPLPSK | -26 | -38 | 2 | 0 | 13 | 2239.2092 | 2322.6755 | 2588.8176 | 3575.7627 | 3538.3818 | 3867.6433 |
| Tmsb4x | EKNPLPSK | -31 | -38 | 1 | 0 | 8 | 1452.3323 | 1448.8685 | 1371.8353 | 1877.7007 | 2105.6777 | 2003.5068 |
| Ddx56 | EKPDVVVGTPSR | -133 | -144 | 1 | 0 | 12 | 819.4855 | 418.43802 | 430.06885 | 1670.0997 | 1454.0121 | 141.76042 |
| Ddx56 | VLPAEQPFLGK | -389 | -399 | 0 | 0 | 11 | 1161.9545 | 982.53845 | 867.762 | NaN | 615.688 | 952.0906 |
| Prpsap2 | FNIPVDNLR | -156 | -164 | 0 | 0 | 9 | 701.86615 | 877.6005 | 964.69836 | 683.73987 | 909.9103 | 914.01154 |
| Prpsap2 | YIQEEIPDYR | -172 | -181 | 0 | 0 | 10 | 485.35764 | NaN | 644.9833 | 129.04173 | 428.86444 | 774.3496 |
| Prpsap2 | SIIGVIPYFPYSK | -103 | -115 | 0 | 0 | 13 | 1066.2986 | NaN | NaN | 706.5185 | 716.7427 | 729.5053 |
| Ppp1r8 | TIEEGNLDIQRPK | -181 | -193 | 1 | 0 | 13 | 690.4927 | 752.6791 | 665.1569 | NaN | NaN | NaN |
| Ppp1r8 | EKPQTLPSAVK | -128 | -138 | 1 | 0.000712251 | 11 | 521.101 | NaN | NaN | NaN | NaN | NaN |
| Caprin1 | LNQDQLDAVSK | -86 | -96 | 0 | 0 | 11 | 1957.2805 | 2388.4863 | 2364.3289 | 2051.2644 | 2088.8672 | 2238.6206 |
| Caprin1 | QEVTNNLEFAK | -98 | -108 | 0 | 0 | 11 | 1648.2562 | 1529.6691 | 1725.8041 | 1274.081 | 1296.9297 | 1220.7922 |
| Caprin1 | YQEVTNNLEFAK | -97 | -108 | 0 | 0 | 12 | 1161.9816 | 1249.9347 | 558.5713 | 1259.5977 | 1396.9863 | 1319.6223 |
| Caprin1 | SLTPVAQSDPLVR | -341 | -353 | 0 | 0 | 13 | 1151.4065 | 1225.977 | 1355.4464 | 4572.6934 | 2318.4387 | 4114.2095 |
| Caprin1 | SQPLYQPSHATEQRPQK | -443 | -459 | 1 | 0 | 17 | 1017.2708 | 1239.3612 | 1491.8223 | 985.5164 | 1453.8674 | 1299.5859 |
| Caprin1 | SQPLYQPSHATEQRPQK | -443 | -459 | 1 | 0 | 17 | 1017.2708 | 1239.3612 | 1491.8223 | 985.5164 | 1453.8674 | 1299.5859 |
| Caprin1 | EKPVCGTTYK | -220 | -229 | 1 | 0 | 10 | 456.186 | 775.6238 | 616.39136 | 511.87668 | 540.8705 | 481.80066 |
| Caprin1 | NAPVPPANEPETLK | -521 | -534 | 0 | 0.00025015 | 14 | 416.80814 | 775.23395 | 508.2298 | 758.82715 | 640.7459 | 1131.8232 |

| | | | | | | | | | | | | |
|---|---|---|---|---|---|---|---|---|---|---|---|---|
| Caprin1 | GKLDDYQER | -71 | -79 | 1 | 0 | 9 | 209.78181 | 1147.4781 | 106.46853 | 1570.2642 | 1304.5273 | 1216.8015 |
| Caprin1 | LDDYQER | -73 | -79 | 0 | 0 | 7 | 2281.0378 | 2469.724 | 610.59296 | NaN | 3220.4712 | 2949.2783 |
| Caprin1 | LNEQYEHASIH | -201 | -211 | 0 | 0 | 11 | 1186.1346 | 937.9753 | 1055.8296 | 1228.326 | NaN | NaN |
| Caprin1 | EVTNNLEFAK | -99 | -108 | 0 | 0 | 10 | 663.9232 | 1093.392 | 1070.9694 | 1442.16 | NaN | NaN |
| Caprin1 | LGDDDVR | -157 | -163 | 0 | 0.000431655 | 7 | 642.38446 | 656.5307 | 430.55035 | 214.9437 | NaN | NaN |
| Tpm4 | KLVILEGELK | -132 | -141 | 1 | 0 | 10 | 24255.58 | 18594.926 | 24107.713 | 32936.26 | 28074.807 | 21868.967 |
| Tpm4 | EKTIDDLEEK | -214 | -223 | 1 | 0 | 10 | 815.9964 | 900.939 | 741.8067 | 1201.3845 | 1265.2852 | 1191.6697 |
| Tpm4 | YSEKEDKYEEEIK | -178 | -190 | 2 | 0 | 13 | 803.0761 | 892.1627 | 1005.65015 | 864.5276 | 1457.2833 | 1265.4757 |
| Tpm4 | YSEKEDKYEEEIK | -178 | -190 | 2 | 0 | 13 | 803.0761 | 892.1627 | 1005.65015 | 864.5276 | 1457.2833 | 1265.4757 |
| Tpm4 | HITDEADR | -117 | -124 | 0 | 0 | 8 | 685.5441 | 741.6467 | 716.22363 | 646.5014 | 716.2475 | 706.1688 |
| Tpm4 | IQLLEEELDR | -56 | -65 | 0 | 0 | 10 | 419.45004 | 925.7634 | 713.2891 | 1325.7626 | 1476.9487 | 831.4729 |
| Tpm4 | QQQADDAEDR | -18 | -27 | 0 | 0.005668207 | 10 | 428.193 | NaN | 282.6397 | NaN | NaN | NaN |
| Cyfip2 | ELFDLALR | -388 | -395 | 0 | 0.00111006 | 8 | 1048.0377 | 1098.1584 | 1229.0715 | 871.0886 | 1192.3961 | 1290.436 |
| Cyfip2 | SVETDSSTVEHVR | -1227 | -1239 | 0 | 0 | 13 | 955.5548 | 1125.9958 | 1024.2445 | 824.1733 | 1095.2921 | 1134.7274 |
| Cyfip2 | QSFFFTHL | -597 | -604 | 0 | 0.00025015 | 8 | 348.1848 | NaN | 103.092995 | NaN | 472.48373 | 431.01062 |
| Cyfip2 | NYGVIIPYPPSNR | -732 | -744 | 0 | 0.000230707 | 13 | 1546.7874 | 1169.7375 | 1919.409 | NaN | NaN | NaN |
| Cyfip2 | SYLQDPIWR | -1101 | -1109 | 0 | 0 | 9 | 1148.2089 | 778.558 | 1162.0701 | NaN | NaN | NaN |
| Cyfip2 | LQDPIWR | -1103 | -1109 | 0 | 0.002810811 | 7 | 237.28693 | NaN | NaN | 601.59705 | NaN | 676.7538 |
| Rps20 | LIDLHSPSEIVK | -88 | -99 | 0 | 0 | 12 | 7255.756 | 8875.919 | 8693.03 | 7010.558 | 8489.925 | 8739.08 |
| Rps20 | LIDLHSPSEIVK | -88 | -99 | 0 | 0 | 12 | 7255.756 | 8875.919 | 8693.03 | 7010.558 | 8489.925 | 8739.08 |
| Rps20 | TPVEPEVAIH | -9 | -18 | 0 | 0.000591124 | 10 | 5124.0923 | 181.21301 | 5490.656 | 6157.951 | 192.53346 | 666.03046 |
| Rps20 | EKVCADLIR | -33 | -41 | 1 | 0.000550307 | 9 | 3924.5056 | 1045.5775 | 2871.522 | 4498.3203 | 4035.4126 | 4050.2778 |
| Rps20 | TPVEPEVAIHR | -9 | -19 | 0 | 0 | 11 | 3405.4844 | 3751.9255 | 3335.51 | 831.279 | 1177.464 | 893.05176 |
| Rps20 | TPVEPEVAIHR | -9 | -19 | 0 | 0 | 11 | 3405.4844 | 3751.9255 | 3335.51 | 831.279 | 1177.464 | 893.05176 |
| Rps20 | HSPSEIVK | -92 | -99 | 0 | 0.003900536 | 8 | 2703.3184 | 2656.6099 | 2627.6787 | 2496.7212 | 2412.0974 | 2506.0298 |
| Top2b | HLTYNDFINK | -701 | -710 | 0 | 0 | 10 | 1890.8107 | 1340.2826 | 1344.4431 | 1201.8986 | 206.24484 | 1863.2145 |
| Top2b | SIPSLVDGFKPGQR | -723 | -736 | 1 | 0 | 14 | 1748.2095 | 211.80716 | 2048.5964 | 2099.673 | 2322.8413 | 2058.5007 |
| Top2b | KYETVQDILK | -1020 | -1029 | 1 | 0 | 10 | 1049.3214 | 1187.1058 | 1245.3973 | 1252.7577 | 1354.7799 | 1441.3082 |
| Top2b | NTVEITELPVR | -928 | -938 | 0 | 0 | 11 | 1036.562 | 959.3062 | 932.841 | 1061.356 | 1603.9102 | 1366.6388 |
| Top2b | LLFPAVDDNLLK | -825 | -836 | 0 | 0 | 12 | 789.9687 | 797.6027 | 789.3087 | 1065.272 | 1007.7473 | 984.75305 |
| Top2b | HVDYVVDQVVSK | -334 | -345 | 0 | 0 | 12 | 756.3396 | 720.59784 | 712.4171 | 1158.8967 | 1032.9619 | 1334.8088 |
| Top2b | GFLEEFITPIVK | -577 | -588 | 0 | 0 | 12 | 232.71378 | 272.4752 | 205.9471 | 317.79395 | 390.145 | 307.6595 |
| Top2b | EKVEELIK | -1145 | -1152 | 1 | 0.00078661 | 8 | 2890.5793 | 2685.5532 | NaN | 3036.0808 | 2974.5752 | 3636.6938 |
| Top2b | SIPEFDEWKK | -600 | -609 | 1 | 0 | 10 | 1654.7936 | 1251.2789 | 1077.2252 | 1990.9683 | 263.40585 | NaN |
| Top2b | HSLECTLILTEGDSAK | -460 | -475 | 0 | 0.000947459 | 16 | 527.45935 | 650.57886 | 681.48663 | NaN | NaN | 942.8781 |
| Top2b | LHGLPEQF | -686 | -693 | 0 | 0.007987481 | 8 | 1681.0542 | NaN | NaN | NaN | NaN | NaN |
| Kcnab2 | GKYDSGIPPYSR | -253 | -264 | 1 | 0 | 12 | 1263.1343 | 1305.8091 | 848.32996 | 1719.7389 | 1381.1206 | 1914.1173 |
| Kcnab2 | EKVEVQLPELFHK | -223 | -235 | 1 | 0 | 13 | 620.7453 | 716.765 | 334.5613 | 805.0961 | 261.76862 | 1022.4798 |
| Kcnab2 | LKELQAIAER | -289 | -298 | 1 | 0 | 10 | 610.9277 | 581.97546 | 621.2244 | 716.824 | 643.4043 | NaN |
| Rfc2 | ESGCDPSESGAQEPSPVPSK | -5 | -24 | 0 | 0 | 20 | 461.29248 | 575.1365 | 595.77954 | 637.27576 | 797.6525 | 765.1483 |
| Rfc2 | TAGHYELPWVEK | -25 | -36 | 0 | 0 | 12 | 1088.4109 | 3325.48 | 308.8242 | NaN | 3899.8987 | 154.017 |
| Rfc2 | EKVPYTDDGLEAIIF | -204 | -218 | 1 | 0.00383302 | 15 | 558.2172 | NaN | NaN | NaN | NaN | NaN |
| Pgrmc2 | EKYDYVGR | -186 | -193 | 1 | 0.000230707 | 8 | 1136.405 | 1099.7373 | 769.3673 | NaN | 753.71655 | 915.657 |
| Csnk2b | YQQGDFGYCPR | -101 | -111 | 0 | 0 | 11 | 1206.8702 | 1422.7224 | 1320.893 | 1312.0065 | 1456.4023 | 1434.9501 |
| Csnk2b | GLNEQVPHYR | -38 | -47 | 0 | 0 | 10 | 752.5103 | 2255.1292 | 2245.7988 | 2913.46 | 3041.076 | 574.02765 |
| Csnk2b | EKYQQGDFGYCPR | -99 | -111 | 1 | 0 | 13 | 483.3131 | 499.08765 | 653.1292 | 516.4878 | 350.5071 | 664.6251 |
| Csnk2b | GNEFFCEVDEDYIQDK | -18 | -33 | 0 | 0 | 16 | 400.3956 | 341.8834 | 423.70612 | 401.26938 | 526.41693 | 445.725 |
| Csnk2b | NEQVPHYR | -40 | -47 | 0 | 0 | 8 | 823.72955 | 625.2645 | 377.06644 | NaN | 1294.7157 | 565.85297 |
| Atp5me | VPPVQVSPLIK | -2 | -12 | 0 | 0 | 11 | 1835.3293 | 1885.6481 | 1869.3986 | 1703.3877 | 1810.026 | 1816.8253 |
| Atp5me | ELAEAQDDSILK | -60 | -71 | 0 | 0 | 12 | 1401.7086 | 1436.783 | 1451.2583 | 826.5736 | 1556.2295 | 1371.8596 |
| Atp5me | YSYLKPR | -30 | -36 | 1 | 0.002502116 | 7 | 697.1244 | 468.0215 | 353.76166 | 241.092 | NaN | NaN |
| Lamtor3 | VANDSAPEHALRPGFLSTF | -35 | -53 | 1 | 0.00025015 | 19 | 231.83485 | 311.14417 | 277.04752 | 528.0876 | 335.4858 | 453.42526 |
| Lamtor3 | ELAPLFEELIK | -109 | -119 | 0 | 0 | 11 | 166.02429 | 236.63596 | 226.05333 | 274.7545 | 470.05746 | 333.27527 |
| Il2ra | ELCLYDPPEVPNATFK | -22 | -37 | 0 | 0 | 16 | 737.91785 | 1068.0444 | 763.2225 | 868.4752 | 1226.5587 | 1146.3096 |
| Il2ra | TGWTQPQLTCVDER | -171 | -184 | 0 | 0.000230707 | 14 | 4861.803 | 2707.5686 | 3394.4297 | 4026.7964 | NaN | 4401.423 |
| Il2ra | IYHFVEGQ | -135 | -142 | 0 | 0.001868731 | 8 | 1918.8726 | 1947.4799 | 1996.7239 | 1641.973 | NaN | 1822.8754 |
| Il2ra | SVHYECIPGYK | -143 | -153 | 0 | 0 | 11 | 632.88354 | 1625.294 | 1953.2507 | 327.58295 | 561.49335 | NaN |
| Il2ra | FLASEESQGSR | -189 | -199 | 0 | 0 | 11 | 237.60986 | 288.66653 | 307.1686 | NaN | NaN | 418.57465 |
| Unc13d | HEFPEVAPR | -957 | -965 | 0 | 0 | 9 | 8025.9517 | 7708.6475 | 726.93243 | 6572.524 | 938.20715 | 6346.2827 |
| Unc13d | HTIPEEETHR | -162 | -171 | 0 | 0 | 10 | 355.22345 | 453.75018 | 451.9046 | 436.55313 | 516.4494 | 690.85297 |
| Unc13d | ELCQLGPVPSDSR | -564 | -576 | 0 | 0 | 13 | 626.91675 | 814.7549 | 829.3706 | NaN | 851.0979 | 867.2791 |
| Unc13d | GQDDFLGNVVLR | -237 | -248 | 0 | 0 | 12 | 529.6755 | 637.93805 | 598.5835 | NaN | 393.15924 | 810.0405 |
| Otud6b | ELDDEEQLVR | -8 | -17 | 0 | 0 | 10 | 127.86831 | NaN | 2185.8264 | NaN | NaN | NaN |
| Ncapd3 | ELDDTVSLEHQK | -427 | -438 | 0 | 0.001463897 | 12 | 666.0956 | NaN | 358.1893 | NaN | NaN | 392.21115 |
| Ncapd3 | SVISHTDTER | -700 | -709 | 0 | 0.000431655 | 10 | 839.3813 | NaN | NaN | NaN | NaN | NaN |
| Ncapd3 | THNNDEQTLPGER | -523 | -535 | 0 | 0.000394361 | 13 | 561.3494 | NaN | NaN | NaN | NaN | NaN |
| Gmps | ESQSVEEALKK | -279 | -289 | 1 | 0 | 11 | 1095.776 | 951.2721 | 895.614 | 1288.1409 | 1053.365 | 948.7619 |
| Gmps | ELDLPEELVSR | -425 | -435 | 0 | 0 | 11 | 908.1348 | 1019.77637 | 990.8191 | 1219.6144 | 1436.9598 | 1400.1984 |
| Gmps | TTLPISDEDR | -308 | -317 | 0 | 0.001092529 | 10 | 677.4911 | 763.959 | 675.77325 | 811.41064 | 603.81036 | 696.3325 |
| Gmps | NAFLLPIK | -508 | -515 | 0 | 0 | 8 | 596.7229 | 465.19473 | 636.8188 | 415.21143 | 366.6568 | 482.76865 |
| Gmps | YDLTSKPPGTTEWE | -680 | -693 | 1 | 0 | 14 | 521.9854 | 553.9311 | 372.6273 | 358.34277 | 498.8968 | 717.5885 |
| Gmps | SEIFPLETPAFAIK | -52 | -65 | 0 | 0 | 14 | 511.73154 | 583.2685 | 553.66626 | 761.4188 | 711.4452 | 701.1086 |
| Gmps | HPFPGPGLAIR | -436 | -446 | 0 | 0 | 11 | 1813.5956 | 1920.3389 | 2049.033 | 2064.546 | 2522.7246 | NaN |
| Gmps | HFDRDPLQK | -619 | -627 | 1 | 0.00025015 | 9 | 1610.5972 | 2177.2178 | 2071.0063 | 952.5716 | NaN | 2143.737 |
| Gmps | VICAEEPYICK | -447 | -457 | 0 | 0 | 11 | 854.8593 | 1866.7362 | 2091.5574 | 3724.8542 | NaN | 1859.7085 |
| Gmps | SGNIVAGIANESK | -170 | -182 | 0 | 0.000431655 | 13 | 1126.9502 | 796.9518 | 1167.9167 | NaN | NaN | NaN |
| Gmps | THHNDTELIR | -390 | -399 | 0 | 0 | 10 | 664.29584 | NaN | 1405.2051 | 1264.1075 | NaN | NaN |
| Gmps | VIEPLKDFHKD | -407 | -417 | 2 | 0.000550307 | 11 | 1400.6409 | 171.97134 | NaN | NaN | NaN | NaN |

| | | | | | | | | | | | | |
|---|---|---|---|---|---|---|---|---|---|---|---|---|
| Tbl3 | ELDNLLHEK | -656 | -664 | 0 | 0 | 9 | 2038.6332 | 2137.0554 | 783.77496 | 1735.2802 | 394.4343 | 795.91205 |
| Tbl3 | FLGPSDSHIVVA | -342 | -353 | 0 | 0 | 12 | 1333.5217 | 1248.4666 | 1414.2596 | 1246.593 | 1334.5516 | 1317.1313 |
| Tbl3 | HCSQLDDHAITGGSDSR | -612 | -628 | 0 | 0.000431655 | 17 | 378.4688 | 521.5652 | 533.1242 | 628.8565 | 539.332 | 642.2136 |
| Tbl3 | GSLEALLPYTER | -755 | -766 | 0 | 0 | 12 | 546.6869 | NaN | NaN | 1100.4941 | 681.3404 | 923.3517 |
| Tbl3 | GSQLLSSGSDGLLK | -575 | -588 | 0 | 0.00025015 | 14 | 1322.0708 | NaN | NaN | NaN | NaN | NaN |
| Ppp2ca | ELDQWIEQLNECK | -9 | -21 | 0 | 0 | 13 | 1515.092 | 1893.8232 | NaN | 2817.566 | 3420.0237 | 2936.4148 |
| Ppp2cb | ELDQWVEQLNECK | -9 | -21 | 0 | 0 | 13 | 1258.5111 | 1286.5442 | 1337.6583 | 1066.9713 | 828.50964 | 885.21814 |
| Eps15 | IDPFGGDPFK | -639 | -648 | 0 | 0.00051306 | 10 | 4900.2285 | 3791.1062 | NaN | 458.9302 | NaN | NaN |
| Eps15 | ELDTLNNEIVDLQR | -334 | -347 | 0 | 0.00025015 | 14 | 1038.727 | NaN | NaN | NaN | 953.6343 | NaN |
| Eps15 | NITGSSPVADFSAIK | -319 | -333 | 0 | 0.00830926 | 15 | 518.90546 | 1136.9202 | NaN | NaN | NaN | NaN |
| Nup62 | ELEDLLSPLEESVK | -406 | -419 | 0 | 0 | 14 | 327.3308 | 364.08493 | 285.1479 | 342.2405 | 264.28778 | NaN |
| Pdcl3 | ELEEEEAEKEEQLLQQ | -28 | -43 | 1 | 0.00025015 | 16 | 229.54448 | NaN | 696.1016 | 173.86938 | 850.0812 | 857.2227 |
| Alyref | SGFGGGAGVETGGK | -93 | -106 | 0 | 0 | 14 | 3662.155 | 3648.9304 | 3540.653 | 3562.6 | 674.8192 | 3472.7017 |
| Alyref | SGFGGGAGVETGGK | -93 | -106 | 0 | 0 | 14 | 3662.155 | 3648.9304 | 3540.653 | 3562.6 | 674.8192 | 3472.7017 |
| Alyref | QLPDKWQHDLFD | -81 | -92 | 1 | 0 | 12 | 2303.489 | 2282.97 | 2289.8398 | 969.38495 | 1021.7963 | 1167.1036 |
| Alyref | LVTSQIDTQR | -179 | -188 | 0 | 0 | 10 | 469.06476 | NaN | NaN | 476.17087 | NaN | NaN |
| Rfc4 | ELFGPELFR | -94 | -102 | 0 | 0 | 9 | 655.742 | 311.33716 | 921.759 | 641.14325 | 625.87146 | 690.5655 |
| Rfc4 | SLEGADLPNLLFYGPPGTGK | -65 | -84 | 0 | 0.001092529 | 20 | 330.33353 | 275.33572 | 229.31607 | 419.65475 | 410.55484 | 338.5657 |
| Acad9 | ELFLGNIEQK | -46 | -55 | 0 | 0 | 10 | 486.48462 | 533.12604 | 485.67615 | 699.6105 | 672.6598 | 779.77435 |
| Acad9 | FFTEEVDSR | -81 | -89 | 0 | 0 | 9 | 2538.0327 | NaN | 3204.7576 | 3119.1064 | 2898.4595 | NaN |
| Acad9 | VPVENVLGEVGGGFK | -284 | -298 | 0 | 0 | 15 | 384.4503 | 904.1947 | 951.0338 | NaN | NaN | 1208.7108 |
| Setd3 | ELFLWVPR | -118 | -125 | 0 | 0.000347792 | 8 | 496.6135 | 499.26044 | 491.03955 | 476.4164 | NaN | 83.40821 |
| Setd3 | YEESDLGLLEGGVGDSR | -502 | -518 | 0 | 0.003360691 | 17 | 426.06158 | NaN | 549.69366 | NaN | NaN | NaN |
| Setd3 | GNAEFPVSWDNEVK | -412 | -425 | 0 | 0 | 14 | 407.4157 | NaN | NaN | 1828.9651 | NaN | NaN |
| Dhx8 | ELFPVLCQPDNPSAR | -117 | -131 | 0 | 0 | 15 | 1051.211 | 1216.0994 | 1297.6737 | 1170.2325 | 2099.1191 | 1845.97 |
| Fth1 | ELGDHVTNLR | -148 | -157 | 0 | 0 | 10 | 6706.658 | 6955.2847 | 6820.734 | 5654.893 | 5043.119 | 4913.455 |
| Fth1 | SVNQSLLELHK | -110 | -120 | 0 | 0 | 11 | 1505.0282 | NaN | 1698.2076 | 1904.3337 | 1785.2446 | 1785.1395 |
| Fth1 | LATDKNDPHLCDFIETY | -121 | -137 | 1 | 0.000431655 | 17 | 418.40924 | NaN | 393.76532 | 223.74261 | NaN | 275.64975 |
| Diaph1 | ITPAEELDFR | -313 | -322 | 0 | 0 | 10 | 2299.4734 | 2662.7163 | 2732.8113 | 3096.2898 | 1026.2329 | 1398.1926 |
| Diaph1 | LIEECVSQIVLHK | -424 | -436 | 0 | 0 | 13 | 722.1323 | 933.5088 | 778.11115 | 809.0681 | 1688.2521 | 1479.9595 |
| Diaph1 | KSEGVPTILEEAK | -1236 | -1248 | 1 | 0 | 13 | 663.7525 | 590.95197 | 734.5413 | 804.68616 | 755.54236 | 442.7978 |
| Diaph1 | ELGDYFVFDPK | -1100 | -1110 | 0 | 0 | 11 | 651.7608 | 424.13696 | 801.3508 | 141.44722 | 1090.4734 | 1025.9463 |
| Diaph1 | FQPLLDGLK | -286 | -294 | 0 | 0 | 9 | 468.8715 | 523.7531 | 520.85693 | 636.854 | 580.4301 | 578.3306 |
| Diaph1 | KQDLEAEVSK | -525 | -534 | 1 | 0 | 10 | 369.67868 | 456.93677 | 569.8271 | 2988.4983 | 981.1558 | 783.46405 |
| Diaph1 | HLQIDIER | -447 | -454 | 0 | 0 | 8 | 1321.6783 | 1304.5996 | NaN | 1148.8701 | 1156.9015 | 1029.1877 |
| Diaph1 | VFDEQGDEDFFDLK | -355 | -368 | 0 | 0 | 14 | 695.46387 | 819.3658 | 820.2694 | 607.0866 | NaN | NaN |
| Diaph1 | LQDLQGEKDALDSEK | -504 | -518 | 1 | 0.000550307 | 15 | 635.06757 | 853.6434 | 694.62537 | NaN | NaN | NaN |
| Ipo9 | ELGENLDQILR | -777 | -787 | 0 | 0 | 11 | 793.15375 | 906.7408 | 1175.1599 | 1181.9349 | 818.95807 | 1336.1089 |
| Ifitm2 | ELGEPSNSAVVR | -30 | -41 | 0 | 0.001023236 | 12 | 1863.1117 | NaN | NaN | NaN | NaN | NaN |
| Armc10 | ELGGIPIVGNK | -99 | -109 | 0 | 0.00051306 | 11 | 463.6255 | NaN | 219.62747 | 222.64328 | NaN | 283.33948 |
| Casp8 | FLCLDYIPHK | -24 | -33 | 0 | 0 | 10 | 992.99054 | 1264.4017 | 1113.7811 | 564.29987 | 1650.4144 | 572.7059 |
| Casp8 | KQETIEDAQK | -34 | -43 | 1 | 0.001092529 | 10 | 808.9405 | 1189.744 | 933.26135 | 406.80334 | 907.73785 | 1414.1763 |
| Casp8 | LSEEVSELELR | -108 | -118 | 0 | 0 | 11 | 707.50336 | 1053.5934 | 922.67236 | 643.46796 | 1057.0549 | 937.88916 |
| Casp8 | ELRDPDNAQISPY | -89 | -101 | 1 | 0.000394361 | 13 | 974.8142 | 753.8039 | NaN | 1289.878 | 1647.8657 | 712.3775 |
| Casp8 | FLLNNEIPK | -122 | -130 | 0 | 0 | 9 | 749.4663 | 794.2142 | 787.6064 | NaN | NaN | NaN |
| Casp8 | IINNHDFSK | -240 | -248 | 0 | 0 | 9 | 942.95306 | 991.5243 | NaN | NaN | NaN | NaN |
| Casp8 | ELGSEDLAALK | -13 | -23 | 0 | 0.002434616 | 11 | 1396.9548 | NaN | NaN | NaN | NaN | NaN |
| Blvra | FGFPAFSGISR | -161 | -171 | 0 | 0 | 11 | 1779.4346 | 2246.5374 | 2118.826 | 2523.4443 | 3202.1514 | 2838.6377 |
| Blvra | TESSSHEDYIR | -74 | -84 | 0 | 0 | 11 | 1030.6891 | 1081.5999 | 988.6646 | 799.1135 | 968.55225 | 890.6421 |
| Blvra | SGSLEEVPNVGVNK | -234 | -247 | 0 | 0 | 14 | 850.5252 | 1112.8892 | 1152.9438 | 1227.3759 | 1344.5323 | 1257.2344 |
| Blvra | ELGSLDNVR | -46 | -54 | 0 | 0 | 9 | 761.2342 | 924.90454 | 617.78265 | 859.6176 | 732.9228 | 210.04013 |
| Blvra | QISLEDALR | -55 | -63 | 0 | 0.00122419 | 9 | 1020.863 | 863.5832 | 974.781 | 783.6093 | NaN | NaN |
| Blvra | FGVVVVGVGR | -9 | -18 | 0 | 0 | 10 | 974.94965 | 1060.7927 | NaN | NaN | NaN | NaN |
| Mri1 | YVAAPSSSCDLHLETGK | -277 | -293 | 0 | 0 | 17 | 1328.3981 | 1580.2845 | 1726.5492 | 893.3394 | 1742.3109 | 1219.353 |
| Mri1 | HLLEQTNPR | -150 | -158 | 0 | 0 | 9 | 1113.0165 | 1169.2129 | 974.79584 | 357.3445 | 1130.7943 | 1138.5605 |
| Mri1 | ELGVFAPEELR | -340 | -350 | 0 | 0 | 11 | 802.5746 | 677.4798 | 601.09674 | 669.6903 | 971.93274 | NaN |
| Mri1 | GVSAVVVGADR | -239 | -249 | 0 | 0.009242859 | 11 | 262.94418 | NaN | NaN | NaN | NaN | NaN |
| Ndrg1 | ELHDVDLAEVKPLVEK | -4 | -19 | 1 | 0.000230707 | 16 | 548.1858 | 690.8302 | 583.7307 | 214.4862 | 647.91 | 689.5699 |
| Ndrg1 | ELHDVDLAEVKPLVEK | -4 | -19 | 1 | 0.000230707 | 16 | 548.1858 | 690.8302 | 583.7307 | 214.4862 | 647.91 | 689.5699 |
| Zc3hc1 | ELIDEGIVPEEGGTEPK | -34 | -50 | 0 | 0 | 17 | 749.79004 | 1004.3308 | 975.15625 | 931.5949 | 972.16113 | 897.3636 |
| Zc3hc1 | SLQPTFDFGR | -127 | -136 | 0 | 0 | 10 | 348.43668 | 383.82587 | NaN | NaN | NaN | 447.82477 |
| Zc3hc1 | FCFWPDSPSPDR | -155 | -166 | 0 | 0 | 12 | 580.0688 | 613.46063 | NaN | NaN | NaN | NaN |
| Bag4 | ELLELDSVETGGQDSVR | -419 | -435 | 0 | 0 | 17 | 426.04367 | 1099.9183 | 950.7103 | 1146.6775 | 871.1824 | 975.4905 |
| Isg15 | NCSEPLSILVR | -75 | -85 | 0 | 0 | 11 | 928.0405 | 707.1642 | 383.5778 | 1413.1583 | 699.7286 | 1302.8138 |
| Isg15 | ELLGEYGLKPQ | -133 | -143 | 1 | 0.00025015 | 11 | 752.0163 | 977.9501 | 791.67737 | 1234.138 | 1580.8729 | 1459.1174 |
| Isg15 | IGVPAFQQR | -36 | -44 | 0 | 0 | 9 | 1207.0747 | 1083.7205 | 1221.0176 | 1027.9677 | 848.27124 | NaN |
| Isg15 | GHSNIYEVF | -89 | -97 | 0 | 0.005758933 | 9 | 1189.7522 | 1073.457 | 1092.4581 | 1557.0044 | 978.2771 | NaN |
| Isg15 | EQVHEDQFWLS | -113 | -123 | 0 | 0.000230707 | 11 | 710.9636 | 634.623 | NaN | 904.65625 | 845.646 | 408.50165 |
| Tnfaip8 | ELLHEIIQR | -137 | -145 | 0 | 0.00025015 | 9 | 3022.7864 | 2607.514 | NaN | NaN | NaN | NaN |
| Cnot6l | ELLLNDNYLR | -106 | -115 | 0 | 0 | 10 | 507.43942 | NaN | 489.5181 | NaN | NaN | NaN |
| Rps9 | KIEDFLER | -101 | -108 | 1 | 0.00025015 | 8 | 6144.147 | 5754.855 | 6147.8013 | 11411.971 | 11500.997 | 11308.103 |
| Rps9 | VVNIPSFIVR | -141 | -150 | 0 | 0 | 10 | 5873.0884 | 6347.09 | 6507.7095 | 7447.4844 | 8682.985 | 8608.159 |
| Rps9 | HIDFSLR | -156 | -162 | 0 | 0 | 7 | 3915.6777 | 3983.06 | 4368.513 | 4329.899 | 4946.872 | 4744.37 |
| Rps9 | SRLDQELK | -23 | -30 | 1 | 0.00122419 | 8 | 3637.7424 | 3565.0596 | 3841.9934 | 449.05112 | 3349.6177 | 3375.0193 |
| Rps9 | ELLTLDEKDPR | -59 | -69 | 1 | 0 | 11 | 2811.8467 | 2972.0947 | 3150.1426 | 3804.0974 | 3425.6992 | 3344.4355 |
| Rps9 | LIGEYGLR | -31 | -38 | 0 | 0 | 8 | 1960.46 | 1747.5464 | 1788.3026 | 885.18494 | 699.7919 | 878.8117 |
| Rps9 | QVVNIPSFIVR | -140 | -150 | 0 | 0 | 11 | 1954.2457 | 2242.9575 | 2165.96 | 201.3076 | 171.50082 | 163.11829 |
| Rps9 | LDYILGLK | -94 | -101 | 0 | 0 | 8 | 1471.3348 | 1595.8137 | 1610.7291 | 940.9536 | 951.98535 | 1069.8152 |

| | | | | | | | | | | | | |
|---|---|---|---|---|---|---|---|---|---|---|---|---|
| Rps9 | LFEGNALLR | -71 | -79 | 0 | 0 | 9 | 1322.5483 | 1199.6569 | 1286.8154 | 1289.2278 | 1201.2592 | 1209.7394 |
| Rps9 | KQVVNIPSFIVR | -139 | -150 | 1 | 0 | 12 | 1027.7748 | 1244.7954 | 1224.8524 | 897.9949 | 1111.4795 | 1132.2969 |
| Rps9 | KQVVNIPSFIVR | -139 | -150 | 1 | 0 | 12 | 1027.7748 | 1244.7954 | 1224.8524 | 897.9949 | 1111.4795 | 1132.2969 |
| Rps9 | IEDFLER | -102 | -108 | 0 | 0.000471409 | 7 | 3062.937 | 2529.2935 | 2528.2488 | 1380.543 | NaN | 963.7987 |
| Rps9 | LTLDEKDPR | -61 | -69 | 1 | 0.00025015 | 9 | 2513.5203 | 1917.3844 | 1667.1985 | 1077.5151 | NaN | 852.8811 |
| Rps9 | MKLDYILGLK | -92 | -101 | 1 | 0 | 10 | 844.5036 | 809.1481 | 783.13416 | NaN | 147.61418 | 629.3724 |
| Rps9 | RLDQELK | -24 | -30 | 1 | 0.003393785 | 7 | 1492.3838 | NaN | NaN | 4274.585 | 4398.879 | 4266.0024 |
| Rps9 | SPYGGGRPGR | -163 | -172 | 1 | 0.001636916 | 10 | 178.31343 | NaN | 1139.1012 | 781.38306 | NaN | 1343.6047 |
| Rps9 | KTYVTPR | -11 | -17 | 1 | 0.001463897 | 7 | 2132.335 | 2171.6514 | NaN | NaN | NaN | 1634.2632 |
| Rps9 | IGVLDEGK | -84 | -91 | 0 | 0.002168954 | 8 | 1353.9325 | NaN | 1192.7192 | NaN | NaN | 855.6588 |
| Lnpep | ELNSYEDFLDAR | -504 | -515 | 0 | 0 | 12 | 1619.3418 | 1134.9105 | 2138.0857 | 2105.86 | 1887.3574 | 2587.818 |
| Lnpep | SSLDGDIIR | -912 | -920 | 0 | 0 | 9 | 1349.159 | 1789.6583 | 927.78925 | 654.4993 | 817.083 | 987.39935 |
| Lnpep | NPYVLSDKDR | -719 | -728 | 1 | 0 | 10 | 1287.9156 | 1201.5052 | 1242.2465 | 1162.5406 | 1490.9551 | 1105.1564 |
| Lnpep | SDDLWDSFNEVTDK | -586 | -599 | 0 | 0.00025015 | 14 | 606.04047 | 674.0355 | 764.9326 | 677.2064 | 756.7891 | 781.54004 |
| Lnpep | YFEIQYPLK | -407 | -415 | 0 | 0 | 9 | 274.27438 | 3062.59 | 412.3029 | 2544.2185 | 3499.2974 | 3284.0935 |
| Lnpep | FENQSEATLK | -987 | -996 | 0 | 0 | 10 | 641.9119 | 281.25256 | 601.82544 | 975.15 | 463.02347 | NaN |
| Lnpep | IGQVHHALDTTIK | -387 | -399 | 0 | 0.000550307 | 13 | 734.80304 | NaN | NaN | 1269.0967 | 706.1721 | 1154.3254 |
| Lnpep | SSVPAEEGLIQDEFSESVK | -335 | -353 | 0 | 0 | 19 | 674.92896 | NaN | 800.6704 | NaN | 344.06595 | 890.81335 |
| Lnpep | ENWGLLTFR | -431 | -439 | 0 | 0 | 9 | 279.08646 | NaN | 301.1493 | NaN | 349.15982 | NaN |
| Lnpep | LGHMDLSSR | -782 | -790 | 0 | 0 | 9 | 158.29771 | 335.63766 | NaN | NaN | 840.0251 | NaN |
| Lnpep | SAFPCFDEPAFK | -301 | -312 | 0 | 0 | 12 | 623.0703 | 566.363 | NaN | NaN | NaN | NaN |
| Lnpep | HIPISYVTDGR | -653 | -663 | 0 | 0.002972566 | 11 | 1208.8466 | NaN | NaN | NaN | NaN | NaN |
| Lnpep | VFPWAQIR | -160 | -167 | 0 | 0.007987481 | 8 | 1084.7089 | NaN | NaN | NaN | NaN | NaN |
| Lnpep | SALLEFACAH | -821 | -830 | 0 | 0.006384543 | 10 | 795.19543 | NaN | NaN | NaN | NaN | NaN |
| Arhgap30 | ELPDPLLTYR | -100 | -109 | 0 | 0 | 10 | 780.0695 | 1014.0779 | 1050.1451 | 972.9818 | 1174.5895 | 1078.153 |
| Ppp6r1 | SQDPTHPSAPQEVTDSSK | -750 | -767 | 0 | 0 | 18 | 552.34894 | 582.32776 | 558.6857 | 616.1152 | 637.0078 | 662.8335 |
| Ppp6r1 | GFNDEEFGEQEESVNAPFDK | -570 | -589 | 0 | 0 | 20 | 326.8588 | 337.94293 | 377.72348 | 364.57104 | 388.2618 | 390.92972 |
| Ppp6r1 | KKDDFVDLLLR | -145 | -155 | 2 | 0 | 11 | 303.7543 | 400.1229 | 693.0316 | 274.07922 | 366.3774 | 468.67706 |
| Ppp6r1 | KKDDFVDLLLR | -145 | -155 | 2 | 0 | 11 | 303.7543 | 400.1229 | 693.0316 | 274.07922 | 366.3774 | 468.67706 |
| Ppp6r1 | QDVFNWLNEEK | -180 | -190 | 0 | 0 | 11 | 1287.6212 | 757.4407 | NaN | 2768.7876 | 3765.7224 | 2927.9531 |
| Ppp6r1 | NAEQGPNAEQLGQLLK | -477 | -492 | 0 | 0.00025015 | 16 | 746.1534 | 895.5202 | 966.74036 | NaN | 1325.396 | NaN |
| Ppp6r1 | SAPSSLDSATR | -793 | -803 | 0 | 0.002068213 | 11 | 1142.8066 | NaN | 1221.1338 | 1454.0266 | NaN | NaN |
| Ppp6r1 | ELPEEQQQR | -493 | -501 | 0 | 0.000394361 | 9 | 210.0658 | 1072.2955 | NaN | 94.41881 | NaN | NaN |
| Hpf1 | IIEDHLASR | -328 | -336 | 0 | 0 | 9 | 1478.1406 | 1272.6306 | 1049.551 | 790.89545 | 928.3431 | 924.6013 |
| Hpf1 | ELPETDADLKR | -240 | -250 | 1 | 0.000712251 | 11 | 1791.2325 | NaN | 1969.6759 | NaN | NaN | NaN |
| Csde1 | VTLLEGDHVR | -308 | -317 | 0 | 0 | 10 | 1695.5809 | 1738.0988 | 1900.6445 | 1602.2877 | 1837.4229 | 1798.9456 |
| Csde1 | IKPEIHPEER | -90 | -99 | 1 | 0 | 10 | 1016.5136 | 1042.6992 | 125.28146 | 782.54553 | 1067.8467 | 1009.594 |
| Csde1 | IKPEIHPEER | -90 | -99 | 1 | 0 | 10 | 1016.5136 | 1042.6992 | 125.28146 | 782.54553 | 1067.8467 | 1009.594 |
| Csde1 | NQNDPLPGR | -278 | -286 | 0 | 0 | 9 | 1292.6997 | 285.79715 | 1140.2954 | 357.28842 | NaN | 913.0527 |
| Csde1 | SNTFQFTNEAR | -338 | -348 | 0 | 0 | 11 | 892.33966 | 726.5299 | 810.92255 | NaN | NaN | 693.1753 |
| Csde1 | INFVIDNNK | -158 | -166 | 0 | 0.000550307 | 9 | 304.18893 | 1128.2587 | 344.83533 | NaN | 1246.7256 | NaN |
| Csde1 | ELPFGDKDTK | -296 | -305 | 1 | 0.00025015 | 10 | 933.4363 | NaN | NaN | 564.3729 | NaN | 1391.4811 |
| Csde1 | SFDPNLLHNN | -2 | -11 | 0 | 0.001610812 | 10 | 1286.1594 | NaN | 89.38702 | NaN | NaN | NaN |
| Zc3h15 | LVNDDDEEADDTR | -301 | -313 | 0 | 0 | 13 | 473.64926 | 548.6537 | 589.9468 | 571.4065 | 781.5342 | 793.3202 |
| Zc3h15 | ELQELNELFKPVVA | -77 | -90 | 1 | 0.00025015 | 14 | 1157.1254 | 849.759 | NaN | 785.0963 | 797.6625 | 1026.6631 |
| Zc3h15 | FSTYAPDKDENK | -355 | -366 | 1 | 0 | 12 | 1248.4584 | 1613.2402 | 1397.7766 | NaN | NaN | 1558.3035 |
| Zc3h15 | ELQELNELFKPVVAA | -77 | -91 | 1 | 0.000394361 | 15 | 394.25644 | NaN | NaN | NaN | 525.67694 | NaN |
| Nup88 | ELQLIPDQLR | -676 | -685 | 0 | 0 | 10 | 598.63324 | 612.1677 | 547.9354 | 607.32874 | 129.10066 | 633.63745 |
| Nup88 | FLGSDEEDKDSLQEL | -428 | -442 | 1 | 0.000712251 | 15 | 890.88324 | NaN | 746.604 | 579.316 | 2352.8154 | 2423.7146 |
| Rnf40 | ELSSGTEVPGCQEGLTR | -117 | -133 | 0 | 0 | 17 | 1170.9335 | 1662.376 | NaN | 1459.1368 | 646.74774 | NaN |
| Rnf40 | EPLPVQFR | -148 | -155 | 0 | 0.000907206 | 8 | 533.31573 | NaN | NaN | NaN | NaN | NaN |
| Gsk3b | ELSSNPPLATILIPPHAR | -366 | -383 | 0 | 0 | 18 | 413.17722 | 490.53693 | 553.5473 | NaN | 682.67487 | 715.698 |
| Gsk3b | NLLLDPDTAVLK | -186 | -197 | 0 | 0.001023236 | 12 | 963.1257 | NaN | 1028.7743 | NaN | 1194.8202 | 1276.824 |
| Rbbp7 | WLPEVTKPEGK | -48 | -58 | 1 | 0 | 11 | 3121.2222 | 3203.4197 | 3463.5942 | 3673.2346 | 3134.9636 | 3109.5696 |
| Rbbp7 | HPAKPDPSGECNPDLR | -156 | -171 | 1 | 0 | 16 | 1595.01 | 1737.7039 | 1712.287 | 1751.2883 | 1944.9713 | 1870.8307 |
| Rbbp7 | HPAKPDPSGECNPDLR | -156 | -171 | 1 | 0 | 16 | 1595.01 | 1737.7039 | 1712.287 | 1751.2883 | 1944.9713 | 1870.8307 |
| Rbbp7 | VHIPNDDAQFDAS | -83 | -95 | 0 | 0.001023236 | 13 | 1039.7338 | 1049.6471 | 1046.9744 | 1203.9354 | 765.3526 | 730.0365 |
| Rbbp7 | SNTTSKPSHLVDAHTA | -258 | -273 | 1 | 0 | 16 | 748.10645 | 953.42926 | 1021.3577 | 732.2908 | 799.13934 | 830.56195 |
| Rbbp7 | SNTTSKPSHLVDAHTA | -258 | -273 | 1 | 0 | 16 | 748.10645 | 953.42926 | 1021.3577 | 732.2908 | 799.13934 | 830.56195 |
| Rbbp7 | SNTTSKPSHLVDAH | -258 | -271 | 1 | 0 | 14 | 700.37585 | 566.3884 | 185.06915 | 972.2653 | 898.9875 | 654.4907 |
| Rbbp7 | TSDEQNHLVVAR | -71 | -82 | 0 | 0 | 12 | 680.2448 | 1397.9398 | 952.40125 | 543.7513 | 619.99866 | 200.92072 |
| Rbbp7 | IGEEQSAEDAEDGPPELLFIH | -349 | -369 | 0 | 0 | 21 | 450.54865 | 571.288 | 593.7919 | 388.64108 | 457.55493 | 494.54645 |
| Rbbp7 | GTHTSDEQNHLVVAR | -68 | -82 | 0 | 0 | 15 | 443.86603 | 562.70374 | 276.42117 | 480.71683 | 563.08923 | 539.50024 |
| Rbbp7 | SDDHTVCLWDINAGPK | -196 | -211 | 0 | 0 | 16 | 438.17938 | 594.4291 | 591.47723 | 602.3598 | 636.41504 | 636.81586 |
| Rbbp7 | LVLGTHTSDEQNHLVVAR | -65 | -82 | 0 | 0 | 18 | 399.08353 | 438.54028 | 223.09418 | 416.04907 | 697.172 | 616.93243 |
| Rbbp7 | SNTTSKPSHLVDAHTAEVN | -258 | -276 | 1 | 0.000394361 | 19 | 341.7186 | 229.98746 | 693.9994 | 945.86865 | 462.48517 | 763.5614 |
| Rbbp7 | SNTTSKPSHLVDAHTAEVN | -258 | -276 | 1 | 0.000394361 | 19 | 341.7186 | 229.98746 | 693.9994 | 945.86865 | 462.48517 | 763.5614 |
| Rbbp7 | LWDINAGPK | -203 | -211 | 0 | 0.000712251 | 9 | 2969.8367 | 3141.4724 | NaN | 3451.1445 | 318.3343 | 3453.2236 |
| Rbbp7 | SASDDHTVCLWDINAGPK | -194 | -211 | 0 | 0 | 18 | 828.3639 | 999.2335 | NaN | 1067.912 | 1258.5305 | 1263.1176 |
| Rbbp7 | SASDDHTVCLWDINAGPK | -194 | -211 | 0 | 0 | 18 | 828.3639 | 999.2335 | NaN | 1067.912 | 1258.5305 | 1263.1176 |
| Rbbp7 | EMFEDTVEER | -5 | -14 | 0 | 0 | 10 | 385.36337 | 746.7626 | 958.33746 | 319.30325 | 484.4447 | NaN |
| Rbbp7 | HTSDEQNHLVVAR | -70 | -82 | 0 | 0 | 13 | 3967.166 | 3352.08 | 4018.5266 | NaN | 4001.0645 | NaN |
| Rbbp7 | HTSDEQNHLVVAR | -70 | -82 | 0 | 0 | 13 | 3967.166 | 3352.08 | 4018.5266 | NaN | 4001.0645 | NaN |
| Rbbp7 | WDINAGPK | -204 | -211 | 0 | 0.00051306 | 8 | 2284.076 | NaN | NaN | 3921.9814 | NaN | NaN |
| Eif3l | RVSSDVIDQK | -78 | -87 | 1 | 0.000550307 | 10 | 2875.8066 | 3001.6438 | 2884.9302 | 3022.6636 | 3187.5422 | 3278.8875 |
| Eif3l | TVSDLIDQK | -62 | -70 | 0 | 0 | 9 | 1633.2698 | 1747.6514 | 1689.0854 | 908.06 | 1524.28 | 1710.687 |
| Eif3l | FLSPVVPNYDNVHPN | -420 | -434 | 0 | 0.000230707 | 15 | 1358.3945 | 1883.4097 | 1560.1644 | 1471.9104 | 2432.737 | 2243.2402 |
| Eif3l | TYQVIPEVIK | -44 | -53 | 0 | 0 | 10 | 1215.4901 | 1714.0614 | 1927.8386 | 2753.723 | 3088.3699 | 3571.9473 |

| | | | | | | | | | | | | |
|---|---|---|---|---|---|---|---|---|---|---|---|---|
| Eif3l | YGDFFIR | -539 | -545 | 0 | 0.001811524 | 7 | 1152.8827 | 1205.9619 | 1300.4783 | 846.51886 | 877.6509 | 979.88477 |
| Eif3l | LHSLLGDYYQAIK | -280 | -292 | 0 | 0 | 13 | 1048.648 | 1420.0298 | 1455.8925 | 1516.3873 | 1714.2391 | 1592.0414 |
| Eif3l | LHSLLGDYYQAIK | -280 | -292 | 0 | 0 | 13 | 1048.648 | 1420.0298 | 1455.8925 | 1516.3873 | 1714.2391 | 1592.0414 |
| Eif3l | GFLDLTEQEFR | -477 | -487 | 0 | 0 | 11 | 939.8831 | 828.7181 | 931.8823 | 1307.6621 | 1512.1174 | 1386.3738 |
| Eif3l | LAGFLDLTEQEFR | -475 | -487 | 0 | 0 | 13 | 503.01474 | 459.48618 | 581.94556 | 499.90967 | 533.4702 | 490.03342 |
| Eif3l | FLSPVVPNYDNVHPNYHK | -420 | -437 | 0 | 0 | 18 | 407.37766 | 641.5198 | 632.87 | 467.1239 | 475.3528 | 725.4713 |
| Eif3l | FLSPVVPNYDNVHPNYHK | -420 | -437 | 0 | 0 | 18 | 407.37766 | 641.5198 | 632.87 | 467.1239 | 475.3528 | 725.4713 |
| Eif3l | GGDPESVAGEYGR | -250 | -262 | 0 | 0 | 13 | 360.88937 | 1458.9592 | 1443.189 | 340.48047 | 377.28766 | 1620.6409 |
| Eif3l | TSGGDPESVAGEYGR | -248 | -262 | 0 | 0 | 15 | 2909.2258 | 1143.5385 | 3298.6545 | NaN | 262.88562 | 640.4525 |
| Eif3l | VSGGPSLEQR | -144 | -153 | 0 | 0 | 10 | 2438.5566 | 1955.7954 | 1466.5044 | NaN | NaN | 576.7202 |
| Eif3l | LHSLLGDYYQA | -280 | -290 | 0 | 0 | 11 | 682.3967 | 525.92194 | 442.36533 | 814.35205 | NaN | NaN |
| Eif3l | ENIELNKK | -295 | -302 | 1 | 0.00111006 | 8 | 492.9412 | NaN | 284.12146 | NaN | 588.4506 | 1106.8223 |
| Eif3l | VLENIELNKK | -293 | -302 | 1 | 0 | 10 | 1121.9442 | 1199.8094 | NaN | 887.87976 | NaN | NaN |
| Eif3l | SEVDFYIDK | -517 | -525 | 0 | 0 | 9 | 373.28506 | 451.08002 | NaN | NaN | NaN | NaN |
| Rgs10 | ENLLEDPEGVQR | -43 | -54 | 0 | 0 | 12 | 689.6968 | 378.1193 | 567.6669 | 1114.12 | 666.7163 | 476.8293 |
| Pycard | ENLSGDELKK | -13 | -22 | 1 | 0.00025015 | 10 | 873.80566 | 1006.93713 | 1120.0765 | 1063.3821 | 1021.70197 | 1108.3832 |
| Pycard | TGHFVDQHR | -109 | -117 | 0 | 0 | 9 | 811.24835 | 838.1789 | 410.38718 | 750.36774 | 854.10864 | 912.65283 |
| Pycard | TGHFVDQHR | -109 | -117 | 0 | 0 | 9 | 811.24835 | 838.1789 | 410.38718 | 750.36774 | 854.10864 | 912.65283 |
| Pycard | SFVPSWNLTCK | -162 | -172 | 0 | 0 | 11 | 513.8541 | 286.58704 | 1218.1671 | NaN | 1959.1583 | 1917.4275 |
| Asf1a | ENPPVKPDFSK | -124 | -134 | 1 | 0 | 11 | 1039.8975 | 811.5519 | 340.77887 | 1046.4424 | 938.2579 | 976.29364 |
| Asf1b | ENPSPFHSPFR | -13 | -23 | 0 | 0 | 11 | 1056.5093 | 1048.9761 | 1257.4235 | NaN | NaN | 560.9468 |
| Asf1b | NEYPDPELR | -115 | -123 | 0 | 0.00719662 | 9 | 530.3165 | NaN | NaN | NaN | NaN | NaN |
| Tes | TVTYEWAPPVQNQALAR | -113 | -129 | 0 | 0 | 17 | 994.29565 | 936.6903 | 929.2977 | 1061.7898 | 1271.2247 | 1323.1652 |
| Tes | QLPAHDQDPSK | -158 | -168 | 0 | 0 | 11 | 955.56616 | 963.97925 | 1123.3324 | 952.66125 | 746.2501 | 901.2493 |
| Tes | GCHNAIDPEVQR | -365 | -376 | 0 | 0 | 12 | 577.67444 | 576.0875 | NaN | NaN | 469.711 | NaN |
| Tes | ENQNWHLK | -319 | -326 | 0 | 0.006100831 | 8 | 3871.6765 | NaN | NaN | NaN | 1534.7509 | NaN |
| Rheb | FVDSYDPTIENTFTK | -31 | -45 | 0 | 0 | 15 | 802.8496 | 836.79297 | 636.7603 | 1831.8809 | 775.5342 | 818.5526 |
| Rheb | ENQTAVDVFK | -152 | -161 | 0 | 0 | 10 | 3615.763 | NaN | 2343.1567 | 1353.8796 | NaN | NaN |
| Ugp2 | ENSLGINVPR | -380 | -389 | 0 | 0.00025015 | 10 | 2091.139 | 240.9148 | 2562.1858 | NaN | NaN | NaN |
| Ipo11 | ENSPCVTPELLR | -676 | -687 | 0 | 0 | 12 | 1134.4574 | 874.84174 | 889.0028 | 763.03534 | 516.8263 | 849.1188 |
| Ipo11 | NFEDSSPETLEAHK | -338 | -351 | 0 | 0.000431655 | 14 | 887.30756 | NaN | 929.9855 | NaN | NaN | NaN |
| Dhx30 | ENYLEENLLYAPSLR | -982 | -996 | 0 | 0 | 15 | 440.68216 | NaN | NaN | 1530.766 | 2282.7827 | 2176.9175 |
| Prpf40a | EPAFEDITLESER | -773 | -785 | 0 | 0 | 13 | 1007.54364 | 1121.2501 | 1134.6335 | 1477.4666 | 1680.6 | 1737.6663 |
| Prpf40a | LAFNSLLEK | -715 | -723 | 0 | 0.000347792 | 9 | 255.00386 | 197.90308 | NaN | 237.12173 | 225.22134 | 247.19379 |
| Ndufs8 | THEELLYNK | -179 | -187 | 0 | 0 | 9 | 1844.4673 | 2173.5554 | 2159.7388 | 1026.2023 | 1649.1029 | 1889.08 |
| Ndufs8 | EPATINYPFEK | -80 | -90 | 0 | 0 | 11 | 1489.2911 | NaN | 1751.0403 | 1885.064 | NaN | NaN |
| Sap18 | KEPEKPIDR | -14 | -22 | 2 | 0.00201047 | 9 | 867.8878 | 806.0124 | 735.6691 | 664.8596 | 1022.7661 | 1006.6525 |
| Sap18 | EPEKPIDR | -15 | -22 | 1 | 0.000550307 | 8 | 431.88144 | NaN | NaN | NaN | NaN | 564.3845 |
| Hspbp1 | EPELGLEELLR | -310 | -320 | 0 | 0 | 11 | 424.1311 | 370.39508 | 558.09784 | 509.7182 | 565.9485 | 441.50525 |
| Hspbp1 | NLLVGHPEHK | -254 | -263 | 0 | 0 | 10 | 3097.3608 | 3199.9353 | NaN | 1949.0854 | 2924.2935 | 2710.3215 |
| Hspbp1 | TEHSPFHEHVLGAL | -281 | -294 | 0 | 0.00025015 | 14 | 532.35046 | NaN | 514.3166 | 936.0921 | 5223.298 | 3435.0808 |
| Sucla2 | EPIDIVEGIK | -206 | -215 | 0 | 0 | 10 | 3556.0588 | 3042.1292 | 3381.2175 | 2040.4572 | 2570.1963 | 963.0347 |
| Sucla2 | LHGGTPANFLDVGGGATVQ | -337 | -355 | 0 | 0.00025015 | 19 | 605.9454 | 778.87177 | 721.2344 | 629.1944 | 774.9591 | 750.8281 |
| Sucla2 | LHGGTPANFLDVGGGA | -337 | -352 | 0 | 0 | 16 | 3266.438 | 3333.5566 | 3048.3018 | NaN | NaN | 2029.3345 |
| Sucla2 | LHGGTPANFLDVGGGAT | -337 | -353 | 0 | 0.003003218 | 17 | 626.69995 | NaN | NaN | NaN | NaN | NaN |
| Ddx18 | LILRPEELGFLR | -518 | -529 | 1 | 0 | 12 | 1195.0571 | 1042.4077 | 1183.4801 | 562.29865 | 970.08105 | 953.8652 |
| Ddx18 | LILRPEELGFLR | -518 | -529 | 1 | 0 | 12 | 1195.0571 | 1042.4077 | 1183.4801 | 562.29865 | 970.08105 | 953.8652 |
| Ddx18 | SIRPLLEGR | -199 | -207 | 1 | 0.001708008 | 9 | 9928.102 | NaN | 10405.932 | 9974.4375 | 9439.475 | 9851.704 |
| Ddx18 | KADDSEVPSLPLGLT | -147 | -161 | 1 | 0.000230707 | 15 | 609.8116 | 672.5781 | 601.9661 | NaN | 754.88983 | 382.6855 |
| Ddx18 | EPLYVGVDDDKEVATVD | -372 | -388 | 1 | 0.005363933 | 17 | 653.3622 | NaN | NaN | 969.9268 | NaN | 987.23663 |
| Ddx18 | NGTGVLILSPTR | -240 | -251 | 0 | 0 | 12 | 525.7681 | NaN | 293.56537 | NaN | NaN | NaN |
| Ddx18 | NLQCLVIDEADR | -316 | -327 | 0 | 0 | 12 | 1637.4929 | NaN | NaN | NaN | NaN | NaN |
| Ddx18 | KVEDLAR | -360 | -366 | 1 | 0.001092529 | 7 | 395.7012 | NaN | NaN | NaN | NaN | NaN |
| Ddx18 | GINIIVATPGR | -291 | -301 | 0 | 0.002099344 | 11 | 358.5664 | NaN | NaN | NaN | NaN | NaN |
| Hmga1 | KEPSEVPTPK | -46 | -55 | 1 | 0 | 10 | 578.2526 | 628.1726 | 770.69977 | 725.60266 | 653.61676 | 749.5316 |
| Hmga1 | SQKEPSEVPTPK | -44 | -55 | 1 | 0 | 12 | 1016.268 | 1005.9767 | 1262.6161 | NaN | 1087.75 | 1102.2113 |
| Hmga1 | KQPPVSPGTALVGSQ | -31 | -45 | 1 | 0.001055699 | 15 | 3672.076 | NaN | 821.9059 | NaN | NaN | NaN |
| Hmga1 | EPSEVPTPK | -47 | -55 | 0 | 0.001392869 | 9 | 980.1448 | NaN | NaN | NaN | NaN | NaN |
| Hmga1 | KQPPVSPGTALVG | -31 | -43 | 1 | 0.006677519 | 13 | 721.42487 | NaN | NaN | NaN | NaN | NaN |
| Larp4 | REIPETTPVEEVK | -200 | -212 | 1 | 0.001463897 | 13 | 878.19275 | 980.46686 | 255.57265 | 1266.2736 | 1238.9728 | 1345.2601 |
| Larp4 | EPSPVLVQPLR | -624 | -634 | 0 | 0 | 11 | 480.42175 | NaN | 503.3334 | 469.72406 | 437.83978 | 513.7197 |
| Trmt112 | INPVEFNPEFVAR | -32 | -44 | 0 | 0 | 13 | 1864.8654 | 2473.2695 | 1268.1505 | 2414.1345 | 2220.9136 | 3113.1345 |
| Trmt112 | EVPKEPTEGYEHDETFLR | -65 | -82 | 1 | 0 | 18 | 1039.3326 | 1238.8881 | 1211.5087 | 1076.1372 | 1443.5386 | 1421.9846 |
| Trmt112 | EPTEGYEHDETFLR | -69 | -82 | 0 | 0 | 14 | 687.8131 | 916.1773 | 803.798 | 335.71628 | 346.55914 | 261.22858 |
| Trmt112 | NLAEVPKEPTEGYEHDETFLR | -62 | -82 | 1 | 0 | 21 | 502.5047 | 399.34674 | 631.2847 | 486.36337 | 640.04407 | 689.53876 |
| Mndal | EQAQLPEPLGTDIK | -204 | -217 | 0 | 0 | 14 | 560.4707 | 665.6161 | 819.816 | 960.84924 | 722.9247 | 834.8257 |
| Mndal | KLQEDYDR | -37 | -44 | 1 | 0.004595494 | 8 | 1255.2789 | NaN | NaN | NaN | 1784.7765 | NaN |
| Add3 | SFEDDSAPLSPLK | -414 | -426 | 0 | 0 | 13 | 8466.292 | 9223.581 | 9687.009 | 10881.773 | 11519.2 | 11666.322 |
| Add3 | INESDPEYLR | -28 | -37 | 0 | 0 | 10 | 2624.2974 | 2737.5554 | 1297.044 | 720.20215 | 1820.8364 | 1943.0825 |
| Add3 | EQDHIIIIPR | -168 | -177 | 0 | 0 | 10 | 562.9401 | 592.3868 | 807.73834 | 682.83844 | 1471.759 | 715.5272 |
| Add3 | EQDHIIIIPR | -168 | -177 | 0 | 0 | 10 | 562.9401 | 592.3868 | 807.73834 | 682.83844 | 1471.759 | 715.5272 |
| Add3 | GHNPSGLLALQ | -82 | -92 | 0 | 0.002168954 | 11 | 464.76666 | NaN | NaN | NaN | NaN | NaN |
| Tmx3 | EQESSSGGSLAPTVQEPK | -431 | -448 | 0 | 0 | 18 | 937.7709 | 293.29187 | 866.8886 | 370.81998 | 411.09885 | 243.41403 |
| Tmx3 | SILDGTVPAQGGDSIFQR | -341 | -358 | 0 | 0.00025015 | 18 | 589.42993 | 651.6674 | 643.3313 | 780.2798 | 1307.1133 | 1488.5007 |
| Tmx3 | SEDVVPEYVTLK | -187 | -198 | 0 | 0 | 12 | 1205.2732 | 1231.9434 | 940.484 | NaN | 681.3794 | 208.31401 |
| Tmx3 | FVYIGGESPLK | -157 | -167 | 0 | 0 | 11 | 4363.1836 | NaN | 5182.298 | 4301.7344 | NaN | 7485.578 |
| Tmx3 | KLEPIWNEVGLE | -61 | -72 | 1 | 0.000394361 | 12 | 524.4274 | NaN | 405.9739 | 364.66205 | NaN | 576.225 |
| Tmx3 | VSGALIRPLPSQQ | -132 | -144 | 1 | 0.00383302 | 13 | 1223.6615 | NaN | 2297.8325 | 705.3296 | NaN | NaN |

| | | | | | | | | | | | | |
|---|---|---|---|---|---|---|---|---|---|---|---|---|
| Tmx3 | FVEDLNESFK | -30 | -39 | 0 | 0.001392869 | 10 | 686.1851 | NaN | NaN | 949.1375 | NaN | NaN |
| Rabl3 | EQFADNQIPLLVIGTK | -133 | -148 | 0 | 0 | 16 | 65.034 | NaN | NaN | NaN | NaN | NaN |
| Pex19 | EQFEAETPTDSEATQR | -230 | -245 | 0 | 0 | 16 | 445.1388 | 488.21503 | 490.1902 | 360.0047 | 511.92838 | 551.1472 |
| Mak16 | EQIDENLIYWPR | -92 | -103 | 0 | 0 | 12 | 1253.4296 | NaN | 1297.982 | 1460.8683 | 1617.578 | 1826.3575 |
| Rad50 | EQISPLETALEK | -910 | -921 | 0 | 0 | 12 | 1555.0518 | 1495.4498 | 1547.0798 | 1572.9436 | 1157.0737 | 1306.9283 |
| Rad50 | LEENIDTIKR | -1060 | -1069 | 1 | 0 | 10 | 765.2038 | 820.2187 | 797.706 | 446.06625 | 1089.6796 | 852.1209 |
| Rad50 | LDNEIKAL | -268 | -275 | 1 | 0.002099344 | 8 | 3502.6372 | 3397.778 | 3770.9158 | 1964.1599 | 1736.323 | NaN |
| Rad50 | SELQQLEGSSDR | -462 | -473 | 0 | 0 | 12 | 645.53577 | NaN | 527.3487 | NaN | 798.4695 | NaN |
| Rpl17 | EQIVPKPEEEVA | -154 | -165 | 1 | 0.00025015 | 12 | 3688.2153 | 3887.713 | 4098.041 | 3021.4368 | 3041.2927 | 3200.7786 |
| Rpl17 | EQIVPKPEEEVA | -154 | -165 | 1 | 0.00025015 | 12 | 3688.2153 | 3887.713 | 4098.041 | 3021.4368 | 3041.2927 | 3200.7786 |
| Rpl17 | YSLDPENPTK | -4 | -13 | 0 | 0 | 10 | 3595.8943 | 3744.003 | 3923.0347 | 3068.6038 | 417.6423 | 3170.278 |
| Rpl17 | YSLDPENPTK | -4 | -13 | 0 | 0 | 10 | 3595.8943 | 3744.003 | 3923.0347 | 3068.6038 | 417.6423 | 3170.278 |
| Rpl17 | EQIVPKPEEEVAQK | -154 | -167 | 1 | 0 | 14 | 2768.4763 | 2963.6057 | 3106.6523 | 5437.2153 | 64.3185 | 5889.0957 |
| Rpl17 | EQIVPKPEEEVAQ | -154 | -166 | 1 | 0 | 13 | 1805.585 | 1733.6519 | 1755.3765 | 148.36026 | 1699.6736 | 1674.6716 |
| Rpl17 | LKDVTLK | -48 | -54 | 1 | 0.009002904 | 7 | 1192.6877 | 1055.8348 | 1043.1062 | 840.4212 | 722.07294 | 883.7906 |
| Rpl17 | KNAESNAELK | -96 | -105 | 1 | 0.001463897 | 10 | 836.34247 | 568.0106 | 822.76276 | 719.4375 | 751.14703 | 665.3701 |
| Rpl17 | GLDVDSLVIEHIQ | -106 | -118 | 0 | 0 | 13 | 512.05054 | 452.2979 | 497.14383 | 766.63 | 714.02094 | NaN |
| Rpl17 | YSLDPENPTKS | -4 | -14 | 1 | 0 | 11 | 338.76328 | NaN | 572.4298 | 334.76648 | 903.4586 | 532.6405 |
| Rpl17 | KSAEFLLH | -86 | -93 | 1 | 0.002434616 | 8 | 2156.316 | 1856.2842 | 1763.7932 | NaN | NaN | NaN |
| Golga5 | EQLQDLQDQIAK | -510 | -521 | 0 | 0 | 12 | 2137.6458 | NaN | 1900.5177 | NaN | NaN | NaN |
| Hsdl2 | LQLQEESQLQK | -299 | -309 | 0 | 0 | 11 | 930.29047 | 545.9513 | 1012.147 | 895.6004 | NaN | NaN |
| Hsdl2 | EQPQLQQQPR | -351 | -360 | 0 | 0.00025015 | 10 | 735.11487 | NaN | 592.48975 | NaN | NaN | 730.7074 |
| Hsdl2 | FIIDENILK | -242 | -250 | 0 | 0.001636916 | 9 | 454.78864 | NaN | NaN | 490.91278 | NaN | 377.67828 |
| Thrap3 | YKDDPVDLR | -707 | -715 | 1 | 0.00025015 | 9 | 4037.0244 | 4110.711 | 3888.519 | 4641.7715 | 4985.3135 | 4956.2017 |
| Thrap3 | SREEEWDPEYTPK | -860 | -872 | 1 | 0 | 13 | 2928.0593 | 2547.9526 | 1857.5426 | 1930.5156 | 1190.2284 | 569.8369 |
| Thrap3 | KAEEMEDEPFTER | -481 | -493 | 1 | 0 | 13 | 1786.6984 | 1679.293 | 1769.567 | 1743.4381 | 1959.1016 | 2041.7982 |
| Thrap3 | LRDDFEK | -411 | -417 | 1 | 0.00272025 | 7 | 1785.6079 | 1695.6007 | 1675.4467 | 241.38138 | 1881.2142 | 2327.1104 |
| Thrap3 | KEPEFDDEPK | -439 | -448 | 1 | 0 | 10 | 1733.2728 | 1566.7329 | 1519.3303 | 723.50195 | 1679.9768 | 1759.9714 |
| Thrap3 | TDSEKPFR | -394 | -401 | 1 | 0.00122419 | 8 | 1431.7212 | 469.38907 | 1505.865 | 1544.4167 | 547.09863 | 736.5775 |
| Thrap3 | GSESSKPWPDATTY | -216 | -229 | 1 | 0 | 14 | 1020.7785 | 175.03842 | 1078.4564 | 1006.1697 | 782.04407 | 1637.4624 |
| Thrap3 | HIVTIVHHVK | -627 | -636 | 0 | 0.000230707 | 10 | 532.8911 | 776.44336 | 773.2932 | 717.1962 | 638.2577 | 413.9162 |
| Thrap3 | HIVTIVHHVK | -627 | -636 | 0 | 0.000230707 | 10 | 532.8911 | 776.44336 | 773.2932 | 717.1962 | 638.2577 | 413.9162 |
| Thrap3 | EYPEEAEEREESTSGFDK | -791 | -808 | 1 | 0 | 18 | 411.99515 | 417.99765 | 382.1933 | 539.3313 | 606.21313 | 526.0937 |
| Thrap3 | SPLQSVVVR | -253 | -261 | 0 | 0 | 9 | 968.6169 | 943.14075 | 1039.6963 | NaN | 1159.7827 | 892.3623 |
| Thrap3 | WESLHTGK | -469 | -476 | 0 | 0 | 8 | 826.4113 | NaN | 471.34586 | 1279.1033 | 993.51465 | 721.6833 |
| Thrap3 | EQTFSGGTSQDIK | -203 | -215 | 0 | 0.000230707 | 13 | 1606.9442 | 2049.399 | 3688.4844 | NaN | 2169.6138 | NaN |
| Thrap3 | HGLTHEELK | -686 | -694 | 0 | 0.00111006 | 9 | 1162.1572 | 1021.2645 | 150.19919 | NaN | NaN | NaN |
| Thrap3 | FSGEEGEIEDDESGTENR | -923 | -940 | 0 | 0.000471409 | 18 | 223.17944 | NaN | NaN | 471.02023 | 419.17618 | NaN |
| Thrap3 | SGKWESLHTGK | -466 | -476 | 1 | 0.005459848 | 11 | 1488.2572 | NaN | NaN | NaN | NaN | NaN |
| Mesd | RPSAPIDFSK | -77 | -86 | 1 | 0.000431655 | 10 | 2123.3513 | 1729.072 | 804.8085 | 2137.0977 | 2249.7393 | 2285.3909 |
| Mesd | EQWEKDDDIEEGDLPEHK | -59 | -76 | 1 | 0 | 18 | 511.3507 | 545.5925 | 588.5735 | 563.8249 | 216.4685 | 694.1593 |
| Mesd | LDPGKPESILK | -87 | -97 | 1 | 0.000347792 | 11 | 2081.25 | NaN | NaN | 1458.8282 | NaN | NaN |
| Frg1 | EVDEGPSPPEQFTAVK | -86 | -101 | 0 | 0 | 16 | 1383.9498 | 570.22034 | 2032.0161 | 1958.2452 | 2449.5454 | 992.62823 |
| Frg1 | EQWEPVFQDGK | -136 | -146 | 0 | 0.00025015 | 11 | 2157.6655 | 2929.0752 | 2808.5933 | 3149.2292 | NaN | NaN |
| Nt5c | RFPEEPHVPLEQR | -31 | -43 | 1 | 0 | 13 | 3470.679 | 3897.4753 | 4086.6484 | 1783.5736 | 3994.3032 | 4180.038 |
| Nt5c | RFPEEPHVPLEQR | -31 | -43 | 1 | 0 | 13 | 3470.679 | 3897.4753 | 4086.6484 | 1783.5736 | 3994.3032 | 4180.038 |
| Nt5c | EQYGALRPDLAEK | -50 | -62 | 1 | 0.00025015 | 13 | 1800.4492 | 1829.0264 | 2312.4128 | 260.16663 | 1925.2267 | 2000.8772 |
| Nt5c | WVEQNLGPEFVER | -117 | -129 | 0 | 0 | 13 | 1627.5331 | 2110.2305 | 2165.8245 | 1651.3523 | 2890.368 | 2643.4768 |
| Nt5c | NEQYGALRPDLAEK | -49 | -62 | 1 | 0 | 14 | 1619.1044 | 1846.5266 | 1847.242 | 2415.8477 | 2534.5776 | 2340.6511 |
| Nt5c | YDHCVGEK | -107 | -114 | 0 | 0.00111006 | 8 | 285.26562 | 287.79437 | 268.1846 | NaN | 213.0911 | 158.98776 |
| Cstb | VFQPLPHENKPLTLS | -69 | -83 | 1 | 0.00025015 | 15 | 3513.3267 | 3050.35 | 3256.6106 | 3748.1653 | 3473.357 | 3818.1284 |
| Cstb | ERHDELSYF | -90 | -98 | 1 | 0.00111006 | 9 | 938.22784 | 964.59454 | 407.4095 | NaN | 1063.7203 | 1047.2072 |
| Cstb | VFQPLPHENKPLTL | -69 | -82 | 1 | 0 | 14 | 545.4747 | 744.9761 | 822.12164 | 691.1409 | NaN | 830.56055 |
| Cstb | VFQPLPHENKPLTL | -69 | -82 | 1 | 0 | 14 | 545.4747 | 744.9761 | 822.12164 | 691.1409 | NaN | 830.56055 |
| Akt2 | ERPEAPDQTLPPLNNF | -40 | -55 | 1 | 0 | 16 | 687.18195 | 836.42084 | 900.88666 | 830.2829 | 1060.6307 | 1126.825 |
| Akt2 | LPFYNQDHER | -348 | -357 | 0 | 0 | 10 | 619.04767 | 777.63007 | NaN | 671.23004 | 488.19583 | 453.80936 |
| Dpy30 | ERPPNPIEFLAS | -75 | -86 | 1 | 0 | 12 | 4533.994 | 4484.303 | 4650.142 | 5342.0015 | 6382.25 | 6325.105 |
| Dpy30 | VAENPHSEYGLTDSVER | -13 | -29 | 0 | 0 | 17 | 486.3371 | 857.12836 | 688.81934 | 524.7845 | 580.6388 | 637.2598 |
| Dpy30 | VAENPHSEYGLTDSVER | -13 | -29 | 0 | 0 | 17 | 486.3371 | 857.12836 | 688.81934 | 524.7845 | 580.6388 | 637.2598 |
| Pf4 | TISSGIHLK | -49 | -57 | 0 | 0 | 9 | 3278.1887 | 3163.4832 | 2197.884 | 1976.8784 | 2187.2412 | 3039.1216 |
| Pf4 | VTSAGPEESDGDLSCVCVK | -30 | -48 | 0 | 0 | 19 | 350.37802 | 439.25824 | 483.8463 | 504.97314 | 565.55536 | 538.26605 |
| Pf4 | SAGPEESDGDLSCVCVK | -32 | -48 | 0 | 0 | 17 | 263.0318 | 259.14975 | 283.85562 | 458.27386 | 436.54156 | 482.00714 |
| Pf4 | ESDGDLSCVCVK | -37 | -48 | 0 | 0 | 12 | 889.4181 | 667.14185 | 876.2087 | 1714.9697 | NaN | NaN |
| Otub1 | GEGGTTNPHVFPEGSEPK | -239 | -256 | 0 | 0.00025015 | 18 | 1638.6412 | 2149.5232 | 1788.4149 | 1839.2661 | 2026.3158 | 2649.3691 |
| Otub1 | GEGGTTNPHVFPEGSEPK | -239 | -256 | 0 | 0.00025015 | 18 | 1638.6412 | 2149.5232 | 1788.4149 | 1839.2661 | 2026.3158 | 2649.3691 |
| Otub1 | IQQEIAVQNPLVSER | -37 | -51 | 0 | 0 | 15 | 974.30304 | 1314.6987 | 1182.044 | 1303.5709 | 1301.4879 | 1403.9933 |
| Otub1 | TRPDGNCFYR | -85 | -94 | 1 | 0.002099344 | 10 | 4353.3135 | 4644.148 | 4733.846 | NaN | 5662.913 | 470.27054 |
| Otub1 | YRPGHYDILYK | -261 | -271 | 1 | 0 | 11 | 669.76044 | 943.3766 | 1233.3649 | NaN | 1235.1791 | 1789.7393 |
| Otub1 | ESDHIHIIAL | -214 | -223 | 0 | 0.001868731 | 10 | 1197.133 | 1470.6733 | 1421.4991 | NaN | 1962.7369 | NaN |
| Otub1 | FFEHFIEGGR | -189 | -198 | 0 | 0 | 10 | 977.466 | 1143.8867 | NaN | 1325.849 | 313.63745 | NaN |
| Otub1 | YKEYAEDDNIYQQK | -58 | -71 | 1 | 0.00025015 | 14 | 227.81512 | 190.21399 | NaN | NaN | NaN | NaN |
| Psmc2 | EVVETPLLHPER | -189 | -200 | 0 | 0 | 12 | 2074.735 | 2561.647 | 2563.8323 | 2170.7864 | 2623.249 | 524.6607 |
| Psmc2 | EVVETPLLHPER | -189 | -200 | 0 | 0 | 12 | 2074.735 | 2561.647 | 2563.8323 | 2170.7864 | 2623.249 | 524.6607 |
| Psmc2 | KIEFSLPDLEGR | -340 | -351 | 1 | 0 | 12 | 1284.18 | 1319.7913 | 1332.574 | 3738.502 | 1945.9648 | 2032.6671 |
| Psmc2 | FDDGAGGDNEVQR | -285 | -297 | 0 | 0 | 13 | 1098.3282 | 1123.0264 | 1162.2131 | 1172.5353 | 359.9753 | 1212.2538 |
| Psmc2 | QVEDDIQQLLK | -47 | -57 | 0 | 0 | 11 | 926.8516 | 965.01086 | 999.4221 | 1099.4335 | 967.2601 | 1123.2943 |
| Psmc2 | KINELTGIK | -58 | -66 | 1 | 0 | 9 | 796.5967 | 791.1896 | 810.16907 | 746.6018 | 875.14716 | 824.84753 |
| Psmc2 | KIINADSEDPK | -100 | -110 | 1 | 0 | 11 | 746.4756 | 814.2037 | 967.8535 | 822.4987 | 751.6633 | 746.28577 |

| | | | | | | | | | | | | |
|---|---|---|---|---|---|---|---|---|---|---|---|---|
| Psmc2 | FDEIDAIGGAR | -274 | -284 | 0 | 0 | 11 | 628.9027 | 456.76874 | 440.56985 | 660.91394 | 107.96874 | 978.7445 |
| Psmc2 | YQIHIPLPPK | -147 | -156 | 0 | 0 | 10 | 581.4458 | 1125.0337 | 1068.5068 | 1540.3032 | 1063.481 | 962.23846 |
| Psmc2 | ESDTGLAPPALWDLAADK | -67 | -84 | 0 | 0.000431655 | 18 | 470.0244 | 434.4521 | 496.74106 | 392.2436 | 461.15698 | 528.239 |
| Psmc2 | QTLQSEQPLQVAR | -85 | -97 | 0 | 0.000230707 | 13 | 2253.1016 | 2474.7388 | 3423.9812 | NaN | 1458.2573 | 1214.9233 |
| Psmc2 | QVEEKPDVTYSDVGGCK | -165 | -181 | 1 | 0 | 17 | 383.7536 | 413.6356 | 442.0834 | 509.73416 | 553.8123 | NaN |
| Psmc2 | VIGSELVQK | -240 | -248 | 0 | 0.00294445 | 9 | 1163.6008 | NaN | NaN | NaN | 857.7375 | 463.4679 |
| Psmc2 | RFDDGAGGDNEVQR | -284 | -297 | 1 | 0.002099344 | 14 | 530.5116 | NaN | 612.1449 | 569.24976 | NaN | NaN |
| Nvl | ESLPLDLSDDQSNSK | -183 | -197 | 0 | 0 | 15 | 524.61536 | 603.62744 | 586.2426 | 190.03589 | 464.9706 | 750.7235 |
| Nvl | NRPDSLDPALR | -410 | -420 | 1 | 0 | 11 | 1903.2755 | 2181.606 | 2262.7295 | NaN | 1905.1918 | 2099.908 |
| Nvl | FEDVGGNDATLK | -263 | -274 | 0 | 0 | 12 | 728.1714 | 913.4258 | NaN | 661.43646 | 903.56415 | NaN |
| Nvl | TLFVGLPPPADR | -742 | -753 | 0 | 0 | 12 | 179.07867 | 652.7381 | 775.84485 | 338.32825 | NaN | NaN |
| Nvl | ESEGGWFIDK | -167 | -176 | 0 | 0 | 10 | 712.2584 | NaN | 545.4153 | NaN | 835.5542 | NaN |
| Smpd4 | ESFTPTEEHVLVVR | -317 | -330 | 0 | 0 | 14 | 1943.9503 | 2482.463 | 2395.243 | 2378.3284 | 3189.8628 | 3026.2344 |
| Smpd4 | SISDQYVESPTGR | -566 | -578 | 0 | 0 | 13 | 786.9772 | NaN | NaN | NaN | NaN | NaN |
| Ptpn1 | LHQEDNDYINAS | -59 | -70 | 0 | 0 | 12 | 1174.9607 | 994.8285 | 1015.123 | 1249.7495 | 601.2097 | 1287.9332 |
| Ptpn1 | ESGSLSLEHGPIVVH | -200 | -214 | 0 | 0 | 15 | 675.6103 | 608.7649 | 729.258 | 681.87256 | 815.7919 | 784.7279 |
| Rab14 | RDVTYEEAK | -132 | -140 | 1 | 0.000431655 | 9 | 772.83936 | 678.34174 | 943.36566 | 611.54004 | 984.2999 | 1053.9158 |
| Rab14 | TGENVEDAFLEAAK | -157 | -170 | 0 | 0 | 14 | 645.61224 | 1121.2294 | 1304.3954 | 1313.3116 | 1799.4857 | 1662.0675 |
| Rab14 | ESGVQHKPSAPQGGR | -187 | -201 | 1 | 0 | 15 | 364.92758 | 435.6284 | 455.37415 | 184.46808 | 303.66278 | 369.8492 |
| Cltb | LTQEPESIR | -96 | -104 | 0 | 0 | 9 | 925.9802 | 532.79016 | 852.8663 | 1020.5499 | 1214.5901 | 1356.3253 |
| Cltb | ESKEETPGTEWEK | -182 | -194 | 1 | 0 | 13 | 1552.6387 | 96.510056 | NaN | 874.0555 | 824.9047 | 718.15515 |
| Cltb | SKEETPGTEWEK | -183 | -194 | 1 | 0.002199523 | 12 | 794.7744 | NaN | NaN | NaN | NaN | NaN |
| U2surp | KFQDELESGK | -853 | -862 | 1 | 0 | 10 | 1021.2028 | 1265.0237 | 1913.2087 | 2318.766 | 1777.2245 | 1748.7856 |
| U2surp | SEEQETEAFVEEPSKK | -516 | -531 | 1 | 0 | 16 | 707.0446 | 219.73717 | 758.6117 | 615.5729 | 815.4994 | 771.48267 |
| U2surp | NPPNQSSNERPPSLLVIE | -169 | -186 | 1 | 0 | 18 | 179.23422 | 756.651 | 615.1827 | 673.17163 | 642.8066 | 717.954 |
| U2surp | ESLCDSPHQNLSR | -62 | -74 | 0 | 0 | 13 | 1409.0256 | NaN | 1195.0492 | 569.2361 | 2158.9902 | 1280.383 |
| U2surp | FLFENQTPAHVY | -461 | -472 | 0 | 0 | 12 | 651.2887 | 886.8604 | 930.33484 | NaN | 1047.3684 | 372.651 |
| U2surp | SLDDDLDGVPLDATEDSK | -731 | -748 | 0 | 0 | 18 | 366.51965 | 374.16794 | NaN | 498.7628 | 575.43365 | NaN |
| Prkar1a | ESLDKWER | -257 | -264 | 1 | 0.000230707 | 8 | 135.46661 | 1128.9268 | 1874.346 | 1813.1322 | 191.69434 | 2429.563 |
| Prkar1a | VYTEEDAASYVR | -104 | -115 | 0 | 0 | 12 | 2103.2253 | 1925.5153 | 2013.7085 | NaN | 734.69684 | 937.2278 |
| Prkar1a | LTVADALEPVQFEDGQK | -265 | -281 | 0 | 0 | 17 | 1654.4594 | 1905.5562 | 1663.1721 | 954.0551 | 1854.7129 | NaN |
| Prkar1a | VLGPCSDILK | -358 | -367 | 0 | 0 | 10 | 796.308 | 910.8782 | 873.59406 | 537.0545 | NaN | 955.08594 |
| Prkar1a | SENEEFVEVGR | -307 | -317 | 0 | 0 | 11 | 1056.0111 | 429.12823 | 1226.597 | 1254.4822 | NaN | NaN |
| Hspa13 | ESLELADSNPQNTIYDAK | -83 | -100 | 0 | 0 | 18 | 1161.8284 | 1078.8235 | NaN | NaN | NaN | 1723.462 |
| As3mt | ESLQNVHEDVSSR | -46 | -58 | 0 | 0 | 13 | 1202.1816 | NaN | NaN | NaN | 1493.0137 | 1601.3798 |
| Anapc4 | ESPLLFPYYPR | -509 | -519 | 0 | 0 | 11 | 1431.5419 | 2914.2415 | NaN | 3841.2192 | NaN | 4106.936 |
| Rad23a | YIQVTPQEK | -313 | -321 | 0 | 0.000230707 | 9 | 579.4575 | 823.69305 | 893.2342 | 979.2472 | 692.31006 | 451.49896 |
| Rad23a | ESQAPEQPATEAAGENPLEFLR | -215 | -236 | 0 | 0.002880184 | 22 | 200.9633 | NaN | 200.26039 | 107.17797 | 231.36375 | 192.62439 |
| Evi2b | ESSEHINDTSNLK | -294 | -306 | 0 | 0 | 13 | 989.2237 | NaN | NaN | NaN | NaN | 1393.5834 |
| Elob | LYKDDQLLDDGK | -44 | -55 | 1 | 0 | 12 | 2422.8513 | 2350.882 | 2783.3975 | 642.9812 | 3279.6265 | 3447.9998 |
| Elob | ESSTVFELK | -20 | -28 | 0 | 0 | 9 | 720.6163 | NaN | 502.7239 | 911.2735 | 806.74835 | 1144.2003 |
| Elob | RADDTFEALR | -80 | -89 | 1 | 0 | 10 | 1132.3035 | 557.6013 | 1575.0787 | NaN | NaN | NaN |
| Ubap2 | QVIFDKNDSDFEAK | -43 | -56 | 1 | 0 | 14 | 520.26434 | 524.19574 | NaN | 649.0996 | 676.1881 | 681.9658 |
| Ubap2 | LEGNSDTTSWETVGGK | -90 | -105 | 0 | 0 | 16 | 520.7132 | 1429.499 | 2000.8658 | NaN | 2733.9182 | NaN |
| Ubap2 | ESTAGDGPSTVSR | -468 | -480 | 0 | 0 | 13 | 5903.1787 | NaN | NaN | NaN | NaN | NaN |
| Camk2d | ESTESSNTTIEDEDVK | -329 | -344 | 0 | 0 | 16 | 709.0305 | 1287.9589 | 1090.2094 | 477.44135 | 516.4697 | 521.92596 |
| Thoc7 | ESVEDKLELR | -154 | -163 | 1 | 0 | 10 | 4327.6543 | NaN | NaN | NaN | 1340.3785 | 4913.712 |
| Mta3 | QYESLPATHIR | -96 | -106 | 0 | 0 | 11 | 1959.6559 | 2006.4203 | 1979.7009 | 1910.7328 | 1368.0703 | 1047.3306 |
| Mta3 | ETEEESETPVEADLTEK | -65 | -81 | 0 | 0 | 17 | 465.727 | 323.76727 | 256.44583 | 362.78482 | 278.4108 | 420.71555 |
| Mta3 | VWDPNSPLTDR | -180 | -190 | 0 | 0 | 11 | 1058.427 | 1624.1415 | NaN | 1328.0978 | NaN | 1670.6558 |
| Dnpep | GGAYEEFIFAPR | -249 | -260 | 0 | 0 | 12 | 717.17926 | 914.39014 | 871.7038 | 431.61014 | 1056.8021 | 478.93552 |
| Dnpep | GFFELFPSVSR | -458 | -468 | 0 | 0 | 11 | 492.4421 | 77.77991 | 457.8209 | 493.04358 | 221.85178 | 613.92316 |
| Dnpep | ETEGWDIVPENK | -50 | -61 | 0 | 0 | 12 | 123.877945 | NaN | 791.4458 | NaN | NaN | 12889.698 |
| Thoc2 | VVVPAEWIK | -6 | -14 | 0 | 0.000591124 | 9 | 456.3328 | 578.08887 | 1150.6843 | 571.2771 | 651.9031 | 602.9447 |
| Thoc2 | FYQEPSGETPSSLYR | -254 | -268 | 0 | 0 | 15 | 850.5322 | 684.7104 | 1366.7229 | NaN | NaN | NaN |
| Thoc2 | ETGEYTHIR | -1113 | -1121 | 0 | 0 | 9 | 614.8203 | NaN | NaN | 1071.2705 | NaN | NaN |
| Coro1c | NILDSKPAANK | -408 | -418 | 1 | 0 | 11 | 1776.177 | 1563.5074 | 3459.976 | 2497.225 | 3240.2502 | 3564.1157 |
| Coro1c | VTWDSSFCAVNPR | -32 | -44 | 0 | 0 | 13 | 716.4002 | 783.1387 | 438.05807 | 652.4994 | 532.6902 | 621.57404 |
| Coro1c | NDQCYDDIR | -20 | -28 | 0 | 0 | 9 | 681.93677 | 751.9969 | 771.918 | 754.924 | 89.5107 | 757.85913 |
| Coro1c | QLALWNPK | -244 | -251 | 0 | 0 | 8 | 574.6017 | 1180.9891 | 998.3313 | 418.1231 | 956.18024 | 905.1986 |
| Coro1c | NADPILISLK | -382 | -391 | 0 | 0.00025015 | 10 | 449.16147 | 389.78387 | 404.0573 | 860.9845 | 744.852 | 261.76678 |
| Coro1c | ETICSQDER | -453 | -461 | 0 | 0 | 9 | 336.2179 | 872.57526 | 282.02396 | 1355.937 | 1099.1263 | 1234.9186 |
| Znf207 | LIHPDEDISLEER | -380 | -392 | 0 | 0 | 13 | 1647.1172 | 820.0164 | 1122.8478 | 2226.2373 | 1531.6465 | 1224.9974 |
| Znf207 | LIHPDEDISLEER | -380 | -392 | 0 | 0 | 13 | 1647.1172 | 820.0164 | 1122.8478 | 2226.2373 | 1531.6465 | 1224.9974 |
| Znf207 | LYTGPGLAIH | -44 | -53 | 0 | 0 | 10 | 1056.217 | 1023.5649 | 944.3015 | 756.91345 | 672.56116 | 680.43634 |
| Znf207 | ETIDAVPNAIPGR | -60 | -72 | 0 | 0 | 13 | 1383.9009 | 1063.5762 | 1263.3342 | NaN | NaN | NaN |
| Tmsb10 | ETIEQEKR | -33 | -40 | 1 | 0.00149795 | 8 | 2455.4758 | 2198.1377 | 2776.22 | 2783.376 | 374.28732 | 2889.4453 |
| Tmsb10 | KETIEQEKR | -32 | -40 | 2 | 0.005821388 | 9 | 279.41577 | NaN | NaN | 342.20767 | 276.4276 | 269.8823 |
| Arcn1 | NTLEWCLPVIDAK | -436 | -448 | 0 | 0 | 13 | 1245.8923 | 1014.026 | 1167.0999 | 294.0426 | 1461.98 | 410.34647 |
| Arcn1 | NSNILEDLETLR | -73 | -84 | 0 | 0 | 12 | 567.25653 | 958.409 | 893.6761 | 1212.6782 | 1320.1558 | 1350.4155 |
| Arcn1 | VAPAPARPSGPSK | -212 | -224 | 1 | 0 | 13 | 528.61127 | 547.2555 | 600.94794 | 383.56177 | 630.42596 | 623.7247 |
| Arcn1 | ISDDKFGR | -305 | -312 | 1 | 0.00025015 | 8 | 593.6893 | 1094.8508 | 1067.2588 | NaN | 1940.5139 | 1267.9844 |
| Arcn1 | GKEVDNFVDK | -232 | -241 | 1 | 0 | 10 | 7165.821 | NaN | 663.7828 | NaN | 7045.168 | 6621.948 |
| Arcn1 | VTQVDGNSPVR | -486 | -496 | 0 | 0 | 11 | 2786.8137 | 754.7041 | NaN | 375.42688 | NaN | 4519.448 |
| Arcn1 | ETIIETDKPK | -202 | -211 | 1 | 0.000550307 | 10 | 686.15674 | NaN | NaN | NaN | 1614.328 | 994.3052 |
| Arcn1 | SFPVNSDVGVLK | -352 | -363 | 0 | 0 | 12 | 352.13947 | 327.5372 | 1680.9373 | NaN | NaN | NaN |
| Lzic | LKQNLEEQLDR | -12 | -22 | 1 | 0 | 11 | 7245.8926 | 7875.139 | 7323.4673 | 5604.342 | 5419.308 | 5533.991 |
| Lzic | ETLEQLSEFNDSLKK | -48 | -62 | 1 | 0 | 15 | 2925.939 | 4135.34 | 4010.4963 | NaN | NaN | NaN |

| | | | | | | | | | | | | |
|---|---|---|---|---|---|---|---|---|---|---|---|---|
| Lzic | QNLEEQLDR | -14 | -22 | 0 | 0 | 9 | 413.6459 | 388.62048 | NaN | NaN | NaN | NaN |
| Nol11 | ETLIPIYSSDTEQNQR | -229 | -244 | 0 | 0 | 16 | 1656.8439 | 2033.3884 | 2014.3801 | 2091.7554 | 2554.9832 | 2576.8145 |
| Nol11 | YSEPFLLPHLK | -600 | -610 | 0 | 0 | 11 | 1069.9525 | 537.7738 | 1248.2289 | 1400.2467 | 1023.9319 | 843.4823 |
| Nol11 | HIEVELR | -411 | -417 | 0 | 0 | 7 | 824.7907 | 1104.1366 | 1312.4325 | 1089.3556 | 1510.2915 | 1475.8073 |
| Nol11 | SVFDYSDSTQDEK | -523 | -535 | 0 | 0 | 13 | 641.976 | 638.93835 | 614.155 | 417.04333 | NaN | NaN |
| Nol11 | LQPEVPGFK | -390 | -398 | 0 | 0.00025015 | 9 | 1010.1683 | NaN | 968.3468 | NaN | NaN | NaN |
| Pcbp1 | ETLSQSPQGR | -168 | -177 | 0 | 0 | 10 | 12817.117 | 12886.717 | 12667.2 | 14410.371 | 13793.27 | 13653.544 |
| Pcbp1 | SIQGQHTISPLDLAK | -223 | -237 | 0 | 0 | 15 | 4118.253 | 4250.904 | 4453.3315 | 2300.2327 | 2591.699 | 2525.0256 |
| Pcbp1 | SIQGQHTISPLDLAK | -223 | -237 | 0 | 0 | 15 | 4118.253 | 4250.904 | 4453.3315 | 2300.2327 | 2591.699 | 2525.0256 |
| Pcbp1 | QGANINEIR | -298 | -306 | 0 | 0 | 9 | 3597.3677 | 3691.5793 | 318.10712 | 3300.296 | 3499.3406 | 4809.417 |
| Pcbp1 | IITLTGPTNAIFK | -58 | -70 | 0 | 0 | 13 | 3346.0925 | 4053.7869 | 3933.222 | 4156.5566 | 4821.737 | 4937.526 |
| Pcbp1 | IANPVEGSSGR | -315 | -325 | 0 | 0 | 11 | 3249.2495 | 365.27066 | 3228.7695 | 2892.8264 | 455.63412 | 3366.884 |
| Pcbp1 | LVVPATQCGSLIGK | -102 | -115 | 0 | 0 | 14 | 2796.5154 | 3036.1602 | 3016.455 | 2503.706 | 1788.4203 | 1568.9484 |
| Pcbp1 | HTISPLDLAK | -228 | -237 | 0 | 0 | 10 | 2453.3643 | 2347.8477 | 299.68243 | 2523.564 | 3162.5361 | 3214.2559 |
| Pcbp1 | GQHTISPLDLAK | -226 | -237 | 0 | 0 | 12 | 1800.3933 | 1843.4878 | 1940.042 | 687.4259 | 576.66626 | 1832.9 |
| Pcbp1 | GQHTISPLDLAK | -226 | -237 | 0 | 0 | 12 | 1800.3933 | 1843.4878 | 1940.042 | 687.4259 | 576.66626 | 1832.9 |
| Pcbp1 | GQHTISPLDLAK | -226 | -237 | 0 | 0 | 12 | 1800.3933 | 1843.4878 | 1940.042 | 687.4259 | 576.66626 | 1832.9 |
| Pcbp1 | HGGTGFAGIDSSSPEVK | -252 | -268 | 0 | 0.000230707 | 17 | 1588.1499 | 1533.6606 | 698.6669 | 629.3125 | 907.0844 | 1043.7375 |
| Pcbp1 | GVPQSVTECVK | -150 | -160 | 0 | 0 | 11 | 1535.1438 | 2701.6274 | 2647.5784 | 3042.9226 | 1634.0944 | 1526.6964 |
| Pcbp1 | TGPTNAIFK | -62 | -70 | 0 | 0 | 9 | 1362.0187 | 1468.3427 | 1597.0706 | 1627.3977 | 1857.9983 | 1926.315 |
| Pcbp1 | GGTGFAGIDSSSPEVK | -253 | -268 | 0 | 0 | 16 | 917.7827 | 926.83453 | 1002.0807 | 1256.5228 | 1400.8599 | 1314.0685 |
| Pcbp1 | TIPYQPMPASSPVICAGGQDR | -180 | -200 | 0 | 0 | 21 | 523.59674 | 478.37497 | 611.33044 | 616.021 | 766.324 | 704.3746 |
| Pcbp1 | TTHELTIPNNLIGCIIGR | -280 | -297 | 0 | 0 | 18 | 404.52158 | 362.94064 | 363.97113 | 422.89215 | 540.52704 | 486.32016 |
| Pcbp1 | RLVVPATQCGSLIGK | -101 | -115 | 1 | 0.00025015 | 15 | 1443.4453 | 741.78033 | 356.57452 | NaN | 625.07556 | NaN |
| Pcbp1 | TIPNNLIGCIIGR | -285 | -297 | 0 | 0 | 13 | 1089.9137 | NaN | NaN | 1259.352 | 1115.3142 | 1099.976 |
| Snrpd3 | VLHEAEGHIVTC | -9 | -20 | 0 | 0.000230707 | 12 | 4295.033 | 4350.6685 | 4144.725 | 1890.6117 | 4429.7163 | 4190.9893 |
| Snrpd3 | VLHEAEGHIVTC | -9 | -20 | 0 | 0.000230707 | 12 | 4295.033 | 4350.6685 | 4144.725 | 1890.6117 | 4429.7163 | 4190.9893 |
| Snrpd3 | QLEQVYIR | -57 | -64 | 0 | 0 | 8 | 1623.5713 | 1511.3927 | 1548.6489 | 1859.645 | 1618.3914 | 1613.0553 |
| Snrpd3 | FLILPDMLK | -70 | -78 | 0 | 0 | 9 | 1226.1233 | 1847.4528 | 1068.998 | 710.4558 | 1035.9655 | 1075.412 |
| Snrpd3 | FLILPDMLK | -70 | -78 | 0 | 0 | 9 | 1226.1233 | 1847.4528 | 1068.998 | 710.4558 | 1035.9655 | 1075.412 |
| Snrpd3 | TCETNTGEVYR | -19 | -29 | 0 | 0 | 11 | 1194.0865 | 2289.8286 | 1320.3049 | 1355.835 | 1541.2644 | 114.40663 |
| Snrpd3 | VAQLEQVYIR | -55 | -64 | 0 | 0 | 10 | 1131.7576 | 1303.9216 | 1406.6877 | 1380.9805 | 919.9656 | 1109.9004 |
| Snrpd3 | ETNTGEVYR | -21 | -29 | 0 | 0.00025015 | 9 | 1784.8917 | 1843.4208 | 1712.6332 | 6573.263 | NaN | 1224.0476 |
| Snrpd3 | VLHEAEGHIV | -9 | -18 | 0 | 0.000627943 | 10 | 2971.1653 | 270.32098 | NaN | 2138.5312 | NaN | 1331.2618 |
| Snrpd3 | IVTCETNTGEVYR | -17 | -29 | 0 | 0 | 13 | 800.0966 | 722.11456 | 641.06226 | NaN | NaN | NaN |
| Snrpd3 | MSNITVTYR | -43 | -51 | 0 | 0.007374243 | 9 | 481.67453 | NaN | NaN | 352.1234 | NaN | NaN |
| Tsfm | SVGSLDDEPGGETETR | -266 | -281 | 0 | 0 | 16 | 1499.0026 | 1292.3445 | 920.9528 | 1027.0559 | 1215.47 | 1069.9775 |
| Tsfm | ETPEQIANLEEVGR | -240 | -253 | 0 | 0 | 14 | 3056.3813 | NaN | NaN | 2653.7021 | NaN | NaN |
| Ado | ETPQADDFWCEGEPYPGPK | -235 | -253 | 0 | 0 | 19 | 246.93413 | 299.29648 | NaN | NaN | 344.99738 | 342.3661 |
| Ndufaf2 | TFQPGSWTPEDGKR | -152 | -165 | 1 | 0 | 14 | 907.24634 | 133.17207 | 1081.2194 | 2054.954 | 2362.5847 | 2528.3945 |
| Ndufaf2 | ETSEELLPSPTATQVK | -114 | -129 | 0 | 0 | 16 | 922.8236 | NaN | NaN | 1500.3212 | 1304.2992 | NaN |
| Ndufaf2 | GHASAPYFGR | -130 | -139 | 0 | 0 | 10 | 791.9668 | NaN | NaN | 485.89484 | NaN | NaN |
| Spr | SYAPGPLDNDMQQLAR | -196 | -211 | 0 | 0 | 16 | 651.02246 | 834.5229 | 857.1275 | 802.2212 | 918.5084 | 908.1242 |
| Spr | QLKEELGAQQPDLK | -50 | -63 | 1 | 0 | 14 | 428.00845 | 492.8759 | 479.37885 | 515.0984 | 640.371 | NaN |
| Spr | ETSKDPELR | -212 | -220 | 1 | 0.00078661 | 9 | 789.50916 | NaN | NaN | NaN | 657.9272 | NaN |
| Spr | SDGALVDCGTSAQK | -228 | -241 | 0 | 0.00025015 | 14 | 1097.8705 | NaN | NaN | NaN | NaN | NaN |
| Prrc2c | ETVEKPEEKPR | -933 | -943 | 2 | 0 | 11 | 965.844 | 608.94336 | 652.10846 | 341.56195 | 494.33063 | 900.8648 |
| Prrc2c | ETTQVVQEAEPESGAQPR | -616 | -633 | 0 | 0 | 18 | 448.12845 | 518.3013 | 391.30148 | 501.94254 | 221.46408 | 543.6624 |
| Prrc2c | SLEDVKPR | -853 | -860 | 1 | 0 | 8 | 1041.5709 | 1258.3839 | 891.37024 | NaN | 1301.1981 | 2389.6184 |
| Prrc2c | KLPDLSLVENK | -2052 | -2062 | 1 | 0.000230707 | 11 | 1234.8293 | NaN | NaN | NaN | 1023.8895 | 1281.7418 |
| Prrc2c | SSSQIPAQPPVTK | -376 | -388 | 0 | 0 | 13 | 902.5407 | 840.33527 | 1158.4794 | NaN | NaN | NaN |
| Prrc2c | IDNRPLEKPYIR | -1286 | -1297 | 2 | 0.000230707 | 12 | 581.5501 | NaN | 682.0358 | NaN | 627.4275 | NaN |
| Prrc2c | LIDTHLLQAR | -2490 | -2499 | 0 | 0 | 10 | 1283.4471 | NaN | 1189.3538 | NaN | NaN | NaN |
| Atad1 | EVAQETDGFSGSDLK | -286 | -300 | 0 | 0 | 15 | 3464.1567 | 3481.6487 | 392.72733 | 1439.9509 | NaN | NaN |
| Atad1 | FINLQPSTLTDK | -154 | -165 | 0 | 0 | 12 | 697.50507 | 898.522 | 457.03525 | NaN | 805.4715 | NaN |
| Atad1 | TDKWYGESQK | -163 | -172 | 1 | 0.00149795 | 10 | 937.4542 | 925.42535 | NaN | NaN | NaN | NaN |
| Edc4 | SIDIEDATQHLR | -6 | -17 | 0 | 0 | 12 | 924.8323 | 996.83673 | 875.6588 | 904.60297 | 1089.589 | 940.95764 |
| Edc4 | SLLPGLLPGPADK | -683 | -695 | 0 | 0 | 13 | 2592.7556 | 2945.0127 | NaN | 2977.1323 | 3630.0405 | 3325.043 |
| Edc4 | KQDPEVPFWR | -359 | -368 | 1 | 0.000347792 | 10 | 718.8839 | 618.6595 | 503.71176 | 1260.9529 | 1033.3322 | NaN |
| Edc4 | EVATPDSQVWPTAPDITR | -823 | -840 | 0 | 0 | 18 | 336.78842 | 1780.4655 | 2201.4414 | NaN | NaN | 3318.312 |
| Edc4 | VVLPAFEK | -1126 | -1133 | 0 | 0.00142999 | 8 | 571.4828 | 790.00916 | 1773.8016 | NaN | NaN | NaN |
| Edc4 | GHSTCLSEGALSPDGTVL | -295 | -312 | 0 | 0.001868731 | 18 | 462.06137 | NaN | NaN | NaN | NaN | 1208.5406 |
| Edc4 | FQPQLNPDVVAPL | -523 | -535 | 0 | 0.007523243 | 13 | 428.26324 | NaN | NaN | NaN | 675.352 | NaN |
| Rpl36 | EVCGFAPYER | -46 | -55 | 0 | 0 | 10 | 1924.8018 | 1795.3817 | 1866.7366 | 1374.0211 | 783.4111 | 809.19495 |
| Rpl36 | VCGFAPYER | -47 | -55 | 0 | 0 | 9 | 597.3251 | 764.95984 | 142.07248 | 351.3459 | 795.42456 | 823.03345 |
| Rpl36 | KREELSNVL | -86 | -94 | 2 | 0.00453359 | 9 | 4227.1826 | 3051.6458 | 2974.1592 | NaN | 2830.4106 | 2511.0154 |
| Nop16 | EVDTEERPR | -77 | -85 | 1 | 0.000471409 | 9 | 885.6469 | 912.43567 | 326.53522 | 1233.0068 | 1438.1632 | 595.33417 |
| Nop16 | NYYQDTPK | -139 | -146 | 0 | 0.00078661 | 8 | 1697.6466 | NaN | NaN | NaN | NaN | NaN |
| Smarca4 | EVEAQLPEKVEYVIK | -983 | -997 | 1 | 0 | 15 | 545.78564 | 478.03958 | 431.96246 | 536.7617 | 625.7386 | 696.10114 |
| Smarca4 | KIPDPDSDDVSEVDAR | -689 | -704 | 1 | 0.00025015 | 16 | 946.77295 | NaN | 976.5388 | 799.6819 | 638.21185 | 3080.6875 |
| Smarca4 | ITPIQKPR | -352 | -359 | 1 | 0.008736733 | 8 | 474.80737 | 363.3822 | 395.33957 | NaN | 348.5409 | 331.44116 |
| Cul1 | EVELKPDTLIK | -660 | -670 | 1 | 0 | 11 | 1933.5354 | 1787.7542 | 1601.9495 | 2725.7922 | 2131.9358 | 1891.7643 |
| NaN | EVHSGVSTDPQAYK | (51);(51) | (64);(64) | 0 | 0 | 14 | 1610.3214 | 1677.0593 | 1708.9446 | 1684.0743 | 1800.7383 | 1844.6674 |
| NaN | EVHSGVSTDPQAYK | (51);(51) | (64);(64) | 0 | 0 | 14 | 1610.3214 | 1677.0593 | 1708.9446 | 1684.0743 | 1800.7383 | 1844.6674 |
| NaN | GFFPDHVELS | (35);(35) | (44);(44) | 0 | 0.001055699 | 10 | 1439.4424 | 1448.1029 | 1476.6353 | 1307.8596 | 1734.921 | 1790.0438 |
| Nxn | GIPTLIVLDPQGEVITR | -274 | -290 | 0 | 0 | 17 | 429.53995 | NaN | NaN | NaN | 493.3321 | NaN |
| Nxn | EVIAGPLLR | -172 | -180 | 0 | 0.002265552 | 9 | 782.4774 | NaN | NaN | NaN | NaN | NaN |
| Tbc1d10a | EVIELPVTER | -350 | -359 | 0 | 0 | 10 | 791.04694 | 811.6803 | 1003.2693 | 1183.5437 | NaN | 1085.5088 |

| | | | | | | | | | | | | |
|---|---|---|---|---|---|---|---|---|---|---|---|---|
| Tbc1d10a | TLPWSSVLR | -282 | -290 | 0 | 0.006384543 | 9 | 447.66003 | NaN | NaN | NaN | NaN | NaN |
| Tbc1d10a | VGLVLLK | -306 | -312 | 0 | 0.00142999 | 7 | 39.282036 | NaN | NaN | NaN | NaN | NaN |
| Rpl15 | EVILIDPFHK | -131 | -140 | 0 | 0.00025015 | 10 | 5078.2466 | 5777.5835 | 5881.853 | 138.23799 | 5793.5513 | 5693.8086 |
| Rpl15 | YIQELWR | -6 | -12 | 0 | 0 | 7 | 2428.2156 | 286.36703 | 2242.0164 | 1757.7651 | 692.63684 | 786.71606 |
| Rpl15 | WITKPVHK | -150 | -157 | 1 | 0 | 8 | 2031.195 | 1959.66 | 2137.676 | 2702.201 | 2860.5474 | 2725.972 |
| Rpl15 | NSYWVGEDSTYK | -117 | -128 | 0 | 0 | 12 | 1712.7802 | 180.10426 | 2356.4807 | 120.56169 | 511.97473 | 1808.3508 |
| Rpl15 | NSYWVGEDSTYK | -117 | -128 | 0 | 0 | 12 | 1712.7802 | 180.10426 | 2356.4807 | 120.56169 | 511.97473 | 1808.3508 |
| Rpl15 | NPDTQWITKPVHK | -145 | -157 | 1 | 0 | 13 | 1398.9111 | 1544.2845 | 1259.8938 | NaN | 514.2817 | 592.99634 |
| Rpl15 | NPDTQWITKPVHK | -145 | -157 | 1 | 0 | 13 | 1398.9111 | 1544.2845 | 1259.8938 | NaN | 514.2817 | 592.99634 |
| Rpl15 | WVGEDSTYK | -120 | -128 | 0 | 0 | 9 | 119.18968 | 3474.7124 | 803.7966 | NaN | 3321.5876 | 3334.1782 |
| Rpl15 | RNPDTQWITKPVHK | -144 | -157 | 2 | 0 | 14 | 1071.5137 | 1590.493 | 1476.414 | NaN | NaN | NaN |
| Rpl15 | RNPDTQWITKPVHK | -144 | -157 | 2 | 0 | 14 | 1071.5137 | 1590.493 | 1476.414 | NaN | NaN | NaN |
| Rpl15 | RNPDTQWITKPVHK | -144 | -157 | 2 | 0 | 14 | 1071.5137 | 1590.493 | 1476.414 | NaN | NaN | NaN |
| Rpl15 | RNPDTQWITK | -144 | -153 | 1 | 0.000627943 | 10 | 1649.2921 | 2167.904 | 2027.6278 | NaN | NaN | NaN |
| Rpl15 | NTLQLHR | -196 | -202 | 0 | 0.000627943 | 7 | 1217.9347 | 1294.1453 | 1177.8167 | NaN | NaN | NaN |
| Rpl15 | NPDTQWITK | -145 | -153 | 0 | 0 | 9 | 970.8436 | 1644.7891 | 1551.849 | NaN | NaN | NaN |
| Rpl15 | FHHTIGGSR | -180 | -188 | 0 | 0 | 9 | 363.737 | 956.5833 | 1144.506 | NaN | NaN | NaN |
| Rpl15 | FHHTIGGS | -180 | -187 | 0 | 0.00272025 | 8 | 716.7117 | NaN | 816.29584 | NaN | NaN | NaN |
| Ppp4r3a | EVIPISDPELK | -248 | -258 | 0 | 0 | 11 | 696.39453 | 688.1797 | NaN | 861.05585 | NaN | NaN |
| Ppp4r3a | LTDEATDEEK | -318 | -327 | 0 | 0.000298805 | 10 | 680.39435 | NaN | NaN | 263.71204 | NaN | NaN |
| Timm9 | LTETCFLDCVK | -24 | -34 | 0 | 0 | 11 | 859.6583 | 822.7334 | 905.88324 | 1297.6263 | 1871.8107 | 1750.5103 |
| Timm9 | EVKPEEVTCSEHCLQK | -40 | -55 | 1 | 0 | 16 | 650.5166 | 705.16113 | 2223.6096 | 1338.5316 | 1268.975 | 2622.8135 |
| Timm9 | EVKPEEVTCSEHCLQK | -40 | -55 | 1 | 0 | 16 | 650.5166 | 705.16113 | 2223.6096 | 1338.5316 | 1268.975 | 2622.8135 |
| Ndufs1 | LIGSPVDLTYR | -431 | -441 | 0 | 0 | 11 | 1788.351 | 1024.428 | 1049.5869 | 1312.4127 | 2355.7168 | 1989.9528 |
| Ndufs1 | FASEIAGVDDLGTTGR | -185 | -200 | 0 | 0 | 16 | 929.95935 | 313.91223 | 846.4861 | 673.0145 | 710.8281 | 160.93591 |
| Ndufs1 | YDHLGDSPK | -442 | -450 | 0 | 0 | 9 | 750.55304 | 578.49426 | 647.1432 | 1016.57574 | 941.90546 | 1311.8126 |
| Ndufs1 | FEAPLFNAR | -409 | -417 | 0 | 0 | 9 | 630.4768 | 335.90427 | 521.83795 | 750.68756 | 868.86255 | 1008.15344 |
| Ndufs1 | SEIAGITLPYDTLDQVR | -627 | -643 | 0 | 0 | 17 | 409.47693 | 437.35248 | 308.9248 | 549.03424 | 494.76324 | 502.04227 |
| Ndufs1 | LVNQEVLADPLVPPQLTIK | -674 | -692 | 0 | 0 | 19 | 346.21432 | 410.90164 | 343.45547 | 438.13525 | 586.3377 | 462.41757 |
| Ndufs1 | VDSDNLCTEEIFPTEGAGTDLR | -361 | -382 | 0 | 0 | 22 | 184.5148 | 223.15683 | 310.6346 | 392.99326 | 411.15634 | 410.1222 |
| Ndufs1 | VALIGSPVDLTYR | -429 | -441 | 0 | 0 | 13 | 621.4058 | 418.6149 | 322.98856 | 1658.14 | NaN | NaN |
| Ndufs1 | SATYVNTEGR | -593 | -602 | 0 | 0 | 10 | 1008.0751 | 1198.8425 | NaN | 1131.59 | NaN | NaN |
| Ndufs1 | VAVTPPGLAR | -608 | -617 | 0 | 0 | 10 | 764.4014 | 647.8136 | NaN | 718.21295 | NaN | NaN |
| Ndufs1 | EVLADPLVPPQLTIK | -678 | -692 | 0 | 0.00025015 | 15 | 304.92764 | NaN | NaN | NaN | 350.34708 | 337.42004 |
| Ndufs1 | LSVAGNCR | -69 | -76 | 0 | 0.00930008 | 8 | 1165.6793 | NaN | NaN | NaN | NaN | NaN |
| Pck2 | FDSEGQLR | -335 | -342 | 0 | 0 | 8 | 2699.5627 | 529.4223 | 484.17258 | 626.828 | 1783.2285 | 1903.1014 |
| Pck2 | PALPGWK | -316 | -322 | 0 | 0 | 7 | 592.05334 | 449.91473 | 484.36963 | 426.40826 | 251.64923 | 381.0637 |
| Pck2 | TLIGHVPDQR | -235 | -244 | 0 | 0 | 10 | 342.49976 | 1288.7808 | 390.90955 | 1300.0394 | 1242.4137 | 1092.9629 |
| Pck2 | EVLAELEALEGR | -625 | -636 | 0 | 0 | 12 | 1010.3847 | NaN | 122.03459 | 1152.9846 | 732.545 | 755.2384 |
| Pck2 | FLWPGFGENAR | -543 | -553 | 0 | 0 | 11 | 785.02264 | 1010.74927 | 331.0845 | NaN | 811.4221 | 814.5151 |
| Pck2 | VLDWICR | -554 | -560 | 0 | 0.003814256 | 7 | 685.82983 | 809.65405 | 814.9302 | NaN | 1003.16345 | 1069.5674 |
| Pck2 | RLEGEDSAQETPIGLVPK | -561 | -578 | 1 | 0 | 18 | 545.04236 | NaN | 1099.6919 | 806.7722 | 1347.2399 | 739.3904 |
| Pck2 | SAHLCQPEGIHIC | -51 | -63 | 0 | 0 | 13 | 418.9294 | 465.2216 | 466.09164 | NaN | NaN | 406.07117 |
| Stk39 | EVLEGLDYLHR | -187 | -197 | 0 | 0 | 11 | 1537.5029 | 1598.8601 | 1666.0615 | 1787.4613 | 2100.1646 | NaN |
| Ppp1r18 | EVLESILSPGEPGDQK | -217 | -232 | 0 | 0 | 16 | 561.96356 | 517.3786 | 470.45837 | 901.4878 | 571.4684 | 421.65564 |
| Cwc27 | YDNTIFHR | -49 | -56 | 0 | 0.000627943 | 8 | 1362.7047 | 1105.0483 | 1049.8912 | 1134.8146 | 462.10077 | NaN |
| Cwc27 | EVLFNPFDDITPR | -165 | -177 | 0 | 0 | 13 | 287.4557 | NaN | 377.1696 | NaN | 328.0151 | 374.6304 |
| Cwc27 | SFGEEAEEEEEEVNR | -206 | -220 | 0 | 0 | 15 | 346.8988 | 239.13586 | NaN | NaN | NaN | NaN |
| Pfdn2 | EVLPALEGNK | -85 | -94 | 0 | 0 | 10 | 2476.2173 | 2637.0986 | NaN | 538.42413 | 2828.9155 | 3301.1313 |
| Aup1 | EVLPHVPLNVIQR | -306 | -318 | 0 | 0 | 13 | 952.38885 | NaN | NaN | NaN | 434.35327 | 588.46747 |
| Aup1 | RVELVESLKK | -135 | -144 | 2 | 0 | 10 | 310.53705 | NaN | NaN | NaN | 365.4413 | 324.85974 |
| Atad2 | LDSIDPALR | -232 | -240 | 0 | 0 | 9 | 2564.7593 | 2032.4941 | 2163.405 | 277.48727 | 1415.4031 | 1418.4379 |
| Atad2 | VFPHVEVGTNK | -378 | -388 | 0 | 0 | 11 | 1744.4692 | 7093.7324 | NaN | 11629.917 | 678.9121 | 638.4246 |
| Atad2 | NVQLPDKEER | -569 | -578 | 1 | 0 | 10 | 1485.5676 | 1537.8715 | 1623.5111 | NaN | 1786.4878 | NaN |
| Atad2 | EVLPVAPPPEPR | -606 | -617 | 0 | 0 | 12 | 560.87805 | 691.7867 | 150.09724 | 698.44824 | NaN | NaN |
| Atad2 | FDEIDGLAPVR | -185 | -195 | 0 | 0 | 11 | 831.97034 | 623.1739 | 580.1627 | NaN | NaN | NaN |
| Gcsh | EVNEALAENPGLVNK | -119 | -133 | 0 | 0 | 15 | 661.6856 | 567.05396 | 734.52924 | NaN | 776.537 | 735.62836 |
| Gcsh | HEWITTEEGIGTVGIS | -54 | -69 | 0 | 0 | 16 | 657.1483 | NaN | 935.8794 | 690.8029 | 983.35925 | 922.95074 |
| Arih1 | LVEPGLGVGGER | -47 | -58 | 0 | 0 | 12 | 1004.6125 | 1209.66 | 1099.8313 | 587.8874 | 951.6347 | 108.4261 |
| Arih1 | EVNEVIQNPATITR | -117 | -130 | 0 | 0 | 14 | 896.28076 | 874.3899 | 842.841 | 590.28564 | 720.4974 | NaN |
| Clgn | EVNFQDGIDCGGAYIK | -142 | -157 | 0 | 0.00025015 | 16 | 744.97345 | 719.86316 | 775.275 | NaN | 661.82935 | 659.0515 |
| Npc2 | VPFPIPEPDGCK | -83 | -94 | 0 | 0 | 12 | 2109.6296 | 2307.8464 | 2313.4404 | 2471.9053 | 2961.6875 | 3141.7163 |
| Npc2 | EVNVSPCPTDPCQLHK | -36 | -51 | 0 | 0 | 16 | 1669.1746 | 1890.6897 | 1820.169 | 1992.6381 | 2270.1326 | 2178.1458 |
| Npc2 | EVNVSPCPTDPCQLHK | -36 | -51 | 0 | 0 | 16 | 1669.1746 | 1890.6897 | 1820.169 | 1992.6381 | 2270.1326 | 2178.1458 |
| Npc2 | SGINCPIQK | -95 | -103 | 0 | 0 | 9 | 1356.0864 | 1378.3 | 1289.0109 | 1556.0272 | 1400.2881 | 1443.4839 |
| Npc2 | VHGILEGIR | -74 | -82 | 0 | 0 | 9 | 1176.9307 | 1173.6882 | 1275.1967 | 904.6936 | 913.6379 | 923.31946 |
| Npc2 | EVNVSPCPTDPCQLHKG | -36 | -52 | 1 | 0.001392869 | 17 | 605.65295 | 526.2391 | 542.57117 | 613.18884 | 616.08594 | 627.228 |
| Usp39 | HCPYLDTINR | -103 | -112 | 0 | 0 | 10 | 1389.7397 | 1276.7377 | 1308.7349 | 1471.303 | 2302.3154 | 1016.198 |
| Usp39 | SLEDITYVLKPTFTK | -184 | -198 | 1 | 0 | 15 | 674.4344 | 611.9505 | 442.82465 | 431.86432 | 380.13382 | 524.05176 |
| Usp39 | EVPAPALPVVR | -60 | -70 | 0 | 0.00025015 | 11 | 316.88162 | 560.80115 | 214.0302 | 287.10135 | 415.7104 | 326.63043 |
| Usp39 | IVHDGKPSEGSYR | -502 | -514 | 1 | 0.001023236 | 13 | 281.1199 | 607.4731 | 431.37677 | 425.4325 | 643.85376 | 206.1326 |
| Usp39 | KNPTIVNFPITNVDLR | -465 | -480 | 1 | 0 | 16 | 249.02263 | 293.7538 | 263.739 | NaN | 426.63455 | 371.43524 |
| Usp39 | NPTIVNFPITNVDLR | -466 | -480 | 0 | 0 | 15 | 87.40642 | 387.46216 | NaN | 376.2863 | 386.56662 | 528.25366 |
| Csrp1 | EVQCEGNSFHK | -22 | -32 | 0 | 0 | 11 | 489.79285 | 481.8288 | 709.3549 | 454.63965 | 820.5037 | 870.4539 |
| Csrp1 | NLDSTTVAVHGEEIYCK | -43 | -59 | 0 | 0 | 17 | 288.50296 | 309.93497 | 271.2202 | 454.08826 | 526.5047 | 482.7218 |
| Csrp1 | TVAVHGEEIYCK | -48 | -59 | 0 | 0 | 12 | 385.82935 | 740.3556 | 500.40167 | NaN | NaN | NaN |
| Diablo | EVSDEGADQEEEAYLRED | -220 | -237 | 1 | 0 | 18 | 272.7199 | NaN | 294.0333 | 408.04288 | 335.22644 | 466.75238 |
| Atp13a1 | EVTPVSSIPIETHR | -553 | -566 | 0 | 0 | 14 | 721.8861 | 1523.3983 | 1005.17896 | 612.29083 | 2505.6477 | 1172.4824 |

| | | | | | | | | | | | | |
|---|---|---|---|---|---|---|---|---|---|---|---|---|
| Atp13a1 | SALLSPEEPPASHR | -914 | -927 | 0 | 0.001092529 | 14 | 1028.9647 | 805.2379 | NaN | NaN | NaN | NaN |
| Kif23 | EVTQEVEVARPVDK | -431 | -444 | 1 | 0.000627943 | 14 | 2735.68 | NaN | NaN | NaN | 3281.5159 | 3212.5085 |
| Dek | EVYENYPAYDLTER | -354 | -367 | 0 | 0 | 14 | 1939.2998 | 2708.8699 | 2657.6492 | 2651.0972 | 3169.16 | 3079.7085 |
| Dek | NVGQFSGFPFEK | -130 | -141 | 0 | 0 | 12 | 1198.4956 | 1473.3169 | 1894.5736 | 1642.775 | 1873.2075 | 1740.7345 |
| Dek | KPPTDEELKETVK | -323 | -335 | 2 | 0 | 13 | 1099.3544 | 1184.3807 | 1317.5503 | 1273.15 | 1538.8727 | 1319.4331 |
| Dek | SGVNSELVK | -173 | -181 | 0 | 0 | 9 | 671.921 | 1834.6284 | 1601.318 | 285.0427 | 1439.5771 | 1496.6611 |
| Dek | SICEVLDLER | -163 | -172 | 0 | 0 | 10 | 576.2455 | 247.59126 | 623.14197 | 626.6839 | 586.7135 | 567.368 |
| Dek | KPPTDEELK | -323 | -331 | 1 | 0 | 9 | 336.0273 | 1006.9207 | 1245.4058 | 721.2194 | 745.4192 | NaN |
| Dek | LLYNRPGTVS | -116 | -125 | 1 | 0.000230707 | 10 | 1418.9612 | NaN | NaN | 1037.2443 | NaN | 1010.19525 |
| Dek | IHFFLSK | -98 | -104 | 0 | 0 | 7 | 726.7206 | 619.696 | 619.2398 | NaN | NaN | NaN |
| Colgalt1 | EVYGFLPVPLR | -281 | -291 | 0 | 0 | 11 | 269.7793 | NaN | NaN | 158.58173 | NaN | NaN |
| Colgalt1 | TPAYIPIR | -209 | -216 | 0 | 0.003745449 | 8 | 235.61473 | 152.24017 | NaN | NaN | NaN | NaN |
| Ndufs5 | IEFDDFEECLLR | -58 | -69 | 0 | 0 | 12 | 1492.8715 | 1701.0693 | 1588.4506 | 1403.3723 | 1925.6825 | 1686.6451 |
| Ndufs5 | EWIECAHGIGGTR | -39 | -51 | 0 | 0 | 13 | 571.27527 | 630.19336 | 499.88617 | 521.0442 | 619.09796 | 605.53723 |
| Ndufs5 | PFLDIQK | -2 | -8 | 0 | 0.000712251 | 7 | 844.4314 | NaN | NaN | 600.13184 | 399.50394 | 494.22864 |
| Ckap5 | YSDTDIEPFLK | -1860 | -1870 | 0 | 0 | 11 | 1082.2513 | 1079.3105 | 1205.3431 | 448.42087 | 551.4725 | 1106.5105 |
| Ckap5 | IEDLEEGQQVIR | -1655 | -1666 | 0 | 0 | 12 | 804.0239 | 1057.826 | 1152.9438 | 1141.8195 | 1344.5323 | 1257.3677 |
| Ckap5 | EWLEGEDLSEELKK | -976 | -989 | 1 | 0 | 14 | 374.716 | 648.55237 | 581.1581 | 719.2291 | 670.445 | 264.48376 |
| Ckap5 | LDDIFEPVLIPEPK | -1510 | -1523 | 0 | 0 | 14 | 146.86925 | 247.34695 | 205.23952 | 346.3451 | 258.92984 | 348.86353 |
| Ckap5 | LEAGDYADLVK | -312 | -322 | 0 | 0.001392869 | 11 | 1963.958 | 1955.7689 | 1782.982 | 1709.9452 | NaN | NaN |
| Ckap5 | SSTTNIDDLKK | -2012 | -2022 | 1 | 0 | 11 | 326.38422 | 625.9809 | 815.98975 | NaN | 675.5292 | NaN |
| Ckap5 | LLPDTINSINLDR | -1719 | -1731 | 0 | 0 | 13 | 1148.7808 | 1985.5012 | 1088.7332 | NaN | NaN | NaN |
| Ckap5 | IGELPTALK | -898 | -906 | 0 | 0 | 9 | 307.0819 | NaN | NaN | NaN | NaN | NaN |
| Rtf1 | IDLQVPSSESK | -673 | -683 | 0 | 0 | 11 | 2442.538 | 2381.9114 | 2554.1287 | 983.60004 | 1550.5787 | 1683.6738 |
| Rtf1 | QIQDQLNELEER | -539 | -550 | 0 | 0 | 12 | 2724.838 | 2994.1982 | NaN | NaN | 3003.7495 | 2876.0789 |
| Rtf1 | FNDQDIEEIVK | -492 | -502 | 0 | 0 | 11 | 881.7034 | 853.3519 | 825.6249 | 1341.4697 | NaN | NaN |
| Rtf1 | YGSGVLPDAPK | -629 | -639 | 0 | 0.00025015 | 11 | 931.6228 | 777.164 | NaN | NaN | NaN | 1014.38574 |
| Rtf1 | EWNIVESEK | -574 | -582 | 0 | 0.00025015 | 9 | 190.98682 | 606.5996 | NaN | NaN | NaN | NaN |
| Necap2 | EWQLDQPSWSGR | -34 | -45 | 0 | 0 | 12 | 1300.2233 | 1674.8018 | 1607.3824 | 2008.8458 | 2295.7937 | 311.08063 |
| Rpf1 | EWVHKPR | -332 | -338 | 1 | 0.00025015 | 7 | 427.12186 | NaN | NaN | NaN | NaN | NaN |
| Supt5h | NIDNVVLDEDR | -107 | -117 | 0 | 0 | 11 | 1025.5112 | 1338.7231 | 1318.3699 | 831.07965 | 1721.1782 | 776.7782 |
| Supt5h | TPAQSGAWDPNNPNTPSR | -808 | -825 | 0 | 0 | 18 | 979.41656 | 1108.9124 | 1124.9153 | 1028.182 | 254.16444 | 1269.6721 |
| Supt5h | TPHYGSQTPLHDGSR | -793 | -807 | 0 | 0 | 15 | 587.2173 | 492.9278 | 350.16437 | 231.67133 | 856.2638 | 1241.492 |
| Supt5h | TPHYGSQTPLHDGSR | -793 | -807 | 0 | 0 | 15 | 587.2173 | 492.9278 | 350.16437 | 231.67133 | 856.2638 | 1241.492 |
| Supt5h | YVEPSQNTISLK | -293 | -304 | 0 | 0 | 12 | 645.28076 | 683.88007 | NaN | 662.04224 | 956.4222 | 658.7404 |
| Supt5h | VIDGPHSGR | -600 | -608 | 0 | 0 | 9 | 1698.7596 | 1723.4591 | NaN | NaN | 354.6656 | 606.5916 |
| Supt5h | SIDGEDGIIR | -1051 | -1060 | 0 | 0.00025015 | 10 | 215.24127 | NaN | NaN | 254.9302 | 739.4268 | 1124.5358 |
| Supt5h | QAIEGVGNLR | -235 | -244 | 0 | 0.00025015 | 10 | 806.36975 | NaN | NaN | NaN | NaN | 259.81952 |
| Supt5h | EYDEEEEEEDDDRPPK | -51 | -66 | 1 | 0.003102268 | 16 | 415.32578 | NaN | 191.35274 | NaN | NaN | NaN |
| Supt5h | LTTVDSQRPGGMTS | -744 | -757 | 1 | 0.002595029 | 14 | 1073.0966 | NaN | NaN | NaN | NaN | NaN |
| Hba | IGGHGAEYGAE | -18 | -28 | 0 | 0.001092529 | 11 | 2221.59 | 2163.3318 | 2191.3945 | 1953.4397 | 816.72656 | 1872.1005 |
| Hba | EYGAEALER | -24 | -32 | 0 | 0 | 9 | 1389.9042 | 1321.464 | 1213.9183 | 1297.0245 | 485.24567 | 1084.5131 |
| Hba | IGGHGAEYGAEALER | -18 | -32 | 0 | 0 | 15 | 1147.7094 | 1385.7175 | 1349.0746 | 1790.0333 | 2654.1367 | 2725.9492 |
| Hba | IGGHGAEYGAEALER | -18 | -32 | 0 | 0 | 15 | 1147.7094 | 1385.7175 | 1349.0746 | 1790.0333 | 2654.1367 | 2725.9492 |
| Hba | TYFPHFDVSHGS | -42 | -53 | 0 | 0 | 12 | 928.02484 | 1273.7427 | 1260.5452 | 955.20953 | 105.61769 | 1183.2701 |
| Hba | TYFPHFDVSHGSAQ | -42 | -55 | 0 | 0 | 14 | 890.2074 | 3761.6965 | 4051.8877 | 3406.8857 | 1124.2018 | 4091.3386 |
| Hba | TYFPHFDVSHGSAQ | -42 | -55 | 0 | 0 | 14 | 890.2074 | 3761.6965 | 4051.8877 | 3406.8857 | 1124.2018 | 4091.3386 |
| Hba | IGGHGAEYGAEAL | -18 | -30 | 0 | 0.000550307 | 13 | 739.1689 | 1176.5566 | 1154.7556 | 398.63547 | 858.25134 | 813.57324 |
| Hba | TYFPHFDVSHGSAQVK | -42 | -57 | 0 | 0 | 16 | 545.4315 | 585.746 | 700.979 | 399.42358 | 1542.5969 | 1356.0836 |
| Hba | VHASLDK | -122 | -128 | 0 | 0.002068213 | 7 | 456.85028 | 566.06934 | 323.49677 | 646.9588 | 422.98917 | 355.27747 |
| Hba | SHHPADFTPAVHASLDK | -112 | -128 | 0 | 0 | 17 | 290.5205 | 1965.7921 | 1694.4521 | 1276.2533 | 1177.046 | 1616.1652 |
| Hba | TYFPHFDVSHGSA | -42 | -54 | 0 | 0.00025015 | 13 | 940.155 | NaN | NaN | 414.30872 | NaN | NaN |
| Rbm22 | YVGGLGDTITETDLR | -235 | -249 | 0 | 0 | 15 | 1309.1876 | 1583.1821 | 1470.1239 | 1583.7202 | 1827.2971 | 1870.926 |
| Rbm22 | TCLGENPYIR | -26 | -35 | 0 | 0 | 10 | 883.7631 | 562.1079 | 728.72455 | 567.3943 | 266.49738 | 791.5863 |
| Rbm22 | YYGINDPVADK | -202 | -212 | 0 | 0 | 11 | 1258.6459 | 1542.5684 | NaN | NaN | NaN | 1288.4858 |
| Rbm22 | EYGLPIQVR | -89 | -97 | 0 | 0.003299042 | 9 | 429.55832 | 692.3575 | NaN | NaN | NaN | NaN |
| Chchd3 | EYHPVCADLQTK | -178 | -189 | 0 | 0 | 12 | 1318.2555 | 719.09906 | 355.08362 | 224.60338 | 1030.9227 | 987.0439 |
| Vrk1 | EYTEKPLYQNLR | -310 | -321 | 1 | 0 | 12 | 1080.4028 | 1166.766 | 1289.687 | 1337.6179 | 1517.5941 | 1416.815 |
| Vrk1 | EYTEKPLYQNLR | -310 | -321 | 1 | 0 | 12 | 1080.4028 | 1166.766 | 1289.687 | 1337.6179 | 1517.5941 | 1416.815 |
| Vrk1 | VEPSDNGPLFTEL | -72 | -84 | 0 | 0 | 13 | 785.29224 | 727.36523 | 727.9643 | 751.4568 | 831.1478 | 889.0875 |
| Vrk1 | NPDQVYLVDYGLAYR | -189 | -203 | 0 | 0 | 15 | 757.65466 | 598.99316 | 798.6525 | 806.291 | 673.71136 | 729.4864 |
| Vrk1 | LLEYTEKPLYQNLR | -308 | -321 | 1 | 0 | 14 | 528.62744 | 310.82422 | 378.37952 | 478.54462 | 474.98694 | 577.8823 |
| Vrk1 | EYIHEHEYVHGDIK | -166 | -179 | 0 | 0 | 14 | 380.77515 | 334.38388 | 506.50372 | 652.5817 | 607.4435 | 307.4246 |
| Vrk1 | EYIHEHEYVHGDIK | -166 | -179 | 0 | 0 | 14 | 380.77515 | 334.38388 | 506.50372 | 652.5817 | 607.4435 | 307.4246 |
| Vrk1 | LGLPIGQGGFGC | -39 | -50 | 0 | 0.00025015 | 12 | 418.60095 | 442.5068 | NaN | 306.01492 | 468.9675 | 471.64908 |
| Vrk1 | NLLLSHK | -182 | -188 | 0 | 0.009416377 | 7 | 952.9666 | 819.42413 | NaN | NaN | NaN | NaN |
| Adpgk | LGPAPVPVGPLSPESR | -41 | -56 | 0 | 0 | 16 | 654.861 | 734.66223 | 725.1272 | 762.7084 | 597.324 | 625.91895 |
| Adpgk | EYLAGEEWGPFK | -211 | -222 | 0 | 0 | 12 | 530.5213 | 641.69507 | 926.99646 | 648.80286 | NaN | 911.079 |
| Adpgk | LLEPELPGTALR | -22 | -33 | 0 | 0 | 12 | 210.60265 | 351.30023 | 341.55634 | NaN | 650.06274 | 515.7514 |
| Adpgk | IVLNPDKPVVEWHR | -444 | -457 | 1 | 0 | 14 | 181.44345 | 961.02155 | 928.2091 | NaN | 904.0182 | 879.5026 |
| Adpgk | HFTPVLVCK | -463 | -471 | 0 | 0.000431655 | 9 | 1364.0747 | 1120.1194 | NaN | NaN | 1460.0875 | 1248.4491 |
| Adpgk | VLLCGPIGPK | -175 | -184 | 0 | 0.000471409 | 10 | 634.61053 | NaN | NaN | NaN | NaN | NaN |
| Cycs | TGPNLHGLFGR | -29 | -39 | 0 | 0 | 11 | 7398.248 | 7584.7676 | 7456.63 | 1128.2551 | 3541.107 | 3615.1934 |
| Cycs | TGPNLHGLFGR | -29 | -39 | 0 | 0 | 11 | 7398.248 | 7584.7676 | 7456.63 | 1128.2551 | 3541.107 | 3615.1934 |
| Cycs | TGPNLHGLF | -29 | -37 | 0 | 0.000591124 | 9 | 669.06256 | NaN | 709.4329 | 666.6301 | 684.1049 | 673.5957 |
| Arsb | EYMEPYGFIQDK | -251 | -262 | 0 | 0.00025015 | 12 | 208.74101 | NaN | NaN | NaN | NaN | NaN |
| Qki | EYPDFNFVGR | -92 | -101 | 0 | 0 | 10 | 8957.144 | 8834.001 | 495.63672 | 4294.6816 | 4325.546 | NaN |
| Qki | LLDEEISR | -44 | -51 | 0 | 0.007493591 | 8 | 555.21796 | NaN | NaN | 681.2179 | NaN | NaN |

| | | | | | | | | | | | | |
|---|---|---|---|---|---|---|---|---|---|---|---|---|
| Agps | EYVDPSNIFGNR | -631 | -642 | 0 | 0 | 12 | 455.84872 | 1462.1163 | 1598.931 | 525.4535 | 1000.03174 | 670.2361 |
| Lat | QPDLLPIPR | -78 | -86 | 0 | 0 | 9 | 3870.1208 | 3434.1353 | 3535.213 | 3312.7073 | 4194.7573 | 4025.8438 |
| Lat | EYVNVSPEQQPVTR | -194 | -207 | 0 | 0 | 14 | 1891.1348 | 1747.7319 | 2127.779 | 2309.2703 | NaN | 1706.8793 |
| Lat | SPQPLGGSHR | -87 | -96 | 0 | 0 | 10 | 1167.4167 | NaN | NaN | 1636.5938 | NaN | NaN |
| Dnajb1 | KVPGEGLPLPK | -296 | -306 | 1 | 0 | 11 | 712.2728 | 722.66113 | 719.76935 | 944.0121 | 796.3644 | 828.4568 |
| Dnajb1 | NPFDTFFGQR | -110 | -119 | 0 | 0.00078661 | 10 | 669.8165 | 680.3993 | 701.10956 | NaN | NaN | NaN |
| Dnajb1 | FAEFFGGR | -102 | -109 | 0 | 0.000298805 | 8 | 467.38046 | NaN | NaN | 429.86282 | NaN | 429.20184 |
| Srek1 | FAFVEFADQNSVPR | -111 | -124 | 0 | 0 | 14 | 576.71625 | NaN | NaN | 818.2269 | 855.3198 | 1027.1218 |
| Ap3d1 | FAGELNPVAPK | -600 | -610 | 0 | 0 | 11 | 598.26514 | 459.2254 | 502.5796 | 461.00974 | 450.8211 | 238.69475 |
| Ap3d1 | LFGALTPLEPR | -244 | -254 | 0 | 0 | 11 | 2643.5024 | 2846.4746 | 3078.3188 | NaN | 2650.8027 | 2799.434 |
| Rps16 | LLEPVLLLGK | -51 | -60 | 0 | 0 | 10 | 6154.093 | 7215.477 | 7297.4854 | 5586.897 | 6504.9346 | 6285.078 |
| Rps16 | GRPLEMIEPR | -36 | -45 | 1 | 0 | 10 | 4184.3545 | 4436.074 | 4348.855 | 5001.291 | 6981.8926 | 6259.108 |
| Rps16 | GRPLEMIEPR | -36 | -45 | 1 | 0 | 10 | 4184.3545 | 4436.074 | 4348.855 | 5001.291 | 6981.8926 | 6259.108 |
| Rps16 | KLLEPVLLLGK | -50 | -60 | 1 | 0 | 11 | 564.5107 | 602.8881 | 624.5124 | 555.444 | 596.9922 | 608.55035 |
| Rps16 | LLEPVLLLGKE | -51 | -61 | 1 | 0 | 11 | 228.41328 | 240.50412 | 216.45004 | 94.62484 | 334.68246 | 336.62546 |
| Rps16 | TLLVADPR | -118 | -125 | 0 | 0.003461185 | 8 | 1738.5143 | 1000.7909 | 888.52234 | 766.13226 | NaN | NaN |
| Rps16 | FAGVDIR | -63 | -69 | 0 | 0.00204437 | 7 | 634.0203 | NaN | 487.79855 | NaN | NaN | 380.3952 |
| Hsd17b4 | FAKPVYPGQTL | -562 | -572 | 1 | 0.00272025 | 11 | 1747.3274 | NaN | NaN | NaN | NaN | NaN |
| Pdha1 | LEEGPPVTTVLTR | -46 | -58 | 0 | 0 | 13 | 2020.2996 | 2482.6072 | 2490.3237 | 2957.3699 | 2136.732 | 2098.5815 |
| Pdha1 | RGDFIPGLR | -245 | -253 | 1 | 0 | 9 | 1900.6774 | 1832.7478 | 1839.6716 | 1858.6398 | 1414.1749 | 1250.5831 |
| Pdha1 | FANDATFEIK | -30 | -39 | 0 | 0 | 10 | 1112.9717 | 114.21922 | 1204.9753 | 66.9401 | 1160.853 | 1349.3606 |
| Pdha1 | TREEIQEVR | -303 | -311 | 1 | 0.00025015 | 9 | 950.6521 | 933.57794 | 888.5214 | 1178.4165 | 1497.6505 | 1315.7483 |
| Uba6 | VCPATESIYSDEFYTK | -545 | -560 | 0 | 0 | 16 | 701.635 | 95.67313 | 851.2139 | 745.7787 | 666.33234 | 787.1394 |
| Uba6 | FAPDADGDEDLPGPPVR | -1028 | -1044 | 0 | 0 | 17 | 443.32367 | 400.8949 | 318.54706 | 461.26016 | 212.94604 | 702.3071 |
| Uba6 | GNPGHEEFLPR | -429 | -439 | 0 | 0 | 11 | 967.1337 | 887.5767 | NaN | NaN | 906.36505 | 717.1111 |
| Uba6 | GDRYDAIR | -440 | -447 | 1 | 0.00142999 | 8 | 1794.8567 | NaN | NaN | NaN | NaN | 1680.4536 |
| Uba6 | SLEIDDGLYSR | -36 | -46 | 0 | 0 | 11 | 252.24036 | NaN | NaN | NaN | NaN | 258.19818 |
| Washc5 | FAPVDESITFIGR | -811 | -823 | 0 | 0 | 13 | 876.01685 | NaN | NaN | 2317.795 | NaN | NaN |
| Snf8 | FAQDVSQDDLIR | -135 | -146 | 0 | 0 | 12 | 1559.05 | 2349.4824 | 2207.3533 | NaN | 1804.452 | 1895.396 |
| Snf8 | QVLEHLLK | -211 | -218 | 0 | 0 | 8 | 811.7749 | 633.90155 | 617.9482 | NaN | 367.1291 | 507.1251 |
| Snf8 | IGVDPLASGK | -75 | -84 | 0 | 0 | 10 | 2940.5427 | 879.74194 | 3057.1665 | NaN | 1596.1 | NaN |
| Rif1 | LTDESPIQENLTEK | -1453 | -1466 | 0 | 0 | 14 | 392.828 | NaN | 707.69727 | 742.31226 | NaN | NaN |
| Rif1 | FASDLIEPVTLDTPLSK | -2347 | -2363 | 0 | 0.00025015 | 17 | 381.89444 | 455.1798 | 436.12756 | NaN | NaN | NaN |
| Bcl2 | FATVVEELFR | -127 | -136 | 0 | 0 | 10 | 581.7282 | 314.3528 | 609.5548 | 768.3472 | 550.6185 | 676.8648 |
| Raver1 | LLLNPLLHGGASGK | -345 | -358 | 0 | 0 | 14 | 1702.9805 | 1921.9158 | 2124.2573 | 1236.7285 | 1578.2087 | 1654.1627 |
| Raver1 | SDLLGKPLGPR | -191 | -201 | 1 | 0 | 11 | 639.6933 | 652.39636 | 670.9041 | 865.1465 | 817.1331 | 793.7758 |
| Raver1 | RAPEQELPPLDPEEIRK | -30 | -46 | 2 | 0 | 17 | 590.1226 | 830.33673 | 1013.8792 | 1027.827 | 1158.1779 | 904.0106 |
| Raver1 | FCAPGPPGR | -296 | -304 | 0 | 0 | 9 | 1580.3049 | NaN | NaN | 1431.6237 | NaN | NaN |
| Stk38 | FCCEWEHR | -360 | -367 | 0 | 0 | 8 | 270.33124 | 315.09567 | 253.15057 | NaN | 252.84468 | 269.92703 |
| Isyna1 | FCEVVPGR | -234 | -241 | 0 | 0 | 8 | 917.959 | 897.89923 | 817.2997 | 1249.4685 | 578.90027 | 618.8262 |
| Isyna1 | HVLYAPGERPDHCVVIK | -368 | -384 | 1 | 0 | 17 | 530.80835 | 431.8078 | 404.10202 | 137.8944 | 799.36426 | 564.90594 |
| Isyna1 | HVLYAPGERPDHCVVIK | -368 | -384 | 1 | 0 | 17 | 530.80835 | 431.8078 | 404.10202 | 137.8944 | 799.36426 | 564.90594 |
| Isyna1 | VSFCTDSDPEPQGFHTVL | -440 | -457 | 0 | 0 | 18 | 200.81082 | 264.29645 | 323.25946 | 319.12656 | 351.94556 | 277.50116 |
| Isyna1 | SCIENIFR | -484 | -491 | 0 | 0 | 8 | 2609.4067 | 666.28296 | NaN | 859.1299 | 874.8593 | 695.396 |
| Isyna1 | VDSPDVVYSPETIEAR | -9 | -24 | 0 | 0 | 16 | 1055.6168 | 1206.8765 | 1017.1459 | 1409.4631 | 1981.9453 | NaN |
| Isyna1 | YVPYVGDSK | -385 | -393 | 0 | 0 | 9 | 1106.6735 | NaN | NaN | NaN | NaN | NaN |
| Nsa2 | FCLPILGVK | -192 | -200 | 0 | 0 | 9 | 616.15576 | 750.74536 | 116.16016 | 823.4689 | 945.81274 | 1006.7294 |
| Acadvl | VLDKPETLSSDASTR | -49 | -63 | 1 | 0.000230707 | 15 | 1645.2625 | 2761.226 | 2780.0476 | 2730.4114 | 2228.3547 | 2578.9465 |
| Acadvl | SLSGIVHPELSR | -521 | -532 | 0 | 0 | 12 | 1448.7104 | 1210.7358 | 1475.75 | 476.37033 | 807.73883 | 762.4683 |
| Acadvl | VEDDTLQGLK | -129 | -138 | 0 | 0 | 10 | 1360.1565 | 1509.3103 | 1076.9839 | 1196.5803 | 1057.5565 | 1040.0011 |
| Acadvl | SSPQHQELFR | -624 | -633 | 0 | 0 | 10 | 1115.558 | 5082.777 | 1213.3518 | 6850.946 | 5258.965 | 1467.1234 |
| Acadvl | FCLTEPSSGSDVASIR | -215 | -230 | 0 | 0 | 16 | 2500.4868 | 3087.4968 | NaN | 3400.908 | 3933.8547 | 3778.4075 |
| Acadvl | VPSENVLGEVGDGFK | -318 | -332 | 0 | 0 | 15 | 711.9733 | NaN | 894.834 | 925.3922 | 1300.1062 | 1288.5973 |
| Acadvl | SFGGVTHGLPEK | -288 | -299 | 0 | 0.000230707 | 12 | 1997.3046 | 1738.1276 | 1841.1833 | NaN | NaN | NaN |
| Hat1 | SFIDVDDER | -187 | -195 | 0 | 0 | 9 | 4536.3706 | 5763.617 | 2994.126 | 10925.479 | 7902.58 | 5438.1455 |
| Hat1 | FQELVEDYR | -399 | -407 | 0 | 0 | 9 | 1980.6261 | 2230.2505 | 2511.134 | 943.569 | 884.5769 | 981.3309 |
| Hat1 | FPEDLENDIR | -37 | -46 | 0 | 0 | 10 | 1577.9548 | 1617.078 | 1220.2426 | 1407.2135 | 1558.1481 | 1639.3097 |
| Hat1 | FQELVEDYRR | -399 | -408 | 1 | 0.00025015 | 10 | 1082.6536 | 1155.6875 | 1243.7975 | 439.1841 | 1323.5098 | 774.774 |
| Hat1 | FCQFLPSFSR | -290 | -299 | 0 | 0 | 10 | 968.3247 | 1135.5876 | 1038.9082 | 1045.2039 | 1013.2963 | 1280.744 |
| Hat1 | LFGDDETAFGYK | -56 | -67 | 0 | 0 | 12 | 946.34827 | 944.4557 | 1095.8118 | 1072.661 | 1017.0339 | 1143.8297 |
| Hat1 | TNDFLSLLEK | -120 | -129 | 0 | 0 | 10 | 732.8541 | 1006.2694 | 922.44434 | 977.9076 | 998.8697 | 1267.8906 |
| Hat1 | TFFPEYTHQ | -47 | -55 | 0 | 0 | 9 | 336.3984 | 517.00287 | 553.98755 | 408.33307 | NaN | 418.40256 |
| Hat1 | QFLPSFSR | -292 | -299 | 0 | 0 | 8 | 249.17549 | NaN | 163.78134 | 1087.7255 | NaN | 1102.4346 |
| Hat1 | LRDFVLVK | -282 | -289 | 1 | 0.00272025 | 8 | 792.88116 | 816.25653 | NaN | 523.7312 | NaN | NaN |
| Hat1 | QLLETVHR | -252 | -259 | 0 | 0 | 8 | 832.4982 | NaN | 1185.9264 | NaN | NaN | NaN |
| Hat1 | VQHEQLEER | -390 | -398 | 0 | 0.002534621 | 9 | 386.21326 | NaN | NaN | NaN | NaN | NaN |
| Trmt61a | FCSFSPCIEQVQR | -203 | -215 | 0 | 0 | 13 | 142.44232 | NaN | NaN | 293.63184 | 458.52405 | 481.05206 |
| Themis | FCWPFNVK | -453 | -460 | 0 | 0 | 8 | 3429.521 | 4895.5264 | 4679.006 | 3533.3052 | 631.6759 | 6899.164 |
| Themis | RHFLIPISYK | -331 | -340 | 1 | 0 | 10 | 106.17186 | 916.2754 | 1002.50464 | 431.13226 | 937.8705 | 601.8781 |
| Themis | RHFLIPISYK | -331 | -340 | 1 | 0 | 10 | 106.17186 | 916.2754 | 1002.50464 | 431.13226 | 937.8705 | 601.8781 |
| Themis | HSETTEVVFEGTR | -394 | -406 | 0 | 0 | 13 | 1971.8191 | 1210.8503 | 2384.672 | NaN | 256.72034 | 1880.2571 |
| Themis | GGFVEVIHDK | -432 | -441 | 0 | 0 | 10 | 1332.6274 | NaN | 561.6124 | 1842.6893 | 1977.1807 | 2022.3733 |
| Themis | KLPSDESGQDSR | -597 | -608 | 1 | 0 | 12 | 664.5385 | 666.8061 | NaN | NaN | 765.153 | 779.6196 |
| Themis | HGPLQPQAPL | -627 | -636 | 0 | 0.002752526 | 10 | 419.75287 | NaN | NaN | NaN | NaN | NaN |
| Cpox | FGLFTPGSR | -394 | -402 | 0 | 0.000298805 | 9 | 2074.8735 | 2102.4407 | 2443.9143 | 2006.1111 | 274.12714 | 1862.1948 |
| Cpox | FDDLDSPSKEEAFR | -328 | -341 | 1 | 0 | 14 | 927.1244 | 563.5016 | 509.81076 | 532.3583 | 894.8079 | 684.06586 |
| Ddx46 | VTYVVLDEADR | -523 | -533 | 0 | 0 | 11 | 5502.4976 | 5604.8696 | 6340.739 | 4011.9053 | 4645.782 | 5427.32 |
| Ddx46 | NFYVEVPELAK | -334 | -344 | 0 | 0 | 11 | 2108.7551 | 1200.2172 | 597.9387 | 1902.199 | 2151.125 | 283.58026 |

| | | | | | | | | | | | | |
|---|---|---|---|---|---|---|---|---|---|---|---|---|
| Ddx46 | NFYVEVPELAK | -334 | -344 | 0 | 0 | 11 | 2108.7551 | 1200.2172 | 597.9387 | 1902.199 | 2151.125 | 283.58026 |
| Ddx46 | HGYEKPTPIQTQ | -390 | -401 | 1 | 0 | 12 | 898.28754 | 967.95166 | 961.59015 | 1467.5391 | 1375.1332 | 986.70056 |
| Ddx46 | VDKQEHADGLLK | -626 | -637 | 1 | 0 | 12 | 796.6404 | 899.60986 | 906.0275 | 880.55707 | 1018.89984 | 1022.7451 |
| Ddx46 | VDKQEHADGLLK | -626 | -637 | 1 | 0 | 12 | 796.6404 | 899.60986 | 906.0275 | 880.55707 | 1018.89984 | 1022.7451 |
| Ddx46 | KLLEPVDHGK | -316 | -325 | 1 | 0.000471409 | 10 | 611.8644 | 328.68488 | 517.5108 | 493.36807 | 341.67395 | 489.26276 |
| Ddx46 | KLLEPVDHGK | -316 | -325 | 1 | 0.000471409 | 10 | 611.8644 | 328.68488 | 517.5108 | 493.36807 | 341.67395 | 489.26276 |
| Ddx46 | LNYVPLEK | -909 | -916 | 0 | 0.00025015 | 8 | 483.8653 | 697.4614 | 480.0648 | NaN | NaN | 837.4611 |
| Ddx46 | LLEPVDHGK | -317 | -325 | 0 | 0 | 9 | 803.63153 | 687.65906 | NaN | 849.86676 | NaN | NaN |
| Ddx46 | FDETEQALANER | -780 | -791 | 0 | 0 | 12 | 610.12695 | 753.4223 | 610.4478 | NaN | NaN | NaN |
| Ddx46 | YEEELEINDFPQTAR | -933 | -947 | 0 | 0.001868731 | 15 | 542.5455 | NaN | NaN | 504.80087 | 913.308 | NaN |
| Ddx46 | SKPIEVQVGGR | -576 | -586 | 1 | 0 | 11 | 647.7364 | NaN | NaN | NaN | NaN | NaN |
| Ppp2r5c | FLESPDFQPNIAK | -144 | -156 | 0 | 0 | 13 | 624.68634 | 859.10754 | 765.968 | 1571.8246 | 491.13297 | 570.3845 |
| Ppp2r5c | FDFVSDPLSDLK | -53 | -64 | 0 | 0 | 12 | 491.9733 | 236.23065 | 266.53445 | 416.02795 | 345.1329 | 440.3739 |
| Ppp2r5c | VLLPLHK | -250 | -256 | 0 | 0.004595494 | 7 | 503.15637 | NaN | 412.14328 | 404.04373 | NaN | NaN |
| Sumo2 | TENNDHINLK | -12 | -21 | 0 | 0 | 10 | 5397.021 | 6101.2925 | 1450.8513 | 5612.4194 | 5635.9478 | 6184.6416 |
| Sumo2 | TENNDHINLK | -12 | -21 | 0 | 0 | 10 | 5397.021 | 6101.2925 | 1450.8513 | 5612.4194 | 5635.9478 | 6184.6416 |
| Sumo2 | NNDHINLK | -14 | -21 | 0 | 0 | 8 | 921.13025 | 873.7417 | 858.96875 | NaN | NaN | 1097.7343 |
| Rtcb | VVDEIFNEYAAK | -233 | -244 | 0 | 0 | 12 | 1406.1909 | 457.80804 | 3860.904 | 1858.2936 | 2387.147 | 2292.2102 |
| Rtcb | NYNDELQFLDK | -4 | -14 | 0 | 0 | 11 | 1186.7761 | 1301.8477 | 224.09944 | 1405.657 | 114.97764 | 254.05873 |
| Rtcb | FDHIPVGVGSK | -148 | -158 | 0 | 0 | 11 | 806.20807 | 1503.3257 | 1616.1948 | 1543.1517 | 1610.5206 | 1563.296 |
| Rtcb | IASPEGQDYLK | -298 | -308 | 0 | 0 | 11 | 660.6045 | 747.01984 | 703.2593 | 647.779 | 842.3793 | NaN |
| Rtcb | TNLDESDVQPVK | -129 | -140 | 0 | 0 | 12 | 1661.1077 | 486.33487 | 868.5033 | NaN | NaN | NaN |
| Rtcb | SPEGQDYLK | -300 | -308 | 0 | 0 | 9 | 896.71765 | 533.4596 | NaN | 1000.0185 | NaN | NaN |
| Rtcb | SIGLPDVHSGY | -82 | -92 | 0 | 0 | 11 | 1606.269 | 298.66327 | NaN | NaN | NaN | NaN |
| Skic8 | IVSVGDDQEIHVY | -289 | -301 | 0 | 0 | 13 | 6505.937 | 7542.826 | 11081.246 | 5893.2676 | 11190.539 | 12019.349 |
| Skic8 | FDHQDQVWGVK | -271 | -281 | 0 | 0 | 11 | 2079.541 | 2250.822 | 2379.155 | 2161.3748 | 2151.0198 | 2120.4792 |
| Skic8 | FDHQDQVWGVK | -271 | -281 | 0 | 0 | 11 | 2079.541 | 2250.822 | 2379.155 | 2161.3748 | 2151.0198 | 2120.4792 |
| Skic8 | SSLDAHIR | -82 | -89 | 0 | 0 | 8 | 455.91962 | 825.9336 | NaN | 1072.4459 | 940.0265 | 1248.3643 |
| Skic8 | SSSLDAHIR | -81 | -89 | 0 | 0.00272025 | 9 | 587.25183 | NaN | NaN | NaN | 386.11667 | NaN |
| Hdhd5 | GFQNVVTIDELR | -150 | -161 | 0 | 0 | 12 | 826.0313 | 394.07077 | NaN | 581.0614 | 603.8015 | 525.62665 |
| Hdhd5 | FDIDGVLVR | -51 | -59 | 0 | 0 | 9 | 1485.2157 | 1246.9662 | 1679.6287 | NaN | NaN | NaN |
| Hm13 | LVFPQDLLEK | -242 | -251 | 0 | 0 | 10 | 2977.8127 | 2935.4036 | 3270.6836 | 3179.7483 | 3694.5515 | 3567.4722 |
| Hm13 | FDISLKK | -279 | -285 | 1 | 0 | 7 | 1542.1876 | 313.50183 | 356.46194 | 1801.7703 | 1442.2336 | 1512.2307 |
| Hm13 | HAQPALLYLVPA | -314 | -325 | 0 | 0 | 12 | 1194.068 | 1354.8436 | 1447.5466 | 1259.2815 | 1476.347 | 1478.1805 |
| Hm13 | FFPANFPNR | -128 | -136 | 0 | 0 | 9 | 1161.4839 | 1411.7505 | 1915.9108 | 1513.8372 | 765.92267 | 1649.5522 |
| Hm13 | SYEESNPKDPAAETESK | -345 | -361 | 1 | 0.00078661 | 17 | 448.97623 | 480.6917 | 601.66 | 448.1744 | 827.8268 | 715.6332 |
| Ncapg2 | FDLVQEVFQK | -946 | -955 | 0 | 0 | 10 | 149.2133 | 1466.7997 | 1514.159 | 2115.5571 | 2971.2546 | 2564.987 |
| Snrpa | FDNEVQAGAAR | -256 | -266 | 0 | 0 | 11 | 821.19006 | 121.22678 | 75.92638 | NaN | 531.524 | 462.84964 |
| Snrpa | HDIAFVEFDNEVQAGAAR | -249 | -266 | 0 | 0.000230707 | 18 | 1144.684 | 918.3922 | NaN | NaN | 981.1918 | NaN |
| Snrpa | TMPVPETR | -6 | -13 | 0 | 0.006384543 | 8 | 881.04315 | NaN | NaN | NaN | NaN | NaN |
| Seh1l | TAAVWEEIVGESNDK | -82 | -96 | 0 | 0 | 15 | 1538.5486 | 1434.5145 | 1856.1228 | 318.0323 | 2129.7588 | 1926.8082 |
| Seh1l | FDNHNSQVWR | -274 | -283 | 0 | 0 | 10 | 703.51434 | 784.6716 | 579.6725 | 857.1345 | 491.21927 | 1246.5703 |
| Seh1l | SFDFHGR | -19 | -25 | 0 | 0 | 7 | 645.53735 | 699.4431 | 633.29944 | 767.41705 | NaN | NaN |
| Seh1l | IFEYNENTR | -199 | -207 | 0 | 0.000230707 | 9 | 276.96472 | 5013.8247 | NaN | 5506.661 | NaN | NaN |
| Tmed5 | LEDILESINSIK | -153 | -164 | 0 | 0 | 12 | 3078.6665 | 3980.8613 | 3122.541 | 260.05603 | 3272.7861 | 3198.897 |
| Tmed5 | FDNTFSTISEK | -108 | -118 | 0 | 0 | 11 | 1035.9406 | 481.4508 | 1007.03064 | NaN | NaN | NaN |
| Ptpn11 | FDSLTDLVEHYK | -187 | -198 | 0 | 0 | 12 | 640.0776 | NaN | NaN | NaN | 477.45325 | 379.87537 |
| Smarcc2 | FDYNSVHAIER | -433 | -443 | 0 | 0 | 11 | 1534.3202 | 516.1485 | 1589.3857 | 2175.4485 | 766.2831 | 1029.7598 |
| Smarcc2 | VEDAPTPEKPR | -225 | -235 | 1 | 0 | 11 | 1263.3192 | 1152.3137 | 1200.1276 | 582.35297 | 1593.4324 | 1542.5793 |
| Ppig | TDAASKPFAEVR | -159 | -170 | 1 | 0 | 12 | 1036.9604 | 1250.8192 | 2034.9513 | 2309.6711 | 1695.3329 | 1532.3727 |
| Ppig | FEDESFAVK | -95 | -103 | 0 | 0.001868731 | 9 | 747.83105 | NaN | 215.86696 | NaN | NaN | NaN |
| Gnai2 | GNLQIDFADPQR | -89 | -100 | 0 | 0 | 12 | 1692.16 | 1617.9485 | 1633.5177 | 2139.6602 | 348.2132 | 532.6013 |
| Gnai2 | IIHEDGYSEEECR | -55 | -67 | 0 | 0 | 13 | 1048.2615 | 965.62665 | 936.536 | 788.48364 | 860.77466 | 977.3644 |
| Gnai2 | IIHEDGYSEEECR | -55 | -67 | 0 | 0 | 13 | 1048.2615 | 965.62665 | 936.536 | 788.48364 | 860.77466 | 977.3644 |
| Gnai2 | IAQSDYIPTQQDVLR | -163 | -177 | 0 | 0.00025015 | 15 | 986.9726 | 1219.2264 | 1173.4998 | 1394.2258 | 1492.5104 | 1538.087 |
| Gnai2 | KIIHEDGYSEEECR | -54 | -67 | 1 | 0 | 14 | 572.66394 | 551.9305 | 506.75116 | 520.2218 | 969.9077 | 917.6524 |
| Gnai2 | SKFEDLNK | -306 | -313 | 1 | 0.00025015 | 8 | 177.74847 | 1300.6229 | 158.72025 | 1413.7827 | 1615.7352 | 1611.5911 |
| Gnai2 | LSAYDLVLAEDEEMNR | -228 | -243 | 0 | 0 | 16 | 1621.3864 | 1822.89 | NaN | 1652.7291 | 1955.9122 | 1948.1665 |
| Gnai2 | SKFEDLNKR | -306 | -314 | 2 | 0.00051306 | 9 | 555.0954 | 528.5094 | 416.03906 | NaN | 646.03326 | 421.92712 |
| Gnai2 | SCAAEEQGMLPEDLSGVIR | -111 | -129 | 0 | 0 | 19 | 430.44894 | NaN | 576.43097 | 282.3983 | 946.80365 | 457.3753 |
| Gnai2 | RLWADHGVQAC | -130 | -140 | 1 | 0 | 11 | 2692.459 | 3212.8047 | 3475.3616 | 4333.4434 | NaN | NaN |
| Gnai2 | LWADHGVQAC | -131 | -140 | 0 | 0 | 10 | 1033.9569 | NaN | NaN | 725.59625 | 1992.2452 | NaN |
| Tm9sf3 | YLDPSFFQHR | -209 | -218 | 0 | 0 | 10 | 2952.8555 | 163.63957 | 3325.9253 | 2485.4116 | 2639.1257 | 2380.6096 |
| Tm9sf3 | SISHYHETLGEAL | -70 | -82 | 0 | 0 | 13 | 1088.2847 | 6012.689 | 4937.9546 | 7792.0522 | 7820.6826 | 6934.248 |
| Tm9sf3 | NLSGQPNFPCR | -417 | -427 | 0 | 0 | 11 | 424.2102 | 1316.5052 | 1305.5448 | 1242.9979 | 609.6009 | 770.1538 |
| Tm9sf3 | FEDRFDK | -202 | -208 | 1 | 0.003521747 | 7 | 830.36084 | NaN | NaN | NaN | 930.94604 | NaN |
| Tm9sf3 | SLPFCVGSK | -60 | -68 | 0 | 0.009955823 | 9 | 1114.6976 | NaN | NaN | NaN | NaN | NaN |
| Mrps11 | FEDVPIAHIK | -77 | -86 | 0 | 0 | 10 | 1195.8104 | 1140.6835 | 1114.252 | 901.17084 | 981.9601 | 950.0836 |
| Tax1bp1 | FEEHVESHWK | -771 | -780 | 0 | 0 | 10 | 1458.2457 | 1196.97 | NaN | NaN | NaN | 1514.0691 |
| Cc2d1b | FEFHYPNSDQAQK | -718 | -730 | 0 | 0 | 13 | 655.00104 | 558.1855 | 502.006 | 545.177 | 669.5262 | 621.5134 |
| Fnbp4 | LAPLLPEGVKEEEER | -330 | -344 | 1 | 0 | 15 | 1393.42 | 1589.6501 | 1607.319 | 1600.6045 | 2123.9448 | 2199.979 |
| Fnbp4 | FEFLGINR | -549 | -556 | 0 | 0.000230707 | 8 | 440.39798 | 373.37598 | 391.0357 | 259.77097 | 302.2706 | NaN |
| Pbdc1 | KVDDQIYSEFR | -66 | -76 | 1 | 0 | 11 | 989.9575 | 1355.542 | 2081.2952 | 1680.3907 | 80.98956 | 2154.6228 |
| Pbdc1 | LDCSQGYTEENTIFAPR | -123 | -139 | 0 | 0.000230707 | 17 | 524.8343 | 804.84686 | 507.67465 | 282.93665 | 665.3949 | 906.9091 |
| Pbdc1 | VDDQIYSEFR | -67 | -76 | 0 | 0 | 10 | 570.8421 | 2621.0813 | 2541.0842 | 3492.705 | 3248.5566 | NaN |
| Pbdc1 | FEGIVEDYNYGTLLR | -108 | -122 | 0 | 0 | 15 | 624.5895 | NaN | 825.7155 | NaN | NaN | 1016.8855 |
| Bola2 | LRQDLEAEHVEVEDTTLNR | -12 | -30 | 1 | 0.000431655 | 19 | 2226.5217 | 2166.989 | 2237.9868 | 2372.1553 | 1774.5817 | 1797.2839 |
| Bola2 | QDLEAEHVEVEDTTLNR | -14 | -30 | 0 | 0.000298805 | 17 | 252.16882 | 313.95825 | 310.43356 | 239.67549 | 289.86212 | 296.37137 |

| | | | | | | | | | | | | |
|---|---|---|---|---|---|---|---|---|---|---|---|---|
| Bola2 | HVEVEDTTLNR | -20 | -30 | 0 | 0 | 11 | 147.95633 | 1795.2468 | 2265.384 | 1058.5378 | 1191.6937 | 823.36127 |
| Bola2 | FEGKPLLQR | -44 | -52 | 1 | 0 | 9 | 2229.5874 | 2421.7107 | 2349.4653 | NaN | 2485.429 | 2368.7537 |
| Bola2 | TLTPEQWTR | -74 | -82 | 0 | 0 | 9 | 1547.7711 | 2006.9028 | 2263.8945 | NaN | 779.6646 | 1943.1777 |
| Bola2 | FEGKPLLQ | -44 | -51 | 1 | 0.007819401 | 8 | 405.59964 | NaN | NaN | NaN | NaN | NaN |
| Snrpb2 | HDIAFVEFENDGQAGAAR | -187 | -204 | 0 | 0.000230707 | 18 | 773.9645 | 846.114 | 461.13974 | 787.2069 | 1139.1042 | 670.1615 |
| Snrpb2 | FENDGQAGAAR | -194 | -204 | 0 | 0 | 11 | 627.0114 | NaN | NaN | 445.53693 | NaN | NaN |
| Snrpb2 | KELGSSTNALR | -57 | -67 | 1 | 0.003102268 | 11 | 4535.117 | NaN | NaN | NaN | NaN | NaN |
| Cox7a2 | FENKVPEK | -24 | -31 | 1 | 0.00025015 | 8 | 1178.8167 | 1167.7091 | 1217.0228 | 1184.9874 | 1454.3158 | 1438.1544 |
| Cox7a2 | LFQEDNGMPVHLK | -34 | -46 | 0 | 0 | 13 | 811.0002 | 544.79193 | 636.80005 | 530.7084 | 570.69183 | 841.22363 |
| Cox7a2 | GGASDALLYR | -47 | -56 | 0 | 0 | 10 | 5474.987 | 4199.212 | 1147.0165 | 1541.6464 | NaN | 1192.5992 |
| Nol10 | FENPDFQVDEESEEFR | -489 | -504 | 0 | 0.00025015 | 16 | 338.72104 | 430.07086 | 327.4941 | 686.68164 | 592.3111 | 462.96286 |
| Nol10 | IVFLHNDR | -110 | -117 | 0 | 0 | 8 | 320.77225 | 472.07828 | 1257.2931 | NaN | 1039.7207 | 1224.3046 |
| Nol10 | YIPVLGPAPR | -340 | -349 | 0 | 0.001291103 | 10 | 508.86777 | 295.44305 | 318.29874 | NaN | 1234.3787 | NaN |
| Nol10 | SLPEWLSDR | -19 | -27 | 0 | 0.009272078 | 9 | 550.3476 | NaN | NaN | 835.2748 | 407.41318 | NaN |
| Poldip2 | TSTDQVPIQHELFER | -185 | -199 | 0 | 0.00025015 | 15 | 792.83044 | 1078.9304 | 1095.092 | 981.2698 | 1271.5648 | 1397.7416 |
| Poldip2 | FERPDGSHFDVR | -336 | -347 | 1 | 0 | 12 | 886.86316 | 978.23785 | 1027.5193 | NaN | 863.22217 | 1042.6266 |
| Poldip2 | NHPWLELSDVHR | -224 | -235 | 0 | 0 | 12 | 177.11133 | NaN | 758.49316 | 254.49834 | 857.6432 | 1930.8853 |
| Poldip2 | GVVLFPWQAR | -92 | -101 | 0 | 0 | 10 | 374.44946 | 377.31055 | 483.3508 | NaN | NaN | NaN |
| Atp5pf | FEVIDKPQS | -100 | -108 | 1 | 0.00201047 | 9 | 2308.959 | 2586.6418 | 2487.367 | 2416.8203 | 1924.378 | 1961.3423 |
| Atp5pf | QASGGPVDIGPEYQQDLDR | -55 | -73 | 0 | 0 | 19 | 1543.1544 | 1989.7216 | 1898.1621 | 1727.1816 | 2055.239 | 125.50676 |
| Atp5pf | GEMDTFPTFK | -85 | -94 | 0 | 0 | 10 | 630.5398 | 624.0105 | 640.9586 | 73.11739 | 646.7173 | 693.25525 |
| Atp5pf | GEMDTFPTFK | -85 | -94 | 0 | 0 | 10 | 630.5398 | 624.0105 | 640.9586 | 73.11739 | 646.7173 | 693.25525 |
| Ctps1 | FEVNPVLK | -482 | -489 | 0 | 0 | 8 | 832.88086 | 819.866 | 912.8454 | 544.26337 | 910.43317 | 795.21533 |
| Ctps1 | RLDLPIER | -269 | -276 | 1 | 0 | 8 | 819.1592 | 876.645 | 1039.1514 | 837.0877 | 936.1887 | 930.5536 |
| Ctps1 | HRFEVNPVLK | -480 | -489 | 1 | 0 | 10 | 755.37054 | 686.4199 | 807.8225 | 676.8906 | 991.5346 | 656.57825 |
| Ctps1 | PIKPSPPYFGLL | -533 | -544 | 1 | 0 | 12 | 603.9446 | 662.4816 | 701.4118 | 795.25867 | 553.4096 | 772.683 |
| Ctps1 | YIDSTDLEPSTLQEEPVR | -336 | -353 | 0 | 0 | 18 | 336.56433 | 483.82092 | 414.1967 | 336.37836 | 481.2556 | 475.29404 |
| Ctps1 | YHEAWQK | -354 | -360 | 0 | 0 | 7 | 113.2251 | 791.2831 | 791.48456 | 249.85767 | 787.31256 | 413.02393 |
| Ctps1 | LVPQPSSTGEQK | -182 | -193 | 0 | 0.000712251 | 12 | 1684.9402 | NaN | NaN | 1580.4475 | NaN | NaN |
| Ctps1 | GVLVPGGFGVR | -366 | -376 | 0 | 0.000591124 | 11 | 489.1051 | NaN | NaN | NaN | NaN | NaN |
| Txndc5 | NLAPTWEELSK | -340 | -350 | 0 | 0 | 11 | 995.4226 | 710.4094 | 924.685 | 1322.0474 | 579.98474 | 731.50024 |
| Txndc5 | FYAPWCGH | -330 | -337 | 0 | 0.002329279 | 8 | 806.23846 | 657.86096 | 766.9415 | 519.20215 | 310.615 | 238.67784 |
| Txndc5 | FFAPWCGH | (70)(198) | (77)(205) | 0 | 0.001291103 | 8 | 403.9455 | 408.4527 | 497.05972 | NaN | NaN | 342.0125 |
| Txndc5 | FFKPGQEAVK | -127 | -136 | 1 | 0.001708008 | 10 | 1540.4666 | 1453.3282 | NaN | 2007.5983 | NaN | NaN |
| Cmpk1 | NQDNLQGWNK | -97 | -106 | 0 | 0 | 10 | 1525.318 | 1293.4944 | 1186.099 | 978.5257 | 834.07367 | 1108.9537 |
| Cmpk1 | KNPDSQYGELIEK | -43 | -55 | 1 | 0 | 13 | 1506.818 | 1709.909 | 1750.0242 | 1887.2618 | 1869.6149 | 1880.4188 |
| Cmpk1 | FFDCNNEICIER | -119 | -130 | 0 | 0 | 12 | 411.55157 | 547.9999 | NaN | 476.2066 | 491.632 | 436.44598 |
| Cmpk1 | SVDEVFGEVVK | -180 | -190 | 0 | 0.000431655 | 11 | 480.5684 | NaN | NaN | 311.8995 | 311.5234 | NaN |
| Cmpk1 | GYTHLSAGELLR | -28 | -39 | 0 | 0 | 12 | 435.90616 | NaN | NaN | NaN | 1740.9111 | 1792.282 |
| Cmpk1 | NPDSQYGELIEK | -44 | -55 | 0 | 0 | 12 | 301.72958 | 281.2046 | NaN | NaN | NaN | NaN |
| Ssr4 | FFDEESYSLLR | -105 | -115 | 0 | 0 | 11 | 1221.8691 | 484.7046 | 1401.1855 | NaN | 693.9942 | 477.45776 |
| Ssr4 | SAHAGTYEVR | -95 | -104 | 0 | 0 | 10 | 1605.1112 | NaN | NaN | 1384.8445 | 1551.049 | 1288.6437 |
| Atp6v0d1 | FFEHEVK | -294 | -300 | 0 | 0 | 7 | 1883.1902 | NaN | 1806.1294 | NaN | NaN | 350.67984 |
| Xrcc5 | FFLPFPIDK | -163 | -171 | 0 | 0 | 9 | 761.4663 | 970.4036 | 499.59427 | 920.05646 | 1541.7754 | 1616.0526 |
| Xrcc5 | YGTDGTDNALAGK | -59 | -71 | 0 | 0 | 13 | 713.346 | 912.8235 | 790.9725 | 1093.6042 | 773.97546 | 756.9163 |
| Xrcc5 | TLFPLTEVIK | -535 | -544 | 0 | 0 | 10 | 439.6457 | 436.85455 | 44.379128 | 503.7376 | 549.46875 | 637.5387 |
| Xrcc5 | FNSFLEALR | -656 | -664 | 0 | 0.000907206 | 9 | 743.1159 | NaN | NaN | NaN | NaN | NaN |
| Cpt2 | FFNEVFR | -383 | -389 | 0 | 0.001055699 | 7 | 504.93625 | NaN | NaN | NaN | 673.36743 | 674.88306 |
| Msmo1 | FFNIPYDWER | -122 | -131 | 0 | 0.001776473 | 10 | 631.3263 | NaN | NaN | 1091.4677 | NaN | NaN |
| Gpam | FFNPSIPSLGLR | -78 | -89 | 0 | 0.00025015 | 12 | 254.83904 | NaN | NaN | 496.92474 | 327.48364 | NaN |
| Pepd | FFNQEVLQR | -434 | -442 | 0 | 0 | 9 | 1384.803 | 1268.6097 | 1175.6174 | 556.2246 | 1692.1062 | 1288.4263 |
| Pepd | IHLEELAR | -336 | -343 | 0 | 0.00025015 | 8 | 924.78784 | 929.1139 | 803.6155 | NaN | 1001.1133 | 967.12976 |
| N6amt1 | FFPLAPELLSPR | -155 | -166 | 0 | 0 | 12 | 474.36243 | 329.6887 | 671.16205 | 584.2333 | 467.79193 | 562.11615 |
| Ddt | SIGVVGTAEQNR | -64 | -75 | 0 | 0 | 12 | 1702.8864 | 1362.7461 | 1212.6088 | 2636.7573 | 2232.5242 | 2067.5159 |
| Ddt | TILDKPEDR | -29 | -37 | 1 | 0 | 9 | 1154.7019 | 1147.6632 | 1165.212 | 916.06683 | 790.49835 | 877.18396 |
| Ddt | VSSIGVVGTAEQNR | -62 | -75 | 0 | 0 | 14 | 851.46387 | 944.32654 | 815.3753 | 1132.2251 | 1231.6298 | 1375.0416 |
| Ddt | THSASFFK | -76 | -83 | 0 | 0.000471409 | 8 | 853.13074 | NaN | 1251.0897 | 329.22382 | 1196.3451 | 1348.4114 |
| Ddt | PFVELETNLPASR | -2 | -14 | 0 | 0 | 13 | 785.63763 | 807.4125 | 751.8985 | 727.1575 | NaN | 738.212 |
| Ddt | FFPLEAWQIGK | -100 | -110 | 0 | 0 | 11 | 309.77866 | NaN | 581.86456 | 667.23627 | 404.8642 | 698.7328 |
| Ddt | FLTEELSLDQDR | -84 | -95 | 0 | 0 | 12 | 1086.5181 | 1250.7133 | 1301.6425 | 1154.9116 | NaN | NaN |
| Ddt | IPAGLENR | -15 | -22 | 0 | 0.001636916 | 8 | 1546.2393 | 1230.2954 | 979.7618 | NaN | NaN | NaN |
| Ddt | STEPCAHLL | -53 | -61 | 0 | 0.001636916 | 9 | 1212.9004 | 1133.7765 | 1223.0499 | NaN | NaN | NaN |
| Ddt | TATILDKPEDR | -27 | -37 | 1 | 0 | 11 | 822.2717 | NaN | NaN | 1319.7236 | NaN | 790.58966 |
| Ddt | PAGLENR | -16 | -22 | 0 | 0.002068213 | 7 | 5927.309 | 144.82445 | NaN | NaN | NaN | NaN |
| Scamp3 | SGELDNPFQDPAVIQHR | -14 | -30 | 0 | 0 | 17 | 815.0633 | 913.82135 | 1008.51276 | 1113.8175 | 1129.3281 | 1143.6805 |
| Scamp3 | FFQDISMEIPQEFQK | -150 | -164 | 0 | 0 | 15 | 590.9128 | 518.15564 | NaN | 423.23434 | 484.46603 | 322.818 |
| S100a13 | TLDVNQDSELR | -62 | -72 | 0 | 0 | 11 | 1714.1956 | NaN | 1864.1604 | 2062.353 | 2152.2122 | 1963.1617 |
| S100a13 | QLPHLLK | -45 | -51 | 0 | 0.003461185 | 7 | 837.06445 | 397.355 | 674.1791 | NaN | NaN | NaN |
| S100a13 | FFTFAGR | -20 | -26 | 0 | 0.000755556 | 7 | 986.9392 | NaN | NaN | NaN | NaN | NaN |
| Tbx21 | FFYPEPGAQDPTDR | -34 | -47 | 0 | 0.002296977 | 14 | 650.37396 | NaN | 765.6742 | 764.2066 | NaN | NaN |
| Rasa3 | FGDEFLGELR | -238 | -247 | 0 | 0 | 10 | 8203.307 | 9469.682 | 9111.394 | 10190.813 | 11513.059 | 10068.066 |
| Rasa3 | NFLDLISSSGR | -556 | -566 | 0 | 0 | 11 | 780.32635 | 827.5197 | 63.39421 | 695.38904 | 1237.498 | 1034.5964 |
| Rasa3 | RFQDDLDVR | -466 | -474 | 1 | 0.002466863 | 9 | 1381.2136 | NaN | NaN | 2771.7117 | 2904.441 | 2851.356 |
| Rasa3 | LSEVITDTGVVCHK | -133 | -146 | 0 | 0 | 14 | 1307.3538 | NaN | NaN | NaN | 1721.4691 | NaN |
| Gramd4 | FQPEENTVETEEPLSAR | -238 | -254 | 0 | 0 | 17 | 461.8909 | 590.0708 | 635.4313 | 958.946 | 785.98035 | 688.96295 |
| Gramd4 | FGEYIEDFR | -229 | -237 | 0 | 0 | 9 | 104.53796 | 334.28473 | 337.47287 | 552.0752 | 510.34277 | NaN |
| Gramd4 | HFLQEELR | -143 | -150 | 0 | 0 | 8 | 696.6801 | 792.58856 | NaN | 920.69385 | 729.86035 | NaN |
| Gramd4 | VEPAKEDLTVSEK | -341 | -353 | 1 | 0.000471409 | 13 | 1271.4127 | NaN | NaN | NaN | NaN | 2158.7107 |

| | | | | | | | | | | | | |
|---|---|---|---|---|---|---|---|---|---|---|---|---|
| Tardbp | VAQSLCGEDLIIK | -239 | -251 | 0 | 0 | 13 | 868.0891 | 803.17847 | 884.964 | 1286.0464 | 1640.2245 | 1485.6528 |
| Tardbp | SLCGEDLIIK | -242 | -251 | 0 | 0 | 10 | 787.3308 | 1040.5781 | 483.479 | 1127.1531 | 967.923 | 1040.1278 |
| Tardbp | GEVVDVFIPKPFR | -215 | -227 | 1 | 0 | 13 | 697.9116 | 864.10345 | 886.1185 | 280.31616 | 250.20914 | 375.93265 |
| Tardbp | FGGNPGGFGNQGGFGNSR | -276 | -293 | 0 | 0 | 18 | 736.72736 | 533.10004 | 880.6945 | 470.94064 | NaN | 414.42896 |
| Tardbp | YGEVVDVFIPK | -214 | -224 | 0 | 0 | 11 | 1280.3417 | NaN | 1347.1992 | 1088.2393 | 1040.6466 | NaN |
| Tardbp | QSPDEPLR | -182 | -189 | 0 | 0.000431655 | 8 | 791.74756 | 1084.6525 | 1340.1176 | 690.3126 | NaN | NaN |
| Tardbp | YGEVVDVFIPKPFR | -214 | -227 | 1 | 0 | 14 | 542.1702 | NaN | NaN | 822.07416 | 1270.7734 | 910.0979 |
| Tardbp | LVEGILHAPDAGWGN | -56 | -70 | 0 | 0 | 15 | 1347.8582 | 1691.3629 | NaN | NaN | 1548.4778 | NaN |
| Tardbp | LVEGILHAPDAGWGNL | -56 | -71 | 0 | 0 | 16 | 398.07608 | NaN | 587.0093 | NaN | 215.1788 | NaN |
| Tardbp | LCGEDLIIK | -243 | -251 | 0 | 0 | 9 | 3086.978 | NaN | NaN | 3768.883 | NaN | NaN |
| Tardbp | GEVVDVFIPK | -215 | -224 | 0 | 0.00051306 | 10 | 1378.4935 | 1184.8057 | NaN | NaN | NaN | NaN |
| Tardbp | VYVVNYPK | -72 | -79 | 0 | 0.001291103 | 8 | 781.31744 | NaN | NaN | NaN | NaN | NaN |
| Pip4k2a | FGIDDQDFQNSLTR | -105 | -118 | 0 | 0 | 14 | 3015.311 | 2023.3506 | 2367.87 | NaN | 3500.1597 | 4969.889 |
| Prkch | FGIDNFEFIR | -350 | -359 | 0 | 0 | 10 | 301.9106 | NaN | NaN | 575.30646 | 386.86563 | 425.69577 |
| Usp36 | FGNQEDAHEFLR | -203 | -214 | 0 | 0 | 12 | 1258.2805 | NaN | 1420.188 | NaN | NaN | NaN |
| Elac2 | VVHIAPESVLIDSR | -336 | -349 | 0 | 0 | 14 | 584.3388 | 437.75827 | 525.6117 | NaN | 673.26666 | 509.7263 |
| Elac2 | RLDNIFLTR | -96 | -104 | 1 | 0.00025015 | 9 | 17841.672 | NaN | NaN | 15286.469 | 20258.428 | 18597.46 |
| Elac2 | FGPDTQHLIL | -357 | -366 | 0 | 0.000591124 | 10 | 485.4562 | NaN | NaN | NaN | 424.28522 | 354.67673 |
| Arhgap25 | FGPHLVPILVEK | -172 | -183 | 0 | 0 | 12 | 221.33371 | 211.26567 | 411.99368 | 562.452 | NaN | 276.59174 |
| Arhgap25 | NSGEDDLDSLQR | -545 | -556 | 0 | 0 | 12 | 742.6128 | NaN | 761.9972 | 613.4015 | NaN | NaN |
| Baz1a | FGPSFPSR | -1279 | -1286 | 0 | 0.003681885 | 8 | 567.9378 | 385.58554 | 587.5451 | NaN | NaN | NaN |
| Scfd1 | SLLDVISDPDAGTPEDK | -434 | -450 | 0 | 0.000230707 | 17 | 1082.1167 | 700.1281 | 727.90063 | 737.7794 | 1017.14124 | 709.808 |
| Scfd1 | FGQDIISPLLSVK | -48 | -60 | 0 | 0 | 13 | 510.96127 | 363.07336 | 322.7394 | 450.24152 | 520.69586 | 918.9589 |
| Ptpa | KFGSLLPIHPVTSG | -310 | -323 | 1 | 0.00025015 | 14 | 1481.7964 | 1714.7009 | 1672.7549 | 1823.2855 | 1211.0612 | 1334.9197 |
| Ptpa | LIDHPHLEPR | -227 | -236 | 0 | 0 | 10 | 1130.8469 | 1255.7075 | 936.2318 | 955.3989 | 1231.9384 | 1050.2511 |
| Ptpa | LIDHPHLEPR | -227 | -236 | 0 | 0 | 10 | 1130.8469 | 1255.7075 | 936.2318 | 955.3989 | 1231.9384 | 1050.2511 |
| Ptpa | WIDETPPVDQPSR | -87 | -99 | 0 | 0 | 13 | 823.08 | 230.53275 | 916.3537 | 444.67276 | 1269.458 | 1293.0818 |
| Ptpa | FGSLLPIHPVTSG | -311 | -323 | 0 | 0 | 13 | 763.5221 | 901.0203 | 946.66595 | 919.24976 | 1114.332 | 915.0435 |
| Fyb1 | FGTTKPPLAAKPT | -57 | -69 | 2 | 0.000431655 | 13 | 1305.3278 | 1210.2316 | 1575.4562 | 1455.8353 | 1455.6826 | 1395.2516 |
| Fyb1 | FKYDGEIR | -735 | -742 | 1 | 0 | 8 | 2870.4783 | NaN | NaN | 1165.2589 | 862.3037 | 1363.1454 |
| Fyb1 | FNTGSNPTEEAATSSR | -4 | -19 | 0 | 0 | 16 | 607.6297 | 407.4314 | 343.1883 | NaN | NaN | 320.8993 |
| Fdxr | LGVIPNTEGR | -380 | -389 | 0 | 0 | 10 | 5743.6255 | 5409.1836 | NaN | 5099.6255 | 4888.795 | 4586.539 |
| Fdxr | SRPIDPSVPFDPK | -367 | -379 | 1 | 0 | 13 | 1317.2793 | 1465.2795 | 277.4378 | 843.21686 | NaN | 1146.308 |
| Fdxr | FGVAPDHPEVK | -83 | -93 | 0 | 0 | 11 | 3835.499 | NaN | NaN | 3795.8118 | 264.65543 | 2998.3276 |
| Abracl | FGVLFQDDR | -30 | -38 | 0 | 0 | 9 | 1485.2253 | 528.9522 | 1162.7654 | 911.0084 | 1146.2421 | 1144.7438 |
| Mrpl15 | ILKPTDENLLK | -281 | -291 | 1 | 0 | 11 | 1105.8206 | 1235.1719 | 1888.6792 | 1188.7805 | 1774.1707 | 2277.24 |
| Mrpl15 | FGYVLPDITK | -239 | -248 | 0 | 0 | 10 | 1227.3578 | 1429.6462 | NaN | NaN | 909.7709 | NaN |
| Pmpcb | LCTDVTESEVAR | -388 | -399 | 0 | 0 | 12 | 952.59674 | 1001.0564 | 939.7191 | 884.00336 | 1102.5986 | 1273.9045 |
| Pmpcb | RIPIPELEAR | -432 | -441 | 1 | 0.001392869 | 10 | 778.2167 | 994.0967 | 1031.9308 | 574.4007 | 1562.3779 | 1503.2604 |
| Pmpcb | FHFGDSLCSHK | -258 | -268 | 0 | 0 | 11 | 338.20834 | 559.7621 | NaN | 635.4532 | 654.91736 | 374.43298 |
| Pmpcb | TILGPTENIK | -210 | -219 | 0 | 0.001392869 | 10 | 821.2727 | NaN | NaN | NaN | 1056.9517 | NaN |
| Pmpcb | LDGSTPICEDIGR | -412 | -424 | 0 | 0.000230707 | 13 | 1057.1941 | NaN | NaN | NaN | NaN | NaN |
| Pak1ip1 | SGHQETWTPVADFTHH | -24 | -39 | 0 | 0 | 16 | 10091.192 | 3618.4954 | 6219.8174 | 6290.183 | 5575.7295 | 1212.7249 |
| Pak1ip1 | SGHQETWTPVADFTHH | -24 | -39 | 0 | 0 | 16 | 10091.192 | 3618.4954 | 6219.8174 | 6290.183 | 5575.7295 | 1212.7249 |
| Pak1ip1 | SDSVLAVAGDEEVVR | -214 | -228 | 0 | 0 | 15 | 675.06976 | 713.1386 | 748.36536 | 616.7082 | 402.98752 | 487.7984 |
| Pak1ip1 | SGDTVQEETSEPNSEK | -331 | -346 | 0 | 0 | 16 | 316.0481 | 369.80862 | NaN | 340.3253 | 409.57895 | 372.9063 |
| Pak1ip1 | FHGNQHLISGAEDGHIC | -90 | -106 | 0 | 0.00111006 | 17 | 596.1644 | NaN | NaN | NaN | NaN | NaN |
| Mrps30 | FHLIPDR | -293 | -299 | 0 | 0 | 7 | 694.9602 | 987.36316 | 1333.0787 | 841.63947 | 1267.6035 | 1258.896 |
| Cox6a1 | HEEHERPPFVAYPH | -65 | -78 | 1 | 0 | 14 | 934.97626 | 1160.1923 | 1250.7764 | 1204.8113 | 1501.7338 | 3231.8062 |
| Cox6a1 | HEEHERPPFVAYPH | -65 | -78 | 1 | 0 | 14 | 934.97626 | 1160.1923 | 1250.7764 | 1204.8113 | 1501.7338 | 3231.8062 |
| Cox6a1 | HEEHERPPFVAYPH | -65 | -78 | 1 | 0 | 14 | 934.97626 | 1160.1923 | 1250.7764 | 1204.8113 | 1501.7338 | 3231.8062 |
| Cox6a1 | FHNPHVNPLPTGYEDE | -96 | -111 | 0 | 0 | 16 | 931.6372 | 1041.5764 | 1063.4818 | 872.9307 | 1336.5406 | 1220.2899 |
| Cox6a1 | FHNPHVNPLPTGYEDE | -96 | -111 | 0 | 0 | 16 | 931.6372 | 1041.5764 | 1063.4818 | 872.9307 | 1336.5406 | 1220.2899 |
| Cox6a1 | HNPHVNPLPTGYEDE | -97 | -111 | 0 | 0.00025015 | 15 | 484.8344 | 2006.1993 | 1690.1635 | 414.26013 | 1369.3846 | 2028.6857 |
| Cox6a1 | SRHEEHERPPFVAYPH | -63 | -78 | 2 | 0.001055699 | 16 | 855.78455 | 1445.2605 | NaN | 616.2465 | 1465.63 | 1383.6384 |
| Cox6a1 | TKPFPWGDGNHTL | -83 | -95 | 1 | 0.000394361 | 13 | 1796.0736 | 2105.8792 | 3266.2114 | 3055.8674 | NaN | NaN |
| Cox6a1 | TKPFPWGDGNHTL | -83 | -95 | 1 | 0.000394361 | 13 | 1796.0736 | 2105.8792 | 3266.2114 | 3055.8674 | NaN | NaN |
| Cbr1 | FHQLDIDNPQSIR | -59 | -71 | 0 | 0 | 13 | 697.9378 | 697.532 | 1212.4138 | 1175.9655 | 779.27216 | 1691.5953 |
| Igf2r | FICNDDIYPGAPK | -1033 | -1045 | 0 | 0 | 13 | 882.2084 | 838.96716 | 1030.6495 | 1027.9423 | 903.9958 | 815.7986 |
| Igf2r | FLHQDIDSTR | -1046 | -1055 | 0 | 0 | 10 | 1617.5317 | 1789.8768 | 2000.2096 | 487.23297 | 2165.9265 | NaN |
| Igf2r | GNQAFDVGRPK | -234 | -244 | 1 | 0.000230707 | 11 | 377.73196 | 353.1535 | 163.22557 | NaN | 395.27048 | 373.18738 |
| Igf2r | SYDECVLEGR | -1906 | -1915 | 0 | 0 | 10 | 769.10046 | NaN | 745.68115 | 765.4151 | NaN | NaN |
| Igf2r | IFFHCDPLVK | -2237 | -2246 | 0 | 0.000230707 | 10 | 610.0559 | NaN | 634.42896 | NaN | 598.90814 | NaN |
| Igf2r | VPVDGPPIDIGR | -1708 | -1719 | 0 | 0 | 12 | 400.51596 | 566.7612 | NaN | 855.00446 | NaN | NaN |
| Timm8a1 | HFIEVETQK | -21 | -29 | 0 | 0.00025015 | 9 | 1292.1638 | 797.0998 | 777.38855 | 1217.0116 | 637.1358 | 644.2009 |
| Timm8a1 | GAVDPQLQHFIEVETQK | -13 | -29 | 0 | 0 | 17 | 450.37817 | 486.66153 | 523.88385 | 510.20197 | 666.24915 | 692.8841 |
| Slfn1 | FIEVHDSGVVCK | -283 | -294 | 0 | 0 | 12 | 2129.2942 | 2613.8718 | 512.17194 | 2586.9194 | 2712.4766 | 2880.6045 |
| Slfn1 | NILPSRPLDFK | -81 | -91 | 1 | 0 | 11 | 1741.6941 | 2197.427 | 2273.5261 | 136.98532 | 3041.6309 | 2865.486 |
| Slfn1 | HGIGEDLETSFK | -69 | -80 | 0 | 0 | 12 | 1606.729 | 478.77417 | 1598.1324 | 1792.2559 | 1232.871 | 1323.5177 |
| Slfn1 | NAGGITLGTESR | -14 | -25 | 0 | 0 | 12 | 887.4781 | 1186.0305 | 560.43744 | 668.5095 | 1219.4856 | 877.44434 |
| Slfn1 | PSDRPGDDTQEEGHIQ | -156 | -171 | 1 | 0.000431655 | 16 | 705.30084 | 757.8365 | 757.13794 | 204.7422 | 847.831 | 901.9358 |
| Slfn1 | SQSPDVSVGKPATIA | -104 | -118 | 1 | 0.00025015 | 15 | 2477.4434 | 3210.6526 | 3189.5994 | 3515.682 | NaN | 4082.5762 |
| Bcat1 | FIGTEPSLGVK | -166 | -176 | 0 | 0 | 11 | 799.04297 | 1370.5089 | 2018.0012 | 443.3765 | 5793.1084 | 1714.0953 |
| Bcat1 | LTDIQYGR | -369 | -376 | 0 | 0 | 8 | 3357.426 | 3920.9421 | NaN | 2672.57 | 2175.9924 | 1263.6886 |
| Bcat1 | VVCPVSDILYK | -336 | -346 | 0 | 0 | 11 | 784.7529 | 656.97205 | 167.35217 | 616.02936 | NaN | NaN |
| Slc25a1 | FIHDQTSSNPK | -150 | -160 | 0 | 0 | 11 | 731.2971 | 860.9487 | NaN | 481.41245 | 950.6616 | 546.53613 |
| Slc25a1 | ITFPTEYVK | -42 | -50 | 0 | 0 | 9 | 1403.6586 | NaN | NaN | NaN | NaN | NaN |
| Ldhb | FIIPQIVK | -120 | -127 | 0 | 0.003299042 | 8 | 546.15063 | NaN | NaN | NaN | NaN | 656.8234 |

| | | | | | | | | | | | | |
|---|---|---|---|---|---|---|---|---|---|---|---|---|
| Septin9 | SITHDIEEK | -342 | -350 | 0 | 0 | 9 | 3799.1204 | 3883.338 | 4352.8867 | 3063.705 | 3901.5774 | 3786.1433 |
| Septin9 | VENTTHCEFAYLR | -523 | -535 | 0 | 0 | 13 | 1317.7954 | 871.42737 | 875.0348 | 769.10675 | 831.42834 | 954.94977 |
| Septin9 | VENTTHCEFAYLR | -523 | -535 | 0 | 0 | 13 | 1317.7954 | 871.42737 | 875.0348 | 769.10675 | 831.42834 | 954.94977 |
| Septin9 | KEFDEDAEDR | -474 | -483 | 1 | 0 | 10 | 731.1446 | 772.3185 | 872.51855 | 865.58014 | 1218.9564 | 1421.7754 |
| Septin9 | VVNIVPVIAK | -434 | -443 | 0 | 0 | 10 | 470.43433 | 465.2198 | 490.91125 | 98.97311 | 335.32434 | 316.328 |
| Septin9 | NTTHCEFAYLR | -525 | -535 | 0 | 0 | 11 | 950.783 | 1108.43 | 934.64557 | 447.02106 | NaN | 1498.4417 |
| Septin9 | IHFEAYR | -553 | -559 | 0 | 0.001392869 | 7 | 1610.9631 | 1272.131 | 1302.8705 | NaN | NaN | NaN |
| Septin9 | FINDQYEK | -380 | -387 | 0 | 0 | 8 | 826.8742 | 1184.3516 | 1302.8353 | NaN | NaN | NaN |
| Septin9 | RTEITIVKPQESVLR | -150 | -164 | 2 | 0.001743443 | 15 | 212.5556 | NaN | NaN | NaN | NaN | NaN |
| Tapbp | SHFFPAEGLEVK | -321 | -332 | 0 | 0 | 12 | 2384.084 | 2957.9192 | 2871.8926 | 2540.893 | 3545.3442 | 3712.4993 |
| Tapbp | VYHSSLPASGR | -387 | -397 | 0 | 0 | 11 | 1592.2517 | 3047.583 | 2513.572 | 2742.0127 | 1737.3263 | 1804.1342 |
| Tapbp | VYHSSLPASGR | -387 | -397 | 0 | 0 | 11 | 1592.2517 | 3047.583 | 2513.572 | 2742.0127 | 1737.3263 | 1804.1342 |
| Tapbp | GHLQLPPVTAK | -367 | -377 | 0 | 0 | 11 | 1285.5288 | 1502.4338 | 1100.1423 | 1092.1143 | 1073.6699 | 1084.6224 |
| Tapbp | KPATLLLR | -44 | -51 | 1 | 0.000230707 | 8 | 645.1942 | 757.3409 | 754.4668 | 740.0384 | 769.9184 | 807.60297 |
| Tapbp | GPEAIECWFVEDAGGGGLSK | -24 | -43 | 0 | 0 | 20 | 366.38318 | 342.7308 | 472.20706 | 559.8705 | 642.8883 | 633.9577 |
| Tapbp | GPPPRPDLDPK | -56 | -66 | 1 | 0 | 11 | 2496.0781 | 2552.734 | NaN | 2147.6726 | 2968.5125 | 3011.045 |
| Tapbp | FIPFPASAK | -99 | -107 | 0 | 0.000230707 | 9 | 2119.4539 | 1794.1406 | 1612.8369 | 1340.5304 | NaN | 149.23586 |
| Tapbp | FVEDAGGGGLSK | -32 | -43 | 0 | 0 | 12 | 1268.6829 | 300.96918 | 1544.4026 | NaN | 1108.5297 | 179.67082 |
| Tapbp | FWLPAVK | -259 | -265 | 0 | 0 | 7 | 672.1441 | 660.24615 | 653.0875 | NaN | 446.00958 | NaN |
| Tapbp | VSLTPAPVVW | -297 | -306 | 0 | 0 | 10 | 422.12637 | NaN | NaN | 563.26526 | 730.07385 | 638.8788 |
| Tapbp | SLSPEQNCPR | -111 | -120 | 0 | 0 | 10 | 594.3652 | NaN | NaN | 2448.6208 | 6942.409 | NaN |
| Tapbp | SLLRPQPEPLR | -140 | -150 | 1 | 0.001463897 | 11 | 413.99292 | 490.97397 | NaN | NaN | NaN | 605.5775 |
| Tapbp | VSLELTVHK | -285 | -293 | 0 | 0.00025015 | 9 | 1199.681 | NaN | 785.0422 | NaN | NaN | NaN |
| Tapbp | TWLSTIR | -350 | -356 | 0 | 0.003966045 | 7 | 646.82025 | NaN | NaN | NaN | NaN | NaN |
| Nfu1 | FIPGKPVLETR | -72 | -82 | 1 | 0 | 11 | 825.5189 | 795.61053 | 756.29565 | 676.5006 | 720.7237 | NaN |
| Nfu1 | SVFFGPDFITVTK | -109 | -121 | 0 | 0 | 13 | 478.3066 | 465.3894 | 458.919 | NaN | 486.00543 | 408.01398 |
| Cops8 | VGLPVEEAVK | -137 | -146 | 0 | 0 | 10 | 803.39777 | 795.4997 | 886.3932 | 731.035 | 667.506 | 836.9706 |
| Cops8 | FIPLSEPAPVPPIPNEQQLAR | -179 | -199 | 0 | 0 | 21 | 387.17838 | 404.35135 | 388.56708 | NaN | 453.74146 | 373.12515 |
| Eml4 | NVSFTHNDSHLISTGGK | -850 | -866 | 0 | 0 | 17 | 776.5476 | 968.3916 | 834.5282 | 724.2828 | 613.9683 | 744.07294 |
| Eml4 | FIPSDVDNYDDIR | -250 | -262 | 0 | 0 | 13 | 350.5803 | 686.7938 | 754.4016 | NaN | 1853.2703 | 388.82068 |
| Taf10 | FISDIANDALQHCK | -162 | -175 | 0 | 0 | 14 | 451.06897 | 577.3027 | 506.5614 | 755.4263 | 926.6299 | 833.9021 |
| Kpna4 | SHFPALLTHPK | -329 | -339 | 0 | 0 | 11 | 1411.053 | 1069.1737 | 1061.7291 | 1044.9961 | 1310.9188 | 1168.7084 |
| Kpna4 | SHFPALLTHPK | -329 | -339 | 0 | 0 | 11 | 1411.053 | 1069.1737 | 1061.7291 | 1044.9961 | 1310.9188 | 1168.7084 |
| Kpna4 | SGILPILVH | -113 | -121 | 0 | 0.000826806 | 9 | 1035.6294 | 1085.1068 | 1150.8975 | 819.2519 | 1130.6096 | 1144.8545 |
| Kpna4 | SGILPILVHC | -113 | -122 | 0 | 0 | 10 | 753.0017 | 716.74603 | 953.6798 | 768.9218 | 889.43835 | 758.2685 |
| Kpna4 | NVPQEDICEDSDIDGDYR | -50 | -67 | 0 | 0 | 18 | 265.6511 | 522.5855 | 430.91922 | 388.62238 | 47.631306 | 599.5527 |
| Kpna4 | ISPSIPITFLR | -208 | -218 | 0 | 0 | 11 | 452.0658 | 504.0436 | 499.46603 | 489.15128 | 479.6554 | NaN |
| Kpna4 | QVVLDGLSNILK | -429 | -440 | 0 | 0 | 12 | 443.29385 | 443.09332 | 496.41827 | NaN | NaN | NaN |
| Kpna4 | FISPSIPITFLR | -207 | -218 | 0 | 0.001392869 | 12 | 332.15497 | NaN | NaN | 79.90744 | 319.57803 | NaN |
| Kpna4 | LSFISPSIPITFLR | -205 | -218 | 0 | 0.001636916 | 14 | 242.71933 | NaN | NaN | NaN | NaN | NaN |
| Hnrnpdl | VFVGGLSPDTSEEQIK | -116 | -131 | 0 | 0 | 16 | 2013.86 | 2567.7551 | 2417.2876 | 1867.184 | 2540.126 | 2456.6768 |
| Hnrnpdl | YTDEEPVKK | -162 | -170 | 1 | 0 | 9 | 1277.6497 | 860.76984 | 1594.5088 | 1185.7628 | 1490.2378 | 1424.8026 |
| Hnrnpdl | KDLTEYLSR | -43 | -51 | 1 | 0.00078661 | 9 | 975.9009 | 983.7943 | 1649.4784 | 1231.0625 | 1642.4691 | 1595.4362 |
| Hnrnpdl | ITYTDEEPVKK | -160 | -170 | 1 | 0 | 11 | 601.0459 | 551.44305 | 534.09283 | 353.90588 | 708.99097 | 663.88336 |
| Hnrnpdl | GLSPDTSEEQIK | -120 | -131 | 0 | 0 | 12 | 1201.3032 | 1240.7041 | 1526.4467 | 2224.4248 | 611.1195 | NaN |
| Hnrnpdl | GEIENIELPMDTK | -138 | -150 | 0 | 0 | 13 | 306.9707 | NaN | 1164.4373 | 788.7248 | 881.50165 | 1029.191 |
| Hnrnpdl | FKDAASVDK | -77 | -85 | 1 | 0.00025015 | 9 | 1099.2437 | NaN | 1674.0657 | 756.3374 | NaN | 1030.1138 |
| Ubr5 | FKDEPGEGSGVAR | -2275 | -2287 | 1 | 0 | 13 | 485.3354 | 611.3554 | 751.26025 | 660.5743 | 600.4644 | 424.22067 |
| Tmem30a | FKDTIKPVNWHK | -229 | -240 | 2 | 0 | 12 | 533.9953 | NaN | NaN | NaN | NaN | NaN |
| Vapa | PHSVSLNDTETR | -162 | -173 | 0 | 0 | 12 | 2052.7454 | 2245.8472 | 2341.6562 | 1261.4355 | 2005.8379 | 1630.79 |
| Vapa | PHSVSLNDTETR | -162 | -173 | 0 | 0 | 12 | 2052.7454 | 2245.8472 | 2341.6562 | 1261.4355 | 2005.8379 | 1630.79 |
| Vapa | HEQILVLDPPSDLK | -11 | -24 | 0 | 0 | 14 | 1337.5709 | 1767.382 | 1747.9617 | 1882.3131 | 2639.2903 | 2295.9224 |
| Vapa | FKGPFTDVVTT | -25 | -35 | 1 | 0.000394361 | 11 | 1665.4567 | 1796.2544 | 1739.9282 | 1940.638 | NaN | 2001.2056 |
| Vapa | TIFAPPNISDMEAVWK | -99 | -114 | 0 | 0.00025015 | 16 | 242.38713 | NaN | NaN | 437.71725 | 542.4652 | NaN |
| Pom121 | SLPAPSSNPLLESLK | -563 | -577 | 0 | 0 | 15 | 1807.9172 | NaN | NaN | NaN | NaN | 2487.272 |
| Pom121 | FKPIFPATPK | -701 | -710 | 1 | 0.00555405 | 10 | 245.49347 | NaN | NaN | NaN | NaN | NaN |
| Arhgap1 | YLSELSEHVK | -191 | -200 | 0 | 0 | 10 | 1493.6746 | 1599.2686 | 1274.5242 | 1302.3652 | 1386.606 | 1149.1367 |
| Arhgap1 | SPGQDPIPIVLR | -253 | -264 | 0 | 0 | 12 | 865.8157 | 1109.0701 | 1016.1109 | 1050.5187 | 1338.7512 | 1282.5288 |
| Arhgap1 | KWDDPYYDIAR | -59 | -69 | 1 | 0 | 11 | 562.3003 | 738.61316 | 388.58374 | 825.90216 | 1112.5192 | 909.4775 |
| Arhgap1 | FLLDHQGELFPSTDAQ | -422 | -437 | 0 | 0.00025015 | 16 | 371.55597 | 605.52795 | 564.1593 | 804.8621 | 975.4391 | 920.14185 |
| Arhgap1 | FLLDHQGELFPSTDAQGV | -422 | -439 | 0 | 0 | 18 | 839.2082 | 773.69916 | NaN | 964.658 | 1616.9116 | 1344.1404 |
| Arhgap1 | FKPLISFK | -174 | -181 | 1 | 0 | 8 | 743.36725 | 710.4673 | 731.47375 | NaN | 449.89777 | 470.10324 |
| Itih4 | FKPTLSQQQK | -215 | -224 | 1 | 0 | 10 | 1837.5675 | 2386.0996 | 2510.8403 | 824.0305 | 1996.5446 | 1983.637 |
| Itih4 | VRPQQLVK | -163 | -170 | 1 | 0.00555405 | 8 | 1169.9802 | 1170.9432 | 1050.5 | 650.88684 | 883.5104 | 804.90845 |
| Itih4 | RIYEDSDSALQLQ | -439 | -451 | 1 | 0.000627943 | 13 | 751.09155 | 853.7965 | NaN | NaN | 1079.8898 | 1007.114 |
| Itih4 | NVIFVIDK | -274 | -281 | 0 | 0 | 8 | 694.56836 | NaN | NaN | 588.4908 | 619.6894 | NaN |
| Itih4 | RIYEDSDSAL | -439 | -448 | 1 | 0.007224385 | 10 | 961.5896 | NaN | 918.99646 | NaN | NaN | NaN |
| Acot13 | LGTLHGGLTAT | -52 | -62 | 0 | 0 | 11 | 1352.8997 | 1468.7872 | 1845.785 | 530.81537 | 414.5195 | 1615.376 |
| Acot13 | FKVPGFDR | -16 | -23 | 1 | 0 | 8 | 1207.5669 | 1101.3801 | 1202.6665 | NaN | 734.8628 | 1052.3508 |
| Hacd3 | SIPVFNESGR | -300 | -309 | 0 | 0.00025015 | 10 | 1186.989 | 432.30142 | 995.3789 | 134.57327 | 660.0142 | 1069.775 |
| Hacd3 | FSFTLPYPVK | -310 | -319 | 0 | 0 | 10 | 984.5183 | 1091.2992 | 1141.5461 | 1251.4343 | 1354.7075 | 1383.3019 |
| Hacd3 | FLAPDFDR | -102 | -109 | 0 | 0 | 8 | 394.6513 | 485.4333 | 571.38165 | 333.98077 | 617.2692 | 456.60776 |
| Hacd3 | HLEFLDLVKPEPAYR | -58 | -72 | 1 | 0 | 15 | 578.3048 | 800.4626 | 560.00806 | 832.6816 | NaN | 1086.4075 |
| Hacd3 | FLDLVKPEPAYR | -61 | -72 | 1 | 0 | 12 | 386.16162 | 403.8029 | 564.5709 | NaN | NaN | NaN |
| Ubash3a | LAGEALLDSGVR | -426 | -437 | 0 | 0 | 12 | 361.30106 | 436.0462 | 476.07816 | NaN | 953.0302 | 741.021 |
| Ubash3a | VDLDYRPALPR | -502 | -512 | 1 | 0.001392869 | 11 | 5224.7793 | 5942.17 | NaN | 6101.7827 | NaN | 360.41162 |
| Ubash3a | FLCPTGPLLEK | -77 | -87 | 0 | 0 | 11 | 1233.6757 | NaN | 1304.022 | 951.923 | NaN | 683.88074 |
| Cfb | FLCTGGVDPYADPNTCK | -677 | -693 | 0 | 0 | 17 | 537.9626 | 616.04224 | 586.5429 | 940.7433 | 1060.5804 | 1020.4708 |

| | | | | | | | | | | | | |
|---|---|---|---|---|---|---|---|---|---|---|---|---|
| Coa3 | FLDELEDEAK | -84 | -93 | 0 | 0 | 10 | 2653.095 | NaN | 4585.357 | NaN | 5295.0825 | 5102.238 |
| Utp20 | FLDESLGEQVVK | -2544 | -2555 | 0 | 0 | 12 | 2857.6982 | 672.6764 | 1869.1342 | 748.6781 | 1845.7194 | 2241.4934 |
| Utp20 | LGTLLPVIEK | -2403 | -2412 | 0 | 0 | 10 | 419.81805 | 481.8282 | 120.73197 | 489.06473 | 513.05505 | 542.2526 |
| Utp20 | LVLPHVPAIL | -1298 | -1307 | 0 | 0 | 10 | 403.0168 | 417.81863 | 340.4078 | 399.12482 | NaN | 482.00418 |
| Utp20 | HLDDINFDVR | -1439 | -1448 | 0 | 0 | 10 | 2028.6313 | NaN | NaN | 1728.9742 | NaN | 1688.7568 |
| Utp20 | LLAENVSLPETLIK | -373 | -386 | 0 | 0 | 14 | 299.50644 | NaN | NaN | NaN | 455.0433 | 451.39502 |
| Utp20 | LAYAEEDIYDTSR | -2208 | -2220 | 0 | 0 | 13 | 782.0886 | NaN | NaN | NaN | 946.0627 | NaN |
| Utp20 | HLFLLLK | -2158 | -2164 | 0 | 0.001776473 | 7 | 384.52576 | NaN | NaN | 186.99625 | NaN | NaN |
| Utp20 | LGSESQYSPTPLLK | -1192 | -1205 | 0 | 0 | 14 | 5875.1187 | NaN | NaN | NaN | NaN | NaN |
| Sltm | FLDFSHR | -766 | -772 | 0 | 0.005394768 | 7 | 866.041 | NaN | NaN | NaN | 665.4629 | 472.5398 |
| Sltm | ITDSEASKPK | -271 | -280 | 1 | 0 | 10 | 216.30606 | NaN | 237.2227 | NaN | NaN | NaN |
| Dcun1d1 | FLDLWNK | -192 | -198 | 0 | 0.00025015 | 7 | 652.06744 | 579.1782 | 651.79224 | 568.0167 | 748.0954 | 777.5395 |
| Med23 | FLELLPVSK | -974 | -982 | 0 | 0 | 9 | 473.23285 | 382.8122 | 383.8414 | 324.00824 | 422.25146 | 420.03705 |
| Uchl3 | FLENYDAIR | -137 | -145 | 0 | 0 | 9 | 3881.8772 | 1451.4093 | 4062.8894 | 1261.6036 | 1512.1627 | 1685.385 |
| Uchl3 | TSDETLLEDAIEVCK | -196 | -210 | 0 | 0.000230707 | 15 | 1032.2054 | 1568.2959 | 624.39746 | 567.06213 | 538.9397 | 525.78375 |
| Uchl3 | VHVDGHLYELDGR | -174 | -186 | 0 | 0 | 13 | 764.79065 | 853.38074 | 876.7114 | 1740.88 | 907.504 | 1000.44763 |
| Uchl3 | VHVDGHLYELDGR | -174 | -186 | 0 | 0 | 13 | 764.79065 | 853.38074 | 876.7114 | 1740.88 | 907.504 | 1000.44763 |
| Uchl3 | TSDETLLEDAIEVCKK | -196 | -211 | 1 | 0.00025015 | 16 | 650.4784 | 720.78204 | 735.88257 | NaN | NaN | NaN |
| Ubtf | FREDHPDLIQ | -179 | -188 | 1 | 0.000550307 | 10 | 1009.5386 | 3981.5488 | 3899.0461 | 2183.9414 | 1335.78 | 1267.9592 |
| Ubtf | QLQEERPELSESELTR | -424 | -439 | 1 | 0 | 16 | 594.22156 | 562.343 | 218.58076 | 574.04755 | 341.80472 | 660.2394 |
| Ubtf | FLESLPEEEQQR | -361 | -372 | 0 | 0 | 12 | 1182.7681 | NaN | 1241.3527 | 1425.0768 | 1190.4044 | 1407.1237 |
| Ubtf | SVIGDYLAR | -495 | -503 | 0 | 0.00078661 | 9 | 914.4164 | 1002.00287 | 1056.6362 | 831.9839 | 1188.5358 | NaN |
| Ubtf | FREDHPDLIQNA | -179 | -190 | 1 | 0 | 12 | 1839.1869 | 336.71973 | 1075.5468 | NaN | 1377.5245 | NaN |
| Pold2 | FLGTSGQNVSDIFR | -341 | -354 | 0 | 0 | 14 | 1295.6425 | 3180.1519 | 3156.1768 | 2653.132 | 3972.3418 | NaN |
| Pold2 | HISPTAPDTLGCYPFYK | -373 | -389 | 0 | 0.000230707 | 17 | 309.38525 | 373.2944 | 452.01004 | NaN | 358.98666 | 345.804 |
| Oga | FLIGDQEPWAFR | -671 | -682 | 0 | 0 | 12 | 2649.8591 | NaN | 4492.8726 | NaN | NaN | NaN |
| Ddx51 | FLIIDEADR | -339 | -347 | 0 | 0 | 9 | 204.9713 | 522.0743 | 487.7203 | NaN | NaN | 1031.1565 |
| Pacs1 | FLIIPLGSHPVAK | -657 | -669 | 0 | 0 | 13 | 297.28165 | 369.81845 | 352.52417 | 337.5752 | 447.391 | 464.6901 |
| Pacs1 | SLSSQPIDHEGIK | -250 | -262 | 0 | 0 | 13 | 518.0199 | 538.88214 | 661.22864 | NaN | 469.57135 | 622.86475 |
| Mark2 | SSELEGDTITLKPR | -365 | -378 | 1 | 0 | 14 | 871.6574 | 816.9268 | 752.0565 | 1036.1923 | 666.5863 | 691.8766 |
| Mark2 | FLILNPSK | -284 | -291 | 0 | 0 | 8 | 647.98517 | 304.7858 | 173.37743 | 432.99173 | 516.8077 | 494.93622 |
| Parg | FLINPELIVSR | -752 | -762 | 0 | 0 | 11 | 617.11847 | NaN | NaN | NaN | NaN | NaN |
| Nipsnap3b | WWNESADSR | -198 | -206 | 0 | 0 | 9 | 667.9149 | 663.1653 | 604.085 | 771.6596 | 751.6958 | 660.35266 |
| Nipsnap3b | FLIPNLAFIDK | -113 | -123 | 0 | 0.00025015 | 11 | 201.85538 | 117.848434 | NaN | NaN | NaN | NaN |
| Nipsnap3b | YLVPWCK | -130 | -136 | 0 | 0.001055699 | 7 | 48.9286 | NaN | NaN | NaN | NaN | NaN |
| Ankrd17 | FLLEAGADQEHK | -415 | -426 | 0 | 0 | 12 | 6061.682 | 6937.155 | NaN | NaN | NaN | NaN |
| Baz2b | FLLPVNLK | -2040 | -2047 | 0 | 0.000431655 | 8 | 5562.9966 | 6487.438 | NaN | 5054.3677 | 6061.0044 | 6135.229 |
| Cfdp1 | VTSPATPLPAGSGIK | -201 | -215 | 0 | 0 | 15 | 1595.3684 | 798.48145 | 1672.0775 | 1334.3751 | 1101.5249 | 1282.3422 |
| Cfdp1 | VAEETEEISSNKPLVK | -131 | -146 | 1 | 0.00051306 | 16 | 762.6968 | 883.18243 | 924.3129 | 1001.8995 | 950.70337 | 947.3111 |
| Cfdp1 | KVFDFAGEEVR | -163 | -173 | 1 | 0 | 11 | 573.2677 | 785.4971 | 780.4293 | 937.36176 | 983.2365 | 969.10803 |
| Cfdp1 | FLNDVGPK | -113 | -120 | 0 | 0.000591124 | 8 | 707.19055 | 840.66895 | 631.69763 | 332.5513 | 659.8477 | NaN |
| Ndufa9 | LFGLSPFEPWTTK | -304 | -316 | 0 | 0 | 13 | 718.32086 | NaN | NaN | 739.72375 | 1022.86224 | 689.6613 |
| Ndufa9 | NFDFEDVFVNIPR | -138 | -150 | 0 | 0 | 13 | 138.1579 | NaN | NaN | 331.22055 | 349.42188 | 494.9918 |
| Ndufa9 | TFLEWDAR | -104 | -111 | 0 | 0.001291103 | 8 | 693.0531 | NaN | 5476.6636 | 2360.672 | NaN | NaN |
| Ndufa9 | FLNHFANYR | -213 | -221 | 0 | 0 | 9 | 3374.4216 | 1182.9833 | NaN | NaN | NaN | NaN |
| Alg3 | FLPETIFLHR | -278 | -287 | 0 | 0 | 10 | 419.35138 | 906.2817 | 940.6602 | 649.22784 | 1156.8236 | 1090.4591 |
| Hspe1 | FLPLFDR | -9 | -15 | 0 | 0 | 7 | 7715.111 | 8487.25 | 8347.145 | 7341.206 | 7359.624 | 7903.489 |
| Hspe1 | VVLDDKDYFLFR | -81 | -92 | 1 | 0 | 12 | 4060.5698 | 4715.2393 | 4801.0454 | 1343.0831 | 7222.5693 | 6907.5522 |
| Hspe1 | VVLDDKDYFLFR | -81 | -92 | 1 | 0 | 12 | 4060.5698 | 4715.2393 | 4801.0454 | 1343.0831 | 7222.5693 | 6907.5522 |
| Hspe1 | KFLPLFDR | -8 | -15 | 1 | 0.001392869 | 8 | 1980.9508 | 2193.7705 | 2267.0735 | 1718.9565 | 1698.0079 | 1788.0316 |
| Hspe1 | GGIMLPEK | -29 | -36 | 0 | 0.000471409 | 8 | 1228.2898 | 1118.6796 | 1208.0682 | 830.5484 | 821.76886 | 353.2036 |
| Hspe1 | VLLPEYGGTK | -71 | -80 | 0 | 0 | 10 | 802.343 | 739.74426 | 6122.6143 | 245.282 | 5099.421 | 4827.0576 |
| Hspe1 | SGEIEPVSVK | -57 | -66 | 0 | 0.000431655 | 10 | 778.38556 | 6153.2627 | 273.56415 | 4635.4487 | 3546.716 | 639.25836 |
| Hspe1 | RDSDILGK | -92 | -99 | 1 | 0.005605646 | 8 | 478.57706 | 475.82465 | 362.1672 | NaN | 742.5486 | 623.8323 |
| Hspe1 | QATVVAVGSGGK | -43 | -54 | 0 | 0 | 12 | 1221.5555 | NaN | NaN | 1102.9663 | 250.88477 | 1264.6101 |
| Ppm1h | SQDEVECSADHIAR | -50 | -63 | 0 | 0 | 14 | 276.6507 | 354.51706 | 360.19055 | 416.74976 | 421.54517 | 479.44177 |
| Ppm1h | FLPNCDPDDPHR | -454 | -465 | 0 | 0.000869338 | 12 | 1406.2383 | NaN | NaN | NaN | NaN | 971.32306 |
| Fkbp3 | FLQDHGSDSFLAEHK | -28 | -42 | 0 | 0 | 15 | 567.0313 | 639.13885 | 757.03876 | 604.6673 | 800.3718 | 98.03023 |
| Fkbp3 | FLQDHGSDSFLAEHK | -28 | -42 | 0 | 0 | 15 | 567.0313 | 639.13885 | 757.03876 | 604.6673 | 800.3718 | 98.03023 |
| Fkbp3 | SEETLDEGPPK | -100 | -110 | 0 | 0 | 11 | 2105.6553 | 2333.9724 | 2212.7178 | 2508.1543 | 2342.3254 | 2330.122 |
| Fkbp3 | LEIEPEWAYGK | -190 | -200 | 0 | 0 | 11 | 1185.8904 | 733.0129 | 759.06055 | 749.6858 | 857.8552 | 976.49023 |
| Fkbp3 | KGDVVHCW | -127 | -134 | 1 | 0 | 8 | 2452.187 | 2245.0254 | 2544.0178 | NaN | 2123.0825 | 2291.6006 |
| Fkbp3 | QDHGSDSFLAEHK | -30 | -42 | 0 | 0 | 13 | 655.9562 | 776.0407 | 835.43024 | 528.63477 | NaN | 797.4823 |
| Fkbp3 | FLQDHGSDSFLAE | -28 | -40 | 0 | 0.00078661 | 13 | 850.1987 | 971.3623 | 899.07825 | NaN | 583.10944 | NaN |
| Ccdc88b | FLQEQENQHQR | -910 | -920 | 0 | 0 | 11 | 802.38824 | 928.55334 | 1014.51624 | 870.14557 | 1580.3623 | 1488.0195 |
| Ccdc88b | SLGGPLETELPEGR | -1353 | -1366 | 0 | 0.000230707 | 14 | 568.2107 | NaN | NaN | NaN | NaN | NaN |
| Sf3b4 | FLSEEDADYAIK | -61 | -72 | 0 | 0 | 12 | 1289.8387 | 1386.0724 | 416.39038 | 1741.8345 | 1648.0828 | 582.0676 |
| Sf3b4 | IFIGNLDPEIDEK | -102 | -114 | 0 | 0 | 13 | 1137.5895 | 1233.9567 | 1261.2539 | 1562.4072 | 1726.5256 | 1554.2325 |
| Xpo7 | FLSHLVDGVK | -303 | -312 | 0 | 0 | 10 | 683.67444 | 499.85184 | 693.2783 | 578.2095 | 529.12695 | 560.1112 |
| Xpo7 | SNYQLGELVK | -339 | -348 | 0 | 0 | 10 | 272.43146 | 352.3751 | 588.76855 | NaN | NaN | NaN |
| Syngr2 | FLSQPQVVTR | -20 | -29 | 0 | 0 | 10 | 957.84015 | 888.5946 | 894.55927 | 678.8732 | 579.5069 | 184.94592 |
| Ssr1 | FLVGFTNK | -103 | -110 | 0 | 0 | 8 | 3078.0388 | NaN | 3143.6758 | 2681.8289 | 1839.2968 | 1735.6174 |
| Ssr1 | GEDFPANNIVK | -92 | -102 | 0 | 0 | 11 | 1070.898 | 1245.483 | 1087.3326 | 1323.8323 | NaN | 1317.9546 |
| Gpx1 | YIIWSPVCR | -147 | -155 | 0 | 0 | 9 | 1046.7413 | 1631.3329 | 1618.0072 | 815.70557 | 1726.4446 | 1375.0642 |
| Gpx1 | FLVGPDGVPVR | -165 | -175 | 0 | 0 | 11 | 849.38617 | 869.64557 | 711.30743 | 677.182 | NaN | 995.35443 |
| Rpl32 | SYCAEIAHNVSSK | -94 | -106 | 0 | 0 | 13 | 899.2828 | 946.4881 | 757.1731 | 952.34656 | 841.5632 | 808.76953 |
| Rpl32 | FLVHNVK | -77 | -83 | 0 | 0 | 7 | 1868.299 | 4551.6567 | 4250.6504 | NaN | NaN | NaN |
| Endog | FLVPIESIER | -260 | -269 | 0 | 0.00122419 | 10 | 681.4834 | NaN | NaN | NaN | 1464.5725 | 1458.6885 |

| | | | | | | | | | | | | |
|---|---|---|---|---|---|---|---|---|---|---|---|---|
| Endog | TGPLFLPR | -194 | -201 | 0 | 0.007969168 | 8 | 469.54727 | NaN | NaN | NaN | NaN | NaN |
| Rngtt | FNAQPVGDCDFNIR | -367 | -380 | 0 | 0 | 14 | 995.204 | 1219.5767 | 1216.3715 | NaN | NaN | NaN |
| Clptm1 | SVAHLPWR | -535 | -542 | 0 | 0 | 8 | 4032.847 | 4554.332 | 5032.0195 | 4118.912 | 412.24792 | 5084.1426 |
| Clptm1 | FNDYWNLQK | -281 | -289 | 0 | 0 | 9 | 949.7997 | 773.3243 | 397.0214 | 763.9963 | 641.8003 | 279.5452 |
| Clptm1 | TAAGALTPAPSTAVSGEDASTVPK | -616 | -639 | 0 | 0.00122419 | 24 | 502.1001 | 522.5043 | 307.00894 | 381.7027 | 449.55887 | 438.914 |
| Clptm1 | GALTPAPSTAVSGEDASTVPK | -619 | -639 | 0 | 0.000907206 | 21 | 1039.8152 | 1277.2678 | NaN | NaN | 796.15936 | 837.45984 |
| Lsm14a | SFFDNISCDDNR | -367 | -378 | 0 | 0 | 12 | 490.1827 | 334.5687 | 495.9224 | 505.8553 | 162.91194 | 653.0336 |
| Lsm14a | NSEGNADEEDPLGPNCYYDK | -345 | -364 | 0 | 0.001776473 | 20 | 195.2223 | 243.72375 | NaN | 246.6619 | 40.473995 | 301.02417 |
| Lsm14a | FNKEEIDR | -304 | -311 | 1 | 0.000907206 | 8 | 1119.8646 | NaN | 1512.8599 | 1048.2988 | 604.35626 | NaN |
| Lsm14a | SVKPQLAQGR | -172 | -181 | 1 | 0.00383302 | 10 | 324.71204 | NaN | NaN | NaN | NaN | NaN |
| Ints8 | FNQLFLRPNK | -403 | -412 | 1 | 0.00078661 | 10 | 284.72995 | NaN | NaN | NaN | NaN | NaN |
| Vps4b | NSLTEADFQELGR | -319 | -331 | 0 | 0 | 13 | 1713.1298 | NaN | 1859.1973 | 1042.8269 | 1572.1355 | 1383.152 |
| Vps4b | IYIPLPEAH | -296 | -304 | 0 | 0.000869338 | 9 | 760.8691 | 776.14923 | 724.0152 | 166.79439 | NaN | NaN |
| Vps4b | IYIPLPEAHAR | -296 | -306 | 0 | 0.003360691 | 11 | 504.8698 | 475.42365 | NaN | NaN | NaN | NaN |
| Vps25 | GNLEWLDK | -85 | -92 | 0 | 0.000230707 | 8 | 1496.4581 | 1466.8546 | 172.83096 | 1511.5854 | 1054.386 | 1071.3574 |
| Vps25 | FPPFFTLQPNVDTR | -13 | -26 | 0 | 0 | 14 | 493.25354 | 491.14127 | NaN | 107.74528 | 1189.3112 | 104.81079 |
| Aurkb | FPSSVPSGAQDLISK | -297 | -311 | 0 | 0 | 15 | 844.6551 | 913.07153 | 1010.55237 | 671.6997 | 817.18384 | 366.28894 |
| Tmprss9 | FPWQVSLR | -216 | -223 | 0 | 0.000471409 | 8 | 1915.9559 | 2168.2043 | 614.00934 | NaN | 2781.4177 | 3046.8682 |
| Mta2 | TLLADQGEIR | -139 | -148 | 0 | 0 | 10 | 8785.834 | 9525.737 | 12378.867 | 3898.6208 | 12839.694 | 8965.244 |
| Mta2 | VWDPDNPLTDR | -178 | -188 | 0 | 0 | 11 | 1671.796 | 1934.8969 | 1960.0591 | 125.69347 | 2060.0151 | 2073.5098 |
| Mta2 | QFESLPATHIR | -93 | -103 | 0 | 0 | 11 | 1629.02 | 1696.7686 | 1581.7529 | 1740.4388 | 1002.01733 | 903.7106 |
| Mta2 | FQAEIPDR | -153 | -160 | 0 | 0 | 8 | 1119.0187 | 1235.264 | 913.1737 | 153.31186 | 1093.6609 | 912.93317 |
| Mta2 | STLVPQGGPVLCR | -250 | -262 | 0 | 0 | 13 | 951.645 | 1045.6635 | 638.3928 | 841.9386 | 920.8189 | 1027.5769 |
| Mta2 | TVRPPVPLPASSHPASTNEPIVLED | -644 | -668 | 1 | 0 | 25 | 612.75806 | 562.2077 | 597.77136 | 589.26855 | 208.55496 | 625.0212 |
| Mta2 | SSSNPYLVR | -17 | -25 | 0 | 0 | 9 | 859.3866 | 751.11 | 706.27 | NaN | NaN | NaN |
| Mta2 | RPYAPINAN | -481 | -489 | 1 | 0.004439092 | 9 | 1200.1692 | NaN | NaN | 2747.706 | NaN | NaN |
| Tap2 | RPEKPVLQGL | -479 | -488 | 2 | 0.003681885 | 10 | 2110.8823 | 1916.7511 | 2942.6917 | 2817.1794 | 226.83006 | 2061.6362 |
| Tap2 | FQDVSFSYPR | -469 | -478 | 0 | 0 | 10 | 1632.3295 | 1563.8223 | 1599.0149 | 2240.6792 | 487.53763 | 2207.5452 |
| Tap2 | LHTVQNADQVLVLK | -662 | -675 | 0 | 0 | 14 | 1160.4519 | 376.98483 | 406.925 | 533.58026 | 391.55728 | 504.60446 |
| Tap2 | VEFQDVSFSYPR | -467 | -478 | 0 | 0 | 12 | 943.72253 | 1251.979 | 1122.5691 | 1289.21 | 1754.7399 | 870.7948 |
| Tap2 | GAEEQEVSHYK | -345 | -355 | 0 | 0 | 11 | 777.03986 | 788.2153 | 904.64996 | 624.73096 | 849.48413 | 866.4004 |
| Tap2 | LYQEEVGQYVR | -413 | -423 | 0 | 0 | 11 | 252.76445 | 1878.6392 | 1958.4961 | 1511.7106 | 1708.5367 | 1687.951 |
| Tap2 | LVEHDQLR | -679 | -686 | 0 | 0 | 8 | 13990.809 | 379.38196 | NaN | 468.20245 | 564.98926 | 345.80173 |
| Tap2 | KPNLPQPGILAPPWLEGR | -449 | -466 | 1 | 0 | 18 | 1985.2306 | 860.0549 | NaN | 1813.922 | 2338.5947 | 2692.7795 |
| Tap2 | NCGVQQILAGEVTR | -392 | -405 | 0 | 0 | 14 | 2435.5354 | NaN | 3007.3435 | NaN | 1850.8584 | 454.0138 |
| Tap2 | QILAGEVTR | -397 | -405 | 0 | 0 | 9 | 1509.0978 | NaN | NaN | 1583.5144 | 1380.6388 | 1303.3699 |
| Tap2 | RPEKPVLQ | -479 | -486 | 2 | 0.008736733 | 8 | 523.95575 | NaN | 283.07547 | NaN | NaN | NaN |
| Acp5 | FQETFEDVFSDR | -88 | -99 | 0 | 0 | 12 | 426.32205 | 8886.069 | 6958.625 | 7955.336 | 6212.655 | 4704.187 |
| Larp1 | FQHPSHELLK | -849 | -858 | 0 | 0 | 10 | 4670.7866 | 3439.1675 | 2891.2537 | 1104.6777 | 4639.579 | 5108.611 |
| Larp1 | LFGAPEPSTIAR | -731 | -742 | 0 | 0 | 12 | 1375.6055 | 130.37479 | 1529.5468 | 1265.0635 | 516.8722 | 698.9436 |
| Larp1 | LNCPEFVPR | -477 | -485 | 0 | 0 | 9 | 1278.9982 | 1251.997 | 193.20569 | 521.5985 | 291.2188 | 1088.6495 |
| Larp1 | SLPTTVPESPNYR | -743 | -755 | 0 | 0 | 13 | 1165.0714 | 1429.082 | 1338.7037 | 1615.8848 | 1530.1663 | 1710.1133 |
| Larp1 | YGLECLFR | -926 | -933 | 0 | 0 | 8 | 909.3013 | 980.6526 | 967.3206 | 918.2669 | 1128.7677 | 1134.9229 |
| Larp1 | SEEDSDYEIDDRDVNK | -604 | -619 | 1 | 0.001092529 | 16 | 592.9064 | 473.32755 | 469.32886 | 618.4347 | 595.50854 | 434.01276 |
| Larp1 | HSSNPPLESHVG | -799 | -810 | 0 | 0 | 12 | 496.65573 | 973.3526 | 7775.953 | 757.71094 | 3733.674 | 8132.1646 |
| Larp1 | SVAPPTPAWQPETK | -273 | -286 | 0 | 0 | 14 | 1648.9116 | 1729.67 | 1485.373 | NaN | NaN | NaN |
| Larp1 | FSVDNLER | -390 | -397 | 0 | 0.002697088 | 8 | 1168.2589 | 815.3974 | NaN | NaN | NaN | 566.1131 |
| Larp1 | VEPAWHDQDETSSVK | -287 | -301 | 0 | 0 | 15 | 708.7945 | 692.8619 | 640.3175 | NaN | NaN | NaN |
| Larp1 | FQQVPTDALANK | -719 | -730 | 0 | 0 | 12 | 7252.5693 | NaN | NaN | NaN | NaN | NaN |
| Derl1 | NFLSTPQFLYR | -198 | -208 | 0 | 0 | 11 | 659.6531 | 754.96814 | 715.7172 | 702.6569 | 729.0403 | 780.09564 |
| Derl1 | FQIWRPFTAT | -50 | -59 | 1 | 0.000230707 | 10 | 605.75653 | 803.4719 | 443.5073 | NaN | 165.01321 | 1002.5797 |
| Ufd1 | FQPQSPDFLDITNPK | -125 | -139 | 0 | 0 | 15 | 1391.9303 | 1297.1659 | 1131.0851 | 1376.8964 | 2067.385 | 2016.7479 |
| Ufd1 | GVEPSPSPIKPGDIK | -241 | -255 | 1 | 0 | 15 | 758.9778 | 719.54803 | 1128.9716 | 1382.9767 | 1267.7089 | 948.8927 |
| Sf1 | FQRPGDPQSAQDK | -294 | -306 | 1 | 0 | 13 | 1793.8625 | 1938.8698 | 2712.433 | 1548.2987 | 2454.6428 | 2210.09 |
| Sf1 | FQRPGDPQSAQDK | -294 | -306 | 1 | 0 | 13 | 1793.8625 | 1938.8698 | 2712.433 | 1548.2987 | 2454.6428 | 2210.09 |
| Sf1 | VALNPDFKPPADYKPPATR | -117 | -135 | 2 | 0 | 19 | 1628.2764 | 1993.7052 | 1840.8009 | 1921.2704 | 2277.7776 | 2145.7532 |
| Sf1 | GIETPEDQNDLRK | -229 | -241 | 1 | 0 | 13 | 761.83813 | 702.3971 | 655.86707 | 768.79694 | 863.30817 | 798.16455 |
| Sf1 | TVIPGMPTVIPPGLTR | -31 | -46 | 0 | 0 | 16 | 440.8395 | 517.51575 | 560.29645 | 389.34274 | 645.6839 | 678.8656 |
| Sf1 | QGIETPEDQNDLR | -228 | -240 | 0 | 0 | 13 | 171.80869 | 2965.4792 | 2904.9155 | 2150.3245 | 2747.174 | 2661.783 |
| Sf1 | GIETPEDQNDLR | -229 | -240 | 0 | 0 | 12 | 861.27356 | 639.4234 | NaN | 1049.1976 | 1090.0752 | 1143.0527 |
| Sf1 | FVGLLIGPR | -152 | -160 | 0 | 0 | 9 | 263.7411 | 253.51039 | 313.68018 | NaN | 295.26544 | 292.09467 |
| Sf1 | LQIEDLTR | -56 | -63 | 0 | 0 | 8 | 1503.7286 | NaN | NaN | 1474.3368 | 1723.625 | 1622.5986 |
| Sf1 | ILRPWQSSETR | -261 | -271 | 1 | 0 | 11 | 604.5068 | 941.62695 | 490.02533 | 3692.4565 | NaN | NaN |
| Sf1 | FQRPGDPQSAQD | -294 | -305 | 1 | 0.001023236 | 12 | 3678.2295 | NaN | NaN | NaN | NaN | NaN |
| Rab11a | FQTILTEIYR | -165 | -174 | 0 | 0 | 10 | 1154.6779 | 970.8782 | 854.48114 | 890.0482 | 873.7735 | 508.8288 |
| Itgb3 | YCECDDFSCVR | -545 | -555 | 0 | 0 | 11 | 373.4165 | 361.25513 | 331.9653 | 399.63043 | 421.29425 | 410.7564 |
| Itgb3 | WDTANNPLYK | -764 | -773 | 0 | 0.00078661 | 10 | 1606.0686 | 1975.2874 | 960.2313 | 2572.9182 | 978.1469 | NaN |
| Itgb3 | FQYYEDTSGR | -692 | -701 | 0 | 0.001811524 | 10 | 785.24695 | 668.5669 | 1343.4958 | NaN | 1611.5674 | 1640.816 |
| Mapk1 | VADPDHDHTGFLT | -171 | -183 | 0 | 0 | 13 | 1309.7871 | 13003.293 | 13284.091 | 13281.526 | 13562.254 | 13822.883 |
| Mapk1 | HENIIGINDIIR | -78 | -89 | 0 | 0 | 12 | 687.4496 | 640.261 | 672.32306 | 1026.8287 | 1124.9773 | 1025.7119 |
| Mapk1 | GSPSQEDLNCIINLK | -243 | -257 | 0 | 0 | 15 | 652.0197 | 850.9244 | 1068.2858 | 1113.8345 | 1489.229 | 1303.0808 |
| Mapk1 | FRHENIIGINDIIR | -76 | -89 | 1 | 0 | 14 | 91.45762 | 197.38274 | 235.6565 | NaN | 376.90598 | 316.35785 |
| Mapk1 | LEQYYDPSDEPIAEAPFK | -311 | -328 | 0 | 0.002168954 | 18 | 293.76343 | 407.1358 | 512.8396 | NaN | NaN | 449.2514 |
| Mapk1 | GQVFDVGPR | -14 | -22 | 0 | 0 | 9 | 799.9544 | 648.91 | 767.56396 | NaN | NaN | NaN |
| Mrpl45 | SLEPAQVVHVR | -188 | -198 | 0 | 0 | 11 | 739.8804 | 1246.5083 | 1222.9354 | 1186.4943 | 1324.8308 | 1248.1825 |
| Mrpl45 | FRPPTPEPK | -41 | -49 | 1 | 0.001636916 | 9 | 1053.9294 | NaN | NaN | NaN | 1182.8684 | 1084.4279 |
| Tpp1 | FRPSFPASSPYVT | -373 | -385 | 1 | 0.000431655 | 13 | 840.8491 | 843.22455 | 941.6582 | NaN | 1308.4241 | 1315.5569 |
| Tpp1 | TWVYSSPGR | -288 | -296 | 0 | 0.001708008 | 9 | 347.34888 | NaN | NaN | NaN | NaN | NaN |

| | | | | | | | | | | | | |
|---|---|---|---|---|---|---|---|---|---|---|---|---|
| Gbf1 | VLFPLLTK | -1594 | -1601 | 0 | 0 | 8 | 405.60986 | NaN | 396.73947 | 77.55687 | 502.16373 | 471.6214 |
| Gbf1 | FSCLLPDPR | -683 | -691 | 0 | 0.000907206 | 9 | 920.00714 | NaN | 919.0779 | NaN | NaN | NaN |
| Msh6 | SVLDGDPSENYSR | -524 | -536 | 0 | 0 | 13 | 1295.8867 | NaN | 899.4828 | 1273.6565 | 1064.7753 | 1023.74255 |
| Msh6 | RWDTAFDHEK | -908 | -917 | 1 | 0 | 10 | 546.0138 | 498.38684 | 532.98846 | NaN | 569.22723 | 668.5264 |
| Msh6 | YQLEIPENFATR | -974 | -985 | 0 | 0.000471409 | 12 | 572.2998 | 502.83893 | NaN | NaN | NaN | NaN |
| Msh6 | SIVYWGIGR | -963 | -971 | 0 | 0 | 9 | 1100.7578 | NaN | NaN | NaN | NaN | NaN |
| Msh6 | FSDSLVQK | -468 | -475 | 0 | 0.00025015 | 8 | 951.30743 | NaN | NaN | NaN | NaN | NaN |
| Msh6 | NLPEEYELK | -986 | -994 | 0 | 0.001023236 | 9 | 904.4503 | NaN | NaN | NaN | NaN | NaN |
| Pdcd6 | LSDQFHDILIR | -126 | -136 | 0 | 0 | 11 | 1449.4083 | 1717.5488 | 1834.3813 | 627.1627 | 2312.8909 | 2321.9785 |
| Pdcd6 | LSDQFHDILIR | -126 | -136 | 0 | 0 | 11 | 1449.4083 | 1717.5488 | 1834.3813 | 627.1627 | 2312.8909 | 2321.9785 |
| Pdcd6 | YITDWQNVFR | -91 | -100 | 0 | 0 | 10 | 981.29016 | 515.4128 | 1533.9744 | 961.5554 | 92.64858 | 504.35303 |
| Pdcd6 | SGVISDNELQQAL | -40 | -52 | 0 | 0.002566359 | 13 | 954.7558 | 296.1787 | 680.8233 | 1157.7656 | 510.1667 | 787.67554 |
| Pdcd6 | RYDTDQDGWIQ | -167 | -177 | 1 | 0 | 11 | 508.5507 | 538.5997 | 403.4857 | 412.9117 | 446.781 | 273.38956 |
| Pdcd6 | SDQFHDILIR | -127 | -136 | 0 | 0 | 10 | 496.18512 | 1056.9265 | 663.3786 | 796.5409 | 381.64877 | 570.35205 |
| Pdcd6 | FSEFTGVWK | -82 | -90 | 0 | 0 | 9 | 414.7806 | 778.64966 | 425.15207 | 483.52206 | 715.26074 | 632.2675 |
| Pdcd6 | GTWTPFNPVTVR | -55 | -66 | 0 | 0 | 12 | 294.0283 | 819.39136 | 886.32635 | 1031.9044 | 1018.988 | 296.18677 |
| Prpsap1 | FSFPVDNLR | -144 | -152 | 0 | 0 | 9 | 2410.6465 | 3169.3467 | 3326.8167 | 2942.1106 | 3695.4119 | 3619.5125 |
| Prpsap1 | YIQEEIPNYR | -160 | -169 | 0 | 0 | 10 | 1081.1259 | 840.3656 | 1315.2559 | 1552.9225 | 1498.2006 | 853.00244 |
| Prpsap1 | NIIGVIPYFPYSK | -91 | -103 | 0 | 0 | 13 | 420.4536 | NaN | 488.41122 | 395.04086 | 539.684 | 466.7776 |
| Ilk | FSFQCPGR | -342 | -349 | 0 | 0 | 8 | 2206.6724 | 2289.4277 | 2397.776 | 1836.3597 | NaN | NaN |
| Ilk | TLEPLIPR | -310 | -317 | 0 | 0.00025015 | 8 | 572.03625 | NaN | 613.3103 | 607.5618 | NaN | NaN |
| Oas3 | VLDPADPTWNVGQGDWK | -749 | -765 | 0 | 0 | 17 | 2072.7703 | 2199.2537 | 2373.983 | 1284.253 | 1329.2135 | 1382.9504 |
| Oas3 | PVILDPADPTGNLGHNA | -1087 | -1103 | 0 | 0.00025015 | 17 | 1658.6533 | 485.3302 | 937.80493 | 393.17273 | 528.05817 | 912.44037 |
| Oas3 | FSLAEGLR | -700 | -707 | 0 | 0.001023236 | 8 | 572.0868 | 636.86383 | 537.57886 | 203.25598 | NaN | 455.02994 |
| Oas3 | HWCQNPVPGLSLQ | -558 | -570 | 0 | 0.000627943 | 13 | 1197.5406 | NaN | NaN | NaN | 1296.1951 | 1127.5337 |
| Oas3 | LDCFQSFGDQK | -80 | -90 | 0 | 0.008828666 | 11 | 598.73376 | NaN | NaN | NaN | NaN | NaN |
| Celf2 | FSPFGQIEECR | -152 | -162 | 0 | 0 | 11 | 3177.9504 | 3380.1868 | 3543.9177 | 388.06277 | 1721.1444 | 1836.3441 |
| Akap13 | FSPGVEAPESR | -1478 | -1488 | 0 | 0.001636916 | 11 | 2153.96 | NaN | 2145.2053 | 2509.54 | NaN | NaN |
| Pkn1 | FSPNASPGAEIR | -535 | -546 | 0 | 0.009657961 | 12 | 816.99005 | NaN | NaN | NaN | NaN | NaN |
| Pkn1 | RLGELPADHPK | -277 | -287 | 1 | 0.00025015 | 11 | 457.20016 | NaN | NaN | NaN | NaN | NaN |
| Elavl1 | SQLYHSPAR | -197 | -205 | 0 | 0 | 9 | 3074.4502 | 3015.4866 | 3635.074 | 1411.2612 | 1475.6377 | 1935.0939 |
| Elavl1 | SLFSSIGEVESAK | -38 | -50 | 0 | 0.000230707 | 13 | 1528.3794 | 1839.7019 | 442.3274 | 1976.435 | 2328.0107 | 559.4049 |
| Elavl1 | YARPSSEVIK | -95 | -104 | 1 | 0.00111006 | 10 | 1012.2167 | 1053.5421 | 1175.7777 | 1373.4891 | 1207.8605 | 1222.4393 |
| Elavl1 | YLPQNMTQEELR | -26 | -37 | 0 | 0 | 12 | 647.1931 | 282.11658 | 634.20825 | 352.16827 | 648.8366 | 768.48517 |
| Elavl1 | LIVNYLPQNMTQEELR | -22 | -37 | 0 | 0 | 16 | 359.04034 | 403.721 | 374.32132 | 482.5756 | 721.1754 | 595.8492 |
| Elavl1 | VLVDQTTGLSR | -137 | -147 | 0 | 0.00025015 | 11 | 137.90594 | 509.4444 | 212.30379 | 1650.5193 | 2185.787 | 2188.1367 |
| Elavl1 | FSSIGEVESAK | -40 | -50 | 0 | 0 | 11 | 701.0508 | NaN | 699.33514 | NaN | 358.76392 | 854.16125 |
| Elavl1 | ILQVSFK | -314 | -320 | 0 | 0.007343498 | 7 | 557.734 | NaN | 471.61487 | NaN | NaN | NaN |
| Trip12 | VEPVGHAPLLAL | -1240 | -1251 | 0 | 0 | 12 | 961.93823 | 1252.6276 | 1146.5199 | 1548.1792 | 1549.8796 | 1526.5215 |
| Trip12 | SLNPPLTIVR | -1964 | -1973 | 0 | 0 | 10 | 687.6949 | 739.1319 | 756.1032 | 627.93 | 594.419 | 673.31915 |
| Trip12 | FSSPLPGEEPVGR | -1227 | -1239 | 0 | 0 | 13 | 503.013 | 666.8676 | 777.9702 | 676.88446 | 1189.7803 | 316.68518 |
| Trip12 | TCPFFFPFDTR | -1570 | -1580 | 0 | 0 | 11 | 328.47552 | 286.24426 | 297.0315 | 421.5903 | 470.6123 | 379.01895 |
| Trip12 | LVDLPLGLPFYK | -1746 | -1757 | 0 | 0.000431655 | 12 | 175.46811 | NaN | NaN | 640.7458 | 356.82565 | 165.39336 |
| Trip12 | THTIWYKPVR | -1426 | -1435 | 1 | 0.00272025 | 10 | 354.92325 | NaN | NaN | NaN | NaN | NaN |
| Tpd52 | SAPEALTEEEQEELRR | -65 | -80 | 1 | 0.000471409 | 16 | 834.2226 | NaN | NaN | 779.8667 | 1223.4421 | 1004.488 |
| Tpd52 | FSSVGSVITK | -153 | -162 | 0 | 0 | 10 | 532.3387 | NaN | NaN | NaN | 695.20337 | NaN |
| Cdc42 | QKPITPETAEKL | -134 | -145 | 2 | 0.001610812 | 12 | 1959.9615 | 2010.5901 | 2188.168 | 2927.609 | 2733.9897 | 2614.037 |
| Cdc42 | SVVSPSSFENVK | -83 | -94 | 0 | 0 | 12 | 1587.257 | 1694.3873 | 1707.935 | 2394.7495 | 2308.5042 | 2122.7607 |
| Cdc42 | PEITHHCPK | -99 | -107 | 0 | 0 | 9 | 1491.4507 | 1371.366 | 1162.1111 | 1613.1222 | 1982.1307 | 1093.6444 |
| Cdc42 | FSVVSPSSFENVK | -82 | -94 | 0 | 0 | 13 | 1398.5555 | 1605.1464 | 1463.5157 | 1654.0967 | 1919.4128 | 1805.7778 |
| Cdc42 | WVPEITHHCPK | -97 | -107 | 0 | 0 | 11 | 1252.5869 | 1268.7551 | 1696.8456 | 2154.213 | 1976.742 | 1870.4111 |
| Cdc42 | WVPEITHHCPK | -97 | -107 | 0 | 0 | 11 | 1252.5869 | 1268.7551 | 1696.8456 | 2154.213 | 1976.742 | 1870.4111 |
| Cdc42 | QKPITPETAEK | -134 | -144 | 1 | 0 | 11 | 1164.0668 | 1135.3596 | 1764.026 | 1909.0797 | 481.93768 | 919.8936 |
| Cdc42 | LRPLSYPQTDVFLVC | -67 | -81 | 1 | 0.002595029 | 15 | 821.66583 | 822.1853 | 564.5206 | 388.29398 | 1173.2942 | 1101.1411 |
| Cdc42 | GLFDTAGQEDYDR | -54 | -66 | 0 | 0 | 13 | 706.80963 | 760.1329 | 762.4586 | 811.6419 | 1037.2983 | 1108.2556 |
| Cdc42 | TLGLFDTAGQEDYDR | -52 | -66 | 0 | 0 | 15 | 368.80118 | 727.5213 | 680.1234 | 720.89624 | 820.3596 | 782.00024 |
| Cdc42 | NVFDEAILA | -167 | -175 | 0 | 0.00437931 | 9 | 1947.0905 | NaN | NaN | 2887.1475 | NaN | NaN |
| Gsr | FSWHVIK | -116 | -122 | 0 | 0.001536401 | 7 | 1928.5035 | 1622.1691 | 1519.4863 | 1788.0967 | 1867.2864 | 1733.942 |
| Gsr | SHPPIGTVGLTEDEAVHK | -395 | -412 | 0 | 0.00025015 | 18 | 1507.5774 | 801.32196 | 1726.6766 | 1818.1625 | 2674.4026 | 2535.5757 |
| Gsr | SHPPIGTVGLTEDEAVHK | -395 | -412 | 0 | 0.00025015 | 18 | 1507.5774 | 801.32196 | 1726.6766 | 1818.1625 | 2674.4026 | 2535.5757 |
| Gsr | SHIEIIHGY | -143 | -151 | 0 | 0.000431655 | 9 | 1398.6003 | 1487.1863 | 1495.313 | 1789.4036 | 1726.8838 | 1690.908 |
| Gsr | KFTAPHIL | -168 | -175 | 1 | 0.000298805 | 8 | 1029.3079 | 786.07837 | 662.06464 | 314.7885 | 645.92883 | 652.8024 |
| Gsr | SDGFFQLEDLPSR | -199 | -211 | 0 | 0 | 13 | 1012.8845 | 1056.0194 | 1084.3613 | 1158.0109 | 1330.6759 | 365.5312 |
| Gsr | VDEFQNTNVK | -337 | -346 | 0 | 0 | 10 | 807.60803 | 898.2305 | 2404.0413 | 829.7422 | 1701.5293 | 1063.8179 |
| Gsr | GHILVDEFQNTNVK | -333 | -346 | 0 | 0 | 14 | 608.0297 | 1411.0366 | 1674.292 | 1395.6991 | 2514.4192 | 2142.8235 |
| Gsr | SSEELVTLR | -492 | -500 | 0 | 0.00025015 | 9 | 537.09393 | 479.16406 | 873.6468 | NaN | NaN | 704.1257 |
| Gsr | FTAPHIL | -169 | -175 | 0 | 0.005639034 | 7 | 2747.2988 | NaN | 1034.223 | NaN | NaN | NaN |
| Ddx52 | FTEDDKPLLR | -517 | -526 | 1 | 0 | 10 | 1427.8312 | 1207.982 | 765.30853 | 711.3886 | 1819.2698 | 1751.3381 |
| Rela | FTGPGWEAR | -228 | -236 | 0 | 0 | 9 | 645.99976 | 712.23376 | 668.7646 | 902.016 | 877.6902 | 725.2223 |
| Rela | LVTGSQRPPDPAPTPLGTS | -502 | -520 | 1 | 0 | 19 | 307.6935 | 119.81356 | 413.70712 | 386.71164 | 392.08972 | 386.85345 |
| Prep | LINIDFTDPDESK | -313 | -325 | 0 | 0 | 13 | 3049.7551 | 2572.845 | 3483.0513 | 754.1619 | 986.6651 | 1011.9116 |
| Prep | SIWEGCDPVNR | -250 | -260 | 0 | 0 | 11 | 1692.5934 | 1407.0175 | 1510.6053 | 1254.3674 | 255.34418 | 1120.5424 |
| Prep | LDPNTLSDDGTVALR | -114 | -128 | 0 | 0 | 15 | 1233.3239 | 2552.1938 | 1908.4215 | 1980.4945 | 1879.1431 | 2615.545 |
| Prep | QHFEWLLK | -606 | -613 | 0 | 0 | 8 | 942.30176 | 1295.27 | 1436.0317 | 1059.0627 | 1425.5576 | 1242.4836 |
| Prep | ITVPFLEQCPIR | -49 | -60 | 0 | 0 | 12 | 917.53937 | 985.7762 | 836.3804 | 896.5158 | 1053.3464 | 976.062 |
| Prep | SWLEDPDSEQTK | -29 | -40 | 0 | 0 | 12 | 547.8005 | 561.9681 | 679.67206 | 1123.7341 | 870.2003 | 929.91016 |
| Prep | ICDPYSWLEDPDSEQTK | -24 | -40 | 0 | 0 | 17 | 525.11145 | 518.8073 | 588.33527 | 400.06165 | 496.28372 | 179.78976 |
| Prep | NILQLHDLTTGALLK | -359 | -373 | 0 | 0 | 15 | 409.4293 | 494.99463 | 436.43515 | 684.7601 | 865.792 | 787.6354 |

| | | | | | | | | | | | | |
|---|---|---|---|---|---|---|---|---|---|---|---|---|
| Prep | SDGTETSTNLHQK | -197 | -209 | 0 | 0 | 13 | 366.08063 | 373.22327 | 383.3789 | 709.32904 | 358.17706 | 562.41144 |
| Prep | TSFLSPGVIYHCDLTK | -399 | -414 | 0 | 0.00025015 | 16 | 953.54193 | 1165.7966 | 1109.5201 | NaN | 1395.376 | 1376.7039 |
| Prep | LDGSHPAFLY | -462 | -471 | 0 | 0.001291103 | 10 | 1098.5414 | 1029.1564 | 1163.5068 | NaN | 1399.8698 | NaN |
| Prep | VVPLHSLK | -644 | -651 | 0 | 0.00272025 | 8 | 599.8055 | 528.6426 | 450.6382 | 461.21887 | NaN | NaN |
| Prep | YSPLHNVK | -614 | -621 | 0 | 0.007493591 | 8 | 949.0916 | NaN | 1024.9263 | NaN | 1041.2723 | NaN |
| Prep | VFLDPNTLSDDGTVALR | -112 | -128 | 0 | 0.00025015 | 17 | 696.5148 | 777.31616 | NaN | NaN | 1076.0995 | NaN |
| Prep | FTIGHAW | -589 | -595 | 0 | 0.002068213 | 7 | 629.6336 | NaN | NaN | NaN | NaN | NaN |
| Nfatc2 | FTLPVPGYEGYR | -153 | -164 | 0 | 0 | 12 | 233.09073 | 665.3189 | 634.75085 | 747.9735 | NaN | 2276.6409 |
| Nfatc2 | NSAPESILLVPPTWPK | -364 | -379 | 0 | 0 | 16 | 234.69727 | NaN | NaN | NaN | 706.3955 | NaN |
| Wdr43 | LHQEYVPSAHLSGT | -48 | -61 | 0 | 0 | 14 | 2828.0388 | 2907.0266 | 3090.4788 | 3906.6829 | 3967.2239 | 4114.859 |
| Wdr43 | VLVPGIPGHHAPIK | -402 | -415 | 0 | 0 | 14 | 987.624 | 1031.069 | 1076.0474 | 565.27325 | 640.55023 | 723.3729 |
| Wdr43 | VLVPGIPGHHAPIK | -402 | -415 | 0 | 0 | 14 | 987.624 | 1031.069 | 1076.0474 | 565.27325 | 640.55023 | 723.3729 |
| Wdr43 | TLSENKEEPVK | -275 | -285 | 1 | 0 | 11 | 658.8656 | 823.6122 | 939.0802 | 482.89206 | 1276.9019 | 825.30597 |
| Wdr43 | VVIPLLQELTK | -499 | -509 | 0 | 0 | 11 | 559.3111 | 533.6147 | 560.66724 | 582.4202 | 626.9478 | 803.394 |
| Wdr43 | VTPKPIPILAA | -327 | -337 | 1 | 0.000431655 | 11 | 454.5716 | 456.26028 | 482.31448 | 272.1976 | NaN | NaN |
| Wdr43 | LHQEYVPSAHLS | -48 | -59 | 0 | 0 | 12 | 1198.8398 | 1286.314 | NaN | 2523.1719 | NaN | NaN |
| Wdr43 | WFQPTIER | -353 | -360 | 0 | 0 | 8 | 615.6108 | NaN | NaN | 452.58615 | 648.417 | NaN |
| Wdr43 | FTTIRPNESQPSDGITGL | -216 | -233 | 1 | 0.001023236 | 18 | 615.07025 | NaN | NaN | 642.5773 | NaN | NaN |
| Clasp2 | FTVDQTQTPSLK | -772 | -783 | 0 | 0 | 12 | 109.22522 | 522.22546 | 761.6952 | NaN | NaN | NaN |
| Emc1 | FYLVDAEQGR | -675 | -684 | 0 | 0 | 10 | 1635.9753 | 1911.9434 | 1796.3469 | 3485.7195 | 369.17075 | 1443.0127 |
| Emc1 | LYLVETGR | -378 | -385 | 0 | 0 | 8 | 680.1688 | 484.58932 | 565.99164 | 621.6302 | 638.7768 | 500.16492 |
| Emc1 | FVFNPIFGK | -606 | -614 | 0 | 0 | 9 | 461.5963 | 512.4067 | 552.9315 | 388.38684 | 607.4695 | 574.6572 |
| Emc1 | HLLIGLPSGAIL | -864 | -875 | 0 | 0 | 12 | 312.47443 | 266.5459 | 294.36398 | 322.79877 | 377.52988 | 361.2887 |
| Txn2 | TTFNVQDGPDFQDR | -60 | -73 | 0 | 0 | 14 | 1366.8785 | 1478.3942 | 1560.4784 | 3592.9739 | 1504.3416 | 1588.3982 |
| Txn2 | VVNSETPVVVDFHAQ | -74 | -88 | 0 | 0 | 15 | 974.06934 | 690.46423 | 1044.784 | 1777.7521 | 1669.7706 | 1080.9515 |
| Txn2 | FVGIKDEDQLEAF | -148 | -160 | 1 | 0.00078661 | 13 | 693.0425 | 698.7942 | 956.992 | NaN | 462.1387 | 1260.8815 |
| Edc3 | FVIPLHSA | -501 | -508 | 0 | 0.001291103 | 8 | 596.2506 | NaN | 458.42426 | NaN | NaN | NaN |
| Eif5 | SDNKDDDIDIDAI | -417 | -429 | 1 | 0.000550307 | 13 | 736.17377 | 1813.9956 | 752.7004 | 717.8041 | 1221.9982 | 780.4937 |
| Eif5 | FVLCPECENPETDLHVNPK | -96 | -114 | 0 | 0 | 19 | 474.53476 | 351.6943 | 557.26416 | 514.82336 | 594.0887 | 590.47876 |
| Eif5 | VNILFDFVK | -238 | -246 | 0 | 0 | 9 | 205.77046 | 406.54834 | 210.05057 | 365.2594 | 194.98021 | 372.51016 |
| Rcn2 | RDPTANEDPEWILVEK | -220 | -235 | 1 | 0.000230707 | 16 | 508.36874 | 601.0053 | 624.4728 | NaN | 890.1708 | 1074.1929 |
| Rcn2 | RDPTANEDPEWILVEK | -220 | -235 | 1 | 0.000230707 | 16 | 508.36874 | 601.0053 | 624.4728 | NaN | 890.1708 | 1074.1929 |
| Rcn2 | FVNDYDKDNDGR | -238 | -249 | 1 | 0 | 12 | 993.9792 | 968.7217 | 1058.9062 | NaN | NaN | NaN |
| Ap2a2 | FVNLFPEVK | -547 | -555 | 0 | 0 | 9 | 350.6033 | 505.9421 | 549.5798 | 640.87067 | 637.7885 | 613.79285 |
| Fau | FVNVVPTFGK | -116 | -125 | 0 | 0 | 10 | 1347.9617 | 1564.6556 | 1594.0693 | 1270.6653 | 1545.8727 | 1398.5164 |
| Fau | RFVNVVPTFGK | -115 | -125 | 1 | 0 | 11 | 852.7827 | 837.15704 | 827.9476 | NaN | NaN | NaN |
| Fau | VNVVPTFGK | -117 | -125 | 0 | 0 | 9 | 675.80414 | NaN | NaN | NaN | NaN | NaN |
| Tomm40 | LALGGELVYHR | -229 | -239 | 0 | 0 | 11 | 3200.6042 | 3262.8315 | 3310.852 | 1885.2878 | 1455.2944 | 1770.3165 |
| Tomm40 | KLPPLPLTL | -331 | -339 | 1 | 0.002199523 | 9 | 1328.3448 | 1616.0377 | 284.8145 | 1113.5292 | 1563.2705 | 1656.5223 |
| Tomm40 | FVNWQVDGEYR | -185 | -195 | 0 | 0 | 11 | 1004.5702 | 1057.913 | 94.65761 | 1210.0425 | 1327.9105 | 1215.0569 |
| Acy1 | LHEDIFLR | -379 | -386 | 0 | 0 | 8 | 758.79614 | 859.1644 | 1066.448 | 1188.6614 | 488.27344 | 1155.8611 |
| Acy1 | TPVLLHDHNER | -368 | -378 | 0 | 0.000627943 | 11 | 1104.0793 | NaN | NaN | NaN | 1213.1221 | 576.06525 |
| Acy1 | FVPDEEVGGHK | -143 | -153 | 0 | 0.009447908 | 11 | 2077.6714 | 2116.3845 | NaN | NaN | NaN | NaN |
| Stoml2 | FVPQQEAWVVER | -40 | -51 | 0 | 0 | 12 | 738.07056 | 725.952 | 512.13934 | 1040.6998 | 781.4983 | NaN |
| Stoml2 | ILEPGLNVLIPVLDR | -58 | -72 | 0 | 0 | 15 | 288.30594 | NaN | NaN | NaN | 294.26077 | NaN |
| Ints1 | LPLFDSVR | -1038 | -1045 | 0 | 0.004628201 | 8 | 604.6153 | 599.9666 | 570.66864 | 600.7568 | 656.73096 | 615.6472 |
| Ints1 | FVQFIHK | -1964 | -1970 | 0 | 0 | 7 | 1097.7313 | NaN | NaN | NaN | 1499.9773 | 1518.7957 |
| Psme3ip1 | FVSEAELDER | -15 | -24 | 0 | 0 | 10 | 976.1706 | 866.85504 | 830.24384 | NaN | NaN | 826.9935 |
| Mpp1 | FVTGDIIQIINK | -186 | -197 | 0 | 0 | 12 | 277.26324 | 268.99448 | 336.58215 | 226.25414 | NaN | 220.52179 |
| Ndufb6 | FWDNFLR | -46 | -52 | 0 | 0 | 7 | 681.01917 | NaN | 129.9538 | 713.87134 | 951.7749 | 870.54535 |
| Rer1 | SEGDSVGDSVHGKPSVVYR | -2 | -20 | 1 | 0 | 19 | 1064.2505 | 263.2467 | 580.70825 | 637.5675 | 693.45465 | 886.89166 |
| Rer1 | SEGDSVGDSVHGKPSVVYR | -2 | -20 | 1 | 0 | 19 | 1064.2505 | 263.2467 | 580.70825 | 637.5675 | 693.45465 | 886.89166 |
| Rer1 | SEGDSVGDSVHGK | -2 | -14 | 0 | 0 | 13 | 932.3786 | 943.98114 | 846.48413 | 697.59973 | 785.8661 | 874.82294 |
| Rer1 | VDPSLMEDSDDGPSLPTK | -87 | -104 | 0 | 0 | 18 | 1099.7223 | NaN | 1092.0979 | 805.2226 | 1376.0503 | 695.4617 |
| Rer1 | FWHAATK | -121 | -127 | 0 | 0.00111006 | 7 | 1949.6614 | NaN | 1678.7203 | 1747.7405 | 1422.4331 | NaN |
| Las1l | FWQPLLR | -365 | -371 | 0 | 0.00204437 | 7 | 696.7821 | 771.0347 | 803.59534 | NaN | NaN | NaN |
| Las1l | SQFWQPLLR | -363 | -371 | 0 | 0.002265552 | 9 | 659.109 | NaN | 1152.8309 | NaN | NaN | NaN |
| Ppp6c | KYLPENDLKR | -17 | -26 | 2 | 0 | 10 | 788.25476 | 767.2384 | 622.86444 | 646.988 | 669.77386 | 758.3133 |
| Ppp6c | FYDLCELFR | -58 | -66 | 0 | 0 | 9 | 408.95047 | 320.08582 | 448.5938 | 540.9083 | 527.0191 | 463.62662 |
| Ppp6c | VTNEFVHIN | -220 | -228 | 0 | 0.001575548 | 9 | 3494.7603 | NaN | 950.4665 | NaN | NaN | 1479.604 |
| Galns | FYEEFPINR | -187 | -195 | 0 | 0 | 9 | 634.7852 | NaN | 530.0998 | 740.7889 | NaN | NaN |
| Csnk2a2 | FYHPAQEYNVR | -182 | -192 | 0 | 0 | 11 | 247.86661 | 805.26385 | 700.8871 | 219.3689 | 907.72766 | 596.1603 |
| Gar1 | VPYFNAPVYLENK | -106 | -118 | 0 | 0 | 13 | 1393.7904 | 1431.0658 | 544.16156 | 558.91174 | 1108.9949 | 979.8168 |
| Gar1 | TTEENKVPYFNAPVYLENK | -100 | -118 | 1 | 0.00025015 | 19 | 653.46106 | 665.57306 | 686.4762 | 478.53723 | 514.53766 | 574.4868 |
| Gar1 | TTEENKVPYFNAPVYLENK | -100 | -118 | 1 | 0.00025015 | 19 | 653.46106 | 665.57306 | 686.4762 | 478.53723 | 514.53766 | 574.4868 |
| Gar1 | FYIDPYK | -155 | -161 | 0 | 0 | 7 | 498.7589 | 779.76 | 732.82526 | 602.0933 | 867.0094 | 829.6718 |
| Gar1 | TEENKVPYFNAPVYLENK | -101 | -118 | 1 | 0 | 18 | 333.478 | 382.9636 | 543.15967 | 369.8179 | 315.59222 | 2319.3008 |
| Gar1 | VDEIFGQLR | -124 | -132 | 0 | 0 | 9 | 1840.1797 | 1687.7961 | 456.40857 | 1692.3182 | NaN | 1746.5668 |
| Atp5mg | FYIGEIIGK | -87 | -95 | 0 | 0 | 9 | 2130.7346 | 2544.2754 | 2586.2249 | 2505.331 | 3022.8154 | 3115.2473 |
| Sec23ip | SVYWEEEPTEVR | -326 | -337 | 0 | 0 | 12 | 912.6439 | 748.8589 | 713.29614 | 964.0585 | 452.7508 | 340.35034 |
| Sec23ip | SIIECVDDFR | -459 | -468 | 0 | 0 | 10 | 449.12558 | NaN | 425.63202 | 370.26956 | 409.2821 | 312.46268 |
| Sec23ip | FYNSPTYCQAIVEK | -537 | -550 | 0 | 0.001392869 | 14 | 905.7266 | NaN | NaN | NaN | NaN | NaN |
| Scamp2 | FYQDFSTEIPADYQR | -133 | -147 | 0 | 0 | 15 | 462.85635 | 847.3261 | 943.70416 | NaN | NaN | NaN |
| Oxct1 | FYTDPVEAVK | -42 | -51 | 0 | 0 | 10 | 1340.0238 | 1272.9038 | 1431.1 | 206.24832 | 955.7087 | 1177.4966 |
| Oxct1 | GYGTLVQEGGSPIK | -160 | -173 | 0 | 0 | 14 | 1116.3401 | 909.1875 | 979.52576 | 2322.29 | 258.04028 | 1157.8994 |
| Oxct1 | YVGENAEFER | -115 | -124 | 0 | 0 | 10 | 708.3785 | 684.3046 | 870.80164 | 787.9348 | 963.1028 | 1128.2142 |
| Oxct1 | SYVGENAEFER | -114 | -124 | 0 | 0 | 11 | 444.53262 | 619.734 | 602.96655 | 505.18994 | 231.4425 | 1219.6577 |
| Oxct1 | GVFDVDKK | -474 | -481 | 1 | 0.000591124 | 8 | 1676.556 | 575.6346 | NaN | 1292.5973 | 503.32123 | NaN |

| | | | | | | | | | | | | |
|---|---|---|---|---|---|---|---|---|---|---|---|---|
| Oxct1 | VGENAEFER | -116 | -124 | 0 | 0.000550307 | 9 | 673.5951 | 990.56714 | NaN | 817.248 | NaN | NaN |
| Bcap31 | LKDELASTK | -190 | -198 | 1 | 0 | 9 | 978.03986 | 906.06506 | 1029.6804 | 1369.5243 | 860.3415 | 1016.6108 |
| Bcap31 | KLKDELASTK | -189 | -198 | 2 | 0.00025015 | 10 | 619.0149 | 767.01385 | 785.329 | 520.5048 | 532.436 | 828.18274 |
| Bcap31 | GAAEDGDKLDIGNTE | -160 | -174 | 1 | 0 | 15 | 636.6732 | 708.8712 | 861.9767 | 598.2313 | NaN | 950.99115 |
| Bcap31 | YMEENDQLKK | -150 | -159 | 1 | 0 | 10 | 134.4151 | 676.897 | 716.433 | NaN | 684.7587 | NaN |
| Bcap31 | LLEEHAK | -225 | -231 | 0 | 0.007880195 | 7 | 962.6549 | NaN | 1002.001 | NaN | NaN | 1069.0509 |
| Bcap31 | LLCIPFISPK | -21 | -30 | 0 | 0 | 10 | 418.32333 | 427.7635 | 622.3965 | NaN | NaN | NaN |
| Ube2c | LSLEFPSGYPYNAPTVK | -81 | -97 | 0 | 0 | 17 | 500.36157 | 198.42581 | 489.44086 | 501.64005 | 651.46924 | 443.19647 |
| Ube2c | GAAGTVYEDLR | -68 | -78 | 0 | 0 | 11 | 1006.23267 | NaN | 450.7181 | 1021.6601 | 747.9418 | NaN |
| Ube2c | GISAFPESDNLFK | -49 | -61 | 0 | 0 | 13 | 1065.8306 | 1482.1884 | NaN | NaN | NaN | 618.64166 |
| Limd2 | GNYDEGFGR | -101 | -109 | 0 | 0 | 9 | 622.62823 | 616.2894 | 592.87555 | 570.4015 | 477.60175 | 486.41705 |
| Limd2 | GAAQATPSHEAK | -6 | -17 | 0 | 0 | 12 | 455.47827 | NaN | NaN | NaN | NaN | 365.2102 |
| Mtpn | KPLHYAADCGQLEIL | -37 | -51 | 1 | 0.000298805 | 15 | 710.14484 | 1160.9624 | 1199.3718 | 1013.3867 | 1185.9531 | 1313.8411 |
| Mtpn | KPLHYAADCGQLEIL | -37 | -51 | 1 | 0.000298805 | 15 | 710.14484 | 1160.9624 | 1199.3718 | 1013.3867 | 1185.9531 | 1313.8411 |
| Mtpn | GADINAPDKHHITPLLSA | -58 | -75 | 1 | 0.00025015 | 18 | 634.34467 | 699.36945 | 800.5867 | 838.1757 | 771.3913 | 768.8248 |
| Mtpn | GADINAPDKHHITPLLSA | -58 | -75 | 1 | 0.00025015 | 18 | 634.34467 | 699.36945 | 800.5867 | 838.1757 | 771.3913 | 768.8248 |
| Mtpn | GADINAPDKHHITPLLS | -58 | -74 | 1 | 0.000907206 | 17 | 913.27277 | 724.9732 | NaN | 227.05904 | 156.15614 | 226.15782 |
| Mtpn | GADINAPDKHHITPLLS | -58 | -74 | 1 | 0.000907206 | 17 | 913.27277 | 724.9732 | NaN | 227.05904 | 156.15614 | 226.15782 |
| Mtpn | HHITPLLSA | -67 | -75 | 0 | 0.000907206 | 9 | 1421.2274 | 1927.9642 | NaN | NaN | 1298.4121 | NaN |
| Aimp1 | IWEQIQPDLHTN | -268 | -279 | 0 | 0 | 12 | 2509.6077 | 923.48145 | 805.89667 | 1397.3615 | 1667.7806 | 1624.4856 |
| Aimp1 | YVEEVDVGEAAPR | -173 | -185 | 0 | 0 | 13 | 1233.8562 | 1337.7709 | 1420.302 | 1545.4097 | 2023.605 | 1452.5839 |
| Aimp1 | STDSKPIDASR | -142 | -152 | 1 | 0 | 11 | 1025.4579 | 1442.815 | 1609.9443 | 716.5155 | 704.94684 | 1134.4957 |
| Aimp1 | GAEADQIIEYLK | -15 | -26 | 0 | 0 | 12 | 949.1048 | 1196.3906 | 868.68726 | 1009.17554 | 1364.7236 | 1318.5258 |
| Aimp1 | ITFDAFPGEPDKELNPK | -249 | -265 | 1 | 0 | 17 | 412.9904 | 1101.9725 | 541.92365 | 415.22397 | 421.11697 | 998.0211 |
| Aimp1 | ITFDAFPGEPDKELNPK | -249 | -265 | 1 | 0 | 17 | 412.9904 | 1101.9725 | 541.92365 | 415.22397 | 421.11697 | 998.0211 |
| Aimp1 | ITFDAFPGEPDK | -249 | -260 | 0 | 0 | 12 | 775.32605 | 884.042 | 896.0346 | 1655.0214 | 1844.9689 | 2775.1792 |
| Aimp1 | IWEQIQPDLHTNAECVAT | -268 | -285 | 0 | 0 | 18 | 636.9947 | 699.6932 | 804.52094 | 355.11765 | 1091.6454 | 1171.6338 |
| Aimp1 | VEILAPPNGSVPGDR | -234 | -248 | 0 | 0.00025015 | 15 | 1400.9948 | 988.2481 | 1326.291 | NaN | 1727.8357 | 2024.6732 |
| Aimp1 | VNHVPLEQMQNR | -192 | -203 | 0 | 0.001055699 | 12 | 523.8264 | NaN | NaN | NaN | NaN | NaN |
| Ltn1 | GAEDIDSLVDSDFR | -53 | -66 | 0 | 0 | 14 | 447.29147 | 488.71057 | 513.3374 | 432.04205 | 433.3826 | 289.33777 |
| Nufip2 | GAEIIPSGTEHPVFPK | -546 | -561 | 0 | 0.000230707 | 16 | 872.95496 | 866.4168 | 914.7088 | 1032.5532 | 919.7182 | 978.35596 |
| Aacs | GIADVPEWFR | -86 | -95 | 0 | 0 | 10 | 856.1579 | 773.95087 | 972.80414 | 571.13477 | 1115.4192 | 1075.6244 |
| Aacs | GAFSNPETLDLYR | -652 | -664 | 0 | 0 | 13 | 2731.314 | NaN | 2464.8596 | 1721.5604 | 1759.7819 | 1542.4524 |
| Aacs | VVLIPYVLPR | -236 | -245 | 0 | 0 | 10 | 286.8299 | 520.6849 | 339.05212 | NaN | NaN | NaN |
| Ttc4 | GALCHLELK | -157 | -165 | 0 | 0.006654467 | 9 | 347.26672 | NaN | NaN | NaN | NaN | NaN |
| Adrm1 | GALFPSLVPGSR | -5 | -16 | 0 | 0 | 12 | 2598.7854 | 3245.4282 | 3109.8384 | 2895.468 | 3769.412 | 3829.431 |
| Adrm1 | LFPSLVPGSR | -7 | -16 | 0 | 0 | 10 | 1110.9675 | 1014.17053 | 1001.52075 | 953.36194 | 123.84501 | 990.89197 |
| Adrm1 | SQSAAVTPSSSTSSAR | -211 | -226 | 0 | 0 | 16 | 484.00208 | 697.5144 | 262.98337 | 516.5954 | 459.24274 | 684.62494 |
| Adrm1 | TTSGALFPSLVPGSR | -2 | -16 | 0 | 0 | 15 | 401.7663 | 235.66199 | 484.83307 | 482.64478 | 692.3412 | 547.59796 |
| Adrm1 | TSGTVEDDLIIFPDDCEFK | -65 | -83 | 0 | 0 | 19 | 193.57655 | 212.59679 | 181.76324 | 71.51232 | 240.49802 | NaN |
| Polr1d | GALPAVEPFQK | -88 | -98 | 0 | 0.00204437 | 11 | 170.11276 | NaN | NaN | NaN | 2057.738 | NaN |
| Nop53 | GASYNPTFEDHQALLR | -244 | -259 | 0 | 0 | 16 | 664.32654 | 1051.2964 | 794.3594 | 505.8593 | 1717.7001 | 1584.1313 |
| Tapbpl | GATLEPEELLR | -257 | -267 | 0 | 0 | 11 | 774.678 | 850.7341 | 1750.3286 | 1617.0264 | 1646.3774 | NaN |
| Tapbpl | TDPPTAPTTAER | -21 | -32 | 0 | 0 | 12 | 2888.4465 | NaN | 344.35916 | NaN | NaN | NaN |
| Rcc1 | SWVGFSGGQHHTVC | -295 | -308 | 0 | 0 | 14 | 2063.0762 | 342.35663 | 876.1594 | 1099.0144 | 1411.0623 | 771.8244 |
| Rcc1 | SFGCNDEGALGR | -90 | -101 | 0 | 0 | 12 | 1001.1062 | 995.72296 | 1161.9753 | 1207.7222 | 966.53143 | 1152.705 |
| Rcc1 | GLGESVLER | -49 | -57 | 0 | 0 | 9 | 933.2017 | 829.32935 | 852.5428 | 1215.5471 | 597.3706 | 605.90155 |
| Rcc1 | GQGDVGQLGLGESVLER | -41 | -57 | 0 | 0 | 17 | 658.18427 | 902.3 | 774.2384 | 1194.869 | 1556.6587 | 1156.0793 |
| Rcc1 | VVQVSAGDSHTAALTEDGR | -121 | -139 | 0 | 0.00025015 | 19 | 377.4821 | 519.4797 | 424.82138 | 279.88162 | 435.9152 | 363.53952 |
| Rcc1 | SHNTEPGLVL | -29 | -38 | 0 | 0.000471409 | 10 | 1466.6978 | 1336.8314 | 1535.7145 | 907.79175 | NaN | 789.6497 |
| Rcc1 | LGLGEGAEEK | -326 | -335 | 0 | 0 | 10 | 1019.73254 | 1167.5383 | 1205.5393 | NaN | 656.85724 | 1064.1044 |
| Rcc1 | KKPALVPLLQDVVQA | -58 | -72 | 2 | 0.001636916 | 15 | 851.30035 | 920.3395 | 507.3346 | 769.0477 | NaN | 1102.756 |
| Rcc1 | SHNTEPGLVLTL | -29 | -40 | 0 | 0 | 12 | 464.98914 | NaN | 264.32483 | 724.6092 | 235.69395 | 785.52203 |
| Rcc1 | GCNDEGALGR | -92 | -101 | 0 | 0 | 10 | 2524.6628 | 2567.572 | NaN | 344.60278 | NaN | NaN |
| Rcc1 | SGGQHTVLLVK | -406 | -416 | 0 | 0.000431655 | 11 | 2314.582 | 2156.2976 | NaN | 1835.5074 | NaN | NaN |
| Itgb7 | GCPAQELEEPR | -90 | -100 | 0 | 0 | 11 | 591.8334 | 5277.5464 | 5582.734 | NaN | NaN | 7619.0337 |
| Psap | GCSFLPDPYQK | -472 | -482 | 0 | 0 | 11 | 2143.1533 | 2098.707 | 2250.6301 | 268.21423 | 2696.385 | 2514.7297 |
| Psap | LVSDVQTAVK | -203 | -212 | 0 | 0.000230707 | 10 | 2108.2097 | 3330.0854 | 2661.276 | 1787.3506 | 3585.7727 | 3661.2766 |
| Psap | TCEWIHDSSLSAS | -93 | -105 | 0 | 0.00025015 | 13 | 1038.708 | 1051.9037 | 1030.7161 | 777.3811 | 624.28503 | 947.2826 |
| Psap | VEVMDPGFVCSK | -500 | -511 | 0 | 0.00025015 | 12 | 894.12616 | 2491.6807 | 1989.9176 | 3637.9797 | 159.77747 | 3125.3633 |
| Psap | TVVTEAGNLLK | -68 | -78 | 0 | 0 | 11 | 568.2106 | 276.78833 | 1549.7023 | 1650.0067 | 2138.5588 | 475.2765 |
| Psap | LGPGVSDICK | -232 | -241 | 0 | 0.000627943 | 10 | 5240.599 | 5396.969 | 5790.749 | 6154.4375 | NaN | 6400.731 |
| Psap | KLVLYLEHNLEK | -447 | -458 | 1 | 0 | 12 | 914.90735 | 893.00775 | NaN | 921.231 | 1719.8196 | 810.48035 |
| Psap | TCEWIHDSSLSASCK | -93 | -107 | 0 | 0 | 15 | 608.56116 | NaN | 707.86676 | 1247.7032 | 1437.6335 | 1423.275 |
| Psap | GFVDHVK | -220 | -226 | 0 | 0.003360691 | 7 | 1655.6908 | 1785.0688 | 1606.4763 | NaN | NaN | NaN |
| Snx3 | GDDGIFDDNFIEER | -105 | -118 | 0 | 0 | 14 | 597.95715 | 646.5889 | 735.0584 | 757.55005 | 1060.1542 | 1009.8055 |
| Snx3 | YSDFEWLR | -71 | -78 | 0 | 0 | 8 | 531.90674 | 524.2266 | 563.761 | 674.7207 | 701.571 | 676.81537 |
| Snx3 | VAGHPLAQNER | -129 | -139 | 0 | 0 | 11 | 384.40854 | 428.6491 | 436.59753 | 322.45792 | 127.01749 | 481.78107 |
| Eif1 | NLHSFDPFADASK | -6 | -18 | 0 | 0 | 13 | 435.59384 | 448.39853 | 544.6309 | 321.0876 | 279.45557 | 255.43855 |
| Eif1 | SAIQNLHSFDPFADASK | -2 | -18 | 0 | 0.00025015 | 17 | 413.89636 | 369.23926 | 363.30185 | 345.88153 | 541.1511 | 628.0752 |
| Eif1 | LHSFDPFADASK | -7 | -18 | 0 | 0.00025015 | 12 | 933.6788 | 1268.4236 | 1042.3445 | 1163.0936 | 1584.7339 | NaN |
| Vps29 | GDFDENLNYPEQK | -61 | -73 | 0 | 0 | 13 | 1123.306 | 1221.9884 | 1152.8562 | 1365.1128 | 1612.146 | 1565.4872 |
| Vps29 | TLAGDVHIVR | -51 | -60 | 0 | 0 | 10 | 854.60956 | 909.579 | 929.8606 | 756.27386 | 822.09265 | 502.78064 |
| Vps29 | VYQLIGDDVK | -164 | -173 | 0 | 0 | 10 | 1789.6606 | 1258.0562 | 1898.7815 | 1571.2351 | 1104.9531 | NaN |
| Vps29 | GDLHIPHR | -7 | -14 | 0 | 0.001776473 | 8 | 546.9546 | NaN | NaN | NaN | 965.1226 | NaN |
| Eif1ad | GDFLIVDPIEEGEK | -66 | -79 | 0 | 0.000230707 | 14 | 874.5342 | NaN | 1564.0066 | 1210.8888 | 1413.7789 | 1196.5742 |
| Eif1ad | RGDFLIVDPIEEGEK | -65 | -79 | 1 | 0 | 15 | 572.45734 | 769.9721 | 776.8054 | NaN | 206.83516 | 843.6956 |
| Eif1ad | RGDFLIVDPIEEGEK | -65 | -79 | 1 | 0 | 15 | 572.45734 | 769.9721 | 776.8054 | NaN | 206.83516 | 843.6956 |

| | | | | | | | | | | | | |
|---|---|---|---|---|---|---|---|---|---|---|---|---|
| Lsm3 | QTTNTVEEPLDLIR | -9 | -22 | 0 | 0 | 14 | 882.68665 | 1020.25507 | 1017.89624 | 1106.2787 | 1337.7517 | 1362.4415 |
| Lsm3 | GDGVVLVAPPLR | -89 | -100 | 0 | 0.00025015 | 12 | 831.7521 | 766.6444 | 732.2052 | 1213.2949 | 1306.3605 | 1198.458 |
| Lsm3 | LHAYDQHLN | -43 | -51 | 0 | 0.000230707 | 9 | 629.7205 | 1117.6764 | 1178.706 | 292.38763 | NaN | 909.65796 |
| Lsm3 | TVEEPLDLIR | -13 | -22 | 0 | 0 | 10 | 299.00647 | 724.2793 | 649.83984 | NaN | 570.9319 | 668.9939 |
| Mov10 | GDHLFALL | -395 | -402 | 0 | 0.00504706 | 8 | 347.3871 | 248.73367 | NaN | NaN | 561.5705 | 490.99936 |
| Mov10 | GTYYPPILLR | -293 | -302 | 0 | 0 | 10 | 332.4511 | NaN | NaN | 749.51324 | NaN | 470.2015 |
| Dpp4 | WVSDFEYLYK | -60 | -69 | 0 | 0 | 10 | 885.6721 | NaN | NaN | 1134.9956 | 333.82742 | NaN |
| Dpp4 | GDHYLCDVVWAT | -290 | -301 | 0 | 0.001291103 | 12 | 423.33014 | 467.95508 | 362.4642 | NaN | NaN | NaN |
| Pgrmc1 | GDQPGASGDNDDDEPPPLPR | -48 | -67 | 0 | 0.000230707 | 20 | 716.96967 | 632.65405 | 308.4344 | 391.1158 | 419.1305 | 353.83786 |
| Pgrmc1 | KFYGPEGPYGVF | -105 | -116 | 1 | 0.001463897 | 12 | 782.67566 | NaN | NaN | 666.63727 | 1408.588 | NaN |
| Pgrmc1 | TLSDWDSQFTFK | -152 | -163 | 0 | 0.00025015 | 12 | 663.6085 | 510.34235 | NaN | NaN | NaN | NaN |
| Srprb | TLIDLPGHESLR | -112 | -123 | 0 | 0 | 12 | 776.45496 | 992.68243 | 941.31476 | 1139.1907 | 1327.3121 | 1258.1543 |
| Srprb | GDTGSADIQDLEK | -251 | -263 | 0 | 0 | 13 | 2570.4045 | 2849.9834 | NaN | NaN | NaN | NaN |
| Immt | SVPAPAVQHEDTIK | -156 | -169 | 0 | 0 | 14 | 6079.5435 | 5805.602 | 5412.04 | 4727.4165 | 6057.935 | 6698.497 |
| Immt | LSEQELEFR | -516 | -524 | 0 | 0 | 9 | 1214.2716 | 1308.6017 | 1300.8558 | 355.47278 | 409.8024 | 1295.2996 |
| Immt | YEFEQGLSEK | -506 | -515 | 0 | 0 | 10 | 963.02344 | 2365.6995 | 1126.5923 | 217.49605 | 903.76276 | 1203.5544 |
| Immt | IEHGDLELAAK | -697 | -707 | 0 | 0 | 11 | 800.77356 | 1057.805 | 959.96265 | 488.5796 | 771.44226 | 996.78467 |
| Immt | VLGSAPYTVPLPK | -89 | -101 | 0 | 0 | 13 | 2680.4314 | NaN | 1199.1449 | 1171.8871 | 2254.413 | 1582.7161 |
| Immt | GDTLSVPAPAVQHEDTIK | -152 | -169 | 0 | 0.001392869 | 18 | 500.19104 | 639.7863 | 652.9489 | 690.7544 | 715.188 | NaN |
| Immt | QLKPPAELYPEDINTF | -672 | -687 | 1 | 0.000826806 | 16 | 983.9521 | 1010.19366 | 1039.6394 | NaN | 1156.6187 | NaN |
| Immt | GSAPYTVPLPK | -91 | -101 | 0 | 0.005459848 | 11 | 3542.6643 | 3791.0784 | NaN | NaN | NaN | 3320.1213 |
| Immt | GVYSEETLR | -623 | -631 | 0 | 0.008498216 | 9 | 3832.692 | 711.7564 | NaN | NaN | NaN | NaN |
| Immt | QHIELALEK | -427 | -435 | 0 | 0.000627943 | 9 | 1265.966 | NaN | NaN | 1162.1962 | NaN | NaN |
| Immt | SLEDALNR | -222 | -229 | 0 | 0.001610812 | 8 | 1142.7955 | 1367.5631 | NaN | NaN | NaN | NaN |
| Immt | VVSQYHELVVQ | -353 | -363 | 0 | 0.003620005 | 11 | 1052.3535 | NaN | NaN | NaN | NaN | NaN |
| Sh3bgrl | GDYDAFFEAR | -77 | -86 | 0 | 0.000471409 | 10 | 581.41345 | NaN | 712.57904 | NaN | NaN | NaN |
| Sdf2l1 | GEDGEGDDLDLWTVR | -134 | -148 | 0 | 0 | 15 | 818.8884 | 1080.7397 | 1013.7641 | 1200.8616 | 1599.8942 | 1565.6989 |
| Dnph1 | GEEAAGGDQFIHER | -55 | -68 | 0 | 0 | 14 | 12557.703 | NaN | 11881.737 | NaN | NaN | NaN |
| Cox17 | GEEHCGHLIEAHK | -41 | -53 | 0 | 0 | 13 | 789.48737 | 920.5688 | 533.0047 | 794.0462 | 936.5853 | 1060.5145 |
| Cox17 | GEEHCGHLIEAHK | -41 | -53 | 0 | 0 | 13 | 789.48737 | 920.5688 | 533.0047 | 794.0462 | 936.5853 | 1060.5145 |
| Cox17 | PGLAAASPAPPEAQEK | -2 | -17 | 0 | 0 | 16 | 1475.3909 | 1517.1605 | NaN | 3724.5842 | 3627.4944 | 3763.7168 |
| Ubxn1 | LYVELHR | -241 | -247 | 0 | 0.000471409 | 7 | 895.62225 | 486.29114 | NaN | 770.79486 | 1135.5862 | 1091.7096 |
| Ubxn1 | GESRPILTEEER | -80 | -91 | 1 | 0.000550307 | 12 | 888.7621 | 968.7381 | NaN | NaN | NaN | 779.61755 |
| Ubxn1 | GESRPILTEEER | -80 | -91 | 1 | 0.000550307 | 12 | 888.7621 | 968.7381 | NaN | NaN | NaN | 779.61755 |
| Ubxn1 | GEEPGQDQDPVQLLSGFPR | -248 | -266 | 0 | 0.00025015 | 19 | 657.0955 | NaN | NaN | NaN | 919.7769 | 974.3873 |
| Ahsa1 | LKETFLTSPEELYR | -211 | -224 | 1 | 0 | 14 | 837.4022 | 445.37637 | 722.9809 | 1166.6798 | 1485.335 | 1516.3748 |
| Ahsa1 | THAPAALEADR | -236 | -246 | 0 | 0 | 11 | 4453.4316 | 2119.2932 | 4501.007 | NaN | 5372.6807 | 5447.14 |
| Ahsa1 | PAPEEER | -308 | -314 | 0 | 0.000431655 | 7 | 1492.8057 | 1679.7144 | 1503.46 | 1904.2021 | 1525.5602 | NaN |
| Ahsa1 | GEFTDLVPEK | -259 | -268 | 0 | 0 | 10 | 1066.6676 | 1124.1986 | 1066.1287 | NaN | 1158.3711 | 634.63824 |
| Ahsa1 | SVDPVGQPALK | -172 | -182 | 0 | 0 | 11 | 760.2725 | 816.973 | 971.554 | 1054.7423 | NaN | 803.8149 |
| Ahsa1 | GHVEIPNLSDENSVDEVEIS | -101 | -120 | 0 | 0.000471409 | 20 | 449.13052 | NaN | NaN | 190.18536 | NaN | 794.92163 |
| Ahsa1 | GHVEIPNLSDEN | -101 | -112 | 0 | 0.000591124 | 12 | 2517.52 | NaN | NaN | NaN | NaN | NaN |
| Srp54 | GEHIDDFEPFK | -276 | -286 | 0 | 0 | 11 | 1838.9648 | 2382.1846 | 2275.056 | 274.98935 | 1834.1482 | 1397.5615 |
| Srp54 | KLDDNEALIEK | -311 | -321 | 1 | 0 | 11 | 847.4095 | 1034.8036 | 907.3306 | 1399.5774 | 1850.204 | 1978.3556 |
| Ppan | YNPDSQELDFR | -185 | -195 | 0 | 0 | 11 | 808.72205 | 741.4413 | 796.6707 | 710.6942 | 668.7227 | 805.6959 |
| Ppan | GEHNTTELPQAVAGR | -246 | -260 | 0 | 0 | 15 | 643.1537 | 556.72266 | 73.11785 | 384.8455 | 727.3206 | NaN |
| Ppan | GEHNTTELPQAVAGR | -246 | -260 | 0 | 0 | 15 | 643.1537 | 556.72266 | 73.11785 | 384.8455 | 727.3206 | NaN |
| Vamp8 | GENLDHLR | -38 | -45 | 0 | 0 | 8 | 6983.66 | 6082.4146 | 4939.529 | 5060.871 | 4885.783 | 5306.5054 |
| Selenot | LNDVPVWSK | -153 | -161 | 0 | 0.000550307 | 9 | 1023.4487 | 473.68643 | 385.4892 | NaN | NaN | NaN |
| Selenot | GENYLPQPIYR | -73 | -83 | 0 | 0.001392869 | 11 | 2186.1284 | 764.5148 | NaN | NaN | NaN | NaN |
| Prpf4 | SLDSDEPVADIEGHTVR | -308 | -324 | 0 | 0 | 17 | 823.3254 | 803.31964 | 843.05005 | 886.2875 | 1391.8723 | 1170.3047 |
| Prpf4 | THPGWSPLK | -477 | -485 | 0 | 0 | 9 | 778.1306 | 635.5008 | 1006.034 | 633.77454 | 756.25903 | 376.08896 |
| Prpf4 | GEPITLFGEGPAER | -115 | -128 | 0 | 0 | 14 | 529.98553 | 261.70587 | 830.0167 | 548.59283 | 1462.4363 | 1042.0879 |
| Prpf4 | TKAPDDLVAPVVK | -13 | -25 | 1 | 0.00051306 | 13 | 14142.853 | 14674.289 | NaN | 11673.555 | 14573.555 | NaN |
| Prpf4 | GHNTNVGAIVFHPK | -270 | -283 | 0 | 0 | 14 | 216.65091 | 430.20645 | NaN | NaN | 314.33398 | NaN |
| Nup160 | NLDLSWNELK | -452 | -461 | 0 | 0 | 10 | 1344.6686 | 1632.5134 | 277.3093 | NaN | 1052.952 | NaN |
| Nup160 | ILDKLEDYQQK | -1378 | -1388 | 1 | 0 | 11 | 2721.8982 | NaN | 3288.2659 | NaN | 3505.8008 | NaN |
| Nup160 | GFDPAQPLNVR | -656 | -666 | 0 | 0 | 11 | 591.76953 | NaN | NaN | 1074.0485 | 854.3429 | NaN |
| Nup160 | WHDAENQTIVK | -376 | -386 | 0 | 0 | 11 | 510.10468 | 608.8014 | NaN | 442.31384 | NaN | NaN |
| Lsm8 | GFDQTINLILDESHER | -29 | -44 | 0 | 0 | 16 | 1244.018 | 1276.5203 | 493.30447 | 1477.6237 | 1168.5475 | 1823.5347 |
| Lsm8 | LILDESHER | -36 | -44 | 0 | 0 | 9 | 1189.0056 | 1108.0519 | 984.82684 | 476.4105 | 458.95673 | 744.2675 |
| Lsm8 | TSALENYINR | -2 | -11 | 0 | 0 | 10 | 789.5977 | 908.9213 | 297.6139 | 1218.9016 | 1204.657 | 242.24321 |
| Lsm8 | INLILDESHER | -34 | -44 | 0 | 0.000591124 | 11 | 5926.803 | NaN | NaN | NaN | NaN | NaN |
| Glipr2 | GFFEENVPPPKK | -143 | -154 | 1 | 0 | 12 | 1193.8196 | 379.33594 | 2691.728 | 574.15704 | 1626.867 | 2672.2803 |
| Glipr2 | SYNFQQPGFTSGTGHFT | -89 | -105 | 0 | 0 | 17 | 669.2312 | 886.0803 | 859.92065 | 731.617 | 776.9414 | 992.4674 |
| Glipr2 | GFFEENVPPPK | -143 | -153 | 0 | 0 | 11 | 590.05524 | 857.57336 | 869.02423 | 1800.0073 | 942.1279 | 1305.6843 |
| Mtnd5 | STLLGFFPSIIHR | -523 | -535 | 0 | 0 | 13 | 1563.9944 | 1765.3126 | 1981.9941 | 1274.3447 | 1409.783 | 1367.2644 |
| Mtnd5 | GFFPSIIHR | -527 | -535 | 0 | 0 | 9 | 372.2916 | 707.19684 | 738.8616 | 940.647 | 1177.3671 | NaN |
| Etf1 | YFDEISQDTGK | -290 | -300 | 0 | 0 | 11 | 1381.4777 | 1420.3164 | 1441.6416 | 1739.2675 | 1153.2483 | 1360.6997 |
| Etf1 | YFDEISQDTGK | -290 | -300 | 0 | 0 | 11 | 1381.4777 | 1420.3164 | 1441.6416 | 1739.2675 | 1153.2483 | 1360.6997 |
| Etf1 | TELSQSDMFDQR | -234 | -245 | 0 | 0.000230707 | 12 | 766.2326 | 791.9125 | 498.93988 | 588.7958 | 550.3287 | 580.4185 |
| Etf1 | ILYLTPEQEKDK | -343 | -354 | 1 | 0 | 12 | 678.27216 | 499.10327 | 627.10693 | 690.1748 | 709.0938 | 626.9304 |
| Etf1 | KVNIDFEPFKPINTS | -109 | -123 | 2 | 0.00051306 | 15 | 641.1711 | 698.75757 | 714.7787 | 671.82825 | 798.0415 | 818.5516 |
| Etf1 | YCFGVEDTLK | -301 | -310 | 0 | 0 | 10 | 606.1438 | 742.8158 | 771.66876 | 774.9504 | 936.4208 | 997.5641 |
| Etf1 | ILYLTPEQEK | -343 | -352 | 0 | 0.00025015 | 10 | 400.7887 | 1407.7167 | 1365.4888 | 1386.2012 | 1050.4546 | 1148.5471 |
| Etf1 | GFGGIGGILR | -405 | -414 | 0 | 0 | 10 | 572.6867 | 576.48157 | 680.2328 | NaN | NaN | NaN |
| Etf1 | SQEGSQFVK | -396 | -404 | 0 | 0.007493591 | 9 | 3499.942 | NaN | NaN | NaN | NaN | NaN |
| Hdgfl3 | IDELPEGAVKPPANK | -29 | -43 | 1 | 0 | 15 | 2711.2725 | NaN | NaN | 2831.544 | 2822.2476 | 2659.0537 |

| | | | | | | | | | | | | |
|---|---|---|---|---|---|---|---|---|---|---|---|---|
| Hdgfl3 | GLWEIENNPGVK | -84 | -95 | 0 | 0 | 12 | 1784.8818 | 2083.8723 | 2141.1458 | 3676.828 | NaN | NaN |
| Hdgfl3 | GFNEGLWEIENNPGVK | -80 | -95 | 0 | 0 | 16 | 454.15167 | NaN | 970.5522 | 374.7043 | 564.3008 | NaN |
| Dhodh | GFNTVTDAIGVDHR | -381 | -394 | 0 | 0 | 14 | 939.3143 | 622.2012 | 646.455 | 695.97327 | 262.23285 | 776.8764 |
| Dhodh | LLDPESAHR | -48 | -56 | 0 | 0.004255025 | 9 | 1318.8265 | NaN | NaN | NaN | NaN | NaN |
| Padi2 | GFPVVLDSPR | -372 | -381 | 0 | 0.001291103 | 10 | 946.268 | 824.7384 | NaN | NaN | NaN | NaN |
| Sqstm1 | LIFPNPFGHL | -166 | -175 | 0 | 0 | 10 | 650.3731 | 526.9048 | 587.45764 | 81.18135 | 716.3246 | 800.42816 |
| Sqstm1 | VAVLFPTLRPGGF | -51 | -63 | 1 | 0 | 13 | 545.3832 | 666.2191 | 573.2358 | 639.4661 | 453.53696 | 540.26263 |
| Sqstm1 | LTPTTPESSSTGTEDK | -268 | -283 | 0 | 0.000230707 | 16 | 322.4082 | 399.0178 | 407.1606 | 369.6134 | 400.9761 | 433.7539 |
| Sqstm1 | GFSDEGGWLTR | -407 | -417 | 0 | 0 | 11 | 343.60223 | 714.23145 | NaN | 745.767 | 937.41864 | 893.6349 |
| Cbx5 | LTWHAYPEDAENK | -172 | -184 | 0 | 0 | 13 | 2016.4313 | 2304.6096 | 2296.7524 | 1916.0936 | 1837.9658 | 1817.0259 |
| Cbx5 | LTWHAYPEDAENK | -172 | -184 | 0 | 0 | 13 | 2016.4313 | 2304.6096 | 2296.7524 | 1916.0936 | 1837.9658 | 1817.0259 |
| Cbx5 | GFSEEHNTWEPEK | -43 | -55 | 0 | 0 | 13 | 1213.2377 | 1315.9369 | 1299.3154 | 870.60126 | 978.85315 | 949.46826 |
| Cbx5 | WKDTDEADLVLAK | -142 | -154 | 1 | 0 | 13 | 1120.3438 | 1204.1747 | 1241.7294 | 1404.3 | 1574.627 | 1303.6654 |
| Cbx5 | SEEHNTWEPEK | -45 | -55 | 0 | 0 | 11 | 837.9741 | 797.45917 | 846.1696 | 1100.7169 | NaN | 1267.4417 |
| Cbx5 | LTWHAYPE | -172 | -179 | 0 | 0.00078661 | 8 | 959.3517 | 1561.3187 | NaN | NaN | NaN | NaN |
| Rpl13 | SGPIRPIVR | -52 | -60 | 1 | 0.003166358 | 9 | 2519.3987 | 2739.9404 | 2729.27 | 2459.997 | 3457.6184 | 3062.743 |
| Rpl13 | TGPVMPIR | -151 | -158 | 0 | 0.000550307 | 8 | 2376.486 | 2851.2617 | 751.53894 | 1726.9037 | 2233.9207 | 3283.447 |
| Rpl13 | TGPVMPIR | -151 | -158 | 0 | 0.000550307 | 8 | 2376.486 | 2851.2617 | 751.53894 | 1726.9037 | 2233.9207 | 3283.447 |
| Rpl13 | LTGPVMPIR | -150 | -158 | 0 | 0.00025015 | 9 | 1520.2874 | 1681.6204 | 1629.7886 | 1196.5117 | 696.21094 | 913.0316 |
| Rpl13 | TQLTGPVMPIR | -148 | -158 | 0 | 0 | 11 | 1122.67 | 1410.7966 | 1439.9457 | 1436.3512 | 1460.4149 | 1394.668 |
| Rpl13 | TQLTGPVMPIR | -148 | -158 | 0 | 0 | 11 | 1122.67 | 1410.7966 | 1439.9457 | 1436.3512 | 1460.4149 | 1394.668 |
| Rpl13 | GFSLEELR | -75 | -82 | 0 | 0.005946382 | 8 | 1010.4731 | 141.12387 | 106.231064 | 101.03066 | NaN | NaN |
| Rpl13 | VDTWFNQPAR | -22 | -31 | 0 | 0 | 10 | 1046.4761 | 721.7953 | 658.8047 | NaN | NaN | NaN |
| Rpl13 | PASGPIRPIVR | -50 | -60 | 1 | 0.004595494 | 11 | 817.00543 | 875.0616 | 835.03485 | NaN | NaN | NaN |
| Rpl13 | NGMILKPH | -6 | -13 | 1 | 0.005015524 | 8 | 4384.04 | NaN | NaN | NaN | 3798.226 | NaN |
| Rpl13 | STESLQANVQR | -106 | -116 | 0 | 0 | 11 | 560.1277 | NaN | 1526.846 | NaN | NaN | NaN |
| Lasp1 | YHEEFEK | -122 | -128 | 0 | 0 | 7 | 1005.80493 | 1063.5798 | 1444.748 | 343.93985 | 1151.4769 | 1536.6179 |
| Lasp1 | SVVADTPELQR | -99 | -109 | 0 | 0.000230707 | 11 | 1962.3613 | 2075.2522 | 2079.315 | NaN | NaN | 827.51117 |
| Lasp1 | RPTEQQQPQPH | -154 | -164 | 1 | 0 | 11 | 315.45837 | NaN | NaN | 324.49304 | 379.4887 | 336.71503 |
| Lasp1 | GFSVVADTPELQR | -97 | -109 | 0 | 0 | 13 | 656.55066 | 134.19633 | NaN | NaN | 812.9364 | NaN |
| Slc16a3 | GGAVVDEGPTGIK | -2 | -14 | 0 | 0 | 13 | 1519.361 | 3247.2292 | 3269.8748 | 4389.953 | 4350.2153 | 2264.1394 |
| Slc16a3 | LHKPPVDVGVDSR | -434 | -446 | 1 | 0 | 13 | 811.3146 | 1309.284 | NaN | 549.7001 | 900.52405 | 1405.7094 |
| Slc16a3 | LHKPPVDVGVDSR | -434 | -446 | 1 | 0 | 13 | 811.3146 | 1309.284 | NaN | 549.7001 | 900.52405 | 1405.7094 |
| Slc16a3 | KRPEVTEPEEVASEEK | -418 | -433 | 2 | 0 | 16 | 545.64557 | NaN | 433.3069 | 489.35498 | 750.5287 | 682.5206 |
| Klhdc4 | GGEFASPDGEQFYHYK | -138 | -153 | 0 | 0 | 16 | 488.34528 | 519.9966 | 535.688 | 803.2706 | 817.52515 | 691.8318 |
| Klhdc4 | INPSGVKPTAR | -286 | -296 | 1 | 0.007405213 | 11 | 660.37854 | NaN | NaN | NaN | NaN | 140.1261 |
| Klhdc4 | QVLTPSGLGVQPSPK | -400 | -414 | 0 | 0 | 15 | 619.3091 | NaN | NaN | NaN | NaN | 540.0058 |
| Gm2a | TSVPLTAPQK | -72 | -81 | 0 | 0 | 10 | 5742.5225 | 5348.953 | 5035.2803 | 3018.9832 | 773.72565 | 1667.3229 |
| Gm2a | GGFSWDNCDEGKDPAVIK | -32 | -49 | 1 | 0 | 18 | 2438.752 | 2918.1794 | 2156.2905 | 344.03043 | 1954.7489 | 5114.9834 |
| Gm2a | GLPCHCPFK | -133 | -141 | 0 | 0 | 9 | 917.5509 | 714.1498 | 953.147 | 871.55853 | 935.44336 | 829.78094 |
| Gm2a | SWDNCDEGKDPAVIK | -35 | -49 | 1 | 0.002466863 | 15 | 526.1114 | 393.35947 | 560.7963 | NaN | 650.6095 | 703.9593 |
| Atrx | TETQNALKEEEER | -1476 | -1488 | 1 | 0 | 13 | 547.86365 | 640.89484 | 610.5913 | 325.03955 | 684.67896 | 704.0591 |
| Atrx | SVPATVDDDDDDNDPENR | -1232 | -1249 | 0 | 0 | 18 | 447.12234 | 442.591 | 439.12265 | 353.23114 | 480.01917 | 450.65445 |
| Atrx | SNQQLEDLINQGR | -2303 | -2315 | 0 | 0 | 13 | 693.44446 | 651.87305 | NaN | 887.32184 | NaN | 687.9336 |
| Atrx | GGGDGNVDDTGNNPSVSLK | -1947 | -1965 | 0 | 0 | 19 | 434.04126 | 101.34955 | 309.70813 | NaN | 652.0725 | NaN |
| Ostc | GGLGFIILDR | -97 | -106 | 0 | 0 | 10 | 455.7739 | 649.96606 | 720.0908 | 450.13205 | 513.7641 | 502.57593 |
| Ca2 | NVEFDDSQDNAVLK | -67 | -80 | 0 | 0 | 14 | 1416.3793 | NaN | NaN | 1543.4211 | NaN | NaN |
| Ca2 | GGPLSDSYR | -81 | -89 | 0 | 0.003521747 | 9 | 797.174 | NaN | NaN | NaN | NaN | NaN |
| Dnaja3 | GGPSVDPEELFR | -176 | -187 | 0 | 0.00025015 | 12 | 910.9746 | 166.44278 | 999.2777 | 918.37225 | 847.9534 | 656.56866 |
| Dnaja3 | VTIPVPAGVEDGQTVR | -304 | -319 | 0 | 0 | 16 | 1992.2848 | 1078.5438 | 366.84164 | NaN | NaN | NaN |
| B4galnt1 | VTIVIADDSDKPER | -306 | -319 | 1 | 0 | 14 | 1114.4066 | 962.3486 | 1221.1781 | 1236.597 | 1107.0321 | 993.20496 |
| B4galnt1 | TPLDLVGGAVR | -377 | -387 | 0 | 0 | 11 | 1007.844 | 1016.71643 | 1061.2769 | 1089.1549 | 1262.3423 | 1242.9904 |
| B4galnt1 | GGSLPLPFLR | -86 | -95 | 0 | 0 | 10 | 1000.7839 | 1238.5408 | 1336.0527 | 990.9967 | 1237.4468 | 1349.4307 |
| B4galnt1 | LDLLDLAPEPR | -49 | -59 | 0 | 0 | 11 | 915.4243 | 976.7797 | 1008.1377 | 327.76056 | 1331.3324 | 1276.3247 |
| B4galnt1 | YAHIPVR | -60 | -66 | 0 | 0.000907206 | 7 | 1183.1971 | 1276.7455 | 1367.6678 | NaN | 1133.8942 | 1002.5268 |
| B4galnt1 | NSPLQYPLQGVEVQPLR | -142 | -158 | 0 | 0 | 17 | 213.04234 | NaN | 271.32806 | NaN | NaN | NaN |
| Cnp | LDEDLAGYCR | -103 | -112 | 0 | 0 | 10 | 3370.84 | 2982.518 | 3160.5562 | 359.4578 | 2332.6055 | 2232.6672 |
| Cnp | GGSQGEAVGELPR | -356 | -368 | 0 | 0 | 13 | 820.9584 | 949.2591 | 1022.25354 | 530.54315 | 868.8955 | 1021.19806 |
| Cnp | HFISGDEPK | -224 | -232 | 0 | 0 | 9 | 598.74023 | NaN | 758.09393 | NaN | 753.71655 | 585.0618 |
| Cnp | VLDDTNHER | -119 | -127 | 0 | 0 | 9 | 216.91629 | 863.6713 | NaN | NaN | 153.79144 | NaN |
| Cnp | VLVLDDTNHER | -117 | -127 | 0 | 0 | 11 | 642.5064 | NaN | NaN | NaN | NaN | NaN |
| Hnrnph2 | GMTLPVDFQGR | -140 | -150 | 0 | 0 | 11 | 917.6342 | NaN | 196.14496 | 427.6658 | NaN | 713.974 |
| Hnrnph2 | GGYDDYGGYNDGYGFGSDR | -241 | -259 | 0 | 0 | 19 | 221.33195 | 283.78217 | 324.12772 | NaN | NaN | NaN |
| Keap1 | GGYDGQDQLNSVER | -523 | -536 | 0 | 0 | 14 | 523.7142 | 649.5517 | NaN | 593.2184 | NaN | NaN |
| Srpk1 | VLQGLDYLHTK | -196 | -206 | 0 | 0 | 11 | 1693.7695 | 1377.8708 | 1648.4352 | 1187.9479 | 978.4009 | 996.70605 |
| Srpk1 | NFLINPLEPK | -470 | -479 | 0 | 0 | 10 | 805.74524 | 806.01416 | 803.8797 | 534.545 | 579.72125 | 667.37854 |
| Srpk1 | LEESNSIGQDQTLTER | -330 | -345 | 0 | 0 | 16 | 3749.238 | 1257.0885 | 4290.799 | NaN | 5845.267 | 4827.506 |
| Srpk1 | HKEDLHNANDCDVHTL | -376 | -391 | 1 | 0.00025015 | 16 | 215.46854 | 343.97488 | 324.7803 | NaN | 325.25742 | 242.94168 |
| Srpk1 | SNYQGLPLPCVK | -179 | -190 | 0 | 0 | 12 | 829.6233 | NaN | 905.31476 | 216.64803 | 799.8063 | NaN |
| Srpk1 | GGYHLVK | -65 | -71 | 0 | 0.005605646 | 7 | 570.8084 | NaN | NaN | NaN | NaN | NaN |
| Plpbp | GHDLSQGPNPDFQR | -187 | -200 | 0 | 0 | 14 | 3635.953 | NaN | 1367.2197 | 1177.0961 | 1091.0017 | 1083.6005 |
| Plpbp | IGSTIFGER | -242 | -250 | 0 | 0.001868731 | 9 | 2828.1284 | 265.4189 | 700.0808 | 318.30444 | NaN | 206.24805 |
| Plpbp | WHFIGHLQ | -91 | -98 | 0 | 0.000431655 | 8 | 725.47424 | NaN | NaN | 984.0096 | NaN | NaN |
| Arhgef2 | GHDRLDLPVTVR | -904 | -915 | 1 | 0.000230707 | 12 | 290.02997 | NaN | NaN | NaN | 233.26666 | 300.91827 |
| Arhgef2 | LDLPVTVR | -908 | -915 | 0 | 0.001868731 | 8 | 473.81067 | NaN | NaN | NaN | NaN | NaN |
| Nck1 | GHEGDFLIR | -300 | -308 | 0 | 0 | 9 | 910.09094 | 998.9342 | 845.57025 | NaN | 1252.5206 | 1236.9167 |
| Gmpr2 | GHIISDGGCSCPGDVAK | -214 | -230 | 0 | 0.00025015 | 17 | 406.78162 | 426.24536 | 497.22696 | 435.37326 | 567.1051 | 588.71313 |
| Gmpr2 | GHSESGGELIER | -247 | -258 | 0 | 0 | 12 | 620.58716 | NaN | 526.5699 | NaN | NaN | NaN |

| | | | | | | | | | | | | |
|---|---|---|---|---|---|---|---|---|---|---|---|---|
| Skic2 | TTGHILGYK | -109 | -117 | 0 | 0.00111006 | 9 | 1914.3387 | 1855.2411 | 1768.0399 | 1703.7673 | 1521.1323 | 1523.8055 |
| Skic2 | GHILGYK | -111 | -117 | 0 | 0.000712251 | 7 | 3708.066 | NaN | NaN | NaN | NaN | NaN |
| Rps29 | GHQQLYWSHPR | -2 | -12 | 0 | 0 | 11 | 2552.832 | 2827.1914 | 3025.9268 | 1478.0144 | 2070.8 | 867.97363 |
| Rps29 | GHQQLYWSHPR | -2 | -12 | 0 | 0 | 11 | 2552.832 | 2827.1914 | 3025.9268 | 1478.0144 | 2070.8 | 867.97363 |
| Rps29 | LYWSHPR | -6 | -12 | 0 | 0.000230707 | 7 | 1836.4409 | 1887.3624 | 1303.9297 | 1197.5538 | 1224.4967 | 1101.8085 |
| Mtx2 | LGTQPYFFNK | -197 | -206 | 0 | 0 | 10 | 2664.2588 | 3608.102 | 2607.805 | 4416.0044 | 3522.1577 | 3738.5789 |
| Mtx2 | IEQHYFEDWGK | -249 | -259 | 0 | 0 | 11 | 1063.8542 | NaN | NaN | 1100.1885 | NaN | NaN |
| Mtx2 | GHSLSDGLDEVQK | -96 | -108 | 0 | 0.000431655 | 13 | 578.0723 | NaN | NaN | NaN | NaN | 995.8612 |
| Rps26 | GHVQPIR | -16 | -22 | 0 | 0 | 7 | 1089.2972 | 798.9956 | 1236.3031 | 727.85803 | 281.93283 | 604.8437 |
| Smc4 | SLVHDLFQK | -572 | -580 | 0 | 0 | 9 | 1379.7744 | 1792.9551 | 1832.6498 | 1724.6741 | 738.01746 | 1624.5249 |
| Smc4 | VIQENEHALQK | -997 | -1007 | 0 | 0 | 11 | 765.3059 | 769.22687 | 802.79614 | 665.4273 | 859.47253 | 987.26794 |
| Smc4 | LGDLGAIDEK | -617 | -626 | 0 | 0 | 10 | 600.9635 | 638.77435 | 594.2212 | 652.0149 | 726.57117 | 729.9587 |
| Smc4 | LTQEEINLK | -563 | -571 | 0 | 0.000431655 | 9 | 1209.7422 | 578.77783 | 724.3959 | NaN | 1204.2725 | 1171.5479 |
| Smc4 | VQHEEAVVK | -796 | -804 | 0 | 0.001023236 | 9 | 965.3338 | 172.63351 | 114.7014 | NaN | 770.44556 | 822.9647 |
| Smc4 | GIDLDHNR | -215 | -222 | 0 | 0 | 8 | 719.9462 | NaN | 513.08 | 499.40143 | 661.461 | NaN |
| Smc4 | QLDECASAITK | -910 | -920 | 0 | 0.001636916 | 11 | 1254.6022 | NaN | NaN | NaN | 830.9181 | 594.06647 |
| Gne | TLVLFPNIDAGSK | -247 | -259 | 0 | 0 | 13 | 643.7397 | 4989.615 | 4926.6094 | NaN | 3339.6536 | NaN |
| Gne | GIEHHPNFR | -269 | -277 | 0 | 0.007224385 | 9 | 286.42953 | NaN | NaN | NaN | NaN | 279.39325 |
| Sh3bp1 | GIFPGLAPQEK | -479 | -489 | 0 | 0.000431655 | 11 | 927.7862 | 1330.2616 | 471.69754 | 1469.0985 | 1379.4384 | 182.07474 |
| Sh3bp1 | SLTSLDTALAELR | -241 | -253 | 0 | 0 | 13 | 718.4554 | NaN | NaN | 454.44803 | NaN | NaN |
| Spag9 | GIFPLPAGDGLLTPDTQK | -205 | -222 | 0 | 0 | 18 | 1112.7006 | 1023.6792 | 1499.793 | 1226.7942 | 1377.0168 | 1400.7847 |
| Adprh | GILDGNSAPVFPQPFGVK | -240 | -257 | 0 | 0 | 18 | 367.86853 | 366.68066 | 305.31006 | 483.39862 | 559.00543 | 469.57178 |
| Adprh | SPWQWGK | -192 | -198 | 0 | 0 | 7 | 1241.314 | NaN | 1281.4297 | 1345.6128 | 1470.2358 | 1482.8713 |
| Adprh | SAPVFPQPFGVK | -246 | -257 | 0 | 0.005394768 | 12 | 460.4604 | 424.87665 | 385.47018 | NaN | NaN | NaN |
| Fyn | TEDDLSFHK | -97 | -105 | 0 | 0 | 9 | 8752.252 | 8810.764 | 9357.12 | 546.4261 | 955.37854 | 9758.58 |
| Fyn | NSSEGDWWEAR | -113 | -123 | 0 | 0 | 11 | 695.7728 | 626.6684 | 791.6721 | 785.9982 | 700.7469 | 732.48157 |
| Fyn | YGTDPTPQHYPSFGVT | -30 | -45 | 0 | 0.000230707 | 16 | 503.7491 | 636.26227 | 634.09546 | 425.57755 | 702.8272 | 463.90622 |
| Fyn | GILLTELVTK | -453 | -462 | 0 | 0 | 10 | 84.791374 | 464.3622 | 467.0287 | 471.0904 | 533.3228 | 517.6636 |
| Nsf | GILLYGPPGCGK | -255 | -266 | 0 | 0 | 12 | 2172.0188 | 2993.2664 | 4121.7393 | 149.45844 | 5837.2715 | 5410.141 |
| Nsf | LLDYVPIGPR | -608 | -617 | 0 | 0 | 10 | 973.9372 | 544.42926 | 704.24286 | 673.32385 | 835.15027 | 572.4467 |
| Nsf | QSIINPDWNFEK | -206 | -217 | 0 | 0 | 12 | 785.05457 | 815.5621 | 800.5961 | 1074.6405 | 4134.314 | 5621.287 |
| Nsf | IAEESNFPFIK | -556 | -566 | 0 | 0 | 11 | 563.59155 | 610.6295 | 507.04584 | 586.80396 | 748.891 | 238.41042 |
| Nsf | NFSGAELEGLVR | -435 | -446 | 0 | 0 | 12 | 513.49493 | 920.93585 | 257.9637 | 391.50842 | 1052.6635 | 917.2663 |
| Nsf | YVGESEANIR | -294 | -303 | 0 | 0 | 10 | 823.6194 | 806.9725 | NaN | 843.77673 | 619.3636 | 906.72845 |
| Nsf | TNRPDLIDEALLRPGR | -373 | -388 | 2 | 0.000230707 | 16 | 443.30185 | 286.2226 | 515.3075 | NaN | 621.7548 | 615.93 |
| Nsf | LSCVVVDDIER | -597 | -607 | 0 | 0 | 11 | 3784.8638 | 7229.071 | 7593.876 | NaN | 6782.703 | NaN |
| Nsf | SQLSCVVVDDIER | -595 | -607 | 0 | 0 | 13 | 459.04987 | 1151.7614 | 1036.9224 | 488.02948 | NaN | NaN |
| Nsf | TIEIDFLQK | -96 | -104 | 0 | 0.00025015 | 9 | 720.7443 | NaN | NaN | 654.5227 | NaN | NaN |
| Yipf5 | TLTVLHPLR | -107 | -115 | 0 | 0 | 9 | 483.9553 | 519.1399 | 584.8538 | 320.18033 | 486.25995 | 447.06207 |
| Cyth4 | GINSGSDLPEEQLR | -219 | -232 | 0 | 0.00025015 | 14 | 12207.874 | 10110.742 | 4127.322 | 2534.5347 | NaN | 6106.2397 |
| Mbnl1 | GIPQAVLPPLPK | -269 | -280 | 0 | 0 | 12 | 999.09186 | 1212.1621 | 1135.8445 | 1191.2806 | 1215.5063 | 1255.8293 |
| Mbnl1 | GTCSRPDTECK | -25 | -35 | 1 | 0.001463897 | 11 | 879.07587 | 544.2336 | 1004.47736 | 1277.3047 | 1075.6531 | 1645.6069 |
| Cox5a | LNDFASAVR | -95 | -103 | 0 | 0 | 9 | 3680.0105 | 3531.3699 | 594.2387 | 2419.5886 | 1649.7764 | 1817.0917 |
| Cox5a | SHGSHETDEEFDAR | -38 | -51 | 0 | 0 | 14 | 1419.2953 | 1575.605 | 1711.7083 | 1412.2291 | 1860.7577 | 1840.655 |
| Cox5a | SHGSHETDEEFDAR | -38 | -51 | 0 | 0 | 14 | 1419.2953 | 1575.605 | 1711.7083 | 1412.2291 | 1860.7577 | 1840.655 |
| Cox5a | GMNTLVGYDLVPEPK | -69 | -83 | 0 | 0 | 15 | 1086.1321 | 1330.8984 | 1261.4216 | 1402.9716 | 1621.0227 | 1630.9104 |
| Cox5a | VGYDLVPEPK | -74 | -83 | 0 | 0 | 10 | 643.1815 | 636.7905 | 684.6278 | 352.68918 | 618.8213 | 597.6675 |
| Cox5a | LNELGISTPEELGLDKV | -130 | -146 | 1 | 0 | 17 | 594.62946 | 463.93683 | 745.5061 | 693.98065 | 727.47174 | 526.3086 |
| Cox5a | GSHETDEEFDAR | -40 | -51 | 0 | 0 | 12 | 1307.647 | NaN | 1251.2776 | 1935.5286 | 837.541 | 568.096 |
| Cox5a | GISTPEELGLDKV | -134 | -146 | 1 | 0 | 13 | 1307.1951 | 1687.9326 | 1632.993 | 1423.8369 | NaN | 1433.1903 |
| Cox5a | LVGYDLVPEPK | -73 | -83 | 0 | 0 | 11 | 785.0114 | 889.03485 | NaN | NaN | 1559.7874 | 1840.6025 |
| Cox5a | TLVGYDLVPEPK | -72 | -83 | 0 | 0.00025015 | 12 | 1069.4188 | NaN | NaN | NaN | NaN | 1513.0824 |
| Psma4 | LLDEVFFSEK | -55 | -64 | 0 | 0 | 10 | 1404.3037 | 1732.0728 | 369.38657 | 1785.8701 | 2327.5225 | 2766.7722 |
| Psma4 | LYQSDPSGNYGGWK | -147 | -160 | 0 | 0 | 14 | 1302.9929 | 1353.0737 | 1390.4907 | 1345.4008 | 1527.1312 | 1396.0592 |
| Psma4 | QSDPSGNYGGWK | -149 | -160 | 0 | 0 | 12 | 637.1284 | 919.1931 | 895.619 | 497.02008 | 270.85745 | 323.05032 |
| Psma4 | TTIFSPEGR | -9 | -17 | 0 | 0 | 9 | 5812.508 | 2289.818 | 3050.6462 | 1367.3744 | 5539.6074 | NaN |
| Psma4 | YQSDPSGNYGGWK | -148 | -160 | 0 | 0 | 13 | 1015.99097 | 1053.686 | 1040.2559 | 503.7657 | 151.10649 | NaN |
| Psma4 | RPFGVSLL | -128 | -135 | 1 | 0.00025015 | 8 | 1869.3894 | NaN | 1898.7894 | 1476.2155 | 1102.1176 | NaN |
| Psma4 | GITSDANVLTNELR | -78 | -91 | 0 | 0 | 14 | 1186.3779 | 799.98193 | NaN | NaN | 1447.1937 | NaN |
| Mad2l1 | GIYPSETFTR | -36 | -45 | 0 | 0 | 10 | 2583.5688 | 2666.8145 | 1217.4021 | 1216.0886 | 1692.9376 | 1751.3038 |
| Mad2l1 | NIESGEVLER | -90 | -99 | 0 | 0 | 10 | 1703.1035 | NaN | 1720.9469 | 2504.172 | 2253.4312 | NaN |
| Rpl21 | KGDIVDIK | -36 | -43 | 1 | 0.000471409 | 8 | 2798.284 | 2364.622 | 2133.148 | 2006.4379 | 1395.768 | 1386.2367 |
| Rpl21 | KKGDIVDIK | -35 | -43 | 2 | 0 | 9 | 1844.8881 | 1548.0404 | 1676.7876 | 2128.5254 | 1758.4304 | 1646.6002 |
| Rpl21 | GKEPELLEPIPYEFMA | -145 | -160 | 1 | 0.001092529 | 16 | 755.0725 | 741.3429 | 753.2838 | 1088.3192 | 135.66275 | 1208.5261 |
| Rpl21 | GKEPELLEPIPYEFMA | -145 | -160 | 1 | 0.001092529 | 16 | 755.0725 | 741.3429 | 753.2838 | 1088.3192 | 135.66275 | 1208.5261 |
| Rpl21 | QHAVGIIVNK | -69 | -78 | 0 | 0.001055699 | 10 | 304.5479 | 156.20193 | 216.61388 | 202.82066 | 296.60373 | 256.12262 |
| Rpl21 | NVTQHAVGIIVNK | -66 | -78 | 0 | 0 | 13 | 1360.9056 | NaN | NaN | 1017.12585 | NaN | NaN |
| Idh2 | LVPGWTKPITIGR | -160 | -172 | 1 | 0 | 13 | 605.32916 | 810.64856 | 845.025 | 540.99805 | 811.9174 | 856.1877 |
| Idh2 | GKLDGNQDLIR | -383 | -393 | 1 | 0 | 11 | 490.28064 | 1098.1499 | 1107.8662 | 951.52167 | 1325.0039 | 1249.5364 |
| Idh2 | YFDLGLPNR | -81 | -89 | 0 | 0 | 9 | 2402.9265 | NaN | 2570.0457 | 2064.7432 | 2165.925 | 177.63818 |
| Idh2 | LVPGWTKPITIG | -160 | -171 | 1 | 0 | 12 | 1207.3589 | NaN | 992.4887 | 528.0649 | 946.5093 | 762.6586 |
| Idh2 | LILPHVDVQ | -70 | -78 | 0 | 0 | 9 | 501.65067 | 1725.54 | 1646.9636 | 1531.9175 | 1662.8425 | NaN |
| Idh2 | LDGNQDLIR | -385 | -393 | 0 | 0 | 9 | 1653.5504 | 1591.9764 | 1526.1565 | 759.0191 | NaN | NaN |
| Idh2 | LNEHFLNTT | -427 | -435 | 0 | 0.004255025 | 9 | 799.58154 | 688.75696 | NaN | NaN | NaN | 1142.3298 |
| Gbp3 | GKLFGVGLK | -605 | -613 | 1 | 0.00111006 | 9 | 1548.0131 | 808.6584 | 1647.89 | 613.4781 | 695.9763 | 1706.9843 |
| Cbx1 | KVEEVLEEEEEEYVVE | -9 | -24 | 1 | 0 | 16 | 884.4793 | 779.1169 | 879.4014 | 649.70087 | 686.69354 | 621.5974 |
| Cbx1 | NSDEADLVPAK | -140 | -150 | 0 | 0 | 11 | 366.61386 | 1338.3048 | 1224.1221 | 1144.8315 | 1693.8867 | 872.54266 |
| Cbx1 | GKVEYLLK | -34 | -41 | 1 | 0 | 8 | 754.79297 | 860.0999 | 773.8286 | NaN | NaN | NaN |

| | | | | | | | | | | | | |
|---|---|---|---|---|---|---|---|---|---|---|---|---|
| Cbx1 | KVEEVLEEEEEEYVVEK | -9 | -25 | 1 | 0.00122419 | 17 | 367.2144 | NaN | NaN | NaN | 297.63712 | 229.86655 |
| Sh3glb1 | GKVPITYLELLN | -354 | -365 | 1 | 0 | 12 | 479.19287 | NaN | NaN | 511.84137 | 339.85773 | 599.8138 |
| Rasal3 | WFPVLGAPAGAVLR | -429 | -442 | 0 | 0 | 14 | 1337.723 | 840.4868 | 860.03674 | 109.15758 | 2267.5479 | 415.27548 |
| Rasal3 | NVAPLEPKPNPK | -118 | -129 | 1 | 0.000298805 | 12 | 621.2579 | 623.5076 | 685.1251 | 538.24255 | 622.7868 | 552.49585 |
| Rasal3 | GLAPEHPAPGPAR | -650 | -662 | 0 | 0 | 13 | 1706.5796 | 1625.5441 | 1462.4589 | NaN | 1052.5994 | 1094.9755 |
| Acot2 | NATLSVEPEGR | -43 | -53 | 0 | 0.001055699 | 11 | 1145.7133 | NaN | 1056.2916 | 603.7893 | NaN | NaN |
| Dhx29 | SLEPTSAEDLAPVR | -536 | -549 | 0 | 0 | 14 | 799.09186 | 519.9348 | 801.7861 | 371.28384 | NaN | 368.15054 |
| Dhx29 | GLEEEEKFDPNQR | -249 | -261 | 1 | 0 | 13 | 557.7905 | 345.02615 | NaN | 295.62296 | NaN | 373.01343 |
| Dhx29 | SPFITPIGR | -1108 | -1116 | 0 | 0 | 9 | 270.11923 | NaN | NaN | NaN | 507.96283 | 478.84814 |
| Dpm3 | GLELPFPCR | -27 | -35 | 0 | 0 | 9 | 1426.5461 | 1639.1687 | 499.8786 | 789.004 | 1368.0481 | 1103.1239 |
| Sun1 | GLETEYQEEGQPLGR | -846 | -860 | 0 | 0.00111006 | 15 | 511.4337 | NaN | 496.11047 | NaN | 728.39886 | NaN |
| Prdx2 | GWKPGSDTIKPNVDDSK | -175 | -191 | 2 | 0.000431655 | 17 | 1627.9255 | 1894.6663 | 1902.988 | 1374.0361 | 1697.0183 | 1665.6361 |
| Prdx2 | GWKPGSDTIKPNVDDSK | -175 | -191 | 2 | 0.000431655 | 17 | 1627.9255 | 1894.6663 | 1902.988 | 1374.0361 | 1697.0183 | 1665.6361 |
| Prdx2 | SDHAEDFR | -59 | -66 | 0 | 0.000394361 | 8 | 1574.6667 | 1651.7882 | 1709.6106 | 1371.3062 | 1749.5317 | 1737.7637 |
| Prdx2 | KNDEGIAYR | -119 | -127 | 1 | 0 | 9 | 1229.994 | 747.89197 | 516.15076 | 503.97562 | 1625.1836 | 1593.451 |
| Prdx2 | KEGGLGPLNIPLLA | -92 | -105 | 1 | 0.000712251 | 14 | 515.9107 | 434.3489 | 435.42416 | 382.96527 | 325.32233 | 512.3397 |
| Prdx2 | KEGGLGPLNIPLLADVTK | -92 | -109 | 1 | 0 | 18 | 456.42557 | 484.90833 | 451.29788 | 558.8119 | 644.26263 | 655.05383 |
| Prdx2 | KEGGLGPLNIPLLADVTK | -92 | -109 | 1 | 0 | 18 | 456.42557 | 484.90833 | 451.29788 | 558.8119 | 644.26263 | 655.05383 |
| Prdx2 | SDHAEDFRK | -59 | -67 | 1 | 0.000431655 | 9 | 229.60782 | 469.70993 | 616.2256 | 304.40875 | 233.99301 | 653.36896 |
| Prdx2 | QITVNDLPVGR | -140 | -150 | 0 | 0 | 11 | 8657.978 | NaN | NaN | 6701.2207 | 7559.552 | 7427.992 |
| Prdx2 | SLSQNYGVLK | -110 | -119 | 0 | 0 | 10 | 866.17957 | NaN | 632.73737 | NaN | 794.1618 | 627.36926 |
| Prdx2 | GLFIIDAK | -128 | -135 | 0 | 0.006472924 | 8 | 511.99796 | 486.8189 | NaN | NaN | NaN | NaN |
| Crkl | YDFPGNDAEDLPFK | -132 | -145 | 0 | 0 | 14 | 271.78876 | 622.0673 | 429.9005 | 776.26337 | 846.32904 | 1081.7277 |
| Crkl | GLFPFTHVK | -284 | -292 | 0 | 0.000591124 | 9 | 1404.6404 | NaN | 534.43713 | 1492.8186 | 1668.5848 | 720.118 |
| Crkl | YLDTTTLIEPAPR | -92 | -104 | 0 | 0 | 13 | 1065.7135 | NaN | NaN | 2117.8535 | 2510.091 | 2124.7192 |
| Crkl | STPGAAINPLPSTQNGPVFAK | -221 | -241 | 0 | 0.00383302 | 21 | 504.99268 | 577.34564 | 526.26404 | NaN | NaN | 640.38806 |
| Brd2 | GLHDYHDIIK | -381 | -390 | 0 | 0 | 10 | 595.5647 | 1283.8362 | 1103.3269 | 1161.5675 | NaN | NaN |
| Fga | GLIDEANQDFTNR | -73 | -85 | 0 | 0 | 13 | 2689.3528 | 1727.7014 | 3514.7449 | 1464.3514 | 572.509 | 2079.5657 |
| Fga | TDTEDKGEFLSEGGGVR | -20 | -36 | 1 | 0 | 17 | 309.1688 | 355.49078 | 394.0994 | 721.54016 | 462.7511 | 465.46893 |
| Fga | NSLFDFQR | -91 | -98 | 0 | 0 | 8 | 793.3999 | 1055.334 | NaN | 867.381 | NaN | 982.33356 |
| Ube2l3 | VNDPQPEHPLR | -112 | -122 | 0 | 0 | 11 | 2203.5833 | 2373.756 | 2494.576 | 99.48317 | 2491.4187 | 2365.2449 |
| Ube2l3 | VNDPQPEHPLR | -112 | -122 | 0 | 0 | 11 | 2203.5833 | 2373.756 | 2494.576 | 99.48317 | 2491.4187 | 2365.2449 |
| Ube2l3 | INFPAEYPFKPPK | -55 | -67 | 1 | 0 | 13 | 1968.0056 | 2601.515 | 2399.8599 | 2277.043 | 2727.151 | 2541.9448 |
| Ube2l3 | INFPAEYPFKPPK | -55 | -67 | 1 | 0 | 13 | 1968.0056 | 2601.515 | 2399.8599 | 2277.043 | 2727.151 | 2541.9448 |
| Ube2l3 | TWQGLIVPDNPPYDK | -34 | -48 | 0 | 0 | 15 | 1680.7794 | 1944.0647 | 2020.8752 | 2031.8348 | 2901.811 | 2686.6882 |
| Ube2l3 | IALVNDPQPEHPLR | -109 | -122 | 0 | 0.00025015 | 14 | 1316.0571 | 1490.4764 | 1502.2202 | 1481.9436 | 1577.4281 | 1607.2064 |
| Ube2l3 | IALVNDPQPEHPLR | -109 | -122 | 0 | 0.00025015 | 14 | 1316.0571 | 1490.4764 | 1502.2202 | 1481.9436 | 1577.4281 | 1607.2064 |
| Ube2l3 | LVNDPQPEHPLR | -111 | -122 | 0 | 0.000471409 | 12 | 875.8754 | 941.73755 | 936.099 | 681.8693 | 2482.5073 | 753.2858 |
| Ube2l3 | IEINFPAEYPFKPPK | -53 | -67 | 1 | 0 | 15 | 460.30225 | 233.26791 | 378.4632 | 557.9792 | 775.18866 | 804.20715 |
| Ube2l3 | IYHPNIDEK | -74 | -82 | 0 | 0 | 9 | 132.47926 | 3691.182 | 3659.0928 | 3763.472 | 3645.6096 | 3719.8704 |
| Ube2l3 | NWKPATK | -94 | -100 | 1 | 0.005946382 | 7 | 1424.7064 | NaN | 2225.7544 | NaN | 1352.8268 | NaN |
| Ube2l3 | GLIVPDNPPYDK | -37 | -48 | 0 | 0.00122419 | 12 | 18240.658 | NaN | NaN | NaN | 16628.22 | NaN |
| Golga7 | GLLLTDPIER | -109 | -118 | 0 | 0.00025015 | 10 | 840.2597 | 903.02875 | 774.2208 | 682.9337 | NaN | NaN |
| Snrpa1 | GLLQSGQIPGR | -222 | -232 | 0 | 0 | 11 | 2678.7737 | 4057.8506 | 4241.218 | 3197.8223 | 2592.9524 | 2510.6533 |
| Snrpa1 | TFNPGAGLPTDK | -180 | -191 | 0 | 0 | 12 | 1275.4963 | 1202.5209 | 1169.6316 | 1070.3384 | 807.6354 | 741.28564 |
| Snrpa1 | KLDGFPLLR | -56 | -64 | 1 | 0.00111006 | 9 | 1062.9343 | 1080.7986 | 1067.4706 | 502.72018 | 833.00543 | 762.9611 |
| Snrpa1 | TFNPGAGLPTDKK | -180 | -192 | 1 | 0 | 13 | 829.6075 | 717.4194 | 861.41345 | 295.36145 | 845.8299 | 961.0145 |
| Snrpa1 | LDQFDAIDFSDNEIR | -41 | -55 | 0 | 0 | 15 | 497.68027 | 717.43195 | 647.7601 | 736.5531 | 677.769 | 705.86395 |
| Snrpa1 | SLVELGDLDPLASLK | -99 | -113 | 0 | 0 | 15 | 381.26923 | 364.04123 | 392.31104 | 439.4626 | 514.0032 | 509.91333 |
| Snrpa1 | LDGFPLLR | -57 | -64 | 0 | 0 | 8 | 357.59656 | 631.2242 | 410.57288 | 259.4734 | 618.56305 | 402.8544 |
| Snrpa1 | NSLVELGDLDPLASLK | -98 | -113 | 0 | 0 | 16 | 220.10492 | 212.64949 | 227.6686 | 279.40652 | 212.75525 | 285.44815 |
| Ak4 | GLLVPDHVITR | -61 | -71 | 0 | 0 | 11 | 512.2185 | 667.10034 | 646.8277 | NaN | 728.9381 | 2401.912 |
| Mpdu1 | GLLVPILLPEK | -11 | -21 | 0 | 0.00025015 | 11 | 372.16223 | 420.06503 | 426.64633 | 257.91995 | 327.99283 | 345.86728 |
| Mpdu1 | WDLLHVPCLK | -30 | -39 | 0 | 0 | 10 | 196.5746 | 756.4392 | 453.7312 | 773.37476 | 809.7855 | 737.0783 |
| Mpdu1 | LVPILLPEK | -13 | -21 | 0 | 0.00025015 | 9 | 493.89276 | 666.04126 | 557.9961 | 837.9936 | 773.47754 | NaN |
| Mpdu1 | VLFYWNAK | -231 | -238 | 0 | 0.009955823 | 8 | 970.6467 | 901.24927 | NaN | 1288.8262 | NaN | NaN |
| Bzw2 | LLELFPVNR | -216 | -224 | 0 | 0 | 9 | 982.77435 | 1025.5358 | 1076.6315 | 969.228 | 1273.3175 | 1182.7421 |
| Bzw2 | SSQGQSELVLLQK | -333 | -345 | 0 | 0 | 13 | 5306.562 | 4870.9536 | NaN | NaN | NaN | NaN |
| Bzw2 | GLNEAGDDLEAVAK | -37 | -50 | 0 | 0 | 14 | 3164.4585 | NaN | NaN | NaN | NaN | NaN |
| Aarsd1 | VVSCSPAELQTDASGGK | -18 | -34 | 0 | 0.001023236 | 17 | 2283.6328 | 2745.7402 | NaN | 595.22943 | 984.19305 | 832.396 |
| Aarsd1 | GLPDDHAGPIR | -187 | -197 | 0 | 0 | 11 | 3500.282 | 3991.6152 | 3280.9749 | NaN | NaN | NaN |
| Xpo5 | SCDPAVEDPCGLNR | -705 | -718 | 0 | 0 | 14 | 397.4564 | 426.72806 | 378.50513 | 490.09787 | 539.7755 | 426.2619 |
| Xpo5 | VVGYTPSGNPIFR | -750 | -762 | 0 | 0 | 13 | 594.87225 | 722.99457 | 698.3611 | 708.431 | 1203.6887 | NaN |
| Xpo5 | GLPQPLLEFNDHPVYR | -815 | -830 | 0 | 0 | 16 | 273.70767 | NaN | 426.56873 | 294.2983 | 340.34515 | 396.8919 |
| Xpo5 | WHVINQR | -931 | -937 | 0 | 0.001055699 | 7 | 9952.671 | NaN | NaN | NaN | NaN | NaN |
| Gnpda1 | LVDPLYSIK | -265 | -273 | 0 | 0 | 9 | 758.46545 | 1365.9719 | 978.2925 | 1233.0695 | 1557.276 | 1473.5413 |
| Gnpda1 | GLPTGSTPLGCYQK | -38 | -51 | 0 | 0 | 14 | 2176.775 | 1723.3823 | NaN | 285.78387 | NaN | NaN |
| Gnpda1 | HIDIHPENTHILDGN | -96 | -110 | 0 | 0.001370415 | 15 | 605.6252 | NaN | NaN | NaN | 1397.5859 | 714.0674 |
| Cdc34 | GLQEEPVEGFR | -19 | -29 | 0 | 0.004628201 | 11 | 2107.1802 | 3374.9119 | NaN | NaN | NaN | NaN |
| Skap1 | GLQNENLSPGAQDQR | -24 | -38 | 0 | 0 | 15 | 600.6366 | 439.40115 | 354.74286 | 846.7581 | 557.3204 | 553.9677 |
| Lypla2 | SCWLPLHR | -146 | -153 | 0 | 0 | 8 | 1910.2664 | 2300.68 | 2169.1213 | 1791.4484 | 2401.1753 | 2439.18 |
| Lypla2 | GLSPDAPEDEAGIK | -80 | -93 | 0 | 0 | 14 | 894.1118 | 1320.6475 | 1353.9521 | 1585.0681 | 2021.2285 | 1015.2811 |
| Lypla2 | GLSPDAPEDEAGIKK | -80 | -94 | 1 | 0 | 15 | 771.99677 | 552.14685 | 1072.5564 | 738.70374 | 875.0002 | 1537.8146 |
| Lypla2 | YICPHAPR | -54 | -61 | 0 | 0 | 8 | 1318.8591 | NaN | NaN | 1079.7601 | 1017.5854 | 1083.28 |
| Slc25a10 | GLTGGFVGTPADLVNVR | -109 | -125 | 0 | 0 | 17 | 324.99588 | NaN | NaN | NaN | NaN | NaN |
| Eefsec | LVDCPGHASLIR | -76 | -87 | 0 | 0 | 12 | 10681.204 | 11262.581 | 11499.867 | 10733.249 | 11377.609 | 11115.206 |
| Eefsec | GLVCAPESLHTVHAA | -287 | -301 | 0 | 0 | 15 | 706.5975 | 171.12546 | 430.42728 | 333.20346 | 613.5017 | 507.67383 |
| Eefsec | LAFHGVLLQ | -431 | -439 | 0 | 0 | 9 | 522.09296 | 598.0934 | 797.4741 | 1234.387 | 1046.0796 | 351.1115 |

| | | | | | | | | | | | | |
|---|---|---|---|---|---|---|---|---|---|---|---|---|
| Eefsec | NYIESFLPALR | -445 | -455 | 0 | 0 | 11 | 337.48877 | 252.17348 | 467.28433 | 490.27576 | 523.4447 | 356.6673 |
| Lgals3 | QSAFPFESGKPFK | -201 | -213 | 1 | 0 | 13 | 4724.9756 | 3942.4521 | 4882.204 | 5201.7314 | 1256.456 | 3815.504 |
| Lgals3 | VLVEADHFK | -216 | -224 | 0 | 0 | 9 | 1556.3754 | 1587.5245 | 1773.01 | 1997.5222 | 1939.6609 | 1881.2048 |
| Lgals3 | GNDVAFHFNPR | -166 | -176 | 0 | 0 | 11 | 749.50024 | 1872.5968 | 2319.2534 | 2340.6338 | 376.2045 | 2208.5867 |
| Rap1gds1 | GNICYDSHEGR | -114 | -124 | 0 | 0 | 11 | 799.35095 | 716.99744 | 500.18445 | NaN | 763.6245 | 815.0377 |
| Rap1gds1 | VDSNKEDDVAELK | -278 | -290 | 1 | 0 | 13 | 758.2598 | 977.009 | 857.5808 | 564.3615 | NaN | 451.83008 |
| Rap1gds1 | KVDSNKEDDVAELK | -277 | -290 | 2 | 0 | 14 | 258.55228 | NaN | 282.5712 | NaN | 484.70444 | 494.03238 |
| Acot7 | GNVHGGTILK | -70 | -79 | 0 | 0.00025015 | 10 | 1567.6848 | 1631.5239 | 1057.1523 | 1047.1615 | 1416.168 | 1715.8325 |
| Acot7 | SVDAINFHDK | -275 | -284 | 0 | 0 | 10 | 1308.0015 | 1392.1487 | 851.6766 | 1438.1273 | 1638.6051 | 1527.1276 |
| Acot7 | VLEVPPIVYLR | -174 | -184 | 0 | 0 | 11 | 1229.2256 | 1293.0177 | 1200.8561 | 1706.7432 | 1884.0128 | 277.28854 |
| Acot7 | VDADPVVDNSQK | -311 | -322 | 0 | 0 | 12 | 1024.4429 | 1087.0226 | 1050.3868 | 772.9804 | 780.7828 | 720.4908 |
| Poldip3 | TSPPIGTVTPALK | -132 | -144 | 0 | 0.00025015 | 13 | 719.04865 | NaN | 941.2143 | 534.0642 | NaN | NaN |
| Poldip3 | GNVITSDQPILLR | -352 | -364 | 0 | 0 | 13 | 234.96646 | NaN | 238.67422 | 239.40561 | NaN | NaN |
| Poldip3 | RGNPASSNPPAEVDPDTVLR | -380 | -399 | 1 | 0.001392869 | 20 | 891.30756 | NaN | NaN | NaN | NaN | NaN |
| Ube2b;Ube2a | GPEGTPFEDGTFK | (42);(42) | (54);(54) | 0 | 0.000550307 | 13 | 875.8663 | 891.5019 | 967.8695 | 867.47974 | 380.59628 | 872.32745 |
| Ube2b;Ube2a | NAVIFGPEGTPFEDGTFK | (37);(37) | (54);(54) | 0 | 0.00201047 | 18 | 1166.6057 | NaN | 1137.0422 | 2364.0781 | 2594.5469 | 2166.6477 |
| Ube2b;Ube2a | VIFGPEGTPFEDGTFK | (39);(39) | (54);(54) | 0 | 0.00025015 | 16 | 223.49438 | 248.53313 | 283.78644 | NaN | 301.20612 | 334.86383 |
| Pitpnb | IAPEGSLVFHEK | -75 | -86 | 0 | 0 | 12 | 2385.9204 | 2417.5737 | NaN | 2736.2534 | NaN | 2146.403 |
| Rtn4 | GPLPAAPPTAPER | -90 | -102 | 0 | 0 | 13 | 374.71918 | 614.97174 | 207.6575 | 1157.2744 | 480.29263 | 605.0998 |
| Rtn4 | HQAQIDHYLGL | -1128 | -1138 | 0 | 0.000627943 | 11 | 927.15173 | NaN | NaN | NaN | 1208.3134 | NaN |
| Urod | GPSFPEPLREER | -105 | -116 | 1 | 0.005605646 | 12 | 308.9588 | NaN | NaN | NaN | 414.64203 | 314.5573 |
| Memo1 | SDFCHWGQR | -188 | -196 | 0 | 0 | 9 | 1072.8955 | 161.15727 | 635.1425 | 1571.541 | 836.8673 | 848.91974 |
| Memo1 | GPSHHVPLSR | -78 | -87 | 0 | 0 | 10 | 558.92096 | 467.24567 | 322.8054 | NaN | 108.736664 | 653.66254 |
| Memo1 | RTPLYDLR | -97 | -104 | 1 | 0 | 8 | 157.24988 | 2847.7812 | 3148.2864 | NaN | NaN | 2616.13 |
| Memo1 | SIEHLDK | -212 | -218 | 0 | 0.00149795 | 7 | 707.5932 | NaN | NaN | 469.85303 | NaN | NaN |
| Aldh18a1 | IIDLIIPR | -551 | -558 | 0 | 0 | 8 | 1612.3463 | 1973.8684 | 1926.4126 | 186.67154 | 3113.84 | 2544.263 |
| Aldh18a1 | VSGHVITDIVEGK | -335 | -347 | 0 | 0 | 13 | 1414.4167 | 1574.7095 | 1666.2628 | 1632.5808 | 1657.529 | 3285.8347 |
| Aldh18a1 | NLDFHDEQK | -191 | -199 | 0 | 0 | 9 | 1274.8073 | 1400.0902 | 1314.636 | 1292.0032 | 1892.1163 | 1366.0721 |
| Aldh18a1 | YGSSHTDVIVTENEK | -691 | -705 | 0 | 0 | 15 | 780.38965 | 802.25366 | 914.93774 | 464.67407 | 688.3431 | 612.2884 |
| Aldh18a1 | YGSSHTDVIVTENEK | -691 | -705 | 0 | 0 | 15 | 780.38965 | 802.25366 | 914.93774 | 464.67407 | 688.3431 | 612.2884 |
| Aldh18a1 | YLHENLPVPQR | -782 | -792 | 0 | 0 | 11 | 1179.4004 | 1299.123 | 1265.2477 | 1525.0479 | 1576.7427 | 1558.6113 |
| Aldh18a1 | YLHENLPVPQR | -782 | -792 | 0 | 0 | 11 | 1179.4004 | 1299.123 | 1265.2477 | 1525.0479 | 1576.7427 | 1558.6113 |
| Aldh18a1 | SDVEGLFDSPPGSDDAK | -266 | -282 | 0 | 0 | 17 | 862.45874 | 960.752 | 927.62 | 946.71857 | 861.95746 | 951.18665 |
| Aldh18a1 | GPVGLEGLLTTK | -750 | -761 | 0 | 0 | 12 | 700.3008 | 706.4348 | 561.30725 | 918.57983 | 1119 | 1130.2993 |
| Aldh18a1 | VVDSVQEAIDHIHK | -677 | -690 | 0 | 0 | 14 | 632.341 | 260.06757 | 847.0242 | 652.4879 | 708.889 | 773.88214 |
| Aldh18a1 | SNIPFITVPLSR | -44 | -55 | 0 | 0 | 12 | 516.4555 | 857.82263 | 816.9035 | 732.82733 | 908.45245 | 929.2138 |
| Aldh18a1 | IEVVDSVQEAIDHIHK | -675 | -690 | 0 | 0 | 16 | 514.4465 | 741.5723 | 751.88324 | 827.9073 | 897.14844 | 1054.4132 |
| Aldh18a1 | NALETLLIHR | -613 | -622 | 0 | 0.000230707 | 10 | 488.07227 | 570.5115 | 739.83997 | 482.20203 | 702.5488 | 570.473 |
| Aldh18a1 | LADLLTDQR | -392 | -400 | 0 | 0.000471409 | 9 | 444.24704 | 560.49817 | 479.44324 | NaN | 780.8417 | NaN |
| Dcaf13 | TVYTGIDHHWK | -157 | -167 | 0 | 0 | 11 | 919.3053 | 1095.5804 | 816.40845 | 961.153 | NaN | 965.3973 |
| Dcaf13 | GQQVDIWDEQR | -175 | -185 | 0 | 0 | 11 | 1199.7107 | NaN | NaN | NaN | NaN | 497.5089 |
| Dcaf13 | NYDPTLHPF | -27 | -35 | 0 | 0.002265552 | 9 | 2784.2424 | NaN | NaN | NaN | NaN | NaN |
| Plaa | ILSLICNNSSEKPTAQ | -579 | -594 | 1 | 0 | 16 | 1217.266 | 1187.1353 | 1149.5308 | 1375.212 | 1216.4104 | 1190.2313 |
| Plaa | KLTEDDLVLLEK | -567 | -578 | 1 | 0 | 12 | 746.374 | 932.43146 | 856.96063 | 1155.1475 | 1128.7981 | 1086.2893 |
| Plaa | TFLGHEDCVR | -186 | -195 | 0 | 0 | 10 | 723.341 | 868.8075 | 792.952 | 860.6266 | 241.0132 | 1049.8073 |
| Plaa | LWAPDSPNR | -45 | -53 | 0 | 0 | 9 | 718.2421 | 1016.9243 | 1477.9011 | 576.405 | 653.58655 | 772.5833 |
| Plaa | TGDLGDINAEQLPGR | -326 | -340 | 0 | 0 | 15 | 554.2138 | 545.1339 | 558.65753 | 693.1548 | 1075.0529 | 1150.9473 |
| Plaa | HPNVNENFCNEK | -623 | -634 | 0 | 0 | 12 | 434.8888 | 543.35767 | 731.3928 | 637.07007 | 1005.6798 | 170.06262 |
| Plaa | YVPGTSGPSNTVQTADPFTGAGR | -477 | -499 | 0 | 0 | 23 | 434.8263 | 682.99304 | 687.04034 | 693.9089 | 909.62036 | 769.83075 |
| Plaa | IGDVVGSSGANQQTSGK | -377 | -393 | 0 | 0 | 17 | 528.5001 | NaN | 587.2757 | 647.7081 | 307.47803 | 501.7801 |
| Plaa | GQTLGLGNTSFSDPFTGGGR | -457 | -476 | 0 | 0.00111006 | 20 | 518.3611 | NaN | 671.5067 | NaN | NaN | 603.1117 |
| Smc6 | GQTTLPFRPVTQEEDDSAS | -1079 | -1097 | 1 | 0.000907206 | 19 | 347.12326 | 434.11728 | 382.66742 | 485.72223 | 473.91046 | 598.19275 |
| Rida | GQVGLDPSSGQLVPGGVVEEAK | -35 | -56 | 0 | 0 | 22 | 329.73395 | 364.25922 | 354.39355 | NaN | NaN | NaN |
| Gss | GRFEDVSER | -237 | -245 | 1 | 0 | 9 | 883.36554 | NaN | 864.622 | NaN | NaN | 1074.2394 |
| Fen1 | YPVPENWLHK | -266 | -275 | 0 | 0 | 10 | 3100.7551 | 4567.038 | 4525.6704 | 4496.184 | 978.21094 | 1014.0139 |
| Fen1 | KLPIQEFHLSR | -199 | -209 | 1 | 0 | 11 | 1818.6141 | 2057.4617 | 2099.581 | 1676.9072 | 2095.2852 | 2018.8835 |
| Fen1 | KLPIQEFHLSR | -199 | -209 | 1 | 0 | 11 | 1818.6141 | 2057.4617 | 2099.581 | 1676.9072 | 2095.2852 | 2018.8835 |
| Fen1 | KLPIQEFHLS | -199 | -208 | 1 | 0.00025015 | 10 | 1085.2715 | 773.16534 | 1105.6472 | 1636.5743 | 1264.1921 | 1323.0292 |
| Fen1 | KWSEPNEEELVK | -295 | -306 | 1 | 0 | 12 | 1060.3184 | 1154.8203 | 1296.0674 | 1257.9777 | 1525.444 | 1503.3241 |
| Fen1 | GRLDDFFK | -336 | -343 | 1 | 0.00025015 | 8 | 3618.726 | 3438.8193 | 3620.329 | 3289.7424 | 2775.7612 | NaN |
| Tomm34 | LKLPPIPVVPVSAQ | -142 | -155 | 1 | 0 | 14 | 2291.457 | 2594.0913 | 2613.6533 | 2863.4207 | 3304.495 | 3705.7212 |
| Tomm34 | WNSLPSDNHK | -158 | -167 | 0 | 0 | 10 | 1545.8951 | 1296.7407 | 1247.4205 | 1573.0356 | NaN | 961.0873 |
| Tomm34 | GSADPEEESVLYSNR | -43 | -57 | 0 | 0 | 15 | 1123.2739 | 921.00415 | 933.19653 | NaN | 286.66397 | 1077.5853 |
| Gsto1 | GSAPPGPVPEGQIR | -12 | -25 | 0 | 0 | 14 | 1185.4952 | 1080.5201 | 1088.4547 | 1201.0228 | 1344.2295 | 1281.7375 |
| Gsto1 | NPLGLVPVLENSQGHLVTE | -67 | -85 | 0 | 0 | 19 | 572.1426 | 873.6073 | 722.7207 | 494.38055 | 637.9347 | 692.76575 |
| Gsto1 | LFPDDPYKK | -102 | -110 | 1 | 0.00051306 | 9 | 845.6515 | 652.3093 | 599.68976 | 827.1927 | NaN | NaN |
| Chchd4 | GSDCIDQFR | -84 | -92 | 0 | 0 | 9 | 660.4207 | 469.7177 | 635.6624 | NaN | 293.02032 | 545.12994 |
| Eri1 | GSDFSDPVYK | -54 | -63 | 0 | 0 | 10 | 506.40936 | NaN | 332.29053 | NaN | NaN | NaN |
| Abce1 | KSGNYFFLDD | -590 | -599 | 1 | 0.000431655 | 10 | 1620.7714 | 2765.7893 | 2822.5864 | 1992.1702 | 1687.2496 | 1823.7997 |
| Abce1 | LDGYVPTENLR | -313 | -323 | 0 | 0 | 11 | 1020.86194 | 2816.9897 | 3339.964 | 3999.5425 | 3725.6755 | 2750.4956 |
| Abce1 | SIVNLPSNLEK | -71 | -81 | 0 | 0 | 11 | 720.45233 | 1018.40405 | 914.08215 | 826.75336 | 906.1185 | 850.4987 |
| Abce1 | LKPDEGGEVPVLNVS | -396 | -410 | 1 | 0 | 15 | 636.1109 | 725.7731 | 481.9506 | 1039.3907 | 1163.5494 | 1265.8226 |
| Abce1 | VIVFDGVPSK | -532 | -541 | 0 | 0 | 10 | 534.97925 | 517.6767 | 539.14386 | 695.9711 | 441.53226 | 365.0696 |
| Abce1 | GSELQNYFTK | -149 | -158 | 0 | 0 | 10 | 280.33603 | 1227.9506 | 1311.4182 | 1122.5323 | 1303.5516 | 1589.7689 |
| Abce1 | LIDEPSAYLDSEQR | -484 | -497 | 0 | 0 | 14 | 2944.3018 | 2243.7874 | 3784.8767 | NaN | 4093.4146 | 3950.3323 |
| Abce1 | NVEDLSGGELQR | -213 | -224 | 0 | 0 | 12 | 2241.7993 | 2580.5059 | 2454.4563 | 2774.6985 | NaN | NaN |
| Prrc2a | GSETGSETHESDLAPSDK | -1103 | -1120 | 0 | 0.000947459 | 18 | 352.30447 | 356.5386 | 389.61435 | NaN | 538.77246 | 453.71417 |
| Spn | VPDEEATTTSGAGGNK | -303 | -318 | 0 | 0 | 16 | 3064.4446 | 1670.8628 | 3950.6992 | 5688.6636 | 5951.5156 | 2566.4297 |

| | | | | | | | | | | | | |
|---|---|---|---|---|---|---|---|---|---|---|---|---|
| Spn | GSEVLETEGSGQRPTLTTF | -319 | -337 | 1 | 0.00025015 | 19 | 959.4443 | 717.85675 | 665.61285 | 766.65656 | 960.9564 | 857.146 |
| Spn | QGSLVLEELKPGSGPNLK | -345 | -362 | 1 | 0 | 18 | 641.85004 | 780.0748 | 705.0958 | 700.71515 | 963.14435 | 828.77997 |
| Spn | GSEVLETEGSGQRPTLT | -319 | -335 | 1 | 0.00122419 | 17 | 2462.8386 | 2497.9104 | 2721.5896 | NaN | 480.14853 | NaN |
| Exosc6 | GSGPAGAGGEAPAALR | -78 | -93 | 0 | 0 | 16 | 1325.1772 | 1556.0634 | 1557.9849 | 1353.6388 | 2282.9531 | 2197.1055 |
| Exosc6 | LGLEGCQR | -244 | -251 | 0 | 0.001291103 | 8 | 1260.3282 | 1429.8977 | 1902.639 | 1331.8335 | 925.28174 | 143.50854 |
| Exosc6 | IRGPEESQPPQLY | -8 | -20 | 1 | 0 | 13 | 807.65735 | 1125.952 | 1236.2853 | 1796.4888 | 1917.6724 | 1462.7108 |
| Exosc6 | GSGEGGQTESWTDAVR | -228 | -243 | 0 | 0 | 16 | 460.5051 | 501.51883 | 571.31287 | 515.3937 | 854.1679 | 701.1812 |
| Jpt2 | GSGIFDESTPVQTR | -68 | -81 | 0 | 0 | 14 | 1613.5763 | 1240.938 | 1417.813 | 999.8556 | NaN | 1159.958 |
| Ndufs2 | VLFGEITR | -172 | -179 | 0 | 0 | 8 | 1430.2311 | 1466.0787 | 1466.4703 | 1309.0659 | 976.9367 | 985.23944 |
| Ndufs2 | LLNIQPPPR | -158 | -166 | 0 | 0 | 9 | 578.7312 | 272.80637 | 315.3261 | 780.79016 | 296.48666 | 798.0033 |
| Ndufs2 | GSGIQWDLR | -295 | -303 | 0 | 0 | 9 | 877.1568 | 1015.0165 | 812.67645 | 880.60486 | NaN | 1119.4723 |
| Ube2d3;Ube2 | TIHFPTDYPFKPPK | (53);(53) | (66);(66) | 1 | 0 | 14 | 1520.8528 | 2077.935 | 2581.1445 | 2154.033 | 2646.8533 | 2267.4695 |
| Ube2d3;Ube2 | TIHFPTDYPFKPPK | (53);(53) | (66);(66) | 1 | 0 | 14 | 1520.8528 | 2077.935 | 2581.1445 | 2154.033 | 2646.8533 | 2267.4695 |
| Ube2d3;Ube2 | IHFPTDYPFKPPK | (54);(54) | (66);(66) | 1 | 0 | 13 | 1283.4907 | 3091.131 | 2669.4004 | 2169.41 | 2877.3074 | 2827.3496 |
| Ube2d3;Ube2 | IHFPTDYPFKPPK | (54);(54) | (66);(66) | 1 | 0 | 13 | 1283.4907 | 3091.131 | 2669.4004 | 2169.41 | 2877.3074 | 2827.3496 |
| Ube2d3;Ube2 | SLLCDPNPDDPLVPEIAR | (108);(108) | (125);(125) | 0 | 0 | 18 | 863.40344 | 863.18445 | 747.7176 | 909.36035 | 1145.9564 | 1050.4668 |
| Atp5mc2;Atp5 | GSLIIGYAR | (101);(96);(91) | (109);(104);(99) | 0 | 0.003360691 | 9 | 1174.7524 | 850.6063 | NaN | 747.19775 | NaN | 500.31693 |
| Vps36 | NISEAFEDLSK | -173 | -183 | 0 | 0 | 11 | 607.54816 | 728.72925 | 703.1681 | 1051.5378 | 538.3197 | 1047.7019 |
| Vps36 | GSLTSEEFAK | -335 | -344 | 0 | 0 | 10 | 639.7698 | 573.6351 | 622.2145 | NaN | 553.36755 | 490.7376 |
| Tmem33 | LSTNQQNILK | -149 | -158 | 0 | 0 | 10 | 1448.3452 | 816.81177 | 706.0223 | 1497.7136 | 2108.1758 | 1264.188 |
| Tmem33 | PSCPLFVR | -222 | -229 | 0 | 0.001291103 | 8 | 1739.4937 | NaN | NaN | NaN | 932.2235 | NaN |
| Tmem33 | GSNSLPLLR | -135 | -143 | 0 | 0.00111006 | 9 | 856.0677 | NaN | NaN | NaN | NaN | NaN |
| Bri3bp | LEHLENQVR | -226 | -234 | 0 | 0.001023236 | 9 | 3256.739 | 2292.2246 | 2429.383 | 3585.8645 | 1750.026 | 2672.1885 |
| Bri3bp | GSPGGLCSPSVEEK | -212 | -225 | 0 | 0 | 14 | 964.24915 | 1033.1307 | 876.0781 | NaN | NaN | NaN |
| Zfr | GSPGLLDPCEKDPFDTL | -983 | -999 | 1 | 0.000471409 | 17 | 377.82996 | NaN | 471.3545 | 479.638 | NaN | 454.11407 |
| Lap3 | GSPNATEAPLVFVGK | -268 | -282 | 0 | 0 | 15 | 595.2954 | 327.1671 | 644.2099 | 879.9179 | 656.25006 | 746.4162 |
| Lap3 | TNKDEIPYLR | -487 | -496 | 1 | 0 | 10 | 520.39026 | 709.55316 | 1458.4543 | 565.652 | 1443.9373 | 3100.8286 |
| Lap3 | LNISGPPLK | -71 | -79 | 0 | 0 | 9 | 741.0303 | 521.27106 | 598.1077 | 592.551 | NaN | NaN |
| Lap3 | TIQVDNTDAEGR | -357 | -368 | 0 | 0.000230707 | 12 | 723.56885 | 3922.5498 | NaN | 2892.4329 | NaN | NaN |
| Pelp1 | NINLLGDGPLR | -356 | -366 | 0 | 0 | 11 | 813.89343 | 1083.4733 | 122.084595 | NaN | NaN | 246.11177 |
| Pelp1 | GSSDGGLQSGKPSAPK | -481 | -496 | 1 | 0 | 16 | 219.6435 | 608.01465 | 757.84247 | NaN | 582.69434 | NaN |
| Pelp1 | HASQLPTLFR | -161 | -170 | 0 | 0 | 10 | 4151.1113 | 4726.2754 | NaN | 3860.641 | NaN | NaN |
| Nup43 | HWDASTDAPEK | -288 | -298 | 0 | 0 | 11 | 1720.9299 | 1999.8457 | 2267.7175 | 1783.0345 | 2004.1946 | NaN |
| Nup43 | VPLHCVDR | -218 | -225 | 0 | 0.000869338 | 8 | 744.1863 | 396.1373 | NaN | 445.65576 | 364.59436 | 370.97333 |
| Nup43 | GTDAEAIYVTR | -366 | -376 | 0 | 0.00204437 | 11 | 411.3294 | NaN | 351.3622 | NaN | NaN | NaN |
| Cblb | GTEPIIVDPFDPR | -417 | -429 | 0 | 0 | 13 | 1387.224 | 1363.241 | 1446.9369 | 1571.44 | 1541.0413 | 1592.8217 |
| Cblb | LFQPWGSILR | -239 | -248 | 0 | 0 | 10 | 531.71173 | NaN | NaN | 451.30914 | 248.23647 | 701.94617 |
| Ehmt2 | GTFLECHPDFR | -557 | -567 | 0 | 0.004350979 | 11 | 1777.2877 | NaN | NaN | NaN | NaN | NaN |
| Emb | GTFNIHVPK | -154 | -162 | 0 | 0 | 9 | 4033.0012 | 1311.8252 | 1619.2363 | 936.0476 | 832.66656 | 1681.1755 |
| Emb | HLLEEDGGSYWCR | -229 | -241 | 0 | 0 | 13 | 1248.9968 | 1509.3345 | 841.91 | 173.26918 | 1827.1292 | 1253.8323 |
| Emb | NDPDAGKEFEQIEQLK | -293 | -308 | 1 | 0 | 16 | 609.9257 | 904.4648 | 764.59674 | NaN | 1192.0249 | 1062.0946 |
| Emb | SLIAYVGDSTVLK | -169 | -181 | 0 | 0 | 13 | 417.0986 | NaN | NaN | 682.05676 | 531.59955 | NaN |
| Lsm2 | GTLHSVDQYLNIK | -27 | -39 | 0 | 0 | 13 | 944.1789 | 1167.9451 | 1084.1937 | 1097.2947 | 1319.9238 | 1277.5614 |
| Lsm2 | GTLHSVDQYLNIK | -27 | -39 | 0 | 0 | 13 | 944.1789 | 1167.9451 | 1084.1937 | 1097.2947 | 1319.9238 | 1277.5614 |
| Lsm2 | YVQLPADEVDTQLLQDAAR | -69 | -87 | 0 | 0 | 19 | 491.06378 | 503.7092 | 516.7068 | 575.0944 | 701.4911 | 677.37897 |
| Lsm2 | LTDISVTDPEKYPH | -40 | -53 | 1 | 0 | 14 | 824.3938 | 933.23456 | 930.71423 | 835.86926 | NaN | NaN |
| Lamtor5 | GTLSDEHAGVISVL | -35 | -48 | 0 | 0.000230707 | 14 | 476.27234 | 781.5039 | 632.06244 | 707.39465 | 575.9795 | 1187.0732 |
| Gatd3 | NHDAAIFPGGFGAAK | -125 | -139 | 0 | 0 | 15 | 1601.6824 | 1036.0928 | 1486.9408 | 1397.7384 | 511.73566 | 1119.7345 |
| Gatd3 | GVEVTVGHEQEEGGK | -187 | -201 | 0 | 0 | 15 | 1439.1965 | 1199.0255 | 1152.2532 | 1511.9049 | 1162.5562 | 1447.2651 |
| Gatd3 | IFAPDVPQMHVIDHTK | -77 | -92 | 0 | 0 | 16 | 273.5399 | NaN | NaN | NaN | 661.7896 | 681.5856 |
| Agpat3 | GVFPGEQFKPAR | -292 | -303 | 1 | 0 | 12 | 5234.809 | 4563.8574 | 5573.8916 | 4821.0576 | 1480.7917 | 3656.8918 |
| Agpat3 | RFPLEDIPADETSAAQ | -257 | -272 | 1 | 0 | 16 | 1060.9773 | NaN | 481.07904 | 1478.639 | NaN | NaN |
| Pmpca | YLVGAEPAWGAPGTVDVDR | -269 | -287 | 0 | 0 | 19 | 997.76373 | 1152.9437 | 1133.758 | 897.99725 | 1113.9176 | 1077.7557 |
| Pmpca | GVGVEHEHLVECAR | -254 | -267 | 0 | 0 | 14 | 3504.033 | 3610.3806 | 3670.2898 | NaN | 4252.7285 | 4054.1318 |
| Exosc3 | NRPNVQVGDLIYGQ | -152 | -165 | 1 | 0.00025015 | 14 | 488.5859 | 657.61273 | 502.18625 | 1053.7424 | 471.06454 | 122.369064 |
| Exosc3 | GVIGQDGLLFK | -190 | -200 | 0 | 0 | 11 | 417.12277 | 540.89575 | 565.9007 | 576.46765 | NaN | NaN |
| Prdx5 | VGDAIPSVEVFEGEPGKK | -54 | -71 | 1 | 0 | 18 | 1057.4188 | 986.56464 | 1187.7073 | 693.32196 | 1011.81744 | 1028.2051 |
| Prdx5 | VGDAIPSVEVFEGEPGKK | -54 | -71 | 1 | 0 | 18 | 1057.4188 | 986.56464 | 1187.7073 | 693.32196 | 1011.81744 | 1028.2051 |
| Prdx5 | VGDAIPSVEVFEGEPGK | -54 | -70 | 0 | 0 | 17 | 767.8579 | 956.41394 | 929.3251 | 1220.2158 | 1211.5897 | 1223.0435 |
| Prdx5 | THLPGFVEQAGAL | -99 | -111 | 0 | 0 | 13 | 763.27936 | 692.1419 | 1410.3864 | 1180.8422 | 1286.2733 | 1252.9675 |
| Prdx5 | VNLAELFK | -72 | -79 | 0 | 0.001636916 | 8 | 543.6862 | 562.1265 | NaN | 425.9821 | 590.6079 | 660.8594 |
| Prdx5 | GVLFGVPGAFTPGCSK | -83 | -98 | 0 | 0 | 16 | 505.59818 | 579.188 | 620.7908 | NaN | 698.2907 | 755.4896 |
| Prdx5 | THLPGFVEQ | -99 | -107 | 0 | 0.000947459 | 9 | 2753.8647 | 96.99247 | 3022.779 | 2586.1123 | NaN | NaN |
| Prdx5 | GVPGAFTPGCSK | -87 | -98 | 0 | 0 | 12 | 1591.6987 | 2356.1375 | 319.39514 | 559.47687 | NaN | NaN |
| Prdx5 | THLPGFVEQAGALK | -99 | -112 | 0 | 0 | 14 | 344.20303 | 183.72287 | 479.46982 | NaN | NaN | NaN |
| Prdx5 | LLADPTGAFGK | -145 | -155 | 0 | 0.00025015 | 11 | 320.93436 | 366.2562 | NaN | 1723.1565 | NaN | NaN |
| Nbeal2 | RPEEAGAEVPLLPLDAL | -47 | -63 | 1 | 0.000627943 | 17 | 254.38358 | 165.13324 | 157.42964 | 301.00824 | 214.89818 | 311.05804 |
| Nbeal2 | GVLLPLLER | -892 | -900 | 0 | 0.00025015 | 9 | 209.01099 | 389.3406 | 310.22156 | 375.0979 | 324.37387 | 370.2125 |
| Nbeal2 | LFEGVCNLLDR | -1545 | -1555 | 0 | 0 | 11 | 764.3465 | NaN | 618.66 | 687.4627 | 927.9733 | 840.6338 |
| Nbeal2 | LIVVGAGQPSEVR | -2699 | -2711 | 0 | 0 | 13 | 605.462 | 486.2339 | 717.6904 | 671.79944 | NaN | 200.25836 |
| Cpsf6 | SLQDCLHGIESK | -472 | -483 | 0 | 0 | 12 | 2309.6348 | 1906.412 | 1656.8009 | 2552.9407 | 1778.6619 | 2487.8215 |
| Cpsf6 | GVNDILEIK | -105 | -113 | 0 | 0 | 9 | 1068.9991 | 3078.7437 | 924.3935 | 1043.4843 | 2617.4255 | 3595.0513 |
| Cpsf6 | TPLSEAEFEEIMNR | -407 | -420 | 0 | 0 | 14 | 418.32806 | NaN | NaN | NaN | 378.12106 | 262.6117 |
| Nubp2 | GVPFLGSVPLDSQLTR | -223 | -238 | 0 | 0 | 16 | 798.7766 | 729.3907 | 649.8097 | 813.5123 | NaN | NaN |
| Ly6c2 | GVPIKDPNIR | -81 | -90 | 1 | 0.000394361 | 10 | 2523.698 | 2889.4487 | 2887.583 | 8336.191 | 8170.12 | 7943.854 |
| Ly6c2 | QCLSFCPAGVPIKDPN | -73 | -88 | 1 | 0 | 16 | 1207.4836 | NaN | 1864.297 | 1384.911 | 2018.1097 | 1151.3594 |
| Ly6c2 | PIKDPNIR | -83 | -90 | 1 | 0.00266297 | 8 | 174.78381 | NaN | NaN | 1558.7314 | 1895.0876 | 1988.2952 |
| Gadd45gip1 | GVPPASLWPTPEQLR | -69 | -83 | 0 | 0.00025015 | 15 | 573.0472 | 769.84454 | 625.59344 | NaN | NaN | 581.1042 |

| | | | | | | | | | | | | |
|---|---|---|---|---|---|---|---|---|---|---|---|---|
| Ndufa12 | GVQQVTGHGGLR | -10 | -21 | 0 | 0 | 12 | 1651.605 | 1561.4347 | 1877.1409 | 145.18118 | 2339.292 | 1504.782 |
| Fkbp1a | GVQVETISPGDGR | -2 | -14 | 0 | 0 | 13 | 1742.8882 | 1939.6171 | 2573.041 | 3604.3428 | 2960.439 | 2966.9163 |
| Fkbp1a | VFDVELLK | -99 | -106 | 0 | 0 | 8 | 401.87225 | 259.93292 | 468.59818 | 312.34488 | 283.31757 | 144.98267 |
| Dnm1l | NVPSAGGGIGDGGQEPTTGNWR | -569 | -590 | 0 | 0.00025015 | 22 | 461.78757 | 371.11102 | 405.3729 | 468.45438 | 448.122 | 479.47968 |
| Dnm1l | NLLDVPVPVAR | -624 | -634 | 0 | 0 | 11 | 398.39136 | 590.87463 | 479.7024 | 384.74005 | 329.20416 | 517.0935 |
| Dnm1l | GVSPEPIHLK | -124 | -133 | 0 | 0.000230707 | 10 | 1298.1648 | 1346.0397 | 669.60114 | NaN | NaN | 839.8955 |
| Dnm1l | LGIIGVVNR | -239 | -247 | 0 | 0 | 9 | 718.88464 | 828.20245 | 732.65356 | NaN | NaN | NaN |
| Dnm1l | RLEEPSLR | -423 | -430 | 1 | 0.002099344 | 8 | 828.78394 | NaN | NaN | NaN | 650.54596 | NaN |
| Ctsc | NVQGVNYVSPVR | -238 | -249 | 0 | 0 | 12 | 1578.0404 | 1900.4185 | 1806.1799 | 2339.4973 | 2590.9827 | 2568.537 |
| Ctsc | ILNLPESWDWR | -227 | -237 | 0 | 0 | 11 | 470.84 | 156.96187 | 326.2727 | 473.91833 | 488.06708 | 445.1006 |
| Ctsc | NSWGSNWGESGYFR | -426 | -439 | 0 | 0 | 14 | 171.24623 | 522.7018 | 534.84705 | 525.6538 | 579.2305 | 584.94403 |
| Ctsc | LPESWDWR | -230 | -237 | 0 | 0 | 8 | 527.41144 | 729.62024 | NaN | 304.39432 | 302.9349 | 848.77423 |
| Ctsc | VHDVLGR | -126 | -132 | 0 | 0.001846793 | 7 | 3829.6555 | 4028.0388 | NaN | NaN | NaN | NaN |
| Ctsc | GVVEESCFPYTAK | -314 | -326 | 0 | 0 | 13 | 908.50024 | NaN | 2118.3284 | NaN | NaN | NaN |
| Osbp | TSKPFNPLLGETF | -489 | -501 | 1 | 0 | 13 | 476.63266 | 503.0257 | 917.66986 | 636.8033 | 811.34265 | 871.69305 |
| Osbp | GVVGPGPAAIAAPGGGGAGPPAVGGGGGR | -8 | -36 | 0 | 0.00078661 | 29 | 398.30835 | 411.42435 | 474.6874 | 474.71948 | 583.20337 | 484.07324 |
| Osbp | KVTGEVTDPSGK | -624 | -635 | 1 | 0.00025015 | 12 | 1321.8735 | 1097.5645 | NaN | 998.71014 | 1309.265 | 1341.4984 |
| Aldoc | GVVPLAGTDGETTTQ | -112 | -126 | 0 | 0 | 15 | 211.36807 | 1092.7072 | 210.17952 | 197.25949 | 155.04915 | 186.21028 |
| Aldoc | GVVPLAGTDGETTTQ | -112 | -126 | 0 | 0 | 15 | 211.36807 | 1092.7072 | 210.17952 | 197.25949 | 155.04915 | 186.21028 |
| Aldoc | VDKGVVPLAGTDGETTTQ | -109 | -126 | 1 | 0 | 18 | 594.12024 | 556.5687 | 494.2779 | 1128.1528 | 699.1186 | 711.9565 |
| Znf330 | GVWDHGGR | -135 | -142 | 0 | 0.002265552 | 8 | 686.00616 | NaN | NaN | NaN | NaN | 713.39526 |
| Ythdc1 | GVWSTLPVNEK | -376 | -386 | 0 | 0 | 11 | 1273.6991 | NaN | NaN | NaN | NaN | NaN |
| Ythdc1 | SAHLTNPWNEHKPVK | -459 | -473 | 1 | 0 | 15 | 424.8133 | NaN | NaN | NaN | NaN | NaN |
| Pik3cd | LCDIQPFLPVLR | -89 | -100 | 0 | 0 | 12 | 544.0222 | 479.79675 | 488.65805 | 581.60077 | 727.39844 | 597.483 |
| Pik3cd | GVYLNFPVSR | -29 | -38 | 0 | 0 | 10 | 332.02515 | 1868.9508 | 1797.5667 | 1535.0923 | 1519.6721 | 1676.2693 |
| Nipsnap1 | GWDENVYYTVPLVR | -255 | -268 | 0 | 0 | 14 | 363.36838 | 414.23233 | NaN | NaN | 387.886 | NaN |
| Prmt3 | GWEDEDADTEPAHGR | -29 | -43 | 0 | 0.001291103 | 15 | 454.17172 | NaN | 403.03662 | 48.023804 | 658.15 | 692.1918 |
| Sod2 | NVRPDYLK | -195 | -202 | 1 | 0.00025015 | 8 | 2923.893 | 2993.9358 | 2837.0474 | 2960.2666 | 3086.7786 | 3004.3171 |
| Sod2 | YHEALAK | -69 | -75 | 0 | 0.001636916 | 7 | 1263.492 | 82.80767 | 1398.98 | 1225.8999 | 1372.445 | 1344.1057 |
| Sod2 | GWGWLGFNK | -146 | -154 | 0 | 0 | 9 | 426.46118 | 349.632 | 442.89056 | 460.2064 | 445.9981 | 495.01645 |
| Cbfb | SKFENEEFFR | -10 | -19 | 1 | 0 | 10 | 2160.7832 | 2272.496 | 2441.1711 | 845.0865 | 3474.1406 | 3287.7395 |
| Cbfb | GWIDLHR | -112 | -118 | 0 | 0 | 7 | 2073.8953 | 1956.7578 | 2060.7058 | 1996.9102 | 1932.4413 | 1820.975 |
| Cbfb | RQQDPSPGSNLGGGDDLK | -168 | -185 | 1 | 0 | 18 | 501.03326 | NaN | 808.4649 | 673.137 | 501.84027 | 464.3623 |
| Cbfb | RQQDPSPGSNLGGGDDLKL | -168 | -186 | 2 | 0.00111006 | 19 | 2582.585 | 461.35135 | NaN | 1703.9486 | NaN | 1609.7507 |
| Prdx4 | LEDSGHTLR | -195 | -203 | 0 | 0 | 9 | 1089.2457 | 1002.7105 | 1434.8678 | 755.6187 | 1035.2891 | 1236.8204 |
| Prdx4 | GWKPGSETIIPDPAGK | -251 | -266 | 1 | 0 | 16 | 765.07245 | 832.0321 | 805.1981 | 678.42395 | 619.06287 | 799.5414 |
| Rpp38 | GWTPVHVR | -98 | -105 | 0 | 0.00111006 | 8 | 670.7169 | 1283.8804 | NaN | NaN | 1229.6788 | 1334.757 |
| Galk1 | SLETSLVPLSDPK | -205 | -217 | 0 | 0 | 13 | 1390.4905 | 1600.5199 | 1668.3914 | 1881.1989 | 2000.4756 | 2111.2874 |
| Galk1 | HSLGSSEYPVR | -229 | -239 | 0 | 0 | 11 | 882.02466 | 722.9187 | 486.33743 | 332.49353 | 588.5272 | 865.79895 |
| Galk1 | GYALLIDCR | -196 | -204 | 0 | 0 | 9 | 819.3165 | 844.594 | 954.3116 | 1071.1897 | 1181.8065 | 1040.5533 |
| Galk1 | VEELLAEAR | -9 | -17 | 0 | 0 | 9 | 1039.7692 | NaN | 1098.7853 | 727.0975 | 1296.1575 | 1721.9717 |
| Galk1 | HVVSEIR | -280 | -286 | 0 | 0 | 7 | 963.92285 | 920.9202 | 847.0682 | NaN | NaN | NaN |
| Galk1 | PGVYGSR | -336 | -342 | 0 | 0.008736733 | 7 | 224.18967 | NaN | NaN | NaN | NaN | NaN |
| Prp4k | VNIGEVLDKR | -677 | -686 | 1 | 0 | 10 | 653.2856 | 721.9282 | 810.3211 | NaN | 738.12683 | 675.85376 |
| Prp4k | GYESGSEEEGEIHEK | -140 | -154 | 0 | 0 | 15 | 262.39728 | 297.88202 | 309.79562 | 368.17667 | 286.04153 | NaN |
| Prp4k | VNIGEVLDK | -677 | -685 | 0 | 0.002068213 | 9 | 1308.876 | 1361.0544 | NaN | NaN | NaN | NaN |
| Fbln1 | QVRPIVGPF | -664 | -672 | 1 | 0.001392869 | 9 | 742.3397 | 816.2872 | 773.97644 | 666.18146 | 662.31116 | 709.89264 |
| Fbln1 | GYHLNEEGTR | -346 | -355 | 0 | 0 | 10 | 605.46155 | 1045.9226 | 996.1679 | 654.25885 | 1154.8888 | 1223.3131 |
| Fbln1 | QCGLVFR | -135 | -141 | 0 | 0.00025015 | 7 | 426.15546 | 424.226 | 594.4884 | 510.2051 | 391.49686 | NaN |
| Tyms | GYNPHPTIK | -294 | -302 | 0 | 0 | 9 | 2819.5378 | 772.6128 | NaN | NaN | 1956.7242 | 2845.9211 |
| Ap1g1 | GYPAHFGQLECLK | -60 | -72 | 0 | 0 | 13 | 1095.5234 | 1429.0278 | 1476.3115 | 1099.728 | 504.87482 | 921.01697 |
| Ap1g1 | TVQTDHNAVQR | -335 | -345 | 0 | 0 | 11 | 724.0794 | 669.3641 | 774.6944 | NaN | 924.66956 | 710.5977 |
| Snx9 | GYQGETDLNDAITEAGK | -453 | -469 | 0 | 0.00025015 | 17 | 793.8222 | 323.23987 | 1017.6771 | NaN | NaN | 503.86533 |
| Stat4 | GYVPSVFIPISTIR | -693 | -706 | 0 | 0.00383302 | 14 | 255.88608 | NaN | NaN | NaN | NaN | NaN |
| Scaf8 | HAQPPPLPVQK | -1219 | -1229 | 0 | 0 | 11 | 3318.2678 | 3186.2993 | NaN | 4397.6313 | 4351.246 | 4082.4668 |
| FAM120A | HCPSAVVPVELQK | -13 | -25 | 0 | 0 | 13 | 1226.489 | 1345.8949 | 1334.092 | 1297.2433 | 1535.6256 | 1478.7927 |
| FAM120A | NIHDTSDLDAIAK | -292 | -304 | 0 | 0 | 13 | 996.9573 | 1100.5076 | 1069.4418 | 1458.7969 | 1662.8005 | 537.13324 |
| FAM120A | NLTEQNSYSNIPHEGK | -409 | -424 | 0 | 0 | 16 | 438.7646 | 868.8156 | 724.63495 | 559.85956 | 573.44275 | 647.65985 |
| FAM120A | HTPLYER | -425 | -431 | 0 | 0 | 7 | 2208.6655 | 2201.6147 | 2103.225 | NaN | NaN | 2253.0422 |
| Hmbs | HDPETLLR | -239 | -246 | 0 | 0 | 8 | 700.47205 | 1898.6635 | 1271.679 | NaN | 1127.9092 | 1102.5547 |
| Hmbs | KFPHLEFK | -157 | -164 | 1 | 0.004658491 | 8 | 372.29285 | NaN | 426.06787 | NaN | NaN | NaN |
| Git2 | SKDNVELILR | -364 | -373 | 1 | 0 | 10 | 786.8664 | 937.0557 | 871.59406 | 1083.003 | 1344.308 | 489.80496 |
| Git2 | HDSYIPCSER | -615 | -624 | 0 | 0 | 10 | 933.46606 | 815.57556 | 860.7833 | 950.90607 | NaN | 883.299 |
| Pcid2 | HETFFIR | -301 | -307 | 0 | 0.000230707 | 7 | 1159.677 | 958.5066 | 643.63385 | 1681.1656 | 941.8004 | 765.34033 |
| Tor1aip1 | ISHLVLPVQPENTLK | -576 | -590 | 0 | 0 | 15 | 883.3884 | 778.69904 | 804.6575 | 457.2194 | 656.06165 | 598.1452 |
| Tor1aip1 | ISHLVLPVQPENTLK | -576 | -590 | 0 | 0 | 15 | 883.3884 | 778.69904 | 804.6575 | 457.2194 | 656.06165 | 598.1452 |
| Tor1aip1 | HEVDQELTDGFK | -471 | -482 | 0 | 0 | 12 | 1976.1237 | NaN | 1340.18 | 1375.5963 | 1362.4497 | 1302.1876 |
| Tor1aip1 | TSGCQENPQEWVDR | -293 | -306 | 0 | 0 | 14 | 558.36304 | 473.36462 | 507.96466 | NaN | 490.37302 | 600.53735 |
| Tor1aip1 | SAIHHPNEPSVK | -336 | -347 | 0 | 0 | 12 | 472.2568 | 256.60257 | 525.19855 | 1284.112 | 610.8641 | NaN |
| Tor1aip1 | LEQHSQQPQLSPATSGR | -130 | -146 | 0 | 0.000230707 | 17 | 2377.7393 | 3029.5938 | 3090.2803 | NaN | NaN | 3348.289 |
| Tor1aip1 | IHHPNEPSVK | -338 | -347 | 0 | 0.003037016 | 10 | 1197.1836 | NaN | NaN | NaN | 1917.5502 | 1868.4971 |
| Tor1aip1 | SNFGNQSPSTSRPQ | -322 | -335 | 1 | 0.00025015 | 14 | 421.27124 | 561.49243 | 665.2804 | NaN | NaN | NaN |
| Dstn | LWAPEQAPLK | -103 | -112 | 0 | 0 | 10 | 966.04016 | 1090.177 | 1121.2019 | NaN | 1197.048 | 864.85895 |
| Dstn | WAPEQAPLK | -104 | -112 | 0 | 0.00025015 | 9 | 208.19246 | NaN | NaN | 882.587 | NaN | NaN |
| Dstn | HEYQANGPEDLNR | -133 | -145 | 0 | 0.00635324 | 13 | 553.8439 | NaN | NaN | NaN | NaN | NaN |
| Wdr46 | VPAELICLNPR | -508 | -518 | 0 | 0 | 11 | 1270.4735 | 1460.047 | 1537.3999 | 1724.9172 | 1565.8619 | 2290.2327 |
| Wdr46 | HFDLNLR | -187 | -193 | 0 | 0 | 7 | 844.70447 | 954.24225 | 756.5769 | 1041.0459 | 843.8852 | 840.2942 |
| Wdr46 | LAEEPGFLVGEDGEDTAK | -152 | -169 | 0 | 0 | 18 | 587.78644 | 742.92725 | NaN | NaN | 1046.9346 | 1014.6158 |

| | | | | | | | | | | | | |
|---|---|---|---|---|---|---|---|---|---|---|---|---|
| Wdr46 | TLSGAQDPFPGPVPAPLEEAR | -89 | -109 | 0 | 0 | 21 | 419.64487 | 315.80844 | NaN | 675.59906 | NaN | 439.76163 |
| Rpl36a | HFELGGDK | -90 | -97 | 0 | 0 | 8 | 2978.5706 | 3015.693 | 3147.9963 | 2441.2058 | 2596.1199 | 2582.1292 |
| Rpl36a | LECVEPNCR | -70 | -78 | 0 | 0 | 9 | 2468.895 | 2463.448 | 1996.9033 | 2004.4565 | 1367.7507 | 1768.6995 |
| Rpl36a | HFELGGDKK | -90 | -98 | 1 | 0.003620005 | 9 | 2396.0037 | NaN | NaN | 2345.2927 | 2266.6426 | NaN |
| Rpl36a | HFELGGDKK | -90 | -98 | 1 | 0.003620005 | 9 | 2396.0037 | NaN | NaN | 2345.2927 | 2266.6426 | NaN |
| Ndufv1 | HFRPELEDR | -441 | -449 | 1 | 0 | 9 | 1426.1511 | 1860.2877 | 1702.3466 | 1853.7336 | 2032.6208 | 2383.4026 |
| Ndufv1 | KTSFGSLKDEDR | -29 | -40 | 2 | 0.00025015 | 12 | 329.53192 | 647.5713 | NaN | 470.30206 | 762.30646 | 846.0854 |
| Ndufv1 | LKPPFPADVGVF | -225 | -236 | 1 | 0.00025015 | 12 | 1321.2476 | NaN | 1828.2025 | NaN | 1978.6464 | 2158.8098 |
| Nploc4 | SISQNPFPIENR | -465 | -476 | 0 | 0 | 12 | 3443.8582 | 2627.2847 | 2690.8394 | NaN | 959.7637 | 762.8884 |
| Nploc4 | SSEQYVPDVFYK | -416 | -427 | 0 | 0 | 12 | 644.91675 | 598.9178 | 357.674 | 554.1243 | 725.8188 | NaN |
| Nploc4 | HGDLLFLFPS | -71 | -80 | 0 | 0.001868731 | 10 | 286.8589 | NaN | 519.658 | NaN | 1161.148 | 371.4329 |
| Pgm1 | HGFFVNPSDSVAVIA | -300 | -314 | 0 | 0.00025015 | 15 | 3279.36 | 782.2968 | 1082.3474 | 3785.115 | 5133.19 | 5027.896 |
| Pgm1 | TIEEYAICPDLK | -153 | -164 | 0 | 0 | 12 | 1823.4366 | 1137.2126 | 1903.3959 | 1263.4081 | 1532.7631 | 1802.2621 |
| Pgm1 | QEATLVVGGDGR | -53 | -64 | 0 | 0 | 12 | 4527.378 | 1688.394 | 4524.376 | 4543.037 | 4941.9355 | NaN |
| Pgm1 | IALYETPTGWK | -350 | -360 | 0 | 0 | 11 | 2337.5996 | 3531.1426 | 3114.2612 | NaN | 3070.1082 | 3025.7102 |
| Pgm1 | KTIEEYAICPDLK | -152 | -164 | 1 | 0 | 13 | 267.2472 | NaN | 465.1321 | 515.4412 | 512.9188 | 613.02246 |
| Pgm1 | SIVSTVEPALR | -42 | -52 | 0 | 0 | 11 | 783.1386 | 795.0953 | 718.759 | NaN | NaN | 875.09186 |
| Pgm1 | LSGTGSAGATIR | -504 | -515 | 0 | 0.00025015 | 12 | 312.43613 | 494.29257 | 1403.6522 | NaN | NaN | NaN |
| Pgm1 | VDLGVLGK | -165 | -172 | 0 | 0.00272025 | 8 | 598.9825 | NaN | NaN | NaN | NaN | NaN |
| Cyc1 | LSDYFPKPYPNPEAAR | -171 | -186 | 1 | 0.00025015 | 16 | 1529.8613 | 2029.7688 | 1852.1454 | 1712.6001 | 2388.6777 | 2234.827 |
| Cyc1 | SDLELHPPSYPWSHR | -85 | -99 | 0 | 0 | 15 | 1475.036 | 1765.6985 | 1798.331 | 1443.285 | 1792.0541 | 1996.9974 |
| Cyc1 | SDLELHPPSYPWSHR | -85 | -99 | 0 | 0 | 15 | 1475.036 | 1765.6985 | 1798.331 | 1443.285 | 1792.0541 | 1996.9974 |
| Cyc1 | LTGYCEPPTGVSLR | -215 | -228 | 0 | 0 | 14 | 1210.7992 | 1144.193 | 1039.6715 | 1246.8157 | 1177.9398 | 1168.1553 |
| Cyc1 | WASEPEHDHR | -276 | -285 | 0 | 0 | 10 | 106.99438 | 402.387 | 322.60733 | 517.5707 | 313.89398 | 701.86847 |
| Cyc1 | HGGEDYVFSL | -205 | -214 | 0 | 0.00025015 | 10 | 2561.4229 | 3682.5789 | 2399.5908 | NaN | 310.4224 | 690.7618 |
| Cyc1 | LSDYFPKPYPNPEAA | -171 | -185 | 1 | 0.00025015 | 15 | 2068.3306 | NaN | 2016.7139 | 2044.4572 | 1802.3821 | 1925.7916 |
| Cyc1 | SDYFPKPYPNPEAAR | -172 | -186 | 1 | 0.00025015 | 15 | 365.48776 | 1044.0039 | 918.61414 | 1535.6053 | 1534.8087 | NaN |
| Cyc1 | RWASEPEHDHR | -275 | -285 | 1 | 0 | 11 | 540.6052 | NaN | 230.54683 | NaN | 473.20715 | NaN |
| Apip | HGNEIYIAPSGVQK | -51 | -64 | 0 | 0 | 14 | 547.1706 | 312.4247 | 523.03973 | 560.1278 | 519.592 | 565.01 |
| Apeh | TVHTEWTQR | -45 | -53 | 0 | 0 | 9 | 1855.1327 | 770.2319 | 1266.1729 | 1262.2808 | 1297.947 | 1237.9395 |
| Apeh | QVLQEEHFDAR | -569 | -579 | 0 | 0 | 11 | 1639.1125 | 1763.7639 | 2218.4949 | 4194.616 | 576.74194 | 3012.8171 |
| Apeh | SLPGNVGHQDVK | -550 | -561 | 0 | 0 | 12 | 798.5345 | 880.6307 | 992.6669 | 667.5372 | 666.87134 | 813.2492 |
| Apeh | HGPVYEDDCFGCL | -141 | -153 | 0 | 0 | 13 | 436.58142 | 516.4768 | 542.86847 | 497.9498 | 587.77795 | 657.90393 |
| Apeh | RVVFDSVQR | -375 | -383 | 1 | 0 | 9 | 1416.8817 | 250.61751 | NaN | 1375.2429 | 1575.0873 | 1129.3356 |
| Fam162a | HGVPLHKPTDFEK | -57 | -69 | 1 | 0 | 13 | 1814.5039 | 1046.2019 | 1717.0979 | 591.35913 | 1953.6287 | 2310.5889 |
| Fam162a | HGVPLHKPTDFEK | -57 | -69 | 1 | 0 | 13 | 1814.5039 | 1046.2019 | 1717.0979 | 591.35913 | 1953.6287 | 2310.5889 |
| Phf5a | ICDECNYGSYQGR | -45 | -57 | 0 | 0 | 13 | 514.1842 | 558.05115 | 554.1404 | 792.87115 | 796.8834 | 66.33836 |
| Phf5a | HHPDLIFCR | -4 | -12 | 0 | 0.000907206 | 9 | 1145.0466 | NaN | NaN | 3492.9207 | 641.9463 | 1386.4932 |
| Crlf3 | LIEHGVNTADDLVR | -98 | -111 | 0 | 0 | 14 | 906.95026 | 998.9727 | 974.0443 | 664.14746 | 655.9513 | 601.17163 |
| Crlf3 | LIEHGVNTADDLVR | -98 | -111 | 0 | 0 | 14 | 906.95026 | 998.9727 | 974.0443 | 664.14746 | 655.9513 | 601.17163 |
| Crlf3 | LDDSILNIVK | -159 | -168 | 0 | 0 | 10 | 826.3156 | 799.37994 | 915.69586 | 956.5334 | 899.7902 | 1026.2028 |
| Crlf3 | HIDPNVDYQFR | -238 | -248 | 0 | 0 | 11 | 573.663 | 1618.5281 | 1752.5132 | 1907.6234 | 1930.9421 | 1953.5518 |
| Crlf3 | SFFYPGWK | -431 | -438 | 0 | 0 | 8 | 471.75217 | 569.9267 | 570.89215 | 342.8484 | 488.12872 | 569.48926 |
| Crlf3 | SRPPVQIEELIEKPGGIIVR | -179 | -198 | 2 | 0.00025015 | 20 | 389.96973 | 383.00256 | 388.35468 | 511.6762 | 677.3764 | 574.4599 |
| Prkd2 | HIQIGLTR | -48 | -55 | 0 | 0.000347792 | 8 | 544.49243 | NaN | NaN | NaN | 675.7806 | 672.3353 |
| Prkd2 | SFYTGRPIELDK | -245 | -256 | 1 | 0.000471409 | 12 | 1309.2651 | NaN | NaN | NaN | NaN | 564.6037 |
| Swi5 | HISLLHEYNDIK | -47 | -58 | 0 | 0.001023236 | 12 | 1052.0045 | NaN | NaN | 2746.1934 | 947.3717 | NaN |
| Idh3g | QAIQDIIR | -374 | -381 | 0 | 0.001392869 | 8 | 1621.3232 | 1226.2832 | NaN | 1110.7397 | 893.9835 | NaN |
| Cdc73 | NVHLSHPVYVR | -66 | -76 | 0 | 0 | 11 | 979.37775 | 575.6996 | 503.06415 | 800.8539 | 758.5432 | 697.8556 |
| Cdc73 | HLDRPVFLR | -505 | -513 | 1 | 0 | 9 | 519.89795 | 502.85587 | 672.4265 | 410.91788 | 516.7255 | 478.78055 |
| Rfc3 | NLVQCGDFPHLL | -28 | -39 | 0 | 0 | 12 | 1811.5178 | NaN | NaN | 658.8768 | 1798.4957 | 1233.4775 |
| Rfc3 | HLEVNPSDAGNSDR | -92 | -105 | 0 | 0.003490217 | 14 | 753.7989 | NaN | NaN | NaN | NaN | NaN |
| Gp1bb | HLGANPWR | -85 | -92 | 0 | 0 | 8 | 803.56244 | NaN | NaN | 818.6842 | NaN | NaN |
| Suclg1 | LIGPNCPGVINPGECK | -167 | -182 | 0 | 0 | 16 | 1485.527 | 2227.4854 | 1520.8561 | 1424.2247 | 1627.8873 | 1377.3805 |
| Suclg1 | HLGLPVFNTVK | -95 | -105 | 0 | 0 | 11 | 1000.2508 | 121.656746 | 233.68059 | NaN | 1257.02 | 1302.9445 |
| Suclg1 | HLGLPVFNTV | -95 | -104 | 0 | 0.000230707 | 10 | 114.19015 | 801.0015 | 915.9156 | NaN | 1018.40155 | 373.15198 |
| Suclg1 | HLGLPVFNT | -95 | -103 | 0 | 0 | 9 | 278.4422 | 204.11406 | 202.60268 | 456.07843 | NaN | NaN |
| Atp5f1c | TEEKPIFSLN | -198 | -207 | 1 | 0.001868731 | 10 | 1757.0238 | 2174.9338 | 586.19116 | 1455.5956 | 1503.9703 | 1623.4783 |
| Atp5f1c | KPPTFGDASVIAL | -159 | -171 | 1 | 0.000471409 | 13 | 1599.3203 | 1867.2397 | 1816.4369 | 2115.322 | 2482.2195 | 2422.544 |
| Atp5f1c | TEEKPIFSL | -198 | -206 | 1 | 0.002697088 | 9 | 1148.47 | 1258.3513 | NaN | 1094.1045 | 1304.2621 | 1291.2267 |
| Atp5f1c | HLIIGVSSDR | -91 | -100 | 0 | 0 | 10 | 1097.2104 | 900.57837 | NaN | 719.844 | 937.2632 | 630.44617 |
| Atp5f1c | THSDQFLVSFK | -144 | -154 | 0 | 0.000394361 | 11 | 3422.5413 | NaN | NaN | 5549.849 | NaN | NaN |
| Wdr18 | SVSTDGSVLLSGSHDESVR | -276 | -294 | 0 | 0 | 19 | 200.58206 | 265.16974 | 246.11807 | 463.79562 | 683.9085 | 709.71027 |
| Wdr18 | HLLGAEHGDEAQGGGL | -342 | -357 | 0 | 0.007880195 | 16 | 1302.3977 | NaN | NaN | 949.95044 | NaN | NaN |
| Ezh2 | SDLDFPAQVIPLK | -87 | -99 | 0 | 0 | 13 | 516.90015 | 1045.6382 | NaN | NaN | 1389.4534 | 1540.1188 |
| Thoc6 | HLLSAGDGEVK | -88 | -98 | 0 | 0 | 11 | 1997.5339 | 1924.38 | 1829.7548 | 1467.1274 | 1670.5399 | 1493.3646 |
| Thoc6 | SSTPTTVFPIR | -248 | -258 | 0 | 0 | 11 | 1046.561 | 897.0654 | 1101.4028 | NaN | 435.70068 | 558.22156 |
| Thoc6 | SPEVLSGGEDGAVR | -180 | -193 | 0 | 0 | 14 | 845.04663 | NaN | 1011.60974 | 372.13467 | 698.71655 | 712.38336 |
| Nup85 | LLTDALPLLEQK | -577 | -588 | 0 | 0 | 12 | 660.3084 | 706.7159 | 807.14136 | 704.56683 | 798.5946 | 774.107 |
| Nup85 | HLQDNTFAANPR | -268 | -279 | 0 | 0 | 12 | 860.6319 | 1030.6525 | NaN | 469.49518 | 1476.9907 | 1739.5396 |
| Nup85 | VPSCPFIYIIR | -48 | -58 | 0 | 0 | 11 | 665.7432 | 600.8441 | 839.93164 | NaN | 772.17065 | 818.6701 |
| Nup85 | IEEELSGK | -85 | -92 | 0 | 0.00111006 | 8 | 1456.6376 | NaN | NaN | NaN | 989.54126 | NaN |
| Nup85 | KVPSCPFIYIIR | -47 | -58 | 1 | 0.001868731 | 12 | 658.45074 | NaN | NaN | NaN | NaN | NaN |
| Lmf2 | HLSAEPGLR | -289 | -297 | 0 | 0.00078661 | 9 | 620.6856 | 551.6082 | 551.9728 | NaN | NaN | 488.0012 |
| Lmf2 | LWETPTLLWEAPR | -57 | -69 | 0 | 0 | 13 | 590.92505 | 319.50864 | 441.68762 | NaN | NaN | NaN |
| Afg3l2 | VGQISFDLPR | -669 | -678 | 0 | 0 | 10 | 2917.616 | 3465.9814 | 3392.1104 | 3084.0015 | 3692.6572 | 3749.962 |
| Afg3l2 | VSEEIFFGR | -632 | -640 | 0 | 0.000230707 | 9 | 557.3298 | 907.29016 | 930.9682 | 897.8766 | 1007.36993 | 912.80396 |
| Afg3l2 | HLSDAINEK | -534 | -542 | 0 | 0 | 9 | 4181.454 | NaN | NaN | 3007.9712 | 3579.809 | 2793.2964 |

| | | | | | | | | | | | | |
|---|---|---|---|---|---|---|---|---|---|---|---|---|
| Afg3l2 | VLKDEIDVK | -297 | -305 | 1 | 0.001392869 | 9 | 633.372 | NaN | NaN | NaN | NaN | NaN |
| Ak3 | REDDKPETVIK | -161 | -171 | 2 | 0.00078661 | 11 | 1068.8655 | NaN | 1072.2036 | 1565.479 | 570.1721 | 991.03485 |
| Ak3 | TVGIDDLTGEPLIQR | -147 | -161 | 0 | 0 | 15 | 1256.0907 | 1164.4825 | 1148.8792 | NaN | 321.15112 | NaN |
| Ak3 | HLSSGDLLR | -35 | -43 | 0 | 0.000591124 | 9 | 641.26666 | NaN | NaN | NaN | 567.15466 | NaN |
| Ebp | TVPLHPYWPR | -5 | -14 | 0 | 0 | 10 | 8384.756 | 7331.5938 | 8447.1875 | 7285.2607 | 6456.637 | 6080.1855 |
| Ebp | TTNTVPLHPYWPR | -2 | -14 | 0 | 0 | 13 | 552.3596 | 637.3083 | 1350.811 | 889.12396 | 1022.26355 | 957.68506 |
| Ebp | HLTSAQSVLDSK | -210 | -221 | 0 | 0 | 12 | 5149.3613 | NaN | NaN | 5995.699 | NaN | 5333.8257 |
| Cisd1 | HNEETGDNVGPLIIK | -90 | -104 | 0 | 0 | 15 | 1436.1116 | 1613.8478 | 1585.0322 | 1550.9857 | 1797.7052 | 1834.1671 |
| Cisd1 | KFPFCDGAHIK | -79 | -89 | 1 | 0 | 11 | 609.25446 | 799.0984 | 812.80634 | 858.5487 | 734.7853 | 934.87915 |
| Cisd1 | KFPFCDGAHIK | -79 | -89 | 1 | 0 | 11 | 609.25446 | 799.0984 | 812.80634 | 858.5487 | 734.7853 | 934.87915 |
| Cisd2 | VQLPAYLK | -12 | -19 | 0 | 0.000394361 | 8 | 765.26495 | 260.10837 | 322.879 | 712.58795 | 685.6773 | 686.6011 |
| Cisd2 | VVNEINIEDLCLTK | -82 | -95 | 0 | 0 | 14 | 461.2557 | 501.82147 | 481.56097 | 542.47107 | 735.1583 | 809.3977 |
| Cisd2 | HNELTGDNVGPL | -117 | -128 | 0 | 0.002068213 | 12 | 1158.5316 | NaN | 1262.1654 | 1641.9678 | 1218.7592 | 1268.5192 |
| Cisd2 | HNELTGDNVGPLIL | -117 | -130 | 0 | 0 | 14 | 573.0433 | 354.112 | NaN | 479.3946 | 930.1309 | 814.3561 |
| Msra | HNFGPITTDIR | -184 | -194 | 0 | 0 | 11 | 739.0886 | 1476.1027 | 972.10315 | 1371.2675 | 2079.6438 | 2124.5693 |
| Dgcr8 | HNSGVPVYLHR | -312 | -322 | 0 | 0 | 11 | 484.62967 | 431.48193 | 722.5594 | 612.1318 | 419.36224 | 208.78673 |
| Gc | HQPQEFPTYVEPTNDEICEAFR | -128 | -149 | 0 | 0 | 22 | 739.8235 | 686.38794 | 730.93176 | 902.29144 | 899.28656 | 932.20624 |
| Gc | YSENTFTEYK | -410 | -419 | 0 | 0 | 10 | 3247.1624 | 2799.6196 | 1899.1593 | 2146.5083 | NaN | 1249.2909 |
| Gc | YVEPTNDEICEAFR | -136 | -149 | 0 | 0.000230707 | 14 | 307.94226 | 1282.438 | 1387.8538 | 797.00525 | 939.27057 | NaN |
| Gc | YSENTFTEYKK | -410 | -420 | 1 | 0 | 11 | 1942.1923 | 2388.6921 | NaN | NaN | NaN | NaN |
| Taok3 | HQTELENQLEYNK | -679 | -691 | 0 | 0.00025015 | 13 | 265.61533 | 323.4464 | NaN | NaN | 378.90057 | 350.20755 |
| Gtf2f2 | TWELKPEYR | -232 | -240 | 1 | 0 | 9 | 1529.476 | 1610.4043 | 1543.4678 | 1108.2383 | 1310.4779 | 1082.8119 |
| Gtf2f2 | NTWELKPEYR | -231 | -240 | 1 | 0 | 10 | 1091.925 | 1040.0258 | 1086.7123 | 664.39386 | 276.3897 | 366.00513 |
| Gtf2f2 | QNTGVWLVK | -14 | -22 | 0 | 0.000230707 | 9 | 797.3561 | NaN | 751.7561 | NaN | NaN | NaN |
| Gtf2f2 | LSLEGIVVQR | -104 | -113 | 0 | 0.00078661 | 10 | 1088.458 | NaN | NaN | NaN | NaN | NaN |
| Gtf2f2 | HQYYNLK | -194 | -200 | 0 | 0.002910424 | 7 | 869.1131 | NaN | NaN | NaN | NaN | NaN |
| Rassf2 | SELLLHLK | -21 | -28 | 0 | 0.001055699 | 8 | 613.92456 | 307.55365 | 501.06052 | 719.5127 | 273.28256 | 461.46274 |
| Rassf2 | HREEEDEFIVE | -43 | -53 | 1 | 0.00025015 | 11 | 1058.7843 | 1046.6522 | 1104.736 | NaN | 1205.5508 | 518.4453 |
| Man1a1 | SHLSGDPVFAEK | -363 | -374 | 0 | 0.00051306 | 12 | 736.5457 | NaN | NaN | 1053.5073 | 885.0327 | 601.3214 |
| Man1a1 | YLSGEEIFR | -296 | -304 | 0 | 0 | 9 | 306.02765 | NaN | NaN | 747.98706 | 1226.1848 | NaN |
| Man1a1 | HREEGAPGDPGAGLEDNLAR | -106 | -125 | 1 | 0 | 20 | 345.32474 | NaN | NaN | NaN | NaN | NaN |
| Pex3 | HSLSLLDLEQK | -198 | -208 | 0 | 0.000471409 | 11 | 599.14185 | NaN | NaN | NaN | 713.94366 | 573.7245 |
| Acot9 | NIIHELFLT | -259 | -267 | 0 | 0.004439092 | 9 | 690.6047 | NaN | NaN | NaN | 814.09796 | NaN |
| Hmgcs1 | LEDTYFDRDVEK | -306 | -317 | 1 | 0 | 12 | 1910.0726 | 2637.5422 | 1287.2948 | 1745.6099 | 1525.4199 | 1532.7937 |
| Hmgcs1 | LIGPNAPLIFDR | -180 | -191 | 0 | 0 | 12 | 486.30472 | 315.16513 | 588.9148 | 722.11115 | 822.1619 | 820.1507 |
| Hmgcs1 | LLIGPNAPLIFDR | -179 | -191 | 0 | 0 | 13 | 394.3332 | 1254.1559 | 1302.108 | 662.0866 | 1647.5742 | 323.93198 |
| Hmgcs1 | VTQDATPGSALDK | -388 | -400 | 0 | 0 | 13 | 1260.3129 | NaN | 835.48004 | 684.2678 | 1263.1311 | 1297.4429 |
| Hmgcs1 | HSLSYDCIGR | -80 | -89 | 0 | 0 | 10 | 2792.008 | NaN | 2593.7898 | 3243.2969 | 2868.8372 | NaN |
| Hmgcs1 | LEDTYFDR | -306 | -313 | 0 | 0 | 8 | 513.7242 | 236.93549 | NaN | NaN | NaN | NaN |
| Ik | HSYFEKPK | -381 | -388 | 1 | 0.00025015 | 8 | 262.54462 | 635.94727 | 627.3072 | 408.44986 | 435.74603 | 529.98395 |
| Ik | HSYFEKPK | -381 | -388 | 1 | 0.00025015 | 8 | 262.54462 | 635.94727 | 627.3072 | 408.44986 | 435.74603 | 529.98395 |
| Ik | KDEDPENKIEFK | -198 | -209 | 2 | 0 | 12 | 498.5657 | 108.719536 | 669.7679 | 482.46573 | 815.00214 | 810.55 |
| Plp2 | HTAAPTDPTDGP | -141 | -152 | 0 | 0 | 12 | 545.27026 | 1074.0205 | 1023.4108 | NaN | NaN | NaN |
| Vac14 | SSPAGQTDDPGAPDGPDLR | -471 | -489 | 0 | 0 | 19 | 765.5076 | 642.8714 | 552.28613 | 611.8351 | 569.83307 | 473.50653 |
| Vac14 | HTDSLFPILLQ | -439 | -449 | 0 | 0.00025015 | 11 | 352.80267 | 370.29932 | 387.40405 | 344.65985 | 97.98276 | 463.11496 |
| Vac14 | VNHSELQVPTSGR | -490 | -502 | 0 | 0 | 13 | 923.1209 | NaN | NaN | NaN | NaN | NaN |
| Ap2s1 | HTNFVEFR | -46 | -53 | 0 | 0 | 8 | 1102.0293 | 1078.0219 | 463.5274 | 1815.4174 | 1183.132 | 1081.1422 |
| Srp9 | HVDGNLCIK | -33 | -41 | 0 | 0 | 9 | 631.0366 | 1694.8501 | 782.8542 | NaN | 1154.4121 | 1032.0874 |
| Srp9 | VTDDLVCLVYR | -42 | -52 | 0 | 0 | 11 | 709.40607 | NaN | NaN | 227.7683 | 1175.9774 | 562.55066 |
| Map2k1 | VGELKDDDFEK | -60 | -70 | 1 | 0 | 11 | 1307.2294 | 1339.1927 | 1409.4816 | 1567.5258 | 1720.9794 | 1728.3239 |
| Map2k1 | KLEELELDEQQR | -36 | -47 | 1 | 0 | 12 | 741.2772 | 765.6658 | 869.86487 | 832.25165 | 821.44867 | 800.15576 |
| Map2k1 | LEELELDEQQR | -37 | -47 | 0 | 0 | 11 | 3733.336 | NaN | 4124.328 | 4779.064 | 5658.006 | 5490.2524 |
| Map2k1 | IPEQILGK | -161 | -168 | 0 | 0.004031417 | 8 | 4969.3203 | 3547.6443 | NaN | 5770.4517 | NaN | NaN |
| Map2k1 | HVEGDAAETPPRPR | -278 | -291 | 1 | 0 | 14 | 568.0818 | NaN | NaN | NaN | 932.23047 | 385.55774 |
| Ntpcr | HVFLTGPPGVGK | -4 | -15 | 0 | 0 | 12 | 1510.6035 | 1621.8387 | 1791.2688 | 2791.903 | 2572.655 | 2588.924 |
| Rbm12 | HVIDFFK | -445 | -451 | 0 | 0.001610812 | 7 | 1992.3907 | 2276.1033 | 2493.1733 | 1693.9656 | 1884.1082 | 87.37396 |
| Ca1 | HVIFDDSSNQSVLK | -68 | -81 | 0 | 0 | 14 | 559.4275 | 747.36414 | 707.83264 | NaN | 410.98325 | 564.4745 |
| Mthfd2 | LVGDVDFEGVK | -289 | -299 | 0 | 0 | 11 | 1986.6715 | 2154.1804 | 2144.3608 | 1002.7342 | 626.65717 | 541.4161 |
| Mthfd2 | VLRPEELEVF | -331 | -340 | 1 | 0.00025015 | 10 | 1140.6685 | 1148.6256 | 1382.6383 | 1483.2178 | 2119.3274 | 1757.9246 |
| Mthfd2 | SVSEEELLNSIR | -107 | -118 | 0 | 0 | 12 | 1020.7203 | 1291.519 | 1069.3147 | 384.04752 | 1819.8413 | 214.16785 |
| Mthfd2 | LVQLPLPEHIDER | -130 | -142 | 0 | 0 | 13 | 805.46747 | 1032.1841 | 944.147 | 1195.6047 | 1381.541 | 1294.2245 |
| Mthfd2 | LVQLPLPEHIDER | -130 | -142 | 0 | 0 | 13 | 805.46747 | 1032.1841 | 944.147 | 1195.6047 | 1381.541 | 1294.2245 |
| Mthfd2 | RVQDPVTAKPK | -278 | -288 | 2 | 0 | 11 | 525.5519 | 613.1614 | 715.8413 | 592.5036 | 784.1043 | 805.30756 |
| Mthfd2 | RVQDPVTAKPK | -278 | -288 | 2 | 0 | 11 | 525.5519 | 613.1614 | 715.8413 | 592.5036 | 784.1043 | 805.30756 |
| Mthfd2 | QEVQQEVEEWVASGNK | -51 | -66 | 0 | 0 | 16 | 448.38904 | NaN | 369.70718 | 386.44846 | 530.6139 | 433.2357 |
| Mthfd2 | HVINVGR | -158 | -164 | 0 | 0.001055699 | 7 | 1064.801 | 1074.9038 | 1236.3383 | 1875.9431 | NaN | NaN |
| Mthfd2 | VCNAVSPDKDVDGFHVINVGR | -144 | -164 | 1 | 0 | 21 | 410.10388 | 431.15247 | NaN | 575.63043 | NaN | 577.0372 |
| Mthfd2 | KVCNAVSPDKDVDGF | -143 | -157 | 2 | 0.00122419 | 15 | 372.45084 | 802.34064 | 519.64734 | NaN | NaN | 506.63788 |
| Smarcd2 | IITDVIGNPEEERR | -481 | -494 | 1 | 0 | 14 | 433.91953 | 365.78186 | 1101.9557 | 731.46173 | 1049.2203 | 357.59177 |
| A2m | TEVSNNHVLIY | -1408 | -1418 | 0 | 0 | 11 | 2190.8555 | 382.3028 | 2305.3142 | 661.5785 | 502.17847 | 2197.6804 |
| A2m | TEVSNNHVLIY | -1408 | -1418 | 0 | 0 | 11 | 2190.8555 | 382.3028 | 2305.3142 | 661.5785 | 502.17847 | 2197.6804 |
| A2m | MVSGFIPLKPTVK | -1385 | -1397 | 1 | 0 | 13 | 1884.4252 | 2062.5923 | 2022.2006 | 2684.9033 | 3138.9443 | 2956.9966 |
| A2m | MVSGFIPLKPTVK | -1385 | -1397 | 1 | 0 | 13 | 1884.4252 | 2062.5923 | 2022.2006 | 2684.9033 | 3138.9443 | 2956.9966 |
| A2m | HVLIYLDK | -1414 | -1421 | 0 | 0.00051306 | 8 | 1583.8301 | 858.3126 | 918.757 | 1034.8326 | 988.333 | 988.80945 |
| A2m | LNTGYQR | -1008 | -1014 | 0 | 0 | 7 | 1542.8497 | 1251.4425 | 1288.8945 | 296.3248 | 3329.4114 | 2142.9053 |
| A2m | TEVSNNHVLIYLDK | -1408 | -1421 | 0 | 0 | 14 | 692.90826 | 809.5692 | 814.2044 | 2055.0928 | 2519.9094 | 2370.581 |
| A2m | VSGFIPLKPTVK | -1386 | -1397 | 1 | 0.00025015 | 12 | 848.4276 | NaN | NaN | 723.23346 | 843.532 | NaN |
| Nup50 | HVNTNPLCDLTPIFK | -173 | -187 | 0 | 0.00051306 | 15 | 471.91418 | NaN | NaN | 433.28653 | 634.3318 | 1050.9082 |

| | | | | | | | | | | | | |
|---|---|---|---|---|---|---|---|---|---|---|---|---|
| Txndc17 | SWCPDCVEAEPVIR | -41 | -54 | 0 | 0 | 14 | 1366.6997 | 1440.6525 | 1677.1473 | 1372.4083 | 211.7821 | 1481.896 |
| Txndc17 | HVTEDCVFIY | -59 | -68 | 0 | 0 | 10 | 814.01483 | 758.0259 | 829.8994 | 817.5974 | NaN | NaN |
| Txndc17 | HVTEDCVFIY | -59 | -68 | 0 | 0 | 10 | 814.01483 | 758.0259 | 829.8994 | 817.5974 | NaN | NaN |
| Ppif | HVVFGHVK | -167 | -174 | 0 | 0 | 8 | 853.81836 | 592.0853 | 509.13162 | 453.55847 | 699.44604 | 529.0482 |
| Sh2d2a | HVVLGEDSAHAQ | -172 | -183 | 0 | 0.00025015 | 12 | 691.23267 | 600.77765 | 732.5212 | NaN | 5046.516 | 915.42914 |
| Hexb | LVQEPVAVLK | -131 | -140 | 0 | 0.00025015 | 10 | 314.5676 | 406.7868 | 355.3728 | 267.58612 | 361.86554 | 357.8355 |
| Hexb | HVYTPNDVR | -243 | -251 | 0 | 0.001868731 | 9 | 1071.9547 | NaN | 2917.629 | NaN | 790.83215 | 686.221 |
| Hexb | VIPEFDTPGHTQ | -264 | -275 | 0 | 0.000591124 | 12 | 699.7489 | NaN | 832.35333 | 812.09125 | NaN | NaN |
| Hexb | LQPALWPFPR | -31 | -40 | 0 | 0 | 10 | 377.60468 | 520.2032 | 443.29614 | NaN | NaN | NaN |
| Tecr | HYEVEIR | -3 | -9 | 0 | 0 | 7 | 2411.3257 | 2525.2659 | 605.66113 | 1924.0924 | 3025.5334 | 2764.1323 |
| Tecr | SLKDEDVLQK | -58 | -67 | 1 | 0.00025015 | 10 | 1068.5995 | 1322.2098 | 1013.5843 | 1001.9226 | 538.8606 | 970.1802 |
| Tecr | LCFLDKVEPQ | -17 | -26 | 1 | 0.00051306 | 10 | 724.466 | NaN | 510.73264 | NaN | 693.9418 | 760.4765 |
| Tecr | KIPYPTK | -227 | -233 | 1 | 0.00713295 | 7 | 440.44128 | NaN | NaN | NaN | NaN | NaN |
| Tbc1d15 | IDVEDILCK | -581 | -589 | 0 | 0 | 9 | 1418.2545 | 1161.0195 | 1403.1553 | 968.85986 | 1036.0085 | 785.24725 |
| Tbc1d15 | HYGFNEILK | -564 | -572 | 0 | 0.00025015 | 9 | 753.57806 | 1008.2982 | 616.46075 | 550.5604 | 901.024 | 795.8138 |
| Atp5pb | HYLFDVQR | -164 | -171 | 0 | 0.00025015 | 8 | 2285.4226 | 2139.9902 | 2456.0547 | 123.02318 | 1862.4343 | 2031.2712 |
| Atp5pb | PLPPLPEYGGK | -43 | -53 | 0 | 0 | 11 | 2197.6707 | 2296.0117 | 2541.5642 | 2667.6118 | 2648.0981 | 2366.907 |
| Atp5pb | YGASFGEFIDK | -116 | -126 | 0 | 0 | 11 | 802.3819 | 583.8234 | 772.2252 | 586.49976 | 576.78357 | 552.16504 |
| Atp5pb | RKEEEHMIDWVEK | -209 | -221 | 2 | 0.001092529 | 13 | 505.4234 | NaN | NaN | 614.29407 | 569.20605 | 546.0277 |
| Atp5pb | YGASFGEFIDKL | -116 | -127 | 1 | 0 | 12 | 417.26724 | NaN | NaN | 474.79755 | 600.53503 | 518.24805 |
| Atp5pb | QIQDAIDMEK | -145 | -154 | 0 | 0 | 10 | 4360.386 | 4033.0398 | NaN | 1271.8568 | NaN | NaN |
| Gbp4 | NKENLEQLR | -527 | -535 | 1 | 0 | 9 | 734.63513 | 1014.56226 | 1421.0402 | 1766.6385 | 1828.4695 | 1250.3677 |
| Gbp4 | HYVTELTELIR | -149 | -159 | 0 | 0 | 11 | 756.8286 | 1255.5544 | 1317.6082 | NaN | 1510.8734 | 1512.4263 |
| Acot8 | HYWVPTSQR | -46 | -54 | 0 | 0 | 9 | 512.9252 | NaN | NaN | 1414.233 | NaN | NaN |
| Pals2 | IAAPELETLR | -469 | -478 | 0 | 0.001092529 | 10 | 848.35535 | NaN | NaN | NaN | NaN | NaN |
| Map2k2 | VGELKDDDFER | -64 | -74 | 1 | 0 | 11 | 867.57465 | 927.18787 | 570.3237 | 1030.0814 | 1309.5151 | 1143.5294 |
| Map2k2 | KLEELDLDEQQR | -40 | -51 | 1 | 0 | 12 | 654.1094 | 739.5684 | 787.2374 | 1271.2104 | 1084.8066 | 1128.0878 |
| Map2k2 | IAEGPSPTSEGASEANLVDLQK | -18 | -39 | 0 | 0.000627943 | 22 | 421.80768 | 397.30515 | 375.56927 | 476.01868 | 231.98862 | 516.38934 |
| Cox6b1 | NCWQNYLDFHR | -29 | -39 | 0 | 0 | 11 | 1323.0348 | 1626.354 | 1708.8066 | 1720.5837 | 2166.519 | 2174.7788 |
| Cox6b1 | NCWQNYLDFHR | -29 | -39 | 0 | 0 | 11 | 1323.0348 | 1626.354 | 1708.8066 | 1720.5837 | 2166.519 | 2174.7788 |
| Cox6b1 | IAEGTFPGKI | -77 | -86 | 1 | 0.00051306 | 10 | 1608.1459 | NaN | NaN | 1409.5931 | 1407.7594 | 374.94806 |
| Cox6b1 | IAEGTFPGK | -77 | -85 | 0 | 0.007546675 | 9 | 737.89825 | NaN | NaN | 510.31757 | NaN | 1338.4418 |
| Pdcd2 | IALEPEQILR | -247 | -256 | 0 | 0.000627943 | 10 | 1058.6052 | 1166.7417 | 1097.5471 | NaN | 1554.1588 | 1450.4667 |
| Ndufa13 | LLIEDLEAR | -60 | -68 | 0 | 0 | 9 | 1007.41 | 951.1221 | 1043.3191 | 900.11676 | 1079.8989 | 999.6864 |
| Ndufa13 | IALMPLFQAEK | -69 | -79 | 0 | 0 | 11 | 400.2501 | 313.86743 | 241.43776 | NaN | 393.31985 | NaN |
| Ndufs6 | ITHTGQVYDEK | -31 | -41 | 0 | 0 | 11 | 600.78424 | 642.31335 | 561.1051 | 417.19116 | 947.6482 | 669.14056 |
| Ndufs6 | IAQQPVNEVEHR | -64 | -75 | 0 | 0.000394361 | 12 | 764.69745 | 212.94202 | 878.1444 | NaN | NaN | NaN |
| Ndufs6 | IAQQPVNEVEHR | -64 | -75 | 0 | 0.000394361 | 12 | 764.69745 | 212.94202 | 878.1444 | NaN | NaN | NaN |
| Ndufs6 | LIAQQPVNEVEHR | -63 | -75 | 0 | 0 | 13 | 314.38452 | NaN | NaN | NaN | NaN | NaN |
| Rprd1b | IASLPQEVQDVSLLEK | -201 | -216 | 0 | 0 | 16 | 582.46497 | 702.6521 | 637.49304 | NaN | NaN | NaN |
| Setd7 | SGPEAPEWYQVELK | -345 | -358 | 0 | 0.000471409 | 14 | 577.15625 | NaN | 549.09485 | NaN | NaN | NaN |
| Setd7 | IAYVYPDQR | -144 | -152 | 0 | 0 | 9 | 4174.2627 | NaN | NaN | NaN | NaN | NaN |
| Eif3k | ICSQEESIKPK | -189 | -199 | 1 | 0 | 11 | 945.16156 | 767.51215 | 677.35547 | 720.5768 | 1049.0352 | 1005.72064 |
| Eif3k | SQEESIKPK | -191 | -199 | 1 | 0 | 9 | 587.3191 | 475.22186 | 859.0715 | 493.3947 | 777.1669 | 598.44183 |
| Eif3k | YNPENLATLER | -21 | -31 | 0 | 0 | 11 | 1736.9684 | 2636.6267 | NaN | 1530.1838 | 3191.1042 | 3038.34 |
| Atp2b4 | VVGIEDPVRPEVPDAIAK | -668 | -685 | 1 | 0.00025015 | 18 | 241.52478 | 274.5563 | NaN | NaN | NaN | NaN |
| Tnfaip8l2 | IDETSSEVLDELYR | -31 | -44 | 0 | 0 | 14 | 930.45496 | 893.31024 | 1101.6453 | 1655.4725 | 2149.916 | 1935.7981 |
| Tnfaip8l2 | LYGPDFTQHLGK | -159 | -170 | 0 | 0 | 12 | 641.1361 | 873.7066 | 662.39777 | 583.72205 | 578.3043 | 498.20615 |
| Ddi2 | IDFSSIAVPGTSNPQQR | -94 | -110 | 0 | 0 | 17 | 2014.7777 | 2559.1301 | 2393.3833 | 2081.9082 | 1380.8586 | 2760.128 |
| Ddi2 | TFLPEGELPECAR | -351 | -363 | 0 | 0 | 13 | 340.0613 | NaN | 559.8656 | 443.165 | 679.8275 | 592.9806 |
| Mrps22 | IDGLLIDQIQR | -280 | -290 | 0 | 0 | 11 | 628.17316 | NaN | 800.76105 | NaN | 675.30817 | 659.22687 |
| Mrps22 | VAQFEPDSAEYIK | -231 | -243 | 0 | 0.006788754 | 13 | 665.81256 | NaN | NaN | 565.28735 | 906.9726 | NaN |
| Mrps22 | ISYNIPHR | -160 | -167 | 0 | 0.00450079 | 8 | 1561.5239 | 1332.8422 | NaN | NaN | NaN | NaN |
| Opa1 | IDQLQEELLHTQ | -215 | -226 | 0 | 0 | 12 | 848.2419 | 1259.1478 | 1245.6611 | 2566.381 | 2710.8909 | 2771.8716 |
| Sar1b | IDRPEAISEER | -136 | -146 | 1 | 0 | 11 | 1142.471 | 1315.891 | 1756.1113 | 2086.6377 | 1405.7932 | 1365.1089 |
| Rpl38 | KIEEIKDFLLT | -4 | -14 | 2 | 0.001291103 | 11 | 2760.496 | 2470.8394 | 2926.9822 | 2638.1826 | 5515.0127 | 4309.202 |
| Rpl38 | QSLPPGLAVK | -58 | -67 | 0 | 0 | 10 | 1936.1606 | 1131.6188 | 749.77124 | 2403.421 | 1011.9285 | 1020.853 |
| Rpl38 | KIEEIKDFLLTA | -4 | -15 | 2 | 0.000712251 | 12 | 731.7133 | 759.16174 | 611.2206 | 1254.4691 | 1531.6744 | 1585.0422 |
| Rpl38 | KIEEIKDFLL | -4 | -13 | 2 | 0.001636916 | 10 | 187.19366 | 930.46985 | 962.89594 | 131.98195 | 1718.0355 | 1608.875 |
| Rpl38 | IEEIKDFLLT | -5 | -14 | 1 | 0 | 10 | 670.4519 | 650.2174 | NaN | 849.54877 | 865.473 | 925.8314 |
| Rpl38 | TLVITDKEK | -44 | -52 | 1 | 0 | 9 | 567.94775 | NaN | 424.13443 | 638.6162 | 443.02448 | 500.36697 |
| Rpl38 | KQSLPPGLAVK | -57 | -67 | 1 | 0 | 11 | 578.6171 | NaN | NaN | 576.0114 | 212.52322 | 381.56622 |
| Rpl38 | KIEEIKDFLLTAR | -4 | -16 | 2 | 0 | 13 | 732.6695 | 411.17322 | 420.70923 | NaN | NaN | NaN |
| Faim | IEFEHGTTSGK | -19 | -29 | 0 | 0.00025015 | 11 | 676.1471 | 1314.8593 | 1318.108 | NaN | 1924.0496 | 1730.0415 |
| Polr2h | LHCESESFK | -28 | -36 | 0 | 0 | 9 | 715.5153 | 636.9895 | 1486.3336 | 822.8479 | 841.1429 | 780.86755 |
| Polr2h | IEGDETSTEAATR | -99 | -111 | 0 | 0 | 13 | 104.83532 | 894.56396 | 937.9464 | 1035.9589 | 1048.9741 | 961.49927 |
| Polr2h | IYPVDLGDKFR | -47 | -57 | 1 | 0 | 11 | 2528.992 | NaN | 2810.5935 | 3816.6223 | 2719.3877 | 2621.0676 |
| Polr2h | STLYEDGTLDDGEYNPTDDRPSR | -62 | -84 | 1 | 0.001291103 | 23 | 478.77774 | 570.3358 | 451.94263 | NaN | 490.18304 | NaN |
| Polr2h | IYPVDLGDK | -47 | -55 | 0 | 0.004690496 | 9 | 3184.0806 | NaN | NaN | NaN | NaN | NaN |
| Nudcd1 | IEKEELDLK | -185 | -193 | 1 | 0 | 9 | 309.10867 | 650.5376 | 477.11572 | 984.45667 | 617.995 | 660.30273 |
| Nudcd1 | LYSSIDHEGSTW | -324 | -335 | 0 | 0.000431655 | 12 | 2710.2205 | 69.389145 | NaN | NaN | NaN | NaN |
| Nudcd1 | LYVIGTGDR | -135 | -143 | 0 | 0.004412728 | 9 | 2353.146 | NaN | NaN | NaN | NaN | NaN |
| Ifit2 | IENPALDCEEGWAR | -133 | -146 | 0 | 0 | 14 | 689.1905 | 832.22974 | 662.99426 | 609.119 | 925.50507 | 507.4725 |
| Csnk1a1 | QGGVGIPHIR | -68 | -77 | 0 | 0.00111006 | 10 | 1181.9792 | NaN | NaN | NaN | NaN | 243.35895 |
| Csnk1a1 | IEYVHTK | -124 | -130 | 0 | 0.003360691 | 7 | 888.96094 | NaN | NaN | NaN | NaN | NaN |
| Fuca2 | IFEERLR | -342 | -348 | 1 | 0.00051306 | 7 | 2276.5532 | NaN | 2450.4712 | 2695.4805 | 2877.296 | 2431.913 |
| Hdac2 | IFKPIISK | -241 | -248 | 1 | 0.001055699 | 8 | 279.5526 | 249.40938 | 226.79675 | 270.37488 | 198.4214 | 237.25775 |
| Hdac2 | YHSDEYIK | -68 | -75 | 0 | 0.001846793 | 8 | 611.6065 | NaN | NaN | 810.17676 | NaN | NaN |

| | | | | | | | | | | | | |
|---|---|---|---|---|---|---|---|---|---|---|---|---|
| Cul2 | IFLESHVR | -63 | -70 | 0 | 0.000627943 | 8 | 342.0244 | NaN | 3689.964 | 3759.7544 | 4817.1855 | 4075.831 |
| Cul2 | LKDEEIR | -243 | -249 | 1 | 0.00416551 | 7 | 1999.5249 | NaN | NaN | NaN | NaN | NaN |
| Rbm24;Rbm3 | IFVGGLPYHTT | (13);(34) | (23);(44) | 0 | 0.00025015 | 11 | 175.99887 | NaN | 430.48053 | NaN | 821.1466 | NaN |
| Bcl2l11 | IGDEFNETYTR | -153 | -163 | 0 | 0 | 11 | 676.1531 | 537.5931 | 824.0622 | 771.4575 | 606.9534 | 562.534 |
| Mrpl39 | IGDFIDVSEGPLIPR | -254 | -268 | 0 | 0 | 15 | 190.88326 | 114.73361 | 538.32855 | 601.00183 | NaN | 648.9961 |
| Clk3 | IGDWLQER | -148 | -155 | 0 | 0 | 8 | 1302.2021 | NaN | NaN | NaN | NaN | NaN |
| Nt5c3b | IGFLNDKVEER | -250 | -260 | 1 | 0 | 11 | 1110.064 | 1095.4446 | 1192.9692 | 968.55005 | 933.6006 | 841.0572 |
| Nt5c3b | VFHPNIHIV | -172 | -180 | 0 | 0.00025015 | 9 | 862.05615 | 1017.95667 | 695.5976 | NaN | 1022.1408 | NaN |
| Ep400 | IGLDWLAK | -1096 | -1103 | 0 | 0 | 8 | 273.62537 | 255.14291 | 292.32315 | 183.28804 | NaN | NaN |
| Dhps | IGNLLVPNDNYCK | -166 | -178 | 0 | 0.001023236 | 13 | 1622.9421 | NaN | 824.9032 | 844.54047 | NaN | NaN |
| Sf3a2 | MDFQHRPGGK | -1 | -10 | 1 | 0 | 10 | 553.41437 | 114.67973 | 495.28467 | 437.89868 | 552.1807 | 1138.6511 |
| Sf3a2 | QIDYPEIAEGIMPR | -135 | -148 | 0 | 0 | 14 | 526.4256 | 641.4792 | 749.20435 | NaN | 994.7803 | 970.1973 |
| Sf3a2 | IGRPGYK | -112 | -118 | 1 | 0.002099344 | 7 | 345.0209 | 184.46495 | NaN | 525.68585 | NaN | NaN |
| Pdpk1 | LDHPFFVK | -140 | -147 | 0 | 0.001291103 | 8 | 5783.18 | NaN | NaN | 9046.641 | NaN | NaN |
| Pdpk1 | IGSFDETCTR | -177 | -186 | 0 | 0.002329279 | 10 | 750.68494 | NaN | 636.3131 | NaN | NaN | NaN |
| Tprkb | IGTLLDAIICR | -158 | -168 | 0 | 0 | 11 | 341.1108 | 445.6277 | 350.49106 | 403.8595 | NaN | 427.53616 |
| Pmm2 | IGVVGGSDFEK | -37 | -47 | 0 | 0 | 11 | 1159.611 | 1274.8286 | 507.02765 | 1169.5201 | 1016.694 | 1189.3298 |
| Pmm2 | LQEQLGNDVVEKY | -48 | -60 | 1 | 0 | 13 | 766.15924 | 440.82693 | 506.9074 | 799.5857 | 425.62708 | 759.5054 |
| Pmm2 | LFDMDGTLTAPR | -6 | -17 | 0 | 0 | 12 | 632.52075 | 953.27264 | 1073.9941 | 1432.3053 | 1481.4773 | 1455.1409 |
| Tasor | IHEEEESVLCK | -1191 | -1201 | 0 | 0.001392869 | 11 | 2549.2563 | NaN | NaN | NaN | NaN | 1771.5127 |
| Eed | KIHFPDFSTR | -293 | -302 | 1 | 0.000947459 | 10 | 674.4894 | 774.02014 | 892.03546 | 789.372 | 855.03375 | 825.83954 |
| Eed | VWDLEVEDPHK | -388 | -398 | 0 | 0 | 11 | 3238.9463 | 2683.575 | NaN | 2763.8892 | 429.4081 | 1615.1442 |
| Eed | WLGDLILSK | -314 | -322 | 0 | 0 | 9 | 538.9729 | 587.9387 | NaN | 573.39355 | 627.0622 | 665.6991 |
| Eed | IHFPDFSTR | -294 | -302 | 0 | 0 | 9 | 885.77264 | 764.3528 | 924.2896 | NaN | 1147.223 | NaN |
| Eed | IKPSESNVTIL | -343 | -353 | 1 | 0.000394361 | 11 | 4381.462 | NaN | NaN | NaN | 4811.531 | 4681.7783 |
| Rpl31 | VTYVPVTTFK | -106 | -115 | 0 | 0 | 10 | 3342.4082 | 3531.9043 | 3805.7703 | 2457.2 | 1838.1355 | 1783.7864 |
| Rpl31 | TYVPVTTFK | -107 | -115 | 0 | 0 | 9 | 1629.7399 | 1650.9387 | 1910.4789 | 1543.5181 | 1912.2136 | 2265.4446 |
| Rpl31 | TLVTYVPVTTFK | -104 | -115 | 0 | 0 | 12 | 622.30005 | 526.145 | 538.5215 | 623.1895 | 571.71954 | 565.2129 |
| Rpl31 | NEDEDSPNKLY | -93 | -103 | 1 | 0 | 11 | 549.4971 | 587.17145 | 493.7351 | 1040.0369 | 872.6824 | 979.6366 |
| Rpl31 | SAINEVVTR | -15 | -23 | 0 | 0 | 9 | 584.34906 | NaN | 260.62442 | NaN | 361.067 | NaN |
| Rpl31 | IHGVGFK | -33 | -39 | 0 | 0.00025015 | 7 | 952.977 | NaN | NaN | NaN | NaN | NaN |
| NaN | IHTDLGALEK | -483 | -492 | 0 | 0 | 10 | 1163.9362 | 895.8507 | NaN | 1036.7845 | NaN | NaN |
| Apaf1 | IHVSPDFR | -1224 | -1231 | 0 | 0 | 8 | 2125.9595 | 1172.019 | 1524.2516 | NaN | 2358.5244 | 2528.7761 |
| Wdr82 | SSDDDSIVLYDCQEGKPK | -39 | -56 | 1 | 0.000755556 | 18 | 429.9047 | NaN | 107.73675 | NaN | NaN | 450.65536 |
| Wdr82 | IHVWNGESGIK | -263 | -273 | 0 | 0 | 11 | 134.41637 | 967.67725 | 977.43896 | NaN | NaN | NaN |
| Ccdc117 | IIDEDEEVESDR | -154 | -165 | 0 | 0.00025015 | 12 | 507.298 | 412.23907 | NaN | NaN | NaN | NaN |
| Ints4 | IIGDHFSDQDPR | -200 | -211 | 0 | 0.000394361 | 12 | 2122.3645 | 1945.5831 | NaN | NaN | 1452.0111 | 1411.3582 |
| Gnai3 | IIHEDGYSEDECK | -55 | -67 | 0 | 0 | 13 | 1194.6481 | 523.3264 | 629.3873 | 740.54047 | 791.87823 | 744.70044 |
| Gnai3 | IIHEDGYSEDECK | -55 | -67 | 0 | 0 | 13 | 1194.6481 | 523.3264 | 629.3873 | 740.54047 | 791.87823 | 744.70044 |
| Gnai3 | KIIHEDGYSEDECK | -54 | -67 | 1 | 0.00025015 | 14 | 269.99478 | 337.77823 | 349.5047 | 462.43243 | 519.11127 | 496.43735 |
| Gnai3 | ISQTNYIPTQQDVLR | -162 | -176 | 0 | 0 | 15 | 1140.4254 | 1326.7776 | NaN | 1586.6829 | 1480.5913 | 1115.4758 |
| Rpa3 | YIDRPVCFVGK | -20 | -30 | 1 | 0.000230707 | 11 | 1031.753 | 1155.7886 | 1277.2888 | 1216.7339 | 936.6131 | 1328.4081 |
| Rpa3 | IINELPQFFPVGLPQHE | -105 | -121 | 0 | 0 | 17 | 266.21094 | 273.1562 | NaN | 216.95131 | 397.26437 | 420.69135 |
| Gmds | VAFDELVR | -349 | -356 | 0 | 0 | 8 | 731.4288 | 877.0328 | 953.886 | 883.44574 | 894.2661 | 111.18053 |
| Gmds | IINEVKPTEIYNL | -96 | -108 | 1 | 0.003620005 | 13 | 574.0719 | NaN | NaN | 766.69366 | NaN | NaN |
| Manf | IINEVSKPLAHHIPVEK | -85 | -101 | 1 | 0 | 17 | 320.6297 | NaN | 548.0965 | 1560.764 | 2499.923 | 581.80035 |
| Manf | TIEEELIK | -52 | -59 | 0 | 0.001291103 | 8 | 12732.451 | 18506.012 | 10407.581 | NaN | 15552.836 | NaN |
| Manf | IINEVSKPL | -85 | -93 | 1 | 0.002466863 | 9 | 817.3133 | 733.484 | NaN | 513.9259 | NaN | NaN |
| Manf | IINEVSKPLAH | -85 | -95 | 1 | 0 | 11 | 571.1519 | NaN | NaN | 1260.0615 | NaN | NaN |
| Ergic1 | LTSNPLASHDYILK | -177 | -190 | 0 | 0 | 14 | 1086.8575 | 1032.8488 | 818.32086 | 693.3206 | 867.2014 | 685.02216 |
| Ergic1 | IIPAIWFR | -225 | -232 | 0 | 0.001610812 | 8 | 553.59546 | 712.7157 | NaN | NaN | 212.67848 | 271.30118 |
| Ephx1 | IIPLLTDPK | -160 | -168 | 0 | 0.000298805 | 9 | 18762.965 | NaN | 18912.531 | 19036.668 | 20425.162 | 15314.925 |
| Ephx1 | SLTPLLGQR | -259 | -267 | 0 | 0.000550307 | 9 | 955.5494 | 1018.75476 | 1197.2618 | NaN | NaN | NaN |
| Jchain | IIPSTEDPNEDIVER | -43 | -57 | 0 | 0 | 15 | 583.05554 | NaN | 616.3461 | 426.1755 | 531.5747 | 640.4061 |
| Rab18 | ILIIGESGVGK | -11 | -21 | 0 | 0 | 11 | 1006.9153 | 1096.4327 | 1158.6035 | 819.903 | 998.0875 | 957.91113 |
| Rab18 | IIQTPGLWESENQNK | -169 | -183 | 0 | 0 | 15 | 913.0031 | 911.73596 | 1098.4661 | 1267.324 | 1328.6733 | 1683.4949 |
| Rab18 | TCDGVQCAFEELVEK | -154 | -168 | 0 | 0 | 15 | 269.19647 | 391.83417 | 396.08292 | 411.38196 | 518.00964 | 521.98596 |
| Afg2a | IIYVPLPDAATR | -791 | -802 | 0 | 0 | 12 | 6769.855 | 7234.3804 | 6579.5796 | 7170.017 | 4633.804 | 3506.4094 |
| Rpl19 | IKDGLIIR | -45 | -52 | 1 | 0.001055699 | 8 | 4202.9043 | 4171.8633 | 4109.2666 | 5801.699 | 5257.6167 | 5435.2812 |
| Rpl19 | KVWLDPNETNEIA | -21 | -33 | 1 | 0.00025015 | 13 | 1475.9464 | 1299.3062 | 1221.0472 | 1745.1154 | 1736.4208 | 86.35112 |
| Rpl19 | VWLDPNETNEIANAN | -22 | -36 | 0 | 0 | 15 | 1229.3987 | 1318.1465 | 1279.3252 | 1345.9498 | 1515.9473 | 1517.0918 |
| Rpl19 | KVWLDPNETNEIANAN | -21 | -36 | 1 | 0.000431655 | 16 | 231.5976 | 1348.9958 | 1314.8931 | 1920.122 | 1766.7867 | 1724.5598 |
| Rpl19 | VWLDPNETNEIA | -22 | -33 | 0 | 0 | 12 | 1029.0541 | 1127.4769 | 1004.6843 | 1178.4563 | NaN | 1519.9053 |
| Rpl19 | KVWLDPNETNEIAN | -21 | -34 | 1 | 0.00025015 | 14 | 1646.3865 | NaN | 2060.829 | NaN | NaN | 573.00586 |
| Rpl19 | VWLDPNETNEIAN | -22 | -34 | 0 | 0.00025015 | 13 | 767.7148 | 943.5214 | 911.3422 | NaN | NaN | NaN |
| Rpl19 | LLADQAEAR | -154 | -162 | 0 | 0.00635324 | 9 | 269.16745 | 342.485 | NaN | NaN | NaN | NaN |
| Cd82 | IKEEDNQLIVK | -179 | -189 | 1 | 0 | 11 | 1610.622 | 1636.1285 | 1658.8494 | 1706.4869 | 1701.6312 | 1668.6517 |
| Ctnnbl1 | IKFPDNPEK | -101 | -109 | 1 | 0.000627943 | 9 | 940.5705 | 646.7613 | 884.52185 | NaN | 1080.7264 | 842.96625 |
| Micos13 | VYDQELLGPSDK | -25 | -36 | 0 | 0 | 12 | 999.012 | 1229.3523 | 1305.2106 | 1326.6814 | 1781.2246 | 1669.3391 |
| Micos13 | IKFPNFR | -76 | -82 | 1 | 0.00078661 | 7 | 467.489 | NaN | NaN | 437.18158 | NaN | 424.69696 |
| Fmr1 | LEFAEDVIQVPR | -279 | -290 | 0 | 0 | 12 | 861.3181 | 964.8667 | 903.91425 | 962.72363 | 1048.3914 | 1141.9445 |
| Fmr1 | IKLEVPEDLR | -129 | -138 | 1 | 0 | 10 | 1780.153 | NaN | 1378.4512 | NaN | NaN | 2271.4966 |
| Sdhb | RIDTDLSK | -118 | -125 | 1 | 0.00078661 | 8 | 1211.8232 | 1259.5776 | 1279.2942 | 1189.0496 | 1104.1588 | 1406.3218 |
| Sdhb | IKNEVDSTLTFR | -81 | -92 | 1 | 0 | 12 | 1136.9165 | 1179.1095 | 1207.7573 | 1312.3727 | 1379.3506 | 1153.9431 |
| Sdhb | LQDPFSVYR | -236 | -244 | 0 | 0 | 9 | 375.98727 | 1505.6373 | 1358.3494 | 429.44147 | 450.90268 | 1535.0806 |
| Sdhb | NEVDSTLTFR | -83 | -92 | 0 | 0 | 10 | 2087.5435 | 1285.517 | 1157.2634 | 1101.441 | NaN | 787.4769 |
| Klf13 | ILADLNQQAPAPAPAER | -66 | -82 | 0 | 0 | 17 | 986.9244 | 1198.5149 | 988.7666 | 925 | 943.73047 | 781.5404 |
| Elp3 | SYSPSDLIELVAR | -339 | -351 | 0 | 0 | 13 | 524.91626 | 301.36163 | 771.7615 | 459.90424 | 448.1148 | 524.78876 |

| | | | | | | | | | | | | |
|---|---|---|---|---|---|---|---|---|---|---|---|---|
| Elp3 | ILALVPPWTR | -352 | -361 | 0 | 0 | 10 | 453.53766 | 477.8612 | 550.89233 | 454.7687 | 456.71747 | 512.66327 |
| Elp3 | QLVEAHEQGK | -25 | -34 | 0 | 0.001868731 | 10 | 1858.4272 | NaN | NaN | NaN | NaN | NaN |
| Thoc1 | ILAPYLEIK | -587 | -595 | 0 | 0 | 9 | 470.33047 | 563.1935 | 560.1925 | 510.43997 | 457.7345 | 545.3118 |
| Thoc1 | NTNEQESTLGQK | -190 | -201 | 0 | 0 | 12 | 450.13522 | 556.66644 | 413.48853 | NaN | 794.7902 | NaN |
| Thoc1 | SIPIDYNLYR | -227 | -236 | 0 | 0 | 10 | 4470.2812 | NaN | 165.88736 | 5518.798 | NaN | NaN |
| Ubr2 | ISKPGDAPHNSPSIL | -956 | -970 | 1 | 0.005605646 | 15 | 223.63843 | 316.70648 | 238.7214 | NaN | NaN | NaN |
| Ubr2 | SLLECSAEEIAGR | -15 | -27 | 0 | 0 | 13 | 434.5422 | NaN | 936.96155 | NaN | NaN | NaN |
| Ubr2 | ILCQEEQEVTVGSR | -1109 | -1122 | 0 | 0 | 14 | 2350.4312 | NaN | NaN | NaN | NaN | NaN |
| Cdc20 | ILDAPEIR | -175 | -182 | 0 | 0.00111006 | 8 | 804.1605 | NaN | NaN | NaN | NaN | NaN |
| Mrps2 | ILDTPLQHSDFFNVK | -59 | -73 | 0 | 0 | 15 | 931.3478 | 920.501 | 1176.6918 | 685.5823 | 763.89606 | 859.4115 |
| Mrps2 | VTGAPVPAVSEPQDGDDFQSR | -38 | -58 | 0 | 0 | 21 | 514.15625 | 1057.3914 | 1351.3466 | 229.7097 | 1306.1179 | 1118.6532 |
| Tpx2 | KLEEEEEGSAPATSR | -168 | -182 | 1 | 0.000627943 | 15 | 1330.7327 | 1235.5818 | 105.539825 | 982.5055 | 1403.2675 | 1377.1229 |
| Tpx2 | ILEDVVGVPEKK | -466 | -477 | 1 | 0 | 12 | 798.8863 | 799.6979 | 881.75214 | NaN | 1415.6393 | 1031.9962 |
| Tpx2 | RAPVKPPTQPVGFD | -421 | -434 | 2 | 0.00025015 | 14 | 438.49255 | 414.84235 | NaN | 485.4398 | 436.77838 | NaN |
| Tpx2 | KLQQLEEVR | -684 | -692 | 1 | 0.001291103 | 9 | 1026.245 | NaN | NaN | NaN | 1095.4712 | NaN |
| Arl1 | ILILGLDGAGK | -20 | -30 | 0 | 0 | 11 | 129.9425 | 161.91626 | 189.71384 | 135.07364 | 182.87373 | 230.5413 |
| Arl1 | LEEEELRK | -111 | -118 | 1 | 0.003299042 | 8 | 1754.9456 | NaN | 1788.3937 | NaN | NaN | NaN |
| Rftn1 | ILLEHIFR | -95 | -102 | 0 | 0.00025015 | 8 | 1995.1803 | 1907.282 | 2353.351 | NaN | NaN | NaN |
| Tbc1d13 | ILLNYLPLER | -47 | -56 | 0 | 0 | 10 | 277.4519 | 359.52502 | 668.1965 | NaN | 538.22705 | 626.5534 |
| Dpm1 | ILLRPGASDLTGSFR | -174 | -188 | 1 | 0 | 15 | 551.524 | 738.6342 | 683.4764 | 462.92294 | 702.2243 | 628.7616 |
| Dpm1 | ILLRPGASDLTGSFR | -174 | -188 | 1 | 0 | 15 | 551.524 | 738.6342 | 683.4764 | 462.92294 | 702.2243 | 628.7616 |
| Ak6 | YVNVGDLAR | -31 | -39 | 0 | 0 | 9 | 745.3743 | 784.17645 | 1130.1512 | 763.0261 | 969.63513 | 981.9995 |
| Ak6 | ILLTGTPGVGK | -6 | -16 | 0 | 0.00450079 | 11 | 578.9437 | NaN | 592.0899 | NaN | NaN | NaN |
| Creb1 | TIAESEDSQESVDSVTDSQK | -90 | -109 | 0 | 0 | 20 | 263.9105 | 422.32556 | 331.00375 | 318.25546 | 430.59695 | 321.29013 |
| Creb1 | ILNDLSSDAPGVPR | -123 | -136 | 0 | 0 | 14 | 1477.4772 | 1900.3562 | 1935.9949 | 2607.747 | 3360.3691 | NaN |
| Tex47 | ILPEQIIK | -208 | -215 | 0 | 0 | 8 | 1277.165 | 848.0129 | 903.4105 | 780.0069 | NaN | 815.4004 |
| Trpv2 | ILPLLQIDR | -131 | -139 | 0 | 0.00142999 | 9 | 426.75958 | 206.21918 | NaN | 258.151 | NaN | NaN |
| Emc2 | YTQGGLENLELSR | -200 | -212 | 0 | 0 | 13 | 597.51874 | 755.0457 | 817.5426 | 1646.7557 | 1797.8052 | 1051.0873 |
| Emc2 | ILQEDPTNTAAR | -113 | -124 | 0 | 0 | 12 | 796.1889 | NaN | NaN | NaN | 2297.9773 | NaN |
| Napg | INEGLEHLAK | -6 | -15 | 0 | 0 | 10 | 1042.1886 | 1496.1654 | 948.23816 | 409.5416 | 2047.4591 | 2233.5283 |
| Mob3a | INNEDIFPTNVGTPFPK | -130 | -146 | 0 | 0.000627943 | 17 | 2638.3916 | NaN | NaN | NaN | NaN | NaN |
| Lcp2 | SLAPLDREPFIL | -246 | -257 | 1 | 0 | 12 | 567.4534 | 526.9791 | 612.463 | 643.9642 | 725.9029 | 735.6257 |
| Lcp2 | INQDGTFLVR | -439 | -448 | 0 | 0 | 10 | 239.2472 | 307.77225 | 344.13037 | 1107.0466 | 898.47455 | 780.8186 |
| Lcp2 | NSDNLADYFR | -16 | -25 | 0 | 0 | 10 | 864.13434 | 897.3335 | 935.2618 | NaN | 1149.4543 | 1100.1549 |
| Lcp2 | PALPPLPTGR | -195 | -204 | 0 | 0 | 10 | 323.5477 | NaN | 259.72034 | 483.07733 | 693.9988 | 237.11612 |
| Lcp2 | KPPFSDKPSAPLGR | -260 | -273 | 2 | 0.000230707 | 14 | 525.06335 | NaN | 575.639 | 266.36166 | 484.50073 | NaN |
| Ptcd3 | IPALQILR | -350 | -357 | 0 | 0.002810811 | 8 | 1175.5408 | 1218.6924 | 396.90625 | 1223.7937 | 415.35004 | NaN |
| Ptcd3 | VEGSDITGIEEIVIPK | -47 | -62 | 0 | 0 | 16 | 242.53424 | NaN | NaN | NaN | 313.31177 | NaN |
| Lsm4 | IPDEIIDMVR | -71 | -80 | 0 | 0.000230707 | 10 | 374.15646 | 1144.7734 | 994.86926 | NaN | NaN | NaN |
| Actl6a | STGLILDSGATHTT | -165 | -178 | 0 | 0 | 14 | 1502.8799 | NaN | NaN | NaN | NaN | NaN |
| Actl6a | VDFPTAIGVVLER | -35 | -47 | 0 | 0 | 13 | 494.96793 | NaN | NaN | NaN | NaN | NaN |
| Gigyf2 | IPFDLLEK | -58 | -65 | 0 | 0.000550307 | 8 | 682.03723 | NaN | 697.29895 | 751.8709 | 963.83606 | 84.247284 |
| Rpl7l1 | TVPLTDNTVIEEHLGR | -164 | -179 | 0 | 0 | 16 | 623.8269 | 810.8383 | 1062.8589 | 791.48395 | 1164.0625 | 955.17163 |
| Rpl7l1 | IPLVPENLLK | -10 | -19 | 0 | 0 | 10 | 435.2198 | 416.79718 | 411.0208 | NaN | 337.1632 | 396.70383 |
| Rpl7l1 | LEDLIHEIAFPGK | -185 | -197 | 0 | 0 | 13 | 349.65283 | 389.72565 | 954.1684 | 795.8637 | NaN | 427.89264 |
| Gorasp2 | ISLPGQMTGTPITPLK | -213 | -228 | 0 | 0 | 16 | 2678.3135 | 2677.567 | 2749.8293 | 2482.8516 | 313.6717 | 2376.7307 |
| Gorasp2 | VSDCTPAVEKPVSDADASEPS | -431 | -451 | 1 | 0.001291103 | 21 | 2165.7737 | 4360.268 | 1781.3025 | 3293.0894 | 3781.4734 | 3572.2195 |
| Gorasp2 | IPTRPFEEGKK | -202 | -212 | 2 | 0 | 11 | 2314.872 | 718.25586 | NaN | 1369.7833 | NaN | NaN |
| Rab2a | IQEGVFDINNEANGIK | -171 | -186 | 0 | 0 | 16 | 921.1726 | 872.923 | 1156.7659 | 919.25006 | 935.1222 | 1374.7853 |
| Impdh1 | NLIDAGVDGLR | -312 | -322 | 0 | 0 | 11 | 1113.275 | 486.99356 | NaN | 869.43677 | 1030.6575 | 1049.69 |
| Impdh1 | IQHGFSGIPITAT | -135 | -147 | 0 | 0 | 13 | 1931.393 | 170.84367 | NaN | NaN | NaN | NaN |
| Parp14 | SIAFPAIGTGNLR | -1138 | -1150 | 0 | 0 | 13 | 0 | 3116.7295 | 363.9037 | NaN | NaN | NaN |
| Parp14 | IQNPALWR | -1667 | -1674 | 0 | 0 | 8 | 593.1688 | NaN | 599.96484 | NaN | NaN | NaN |
| Wdr55 | IRDTPEDIVLEAPASGL | -29 | -45 | 1 | 0 | 17 | 250.35313 | 220.33774 | 262.06415 | 327.75775 | 419.99658 | 637.1629 |
| Wdr55 | VVGTVGQHAGEPVEAL | -287 | -302 | 0 | 0 | 16 | 465.33798 | NaN | 547.82916 | NaN | 607.1618 | 671.5895 |
| Ifit1 | LLFENNDIPDLEVR | -25 | -38 | 0 | 0 | 14 | 1624.5071 | 1978.2632 | 1931.9381 | 1349.9108 | 2252.2573 | 2400.1428 |
| Ifit1 | ISEQVQFLDIK | -39 | -49 | 0 | 0 | 11 | 1209.698 | 1100.2518 | 2952.658 | 3519.4617 | NaN | 364.09485 |
| Lrch3 | ISHEVSPVKPIAVR | -414 | -427 | 1 | 0.00122419 | 14 | 4227.91 | 3332.3196 | 3048.9084 | NaN | NaN | NaN |
| Mtnd4 | ISIILDPLTK | -246 | -255 | 0 | 0 | 10 | 578.0852 | 624.6227 | 572.3503 | 456.96185 | 455.16348 | 581.59393 |
| Naa50 | ISNESAIDFYR | -115 | -125 | 0 | 0 | 11 | 1321.6753 | 1387.5923 | 1390.4581 | 1456.368 | 1510.2976 | 1484.2112 |
| Naa50 | TLGCLAPYR | -76 | -84 | 0 | 0.000550307 | 9 | 1023.575 | 882.01953 | 303.84256 | 637.8764 | 783.44214 | 1017.47406 |
| Naa50 | KFGFEIIETK | -126 | -135 | 1 | 0 | 10 | 1068.075 | NaN | NaN | NaN | NaN | NaN |
| Naa50 | LNQVIFPVSYNDK | -22 | -34 | 0 | 0 | 13 | 607.38116 | NaN | NaN | NaN | NaN | NaN |
| Lum | ISNIPDEYFK | -217 | -226 | 0 | 0 | 10 | 1092.6805 | 1450.8285 | 944.8785 | 654.94476 | 407.43784 | 1349.9213 |
| Polr2b | TVTLPENEDELESTNR | -870 | -885 | 0 | 0 | 16 | 1209.2429 | 1026.2472 | 798.6491 | 416.42322 | 636.98145 | 708.40295 |
| Polr2b | KGFDQEEVFEKPTR | -821 | -834 | 2 | 0 | 14 | 268.90433 | NaN | 1528.8013 | 1394.8401 | 1684.1276 | 1769.5771 |
| Polr2b | YYPQKPLVTTR | -752 | -762 | 1 | 0 | 11 | 859.22815 | NaN | 903.11566 | NaN | 984.8046 | 924.0566 |
| Polr2b | LFEASDPYQVH | -1107 | -1117 | 0 | 0.002697088 | 11 | 1293.761 | 1151.7087 | NaN | 1812.4772 | NaN | NaN |
| Polr2b | ISNLLSDYGYHLR | -1011 | -1023 | 0 | 0 | 13 | 1451.7294 | NaN | NaN | NaN | 2243.4487 | NaN |
| Polr2b | SAPLYVDITK | -121 | -130 | 0 | 0.00025015 | 10 | 659.99585 | NaN | NaN | NaN | NaN | NaN |
| Dlat | LQPHEFQGGTF | -548 | -558 | 0 | 0 | 11 | 2285.4673 | 2309.4785 | 2527.102 | 2313.4272 | 2164.0989 | 2183.509 |
| Dlat | ISVNDFIIK | -469 | -477 | 0 | 0 | 9 | 535.84186 | 633.5635 | 602.01263 | 424.69534 | 603.759 | 572.35315 |
| Dlat | TDIPISNIR | -416 | -424 | 0 | 0.00051306 | 9 | 1123.3635 | NaN | 1665.542 | 659.80884 | 1168.615 | 1629.8237 |
| Dlat | VAPAPAGVFTDIPISNIR | -407 | -424 | 0 | 0 | 18 | 537.21124 | 440.30524 | 523.82416 | NaN | 732.0343 | 317.391 |
| Prim1 | ISVPIDFHK | -329 | -337 | 0 | 0 | 9 | 935.82306 | NaN | 1159.9868 | 1148.3365 | NaN | 1249.5566 |
| Mycbp2 | ITFGFISCPICK | -4539 | -4550 | 0 | 0.00025015 | 12 | 871.30835 | NaN | NaN | NaN | 849.2803 | NaN |
| Prpf38b | TYHEVVDEIYFK | -82 | -93 | 0 | 0 | 12 | 338.97815 | 329.51044 | 439.18933 | 643.736 | 524.5635 | 356.9729 |
| Prpf38b | ITHTDSPYIR | -150 | -159 | 0 | 0.000230707 | 10 | 1299.5492 | NaN | NaN | NaN | 1169.8662 | NaN |

| | | | | | | | | | | | | |
|---|---|---|---|---|---|---|---|---|---|---|---|---|
| Tgs1 | ITNNIVYFLPR | -789 | -799 | 0 | 0.00025015 | 11 | 498.88675 | NaN | NaN | NaN | NaN | NaN |
| Utp15 | NYHVVSGADDYTVK | -134 | -147 | 0 | 0 | 14 | 811.03284 | 969.3567 | 957.8736 | 1008.5055 | 371.32428 | 736.42053 |
| Utp15 | NDSDPVPEHVPAELPEEK | -497 | -514 | 0 | 0 | 18 | 491.4695 | 428.96463 | 360.9244 | 503.9611 | 694.48615 | 948.906 |
| Utp15 | ITQDTLYWNNYK | -19 | -30 | 0 | 0 | 12 | 861.929 | 107.62693 | 817.99603 | NaN | 899.9998 | 722.7008 |
| Pzp | IVEPEGIEK | -916 | -924 | 0 | 0.000298805 | 9 | 2658.1614 | 2694.6812 | 2568.1555 | 1295.7415 | 1925.7878 | 1801.2784 |
| Pzp | KMVSGFIPMK | -1403 | -1412 | 1 | 0.002595029 | 10 | 1583.1359 | 1662.9857 | 1809.9996 | 1866.6382 | 1663.791 | 1550.4857 |
| Polr1b | QGEDNLVFGVKPGDPR | -787 | -802 | 1 | 0.00025015 | 16 | 303.9356 | NaN | 282.96286 | 502.2446 | 389.4475 | 829.56287 |
| Polr1b | IVIEEGCHSQK | -288 | -298 | 0 | 0.002068213 | 11 | 2489.2947 | NaN | NaN | NaN | NaN | NaN |
| Gbe1 | IVLDSDAAEYGGHQR | -648 | -662 | 0 | 0 | 15 | 660.66437 | 522.4876 | 557.63196 | 589.28625 | 551.1785 | 656.0802 |
| Ppat | IVLIDDSIVR | -385 | -394 | 0 | 0.00025015 | 10 | 1116.5676 | 1120.7198 | 251.24753 | 946.2778 | 1240.1179 | 330.77258 |
| Ppat | LYDSNLGIGHTR | -82 | -93 | 0 | 0 | 12 | 1013.73096 | 983.5331 | 1175.1705 | 2277.4868 | 1089.9731 | 1052.6377 |
| Uck2 | TVQIPVYDFVSH | -106 | -117 | 0 | 0.000230707 | 12 | 1223.1525 | 589.73663 | 736.1947 | 891.1612 | 1222.1672 | 1086.8146 |
| Uck2 | IVQLLGQNEVDYHQK | -40 | -54 | 0 | 0 | 15 | 679.59375 | NaN | NaN | 1017.48956 | 1367.8538 | 1048.2643 |
| Snx12 | IVVPPLPGK | -88 | -96 | 0 | 0 | 9 | 117.079445 | 414.5229 | 112.494484 | NaN | NaN | NaN |
| Tbc1d10c | IWDAFLSEGAK | -270 | -280 | 0 | 0 | 11 | 716.053 | 412.47452 | 1419.9912 | 1109.4305 | 1361.206 | 659.2033 |
| Tbc1d10c | SLPFPTVLR | -261 | -269 | 0 | 0.00051306 | 9 | 448.81244 | 542.649 | 484.56888 | 409.8082 | 577.68506 | NaN |
| Tbc1d10c | YGFIGGNSGELR | -38 | -49 | 0 | 0 | 12 | 540.5496 | 676.53796 | 249.7918 | NaN | NaN | NaN |
| Coro7 | TECDIQDVEFAR | -785 | -796 | 0 | 0.00025015 | 12 | 679.15125 | 584.6289 | 659.362 | 480.5618 | 503.89703 | 653.0572 |
| Coro7 | IWDLQTGAER | -618 | -627 | 0 | 0 | 10 | 281.37976 | 633.60846 | 267.31747 | 430.8692 | 498.64313 | 460.53552 |
| Coro7 | SSPLPLQEGPGPEGGR | -665 | -680 | 0 | 0 | 16 | 1077.9894 | NaN | 1326.7507 | 1413.5397 | NaN | 1854.1194 |
| Coro7 | LVWDPFDPHR | -547 | -556 | 0 | 0.00025015 | 10 | 340.8416 | 385.19098 | NaN | 680.38654 | 571.3519 | NaN |
| Coro7 | QLSDTAIIPISHHVPR | -332 | -347 | 0 | 0 | 16 | 311.36676 | NaN | 263.0168 | 318.70267 | NaN | NaN |
| Il4r | IWWDQIPTPAR | -262 | -272 | 0 | 0 | 11 | 1781.9978 | NaN | 1825.5903 | NaN | 4440.504 | NaN |
| Il4r | LGLELGLK | -674 | -681 | 0 | 0.006100831 | 8 | 604.8883 | NaN | 678.96606 | NaN | NaN | NaN |
| Tex10 | IYDPQEGAVVVLPAESQQR | -598 | -616 | 0 | 0 | 19 | 244.31586 | 356.17642 | 287.8959 | NaN | 291.44965 | 259.83826 |
| Tex10 | THITEGIQEDSLK | -150 | -162 | 0 | 0.000627943 | 13 | 2594.969 | NaN | 2778.743 | NaN | 277.6535 | 3126.8613 |
| Rpl24 | IYPGHGR | -13 | -19 | 0 | 0.000230707 | 7 | 1454.0117 | 1162.3123 | NaN | 1235.2964 | 1126.4185 | 1221.6736 |
| Rpl24 | VFQFLNAK | -28 | -35 | 0 | 0.00025015 | 8 | 942.8181 | 957.89215 | 594.44073 | NaN | NaN | NaN |
| Rpl24 | QINWTVLYR | -48 | -56 | 0 | 0 | 9 | 507.0997 | 510.73975 | 520.8786 | NaN | NaN | NaN |
| Uqcrq | IYTWGNQEFEQSK | -59 | -71 | 0 | 0 | 13 | 3879.5278 | NaN | 2907.9536 | 1548.907 | 4605.7896 | 1715.5288 |
| Uqcrq | SLSPFEQR | -18 | -25 | 0 | 0.00025015 | 8 | 1513.7311 | 1201.6525 | 1281.5316 | 1610.1187 | 890.0334 | NaN |
| Rnaset2a;Rnas | IYWPDVIHR | (99);(99) | (107);(107) | 0 | 0 | 9 | 400.12775 | 779.271 | 577.7904 | 582.94446 | 841.97815 | 778.80695 |
| Elf1 | KAEDDLNEDAEK | -566 | -577 | 1 | 0 | 12 | 396.02875 | 327.79312 | 438.51 | 439.35443 | NaN | 267.8404 |
| Ndufa2 | KAHPNLPILIR | -46 | -56 | 1 | 0.00111006 | 11 | 574.60236 | 433.42694 | 448.34018 | 73.195305 | 689.6873 | NaN |
| Ndufa2 | TVSLNNLSADEVTR | -76 | -89 | 0 | 0 | 14 | 557.4622 | 705.9573 | 1317.6642 | 269.45697 | 1281.2672 | NaN |
| Safb | SKDADDQKPGPSER | -558 | -571 | 2 | 0.000907206 | 14 | 195.58925 | 199.83563 | 197.65018 | 243.87436 | 281.97925 | 257.54974 |
| Safb | RPYEVDGR | -733 | -740 | 1 | 0.002296977 | 8 | 1012.6128 | 1053.4901 | NaN | NaN | 1562.8833 | 1532.9358 |
| Safb | RDDAYWPEAK | -741 | -750 | 1 | 0.00025015 | 10 | 1188.8849 | NaN | NaN | 909.9375 | NaN | 670.6844 |
| Safb | KAIEDEGGNPDEIEVT | -63 | -78 | 1 | 0.003490217 | 16 | 260.15125 | NaN | NaN | NaN | NaN | NaN |
| Dnttip2 | KAPELTDELK | -651 | -660 | 1 | 0 | 10 | 1356.4102 | 873.8392 | 736.91626 | 843.4646 | 1048.3899 | 915.65344 |
| Dnttip2 | SVLEDQEPIVR | -87 | -97 | 0 | 0 | 11 | 1153.6394 | 1026.748 | 1172.2212 | 1764.865 | 1199.9944 | 1355.3411 |
| Dnttip2 | SSQSGVPLECDTKPCK | -413 | -428 | 1 | 0.000591124 | 16 | 504.13596 | 334.48813 | 452.5239 | 450.6222 | 85.15876 | 726.75806 |
| Znf622 | KAPFVPTDECGR | -148 | -159 | 1 | 0 | 12 | 480.1847 | 629.72546 | 551.723 | NaN | NaN | 456.76965 |
| Gimap8 | KEDLGDGDLHDY | -604 | -615 | 1 | 0.00025015 | 12 | 871.45044 | 787.6326 | 809.32214 | 476.95068 | 558.40814 | 447.45404 |
| Gimap8 | NYDEHSNSWVGQLK | -664 | -677 | 0 | 0.000471409 | 14 | 840.341 | 876.87665 | NaN | 624.2232 | NaN | NaN |
| Mlec | LQPHPGLEK | -220 | -228 | 0 | 0.00025015 | 9 | 781.16766 | 875.5135 | 1144.3083 | NaN | 679.9913 | 1089.2069 |
| Mlec | KEEEEEEEYDEGSNL | -229 | -244 | 1 | 0.00111006 | 16 | 260.2555 | 256.88626 | NaN | 296.97174 | 286.89645 | 281.42102 |
| Mlec | YILAGTVDDVPK | -208 | -219 | 0 | 0 | 12 | 3629.6626 | NaN | 1988.8271 | 2597.465 | NaN | NaN |
| Ppie | KFDDENFILK | -218 | -227 | 1 | 0.00025015 | 10 | 782.51215 | 1408.977 | 1619.9915 | 1363.4825 | 837.86426 | 1503.2948 |
| Ppie | PVWSDDDWLK | -94 | -103 | 0 | 0 | 10 | 152.28981 | 284.54556 | 298.30896 | 157.43707 | 622.76825 | 446.0824 |
| Ppil3 | KFEDEYSEYLK | -70 | -80 | 1 | 0 | 11 | 1005.38745 | 992.1783 | 450.519 | NaN | 1668.2549 | 1663.3304 |
| Ifng | KFEVNNPQVQR | -118 | -128 | 1 | 0 | 11 | 1059.3628 | 1139.6105 | 1202.87 | 1077.9865 | 1187.5485 | 1218.2709 |
| Ifng | VVHQLLPESSLR | -137 | -148 | 0 | 0 | 12 | 1869.6213 | 1631.509 | 802.5472 | NaN | 4457.5176 | NaN |
| Cox6c | SSGALLPKPQMR | -2 | -13 | 1 | 0 | 12 | 393.6905 | NaN | 950.1979 | 1008.0557 | 1022.87286 | 1218.0137 |
| Cox6c | KFGVAEPR | -39 | -46 | 1 | 0.002780285 | 8 | 959.6106 | 1052.4086 | 1078.7773 | 1800.625 | NaN | NaN |
| Fis1 | KFQSEQAAGSVSK | -20 | -32 | 1 | 0 | 13 | 298.85397 | 358.95306 | 344.4477 | 354.94556 | 468.8767 | 428.04163 |
| Fis1 | SKYNEDIR | -45 | -52 | 1 | 0 | 8 | 531.6192 | NaN | 699.2777 | NaN | NaN | 2225.3972 |
| Sec62 | KGEDALFTTR | -66 | -75 | 1 | 0 | 10 | 2009.794 | 2503.7256 | 2570.3418 | NaN | 1185.4496 | 1202.2123 |
| Dnajb11 | KQYDTYGEEGLK | -84 | -95 | 1 | 0 | 12 | 977.03845 | 951.1462 | 906.6047 | 1257.9966 | 1505.316 | 1194.3906 |
| Dnajb11 | KGEGLPNFDNNNIK | -302 | -315 | 1 | 0 | 14 | 521.13885 | NaN | 1046.2681 | NaN | 950.7787 | NaN |
| Dnajb11 | KGEGLPNFDNNNIK | -302 | -315 | 1 | 0 | 14 | 521.13885 | NaN | 1046.2681 | NaN | 950.7787 | NaN |
| Eif3j1;Eif3j2 | LEIDDLKK | (194);(196) | (201);(203) | 1 | 0 | 8 | 1566.1476 | 1852.6278 | 1509.0343 | 1753.9607 | 1626.442 | 1856.5314 |
| Eif3j1;Eif3j2 | KGVVPGGGLK | (227);(229) | (236);(238) | 1 | 0 | 10 | 1351.417 | 1081.6162 | 1077.637 | 881.3044 | 870.34595 | 1077.8739 |
| Eif3j1;Eif3j2 | VLTPEEQLADKL | (110);(112) | (121);(123) | 1 | 0 | 12 | 1204.6997 | 633.86615 | 454.32153 | 2746.447 | 2941.2803 | 2101.3154 |
| Eif3j1;Eif3j2 | KLQEESDLELAK | (125);(127) | (136);(138) | 1 | 0 | 12 | 677.1042 | 644.28564 | 524.9177 | 1164.0785 | 1085.9072 | 1326.3197 |
| Eif3j1;Eif3j2 | RLEEPEESK | (101);(103) | (109);(111) | 1 | 0 | 9 | 657.01263 | 246.10912 | 990.36926 | 672.04193 | 882.90173 | 859.8951 |
| Eif3j1;Eif3j2 | WDDDDDENKEEAEVKPEVK | (55);(57) | (73);(75) | 2 | 0 | 19 | 351.55612 | 477.1685 | 374.37326 | 483.1344 | 510.92746 | 524.0129 |
| Eif3j1;Eif3j2 | VLTPEEQLADK | (110);(112) | (120);(122) | 0 | 0 | 11 | 3075.4744 | 3380.3784 | 2100.3699 | 5662.4 | NaN | 4156.365 |
| Eif3j1;Eif3j2 | LQEESDLELAK | (126);(128) | (136);(138) | 0 | 0 | 11 | 1127.7227 | 837.9941 | 985.7481 | NaN | NaN | NaN |
| Slc39a8 | KIDNYVEK | -191 | -198 | 1 | 0 | 8 | 7690.0986 | 8119.5312 | 8021.596 | 819.2445 | 1857.3461 | 3624.1772 |
| Dgka | NIPSTHGGSNLWGDTK | -596 | -611 | 0 | 0 | 16 | 985.2607 | NaN | 950.8975 | 897.09143 | 1388.577 | 1482.4766 |
| Dgka | KIITDPDILK | -629 | -638 | 1 | 0.001370415 | 10 | 468.52344 | NaN | NaN | NaN | 487.48364 | NaN |
| Hmcn1 | KILETSLK | -65 | -72 | 1 | 0.000550307 | 8 | 653.0127 | 604.08453 | NaN | 1044.8701 | 791.5204 | 842.625 |
| Plgrkt | KKPAFLVPIVPL | -76 | -87 | 2 | 0.000471409 | 12 | 443.22067 | 371.91812 | 437.95746 | 401.90497 | 498.02985 | 474.20212 |
| H1-1 | KKPAGPSVSEL | -35 | -45 | 2 | 0.000591124 | 11 | 2134.9775 | 2058.878 | 2084.1062 | 1295.7339 | 1444.3154 | 1337.1627 |
| H1-1 | KKPAGPSVSELIVQ | -35 | -48 | 2 | 0.00111006 | 14 | 1407.3296 | 1521.6819 | 1663.7378 | 973.73663 | 1047.9886 | 1037.9497 |
| H1-1 | KKPAGPSVSELIVQ | -35 | -48 | 2 | 0.00111006 | 14 | 1407.3296 | 1521.6819 | 1663.7378 | 973.73663 | 1047.9886 | 1037.9497 |
| H1-1 | KKPAGPSVSELIVQA | -35 | -49 | 2 | 0.00659934 | 15 | 1361.7662 | 1628.696 | 100.22238 | NaN | 1009.012 | 377.43124 |

| | | | | | | | | | | | | |
|---|---|---|---|---|---|---|---|---|---|---|---|---|
| Cd2 | RKDEELEIK | -238 | -246 | 2 | 0.000627943 | 9 | 706.47485 | 243.56181 | 1370.8887 | 805.45245 | 806.52344 | 743.5221 |
| Cd2 | RKDEELEIK | -238 | -246 | 2 | 0.000627943 | 9 | 706.47485 | 243.56181 | 1370.8887 | 805.45245 | 806.52344 | 743.5221 |
| Cd2 | WTNLNAPFK | -171 | -179 | 0 | 0 | 9 | 727.34827 | 821.62573 | 937.7071 | 825.16565 | 828.6798 | 988.6395 |
| Cd2 | KKPPFLISE | -67 | -75 | 2 | 0.004595494 | 9 | 551.5973 | NaN | 737.7046 | 411.88284 | 370.96625 | NaN |
| Ndufs7 | KLDDLINWAR | -77 | -86 | 1 | 0.000627943 | 10 | 2901.3647 | 3624.4133 | NaN | NaN | 5955.962 | 679.8085 |
| Ndufs7 | LDDLINWAR | -78 | -86 | 0 | 0 | 9 | 994.9138 | 1089.8951 | NaN | NaN | NaN | NaN |
| Zranb2 | VSDGDWICPDKK | -8 | -19 | 1 | 0 | 12 | 8653.099 | 7608.3037 | 10041.91 | NaN | 5706.0425 | 6129.785 |
| Zranb2 | KLDEDEDEDDADLSK | -168 | -182 | 1 | 0.003681885 | 15 | 406.64377 | NaN | 424.56253 | 376.8114 | 349.50928 | 423.0433 |
| Zranb2 | TGYGGGFNER | -100 | -109 | 0 | 0.001092529 | 10 | 733.5558 | NaN | 397.05487 | NaN | NaN | NaN |
| Scp2 | KLEEEGEQFVK | -443 | -453 | 1 | 0 | 11 | 1693.2434 | 1988.1079 | 2346.128 | 2049.6907 | 2057.6714 | 2091.0398 |
| Nxf1 | KLEELWLDR | -315 | -323 | 1 | 0 | 9 | 1313.0342 | 1652.2726 | NaN | NaN | 2105.9001 | 1745.0057 |
| Nxf1 | SIPYNPQNPVR | -418 | -428 | 0 | 0.002595029 | 11 | 162.86888 | NaN | NaN | NaN | NaN | NaN |
| Pyhin1 | SSETLQSIIQSPK | -166 | -178 | 0 | 0 | 13 | 2226.7234 | 2183.2314 | 2264.2822 | 1376.2178 | 1871.1312 | 1879.4626 |
| Pyhin1 | RPSSSILDPPVSSGTAS | -182 | -198 | 1 | 0 | 17 | 1574.5532 | 2457.51 | 2081.6628 | 1534.755 | 3230.3965 | 3344.7773 |
| Pyhin1 | KVTEPFSYDVTEEK | -241 | -254 | 1 | 0 | 14 | 794.387 | 833.71387 | 995.1749 | 1029.484 | 1196.3466 | 1146.2037 |
| Pyhin1 | RPSSSILDPPVSSGTASSS | -182 | -200 | 1 | 0 | 19 | 675.958 | 909.79626 | 1106.069 | 616.0077 | 780.24506 | 98.17722 |
| Pyhin1 | KLIPDEEK | -395 | -402 | 1 | 0 | 8 | 1140.5206 | 1332.7291 | NaN | 877.73004 | 932.33716 | NaN |
| Pyhin1 | RPSSSILDPPVSSGTASSSAQ | -182 | -202 | 1 | 0.003681885 | 21 | 226.01335 | 272.15378 | 241.10568 | NaN | NaN | NaN |
| Tcea1 | LLDGPSTDKDPEEK | -76 | -89 | 1 | 0 | 14 | 1184.0452 | 1390.747 | 1434.9064 | 1512.3806 | 1531.8298 | 1594.7142 |
| Tcea1 | LLDGPSTDKDPEEK | -76 | -89 | 1 | 0 | 14 | 1184.0452 | 1390.747 | 1434.9064 | 1512.3806 | 1531.8298 | 1594.7142 |
| Tcea1 | KLLDGPSTDKDPEEK | -75 | -89 | 2 | 0.000826806 | 15 | 1033.6051 | 1173.8438 | 1178.6736 | 1211.108 | 1374.6249 | 1111.1506 |
| Tcea1 | KQSTDEEVTSL | -55 | -65 | 1 | 0.00025015 | 11 | 1391.8506 | 1360.9717 | 1486.5781 | NaN | NaN | NaN |
| Psme3 | KLLELDSFLK | -37 | -46 | 1 | 0 | 10 | 1760.2235 | 1637.0211 | 766.5681 | 1278.935 | 1359.6327 | 1512.8949 |
| Psme3 | TVESEAASYLDQISR | -167 | -181 | 0 | 0 | 15 | 1104.8083 | 1034.4397 | 1249.2937 | 1773.3718 | 2272.2544 | 2191.3738 |
| Psme3 | RLDECEEAFQGT | -88 | -99 | 1 | 0.000550307 | 12 | 879.1482 | 890.7704 | 816.8857 | 2282.5618 | 1310.2107 | 732.92706 |
| Psme3 | RLDECEEAFQGTK | -88 | -100 | 1 | 0.00051306 | 13 | 796.9181 | 567.3286 | 864.84515 | 666.1706 | 650.8681 | 666.1951 |
| Psme3 | QQLVDIIEK | -113 | -121 | 0 | 0 | 9 | 604.34235 | 588.369 | 912.0842 | 2317.2817 | 606.8638 | 1105.0681 |
| Psme3 | LLELDSFLK | -38 | -46 | 0 | 0 | 9 | 467.98392 | 466.62207 | 486.39243 | 467.15894 | 634.29266 | 589.01117 |
| Psme3 | MWVQLLIPR | -138 | -146 | 0 | 0 | 9 | 343.0956 | 252.31232 | 192.69466 | 331.8355 | 391.7068 | 551.7767 |
| Psme3 | YPHVEDYR | -196 | -203 | 0 | 0 | 8 | 1140.3136 | 995.8508 | 964.6251 | NaN | NaN | 245.60715 |
| Gpaa1 | KLPFPDETHER | -111 | -121 | 1 | 0 | 11 | 500.76834 | 531.4796 | 638.26624 | 614.15106 | 617.7827 | 733.2636 |
| Fkbp2 | LVIPSELGYGER | -102 | -113 | 0 | 0 | 12 | 403.26254 | NaN | 542.7523 | 420.89832 | 540.9832 | 512.2661 |
| Fkbp2 | KLQIGVK | -28 | -34 | 1 | 0.00635324 | 7 | 382.70026 | 403.8029 | 295.75555 | NaN | NaN | NaN |
| Fkbp2 | VFEVELLK | -126 | -133 | 0 | 0.003102268 | 8 | 580.20825 | NaN | NaN | NaN | NaN | NaN |
| Dpp9 | KLSGPDDDPLHK | -575 | -586 | 1 | 0.000431655 | 12 | 758.5029 | 695.95215 | 294.95706 | NaN | 508.16895 | 786.1562 |
| Ndufa4 | LALFNPDVSWDR | -36 | -47 | 0 | 0 | 12 | 1869.4419 | 1912.1774 | 2128.343 | 1976.556 | 2593.9998 | 2454.795 |
| Ndufa4 | KNNPEPWNK | -48 | -56 | 1 | 0.001463897 | 9 | 1215.6476 | 505.84195 | 1363.9359 | 351.0169 | 1510.1404 | 1748.9678 |
| Ndufa4 | LFNPDVSWDR | -38 | -47 | 0 | 0 | 10 | 1126.5012 | 1412.9071 | 1719.8461 | 230.85568 | 190.82675 | 3049.6572 |
| Ndufa4 | LGPNEQYK | -57 | -64 | 0 | 0.001636916 | 8 | 3719.9834 | NaN | NaN | 1932.343 | 1628.041 | NaN |
| Asl | KNPDSLELIR | -288 | -297 | 1 | 0 | 10 | 477.82867 | 851.12366 | 800.33997 | 1190.1466 | 527.744 | 665.54785 |
| Idi1 | NVTLNPDPNEIK | -158 | -169 | 0 | 0 | 12 | 1028.1809 | 1554.9089 | 1876.607 | 1049.4641 | 753.18164 | 1984.5641 |
| Idi1 | KNVTLNPDPNEIK | -157 | -169 | 1 | 0 | 13 | 630.66864 | 615.5011 | 621.2978 | 661.4331 | 612.5871 | 612.0333 |
| Idi1 | NCHLNENIDK | -38 | -47 | 0 | 0 | 10 | 221.67328 | 378.0755 | 215.85199 | 2117.3699 | 818.47894 | 824.3785 |
| Idi1 | SNPGELEENNAIGVK | -91 | -105 | 0 | 0 | 15 | 560.70325 | NaN | NaN | 661.07086 | 954.35455 | 601.0235 |
| Sec63 | SPLLQLPHIE | -382 | -391 | 0 | 0 | 10 | 891.90216 | 835.57684 | 242.30264 | 560.67957 | 523.83997 | 466.57565 |
| Sec63 | KNWEEFGNPDGPQAT | -159 | -173 | 1 | 0 | 15 | 566.8871 | 638.98016 | 723.6672 | 621.3599 | 801.687 | 742.96936 |
| Sec63 | TLKDTEEVELK | -672 | -682 | 1 | 0.00025015 | 11 | 428.8529 | NaN | NaN | NaN | NaN | NaN |
| Sec63 | YSGDQILIR | -218 | -226 | 0 | 0.000431655 | 9 | 274.13684 | NaN | NaN | NaN | NaN | NaN |
| Ppp6r3 | SHIDTLLER | -11 | -19 | 0 | 0 | 9 | 1976.04 | 2093.1462 | 2070.4521 | 3740.19 | 2906.6426 | 2979.839 |
| Ppp6r3 | KPEQIVDFLK | -135 | -144 | 1 | 0 | 10 | 789.525 | 742.149 | 824.2893 | 786.36725 | 921.8443 | 850.29425 |
| Ppp6r3 | LVEIVHPSQEEDR | -195 | -207 | 0 | 0 | 13 | 338.60614 | 914.09265 | 813.5108 | 913.1017 | 1023.0004 | 904.85126 |
| Ppp6r3 | LLTCIEPPQPR | -169 | -179 | 0 | 0 | 11 | 658.7548 | 967.3316 | NaN | 1091.4938 | 1594.8152 | 1359.5356 |
| Ppp6r3 | QDVLNWLNEER | -180 | -190 | 0 | 0 | 11 | 508.08282 | NaN | 540.7426 | NaN | 522.4337 | NaN |
| Cd8a | KPQAPELR | -28 | -35 | 1 | 0.000471409 | 8 | 1187.7969 | 2442.3894 | 1445.6896 | 2617.3652 | 3168.437 | 3004.8086 |
| Mrps7 | NCEPVIGLVPILK | -151 | -163 | 0 | 0 | 13 | 457.25363 | 490.22495 | 642.9235 | 483.21405 | 569.9658 | 542.2191 |
| Mrps7 | KPVAELTEEEKYDQELK | -53 | -69 | 2 | 0.00025015 | 17 | 483.50293 | NaN | NaN | 600.366 | 689.71185 | 726.881 |
| Xrcc6 | LGSLADEFK | -516 | -524 | 0 | 0 | 9 | 468.0307 | 400.93176 | 360.51733 | 503.1326 | 538.39905 | 615.0405 |
| Xrcc6 | KVPFTEK | -443 | -449 | 1 | 0.000431655 | 7 | 790.2136 | 704.4064 | NaN | 490.29456 | NaN | NaN |
| Xrcc6 | TGSLLLPSDTK | -305 | -315 | 0 | 0.00266297 | 11 | 2422.6558 | NaN | NaN | 2806.2756 | NaN | NaN |
| Xrcc6 | KQELLDALIR | -594 | -603 | 1 | 0.00201047 | 10 | 302.1588 | NaN | 259.58917 | NaN | NaN | NaN |
| Gdi1 | TDDYLDQPCLETINR | -194 | -208 | 0 | 0 | 15 | 357.4729 | 516.51855 | 544.0825 | 679.6053 | 608.907 | 491.8485 |
| Gdi1 | SITPLEELYK | -45 | -54 | 0 | 0 | 10 | 328.40152 | 152.39638 | 619.59686 | 334.92917 | 627.05115 | NaN |
| Gdi1 | KQNDVFGEADQ | -437 | -447 | 1 | 0.001392869 | 11 | 1691.7971 | NaN | NaN | 984.39996 | 591.74927 | NaN |
| Gdi1 | QLICDPSYIPDR | -279 | -290 | 0 | 0.001743443 | 12 | 405.32977 | NaN | NaN | 831.4727 | NaN | NaN |
| Polr2d | KQQNESAEDEQELSEVF | -45 | -61 | 1 | 0.000947459 | 17 | 472.6196 | 435.49933 | 467.8918 | 563.26526 | 621.06116 | 657.9227 |
| Lsm6 | QIIGRPVVVK | -14 | -23 | 1 | 0 | 10 | 707.99 | 807.9355 | 858.4855 | 641.3966 | 1003.96533 | 887.1859 |
| Lsm6 | KQTPSDFLK | -5 | -13 | 1 | 0.001055699 | 9 | 2822.054 | NaN | 3019.6511 | 2806.992 | 3319.12 | 3186.4292 |
| Lsm6 | YGDAFIR | -60 | -66 | 0 | 0 | 7 | 620.1121 | 532.8645 | 486.77356 | 481.91748 | 442.5269 | NaN |
| Lsm6 | QTPSDFLK | -6 | -13 | 0 | 0.00025015 | 8 | 717.33966 | 1218.9484 | 552.6687 | NaN | NaN | NaN |
| Rpl28 | NCSSFLIK | -12 | -19 | 0 | 0 | 8 | 2848.041 | 107.81986 | 2866.3447 | 2261.807 | 2035.992 | 2093.5264 |
| Rpl28 | TVGVEPAADGK | -48 | -58 | 0 | 0 | 11 | 2385.7427 | 1335.056 | 1546.1013 | NaN | 2170.828 | 2315.6782 |
| Rpl28 | STEPNNLK | -26 | -33 | 0 | 0.00122419 | 8 | 1139.221 | NaN | 1252.0851 | NaN | 1193.0864 | 1234.3346 |
| Rpl28 | KYRPDLR | -101 | -107 | 2 | 0.000947459 | 7 | 617.24365 | 375.93542 | NaN | NaN | 560.1498 | NaN |
| Rpl28 | KYRPDLR | -101 | -107 | 2 | 0.000947459 | 7 | 617.24365 | 375.93542 | NaN | NaN | 560.1498 | NaN |
| Rpl28 | QTYSTEPNNLK | -23 | -33 | 0 | 0 | 11 | 5621.8726 | 418.89734 | NaN | NaN | NaN | NaN |
| Rpl28 | KQTYSTEPNNLK | -22 | -33 | 1 | 0.00025015 | 12 | 317.11337 | NaN | NaN | NaN | 484.54684 | NaN |
| Raly | KSDVETIFSK | -35 | -44 | 1 | 0 | 10 | 1467.7496 | 538.2204 | 1381.2056 | 1700.5898 | 589.91113 | 1288.4225 |
| Raly | SNIDALLGR | -199 | -207 | 0 | 0 | 9 | 1393.334 | 849.06 | NaN | 349.65332 | 2003.0885 | 2417.8496 |

| | | | | | | | | | | | | |
|---|---|---|---|---|---|---|---|---|---|---|---|---|
| Raly | LPAPQEDTASEAGTPQGEVQTR | -261 | -282 | 0 | 0.001392869 | 22 | 373.51968 | NaN | 323.28415 | 217.30748 | 506.29538 | NaN |
| Raly | LEQIAEEQK | -208 | -216 | 0 | 0.008890469 | 9 | 1594.5515 | NaN | NaN | NaN | NaN | NaN |
| Tsnax | KSFQQELDAR | -40 | -49 | 1 | 0.001190916 | 10 | 313.84232 | 609.76556 | NaN | NaN | NaN | 799.48303 |
| Tsnax | SFIGNTGPYEVSK | -229 | -241 | 0 | 0 | 13 | 2333.459 | 1952.0295 | NaN | NaN | NaN | NaN |
| Ptk2b | LIGSVDDLLPSLPASSR | -920 | -936 | 0 | 0 | 17 | 416.62152 | 524.57355 | 398.65063 | 390.92773 | 572.0738 | 632.5451 |
| Ptk2b | KSNFELLEK | -198 | -206 | 1 | 0.003681885 | 9 | 1937.158 | 1934.0371 | NaN | 2184.6611 | 116.958916 | NaN |
| Ptk2b | SDEIHWLHPQ | -99 | -108 | 0 | 0.00078661 | 10 | 2505.8713 | NaN | NaN | NaN | NaN | NaN |
| Aldh2 | TEQGPQVDETQFK | -358 | -370 | 0 | 0 | 13 | 4527.3945 | 3745.7542 | 3482.1973 | 4023.9937 | 762.02704 | 872.9494 |
| Aldh2 | KTFPTVNPSTGEVICQ | -54 | -69 | 1 | 0.00078661 | 16 | 244.12157 | 374.40735 | 603.99536 | 404.58295 | 1582.5243 | 1540.6797 |
| Aldh2 | VVGNPFDSR | -349 | -357 | 0 | 0.00025015 | 9 | 656.4535 | NaN | 536.57544 | 470.50797 | 896.0875 | 623.83246 |
| Napa | KYEELFPAFSDSR | -227 | -239 | 1 | 0.000230707 | 13 | 1252.6625 | 1469.1318 | 896.1405 | 197.06102 | 3866.6619 | 2886.4133 |
| Napa | KTIQGDEEDLR | -285 | -295 | 1 | 0 | 11 | 848.1582 | 888.85205 | 950.57904 | 926.10736 | 964.91705 | 847.9197 |
| Napa | NSQSFFSGLFGGSSK | -23 | -37 | 0 | 0 | 15 | 579.0076 | 641.1062 | 1103.4133 | 754.6684 | 1385.0525 | 1025.8068 |
| Napa | TIQGDEEDLR | -286 | -295 | 0 | 0 | 10 | 950.72876 | 1246.0737 | 1141.9712 | 681.698 | NaN | 977.4473 |
| Napa | YEELFPAFSDSR | -228 | -239 | 0 | 0 | 12 | 345.2132 | NaN | 1317.4503 | 1029.9187 | 931.1396 | NaN |
| Chmp2a | KTPEELLR | -9 | -16 | 1 | 0.00051306 | 8 | 840.91675 | 889.25574 | 872.7834 | 161.67372 | 1846.567 | 1564.5315 |
| Rab10 | NIDEHANEDVER | -106 | -117 | 0 | 0 | 12 | 1913.8585 | 2179.9634 | 2263.4438 | 1800.1057 | 2453.6013 | 2590.9976 |
| Rab10 | KTYDLLFK | -4 | -11 | 1 | 0 | 8 | 1325.1445 | 1510.3215 | 1814.6443 | 1388.6935 | 1777.6711 | 1881.8345 |
| Rab10 | KTPVKEPNSENVDIS | -173 | -187 | 2 | 0 | 15 | 842.3125 | 930.12714 | 994.57684 | 1145.0387 | 1294.8925 | 1197.2135 |
| Rab10 | KTPVKEPNSENVDISSGGGVT | -173 | -193 | 2 | 0.00713295 | 21 | 598.3468 | 719.70624 | 534.0275 | NaN | NaN | NaN |
| H1-4 | TSGPPVSELITK | -35 | -46 | 0 | 0 | 12 | 14221.326 | 14933.899 | 15287.15 | 13697.468 | 16355.849 | 16249.601 |
| H1-4 | TSGPPVSELITK | -35 | -46 | 0 | 0 | 12 | 14221.326 | 14933.899 | 15287.15 | 13697.468 | 16355.849 | 16249.601 |
| H1-4 | KTSGPPVSELITK | -34 | -46 | 1 | 0 | 13 | 5584.913 | 5134.028 | 5632.499 | 3327.4639 | 3640.475 | 3598.6133 |
| H1-4 | KTSGPPVSELITK | -34 | -46 | 1 | 0 | 13 | 5584.913 | 5134.028 | 5632.499 | 3327.4639 | 3640.475 | 3598.6133 |
| H1-4 | SETAPAAPAAPAPAEK | -2 | -17 | 0 | 0 | 16 | 3546.8044 | 3744.0193 | 3201.589 | 2625.3977 | 1939.2856 | 1863.2377 |
| H1-4 | SETAPAAPAAPAPAEK | -2 | -17 | 0 | 0 | 16 | 3546.8044 | 3744.0193 | 3201.589 | 2625.3977 | 1939.2856 | 1863.2377 |
| H1-4 | KTSGPPVSEL | -34 | -43 | 1 | 0.00025015 | 10 | 2512.557 | 4399.701 | 2459.48 | 2186.4702 | 3171.9956 | 3362.0352 |
| H1-4 | SETAPAAPAAPAPAEKTPVK | -2 | -21 | 1 | 0 | 20 | 352.18958 | 251.26532 | 308.36557 | 398.3769 | 382.02008 | 379.04605 |
| H1-4 | TAPAAPAAPAPAEK | -4 | -17 | 0 | 0 | 14 | 3740.436 | 3281.685 | 3247.3054 | 3264.242 | NaN | 3660.3774 |
| H1-4 | KTSGPPVSELIT | -34 | -45 | 1 | 0.00078661 | 12 | 14099.613 | 15014.37 | 15528.963 | NaN | 16133.59 | NaN |
| H1-4 | TSGPPVSELIT | -35 | -45 | 0 | 0 | 11 | 3029.1526 | NaN | NaN | 4528.594 | NaN | NaN |
| Tmx1 | KVDVTEQTGLSGR | -82 | -94 | 1 | 0 | 13 | 964.78595 | 965.0513 | 951.9959 | 1013.8924 | 1366.0955 | 2010.0859 |
| Tmx1 | VDVTEQTGLSGR | -83 | -94 | 0 | 0 | 12 | 2503.7224 | 2604.2314 | NaN | 1947.099 | NaN | NaN |
| Cbx3 | KVEEAEPEEFVVE | -21 | -33 | 1 | 0.00025015 | 13 | 4639.678 | 5023.8545 | 5075.6143 | 6047.6113 | 5216.5815 | 5220.9062 |
| Cbx3 | LTWHSCPEDEAQ | -172 | -183 | 0 | 0.00111006 | 12 | 3890.5063 | 4077.8918 | 4042.5117 | 2235.2053 | 2091.5354 | 2025.8938 |
| Cbx3 | WKDSDEADLVLAK | -142 | -154 | 1 | 0 | 13 | 2817.9705 | 2115.7427 | 2170.0032 | 3177.2446 | 2227.3862 | 2029.3016 |
| Cbx3 | KVEEAEPEEFVVEK | -21 | -34 | 1 | 0 | 14 | 1180.8556 | 595.18115 | 1488.6837 | 1867.2545 | 2144.3508 | 2012.9388 |
| Cbx3 | SLSDSESDDSK | -93 | -103 | 0 | 0 | 11 | 521.5981 | NaN | 534.1835 | 333.11594 | 621.2418 | NaN |
| Tpd52l2 | KVEEEIVTLR | -55 | -64 | 1 | 0.00025015 | 10 | 577.6199 | NaN | NaN | 692.26416 | 693.8197 | 699.41833 |
| Tpd52l2 | TPVVEGLTEGEEEELRAE | -35 | -52 | 1 | 0.001776473 | 18 | 519.9057 | 607.63403 | 423.69635 | NaN | NaN | NaN |
| Bud31 | VESLWPIFR | -43 | -51 | 0 | 0.00025015 | 9 | 644.65955 | 703.17426 | 792.4603 | 867.781 | 1005.5843 | 1160.104 |
| Bud31 | KVESLWPIFR | -42 | -51 | 1 | 0 | 10 | 439.71866 | 496.53192 | 464.22327 | 637.3502 | 833.39166 | 726.7837 |
| Cab39 | VADLQLIDFEGK | -84 | -95 | 0 | 0 | 12 | 383.0557 | 568.05176 | 746.07434 | 896.9559 | 1000.02527 | 1124.6265 |
| Cab39 | TQPILDILLK | -288 | -297 | 0 | 0.000394361 | 10 | 171.28813 | 341.02634 | 69.051384 | 257.20032 | 170.35939 | 391.28055 |
| Cab39 | VADLQLIDFEGKK | -84 | -96 | 1 | 0 | 13 | 784.5419 | NaN | NaN | 1421.1392 | 1268.2229 | NaN |
| Cab39 | QFYDFFR | -165 | -171 | 0 | 0.002168954 | 7 | 433.70496 | 443.43988 | 448.41058 | NaN | NaN | NaN |
| Cab39 | KVFVANPNK | -279 | -287 | 1 | 0.002752526 | 9 | 231.92279 | NaN | NaN | NaN | NaN | NaN |
| Adprs | VGELLDQDVVSR | -255 | -266 | 0 | 0 | 12 | 7132.54 | 7564.0225 | 7936.1978 | 3560.3223 | 7166.2974 | 7180.7964 |
| Adprs | SHVESLEPDPGTPGSAR | -59 | -75 | 0 | 0 | 17 | 603.0175 | 681.23224 | 366.1555 | 799.9464 | 676.30554 | 899.4217 |
| Adprs | KVGELLDQDVVSR | -254 | -266 | 1 | 0.000431655 | 13 | 368.0781 | 330.03598 | 377.93356 | NaN | NaN | 425.04657 |
| Psmb5 | KVIEINPY | -92 | -99 | 1 | 0.004439092 | 8 | 576.3744 | NaN | NaN | NaN | NaN | NaN |
| Sec22b | LEDLHSEFDEQHGK | -88 | -101 | 0 | 0 | 14 | 1150.1904 | 731.8833 | 1163.9232 | 616.7876 | 783.2536 | 1006.8955 |
| Sec22b | LEDLHSEFDEQHGK | -88 | -101 | 0 | 0 | 14 | 1150.1904 | 731.8833 | 1163.9232 | 616.7876 | 783.2536 | 1006.8955 |
| Sec22b | KVPTVSRPY | -102 | -110 | 2 | 0.00111006 | 9 | 879.3009 | 1025.1664 | 1166.8428 | 1090.5948 | 1118.6282 | 1087.4163 |
| Arrb2 | KVQFAPETPGPQPSAE | -171 | -186 | 1 | 0.002068213 | 16 | 679.0558 | NaN | NaN | NaN | NaN | NaN |
| Mtarc2 | KVWIYPIK | -59 | -66 | 1 | 0 | 8 | 417.10162 | 377.68076 | 500.15945 | 341.29825 | 538.66864 | 534.0679 |
| Sf3b5 | WEWLVNQHR | -29 | -37 | 0 | 0 | 9 | 959.4896 | 1095.2709 | 1124.4868 | 1019.8473 | 1913.6823 | 1350.9984 |
| Sf3b5 | KWEWLVNQHR | -28 | -37 | 1 | 0 | 10 | 512.7367 | 370.34897 | 1070.8591 | 210.7155 | 1112.9049 | 1543.5741 |
| Sf3b5 | MLQPSGPPADKPEEN | -72 | -86 | 1 | 0.001055699 | 15 | 2522.6455 | NaN | NaN | 1528.4899 | 2928.5886 | NaN |
| Selenof | KWNTDSVEEFLSEK | -145 | -158 | 1 | 0.003003218 | 14 | 261.68594 | 577.9369 | 749.19196 | 964.0918 | 692.96436 | 487.77286 |
| Selenof | NLLPLDPVCR | -59 | -68 | 0 | 0.000907206 | 10 | 101.680855 | 1010.7775 | 1037.8542 | 456.51236 | 244.25912 | 1024.8041 |
| Selenof | WNTDSVEEFLSEK | -146 | -158 | 0 | 0.003299042 | 13 | 342.0722 | 463.61465 | NaN | NaN | NaN | NaN |
| Atp6v1h | KYDDEDISEDIK | -325 | -336 | 1 | 0 | 12 | 1124.4576 | 1353.3641 | 1270.9141 | 1162.1993 | NaN | 1236.7422 |
| Cks1b | KYDDEEFEYR | -11 | -20 | 1 | 0 | 10 | 679.5632 | 749.3665 | 743.0865 | 545.59906 | 747.0693 | 745.1784 |
| Cks2 | YFDEHYEYR | -12 | -20 | 0 | 0 | 9 | 548.32556 | 615.9178 | 614.6092 | 866.0317 | 909.4595 | 916.99664 |
| Cks2 | KYFDEHYEYR | -11 | -20 | 1 | 0 | 10 | 3186.0344 | 2779.4824 | NaN | 400.12177 | 1650.3833 | 1685.5648 |
| Nceh1 | KYFLQPEVLDK | -167 | -177 | 1 | 0.000230707 | 11 | 365.7354 | 1051.823 | NaN | 3350.9807 | 4584.5176 | 3148.9375 |
| Nceh1 | TCEHDVLR | -344 | -351 | 0 | 0.002329279 | 8 | 1397.7511 | NaN | NaN | NaN | NaN | 1123.8405 |
| Rpl27a | TGVAPIIDVVR | -95 | -105 | 0 | 0 | 11 | 7519.5996 | 8754.558 | 8778.574 | 8354.744 | 10422.738 | 10800.305 |
| Rpl27a | YHPGYFGK | -48 | -55 | 0 | 0 | 8 | 1990.7314 | 1998.9473 | 2079.15 | 366.64404 | 1542.7108 | 1642.7015 |
| Rpl27a | NQSFCPTVNLDK | -66 | -77 | 0 | 0 | 12 | 1329.2747 | 1584.5333 | 1603.6232 | 1413.953 | 1404.3146 | 1488.3776 |
| Rpl27a | KYHPGYFGK | -47 | -55 | 1 | 0 | 9 | 675.9967 | 1380.0787 | 1937.7855 | 372.40158 | 3525.8623 | 1200.5298 |
| Rpl27a | KYHPGYFGK | -47 | -55 | 1 | 0 | 9 | 675.9967 | 1380.0787 | 1937.7855 | 372.40158 | 3525.8623 | 1200.5298 |
| Rpl27a | LWTLVSEQTR | -78 | -87 | 0 | 0 | 10 | 506.6578 | 541.4309 | 557.40436 | NaN | NaN | NaN |
| C4b | KYVLPNFEVK | -233 | -242 | 1 | 0 | 10 | 834.3178 | 742.7785 | 780.352 | 834.7975 | 493.06573 | 768.47784 |
| C4b | YVLPNFEVK | -234 | -242 | 0 | 0 | 9 | 1100.8083 | 1269.4045 | NaN | 1322.9973 | 1494.1208 | 1498.517 |
| C4b | SGFHALR | -1486 | -1492 | 0 | 0.005363933 | 7 | 785.3201 | 551.41394 | 518.22815 | NaN | NaN | NaN |
| Dctn2 | WSPVASTLPELVQR | -320 | -333 | 0 | 0 | 14 | 1178.1729 | 95.476524 | 1020.12854 | 836.6949 | 1422.9176 | 1233.8119 |

| | | | | | | | | | | | | |
|---|---|---|---|---|---|---|---|---|---|---|---|---|
| Dctn2 | LADPDGALAK | -166 | -175 | 0 | 0 | 10 | 551.6781 | 468.0772 | 386.95444 | NaN | 852.8229 | 567.535 |
| Dctn2 | VHQLYETIQR | -310 | -319 | 0 | 0 | 10 | 1387.3123 | 1447.9813 | 1536.7542 | 1460.8776 | NaN | NaN |
| Dctn2 | LLHEVQELTTEVEK | -106 | -119 | 0 | 0.000431655 | 14 | 835.0145 | NaN | NaN | 1101.545 | NaN | NaN |
| Mrpl43 | LADQSGLDVIR | -104 | -114 | 0 | 0 | 11 | 1527.397 | 1650.0261 | 513.8919 | 456.97324 | 2425.3447 | NaN |
| Arglu1 | RVEELVAK | -131 | -138 | 1 | 0.00025015 | 8 | 742.633 | 351.72964 | 901.439 | 1129.7397 | 952.0116 | 782.2378 |
| Arglu1 | QLLEELER | -169 | -176 | 0 | 0.002810811 | 8 | 1031.6697 | NaN | 1368.3605 | NaN | NaN | NaN |
| Arglu1 | LAEEQLR | -217 | -223 | 0 | 0.00719662 | 7 | 1677.8688 | NaN | NaN | NaN | NaN | NaN |
| Ahcyl2 | LAESGFPVFAWK | -273 | -284 | 0 | 0 | 12 | 191.46506 | 424.64545 | 429.6093 | NaN | NaN | NaN |
| Rcsd1 | LAFDPAALLPGASPK | -93 | -107 | 0 | 0 | 15 | 944.50366 | 852.9294 | 1014.1112 | 832.6692 | 834.3104 | 852.55457 |
| Rcsd1 | NTCNSTEKPEELVR | -242 | -255 | 1 | 0.003003218 | 14 | 414.80795 | NaN | NaN | NaN | 401.6333 | NaN |
| Sdr39u1 | LAGENILNPLR | -58 | -68 | 0 | 0 | 11 | 620.6506 | 456.94385 | 804.3847 | 484.51126 | 650.2533 | 293.20377 |
| Sdr39u1 | LGLGGPIGSGR | -177 | -187 | 0 | 0 | 11 | 582.3143 | 522.0338 | 311.95477 | 430.2052 | 188.9001 | 455.6522 |
| Sdr39u1 | PAFIPVPSTVVR | -240 | -251 | 0 | 0.000550307 | 12 | 1468.1672 | 2479.3374 | 1194.6832 | NaN | NaN | NaN |
| Sdr39u1 | LPGESTR | -145 | -151 | 0 | 0.009242859 | 7 | 499.10986 | NaN | NaN | NaN | NaN | NaN |
| Rab21 | NLAIWDTAGQER | -67 | -78 | 0 | 0 | 12 | 1806.5531 | 1954.5159 | 2463.9893 | 1729.6057 | 2120.4902 | 1966.5096 |
| Rab21 | RGVQIIDDEPQAQ | -201 | -213 | 1 | 0 | 13 | 881.74963 | 856.27924 | 953.2528 | 410.46255 | 777.74384 | 751.8984 |
| Rab21 | VYDVTDEDSFQK | -96 | -107 | 0 | 0 | 12 | 810.6069 | 439.95508 | 432.88577 | 648.2634 | NaN | NaN |
| Pitpna | LAPEGALNIHEK | -76 | -87 | 0 | 0 | 12 | 1040.8759 | 1090.1794 | 1081.4817 | 1437.093 | 1423.505 | 1371.0001 |
| Pitpna | VNEPYEKDDGEKGQYTHK | -44 | -61 | 2 | 0.000230707 | 18 | 472.51758 | 572.27716 | 636.0008 | 595.8353 | 686.54803 | 839.8165 |
| Pitpna | TWHKPDLGTQENVHK | -114 | -128 | 1 | 0.00025015 | 15 | 1591.1669 | NaN | 721.793 | NaN | 1066.1033 | 1079.9486 |
| Ppil1 | QFEDELHPDLK | -81 | -91 | 0 | 0 | 11 | 1129.7871 | 1090.3547 | 1119.9312 | 1278.3572 | 1243.5613 | 1117.8234 |
| Ppil1 | SQDRPVDDVK | -149 | -158 | 1 | 0.000230707 | 10 | 904.1531 | 976.7298 | 1230.829 | 540.10065 | 640.4886 | 1613.0104 |
| Ppil1 | LAPTQWLDGK | -116 | -125 | 0 | 0.00025015 | 10 | 652.9972 | NaN | NaN | 1212.639 | NaN | 588.1522 |
| Ppil1 | TNSQDRPVDDVK | -147 | -158 | 1 | 0 | 12 | 482.10947 | 547.16296 | 930.8827 | NaN | NaN | NaN |
| Birc6 | VIFVDDYAVGCR | -129 | -140 | 0 | 0 | 12 | 345.91953 | 1055.053 | 1340.0616 | NaN | 447.6791 | NaN |
| Birc6 | LHTGLLLVQ | -2090 | -2098 | 0 | 0.00688377 | 9 | 400.3853 | 402.26495 | NaN | NaN | NaN | NaN |
| Birc6 | LATSPAQLPSADGADR | -375 | -390 | 0 | 0.008653659 | 16 | 440.45892 | NaN | NaN | NaN | NaN | NaN |
| Slamf7 | LCEDAATDLTSLR | -211 | -223 | 0 | 0 | 13 | 1352.0792 | 1017.3139 | 707.0554 | 2968.8088 | 796.3729 | 2560.3232 |
| Slamf7 | SFSTPVFPQK | -201 | -210 | 0 | 0 | 10 | 1001.42554 | 556.9537 | 397.9803 | NaN | 1155.3231 | NaN |
| Chchd2 | LCEGFNEVLR | -135 | -144 | 0 | 0 | 10 | 1494.2649 | 1928.1857 | 1913.76 | 1345.9385 | 1125.3794 | 2078.9182 |
| Chchd2 | QFLECAQNQSDVK | -122 | -134 | 0 | 0 | 13 | 1166.3516 | 795.0512 | 743.4908 | 1439.4519 | 1498.9243 | 1363.3754 |
| Chchd2 | RAPAAQPPAAAAPS | -28 | -41 | 1 | 0.002566359 | 14 | 1060.858 | 1088.5869 | NaN | 1192.2368 | NaN | NaN |
| Exosc1 | LCNLEEGSPGSGTYTR | -14 | -29 | 0 | 0 | 16 | 583.3943 | 573.9731 | 583.4283 | 733.5414 | 647.05817 | 625.9023 |
| Inhca | LCQLCAGK | -184 | -191 | 0 | 0 | 8 | 4688.992 | 4634.85 | 4847.9697 | 3732.459 | 4410.2705 | 5512.138 |
| Denr | LDADYPLR | -23 | -30 | 0 | 0 | 8 | 1352.7806 | 1007.60565 | 1055.964 | 1496.0308 | 1186.3542 | NaN |
| Cdc26 | LDDIEEFESIR | -13 | -23 | 0 | 0 | 11 | 375.39816 | NaN | 487.48944 | 519.2691 | 261.8424 | 480.10547 |
| Cryzl1 | LDEPIPLYEAK | -319 | -329 | 0 | 0.000471409 | 11 | 584.194 | NaN | 394.51233 | 1059.8771 | NaN | 1238.9451 |
| Ggct | LDFGNFQGK | -51 | -59 | 0 | 0 | 9 | 784.43744 | 855.1767 | 843.9918 | 839.7417 | 898.3118 | 547.3074 |
| Gclc | LDFLIPLSK | -457 | -465 | 0 | 0 | 9 | 238.94171 | 323.18518 | 336.3608 | 348.44193 | 314.752 | 447.17593 |
| Gclc | VVINVPIFK | -201 | -209 | 0 | 0 | 9 | 2922.3887 | 2822.3508 | 5111.953 | NaN | NaN | 1221.9677 |
| Cfap54 | LDFPQTSLL | -467 | -475 | 0 | 0.001392869 | 9 | 4197.636 | 4649.927 | 4906.876 | 3994.9636 | NaN | 5455.7847 |
| Sipa1 | LDGAPGQAVGEVVAR | -656 | -670 | 0 | 0 | 15 | 2736.011 | 2202.2354 | 230.95898 | NaN | 658.41187 | 419.22 |
| Nup54 | LDIISEDISELQK | -354 | -366 | 0 | 0 | 13 | 1995.2378 | 2080.0515 | 2005.156 | 1976.0271 | 516.6216 | 2033.837 |
| Nup54 | LVEHGLNETIHSR | -493 | -505 | 0 | 0 | 13 | 884.2423 | 1035.4 | 1121.8868 | 375.66418 | 737.251 | 581.0017 |
| Nup54 | LVEHGLNETIHSR | -493 | -505 | 0 | 0 | 13 | 884.2423 | 1035.4 | 1121.8868 | 375.66418 | 737.251 | 581.0017 |
| Nup54 | SQQQQLVESLHK | -207 | -218 | 0 | 0 | 12 | 540.07465 | 716.20795 | 655.511 | 277.30023 | 1008.75525 | 946.61725 |
| Nup54 | LDTIQGELNAPTQFK | -417 | -431 | 0 | 0 | 15 | 462.8019 | 237.30244 | 480.49588 | 588.5271 | 636.4241 | 622.047 |
| Nup54 | VQLDTIQGELNAPTQFK | -415 | -431 | 0 | 0.00122419 | 17 | 560.3088 | NaN | 599.2703 | NaN | NaN | NaN |
| Nae1 | SHPDNALEDLR | -174 | -184 | 0 | 0 | 11 | 1605.8801 | NaN | 1493.8053 | 2231.494 | 2564.5615 | 1620.3003 |
| Nae1 | LDKPFPELR | -185 | -193 | 1 | 0 | 9 | 583.56506 | NaN | 440.98358 | 633.12823 | 806.2943 | NaN |
| Nae1 | WNSQIPLLICR | -144 | -154 | 0 | 0 | 11 | 1217.628 | 1476.1481 | 1570.396 | NaN | NaN | NaN |
| Hemk1 | LDLVVSNPPYIFR | -235 | -247 | 0 | 0.000550307 | 13 | 357.5972 | 297.86874 | NaN | NaN | NaN | NaN |
| Ndrg3 | LDPINTTLLK | -277 | -286 | 0 | 0 | 10 | 1266.94 | 1113.177 | 1165.1638 | 736.9455 | 597.9786 | 487.98517 |
| Acyp1 | LDYSDFQIVK | -90 | -99 | 0 | 0 | 10 | 514.5172 | 1044.0842 | 317.76514 | 477.2894 | 1361.8127 | 422.3767 |
| Gtf2e2 | LEDIEEGLPNSQK | -169 | -181 | 0 | 0.000627943 | 13 | 1518.46 | NaN | NaN | 1920.7891 | NaN | 2134.338 |
| Senp3 | LEDIFQQEFSTPSR | -337 | -350 | 0 | 0 | 14 | 506.94016 | 434.80524 | NaN | 596.61145 | 644.45825 | 780.77185 |
| Senp3 | SGGGFGPDPGSGTTVPTR | -42 | -59 | 0 | 0 | 18 | 347.47815 | 755.3043 | 790.8196 | 806.216 | 792.43494 | NaN |
| Senp3 | NHLSPQEGGATPQVPSPC | -160 | -177 | 0 | 0.005605646 | 18 | 823.2935 | NaN | NaN | NaN | NaN | NaN |
| Noc2l | LEDLNFPEIK | -641 | -650 | 0 | 0 | 10 | 9753.111 | 10731.845 | 11191.578 | NaN | 8117.5933 | 8381.009 |
| Noc2l | LLPSSSPLWGK | -236 | -246 | 0 | 0 | 11 | 177.48853 | 202.99544 | 1000.8047 | NaN | 1217.6223 | 1135.8617 |
| Cnpy4 | LEEALENLCER | -92 | -102 | 0 | 0 | 11 | 708.80286 | 666.2003 | 736.2996 | 1231.214 | 2767.279 | 951.05896 |
| Cfap45 | LEELRATGLPEK | -522 | -533 | 1 | 0.00025015 | 12 | 3691.332 | NaN | NaN | 8007.3184 | NaN | 7808.824 |
| Psmc5 | LEGGSGGDSEVQR | -259 | -271 | 0 | 0 | 13 | 1462.7406 | 1522.9927 | 1717.4946 | 888.77936 | 1165.9801 | 1289.0098 |
| Psmc5 | VSGSELVQK | -214 | -222 | 0 | 0 | 9 | 1335.0624 | 1256.4495 | 1197.9382 | 288.63507 | 865.8045 | 1039.0975 |
| Psmc5 | NIDINDVTPNCR | -102 | -113 | 0 | 0 | 12 | 803.94116 | 691.3696 | 742.96136 | 840.06067 | 679.6992 | 902.8858 |
| Psmc5 | QEQGSYVGEVVR | -67 | -78 | 0 | 0 | 12 | 620.17365 | 1199.2383 | 1147.4392 | 2112.015 | 2506.1213 | 2268.9238 |
| Psmc5 | NQLDGFEATK | -278 | -287 | 0 | 0.009362332 | 10 | 4343.902 | NaN | 3267.8535 | NaN | NaN | NaN |
| Rab8a | LEGNSPQGSSHGVK | -177 | -190 | 0 | 0 | 14 | 199.23474 | 307.20322 | 367.8494 | 403.61365 | 374.09848 | 306.83017 |
| Rab8a | NIEEHASADVEK | -105 | -116 | 0 | 0 | 12 | 1626.1305 | NaN | NaN | 1634.8071 | 2085.9045 | 2175.0735 |
| Rab8a | TIELDGKR | -49 | -56 | 1 | 0.005668207 | 8 | 7740.181 | NaN | NaN | 7821.4194 | NaN | NaN |
| Wdr4 | LEGSVGTDDSFR | -349 | -360 | 0 | 0 | 12 | 807.8023 | 569.4439 | NaN | 335.2837 | 756.056 | 874.07385 |
| Ndufa11 | LEGWELFPTPK | -130 | -140 | 0 | 0 | 11 | 1064.9604 | 265.31808 | 1240.4231 | 959.0567 | 1179.4708 | 255.25972 |
| Ndufa11 | SLNPADSTLEAVAR | -42 | -55 | 0 | 0 | 14 | 6378.002 | 6026.9707 | NaN | NaN | 5860.2046 | 5726.508 |
| Polr2g | LEHEILLHPR | -7 | -16 | 0 | 0 | 10 | 966.7558 | 1075.2502 | 885.40784 | NaN | NaN | 1374.2762 |
| Polr2g | VTTIDNIGAGVIQPGR | -48 | -63 | 0 | 0 | 16 | 412.17627 | 458.67532 | NaN | NaN | NaN | NaN |
| Snx1 | LQEVECEEQR | -313 | -322 | 0 | 0 | 10 | 976.6478 | 1000.48145 | 316.7848 | 957.1347 | 886.683 | 830.54706 |
| Snx1 | LEKEELPR | -265 | -272 | 1 | 0.001291103 | 8 | 408.3067 | 750.2533 | NaN | NaN | NaN | NaN |
| Nub1 | WLQDADPENNSR | -474 | -485 | 0 | 0 | 12 | 767.20056 | 813.06934 | 762.40094 | 878.33563 | 379.4102 | 693.0427 |

| | | | | | | | | | | | | |
|---|---|---|---|---|---|---|---|---|---|---|---|---|
| Nub1 | LEQLECLDDAEK | -293 | -304 | 0 | 0 | 12 | 165.2352 | 178.74445 | 202.79712 | 1725.0569 | 1548.7141 | 1573.0513 |
| Nub1 | NLQLEEEEQNEAELK | -169 | -183 | 0 | 0 | 15 | 4532.456 | 5761.826 | 5947.178 | 6203.2964 | NaN | NaN |
| Pgp | LFDCDGVLWR | -32 | -41 | 0 | 0 | 10 | 553.6342 | 572.86017 | 647.773 | 488.29868 | 646.58014 | 104.98801 |
| Rab27a | LFDLTNEQSFLNVR | -98 | -111 | 0 | 0 | 14 | 479.49072 | 400.32083 | 459.41226 | 377.33575 | 491.7649 | 462.5127 |
| Ppp2r2a | YRDPTTVTTL | -143 | -152 | 1 | 0.000627943 | 10 | 4471.2954 | 123.62071 | 9521.101 | 5678.096 | 6308.922 | NaN |
| Slc38a7 | LFIPDIGK | -387 | -394 | 0 | 0.001055699 | 8 | 13784.903 | 15290.182 | 15416.894 | 6043.2354 | 6152.335 | 6612.4365 |
| Ly75 | LFLEDIVNR | -1440 | -1448 | 0 | 0 | 9 | 235.71149 | NaN | 243.32826 | NaN | 424.6856 | 413.95776 |
| Ctdp1 | SGLHPTNFPVEK | -632 | -643 | 0 | 0 | 12 | 2008.8259 | 1985.4689 | 1900.9778 | 3121.9824 | 3124.978 | 2740.1384 |
| Ctdp1 | LFPLIDDDTR | -715 | -724 | 0 | 0 | 10 | 573.66394 | 549.5029 | 681.1514 | 563.0541 | 673.68677 | 796.94775 |
| Tcf25 | LFQEWEAHR | -402 | -410 | 0 | 0 | 9 | 464.7819 | 358.77408 | 1165.2594 | NaN | 497.85544 | 677.5018 |
| Cept1 | LFQLPTPPLSR | -35 | -45 | 0 | 0 | 11 | 2202.1858 | 2408.2986 | 2445.1897 | 1666.9215 | 556.12476 | 1935.4757 |
| Etnk1 | LFTDGITNK | -53 | -61 | 0 | 0.001023236 | 9 | 517.35126 | 435.45868 | 810.2448 | 676.9272 | 884.97394 | 769.8527 |
| Etnk1 | TELLVDRDEEVK | -84 | -95 | 1 | 0.00025015 | 12 | 769.7226 | 821.1096 | 737.97845 | NaN | 777.2272 | 848.46136 |
| Wdhd1 | SDNPDISDETDIIK | -1024 | -1037 | 0 | 0 | 14 | 796.3979 | 785.54016 | 879.6005 | NaN | 1214.0873 | NaN |
| Wdhd1 | LFTIGGVQK | -549 | -557 | 0 | 0.001868731 | 9 | 324.4867 | NaN | NaN | NaN | NaN | NaN |
| Nup37 | TSDLQDKNEYK | -108 | -118 | 1 | 0 | 11 | 2143.1807 | 1795.314 | 2020.4591 | 1204.2026 | 2521.315 | 2766.007 |
| Nup37 | WHPEETFK | -173 | -180 | 0 | 0 | 8 | 792.86383 | 766.28253 | 862.05927 | 1023.1652 | 1130.2114 | 1052.011 |
| Nup37 | TFQEEETDIEGIQYK | -50 | -64 | 0 | 0 | 15 | 439.87708 | 683.1018 | 514.88165 | 537.0476 | 460.21066 | 456.9442 |
| Nup37 | LFTSDLQDKNEYK | -106 | -118 | 1 | 0 | 13 | 1644.2338 | 1837.1912 | 1786.2913 | NaN | NaN | NaN |
| Nup37 | TLPLCAVGGDHK | -307 | -318 | 0 | 0 | 12 | 854.1881 | 772.481 | NaN | 668.47424 | NaN | NaN |
| Gphn | STGNELLNPEDDLLPGK | -538 | -554 | 0 | 0 | 17 | 617.27814 | NaN | 751.9363 | 866.36035 | NaN | 1298.6437 |
| Gphn | TEQYVELHK | -749 | -757 | 0 | 0 | 9 | 2095.294 | NaN | NaN | NaN | 1584.9247 | 1691.6791 |
| Gphn | LFVVPALR | -670 | -677 | 0 | 0.002780285 | 8 | 435.19562 | NaN | 392.5995 | 417.09167 | NaN | NaN |
| Gphn | NLFVVPALR | -669 | -677 | 0 | 0.000471409 | 9 | 166.56029 | 329.33597 | NaN | 109.90151 | NaN | NaN |
| Ubxn7 | LGDFPYVSILDPR | -200 | -212 | 0 | 0 | 13 | 930.72986 | 854.1828 | 743.79486 | 953.4109 | 914.0901 | 1091.0039 |
| Naa40 | LGDPLEAFPVFK | -35 | -46 | 0 | 0 | 12 | 617.21436 | 627.1792 | 630.16595 | 690.25146 | 777.3598 | 832.8998 |
| Tma16 | LGELELSGPTGATTDGK | -183 | -199 | 0 | 0 | 17 | 475.67398 | 632.37256 | 558.05865 | NaN | NaN | NaN |
| Atox1 | VCIDSEHSSDTLLAT | -40 | -54 | 0 | 0.00025015 | 15 | 1879.2134 | 352.72827 | 2098.4163 | 1693.6268 | 772.19586 | 1020.2473 |
| Atox1 | LGGVEFNIDLPNKK | -26 | -39 | 1 | 0 | 14 | 1640.133 | 2344.7507 | 2192.9988 | 2967.6736 | 3493.1865 | 3542.5242 |
| Atox1 | LGGVEFNIDLPNKK | -26 | -39 | 1 | 0 | 14 | 1640.133 | 2344.7507 | 2192.9988 | 2967.6736 | 3493.1865 | 3542.5242 |
| Niban1 | LGILPEGR | -778 | -785 | 0 | 0.00025015 | 8 | 694.39166 | 540.6798 | 591.9341 | NaN | NaN | NaN |
| Niban1 | TKPPLEPGTVLYEAE | -61 | -75 | 1 | 0.001023236 | 15 | 2776.0112 | NaN | NaN | NaN | 4054.4897 | NaN |
| Polr1c | LGLIPILADPR | -110 | -120 | 0 | 0 | 11 | 573.5367 | 669.34845 | 668.4614 | 591.34393 | 144.80128 | 688.40875 |
| Arf5 | LGLQHLR | -143 | -149 | 0 | 0 | 7 | 951.2329 | 824.4121 | 901.2545 | 534.8031 | 598.61145 | 593.4514 |
| Lrch1 | LGSLPEEIGQLK | -167 | -178 | 0 | 0 | 12 | 335.61496 | 303.8075 | 625.44995 | NaN | 1079.2275 | 1104.9734 |
| Lrch1 | VLPPELVDLPLVK | -215 | -227 | 0 | 0.002265552 | 13 | 304.49875 | NaN | NaN | NaN | NaN | 544.8716 |
| Ikbkb | LGTGGFGNVIR | -21 | -31 | 0 | 0 | 11 | 1401.7589 | 1200.4222 | 1354.7098 | NaN | NaN | 1342.5298 |
| Pde12 | LGVEFGDPASSVFR | -182 | -195 | 0 | 0 | 14 | 317.7451 | NaN | 709.8664 | 797.91064 | 808.9032 | 383.60303 |
| Utp4 | TASPSGSQLLVGCEDGSVK | -108 | -126 | 0 | 0 | 19 | 2734.4521 | 3049.088 | 2977.6716 | 2139.5356 | 2697.988 | 2703.9402 |
| Utp4 | LHCTVPAYNFPVTAL | -518 | -532 | 0 | 0.000298805 | 15 | 1068.8549 | 557.64343 | 1010.77856 | 1143.3208 | 1698.4233 | 1758.2839 |
| Utp4 | NADHLLHLK | -378 | -386 | 0 | 0 | 9 | 502.41742 | 708.7454 | 565.86426 | 664.42865 | 420.48776 | 680.13574 |
| Vbp1 | QPGNETADTVLK | -39 | -50 | 0 | 0 | 12 | 990.51904 | 1098.9509 | 969.2091 | 1088.5369 | 1114.5298 | 152.77621 |
| Vbp1 | LHLGIPEAVFVE | -20 | -31 | 0 | 0.002910424 | 12 | 378.50946 | 404.81952 | 282.84818 | NaN | NaN | NaN |
| Ndufc2 | TYAEILEPFHPVR | -108 | -120 | 0 | 0 | 13 | 3435.4211 | 4590.2695 | 4231.8125 | 4445.106 | 5706.92 | 5904.119 |
| Ndufc2 | LHPEDFPEK | -96 | -104 | 0 | 0 | 9 | 775.9921 | NaN | NaN | NaN | NaN | NaN |
| Bacc1 | LHPVSDSSPAGAK | -29 | -41 | 0 | 0 | 13 | 871.1142 | 721.7046 | 904.1729 | 678.52106 | 920.555 | 998.23175 |
| Mtatp6 | LHSFQHWLVK | -42 | -51 | 0 | 0 | 10 | 951.08954 | 783.18176 | 1733.3104 | 3564.3384 | 2230.7588 | 1195.6958 |
| Mtatp6 | LHSFQHW | -42 | -48 | 0 | 0.00111006 | 7 | 876.6914 | 1205.9921 | 1033.1923 | 641.3954 | 409.2841 | 460.04507 |
| Mrps10 | LHVDVPK | -9 | -15 | 0 | 0 | 7 | 2187.9836 | NaN | NaN | 2302.5647 | NaN | NaN |
| Suox | LHVVGAPGGQSL | -233 | -244 | 0 | 0.00383302 | 12 | 1813.8721 | NaN | NaN | NaN | NaN | NaN |
| Gsta3 | LHYFDGR | -7 | -13 | 0 | 0.00025015 | 7 | 519.7085 | 1237.6189 | 1507.5463 | 788.0934 | NaN | 703.3336 |
| Tm9sf4 | LIADNLPVATR | -133 | -143 | 0 | 0 | 11 | 788.37805 | 616.6268 | 587.9693 | 1044.6112 | 842.3132 | 791.9305 |
| Tm9sf4 | SSCTLPEGANSLPQEIDPTK | -226 | -245 | 0 | 0.000230707 | 20 | 1550.4506 | 1222.5201 | NaN | 3437.9639 | 3088.9082 | 3352.0981 |
| Tm9sf4 | SVHWEESDIK | -254 | -263 | 0 | 0 | 10 | 1007.4889 | 1032.367 | NaN | 917.8627 | 800.3587 | NaN |
| Srpra | LIDDVHR | -82 | -88 | 0 | 0.002068213 | 7 | 364.0423 | NaN | NaN | 4747.8013 | 1570.4882 | NaN |
| Srpra | NQGFDVVLVDTAGR | -509 | -522 | 0 | 0 | 14 | 686.3695 | NaN | 720.1368 | NaN | NaN | NaN |
| Rrp1b | SACVPQESVEER | -224 | -235 | 0 | 0 | 12 | 956.5109 | NaN | 1486.2908 | 1345.3816 | 1260.2185 | 1189.4486 |
| Rrp1b | LIDPFCR | -192 | -198 | 0 | 0.005698678 | 7 | 385.2749 | NaN | NaN | NaN | NaN | NaN |
| Srsf6 | TNEGVIEFR | -146 | -154 | 0 | 0 | 9 | 685.2437 | 2586.5376 | 2664.4485 | 1721.0156 | 1974.9961 | 1850.732 |
| Srsf6 | LIEDKPR | -178 | -184 | 1 | 0.002265552 | 7 | 1142.0697 | 1037.2167 | NaN | 455.6336 | 1606.2383 | 1612.9767 |
| Dhfr | LLPEYPGVLSEVQEEK | -159 | -174 | 0 | 0 | 16 | 2136.7769 | NaN | 3369.7373 | 345.0798 | 971.1381 | 933.495 |
| Dhfr | NGDLPWPPLR | -20 | -29 | 0 | 0 | 10 | 1184.3896 | 463.3693 | 1251.0992 | 1143.2946 | 1218.3528 | NaN |
| Dhfr | LIEQPELASK | -100 | -109 | 0 | 0.000627943 | 10 | 1235.2522 | 1428.9064 | 1555.1543 | NaN | 1581.8207 | NaN |
| Dhfr | VRPLNCIVAV | -2 | -11 | 1 | 0.000907206 | 10 | 729.37836 | NaN | NaN | 664.63885 | 774.3275 | 747.8598 |
| Sco1 | SIGKPLLGGPF | -113 | -123 | 1 | 0.001392869 | 11 | 2120.54 | NaN | 1635.6234 | 1740.2363 | 1836.7954 | NaN |
| Sco1 | LIGPDGEFLDYFGQNK | -249 | -264 | 0 | 0 | 16 | 124.95874 | NaN | NaN | NaN | 291.47427 | 403.4528 |
| Sec61g | LIHIPINNIIVGG | -56 | -68 | 0 | 0 | 13 | 1394.3674 | 1289.7227 | 1116.7676 | 1409.3302 | 1554.233 | 1613.6427 |
| Lyrm4 | LIIENQEKPR | -81 | -90 | 1 | 0 | 10 | 379.73264 | 486.88354 | 475.94238 | 526.7211 | NaN | 460.34894 |
| Scyl2 | LILPELGPVFK | -408 | -418 | 0 | 0 | 11 | 262.4136 | 354.29453 | NaN | 327.92972 | 328.81033 | 373.936 |
| Inip | LILPVLPR | -93 | -100 | 0 | 0 | 8 | 280.03894 | 307.83112 | 316.32245 | NaN | 259.2878 | 276.65747 |
| Utp18 | LTLEDDKPAAER | -72 | -83 | 1 | 0 | 12 | 1010.4364 | 587.3244 | 888.37463 | 651.4826 | 969.6643 | 814.92255 |
| Utp18 | NQDSCLQQTNPKPIK | -449 | -463 | 1 | 0.00142999 | 15 | 640.26624 | 575.2576 | 974.8658 | 498.52744 | 509.30493 | 637.34985 |
| Utp18 | VFGDVEDDEDALLQR | -89 | -103 | 0 | 0 | 15 | 575.7822 | 503.13782 | 526.072 | 865.0864 | 543.6935 | 531.30383 |
| Utp18 | LIPVHQVR | -326 | -333 | 0 | 0.005363933 | 8 | 679.3978 | 527.0255 | 409.5414 | 355.43155 | NaN | NaN |
| Iigp1 | TVFGVDETSLQR | -318 | -329 | 0 | 0 | 12 | 672.60864 | 850.13885 | 1091.5355 | 2011.4819 | 2179.6118 | 921.694 |
| Iigp1 | SPAVFKPTDEETIQER | -347 | -362 | 1 | 0.00025015 | 16 | 527.85864 | 575.54395 | 635.01874 | 660.53143 | 686.7239 | 703.2421 |
| Iigp1 | SDENNDLPSSFTGYFK | -10 | -25 | 0 | 0 | 16 | 386.8429 | 411.60336 | 387.6997 | 559.2208 | 496.63132 | 564.79553 |
| Iigp1 | VFGVDETSLQR | -319 | -329 | 0 | 0 | 11 | 4110.444 | NaN | 4267.0024 | 2750.8862 | 1087.4092 | 922.80255 |

| | | | | | | | | | | | | |
|---|---|---|---|---|---|---|---|---|---|---|---|---|
| Iigp1 | LISDLPIYK | -247 | -255 | 0 | 0.00025015 | 9 | 459.27408 | NaN | 452.21332 | NaN | 172.09166 | 223.73593 |
| Iigp1 | LTNSAISDALK | -53 | -63 | 0 | 0 | 11 | 3763.7803 | NaN | NaN | NaN | NaN | NaN |
| Gsk3a | SFAEPGGGGGGGGGPGGSASGPGGTGGGK | -21 | -50 | 0 | 0.000712251 | 30 | 280.17282 | 267.9353 | 401.85474 | 283.13666 | 463.09625 | 355.66287 |
| Gsk3a | LITPIIYIK | -214 | -222 | 0 | 0 | 9 | 265.5361 | 278.8947 | 297.71 | 239.01152 | 351.54294 | 341.3884 |
| Gsk3a | SQEVAYTDIK | -114 | -123 | 0 | 0.00266297 | 10 | 869.4009 | NaN | 887.9911 | NaN | NaN | NaN |
| Was | QLPPPPAPINEER | -160 | -172 | 0 | 0 | 13 | 1402.1093 | 1517.1353 | 1574.255 | 2350.5635 | 3120.5784 | 3044.2468 |
| Was | NIPSNLLQDHENQR | -23 | -36 | 0 | 0 | 14 | 482.89966 | 884.2445 | NaN | 807.4823 | NaN | NaN |
| Was | LIYDFIEDQGGLEAVR | -291 | -306 | 0 | 0 | 16 | 263.3082 | NaN | 314.00076 | NaN | NaN | NaN |
| Atl2 | NLVPLLLAPENLVEK | -326 | -340 | 0 | 0.000394361 | 15 | 219.1738 | 317.98035 | 229.13486 | 508.7978 | 274.7632 | 407.1116 |
| Atl2 | LKDIDEDFKR | -313 | -322 | 2 | 0 | 10 | 360.59677 | NaN | 382.80197 | NaN | 585.13245 | 417.46027 |
| Ufc1 | LKEEYQSLIR | -32 | -41 | 1 | 0 | 10 | 1288.5548 | 1152.1105 | 1141.4098 | 379.28336 | 1200.2693 | 1199.004 |
| Ufc1 | VVSEIPVLK | -9 | -17 | 0 | 0 | 9 | 554.84796 | 504.01736 | NaN | 521.94946 | 398.84454 | 412.14914 |
| Cd3d | LKNEQLYQPLR | -143 | -153 | 1 | 0 | 11 | 566.9084 | 460.0315 | 461.22107 | 638.5824 | 471.0739 | 448.46832 |
| Cd3d | LKNEQLYQPLR | -143 | -153 | 1 | 0 | 11 | 566.9084 | 460.0315 | 461.22107 | 638.5824 | 471.0739 | 448.46832 |
| Cd3d | NEQLYQPLR | -145 | -153 | 0 | 0.00051306 | 9 | 4875.688 | 4654.9473 | 242.80156 | 4273.512 | 2831.3965 | NaN |
| Sil1 | LLALPEHDAR | -389 | -398 | 0 | 0 | 10 | 771.7179 | NaN | NaN | NaN | NaN | NaN |
| Nudt16 | LLDPGIIAK | -180 | -188 | 0 | 0.00025015 | 9 | 611.00085 | 626.6263 | 240.9775 | 252.08215 | 165.71341 | 336.84485 |
| Alg8 | LLDPSQIPR | -305 | -313 | 0 | 0 | 9 | 403.4517 | 358.0079 | 520.10706 | 523.5335 | 482.7573 | 461.3065 |
| Ap3b1 | LLDSITVPVAR | -487 | -497 | 0 | 0 | 11 | 220.1434 | 351.1082 | 303.40378 | 402.45572 | 363.54626 | 425.8667 |
| Sympk | LLDSSLEGPLPK | -1235 | -1246 | 0 | 0 | 12 | 772.1561 | 816.1007 | NaN | 685.3515 | 1039.3441 | 933.26105 |
| Paf1 | LLEEEIQAPTSSK | -107 | -119 | 0 | 0 | 13 | 601.74414 | 557.9191 | 514.1463 | 889.37604 | 689.2077 | 689.944 |
| Stag2 | LLEEPLSGEEALTDR | -502 | -516 | 0 | 0 | 15 | 531.25323 | 863.0191 | NaN | 737.7201 | 876.16943 | 783.8263 |
| Akap8 | SFAPEPCTDNSDSLIAK | -79 | -95 | 0 | 0 | 17 | 597.0977 | 638.22406 | 648.494 | 713.5932 | 832.5368 | 841.2088 |
| Akap8 | LLEEQTCEAASETR | -620 | -633 | 0 | 0 | 14 | 926.6841 | NaN | NaN | 320.38397 | 150.5444 | NaN |
| Pafah1b2 | LLEETPEEK | -216 | -224 | 0 | 0.00025015 | 9 | 993.78986 | 340.4382 | 1175.5728 | 1049.9133 | 1084.4404 | 499.0545 |
| Pafah1b2 | NFGIGGDTTR | -70 | -79 | 0 | 0 | 10 | 889.1362 | 976.43713 | 929.0913 | 656.3203 | NaN | NaN |
| Pafah1b2 | SQGDSNPAAIPHAAEDIQGDDR | -2 | -23 | 0 | 0.000627943 | 22 | 536.1833 | 507.61362 | 523.7765 | NaN | NaN | NaN |
| P4ha1 | LLELDPEHQR | -231 | -240 | 0 | 0 | 10 | 1469.127 | 1467.122 | 990.7045 | 81.41062 | 1001.1021 | 783.1376 |
| P4ha1 | LQDTYNLDTNTISK | -137 | -150 | 0 | 0 | 14 | 287.0173 | 623.4986 | 630.8712 | 514.20605 | 734.05865 | 516.57874 |
| P4ha1 | NYGVGGQYEPHFDFAR | -419 | -434 | 0 | 0 | 16 | 554.2389 | NaN | 692.86646 | 1005.0245 | 1324.886 | 1305.8318 |
| Irf2bpl | LLGDLLPEAVR | -452 | -462 | 0 | 0.00051306 | 11 | 332.1072 | NaN | NaN | 548.9345 | NaN | NaN |
| Lamtor1 | LLLDPSSTPTK | -21 | -31 | 0 | 0 | 11 | 285.80252 | NaN | 320.36847 | 346.52576 | NaN | NaN |
| Dnajc8 | SILVHPDKNQDDADR | -81 | -95 | 1 | 0.000471409 | 15 | 992.2736 | 689.1571 | 438.78314 | 185.49352 | 647.39075 | 581.1226 |
| Dnajc8 | SILVHPDKNQDDADR | -81 | -95 | 1 | 0.000471409 | 15 | 992.2736 | 689.1571 | 438.78314 | 185.49352 | 647.39075 | 581.1226 |
| Dnajc8 | NVEEDDPELFK | -149 | -159 | 0 | 0 | 11 | 958.8613 | 1412.2306 | 909.7794 | 686.69794 | 1458.185 | 1457.141 |
| Dnajc8 | QIDPEVTDEEIK | -63 | -74 | 0 | 0.000230707 | 12 | 1409.5222 | 814.8457 | 714.69543 | 2416.1167 | 1369.9847 | NaN |
| Dnajc8 | LLLDQEQK | -109 | -116 | 0 | 0.000230707 | 8 | 1118.1543 | 1153.473 | NaN | NaN | NaN | NaN |
| Rcl1 | LLLEEIYR | -282 | -289 | 0 | 0 | 8 | 250.45094 | 352.82898 | 325.42453 | 257.83563 | 424.88577 | 682.6059 |
| Rcl1 | SVEHDCSVLR | -84 | -93 | 0 | 0.000627943 | 10 | 928.5088 | NaN | 912.68134 | NaN | 1169.9269 | 1153.2697 |
| Ints2 | LLLPCLVR | -37 | -44 | 0 | 0 | 8 | 312.6304 | 414.07254 | 415.45755 | 215.4734 | 542.71204 | 386.7212 |
| Lig3 | LLLPGVIK | -314 | -321 | 0 | 0.003681885 | 8 | 440.70007 | 410.61694 | 394.39944 | 335.95798 | 235.05328 | 276.68555 |
| Orc3 | LLLTPQFPFK | -298 | -307 | 0 | 0 | 10 | 526.4748 | 531.9788 | 558.9764 | 506.82733 | 677.19385 | 733.9441 |
| Stat6 | LLNEPDGTFLLR | -551 | -562 | 0 | 0 | 12 | 1630.1127 | 1955.1116 | 1545.1719 | 1662.6548 | 2722.7324 | 1962.5901 |
| Stat6 | LYVDFPQR | -18 | -25 | 0 | 0.001055699 | 8 | 231.98524 | 355.46625 | 282.8865 | 235.72563 | 271.5634 | 946.3013 |
| Dennd2d | LLPAGPGPQLPK | -141 | -152 | 0 | 0 | 12 | 886.3589 | 870.0791 | 787.2471 | 1019.56586 | 763.09326 | 771.0129 |
| Ifrd2 | LLPDFLLER | -107 | -115 | 0 | 0 | 9 | 1650.024 | 1740.691 | NaN | 440.68854 | 3590.638 | 2386.3408 |
| Ercc6l | LLPLQEEIYR | -374 | -383 | 0 | 0 | 10 | 214.34811 | 384.61862 | NaN | 312.06372 | 593.7869 | NaN |
| Ncdn | LLSTSPALQGTPASR | -579 | -593 | 0 | 0 | 15 | 393.92084 | NaN | 808.0963 | 648.53357 | 687.50085 | NaN |
| Fgg | LLTHETSIR | -125 | -133 | 0 | 0.000431655 | 9 | 961.42084 | 1048.9609 | NaN | NaN | 1222.0679 | 686.0975 |
| Fgg | TLEDILFR | -67 | -74 | 0 | 0.001392869 | 8 | 507.3821 | 73.06609 | NaN | 488.43527 | NaN | 641.13617 |
| Mrps9 | LLTLPCGPAEEEFVQR | -222 | -237 | 0 | 0 | 16 | 3015.9536 | 3432.6946 | 3163.0745 | 787.0954 | 3541.8013 | 3536.4084 |
| Mrps9 | LSDEDYAQFIR | -207 | -217 | 0 | 0 | 11 | 831.3302 | 790.93976 | 686.16925 | 303.84195 | NaN | 908.72363 |
| Cdk9 | LLVLDPAQR | -295 | -303 | 0 | 0 | 9 | 256.07596 | 122.308525 | 254.62085 | 728.6679 | 274.88333 | 227.4664 |
| Cr1l | SQCQSDGSWNPLLAK | -383 | -397 | 0 | 0 | 15 | 646.9523 | 659.4339 | 679.3264 | 325.04553 | 767.3736 | 754.383 |
| Cr1l | LLYECLPGYIK | -111 | -121 | 0 | 0 | 11 | 421.01266 | 659.1288 | 476.45428 | 873.0331 | 761.3006 | 475.61774 |
| Cr1l | TPSDPENGLVHVH | -148 | -160 | 0 | 0.00149795 | 13 | 1132.2068 | NaN | NaN | 2836.6926 | NaN | NaN |
| Rwdd1 | LNEIVDQIK | -111 | -119 | 0 | 0 | 9 | 887.0307 | 867.9527 | 526.34753 | 1030.6548 | 828.3051 | 1219.1466 |
| Rwdd1 | VTSEAGENDETVQTTLK | -37 | -53 | 0 | 0.001708008 | 17 | 205.15428 | 268.3154 | 245.80038 | NaN | NaN | NaN |
| Gimap9 | LNHCLEELK | -211 | -219 | 0 | 0 | 9 | 3002.7527 | 3454.912 | 196.23444 | 1918.634 | 1225.3622 | 1803.55 |
| Gimap9 | VVVDTPGLFDTK | -60 | -71 | 0 | 0 | 12 | 620.0495 | 766.90094 | 616.4339 | NaN | 1259.9609 | 1227.5155 |
| Gimap9 | LNHCLEELKETYAQ | -211 | -224 | 1 | 0.00051306 | 14 | 447.95755 | 438.2205 | 211.17145 | 505.79395 | NaN | 695.5037 |
| Gimap9 | SLDNFVSDAGEK | -141 | -152 | 0 | 0 | 12 | 663.4868 | 862.49603 | 798.4536 | NaN | NaN | 597.7154 |
| Gimap9 | NLVVVDTPGLFDTK | -58 | -71 | 0 | 0 | 14 | 863.3197 | 879.8107 | 645.0904 | NaN | NaN | NaN |
| Gimap9 | TPGLFDTK | -64 | -71 | 0 | 0.00025015 | 8 | 261.10773 | NaN | 317.35995 | NaN | NaN | NaN |
| Ppp1ca | LNLDSIIGR | -7 | -15 | 0 | 0 | 9 | 1555.6473 | 2335.8472 | 1858.9452 | 2144.935 | 2278.6064 | 2347.1257 |
| Ppp1ca | NVQLTENEIR | -27 | -36 | 0 | 0 | 10 | 1507.8691 | 1560.1456 | 1686.3098 | 4848.123 | 4383.687 | 4809.644 |
| Ubl4a | VDSPATPIWQLISK | -88 | -101 | 0 | 0 | 14 | 2251.852 | 2246.6582 | 2114.8372 | 1843.1493 | 1807.1521 | 1752.5801 |
| Ubl4a | LNLVVKPLEK | -67 | -76 | 1 | 0 | 10 | 414.48413 | 436.6853 | 520.9922 | 561.92035 | 579.34753 | 625.7022 |
| Ubl4a | LVDSPATPIWQLISK | -87 | -101 | 0 | 0 | 15 | 330.71625 | NaN | NaN | NaN | 441.2962 | 260.55267 |
| Usp9x | LNNEIDWLK | -1400 | -1408 | 0 | 0.002329279 | 9 | 768.1326 | 608.6914 | 645.4628 | NaN | NaN | NaN |
| Map1s | LNPPAQLPASEGLR | -188 | -201 | 0 | 0 | 14 | 771.16095 | 792.0789 | 760.26764 | 1051.6525 | 796.3143 | 920.1456 |
| Bst2 | LNQELENLR | -131 | -139 | 0 | 0 | 9 | 1385.1208 | 621.8012 | 1385.7742 | 1227.0533 | 1027.1536 | 1157.2781 |
| Ap2m1 | LNYSDHDVIK | -411 | -420 | 0 | 0 | 10 | 2100.5168 | 1305.6942 | 1761.2898 | 997.69366 | 1536.3497 | 1547.8402 |
| Ap2m1 | NFEVPFAPSGLK | -389 | -400 | 0 | 0 | 12 | 320.67435 | 427.12936 | 557.35345 | 492.76758 | 363.37796 | 295.07446 |
| Flii | YTVEDTHYVTR | -610 | -620 | 0 | 0 | 11 | 1964.2365 | 1910.8157 | 1013.2524 | 1149.5034 | 1428.8163 | 1016.92 |
| Flii | LVTLPEAIHF | -352 | -361 | 0 | 0 | 10 | 2481.9744 | 2424.6248 | 3045.9421 | 1744.0038 | 1545.6053 | NaN |
| Flii | VPECLYTLPSLR | -238 | -249 | 0 | 0 | 12 | 1388.6036 | NaN | 1537.0852 | 1084.0596 | 2424.2126 | 2146.3276 |
| Flii | LPEEEFGHFYTQ | -920 | -931 | 0 | 0 | 12 | 133.10158 | NaN | 343.23676 | 485.3519 | 565.468 | 484.89755 |

| | | | | | | | | | | | | |
|---|---|---|---|---|---|---|---|---|---|---|---|---|
| Flii | RWDQGLEKPR | -475 | -484 | 2 | 0.001392869 | 10 | 754.7715 | 879.90985 | 977.1767 | NaN | NaN | NaN |
| Uba3 | LPEHCIEYVR | -245 | -254 | 0 | 0 | 10 | 2222.1226 | 2355.996 | 2644.108 | 3015.297 | 3082.3591 | 2879.0215 |
| Uba3 | VLVIGAGGLGCELLK | -72 | -86 | 0 | 0 | 15 | 574.5465 | NaN | 376.64944 | NaN | 566.46326 | 555.3242 |
| Uba3 | SGPFTHPDFEPSTESL | -48 | -63 | 0 | 0.001055699 | 16 | 162.49205 | 268.23758 | NaN | NaN | NaN | NaN |
| Pnn | LPEVSVEPVLTVH | -545 | -557 | 0 | 0.000394361 | 13 | 653.7463 | 257.2585 | 633.06104 | 491.26965 | 583.0439 | 1031.5717 |
| Pnn | RIEFAEQINK | -270 | -279 | 1 | 0 | 10 | 332.56036 | 903.76044 | 808.4452 | 1081.5404 | 1123.9768 | 1030.6959 |
| Pnn | NEEQKAEQEEGK | -299 | -310 | 1 | 0.001291103 | 12 | 270.33472 | 204.00578 | 355.89767 | NaN | 250.53996 | 304.71106 |
| Pnn | QLQEEWNEHNAK | -216 | -227 | 0 | 0.000471409 | 12 | 635.1461 | NaN | NaN | NaN | NaN | 780.61255 |
| Nudt21 | RTVEGVLIVH | -78 | -87 | 1 | 0.001023236 | 10 | 659.87085 | 742.06586 | 837.3054 | 1045.4841 | 1150.7937 | 1125.7683 |
| Nudt21 | LPGGELNPGEDEVEGLKR | -106 | -123 | 1 | 0 | 18 | 651.7634 | 873.9531 | 847.16876 | 636.4972 | 837.6337 | 774.6822 |
| Nudt21 | LPGGELNPGEDEVEGLKR | -106 | -123 | 1 | 0 | 18 | 651.7634 | 873.9531 | 847.16876 | 636.4972 | 837.6337 | 774.6822 |
| Nudt21 | TINLYPLTNYTFGTK | -36 | -50 | 0 | 0.000230707 | 15 | 1393.4275 | NaN | 1544.3204 | 804.7741 | 660.98975 | 1014.3363 |
| Ssu72 | VYDQVVEDLNSR | -115 | -126 | 0 | 0 | 12 | 1229.4618 | NaN | 1021.95746 | NaN | 1293.8429 | 946.0535 |
| Ssu72 | LPGPAPDKPNVY | -44 | -55 | 1 | 0.00025015 | 12 | 946.96735 | NaN | NaN | NaN | NaN | NaN |
| Abcb1b;Abcb1 | LPHQFDTLVGER | (515);(512) | (526);(523) | 0 | 0.00122419 | 12 | 358.5351 | 491.29712 | 468.10043 | NaN | NaN | NaN |
| Gatd1 | SLTGPSVYELIR | -147 | -158 | 0 | 0 | 12 | 668.8112 | 677.5752 | 439.7052 | NaN | 713.40564 | 1007.5142 |
| Gatd1 | LPLIVEDFVK | -165 | -174 | 0 | 0 | 10 | 345.8941 | 370.78036 | NaN | 474.74713 | 503.00595 | 509.30798 |
| Leo1 | LPNFLSVEPR | -379 | -388 | 0 | 0 | 10 | 220.38672 | 948.6465 | 435.039 | 703.6526 | 443.02798 | 467.79514 |
| Cyb561a3 | LPSSWVGPR | -68 | -76 | 0 | 0 | 9 | 2480.9092 | NaN | 1391.5626 | NaN | 1143.1514 | NaN |
| Mms19 | LPTSVLLPYK | -985 | -994 | 0 | 0.00025015 | 10 | 359.63104 | 343.83676 | 440.33496 | NaN | NaN | NaN |
| Mms19 | TDPSTQLQLVGIR | -454 | -466 | 0 | 0 | 13 | 474.8441 | NaN | NaN | NaN | 974.78973 | NaN |
| Mms19 | SAVSTHPSIVK | -561 | -571 | 0 | 0 | 11 | 1295.8536 | NaN | NaN | NaN | NaN | NaN |
| Babam2 | SVSFEDTEATQVYPK | -204 | -218 | 0 | 0 | 15 | 87.81958 | 1266.307 | 1219.3816 | 688.5907 | 718.60394 | 711.4521 |
| Babam2 | LPVDFSNIPTYLLK | -177 | -190 | 0 | 0 | 14 | 199.68655 | 323.63678 | 219.46632 | NaN | NaN | NaN |
| Babam2 | VGLDATNCLR | -27 | -36 | 0 | 0.00025015 | 10 | 1120.0303 | NaN | NaN | NaN | 2347.3308 | NaN |
| Babam2 | NNWTGEFSAR | -163 | -172 | 0 | 0 | 10 | 558.1463 | NaN | NaN | 694.18616 | NaN | NaN |
| Eif4a2 | LQAEAPHIVVGTPGR | -148 | -162 | 0 | 0.003235556 | 15 | 372.2258 | NaN | NaN | NaN | NaN | NaN |
| Ergic2 | LQEEHSLQDVIFK | -134 | -146 | 0 | 0.000627943 | 13 | 543.2109 | NaN | NaN | NaN | 497.3019 | NaN |
| Eml3 | LQEIGLGAFER | -351 | -361 | 0 | 0.001291103 | 11 | 536.1323 | NaN | 840.577 | 4012.7864 | NaN | NaN |
| Cd5 | LQLVPGHEGLR | -159 | -169 | 0 | 0 | 11 | 1878.9138 | 1787.7405 | 1923.0771 | 1257.1921 | 1843.0808 | NaN |
| Cd5 | TVICSDFQPK | -264 | -273 | 0 | 0 | 10 | 489.68692 | 1401.0502 | NaN | NaN | NaN | NaN |
| Eps15l1 | LQQEETQLEQSIQAGR | -506 | -521 | 0 | 0 | 16 | 459.01367 | 458.62766 | 516.83704 | 469.17752 | 374.2652 | 513.0655 |
| Eps15l1 | LSQLQESHLEAHR | -545 | -557 | 0 | 0 | 13 | 222.69446 | 652.95734 | 921.0893 | NaN | NaN | NaN |
| Eps15l1 | YSLEQDIR | -399 | -406 | 0 | 0.001392869 | 8 | 247.84149 | NaN | NaN | NaN | 493.55182 | NaN |
| Nt5c3a | SAGIGDVLEEVIR | -198 | -210 | 0 | 0 | 13 | 377.21808 | 447.20905 | 568.1688 | 580.10486 | 497.67114 | 431.81503 |
| Nt5c3a | LQQHGIPVFIF | -187 | -197 | 0 | 0.000989247 | 11 | 371.0614 | NaN | NaN | NaN | NaN | NaN |
| Cul5 | LQTELVEILK | -734 | -743 | 0 | 0.003521747 | 10 | 3395.6665 | NaN | NaN | 4809.307 | 4450.675 | 4661.6865 |
| Tex30 | LRDEDLFR | -143 | -150 | 1 | 0.00078661 | 8 | 1911.5342 | 1992.0979 | 515.5149 | 471.53796 | 1104.6763 | 1100.8441 |
| Dbt | TFPTPIAKPPVFTGK | -236 | -250 | 1 | 0 | 15 | 569.69214 | 694.9262 | 991.1115 | 689.64087 | 1018.6019 | 830.5481 |
| Dbt | LREELKPVAL | -290 | -299 | 2 | 0.00025015 | 10 | 389.38965 | 428.4043 | NaN | 531.98895 | 776.23584 | 698.3081 |
| Fsaf1 | LREELQSAPAAPSEVPVTTA | -83 | -102 | 1 | 0.00025015 | 20 | 302.18167 | 341.10388 | 374.6796 | 380.16965 | 423.43155 | 351.20444 |
| Fsaf1 | LREELQSAPAAPSEVPVTT | -83 | -101 | 1 | 0.00025015 | 19 | 240.28143 | 244.79065 | 228.0547 | NaN | 281.37717 | NaN |
| Ncaph | LSAAGDHEEFR | -374 | -384 | 0 | 0 | 11 | 1617.7156 | 2468.1936 | 2355.3423 | NaN | NaN | NaN |
| Ncaph | TPVLEDFPQNDDEKER | -46 | -61 | 1 | 0 | 16 | 577.56604 | NaN | NaN | NaN | 895.82654 | 1178.3994 |
| Hexa | LSAPWYLNR | -415 | -423 | 0 | 0.001291103 | 9 | 207.1147 | NaN | NaN | 547.1655 | 882.90405 | 979.5931 |
| Commd1 | LSEVEESINR | -174 | -183 | 0 | 0.00111006 | 10 | 1701.4905 | NaN | NaN | NaN | NaN | NaN |
| Snrpd1 | NREPVQLETL | -49 | -58 | 1 | 0 | 10 | 4813.0215 | 779.1033 | 4999.581 | 5150.6445 | 8056.6597 | 7376.103 |
| Snrpd1 | SHETVTIELK | -11 | -20 | 0 | 0 | 10 | 2217.4268 | 2417.9858 | 1937.2134 | 2137.6975 | 1422.3813 | 1104.9633 |
| Snrpd1 | LSHETVTIELK | -10 | -20 | 0 | 0 | 11 | 1510.3391 | 1740.591 | 1780.5122 | 1791.2058 | 2179.841 | 2663.605 |
| Snrpd1 | LSHETVTIELK | -10 | -20 | 0 | 0 | 11 | 1510.3391 | 1740.591 | 1780.5122 | 1791.2058 | 2179.841 | 2663.605 |
| Snrpd1 | SLPLDTLLVDVEPK | -73 | -86 | 0 | 0 | 14 | 528.86005 | 475.1157 | 450.87527 | 446.62418 | 461.918 | 500.93512 |
| Snrpd1 | NREPVQLE | -49 | -56 | 1 | 0.002780285 | 8 | 864.4705 | NaN | NaN | NaN | NaN | NaN |
| Snrpd1 | LSHETVTIE | -10 | -18 | 0 | 0.007343498 | 9 | 486.46774 | NaN | NaN | NaN | NaN | NaN |
| Gsdma2;Gsdm | LSHLLGK | (284);(296);(287) | (290);(302);(293) | 0 | 0.00078661 | 7 | 371.17178 | NaN | NaN | NaN | NaN | NaN |
| Maz | LSHSDEKPYQCPVCQ | -299 | -313 | 1 | 0.005946382 | 15 | 605.58234 | 555.58875 | 389.8442 | NaN | NaN | NaN |
| Ndufaf3 | LSPADDELYQR | -35 | -45 | 0 | 0 | 11 | 387.71634 | NaN | NaN | 890.63983 | 344.63284 | NaN |
| Yme1l1 | LSPETQSAIEQEIR | -651 | -664 | 0 | 0 | 14 | 637.62335 | 652.0638 | 436.21658 | 493.18542 | 799.0156 | 743.72473 |
| Atp5po | LVRPPVQVY | -27 | -35 | 1 | 0.00111006 | 9 | 4980.5537 | 5758.9404 | 6167.3394 | 5978.979 | 6607.061 | 6529.7764 |
| Atp5po | SFLSPNQILK | -163 | -172 | 0 | 0 | 10 | 2443.9434 | 2703.2156 | 2749.4387 | 2597.0933 | 2715.1387 | 2482.35 |
| Atp5po | TASPLDDAVLSELK | -145 | -158 | 0 | 0 | 14 | 1263.4701 | 1454.1388 | 1435.128 | 1811.1647 | 2251.3184 | 2194.4324 |
| Atp5po | SPLDDAVLSELK | -147 | -158 | 0 | 0 | 12 | 821.9988 | 843.53094 | 224.76083 | 926.1915 | 1132.5535 | 1153.1388 |
| Atp5po | LSPNQILK | -165 | -172 | 0 | 0 | 8 | 669.39105 | NaN | NaN | 1499.1171 | 1206.9644 | 1032.5924 |
| Eif2b2 | LSPQFPSEEDSFHKF | -274 | -288 | 1 | 0.000471409 | 15 | 517.3054 | 652.9953 | 409.37146 | 696.7054 | 935.36346 | 918.6896 |
| Eif2b2 | SDESDQQESLHK | -100 | -111 | 0 | 0 | 12 | 274.08823 | 270.73605 | 278.60544 | 360.56488 | 275.92566 | 303.64883 |
| Cpsf1 | NVLDGELLNR | -1395 | -1404 | 0 | 0 | 10 | 5680.4087 | 5876.8423 | 1104.5874 | 588.40533 | 490.0985 | 6144.3516 |
| Cpsf1 | LSVVEYDPGTHDLK | -103 | -116 | 0 | 0 | 14 | 1018.52075 | 2969.6814 | 4347.211 | 1108.6448 | 474.56097 | 2459.3 |
| Cpsf1 | SLHYFEEPELR | -119 | -129 | 0 | 0 | 11 | 507.84464 | 601.9911 | 612.61554 | 413.78177 | 776.7893 | 736.7613 |
| Cpsf1 | LVVLPFR | -159 | -165 | 0 | 0 | 7 | 483.3246 | 471.0121 | 475.11478 | 332.34488 | NaN | NaN |
| Mapk14 | LVGTPGAELLK | -238 | -248 | 0 | 0 | 11 | 859.3152 | 390.30518 | 414.6425 | NaN | 481.36234 | 400.30328 |
| Mapk14 | LTDDHVQFL | -122 | -130 | 0 | 0.002534621 | 9 | 595.0928 | NaN | NaN | NaN | NaN | NaN |
| Gpd1l | LTDIINNDHENVK | -52 | -64 | 0 | 0.000230707 | 13 | 448.9736 | 401.4267 | NaN | 858.87616 | 803.9123 | 767.595 |
| Nr2c2ap | LTFEDTTDFFGR | -115 | -126 | 0 | 0.007987481 | 12 | 169.04773 | NaN | NaN | NaN | NaN | NaN |
| Snrpc | LTHDSPSVR | -13 | -21 | 0 | 0 | 9 | 1199.88 | 1072.6805 | 1777.9943 | 1129.6367 | 987.5804 | 1305.6329 |
| Psmb6 | QVLLGDQIPK | -220 | -229 | 0 | 0 | 10 | 983.7117 | 1002.0154 | 1002.26855 | 421.3369 | 837.6746 | 884.54553 |
| Psmb6 | LTPIHDHIFCCR | -67 | -78 | 0 | 0 | 12 | 602.1064 | 660.35504 | 879.50574 | 120.98542 | 786.2726 | 815.53375 |
| Psmb6 | LTPIHDHIFCCR | -67 | -78 | 0 | 0 | 12 | 602.1064 | 660.35504 | 879.50574 | 120.98542 | 786.2726 | 815.53375 |
| Psmb6 | LTPIHDHIF | -67 | -75 | 0 | 0.00025015 | 9 | 733.3864 | 898.5929 | 937.003 | NaN | NaN | NaN |
| Psmb6 | TTTGSYIANR | -53 | -62 | 0 | 0.001392869 | 10 | 2349.0537 | 2651.0032 | NaN | NaN | NaN | NaN |

| | | | | | | | | | | | | |
|---|---|---|---|---|---|---|---|---|---|---|---|---|
| Psmb6 | LTPIHDHIFCC | -67 | -77 | 0 | 0.00078661 | 11 | 1194.1903 | NaN | NaN | 4146.5586 | NaN | NaN |
| Cd8b | QCPFPHPETQK | -157 | -167 | 0 | 0 | 11 | 1015.6545 | 994.36145 | 978.2374 | 1066.2795 | 973.43494 | 962.9589 |
| Cd8b | LTSIYWLR | -49 | -56 | 0 | 0 | 8 | 872.9284 | 935.57117 | 913.4888 | 1253.0748 | NaN | 1429.1658 |
| Cd8b | NIILESSDSR | -89 | -98 | 0 | 0.00025015 | 10 | 2976.056 | 3035.8123 | 3189.1575 | 3041.3162 | NaN | NaN |
| Adam10 | TDHLFFK | -229 | -235 | 0 | 0.000712251 | 7 | 756.7783 | 1469.905 | 1177.2388 | 1337.0306 | 828.9047 | 689.2184 |
| Adam10 | TITLQPGSPCNDFR | -624 | -637 | 0 | 0 | 14 | 1113.1731 | 1009.81287 | 1123.5587 | NaN | 1074.0449 | 1482.1808 |
| Adam10 | LVDADGPLAR | -648 | -657 | 0 | 0 | 10 | 1075.1613 | 922.74817 | 1178.9894 | NaN | NaN | 641.0532 |
| Txnl4a | LVDITEVPDFNK | -60 | -71 | 0 | 0 | 12 | 2314.5132 | 1893.6674 | 839.6784 | 2501.7058 | 1087.2539 | NaN |
| Ccna2 | YFLHLQPANCK | -308 | -318 | 0 | 0 | 11 | 1070.4685 | 509.97156 | 1344.9594 | 755.69385 | 1141.0028 | 729.86865 |
| Ccna2 | LVDLHQTYLK | -381 | -390 | 0 | 0 | 10 | 247.11148 | NaN | NaN | 982.13855 | 1346.9285 | 391.48047 |
| Prkag1 | SHYFEGVLK | -269 | -277 | 0 | 0 | 9 | 3489.0947 | 3488.1653 | 3265.587 | 795.7945 | 3332.7378 | 3792.4246 |
| Prkag1 | VSALPVVDEK | -224 | -233 | 0 | 0 | 10 | 1297.3812 | 716.3831 | 1547.9989 | 786.6406 | 1849.5704 | 853.747 |
| Prkag1 | LVEAEVHR | -291 | -298 | 0 | 0.006735683 | 8 | 686.01697 | 348.4453 | NaN | NaN | NaN | NaN |
| Gapvd1 | SEDIPNKIEDLR | -775 | -786 | 1 | 0 | 12 | 848.8937 | 288.3167 | 562.6638 | 1193.708 | 219.67285 | 1316.4778 |
| Gapvd1 | VEDEDHLETDPNK | -209 | -221 | 0 | 0.00025015 | 13 | 272.46634 | NaN | 303.7921 | NaN | 346.33228 | 337.86993 |
| Gapvd1 | LVEDEDHLETDPNK | -208 | -221 | 0 | 0.00025015 | 14 | 386.15012 | NaN | NaN | NaN | NaN | NaN |
| Luc7l3 | WDHESVCK | -29 | -36 | 0 | 0 | 8 | 671.80786 | 681.9599 | 728.6192 | 210.85626 | 742.5992 | 607.4591 |
| Luc7l3 | LVEQLKEER | -161 | -169 | 1 | 0 | 9 | 750.3028 | 654.10474 | NaN | 982.19684 | 983.406 | 1025.8958 |
| Ighm | LVESGFTTDPVTIENK | -155 | -170 | 0 | 0.00025015 | 16 | 877.7836 | 139.26854 | 289.08206 | 915.56287 | 602.4216 | 746.06445 |
| Ighm | VVGHEALPHLVTER | -415 | -428 | 0 | 0 | 14 | 1240.0519 | 1563.0449 | NaN | NaN | 916.90533 | 720.2542 |
| Acap1 | LVETEVSELETR | -23 | -34 | 0 | 0 | 12 | 4068.333 | 3962.3435 | 3937.9185 | NaN | 3515.1763 | NaN |
| Acap1 | TLDSWEPELVK | -461 | -471 | 0 | 0 | 11 | 1111.7986 | NaN | 1471.0027 | NaN | 971.93835 | 1439.6494 |
| Atxn10 | NLTEDNSQNQDVIAK | -424 | -438 | 0 | 0.000591124 | 15 | 678.71985 | 838.1101 | 864.95416 | 1315.0833 | 1050.6907 | 960.3805 |
| Atxn10 | LVGEEQLTK | -254 | -262 | 0 | 0.000394361 | 9 | 9215.22 | 910.78345 | 890.1019 | NaN | NaN | NaN |
| Atxn10 | NEESQSIVWVH | -146 | -156 | 0 | 0.00025015 | 11 | 375.27527 | 1344.8029 | NaN | NaN | NaN | NaN |
| Pdlim5 | LVGPAPWGFR | -8 | -17 | 0 | 0 | 10 | 247.17325 | 1149.0815 | 1028.046 | 963.4154 | 711.0798 | 684.67725 |
| Plin3 | LVGPFAPGITEK | -421 | -432 | 0 | 0 | 12 | 627.82404 | NaN | 389.04166 | NaN | NaN | NaN |
| Snrpe | VLDDAEEIHSK | -57 | -67 | 0 | 0 | 11 | 4795.704 | 6914.2515 | 5014.533 | 4993.556 | 4908.5513 | 5081.9263 |
| Snrpe | VQPINLIFR | -15 | -23 | 0 | 0 | 9 | 1521.659 | 319.34146 | 1790.9073 | 1315.8618 | 1716.6654 | 1785.0677 |
| Snrpe | LVLDDAEEIHSK | -56 | -67 | 0 | 0 | 12 | 1002.4809 | 1500.3566 | 1207.2587 | 1145.226 | 894.6603 | 1296.0237 |
| Snrpe | VMVQPINLIFR | -13 | -23 | 0 | 0 | 11 | 936.4186 | 1050.0477 | 979.3269 | 830.4761 | 910.9103 | 599.80066 |
| Snrpe | VMVQPINLIFR | -13 | -23 | 0 | 0 | 11 | 936.4186 | 1050.0477 | 979.3269 | 830.4761 | 910.9103 | 599.80066 |
| Snrpe | NLVLDDAEEIHSK | -55 | -67 | 0 | 0 | 13 | 735.7794 | 438.9442 | 545.04614 | 472.65747 | 787.50287 | 643.02423 |
| Vps26a | SNTHEFVNLVK | -83 | -93 | 0 | 0.00025015 | 11 | 537.5368 | NaN | NaN | NaN | NaN | 756.202 |
| Vps26a | LVLVDEEDRR | -276 | -285 | 1 | 0.00111006 | 10 | 357.0668 | NaN | NaN | NaN | NaN | 977.1937 |
| Nln | LVNTGLLTLR | -568 | -577 | 0 | 0.00025015 | 10 | 263.9049 | 296.9317 | 281.48566 | NaN | NaN | NaN |
| Nln | WDLSPEQIR | -50 | -58 | 0 | 0.000298805 | 9 | 735.07855 | 766.19165 | NaN | NaN | NaN | NaN |
| Mrpl23 | LVRPGTAQPEDTVQFR | -28 | -43 | 1 | 0.00111006 | 16 | 955.76117 | 621.7258 | 735.49365 | 447.15845 | 479.35733 | 589.37506 |
| Mrpl23 | TFTFPDLFPEKDPR | -104 | -117 | 1 | 0.00025015 | 14 | 717.63635 | 667.1312 | 450.208 | NaN | 593.70386 | NaN |
| Lman1 | LVSGIQHPGSAGVY | -424 | -437 | 0 | 0 | 14 | 914.42017 | 844.6677 | 775.9043 | 1219.679 | 1101.9817 | 996.316 |
| Lman1 | YQEEFEHFQQELDK | -297 | -310 | 0 | 0 | 14 | 540.1409 | 672.6607 | 755.1823 | 671.7112 | 745.8535 | 722.02527 |
| Slc30a7 | LVVAPDADAR | -342 | -351 | 0 | 0 | 10 | 1069.9614 | NaN | NaN | NaN | NaN | NaN |
| Mccc2 | LWDDGIIDPVDTR | -526 | -538 | 0 | 0 | 13 | 3850.3054 | 4477.249 | 3033.4373 | NaN | 4907.102 | 229.08447 |
| Nle1 | LWDPSTGLQVGR | -182 | -193 | 0 | 0 | 12 | 1004.8591 | 986.1856 | 1024.3928 | 790.73126 | 984.7744 | 937.97705 |
| Nle1 | YLASGSGDTTVR | -128 | -139 | 0 | 0 | 12 | 1532.9489 | NaN | NaN | NaN | NaN | NaN |
| Paip2 | LWEEEFIER | -51 | -59 | 0 | 0 | 9 | 346.6131 | 259.40756 | 486.90393 | NaN | NaN | 867.4442 |
| H6pd | LWLVDER | -614 | -620 | 0 | 0.003299042 | 7 | 796.6404 | NaN | NaN | 640.70013 | 465.17426 | NaN |
| Tle3 | TIWDLASPTPR | -557 | -567 | 0 | 0 | 11 | 552.50244 | 1503.3563 | NaN | 1371.0913 | NaN | 1355.8722 |
| Tle3 | NDAPTPGTSTTPGLR | -324 | -338 | 0 | 0 | 15 | 581.99725 | 472.5702 | 456.11285 | NaN | NaN | NaN |
| Apoo | LYDWGLR | -166 | -172 | 0 | 0.00051306 | 7 | 730.4762 | 766.1544 | 862.24286 | 844.878 | 923.80084 | 787.77637 |
| Slamf1 | LYEQVSPPEIK | -138 | -148 | 0 | 0 | 11 | 2302.9404 | 2396.2625 | 3015.7153 | 1454.8845 | 420.61084 | NaN |
| Cfap20 | LYLPVQNK | -183 | -190 | 0 | 0 | 8 | 510.31216 | 510.86844 | 514.958 | 325.35864 | 533.2642 | 439.56256 |
| Pigt | LYNSHPYR | -376 | -383 | 0 | 0.000550307 | 8 | 274.48724 | NaN | NaN | 1159.721 | NaN | NaN |
| Pigt | RPPDNEALPVPFLHAQ | -344 | -359 | 1 | 0 | 16 | 7920.479 | NaN | NaN | NaN | NaN | NaN |
| Mdp1 | WVDTHVDPPFHK | -20 | -31 | 0 | 0 | 12 | 892.95044 | 1006.34735 | 1028.988 | 967.7893 | 1077.9221 | 1258.2506 |
| Mdp1 | LYPEVPEVLGR | -47 | -57 | 0 | 0 | 11 | 633.8728 | 456.13388 | 700.9274 | NaN | NaN | 546.8982 |
| Mdp1 | NQLLELFDLGK | -78 | -88 | 0 | 0 | 11 | 450.32712 | 385.349 | 414.53772 | NaN | NaN | NaN |
| Mrpl55 | LYPVLLVK | -50 | -57 | 0 | 0 | 8 | 512.8067 | 507.182 | 464.59985 | 405.65182 | 347.51953 | 351.21198 |
| Cnot10 | LYQFIEPFEEK | -136 | -146 | 0 | 0 | 11 | 382.11832 | 328.0419 | 284.134 | NaN | NaN | NaN |
| Trex1 | LYWQAPTDSHTAEGDVL | -186 | -202 | 0 | 0.00025015 | 17 | 409.3677 | 338.60168 | NaN | NaN | NaN | NaN |
| Nubp1 | TTPQEVALQDVR | -199 | -210 | 0 | 0 | 12 | 937.7208 | 1558.6748 | 1495.3969 | NaN | 1099.515 | NaN |
| Nubp1 | SSPDDAVIWR | -136 | -145 | 0 | 0 | 10 | 3917.9832 | 810.08466 | 3481.2085 | NaN | NaN | NaN |
| Nubp1 | MEEAPHGCPGADSAQAGR | -1 | -18 | 0 | 0.004690496 | 18 | 263.65594 | NaN | NaN | 159.78954 | NaN | 350.3114 |
| Lyar | MKLPEQPEEGEAK | -296 | -308 | 1 | 0.00266297 | 13 | 11933.097 | NaN | 11840.74 | 3891.7134 | 4956.349 | 3766.0771 |
| Rae1 | NAAEELKPR | -357 | -365 | 1 | 0 | 9 | 235.79529 | NaN | 1823.8013 | 1152.9685 | 1359.3728 | 1409.833 |
| Rae1 | QLENQPSEFR | -196 | -205 | 0 | 0.005459848 | 10 | 2108.5017 | NaN | NaN | NaN | NaN | NaN |
| Dtymk | NAAQRPLGELWK | -201 | -212 | 1 | 0 | 12 | 995.60895 | 924.38245 | 995.6699 | 799.4134 | 775.8798 | 880.31085 |
| Sf3b6 | NACDHLSGF | -72 | -80 | 0 | 0 | 9 | 701.1096 | 698.9307 | 650.4027 | 592.78937 | 623.22656 | 599.0386 |
| Sf3b6 | VVYEDIFDAK | -62 | -71 | 0 | 0 | 10 | 431.93747 | 936.41846 | 514.68207 | 614.6647 | 1256.6847 | 579.86914 |
| NaN | NAEDCLYELPENIR | -141 | -154 | 0 | 0 | 14 | 436.5625 | 429.02573 | 424.92535 | 289.57828 | 739.6155 | 774.6947 |
| Timm17b | NAPPFLEDPSQLTPK | -143 | -157 | 0 | 0 | 15 | 583.4634 | 573.48663 | 525.3532 | 468.521 | 443.83386 | 323.55026 |
| Ivns1abp | NCDVFDPVTK | -481 | -490 | 0 | 0 | 10 | 893.2747 | NaN | NaN | 389.95285 | 1546.4053 | NaN |
| Prps2 | NCIIVSPDAGGAK | -164 | -176 | 0 | 0 | 13 | 1291.2828 | 1243.6027 | 1224.1824 | 1292.3857 | NaN | 1464.2393 |
| Pon2 | NDHYFSDPFLK | -181 | -191 | 0 | 0 | 11 | 96.88076 | 594.043 | 574.35803 | 310.09998 | 669.60693 | NaN |
| Bcap29 | NDNAEEHLLEAENKK | -164 | -178 | 1 | 0 | 15 | 1726.2455 | 531.60535 | 1342.2977 | 251.86755 | 566.15656 | 752.9837 |
| Bcap29 | NDNAEEHLLEAENKK | -164 | -178 | 1 | 0 | 15 | 1726.2455 | 531.60535 | 1342.2977 | 251.86755 | 566.15656 | 752.9837 |
| Bcap29 | NDNAEEHLLEAENK | -164 | -177 | 0 | 0 | 14 | 391.4008 | 717.00287 | 463.3598 | 891.7463 | NaN | NaN |
| Hnrnpll | VSVSPVVHVR | -121 | -130 | 0 | 0 | 10 | 1268.9266 | 1058.2388 | 1313.5431 | 1067.4834 | 1478.4104 | 1527.6632 |

| | | | | | | | | | | | | |
|---|---|---|---|---|---|---|---|---|---|---|---|---|
| Hnrnpll | NDNDSWDYTKPYLGR | -301 | -315 | 1 | 0 | 15 | 1818.7717 | 1972.8722 | 2131.202 | NaN | 2684.0586 | 2696.8076 |
| Cyp51a1 | NEDLNAEEVYGR | -122 | -133 | 0 | 0 | 12 | 4482.928 | 705.2065 | 1779.6667 | 2280.0854 | 1654.0021 | 1522.9108 |
| Cyp51a1 | WLPLPSFR | -244 | -251 | 0 | 0.00025015 | 8 | 498.57047 | NaN | 560.2869 | 416.58392 | 417.8738 | 449.42303 |
| Picalm | SSGDVHLPIASDVSTF | -443 | -458 | 0 | 0 | 16 | 3249.012 | 1400.4819 | 992.5598 | 3756.4888 | 3767.435 | 3442.5369 |
| Picalm | NEGIINLLEK | -212 | -221 | 0 | 0 | 10 | 961.59937 | 961.3499 | 979.99243 | 670.3068 | 897.7812 | 690.7158 |
| Picalm | VAEQVGIDR | -253 | -261 | 0 | 0.00025015 | 9 | 2484.4343 | 2125.864 | 2232.7983 | NaN | 2121.8762 | NaN |
| Picalm | QAALEEEQAR | -325 | -334 | 0 | 0 | 10 | 590.5948 | NaN | NaN | NaN | 651.22284 | 762.9471 |
| Pola1 | VVEPTSDGNYITK | -1063 | -1075 | 0 | 0.000431655 | 13 | 2193.9417 | 2344.5767 | 2481.2024 | 528.75366 | 2748.555 | 3450.8499 |
| Pola1 | NEPLLTPIPLK | -220 | -230 | 0 | 0 | 11 | 729.7382 | 719.38306 | 623.10596 | 654.8835 | 567.9608 | 570.0183 |
| Pola1 | TWDQETEPVER | -278 | -288 | 0 | 0 | 11 | 686.8407 | 928.83716 | NaN | NaN | 788.3429 | 714.49084 |
| Pola1 | YENNYIVPDK | -801 | -810 | 0 | 0.001392869 | 10 | 415.5011 | NaN | NaN | NaN | 520.53156 | NaN |
| M6pr | SFESTVGQGSDTYSYIFR | -60 | -77 | 0 | 0 | 18 | 626.8342 | 612.49414 | 634.81055 | 901.0245 | 1069.4974 | 1088.472 |
| M6pr | NFNPVSEER | -154 | -162 | 0 | 0 | 9 | 1274.9443 | 1210.3425 | 1037.2987 | NaN | 909.43774 | NaN |
| M6pr | SCDLVGEK | -32 | -39 | 0 | 0 | 8 | 1299.307 | 269.75778 | 1025.6594 | NaN | NaN | NaN |
| M6pr | SFESTVGQGSDTYSY | -60 | -74 | 0 | 0.001708008 | 15 | 731.8048 | 770.0135 | 661.1967 | NaN | NaN | NaN |
| M6pr | NEVALLER | -45 | -52 | 0 | 0.00078661 | 8 | 278.23218 | NaN | NaN | NaN | NaN | NaN |
| Lipa | NFASGPLLQLGR | -205 | -216 | 0 | 0 | 12 | 840.4939 | 860.4233 | 940.42957 | NaN | 142.9928 | NaN |
| Atp6v1b2 | QIYPPINVLPSLSR | -387 | -400 | 0 | 0.00025015 | 14 | 503.90857 | 516.5007 | 340.95206 | 146.7652 | 243.26477 | 413.00543 |
| Atp6v1b2 | VFEGTSGIDAK | -98 | -108 | 0 | 0.000431655 | 11 | 735.3331 | 757.8731 | 316.59894 | NaN | NaN | 259.7338 |
| Atp6v1b2 | TSCEFTGDILR | -110 | -120 | 0 | 0 | 11 | 827.9119 | 1142.996 | 1173.2665 | NaN | NaN | NaN |
| Ints9 | NFLPLPLVQSPR | -37 | -48 | 0 | 0 | 12 | 914.6176 | 475.9426 | 740.756 | 860.40344 | 524.7704 | 633.7376 |
| Ints9 | SIHGDFSNDFR | -378 | -388 | 0 | 0 | 11 | 1358.9286 | 246.39926 | NaN | 1401.514 | NaN | NaN |
| Klhl9 | NFPALLNTGEFLK | -180 | -192 | 0 | 0 | 13 | 486.06732 | NaN | NaN | 470.55835 | 340.41388 | 735.04944 |
| Mios | NHILAGNEDPQLK | -412 | -424 | 0 | 0.00897527 | 13 | 234.36621 | NaN | 1059.1259 | NaN | NaN | NaN |
| Gaa | NHNDLNSVPQEPYR | -673 | -686 | 0 | 0 | 14 | 1350.5676 | 2039.6324 | 2042.1527 | NaN | 2472.1572 | 3049.5261 |
| Sdad1 | NHTVLDPDLR | -88 | -97 | 0 | 0.000230707 | 10 | 2397.142 | NaN | 1871.1398 | 537.6552 | NaN | NaN |
| Rab8b | NIEEHASSDVER | -105 | -116 | 0 | 0 | 12 | 1146.3885 | 1387.7751 | 1483.0862 | 1516.9132 | 1884.102 | 1333.0355 |
| Rab8b | NIEEHASSDVER | -105 | -116 | 0 | 0 | 12 | 1146.3885 | 1387.7751 | 1483.0862 | 1516.9132 | 1884.102 | 1333.0355 |
| Krr1 | NIHPIYNIK | -211 | -219 | 0 | 0 | 9 | 700.7239 | 754.7074 | 707.8657 | 344.1264 | 595.5752 | 446.5392 |
| Lpxn | NIQEPNVYSEVQEPK | -65 | -79 | 0 | 0 | 15 | 6328.372 | 8832.574 | 8176.571 | 246.9439 | 7205.9053 | 7472.5356 |
| Prkar2a | NISHYEEQLVK | -378 | -388 | 0 | 0 | 11 | 3428.5273 | NaN | 4753.1846 | 3764.1108 | 3712.8677 | 937.919 |
| Chmp5 | NKDGVLVDEFGLPQIPAS | -202 | -219 | 1 | 0 | 18 | 250.92033 | 378.84232 | 305.7729 | 580.62103 | 745.07764 | 719.43994 |
| Ccdc47 | NKDPITIVDVPAHLQ | -116 | -130 | 1 | 0 | 15 | 2275.5684 | 2209.0994 | NaN | 2133.9883 | 2382.281 | 2298.1848 |
| Ccdc47 | RQDLLNVL | -227 | -234 | 1 | 0.001055699 | 8 | 426.12552 | 565.1624 | 512.9412 | NaN | NaN | NaN |
| Tmprss13 | NKPGVYTK | -515 | -522 | 1 | 0 | 8 | 5371.652 | 4163.024 | 5672.0835 | 5770.3286 | 5185.2764 | 5649.4673 |
| Krt7 | NKYEEEINRR | -172 | -181 | 2 | 0.002595029 | 10 | 348.07605 | NaN | 744.8238 | NaN | 425.77213 | NaN |
| Plg | PNKPGVYVR | -789 | -797 | 1 | 0.00025015 | 9 | 2423.2375 | 1519.8782 | 1548.6847 | 1702.7792 | 1381.5895 | 81.78608 |
| Plg | PNKPGVYVR | -789 | -797 | 1 | 0.00025015 | 9 | 2423.2375 | 1519.8782 | 1548.6847 | 1702.7792 | 1381.5895 | 81.78608 |
| Plg | TPENFPCK | -310 | -317 | 0 | 0 | 8 | 1163.8364 | 1104.7593 | 1333.4497 | 555.92725 | 1372.1249 | 635.9889 |
| Plg | SCQGDSGGPLVCFEK | -757 | -771 | 0 | 0 | 15 | 840.4021 | 791.85803 | 822.9019 | 1033.2272 | 1162.6693 | 1153.205 |
| Plg | NPDGEPRPW | -236 | -244 | 1 | 0 | 9 | 732.0623 | 796.81647 | 938.5685 | 698.08984 | 926.9052 | 1298.876 |
| Plg | NLEENYCR | -318 | -325 | 0 | 0 | 8 | 187.69977 | 835.2998 | 752.0336 | 938.9843 | 495.68503 | 803.6214 |
| Plg | PNKPGVY | -789 | -795 | 1 | 0.00622886 | 7 | 384.0358 | NaN | NaN | 766.0804 | NaN | NaN |
| Trnt1 | YQGEHGLLK | -368 | -376 | 0 | 0 | 9 | 1835.1259 | 422.01703 | 626.23334 | 575.68494 | 301.35718 | 403.78693 |
| Trnt1 | NLGLFIVK | -321 | -328 | 0 | 0.002972566 | 8 | 119.489555 | 106.36298 | 135.44133 | 93.84122 | 181.5755 | NaN |
| Rasa2 | NLLPYLGPNK | -48 | -57 | 0 | 0 | 10 | 312.17548 | 174.03915 | 765.32697 | 615.03674 | NaN | NaN |
| Cox7c | SHYEEGPGK | -17 | -25 | 0 | 0 | 9 | 651.35474 | 485.12604 | 505.71408 | 443.33685 | 487.5126 | 603.83246 |
| Cox7c | NLPFSVENK | -26 | -34 | 0 | 0 | 9 | 6265.0674 | NaN | NaN | 6787.66 | 3608.0884 | NaN |
| Ambp | NLPIVQGPCR | -287 | -296 | 0 | 0 | 10 | 4424.7026 | 556.2868 | 4862.145 | 3460.9167 | 3997.2395 | 3980.3838 |
| Pus10 | NLPQTPWIIDGER | -298 | -310 | 0 | 0 | 13 | 429.5354 | NaN | NaN | 813.79364 | 1040.0309 | 737.3037 |
| Cnot9 | NLQEGQVTDPR | -282 | -292 | 0 | 0 | 11 | 1871.0625 | 1674.5376 | 2365.4807 | 1651.0953 | NaN | 1427.73 |
| Ndufv3 | SVLKEPEPTDTTTYK | -54 | -68 | 1 | 0.00025015 | 15 | 842.91296 | 1062.645 | NaN | 320.68875 | 455.44153 | 459.05438 |
| Ndufv3 | NLQHHDYNTY | -69 | -78 | 0 | 0 | 10 | 786.77997 | 354.01953 | 685.1146 | NaN | NaN | NaN |
| Nit2 | NLTTGPAHWELLQR | -180 | -193 | 0 | 0 | 14 | 1082.4022 | 3439.6077 | 5130.97 | 4957.9775 | 1359.9805 | 968.167 |
| Nit2 | TTGPAHWELLQR | -182 | -193 | 0 | 0 | 12 | 450.78973 | 585.01886 | 673.21173 | 366.0374 | 599.68976 | NaN |
| Nit2 | SVFGPDGSLLVK | -98 | -109 | 0 | 0.000431655 | 12 | 2844.7764 | 3228.0342 | NaN | NaN | NaN | NaN |
| Cd3g | NLVQVDGSR | -29 | -37 | 0 | 0.001463897 | 9 | 538.1047 | NaN | NaN | NaN | NaN | NaN |
| Atp6ap1 | NLWAPVADTHEGHIT | -49 | -63 | 0 | 0 | 15 | 431.7244 | 556.4722 | 492.33725 | 558.1552 | 573.90155 | 524.775 |
| Mrpl32 | NNIDICPECGHLK | -104 | -116 | 0 | 0 | 13 | 514.8596 | 489.66602 | 933.8203 | 879.42993 | 1346.4534 | 1156.6188 |
| Eci1 | NPAHYAEYWK | -104 | -113 | 0 | 0 | 10 | 1490.5912 | 1460.8438 | 1526.2214 | 446.162 | 690.3266 | 482.00656 |
| Eci1 | VGVVDEVVPEDQVHSK | -207 | -222 | 0 | 0 | 16 | 538.2423 | 888.6603 | 785.205 | 1559.8074 | 914.4631 | 1115.9368 |
| Eci1 | WLAIPDHSR | -230 | -238 | 0 | 0 | 9 | 755.35645 | 664.0495 | NaN | 746.9031 | 827.6626 | 774.57104 |
| Tram1 | NPPLLSHEFLLQ | -10 | -21 | 0 | 0 | 12 | 1926.8496 | 768.61926 | 2139.8743 | 2141.1816 | 2697.5298 | 2311.7566 |
| Iws1 | NQEDLEEEKNETQLK | -439 | -453 | 1 | 0.000298805 | 15 | 229.4034 | 509.54727 | 501.44046 | 486.41898 | 571.6897 | 616.3094 |
| Ctla2a | NQFSDLTPEEFK | -89 | -100 | 0 | 0 | 12 | 1707.1857 | 719.88947 | 2210.3545 | 464.7354 | NaN | NaN |
| Cisd3 | NQPFCDGSHFFQR | -77 | -89 | 0 | 0 | 13 | 376.9288 | NaN | 508.87256 | NaN | NaN | 293.1694 |
| Gmppa | NQQDGQLEESPGSWPGAGTIR | -201 | -221 | 0 | 0 | 21 | 370.20248 | 419.1331 | 369.0034 | 511.7643 | 600.7233 | 1071.8953 |
| Gmppa | VEGTPNDPNPNDPR | -353 | -366 | 0 | 0.00078661 | 14 | 5968.4507 | NaN | 5370.9785 | NaN | 2779.34 | 2552.4983 |
| Ncstn | SNSYGPEFAHCK | -148 | -159 | 0 | 0 | 12 | 686.4369 | NaN | 321.7283 | 549.448 | 463.7628 | 515.5586 |
| Ncstn | SGAGVPEVVLR | -403 | -413 | 0 | 0 | 11 | 798.7441 | 808.04877 | 649.32153 | 510.7594 | NaN | NaN |
| Ncstn | VVEKEEDLK | -74 | -82 | 1 | 0 | 9 | 770.0872 | 782.1714 | 784.23236 | 768.58636 | NaN | NaN |
| Ncstn | NQVEDLLATLEK | -391 | -402 | 0 | 0.001743443 | 12 | 454.1298 | NaN | NaN | NaN | 284.0137 | 332.26245 |
| Ncstn | SFFWNVAPGAESA | -285 | -297 | 0 | 0.000550307 | 13 | 236.16095 | NaN | NaN | NaN | 384.12204 | NaN |
| Rpe | NSDLANLGAECLR | -13 | -25 | 0 | 0 | 13 | 1036.9757 | 1117.2971 | 700.95154 | 615.3652 | 938.1745 | 272.4985 |
| Pacsin1 | NSLLNEDLEK | -100 | -109 | 0 | 0 | 10 | 1300.5978 | NaN | 709.8714 | 1670.9738 | NaN | 229.09148 |
| Ctsz | NSWGEPWGEK | -264 | -273 | 0 | 0 | 10 | 453.53455 | 445.0743 | 522.5283 | 324.21033 | 489.33917 | NaN |
| Ctsz | VGDYGSLSGR | -192 | -201 | 0 | 0 | 10 | 167.78772 | NaN | NaN | 648.1348 | 414.9198 | 791.5899 |
| Gmfb | NTEDLTEEWLR | -125 | -135 | 0 | 0 | 11 | 1337.4902 | 430.46548 | 1587.0055 | 550.57874 | 1385.2517 | 365.88232 |

| | | | | | | | | | | | | |
|---|---|---|---|---|---|---|---|---|---|---|---|---|
| Nmt1 | NTHGPVEPDKDNIR | -133 | -146 | 1 | 0 | 14 | 1722.6436 | 874.3567 | 1582.0884 | 661.791 | 1319.2832 | 1353.6692 |
| Nmt1 | NTHGPVEPDKDNIR | -133 | -146 | 1 | 0 | 14 | 1722.6436 | 874.3567 | 1582.0884 | 661.791 | 1319.2832 | 1353.6692 |
| Nmt1 | SYQFWDTQPVPK | -116 | -127 | 0 | 0 | 12 | 2551.4365 | 4133.807 | 4050.5781 | 5440.1685 | NaN | 6331.9136 |
| Utp23 | NTIVLDKPSPR | -143 | -153 | 1 | 0 | 11 | 5582.686 | 5516.9414 | 4675.242 | 6046.661 | 6265.203 | 5930.179 |
| Sdhc | NTSSNRPLSPHLTIY | -45 | -59 | 1 | 0 | 15 | 991.3557 | 1208.7437 | 1334.8248 | 953.4831 | 1449.289 | 1337.7417 |
| Dimt1 | NTVIPEDFSIADK | -263 | -275 | 0 | 0.00025015 | 13 | 375.6552 | NaN | NaN | NaN | 536.7104 | 543.7977 |
| Aldh3a2 | NVDEAINFINDR | -344 | -355 | 0 | 0 | 12 | 363.5598 | 476.82263 | 1030.2959 | 611.7844 | 2057.886 | 1997.6681 |
| Pcif1 | NVEDTFSWLR | -314 | -323 | 0 | 0 | 10 | 387.5624 | NaN | 397.24045 | NaN | NaN | 254.29898 |
| Degs1 | NVGYHNEHHDFPNVPGK | -255 | -271 | 0 | 0 | 17 | 593.76965 | NaN | NaN | 463.24893 | 646.73895 | 700.66254 |
| Glrx5 | NVLDDPELR | -85 | -93 | 0 | 0 | 9 | 839.3765 | 208.1397 | 1112.9602 | 350.42065 | 428.73346 | 1434.9458 |
| Glrx5 | SALVDEKDQDSK | -141 | -152 | 1 | 0 | 12 | 1470.5801 | 1670.2797 | 2089.0027 | NaN | 1856.6804 | 2062.812 |
| Birc5 | NWPFLEDCACTPER | -24 | -37 | 0 | 0 | 14 | 466.41208 | 547.6941 | 618.8992 | 549.7732 | 723.27515 | 743.8684 |
| Vrk3 | NYEEKPPYATLR | -418 | -429 | 1 | 0 | 12 | 1536.6149 | NaN | 1534.243 | NaN | 1683.9872 | NaN |
| Pam16 | NYEHLFK | -76 | -82 | 0 | 0 | 7 | 2478.2307 | 3446.2708 | 2168.4683 | 1599.6733 | 1620.6971 | 1558.0775 |
| Mcat | NYLFPDCR | -227 | -234 | 0 | 0.001291103 | 8 | 3125.4736 | NaN | NaN | NaN | NaN | NaN |
| Rbm27 | NYLPPLEPVKPEPK | -76 | -89 | 1 | 0.000230707 | 14 | 674.86096 | NaN | 804.3165 | 664.176 | NaN | 1155.6842 |
| Rbm27 | VFQEPAEEER | -102 | -111 | 0 | 0.00025015 | 10 | 861.633 | NaN | 1392.8029 | NaN | NaN | NaN |
| Ccdc12 | NYVPEDEDLKR | -61 | -71 | 1 | 0 | 11 | 3057.0588 | 2730.1116 | 2803.545 | 3567.7744 | NaN | NaN |
| H2bc7;H2bc9; | PEPAKSAPAPK | (2);(2);(2);(2) | (12);(12);(12);(12) | 1 | 0 | 11 | 343.50806 | 341.83438 | 397.22443 | 571.1663 | 559.0014 | 591.0115 |
| Arpp19 | QLPAAAPDKTEVTGDHIPTPQDLPQR | -76 | -101 | 1 | 0 | 26 | 1391.7051 | 1731.0903 | 1673.9998 | 1944.6448 | 2214.1948 | 2327.5056 |
| Arpp19 | PGGSDFLR | -43 | -50 | 0 | 0.004321969 | 8 | 714.7774 | NaN | NaN | NaN | NaN | NaN |
| Sec61b | PGPTPSGTNVGSSGR | -2 | -16 | 0 | 0 | 15 | 1813.253 | 1842.2782 | 1866.9978 | 830.2726 | 1464.5181 | 1409.2615 |
| Sec61b | RFYTEDSPGLK | -57 | -67 | 1 | 0 | 11 | 1593.3877 | 1692.9597 | 1624.4921 | 357.49237 | 1831.576 | 1913.7378 |
| Clpp | PLIPIVVEQTGR | -53 | -64 | 0 | 0 | 12 | 1042.1278 | 947.73846 | 1171.4955 | 279.79565 | 900.8173 | 933.5142 |
| Clpp | VLVHPPQDGEDEPELVQK | -240 | -257 | 0 | 0.000550307 | 18 | 660.369 | 752.782 | 803.51807 | 938.476 | 833.66187 | 1228.1533 |
| Tf | PSHAVVAR | -266 | -273 | 0 | 0 | 8 | 1829.5881 | 1261.105 | NaN | 3381.463 | 504.041 | 343.2141 |
| Polr2c | PYANQPTVR | -2 | -10 | 0 | 0.000230707 | 9 | 1284.6162 | 1203.8967 | 934.6318 | 1169.848 | NaN | NaN |
| Polr2c | SSVLHDEFIAHR | -56 | -67 | 0 | 0 | 12 | 602.9287 | NaN | 612.48865 | NaN | 603.9582 | 190.5304 |
| Rrs1 | QAGPSPEAELR | -52 | -62 | 0 | 0 | 11 | 1143.1724 | 2272.0945 | NaN | 2457.6982 | NaN | 777.3061 |
| Morc3 | QDFDYTNEYR | -359 | -368 | 0 | 0 | 10 | 433.70187 | NaN | NaN | 498.95154 | NaN | NaN |
| Traf3ip3 | QLLENEHQELQTK | -366 | -378 | 0 | 0 | 13 | 2083.1597 | 1768.8213 | 660.42206 | 2139.567 | 2130.2717 | 2238.6025 |
| Traf3ip3 | QDLLPALIR | -220 | -228 | 0 | 0 | 9 | 430.80023 | 390.69608 | 635.59766 | 705.2262 | 764.669 | 624.1632 |
| Mvp | QDPFPLYPGELLEK | -94 | -107 | 0 | 0 | 14 | 446.1414 | 1140.3876 | NaN | 279.14453 | 818.2355 | 517.2214 |
| Golga2 | QELQEAQER | -673 | -681 | 0 | 0.001776473 | 9 | 1733.3696 | NaN | NaN | NaN | NaN | NaN |
| Sin3a | QEQATEENAGVPVGPHLSL | -762 | -780 | 0 | 0.00025015 | 19 | 735.9426 | 751.8939 | 980.98346 | 828.034 | 668.16925 | 838.75854 |
| Slc2a1 | TFDEIASGFR | -459 | -468 | 0 | 0 | 10 | 580.7163 | 928.39606 | 868.45715 | 748.0333 | 342.85464 | 1039.2587 |
| Slc2a1 | QFGYNTGVINAPQK | -25 | -38 | 0 | 0 | 14 | 486.16843 | 547.436 | 552.2713 | 658.803 | 735.145 | 803.4574 |
| Micos10 | QHDFQAPYLLHGK | -59 | -71 | 0 | 0 | 13 | 440.88293 | 475.2743 | 557.2186 | NaN | NaN | NaN |
| Cox7b | QIGIEWNMSPVGR | -59 | -71 | 0 | 0 | 13 | 805.6859 | NaN | NaN | 1108.7433 | NaN | NaN |
| Rimbp3 | QLADHLR | -74 | -80 | 0 | 0.00025015 | 7 | 2730.0132 | 775.4135 | 4249.5195 | 1647.802 | 3744.752 | 913.7898 |
| Lsm7 | QLGLVVCR | -70 | -77 | 0 | 0 | 8 | 1340.9607 | 1293.8212 | 1542.5682 | 1399.5938 | 1252.9446 | 1343.7897 |
| Tubd1 | QLLVKPLDMIVGK | -400 | -412 | 1 | 0.001023236 | 13 | 365.75183 | 421.68567 | 387.86584 | 444.58176 | 442.69904 | 488.37064 |
| Aatf | QLPQPDVFPVFK | -220 | -231 | 0 | 0 | 12 | 447.5183 | 386.12543 | 403.0788 | 337.31445 | 465.72525 | 306.2014 |
| Aatf | VIDRFDEGEEEKDDLAVS | -36 | -53 | 2 | 0.007523243 | 18 | 432.78696 | NaN | 477.30573 | NaN | NaN | NaN |
| Stx8 | QNLLDDLVTR | -80 | -89 | 0 | 0 | 10 | 311.0268 | 245.26376 | 262.18866 | 758.086 | 461.27216 | 558.3049 |
| Hibch | TVEQIIENLR | -275 | -284 | 0 | 0 | 10 | 449.84027 | 605.4436 | 580.50543 | 676.0488 | 887.99567 | 767.0997 |
| Hibch | QNLTQDLFR | -110 | -118 | 0 | 0 | 9 | 1429.3582 | 1290.7899 | 1585.712 | NaN | NaN | 1386.4089 |
| Stk4 | VINTEDEEEEGTMK | -369 | -382 | 0 | 0 | 14 | 1238.2373 | NaN | 640.91846 | 493.08386 | 959.0457 | NaN |
| Pus7 | QQLEELQLFK | -176 | -185 | 0 | 0.000471409 | 10 | 541.982 | NaN | NaN | NaN | 456.4068 | NaN |
| Pus7 | SSQDGVENDGLHR | -39 | -51 | 0 | 0 | 13 | 322.0556 | 422.2722 | NaN | NaN | NaN | NaN |
| Rbm5 | QQLLPELVR | -596 | -604 | 0 | 0.001536401 | 9 | 367.49838 | NaN | NaN | NaN | NaN | NaN |
| Ikbkg | QQLQQAEEALVAK | -258 | -270 | 0 | 0.000394361 | 13 | 3078.6577 | NaN | NaN | NaN | NaN | NaN |
| Snx2 | QQQFENLDQQLR | -310 | -321 | 0 | 0 | 12 | 1468.7432 | 2430.2646 | 2293.2917 | 1570.1298 | 1862.5096 | 1486.9474 |
| Snx2 | SEPSPAVTPVTPTTLIAPR | -94 | -112 | 0 | 0 | 19 | 827.8903 | 902.251 | 1151.4404 | 1186.9706 | 1396.6794 | 1318.481 |
| Snx2 | SSEPSPAVTPVTPTTLIAPR | -93 | -112 | 0 | 0.003745449 | 20 | 280.39233 | 309.7474 | 334.3715 | 390.37146 | 407.6417 | 366.42972 |
| Snx2 | SISAPVIFDR | -117 | -126 | 0 | 0 | 10 | 647.19556 | 669.77496 | 237.75919 | NaN | 576.14905 | 706.6759 |
| Ssr3 | QQSEEDLLLQDFSR | -9 | -22 | 0 | 0 | 14 | 671.75006 | 1894.9751 | NaN | 1206.6545 | 1557.201 | 1246.942 |
| ;;Igh-VJ558; | QSGPELVKPGASVK | (6);(6);(25);(6) | (19);(19);(38);(19) | 1 | 0 | 14 | 3515.3157 | NaN | 434.33878 | NaN | 3691.4148 | 1686.6731 |
| Ptprv;Mcl1 | QSLEIISR | (477);(158) | (484);(165) | 0 | 0.000471409 | 8 | 265.3853 | 316.54306 | NaN | NaN | NaN | NaN |
| Bcat2 | QSLLDLAR | -300 | -307 | 0 | 0.001868731 | 8 | 3658.2544 | 2801.7507 | 1596.7479 | NaN | NaN | NaN |
| Thumpd1 | QSVESGANNVVFIR | -118 | -131 | 0 | 0 | 14 | 781.9932 | 875.76697 | 730.4268 | 837.31494 | 805.5459 | 703.19403 |
| Thumpd1 | SIEELGQTEPGSETQAGSEGDAKPEPESQVSEVPK | -309 | -343 | 1 | 0.00025015 | 35 | 482.29553 | 514.19824 | 558.08856 | NaN | NaN | NaN |
| Isoc1 | QTALELVGR | -217 | -225 | 0 | 0.000907206 | 9 | 610.1602 | 666.9013 | 792.9485 | NaN | NaN | NaN |
| Arhgef6 | SEPIQAWEGDDIK | -432 | -444 | 0 | 0 | 13 | 2150.6343 | 1899.1915 | 2202.2585 | NaN | 668.0534 | NaN |
| Arhgef6 | TVCDPEEFLK | -23 | -32 | 0 | 0 | 10 | 339.9784 | 815.04846 | 626.83093 | NaN | NaN | NaN |
| Arhgef6 | QTNEDELSVCK | -172 | -182 | 0 | 0.00122419 | 11 | 496.04956 | 826.5405 | NaN | NaN | NaN | NaN |
| Mvd | QVAPVLLSDELK | -345 | -356 | 0 | 0 | 12 | 372.28366 | 419.40778 | 343.3067 | NaN | NaN | NaN |
| Pycr3 | QVLASAPTDNNLCHFR | -37 | -52 | 0 | 0 | 16 | 430.96558 | NaN | 385.42944 | 461.92557 | 637.0852 | 350.943 |
| Vapb | SLTSPLDDTEVKK | -156 | -168 | 1 | 0 | 13 | 683.2055 | 639.27936 | 779.8365 | 642.27185 | 600.43286 | 803.54126 |
| Vapb | VEQVLSLEPQHELK | -4 | -17 | 0 | 0 | 14 | 425.40143 | 576.32434 | 500.38776 | 553.5933 | 765.6571 | 706.45605 |
| Vapb | QVLSLEPQHELK | -6 | -17 | 0 | 0 | 12 | 1080.19 | 1249.2537 | NaN | NaN | NaN | NaN |
| Stim2 | QYAEQELEQVR | -319 | -329 | 0 | 0.000230707 | 11 | 320.50858 | NaN | NaN | 782.125 | NaN | NaN |
| Mbd2 | RDPVPFPSGSSGPGPR | -127 | -142 | 1 | 0 | 16 | 400.53955 | 537.8016 | 201.86974 | NaN | NaN | NaN |
| Max | RDSVPSLQGEK | -47 | -57 | 1 | 0.000230707 | 11 | 1715.2954 | 1277.1704 | 1672.3812 | 1666.0167 | 1547.1492 | 1828.3329 |
| Myef2 | RDTHPGEPEKPPR | -25 | -37 | 2 | 0.000550307 | 13 | 286.28033 | NaN | 274.88898 | NaN | 321.06876 | NaN |
| Pgam5 | REPLSLINLK | -64 | -73 | 1 | 0.000431655 | 10 | 565.68677 | 595.06537 | 593.266 | NaN | 507.82288 | NaN |
| Cul3 | REVVTEHLINK | -78 | -88 | 1 | 0 | 11 | 427.02475 | 447.1553 | 489.51703 | 189.86002 | 671.30457 | 638.8119 |
| Sumo1 | VIGQDSSEIHFK | -26 | -37 | 0 | 0 | 12 | 2812.246 | 3137.6248 | 3165.7615 | 2703.3325 | 3305.213 | 3239.8755 |

| | | | | | | | | | | | | |
|---|---|---|---|---|---|---|---|---|---|---|---|---|
| Sumo1 | SDQEAKPSTEDLGDKK | -2 | -17 | 2 | 0 | 16 | 637.34015 | 634.6158 | 598.92053 | 1991.436 | 679.0374 | 764.60254 |
| Sumo1 | SDQEAKPSTEDLGDKK | -2 | -17 | 2 | 0 | 16 | 637.34015 | 634.6158 | 598.92053 | 1991.436 | 679.0374 | 764.60254 |
| Sumo1 | RFLFEGQR | -63 | -70 | 1 | 0.002099344 | 8 | 2780.8374 | 3833.5261 | 691.1786 | NaN | NaN | NaN |
| Usp25 | RFPLVDVL | -445 | -452 | 1 | 0.003681885 | 8 | 95.411224 | 119.0848 | NaN | 462.66046 | 401.87762 | 361.55472 |
| Clns1a | RFVPSDK | -112 | -118 | 1 | 0.002972566 | 7 | 1975.5438 | NaN | 1879.585 | NaN | NaN | 571.68384 |
| Clns1a | TTPTVAGQFEDADVDH | -221 | -236 | 0 | 0.000431655 | 16 | 373.61115 | NaN | NaN | NaN | NaN | NaN |
| Tm9sf1 | RGDDGGFFPR | -221 | -230 | 1 | 0.002595029 | 10 | 13113.61 | 307.6096 | 9641.842 | 8788.283 | 8455.799 | 7563.754 |
| Ppih | RGPFADENFK | -92 | -101 | 1 | 0 | 10 | 581.76355 | 172.06079 | 1844.2721 | 741.8894 | 2105.368 | 696.38605 |
| Dbr1 | RGSGPVDLLLCC | -26 | -37 | 1 | 0.00025015 | 12 | 468.58585 | NaN | NaN | 539.719 | 655.47205 | 875.5714 |
| Dbr1 | RGSGPVDLLL | -26 | -35 | 1 | 0.003461185 | 10 | 194.74754 | 290.07498 | 469.88394 | NaN | NaN | NaN |
| Psmd9 | SGEDVPHR | -2 | -9 | 0 | 0 | 8 | 1505.6648 | 576.3303 | 577.35455 | 767.4612 | 1226.4808 | 1299.1539 |
| Psmd9 | RKEEIEAEIK | -29 | -38 | 2 | 0.00025015 | 10 | 1021.8025 | 881.05414 | 891.3836 | 1319.8337 | 860.6873 | 901.2552 |
| Psmd9 | RKEEIEAEIK | -29 | -38 | 2 | 0.00025015 | 10 | 1021.8025 | 881.05414 | 891.3836 | 1319.8337 | 860.6873 | 901.2552 |
| RtcA | RLDPINLTDR | -177 | -186 | 1 | 0.00025015 | 10 | 1123.1937 | 1935.4453 | 2087.0874 | NaN | 1669.4202 | 1606.4624 |
| Saraf | TVVSCEGYESSEDQYVLR | -111 | -128 | 0 | 0 | 18 | 238.76302 | 259.21243 | 318.86792 | 328.9806 | 425.4463 | 414.3032 |
| Saraf | RLDPIPQLK | -58 | -66 | 1 | 0.00025015 | 9 | 439.42352 | 528.7034 | NaN | NaN | NaN | 711.8764 |
| Grhpr | SVGVDHLALDEIK | -82 | -94 | 0 | 0 | 13 | 830.0325 | 1175.9576 | 1001.5379 | 895.2511 | 1139.7094 | 1479.4333 |
| Grhpr | RLPEAIEEVK | -125 | -134 | 1 | 0.001291103 | 10 | 1850.2749 | NaN | 1171.9116 | NaN | 2645.5962 | 2399.574 |
| Grhpr | SKDFFQK | -227 | -233 | 1 | 0 | 7 | 887.3542 | NaN | NaN | 736.83514 | NaN | NaN |
| Rcor1 | RLPEVIQK | -368 | -375 | 1 | 0.00272025 | 8 | 502.59833 | NaN | NaN | 494.5688 | 544.93945 | NaN |
| Rcor1 | VWSPNQSLSEAK | -131 | -142 | 0 | 0.001291103 | 12 | 825.68695 | NaN | NaN | NaN | NaN | NaN |
| Dhx36 | RLPVEPHIGK | -688 | -697 | 1 | 0.001636916 | 10 | 360.40582 | NaN | 297.02463 | NaN | NaN | 295.50385 |
| Dnajb6 | RNPDDVFR | -95 | -102 | 1 | 0.00025015 | 8 | 1271.6627 | 1387.7279 | 1360.9514 | 1310.5541 | 1426.4961 | 1475.4136 |
| Papss1 | RNPEFFEHR | -339 | -347 | 1 | 0.00025015 | 9 | 537.0116 | 392.09558 | 294.8522 | 435.26227 | 332.14655 | 549.253 |
| Znf593 | RPDLDEIHR | -25 | -33 | 1 | 0 | 9 | 1127.3278 | 962.014 | 947.93536 | 195.6553 | 879.11096 | 934.10785 |
| Mat2b | RPDVVESQPDAASQL | -98 | -112 | 1 | 0.00025015 | 15 | 710.09326 | 779.5148 | 717.5222 | 999.3997 | 484.74854 | 490.31305 |
| Ctdspl;Ctdsp1 | RPHVDEFLQR | (148);(132) | (157);(141) | 1 | 0.000591124 | 10 | 677.4448 | NaN | NaN | 484.4854 | NaN | NaN |
| Mrpl44 | SAEFPVPLPVLR | -181 | -192 | 0 | 0 | 12 | 382.99445 | 359.95197 | 422.7427 | 332.6738 | 402.77545 | 696.36017 |
| Mrpl44 | RPWDYSKPK | -312 | -320 | 2 | 0.00635324 | 9 | 246.28746 | NaN | NaN | NaN | NaN | NaN |
| Dcaf8 | RQPVLDFESGHK | -219 | -230 | 1 | 0 | 12 | 1169.7119 | 1066.8912 | 981.5719 | 1398.8741 | 1352.09 | 1291.3359 |
| Dcaf8 | SLSLTGDDGGPNR | -48 | -60 | 0 | 0.00025015 | 13 | 1752.3094 | NaN | NaN | NaN | NaN | NaN |
| Casp6 | RVDFCKDPDAIGK | -242 | -254 | 2 | 0.000431655 | 13 | 910.96686 | NaN | 1495.6539 | 768.2704 | 958.4846 | 1026.7026 |
| Kin | RVEGIQYEDISK | -378 | -389 | 1 | 0 | 12 | 542.2333 | 475.26202 | 387.37958 | 585.91705 | 619.94543 | 510.0773 |
| Them6 | RVEPPELPEDLK | -171 | -182 | 1 | 0.000431655 | 12 | 462.5223 | NaN | NaN | 1005.7612 | NaN | NaN |
| Sft2d1 | RVLSGQDDEEQGLTAQ | -6 | -21 | 1 | 0.00025015 | 16 | 542.5651 | 449.19284 | 529.64496 | 1245.3667 | 1746.0792 | 611.6568 |
| Ptpn18 | RVPADQNSSGPDAYEEVTD | -406 | -424 | 1 | 0.000471409 | 19 | 1356.0625 | 1415.3058 | 1369.4742 | 1154.6982 | 1231.8047 | 1237.1924 |
| Cwc15 | WDDDVVFK | -188 | -195 | 0 | 0 | 8 | 583.4149 | 539.7095 | 589.02826 | 550.58295 | 643.22974 | 708.65 |
| Cwc15 | RWDDDVVFK | -187 | -195 | 1 | 0.00025015 | 9 | 972.5117 | 949.0023 | 986.6298 | NaN | 840.53894 | 1082.1324 |
| Mtco1 | RYSDYPDAYTTW | -439 | -450 | 1 | 0 | 12 | 760.09735 | 418.6669 | 880.69 | 604.4172 | 620.42615 | 625.7598 |
| Mtco1 | RYSDYPDAYTT | -439 | -449 | 1 | 0.001055699 | 11 | 1232.3307 | 1452.5121 | 1821.2522 | 755.02686 | 1030.2006 | NaN |
| Mtco1 | TMIIAIPTGVK | -309 | -319 | 0 | 0.002199523 | 11 | 998.1925 | 1589.643 | NaN | NaN | NaN | 1523.3892 |
| Samsn1 | SAAESLLDEETTVEHEK | -297 | -313 | 0 | 0 | 17 | 952.7744 | NaN | 1150.3257 | NaN | 215.58597 | 1351.8035 |
| Ppil4 | SAEEVEEIKAEK | -204 | -215 | 1 | 0 | 12 | 216.09811 | 629.14856 | 583.1225 | NaN | 648.57214 | 577.5827 |
| Ncln | SAGAVVIILPR | -104 | -114 | 0 | 0 | 11 | 250.3081 | NaN | NaN | 460.937 | 320.5623 | 333.9578 |
| Napsa | SAILDTGTSLITGPSEEIR | -274 | -292 | 0 | 0.00111006 | 19 | 315.91522 | NaN | NaN | 459.86633 | NaN | 575.5258 |
| Rnf114 | SAKPASETDPLSR | -15 | -27 | 1 | 0.00025015 | 13 | 2584.5962 | 2404.7349 | 2220.2402 | 2922.8027 | 4282.055 | 3598.4219 |
| Rab4b | SALTGENVEEAFLK | -151 | -164 | 0 | 0 | 14 | 8374.27 | 8461.918 | 8763.242 | NaN | NaN | NaN |
| Ddx31 | SAPSSPAEYIHR | -468 | -479 | 0 | 0.00450079 | 12 | 264.39554 | NaN | NaN | NaN | NaN | NaN |
| Ythdf1 | SAPSVESHPVLEK | -358 | -370 | 0 | 0 | 13 | 619.7784 | 572.43274 | 651.80176 | NaN | 594.75745 | NaN |
| Eif4ebp2 | SASAGGSHQPSQSR | -2 | -15 | 0 | 0.00025015 | 14 | 152.56425 | 311.95728 | 332.06705 | 228.83138 | 295.67816 | 251.27573 |
| Vps28 | SASDELDDSQVR | -191 | -202 | 0 | 0 | 12 | 815.59515 | 719.35706 | 964.018 | 799.4483 | 918.0703 | 1130.5392 |
| Ist1 | SASEDIDFDDLSR | -341 | -353 | 0 | 0 | 13 | 471.13025 | 383.79855 | 465.87057 | NaN | NaN | 407.5342 |
| Slc7a1 | TFDELIGKPIGEFSR | -135 | -149 | 1 | 0 | 15 | 602.009 | 863.26056 | 855.1985 | 716.5434 | 448.9684 | 343.16086 |
| Slc7a1 | SATFDELIGKPIGEFSR | -133 | -149 | 1 | 0 | 17 | 468.26965 | 538.3138 | NaN | 649.5255 | 937.36334 | 729.9598 |
| Slc7a1 | VGFDCIATTGEEVKNPQK | -260 | -277 | 1 | 0.00025015 | 18 | 280.25317 | 465.71353 | 396.50894 | NaN | 369.81638 | NaN |
| Ngp | SATPPSSQNPATNIPLQFR | -52 | -70 | 0 | 0 | 19 | 615.5293 | 799.8474 | 877.29333 | 1043.972 | 1665.9391 | 731.0287 |
| Mt2 | SCCSCCPVGCAK | -32 | -43 | 0 | 0 | 12 | 1516.669 | 1607.7643 | 1595.9586 | 2085.7874 | 2639.0876 | 2557.9983 |
| Mt1 | SCCSCCPVGCSK | -32 | -43 | 0 | 0 | 12 | 529.9089 | 914.86584 | 563.72943 | 485.57816 | 648.19183 | 672.02374 |
| Ttc27 | SCEFSPAPTPQECLAK | -302 | -317 | 0 | 0 | 16 | 967.49396 | 984.17175 | 1111.624 | 1276.8098 | 1527.9128 | 1342.5404 |
| Serinc1 | SDGSLDDGDGIHR | -361 | -373 | 0 | 0 | 13 | 729.4861 | 730.8085 | 712.59326 | 289.24445 | NaN | 524.1705 |
| Rgs3 | SDLYLDLINQK | -950 | -960 | 0 | 0 | 11 | 494.55072 | 461.83136 | 456.79407 | 382.61703 | 933.1833 | NaN |
| Rbm28 | SDQLEELFSQVGPVK | -17 | -31 | 0 | 0 | 15 | 934.77374 | 655.72565 | 1076.8972 | 751.76624 | 716.6743 | 509.21445 |
| Rock2 | SIAEEQYSDLEK | -930 | -941 | 0 | 0 | 12 | 1175.9623 | 1127.7383 | 1182.284 | NaN | NaN | NaN |
| Unc119b | SDSFYFVDNK | -227 | -236 | 0 | 0.000869338 | 10 | 655.69446 | NaN | NaN | NaN | NaN | NaN |
| Ykt6 | SDSLAGVVIADSEYPSR | -72 | -88 | 0 | 0 | 17 | 471.06158 | 620.0863 | 548.18567 | 1325.8364 | 622.5923 | 1361.8422 |
| Tnfrsf26 | SECAPCDSEHFIDHK | -59 | -73 | 0 | 0 | 15 | 603.45435 | 628.86505 | 615.6968 | NaN | 330.31506 | 190.19548 |
| Cdipt | SVISVIHLITAAR | -187 | -199 | 0 | 0 | 13 | 1243.2808 | 1409.7153 | 1418.0938 | 2105.6445 | 2200.0183 | 1916.6859 |
| Cdipt | SEGPLVGSVGLFR | -161 | -173 | 0 | 0 | 13 | 564.6707 | 771.37195 | 701.2411 | 655.7562 | 660.49976 | 560.00366 |
| Cdipt | SVISVIHLIT | -187 | -196 | 0 | 0 | 10 | 323.26807 | 413.70825 | 792.03827 | 648.58246 | 794.4451 | 818.8578 |
| LRWD1 | SEHLDPNLLGR | -35 | -45 | 0 | 0.002099344 | 11 | 540.6578 | NaN | NaN | NaN | NaN | NaN |
| Cmss1 | SEPKPGTPEDLQK | -82 | -94 | 1 | 0.00025015 | 13 | 911.29626 | 1149.6316 | NaN | 1074.4374 | 1382.3812 | 1195.461 |
| Bid | SESQEEIIHNIAR | -76 | -88 | 0 | 0 | 13 | 497.8431 | NaN | 691.4408 | NaN | 453.62097 | NaN |
| Wdr89 | SETHSDDITQVR | -165 | -176 | 0 | 0 | 12 | 655.2354 | 716.6152 | NaN | 505.95197 | 1354.2266 | 685.6582 |
| Anxa4 | SETSGSFEDALLAIVK | -226 | -241 | 0 | 0 | 16 | 314.31546 | NaN | NaN | 456.995 | 553.339 | 491.75748 |
| Sigmar1 | SEVFYPGETVVHGPGEATAL | -143 | -162 | 0 | 0.00025015 | 20 | 574.1752 | 743.0579 | 560.79047 | 699.8114 | 953.0901 | 1019.83765 |
| Sigmar1 | SEVFYPGETVVHGPGEAT | -143 | -160 | 0 | 0.000230707 | 18 | 572.6588 | 728.2284 | 819.8337 | NaN | NaN | 1036.5948 |
| Emc7 | SEVPGAAAEGPGGSGVGLGDR | -23 | -43 | 0 | 0 | 21 | 1670.4891 | 1540.5005 | NaN | NaN | NaN | 1849.7334 |
| Rbl1 | SFAPSTPLTGR | -364 | -374 | 0 | 0 | 11 | 561.3959 | 850.9601 | 912.3954 | 815.5654 | 763.57227 | 854.81 |

| | | | | | | | | | | | | |
|---|---|---|---|---|---|---|---|---|---|---|---|---|
| Mmut | SFDEALGLPTVK | -387 | -398 | 0 | 0 | 12 | 2836.1826 | 3400.022 | 3354.0464 | 2635.079 | 3526.9534 | NaN |
| Samm50 | VTGQFPWSSLR | -195 | -205 | 0 | 0 | 11 | 861.4712 | 878.46985 | 648.60095 | 675.3258 | 536.38043 | 446.53104 |
| Samm50 | SFFKPQPGNFER | -175 | -186 | 1 | 0 | 12 | 715.31757 | 681.6723 | 347.72592 | 806.617 | 728.7545 | 1328.1382 |
| Samm50 | WSYGAGVVLR | -424 | -433 | 0 | 0.001023236 | 10 | 718.31384 | 507.91824 | 422.46317 | 991.50964 | NaN | 1165.2534 |
| Ociad1 | SSFGTSPAADNIEK | -142 | -155 | 0 | 0 | 14 | 439.5567 | 679.2316 | 650.437 | 502.6369 | 731.5919 | 688.81525 |
| Ociad1 | SFGTSPAADNIEK | -143 | -155 | 0 | 0.005668207 | 13 | 1149.845 | NaN | NaN | 757.0639 | NaN | NaN |
| Trmt6 | SFLSGLYEFPLNK | -250 | -262 | 0 | 0 | 13 | 60.919296 | 362.61584 | 214.73718 | 368.2523 | 296.5631 | 313.62378 |
| Paxbp1 | SFQDEEEENEEVFK | -110 | -123 | 0 | 0.000230707 | 14 | 170.13187 | 613.5914 | 458.4105 | 332.71997 | 273.74643 | NaN |
| Tmem87a | SFSGDFTHR | -136 | -144 | 0 | 0 | 9 | 673.2615 | 908.3732 | 873.13916 | 438.86935 | NaN | 769.717 |
| Slc38a2 | SFVCHPAVLPIYEELK | -299 | -314 | 0 | 0 | 16 | 2701.1343 | 3228.511 | 3324.9983 | 3347.3164 | 4164.3804 | 3576.984 |
| Slc38a2 | SHYADVDPENQNFLLE | -39 | -54 | 0 | 0 | 16 | 684.4563 | 686.44806 | 2372.3835 | 2975.1802 | 1247.3586 | NaN |
| Nipsnap2 | SFWNEPVPR | -168 | -176 | 0 | 0 | 9 | 1327.6202 | 1215.8514 | NaN | 1189.6381 | 319.2127 | 1211.2418 |
| Gpatch11 | SGDGIVEPIPLNVK | -91 | -104 | 0 | 0.000347792 | 14 | 372.56723 | 1113.2844 | 705.36035 | NaN | 1544.2137 | NaN |
| Col1a1 | SGEYWIDPNQGCNLDAIK | -1260 | -1277 | 0 | 0 | 18 | 356.52954 | 512.6116 | 470.8738 | 165.0741 | 524.89264 | 550.9081 |
| Gid8 | SGIEPSVDLETLDER | -53 | -67 | 0 | 0 | 15 | 1410.6196 | NaN | NaN | 3887.652 | 3863.5193 | 2265.0908 |
| Cox15 | SGLEEKPESYDIPR | -207 | -220 | 1 | 0.00025015 | 14 | 554.94305 | NaN | 768.64996 | 872.01086 | 1125.5764 | NaN |
| Haus4 | SGPGQETELPPLLGLER | -128 | -144 | 0 | 0 | 17 | 394.19568 | NaN | 414.02118 | 656.8462 | 611.0416 | 966.46094 |
| Foxo1 | SGQEGPGDSPGSQFSK | -276 | -291 | 0 | 0 | 16 | 732.60266 | NaN | NaN | NaN | NaN | NaN |
| Casp3 | SLDSTFHAK | -251 | -259 | 0 | 0 | 9 | 3640.3657 | NaN | 427.62283 | NaN | 3583.1328 | NaN |
| Casp3 | STAPGYYSWR | -198 | -207 | 0 | 0.00025015 | 10 | 1078.2433 | NaN | NaN | NaN | NaN | NaN |
| Casp3 | SGTDVDAANLR | -65 | -75 | 0 | 0 | 11 | 247.9626 | NaN | NaN | NaN | NaN | NaN |
| Capns1 | SHYSNIEANESEEVR | -85 | -99 | 0 | 0 | 15 | 895.4149 | 1002.31274 | 842.4337 | 613.6285 | 951.74805 | 953.9391 |
| Capns1 | SGTIGSHELPGAF | -186 | -198 | 0 | 0 | 13 | 544.81036 | 820.0988 | 730.1471 | 1195.133 | 1140.7148 | 1174.711 |
| Rcn1 | TFDQLSPDESK | -65 | -75 | 0 | 0 | 11 | 4904.2437 | 5191.0503 | 163.35384 | 5932.1475 | 928.8679 | 5129.9346 |
| Rcn1 | SHEDNGPEPDWVLSER | -228 | -243 | 0 | 0.000347792 | 16 | 428.09644 | 654.59015 | 372.66953 | 424.9651 | 553.61456 | NaN |
| Ccdc124 | SHLELPLEENLNR | -117 | -129 | 0 | 0 | 13 | 904.9474 | 986.5707 | 1077.1477 | 869.0491 | 1292.8542 | 1191.8146 |
| Ccdc124 | SVAEEADRHPER | -150 | -161 | 1 | 0.00383302 | 12 | 389.78012 | NaN | NaN | 647.7812 | 410.9882 | NaN |
| Lpp | SIASPPVSTPVTGHK | -149 | -163 | 0 | 0 | 15 | 822.3878 | NaN | 862.91504 | 533.73737 | NaN | 708.6656 |
| Zmpste24 | SIDFPLTK | -244 | -251 | 0 | 0.00025015 | 8 | 1385.888 | 177.41006 | 1317.9847 | 1115.9089 | 1236.814 | 710.85956 |
| Icos | SIFDPPPFQER | -112 | -122 | 0 | 0 | 11 | 1603.5533 | 1152.7802 | 709.1365 | 618.06146 | 1483.0051 | 1533.0975 |
| Scrn2 | SIGTDISAEHPELR | -196 | -209 | 0 | 0 | 14 | 377.45706 | 483.8282 | 474.8418 | 514.94385 | NaN | 403.89795 |
| Naa20 | SIIPLPHPVRPEDIE | -164 | -178 | 1 | 0.000471409 | 15 | 1293.6017 | 1307.8411 | NaN | 1359.1422 | 592.27295 | 1702.073 |
| Cdca8 | SIKEEEEEEEEGGGGGGR | -122 | -139 | 1 | 0.00437931 | 18 | 316.73102 | 278.16565 | 248.06256 | NaN | NaN | NaN |
| Tmem115 | SITISLPGTDPQDAER | -267 | -282 | 0 | 0.00025015 | 16 | 1844.8257 | NaN | NaN | 2274.6165 | 2276.445 | NaN |
| Nup35 | SPLLAGGSPPQPVVPAHK | -66 | -83 | 0 | 0 | 18 | 314.3424 | 373.91467 | 341.3062 | 482.68982 | 402.8698 | 417.75293 |
| Nup35 | SIYDDISSPGLGSTPLTSR | -93 | -111 | 0 | 0.000550307 | 19 | 1107.714 | 1228.4683 | 1306.3213 | 1822.0188 | 2172.8303 | NaN |
| Pdcd10 | SKIPDEINDR | -115 | -124 | 1 | 0.00025015 | 10 | 305.10083 | 497.91006 | 481.03833 | 463.8663 | 584.11066 | 478.64648 |
| Nob1 | SKPLQEAVDR | -134 | -143 | 1 | 0 | 10 | 2748.1978 | 2914.359 | NaN | NaN | NaN | 2848.0598 |
| Nup214 | TLELISTEGER | -416 | -426 | 0 | 0 | 11 | 6798.7866 | 546.52936 | 5816.665 | NaN | 4088.3474 | NaN |
| Nup214 | SLAGETLGSFSGLR | -1337 | -1350 | 0 | 0.00051306 | 14 | 1661.0411 | 2127.9595 | NaN | NaN | 2782.2673 | NaN |
| Tmem167a | SLAPSILDR | -26 | -34 | 0 | 0 | 9 | 1239.4642 | 2360.287 | 1205.8295 | 1513.228 | 1209.1733 | 1168.8923 |
| Nup188 | SLDAPSWPGVYR | -1387 | -1398 | 0 | 0 | 12 | 162.01875 | 952.0704 | 792.3372 | 1678.2227 | NaN | NaN |
| Nup188 | VKPIIDLVHK | -565 | -574 | 1 | 0.000431655 | 10 | 400.19095 | NaN | NaN | 218.40797 | NaN | NaN |
| Sec11a | VGEIVVFR | -79 | -86 | 0 | 0 | 8 | 1140.8203 | 831.1704 | 946.20435 | 757.1028 | 595.1584 | 589.7742 |
| Sec11a | SLDFLDDVR | -3 | -11 | 0 | 0.00025015 | 9 | 294.29477 | 286.57178 | 251.75598 | 556.3798 | 257.99786 | 512.7079 |
| Atp2b1 | SLDKDPLLLSGT | -253 | -264 | 1 | 0.00025015 | 12 | 727.4471 | 728.7386 | 828.0954 | 638.21674 | 748.5694 | NaN |
| Iqgap2 | SLDLLPYGLR | -1065 | -1074 | 0 | 0 | 10 | 277.70065 | 355.98624 | NaN | NaN | 310.168 | NaN |
| Fkbp15 | SLEEELTDLR | -722 | -731 | 0 | 0 | 10 | 1284.3486 | 1067.5476 | 160.77917 | 1376.5269 | 1562.8566 | 1351.1923 |
| Pxn | SLGSNLSELDR | -137 | -147 | 0 | 0 | 11 | 1172.8386 | NaN | 1619.7133 | NaN | 1032.9039 | 792.7476 |
| Lsm1 | SLIEDIDKK | -9 | -17 | 1 | 0 | 9 | 1190.545 | 471.1443 | 854.19714 | 854.2226 | 798.1553 | 546.33496 |
| Churc1 | SLKEEDGEEIVTY | -45 | -57 | 1 | 0.00025015 | 13 | 1340.3123 | 1900.278 | 1898.2355 | 1369.21 | 1860.6003 | NaN |
| Agfg1 | SLLGESAPALHLNK | -162 | -175 | 0 | 0 | 14 | 997.38214 | 1072.8362 | 1103.5181 | 1096.1 | 1204.5076 | 1287.141 |
| Agfg1 | SSAIPDFRDPQK | -101 | -112 | 1 | 0 | 12 | 968.47296 | 870.5164 | 976.3884 | 963.51 | 1046.7688 | NaN |
| Cyp20a1 | SLLGYQSGGGSAGEDHVR | -106 | -123 | 0 | 0 | 18 | 735.8965 | 660.8541 | 428.31274 | 668.17725 | 514.41583 | 397.09784 |
| Tmem214 | SLLLEHLLK | -415 | -423 | 0 | 0 | 9 | 314.91296 | 355.11157 | 381.381 | NaN | 127.93752 | 614.722 |
| Plek | SLLSEGYLQPAGDLSK | -183 | -198 | 0 | 0 | 16 | 4379.548 | 4827.2905 | 5115.8423 | 4212.5376 | 5178.489 | 5226.5674 |
| Polr1e | SLPIDEVHR | -325 | -333 | 0 | 0.000869338 | 9 | 4479.8677 | 2132.8936 | 457.04486 | 669.29254 | NaN | NaN |
| Atp2a3 | SLPVILLDEALK | -974 | -985 | 0 | 0 | 12 | 291.9894 | 352.9606 | 326.2778 | 394.4105 | 394.88687 | 336.3711 |
| Atp2a3 | YGPNELPTEEGK | -36 | -47 | 0 | 0 | 12 | 2141.5088 | 416.10223 | NaN | NaN | 4436.8804 | 4387.9414 |
| Ube2r2 | SLQEEPVEGFR | -19 | -29 | 0 | 0 | 11 | 772.8863 | NaN | NaN | NaN | NaN | NaN |
| Def6 | SLQHAQQQLEEVR | -504 | -516 | 0 | 0 | 13 | 936.6532 | 927.54 | 987.37866 | 1759.6647 | 1799.7344 | 410.76266 |
| Wdr70 | SLQPCECHQIK | -226 | -236 | 0 | 0.00078661 | 11 | 2269.354 | NaN | NaN | 2183.3008 | 2570.6406 | 2686.48 |
| Erbin | SLQQLPETIGSLK | -264 | -276 | 0 | 0 | 13 | 446.84003 | 512.55945 | 404.95392 | 497.65662 | 275.2768 | 419.63266 |
| Ring1;Rnf2 | SLRPDPNFDAL | (96);(99) | (106);(109) | 1 | 0 | 11 | 458.492 | 616.5643 | 762.34924 | 792.7506 | 649.98505 | 686.6494 |
| Serpinf1 | VSNFGYDLYR | -57 | -66 | 0 | 0 | 10 | 905.4874 | 842.21875 | 268.1115 | 832.13837 | 316.167 | 1049.8137 |
| Serpinf1 | SLTSEFIHDIDR | -291 | -302 | 0 | 0.000591124 | 12 | 875.9421 | 874.22174 | 903.5397 | NaN | 933.92694 | 962.4317 |
| Serpinf1 | YGLDSDLNCK | -252 | -261 | 0 | 0 | 10 | 622.99896 | 574.5178 | 733.56714 | NaN | 699.8105 | 553.1862 |
| Mrpl46 | SLYSDHELR | -80 | -88 | 0 | 0 | 9 | 1238.1763 | 1280.1772 | 1025.4149 | 1618.3994 | 1301.4177 | 1631.4283 |
| Npc1 | SNLQLPLQFLSR | -85 | -96 | 0 | 0 | 12 | 467.82318 | 531.3728 | 500.65076 | 591.7538 | 576.82837 | 539.61615 |
| Bag1 | SNPEEEVELKK | -232 | -242 | 1 | 0 | 11 | 5378.2363 | NaN | 4645.2847 | 5014.587 | 5285.5312 | NaN |
| Ndufaf7 | SPAFQLVELGPGR | -110 | -122 | 0 | 0 | 13 | 1422.7728 | 1272.6245 | 289.96848 | 1309.9685 | 2042.033 | NaN |
| Fam3c | SPFEQHIK | -194 | -201 | 0 | 0 | 8 | 1952.8765 | 1062.521 | 1397.2249 | NaN | 303.76086 | 1219.854 |
| Acp1 | SPIAEAVFR | -20 | -28 | 0 | 0 | 9 | 2570.2627 | 1479.6776 | 1701.5193 | 1595.4093 | 1257.78 | 1049.7543 |
| Hsd11b1 | SPLTPILLGNPGR | -261 | -273 | 0 | 0 | 13 | 562.84784 | 609.7891 | 584.47156 | 762.7235 | 753.529 | 535.8764 |
| Ranbp3 | SPSESAEETHTLEEK | -146 | -160 | 0 | 0 | 15 | 326.43372 | 454.73825 | 484.95767 | 470.5385 | 575.73486 | 725.6255 |
| Ranbp3 | YISSSADNATHSADNSTK | -227 | -244 | 0 | 0.000471409 | 18 | 169.49896 | 210.93016 | 271.67966 | NaN | 365.79758 | 364.16916 |
| Dcp1a | SQLQDTLIHLIK | -565 | -576 | 0 | 0 | 12 | 553.0802 | 303.5485 | 547.60126 | NaN | 375.25055 | 397.19925 |
| Brix1 | SQPFVDHVFTF | -198 | -208 | 0 | 0 | 11 | 784.3474 | 663.5351 | 733.5947 | 876.8013 | 902.7955 | 1082.5333 |

| | | | | | | | | | | | | |
|---|---|---|---|---|---|---|---|---|---|---|---|---|
| Brix1 | TILPHDPTAD | -296 | -305 | 0 | 0.00025015 | 10 | 652.7652 | 808.84283 | 886.43274 | 649.2393 | 386.1741 | 980.2049 |
| Wbp4 | STNPYGEWQEIK | -291 | -302 | 0 | 0 | 12 | 759.14294 | NaN | 904.7044 | NaN | 1454.6329 | 408.95267 |
| Wbp4 | SQWEKPEGFQGNLK | -149 | -162 | 1 | 0 | 14 | 952.4337 | 921.80865 | NaN | NaN | NaN | NaN |
| Prpf38a | VCDIILPR | -157 | -164 | 0 | 0 | 8 | 1706.8716 | 1711.7915 | 1723.7015 | 1464.8394 | 1719.9425 | 1852.6598 |
| Prpf38a | SSEEEEEEDEKLER | -193 | -206 | 1 | 0.001291103 | 14 | 323.7671 | NaN | NaN | 257.63397 | NaN | NaN |
| Ablim1 | VDNEILDYK | -420 | -428 | 0 | 0 | 9 | 945.72754 | 702.3959 | 1075.5941 | 290.55167 | NaN | 952.9911 |
| Ablim1 | SSGREEDEEELLR | -670 | -682 | 1 | 0 | 13 | 481.38824 | NaN | 652.68146 | NaN | 727.8162 | 406.817 |
| Ints10 | SSHIVDPWER | -249 | -258 | 0 | 0 | 10 | 1512.4484 | NaN | NaN | NaN | NaN | NaN |
| Edrf1 | SSKAYYILSDA | -692 | -702 | 1 | 0.000591124 | 11 | 1138.1941 | 441.82593 | 1051.8658 | 1376.2678 | 1071.097 | 151.87645 |
| Pigu | SSLAEFISER | -22 | -31 | 0 | 0 | 10 | 1431.1039 | 1301.239 | 491.28497 | 2006.3842 | 1732.095 | NaN |
| Slc38a1 | SSLFELAK | -375 | -382 | 0 | 0.006195994 | 8 | 566.0717 | 690.5429 | NaN | NaN | NaN | NaN |
| Qtrt2 | SSVTAELPEDKPR | -237 | -249 | 1 | 0 | 13 | 528.264 | 558.48315 | 261.86737 | NaN | 482.1832 | 431.74203 |
| Suv39h1 | SSWNQLQDLCR | -14 | -24 | 0 | 0 | 11 | 1450.3793 | 767.14984 | 1159.24 | 1306.5713 | 1097.5774 | 970.10486 |
| Nmd3 | STFSVEIVPICK | -240 | -251 | 0 | 0 | 12 | 360.30286 | 517.32367 | NaN | 683.9455 | NaN | NaN |
| Pno1 | STGTEDGFTPVTHR | -5 | -18 | 0 | 0 | 14 | 1190.269 | 1020.97314 | 1204.4482 | NaN | NaN | NaN |
| Kifc1 | STLSTQLEEQER | -243 | -254 | 0 | 0 | 12 | 4237.9062 | NaN | NaN | 1037.984 | 4155.431 | NaN |
| Hmgb3 | STNPGISIGDVAK | -113 | -125 | 0 | 0 | 13 | 2135.6284 | 938.6197 | 1528.0061 | 1478.7074 | 1416.4906 | 1381.6804 |
| Toe1 | STNPVPGDGLHR | -429 | -440 | 0 | 0.002329279 | 12 | 679.36914 | NaN | 512.6043 | NaN | 347.8287 | 219.39212 |
| Toe1 | VAEGEAGDQPGSR | -385 | -397 | 0 | 0 | 13 | 630.808 | NaN | NaN | 342.99673 | NaN | NaN |
| Ntan1 | STSPLAEPPHFVEHIR | -254 | -269 | 0 | 0 | 16 | 742.23157 | NaN | 639.3293 | 318.468 | 1269.1588 | 580.0925 |
| Strn | STTNPNVDPYDAYDPSVLR | -544 | -562 | 0 | 0 | 19 | 371.87207 | 163.20685 | 360.08734 | 333.57523 | 523.0062 | 407.49487 |
| Mcm3ap | STVFPLDGVVR | -889 | -899 | 0 | 0 | 11 | 461.89407 | 379.32462 | 517.8936 | 501.8335 | NaN | NaN |
| Vps26b | STYPELNSSIK | -153 | -163 | 0 | 0.003299042 | 11 | 575.4452 | NaN | NaN | NaN | NaN | NaN |
| Wdr76 | SVALHPSEVR | -317 | -326 | 0 | 0.00078661 | 10 | 572.1487 | NaN | NaN | NaN | NaN | NaN |
| Cstf1 | SVCAPSEQLLHLIK | -42 | -55 | 0 | 0 | 14 | 720.41815 | 791.5871 | 732.106 | 638.7309 | NaN | 692.43866 |
| Pef1 | SVDADHSGYISLK | -116 | -128 | 0 | 0 | 13 | 542.5581 | 1890.5752 | 2819.4355 | 397.15622 | 4078.3442 | 3911.6992 |
| Trappc11 | SVDLFLGSESGR | -686 | -697 | 0 | 0.000230707 | 12 | 1600.4474 | 1796.3732 | NaN | 2009.1124 | NaN | NaN |
| Esyt2 | SVDLPGLLGQLAR | -30 | -42 | 0 | 0 | 13 | 363.89694 | 342.65802 | NaN | 483.88812 | 462.55573 | 486.2367 |
| Esyt2 | VDVGQQPLR | -164 | -172 | 0 | 0 | 9 | 727.1274 | 812.035 | 743.3888 | NaN | NaN | NaN |
| Cd3e | SVELTCPLDSDENLK | -37 | -51 | 0 | 0 | 15 | 865.6783 | NaN | 936.80994 | 1027.4564 | 1071.976 | 1068.2625 |
| Cyb561d2 | SVETESHIYR | -4 | -13 | 0 | 0 | 10 | 268.72842 | NaN | NaN | NaN | NaN | NaN |
| Castor1 | SVIAPLAEHHVSVL | -99 | -112 | 0 | 0.000627943 | 14 | 498.49402 | NaN | 767.6361 | NaN | NaN | 628.2593 |
| Helb | SVNDDVDTDEESAQLR | -985 | -1000 | 0 | 0.001776473 | 16 | 383.4624 | 518.81006 | 438.5162 | NaN | NaN | NaN |
| Lilrb4a | SVNPWDTQVPLETR | -65 | -78 | 0 | 0 | 14 | 1221.9315 | 1596.4849 | 1553.3882 | 610.6195 | 892.3345 | 610.3576 |
| Gemin6 | SVQTVETISEGDHR | -63 | -76 | 0 | 0 | 14 | 2456.2107 | 1644.2568 | 2729.4465 | 3733.958 | 2566.6404 | 1043.2878 |
| Dtd1 | SYRPELIR | -116 | -123 | 1 | 0.001636916 | 8 | 672.48157 | 1118.0863 | 812.9735 | 893.212 | 814.819 | 952.911 |
| Dtd1 | SVTVGGEQISAIGR | -12 | -25 | 0 | 0 | 14 | 812.2941 | NaN | 781.92975 | 1123.5369 | 747.8401 | 836.54297 |
| Dtd1 | YNSFLEQLR | -106 | -114 | 0 | 0 | 9 | 1135.8282 | 483.48904 | 2726.2063 | NaN | NaN | NaN |
| Lta | SVYPGLQGPWVR | -151 | -162 | 0 | 0 | 12 | 3205.6526 | 3385.5378 | 6135.3394 | 7243.8237 | 3798.3037 | 4238.0786 |
| Ldah | SYLLFKPCPEVIK | -195 | -207 | 1 | 0 | 13 | 1059.7198 | NaN | NaN | 723.38416 | 1383.429 | 1167.034 |
| Ric8a | SYPHEVQFFDLR | -155 | -166 | 0 | 0.000550307 | 12 | 367.0933 | NaN | NaN | NaN | NaN | NaN |
| Mrps18b | SYYVPQVEPR | -162 | -171 | 0 | 0 | 10 | 4354.757 | NaN | NaN | 4727.3916 | 3212.5945 | 3003.147 |
| Mrps18b | YLDSEEYHNR | -56 | -65 | 0 | 0 | 10 | 362.45575 | NaN | NaN | 149.7051 | 1683.2737 | 249.9647 |
| Ptms | TAEEEDEADPKR | -81 | -92 | 1 | 0 | 12 | 736.5706 | 536.2323 | 290.55478 | 299.24124 | 467.13937 | NaN |
| Sdf4 | TAEHFQEAVK | -130 | -139 | 0 | 0 | 10 | 2123.3218 | NaN | NaN | NaN | NaN | NaN |
| Meaf6 | TAPPQIPDTR | -7 | -16 | 0 | 0 | 10 | 7834.1147 | NaN | 3844.7246 | 6788.939 | 2903.2175 | 6479.6284 |
| Armt1 | TASDPHWLTTGR | -418 | -429 | 0 | 0 | 12 | 2487.7122 | 2744.5505 | 2556.6555 | 1005.2737 | 943.8239 | 839.1555 |
| Nit1 | VVDPWGTVVAR | -270 | -280 | 0 | 0 | 11 | 685.8687 | 829.6416 | 948.1571 | 708.7652 | 1240.9229 | 771.7189 |
| Nit1 | TCAELVQEAAR | -62 | -72 | 0 | 0 | 11 | 1781.0969 | NaN | NaN | NaN | NaN | NaN |
| Il2rb | TCDFHPFDNLR | -122 | -132 | 0 | 0 | 11 | 534.11475 | 557.11664 | 302.71497 | 645.6865 | 874.4138 | 551.7241 |
| Rbbp6 | TDTLFVLPSR | -972 | -981 | 0 | 0 | 10 | 1081.7732 | 1173.7885 | 1247.949 | NaN | NaN | NaN |
| Elp2 | TEDGSPSNELVSGGSDNR | -69 | -86 | 0 | 0 | 18 | 753.1215 | 661.9078 | 573.2775 | 879.1427 | 650.9226 | 860.8515 |
| Slfn2 | TEFQYEETFPFTR | -218 | -230 | 0 | 0.00025015 | 13 | 375.22052 | NaN | 465.4105 | NaN | 324.90216 | NaN |
| Atg4b | TEKDEILSDVASR | -37 | -49 | 1 | 0 | 13 | 1303.0996 | 1581.2181 | 1096.0316 | 1352.3246 | 1283.0758 | 1362.9794 |
| Ube3c | TELPFVVPFEER | -687 | -698 | 0 | 0 | 12 | 261.65997 | 493.78427 | 521.71893 | 285.32993 | 305.9105 | NaN |
| Cdc23 | TELQLIEEALQK | -239 | -250 | 0 | 0 | 12 | 622.7619 | 553.6526 | 565.5868 | 552.114 | 497.41125 | 746.44025 |
| Sumo3 | TENDHINLK | -12 | -20 | 0 | 0 | 9 | 824.7124 | 1094.1359 | 1176.4507 | NaN | 1081.0737 | NaN |
| Abhd12 | TEPVTLEHER | -5 | -14 | 0 | 0.00025015 | 10 | 1341.149 | NaN | NaN | 1031.5214 | NaN | NaN |
| Tfip11 | TEQEIIQSDR | -332 | -341 | 0 | 0.000230707 | 10 | 767.21185 | 737.0711 | 905.9347 | NaN | NaN | 995.4497 |
| Nek9 | TFDATNPLNLCVK | -250 | -262 | 0 | 0.003003218 | 13 | 406.23688 | NaN | NaN | NaN | NaN | 967.368 |
| Serpinb1a | TFHFDSVEDIHSR | -57 | -69 | 0 | 0 | 13 | 1672.2825 | NaN | 1312.5391 | NaN | 1484.591 | NaN |
| Mcrip1 | TFKPIDLSDLK | -77 | -87 | 1 | 0 | 11 | 535.01794 | 583.9626 | 556.14624 | 556.7532 | 647.3833 | 649.88403 |
| Mtatp8 | VSSQTFPLAPSPK | -28 | -40 | 0 | 0 | 13 | 6495.7705 | 369.19772 | 1556.938 | 8022.592 | 7761.3237 | 5979.066 |
| Mtatp8 | TFPLAPSPK | -32 | -40 | 0 | 0.000431655 | 9 | 1257.9253 | 1078.8599 | 1184.3564 | NaN | NaN | NaN |
| Yy1 | TGDRPYVCPFDGCNK | -378 | -392 | 1 | 0 | 15 | 268.98407 | 222.60861 | NaN | 101.221565 | 412.79883 | 521.96576 |
| Ncf1 | TGDVVDVVEK | -179 | -188 | 0 | 0 | 10 | 2164.5012 | NaN | 2853.9102 | NaN | 3802.6384 | NaN |
| Rpl22l1 | TGNLGNVVHIE | -48 | -58 | 0 | 0.00025015 | 11 | 1513.0658 | 1237.7235 | 1290.3951 | 1981.4689 | 1069.0713 | 1399.0889 |
| Rpl22l1 | VVASDKETYELR | -96 | -107 | 1 | 0 | 12 | 1219.5493 | 1034.9476 | 939.77386 | 900.0226 | 869.3361 | 908.43 |
| Rpl22l1 | TGNLGNVVHIER | -48 | -59 | 0 | 0 | 12 | 959.61505 | 158.57568 | 514.0805 | NaN | NaN | NaN |
| Rps28 | TGSQGQCTQVR | -21 | -31 | 0 | 0 | 11 | 859.59436 | 729.5022 | 709.5733 | 859.86676 | 765.42676 | 1043.4497 |
| Rps28 | VEFMDDTSR | -32 | -40 | 0 | 0 | 9 | 644.4415 | 537.14075 | 661.84937 | 347.3942 | 941.489 | 726.94135 |
| Bdh1 | WDDLPEVVR | -261 | -269 | 0 | 0 | 9 | 1104.6349 | 119.700134 | 1292.0267 | 1822.9126 | 2068.7747 | 140.95428 |
| Bdh1 | THFPGAISDK | -330 | -339 | 0 | 0 | 10 | 458.2779 | 12682.276 | 8956.8125 | 9784.594 | 4616.2393 | 5550.2896 |
| Dph5 | THSDLILR | -90 | -97 | 0 | 0.001370415 | 8 | 2042.1974 | NaN | 882.4348 | NaN | NaN | NaN |
| Ghitm | TILGGPLLLR | -209 | -218 | 0 | 0 | 10 | 374.0075 | 367.56375 | 346.74063 | 312.73605 | 426.8234 | 391.3857 |
| Commd2 | TILNELAPR | -112 | -120 | 0 | 0.00149795 | 9 | 1423.6738 | NaN | NaN | NaN | 1879.8689 | NaN |
| Kdm3a | TIQDGDSDELTIK | -1151 | -1163 | 0 | 0.00111006 | 13 | 1239.719 | NaN | NaN | NaN | NaN | NaN |
| Htra2 | TKEPLPTLPLGR | -236 | -247 | 1 | 0.00025015 | 12 | 842.7957 | 672.9669 | NaN | 680.4556 | NaN | 545.5863 |

| | | | | | | | | | | | | |
|---|---|---|---|---|---|---|---|---|---|---|---|---|
| Rbm34 | TKIPVNPEEER | -174 | -184 | 1 | 0.000230707 | 11 | 879.22644 | 1042.7072 | NaN | NaN | NaN | NaN |
| Dhrs7b | TLAPGLFFR | -300 | -308 | 0 | 0 | 9 | 508.47916 | 573.8762 | 541.06683 | 406.64014 | 468.9111 | 655.387 |
| Cpsf2 | TLDDVDAAFDK | -115 | -125 | 0 | 0 | 11 | 97.65392 | NaN | 666.0403 | NaN | NaN | NaN |
| Cfap100 | TLDELNVTLK | -416 | -425 | 0 | 0.00149795 | 10 | 1425.643 | NaN | NaN | NaN | NaN | NaN |
| Exosc5 | TLEVILRPK | -75 | -83 | 1 | 0 | 9 | 360.63834 | 358.20023 | NaN | 597.9222 | 688.8921 | 678.816 |
| Rbm7 | TLFVGNLETK | -11 | -20 | 0 | 0.00122419 | 10 | 656.6983 | 898.371 | 1043.6919 | NaN | NaN | 450.38724 |
| Hgh1 | TLLAGQPELLR | -61 | -71 | 0 | 0 | 11 | 1038.413 | 1051.1659 | 1070.7535 | NaN | NaN | NaN |
| Stx7 | TLNQLGTPQDSPELR | -35 | -49 | 0 | 0 | 15 | 497.01794 | NaN | NaN | 1138.6094 | 727.3491 | NaN |
| Wdr5 | TLPAHSDPVSAVHFNR | -166 | -181 | 0 | 0 | 16 | 872.5066 | 370.9626 | 867.36145 | 917.84424 | 1261.2633 | 1190.4325 |
| Dennd10 | TLPALVWHR | -199 | -207 | 0 | 0.002296977 | 9 | 259.45944 | NaN | NaN | 174.84982 | NaN | 401.8326 |
| Ndufs4 | TLTGVPEEHIK | -63 | -73 | 0 | 0 | 11 | 730.91956 | 901.24554 | NaN | 995.4166 | 873.05634 | 762.0442 |
| Rnaseh2c | TLWGLETVPGPDAK | -132 | -145 | 0 | 0 | 14 | 423.7086 | 372.9267 | NaN | 1397.039 | 1771.4579 | 1158.6859 |
| Rabggtb | TLYDSVHVINVDK | -114 | -126 | 0 | 0 | 13 | 528.019 | 619.0098 | 641.87036 | 713.8831 | 578.2621 | NaN |
| Siae | TMNFTLR | -105 | -111 | 0 | 0.006788754 | 7 | 5671.041 | NaN | NaN | NaN | NaN | 4025.2473 |
| Smg1 | TNTEIALPGEVSAR | -2101 | -2114 | 0 | 0.000471409 | 14 | 853.9585 | NaN | NaN | NaN | NaN | NaN |
| Etv6 | TPAESVHHNPPTIE | -165 | -178 | 0 | 0.000230707 | 14 | 1110.2438 | NaN | 1165.2583 | NaN | NaN | NaN |
| Anxa1 | TPAQFDADELR | -114 | -124 | 0 | 0 | 11 | 1345.55 | NaN | NaN | 1301.1208 | NaN | 1348.5364 |
| Nup58 | TPPGLQHENTAPADYFR | -319 | -335 | 0 | 0 | 17 | 1007.685 | NaN | 1198.7708 | NaN | 1135.5856 | 1198.6686 |
| Faf1 | TPSGEFLER | -579 | -587 | 0 | 0.000471409 | 9 | 4193.204 | 543.7793 | 3673.9248 | NaN | 4238.6304 | 3895.1667 |
| Galk2 | TQILTPNTQDELTFK | -313 | -327 | 0 | 0.00025015 | 15 | 1094.8778 | 1330.121 | 1435.5911 | 1302.4609 | 1554.2076 | 1641.775 |
| Cuta | TQSSLVPALTEFVR | -123 | -136 | 0 | 0 | 14 | 1006.39124 | 1038.5165 | 1169.031 | 1027.4379 | 1309.0892 | 1343.92 |
| Impa2 | TSAADLVTETDHR | -50 | -62 | 0 | 0 | 13 | 1064.879 | 1876.5675 | NaN | 1465.4071 | NaN | 1513.8319 |
| Mrpl1 | TSDEKPVDDIEK | -65 | -76 | 1 | 0 | 12 | 561.3083 | 612.33716 | 707.3388 | 688.39777 | 736.6105 | 654.2421 |
| Eif1ax | TSDIILVGLR | -73 | -82 | 0 | 0 | 10 | 1918.971 | 2073.579 | 1948.3948 | 1582.4154 | 1445.6733 | 1511.9784 |
| Rassf5 | TSFYLPLDAIK | -273 | -283 | 0 | 0 | 11 | 537.64526 | 456.5774 | 546.65466 | 760.8648 | 762.7146 | 896.7724 |
| Eif2d | TSGGNPILFELEK | -65 | -77 | 0 | 0 | 13 | 295.9902 | NaN | NaN | 17529.7 | 1769.7322 | NaN |
| Kif15 | TSLLNLVDLAGSER | -258 | -271 | 0 | 0 | 14 | 683.44293 | 462.3606 | 415.22168 | NaN | 559.78326 | NaN |
| Ahctf1 | TWIIEEQPNSAANLR | -567 | -581 | 0 | 0 | 15 | 483.81778 | 492.05072 | 489.99283 | NaN | 531.436 | 737.11456 |
| Ahctf1 | TSPHHILDIL | -440 | -449 | 0 | 0.00078661 | 10 | 393.9394 | 203.88916 | NaN | NaN | 443.22815 | NaN |
| Dgat1 | TSVGDGYWDLR | -66 | -76 | 0 | 0.000394361 | 11 | 1057.6356 | NaN | NaN | NaN | NaN | NaN |
| Ube2j1 | TSYGAQNPSGAPLPQPTQPAPK | -225 | -246 | 0 | 0 | 22 | 273.65887 | 299.4226 | 273.2003 | 309.53275 | 275.04327 | 282.35016 |
| Tbrg4 | TTALLEHPEYK | -458 | -468 | 0 | 0 | 11 | 3211.1033 | NaN | NaN | NaN | NaN | 593.20807 |
| Tnfrsf8 | TTCAGDLSHYPGEAAR | -27 | -42 | 0 | 0 | 16 | 326.6933 | 368.01364 | 338.00644 | 406.55066 | 526.5521 | 466.2067 |
| Bcl7b | TTEEEEDSGAPPLKR | -173 | -187 | 1 | 0 | 15 | 692.3122 | 791.3057 | 867.8341 | 985.6937 | 1224.513 | 1181.2716 |
| Hmgn1 | TTELPAENGETENQSPASEEEK | -70 | -91 | 0 | 0.000431655 | 22 | 191.02394 | 408.36813 | 413.17584 | 216.37105 | 359.04816 | 277.26328 |
| Sp2 | TTEPTPTQVYIR | -345 | -356 | 0 | 0.002752526 | 12 | 951.97455 | 852.101 | NaN | NaN | NaN | NaN |
| Parvg | TTLLPEHIVVR | -56 | -66 | 0 | 0 | 11 | 405.5082 | 360.0543 | 375.8167 | 502.11133 | 451.5905 | 453.03284 |
| Ndfip1 | TTLPSYDEAER | -71 | -81 | 0 | 0 | 11 | 1153.1156 | 896.3529 | 1171.677 | 959.0751 | NaN | 944.51904 |
| Junb | TTPTPPGQYFYPR | -101 | -113 | 0 | 0 | 13 | 1459.4136 | NaN | NaN | NaN | 1546.3092 | NaN |
| Camk4 | TVCGTPGYCAPEILR | -196 | -210 | 0 | 0 | 15 | 2915.4373 | 2000.3984 | 165.80849 | 509.64838 | 1053.9113 | 1404.4521 |
| Cpt1a | TVDREELKPIR | -285 | -295 | 2 | 0 | 11 | 385.031 | NaN | NaN | 358.47562 | 516.2468 | 467.03342 |
| Cpt1a | YLAVDSPFLK | -666 | -675 | 0 | 0.00025015 | 10 | 402.33286 | NaN | NaN | NaN | NaN | NaN |
| Mpc2 | YSLVIIPK | -85 | -92 | 0 | 0.002168954 | 8 | 375.5644 | 408.6477 | 406.80234 | 176.43224 | NaN | 143.93408 |
| Mpc2 | TVFFWAPIMK | -40 | -49 | 0 | 0 | 10 | 556.9153 | NaN | 647.4872 | NaN | 763.56433 | NaN |
| Hspbap1 | TVGEEGEPVKPFTPEK | -17 | -32 | 1 | 0.001463897 | 16 | 485.41296 | 734.4434 | 653.7271 | 380.47546 | 526.1609 | 667.7094 |
| Cd47 | TVSWFSPNEK | -131 | -140 | 0 | 0.00025015 | 10 | 4187.519 | 4360.6177 | 4485.946 | 3774.086 | 65.98036 | 3266.7295 |
| Cd47 | VSDLINGIASLK | -88 | -99 | 0 | 0 | 12 | 219.74895 | NaN | NaN | 381.63486 | NaN | NaN |
| Rabepk | TWSTPEVTGSPPSPR | -133 | -147 | 0 | 0 | 15 | 5755.506 | NaN | 453.3226 | 2624.06 | NaN | NaN |
| Prf1 | TYQDQYNFNSDTVECR | -161 | -176 | 0 | 0 | 16 | 314.48395 | 345.2467 | 328.52597 | 373.65567 | 418.55197 | 429.19223 |
| Osgep | TYVTAPGTGFLPGDTAR | -31 | -47 | 0 | 0 | 17 | 651.7544 | 713.75494 | 756.08264 | 1216.745 | 1178.465 | 943.6438 |
| Kptn | VAEGPCPLLEDSFTR | -6 | -20 | 0 | 0.00051306 | 15 | 817.1044 | NaN | NaN | NaN | NaN | NaN |
| Serpinf2 | VAHFPFK | -287 | -293 | 0 | 0.000826806 | 7 | 1425.3007 | 99.465004 | 1170.6985 | 1323.5803 | NaN | NaN |
| Skic3 | VALVDFLDGK | -682 | -691 | 0 | 0 | 10 | 197.38373 | 377.5794 | NaN | NaN | 226.23717 | NaN |
| Tmx2 | VANAILFFR | -99 | -107 | 0 | 0 | 9 | 395.89597 | 457.1779 | 616.41864 | 396.5951 | 555.8701 | 549.0277 |
| Acat2 | VAPEEVSEVIFGHVL | -47 | -61 | 0 | 0 | 15 | 273.8466 | 258.9309 | 172.63818 | 208.00815 | 143.0732 | 335.50397 |
| Acat2 | VAPEEVSEVIFGH | -47 | -59 | 0 | 0.00272025 | 13 | 445.91837 | NaN | NaN | NaN | NaN | NaN |
| Cmtr1 | VSWFPECTTEIPDSR | -154 | -168 | 0 | 0 | 15 | 420.02966 | 368.66943 | 472.46704 | 608.2515 | 368.27408 | 364.51752 |
| Cmtr1 | VCLFKPITSR | -426 | -435 | 1 | 0 | 10 | 836.7112 | 450.85303 | 825.1823 | NaN | 934.5843 | 403.13525 |
| Dpf2 | VDDDSLGEFPVSNSR | -138 | -152 | 0 | 0 | 15 | 1306.0099 | 2053.9998 | 2087.6104 | NaN | NaN | NaN |
| Emc8 | VDFDNHLDDIR | -181 | -191 | 0 | 0 | 11 | 2859.452 | 5893.2393 | 4060.9827 | NaN | 3222.3914 | 3183.3955 |
| Rfc5 | VDFPSSVR | -290 | -297 | 0 | 0.00142999 | 8 | 1632.5894 | 1631.7725 | NaN | NaN | NaN | NaN |
| Nsd3 | VDTAPEKPR | -231 | -239 | 1 | 0 | 9 | 1031.8104 | NaN | NaN | NaN | 1229.3121 | 441.53635 |
| Timm21 | VEENPGSGQFEFR | -210 | -222 | 0 | 0 | 13 | 2379.823 | 2412.1855 | NaN | NaN | 1807.4153 | NaN |
| Fam241a | VEEPEHGASPR | -30 | -40 | 0 | 0.00025015 | 11 | 530.75433 | 611.4515 | NaN | NaN | NaN | 556.2286 |
| H2-Aa;H2-Aa | VEHWGLEEPVLK | (192);(192);(192);(169) | (203);(203);(203);(18( | 0 | 0.00025015 | 12 | 534.20233 | NaN | 398.9862 | NaN | 592.79236 | NaN |
| Shcbp1 | VEVTNVDFPSLK | -128 | -139 | 0 | 0 | 12 | 1048.7732 | NaN | NaN | NaN | NaN | NaN |
| Inpp1 | VFGEESNEFTNDLGEK | -76 | -91 | 0 | 0 | 16 | 470.56805 | 509.67258 | 555.46765 | 596.3053 | 678.2224 | 697.9155 |
| Dpagt1 | VFGPIHER | -372 | -379 | 0 | 0 | 8 | 760.3321 | 811.7331 | 857.1914 | 578.6731 | 853.97046 | 610.3258 |
| Wdr37 | VFTVGDNVVSGSDDR | -376 | -390 | 0 | 0 | 15 | 5573.738 | 6349.2236 | 6510.1846 | 2731.6135 | 3057.6016 | 2875.6494 |
| Pnpo | VFYWEPLNR | -130 | -138 | 0 | 0 | 9 | 376.48947 | 257.74216 | 363.6088 | 731.6209 | 71.20219 | 413.6175 |
| Pbrm1 | VGDCVFIK | -1160 | -1167 | 0 | 0.00111006 | 8 | 1290.4121 | NaN | NaN | NaN | NaN | NaN |
| Oxnad1 | VGGEFFFDPQPTDAPR | -154 | -169 | 0 | 0.000298805 | 16 | 2127.752 | NaN | NaN | NaN | NaN | NaN |
| Stub1 | VGHFDPVTR | -265 | -273 | 0 | 0.000550307 | 9 | 1261.5582 | NaN | NaN | 1065.2047 | 327.0618 | NaN |
| Imp3 | VGPDVVTDPAFLVTR | -138 | -152 | 0 | 0 | 15 | 1493.0535 | 147.5773 | 1748.4677 | 1403.5642 | 1930.7023 | 1800.9724 |
| Ifngr1 | VGPPGLEIR | -133 | -141 | 0 | 0.00025015 | 9 | 1592.0157 | 1523.5709 | 1642.6469 | 1543.0021 | 1429.9003 | NaN |
| Polb | VHFITDTLSK | -221 | -230 | 0 | 0 | 10 | 828.82605 | 431.81143 | NaN | 1102.8589 | 1147.2277 | 1510.9609 |
| Polb | YFTGSDIFNK | -271 | -280 | 0 | 0 | 10 | 1211.2125 | 1155.4999 | 328.5166 | NaN | NaN | NaN |
| Phpt1 | VHLAEPSGDPAK | -26 | -37 | 0 | 0 | 12 | 1187.3784 | 971.75415 | 1842.2106 | 425.13824 | NaN | 1380.4869 |

| | | | | | | | | | | | | |
|---|---|---|---|---|---|---|---|---|---|---|---|---|
| Sephs1 | VHQWLDIPEK | -211 | -220 | 0 | 0 | 10 | 2091.992 | 501.03665 | 585.159 | 957.8076 | 1378.0688 | 1058.6498 |
| Trappc2l | VHTSLDVVDEK | -32 | -42 | 0 | 0 | 11 | 1269.1161 | NaN | NaN | NaN | 1364.7522 | NaN |
| Actg1 | VIDNGSGMCK | -9 | -18 | 0 | 0 | 10 | 544.937 | 245.67511 | 306.44595 | 229.39172 | 258.79672 | 300.68112 |
| Echdc1 | VIELENWTEGK | -105 | -115 | 0 | 0 | 11 | 797.0655 | 609.75336 | 558.81573 | 868.3917 | 1321.4302 | 1131.0215 |
| Pdp1 | VIESGPDQLNDNEYTK | -368 | -383 | 0 | 0.00111006 | 16 | 799.4409 | NaN | NaN | NaN | 897.1857 | NaN |
| Herc4 | VIFVGEDAVDAGGVR | -740 | -754 | 0 | 0 | 15 | 879.81696 | 789.54193 | 534.5621 | NaN | 859.109 | 1308.8328 |
| Herc4 | VLEILHR | -596 | -602 | 0 | 0.00719662 | 7 | 501.52655 | NaN | NaN | NaN | NaN | NaN |
| Znf638 | VILDEKEEDEFNVK | -1303 | -1316 | 1 | 0 | 14 | 753.3066 | 871.68945 | NaN | 686.95605 | 724.9276 | 730.8651 |
| Tradd | VILSEAYTDPK | -26 | -36 | 0 | 0 | 11 | 438.5216 | NaN | NaN | NaN | NaN | NaN |
| Fam98b | VKDILCR | -178 | -184 | 1 | 0.003333688 | 7 | 1710.0983 | NaN | NaN | 2050.472 | NaN | NaN |
| Myo18a | VKDQEEELDEQAGSIQ | -1547 | -1562 | 1 | 0.002296977 | 16 | 1229.0303 | 1277.5161 | 955.63824 | 586.62067 | 1693.7458 | NaN |
| Myo18a | VPPPPPALR | -211 | -219 | 0 | 0.00204437 | 9 | 10509.278 | NaN | 9454.865 | NaN | 7805.818 | 9618.104 |
| Rala | VKEDENVPFLLVGNK | -114 | -128 | 1 | 0 | 15 | 325.7094 | 407.34705 | NaN | 411.98813 | 407.83734 | 417.5043 |
| Nudt16l1 | VLGLGLGGLR | -81 | -90 | 0 | 0.000591124 | 10 | 171.53651 | 235.5921 | NaN | NaN | NaN | NaN |
| Gstt2 | VLGPLIGVQVPQEK | -118 | -131 | 0 | 0 | 14 | 399.4134 | 477.13916 | 484.87613 | 449.7404 | 655.1488 | 659.68475 |
| Gstt2 | YQVADHWYPADLQAR | -78 | -92 | 0 | 0 | 15 | 882.9535 | NaN | NaN | NaN | 965.08167 | 1172.9199 |
| Utp25 | VLIVVPFR | -371 | -378 | 0 | 0.00272025 | 8 | 339.91827 | NaN | NaN | 341.17505 | 369.44128 | NaN |
| Man2a1 | VLLAPLGDDFR | -417 | -427 | 0 | 0 | 11 | 552.84015 | 107.06215 | 473.04663 | 881.31964 | 1114.248 | 1135.9084 |
| Man2a1 | YLVVYNPFEQER | -649 | -660 | 0 | 0.00111006 | 12 | 2047.2992 | NaN | 2154.626 | 649.68695 | 1880.0138 | 2920.364 |
| Srp14 | VLLESEQFLTELTR | -2 | -15 | 0 | 0 | 14 | 428.22055 | 431.6386 | 442.81256 | 584.1323 | 614.942 | 533.0499 |
| Psmg3 | VLLGQDEPLIHVF | -66 | -78 | 0 | 0 | 13 | 478.345 | 444.46487 | 425.78906 | 506.00934 | 500.54523 | 486.8821 |
| Gtf2a2 | VLNDVEFR | -75 | -82 | 0 | 0 | 8 | 2848.065 | NaN | 2240.2607 | NaN | NaN | NaN |
| Slc35c1 | VLPAVDHSIWR | -218 | -228 | 0 | 0 | 11 | 853.4426 | 1037.3549 | 998.6898 | 1324.2289 | 556.6723 | 1566.3098 |
| Rpl35 | VLTVINQTQK | -57 | -66 | 0 | 0 | 10 | 1597.4032 | 1665.5468 | 1442.889 | 1115.6526 | 1143.5743 | 1232.0664 |
| Rpl35 | YKPLDLRPK | -78 | -86 | 2 | 0.000431655 | 9 | 564.6217 | 454.19815 | 433.62003 | NaN | NaN | NaN |
| Grap | VNYGDQVQHFK | -99 | -109 | 0 | 0 | 11 | 1241.5973 | 1101.0112 | 807.34827 | NaN | 1552.1245 | NaN |
| Emc3 | VPFPLTLR | -140 | -147 | 0 | 0 | 8 | 3440.3335 | 757.67676 | 3663.482 | 3265.389 | 3722.9954 | 3812.985 |
| Lamp2 | VPLDVIFK | -141 | -148 | 0 | 0 | 8 | 920.1698 | 1079.2648 | 1141.4624 | 1145.6046 | 1391.8652 | 1337.6359 |
| Fancd2 | VPQFLFESENR | -155 | -165 | 0 | 0 | 11 | 452.8545 | 520.12555 | NaN | NaN | NaN | NaN |
| Ptpn2 | VQDTVEESSESILR | -332 | -345 | 0 | 0 | 14 | 964.75696 | NaN | 529.0528 | 1438.089 | 1566.0862 | NaN |
| Dguok | VQLEPIPGR | -121 | -129 | 0 | 0.000394361 | 9 | 293.5105 | 222.34785 | 260.04517 | NaN | NaN | NaN |
| Apoa1 | VQPYLDEFQK | -120 | -129 | 0 | 0 | 10 | 5500.295 | 5352.594 | 5730.578 | 4077.4814 | 4900.801 | 4680.253 |
| Apoa1 | VQPYLDEFQK | -120 | -129 | 0 | 0 | 10 | 5500.295 | 5352.594 | 5730.578 | 4077.4814 | 4900.801 | 4680.253 |
| Txndc12 | VRPEIINESGNPSYK | -122 | -136 | 1 | 0.00025015 | 15 | 1075.6179 | 1281.6495 | 1492.3687 | 1256.1378 | 1305.7228 | 1363.0499 |
| Efl1 | VSEPIIPFR | -685 | -693 | 0 | 0 | 9 | 393.04645 | 435.8647 | 329.83066 | 513.5897 | 442.80206 | 764.67456 |
| Alkbh5 | VSEPVLSLPVR | -240 | -250 | 0 | 0 | 11 | 483.78094 | 836.4826 | 677.7964 | 542.6939 | 617.5634 | 699.4751 |
| Supt4h1a;Supt | VSNFKPGVY | (77);(77) | (85);(85) | 1 | 0.00111006 | 9 | 1169.131 | 794.9663 | 704.81665 | 584.8367 | NaN | 554.3411 |
| Ifi35 | VSPYVSGEIQK | -211 | -221 | 0 | 0.000230707 | 11 | 594.21936 | 655.7904 | 737.4112 | 1068.585 | NaN | NaN |
| H2-Q8 | VSWPGLVEPR | -33 | -42 | 0 | 0 | 10 | 6265.981 | NaN | 6267.538 | 6462.2705 | 4748.425 | 7239.7256 |
| Cggbp1 | VTEFGGELHEDGGK | -27 | -40 | 0 | 0 | 14 | 409.771 | 496.0317 | 1608.846 | NaN | 877.89307 | 1529.1003 |
| Ngdn | VTGSLSENDPLR | -118 | -129 | 0 | 0.001392869 | 12 | 410.7992 | NaN | NaN | NaN | NaN | NaN |
| Eea1 | VTNSTELQHQLEK | -473 | -485 | 0 | 0.000431655 | 13 | 433.53998 | NaN | NaN | NaN | NaN | NaN |
| Ndufa6 | VVDLLVIK | -81 | -88 | 0 | 0.00111006 | 8 | 401.68332 | 509.52676 | 67.6965 | 569.41833 | 543.9603 | 594.6516 |
| Mettl13 | VVDNPAVKPSR | -286 | -296 | 1 | 0.000550307 | 11 | 138.91844 | NaN | NaN | NaN | NaN | NaN |
| Pcca | VVEEAPSIFLDPETR | -295 | -309 | 0 | 0 | 15 | 492.22052 | 589.495 | 480.20398 | 279.84662 | 1082.2626 | 1119.7008 |
| Polr1f | VVGELGDIYDDQGHIH | -98 | -113 | 0 | 0 | 16 | 997.02625 | 1144.3976 | NaN | NaN | 822.6804 | NaN |
| Kgd4 | VVKPHAPLIK | -14 | -23 | 1 | 0.001392869 | 10 | 325.61935 | 389.50665 | 365.1917 | NaN | NaN | NaN |
| Mpo | VVLEGGIDPILR | -511 | -522 | 0 | 0 | 12 | 1254.4609 | 1489.1233 | 1544.2041 | 994.63153 | 1384.7545 | 1239.7742 |
| Trip13 | VVLLHGPPGTGK | -174 | -185 | 0 | 0 | 12 | 575.10114 | 635.7739 | 705.8463 | NaN | NaN | NaN |
| Ppwd1 | VVPTPWLDNK | -596 | -605 | 0 | 0 | 10 | 434.21658 | 323.34668 | 544.05505 | 445.3831 | 211.89459 | 409.51965 |
| Tiprl | VVPTTDHIDTEK | -129 | -140 | 0 | 0 | 12 | 1041.0149 | 1593.6267 | 1445.9034 | 1230.3958 | 2481.719 | 1130.7242 |
| Coil | VVTDLISLIR | -37 | -46 | 0 | 0 | 10 | 237.7581 | 252.10844 | NaN | 263.1164 | 271.64908 | 327.14957 |
| Enoph1 | VVVRPGNAGLTDDEK | -226 | -240 | 1 | 0 | 15 | 3171.1868 | 3291.1147 | 3119.7693 | 2747.947 | 2649.0247 | 2640.4202 |
| Lpcat3 | VWLFETTPR | -321 | -329 | 0 | 0.000230707 | 9 | 1579.3097 | 1540.9991 | 1593.3824 | 707.12976 | 1685.2379 | 1376.2184 |
| Ankrd13a | VWNNDYEQLEK | -17 | -27 | 0 | 0 | 11 | 3044.377 | 2176.7693 | 2597.9954 | NaN | 2879.274 | NaN |
| Brcc3;C6.1al | VWPSHVDVR | (129);(129) | (137);(137) | 0 | 0 | 9 | 494.7256 | 1194.8844 | 1338.9438 | 567.047 | NaN | 373.5555 |
| Mrpl13 | VWTPPDDFR | -169 | -177 | 0 | 0.000431655 | 9 | 1049.5051 | 761.54083 | NaN | 2085.163 | NaN | NaN |
| Ndufa10 | YNQGYIR | -174 | -180 | 0 | 0.00025015 | 7 | 1126.02 | 1273.5005 | NaN | 1097.5886 | NaN | 1888.0906 |
| Ndufa10 | VYDSWEAEDPTK | -257 | -268 | 0 | 0 | 12 | 679.9221 | 572.0082 | 509.14688 | 425.90433 | NaN | NaN |
| Mrpl28 | VYVEELVQR | -229 | -237 | 0 | 0.002534621 | 9 | 1581.1108 | NaN | NaN | NaN | NaN | NaN |
| Eif2b5 | WCHPTSPNVVR | -107 | -117 | 0 | 0.000394361 | 11 | 833.5741 | 850.51764 | NaN | 1052.6805 | 1187.1017 | NaN |
| Clu | WEECKPCLK | -109 | -117 | 1 | 0 | 9 | 393.4535 | 716.2782 | 217.38298 | 737.57043 | 254.88002 | 849.5029 |
| Gclm | WEELENLVQSK | -147 | -157 | 0 | 0 | 11 | 248.98784 | 458.33395 | NaN | NaN | 435.9795 | NaN |
| Chia | WEGYTGENSPLYK | -218 | -230 | 0 | 0.00078661 | 13 | 2057.5352 | NaN | 1563.2231 | NaN | NaN | NaN |
| Mrpl9 | WELNPEIVAR | -179 | -188 | 0 | 0 | 10 | 293.7179 | 1103.5315 | 409.8551 | 822.55237 | 837.4739 | 413.66156 |
| Cdc16 | WFLDALEK | -424 | -431 | 0 | 0.00025015 | 8 | 442.00403 | 499.7819 | NaN | 439.70316 | 762.1776 | 611.75616 |
| Wls | WLAPWGPNK | -62 | -70 | 0 | 0.000298805 | 9 | 348.0215 | 359.37512 | NaN | NaN | 292.0551 | NaN |
| Spcs1 | WLPVQDLGTEDKK | -136 | -148 | 1 | 0 | 13 | 919.00684 | 1176.5049 | 957.8541 | 1437.5677 | 1834.9491 | 1539.3778 |
| Slc33a1 | WLVDPLTVK | -470 | -478 | 0 | 0 | 9 | 893.96173 | 579.03595 | 762.63556 | NaN | 807.17126 | NaN |
| Acad10 | WLVPLLEGR | -779 | -787 | 0 | 0.000471409 | 9 | 277.59088 | 329.27658 | NaN | NaN | 332.6686 | 461.66028 |
| Ndufb11 | WQEDPEPEDENVYAK | -53 | -67 | 0 | 0 | 15 | 663.09314 | 713.16455 | 772.91693 | 897.9877 | 1038.5592 | 1024.2396 |
| Ube2g2 | WSPVQSVEK | -110 | -118 | 0 | 0 | 9 | 806.29456 | 993.00555 | NaN | 577.4894 | 699.5749 | 718.4746 |
| Nkg7 | WSQTPFSQVQTF | -117 | -128 | 0 | 0.000230707 | 12 | 792.42163 | 973.9056 | 1408.6115 | 1049.3586 | 1412.0983 | 1295.9703 |
| Arl6ip1 | WTTEQQQR | -97 | -104 | 0 | 0 | 8 | 4786.143 | 454.75266 | 530.9341 | 960.68884 | 3877.2263 | 4208.632 |
| Cxcr3 | WVFGPGLCK | -116 | -124 | 0 | 0 | 9 | 580.4293 | 628.09247 | 401.44824 | NaN | NaN | NaN |
| Galnt7 | WYAPLVAPISK | -310 | -320 | 0 | 0 | 11 | 10257.811 | 10636.655 | 9400.852 | 8809.979 | 8917.533 | 8789.081 |
| Eef1akmt1 | WYSQDTALR | -54 | -62 | 0 | 0.000947459 | 9 | 417.4646 | 1370.4965 | NaN | NaN | 1006.20917 | NaN |
| Pclaf | YAGGNPVCVRPTPK | -46 | -59 | 1 | 0 | 14 | 2936.8796 | 1847.4912 | 3084.845 | 3102.0195 | 3020.8054 | 1174.0481 |

| | | | | | | | | | | | | |
|---|---|---|---|---|---|---|---|---|---|---|---|---|
| Fgb | YCGLPGEYWLGNDK | -305 | -318 | 0 | 0.000394361 | 14 | 781.8061 | 778.7765 | NaN | NaN | 1291.8221 | 1300.2468 |
| Wdr74 | YCPGGEGTFR | -82 | -91 | 0 | 0 | 10 | 523.9337 | 703.62976 | 709.83044 | NaN | NaN | NaN |
| Timm44 | YDESDNVLIR | -265 | -274 | 0 | 0 | 10 | 792.61163 | 898.3978 | 945.4151 | 463.38565 | 1177.9712 | 705.6006 |
| Cnn3 | YDQQAEEDLR | -24 | -33 | 0 | 0 | 10 | 1553.0643 | 1637.2305 | 1314.9727 | NaN | NaN | NaN |
| H2-Ab1 | YDSDVGEHR | -67 | -75 | 0 | 0 | 9 | 743.2418 | 482.2209 | 833.48126 | 416.76453 | 780.71063 | 701.9066 |
| Sdhd | YFNYHDVGICR | -141 | -151 | 0 | 0 | 11 | 4913.1265 | 386.34686 | 1590.0563 | 12416.312 | 1598.5956 | 2164.3784 |
| Sdhd | YFNYHDVGICR | -141 | -151 | 0 | 0 | 11 | 4913.1265 | 386.34686 | 1590.0563 | 12416.312 | 1598.5956 | 2164.3784 |
| Sdhd | YVHGDTLPK | -114 | -122 | 0 | 0.000431655 | 9 | 2058.6284 | 1498.7802 | 2046.7539 | 1460.3779 | 1618.8286 | 1825.9518 |
| Cd44 | YGFIEGNVVIPR | -81 | -92 | 0 | 0 | 12 | 383.8414 | 735.3184 | 698.6242 | 287.1955 | 501.47308 | NaN |
| Rad51 | YGLSGSDVLDNVAYAR | -178 | -193 | 0 | 0 | 16 | 886.9825 | 1460.421 | 1214.3448 | 1826.9868 | 2147.6577 | 1497.614 |
| Glb1 | YHLGNDVILF | -210 | -219 | 0 | 0.001055699 | 10 | 382.42175 | 366.43024 | 536.2117 | 607.86926 | 570.5531 | 603.95715 |
| Pbk | YIGTEPWKPK | -205 | -214 | 1 | 0.00051306 | 10 | 211.7098 | 310.62195 | NaN | NaN | 1020.8247 | NaN |
| Ctss | YIQLPFGDEDALK | -236 | -248 | 0 | 0 | 13 | 861.84827 | 1711.3438 | 1509.6287 | 1401.9431 | 2039.9185 | 1772.79 |
| Timm10 | YLDIHER | -58 | -64 | 0 | 0 | 7 | 2411.3257 | 2572.8708 | 624.0477 | 2040.4484 | 827.7904 | 482.43958 |
| Dctn3 | YLDPEYIDR | -67 | -75 | 0 | 0 | 9 | 614.0998 | 895.69257 | 890.91583 | 639.0777 | 860.2193 | 469.50742 |
| Hells | YLIVDEGHR | -333 | -341 | 0 | 0 | 9 | 1186.0558 | NaN | 1411.2141 | NaN | NaN | 1990.3721 |
| Pex14 | YLLPLILGGR | -130 | -139 | 0 | 0 | 10 | 237.65706 | NaN | 254.22195 | 283.44116 | 264.3999 | 360.3666 |
| Rsu1 | YLSDNDFEILPPDIGK | -140 | -155 | 0 | 0 | 16 | 746.0561 | NaN | NaN | 528.4244 | 1182.7058 | 708.8681 |
| Cpsf4 | YLVGFCPEGPSCK | -119 | -131 | 0 | 0 | 13 | 167.23631 | NaN | 509.57355 | NaN | 483.9894 | NaN |
| Exosc2 | YNGEVGDIVVGR | -76 | -87 | 0 | 0 | 12 | 3874.6765 | 3787.3606 | 3788.8062 | 4698.147 | 1285.3656 | NaN |
| Higd1a | YQEFWANPK | -84 | -92 | 0 | 0 | 9 | 1279.8772 | 1570.382 | 1726.1458 | 1539.3134 | 1822.6272 | 1660.7401 |
| Irgm2 | YQQDDWVLAR | -330 | -339 | 0 | 0 | 10 | 2505.9734 | 344.01578 | NaN | NaN | 187.54222 | NaN |
| Nusap1 | YSDLQNLAK | -14 | -22 | 0 | 0.000347792 | 9 | 370.7753 | NaN | NaN | NaN | NaN | NaN |
| Ctdspl2 | YSEVDDSLPSGGEKPSK | -27 | -43 | 1 | 0 | 17 | 565.87976 | 625.4904 | NaN | 386.66702 | 533.9424 | 506.43082 |
| Capg | YSPNTQVEILPQGR | -326 | -339 | 0 | 0.00025015 | 14 | 360.5011 | NaN | NaN | NaN | NaN | NaN |
| Wtap | YTDLNSNDVTGLR | -44 | -56 | 0 | 0 | 13 | 154.18849 | 412.81198 | NaN | 897.9995 | 439.74426 | NaN |
| Rexo4 | YVKPTEPVTDYR | -289 | -300 | 1 | 0 | 12 | 821.8683 | 864.9695 | 998.57367 | 562.7596 | 390.7558 | 618.74426 |
| Lsm12 | YVSEVEIINDR | -62 | -72 | 0 | 0 | 11 | 2696.5754 | 2245.9263 | 2376.5298 | 453.4322 | 199.27808 | NaN |
| Enpp4 | YWEEPDVSGHK | -187 | -197 | 0 | 0 | 11 | 949.43176 | NaN | NaN | NaN | 970.14264 | 719.60516 |
| Nop10 | YYLNEQGDR | -5 | -13 | 0 | 0 | 9 | 1302.734 | 1198.9695 | 1171.9404 | 1409.0914 | 1067.0306 | 1139.6027 |
| Hacd2 | YYSIERPLK | -74 | -82 | 1 | 0 | 9 | 31665.836 | 26985.164 | 31975.191 | 27077.93 | 30331.115 | 38144.973 |
| Tbcel | YYVDVPQEEVPFR | -307 | -319 | 0 | 0 | 13 | 3182.939 | NaN | 2389.311 | 2854.7273 | 3986.1577 | 3477.5417 |
| Naca | EESEEEEVDETGVEVK | (2136);(164) | (2151);(179) | 0 | 0.000947459 | 16 | NaN | 943.25885 | 830.73126 | 2266.335 | 1366.8069 | 481.6342 |
| Naca | EIDEEPVSK | (2030);(58) | (2038);(66) | 0 | 0.006677519 | 9 | NaN | 1205.8391 | NaN | NaN | NaN | NaN |
| Mcm6 | AAAEPGAGSQHPEVR | -5 | -19 | 0 | 0 | 15 | NaN | 758.6723 | 709.04034 | 772.284 | 765.2646 | 649.5759 |
| Mcm6 | QFAEELIRPER | -48 | -58 | 1 | 0 | 11 | NaN | 787.40295 | 1675.2 | 823.8362 | 2894.1982 | 2232.23 |
| Mcm6 | QFAEELIRPER | -48 | -58 | 1 | 0 | 11 | NaN | 787.40295 | 1675.2 | 823.8362 | 2894.1982 | 2232.23 |
| Mcm6 | VDECNEVTDYAIAR | -537 | -550 | 0 | 0 | 14 | NaN | 390.39868 | 612.8251 | 299.26273 | 601.7251 | 494.98477 |
| Mcm6 | ANPVSGHYDR | -503 | -512 | 0 | 0.000471409 | 10 | NaN | 108.62066 | 3730.2654 | 1593.6272 | NaN | 2603.2605 |
| Mcm6 | LGFAEYCR | -715 | -722 | 0 | 0 | 8 | NaN | 774.58923 | NaN | 934.6257 | 851.8143 | NaN |
| Mcm6 | CDFTGALIVVPDVSK | -242 | -256 | 0 | 0 | 15 | NaN | 451.8961 | NaN | NaN | 332.61813 | 287.45847 |
| Mcm6 | YLQFAEELIRPER | -46 | -58 | 1 | 0.004957095 | 13 | NaN | 1375.925 | 1201.8162 | NaN | NaN | NaN |
| Mcm6 | VETPDVNLDQEEEIQ | -659 | -673 | 0 | 0.00078661 | 15 | NaN | 2005.7555 | NaN | NaN | NaN | NaN |
| Mcm6 | VVPYLCR | -86 | -92 | 0 | 0.005270854 | 7 | NaN | 780.3054 | NaN | NaN | NaN | NaN |
| H1-3 | PAAPAAPAPVEK | -6 | -17 | 0 | 0.00078661 | 12 | NaN | 510.8079 | 510.30728 | 570.08295 | 606.22656 | 593.2275 |
| Mcm3 | ASEDESDLEDEEEKSQEDTEQKR | -667 | -689 | 2 | 0 | 23 | NaN | 270.254 | 288.98138 | 105.88504 | 189.27927 | 360.84503 |
| Mcm3 | ELENGSHIR | -328 | -336 | 0 | 0 | 9 | NaN | 1156.7749 | 1528.8445 | NaN | 1842.5817 | NaN |
| Mcm3 | ISDNQYR | -40 | -46 | 0 | 0.005605646 | 7 | NaN | 407.82196 | NaN | NaN | 678.5408 | 137.64793 |
| Mcm3 | IQVVGTYR | -235 | -242 | 0 | 0.004350979 | 8 | NaN | 1169.7578 | 1184.054 | NaN | NaN | NaN |
| Mcm3 | ASEDESDLEDEEEKSQ | -667 | -682 | 1 | 0 | 16 | NaN | 454.57275 | 378.92688 | NaN | NaN | NaN |
| Mcm3 | LIVSVNDLR | -47 | -55 | 0 | 0.002910424 | 9 | NaN | 281.78522 | 378.78174 | NaN | NaN | NaN |
| Lcp1 | EGICAIGGTSEQSSVGTQ | -98 | -115 | 0 | 0 | 18 | NaN | 774.2951 | 745.0781 | 749.66 | 921.72156 | 941.9236 |
| Lcp1 | LVGIAGQDLNEGNR | -475 | -488 | 0 | 0 | 14 | NaN | 1386.2795 | NaN | 2114.6665 | 2550.23 | 2474.125 |
| Lcp1 | PEDLVEVNPK | -601 | -610 | 0 | 0.000550307 | 10 | NaN | 1104.4475 | NaN | 2105.3796 | 1769.4082 | 3228.4426 |
| Lcp1 | VVNIGAEDLK | -208 | -217 | 0 | 0 | 10 | NaN | 1347.1759 | NaN | 1009.37305 | NaN | 671.4506 |
| Psmb8 | AADCQYWER | -121 | -129 | 0 | 0 | 9 | NaN | 1034.1746 | 988.51306 | NaN | 1079.2516 | 1028.2712 |
| Psmb8 | FQHGVIVAVDSR | -80 | -91 | 0 | 0.000230707 | 12 | NaN | 220.86673 | 732.42896 | NaN | 771.8756 | 684.2187 |
| Psmb8 | DCQYWER | -123 | -129 | 0 | 0.001291103 | 7 | NaN | 713.9459 | NaN | 610.8906 | NaN | NaN |
| Apex1 | VSYGIGEEEHDQEGR | -141 | -155 | 0 | 0 | 15 | NaN | 1762.3945 | 1979.0596 | 1904.3899 | 2145.4539 | 2069.7275 |
| Apex1 | YGIGEEEHDQEGR | -143 | -155 | 0 | 0 | 13 | NaN | 639.38116 | 587.4059 | 508.15274 | 627.68243 | 707.72925 |
| Apex1 | EKEAAGEGPVLYEDPPDQK | -33 | -51 | 1 | 0.000471409 | 19 | NaN | 428.74832 | 399.9676 | 570.2447 | 645.4839 | 587.1205 |
| Apex1 | GPVLYEDPPDQK | -40 | -51 | 0 | 0.001264841 | 12 | NaN | 2313.2603 | 2032.577 | 2030.7712 | NaN | 2231.3513 |
| Apex1 | GDLNVAHEEIDLR | -208 | -220 | 0 | 0 | 13 | NaN | 1385.1248 | NaN | 1805.112 | 1868.1411 | 921.48584 |
| Apex1 | GIGEEEHDQEGR | -144 | -155 | 0 | 0 | 12 | NaN | 452.55157 | 460.00937 | NaN | 520.3055 | 590.0997 |
| Apex1 | AADDGEEPKSEPETK | -9 | -23 | 1 | 0.00025015 | 15 | NaN | 279.38086 | NaN | NaN | 376.92084 | NaN |
| Nap1l1 | TGCQIDWK | -256 | -263 | 0 | 0 | 8 | NaN | 191.7792 | 467.83713 | NaN | 480.08853 | 543.14764 |
| Nap1l1 | GCTGCQIDWK | -254 | -263 | 0 | 0.000431655 | 10 | NaN | 340.1229 | NaN | NaN | NaN | 716.61755 |
| Vdac3 | AADFQLHTH | -175 | -183 | 0 | 0.00111006 | 9 | NaN | 594.5946 | 676.47076 | NaN | NaN | NaN |
| Hspa9 | SSGGLSKDDIENMVK | -549 | -563 | 1 | 0 | 15 | NaN | 1158.0297 | 1205.3643 | 694.5397 | 2070.9805 | 2254.3657 |
| Hspa9 | DIGEVILVGGMTR | -379 | -391 | 0 | 0 | 13 | NaN | 1047.0482 | 1046.7297 | 1142.2855 | 1274.053 | 1250.4812 |
| Hspa9 | QFEGIVTDLIK | -350 | -360 | 0 | 0 | 11 | NaN | 1135.4763 | 1316.219 | 970.9766 | 208.68594 | 1063.3657 |
| Hspa9 | LGGEDFDQALLR | -273 | -284 | 0 | 0 | 12 | NaN | 1059.7024 | 704.46375 | 1391.4211 | 1477.7596 | 1287.2355 |
| Hspa9 | YAEEDRR | -568 | -574 | 1 | 0.006677519 | 7 | NaN | 749.9779 | 651.2336 | 511.00687 | 715.4958 | 699.78815 |
| Hspa9 | RYDDPEVQKD | -127 | -136 | 2 | 0.001055699 | 10 | NaN | 3665.5854 | 3876.3086 | 1410.9825 | 2342.4485 | NaN |
| Hspa9 | VEAVNMAEGIIHDTETK | -579 | -595 | 0 | 0 | 17 | NaN | 815.2607 | 781.39856 | NaN | NaN | 874.91516 |
| Hspa9 | EGIIHDTETK | -586 | -595 | 0 | 0 | 10 | NaN | 917.7954 | NaN | 2879.3928 | 2836.959 | NaN |
| Hspa9 | NGDTFLGGEDFDQALLR | -268 | -284 | 0 | 0 | 17 | NaN | 575.99054 | NaN | NaN | 893.98395 | 671.016 |
| Hspa9 | NMAEGIIHDTETK | -583 | -595 | 0 | 0 | 13 | NaN | 1465.5425 | 1267.6085 | NaN | NaN | NaN |

| | | | | | | | | | | | | |
|---|---|---|---|---|---|---|---|---|---|---|---|---|
| Hspa9 | VLENAEGAR | -77 | -85 | 0 | 0 | 9 | NaN | 410.9127 | NaN | 1197.5713 | NaN | NaN |
| Stat5b | AADGYVKPQIK | -695 | -705 | 1 | 0.001023236 | 11 | NaN | 877.0741 | NaN | 472.3896 | NaN | NaN |
| Spr | AADLGTEAGVQR | -68 | -79 | 0 | 0.001392869 | 12 | NaN | 575.3498 | NaN | NaN | NaN | NaN |
| Ybx1 | KENQGDETQGQQPPQR | -262 | -277 | 1 | 0 | 16 | NaN | 401.4797 | 526.92993 | 355.85657 | 529.01227 | 606.93585 |
| Ybx1 | RPENPKPQDGK | -289 | -299 | 2 | 0.001868731 | 11 | NaN | 413.5733 | 463.7336 | 307.1235 | NaN | 345.1091 |
| Ybx1 | EDGNEEDKENQGDETQGQQPPQ | -255 | -276 | 1 | 0.000471409 | 22 | NaN | 275.94415 | NaN | NaN | 268.05832 | NaN |
| Stxbp2 | KADTPSLGEGPEK | -213 | -225 | 1 | 0.001392869 | 13 | NaN | 1605.802 | NaN | NaN | 1784.0432 | NaN |
| Stxbp2 | KVTELLK | -302 | -308 | 1 | 0.00025015 | 7 | NaN | 270.03833 | NaN | NaN | NaN | NaN |
| Pkm | LNFSHGTHEYHAETIK | -74 | -89 | 0 | 0 | 16 | NaN | 636.0616 | 812.20795 | 808.4943 | 191.13763 | 1163.1514 |
| Pkm | SGETAKGDYPLEAVR | -362 | -376 | 1 | 0 | 15 | NaN | 572.38007 | 753.02905 | 897.8811 | 808.7068 | 755.93524 |
| Pkm | TESFASDPILYR | -95 | -106 | 0 | 0 | 12 | NaN | 6118.4453 | 6777.011 | 3704.122 | 4343.6084 | 4174.6777 |
| Pkm | GTAEVELKK | -128 | -136 | 1 | 0 | 9 | NaN | 740.3289 | 869.84796 | 363.30658 | 714.83405 | 858.81726 |
| Pkm | RLAPITSDPTEAAAVG | -400 | -415 | 1 | 0.000712251 | 16 | NaN | 589.37683 | 621.67114 | 557.785 | 967.483 | 462.84756 |
| Pkm | WAEDVDLR | -482 | -489 | 0 | 0 | 8 | NaN | 474.78668 | NaN | 6352.3833 | 901.80066 | 185.74011 |
| Pkm | EATESFASDPILYRPVA | -93 | -109 | 1 | 0.000712251 | 17 | NaN | 363.95996 | 724.66016 | NaN | 428.6389 | 434.52216 |
| Pdcd6ip | KFTDLFEK | -350 | -357 | 1 | 0.00025015 | 8 | NaN | 227.12622 | 489.6466 | 935.9308 | 907.9627 | 876.7055 |
| Pdcd6ip | DNDFIYHDR | -314 | -322 | 0 | 0 | 9 | NaN | 946.7303 | NaN | 913.87463 | 731.6936 | NaN |
| Pdcd6ip | SALGRPLDK | -52 | -60 | 1 | 0.002880184 | 9 | NaN | 3509.6677 | NaN | NaN | 3958.7705 | NaN |
| Pdcd6ip | DEEEATDNDLR | -459 | -469 | 0 | 0.000627943 | 11 | NaN | 1130.3395 | NaN | NaN | NaN | 1763.8241 |
| Pdcd6ip | ADYFGDAFK | -221 | -229 | 0 | 0.001811524 | 9 | NaN | 406.1543 | 337.0152 | NaN | NaN | NaN |
| Cct2 | AIEHADFAGVER | -311 | -322 | 0 | 0 | 12 | NaN | 3659.7385 | 3994.337 | 4185.6206 | 4476.22 | 4419.4937 |
| Cct2 | NAGYDSADLVAQLR | -453 | -466 | 0 | 0 | 14 | NaN | 2453.5876 | 2679.3489 | 671.6307 | 4697.4907 | 3162.3804 |
| Cct2 | AAEAAEVILR | -507 | -516 | 0 | 0 | 10 | NaN | 891.39154 | 222.66489 | 899.85876 | 790.09326 | 191.6908 |
| Cct2 | IEHADFAGVER | -312 | -322 | 0 | 0 | 11 | NaN | 214.63763 | 1957.177 | 1789.341 | 1834.1049 | 1227.9707 |
| Cct2 | IGVNQPK | -224 | -230 | 0 | 0.005270854 | 7 | NaN | 461.3483 | 455.99854 | 778.2373 | NaN | 334.75223 |
| Cct2 | SLAPVNIFK | -5 | -13 | 0 | 0.00204437 | 9 | NaN | 7427.494 | 5640.1514 | NaN | NaN | NaN |
| Cct2 | VAEIEHAEK | -264 | -272 | 0 | 0.001811524 | 9 | NaN | 2852.0884 | NaN | NaN | NaN | NaN |
| Cct2 | SYLDEGFLLDKK | -212 | -223 | 1 | 0.004824802 | 12 | NaN | 2035.0681 | NaN | NaN | NaN | NaN |
| Dhx9 | RFESILPR | -476 | -483 | 1 | 0.00122419 | 8 | NaN | 823.4513 | 725.2476 | 539.9301 | 820.14276 | 830.2193 |
| Dhx9 | DFVNYLVR | -64 | -71 | 0 | 0.001904399 | 8 | NaN | 789.8677 | NaN | NaN | NaN | NaN |
| Hspa4 | AFSDPFVEAEK | -74 | -84 | 0 | 0.002329279 | 11 | NaN | 987.93317 | 718.4112 | 999.7566 | 860.58856 | NaN |
| Hspa4 | KQDLPALEEKPR | -185 | -196 | 2 | 0.000230707 | 12 | NaN | 602.90063 | NaN | 996.7087 | 818.23456 | 839.14813 |
| Hspa4 | HAEQNGPVDGQGDNPGSQAAEHGADTAVPSDGD | -800 | -833 | 0 | 0 | 34 | NaN | 1315.8455 | NaN | NaN | NaN | NaN |
| Hspa4 | VDQEEPHTEEQQQPQTPAENK | -524 | -545 | 0 | 0.00025015 | 22 | NaN | 585.25476 | 668.78186 | NaN | NaN | NaN |
| Stat1 | LAQWLEK | -34 | -40 | 0 | 0.001055699 | 7 | NaN | 1442.2506 | 159.24771 | 126.53499 | 1466.3326 | 1206.725 |
| Stat1 | TLINDELVEWK | -230 | -240 | 0 | 0 | 11 | NaN | 577.2232 | 809.9231 | 1093.6731 | 1361.4055 | 1289.8431 |
| Stat1 | KILENAQR | -114 | -121 | 1 | 0 | 8 | NaN | 476.06305 | 554.0178 | 912.23987 | 463.93774 | 510.90204 |
| Stat1 | LKDQQPGTFLLR | -591 | -602 | 1 | 0 | 12 | NaN | 668.9421 | 708.57135 | 561.04584 | NaN | NaN |
| Stat1 | QNTLINDELVEWK | -228 | -240 | 0 | 0 | 13 | NaN | 1318.2065 | NaN | NaN | 1499.0573 | 1584.9073 |
| Stat1 | FDKDVNEK | -364 | -371 | 1 | 0.001941082 | 8 | NaN | 619.0682 | 774.10754 | 618.17145 | NaN | NaN |
| Plrg1 | LWDLVAGK | -355 | -362 | 0 | 0 | 8 | NaN | 409.31805 | NaN | 468.5418 | 450.04187 | 487.75916 |
| Plrg1 | IITGSHDTTIR | -344 | -354 | 0 | 0.00201047 | 11 | NaN | 499.37793 | 540.64185 | NaN | 1000.6859 | NaN |
| Plrg1 | THPVSWKPEIIK | -498 | -509 | 1 | 0.003461185 | 12 | NaN | 749.9189 | NaN | NaN | NaN | NaN |
| Nup93 | FDEYHSGHIDR | -699 | -709 | 0 | 0 | 11 | NaN | 152.70663 | 207.57501 | 785.898 | 455.89786 | 385.6423 |
| Ganab | GPSVFDVFR | -369 | -377 | 0 | 0 | 9 | NaN | 549.784 | 592.2969 | 539.65814 | 630.6372 | 571.97253 |
| Ganab | KDAVHYGGWEHR | -558 | -569 | 1 | 0 | 12 | NaN | 512.4128 | 488.5693 | 568.54395 | 300.05588 | 576.6507 |
| Ganab | DDNSVELTVAEGPYK | -134 | -148 | 0 | 0 | 15 | NaN | 362.3257 | 775.3826 | 309.9631 | 472.10767 | 454.97943 |
| Ganab | SGGIERPFVL | -591 | -600 | 1 | 0 | 10 | NaN | 1350.179 | 350.6341 | 1226.9396 | 1274.7137 | NaN |
| Ganab | WNYRDEADVL | -401 | -410 | 1 | 0.000712251 | 10 | NaN | 810.7057 | 868.80963 | NaN | 823.9991 | 730.42596 |
| Ganab | AAETWVDISSNTAGK | -318 | -332 | 0 | 0 | 15 | NaN | 1604.7772 | NaN | 1508.2495 | NaN | 1365.8274 |
| Ganab | AEWDHLK | -621 | -627 | 0 | 0.003620005 | 7 | NaN | 2158.5286 | NaN | NaN | NaN | NaN |
| Snrnp200 | FNDPHVK | -1884 | -1890 | 0 | 0 | 7 | NaN | 1116.3951 | 1116.07 | 938.1875 | 1349.6589 | 1101.9934 |
| Snrnp200 | DTLGLFLR | -746 | -753 | 0 | 0 | 8 | NaN | 892.313 | 737.1337 | 766.5718 | 761.5006 | 496.0665 |
| Snrnp200 | FYDDAIVSQK | -275 | -284 | 0 | 0.001392869 | 10 | NaN | 3811.035 | 804.6743 | 2811.1963 | NaN | 807.83527 |
| Snrnp200 | KADEVLEIL | -285 | -293 | 1 | 0.00025015 | 9 | NaN | 591.9187 | NaN | 581.072 | 567.7541 | 607.09717 |
| Snrnp200 | LETDENLLLCAPTGAGK | -493 | -509 | 0 | 0.000627943 | 17 | NaN | 218.84297 | NaN | 372.58826 | 878.6576 | 776.868 |
| Snrnp200 | HLSDHLSELVEQ | -1780 | -1791 | 0 | 0.00149795 | 12 | NaN | 7937.065 | 5669.9336 | NaN | 1766.1229 | NaN |
| Snrnp200 | VVLLTGETSTDLK | -1405 | -1417 | 0 | 0.007819401 | 13 | NaN | 2789.4277 | NaN | 168.97237 | 2038.0311 | NaN |
| Snrnp200 | AELQSDTEEILSK | -1906 | -1918 | 0 | 0 | 13 | NaN | 1345.9878 | NaN | NaN | 1138.9309 | 1388.0339 |
| Snrnp200 | AIFEIVLNR | -1094 | -1102 | 0 | 0 | 9 | NaN | 2534.8281 | NaN | NaN | 2452.801 | NaN |
| Snrnp200 | RLVEQLFSS | -1630 | -1638 | 1 | 0.003745449 | 9 | NaN | 1848.3806 | 1866.6786 | NaN | NaN | NaN |
| Tln1 | EQGVEEHETLLLR | -182 | -194 | 0 | 0 | 13 | NaN | 448.50253 | 474.17038 | 443.1938 | 446.37192 | 312.0543 |
| Tln1 | SNTSPEELGPLANQL | -1875 | -1889 | 0 | 0.000230707 | 15 | NaN | 2080.1184 | 1297.3019 | NaN | 3125.3235 | 3529.2686 |
| Tln1 | AVAAENEEIGAHIK | -1904 | -1917 | 0 | 0 | 14 | NaN | 806.6471 | NaN | 632.8063 | 807.70844 | 829.8883 |
| Tln1 | TLAESALQLLYTAK | -1767 | -1780 | 0 | 0 | 14 | NaN | 571.28534 | NaN | 337.20627 | 902.8628 | 890.2921 |
| Tln1 | EADESLNFEEQILEAAK | -2334 | -2350 | 0 | 0 | 17 | NaN | 282.97693 | 467.1896 | 526.162 | NaN | 655.21136 |
| Tln1 | CVSCLPGQR | -1199 | -1207 | 0 | 0 | 9 | NaN | 867.67883 | 648.2208 | NaN | NaN | 227.31975 |
| Tln1 | DELNWLDHGR | -169 | -178 | 0 | 0.00025015 | 10 | NaN | 1054.1194 | 1027.1368 | NaN | NaN | NaN |
| Tln1 | ASQAATEDGQLLR | -753 | -765 | 0 | 0.009688493 | 13 | NaN | 808.7004 | 618.29974 | NaN | NaN | NaN |
| Tln1 | SAQPASAEPR | -625 | -634 | 0 | 0.008653659 | 10 | NaN | 1034.2454 | NaN | NaN | NaN | NaN |
| Tln1 | ALGDLISATK | -2090 | -2099 | 0 | 0.001092529 | 10 | NaN | 288.93674 | NaN | NaN | NaN | NaN |
| Dld | HAHPTLSEAFR | -485 | -495 | 0 | 0 | 11 | NaN | 1121.9943 | 990.4655 | 579.6333 | 579.9882 | 1020.1621 |
| Dld | NETLGGTCLNVGCIPSK | -73 | -89 | 0 | 0 | 17 | NaN | 419.0541 | 532.03845 | 1252.6776 | 1344.649 | 1391.3385 |
| Dld | KADGSTQVIDTK | -166 | -177 | 1 | 0.000230707 | 12 | NaN | 670.9657 | 629.00275 | NaN | 575.1279 | 580.2529 |
| Srm | IANLPLCSHPNPR | -83 | -95 | 0 | 0 | 13 | NaN | 2147.8757 | 2431.4414 | 578.7724 | 1914.4404 | 2017.4857 |
| Srm | HVGDGFEFMK | -152 | -161 | 0 | 0.000431655 | 10 | NaN | 683.7807 | 715.6417 | 165.60652 | 756.0954 | 679.1095 |
| Srm | ETCSLWPGQAL | -23 | -33 | 0 | 0.00078661 | 11 | NaN | 86.02806 | 233.4056 | 192.23169 | 336.65732 | 242.7098 |
| Srm | KVLIIGGGDGGVLR | -96 | -109 | 1 | 0.000431655 | 14 | NaN | 752.06067 | 616.6064 | NaN | 612.7729 | NaN |

| | | | | | | | | | | | | |
|---|---|---|---|---|---|---|---|---|---|---|---|---|
| Srm | WLHLDLIK | -211 | -218 | 0 | 0 | 8 | NaN | 593.9981 | NaN | 514.4305 | NaN | 495.2185 |
| Srm | SVVQCEIDEDVIEVSK | -119 | -134 | 0 | 0.005112649 | 16 | NaN | 1129.7065 | NaN | NaN | NaN | NaN |
| Dnajc2 | NWSEDDLQLLIK | -453 | -464 | 0 | 0.00025015 | 12 | NaN | 1867.7638 | 1824.3678 | NaN | 1986.7529 | NaN |
| Dnajc2 | SVCQVEPVGR | -24 | -33 | 0 | 0.003461185 | 10 | NaN | 802.6044 | NaN | NaN | NaN | 658.31506 |
| Gart | GGGLLENIPR | -683 | -692 | 0 | 0 | 10 | NaN | 2995.7778 | 3396.3704 | NaN | 3055.8352 | 2619.1072 |
| Gart | VLEFNCR | -286 | -292 | 0 | 0.000947459 | 7 | NaN | 936.8165 | 937.2979 | 858.31726 | 832.60767 | NaN |
| Gart | AFTNPEDACSFIT | -126 | -138 | 0 | 0.003521747 | 13 | NaN | 426.91 | 489.43173 | NaN | NaN | NaN |
| Nolc1 | QNETADEATTPQAK | -591 | -604 | 0 | 0.00025015 | 14 | NaN | 1163.5417 | 804.7223 | NaN | NaN | 1602.2684 |
| Nolc1 | VKLETPNTFPK | -606 | -616 | 1 | 0.000627943 | 11 | NaN | 604.1112 | 647.1931 | NaN | NaN | NaN |
| Nolc1 | KAAGGAVSTPAPGK | -496 | -509 | 1 | 0.002629273 | 14 | NaN | 413.59537 | NaN | NaN | NaN | NaN |
| Uqcrc1 | SRTDLTDYLNR | -212 | -222 | 1 | 0.000431655 | 11 | NaN | 851.0497 | NaN | 146.3264 | 675.7592 | 730.41144 |
| Uqcrc1 | RIPLAEWESR | -423 | -432 | 1 | 0.000627943 | 10 | NaN | 7952.825 | NaN | NaN | NaN | NaN |
| Ilf2 | RNQDLAPNSAEQA | -60 | -72 | 1 | 0.00025015 | 13 | NaN | 1361.9891 | NaN | NaN | NaN | NaN |
| Rrbp1 | AAGPLESSGKEEITQLK | -1529 | -1545 | 1 | 0 | 17 | NaN | 624.61505 | 695.69507 | NaN | NaN | NaN |
| Rrbp1 | VGAAEEELHK | -1359 | -1368 | 0 | 0.009388464 | 10 | NaN | 219.25179 | NaN | NaN | NaN | NaN |
| Eef1a1 | MDSTEPPYSQK | -155 | -165 | 0 | 0 | 11 | NaN | 7248.169 | 7739.9966 | 7070.357 | 8050.1455 | 7837.546 |
| Eef1a1 | KMDSTEPPYSQK | -154 | -165 | 1 | 0 | 12 | NaN | 772.09717 | 834.46454 | 770.81445 | 1373.8427 | 1250.0421 |
| Rplp1 | ILHDDEVTVTEDK | -15 | -27 | 0 | 0 | 13 | NaN | 2803.5842 | 2770.2192 | 2355.148 | 2604.7153 | 2530.5671 |
| Mogs | AAHANPPTLLLPVVH | -504 | -518 | 0 | 0.00025015 | 15 | NaN | 682.80945 | 181.12529 | NaN | 810.8833 | 773.75446 |
| Mogs | TDGQEVLLPEIGAK | -210 | -223 | 0 | 0.003900536 | 14 | NaN | 397.9364 | NaN | NaN | NaN | 706.82825 |
| Ddx21 | TFHHVYSGK | -288 | -296 | 0 | 0.00025015 | 9 | NaN | 423.17438 | 217.23184 | 227.57275 | 188.55383 | NaN |
| Ddx21 | NFLFPIQAK | -279 | -287 | 0 | 0 | 9 | NaN | 544.8991 | 621.8757 | 780.8372 | NaN | 1274.4108 |
| Ddx21 | TYEQVDLIGK | -465 | -474 | 0 | 0 | 10 | NaN | 1657.0936 | 1659.7961 | NaN | 1786.4354 | NaN |
| Ddx21 | DAQSLHGDIPQK | -535 | -546 | 0 | 0.00078661 | 12 | NaN | 1047.3369 | 823.10095 | 1286.9666 | NaN | NaN |
| Ddx21 | SATCPHWVFNVAK | -447 | -459 | 0 | 0 | 13 | NaN | 658.3932 | 685.56476 | NaN | NaN | NaN |
| Ddx21 | LFSATCPHWVFNVAK | -445 | -459 | 0 | 0.002296977 | 15 | NaN | 883.17035 | NaN | NaN | NaN | NaN |
| Cap1 | VQAFDSLLANPVAEYLK | -43 | -59 | 0 | 0.000230707 | 17 | NaN | 178.56116 | NaN | 216.45195 | 286.89978 | 277.46463 |
| Vta1 | AALAPLPPLPAQFK | -2 | -15 | 0 | 0 | 14 | NaN | 275.7843 | 286.0911 | 267.4731 | 384.78925 | 378.70334 |
| Rplp2 | KILDSVGIEADDDRLNK | -25 | -41 | 2 | 0.000230707 | 17 | NaN | 2615.7368 | 2107.708 | 1716.7281 | 2350.7522 | 2191.5212 |
| Rplp2 | DSVGIEADDDRLNK | -28 | -41 | 1 | 0 | 14 | NaN | 1200.2328 | 850.45465 | 379.65515 | 295.55853 | 395.78497 |
| Mcm5 | QPAEHLQLLEEAAK | -91 | -104 | 0 | 0 | 14 | NaN | 2933.3262 | 3118.276 | NaN | NaN | NaN |
| Mcm5 | ALPNIYELISK | -327 | -337 | 0 | 0.00111006 | 11 | NaN | 572.5982 | NaN | 1004.1069 | 1175.354 | NaN |
| Mcm5 | CPLDPYFIMPDK | -207 | -218 | 0 | 0 | 12 | NaN | 201.51472 | 413.61826 | NaN | 456.2217 | NaN |
| Mcm5 | GGVVCIDEFDK | -439 | -449 | 0 | 0 | 11 | NaN | 171.74268 | NaN | NaN | 1115.8026 | 1028.8705 |
| Mcm5 | VSPQEEEEFRR | -314 | -324 | 1 | 0.003998192 | 11 | NaN | 896.89056 | NaN | NaN | NaN | NaN |
| Mcm5 | SFAGSVSPQEEEEFR | -309 | -323 | 0 | 0.001092529 | 15 | NaN | 165.53639 | NaN | NaN | NaN | NaN |
| Septin1 | GLRPLDVAFL | -147 | -156 | 1 | 0.002534621 | 10 | NaN | 423.92584 | 325.60504 | NaN | 331.203 | 769.5636 |
| Eif4g1 | TLAFEELRR | -1437 | -1445 | 1 | 0 | 9 | NaN | 745.5559 | 809.64777 | 1796.6462 | 3515.6714 | 2111.874 |
| Eif4g1 | EFEKDKDDDEVFEK | -854 | -867 | 2 | 0.001392869 | 14 | NaN | 439.91437 | 429.2779 | NaN | 506.71515 | NaN |
| Cad | LPGLIDVH | -1464 | -1471 | 0 | 0.000471409 | 8 | NaN | 1142.9441 | 1169.1631 | 887.05646 | 866.1566 | 967.024 |
| Cad | LYLNETFSELR | -1557 | -1567 | 0 | 0 | 11 | NaN | 583.1402 | 895.7866 | 540.55383 | 724.53845 | 972.48694 |
| Cad | PGLQGVDTR | -114 | -122 | 0 | 0.003620005 | 9 | NaN | 1029.9414 | NaN | 948.4584 | 1701.7125 | 1691.0426 |
| Cad | ELSDLESAR | -1067 | -1075 | 0 | 0.005112649 | 9 | NaN | 1528.5637 | 2632.544 | NaN | 1206.7655 | NaN |
| Cad | GIPSDEEDEFGLSK | -61 | -74 | 0 | 0 | 14 | NaN | 1040.168 | NaN | 445.2188 | NaN | 1039.8782 |
| Cad | LVTPPQDITPK | -1167 | -1177 | 0 | 0.001708008 | 11 | NaN | 1330.0964 | 1581.4528 | NaN | NaN | NaN |
| Cad | SGDATLVTPPQDITPK | -1162 | -1177 | 0 | 0.000550307 | 16 | NaN | 762.48645 | NaN | NaN | 1072.7003 | NaN |
| Cad | ACLLTQYR | -2091 | -2098 | 0 | 0.002972566 | 8 | NaN | 761.42554 | 1218.2505 | NaN | NaN | NaN |
| Cad | DTIGISQPQWR | -1056 | -1066 | 0 | 0.002780285 | 11 | NaN | 421.87854 | NaN | NaN | NaN | NaN |
| Lmnb1 | AANAGVTASPPTDLIWK | -501 | -517 | 0 | 0 | 17 | NaN | 1297.2222 | 1093.8524 | 1964.6932 | 1397.2461 | 1423.0337 |
| Lmnb1 | NTSEQDQPMGGWEMIR | -459 | -474 | 0 | 0 | 16 | NaN | 355.6676 | 450.27832 | 427.18423 | 492.3964 | 493.91275 |
| Lmnb1 | CQSLTEDLEFR | -199 | -209 | 0 | 0 | 11 | NaN | 188.11179 | 2580.0308 | 713.1793 | 4790.698 | 7790.8145 |
| Lmnb1 | SDYEQLLDVK | -359 | -368 | 0 | 0 | 10 | NaN | 1876.8464 | 1996.875 | NaN | NaN | NaN |
| Lmnb1 | IEEIDVDGK | -445 | -453 | 0 | 0.006654467 | 9 | NaN | 3004.7412 | NaN | NaN | NaN | NaN |
| Anxa6 | KSEISGDLAR | -418 | -427 | 1 | 0 | 10 | NaN | 792.3538 | 1088.921 | 1134.2924 | 1457.759 | 1497.6187 |
| Anxa6 | VCAANDFNPDADAK | -357 | -370 | 0 | 0 | 14 | NaN | 2308.3936 | 2252.6782 | NaN | 2602.2205 | 2576.4058 |
| Anxa6 | IRGELSGDFEK | -230 | -240 | 1 | 0 | 11 | NaN | 1921.3546 | NaN | 1467.8562 | 1834.6904 | 2414.5632 |
| Anxa6 | KGFGSDKESILELITSR | -34 | -50 | 2 | 0.001291103 | 17 | NaN | 215.2398 | NaN | 650.5799 | 437.9843 | 405.02997 |
| Anxa6 | DEATIIDIVTHR | -382 | -393 | 0 | 0 | 12 | NaN | 641.60767 | NaN | 870.1638 | 516.7378 | NaN |
| Anxa6 | AANDFNPDADAK | -359 | -370 | 0 | 0 | 12 | NaN | 895.1588 | 1748.7506 | NaN | NaN | NaN |
| Nudt16l1 | AANPGQLFGR | -34 | -43 | 0 | 0 | 10 | NaN | 873.201 | NaN | NaN | 467.87488 | NaN |
| Nudt16l1 | DYLSSHLTEGPHR | -95 | -107 | 0 | 0 | 13 | NaN | 460.61942 | NaN | NaN | NaN | NaN |
| Tpi1 | AYEPVWAIGTGK | -164 | -175 | 0 | 0 | 12 | NaN | 1806.8832 | 365.3312 | 509.3487 | 833.718 | 477.53198 |
| Tpi1 | GATWVVLGHSER | -88 | -99 | 0 | 0 | 12 | NaN | 1939.6096 | 1310.3949 | 3029.8662 | 3594.086 | 1946.963 |
| Tpi1 | GASLKPEFVDIINAK | -234 | -248 | 1 | 0 | 15 | NaN | 421.77365 | 667.9838 | NaN | 617.72797 | 484.19177 |
| Tpi1 | KVIADNVK | -149 | -156 | 1 | 0 | 8 | NaN | 276.46 | NaN | 635.4563 | 509.76816 | 617.05835 |
| Tpi1 | IIYGGSVTGATCK | -207 | -219 | 0 | 0 | 13 | NaN | 68.04303 | NaN | 317.96542 | 2303.054 | 499.82153 |
| Tpi1 | TPQQAQEVHEK | -178 | -188 | 0 | 0 | 11 | NaN | 380.97888 | NaN | NaN | NaN | NaN |
| Vim | LQEEMLQR | -189 | -196 | 0 | 0 | 8 | NaN | 472.10565 | 2275.7576 | 5095.176 | 4322.097 | 2495.3372 |
| Vim | SLYSSSPGGAYVT | -51 | -63 | 0 | 0.00025015 | 13 | NaN | 220.18448 | 258.88474 | 241.83495 | 230.34604 | 325.8743 |
| Vim | SLADAINTEFK | -87 | -97 | 0 | 0.000907206 | 11 | NaN | 2822.9353 | NaN | NaN | NaN | 2649.1636 |
| Vim | LHDEEIQELQAQIQ | -237 | -250 | 0 | 0.002199523 | 14 | NaN | 559.8933 | NaN | NaN | NaN | NaN |
| Vim | KVESLQEEIAF | -223 | -233 | 1 | 0.006472924 | 11 | NaN | 169.44174 | NaN | NaN | NaN | NaN |
| Faf2 | LHGDDHQDSDEFCR | -177 | -190 | 0 | 0 | 14 | NaN | 608.3071 | 545.82385 | 362.4311 | 151.85948 | 537.4539 |
| Me2 | VESGLSEEEAQR | -328 | -339 | 0 | 0 | 12 | NaN | 625.3285 | NaN | 577.1708 | 620.8047 | NaN |
| Me2 | FTHGVIK | -399 | -405 | 0 | 0.001811524 | 7 | NaN | 482.44666 | NaN | NaN | NaN | NaN |
| Ppa1 | INVDDPDAANYK | -159 | -170 | 0 | 0 | 12 | NaN | 598.60803 | 852.9468 | 1284.7831 | 810.3267 | 755.6953 |
| Ppa1 | LKPGYLEAT | -178 | -186 | 1 | 0.00830926 | 9 | NaN | 3643.9216 | NaN | 1536.4086 | 3590.477 | 3286.5537 |
| Ppa1 | AAPFTLEYR | -10 | -18 | 0 | 0.00111006 | 9 | NaN | 473.5302 | 756.5112 | NaN | 663.3808 | NaN |

| | | | | | | | | | | | | |
|---|---|---|---|---|---|---|---|---|---|---|---|---|
| Ppa1 | YKVPDGKPENEFAF | -193 | -206 | 2 | 0.004199202 | 14 | NaN | 1479.1307 | 2968.3909 | NaN | NaN | NaN |
| Ppa1 | TKDPLNPIK | -62 | -70 | 1 | 0.004439092 | 9 | NaN | 617.12366 | 483.7162 | NaN | NaN | NaN |
| Uqcrc2 | IIENLHDVAYK | -173 | -183 | 0 | 0 | 11 | NaN | 2530.9312 | 2811.7202 | 2412.1587 | 2710.4902 | 2696.623 |
| Wars1 | AGCPPGNPTAGR | -64 | -75 | 0 | 0 | 12 | NaN | 513.4136 | NaN | NaN | 358.4548 | 566.9153 |
| Wars1 | GIFGFTDSDCIGK | -269 | -281 | 0 | 0.00025015 | 13 | NaN | 554.3789 | NaN | NaN | NaN | NaN |
| Wars1 | GIFFSHR | -139 | -145 | 0 | 0.004923919 | 7 | NaN | 468.66382 | NaN | NaN | NaN | NaN |
| Eif2a | CDPVFDFGTGPR | -306 | -317 | 0 | 0 | 12 | NaN | 775.10565 | 258.17764 | NaN | 996.3346 | NaN |
| Tcp1 | IHPTSVISGYR | -112 | -122 | 0 | 0 | 11 | NaN | 967.1968 | 788.76056 | NaN | 890.8751 | 688.0974 |
| Tcp1 | IHPTSVISGY | -112 | -121 | 0 | 0.000431655 | 10 | NaN | 4668.4575 | NaN | 204.80777 | NaN | NaN |
| Hnrnpd | VGGLSPDTPEEK | -186 | -197 | 0 | 0 | 12 | NaN | 498.6684 | NaN | NaN | 3414.1348 | 681.5303 |
| Hnrnpd | SIELPMDNK | -210 | -218 | 0 | 0.00025015 | 9 | NaN | 867.8962 | NaN | 688.987 | NaN | NaN |
| Hnrnpd | SIELPMDNK | -210 | -218 | 0 | 0.00025015 | 9 | NaN | 867.8962 | NaN | 688.987 | NaN | NaN |
| Dpp3 | LEEEDKDLYIR | -427 | -437 | 1 | 0 | 11 | NaN | 636.4247 | 649.95795 | 531.30554 | 479.3494 | 630.90247 |
| Dpp3 | VVEHLEK | -258 | -264 | 0 | 0.000230707 | 7 | NaN | 1098.5408 | NaN | 771.6356 | NaN | NaN |
| Dpp3 | GIPAGINIPNYDDLR | -385 | -399 | 0 | 0 | 15 | NaN | 531.004 | NaN | NaN | NaN | 782.91846 |
| Slc16a1 | EGKEDEASTDVDEKPK | -454 | -469 | 2 | 0 | 16 | NaN | 779.50366 | 776.98206 | 924.96735 | 907.9996 | 702.7929 |
| Slc16a1 | AAQSPQQHSSGDPTEEESPV | -474 | -493 | 0 | 0.000907206 | 20 | NaN | 753.18744 | 872.435 | 397.7945 | 669.4094 | 787.00964 |
| Igbp1 | LPELFETSK | -13 | -21 | 0 | 0.000712251 | 9 | NaN | 468.82977 | NaN | NaN | NaN | 519.3205 |
| Esd | LQEGYDHSYYFIA | -254 | -266 | 0 | 0.001868731 | 13 | NaN | 747.89886 | 782.3834 | 657.568 | 711.2227 | NaN |
| Esd | SYVTEELPQLINANFPVDPQR | -122 | -142 | 0 | 0.00078661 | 21 | NaN | 315.77734 | NaN | 345.48148 | 389.85532 | 388.81985 |
| Esd | KAFSGYLGPDESK | -186 | -198 | 1 | 0 | 13 | NaN | 471.05182 | 519.16583 | NaN | NaN | 1066.7039 |
| Esd | ELPQLINANFPVDPQR | -127 | -142 | 0 | 0.000550307 | 16 | NaN | 378.85553 | 262.99985 | NaN | NaN | NaN |
| Nosip | ALPSFWIPSLTPEAK | -158 | -172 | 0 | 0.00201047 | 15 | NaN | 119.84894 | NaN | NaN | NaN | NaN |
| Got1 | AATLSDPELFK | -308 | -318 | 0 | 0.001291103 | 11 | NaN | 668.2895 | 512.341 | 788.2926 | 889.7825 | 808.2359 |
| Got1 | QVEYLVNEK | -370 | -378 | 0 | 0.001055699 | 9 | NaN | 853.0453 | 1103.2802 | 1235.989 | 1465.5305 | NaN |
| Vcp | AATNRPNSIDPALR | -345 | -358 | 1 | 0.000826806 | 14 | NaN | 1075.4487 | 310.81094 | NaN | 2976.907 | 1009.44244 |
| Vcp | CIVAPDTVIHCEGEPIKR | -174 | -191 | 1 | 0 | 18 | NaN | 525.47375 | 482.36362 | NaN | 573.5668 | 555.60474 |
| Vcp | WEDIGGLEDVKR | -476 | -487 | 1 | 0 | 12 | NaN | 4118.506 | NaN | 347.3346 | 3212.2883 | 2859.6074 |
| Vcp | KAFEEAEK | -288 | -295 | 1 | 0.001463897 | 8 | NaN | 3556.3135 | NaN | 3814.8079 | 4017.4583 | NaN |
| Vcp | GDDLSTAILK | -9 | -18 | 0 | 0.000431655 | 10 | NaN | 955.2891 | 567.0348 | NaN | NaN | NaN |
| Vcp | RGDTVLLK | -53 | -60 | 1 | 0.009745776 | 8 | NaN | 410.07907 | NaN | NaN | 746.58905 | NaN |
| Vcp | SLNEVGYDDIGGCR | -197 | -210 | 0 | 0.00078661 | 14 | NaN | 3735.2458 | NaN | NaN | NaN | NaN |
| Vcp | ESIESEIRR | -701 | -709 | 1 | 0.00025015 | 9 | NaN | 1822.4786 | NaN | NaN | NaN | NaN |
| Vcp | LADDVDLEQVANETHGHVGADLA | -390 | -412 | 0 | 0.001636916 | 23 | NaN | 212.0085 | NaN | NaN | NaN | NaN |
| Erap1 | YQSSLSSTEK | -778 | -787 | 0 | 0 | 10 | NaN | 12116.258 | 12968.701 | NaN | 13030.324 | 14499.443 |
| Erap1 | TKTDVLILPE | -573 | -582 | 1 | 0.003166358 | 10 | NaN | 483.92105 | 887.4315 | NaN | NaN | 820.9545 |
| Erap1 | FKDFLLK | -688 | -694 | 1 | 0 | 7 | NaN | 279.20322 | 463.33826 | NaN | NaN | NaN |
| Dync1h1 | TKPVTGNLRPEEALQAL | -1262 | -1278 | 2 | 0.001463897 | 17 | NaN | 2931.6973 | 2254.9253 | 3041.5269 | 3803.6091 | 4823.8936 |
| Dync1h1 | VAAPDVVVPTLDTVR | -2560 | -2574 | 0 | 0 | 15 | NaN | 511.7882 | NaN | 614.7325 | 551.10834 | 626.0861 |
| Dync1h1 | TLPVYLNFTR | -4604 | -4613 | 0 | 0.001610812 | 10 | NaN | 117.046234 | NaN | 1041.4778 | 1300.1713 | 1415.347 |
| Dync1h1 | LAETVFNFQEK | -834 | -844 | 0 | 0 | 11 | NaN | 4312.4 | 2225.009 | NaN | NaN | NaN |
| Dync1h1 | IFVFEPPPGVK | -4142 | -4152 | 0 | 0.00078661 | 11 | NaN | 1555.7369 | 453.49258 | NaN | NaN | NaN |
| Dync1h1 | EFGPVVIDYGK | -1099 | -1109 | 0 | 0.00025015 | 11 | NaN | 1301.8612 | 380.60858 | NaN | NaN | NaN |
| Dync1h1 | EIFDDWAR | -695 | -702 | 0 | 0.009657961 | 8 | NaN | 508.88586 | 699.69476 | NaN | NaN | NaN |
| Dync1h1 | TLINELVK | -4448 | -4455 | 0 | 0.002845514 | 8 | NaN | 2348.899 | NaN | NaN | NaN | NaN |
| Sri | ITFDDYIACCVK | -154 | -165 | 0 | 0 | 12 | NaN | 1124.1814 | 284.34076 | 1379.2963 | 1566.7228 | 477.9527 |
| Sri | GIAGGYKPFNLETCR | -62 | -76 | 1 | 0 | 15 | NaN | 953.749 | 945.2795 | NaN | 1196.3733 | 991.6035 |
| Acat1 | LEDLIVK | -172 | -178 | 0 | 0.000230707 | 7 | NaN | 739.05914 | 646.05237 | NaN | NaN | NaN |
| Rrp12 | KFGFELVK | -1015 | -1022 | 1 | 0.00025015 | 8 | NaN | 1031.2272 | 879.8637 | 807.60315 | 877.7535 | 900.2539 |
| Kpnb1 | LLETTDRPDGHQNNLR | -510 | -525 | 1 | 0 | 16 | NaN | 2683.64 | 2987.5347 | 2102.7214 | 4145.4023 | 3784.8276 |
| Kpnb1 | EHIKNPDWR | -373 | -381 | 1 | 0 | 9 | NaN | 667.99396 | 931.6053 | 500.85382 | 2171.3618 | 1041.7607 |
| Kpnb1 | EAAEQGRPPEHTSK | -299 | -312 | 1 | 0 | 14 | NaN | 391.91 | 736.9764 | 429.3118 | 666.8458 | 805.0255 |
| Kpnb1 | ICELLPEAAIN | -435 | -445 | 0 | 0.00111006 | 11 | NaN | 297.97086 | 283.80527 | 335.57602 | 267.57434 | NaN |
| Kpnb1 | KVQHQDALQIS | -594 | -604 | 1 | 0.000471409 | 11 | NaN | 1127.3043 | 1009.6704 | NaN | 3253.6428 | NaN |
| Kpnb1 | DCYPAVQK | -542 | -549 | 0 | 0.001291103 | 8 | NaN | 601.09424 | NaN | NaN | NaN | NaN |
| Stip1 | PQNHVLYSNR | -35 | -44 | 0 | 0.000591124 | 10 | NaN | 6567.1973 | 5683.313 | 366.32452 | 456.6632 | 8362.354 |
| Stip1 | LDPQNHVLYSN | -33 | -43 | 0 | 0 | 11 | NaN | 2148.5813 | 2642.9478 | 2767.5928 | 2599.0798 | 2401.5444 |
| Stip1 | TYEEGLKHEANNLQLK | -94 | -109 | 1 | 0 | 16 | NaN | 801.32196 | 694.9815 | 731.3296 | 1049.5466 | 1007.4974 |
| Stip1 | FMNPFNLPNLYQK | -124 | -136 | 0 | 0 | 13 | NaN | 803.8417 | 574.17694 | 820.42474 | 835.24963 | 1013.81177 |
| Stip1 | AAVHFEK | -266 | -272 | 0 | 0.000230707 | 7 | NaN | 2983.6409 | 3170.2925 | 3158.7202 | 2039.7933 | 2380.7974 |
| Stip1 | DLKPDWGK | -66 | -73 | 1 | 0 | 8 | NaN | 993.31323 | 1233.4125 | 1881.3191 | 1660.9613 | NaN |
| Stip1 | RTYEEGLK | -93 | -100 | 1 | 0.002845514 | 8 | NaN | 1442.233 | NaN | 384.777 | 707.79645 | NaN |
| Stip1 | EAADGYQR | -463 | -470 | 0 | 0.002265552 | 8 | NaN | 366.54196 | 216.97156 | 232.11066 | NaN | NaN |
| Ly6a | AAVIVDSQTR | -58 | -67 | 0 | 0.002068213 | 10 | NaN | 959.12177 | NaN | NaN | NaN | NaN |
| Hsp90ab1 | SLEDPQTHSNR | -669 | -679 | 0 | 0 | 11 | NaN | 5443.865 | 6268.35 | 3539.0864 | 5513.1704 | 5366.2256 |
| Hsp90ab1 | SLTNDWEDHLAVK | -307 | -319 | 0 | 0 | 13 | NaN | 5304.401 | 5311.8315 | 3384.3643 | 3808.8477 | 3801.0098 |
| Hsp90ab1 | RAPFDLFENKK | -338 | -348 | 2 | 0 | 11 | NaN | 4837.727 | 4931.78 | 1631.2612 | 2613.1523 | 2671.1394 |
| Hsp90ab1 | ADHGEPIGR | -169 | -177 | 0 | 0 | 9 | NaN | 3773.4617 | 620.57886 | 4077.366 | 4161.2915 | 4451.43 |
| Hsp90ab1 | FSELAEDKENYKK | -416 | -428 | 2 | 0 | 13 | NaN | 816.4855 | 910.55273 | 625.3216 | 894.7237 | 1303.6432 |
| Hsp90ab1 | LSSGFSLEDPQTHSNR | -664 | -679 | 0 | 0 | 16 | NaN | 770.51025 | 800.2434 | 1042.2959 | 732.72375 | 729.96106 |
| Hsp90ab1 | LGIHEDSTNRR | -439 | -449 | 1 | 0 | 11 | NaN | 554.14825 | 576.6675 | 1951.1781 | 651.1079 | 632.05676 |
| Hsp90ab1 | PPLEGDEDASR | -709 | -719 | 0 | 0.002199523 | 11 | NaN | 4010.5945 | 4749.7188 | 1326.2234 | 931.17395 | 7753.2065 |
| Hsp90ab1 | EQVANSAFVER | -492 | -502 | 0 | 0.00025015 | 11 | NaN | 3436.1462 | 3473.1768 | 3706.3728 | 1821.0604 | 976.927 |
| Hsp90ab1 | LELFSELAEDKENYKK | -413 | -428 | 2 | 0 | 16 | NaN | 452.6071 | 409.95575 | 164.66202 | 947.05945 | 889.4998 |
| Hsp90ab1 | AVPDEIPPLEGDEDASR | -703 | -719 | 0 | 0 | 17 | NaN | 439.6196 | 585.8205 | 1135.9065 | 1295.728 | 1266.3774 |
| Hsp90ab1 | EISDDEAEEEKGEKE | -224 | -238 | 2 | 0.00051306 | 15 | NaN | 808.8364 | 791.3695 | NaN | 1373.7764 | 1234.6971 |
| Hsp90ab1 | KHLEINPDHPIVET | -624 | -637 | 1 | 0.000230707 | 14 | NaN | 1359.4185 | 1624.2045 | NaN | NaN | NaN |
| Hsp90ab1 | YHTSQSGDEMTSL | -457 | -469 | 0 | 0 | 13 | NaN | 521.7813 | 700.92065 | NaN | NaN | NaN |

| | | | | | | | | | | | | |
|---|---|---|---|---|---|---|---|---|---|---|---|---|
| Hsp90ab1 | PEEVHHGEEEVE | -2 | -13 | 0 | 0.000230707 | 12 | NaN | 356.5149 | 293.38745 | NaN | NaN | NaN |
| Snx5 | LSSHPVLSK | -150 | -158 | 0 | 0.00078661 | 9 | NaN | 1205.3623 | 1164.5226 | NaN | NaN | 1362.6226 |
| Eno1 | GNPEVILPVPAFNVINGGSHAGNK | -139 | -162 | 0 | 0 | 24 | NaN | 849.53204 | 935.9304 | 667.91614 | 722.99884 | 827.1282 |
| Eno1 | SQAVEHINK | -63 | -71 | 0 | 0 | 9 | NaN | 1943.8936 | 2298.559 | 1261.9049 | 2913.0513 | 3049.0916 |
| Eno1 | IEDPFDQDDWGAWQK | -292 | -306 | 0 | 0 | 15 | NaN | 343.40744 | 323.28195 | 152.55688 | 605.19696 | 599.36646 |
| Eno1 | IEEELGSK | -413 | -420 | 0 | 0.006735683 | 8 | NaN | 1171.7141 | NaN | NaN | NaN | 976.3336 |
| Eno1 | NVGDEGGFAPNILENK | -206 | -221 | 0 | 0.000431655 | 16 | NaN | 740.5169 | 472.92673 | NaN | NaN | NaN |
| Rpl5 | AVDGGLSIPHSTK | -166 | -178 | 0 | 0 | 13 | NaN | 10908.067 | 10477.96 | 8031.4907 | 13665.507 | 9097.761 |
| Rpl5 | LKGAVDGGLSIPHSTK | -163 | -178 | 1 | 0 | 16 | NaN | 296.22043 | NaN | 810.0762 | 558.4304 | 809.74243 |
| Slc25a5 | LTSYPFDTVR | -226 | -235 | 0 | 0.00025015 | 10 | NaN | 2642.2388 | 2537.8323 | NaN | NaN | NaN |
| Rars1 | FPEILQK | -581 | -587 | 0 | 0.004321969 | 7 | NaN | 2210.132 | 2086.2285 | 1951.6556 | 1984.9641 | 1854.4414 |
| Rars1 | LNDYIFSFDK | -513 | -522 | 0 | 0 | 10 | NaN | 237.3634 | 264.1624 | 186.71364 | 489.1067 | 567.6304 |
| Rars1 | KIVFVPGCSVPL | -362 | -373 | 1 | 0.002099344 | 12 | NaN | 1801.9713 | 521.79895 | 2683.551 | NaN | 258.46924 |
| Rars1 | DILGIKPVQR | -650 | -659 | 1 | 0 | 10 | NaN | 661.796 | NaN | 448.92755 | 717.3881 | 507.44217 |
| Rars1 | RFDADEEFK | -279 | -287 | 1 | 0.000431655 | 9 | NaN | 920.9234 | NaN | NaN | 1167.0255 | 1196.5804 |
| Rars1 | NCGCLEASPSLEQLREENLK | -31 | -50 | 1 | 0.00051306 | 20 | NaN | 177.54042 | NaN | NaN | 403.18933 | NaN |
| Gars1 | EHTFHVR | -580 | -586 | 0 | 0.00025015 | 7 | NaN | 1098.2485 | 1958.8738 | 194.62929 | 622.8589 | 1170.6282 |
| Gars1 | VSELPNVVR | -689 | -697 | 0 | 0 | 9 | NaN | 820.1634 | 788.4693 | 709.80347 | NaN | 349.11267 |
| Gars1 | EDKAPQVDVDR | -73 | -83 | 1 | 0.000591124 | 11 | NaN | 999.7677 | NaN | NaN | 1135.7329 | NaN |
| Gars1 | SQNQEFMPFVK | -612 | -622 | 0 | 0.00111006 | 11 | NaN | 910.6927 | NaN | NaN | NaN | NaN |
| C3 | IWDVVEK | -615 | -621 | 0 | 0.002168954 | 7 | NaN | 841.7109 | NaN | NaN | 898.6894 | NaN |
| Paics | FKDDANNDPQWSEEQLIAAK | -130 | -149 | 1 | 0 | 20 | NaN | 346.96973 | 440.11102 | 371.76404 | 381.94135 | 441.07602 |
| Paics | FGIPCELR | -291 | -298 | 0 | 0.00149795 | 8 | NaN | 2883.9883 | NaN | 1771.2639 | NaN | 1046.9409 |
| Aco2 | FNPETDFLTGK | -507 | -517 | 0 | 0 | 11 | NaN | 1636.8303 | 1067.655 | 953.9837 | 1079.1338 | 937.7846 |
| Aco2 | GLLPLTFADPSDYNK | -703 | -717 | 0 | 0 | 15 | NaN | 310.9713 | 451.4359 | 527.4799 | 507.51752 | 531.95374 |
| Aco2 | LLEPFDKWDGK | -567 | -577 | 1 | 0.00025015 | 11 | NaN | 1520.636 | 1589.4094 | NaN | 2115.225 | 1236.2223 |
| Rbm24;Rbm3[illegible] | ACKDPNPIIDGR | (68);(89) | (79);(100) | 1 | 0 | 12 | NaN | 2785.14 | 2912.5994 | NaN | 2081.713 | 2653.1565 |
| Sec23b | VNPSQTHNNLYAW | -724 | -736 | 0 | 0.001904399 | 13 | NaN | 1409.8254 | NaN | NaN | 1527.0498 | 1069.4185 |
| Sec23b | ACLLTPLK | -39 | -46 | 0 | 0.000712251 | 8 | NaN | 671.82446 | NaN | NaN | NaN | NaN |
| Cct3 | KIGDEYFTFITDCKDPK | -354 | -370 | 2 | 0.000230707 | 17 | NaN | 746.6845 | 833.98724 | 1167.5809 | 1563.2285 | 1657.756 |
| Cct3 | RIDDIVSGHK | -518 | -527 | 1 | 0.001092529 | 10 | NaN | 557.5231 | 521.6316 | 792.3123 | 1572.4613 | 713.96735 |
| Itgb2 | TVLPFVNTHPEK | -176 | -187 | 0 | 0 | 12 | NaN | 4008.1123 | 3770.5151 | 4207.3145 | 4714.178 | 4875.684 |
| Itgb2 | LSESNIQPIFAVTK | -312 | -325 | 0 | 0 | 14 | NaN | 584.7354 | 638.54486 | 907.5478 | 996.62134 | 969.0096 |
| Prkcsh | ESLQQLAEVTR | -125 | -135 | 0 | 0 | 11 | NaN | 2650.1226 | 2706.5815 | 2599.2317 | 2905.547 | 2977.289 |
| Prkcsh | KLWEEQQAAAK | -196 | -206 | 1 | 0 | 11 | NaN | 1381.3096 | 1557.9481 | 1510.4073 | 1677.2888 | 1732.8058 |
| Prkcsh | YEQGTGCWQGPNR | -458 | -470 | 0 | 0 | 13 | NaN | 468.83707 | NaN | NaN | NaN | NaN |
| Pclaf | ACPLQPDHRDDENE | -97 | -110 | 1 | 0 | 14 | NaN | 753.60223 | 742.0172 | 701.5059 | 296.67328 | 674.0235 |
| Hnrnpm | KACQIFVR | -650 | -657 | 1 | 0 | 8 | NaN | 734.791 | 569.4348 | NaN | 430.6306 | 397.46832 |
| Hnrnpm | GPGGQPIDANHLSK | -308 | -321 | 0 | 0.000431655 | 14 | NaN | 500.77875 | NaN | 628.8763 | 321.25 | 376.42593 |
| Hnrnpm | VGSEIER | -429 | -435 | 0 | 0.003133457 | 7 | NaN | 1481.8254 | NaN | 1025.5691 | NaN | 806.7577 |
| Hnrnpm | AGSFGGAGGHAPGVAR | -634 | -649 | 0 | 0 | 16 | NaN | 1066.396 | 1159.1177 | NaN | NaN | NaN |
| Hnrnpm | GCAVVEFK | -112 | -119 | 0 | 0.006100831 | 8 | NaN | 931.1161 | NaN | NaN | NaN | NaN |
| Rps3a | NCLTNFHGMDLTR | -95 | -107 | 0 | 0 | 13 | NaN | 185.93083 | 605.23486 | 2138.6934 | 2456.9973 | 2380.9841 |
| Rps3a | KLIPDSIGK | -187 | -195 | 1 | 0.001463897 | 9 | NaN | 280.00827 | 329.1444 | NaN | NaN | NaN |
| Fasn | VSVHIIEGDHR | -2465 | -2475 | 0 | 0 | 11 | NaN | 1633.0687 | 1173.752 | 911.4767 | 921.55896 | 1065.7461 |
| Fasn | EVLAGEGHLYSR | -1239 | -1250 | 0 | 0 | 12 | NaN | 2987.4417 | 2249.3206 | 879.7435 | 1039.2487 | 956.35376 |
| Fasn | FPQLDDTSFANSR | -1705 | -1717 | 0 | 0 | 13 | NaN | 1880.367 | 559.6618 | 1272.2782 | 2315.3933 | 848.0105 |
| Fasn | GSSQQSLQLPTR | -1041 | -1052 | 0 | 0 | 12 | NaN | 922.6535 | 727.2384 | 627.6371 | 736.8434 | 597.4857 |
| Fasn | RQQEQLVPTLEKF | -1106 | -1118 | 2 | 0.000550307 | 13 | NaN | 514.4514 | 461.36035 | 554.72906 | 782.3628 | 694.6357 |
| Fasn | QFLDVEHSK | -2367 | -2375 | 0 | 0 | 9 | NaN | 2407.2817 | 2102.9497 | NaN | 2554.958 | 2211.7922 |
| Fasn | LPESENLQEFWAN | -13 | -25 | 0 | 0.000230707 | 13 | NaN | 547.04205 | 684.433 | NaN | 385.88495 | 975.17993 |
| Fasn | EHDLVLPMR | -2163 | -2171 | 0 | 0.00025015 | 9 | NaN | 947.3379 | NaN | 6010.4976 | 415.26367 | NaN |
| Fasn | FFGVHPK | -64 | -70 | 0 | 0.00272025 | 7 | NaN | 528.49335 | 471.08905 | 621.29834 | NaN | NaN |
| Fasn | FHNPNPEIPALLDGR | -359 | -373 | 0 | 0 | 15 | NaN | 422.70578 | 1723.2015 | NaN | NaN | 3108.9294 |
| Fasn | HFQLEQDKPK | -1509 | -1518 | 1 | 0.000230707 | 10 | NaN | 1497.6136 | 1760.304 | NaN | NaN | NaN |
| Fasn | GEDLGADYNLSQVCDGK | -2448 | -2464 | 0 | 0 | 17 | NaN | 234.06082 | NaN | 252.20273 | NaN | NaN |
| Cct8 | LYSVHQEGNK | -467 | -476 | 0 | 0 | 10 | NaN | 1389.8662 | 1400.7767 | 696.349 | 2012.1068 | 2033.7913 |
| Cct8 | GSTDNLMDDIER | -379 | -390 | 0 | 0 | 12 | NaN | 656.12665 | 533.6266 | 1084.8986 | 962.0515 | 707.9555 |
| Cct8 | RDVDEVSSLLR | -155 | -165 | 1 | 0 | 11 | NaN | 2617.3247 | NaN | 1478.6848 | 842.6566 | 790.02734 |
| Cct8 | AHEILPELVC | -139 | -148 | 0 | 0.00676572 | 10 | NaN | 820.72296 | 816.9303 | NaN | NaN | 315.28653 |
| Cct8 | TAEELMNFSK | -261 | -270 | 0 | 0 | 10 | NaN | 869.3497 | NaN | NaN | 1851.7054 | NaN |
| Cct8 | KAHEILPELVC | -138 | -148 | 1 | 0.006472924 | 11 | NaN | 3611.3489 | NaN | NaN | NaN | NaN |
| Atp1a1 | DMTSEELDDILR | -672 | -683 | 0 | 0 | 12 | NaN | 625.2293 | 915.09485 | 903.10297 | 903.9317 | 617.2977 |
| Atp1a1 | KYGTDLSR | -54 | -61 | 1 | 0 | 8 | NaN | 402.65805 | 961.9251 | 830.76636 | 828.7661 | 1030.1031 |
| Atp1a1 | WADLVICK | -931 | -938 | 0 | 0 | 8 | NaN | 613.94415 | 306.5575 | NaN | 625.07324 | 672.54767 |
| Atp1a1 | TSEELDDILR | -674 | -683 | 0 | 0 | 10 | NaN | 493.9792 | 1097.1998 | NaN | 1153.7272 | 889.31854 |
| Atp1a1 | RPGGWVEKE | -1012 | -1020 | 2 | 0.005605646 | 9 | NaN | 454.61438 | NaN | NaN | NaN | NaN |
| Atp1a1 | EQPLDEELKDAFQNA | -527 | -541 | 1 | 0.00025015 | 15 | NaN | 382.59454 | NaN | NaN | NaN | NaN |
| Rps3 | GGKPEPPAMPQPVPTA | -228 | -243 | 1 | 0 | 16 | NaN | 3219.4673 | 4580.8984 | 5596.0347 | 3823.6714 | 5453.6743 |
| Rps3 | KGGKPEPPAMPQPVPTA | -227 | -243 | 2 | 0.00078661 | 17 | NaN | 2059.3987 | 2148.1904 | 1925.0695 | 2006.9116 | 1959.1193 |
| Rps3 | DEILPTTPISEQ | -215 | -226 | 0 | 0.00025015 | 12 | NaN | 708.45807 | 518.6429 | NaN | 159.18364 | NaN |
| Atg3 | ADAGGEDAILQTR | -186 | -198 | 0 | 0 | 13 | NaN | 723.3684 | NaN | NaN | 2157.7617 | NaN |
| Api5 | PDEVLTK | -163 | -169 | 0 | 0 | 7 | NaN | 521.0403 | 769.61 | 760.463 | 645.5143 | 776.1048 |
| Snd1 | VDYIRPASPATETVPAFSER | -419 | -438 | 1 | 0 | 20 | NaN | 513.3065 | 539.3142 | 606.4843 | 723.77124 | 579.2318 |
| Snd1 | ETDGSETPEPFAAEAK | -234 | -249 | 0 | 0 | 16 | NaN | 338.1159 | 836.66266 | 826.0103 | 756.40326 | 386.7636 |
| Snd1 | HFVDSHHQKPVNAIIE | -185 | -200 | 1 | 0 | 16 | NaN | 305.12442 | 471.53903 | NaN | NaN | NaN |
| Mcm4 | RLHGLDEEAEQK | -427 | -438 | 1 | 0 | 12 | NaN | 343.78015 | 343.82492 | 954.2277 | 649.81146 | 595.5444 |
| Mcm4 | LDPQDEAYDRR | -650 | -660 | 1 | 0 | 11 | NaN | 1005.738 | 1086.2933 | 1273.4124 | 1029.2294 | 951.4032 |

| | | | | | | | | | | | | |
|---|---|---|---|---|---|---|---|---|---|---|---|---|
| Mcm4 | LCGDPGTSK | -507 | -515 | 0 | 0 | 9 | NaN | 593.6262 | NaN | 1002.98267 | 655.901 | 913.2804 |
| Mcm4 | KPDIYER | -454 | -460 | 1 | 0.005112649 | 7 | NaN | 1869.4899 | NaN | 1192.8108 | NaN | 1485.4985 |
| Mcm4 | MLDPQDEAYDR | -649 | -659 | 0 | 0.000431655 | 11 | NaN | 488.7983 | 740.2723 | NaN | 1106.0674 | NaN |
| Samhd1 | FHPLLIR | -160 | -166 | 0 | 0.000627943 | 7 | NaN | 1809.4187 | 2085.708 | 1641.9587 | 646.041 | 882.79895 |
| Samhd1 | PGASHNR | -190 | -196 | 0 | 0.002401448 | 7 | NaN | 423.86768 | 798.92474 | NaN | 716.71094 | 546.5235 |
| Ppia | KITISDCGQL | -155 | -164 | 1 | 0.000627943 | 10 | NaN | 5239.037 | 5984.0894 | NaN | NaN | NaN |
| Ppia | ITADDEPLGR | -10 | -19 | 0 | 0.00025015 | 10 | NaN | 2696.1333 | NaN | NaN | NaN | NaN |
| Psme2 | LYHIISSNLEK | -216 | -226 | 0 | 0 | 11 | NaN | 2694.364 | 2653.726 | 2534.0718 | 2491.2603 | NaN |
| Psme2 | YAELYHIISSNLEK | -213 | -226 | 0 | 0 | 14 | NaN | 1244.8208 | 1167.7881 | 1839.2911 | 2248.1206 | 2205.8735 |
| Psme2 | LCTFLPR | -32 | -38 | 0 | 0.002534621 | 7 | NaN | 1920.6927 | 118.44995 | NaN | NaN | NaN |
| Psme2 | RAPLDIPIPDPPPK | -61 | -74 | 1 | 0.00051306 | 14 | NaN | 435.1706 | 494.88794 | NaN | NaN | NaN |
| Aldoa | PHPYPALTPEQK | -2 | -13 | 0 | 0 | 12 | NaN | 11841.2295 | 11183.872 | 11134.3955 | 12219.063 | 11913.5205 |
| Dars1 | YPLAVRPF | -394 | -401 | 1 | 0 | 8 | NaN | 825.9565 | 1021.1943 | 981.1051 | 1425.8317 | 1214.113 |
| Dars1 | KFLEPTLR | -338 | -345 | 1 | 0.004595494 | 8 | NaN | 463.22858 | 342.93796 | NaN | NaN | NaN |
| Dars1 | DVELHVQK | -134 | -141 | 0 | 0.003745449 | 8 | NaN | 1426.3774 | NaN | NaN | NaN | NaN |
| Dars1 | QFPCEPFK | -331 | -338 | 0 | 0.00676572 | 8 | NaN | 1330.1722 | NaN | NaN | NaN | NaN |
| Commd3 | ADEAALDHPYLK | -41 | -52 | 0 | 0 | 12 | NaN | 1436.3525 | NaN | 1781.6747 | 714.2179 | 1318.822 |
| Commd3 | NSLETLLGSIGR | -109 | -120 | 0 | 0 | 12 | NaN | 386.25647 | NaN | 461.56357 | 412.6409 | 411.3179 |
| Pdcd5 | NLALVKPEK | -58 | -66 | 1 | 0 | 9 | NaN | 2438.6753 | 2682.1172 | 2287.9243 | 1837.6678 | 2039.0719 |
| Pdcd5 | LSNLALVKPEK | -56 | -66 | 1 | 0.00025015 | 11 | NaN | 606.54736 | 717.2145 | NaN | NaN | 2128.7383 |
| Ipo7 | EDDRPELPWWK | -249 | -259 | 1 | 0 | 11 | NaN | 912.38983 | 926.84375 | 334.73502 | 949.366 | 832.12445 |
| Ipo7 | IIDDEDNPVDEYQIFK | -948 | -963 | 0 | 0.000627943 | 16 | NaN | 187.28009 | 363.05124 | NaN | NaN | NaN |
| Prdx1 | HGEVCPAGWKPGSDTIKPDVNK | -169 | -190 | 2 | 0 | 22 | NaN | 203.29172 | 696.7845 | 5564.3174 | 5649.3496 | 4545.9272 |
| Prdx1 | QGGLGPMNIPLISDPK | -94 | -109 | 0 | 0 | 16 | NaN | 680.6803 | NaN | 509.6705 | 848.3883 | 1137.8132 |
| Ldha | ADELALVDVMEDK | -45 | -57 | 0 | 0 | 13 | NaN | 1295.3624 | NaN | 1474.616 | NaN | 1473.2354 |
| Slc25a20 | VFVGHPLDTVK | -25 | -35 | 0 | 0 | 11 | NaN | 818.8777 | 1131.99 | NaN | NaN | NaN |
| Myh9 | DELADEIANSSGK | -1704 | -1716 | 0 | 0 | 13 | NaN | 1870.7555 | 2209.527 | 1131.559 | 154.22571 | 2178.796 |
| Myh9 | KLEMDLK | -1614 | -1620 | 1 | 0.001636916 | 7 | NaN | 1640.8655 | 1815.5189 | 1514.4673 | 1682.7025 | 2048.744 |
| Myh9 | EQLEEEEEAKR | -1343 | -1353 | 1 | 0 | 11 | NaN | 1344.2202 | 5412.1553 | 3732.554 | 5197.677 | 5019.879 |
| Myh9 | EMEAELEDERK | -1593 | -1603 | 1 | 0 | 11 | NaN | 1176.6028 | 979.0192 | 1037.5557 | 1577.3992 | 1481.8413 |
| Myh9 | QAQQERDELADEIANSSGK | -1698 | -1716 | 1 | 0 | 19 | NaN | 514.7532 | 728.2595 | 871.91437 | 700.06464 | 676.64026 |
| Myh9 | HEAMITDLEER | -1025 | -1035 | 0 | 0 | 11 | NaN | 730.8906 | 1071.7747 | 1119.0979 | 1187.899 | 1229.1257 |
| Myh9 | QLEEAEEEAQRA | -1878 | -1889 | 1 | 0 | 12 | NaN | 2080.1128 | 1715.766 | 184.06671 | 1163.2241 | 1479.1945 |
| Myh9 | LKDVLLQVEDER | -1844 | -1855 | 1 | 0 | 12 | NaN | 1016.33026 | 456.93314 | 2063.531 | NaN | 1492.8016 |
| Myh9 | LEGDSTDLSDQIAELQ | -1053 | -1068 | 0 | 0.00025015 | 16 | NaN | 715.7458 | 883.9055 | 372.51212 | 661.0481 | 955.495 |
| Myh9 | TDLLLEPYNKY | -290 | -300 | 1 | 0 | 11 | NaN | 469.23932 | 712.9345 | 540.9884 | 694.23987 | 654.17706 |
| Myh9 | ETQISELQEDLESER | -1110 | -1124 | 0 | 0 | 15 | NaN | 427.23816 | 836.6791 | 716.3167 | 958.94794 | 772.63513 |
| Myh9 | QTLENER | -1220 | -1226 | 0 | 0.00383302 | 7 | NaN | 1114.884 | NaN | 958.1633 | 665.6135 | 1137.8645 |
| Myh9 | GVLQLGNIAFK | -345 | -355 | 0 | 0 | 11 | NaN | 429.7292 | NaN | 715.16956 | 755.64655 | 655.9061 |
| Myh9 | EMEAELEDER | -1593 | -1602 | 0 | 0 | 10 | NaN | 477.3109 | NaN | 609.8196 | 455.4649 | NaN |
| Myh9 | NMDPLNDNIATL | -588 | -599 | 0 | 0.005605646 | 12 | NaN | 417.3988 | 535.88055 | NaN | NaN | 456.39902 |
| Myh9 | RQLEEAEEEAQ | -1877 | -1887 | 1 | 0.000471409 | 11 | NaN | 472.27573 | 510.04535 | NaN | NaN | NaN |
| Myh9 | ADKYLYVDK | -6 | -14 | 1 | 0.00051306 | 9 | NaN | 470.77454 | NaN | NaN | 1467.2842 | NaN |
| Myh9 | KLEGDSTDLSDQIAELQAQ | -1052 | -1070 | 1 | 0.001392869 | 19 | NaN | 995.23505 | NaN | NaN | NaN | NaN |
| Myh9 | EELEEEQGNTELINDR | -1736 | -1751 | 0 | 0.00204437 | 16 | NaN | 571.69116 | NaN | NaN | NaN | NaN |
| Gsr | ADFDNTVAIHPTSSEELVTLR | -480 | -500 | 0 | 0 | 21 | NaN | 280.56198 | NaN | NaN | 291.26138 | 686.412 |
| Gsr | ADFDNTVAIHPT | -480 | -491 | 0 | 0 | 12 | NaN | 709.42004 | 646.50397 | NaN | NaN | NaN |
| Slc3a2 | LQDGVDGFQFR | -232 | -242 | 0 | 0 | 11 | NaN | 944.6984 | 991.4243 | 1003.6765 | 1067.2363 | 1024.9962 |
| Slc3a2 | IGDLQAFVGR | -120 | -129 | 0 | 0.000869338 | 10 | NaN | 340.1098 | NaN | 652.9639 | 969.72833 | 1044.2731 |
| Slc3a2 | DFIPDHLLR | -337 | -345 | 0 | 0.00025015 | 9 | NaN | 416.3154 | 390.4904 | NaN | NaN | NaN |
| Vars1 | LPALEQGPGGL | -57 | -67 | 0 | 0.002566359 | 11 | NaN | 1477.124 | 1551.8198 | 1306.8159 | NaN | 1580.4585 |
| Vars1 | QGFFKPEYGRPS | -315 | -326 | 2 | 0 | 12 | NaN | 492.81406 | 603.599 | 729.73083 | NaN | 787.31824 |
| Vars1 | VPPPFLGLPR | -623 | -632 | 0 | 0.00025015 | 10 | NaN | 410.98907 | NaN | 29.64568 | 233.60909 | 368.40125 |
| Vars1 | GPGQDPQAALGALGK | -123 | -137 | 0 | 0 | 15 | NaN | 1191.9001 | 2946.189 | NaN | NaN | NaN |
| Vars1 | GETTLWNPGCDHAGIAT | -371 | -387 | 0 | 0.006820063 | 17 | NaN | 1179.6321 | NaN | NaN | NaN | NaN |
| Vars1 | RYGEAGDGPGWGGPHPR | -23 | -39 | 1 | 0.000989247 | 17 | NaN | 235.11496 | NaN | NaN | NaN | NaN |
| Icam1 | RTFDLPATIPK | -212 | -222 | 1 | 0 | 11 | NaN | 1137.8848 | NaN | 1165.1671 | 1155.116 | 1165.3153 |
| Ppp1cb | ADGELNVDSLITR | -2 | -14 | 0 | 0 | 13 | NaN | 773.9578 | 696.1544 | 1497.7953 | 1914.879 | 1955.9191 |
| Mrpl24 | DTSVEDALER | -176 | -185 | 0 | 0.00204437 | 10 | NaN | 2500.6077 | NaN | NaN | NaN | NaN |
| Rps8 | ELEFYLR | -194 | -200 | 0 | 0.007791272 | 7 | NaN | 733.81177 | 674.9965 | 684.4896 | 739.28754 | 638.5203 |
| Rps8 | ASNNELVR | -85 | -92 | 0 | 0.006078721 | 8 | NaN | 1610.3629 | NaN | NaN | NaN | 1397.2979 |
| Hnrnpk | GSYGDLGGPIITTQ | -378 | -391 | 0 | 0 | 14 | NaN | 1086.3854 | 1086.4359 | 1030.1305 | 221.2301 | 228.04102 |
| Hnrnpk | METEQPEETFPNTETNGEFGK | -1 | -21 | 0 | 0 | 21 | NaN | 170.8446 | NaN | 260.3871 | 352.20566 | 284.95663 |
| Hnrnpk | VLIGGKPDR | -193 | -201 | 1 | 0.002365814 | 9 | NaN | 734.46826 | NaN | 784.61725 | 909.3526 | NaN |
| Hnrnpk | RPAEDMEEEQAF | -22 | -33 | 1 | 0.001811524 | 12 | NaN | 678.7187 | 578.6669 | NaN | 215.30095 | NaN |
| Hnrnpk | SISADIETIGEILKK | -89 | -103 | 1 | 0.000431655 | 15 | NaN | 649.2341 | 673.91376 | NaN | NaN | NaN |
| Glg1 | VAELSSDDFHLDR | -294 | -306 | 0 | 0 | 13 | NaN | 1551.2224 | 1341.0212 | NaN | 1111.9639 | 1527.857 |
| Npm1 | GQHLVAVEEDAESEDEDEEDVK | -113 | -134 | 0 | 0.00111006 | 22 | NaN | 347.72287 | 328.2422 | 232.32639 | 198.75003 | 271.9668 |
| Npm1 | ADKDYHFKVD | -25 | -34 | 2 | 0.000591124 | 10 | NaN | 713.53735 | 1394.3793 | 1303.8856 | 954.4399 | 1071.5388 |
| Npm1 | HLVAVEEDAESEDEDEEDVK | -115 | -134 | 0 | 0.001291103 | 20 | NaN | 354.32526 | 542.28815 | 237.74136 | 285.0978 | 289.60645 |
| Npm1 | MSVQPTVSLGGFEITPPVVLR | -81 | -101 | 0 | 0.000907206 | 21 | NaN | 276.30295 | 307.51456 | 312.45392 | 339.57596 | 339.35126 |
| Npm1 | PSSVEDIK | -239 | -246 | 0 | 0.00122419 | 8 | NaN | 882.6704 | 1015.018 | NaN | 945.17065 | 807.2132 |
| Npm1 | VKLDEDDEDDDEDDEDDEDDDDDDFDEEETEEK | -156 | -188 | 1 | 0.00111006 | 33 | NaN | 458.49396 | 446.11487 | 425.19113 | NaN | NaN |
| Rbbp4 | AENIYNDEDPEGSVDPEGQGS | -405 | -425 | 0 | 0.00025015 | 21 | NaN | 357.73895 | 338.46097 | 341.59735 | 403.59445 | 419.74396 |
| Tmsb10 | ADKPDMGEIASFDK | -2 | -15 | 1 | 0 | 14 | NaN | 656.04114 | 719.469 | 885.3776 | 238.5409 | 956.25183 |
| Ranbp3 | ADLANEEKPAVAPSVF | -2 | -17 | 1 | 0.000431655 | 16 | NaN | 780.104 | 863.6184 | 896.553 | 889.76337 | 771.443 |
| Orc6 | ADLDLSRPLFTTA | -131 | -143 | 1 | 0.001092529 | 13 | NaN | 761.15027 | NaN | NaN | 1277.2725 | NaN |
| Capza2 | EATDPRPYEAENAIESWR | -104 | -121 | 1 | 0 | 18 | NaN | 794.27203 | NaN | 837.9136 | 639.7761 | 648.6427 |

| | | | | | | | | | | | |
|---|---|---|---|---|---|---|---|---|---|---|---|
| Capza2 | RTSVETALR | -121 | -129 | 1 | 0.00272025 | 9 | NaN | 365.49783 | NaN | 638.5307 | 676.9404 | 628.0398 |
| Spcs3 | GHVSVPFPDTYEITK | -164 | -178 | 0 | 0 | 15 | NaN | 4518.517 | NaN | NaN | NaN | NaN |
| Efhd2 | FEEEIKAEQEER | -207 | -218 | 1 | 0 | 12 | NaN | 822.584 | 710.0482 | 1161.5989 | 516.6773 | 895.80414 |
| Efhd2 | DLNQGIGEPQSPSR | -64 | -77 | 0 | 0 | 14 | NaN | 898.4263 | NaN | 627.0441 | NaN | NaN |
| Efhd2 | GSADDELSAK | -49 | -58 | 0 | 0.000907206 | 10 | NaN | 569.8555 | NaN | NaN | 412.01672 | NaN |
| Adsl | LEHLLDPSSFTGR | -440 | -452 | 0 | 0 | 13 | NaN | 1906.5247 | 1956.6714 | 1624.9104 | 2125.1216 | 1510.4574 |
| Adsl | LFEGDHQK | -211 | -218 | 0 | 0.003003218 | 8 | NaN | 1258.4446 | NaN | NaN | 2153.39 | NaN |
| Mical1 | FCVDGHFFHR | -696 | -705 | 0 | 0 | 10 | NaN | 331.28992 | NaN | 342.4815 | 354.01495 | 370.50375 |
| Mical1 | FCTGTLDHISIR | -147 | -158 | 0 | 0 | 12 | NaN | 499.9705 | 524.3582 | NaN | NaN | NaN |
| Ndufs3 | SLADLTAVDVPTR | -109 | -121 | 0 | 0.00122419 | 13 | NaN | 1667.9377 | NaN | NaN | NaN | NaN |
| Cct6a | VVDSILAIR | -172 | -180 | 0 | 0.00078661 | 9 | NaN | 442.2155 | 424.93692 | 161.46573 | NaN | 164.90836 |
| Glod4 | ADPDGHEICFVGDEAFR | -246 | -262 | 0 | 0 | 17 | NaN | 341.26212 | 464.752 | NaN | 1331.86 | 1173.8403 |
| Glod4 | HEEFEEGCK | -34 | -42 | 0 | 0.000230707 | 9 | NaN | 1130.7936 | 1337.088 | NaN | 1284.1718 | 1278.2444 |
| Uba2 | NTPSEPIHCIVW | -177 | -188 | 0 | 0 | 12 | NaN | 729.7851 | 711.0608 | 663.3571 | 784.2274 | 836.3955 |
| Uba2 | GVTECYECHPKPTQR | -154 | -168 | 1 | 0 | 15 | NaN | 402.77982 | NaN | NaN | NaN | NaN |
| Stk24 | SQCDSPYVTK | -75 | -84 | 0 | 0.00025015 | 10 | NaN | 421.97458 | NaN | 471.83505 | 324.4334 | NaN |
| Pabpn1 | TNRPGIS | -245 | -251 | 1 | 0.008890469 | 7 | NaN | 746.6129 | NaN | 386.49686 | NaN | NaN |
| Arhgdia | EFLTPMEEAPK | -157 | -167 | 0 | 0 | 11 | NaN | 453.67233 | 870.3716 | NaN | NaN | NaN |
| Rps6ka3 | YVDESGNPESIR | -547 | -558 | 0 | 0 | 12 | NaN | 3530.572 | NaN | NaN | NaN | 2606.5525 |
| Copb2 | QLGSSSPNFTLEGHEK | -171 | -186 | 0 | 0 | 16 | NaN | 3833.8743 | 1175.8088 | 3020.3474 | 759.03796 | 4939.4727 |
| Copb2 | TCVQTLEGHAQ | -220 | -230 | 0 | 0 | 11 | NaN | 1337.3875 | NaN | 1279.0737 | 740.7645 | 437.33658 |
| Copb2 | VWQLGSSSPNFTLEGHEK | -169 | -186 | 0 | 0.003900536 | 18 | NaN | 1209.1149 | NaN | NaN | NaN | NaN |
| Cpsf1 | LISPVSWEAIPNAR | -1076 | -1089 | 0 | 0.00025015 | 14 | NaN | 2314.9802 | 2378.3555 | NaN | NaN | 2617.8057 |
| Cpsf1 | ADPTHWCLLVR | -777 | -787 | 0 | 0.00025015 | 11 | NaN | 375.3013 | 377.8989 | NaN | NaN | NaN |
| Coro1a | WDSGFCAVNPK | -35 | -45 | 0 | 0 | 11 | NaN | 1410.6923 | 1410.9532 | 1711.3965 | 2137.9573 | 2112.5881 |
| Coro1a | DRPHEGTRPVHAVFVSEGK | -215 | -233 | 2 | 0.00025015 | 19 | NaN | 1306.3799 | NaN | 753.53253 | 402.13913 | 1388.9989 |
| Coro1a | PEPSGTPSSDTVSR | -419 | -432 | 0 | 0 | 14 | NaN | 867.65955 | 660.66125 | NaN | 1706.8577 | 1535.821 |
| Coro1a | DRPHEGTRPVHAVF | -215 | -228 | 2 | 0.002629273 | 14 | NaN | 303.7329 | NaN | 3616.6504 | 1324.23 | 419.70978 |
| Calm1;Calm2;( | EAFSLFDKDGDGTIT | (15);(15);(15) | (29);(29);(29) | 1 | 0.000471409 | 15 | NaN | 441.49915 | NaN | NaN | 435.50836 | NaN |
| Nup133 | WELDDSSEK | -296 | -304 | 0 | 0.004439092 | 9 | NaN | 716.64276 | NaN | NaN | NaN | NaN |
| Cdk6 | GALNHPYFQDLER | -293 | -305 | 0 | 0 | 13 | NaN | 16490.13 | 1543.5391 | 1185.0741 | 1477.2466 | 1474.7383 |
| Cdk6 | GLDFLHSH | -132 | -139 | 0 | 0.000431655 | 8 | NaN | 623.7425 | 643.59247 | 754.58746 | 918.0011 | 732.8792 |
| Cdk6 | FVTDIDELGK | -265 | -274 | 0 | 0.000431655 | 10 | NaN | 1162.8358 | 1394.3795 | NaN | 134.29872 | 1236.0339 |
| Dph5 | LLEIVQNHR | -197 | -205 | 0 | 0.00204437 | 9 | NaN | 490.34604 | NaN | 240.05095 | NaN | NaN |
| Dph5 | ADREEVEQEADNIFK | -56 | -70 | 1 | 0.002629273 | 15 | NaN | 394.1747 | NaN | NaN | NaN | 771.6954 |
| Dph5 | EALEEFYGR | -43 | -51 | 0 | 0.00025015 | 9 | NaN | 1164.3505 | NaN | NaN | NaN | NaN |
| Rab1a | VYDVTDQESFNNVK | -90 | -103 | 0 | 0.000431655 | 14 | NaN | 1105.0652 | 1046.9802 | NaN | 693.86847 | 539.55963 |
| Pfkp | KFVSDDSICVL | -709 | -719 | 1 | 0 | 11 | NaN | 1724.7427 | 1673.1628 | NaN | 1607.3307 | 1435.4164 |
| Pfkp | GAIDMQNKPITSEK | -273 | -286 | 1 | 0.00111006 | 14 | NaN | 769.9923 | 902.92773 | NaN | NaN | 565.5628 |
| Pfkp | DLLFQPVAELKK | -725 | -736 | 1 | 0.00025015 | 12 | NaN | 528.0417 | NaN | NaN | 173.29816 | 439.5313 |
| Pfkp | KVTDFEHR | -736 | -743 | 1 | 0.000431655 | 8 | NaN | 545.78955 | NaN | NaN | 274.64105 | NaN |
| Pfkp | SIGADTALNTITDTCDR | -548 | -564 | 0 | 0 | 17 | NaN | 475.4392 | NaN | NaN | 616.1044 | NaN |
| Nucb1 | LSQELDFVSHNVR | -84 | -96 | 0 | 0 | 13 | NaN | 2366.2646 | NaN | 1750.5244 | 1621.6848 | 1590.4103 |
| Nucb1 | EFGDTGEGWK | -324 | -333 | 0 | 0 | 10 | NaN | 614.4442 | 594.46375 | NaN | NaN | NaN |
| Tars1 | ELNPGDGAFYGPK | -530 | -542 | 0 | 0.000394361 | 13 | NaN | 1503.4291 | 1448.2645 | 1463.8158 | 1906.4114 | 1881.1333 |
| Tars1 | LNPWPEYINTR | -46 | -56 | 0 | 0 | 11 | NaN | 960.1895 | 846.66064 | 800.0174 | 978.34143 | 302.74783 |
| Tars1 | ELSPGSCFFLPK | -336 | -347 | 0 | 0 | 12 | NaN | 313.62848 | 604.069 | 773.3003 | 952.2053 | 712.35724 |
| Tars1 | SPGSCFFLPK | -338 | -347 | 0 | 0.006735683 | 10 | NaN | 229.15756 | NaN | NaN | NaN | NaN |
| Psmc4 | KIEFPLPDR | -330 | -338 | 1 | 0 | 9 | NaN | 1075.7969 | 1958.7924 | 484.04288 | 1382.7844 | 1497.0787 |
| Psmc4 | NLSEEVDLEDYVARPDK | -353 | -369 | 1 | 0.00025015 | 17 | NaN | 832.6902 | NaN | 2620.4514 | 915.9471 | 1660.5322 |
| Psmc4 | EAVELPLTHF | -179 | -188 | 0 | 0.001092529 | 10 | NaN | 595.9826 | NaN | 530.7339 | NaN | 696.98895 |
| Psmc4 | STTGSNYYVR | -105 | -114 | 0 | 0.004983786 | 10 | NaN | 797.79724 | 928.7007 | NaN | NaN | NaN |
| Septin7 | ADTLTPEECQQFK | -195 | -207 | 0 | 0 | 13 | NaN | 1388.6176 | 1282.5245 | NaN | NaN | NaN |
| Sptan1 | HQAFEAELHAN | -1592 | -1602 | 0 | 0 | 11 | NaN | 782.0646 | 699.28656 | NaN | 1227.0205 | 1249.0497 |
| Sptan1 | VLETAEDIQER | -8 | -18 | 0 | 0.000230707 | 11 | NaN | 4511.974 | 659.2672 | NaN | NaN | NaN |
| Sptan1 | EELITNWEQIR | -337 | -347 | 0 | 0 | 11 | NaN | 3019.2122 | NaN | NaN | NaN | 3701.1838 |
| Sptan1 | GVIDMGNSLIER | -1608 | -1619 | 0 | 0 | 12 | NaN | 809.7326 | NaN | 1574.3429 | NaN | NaN |
| Sptan1 | LSDDNTIGQEEIQQR | -1830 | -1844 | 0 | 0 | 15 | NaN | 521.1722 | 594.069 | NaN | NaN | NaN |
| Sptan1 | DVEDEETWIR | -792 | -801 | 0 | 0.00025015 | 10 | NaN | 170.99173 | NaN | NaN | NaN | 332.3108 |
| Sptan1 | VNSLGETAQR | -1289 | -1298 | 0 | 0 | 10 | NaN | 1967.3652 | NaN | NaN | NaN | NaN |
| Sptan1 | SLQQLAEER | -1217 | -1225 | 0 | 0 | 9 | NaN | 420.71024 | NaN | NaN | NaN | NaN |
| Gbp2 | TLCGGIIVNGPR | -279 | -290 | 0 | 0 | 12 | NaN | 7637.4785 | 8071.7607 | NaN | 5806.9023 | 6536.002 |
| Gbp2 | LKNYFDSK | -459 | -466 | 1 | 0.000230707 | 8 | NaN | 1007.5587 | NaN | 650.20636 | 751.306 | 599.9382 |
| Erh | ADYESVNECMEGVCK | -20 | -34 | 0 | 0 | 15 | NaN | 387.93207 | 402.60132 | 822.1075 | 821.7761 | 822.80597 |
| Erh | RMNPNSPSITYDIS | -42 | -55 | 1 | 0 | 14 | NaN | 615.3609 | NaN | NaN | 409.43216 | 501.54086 |
| Rmdn1 | ADYLYESGETEK | -85 | -96 | 0 | 0.00122419 | 12 | NaN | 326.5093 | NaN | NaN | 402.29108 | NaN |
| Gc | NDEICEAFR | -141 | -149 | 0 | 0 | 9 | NaN | 290.3247 | NaN | NaN | 486.011 | 524.20764 |
| Gc | ADYSENTFTEYK | -408 | -419 | 0 | 0 | 12 | NaN | 4583.8086 | NaN | NaN | NaN | NaN |
| Gcn1 | LVLPSPDTDEK | -1111 | -1121 | 0 | 0.00025015 | 11 | NaN | 990.7119 | 1628.7838 | 1851.6492 | 1076.5791 | NaN |
| Gcn1 | VISESPPDQWEAR | -1222 | -1234 | 0 | 0 | 13 | NaN | 1130.5895 | NaN | NaN | 830.5948 | 290.34027 |
| Gcn1 | SEDALCTVLR | -535 | -544 | 0 | 0 | 10 | NaN | 769.16797 | NaN | 1302.9972 | NaN | NaN |
| Tkt | MEGYHKPDQQK | -1 | -11 | 1 | 0 | 11 | NaN | 333.3166 | 832.63336 | 554.6917 | 819.00244 | 783.458 |
| Tkt | ISSDLDGHPVPK | -103 | -114 | 0 | 0 | 12 | NaN | 1550.4496 | 5598.6274 | 5769.725 | 6208.0615 | 5880.5703 |
| Tkt | IIVDGHSVEELCK | -214 | -226 | 0 | 0 | 13 | NaN | 378.42856 | 891.18054 | 480.60403 | 349.15866 | 519.2398 |
| Tkt | SKDDQVTVIGAGVTLHEA | -498 | -515 | 1 | 0.00111006 | 18 | NaN | 507.0233 | NaN | NaN | 585.55646 | NaN |
| Tkt | ILATPPQEDAPSVDIANIR | -284 | -302 | 0 | 0.000230707 | 19 | NaN | 307.2633 | NaN | NaN | 736.3124 | NaN |
| Tkt | FIECYIAEQN | -359 | -368 | 0 | 0.00025015 | 10 | NaN | 288.38162 | NaN | NaN | NaN | NaN |
| Zbp1 | HPSYSFSDSPPEVSTT | -265 | -280 | 0 | 0.000431655 | 16 | NaN | 1338.469 | 1286.4636 | 241.74413 | 1456.8754 | NaN |

| | | | | | | | | | | | | |
|---|---|---|---|---|---|---|---|---|---|---|---|---|
| Vdac2 | TGDFQLHTN | -187 | -195 | 0 | 0.00025015 | 9 | NaN | 845.9081 | 1059.5831 | NaN | 1560.8954 | 720.23944 |
| Ptma | VAEDDEDDDVDTK | -91 | -103 | 0 | 0 | 13 | NaN | 8321.947 | 8359.123 | 6159.997 | 7768.763 | 1533.2742 |
| Ptma | DTSSEITTK | -7 | -15 | 0 | 0.002972566 | 9 | NaN | 1404.0977 | 1349.2726 | 2683.5747 | NaN | 3246.2295 |
| Notch2 | AEDEALLSEDDPIDR | -1773 | -1787 | 0 | 0.00025015 | 15 | NaN | 4609.993 | 4706.609 | NaN | 462.09082 | NaN |
| Surf4 | AEDFADQFLR | -10 | -19 | 0 | 0 | 10 | NaN | 936.6784 | 224.2202 | 1027.02 | 906.80975 | 1145.0853 |
| Surf4 | EDFADQFLR | -11 | -19 | 0 | 0 | 9 | NaN | 497.8708 | 458.8173 | 400.312 | 561.33734 | 554.9154 |
| Copa | KETFDPEKETIPDIDPNAK | -775 | -793 | 2 | 0 | 19 | NaN | 569.1136 | 525.47186 | 541.1094 | 612.71625 | 488.81125 |
| Copa | AEDGFFVPPTK | -902 | -912 | 0 | 0 | 11 | NaN | 435.3818 | 161.5054 | 1842.3291 | 567.20447 | 1191.5275 |
| Copa | RLLELGPKPEVA | -1127 | -1138 | 2 | 0.000591124 | 12 | NaN | 540.7618 | 593.32495 | 575.27405 | 663.0873 | NaN |
| Copa | DTNWPLLTVSK | -805 | -815 | 0 | 0.000394361 | 11 | NaN | 757.9179 | 662.8352 | NaN | NaN | NaN |
| Copa | SVPLLVVDNK | -1035 | -1044 | 0 | 0 | 10 | NaN | 592.8436 | 620.9251 | NaN | NaN | NaN |
| Copa | LDALCNIHENIR | -518 | -529 | 0 | 0.00204437 | 12 | NaN | 316.306 | 905.2494 | NaN | NaN | NaN |
| Hdlbp | AEDKDQDLITIIGK | -766 | -779 | 1 | 0 | 14 | NaN | 799.487 | 926.8005 | 636.83215 | 797.85065 | 640.24634 |
| Hdlbp | VSEDVPLDHR | -1128 | -1137 | 0 | 0 | 10 | NaN | 1060.6249 | NaN | 326.30432 | 981.04724 | 320.8507 |
| Hdlbp | LHNSLIGTK | -664 | -672 | 0 | 0 | 9 | NaN | 545.84985 | 583.57227 | NaN | NaN | NaN |
| Hdlbp | SYSISVPIFK | -581 | -590 | 0 | 0.009388464 | 10 | NaN | 788.19165 | NaN | NaN | NaN | NaN |
| Rtcb | RNLDFQDVLDK | -441 | -451 | 1 | 0.000230707 | 11 | NaN | 696.30115 | 1249.587 | 1072.5422 | 887.2195 | 827.96606 |
| Rtcb | AEDKEHCEEYGR | -187 | -198 | 1 | 0.00051306 | 12 | NaN | 453.84256 | 451.6995 | 459.83844 | 356.2693 | 426.72983 |
| Rtcb | SIGLPDVHSGYGF | -82 | -94 | 0 | 0.00025015 | 13 | NaN | 911.4251 | NaN | 1277.7781 | 1221.6688 | NaN |
| Rtcb | LVMEEAPESYK | -466 | -476 | 0 | 0.00051306 | 11 | NaN | 1324.4967 | NaN | 1403.9803 | NaN | NaN |
| Rtcb | DKEHCEEYGR | -189 | -198 | 1 | 0.005242018 | 10 | NaN | 445.71884 | 361.50354 | NaN | NaN | NaN |
| Arfgap1 | LETQDDYEPSWSLQDK | -87 | -102 | 0 | 0 | 16 | NaN | 908.60834 | NaN | NaN | NaN | NaN |
| Arfgap1 | NSNSDGWESWEGASGEGR | -376 | -393 | 0 | 0.00025015 | 18 | NaN | 521.81323 | NaN | NaN | NaN | NaN |
| Churc1 | AEDTISILPDDPR | -94 | -106 | 0 | 0 | 13 | NaN | 626.9013 | 187.89043 | 773.02356 | NaN | NaN |
| Hsph1 | RGPFELEAF | -437 | -445 | 1 | 0.008014596 | 9 | NaN | 1199.0695 | NaN | 1198.7986 | 1464.2913 | 1479.1353 |
| Hsph1 | KPVTDCVISVPSFFTD | -135 | -150 | 1 | 0.00078661 | 16 | NaN | 355.842 | 419.71866 | 428.6161 | 506.6266 | NaN |
| Hsph1 | YNHIDESEMK | -741 | -750 | 0 | 0 | 10 | NaN | 258.6025 | NaN | 413.12183 | 304.5318 | 432.4489 |
| Atp5f1d | ASHVPTLQVLR | -71 | -81 | 0 | 0 | 11 | NaN | 749.8134 | 693.13477 | 635.7374 | 729.96 | NaN |
| Atp5f1d | VLRPGLVVVHTEDGTTTK | -79 | -96 | 1 | 0 | 18 | NaN | 420.53546 | NaN | NaN | NaN | 479.94647 |
| Atp5f1d | AEEAVTLDMLDLGAAR | -116 | -131 | 0 | 0.001055699 | 16 | NaN | 593.3748 | NaN | NaN | NaN | NaN |
| Tmed2 | AEECFFER | -29 | -36 | 0 | 0.006568856 | 8 | NaN | 130.6662 | NaN | NaN | 450.5224 | NaN |
| Canx | IADPDAVKPDDWDEDAPSK | -294 | -312 | 1 | 0 | 19 | NaN | 590.81116 | 592.19366 | 564.08417 | 661.664 | 643.7696 |
| Canx | VDQSVVNSGNLLNDMTPPVNPSR | -253 | -275 | 0 | 0 | 23 | NaN | 97.76054 | 306.20508 | 96.577065 | 266.69824 | 128.2648 |
| Canx | VVDDWANDGWGLKK | -447 | -460 | 1 | 0 | 14 | NaN | 778.2366 | 784.07446 | 695.8686 | 850.85364 | NaN |
| Canx | SKSDASTPPSPK | -47 | -58 | 1 | 0.00025015 | 12 | NaN | 694.929 | NaN | 633.7615 | 830.4573 | 780.51025 |
| Canx | WEVDEMK | -105 | -111 | 0 | 0.003393785 | 7 | NaN | 322.84186 | NaN | NaN | NaN | NaN |
| Canx | APVPTGEVYFAD | -63 | -74 | 0 | 0.00122419 | 12 | NaN | 224.35385 | NaN | NaN | NaN | NaN |
| P4ha1 | AEEDKLEQIKK | -48 | -58 | 2 | 0 | 11 | NaN | 385.1031 | 195.33043 | NaN | 471.7463 | 301.67654 |
| P4ha1 | KDEPDAFR | -435 | -442 | 1 | 0.008919338 | 8 | NaN | 2019.8813 | NaN | NaN | NaN | NaN |
| Cdc37 | EVAESDGQVELER | -72 | -84 | 0 | 0 | 13 | NaN | 952.2738 | 389.23557 | 964.85944 | 952.9737 | 960.18555 |
| Cdc37 | YLSDNVHLVCEETANYL | -175 | -191 | 0 | 0.00111006 | 17 | NaN | 443.2492 | NaN | 475.81003 | 728.6882 | 516.1514 |
| Dbnl | TRQEWESAGQQAPHPR | -251 | -266 | 1 | 0 | 16 | NaN | 488.31238 | NaN | 624.19037 | 811.58606 | 728.4415 |
| Dbnl | EPTAPVSR | -308 | -315 | 0 | 0 | 8 | NaN | 1132.714 | NaN | 1816.3802 | NaN | NaN |
| Dbnl | VIDEGWWR | -409 | -416 | 0 | 0.00078661 | 8 | NaN | 477.4088 | NaN | NaN | NaN | NaN |
| Clic1 | NSNPALNDNLEK | -120 | -131 | 0 | 0 | 12 | NaN | 5043.9463 | 4848.5234 | 2989.6892 | 2433.5886 | 39.680405 |
| Clic1 | KFLDGNELTLADCNLLPK | -166 | -183 | 1 | 0 | 18 | NaN | 470.45908 | NaN | 358.19608 | 368.8093 | 427.85266 |
| Clic1 | KFLDGNELTLA | -166 | -176 | 1 | 0.00149795 | 11 | NaN | 13589.803 | NaN | NaN | NaN | NaN |
| Ipo5 | AFDEVYPALPSDVQTAIK | -73 | -90 | 0 | 0.00025015 | 18 | NaN | 433.96442 | 491.172 | 613.88654 | 722.20264 | 575.0063 |
| Ipo5 | QQQHYLDVIK | -165 | -174 | 0 | 0.00078661 | 10 | NaN | 1084.4482 | NaN | 747.82367 | 1318.2969 | NaN |
| Ipo5 | AKLEEHFK | -799 | -806 | 1 | 0.00272025 | 8 | NaN | 673.0498 | 719.04254 | NaN | NaN | NaN |
| Ipo5 | IDEDGNNQWPEGLK | -121 | -134 | 0 | 0.00025015 | 14 | NaN | 429.3505 | NaN | NaN | NaN | 1042.8994 |
| Ipo5 | LMVPLLK | -706 | -712 | 0 | 0.000431655 | 7 | NaN | 310.25958 | 478.84198 | NaN | NaN | NaN |
| Pa2g4 | HLCAEAALR | -147 | -155 | 0 | 0 | 9 | NaN | 276.45685 | 1406.9758 | 1136.9016 | 2085.516 | 2099.5173 |
| Pa2g4 | KSDQDYILK | -93 | -101 | 1 | 0.000471409 | 9 | NaN | 4497.609 | NaN | 4532.0615 | 3010.5542 | 2955.5142 |
| Twf2 | ETIELVHTEPTNVA | -210 | -223 | 0 | 0 | 14 | NaN | 2543.473 | 1997.1436 | 488.58588 | 236.65848 | 499.08585 |
| Twf2 | ETIELVHTEPTNVAQLPSR | -210 | -228 | 0 | 0 | 19 | NaN | 700.92944 | 623.66174 | 543.431 | 591.09033 | 546.553 |
| Twf2 | AFPLQPEAQR | -179 | -188 | 0 | 0 | 10 | NaN | 1186.1174 | NaN | 1115.2992 | 1347.1196 | 1323.324 |
| Twf2 | KEFGGGHIKDELFGTVK | -110 | -126 | 2 | 0 | 17 | NaN | 882.83795 | 644.1848 | NaN | NaN | 741.06464 |
| Dhx15 | LTGQEEIDEACKR | -356 | -368 | 1 | 0 | 13 | NaN | 10491.066 | 11595.573 | 7595.187 | 9926.008 | 10522.056 |
| Dhx15 | TLATDILMGVLK | -266 | -277 | 0 | 0 | 12 | NaN | 359.91595 | 285.4933 | 465.07782 | 696.47644 | 527.9164 |
| Dhx15 | GVIILDEAHER | -255 | -265 | 0 | 0 | 11 | NaN | 401.586 | NaN | 1336.5026 | 967.3567 | 1036.8146 |
| Dhx15 | CINPFTNLPHTPR | -114 | -126 | 0 | 0 | 13 | NaN | 2609.9858 | NaN | 2666.0398 | NaN | NaN |
| Dhx15 | HRLDLGEDYPSGK | -5 | -17 | 1 | 0.00025015 | 13 | NaN | 208.06998 | NaN | NaN | 421.91302 | NaN |
| Fip1l1 | KPGADLSDYFNY | -166 | -177 | 1 | 0.002534621 | 12 | NaN | 10924.6 | 7734.8813 | 12740.4375 | NaN | 12937.262 |
| Tnpo1 | AGDEEAEVR | -232 | -240 | 0 | 0 | 9 | NaN | 517.2613 | NaN | 434.91962 | NaN | NaN |
| Ddb1 | FVYQDPQGR | -180 | -188 | 0 | 0 | 9 | NaN | 1952.977 | 2327.3633 | 1858.3499 | NaN | 1256.0333 |
| Ddb1 | QSGESIDIITR | -93 | -103 | 0 | 0 | 11 | NaN | 1460.7649 | NaN | 1563.235 | 1392.4014 | 1777.7102 |
| Ddb1 | ETGIIGIIDPECR | -117 | -129 | 0 | 0 | 13 | NaN | 508.34088 | NaN | NaN | 1286.5233 | 402.18314 |
| Ddb1 | TYLDNGVVFVGSR | -315 | -327 | 0 | 0 | 13 | NaN | 2074.9824 | NaN | NaN | 2353.2002 | NaN |
| Flna | VAQPSITDNKDGTVTVR | -1815 | -1831 | 1 | 0 | 17 | NaN | 1996.2555 | 2123.8503 | 1913.807 | 1981.4298 | 1911.8617 |
| Flna | VHSPSGALEECYVT | -2368 | -2381 | 0 | 0.00025015 | 14 | NaN | 1545.3019 | 490.14294 | 519.9177 | 1517.6558 | 1112.6736 |
| Flna | YCPTEPGNYIINIK | -2106 | -2119 | 0 | 0 | 14 | NaN | 771.65955 | 830.10425 | 744.18427 | 703.47754 | 983.6699 |
| Flna | VHGPGIQSGTTNKPNK | -1360 | -1375 | 1 | 0.00025015 | 16 | NaN | 469.68817 | 2868.1743 | 718.523 | 427.08868 | 611.7482 |
| Flna | TVPQHATSGPGPADVSK | -2543 | -2559 | 0 | 0 | 17 | NaN | 299.6477 | 1519.9497 | 1261.0774 | 265.25818 | 175.4666 |
| Flna | YIPVQQGPVGVNVTYGGDHIPK | -937 | -958 | 0 | 0 | 22 | NaN | 640.453 | NaN | 888.1365 | 958.78046 | 922.4823 |
| Flna | VQDNEGCSVEATVK | -711 | -724 | 0 | 0 | 14 | NaN | 361.27603 | NaN | 1079.6454 | 295.4555 | 1022.85974 |
| Flna | GDEHIPGSPYR | -2633 | -2643 | 0 | 0.000298805 | 11 | NaN | 2258.2363 | 2690.5369 | 3017.5935 | NaN | NaN |
| Flna | WGGQNIGR | -556 | -563 | 0 | 0.00122419 | 8 | NaN | 703.9686 | 720.9119 | NaN | NaN | 574.7659 |

| | | | | | | | | | | | | |
|---|---|---|---|---|---|---|---|---|---|---|---|---|
| Flna | LAVQITDPEGKPK | -1580 | -1592 | 1 | 0 | 13 | NaN | 508.2763 | 761.6162 | NaN | NaN | NaN |
| Flna | VQITDPEGKPK | -1582 | -1592 | 1 | 0 | 11 | NaN | 526.595 | NaN | NaN | NaN | NaN |
| Flna | MDCQECPEGYR | -2474 | -2484 | 0 | 0.005605646 | 11 | NaN | 263.71558 | NaN | NaN | NaN | NaN |
| Ppp5c | AEGYEVAHGGR | -431 | -441 | 0 | 0 | 11 | NaN | 1060.3823 | 1007.92706 | 245.20634 | 1987.5895 | 2172.721 |
| Ppp5c | LEENQLDYIIR | -415 | -425 | 0 | 0 | 11 | NaN | 415.45242 | 644.6576 | 761.15393 | 1023.3284 | 1042.3025 |
| Ppp5c | GYEVAHGGR | -433 | -441 | 0 | 0 | 9 | NaN | 292.72916 | NaN | 596.3447 | NaN | NaN |
| Ppp5c | IFNGDFVDR | -267 | -275 | 0 | 0.001636916 | 9 | NaN | 871.6109 | NaN | NaN | NaN | NaN |
| Tmbim6 | AEHGDKDYIWHC | -196 | -207 | 1 | 0 | 12 | NaN | 468.80963 | 554.2461 | NaN | 373.2981 | 324.1613 |
| Tmbim6 | INFDALLK | -8 | -15 | 0 | 0 | 8 | NaN | 379.19794 | NaN | 1241.2261 | 1144.1091 | 1624.5524 |
| Tgtp2;Tgtp1 | KIDSDLDNEQK | (171);(171) | (181);(181) | 1 | 0 | 11 | NaN | 761.23425 | 859.7571 | NaN | 1791.5034 | 918.55664 |
| Tgtp2;Tgtp1 | FPHLFAEHNDESLEDK | (333);(333) | (348);(348) | 0 | 0 | 16 | NaN | 141.95561 | NaN | 3626.3943 | 519.15717 | 446.9215 |
| Tgtp2;Tgtp1 | FPHLFAEHNDESLEDK | (333);(333) | (348);(348) | 0 | 0 | 16 | NaN | 141.95561 | NaN | 3626.3943 | 519.15717 | 446.9215 |
| Psph | LAEHPPHLTPGIR | -80 | -92 | 0 | 0 | 13 | NaN | 1932.1111 | 1720.4222 | 2355.2593 | 3086.4714 | 2968.6023 |
| Psph | AEHPPHLTPGIR | -81 | -92 | 0 | 0 | 12 | NaN | 646.8245 | 727.548 | 821.1369 | 814.82666 | 758.87244 |
| Uchl5 | TPAKEEDAFHFVS | -155 | -167 | 1 | 0.00025015 | 13 | NaN | 1717.9585 | 2677.6826 | 2236.8848 | 1846.0444 | 667.94385 |
| Uchl5 | ESDPGVFTELIK | -12 | -23 | 0 | 0 | 12 | NaN | 4282.5117 | 4311.43 | 543.2041 | NaN | 135.5066 |
| Mthfd1l | LQPLSPVPSDIEIS | -352 | -365 | 0 | 0 | 14 | NaN | 654.8111 | 1025.0845 | 609.57104 | 1527.3845 | 1211.6007 |
| Mthfd1l | LVADGCCNLQK | -772 | -782 | 0 | 0 | 11 | NaN | 1318.0029 | 1315.6049 | 1717.8704 | NaN | 1482.0304 |
| Mthfd1l | VQLSLLER | -397 | -404 | 0 | 0.00025015 | 8 | NaN | 268.9634 | 386.96674 | NaN | NaN | NaN |
| Eef2 | ARPFPDGLAEDIDKGEVSAR | -606 | -625 | 2 | 0.000230707 | 20 | NaN | 10226.626 | 10140.84 | 11531.882 | 11779.682 | 15481.523 |
| Eef2 | FDVHDVTLHAD | -702 | -712 | 0 | 0.00025015 | 11 | NaN | 5170.8643 | 5184.744 | 798.0894 | 3443.7114 | 3044.1511 |
| Eef2 | CELLYEGPPDDEAAMGIK | -369 | -386 | 0 | 0 | 18 | NaN | 2020.5867 | 1710.497 | 148.18707 | 2558.5017 | 2305.6887 |
| Eef2 | EESQVAGTPMFVVK | -772 | -785 | 0 | 0 | 14 | NaN | 1650.9746 | 1773.0336 | 2446.7966 | 2577.942 | 2320.505 |
| Eef2 | YLAEKYEWDVAEAR | -634 | -647 | 1 | 0 | 14 | NaN | 1197.4467 | 1037.4662 | 1117.0873 | 1153.9438 | 1177.4324 |
| Eef2 | HLEICLK | -553 | -559 | 0 | 0.00025015 | 7 | NaN | 919.63617 | 918.37396 | 1724.0217 | 1191.404 | 1186.0283 |
| Eef2 | LTDSLVCK | -35 | -42 | 0 | 0 | 8 | NaN | 648.4585 | 910.68805 | 1134.688 | 1221.6565 | 556.72736 |
| Eef2 | AYLPVNESFGFT | -786 | -797 | 0 | 0.000394361 | 12 | NaN | 559.28644 | 582.527 | 833.672 | 953.4485 | 776.6435 |
| Eef2 | QLILDPIFK | -291 | -299 | 0 | 0 | 9 | NaN | 521.86255 | 479.23895 | 444.1246 | 135.27124 | 455.9306 |
| Eef2 | VTDGALVVVDCVSGVCVQ | -121 | -138 | 0 | 0 | 18 | NaN | 730.55896 | NaN | 1071.3809 | 83.02104 | 1084.4667 |
| Eef2 | RWLPAGDAL | -342 | -350 | 1 | 0.00294445 | 9 | NaN | 745.3639 | 367.43274 | 381.91058 | NaN | NaN |
| Eef2 | GVQYLNEIKDS | -668 | -678 | 1 | 0.000947459 | 11 | NaN | 965.3929 | 969.8227 | NaN | NaN | NaN |
| Eef2 | QVAGTPMFVVK | -775 | -785 | 0 | 0.000755556 | 11 | NaN | 847.0076 | 1060.0991 | NaN | NaN | NaN |
| Eef2 | LSENDLNFIK | -81 | -90 | 0 | 0.000591124 | 10 | NaN | 577.69586 | NaN | NaN | 520.8288 | NaN |
| Eef2 | PFPDGLAEDIDKGEVS | -608 | -623 | 1 | 0.00272025 | 16 | NaN | 480.07742 | NaN | NaN | NaN | NaN |
| Got2 | CFTGLKPEQVER | -382 | -393 | 1 | 0 | 12 | NaN | 1601.1348 | 1620.2151 | 1182.7437 | 1357.9816 | 505.06674 |
| Got2 | NLDKEYLPIGGLAE | -91 | -104 | 1 | 0.00025015 | 14 | NaN | 1881.5962 | 448.93747 | NaN | 821.1699 | NaN |
| Got2 | SRDVFLPK | -152 | -159 | 1 | 0.001023236 | 8 | NaN | 1391.7484 | NaN | 1197.8943 | NaN | 2073.606 |
| Got2 | IPEQSVLLLHAC | -201 | -212 | 0 | 0 | 12 | NaN | 1102.6163 | 1044.4286 | NaN | 879.13226 | NaN |
| Got2 | ASAELALGENNEVLK | -108 | -122 | 0 | 0 | 15 | NaN | 3985.419 | 4243.472 | NaN | NaN | NaN |
| Iars2 | SLGNVINPDTIISGGK | -668 | -683 | 0 | 0 | 16 | NaN | 1509.8174 | 1488.9536 | NaN | 2449.819 | 2531.4583 |
| Iars2 | NIVHSYPCDWR | -458 | -468 | 0 | 0.00111006 | 11 | NaN | 853.1234 | 742.91394 | 903.3293 | 735.1124 | NaN |
| Iars2 | VWGVPIPVFHHK | -527 | -538 | 0 | 0 | 12 | NaN | 746.3279 | 688.2777 | 327.6446 | 300.55676 | NaN |
| Sf1 | VALNPDFKPPADYKPPATR | -117 | -135 | 2 | 0 | 19 | NaN | 1993.7052 | 1840.8009 | 1921.2704 | 2277.7776 | 2145.7532 |
| Sf1 | VALNPDFKPPADYKPPATR | -117 | -135 | 2 | 0 | 19 | NaN | 1993.7052 | 1840.8009 | 1921.2704 | 2277.7776 | 2145.7532 |
| Sf1 | TVIPGMPTVIPPGLTR | -31 | -46 | 0 | 0 | 16 | NaN | 517.51575 | 560.29645 | 389.34274 | 645.6839 | 678.8656 |
| Sf1 | AELGEAPVPASVGSTSGPATTPLASAPR | -318 | -345 | 0 | 0 | 28 | NaN | 191.19008 | 233.36258 | 251.61368 | 295.14487 | 274.21396 |
| Sf1 | MVALNPDFKPPADYKPPATR | -116 | -135 | 2 | 0.008014596 | 20 | NaN | 150.5766 | NaN | 277.11243 | NaN | 504.98364 |
| Eif2s2 | EVEPEPTEEKDVDADEEDSR | -43 | -62 | 1 | 0.00025015 | 20 | NaN | 695.8289 | 748.9324 | 634.6947 | 1149.5663 | 1089.265 |
| Eif2s2 | GTSGSIDGNNQLVIK | -243 | -257 | 0 | 0.00201047 | 15 | NaN | 1623.4996 | NaN | NaN | NaN | NaN |
| Aldh4a1 | AELIDFFR | -163 | -170 | 0 | 0.002068213 | 8 | NaN | 524.29626 | 525.72894 | 74.17956 | 546.4337 | 590.258 |
| Umps | RHEFLIF | -304 | -310 | 1 | 0.006100831 | 7 | NaN | 1254.9753 | 1331.8433 | 1999.0977 | 1345.4741 | NaN |
| Umps | KFADIGNTVK | -314 | -323 | 1 | 0 | 10 | NaN | 1226.969 | 1288.5164 | NaN | 2066.7131 | 2575.3396 |
| Umps | GLSSPVYIDLR | -31 | -41 | 0 | 0 | 11 | NaN | 1118.2432 | NaN | 1668.5844 | 2687.0444 | 1971.9391 |
| Tubb4b | INVYYNEATGGK | -47 | -58 | 0 | 0 | 12 | NaN | 1057.9631 | 1028.7478 | NaN | NaN | 1108.5247 |
| Nasp | NSQYDEAVAQFGK | -583 | -595 | 0 | 0 | 13 | NaN | 815.6318 | 569.4602 | 924.5886 | NaN | 868.33746 |
| Nasp | CVTDISHLVR | -694 | -703 | 0 | 0 | 10 | NaN | 452.15128 | 941.855 | NaN | 664.58594 | 1007.50726 |
| Nasp | EAEGSFTEYEKEIEELK | -613 | -629 | 1 | 0.00078661 | 17 | NaN | 259.67947 | 395.70297 | NaN | NaN | NaN |
| Cct4 | AENAGLNPISTVTELR | -466 | -481 | 0 | 0.00025015 | 16 | NaN | 2118.929 | 1980.5308 | 2169.0134 | 2911.5806 | 3007.7576 |
| Cct4 | GAPTAGPGSR | -10 | -19 | 0 | 0.006078721 | 10 | NaN | 354.31384 | 502.81512 | 622.99744 | NaN | 832.50977 |
| Gvin1 | ELSDEELHEK | -1885 | -1894 | 0 | 0 | 10 | NaN | 1208.2761 | 1495.1606 | 662.10425 | NaN | 1032.0969 |
| Gvin1 | SQYENQLLER | -1864 | -1873 | 0 | 0 | 10 | NaN | 833.1611 | 2243.0798 | NaN | 944.55237 | 1248.7117 |
| Gvin1 | LNISLGDIK | -2184 | -2192 | 0 | 0 | 9 | NaN | 107.48307 | 670.17426 | NaN | 388.42715 | 1303.3146 |
| Gvin1 | WVAAIFDK | -1471 | -1478 | 0 | 0 | 8 | NaN | 1131.0206 | NaN | 1533.4204 | 1020.52234 | NaN |
| Gvin1 | QFSLVGPEHGTEIK | -216 | -229 | 0 | 0 | 14 | NaN | 384.55505 | 472.72086 | NaN | NaN | NaN |
| Arpc4 | KPVEGYDISFL | -107 | -117 | 1 | 0.00111006 | 11 | NaN | 338.5974 | 519.52655 | 281.43518 | 311.81845 | 718.2625 |
| Arpc4 | RKPVEGYDISF | -106 | -116 | 2 | 0.004595494 | 11 | NaN | 905.3534 | 888.2795 | NaN | NaN | NaN |
| Smarcc1 | ENEENKELTDTCK | -833 | -845 | 1 | 0 | 13 | NaN | 860.216 | 847.3973 | NaN | 982.012 | 1047.6923 |
| Kpna3 | INPSIPITFLR | -208 | -218 | 0 | 0.00025015 | 11 | NaN | 1187.113 | NaN | 934.8909 | 1006.17456 | 534.79205 |
| Kpna3 | DVLSHFPNLLSHPK | -326 | -339 | 0 | 0.00051306 | 14 | NaN | 968.2596 | NaN | NaN | 578.2238 | 588.081 |
| Lrrfip1 | EVEERPDKDFAEK | -51 | -63 | 2 | 0 | 13 | NaN | 897.74133 | 1078.509 | 1054.3322 | 1026.6869 | 977.6523 |
| Lrrfip1 | AENQRPAEDSALSPGPLAGAK | -290 | -310 | 1 | 0 | 21 | NaN | 530.90155 | 639.2051 | 970.41833 | 1053.6827 | 875.66174 |
| Lrrfip1 | EPVEDPQSGSSGK | -616 | -628 | 0 | 0 | 13 | NaN | 282.18585 | 284.5613 | 399.23135 | NaN | NaN |
| Nap1l4 | DFEIGHFFR | -316 | -324 | 0 | 0.001811524 | 9 | NaN | 242.38396 | NaN | NaN | NaN | NaN |
| Uba1 | SLVLQRPQTWGDCVTW | -694 | -709 | 1 | 0.001743443 | 16 | NaN | 564.9831 | 939.56854 | 899.0857 | 1210.4633 | 1158.9935 |
| Uba1 | FKQPAENVNQYLTDSK | -656 | -671 | 1 | 0.000550307 | 16 | NaN | 502.3872 | 373.40668 | 354.74915 | 491.5955 | 533.0042 |
| Uba1 | GCGEGGEVVVTDMDTIEK | -493 | -510 | 0 | 0 | 18 | NaN | 986.85425 | NaN | 2218.3386 | 1163.857 | 1045.7944 |
| Uba1 | LDQPMTEIVSR | -1011 | -1021 | 0 | 0 | 11 | NaN | 2073.9243 | NaN | NaN | 2215.383 | 781.7868 |
| Uba1 | NEEDATELVGLA | -351 | -362 | 0 | 0.000627943 | 12 | NaN | 112.03233 | NaN | NaN | 219.87395 | NaN |

| | | | | | | | | | | | | |
|---|---|---|---|---|---|---|---|---|---|---|---|---|
| Mybbp1a | VGTFLEGCQDDPKR | -354 | -367 | 1 | 0 | 14 | NaN | 1009.4659 | 1408.1217 | 2781.5303 | 1186.8589 | 1530.4509 |
| Mybbp1a | SEVPESMFQEILPK | -184 | -197 | 0 | 0 | 14 | NaN | 2177.9348 | 794.7979 | 2406.8792 | 415.16995 | 2822.7007 |
| Mybbp1a | LVSLVDHLHLEK | -454 | -465 | 0 | 0 | 12 | NaN | 1162.4141 | 1137.6079 | NaN | 294.3565 | 995.6843 |
| Mybbp1a | LVSLVDHLHLEK | -454 | -465 | 0 | 0 | 12 | NaN | 1162.4141 | 1137.6079 | NaN | 294.3565 | 995.6843 |
| Mybbp1a | LDLDFVTR | -964 | -971 | 0 | 0 | 8 | NaN | 939.2461 | NaN | NaN | 612.696 | 802.066 |
| Fam107b | GLAPQNKPELQ | -44 | -54 | 1 | 0.00078661 | 11 | NaN | 2006.3591 | 2002.7462 | 1047.0686 | 2190.3577 | 2219.835 |
| Fam107b | LEQLELEK | -91 | -98 | 0 | 0.00555405 | 8 | NaN | 1294.6934 | NaN | 1557.4407 | NaN | NaN |
| Actr3 | DREVGIPPEQSLETAK | -210 | -225 | 1 | 0.00025015 | 16 | NaN | 1515.4865 | 1682.2264 | 4062.5264 | 4096.9385 | 2323.3186 |
| Actr3 | KDYEEIGPSIC | -398 | -408 | 1 | 0 | 11 | NaN | 1124.1141 | 1148.934 | 1243.2124 | 1560.3562 | 462.45636 |
| Actr3 | EFNKYDTDGSK | -241 | -251 | 1 | 0 | 11 | NaN | 654.7435 | 731.58673 | 667.70667 | 897.91016 | 951.583 |
| Actr3 | LSEELSGGR | -349 | -357 | 0 | 0 | 9 | NaN | 471.47208 | 278.80487 | 2139.9587 | 378.0839 | 326.02768 |
| Actr3 | LGYAGNTEPQFIIPSCIAIK | -19 | -38 | 0 | 0 | 20 | NaN | 263.5244 | 379.65466 | 433.1606 | 509.4475 | 513.0997 |
| Actr3 | KEFSIDVGYER | -265 | -275 | 1 | 0.000550307 | 11 | NaN | 1471.4175 | NaN | NaN | 1399.8005 | NaN |
| Hyou1 | GQAGPEGAAPAPEEDKKPK | -670 | -688 | 2 | 0 | 19 | NaN | 431.63824 | 493.35757 | NaN | 480.30502 | 355.23523 |
| Hyou1 | LDQIEQVILVGGATR | -374 | -388 | 0 | 0 | 15 | NaN | 189.66974 | 159.3647 | NaN | 172.91312 | 426.84683 |
| Csk | LVEHYTTDADGLCTR | -152 | -166 | 0 | 0.000947459 | 15 | NaN | 261.1988 | NaN | 299.68692 | NaN | NaN |
| Stim1 | LTEPQLGLGSQR | -503 | -514 | 0 | 0 | 12 | NaN | 2292.331 | 2878.9106 | 3683.7415 | NaN | NaN |
| Stim1 | AEQSLHDLQER | -254 | -264 | 0 | 0 | 11 | NaN | 983.9011 | 781.3476 | NaN | NaN | NaN |
| Ubqln1 | DQDTLSQHGIHDGLTVH | -78 | -94 | 0 | 0 | 17 | NaN | 486.67938 | NaN | NaN | 605.15326 | 501.52148 |
| Smc1a | IAFGGHQR | -628 | -635 | 0 | 0.00111006 | 8 | NaN | 1102.8392 | 1010.0219 | 1087.7789 | 935.71954 | 992.9973 |
| Smc1a | RLEFENQK | -807 | -814 | 1 | 0.001743443 | 8 | NaN | 4032.3892 | 4392.884 | 1502.7666 | 2264.8906 | NaN |
| Smc1a | ALVCDNVEDAR | -616 | -626 | 0 | 0 | 11 | NaN | 2195.1382 | 2229.9512 | 2229.7075 | NaN | NaN |
| Smc1a | SLIGVYPEQGDCVISK | -1199 | -1214 | 0 | 0 | 16 | NaN | 362.81168 | 386.56824 | NaN | NaN | NaN |
| Dis3 | TANPELIDR | -202 | -210 | 0 | 0 | 9 | NaN | 2244.444 | NaN | NaN | NaN | NaN |
| Dis3 | DVSHFIRPGNALDQESAR | -508 | -525 | 1 | 0.00025015 | 18 | NaN | 436.5211 | NaN | NaN | NaN | NaN |
| Tpr | FHNELNAHIK | -349 | -358 | 0 | 0 | 10 | NaN | 4315.913 | 4196.4575 | 1080.5966 | 4820.974 | 5467.151 |
| Tpr | ETPLFLAHEEESGGR | -2263 | -2277 | 0 | 0 | 15 | NaN | 926.2842 | 427.59222 | NaN | 774.75714 | 1110.0706 |
| Tpr | AESQLLECK | -1194 | -1202 | 0 | 0.001023236 | 9 | NaN | 3929.2378 | NaN | NaN | NaN | NaN |
| Fkbp4 | TGWLLDGTK | -58 | -66 | 0 | 0.001636916 | 9 | NaN | 406.89185 | NaN | NaN | 384.65518 | NaN |
| Fkbp4 | AEVAAGDHPTDAEMK | -427 | -441 | 0 | 0 | 15 | NaN | 223.50418 | 722.0708 | NaN | NaN | NaN |
| F5 | DIHSGLIGPLLICR | -1699 | -1712 | 0 | 0 | 14 | NaN | 197.68332 | 309.3971 | 466.54443 | 724.5076 | 712.53125 |
| Dynlrb1 | AEVEETLKR | -2 | -10 | 1 | 0.00078661 | 9 | NaN | 3254.173 | 556.40564 | NaN | 1051.2683 | 6621.3965 |
| Bzw1 | GLTETGTDLEAVAK | -39 | -52 | 0 | 0 | 14 | NaN | 2690.0554 | 2727.5322 | NaN | 3920.025 | 2968.1252 |
| Bzw1 | FDPTQFQDCIIQ | -27 | -38 | 0 | 0 | 12 | NaN | 1010.1194 | NaN | NaN | 648.35315 | NaN |
| Rpn1 | DIPAYSQDTFK | -206 | -216 | 0 | 0 | 11 | NaN | 1920.1204 | 2064.0469 | 1872.054 | 1596.0226 | 1664.8248 |
| Rpn1 | EGNHYFYSPYPTK | -158 | -170 | 0 | 0 | 13 | NaN | 922.13794 | 229.53258 | 1212.2781 | 825.699 | 812.1293 |
| Rpn1 | FYSPYPTK | -163 | -170 | 0 | 0 | 8 | NaN | 909.62286 | 1163.5112 | NaN | 860.1517 | NaN |
| Rpn1 | NIQVDSPYDISR | -381 | -392 | 0 | 0 | 12 | NaN | 686.8506 | 2205.6047 | 550.9298 | NaN | NaN |
| Psmd6 | KGDLLLNR | -372 | -379 | 1 | 0.002265552 | 8 | NaN | 1172.6506 | 950.6805 | 769.98413 | 1067.8339 | 905.99225 |
| Psmd6 | GVDFIDQELSR | -326 | -336 | 0 | 0.004133093 | 11 | NaN | 700.0274 | 757.1542 | NaN | 1443.0713 | NaN |
| Psmd6 | AEYLCQIGDK | -108 | -117 | 0 | 0 | 10 | NaN | 930.50146 | NaN | NaN | 1961.2091 | NaN |
| Eif3e | RVLVPATDR | -155 | -163 | 1 | 0.003299042 | 9 | NaN | 5759.045 | 5384.801 | 2887.3914 | 3974.5166 | 5143.198 |
| Eif3e | LKETIDNNSVSSPLQSL | -192 | -208 | 1 | 0 | 17 | NaN | 793.47156 | 746.61115 | 1298.4673 | 870.7574 | 1447.1394 |
| Eif3e | LKETIDNN | -192 | -199 | 1 | 0.00294445 | 8 | NaN | 892.89246 | NaN | NaN | 1626.8796 | 1397.3846 |
| Eif3e | GRDNIIDLF | -228 | -236 | 1 | 0.00272025 | 9 | NaN | 681.47797 | NaN | 592.3358 | 731.4251 | NaN |
| Rangap1 | AFAINPLLR | -231 | -239 | 0 | 0 | 9 | NaN | 11964.301 | 12039.651 | 10479.175 | NaN | NaN |
| Rangap1 | GSGEEPATPSR | -404 | -414 | 0 | 0 | 11 | NaN | 241.04355 | NaN | 3972.3167 | 4833.679 | NaN |
| Ddx27 | LIDHLHNCPSFHLS | -316 | -329 | 0 | 0 | 14 | NaN | 1874.3646 | 1341.8359 | NaN | NaN | 2042.37 |
| Ddx27 | SNTDVAPFLR | -393 | -402 | 0 | 0 | 10 | NaN | 1073.2977 | 1978.7639 | NaN | NaN | NaN |
| Ywhae | HLIPAANTGESK | -107 | -118 | 0 | 0 | 12 | NaN | 5353.7646 | 5423.2344 | 3995.8098 | 4783.1455 | 5071.0796 |
| Ywhae | LAEFATGNDR | -132 | -141 | 0 | 0 | 10 | NaN | 1140.7915 | NaN | 1148.133 | 1133.4427 | 1146.6022 |
| Ywhab | KAVTEQGHELSNEER | -29 | -43 | 1 | 0 | 15 | NaN | 966.83856 | 3114.5513 | 916.8196 | 883.78687 | 2530.4363 |
| Ywhaz | KSVTEQGAELSNEER | -27 | -41 | 1 | 0 | 15 | NaN | 2442.2383 | 1957.4429 | 2729.402 | 3662.0928 | 3386.8665 |
| Ywhaz | KGIVDQSQQAYQE | -139 | -151 | 1 | 0.00025015 | 13 | NaN | 6815.1743 | 7176.9575 | 4654.8574 | 6797.8433 | 6921.1245 |
| Ywhaz | PNASQPESK | -107 | -115 | 0 | 0.002780285 | 9 | NaN | 575.5416 | 559.7402 | 749.2878 | 730.5733 | 1432.5575 |
| Cct5 | LLQNGLPAVR | -314 | -323 | 0 | 0.001463897 | 10 | NaN | 597.9618 | 613.16125 | NaN | 607.5191 | NaN |
| Uqcr10 | AFDQGADAIYEHIN | -35 | -48 | 0 | 0.000394361 | 14 | NaN | 638.3276 | 645.24347 | NaN | NaN | 686.497 |
| Eif3i | DPSQIDSNEPYMK | -129 | -141 | 0 | 0.00025015 | 13 | NaN | 652.7627 | 655.3652 | NaN | 867.0354 | 555.793 |
| Erp44 | SLDEVTNLDR | -149 | -158 | 0 | 0 | 10 | NaN | 1259.0676 | 1070.6096 | NaN | 779.9472 | 1577.5334 |
| Erp44 | KTPADCPVIAIDSFR | -313 | -327 | 1 | 0.00025015 | 15 | NaN | 1197.1859 | NaN | 882.949 | 2344.4219 | 1767.7366 |
| Erp44 | GTINFLHADCDK | -292 | -303 | 0 | 0 | 12 | NaN | 408.4439 | NaN | 584.7083 | 527.18066 | 580.33215 |
| Erp44 | FLHADCDK | -296 | -303 | 0 | 0.00025015 | 8 | NaN | 451.70993 | 414.3462 | 328.13205 | NaN | NaN |
| Ilf3 | AFGQLHK | -326 | -332 | 0 | 0.000431655 | 7 | NaN | 2822.9438 | NaN | 3111.767 | 2716.7527 | NaN |
| Ilf3 | IHLTSPVVR | -169 | -177 | 0 | 0.007288197 | 9 | NaN | 377.24203 | NaN | NaN | NaN | NaN |
| Cul4b | AFGSTIVINPEKDK | -557 | -570 | 1 | 0.00025015 | 14 | NaN | 534.16724 | NaN | NaN | NaN | 877.41205 |
| Sfxn1 | YAYDSAFHPDTGEK | -73 | -86 | 0 | 0 | 14 | NaN | 432.48486 | NaN | 979.90497 | 550.9784 | 557.20856 |
| Actn1 | MLDAEDIVGTARPDEK | -221 | -236 | 1 | 0 | 16 | NaN | 1447.9663 | 1297.1506 | 887.41235 | 978.7892 | 464.37695 |
| Actn1 | TIEEIQGLTTAHEQFK | -540 | -555 | 0 | 0 | 16 | NaN | 302.84686 | 405.83423 | 540.9193 | 786.76746 | 777.9147 |
| Actn1 | LEDQLSHLR | -664 | -672 | 0 | 0 | 9 | NaN | 2645.2312 | 2632.6287 | NaN | 1793.4143 | 1665.228 |
| Actn1 | ICDQWDNLGALTQK | -479 | -492 | 0 | 0.000230707 | 14 | NaN | 2711.9673 | NaN | 3146.2827 | NaN | 3844.68 |
| Actn1 | IENIEEDFRDGLK | -52 | -64 | 1 | 0.00025015 | 13 | NaN | 1234.0834 | NaN | NaN | 1949.3489 | 186.06073 |
| Actn1 | SDINNAWGCLEQAEK | -362 | -376 | 0 | 0 | 15 | NaN | 93.81389 | NaN | 126.26246 | NaN | 534.4343 |
| Actn1 | NTAFDVAER | -206 | -214 | 0 | 0.001463897 | 9 | NaN | 808.4557 | NaN | NaN | NaN | NaN |
| Actn1 | ILAGDKNYITEDELR | -835 | -849 | 1 | 0.004658491 | 15 | NaN | 782.6145 | NaN | NaN | NaN | NaN |
| Actn1 | NIEEDFRDGLK | -54 | -64 | 1 | 0.00025015 | 11 | NaN | 368.01126 | NaN | NaN | NaN | NaN |
| Lck | ITFPGLHDLVR | -197 | -207 | 0 | 0.000550307 | 11 | NaN | 502.95062 | 422.87015 | 471.32962 | 2240.924 | 482.88083 |
| Lck | NILVSDTLSCK | -369 | -379 | 0 | 0 | 11 | NaN | 10698.881 | 12313.333 | 8822.008 | 6524.524 | NaN |
| Snrnp70 | HHNQPYCGIAPYIR | -33 | -46 | 0 | 0 | 14 | NaN | 932.0005 | 811.6365 | 2090.8672 | 1006.5403 | 1199.7051 |

| | | | | | | | | | | | | |
|---|---|---|---|---|---|---|---|---|---|---|---|---|
| Snrnp70 | PGPSPLPHR | -223 | -231 | 0 | 0.007987481 | 9 | NaN | 480.73703 | NaN | NaN | NaN | NaN |
| Syncrip | SENFQTLLDAGLPQK | -24 | -38 | 0 | 0 | 15 | NaN | 130.27615 | 454.69897 | 806.2469 | 1056.2128 | 875.7994 |
| Syncrip | KYGGPPPDSVYSGQQPSVGTEIFVGK | -143 | -168 | 1 | 0.000230707 | 26 | NaN | 658.3431 | NaN | 791.032 | 541.6589 | 490.4255 |
| Syncrip | GYDYHNYR | -471 | -478 | 0 | 0.000550307 | 8 | NaN | 7522.258 | 8566.73 | 7158.5186 | NaN | NaN |
| Dnm2 | KVPVGDQPPDIEYQIK | -142 | -157 | 1 | 0 | 16 | NaN | 886.2707 | 639.5902 | 714.1172 | 492.70493 | 752.56134 |
| Dnm2 | KVPVGDQPPDIEYQIK | -142 | -157 | 1 | 0 | 16 | NaN | 886.2707 | 639.5902 | 714.1172 | 492.70493 | 752.56134 |
| Dnm2 | NDPFSAPPQIPSR | -822 | -834 | 0 | 0 | 13 | NaN | 2573.2612 | 2592.3142 | 1429.4818 | NaN | NaN |
| Dnm2 | AGVYPEKDQAENEDGAQEN | -623 | -641 | 1 | 0.000550307 | 19 | NaN | 302.66086 | NaN | 172.49289 | NaN | 77.75952 |
| Ide | EALDDVTLPR | -702 | -711 | 0 | 0 | 10 | NaN | 2822.4102 | NaN | 2606.4888 | NaN | 2509.4463 |
| Ide | TKPLLPSQLVR | -755 | -765 | 1 | 0.00078661 | 11 | NaN | 853.5927 | NaN | 494.32407 | 502.8508 | NaN |
| Ide | YTLETRPNQEGIDVR | -224 | -238 | 1 | 0.001190916 | 15 | NaN | 655.0206 | NaN | NaN | 772.72327 | 829.16473 |
| Tubb5 | EDFGEEAEEEA | -434 | -444 | 0 | 0 | 11 | NaN | 333.41556 | 263.89435 | NaN | NaN | NaN |
| Sec31a | AFLPDNTNQPNIVQLR | -767 | -782 | 0 | 0 | 16 | NaN | 443.37772 | 436.4705 | 170.91849 | 557.4299 | 601.3905 |
| Hnrnpu | GYFEYIEENK | -232 | -241 | 0 | 0 | 10 | NaN | 1121.7859 | 1187.4911 | 1025.0881 | 277.6497 | 940.4167 |
| Hnrnpu | SPQPPVEEEDEHFDDTVVCL | -247 | -266 | 0 | 0.00025015 | 20 | NaN | 390.1185 | 382.38925 | 497.4503 | 632.5261 | 590.43823 |
| Hnrnpu | KVSELKEELK | -12 | -21 | 2 | 0.00025015 | 10 | NaN | 323.47888 | 364.07242 | 623.1066 | 384.8827 | 396.01605 |
| Hnrnpu | TCNCETEDYGEKFDENDVITCF | -364 | -385 | 1 | 0.003003218 | 22 | NaN | 195.35002 | 219.20528 | 194.72388 | 187.85489 | 186.60974 |
| Hnrnpu | GNFTLPEVAECF | -614 | -625 | 0 | 0 | 12 | NaN | 569.70557 | NaN | 558.93555 | 738.2899 | 845.6156 |
| Rpa1 | LETYNDESR | -587 | -595 | 0 | 0 | 9 | NaN | 1022.0647 | 2253.643 | 167.81754 | 143.93832 | 1446.7776 |
| Rpa1 | STVIVNPDIPEAYK | -406 | -419 | 0 | 0 | 14 | NaN | 3829.6675 | NaN | NaN | 4660.93 | 2434.013 |
| Rpa1 | AKDALVDIIGICK | -321 | -333 | 1 | 0 | 13 | NaN | 1188.8013 | NaN | NaN | 1932.2802 | 1927.6187 |
| Actr2 | SNFPEHIFPALVGR | -25 | -38 | 0 | 0 | 14 | NaN | 1477.59 | 1660.1713 | 878.8754 | 1630.4806 | 1268.5475 |
| Actr2 | WDYTFGPEK | -89 | -97 | 0 | 0 | 9 | NaN | 491.2816 | 696.38745 | 617.04346 | 674.65784 | 746.78827 |
| Aimp1 | AFPGEPDKELNPK | -253 | -265 | 1 | 0 | 13 | NaN | 1516.3743 | 1991.2025 | NaN | 2019.997 | NaN |
| Gzma | HYNFHPVIGLN | -183 | -193 | 0 | 0.000230707 | 11 | NaN | 2099.857 | 2198.3467 | 611.6161 | 2832.0425 | 2583.8953 |
| Prdx3 | AFQFVETHGEVCPA | -219 | -232 | 0 | 0.00078661 | 14 | NaN | 380.06384 | NaN | NaN | NaN | NaN |
| Uggt1 | MINNPSQEISDSSTPIFR | -795 | -812 | 0 | 0 | 18 | NaN | 327.56848 | 328.08524 | 414.89673 | 462.73178 | 376.7887 |
| Uggt1 | SEIGHEEFSNIHHQLISK | -200 | -217 | 0 | 0.000471409 | 18 | NaN | 462.47858 | NaN | 435.92108 | 583.5411 | 239.00928 |
| Uggt1 | FLSPLQQNLLK | -109 | -119 | 0 | 0 | 11 | NaN | 2449.0876 | NaN | 3043.0564 | 3555.3337 | NaN |
| Pes1 | ERYPTFIDALR | -122 | -132 | 1 | 0.000550307 | 11 | NaN | 1452.5314 | NaN | 805.1407 | NaN | NaN |
| Pes1 | LAQEEESEAKR | -520 | -530 | 1 | 0.000431655 | 11 | NaN | 732.4014 | 85.32976 | NaN | NaN | NaN |
| Pes1 | YPTFIDALR | -124 | -132 | 0 | 0.001023236 | 9 | NaN | 1557.8118 | NaN | NaN | NaN | NaN |
| Sfpq | VGNLPADITEDEFK | -293 | -306 | 0 | 0 | 14 | NaN | 2008.0739 | 1934.3768 | 2352.7158 | 2098.3071 | 1955.8647 |
| Sfpq | KAELDDTPMR | -341 | -350 | 1 | 0 | 10 | NaN | 417.96695 | NaN | 307.3778 | 3840.1497 | 3735.5662 |
| Sfpq | NMGDPYGSGGQK | -611 | -622 | 0 | 0.000907206 | 12 | NaN | 499.40564 | NaN | 255.05284 | NaN | NaN |
| Mars1 | LQQHPVLPVPGER | -254 | -266 | 0 | 0 | 13 | NaN | 941.25397 | 956.0083 | 675.1769 | 1563.585 | 1419.2194 |
| Mars1 | LENDQIENLR | -807 | -816 | 0 | 0 | 10 | NaN | 1268.1147 | NaN | 1454.3132 | 1314.6302 | 1494.8549 |
| Mars1 | WELQHYHQLLEK | -689 | -700 | 0 | 0 | 12 | NaN | 1262.2811 | 979.8696 | NaN | 960.94885 | 1078.7867 |
| Mars1 | AFSWTDLLIK | -634 | -643 | 0 | 0 | 10 | NaN | 274.93256 | 226.42651 | NaN | 331.54022 | 238.53809 |
| Mars1 | SSQHLFLDLPK | -448 | -458 | 0 | 0 | 11 | NaN | 7389.5127 | 7627.5293 | NaN | NaN | 8560.968 |
| Mars1 | RLEDWLGK | -462 | -469 | 1 | 0.00051306 | 8 | NaN | 2913.9487 | 2681.6838 | NaN | NaN | 3110.329 |
| Mars1 | FFGGCVPEMALTPDDR | -666 | -681 | 0 | 0.001023236 | 16 | NaN | 384.2432 | NaN | NaN | NaN | NaN |
| Srsf7 | GHYAYDCHR | -142 | -150 | 0 | 0.00111006 | 9 | NaN | 292.52554 | 198.01282 | 457.77542 | NaN | NaN |
| Prpf31 | QVKPLPAPLDGQ | -339 | -350 | 1 | 0.001291103 | 12 | NaN | 1881.5 | 1979.6111 | NaN | 2140.9697 | 2043.2789 |
| Prpf31 | IYEYVESR | -204 | -211 | 0 | 0 | 8 | NaN | 861.5936 | 1469.835 | NaN | NaN | 1706.9637 |
| Rpl7a | AGVNTVTTLVENK | -138 | -150 | 0 | 0 | 13 | NaN | 2351.851 | 1072.607 | 1346.8529 | NaN | NaN |
| Rpl7a | RHWGGNVLGPK | -235 | -245 | 1 | 0.001575548 | 11 | NaN | 447.88632 | NaN | NaN | NaN | NaN |
| Rpl7a | TVTTLVENKK | -142 | -151 | 1 | 0.00111006 | 10 | NaN | 302.5283 | NaN | NaN | NaN | NaN |
| Rps2 | LSIVPVR | -160 | -166 | 0 | 0.007105559 | 7 | NaN | 589.768 | NaN | NaN | NaN | 218.65804 |
| Rps2 | KIGKPHTVPCK | -173 | -183 | 2 | 0.00025015 | 11 | NaN | 255.73605 | 255.18117 | NaN | NaN | NaN |
| Acadl | IFSSEHDIFR | -52 | -61 | 0 | 0 | 10 | NaN | 3034.8777 | 3184.8647 | 1747.21 | 1674.3787 | 1707.0868 |
| Acadl | AFVDSCLQLHETK | -346 | -358 | 0 | 0 | 13 | NaN | 769.2714 | 880.6122 | 1093.1385 | 445.45792 | 1497.17 |
| Acadl | AQDTAELFFEDVR | -255 | -267 | 0 | 0 | 13 | NaN | 720.25323 | 796.53973 | 789.4246 | 1081.1584 | 983.7732 |
| Acadl | KFFQEEVIPHHTEWEK | -66 | -81 | 1 | 0 | 16 | NaN | 622.1359 | 496.27173 | 7616.3257 | 921.11523 | 664.4708 |
| Acadl | KFFQEEVIPH | -66 | -75 | 1 | 0.001055699 | 10 | NaN | 972.4212 | NaN | 1982.5979 | 1499.6411 | 1456.0837 |
| C1qbp | EHQEYITFLEDLK | -260 | -272 | 0 | 0 | 13 | NaN | 427.29266 | 576.74585 | 776.74805 | 899.46875 | 782.89496 |
| C1qbp | GVDNTFADELVEL | -243 | -255 | 0 | 0.008947895 | 13 | NaN | 282.04562 | NaN | 304.10605 | 280.54092 | 100.3976 |
| C1qbp | LTDEIKEEK | -83 | -91 | 1 | 0.00025015 | 9 | NaN | 285.71024 | NaN | 259.3672 | NaN | NaN |
| Ddx49 | HAIEEQIK | -361 | -368 | 0 | 0.003780189 | 8 | NaN | 2751.2876 | NaN | NaN | 3036.279 | NaN |
| Atp5mk | AGAESDGQFQFTGIK | -2 | -16 | 0 | 0 | 15 | NaN | 673.92255 | 913.8204 | 833.65924 | 1420.5824 | 1414.755 |
| Rplp0 | IAPCEVTVPAQNTGLGPEK | -116 | -134 | 0 | 0 | 19 | NaN | 307.93353 | 208.16394 | 684.96295 | 528.9726 | 572.4243 |
| Taldo1 | WLHNEDQMAVEK | -296 | -307 | 0 | 0 | 12 | NaN | 503.6396 | 573.4712 | 457.2211 | 472.28552 | 472.20447 |
| Taldo1 | WLHNEDQMAVEK | -296 | -307 | 0 | 0 | 12 | NaN | 503.6396 | 573.4712 | 457.2211 | 472.28552 | 472.20447 |
| Taldo1 | STWEGIQAGK | -145 | -154 | 0 | 0.00025015 | 10 | NaN | 80.72862 | NaN | 761.2016 | NaN | NaN |
| Txnl1 | QHLENDPGSNEDADIPK | -105 | -121 | 0 | 0 | 17 | NaN | 874.2398 | 865.26324 | 919.9006 | 1023.4694 | 967.9746 |
| Txnl1 | VGVKPVGSDPDFQPELSGAGSR | -2 | -23 | 1 | 0 | 22 | NaN | 813.5239 | 863.49347 | 1853.4048 | 1012.3716 | 812.4974 |
| Txnl1 | AGCECLNESDEHGFDNCLR | -133 | -151 | 0 | 0.000907206 | 19 | NaN | 1208.6556 | NaN | NaN | 402.0809 | NaN |
| Actb | DDDIAALVVDNGSGMCK | -2 | -18 | 0 | 0 | 17 | NaN | 317.93256 | 348.55643 | 349.64142 | 500.39026 | 336.76212 |
| Hadh | GHTVVLVDQTEDILAK | -50 | -65 | 0 | 0 | 16 | NaN | 715.42114 | 732.0994 | 773.56964 | 881.1814 | 834.7941 |
| Acap1 | AGDETYLDIFR | -707 | -717 | 0 | 0 | 11 | NaN | 2885.3564 | 2935.4248 | NaN | 3103.7375 | 3229.55 |
| Acap1 | ELGGEEPEPSLK | -249 | -260 | 0 | 0 | 12 | NaN | 1617.9705 | NaN | 490.45602 | NaN | NaN |
| Fermt3 | LFFGPQHRPVIL | -91 | -102 | 1 | 0 | 12 | NaN | 487.53217 | 463.7682 | 524.17725 | 686.6527 | 615.8685 |
| Fermt3 | LFFGPQHRPVIL | -91 | -102 | 1 | 0 | 12 | NaN | 487.53217 | 463.7682 | 524.17725 | 686.6527 | 615.8685 |
| Fermt3 | GESHIGGVLLK | -38 | -48 | 0 | 0 | 11 | NaN | 542.4571 | NaN | 540.21936 | 576.37726 | 796.68317 |
| Fermt3 | SFFDLDPK | -255 | -262 | 0 | 0.00383302 | 8 | NaN | 579.143 | NaN | 633.17255 | NaN | 719.6914 |
| Ybx3 | SPAASGAPQAPAPAAL | -33 | -48 | 0 | 0.001392869 | 16 | NaN | 715.41925 | NaN | NaN | NaN | NaN |
| Xpo1 | DLLGLCEQK | -523 | -531 | 0 | 0 | 9 | NaN | 1981.9802 | NaN | NaN | 1826.6051 | 1940.7587 |
| Xpo1 | QLLDFSQK | -15 | -22 | 0 | 0.003102268 | 8 | NaN | 794.9904 | NaN | NaN | 610.7324 | 470.22003 |

| | | | | | | | | | | | | |
|---|---|---|---|---|---|---|---|---|---|---|---|---|
| Sdha | SIAESCRPGDKVPSIK | -470 | -485 | 2 | 0.001055699 | 16 | NaN | 540.5812 | 573.9579 | 486.8829 | 1148.0044 | 1166.4548 |
| Sdha | KPFGEHWR | -616 | -623 | 1 | 0 | 8 | NaN | 1353.6534 | 215.3353 | NaN | 1411.0529 | NaN |
| Sdha | SVLQEGCEK | -530 | -538 | 0 | 0.001636916 | 9 | NaN | 1518.5096 | NaN | 1485.5687 | NaN | NaN |
| Sdha | AGEESVMNLDK | -488 | -498 | 0 | 0 | 11 | NaN | 552.42267 | NaN | 738.00446 | NaN | NaN |
| Sdha | PFGEHWR | -617 | -623 | 0 | 0.007880195 | 7 | NaN | 2275.27 | NaN | NaN | NaN | NaN |
| Copb1 | EAELLEPLMPAIR | -129 | -141 | 0 | 0 | 13 | NaN | 1414.8447 | 1057.0662 | 3912.1357 | 1688.0449 | 1773.2065 |
| Copb1 | VLSTPDLEVR | -328 | -337 | 0 | 0 | 10 | NaN | 4418.905 | 578.5353 | 1054.4454 | 3722.9688 | 3714.1821 |
| Copb1 | WAEFEWENK | -844 | -852 | 0 | 0 | 9 | NaN | 613.2281 | NaN | 739.2148 | 816.41785 | 794.39685 |
| Copb1 | LVEKPSPLTLAPHDF | -768 | -782 | 1 | 0 | 15 | NaN | 446.4617 | 357.62582 | NaN | NaN | NaN |
| Copb1 | VCHANPSER | -234 | -242 | 0 | 0.007343498 | 9 | NaN | 904.20905 | NaN | NaN | NaN | NaN |
| Copb1 | AGELKPEEEITVGPVQK | -497 | -513 | 1 | 0 | 17 | NaN | 634.06067 | NaN | NaN | NaN | NaN |
| Ssrp1 | GWNWGTVK | -106 | -113 | 0 | 0.000431655 | 8 | NaN | 3558.5667 | 4621.3364 | NaN | NaN | NaN |
| Numa1 | HLCQQLQAEQAAAEK | -1361 | -1375 | 0 | 0.000230707 | 15 | NaN | 2676.0125 | 2261.8289 | 1411.0112 | 1652.6351 | 2649.6943 |
| Numa1 | LENELSTALQDK | -360 | -371 | 0 | 0 | 12 | NaN | 490.80023 | 522.39923 | 161.61888 | 262.20752 | 888.54114 |
| Numa1 | SLDLSLEEGTPCSVASK | -1706 | -1722 | 0 | 0.000230707 | 17 | NaN | 1493.9032 | 1569.7046 | NaN | 1135.4089 | NaN |
| Numa1 | LQNTLNEQR | -1021 | -1029 | 0 | 0.001868731 | 9 | NaN | 1477.2771 | 1724.9152 | NaN | NaN | NaN |
| Numa1 | SEELSPPSHQTK | -163 | -174 | 0 | 0.00025015 | 12 | NaN | 1367.1332 | NaN | NaN | 1532.4169 | NaN |
| Ly6e | GFGNVNLGYTLNK | -62 | -74 | 0 | 0 | 13 | NaN | 453.36505 | NaN | 1286.8337 | 1316.9165 | 1246.3163 |
| Nat10 | KDLPPLLL | -689 | -696 | 1 | 0.001291103 | 8 | NaN | 352.8387 | 305.07446 | 386.20465 | 372.4839 | 319.15134 |
| Cyb5a | STWVILHHK | -25 | -33 | 0 | 0 | 9 | NaN | 4656.1685 | NaN | 694.16864 | 8590.342 | 7107.8286 |
| Ppp2r1a | LTQDQDVDVK | -567 | -576 | 0 | 0.00025015 | 10 | NaN | 3766.6516 | 3622.9873 | 1907.8737 | NaN | NaN |
| Eif2s3x | LDDPSCPRPECYR | -91 | -103 | 1 | 0 | 13 | NaN | 1467.8479 | 1532.4556 | 1774.5839 | 1858.3224 | 1813.99 |
| Eif2s3x | LDDPSCPRPECY | -91 | -102 | 1 | 0 | 12 | NaN | 401.6314 | NaN | 2715.9756 | NaN | NaN |
| Nek9 | AGGGAAEQEELHYIPIR | -40 | -56 | 0 | 0.000230707 | 17 | NaN | 692.95465 | 725.2834 | NaN | 1081.8534 | 895.2097 |
| Ttll12 | EFIPQFEK | -525 | -532 | 0 | 0.002566359 | 8 | NaN | 1021.7637 | 928.98773 | NaN | NaN | NaN |
| Ttll12 | ALDDLDDYEK | -496 | -505 | 0 | 0 | 10 | NaN | 421.2341 | NaN | 537.6729 | NaN | NaN |
| Ahcy | TGCVDIILGR | -276 | -285 | 0 | 0 | 10 | NaN | 3126.9817 | 1727.4946 | NaN | NaN | 4345.5273 |
| Ahcy | AGIPVFAWKG | -95 | -104 | 1 | 0.002595029 | 10 | NaN | 1183.631 | 940.2286 | NaN | NaN | 1732.6722 |
| Ahcy | SKFDNLYGCRE | -187 | -197 | 2 | 0.00122419 | 11 | NaN | 1803.7997 | NaN | NaN | NaN | NaN |
| Ahcy | ETTTGVHNLYK | -156 | -166 | 0 | 0.003900536 | 11 | NaN | 428.0208 | NaN | NaN | NaN | NaN |
| Gsn | DSQEEEKTEALTS | -712 | -724 | 1 | 0.00025015 | 13 | NaN | 861.9069 | NaN | 477.09814 | 413.81653 | 380.12512 |
| Gsn | TVQLDDYLNGR | -130 | -140 | 0 | 0.00025015 | 11 | NaN | 522.9654 | NaN | 2645.776 | NaN | NaN |
| Gsn | QVSVLPEGGETPLFK | -374 | -388 | 0 | 0.00025015 | 15 | NaN | 82.76301 | NaN | NaN | NaN | NaN |
| Nifk | VEFESEDVAK | -93 | -102 | 0 | 0 | 10 | NaN | 1023.621 | 990.58044 | 1289.4963 | 1376.671 | 1116.9216 |
| Nifk | KDDEIILK | -279 | -286 | 1 | 0.009031675 | 8 | NaN | 1387.6527 | 1730.013 | NaN | NaN | 2428.2449 |
| Nifk | SSVDSQGPTPVCTPTFLER | -249 | -267 | 0 | 0.00223472 | 19 | NaN | 315.94107 | NaN | NaN | NaN | NaN |
| Sigmar1 | AGLDHELAFSR | -50 | -60 | 0 | 0.005394768 | 11 | NaN | 2212.8208 | 2154.3882 | NaN | NaN | NaN |
| Dnajc11 | GIRPGLTTVL | -249 | -258 | 1 | 0.006788754 | 10 | NaN | 459.17673 | 418.00488 | 369.99744 | NaN | NaN |
| Pdhb | SAGLQPVPIVFR | -134 | -145 | 0 | 0 | 12 | NaN | 313.29395 | 381.43665 | 382.79156 | 435.60846 | 415.35922 |
| Pdhb | VFLLGEEVAQYDGAYK | -53 | -68 | 0 | 0.000431655 | 16 | NaN | 560.9725 | NaN | NaN | NaN | NaN |
| Ddx54 | HENPDIIIATPGR | -213 | -225 | 0 | 0 | 13 | NaN | 1033.7443 | 1527.7415 | 1336.8186 | 1917.9308 | 1956.8674 |
| Ddx54 | ALPSFPTSECVSDVEPDTR | -63 | -81 | 0 | 0.000230707 | 19 | NaN | 280.7939 | NaN | 432.74377 | 639.866 | 521.4079 |
| Ddx54 | SRPAPSPESIK | -523 | -533 | 1 | 0.000431655 | 11 | NaN | 1874.128 | NaN | NaN | 1848.6256 | 518.6125 |
| Plec | LEAQAGTGHIIDPATSAR | -3202 | -3219 | 0 | 0.000431655 | 18 | NaN | 1180.7646 | 405.78708 | 2136.8071 | 1184.4408 | 2456.4773 |
| Plec | KQEELQHLEQQR | -2714 | -2725 | 1 | 0 | 12 | NaN | 1044.469 | 719.88074 | NaN | 882.54785 | 892.76196 |
| Plec | RLLDPEDVDVPQPDEK | -382 | -397 | 1 | 0 | 16 | NaN | 768.9224 | 483.45316 | 476.84717 | 544.12585 | 489.96252 |
| Plec | AGLVGPELHEK | -3228 | -3238 | 0 | 0 | 11 | NaN | 2984.654 | 3701.5325 | 3624.9648 | 3117.9038 | NaN |
| Plec | KGLVGPELHDR | -3894 | -3904 | 1 | 0 | 11 | NaN | 903.94293 | 641.4947 | NaN | 955.27704 | 1001.4167 |
| Plec | RLEDLLQDAQDEK | -904 | -916 | 1 | 0.00149795 | 13 | NaN | 481.072 | NaN | 456.71207 | 576.6944 | 141.50359 |
| Plec | LEQLFQDEVAK | -2661 | -2671 | 0 | 0.000431655 | 11 | NaN | 2159.5247 | 1284.2922 | NaN | NaN | 5186.5347 |
| Plec | GLLLKPTNEK | -2844 | -2853 | 1 | 0 | 10 | NaN | 336.5221 | NaN | NaN | 398.39163 | 400.42493 |
| Plec | SVEAEPEYTFEGLR | -2781 | -2794 | 0 | 0.001868731 | 14 | NaN | 1496.8665 | NaN | NaN | NaN | NaN |
| Pcna | HVSLVQLTLR | -44 | -53 | 0 | 0.000394361 | 10 | NaN | 503.0923 | NaN | NaN | NaN | 701.2336 |
| Hsp90b1 | FQSSHHSTDITSLDQYVER | -512 | -530 | 0 | 0 | 19 | NaN | 317.64267 | 323.42368 | 588.9941 | 575.74634 | 531.6589 |
| Hsp90b1 | STDITSLDQYVER | -518 | -530 | 0 | 0 | 13 | NaN | 1782.1958 | 1615.2856 | 1420.5248 | 367.4206 | NaN |
| Hsp90b1 | VLKEEASDYLELDTIK | -250 | -265 | 1 | 0 | 16 | NaN | 358.6849 | 285.79108 | 648.39154 | 568.66943 | NaN |
| Hsp90b1 | VFITDDFHDMMPK | -416 | -428 | 0 | 0.000550307 | 13 | NaN | 629.2963 | 587.46106 | NaN | NaN | 496.04657 |
| Hsp90b1 | FQSSHHSTDITSL | -512 | -524 | 0 | 0.006654467 | 13 | NaN | 603.8524 | 525.5689 | NaN | NaN | NaN |
| Hsp90b1 | AVEEEEEEKKPK | -311 | -322 | 2 | 0.001023236 | 12 | NaN | 213.52571 | 196.06894 | NaN | NaN | NaN |
| Hsp90b1 | AGNEELTVK | -127 | -135 | 0 | 0 | 9 | NaN | 2756.9253 | NaN | NaN | NaN | NaN |
| Top1 | KIEPPGLFR | -356 | -364 | 1 | 0.00025015 | 9 | NaN | 247.15785 | NaN | NaN | 418.031 | 791.41003 |
| Top1 | SWTENIQGSIK | -417 | -427 | 0 | 0 | 11 | NaN | 476.787 | 409.69144 | NaN | NaN | NaN |
| Top1 | VEHINLHPELDGQ | -511 | -523 | 0 | 0.00025015 | 13 | NaN | 1049.7025 | NaN | NaN | NaN | NaN |
| Rpl7 | MEAVPEKK | -1 | -8 | 1 | 0 | 8 | NaN | 995.5438 | NaN | 1099.9718 | NaN | 1651.1279 |
| Rpl22 | ITVTSEVPFSK | -70 | -80 | 0 | 0 | 11 | NaN | 565.0925 | 632.0355 | NaN | NaN | NaN |
| Rpl22 | FEQFLQER | -39 | -46 | 0 | 0.009745776 | 8 | NaN | 968.5584 | NaN | NaN | NaN | NaN |
| Aarsd1 | DRLPVSVR | -164 | -171 | 1 | 0 | 8 | NaN | 546.2454 | 629.8589 | 559.3842 | 1052.3556 | 872.7799 |
| Aarsd1 | AGPAEAVETLGPR | -356 | -368 | 0 | 0 | 13 | NaN | 4191.883 | NaN | NaN | NaN | 300.73386 |
| Aarsd1 | VADLLFGLK | -118 | -126 | 0 | 0.00830926 | 9 | NaN | 258.3185 | NaN | NaN | NaN | NaN |
| Arpc5 | FVDEEDGGDGQAGPDEGEVDSCLR | -24 | -47 | 0 | 0 | 24 | NaN | 373.28644 | 330.1364 | 323.26685 | 399.82474 | 358.70746 |
| Arpc5 | GFESPSDNSSAVLLQ | -113 | -127 | 0 | 0.00025015 | 15 | NaN | 1657.0398 | NaN | NaN | NaN | NaN |
| Sart3 | TEGTLEDWDLAIQK | -556 | -569 | 0 | 0.00025015 | 14 | NaN | 602.6117 | NaN | NaN | 741.0601 | 177.18726 |
| Ddx6 | EYHSEPAEDEKP | -472 | -483 | 1 | 0.00025015 | 12 | NaN | 1804.7158 | 2137.9443 | 400.61417 | 1302.6376 | 1383.5756 |
| Ddx6 | YHSEPAEDEKP | -473 | -483 | 1 | 0 | 11 | NaN | 750.55426 | 810.46234 | 641.18146 | 1035.2673 | 302.61478 |
| Sod1 | QYGDNTQGCTSAGPHFNPHSK | -50 | -70 | 0 | 0 | 21 | NaN | 2416.7551 | 2845.526 | 655.79205 | 568.0916 | 2870.185 |
| Sod1 | GEHSIIGR | -109 | -116 | 0 | 0.003133457 | 8 | NaN | 3129.6 | 3173.1006 | NaN | NaN | 1833.1346 |
| Sod1 | GDNTQGCTSAGPHFNPHSK | -52 | -70 | 0 | 0.00025015 | 19 | NaN | 376.25577 | 342.13617 | NaN | NaN | NaN |
| Pgd | LVPLLDTGDIIIDGGNSEYR | -88 | -107 | 0 | 0 | 20 | NaN | 230.63254 | 206.50604 | 252.3853 | 320.64984 | 266.07504 |

| | | | | | | | | | | | | |
|---|---|---|---|---|---|---|---|---|---|---|---|---|
| Pgd | LDTGDIIIDGGNSEYR | -92 | -107 | 0 | 0 | 16 | NaN | 192.4325 | 256.70462 | 756.5777 | 821.9025 | 706.1331 |
| Arid1a | VDENHSEFTLY | -2242 | -2252 | 0 | 0.001636916 | 11 | NaN | 899.48627 | NaN | NaN | NaN | NaN |
| Slc25a12 | SHMLTPFVEENLVSAA | -191 | -206 | 0 | 0 | 16 | NaN | 357.90717 | 394.72955 | 499.9308 | 559.62317 | 752.1655 |
| Slc25a12 | IAPLAEGALPYNLAELQR | -292 | -309 | 0 | 0 | 18 | NaN | 252.44904 | 378.78503 | 500.24918 | 475.79758 | 477.74246 |
| Slc25a12 | AGQTTYSGVVDCFR | -552 | -565 | 0 | 0 | 14 | NaN | 1324.571 | NaN | 1159.6539 | 1393.4005 | 1256.521 |
| Slc25a12 | RDGSIPLPAEIL | -420 | -431 | 1 | 0 | 12 | NaN | 298.1528 | 310.65033 | NaN | 601.0792 | NaN |
| Acaca | LLLEDLVK | -2221 | -2228 | 0 | 0.003299042 | 8 | NaN | 818.1415 | NaN | NaN | NaN | NaN |
| Clic4 | DEFTNTCPSDK | -228 | -238 | 0 | 0.00122419 | 11 | NaN | 362.35236 | NaN | NaN | NaN | NaN |
| Clic4 | IEEFLEEVLCPPK | -91 | -103 | 0 | 0.000431655 | 13 | NaN | 344.39517 | NaN | NaN | NaN | NaN |
| Arhgap45 | LKECVLQDDLLEAR | -132 | -145 | 1 | 0 | 14 | NaN | 844.9692 | 878.05176 | 514.82886 | 1322.1886 | 1213.532 |
| Arhgap45 | RFAEGLEK | -124 | -131 | 1 | 0.002068213 | 8 | NaN | 779.9339 | NaN | 835.36096 | 742.19055 | NaN |
| Arhgap45 | RPLAHECLGEALR | -146 | -158 | 1 | 0.001291103 | 13 | NaN | 2539.9763 | NaN | NaN | NaN | 1411.1229 |
| Arhgap45 | STPDGVPFIVK | -766 | -776 | 0 | 0 | 11 | NaN | 1007.13635 | 733.823 | NaN | NaN | NaN |
| Eef1d | VGTDLLEEEITKF | -254 | -266 | 1 | 0 | 13 | NaN | 558.0296 | 592.0411 | 581.72015 | NaN | 314.29706 |
| Eef1d | RQVEPPTK | -136 | -143 | 1 | 0.000230707 | 8 | NaN | 384.89206 | NaN | 540.6039 | 631.59894 | 359.11584 |
| Nop2 | LAPQEHER | -365 | -372 | 0 | 0.007009655 | 8 | NaN | 392.1184 | NaN | NaN | 566.5339 | 684.7023 |
| Fmnl1 | VFTELNDEK | -654 | -662 | 0 | 0.000550307 | 9 | NaN | 120.00413 | 320.55768 | 2498.7651 | NaN | 570.9316 |
| Lman2 | GTGDLSDNHDIISIK | -260 | -274 | 0 | 0.000471409 | 15 | NaN | 2966.5188 | 2972.8835 | NaN | NaN | NaN |
| Baz1b | KDHSNLPDDDYCPR | -943 | -956 | 1 | 0 | 14 | NaN | 621.95276 | 547.3991 | 828.4664 | 465.46332 | 446.66687 |
| Baz1b | ELDELLSCLHPQGIR | -999 | -1013 | 0 | 0 | 15 | NaN | 533.297 | 412.35638 | 396.68732 | 559.35583 | 614.04236 |
| Baz1b | PGLFIEK | -921 | -927 | 0 | 0.005605646 | 7 | NaN | 540.65405 | NaN | 155.51566 | NaN | 321.09674 |
| Baz1b | WFLCDSQK | -991 | -998 | 0 | 0.001610812 | 8 | NaN | 452.98245 | 598.3991 | NaN | 314.41965 | NaN |
| Baz1b | DGNQELLNFLR | -1046 | -1056 | 0 | 0.003166358 | 11 | NaN | 707.7786 | 229.60303 | NaN | NaN | NaN |
| Phgdh | LQDCEGLIVR | -45 | -54 | 0 | 0.00025015 | 10 | NaN | 786.96893 | 789.3827 | NaN | 650.28503 | 874.7765 |
| Phgdh | QDGGLQVVEK | -24 | -33 | 0 | 0.009272078 | 10 | NaN | 893.687 | NaN | NaN | NaN | NaN |
| Phgdh | SLDPCCR | -14 | -20 | 0 | 0.006677519 | 7 | NaN | 676.14124 | NaN | NaN | NaN | NaN |
| Phgdh | GQPLLVFR | -462 | -469 | 0 | 0.003360691 | 8 | NaN | 527.18 | NaN | NaN | NaN | NaN |
| Phgdh | RGGIVDEGALLR | -236 | -247 | 1 | 0.001743443 | 12 | NaN | 523.7744 | NaN | NaN | NaN | NaN |
| Actn4 | RDHALLEEQSK | -634 | -644 | 1 | 0 | 11 | NaN | 928.1169 | 405.9387 | 882.00494 | 1637.7327 | 1430.2625 |
| Actn4 | ICDQWDNLGSLTHSR | -499 | -513 | 0 | 0 | 15 | NaN | 537.5513 | 521.20306 | 806.1993 | 995.4725 | 990.44147 |
| Actn4 | TIPWLEDR | -313 | -320 | 0 | 0.001868731 | 8 | NaN | 2190.7998 | 2302.3765 | 2228.315 | 2049.0234 | 2100.8994 |
| Actn4 | DHGGALGPEEFK | -781 | -792 | 0 | 0 | 12 | NaN | 956.9765 | 980.663 | 1136.2726 | 622.7149 | 1102.2534 |
| Actn4 | MAPYQGPDAAPGALDYK | -884 | -900 | 0 | 0 | 17 | NaN | 644.8504 | 751.8406 | 658.6945 | 580.3791 | 876.0499 |
| Actn4 | IENIDEDFRDGLK | -72 | -84 | 1 | 0 | 13 | NaN | 600.4741 | 493.3702 | 781.16284 | 961.2455 | 934.01117 |
| Actn4 | DYETATLSDIK | -441 | -451 | 0 | 0 | 11 | NaN | 564.5138 | NaN | 706.5784 | 468.5289 | 796.0273 |
| Actn4 | IENIDEDFR | -72 | -80 | 0 | 0 | 9 | NaN | 312.86536 | 515.1013 | NaN | 560.83966 | 733.68933 |
| Actn4 | KDDPVTNLNNAF | -218 | -229 | 1 | 0.000550307 | 12 | NaN | 1192.6417 | 1197.0499 | NaN | NaN | 1498.581 |
| Actn4 | AGTQIENIDEDFRDGLK | -68 | -84 | 1 | 0.000298805 | 17 | NaN | 447.3586 | 2470.1301 | NaN | 1313.3359 | NaN |
| Actn4 | DIVNTARPDEK | -246 | -256 | 1 | 0.00025015 | 11 | NaN | 1603.6204 | 1007.8854 | NaN | NaN | NaN |
| Actn4 | NELDYYDSHNVNTR | -482 | -495 | 0 | 0 | 14 | NaN | 471.17746 | 165.28061 | NaN | NaN | NaN |
| Actn4 | ASFNHFDKDHGGALGPEEFK | -773 | -792 | 1 | 0.00025015 | 20 | NaN | 529.23987 | NaN | NaN | NaN | NaN |
| Pgk1 | SHLGRPDGVPMPDK | -62 | -75 | 1 | 0.00025015 | 14 | NaN | 745.8015 | 792.5867 | 843.1508 | 975.63885 | 1017.12897 |
| Pgk1 | SHLGRPDGVPMPDKYSLEPVAAELK | -62 | -86 | 2 | 0 | 25 | NaN | 741.02527 | NaN | 1761.4031 | 1022.8556 | 1433.913 |
| Pgk1 | SHLGRPDGVPMPDKYSLEPVAAELK | -62 | -86 | 2 | 0 | 25 | NaN | 741.02527 | NaN | 1761.4031 | 1022.8556 | 1433.913 |
| Pgk1 | FCLDNGAK | -49 | -56 | 0 | 0 | 8 | NaN | 913.74005 | 769.3197 | 623.45514 | NaN | NaN |
| Pgk1 | SGIPAGWMGLDCGTESSKK | -305 | -323 | 1 | 0 | 19 | NaN | 196.376 | NaN | NaN | 214.90656 | 503.2648 |
| Prkcb | SLNPEWNETFR | -218 | -228 | 0 | 0.005884925 | 11 | NaN | 983.7983 | NaN | NaN | NaN | NaN |
| Fto | DPDTWHVGFK | -263 | -272 | 0 | 0 | 10 | NaN | 536.75854 | 298.10837 | NaN | 808.38904 | 1156.8136 |
| Macroh2a1 | SGGVLPNIHPELLAK | -102 | -116 | 0 | 0 | 15 | NaN | 740.51825 | 3184.8157 | 3790.8982 | 3961.551 | 3849.7854 |
| Macroh2a1 | CNSPVWGADKCEELLEK | -273 | -289 | 1 | 0 | 17 | NaN | 551.3534 | 527.4742 | 613.001 | 243.98062 | 354.2528 |
| Macroh2a1 | NSPVWGADKCEELLEK | -274 | -289 | 1 | 0 | 16 | NaN | 787.3749 | 752.8778 | 671.0392 | NaN | 856.3663 |
| Macroh2a1 | GGVLPNIHPELLAK | -103 | -116 | 0 | 0 | 14 | NaN | 669.8563 | 411.3063 | NaN | 1627.7161 | 590.6446 |
| Macroh2a1 | GKLEAIITPPPAK | -122 | -134 | 1 | 0.00111006 | 13 | NaN | 1292.2539 | 1791.5631 | NaN | NaN | NaN |
| Metap1 | TVEGDVNTDPWAGYR | -68 | -82 | 0 | 0 | 15 | NaN | 824.2598 | 679.88727 | NaN | 588.6766 | NaN |
| Prmt5 | EIGADLPSNHVIDR | -207 | -220 | 0 | 0.00025015 | 14 | NaN | 2891.373 | 3208.4202 | 2878.362 | 2969.163 | 2967.1963 |
| Sgpl1 | GSNDFDIYR | -454 | -462 | 0 | 0 | 9 | NaN | 617.9034 | NaN | 496.68384 | NaN | NaN |
| Cebpz | ELLITDLLPDSR | -291 | -302 | 0 | 0 | 12 | NaN | 397.55795 | 463.05148 | 402.52264 | 397.71637 | 419.72214 |
| Rpl13a | KVLDGIPPPYDK | -103 | -114 | 1 | 0 | 12 | NaN | 411.8557 | NaN | 1794.9027 | 1072.8417 | 1228.462 |
| Ap1m1 | AHFGLPSVEAEDK | -358 | -370 | 0 | 0 | 13 | NaN | 1400.141 | 1826.7819 | 2411.4023 | 2323.2969 | 432.46063 |
| Atp5f1b | PIEEAVAK | -512 | -519 | 0 | 0.000550307 | 8 | NaN | 963.4498 | 928.4984 | 1180.2255 | 1849.1664 | 1692.5419 |
| Atp5f1b | IGEPIDER | -148 | -155 | 0 | 0.000230707 | 8 | NaN | 637.3406 | 834.02844 | 960.49097 | 1073.3059 | 978.1644 |
| Atp5f1b | PNIVGNEHYDVAR | -410 | -422 | 0 | 0.00025015 | 13 | NaN | 318.04016 | 715.33936 | 1342.7932 | 237.24176 | 369.06137 |
| Atp5f1b | MVGPIEEAVAK | -509 | -519 | 0 | 0.001708008 | 11 | NaN | 294.42523 | 610.8649 | NaN | 862.1043 | 665.59705 |
| Atp5f1b | TREGNDLYHE | -240 | -249 | 1 | 0.001023236 | 10 | NaN | 1307.9437 | 1150.0353 | 1406.0093 | 1983.5128 | NaN |
| Atp5f1b | MIESGVINLK | -250 | -259 | 0 | 0.000230707 | 10 | NaN | 283.8452 | 945.68427 | NaN | 247.57036 | 973.23956 |
| Atp5f1b | IMNVIGEPIDER | -144 | -155 | 0 | 0 | 12 | NaN | 1397.1594 | 1631.8088 | NaN | NaN | NaN |
| Atp5f1b | SVEQEILVTGIK | -177 | -188 | 0 | 0 | 12 | NaN | 606.9705 | NaN | 420.8085 | 382.43378 | NaN |
| Psmc5 | AHHTDCTFIR | -204 | -213 | 0 | 0 | 10 | NaN | 503.9172 | 2769.9272 | 1357.7897 | 2404.246 | 2347.6526 |
| Psmc5 | AHHTDCTFIR | -204 | -213 | 0 | 0 | 10 | NaN | 503.9172 | 2769.9272 | 1357.7897 | 2404.246 | 2347.6526 |
| Psmc5 | NDSYTLHK | -118 | -125 | 0 | 0 | 8 | NaN | 1910.3978 | 987.2196 | NaN | 2176.8289 | 2422.8545 |
| Psmc5 | AVAHHTDCTFIR | -202 | -213 | 0 | 0 | 12 | NaN | 867.99493 | 1084.8334 | NaN | 1201.5942 | 1190.1205 |
| Psmc5 | EVIELPVK | -163 | -170 | 0 | 0.007819401 | 8 | NaN | 281.80566 | NaN | NaN | NaN | NaN |
| Tgm2 | VDLFPTDIGLHK | -651 | -662 | 0 | 0 | 12 | NaN | 1771.511 | 2082.6907 | 2363.5195 | 4261.1445 | 572.5576 |
| Tgm2 | LCTPANAPIGLYR | -104 | -116 | 0 | 0 | 13 | NaN | 1008.8306 | 1055.1458 | 1124.287 | 1259.9877 | NaN |
| Tgm2 | ARVDLFPTDIGLHK | -649 | -662 | 1 | 0 | 14 | NaN | 482.0474 | NaN | NaN | 808.76996 | 531.47974 |
| Osbpl11 | AHLHPDHLVEVR | -215 | -226 | 0 | 0.00025015 | 12 | NaN | 489.17868 | NaN | NaN | NaN | NaN |
| Cnpy2 | RTDLCDHALHR | -167 | -177 | 1 | 0 | 11 | NaN | 669.16595 | 979.5305 | 510.21582 | 561.3763 | 878.30804 |
| Cnpy2 | ELLEEVCDR | -80 | -88 | 0 | 0 | 9 | NaN | 137.76729 | 1295.0664 | NaN | 1089.5844 | 938.0316 |

| | | | | | | | | | | | | |
|---|---|---|---|---|---|---|---|---|---|---|---|---|
| Cnpy2 | IDSDISGTLK | -125 | -134 | 0 | 0.00025015 | 10 | NaN | 1223.8383 | 1067.0564 | NaN | 1024.2268 | NaN |
| Nampt | YDGHLPIEVK | -108 | -117 | 0 | 0 | 10 | NaN | 32738.92 | 21263.05 | 15195.944 | 36799.504 | 33188.016 |
| Nampt | VFKDPVADPNKR | -413 | -424 | 2 | 0 | 12 | NaN | 860.56213 | NaN | NaN | 191.48923 | 190.34026 |
| Cand1 | KHTVDDGLDIR | -1072 | -1082 | 1 | 0 | 11 | NaN | 1294.2203 | 4922.9077 | 1079.6906 | 3761.731 | 3634.5164 |
| Cand1 | VIRPLDQPSSFDATPYIK | -549 | -566 | 1 | 0 | 18 | NaN | 622.4502 | 634.22125 | NaN | 805.61053 | 968.6211 |
| Cand1 | VVTGTNNLGYMDLLR | -763 | -777 | 0 | 0 | 15 | NaN | 349.52847 | NaN | NaN | 378.65924 | NaN |
| Polr2a | YPETTEGGRPK | -43 | -53 | 1 | 0 | 11 | NaN | 1084.7684 | 1186.2186 | 361.2222 | 858.22046 | 963.5868 |
| Polr2a | YSPTSPTYSPTSPK | (1909)(1923) | (1922)(1936) | 0 | 0 | 14 | NaN | 743.238 | 911.73126 | 564.7657 | 817.11035 | 865.8876 |
| Polr2a | VVLPCNLLR | -977 | -985 | 0 | 0 | 9 | NaN | 318.76776 | 469.90018 | 351.0652 | NaN | 592.49146 |
| Polr2a | AKQDVIEVIEK | -709 | -719 | 1 | 0 | 11 | NaN | 1085.9664 | NaN | 1173.657 | 1209.4329 | NaN |
| Polr2a | SVTEGGIKYPETTEGGRPK | -35 | -53 | 2 | 0.001291103 | 19 | NaN | 372.5406 | NaN | NaN | 510.93417 | 535.5692 |
| Polr2a | YTPTSPSYSPSSPEYTPASPK | -1839 | -1859 | 0 | 0.00111006 | 21 | NaN | 380.25122 | 266.4654 | NaN | NaN | NaN |
| Polr2a | SFDGSYVNYR | -1387 | -1396 | 0 | 0.006654467 | 10 | NaN | 460.37833 | NaN | NaN | NaN | NaN |
| Ncapd2 | LRDPAQQVR | -1091 | -1099 | 1 | 0.001610812 | 9 | NaN | 8460.629 | 8135.254 | NaN | 7368.412 | 7673.892 |
| Ncapd2 | AHPDPLWIPFK | -848 | -858 | 0 | 0 | 11 | NaN | 1716.763 | 245.81488 | NaN | 3081.2036 | 3645.943 |
| Ncapd2 | AIIDEFEQK | -1270 | -1278 | 0 | 0.001811524 | 9 | NaN | 731.55634 | NaN | 857.4725 | NaN | NaN |
| Nmi | VSAHPVPLNTGVR | -136 | -148 | 0 | 0.00078661 | 13 | NaN | 460.1591 | NaN | 481.5387 | NaN | 406.4859 |
| Dock8 | WIADLPSMQLNR | -1316 | -1327 | 0 | 0 | 12 | NaN | 3930.1514 | NaN | 2860.7156 | NaN | 2269.2842 |
| Dock8 | AVSSVHTQDNHLEK | -729 | -742 | 0 | 0 | 14 | NaN | 424.61002 | 401.36023 | NaN | NaN | 456.9352 |
| Dock8 | ISENFHCDLNSDQFK | -300 | -314 | 0 | 0.000989247 | 15 | NaN | 228.38397 | NaN | 457.39294 | 478.72986 | NaN |
| Dock8 | EKVPLQNPPIK | -704 | -714 | 1 | 0.000230707 | 11 | NaN | 729.866 | NaN | 686.6822 | NaN | NaN |
| Dock8 | ESQVSFPIR | -751 | -759 | 0 | 0.002534621 | 9 | NaN | 1228.5878 | NaN | NaN | NaN | NaN |
| Dock8 | SGGPTTVVPDPR | -869 | -880 | 0 | 0.005176406 | 12 | NaN | 601.7107 | NaN | NaN | NaN | NaN |
| Dock8 | VPLQNPPIK | -706 | -714 | 0 | 0.000947459 | 9 | NaN | 106.15299 | NaN | NaN | NaN | NaN |
| Irf4 | ELPEHVTTPDYHR | -429 | -441 | 0 | 0 | 13 | NaN | 1131.5944 | 928.9425 | 1198.0432 | 586.0161 | 427.36493 |
| Irf4 | DYAPDQSHPEIPYQCPVTFGPR | -180 | -201 | 0 | 0.00204437 | 22 | NaN | 237.95197 | NaN | NaN | NaN | NaN |
| Pdia4 | AIADEEDYATEVK | -444 | -456 | 0 | 0 | 13 | NaN | 1533.1553 | NaN | NaN | NaN | NaN |
| Mdh1 | GNHSSTQYPDVNHAK | -185 | -199 | 0 | 0 | 15 | NaN | 1789.8663 | 2013.6013 | 1857.2352 | 1946.3481 | 2079.9639 |
| Mdh1 | HSSTQYPDVNHAK | -187 | -199 | 0 | 0 | 13 | NaN | 540.04517 | 462.439 | 671.51807 | 677.4541 | 807.2848 |
| Mdh1 | GEFITTVQQR | -221 | -230 | 0 | 0 | 10 | NaN | 3769.2 | 3983.139 | 4261.1294 | 4859.789 | 4534.1797 |
| Mdh1 | DKEEIAFK | -72 | -79 | 1 | 0 | 8 | NaN | 2931.508 | 2952.6719 | 2360.3362 | 3391.2026 | 2864.5767 |
| Mdh1 | TEEKETAFEFLSSA | -321 | -334 | 1 | 0.00111006 | 14 | NaN | 735.5765 | 518.40753 | 427.10397 | NaN | 540.1138 |
| Psmb1 | SGFHGDCLTLTK | -82 | -93 | 0 | 0 | 12 | NaN | 2342.2273 | 3089.2942 | 5540.29 | 967.2776 | 5391.5747 |
| Psmb1 | NVEHVPLTLDR | -187 | -197 | 0 | 0.001023236 | 11 | NaN | 1019.6846 | 1140.7678 | 767.36127 | 1423.238 | NaN |
| Mthfd1 | LSLSHNPEQK | -869 | -878 | 0 | 0 | 10 | NaN | 2717.3667 | 2324.2688 | 2909.8071 | 1236.364 | 2692.616 |
| Mthfd1 | TDPTTLTDDEINRF | -505 | -518 | 1 | 0 | 14 | NaN | 710.97186 | 704.73956 | 254.42656 | 530.77924 | 1145.6495 |
| Mthfd1 | AYTEEDLDLVEK | -722 | -733 | 0 | 0 | 12 | NaN | 524.1275 | 883.1604 | 931.09985 | 1177.3412 | 542.0159 |
| Mthfd1 | TEEDLDLVEK | -724 | -733 | 0 | 0 | 10 | NaN | 794.5563 | NaN | 926.24066 | NaN | NaN |
| Qdpr | MTDSFTEQADQVTADVGK | -53 | -70 | 0 | 0 | 18 | NaN | 602.5507 | 705.3991 | 652.0053 | 811.3148 | 200.83649 |
| Qdpr | TDSFTEQADQVTADVGK | -54 | -70 | 0 | 0 | 17 | NaN | 1836.0457 | NaN | 1084.2478 | 1350.0812 | 992.1477 |
| Gabpa | DCISWVGDEGEFK | -337 | -349 | 0 | 0 | 13 | NaN | 2202.0115 | NaN | NaN | 1744.8556 | 817.9493 |
| St13 | AIEINPDSAQPYK | -173 | -185 | 0 | 0 | 13 | NaN | 820.3316 | 983.61993 | NaN | NaN | NaN |
| St13 | ALNDGELQK | -123 | -131 | 0 | 0 | 9 | NaN | 313.51477 | 347.39777 | NaN | NaN | NaN |
| Set | FTDHSDAGADELGEVIK | -205 | -221 | 0 | 0 | 17 | NaN | 1597.7021 | 1514.6342 | 1500.626 | 1644.7804 | 1679.1033 |
| Set | RQPFFQK | -76 | -82 | 1 | 0.001846793 | 7 | NaN | 1444.3767 | 1397.0651 | 978.5263 | 1423.2787 | 1431.0251 |
| Set | SKEFHLNESGDPSSK | -152 | -166 | 1 | 0 | 15 | NaN | 1033.465 | 599.4776 | 977.6622 | 1677.4642 | 1079.7859 |
| Set | YFDENPYFENK | -139 | -149 | 0 | 0 | 11 | NaN | 970.03534 | NaN | 1052.4225 | 837.1346 | 1211.6307 |
| Set | TRVEVTEFEDIK | -120 | -131 | 1 | 0.00025015 | 12 | NaN | 761.4587 | 714.61914 | NaN | 1809.1786 | 884.3445 |
| Set | QHEEPESFFT | -194 | -203 | 0 | 0.005917757 | 10 | NaN | 516.66956 | 700.08636 | NaN | NaN | NaN |
| Set | HIDEVQNEIDR | -46 | -56 | 0 | 0.00025015 | 11 | NaN | 2897.1191 | NaN | NaN | NaN | NaN |
| Gosr1 | AIEIEQLLAR | -71 | -80 | 0 | 0 | 10 | NaN | 702.9013 | NaN | 848.1738 | 683.613 | 895.29083 |
| Gosr1 | NYWEDLRK | -7 | -14 | 1 | 0.000712251 | 8 | NaN | 1086.9103 | 1445.633 | NaN | NaN | 911.6112 |
| Oxsr1 | REVLEGLEYLHK | -128 | -139 | 1 | 0 | 12 | NaN | 364.25922 | 679.6356 | 231.14519 | 621.11804 | 648.05444 |
| Oxsr1 | EVLEGLEYLHK | -129 | -139 | 0 | 0 | 11 | NaN | 1740.4739 | NaN | 2193.1106 | 98.67355 | 2284.5847 |
| Fxr1 | SELSNPSETESER | -432 | -444 | 0 | 0 | 13 | NaN | 1052.6343 | 1094.1875 | 1250.3824 | 1446.8749 | 1364.8 |
| Fxr1 | FVEDFIQVPR | -281 | -290 | 0 | 0.00025015 | 10 | NaN | 307.86856 | NaN | NaN | NaN | NaN |
| Hnrnpab | FKEEDPVKK | -207 | -215 | 2 | 0 | 9 | NaN | 4204.509 | 4885.9697 | 4201.437 | 5105.749 | 5316.9214 |
| Hnrnpab | VGGLSWDTSK | -79 | -88 | 0 | 0.00025015 | 10 | NaN | 2344.7786 | NaN | 1469.3826 | 1330.0975 | 1096.5862 |
| Hnrnpab | KIFVGGLNPEATEEK | -160 | -174 | 1 | 0 | 15 | NaN | 2322.973 | 2231.674 | NaN | 2460.5427 | 1770.7932 |
| Hnrnpab | IFVGGLNPEATEE | -161 | -173 | 0 | 0 | 13 | NaN | 218.58073 | 805.35596 | NaN | NaN | NaN |
| Otulin | HYNIPVR | -339 | -345 | 0 | 0.000230707 | 7 | NaN | 1197.3654 | 1256.415 | 540.5066 | 1079.6793 | NaN |
| Reep5 | IIRPIFL | -136 | -142 | 1 | 0.002068213 | 7 | NaN | 131.4206 | 371.62888 | 425.06952 | 560.0106 | 517.02094 |
| Reep5 | AIESPNKDDDTQWLTYW | -70 | -86 | 1 | 0.003003218 | 17 | NaN | 412.38007 | 258.04486 | NaN | 394.1128 | NaN |
| Jak1 | VSLVDGYFR | -406 | -414 | 0 | 0.000230707 | 9 | NaN | 701.93677 | 873.0127 | NaN | 1164.5834 | NaN |
| Dhx30 | AIFQQPPLGVR | -738 | -748 | 0 | 0.00713295 | 11 | NaN | 2976.553 | NaN | NaN | NaN | NaN |
| Dlat | VEKPQDIEAFK | -166 | -176 | 1 | 0 | 11 | NaN | 2295.282 | 2275.521 | 2535.7957 | 2419.3613 | 1795.6321 |
| Dlat | YLEKPITMLL | -633 | -642 | 1 | 0.000431655 | 10 | NaN | 915.48285 | 1288.0216 | 274.50143 | 607.9063 | NaN |
| Dlat | AIGASEDKLIPADNEK | -584 | -599 | 1 | 0.00149795 | 16 | NaN | 609.20233 | NaN | 321.1597 | 775.46063 | 975.1824 |
| Dlat | DVPLGAPLCIIVEK | -282 | -295 | 0 | 0 | 14 | NaN | 212.99411 | 2068.8745 | 348.62955 | NaN | NaN |
| Ppm1g | ETPSQENGPTAK | -198 | -209 | 0 | 0.002595029 | 12 | NaN | 662.10754 | NaN | NaN | NaN | NaN |
| Ppm1g | LTTEEVIK | -105 | -112 | 0 | 0.002910424 | 8 | NaN | 403.80676 | NaN | NaN | NaN | NaN |
| Asns | VEPFLPGHYEVL | -177 | -188 | 0 | 0.000394361 | 12 | NaN | 1145.538 | 1304.5466 | 1528.14 | 1523.8269 | 1528.9323 |
| Asns | ILFDNAIK | -237 | -244 | 0 | 0.003360691 | 8 | NaN | 932.3904 | NaN | 1058.4557 | 926.3606 | 1188.5286 |
| Asns | AIGMEDSPDLLAAR | -287 | -300 | 0 | 0 | 14 | NaN | 485.80945 | NaN | 962.56134 | 704.23846 | 743.5199 |
| Asns | EILWRPK | -461 | -467 | 1 | 0.00111006 | 7 | NaN | 8793.042 | NaN | 15010.08 | 1874.0828 | NaN |
| Asns | DTYGVRPL | -138 | -145 | 1 | 0.005363933 | 8 | NaN | 2714.464 | NaN | NaN | NaN | NaN |
| Kpna2 | FPSLLTNPK | -340 | -348 | 0 | 0.001190916 | 9 | NaN | 1288.1461 | 1325.6119 | 1085.4135 | 628.98 | 1239.2213 |
| Kpna2 | PSLLTNPK | -341 | -348 | 0 | 0.003490217 | 8 | NaN | 422.02344 | 407.29697 | 380.94238 | 619.76263 | 675.5478 |

| | | | | | | | | | | | | |
|---|---|---|---|---|---|---|---|---|---|---|---|---|
| Kpna2 | LLHHNDPEVL | -259 | -268 | 0 | 0.002697088 | 10 | NaN | 2728.0674 | 2611.4756 | NaN | NaN | NaN |
| Kpna2 | LTDGPNER | -278 | -285 | 0 | 0.000826806 | 8 | NaN | 744.24615 | NaN | NaN | NaN | NaN |
| Impdh2 | EDLVVAPAGVTLK | -183 | -195 | 0 | 0 | 13 | NaN | 1064.8966 | 1150.7188 | 859.8187 | NaN | 873.6933 |
| Impdh2 | FGVPVIADGGIQNVGHIA | -357 | -374 | 0 | 0 | 18 | NaN | 241.86588 | NaN | 574.9593 | 645.1725 | 623.59296 |
| Rps7 | AQQNNVEHKVETF | -161 | -173 | 1 | 0 | 13 | NaN | 565.54517 | 598.3643 | 528.4972 | 756.27185 | 778.2068 |
| Cstf3 | GAVVPPVHDIYR | -698 | -709 | 0 | 0 | 12 | NaN | 255.81256 | 465.79956 | 838.8148 | 724.402 | 712.58984 |
| Rcc2 | NWDLIGR | -111 | -117 | 0 | 0.002099344 | 7 | NaN | 2252.1265 | NaN | NaN | 1203.7782 | NaN |
| Prmt1 | AIKEPLVDVVDPK | -231 | -243 | 1 | 0 | 13 | NaN | 162.33447 | 600.3715 | 481.08447 | 485.5396 | 524.0187 |
| Prmt1 | VIGIECSSISDYAVK | -114 | -128 | 0 | 0 | 15 | NaN | 325.4184 | NaN | 872.09906 | 392.22284 | 743.4727 |
| Prmt1 | ANKLDHVVTIIK | -132 | -143 | 1 | 0.001023236 | 12 | NaN | 1142.0868 | 977.5848 | NaN | NaN | 1263.8784 |
| Srp68 | RVEEISPNIR | -235 | -244 | 1 | 0.000627943 | 10 | NaN | 4086.7754 | NaN | 4759.277 | 4347.0776 | 4951.596 |
| Zwint | EDTEQSPDALASEDASR | -76 | -92 | 0 | 0.001610812 | 17 | NaN | 1213.0819 | NaN | 206.18787 | NaN | NaN |
| Fbl | DQIHIKPGAK | -159 | -168 | 1 | 0.00025015 | 10 | NaN | 537.95154 | 822.14526 | 599.6177 | 931.3888 | 163.8069 |
| Copz1 | ESDPQQVVHR | -136 | -145 | 0 | 0.000230707 | 10 | NaN | 1870.5923 | NaN | 1594.938 | 2390.2563 | 2042.4064 |
| Lta4h | QWLTPEQTSGK | -117 | -127 | 0 | 0 | 11 | NaN | 3020.3828 | 3246.7385 | 500.73672 | 196.63585 | NaN |
| Lta4h | TFGESHPFTK | -356 | -365 | 0 | 0 | 10 | NaN | 5186.158 | 12810.3545 | NaN | NaN | 9574.371 |
| Lta4h | EGHTVYLER | -319 | -327 | 0 | 0.003003218 | 9 | NaN | 793.6164 | NaN | NaN | 1186.4772 | NaN |
| Pds5b | DHLLNFVR | -387 | -394 | 0 | 0 | 8 | NaN | 1774.7677 | NaN | NaN | NaN | NaN |
| Iqgap1 | KLPYDVTPEQALSHEEVK | -1111 | -1128 | 1 | 0 | 18 | NaN | 875.27985 | 848.4695 | 279.979 | 195.54585 | 1145.4971 |
| Iqgap1 | EQLWLANEGLITK | -739 | -751 | 0 | 0 | 13 | NaN | 997.1856 | 910.1875 | 1282.344 | 1784.3435 | 1789.5126 |
| Iqgap1 | LGEDLPPTTELEEGLR | -58 | -73 | 0 | 0.000431655 | 16 | NaN | 902.6502 | 951.0532 | 1215.6166 | 1487.6228 | 1424.4448 |
| Iqgap1 | EIGPTEEVGDFEVK | -1591 | -1604 | 0 | 0 | 14 | NaN | 877.68787 | 1405.4812 | 1186.1211 | 1485.4033 | 2304.2034 |
| Iqgap1 | SSSVTGLTNIEEENCQR | -480 | -496 | 0 | 0 | 17 | NaN | 331.12537 | 396.22028 | 525.138 | 525.203 | 655.8461 |
| Iqgap1 | TCLDNLASK | -1533 | -1541 | 0 | 0 | 9 | NaN | 1084.0842 | 1075.1122 | NaN | 1606.0442 | 1972.0128 |
| Iqgap1 | EEVQAGVDAANSAAQQYQR | -369 | -387 | 0 | 0 | 19 | NaN | 431.5363 | NaN | 282.14584 | 557.96826 | 543.3225 |
| Iqgap1 | QLSSSVTGLTNIEEENCQR | -478 | -496 | 0 | 0 | 19 | NaN | 280.6532 | 205.86444 | 222.33647 | 375.54672 | NaN |
| Iqgap1 | VAQHYQDTLIR | -575 | -585 | 0 | 0.001463897 | 11 | NaN | 949.90063 | 518.17975 | NaN | NaN | 620.67804 |
| Iqgap1 | QNVAYEYLCHLEEAK | -37 | -51 | 0 | 0 | 15 | NaN | 483.79108 | NaN | NaN | NaN | 667.9902 |
| Iqgap1 | KLTELGTVDPK | -1465 | -1475 | 1 | 0 | 11 | NaN | 433.61346 | NaN | NaN | 345.39584 | NaN |
| Iqgap1 | DQSDQDFQEELDLMK | -877 | -891 | 0 | 0.002099344 | 15 | NaN | 386.8777 | NaN | 580.7188 | NaN | NaN |
| Khsrp | MILIQDGSQNTNVDKPLR | -268 | -285 | 1 | 0.000230707 | 18 | NaN | 19895.043 | 21166.898 | 24536.943 | 27666.062 | 26896.59 |
| Khsrp | VNNNTPDFGFGGQK | -97 | -110 | 0 | 0 | 14 | NaN | 1272.2969 | 1436.7963 | 1370.3527 | NaN | 1369.003 |
| Khsrp | KLASQGDSIGSQLGPIHPPPR | -123 | -143 | 1 | 0 | 21 | NaN | 874.6987 | NaN | 712.22516 | 859.93646 | 1647.5594 |
| Khsrp | GAGGGPPPGPPGAGDR | -25 | -40 | 0 | 0.000230707 | 16 | NaN | 1226.2778 | NaN | 1214.8629 | NaN | NaN |
| Nmt1 | GPVEPDKDNIR | -136 | -146 | 1 | 0 | 11 | NaN | 728.7775 | 569.2024 | 389.5888 | 903.0428 | 914.324 |
| Nmt1 | AIPANIHIYDTEK | -228 | -240 | 0 | 0 | 13 | NaN | 533.6635 | NaN | 1053.4492 | 737.0439 | 1328.4839 |
| Psmb6 | AIQESGVER | -211 | -219 | 0 | 0.001636916 | 9 | NaN | 918.162 | NaN | 701.19574 | NaN | NaN |
| Zpr1 | IPELDFEIPAFSQK | -127 | -140 | 0 | 0 | 14 | NaN | 591.66266 | 724.0565 | 701.788 | NaN | 1124.6611 |
| Nop58 | SQMDGLIPGVEPR | -121 | -133 | 0 | 0 | 13 | NaN | 1641.7773 | 2112.6982 | 1005.7225 | NaN | 1125.893 |
| Nop58 | ELNNYIMR | -175 | -182 | 0 | 0.007546675 | 8 | NaN | 1031.0427 | 660.0699 | 578.9387 | NaN | NaN |
| Ddx39b | AITFVSDENDAK | -387 | -398 | 0 | 0.00025015 | 12 | NaN | 3367.1296 | 1384.4952 | NaN | NaN | NaN |
| Rpl14 | AIVDVIDQNR | -26 | -35 | 0 | 0 | 10 | NaN | 0 | 1530.2115 | 2664.9019 | 1628.6844 | 1544.7617 |
| Rpl14 | QLTDFILK | -56 | -63 | 0 | 0.005758933 | 8 | NaN | 526.2541 | 494.13025 | 528.8967 | NaN | 488.36307 |
| Rpl14 | VDGPCTR | -38 | -44 | 0 | 0.006735683 | 7 | NaN | 450.2282 | NaN | 651.9571 | 652.03955 | 714.41125 |
| Polr2g | TIDNIGAGVIQPGR | -50 | -63 | 0 | 0 | 14 | NaN | 615.42773 | 606.8571 | 653.8069 | 458.87088 | 479.7155 |
| Polr2g | AIVFRPFK | -74 | -81 | 1 | 0.006100831 | 8 | NaN | 659.35394 | 543.9554 | NaN | 643.05035 | NaN |
| Bin2 | NDLLWEDYEEK | -106 | -116 | 0 | 0 | 11 | NaN | 741.9115 | 758.8734 | 586.0673 | NaN | 150.6326 |
| Hnrnpul1 | NCAVEFNFGQR | -377 | -387 | 0 | 0.00025015 | 11 | NaN | 416.93533 | NaN | NaN | NaN | NaN |
| Sod2 | AIWNVINWENVTER | -203 | -216 | 0 | 0 | 14 | NaN | 510.42752 | NaN | NaN | NaN | NaN |
| Psmc1 | RVVGSELIQK | -249 | -258 | 1 | 0.00428947 | 10 | NaN | 2511.2537 | NaN | NaN | 3174.8135 | 3348.1233 |
| Psmc1 | RIETLDPALIRPGR | -333 | -346 | 2 | 0.00111006 | 14 | NaN | 267.4401 | NaN | NaN | NaN | 530.5102 |
| Psmc1 | NQLDGFDSR | -314 | -322 | 0 | 0.002199523 | 9 | NaN | 544.26843 | NaN | NaN | NaN | NaN |
| Eef1g | KAAAPAPEEEMDECEQAL | -253 | -270 | 1 | 0 | 18 | NaN | 690.7512 | 772.5567 | 814.0033 | 879.6253 | 589.99945 |
| Eef1g | CTLLWLYK | -166 | -173 | 0 | 0 | 8 | NaN | 708.299 | 746.4696 | 627.90686 | 320.6115 | 798.03534 |
| Eef1g | EGTFQHVGK | -420 | -428 | 0 | 0.001610812 | 9 | NaN | 1531.0166 | 1884.051 | 578.337 | 289.30865 | NaN |
| Eef1g | EYFSWEGTFQ | -415 | -424 | 0 | 0 | 10 | NaN | 101.42584 | 166.90553 | NaN | 195.15292 | 162.74103 |
| Eef1g | VLSAPPHFHFGQTN | -31 | -44 | 0 | 0.00383302 | 14 | NaN | 645.613 | NaN | NaN | 1169.2809 | NaN |
| Eef1g | VPAFEGDDGFCVFE | -58 | -71 | 0 | 0.005758933 | 14 | NaN | 357.38892 | 338.41254 | NaN | NaN | NaN |
| Ezr | THNDIIHNENMR | -548 | -559 | 0 | 0 | 12 | NaN | 483.0938 | 79.49807 | NaN | 746.80164 | 764.18414 |
| Ezr | ITEAEKNER | -518 | -526 | 1 | 0.002329279 | 9 | NaN | 371.30035 | 753.09125 | 338.54102 | NaN | 319.29752 |
| Hspa5 | ATNGDTHLGGEDFDQR | -247 | -262 | 0 | 0 | 16 | NaN | 3840.5483 | 470.31717 | 488.2286 | 529.6325 | 5156.1865 |
| Hspa5 | KIEWLESHQDADIEDFK | -602 | -618 | 1 | 0 | 17 | NaN | 1429.4335 | 1880.99 | 1920.9169 | 1321.9895 | 460.26282 |
| Hspa5 | FEGEDFSETLTR | -314 | -325 | 0 | 0 | 12 | NaN | 1533.253 | 1382.9341 | 1475.8008 | 136.87254 | 1509.7694 |
| Hspa5 | VTHAVVTVPAYFNDAQR | -166 | -182 | 0 | 0 | 17 | NaN | 202.51358 | 539.6668 | NaN | 1132.6163 | NaN |
| Hspa5 | ESHQDADIEDFK | -607 | -618 | 0 | 0 | 12 | NaN | 457.6849 | NaN | 412.4137 | 870.55676 | 703.1299 |
| Hspa5 | KNQLTSNPENTVFDAK | -82 | -97 | 1 | 0.000431655 | 16 | NaN | 281.7269 | 376.0286 | NaN | 736.08167 | 141.1639 |
| Hspa5 | GIDLGTTYSCVGVFK | -33 | -47 | 0 | 0 | 15 | NaN | 541.02625 | NaN | 1089.0764 | NaN | 762.39636 |
| Hspa5 | VTHAVVTVPAYFNDA | -166 | -180 | 0 | 0.00025015 | 15 | NaN | 1259.9604 | 1134.8406 | NaN | NaN | NaN |
| Hspa5 | IQVDIGGGQTK | -129 | -139 | 0 | 0.004658491 | 11 | NaN | 1093.3123 | NaN | NaN | NaN | NaN |
| Ncl | KVTLDWAKPK | -636 | -645 | 2 | 0.00051306 | 10 | NaN | 1189.423 | 1415.1665 | 1034.5558 | 1415.2927 | 1151.1235 |
| Ncl | KEAMEDGEIDGNK | -624 | -636 | 1 | 0 | 13 | NaN | 417.12988 | 482.76813 | 536.64264 | 633.5802 | 574.8558 |
| Ncl | ESFEGSVR | -587 | -594 | 0 | 0 | 8 | NaN | 1240.4064 | NaN | 871.48315 | NaN | NaN |
| Ncl | LFIGNLNPNK | -311 | -320 | 0 | 0.00025015 | 10 | NaN | 687.5718 | NaN | NaN | NaN | 705.62805 |
| Cs | GYTDPQFTELMR | -245 | -256 | 0 | 0 | 12 | NaN | 495.05423 | 714.58545 | 302.86432 | 1364.1876 | 1325.8073 |
| Cs | LDWSHNFTNML | -234 | -244 | 0 | 0.00025015 | 11 | NaN | 121.43327 | 498.609 | 175.09784 | 445.3235 | NaN |
| Cs | LDWSHNFTN | -234 | -242 | 0 | 0.009985598 | 9 | NaN | 259.40195 | NaN | 3468.6255 | NaN | NaN |
| Cs | AQLIWSR | -434 | -440 | 0 | 0.001392869 | 7 | NaN | 246.61327 | NaN | NaN | 90.610085 | NaN |
| Eif3b | AQSEEETATSPAASPTPQSAER | -66 | -87 | 0 | 0.00025015 | 22 | NaN | 158.61018 | 172.65486 | 239.96268 | 405.35776 | 293.11475 |

| | | | | | | | | | | | | |
|---|---|---|---|---|---|---|---|---|---|---|---|---|
| Eif3b | DRPQEADGIDSVIVVD | -164 | -179 | 1 | 0 | 16 | NaN | 703.10895 | 1037.3723 | 997.1056 | NaN | 1194.3271 |
| Eif3b | AKPAAQSEEETATSPAASPTPQ | -62 | -83 | 1 | 0.005758933 | 22 | NaN | 284.52646 | 394.26538 | NaN | NaN | NaN |
| Eif3b | YASPAHAVDAVK | -226 | -237 | 0 | 0.000471409 | 12 | NaN | 850.0087 | NaN | NaN | NaN | NaN |
| Eif3b | TSIFWNDVKDPVSIEER | -305 | -321 | 1 | 0 | 17 | NaN | 792.2603 | NaN | NaN | NaN | NaN |
| Dlst | KNELAIEDMDGGTF | -357 | -370 | 1 | 0.001055699 | 14 | NaN | 345.41513 | 388.70575 | NaN | NaN | NaN |
| Rrp9 | KTEEEEEEELEETAQEK | -62 | -78 | 1 | 0.001610812 | 17 | NaN | 510.66058 | NaN | NaN | 502.68448 | 566.7896 |
| Serbp1 | TENKENEVEEVKEEGPK | -269 | -285 | 2 | 0 | 17 | NaN | 2640.2249 | 2939.5117 | 2658.3794 | 3422.3186 | 3372.5767 |
| Serbp1 | DKSSASAPDVDDPEAFPALA | -388 | -407 | 1 | 0 | 20 | NaN | 139.08197 | 126.06947 | 645.9543 | 811.2036 | 352.94983 |
| Serbp1 | SEEAHAEDSVMDHHFR | -329 | -344 | 0 | 0 | 16 | NaN | 257.6791 | NaN | 504.71872 | 724.22003 | 710.7787 |
| Serbp1 | KEETQPPVAL | -92 | -101 | 1 | 0.001811524 | 10 | NaN | 730.2566 | 739.4997 | 569.0107 | NaN | NaN |
| Serbp1 | NVTEETPEGEEHPVADTENK | -253 | -272 | 0 | 0.001776473 | 20 | NaN | 188.51003 | NaN | 220.10631 | NaN | 247.7969 |
| Vdac1 | GTEITVEDQLAR | -95 | -106 | 0 | 0 | 12 | NaN | 3020.6465 | 2649.89 | NaN | NaN | NaN |
| Vdac1 | AKYQVDPDACFSAK | -236 | -249 | 1 | 0.008680443 | 14 | NaN | 1048.9545 | NaN | NaN | NaN | NaN |
| H2-K1 | HKWEQAGEAERL | -166 | -177 | 2 | 0 | 12 | NaN | 752.05206 | 840.0666 | 442.58145 | 894.96576 | 887.1707 |
| Arpc5l | AVGGLGSIIR | -136 | -145 | 0 | 0 | 10 | NaN | 661.5584 | NaN | NaN | 475.61267 | 446.5797 |
| Xrn2 | ALLPFVDER | -648 | -656 | 0 | 0.00025015 | 9 | NaN | 1271.6992 | 1347.524 | 1279.8755 | 1560.6813 | 1469.314 |
| Xrn2 | NSSPSISPNTSFASDGSPSPLGGIK | -469 | -493 | 0 | 0.00025015 | 25 | NaN | 401.00305 | 308.38766 | 556.5127 | 467.14032 | 668.34467 |
| Xrn2 | ALCNQFGHEVK | -264 | -274 | 0 | 0 | 11 | NaN | 7118.7383 | NaN | 7380.6777 | 6888.1084 | 7042.5186 |
| Xrn2 | GLATHEPNFTIIR | -240 | -252 | 0 | 0.000230707 | 13 | NaN | 686.54407 | 400.11218 | NaN | 603.8072 | 1079.2836 |
| Xrn2 | QWKPQLGFNR | -794 | -803 | 1 | 0.002810811 | 10 | NaN | 557.5458 | NaN | NaN | NaN | NaN |
| Rnh1 | ASLQELDLSSNK | -222 | -233 | 0 | 0.00078661 | 12 | NaN | 7520.3667 | 8314.832 | NaN | 7769.2056 | NaN |
| Rnh1 | DLCDVVASK | -213 | -221 | 0 | 0.005080642 | 9 | NaN | 5176.9897 | NaN | 383.12433 | NaN | NaN |
| Rnh1 | VLSNNDLHEPGVR | -171 | -183 | 0 | 0 | 13 | NaN | 825.37286 | 1085.3971 | NaN | NaN | NaN |
| Rnh1 | SNELKDEGAR | -288 | -297 | 1 | 0.00051306 | 10 | NaN | 74.45954 | 1537.0531 | NaN | NaN | NaN |
| Prpf8 | DYNFNLKPVK | -519 | -528 | 1 | 0 | 10 | NaN | 564.48 | 597.5397 | 737.8399 | 879.2721 | 812.09424 |
| Prpf8 | DEDWNEFNDINK | -290 | -301 | 0 | 0 | 12 | NaN | 182.15616 | 918.2656 | NaN | 702.4405 | 715.9613 |
| Prpf8 | GHVLIPQSDLR | -1331 | -1341 | 0 | 0.00025015 | 11 | NaN | 1279.6951 | NaN | 1153.8828 | 1322.9573 | NaN |
| Prpf8 | YNFNLKPVK | -520 | -528 | 1 | 0.000230707 | 9 | NaN | 1086.3258 | 783.73395 | NaN | NaN | NaN |
| Prpf8 | WEGLFWEK | -1498 | -1505 | 0 | 0.005301614 | 8 | NaN | 389.30493 | NaN | NaN | NaN | 398.49045 |
| Srsf6 | ALDKLDGTEINGR | -162 | -174 | 1 | 0 | 13 | NaN | 1036.4259 | 683.6293 | 985.1683 | 354.1108 | 1244.9578 |
| Ccar2 | ESDSPLKEDGLLPK | -609 | -622 | 1 | 0.00025015 | 14 | NaN | 2317.0176 | NaN | 1183.0829 | 2722.4429 | 2083.2793 |
| Ccar2 | EQAPDVSEQADTSK | -465 | -478 | 0 | 0 | 14 | NaN | 1231.6719 | NaN | NaN | 1033.8428 | NaN |
| Ccar2 | VVSPPEPEKEEAAKED | -566 | -581 | 2 | 0.001463897 | 16 | NaN | 1142.4342 | 770.14374 | NaN | NaN | NaN |
| Ccar2 | WCGYLHR | -715 | -721 | 0 | 0.001636916 | 7 | NaN | 775.1657 | 1152.4738 | NaN | NaN | NaN |
| Rabggta | LLPQLHPQPDSGPQGR | -187 | -202 | 0 | 0.000550307 | 16 | NaN | 649.32886 | NaN | 482.3038 | 292.264 | 288.80853 |
| Rabggta | ALDPLLYEK | -394 | -402 | 0 | 0 | 9 | NaN | 3469.2666 | NaN | 1835.9227 | NaN | NaN |
| Nup210 | KEIVVGTPQK | -1547 | -1556 | 1 | 0 | 10 | NaN | 1036.2546 | 484.4603 | 888.0781 | 480.5255 | 459.80548 |
| Nup210 | VSDIQELYVR | -975 | -984 | 0 | 0 | 10 | NaN | 923.7439 | 1101.9153 | 726.4756 | 889.3792 | 302.37128 |
| Nup210 | GAVESLENLEVK | -1713 | -1724 | 0 | 0 | 12 | NaN | 1504.9629 | 1921.1881 | NaN | 2987.141 | 1748.1884 |
| Nup210 | VGAEVPFSPGLY | -1681 | -1692 | 0 | 0 | 12 | NaN | 366.77332 | NaN | 363.37112 | 405.521 | 427.09485 |
| Nup210 | VGAEVPFSPGLYAN | -1681 | -1694 | 0 | 0 | 14 | NaN | 576.4818 | NaN | 881.9817 | 805.84296 | NaN |
| Rps17 | NLQVTQPTVGMNFK | -116 | -129 | 0 | 0.000550307 | 14 | NaN | 967.30615 | NaN | 906.81024 | NaN | NaN |
| Rps17 | NLQVTQPTVGMNFK | -116 | -129 | 0 | 0.000550307 | 14 | NaN | 967.30615 | NaN | 906.81024 | NaN | NaN |
| Rps17 | EIIEVDPDTK | -94 | -103 | 0 | 0.001636916 | 10 | NaN | 1698.8483 | NaN | NaN | NaN | NaN |
| Sptbn1 | ALDTGWNELHK | -1146 | -1156 | 0 | 0 | 11 | NaN | 2536.6648 | 2354.1318 | 176.5271 | 1323.223 | 830.10236 |
| Sptbn1 | EIGQSVDEVEK | -2044 | -2054 | 0 | 0 | 11 | NaN | 2775.551 | 1814.0901 | NaN | 3103.5178 | 1761.7748 |
| Sptbn1 | KLPEELGR | -1827 | -1834 | 1 | 0.001023236 | 8 | NaN | 1407.5732 | 1219.0623 | NaN | 1230.3977 | 1091.7914 |
| Sptbn1 | ISEKPETEAVVK | -1341 | -1352 | 1 | 0 | 12 | NaN | 923.6501 | 674.8819 | NaN | NaN | NaN |
| Cndp2 | EGGSIPVTLTFQEATGK | -414 | -430 | 0 | 0 | 17 | NaN | 1716.167 | 1506.908 | 2087.3406 | NaN | 2069.1104 |
| Cndp2 | EESGSEGLDELIFAQK | -166 | -181 | 0 | 0.000550307 | 16 | NaN | 888.7784 | NaN | NaN | NaN | 834.84265 |
| Cndp2 | CISDNYWLGK | -192 | -201 | 0 | 0 | 10 | NaN | 314.56046 | 344.78284 | NaN | NaN | NaN |
| Ppp4r3a | ALEDVDYVQTFK | -614 | -625 | 0 | 0 | 12 | NaN | 1927.2539 | 1985.5522 | NaN | NaN | NaN |
| Osbpl8 | ALEENPYFR | -453 | -461 | 0 | 0.005946382 | 9 | NaN | 155.62039 | 139.93506 | NaN | NaN | 1600.4482 |
| Osbpl8 | LSDYYYHAD | -439 | -447 | 0 | 0.007987481 | 9 | NaN | 228.44467 | NaN | NaN | NaN | NaN |
| Krt10 | ALEEQLQQIR | -411 | -420 | 0 | 0.003235556 | 10 | NaN | 1308.1776 | NaN | NaN | 590.8915 | 1030.5608 |
| Cltc | VLIEHIGNLDR | -1084 | -1094 | 0 | 0 | 11 | NaN | 1293.4834 | 1533.1094 | 366.1754 | 1692.3226 | 1539.613 |
| Cltc | NASGNWEELVK | -1145 | -1155 | 0 | 0 | 11 | NaN | 2169.4956 | 2216.6238 | 2557.287 | 2359.7793 | 2184.8315 |
| Cltc | GQFSTDELVAEVEK | -838 | -851 | 0 | 0 | 14 | NaN | 1717.2664 | 1570.1935 | 1381.7491 | 963.18665 | 739.8215 |
| Cltc | LLYNNVSNFGR | -1216 | -1226 | 0 | 0 | 11 | NaN | 1026.976 | 1175.8949 | 1449.3247 | 811.8846 | 1345.0884 |
| Cltc | RPLIDQVVQ | -968 | -976 | 1 | 0.004224523 | 9 | NaN | 786.1989 | 198.56113 | 1453.7869 | 2615.4775 | 2356.7712 |
| Cltc | LESNPYR | -961 | -967 | 0 | 0.001610812 | 7 | NaN | 1155.8501 | 509.15854 | 1043.3147 | 696.69495 | NaN |
| Cltc | ANASGNWEELVK | -1144 | -1155 | 0 | 0.001392869 | 12 | NaN | 869.1305 | 953.0314 | NaN | 744.0482 | 1462.4244 |
| Cltc | LDVDCSEDVIK | -820 | -830 | 0 | 0 | 11 | NaN | 852.945 | 717.30695 | NaN | 384.3012 | 850.27374 |
| Cltc | ISGETIFVTAPHEATAGIIGVNR | -298 | -320 | 0 | 0 | 23 | NaN | 520.8344 | NaN | 500.07257 | 610.49854 | 213.24973 |
| Cltc | NLAGAEELFAR | -356 | -366 | 0 | 0.00025015 | 11 | NaN | 317.96863 | NaN | NaN | 319.6984 | 261.06155 |
| Cltc | KLECSEELGDLVK | -456 | -468 | 1 | 0.001392869 | 13 | NaN | 525.54034 | 574.8966 | NaN | NaN | NaN |
| Cltc | FITEEDYQALR | -1471 | -1481 | 0 | 0.002629273 | 11 | NaN | 1895.2208 | NaN | NaN | NaN | NaN |
| Cltc | VDKLDASESLR | -1610 | -1620 | 1 | 0 | 11 | NaN | 461.66937 | NaN | NaN | NaN | NaN |
| Prpf6 | EKEEIEKY | -124 | -131 | 2 | 0.002629273 | 8 | NaN | 1168.1997 | 1089.412 | NaN | 1080.2014 | 1131.6339 |
| Prpf6 | ALEHVPNSVR | -396 | -405 | 0 | 0 | 10 | NaN | 6170.908 | NaN | 4094.5095 | NaN | NaN |
| Prpf6 | AVELEEPEDAR | -410 | -420 | 0 | 0 | 11 | NaN | 546.8269 | 1614.8306 | NaN | NaN | NaN |
| Prpf6 | LESENNEYER | -651 | -660 | 0 | 0.000230707 | 10 | NaN | 96.90971 | NaN | NaN | NaN | NaN |
| Rdx | IQNWHEEHR | -172 | -180 | 0 | 0 | 9 | NaN | 563.8528 | 505.0623 | 854.15753 | 459.0515 | 1184.643 |
| Rdx | EEEATEWQHK | -439 | -448 | 0 | 0 | 10 | NaN | 566.0379 | 796.02203 | 756.1219 | 818.51697 | 690.05316 |
| Pcmt1 | SGGASHSELIHNLR | -5 | -18 | 0 | 0 | 14 | NaN | 563.42365 | 689.2246 | NaN | NaN | NaN |
| Chmp4b | KQEFLEK | -39 | -45 | 1 | 0.007523243 | 7 | NaN | 1241.186 | NaN | 1220.8032 | NaN | NaN |
| Chmp4b | LRDTEEMLSK | -29 | -38 | 1 | 0.00025015 | 10 | NaN | 1002.0232 | 1339.1487 | NaN | NaN | NaN |
| Chmp4b | ALENANTNTEVLK | -95 | -107 | 0 | 0.00142999 | 13 | NaN | 322.078 | NaN | NaN | NaN | NaN |

| | | | | | | | | | | | | |
|---|---|---|---|---|---|---|---|---|---|---|---|---|
| Itih3 | YHFVTPLTSM | -592 | -601 | 0 | 0.003620005 | 10 | NaN | 494.58206 | NaN | NaN | NaN | NaN |
| Ubac2 | ALEPIFSSSEPTSEAR | -217 | -232 | 0 | 0 | 16 | NaN | 1461.1067 | 1699.3795 | 589.74615 | 1628.1636 | NaN |
| Ubac2 | HDLQIWR | -50 | -56 | 0 | 0 | 7 | NaN | 746.4304 | 774.01056 | NaN | 387.99213 | 826.391 |
| Nans | HLEFSHDQYK | -87 | -96 | 0 | 0 | 10 | NaN | 1223.9158 | 1436.543 | 1332.309 | 1792.3712 | 1682.3861 |
| Kif11 | LNLVDLAGSENIGR | -260 | -273 | 0 | 0 | 14 | NaN | 694.9713 | NaN | NaN | NaN | NaN |
| Emc1 | LFVFNPIFGK | -605 | -614 | 0 | 0 | 10 | NaN | 332.91647 | NaN | 247.09012 | 283.70874 | 285.91208 |
| Emc1 | ALETEWELR | -250 | -258 | 0 | 0.000230707 | 9 | NaN | 2956.3567 | NaN | NaN | 2481.0115 | NaN |
| Emc1 | VFNPIFGK | -607 | -614 | 0 | 0.00897527 | 8 | NaN | 538.2616 | NaN | NaN | NaN | 510.10004 |
| Ogt | ASSVGNVADSTEPTKR | -2 | -17 | 1 | 0 | 16 | NaN | 1065.7722 | 945.05975 | 1435.8135 | 2072.5437 | 1990.5522 |
| Ogt | QIVCDWTDYDER | -474 | -485 | 0 | 0.000230707 | 12 | NaN | 154.37773 | 247.58145 | 226.3583 | NaN | 272.51727 |
| Ogt | LKPDFPDAYCNL | -458 | -469 | 1 | 0.00149795 | 12 | NaN | 1019.10504 | NaN | NaN | NaN | 826.7021 |
| Ogt | ALSPDDGTNFR | -592 | -602 | 0 | 0.001636916 | 11 | NaN | 3758.8179 | NaN | NaN | NaN | NaN |
| Ogt | KALEVFPEFAAA | -352 | -363 | 1 | 0.00078661 | 12 | NaN | 376.7032 | NaN | NaN | NaN | NaN |
| Psmd2 | SCVNYVPEPENSALLR | -235 | -250 | 0 | 0 | 16 | NaN | 1117.9169 | 938.81854 | 905.4202 | 656.4939 | 824.36536 |
| Psmd2 | DPNNLFMVR | -755 | -763 | 0 | 0 | 9 | NaN | 601.00604 | 765.52045 | 683.4518 | 762.7665 | 745.9421 |
| Psmd2 | DTSLYRPALEELR | -67 | -79 | 1 | 0 | 13 | NaN | 567.73285 | 527.9014 | NaN | 856.93805 | 2029.5918 |
| Psmd2 | VTFDEELRPLPVSVR | -831 | -845 | 1 | 0 | 15 | NaN | 511.40402 | NaN | 721.26447 | 816.4137 | 891.48303 |
| Psmd2 | QTHTTPVLLAHGER | -866 | -879 | 0 | 0.000230707 | 14 | NaN | 6636.823 | NaN | NaN | 7271.3296 | NaN |
| Grb2 | HDGAFLIR | -79 | -86 | 0 | 0 | 8 | NaN | 2330.284 | 1924.0577 | 2070.1775 | 1790.3359 | 2047.3987 |
| Grb2 | ESESAPGDFSLSVK | -87 | -100 | 0 | 0 | 14 | NaN | 526.20984 | 645.52264 | 1238.4279 | 1278.5076 | 154.19833 |
| Grb2 | VLNEECDQNWYK | -27 | -38 | 0 | 0 | 12 | NaN | 242.10286 | 257.71802 | NaN | NaN | NaN |
| Atic | DGQVIGIGAGQQSR | -438 | -451 | 0 | 0 | 14 | NaN | 745.128 | 1098.2833 | 1348.5691 | 1105.6056 | 979.8089 |
| Atic | SVSSDAFFPFR | -535 | -545 | 0 | 0 | 11 | NaN | 1436.1511 | 723.2572 | 1952.3389 | 54.8311 | NaN |
| Atic | DVSELTGFPEMLGGR | -50 | -64 | 0 | 0 | 15 | NaN | 1077.3131 | 900.1595 | 916.0186 | 978.55725 | NaN |
| Atic | IGEGEDLVK | -499 | -507 | 0 | 0.00025015 | 9 | NaN | 696.4109 | 294.4636 | NaN | 1198.1316 | 865.0813 |
| Atic | GISQMPLR | -200 | -207 | 0 | 0.006100831 | 8 | NaN | 2002.6814 | NaN | NaN | NaN | NaN |
| Abcf2 | VWLEEELK | -270 | -277 | 0 | 0.00025015 | 8 | NaN | 792.1786 | 122.57007 | 704.70325 | 135.48233 | 70.940605 |
| Abcf2 | EVPIPEHIDIYHLTR | -140 | -154 | 0 | 0 | 15 | NaN | 394.44125 | 4785.7407 | 473.8694 | 497.44894 | 378.1134 |
| Abcf2 | ERLEELDADK | -199 | -208 | 1 | 0.000431655 | 10 | NaN | 2007.6641 | 2566.463 | 1248.2499 | 1326.3007 | NaN |
| Abcf2 | LVDEEPQLTK | -614 | -623 | 0 | 0 | 10 | NaN | 1099.09 | 540.1717 | 3909.8877 | NaN | NaN |
| Abcf2 | LAHEDAECEK | -184 | -193 | 0 | 0.00676572 | 10 | NaN | 504.8072 | 515.37494 | NaN | NaN | NaN |
| Bysl | ALFKPGAWFK | -278 | -287 | 1 | 0.000755556 | 10 | NaN | 3483.3179 | 3420.7542 | NaN | 2548.0217 | NaN |
| Ndufa4 | FNPDVSWDR | -39 | -47 | 0 | 0 | 9 | NaN | 728.3743 | 731.6226 | 444.028 | 268.14798 | 468.0907 |
| Ndufa4 | ALFNPDVSWDR | -37 | -47 | 0 | 0 | 11 | NaN | 538.00696 | 761.425 | NaN | 955.2269 | 158.46707 |
| Ftl1 | LVNLHLR | -20 | -26 | 0 | 0.007343498 | 7 | NaN | 1049.8026 | 1095.4933 | 235.50664 | NaN | 953.7596 |
| Ftl1 | FQDVQKPSQDEWGK | -79 | -92 | 1 | 0.002595029 | 14 | NaN | 811.99304 | 719.45557 | NaN | NaN | NaN |
| Zap70 | WYAPECINFR | -504 | -513 | 0 | 0 | 10 | NaN | 539.7843 | 584.1438 | NaN | NaN | 483.78006 |
| Zap70 | ALGADDSYYTAR | -484 | -495 | 0 | 0 | 12 | NaN | 2590.8965 | NaN | NaN | NaN | NaN |
| Tbcd | SDTAWDAELPVVR | -1159 | -1171 | 0 | 0 | 13 | NaN | 2082.431 | 2592.069 | 1608.5311 | 1842.9293 | 2233.2727 |
| Tbcd | HFDSPPIPHVPHR | -933 | -945 | 0 | 0.001392869 | 13 | NaN | 477.13724 | NaN | 3861.9104 | NaN | NaN |
| Dcps | TITLPYLESQ | -159 | -168 | 0 | 0.001291103 | 10 | NaN | 6243.6807 | NaN | 900.5695 | 5198.7266 | 5670.336 |
| Dcps | DKIIFLHGK | -58 | -66 | 1 | 0.008768524 | 9 | NaN | 236.13332 | NaN | NaN | NaN | NaN |
| Rtn3 | TQIDHYVGIA | -931 | -940 | 0 | 0.003681885 | 10 | NaN | 473.4164 | 867.3666 | NaN | 287.5775 | NaN |
| Rtn3 | SSSSGAEPSALGGGGGSPGACPALGAK | -15 | -41 | 0 | 0.007374243 | 27 | NaN | 299.54358 | 280.42255 | 323.21088 | NaN | NaN |
| Dnajc7 | ALGIDPNNIK | -282 | -291 | 0 | 0.00025015 | 10 | NaN | 1457.4539 | NaN | 1204.9585 | NaN | 1208.8176 |
| Lrrc47 | ELSPDIAHLA | -195 | -204 | 0 | 0.000431655 | 10 | NaN | 683.85913 | NaN | 798.7298 | 728.775 | 515.1171 |
| Lrrc47 | DGQCPLVVEQVR | -541 | -552 | 0 | 0 | 12 | NaN | 1725.6696 | 1743.2693 | NaN | NaN | 1367.9202 |
| Cnot3 | VSEGVEQFEDIWQK | -18 | -31 | 0 | 0 | 14 | NaN | 2806.9058 | 2611.7554 | 2535.7578 | 3588.1108 | 3094.9453 |
| Cnot3 | ALGPTSGPHNPAPSTSK | -452 | -468 | 0 | 0 | 17 | NaN | 1419.6893 | NaN | 1236.6981 | 594.1014 | NaN |
| Cnot3 | APEPLSSLK | -534 | -542 | 0 | 0.003299042 | 9 | NaN | 926.27655 | NaN | 725.2683 | NaN | NaN |
| Ptbp1 | PPSVSEDDLK | -486 | -495 | 0 | 0 | 10 | NaN | 794.16675 | 608.44586 | 986.3923 | 1181.8552 | 1233.2988 |
| Ptbp1 | LHLSNIPPSVSEDDLK | -480 | -495 | 0 | 0 | 16 | NaN | 703.73486 | 588.1969 | 1245.9167 | 1136.4381 | 1018.1721 |
| Top2a | IPNFDVR | -863 | -869 | 0 | 0.002780285 | 7 | NaN | 1672.0187 | 1794.0531 | 1546.9601 | 1297.0918 | 102.1631 |
| Top2a | IHHNWPSLLR | -556 | -565 | 0 | 0 | 10 | NaN | 1449.5709 | 1334.5872 | 682.7784 | 503.8472 | 1190.2825 |
| Top2a | NEKEQELNTLK | -1146 | -1156 | 1 | 0 | 11 | NaN | 1038.6179 | 1713.8546 | 4386.7705 | 3029.5186 | 3541.0364 |
| Top2a | LLDGEEPLPMLPSYK | -878 | -892 | 0 | 0 | 15 | NaN | 729.22864 | 674.37396 | 993.07086 | 1119.2488 | 1420.3541 |
| Top2a | KKYDTVLDILR | -1009 | -1019 | 2 | 0.00122419 | 11 | NaN | 457.3728 | NaN | 274.15656 | 491.80032 | 496.76068 |
| Top2a | SVVDLESDVK | -1373 | -1382 | 0 | 0 | 10 | NaN | 557.2394 | NaN | 1595.7349 | NaN | NaN |
| Top2a | STTIEISELPIR | -917 | -928 | 0 | 0 | 12 | NaN | 436.9369 | 1250.8064 | NaN | NaN | NaN |
| Nop56 | EELGYNCQTGGVIAEILR | -106 | -123 | 0 | 0 | 18 | NaN | 180.59676 | NaN | 368.17648 | 443.44696 | 453.97336 |
| Nop56 | VEECVLNLGK | -32 | -41 | 0 | 0.000431655 | 10 | NaN | 16487.9 | 10938.8125 | NaN | NaN | 14033.527 |
| Nop56 | QVALENANAVSEGVVHEDLR | -58 | -77 | 0 | 0.002972566 | 20 | NaN | 376.93298 | NaN | NaN | NaN | NaN |
| Rrm1 | ALKEEEEKER | -763 | -772 | 2 | 0.000431655 | 10 | NaN | 371.78748 | 368.35522 | 303.0755 | 340.35175 | 385.87405 |
| Rrm1 | YPFESPEAQLLNK | -517 | -529 | 0 | 0 | 13 | NaN | 807.2524 | 969.86145 | 2896.695 | 3252.3877 | 1223.8438 |
| Rrm1 | NIHIAEPNYGK | -709 | -719 | 0 | 0 | 11 | NaN | 736.0626 | 738.2622 | 1749.7246 | 758.4922 | 490.95944 |
| Rrm1 | VLSGEFQIVNPH | -629 | -640 | 0 | 0.00025015 | 12 | NaN | 1416.4121 | NaN | NaN | 2553.208 | 2683.8528 |
| Rrm1 | WFTHASPT | -197 | -204 | 0 | 0.002910424 | 8 | NaN | 1025.8961 | NaN | 989.10986 | NaN | NaN |
| Rrm1 | RVETNQDWSL | -341 | -350 | 1 | 0.007405213 | 10 | NaN | 979.976 | NaN | NaN | NaN | NaN |
| Gltp | DENHPNLIR | -117 | -125 | 0 | 0 | 9 | NaN | 735.25183 | NaN | 971.1723 | 506.59436 | 1001.1065 |
| Eif3d | KLGDDIDLIVR | -364 | -374 | 1 | 0 | 11 | NaN | 747.1518 | 841.0042 | 625.2706 | 1270.9557 | 1144.6698 |
| Eif3d | TVSETANEPPQDEGNSFNSPR | -290 | -310 | 0 | 0.00111006 | 21 | NaN | 641.8236 | 643.07404 | 530.11035 | 720.418 | 693.582 |
| Eif3d | VTTTDDPVIR | -227 | -236 | 0 | 0.001150701 | 10 | NaN | 348.58905 | NaN | 590.6291 | NaN | 450.16638 |
| Eif3d | ANEPPQDEGNSFNSPR | -295 | -310 | 0 | 0.002068213 | 16 | NaN | 904.70154 | NaN | NaN | 948.38513 | NaN |
| Eif3d | TTTDDPVIR | -228 | -236 | 0 | 0 | 9 | NaN | 772.33466 | NaN | 965.03107 | NaN | NaN |
| Eif3h | EKDFSPEALK | -179 | -188 | 1 | 0.00111006 | 10 | NaN | 721.96533 | NaN | NaN | 2486.6404 | 2238.4844 |
| Psma7 | ILNPEEIEKY | -219 | -228 | 1 | 0 | 10 | NaN | 2565.9617 | 437.1703 | 2388.399 | 2542.5378 | 2616.552 |
| Xpot | LQNWQTTR | -436 | -443 | 0 | 0.008680443 | 8 | NaN | 1887.7736 | NaN | NaN | NaN | NaN |
| Gimap4 | ELYREELER | -242 | -250 | 1 | 0.00111006 | 9 | NaN | 1812.2422 | 1924.8829 | 2778.9421 | 2532.142 | 2575.669 |

| | | | | | | | | | | | | |
|---|---|---|---|---|---|---|---|---|---|---|---|---|
| Gimap4 | STGNSILGEK | -45 | -54 | 0 | 0 | 10 | NaN | 775.1161 | 813.1463 | 657.4373 | 623.8752 | 838.5272 |
| Gimap4 | DLRDELER | -265 | -272 | 1 | 0.00051306 | 8 | NaN | 5532.966 | 5980.611 | NaN | 6378.96 | NaN |
| Hnrnpul2 | FPTLWSGAR | -272 | -280 | 0 | 0.000627943 | 9 | NaN | 1929.2028 | 2127.1963 | 1769.0645 | 2264.2397 | 2174.2783 |
| Hnrnpul2 | ANFETEEVELSFSK | -363 | -376 | 0 | 0 | 14 | NaN | 420.81476 | 403.06702 | 392.92203 | NaN | NaN |
| Hnrnpul2 | NCVVELNFGQK | -405 | -415 | 0 | 0.001291103 | 11 | NaN | 1938.2915 | NaN | 2002.6702 | NaN | NaN |
| Hnrnpul2 | KVVVVVPNEEDWKR | -556 | -569 | 2 | 0.002068213 | 14 | NaN | 241.3933 | NaN | 342.38382 | NaN | NaN |
| Gpi | HWMDQHFLK | -301 | -309 | 0 | 0 | 9 | NaN | 521.19293 | 848.4741 | 2794.0166 | 6300.82 | 6417.4717 |
| Gpi | HWMDQHFLK | -301 | -309 | 0 | 0 | 9 | NaN | 521.19293 | 848.4741 | 2794.0166 | 6300.82 | 6417.4717 |
| Gpi | KLLEWHR | -12 | -18 | 1 | 0.002329279 | 7 | NaN | 1966.6442 | NaN | 1790.3876 | 2033.3195 | 1757.6545 |
| Gpi | KIEPELEGSSAVTS | -524 | -537 | 1 | 0.001846793 | 14 | NaN | 1250.6702 | NaN | 1166.0566 | NaN | NaN |
| Gpi | LRELFEADPER | -26 | -36 | 1 | 0 | 11 | NaN | 680.9151 | NaN | NaN | NaN | 331.22897 |
| Tbl3 | VLLPEQPAPALGVK | -243 | -256 | 0 | 0 | 14 | NaN | 418.5625 | 444.2407 | 380.44882 | 496.13925 | NaN |
| Tbl3 | HEAPEELLAYDGVR | -741 | -754 | 0 | 0.000627943 | 14 | NaN | 960.5713 | NaN | NaN | NaN | 313.86954 |
| Tbl3 | ALLPYTER | -759 | -766 | 0 | 0.002168954 | 8 | NaN | 998.22003 | NaN | NaN | NaN | NaN |
| Shmt2 | LLKDPETSQR | -472 | -481 | 1 | 0 | 10 | NaN | 3815.2395 | 3659.19 | 3355.889 | 4866.2573 | 4852.1216 |
| Shmt2 | IMGLDLPDGGHLTHGY | -161 | -176 | 0 | 0.00025015 | 16 | NaN | 553.1973 | NaN | 282.5462 | 469.80923 | 822.84344 |
| Hsp90aa1 | SLTNDWEEHLAVK | -316 | -328 | 0 | 0 | 13 | NaN | 2733.851 | 2802.5757 | 1674.5431 | 2256.3994 | 2343.3538 |
| Hsp90aa1 | LLSSGFSLEDPQTHANR | -672 | -688 | 0 | 0 | 17 | NaN | 1206.2711 | 1156.7208 | 1405.9403 | 575.90125 | 479.10266 |
| Hsp90aa1 | KHLEINPDHSIIETLR | -633 | -648 | 1 | 0 | 16 | NaN | 247.85152 | 480.81525 | 550.388 | 1568.1277 | 1012.28094 |
| Hsp90aa1 | KHLEINPDHSIIETLR | -633 | -648 | 1 | 0 | 16 | NaN | 247.85152 | 480.81525 | 550.388 | 1568.1277 | 1012.28094 |
| Hsp90aa1 | KHSQFIGYPITLF | -209 | -221 | 1 | 0.00078661 | 13 | NaN | 460.90976 | NaN | 570.9198 | 756.5309 | 308.22928 |
| Hsp90aa1 | KHGLEVIYM | -514 | -522 | 1 | 0.005394768 | 9 | NaN | 1362.0022 | NaN | NaN | 1158.4762 | NaN |
| Hsp90aa1 | KCLELFTEL | -420 | -428 | 1 | 0.00450079 | 9 | NaN | 658.47516 | NaN | NaN | NaN | 730.09717 |
| Hsp90aa1 | HLEINPDHSIIET | -634 | -646 | 0 | 0.001392869 | 13 | NaN | 917.8498 | NaN | NaN | NaN | NaN |
| Hsp90aa1 | NDWEEHLAVK | -319 | -328 | 0 | 0.00025015 | 10 | NaN | 524.93726 | NaN | NaN | NaN | NaN |
| Gtpbp1 | REEGGPCGVPAAGGPPTGDEASSL | -602 | -625 | 1 | 0.000826806 | 24 | NaN | 530.328 | NaN | 650.71515 | NaN | NaN |
| Gtf2i | PEGVPFR | -775 | -781 | 0 | 0.00437931 | 7 | NaN | 832.5784 | 823.4863 | NaN | 749.6678 | 617.6485 |
| Gtf2i | STVVPVPYEK | -131 | -140 | 0 | 0 | 10 | NaN | 683.43207 | 1333.7378 | 919.6332 | 1238.2094 | NaN |
| Trap1 | ALPAGQLTSLPDYASR | -476 | -491 | 0 | 0 | 16 | NaN | 945.10706 | NaN | 1965.3923 | 1810.119 | 1907.5178 |
| Il16 | SLAGGADLENK | -1120 | -1130 | 0 | 0.000394361 | 11 | NaN | 1795.8337 | NaN | 463.96112 | NaN | NaN |
| Il16 | ALSPDPLLR | -928 | -936 | 0 | 0.001708008 | 9 | NaN | 1247.7286 | NaN | NaN | NaN | NaN |
| Pgam1 | SYDVPPPPMEPDHPFYSN | -118 | -135 | 0 | 0 | 18 | NaN | 456.86282 | 538.95593 | 673.03033 | 548.5002 | 574.7627 |
| Pgam1 | RYADLTEDQLPSCESLK | -141 | -157 | 1 | 0.000591124 | 17 | NaN | 301.41956 | NaN | NaN | 1351.3646 | NaN |
| Pgam1 | HLEGLSEEAIME | -196 | -207 | 0 | 0.004224523 | 12 | NaN | 1456.5234 | NaN | NaN | NaN | NaN |
| Pgam1 | HLEGLSEEAIME | -196 | -207 | 0 | 0.004224523 | 12 | NaN | 1456.5234 | NaN | NaN | NaN | NaN |
| Pgam1 | HLEGLSEEAIMEL | -196 | -208 | 0 | 0.001291103 | 13 | NaN | 282.48523 | NaN | NaN | NaN | NaN |
| Utp14a | NKQPVTDPLLTPVEK | -195 | -209 | 1 | 0 | 15 | NaN | 514.29926 | 511.59418 | 452.42175 | NaN | 566.8432 |
| Utp14a | LLEAIGSLSGK | -46 | -56 | 0 | 0.001291103 | 11 | NaN | 492.80212 | NaN | NaN | 331.78638 | NaN |
| Bcas2 | RYELPAPSSGQK | -86 | -97 | 1 | 0 | 12 | NaN | 964.2965 | 1077.299 | 808.7099 | 1097.8796 | 1207.498 |
| Cse1l | FQSGDFHVINGVLR | -138 | -151 | 0 | 0 | 14 | NaN | 1404.0017 | 1279.136 | 1479.717 | 1809.9141 | 1982.8965 |
| Cse1l | QEIFDGIQPK | -807 | -816 | 0 | 0.003490217 | 10 | NaN | 537.4273 | 481.5561 | NaN | NaN | NaN |
| Igtp | ENLQQAQVR | -225 | -233 | 0 | 0 | 9 | NaN | 656.1104 | 621.3112 | NaN | NaN | NaN |
| Ppp2r1b | ANDPNYLHR | -502 | -510 | 0 | 0 | 9 | NaN | 624.7671 | 920.3751 | 977.2069 | 930.5969 | 839.6199 |
| Ppp2r1b | ELCENLPAEGR | -320 | -330 | 0 | 0 | 11 | NaN | 124.36192 | NaN | 1568.7191 | NaN | 441.11795 |
| Mvp | ALQPLEEGEGEEK | -336 | -348 | 0 | 0.00111006 | 13 | NaN | 477.69904 | 603.08167 | NaN | NaN | NaN |
| Gatad2b | ALQQEQEIEQR | -469 | -479 | 0 | 0 | 11 | NaN | 806.67633 | 2371.6855 | 4266.1655 | 1399.7612 | 1475.251 |
| Psmc6 | ATNRPDTLDPALLRPGR | -278 | -294 | 2 | 0 | 17 | NaN | 1101.4093 | 80.76836 | 574.3933 | 843.09784 | 890.35504 |
| Psmc6 | DHQPCIIF | -224 | -231 | 0 | 0.002434616 | 8 | NaN | 762.79443 | 509.7115 | 543.8183 | 641.5869 | NaN |
| Psmc6 | FSEGTSADR | -243 | -251 | 0 | 0 | 9 | NaN | 611.82104 | 511.93546 | NaN | NaN | NaN |
| Shmt1 | SDALTELGYK | -333 | -342 | 0 | 0 | 10 | NaN | 1269.9856 | NaN | 2739.8413 | 1990.655 | 1789.0952 |
| Uhrf1 | IEEVFHVEPQLQR | -32 | -44 | 0 | 0 | 13 | NaN | 889.9846 | 1130.2766 | 1028.3011 | 550.67334 | 942.69226 |
| Itga2b | VPIENIISSYRPGTL | -228 | -242 | 1 | 0.00025015 | 15 | NaN | 482.6004 | NaN | 542.60095 | 574.45386 | 610.4705 |
| Itga2b | DSYYQPLHR | -301 | -309 | 0 | 0.00111006 | 9 | NaN | 250.08107 | NaN | NaN | NaN | NaN |
| Itpa | LKPEGLHQLLA | -95 | -105 | 1 | 0 | 11 | NaN | 4452.1416 | 4648.9663 | NaN | NaN | 5214.1475 |
| Psmb10 | GAGVAADTEMTTR | -84 | -96 | 0 | 0 | 13 | NaN | 1909.3081 | NaN | NaN | 456.23322 | 2224.5962 |
| Psmb10 | STPTEPVQR | -229 | -237 | 0 | 0.00025015 | 9 | NaN | 1016.0501 | NaN | 711.39874 | 591.077 | NaN |
| Cdk2 | KELNHPNIVK | -56 | -65 | 1 | 0.000591124 | 10 | NaN | 310.11743 | NaN | 867.00085 | 696.3378 | NaN |
| Eif3k | TNLPHTDFTLCK | -75 | -86 | 0 | 0 | 12 | NaN | 906.43604 | 885.3957 | 743.14636 | 691.9665 | 834.82935 |
| Eif3k | IDQAHQEERPIR | -89 | -100 | 1 | 0.00025015 | 12 | NaN | 575.2012 | 386.91553 | 401.84235 | 505.1949 | 505.9503 |
| Eif3k | FICHVVGITY | -141 | -150 | 0 | 0.00272025 | 10 | NaN | 1119.9598 | NaN | 1997.7369 | 2735.5537 | 2448.523 |
| Eif3k | ALTNLPHTDFTLCK | -73 | -86 | 0 | 0 | 14 | NaN | 585.02765 | 341.095 | NaN | 845.4199 | NaN |
| Msn | AQQELEEQTR | -361 | -370 | 0 | 0 | 10 | NaN | 1022.5496 | 4654.8584 | 3579.6882 | 4177.045 | 4712.5576 |
| Msn | ESEAVEWQQK | -439 | -448 | 0 | 0 | 10 | NaN | 494.33978 | 447.4931 | 407.6252 | 467.75623 | 404.32394 |
| Msn | TQEQLASEMAELTAR | -413 | -427 | 0 | 0 | 15 | NaN | 1408.229 | 1484.4674 | 2871.246 | 3751.6343 | NaN |
| Msn | LIQDITQR | -93 | -100 | 0 | 0.004439092 | 8 | NaN | 1378.6417 | 1188.291 | NaN | NaN | NaN |
| Elmo1 | TTSPAQNAQQLHER | -78 | -91 | 0 | 0 | 14 | NaN | 432.91803 | 426.95782 | NaN | 358.0353 | 478.40067 |
| Elmo1 | VLHYGDLEESPQGEVPHDSLQDK | -585 | -607 | 0 | 0.001291103 | 23 | NaN | 401.6632 | NaN | NaN | NaN | NaN |
| Dock11 | EDQIEYHEGLK | -2007 | -2017 | 0 | 0.00078661 | 11 | NaN | 971.9892 | 680.459 | NaN | NaN | NaN |
| Dock11 | ALVAEFLHR | -1647 | -1655 | 0 | 0.001636916 | 9 | NaN | 702.0974 | NaN | NaN | NaN | NaN |
| Dock11 | YEDFSGDFR | -119 | -127 | 0 | 0.00025015 | 9 | NaN | 193.8237 | NaN | NaN | NaN | NaN |
| Map4k1 | SCWLDDSSTEHK | -703 | -714 | 0 | 0.000230707 | 12 | NaN | 323.56808 | NaN | 317.92203 | NaN | NaN |
| Ubr4 | ALVEGENDEQSSPDQVSAAK | -215 | -234 | 0 | 0.000591124 | 20 | NaN | 1931.1263 | 2210.5217 | 2558.6914 | NaN | 2962.5134 |
| Ubr4 | SIAVTRPNNLVHFT | -2577 | -2590 | 1 | 0.00078661 | 14 | NaN | 2112.6477 | 1398.3986 | NaN | NaN | NaN |
| Ubr4 | ELASPVSPELR | -175 | -185 | 0 | 0 | 11 | NaN | 1192.7305 | NaN | 233.39136 | NaN | NaN |
| Ubr4 | HLILLIK | -128 | -134 | 0 | 0.006472924 | 7 | NaN | 298.46973 | NaN | NaN | 344.46194 | NaN |
| Ubr4 | DSDQLEVIQENR | -1477 | -1488 | 0 | 0 | 12 | NaN | 1814.1503 | NaN | NaN | NaN | NaN |
| Upf1 | SSVGAPVEVTHNF | -403 | -415 | 0 | 0.00025015 | 13 | NaN | 1716.0499 | NaN | 1895.1786 | NaN | NaN |
| Phb1 | PIIFDCR | -64 | -70 | 0 | 0.00078661 | 7 | NaN | 1695.3673 | NaN | NaN | NaN | NaN |

| | | | | | | | | | | | | |
|---|---|---|---|---|---|---|---|---|---|---|---|---|
| Phb1 | FRPVASQLPR | -96 | -105 | 1 | 0.00122419 | 10 | NaN | 477.62622 | NaN | NaN | NaN | NaN |
| Pabpc1 | AYVNFQQPADAER | -55 | -67 | 0 | 0 | 13 | NaN | 1234.1969 | NaN | 1918.4464 | 952.99634 | 786.3897 |
| Snx1 | AMLGSSEDNTALSR | -352 | -365 | 0 | 0.000431655 | 14 | NaN | 576.93756 | NaN | NaN | 816.2082 | NaN |
| Satb1 | DCKEEHAEFVLVR | -90 | -102 | 1 | 0 | 13 | NaN | 488.9142 | 570.92694 | NaN | 736.3118 | 594.39 |
| Satb1 | TLWENLSMIR | -544 | -553 | 0 | 0.00078661 | 10 | NaN | 1549.6847 | NaN | NaN | NaN | 1762.5962 |
| Myo1g | AVLDEACSTAGPITDR | -472 | -487 | 0 | 0 | 16 | NaN | 719.7608 | 638.486 | 580.854 | NaN | 870.5212 |
| Myo1g | AMWPDGQQDITEVTK | -563 | -577 | 0 | 0.001291103 | 15 | NaN | 1127.4125 | NaN | 612.764 | 1266.4386 | NaN |
| Myo1g | HLGNIEFVETEENGPQK | -287 | -303 | 0 | 0.000550307 | 17 | NaN | 424.3255 | NaN | NaN | 263.68277 | 611.4464 |
| Myo1g | ELVEQPHR | -461 | -468 | 0 | 0.00025015 | 8 | NaN | 1449.0133 | 2333.597 | NaN | NaN | NaN |
| Myo1g | DYLSSDTDNPTASHLF | -844 | -859 | 0 | 0.001636916 | 16 | NaN | 306.2202 | NaN | NaN | NaN | 249.97723 |
| Myo1g | GVHNALDSDEK | -246 | -256 | 0 | 0.000712251 | 11 | NaN | 894.6489 | NaN | NaN | NaN | NaN |
| Myo1g | LISVEPRPEQPEPDFQSSR | -996 | -1014 | 1 | 0.00025015 | 19 | NaN | 421.18262 | NaN | NaN | NaN | NaN |
| Pfkl | ANAPDSACVIGLR | -701 | -713 | 0 | 0 | 13 | NaN | 349.864 | NaN | NaN | 787.1089 | NaN |
| Tbc1d15 | REPVSLEEWNK | -297 | -307 | 1 | 0 | 11 | NaN | 999.2195 | 874.0406 | 2250.346 | 2807.7524 | 2943.6187 |
| Tbc1d15 | QQEEPGFEVITR | -274 | -285 | 0 | 0.00025015 | 12 | NaN | 928.0951 | NaN | 413.68814 | 429.66086 | 826.62665 |
| Tbc1d15 | INQQEEPGFEVITR | -272 | -285 | 0 | 0.00025015 | 14 | NaN | 1046.793 | 904.03357 | NaN | NaN | NaN |
| Tbc1d15 | ANDQDSLISGILR | -27 | -39 | 0 | 0.001708008 | 13 | NaN | 601.96515 | NaN | NaN | 375.8156 | NaN |
| Pdia3 | DLFSDGHSEFLK | -162 | -173 | 0 | 0 | 12 | NaN | 6093.019 | 6006.2744 | 1119.376 | 888.3793 | 7909.7134 |
| Pdia3 | FISDKDASVVGFFR | -148 | -161 | 1 | 0 | 14 | NaN | 509.04028 | 407.76666 | 477.88055 | 594.9037 | 529.0777 |
| Pdia3 | MTEDNKDLIQGK | -247 | -258 | 1 | 0 | 12 | NaN | 1505.9489 | NaN | 1090.956 | 1452.0126 | 1564.1029 |
| Pdia3 | PIPESNEGPVK | -369 | -379 | 0 | 0 | 11 | NaN | 741.83545 | 935.6213 | 150.65613 | 1068.8566 | 234.54813 |
| Pdia3 | RDLFSDGHSEFLK | -161 | -173 | 1 | 0 | 13 | NaN | 614.66547 | 646.3663 | 1157.069 | 1781.0684 | 1652.6646 |
| Pdia3 | FIQDSIFGLCPH | -235 | -246 | 0 | 0 | 12 | NaN | 318.75616 | 728.92194 | 561.03033 | 569.9694 | NaN |
| Ptbp3 | INYYTPVTPHLR | -82 | -93 | 0 | 0 | 12 | NaN | 1041.2656 | NaN | 518.2599 | 1294.6516 | 848.3343 |
| Ptbp3 | NEGTLLPGQSPVLR | -141 | -154 | 0 | 0 | 14 | NaN | 375.7039 | NaN | 402.93262 | 834.0418 | 770.0265 |
| Psat1 | EFDFVPDVK | -161 | -169 | 0 | 0 | 9 | NaN | 7068.344 | 260.85977 | NaN | NaN | 6036.5293 |
| Psat1 | SQGFYVCPVER | -285 | -295 | 0 | 0 | 11 | NaN | 875.96533 | NaN | NaN | 549.29297 | NaN |
| Psat1 | SAVPLNLIGLK | -84 | -94 | 0 | 0.001023236 | 11 | NaN | 403.7807 | NaN | 402.2138 | NaN | NaN |
| Iap | ANFTLVQLDR | -252 | -261 | 0 | 0 | 10 | NaN | 335.88492 | 509.9991 | NaN | NaN | NaN |
| Lap3 | ANKPGDVVR | -343 | -351 | 1 | 0.000550307 | 9 | NaN | 876.568 | NaN | 226.69394 | 993.18225 | 1243.5818 |
| Lap3 | SAGVDDQENWHEGK | -105 | -118 | 0 | 0 | 14 | NaN | 532.9687 | 402.72205 | NaN | 455.21008 | 464.53937 |
| Lap3 | GSEEPPVFLEIH | -254 | -265 | 0 | 0 | 12 | NaN | 689.0675 | 714.2316 | NaN | NaN | 1068.9768 |
| Agfg1 | APVGSVVSVPSHSSASSDK | -355 | -373 | 0 | 0 | 19 | NaN | 1019.99 | 1026.5658 | 878.8128 | 930.711 | 928.73566 |
| Agfg1 | ANLDNIFSAGQGGDQGSGFGTTGK | -331 | -354 | 0 | 0.00025015 | 24 | NaN | 193.00963 | 362.115 | 367.6437 | 478.49646 | 420.9586 |
| Pdcd11 | FLPISHPR | -1114 | -1121 | 0 | 0.005270854 | 8 | NaN | 977.51776 | 847.86304 | NaN | NaN | 741.7013 |
| Pdcd11 | ANLEESFPR | -2 | -10 | 0 | 0 | 9 | NaN | 389.94672 | NaN | 308.96335 | NaN | 1435.6301 |
| Pdcd11 | ILDDVPVGTSPTTTLK | -1079 | -1094 | 0 | 0.00025015 | 16 | NaN | 400.79044 | NaN | NaN | 412.93185 | NaN |
| Xrcc6 | NVNTGSLLLPSDTK | -302 | -315 | 0 | 0 | 14 | NaN | 3621.181 | 3462.197 | NaN | NaN | 4291.841 |
| Xrcc6 | ANLFSDVQLK | -149 | -158 | 0 | 0.001708008 | 10 | NaN | 436.72824 | NaN | NaN | NaN | 467.9933 |
| Sae1 | EALEVDWSGEK | -222 | -232 | 0 | 0 | 11 | NaN | 760.7988 | 703.58264 | 639.916 | 1274.641 | 1226.2957 |
| Sae1 | VLFCPVKE | -215 | -222 | 1 | 0.003681885 | 8 | NaN | 1001.1474 | NaN | NaN | NaN | NaN |
| Ncbp1 | KDGALEEQIER | -684 | -694 | 1 | 0 | 11 | NaN | 1984.2434 | 775.5221 | 5853.1157 | 6679.474 | 421.54224 |
| Ncbp1 | FVIEENLHCIIK | -312 | -323 | 0 | 0 | 12 | NaN | 711.86017 | 685.69354 | 796.5021 | 701.21643 | 908.0396 |
| Ncbp1 | DGALEEQIER | -685 | -694 | 0 | 0 | 10 | NaN | 1350.3497 | 1056.977 | 2564.7231 | 2512.6782 | NaN |
| Aimp2 | ATNTLDLNSVLGK | -102 | -114 | 0 | 0.001291103 | 13 | NaN | 579.59973 | NaN | NaN | NaN | NaN |
| Tra2b | RPHTPTPGIY | -198 | -207 | 1 | 0.002780285 | 10 | NaN | 4274.33 | 4124.8433 | 3428.214 | NaN | NaN |
| Rps23 | KGHAVGDIPGVR | -108 | -119 | 1 | 0 | 12 | NaN | 2032.2485 | 1963.8018 | 2868.7817 | NaN | NaN |
| Rps23 | ITAFVPNDGCLN | -81 | -92 | 0 | 0.006978206 | 12 | NaN | 3235.4626 | NaN | NaN | NaN | NaN |
| Nup107 | VDSLVEQEIR | -458 | -467 | 0 | 0 | 10 | NaN | 2141.0562 | NaN | NaN | NaN | 2626.6572 |
| Nup107 | HQCLELAK | -620 | -627 | 0 | 0.001392869 | 8 | NaN | 1219.8469 | NaN | NaN | NaN | NaN |
| Nup107 | QKLPLDDLDREDEVR | -332 | -346 | 2 | 0 | 15 | NaN | 1093.011 | NaN | NaN | NaN | NaN |
| Smarca5 | FITDNTVEER | -606 | -615 | 0 | 0.000471409 | 10 | NaN | 1237.1746 | 1937.2485 | NaN | NaN | 1028.8641 |
| Serpinc1 | QVLELPFK | -301 | -308 | 0 | 0 | 8 | NaN | 866.743 | 805.07513 | 1023.8915 | 818.8601 | 986.83215 |
| Serpinc1 | LGACNDTLK | -125 | -133 | 0 | 0.00025015 | 9 | NaN | 2328.8723 | NaN | 1936.1792 | 1493.8516 | 1966.9087 |
| Zcchc9 | ANVDPALGEFPFAK | -172 | -185 | 0 | 0 | 14 | NaN | 364.57422 | NaN | 1673.2007 | 1428.3649 | 1838.0496 |
| Acly | SGGMSNELNNIISR | -653 | -666 | 0 | 0 | 14 | NaN | 526.86426 | 422.35574 | 505.4321 | 1116.0215 | 1021.26044 |
| Acly | KEILIPVFK | -536 | -544 | 1 | 0 | 9 | NaN | 207.61398 | 274.8804 | 332.04318 | 370.382 | 355.36554 |
| Acly | VDATADYICK | -221 | -230 | 0 | 0 | 10 | NaN | 1189.0076 | 1114.3408 | 369.92285 | 1208.2783 | NaN |
| Acly | KHPEVDVL | -552 | -559 | 1 | 0.00897527 | 8 | NaN | 335.05972 | NaN | NaN | NaN | NaN |
| Ada | SLFPGHVEAYEGAVK | -192 | -206 | 0 | 0 | 15 | NaN | 301.9479 | 566.33527 | 666.3233 | 800.9409 | 582.8906 |
| Mrpl12 | APASEAAEEEDVPK | -113 | -126 | 0 | 0.00025015 | 14 | NaN | 4365.178 | 2351.5513 | NaN | 7118.284 | 7450.8804 |
| Rpl23a | PDYDALDVANK | -142 | -152 | 0 | 0 | 11 | NaN | 160.42831 | 370.92285 | 515.6496 | 255.52763 | 216.03398 |
| Rpl23a | VNTLIRPDGEKK | -124 | -135 | 2 | 0.002595029 | 12 | NaN | 490.66095 | NaN | NaN | NaN | NaN |
| Anp32e | IKDLSTVEAL | -100 | -109 | 1 | 0.003003218 | 10 | NaN | 6716.4014 | 6981.542 | NaN | 7726.583 | 7480.0054 |
| Cfl1 | PENAPLK | -106 | -112 | 0 | 0 | 7 | NaN | 2070.2283 | 2015.6918 | 2194.6824 | 3525.3086 | 3268.852 |
| Cfl1 | APENAPLK | -105 | -112 | 0 | 0 | 8 | NaN | 1154.46 | 965.4719 | 1345.3832 | 428.10333 | 1215.3544 |
| Cfl1 | LTGIKHELQAN | -128 | -138 | 1 | 0.00025015 | 11 | NaN | 678.4619 | 875.01105 | NaN | 380.81854 | 442.84998 |
| Raver1 | IPLNPYLNLH | -510 | -519 | 0 | 0 | 10 | NaN | 371.91183 | NaN | 424.3415 | 502.7265 | 496.84644 |
| Raver1 | GKPQPLLPPLLGPSGGDR | -439 | -456 | 1 | 0 | 18 | NaN | 357.6717 | 489.49786 | NaN | 441.9809 | 333.85907 |
| Raver1 | EVHDLLSDYELK | -74 | -85 | 0 | 0.001023236 | 12 | NaN | 1680.1799 | NaN | 834.51324 | NaN | NaN |
| Raver1 | FCQLACGQDGQLK | -250 | -262 | 0 | 0 | 13 | NaN | 835.958 | NaN | NaN | 921.5767 | NaN |
| Raver1 | SAGGGLAPELPPR | -425 | -437 | 0 | 0.001636916 | 13 | NaN | 627.13306 | NaN | NaN | NaN | 653.1288 |
| Raver1 | APEQELPPLDPEEIR | -31 | -45 | 0 | 0.003867596 | 15 | NaN | 1291.0958 | NaN | NaN | NaN | NaN |
| Psmd12 | CTYVEEITDLPVK | -109 | -121 | 0 | 0 | 13 | NaN | 272.537 | 1151.7704 | NaN | NaN | NaN |
| Tmem43 | SDPNYGVHLPAVK | -102 | -114 | 0 | 0 | 13 | NaN | 1425.6165 | NaN | 1237.8558 | 1730.1415 | 1997.5424 |
| Tmem43 | VVSPDSIHSVAPENEGR | -71 | -87 | 0 | 0.001291103 | 17 | NaN | 347.11618 | NaN | 653.64264 | 601.57465 | NaN |
| Hspd1 | NAGVEGSLIVEK | -482 | -493 | 0 | 0 | 12 | NaN | 1578.7286 | 433.91995 | 1688.3186 | 672.48926 | 682.8449 |
| Hspd1 | LVQDVANNTNEEAGDGTTTA | -97 | -116 | 0 | 0.00025015 | 20 | NaN | 511.49622 | 489.06104 | 547.67395 | 753.19507 | 396.2702 |

| | | | | | | | | | | | | |
|---|---|---|---|---|---|---|---|---|---|---|---|---|
| Hspd1 | PGFGDNR | -303 | -309 | 0 | 0 | 7 | NaN | 899.31964 | NaN | 70.89336 | NaN | NaN |
| Hspd1 | DIGNIISDAMK | -181 | -191 | 0 | 0.001291103 | 11 | NaN | 647.55927 | NaN | NaN | NaN | 633.3779 |
| Hspd1 | QSKPVTTPEEIA | -158 | -169 | 1 | 0.003393785 | 12 | NaN | 3992.0437 | NaN | NaN | NaN | NaN |
| Hspd1 | KGANPVEIR | -133 | -141 | 1 | 0.000826806 | 9 | NaN | 1388.1782 | NaN | NaN | NaN | NaN |
| Mki67 | KVDQQVQDNENAPQR | -853 | -867 | 1 | 0.001291103 | 15 | NaN | 209.5788 | NaN | NaN | 267.32672 | 331.97443 |
| Mki67 | TLKEDEDIVCTK | -3160 | -3171 | 1 | 0 | 12 | NaN | 1046.7175 | NaN | NaN | 1786.5068 | NaN |
| Mki67 | KEDEDIVCTK | -3162 | -3171 | 1 | 0.00111006 | 10 | NaN | 881.3806 | NaN | NaN | 167.97389 | NaN |
| Mki67 | QKIDPVASVPVSK | -2712 | -2724 | 1 | 0.003360691 | 13 | NaN | 1092.6161 | NaN | NaN | NaN | NaN |
| Mki67 | NTLKEPVGDSINVE | -2691 | -2704 | 1 | 0 | 14 | NaN | 960.4517 | NaN | NaN | NaN | NaN |
| Mki67 | ESTPQTLDSSASR | -1764 | -1776 | 0 | 0.000550307 | 13 | NaN | 914.55066 | NaN | NaN | NaN | NaN |
| Mki67 | VSFGGHLRPE | -466 | -475 | 1 | 0 | 10 | NaN | 716.8651 | NaN | NaN | NaN | NaN |
| Eif4a1 | LQMEAPHIIVGTPGR | -147 | -161 | 0 | 0 | 15 | NaN | 1161.5322 | 1285.1143 | 1118.0997 | 1356.303 | 1342.8134 |
| Eif4a1 | LQMEAPHIIVGTPGR | -147 | -161 | 0 | 0 | 15 | NaN | 1161.5322 | 1285.1143 | 1118.0997 | 1356.303 | 1342.8134 |
| Eif4a1 | QMEAPHIIVGTPGR | -148 | -161 | 0 | 0 | 14 | NaN | 968.2677 | 1047.4728 | 1378.8909 | 1372.5417 | 1382.568 |
| Eif4a1 | SCHACIGGTNVR | -130 | -141 | 0 | 0 | 12 | NaN | 5299.299 | 5892.904 | 3646.1594 | 387.01648 | 1367.7959 |
| Idh3a | GLFDHAAK | -319 | -326 | 0 | 0 | 8 | NaN | 794.62744 | 410.53687 | NaN | 1281.2606 | 1269.4506 |
| Rpl4 | IEEVPELPLVVEDKVEGYKK | -144 | -163 | 2 | 0.00025015 | 20 | NaN | 307.55957 | 262.14014 | NaN | 414.22064 | 366.32504 |
| Srrm1 | KETESEAEDDNLDDLER | -911 | -927 | 1 | 0 | 17 | NaN | 856.3045 | 944.8233 | 985.87384 | 934.07324 | 955.58887 |
| Psme1 | KQPHVGDYR | -190 | -198 | 1 | 0.000907206 | 9 | NaN | 5100.7437 | 5448.454 | 5127.59 | 3168.077 | 6693.7256 |
| Psme1 | LVHELDEAEYQEIR | -200 | -213 | 0 | 0 | 14 | NaN | 873.67084 | 707.1322 | 814.6284 | 771.8621 | 835.8181 |
| Nckap1l | EFVTPDTDIK | -945 | -954 | 0 | 0.000627943 | 10 | NaN | 3003.1785 | 2438.2563 | NaN | NaN | NaN |
| Iars1 | STACGTPLSNFESNQNYK | -182 | -199 | 0 | 0.000347792 | 18 | NaN | 682.00745 | 693.5241 | 650.4639 | 529.6844 | 577.0144 |
| Iars1 | ILEFWSK | -18 | -24 | 0 | 0.006624942 | 7 | NaN | 908.4711 | NaN | 857.60535 | 792.83624 | NaN |
| Iars1 | SVAELEELSGTK | -483 | -494 | 0 | 0.003037016 | 12 | NaN | 820.1507 | NaN | NaN | 899.7245 | NaN |
| Leo1 | APLQGDHNHLFIR | -463 | -475 | 0 | 0 | 13 | NaN | 658.3175 | 647.25055 | 460.13196 | 671.0624 | 696.3607 |
| Chd4 | GIINHGYAR | -1743 | -1751 | 0 | 0.00025015 | 9 | NaN | 2042.4785 | 1886.3374 | 1406.2532 | 1476.5007 | 1686.1053 |
| Chd4 | NDKDKPLPPLLA | -1387 | -1398 | 2 | 0.000431655 | 12 | NaN | 464.90628 | NaN | NaN | 786.42816 | 689.50977 |
| Chd4 | SIDWACLIVDEAHR | -857 | -870 | 0 | 0.000431655 | 14 | NaN | 473.74344 | NaN | NaN | NaN | NaN |
| Wdr12 | LWEWNVEK | -169 | -176 | 0 | 0.00111006 | 8 | NaN | 1158.8518 | NaN | NaN | 531.73315 | NaN |
| Rpl18a | NFGIWLR | -77 | -83 | 0 | 0.004350979 | 7 | NaN | 973.4238 | NaN | NaN | 411.42218 | 1396.7839 |
| Crkl | VIIEKPEEQWWSAR | -151 | -164 | 1 | 0 | 14 | NaN | 1413.6952 | 1756.0374 | NaN | NaN | 2806.043 |
| Crkl | APNLPTAEENLEYVR | -115 | -129 | 0 | 0.000230707 | 15 | NaN | 740.7728 | 1028.3369 | NaN | 1027.985 | NaN |
| Cope | SIYFHDQNPDAALR | -125 | -138 | 0 | 0 | 14 | NaN | 5263.7905 | 4771.249 | 4139.883 | 4880.008 | 5021.9897 |
| Cope | IYFHDQNPDAALR | -126 | -138 | 0 | 0.00025015 | 13 | NaN | 271.82526 | NaN | NaN | NaN | NaN |
| Aifm1 | LNDGSQITFEK | -244 | -254 | 0 | 0 | 11 | NaN | 1041.0583 | 1314.5995 | 1432.7094 | 1864.4512 | 1440.8119 |
| Tcea1 | APSTSDSVR | -133 | -141 | 0 | 0.005015524 | 9 | NaN | 450.26538 | 503.22018 | NaN | NaN | NaN |
| Rrm2 | SLLIDTYIKDPK | -178 | -189 | 1 | 0 | 12 | NaN | 1856.8186 | NaN | 1639.2806 | 1703.512 | 1767.4315 |
| Cars1 | LATEPLEQAVR | -244 | -254 | 0 | 0 | 11 | NaN | 1251.7975 | 1540.6963 | 1110.2997 | 1278.7212 | 1398.4728 |
| Cars1 | FWEEEFHK | -301 | -308 | 0 | 0 | 8 | NaN | 945.4785 | NaN | 925.25726 | 1000.6841 | 964.3826 |
| Cars1 | APVDITGQFEK | -555 | -565 | 0 | 0.001868731 | 11 | NaN | 888.5334 | NaN | 1260.8225 | 3245.6948 | NaN |
| Acin1 | KVTLGDTLTR | -969 | -978 | 1 | 0 | 10 | NaN | 1100.6642 | 1087.394 | 299.6454 | 1256.7087 | 1137.0282 |
| Acin1 | LLDDLFR | -1209 | -1215 | 0 | 0.000907206 | 7 | NaN | 859.4023 | NaN | 650.37646 | 778.3721 | 834.21765 |
| Acin1 | VTLGDTLTR | -970 | -978 | 0 | 0.00025015 | 9 | NaN | 1094.504 | 1109.7528 | NaN | 1105.0756 | NaN |
| Acin1 | APVVLQPEQIVSEEETPPPLLTK | -402 | -424 | 0 | 0.00659934 | 23 | NaN | 244.16539 | NaN | NaN | 258.93695 | 242.64778 |
| Yars1 | TSYYENVIK | -94 | -102 | 0 | 0 | 9 | NaN | 1692.0385 | NaN | 2952.1958 | 1683.2136 | 1500.1495 |
| Yars1 | IDVGEAEPR | -392 | -400 | 0 | 0.009897819 | 9 | NaN | 530.8514 | NaN | NaN | 350.48105 | 609.6678 |
| Hadhb | VGSPPLEKF | -410 | -418 | 1 | 0 | 9 | NaN | 9034.722 | 15538.188 | 12805.04 | 13429.558 | 15082.343 |
| Esyt1 | RVELGEKPLR | -173 | -182 | 2 | 0.00025015 | 10 | NaN | 1247.5146 | 1304.493 | 1275.9629 | 1416.764 | 1461.05 |
| Esyt1 | DKDPDKDDFLGR | -394 | -405 | 2 | 0 | 12 | NaN | 460.9512 | 626.7742 | 643.3018 | 675.3391 | 715.856 |
| Esyt1 | DKDPDKDDFLGR | -394 | -405 | 2 | 0 | 12 | NaN | 460.9512 | 626.7742 | 643.3018 | 675.3391 | 715.856 |
| Esyt1 | ATYSTNSPVWEEAFR | -506 | -520 | 0 | 0.000550307 | 15 | NaN | 4537.422 | 4777.0854 | NaN | 8056.6323 | 7705.2793 |
| Esyt1 | LTLEDVPSGR | -738 | -747 | 0 | 0 | 10 | NaN | 1364.485 | NaN | NaN | 1070.1667 | NaN |
| Esyt1 | NSPVWEEAFR | -511 | -520 | 0 | 0 | 10 | NaN | 1183.6116 | NaN | NaN | 1283.1743 | NaN |
| Esyt1 | SPVWEEAFR | -512 | -520 | 0 | 0.002329279 | 9 | NaN | 997.3527 | 1006.083 | NaN | NaN | NaN |
| Nsun2 | YALDPSFPR | -536 | -544 | 0 | 0 | 9 | NaN | 392.5606 | NaN | NaN | 758.4221 | 1039.0803 |
| Nsun2 | EILHCLK | -89 | -95 | 0 | 0.006654467 | 7 | NaN | 1319.8646 | NaN | NaN | 728.5775 | NaN |
| Nsun2 | KNIDVWK | -279 | -285 | 1 | 0.009031675 | 7 | NaN | 780.7339 | 403.36548 | NaN | NaN | NaN |
| Nsun2 | TNENAASPEQPGDEDAK | -717 | -733 | 0 | 0.00025015 | 17 | NaN | 359.84354 | NaN | NaN | NaN | 145.55081 |
| Kri1 | AQEEVDYLEWLK | -202 | -213 | 0 | 0 | 12 | NaN | 414.69464 | 629.4541 | 613.1798 | 615.4384 | 649.2048 |
| Ppid | TVPTPHLDGK | -136 | -145 | 0 | 0.002780285 | 10 | NaN | 692.7879 | 423.31287 | NaN | NaN | NaN |
| Ppid | LFADIVPK | -36 | -43 | 0 | 0.002168954 | 8 | NaN | 546.2696 | NaN | 472.80563 | NaN | NaN |
| Ppid | VFFDVDIGGER | -18 | -28 | 0 | 0.004983786 | 11 | NaN | 470.21262 | 1908.8832 | NaN | NaN | NaN |
| Scin | AQELQHPEFAR | -2 | -12 | 0 | 0 | 11 | NaN | 3524.0747 | NaN | NaN | 4130.986 | 4218.6206 |
| Hdhd5 | AQELSDLLR | -100 | -108 | 0 | 0.00025015 | 9 | NaN | 992.228 | NaN | NaN | 541.62537 | NaN |
| Sf3a1 | QQQQQATQQQLPQK | -118 | -131 | 0 | 0 | 14 | NaN | 340.6615 | NaN | 339.8329 | 537.8963 | 537.58453 |
| Sf3a1 | ITLQEQIEAIHK | -518 | -529 | 0 | 0 | 12 | NaN | 406.18906 | NaN | NaN | 294.97427 | 347.26266 |
| Sf3a1 | IGPSKPNEIPQQPPPPSSAT | -543 | -562 | 1 | 0.003681885 | 20 | NaN | 771.594 | 548.16815 | NaN | NaN | NaN |
| Sf3a1 | AQEPSAAIPK | -106 | -115 | 0 | 0.000471409 | 10 | NaN | 1284.4365 | NaN | NaN | NaN | NaN |
| Diablo | AQEVSDEGADQEEEAYLRED | -218 | -237 | 1 | 0.006164361 | 20 | NaN | 251.79158 | NaN | NaN | NaN | 292.1691 |
| Echs1 | LVEEAIQCAEK | -218 | -228 | 0 | 0 | 11 | NaN | 4456.626 | 4606.187 | 3947.4016 | 4153.754 | 1762.0217 |
| Atp5f1a | LETELFYK | -395 | -402 | 0 | 0.00025015 | 8 | NaN | 1362.4589 | 599.9856 | 1555.7993 | 778.2831 | 1620.0228 |
| Atp5f1a | EAYPGDVFYLHSR | -335 | -347 | 0 | 0 | 13 | NaN | 1282.8873 | 1684.3156 | 1835.23 | 2218.1624 | 2115.8872 |
| Atp5f1a | KLELAQYR | -434 | -441 | 1 | 0.00025015 | 8 | NaN | 373.49857 | 771.05164 | 1710.3877 | 1644.6848 | 1668.7842 |
| Atp5f1a | GIRPAINVGLS | -403 | -413 | 1 | 0.001392869 | 11 | NaN | 800.28406 | 724.8838 | 1569.8464 | NaN | NaN |
| Gapdh | AQNIIPASTGAAK | -201 | -213 | 0 | 0 | 13 | NaN | 1571.2454 | 1613.8889 | 2165.917 | 1522.8073 | 1587.5958 |
| Gapdh | ERDPTNIK | -77 | -84 | 1 | 0.005605646 | 8 | NaN | 419.17014 | 821.9939 | 429.6739 | 659.3032 | 728.4049 |
| Gapdh | LEKPAKY | -247 | -253 | 2 | 0.000627943 | 7 | NaN | 217.24829 | NaN | 2009.0726 | 1802.1152 | NaN |
| Gapdh | QASEGPLK | -262 | -269 | 0 | 0.001392869 | 8 | NaN | 254.57967 | 809.34296 | NaN | NaN | NaN |

| | | | | | | | | | | | | |
|---|---|---|---|---|---|---|---|---|---|---|---|---|
| Sp110 | RDNSFITER | -38 | -46 | 1 | 0.002265552 | 9 | NaN | 2732.238 | NaN | 1956.5132 | 2758.7542 | 175.93701 |
| Sp110 | VVHNILTSL | -64 | -72 | 0 | 0.00122419 | 9 | NaN | 2032.0887 | NaN | 255.12157 | NaN | 1656.2666 |
| Sp110 | GVASPGHGVQEK | -274 | -285 | 0 | 0.00111006 | 12 | NaN | 1183.1467 | NaN | NaN | NaN | NaN |
| Sp110 | DDSPAANDLEMAR | -224 | -236 | 0 | 0.005270854 | 13 | NaN | 513.94037 | NaN | NaN | NaN | NaN |
| Fkbp8 | GQPTVEEAEQPGALAR | -49 | -64 | 0 | 0.00025015 | 16 | NaN | 984.34753 | 1023.18634 | 1639.6733 | NaN | NaN |
| Stat5a | WIESQPWDAIDLDNPQDR | -37 | -54 | 0 | 0.001392869 | 18 | NaN | 746.2776 | 650.65045 | 640.294 | 786.61194 | 684.21265 |
| Stat5a | TLSLPVVVIVHGSQDHNATAT | -461 | -481 | 0 | 0 | 21 | NaN | 410.24066 | NaN | NaN | NaN | NaN |
| Psmd11 | LRDDPIISTHLAK | -313 | -325 | 1 | 0 | 13 | NaN | 779.2256 | 785.8817 | 701.7285 | 814.5907 | 734.84393 |
| Psmd11 | AQSLLSTDR | -12 | -20 | 0 | 0.002068213 | 9 | NaN | 586.32056 | 282.3509 | NaN | NaN | NaN |
| Rpl12 | HDIIDDINSGAVECPAS | -149 | -165 | 0 | 0 | 17 | NaN | 341.54892 | NaN | NaN | 524.38416 | 554.37683 |
| Grwd1 | IKPIFSFA | -204 | -211 | 1 | 0.002502116 | 8 | NaN | 707.6576 | NaN | 533.3076 | 673.2389 | NaN |
| Stat3 | IGGPPNICLDR | -252 | -262 | 0 | 0 | 11 | NaN | 1500.7952 | 1965.0947 | NaN | 2402.708 | 1694.7113 |
| Hspa8 | SQIHDIVLVGGST | -329 | -341 | 0 | 0 | 13 | NaN | 2781.779 | 3144.2065 | 4018.0642 | 2752.853 | 2247.1294 |
| Hspa8 | HWPFMVVNDAGRPK | -89 | -102 | 1 | 0 | 14 | NaN | 1226.919 | 966.59827 | 957.7147 | 1102.7919 | 1112.7008 |
| Hspa8 | HWPFMVVNDAGRPK | -89 | -102 | 1 | 0 | 14 | NaN | 1226.919 | 966.59827 | 957.7147 | 1102.7919 | 1112.7008 |
| Hspa8 | CNEIISWLDK | -574 | -583 | 0 | 0 | 10 | NaN | 1204.1647 | 1263.192 | 1269.5618 | 1494.2542 | 1207.9254 |
| Hspa8 | SFYPEEVSSMVLTK | -113 | -126 | 0 | 0 | 14 | NaN | 1031.2952 | 1036.3208 | 1331.1309 | 1241.6823 | 820.3951 |
| Hspa8 | DKSQIHDIVLVGGSTR | -327 | -342 | 1 | 0 | 16 | NaN | 425.82858 | 548.8624 | NaN | NaN | NaN |
| Hspa8 | PGGFPGGGAPPSGGASSGPTIEEVD | -622 | -646 | 0 | 0.00025015 | 25 | NaN | 451.88434 | 517.9847 | NaN | NaN | 399.68176 |
| Fkbp3 | ARLEIEPEWAYGK | -188 | -200 | 1 | 0 | 13 | NaN | 1193.8655 | NaN | 1088.8035 | NaN | 731.1412 |
| Fkbp3 | NHLFESK | -64 | -70 | 0 | 0.000431655 | 7 | NaN | 428.8255 | NaN | 1955.69 | NaN | NaN |
| Ndufa8 | TDRPLPENPYHSR | -133 | -145 | 1 | 0 | 13 | NaN | 1391.7689 | 1458.7312 | 1278.503 | 1644.9801 | 1797.9169 |
| Ndufa8 | FDQCVLDK | -107 | -114 | 0 | 0 | 8 | NaN | 485.10205 | 541.5073 | 805.50085 | NaN | NaN |
| Kif2a | LIDIGNSCR | -412 | -420 | 0 | 0.001811524 | 9 | NaN | 812.9127 | 985.3178 | NaN | 974.0476 | NaN |
| Ptgr3 | DCPVPLPGDGDLLVR | -54 | -68 | 0 | 0 | 15 | NaN | 971.509 | 927.53815 | 454.14923 | NaN | 1139.934 |
| Ptgr3 | ASDINYSAGR | -76 | -85 | 0 | 0.001055699 | 10 | NaN | 401.20383 | NaN | NaN | NaN | NaN |
| Capza1 | IESHQFQPK | -158 | -166 | 0 | 0 | 9 | NaN | 2904.4114 | 2995.704 | 2308.7366 | 3152.1917 | 2824.5042 |
| Capza1 | GYDDQVLITEHGDLGNSR | -69 | -86 | 0 | 0 | 18 | NaN | 483.3595 | 527.18567 | 430.01425 | 595.28894 | 422.4124 |
| Capza1 | IIESAENEYQTA | -231 | -242 | 0 | 0 | 12 | NaN | 205.77998 | NaN | 154.2149 | 172.00359 | 201.66988 |
| Cops6 | NISDHWIR | -47 | -54 | 0 | 0.00025015 | 8 | NaN | 1661.9564 | 1247.8907 | NaN | 2358.0596 | 919.95386 |
| Cops6 | EAGEVPFNHEILR | -243 | -255 | 0 | 0 | 13 | NaN | 2378.178 | NaN | 2804.7834 | NaN | 2697.6104 |
| Gapvd1 | ASEEQLQDAQLAIER | -1285 | -1299 | 0 | 0 | 15 | NaN | 5870.0806 | 1149.1609 | 1717.0646 | 3184.2322 | NaN |
| Sf3a3 | WQPDTEEEYEDSSGNVVNKK | -471 | -490 | 1 | 0 | 20 | NaN | 673.54224 | 612.8506 | 632.1946 | 773.6264 | 728.2063 |
| Sf3a3 | TYEDLKR | -491 | -497 | 1 | 0.008222027 | 7 | NaN | 2932.3484 | 1444.2208 | NaN | 1131.4114 | 1768.2322 |
| Sf3a3 | ASEKLDYITYL | -154 | -164 | 1 | 0.001636916 | 11 | NaN | 839.0863 | NaN | 666.6824 | 649.09656 | NaN |
| Sf3a3 | QDQNELFGK | -204 | -212 | 0 | 0.00051306 | 9 | NaN | 747.13184 | NaN | 431.44867 | 812.354 | NaN |
| Sf3a3 | EELNAISGPNEFAEFYNR | -70 | -87 | 0 | 0.002296977 | 18 | NaN | 463.38507 | NaN | NaN | NaN | NaN |
| Ranbp1 | ASENDLPEWK | -59 | -68 | 0 | 0 | 10 | NaN | 1423.3928 | 424.72968 | 105.71017 | 1556.4167 | 286.62396 |
| Ranbp1 | MELKPNAGSDR | -108 | -118 | 1 | 0 | 11 | NaN | 1050.9143 | 1190.7501 | 1115.9314 | 1156.0953 | 1487.6505 |
| Uqcrfs1 | ASEPPVLDVK | -37 | -46 | 0 | 0.00223472 | 10 | NaN | 1058.9252 | NaN | NaN | NaN | NaN |
| Itgal | GSLLACDPGLSR | -103 | -114 | 0 | 0 | 12 | NaN | 4151.8467 | NaN | NaN | 1607.5487 | 1603.9714 |
| Itgal | SELQGDPGYPLGR | -506 | -518 | 0 | 0.00025015 | 13 | NaN | 416.89822 | 419.83765 | NaN | 963.7223 | NaN |
| Itgal | VDLIHEK | -1078 | -1084 | 0 | 0.000298805 | 7 | NaN | 368.87244 | NaN | 371.37653 | NaN | NaN |
| Hddc2 | QITQLLPEDLR | -103 | -113 | 0 | 0.007343498 | 11 | NaN | 604.6085 | NaN | 688.4211 | NaN | NaN |
| Rad23b | QNPSLLPALLQQIGR | -294 | -308 | 0 | 0.002068213 | 15 | NaN | 606.2632 | NaN | NaN | 658.7661 | NaN |
| Txnrd2 | ASFVDEHTVR | -159 | -168 | 0 | 0 | 10 | NaN | 1830.8959 | NaN | NaN | NaN | NaN |
| Tnpo3 | TSFYELPTDSHASLR | -73 | -87 | 0 | 0 | 15 | NaN | 1106.6696 | 1411.5083 | 1122.0812 | 1518.3157 | 1303.8682 |
| Tnpo3 | EISDQLLQIR | -45 | -54 | 0 | 0 | 10 | NaN | 659.40656 | 647.0878 | 676.59393 | 715.8512 | 825.3408 |
| Tnpo3 | DSLLTHIQNLK | -88 | -98 | 0 | 0 | 11 | NaN | 1347.5569 | NaN | 1877.4083 | 1864.8816 | 1729.2571 |
| Tnpo3 | LGEHLYK | -349 | -355 | 0 | 0.001868731 | 7 | NaN | 359.47543 | NaN | 597.00433 | NaN | NaN |
| Tnpo3 | TVLPEEVHSR | -148 | -157 | 0 | 0.00142999 | 10 | NaN | 1522.8672 | NaN | NaN | NaN | NaN |
| Dtx3l | ASGGGECSVQPVGPSAPDTYEVK | -34 | -56 | 0 | 0.00025015 | 23 | NaN | 264.48035 | NaN | 302.89078 | 340.90527 | 420.93347 |
| Cox5b | SGGGVPTDEEQATGLER | -32 | -48 | 0 | 0.00025015 | 17 | NaN | 1385.9236 | 1971.3923 | 752.3696 | 833.81726 | 759.91583 |
| Cox5b | PNLVPSISNK | -76 | -85 | 0 | 0 | 10 | NaN | 2045.6097 | NaN | NaN | 780.43854 | 2739.3835 |
| Exosc10 | HGADSDIEWLQK | -366 | -377 | 0 | 0 | 12 | NaN | 778.7005 | 655.79803 | 535.94214 | 892.8139 | 874.0752 |
| Exosc10 | HQQPVLPAGLQVPK | -137 | -150 | 0 | 0.006677519 | 14 | NaN | 2145.2493 | NaN | NaN | 3743.5432 | NaN |
| Ruvbl1 | GLGLDESGLAK | -23 | -33 | 0 | 0 | 11 | NaN | 980.859 | 516.0171 | 728.09314 | 545.63776 | 858.5173 |
| Ruvbl1 | EEALNHLGEIGTK | -388 | -400 | 0 | 0 | 13 | NaN | 578.9368 | 1674.7657 | 836.33777 | 2128.3564 | 1407.2559 |
| Ruvbl1 | EIIQDVTLHDLDVANAR | -233 | -249 | 0 | 0.008528992 | 17 | NaN | 868.13153 | NaN | NaN | NaN | NaN |
| Ruvbl1 | EIIQDVTLHDLDVA | -233 | -246 | 0 | 0.003681885 | 14 | NaN | 861.87177 | NaN | NaN | NaN | NaN |
| Clip1 | YGLFAPVHK | -268 | -276 | 0 | 0.000550307 | 9 | NaN | 495.1719 | 330.5658 | NaN | NaN | NaN |
| Akr1b1 | AYSPLGSPDRPWAKPEDPSLLEDPR | -209 | -233 | 2 | 0 | 25 | NaN | 519.0342 | 611.2783 | 575.4821 | 693.7005 | 733.15454 |
| Akr1b1 | VNQIECHPYLTQEK | -182 | -195 | 0 | 0.00025015 | 14 | NaN | 506.3415 | NaN | NaN | 572.6992 | 550.2581 |
| Akr1b1 | VNQIECHPYLTQEK | -182 | -195 | 0 | 0.00025015 | 14 | NaN | 506.3415 | NaN | NaN | 572.6992 | 550.2581 |
| Akr1b1 | YKPAVNQIECHPYLTQEK | -178 | -195 | 1 | 0.00204437 | 18 | NaN | 413.83463 | 459.76282 | NaN | NaN | NaN |
| Arpc2 | VFKDDDDVVIGK | -168 | -179 | 1 | 0.000230707 | 12 | NaN | 3539.292 | 3915.9255 | 669.17694 | 4110.501 | 4225.222 |
| Polr2b | DDDGLIAPGVR | -849 | -859 | 0 | 0 | 11 | NaN | 638.002 | 830.59155 | NaN | NaN | NaN |
| Polr2b | ASIIPFPDHNQSPR | -708 | -721 | 0 | 0 | 14 | NaN | 482.3732 | 646.8908 | NaN | NaN | NaN |
| Rpn2 | VDLSTVDKDQSIAPK | -378 | -392 | 1 | 0 | 15 | NaN | 553.051 | 526.3623 | 504.40353 | 780.7763 | 632.6416 |
| Rpn2 | YHVPVVVVPEGSTSDTQEQAILR | -267 | -289 | 0 | 0 | 23 | NaN | 790.2365 | 817.83167 | 925.2641 | 1129.882 | 1098.9912 |
| Rpn2 | NVDLSTVDKDQSIAPK | -377 | -392 | 1 | 0 | 16 | NaN | 479.48102 | 622.64307 | 601.72485 | 681.22253 | 582.8046 |
| Rpn2 | YHVPVVVVPEGSTS | -267 | -280 | 0 | 0 | 14 | NaN | 1165.6388 | 972.1645 | NaN | 980.9569 | 793.64484 |
| Rpn2 | PTHYLTK | -25 | -31 | 0 | 0.00149795 | 7 | NaN | 977.76465 | NaN | 728.4956 | 1255.6482 | NaN |
| Rpn2 | SHLSQQADLR | -169 | -178 | 0 | 0.000550307 | 10 | NaN | 1609.9642 | NaN | 1113.6456 | NaN | NaN |
| Rpn2 | GTFIADSHQNFAL | -404 | -416 | 0 | 0.000431655 | 13 | NaN | 902.4093 | NaN | NaN | 412.21082 | NaN |
| Sil1 | ASLELQEGEDDGYFR | -437 | -451 | 0 | 0.001811524 | 15 | NaN | 866.8742 | NaN | NaN | NaN | NaN |
| Psmd1 | LLDEEEPQLK | -11 | -20 | 0 | 0.00025015 | 10 | NaN | 5361.5337 | 6555.0083 | 4591.351 | 2512.7407 | 591.5276 |
| Psmd1 | DKDPILR | -573 | -579 | 1 | 0 | 7 | NaN | 997.4273 | 699.65326 | 921.321 | 1077.0922 | 1056.3789 |

| | | | | | | | | | | | | |
|---|---|---|---|---|---|---|---|---|---|---|---|---|
| Psmd1 | SESYNPHVR | -641 | -649 | 0 | 0 | 9 | NaN | 5052.7646 | 5609.1533 | NaN | 4890.1724 | 5180.7705 |
| Psmd1 | TNDPVNYVR | -676 | -684 | 0 | 0 | 9 | NaN | 1180.5261 | 2884.8547 | NaN | 2361.2368 | 3162.3982 |
| Psmd1 | DNLEWLAR | -388 | -395 | 0 | 0 | 8 | NaN | 759.22485 | NaN | NaN | NaN | NaN |
| Rad21 | LLEPEQSTSNLNEK | -192 | -205 | 0 | 0.000471409 | 14 | NaN | 636.2708 | 533.4006 | NaN | NaN | NaN |
| Atad3a | DKWSNFDPTGLER | -44 | -56 | 1 | 0 | 13 | NaN | 1377.507 | 1328.8453 | NaN | 978.9992 | NaN |
| Atad3a | GEGTGPPLPLPPAQPGAEGGGDR | -12 | -34 | 0 | 0.00078661 | 23 | NaN | 679.94336 | NaN | 585.4554 | 660.70825 | NaN |
| Atad3a | QTILESIR | -219 | -226 | 0 | 0.004658491 | 8 | NaN | 625.96094 | NaN | 396.11194 | NaN | 4135.561 |
| Acadvl | ASNTSEVYFDGVK | -305 | -317 | 0 | 0 | 13 | NaN | 935.50354 | 719.2636 | NaN | 1094.1307 | 940.3266 |
| Acadvl | SEEQAQFLK | -96 | -104 | 0 | 0 | 9 | NaN | 890.03894 | NaN | NaN | 1511.9729 | NaN |
| Clint1 | ASPDQNASTHTPQSSAKPSVPSSK | -306 | -329 | 1 | 0 | 24 | NaN | 437.0256 | NaN | 383.48734 | 468.21674 | NaN |
| Clint1 | ENYHFVDEHGK | -120 | -130 | 0 | 0 | 11 | NaN | 350.4626 | NaN | NaN | 4978.108 | 5159.1587 |
| Tsc22d4 | SFTGGLGQLAGPGK | -165 | -178 | 0 | 0.00025015 | 14 | NaN | 2051.35 | NaN | NaN | NaN | 1819.467 |
| Hnrnpa0 | VYFQSHDAADK | -144 | -154 | 0 | 0 | 11 | NaN | 703.52496 | 540.48694 | 650.105 | 309.76498 | 1155.4028 |
| Hnrnpa0 | TDCVVVVNPQTK | -34 | -45 | 0 | 0 | 12 | NaN | 3133.562 | 3151.3118 | 2070.7068 | NaN | NaN |
| Hnrnpa0 | LTDCVVVVNPQTK | -33 | -45 | 0 | 0 | 13 | NaN | 682.312 | 519.40765 | NaN | NaN | 516.82666 |
| Atp5po | ASPLDDAVLSELK | -146 | -158 | 0 | 0 | 13 | NaN | 1619.1023 | 1678.4622 | 2155.6658 | 2372.623 | 2591.5208 |
| Atp5po | GEVPCTVTTASPLDDAVLSELK | -137 | -158 | 0 | 0 | 22 | NaN | 237.24286 | NaN | 487.65417 | 293.1055 | 488.42993 |
| Wdr46 | ASPPSLEQPYLTHR | -431 | -444 | 0 | 0 | 14 | NaN | 818.89923 | 768.6454 | NaN | 750.73865 | 789.081 |
| Wdr46 | QDPFPGPVPAPLEEAR | -94 | -109 | 0 | 0.00025015 | 16 | NaN | 677.46313 | 811.26794 | NaN | NaN | NaN |
| Wdr46 | EEPGFLVGEDGEDTAK | -154 | -169 | 0 | 0 | 16 | NaN | 1035.6847 | NaN | NaN | NaN | NaN |
| Hnrnpl | NYDDPHKTPASPVVHIR | -88 | -104 | 1 | 0.00149795 | 17 | NaN | 351.1993 | 167.60072 | NaN | 834.4202 | 400.02597 |
| Hnrnpl | VLYTICNPCGPVQR | -207 | -220 | 0 | 0.000347792 | 14 | NaN | 812.3323 | 410.24118 | NaN | NaN | NaN |
| Hnrnpl | NNNFMFGQK | -437 | -445 | 0 | 0.002972566 | 9 | NaN | 734.1338 | NaN | NaN | NaN | NaN |
| Hnrnpl | DQDTWDYTNPNLSGQGDPGSNPNKR | -276 | -300 | 1 | 0.000230707 | 25 | NaN | 340.28223 | 351.46964 | NaN | NaN | NaN |
| Zc3hav1 | FVLLETGDQAGITR | -49 | -62 | 0 | 0 | 14 | NaN | 1163.7112 | NaN | 862.3707 | 898.6826 | 701.8436 |
| Zc3hav1 | TCDSYPIR | -670 | -677 | 0 | 0.001636916 | 8 | NaN | 1196.405 | NaN | NaN | 877.0379 | NaN |
| Zc3hav1 | NHELSGLNQEELAVL | -123 | -137 | 0 | 0 | 15 | NaN | 294.5859 | NaN | NaN | 1105.462 | NaN |
| Smarcb1 | GQLSWHQK | -317 | -324 | 0 | 0.009897819 | 8 | NaN | 576.3077 | 1904.4465 | NaN | NaN | NaN |
| Eci2 | ASQQDFENALNQVK | -36 | -49 | 0 | 0 | 14 | NaN | 618.1643 | NaN | NaN | 687.6442 | 417.066 |
| Psmd8 | ASQQQKPEDSTIPSTELAK | -322 | -340 | 1 | 0.00025015 | 19 | NaN | 594.37195 | 450.60883 | 333.33255 | NaN | 397.59537 |
| Arpc1b | WVSHDSTVCLVDADKK | -219 | -234 | 1 | 0.00025015 | 16 | NaN | 788.6231 | 950.4599 | 415.03732 | 920.49695 | 1098.6764 |
| Arpc1b | PAPTPWGSK | -180 | -188 | 0 | 0 | 9 | NaN | 1429.8102 | 1227.301 | 1049.6708 | NaN | 1436.1338 |
| Arpc1b | EVEERPAPTPW | -175 | -185 | 1 | 0.000869338 | 11 | NaN | 822.38794 | 1295.2928 | 988.7749 | 607.85223 | NaN |
| Eif4g2 | LEVDIPLVK | -631 | -639 | 0 | 0.00025015 | 9 | NaN | 899.6522 | 1022.94006 | 1498.871 | 1870.1157 | 1951.0823 |
| Rpl3 | GIVGYVETPR | -88 | -97 | 0 | 0.00025015 | 10 | NaN | 1572.1376 | 1752.3 | 1488.6871 | 1226.7115 | 1160.3896 |
| Rpl3 | DEMIDVIGVTK | -214 | -224 | 0 | 0 | 11 | NaN | 1101.9921 | 587.12537 | 1196.3613 | NaN | 1343.8313 |
| Rpl3 | VHYGEVTNDFIMLK | -321 | -334 | 0 | 0 | 14 | NaN | 1360.7649 | NaN | 921.0713 | NaN | 2090.338 |
| Rpl3 | SINPLGGFVHYGEVT | -313 | -327 | 0 | 0 | 15 | NaN | 857.14514 | NaN | NaN | NaN | NaN |
| Sh3glb1 | ASTGSLVITCPSNLNDLK | -285 | -302 | 0 | 0.005242018 | 18 | NaN | 222.98068 | NaN | NaN | NaN | NaN |
| Psmd10 | ASTNIQDTEGNTPLHLA | -163 | -179 | 0 | 0 | 17 | NaN | 639.6774 | 743.9881 | NaN | NaN | NaN |
| Pepd | ASTVRPSFSLGNETLK | -2 | -17 | 1 | 0.000550307 | 16 | NaN | 823.0431 | NaN | 915.3507 | 1109.9377 | 1002.3612 |
| Mif | PMFIVNTNVPR | -2 | -12 | 0 | 0 | 11 | NaN | 371.11694 | 683.91895 | 1985.6246 | 2642.8374 | 2584.9524 |
| Stk10 | GGLELPSSVTPSHSK | -467 | -481 | 0 | 0.000230707 | 15 | NaN | 821.6701 | NaN | NaN | 1339.7338 | 1166.6788 |
| Stk10 | IALDKNPETRPSAA | -273 | -286 | 2 | 0.003069674 | 14 | NaN | 762.5363 | NaN | NaN | NaN | NaN |
| Hp1bp3 | GSALDPEPQVK | -248 | -258 | 0 | 0.000347792 | 11 | NaN | 2162.0227 | 1700.5186 | 2090.3289 | NaN | NaN |
| Sf3b1 | IYNDDKNTYIR | -1287 | -1297 | 1 | 0 | 11 | NaN | 966.39197 | 1079.1743 | 791.34375 | 853.6674 | 841.8422 |
| Sf3b1 | DQAETPGHTPSLR | -219 | -231 | 0 | 0 | 13 | NaN | 1364.5907 | 1467.2578 | 741.4299 | 913.32263 | 631.9943 |
| Sf3b1 | IADREDEYKK | -112 | -121 | 2 | 0.00111006 | 10 | NaN | 430.5093 | 512.2048 | 684.196 | 606.3163 | 447.124 |
| Sf3b1 | LSSWDQAETPGHTPSLR | -215 | -231 | 0 | 0 | 17 | NaN | 399.95792 | 355.32144 | 447.21298 | 416.70007 | 527.5432 |
| Sf3b1 | EILPPFFK | -809 | -816 | 0 | 0.003333688 | 8 | NaN | 2095.8628 | 2141.8845 | NaN | 2018.3481 | 1937.796 |
| Sf3b1 | IVDDLKDEAEQYRK | -853 | -866 | 2 | 0.00025015 | 14 | NaN | 957.83826 | 1160.1279 | NaN | 904.5382 | 827.87744 |
| Apoh | ATFGCHETY | -230 | -238 | 0 | 0.00025015 | 9 | NaN | 626.47394 | 580.6389 | 553.11005 | 459.25345 | 472.25284 |
| Pycr2 | HFLESGGFR | -243 | -251 | 0 | 0 | 9 | NaN | 1444.9531 | 1077.9209 | 2050.4485 | 1705.6582 | 1492.1968 |
| Pycr2 | ATGTHALVEDGK | -136 | -147 | 0 | 0 | 12 | NaN | 577.45807 | 624.7352 | 567.82214 | 430.40436 | 498.13644 |
| Qars1 | VPNFPADETK | -592 | -601 | 0 | 0 | 10 | NaN | 644.883 | 1010.54877 | 456.69415 | 1333.1028 | 992.0348 |
| Qars1 | ATGVLLYDLVSR | -51 | -62 | 0 | 0 | 12 | NaN | 112.1228 | NaN | NaN | 323.33566 | 137.3614 |
| Rad50 | ATHLELDGFER | -375 | -385 | 0 | 0 | 11 | NaN | 1492.9633 | 1518.148 | 1457.3295 | 1048.3413 | 1517.5366 |
| Rab7a | IGADFLTK | -41 | -48 | 0 | 0 | 8 | NaN | 793.81946 | 901.6709 | 918.5903 | 580.34503 | 729.2947 |
| Vav1 | DTTLQFPYKEPER | -758 | -770 | 1 | 0.00025015 | 13 | NaN | 179.9619 | 653.44745 | NaN | NaN | 487.6063 |
| Vav1 | SWTPIAQNK | -116 | -124 | 0 | 0.003069674 | 9 | NaN | 1506.1908 | NaN | NaN | NaN | NaN |
| Glrx | TQEILSQLPFK | -29 | -39 | 0 | 0 | 11 | NaN | 604.19165 | 591.8299 | 500.2616 | 660.59094 | NaN |
| Pnkp | LQEHLDPALQR | -504 | -514 | 0 | 0 | 11 | NaN | 3506.7192 | 3392.5173 | NaN | NaN | NaN |
| Tcof1 | EKPDGELLGIK | -1250 | -1260 | 1 | 0.001636916 | 11 | NaN | 582.7576 | 810.37134 | NaN | NaN | NaN |
| Tcof1 | TELLEQER | -1164 | -1171 | 0 | 0.00025015 | 8 | NaN | 312.08752 | NaN | NaN | NaN | 239.33926 |
| Tcof1 | GNVAPTPAKPAR | -243 | -254 | 1 | 0.001023236 | 12 | NaN | 219.00499 | NaN | NaN | 346.57376 | NaN |
| Tcof1 | GKPPVPNSTVSAR | -715 | -727 | 1 | 0 | 13 | NaN | 668.07465 | NaN | NaN | NaN | NaN |
| Ap1b1 | SAEPELQYVALR | -293 | -304 | 0 | 0 | 12 | NaN | 523.02356 | 1324.7476 | 1233.9852 | NaN | 1368.3683 |
| Ap1b1 | LCENLDSLDEPEAR | -432 | -445 | 0 | 0 | 14 | NaN | 563.41254 | 511.2748 | 408.87177 | NaN | NaN |
| Ap1b1 | SEIAESHPSSNLLDLNPQSINK | -185 | -206 | 0 | 0 | 22 | NaN | 301.81305 | 600.461 | NaN | 542.3848 | NaN |
| Abcf1 | QQPPEPEWIGDGEGTSPADK | -7 | -26 | 0 | 0 | 20 | NaN | 834.1299 | NaN | 778.40607 | 776.09814 | 827.8958 |
| Abcf1 | SKPAAADSEGEEEEEDTAK | -187 | -205 | 1 | 0.008800311 | 19 | NaN | 361.78387 | NaN | NaN | 615.9752 | 380.05072 |
| Abcf1 | AVAEEPPGLR | -163 | -172 | 0 | 0.001291103 | 10 | NaN | 436.41144 | 745.7355 | NaN | NaN | NaN |
| Psma2 | LFQSDPSGAYFAWK | -146 | -159 | 0 | 0 | 14 | NaN | 462.0244 | 499.20917 | NaN | NaN | NaN |
| Septin11 | FESDPATHNEPGVR | -66 | -79 | 0 | 0 | 14 | NaN | 4171.109 | 3891.9226 | 4184.3916 | 3863.4785 | 3808.453 |
| Septin11 | KFESDPATHNEPGVR | -65 | -79 | 1 | 0 | 15 | NaN | 810.50995 | 800.9845 | 853.3942 | 931.5767 | 944.3238 |
| Septin11 | AVAVGRPSNEELR | -2 | -14 | 1 | 0 | 13 | NaN | 2525.0735 | 2543.1482 | 3028.9797 | 2940.8027 | 3001.5237 |
| Psmd14 | HLEEHVDVL | -283 | -291 | 0 | 0.000712251 | 9 | NaN | 982.9932 | 1543.6233 | 3772.2268 | 1840.0199 | 1100.2048 |
| Psmd14 | AVEEEDKMTPEQLAIK | -258 | -273 | 1 | 0.00051306 | 16 | NaN | 821.2264 | 718.572 | 809.68085 | 800.66626 | 771.0588 |

| | | | | | | | | | | | | |
|---|---|---|---|---|---|---|---|---|---|---|---|---|
| Psmd14 | MLGEFVDDYTVR | -57 | -68 | 0 | 0 | 12 | NaN | 2475.0232 | NaN | 2672.2483 | 2400.3894 | 2683.9944 |
| Ndufa5 | KYTEQITNEK | -46 | -55 | 1 | 0 | 10 | NaN | 633.5509 | NaN | 525.1384 | 773.4938 | 760.999 |
| Ddx19a | SNLVDNTNQVEVLQR | -67 | -81 | 0 | 0 | 15 | NaN | 1052.547 | NaN | 762.6286 | 1555.1536 | NaN |
| Ddx19a | VVPDPNIIK | -287 | -295 | 0 | 0.009272078 | 9 | NaN | 871.53564 | 86.4913 | NaN | NaN | NaN |
| Ddx19a | TSTTAEKTEEEEKEDR | -40 | -55 | 2 | 0.002910424 | 16 | NaN | 199.33022 | NaN | NaN | NaN | NaN |
| Thbs1 | GTSQNDPNWVVR | -969 | -980 | 0 | 0 | 12 | NaN | 926.7832 | 1151.3837 | 2110.2517 | 1537.1831 | 1285.8118 |
| Thbs1 | QVTQSYWDTNPTR | -1042 | -1054 | 0 | 0 | 13 | NaN | 9045.823 | NaN | 1784.401 | NaN | NaN |
| Thbs1 | ELDVPIQSIFTR | -178 | -189 | 0 | 0.000230707 | 12 | NaN | 915.8352 | 1045.6056 | NaN | NaN | NaN |
| Rpl10a | KNWQNVR | -196 | -202 | 1 | 0.003745449 | 7 | NaN | 952.52454 | NaN | NaN | 1204.4121 | 604.89105 |
| Rpl10a | ILGPGLNKA | -123 | -131 | 1 | 0.002199523 | 9 | NaN | 292.26495 | NaN | 218.08511 | NaN | 251.75415 |
| Rpl10a | FPSLLTHNEN | -134 | -143 | 0 | 0.00025015 | 10 | NaN | 731.142 | 1403.7443 | NaN | NaN | NaN |
| Rpl10a | FPSLLTHNENMVAK | -134 | -147 | 0 | 0 | 14 | NaN | 512.6321 | 719.58167 | NaN | NaN | NaN |
| Rpl6 | YEITEQR | -248 | -254 | 0 | 0.00111006 | 7 | NaN | 919.0875 | 410.3623 | NaN | NaN | NaN |
| Lgals3bp | STETIASHEDIER | -329 | -341 | 0 | 0 | 13 | NaN | 1951.5836 | 931.903 | 1167.9808 | 1461.4816 | 1460.9573 |
| Lgals3bp | LYQDHQALFQR | -365 | -375 | 0 | 0 | 11 | NaN | 632.7106 | NaN | 339.40314 | NaN | NaN |
| Tufm | EEIDNAPEER | -93 | -102 | 0 | 0 | 10 | NaN | 1325.3263 | 1328.6472 | 766.3048 | 1135.8083 | 1247.9022 |
| Tufm | EHLLLAK | -163 | -169 | 0 | 0.00111006 | 7 | NaN | 901.1486 | NaN | 1167.2418 | 807.97534 | 633.7087 |
| Tufm | TVVTGIEMFHK | -301 | -311 | 0 | 0.004658491 | 11 | NaN | 765.1089 | 745.5904 | 396.1457 | NaN | NaN |
| Tufm | VKPGSIQPHQK | -341 | -351 | 1 | 0 | 11 | NaN | 261.8406 | NaN | 256.28094 | NaN | NaN |
| Psmb9 | YREDLLAHL | -110 | -118 | 1 | 0.00201047 | 9 | NaN | 1460.9456 | 1510.0822 | 761.2963 | 1309.2684 | 1471.896 |
| Psma3 | DIREEAEKY | -231 | -239 | 2 | 0.002401448 | 9 | NaN | 1201.6244 | 959.8726 | 2074.6619 | 2141.76 | 1757.1812 |
| Psma3 | FGYNIPLK | -103 | -110 | 0 | 0 | 8 | NaN | 408.0831 | 374.2041 | NaN | 218.15648 | NaN |
| Psma3 | DIREEAEK | -231 | -238 | 1 | 0.009541249 | 8 | NaN | 1313.1125 | NaN | NaN | 1926.446 | NaN |
| Psmd5 | YPVFLEK | -293 | -299 | 0 | 0.004321969 | 7 | NaN | 507.32556 | NaN | 598.9898 | 328.69138 | 463.74774 |
| Psmd5 | IVENSEAVTEILNNAELLK | -110 | -128 | 0 | 0 | 19 | NaN | 252.26706 | NaN | 333.70764 | 271.51886 | 241.2745 |
| Psmd5 | ELTGEDVLVR | -215 | -224 | 0 | 0.000550307 | 10 | NaN | 742.24426 | NaN | NaN | 376.86264 | NaN |
| Psmd5 | VFTAIADQPWAQR | -417 | -429 | 0 | 0.000230707 | 13 | NaN | 426.52896 | 242.47772 | NaN | NaN | NaN |
| Sh3bgrl3 | LVDISQDNALRDEMR | -37 | -51 | 1 | 0.006100831 | 15 | NaN | 319.51566 | NaN | 798.77264 | NaN | NaN |
| Usp9x | TGETGVEETILEGHLGVTK | -1417 | -1435 | 0 | 0 | 19 | NaN | 469.26294 | NaN | 602.97217 | 792.8926 | 767.6915 |
| Usp9x | AVEWLGDELER | -2418 | -2428 | 0 | 0.002068213 | 11 | NaN | 335.834 | NaN | NaN | NaN | NaN |
| Aars1 | LADGGRPDNTGR | -309 | -320 | 1 | 0 | 12 | NaN | 453.45837 | 476.93976 | 506.4843 | 541.3698 | 461.2592 |
| Aars1 | LFTVDNEAGK | -889 | -898 | 0 | 0 | 10 | NaN | 392.0024 | 572.2177 | 2615.5718 | 613.11194 | 592.32623 |
| Aars1 | HNDLDDVGKDVYHHT | -82 | -96 | 1 | 0 | 15 | NaN | 389.38104 | NaN | 697.4961 | 731.50146 | 722.35266 |
| Aars1 | FGGDEAAGLEPDLECR | -138 | -153 | 0 | 0.000230707 | 16 | NaN | 1226.0978 | NaN | NaN | 1618.5374 | NaN |
| Wdr3 | DLIHSPQGELK | -368 | -378 | 0 | 0 | 11 | NaN | 1282.449 | 1287.1108 | NaN | NaN | NaN |
| Wdr3 | AVFGLIGSQK | -14 | -23 | 0 | 0 | 10 | NaN | 494.9146 | NaN | NaN | 438.6614 | NaN |
| Wdr3 | FEHIQTLEGHHQ | -664 | -675 | 0 | 0.009745776 | 12 | NaN | 10601.248 | NaN | NaN | NaN | NaN |
| Msh2 | AVFVTPLIDLR | -434 | -444 | 0 | 0 | 11 | NaN | 586.3932 | 580.0622 | 535.61914 | 654.2462 | 638.47845 |
| Msh2 | FTNSELSSLNEEYTK | -551 | -565 | 0 | 0.00025015 | 15 | NaN | 2175.23 | 524.3657 | 601.437 | 1449.3093 | NaN |
| Msh2 | LDSSAQFGYYFR | -513 | -524 | 0 | 0 | 12 | NaN | 468.85345 | 388.15024 | NaN | 778.5619 | NaN |
| Sf3b3 | VLDPVTGDLSDTR | -678 | -690 | 0 | 0 | 13 | NaN | 2361.8833 | NaN | 1342.1608 | 1083.3196 | 620.9914 |
| Sf3b3 | RNENQLIIF | -1016 | -1024 | 1 | 0.004824802 | 9 | NaN | 1346.1227 | 1314.474 | 471.577 | 1220.8264 | NaN |
| Tap1 | LVPEVPLLR | -31 | -39 | 0 | 0.005363933 | 9 | NaN | 597.4045 | 581.9674 | 601.59326 | 626.13855 | NaN |
| Tap1 | DVSFAYPNQPK | -483 | -493 | 0 | 0 | 11 | NaN | 468.30966 | 2799.348 | NaN | NaN | NaN |
| Chchd2 | AVGSPAAAPR | -42 | -51 | 0 | 0.000907206 | 10 | NaN | 537.8704 | 548.98785 | NaN | NaN | NaN |
| Pex11b | AVHLSDVVLR | -74 | -83 | 0 | 0.00025015 | 10 | NaN | 1429.0814 | 1500.6497 | 1568.7225 | NaN | NaN |
| Pex11b | TVGPGGPGTPGGSLPQLALK | -170 | -189 | 0 | 0.00025015 | 20 | NaN | 5348.0093 | NaN | NaN | NaN | NaN |
| Wdr1 | IKDIAWTEDSKR | -105 | -116 | 2 | 0 | 12 | NaN | 1084.0438 | NaN | 717.76715 | 852.66455 | 841.3138 |
| Wdr1 | SILASTLKDEGK | -473 | -484 | 1 | 0 | 12 | NaN | 646.65533 | NaN | 1412.5591 | 1143.3267 | 839.2428 |
| Wdr1 | VVAVHPGGDTVAVGGTDGNVR | -450 | -470 | 0 | 0.00383302 | 21 | NaN | 746.40137 | NaN | NaN | NaN | NaN |
| Wdr1 | DIYTEHAHQVVVAK | -52 | -65 | 0 | 0.00025015 | 14 | NaN | 265.40073 | NaN | NaN | NaN | NaN |
| Acsl4 | HIIYVDNK | -224 | -231 | 0 | 0.00111006 | 8 | NaN | 980.0132 | NaN | 699.90186 | 504.9715 | NaN |
| Cops4 | QNESTNEQLQIHYK | -175 | -188 | 0 | 0 | 14 | NaN | 1376.5729 | 1374.2872 | 1204.0216 | 1621.2543 | 1426.2059 |
| Cops4 | NAAQVLVGIPLETGQK | -122 | -137 | 0 | 0 | 16 | NaN | 507.62405 | NaN | 661.5056 | 643.2825 | 641.3833 |
| Prps1 | KFSNQETCVEIGESVR | -34 | -49 | 1 | 0.00025015 | 16 | NaN | 375.53427 | 450.49506 | 519.0092 | 609.73315 | 530.963 |
| Rnps1 | AVLAPWPRPPPR | -237 | -248 | 1 | 0.004224523 | 12 | NaN | 705.3228 | 800.4087 | NaN | NaN | NaN |
| Rac1 | KLTPITYPQ | -133 | -141 | 1 | 0.001811524 | 9 | NaN | 1504.8915 | NaN | 784.16864 | 1286.6648 | 1220.6631 |
| Rac2 | SLVSPASYENVR | -83 | -94 | 0 | 0.000230707 | 12 | NaN | 1065.5808 | 947.8244 | 2108.6777 | NaN | 2915.4387 |
| Rac2 | AVLCPQPTRQQK | -175 | -186 | 1 | 0.008593015 | 12 | NaN | 681.9945 | NaN | NaN | 451.8164 | 341.66516 |
| Cmpk2 | STNFPVIVDR | -322 | -331 | 0 | 0 | 10 | NaN | 426.2417 | 517.4929 | NaN | NaN | NaN |
| Ipo4 | LFPVLLNNAR | -901 | -910 | 0 | 0 | 10 | NaN | 539.8179 | 658.57837 | 439.28558 | 634.7541 | 563.37103 |
| Ipo4 | LLYPLLQIVCK | -391 | -401 | 0 | 0 | 11 | NaN | 243.51651 | 304.88486 | 409.45435 | 448.0039 | 508.23566 |
| Ipo4 | RTLIPLDEVK | -211 | -220 | 1 | 0.00025015 | 10 | NaN | 399.67252 | NaN | 459.63693 | 483.75842 | 506.55438 |
| Ipo4 | AVLESLTGVLR | -772 | -782 | 0 | 0 | 11 | NaN | 298.15683 | NaN | NaN | 310.38184 | 407.69943 |
| Ipo4 | TLIPLDEVK | -212 | -220 | 0 | 0.00122419 | 9 | NaN | 245.02899 | 421.24246 | NaN | NaN | NaN |
| Gemin5 | EEPLFNFR | -672 | -679 | 0 | 0.003037016 | 8 | NaN | 633.942 | 607.4729 | 650.9844 | 701.94916 | NaN |
| Gemin5 | SNPESPITITEPYR | -620 | -633 | 0 | 0.006677519 | 14 | NaN | 1625.009 | NaN | NaN | NaN | NaN |
| Calr | HEQNIDCGGGYVK | -99 | -111 | 0 | 0 | 13 | NaN | 6192.355 | 6605.843 | 6502.2856 | 7460.454 | 7449.5933 |
| Calr | IKDPDAAKPEDWDER | -208 | -222 | 2 | 0 | 15 | NaN | 4348.2236 | 4515.414 | 4287.6743 | 5105.2847 | 5050.4614 |
| Calr | WEPPVIQNPEYK | -261 | -272 | 0 | 0 | 12 | NaN | 1776.7737 | 1584.5973 | 2472.708 | 1381.0844 | 2341.7507 |
| Calr | PSGLDQK | -114 | -120 | 0 | 0.003299042 | 7 | NaN | 1523.9164 | 1494.5016 | 2158.5042 | 2820.3484 | 3008.0552 |
| Calr | KFEPFSNK | -80 | -87 | 1 | 0.00111006 | 8 | NaN | 1190.243 | 345.6316 | 2564.2302 | 1485.536 | 1451.5256 |
| Calr | DDEFTHLYTL | -165 | -174 | 0 | 0 | 10 | NaN | 167.8288 | 648.91486 | 609.6535 | 258.6928 | 749.3681 |
| Calr | IKDPDAAKPE | -208 | -217 | 2 | 0.000230707 | 10 | NaN | 2092.9436 | 1926.8129 | NaN | 1434.8997 | 1464.8003 |
| Calr | IDDPTDSKPEDWDKPE | -225 | -240 | 2 | 0.001055699 | 16 | NaN | 349.45557 | NaN | 304.42664 | NaN | 271.0703 |
| Calr | QIDNPDYK | -279 | -286 | 0 | 0.001055699 | 8 | NaN | 1533.6229 | NaN | 1869.9113 | NaN | NaN |
| Gmppa | AVLGPNVSIGK | -299 | -309 | 0 | 0.002502116 | 11 | NaN | 708.39166 | NaN | NaN | NaN | NaN |
| Hibch | AVLIDKDQTPK | -347 | -357 | 1 | 0.00078661 | 11 | NaN | 385.8901 | NaN | NaN | NaN | 505.12042 |

| | | | | | | | | | | | |
|---|---|---|---|---|---|---|---|---|---|---|---|
| Tubg1;Tubg2 | AVLLDLEPR | (64);(64) | (72);(72) | 0 | 0 | 9 | NaN | 925.01935 | 1011.54926 | 756.67786 | 728.56476 | NaN |
| Rhog | NLNLWDTAGQEEYDR | -52 | -66 | 0 | 0.000471409 | 15 | NaN | 1163.1855 | 790.4944 | 1500.4934 | 1705.5662 | 1692.1974 |
| Rhog | AVLNPTPIKR | -175 | -184 | 1 | 0 | 10 | NaN | 335.20142 | NaN | 234.59827 | NaN | 891.86694 |
| Noc4l | LEELLAHPTFQVK | -103 | -115 | 0 | 0 | 13 | NaN | 1932.2308 | 1508.4404 | NaN | 2784.9797 | 3126.681 |
| Noc4l | EKPQWESHYLFPR | -135 | -147 | 1 | 0.000431655 | 13 | NaN | 812.7801 | 422.2242 | NaN | 1393.5511 | 834.7183 |
| Pola1 | NEPLLTPIPLKR | -220 | -231 | 1 | 0.000907206 | 12 | NaN | 1566.5096 | 1521.0963 | 1134.0326 | 1133.5499 | 867.7888 |
| Pola1 | AVLRPEYSDK | -1381 | -1390 | 1 | 0.001291103 | 10 | NaN | 2537.8872 | NaN | NaN | 1805.8348 | 1700.5076 |
| Pola1 | IDPDILVGH | -633 | -641 | 0 | 0.000989247 | 9 | NaN | 618.8547 | 644.468 | NaN | NaN | NaN |
| Pola1 | NPQLLNQPISWCK | -507 | -519 | 0 | 0 | 13 | NaN | 1322.4517 | NaN | NaN | NaN | NaN |
| Zc3h4 | AVNIPLDPLPGHPLR | -943 | -957 | 0 | 0.00025015 | 15 | NaN | 176.42209 | NaN | 240.12103 | NaN | NaN |
| Aip | VLELDPALAPVVSR | -291 | -304 | 0 | 0 | 14 | NaN | 341.3278 | 338.59192 | NaN | NaN | NaN |
| G3bp1 | PGDVEPR | -323 | -329 | 0 | 0 | 7 | NaN | 1019.05383 | NaN | 650.7809 | 971.2429 | 1751.9814 |
| G3bp1 | HPDSHQLFIG | -334 | -343 | 0 | 0.007546675 | 10 | NaN | 4673.5454 | 1773.4658 | NaN | 1238.857 | NaN |
| G3bp1 | VPASQPRPESKPD | -275 | -287 | 2 | 0.00111006 | 13 | NaN | 1348.3802 | 1365.0839 | 1250.5947 | NaN | NaN |
| G3bp1 | EAAPDDVQK | -220 | -228 | 0 | 0 | 9 | NaN | 1240.0435 | NaN | 525.2576 | NaN | NaN |
| Cnot1 | LAEVGQYEQVK | -460 | -470 | 0 | 0 | 11 | NaN | 1236.839 | 986.57837 | 451.38513 | 1210.8094 | 701.02954 |
| Cnot1 | SGADADLLLR | -1673 | -1682 | 0 | 0 | 10 | NaN | 1274.0948 | 849.5535 | NaN | 787.858 | 788.67346 |
| Cnot1 | NKPILHTDLDVK | -1206 | -1217 | 1 | 0 | 12 | NaN | 444.252 | NaN | 489.6631 | 466.72495 | NaN |
| Cnot1 | EFQQLPYHR | -1983 | -1991 | 0 | 0.00713295 | 9 | NaN | 1489.4833 | 2364.9194 | NaN | NaN | NaN |
| Cnot1 | GLHNVVYGIQR | -373 | -383 | 0 | 0.00122419 | 11 | NaN | 969.16785 | NaN | NaN | NaN | NaN |
| Nol6 | YVGGFLDALIQNPK | -642 | -655 | 0 | 0 | 14 | NaN | 236.06868 | 86.358765 | 95.30165 | 518.6806 | 455.63773 |
| Nol6 | VIDLGPEADKPEAADFR | -570 | -586 | 1 | 0 | 17 | NaN | 462.84717 | NaN | NaN | 329.3117 | NaN |
| Nol6 | VPLHQVPYAVK | -148 | -158 | 0 | 0.00142999 | 11 | NaN | 1059.0706 | NaN | NaN | NaN | NaN |
| Mios | AVQGVTVDPYFHDR | -226 | -239 | 0 | 0 | 14 | NaN | 1711.431 | NaN | NaN | NaN | 2568.142 |
| Eif4b | VDVADQAQDKDRDDR | -168 | -182 | 2 | 0 | 15 | NaN | 370.31122 | 565.6511 | 498.45917 | 868.77527 | 640.2798 |
| Eif4b | VDVADQAQDK | -168 | -177 | 0 | 0.003102268 | 10 | NaN | 791.40015 | 1156.8987 | NaN | NaN | NaN |
| Krt19 | AVSDGLLSGNEK | -73 | -84 | 0 | 0.001392869 | 12 | NaN | 2610.126 | NaN | 2581.3506 | NaN | NaN |
| Eif2s1 | CYGYEGIDAVK | -199 | -209 | 0 | 0 | 11 | NaN | 3811.701 | 4301.7485 | NaN | 4972.6484 | 5125.4414 |
| Eif2s1 | TKDEQLESLFQR | -122 | -133 | 1 | 0 | 12 | NaN | 732.27075 | 765.3209 | NaN | 718.73346 | 676.12506 |
| Eif2s1 | NECVVVIR | -68 | -75 | 0 | 0 | 8 | NaN | 1632.3494 | 723.49976 | NaN | NaN | NaN |
| Smg8 | AVSVGEAGGAGDPGAAAGDSLR | -96 | -117 | 0 | 0 | 22 | NaN | 1169.3414 | 1558.0602 | 1104.8583 | 1002.31714 | 1262.0635 |
| Coro1b | NDQCYEDIR | -22 | -30 | 0 | 0 | 9 | NaN | 2007.7361 | 1906.9103 | 2222.475 | 1949.7803 | 2035.1652 |
| Coro1b | LDSLHPDLIYNVS | -171 | -183 | 0 | 0.000230707 | 13 | NaN | 1002.77026 | 1147.689 | 876.2394 | 1183.9124 | 774.1559 |
| Coro1b | VTWDSTFCAVNPK | -34 | -46 | 0 | 0 | 13 | NaN | 937.63654 | 1077.884 | 731.83765 | 581.2982 | 631.45905 |
| Itgb1 | SHYYDYPSIAHLVQK | -311 | -325 | 0 | 0.00025015 | 15 | NaN | 2920.789 | 849.3383 | NaN | 168.93047 | 825.4217 |
| Itgb1 | KGCQPSDIENPR | -73 | -84 | 1 | 0 | 12 | NaN | 647.2379 | NaN | NaN | 840.9895 | NaN |
| Snrnp40 | DVRPFAPK | -264 | -271 | 1 | 0.001811524 | 8 | NaN | 947.72034 | NaN | 870.5803 | NaN | NaN |
| Coro7 | AVTTPTCGNHIK | -28 | -39 | 0 | 0 | 12 | NaN | 609.4784 | NaN | NaN | 1712.9696 | 1182.3284 |
| H4c1 | TVTAMDVVYALK | -81 | -92 | 0 | 0 | 12 | NaN | 3726.289 | 4449.667 | 3812.872 | 1475.4908 | 4309.8525 |
| H4c1 | ITKPAIR | -30 | -36 | 1 | 0.000947459 | 7 | NaN | 1005.24695 | 1023.4331 | 901.2106 | 1194.9277 | 1137.0604 |
| H4c1 | SGLIYEETR | -48 | -56 | 0 | 0.001055699 | 9 | NaN | 747.556 | 928.24945 | 87.98962 | 925.7996 | 1364.5573 |
| Adk | HKELFDELVK | -61 | -70 | 1 | 0 | 10 | NaN | 3580.8516 | 1198.6862 | 2567.8928 | 3671.4688 | 2663.5803 |
| Adk | FKVEYHAGGSTQ | -72 | -83 | 1 | 0.00051306 | 12 | NaN | 511.97388 | 259.45325 | NaN | NaN | NaN |
| Glud1 | GFIGPGIDVPAPDMSTGER | -213 | -231 | 0 | 0 | 19 | NaN | 921.38165 | 901.6222 | 779.61383 | 1049.5502 | 1031.5239 |
| Glud1 | DDGSWEVIEGY | -125 | -135 | 0 | 0 | 11 | NaN | 232.55898 | 284.9915 | 311.1785 | 293.71783 | 304.0994 |
| Glud1 | GVGESDGSIWNPDGIDPK | -329 | -346 | 0 | 0 | 18 | NaN | 1508.7037 | NaN | 1912.7411 | 1916.2418 | 1978.5457 |
| Glud1 | AVVDVPFGGAK | -173 | -183 | 0 | 0 | 11 | NaN | 3399.9485 | 3530.2263 | NaN | 4371.4883 | NaN |
| Glud1 | REDDPNFFK | -60 | -68 | 1 | 0 | 9 | NaN | 1232.3303 | 1265.1078 | 1093.725 | NaN | NaN |
| Glud1 | IIAEGANGPTTPEADK | -400 | -415 | 0 | 0 | 16 | NaN | 680.95544 | NaN | 542.30096 | 942.3943 | NaN |
| Glud1 | YSTDVSVDEVK | -152 | -162 | 0 | 0 | 11 | NaN | 595.1747 | NaN | NaN | 1123.5488 | 936.211 |
| Slc25a3 | SCGLTHTAVVPLDLVK | -70 | -85 | 0 | 0 | 16 | NaN | 2900.6243 | 2964.7607 | 2941.2756 | 3562.6104 | 3684.2002 |
| Slc25a3 | LPRPPPPEMPESLK | -337 | -350 | 1 | 0 | 14 | NaN | 1003.72 | 271.0044 | 549.17847 | 482.0347 | 854.4066 |
| Slc25a3 | GLTHTAVVPLDLVK | -72 | -85 | 0 | 0 | 14 | NaN | 656.49994 | 645.7615 | 738.91864 | 955.7894 | 1037.7485 |
| Pfas | SQLGEHPPDLDLPENLVR | -872 | -889 | 0 | 0 | 18 | NaN | 548.71814 | 407.72522 | 582.672 | 1183.9906 | 874.59973 |
| Pfas | FLRPEDSTRPSCF | -283 | -295 | 2 | 0.001023236 | 13 | NaN | 352.09393 | 433.67166 | NaN | 281.03918 | 616.54663 |
| Pfas | VEPGPALMLR | -1210 | -1219 | 0 | 0 | 10 | NaN | 2544.5728 | 2582.8088 | NaN | NaN | NaN |
| Pfas | FLRPEDSTR | -283 | -291 | 1 | 0.001092529 | 9 | NaN | 514.75275 | 634.23883 | NaN | NaN | NaN |
| Pfas | PSDRDFLSR | -569 | -577 | 1 | 0.002265552 | 9 | NaN | 179.8182 | NaN | NaN | NaN | NaN |
| Sars1 | VSCSNCTDYQAR | -393 | -404 | 0 | 0 | 12 | NaN | 5060.669 | 4126.3213 | 2907.1538 | 3868.5403 | 4712.218 |
| Sars1 | VKPAPIDQEPSK | -471 | -482 | 1 | 0.00025015 | 12 | NaN | 834.6324 | NaN | 784.142 | 432.50027 | 451.648 |
| Sars1 | VKLEAER | -122 | -128 | 1 | 0.00930008 | 7 | NaN | 1048.3442 | NaN | 1056.9244 | 1196.441 | NaN |
| Cyrib | FFLDFENAQPTESEK | -19 | -33 | 0 | 0 | 15 | NaN | 567.0637 | 655.35315 | 433.64297 | 909.40625 | NaN |
| Cyria | AWNAVCPLVVR | -78 | -88 | 0 | 0 | 11 | NaN | 1303.4475 | NaN | NaN | 1455.0278 | NaN |
| Tbl1x | NTQSGSLVHSYR | -483 | -494 | 0 | 0 | 12 | NaN | 886.6049 | 866.7225 | NaN | 500.91406 | 333.411 |
| Tspo | PSWHPPR | -40 | -46 | 0 | 0.002780285 | 7 | NaN | 1195.7954 | NaN | 232.45023 | 1453.7588 | 1060.5074 |
| Gnai2 | KIIHEDGYSEEECR | -54 | -67 | 1 | 0 | 14 | NaN | 551.9305 | 506.75116 | 520.2218 | 969.9077 | 917.6524 |
| Gnai2 | AYDLVLAEDEEMNR | -230 | -243 | 0 | 0.000947459 | 14 | NaN | 2414.8757 | 2483.9604 | NaN | NaN | NaN |
| Dnaja1 | AYEDDEHHPR | -380 | -389 | 0 | 0.00025015 | 10 | NaN | 224.1699 | 387.37347 | 166.85237 | 436.87665 | 401.86685 |
| Dnaja1 | SVTLEDLYNGATR | -112 | -124 | 0 | 0 | 13 | NaN | 1093.2383 | 1127.0378 | 1006.78595 | 1144.0599 | 1140.3414 |
| Glo1 | RFEELGVK | -141 | -148 | 1 | 0.001392869 | 8 | NaN | 354.5872 | NaN | NaN | 1121.5994 | NaN |
| Glo1 | GFGHIGIAVPDVYSAC | -124 | -139 | 0 | 0.003393785 | 16 | NaN | 402.46967 | NaN | NaN | NaN | NaN |
| Spcs2 | FFDHSGTLVMD | -196 | -206 | 0 | 0.007374243 | 11 | NaN | 494.13995 | 616.5949 | NaN | NaN | NaN |
| Kars1 | KVTGEDPYPHK | -86 | -96 | 1 | 0 | 11 | NaN | 469.4934 | 559.5447 | 897.0333 | 242.64218 | 582.9712 |
| Kars1 | RGDIIGVEGNPGK | -176 | -188 | 1 | 0 | 13 | NaN | 711.2219 | NaN | 982.5536 | NaN | NaN |
| Bclaf1 | TITPQNAPR | -303 | -311 | 0 | 0.008919338 | 9 | NaN | 615.7066 | NaN | NaN | NaN | NaN |
| Snu13 | KLLDLVQQ | -21 | -28 | 1 | 0.00383302 | 8 | NaN | 2345.8257 | 577.11005 | 1435.2549 | 874.792 | NaN |
| Snu13 | HLPLLCEDK | -68 | -76 | 0 | 0 | 9 | NaN | 365.49106 | NaN | 512.4109 | 335.0692 | 435.77005 |
| Ptprc | YIDGFKEPR | -701 | -709 | 1 | 0 | 9 | NaN | 1374.5067 | 1569.7421 | 1818.0714 | NaN | 1447.2476 |

| | | | | | | | | | | | | |
|---|---|---|---|---|---|---|---|---|---|---|---|---|
| Ptprc | LFLAEFQSIPR | -641 | -651 | 0 | 0 | 11 | NaN | 246.46193 | 603.9606 | NaN | 1005.44196 | 1287.5775 |
| Ptprc | TLVSWPEPVSKPESA | -392 | -406 | 1 | 0.000591124 | 15 | NaN | 6163.881 | 7601.867 | NaN | NaN | NaN |
| Ptprc | TIDLYVPPGTDK | -268 | -279 | 0 | 0.00025015 | 12 | NaN | 302.13428 | NaN | NaN | NaN | 326.84674 |
| Alb | ECCHGDLLEC | -268 | -277 | 0 | 0 | 10 | NaN | 2075.685 | 2048.8281 | 2179.2407 | 2313.4846 | 2122.6116 |
| Alb | KECCHGDLLECADDR | -267 | -281 | 1 | 0 | 15 | NaN | 475.25558 | 503.21146 | 736.3922 | 954.338 | 1036.4586 |
| Alb | KECCHGDLLECADDR | -267 | -281 | 1 | 0 | 15 | NaN | 475.25558 | 503.21146 | 736.3922 | 954.338 | 1036.4586 |
| Psma6 | KKFDWTFEQ | -185 | -193 | 2 | 0.000591124 | 9 | NaN | 693.9861 | 1046.993 | 767.6886 | 1213.5367 | 731.92285 |
| Psma6 | GKDCAVIVTQK | -44 | -54 | 1 | 0.00025015 | 11 | NaN | 2723.3542 | 3682.547 | 2362.4539 | 269.8878 | NaN |
| Psma6 | MILIGIDEEQGPQVYK | -138 | -153 | 0 | 0.005946382 | 16 | NaN | 1241.5144 | NaN | 2415.5195 | NaN | NaN |
| Psma6 | LYQVEYAFK | -22 | -30 | 0 | 0.000471409 | 9 | NaN | 1555.278 | NaN | NaN | NaN | NaN |
| Lsp1 | SSELDEDEGFGDWSQKPEPR | -77 | -96 | 1 | 0 | 20 | NaN | 648.5692 | 823.05084 | 910.1064 | 726.3482 | 709.44946 |
| Lsp1 | REPDPEDAVGGSGEAEEHLIR | -141 | -161 | 1 | 0 | 21 | NaN | 533.6259 | NaN | 530.3417 | 1090.0641 | 878.8727 |
| Tfrc | LELSQNQNVK | -367 | -376 | 0 | 0 | 10 | NaN | 2468.284 | 1027.6971 | 3470.615 | 1830.4462 | 2327.4673 |
| Tfrc | ETEDVPTSSR | -114 | -123 | 0 | 0 | 10 | NaN | 534.6165 | 578.1774 | 846.06244 | NaN | 411.52313 |
| Tfrc | FSKPTEVSGK | -224 | -233 | 1 | 0 | 10 | NaN | 3063.679 | 2366.453 | 4700.298 | NaN | NaN |
| Tfrc | VEYHFLSPYVSPR | -684 | -696 | 0 | 0 | 13 | NaN | 973.012 | 211.74467 | NaN | NaN | NaN |
| Igkc | CFLNNFYPK | -27 | -35 | 0 | 0 | 9 | NaN | 702.1526 | 678.19305 | 790.51294 | 580.3413 | 799.6437 |
| Rhoa | MKQEPVKPEEGR | -134 | -145 | 2 | 0 | 12 | NaN | 483.9399 | NaN | 629.0545 | 626.83307 | 679.7991 |
| Dock2 | EDGTTLHDGYHELVVLK | -537 | -553 | 0 | 0.000394361 | 17 | NaN | 780.40985 | 534.84045 | 483.44293 | 481.2027 | 496.83258 |
| Dock2 | LGFPEIIMPGDVR | -411 | -423 | 0 | 0 | 13 | NaN | 535.0552 | 1049.069 | 634.4055 | 766.9026 | 785.54 |
| Dock2 | ISSSPTHSLYVF | -215 | -226 | 0 | 0.00078661 | 12 | NaN | 2483.6948 | NaN | 1461.2695 | 2069.0605 | NaN |
| Dock2 | NKPVPDQIINFY | -1422 | -1433 | 1 | 0.002068213 | 12 | NaN | 1197.4604 | NaN | 891.86273 | 2422.0203 | NaN |
| Dock2 | LHTWLLK | -1245 | -1251 | 0 | 0.00025015 | 7 | NaN | 959.5608 | 1035.7391 | NaN | 426.72964 | NaN |
| Gbp5 | TILQTDQVLTDKEIQSK | -473 | -489 | 1 | 0.00025015 | 17 | NaN | 201.01857 | 266.85724 | 195.00526 | 363.65823 | 531.66943 |
| Cnbp | SSLPDICYR | -48 | -56 | 0 | 0.000550307 | 9 | NaN | 620.73395 | 1152.7747 | 398.89694 | 556.65985 | NaN |
| Dpp4 | FRPAEPHFTSDGSSFYK | -351 | -367 | 1 | 0.002265552 | 17 | NaN | 761.96606 | 440.3084 | NaN | NaN | 296.61118 |
| Dpp4 | CGIAVAPVSR | -643 | -652 | 0 | 0.00051306 | 10 | NaN | 1640.7014 | NaN | 213.1459 | NaN | NaN |
| Dpp4 | HICHFPK | -377 | -383 | 0 | 0.008890469 | 7 | NaN | 230.529 | 253.54797 | NaN | NaN | NaN |
| Srrt | ISHGEVLEWQK | -653 | -663 | 0 | 0 | 11 | NaN | 978.2629 | 1534.442 | 1301.9535 | 2203.573 | 2125.2563 |
| Srrt | LLSLDDSVDETEAVKR | -156 | -171 | 1 | 0 | 16 | NaN | 633.7294 | NaN | 786.742 | NaN | 921.8907 |
| Srrt | LSEPQPER | -451 | -458 | 0 | 0.001708008 | 8 | NaN | 2676.8257 | 2802.7969 | NaN | NaN | NaN |
| Srrt | WLCPLSGK | -712 | -719 | 0 | 0 | 8 | NaN | 467.45963 | 613.3534 | NaN | NaN | NaN |
| Srrt | ECELSPGVNR | -488 | -497 | 0 | 0.00025015 | 10 | NaN | 972.6744 | NaN | NaN | NaN | NaN |
| Eif4a3 | VLDEADEMLNK | -185 | -195 | 0 | 0 | 11 | NaN | 550.3827 | 554.42206 | 844.98206 | 626.86523 | 655.3441 |
| Eif4a3 | KLDYGQHVVAGTPGR | -152 | -166 | 1 | 0 | 15 | NaN | 621.99664 | 668.2124 | NaN | 455.73547 | 345.12595 |
| Eif4a3 | KLDYGQHVVAGTPGR | -152 | -166 | 1 | 0 | 15 | NaN | 621.99664 | 668.2124 | NaN | 455.73547 | 345.12595 |
| Bsg | RKPDQTLDEDDPGAAPLK | -353 | -370 | 2 | 0.00025015 | 18 | NaN | 470.63373 | 767.4914 | 419.13034 | 569.1868 | 560.3288 |
| Timm13 | MDSGFGSDFGGTGGGK | -1 | -16 | 0 | 0.00078661 | 16 | NaN | 329.95398 | 726.7384 | NaN | NaN | NaN |
| Gzmb | TQQVIPMVK | -85 | -93 | 0 | 0.000230707 | 9 | NaN | 2882.9521 | 3086.2288 | 3129.1296 | 3248.2837 | 3445.5461 |
| Gzmb | TFSNDIMLLK | -104 | -113 | 0 | 0 | 10 | NaN | 226.82524 | 958.0323 | NaN | 690.5866 | 1087.713 |
| Ube2i | EEDKDWRPAITIK | -98 | -110 | 2 | 0 | 13 | NaN | 1829.9257 | 1765.5785 | 1352.8379 | 1403.4711 | 1505.7646 |
| Ube2i | LSILEEDKDWR | -94 | -104 | 1 | 0 | 11 | NaN | 2831.4026 | NaN | 3307.1504 | 1934.44 | 2704.7246 |
| Ube2i | LSILEEDKDWRPAITIK | -94 | -110 | 2 | 0 | 17 | NaN | 309.13174 | NaN | 385.84375 | 471.77066 | 478.9923 |
| Tpt1 | ISHDELFSDIYK | -8 | -19 | 0 | 0 | 12 | NaN | 5912.5293 | 5473.9917 | 8692.976 | 4228.8184 | 3616.2708 |
| Tpt1 | TEGAIDDSLIGGNAS | -39 | -53 | 0 | 0.000230707 | 15 | NaN | 1343.4691 | 1266.2947 | 159.28142 | 157.40436 | 992.78925 |
| Tpt1 | TEGAIDDSLIGGN | -39 | -51 | 0 | 0 | 13 | NaN | 276.12674 | 331.69785 | 491.00613 | 499.8471 | 440.43463 |
| Tpt1 | HILANFNNY | -124 | -132 | 0 | 0.000712251 | 9 | NaN | 1443.7018 | 777.5314 | 1033.4647 | 986.78424 | 914.04846 |
| Tpt1 | MTGAAEQIK | -115 | -123 | 0 | 0.000230707 | 9 | NaN | 816.07635 | 570.8938 | 1134.6455 | 904.64954 | 940.3788 |
| Tpt1 | DLISHDELFSDIY | -6 | -18 | 0 | 0 | 13 | NaN | 287.30902 | 386.68484 | 402.79684 | 428.02573 | 374.04877 |
| Tpt1 | LISHDELFSDIYK | -7 | -19 | 0 | 0 | 13 | NaN | 926.23816 | 709.6465 | NaN | 1635.41 | 1079.7557 |
| Lgals1 | FNAHGDANTIVC | -50 | -61 | 0 | 0 | 12 | NaN | 972.7826 | NaN | 1411.9198 | 964.63586 | 992.64954 |
| Lgals1 | LNLKPGECLK | -10 | -19 | 1 | 0 | 10 | NaN | 435.4995 | NaN | 646.8259 | 215.56367 | 402.82532 |
| Ppp2cb | QLNENQVR | -22 | -29 | 0 | 0.001392869 | 8 | NaN | 1673.2117 | 2068.409 | NaN | 1972.4061 | NaN |
| Rpl11 | KNNFSDTGNFGF | -96 | -107 | 1 | 0.001392869 | 12 | NaN | 670.4813 | 684.8684 | NaN | 822.59875 | NaN |
| Pebp1 | FREWHHFLVVN | -81 | -91 | 1 | 0 | 11 | NaN | 842.7123 | 744.6613 | 586.8032 | 1243.4573 | 982.965 |
| Hat1 | VDENFDCVEADDVEGK | -92 | -107 | 0 | 0.000591124 | 16 | NaN | 646.51465 | 602.15845 | 628.25073 | 800.04376 | 825.56665 |
| Hat1 | CLRPEELTNQ | -373 | -382 | 1 | 0.000230707 | 10 | NaN | 1525.4144 | NaN | 947.9095 | 1273.7933 | 1631.5962 |
| Hat1 | GHGAQLLETVHR | -248 | -259 | 0 | 0 | 12 | NaN | 398.6175 | 402.11395 | NaN | 390.27118 | 432.9751 |
| Oxa1l | CLVFPLIVK | -148 | -156 | 0 | 0.000712251 | 9 | NaN | 394.93253 | 436.39252 | 457.93906 | 676.0058 | 606.7516 |
| Dnmt1 | TQLGPEEPCEQEDDNR | -224 | -239 | 0 | 0 | 16 | NaN | 1649.2805 | 1707.4473 | 1434.9735 | 1815.7211 | 1504.881 |
| Dnmt1 | CPECGQHLDDPNLK | -359 | -372 | 0 | 0.00111006 | 14 | NaN | 598.64417 | 824.7275 | 827.4097 | 861.9403 | 734.46783 |
| Dnmt1 | FTEDSLLR | -551 | -558 | 0 | 0 | 8 | NaN | 131.7518 | 1214.7604 | 648.13446 | 901.0002 | 715.11395 |
| Dnmt1 | LGPEEPCEQEDDNR | -226 | -239 | 0 | 0 | 14 | NaN | 89.86159 | 722.4756 | 831.92615 | 951.5995 | 926.4625 |
| Dnmt1 | HNHWAGLY | -1509 | -1516 | 0 | 0.007493591 | 8 | NaN | 1051.3903 | 1225.8636 | NaN | 1128.1143 | 1147.1775 |
| Dnmt1 | STLIPWCLPHTGNR | -1495 | -1508 | 0 | 0.008075282 | 14 | NaN | 876.61597 | 948.22955 | NaN | 615.6569 | NaN |
| Dnmt1 | KYSDYIK | -979 | -985 | 1 | 0 | 7 | NaN | 361.73505 | NaN | NaN | NaN | NaN |
| Dnmt1 | EESEEGNSAESAAEER | -180 | -195 | 0 | 0.002099344 | 16 | NaN | 246.95557 | NaN | NaN | NaN | NaN |
| Rps11 | VHLSPCFR | -111 | -118 | 0 | 0.00025015 | 8 | NaN | 3028.3716 | 3204.986 | 3470.815 | 3149.4744 | 3097.778 |
| Rps11 | QIGDIVTVGECRPLSK | -121 | -136 | 1 | 0.001392869 | 16 | NaN | 178.6417 | 339.50565 | 280.001 | 492.43924 | 345.46585 |
| Rps11 | EAIEGTYIDK | -49 | -58 | 0 | 0 | 10 | NaN | 672.2403 | 801.3767 | 721.8671 | NaN | NaN |
| Pfn1 | DSLLQDGEFTMDLR | -76 | -89 | 0 | 0 | 14 | NaN | 4654.676 | 4558.7593 | 1769.891 | 9913.92 | 9708.3125 |
| Pfn1 | LLQDGEFTMDLR | -78 | -89 | 0 | 0 | 12 | NaN | 2178.4104 | NaN | 1280.2 | 1626.5198 | 509.2368 |
| Tagln2 | PEGQAPVK | -71 | -78 | 0 | 0.003998192 | 8 | NaN | 519.9182 | 929.95605 | 689.23413 | 950.0295 | 830.23444 |
| Pml | NHSHLHCDIGEEIQ | -225 | -238 | 0 | 0.00025015 | 14 | NaN | 1656.4114 | 2207.5088 | NaN | 1403.9401 | 1704.0359 |
| Pml | LATSSPEQSWPSTFK | -511 | -525 | 0 | 0.000298805 | 15 | NaN | 1705.6968 | 1220.8755 | NaN | NaN | NaN |
| Pml | LLPCLHTL | -74 | -81 | 0 | 0.003490217 | 8 | NaN | 967.91016 | NaN | 930.2953 | NaN | NaN |
| Pml | LSCLEAVLQR | -320 | -329 | 0 | 0.00078661 | 10 | NaN | 556.4775 | NaN | NaN | NaN | NaN |
| Ptges3 | WLSVDFNNWK | -98 | -107 | 0 | 0 | 10 | NaN | 1434.2211 | 103.72403 | 1990.6345 | 2419.5078 | 2445.265 |

| | | | | | | | | | | | | |
|---|---|---|---|---|---|---|---|---|---|---|---|---|
| Ptges3 | HLNEIDLF | -49 | -56 | 0 | 0.002972566 | 8 | NaN | 749.64795 | 617.5761 | 663.0472 | NaN | 791.0247 |
| Ptges3 | HLNEIDLFH | -49 | -57 | 0 | 0.000550307 | 9 | NaN | 821.2982 | 629.1313 | 372.27194 | NaN | NaN |
| Ptges3 | GESGQSWPR | -80 | -88 | 0 | 0 | 9 | NaN | 797.1327 | 618.04626 | 771.6636 | NaN | NaN |
| Pstpip1 | CVKDDELYEEVR | -242 | -253 | 1 | 0 | 12 | NaN | 389.94504 | NaN | NaN | 546.9504 | 538.79205 |
| Strap | KYDYNSGEELESYK | -249 | -262 | 1 | 0.000431655 | 14 | NaN | 781.48413 | 565.1249 | 916.71045 | 792.4351 | 820.56116 |
| Strap | GHFGPIHCV | -263 | -271 | 0 | 0.00111006 | 9 | NaN | 524.99854 | NaN | 383.78528 | 843.5141 | 773.46204 |
| Strap | YDYNSGEELESYK | -250 | -262 | 0 | 0 | 13 | NaN | 478.58078 | 469.14722 | NaN | 367.31882 | 326.78143 |
| Xab2 | FYEDNGQLDDAR | -404 | -415 | 0 | 0 | 12 | NaN | 398.1127 | 448.74496 | NaN | 457.5307 | 408.81763 |
| Xab2 | GREEEDDVDLELR | -318 | -330 | 1 | 0.001776473 | 13 | NaN | 569.93634 | NaN | NaN | 602.36945 | NaN |
| Rae1 | GHEFYNPQK | -341 | -349 | 0 | 0 | 9 | NaN | 2035.941 | NaN | NaN | 221.74301 | NaN |
| Rae1 | CWEVQDSGQTIPK | -68 | -80 | 0 | 0.002265552 | 13 | NaN | 1827.9758 | NaN | NaN | NaN | NaN |
| Slc7a5 | GIFVTPTGVLK | -68 | -78 | 0 | 0 | 11 | NaN | 580.9306 | 636.01794 | NaN | NaN | NaN |
| Eif4h | TVATPLNQVANPNSAIFGGAR | -217 | -237 | 0 | 0 | 21 | NaN | 571.3281 | 523.4764 | 582.50354 | 986.9455 | 833.4572 |
| Eif4h | CYVEFDEVDSLK | -85 | -96 | 0 | 0 | 12 | NaN | 567.37524 | NaN | 718.2386 | 898.51697 | NaN |
| Srsf5 | DADDAVYELDGK | -49 | -60 | 0 | 0 | 12 | NaN | 1361.3601 | 1764.9531 | 1499.3054 | 1352.5293 | 1410.666 |
| Safb | SKDADDQKPGPSER | -558 | -571 | 2 | 0.000907206 | 14 | NaN | 199.83563 | 197.65018 | 243.87436 | 281.97925 | 257.54974 |
| Safb | DADDQKPGPSER | -560 | -571 | 1 | 0 | 12 | NaN | 103.9077 | 277.41888 | 222.87798 | NaN | NaN |
| Srsf2 | SYGRPPPDVEGMTSL | -2 | -16 | 1 | 0.003620005 | 15 | NaN | 480.52017 | 684.5675 | 393.36743 | 387.66318 | 664.36505 |
| Nudc | VVTVHLEK | -241 | -248 | 0 | 0.000627943 | 8 | NaN | 1026.283 | 1275.5093 | 979.1275 | 1118.2974 | 667.10895 |
| Dck | LKDAEKPVLFFER | -116 | -128 | 2 | 0 | 13 | NaN | 546.335 | 538.3675 | 455.132 | 734.6274 | 704.2533 |
| Dck | LKDAEKPVLFFER | -116 | -128 | 2 | 0 | 13 | NaN | 546.335 | 538.3675 | 455.132 | 734.6274 | 704.2533 |
| Rps21 | MGESDDSILR | -62 | -71 | 0 | 0 | 10 | NaN | 824.8371 | 1458.2042 | 1668.6685 | 1862.6316 | 411.43945 |
| Akr1a1 | KHHPEDVEPALR | -85 | -96 | 1 | 0.00025015 | 12 | NaN | 216.03828 | NaN | 910.84076 | NaN | NaN |
| Eif3c | GTTEEICQIYLR | -492 | -503 | 0 | 0 | 12 | NaN | 1074.2004 | 1102.5796 | 1211.3628 | 1346.7152 | 1388.7136 |
| Eif3c | RFEELTNLIR | -56 | -65 | 1 | 0.000591124 | 10 | NaN | 568.4062 | NaN | 752.28204 | 337.59985 | NaN |
| Eif3c | RLDEEEEDNEGGEWER | -286 | -301 | 1 | 0.000230707 | 16 | NaN | 331.05405 | NaN | NaN | 387.88263 | 384.35208 |
| Eif3c | QGTYGGYFR | -876 | -884 | 0 | 0.002329279 | 9 | NaN | 631.9768 | NaN | NaN | 304.7794 | NaN |
| Nudt9 | EFGEEALNSLQK | -230 | -241 | 0 | 0.005698678 | 12 | NaN | 1571.4108 | NaN | NaN | NaN | NaN |
| Cat | DAILFPSFIHSQK | -157 | -169 | 0 | 0 | 13 | NaN | 2087.4258 | 2329.6775 | 1660.9564 | 1818.9 | 1727.9908 |
| Ppp1r10 | PLVLPSPLVIPGSNSQER | -466 | -483 | 0 | 0 | 18 | NaN | 294.43887 | NaN | 273.69794 | 280.53992 | 308.35785 |
| Psmg1 | TPESTCSLSSPFLR | -178 | -191 | 0 | 0 | 14 | NaN | 1146.9861 | 736.9756 | 835.5757 | 244.28955 | 1071.818 |
| Psmg1 | NIPESTEILK | -266 | -275 | 0 | 0.003490217 | 10 | NaN | 1371.773 | NaN | NaN | NaN | NaN |
| P4hb | QLDKDGVVLFK | -199 | -209 | 1 | 0 | 11 | NaN | 764.38446 | 962.08905 | 1868.9783 | 1552.9076 | 1348.5612 |
| P4hb | LITLEEEMTK | -319 | -328 | 0 | 0 | 10 | NaN | 161.27127 | 1536.1223 | 823.7952 | NaN | NaN |
| Smyd5 | KGETIFIER | -44 | -52 | 1 | 0 | 9 | NaN | 639.3061 | 677.0171 | 304.96094 | 885.0365 | NaN |
| Smyd5 | LTGKPSQILPHPELC | -87 | -101 | 1 | 0.003003218 | 15 | NaN | 542.6934 | NaN | NaN | 873.05383 | NaN |
| Prkar1a | DALEPVQFEDGQK | -269 | -281 | 0 | 0 | 13 | NaN | 1033.6237 | 994.9567 | 1007.1343 | 1157.5913 | 1206.5844 |
| Prkar1a | TVADALEPVQFEDGQK | -266 | -281 | 0 | 0 | 16 | NaN | 692.60675 | 532.2562 | 657.95715 | 984.89453 | 978.5966 |
| Ppp6r1 | DVPQINDALGADESLLNR | -88 | -105 | 0 | 0 | 18 | NaN | 688.46796 | NaN | 928.0285 | 1266.2191 | NaN |
| Ppp6r1 | DALGADESLLNR | -94 | -105 | 0 | 0 | 12 | NaN | 536.11084 | 943.90265 | NaN | NaN | 838.1792 |
| Ppp6r1 | SIQIPSSPPAHGSPQLR | -733 | -749 | 0 | 0.002296977 | 17 | NaN | 658.156 | NaN | NaN | NaN | NaN |
| Kpna4 | DALSHFPALLTHPK | -326 | -339 | 0 | 0 | 14 | NaN | 479.76773 | 427.73212 | NaN | 493.68417 | 456.4579 |
| Ctbp1 | FYDPYLSDGIER | -202 | -213 | 0 | 0 | 12 | NaN | 1310.5742 | NaN | 337.08374 | 1753.3167 | 1846.0739 |
| Mier1 | VHDFDDER | -30 | -37 | 0 | 0.001291103 | 8 | NaN | 297.77588 | 325.76987 | NaN | 347.63422 | 343.5556 |
| Arhgdib | GPRPEEYEFLTPVEEAPK | -146 | -163 | 1 | 0 | 18 | NaN | 243.20601 | 523.39435 | 568.92456 | 702.1496 | 897.8764 |
| Arhgdib | KEGIEYR | -101 | -107 | 1 | 0.000394361 | 7 | NaN | 4025.795 | NaN | 163.10518 | 3682.3816 | NaN |
| Prrc2c | KEEEPIFTR | -599 | -607 | 1 | 0 | 9 | NaN | 1759.3163 | 1271.3514 | 1680.9325 | 958.53516 | 1228.7363 |
| Prrc2c | DAQQVEPEGQEKPSPAVVR | -2082 | -2100 | 1 | 0.001708008 | 19 | NaN | 550.2342 | 474.00076 | NaN | NaN | 497.75482 |
| Prrc2c | QLGQDESTAITSEQNDILK | -198 | -216 | 0 | 0 | 19 | NaN | 245.4159 | NaN | NaN | 348.56296 | 270.5683 |
| Prrc2c | KAWENSPNLR | -2207 | -2216 | 1 | 0.002566359 | 10 | NaN | 747.1763 | NaN | NaN | NaN | 8790.482 |
| Gfpt1 | WATHGEPNPVNSHPQR | -96 | -111 | 0 | 0 | 16 | NaN | 809.6174 | 177.66505 | 662.5796 | 652.73004 | 682.53033 |
| Gfpt1 | LSTDHIPILYR | -217 | -227 | 0 | 0 | 11 | NaN | 1201.9083 | NaN | 291.7907 | 818.0497 | NaN |
| Gfpt1 | RLPDLIK | -528 | -534 | 1 | 0.009416377 | 7 | NaN | 1819.305 | 1857.3865 | NaN | NaN | NaN |
| Isoc2a | IIKEPVPDSGLL | -183 | -194 | 1 | 0.001023236 | 12 | NaN | 2971.7185 | 1414.4202 | NaN | 429.6805 | 597.1057 |
| Nt5c2 | DAVDWVHYK | -203 | -211 | 0 | 0 | 9 | NaN | 784.89246 | 900.09204 | 213.02469 | 1227.4452 | 949.5581 |
| Atl3 | EQHSFELEER | -35 | -44 | 0 | 0 | 10 | NaN | 1429.362 | 1374.0288 | 792.74255 | 302.83286 | 833.5045 |
| Atl3 | DAVVGRPPADKK | -528 | -539 | 2 | 0.001636916 | 12 | NaN | 113.893135 | 612.20764 | 319.11227 | 441.32196 | 1190.8334 |
| Atl3 | VLLQDHIR | -49 | -56 | 0 | 0.002099344 | 8 | NaN | 1836.3345 | NaN | NaN | 1432.3848 | NaN |
| Ero1a | DCAVKPCHSDEVPDGIK | -98 | -114 | 1 | 0.00025015 | 17 | NaN | 431.5464 | 472.77353 | 541.0651 | 530.42426 | 504.69318 |
| Ero1a | SFFAGDKNEAHK | -358 | -369 | 1 | 0 | 12 | NaN | 390.12558 | 502.01706 | NaN | NaN | 513.9703 |
| Serinc3 | KIPGFCEGGFQIK | -67 | -79 | 1 | 0.002780285 | 13 | NaN | 399.23633 | NaN | NaN | NaN | NaN |
| Smc3 | LFYHIVDSDEVSTK | -558 | -571 | 0 | 0 | 14 | NaN | 1411.816 | 1178.7559 | NaN | NaN | 794.8214 |
| Smc3 | VLSDEFSHLRPEQR | -48 | -61 | 1 | 0.00025015 | 14 | NaN | 694.45905 | 612.4683 | NaN | 493.974 | 702.5917 |
| Smc3 | TFRPELLESADK | -1174 | -1185 | 1 | 0.00025015 | 12 | NaN | 1245.928 | NaN | 3293.9016 | NaN | 1157.8994 |
| Smc3 | KAEEELGELEAK | -684 | -695 | 1 | 0.000431655 | 12 | NaN | 813.5335 | 1339.8875 | NaN | NaN | NaN |
| Smc3 | NFSEVFQK | -1039 | -1046 | 0 | 0.00025015 | 8 | NaN | 439.9441 | NaN | NaN | NaN | NaN |
| Itm2b | SSEEALIVPPDAVAVDCK | -22 | -39 | 0 | 0.001392869 | 18 | NaN | 698.5474 | 1674.5116 | NaN | NaN | NaN |
| Hpcal1 | LSLEEFIK | -164 | -171 | 0 | 0.002199523 | 8 | NaN | 816.87256 | 992.251 | 318.32477 | 1124.5664 | 1116.2018 |
| Tnfrsf9 | SSIGGQPNCNICR | -53 | -65 | 0 | 0 | 13 | NaN | 704.6525 | 715.6751 | 720.6821 | 961.63873 | 896.6001 |
| Tnfrsf9 | VQNSCDNCQPGTFCR | -24 | -38 | 0 | 0 | 15 | NaN | 210.80731 | 230.30255 | 356.99963 | 319.5668 | 367.14102 |
| Tnfrsf9 | DCRPGQELTK | -104 | -113 | 1 | 0.00025015 | 10 | NaN | 1139.6726 | 1041.6886 | NaN | 1295.1469 | 1150.9166 |
| Tnfrsf9 | KYNPVCK | -39 | -45 | 1 | 0.00111006 | 7 | NaN | 294.67682 | NaN | 551.8527 | 387.2961 | 367.37363 |
| Magoh;Magol | GKFGHEFLEF | (15);(15) | (24);(24) | 1 | 0.006677519 | 10 | NaN | 318.87317 | 506.88052 | NaN | NaN | 657.767 |
| Psma1 | ILHVDNHIGIS | -63 | -73 | 0 | 0.000230707 | 11 | NaN | 1572.8064 | NaN | NaN | 2198.8484 | 1944.9471 |
| Mesd | DDDIEEGDLPEHK | -64 | -76 | 0 | 0 | 13 | NaN | 1989.3115 | 2094.9448 | 2388.215 | 1456.4097 | 1760.0676 |
| Usp14 | LEAIEDDSGR | -215 | -224 | 0 | 0 | 10 | NaN | 675.91583 | 1089.3472 | NaN | NaN | 477.33875 |
| Usp14 | ETDSSSAPAVTPSKK | -225 | -239 | 1 | 0.00111006 | 15 | NaN | 332.2793 | NaN | NaN | NaN | NaN |
| Hnrnpa2b1 | EESGKPGAHVTVK | -100 | -112 | 1 | 0 | 13 | NaN | 552.7358 | 728.75836 | 511.47726 | 596.2703 | 627.86884 |

| | | | | | | | | | | | | |
|---|---|---|---|---|---|---|---|---|---|---|---|---|
| Hnrnpa2b1 | YHTINGHNAEVR | -174 | -185 | 0 | 0 | 12 | NaN | 580.66986 | 638.9182 | 300.8222 | 1234.3539 | 697.95044 |
| Hnrnpa2b1 | GGLSFETTEESLR | -26 | -38 | 0 | 0 | 13 | NaN | 373.5006 | 440.59262 | NaN | NaN | 476.87122 |
| Hnrnpa2b1 | GNYNQQPSNYGPMK | -304 | -317 | 0 | 0 | 14 | NaN | 1322.3624 | 1471.9628 | NaN | NaN | NaN |
| Hnrnpa2b1 | GNYNQQPSNYGPMK | -304 | -317 | 0 | 0 | 14 | NaN | 1322.3624 | 1471.9628 | NaN | NaN | NaN |
| Hnrnpa2b1 | GGYGGGGPGYGNQGGGYGGGYDNYGG | -267 | -292 | 0 | 0 | 26 | NaN | 243.35762 | 200.95108 | NaN | NaN | NaN |
| Rbbp7 | WLPEVTKPEGK | -48 | -58 | 1 | 0 | 11 | NaN | 3203.4197 | 3463.5942 | 3673.2346 | 3134.9636 | 3109.5696 |
| Rbbp7 | WLPEVTKPEGK | -48 | -58 | 1 | 0 | 11 | NaN | 3203.4197 | 3463.5942 | 3673.2346 | 3134.9636 | 3109.5696 |
| Rbbp7 | SDDHTVCLWDINAGPK | -196 | -211 | 0 | 0 | 16 | NaN | 594.4291 | 591.47723 | 602.3598 | 636.41504 | 636.81586 |
| Rbbp7 | SNTTSKPSHLVDAH | -258 | -271 | 1 | 0 | 14 | NaN | 566.3884 | 185.06915 | 972.2653 | 898.9875 | 654.4907 |
| Rbbp7 | LVLGTHTSDEQNHLVVAR | -65 | -82 | 0 | 0 | 18 | NaN | 438.54028 | 223.09418 | 416.04907 | 697.172 | 616.93243 |
| Rbbp7 | DDHTVCLWDINAGPK | -197 | -211 | 0 | 0 | 15 | NaN | 399.19028 | 728.99554 | 438.37567 | 159.66121 | 488.51614 |
| Rbbp7 | PSHLVDAHTAEVN | -264 | -276 | 0 | 0 | 13 | NaN | 2721.7183 | NaN | 3480.495 | 3613.584 | 3760.3552 |
| Rbbp7 | LHTFESHKDEIFQVH | -309 | -323 | 1 | 0.000431655 | 15 | NaN | 1084.961 | 1333.1323 | NaN | 60.652847 | NaN |
| Rbbp7 | LHTFESHKDEIFQVH | -309 | -323 | 1 | 0.000431655 | 15 | NaN | 1084.961 | 1333.1323 | NaN | 60.652847 | NaN |
| Suclg2 | LNLQEYQSK | -39 | -47 | 0 | 0 | 9 | NaN | 706.64124 | NaN | 520.34766 | 801.0449 | NaN |
| Sub1 | KQAVPEKPVK | -29 | -38 | 2 | 0.00025015 | 10 | NaN | 293.9525 | 247.14339 | 240.21843 | 287.96722 | 327.51337 |
| Sub1 | KEQISDIDDAVR | -114 | -125 | 1 | 0 | 12 | NaN | 2221.0315 | NaN | 948.4574 | 1707.9816 | 1367.647 |
| Calu | SFDQLTPEESK | -60 | -70 | 0 | 0 | 11 | NaN | 566.83673 | 1573.5862 | NaN | NaN | NaN |
| Calu | VHNDAQNFDYDHDAFLGAE | -38 | -56 | 0 | 0.001610812 | 19 | NaN | 458.26743 | 271.85138 | NaN | NaN | NaN |
| Calu | VHNDAQNFDYDHDAFLGAE | -38 | -56 | 0 | 0.001610812 | 19 | NaN | 458.26743 | 271.85138 | NaN | NaN | NaN |
| Skp1 | GKTPEEIR | -129 | -136 | 1 | 0.000627943 | 8 | NaN | 117.231026 | 435.8008 | NaN | NaN | NaN |
| Szrd1 | DDSLPTGPPPQIR | -49 | -61 | 0 | 0 | 13 | NaN | 335.5041 | 1674.4912 | 1741.2031 | 1866.4861 | 1889.4111 |
| Bub3 | MHDLNTDQENLVGTHDAPIR | -81 | -100 | 0 | 0 | 20 | NaN | 690.976 | 724.43604 | NaN | 339.3032 | 413.74707 |
| Bub3 | VAVEYLDPSPEVQK | -203 | -216 | 0 | 0.00051306 | 14 | NaN | 362.61282 | NaN | NaN | NaN | NaN |
| Psmc3 | DSYLILETLPTEYDSR | -159 | -174 | 0 | 0 | 16 | NaN | 1227.9203 | 1326.4576 | 1785.9597 | 2487.4578 | 2399.7727 |
| Psmc3 | TNRVDILDPALLR | -335 | -347 | 1 | 0 | 13 | NaN | 782.20526 | 475.74512 | 549.7326 | 595.142 | 577.3086 |
| Psmc3 | NVSPDVNYEELAR | -377 | -389 | 0 | 0 | 13 | NaN | 1685.5521 | NaN | 1288.8362 | 1470.3287 | NaN |
| Psmc3 | NQLDGFQPNTQVK | -318 | -330 | 0 | 0.00111006 | 13 | NaN | 759.6124 | 787.5749 | 548.69135 | NaN | NaN |
| Psmc3 | STEEIVQR | -40 | -47 | 0 | 0.00025015 | 8 | NaN | 3865.7483 | NaN | NaN | NaN | 3622.144 |
| Ndufaf1 | DEAIEHFR | -89 | -96 | 0 | 0 | 8 | NaN | 628.9283 | NaN | NaN | 1233.5897 | 1587.9619 |
| Mtrex | DLDVEGTDEPIFGK | -63 | -76 | 0 | 0 | 14 | NaN | 1363.246 | 1393.3395 | 1231.3202 | 1460.9924 | 1265.7784 |
| Mtrex | GIGIHHGGLLPILK | -449 | -462 | 0 | 0 | 14 | NaN | 539.4637 | 444.65482 | 277.09436 | 305.3212 | 501.2307 |
| Mtrex | GIGIHHGGLLPILK | -449 | -462 | 0 | 0 | 14 | NaN | 539.4637 | 444.65482 | 277.09436 | 305.3212 | 501.2307 |
| Dctn1 | DEGHGIFVR | -82 | -90 | 0 | 0.00025015 | 9 | NaN | 688.511 | 1187.2368 | NaN | 1322.9604 | NaN |
| Dctn1 | STHTHVVDITR | -1166 | -1176 | 0 | 0.000431655 | 11 | NaN | 1410.6888 | NaN | NaN | 911.8927 | NaN |
| Mtco2 | MLISSEDVLHSW | -152 | -163 | 0 | 0 | 12 | NaN | 440.9084 | NaN | 322.13998 | 403.31866 | 218.0045 |
| Cwc22 | NILHESEIDKR | -319 | -329 | 1 | 0 | 11 | NaN | 164.72185 | 883.32874 | 753.05865 | 798.6646 | 860.50653 |
| Cwc22 | KKDELDPLLTR | -116 | -126 | 2 | 0 | 11 | NaN | 851.53107 | 777.7459 | 746.2073 | 820.068 | 899.029 |
| Polr2e | FGDKPSEGRPR | -44 | -54 | 2 | 0 | 11 | NaN | 468.93353 | 675.1898 | NaN | 831.43713 | 645.15955 |
| Polr2e | DELDQTLEEFK | -31 | -41 | 0 | 0 | 11 | NaN | 202.8377 | NaN | NaN | NaN | NaN |
| Parp1 | SATLGLVDIVK | -553 | -563 | 0 | 0 | 11 | NaN | 505.62973 | 234.56519 | 408.2682 | 885.0537 | 320.44757 |
| Parp1 | LGLVDIVK | -556 | -563 | 0 | 0 | 8 | NaN | 385.21286 | NaN | 297.50256 | 247.95282 | 270.74033 |
| Parp1 | QQVPSGESAILDR | -270 | -282 | 0 | 0 | 13 | NaN | 369.83493 | NaN | NaN | 522.3621 | 510.49237 |
| Eftud2 | HHWQIVPGDPLDK | -901 | -913 | 0 | 0 | 13 | NaN | 863.00916 | NaN | 2492.2183 | 1133.0292 | NaN |
| Eftud2 | TIVQEEDTQPLTEPIIKPVK | -78 | -97 | 1 | 0.00051306 | 20 | NaN | 395.90213 | 401.32965 | NaN | 185.86147 | NaN |
| Eftud2 | PGDPLDK | -907 | -913 | 0 | 0.006788754 | 7 | NaN | 215.33508 | NaN | 590.6728 | 442.69714 | 180.81538 |
| Eftud2 | VLSGTIHAGQPVK | -495 | -507 | 0 | 0.00149795 | 13 | NaN | 435.80338 | NaN | NaN | NaN | 334.54953 |
| Eif5b | KQDFDENDILR | -50 | -60 | 1 | 0.000230707 | 11 | NaN | 840.06195 | 781.9421 | NaN | 907.8634 | 703.2892 |
| Eif5b | RPEEEVLTLR | -276 | -285 | 1 | 0.00272025 | 10 | NaN | 964.6388 | NaN | 511.95288 | 1648.7701 | NaN |
| Eif5b | HIAVFPCK | -1082 | -1089 | 0 | 0 | 8 | NaN | 1322.6283 | 1474.0607 | NaN | NaN | NaN |
| Rps19 | VPEWVDTVK | -30 | -38 | 0 | 0 | 9 | NaN | 1941.2416 | 2061.0088 | 2085.6973 | 2574.5828 | 2530.0261 |
| Rps19 | HKELAPYDENWFYTR | -42 | -56 | 1 | 0 | 15 | NaN | 637.1181 | NaN | NaN | 709.8447 | 710.66644 |
| Rps19 | YDENWFYTR | -48 | -56 | 0 | 0 | 9 | NaN | 67.330414 | NaN | NaN | 358.25592 | 389.67944 |
| Rack1 | FSPNSSNPIIVSC | -156 | -168 | 0 | 0 | 13 | NaN | 5576.81 | 5687.252 | 4082.3918 | 231.20468 | 3160.9878 |
| Rack1 | VQDESHSEWVSCVR | -142 | -155 | 0 | 0 | 14 | NaN | 1236.6418 | 1061.769 | 645.318 | 716.96796 | 757.4452 |
| Rack1 | TVQDESHSEWVSCVR | -141 | -155 | 0 | 0 | 15 | NaN | 1269.3804 | 1350.5234 | 1557.9727 | 1547.1772 | 1438.3712 |
| Rack1 | LSGSWDGTLR | -79 | -88 | 0 | 0.000591124 | 10 | NaN | 1327.0541 | NaN | 1073.2715 | 693.71783 | NaN |
| Rack1 | YTDNLVR | -302 | -308 | 0 | 0.000471409 | 7 | NaN | 839.7369 | NaN | 858.0958 | 444.41473 | NaN |
| Rack1 | VWNLANCK | -176 | -183 | 0 | 0 | 8 | NaN | 4051.039 | NaN | NaN | NaN | 2089.7424 |
| Anxa11 | SLYHDITGDTSGDYR | -478 | -492 | 0 | 0 | 15 | NaN | 857.08276 | 806.2224 | 624.28015 | 928.5062 | 886.9554 |
| Rpl35a | TVTPGGKPNK | -57 | -66 | 1 | 0.00111006 | 10 | NaN | 1553.0153 | 1474.8676 | 824.27124 | 938.0639 | 1228.1151 |
| Capzb | DETVSDCSPHIANIGR | -200 | -215 | 0 | 0.00025015 | 16 | NaN | 122.670364 | NaN | 503.2543 | 602.01697 | NaN |
| Capzb | LWLQTNK | -175 | -181 | 0 | 0.001868731 | 7 | NaN | 417.51318 | NaN | 511.92944 | NaN | NaN |
| Supt16h | LQLQEELLK | -306 | -314 | 0 | 0 | 9 | NaN | 74.92233 | 1022.48303 | 353.4227 | 497.17874 | 465.12674 |
| Supt16h | QKPELLNK | -336 | -343 | 1 | 0.007819401 | 8 | NaN | 4525.59 | 1113.2046 | NaN | NaN | NaN |
| Lrpprc | MVAGLDTIGLSK | -1163 | -1174 | 0 | 0 | 12 | NaN | 551.43225 | 729.5792 | 681.10736 | 757.5172 | 691.3834 |
| Lrpprc | QLHEAIVTL | -806 | -814 | 0 | 0.001776473 | 9 | NaN | 1259.0872 | 1369.9127 | 1458.5472 | NaN | NaN |
| Lrpprc | LKAEGQPVGSALK | -670 | -682 | 1 | 0 | 13 | NaN | 1115.8658 | NaN | NaN | 793.97986 | 1130.4653 |
| Otulinl | DFAVCYPEEPLR | -322 | -333 | 0 | 0 | 12 | NaN | 1733.3198 | 1770.4407 | NaN | NaN | NaN |
| Sltm | DFDGDDLLETGK | -256 | -267 | 0 | 0 | 12 | NaN | 831.3391 | NaN | 917.9059 | 975.96875 | 980.41205 |
| Pdlim1 | IWSPLVTEEGK | -88 | -98 | 0 | 0 | 11 | NaN | 1249.5913 | NaN | 1657.5326 | 1723.9232 | 1735.9379 |
| Pdlim1 | NISNFNNAVESK | -153 | -164 | 0 | 0 | 12 | NaN | 1145.8856 | 1150.839 | 997.95514 | NaN | 891.1408 |
| Pdlim1 | VEDQIYCEK | -299 | -307 | 0 | 0 | 9 | NaN | 378.38498 | NaN | 777.744 | NaN | 378.11566 |
| Matr3 | VDYYTTTPALVFGK | -451 | -464 | 0 | 0 | 14 | NaN | 516.26807 | NaN | 565.21246 | 660.538 | 598.94403 |
| Pdlim2 | LLQEALEAEER | -241 | -251 | 0 | 0.000591124 | 11 | NaN | 332.85263 | 1571.9677 | NaN | NaN | 721.231 |
| Psip1 | IDNLDVNR | -363 | -370 | 0 | 0.000550307 | 8 | NaN | 1453.3569 | 444.6478 | 1164.674 | NaN | NaN |
| Tbcb | TDFKPGYWVGVR | -185 | -196 | 1 | 0.001868731 | 12 | NaN | 671.3192 | 311.27298 | NaN | 342.46744 | NaN |
| Ebna1bp2 | KLDFLEGDQKPVER | -209 | -222 | 2 | 0.002068213 | 14 | NaN | 598.83295 | 593.3876 | NaN | 807.9518 | NaN |

| | | | | | | | | | | | | |
|---|---|---|---|---|---|---|---|---|---|---|---|---|
| Rab3gap2 | DFLFPHLR | -19 | -26 | 0 | 0.00078661 | 8 | NaN | 652.04425 | 1111.8076 | NaN | NaN | NaN |
| Ap2a1 | FINLFPETK | -548 | -556 | 0 | 0 | 9 | NaN | 820.15814 | 893.2323 | 694.04346 | 254.60475 | 271.69635 |
| Rpl9 | TICSHVQNMIK | -72 | -82 | 0 | 0 | 11 | NaN | 814.4303 | 793.07367 | 905.95715 | 984.11224 | 303.11224 |
| Rpl9 | TICSHVQNMIK | -72 | -82 | 0 | 0 | 11 | NaN | 814.4303 | 793.07367 | 905.95715 | 984.11224 | 303.11224 |
| Rpl9 | KFLDGIYVS | -174 | -182 | 1 | 0.001776473 | 9 | NaN | 1249.6493 | NaN | 551.4363 | 1455.2435 | 1568.7366 |
| Itih2 | VQFELHYQ | -177 | -184 | 0 | 0.00078661 | 8 | NaN | 1084.9692 | NaN | NaN | NaN | NaN |
| Mfap1a;Mfap1 | EDSSEEEEEEIDDEEIER | (130);(130) | (147);(147) | 0 | 0.00025015 | 18 | NaN | 197.4231 | 275.54962 | NaN | 361.67212 | 224.98581 |
| Mfap1a;Mfap1 | QPPIQSTAGAVPVR | (10);(10) | (23);(23) | 0 | 0.002265552 | 14 | NaN | 286.9151 | 230.6174 | NaN | NaN | NaN |
| Anapc1 | LDSYLDHYYR | -799 | -808 | 0 | 0 | 10 | NaN | 3051.7356 | NaN | NaN | 1790.2388 | 2360.5388 |
| Psmd13 | LYENFISEFEHR | -59 | -70 | 0 | 0 | 12 | NaN | 1020.36676 | 153.34659 | 936.05347 | 1031.064 | 703.89764 |
| Psmd13 | PVSEQQER | -189 | -196 | 0 | 0.003745449 | 8 | NaN | 741.4229 | NaN | NaN | 989.0503 | NaN |
| Txnrd1 | YLGIPGDKEYCIS | -281 | -293 | 1 | 0.000230707 | 13 | NaN | 664.84937 | 751.4738 | 2022.8975 | 844.4206 | 775.1387 |
| Ctnnbl1 | DGEEEEEEEEPLDESSVK | -66 | -83 | 0 | 0.000627943 | 18 | NaN | 223.23079 | 323.274 | NaN | NaN | NaN |
| Adss2 | LDGETIPHFPANQEVLNK | -380 | -397 | 0 | 0 | 18 | NaN | 710.8279 | 728.98126 | 307.45718 | 891.20374 | NaN |
| Adss2 | KLDGETIPHFPANQEVLNK | -379 | -397 | 1 | 0.001023236 | 19 | NaN | 527.0307 | 398.78293 | NaN | 395.12772 | 549.4157 |
| Pak2 | DGFPSGTPALNTK | -148 | -160 | 0 | 0.00111006 | 13 | NaN | 1797.6162 | 7321.7334 | NaN | NaN | NaN |
| Pak2 | KFYDSNTVK | -128 | -136 | 1 | 0.002780285 | 9 | NaN | 1249.7975 | NaN | NaN | 1778.0507 | NaN |
| Hdac1 | DGIDDESYEAIFKPV | -230 | -244 | 1 | 0.00025015 | 15 | NaN | 288.50467 | 260.5028 | 270.63593 | 88.81408 | 414.77728 |
| Hdac2 | DGIDDESYGQIFKPIISK | -231 | -248 | 1 | 0.000550307 | 18 | NaN | 443.71094 | 520.61176 | 661.10175 | 661.216 | 927.52124 |
| Psmb2 | VIDKDGIHNL | -182 | -191 | 1 | 0.006472924 | 10 | NaN | 1223.6195 | 1010.896 | NaN | NaN | NaN |
| Usp5 | GLQPGEEELPDIAPPLVTPDEPK | -606 | -628 | 0 | 0.00111006 | 23 | NaN | 398.9352 | 341.48938 | 478.02414 | 576.3915 | 574.32526 |
| Usp5 | SISESVPVGPK | -783 | -793 | 0 | 0.000550307 | 11 | NaN | 1136.3662 | 1343.1829 | NaN | NaN | NaN |
| Tsr1 | VVLSGHPFK | -696 | -704 | 0 | 0.00383302 | 9 | NaN | 4766.8555 | NaN | NaN | NaN | NaN |
| Fubp1 | VPDGMVGFIIGR | -103 | -114 | 0 | 0 | 12 | NaN | 190.47325 | 236.28772 | 572.2126 | 632.5767 | 671.8205 |
| Gmps | HFDRDPLQK | -619 | -627 | 1 | 0.00025015 | 9 | NaN | 2177.2178 | 2071.0063 | 952.5716 | NaN | 2143.737 |
| Gmps | VIEPLKDFHKDEVR | -407 | -420 | 2 | 0.000230707 | 14 | NaN | 367.74124 | NaN | 320.2125 | 754.71594 | NaN |
| Gmps | DGSHHYEGAVVIL | -19 | -31 | 0 | 0.000591124 | 13 | NaN | 9155.716 | NaN | 7644.773 | NaN | NaN |
| Gmps | VIEPLKDFHKD | -407 | -417 | 2 | 0.000550307 | 11 | NaN | 171.97134 | NaN | NaN | NaN | NaN |
| Ubap2l | IDLAVLLGK | -302 | -310 | 0 | 0 | 9 | NaN | 140.38077 | 297.1468 | 308.0653 | 165.96468 | 146.76355 |
| Ubap2l | NNTGHFEPDDGTR | -223 | -235 | 0 | 0 | 13 | NaN | 597.86633 | 889.66406 | 835.01044 | NaN | 885.65955 |
| Ubap2l | NNTGHFEPDDGTR | -223 | -235 | 0 | 0 | 13 | NaN | 597.86633 | 889.66406 | 835.01044 | NaN | 885.65955 |
| Ubap2l | TEEWGTEDWNEDLSETK | -261 | -277 | 0 | 0.000471409 | 17 | NaN | 152.16272 | 106.83395 | NaN | 377.12463 | NaN |
| Ptpn6 | QISPLDNGDLVR | -396 | -407 | 0 | 0 | 12 | NaN | 10473.587 | 10361.205 | 11088.432 | 8633.99 | NaN |
| Ptpn6 | WYHGHIS | -110 | -116 | 0 | 0.000230707 | 7 | NaN | 875.82837 | 845.9136 | 738.6395 | NaN | 1143.2258 |
| Ptpn6 | IAQFIETTK | -512 | -520 | 0 | 0 | 9 | NaN | 1049.8726 | NaN | NaN | NaN | 551.9473 |
| Ddx5 | DGWPAMGIHGDK | -364 | -375 | 0 | 0 | 12 | NaN | 1027.3734 | 967.463 | 813.42944 | 524.27484 | 945.8303 |
| Ddx5 | YQEHPDLAR | -59 | -67 | 0 | 0 | 9 | NaN | 5297.171 | NaN | 6772.142 | 11880.8545 | 257.49945 |
| Ddx5 | ILQIVDVCHDVEKDEK | -313 | -328 | 1 | 0 | 16 | NaN | 409.64587 | NaN | NaN | 118.06566 | NaN |
| Ddx5 | TFFTPNNIK | -443 | -451 | 0 | 0.008129024 | 9 | NaN | 2163.078 | NaN | NaN | NaN | NaN |
| Pgls | LVPFDHAESTYGLYR | -82 | -96 | 0 | 0 | 15 | NaN | 526.0225 | 536.5467 | 485.53604 | 1109.3392 | 1039.1753 |
| Aqr | SYLCLLPTLPK | -382 | -392 | 0 | 0 | 11 | NaN | 126.742905 | NaN | NaN | 463.49203 | NaN |
| Nup155 | SVPSLPEISTIR | -74 | -85 | 0 | 0 | 12 | NaN | 514.0985 | NaN | 1062.954 | 1011.5545 | 1089.6716 |
| Nup155 | SAIDELKVDK | -462 | -471 | 1 | 0 | 10 | NaN | 888.136 | 808.8397 | 934.89185 | NaN | NaN |
| Eif3m | YIPTELDQVR | -139 | -148 | 0 | 0.00025015 | 10 | NaN | 1009.43085 | 1117.6125 | 1822.3894 | 2183.4512 | 1818.9017 |
| Eif3m | PTELDQVR | -141 | -148 | 0 | 0.003003218 | 8 | NaN | 1020.1186 | NaN | 596.254 | 1692.7959 | 1860.184 |
| Eif3m | DHLLTLKPVK | -221 | -230 | 1 | 0.000989247 | 10 | NaN | 487.90698 | 406.64078 | 301.20563 | 636.78296 | NaN |
| Eif3m | GAIQYIPTELDQVR | -135 | -148 | 0 | 0.00025015 | 14 | NaN | 586.4147 | NaN | NaN | 626.58203 | 1486.187 |
| Eif3m | LLGSYTEDNASQAR | -188 | -201 | 0 | 0 | 14 | NaN | 767.6095 | 570.08344 | NaN | NaN | NaN |
| Nol10 | DHQYGLPIK | -268 | -276 | 0 | 0.00111006 | 9 | NaN | 942.82745 | 1021.9568 | 918.697 | 416.10535 | 1234.9894 |
| Trim28 | MNDAFGDTK | -796 | -804 | 0 | 0.003620005 | 9 | NaN | 216.44292 | NaN | NaN | 113.29972 | 600.40576 |
| Trim28 | DHQYQFLED | -240 | -248 | 0 | 0.000471409 | 9 | NaN | 435.06308 | 491.9969 | NaN | NaN | NaN |
| Jpt2 | DHVLLCEGEDSK | -113 | -124 | 0 | 0.001055699 | 12 | NaN | 885.5003 | NaN | NaN | 414.4296 | NaN |
| Supt6h | SEEFEDLDEIVAR | -1402 | -1414 | 0 | 0 | 13 | NaN | 609.7782 | NaN | 817.94775 | NaN | 932.70105 |
| Supt6h | GALEEILENPER | -1087 | -1098 | 0 | 0.000591124 | 12 | NaN | 1611.6401 | NaN | NaN | NaN | NaN |
| Ctsd | GYLSQDTVSVPCK | -149 | -161 | 0 | 0 | 13 | NaN | 957.8599 | 946.7255 | 1051.4474 | 1151.9838 | 1139.3395 |
| Ctsd | DIACWVHHK | -114 | -122 | 0 | 0 | 9 | NaN | 720.9385 | 995.0217 | 1273.6089 | 1530.3857 | 1478.2996 |
| Ctsd | TTEPVSELLK | -63 | -72 | 0 | 0.00025015 | 10 | NaN | 1684.1318 | NaN | 1654.2913 | NaN | 1630.9528 |
| Puf60 | WKPPQGTESIK | -38 | -48 | 1 | 0 | 11 | NaN | 694.4836 | 2181.6243 | 2042.9918 | 2263.889 | 2612.836 |
| Clta | LQSEPESIR | -102 | -110 | 0 | 0.006677519 | 9 | NaN | 1483.9514 | 552.2187 | NaN | NaN | NaN |
| Sms | HSTLDFK | -6 | -12 | 0 | 0 | 7 | NaN | 343.82413 | 443.5198 | 850.6179 | NaN | 465.65945 |
| Sms | SLYEEQLGR | -326 | -334 | 0 | 0 | 9 | NaN | 721.9315 | NaN | NaN | 1341.1084 | NaN |
| Cbr1 | FHQLDIDNPQSIR | -59 | -71 | 0 | 0 | 13 | NaN | 697.532 | 1212.4138 | 1175.9655 | 779.27216 | 1691.5953 |
| Cbr1 | DIDNPQSIR | -63 | -71 | 0 | 0 | 9 | NaN | 5655.2485 | NaN | NaN | NaN | 5841.106 |
| Cbr1 | VNDDTPFHIQ | -97 | -106 | 0 | 0.00111006 | 10 | NaN | 891.77496 | NaN | NaN | NaN | NaN |
| Mrps6 | SAVENILEHLAR | -73 | -84 | 0 | 0 | 12 | NaN | 1363.3308 | NaN | 598.0492 | 879.34937 | 1061.1204 |
| Lonp1 | LALLDNHSSEFNVTR | -434 | -448 | 0 | 0 | 15 | NaN | 671.4903 | NaN | NaN | NaN | NaN |
| Cstf2 | DIFSEVGPVVSFR | -34 | -46 | 0 | 0.003745449 | 13 | NaN | 618.23047 | NaN | NaN | NaN | NaN |
| Cstf2 | LCVQNSPQEAR | -149 | -159 | 0 | 0.001868731 | 11 | NaN | 406.60083 | NaN | NaN | NaN | NaN |
| Grap2 | YLQHFHQDR | -218 | -226 | 0 | 0 | 9 | NaN | 1491.579 | 338.40112 | NaN | 1503.9478 | 487.89087 |
| Grap2 | EEDELGFR | -278 | -285 | 0 | 0.001392869 | 8 | NaN | 500.5169 | NaN | NaN | NaN | NaN |
| Son | ETLLHPGEEPR | -1533 | -1543 | 0 | 0 | 11 | NaN | 1143.4747 | 1368.8088 | 1679.8865 | 1036.0541 | 1422.9681 |
| Son | TVGPVGEASEEK | -1583 | -1594 | 0 | 0.00025015 | 12 | NaN | 1322.059 | 1415.6277 | 1565.1906 | NaN | 615.26074 |
| Son | LGHDPYR | (925)(946) | (931)(952) | 0 | 0.001023236 | 7 | NaN | 371.6728 | 258.5737 | 576.4997 | 580.8469 | NaN |
| Son | TDPTDEVPTKK | -97 | -107 | 1 | 0 | 11 | NaN | 789.4338 | 949.69977 | NaN | NaN | 1294.5574 |
| Son | PNEEIVQK | -57 | -64 | 0 | 0.00111006 | 8 | NaN | 703.01447 | NaN | NaN | 998.54456 | NaN |
| Son | LALELDTVGTSK | -1626 | -1637 | 0 | 0.000230707 | 12 | NaN | 88.02118 | 171.36183 | NaN | NaN | NaN |
| Usp47 | EVDEYPEHIK | -574 | -583 | 0 | 0 | 10 | NaN | 4204.322 | 4498.781 | 2134.1243 | NaN | NaN |
| Nsdhl | DIHQGFDNPR | -57 | -66 | 0 | 0.001610812 | 10 | NaN | 1584.0808 | NaN | NaN | NaN | NaN |

| | | | | | | | | | | | | |
|---|---|---|---|---|---|---|---|---|---|---|---|---|
| Nsdhl | DPQLVPILIDAAR | -197 | -209 | 0 | 0.002752526 | 13 | NaN | 445.8699 | NaN | NaN | NaN | NaN |
| Stk38 | DIKPDNLLLDSK | -212 | -223 | 1 | 0 | 12 | NaN | 1828.5447 | 1831.3866 | 2390.1096 | NaN | 2454.786 |
| Man2a1 | LLAENNEIISNIR | -61 | -73 | 0 | 0 | 13 | NaN | 4160.963 | 2720.9053 | NaN | 3496.1304 | 4906.1436 |
| Man2a1 | DIPHTCGPDPK | -354 | -364 | 0 | 0.008593015 | 11 | NaN | 530.38324 | NaN | NaN | NaN | NaN |
| Akap13 | DIPISELLNQEK | -1094 | -1105 | 0 | 0 | 12 | NaN | 1188.2461 | NaN | 557.08875 | 585.59546 | 807.3613 |
| Hcfc1 | STPVPGPSVPPPEELQVSPGPR | -1499 | -1520 | 0 | 0.00111006 | 22 | NaN | 404.81244 | 433.7223 | NaN | 237.9379 | NaN |
| Parn | SLELDPCTGFQR | -192 | -203 | 0 | 0 | 12 | NaN | 1539.7714 | 1314.9639 | 1842.0414 | 601.26917 | 1234.117 |
| Parn | FIDQVIEK | -173 | -180 | 0 | 0 | 8 | NaN | 319.04742 | NaN | 213.96362 | 427.35577 | NaN |
| Dut | HFIDVGAGVIDEDYR | -90 | -104 | 0 | 0 | 15 | NaN | 1210.9724 | 2586.2598 | 750.4803 | 880.8439 | 1416.3237 |
| Dut | HFIDVGAGVIDEDYR | -90 | -104 | 0 | 0 | 15 | NaN | 1210.9724 | 2586.2598 | 750.4803 | 880.8439 | 1416.3237 |
| Gnl3 | KLEVSPGDEQSNVETR | -91 | -106 | 1 | 0 | 16 | NaN | 747.56006 | 744.33636 | 939.372 | 911.9203 | 1057.5083 |
| Gnl3 | GFNLEELEK | -421 | -429 | 0 | 0 | 9 | NaN | 1184.2006 | 697.1503 | NaN | 1357.9136 | 1250.2698 |
| Ube2m | LFLEPNPEDPLNK | -135 | -147 | 0 | 0.000431655 | 13 | NaN | 675.6726 | 169.59323 | NaN | 3736.509 | NaN |
| Rab5c | VYDITNTDTFAR | -100 | -111 | 0 | 0 | 12 | NaN | 1093.5288 | 1103.8375 | 430.51526 | NaN | NaN |
| Cpne3 | NFSPIINHVAR | -396 | -406 | 0 | 0 | 11 | NaN | 1776.4777 | NaN | NaN | 2223.2627 | NaN |
| Irf3 | SSLPHSQENLPK | -130 | -141 | 0 | 0 | 12 | NaN | 529.5918 | 1249.4377 | NaN | 583.029 | NaN |
| Ash2l | YHVETEVDGR | -607 | -616 | 0 | 0 | 10 | NaN | 1148.4447 | NaN | 1142.4705 | 1292.8961 | NaN |
| Ankrd44 | LETADNPEVVDVK | -654 | -666 | 0 | 0 | 13 | NaN | 587.9825 | 674.3139 | 837.959 | 680.7684 | 784.65173 |
| Ankrd44 | DKEGYNSIHYA | -529 | -539 | 1 | 0.00025015 | 11 | NaN | 107.26807 | 160.81525 | 907.39404 | 621.6514 | 811.2848 |
| Ankrd44 | SLVDLDIRDEK | -589 | -599 | 1 | 0.00025015 | 11 | NaN | 317.11658 | 1166.3262 | NaN | 1699.637 | 1971.8035 |
| Ankrd44 | VTTGANVNETDDWGR | -458 | -472 | 0 | 0 | 15 | NaN | 687.4754 | 764.8193 | NaN | NaN | 878.595 |
| Ankrd44 | VVVEELLAK | -956 | -964 | 0 | 0.00111006 | 9 | NaN | 617.23505 | NaN | 923.48456 | NaN | 522.3172 |
| Ankrd44 | GFEIDTPDTFGR | -395 | -406 | 0 | 0 | 12 | NaN | 746.43195 | 651.4928 | NaN | NaN | NaN |
| Ankrd44 | FIGNPFTPLHC | -800 | -810 | 0 | 0.008736733 | 11 | NaN | 354.90704 | NaN | NaN | NaN | NaN |
| Naa15 | LFHSVCESK | -715 | -723 | 0 | 0 | 9 | NaN | 959.5525 | 5146.6123 | 859.2582 | 1311.5245 | 1272.7886 |
| Naa15 | RLPLNFLSGE | -295 | -304 | 1 | 0.003201138 | 10 | NaN | 535.8065 | 1956.7007 | NaN | 2235.0012 | 523.57275 |
| Naa15 | EALCDGSLGDCK | -806 | -817 | 0 | 0 | 12 | NaN | 426.46582 | NaN | 446.65872 | 605.17194 | 525.00085 |
| Naa15 | WDKDNLQILR | -109 | -118 | 1 | 0 | 10 | NaN | 451.59406 | NaN | 1465.7795 | NaN | NaN |
| Naa15 | NFNETFLK | -748 | -755 | 0 | 0.00025015 | 8 | NaN | 893.7629 | NaN | NaN | 1037.9941 | NaN |
| Csde1 | DKEAEDGIIAYDDCGVK | -451 | -467 | 1 | 0 | 17 | NaN | 296.92264 | NaN | 295.99475 | 603.9528 | 612.1675 |
| Csde1 | QRPGQQIATC | -497 | -506 | 1 | 0.002534621 | 10 | NaN | 793.1728 | NaN | NaN | NaN | NaN |
| Csde1 | DVEGSTSPQIGDKVEF | -476 | -491 | 1 | 0.004983786 | 16 | NaN | 633.877 | NaN | NaN | NaN | NaN |
| Wdr75 | DKETNNNTDLGEDAIHQL | -791 | -808 | 1 | 0.000471409 | 18 | NaN | 462.26318 | 626.8689 | 954.59515 | 612.0934 | 491.73288 |
| Wdr75 | EYINDEGLTQTELTK | -377 | -391 | 0 | 0 | 15 | NaN | 330.18817 | 387.10056 | 712.3355 | 857.0554 | 914.7254 |
| Nono | PVTVEPMDQLDDEEGLPEK | -223 | -241 | 0 | 0.001055699 | 19 | NaN | 3410.5222 | 3713.462 | 3496.4858 | NaN | 3755.1558 |
| Nono | TFPRPVTVEPMDQLDDEEGLPEK | -219 | -241 | 1 | 0.00025015 | 23 | NaN | 347.85107 | NaN | 772.5981 | 3907.405 | 31.894033 |
| Alad | DKHPELPLAVYQ | -266 | -277 | 1 | 0.002972566 | 12 | NaN | 392.4392 | NaN | NaN | NaN | NaN |
| Usp19 | KEPTKDELLLDWR | -47 | -59 | 2 | 0 | 13 | NaN | 728.497 | NaN | NaN | 787.2513 | 657.90924 |
| Uso1 | EQDLQLEELK | -661 | -670 | 0 | 0 | 10 | NaN | 2336.3147 | 1638.3691 | 2084.0752 | NaN | NaN |
| Uso1 | DKLDLEVTDSKK | -893 | -904 | 2 | 0.00266297 | 12 | NaN | 4960.6846 | NaN | NaN | NaN | NaN |
| Rps4x | VNDTIQIDLETGK | -156 | -168 | 0 | 0 | 13 | NaN | 1370.1393 | 1097.0404 | 1295.7827 | 651.6806 | 761.4466 |
| Rps4x | MVTGGANLGR | -182 | -191 | 0 | 0 | 10 | NaN | 388.94156 | NaN | 936.5697 | NaN | NaN |
| Rps4x | LRECLPLIIF | -38 | -47 | 1 | 0.003037016 | 10 | NaN | 235.05334 | NaN | NaN | 180.10873 | NaN |
| Ppib | DKPLKDVIIVDSGK | -191 | -204 | 2 | 0 | 14 | NaN | 703.8707 | 869.67096 | 968.69543 | 833.3168 | 822.76465 |
| Elp1 | DKPNQQDVVFFEK | -258 | -270 | 1 | 0 | 13 | NaN | 1018.4145 | 1193.9601 | 380.6853 | 591.3764 | 448.9794 |
| Elp1 | TIGGEECVLGLTDR | -615 | -628 | 0 | 0 | 14 | NaN | 742.10693 | 563.63293 | NaN | 1164.8823 | 953.24634 |
| Elp1 | INLNLIHDHNPK | -751 | -762 | 0 | 0 | 12 | NaN | 499.92593 | 546.15155 | NaN | 452.2984 | NaN |
| Hmgb2 | KKHPDSSVNFAEF | -29 | -41 | 2 | 0.00122419 | 13 | NaN | 1700.4716 | 2483.9587 | NaN | 2984.6523 | 2766.4453 |
| Hmgb2 | KHPDSSVNFAE | -30 | -40 | 1 | 0.001291103 | 11 | NaN | 1140.6578 | NaN | 558.8785 | 1178.834 | 1256.7537 |
| Hmgb2 | HPDSSVNFAEF | -31 | -41 | 0 | 0.007546675 | 11 | NaN | 454.202 | NaN | NaN | 678.7841 | 630.68823 |
| Hmgb2 | KHPDSSVNFAEFSK | -30 | -43 | 1 | 0 | 14 | NaN | 331.40588 | 385.83972 | NaN | NaN | NaN |
| Hmgb2 | KNEPEDEEEEEEEEE | -184 | -199 | 1 | 0.003299042 | 16 | NaN | 746.5776 | 700.86536 | NaN | NaN | NaN |
| Hmgb2 | KHPDSSVNFAEFS | -30 | -42 | 1 | 0.001868731 | 13 | NaN | 234.22482 | NaN | 528.54193 | NaN | NaN |
| Hmgb2 | KKNEPEDEEEEEEE | -183 | -197 | 2 | 0.00272025 | 15 | NaN | 770.04205 | NaN | NaN | NaN | NaN |
| Sephs1 | ESFNPETYELDK | -5 | -16 | 0 | 0 | 12 | NaN | 619.8898 | 814.6609 | 205.70743 | 923.7775 | 731.814 |
| Sephs1 | DKVIPLIIQ | -138 | -146 | 1 | 0.00416551 | 9 | NaN | 427.47278 | 486.0972 | 429.11133 | 504.41116 | NaN |
| Ruvbl2 | IGLETSLR | -393 | -400 | 0 | 0.004133093 | 8 | NaN | 1523.881 | 1234.6237 | 1894.6693 | 1357.0542 | 1688.3003 |
| Ruvbl2 | GLDDALEPR | -32 | -40 | 0 | 0 | 9 | NaN | 1248.8071 | 459.22827 | 1294.1633 | 1124.9296 | 1173.9401 |
| Ruvbl2 | IDRPATGTGSK | -147 | -157 | 1 | 0.002566359 | 11 | NaN | 908.85974 | 104.93273 | 937.51154 | 1140.7731 | 1143.345 |
| Ruvbl2 | FSGDTGEIK | -261 | -269 | 0 | 0 | 9 | NaN | 197.82416 | NaN | NaN | NaN | NaN |
| Ube2n | IYHPNVDK | -75 | -82 | 0 | 0.002752526 | 8 | NaN | 4122.222 | 4250.568 | 2227.3862 | 4933.32 | 4658.217 |
| Dctn2 | DLAVLDQVEAR | -273 | -283 | 0 | 0.007791272 | 11 | NaN | 507.63104 | NaN | NaN | NaN | NaN |
| Ctps2 | DLCFVGEDVDGKR | -496 | -508 | 1 | 0.00025015 | 13 | NaN | 184.28548 | NaN | 810.954 | NaN | 246.0625 |
| Dis3l2 | FADVIVHR | -704 | -711 | 0 | 0.00719662 | 8 | NaN | 1072.3599 | NaN | NaN | NaN | NaN |
| Dis3l2 | DLDDALACR | -387 | -395 | 0 | 0.002329279 | 9 | NaN | 668.2191 | NaN | NaN | NaN | NaN |
| Ik | KDEDPENKIEFK | -198 | -209 | 2 | 0 | 12 | NaN | 108.719536 | 669.7679 | 482.46573 | 815.00214 | 810.55 |
| Ik | DLDDEYADTDIPTTLIR | -242 | -258 | 0 | 0.001636916 | 17 | NaN | 268.7904 | NaN | NaN | NaN | 434.6559 |
| Nt5dc1 | DLDHTLCR | -16 | -23 | 0 | 0.00272025 | 8 | NaN | 15442.344 | NaN | NaN | NaN | 15679.28 |
| Rbm39 | SFLDSDELER | -337 | -346 | 0 | 0 | 10 | NaN | 797.87805 | 776.58344 | 922.9684 | 874.1515 | 965.468 |
| Cltb | DLEEWNQR | -135 | -142 | 0 | 0.003360691 | 8 | NaN | 1254.9742 | 1212.9998 | NaN | NaN | NaN |
| Ndrg3 | TYHDIGLNHK | -59 | -68 | 0 | 0 | 10 | NaN | 4122.9062 | 1482.9093 | NaN | NaN | NaN |
| Ndrg3 | DLEIERPILGQNDNR | -234 | -248 | 1 | 0.000230707 | 15 | NaN | 923.3132 | NaN | NaN | NaN | 829.9715 |
| Ogdh | SSPYPTDVAR | -457 | -466 | 0 | 0.000471409 | 10 | NaN | 849.6361 | 809.5359 | 620.0927 | NaN | NaN |
| Ogdh | VIPENGPAAQDPHK | -884 | -897 | 0 | 0 | 14 | NaN | 1144.053 | NaN | NaN | 1429.0657 | NaN |
| Ogdh | VSQGVVNQPEYEEEISK | -545 | -561 | 0 | 0.002810811 | 17 | NaN | 408.51074 | NaN | NaN | NaN | NaN |
| Eed | DLEVEDPHK | -390 | -398 | 0 | 0.006472924 | 9 | NaN | 260.6603 | NaN | NaN | NaN | NaN |
| Plekho2 | DLGELNQETPGLVQK | -468 | -482 | 0 | 0.000907206 | 15 | NaN | 852.5432 | 827.2597 | 925.7051 | NaN | NaN |
| Gdi2 | ICDPSYVK | -281 | -288 | 0 | 0.000298805 | 8 | NaN | 758.5994 | NaN | 312.74094 | 844.1691 | 575.28595 |

| | | | | | | | | | | | | |
|---|---|---|---|---|---|---|---|---|---|---|---|---|
| Gdi2 | TGSEFDFEEMKR | -425 | -436 | 1 | 0.000627943 | 12 | NaN | 1415.6827 | NaN | 1907.8987 | 1974.8761 | NaN |
| Gdi2 | DLGTDSQIFISR | -391 | -402 | 0 | 0.00025015 | 12 | NaN | 1281.8594 | NaN | NaN | NaN | NaN |
| Ctcf | FHCPHCDTVIAR | -437 | -448 | 0 | 0 | 12 | NaN | 161.75946 | NaN | 520.7913 | 691.1587 | NaN |
| Ctcf | THSGEKPYECY | -400 | -410 | 1 | 0.005363933 | 11 | NaN | 497.98965 | NaN | 517.2026 | NaN | NaN |
| Ndufs1 | SEIAGVDDLGTTGR | -187 | -200 | 0 | 0 | 14 | NaN | 1415.0369 | 1464.7156 | 1361.8778 | 1456.5321 | 1534.9796 |
| Ndufs1 | HSFCEVLK | -460 | -467 | 0 | 0.00111006 | 8 | NaN | 3646.126 | 2957.2542 | NaN | NaN | 1082.163 |
| Ndufs1 | DLGYKPGVEAIR | -527 | -538 | 1 | 0.003299042 | 12 | NaN | 2540.6113 | NaN | NaN | NaN | NaN |
| Ndufs1 | IFPTEGAGTDLR | -371 | -382 | 0 | 0.001941082 | 12 | NaN | 1339.6063 | NaN | NaN | NaN | NaN |
| Ndufs1 | SWLHNDLK | -421 | -428 | 0 | 0 | 8 | NaN | 832.7372 | NaN | NaN | NaN | NaN |
| Rnf213 | DLHDLGSLVEGK | -393 | -404 | 0 | 0 | 12 | NaN | 802.8703 | 1450.9009 | 2284.4395 | NaN | 387.80588 |
| Rnf213 | VLDCLQGISYR | -761 | -771 | 0 | 0 | 11 | NaN | 235.65587 | NaN | NaN | 63.917 | 289.2892 |
| Nsfl1c | SGQQIVGPPR | -102 | -111 | 0 | 0 | 10 | NaN | 1397.7822 | NaN | 1173.1519 | 353.9978 | 822.756 |
| Nsfl1c | GFSLDNGDLR | -190 | -199 | 0 | 0.00025015 | 10 | NaN | 887.8981 | NaN | 1187.263 | NaN | 1155.5758 |
| Nsfl1c | HSGQDVHVVLK | -175 | -185 | 0 | 0.00078661 | 11 | NaN | 309.32965 | 468.51566 | NaN | NaN | NaN |
| Lars1 | TNSLELEHIEVK | -1030 | -1041 | 0 | 0 | 12 | NaN | 1367.2898 | 1438.568 | 1484.6371 | 428.34482 | 1516.9622 |
| Lars1 | SAKPGDIVVR | -130 | -139 | 1 | 0.002972566 | 10 | NaN | 1500.5455 | NaN | NaN | NaN | NaN |
| Lars1 | RTEPGVPVSLVNPQPSSGHFSTK | -1064 | -1086 | 1 | 0.00111006 | 23 | NaN | 566.5618 | NaN | NaN | NaN | NaN |
| Lars1 | TWVEWVK | -771 | -777 | 0 | 0.00272025 | 7 | NaN | 505.09488 | NaN | NaN | NaN | NaN |
| Tpp2 | KYSDPGPVYDCL | -187 | -198 | 1 | 0 | 12 | NaN | 2963.7922 | 2626.0254 | 2774.152 | 891.60803 | NaN |
| Tpp2 | KIPANWTNPLGK | -98 | -109 | 1 | 0 | 12 | NaN | 940.9557 | 882.5062 | NaN | 1082.2506 | 995.6481 |
| Tpp2 | SLSETYWETTK | -1175 | -1185 | 0 | 0.00223472 | 11 | NaN | 378.32578 | 3120.972 | NaN | NaN | NaN |
| Grk6 | GVDLEPTDQDFYQK | -488 | -501 | 0 | 0.000230707 | 14 | NaN | 230.98383 | 197.84204 | 287.73807 | NaN | NaN |
| Camk4 | DLKPENLLYATPAPDAPLK | -160 | -178 | 1 | 0 | 19 | NaN | 251.42381 | 155.86633 | 176.88257 | 315.6667 | 58.6241 |
| Ripk3 | TSSDPVAGTPQIPHTLPFR | -378 | -396 | 0 | 0.00025015 | 19 | NaN | 1322.841 | NaN | NaN | 1201.7524 | 1268.2122 |
| Sart1 | SVDDLAQQKPR | -321 | -331 | 1 | 0.003270762 | 11 | NaN | 401.52197 | NaN | NaN | NaN | NaN |
| Ppil2 | TDNTHIVAIR | -111 | -120 | 0 | 0 | 10 | NaN | 3847.086 | 4526.1685 | 4343.9565 | NaN | NaN |
| Esyt2 | DLPAWVHFPDTER | -106 | -118 | 0 | 0 | 13 | NaN | 469.5742 | 481.98737 | 501.0838 | 336.6626 | 554.0375 |
| Esyt2 | WYDLTEDGTRPQVIT | -831 | -845 | 1 | 0 | 15 | NaN | 459.71555 | NaN | NaN | 732.1085 | 794.94214 |
| Pdcd4 | DSVWGSGGGQQPVNHLVK | -312 | -329 | 0 | 0.001708008 | 18 | NaN | 1437.6523 | NaN | NaN | NaN | NaN |
| Arl8b | DLPNALDEK | -133 | -141 | 0 | 0.000431655 | 9 | NaN | 898.6771 | 948.71814 | NaN | 831.54 | NaN |
| Ddx3x | IYADGPGEALR | -242 | -252 | 0 | 0 | 11 | NaN | 303.5003 | 8086.5356 | 3919.1313 | 3854.4568 | 3894.7805 |
| Ddx3x | VVYGGAEIGQQIR | -299 | -311 | 0 | 0 | 13 | NaN | 3016.9033 | 2984.1672 | 1291.2659 | NaN | 3844.1255 |
| Gtpbp4 | KVNEVLNR | -352 | -359 | 1 | 0 | 8 | NaN | 602.0638 | 741.55164 | 444.07535 | 772.5016 | 774.18823 |
| Gtpbp4 | EQLELFQNIRPL | -267 | -278 | 1 | 0 | 12 | NaN | 607.0459 | NaN | 432.50406 | 479.69754 | 458.55997 |
| Gtpbp4 | RIAELSEEDQK | -295 | -305 | 1 | 0.002068213 | 11 | NaN | 2333.4321 | NaN | 1235.7756 | NaN | 1790.8635 |
| Gtpbp4 | TLLLCGYPNVGK | -170 | -181 | 0 | 0 | 12 | NaN | 146.04327 | NaN | NaN | NaN | NaN |
| Ngdn | LVPVHYDETEAER | -176 | -188 | 0 | 0 | 13 | NaN | 847.4185 | NaN | 921.55597 | 732.7762 | 404.45386 |
| Ngdn | DLSHLILDK | -66 | -74 | 0 | 0.002502116 | 9 | NaN | 528.9303 | 506.56485 | NaN | 701.0077 | 735.1527 |
| Flii | DLSHNQLTECPR | -110 | -121 | 0 | 0 | 12 | NaN | 1097.7947 | 1538.4634 | 1170.1952 | 1344.2997 | NaN |
| Flii | SNLSDVDLSCNDLTR | -223 | -237 | 0 | 0.00272025 | 15 | NaN | 602.96185 | 592.78973 | 636.26904 | NaN | NaN |
| Flii | LPEEEFGHFY | -920 | -929 | 0 | 0.001023236 | 10 | NaN | 483.67642 | 575.9178 | NaN | NaN | NaN |
| Lig1 | NQEDNTGKYPDIISR | -588 | -602 | 1 | 0 | 15 | NaN | 1484.5238 | 1560.8889 | 1910.3456 | 2793.9797 | 2717.7654 |
| Lig1 | LKPQEEEQSKPPAR | -209 | -222 | 2 | 0 | 14 | NaN | 399.833 | 539.42535 | 543.2788 | 160.27742 | 583.0057 |
| Lig1 | DLSLSPIYPAAR | -846 | -857 | 0 | 0 | 12 | NaN | 453.9708 | 472.3283 | 502.0346 | 470.53345 | 416.4895 |
| Lig1 | YDEESEELQAICK | -781 | -793 | 0 | 0 | 13 | NaN | 875.8619 | 450.5359 | NaN | NaN | NaN |
| Irgm1 | SQNFYTLR | -338 | -345 | 0 | 0.000471409 | 8 | NaN | 955.2928 | NaN | NaN | NaN | NaN |
| Hnrnpdl | KVFVGGLSPDTSEEQIK | -115 | -131 | 1 | 0.002629273 | 17 | NaN | 697.1953 | 748.71277 | NaN | 1454.0148 | NaN |
| Hnrnpdl | DLTEYLSR | -44 | -51 | 0 | 0.000755556 | 8 | NaN | 1243.1504 | NaN | NaN | 1107.5894 | NaN |
| Pgm2 | WDNGAQIISPHDR | -186 | -198 | 0 | 0 | 13 | NaN | 836.96216 | 916.9031 | 1328.7173 | NaN | 569.23425 |
| Pgm2 | IVLANDPDADR | -325 | -335 | 0 | 0 | 11 | NaN | 3488.3237 | NaN | NaN | NaN | NaN |
| Anxa5 | LEDDVVGDTSGYYQR | -135 | -149 | 0 | 0.003461185 | 15 | NaN | 455.11053 | NaN | 176.15137 | NaN | NaN |
| Eif3a | RANEFLEVGK | -14 | -23 | 1 | 0.000591124 | 10 | NaN | 637.9827 | 1237.0343 | 1118.6782 | 811.34973 | 306.95807 |
| Eif3a | FCLQYTR | -184 | -190 | 0 | 0 | 7 | NaN | 574.49005 | 686.92773 | 876.35645 | 1135.344 | 94.45312 |
| Eif3a | EDAPVGPHLQ | -503 | -512 | 0 | 0.00025015 | 10 | NaN | 674.09595 | 903.76294 | 488.44562 | NaN | 416.09036 |
| Eif3a | HHNQSTAINL | -211 | -220 | 0 | 0.001868731 | 10 | NaN | 3150.641 | 1129.0417 | 794.1139 | NaN | NaN |
| Eif3a | NADDDRPPR | -991 | -999 | 1 | 0.002199523 | 9 | NaN | 573.4002 | NaN | NaN | 630.11115 | 617.3455 |
| Eif3a | NETDEDGWTTVR | -1332 | -1343 | 0 | 0 | 12 | NaN | 686.1092 | NaN | NaN | NaN | NaN |
| Ddi2 | GSQTTFLPEGELPECAR | -347 | -363 | 0 | 0 | 17 | NaN | 2584.8855 | 2371.5005 | NaN | 3453.0227 | 3457.818 |
| Ddi2 | DMLLANPHELSLLK | -140 | -153 | 0 | 0 | 14 | NaN | 734.1269 | NaN | 747.9231 | 1004.031 | 991.25574 |
| Map4 | EFNNVTPLSEEEVTSVK | -498 | -514 | 0 | 0.005789881 | 17 | NaN | 1152.4788 | 1221.788 | NaN | 2048.5044 | NaN |
| Map4 | LAPPEDVLLTK | -359 | -369 | 0 | 0 | 11 | NaN | 569.23285 | NaN | 785.7829 | NaN | 520.0628 |
| Cip2a | DNAPALNIEDLIEK | -592 | -605 | 0 | 0.00025015 | 14 | NaN | 499.0034 | NaN | NaN | NaN | NaN |
| Fgg | DNCCILDER | -31 | -39 | 0 | 0 | 9 | NaN | 437.48267 | 411.09125 | NaN | 521.6978 | 493.90964 |
| Fgg | VGPESDKYR | -292 | -300 | 1 | 0.001636916 | 9 | NaN | 389.32532 | 1585.924 | NaN | 182.72821 | NaN |
| Tfdp1 | DNHILPNESAYDQK | -150 | -163 | 0 | 0 | 14 | NaN | 1469.4347 | 1482.4718 | 1275.5499 | 1179.939 | 1072.3354 |
| Tbcel | DNNLQDWTEIR | -183 | -193 | 0 | 0.000471409 | 11 | NaN | 452.62177 | NaN | NaN | NaN | NaN |
| Lrrc59 | VTGLHQQPLCTSVNTIYDN | -268 | -286 | 0 | 0.000712251 | 19 | NaN | 303.08572 | 579.5532 | 617.836 | 633.7405 | 656.4498 |
| Lrrc59 | LVNLQHLDLLNNR | -84 | -96 | 0 | 0 | 13 | NaN | 296.05396 | 241.56564 | 370.67 | 440.5125 | 281.4684 |
| Cdk7 | VPFLPGDSDLDQLTR | -210 | -224 | 0 | 0 | 15 | NaN | 962.2426 | 923.74506 | 736.0182 | 980.63696 | 261.73123 |
| Cdk7 | DNSLVLTPSHIK | -104 | -115 | 0 | 0.00025015 | 12 | NaN | 440.83673 | 614.9092 | 692.85315 | 1013.1261 | 731.4183 |
| Cdk7 | GLFLFNPCTR | -274 | -283 | 0 | 0.00111006 | 10 | NaN | 544.0697 | 693.7928 | 662.31726 | NaN | NaN |
| Ube2e1;Ube2e | DNWSPALTISK | (137);(151);(145) | (147);(161);(155) | 0 | 0 | 11 | NaN | 1632.6152 | NaN | NaN | NaN | 2049.3 |
| Ogfr | GELGLEHYQAPLVR | -211 | -224 | 0 | 0 | 14 | NaN | 3284.2114 | 3205.1143 | 2751.67 | 2898.3418 | 2145.0542 |
| Ogfr | GELGLEHYQAPLVR | -211 | -224 | 0 | 0 | 14 | NaN | 3284.2114 | 3205.1143 | 2751.67 | 2898.3418 | 2145.0542 |
| Ogfr | QEQVPGEPDPQGVSEVEK | -375 | -392 | 0 | 0.00111006 | 18 | NaN | 991.8298 | 1111.6658 | 1094.1633 | 1479.3525 | 1295.6962 |
| Ogfr | REFVWGPR | -272 | -279 | 1 | 0.00830926 | 8 | NaN | 1315.086 | NaN | NaN | NaN | NaN |
| Emg1 | VSVEYTEK | -208 | -215 | 0 | 0.000591124 | 8 | NaN | 634.8941 | NaN | 773.74713 | NaN | NaN |
| Khdrbs1 | KDDEENYLDLFSHK | -139 | -152 | 1 | 0.00025015 | 14 | NaN | 632.8101 | NaN | NaN | 609.5238 | 700.7851 |

| | | | | | | | | | | | | |
|---|---|---|---|---|---|---|---|---|---|---|---|---|
| Rap1gds1 | DQEVLLQTGR | -102 | -111 | 0 | 0.00025015 | 10 | NaN | 4865.4966 | 4917.2466 | NaN | 4805.603 | 5105.0454 |
| Rap1gds1 | IVIDHLR | -135 | -141 | 0 | 0.007987481 | 7 | NaN | 1288.0658 | NaN | NaN | NaN | 861.09106 |
| Prpf3 | DQTKPTPLILDEQGR | -249 | -263 | 1 | 0 | 15 | NaN | 713.05597 | 628.2344 | 473.58554 | 495.43958 | 603.9684 |
| Prpf3 | LGLTPPPEPK | -466 | -475 | 0 | 0.00025015 | 10 | NaN | 1236.2943 | 1450.1494 | 1562.898 | NaN | NaN |
| Prpf3 | EVFGDDSEISK | -93 | -103 | 0 | 0 | 11 | NaN | 1216.7418 | NaN | 554.0319 | 201.87666 | NaN |
| Eprs1 | EFLWQEGHSAF | -1166 | -1176 | 0 | 0.00025015 | 11 | NaN | 424.5375 | NaN | 286.9551 | 240.88336 | 221.72733 |
| Eprs1 | SLTGIEYKPVSAT | -944 | -956 | 1 | 0.000431655 | 13 | NaN | 1149.9177 | NaN | 1629.8401 | NaN | 945.4052 |
| Eprs1 | RWFGFLEAQ | -149 | -157 | 1 | 0.00450079 | 9 | NaN | 95.95115 | NaN | 722.77075 | 1003.0905 | NaN |
| Eprs1 | HYQVNFK | -223 | -229 | 0 | 0.004439092 | 7 | NaN | 1055.7291 | NaN | NaN | NaN | NaN |
| Xrcc5 | GDLDSGLDHLKPSFPQK | -179 | -195 | 1 | 0 | 17 | NaN | 1468.225 | 2748.4858 | NaN | 503.20328 | 531.0707 |
| Xrcc5 | TEKEEDHISIS | -570 | -580 | 1 | 0.001708008 | 11 | NaN | 8951.529 | NaN | NaN | NaN | NaN |
| Xrcc5 | DQYQNITVCR | -72 | -81 | 0 | 0 | 10 | NaN | 1824.8536 | NaN | NaN | NaN | NaN |
| Rbm25 | LKEFLEDYDDDRDDPK | -491 | -506 | 2 | 0.000471409 | 16 | NaN | 947.27295 | NaN | NaN | 395.9576 | 445.2575 |
| Rbm25 | REEPVEEEEEPEQKPC | -592 | -607 | 2 | 0.002068213 | 16 | NaN | 491.18958 | NaN | NaN | 519.1009 | 489.10928 |
| Rbm25 | NKFEDEDSDDVPR | -691 | -703 | 1 | 0.003299042 | 13 | NaN | 805.037 | NaN | NaN | NaN | NaN |
| Snrpa1 | DRELDLR | -21 | -27 | 1 | 0.000431655 | 7 | NaN | 2278.3381 | NaN | NaN | 1884.7063 | 2300.5627 |
| Snrpa1 | RSGPSDEGEEEIEDDTVTNGS | -235 | -255 | 1 | 0.00383302 | 21 | NaN | 519.5354 | NaN | NaN | 642.6718 | 594.98676 |
| Dkc1 | IMLPGLLR | -315 | -322 | 0 | 0.00025015 | 8 | NaN | 1109.5112 | 1921.6809 | 181.33559 | 1815.9822 | 1889.9415 |
| Dkc1 | DRKPLQEDDVAEIQ | -18 | -31 | 2 | 0.000550307 | 14 | NaN | 1361.6467 | 1236.393 | 1586.7736 | 1748.2717 | 1690.2512 |
| Dkc1 | TAHYTPLPCGSNPLK | -66 | -80 | 0 | 0 | 15 | NaN | 1121.5935 | 392.74286 | NaN | NaN | NaN |
| Nme2 | NIIHGSDSVESAEK | -115 | -128 | 0 | 0 | 14 | NaN | 7957.357 | 7483.6455 | 9902.234 | 9571.254 | 9740.7705 |
| Nme2 | NIIHGSDSVESA | -115 | -126 | 0 | 0.00111006 | 12 | NaN | 2074.0168 | NaN | 1646.7755 | NaN | 762.97644 |
| Nme1 | EHYTDLK | -50 | -56 | 0 | 0.000947459 | 7 | NaN | 2053.256 | NaN | NaN | NaN | 1800.146 |
| Farsb | KIDVPANR | -64 | -71 | 1 | 0 | 8 | NaN | 885.6594 | 933.77625 | 629.22034 | 3092.7458 | 1698.9764 |
| Farsb | YDLLCLEGLAR | -72 | -82 | 0 | 0 | 11 | NaN | 985.3762 | 1185.9324 | NaN | 1571.0457 | 1374.0464 |
| Farsb | GLELDEITSEK | -32 | -42 | 0 | 0 | 11 | NaN | 573.71155 | NaN | 272.343 | NaN | 1069.2538 |
| Try5;Try4 | NNMICVGFLEGGK | (181);(181) | (193);(193) | 0 | 0 | 13 | NaN | 348.39395 | 450.71207 | 1518.2925 | 1398.2175 | 1202.7169 |
| Cbx3 | LTWHSCPEDEAQ | -172 | -183 | 0 | 0.00111006 | 12 | NaN | 4077.8918 | 4042.5117 | 2235.2053 | 2091.5354 | 2025.8938 |
| Cbx3 | DSDEADLVLAK | -144 | -154 | 0 | 0 | 11 | NaN | 1866.0496 | NaN | 2516.092 | 1533.6631 | 1233.6941 |
| Nars1 | YGTCPHGGYGLGLER | -519 | -533 | 0 | 0 | 15 | NaN | 876.2028 | 835.1187 | 986.6543 | 825.6222 | 870.8419 |
| Nars1 | KEDGTFYEF | -418 | -426 | 1 | 0.001392869 | 9 | NaN | 1098.8889 | NaN | NaN | 1330.1613 | 1360.0892 |
| Nars1 | ELNPNFKPPK | -387 | -396 | 1 | 0.001636916 | 10 | NaN | 2308.6382 | 1228.8102 | NaN | NaN | NaN |
| Nars1 | KYGTCPHGGYGL | -518 | -529 | 1 | 0.002466863 | 12 | NaN | 1329.8508 | 1218.1385 | NaN | NaN | NaN |
| Nars1 | TDTINEPILLCR | -439 | -450 | 0 | 0.001392869 | 12 | NaN | 1105.6362 | NaN | 1212.9915 | NaN | NaN |
| Nars1 | INEPILLCR | -442 | -450 | 0 | 0.000550307 | 9 | NaN | 19996.822 | NaN | NaN | NaN | NaN |
| Ndufb7 | HEQHDWDYCEHLDYVK | -82 | -97 | 0 | 0 | 16 | NaN | 450.16705 | 394.0652 | NaN | 678.6016 | 470.5545 |
| Sash3 | EEGSASPTSPDCSLDSPGPEK | -105 | -125 | 0 | 0.002972566 | 21 | NaN | 544.49164 | NaN | 679.0766 | 123.998474 | 308.70236 |
| Usp24 | DSIPSEVDYETR | -1218 | -1229 | 0 | 0.005176406 | 12 | NaN | 721.3099 | NaN | NaN | NaN | NaN |
| Nup98 | NLNLDDIVHIR | -785 | -795 | 0 | 0 | 11 | NaN | 1356.4204 | 1299.1948 | NaN | NaN | NaN |
| Nup98 | TVPSPAPEVQLK | -1095 | -1106 | 0 | 0 | 12 | NaN | 1056.7802 | 1170.3739 | NaN | NaN | NaN |
| Nup98 | SKPVDENNQQDGEDDSLVSR | -638 | -657 | 1 | 0.004224523 | 20 | NaN | 431.42938 | NaN | NaN | NaN | 364.4753 |
| Gls | SSPSEILQELGK | -74 | -85 | 0 | 0.001023236 | 12 | NaN | 530.4016 | NaN | NaN | NaN | 624.41113 |
| Trnt1 | IVDRPGDHDHETLEAIAE | -208 | -225 | 1 | 0.000230707 | 18 | NaN | 613.7094 | 3820.746 | 397.25574 | 286.24524 | 252.98422 |
| Trnt1 | DSSEPLKPYQDFVIDSR | -338 | -354 | 1 | 0.000591124 | 17 | NaN | 754.632 | NaN | NaN | 396.5072 | 1037.7372 |
| Trnt1 | IVDRPGDHDHETLE | -208 | -221 | 1 | 0.00659934 | 14 | NaN | 639.4705 | NaN | NaN | NaN | NaN |
| Sdcbp | FIFEHIIK | -274 | -281 | 0 | 0.002595029 | 8 | NaN | 718.28973 | NaN | NaN | NaN | NaN |
| Smchd1 | GLVPIECFNR | -499 | -508 | 0 | 0.000550307 | 10 | NaN | 1782.0515 | 148.21515 | 1491.8893 | NaN | 1578.5511 |
| Smchd1 | IDFLPHYDTLVK | -115 | -126 | 0 | 0 | 12 | NaN | 847.67285 | NaN | 509.1979 | 1043.6748 | 1055.4711 |
| Smchd1 | IDFLPHYDTL | -115 | -124 | 0 | 0.000230707 | 10 | NaN | 549.9305 | 286.9331 | 586.0548 | 766.7098 | NaN |
| Smchd1 | GNDKEDGCFYFR | -1439 | -1450 | 1 | 0 | 12 | NaN | 61.285324 | NaN | 1151.9979 | 1699.3008 | NaN |
| Hnrnpa3 | IFVGGIKEDTEEYNLR | -128 | -143 | 1 | 0 | 16 | NaN | 11845.832 | 11417.129 | 9006.77 | 15392.79 | 9064.033 |
| Hnrnpa3 | VTFDDHDTVDK | -172 | -182 | 0 | 0 | 11 | NaN | 8917.49 | 8890.168 | 10768.099 | 11152.848 | 11288.888 |
| Hnrnpa3 | EDTEEYNLR | -135 | -143 | 0 | 0 | 9 | NaN | 6150.4956 | 4561.777 | 166.79515 | 196.96744 | 3209.8523 |
| Hnrnpa3 | RGEEGHDPKEPEQLR | -21 | -35 | 2 | 0 | 15 | NaN | 1198.0055 | 1185.4208 | NaN | 685.24774 | 870.55597 |
| Hnrnpa3 | TFDDHDTVDK | -173 | -182 | 0 | 0.000230707 | 10 | NaN | 591.4686 | 673.07904 | 756.06604 | 1091.8203 | 317.11584 |
| Hnrnpa3 | IKEDTEEYNLR | -133 | -143 | 1 | 0.00025015 | 11 | NaN | 2620.1814 | NaN | NaN | 797.3838 | 2813.3262 |
| Hnrnpa3 | GEEGHDPKEPEQL | -22 | -34 | 1 | 0 | 13 | NaN | 659.57416 | NaN | NaN | 1980.5933 | NaN |
| Nusap1 | DTAEVPSLPEQRPEDGNAA | -133 | -151 | 1 | 0.00025015 | 19 | NaN | 743.42834 | 355.30026 | 1375.5703 | 253.80624 | 1570.1826 |
| Nusap1 | GTDSSNFQNQENQENQDPR | -114 | -132 | 0 | 0 | 19 | NaN | 251.34595 | 222.85658 | NaN | NaN | NaN |
| Nusap1 | TVPSAEELDSFK | -2 | -13 | 0 | 0 | 12 | NaN | 1685.4281 | NaN | NaN | NaN | NaN |
| Get3 | LEETLPVIR | -222 | -230 | 0 | 0 | 9 | NaN | 590.5274 | 160.19365 | NaN | 479.65933 | 519.5088 |
| Nceh1 | DTDFDGVEVR | -84 | -93 | 0 | 0.00635324 | 10 | NaN | 1989.0917 | NaN | NaN | NaN | NaN |
| Mtdh | EEAAPPTPAPDDLAQLK | -90 | -106 | 0 | 0.000431655 | 17 | NaN | 949.97845 | 922.1096 | 934.4955 | 1163.8387 | NaN |
| Nhlrc2 | DTENHLIR | -295 | -302 | 0 | 0.000230707 | 8 | NaN | 1523.3754 | 1537.4958 | 671.7964 | NaN | NaN |
| Hmgn5 | EQINEDTEEDGGEKK | -99 | -113 | 1 | 0 | 15 | NaN | 377.23267 | 462.87357 | NaN | NaN | NaN |
| Unc13d | ELEKPVFCLK | -104 | -113 | 1 | 0 | 10 | NaN | 3566.6482 | NaN | NaN | NaN | NaN |
| Unc13d | DTLVPLGELTK | -614 | -624 | 0 | 0.003620005 | 11 | NaN | 584.00684 | NaN | NaN | NaN | NaN |
| Xpnpep1 | TLSLDEVYLIDSGAQYK | -405 | -421 | 0 | 0 | 17 | NaN | 563.06915 | 469.0181 | NaN | NaN | NaN |
| Inpp5d | FWLGDLNYR | -586 | -594 | 0 | 0 | 9 | NaN | 1473.0286 | 1206.0594 | 1823.8922 | 1060.5671 | 1104.3633 |
| Inpp5d | TLPSLESLQR | -230 | -239 | 0 | 0.000298805 | 10 | NaN | 3278.5654 | NaN | 1104.9814 | NaN | 3466.8481 |
| Mrpl40 | DTSLFPFEK | -165 | -173 | 0 | 0.00051306 | 9 | NaN | 271.9824 | 342.95697 | NaN | NaN | NaN |
| Atp6v1f | KDTTINEIEDTFR | -41 | -53 | 1 | 0.00272025 | 13 | NaN | 1054.5956 | NaN | NaN | NaN | NaN |
| Eif6 | NTTDQELQHIR | -75 | -85 | 0 | 0 | 11 | NaN | 9923.213 | 10543.408 | 9284.844 | NaN | 10727.364 |
| Mcm2 | ISHLPLVEELR | -285 | -295 | 0 | 0 | 11 | NaN | 1103.2738 | 1046.7871 | 973.7197 | 1013.03485 | 1004.69525 |
| Mcm2 | TYGVEPLPQEVLK | -709 | -721 | 0 | 0 | 13 | NaN | 1198.9938 | NaN | 980.9663 | 1058.9907 | 836.23975 |
| Mcm2 | VDSHGHNVFK | -207 | -216 | 0 | 0.00025015 | 10 | NaN | 692.87537 | NaN | 1245.254 | 1150.3845 | 1007.6024 |
| Mcm2 | LCGDPGTAK | -521 | -529 | 0 | 0 | 9 | NaN | 1105.683 | NaN | 1134.7378 | NaN | 1286.1998 |
| Mcm2 | VIAAANPIGGR | -626 | -636 | 0 | 0.00025015 | 11 | NaN | 462.6256 | NaN | NaN | 351.82407 | 401.75592 |

| | | | | | | | | | | | | |
|---|---|---|---|---|---|---|---|---|---|---|---|---|
| Ggps1 | VLTLDHPDAVK | -103 | -113 | 0 | 0 | 11 | NaN | 1345.7214 | 1291.6173 | 2045.612 | 2079.023 | 1695.3802 |
| Mcmbp | LSGCPQNSTFTEHLYR | -393 | -408 | 0 | 0 | 16 | NaN | 400.53735 | 462.45386 | 398.2369 | 457.03754 | 399.79486 |
| Mcmbp | KVIEYFK | -31 | -37 | 1 | 0.001023236 | 7 | NaN | 906.8031 | 682.8247 | 330.57043 | 838.3443 | 734.4199 |
| Mcmbp | LQHINPLLPTCLNK | -315 | -328 | 0 | 0 | 14 | NaN | 378.917 | 425.344 | NaN | NaN | 523.24915 |
| Mcmbp | RLETEASSGQQL | -205 | -216 | 1 | 0.007905138 | 12 | NaN | 835.56024 | 1197.9282 | NaN | NaN | NaN |
| Mcmbp | HINPLLPTCLNK | -317 | -328 | 0 | 0.002566359 | 12 | NaN | 5043.48 | NaN | NaN | NaN | NaN |
| G6pdx | EKPQPIPYVY | -475 | -484 | 1 | 0.001868731 | 10 | NaN | 1399.4912 | 1363.0616 | 1627.9694 | 1612.9591 | 1824.6683 |
| G6pdx | NSYVAGQYDDAASYK | -105 | -119 | 0 | 0 | 15 | NaN | 329.8306 | NaN | 441.03156 | 431.44995 | 464.11603 |
| G6pdx | YVAGQYDDAASYK | -107 | -119 | 0 | 0 | 13 | NaN | 760.3549 | NaN | NaN | 255.40671 | NaN |
| Pdia6 | GLYSSSDDVIELTPSNFNR | -19 | -37 | 0 | 0.000471409 | 19 | NaN | 455.46112 | 501.1811 | 619.53735 | 852.25616 | 703.60736 |
| Pdia6 | GSFSEQGINEFLR | -374 | -386 | 0 | 0 | 13 | NaN | 1264.3202 | NaN | 1411.7153 | 1456.3169 | 1448.605 |
| Pdia6 | LYSSSDDVIELTPSNFNR | -20 | -37 | 0 | 0 | 18 | NaN | 932.4575 | NaN | NaN | NaN | 903.2252 |
| Pdia6 | DVVELTDDTFDK | -161 | -172 | 0 | 0.002566359 | 12 | NaN | 1224.0797 | NaN | NaN | NaN | NaN |
| Rpl27 | DVFRDPALKR | -99 | -108 | 2 | 0 | 10 | NaN | 783.83435 | 740.5433 | 499.96036 | 543.20337 | 393.17935 |
| Rpl27 | YSVDIPLDKT | -85 | -94 | 1 | 0.000431655 | 10 | NaN | 692.701 | NaN | 243.43741 | 1315.8042 | 902.55225 |
| Metap2 | NFDVGHVPIR | -386 | -395 | 0 | 0 | 10 | NaN | 514.576 | NaN | 700.9366 | 870.1439 | NaN |
| Evl | SPLQSQPHSR | -352 | -361 | 0 | 0 | 10 | NaN | 336.18933 | 560.38544 | 213.33478 | 83.29876 | NaN |
| Evl | EDESQTEDPSTSPSPGTR | -291 | -308 | 0 | 0.00025015 | 18 | NaN | 303.77063 | NaN | 245.75104 | 323.55472 | 343.45932 |
| Evl | LQDQQVVINY | -54 | -63 | 0 | 0.001941082 | 10 | NaN | 498.80478 | 533.2887 | NaN | NaN | NaN |
| Evl | WVPIKPGQQGFSR | -23 | -35 | 1 | 0 | 13 | NaN | 946.5951 | NaN | NaN | NaN | NaN |
| Pold1 | WGEPEPFLR | -342 | -350 | 0 | 0.001776473 | 9 | NaN | 209.03215 | 3054.9333 | 2127.1204 | NaN | NaN |
| Pold1 | NFDLPYLISR | -398 | -407 | 0 | 0 | 10 | NaN | 245.17888 | 2636.3901 | NaN | NaN | NaN |
| Mrpl50 | DVLDFYNVPVQDK | -126 | -138 | 0 | 0 | 13 | NaN | 312.10925 | 2090.873 | 1989.9357 | NaN | 2118.344 |
| Mcm7 | IAPEIYGHEDVK | -340 | -351 | 0 | 0 | 12 | NaN | 728.35846 | 1026.4712 | 735.81866 | 676.56647 | 746.41174 |
| Mcm7 | WLIQDRPDR | -519 | -527 | 1 | 0.002199523 | 9 | NaN | 603.76404 | NaN | 1540.0863 | 2329.6501 | 1906.7538 |
| Mcm7 | DVLDVYIEHR | -97 | -106 | 0 | 0 | 10 | NaN | 763.1376 | NaN | NaN | 816.7687 | 899.0932 |
| Mcm7 | SQLLSYIDR | -388 | -396 | 0 | 0 | 9 | NaN | 345.00742 | 475.4538 | NaN | NaN | NaN |
| Asl | DVLFPGYTHLQ | -152 | -162 | 0 | 0.00025015 | 11 | NaN | 1673.6443 | 144.03215 | 442.61356 | 1703.2888 | 1719.013 |
| Asl | NPDSLELIR | -289 | -297 | 0 | 0.000230707 | 9 | NaN | 670.9377 | 1455.3586 | 327.27246 | NaN | 2009.6941 |
| Asl | LHPNDEDIHTA | -81 | -91 | 0 | 0.001708008 | 11 | NaN | 8221.708 | NaN | NaN | 1686.4231 | NaN |
| Asl | HLWNVDVQGSK | -33 | -43 | 0 | 0 | 11 | NaN | 783.11237 | 477.87433 | NaN | NaN | NaN |
| Cluh | LGYEEHIPGQTR | -344 | -355 | 0 | 0 | 12 | NaN | 489.49298 | 536.81775 | 424.55994 | 464.1532 | 518.9594 |
| Cluh | DVLNIFPVVK | -962 | -971 | 0 | 0 | 10 | NaN | 154.8957 | NaN | NaN | NaN | NaN |
| Epb41 | SLAAEAENSQHQQQK | -6 | -20 | 0 | 0 | 15 | NaN | 261.85782 | 525.98505 | NaN | NaN | NaN |
| Ppp2r5e | DVPTSEQPELFLK | -53 | -65 | 0 | 0 | 13 | NaN | 1918.8982 | 1816.9835 | NaN | NaN | NaN |
| Letm1 | SALDDNKDGNINIDDLVK | -672 | -689 | 1 | 0 | 18 | NaN | 630.50696 | NaN | 625.4645 | 642.00446 | 504.78552 |
| Letm1 | HYYHGFR | -162 | -168 | 0 | 0.000298805 | 7 | NaN | 431.4432 | 485.82068 | NaN | 391.911 | NaN |
| Letm1 | YLPDTLSPADQLK | -421 | -433 | 0 | 0 | 13 | NaN | 305.2909 | 425.25366 | 518.8843 | NaN | NaN |
| Dnaja2 | EKEHEVFQR | -254 | -262 | 1 | 0.00111006 | 9 | NaN | 765.0142 | 626.79395 | NaN | NaN | 627.093 |
| Dnaja2 | EGSGGGGGMDDIFSH | -76 | -90 | 0 | 0.000947459 | 15 | NaN | 314.72675 | NaN | 313.2426 | NaN | 243.3084 |
| Hk2 | STPDGTEHGEFLAL | -70 | -83 | 0 | 0 | 14 | NaN | 1274.7938 | 1445.3274 | 2968.9272 | 1943.6091 | 1767.5104 |
| Hk2 | VDELSLNPGK | -729 | -738 | 0 | 0 | 10 | NaN | 897.53375 | 1035.3884 | 721.2471 | 739.25696 | 837.5327 |
| Hk2 | KLSHEQLLEVK | -478 | -488 | 1 | 0 | 11 | NaN | 698.1635 | 708.679 | 592.698 | 921.6461 | 915.72205 |
| Hk2 | KLSHEQLLEVK | -478 | -488 | 1 | 0 | 11 | NaN | 698.1635 | 708.679 | 592.698 | 921.6461 | 915.72205 |
| Hk2 | GLESTCDDSIIVK | -808 | -820 | 0 | 0 | 13 | NaN | 921.16925 | NaN | 1957.8235 | 1240.7601 | 605.99384 |
| Hk2 | TELNQNQVQK | -12 | -21 | 0 | 0 | 10 | NaN | 856.3015 | NaN | 968.5817 | 1350.565 | 1200.2992 |
| Hk2 | FTELNQNQVQK | -11 | -21 | 0 | 0 | 11 | NaN | 665.27234 | NaN | NaN | 1040.4833 | 593.65906 |
| Hk2 | EWGAFGDDGTLNDIR | -260 | -274 | 0 | 0 | 15 | NaN | 691.9133 | 427.8545 | NaN | NaN | NaN |
| Hk2 | LSDETLLEISR | -31 | -41 | 0 | 0.000431655 | 11 | NaN | 582.1314 | 345.9177 | NaN | NaN | NaN |
| Hk2 | STPDGTEHGEFL | -70 | -81 | 0 | 0.000394361 | 12 | NaN | 568.4395 | 660.0025 | NaN | NaN | NaN |
| Hk2 | GSGTQLFDHIAECL | -121 | -134 | 0 | 0.001392869 | 14 | NaN | 911.3682 | NaN | NaN | NaN | NaN |
| Dctpp1 | FSPEPTLEDIR | -25 | -35 | 0 | 0 | 11 | NaN | 853.1191 | 760.03253 | NaN | 352.58725 | NaN |
| Pld3 | SLLQSGAQVR | -177 | -186 | 0 | 0 | 10 | NaN | 1444.0884 | NaN | NaN | 603.5701 | NaN |
| Ddx17 | DWVLNEFR | -379 | -386 | 0 | 0 | 8 | NaN | 629.3445 | 685.3144 | 589.92096 | 789.5497 | 787.24066 |
| Ddx17 | FYVEHPEVAR | -55 | -64 | 0 | 0 | 10 | NaN | 444.37857 | 942.2484 | 887.57367 | 1543.7184 | 1494.7167 |
| Ddx17 | GGDVCPKPVFAF | -83 | -94 | 1 | 0 | 12 | NaN | 846.95 | NaN | 631.71405 | 388.47495 | 1044.1602 |
| Ddx17 | TAYQYPPPPPPPPSR | -634 | -649 | 0 | 0.004439092 | 16 | NaN | 1123.9374 | 1556.7798 | NaN | NaN | NaN |
| Ranbp2 | KLPVPLESVK | -734 | -743 | 1 | 0 | 10 | NaN | 413.83313 | 450.85675 | 627.0359 | 691.8346 | 591.1244 |
| Ranbp2 | GENKPVLFGF | -2667 | -2676 | 1 | 0.001392869 | 10 | NaN | 387.99695 | 447.2589 | 395.4756 | NaN | 482.09937 |
| Cd2ap | DYDAVHDDELTIR | -9 | -21 | 0 | 0.000394361 | 13 | NaN | 518.8331 | 652.4439 | 617.46826 | NaN | 465.11847 |
| Cd2ap | TLFPYTGTNEDELTFR | -276 | -291 | 0 | 0.001708008 | 16 | NaN | 768.1992 | NaN | NaN | NaN | NaN |
| Rras2 | VDQAFHELVR | -164 | -173 | 0 | 0 | 10 | NaN | 1764.8289 | NaN | 1750.4653 | 1115.3037 | 1752.0527 |
| Morf4l1 | SASDYEVAPPEYHR | -346 | -359 | 0 | 0 | 14 | NaN | 2027.8945 | 1695.6667 | 2025.5911 | 2360.6545 | 2488.1335 |
| Morf4l1 | LVDDWDLITR | -207 | -216 | 0 | 0 | 10 | NaN | 1000.90753 | 978.1808 | 573.34595 | 892.90424 | 103.743614 |
| Morf4l1 | VYGAPHLLR | -296 | -304 | 0 | 0 | 9 | NaN | 874.87665 | 516.8607 | 908.9842 | 791.76385 | 554.5019 |
| Hnrnpa1 | ESPKEPEQLR | -5 | -14 | 1 | 0.000471409 | 10 | NaN | 2155.8284 | 2459.0251 | 2937.2878 | 4503.788 | 4353.0767 |
| Npepps | NWEELHNR | -835 | -842 | 0 | 0 | 8 | NaN | 6395.953 | 5763.025 | 4924.067 | NaN | NaN |
| Npepps | LNLGTVGFYR | -582 | -591 | 0 | 0.009388464 | 10 | NaN | 6722.479 | NaN | NaN | NaN | NaN |
| F2 | SWGEGCDR | -586 | -593 | 0 | 0.001392869 | 8 | NaN | 116.78416 | NaN | NaN | 457.05798 | NaN |
| Hcls1 | DYKGEVEK | -139 | -146 | 1 | 0.002595029 | 8 | NaN | 841.4821 | 656.74335 | 729.20935 | 762.4758 | 1867.9307 |
| Hcls1 | NEMEAPTTAYK | -213 | -223 | 0 | 0 | 11 | NaN | 779.2509 | 609.32104 | 797.18646 | NaN | NaN |
| Hcls1 | TEHINIH | -49 | -55 | 0 | 0.00555405 | 7 | NaN | 930.8366 | NaN | NaN | NaN | 370.45724 |
| Rps10 | RGEADRDTYR | -119 | -128 | 2 | 0.002910424 | 10 | NaN | 294.9543 | 328.23624 | 307.3634 | 393.08362 | 461.12973 |
| Rps10 | GEADRDTYR | -120 | -128 | 1 | 0.006260252 | 9 | NaN | 710.8614 | NaN | NaN | NaN | 784.94543 |
| Emd | SSSPSSWLTR | -195 | -204 | 0 | 0.00078661 | 10 | NaN | 1535.7407 | 1627.9501 | NaN | NaN | NaN |
| Cox4i1 | EKADWSSLSR | -66 | -75 | 1 | 0.00078661 | 10 | NaN | 339.21887 | NaN | NaN | NaN | NaN |
| Etfdh | NLSIYDGPEQR | -548 | -558 | 0 | 0 | 11 | NaN | 1296.0061 | NaN | 1579.4374 | 1299.127 | 986.61395 |
| Etfdh | SGACLDPAAFK | -114 | -124 | 0 | 0 | 11 | NaN | 1200.724 | 580.5645 | 495.40198 | 453.4163 | NaN |

| | | | | | | | | | | | | |
|---|---|---|---|---|---|---|---|---|---|---|---|---|
| Aatf | LAPVSLLDTDKR | -58 | -69 | 1 | 0 | 12 | NaN | 1785.059 | 1216.109 | NaN | NaN | NaN |
| Aatf | EADPEADPEEATR | -21 | -33 | 0 | 0 | 13 | NaN | 1098.3673 | NaN | NaN | NaN | NaN |
| Ccar1 | EAEDEDDDREEEEVKR | -874 | -889 | 2 | 0 | 16 | NaN | 490.35532 | 552.36865 | 462.0264 | 572.80316 | 520.0518 |
| Ccar1 | YILPDEPAIIVHPN | -718 | -731 | 0 | 0.000712251 | 14 | NaN | 1963.7284 | NaN | 2751.872 | 756.3938 | 3387.0623 |
| Psmd7 | ISVEEVHDDGTPTSK | -139 | -153 | 0 | 0 | 15 | NaN | 960.897 | NaN | 873.79614 | 920.88074 | 908.4919 |
| Psmd7 | LPINHQIIY | -220 | -228 | 0 | 0.002329279 | 9 | NaN | 652.0163 | 528.08655 | 529.8788 | 459.28836 | NaN |
| Psmd7 | LPINHQIIYQ | -220 | -229 | 0 | 0.00051306 | 10 | NaN | 1049.3077 | NaN | NaN | NaN | 1062.8799 |
| Noc2l | EAFLGPILK | -327 | -335 | 0 | 0.004255025 | 9 | NaN | 1446.6078 | NaN | NaN | NaN | NaN |
| Myl12b | GDRFTDEEVDELYR | -131 | -144 | 1 | 0.00051306 | 14 | NaN | 1496.7681 | 1338.3827 | 683.39795 | 1853.5492 | 1878.6201 |
| Edc4 | SSHNTWPVDVSQIK | -274 | -287 | 0 | 0.002697088 | 14 | NaN | 1157.7043 | NaN | 1071.9602 | 1222.3242 | 999.2487 |
| Edc4 | EAFQSVVLPAFEK | -1121 | -1133 | 0 | 0 | 13 | NaN | 463.68372 | NaN | 734.08105 | 749.34534 | 758.2039 |
| Hopx | RSEGLPSECR | -60 | -69 | 1 | 0.00111006 | 10 | NaN | 2237.277 | 953.3297 | 2847.0693 | 837.53467 | 3642.1885 |
| Gale | EAGYSPVVIDNFHNAIR | -23 | -39 | 0 | 0 | 17 | NaN | 2301.3413 | 2449.4429 | 1146.8098 | NaN | NaN |
| Vasp | PAQSEPVR | -295 | -302 | 0 | 0.000431655 | 8 | NaN | 461.45657 | 327.83673 | NaN | 1132.2968 | 701.11804 |
| Vasp | QEEASGGPLAPK | -237 | -248 | 0 | 0 | 12 | NaN | 803.23535 | 720.7645 | NaN | NaN | 1849.387 |
| Alg5 | EVIVVDDGSEDQTSK | -104 | -118 | 0 | 0 | 15 | NaN | 9302.532 | NaN | NaN | NaN | NaN |
| Selenoh | EAPELPVQVNPSKPR | -55 | -69 | 1 | 0 | 15 | NaN | 1304.917 | 876.3596 | 1296.8849 | NaN | NaN |
| Eif3f | EAPNPIHLTVDTGLQHGR | -197 | -214 | 0 | 0 | 18 | NaN | 922.62445 | 827.33057 | 887.3608 | 1136.7792 | 202.55154 |
| Eif3f | IGVDLIMK | -251 | -258 | 0 | 0 | 8 | NaN | 1158.51 | 1110.0912 | 713.8633 | 1046.1558 | 1098.319 |
| Sh3kbp1 | RGVFPDNFVK | -359 | -368 | 1 | 0.006788754 | 10 | NaN | 1260.9178 | NaN | NaN | NaN | 2131.015 |
| Sh3kbp1 | QLLSELDEEK | -680 | -689 | 0 | 0 | 10 | NaN | 1035.1392 | 2323.7168 | NaN | NaN | NaN |
| Mtpn | GADINAPDKHHITPL | -58 | -72 | 1 | 0.00025015 | 15 | NaN | 432.2543 | 344.154 | 314.36108 | 529.60736 | 482.44122 |
| Mtpn | VYEGHVSCVK | -76 | -85 | 0 | 0 | 10 | NaN | 11179.197 | NaN | 1216.2992 | 8486.931 | 5655.39 |
| Mtpn | EATDNQAIK | -106 | -114 | 0 | 0.005176406 | 9 | NaN | 383.59555 | NaN | NaN | NaN | NaN |
| Eef1b | EAVSGPPPADLCHALR | -39 | -54 | 0 | 0 | 16 | NaN | 856.9081 | NaN | 2159.3767 | 2549.183 | 2401.7803 |
| Eef1b | VFEAVSGPPPADLCHALR | -37 | -54 | 0 | 0 | 18 | NaN | 594.6281 | 812.5052 | 272.44077 | 895.3797 | 992.47534 |
| Stt3a | IHEFDPYFNYR | -45 | -55 | 0 | 0 | 11 | NaN | 396.0207 | NaN | NaN | NaN | 223.86775 |
| Rpl23 | YFEDNAGVIVNNK | -97 | -109 | 0 | 0 | 13 | NaN | 1390.9421 | 336.89276 | 2241.5542 | 2604.4407 | 2457.7388 |
| Rpl23 | ISLGLPVGAVINCADNTGAK | -16 | -35 | 0 | 0.00025015 | 20 | NaN | 657.71014 | 277.2322 | 402.31686 | 477.86453 | 500.559 |
| Elac2 | VVHIAPESVLIDSR | -336 | -349 | 0 | 0 | 14 | NaN | 437.75827 | 525.6117 | NaN | 673.26666 | 509.7263 |
| Elac2 | ECPDEGFILPICENDTFK | -306 | -323 | 0 | 0.00025015 | 18 | NaN | 259.14395 | 326.92624 | 234.30139 | 330.36758 | 236.659 |
| Ddx47 | YDVELFQR | -370 | -377 | 0 | 0.00025015 | 8 | NaN | 394.7015 | NaN | NaN | 853.6705 | 917.8397 |
| Ephx1 | EDESIRPF | -44 | -51 | 1 | 0 | 8 | NaN | 664.92944 | NaN | NaN | NaN | NaN |
| Ola1 | KFNTPQQPK | -386 | -394 | 1 | 0.000989247 | 9 | NaN | 318.63992 | 507.6605 | NaN | 652.0885 | 321.5923 |
| Ola1 | GGDGIKPPPIIGR | -6 | -18 | 1 | 0 | 13 | NaN | 2110.3208 | 939.719 | 2208.7107 | NaN | NaN |
| Ola1 | NFPFCTIDPNESR | -51 | -63 | 0 | 0 | 13 | NaN | 1284.9874 | 1152.3031 | NaN | NaN | NaN |
| Rad23a | EDKSPSEESTTTTSPESISGSVPSSGSSGR | -120 | -149 | 1 | 0.00025015 | 30 | NaN | 387.7102 | 338.03986 | NaN | 332.68683 | NaN |
| Rad23a | ILSDDVPIK | -54 | -62 | 0 | 0.000431655 | 9 | NaN | 548.22906 | NaN | NaN | NaN | NaN |
| Pole | EDLDLPNHLVGLK | -137 | -149 | 0 | 0.00025015 | 13 | NaN | 389.14703 | 387.10706 | NaN | NaN | NaN |
| Cyb5r1 | EDLEELQAQYPNR | -224 | -236 | 0 | 0 | 13 | NaN | 517.2394 | 495.4452 | 426.96304 | 590.14453 | NaN |
| Ssb | FSGDLDDQTCR | -235 | -245 | 0 | 0 | 11 | NaN | 2023.5092 | 2326.7231 | 2411.2817 | 71.26615 | 66.60305 |
| Ssb | FKEDYFAK | -184 | -191 | 1 | 0.000591124 | 8 | NaN | 1784.571 | 2007.081 | 1456.3386 | 3932.8157 | 3878.6865 |
| Dock10 | FHVDLNHPAVR | -442 | -452 | 0 | 0 | 11 | NaN | 314.49747 | 245.63095 | NaN | 336.84995 | 365.70212 |
| Dock10 | HATSVDTSFSK | -1254 | -1264 | 0 | 0 | 11 | NaN | 739.4716 | NaN | NaN | 445.28906 | 944.6766 |
| Dock10 | FVFETPFTLSGK | -1977 | -1988 | 0 | 0.002099344 | 12 | NaN | 310.1526 | 411.99915 | NaN | NaN | 398.8137 |
| Dock10 | SDLYYDIHR | -1850 | -1858 | 0 | 0 | 9 | NaN | 730.0906 | NaN | NaN | 1687.1195 | NaN |
| Rer1 | SEGDSVGDSVHGKPSVVYR | -2 | -20 | 1 | 0 | 19 | NaN | 263.2467 | 580.70825 | 637.5675 | 693.45465 | 886.89166 |
| Rer1 | EDSDDGPSLPTK | -93 | -104 | 0 | 0 | 12 | NaN | 473.94113 | NaN | NaN | 2058.5178 | NaN |
| Ly9 | QSVCDPDLPYTCK | -211 | -223 | 0 | 0.00025015 | 13 | NaN | 351.98053 | 277.23697 | 185.31606 | NaN | 382.3819 |
| Ly9 | EDSNTIYCSVQKPK | -619 | -632 | 1 | 0.000230707 | 14 | NaN | 297.39764 | NaN | 381.518 | 252.01675 | 340.15298 |
| Septin6 | TVPLAGHVGFD | -16 | -26 | 0 | 0.000431655 | 11 | NaN | 1009.56354 | NaN | 824.0646 | 711.7617 | 673.11224 |
| Mob1b;Mob1 | HAEATLGSGNLR | (31);(31) | (42);(42) | 0 | 0.005493574 | 12 | NaN | 4964.2754 | 4715.025 | NaN | NaN | 5673.2183 |
| Atp1b3 | SFHQSLAEWK | -8 | -17 | 0 | 0 | 10 | NaN | 2029.7346 | 6175.273 | NaN | 3156.7817 | 3908.7754 |
| Atp1b3 | VKFDSGLNKK | -234 | -243 | 2 | 0.00450079 | 10 | NaN | 662.28766 | 176.05528 | NaN | NaN | NaN |
| Uqcrb | RLPEDLYNDR | -50 | -59 | 1 | 0.000550307 | 10 | NaN | 4961.8716 | 4951.1646 | 5580.8345 | 4298.258 | 4536.237 |
| Snrpd2 | EEEEFNTGPLSVLTQ | -20 | -34 | 0 | 0.001846793 | 15 | NaN | 480.51096 | 524.9424 | 765.5832 | 426.91748 | 828.0022 |
| Psmc2 | KIEFSLPDLEGR | -340 | -351 | 1 | 0 | 12 | NaN | 1319.7913 | 1332.574 | 3738.502 | 1945.9648 | 2032.6671 |
| Psmc2 | IEFSLPDLEGR | -341 | -351 | 0 | 0 | 11 | NaN | 4346.921 | 4000.6074 | 3912.1467 | 886.7261 | 3956.3245 |
| Psmc2 | EEEKDDKPIR | -14 | -23 | 2 | 0.004957095 | 10 | NaN | 624.431 | NaN | NaN | 529.4313 | NaN |
| Tpx2 | VPIKDEEEEKPVVIK | -497 | -511 | 2 | 0.000230707 | 15 | NaN | 550.7317 | 604.67505 | 695.8836 | 622.43524 | 656.8065 |
| Tpx2 | EEEQQKEELAR | -695 | -705 | 1 | 0.00025015 | 11 | NaN | 534.38763 | 664.3917 | NaN | 725.6211 | 717.55414 |
| U2af2 | FTSVFDCQK | -436 | -444 | 0 | 0.000394361 | 9 | NaN | 672.50433 | 721.77374 | NaN | NaN | 335.3821 |
| Mark2 | EEIQDSLVGQR | -340 | -350 | 0 | 0 | 11 | NaN | 2504.7246 | NaN | NaN | NaN | NaN |
| Casp8 | TGVNYDVSNKDDR | -446 | -458 | 1 | 0 | 13 | NaN | 450.78204 | 554.8957 | 614.5598 | 818.15497 | 743.7162 |
| Casp8 | GMLEEGNLSFLK | -52 | -63 | 0 | 0 | 12 | NaN | 590.6197 | NaN | 1550.2325 | 1749.5063 | 440.2389 |
| Casp8 | KLSEEVSELELR | -107 | -118 | 1 | 0 | 12 | NaN | 466.36002 | NaN | 890.1314 | 1002.4552 | 1150.636 |
| Casp8 | EELGSEDLAALK | -12 | -23 | 0 | 0 | 12 | NaN | 661.32025 | 826.82153 | 474.28076 | NaN | NaN |
| Cdc123 | EELTSENNLR | -262 | -271 | 0 | 0.000627943 | 10 | NaN | 580.0786 | 743.91974 | NaN | NaN | 514.2389 |
| Cdc123 | EGDAQEQDSPAFR | -276 | -288 | 0 | 0.004255025 | 13 | NaN | 579.8535 | NaN | NaN | NaN | NaN |
| Dek | KPPTDEELKETVK | -323 | -335 | 2 | 0 | 13 | NaN | 1184.3807 | 1317.5503 | 1273.15 | 1538.8727 | 1319.4331 |
| Dek | LKKPPTDEELK | -321 | -331 | 2 | 0 | 11 | NaN | 849.99146 | 1428.0332 | NaN | NaN | 1005.57623 |
| Dek | LLADANLEEVTMK | -337 | -349 | 0 | 0 | 13 | NaN | 1223.3806 | 342.54712 | NaN | NaN | NaN |
| Dek | EESEEEEEDDEDDDEEDEEEEKEK | -31 | -54 | 1 | 0.001092529 | 24 | NaN | 194.02565 | NaN | NaN | NaN | 256.48245 |
| Cct7 | VDETIKNPR | -516 | -524 | 1 | 0 | 9 | NaN | 2018.2196 | 628.3085 | 2061.5012 | 2621.8687 | NaN |
| Cct7 | VQGGALEESQLVAGVAFK | -200 | -217 | 0 | 0 | 18 | NaN | 461.08286 | 419.19586 | NaN | 319.6115 | 231.74893 |
| Cct7 | SVDETIKNPR | -515 | -524 | 1 | 0 | 10 | NaN | 1942.0232 | NaN | 978.79016 | 1750.2004 | NaN |
| Rcsd1 | GVEGTPWSAEKPR | -227 | -239 | 1 | 0 | 13 | NaN | 1131.1907 | 1152.228 | NaN | 1317.0001 | 1253.9622 |
| Rcsd1 | EETTPGEHTDTGK | -300 | -312 | 0 | 0 | 13 | NaN | 281.86282 | 248.78024 | NaN | 323.76904 | 273.24323 |

| | | | | | | | | | | | | |
|---|---|---|---|---|---|---|---|---|---|---|---|---|
| Ufc1 | NSDNDWFR | -48 | -55 | 0 | 0.00111006 | 8 | NaN | 645.12164 | 704.2993 | 733.6085 | 861.5544 | 848.31335 |
| Ufc1 | EEYQSLIR | -34 | -41 | 0 | 0.00025015 | 8 | NaN | 443.1473 | NaN | 1127.0825 | NaN | NaN |
| Spag7 | EKPPSLGDQESR | -13 | -24 | 1 | 0.00025015 | 12 | NaN | 944.0061 | NaN | NaN | 1210.556 | 1112.0232 |
| Tmem109 | KETSADILTQIGR | -47 | -59 | 1 | 0 | 13 | NaN | 332.4964 | NaN | 1269.7788 | 804.9139 | 756.7465 |
| Septin9 | SVQPTSEER | -325 | -333 | 0 | 0.00025015 | 9 | NaN | 165.05711 | NaN | 645.05176 | NaN | 735.183 |
| Septin9 | IPEGSAVPATDAAPK | -173 | -187 | 0 | 0.00025015 | 15 | NaN | 499.45697 | NaN | 735.3714 | NaN | NaN |
| Septin9 | EFDEDAEDR | -475 | -483 | 0 | 0.00025015 | 9 | NaN | 513.0805 | NaN | NaN | NaN | NaN |
| Rps15a | GYIGEFEIIDDHR | -45 | -57 | 0 | 0 | 13 | NaN | 781.62146 | 717.0285 | 1102.5839 | 358.26147 | 1444.06 |
| Rps15a | FEIIDDHR | -50 | -57 | 0 | 0 | 8 | NaN | 2331.0022 | NaN | NaN | NaN | NaN |
| Ckap5 | EFGFSELNVK | -745 | -754 | 0 | 0 | 10 | NaN | 737.6121 | NaN | NaN | 2516.9148 | 3012.4023 |
| Gclc | EFIANHPDYK | -579 | -588 | 0 | 0.00051306 | 10 | NaN | 1230.8662 | NaN | 1507.1833 | 1656.5468 | NaN |
| Gclc | LVSFDHENR | -55 | -63 | 0 | 0.002068213 | 9 | NaN | 1324.2502 | NaN | NaN | 1340.7261 | NaN |
| Ndufb5 | EFIDHSPK | -177 | -184 | 0 | 0.00025015 | 8 | NaN | 1076.2156 | 1184.7117 | 1334.0502 | 1123.5011 | 1185.2883 |
| Atxn2l | SIPVTSSVVDPGAGSISPASPK | -480 | -501 | 0 | 0.000431655 | 22 | NaN | 358.19647 | 292.9981 | 385.9208 | 420.10785 | 377.12393 |
| Atxn2l | RNVDFNYATK | -194 | -203 | 1 | 0.004350979 | 10 | NaN | 1582.1692 | NaN | NaN | NaN | NaN |
| Atxn2l | GDEKEEGPVTEQ | -645 | -656 | 1 | 0.004065182 | 12 | NaN | 872.68945 | NaN | NaN | NaN | NaN |
| Wbp11 | SILPLLGHGVPR | -181 | -192 | 0 | 0 | 12 | NaN | 735.79224 | 720.8219 | NaN | NaN | NaN |
| Map2k1 | EFQDFVNK | -333 | -340 | 0 | 0.001776473 | 8 | NaN | 1123.4591 | 1206.1298 | 842.6259 | 1037.1777 | 944.745 |
| Map2k1 | LIHLEIKPAIR | -98 | -108 | 1 | 0.000471409 | 11 | NaN | 165.28767 | 117.86719 | NaN | NaN | 233.03902 |
| Gfm1 | TIVAPVPFDFTHK | -528 | -540 | 0 | 0.000394361 | 13 | NaN | 209.9858 | 299.75504 | NaN | 305.80585 | NaN |
| Gfm1 | VHFDPESK | -482 | -489 | 0 | 0.002780285 | 8 | NaN | 1171.4767 | 1080.783 | NaN | NaN | NaN |
| Riox2 | TVGPDKNPFDEAQQK | -375 | -389 | 1 | 0 | 15 | NaN | 1813.9423 | 1927.8441 | NaN | NaN | 2463.5151 |
| Riox2 | TVGPDKNPFDEAQQK | -375 | -389 | 1 | 0 | 15 | NaN | 1813.9423 | 1927.8441 | NaN | NaN | 2463.5151 |
| Riox2 | QIWCGSPIR | -429 | -437 | 0 | 0.002329279 | 9 | NaN | 891.4682 | NaN | NaN | NaN | NaN |
| Riox2 | KLSGFLR | -307 | -313 | 1 | 0.003681885 | 7 | NaN | 220.55493 | NaN | NaN | NaN | NaN |
| Unc45a | EGAIIVDPAR | -280 | -289 | 0 | 0.006735683 | 10 | NaN | 196.5645 | NaN | NaN | NaN | NaN |
| Ecpas | GYFPVGDGVFPHQK | -861 | -874 | 0 | 0 | 14 | NaN | 3661.7195 | 3580.3904 | NaN | 4845.187 | 5264.8555 |
| Ecpas | SENDELSQDVASK | -990 | -1002 | 0 | 0 | 13 | NaN | 548.5394 | NaN | NaN | NaN | NaN |
| Ecpas | FLPPVLLK | -33 | -40 | 0 | 0.003998192 | 8 | NaN | 544.0392 | NaN | NaN | NaN | NaN |
| Ppt1 | ETIPLQESTLYTEDR | -254 | -268 | 0 | 0 | 15 | NaN | 688.9385 | 536.6456 | 753.99866 | 875.13477 | 905.5788 |
| Ppt1 | ISVGGQHQGVFGLPR | -137 | -151 | 0 | 0 | 15 | NaN | 219.48312 | NaN | NaN | 2842.4849 | 302.4435 |
| Ppt1 | EGDHLQISK | -286 | -294 | 0 | 0.000471409 | 9 | NaN | 2889.8108 | NaN | NaN | 2151.3718 | NaN |
| Rps28 | VEFMDDTSR | -32 | -40 | 0 | 0 | 9 | NaN | 537.14075 | 661.84937 | 347.3942 | 941.489 | 726.94135 |
| Rps28 | EGDVLTLLESER | -52 | -63 | 0 | 0 | 12 | NaN | 573.47845 | 647.6851 | 970.94025 | 1133.2819 | 1098.4333 |
| Treml1 | SSDSGLPSDIPHVR | -228 | -241 | 0 | 0.00111006 | 14 | NaN | 4426.366 | 2095.1116 | 3624.3286 | NaN | NaN |
| Ndufb10 | PDSWDKDVYPEPPSR | -2 | -16 | 1 | 0.00025015 | 15 | NaN | 1390.5819 | 1425.2922 | 796.5797 | 751.84015 | 549.3385 |
| Cdk11b | NYVEHDLK | -506 | -513 | 0 | 0 | 8 | NaN | 1612.937 | 1344.0942 | NaN | 558.2574 | 2016.3505 |
| Cdk11b | TFSEYPYNNLR | -664 | -674 | 0 | 0.004983786 | 11 | NaN | 1916.3046 | NaN | 3896.4724 | NaN | NaN |
| Cdk11b | YLPALQGCR | -414 | -422 | 0 | 0.008800311 | 9 | NaN | 1346.1166 | NaN | NaN | 940.1846 | NaN |
| Cdk11b | IWPGYNDLPAVK | -650 | -661 | 0 | 0 | 12 | NaN | 1040.7343 | NaN | 1084.4552 | NaN | NaN |
| Sun2 | SVADEVGLLPQK | -448 | -459 | 0 | 0 | 12 | NaN | 625.0687 | 655.8805 | 3136.0356 | 2135.123 | 2221.447 |
| Dhcr24 | EGLEYIPLR | -317 | -325 | 0 | 0 | 9 | NaN | 2416.1077 | 2395.9883 | NaN | 1387.9169 | NaN |
| Srrm2 | EMPGSNIESSPEVEERPAVL | -1221 | -1240 | 1 | 0 | 20 | NaN | 366.76233 | NaN | 489.75546 | 664.2324 | 740.7328 |
| Spout1 | SVALPGSILDNAQSPELR | -77 | -94 | 0 | 0.00111006 | 18 | NaN | 363.69263 | 441.36325 | NaN | 307.16626 | NaN |
| Smc2 | LAPPEGQTVLDGLEFK | -1056 | -1071 | 0 | 0 | 16 | NaN | 2140.5176 | NaN | 2709.7056 | 3716.314 | 4660.63 |
| Smc2 | KITDGLHGLQEASNK | -354 | -368 | 1 | 0 | 15 | NaN | 1522.0215 | 420.86646 | 1052.9111 | NaN | 896.6347 |
| Smc2 | NCEIEELER | -278 | -286 | 0 | 0 | 9 | NaN | 548.8951 | NaN | 1026.852 | 279.1751 | 665.89136 |
| Smc2 | IVNLQEVLSENEK | -260 | -272 | 0 | 0 | 13 | NaN | 93.22066 | NaN | 151.78664 | 704.4237 | 679.34766 |
| Smc2 | HLFGQPNSAYDFK | -946 | -958 | 0 | 0.00025015 | 13 | NaN | 341.3836 | NaN | NaN | 641.2267 | 538.22705 |
| Drg2 | LVQLILHEYK | -202 | -211 | 0 | 0 | 10 | NaN | 459.28763 | NaN | NaN | NaN | 4835.2305 |
| Mtap | YVDTPFGKPSDALILGK | -33 | -49 | 1 | 0 | 17 | NaN | 1330.7142 | 1230.3616 | 779.927 | NaN | 563.1959 |
| Mtap | KYVDTPFGK | -32 | -40 | 1 | 0.000712251 | 9 | NaN | 2064.887 | NaN | 2925.0413 | NaN | NaN |
| Mtap | GTGLDDPEILEGR | -17 | -29 | 0 | 0 | 13 | NaN | 691.68036 | NaN | NaN | NaN | NaN |
| Idh3g | EVAAHYPQITFD | -238 | -249 | 0 | 0.001023236 | 12 | NaN | 4097.6533 | NaN | 3881.6973 | NaN | NaN |
| Idh3g | HACVPVDFEEVHVS | -79 | -92 | 0 | 0.000394361 | 14 | NaN | 627.057 | 532.2034 | NaN | NaN | NaN |
| Idh3g | SNADEEDIR | -93 | -101 | 0 | 0 | 9 | NaN | 511.34332 | NaN | 530.3464 | NaN | NaN |
| Idh3g | EHESVAGVVESLK | -178 | -190 | 0 | 0 | 13 | NaN | 2911.9622 | NaN | NaN | NaN | NaN |
| Cops2 | SGINPFDSQEAKPYK | -289 | -303 | 1 | 0 | 15 | NaN | 851.4613 | 873.63556 | 800.77625 | NaN | 1075.0426 |
| Cops2 | YEQSLHIK | -218 | -225 | 0 | 0.00025015 | 8 | NaN | 192.43835 | NaN | 645.2253 | 538.8411 | 676.1293 |
| Cops2 | SCQTDDGEDDLKK | -178 | -190 | 1 | 0.002752526 | 13 | NaN | 154.05556 | NaN | NaN | 371.262 | 274.74762 |
| Chordc1 | HNSEKPPEPVKPEVK | -64 | -78 | 2 | 0 | 15 | NaN | 740.1529 | 579.1462 | 414.79742 | 759.0864 | 670.19763 |
| Dhx33 | NLDNAVLIGETGSGK | -80 | -94 | 0 | 0 | 15 | NaN | 1203.6896 | 1169.2434 | NaN | NaN | NaN |
| Plcg1 | GVLDVPACQIAIRPEGK | -874 | -890 | 1 | 0 | 17 | NaN | 473.86795 | NaN | NaN | 909.4936 | 730.17114 |
| Acads | STANLIFEDCR | -237 | -247 | 0 | 0 | 11 | NaN | 2424.1812 | 2298.9126 | NaN | NaN | 1086.7518 |
| Acads | SIALEEISR | -99 | -107 | 0 | 0.00025015 | 9 | NaN | 508.81424 | NaN | 686.71497 | NaN | 490.00125 |
| Plaa | EHLSEPGTR | -341 | -349 | 0 | 0.000431655 | 9 | NaN | 469.86505 | 883.7615 | NaN | NaN | 850.8816 |
| Supt5h | LDSDQNNIHVK | -585 | -595 | 0 | 0 | 11 | NaN | 1244.8009 | 330.7707 | 690.9551 | 692.6367 | 689.9396 |
| Supt5h | EHNFQPGDNVEVC | -415 | -427 | 0 | 0 | 13 | NaN | 409.292 | 449.26913 | NaN | 456.55328 | 468.82385 |
| Supt5h | FEGDTGLIVR | -483 | -492 | 0 | 0 | 10 | NaN | 2650.8403 | NaN | NaN | 1044.1624 | NaN |
| Ilkap | EHNPTQYEER | -259 | -268 | 0 | 0.008680443 | 10 | NaN | 1725.5944 | NaN | NaN | 1437.196 | 907.019 |
| Cmah | LPLTPPNWK | -562 | -570 | 0 | 0.00111006 | 9 | NaN | 551.14307 | 593.59265 | NaN | 659.91644 | NaN |
| Pfdn2 | EHSLVIDTLK | -53 | -62 | 0 | 0.000298805 | 10 | NaN | 1160.4216 | NaN | NaN | 964.65784 | NaN |
| Hmox2 | IHYVGQNEPELLVA | -137 | -150 | 0 | 0.002566359 | 14 | NaN | 903.092 | NaN | 877.84973 | NaN | NaN |
| Immt | ELDSITPDITPGWK | -371 | -384 | 0 | 0 | 14 | NaN | 735.1593 | 1055.4813 | 1327.5754 | 902.3702 | 2076.7183 |
| Immt | YCIEHGDLELAAK | -695 | -707 | 0 | 0 | 13 | NaN | 1270.8898 | NaN | NaN | 1777.087 | 2201.3545 |
| Immt | EIAGATPHIT | -315 | -324 | 0 | 0.000989247 | 10 | NaN | 1357.0249 | 734.1399 | NaN | NaN | NaN |
| Immt | LSTDDLNSLIAH | -395 | -406 | 0 | 0.000947459 | 12 | NaN | 672.2502 | 573.0382 | NaN | NaN | NaN |
| Capn2 | WNDNCPSWNTVDPEVR | -297 | -312 | 0 | 0 | 16 | NaN | 570.2465 | NaN | NaN | NaN | 593.55396 |

| | | | | | | | | | | | | |
|---|---|---|---|---|---|---|---|---|---|---|---|---|
| Capn2 | GAEEVESSGSLQK | -268 | -280 | 0 | 0 | 13 | NaN | 453.47897 | NaN | NaN | NaN | NaN |
| Impa1 | YPCHSFIGEESVAAGEK | -62 | -78 | 0 | 0.000550307 | 17 | NaN | 249.60716 | NaN | 369.7627 | 132.168 | 140.98006 |
| Impa1 | HSFIGEESVAAGEK | -65 | -78 | 0 | 0.001392869 | 14 | NaN | 664.9887 | NaN | 439.80182 | NaN | NaN |
| Eml4 | EIEVPDQYGTIR | -556 | -567 | 0 | 0 | 12 | NaN | 1126.1783 | 1149.0771 | 901.3786 | 1180.5243 | 1170.2635 |
| Hip1r | EIGDLWGHLR | -114 | -123 | 0 | 0.001392869 | 10 | NaN | 673.2481 | 844.8235 | NaN | NaN | NaN |
| Dpm1 | LGGNEIVSFLK | -241 | -251 | 0 | 0 | 11 | NaN | 366.33398 | 429.39615 | 402.39886 | 410.3996 | 1138.4601 |
| Dpm1 | EIIIIDDGSPDGTR | -59 | -72 | 0 | 0 | 14 | NaN | 5120.1665 | 5563.7056 | NaN | 4575.0225 | 4355.525 |
| Arhgef1 | EILHHVNQAVR | -596 | -606 | 0 | 0 | 11 | NaN | 926.6736 | 886.45435 | 975.13385 | 1235.3396 | 1126.0979 |
| Arhgef1 | LLNDLLPFCR | -806 | -815 | 0 | 0 | 10 | NaN | 725.27856 | NaN | 719.6753 | 485.29523 | 650.7 |
| Arhgef1 | GKVELAAECCR | -585 | -595 | 1 | 0 | 11 | NaN | 43.468758 | NaN | 969.4812 | 1471.9353 | 1663.6042 |
| Ppp4c | WSDPEDTTGWGVSPR | -197 | -211 | 0 | 0 | 15 | NaN | 1163.1726 | NaN | 1313.7279 | 1044.3998 | 745.55347 |
| Naa25 | TLLEEGLTHSPSNAQFK | -493 | -509 | 0 | 0 | 17 | NaN | 679.4949 | 538.0877 | 509.91223 | 605.5507 | 568.5071 |
| Pdk1 | EISLLPDNLLR | -93 | -103 | 0 | 0 | 11 | NaN | 353.58673 | NaN | 371.69543 | 939.356 | NaN |
| Bub1b | EISPAENILQEQPDSK | -500 | -515 | 0 | 0.00111006 | 16 | NaN | 480.76678 | NaN | NaN | NaN | NaN |
| Hnrnpf | QSGEAFVELESEDDVK | -53 | -68 | 0 | 0 | 16 | NaN | 461.34637 | NaN | 868.92017 | 954.1969 | 210.78986 |
| Hnrnpf | KHSGPNSADSANDGFVR | -98 | -114 | 1 | 0.001291103 | 17 | NaN | 506.95444 | 239.83307 | NaN | 343.94827 | NaN |
| Hnrnpf | GEAFVELESEDDVK | -55 | -68 | 0 | 0.002401448 | 14 | NaN | 1598.1405 | NaN | NaN | NaN | NaN |
| Tpm4 | RIQLLEEELDR | -55 | -65 | 1 | 0 | 11 | NaN | 512.09937 | 542.00684 | NaN | NaN | NaN |
| Tpm4 | EKAEGDAAALNR | -43 | -54 | 1 | 0.00025015 | 12 | NaN | 1342.956 | NaN | NaN | NaN | NaN |
| Srsf9 | IYVGNLPSDVR | -17 | -27 | 0 | 0 | 11 | NaN | 3000.8062 | NaN | NaN | 924.6453 | NaN |
| Tm9sf4 | EKDVQFEHGYR | -159 | -169 | 1 | 0 | 11 | NaN | 604.5809 | 630.9974 | 563.65576 | 1659.2946 | NaN |
| Hsd17b12 | LNQVSNNIK | -87 | -95 | 0 | 0.001023236 | 9 | NaN | 1410.2068 | NaN | 3166.4678 | NaN | NaN |
| Thrap3 | EKGGFSDADVK | -374 | -384 | 1 | 0 | 11 | NaN | 558.34705 | NaN | 529.2875 | 955.3504 | 1386.728 |
| Rpl21 | EKGTWVQLK | -121 | -129 | 1 | 0.000712251 | 9 | NaN | 466.17288 | 949.3966 | NaN | NaN | NaN |
| Sf3b2 | YGPPPSYPNLK | -622 | -632 | 0 | 0 | 11 | NaN | 9659.414 | 9791.944 | 6579.98 | 5273.826 | 5299.6313 |
| Sf3b2 | SSVPAGMETPELIELR | -735 | -750 | 0 | 0 | 16 | NaN | 609.5968 | 723.96155 | 651.55005 | 845.6819 | 801.675 |
| Sf3b2 | VPPPWLIA | -611 | -618 | 0 | 0.008828666 | 8 | NaN | 167.35112 | NaN | NaN | NaN | NaN |
| Eif3g | EKLPGELEPVQAAQ | -194 | -207 | 1 | 0 | 14 | NaN | 677.89575 | 804.1146 | 1209.3273 | NaN | NaN |
| Tmsb4x | ETQEKNPLPSK | -28 | -38 | 1 | 0 | 11 | NaN | 618.762 | NaN | 1064.7505 | 1115.643 | 1332.1923 |
| Bcat1 | EKPDPDSLVFGAT | -40 | -52 | 1 | 0 | 13 | NaN | 818.6767 | 839.4608 | 858.3515 | 1238.8678 | 1284.7599 |
| Ddx56 | SLLCHLPR | -182 | -189 | 0 | 0 | 8 | NaN | 1920.9412 | 1961.4193 | 376.3779 | NaN | NaN |
| Caprin1 | LWDLLEGK | -212 | -219 | 0 | 0.001392869 | 8 | NaN | 80.02338 | 1030.6538 | 969.4447 | 1318.742 | 1363.769 |
| Caprin1 | SIHLWDLLEGK | -209 | -219 | 0 | 0 | 11 | NaN | 472.49237 | 605.9515 | NaN | 803.63666 | 670.66016 |
| Rps20 | GKTPVEPEVAIHR | -7 | -19 | 1 | 0 | 13 | NaN | 567.9024 | 681.6265 | NaN | NaN | NaN |
| Top2b | HGFLEEFITPIVK | -576 | -588 | 0 | 0 | 13 | NaN | 502.58298 | NaN | 604.3679 | 749.1526 | NaN |
| Top2b | IHHNWPSLLK | -566 | -575 | 0 | 0.002595029 | 10 | NaN | 1707.7113 | NaN | NaN | NaN | NaN |
| Top2b | LPNYDAR | -873 | -879 | 0 | 0.001776473 | 7 | NaN | 404.367 | NaN | NaN | NaN | NaN |
| Kcnab2 | QFNLIPPICEQAEYH | -204 | -218 | 0 | 0.002910424 | 15 | NaN | 399.8456 | 407.67282 | NaN | NaN | NaN |
| Pgrmc2 | GLCSGPGAGEESPAATLPR | -73 | -91 | 0 | 0 | 19 | NaN | 866.4863 | 960.5321 | 1004.22003 | 1079.3429 | 992.56226 |
| Csnk2b | HHHTDGAYFGT | -151 | -161 | 0 | 0 | 11 | NaN | 1240.6216 | 1101.5479 | 2062.9685 | 1320.8733 | 1034.1273 |
| Atp5me | RYSYLKPR | -29 | -36 | 2 | 0.001291103 | 8 | NaN | 91.39494 | 411.291 | NaN | NaN | NaN |
| Pitrm1 | SEIFNNPCFEEEEHFK | -685 | -700 | 0 | 0 | 16 | NaN | 518.6448 | 485.5801 | NaN | NaN | 509.81384 |
| Psmd3 | ELDTVTLEDIK | -62 | -72 | 0 | 0 | 11 | NaN | 4632.3745 | 4467.1 | 239.16272 | 4446.1685 | 255.01732 |
| Psmd3 | ISLADIAQK | -413 | -421 | 0 | 0 | 9 | NaN | 955.9977 | 912.5729 | 525.6718 | 632.77106 | 637.9656 |
| Psmd3 | QTDGTYTLIIR | -383 | -393 | 0 | 0 | 11 | NaN | 2432.6514 | 557.65515 | 1670.3951 | NaN | NaN |
| Rftn1 | ELDYCSSLEHLADQK | -125 | -139 | 0 | 0 | 15 | NaN | 348.51093 | 426.18994 | 555.4335 | 730.85443 | 696.959 |
| Cotl1 | YDGATIVPGDQGADYQ | -31 | -46 | 0 | 0.00025015 | 16 | NaN | 769.1975 | 386.22723 | 467.7569 | NaN | NaN |
| Usp25 | VIDLTGDDKDDLQR | -91 | -104 | 1 | 0 | 14 | NaN | 598.2407 | 683.9186 | 1128.2225 | 1139.6384 | 897.06934 |
| Usp25 | ELEEWDTQLAQR | -700 | -711 | 0 | 0 | 12 | NaN | 665.2191 | 522.0425 | 590.75653 | NaN | 697.83276 |
| Birc5 | ELEGWEPDDNPIEEHRK | -63 | -79 | 1 | 0 | 17 | NaN | 1171.2485 | NaN | NaN | 293.8613 | NaN |
| Birc5 | ELEGWEPDDNPIEEHR | -63 | -78 | 0 | 0.001392869 | 16 | NaN | 334.94244 | NaN | NaN | NaN | NaN |
| Cyfip2 | FDTNFEDR | -42 | -49 | 0 | 0.009416377 | 8 | NaN | 547.8187 | NaN | NaN | NaN | NaN |
| Dhx8 | FYEAPIFTIPGR | -753 | -764 | 0 | 0 | 12 | NaN | 1048.2633 | 1066.9309 | NaN | NaN | NaN |
| Fth1 | YFDRDDVALK | -41 | -50 | 1 | 0.000989247 | 10 | NaN | 1234.3767 | 1272.1512 | NaN | 1293.1882 | NaN |
| Fth1 | SCYFDRDDVALK | -39 | -50 | 1 | 0.001291103 | 12 | NaN | 718.8195 | 889.0272 | NaN | NaN | NaN |
| Ipo9 | LLQHGINADDKR | -866 | -877 | 1 | 0 | 12 | NaN | 1761.448 | NaN | NaN | NaN | NaN |
| Nelfe | ELGQDGEEAEAPGAGDGPPR | -145 | -164 | 0 | 0 | 20 | NaN | 1412.3708 | 551.2163 | NaN | NaN | 631.23846 |
| Nelfe | SLSEQPVVDTATATEQAK | -49 | -66 | 0 | 0.00272025 | 18 | NaN | 518.0628 | NaN | NaN | NaN | 336.9014 |
| Ddost | GFELTFK | -56 | -62 | 0 | 0.009447908 | 7 | NaN | 646.25574 | NaN | NaN | NaN | NaN |
| Stx12 | ELGSLPLPLSASEQR | -83 | -97 | 0 | 0 | 15 | NaN | 332.3487 | NaN | 550.7646 | NaN | NaN |
| Slc7a6 | ELGSPTPTYHLLPK | -5 | -18 | 0 | 0.00025015 | 14 | NaN | 292.60223 | 282.91052 | 372.26328 | 984.6505 | 972.217 |
| Crlf3 | QEWSPWSVPQTGHST | -257 | -271 | 0 | 0 | 15 | NaN | 784.578 | 465.3147 | 939.1932 | 1289.746 | 1245.1147 |
| Crlf3 | ELIEKPGGIIVR | -187 | -198 | 1 | 0 | 12 | NaN | 733.39264 | 789.7567 | 463.53638 | 602.0122 | 550.50165 |
| Tnfaip8 | HQVEYTFDR | -117 | -125 | 0 | 0 | 9 | NaN | 432.10233 | NaN | 281.68738 | 459.03217 | 291.65503 |
| Sucla2 | EPIDIVEGIKK | -206 | -216 | 1 | 0 | 11 | NaN | 283.1937 | 343.0224 | NaN | 196.54506 | 152.75401 |
| Sucla2 | ELLQEAGVSVPK | -62 | -73 | 0 | 0 | 12 | NaN | 2009.4122 | NaN | NaN | 1108.7797 | 513.065 |
| Sucla2 | GGVNIEDVAAENPEAIVK | -188 | -205 | 0 | 0 | 18 | NaN | 405.69894 | NaN | 482.11646 | 922.4859 | NaN |
| Nup188 | ILLIIAASHA | -1109 | -1118 | 0 | 0.008014596 | 10 | NaN | 861.77344 | NaN | 709.99927 | 718.93164 | NaN |
| Nup188 | ELNQIEAELNK | -28 | -38 | 0 | 0.000755556 | 11 | NaN | 1134.2385 | NaN | NaN | 1997.1703 | NaN |
| Lnpep | SYLSEDVFR | -561 | -569 | 0 | 0.007791272 | 9 | NaN | 1833.2405 | 2363.928 | NaN | NaN | 1914.5934 |
| Lnpep | GFPLVTVQR | -615 | -623 | 0 | 0.003360691 | 9 | NaN | 950.6873 | NaN | NaN | NaN | NaN |
| Lnpep | IIAHELAHQ | -461 | -469 | 0 | 0.001023236 | 9 | NaN | 381.18274 | NaN | NaN | NaN | NaN |
| Prep | YVQDSLEGEAR | -101 | -111 | 0 | 0 | 11 | NaN | 263.0297 | 782.3144 | NaN | 349.07477 | 330.0015 |
| Prep | SFLSPGVIYHCDLTK | -400 | -414 | 0 | 0.00025015 | 15 | NaN | 1221.7045 | 1246.6548 | NaN | NaN | NaN |
| Prep | ELPDVLER | -163 | -170 | 0 | 0.008653659 | 8 | NaN | 956.33545 | NaN | NaN | NaN | NaN |
| Zc3h15 | HFLEAIENNK | -182 | -191 | 0 | 0 | 10 | NaN | 1212.2325 | NaN | 601.4048 | 1354.3182 | 1357.1984 |
| Zc3h15 | HALPPGFVLK | -209 | -218 | 0 | 0 | 10 | NaN | 950.7827 | 795.6204 | NaN | NaN | NaN |
| Nup88 | HAQLPVLSDSER | -660 | -671 | 0 | 0 | 12 | NaN | 894.24664 | 848.4408 | NaN | 84.85535 | 1198.4377 |

| | | | | | | | | | | | | |
|---|---|---|---|---|---|---|---|---|---|---|---|---|
| Def6 | KLQELELLQEAQR | -360 | -372 | 1 | 0 | 13 | NaN | 400.72415 | 594.6611 | 599.29694 | 412.68628 | 447.49457 |
| Def6 | ELQQALEGQLR | -391 | -401 | 0 | 0 | 11 | NaN | 561.0568 | 860.05585 | 1090.4971 | NaN | 776.32196 |
| Def6 | LQELELLQEAQR | -361 | -372 | 0 | 0.002434616 | 12 | NaN | 343.00696 | NaN | NaN | NaN | NaN |
| Tbrg4 | TSSVSSPSSGSLAELLGK | -40 | -57 | 0 | 0.002880184 | 18 | NaN | 358.169 | 845.7714 | NaN | NaN | 1539.0171 |
| Tbrg4 | ELQSVEQEVR | -162 | -171 | 0 | 0.00897527 | 10 | NaN | 2029.0148 | NaN | NaN | NaN | NaN |
| Sfxn3 | IQAQNPSIDVVYYNK | -305 | -319 | 0 | 0.001055699 | 15 | NaN | 214.61032 | NaN | 314.07318 | 405.81635 | 295.82135 |
| Tmpo | SSILPITEFSDITR | -305 | -318 | 0 | 0 | 14 | NaN | 495.23373 | 455.19394 | 589.1246 | 646.4911 | 712.3222 |
| Gsk3b | LVLDYVPETVYR | -130 | -141 | 0 | 0 | 12 | NaN | 294.4115 | 408.90466 | 524.549 | 721.1399 | 579.40826 |
| Nudcd1 | ENDSLEISLIK | -339 | -349 | 0 | 0.000230707 | 11 | NaN | 378.02188 | NaN | NaN | NaN | NaN |
| Msh6 | IHNVGSPLK | -822 | -830 | 0 | 0 | 9 | NaN | 749.06647 | 748.2495 | 828.1038 | 1042.5873 | 857.91077 |
| Msh6 | ENEQSLLEYLDK | -943 | -954 | 0 | 0 | 12 | NaN | 885.17676 | NaN | 697.3812 | 971.6763 | 1110.881 |
| Msh6 | VSGLLEEVAGGFTSK | -868 | -882 | 0 | 0.000230707 | 15 | NaN | 453.7814 | 416.6654 | 352.42078 | NaN | NaN |
| Msh6 | RRPDHPEFNPTTL | -383 | -395 | 2 | 0.002434616 | 13 | NaN | 246.31244 | NaN | 569.2863 | 810.66364 | NaN |
| Ap2b1 | TLCENLDSLDEPDAR | -431 | -445 | 0 | 0 | 15 | NaN | 1366.4404 | 1134.2661 | 752.8824 | 137.77933 | NaN |
| Ap2b1 | MVEDQGFLDSLR | -155 | -166 | 0 | 0.000230707 | 12 | NaN | 1038.0676 | 1525.642 | NaN | NaN | 696.7859 |
| Ap2b1 | SGEPEVQYVALR | -293 | -304 | 0 | 0.004224523 | 12 | NaN | 1369.7135 | NaN | NaN | NaN | NaN |
| Ap2b1 | LLSTDPVTAK | -529 | -538 | 0 | 0 | 10 | NaN | 468.22626 | NaN | NaN | NaN | NaN |
| Pycard | LFSFVPSWNLTCK | -160 | -172 | 0 | 0.001868731 | 13 | NaN | 326.37997 | NaN | NaN | NaN | NaN |
| Hmbs | ENPCDAVVFHPK | -117 | -128 | 0 | 0 | 12 | NaN | 902.9834 | 1017.22314 | NaN | NaN | 1232.8441 |
| Pdcd10 | ENPGLTQDIIMK | -54 | -65 | 0 | 0.000907206 | 12 | NaN | 976.2863 | NaN | NaN | 722.9278 | 791.13153 |
| Ugp2 | IQRPPEDSIQPY | -78 | -89 | 1 | 0.000550307 | 12 | NaN | 2969.009 | NaN | NaN | NaN | NaN |
| Ipo11 | VTEDEEPPTEQDKR | -905 | -918 | 1 | 0 | 14 | NaN | 265.58685 | 342.54672 | 432.6855 | 637.4352 | 333.07175 |
| Fmr1 | ENTHFSQPNSTK | -360 | -371 | 0 | 0 | 12 | NaN | 393.01556 | NaN | 410.67078 | NaN | 441.4745 |
| Fmr1 | SIVAGGPQKPEPK | -380 | -392 | 1 | 0.00025015 | 13 | NaN | 228.71722 | 588.01855 | NaN | NaN | NaN |
| Sh3bgrl | ENVPEDSRPSTGYPLPPQIF | -51 | -70 | 1 | 0.00025015 | 20 | NaN | 425.05438 | NaN | NaN | 201.96486 | 497.22247 |
| Hspbp1 | TEHSPFHEHVLGALC | -281 | -295 | 0 | 0.00051306 | 15 | NaN | 1719.7748 | 1710.9056 | NaN | 5088.7104 | 4724.235 |
| Hspbp1 | SLVTDFPQGVR | -296 | -306 | 0 | 0 | 11 | NaN | 1764.4913 | 1739.8613 | NaN | 1139.0394 | NaN |
| Mrps5 | EPEPEPEVPDTK | -396 | -407 | 0 | 0.002068213 | 12 | NaN | 696.21735 | NaN | NaN | NaN | NaN |
| Csnk2a1 | FYHPGQEYNVR | -181 | -191 | 0 | 0 | 11 | NaN | 3788.889 | 2748.0479 | 3582.9707 | 4019.025 | 232.63094 |
| Csnk2a1 | EPFFHGHDNYDQLVR | -230 | -244 | 0 | 0 | 15 | NaN | 1311.5477 | NaN | 1201.6686 | 1772.9377 | 1446.5892 |
| Csnk2a1 | KEPFFHGHDNYDQLVR | -229 | -244 | 1 | 0 | 16 | NaN | 514.10974 | 1063.9033 | 817.28735 | 427.80518 | 600.8844 |
| Inpp5b | EPILDLPLK | -796 | -804 | 0 | 0.000431655 | 9 | NaN | 561.85474 | 526.45135 | 553.7754 | 785.71484 | 769.74347 |
| Rnaseh1 | EPLGEGEELPEPGPK | -101 | -115 | 0 | 0.000712251 | 15 | NaN | 2647.1028 | 2916.0046 | 2948.4897 | 2771.5845 | NaN |
| Kpna6 | EPSPPIDEVINTPGVVDR | -111 | -128 | 0 | 0.00051306 | 18 | NaN | 2136.2583 | 2074.6692 | NaN | NaN | NaN |
| Kpna6 | SKEPSPPIDEVINTPGVVDR | -109 | -128 | 1 | 0 | 20 | NaN | 441.0626 | NaN | NaN | NaN | NaN |
| Mndal | NQKPQPQNQNIPR | -294 | -306 | 1 | 0.001291103 | 13 | NaN | 207.39716 | NaN | NaN | 337.25473 | NaN |
| Add3 | SHLLEGELEEYTK | -555 | -567 | 0 | 0 | 13 | NaN | 961.23596 | 632.73615 | NaN | 1121.718 | 1426.0367 |
| Add3 | LEENHELFSK | -608 | -617 | 0 | 0 | 10 | NaN | 3105.3528 | 3138.4387 | 2331.3342 | NaN | NaN |
| Add3 | KQQGLDDAEQGSLSDDAASVS | -572 | -592 | 1 | 0.001868731 | 21 | NaN | 266.2652 | 385.1916 | NaN | NaN | NaN |
| Larp1 | THFDYQFGYR | -332 | -341 | 0 | 0 | 10 | NaN | 1719.245 | 1768.9597 | 1869.699 | 1338.4811 | 1726.0214 |
| Larp1 | EQFDTLTPEPPVDPNQEVPPGPPR | -695 | -718 | 0 | 0.00025015 | 24 | NaN | 6720.1533 | NaN | 7776.3174 | NaN | 1487.8936 |
| Larp1 | SGPGDGGGEPDGGVR | -36 | -50 | 0 | 0 | 15 | NaN | 722.19 | 699.0607 | NaN | NaN | NaN |
| Txlna | VQDLTAGGITDIGSER | -485 | -500 | 0 | 0 | 16 | NaN | 828.02454 | NaN | NaN | NaN | NaN |
| Rpl17 | IVPKPEEEVAQK | -156 | -167 | 1 | 0 | 12 | NaN | 466.7874 | 490.26746 | NaN | NaN | NaN |
| Rpl17 | SLVIEHIQVNK | -111 | -121 | 0 | 0.00025015 | 11 | NaN | 5073.1714 | NaN | NaN | NaN | NaN |
| Cast | KVEEEVINDQALQAL | -360 | -374 | 1 | 0.000947459 | 15 | NaN | 610.1051 | 565.37146 | 604.7814 | NaN | NaN |
| Cast | SEDLDSCPSTTETSK | -730 | -744 | 0 | 0.000471409 | 15 | NaN | 346.38138 | NaN | 394.32758 | NaN | NaN |
| Tax1bp1 | EQLECQLQTEKDEK | -267 | -280 | 1 | 0.00025015 | 14 | NaN | 845.3547 | 231.79655 | 385.30423 | 1031.4623 | 506.18924 |
| Tax1bp1 | LELAEVEDNYK | -580 | -590 | 0 | 0 | 11 | NaN | 1591.4824 | 1404.3661 | NaN | NaN | 2269.2756 |
| Nomo1 | EQQLAEIETR | -700 | -709 | 0 | 0.001291103 | 10 | NaN | 189.00984 | NaN | NaN | NaN | NaN |
| Ttc5 | EQQLLEFLSR | -264 | -273 | 0 | 0 | 10 | NaN | 520.74786 | NaN | NaN | 2449.537 | 2307.0352 |
| Pds5a | YLVPHNLETEER | -471 | -482 | 0 | 0 | 12 | NaN | 1101.3757 | 1210.1676 | 494.42322 | 1353.3806 | 1067.2511 |
| Frg1 | KREEDEETQLDIV | -32 | -44 | 2 | 0 | 13 | NaN | 1184.7568 | 1349.1212 | NaN | 1628.9933 | 1571.0914 |
| Ctsb | RGENHCGIESEIVAGIPR | -314 | -331 | 1 | 0.000550307 | 18 | NaN | 345.2953 | NaN | 2216.6494 | 447.74072 | 889.44336 |
| Nt5c | GFLANEQYGALRPDLAEK | -45 | -62 | 1 | 0.00025015 | 18 | NaN | 524.4906 | NaN | 498.91452 | 626.38544 | 620.52106 |
| Nt5c | FPEEPHVPLEQR | -32 | -43 | 0 | 0 | 12 | NaN | 518.3223 | NaN | NaN | NaN | 869.94434 |
| Nt5c | HNQHLALPPTR | -169 | -179 | 0 | 0.001055699 | 11 | NaN | 70.356125 | NaN | NaN | NaN | 429.1835 |
| Nt5c | LLSWSDNWR | -182 | -190 | 0 | 0.00025015 | 9 | NaN | 473.63348 | NaN | NaN | NaN | NaN |
| Cstb | VFQPLPHENKPLTLS | -69 | -83 | 1 | 0.00025015 | 15 | NaN | 3050.35 | 3256.6106 | 3748.1653 | 3473.357 | 3818.1284 |
| Cd3e | ERPPPVPNPDYEPIR | -160 | -174 | 1 | 0 | 15 | NaN | 897.5822 | NaN | 718.59045 | 1020.0058 | 959.62415 |
| Cd3e | ERPPPVPNPDYEPIR | -160 | -174 | 1 | 0 | 15 | NaN | 897.5822 | NaN | 718.59045 | 1020.0058 | 959.62415 |
| Ptpn11 | ESAAHDYTLR | -390 | -399 | 0 | 0.000712251 | 10 | NaN | 3459.939 | NaN | NaN | NaN | 5152.0073 |
| Otub1 | LLTSGYLQR | -177 | -185 | 0 | 0 | 9 | NaN | 1716.127 | 1790.2682 | 1029.534 | 1160.4774 | 1067.462 |
| Otub1 | EYAEDDNIYQQK | -60 | -71 | 0 | 0.00025015 | 12 | NaN | 3181.8455 | 2885.2722 | NaN | NaN | NaN |
| Otub1 | SHLEALLDDSK | -99 | -109 | 0 | 0 | 11 | NaN | 593.3777 | NaN | 3484.4666 | NaN | NaN |
| Dhfr | ESDTFFPEIDLGK | -144 | -156 | 0 | 0.008736733 | 13 | NaN | 949.9336 | NaN | NaN | NaN | NaN |
| Mydgf | ESDVPLKSEEFEVTK | -124 | -138 | 1 | 0 | 15 | NaN | 1356.4288 | NaN | 544.3643 | 1279.4081 | 1261.9988 |
| Mydgf | TAVSHRPGAFK | -139 | -149 | 1 | 0.00383302 | 11 | NaN | 429.4252 | NaN | NaN | NaN | NaN |
| Rfc2 | TAGHYELPWVEK | -25 | -36 | 0 | 0 | 12 | NaN | 3325.48 | 308.8242 | NaN | 3899.8987 | 154.017 |
| Ptpn1 | QLELENLTTK | -157 | -166 | 0 | 0 | 10 | NaN | 574.1643 | 383.18387 | 1679.2996 | NaN | NaN |
| Rab14 | KIYQNIQDGSLDLNAA | -171 | -186 | 1 | 0.003360691 | 16 | NaN | 311.6137 | NaN | NaN | NaN | NaN |
| Mta1 | KPLEAVLR | -532 | -539 | 1 | 0.001743443 | 8 | NaN | 498.95972 | NaN | 87.08769 | 636.70514 | 587.28 |
| Mta1 | QVYIPNYNKPNPNQISAS | -353 | -370 | 1 | 0.006946742 | 18 | NaN | 1722.0717 | 1609.0347 | NaN | NaN | NaN |
| Nvl | HFEDAFK | -826 | -832 | 0 | 0.001610812 | 7 | NaN | 1641.6726 | 1282.7675 | NaN | NaN | 1337.9412 |
| Nvl | GVLLHGPPGCGK | -299 | -310 | 0 | 0.000550307 | 12 | NaN | 315.28223 | 546.1158 | NaN | NaN | NaN |
| Mapre1 | INESLQLNLTK | -24 | -34 | 0 | 0.001868731 | 11 | NaN | 703.0978 | NaN | NaN | NaN | NaN |
| Papola | VPNPIVGVK | -636 | -644 | 0 | 0.00025015 | 9 | NaN | 478.5062 | 52.660248 | NaN | 520.15497 | NaN |
| Papola | ESPDREEFR | -426 | -434 | 1 | 0.00504706 | 9 | NaN | 1855.5865 | NaN | NaN | NaN | NaN |

| | | | | | | | | | | | | |
|---|---|---|---|---|---|---|---|---|---|---|---|---|
| Papola | LFEAPNFFQK | -361 | -370 | 0 | 0 | 10 | NaN | 507.66348 | NaN | NaN | NaN | NaN |
| Camk2d | SNKPIHTIILNPHVH | -406 | -420 | 1 | 0.007493591 | 15 | NaN | 506.09372 | NaN | 387.30298 | 379.59204 | 506.96417 |
| Ociad1 | ESYGVTLPHK | -210 | -219 | 0 | 0 | 10 | NaN | 1005.1778 | 1028.884 | 1207.5659 | 1200.7336 | 1139.1415 |
| Ahsa1 | ETFLTSPEELYR | -213 | -224 | 0 | 0 | 12 | NaN | 1309.1144 | 1209.0051 | 1341.7677 | 1541.1411 | 1366.9126 |
| Ahsa1 | VENEEGKCEVTEVNKL | -49 | -64 | 2 | 0.002199523 | 16 | NaN | 499.583 | NaN | 570.51697 | 610.65643 | 565.569 |
| Ahsa1 | FLTSPEELYR | -215 | -224 | 0 | 0.000394361 | 10 | NaN | 1884.66 | NaN | NaN | 2982.6392 | NaN |
| Gm12250 | YFLDTVTEDAK | -391 | -401 | 0 | 0 | 11 | NaN | 1014.6282 | 1244.3165 | 543.2625 | 1665.7842 | 535.5678 |
| Gm12250 | IWLEALK | -281 | -287 | 0 | 0.001636916 | 7 | NaN | 1253.5375 | 1223.197 | NaN | NaN | 1134.1469 |
| Gm12250 | QTIWLEALK | -279 | -287 | 0 | 0.000431655 | 9 | NaN | 551.62744 | NaN | NaN | 686.12756 | 687.0012 |
| Gm12250 | SCIQEGDIQK | -43 | -52 | 0 | 0.00272025 | 10 | NaN | 1511.856 | NaN | NaN | NaN | NaN |
| Nol11 | LIPIYSSDTEQNQR | -231 | -244 | 0 | 0 | 14 | NaN | 756.7427 | 888.0434 | NaN | NaN | NaN |
| Pcbp1 | TIPYQPMPASSPVICAGGQDR | -180 | -200 | 0 | 0 | 21 | NaN | 478.37497 | 611.33044 | 616.021 | 766.324 | 704.3746 |
| Pcbp1 | PQSVTECVK | -152 | -160 | 0 | 0.001055699 | 9 | NaN | 1734.8279 | 1978.254 | 526.06213 | 3932.8157 | 3845.0608 |
| Pcbp1 | TQCGSLIGK | -107 | -115 | 0 | 0 | 9 | NaN | 1415.4948 | 1415.7421 | 1645.0176 | 2095.1904 | 2021.3513 |
| Pcbp1 | LETLSQSPQGR | -167 | -177 | 0 | 0 | 11 | NaN | 4364.3706 | 5245.4307 | 4472.385 | NaN | 5250.4897 |
| Pcbp1 | SSPVICAGGQDR | -189 | -200 | 0 | 0 | 12 | NaN | 1036.0219 | 1127.2977 | 1629.993 | NaN | 1525.5288 |
| Sec16a | ETLSVDPFPLSDQIR | -489 | -503 | 0 | 0.000230707 | 15 | NaN | 365.38342 | 431.22842 | NaN | 516.333 | 464.76016 |
| Sec16a | NQSWGDAVVGER | -879 | -890 | 0 | 0.001868731 | 12 | NaN | 340.63046 | 459.54724 | NaN | NaN | NaN |
| Tut1 | ETPQDWALLH | -701 | -710 | 0 | 0 | 10 | NaN | 1326.4413 | NaN | NaN | NaN | NaN |
| Isca2 | ETTSSIPEAGEGQIR | -37 | -51 | 0 | 0 | 15 | NaN | 958.31415 | 898.50977 | NaN | NaN | NaN |
| Rasa2 | ETWFSLQPIDSNSEVQGK | -134 | -151 | 0 | 0.00111006 | 18 | NaN | 898.4591 | NaN | 518.17004 | NaN | NaN |
| Smarca4 | IQELENLPGSLAGDLR | -382 | -397 | 0 | 0 | 16 | NaN | 250.8176 | NaN | NaN | NaN | 230.05211 |
| Cul1 | KYIEDKDVFQK | -454 | -464 | 2 | 0.002972566 | 11 | NaN | 379.04907 | NaN | NaN | 275.1652 | NaN |
| Hectd1 | GGDIFLDQLAR | -608 | -618 | 0 | 0 | 11 | NaN | 562.29224 | NaN | 361.21198 | 605.8605 | 739.3891 |
| Hectd1 | LVDLPISKPFFK | -2266 | -2277 | 1 | 0.00078661 | 12 | NaN | 449.86514 | 94.63985 | NaN | 356.02225 | NaN |
| Pon2 | EVESVDLPNCHLIK | -33 | -46 | 0 | 0 | 14 | NaN | 1407.1853 | 1215.9437 | NaN | 959.91473 | 933.0441 |
| Pon2 | LFVYHPNHPPASEVLR | -290 | -305 | 0 | 0 | 16 | NaN | 478.18698 | NaN | NaN | 402.59967 | NaN |
| Rpl15 | FHHTIGGSR | -180 | -188 | 0 | 0 | 9 | NaN | 956.5833 | 1144.506 | NaN | NaN | NaN |
| Oat | FIADEIQTGLAR | -260 | -271 | 0 | 0 | 12 | NaN | 1825.1205 | 1908.5582 | 1588.0034 | 2105.5947 | 2115.7563 |
| Stk39 | LASGCDGSEIPDEVK | -532 | -546 | 0 | 0.00078661 | 15 | NaN | 1759.271 | NaN | 285.52292 | 654.9351 | 659.1332 |
| Atad2 | SQPTPSLVLDHK | -969 | -980 | 0 | 0 | 12 | NaN | 917.32104 | 263.63947 | 704.9647 | 661.05896 | 1655.0734 |
| Aldh9a1 | LGDPLLEDTR | -317 | -326 | 0 | 0 | 10 | NaN | 10867.831 | 12248.303 | NaN | 6924.427 | 12676.153 |
| Usp39 | IVHDGKPSEGSYR | -502 | -514 | 1 | 0.001023236 | 13 | NaN | 607.4731 | 431.37677 | 425.4325 | 643.85376 | 206.1326 |
| Usp39 | EYLSEEVQAVHK | -481 | -492 | 0 | 0 | 12 | NaN | 1146.2761 | 1053.3687 | 1189.145 | 1197.192 | 1337.6761 |
| Usp39 | SVLDFDFEK | -113 | -121 | 0 | 0 | 9 | NaN | 1347.2821 | NaN | 1186.9995 | 1298.4733 | 1377.8578 |
| Rpl7l1 | TVPLTDNTVIEEHLGR | -164 | -179 | 0 | 0 | 16 | NaN | 810.8383 | 1062.8589 | 791.48395 | 1164.0625 | 955.17163 |
| Rpl7l1 | EVPDKHPLAFVIR | -80 | -92 | 1 | 0 | 13 | NaN | 528.5206 | 638.2782 | 644.8686 | NaN | 749.32025 |
| Rpl7l1 | TDNTVIEEHLGR | -168 | -179 | 0 | 0 | 12 | NaN | 1058.7131 | NaN | 1497.9016 | NaN | NaN |
| Rpl7l1 | KIPLVPENLLK | -9 | -19 | 1 | 0.000471409 | 11 | NaN | 467.7921 | NaN | NaN | 58.59253 | NaN |
| Trmt10c | EVPEHITEEDLK | -102 | -113 | 0 | 0 | 12 | NaN | 1114.8751 | NaN | 1435.7694 | 1478.5553 | 1427.4988 |
| Acot7 | VDADPVVDNSQK | -311 | -322 | 0 | 0 | 12 | NaN | 1087.0226 | 1050.3868 | 772.9804 | 780.7828 | 720.4908 |
| Acot7 | VLVDADPVVDNSQK | -309 | -322 | 0 | 0 | 14 | NaN | 1471.6837 | NaN | NaN | NaN | NaN |
| Acot7 | EVPPIVYLR | -176 | -184 | 0 | 0.000550307 | 9 | NaN | 500.45218 | NaN | NaN | NaN | NaN |
| Uri1 | EVPSEVSEEPTKR | -506 | -518 | 1 | 0 | 13 | NaN | 690.3862 | 754.7182 | NaN | 717.3513 | 794.6465 |
| Uri1 | NTFDADELWAR | -201 | -211 | 0 | 0 | 11 | NaN | 330.76987 | NaN | NaN | NaN | NaN |
| Urb1 | EVTFSFLPR | -377 | -385 | 0 | 0.001846793 | 9 | NaN | 360.45325 | 391.4476 | 175.43263 | NaN | NaN |
| Ndufs5 | EWIECAHGIGGTR | -39 | -51 | 0 | 0 | 13 | NaN | 630.19336 | 499.88617 | 521.0442 | 619.09796 | 605.53723 |
| Iws1 | EWLSPLPDR | -565 | -573 | 0 | 0 | 9 | NaN | 818.5303 | NaN | 705.23334 | 680.8633 | NaN |
| Cpsf3 | EYEDNDEVHIE | -434 | -444 | 0 | 0 | 11 | NaN | 498.9054 | NaN | NaN | 464.4685 | 583.0429 |
| Cpsf3 | NTVHDIVNR | -224 | -232 | 0 | 0 | 9 | NaN | 1414.2661 | NaN | NaN | NaN | NaN |
| Hba | SHHPADFTPAVHASLDK | -112 | -128 | 0 | 0 | 17 | NaN | 1965.7921 | 1694.4521 | 1276.2533 | 1177.046 | 1616.1652 |
| Hba | LSALSDLHAHK | -81 | -91 | 0 | 0.00713295 | 11 | NaN | 1741.151 | NaN | NaN | 478.68604 | NaN |
| Vrk1 | LADTNSSKPVGSDAPCVVK | -53 | -71 | 1 | 0.000591124 | 19 | NaN | 657.58954 | 675.1163 | 505.15958 | 803.4706 | 696.36786 |
| Vrk1 | RYCPDGVHK | -203 | -211 | 1 | 0.001392869 | 9 | NaN | 192.5485 | 137.52881 | 462.34512 | 266.69403 | 143.62193 |
| Gsto1 | EYLDEAYPEK | -91 | -100 | 0 | 0.000298805 | 10 | NaN | 597.308 | NaN | 327.9513 | NaN | NaN |
| Gsto1 | TWPWFQR | -176 | -182 | 0 | 0.006537474 | 7 | NaN | 653.8569 | NaN | NaN | NaN | NaN |
| Gstt2 | EYLGWHADNIR | -97 | -107 | 0 | 0.005015524 | 11 | NaN | 621.45337 | NaN | NaN | NaN | 1332.2255 |
| Ikbkg | EYLQEQLEQLQR | -320 | -331 | 0 | 0.00025015 | 12 | NaN | 511.15173 | NaN | 220.00708 | 617.16656 | 678.0278 |
| Lat | SILIKPPQITVPR | -49 | -61 | 1 | 0 | 13 | NaN | 582.27246 | 634.95825 | 565.8523 | 677.8531 | 742.3242 |
| Elavl1 | FAANPNQNK | -183 | -191 | 0 | 0.00051306 | 9 | NaN | 419.04013 | 1193.3662 | NaN | NaN | 327.63956 |
| Elavl1 | FGGPVHHQ | -207 | -214 | 0 | 0.006850902 | 8 | NaN | 4767.3926 | 5148.4507 | NaN | NaN | NaN |
| Flnb | FADEHVPGSPFTVK | -2075 | -2088 | 0 | 0 | 14 | NaN | 9826.556 | 10058.317 | 6042.5757 | NaN | NaN |
| Flnb | SGCTINNPAEFIVDPK | -658 | -673 | 0 | 0.00025015 | 16 | NaN | 1283.6028 | 1534.7305 | NaN | NaN | 678.9756 |
| Pdha1 | SSDPPFEVR | -370 | -378 | 0 | 0.00025015 | 9 | NaN | 916.6923 | 1049.2313 | NaN | 1232.121 | 1114.2621 |
| Aspscr1 | FANLPNNAK | -67 | -75 | 0 | 0.002780285 | 9 | NaN | 482.97287 | NaN | NaN | NaN | NaN |
| Isyna1 | HVLYAPGERPDHCVVIK | -368 | -384 | 1 | 0 | 17 | NaN | 431.8078 | 404.10202 | 137.8944 | 799.36426 | 564.90594 |
| Isyna1 | NHLGNNDGQNLSAPLQFR | -333 | -350 | 0 | 0 | 18 | NaN | 477.55508 | 412.18088 | NaN | 412.77475 | 473.948 |
| Isyna1 | RFCEVVPGR | -233 | -241 | 1 | 0.008919338 | 9 | NaN | 2530.3557 | NaN | NaN | NaN | NaN |
| Nsa2 | VCFVGDGFTR | -153 | -162 | 0 | 0 | 10 | NaN | 680.70026 | NaN | NaN | NaN | NaN |
| Tm9sf2 | RPSENLGQVLFGER | -90 | -103 | 1 | 0.000550307 | 14 | NaN | 275.6044 | NaN | NaN | 376.09924 | 244.9417 |
| Tm9sf2 | RPSENLGQVLFGE | -90 | -102 | 1 | 0.008222027 | 13 | NaN | 651.01117 | 891.9958 | NaN | NaN | NaN |
| Hmgb1 | KFKDPNAPK | -88 | -96 | 2 | 0.002329279 | 9 | NaN | 367.84882 | 378.98785 | 259.3143 | 280.6328 | 280.37543 |
| Hmgb1 | NNTAADDKQPYEK | -134 | -146 | 1 | 0 | 13 | NaN | 259.7233 | 849.6865 | 296.06607 | 211.50739 | 583.49365 |
| Hmgb1 | NNTAADDKQPYEK | -134 | -146 | 1 | 0 | 13 | NaN | 259.7233 | 849.6865 | 296.06607 | 211.50739 | 583.49365 |
| Hmgb1 | IKGEHPGLSIGDV | -113 | -125 | 1 | 0.00272025 | 13 | NaN | 277.86017 | NaN | 949.5381 | 550.38086 | NaN |
| Utp20 | FDCLQESLLELHNK | -293 | -306 | 0 | 0 | 14 | NaN | 406.78534 | 422.16388 | NaN | NaN | 269.78098 |
| Usp7 | NYGGHDDDLSVR | -498 | -509 | 0 | 0 | 12 | NaN | 1384.3037 | 1586.6646 | 1924.8535 | 2467.2852 | 2119.0027 |
| Ltv1 | FDDDYDYLQHLK | -62 | -73 | 0 | 0.00025015 | 12 | NaN | 513.3526 | NaN | NaN | NaN | NaN |

| | | | | | | | | | | | | |
|---|---|---|---|---|---|---|---|---|---|---|---|---|
| Fnbp1 | HIPNIFQK | -208 | -215 | 0 | 0.00111006 | 8 | NaN | 1389.8016 | 1105.5635 | 703.2514 | 1262.3867 | 1180.7504 |
| Fnbp1 | FDDEFDDEEPLPAIGTCK | -538 | -555 | 0 | 0.000907206 | 18 | NaN | 278.12027 | 282.58798 | 274.83347 | 298.79703 | 293.61566 |
| Ppie | FDDENFILK | -219 | -227 | 0 | 0 | 9 | NaN | 524.688 | NaN | 316.5652 | 543.3794 | 472.5703 |
| Ppie | VLYVGGLAEEVDDK | -7 | -20 | 0 | 0 | 14 | NaN | 857.66516 | NaN | NaN | NaN | NaN |
| Dlgap4 | FDELYHLK | -899 | -906 | 0 | 0 | 8 | NaN | 7558.8945 | 7809.0557 | NaN | 7744.2915 | 7812.2476 |
| Ddx46 | IVDNVRPDR | -546 | -554 | 1 | 0.004439092 | 9 | NaN | 1208.3466 | NaN | NaN | NaN | 1762.2052 |
| Ddx46 | SCPNHYEDYVHR | -695 | -706 | 0 | 0 | 12 | NaN | 640.71326 | 504.12756 | NaN | NaN | NaN |
| Skic8 | SLDAHIR | -83 | -89 | 0 | 0 | 7 | NaN | 1051.3608 | NaN | 1266.7349 | 941.52325 | 1140.627 |
| Rpa3 | FDLELYNEAVK | -94 | -104 | 0 | 0 | 11 | NaN | 1061.8158 | NaN | 848.8003 | NaN | 809.8832 |
| Acbd3 | FDNSYSLWR | -505 | -513 | 0 | 0.00025015 | 9 | NaN | 728.6184 | 668.58795 | NaN | NaN | NaN |
| Abcb7 | FDNVHFEYIEGQK | -474 | -486 | 0 | 0 | 13 | NaN | 12555.776 | NaN | NaN | 16471.037 | 16109.564 |
| Abcb7 | LSYVWPEDRPDLR | -125 | -137 | 1 | 0.00025015 | 13 | NaN | 247.60866 | NaN | NaN | 100.44284 | 307.164 |
| Pck2 | GYLTEQVNQDLPK | -612 | -624 | 0 | 0.00830926 | 13 | NaN | 267.03644 | NaN | NaN | NaN | NaN |
| Smarcc2 | SDGDPIVDPEKDKEPTEGQEEVLK | -813 | -836 | 2 | 0.006978206 | 24 | NaN | 399.68964 | NaN | NaN | 395.69244 | NaN |
| Cc2d1b | VSPSHLLTEPK | -501 | -511 | 0 | 0 | 11 | NaN | 628.28296 | NaN | NaN | NaN | NaN |
| Fnbp4 | WEFPDGEEEEESQTK | -632 | -646 | 0 | 0 | 15 | NaN | 471.94083 | 457.8366 | 479.79904 | 456.69702 | 411.79285 |
| Bola2 | QDLEAEHVEVEDTTLNR | -14 | -30 | 0 | 0.000298805 | 17 | NaN | 313.95825 | 310.43356 | 239.67549 | 289.86212 | 296.37137 |
| Bola2 | LVNECLAEELPHIH | -55 | -68 | 0 | 0 | 14 | NaN | 1742.1027 | NaN | 953.7219 | 1271.0176 | 1160.2406 |
| Cbfb | RQQDPSPGSNLGGGDDLKL | -168 | -186 | 2 | 0.00111006 | 19 | NaN | 461.35135 | NaN | 1703.9486 | NaN | 1609.7507 |
| Cbfb | FENEEFFR | -12 | -19 | 0 | 0.000627943 | 8 | NaN | 430.73117 | NaN | NaN | NaN | NaN |
| Atp5pf | RQASGGPVDIGPEYQQDLDR | -54 | -73 | 1 | 0 | 20 | NaN | 486.56393 | NaN | NaN | 504.4972 | 199.87338 |
| Ctps1 | HRFEVNPVLK | -480 | -489 | 1 | 0 | 10 | NaN | 686.4199 | 807.8225 | 676.8906 | 991.5346 | 656.57825 |
| Ctps1 | GLSPDLVVCR | -208 | -217 | 0 | 0.001291103 | 10 | NaN | 2532.2842 | 3729.1934 | 4085.0293 | NaN | NaN |
| Ctps1 | YHPEFLSR | -525 | -532 | 0 | 0.002752526 | 8 | NaN | 835.1313 | NaN | 1001.4585 | NaN | NaN |
| Txndc5 | GYPTLLLFR | -380 | -388 | 0 | 0.003681885 | 9 | NaN | 204.54529 | NaN | 161.28958 | 491.66904 | 427.39606 |
| Hells | FFFDPVR | -180 | -186 | 0 | 0.004983786 | 7 | NaN | 1041.6143 | 1047.2756 | NaN | 773.84985 | 957.1732 |
| Hells | FTPEIPTLLYHGT | -276 | -288 | 0 | 0.002595029 | 13 | NaN | 581.25824 | NaN | NaN | 575.8723 | NaN |
| Cmc2 | FFGHCNDLDR | -33 | -42 | 0 | 0 | 10 | NaN | 742.73 | NaN | 786.55444 | 764.5217 | NaN |
| Prrc2a | FGAPDKR | -495 | -501 | 1 | 0.004439092 | 7 | NaN | 5034.186 | NaN | NaN | NaN | NaN |
| Nop14 | HTSADDLNDGFIL | -309 | -321 | 0 | 0.007546675 | 13 | NaN | 627.81366 | NaN | NaN | NaN | NaN |
| Naa50 | FGFEIIETK | -127 | -135 | 0 | 0.00294445 | 9 | NaN | 579.00757 | 553.8725 | NaN | 675.6271 | NaN |
| Tardbp | FTEYETQVK | -152 | -160 | 0 | 0 | 9 | NaN | 1691.3231 | 1773.8229 | 1379.3035 | NaN | NaN |
| Prkch | FGINIPHK | -240 | -247 | 0 | 0.002265552 | 8 | NaN | 844.17194 | NaN | NaN | 632.2413 | 572.4415 |
| Arhgap25 | FGPHLVPILVEK | -172 | -183 | 0 | 0 | 12 | NaN | 211.26567 | 411.99368 | 562.452 | NaN | 276.59174 |
| Arhgap25 | SSVGWDATEDPPLSR | -379 | -393 | 0 | 0.00025015 | 15 | NaN | 549.80286 | 413.9026 | NaN | 1143.894 | 521.60486 |
| Snx9 | FGSAIPIPSLPDK | -301 | -313 | 0 | 0.001291103 | 13 | NaN | 344.04193 | NaN | NaN | NaN | NaN |
| Ptpa | SQLIDHPHLEPR | -225 | -236 | 0 | 0.00025015 | 12 | NaN | 2922.5486 | 2458.5574 | NaN | 4811.5234 | 3322.9004 |
| Ptpa | FPVIQHF | -303 | -309 | 0 | 0.003681885 | 7 | NaN | 664.04065 | 190.78181 | 662.0485 | NaN | NaN |
| Abracl | MNVEHEVNLLVEEIHR | -1 | -16 | 0 | 0.00025015 | 16 | NaN | 328.9158 | NaN | 298.21384 | 697.8114 | 550.531 |
| Prdx6 | PTLSEEEAK | -191 | -199 | 0 | 0.000550307 | 9 | NaN | 1204.5405 | 1196.0626 | 1658.8848 | 204.6254 | NaN |
| Utp25 | FHFFINK | -611 | -617 | 0 | 0 | 7 | NaN | 6017.734 | 5068.149 | NaN | 10976.642 | 11236.374 |
| Cox6a1 | LFHNPHVNPLPTGYEDE | -95 | -111 | 0 | 0 | 17 | NaN | 518.739 | 595.07965 | NaN | 912.6115 | 613.2094 |
| Cox6a1 | LFHNPHVNPLPTGYEDE | -95 | -111 | 0 | 0 | 17 | NaN | 518.739 | 595.07965 | NaN | 912.6115 | 613.2094 |
| Igf2r | VRDGPQWTDGVTVL | -1433 | -1446 | 1 | 0 | 14 | NaN | 711.19965 | NaN | NaN | 117.55255 | 1093.3412 |
| Igf2r | LSSDVCSAHDGSK | -1258 | -1270 | 0 | 0.003521747 | 13 | NaN | 425.30457 | NaN | NaN | NaN | NaN |
| Slfn1 | PSDRPGDDTQEEGHIQ | -156 | -171 | 1 | 0.000431655 | 16 | NaN | 757.8365 | 757.13794 | 204.7422 | 847.831 | 901.9358 |
| Slfn1 | SQSPDVSVGKPA | -104 | -115 | 1 | 0 | 12 | NaN | 3340.0347 | NaN | NaN | NaN | NaN |
| Slc25a1 | SVFGNTPLDVIK | -234 | -245 | 0 | 0 | 12 | NaN | 503.48135 | 476.56708 | 782.10095 | 372.36914 | 739.09717 |
| Cnot10 | NALLLLPEEQQDPK | -467 | -480 | 0 | 0 | 14 | NaN | 2007.4597 | NaN | NaN | 2741.842 | 2889.7476 |
| Cnot10 | NHADQLLQQPK | -554 | -564 | 0 | 0.000230707 | 11 | NaN | 685.2946 | NaN | 1190.5048 | NaN | 906.36066 |
| Cnot10 | FIPAPPSSPLR | -514 | -524 | 0 | 0 | 11 | NaN | 468.0005 | 446.17578 | NaN | NaN | NaN |
| Pacs1 | QVDYWLGHPGER | -821 | -832 | 0 | 0 | 12 | NaN | 675.8652 | 730.00214 | 856.95447 | NaN | 899.43005 |
| Pacs1 | FIPFIGVVK | -743 | -751 | 0 | 0.007791272 | 9 | NaN | 314.16824 | NaN | 182.74399 | NaN | NaN |
| Tapbp | QSGHLQLPPVTAK | -365 | -377 | 0 | 0 | 13 | NaN | 1816.988 | 2420.0522 | NaN | 1741.2683 | NaN |
| Tapbp | SGHLQLPPVTAK | -366 | -377 | 0 | 0 | 12 | NaN | 258.00772 | NaN | NaN | NaN | NaN |
| Cops8 | GVLEQGWQADSTTR | -147 | -160 | 0 | 0 | 14 | NaN | 766.2361 | 893.6812 | 1148.9393 | 1411.3037 | 1475.1288 |
| Eif2d | FITWPLVLEK | -96 | -105 | 0 | 0.001868731 | 10 | NaN | 937.2792 | NaN | NaN | NaN | 1156.5477 |
| Ubr5 | NSLEDLTAEDFR | -2658 | -2669 | 0 | 0 | 12 | NaN | 141.20609 | 421.8495 | 542.6391 | NaN | 397.0138 |
| Arhgap1 | LEQLGIPR | -201 | -208 | 0 | 0 | 8 | NaN | 900.1112 | 837.1482 | 397.07288 | 683.1157 | 659.9238 |
| Thop1 | NFEDVSYESTLK | -55 | -66 | 0 | 0 | 12 | NaN | 630.7493 | 1040.6445 | 849.36 | 1088.9536 | 1230.2769 |
| Thop1 | FLDELAQK | -294 | -301 | 0 | 0.005884925 | 8 | NaN | 382.33115 | NaN | 258.967 | 813.05334 | NaN |
| Pitpna | VENFIHK | -210 | -216 | 0 | 0.001291103 | 7 | NaN | 715.89276 | NaN | NaN | NaN | 506.0375 |
| Pitpna | FKWWGLQNK | -201 | -209 | 1 | 0.004439092 | 9 | NaN | 1840.3053 | NaN | NaN | NaN | NaN |
| Hacd3 | RPLFLAPDFDR | -99 | -109 | 1 | 0.00142999 | 11 | NaN | 1367.1675 | NaN | 939.1527 | 1289.3196 | 1818.7472 |
| Hacd3 | GDNVYEFHLE | -51 | -60 | 0 | 0.005301614 | 10 | NaN | 227.76637 | NaN | NaN | NaN | NaN |
| Bag2 | FLDDLGNAK | -124 | -132 | 0 | 0.000431655 | 9 | NaN | 672.1029 | 655.8496 | NaN | 740.3366 | 665.8794 |
| Ppcs | FLDNFSSGR | -56 | -64 | 0 | 0 | 9 | NaN | 931.10675 | 445.58347 | NaN | 84.583206 | 648.0309 |
| Pzp | FLELTLPYSVVR | -805 | -816 | 0 | 0 | 12 | NaN | 907.53864 | NaN | NaN | 870.1733 | 977.1563 |
| Smarce1 | IAAEIAQAEEQAR | -285 | -297 | 0 | 0.000712251 | 13 | NaN | 826.61 | 735.30975 | NaN | 664.24133 | NaN |
| Huwe1 | LLEEICNLGR | -684 | -693 | 0 | 0 | 10 | NaN | 502.87518 | NaN | 730.24146 | NaN | NaN |
| Huwe1 | IVNQPSSLFGSK | -2256 | -2267 | 0 | 0 | 12 | NaN | 419.07736 | 184.99286 | NaN | NaN | NaN |
| Huwe1 | FLGDEQDQITFVTR | -418 | -431 | 0 | 0 | 14 | NaN | 395.20468 | NaN | NaN | NaN | NaN |
| Ddx51 | GGYQPSDLCVSAPTGSGK | -217 | -234 | 0 | 0.00025015 | 18 | NaN | 1614.8779 | 1639.9408 | 1685.7184 | NaN | 1932.2141 |
| Ddx51 | YAGPEAAGQGDTDAEAGSR | -9 | -27 | 0 | 0.001636916 | 19 | NaN | 392.32178 | NaN | 311.89697 | NaN | NaN |
| Vbp1 | FLLADNLYCK | -104 | -113 | 0 | 0 | 10 | NaN | 896.4395 | 833.74634 | NaN | NaN | NaN |
| Cfdp1 | VFDFAGEEVR | -164 | -173 | 0 | 0 | 10 | NaN | 1216.5159 | 1036.0687 | NaN | 317.7952 | NaN |
| Ndufa9 | SEIEETKPAK | -364 | -373 | 1 | 0 | 10 | NaN | 2050.3765 | 406.60123 | 361.02698 | 2041.1752 | 2366.0938 |
| Ndufa9 | YLLFHLVK | -271 | -278 | 0 | 0.001023236 | 8 | NaN | 399.38184 | 471.6516 | 465.98456 | 591.273 | 735.359 |

| | | | | | | | | | | | | |
|---|---|---|---|---|---|---|---|---|---|---|---|---|
| Ndufa9 | WLSSEIEETKPAK | -361 | -373 | 1 | 0 | 13 | NaN | 622.49945 | 686.6254 | NaN | NaN | NaN |
| Hspe1 | VVLDDKDYFLF | -81 | -91 | 1 | 0.001092529 | 11 | NaN | 949.518 | NaN | 243.24146 | 1230.1924 | 732.1752 |
| Gar1 | TEENKVPYFNAPVYLENK | -101 | -118 | 1 | 0 | 18 | NaN | 382.9636 | 543.15967 | 369.8179 | 315.59222 | 2319.3008 |
| Gar1 | FLPRPPGEK | -168 | -176 | 1 | 0 | 9 | NaN | 3105.1792 | 3339.5852 | 2753.6118 | 2612.2148 | 3457.4045 |
| Gar1 | KFQDQGPPER | -71 | -80 | 1 | 0.000431655 | 10 | NaN | 1218.0884 | 1336.1703 | 1024.3524 | NaN | NaN |
| Sf3b4 | FLQAGPVVNTHMPK | -33 | -46 | 0 | 0.002199523 | 14 | NaN | 2529.8125 | NaN | NaN | NaN | NaN |
| Ccdc88b | KLGVLEEEVR | -1054 | -1063 | 1 | 0.000230707 | 10 | NaN | 715.39185 | NaN | NaN | NaN | NaN |
| Cul3 | NNSGLSFEELYR | -48 | -59 | 0 | 0 | 12 | NaN | 881.4621 | NaN | 854.57544 | 885.2008 | 1298.8239 |
| Cul3 | FLQESFNNDR | -355 | -364 | 0 | 0 | 10 | NaN | 1662.8201 | 1191.7635 | 1743.7966 | NaN | NaN |
| Cul3 | TEQEVETILDK | -404 | -414 | 0 | 0.004690496 | 11 | NaN | 1338.8757 | NaN | NaN | NaN | NaN |
| Xpo7 | TNNPLPLEQR | -74 | -83 | 0 | 0.000907206 | 10 | NaN | 1096.5262 | 948.28174 | 371.5308 | 933.87115 | 981.1911 |
| Dap3 | FLSNCNPEQLER | -375 | -386 | 0 | 0 | 12 | NaN | 1041.9523 | 649.94165 | NaN | NaN | NaN |
| Eif3l | RYQDAIR | -328 | -334 | 1 | 0 | 7 | NaN | 3029.3335 | 3137.1787 | 2637.8962 | 2480.0825 | 2618.401 |
| Eif3l | NFIQYFHK | -54 | -61 | 0 | 0.00025015 | 8 | NaN | 1550.665 | 1734.2639 | 2533.7615 | 1896.8965 | 1790.4026 |
| Eif3l | VLENIELNK | -293 | -301 | 0 | 0 | 9 | NaN | 11158.092 | 12368.051 | 11078.942 | 9551.094 | NaN |
| Gpx1 | TGGEPVSLGSLR | -23 | -34 | 0 | 0 | 12 | NaN | 886.82153 | 834.3706 | NaN | 628.9633 | NaN |
| Xpo6 | FNDALTVVER | -580 | -589 | 0 | 0 | 10 | NaN | 219.32755 | 220.04306 | NaN | NaN | NaN |
| Clptm1 | RAEDYGPVEVISH | -225 | -237 | 1 | 0.001023236 | 13 | NaN | 1643.4095 | 1700.0137 | 1872.1692 | 2156.2075 | 2012.1833 |
| Clptm1 | GIFPCPTFK | -450 | -458 | 0 | 0.001610812 | 9 | NaN | 75.93087 | NaN | NaN | NaN | 801.61127 |
| Lgals9 | FNENAVVR | -270 | -277 | 0 | 0.003133457 | 8 | NaN | 880.9002 | NaN | NaN | NaN | NaN |
| M6pr | FNPVSEER | -155 | -162 | 0 | 0 | 8 | NaN | 1481.162 | NaN | NaN | NaN | NaN |
| Aebp2 | FNSSPDLADHIR | -265 | -276 | 0 | 0 | 12 | NaN | 338.77713 | 1340.0388 | NaN | NaN | NaN |
| Ppif | FPDENFTLK | -124 | -132 | 0 | 0 | 9 | NaN | 1039.7264 | 2001.7454 | 2015.1708 | 1244.4961 | 1633.4905 |
| Vps4b | SSTKPTVNEQDLLK | -419 | -432 | 1 | 0 | 14 | NaN | 2033.7069 | 1997.6196 | 2269.9326 | 2274.7847 | 2393.0842 |
| Vps4b | IVIERPNVK | -124 | -132 | 1 | 0.001291103 | 9 | NaN | 546.9411 | NaN | 694.5186 | 675.13354 | 715.6292 |
| Tap2 | LDGEPLTEYDHHYLHR | -528 | -543 | 0 | 0 | 16 | NaN | 705.43085 | 536.23926 | NaN | 199.71378 | 676.8057 |
| Tap2 | SFLLYQEEVGQYVR | -410 | -423 | 0 | 0 | 14 | NaN | 613.1029 | NaN | 870.518 | 1234.3391 | 1037.0482 |
| Tap2 | QVVLVGQEPVLFSG | -544 | -557 | 0 | 0.002629273 | 14 | NaN | 528.8162 | NaN | 477.7511 | 531.8457 | 526.25195 |
| Tap2 | LHTVQNADQVLVL | -662 | -674 | 0 | 0.00111006 | 13 | NaN | 387.50873 | 414.6867 | NaN | 629.0011 | 632.79584 |
| Nbeal2 | LPPTLLLR | -193 | -200 | 0 | 0.002296977 | 8 | NaN | 695.661 | 638.0272 | 710.3686 | NaN | 742.9163 |
| Nbeal2 | NNICLDLSPGHGLDGR | -855 | -870 | 0 | 0 | 16 | NaN | 871.1458 | 694.1501 | NaN | 526.97046 | NaN |
| Nbeal2 | TVQTSLLGLVR | -1087 | -1097 | 0 | 0.002296977 | 11 | NaN | 218.25087 | NaN | NaN | 210.30559 | 221.32907 |
| Nbeal2 | FQENLQDAER | -183 | -192 | 0 | 0 | 10 | NaN | 1401.0496 | 649.4418 | NaN | NaN | NaN |
| Nbeal2 | LWWLCDSCPASR | -474 | -485 | 0 | 0 | 12 | NaN | 496.9162 | NaN | NaN | 329.8069 | NaN |
| Nbeal2 | QAGWQDVLTR | -1277 | -1286 | 0 | 0.002780285 | 10 | NaN | 3644.0852 | NaN | NaN | NaN | NaN |
| Derl1 | TVAVPLIGK | -22 | -30 | 0 | 0.00383302 | 9 | NaN | 521.1558 | NaN | NaN | NaN | NaN |
| Cab39 | FQNDRTEDEQFNDEK | -309 | -323 | 1 | 0.001636916 | 15 | NaN | 636.53723 | 600.1996 | 509.04288 | 489.62985 | 486.19904 |
| Cab39 | TEDEQFNDEK | -314 | -323 | 0 | 0 | 10 | NaN | 67.51812 | 774.2026 | 757.6413 | 789.2261 | 914.1393 |
| Mug1 | FQVENSNR | -1292 | -1299 | 0 | 0 | 8 | NaN | 1223.26 | 986.95746 | 1153.2611 | 250.75931 | NaN |
| Ubtf | RAEEIWQQ | -487 | -494 | 1 | 0.003681885 | 8 | NaN | 276.10892 | NaN | 481.21765 | NaN | 268.6172 |
| Psmb7 | FRPDMEEEEAK | -185 | -195 | 1 | 0.000230707 | 11 | NaN | 675.275 | NaN | 435.87286 | 980.07245 | 1009.9517 |
| Psmb7 | SIYPHGSTDK | -155 | -164 | 0 | 0.00437931 | 10 | NaN | 1152.9684 | NaN | NaN | NaN | NaN |
| Hpf1 | FSEADVSSDLRK | -28 | -39 | 1 | 0 | 12 | NaN | 3056.2937 | 2363.0278 | 918.2656 | 3420.5989 | 2820.9956 |
| Ilk | LHTLEPLIPR | -308 | -317 | 0 | 0 | 10 | NaN | 509.0915 | 493.00903 | 635.6158 | 497.90878 | 527.2323 |
| Oas3 | SGVLVQPWEGPGLPR | -331 | -345 | 0 | 0 | 15 | NaN | 719.6906 | NaN | NaN | 927.5089 | 966.9821 |
| Prps2 | FSNQETSVEIGESVR | -35 | -49 | 0 | 0 | 15 | NaN | 1093.012 | 1039.6858 | NaN | 824.81256 | 743.3977 |
| Prps2 | SNQETSVEIGESVR | -36 | -49 | 0 | 0 | 14 | NaN | 614.9129 | NaN | NaN | NaN | 697.7206 |
| Celf2 | MFVGQIPR | -42 | -49 | 0 | 0.005459848 | 8 | NaN | 1433.1519 | 718.31 | NaN | NaN | NaN |
| Scarb2 | SFPHFYQADEK | -338 | -348 | 0 | 0.000471409 | 11 | NaN | 1675.0455 | NaN | 1256.2418 | NaN | NaN |
| Trip12 | VDGGAECLVEIR | -1167 | -1178 | 0 | 0.001023236 | 12 | NaN | 788.2298 | NaN | NaN | 609.7471 | NaN |
| Trip12 | TNASADLGSPSLQHSR | -1002 | -1017 | 0 | 0 | 16 | NaN | 261.02716 | 335.05765 | NaN | NaN | NaN |
| Nedd4 | FTADPYLELHNDDTR | -64 | -78 | 0 | 0.000431655 | 15 | NaN | 681.5916 | NaN | NaN | NaN | NaN |
| Elof1 | FTCPFCNHEK | -24 | -33 | 0 | 0 | 10 | NaN | 880.47 | 881.8156 | 935.4884 | 751.46136 | NaN |
| Nfatc2 | YLNPIEEEPIAHK | -38 | -50 | 0 | 0 | 13 | NaN | 375.53802 | NaN | 484.588 | 450.16727 | 449.85092 |
| Nfatc2 | TYHPVPAIK | -676 | -684 | 0 | 0.00025015 | 9 | NaN | 324.10648 | 656.589 | NaN | 746.81506 | NaN |
| Nfatc2 | YDYLNPIEEEPIAHK | -36 | -50 | 0 | 0 | 15 | NaN | 167.00887 | 157.04524 | 396.88843 | NaN | NaN |
| Nrm | FTSLRPLLGGIPESGGPDAR | -28 | -47 | 1 | 0 | 20 | NaN | 2455.7258 | 2318.8196 | NaN | NaN | 1600.4785 |
| Bud13 | FVDERPEEVK | -82 | -91 | 1 | 0.000591124 | 10 | NaN | 272.16034 | 313.56265 | 628.0012 | 347.7604 | 478.26572 |
| Eif5 | FVLCPECENPETDLHVNPK | -96 | -114 | 0 | 0 | 19 | NaN | 351.6943 | 557.26416 | 514.82336 | 594.0887 | 590.47876 |
| Prorsd1 | FVLDSAFLEGGHEK | -120 | -133 | 0 | 0 | 14 | NaN | 755.3016 | 618.0622 | NaN | 77.20099 | 679.32733 |
| Ap2a2 | TTQIGCLLR | -897 | -905 | 0 | 0 | 9 | NaN | 583.92596 | 1108.1754 | 337.4345 | 1206.8488 | 462.95438 |
| Tomm40 | VIHQLSPGLR | -164 | -173 | 0 | 0.001055699 | 10 | NaN | 371.74954 | 479.09875 | NaN | 478.3709 | 470.21268 |
| Tomm40 | SEDGSCGCLPNPGTFEECHR | -69 | -88 | 0 | 0.00111006 | 20 | NaN | 413.10904 | NaN | NaN | NaN | NaN |
| Psme3ip1 | KQQEYEEQFK | -65 | -74 | 1 | 0.000550307 | 10 | NaN | 852.22974 | NaN | NaN | NaN | NaN |
| Mpp1 | IAPTDQGTQTEALQQLQK | -400 | -417 | 0 | 0.002401448 | 18 | NaN | 373.25336 | NaN | NaN | NaN | 1147.2502 |
| Pik3r4 | FWDLVSPER | -1267 | -1275 | 0 | 0.000550307 | 9 | NaN | 705.54376 | NaN | 343.07733 | NaN | 423.66107 |
| Prkcd | FWLDLQPQAK | -104 | -113 | 0 | 0.00201047 | 10 | NaN | 77.85839 | NaN | NaN | NaN | NaN |
| Las1l | YLLDQPVIEHR | -715 | -725 | 0 | 0 | 11 | NaN | 554.3847 | NaN | 410.75565 | 695.61 | 805.6027 |
| Las1l | WGTFPLGR | -688 | -695 | 0 | 0.004224523 | 8 | NaN | 529.2174 | NaN | 545.9143 | 406.0824 | NaN |
| Ppp6c | VWSDPEDVDTWAISPR | -195 | -210 | 0 | 0 | 16 | NaN | 397.42676 | 119.90901 | 472.16144 | 708.89014 | 392.24976 |
| Oxct1 | SFAPEDIHIPK | -253 | -263 | 0 | 0 | 11 | NaN | 475.52612 | 915.693 | 1552.4698 | 1181.5253 | 237.66364 |
| Sec61b | FYTEDSPGLK | -58 | -67 | 0 | 0 | 10 | NaN | 1520.05 | 209.16116 | 1992.6658 | 212.08609 | 1755.76 |
| Bcap31 | YMEENDQLK | -150 | -158 | 0 | 0.002910424 | 9 | NaN | 9390.655 | 7201.5234 | NaN | NaN | NaN |
| Adh5 | GHEGAGIVESVGEGVTK | -66 | -82 | 0 | 0 | 17 | NaN | 727.29175 | 575.39325 | 1630.2358 | 1113.0989 | 1055.8086 |
| Tes | HFCCFDCDHILAGK | -327 | -340 | 0 | 0 | 14 | NaN | 664.7856 | 641.13293 | 496.3529 | 924.7563 | 755.5974 |
| Pus7 | IEEYGLRPVPGDLVLK | -491 | -506 | 1 | 0.00111006 | 16 | NaN | 228.04938 | 296.18994 | 472.1481 | 481.67593 | 500.02603 |
| Pus7 | GCLVVEDNDSVTPHDETK | -11 | -28 | 0 | 0.00025015 | 18 | NaN | 322.04282 | 395.464 | NaN | 422.56348 | 395.4176 |
| Pus7 | GLRPVPGDLVLK | -495 | -506 | 1 | 0.000755556 | 12 | NaN | 650.3308 | NaN | 688.3073 | 773.6893 | NaN |

| | | | | | | | | | | | | |
|---|---|---|---|---|---|---|---|---|---|---|---|---|
| Rcc1 | KKPALVPLLQDVVQA | -58 | -72 | 2 | 0.001636916 | 15 | NaN | 920.3395 | 507.3346 | 769.0477 | NaN | 1102.756 |
| Rcc1 | QLGTGQDEDAWSPVEMTGK | -376 | -394 | 0 | 0.00111006 | 19 | NaN | 539.2711 | NaN | 365.786 | 1117.0392 | 1108.0334 |
| Rcc1 | VPELFANR | -207 | -214 | 0 | 0.001636916 | 8 | NaN | 1147.8943 | 1103.706 | NaN | NaN | 414.13028 |
| Rcc1 | SMVPVQVQLDAPVVK | -162 | -176 | 0 | 0.001392869 | 15 | NaN | 989.66705 | NaN | NaN | NaN | NaN |
| Itgb7 | LTLAPNLDDGWCK | -677 | -689 | 0 | 0.005946382 | 13 | NaN | 1059.9072 | 1242.7059 | NaN | NaN | NaN |
| Psap | GCSFLPDPYQK | -472 | -482 | 0 | 0 | 11 | NaN | 2098.707 | 2250.6301 | 268.21423 | 2696.385 | 2514.7297 |
| Psap | LVLYLEHNLEK | -448 | -458 | 0 | 0 | 11 | NaN | 2368.0227 | 2447.7214 | NaN | 3190.5444 | 3162.5515 |
| Psap | TCSGGSAVLCR | -24 | -34 | 0 | 0.00025015 | 11 | NaN | 599.0069 | NaN | 671.80554 | NaN | NaN |
| Psap | GFVDHVKEDCDR | -220 | -231 | 1 | 0 | 12 | NaN | 283.87534 | NaN | NaN | NaN | 394.07294 |
| Polr2h | GDANNLHGFEVDSR | -127 | -140 | 0 | 0 | 14 | NaN | 908.82983 | NaN | 770.7989 | 249.1075 | 593.40735 |
| Polr2h | LQGDANNLHGF | -125 | -135 | 0 | 0.00025015 | 11 | NaN | 763.3804 | 3619.6267 | NaN | NaN | 820.3574 |
| Eif1 | NLHSFDPFADASK | -6 | -18 | 0 | 0 | 13 | NaN | 448.39853 | 544.6309 | 321.0876 | 279.45557 | 255.43855 |
| Eif1 | SKGDDLLPAGTEDYIHIR | -17 | -34 | 1 | 0.000431655 | 18 | NaN | 3308.5798 | 3261.2368 | NaN | NaN | 3040.5813 |
| Lsm3 | TTNTVEEPLDLIR | -10 | -22 | 0 | 0 | 13 | NaN | 520.5806 | 471.89655 | 833.02185 | NaN | 124.16629 |
| Txndc12 | GDHIHWR | -33 | -39 | 0 | 0.00025015 | 7 | NaN | 300.3617 | 452.5443 | 477.79785 | 1706.349 | 880.14886 |
| Txndc12 | GFGDHIHWR | -31 | -39 | 0 | 0.000230707 | 9 | NaN | 221.87274 | NaN | NaN | 7312.7427 | NaN |
| Sec11a | GDLLFLTNR | -64 | -72 | 0 | 0.00025015 | 9 | NaN | 666.6309 | 682.11084 | 531.18634 | NaN | 614.98456 |
| Emb | YSCVFGEK | -143 | -150 | 0 | 0 | 8 | NaN | 541.68036 | 531.0954 | 637.04114 | 685.9807 | 665.23944 |
| Emb | GDPTDPTFTSLPVR | -33 | -46 | 0 | 0.001150701 | 14 | NaN | 1414.057 | 519.7611 | NaN | NaN | NaN |
| Emb | YVGDSTVLK | -173 | -181 | 0 | 0.000869338 | 9 | NaN | 175.84 | 453.79956 | NaN | NaN | NaN |
| Srprb | VGDGAGGAFQPYLDSLR | -9 | -25 | 0 | 0.00025015 | 17 | NaN | 670.53546 | 697.72546 | NaN | NaN | 928.8481 |
| Ttc27 | GDVLQDHSCYDK | -510 | -521 | 0 | 0.00111006 | 12 | NaN | 611.01605 | NaN | 747.2174 | NaN | NaN |
| Spn | GEEEPLVGSEDEAVETPTSDGPQAK | -363 | -387 | 0 | 0 | 25 | NaN | 254.615 | 264.8589 | 316.28873 | NaN | 334.05716 |
| Cdc23 | GETVEHLEESTAFR | -501 | -514 | 0 | 0.00025015 | 14 | NaN | 1001.58246 | 1404.9132 | NaN | NaN | NaN |
| Agk | GGDGTLQEVVTGVLR | -123 | -137 | 0 | 0.00025015 | 15 | NaN | 434.96143 | NaN | NaN | 432.9087 | 517.52344 |
| Agk | GETVPLDVLQIK | -185 | -196 | 0 | 0 | 12 | NaN | 162.91167 | NaN | 556.4951 | NaN | NaN |
| Luc7l3 | GFCPAELFTNTR | -41 | -52 | 0 | 0.00025015 | 12 | NaN | 172.82402 | NaN | 526.86285 | 471.55762 | 188.43614 |
| Luc7l3 | TEEPDRDER | -238 | -246 | 1 | 0.001392869 | 9 | NaN | 428.05505 | NaN | 700.8402 | NaN | NaN |
| Ngly1 | GFEEGETHLIFPK | -79 | -91 | 0 | 0 | 13 | NaN | 3375.4731 | NaN | NaN | NaN | NaN |
| Glipr2 | SYNFQQPGFTSGTGH | -89 | -103 | 0 | 0.00078661 | 15 | NaN | 246.1704 | NaN | NaN | NaN | NaN |
| Etf1 | VNIDFEPFKPINTS | -110 | -123 | 1 | 0.00078661 | 14 | NaN | 461.58673 | 629.1922 | NaN | NaN | NaN |
| Dhodh | TFLGPPVVAR | -359 | -368 | 0 | 0.002534621 | 10 | NaN | 1664.7723 | 1700.9661 | NaN | 1778.4567 | 2022.164 |
| Padi2 | SLTQENQYFQR | -544 | -554 | 0 | 0 | 11 | NaN | 1047.5112 | NaN | NaN | NaN | NaN |
| Sqstm1 | SAPPEDPNVNFLK | -226 | -238 | 0 | 0.005884925 | 13 | NaN | 3543.6047 | 4388.1006 | NaN | NaN | NaN |
| Sqstm1 | LIFPNPFGHLS | -166 | -176 | 0 | 0.005394768 | 11 | NaN | 592.78796 | NaN | NaN | 743.6789 | NaN |
| Cbx5 | NLDCPELISEF | -56 | -66 | 0 | 0 | 11 | NaN | 166.60254 | 187.46896 | 215.51056 | 255.46916 | 246.29533 |
| Cbx5 | LTWHAYPEDAENKEK | -172 | -186 | 1 | 0.002595029 | 15 | NaN | 2280.8245 | NaN | NaN | NaN | NaN |
| Lasp1 | TQDQISNIK | -113 | -121 | 0 | 0.00025015 | 9 | NaN | 1213.3568 | NaN | 546.6189 | NaN | NaN |
| Slc16a3 | KLHKPPVDVGVDSR | -433 | -446 | 2 | 0.00122419 | 14 | NaN | 368.54056 | NaN | 487.66016 | NaN | 746.3064 |
| Aldh3a2 | GGIPETTELLK | -165 | -175 | 0 | 0.000230707 | 11 | NaN | 3174.5386 | 211.35449 | 3668.8718 | 2246.3145 | 2578.0425 |
| Ostc | VPFLVLECPNLK | -7 | -18 | 0 | 0 | 12 | NaN | 351.2814 | NaN | 988.10864 | 1318.3777 | 523.6841 |
| Acad10 | GGLVISPEGLSPAVR | -650 | -664 | 0 | 0 | 15 | NaN | 383.5328 | 447.8167 | 450.30377 | NaN | 421.4055 |
| Ptprcap | LWGHTCR | -74 | -80 | 0 | 0.00266297 | 7 | NaN | 1379.5416 | NaN | 684.2742 | 842.91833 | NaN |
| B4galnt1 | WVDDDFVFTAR | -354 | -364 | 0 | 0 | 11 | NaN | 412.55133 | 388.25894 | 240.90646 | 522.27966 | NaN |
| B4galnt1 | NAPSLPNPLAL | -29 | -39 | 0 | 0.002099344 | 11 | NaN | 346.60883 | NaN | 323.23978 | NaN | 334.06604 |
| B4galnt1 | IVIADDSDKPER | -308 | -319 | 1 | 0.00025015 | 12 | NaN | 1200.2699 | NaN | NaN | NaN | NaN |
| B4galnt1 | WSPPQGPPR | -40 | -48 | 0 | 0.000907206 | 9 | NaN | 891.3192 | NaN | NaN | NaN | NaN |
| Atox1 | GGVEFNIDLPNKK | -27 | -39 | 1 | 0 | 13 | NaN | 1219.5775 | 857.0384 | 1227.7601 | 1643.0116 | 1233.1207 |
| Atox1 | VCIDSEHSSDTLLATL | -40 | -55 | 0 | 0 | 16 | NaN | 722.4613 | 809.9709 | NaN | 719.1908 | 748.1135 |
| Atox1 | MTCEGCAEAVSR | -10 | -21 | 0 | 0.000230707 | 12 | NaN | 168.55937 | 175.86206 | NaN | NaN | 254.24136 |
| Hnrnph2 | GLPWSCSAEEVMR | -17 | -29 | 0 | 0 | 13 | NaN | 816.8738 | NaN | NaN | 1106.12 | 1040.1442 |
| Keap1 | NFAEQIGCTELHQR | -189 | -202 | 0 | 0 | 14 | NaN | 727.18353 | 592.51855 | 613.3546 | 412.9024 | 771.9168 |
| Nudt2 | GHVDPGENDLETALR | -43 | -57 | 0 | 0 | 15 | NaN | 495.3183 | NaN | NaN | 457.30618 | 471.4083 |
| Rps26 | VFDAYVLPK | -58 | -66 | 0 | 0 | 9 | NaN | 124.66084 | NaN | 197.60524 | NaN | 723.98944 |
| Sh3bp1 | LPQENFNNLR | -391 | -400 | 0 | 0 | 10 | NaN | 984.58923 | 1148.892 | 682.1404 | NaN | 841.68134 |
| Sh3bp1 | LAEEQDVNK | -409 | -417 | 0 | 0.001904399 | 9 | NaN | 2864.3237 | 2053.8108 | NaN | 2494.563 | NaN |
| Sh3bp1 | KVEQCKDEYLAD | -201 | -212 | 2 | 0.002810811 | 12 | NaN | 557.32074 | NaN | 1030.5981 | NaN | NaN |
| Yme1l1 | NIPEAHQDAFK | -201 | -211 | 0 | 0 | 11 | NaN | 1385.1704 | 3942.106 | 2987.2847 | 4411.9985 | 5492.161 |
| Yme1l1 | GILLVGPPGTGK | -316 | -327 | 0 | 0 | 12 | NaN | 832.9386 | NaN | 695.30914 | 767.3543 | 794.13477 |
| Nsf | NIDSNPYDTDK | -106 | -116 | 0 | 0 | 11 | NaN | 490.7755 | 634.7541 | 509.37897 | NaN | 752.552 |
| Ncoa5 | GINFDNPSVQK | -532 | -542 | 0 | 0.00111006 | 11 | NaN | 4456.364 | NaN | NaN | NaN | NaN |
| Cyth4 | YCLCNPGVFR | -171 | -180 | 0 | 0.004923919 | 10 | NaN | 1057.1294 | NaN | NaN | NaN | NaN |
| Isoc1 | GIPVIITEQYPK | -150 | -161 | 0 | 0 | 12 | NaN | 943.6942 | 959.4182 | 917.0234 | 1111.8967 | 851.16223 |
| Isoc1 | YFGDIISVGQR | -131 | -141 | 0 | 0 | 11 | NaN | 414.18237 | 448.469 | 1122.6614 | 505.15622 | 654.45276 |
| Cox5a | GMNTLVGYDLVPEPK | -69 | -83 | 0 | 0 | 15 | NaN | 1330.8984 | 1261.4216 | 1402.9716 | 1621.0227 | 1630.9104 |
| Psma4 | LYIGWDK | -135 | -141 | 0 | 0.001708008 | 7 | NaN | 433.6565 | NaN | 754.0109 | 727.7107 | NaN |
| Polr1b | GKEEDSFFR | -271 | -279 | 1 | 0 | 9 | NaN | 1114.6959 | 186.05316 | NaN | 164.31451 | 197.97377 |
| Polr1b | LVRPVQNLELGR | -619 | -630 | 1 | 0 | 12 | NaN | 396.91452 | NaN | NaN | NaN | NaN |
| Idh2 | LVPGWTKPITIGR | -160 | -172 | 1 | 0 | 13 | NaN | 810.64856 | 845.025 | 540.99805 | 811.9174 | 856.1877 |
| Mbd2 | GKPDLNTTLPIR | -238 | -249 | 1 | 0 | 12 | NaN | 1040.5414 | 1129.1895 | 1119.1417 | 1293.6847 | 1294.1292 |
| Cbx1 | KKVEEVLEEEEEEYVVE | -8 | -24 | 2 | 0.000869338 | 17 | NaN | 962.73816 | NaN | 1055.153 | 1072.1815 | NaN |
| Stk26 | TACPGITDK | -390 | -398 | 0 | 0.002265552 | 9 | NaN | 1327.5481 | NaN | NaN | NaN | NaN |
| Kti12 | GLEEPLPLAEIR | -231 | -242 | 0 | 0 | 12 | NaN | 590.88385 | 625.2211 | 701.1183 | 1014.4536 | 685.2155 |
| Prdx2 | NIPLLADVTK | -100 | -109 | 0 | 0.00078661 | 10 | NaN | 2818.8542 | 2765.2886 | NaN | 1153.5024 | NaN |
| Brd2 | GLHDYHDIIK | -381 | -390 | 0 | 0 | 10 | NaN | 1283.8362 | 1103.3269 | 1161.5675 | NaN | NaN |
| Fga | SGSLDELSER | -451 | -460 | 0 | 0.001291103 | 10 | NaN | 641.04004 | 889.43146 | 199.44186 | NaN | 940.52716 |
| Fga | NPGTTGSDGTGDWGTGSPR | -324 | -342 | 0 | 0.001055699 | 19 | NaN | 364.80267 | NaN | 716.357 | NaN | NaN |
| Fga | PGSDSGNFRPANPNWGVF | -343 | -360 | 1 | 0.001868731 | 18 | NaN | 232.49554 | NaN | NaN | NaN | NaN |

| | | | | | | | | | | | | |
|---|---|---|---|---|---|---|---|---|---|---|---|---|
| Golga7 | IYAPQGLLLTDPIER | -104 | -118 | 0 | 0.007791272 | 15 | NaN | 174.59656 | NaN | NaN | NaN | NaN |
| Skap1 | WDCHGDQPDELSFQR | -299 | -313 | 0 | 0 | 15 | NaN | 338.39014 | 282.74567 | NaN | 150.76202 | 324.6258 |
| Dpysl2 | MVIPGGIDVHTR | -64 | -75 | 0 | 0.000431655 | 12 | NaN | 2394.285 | 2033.67 | NaN | 980.7646 | 1054.4849 |
| Dpysl2 | YDGPVCEVSVTPK | -499 | -511 | 0 | 0 | 13 | NaN | 626.247 | NaN | 1642.3524 | 2010.9117 | 777.76575 |
| Dpysl2 | SRPEEVEAEAVNR | -226 | -238 | 1 | 0 | 13 | NaN | 7800.187 | NaN | NaN | NaN | 8982.981 |
| Dpysl2 | IVLEDGTLHVT | -452 | -462 | 0 | 0 | 11 | NaN | 121.38603 | NaN | NaN | 1412.9341 | 1709.0138 |
| Tmed4 | GLYFHIGETEK | -29 | -39 | 0 | 0 | 11 | NaN | 1461.829 | 725.3215 | 942.8895 | NaN | NaN |
| Lgals3 | GVPAGPLTVPYDLPLPGGVMPR | -122 | -143 | 0 | 0.001291103 | 22 | NaN | 324.85114 | NaN | 292.36993 | 454.38025 | 412.9883 |
| Vps26b | GNHHEFVSL | -81 | -89 | 0 | 0.008161009 | 9 | NaN | 679.2679 | 289.16928 | NaN | NaN | NaN |
| Poldip3 | GNPASSNPPAEVDPDTVLR | -381 | -399 | 0 | 0 | 19 | NaN | 6906.4517 | 6536.888 | 6298.8247 | NaN | NaN |
| Aco1 | GPFLLGIK | -786 | -793 | 0 | 0.00830926 | 8 | NaN | 212.91058 | 309.3997 | NaN | NaN | NaN |
| Aco1 | NIEVPFKPAR | -73 | -82 | 1 | 0.002810811 | 10 | NaN | 5512.9736 | NaN | NaN | NaN | NaN |
| Memo1 | IYGELWK | -109 | -115 | 0 | 0.000947459 | 7 | NaN | 1478.0387 | 2235.4983 | 883.66864 | NaN | NaN |
| Aldh18a1 | VSGHVITDIVEGK | -335 | -347 | 0 | 0 | 13 | NaN | 1574.7095 | 1666.2628 | 1632.5808 | 1657.529 | 3285.8347 |
| Aldh18a1 | VTNLDFHDEQK | -189 | -199 | 0 | 0 | 11 | NaN | 967.58777 | NaN | 3374.0898 | 267.13287 | 3207.8962 |
| Aldh18a1 | GQDHVVSDFSEHGSLK | -766 | -781 | 0 | 0 | 16 | NaN | 884.4508 | NaN | 344.84036 | 560.0849 | 487.15817 |
| Aldh18a1 | LDFHDEQK | -192 | -199 | 0 | 0.005270854 | 8 | NaN | 3299.4438 | NaN | NaN | NaN | 4015.9568 |
| Aldh18a1 | GQDHVVSDF | -766 | -774 | 0 | 0.007987481 | 9 | NaN | 803.95703 | NaN | NaN | NaN | NaN |
| Fen1 | LDPSKYPVPENWLHK | -261 | -275 | 1 | 0 | 15 | NaN | 2799.7935 | 2858.5469 | 764.37463 | 919.1631 | 982.598 |
| Fen1 | WSEPNEEELVK | -296 | -306 | 0 | 0 | 11 | NaN | 1130.0396 | 1241.3362 | 1602.1653 | 575.11633 | 239.20792 |
| Fen1 | YVFDGKPPQLK | -81 | -91 | 1 | 0 | 11 | NaN | 1113.109 | 1127.5253 | 1803.2971 | 2186.6648 | 757.88214 |
| Fen1 | WSEPNEEELVKF | -296 | -307 | 1 | 0 | 12 | NaN | 738.9142 | 795.2169 | 839.44214 | 1133.1273 | 547.0879 |
| Add1 | YHDYHGILVDEEEK | -270 | -283 | 0 | 0.004567621 | 14 | NaN | 587.86786 | NaN | NaN | NaN | NaN |
| Abce1 | QLDLTHLK | -203 | -210 | 0 | 0 | 8 | NaN | 840.9785 | NaN | 930.89594 | 1019.5588 | 1053.0513 |
| Atp5mc2;Atp5 | VFGSLIIGYAR | (99);(94);(89) | (109);(104);(99) | 0 | 0.001392869 | 11 | NaN | 369.3354 | 478.30313 | NaN | NaN | NaN |
| Pelp1 | LHDLVLPL | -549 | -556 | 0 | 0.001092529 | 8 | NaN | 416.02643 | NaN | 349.77185 | 433.8304 | 364.25064 |
| Mycbp2 | GSTPTQISEIIIR | -4236 | -4248 | 0 | 0 | 13 | NaN | 1295.8798 | NaN | 1470.0443 | NaN | NaN |
| Mapk3 | GTAGVVPVVPGEVEVVK | -17 | -33 | 0 | 0 | 17 | NaN | 1182.355 | NaN | 328.33612 | 1128.198 | 1166.7598 |
| Git2 | GTFICDECCSVHR | -27 | -39 | 0 | 0 | 13 | NaN | 475.22958 | NaN | NaN | 900.59985 | 1004.5248 |
| Lsm2 | LTDISVTDPEK | -40 | -50 | 0 | 0.00713295 | 11 | NaN | 4509.7935 | 4573.01 | NaN | NaN | 2680.1824 |
| Stx8 | GTPNPWLCEEPEETR | -119 | -133 | 0 | 0 | 15 | NaN | 521.2279 | NaN | NaN | 712.08075 | 329.6966 |
| Nup85 | GVDYFDYCPELGR | -420 | -432 | 0 | 0 | 13 | NaN | 308.4876 | NaN | NaN | NaN | NaN |
| Plk1 | GVENPLPDRPR | -346 | -356 | 1 | 0.000907206 | 11 | NaN | 757.58545 | 1107.7858 | NaN | 874.8437 | 687.2648 |
| Rpl28 | GVEPAADGK | -50 | -58 | 0 | 0.000947459 | 9 | NaN | 901.2491 | 954.60297 | 1091.6979 | 1445.4441 | 1180.4319 |
| Pithd1 | GVFKPWEER | -53 | -61 | 1 | 0.000431655 | 9 | NaN | 2811.8677 | NaN | NaN | 3982.398 | 124.76494 |
| Agpat3 | RFPLEDIPADETSAA | -257 | -271 | 1 | 0.00025015 | 15 | NaN | 1196.6133 | 1455.4125 | NaN | NaN | NaN |
| Gbp7 | LAGQNHGFNLGTTVR | -58 | -72 | 0 | 0.00223472 | 15 | NaN | 384.6287 | NaN | NaN | NaN | NaN |
| Gbp7 | KVEQDYER | -441 | -448 | 1 | 0.002199523 | 8 | NaN | 305.05612 | NaN | NaN | NaN | NaN |
| Pmpca | SSGATYPNIPLSSPLPGVPK | -33 | -52 | 0 | 0 | 20 | NaN | 570.72986 | 549.30096 | 519.8475 | 574.9417 | 516.5629 |
| Pmpca | LTDEEIEMTR | -175 | -184 | 0 | 0.002099344 | 10 | NaN | 192.53778 | NaN | 321.96652 | NaN | NaN |
| Exosc3 | LLAPDCEIVQELGK | -209 | -222 | 0 | 0 | 14 | NaN | 1445.8999 | NaN | NaN | NaN | 2739.7583 |
| Exosc3 | NRPNVQVGDLIYGQC | -152 | -166 | 1 | 0.002595029 | 15 | NaN | 600.89453 | NaN | NaN | 694.3951 | NaN |
| Srpk1 | HKEDLHNANDCDVHTL | -376 | -391 | 1 | 0.00025015 | 16 | NaN | 343.97488 | 324.7803 | NaN | 325.25742 | 242.94168 |
| Srpk1 | GVIGVVNYPENSNNETLR | -358 | -375 | 0 | 0 | 18 | NaN | 472.53116 | 492.47388 | NaN | 413.1753 | NaN |
| Prdx5 | THLPGFVEQA | -99 | -108 | 0 | 0 | 10 | NaN | 2468.3328 | 2979.2068 | 1668.6543 | 1964.0713 | 2401.9836 |
| Prdx5 | KVNLAELFK | -71 | -79 | 1 | 0.003490217 | 9 | NaN | 1415.858 | NaN | 989.64185 | NaN | 1333.098 |
| Cpsf6 | RELHGQSPVVTPC | -147 | -159 | 1 | 0.00122419 | 13 | NaN | 315.49066 | 616.9282 | NaN | 647.59155 | NaN |
| Nubp2 | PGNLAGVR | -11 | -18 | 0 | 0 | 8 | NaN | 601.2134 | 702.7042 | 550.0113 | 564.50494 | 576.4657 |
| Nubp2 | VVTTPQAVSIGDVR | -162 | -175 | 0 | 0 | 14 | NaN | 251.42116 | 170.36386 | NaN | 1310.9132 | NaN |
| Man2b1 | GVSEPLLETDTGDK | -825 | -838 | 0 | 0.001868731 | 14 | NaN | 709.4756 | 661.9953 | 708.1305 | NaN | NaN |
| Man2b1 | VAWHIDPFGHSR | -191 | -202 | 0 | 0 | 12 | NaN | 451.83298 | 443.09924 | NaN | NaN | NaN |
| Pik3cd | YESYLDCELTK | -620 | -630 | 0 | 0 | 11 | NaN | 1447.2247 | 1427.4489 | 1914.371 | 1363.0316 | 1070.8956 |
| Pik3cd | TYDFVHVIQQGK | -934 | -945 | 0 | 0.000230707 | 12 | NaN | 1048.5898 | 796.2276 | NaN | NaN | NaN |
| Nipsnap1 | SFWNEPQPR | -171 | -179 | 0 | 0 | 9 | NaN | 687.895 | 800.4852 | 854.4691 | 758.85706 | NaN |
| Prdx4 | ISKPAPYWEGT | -84 | -94 | 1 | 0.000591124 | 11 | NaN | 1233.881 | 1651.7683 | NaN | 384.715 | 5794.5386 |
| Cyc1 | LSDYFPKPYPNPEAAR | -171 | -186 | 1 | 0.00025015 | 16 | NaN | 2029.7688 | 1852.1454 | 1712.6001 | 2388.6777 | 2234.827 |
| Cyc1 | HGGEDYVFSLL | -205 | -215 | 0 | 0.002068213 | 11 | NaN | 325.61267 | NaN | NaN | 419.96646 | NaN |
| Cyc1 | GYCEPPTGVSLR | -217 | -228 | 0 | 0.002566359 | 12 | NaN | 698.90076 | NaN | NaN | NaN | NaN |
| Skic3 | GYLNEHLQLK | -987 | -996 | 0 | 0.005884925 | 10 | NaN | 1782.6974 | NaN | NaN | NaN | NaN |
| Tyms | HGELQYLR | -22 | -29 | 0 | 0 | 8 | NaN | 2624.4263 | NaN | NaN | 2880.0088 | 2844.0996 |
| Tyms | LNHIEPLK | -253 | -260 | 0 | 0.00078661 | 8 | NaN | 569.0596 | NaN | NaN | NaN | 503.13882 |
| FAM120A | LPPEFSPLIIK | -631 | -641 | 0 | 0.002068213 | 11 | NaN | 664.22644 | 613.1213 | NaN | NaN | NaN |
| Snrpb2 | MDIRPNHTIYIN | -1 | -12 | 1 | 0.00122419 | 12 | NaN | 1046.0151 | NaN | NaN | 1724.314 | 1441.1346 |
| Snrpb2 | QLQGFPFYGK | -68 | -77 | 0 | 0.000230707 | 10 | NaN | 931.18317 | 960.153 | NaN | NaN | 407.84094 |
| Arhgap29 | HDICDVLK | -739 | -746 | 0 | 0.001392869 | 8 | NaN | 3161.8054 | 3013.0154 | NaN | NaN | NaN |
| Rpusd2 | HEPPVTAEPIHLL | -227 | -239 | 0 | 0.000550307 | 13 | NaN | 545.6663 | 532.96094 | 344.23825 | 708.1893 | NaN |
| Ndufa11 | LNPADSTLEAVAR | -43 | -55 | 0 | 0.00025015 | 13 | NaN | 1065.7429 | 899.3634 | 1265.3322 | 2071.1602 | NaN |
| Ndufa11 | HEVPDGTQCHR | -10 | -20 | 0 | 0.001743443 | 11 | NaN | 303.13135 | NaN | NaN | NaN | 260.00455 |
| Fhl3 | HFHEGCFR | -60 | -67 | 0 | 0.000712251 | 8 | NaN | 300.739 | NaN | NaN | NaN | NaN |
| Rpl26 | HVGIHPSK | -96 | -103 | 0 | 0.004099215 | 8 | NaN | 582.927 | 1411.5073 | 1932.5173 | NaN | 1506.4633 |
| Ndufv1 | KTSFGSLKDEDR | -29 | -40 | 2 | 0.00025015 | 12 | NaN | 647.5713 | NaN | 470.30206 | 762.30646 | 846.0854 |
| Apip | NEYPDSCAVLVR | -181 | -192 | 0 | 0 | 12 | NaN | 434.9337 | 431.9571 | NaN | 672.8138 | NaN |
| Mycbp | HHLGAATPENPEIELLR | -50 | -66 | 0 | 0 | 17 | NaN | 781.7848 | 1813.2827 | 407.7751 | 355.38736 | 3284.4785 |
| Dhx36 | HIVLDEIHER | -323 | -332 | 0 | 0 | 10 | NaN | 729.33594 | 700.9281 | 616.07996 | 441.08707 | 364.8516 |
| Dhx36 | LLDDYQLPEILR | -615 | -626 | 0 | 0.008736733 | 12 | NaN | 2635.9082 | NaN | NaN | NaN | NaN |
| Kdm3a | HLDESHLLQGDK | -177 | -188 | 0 | 0 | 12 | NaN | 747.00226 | NaN | 1534.1205 | NaN | NaN |
| Kdm3a | IVDPALIHVE | -233 | -242 | 0 | 0.009955823 | 10 | NaN | 168.01378 | NaN | NaN | NaN | NaN |
| Ampd2 | WLVQVPR | -516 | -522 | 0 | 0 | 7 | NaN | 774.29895 | 830.14886 | 583.0469 | 482.59427 | 570.08655 |

| | | | | | | | | | | | | |
|---|---|---|---|---|---|---|---|---|---|---|---|---|
| Ampd2 | SAPYEFPEESPIEQLEER | -87 | -104 | 0 | 0.00025015 | 18 | NaN | 1012.25696 | 1081.4722 | NaN | 2369.9841 | 2210.4038 |
| Ampd2 | YFAHIIK | -463 | -469 | 0 | 0.001291103 | 7 | NaN | 1112.9446 | NaN | NaN | NaN | NaN |
| Qtrt1 | SLSEVTEEGVHFR | -116 | -128 | 0 | 0 | 13 | NaN | 952.7849 | 1391.4348 | NaN | NaN | NaN |
| Wdr18 | HLLGAEHGDEAQGGGLR | -342 | -358 | 0 | 0 | 17 | NaN | 448.7821 | 529.3284 | NaN | 888.85315 | 1018.84827 |
| Wdr18 | HLLGAEHGDEAQGGGLR | -342 | -358 | 0 | 0 | 17 | NaN | 448.7821 | 529.3284 | NaN | 888.85315 | 1018.84827 |
| Wdr18 | LSGSHDESVR | -285 | -294 | 0 | 0.00078661 | 10 | NaN | 437.75296 | NaN | 433.50092 | NaN | NaN |
| Sdhc | HLLWDLGK | -134 | -141 | 0 | 0.00025015 | 8 | NaN | 1863.8584 | 1986.3385 | 1836.855 | 2131.249 | 2217.2563 |
| Sdhc | NTSSNRPLSPHLT | -45 | -57 | 1 | 0.000431655 | 13 | NaN | 904.8616 | 919.8682 | 840.6128 | 647.8483 | 833.0526 |
| Tmem260 | HLPGVHFPGDR | -476 | -486 | 0 | 0.000431655 | 11 | NaN | 492.78564 | NaN | NaN | 377.44742 | NaN |
| Cisd2 | HNELTGDNVGPLILK | -117 | -131 | 0 | 0 | 15 | NaN | 797.47144 | 735.13965 | 516.61694 | 573.9786 | 610.6042 |
| Cisd2 | TFPACDGSHNK | -106 | -116 | 0 | 0.009745776 | 11 | NaN | 1174.9458 | 1172.064 | NaN | NaN | NaN |
| Bzw2 | HQKPVLTGQ | -4 | -12 | 1 | 0.001868731 | 9 | NaN | 377.4636 | 388.2904 | 429.81274 | NaN | NaN |
| Nipbl | LSESKPNENQLGESK | -620 | -634 | 1 | 0 | 15 | NaN | 947.33923 | 1542.6805 | NaN | 1916.9768 | 2074.7913 |
| Nipbl | HQLNELGSESAK | -1230 | -1241 | 0 | 0 | 12 | NaN | 1747.8998 | NaN | NaN | 995.71783 | NaN |
| Ankfy1 | HQLPLVVDAICTR | -665 | -677 | 0 | 0 | 13 | NaN | 2590.9111 | NaN | NaN | 1893.4991 | 2123.5588 |
| Rassf2 | LQEEEDREVEK | -290 | -300 | 1 | 0.000907206 | 11 | NaN | 965.6319 | 1101.7938 | NaN | NaN | NaN |
| Pisd | HSPTDWTISHR | -264 | -274 | 0 | 0 | 11 | NaN | 552.7297 | 495.01263 | NaN | 467.18225 | NaN |
| Pgam5 | HSQYHVDGSLEK | -104 | -115 | 0 | 0 | 12 | NaN | 2422.8838 | NaN | NaN | 3314.7004 | NaN |
| Utp6 | RFHPECPK | -170 | -177 | 1 | 0.001463897 | 8 | NaN | 444.23663 | NaN | 105.19514 | NaN | 488.54517 |
| Tnfaip8l2 | HVFDHYSDPDLLAA | -145 | -158 | 0 | 0.000712251 | 14 | NaN | 1564.9669 | NaN | NaN | NaN | NaN |
| Tnfaip8l2 | YGPDFTQHLGK | -160 | -170 | 0 | 0 | 11 | NaN | 780.7811 | NaN | NaN | NaN | NaN |
| Samm50 | HVHFDGLGR | -49 | -57 | 0 | 0.00078661 | 9 | NaN | 201.49173 | 249.95552 | NaN | NaN | 122.92565 |
| Mthfd2 | KVLRPEELEVF | -330 | -340 | 2 | 0.000230707 | 11 | NaN | 847.5443 | 759.765 | 951.7682 | 1032.3402 | 1094.0571 |
| Mthfd2 | VQDPVTAKPK | -279 | -288 | 1 | 0.000431655 | 10 | NaN | 354.01913 | 431.37418 | NaN | NaN | 388.00977 |
| Mthfd2 | VSPDKDVDGFHVINVGR | -148 | -164 | 1 | 0.001392869 | 17 | NaN | 381.04315 | NaN | 1389.6041 | NaN | NaN |
| Chd8 | HVVLGSLPGK | -304 | -313 | 0 | 0.001291103 | 10 | NaN | 446.58456 | NaN | NaN | NaN | NaN |
| Rcn1 | HWILPQDYDHAQAEAR | -265 | -280 | 0 | 0 | 16 | NaN | 100.275314 | NaN | NaN | NaN | NaN |
| Rex1bd | HYDFPSYR | -63 | -70 | 0 | 0.00078661 | 8 | NaN | 1812.4276 | NaN | NaN | 685.2098 | 2022.6669 |
| Tecr | LLETLFVHR | -142 | -150 | 0 | 0 | 9 | NaN | 1363.8907 | NaN | 1469.9702 | 1855.6295 | 1912.2572 |
| Ints7 | HYFSVVPGNVGSSPR | -326 | -340 | 0 | 0 | 15 | NaN | 425.14236 | 599.6389 | 609.3181 | 656.6318 | 457.78012 |
| Zmpste24 | HYSHPPLLER | -456 | -465 | 0 | 0.00142999 | 10 | NaN | 744.2339 | 1139.3608 | 881.3505 | 1000.66437 | 54.14399 |
| Stat4 | LPELNYQVK | -351 | -359 | 0 | 0 | 9 | NaN | 807.4634 | NaN | NaN | 723.1975 | 439.33493 |
| Stat4 | HYSSQPCEVSRPTER | -675 | -689 | 1 | 0.00025015 | 15 | NaN | 363.64243 | NaN | NaN | NaN | NaN |
| Wdhd1 | IAAGSSDFLVK | -109 | -119 | 0 | 0 | 11 | NaN | 557.8472 | 1082.3058 | 985.5464 | 1194.0621 | 354.27737 |
| Wdhd1 | NQVLHGDPLPLTR | -604 | -616 | 0 | 0 | 13 | NaN | 347.18027 | NaN | NaN | 2130.1716 | NaN |
| Rprd1b | LAAELEDR | -244 | -251 | 0 | 0.003966045 | 8 | NaN | 522.6072 | NaN | 886.9506 | NaN | NaN |
| Phf5a | TDLFYER | -96 | -102 | 0 | 0.002265552 | 7 | NaN | 463.82114 | NaN | 291.80698 | NaN | NaN |
| Prkcq | IDDFILHK | -377 | -384 | 0 | 0 | 8 | NaN | 2621.0266 | 2887.3171 | NaN | 1705.256 | 2608.2341 |
| Ndufa10 | IDVPVPEVQSR | -207 | -217 | 0 | 0 | 11 | NaN | 551.488 | 768.46387 | NaN | 291.83813 | 1019.76746 |
| Napa | KYEELFPAFSDSR | -227 | -239 | 1 | 0.000230707 | 13 | NaN | 1469.1318 | 896.1405 | 197.06102 | 3866.6619 | 2886.4133 |
| Napa | IEEACEIYAR | -38 | -47 | 0 | 0 | 10 | NaN | 2459.0747 | 2601.2346 | 2978.7815 | 2416.4958 | 2505.425 |
| Napa | LLEAHEEQNVDSYTEAVK | -247 | -264 | 0 | 0 | 18 | NaN | 654.8163 | NaN | 574.0266 | 702.66034 | 652.81036 |
| Rpl38 | KIEEIKDFLLTAR | -4 | -16 | 2 | 0 | 13 | NaN | 411.17322 | 420.70923 | NaN | NaN | NaN |
| Rpl38 | KIEEIKDF | -4 | -11 | 2 | 0.00201047 | 8 | NaN | 3244.6443 | NaN | NaN | NaN | NaN |
| Terf1 | IFGDPEFYTPLER | -187 | -199 | 0 | 0.000627943 | 13 | NaN | 725.0803 | NaN | NaN | NaN | NaN |
| Zc3h18 | IGEDDREFDKENEVFR | -330 | -345 | 2 | 0.000712251 | 16 | NaN | 386.55432 | NaN | NaN | NaN | 406.4819 |
| Ganc | IGEDWEVR | -889 | -896 | 0 | 0 | 8 | NaN | 1037.5778 | 834.1498 | 422.90695 | NaN | NaN |
| Rab6a | QVSIEEGER | -135 | -143 | 0 | 0.001291103 | 9 | NaN | 640.3158 | NaN | NaN | NaN | 763.26636 |
| Stn1 | IGIGDIIR | -136 | -143 | 0 | 0.00025015 | 8 | NaN | 153.80301 | 398.045 | 346.86057 | 210.89873 | 163.45224 |
| Ca2 | SHHWGYSK | -2 | -9 | 0 | 0 | 8 | NaN | 742.3888 | 780.89746 | NaN | 456.4118 | 786.0399 |
| Ca2 | SHHWGYSK | -2 | -9 | 0 | 0 | 8 | NaN | 742.3888 | 780.89746 | NaN | 456.4118 | 786.0399 |
| Ca2 | IGPASQGLQK | -149 | -158 | 0 | 0.004595494 | 10 | NaN | 650.8051 | 109.53823 | NaN | NaN | NaN |
| Med30 | IGQETVQDIVYR | -36 | -47 | 0 | 0 | 12 | NaN | 511.13458 | NaN | 918.1123 | NaN | 703.8311 |
| Sf3a2 | MDFQHRPGGK | -1 | -10 | 1 | 0 | 10 | NaN | 114.67973 | 495.28467 | 437.89868 | 552.1807 | 1138.6511 |
| Sf3a2 | MDFQHRPGGK | -1 | -10 | 1 | 0 | 10 | NaN | 114.67973 | 495.28467 | 437.89868 | 552.1807 | 1138.6511 |
| Sf3a2 | NHLGSYECK | -49 | -57 | 0 | 0.00051306 | 9 | NaN | 898.9152 | NaN | NaN | NaN | NaN |
| Plpbp | TFGENYVQELLEK | -64 | -76 | 0 | 0 | 13 | NaN | 598.6273 | NaN | 466.30914 | NaN | 696.59924 |
| Cd48 | VYLEENNGALHIS | -84 | -96 | 0 | 0.001291103 | 13 | NaN | 1658.7874 | 1734.2198 | NaN | NaN | NaN |
| Bop1 | IHPVTNRPADK | -219 | -229 | 1 | 0.00025015 | 11 | NaN | 222.16745 | 229.98378 | 190.154 | 254.4802 | 261.40167 |
| Thoc1 | TGGEHVYFAK | -291 | -300 | 0 | 0 | 10 | NaN | 1862.2229 | 1162.0325 | 1008.76495 | 1951.5175 | 1977.0121 |
| Thoc1 | ILIGNEELTR | -431 | -440 | 0 | 0.000431655 | 10 | NaN | 2825.529 | 2940.7747 | NaN | 1424.9741 | 1617.9762 |
| Rfc1 | ILDEDGLLDLIR | -466 | -477 | 0 | 0 | 12 | NaN | 303.49146 | NaN | NaN | 345.70004 | NaN |
| Ptcd2 | ILEDSLDTNLSK | -285 | -296 | 0 | 0 | 12 | NaN | 1057.1216 | NaN | NaN | NaN | NaN |
| Creb1 | SPALPTQPAEEAAR | -257 | -270 | 0 | 0 | 14 | NaN | 927.1979 | NaN | 462.41287 | NaN | 318.67404 |
| Ctsc | NWACFVGK | -133 | -140 | 0 | 0 | 8 | NaN | 1599.8068 | 1674.3419 | 897.03143 | 1603.1171 | 1508.8362 |
| Ctsc | LNLPESWDWR | -228 | -237 | 0 | 0 | 10 | NaN | 306.39975 | 743.8949 | 710.1319 | 344.21255 | 752.17914 |
| Ciapin1 | ILRPGGCLFLK | -86 | -96 | 1 | 0 | 11 | NaN | 735.51056 | 852.1955 | NaN | NaN | NaN |
| Napg | LPEAVQLIEK | -91 | -100 | 0 | 0 | 10 | NaN | 561.6298 | 764.44464 | 832.3997 | 630.26825 | 574.13275 |
| Ppil4 | INETFVDKDFVPYQDIR | -139 | -155 | 1 | 0.005459848 | 17 | NaN | 322.87582 | NaN | NaN | NaN | NaN |
| Flad1 | INPLLDWTYR | -419 | -428 | 0 | 0.000230707 | 10 | NaN | 641.8166 | 1343.4148 | NaN | NaN | 759.1593 |
| Nudcd3 | INTESSLWSLEPGR | -248 | -261 | 0 | 0 | 14 | NaN | 791.2181 | 1396.8898 | 1876.4443 | NaN | NaN |
| Lsm4 | TAQNHPMLVELK | -9 | -20 | 0 | 0 | 12 | NaN | 902.47314 | 841.1843 | 1098.388 | 631.154 | NaN |
| Actl6a | SIPAFFLCK | -145 | -153 | 0 | 0.001291103 | 9 | NaN | 549.74146 | NaN | NaN | NaN | NaN |
| Gorasp2 | IPTRPFEEGK | -202 | -211 | 1 | 0.00025015 | 10 | NaN | 1558.4493 | 2654.0684 | 2093.5728 | 2076.6575 | 2254.1902 |
| Gorasp2 | LYVYNTDTDNCR | -163 | -174 | 0 | 0 | 12 | NaN | 505.4383 | NaN | NaN | NaN | NaN |
| Hvcn1 | IQHLEFSCSEK | -238 | -248 | 0 | 0 | 11 | NaN | 1653.1113 | 660.2112 | 700.5401 | 716.0216 | 1083.1534 |
| Smc4 | LHPVEDNPVETVAVL | -1042 | -1056 | 0 | 0 | 15 | NaN | 1433.0349 | NaN | 1921.3574 | 2611.114 | 2677.67 |
| Smc4 | LKLEQIDGHIS | -1014 | -1024 | 1 | 0 | 11 | NaN | 1145.112 | 957.8835 | NaN | 1801.3914 | 937.8753 |

| | | | | | | | | | | | | |
|---|---|---|---|---|---|---|---|---|---|---|---|---|
| Smc4 | SAAPAKPPGEELDNR | -40 | -54 | 1 | 0.000755556 | 15 | NaN | 1059.0427 | 831.47205 | 1149.5734 | 1292.616 | NaN |
| Smc4 | IQTPENTPR | -682 | -690 | 0 | 0.001636916 | 9 | NaN | 2165.3596 | NaN | 1524.898 | NaN | NaN |
| Smc4 | LHNTIIDINNR | -885 | -895 | 0 | 0.000394361 | 11 | NaN | 2609.111 | NaN | NaN | NaN | NaN |
| Znf638 | ISDVLLVPCR | -703 | -712 | 0 | 0 | 10 | NaN | 4855.5933 | 5100.158 | 4461.8125 | 6002.6904 | NaN |
| Lmnb2 | RVELEQTYQAK | -253 | -263 | 1 | 0.00025015 | 11 | NaN | 833.4979 | 856.9371 | 216.42894 | 1169.1753 | 1258.4797 |
| Lmnb2 | VELEQTYQAK | -254 | -263 | 0 | 0 | 10 | NaN | 2399.5808 | 10190.665 | NaN | NaN | NaN |
| Afg3l2 | YLEHADPLLK | -583 | -592 | 0 | 0 | 10 | NaN | 755.0612 | 702.3185 | NaN | 1490.6503 | 720.7157 |
| Afg3l2 | NLETLQQELGIEGENR | -224 | -239 | 0 | 0.000230707 | 16 | NaN | 329.11545 | NaN | NaN | 486.06696 | 500.46655 |
| Pno1 | IVLADVHVH | -198 | -206 | 0 | 0.002068213 | 9 | NaN | 2972.4304 | NaN | NaN | NaN | NaN |
| Pgm2l1 | IVLATDPDADR | -327 | -337 | 0 | 0.001636916 | 11 | NaN | 2770.6218 | NaN | NaN | NaN | NaN |
| Gbe1 | SPLIPHGSK | -133 | -141 | 0 | 0.008014596 | 9 | NaN | 3455.0222 | NaN | NaN | NaN | NaN |
| Snx12 | RQGLEQFINK | -119 | -128 | 1 | 0.00025015 | 10 | NaN | 655.9315 | 1379.4071 | 1117.0536 | 856.46423 | 777.651 |
| Eps15l1 | IWDLADPEGK | -56 | -65 | 0 | 0.001536401 | 10 | NaN | 1166.2384 | 161.37643 | NaN | 1335.7617 | NaN |
| Ppp2r2a | IWDLNMENRPVETY | -310 | -323 | 1 | 0.000431655 | 14 | NaN | 825.8527 | 273.35876 | 629.3328 | 925.74817 | 529.47003 |
| Ppp2r2a | IWDLNMENRPVETY | -310 | -323 | 1 | 0.000431655 | 14 | NaN | 825.8527 | 273.35876 | 629.3328 | 925.74817 | 529.47003 |
| Ube2g1 | IWHPNVDK | -78 | -85 | 0 | 0.000431655 | 8 | NaN | 744.4845 | 843.60065 | NaN | 4381.844 | NaN |
| Rps24 | IYDSLDYAK | -75 | -83 | 0 | 0 | 9 | NaN | 1803.0483 | 1928.768 | 1323.3318 | 1349.4362 | 832.7138 |
| Rnaset2a;Rnas | IYWPDVIHR | (99);(99) | (107);(107) | 0 | 0 | 9 | NaN | 779.271 | 577.7904 | 582.94446 | 841.97815 | 778.80695 |
| Ddx18 | VPLNQFDFSWSK | -536 | -547 | 0 | 0 | 12 | NaN | 855.81433 | 1059.3424 | 1084.1096 | 1116.657 | 764.8598 |
| Ddx18 | NYIDLPVLAIHGK | -436 | -448 | 0 | 0 | 13 | NaN | 487.51627 | NaN | NaN | NaN | NaN |
| Elf1 | TVHVVQPVQAVPEEATIA | -385 | -402 | 0 | 0 | 18 | NaN | 577.9438 | 466.10034 | NaN | 641.5149 | 583.0051 |
| Elf1 | TVQPSQAPYPTQLFR | -370 | -384 | 0 | 0.00078661 | 15 | NaN | 307.89713 | 1370.2332 | NaN | 2822.7874 | 2605.1582 |
| Ylpm1 | KDAEEEESELGYIPK | -1268 | -1282 | 1 | 0.007791272 | 15 | NaN | 768.7215 | NaN | NaN | 815.2661 | NaN |
| Grhpr | KDLEQGVVGAH | -43 | -53 | 1 | 0 | 11 | NaN | 9846.667 | 32583.004 | NaN | 25680.592 | 28172.518 |
| Polr2j | KEDHTLGNIIK | -37 | -47 | 1 | 0.005946382 | 11 | NaN | 474.26956 | NaN | NaN | NaN | 1444.1971 |
| Mrpl46 | KEELGDYLQPK | -255 | -265 | 1 | 0 | 11 | NaN | 706.1389 | 582.7932 | 444.4164 | 745.3263 | 702.17303 |
| Sap18 | KEPEKPIDR | -14 | -22 | 2 | 0.00201047 | 9 | NaN | 806.0124 | 735.6691 | 664.8596 | 1022.7661 | 1006.6525 |
| Hmga1 | KQPPVSPGTAL | -31 | -41 | 1 | 0 | 11 | NaN | 1912.5924 | 1751.427 | 2399.7666 | 2616.051 | 2443.5188 |
| Ppil3 | TYRPLNDVHIK | -140 | -150 | 1 | 0 | 11 | NaN | 415.54633 | 409.1788 | 491.48923 | 795.8192 | 447.95627 |
| Ppil3 | TYRPLNDVHIK | -140 | -150 | 1 | 0 | 11 | NaN | 415.54633 | 409.1788 | 491.48923 | 795.8192 | 447.95627 |
| Tbl2 | KGEFQEVLR | -259 | -267 | 1 | 0 | 9 | NaN | 583.4194 | 719.07196 | 560.4117 | 780.2206 | NaN |
| Eif3j1;Eif3j2 | NWDDDDDENKEEAEVKPEVK | (54);(56) | (73);(75) | 2 | 0.00025015 | 20 | NaN | 576.87683 | 526.2119 | 388.78198 | 448.4038 | 382.6841 |
| Ccnh | KGYEDDDYVSK | -295 | -305 | 1 | 0 | 11 | NaN | 1043.9604 | 452.44223 | 820.4769 | 936.89905 | 836.54944 |
| Prpf18 | KIEILTPEVNK | -111 | -121 | 1 | 0.00111006 | 11 | NaN | 953.88446 | NaN | NaN | NaN | NaN |
| Ints1 | KLAVAHPVLLL | -1873 | -1883 | 1 | 0.00122419 | 11 | NaN | 358.5993 | 371.02234 | 417.42807 | 340.11285 | 378.95874 |
| Ktn1 | KLDDSYSEAVR | -1216 | -1226 | 1 | 0 | 11 | NaN | 718.4546 | NaN | NaN | NaN | NaN |
| Scp2 | LEEEGEQFVK | -444 | -453 | 0 | 0.000627943 | 10 | NaN | 749.1326 | 592.4436 | NaN | NaN | 749.51 |
| Nxf1 | SNSGLCIVNDELFVR | -522 | -536 | 0 | 0 | 15 | NaN | 1085.0347 | 1153.2257 | NaN | 1388.2054 | 913.38055 |
| Psme3 | SHDGLDGPTYK | -75 | -85 | 0 | 0 | 11 | NaN | 2139.7727 | NaN | 938.94666 | NaN | 12555.587 |
| Psme3 | SYLDQISR | -174 | -181 | 0 | 0.00051306 | 8 | NaN | 853.53314 | NaN | NaN | NaN | 1517.4414 |
| Gpaa1 | KLPFPDETHER | -111 | -121 | 1 | 0 | 11 | NaN | 531.4796 | 638.26624 | 614.15106 | 617.7827 | 733.2636 |
| Fkbp2 | VDHCPIK | -37 | -43 | 0 | 0.006946742 | 7 | NaN | 520.4035 | NaN | NaN | NaN | 1452.0565 |
| Mtarc2 | KLYPSESYLQ | -193 | -202 | 1 | 0.009060424 | 10 | NaN | 1036.4253 | NaN | NaN | NaN | NaN |
| Hmgb3 | KNPEVPVNFAEF | -30 | -41 | 1 | 0 | 12 | NaN | 525.4087 | 375.7988 | 559.5945 | 520.6559 | 4985.0312 |
| Clptm1l | KPSNALDEPVSHWRPR | -159 | -174 | 2 | 0 | 16 | NaN | 738.52423 | 530.1558 | NaN | NaN | 662.7365 |
| Cd8b | KQCPFPHPETQK | -156 | -167 | 1 | 0 | 12 | NaN | 670.21814 | 666.7331 | NaN | 771.2422 | 746.58057 |
| Dnajb11 | QYDTYGEEGLK | -85 | -95 | 0 | 0 | 11 | NaN | 386.9634 | NaN | 298.68912 | NaN | NaN |
| Tsnax | SFQHFIK | -132 | -138 | 0 | 0.00078661 | 7 | NaN | 386.68466 | 399.74695 | NaN | 868.5886 | 573.2166 |
| Mpc1 | KSPEIISGR | -46 | -54 | 1 | 0.001636916 | 9 | NaN | 630.55 | NaN | NaN | NaN | NaN |
| Guk1 | KTDLCPIYIF | -112 | -121 | 1 | 0.000550307 | 10 | NaN | 841.5747 | 271.6761 | NaN | NaN | NaN |
| Dtymk | KTELEDHSVHLL | -60 | -71 | 1 | 0.001868731 | 12 | NaN | 490.5409 | 486.86224 | 436.3137 | 525.97296 | 535.0273 |
| Rab10 | NIDEHANEDVER | -106 | -117 | 0 | 0 | 12 | NaN | 2179.9634 | 2263.4438 | 1800.1057 | 2453.6013 | 2590.9976 |
| Rab10 | TYDLLFK | -5 | -11 | 0 | 0.002168954 | 7 | NaN | 480.90598 | 668.7412 | 671.20844 | 430.69962 | NaN |
| Pbdc1 | RVDVLDPEELK | -83 | -93 | 1 | 0 | 11 | NaN | 746.245 | 812.71436 | 1055.5135 | 1162.5504 | 1193.6691 |
| Mrps31 | KVDQSVPANEISQK | -59 | -72 | 1 | 0.00051306 | 14 | NaN | 232.944 | NaN | 263.7703 | NaN | 342.61157 |
| Mob2 | KVYLEPEHTK | -23 | -32 | 1 | 0.00025015 | 10 | NaN | 410.73273 | NaN | NaN | 913.9276 | NaN |
| Bpnt1 | KWDTCAPEVILH | -245 | -256 | 1 | 0 | 12 | NaN | 1143.6769 | 887.7299 | NaN | 889.8513 | NaN |
| Selenof | LLPLDPVCR | -60 | -68 | 0 | 0.009416377 | 9 | NaN | 698.41223 | NaN | NaN | NaN | NaN |
| Rpl27a | SFCPTVNLDK | -68 | -77 | 0 | 0 | 10 | NaN | 1516.4509 | 904.6534 | 980.59515 | 1248.4757 | 1275.4557 |
| Rpl27a | RNQSFCPTVNLDK | -65 | -77 | 1 | 0.00078661 | 13 | NaN | 3430.5493 | 1276.1079 | NaN | 1305.475 | NaN |
| C4b | YIYGKPVQGVAY | -268 | -279 | 1 | 0.006100831 | 12 | NaN | 1007.08057 | NaN | NaN | NaN | 1576.7056 |
| Pxn | LADLESTTSHISK | -8 | -20 | 0 | 0 | 13 | NaN | 430.9787 | NaN | 658.01636 | 420.89523 | 629.89905 |
| Tradd | LAELEDELCK | -152 | -161 | 0 | 0.003133457 | 10 | NaN | 4190.1973 | 4186.868 | 5564.467 | 10955.743 | 7864.3765 |
| Hikeshi | LAQNPLFWK | -188 | -196 | 0 | 0 | 9 | NaN | 1966.685 | 1856.5435 | 1584.9806 | 1859.1404 | 1797.2229 |
| Exosc1 | SFRPGDIVLAK | -121 | -131 | 1 | 0 | 11 | NaN | 3337.022 | 4014.5781 | NaN | 663.45514 | 2217.8582 |
| Mon2 | LCVQPQLLR | -361 | -369 | 0 | 0.007546675 | 9 | NaN | 634.6778 | NaN | 868.5054 | NaN | NaN |
| Prpf38b | TYHEVVDEIYFK | -82 | -93 | 0 | 0 | 12 | NaN | 329.51044 | 439.18933 | 643.736 | 524.5635 | 356.9729 |
| Prpf38b | QGNVLPLWGNEK | -44 | -55 | 0 | 0 | 12 | NaN | 662.4717 | 1933.1998 | 1396.0195 | NaN | NaN |
| Prpf38b | LDDEEDLDVK | -183 | -192 | 0 | 0.005242018 | 10 | NaN | 650.75433 | NaN | NaN | NaN | NaN |
| Sipa1 | TFSEHQLDLYHGR | -636 | -648 | 0 | 0 | 13 | NaN | 818.56757 | 1042.0419 | 389.59564 | 1082.4287 | 1378.3837 |
| Ttc4 | LDGLSSENPYGAR | -254 | -266 | 0 | 0 | 13 | NaN | 1198.0278 | NaN | 1582.1733 | 1506.8004 | NaN |
| Nup54 | SAPTNTGSTGLLGGTQNK | -42 | -59 | 0 | 0 | 18 | NaN | 489.3041 | 798.08514 | NaN | 981.61224 | 854.6172 |
| Becn1 | LDQEEAQYQR | -220 | -229 | 0 | 0 | 10 | NaN | 1206.6017 | NaN | 1489.9102 | 1523.9841 | 1028.2546 |
| Tsg101 | LDQEVAEVDKNIELLK | -278 | -293 | 1 | 0.00025015 | 16 | NaN | 526.01337 | 1042.3964 | NaN | 1930.6184 | NaN |
| Gimap9 | LDRYTEEEQK | -101 | -110 | 1 | 0.001392869 | 10 | NaN | 1304.3868 | 1345.716 | NaN | NaN | 1519.631 |
| Fancd2 | LEDFPVIVK | -273 | -281 | 0 | 0 | 9 | NaN | 595.8568 | NaN | 263.07397 | 475.301 | 263.15457 |
| Hmgcs1 | TATEHIPSPAK | -488 | -498 | 0 | 0.00383302 | 11 | NaN | 1008.5337 | NaN | NaN | NaN | 1880.9293 |
| Ppp1r9b | SLCQAVDEHLR | -732 | -742 | 0 | 0 | 11 | NaN | 1115.4098 | NaN | NaN | 1546.7742 | 1115.0468 |

| | | | | | | | | | | | | |
|---|---|---|---|---|---|---|---|---|---|---|---|---|
| Ppp1r9b | LELFPVELEK | -492 | -501 | 0 | 0 | 10 | NaN | 282.97458 | NaN | NaN | NaN | NaN |
| Ppp1r8 | LEPHKPQQIPIDSTVSF | -101 | -117 | 1 | 0 | 17 | NaN | 344.57385 | 365.9757 | 685.77026 | 807.1038 | 769.505 |
| Ppp1r8 | LEPHKPQQIPIDSTVSF | -101 | -117 | 1 | 0 | 17 | NaN | 344.57385 | 365.9757 | 685.77026 | 807.1038 | 769.505 |
| Erlin2 | VTKPNIPEAIR | -170 | -180 | 1 | 0 | 11 | NaN | 2188.5046 | 3050.7595 | NaN | 1322.1824 | 1097.9054 |
| Rwdd1 | SQENLEDNDVSDILK | -71 | -85 | 0 | 0 | 15 | NaN | 857.47375 | 883.321 | 869.5307 | 1093.489 | 1116.0817 |
| Rwdd1 | LFHGTPVTIE | -140 | -149 | 0 | 0.004690496 | 10 | NaN | 304.89523 | NaN | NaN | NaN | NaN |
| Bud23 | LFSGPSTSLPK | -204 | -214 | 0 | 0.001846793 | 11 | NaN | 1053.6212 | NaN | NaN | 901.8622 | NaN |
| Tram1 | LFYFSDEK | -238 | -245 | 0 | 0 | 8 | NaN | 357.39612 | 438.17487 | 2970.605 | 2983.7717 | 2370.1785 |
| Grsf1 | LGDEVDDVYLIR | -141 | -152 | 0 | 0.00025015 | 12 | NaN | 834.32947 | NaN | NaN | NaN | NaN |
| Hsd17b10 | TIIENPFLNGEVIR | -239 | -252 | 0 | 0 | 14 | NaN | 498.73862 | 706.895 | NaN | 455.38684 | 498.4934 |
| Hsd17b10 | LGDPAEYAHL | -227 | -236 | 0 | 0.003299042 | 10 | NaN | 1695.5156 | NaN | NaN | NaN | NaN |
| Niban1 | TKPPLEPGTVLYEA | -61 | -74 | 1 | 0.00025015 | 14 | NaN | 296.17493 | 483.20715 | NaN | NaN | NaN |
| Apoe | LGQSTEEIR | -136 | -144 | 0 | 0.000712251 | 9 | NaN | 787.39514 | NaN | NaN | NaN | NaN |
| Lrch1 | TTAPGHDLSDTVR | -79 | -91 | 0 | 0 | 13 | NaN | 5127.9897 | NaN | NaN | NaN | NaN |
| Hsdl1 | LGSRPDLIK | -56 | -64 | 1 | 0 | 9 | NaN | 691.1935 | 634.32556 | 682.995 | 670.658 | 774.37134 |
| Fdxr | SPQQVLPTPDGQR | -313 | -325 | 0 | 0 | 13 | NaN | 2435.73 | NaN | NaN | NaN | NaN |
| Rbm27 | LHDDDDEDEDEYESR | -1042 | -1056 | 0 | 0.000627943 | 15 | NaN | 475.37378 | NaN | 406.99307 | 465.9379 | 542.5414 |
| Exosc4 | LHEDHLEQVLEAA | -205 | -217 | 0 | 0.000394361 | 13 | NaN | 2548.841 | 2494.1824 | 1166.2673 | 2541.3245 | 2877.6057 |
| Dcp1a | LHEPFLLYR | -110 | -118 | 0 | 0 | 9 | NaN | 758.21686 | 668.4679 | NaN | NaN | NaN |
| Wdr43 | SKDTHICLER | -365 | -374 | 1 | 0.000230707 | 10 | NaN | 528.24615 | 436.17957 | NaN | 486.41376 | NaN |
| Birc6 | SYILPWESELK | -1896 | -1906 | 0 | 0 | 11 | NaN | 356.53256 | 818.6235 | 527.99866 | 893.14825 | 496.10965 |
| Trappc1 | LHYYETPTGIK | -71 | -81 | 0 | 0 | 11 | NaN | 1030.6141 | 1014.4834 | NaN | NaN | NaN |
| Srpra | LIDGIVLTK | -579 | -587 | 0 | 0.00025015 | 9 | NaN | 512.923 | NaN | NaN | 388.19708 | 423.9921 |
| Srpra | SLVQILQPQR | -390 | -399 | 0 | 0.001392869 | 10 | NaN | 94.88246 | 104.66323 | NaN | NaN | NaN |
| Por | LIHEGGAHIY | -619 | -628 | 0 | 0.000230707 | 10 | NaN | 402.1966 | 878.32086 | 1267.8431 | NaN | 1145.5046 |
| Por | YLDITNPPR | -374 | -382 | 0 | 0.00111006 | 9 | NaN | 1544.748 | 485.42584 | NaN | 1105.0542 | NaN |
| Rfk | LKDDNFFQVSK | -138 | -148 | 1 | 0 | 11 | NaN | 1553.5869 | NaN | NaN | 1462.0863 | 283.23926 |
| Nfatc2ip | LLDPGEAPVVPVYSGK | -105 | -120 | 0 | 0.001392869 | 16 | NaN | 2498.1155 | NaN | NaN | NaN | NaN |
| Sympk | SPQNLSHAVEEALK | -1260 | -1273 | 0 | 0 | 14 | NaN | 656.2478 | 222.27498 | NaN | 705.84143 | 784.5226 |
| Sympk | RQEHDISLDR | -206 | -215 | 1 | 0.00025015 | 10 | NaN | 485.50104 | 584.7543 | 391.78693 | NaN | 291.95975 |
| Paf1 | YCNSLPDIPFDPK | -35 | -47 | 0 | 0 | 13 | NaN | 424.3906 | 424.2302 | NaN | NaN | 442.23193 |
| Mrpl28 | LLEEKDPVPLFK | -217 | -228 | 1 | 0 | 12 | NaN | 611.5535 | 809.49255 | NaN | NaN | 540.57446 |
| Mrpl28 | LLEEKDPVPLFK | -217 | -228 | 1 | 0 | 12 | NaN | 611.5535 | 809.49255 | NaN | NaN | 540.57446 |
| Ttc39b | LLEFIGFSGNR | -257 | -267 | 0 | 0 | 11 | NaN | 578.06757 | 1607.2924 | 1014.6734 | NaN | 438.04047 |
| Ldah | LLFKPCPEVIK | -197 | -207 | 1 | 0.009688493 | 11 | NaN | 421.25003 | NaN | NaN | NaN | NaN |
| Racgap1 | LLGPVTTPEFQLVK | -571 | -584 | 0 | 0 | 14 | NaN | 341.86963 | NaN | 459.41003 | 387.05566 | NaN |
| Bccip | LLNDTSKPVGLLL | -170 | -182 | 1 | 0.00025015 | 13 | NaN | 419.46204 | 364.151 | 387.68845 | 498.1568 | 416.2832 |
| Kif15 | LLNLVDLAGSER | -260 | -271 | 0 | 0 | 12 | NaN | 246.15352 | NaN | 326.4047 | NaN | 214.97844 |
| Ddx41 | LLQEDSSPLLR | -283 | -293 | 0 | 0.004690496 | 11 | NaN | 359.29367 | NaN | NaN | NaN | NaN |
| Kifc1 | LLQELQGER | -227 | -235 | 0 | 0.005698678 | 9 | NaN | 1549.4388 | NaN | NaN | NaN | 1422.7971 |
| Cdk9 | NPATTNQTEFER | -359 | -370 | 0 | 0 | 12 | NaN | 1059.9022 | 810.61707 | 713.7146 | 790.1679 | 840.4012 |
| Cr1l | SQSQCQSDGSWNPLLAK | -381 | -397 | 0 | 0 | 17 | NaN | 198.21501 | 1372.6725 | NaN | 2446.388 | NaN |
| Thumpd1 | LNEYGDDMYGPEK | -65 | -77 | 0 | 0.002697088 | 13 | NaN | 502.46957 | NaN | NaN | 364.17007 | NaN |
| Ccdc47 | LNQENEHIYNLW | -197 | -208 | 0 | 0.00025015 | 12 | NaN | 631.3144 | NaN | 516.1675 | 385.84283 | 337.5077 |
| Nudt21 | LPGGELNPGEDEVEGLK | -106 | -122 | 0 | 0 | 17 | NaN | 685.397 | 613.6104 | 661.53107 | 505.4926 | 1150.1064 |
| Babam2 | SGCTSLTPGPNCDR | -42 | -55 | 0 | 0 | 14 | NaN | 448.68552 | NaN | NaN | 765.11285 | 420.41232 |
| Gnas | LQEALNLFK | (266);(1005) | (274);(1013) | 0 | 0.00025015 | 9 | NaN | 629.1523 | 378.94415 | 821.56793 | 802.6411 | 929.6676 |
| Chp1 | LRDEELEEIKK | -9 | -19 | 2 | 0 | 11 | NaN | 780.5548 | NaN | 1679.0153 | 714.0419 | 990.1878 |
| Far1 | LRDENPDFR | -68 | -76 | 1 | 0 | 9 | NaN | 2420.9846 | 2939.2656 | 2651.134 | NaN | NaN |
| Mcu | LRDPLQVHLPLR | -333 | -344 | 1 | 0.00025015 | 12 | NaN | 340.10516 | 382.21326 | 442.5065 | 525.57294 | 423.3334 |
| Exoc5 | LSDPSDLPR | -433 | -441 | 0 | 0.00025015 | 9 | NaN | 900.76324 | NaN | NaN | 871.8831 | NaN |
| Hbb-y | LSELHCDKL | -89 | -97 | 1 | 0.001023236 | 9 | NaN | 2098 | NaN | NaN | NaN | NaN |
| Snrpd1 | NGTQVHGTIT | -21 | -30 | 0 | 0.00830926 | 10 | NaN | 1382.2936 | NaN | NaN | NaN | NaN |
| Maz | LSHSDEKPYQCPVCQQ | -299 | -314 | 1 | 0.000347792 | 16 | NaN | 191.81192 | NaN | 356.8285 | 253.7109 | NaN |
| Brix1 | TILPHDPTADVF | -296 | -307 | 0 | 0.000712251 | 12 | NaN | 1022.8959 | 1516.1719 | 1305.781 | 1414.336 | 1147.2322 |
| Brix1 | SQPFVDHVF | -198 | -206 | 0 | 0.002780285 | 9 | NaN | 246.87274 | 585.6088 | 258.435 | 343.2554 | NaN |
| Brix1 | LSNSPHGPSAK | -130 | -140 | 0 | 0.001092529 | 11 | NaN | 303.9433 | NaN | NaN | NaN | 367.37613 |
| Arhgef2 | LSNVDQDVHELEK | -427 | -439 | 0 | 0 | 13 | NaN | 898.7542 | NaN | NaN | NaN | 1279.3737 |
| Eif2b2 | LSPQFPSEEDSFHKF | -274 | -288 | 1 | 0.000471409 | 15 | NaN | 652.9953 | 409.37146 | 696.7054 | 935.36346 | 918.6896 |
| Eif2b2 | SDESDQQESLHKL | -100 | -112 | 1 | 0.000591124 | 13 | NaN | 2226.1265 | NaN | 2043.8855 | 2183.6165 | 2276.92 |
| Xpo4 | LSSQQDPVIYQR | -1097 | -1108 | 0 | 0.00025015 | 12 | NaN | 677.02295 | 1401.4067 | NaN | NaN | 1672.0153 |
| Erbin | LTFIPGFIGSLR | -219 | -230 | 0 | 0.00416551 | 12 | NaN | 335.66696 | NaN | NaN | NaN | NaN |
| Snrpc | YCDTYLTHDSPSVR | -8 | -21 | 0 | 0 | 14 | NaN | 1075.1696 | NaN | 984.2376 | 566.20135 | 555.1911 |
| Lancl1 | LTPEFSHR | -28 | -35 | 0 | 0.009657961 | 8 | NaN | 1319.3623 | 1811.6796 | NaN | NaN | NaN |
| Prkag1 | LVVVDEHDVVK | -299 | -309 | 0 | 0 | 11 | NaN | 1601.945 | 1641.9781 | NaN | NaN | NaN |
| Atxn10 | VEQDSLLTAFR | -123 | -133 | 0 | 0 | 11 | NaN | 433.51624 | NaN | 565.3594 | NaN | 629.9361 |
| Snrpe | LYEQVNMR | -35 | -42 | 0 | 0.00051306 | 8 | NaN | 1574.2798 | NaN | NaN | 519.99097 | 1605.2272 |
| Zcchc8 | SVASSPCPNEFDLPVPEGK | -556 | -574 | 0 | 0 | 19 | NaN | 812.8865 | 315.5754 | 584.2193 | 134.8084 | 375.20737 |
| Zcchc8 | LVNYPGFNISTPR | -366 | -378 | 0 | 0 | 13 | NaN | 1032.2725 | 1052.0073 | NaN | 791.33936 | 736.1217 |
| Prf1 | LVQKPPLHLDFK | -182 | -193 | 1 | 0 | 12 | NaN | 435.90762 | 298.6178 | NaN | 537.60095 | 590.31573 |
| Prf1 | SPHSGFHEVTCELNHGR | -499 | -515 | 0 | 0 | 17 | NaN | 348.45062 | 281.63013 | NaN | 321.2711 | 533.1634 |
| Mrpl23 | SPEPLEEELPQQR | -118 | -130 | 0 | 0 | 13 | NaN | 806.2652 | 853.80566 | 998.43414 | 732.4032 | 618.427 |
| Mrpl23 | YQLGGPQLR | -10 | -18 | 0 | 0.000712251 | 9 | NaN | 744.25134 | 721.7744 | 640.6097 | 854.61084 | 615.3972 |
| Mrpl23 | TFPDLFPEKDPR | -106 | -117 | 1 | 0.003299042 | 12 | NaN | 3164.4814 | NaN | NaN | NaN | NaN |
| Wdr70 | LVTGGYDYDVK | -200 | -210 | 0 | 0.006788754 | 11 | NaN | 755.4134 | NaN | NaN | NaN | NaN |
| Tle3 | SPISQLDCLNR | -521 | -531 | 0 | 0 | 11 | NaN | 968.9116 | 919.4454 | NaN | 612.7812 | 1557.9746 |
| Itk | LYDYQTNDPQELALR | -185 | -199 | 0 | 0.002199523 | 15 | NaN | 744.37024 | NaN | NaN | NaN | NaN |
| Elob | RIVEGILK | -29 | -36 | 1 | 0.005605646 | 8 | NaN | 939.49066 | 921.53754 | 1295.7452 | 1513.1089 | 1477.913 |

| | | | | | | | | | | | | |
|---|---|---|---|---|---|---|---|---|---|---|---|---|
| Cd2 | LYQGETLLNSLPQK | -152 | -165 | 0 | 0 | 14 | NaN | 1222.1869 | 1044.6643 | 988.8263 | 1162.9801 | 1244.3514 |
| Polr2i | MEPDGTYEPGFVGIR | -1 | -15 | 0 | 0.001392869 | 15 | NaN | 193.55185 | NaN | NaN | NaN | NaN |
| Mtco1 | VFSWLATLHGGNIK | -320 | -333 | 0 | 0 | 14 | NaN | 1363.7377 | 1498.4479 | 546.97754 | 1426.828 | 1327.8995 |
| Mtco1 | MIIAIPTGVK | -310 | -319 | 0 | 0.000471409 | 10 | NaN | 1203.7207 | NaN | NaN | NaN | 1089.1056 |
| Mtco1 | RYSDYPDAY | -439 | -447 | 1 | 0.00897527 | 9 | NaN | 357.07663 | NaN | NaN | NaN | NaN |
| Med22 | NDFPSVNEAIDQR | -89 | -101 | 0 | 0 | 13 | NaN | 1325.4066 | NaN | NaN | 1119.4889 | NaN |
| Nqo1 | NFNPIISR | -46 | -53 | 0 | 0 | 8 | NaN | 858.34717 | NaN | 648.4968 | NaN | 492.21756 |
| Med27 | NFQDNLHSVNR | -55 | -65 | 0 | 0 | 11 | NaN | 2180.5234 | 2286.5366 | NaN | 410.31537 | 1649.3611 |
| Gemin6 | NHIPVTEQGDAQR | -112 | -124 | 0 | 0 | 13 | NaN | 503.096 | 530.3532 | NaN | 501.10464 | NaN |
| Vcpip1 | NHYIPLVGIK | -352 | -361 | 0 | 0 | 10 | NaN | 326.40894 | 392.42444 | NaN | 615.8567 | 618.92786 |
| Krr1 | TVPDGWKEPAFSK | -34 | -46 | 1 | 0.002265552 | 13 | NaN | 973.76025 | 953.20074 | NaN | NaN | NaN |
| Cdc16 | NIISPPWDFR | -557 | -566 | 0 | 0 | 10 | NaN | 542.2413 | NaN | 514.48706 | NaN | NaN |
| Uba5 | NINPDVLFEVH | -135 | -145 | 0 | 0 | 11 | NaN | 743.8265 | NaN | NaN | 801.5119 | 781.232 |
| Pef1 | NLFQQYDR | -179 | -186 | 0 | 0 | 8 | NaN | 675.0908 | NaN | 201.39828 | 1001.72656 | 858.0677 |
| Pptc7 | NLGDSGFLVVR | -161 | -171 | 0 | 0 | 11 | NaN | 3656.2578 | NaN | 1877.8794 | 2109.5017 | NaN |
| Synj2bp | NLLHQDAVDLFR | -75 | -86 | 0 | 0 | 12 | NaN | 663.3856 | NaN | 444.14453 | 576.7709 | 541.9048 |
| Vrk2 | NLLLDFTNPDR | -171 | -181 | 0 | 0 | 11 | NaN | 495.44354 | NaN | 555.4443 | 416.06802 | NaN |
| Kat7 | NLPSFGTEEPAYSTR | -63 | -77 | 0 | 0 | 15 | NaN | 738.3357 | 521.25256 | 556.1776 | 502.73685 | 796.3301 |
| Wdr53 | NLQEDETEETEGPQTPGR | -176 | -193 | 0 | 0 | 18 | NaN | 415.43323 | 330.3459 | 405.4983 | NaN | NaN |
| Icos | NLSGGYLHIY | -123 | -132 | 0 | 0.003681885 | 10 | NaN | 572.91034 | 453.90158 | NaN | 569.19885 | 797.63367 |
| Pdcl3 | NLTIDELEWK | -189 | -198 | 0 | 0.00111006 | 10 | NaN | 992.55566 | 1225.6523 | 926.77356 | 1118.8911 | NaN |
| Nit2 | NLTTGPAHWELLQR | -180 | -193 | 0 | 0 | 14 | NaN | 3439.6077 | 5130.97 | 4957.9775 | 1359.9805 | 968.167 |
| Atp6ap1 | TYLDPALELGPR | -70 | -81 | 0 | 0 | 12 | NaN | 1164.5815 | NaN | NaN | 999.0925 | 926.97034 |
| Slu7 | NPDEVSYAGDNFVR | -303 | -316 | 0 | 0.000298805 | 14 | NaN | 463.4999 | 454.80917 | NaN | 426.1728 | 529.3217 |
| Uqcc1 | NPQSILKPHAPTYNDE | -278 | -293 | 1 | 0.000471409 | 16 | NaN | 1198.4956 | 1389.3203 | NaN | NaN | NaN |
| Soat1 | NPTFLDYVRPR | -522 | -532 | 1 | 0.00078661 | 11 | NaN | 2013.5094 | NaN | 1576.9897 | 524.43726 | 1587.1266 |
| Cers5 | NQDKPPTLTK | -131 | -140 | 1 | 0.001055699 | 10 | NaN | 1048.2068 | NaN | NaN | NaN | NaN |
| Ctla2a | RLVWEENKK | -60 | -68 | 2 | 0.000550307 | 9 | NaN | 1241.545 | 968.2336 | 1367.7666 | 567.74396 | 1695.7434 |
| Ctla2a | QFSDLTPEEFK | -90 | -100 | 0 | 0 | 11 | NaN | 1213.6351 | 2017.4178 | NaN | 1035.4639 | 2029.3315 |
| Lsm14a | SAHLPAPAPVGR | -203 | -214 | 0 | 0.000471409 | 12 | NaN | 655.2318 | 695.7741 | 394.24057 | NaN | NaN |
| Mrpl27 | NTEAVDLVTSLPK | -110 | -122 | 0 | 0 | 13 | NaN | 975.7513 | 1157.794 | 1997.6487 | 1591.0793 | 1994.026 |
| Dcaf7 | NTFDHPYPTTK | -63 | -73 | 0 | 0 | 11 | NaN | 1082.3619 | NaN | NaN | 786.2279 | 718.3945 |
| Atp6v0d1 | NVADYYPEYK | -266 | -275 | 0 | 0.000591124 | 10 | NaN | 3291.395 | 595.52325 | NaN | NaN | NaN |
| Trip13 | NVDSNLITWNR | -163 | -173 | 0 | 0 | 11 | NaN | 2501.1008 | NaN | 2484.6853 | 2626.7236 | 250.82076 |
| Dpf2 | NYCDFCLGDSK | -271 | -281 | 0 | 0 | 11 | NaN | 301.07144 | NaN | 393.49927 | NaN | NaN |
| Pam16 | SLQEAQQILNVSK | -55 | -67 | 0 | 0 | 13 | NaN | 1120.775 | NaN | NaN | NaN | NaN |
| Itch | NYEQWQLQR | -357 | -365 | 0 | 0.002910424 | 9 | NaN | 482.59418 | NaN | NaN | NaN | NaN |
| Snx27 | NYTSAVPGTCLTIR | -343 | -356 | 0 | 0 | 14 | NaN | 706.88214 | NaN | NaN | NaN | NaN |
| H2bc7;H2bc9; | PEPAKSAPAPK | (2);(2);(2);(2) | (12);(12);(12);(12) | 1 | 0 | 11 | NaN | 341.83438 | 397.22443 | 571.1663 | 559.0014 | 591.0115 |
| Vwf | PIAPIFIR | -1645 | -1652 | 0 | 0 | 8 | NaN | 328.13666 | 378.58292 | NaN | 284.8433 | NaN |
| Rbm12b2;Rbn | PVLIDPVSR | (351);(351) | (359);(359) | 0 | 0.000947459 | 9 | NaN | 726.0973 | 793.8783 | NaN | NaN | NaN |
| Pcnt | VLESELEDQLVQHR | -1215 | -1228 | 0 | 0.00025015 | 14 | NaN | 2360.0664 | NaN | NaN | NaN | NaN |
| Pcnt | QDWESER | -221 | -227 | 0 | 0.00555405 | 7 | NaN | 97.11068 | NaN | NaN | NaN | NaN |
| Micos10 | SNCQHDFQAPYLLHGK | -56 | -71 | 0 | 0 | 16 | NaN | 916.2376 | 1188.6521 | 1246.5345 | 1102.4648 | 1096.772 |
| Micos10 | SNCQHDFQAPYLLHGK | -56 | -71 | 0 | 0 | 16 | NaN | 916.2376 | 1188.6521 | 1246.5345 | 1102.4648 | 1096.772 |
| Nae1 | SVGQAPESISEK | -367 | -378 | 0 | 0.000550307 | 12 | NaN | 1896.6796 | 874.22766 | 365.5787 | 2216.1152 | NaN |
| Nae1 | QIPSSIEDIFNDDR | -280 | -293 | 0 | 0.00122419 | 14 | NaN | 866.4485 | NaN | NaN | NaN | NaN |
| Rpl35 | QLDDLKVE | -20 | -27 | 1 | 0.006788754 | 8 | NaN | 601.0797 | NaN | NaN | NaN | NaN |
| Morf4l2 | QLFQLPAK | -145 | -152 | 0 | 0 | 8 | NaN | 1397.2118 | NaN | 1303.4708 | 1207.1237 | NaN |
| Qrich1 | QLGAHSPSTLLTTL | -614 | -627 | 0 | 0.002780285 | 14 | NaN | 912.56433 | NaN | 1417.1943 | NaN | NaN |
| Diaph3 | QLSESMIQNLIK | -727 | -738 | 0 | 0.001610812 | 12 | NaN | 8083.1074 | 8283.014 | 7292.9683 | 8062.1597 | 6563.876 |
| Mpo | QNQIVVDEIR | -534 | -543 | 0 | 0 | 10 | NaN | 661.83325 | 433.24496 | 894.6764 | NaN | NaN |
| Ctc1 | QQFQCGELPLLTR | -1172 | -1184 | 0 | 0.005363933 | 13 | NaN | 11628.57 | NaN | 17090.795 | 15059.257 | 13993.656 |
| Gid8 | QQHLIELIR | -106 | -114 | 0 | 0 | 9 | NaN | 1167.5907 | 354.43228 | NaN | 2291.0198 | NaN |
| Ssr3 | SEEDLLLQDFSR | -11 | -22 | 0 | 0 | 12 | NaN | 897.8101 | 415.76035 | 814.8056 | 426.9552 | 783.8893 |
| Tiprl | QSSHQDFSFGPWK | -7 | -19 | 0 | 0.002434616 | 13 | NaN | 292.86792 | 355.8695 | NaN | 336.58652 | 297.22998 |
| Cd3g | QTLLQNEQLYQPLK | -151 | -164 | 0 | 0 | 14 | NaN | 699.2908 | NaN | 450.19446 | NaN | 494.39035 |
| Mtx1 | YNADYDLSAR | -79 | -88 | 0 | 0.002629273 | 10 | NaN | 1355.5775 | NaN | NaN | 1641.2126 | NaN |
| Rbx1 | QVCPLDNR | -92 | -99 | 0 | 0.00025015 | 8 | NaN | 1226.571 | 1221.4369 | 1336.0212 | 764.88513 | 1136.0905 |
| Faf1 | RDVTQLDPNK | -621 | -630 | 1 | 0.001055699 | 10 | NaN | 589.86365 | 544.07184 | 799.14874 | 1355.3993 | NaN |
| Exoc4 | REEEEDFIR | -669 | -677 | 1 | 0.005270854 | 9 | NaN | 687.8383 | NaN | NaN | NaN | NaN |
| Rfc3 | RELYGIGVEK | -57 | -66 | 1 | 0.003201138 | 10 | NaN | 482.84088 | NaN | NaN | 825.99335 | 1041.5325 |
| Sumo1 | SDQEAKPSTEDLGDK | -2 | -16 | 1 | 0.001743443 | 15 | NaN | 422.88858 | 475.21274 | 312.97305 | 251.50815 | 516.2788 |
| RtcA | SCLLGLPLR | -27 | -35 | 0 | 0.001023236 | 9 | NaN | 388.88193 | NaN | 359.8076 | NaN | NaN |
| Ddx3y | RNDYDGIGGR | -101 | -110 | 1 | 0.001743443 | 10 | NaN | 1780.2217 | NaN | 2112.6672 | 1998.7062 | 1892.956 |
| Mtmr3 | SAERPQWDSVLHR | -886 | -898 | 1 | 0 | 13 | NaN | 168.66739 | NaN | 161.68349 | 395.69916 | 381.1808 |
| Slc7a1 | SESQTGFLPVAEK | -455 | -467 | 0 | 0.00142999 | 13 | NaN | 387.90836 | NaN | NaN | NaN | NaN |
| Anln | SDLEVEQNTEAVR | -531 | -543 | 0 | 0.001536401 | 13 | NaN | 3111.3318 | NaN | NaN | NaN | NaN |
| Pak1ip1 | SVLAVAGDEEVVR | -216 | -228 | 0 | 0 | 13 | NaN | 1309.5835 | 2046.5287 | NaN | NaN | NaN |
| Pak1ip1 | YVVSGSKDETIHIY | -54 | -67 | 1 | 0.000298805 | 14 | NaN | 144.02374 | NaN | NaN | NaN | NaN |
| Gns | SDVLVEYQGEGR | -402 | -413 | 0 | 0.001868731 | 12 | NaN | 1243.1952 | NaN | NaN | 1376.7963 | NaN |
| Cdipt | SGLLDAFDGHAAR | -43 | -55 | 0 | 0.000627943 | 13 | NaN | 1015.5405 | 971.03394 | NaN | NaN | 813.4715 |
| Pik3r1 | SEIFSPVLFR | -265 | -274 | 0 | 0 | 10 | NaN | 754.8583 | 524.2882 | 593.51465 | 788.98676 | 305.56113 |
| Ikzf1 | SENSQDAFR | -437 | -445 | 0 | 0 | 9 | NaN | 834.1088 | 801.31903 | 865.9925 | 836.0064 | 1041.6757 |
| Ikzf1 | YQLHKPPSDGPPR | -344 | -356 | 1 | 0 | 13 | NaN | 699.7663 | 828.3028 | 610.47864 | 925.9608 | 798.37274 |
| Ikzf1 | YQLHKPPSDGPPR | -344 | -356 | 1 | 0 | 13 | NaN | 699.7663 | 828.3028 | 610.47864 | 925.9608 | 798.37274 |
| Emc7 | VLVDGEEHVGFLK | -66 | -78 | 0 | 0 | 13 | NaN | 591.4502 | 559.9874 | NaN | 1106.2775 | 574.1361 |
| Clns1a | SFPPPGSADGLR | -6 | -17 | 0 | 0.005821388 | 12 | NaN | 3039.64 | NaN | NaN | NaN | NaN |

| | | | | | | | | | | | | |
|---|---|---|---|---|---|---|---|---|---|---|---|---|
| Foxk1 | SFQEPHFYLR | -138 | -147 | 0 | 0 | 10 | NaN | 1316.0596 | 360.24716 | NaN | 347.9594 | 1641.6692 |
| Trim32 | SFTEEQLRPK | -27 | -36 | 1 | 0.000431655 | 10 | NaN | 610.4244 | 848.0692 | NaN | NaN | NaN |
| Akr1c13 | SGDNDFPVNEQGK | -124 | -136 | 0 | 0.00025015 | 13 | NaN | 940.6819 | 903.38104 | 903.0021 | 925.5803 | 908.09595 |
| Coq9 | SGEEEEDYESEEQLQHR | -81 | -97 | 0 | 0 | 17 | NaN | 210.811 | 227.02943 | 178.82552 | 107.065 | 238.2933 |
| Coq9 | STDFNWYTR | -230 | -238 | 0 | 0 | 9 | NaN | 272.50702 | 192.52756 | NaN | 953.1778 | 820.52893 |
| Ankrd13a | SGFWGWR | -248 | -254 | 0 | 0.000627943 | 7 | NaN | 127.234436 | NaN | NaN | NaN | NaN |
| Parp14 | SNQQLQLAPR | -212 | -221 | 0 | 0 | 10 | NaN | 2106.1953 | 1193.028 | 5059.394 | 6915.188 | NaN |
| Tbce | SIHFPDAEIGCK | -292 | -303 | 0 | 0 | 12 | NaN | 1232.1188 | 1613.3849 | 855.415 | 956.5625 | NaN |
| Micos13 | SKEGWEYLK | -107 | -115 | 1 | 0 | 9 | NaN | 3631.3374 | 675.7122 | NaN | 2855.0007 | 2897.071 |
| Pigk | SLCVSTPGHR | -278 | -287 | 0 | 0.000347792 | 10 | NaN | 125.95018 | 1131.6671 | NaN | 576.718 | 361.06625 |
| Plbd2 | SLLLDAASGQLR | -68 | -79 | 0 | 0 | 12 | NaN | 1487.2668 | 409.66916 | 2431.4243 | 1452.9806 | 1787.2812 |
| Tmem214 | TPGESLHGYR | -211 | -220 | 0 | 0 | 10 | NaN | 1864.6296 | NaN | 1538.5685 | NaN | NaN |
| Pbk | SLNDLIEER | -121 | -129 | 0 | 0.002595029 | 9 | NaN | 750.0365 | NaN | NaN | 650.36475 | NaN |
| Trappc3 | SNLLCGVLR | -128 | -136 | 0 | 0.001392869 | 9 | NaN | 692.2627 | 595.7734 | 431.83255 | 772.36395 | NaN |
| Mtif2 | SNVLPIIIK | -513 | -521 | 0 | 0.000230707 | 9 | NaN | 259.86237 | NaN | 280.34396 | NaN | 240.60947 |
| Urod | SPEACCELTLQPLR | -61 | -74 | 0 | 0.001636916 | 14 | NaN | 1947.456 | NaN | NaN | NaN | NaN |
| Mcrip1 | SPTNSSEIFTPAHEENVR | -21 | -38 | 0 | 0.002265552 | 18 | NaN | 949.5067 | 1896.9049 | NaN | NaN | NaN |
| Ctdspl2 | SPVFNFFSPANK | -146 | -157 | 0 | 0 | 12 | NaN | 295.88577 | NaN | NaN | 773.86115 | NaN |
| Ppm1h | VHDSNIYIKPF | -398 | -408 | 1 | 0.00025015 | 11 | NaN | 637.2835 | NaN | NaN | NaN | NaN |
| Nab2 | SQPVPAVPVSSIPLFK | -114 | -129 | 0 | 0.000826806 | 16 | NaN | 340.51862 | 490.1437 | NaN | 417.67865 | 496.65186 |
| Wnk1 | SQQPLEDLDAQLR | -1787 | -1799 | 0 | 0 | 13 | NaN | 1183.6554 | 714.6551 | NaN | 1349.0988 | 1346.2278 |
| Pdpk1 | SQTEPGSSPGIPSGVSR | -31 | -47 | 0 | 0.001610812 | 17 | NaN | 877.8232 | NaN | NaN | NaN | NaN |
| Abhd11 | SSGQENADLRPLPL | -34 | -47 | 1 | 0 | 14 | NaN | 1581.2078 | 1648.9845 | 1287.9385 | 656.531 | 1221.2604 |
| Plcl2 | SSHNTYLIEDQFR | -440 | -452 | 0 | 0.000431655 | 13 | NaN | 903.5762 | 859.36035 | NaN | NaN | 1295.9946 |
| Smad2 | SSILPFTPPVVK | -2 | -13 | 0 | 0.00025015 | 12 | NaN | 331.33997 | NaN | NaN | 337.7963 | 463.59265 |
| Wasf2 | SSLPAVSDAR | -427 | -436 | 0 | 0.000431655 | 10 | NaN | 700.0865 | 709.7219 | NaN | NaN | NaN |
| Nmd3 | TIPVESDTDDEGAPR | -463 | -477 | 0 | 0.001392869 | 15 | NaN | 572.4231 | NaN | NaN | NaN | NaN |
| Nfyb | STLGFDSYVEPLK | -123 | -135 | 0 | 0 | 13 | NaN | 1435.7836 | NaN | NaN | 1264.5405 | 680.50885 |
| Strn | VISHPTLPISIT | -670 | -681 | 0 | 0.001291103 | 12 | NaN | 1099.4329 | 877.43243 | NaN | 1022.5262 | 1413.7684 |
| Chaf1b | SVDNTVIIWDVSK | -148 | -160 | 0 | 0.005884925 | 13 | NaN | 9667.53 | NaN | NaN | NaN | NaN |
| Pde2a | SVFLLDQNELVAK | -430 | -442 | 0 | 0.000550307 | 13 | NaN | 481.61957 | NaN | NaN | 683.9936 | NaN |
| Rdh11 | SVHPGTVHSELTR | -226 | -238 | 0 | 0 | 13 | NaN | 1676.3856 | 1524.6035 | 1067.7189 | NaN | 1808.0792 |
| Fn1 | SVYEQHESIPLR | -1430 | -1441 | 0 | 0 | 12 | NaN | 684.35657 | NaN | 2848.572 | 584.30133 | 2880.587 |
| Sec23ip | VANQIKEEEEK | -903 | -913 | 1 | 0.000591124 | 11 | NaN | 315.9081 | NaN | NaN | NaN | NaN |
| Ak3 | SWLLDGFPR | -86 | -94 | 0 | 0.000627943 | 9 | NaN | 1451.0222 | 1451.1371 | NaN | NaN | 1419.0835 |
| Cmtr1 | SYKPTLLTAK | -570 | -579 | 1 | 0 | 10 | NaN | 2883.941 | 4779.426 | NaN | NaN | NaN |
| Atp1a2 | TAGDASESALLK | -445 | -456 | 0 | 0 | 12 | NaN | 9198.74 | NaN | 851.9526 | NaN | NaN |
| Rbpj | TALLDADDPVSQLHK | -297 | -311 | 0 | 0 | 15 | NaN | 907.356 | 736.24725 | 756.12976 | 615.67426 | 747.5233 |
| Pih1d1 | TARPDSTQIQPK | -40 | -51 | 1 | 0.005587349 | 12 | NaN | 477.59552 | 399.0576 | 332.37552 | 369.75644 | NaN |
| Rnf20 | TAVEDSGTTVETIK | -23 | -36 | 0 | 0.00025015 | 14 | NaN | 580.67377 | NaN | NaN | NaN | 1296.7515 |
| Pycr3 | TDVLTPAGTTIHGL | -232 | -245 | 0 | 0.001291103 | 14 | NaN | 392.2408 | NaN | NaN | NaN | 1005.2829 |
| Elp2 | WAPDEESGVWLEQVR | -318 | -332 | 0 | 0 | 15 | NaN | 321.50443 | NaN | 349.2624 | NaN | 542.3367 |
| Larp4b | TEDLFENR | -548 | -555 | 0 | 0.001023236 | 8 | NaN | 454.29364 | NaN | NaN | NaN | 735.26465 |
| Atg4b | TEEDFNDWCQQVK | -325 | -337 | 0 | 0 | 13 | NaN | 431.90332 | 326.45468 | 352.07907 | 409.72668 | NaN |
| Haao | TGKPNPDQLLK | -156 | -166 | 1 | 0 | 11 | NaN | 1720.9126 | 756.20087 | 544.86523 | 404.28223 | 2163.7957 |
| Utp11 | THLQTAPELVDR | -153 | -164 | 0 | 0 | 12 | NaN | 1157.169 | NaN | 1516.2009 | 1669.4296 | 1775.7651 |
| Pus10 | THYLDEHHFR | -464 | -473 | 0 | 0 | 10 | NaN | 1632.1003 | NaN | NaN | NaN | 1046.1569 |
| Timm50 | TIALNQVEDVR | -285 | -295 | 0 | 0 | 11 | NaN | 805.64185 | 630.9794 | NaN | 564.67755 | 760.1858 |
| Sh3gl1 | VDAQLDYHR | -219 | -227 | 0 | 0.005698678 | 9 | NaN | 1294.3693 | 1364.738 | NaN | NaN | NaN |
| Lad1 | TLEDEEEQER | -19 | -28 | 0 | 0.000712251 | 10 | NaN | 554.8817 | 394.46613 | NaN | NaN | NaN |
| Lmna | TLEGELHDLR | -157 | -166 | 0 | 0 | 10 | NaN | 573.3015 | 2647.8667 | 2709.4165 | 3025.1636 | 3155.7793 |
| Rbm26 | TLEPICDADPSALAK | -20 | -34 | 0 | 0 | 15 | NaN | 595.8174 | 427.67584 | 413.06366 | 615.9786 | 569.5765 |
| Trmt2a | TNELSNVEFHCGR | -466 | -478 | 0 | 0 | 13 | NaN | 503.00458 | NaN | 16734.734 | NaN | NaN |
| Pnpt1 | TNPEKGEIEDYR | -517 | -528 | 1 | 0 | 12 | NaN | 378.72498 | 433.2073 | NaN | 421.04227 | 471.52844 |
| Ptpn22 | TPESFIVVEEAGEPSPR | -620 | -636 | 0 | 0.001636916 | 17 | NaN | 1632.4563 | NaN | NaN | NaN | NaN |
| Ech1 | TQDAFFQIK | -187 | -195 | 0 | 0 | 9 | NaN | 835.47003 | NaN | 968.0796 | 1299.6891 | 1211.798 |
| Pcyt2 | TQGVSTTDLVGR | -141 | -152 | 0 | 0 | 12 | NaN | 1863.033 | 1558.0067 | 1850.6953 | 2264.359 | NaN |
| Mrfap1 | TQVEASEESALNHLQGAGGAEPR | -69 | -91 | 0 | 0.00025015 | 23 | NaN | 254.29295 | NaN | 266.42706 | NaN | 235.83054 |
| Prmt7 | TSLGEQVIVPPSELER | -203 | -218 | 0 | 0.000230707 | 16 | NaN | 1079.2979 | NaN | NaN | NaN | 1196.9178 |
| Cmtm7 | TTCSSGGALGPGQPSEGLLDR | -10 | -30 | 0 | 0.009985598 | 21 | NaN | 530.70886 | NaN | NaN | NaN | NaN |
| Mtfr1l | TTELQDELSHLR | -125 | -136 | 0 | 0.000907206 | 12 | NaN | 12143.133 | NaN | NaN | NaN | NaN |
| Riox1 | TVLEPGDLLYFPR | -351 | -363 | 0 | 0 | 13 | NaN | 280.71268 | 319.4938 | 425.61496 | 423.86 | 404.04633 |
| Dcaf13 | TVYTGIDHHWK | -157 | -167 | 0 | 0 | 11 | NaN | 1095.5804 | 816.40845 | 961.153 | NaN | 965.3973 |
| Carmil2 | TWLPQEGAEQSHIL | -31 | -44 | 0 | 0.000627943 | 14 | NaN | 1230.5273 | NaN | NaN | NaN | NaN |
| Cd226 | TWPVQQVIWEK | -158 | -168 | 0 | 0.00025015 | 11 | NaN | 902.2255 | NaN | NaN | 800.434 | NaN |
| Hbz | TYFPHFDL | -42 | -49 | 0 | 0.00078661 | 8 | NaN | 700.16516 | 842.1282 | 639.44073 | NaN | 284.0263 |
| Imp3 | VAFVEQGHVR | -128 | -137 | 0 | 0.00078661 | 10 | NaN | 425.22528 | NaN | NaN | 662.27997 | 620.7685 |
| Glb1 | VDGVPQGILDR | -449 | -459 | 0 | 0.000394361 | 11 | NaN | 2021.3135 | NaN | NaN | NaN | 2483.2205 |
| Ube2h | VDLPDKYPFK | -55 | -64 | 1 | 0.000550307 | 10 | NaN | 361.79626 | NaN | 473.6842 | NaN | NaN |
| Mmtag2 | VDSEGPSTAPTAPR | -153 | -166 | 0 | 0.002780285 | 14 | NaN | 3981.0176 | NaN | NaN | NaN | NaN |
| Usp3 | VDTYVQFPLR | -424 | -433 | 0 | 0 | 10 | NaN | 3178.7207 | 293.67618 | NaN | NaN | NaN |
| Vps11 | VELDTHGCGLR | -224 | -234 | 0 | 0 | 11 | NaN | 432.91104 | NaN | NaN | NaN | 2441.2913 |
| Ube2o | VEVVWADNSK | -685 | -694 | 0 | 0 | 10 | NaN | 1606.3835 | NaN | NaN | NaN | NaN |
| Slc29a1 | VFDWLGR | -339 | -345 | 0 | 0.008947895 | 7 | NaN | 409.19434 | NaN | NaN | NaN | NaN |
| Mrpl30 | VFQPKPEDHEK | -45 | -55 | 1 | 0.000230707 | 11 | NaN | 903.1226 | 852.512 | NaN | 1077.1066 | 1153.211 |
| Arl6ip6 | VGDGSPVLPDKR | -76 | -87 | 1 | 0.00051306 | 12 | NaN | 2678.9895 | NaN | NaN | NaN | NaN |
| Sdhd | VVTDYVHGDTLPK | -110 | -122 | 0 | 0 | 13 | NaN | 1294.9296 | 996.23706 | 1011.1686 | NaN | 1366.6213 |
| Sdhd | VHGDTLPK | -115 | -122 | 0 | 0.005176406 | 8 | NaN | 1005.47687 | 1109.3177 | NaN | NaN | NaN |

| | | | | | | | | | | | | |
|---|---|---|---|---|---|---|---|---|---|---|---|---|
| Itga4 | VIELNKDENVAHVF | -957 | -970 | 1 | 0.00111006 | 14 | NaN | 686.7905 | 526.6747 | NaN | 665.15765 | 546.77875 |
| Herc4 | YWADHPTIK | -962 | -970 | 0 | 0.00025015 | 9 | NaN | 264.24 | NaN | NaN | 2213.7578 | 1892.7516 |
| Rpia | VKQENLDLICIPT | -112 | -124 | 1 | 0.00078661 | 13 | NaN | 679.5249 | 1054.2996 | NaN | 599.2517 | 945.2631 |
| Btrc;Fbxw11 | VLEGHEELVR | (501);(438) | (510);(447) | 0 | 0 | 10 | NaN | 712.2942 | NaN | NaN | NaN | NaN |
| Dcakd | VLGDLIFNQPDR | -68 | -79 | 0 | 0.000471409 | 12 | NaN | 476.13593 | NaN | NaN | NaN | NaN |
| Msi2 | VLGQPHHELDSK | -78 | -89 | 0 | 0 | 12 | NaN | 1739.7236 | 1835.2316 | 1105.4299 | NaN | 3081.9165 |
| Tssc4 | VLPVPDYVSHPER | -146 | -158 | 0 | 0 | 13 | NaN | 1185.634 | NaN | NaN | 633.52356 | 703.173 |
| Ifi35 | VLVSGFPAGLR | -135 | -145 | 0 | 0 | 11 | NaN | 779.35187 | 839.86115 | NaN | NaN | NaN |
| Rrp1b | VPDSESEPDTAQR | -402 | -414 | 0 | 0.004658491 | 13 | NaN | 870.237 | NaN | NaN | NaN | NaN |
| Wdr74 | VPIWDQDTQFLPGSQK | -182 | -197 | 0 | 0 | 16 | NaN | 2077.5586 | 1783.7119 | 801.8263 | 2070.4429 | 1906.3157 |
| Wdr74 | VWDLQGSEEPVFR | -157 | -169 | 0 | 0 | 13 | NaN | 3002.9336 | 3029.0686 | 1634.4536 | NaN | 3358.9294 |
| Gtse1 | VPTLSPLTQQPQTPEQR | -394 | -410 | 0 | 0 | 17 | NaN | 385.94794 | 390.84732 | NaN | NaN | 536.5361 |
| Med8 | VPVFSHEVVPDHLR | -97 | -110 | 0 | 0 | 14 | NaN | 955.8555 | 584.65906 | NaN | 1052.1156 | NaN |
| H6pd | VQQDGLWEAGR | -125 | -135 | 0 | 0.00555405 | 11 | NaN | 5671.741 | NaN | NaN | NaN | NaN |
| Sp3 | VQVVDEEGDQQHQEGK | -580 | -595 | 0 | 0 | 16 | NaN | 200.16435 | 272.8658 | 178.99731 | 408.8364 | 372.7215 |
| Mettl1 | VSDYVQDR | -106 | -113 | 0 | 0.00635324 | 8 | NaN | 525.8255 | NaN | NaN | NaN | NaN |
| Pacsin2 | VSELHLEVK | -93 | -101 | 0 | 0.00025015 | 9 | NaN | 295.62415 | 1066.6139 | NaN | NaN | NaN |
| Efl1 | YSPVDFLQR | -529 | -537 | 0 | 0.00111006 | 9 | NaN | 579.7756 | 417.87173 | NaN | 405.45776 | NaN |
| Galm | VSPDGEEGYPGELK | -132 | -145 | 0 | 0 | 14 | NaN | 2773.5842 | NaN | NaN | NaN | NaN |
| Sars2 | VVGEKPAFSFQPR | -191 | -203 | 1 | 0 | 13 | NaN | 920.4226 | 457.15234 | 1579.2311 | 2094.6665 | 1961.7927 |
| Thoc5 | VVLNPGYSSIPPVFR | -567 | -581 | 0 | 0 | 15 | NaN | 907.5917 | 879.9967 | 382.5515 | NaN | 706.059 |
| Sptlc1 | VVVTVEQTEEELQR | -446 | -459 | 0 | 0 | 14 | NaN | 779.07916 | 512.08325 | 598.04095 | 494.15894 | 538.1307 |
| Brap | VWDYAGDNYVHR | -364 | -375 | 0 | 0.001291103 | 12 | NaN | 768.53516 | NaN | NaN | NaN | NaN |
| Uba7 | WAQDQFEGLFR | -601 | -611 | 0 | 0 | 11 | NaN | 517.05884 | NaN | 436.7096 | NaN | NaN |
| Eif2b5 | YCALLLPLLK | -613 | -622 | 0 | 0 | 10 | NaN | 257.83533 | NaN | 410.20712 | 555.7969 | 420.34445 |
| Galnt2 | WYLDNVYPELR | -427 | -437 | 0 | 0.005758933 | 11 | NaN | 339.2812 | NaN | NaN | NaN | NaN |
| Spcs1 | WLPVQDLGTEDKK | -136 | -148 | 1 | 0 | 13 | NaN | 1176.5049 | 957.8541 | 1437.5677 | 1834.9491 | 1539.3778 |
| Lrrc40 | WWDQTDLTK | -78 | -86 | 0 | 0.007905138 | 9 | NaN | 521.8976 | NaN | 629.1595 | NaN | 414.4937 |
| Pacsin1 | YDYDGQEQDELSFK | -391 | -404 | 0 | 0.000230707 | 14 | NaN | 238.61252 | NaN | NaN | NaN | NaN |
| Arpc1a | YFESENDWWVSK | -124 | -135 | 0 | 0 | 12 | NaN | 656.4447 | NaN | NaN | NaN | NaN |
| Tmem254 | YLVDHYHTF | -52 | -60 | 0 | 0 | 9 | NaN | 1080.2517 | 1109.5198 | NaN | NaN | NaN |
| Parp9 | YPASLTQELQDR | -653 | -664 | 0 | 0.001055699 | 12 | NaN | 578.235 | NaN | NaN | NaN | NaN |
| Msmo1 | YVDSLLPENPLQEPFK | -19 | -34 | 0 | 0 | 16 | NaN | 413.0681 | NaN | 478.93503 | 527.24475 | 491.46136 |
| Adk | SLKPNDQILAEDK | -48 | -60 | 1 | 0.000627943 | 13 | NaN | NaN | 470.69757 | 628.18713 | 558.10913 | NaN |
| Adk | AAADEPKPK | -2 | -10 | 1 | 0.008129024 | 9 | NaN | NaN | 1198.5234 | NaN | NaN | NaN |
| Adk | VAQWLIQEPHK | -88 | -98 | 0 | 0 | 11 | NaN | NaN | 284.39008 | NaN | NaN | NaN |
| Naca | SPASDTYIVF | (2086);(114) | (2095);(123) | 0 | 0.005333786 | 10 | NaN | NaN | 1727.614 | 1512.5398 | 2752.937 | 2620.1226 |
| Mcm6 | VSGADGYETEGIR | -270 | -282 | 0 | 0 | 13 | NaN | NaN | 4849.4004 | 2539.9456 | 1669.2427 | 1937.2328 |
| Mcm6 | NINLSAPIMSR | -519 | -529 | 0 | 0 | 11 | NaN | NaN | 889.46783 | 1607.1339 | NaN | NaN |
| Mcm6 | HVAPTNPR | -302 | -309 | 0 | 0.003461185 | 8 | NaN | NaN | 380.84174 | NaN | NaN | 445.72308 |
| Mcm6 | SLFPTIHGNDEVK | -353 | -365 | 0 | 0.009871162 | 13 | NaN | NaN | 1227.1323 | NaN | NaN | NaN |
| Tln1 | AAENEEIGAHIK | -1906 | -1917 | 0 | 0.000550307 | 12 | NaN | NaN | 447.3289 | 273.83228 | NaN | 770.3345 |
| Tln1 | AAAFEDQENETVVVK | -2477 | -2491 | 0 | 0.001023236 | 15 | NaN | NaN | 1273.0087 | NaN | NaN | NaN |
| Hnrnpd | GGTEGGSAEAEGAK | -54 | -67 | 0 | 0.00025015 | 14 | NaN | NaN | 1095.1084 | 1222.0558 | NaN | NaN |
| Hnrnpd | LKLDPITGR | -128 | -136 | 1 | 0.00383302 | 9 | NaN | NaN | 756.74194 | 640.5828 | NaN | NaN |
| Hnrnpd | KYHNVGL | -243 | -249 | 1 | 0.005270854 | 7 | NaN | NaN | 496.1999 | NaN | NaN | NaN |
| Lcp1 | EGICAIGGTSEQSSVGTQH | -98 | -116 | 0 | 0 | 19 | NaN | NaN | 1446.8171 | 1803.3563 | 2259.2986 | 2386.782 |
| Lcp1 | VNHLYSDLSDAL | -413 | -424 | 0 | 0.00051306 | 12 | NaN | NaN | 1308.016 | 936.42444 | 1138.185 | 1324.5627 |
| Lcp1 | EGICAIGGTSEQS | -98 | -110 | 0 | 0 | 13 | NaN | NaN | 203.65778 | 209.64496 | NaN | 249.34885 |
| Lcp1 | CLPLPGYR | -42 | -49 | 0 | 0.001023236 | 8 | NaN | NaN | 1244.3372 | 1549.8418 | 1452.6257 | 1411.9285 |
| Lcp1 | YPALHKPENQDIDWGAL | -374 | -390 | 1 | 0.000431655 | 17 | NaN | NaN | 442.1394 | 824.46265 | 827.18567 | 684.8598 |
| Lcp1 | YTLNILEDIGGGQK | -502 | -515 | 0 | 0 | 14 | NaN | NaN | 310.87842 | 370.7792 | 525.5714 | 374.44894 |
| Lcp1 | YPALHKPENQDIDWGALE | -374 | -391 | 1 | 0.000712251 | 18 | NaN | NaN | 593.3142 | NaN | NaN | NaN |
| Lcp1 | SSVGTQHSYSEEEK | -110 | -123 | 0 | 0.00078661 | 14 | NaN | NaN | 468.47498 | NaN | NaN | NaN |
| Psmb8 | KGPGLYYVDDNGTR | -179 | -192 | 1 | 0 | 14 | NaN | NaN | 478.3274 | 1061.627 | 1030.9998 | 951.55585 |
| Hmgb1 | GEMWNNTAADDKQPYEK | -130 | -146 | 1 | 0.00204437 | 17 | NaN | NaN | 1529.115 | 293.23486 | NaN | NaN |
| Hmgb1 | IKGEHPGLSIG | -113 | -123 | 1 | 0 | 11 | NaN | NaN | 613.9967 | 291.10953 | NaN | NaN |
| Nap1l1 | YAVLYQPLFDK | -106 | -116 | 0 | 0 | 11 | NaN | NaN | 841.6649 | 1232.0632 | 1885.2954 | 1580.2828 |
| Nap1l1 | IYEPTEEECEWKPDEEDEVSEELKEK | -124 | -149 | 2 | 0.00025015 | 26 | NaN | NaN | 532.8219 | 834.16003 | 1048.0182 | 904.15405 |
| Hspa9 | SLGIETLGGVFTK | -444 | -456 | 0 | 0 | 13 | NaN | NaN | 809.56824 | 711.58093 | 555.4419 | 550.0081 |
| Rfc1 | AADSICDGDLVDNQIR | -895 | -910 | 0 | 0.00025015 | 16 | NaN | NaN | 2433.88 | 2201.007 | 2398.405 | NaN |
| Pkm | MADTFLEHMCR | -22 | -32 | 0 | 0 | 11 | NaN | NaN | 519.8714 | 583.61945 | 853.86334 | 398.218 |
| Pkm | DEILEASDGIMVAR | -281 | -294 | 0 | 0 | 14 | NaN | NaN | 3890.1448 | NaN | 3569.3308 | 3827.5698 |
| Pdcd6ip | QTYPSGGEEQAQYCR | -27 | -41 | 0 | 0 | 15 | NaN | NaN | 387.15387 | 658.3796 | NaN | NaN |
| Cct2 | GVMAIEHADFAGVER | -308 | -322 | 0 | 0 | 15 | NaN | NaN | 826.5432 | NaN | 1019.34326 | 503.59528 |
| Cct2 | HGINCFINR | -285 | -293 | 0 | 0 | 9 | NaN | NaN | 3470.2659 | 993.1203 | 2262.5403 | 3046.636 |
| Cct2 | HADFAGVER | -314 | -322 | 0 | 0.00111006 | 9 | NaN | NaN | 1301.7849 | 452.1405 | NaN | NaN |
| Cct2 | VQDDEVGDGTTSVTVL | -90 | -105 | 0 | 0.00025015 | 16 | NaN | NaN | 427.68387 | NaN | NaN | NaN |
| Psmd4 | VGSPVEDNEKDLVK | -113 | -126 | 1 | 0 | 14 | NaN | NaN | 1004.5348 | 873.0836 | 866.77454 | 888.9617 |
| Psmd4 | AAEAGIATPGTEDSDDALLK | -243 | -262 | 0 | 0.001636916 | 20 | NaN | NaN | 366.9955 | NaN | NaN | 446.53854 |
| Psmd4 | VNVDIINFGEEEVNTEK | -136 | -152 | 0 | 0.001636916 | 17 | NaN | NaN | 1155.4196 | NaN | NaN | NaN |
| Dhx9 | VQSDGQIVFIDDWIR | -1077 | -1091 | 0 | 0.001291103 | 15 | NaN | NaN | 660.92694 | NaN | 375.75278 | 331.83707 |
| Smchd1 | PGPPHQLK | -924 | -931 | 0 | 0 | 8 | NaN | NaN | 257.8332 | 449.57715 | 519.6997 | 543.83655 |
| Smchd1 | QVEVVDESDNITAQPK | -950 | -965 | 0 | 0.000471409 | 16 | NaN | NaN | 674.2179 | NaN | NaN | 546.7966 |
| Smchd1 | VITGVQPEHIQYLK | -323 | -336 | 0 | 0.00025015 | 14 | NaN | NaN | 271.98965 | 250.39998 | NaN | NaN |
| Smchd1 | AAEGASDPAGLSEGSGR | -2 | -18 | 0 | 0.004824802 | 17 | NaN | NaN | 2037.3004 | NaN | NaN | NaN |
| Smchd1 | DVHELVLSK | -261 | -269 | 0 | 0.000230707 | 9 | NaN | NaN | 495.72064 | NaN | NaN | NaN |
| Hspa4 | ELTSICSPIISKPK | -775 | -788 | 1 | 0 | 14 | NaN | NaN | 800.79236 | NaN | 686.7322 | NaN |

| | | | | | | | | | | | | |
|---|---|---|---|---|---|---|---|---|---|---|---|---|
| Hspa4 | NKEDQYEHLDAADVT | -719 | -733 | 1 | 0.000550307 | 15 | NaN | NaN | 618.6817 | NaN | NaN | NaN |
| Gvin1 | KREELLSVPK | -206 | -215 | 2 | 0.001291103 | 10 | NaN | NaN | 614.0367 | 460.25845 | 591.3481 | NaN |
| Gvin1 | DKEFYHLR | -1292 | -1299 | 1 | 0.002780285 | 8 | NaN | NaN | 1881.9144 | NaN | NaN | 1487.4563 |
| Gvin1 | AAELEECSNITR | -1648 | -1659 | 0 | 0 | 12 | NaN | NaN | 1599.8096 | 193.7716 | NaN | NaN |
| Gvin1 | EGEEFLTAR | -514 | -522 | 0 | 0 | 9 | NaN | NaN | 1309.4556 | NaN | NaN | 928.1838 |
| Stat1 | AETLQQIR | -267 | -274 | 0 | 0.00383302 | 8 | NaN | NaN | 403.32703 | NaN | NaN | NaN |
| Plrg1 | GPQNATDSYPHK | -69 | -80 | 0 | 0 | 12 | NaN | NaN | 938.35706 | 695.4507 | 608.49414 | 547.33264 |
| Nup93 | KPGVIDKF | -594 | -601 | 2 | 0.001708008 | 8 | NaN | NaN | 5410.9844 | 4926.327 | 5449.418 | 4798.2437 |
| Nup93 | KFESTDPR | -538 | -545 | 1 | 0.002329279 | 8 | NaN | NaN | 737.288 | 1059.3387 | 967.3037 | NaN |
| Nup93 | TSDTKPIINK | -602 | -611 | 1 | 0 | 10 | NaN | NaN | 234.19003 | 293.58133 | 135.27448 | NaN |
| Nup93 | CVSELVIESR | -569 | -578 | 0 | 0.001636916 | 10 | NaN | NaN | 631.4037 | NaN | NaN | NaN |
| Snrnp200 | VKLDFVAPATGGHNY | -2090 | -2104 | 1 | 0.000230707 | 15 | NaN | NaN | 501.55588 | NaN | 583.84216 | 456.3796 |
| Snrnp200 | ETQLPVSFR | -1279 | -1287 | 0 | 0.002780285 | 9 | NaN | NaN | 497.606 | NaN | NaN | NaN |
| Snrnp200 | VVDKDSIR | -2023 | -2030 | 1 | 0.009242859 | 8 | NaN | NaN | 407.60715 | NaN | NaN | NaN |
| Dld | VCHAHPTLSEAFR | -483 | -495 | 0 | 0 | 13 | NaN | NaN | 742.8714 | 723.57434 | 1024.795 | 941.9256 |
| Dld | CLNVGCIPSK | -80 | -89 | 0 | 0.002780285 | 10 | NaN | NaN | 2997.7803 | NaN | 1046.766 | NaN |
| Dld | EQLKEEGIEFK | -407 | -417 | 1 | 0.002534621 | 11 | NaN | NaN | 3997.3376 | NaN | NaN | NaN |
| Srm | HVGDGFEFMK | -152 | -161 | 0 | 0.000431655 | 10 | NaN | NaN | 715.6417 | 165.60652 | 756.0954 | 679.1095 |
| Srm | EIDEDVIEVSK | -124 | -134 | 0 | 0 | 11 | NaN | NaN | 1771.5942 | NaN | NaN | 1975.0197 |
| Srm | EMIANLPLCSHPNPR | -81 | -95 | 0 | 0.00025015 | 15 | NaN | NaN | 525.82166 | NaN | NaN | 523.5177 |
| Gart | RQEEAWVIGSVV | -756 | -767 | 1 | 0.000550307 | 12 | NaN | NaN | 590.6859 | NaN | 423.52997 | 343.50403 |
| Uqcrc1 | TSATESEVTR | -381 | -390 | 0 | 0 | 10 | NaN | NaN | 2614.5613 | NaN | 1248.9725 | 1553.4855 |
| Uqcrc1 | ADIVQNSSLEDSQIEK | -148 | -163 | 0 | 0.000431655 | 16 | NaN | NaN | 1837.9537 | NaN | NaN | NaN |
| Uqcrc1 | SGMFWLR | -473 | -479 | 0 | 0.007987481 | 7 | NaN | NaN | 209.85338 | NaN | NaN | NaN |
| Kpnb1 | STCCEDDIVPHVLPFIK | -356 | -372 | 0 | 0 | 17 | NaN | NaN | 381.87958 | NaN | 1027.9775 | NaN |
| Rrbp1 | AHQNYAEWLQEFK | -1278 | -1290 | 0 | 0.002099344 | 13 | NaN | NaN | 574.2644 | NaN | 326.90396 | 958.182 |
| Rrbp1 | LTAEFEEAQR | -1506 | -1515 | 0 | 0.000230707 | 10 | NaN | NaN | 664.4006 | NaN | NaN | 2409.9578 |
| Eef1a1 | MDSTEPPYSQK | -155 | -165 | 0 | 0 | 11 | NaN | NaN | 7739.9966 | 7070.357 | 8050.1455 | 7837.546 |
| Eef1a1 | LEALDCILPPTRPTDKPLR | -229 | -247 | 2 | 0 | 19 | NaN | NaN | 4434.963 | 5084.0806 | 7353.383 | 7643.312 |
| Eef1a1 | LEALDCILPPTRPTDK | -229 | -244 | 1 | 0 | 16 | NaN | NaN | 1585.9968 | 2193.4202 | 2406.532 | 3342.9004 |
| Eef1a1 | YVTIIDAPGHR | -86 | -96 | 0 | 0 | 11 | NaN | NaN | 2142.445 | 3170.9216 | 2507.3909 | 2324.7708 |
| Eef1a1 | GNVAGDSKNDPPMEAA | -323 | -338 | 1 | 0.001463897 | 16 | NaN | NaN | 697.1956 | 803.4189 | 917.78345 | 770.49304 |
| Rplp1 | GGAAPSTAAAPAEEK | -78 | -92 | 0 | 0 | 15 | NaN | NaN | 1621.6012 | 910.4873 | NaN | NaN |
| Mogs | PSGLDDYPR | -576 | -584 | 0 | 0.004690496 | 9 | NaN | NaN | 630.13135 | 289.25568 | 874.7234 | 854.07434 |
| Rcc2 | AAIITEPEHTK | -80 | -90 | 0 | 0.001023236 | 11 | NaN | NaN | 1978.6125 | NaN | NaN | NaN |
| Ddx21 | QDAQSLHGDIPQK | -534 | -546 | 0 | 0 | 13 | NaN | NaN | 5656.6113 | 3849.588 | 3240.7761 | 4692.6543 |
| Ddx21 | AITVEHLAIK | -480 | -489 | 0 | 0 | 10 | NaN | NaN | 451.80008 | 402.05365 | 461.9704 | 442.93723 |
| Ddx21 | KQEEEPQDDTASTS | -78 | -91 | 1 | 0.001023236 | 14 | NaN | NaN | 352.00607 | 874.64355 | NaN | 1202.5011 |
| Ddx21 | KDSEDNPQTLLF | -435 | -446 | 1 | 0.009955823 | 12 | NaN | NaN | 606.5524 | NaN | NaN | NaN |
| Cap1 | HAEMVHTGLK | -71 | -80 | 0 | 0.000550307 | 10 | NaN | NaN | 597.7179 | 1222.2727 | 841.79645 | 1171.9385 |
| Cap1 | TDGCHAYLSK | -412 | -421 | 0 | 0 | 10 | NaN | NaN | 1507.0286 | 442.27576 | 1680.1864 | 1701.2977 |
| Cap1 | LAAKPGPFVK | -145 | -154 | 1 | 0.001904399 | 10 | NaN | NaN | 582.0995 | NaN | NaN | 512.683 |
| Mcm5 | KYPEHAIR | -702 | -709 | 1 | 0 | 8 | NaN | NaN | 1036.6653 | 424.36765 | 500.4669 | 519.47064 |
| Mcm5 | QKYPEHAIR | -701 | -709 | 1 | 0.000431655 | 9 | NaN | NaN | 2487.202 | NaN | NaN | 1379.223 |
| Septin1 | EVTHDLLYEGYR | -278 | -289 | 0 | 0 | 12 | NaN | NaN | 1692.6978 | NaN | 1701.039 | NaN |
| Septin1 | IRDQLKEEEINIY | -187 | -199 | 2 | 0.00078661 | 13 | NaN | NaN | 816.00494 | 678.9929 | NaN | NaN |
| Septin1 | DQLKEEEINIY | -189 | -199 | 1 | 0.00504706 | 11 | NaN | NaN | 851.8798 | NaN | NaN | NaN |
| Eif4g1 | TIDQIHK | -1005 | -1011 | 0 | 0.001463897 | 7 | NaN | NaN | 1690.5476 | NaN | 1619.9803 | 700.10406 |
| Eif4g1 | VIIRPDDR | -235 | -242 | 1 | 0.004983786 | 8 | NaN | NaN | 624.22906 | 589.20605 | NaN | NaN |
| Cad | AVDWHFEEAVDGECPPQR | -1361 | -1378 | 0 | 0.003681885 | 18 | NaN | NaN | 313.27536 | NaN | NaN | NaN |
| Lmnb1 | VCIEEIDVDGK | -443 | -453 | 0 | 0 | 11 | NaN | NaN | 1523.7233 | NaN | 2001.7142 | 1042.2971 |
| Anxa6 | SLHQAIEGDTSGDFMK | -648 | -663 | 0 | 0 | 16 | NaN | NaN | 7295.973 | 2938.5369 | 2540.0732 | 2611.4458 |
| Anxa6 | LIVNLMRPL | -85 | -93 | 1 | 0 | 9 | NaN | NaN | 679.53955 | 417.25888 | 485.91553 | 498.41144 |
| Anxa6 | ILEIADTPSGDK | -529 | -540 | 0 | 0.000826806 | 12 | NaN | NaN | 2157.8257 | NaN | NaN | NaN |
| Anxa6 | LVQQDVQDLYEAGELK | -176 | -191 | 0 | 0.001055699 | 16 | NaN | NaN | 1501.8127 | NaN | NaN | NaN |
| Elf1 | AANPKDPVEVGQPSEVLR | -352 | -369 | 1 | 0.001610812 | 18 | NaN | NaN | 446.32248 | NaN | NaN | NaN |
| Ftsj3 | QKEQQPTDPPPPPTNLK | -555 | -571 | 1 | 0.001092529 | 17 | NaN | NaN | 563.6609 | 436.5775 | 614.10144 | NaN |
| Ftsj3 | EISIDDEELAQHPATTEDIR | -272 | -291 | 0 | 0.00078661 | 20 | NaN | NaN | 193.86404 | NaN | 326.62347 | 255.39024 |
| Tpi1 | NGAFTGEISPGMIK | -72 | -85 | 0 | 0 | 14 | NaN | NaN | 4730.2993 | 5194.6104 | 853.9885 | 4931.0464 |
| Vim | LQDEIQNMKEEMAR | -365 | -378 | 1 | 0 | 14 | NaN | NaN | 409.80646 | 373.90814 | 569.6821 | 525.70197 |
| Vim | VQSLTCEVDALK | -323 | -334 | 0 | 0.00025015 | 12 | NaN | NaN | 1651.96 | 1500.7643 | 1261.0542 | 2302.3284 |
| Vim | EMEENFALEAA | -346 | -356 | 0 | 0 | 11 | NaN | NaN | 263.58554 | 346.34982 | 252.74963 | 250.52106 |
| Vim | RETNLESLPLVDTHSK | -424 | -439 | 1 | 0.00025015 | 16 | NaN | NaN | 988.0156 | NaN | 683.94403 | 663.1736 |
| Me2 | NTLIQFEDFGNHNAFR | -249 | -264 | 0 | 0.00111006 | 16 | NaN | NaN | 250.8084 | NaN | 545.8377 | 604.2162 |
| Ppa1 | SLPTDVDKWFHQQK | -275 | -288 | 1 | 0.00078661 | 14 | NaN | NaN | 969.3528 | 558.7097 | 1098.8085 | 852.4827 |
| Uqcrc2 | SVLQHLLGAGPHIK | -287 | -300 | 0 | 0.000627943 | 14 | NaN | NaN | 478.89795 | 205.45508 | 550.72095 | 467.78635 |
| Uqcrc2 | QQIDSVADADVVK | -414 | -426 | 0 | 0.007905138 | 13 | NaN | NaN | 529.11664 | NaN | NaN | NaN |
| Wars1 | IDKELINR | -116 | -123 | 1 | 0.006735683 | 8 | NaN | NaN | 1794.4446 | NaN | 1598.7588 | NaN |
| N6amt1 | AAPSVPTPLYGHVGR | -2 | -16 | 0 | 0.00078661 | 15 | NaN | NaN | 604.32764 | NaN | 870.08057 | NaN |
| Tcp1 | VLCELADLQDKEVGDGTTS | -74 | -92 | 1 | 0.002595029 | 19 | NaN | NaN | 75.427666 | NaN | NaN | NaN |
| Ran | HLTGEFEKK | -30 | -38 | 1 | 0 | 9 | NaN | NaN | 378.97092 | 354.63217 | NaN | NaN |
| Dpp3 | LTFLEEEDKDLYIR | -424 | -437 | 1 | 0 | 14 | NaN | NaN | 2484.8223 | 4042.6196 | 5938.84 | 5263.1753 |
| Dpp3 | AGSGIPAGINIPNYDDLR | -382 | -399 | 0 | 0.001743443 | 18 | NaN | NaN | 454.6727 | 626.9787 | NaN | 663.4853 |
| Slc16a1 | FDIFDWR | -169 | -175 | 0 | 0.007905138 | 7 | NaN | NaN | 343.68463 | NaN | NaN | 303.20575 |
| Slc16a1 | SDANTDLIGGSPK | -220 | -232 | 0 | 0 | 13 | NaN | NaN | 676.83514 | NaN | NaN | NaN |
| Ywhae | AASDIAMTELPPTHPIR | -154 | -170 | 0 | 0.00025015 | 17 | NaN | NaN | 361.53342 | 928.0046 | 901.9398 | 1029.8469 |
| Ywhae | EAAENSLVAYK | -143 | -153 | 0 | 0.00025015 | 11 | NaN | NaN | 1068.7332 | NaN | NaN | 2540.1843 |
| Pfdn1 | EVIHNQLLEK | -74 | -83 | 0 | 0 | 10 | NaN | NaN | 658.0018 | 667.6649 | 1174.86 | 1335.777 |

| | | | | | | | | | | | | |
|---|---|---|---|---|---|---|---|---|---|---|---|---|
| Pfdn1 | AASVDLELKK | -2 | -11 | 1 | 0.000550307 | 10 | NaN | NaN | 742.73047 | 163.02022 | NaN | 826.0557 |
| Mrps7 | AATETSSVFADPVISK | -78 | -93 | 0 | 0 | 16 | NaN | NaN | 1038.858 | 668.19324 | 1848.408 | 691.4705 |
| Got1 | KVNLGVGAYR | -33 | -42 | 1 | 0.00025015 | 10 | NaN | NaN | 399.88718 | NaN | NaN | NaN |
| Vcp | LADDVDLEQVANETHGHVGAD | -390 | -410 | 0 | 0 | 21 | NaN | NaN | 849.49805 | 755.26355 | 978.0798 | 906.08325 |
| Vcp | VTWEDIGGLEDVKR | -474 | -487 | 1 | 0.000471409 | 14 | NaN | NaN | 669.6553 | NaN | NaN | 312.94568 |
| Vcp | KNVFIIGAT | -615 | -623 | 1 | 0.00201047 | 9 | NaN | NaN | 878.4989 | NaN | NaN | NaN |
| Dync1h1 | HYLDFINHY | -3173 | -3181 | 0 | 0 | 9 | NaN | NaN | 472.7915 | 507.9596 | 468.47784 | NaN |
| Dync1h1 | ITNQVIYLNPPIEECR | -962 | -977 | 0 | 0.00122419 | 16 | NaN | NaN | 327.2479 | NaN | 243.84424 | 403.25632 |
| Dync1h1 | RVEDVLGK | -662 | -669 | 1 | 0.006537474 | 8 | NaN | NaN | 101.995544 | NaN | 3082.8484 | 5434.903 |
| Dync1h1 | DHLYGTLDPNTR | -2260 | -2271 | 0 | 0.002972566 | 12 | NaN | NaN | 1125.0735 | NaN | NaN | 896.37976 |
| Dync1h1 | GEWSPWQAK | -2541 | -2549 | 0 | 0 | 9 | NaN | NaN | 3360.8025 | NaN | NaN | NaN |
| Dync1h1 | SDVFPGVQYHR | -2160 | -2170 | 0 | 0 | 11 | NaN | NaN | 1494.5745 | NaN | NaN | NaN |
| Dync1h1 | ALEHAFK | -2434 | -2440 | 0 | 0.00122419 | 7 | NaN | NaN | 446.7231 | NaN | NaN | NaN |
| Acat1 | FASEITPITISVK | -228 | -240 | 0 | 0.00025015 | 13 | NaN | NaN | 2077.5764 | NaN | NaN | NaN |
| Rrp12 | GLLPAEYHK | -1023 | -1031 | 0 | 0 | 9 | NaN | NaN | 684.3125 | 657.09827 | 808.4241 | 455.45306 |
| Rrp12 | FWESNSAAHK | -113 | -122 | 0 | 0 | 10 | NaN | NaN | 1436.8297 | NaN | NaN | NaN |
| Rrp12 | EEEEEEEEEPVQSK | -1056 | -1070 | 0 | 0.002972566 | 15 | NaN | NaN | 80.70664 | NaN | NaN | NaN |
| Stip1 | LKDCEECIQLEPTFIK | -414 | -429 | 1 | 0 | 16 | NaN | NaN | 656.03674 | 360.5848 | 400.01614 | NaN |
| Stip1 | LGSMDEEEEAATPPPPPPPK | -187 | -206 | 0 | 0.00111006 | 20 | NaN | NaN | 241.22252 | 440.88358 | NaN | 711.95374 |
| Stip1 | HDSPEDVK | -479 | -486 | 0 | 0.000347792 | 8 | NaN | NaN | 557.96405 | 548.54895 | NaN | NaN |
| Stip1 | IGNSYFKEE | -306 | -314 | 1 | 0.000550307 | 9 | NaN | NaN | 538.4048 | NaN | NaN | 624.12006 |
| Hsp90ab1 | KSLTNDWEDHLAVK | -306 | -319 | 1 | 0 | 14 | NaN | NaN | 1089.1301 | 1278.5544 | 1930.299 | 2025.1257 |
| Hsp90ab1 | GFSLEDPQTHSNR | -667 | -679 | 0 | 0 | 13 | NaN | NaN | 870.0639 | 645.28485 | 6048.8774 | 5934.744 |
| Hsp90ab1 | ALLSSGFSLEDPQTHSNR | -662 | -679 | 0 | 0 | 18 | NaN | NaN | 575.33875 | 670.22327 | 915.88434 | 808.3267 |
| Hsp90ab1 | SGDEMTSLSEYVSR | -462 | -475 | 0 | 0 | 14 | NaN | NaN | 304.62036 | NaN | 654.82056 | 774.4199 |
| Hsp90ab1 | KCLELFSEL | -411 | -419 | 1 | 0.007320451 | 9 | NaN | NaN | 912.66907 | NaN | NaN | NaN |
| Eno1 | HIADLAGNPEVILPVPAFNVINGGSHAGNK | -133 | -162 | 0 | 0 | 30 | NaN | NaN | 743.79407 | 1232.7979 | 1177.3108 | 1061.1272 |
| Mrps15 | EKNEGVPENPSNAVPEK | -238 | -254 | 1 | 0 | 17 | NaN | NaN | 1709.7898 | 142.38168 | 1791.6333 | 1439.0989 |
| Mrps15 | AAVQKPVQPK | -53 | -62 | 1 | 0.001463897 | 10 | NaN | NaN | 589.27905 | NaN | NaN | NaN |
| Eprs1 | AAWQEHLK | -132 | -139 | 0 | 0.00025015 | 8 | NaN | NaN | 1425.7551 | 1564.4867 | 370.45288 | 1329.3812 |
| Eprs1 | STCATAEDSSVLYSR | -742 | -756 | 0 | 0 | 15 | NaN | NaN | 326.55353 | 471.4169 | 398.67184 | 462.98923 |
| Eprs1 | KEENLAEWYSQVITK | -1020 | -1034 | 1 | 0.00025015 | 15 | NaN | NaN | 703.7219 | NaN | 844.37195 | 775.46423 |
| Eprs1 | YEHLITKPVLGK | -626 | -637 | 1 | 0.00025015 | 12 | NaN | NaN | 347.88486 | 626.6099 | NaN | NaN |
| Rpl5 | RYLMEEDEDAYKK | -209 | -221 | 2 | 0 | 13 | NaN | NaN | 799.70245 | NaN | 982.11884 | 228.4907 |
| Rpl5 | DGGLSIPHSTK | -168 | -178 | 0 | 0.000230707 | 11 | NaN | NaN | 889.04407 | NaN | NaN | 771.9369 |
| Rars1 | KIILDHEK | -563 | -570 | 1 | 0.003299042 | 8 | NaN | NaN | 228.93637 | 369.9465 | 602.2531 | NaN |
| Rars1 | LMDLLEEGLKR | -462 | -472 | 1 | 0.000347792 | 11 | NaN | NaN | 650.564 | NaN | NaN | NaN |
| Vps35 | EDGPGIPAEIK | -313 | -323 | 0 | 0 | 11 | NaN | NaN | 2016.8744 | 1037.8834 | 1281.6816 | 1449.4854 |
| Vps35 | HFHNTLEHLR | -769 | -778 | 0 | 0.001776473 | 10 | NaN | NaN | 376.25894 | NaN | NaN | 546.1646 |
| Itgal | MPVSCEELTEGSSLLTK | -837 | -853 | 0 | 0 | 17 | NaN | NaN | 608.09045 | 889.0798 | 784.5704 | 502.93185 |
| Itgal | VGQEPLTSDVR | -388 | -398 | 0 | 0.001023236 | 11 | NaN | NaN | 730.9254 | NaN | 502.64825 | 695.7881 |
| Itgal | HFVSVQK | -287 | -293 | 0 | 0.002068213 | 7 | NaN | NaN | 563.04663 | 378.95157 | 368.30774 | NaN |
| Itgal | GHAVVGAVGAK | -359 | -369 | 0 | 0 | 11 | NaN | NaN | 419.575 | NaN | 420.72327 | 680.04456 |
| Itgal | LFDGSQSLDR | -157 | -166 | 0 | 0.003461185 | 10 | NaN | NaN | 1042.451 | NaN | 967.4595 | NaN |
| Itgal | WTLSNSGEDAYWVR | -801 | -814 | 0 | 0.00025015 | 14 | NaN | NaN | 596.312 | 930.33295 | NaN | NaN |
| C3 | FYHPEKDDGMLSK | -1492 | -1504 | 1 | 0 | 13 | NaN | NaN | 542.5521 | 633.36554 | 324.58313 | 687.76624 |
| C3 | KFISHIK | -1583 | -1589 | 1 | 0 | 7 | NaN | NaN | 1121.05 | NaN | 1511.2296 | NaN |
| Gnb2 | KACGDSTLTQITAGLDPVGR | -23 | -42 | 1 | 0.00025015 | 20 | NaN | NaN | 1331.8912 | NaN | NaN | 718.2535 |
| Gnb2 | ITAGLDPVGR | -33 | -42 | 0 | 0.007701635 | 10 | NaN | NaN | 415.4948 | NaN | NaN | NaN |
| Paics | SWLPQDCTLVDMK | -179 | -191 | 0 | 0 | 13 | NaN | NaN | 710.3359 | 515.2381 | 1131.3264 | 725.9215 |
| Paics | IKAEYEGDGIPTVF | -312 | -325 | 1 | 0.000826806 | 14 | NaN | NaN | 562.4737 | NaN | NaN | NaN |
| Aco2 | GGTGAIVEYHGPGVDSISCT | -259 | -278 | 0 | 0.000471409 | 20 | NaN | NaN | 288.68082 | 551.8134 | 914.02704 | 706.2118 |
| Aco2 | WVIGDENYGEGS | -657 | -669 | 0 | 0.009242859 | 13 | NaN | NaN | 258.78857 | NaN | NaN | NaN |
| Ddx27 | ACIPVGLLGK | -213 | -222 | 0 | 0.002697088 | 10 | NaN | NaN | 821.34 | NaN | NaN | NaN |
| Ddx27 | FKDEQIDIL | -475 | -483 | 1 | 0.009242859 | 9 | NaN | NaN | 362.3738 | NaN | NaN | NaN |
| Cct3 | GVNGETGTLVDMK | -479 | -491 | 0 | 0.00025015 | 13 | NaN | NaN | 1218.7948 | NaN | NaN | 703.2329 |
| Itgb2 | WNNDNPLFK | -749 | -757 | 0 | 0.000591124 | 9 | NaN | NaN | 1133.984 | 1333.1335 | 109.44806 | 1501.6343 |
| Itgb2 | CVCHTSDVPNK | -518 | -528 | 0 | 0.000431655 | 11 | NaN | NaN | 886.53357 | NaN | 981.21625 | NaN |
| Prkcsh | LWEEQQAAAK | -197 | -206 | 0 | 0 | 10 | NaN | NaN | 827.60297 | 1285.7379 | 252.07722 | 1318.7018 |
| Rpa2 | SATLTDEVFR | -56 | -65 | 0 | 0.00266297 | 10 | NaN | NaN | 1084.3108 | NaN | NaN | NaN |
| Hnrnpm | ESPEVAER | -699 | -706 | 0 | 0.003360691 | 8 | NaN | NaN | 832.9292 | 1000.57574 | 1002.352 | 916.2899 |
| Rps3a | APAMFNIR | -35 | -42 | 0 | 0.000627943 | 8 | NaN | NaN | 824.1563 | 1217.8773 | 1275.3938 | 1130.4452 |
| Fasn | RHDGLPGLAVQW | -2042 | -2053 | 1 | 0.000591124 | 12 | NaN | NaN | 1235.6746 | 1896.6237 | 1285.4346 | 844.1877 |
| Fasn | QAHTMDPQLR | -71 | -80 | 0 | 0.000591124 | 10 | NaN | NaN | 541.25995 | 479.22736 | 495.61383 | 727.2942 |
| Fasn | SFDDSGSGYCR | -214 | -224 | 0 | 0 | 11 | NaN | NaN | 368.56396 | 419.7643 | 511.0617 | 507.47006 |
| Fasn | FHNPNPEIPALLDGR | -359 | -373 | 0 | 0 | 15 | NaN | NaN | 1723.2015 | NaN | NaN | 3108.9294 |
| Fasn | VSGGIHISR | -1089 | -1097 | 0 | 0.00122419 | 9 | NaN | NaN | 515.8012 | NaN | NaN | 605.92004 |
| Fasn | EQGVTFPSGEVQEQ | -258 | -271 | 0 | 0.001291103 | 14 | NaN | NaN | 512.9441 | 453.16467 | NaN | NaN |
| Fasn | ENQTPELFQDVNKPK | -1974 | -1988 | 1 | 0.00025015 | 15 | NaN | NaN | 475.6086 | NaN | NaN | NaN |
| Atp1a1 | IHEADTTENQSGVSFDK | -397 | -413 | 0 | 0 | 17 | NaN | NaN | 883.9645 | 130.15955 | 929.6595 | 532.43414 |
| Api5 | VTLSWKPVQK | -439 | -448 | 1 | 0 | 10 | NaN | NaN | 457.61707 | 368.97287 | 383.88913 | NaN |
| Mcm4 | YPDSILEHQIQVRPF | -248 | -262 | 1 | 0.00025015 | 15 | NaN | NaN | 818.2264 | 290.85864 | 1245.7313 | 1715.3988 |
| Mcm4 | FSNKVEAIDVEEAKR | -748 | -762 | 2 | 0.00111006 | 15 | NaN | NaN | 259.8888 | 971.2118 | 700.5314 | 605.61804 |
| Samhd1 | MFEHLVNSNELK | -272 | -283 | 0 | 0 | 12 | NaN | NaN | 1113.1471 | 3096.8975 | 1407.3231 | 1290.6984 |
| Samhd1 | EQIMGPPITPVK | -302 | -313 | 0 | 0 | 12 | NaN | NaN | 1447.7233 | 1119.7565 | 1428.3912 | 1378.2327 |
| Samhd1 | DCHHLGIQ | -351 | -358 | 0 | 0.00025015 | 8 | NaN | NaN | 1184.6863 | 1274.8174 | 1223.3715 | NaN |
| Samhd1 | YLGETQPK | -499 | -506 | 0 | 0.007493591 | 8 | NaN | NaN | 883.5147 | 814.3512 | NaN | NaN |
| Samhd1 | LPQEVAK | -518 | -524 | 0 | 0.006164361 | 7 | NaN | NaN | 393.48267 | NaN | 610.53235 | NaN |

| | | | | | | | | | | | | |
|---|---|---|---|---|---|---|---|---|---|---|---|---|
| Samhd1 | ICEVEYK | -372 | -378 | 0 | 0.003133457 | 7 | NaN | NaN | 1622.3966 | NaN | NaN | NaN |
| Pds5a | IYAPEAPYTSHDK | -99 | -111 | 0 | 0 | 13 | NaN | NaN | 1064.693 | 894.2473 | 960.3372 | 930.30896 |
| Pds5a | AHDPDFTR | -1009 | -1016 | 0 | 0.002780285 | 8 | NaN | NaN | 542.616 | 412.66904 | NaN | NaN |
| Ppia | SIYGEKFEDE | -77 | -86 | 1 | 0.002168954 | 10 | NaN | NaN | 1043.2379 | NaN | NaN | NaN |
| Ppia | VNPTVFFDITADDEPLGR | -2 | -19 | 0 | 0.00025015 | 18 | NaN | NaN | 456.91736 | NaN | NaN | NaN |
| Psme2 | DTHVMDYR | -181 | -188 | 0 | 0.001291103 | 8 | NaN | NaN | 844.9652 | NaN | 186.35268 | NaN |
| Aldoa | GRPFPQVIK | -91 | -99 | 1 | 0.001743443 | 9 | NaN | NaN | 1453.4866 | 446.21237 | 1160.4686 | 1255.7913 |
| Pin1 | ADEEKLPPGWEK | -2 | -13 | 1 | 0 | 12 | NaN | NaN | 1887.6796 | 3110.349 | 3518.31 | 3972.9773 |
| Ipo7 | TIPEEDRHCIR | -82 | -92 | 1 | 0 | 11 | NaN | NaN | 1292.1146 | NaN | NaN | 1443.5726 |
| Prdx1 | FIIDDKGILR | -131 | -140 | 1 | 0.002068213 | 10 | NaN | NaN | 310.6745 | 417.16547 | 466.1045 | 394.90787 |
| Rps5 | KAQCPIVER | -63 | -71 | 1 | 0 | 9 | NaN | NaN | 507.52133 | NaN | 612.20264 | NaN |
| Myh9 | KMEDGVGCLETAEEAK | -1372 | -1387 | 1 | 0 | 16 | NaN | NaN | 1263.9207 | 3132.163 | 1830.2399 | 1305.8148 |
| Myh9 | LHQSSDKFVSELWK | -600 | -613 | 1 | 0.000627943 | 14 | NaN | NaN | 2072.084 | 1132.5717 | 1996.5864 | 314.6356 |
| Myh9 | EQLQAETELCAEAEELR | -887 | -903 | 0 | 0.000550307 | 17 | NaN | NaN | 822.3056 | NaN | NaN | 1222.0131 |
| Myh9 | EEELEEEQGNTELINDR | -1735 | -1751 | 0 | 0.00025015 | 17 | NaN | NaN | 416.6522 | NaN | 584.9372 | NaN |
| Myh9 | SALESQLQDTQELLQEENR | -1304 | -1322 | 0 | 0.002780285 | 19 | NaN | NaN | 207.91696 | NaN | NaN | NaN |
| Gsr | GHILVDEFQNTNVK | -333 | -346 | 0 | 0 | 14 | NaN | NaN | 1674.292 | 1395.6991 | 2514.4192 | 2142.8235 |
| Gsr | VGIQTDEK | -325 | -332 | 0 | 0.002534621 | 8 | NaN | NaN | 312.07315 | NaN | 1221.279 | 1083.3638 |
| Slc3a2 | EEDTSLKLENL | -500 | -510 | 1 | 0.00025015 | 11 | NaN | NaN | 2587.0762 | 2691.946 | 2887.9922 | 2827.5632 |
| Vars1 | QGFFKPEYGRPS | -315 | -326 | 2 | 0 | 12 | NaN | NaN | 603.599 | 729.73083 | NaN | 787.31824 |
| Vars1 | PAHDQNDYEVGQR | -593 | -605 | 0 | 0.000550307 | 13 | NaN | NaN | 495.9923 | 524.50037 | 542.9116 | 644.4173 |
| Vars1 | RYVLDPSAR | -173 | -181 | 1 | 0.00078661 | 9 | NaN | NaN | 478.8692 | 1121.2312 | 1328.2958 | 1229.9198 |
| Hnrnpk | PLEGSEDR | -426 | -433 | 0 | 0.000471409 | 8 | NaN | NaN | 625.40985 | NaN | NaN | 441.86655 |
| Hnrnpk | DRVVLIGGKPDR | -190 | -201 | 2 | 0.00078661 | 12 | NaN | NaN | 1035.3584 | NaN | NaN | NaN |
| Glg1 | FCENTQAGEGR | -319 | -329 | 0 | 0 | 11 | NaN | NaN | 787.218 | 457.27695 | 488.7873 | 899.7175 |
| Atp6v0a2 | VWCPEVDLPGLR | -328 | -339 | 0 | 0.006100831 | 12 | NaN | NaN | 429.56827 | NaN | 496.73358 | NaN |
| Csnk2a1 | KEPFFHGHDNYDQLVR | -229 | -244 | 1 | 0 | 16 | NaN | NaN | 1063.9033 | 817.28735 | 427.80518 | 600.8844 |
| Csnk2a1 | YQTLTDYDIR | -125 | -134 | 0 | 0 | 10 | NaN | NaN | 1244.1912 | 1470.4635 | NaN | NaN |
| Npm1 | ADKDYHF | -25 | -31 | 1 | 0.005605646 | 7 | NaN | NaN | 810.0826 | 993.3727 | 1059.1672 | 1156.9628 |
| Npm1 | GPSSVEDIKAK | -238 | -248 | 1 | 0.000431655 | 11 | NaN | NaN | 672.4454 | NaN | 1525.6417 | 1307.7551 |
| Npm1 | LDEDDEDDDEDDEDDEDDDDDDFDEEETEEKVPVKK | -158 | -193 | 2 | 0.003393785 | 36 | NaN | NaN | 822.2422 | 901.1712 | NaN | NaN |
| Npm1 | LDEDDEDDDEDDEDDEDDDDDDFDEEETEEKVPVKK | -158 | -193 | 2 | 0.003393785 | 36 | NaN | NaN | 822.2422 | 901.1712 | NaN | NaN |
| Rbbp4 | PSHSVDAHTA | -265 | -274 | 0 | 0.007493591 | 10 | NaN | NaN | 390.1654 | NaN | NaN | NaN |
| Adsl | RNAFDLLLPK | -125 | -134 | 1 | 0.000471409 | 10 | NaN | NaN | 1294.5033 | NaN | 1529.396 | 1353.82 |
| Mdh2 | VNVPVIGGHAGK | -192 | -203 | 0 | 0 | 12 | NaN | NaN | 9094.548 | 9700.752 | 7926.169 | 8028.7437 |
| Mdh2 | PVIGGHAGK | -195 | -203 | 0 | 0.001392869 | 9 | NaN | NaN | 1988.0134 | 692.9847 | 2448.4421 | 2172.9365 |
| Hdgfl2 | LEEQPGEQAPR | -572 | -582 | 0 | 0.00025015 | 11 | NaN | NaN | 1056.3992 | NaN | 1018.1448 | NaN |
| Hdgfl2 | ELAEDEPSTDR | -583 | -593 | 0 | 0.001868731 | 11 | NaN | NaN | 1266.901 | NaN | NaN | NaN |
| Phb2 | RLGLDYEER | -123 | -131 | 1 | 0.000230707 | 9 | NaN | NaN | 786.92993 | NaN | 549.64935 | 457.46588 |
| Glod4 | FQTVHFF | -17 | -23 | 0 | 0.005015524 | 7 | NaN | NaN | 566.71906 | 515.95483 | NaN | 416.382 |
| Glod4 | SDEWFATR | -284 | -291 | 0 | 0.000394361 | 8 | NaN | NaN | 338.378 | 451.8813 | NaN | NaN |
| Glod4 | SPSQSDPVLK | -131 | -140 | 0 | 0.004595494 | 10 | NaN | NaN | 393.73517 | NaN | NaN | NaN |
| Uba2 | RKPPVPLDWAEVQ | -276 | -288 | 2 | 0.001868731 | 13 | NaN | NaN | 504.39862 | NaN | 1273.4669 | 1149.9388 |
| Pabpn1 | APPGAPGPGPGSGAPGSQEEEEEPGLVE | -75 | -102 | 0 | 0.001055699 | 28 | NaN | NaN | 250.97 | NaN | 382.03793 | 383.48563 |
| Pabpn1 | FHGCGSVNR | -188 | -196 | 0 | 0.007791272 | 9 | NaN | NaN | 989.28577 | NaN | 893.44775 | NaN |
| Arhgdia | IAAENEEDEHSVNYKPPAQK | -14 | -33 | 1 | 0 | 20 | NaN | NaN | 906.2069 | 1217.5166 | 1359.7676 | 1437.5967 |
| Arhgdia | EYEFLTPMEEAPK | -155 | -167 | 0 | 0 | 13 | NaN | NaN | 1106.7201 | NaN | 408.1268 | 216.34094 |
| Arhgdia | EFLTPMEEAPK | -157 | -167 | 0 | 0 | 11 | NaN | NaN | 870.3716 | NaN | NaN | NaN |
| Copb2 | KSFKPDFGAE | -422 | -431 | 2 | 0.002068213 | 10 | NaN | NaN | 722.69965 | NaN | 689.2429 | NaN |
| Copb2 | QLAELAISK | -672 | -680 | 0 | 0 | 9 | NaN | NaN | 392.85538 | 231.6036 | NaN | NaN |
| Copb2 | GVNCIDYYSGGDKPYLIS | -187 | -204 | 1 | 0.000230707 | 18 | NaN | NaN | 226.69807 | NaN | 1383.0999 | NaN |
| Coro1a | DRPHEGTRPVHAV | -215 | -227 | 2 | 0.000230707 | 13 | NaN | NaN | 1214.3545 | 509.30185 | 369.1738 | 1824.5701 |
| Coro1a | HLEEPLSLQEL | -254 | -264 | 0 | 0.00025015 | 11 | NaN | NaN | 833.5406 | 2028.2051 | 2654.4265 | 2623.7607 |
| Nup133 | HTDLDSDPELDK | -547 | -558 | 0 | 0 | 12 | NaN | NaN | 616.2309 | 1000.9988 | NaN | 977.83923 |
| Cdk6 | VPEPGVPTETIK | -112 | -123 | 0 | 0.002068213 | 12 | NaN | NaN | 1459.7693 | 1380.9152 | NaN | NaN |
| Tnpo2 | ADSVGHHLNQPEYIQK | -546 | -561 | 0 | 0.007819401 | 16 | NaN | NaN | 1268.0151 | NaN | NaN | NaN |
| Pfkp | LEATPETPACVVSLR | -333 | -347 | 0 | 0.00025015 | 15 | NaN | NaN | 2119.8083 | 1864.7523 | 1849.2463 | 1864.6821 |
| Pfkp | GADTALNTITDTCDR | -550 | -564 | 0 | 0 | 15 | NaN | NaN | 930.3331 | 801.03613 | 834.39624 | 369.5196 |
| Pfkp | NVLGHMQQGGAPSPFDR | -666 | -682 | 0 | 0.000471409 | 17 | NaN | NaN | 328.31677 | NaN | NaN | NaN |
| Nucb1 | KVPEQPPELPQLDSQHL | -443 | -459 | 1 | 0 | 17 | NaN | NaN | 918.37756 | 716.41595 | 932.0106 | 740.19727 |
| Nucb1 | QFEHLDPQNQHTF | -136 | -148 | 0 | 0.000431655 | 13 | NaN | NaN | 542.6455 | NaN | NaN | NaN |
| Tars1 | AEHDSILAEK | -65 | -74 | 0 | 0.005522803 | 10 | NaN | NaN | 1060.9216 | NaN | NaN | NaN |
| Psmc4 | IQDEIPALSVSR | -11 | -22 | 0 | 0 | 12 | NaN | NaN | 258.6502 | 592.38934 | 484.74152 | 475.47302 |
| Psmc4 | HSNALVDVLPPEADSSIM | -135 | -152 | 0 | 0 | 18 | NaN | NaN | 399.34842 | NaN | 389.75604 | 444.37024 |
| Psmc4 | KIEFPLPDRR | -330 | -339 | 2 | 0.007546675 | 10 | NaN | NaN | 249.96803 | NaN | NaN | NaN |
| Ldha | RVHPISTMIK | -269 | -278 | 1 | 0 | 10 | NaN | NaN | 571.1889 | 662.44183 | 432.1757 | 469.5753 |
| Scly | ADVDLIVQDLK | -412 | -422 | 0 | 0.000550307 | 11 | NaN | NaN | 714.9381 | NaN | NaN | NaN |
| Rbm8a | ADVLDLHEA | -2 | -10 | 0 | 0.004321969 | 9 | NaN | NaN | 403.5814 | NaN | NaN | NaN |
| Pcna | MPSGEFAR | -139 | -146 | 0 | 0 | 8 | NaN | NaN | 864.613 | NaN | 512.3796 | 516.46594 |
| Dcp1a | ADVVEEETRR | -143 | -152 | 1 | 0.003620005 | 10 | NaN | NaN | 1043.3322 | 478.20853 | 707.8682 | 1391.9121 |
| Sptan1 | VEEGHFAAEDVK | -851 | -862 | 0 | 0 | 12 | NaN | NaN | 6049.835 | 849.7859 | 5586.0156 | 523.45325 |
| Sptan1 | SSEGKPYVTK | -2417 | -2426 | 1 | 0.000712251 | 10 | NaN | NaN | 947.1361 | NaN | 2071.2974 | 1960.5398 |
| Sptan1 | SHPESAEDLK | -1302 | -1311 | 0 | 0 | 10 | NaN | NaN | 422.75076 | NaN | 644.73236 | 225.97545 |
| Sptan1 | LQVASDENYK | -64 | -73 | 0 | 0.00830926 | 10 | NaN | NaN | 580.3006 | NaN | NaN | NaN |
| Gcn1 | HLDQIIPR | -705 | -712 | 0 | 0 | 8 | NaN | NaN | 1421.7125 | 1063.5314 | 894.15 | NaN |
| Gcn1 | NYLPITFGDK | -1761 | -1770 | 0 | 0.005668207 | 10 | NaN | NaN | 2832.6055 | NaN | NaN | NaN |
| Fubp1 | PQQIDYAR | -429 | -436 | 0 | 0 | 8 | NaN | NaN | 2006.8042 | 2947.2236 | 1252.8962 | 1201.2179 |
| Fubp1 | ADYSTVPPPSSGSAGGGGGGVVNDAFK | -2 | -28 | 0 | 0.00025015 | 27 | NaN | NaN | 275.67017 | 402.94366 | 627.4244 | 636.6691 |

| | | | | | | | | | | | | |
|---|---|---|---|---|---|---|---|---|---|---|---|---|
| Fubp1 | MGQAVPAPAGAPPGGQPDYSAA | -589 | -610 | 0 | 0.001023236 | 22 | NaN | NaN | 1023.36847 | NaN | 967.58673 | NaN |
| Cct8 | EDGAISTIVLR | -368 | -378 | 0 | 0 | 11 | NaN | NaN | 1569.5284 | 2060.4375 | 1216.9371 | 1827.1183 |
| Cct8 | KAHEILPELVCCS | -138 | -150 | 1 | 0.00025015 | 13 | NaN | NaN | 1564.444 | 1600.2673 | 1563.3835 | 1765.4708 |
| Cct8 | KETEGDVTSVK | -225 | -235 | 1 | 0.008653659 | 11 | NaN | NaN | 506.05206 | NaN | NaN | 527.8993 |
| Tkt | TVFYPSDGVATEK | -444 | -456 | 0 | 0.007905138 | 13 | NaN | NaN | 1139.4625 | NaN | NaN | 748.51184 |
| Sart1 | AEDESVDDLAQQKPR | -317 | -331 | 1 | 0 | 15 | NaN | NaN | 583.53174 | 1038.9363 | 500.42224 | 1156.0802 |
| Sart1 | STTILDEEPIVNR | -630 | -642 | 0 | 0 | 13 | NaN | NaN | 3659.5654 | 2172.1106 | NaN | NaN |
| Surf4 | LISTFLEDGIR | -33 | -43 | 0 | 0 | 11 | NaN | NaN | 2086.8887 | 1033.1929 | 1114.3947 | 146.34059 |
| Copa | DADSQNPDAPEGK | -399 | -411 | 0 | 0 | 13 | NaN | NaN | 950.72424 | NaN | 1351.9209 | 795.4238 |
| Copa | ERPAYAVHGN | -322 | -331 | 1 | 0.003133457 | 10 | NaN | NaN | 1113.2874 | NaN | 893.4604 | NaN |
| Copa | GFVEAPEGLGEDVLGK | -857 | -872 | 0 | 0 | 16 | NaN | NaN | 643.9719 | NaN | NaN | NaN |
| Rps3 | IHSGDPVNYYVDTAVR | -158 | -173 | 0 | 0.007905138 | 16 | NaN | NaN | 874.22 | NaN | NaN | NaN |
| Hdlbp | IQIPRPDDPSNQIK | -184 | -197 | 1 | 0.00025015 | 14 | NaN | NaN | 1065.0386 | 907.5602 | 950.3624 | 839.9885 |
| Hdlbp | IEGDPQGVQQAK | -483 | -494 | 0 | 0 | 12 | NaN | NaN | 970.42847 | 1388.4639 | NaN | NaN |
| Rtcb | AEDKEHCEEYGR | -187 | -198 | 1 | 0.00051306 | 12 | NaN | NaN | 451.6995 | 459.83844 | 356.2693 | 426.72983 |
| Rps2 | AEDKEWIPVTK | -55 | -65 | 1 | 0.00025015 | 11 | NaN | NaN | 591.6141 | NaN | NaN | 186.24525 |
| Rps2 | SKEVATAIR | -144 | -152 | 1 | 0.001392869 | 9 | NaN | NaN | 280.09714 | 751.3514 | NaN | 433.19623 |
| Rps2 | KIGKPHTVPCK | -173 | -183 | 2 | 0.00025015 | 11 | NaN | NaN | 255.18117 | NaN | NaN | NaN |
| Nup50 | AEDTSEKVEFTAEK | -243 | -256 | 1 | 0.000947459 | 14 | NaN | NaN | 746.9244 | NaN | NaN | NaN |
| Nup50 | LQQESPFSFHGNK | -230 | -242 | 0 | 0 | 13 | NaN | NaN | 460.26065 | NaN | NaN | NaN |
| Hsph1 | SLDQDPVVR | -774 | -782 | 0 | 0.00025015 | 9 | NaN | NaN | 2573.9468 | 2312.2515 | 3123.9658 | 2943.1538 |
| Hsph1 | NHDSEETEGVHEVFSR | -407 | -422 | 0 | 0 | 16 | NaN | NaN | 441.0856 | 578.65344 | 223.84023 | 154.85033 |
| Hsph1 | KPVTDCVISVPSFFTDAER | -135 | -153 | 1 | 0 | 19 | NaN | NaN | 344.74512 | 615.9228 | 603.6274 | 941.6356 |
| Canx | RDEEEEEEKLEEK | -538 | -550 | 2 | 0 | 13 | NaN | NaN | 1114.5674 | 936.4568 | 1385.7059 | 1324.9275 |
| Canx | SVVNSGNLLNDMTPPVNPSR | -256 | -275 | 0 | 0 | 20 | NaN | NaN | 449.62988 | 353.1629 | 698.3798 | 621.20825 |
| Canx | PMIDNPNYK | -373 | -381 | 0 | 0.00078661 | 9 | NaN | NaN | 6454.958 | 6754.8325 | 7865.3477 | 295.8373 |
| P4ha1 | DPEGFVGHPVNAF | -73 | -85 | 0 | 0.002401448 | 13 | NaN | NaN | 556.6368 | NaN | 1120.9924 | NaN |
| P4ha1 | KLLELDPEHQR | -230 | -240 | 1 | 0 | 11 | NaN | NaN | 504.0238 | 489.22855 | NaN | NaN |
| Cdc37 | QFILELAK | -218 | -225 | 0 | 0.000431655 | 8 | NaN | NaN | 444.6398 | 482.60364 | 589.726 | 516.8561 |
| Cdc37 | ESDGQVELER | -75 | -84 | 0 | 0 | 10 | NaN | NaN | 1042.2007 | NaN | NaN | 1539.2617 |
| Cdc37 | NMPWNVDTLSK | -112 | -122 | 0 | 0.002910424 | 11 | NaN | NaN | 942.2127 | NaN | NaN | NaN |
| Ppp4r2 | SGSDCLETQESGGPPSSK | -373 | -390 | 0 | 0.000907206 | 18 | NaN | NaN | 268.5788 | NaN | NaN | NaN |
| Ipo5 | LMVPLLK | -706 | -712 | 0 | 0.000431655 | 7 | NaN | NaN | 478.84198 | NaN | NaN | NaN |
| Ipo5 | RQDEDYDEQVE | -815 | -825 | 1 | 0.003360691 | 11 | NaN | NaN | 343.60422 | NaN | NaN | NaN |
| Psmd7 | SVVDHFNR | -18 | -25 | 0 | 0.00051306 | 8 | NaN | NaN | 1408.7178 | 509.83646 | 1401.539 | 1303.923 |
| Psmd7 | AEEVGVEHLLR | -167 | -177 | 0 | 0 | 11 | NaN | NaN | 194.69998 | NaN | NaN | NaN |
| Pa2g4 | RAFFSEVER | -263 | -271 | 1 | 0.00025015 | 9 | NaN | NaN | 354.37003 | 6966.508 | 6842.2783 | 3015.976 |
| Pa2g4 | LKQHVIDGEK | -190 | -199 | 1 | 0.003393785 | 10 | NaN | NaN | 210.05655 | NaN | NaN | NaN |
| Twf2 | ETIELVHTEPTNVAQLPSR | -210 | -228 | 0 | 0 | 19 | NaN | NaN | 623.66174 | 543.431 | 591.09033 | 546.553 |
| Dhx15 | FDNCPLLTIPGR | -304 | -315 | 0 | 0 | 12 | NaN | NaN | 7636.9497 | 1143.0082 | 11446.502 | 10449.498 |
| Dhx15 | YTPEPERDYLEAAIR | -323 | -337 | 1 | 0.00078661 | 15 | NaN | NaN | 1638.7361 | NaN | 1748.609 | 1094.9502 |
| Dhx15 | EMQDNTYPEILR | -493 | -504 | 0 | 0 | 12 | NaN | NaN | 1204.7981 | 867.92926 | 795.40704 | NaN |
| Dhx15 | TGHYLTVK | -707 | -714 | 0 | 0.00111006 | 8 | NaN | NaN | 934.982 | NaN | 969.7284 | NaN |
| Mrpl4 | AEGAERPAEISEPR | -25 | -38 | 1 | 0.000431655 | 14 | NaN | NaN | 605.9497 | NaN | NaN | 674.4819 |
| Tnpo1 | RFSDQFPLPLK | -879 | -889 | 1 | 0 | 11 | NaN | NaN | 791.5183 | NaN | 673.8472 | 748.2557 |
| Ddb1 | EHASIDLPGIK | -398 | -408 | 0 | 0 | 11 | NaN | NaN | 6019.123 | 7078.141 | 210.35246 | NaN |
| Flna | FADQATPTSPIR | -852 | -863 | 0 | 0 | 12 | NaN | NaN | 967.2787 | 1285.632 | 1293.1897 | 1330.0386 |
| Flna | VHSPSGALEECYVTEIDQDK | -2368 | -2387 | 0 | 0 | 20 | NaN | NaN | 432.91827 | 360.1719 | 917.58563 | 1473.4473 |
| Flna | IANLQTDLSDGLR | -64 | -76 | 0 | 0 | 13 | NaN | NaN | 378.84552 | 639.89496 | 560.9779 | 648.0923 |
| Flna | FNGTHIPGSPFK | -2406 | -2417 | 0 | 0 | 12 | NaN | NaN | 876.1839 | NaN | 2682.1296 | 2791.9688 |
| Flna | TAGAGMGEVEVVIQDPTGQK | -408 | -427 | 0 | 0.001092529 | 20 | NaN | NaN | 309.76492 | 287.69385 | 280.47345 | NaN |
| Flna | VVQEPGDYEVSVK | -2298 | -2310 | 0 | 0 | 13 | NaN | NaN | 1310.3951 | NaN | NaN | 1416.0507 |
| Flna | GLHEGHTFEPAEFIIDTR | -2056 | -2073 | 0 | 0.000230707 | 18 | NaN | NaN | 324.6877 | NaN | NaN | 543.1747 |
| Flna | YDNMHIPGSPLQFYVD | -1846 | -1861 | 0 | 0 | 16 | NaN | NaN | 173.30336 | NaN | NaN | 777.60986 |
| Pfas | VQGDNTSDLDFGAVQR | -471 | -486 | 0 | 0 | 16 | NaN | NaN | 811.62756 | NaN | 927.18115 | NaN |
| Pfas | PCKPGGPVPR | -1054 | -1063 | 1 | 0 | 10 | NaN | NaN | 385.4668 | NaN | 402.2698 | NaN |
| Pfas | AEGRPALEK | -178 | -186 | 1 | 0.009955823 | 9 | NaN | NaN | 390.4416 | NaN | NaN | NaN |
| Tgtp2;Tgtp1 | VGPYISEPPEYWEA | (402);(402) | (415);(415) | 0 | 0.000230707 | 14 | NaN | NaN | 341.98666 | NaN | 533.26483 | 413.62024 |
| Tgtp2;Tgtp1 | TYIEENKLQK | (31);(31) | (40);(40) | 1 | 0.00025015 | 10 | NaN | NaN | 1237.79 | NaN | NaN | NaN |
| Tgtp2;Tgtp1 | HNDESLEDK | (340);(340) | (348);(348) | 0 | 0.004658491 | 9 | NaN | NaN | 672.89746 | NaN | NaN | NaN |
| Nop9 | AEHPIANFPLQR | -360 | -371 | 0 | 0 | 12 | NaN | NaN | 903.1255 | 600.21576 | NaN | 1728.9532 |
| Psph | AMGGALPFK | -51 | -59 | 0 | 0.003966045 | 9 | NaN | NaN | 1810.0897 | NaN | NaN | NaN |
| Stat5b | YYTPVPCEPATAK | -682 | -694 | 0 | 0 | 13 | NaN | NaN | 1421.109 | 400.59836 | 160.94772 | 1948.9077 |
| Morc3 | AEHVVVPIVTF | -140 | -150 | 0 | 0.001392869 | 11 | NaN | NaN | 1129.0493 | NaN | NaN | NaN |
| Acaa1a | AEIVPVTTTVLDDKGDKK | -221 | -238 | 2 | 0 | 18 | NaN | NaN | 556.12244 | NaN | 239.72845 | 409.75848 |
| Ganab | FKNPEPELLVR | -652 | -662 | 1 | 0 | 11 | NaN | NaN | 655.1205 | 2492.911 | 2172.3223 | 2060.0322 |
| Eef2 | WLPAGDALLQ | -343 | -352 | 0 | 0 | 10 | NaN | NaN | 2731.3882 | 2714.0015 | NaN | 4987.2324 |
| Eef2 | RYFDPANGK | -264 | -272 | 1 | 0 | 9 | NaN | NaN | 2267.806 | 2552.125 | NaN | 2525.115 |
| Got2 | RTQLVSNLK | -355 | -363 | 1 | 0.00111006 | 9 | NaN | NaN | 445.45898 | 530.7283 | 554.21 | 609.4273 |
| Sf1 | QGIETPEDQNDLRK | -228 | -241 | 1 | 0.00111006 | 14 | NaN | NaN | 672.4854 | NaN | NaN | NaN |
| Grap2 | SQGGPHPSGTVGEEIRPS | -163 | -180 | 1 | 0.000230707 | 18 | NaN | NaN | 671.0416 | 102.73286 | 430.65396 | 564.49036 |
| Grap2 | ASGEDELSFR | -11 | -20 | 0 | 0 | 10 | NaN | NaN | 907.72516 | 1018.3731 | 1034.3425 | 469.88232 |
| Grap2 | SGEVVEVLDSSNPSWWTGR | -286 | -304 | 0 | 0 | 19 | NaN | NaN | 480.30344 | 414.5784 | 574.70605 | 526.26086 |
| Grap2 | HEDDVQHF | -98 | -105 | 0 | 0.009060424 | 8 | NaN | NaN | 335.0604 | NaN | NaN | NaN |
| Eif2s2 | TEETQPSETK | -33 | -42 | 0 | 0.000627943 | 10 | NaN | NaN | 283.29892 | 304.77896 | 629.71716 | 356.49283 |
| Arid2 | AELLLDPVDFR | -340 | -350 | 0 | 0.00025015 | 11 | NaN | NaN | 398.23148 | 713.77606 | NaN | 713.6667 |
| Umps | PGTHPLASK | -234 | -242 | 0 | 0.001392869 | 9 | NaN | NaN | 1621.0713 | 1435.6746 | NaN | 1291.2572 |
| Sin3a | AELVQLVSPFLGK | -499 | -511 | 0 | 0 | 13 | NaN | NaN | 326.99258 | 162.1545 | 499.67786 | 430.80182 |

| | | | | | | | | | | | | |
|---|---|---|---|---|---|---|---|---|---|---|---|---|
| Nasp | EQYLEAHDR | -559 | -567 | 0 | 0.000550307 | 9 | NaN | NaN | 2971.9912 | NaN | NaN | NaN |
| Nasp | GGREDMDISEPEEKLQETVE | -180 | -199 | 2 | 0.00111006 | 20 | NaN | NaN | 172.57002 | NaN | NaN | NaN |
| Cct4 | SRPVQLSDR | -152 | -160 | 1 | 0.003681885 | 9 | NaN | NaN | 1595.4442 | 1390.9644 | NaN | 2405.6936 |
| Cct4 | ALIAGGGAPEIELALR | -420 | -435 | 0 | 0 | 16 | NaN | NaN | 937.64874 | NaN | NaN | NaN |
| Cct4 | KLGGTIDDCELVEGL | -213 | -227 | 1 | 0.00670918 | 15 | NaN | NaN | 678.9198 | NaN | NaN | NaN |
| Eloc | PSHVLSK | -66 | -72 | 0 | 0.002845514 | 7 | NaN | NaN | 658.8047 | NaN | 851.25165 | 709.4659 |
| Fga | AENPTNPGPGGSGYWR | -296 | -311 | 0 | 0.00025015 | 16 | NaN | NaN | 588.4705 | NaN | 401.03232 | 945.6977 |
| Lrrfip1 | AENQRPAEDSALSPGPLAGAK | -290 | -310 | 1 | 0 | 21 | NaN | NaN | 639.2051 | 970.41833 | 1053.6827 | 875.66174 |
| Mybbp1a | HSLLEHSR | -24 | -31 | 0 | 0 | 8 | NaN | NaN | 134.97392 | 653.51044 | 682.4972 | 735.2579 |
| Mybbp1a | KGFLPETK | -1173 | -1180 | 1 | 0.004255025 | 8 | NaN | NaN | 2386.051 | 2035.856 | 2376.0535 | 2426.3445 |
| Mybbp1a | SRYPVICK | -998 | -1005 | 1 | 0 | 8 | NaN | NaN | 792.6715 | 883.5905 | 893.22925 | 852.1214 |
| Mybbp1a | FLEGCQDDPKR | -357 | -367 | 1 | 0 | 11 | NaN | NaN | 1039.3033 | NaN | 854.01715 | 147.01299 |
| Mybbp1a | QTPDLAENGKPWTYR | -528 | -542 | 1 | 0.000230707 | 15 | NaN | NaN | 843.8732 | NaN | 1071.8403 | 1230.8801 |
| Mybbp1a | AQDASLNVPER | -431 | -441 | 0 | 0.000230707 | 11 | NaN | NaN | 274.12912 | NaN | NaN | NaN |
| Fam107b | AEPDYIEDDNPELIRPQ | -2 | -18 | 1 | 0 | 17 | NaN | NaN | 388.02252 | 479.5586 | 533.57806 | NaN |
| Hyou1 | GQAGPEGAAPAPEEDKKPK | -670 | -688 | 2 | 0 | 19 | NaN | NaN | 493.35757 | NaN | 480.30502 | 355.23523 |
| Hyou1 | LGNTISSLFGGGTSSDAK | -577 | -594 | 0 | 0.00122419 | 18 | NaN | NaN | 1156.3174 | NaN | NaN | NaN |
| Psmd12 | AESDKWQQALK | -258 | -268 | 1 | 0.00025015 | 11 | NaN | NaN | 1334.012 | 523.0882 | 1505.1995 | 1526.6552 |
| Psmd12 | TYVEEITDLPVK | -110 | -121 | 0 | 0 | 12 | NaN | NaN | 616.699 | 596.0797 | NaN | NaN |
| Psmd12 | DNEQSDLVHR | -280 | -289 | 0 | 0 | 10 | NaN | NaN | 306.1279 | NaN | 572.7154 | NaN |
| Psmd12 | WSTLVEDYGVELR | -317 | -329 | 0 | 0 | 13 | NaN | NaN | 79.402306 | 1475.936 | NaN | NaN |
| Smc1a | KYPDANPNPNEQ | -1222 | -1233 | 1 | 0.00025015 | 12 | NaN | NaN | 448.6147 | 191.24878 | 679.5181 | 925.8235 |
| Smc1a | LKEEFYTK | -1189 | -1196 | 1 | 0.001463897 | 8 | NaN | NaN | 2447.8728 | NaN | NaN | NaN |
| Smc1a | LVIDVIR | -593 | -599 | 0 | 0.004595494 | 7 | NaN | NaN | 305.56387 | NaN | NaN | NaN |
| Fkbp4 | KVLQLYPSNK | -378 | -387 | 1 | 0.00025015 | 10 | NaN | NaN | 284.418 | NaN | 181.20337 | NaN |
| F5 | DIHSGLIGPLLICR | -1699 | -1712 | 0 | 0 | 14 | NaN | NaN | 309.3971 | 466.54443 | 724.5076 | 712.53125 |
| Rpn1 | HYFYSPYPTK | -161 | -170 | 0 | 0 | 10 | NaN | NaN | 1759.5848 | 1403.6433 | NaN | 1098.0193 |
| Rpn1 | SFVLALEPELESR | -69 | -81 | 0 | 0 | 13 | NaN | NaN | 597.4833 | NaN | 390.9983 | 198.76865 |
| Rangap1 | LVELDLSDNAFGPDGVR | -114 | -130 | 0 | 0.00025015 | 17 | NaN | NaN | 763.08325 | 575.255 | 580.9697 | 506.0117 |
| Utp3 | AFAKPVPQVDEAETR | -184 | -198 | 1 | 0.000471409 | 15 | NaN | NaN | 388.09973 | NaN | 436.25464 | 463.5447 |
| Ywhaq | TAFDEAIAELDTLNEDSYK | -194 | -212 | 0 | 0 | 19 | NaN | NaN | 242.71225 | NaN | 205.23534 | 177.8199 |
| Ywhaz | LAEVAAGDDKK | -129 | -139 | 1 | 0 | 11 | NaN | NaN | 4421.476 | 3989.5112 | 4511.814 | 2972.5347 |
| Cct5 | SMMDVDHQIAK | -79 | -89 | 0 | 0 | 11 | NaN | NaN | 495.0298 | 498.9236 | 336.91913 | 527.4027 |
| Cct5 | DVDHQIAK | -82 | -89 | 0 | 0 | 8 | NaN | NaN | 1603.6804 | 219.18831 | 416.6344 | NaN |
| Cct5 | DFSHPQMPK | -233 | -241 | 0 | 0.003360691 | 9 | NaN | NaN | 632.4693 | NaN | NaN | NaN |
| Psmd11 | EAFEGYDSIDSPK | -234 | -246 | 0 | 0.00051306 | 13 | NaN | NaN | 861.33813 | NaN | 726.54315 | 777.7529 |
| Psmd11 | AFEGYDSIDSPK | -235 | -246 | 0 | 0.009569685 | 12 | NaN | NaN | 248.14482 | NaN | NaN | NaN |
| Rnpep | AIGDLASAEVGPR | -234 | -246 | 0 | 0 | 13 | NaN | NaN | 22002.26 | 23677.299 | 17460.545 | NaN |
| Stmn1 | NKESKDPADETEAD | -136 | -149 | 2 | 0.000712251 | 14 | NaN | NaN | 318.22827 | 667.06244 | NaN | 404.35007 |
| Pdhb | SKEGIECEVINLR | -257 | -269 | 1 | 0 | 13 | NaN | NaN | 500.8555 | 408.10504 | 345.75162 | 713.17285 |
| Pdhb | DFLIPIGK | -220 | -227 | 0 | 0 | 8 | NaN | NaN | 368.07556 | 341.38397 | 413.96652 | 460.1048 |
| Pdhb | AFELPAEAQSK | -209 | -219 | 0 | 0.005789881 | 11 | NaN | NaN | 1618.5049 | NaN | NaN | NaN |
| Adprh | LAQMGDLEAIDVAR | -42 | -55 | 0 | 0 | 14 | NaN | NaN | 1362.9124 | 915.51434 | 1014.3673 | NaN |
| Adprh | AFFHGGDSDSTAAIA | -301 | -315 | 0 | 0.001776473 | 15 | NaN | NaN | 473.1481 | NaN | NaN | NaN |
| Erp44 | TPADCPVIAIDSFR | -314 | -327 | 0 | 0 | 14 | NaN | NaN | 1051.1855 | 1181.8112 | 1335.3611 | 1730.4161 |
| Erp44 | LHPIFEEASDVIK | -64 | -76 | 0 | 0 | 13 | NaN | NaN | 563.3239 | NaN | 813.6459 | 346.12234 |
| Cul4b | DKVDHIIDTCFLK | -581 | -593 | 1 | 0.00025015 | 13 | NaN | NaN | 379.01044 | NaN | 158.2687 | 195.56178 |
| Cul4b | DKENPNQYNYIA | -959 | -970 | 1 | 0.009002904 | 12 | NaN | NaN | 561.03485 | NaN | NaN | NaN |
| Nrm | YWETTPR | -115 | -121 | 0 | 0.009242859 | 7 | NaN | NaN | 671.2733 | NaN | NaN | NaN |
| Lck | DFDQNQGEVVK | -169 | -179 | 0 | 0 | 11 | NaN | NaN | 195.62967 | NaN | 206.2589 | 1317.4391 |
| Syncrip | WADPIEDPDPEVMAK | -322 | -336 | 0 | 0 | 15 | NaN | NaN | 1390.8252 | 619.9224 | 1312.1617 | 1086.5989 |
| Syncrip | ENQEFYQDTFGQQWK | -609 | -623 | 0 | 0.002434616 | 15 | NaN | NaN | 419.25153 | NaN | NaN | 547.5116 |
| Syncrip | DLFEDELVPLFEK | -172 | -184 | 0 | 0 | 13 | NaN | NaN | 316.15762 | NaN | 531.57495 | NaN |
| Ide | HTKPLLPSQLVR | -754 | -765 | 1 | 0.00025015 | 12 | NaN | NaN | 321.2073 | NaN | 243.16321 | 442.33658 |
| Ide | NVPLPEFPEHPFQEEHLR | -282 | -299 | 0 | 0.00111006 | 18 | NaN | NaN | 84.91177 | NaN | NaN | 2792.2397 |
| Gbp6 | AFLEEGFKK | -554 | -562 | 1 | 0 | 9 | NaN | NaN | 1408.9183 | NaN | NaN | NaN |
| Rragd;Rragc | VDGLSDDHKIETQ | (230);(179) | (242);(191) | 1 | 0.004595494 | 13 | NaN | NaN | 561.4525 | NaN | NaN | NaN |
| Tubb5 | EHGIDPTGTYHGDSDLQLDR | -27 | -46 | 0 | 0 | 20 | NaN | NaN | 479.26913 | 590.56226 | 633.0909 | 852.9442 |
| Nfkb1 | GDFSPTDVHR | -296 | -305 | 0 | 0 | 10 | NaN | NaN | 359.15823 | 600.0481 | 637.85394 | 372.07742 |
| Hnrnpu | VVCLDTYNCDLHFK | -263 | -276 | 0 | 0 | 14 | NaN | NaN | 689.1927 | 548.25696 | 1080.9165 | 959.8122 |
| Hnrnpu | FDENDVITCF | -376 | -385 | 0 | 0.000627943 | 10 | NaN | NaN | 230.23114 | 257.57782 | 285.90918 | 68.3429 |
| Hnrnpu | GQKPWSQHYHQGYY | -787 | -800 | 1 | 0.001463897 | 14 | NaN | NaN | 456.2221 | NaN | NaN | NaN |
| Hnrnpu | CHNCAVEFNFGQK | -426 | -438 | 0 | 0.00078661 | 13 | NaN | NaN | 385.6699 | NaN | NaN | 2248.47 |
| Hnrnpu | RNFILDQTNVSAA | -551 | -563 | 1 | 0.002296977 | 13 | NaN | NaN | 575.7867 | NaN | NaN | NaN |
| Rpl3 | NNASTDYDLSDK | -301 | -312 | 0 | 0 | 12 | NaN | NaN | 1588.9435 | 1395.7756 | 126.88541 | 382.01163 |
| Rpl3 | SINPLGGFVH | -313 | -322 | 0 | 0.00025015 | 10 | NaN | NaN | 647.7157 | 806.80035 | 1355.9412 | 904.6633 |
| Actr2 | MVGDEASELR | -56 | -65 | 0 | 0.00025015 | 10 | NaN | NaN | 1042.7986 | 574.3052 | 1476.8652 | 2080.4067 |
| Krtcap2 | NNLENLVFGK | -56 | -65 | 0 | 0.001055699 | 10 | NaN | NaN | 749.2189 | 549.7522 | 643.6709 | 730.8831 |
| Elp1 | GNLEVVHHR | -715 | -723 | 0 | 0.006078721 | 9 | NaN | NaN | 2569.624 | NaN | NaN | NaN |
| Elp1 | AFPVQLPENALPWDDR | -179 | -194 | 0 | 0.004658491 | 16 | NaN | NaN | 565.9619 | NaN | NaN | NaN |
| Gzma | FHPVIGLNMICAGDLRGGKDSCNGDSGSPLLCDGILF | -186 | -222 | 2 | 0.00383302 | 37 | NaN | NaN | 115.04285 | NaN | NaN | NaN |
| Uggt1 | LSNHIPLR | -521 | -528 | 0 | 0 | 8 | NaN | NaN | 332.57626 | 333.5802 | 354.1037 | 377.73993 |
| Uggt1 | NNFYVDDFAR | -713 | -722 | 0 | 0 | 10 | NaN | NaN | 250.04156 | NaN | 385.86584 | 465.6922 |
| Uggt1 | SVASGHLYER | -1260 | -1269 | 0 | 0.000471409 | 10 | NaN | NaN | 1620.016 | NaN | NaN | 1583.3826 |
| Uggt1 | SFLSVHGK | -149 | -156 | 0 | 0.003360691 | 8 | NaN | NaN | 852.97705 | NaN | NaN | NaN |
| Pes1 | GIYPHEPK | -42 | -49 | 0 | 0 | 8 | NaN | NaN | 665.70557 | 557.5746 | NaN | 493.0806 |
| Sfpq | VSNELLEEAFSQFGPIER | -374 | -391 | 0 | 0 | 18 | NaN | NaN | 393.2195 | 364.5994 | 521.37964 | 514.38965 |
| Sfpq | EDAYHEHQANLLR | -516 | -528 | 0 | 0.00025015 | 13 | NaN | NaN | 875.6167 | NaN | NaN | NaN |

| | | | | | | | | | | | | |
|---|---|---|---|---|---|---|---|---|---|---|---|---|
| Mars1 | ENDQIENLR | -808 | -816 | 0 | 0.00025015 | 9 | NaN | NaN | 2115.1135 | 1848.3914 | 1939.0034 | NaN |
| Mars1 | TVGPEECVVPFLTRPK | -32 | -47 | 1 | 0.004199202 | 16 | NaN | NaN | 1346.8586 | NaN | NaN | NaN |
| Mars1 | QPQPQPPPPEGR | -205 | -216 | 0 | 0.007905138 | 12 | NaN | NaN | 719.8666 | NaN | NaN | NaN |
| Prpf31 | TTQGQQLSDEELER | -173 | -186 | 0 | 0.002434616 | 14 | NaN | NaN | 893.6387 | NaN | NaN | NaN |
| Prpf31 | ASTTQGQQLSDEELER | -171 | -186 | 0 | 0.002780285 | 16 | NaN | NaN | 372.71057 | NaN | NaN | NaN |
| Acadl | SGSDWILNGSK | -191 | -201 | 0 | 0.000431655 | 11 | NaN | NaN | 273.6338 | NaN | NaN | NaN |
| C1qbp | YDHLMDFLADR | -232 | -242 | 0 | 0.001291103 | 11 | NaN | NaN | 1061.3973 | NaN | NaN | NaN |
| C1qbp | STPNFVVEVTK | -161 | -171 | 0 | 0 | 11 | NaN | NaN | 307.29297 | NaN | NaN | NaN |
| C1qbp | STALEHQEYITFLEDLK | -256 | -272 | 0 | 0.00272025 | 17 | NaN | NaN | 294.55676 | NaN | NaN | NaN |
| Cyb5b | QYYIGDVHPSDLKPK | -89 | -103 | 1 | 0 | 15 | NaN | NaN | 1036.8622 | 435.9421 | NaN | NaN |
| Gigyf2 | AGAEASEEVPQTSLSSAR | -363 | -380 | 0 | 0.003166358 | 18 | NaN | NaN | 1867.346 | NaN | NaN | NaN |
| Rplp0 | AGAIAPCEVTVPAQNTGLGPEK | -113 | -134 | 0 | 0 | 22 | NaN | NaN | 1065.3098 | 782.4769 | 1147.8203 | 1149.7083 |
| Rplp0 | VASVPHSIINGYK | -234 | -246 | 0 | 0.00122419 | 13 | NaN | NaN | 557.19946 | NaN | NaN | NaN |
| Ube2k | LVDENFTELR | -31 | -40 | 0 | 0 | 10 | NaN | NaN | 2010.6771 | 1411.415 | 1039.7903 | 678.5123 |
| Taldo1 | KLGGPQEEQIK | -71 | -81 | 1 | 0 | 11 | NaN | NaN | 566.2609 | 489.46042 | 443.27805 | 571.27496 |
| Hadh | TFESLVDFCK | -193 | -202 | 0 | 0 | 10 | NaN | NaN | 976.23 | 867.65515 | NaN | NaN |
| Acap1 | YKDPVTVVVDDLR | -301 | -313 | 1 | 0.000394361 | 13 | NaN | NaN | 152.7455 | NaN | 1661.122 | 1655.6199 |
| Fermt3 | KDEILGIANNR | -565 | -575 | 1 | 0 | 11 | NaN | NaN | 1125.7474 | NaN | 963.5216 | 1140.1289 |
| Fermt3 | QDWSDHAIWWEQK | -57 | -69 | 0 | 0.00025015 | 13 | NaN | NaN | 672.5143 | NaN | NaN | 221.79472 |
| Xpo1 | DFEEYPEHR | -837 | -845 | 0 | 0.00025015 | 9 | NaN | NaN | 1570.9938 | 1819.0563 | 1613.3346 | 1666.3068 |
| Xpo1 | EHGQLLEK | -332 | -339 | 0 | 0.001055699 | 8 | NaN | NaN | 3001.2163 | 1917.8357 | NaN | 2936.9905 |
| Xpo1 | MKDTDSINLYK | -445 | -455 | 1 | 0.004255025 | 11 | NaN | NaN | 2453.4927 | 2404.0696 | NaN | NaN |
| Xpo1 | THLDYVDTEIIMTK | -465 | -478 | 0 | 0.001463897 | 14 | NaN | NaN | 545.898 | NaN | NaN | 1166.0345 |
| Cbr4 | AGELGGNHLAFR | -43 | -54 | 0 | 0.000627943 | 12 | NaN | NaN | 884.1266 | NaN | NaN | NaN |
| Ecpas | ESSCLALNDLLR | -1175 | -1186 | 0 | 0 | 12 | NaN | NaN | 389.71527 | NaN | 776.1771 | NaN |
| Ecpas | LGHAETDEQLQNIISK | -17 | -32 | 0 | 0.002296977 | 16 | NaN | NaN | 636.9998 | NaN | NaN | NaN |
| Numa1 | ASDQLGEQQGRPF | -935 | -947 | 1 | 0 | 13 | NaN | NaN | 739.40546 | 1362.5593 | 1244.4036 | 1172.0083 |
| Numa1 | RELGELGSLR | -1400 | -1409 | 1 | 0.000471409 | 10 | NaN | NaN | 732.06616 | 821.1773 | 1006.7133 | NaN |
| Numa1 | TCYPLESRPTLSLA | -1918 | -1931 | 1 | 0.004350979 | 14 | NaN | NaN | 692.7868 | NaN | NaN | 1960.9764 |
| Numa1 | LESLQQQLQAANDAR | -611 | -625 | 0 | 0.00051306 | 15 | NaN | NaN | 456.26218 | 808.496 | NaN | NaN |
| Bax | MAGETPELTLEQPPQDASTK | -38 | -57 | 0 | 0 | 20 | NaN | NaN | 735.879 | NaN | NaN | NaN |
| Msh6 | LQEGLIPGSQFWDATK | -616 | -631 | 0 | 0.006677519 | 16 | NaN | NaN | 1326.8031 | NaN | 734.4738 | 763.0275 |
| Msh6 | AGFDSDYDQALADIR | -928 | -942 | 0 | 0 | 15 | NaN | NaN | 1465.9232 | NaN | NaN | NaN |
| Eif2s3x | DDPSCPRPECYR | -92 | -103 | 1 | 0.00025015 | 12 | NaN | NaN | 719.7806 | 842.89514 | 821.1372 | 834.69543 |
| Eif2s3x | KLDDPSCPRPECYR | -90 | -103 | 2 | 0.00025015 | 14 | NaN | NaN | 521.83453 | 867.02423 | 806.885 | 184.66144 |
| Hnrnpl | AGGGGGENYDDPHK | -81 | -94 | 0 | 0 | 14 | NaN | NaN | 2542.017 | 393.076 | 908.3066 | 1530.1794 |
| Slc25a12 | QFLQELQLEHAR | -152 | -163 | 0 | 0 | 12 | NaN | NaN | 1559.9498 | 1716.8284 | 1227.1855 | 1132.2222 |
| Slc25a12 | SGNGEVTFENVK | -101 | -112 | 0 | 0.009897819 | 12 | NaN | NaN | 925.0386 | NaN | NaN | NaN |
| Ahcy | KLDEAVAEAHLGK | -389 | -401 | 1 | 0 | 13 | NaN | NaN | 521.5319 | 902.76306 | 879.4555 | 774.5853 |
| Ahcy | IELWTHPDKYPVGVHFLPK | -370 | -388 | 1 | 0.000230707 | 19 | NaN | NaN | 463.381 | 627.77747 | 669.5522 | 751.3206 |
| Ahcy | SKFDNLYGC | -187 | -195 | 1 | 0.000627943 | 9 | NaN | NaN | 364.90475 | 791.83026 | NaN | 599.0737 |
| Ahcy | WLNENAVEK | -310 | -318 | 0 | 0.00025015 | 9 | NaN | NaN | 3347.8523 | 5596.652 | NaN | NaN |
| Ahcy | GETDEEYLWCIEQ | -104 | -116 | 0 | 0 | 13 | NaN | NaN | 508.83014 | NaN | NaN | NaN |
| Sigmar1 | QYAGLDHELAFSR | -48 | -60 | 0 | 0 | 13 | NaN | NaN | 1097.0298 | 1156.1807 | 1172.472 | 1139.1769 |
| Cd6 | AGLHFTPGQGPIHR | -211 | -224 | 0 | 0 | 14 | NaN | NaN | 412.0203 | NaN | 578.3136 | 760.56946 |
| Ube4b | AGLLDESFLR | -703 | -712 | 0 | 0.000712251 | 10 | NaN | NaN | 954.77246 | 650.41205 | 194.72067 | 692.02136 |
| Fdxr | GVRPVSFSDWEK | -452 | -463 | 1 | 0.00025015 | 12 | NaN | NaN | 2266.228 | NaN | 1730.8563 | 2976.4785 |
| Fdxr | AGLLPSGPRPGYVAIQ | -430 | -445 | 1 | 0.00078661 | 16 | NaN | NaN | 414.73615 | 527.4897 | NaN | 770.07635 |
| Sdha | SIAESCRPGDK | -470 | -480 | 1 | 0 | 11 | NaN | NaN | 879.22266 | 1287.919 | 1953.639 | 1520.7106 |
| Sdha | GCLITEGCR | -304 | -312 | 0 | 0.003461185 | 9 | NaN | NaN | 1124.0581 | NaN | NaN | 348.56924 |
| Sdha | HTLSYVDIK | -625 | -633 | 0 | 0.003235556 | 9 | NaN | NaN | 1546.491 | NaN | NaN | NaN |
| Sdha | VDEYDYSKPIQGQQK | -601 | -615 | 1 | 0.00025015 | 15 | NaN | NaN | 473.31592 | NaN | NaN | NaN |
| Plec | LEAQAGTGHIIDPATSAR | -3202 | -3219 | 0 | 0.000431655 | 18 | NaN | NaN | 405.78708 | 2136.8071 | 1184.4408 | 2456.4773 |
| Plec | RLQLEETDHQK | -2337 | -2347 | 1 | 0.000298805 | 11 | NaN | NaN | 438.2325 | NaN | 524.255 | NaN |
| Plec | KVALEEVER | -2150 | -2158 | 1 | 0 | 9 | NaN | NaN | 398.20047 | 340.78772 | NaN | NaN |
| Plec | ALPEPSPAAPTLR | -1197 | -1209 | 0 | 0.001291103 | 13 | NaN | NaN | 961.9017 | NaN | NaN | NaN |
| Rpl7 | KTTHFVEGGDAGNREDQIN | -245 | -263 | 2 | 0.00078661 | 19 | NaN | NaN | 335.81113 | NaN | NaN | NaN |
| Sf3b3 | RNDLDDPER | -275 | -283 | 1 | 0.003966045 | 9 | NaN | NaN | 2343.907 | 3768.9167 | 272.31512 | NaN |
| Sf3b3 | AGNGQWASVIR | -857 | -867 | 0 | 0.002296977 | 11 | NaN | NaN | 315.72797 | NaN | NaN | NaN |
| Rpl22 | SKESYELR | -106 | -113 | 1 | 0.000826806 | 8 | NaN | NaN | 2539.9517 | 2903.704 | 2809.7642 | 3126.1252 |
| Apeh | AGPAGNSVETR | -79 | -89 | 0 | 0.008075282 | 11 | NaN | NaN | 921.7593 | NaN | NaN | NaN |
| Pnp | DYNEIPNFPQSTVQGHAGR | -49 | -67 | 0 | 0.00025015 | 19 | NaN | NaN | 1266.9128 | 1628.1588 | 1976.8219 | 1910.6267 |
| Sf3b4 | VTGQHQGYGFVE | -49 | -60 | 0 | 0.000591124 | 12 | NaN | NaN | 843.55646 | 1077.0382 | NaN | 1136.3405 |
| Sf3b4 | AGPVVNTHMPK | -36 | -46 | 0 | 0.007987481 | 11 | NaN | NaN | 6666.4204 | NaN | NaN | NaN |
| Bhmt | AGPWTPEAAVEHPEAVR | -41 | -57 | 0 | 0 | 17 | NaN | NaN | 867.2068 | 810.4692 | 1092.9614 | 1167.9009 |
| Mrpl11 | AGQAIPGPPLGPILGQR | -26 | -42 | 0 | 0.000431655 | 17 | NaN | NaN | 210.21109 | NaN | NaN | NaN |
| Arid1a | SVGIQDAFPDGSDPTFQK | -1185 | -1202 | 0 | 0 | 18 | NaN | NaN | 1301.5437 | 2095.2815 | 2095.2683 | 1665.3611 |
| Arid1a | KPLDLYR | -1048 | -1054 | 1 | 0.002466863 | 7 | NaN | NaN | 958.79675 | NaN | NaN | NaN |
| Dennd2d | AGQSQNNPGEAVTEPER | -22 | -38 | 0 | 0.004255025 | 17 | NaN | NaN | 345.1885 | NaN | NaN | NaN |
| Acaca | SFGPQEDLLFLR | -1689 | -1700 | 0 | 0 | 12 | NaN | NaN | 411.9267 | 247.20216 | 569.8012 | 241.3812 |
| Acaca | GGSWVVIDPTINPR | -2089 | -2102 | 0 | 0.002810811 | 14 | NaN | NaN | 527.39905 | NaN | 1079.2949 | 1079.1025 |
| Acaca | ILNVPQDLYEK | -277 | -287 | 0 | 0.002466863 | 11 | NaN | NaN | 381.3012 | 568.1684 | NaN | NaN |
| Clic4 | GSDGESIGNCPFSQR | -26 | -40 | 0 | 0 | 15 | NaN | NaN | 911.7273 | 782.3853 | 765.5081 | 748.3186 |
| Ythdf2 | AGSGSTPSEPHPVLEK | -376 | -391 | 0 | 0 | 16 | NaN | NaN | 582.755 | 568.8256 | 501.20837 | 585.90845 |
| Ythdf2 | SASSLLEQRPK | -2 | -12 | 1 | 0 | 11 | NaN | NaN | 3006.2004 | 2227.6226 | 2876.8499 | 2736.1145 |
| Pld3 | FWPAIDDGLR | -339 | -348 | 0 | 0 | 10 | NaN | NaN | 835.33716 | NaN | NaN | NaN |
| Fmnl1 | VFLQYEFTHL | -343 | -352 | 0 | 0 | 10 | NaN | NaN | 600.7169 | 743.13965 | 926.8735 | 394.45126 |
| Fmnl1 | VVEYFGENPK | -965 | -974 | 0 | 0.00149795 | 10 | NaN | NaN | 889.7156 | NaN | NaN | NaN |

| | | | | | | | | | | | | |
|---|---|---|---|---|---|---|---|---|---|---|---|---|
| Fmnl1 | KAEQEVEQWK | -994 | -1003 | 1 | 0.00025015 | 10 | NaN | NaN | 605.19916 | NaN | NaN | NaN |
| Actn4 | AGTQIENIDEDFRDGLK | -68 | -84 | 1 | 0.000298805 | 17 | NaN | NaN | 2470.1301 | NaN | 1313.3359 | NaN |
| Actn4 | KAGTQIENIDEDFR | -67 | -80 | 1 | 0.00025015 | 14 | NaN | NaN | 1711.0707 | NaN | 2141.2688 | 2076.0432 |
| Pgk1 | VASGIPAGWMGLDCGTESSK | -303 | -322 | 0 | 0.00025015 | 20 | NaN | NaN | 628.2429 | 588.75726 | 853.34344 | 893.6667 |
| Pgk1 | WNTEDKVSHVSTGGGASL | -383 | -400 | 1 | 0.001392869 | 18 | NaN | NaN | 496.44666 | NaN | 601.8817 | NaN |
| Pgk1 | VADKIQLIN | -217 | -225 | 1 | 0.006735683 | 9 | NaN | NaN | 420.44763 | NaN | NaN | NaN |
| Mcm3 | SKDVFEQL | -292 | -299 | 1 | 0.003745449 | 8 | NaN | NaN | 1711.2976 | 1794.3876 | NaN | NaN |
| Mcm3 | GCYTSGTFR | -249 | -257 | 0 | 0.00670918 | 9 | NaN | NaN | 618.53284 | NaN | NaN | NaN |
| Prkcb | SFGISELQK | -260 | -268 | 0 | 0 | 9 | NaN | NaN | 58.53753 | NaN | 769.9091 | NaN |
| Fto | NVLNDSDDGDVSLK | -341 | -354 | 0 | 0.000550307 | 14 | NaN | NaN | 575.7026 | NaN | NaN | NaN |
| Fto | WHHDENLVDR | -227 | -236 | 0 | 0 | 10 | NaN | NaN | 470.309 | NaN | NaN | NaN |
| Tmed2 | IDAHAEECFFER | -25 | -36 | 0 | 0.003620005 | 12 | NaN | NaN | 162.8821 | NaN | NaN | NaN |
| Cebpz | LYEHEVSLF | -212 | -220 | 0 | 0.003133457 | 9 | NaN | NaN | 897.2186 | NaN | NaN | 657.7379 |
| Prmt1 | KLDHVVTIIK | -134 | -143 | 1 | 0 | 10 | NaN | NaN | 1753.695 | 1654.499 | 2572.7034 | 2334.4321 |
| Atp5f1b | VGNEHYDVAR | -413 | -422 | 0 | 0.004031417 | 10 | NaN | NaN | 163.15326 | 1632.4808 | NaN | 180.155 |
| Atp5f1b | VVDLLAPYAKGG | -189 | -200 | 1 | 0.003998192 | 12 | NaN | NaN | 1333.0306 | NaN | NaN | NaN |
| Atp5f1b | TREGNDLYHEM | -240 | -250 | 1 | 0.00201047 | 11 | NaN | NaN | 668.1253 | NaN | NaN | NaN |
| Manf | IINEVSKPLAHHIPVEK | -85 | -101 | 1 | 0 | 17 | NaN | NaN | 548.0965 | 1560.764 | 2499.923 | 581.80035 |
| Manf | IINEVSKPLAHHIPVEK | -85 | -101 | 1 | 0 | 17 | NaN | NaN | 548.0965 | 1560.764 | 2499.923 | 581.80035 |
| Manf | AHHIPVEK | -94 | -101 | 0 | 0.00676572 | 8 | NaN | NaN | 652.6501 | NaN | NaN | NaN |
| Psmc5 | HTDCTFIR | -206 | -213 | 0 | 0 | 8 | NaN | NaN | 676.3177 | 807.13965 | 995.6644 | 753.40674 |
| Psmc5 | KIEFPPPNEEAR | -314 | -325 | 1 | 0.00122419 | 12 | NaN | NaN | 553.52277 | 780.04663 | NaN | NaN |
| Tgm2 | EDITHTYK | -437 | -444 | 0 | 0.00635324 | 8 | NaN | NaN | 1040.6102 | NaN | NaN | NaN |
| Cand1 | TVIGELPPASSGSAL | -112 | -126 | 0 | 0.00051306 | 15 | NaN | NaN | 661.7089 | NaN | NaN | NaN |
| Cand1 | LIDPETLLPR | -960 | -969 | 0 | 0.003360691 | 10 | NaN | NaN | 327.92465 | NaN | NaN | NaN |
| Polr2a | LPSDLHPIK | -1000 | -1008 | 0 | 0 | 9 | NaN | NaN | 860.78906 | 379.82422 | 663.33136 | 821.48804 |
| Polr2a | STHPDDEDSGPYK | -607 | -619 | 0 | 0 | 13 | NaN | NaN | 951.8774 | 582.35767 | 990.321 | NaN |
| Polr2a | VPQPAILKPR | -574 | -583 | 1 | 0 | 10 | NaN | NaN | 394.42618 | 313.57773 | NaN | NaN |
| Gpx1 | PLTGGEPVSLGSLR | -21 | -34 | 0 | 0 | 14 | NaN | NaN | 785.16724 | 4878.5225 | 3941.186 | 5110.215 |
| Gpx1 | AHPLFTFLR | -120 | -128 | 0 | 0.000471409 | 9 | NaN | NaN | 183.3898 | NaN | NaN | 1005.23956 |
| Nmi | SENKELDEEIK | -36 | -46 | 1 | 0 | 11 | NaN | NaN | 448.0043 | 291.49857 | 384.09863 | 620.1768 |
| Ppp6c | AHQLVHEGYK | -236 | -245 | 0 | 0.00025015 | 10 | NaN | NaN | 658.4922 | NaN | NaN | 423.29297 |
| Snap23 | AHQVTDESLESTR | -13 | -25 | 0 | 0 | 13 | NaN | NaN | 717.17126 | NaN | NaN | NaN |
| B2m | TVYWDRDM | -112 | -119 | 1 | 0.002099344 | 8 | NaN | NaN | 699.07874 | NaN | NaN | NaN |
| Ptpn1 | QWVSEETCGDEDSLAR | -332 | -347 | 0 | 0.00025015 | 16 | NaN | NaN | 436.32864 | 438.09787 | 173.1508 | 516.06604 |
| Ptpn1 | AHWPSHWKPFLVN | -400 | -412 | 1 | 0.002068213 | 13 | NaN | NaN | 1262.8981 | NaN | NaN | 417.67307 |
| Mthfd1 | GADDIELLPEAQNK | -835 | -848 | 0 | 0 | 14 | NaN | NaN | 724.50555 | 596.6016 | 1016.48083 | 924.67737 |
| Ier3ip1 | SQLMNLIR | -50 | -57 | 0 | 0.00078661 | 8 | NaN | NaN | 96.30671 | NaN | NaN | NaN |
| Qdpr | TFHDWITGNK | -208 | -217 | 0 | 0 | 10 | NaN | NaN | 4015.4253 | NaN | 7175.8774 | 7500.269 |
| Qdpr | VLPVTLDTPMNR | -177 | -188 | 0 | 0 | 12 | NaN | NaN | 8905.232 | 10074.835 | NaN | NaN |
| Set | GEKEQQEAIEHIDEVQNEIDR | -36 | -56 | 1 | 0.001291103 | 21 | NaN | NaN | 402.74976 | 385.42062 | 497.8992 | 598.7692 |
| Set | FDENPYFENK | -140 | -149 | 0 | 0 | 10 | NaN | NaN | 3231.3464 | NaN | NaN | 4867.1147 |
| Oxsr1 | TLQNDPPSLETGVQDK | -239 | -254 | 0 | 0.005917757 | 16 | NaN | NaN | 760.7604 | 409.64185 | NaN | NaN |
| Fxr1 | IEGDNENKLPR | -318 | -328 | 1 | 0 | 11 | NaN | NaN | 445.8883 | 1444.4114 | 1001.74976 | 943.80206 |
| Fxr2 | LEFSEDSVQVPR | -289 | -300 | 0 | 0 | 12 | NaN | NaN | 1934.0935 | 570.26733 | 1251.7205 | NaN |
| Fxr2 | AIELGEETCTFR | -262 | -273 | 0 | 0 | 12 | NaN | NaN | 1616.5188 | NaN | 863.73047 | NaN |
| Hnrnpab | IELPIDPK | -188 | -195 | 0 | 0.002595029 | 8 | NaN | NaN | 626.76733 | 704.8574 | 674.2206 | NaN |
| Hnrnpab | FHTVSGSK | -221 | -228 | 0 | 0.00676572 | 8 | NaN | NaN | 1151.0535 | NaN | 1037.0809 | NaN |
| Hnrnpab | GGHQNNYKPY | -276 | -285 | 1 | 0.001291103 | 10 | NaN | NaN | 691.6013 | NaN | NaN | NaN |
| Jak1 | SLKPESGGNHIADLK | -908 | -922 | 1 | 0.00111006 | 15 | NaN | NaN | 466.25757 | NaN | NaN | NaN |
| Ap3d1 | AIFHEEEPR | -643 | -651 | 0 | 0 | 9 | NaN | NaN | 5183.3315 | NaN | 1681.566 | 1818.1204 |
| Kpna2 | LLHHNDPEVLADSC | -259 | -272 | 0 | 0 | 14 | NaN | NaN | 3395.7927 | NaN | NaN | NaN |
| Mtrex | EYPFILDAFQR | -133 | -143 | 0 | 0 | 11 | NaN | NaN | 4531.8975 | 554.2958 | 1752.0878 | 2128.274 |
| Mtrex | AIGNTELENK | -1011 | -1020 | 0 | 0.003069674 | 10 | NaN | NaN | 664.47656 | NaN | NaN | NaN |
| Ifit1 | AIHEFQETLK | -314 | -323 | 0 | 0 | 10 | NaN | NaN | 3978.5884 | NaN | 1760.0316 | 1646.7999 |
| Pip4k2b | FGIDDQDYQNSVTR | -110 | -123 | 0 | 0 | 14 | NaN | NaN | 383.34872 | 904.5794 | 277.16544 | NaN |
| Rps7 | DLVFPSEIVGK | -132 | -142 | 0 | 0.000394361 | 11 | NaN | NaN | 625.0059 | NaN | NaN | NaN |
| Rps7 | ILEDLVFPSEIVGK | -129 | -142 | 0 | 0.001392869 | 14 | NaN | NaN | 365.20245 | NaN | NaN | NaN |
| Cstf3 | KLEENPYDLDAW | -24 | -35 | 1 | 0.000230707 | 12 | NaN | NaN | 739.3839 | 424.8669 | 770.77325 | NaN |
| Card11 | AILDILEHDRK | -300 | -310 | 1 | 0.00025015 | 11 | NaN | NaN | 218.77417 | NaN | 252.1788 | NaN |
| Vps52 | AILEAPVTEPR | -184 | -194 | 0 | 0 | 11 | NaN | NaN | 903.7474 | 396.86526 | NaN | NaN |
| Fbl | AILGGVDQIHIKPGAK | -153 | -168 | 1 | 0 | 16 | NaN | NaN | 361.19415 | 445.70062 | 475.14035 | 463.9112 |
| Fbl | MKPQEQLTLEPYER | -297 | -310 | 1 | 0 | 14 | NaN | NaN | 486.7409 | 413.78494 | 677.21655 | 514.5715 |
| Lta4h | QHPYLFSQ | -128 | -135 | 0 | 0.001708008 | 8 | NaN | NaN | 1039.5093 | NaN | 901.9247 | NaN |
| Lta4h | FVTPTLLAGDK | -278 | -288 | 0 | 0.000298805 | 11 | NaN | NaN | 948.7517 | NaN | NaN | NaN |
| Pds5b | IEEDFPHIR | -699 | -707 | 0 | 0 | 9 | NaN | NaN | 256.51965 | 859.31323 | 1339.003 | 1262.3511 |
| Iqgap1 | SNQQLENDLNLMDIK | -902 | -916 | 0 | 0 | 15 | NaN | NaN | 3056.918 | NaN | 2949.9387 | 2928.2598 |
| Iqgap1 | LEGVLAEVAQHY | -568 | -579 | 0 | 0.00051306 | 12 | NaN | NaN | 1192.7601 | 1227.5763 | 1483.375 | 1382.408 |
| Iqgap1 | GLAPQIQDLYGK | -163 | -174 | 0 | 0 | 12 | NaN | NaN | 824.98376 | 781.6278 | 177.18414 | 753.20483 |
| Iqgap1 | SVTGLTNIEEENCQR | -482 | -496 | 0 | 0 | 15 | NaN | NaN | 772.49927 | 918.3642 | 1054.4965 | 1094.1135 |
| Iqgap1 | TNIEEENCQR | -487 | -496 | 0 | 0 | 10 | NaN | NaN | 337.8206 | 289.5419 | 466.54398 | 536.44025 |
| Khsrp | LASQGDSIGSQLGPIHPPPR | -124 | -143 | 0 | 0.00025015 | 20 | NaN | NaN | 655.7986 | 487.6258 | 441.27997 | 370.31787 |
| Khsrp | DQGGFGDRNEYGSR | -308 | -321 | 1 | 0 | 14 | NaN | NaN | 425.3514 | NaN | NaN | NaN |
| Khsrp | IQDGSQNTNVDKPLR | -271 | -285 | 1 | 0.004224523 | 15 | NaN | NaN | 489.78583 | 363.23624 | NaN | 439.3822 |
| Khsrp | QTGAFVEISR | -454 | -463 | 0 | 0.001811524 | 10 | NaN | NaN | 742.38293 | NaN | NaN | 877.0984 |
| Edf1 | AIPNNQVLGK | -114 | -123 | 0 | 0.009002904 | 10 | NaN | NaN | 339.81503 | NaN | NaN | NaN |
| Rsl1d1 | VPLPHSIL | -82 | -89 | 0 | 0.001055699 | 8 | NaN | NaN | 1988.9194 | 1158.3896 | 380.16824 | 1004.7987 |
| H2az1;H2az2 | GDEELDSLIKA | (93);(93) | (103);(103) | 1 | 0 | 11 | NaN | NaN | 540.6846 | 957.5676 | 783.79254 | 843.98914 |

| | | | | | | | | | | | | |
|---|---|---|---|---|---|---|---|---|---|---|---|---|
| Ncdn | EAPDGCPDHVLR | -93 | -104 | 0 | 0.00111006 | 12 | NaN | NaN | 979.72754 | NaN | NaN | NaN |
| Ncdn | AISPTTPGPSWPGDALR | -446 | -462 | 0 | 0.005112649 | 17 | NaN | NaN | 796.516 | NaN | NaN | NaN |
| Ythdf3 | AITDGQAGFGNDTLSK | -165 | -180 | 0 | 0.00025015 | 16 | NaN | NaN | 2099.825 | NaN | NaN | 467.2193 |
| Psmc1 | IEFPLPDEK | -351 | -359 | 0 | 0.003133457 | 9 | NaN | NaN | 601.49133 | NaN | NaN | NaN |
| Psmc1 | KAPQETYADIGGLDNQIQEIK | -178 | -198 | 1 | 0.00272025 | 21 | NaN | NaN | 580.9477 | NaN | NaN | NaN |
| Eef1g | PGSEETQTLVR | -404 | -414 | 0 | 0 | 11 | NaN | NaN | 464.31772 | 1443.9438 | 2269.951 | 1197.1284 |
| Eef1g | PPHFHFGQTNR | -35 | -45 | 0 | 0 | 11 | NaN | NaN | 1127.8721 | NaN | 1554.2467 | 233.77191 |
| Ezr | EKEELMLR | -343 | -350 | 1 | 0 | 8 | NaN | NaN | 9790.357 | 9842.979 | 9430.385 | 9782.268 |
| Ezr | ELSEQIEK | -364 | -371 | 0 | 0.002068213 | 8 | NaN | NaN | 773.8881 | NaN | NaN | 1039.8712 |
| Rrp7a | DYEDSVLDPEALR | -158 | -170 | 0 | 0 | 13 | NaN | NaN | 638.0149 | 253.0522 | 454.11206 | NaN |
| Hspa5 | GDTHLGGEDFDQR | -250 | -262 | 0 | 0 | 13 | NaN | NaN | 2403.6653 | 533.05646 | 2667.9006 | 197.86172 |
| Hspa5 | NGDTHLGGEDFDQR | -249 | -262 | 0 | 0 | 14 | NaN | NaN | 1021.6168 | 488.35907 | 1079.1819 | NaN |
| Mapre1 | GKFQDNFEFVQ | -99 | -109 | 1 | 0 | 11 | NaN | NaN | 239.05843 | 662.0581 | 527.265 | 757.60645 |
| Cs | DVLSNLIPK | -35 | -43 | 0 | 0.000550307 | 9 | NaN | NaN | 1049.7496 | 911.78815 | 1267.7391 | 200.26405 |
| Cs | LDWSHNF | -234 | -240 | 0 | 0.002780285 | 7 | NaN | NaN | 961.3017 | 722.5927 | 788.2471 | 707.5389 |
| Eif3b | GFHCESSAHWPIFK | -406 | -419 | 0 | 0.002780285 | 14 | NaN | NaN | 311.12378 | NaN | 622.62177 | 575.31604 |
| Eif3b | GVQLIDFSPCER | -364 | -375 | 0 | 0.00025015 | 12 | NaN | NaN | 881.10077 | 636.9866 | NaN | NaN |
| Rrp9 | AKPAPGPGAAPVAGK | -11 | -25 | 1 | 0.00078661 | 15 | NaN | NaN | 1501.5334 | 2790.1582 | 2571.0278 | 3163.3323 |
| Rrp9 | LYLEQLR | -85 | -91 | 0 | 0.007701635 | 7 | NaN | NaN | 687.3497 | NaN | NaN | NaN |
| Serbp1 | SASAPDVDDPEAFPALA | -391 | -407 | 0 | 0 | 17 | NaN | NaN | 681.0241 | 775.03076 | 1118.3691 | 1063.4327 |
| Serbp1 | EETPEGEEHPVADTENKENEVEEVKEEGPK | -256 | -285 | 2 | 0.001636916 | 30 | NaN | NaN | 460.67203 | 612.43036 | 671.3804 | 674.44916 |
| Serbp1 | LEINFGDLGR | -354 | -363 | 0 | 0.00078661 | 10 | NaN | NaN | 1268.0782 | NaN | NaN | NaN |
| Dbi | DVNTDRPGLLDLK | -39 | -51 | 1 | 0.00223472 | 13 | NaN | NaN | 743.94354 | NaN | NaN | NaN |
| H2-K1 | YMEVGYVDDTEFVR | -43 | -56 | 0 | 0 | 14 | NaN | NaN | 929.96216 | 852.1374 | 1015.74426 | 981.47845 |
| H2-K1 | WEQAGEAERL | -168 | -177 | 1 | 0 | 10 | NaN | NaN | 391.29593 | 678.8314 | 1058.8942 | 1207.6226 |
| H2-K1 | HKWEQAGEAE | -166 | -175 | 1 | 0.00272025 | 10 | NaN | NaN | 2402.8618 | NaN | NaN | NaN |
| Tomm34 | ALCHLVLK | -234 | -241 | 0 | 0.00025015 | 8 | NaN | NaN | 4082.6194 | 3547.7166 | 4043.446 | 4246.183 |
| Rnh1 | DSACQLESLK | -191 | -200 | 0 | 0 | 10 | NaN | NaN | 752.0834 | 871.71735 | 673.2331 | 632.5624 |
| Rnh1 | WTNEVEEQLR | -434 | -443 | 0 | 0 | 10 | NaN | NaN | 3869.7493 | 234.91737 | 219.54628 | NaN |
| Pml | DESLAEPHLEDR | -607 | -618 | 0 | 0 | 12 | NaN | NaN | 946.9909 | 231.03131 | 1046.3859 | 1201.3809 |
| Prpf8 | TTTVHWLESR | -811 | -820 | 0 | 0 | 10 | NaN | NaN | 948.6259 | NaN | 1565.8198 | 1287.4412 |
| Prpf8 | LTDEEWIK | -1971 | -1978 | 0 | 0.005639034 | 8 | NaN | NaN | 1194.4081 | NaN | 1326.4685 | NaN |
| Prpf8 | GMLDPLEVHLL | -1867 | -1877 | 0 | 0 | 11 | NaN | NaN | 760.2417 | NaN | 1108.8994 | NaN |
| Prpf8 | QDIATEAAHPIR | -1075 | -1086 | 0 | 0.007791272 | 12 | NaN | NaN | 2611.7744 | NaN | NaN | NaN |
| Prpf8 | LFPPWIKPA | -935 | -943 | 1 | 0.001708008 | 9 | NaN | NaN | 655.1744 | NaN | NaN | NaN |
| Prpf8 | DFPNIVIK | -1878 | -1885 | 0 | 0.008436628 | 8 | NaN | NaN | 131.6763 | NaN | NaN | NaN |
| Hk1 | SIPDGTEHGDFLAL | -574 | -587 | 0 | 0 | 14 | NaN | NaN | 2373.4336 | 1572.2102 | 2133.7476 | 1653.6848 |
| Hk1 | QIEETLSHFR | -525 | -534 | 0 | 0.000627943 | 10 | NaN | NaN | 1867.7341 | NaN | NaN | NaN |
| Rnf213 | YKEEVSTVELIK | -2969 | -2980 | 1 | 0 | 12 | NaN | NaN | 321.11102 | 591.8942 | 498.24667 | 696.317 |
| Rnf213 | ETHYCLSPDIR | -1184 | -1194 | 0 | 0.000347792 | 11 | NaN | NaN | 1074.2324 | NaN | NaN | 1512.7085 |
| Ccar2 | GRLPQLGEK | -85 | -93 | 1 | 0.005522803 | 9 | NaN | NaN | 637.72003 | NaN | NaN | NaN |
| Nup210 | VLLPFTR | -33 | -39 | 0 | 0.006677519 | 7 | NaN | NaN | 394.8525 | NaN | NaN | NaN |
| Nup210 | GVFQPLPGR | -589 | -597 | 0 | 0.004224523 | 9 | NaN | NaN | 361.061 | NaN | NaN | NaN |
| Rps17 | QEIIEVDPDTK | -93 | -103 | 0 | 0 | 11 | NaN | NaN | 159.08987 | 232.68674 | NaN | 195.51062 |
| Myg1 | AMDLVQEEFLQR | -219 | -230 | 0 | 0 | 12 | NaN | NaN | 903.685 | NaN | 968.5901 | 1028.5989 |
| Sptbn1 | KLLDPEDISVDHPDEK | -249 | -264 | 1 | 0.003745449 | 16 | NaN | NaN | 532.00836 | NaN | 3104.44 | NaN |
| Sptbn1 | LLTQHENIK | -1100 | -1108 | 0 | 0 | 9 | NaN | NaN | 220.83797 | NaN | NaN | NaN |
| Hprt1 | VFIPHGLIMDR | -35 | -45 | 0 | 0 | 11 | NaN | NaN | 692.2811 | 789.79663 | 500.32437 | 928.3939 |
| Cndp2 | LLPVGSADDGAHSQNEK | -434 | -450 | 0 | 0.002810811 | 17 | NaN | NaN | 280.86267 | 149.40251 | NaN | NaN |
| Cltc | SIVNFSQDPDVHFK | -717 | -730 | 0 | 0 | 14 | NaN | NaN | 747.2306 | 534.079 | 755.6977 | 755.773 |
| Cltc | SQDPDVHFK | -722 | -730 | 0 | 0 | 9 | NaN | NaN | 6151.85 | 692.32776 | NaN | NaN |
| Cltc | IHEGCEEPATHN | -866 | -877 | 0 | 0.00111006 | 12 | NaN | NaN | 597.4747 | NaN | NaN | NaN |
| Cltc | LEKHELIEFR | -1499 | -1508 | 1 | 0.002466863 | 10 | NaN | NaN | 566.0187 | NaN | NaN | NaN |
| Cltc | IHEGCEEPATH | -866 | -876 | 0 | 0.008890469 | 11 | NaN | NaN | 563.2001 | NaN | NaN | NaN |
| Itih3 | GHVSFKPSLDQQR | -222 | -234 | 1 | 0.00025015 | 13 | NaN | NaN | 768.07275 | NaN | NaN | NaN |
| Htatsf1 | KLLLFDR | -304 | -310 | 1 | 0.005758933 | 7 | NaN | NaN | 325.08362 | NaN | 314.67047 | 304.27203 |
| Htatsf1 | AWDGTTDYQVEETSR | -347 | -361 | 0 | 0 | 15 | NaN | NaN | 563.13586 | 998.4492 | NaN | NaN |
| Rbm39 | ALEQLNGFELAGR | -307 | -319 | 0 | 0.009928578 | 13 | NaN | NaN | 899.56586 | 684.53625 | NaN | NaN |
| Sugt1 | LLHPIIPEQSTFK | -195 | -207 | 0 | 0 | 13 | NaN | NaN | 445.5121 | 747.9713 | 980.0911 | 941.72723 |
| Nans | LHELNVPFFK | -122 | -131 | 0 | 0.00051306 | 10 | NaN | NaN | 669.38025 | 1015.2367 | 1069.075 | 635.6518 |
| Kif11 | IAVLEEELSK | -425 | -434 | 0 | 0.005884925 | 10 | NaN | NaN | 12622.492 | NaN | NaN | NaN |
| Oat | GLLAKPTHGDIIR | -401 | -413 | 1 | 0.000550307 | 13 | NaN | NaN | 804.7001 | NaN | 345.6432 | 557.887 |
| Oat | LPSDVVTSVR | -363 | -372 | 0 | 0.00025015 | 10 | NaN | NaN | 1474.5934 | 11628.794 | NaN | 9040.893 |
| Psmd2 | SETELKDTYAR | -546 | -556 | 1 | 0 | 11 | NaN | NaN | 1554.1776 | 1317.1355 | 619.28217 | 999.74194 |
| Psmd2 | GLGLNHLGK | -561 | -569 | 0 | 0.001291103 | 9 | NaN | NaN | 477.2082 | 688.6477 | 462.0624 | 470.80753 |
| Psmd2 | FLRPHYGK | -98 | -105 | 1 | 0.002752526 | 8 | NaN | NaN | 542.1552 | NaN | NaN | NaN |
| Atic | DVSELTGFPEMLGGR | -50 | -64 | 0 | 0 | 15 | NaN | NaN | 900.1595 | 916.0186 | 978.55725 | NaN |
| Atic | DQSYKPDENEVR | -368 | -379 | 1 | 0.000431655 | 12 | NaN | NaN | 456.06476 | NaN | 77.2966 | 478.59744 |
| Atic | FEEVPELLTEAEK | -512 | -524 | 0 | 0.000230707 | 13 | NaN | NaN | 2522.984 | NaN | 2549.6074 | NaN |
| Atic | SLFSNIVTK | -398 | -406 | 0 | 0.004439092 | 9 | NaN | NaN | 1436.0027 | NaN | 1055.9315 | NaN |
| Atic | EVPELLTEAEKK | -514 | -525 | 1 | 0.001055699 | 12 | NaN | NaN | 729.12225 | NaN | NaN | NaN |
| Ftl1 | GNHLTNLR | -146 | -153 | 0 | 0.001610812 | 8 | NaN | NaN | 544.56934 | NaN | 1258.0365 | NaN |
| Mrpl41 | ALFQETVAPAIEK | -88 | -100 | 0 | 0 | 13 | NaN | NaN | 1118.3445 | NaN | 1586.7599 | 1580.1111 |
| Raver1 | ALGDSHLR | -286 | -293 | 0 | 0.001291103 | 8 | NaN | NaN | 3193.1587 | 1874.76 | NaN | 1796.8174 |
| Rtn3 | SLEGEVASQVPN | -529 | -540 | 0 | 0.003003218 | 12 | NaN | NaN | 825.73627 | NaN | NaN | NaN |
| Lrrc47 | DLSNNQLTEIPAELADCPK | -210 | -228 | 0 | 0 | 19 | NaN | NaN | 900.79816 | 731.19745 | 964.1064 | NaN |
| Apex1 | SLLPALCDSK | -289 | -298 | 0 | 0.001636916 | 10 | NaN | NaN | 531.05426 | NaN | NaN | 772.3576 |
| Ago2 | ALHDALSGR | -143 | -151 | 0 | 0.005789881 | 9 | NaN | NaN | 755.6974 | 546.47406 | NaN | NaN |

| | | | | | | | | | | | | |
|---|---|---|---|---|---|---|---|---|---|---|---|---|
| Cers2 | NQDRPSLLK | -123 | -131 | 1 | 0.001023236 | 9 | NaN | NaN | 868.8272 | 719.5462 | 896.9204 | 929.8047 |
| Cers2 | LHDASDYLLESAK | -238 | -250 | 0 | 0 | 13 | NaN | NaN | 550.58826 | NaN | NaN | NaN |
| Dars1 | EKYDTDFYVLDK | -382 | -393 | 1 | 0 | 12 | NaN | NaN | 989.3779 | 1773.234 | 537.5252 | 768.0124 |
| Ptbp1 | KHQSVQLPR | -434 | -442 | 1 | 0.00078661 | 9 | NaN | NaN | 1376.8586 | 691.6298 | 1134.6903 | 1429.0834 |
| Plcl2 | ALIENADAVYEK | -1014 | -1025 | 0 | 0 | 12 | NaN | NaN | 4294.565 | NaN | 4295.6255 | 1334.1641 |
| Top2a | LDMPLWYLTK | -1126 | -1135 | 0 | 0 | 10 | NaN | NaN | 188.00336 | 719.89746 | 338.45578 | 314.83643 |
| Top2a | VYDEDVGINYR | -62 | -72 | 0 | 0 | 11 | NaN | NaN | 1257.7285 | 2512.908 | 3115.659 | 3488.957 |
| Top2a | KNYEDEDSLK | -519 | -528 | 1 | 0 | 10 | NaN | NaN | 1251.5939 | 853.2597 | 1012.18414 | NaN |
| Top2a | FIHHNWPSLLR | -555 | -565 | 0 | 0.000471409 | 11 | NaN | NaN | 871.12317 | 2981.7483 | NaN | NaN |
| Rps12 | NCDEPMYVK | -55 | -63 | 0 | 0.004439092 | 9 | NaN | NaN | 328.48038 | NaN | 401.17014 | NaN |
| Eif3d | LSVENAWGILR | -482 | -492 | 0 | 0.008736733 | 11 | NaN | NaN | 460.8492 | NaN | 489.09753 | NaN |
| Eif3h | GEPPLPEEDLSK | -292 | -303 | 0 | 0.008129024 | 12 | NaN | NaN | 330.65402 | NaN | NaN | NaN |
| Msn | KTQEQLASEMAELT | -412 | -425 | 1 | 0.00025015 | 14 | NaN | NaN | 669.8354 | NaN | 894.58124 | 1067.767 |
| Msn | KGSELWLGVD | -212 | -221 | 1 | 0.003745449 | 10 | NaN | NaN | 1175.7615 | NaN | NaN | NaN |
| Msn | TANDMIHAE | -539 | -547 | 0 | 0.00025015 | 9 | NaN | NaN | 289.9769 | NaN | NaN | NaN |
| Gimap4 | FFQEVMHEF | -174 | -182 | 0 | 0.00025015 | 9 | NaN | NaN | 390.6183 | 380.41113 | 357.68707 | 420.4143 |
| Gimap4 | DDLEDTDIHEY | -157 | -167 | 0 | 0.000550307 | 11 | NaN | NaN | 242.64726 | NaN | NaN | 260.81003 |
| Xrn2 | SDASAPGEGEHK | -196 | -207 | 0 | 0.001392869 | 12 | NaN | NaN | 502.57147 | NaN | 588.505 | 585.81067 |
| Gpi | PVQTQHPIR | -409 | -417 | 0 | 0.002752526 | 9 | NaN | NaN | 465.27298 | NaN | 419.1313 | NaN |
| Gpi | IDGTHIAK | -187 | -194 | 0 | 0.00897527 | 8 | NaN | NaN | 614.59094 | NaN | NaN | NaN |
| Ube2n | YHPNVDK | -76 | -82 | 0 | 0.002199523 | 7 | NaN | NaN | 1400.2255 | NaN | NaN | 2255.4666 |
| Hsp90aa1 | TSASGDEMVSLK | -468 | -479 | 0 | 0 | 12 | NaN | NaN | 2846.9773 | 2896.2126 | 3363.3145 | 3309.7969 |
| Hsp90aa1 | ELHINLIPSK | -75 | -84 | 0 | 0.00025015 | 10 | NaN | NaN | 1504.6456 | 1950.6437 | 2171.6323 | 1652.2589 |
| Hsp90aa1 | APFDLFENR | -348 | -356 | 0 | 0 | 9 | NaN | NaN | 1413.6917 | 862.1896 | 2379.928 | 222.90613 |
| Hsp90aa1 | IEPIDEYCVQQLK | -523 | -535 | 0 | 0 | 13 | NaN | NaN | 378.50656 | 809.628 | 721.701 | 869.4045 |
| Hsp90aa1 | GIHEDSQNR | -449 | -457 | 0 | 0.002534621 | 9 | NaN | NaN | 609.28705 | 821.2395 | 151.83154 | NaN |
| Hsp90aa1 | NPDDITNEEYGEFY | -301 | -314 | 0 | 0.00025015 | 14 | NaN | NaN | 501.2334 | 628.8215 | NaN | 454.67963 |
| Hsp90aa1 | TDTGEPMGR | -174 | -182 | 0 | 0 | 9 | NaN | NaN | 400.47382 | 602.01355 | 368.93756 | NaN |
| Hsp90aa1 | EKVTVITK | -146 | -153 | 1 | 0.000394361 | 8 | NaN | NaN | 301.4568 | 299.53238 | 268.43503 | NaN |
| Lrpap1 | HVESIGDPEHISR | -310 | -322 | 0 | 0 | 13 | NaN | NaN | 740.17004 | 605.6561 | 701.8543 | 657.64514 |
| Urb1 | DTGVFEHTWR | -667 | -676 | 0 | 0 | 10 | NaN | NaN | 949.06494 | NaN | 1360.2424 | NaN |
| Urb1 | ALNLDSIPVK | -443 | -452 | 0 | 0.003360691 | 10 | NaN | NaN | 285.40952 | NaN | NaN | NaN |
| Clic1 | KFLDGNELTLADCN | -166 | -179 | 1 | 0 | 14 | NaN | NaN | 1177.5908 | NaN | 1240.5115 | NaN |
| Gtpbp1 | LVLVSPTSEQYDSLLR | -65 | -80 | 0 | 0 | 16 | NaN | NaN | 839.1675 | 897.0711 | NaN | 880.44525 |
| Pgam1 | HGESAWNLENR | -11 | -21 | 0 | 0 | 11 | NaN | NaN | 131.30453 | 6202.1694 | 7089.5933 | 6899.901 |
| Tap1 | YLGGQLVIR | -404 | -412 | 0 | 0.000627943 | 9 | NaN | NaN | 814.95874 | NaN | 1246.8027 | NaN |
| Tap1 | QDVSFAYPNQPK | -482 | -493 | 0 | 0.000431655 | 12 | NaN | NaN | 891.5088 | NaN | NaN | NaN |
| Tap1 | EGSVGEQGTHL | -693 | -703 | 0 | 0.001636916 | 11 | NaN | NaN | 649.757 | NaN | NaN | NaN |
| Utp14a | TLEELEELGKED | -509 | -520 | 1 | 0.006624942 | 12 | NaN | NaN | 2923.9993 | NaN | NaN | NaN |
| Bcas2 | FDQGYEAPGVR | -19 | -29 | 0 | 0 | 11 | NaN | NaN | 1217.447 | 2249.955 | 2409.9038 | 2627.9937 |
| Cse1l | IVEDEPNKICEADR | -76 | -89 | 1 | 0 | 14 | NaN | NaN | 3687.4717 | NaN | 3933.5002 | 3830.8594 |
| Cse1l | DEEAFEDNSEEYIR | -358 | -371 | 0 | 0 | 14 | NaN | NaN | 513.4368 | 818.3237 | 768.5404 | 652.0467 |
| Cse1l | KYDEEFQR | -281 | -288 | 1 | 0.002199523 | 8 | NaN | NaN | 1894.4028 | NaN | NaN | 1828.2158 |
| Cse1l | RDLEGSDIDTR | -372 | -382 | 1 | 0.00272025 | 11 | NaN | NaN | 1188.8252 | NaN | NaN | 2242.2983 |
| Selenoh | ALQLEAPELPVQVNPSKPR | -51 | -69 | 1 | 0 | 19 | NaN | NaN | 348.70184 | 346.9016 | 302.09406 | 391.37238 |
| Mvp | HYCIVANPVSR | -57 | -67 | 0 | 0.009657961 | 11 | NaN | NaN | 3213.8816 | NaN | NaN | NaN |
| Copg1 | YTLEPSEKPFDLK | -565 | -577 | 1 | 0.00025015 | 13 | NaN | NaN | 597.43463 | 884.85126 | 1087.7605 | 1044.3184 |
| Copg1 | SVPLATTPMAEQRPESTAT | -578 | -596 | 1 | 0.000826806 | 19 | NaN | NaN | 466.40976 | 439.58264 | NaN | 393.23206 |
| Copg1 | SSPEPVALTESETEYVIR | -632 | -649 | 0 | 0.000550307 | 18 | NaN | NaN | 387.76645 | NaN | 442.56415 | 425.70374 |
| Copg1 | EAASAIVNLPGCSAK | -269 | -283 | 0 | 0.00635324 | 15 | NaN | NaN | 3235.145 | NaN | NaN | NaN |
| Psmc6 | NQMDGFDTLHR | -262 | -272 | 0 | 0.00383302 | 11 | NaN | NaN | 334.4295 | NaN | NaN | NaN |
| Lypla1 | ALSCWLPLR | -141 | -149 | 0 | 0 | 9 | NaN | NaN | 631.0771 | 702.38214 | 916.3691 | 562.2932 |
| Uhrf1 | EDGHTLFDYDVR | -53 | -64 | 0 | 0.002845514 | 12 | NaN | NaN | 1151.1764 | NaN | NaN | NaN |
| Ubr4 | ILGPAESDEFLAR | -1336 | -1348 | 0 | 0 | 13 | NaN | NaN | 963.6822 | 979.65204 | 1154.5104 | NaN |
| Ubr4 | ALSPTFYFLLPSSK | -2030 | -2043 | 0 | 0 | 14 | NaN | NaN | 319.3789 | NaN | NaN | 340.40698 |
| Rbm10 | NSFQPISALR | -671 | -680 | 0 | 0.009242859 | 10 | NaN | NaN | 576.9602 | NaN | NaN | NaN |
| Scfd1 | ALTDAGCNLSPLQYIK | -476 | -491 | 0 | 0 | 16 | NaN | NaN | 427.36823 | NaN | NaN | 444.90356 |
| Rab4b | ALTGENVEEAFLK | -152 | -164 | 0 | 0 | 13 | NaN | NaN | 619.5157 | NaN | 1043.5001 | NaN |
| Cdk2 | LLDVIHTENK | -66 | -75 | 0 | 0 | 10 | NaN | NaN | 1015.81165 | 529.3331 | 393.00876 | 676.44037 |
| Eif3k | IDQAHQEERPIR | -89 | -100 | 1 | 0.00025015 | 12 | NaN | NaN | 386.91553 | 401.84235 | 505.1949 | 505.9503 |
| Eif3k | LYQFNPAFFQTT | -53 | -64 | 0 | 0.003745449 | 12 | NaN | NaN | 395.3006 | 404.36646 | 378.7224 | 337.2365 |
| Eif3k | ALTNLPHTDFTLC | -73 | -85 | 0 | 0 | 13 | NaN | NaN | 694.4377 | NaN | 1341.8082 | 178.55087 |
| Eif3k | ALTNLPHTDFTLCK | -73 | -86 | 0 | 0 | 14 | NaN | NaN | 341.095 | NaN | 845.4199 | NaN |
| Eipr1 | ALTPQTAETDAIR | -17 | -29 | 0 | 0 | 13 | NaN | NaN | 1897.8828 | 789.69635 | 878.3223 | 998.775 |
| Elmo1 | DFQSRPILELK | -527 | -537 | 1 | 0 | 11 | NaN | NaN | 957.1802 | NaN | 1728.2865 | 1546.207 |
| Xpo7 | ALVEFTNSPDCLSK | -33 | -46 | 0 | 0.007609801 | 14 | NaN | NaN | 1113.8993 | NaN | NaN | NaN |
| Xpo7 | SDPNNYHEFCR | -322 | -332 | 0 | 0.00149795 | 11 | NaN | NaN | 285.93332 | NaN | NaN | NaN |
| Upf1 | EHQGIGFLNDPR | -852 | -863 | 0 | 0 | 12 | NaN | NaN | 1830.0035 | 1454.5698 | 3850.3784 | 2227.846 |
| Upf1 | ANEHQGIGFLNDPR | -850 | -863 | 0 | 0.00025015 | 14 | NaN | NaN | 970.1857 | 4518.0386 | NaN | NaN |
| Srpra | ALVGNEAVDQLVK | -551 | -563 | 0 | 0 | 13 | NaN | NaN | 1088.2515 | NaN | 1544.6287 | NaN |
| Gna13 | ALWEDSGIQNAYDR | -151 | -164 | 0 | 0.00111006 | 14 | NaN | NaN | 371.94287 | NaN | NaN | NaN |
| Pdia4 | DATANDITNDQYK | -577 | -589 | 0 | 0 | 13 | NaN | NaN | 126.67043 | NaN | 458.4001 | 550.3312 |
| Zc3hav1 | SFFHSCPR | -730 | -737 | 0 | 0.00504706 | 8 | NaN | NaN | 1576.2262 | NaN | 1541.8638 | NaN |
| Zc3hav1 | AMPGETTTPVQGSNR | -520 | -534 | 0 | 0.009388464 | 15 | NaN | NaN | 1000.22687 | NaN | NaN | NaN |
| Hmgn5 | AMPVPFTPELKPK | -23 | -35 | 1 | 0.00025015 | 13 | NaN | NaN | 551.4992 | NaN | NaN | NaN |
| Satb1 | MDHLNEATQGK | -1 | -11 | 0 | 0 | 11 | NaN | NaN | 2319.3643 | 373.0917 | 3425.7563 | 1177.8799 |
| Satb1 | LEDLPPEQWSHTTVR | -176 | -190 | 0 | 0.002466863 | 15 | NaN | NaN | 2016.254 | NaN | 1630.1462 | 2490.4324 |
| Satb1 | GSTDVNADLKD | -754 | -764 | 1 | 0.004224523 | 11 | NaN | NaN | 766.2292 | 4107.0366 | NaN | 463.44714 |

| | | | | | | | | | | | | |
|---|---|---|---|---|---|---|---|---|---|---|---|---|
| Yme1l1 | ANAPCVIFIDELDSVGGK | -371 | -388 | 0 | 0.000431655 | 18 | NaN | NaN | 125.493416 | 253.24171 | NaN | 118.67668 |
| Abcf1 | ETPTEYLQR | -697 | -705 | 0 | 0.009388464 | 9 | NaN | NaN | 619.32935 | NaN | NaN | NaN |
| Eif3a | GPAEESSSWR | -1212 | -1221 | 0 | 0 | 10 | NaN | NaN | 6493.786 | 6502.6924 | 7750.581 | 7171.071 |
| Eif3a | IHEPIMLK | -46 | -53 | 0 | 0.001463897 | 8 | NaN | NaN | 1287.2192 | 921.35583 | 1371.8082 | NaN |
| Eif3a | ANEFLEVGK | -15 | -23 | 0 | 0.002972566 | 9 | NaN | NaN | 830.54205 | 1047.5874 | NaN | 7462.347 |
| Eif3a | NICQQVNIK | -76 | -84 | 0 | 0.000230707 | 9 | NaN | NaN | 460.1851 | 669.1214 | NaN | 329.71088 |
| Eif3a | RGPAEESSSWR | -1211 | -1221 | 1 | 0.00272025 | 11 | NaN | NaN | 991.3072 | 15443.285 | NaN | NaN |
| Eif3a | RGLDDDRGPR | -960 | -969 | 2 | 0.00111006 | 10 | NaN | NaN | 629.013 | NaN | 1452.9661 | NaN |
| Psat1 | LYNAVTTEDVEK | -345 | -356 | 0 | 0 | 12 | NaN | NaN | 719.67145 | 1541.2551 | 1111.516 | 1087.5532 |
| Psat1 | ASLYNAVTTEDVEK | -343 | -356 | 0 | 0 | 14 | NaN | NaN | 598.84283 | NaN | 1110.6669 | NaN |
| Agfg1 | TFTQQEIEFLQK | -72 | -83 | 0 | 0 | 12 | NaN | NaN | 1691.2297 | 1761.88 | 194.5552 | 481.7972 |
| Septin2 | YLHDESGLNR | -129 | -138 | 0 | 0 | 10 | NaN | NaN | 738.2668 | 722.33575 | 806.9116 | 1261.4152 |
| Septin2 | SVVGSNQLIEAK | -238 | -249 | 0 | 0.00111006 | 12 | NaN | NaN | 680.4669 | NaN | NaN | NaN |
| Ncbp1 | KDVPNPNQVDDDDEGFR | -522 | -538 | 1 | 0.00078661 | 17 | NaN | NaN | 706.7334 | 609.10406 | 391.91263 | 1048.5992 |
| Aimp2 | ANPASPPLSLLVLHR | -126 | -140 | 0 | 0 | 15 | NaN | NaN | 684.50543 | 717.34326 | 953.798 | 1027.1953 |
| Aimp2 | TLDLNSVLGK | -105 | -114 | 0 | 0.000347792 | 10 | NaN | NaN | 594.42413 | 421.86603 | 309.08524 | 196.77426 |
| Tra2b | DDRDYYSR | -231 | -238 | 1 | 0.002068213 | 8 | NaN | NaN | 8491.423 | NaN | 8332.249 | NaN |
| Imp4 | ANQDDYISFR | -221 | -230 | 0 | 0.00622886 | 10 | NaN | NaN | 1915.2883 | NaN | NaN | NaN |
| Nup107 | VIDWLVFDPAQR | -675 | -686 | 0 | 0 | 12 | NaN | NaN | 156.29956 | 172.38733 | 413.97327 | 294.75583 |
| Nup107 | FLEGVTEFEQR | -609 | -619 | 0 | 0 | 11 | NaN | NaN | 410.42535 | 673.8916 | NaN | 458.35696 |
| Smarca5 | EAEPLNDEELEEKEK | -821 | -835 | 1 | 0 | 15 | NaN | NaN | 551.0032 | 384.25803 | 432.6946 | 432.70908 |
| Smarca5 | IDEAEPLNDEELEEKEK | -819 | -835 | 1 | 0.002972566 | 17 | NaN | NaN | 435.00598 | NaN | NaN | NaN |
| Serpinc1 | ANRPFLVL | -437 | -444 | 1 | 0.004224523 | 8 | NaN | NaN | 1793.7556 | 1577.7604 | NaN | NaN |
| Acly | HLLVHAPEDK | -168 | -177 | 0 | 0.000431655 | 10 | NaN | NaN | 555.3318 | 1217.1064 | 1338.9314 | 1030.2999 |
| Acly | YLEINPLVVTK | -199 | -209 | 0 | 0.00294445 | 11 | NaN | NaN | 376.44528 | NaN | NaN | NaN |
| Ada | DKPLSLPGFLAK | -53 | -64 | 1 | 0.000550307 | 12 | NaN | NaN | 723.1501 | NaN | 590.7181 | NaN |
| Gtf2b | VIDVGSEWR | -45 | -53 | 0 | 0.00025015 | 9 | NaN | NaN | 1432.196 | NaN | 581.6126 | 1602.2207 |
| Gtf2b | FDTPVDKLPQL | -306 | -316 | 1 | 0.001092529 | 11 | NaN | NaN | 469.55908 | 400.70682 | NaN | NaN |
| Anp32e | EEEEEEGLRGEK | -235 | -246 | 1 | 0.00025015 | 12 | NaN | NaN | 866.84326 | NaN | NaN | NaN |
| Cfl1 | TVDDPYTTFVK | -63 | -73 | 0 | 0.001636916 | 11 | NaN | NaN | 856.9639 | 630.1232 | 782.8116 | NaN |
| Cfl1 | NIILEEGKEILVGDVGQT | -46 | -63 | 1 | 0.00122419 | 18 | NaN | NaN | 384.66638 | 574.14343 | 587.75433 | NaN |
| Bcl11b | LLNPFQPSPK | -391 | -400 | 0 | 0.00142999 | 10 | NaN | NaN | 357.5643 | NaN | 682.4461 | 121.14155 |
| Bcl11b | APGAEPFPALFPR | -645 | -657 | 0 | 0.002199523 | 13 | NaN | NaN | 3583.0164 | NaN | 5655.2725 | NaN |
| Mki67 | EHFETPNPK | -977 | -985 | 0 | 0.001392869 | 9 | NaN | NaN | 740.3463 | NaN | 1135.9305 | NaN |
| Mki67 | TSKPAAEILIKPQEEK | -3107 | -3122 | 2 | 0.000230707 | 16 | NaN | NaN | 600.8573 | NaN | NaN | 522.92584 |
| Mki67 | GDVASPADTPEHSSSK | -369 | -384 | 0 | 0.003428289 | 16 | NaN | NaN | 211.98662 | NaN | NaN | NaN |
| Eif4a1 | VKKEELTLEGIR | -236 | -247 | 2 | 0 | 12 | NaN | NaN | 709.75934 | 928.13947 | 1061.3624 | 1081.4248 |
| Eif4a1 | DQIYDIFQK | -194 | -202 | 0 | 0.00025015 | 9 | NaN | NaN | 1476.6229 | NaN | NaN | 1309.477 |
| Eif4a1 | ATMPSDVLEVTK | -214 | -225 | 0 | 0.001392869 | 12 | NaN | NaN | 2104.5938 | NaN | NaN | NaN |
| Idh3a | HMGLFDHAAK | -317 | -326 | 0 | 0.00111006 | 10 | NaN | NaN | 472.43542 | NaN | NaN | NaN |
| Rpl4 | KSNYNLPMHK | -274 | -283 | 1 | 0.001463897 | 10 | NaN | NaN | 310.92532 | NaN | NaN | 727.73114 |
| Iars1 | HNCFQECLK | -25 | -33 | 0 | 0 | 9 | NaN | NaN | 1125.469 | NaN | 1200.9819 | 1264.7384 |
| Iars1 | DADKNIIR | -372 | -379 | 1 | 0.001708008 | 8 | NaN | NaN | 3214.8472 | NaN | 3425.541 | NaN |
| Iars1 | YKPLFDYFIK | -289 | -298 | 1 | 0.000230707 | 10 | NaN | NaN | 463.3522 | NaN | 729.91296 | NaN |
| Chd4 | KLERPPETPTVDPTVK | -689 | -704 | 2 | 0.000712251 | 16 | NaN | NaN | 306.14423 | NaN | 837.7742 | 838.83636 |
| Chd4 | VQEFEHVNGR | -1499 | -1508 | 0 | 0.005394768 | 10 | NaN | NaN | 1196.6372 | NaN | NaN | NaN |
| Wdr12 | SVDWTDTGLLLSGGADNK | -391 | -408 | 0 | 0.00051306 | 18 | NaN | NaN | 3166.8877 | 2684.8557 | NaN | NaN |
| Crkl | VIIEKPEEQWWSAR | -151 | -164 | 1 | 0 | 14 | NaN | NaN | 1756.0374 | NaN | NaN | 2806.043 |
| Ass1 | KQHGIPIPVTPK | -165 | -176 | 1 | 0.000826806 | 12 | NaN | NaN | 262.53146 | 189.96333 | NaN | 182.57564 |
| Irf4 | DYNREEDAALFK | -61 | -72 | 1 | 0 | 12 | NaN | NaN | 361.8456 | 557.4114 | 1132.9279 | 1165.5026 |
| Irf4 | GYELPEHVTTPDYHR | -427 | -441 | 0 | 0 | 15 | NaN | NaN | 253.23122 | NaN | NaN | NaN |
| Cdk4 | APPPGLPVETIK | -107 | -118 | 0 | 0.008947895 | 12 | NaN | NaN | 797.4906 | 617.3805 | NaN | NaN |
| Gars1 | YPLFEGQETGK | -713 | -723 | 0 | 0 | 11 | NaN | NaN | 1941.0022 | 439.82712 | 343.49692 | 1909.7354 |
| Gars1 | VPLVAEKPLKEPK | -468 | -480 | 2 | 0 | 13 | NaN | NaN | 559.25116 | NaN | 530.83765 | NaN |
| Gars1 | TSYGWIEIVGCADR | -441 | -454 | 0 | 0.00025015 | 14 | NaN | NaN | 335.64047 | NaN | 414.57205 | 380.27908 |
| Gars1 | VDPTEKDHPK | -345 | -354 | 1 | 0.00025015 | 10 | NaN | NaN | 84.27854 | 919.1932 | 1070.6483 | NaN |
| Gars1 | YDFGPVGCALK | -138 | -148 | 0 | 0 | 11 | NaN | NaN | 416.04205 | 208.15662 | NaN | NaN |
| Gars1 | KVPLVAEKPLK | -467 | -477 | 2 | 0.00078661 | 11 | NaN | NaN | 273.8736 | 350.22018 | NaN | NaN |
| Asns | EDSPDLLAAR | -291 | -300 | 0 | 0 | 10 | NaN | NaN | 1281.9935 | NaN | 978.0145 | NaN |
| Asns | APSPEKAEEESER | -380 | -392 | 1 | 0.000431655 | 13 | NaN | NaN | 374.2127 | NaN | 576.51996 | NaN |
| Asns | WINATDPSAR | -541 | -550 | 0 | 0 | 10 | NaN | NaN | 686.80975 | NaN | NaN | NaN |
| Eif2a | VNDFNLSPGTQPYK | -170 | -183 | 0 | 0 | 14 | NaN | NaN | 1367.0087 | 510.7978 | NaN | 1069.3352 |
| Apbb1ip | DVLDNLFEK | -205 | -213 | 0 | 0.00025015 | 9 | NaN | NaN | 830.49445 | 183.21587 | 218.80785 | NaN |
| Apbb1ip | VFEDHENVVEVL | -236 | -247 | 0 | 0.00383302 | 12 | NaN | NaN | 1146.9955 | NaN | NaN | NaN |
| Hint1 | QVAQPGGDTIFGK | -9 | -21 | 0 | 0.000230707 | 13 | NaN | NaN | 1545.5891 | 1424.1396 | 976.42706 | 848.3227 |
| Rrm2 | SKAPTNPSVEDEPLLR | -60 | -75 | 1 | 0 | 16 | NaN | NaN | 443.70663 | NaN | 1390.8911 | 1504.3597 |
| Ppil1 | APTQWLDGK | -117 | -125 | 0 | 0.00025015 | 9 | NaN | NaN | 842.1405 | 838.7011 | NaN | NaN |
| Stoml2 | APVPGAQNSSQSR | -322 | -334 | 0 | 0.00025015 | 13 | NaN | NaN | 688.091 | NaN | 821.69073 | NaN |
| Yars1 | VVHPGDLK | -303 | -310 | 0 | 0 | 8 | NaN | NaN | 604.4211 | 885.2224 | 1194.4271 | 632.94324 |
| Zdhhc20 | SIGSNQPFPIKPL | -330 | -342 | 1 | 0.001055699 | 13 | NaN | NaN | 476.9679 | 418.12772 | 537.0499 | 550.7196 |
| Esyt1 | ESLETGSSVDAPPRPY | -607 | -622 | 1 | 0.00025015 | 16 | NaN | NaN | 2061.0513 | NaN | 2751.659 | 1204.0857 |
| Esyt1 | SSLNDEPEALGGPTHPASPVLEVR | -920 | -943 | 0 | 0 | 24 | NaN | NaN | 174.01353 | NaN | 606.88934 | 657.70746 |
| Esyt1 | SQELDVQVK | -528 | -536 | 0 | 0.000230707 | 9 | NaN | NaN | 3306.5928 | 248.26862 | NaN | NaN |
| Esyt1 | GTKPPSPYATIT | -802 | -813 | 1 | 0.00025015 | 12 | NaN | NaN | 756.62317 | 884.52704 | NaN | NaN |
| Nsun2 | LVSETESGNISR | -149 | -160 | 0 | 0.000230707 | 12 | NaN | NaN | 3495.9072 | 2429.0698 | 514.4576 | NaN |
| Nsun2 | WHEVPQGR | -377 | -384 | 0 | 0.001291103 | 8 | NaN | NaN | 1210.5293 | NaN | 2841.9011 | NaN |
| Nsun2 | EILFYDR | -258 | -264 | 0 | 0.003428289 | 7 | NaN | NaN | 647.32605 | 736.137 | NaN | NaN |
| Nsun2 | QTAQDPCVPDSVPGCDAAAAEPSR | -734 | -757 | 0 | 0.00437931 | 24 | NaN | NaN | 162.72379 | NaN | NaN | NaN |

| | | | | | | | | | | | | |
|---|---|---|---|---|---|---|---|---|---|---|---|---|
| Fam98a | AQEGGGSEVFQELK | -134 | -147 | 0 | 0.000907206 | 14 | NaN | NaN | 1156.0336 | 319.87506 | NaN | NaN |
| Ppid | GGDFSNQNGTGGESIYGEK | -84 | -102 | 0 | 0 | 19 | NaN | NaN | 1149.21 | 1389.4257 | NaN | 1413.7246 |
| Ppid | KAQGWQGLK | -313 | -321 | 1 | 0.00025015 | 9 | NaN | NaN | 873.19244 | 406.03278 | NaN | NaN |
| Ddx5 | AQEVDTYR | -70 | -77 | 0 | 0.001291103 | 8 | NaN | NaN | 1323.5691 | NaN | NaN | 222.73953 |
| Ddx5 | WNLDELPKFE | -46 | -55 | 1 | 0.002780285 | 10 | NaN | NaN | 285.67987 | 422.5969 | NaN | NaN |
| Ddx5 | KNFYQEHPDLAR | -56 | -67 | 1 | 0.001092529 | 12 | NaN | NaN | 529.8382 | NaN | NaN | NaN |
| Polr2e | AQFGDKPSEGRPR | -42 | -54 | 2 | 0.002199523 | 13 | NaN | NaN | 413.56177 | NaN | NaN | NaN |
| Echs1 | FLSHWDHITR | -116 | -125 | 0 | 0 | 10 | NaN | NaN | 19898.924 | 609.48535 | 828.4708 | 659.65405 |
| Gsn | VVEHPEFLK | -51 | -59 | 0 | 0 | 9 | NaN | NaN | 534.3091 | NaN | 412.73868 | 892.72363 |
| Gsn | DLGNNIYQWCGSGSNK | -217 | -232 | 0 | 0 | 16 | NaN | NaN | 392.22107 | 478.2685 | NaN | 475.3113 |
| Trap1 | LLQPTLEINPR | -639 | -649 | 0 | 0 | 11 | NaN | NaN | 1002.56 | NaN | NaN | NaN |
| Pola2 | LTSPFEDVFK | -394 | -403 | 0 | 0.00025015 | 10 | NaN | NaN | 633.28546 | NaN | NaN | NaN |
| Gapdh | PASTGAAK | -206 | -213 | 0 | 0.002780285 | 8 | NaN | NaN | 673.0582 | 4116.9707 | 5226.398 | 3610.2148 |
| Gapdh | VHAITATQK | -176 | -184 | 0 | 0 | 9 | NaN | NaN | 607.1049 | 707.359 | 803.198 | 763.8361 |
| Gapdh | RVIISAPSADAPMFV | -116 | -130 | 1 | 0.00025015 | 15 | NaN | NaN | 661.3541 | NaN | NaN | NaN |
| Gapdh | WGEAGAEYVVE | -85 | -95 | 0 | 0.00111006 | 11 | NaN | NaN | 603.77423 | 946.64355 | NaN | 984.9724 |
| Adi1 | NRPADHFDAR | -159 | -168 | 1 | 0.00111006 | 10 | NaN | NaN | 487.50143 | NaN | NaN | 434.9213 |
| Mpst | THEDILENLDAR | -165 | -176 | 0 | 0 | 12 | NaN | NaN | 3415.8406 | NaN | 3464.1543 | NaN |
| Mpst | AQPEHIISEGR | -283 | -293 | 0 | 0 | 11 | NaN | NaN | 2882.8518 | 1860.8926 | NaN | NaN |
| Kdm1a | EAACFPDIISGPQQTQK | -193 | -209 | 0 | 0.002595029 | 17 | NaN | NaN | 1506.9401 | 1249.2003 | NaN | NaN |
| Kdm1a | TLQLWLDNPK | -218 | -227 | 0 | 0 | 10 | NaN | NaN | 1123.8047 | 961.1844 | NaN | NaN |
| Psma1 | IHQIEYAMEAVK | -19 | -30 | 0 | 0.00051306 | 12 | NaN | NaN | 141.15547 | 569.49536 | 204.13885 | NaN |
| Psma1 | AQPSQAAEEPAEK | -244 | -256 | 0 | 0.007819401 | 13 | NaN | NaN | 641.8011 | NaN | NaN | NaN |
| Fkbp8 | QGEYSEAIPILR | -309 | -320 | 0 | 0 | 12 | NaN | NaN | 906.5566 | 1031.2366 | NaN | NaN |
| Atp5f1d | QSELSGAADEAAR | -138 | -150 | 0 | 0 | 13 | NaN | NaN | 1766.0364 | 1756.0232 | 1884.7004 | 1686.5414 |
| Stat3 | VYLYPDIPKEEAFGK | -671 | -685 | 1 | 0.000230707 | 15 | NaN | NaN | 522.1186 | 490.09152 | 629.0137 | 575.73004 |
| Stat3 | FLQESNVLYQH | -71 | -81 | 0 | 0 | 11 | NaN | NaN | 661.1232 | NaN | 1062.8461 | 2063.0513 |
| Hspa8 | QTAEKEEFEHQQK | -585 | -597 | 1 | 0 | 13 | NaN | NaN | 425.8634 | 563.41724 | 214.74072 | 197.5429 |
| Ndufa8 | ARPEPNPVIEGDLKPAK | -146 | -162 | 2 | 0.000230707 | 17 | NaN | NaN | 1921.9619 | 1760.827 | 2337.4285 | 2376.932 |
| Ndufa8 | LGWVRPDLGQL | -115 | -125 | 1 | 0.000712251 | 11 | NaN | NaN | 680.72595 | NaN | NaN | NaN |
| Capza1 | KEASDPQPEDVDGGLK | -103 | -118 | 1 | 0 | 16 | NaN | NaN | 1738.2905 | 2206.8235 | 841.88513 | 1940.7178 |
| Hsp90b1 | SLDQYVER | -523 | -530 | 0 | 0.000431655 | 8 | NaN | NaN | 1752.2283 | 1444.9949 | 1307.1022 | 1323.1816 |
| Hsp90b1 | FDEYGSK | -398 | -404 | 0 | 0.005270854 | 7 | NaN | NaN | 866.3363 | 684.1768 | 552.3679 | 476.19543 |
| Hsp90b1 | MTEAQEDGQSTSELIGQF | -178 | -195 | 0 | 0.000431655 | 18 | NaN | NaN | 328.00183 | 145.10344 | 539.0292 | 416.08636 |
| Cops6 | QVCEIIESPLFLK | -138 | -150 | 0 | 0 | 13 | NaN | NaN | 1010.6212 | NaN | NaN | NaN |
| Dck | ASEDWEVVPEPVAR | -44 | -57 | 0 | 0 | 14 | NaN | NaN | 251.97652 | 492.64456 | 436.39114 | 506.54047 |
| Ranbp1 | NTHADFADECPKPELLAIR | -123 | -141 | 1 | 0 | 19 | NaN | NaN | 1719.4194 | 1792.7136 | 2297.512 | 1135.3552 |
| Tnpo3 | DHEEDFELRK | -810 | -819 | 1 | 0 | 10 | NaN | NaN | 374.20853 | NaN | NaN | 349.1676 |
| Chd1 | ASGEPVPIAEESEELDQK | -1395 | -1412 | 0 | 0.001811524 | 18 | NaN | NaN | 548.77484 | NaN | NaN | NaN |
| Chd1 | NKEPGDIQYLIK | -308 | -319 | 1 | 0 | 12 | NaN | NaN | 526.5376 | NaN | NaN | NaN |
| Exosc10 | SLDELVELNEK | -293 | -303 | 0 | 0 | 11 | NaN | NaN | 541.35254 | 370.3983 | NaN | 894.6174 |
| Ruvbl1 | EEALNHLGEIGTK | -388 | -400 | 0 | 0 | 13 | NaN | NaN | 1674.7657 | 836.33777 | 2128.3564 | 1407.2559 |
| Rpl8 | EGIHTGQFVYCGK | -80 | -92 | 0 | 0 | 13 | NaN | NaN | 610.43274 | 749.0714 | 118.3015 | 822.4023 |
| Acin1 | QIVSEEETPPPLLTK | -410 | -424 | 0 | 0 | 15 | NaN | NaN | 501.05994 | 451.93436 | 531.2426 | NaN |
| Acin1 | ASHALFPEHSGK | -560 | -571 | 0 | 0 | 12 | NaN | NaN | 1968.8618 | NaN | NaN | NaN |
| Acin1 | GVQAGNSDTEGGQPGR | -832 | -847 | 0 | 0 | 16 | NaN | NaN | 513.54333 | NaN | NaN | NaN |
| Arpc2 | SHREPPLELK | -201 | -210 | 1 | 0.002752526 | 10 | NaN | NaN | 649.18774 | 643.47095 | 469.35306 | 605.4959 |
| Polr2b | KGFDQEEVFEKPTR | -821 | -834 | 2 | 0 | 14 | NaN | NaN | 1528.8013 | 1394.8401 | 1684.1276 | 1769.5771 |
| Polr2b | LDDDGLIAPGVR | -848 | -859 | 0 | 0 | 12 | NaN | NaN | 3594.3564 | NaN | NaN | 2453.263 |
| Polr2b | VSGDDVIIGK | -860 | -869 | 0 | 0.001291103 | 10 | NaN | NaN | 1288.1486 | NaN | NaN | NaN |
| Sf3a1 | TVDFQPNEQGNFPPPTTPEELGAR | -284 | -307 | 0 | 0.000627943 | 24 | NaN | NaN | 337.79782 | 340.66785 | 320.2959 | 508.86115 |
| Sf3a1 | LQEQIEAIHK | -520 | -529 | 0 | 0 | 10 | NaN | NaN | 2959.3013 | NaN | NaN | NaN |
| Psmd1 | LDDPQAVSDILEK | -223 | -235 | 0 | 0 | 13 | NaN | NaN | 1891.4799 | 2483.293 | 3004.2732 | 2689.555 |
| Lmnb2 | SDKQGLETEVAELR | -146 | -159 | 1 | 0.00025015 | 14 | NaN | NaN | 1679.5038 | 1620.243 | NaN | 3150.6177 |
| Lmnb2 | VKDLESLFHR | -127 | -136 | 1 | 0.002199523 | 10 | NaN | NaN | 409.87766 | 973.00635 | 328.3961 | NaN |
| Acadvl | GAHQSIGFK | -180 | -188 | 0 | 0.002296977 | 9 | NaN | NaN | 2437.6025 | 2077.3086 | NaN | 2405.2832 |
| Acadvl | EATQAVLDKPETLSSDASTR | -44 | -63 | 1 | 0.000826806 | 20 | NaN | NaN | 905.8533 | NaN | 379.1783 | NaN |
| Acadvl | FFEEVNDPAK | -113 | -122 | 0 | 0 | 10 | NaN | NaN | 1048.3854 | NaN | NaN | NaN |
| Dnmt1 | ASPAGSLPDHVR | -14 | -25 | 0 | 0 | 12 | NaN | NaN | 1792.7192 | 1237.005 | 1002.08716 | 1362.1725 |
| Dnmt1 | LHKEELSEEGYLAK | -68 | -81 | 1 | 0.000230707 | 14 | NaN | NaN | 1643.7178 | 952.2812 | 2230.604 | 2440.0347 |
| Dnmt1 | WYNQEYAR | -872 | -879 | 0 | 0.006788754 | 8 | NaN | NaN | 1860.6431 | NaN | NaN | NaN |
| Dnmt1 | VPALASPAGSLPDHVR | -10 | -25 | 0 | 0.000947459 | 16 | NaN | NaN | 250.50447 | NaN | NaN | NaN |
| Nceh1 | ASPLIAEQEVLEALPK | -324 | -339 | 0 | 0 | 16 | NaN | NaN | 241.02634 | 317.60254 | 480.42047 | 376.91998 |
| Wdr46 | SGAQDPFPGPVPAPLEEAR | -91 | -109 | 0 | 0.000907206 | 19 | NaN | NaN | 1730.529 | NaN | 1540.0094 | 1447.2559 |
| Psmd8 | SQQQKPEDSTIPSTELAK | -323 | -340 | 1 | 0.001463897 | 18 | NaN | NaN | 1142.3796 | NaN | NaN | NaN |
| Cat | ASQRPDVLTTGGGNPIGDK | -20 | -38 | 1 | 0 | 19 | NaN | NaN | 377.76868 | 472.86627 | 488.56674 | 484.0715 |
| Pcbp2 | TIQGQYAIPQPDLTK | -227 | -241 | 0 | 0 | 15 | NaN | NaN | 640.5464 | 968.1586 | 1118.2162 | 1106.895 |
| Ddost | TAVIDHHNYDVS | -139 | -150 | 0 | 0.00272025 | 12 | NaN | NaN | 994.02594 | NaN | NaN | NaN |
| Ddost | TAVIDHHNYDVSDLGQHT | -139 | -156 | 0 | 0.00025015 | 18 | NaN | NaN | 578.5035 | NaN | NaN | NaN |
| Smndc1 | ASSDSFASTQPTHSWK | -59 | -74 | 0 | 0.000471409 | 16 | NaN | NaN | 882.48804 | NaN | NaN | 841.4292 |
| Glrx5 | ASSGGQAEQLDALVK | -32 | -46 | 0 | 0 | 15 | NaN | NaN | 1101.2849 | NaN | 1234.3639 | 1201.3616 |
| Rel | ASSGYNPYVEIIEQPR | -2 | -17 | 0 | 0.009808007 | 16 | NaN | NaN | 547.21375 | NaN | NaN | NaN |
| Phb1 | KAAIISAEGDSK | -208 | -219 | 1 | 0 | 12 | NaN | NaN | 669.4411 | 1029.9426 | 1825.3055 | NaN |
| Pdia3 | RDLFSDGHSEFLK | -161 | -173 | 1 | 0 | 13 | NaN | NaN | 646.3663 | 1157.069 | 1781.0684 | 1652.6646 |
| Pdia3 | KFIQDSIFGLCPH | -234 | -246 | 1 | 0 | 13 | NaN | NaN | 2649.986 | 1586.3593 | 909.7377 | 1344.6206 |
| Pdia3 | TDENFESR | -31 | -38 | 0 | 0 | 8 | NaN | NaN | 452.37875 | 1080.269 | 530.0375 | 709.1513 |
| Pdia3 | FLQEYFDGNLK | -352 | -362 | 0 | 0 | 11 | NaN | NaN | 357.08728 | 452.46484 | 1673.5027 | 1651.9862 |
| Drg1 | ATCPQSELDAETVK | -193 | -206 | 0 | 0 | 14 | NaN | NaN | 1939.0498 | NaN | 2615.9897 | 2497.309 |

| | | | | | | | | | | | | |
|---|---|---|---|---|---|---|---|---|---|---|---|---|
| Drg1 | GQLPDYTSPVVLPY | -300 | -313 | 0 | 0.00025015 | 14 | NaN | NaN | 684.09705 | NaN | 2128.0894 | 2112.5989 |
| Drg1 | DHTLEDEDVIQIVK | -353 | -366 | 0 | 0 | 14 | NaN | NaN | 676.381 | 773.8862 | NaN | NaN |
| Denr | ATDISESSGADCKGDTK | -2 | -18 | 1 | 0.00025015 | 17 | NaN | NaN | 786.66394 | 504.20145 | 368.71692 | 473.91495 |
| Denr | QETGITEGQGPVGEEEEK | -76 | -93 | 0 | 0.000712251 | 18 | NaN | NaN | 660.0405 | 868.0915 | 885.7307 | 924.13153 |
| Denr | EKWPEVDDDSIEDLGEVKK | -180 | -198 | 2 | 0.00025015 | 19 | NaN | NaN | 2711.1646 | NaN | 3200.2036 | 3069.2986 |
| Hp1bp3 | YPSLGLER | -193 | -200 | 0 | 0 | 8 | NaN | NaN | 1699.401 | 1364.2777 | 1607.7747 | NaN |
| Hp1bp3 | YVLENHPGANSNYQ | -367 | -380 | 0 | 0.000550307 | 14 | NaN | NaN | 412.07343 | NaN | NaN | NaN |
| Ubap2l | ATEEWGTEDWNEDLSETK | -260 | -277 | 0 | 0.00025015 | 18 | NaN | NaN | 302.3049 | 247.63431 | 292.7432 | 329.24557 |
| Ubap2l | WEQPQNQNQTQHK | -14 | -26 | 0 | 0 | 13 | NaN | NaN | 767.50104 | NaN | 841.74054 | 798.9035 |
| Ubap2l | KFETAPGAWR | -249 | -258 | 1 | 0.002810811 | 10 | NaN | NaN | 944.6924 | NaN | NaN | NaN |
| Ubap2l | TATEEWGTEDWNEDLSETK | -259 | -277 | 0 | 0.001392869 | 19 | NaN | NaN | 207.85614 | NaN | NaN | NaN |
| Sf3b1 | PPPAGYVPIR | -416 | -425 | 0 | 0.00111006 | 10 | NaN | NaN | 606.57184 | 908.3158 | 1689.7072 | 1393.415 |
| Sf3b1 | LMSPTLEDQER | -539 | -549 | 0 | 0.00149795 | 11 | NaN | NaN | 677.20636 | NaN | NaN | NaN |
| Drg2 | ATEYHLGLLK | -22 | -31 | 0 | 0 | 10 | NaN | NaN | 1375.1416 | NaN | NaN | NaN |
| Eif5a | ATFPMQCSALR | -16 | -26 | 0 | 0.001291103 | 11 | NaN | NaN | 1074.6793 | NaN | NaN | 1053.7565 |
| Rad50 | LIQDQQEQIQHLK | -852 | -864 | 0 | 0.000431655 | 13 | NaN | NaN | 959.7089 | NaN | NaN | NaN |
| Nomo1 | IAPNTPQLADLIAT | -385 | -398 | 0 | 0.007523243 | 14 | NaN | NaN | 567.3309 | NaN | NaN | NaN |
| Myo1g | DEACSTAGPITDR | -475 | -487 | 0 | 0 | 13 | NaN | NaN | 421.37805 | 456.46298 | NaN | NaN |
| Myo1g | DYLSSDTDNPTASHLFAE | -844 | -861 | 0 | 0.002810811 | 18 | NaN | NaN | 201.98941 | NaN | NaN | NaN |
| Mtch2 | ATLITHPFHVITLR | -134 | -147 | 0 | 0 | 14 | NaN | NaN | 324.87036 | 287.72668 | 469.1515 | 366.04996 |
| Pygb | SIGDVVNHDPVVGDR | -626 | -640 | 0 | 0 | 15 | NaN | NaN | 465.58606 | 696.07715 | 683.5696 | 662.4623 |
| Pygb | TCFETFPDK | -325 | -333 | 0 | 0 | 9 | NaN | NaN | 148.65602 | 844.6622 | 506.62027 | 828.3258 |
| Pygb | DFYELEPEKF | -471 | -480 | 1 | 0.00025015 | 10 | NaN | NaN | 967.7699 | NaN | NaN | NaN |
| Pnkp | VVIDNTNPDVPSR | -418 | -430 | 0 | 0.00078661 | 13 | NaN | NaN | 1008.1215 | 680.1559 | NaN | 1457.2428 |
| Hsd17b10 | LGDPAEYAHLV | -227 | -237 | 0 | 0.003461185 | 11 | NaN | NaN | 510.34634 | NaN | NaN | 787.37036 |
| Tcof1 | THWQQTSELGQK | -51 | -62 | 0 | 0 | 12 | NaN | NaN | 1437.1606 | 1445.0721 | 1977.2896 | 1819.4026 |
| Tcof1 | WQQTSELGQK | -53 | -62 | 0 | 0.005884925 | 10 | NaN | NaN | 808.45074 | 1216.2216 | 657.714 | NaN |
| Lsm5 | EDVTEFEITPEGR | -52 | -64 | 0 | 0 | 13 | NaN | NaN | 1196.2943 | NaN | NaN | NaN |
| Psma2 | RYNEDLELEDAIHTA | -177 | -191 | 1 | 0.000947459 | 15 | NaN | NaN | 327.06964 | 143.89204 | NaN | NaN |
| Thbs1 | NALWHTGNTPGQVR | -1078 | -1091 | 0 | 0 | 14 | NaN | NaN | 443.19476 | 903.26843 | 1182.0492 | 837.79895 |
| Thbs1 | EVPDACFNHNGEHR | -594 | -607 | 0 | 0.00111006 | 14 | NaN | NaN | 611.30884 | NaN | NaN | NaN |
| Rpl10a | FPSLLTHNENMVAK | -134 | -147 | 0 | 0 | 14 | NaN | NaN | 719.58167 | NaN | NaN | NaN |
| Rpl6 | HLTDAYFK | -219 | -226 | 0 | 0.001392869 | 8 | NaN | NaN | 3904.5144 | NaN | NaN | NaN |
| Rpl6 | LVTGPLVINR | -182 | -191 | 0 | 0.007987481 | 10 | NaN | NaN | 750.0752 | NaN | NaN | NaN |
| Sdf4 | AVDPDGDGHVSWDEYK | -147 | -162 | 0 | 0.001055699 | 16 | NaN | NaN | 2916.3938 | 2729.0166 | NaN | 1913.1007 |
| Nherf1 | VIPSQEHLDGPLPEPF | -239 | -254 | 0 | 0.001610812 | 16 | NaN | NaN | 996.14044 | 200.73239 | 1662.9541 | 1369.9329 |
| Nherf1 | EALVEPASESPRPALAR | -266 | -282 | 1 | 0.00051306 | 17 | NaN | NaN | 560.2261 | 439.44412 | 460.3977 | 504.62946 |
| Nherf1 | VVDKETDEFFKK | -225 | -236 | 2 | 0 | 12 | NaN | NaN | 546.6257 | 538.09235 | 507.2066 | 556.6681 |
| Lgals3bp | TIASHEDIER | -332 | -341 | 0 | 0 | 10 | NaN | NaN | 1325.817 | NaN | NaN | NaN |
| Tufm | DKPHVNVGTIGHVDHGK | -54 | -70 | 1 | 0.00383302 | 17 | NaN | NaN | 750.27606 | NaN | NaN | NaN |
| Rnf20 | AVEEQIEYLQK | -704 | -714 | 0 | 0.00025015 | 11 | NaN | NaN | 1340.9021 | NaN | NaN | NaN |
| Fyb1 | AVEIDYDSLKR | -554 | -564 | 1 | 0.000550307 | 11 | NaN | NaN | 666.3614 | NaN | NaN | NaN |
| Sh3bgrl3 | VEQDTLQEFLK | -81 | -91 | 0 | 0.00078661 | 11 | NaN | NaN | 1336.9692 | NaN | 1670.2173 | 1464.4238 |
| Aars1 | VIVTEEAIAK | -738 | -747 | 0 | 0.00383302 | 10 | NaN | NaN | 2761.2495 | NaN | NaN | 769.4012 |
| Hcls1 | AVGHEYVADVEK | -98 | -109 | 0 | 0.000230707 | 12 | NaN | NaN | 1201.1361 | 1138.0818 | 1484.2292 | NaN |
| Hcls1 | ISSEVWPPAESHLPPESQPVR | -298 | -318 | 0 | 0.000230707 | 21 | NaN | NaN | 210.09105 | NaN | 551.51636 | NaN |
| Hcls1 | REYPVPSLPTR | -321 | -331 | 1 | 0.006078721 | 11 | NaN | NaN | 1851.965 | NaN | NaN | NaN |
| Flii | NNLELIPESLCR | -327 | -338 | 0 | 0.003393785 | 12 | NaN | NaN | 807.3722 | NaN | NaN | NaN |
| Flii | AVGSGSKDPLAR | -415 | -426 | 1 | 0.00365438 | 12 | NaN | NaN | 704.6765 | NaN | NaN | NaN |
| Flii | KFESLFPGK | -1009 | -1017 | 1 | 0.003299042 | 9 | NaN | NaN | 473.99084 | NaN | NaN | NaN |
| Dis3 | AVHEDIVAVELLPR | -286 | -299 | 0 | 0.00025015 | 14 | NaN | NaN | 383.6477 | NaN | NaN | 418.7027 |
| Wdr1 | SVADGYSENNVFYGHHAK | -517 | -534 | 0 | 0 | 18 | NaN | NaN | 209.60004 | 387.2342 | 516.7281 | 515.1705 |
| Wdr1 | NIDNPAIADIYTEHAHQVVVAK | -44 | -65 | 0 | 0 | 22 | NaN | NaN | 404.03485 | 670.97266 | 840.02563 | 800.01025 |
| Wdr1 | LATGSDDNCAAF | -162 | -173 | 0 | 0 | 12 | NaN | NaN | 363.37457 | 449.45938 | 390.02707 | 419.69748 |
| Wdr1 | LHHVSSL | -576 | -582 | 0 | 0.006100831 | 7 | NaN | NaN | 3043.6177 | 4763.5405 | 2983.018 | NaN |
| Wdr1 | VCIGQIVLLK | -424 | -433 | 0 | 0 | 10 | NaN | NaN | 5803.867 | NaN | NaN | NaN |
| Acsl4 | LFDHAVAK | -82 | -89 | 0 | 0.002810811 | 8 | NaN | NaN | 895.52014 | NaN | NaN | NaN |
| Gale | AVIHFAGLK | -83 | -91 | 0 | 0 | 9 | NaN | NaN | 1082.7256 | NaN | NaN | 880.5767 |
| Pstpip1 | AVILEGEDGWWTVER | -386 | -400 | 0 | 0 | 15 | NaN | NaN | 523.25183 | NaN | 291.2845 | NaN |
| Cert1 | AVITPHDFDECR | -74 | -85 | 0 | 0 | 12 | NaN | NaN | 2792.172 | 1574.9294 | 1409.678 | 1581.9657 |
| Rac2 | HHCPSTPIILVG | -103 | -114 | 0 | 0.000826806 | 12 | NaN | NaN | 1477.4569 | NaN | NaN | NaN |
| Gemin5 | NDSQHLFTEPR | -142 | -152 | 0 | 0 | 11 | NaN | NaN | 488.57764 | NaN | 381.97702 | NaN |
| Calr | TLIVRPDNTYEVK | -173 | -185 | 1 | 0.00025015 | 13 | NaN | NaN | 3903.9697 | 3122.5442 | 3840.3457 | 3817.7317 |
| Calr | IDDPTDSKPEDWDKPEHIPDPDAK | -225 | -248 | 2 | 0 | 24 | NaN | NaN | 2195.1501 | 1902.2278 | 3728.71 | 2779.377 |
| Calr | HEQNIDCGGGY | -99 | -109 | 0 | 0 | 11 | NaN | NaN | 897.11755 | NaN | NaN | 197.03635 |
| Calr | PDAAKPEDWDER | -211 | -222 | 1 | 0.003966045 | 12 | NaN | NaN | 855.5346 | 465.1353 | 904.88165 | NaN |
| Calr | VRPDNTYEVK | -176 | -185 | 1 | 0.002595029 | 10 | NaN | NaN | 586.8431 | NaN | 896.47986 | 1352.4619 |
| Calr | DEAYAEEFGNETWGVTK | -335 | -351 | 0 | 0 | 17 | NaN | NaN | 113.85713 | NaN | NaN | 184.32222 |
| Calr | QFLDGDAWTNR | -26 | -36 | 0 | 0 | 11 | NaN | NaN | 589.21857 | NaN | NaN | NaN |
| Cyb5r3 | DILLRPELEELR | -215 | -226 | 1 | 0 | 12 | NaN | NaN | 1405.0978 | 976.74475 | 1491.7682 | NaN |
| Cyb5r3 | DPNDHTVCYL | -197 | -206 | 0 | 0.009060424 | 10 | NaN | NaN | 159.63605 | NaN | NaN | NaN |
| Tubg1;Tubg2 | NIIQGEVDPTQVHK | (322);(322) | (335);(335) | 0 | 0 | 14 | NaN | NaN | 733.6871 | 582.50134 | 658.379 | 668.6255 |
| Cpne1 | NNLNPTWK | -182 | -189 | 0 | 0.00272025 | 8 | NaN | NaN | 607.3263 | NaN | NaN | NaN |
| Cnot1 | ELKPGNLLK | -1295 | -1303 | 1 | 0.003360691 | 9 | NaN | NaN | 176.15073 | NaN | NaN | NaN |
| Nol6 | LIPHQVVTHL | -619 | -628 | 0 | 0 | 10 | NaN | NaN | 643.6627 | 705.38434 | 1286.7887 | 711.04346 |
| Nol6 | DHVVHLHPLSR | -477 | -487 | 0 | 0 | 11 | NaN | NaN | 505.247 | NaN | 538.90405 | NaN |
| Eif4b | SQSSDTEQPSPTSGGGK | -495 | -511 | 0 | 0 | 17 | NaN | NaN | 372.28568 | 367.57877 | 382.84637 | NaN |
| Eif4b | SLENETLNKE | -445 | -454 | 1 | 0.003069674 | 10 | NaN | NaN | 686.574 | 289.464 | NaN | NaN |

| | | | | | | | | | | | | |
|---|---|---|---|---|---|---|---|---|---|---|---|---|
| Ppp1r12a | LHDTNMELTDLK | -949 | -960 | 0 | 0 | 12 | NaN | NaN | 447.1105 | 320.57703 | 393.21542 | 620.64905 |
| Itgb1 | SHYYDYPSIAHLVQK | -311 | -325 | 0 | 0.00025015 | 15 | NaN | NaN | 849.3383 | NaN | 168.93047 | 825.4217 |
| Itgb1 | LRPEDITQIQPQ | -106 | -117 | 1 | 0.00078661 | 12 | NaN | NaN | 1514.8069 | NaN | NaN | NaN |
| Ywhah | ELETVCNDVLALLDK | -92 | -106 | 0 | 0 | 15 | NaN | NaN | 457.40057 | 601.31775 | 702.5291 | 809.1188 |
| H4c1 | TYTEHAK | -72 | -78 | 0 | 0.007546675 | 7 | NaN | NaN | 1055.212 | 120.371605 | 3896.234 | 3418.8809 |
| Glud1 | GANGPTTPEADKIFLER | -404 | -420 | 1 | 0.00025015 | 17 | NaN | NaN | 701.69336 | 582.58307 | 578.9271 | 468.89777 |
| Glud1 | KGFIGPGIDVPAPDMSTGER | -212 | -231 | 1 | 0 | 20 | NaN | NaN | 675.1585 | 553.6904 | 532.40247 | 908.0329 |
| Glud1 | VTGKPISQGGIHGR | -255 | -268 | 1 | 0 | 14 | NaN | NaN | 513.3282 | 378.46158 | NaN | NaN |
| Glud1 | VTGKPISQGGIHGR | -255 | -268 | 1 | 0 | 14 | NaN | NaN | 513.3282 | 378.46158 | NaN | NaN |
| Eif4e2 | AVVPGPAEHPLQYNYTF | -46 | -62 | 0 | 0.00122419 | 17 | NaN | NaN | 744.186 | NaN | NaN | 1015.3082 |
| Tap2 | GVQQILAGEVTR | -394 | -405 | 0 | 0 | 12 | NaN | NaN | 823.0449 | NaN | 918.2718 | 941.8166 |
| Tap2 | QVVLVGQEPVLFSGSVK | -544 | -560 | 0 | 0.000471409 | 17 | NaN | NaN | 516.091 | 90.209076 | 582.6431 | NaN |
| Tap2 | SFGAEEQEVSHYK | -343 | -355 | 0 | 0 | 13 | NaN | NaN | 1096.014 | NaN | 1245.9886 | NaN |
| Tap2 | AVWAVLSPAGVQEK | -114 | -127 | 0 | 0.000230707 | 14 | NaN | NaN | 423.87357 | NaN | 230.92087 | NaN |
| Tap2 | DAQCEQALQNWR | -637 | -648 | 0 | 0.000230707 | 12 | NaN | NaN | 1568.9512 | NaN | NaN | NaN |
| Tap2 | KPNLPQPGIL | -449 | -458 | 1 | 0.003681885 | 10 | NaN | NaN | 478.7392 | NaN | NaN | NaN |
| Emc1 | AWTSHLWK | -501 | -508 | 0 | 0.00025015 | 8 | NaN | NaN | 1626.2925 | 990.7268 | 1329.3578 | NaN |
| Emc1 | TGEILWR | -71 | -77 | 0 | 0 | 7 | NaN | NaN | 271.32126 | 748.4581 | NaN | NaN |
| Dock11 | LTGLSEISLR | -1789 | -1798 | 0 | 0.000627943 | 10 | NaN | NaN | 707.8573 | NaN | NaN | NaN |
| Usp9x | AYDNLSLSDHLLR | -2146 | -2158 | 0 | 0 | 13 | NaN | NaN | 9836.105 | 10518.241 | 10356.64 | 10516.417 |
| Usp9x | FSEYLLECPSAEVR | -2100 | -2113 | 0 | 0 | 14 | NaN | NaN | 1109.0248 | NaN | 1835.4509 | 1755.1416 |
| Usp9x | DGQLWLCAPQAK | -667 | -678 | 0 | 0 | 12 | NaN | NaN | 4056.9155 | 213.96402 | NaN | NaN |
| Dnaja1 | KILEVHIDK | -206 | -214 | 1 | 0.002136192 | 9 | NaN | NaN | 245.46367 | NaN | NaN | NaN |
| Tmed10 | YEITDQSGGAGGLR | -58 | -71 | 0 | 0 | 14 | NaN | NaN | 1306.7244 | 732.45374 | 761.1271 | 1027.2529 |
| Tmed10 | EITDQSGGAGGLR | -59 | -71 | 0 | 0 | 13 | NaN | NaN | 2882.4941 | NaN | NaN | NaN |
| Cars1 | SKPGEPSWPCPWGK | -409 | -422 | 1 | 0.002265552 | 14 | NaN | NaN | 462.14417 | NaN | 684.2258 | 144.25378 |
| Hspa14 | LIHEPSAALL | -169 | -178 | 0 | 0.002752526 | 10 | NaN | NaN | 5039.066 | NaN | NaN | 2811.3535 |
| Hspa14 | QHTLQAPGR | -427 | -435 | 0 | 0.008947895 | 9 | NaN | NaN | 234.07643 | NaN | NaN | NaN |
| Bclaf1 | ELFDYSPPLHK | -505 | -515 | 0 | 0 | 11 | NaN | NaN | 988.8724 | NaN | 1018.5335 | 443.0703 |
| Prrc2c | AYPVEHSQLETHSK | -821 | -834 | 0 | 0 | 14 | NaN | NaN | 290.24118 | 223.6112 | 339.58667 | 288.6396 |
| Prrc2c | RGPFDDQASGTAGADPVSGN | -1558 | -1577 | 1 | 0.007670622 | 20 | NaN | NaN | 1410.3562 | NaN | NaN | NaN |
| Ranbp9 | AYSDPWNSPVGNQLDPIQR | -580 | -598 | 0 | 0 | 19 | NaN | NaN | 574.82715 | 676.1052 | 587.9701 | 598.17334 |
| Lig1 | CADLSLSPIYPAAR | -844 | -857 | 0 | 0.00266297 | 14 | NaN | NaN | 956.37915 | NaN | NaN | 697.1065 |
| Ankrd44 | CALLILDK | -921 | -928 | 0 | 0 | 8 | NaN | NaN | 1177.5507 | 552.45 | 901.5945 | NaN |
| Alb | YEATLEKCCAE | -377 | -387 | 1 | 0.000230707 | 11 | NaN | NaN | 668.4997 | 297.02322 | 480.51993 | 473.22665 |
| Alb | SLHTLFGD | -89 | -96 | 0 | 0.009745776 | 8 | NaN | NaN | 3254.8381 | NaN | 3364.0437 | NaN |
| Aprt | CDLLHQLR | -140 | -147 | 0 | 0.001291103 | 8 | NaN | NaN | 983.12683 | NaN | NaN | NaN |
| Aprt | SFPDFPIPGVLFR | -15 | -27 | 0 | 0.00272025 | 13 | NaN | NaN | 545.4648 | NaN | NaN | NaN |
| Hk2 | LGLSPLQEDCVATH | -359 | -372 | 0 | 0.00025015 | 14 | NaN | NaN | 958.3368 | NaN | 246.73795 | 3509.4944 |
| Hk2 | RLVPDCDVR | -433 | -441 | 1 | 0 | 9 | NaN | NaN | 1934.9556 | NaN | 2025.743 | NaN |
| Lsp1 | KSLWEQK | -285 | -291 | 1 | 0.007670622 | 7 | NaN | NaN | 1574.9148 | 2120.55 | 2015.0952 | NaN |
| Lsp1 | QQFWGNEGTAEGTEPSQSERPEEK | -97 | -120 | 1 | 0.00122419 | 24 | NaN | NaN | 474.1586 | NaN | NaN | NaN |
| Dock2 | LGFPEIIMPGDVR | -411 | -423 | 0 | 0 | 13 | NaN | NaN | 1049.069 | 634.4055 | 766.9026 | 785.54 |
| Dock2 | FHAHEEATYK | -184 | -193 | 0 | 0 | 10 | NaN | NaN | 189.36108 | 1254.9913 | 558.4181 | NaN |
| Gbp5 | HSFKDEGEFFQK | -370 | -381 | 1 | 0 | 12 | NaN | NaN | 688.95624 | 723.0924 | 323.11615 | NaN |
| Gbp5 | SFKDEGEFFQK | -371 | -381 | 1 | 0.000627943 | 11 | NaN | NaN | 5651.403 | NaN | NaN | 5794.4585 |
| Rpl36 | CGFAPYER | -48 | -55 | 0 | 0.001868731 | 8 | NaN | NaN | 812.2344 | NaN | 924.2095 | 59.545216 |
| Ptprc | FTSWPDHGVPEDPHLLLK | -805 | -822 | 0 | 0 | 18 | NaN | NaN | 641.5241 | 622.53436 | 735.1917 | 689.18774 |
| Ptprc | KTLVSWPEPVSKPESA | -391 | -406 | 2 | 0 | 16 | NaN | NaN | 619.63074 | 938.17395 | 697.0044 | NaN |
| Ptprc | RDPPSDPSPLEAEYQ | -924 | -938 | 1 | 0.001868731 | 15 | NaN | NaN | 199.83183 | 685.07874 | 366.9956 | 574.2596 |
| Ptprc | LEGLEAEGK | -859 | -867 | 0 | 0.00025015 | 9 | NaN | NaN | 954.2713 | NaN | NaN | 1659.2905 |
| Ociad1 | CHEECFWFR | -38 | -46 | 0 | 0.00025015 | 9 | NaN | NaN | 326.0201 | NaN | NaN | NaN |
| Tnfrsf18 | SVVEEPGCGPGK | -22 | -33 | 0 | 0.000431655 | 12 | NaN | NaN | 709.81805 | NaN | NaN | NaN |
| Gzmb | GEIIGGHEVKPHSR | -19 | -32 | 1 | 0 | 14 | NaN | NaN | 940.61584 | 904.22504 | 1138.2012 | 1635.7065 |
| Gzmb | SFLSWIK | -235 | -241 | 0 | 0.002329279 | 7 | NaN | NaN | 78.82566 | 526.0624 | 170.75858 | NaN |
| Lgals1 | EPAFPFQPGSIT | -75 | -86 | 0 | 0 | 12 | NaN | NaN | 1125.0691 | 963.39734 | 671.4209 | 1029.9183 |
| Pebp1 | RVDYAGVTVDELGK | -26 | -39 | 1 | 0.00025015 | 14 | NaN | NaN | 603.8691 | 625.5739 | 1164.1382 | 1080.022 |
| Nutf2 | IQHYYQLFDNDR | -15 | -26 | 0 | 0 | 12 | NaN | NaN | 1328.3423 | 645.4405 | 1731.6545 | 1619.3066 |
| Nutf2 | KLSSLPFQK | -55 | -63 | 1 | 0.00078661 | 9 | NaN | NaN | 334.5864 | NaN | NaN | NaN |
| Ahsg | KCNLLAEK | -218 | -225 | 1 | 0.001392869 | 8 | NaN | NaN | 813.21735 | NaN | NaN | NaN |
| Snrpf | GVEEEEEDGEMRE | -74 | -86 | 1 | 0 | 13 | NaN | NaN | 1440.927 | 1233.3672 | 1599.8691 | 1585.2056 |
| Hectd1 | LVDLPISKPF | -2266 | -2275 | 1 | 0.00078661 | 10 | NaN | NaN | 461.98312 | NaN | 2274.9688 | NaN |
| Hectd1 | GALPDWCEQLTSK | -2063 | -2075 | 0 | 0.000712251 | 13 | NaN | NaN | 934.14294 | NaN | NaN | NaN |
| Micos10 | CQHDFQAPYLLHGK | -58 | -71 | 0 | 0.001055699 | 14 | NaN | NaN | 317.533 | NaN | NaN | NaN |
| Tagln2 | QYDADLEQILIQ | -21 | -32 | 0 | 0 | 12 | NaN | NaN | 842.60376 | 1044.814 | 1220.9464 | 1160.8143 |
| Tagln2 | KLINSLYPEGQAPVK | -64 | -78 | 1 | 0.002199523 | 15 | NaN | NaN | 333.4207 | NaN | NaN | NaN |
| Itgb3 | EQSFTIKPVGFK | -436 | -447 | 1 | 0 | 12 | NaN | NaN | 1615.9116 | NaN | 688.95764 | NaN |
| Itgb3 | CSDETLSQGSPR | -51 | -62 | 0 | 0.000230707 | 12 | NaN | NaN | 1066.4996 | NaN | NaN | 779.1729 |
| Itgb3 | THIALDGR | -279 | -286 | 0 | 0.000755556 | 8 | NaN | NaN | 688.66565 | NaN | NaN | NaN |
| Itgb3 | NEDDCVVR | -684 | -691 | 0 | 0 | 8 | NaN | NaN | 467.5093 | NaN | NaN | NaN |
| Nbeal2 | CSWLVPLVR | -1649 | -1657 | 0 | 0.00025015 | 9 | NaN | NaN | 1377.4491 | NaN | NaN | NaN |
| Eif3i | DPSQIDSNEPYMK | -129 | -141 | 0 | 0.00025015 | 13 | NaN | NaN | 655.3652 | NaN | 867.0354 | 555.793 |
| Eif3i | HLAFEEEFGR | -271 | -280 | 0 | 0 | 10 | NaN | NaN | 619.78516 | 865.0798 | 1028.5312 | 957.868 |
| Ptges3 | DWEDDSDEDMSNFDR | -108 | -122 | 0 | 0.001636916 | 15 | NaN | NaN | 280.2256 | NaN | NaN | NaN |
| Strap | EISGHTSGIK | -138 | -147 | 0 | 0.00025015 | 10 | NaN | NaN | 1269.1047 | 118.23118 | 1192.5278 | 982.41675 |
| Xab2 | SERPDLVFEEE | -10 | -20 | 1 | 0.000712251 | 11 | NaN | NaN | 669.2468 | NaN | NaN | NaN |
| Slc7a5 | EVYGSLPAFLK | -123 | -133 | 0 | 0.001868731 | 11 | NaN | NaN | 1442.8384 | NaN | 1418.7826 | NaN |
| Slc7a5 | DMGQGDASNLQQK | -223 | -235 | 0 | 0.001023236 | 13 | NaN | NaN | 1270.9888 | NaN | NaN | NaN |

| | | | | | | | | | | | | |
|---|---|---|---|---|---|---|---|---|---|---|---|---|
| Cnbp | VSSSLPDICYR | -46 | -56 | 0 | 0 | 11 | NaN | NaN | 1002.54395 | 384.2172 | 1334.3777 | 1129.9065 |
| Cnbp | EQCCYNCGK | -96 | -104 | 0 | 0.00111006 | 9 | NaN | NaN | 274.35107 | 42.863598 | 324.29388 | NaN |
| Srrt | DAAGLECKPR | -405 | -414 | 1 | 0.000471409 | 10 | NaN | NaN | 668.8289 | NaN | 3539.5144 | 3535.2803 |
| Srrt | YHPDEVGK | -201 | -208 | 0 | 0 | 8 | NaN | NaN | 486.47034 | 525.8701 | 576.3939 | NaN |
| Srsf5 | GFGFVEFEDPR | -38 | -48 | 0 | 0 | 11 | NaN | NaN | 500.9025 | NaN | NaN | NaN |
| Nudc | KVEESSWLIEDGK | -228 | -240 | 1 | 0 | 13 | NaN | NaN | 899.51624 | 736.6492 | 1000.4563 | 1223.6864 |
| Nudc | DAEDEEDEEDEKDKGK | -144 | -159 | 2 | 0 | 16 | NaN | NaN | 455.13498 | 1079.7441 | 1022.3589 | 1021.1567 |
| Nudc | DAEDEEDEEDEKDKGK | -144 | -159 | 2 | 0 | 16 | NaN | NaN | 455.13498 | 1079.7441 | 1022.3589 | 1021.1567 |
| Nudc | FMDQHPEMDFSK | -317 | -328 | 0 | 0.00051306 | 12 | NaN | NaN | 585.25635 | 513.0319 | NaN | 675.9258 |
| Ly9 | DAEIEHIIWNCPPK | -72 | -85 | 0 | 0.000394361 | 14 | NaN | NaN | 1492.1074 | NaN | 2250.5918 | NaN |
| Eif3c | VTKPVSGNYGK | -23 | -33 | 1 | 0 | 11 | NaN | NaN | 141.7372 | NaN | NaN | NaN |
| Prpf38a | DAHSIHGTNPQYLVE | -8 | -22 | 0 | 0.001868731 | 15 | NaN | NaN | 1227.4961 | 1314.8627 | NaN | 1731.3486 |
| Prpf38a | LTGTAIDCYK | -109 | -118 | 0 | 0.00025015 | 10 | NaN | NaN | 1378.0878 | NaN | NaN | NaN |
| P4hb | HNQLPLVIEFTEQTAPK | -233 | -249 | 0 | 0 | 17 | NaN | NaN | 789.0011 | 1272.3928 | 1174.7322 | 1135.7224 |
| P4hb | HNQLPLVIEFT | -233 | -243 | 0 | 0.00025015 | 11 | NaN | NaN | 493.03107 | 721.87164 | 820.4328 | NaN |
| P4hb | FEEVAFDEK | -379 | -387 | 0 | 0.000230707 | 9 | NaN | NaN | 315.63824 | 401.409 | NaN | NaN |
| P4hb | IDSDHTDNQR | -293 | -302 | 0 | 0.002296977 | 10 | NaN | NaN | 918.4096 | NaN | NaN | NaN |
| Ttc39b | DALELLRPWAK | -88 | -98 | 1 | 0.001708008 | 11 | NaN | NaN | 662.3076 | NaN | NaN | 762.9568 |
| Ppp6r1 | VSGPLAETNKK | -506 | -516 | 1 | 0.00111006 | 11 | NaN | NaN | 2934.8838 | NaN | NaN | NaN |
| Ppp6r1 | KTDQLVSFLR | -135 | -144 | 1 | 0.001636916 | 10 | NaN | NaN | 949.8349 | NaN | NaN | NaN |
| Nudt5 | DAVSVIPVLQR | -59 | -69 | 0 | 0.002752526 | 11 | NaN | NaN | 896.3808 | 289.5543 | 570.66516 | NaN |
| Atp5f1e | DALKTEFK | -25 | -32 | 1 | 0 | 8 | NaN | NaN | 744.30255 | NaN | NaN | NaN |
| Mia2 | KLTVEENYR | -994 | -1002 | 1 | 0.002752526 | 9 | NaN | NaN | 7974.7983 | NaN | NaN | NaN |
| Kpna4 | VVLDGLSNILK | -430 | -440 | 0 | 0.00025015 | 11 | NaN | NaN | 317.0041 | 304.28244 | NaN | 389.97662 |
| Ikzf3 | SFELLKPPPICLR | -422 | -434 | 1 | 0.000627943 | 13 | NaN | NaN | 1049.6896 | 759.7087 | NaN | 428.83588 |
| Brd4 | DAQEFGADVR | -415 | -424 | 0 | 0 | 10 | NaN | NaN | 1054.9954 | NaN | 755.72266 | NaN |
| Brd4 | GLHDYCDIIK | -387 | -396 | 0 | 0.00025015 | 10 | NaN | NaN | 602.90625 | NaN | NaN | 1270.0887 |
| Arhgdib | ELQEMDKDDESLTK | -33 | -46 | 1 | 0 | 14 | NaN | NaN | 3695.24 | 3117.2825 | 5471.559 | 3906.072 |
| Arhgdib | GDVPVVADPTVPNVTVTR | -53 | -70 | 0 | 0.007670622 | 18 | NaN | NaN | 805.09094 | NaN | NaN | NaN |
| Arhgdib | YVQHTYR | -124 | -130 | 0 | 0.006568856 | 7 | NaN | NaN | 119.62323 | NaN | NaN | NaN |
| Sp100 | SCSLLPQNEER | -212 | -222 | 0 | 0.002265552 | 11 | NaN | NaN | 668.95654 | 3193.786 | NaN | NaN |
| Tpr | FKVESEQQYFEIEK | -120 | -133 | 1 | 0.000431655 | 14 | NaN | NaN | 1405.4518 | NaN | 1340.9617 | 1157.5975 |
| Tpr | FHNELNAH | -349 | -356 | 0 | 0 | 8 | NaN | NaN | 909.6593 | 384.07553 | 487.8591 | NaN |
| Tpr | LLEEDVKR | -1401 | -1408 | 1 | 0.003681885 | 8 | NaN | NaN | 1408.0148 | NaN | NaN | NaN |
| Atl3 | EQASSHIGNSTQAAVR | -512 | -527 | 0 | 0.00111006 | 16 | NaN | NaN | 905.0237 | NaN | 1360.6368 | NaN |
| Ndrg1 | DCGGLPQISQPAK | -288 | -300 | 0 | 0.00272025 | 13 | NaN | NaN | 740.2858 | NaN | NaN | NaN |
| Smc3 | KDQYFLDK | -106 | -113 | 1 | 0 | 8 | NaN | NaN | 10037.468 | NaN | 644.07245 | NaN |
| Smc3 | LDQVEQELNELR | -840 | -851 | 0 | 0.00025015 | 12 | NaN | NaN | 1172.2565 | NaN | NaN | 1624.8044 |
| Smc3 | LHTLEEEKEELAQYQ | -200 | -214 | 1 | 0.008653659 | 15 | NaN | NaN | 913.3965 | NaN | NaN | NaN |
| Magoh;Magol | LEFEFRPDGK | (22);(22) | (31);(31) | 1 | 0.002199523 | 10 | NaN | NaN | 798.6345 | NaN | 541.4934 | 525.632 |
| Hnrnpc | DLDYDFQR | -123 | -130 | 0 | 0.000712251 | 8 | NaN | NaN | 662.8852 | NaN | NaN | NaN |
| Ncl | VTLDWAKPKG | -637 | -646 | 2 | 0.000230707 | 10 | NaN | NaN | 1312.1069 | 555.436 | 1212.3021 | 1238.1791 |
| Ncl | FNSEEDAK | -614 | -621 | 0 | 0 | 8 | NaN | NaN | 425.05502 | NaN | 279.06046 | NaN |
| Ncl | EAMEDGEIDGNKVTLD | -625 | -640 | 1 | 0.001868731 | 16 | NaN | NaN | 550.55237 | NaN | NaN | NaN |
| Ndfip1 | DDFDDTDQLR | -102 | -111 | 0 | 0 | 10 | NaN | NaN | 730.9162 | 793.6097 | 725.6557 | 720.4573 |
| Hnrnpa2b1 | GGGGNFGPGPGSNF | -214 | -227 | 0 | 0 | 14 | NaN | NaN | 109.66415 | 1126.782 | 903.6577 | 951.6668 |
| Hnrnpa2b1 | YHTINGHNAEVR | -174 | -185 | 0 | 0 | 12 | NaN | NaN | 638.9182 | 300.8222 | 1234.3539 | 697.95044 |
| Hnrnpa2b1 | DHDPVDK | -162 | -168 | 0 | 0.006850902 | 7 | NaN | NaN | 551.2616 | NaN | 820.0126 | 623.5786 |
| Rbbp7 | GTHTSDEQNHLVVAR | -68 | -82 | 0 | 0 | 15 | NaN | NaN | 276.42117 | 480.71683 | 563.08923 | 539.50024 |
| Rbbp7 | DDHTVCLWDINAGPK | -197 | -211 | 0 | 0 | 15 | NaN | NaN | 728.99554 | 438.37567 | 159.66121 | 488.51614 |
| Ckap5 | DDIFEPVLIPEPK | -1511 | -1523 | 0 | 0 | 13 | NaN | NaN | 318.36667 | NaN | 1330.4492 | 608.58844 |
| Suclg2 | FDDNAEFR | -263 | -270 | 0 | 0 | 8 | NaN | NaN | 460.5033 | 574.5573 | 556.24 | 588.84784 |
| Tsr1 | DGPPHQVLVVPLHSR | -79 | -93 | 0 | 0 | 15 | NaN | NaN | 763.6576 | NaN | 432.39862 | 551.8224 |
| Sub1 | EYWMDSEGEMKPGR | -87 | -100 | 1 | 0 | 14 | NaN | NaN | 575.76996 | 700.89514 | 808.8537 | 814.2425 |
| Nxn | DDPEGLEFPWGPK | -156 | -168 | 0 | 0 | 13 | NaN | NaN | 95.63003 | 247.53595 | 266.64084 | NaN |
| Skp1 | DFTEEEEAQVR | -144 | -154 | 0 | 0.000712251 | 11 | NaN | NaN | 869.3082 | NaN | NaN | NaN |
| Ptpn6 | DDQVTHIR | -46 | -53 | 0 | 0 | 8 | NaN | NaN | 254.81415 | NaN | 403.4763 | NaN |
| Ptpn6 | DRDGTIIHLK | -88 | -97 | 1 | 0.002972566 | 10 | NaN | NaN | 316.93958 | NaN | NaN | NaN |
| Szrd1 | QPLGPDGSQGFK | -138 | -149 | 0 | 0.001811524 | 12 | NaN | NaN | 657.8845 | NaN | NaN | NaN |
| Tbc1d15 | DDSPTQTLASPNACR | -653 | -667 | 0 | 0.00142999 | 15 | NaN | NaN | 499.04367 | NaN | 200.18411 | 707.38135 |
| Uqcrb | DDTLHETEDVK | -35 | -45 | 0 | 0.00078661 | 11 | NaN | NaN | 150.70221 | 197.53171 | 219.6784 | 1208.6711 |
| Psmc3 | FENLGIQPPK | -215 | -224 | 0 | 0.001392869 | 10 | NaN | NaN | 1625.1318 | NaN | NaN | NaN |
| Ddx39a | NCPHVVVGTPGR | -163 | -174 | 0 | 0.00025015 | 12 | NaN | NaN | 1212.4504 | 288.51346 | 3811.6482 | 3625.149 |
| Ddx39a | LLDYDEDEEPQAPQESTPAPPK | -10 | -31 | 0 | 0.000431655 | 22 | NaN | NaN | 365.3885 | 589.4959 | 767.8966 | 733.0032 |
| Prkcq | KLDNILLDR | -506 | -514 | 1 | 0.00025015 | 9 | NaN | NaN | 1294.7738 | 903.9628 | 829.7246 | 1441.6144 |
| Prkcq | EIDPPFRPK | -646 | -654 | 1 | 0.002329279 | 9 | NaN | NaN | 1280.706 | 991.4616 | NaN | NaN |
| Gmfb | DEELEGVSPDELKDELPER | -46 | -64 | 1 | 0.001055699 | 19 | NaN | NaN | 440.2284 | 568.64307 | 601.97455 | 668.53674 |
| Ranbp2 | DEENASSGADAPSASDTTAK | -2030 | -2049 | 0 | 0.00025015 | 20 | NaN | NaN | 126.14557 | 456.35828 | 315.98535 | 443.22363 |
| Mtco2 | NDLKPGELR | -126 | -134 | 1 | 0 | 9 | NaN | NaN | 1436.7721 | NaN | NaN | NaN |
| Cwc22 | KKDELDPLLTR | -116 | -126 | 2 | 0 | 11 | NaN | NaN | 777.7459 | 746.2073 | 820.068 | 899.029 |
| Parp1 | GGSDDSSKDPIDVNYEK | -779 | -795 | 1 | 0 | 17 | NaN | NaN | 712.42706 | 2102.3772 | 1629.7468 | 1482.1042 |
| Parp1 | VVDRDSEEAEVIR | -802 | -814 | 1 | 0 | 13 | NaN | NaN | 1506.5094 | 333.23767 | 1477.0778 | 1755.124 |
| Parp1 | NELIWNIKDELK | -242 | -253 | 1 | 0 | 12 | NaN | NaN | 378.05026 | 598.2828 | 179.50134 | 578.29596 |
| Eftud2 | ILDAVVAQEPLHR | -790 | -802 | 0 | 0 | 13 | NaN | NaN | 1994.6158 | NaN | 1925.7404 | 1945.623 |
| Eftud2 | LNIRPLLR | -409 | -416 | 1 | 0.002068213 | 8 | NaN | NaN | 433.82693 | NaN | 401.11444 | NaN |
| Snrnp70 | RVNYDTTESK | -109 | -118 | 1 | 0 | 10 | NaN | NaN | 346.39578 | 520.4544 | 324.56534 | 319.92517 |
| Snrnp70 | GYAFIEYEHER | -145 | -155 | 0 | 0 | 11 | NaN | NaN | 899.4572 | NaN | NaN | NaN |
| Snrnp70 | RYDERPGPSPLPHR | -218 | -231 | 2 | 0.00294445 | 14 | NaN | NaN | 429.2846 | NaN | NaN | NaN |

| | | | | | | | | | | | | |
|---|---|---|---|---|---|---|---|---|---|---|---|---|
| Anxa11 | SLYHDITGDTSGDYRK | -478 | -493 | 1 | 0.00204437 | 16 | NaN | NaN | 657.3038 | NaN | NaN | NaN |
| Capzb | SGSGTMNLGGSLTR | -182 | -195 | 0 | 0 | 14 | NaN | NaN | 2234.547 | 751.9473 | 1756.5458 | 312.54373 |
| Capzb | LVEDMENK | -216 | -223 | 0 | 0.00272025 | 8 | NaN | NaN | 1429.8765 | 767.92896 | NaN | NaN |
| Supt16h | VGVDEEIVYAK | -38 | -48 | 0 | 0 | 11 | NaN | NaN | 297.089 | NaN | 530.03455 | 309.88342 |
| Supt16h | DKDAYYR | -6 | -12 | 1 | 0.001392869 | 7 | NaN | NaN | 954.8064 | NaN | NaN | 728.6728 |
| Lrpprc | HDNAEDAWNLK | -723 | -733 | 0 | 0 | 11 | NaN | NaN | 1626.3964 | 1668.5054 | 1728.8774 | 1929.6616 |
| Lrpprc | VYLQNEYK | -170 | -177 | 0 | 0.004628201 | 8 | NaN | NaN | 9238.897 | NaN | NaN | NaN |
| Ctcf | DFEEEQQEGLLSEVNAEK | -227 | -244 | 0 | 0 | 18 | NaN | NaN | 474.22388 | NaN | NaN | 667.3488 |
| G3bp1 | NLPPSGAVPVTGTPPHVV | -256 | -273 | 0 | 0.009416377 | 18 | NaN | NaN | 385.81964 | NaN | NaN | NaN |
| Matr3 | KVDKIEELDQENEAALEN | -715 | -732 | 2 | 0.002434616 | 18 | NaN | NaN | 251.0116 | 585.49866 | 423.9854 | 390.81213 |
| Matr3 | DYEDDRLR | -218 | -225 | 1 | 0.000471409 | 8 | NaN | NaN | 273.97928 | NaN | 1023.761 | 1025.9209 |
| Fkbp5 | VVGEGEDHDIPIGIDK | -189 | -204 | 0 | 0 | 16 | NaN | NaN | 517.7441 | 425.16968 | 728.6572 | 524.6436 |
| Psip1 | KGFNEGLWEIDNNPK | -75 | -89 | 1 | 0.00025015 | 15 | NaN | NaN | 94.76667 | NaN | 3287.8103 | 5029.4453 |
| Psip1 | DFKPGDLIF | -4 | -12 | 1 | 0.00078661 | 9 | NaN | NaN | 729.63806 | NaN | 641.5131 | NaN |
| Aqr | LCLLPTLPK | -384 | -392 | 0 | 0 | 9 | NaN | NaN | 1213.2145 | 371.2106 | 418.8926 | 589.78015 |
| Aqr | DFNDTFLSIEHLK | -713 | -725 | 0 | 0 | 13 | NaN | NaN | 1080.6334 | 878.5888 | 922.54816 | 665.7237 |
| Aqr | HLHIIPTEPFPTSR | -1333 | -1346 | 0 | 0 | 14 | NaN | NaN | 973.90674 | 907.2799 | 1500.8792 | NaN |
| Aqr | VGVPTVDLDAQGR | -1130 | -1142 | 0 | 0.002365814 | 13 | NaN | NaN | 10486.134 | NaN | 205.86337 | NaN |
| Aqr | IVEPHVIPNR | -778 | -787 | 0 | 0.009720262 | 10 | NaN | NaN | 2432.105 | NaN | NaN | NaN |
| Itih2 | HADPDFTKK | -574 | -582 | 1 | 0.004983786 | 9 | NaN | NaN | 235.03827 | NaN | NaN | 193.19676 |
| Camk2g | FTDDYQLFEELGK | -10 | -22 | 0 | 0.000826806 | 13 | NaN | NaN | 371.43915 | 570.8259 | NaN | NaN |
| Nfu1 | DFPTPAAAFR | -85 | -94 | 0 | 0.00450079 | 10 | NaN | NaN | 848.87427 | NaN | NaN | NaN |
| Rbm17 | DFPYEEDSRPR | -191 | -201 | 1 | 0.00025015 | 11 | NaN | NaN | 1610.305 | NaN | 1627.8389 | 1937.4476 |
| Tacc3 | ENQRPLAPVDDAPVVQ | -95 | -110 | 1 | 0.000471409 | 16 | NaN | NaN | 655.63495 | 731.0138 | 574.563 | 822.29004 |
| Mfap1a;Mfap1 | DVFERPSAK | (426);(426) | (434);(434) | 1 | 0.001023236 | 9 | NaN | NaN | 422.7201 | NaN | 457.8953 | 514.5358 |
| Acaa2 | HNFTPLAR | -271 | -278 | 0 | 0.00025015 | 8 | NaN | NaN | 696.39545 | 633.05927 | 501.00586 | 542.4695 |
| Acaa2 | QSIVSGCQEICSK | -97 | -109 | 0 | 0 | 13 | NaN | NaN | 723.39624 | 121.318 | NaN | NaN |
| Prdx6 | DSVEDHLAWSK | -74 | -84 | 0 | 0.00025015 | 11 | NaN | NaN | 879.75946 | NaN | 582.3604 | 667.27264 |
| Prdx6 | PVATPVDWK | -174 | -182 | 0 | 0 | 9 | NaN | NaN | 1491.7461 | NaN | 1741.519 | NaN |
| Anapc1 | SYHPVPTEPLPVPK | -1091 | -1104 | 0 | 0.00025015 | 14 | NaN | NaN | 551.9614 | 475.95355 | 409.23022 | 418.23438 |
| Psmd13 | HRLEELYTK | -23 | -31 | 1 | 0.001023236 | 9 | NaN | NaN | 432.43384 | 633.2426 | 805.7476 | 813.75665 |
| Txnrd1 | FWPLEWTVPSR | -520 | -530 | 0 | 0 | 11 | NaN | NaN | 835.5834 | NaN | 1134.2192 | 1122.0038 |
| Blmh | KTLYNNQPIDFLK | -297 | -309 | 1 | 0 | 13 | NaN | NaN | 537.21014 | 512.035 | 494.38553 | 563.6417 |
| Blmh | ICFVNDPRPQHK | -268 | -279 | 1 | 0 | 12 | NaN | NaN | 615.0711 | 761.34265 | 821.00574 | 719.91724 |
| Ctnnbl1 | LDESVREEADGVHNTL | -204 | -219 | 1 | 0.000591124 | 16 | NaN | NaN | 750.77856 | NaN | 938.2632 | 961.5783 |
| Pak2 | EEDDDDEDAAPPVIAPRPDHTK | -170 | -191 | 1 | 0.00078661 | 22 | NaN | NaN | 715.4597 | NaN | NaN | NaN |
| Gusb | GFDWPLLVK | -356 | -364 | 0 | 0.001463897 | 9 | NaN | NaN | 734.1535 | 585.13354 | NaN | NaN |
| Ubap2 | DGNLANNPYSGDVTK | -868 | -882 | 0 | 0 | 15 | NaN | NaN | 15145.173 | NaN | 15907.688 | NaN |
| Ubap2 | LKPPGLSPFPAAS | -380 | -392 | 1 | 0.008947895 | 13 | NaN | NaN | 600.71594 | NaN | NaN | NaN |
| Ddx19a | IQEHFNK | -454 | -460 | 0 | 0.001291103 | 7 | NaN | NaN | 1787.2373 | 1154.898 | 2151.0483 | 2445.4941 |
| Ddx19a | TTAEKTEEEEKEDR | -42 | -55 | 2 | 0.000230707 | 14 | NaN | NaN | 230.40869 | NaN | 290.23285 | 308.4848 |
| Hadha | QLLLDHANNSSK | -748 | -759 | 0 | 0.00025015 | 12 | NaN | NaN | 879.1576 | 378.6982 | 525.89825 | NaN |
| Rela | NNPFHVPIEEQR | -138 | -149 | 0 | 0.000471409 | 12 | NaN | NaN | 2133.109 | 2368.675 | 1765.1195 | NaN |
| Rela | DGYYEADLCPDR | -97 | -108 | 0 | 0.000627943 | 12 | NaN | NaN | 721.6616 | NaN | 1950.4631 | NaN |
| Rela | EFQYLPDTDDR | -285 | -295 | 0 | 0 | 11 | NaN | NaN | 685.14014 | NaN | NaN | NaN |
| Rela | RTPPYADPSLQAPVR | -253 | -267 | 1 | 0.003360691 | 15 | NaN | NaN | 360.82697 | NaN | NaN | NaN |
| Parp9 | DHHIIENNHILYLGK | -550 | -564 | 0 | 0.005522803 | 15 | NaN | NaN | 641.5188 | NaN | NaN | NaN |
| Ube3a | EETTEYDGGYTR | -766 | -777 | 0 | 0 | 12 | NaN | NaN | 457.58206 | 374.61392 | NaN | 384.6756 |
| Riox2 | DHIVLTVGPDKNPFDEAQQK | -370 | -389 | 1 | 0.00025015 | 20 | NaN | NaN | 3353.6238 | NaN | NaN | NaN |
| Mrpl22 | IVYPPQLPGEPR | -51 | -62 | 0 | 0 | 12 | NaN | NaN | 1165.3807 | 1801.2747 | 1269.9795 | NaN |
| Nol10 | HGTLNVSDTTVGSK | -627 | -640 | 0 | 0 | 14 | NaN | NaN | 681.1161 | 1497.6205 | 1779.9955 | NaN |
| Trim28 | STGPGPMAPPR | -418 | -428 | 0 | 0.005884925 | 11 | NaN | NaN | 677.09717 | NaN | NaN | NaN |
| Supt6h | EFEDLDEIVAR | -1404 | -1414 | 0 | 0.005605646 | 11 | NaN | NaN | 570.4767 | NaN | NaN | NaN |
| Ctsd | NVLPVFDNLMQQK | -208 | -220 | 0 | 0 | 13 | NaN | NaN | 1032.3076 | 270.6552 | 909.42975 | 830.6513 |
| Puf60 | QAFAPFGPIK | -152 | -161 | 0 | 0.00025015 | 10 | NaN | NaN | 1025.2266 | 913.8267 | NaN | NaN |
| Igf2r | LSYTDGDDCGSDKK | -558 | -571 | 1 | 0 | 14 | NaN | NaN | 755.68463 | 963.9975 | 750.3679 | 1135.6191 |
| Igf2r | DIDSLRDPSTQLR | -209 | -221 | 1 | 0.00025015 | 13 | NaN | NaN | 318.4856 | NaN | 1776.1437 | NaN |
| Wdr3 | SLNASLPAPQPVR | -392 | -404 | 0 | 0 | 13 | NaN | NaN | 3592.699 | 3326.498 | 3922.9827 | 4606.3228 |
| Wdr3 | EIDDPEEPEPK | -225 | -235 | 0 | 0.001776473 | 11 | NaN | NaN | 844.7395 | NaN | NaN | NaN |
| Pus10 | DIGFLDDLK | -426 | -434 | 0 | 0 | 9 | NaN | NaN | 347.30786 | 296.91467 | NaN | 235.70375 |
| Lrrc40 | DIHDNQLTSLPSAIR | -111 | -125 | 0 | 0 | 15 | NaN | NaN | 320.5781 | 383.03265 | 445.6999 | 565.0216 |
| Lrrc40 | QATLIPDDLFDATK | -410 | -423 | 0 | 0.005176406 | 14 | NaN | NaN | 554.2129 | NaN | NaN | NaN |
| Son | TSPSGGSEADIEGPLPAK | -1672 | -1689 | 0 | 0.000471409 | 18 | NaN | NaN | 511.38287 | 525.66547 | 544.8214 | NaN |
| Pitrm1 | DIKPILR | -757 | -763 | 1 | 0.000550307 | 7 | NaN | NaN | 2507.382 | 2682.7095 | 147.52744 | NaN |
| Pitrm1 | NNPCFEEEEHFK | -689 | -700 | 0 | 0.003003218 | 12 | NaN | NaN | 359.07166 | NaN | NaN | 663.464 |
| Pitrm1 | SSLLIAGPNSPFYK | -350 | -363 | 0 | 0 | 14 | NaN | NaN | 1801.8265 | NaN | NaN | NaN |
| Rps6ka1 | QVLQHPWITQK | -657 | -667 | 0 | 0 | 11 | NaN | NaN | 509.96332 | 419.15152 | NaN | NaN |
| Glrx3 | SNWPTYPQLYVR | -299 | -310 | 0 | 0 | 12 | NaN | NaN | 1599.6637 | NaN | 2182.2566 | 2108.7134 |
| Rbm25 | IPTAKPELF | -740 | -748 | 1 | 0.002265552 | 9 | NaN | NaN | 1449.1792 | NaN | 1367.7377 | 1427.467 |
| Rbm25 | KLVPLDYGEDDKNATK | -706 | -721 | 2 | 0 | 16 | NaN | NaN | 922.7165 | NaN | 226.57938 | 1359.1324 |
| Hcfc1 | KTCLPGFPGAPCAIK | -1894 | -1908 | 1 | 0.007734155 | 15 | NaN | NaN | 112.806526 | NaN | NaN | NaN |
| Wbp11 | DIPLPGAQPPSILK | -155 | -168 | 0 | 0.00201047 | 14 | NaN | NaN | 753.52637 | NaN | 785.7149 | NaN |
| Ap2b1 | DIPNENELQFQIK | -843 | -855 | 0 | 0 | 13 | NaN | NaN | 1793.3276 | 1789.8292 | 1985.4084 | 2755.9272 |
| Gnl3 | ELSPEQSTAGKPSDGSSALDR | -503 | -523 | 1 | 0.001708008 | 21 | NaN | NaN | 3402.5195 | NaN | 1633.1097 | NaN |
| Ehd1 | FQALKPK | -371 | -377 | 1 | 0.00930008 | 7 | NaN | NaN | 297.5472 | NaN | NaN | NaN |
| Cpne3 | GHQWYEVER | -39 | -47 | 0 | 0.005605646 | 9 | NaN | NaN | 795.8438 | NaN | NaN | NaN |
| Arhgef1 | LEELEEEFCR | -883 | -892 | 0 | 0 | 10 | NaN | NaN | 618.7854 | 1070.1282 | 1402.6427 | 1394.4917 |
| Arhgef1 | LSHLEETQHTISTDEEK | -196 | -212 | 0 | 0.00025015 | 17 | NaN | NaN | 590.3783 | 659.9485 | 565.64825 | 611.5434 |

| | | | | | | | | | | | | |
|---|---|---|---|---|---|---|---|---|---|---|---|---|
| Arhgef1 | QLGGTLSPNLAAPER | -899 | -913 | 0 | 0 | 15 | NaN | NaN | 447.44525 | 605.9685 | 652.6673 | 692.20544 |
| Arhgef1 | DKAIEVHVL | -660 | -668 | 1 | 0.000230707 | 9 | NaN | NaN | 1578.4199 | NaN | 1178.6257 | NaN |
| Irf3 | NSQPISLTSDQYK | -389 | -401 | 0 | 0.003620005 | 13 | NaN | NaN | 328.95963 | NaN | 269.58078 | NaN |
| Cast | SNDTSQTPPGETVPR | -473 | -487 | 0 | 0 | 15 | NaN | NaN | 1416.2465 | 1237.0326 | NaN | NaN |
| Naa15 | LHDNPLTDENKEHEADTA | -568 | -585 | 1 | 0.00111006 | 18 | NaN | NaN | 445.4446 | 506.62735 | 670.0046 | 688.6274 |
| Naa15 | LFNPNDDGKEEPPTTLL | -358 | -374 | 1 | 0.00051306 | 17 | NaN | NaN | 2233.8591 | NaN | 3309.1497 | 3165.7354 |
| Naa15 | LHDNPLTDENKEHEA | -568 | -582 | 1 | 0.001392869 | 15 | NaN | NaN | 862.64844 | NaN | NaN | NaN |
| Srrm2 | DKFSPTQDRPESSTVLK | -1148 | -1164 | 2 | 0.00149795 | 17 | NaN | NaN | 425.40085 | NaN | 439.37564 | 403.8137 |
| Srrm2 | HASSSPESLKPTPAPGSR | -431 | -448 | 1 | 0.001392869 | 18 | NaN | NaN | 332.24924 | NaN | 91.90665 | 559.1021 |
| Nono | EELHNQEVQK | -329 | -338 | 0 | 0.000471409 | 10 | NaN | NaN | 1202.6521 | 963.7831 | 1115.6741 | 1113.1144 |
| Usp4 | VIEPQNEDGTWPR | -214 | -226 | 0 | 0.001055699 | 13 | NaN | NaN | 810.5901 | 1212.7139 | NaN | NaN |
| Tmed9 | LYFHIGETEKK | -38 | -48 | 1 | 0 | 11 | NaN | NaN | 758.26794 | 777.017 | 427.4142 | 530.98413 |
| Rps4x | ITDFIKF | -169 | -175 | 1 | 0.00025015 | 7 | NaN | NaN | 1065.9569 | 1716.1835 | 2020.5396 | 2037.8245 |
| Rps4x | PGSFDVVHVK | -202 | -211 | 0 | 0.005605646 | 10 | NaN | NaN | 1624.4963 | NaN | 2490.4597 | 2230.4807 |
| Rasa3 | SLKPDDLGSLR | -274 | -284 | 1 | 0 | 11 | NaN | NaN | 2804.9712 | 4413.923 | NaN | 2703.3967 |
| Rasa3 | DKQEAAIPLVR | -330 | -340 | 1 | 0 | 11 | NaN | NaN | 340.29114 | 436.31854 | 515.41656 | NaN |
| Ube2f | DKVDEYIK | -174 | -181 | 1 | 0 | 8 | NaN | NaN | 1501.5977 | 1788.9518 | 1622.1235 | 1674.6248 |
| Ruvbl2 | KGTEVQVDDIKR | -417 | -428 | 2 | 0.00025015 | 12 | NaN | NaN | 183.82117 | 317.5948 | 209.50885 | 357.34286 |
| Ruvbl2 | EMETIYDLGTK | -167 | -177 | 0 | 0.000230707 | 11 | NaN | NaN | 1890.5939 | 5855.7725 | NaN | NaN |
| Ruvbl2 | QSPHGIPIDLLDR | -341 | -353 | 0 | 0.000471409 | 13 | NaN | NaN | 614.2837 | NaN | NaN | 857.2916 |
| Ruvbl2 | GLGLDDALEPR | -30 | -40 | 0 | 0.002595029 | 11 | NaN | NaN | 1731.5813 | NaN | NaN | NaN |
| Baz1a | DLDPDVVTEDEDDPGSHK | -725 | -742 | 0 | 0.000826806 | 18 | NaN | NaN | 414.39554 | 282.1882 | 346.67947 | 460.12708 |
| Baz1a | HYDDFFER | -36 | -43 | 0 | 0 | 8 | NaN | NaN | 983.3871 | NaN | NaN | 823.3732 |
| Baz1a | FHFSDKPQADSKPVSSR | -943 | -959 | 2 | 0.000230707 | 17 | NaN | NaN | 369.58612 | NaN | NaN | NaN |
| Hdgf | IDEMPEAAVK | -30 | -39 | 0 | 0.006850902 | 10 | NaN | NaN | 380.96948 | 550.16376 | NaN | NaN |
| Sf3a3 | SSWEELASLGLDR | -250 | -262 | 0 | 0 | 13 | NaN | NaN | 420.17502 | 613.053 | 698.6019 | 656.43616 |
| Sf3a3 | KEELNAISGPNEFAE | -69 | -83 | 1 | 0.000627943 | 15 | NaN | NaN | 2219.2998 | NaN | NaN | NaN |
| Aga | DLHSDWLSR | -153 | -161 | 0 | 0.000471409 | 9 | NaN | NaN | 267.57974 | NaN | NaN | NaN |
| Nsfl1c | FYAGGSER | -94 | -101 | 0 | 0.002752526 | 8 | NaN | NaN | 836.1442 | NaN | NaN | NaN |
| Tpt1 | FKDGLEMEK | -163 | -171 | 1 | 0 | 9 | NaN | NaN | 2686.148 | 2699.4614 | 2985.4841 | 2923.8748 |
| Tpp2 | TDVHFKPGQIR | -650 | -660 | 1 | 0.001291103 | 11 | NaN | NaN | 486.9255 | NaN | NaN | NaN |
| Prkaca | KWETPSQNTAQLDQFDR | -30 | -46 | 1 | 0 | 17 | NaN | NaN | 938.70886 | NaN | 421.84167 | NaN |
| Ripk3 | DLKPSNILLDPELHAK | -143 | -158 | 1 | 0 | 16 | NaN | NaN | 561.6542 | 388.46146 | 317.46014 | 588.55365 |
| Pgls | SLFPDHPLLQER | -157 | -168 | 0 | 0 | 12 | NaN | NaN | 1268.4293 | 1620.4713 | 1813.0822 | 1693.6917 |
| Arl8b | PNALDEK | -135 | -141 | 0 | 0.002329279 | 7 | NaN | NaN | 433.81683 | NaN | 625.5817 | 443.89865 |
| Bop1 | DLQPFPVCQAL | -383 | -393 | 0 | 0.001055699 | 11 | NaN | NaN | 714.0831 | 684.367 | 972.3345 | NaN |
| Bop1 | RWLEVSEEEHQR | -500 | -511 | 1 | 0 | 12 | NaN | NaN | 348.87674 | NaN | NaN | 370.01337 |
| Gtpbp4 | LEELEKEEELR | -450 | -460 | 1 | 0.00025015 | 11 | NaN | NaN | 828.9726 | NaN | NaN | NaN |
| Irgm1 | LLQNIQENIR | -206 | -215 | 0 | 0 | 10 | NaN | NaN | 1499.4583 | NaN | NaN | NaN |
| Emg1 | NVLIEVNPQTR | -115 | -125 | 0 | 0.00111006 | 11 | NaN | NaN | 658.18353 | NaN | NaN | NaN |
| Uqcrfs1 | LSDIPEGK | -156 | -163 | 0 | 0.000471409 | 8 | NaN | NaN | 344.7574 | NaN | NaN | NaN |
| Uqcrfs1 | YCPCHGSHYDASGR | -235 | -248 | 0 | 0 | 14 | NaN | NaN | 308.82513 | NaN | NaN | NaN |
| Ctsc | DPVTGIEYWIIK | -414 | -425 | 0 | 0 | 12 | NaN | NaN | 182.7099 | 395.78226 | 278.73895 | 248.50494 |
| Ndufb8 | GGDPTKEPEPVVHYDI | -171 | -186 | 1 | 0 | 16 | NaN | NaN | 572.3999 | 776.82715 | 1193.4038 | 1110.9861 |
| Cd27 | SSHRPLCSSDCIR | -170 | -182 | 1 | 0 | 13 | NaN | NaN | 860.0093 | NaN | 548.7781 | NaN |
| Farsa | FQPQQHPAR | -263 | -271 | 0 | 0.002168954 | 9 | NaN | NaN | 983.06555 | NaN | 130.40176 | 832.9378 |
| Farsa | GVLPDSGHLHPLLK | -216 | -229 | 0 | 0 | 14 | NaN | NaN | 1521.5638 | NaN | NaN | 489.946 |
| Farsa | KWVEVGNSGVFRPE | -428 | -441 | 2 | 0.008129024 | 14 | NaN | NaN | 543.2629 | NaN | NaN | NaN |
| Ddx3x | GRGDYDGIGGR | -100 | -110 | 1 | 0.00111006 | 11 | NaN | NaN | 459.81384 | NaN | NaN | NaN |
| Dkc1 | NTLVTEAVQAPQLAAE | -425 | -440 | 0 | 0.001636916 | 16 | NaN | NaN | 224.2753 | NaN | 651.12067 | 657.18164 |
| Aarsd1 | SLDDPEVEQVR | -174 | -184 | 0 | 0 | 11 | NaN | NaN | 573.81464 | 375.00494 | NaN | NaN |
| Aarsd1 | TESPLSPGSQVQVR | -83 | -96 | 0 | 0 | 14 | NaN | NaN | 258.9476 | NaN | NaN | 332.37952 |
| Aarsd1 | HFTESPLSPGSQVQVR | -81 | -96 | 0 | 0.002099344 | 16 | NaN | NaN | 4086.968 | NaN | NaN | NaN |
| Nme2 | DRPFFPGLV | -57 | -65 | 1 | 0.002199523 | 9 | NaN | NaN | 530.4228 | NaN | 386.01318 | 464.99222 |
| Farsb | SEVIGDGNQIEVEIPPTR | -338 | -355 | 0 | 0.005270854 | 18 | NaN | NaN | 565.5058 | NaN | NaN | 558.7275 |
| Farsb | DRYDSFIELQ | -134 | -143 | 1 | 0.00025015 | 10 | NaN | NaN | 446.8477 | NaN | 479.2631 | NaN |
| Try5;Try4 | VGFLEGGK | (186);(186) | (193);(193) | 0 | 0.00223472 | 8 | NaN | NaN | 615.6556 | NaN | NaN | NaN |
| Nars1 | SWDSEEILEGYKR | -490 | -502 | 1 | 0.002845514 | 13 | NaN | NaN | 2941.157 | NaN | NaN | NaN |
| Mat2a | KYLDEDTIYHLQPSGR | -234 | -249 | 1 | 0.00051306 | 16 | NaN | NaN | 446.24902 | 516.6278 | 615.1043 | 490.696 |
| Mat2a | VACETVAK | -54 | -61 | 0 | 0.003461185 | 8 | NaN | NaN | 1338.1671 | NaN | NaN | NaN |
| Sash3 | VHTDFTPSPYDHDSLK | -180 | -195 | 0 | 0 | 16 | NaN | NaN | 1966.7354 | 2516.4834 | 2336.8174 | 2214.393 |
| Gls | GGTPPQQQQQQQQQPGASPPAAPGPK | -86 | -111 | 0 | 0.005605646 | 26 | NaN | NaN | 130.17339 | NaN | 266.81668 | NaN |
| Thrap3 | EYPEEAEEREESTSGFDK | -791 | -808 | 1 | 0 | 18 | NaN | NaN | 382.1933 | 539.3313 | 606.21313 | 526.0937 |
| Thrap3 | MDSFDEDLARPS | -570 | -581 | 1 | 0 | 12 | NaN | NaN | 312.6514 | 287.82175 | 289.06226 | 307.1759 |
| Thrap3 | YYLHDDR | -876 | -882 | 0 | 0.00051306 | 7 | NaN | NaN | 1141.3617 | NaN | NaN | NaN |
| Thrap3 | DSRPSQAAGDNQGDEAK | -186 | -202 | 1 | 0.00223472 | 17 | NaN | NaN | 260.79578 | NaN | NaN | NaN |
| Itk | DSSEIHWWR | -208 | -216 | 0 | 0 | 9 | NaN | NaN | 231.28871 | 357.19818 | 319.91336 | 350.9303 |
| Tm9sf2 | DSTEDAQEEFGWK | -336 | -348 | 0 | 0 | 13 | NaN | NaN | 517.1368 | NaN | NaN | NaN |
| Hnrnpa3 | DSVKPGAHLTVK | -115 | -126 | 1 | 0.000230707 | 12 | NaN | NaN | 468.22467 | NaN | NaN | NaN |
| Nusap1 | NQDENQFSTDETEIHVS | -53 | -69 | 0 | 0 | 17 | NaN | NaN | 235.40804 | NaN | 545.8751 | 332.57666 |
| Arpc3 | LYISECLKK | -73 | -81 | 1 | 0.00078661 | 9 | NaN | NaN | 978.6189 | NaN | NaN | NaN |
| Wdr43 | DTHICLER | -367 | -374 | 0 | 0 | 8 | NaN | NaN | 2755.529 | 2686.977 | 2661.8455 | 2659.2722 |
| Wdr43 | GNWFQPTIER | -351 | -360 | 0 | 0.00025015 | 10 | NaN | NaN | 953.91956 | 1128.6741 | 677.92706 | NaN |
| Wdr43 | VYGNWFQPTIER | -349 | -360 | 0 | 0 | 12 | NaN | NaN | 486.63943 | NaN | 573.4868 | NaN |
| Xpnpep1 | GLDYTGISWK | -170 | -179 | 0 | 0 | 10 | NaN | NaN | 1439.1423 | NaN | 1915.5146 | NaN |
| Ppp3r1 | SHFDADEIKR | -13 | -22 | 1 | 0 | 10 | NaN | NaN | 890.04535 | NaN | 315.57642 | NaN |
| Inpp5d | IVVLAKPEHENR | -493 | -504 | 1 | 0.00078661 | 12 | NaN | NaN | 442.09833 | 133.3831 | 507.67673 | 463.3517 |
| Inpp5d | EGENEEGSEGELVVR | -768 | -782 | 0 | 0 | 15 | NaN | NaN | 964.63586 | NaN | 75.66916 | 977.17456 |

| | | | | | | | | | | | | |
|---|---|---|---|---|---|---|---|---|---|---|---|---|
| Eif2b5 | DTTDEGQQLR | -685 | -694 | 0 | 0 | 10 | NaN | NaN | 903.6695 | NaN | NaN | NaN |
| Eif6 | HGLLVPNNTTDQELQHIR | -68 | -85 | 0 | 0 | 18 | NaN | NaN | 2191.5261 | 3141.924 | 3007.8704 | 117.24656 |
| Mcm2 | IAPSIYGHEDIKR | -481 | -493 | 1 | 0 | 13 | NaN | NaN | 259.41278 | 966.98596 | 1002.73267 | 1039.8313 |
| Mcm2 | QDTIEIPEKDLMDK | -855 | -868 | 1 | 0.00078661 | 14 | NaN | NaN | 3792.6475 | 935.71484 | 2563.7485 | 2582.164 |
| G6pdx | VGFQYEGTYK | -499 | -508 | 0 | 0 | 10 | NaN | NaN | 154.26456 | 4931.5874 | 4216.915 | 4214.2305 |
| Snx6 | TFLLEYHNR | -215 | -223 | 0 | 0 | 9 | NaN | NaN | 881.5191 | NaN | 1390.3173 | 1215.5793 |
| Notch1 | DVDECALGANPCEHAGK | -412 | -428 | 0 | 0.000907206 | 17 | NaN | NaN | 666.3451 | 633.9429 | 598.8578 | NaN |
| Iah1 | LCPLLDKK | -213 | -220 | 1 | 0.003681885 | 8 | NaN | NaN | 1480.3544 | NaN | NaN | NaN |
| Npepps | DVFSPIGER | -682 | -690 | 0 | 0 | 9 | NaN | NaN | 836.329 | 1409.2379 | 610.4096 | 1672.6705 |
| Npepps | IYVEAEQVEDDR | -507 | -518 | 0 | 0.000230707 | 12 | NaN | NaN | 1840.2979 | NaN | NaN | NaN |
| Npepps | LGWDPKPGEGHLD | -691 | -703 | 1 | 0.002265552 | 13 | NaN | NaN | 951.35956 | NaN | NaN | NaN |
| Coq3 | DVGCGGGLLTEPLGR | -153 | -167 | 0 | 0 | 15 | NaN | NaN | 710.1769 | NaN | 788.2917 | NaN |
| Evl | VKPAGSVNDVGLDALDLDR | -362 | -380 | 1 | 0 | 19 | NaN | NaN | 288.9447 | NaN | 463.94183 | 416.34436 |
| Evl | NIMNSQEGGPSTQR | -110 | -123 | 0 | 0.009631128 | 14 | NaN | NaN | 911.2511 | NaN | NaN | NaN |
| Nucks1 | VVDYSQFQESDDADEDYGR | -10 | -28 | 0 | 0.00111006 | 19 | NaN | NaN | 179.69258 | NaN | NaN | NaN |
| Pold1 | WSDVISHPPEGQWQR | -290 | -304 | 0 | 0.007105559 | 15 | NaN | NaN | 207.27644 | 484.05133 | NaN | NaN |
| Naa50 | RIEPADAHVL | -141 | -150 | 1 | 0.002296977 | 10 | NaN | NaN | 1629.2123 | NaN | 1289.4695 | 1063.9564 |
| Naa50 | DVLEVGELAK | -38 | -47 | 0 | 0.00025015 | 10 | NaN | NaN | 386.43118 | NaN | 627.11884 | 1142.6362 |
| Park7 | GAEEMETVIPVDVMR | -13 | -27 | 0 | 0.00025015 | 15 | NaN | NaN | 611.1475 | NaN | 1358.2516 | NaN |
| Ppp2r5e | VLIPLHTVR | -267 | -275 | 0 | 0.000591124 | 9 | NaN | NaN | 292.86792 | NaN | NaN | NaN |
| Dnaja2 | ILEVHVDK | -216 | -223 | 0 | 0.007009655 | 8 | NaN | NaN | 1229.8704 | NaN | NaN | NaN |
| Diaph1 | DVQNFPAATDEKDK | -1055 | -1068 | 1 | 0.000907206 | 14 | NaN | NaN | 394.72266 | 511.9604 | NaN | NaN |
| Diaph1 | FPDELAHVE | -1020 | -1028 | 0 | 0.002068213 | 9 | NaN | NaN | 386.87265 | NaN | NaN | NaN |
| Calu | IDDDKDGFVTVDELK | -80 | -94 | 1 | 0.002595029 | 15 | NaN | NaN | 595.16364 | NaN | NaN | 490.69073 |
| Gdi2 | KFVSISDLFVPK | -379 | -390 | 1 | 0.00078661 | 12 | NaN | NaN | 502.62448 | 523.535 | 180.55714 | 649.5154 |
| Gdi2 | MTGSEFDFEEMKR | -424 | -436 | 1 | 0.00025015 | 13 | NaN | NaN | 306.20407 | 309.07907 | 256.12964 | 328.05396 |
| Ddx17 | KWDLSELPK | -42 | -50 | 1 | 0.000591124 | 9 | NaN | NaN | 3108.7441 | 3182.1375 | 3667.3625 | 3275.7664 |
| Ddx17 | LTPYEVDELRR | -65 | -75 | 1 | 0.00025015 | 11 | NaN | NaN | 1940.0532 | 2173.7156 | 2235.531 | 2199.4297 |
| Rab35 | WLHEINQNCDDVCR | -102 | -115 | 0 | 0.00025015 | 14 | NaN | NaN | 1050.0602 | 1829.176 | 242.42249 | 45.044273 |
| Morf4l1 | KYVDTNLQK | -103 | -111 | 1 | 0.001392869 | 9 | NaN | NaN | 547.6085 | NaN | 631.9903 | 544.57947 |
| Hnrnpa1 | TFDDHDSVDK | -152 | -161 | 0 | 0 | 10 | NaN | NaN | 342.04663 | 1153.4921 | 812.9146 | 723.8041 |
| Hnrnpa1 | SPKEPEQLR | -6 | -14 | 1 | 0.00635324 | 9 | NaN | NaN | 4472.339 | NaN | 485.24152 | NaN |
| RTRAF | DYHNPSGFNCKDETEFR | -10 | -26 | 1 | 0.003133457 | 17 | NaN | NaN | 6784.236 | NaN | NaN | NaN |
| RTRAF | DYHNPSGFNCK | -10 | -20 | 0 | 0 | 11 | NaN | NaN | 727.16644 | NaN | NaN | NaN |
| Pura | VSEVKPTYR | -245 | -253 | 1 | 0 | 9 | NaN | NaN | 605.29846 | NaN | NaN | NaN |
| Rps10 | NEGIQYLR | -73 | -80 | 0 | 0.000230707 | 8 | NaN | NaN | 3679.7903 | 2341.7886 | 1794.3481 | 1987.8702 |
| Pfkl | YVFEDPFNIHDLK | -593 | -605 | 0 | 0 | 13 | NaN | NaN | 1715.4994 | 920.65393 | 1480.8771 | 2151.4387 |
| Crnkl1 | FVLVHPAVK | -208 | -216 | 0 | 0 | 9 | NaN | NaN | 406.44308 | NaN | NaN | NaN |
| Trappc5 | EADKLEQANDDAR | -101 | -113 | 1 | 0.00122419 | 13 | NaN | NaN | 540.04895 | NaN | 606.7714 | NaN |
| Sell | EIEYLENTLPK | -72 | -82 | 0 | 0 | 11 | NaN | NaN | 1353.1006 | 792.45276 | 964.3898 | 986.23645 |
| Adprs | EAFDEVDMAHR | -99 | -109 | 0 | 0.001636916 | 11 | NaN | NaN | 703.3014 | NaN | NaN | NaN |
| Vasp | QNGPSPEELEQQK | -130 | -142 | 0 | 0.008045078 | 13 | NaN | NaN | 4608.6387 | NaN | NaN | NaN |
| Sp1 | EAICPEGIAR | -751 | -760 | 0 | 0.00025015 | 10 | NaN | NaN | 789.68225 | 462.28397 | 556.12756 | 754.9579 |
| Eif3f | IGVDLIMK | -251 | -258 | 0 | 0 | 8 | NaN | NaN | 1110.0912 | 713.8633 | 1046.1558 | 1098.319 |
| Rnmt | HYNELQEVGLAK | -141 | -152 | 0 | 0 | 12 | NaN | NaN | 347.11697 | 626.74585 | NaN | 458.26196 |
| Ethe1 | EAVLIDPVLETAHR | -47 | -60 | 0 | 0 | 14 | NaN | NaN | 4812.869 | 2875.5623 | 3413.2698 | 3292.6128 |
| Ethe1 | SQQSASGAPVLLR | -14 | -26 | 0 | 0.00025015 | 13 | NaN | NaN | 4338.939 | NaN | 5455.455 | NaN |
| Zwilch | EAVNHDAAAVVR | -323 | -334 | 0 | 0 | 12 | NaN | NaN | 612.7424 | NaN | 572.8203 | NaN |
| Ndufv1 | SFGSLKDEDR | -31 | -40 | 1 | 0 | 10 | NaN | NaN | 1275.1118 | 1100.2117 | 2152.3232 | 1625.4783 |
| Ndufv1 | EAYEAGLIGK | -175 | -184 | 0 | 0 | 10 | NaN | NaN | 452.95276 | NaN | NaN | 846.8183 |
| Adh5 | EFGASECISPQDFSK | -234 | -248 | 0 | 0 | 15 | NaN | NaN | 642.1541 | 692.9528 | 898.4203 | 850.3889 |
| Elac2 | FGDFPTVPK | -750 | -758 | 0 | 0.004321969 | 9 | NaN | NaN | 3159.4475 | 4196.0376 | NaN | NaN |
| Elac2 | LVVAPTQLR | -573 | -581 | 0 | 0.000394361 | 9 | NaN | NaN | 325.82306 | 300.64465 | NaN | NaN |
| Huwe1 | ECPFVIKPK | -470 | -478 | 1 | 0.00051306 | 9 | NaN | NaN | 280.62198 | 330.2356 | NaN | 533.84534 |
| Huwe1 | TLTSIVHLER | -330 | -339 | 0 | 0 | 10 | NaN | NaN | 2074.16 | NaN | NaN | NaN |
| Itgb7 | ECSEADLSSPDLESGCR | -512 | -528 | 0 | 0.002168954 | 17 | NaN | NaN | 241.58968 | NaN | NaN | NaN |
| Ola1 | QLEYFFTAGPDEVR | -304 | -317 | 0 | 0 | 14 | NaN | NaN | 539.17804 | 457.2557 | 856.39166 | 722.41187 |
| Ola1 | GGDGIKPPPIIGR | -6 | -18 | 1 | 0 | 13 | NaN | NaN | 939.719 | 2208.7107 | NaN | NaN |
| Eed | EDHNQPLFGVQ | -90 | -100 | 0 | 0.00383302 | 11 | NaN | NaN | 1475.6887 | NaN | NaN | NaN |
| Rad23a | SPSEESTTTTSPESISGSVPSSGSSGR | -123 | -149 | 0 | 0.004983786 | 27 | NaN | NaN | 151.14459 | NaN | 222.96431 | 227.03766 |
| Ssb | ITDDQQESLNK | -317 | -327 | 0 | 0 | 11 | NaN | NaN | 177.37863 | 2309.2742 | 1051.4896 | 1955.7375 |
| Ssb | SPSRPLPEVTDEYKN | -92 | -106 | 2 | 0.004658491 | 15 | NaN | NaN | 795.28577 | 500.65936 | NaN | NaN |
| Dock10 | LIKEDQLEYQEELR | -2118 | -2131 | 1 | 0 | 14 | NaN | NaN | 964.11017 | NaN | 1341.6042 | NaN |
| Dock10 | YEQYSGDIR | -127 | -135 | 0 | 0.00272025 | 9 | NaN | NaN | 384.46848 | NaN | NaN | NaN |
| Ppme1 | GIIEEEEEDEEGSESVNKR | -253 | -271 | 1 | 0.00025015 | 19 | NaN | NaN | 242.24973 | 344.2329 | 375.97586 | 401.4968 |
| Ppme1 | YWDGWFR | -297 | -303 | 0 | 0.002068213 | 7 | NaN | NaN | 237.9018 | NaN | NaN | 353.51547 |
| Ube2m | KVGQGYPHDPPK | -81 | -92 | 1 | 0.000230707 | 12 | NaN | NaN | 593.0775 | 268.1955 | 790.4886 | 704.5689 |
| Ube2m | YLFLEPNPEDPLNK | -134 | -147 | 0 | 0.000431655 | 14 | NaN | NaN | 524.6857 | NaN | NaN | NaN |
| Lars1 | EECCPGKPLNVFR | -1052 | -1064 | 1 | 0 | 13 | NaN | NaN | 578.7465 | 724.0423 | 1255.4633 | 1153.8387 |
| Lars1 | EECCPGKPLNVF | -1052 | -1063 | 1 | 0.003003218 | 12 | NaN | NaN | 940.4947 | NaN | NaN | 1610.8422 |
| Lars1 | VNPQPSSGHFSTK | -1074 | -1086 | 0 | 0.00272025 | 13 | NaN | NaN | 2182.825 | NaN | NaN | NaN |
| Atp1b3 | VKFDSGLNK | -234 | -242 | 1 | 0.000550307 | 9 | NaN | NaN | 1104.6328 | NaN | NaN | 1075.3312 |
| Atp1b3 | TFSMSEPQTYK | -94 | -104 | 0 | 0.007987481 | 11 | NaN | NaN | 2512.5947 | NaN | NaN | NaN |
| Sec13 | GHEGPVWQVA | -55 | -64 | 0 | 0.003490217 | 10 | NaN | NaN | 4035.73 | NaN | NaN | NaN |
| Snrpd2 | REEEEFNTGPLSVLTQ | -19 | -34 | 1 | 0 | 16 | NaN | NaN | 3142.3735 | NaN | 2680.4702 | 2884.8855 |
| Tpx2 | QQEEEQQKEELAR | -693 | -705 | 1 | 0.002329279 | 13 | NaN | NaN | 218.31903 | NaN | 138.36488 | 703.2707 |
| Tpx2 | TEDEQFEFHSR | -450 | -460 | 0 | 0 | 11 | NaN | NaN | 285.67166 | NaN | 402.41455 | NaN |
| Degs1 | EEFEWVYTDQPHAAR | -8 | -22 | 0 | 0 | 15 | NaN | NaN | 400.1522 | 677.5807 | 816.5371 | 506.15286 |

| | | | | | | | | | | | | |
|---|---|---|---|---|---|---|---|---|---|---|---|---|
| Casp8 | KFLLNNEIPK | -121 | -130 | 1 | 0.00025015 | 10 | NaN | NaN | 656.2784 | 727.6469 | 1013.85297 | 945.7202 |
| Crtc3 | EENLLNVPKPLPK | -214 | -226 | 1 | 0.00111006 | 13 | NaN | NaN | 397.2661 | NaN | NaN | NaN |
| Gnl1 | EERSFQEYLGK | -148 | -158 | 1 | 0.002595029 | 11 | NaN | NaN | 935.8325 | NaN | 1250.0184 | NaN |
| Nt5c3a | EGYENFFGK | -178 | -186 | 0 | 0.008919338 | 9 | NaN | NaN | 447.85693 | NaN | NaN | NaN |
| Nt5c3a | EESLEVVNSILQK | -317 | -329 | 0 | 0 | 13 | NaN | NaN | 236.46771 | NaN | NaN | NaN |
| Cct7 | KVQGGALEESQLVAGV | -199 | -214 | 1 | 0.001055699 | 16 | NaN | NaN | 808.0159 | 5754.7676 | 1236.4221 | 2080.1023 |
| Cct7 | KVQGGALEESQLVA | -199 | -212 | 1 | 0.001868731 | 14 | NaN | NaN | 467.1499 | NaN | 817.91504 | 823.2184 |
| Cct7 | EGTDSSQGIPQLVSNIS | -11 | -27 | 0 | 0.004595494 | 17 | NaN | NaN | 981.82623 | 1116.665 | NaN | NaN |
| Cct7 | ISNDGATILK | -58 | -67 | 0 | 0.004321969 | 10 | NaN | NaN | 1086.8802 | NaN | NaN | NaN |
| Cct7 | GGAEQFMEETER | -376 | -387 | 0 | 0.003201138 | 12 | NaN | NaN | 658.8721 | NaN | NaN | NaN |
| Atp6v1a | FKDPVKDGEAK | -586 | -596 | 2 | 0.00437931 | 11 | NaN | NaN | 1204.3815 | NaN | NaN | 1225.2423 |
| Mien1 | LENGGFPYEK | -78 | -87 | 0 | 0.003393785 | 10 | NaN | NaN | 841.554 | NaN | NaN | NaN |
| Atp5f1c | EFDEGSIIFNQFK | -179 | -191 | 0 | 0 | 13 | NaN | NaN | 438.65723 | 94.19151 | 629.37976 | 192.57947 |
| Stt3b | EFDPWFNYR | -98 | -106 | 0 | 0.002199523 | 9 | NaN | NaN | 762.5868 | NaN | NaN | NaN |
| Abcf2 | LVSHDFR | -573 | -579 | 0 | 0.004923919 | 7 | NaN | NaN | 5222.1553 | 1169.0366 | NaN | 1324.7422 |
| Ndufb5 | EIPEGYIPEHWEYYK | -100 | -114 | 0 | 0.00025015 | 15 | NaN | NaN | 265.17676 | 730.037 | NaN | 964.9437 |
| S100a10 | KDPLAVDK | -47 | -54 | 1 | 0.006624942 | 8 | NaN | NaN | 913.3308 | NaN | 1009.07367 | NaN |
| Gfm1 | TIVAPVPFDFTHK | -528 | -540 | 0 | 0.000394361 | 13 | NaN | NaN | 299.75504 | NaN | 305.80585 | NaN |
| Gfm1 | VIGVLEPLPPEDYTK | -552 | -566 | 0 | 0 | 15 | NaN | NaN | 464.27347 | NaN | 531.29126 | 218.62695 |
| Gfm1 | YDEIPAGLR | -236 | -244 | 0 | 0.000230707 | 9 | NaN | NaN | 1007.49316 | 253.36578 | NaN | NaN |
| Tm9sf1 | EFVVDDLPIR | -130 | -139 | 0 | 0.00025015 | 10 | NaN | NaN | 866.2276 | NaN | 790.4568 | 1023.3702 |
| Mcm7 | IMCPSQECQTNR | -204 | -215 | 0 | 0.00025015 | 12 | NaN | NaN | 1404.8594 | NaN | 1066.0306 | 1028.2013 |
| Ndufb10 | TPAPSPQTSLPNPITYLTK | -17 | -35 | 0 | 0.008828666 | 19 | NaN | NaN | 945.1555 | 1228.9718 | 1685.3314 | 1535.238 |
| Elp5 | EGFDSDVNSR | -54 | -63 | 0 | 0.002199523 | 10 | NaN | NaN | 597.8285 | NaN | NaN | 742.59644 |
| Smarcc2 | EGGGAVEEEAKEEISEVPK | -777 | -795 | 1 | 0.000627943 | 19 | NaN | NaN | 673.46844 | 806.18445 | 492.70303 | 1248.4569 |
| Smarcc2 | EGGGAVEEEAKEEISEVPK | -777 | -795 | 1 | 0.000627943 | 19 | NaN | NaN | 673.46844 | 806.18445 | 492.70303 | 1248.4569 |
| Ensa | QEEENPAEETGEEK | -5 | -18 | 0 | 0 | 14 | NaN | NaN | 389.06708 | NaN | NaN | NaN |
| Atp1b1 | EGKPCIIIK | -171 | -179 | 1 | 0.009657961 | 9 | NaN | NaN | 1776.6946 | NaN | NaN | NaN |
| Rfc2 | EGNVPNIIIAGPPGTGK | -61 | -77 | 0 | 0 | 17 | NaN | NaN | 529.75165 | NaN | 604.29944 | NaN |
| Smc2 | GGDVFDPHGTLSGGAR | -653 | -668 | 0 | 0 | 16 | NaN | NaN | 386.07956 | 384.80505 | 616.5471 | 385.50934 |
| Mtap | TFYDGSHCSAR | -123 | -133 | 0 | 0 | 11 | NaN | NaN | 890.0829 | NaN | 645.4946 | 593.33966 |
| Chordc1 | HSGVPIFHEGMK | -179 | -190 | 0 | 0.003102268 | 12 | NaN | NaN | 1106.2996 | NaN | NaN | 368.33914 |
| Haus3 | EHLEEVAEK | -469 | -477 | 0 | 0.00078661 | 9 | NaN | NaN | 993.3283 | NaN | NaN | NaN |
| Plaa | SIDVNEGGPSYK | -406 | -417 | 0 | 0.00025015 | 12 | NaN | NaN | 1387.6133 | 551.95996 | 2148.429 | 1231.5745 |
| Pum3 | SADHPTLDK | -284 | -292 | 0 | 0.007493591 | 9 | NaN | NaN | 1402.5863 | NaN | NaN | NaN |
| Smap | LVIGDHK | -56 | -62 | 0 | 0.00450079 | 7 | NaN | NaN | 619.4235 | NaN | NaN | 419.1468 |
| Immt | TSSAEMPTIPLGSAVEAIR | -581 | -599 | 0 | 0 | 19 | NaN | NaN | 751.8894 | NaN | 740.3424 | 881.44025 |
| Immt | QLKPPAELYPEDINTFK | -672 | -688 | 1 | 0.008736733 | 17 | NaN | NaN | 411.6546 | NaN | NaN | NaN |
| Cdc5l | SEHITYLEHSPYEK | -610 | -623 | 0 | 0 | 14 | NaN | NaN | 642.7277 | 754.1348 | 1082.4478 | NaN |
| Tmem30a | EIEIDYTGTEPSSPCNK | -77 | -93 | 0 | 0.000230707 | 17 | NaN | NaN | 89.665016 | 559.71606 | 576.2252 | 557.50085 |
| Impa1 | EKYPCHSFIGEESVAAGEK | -60 | -78 | 1 | 0.00111006 | 19 | NaN | NaN | 817.0582 | 6363.2183 | 479.358 | 1136.3683 |
| Eml4 | LAISEDHVASVK | -52 | -63 | 0 | 0.00111006 | 12 | NaN | NaN | 641.06213 | 134.48239 | NaN | NaN |
| Hnrnpf | TEMDWVLK | -91 | -98 | 0 | 0 | 8 | NaN | NaN | 3736.736 | 1642.8984 | 670.6501 | 2196.932 |
| Hnrnpf | PGPYDRPGTAR | -207 | -217 | 1 | 0.00437931 | 11 | NaN | NaN | 1130.1848 | NaN | 1904.8662 | 2071.9167 |
| Hnrnpf | KHSGPNSADSANDGFVR | -98 | -114 | 1 | 0.001291103 | 17 | NaN | NaN | 239.83307 | NaN | 343.94827 | NaN |
| Hnrnpf | SDGYGFTTDLFGR | -250 | -262 | 0 | 0 | 13 | NaN | NaN | 208.36581 | NaN | 309.32394 | 232.1198 |
| Pwp2 | FDPFELDTSVTPGR | -736 | -749 | 0 | 0 | 14 | NaN | NaN | 555.6203 | NaN | NaN | NaN |
| Ik | EKEEEELMEKPQK | -182 | -194 | 2 | 0.00025015 | 13 | NaN | NaN | 329.46725 | NaN | 718.55896 | NaN |
| Gtf2f2 | EKHQYYNLK | -192 | -200 | 1 | 0.00122419 | 9 | NaN | NaN | 852.78375 | NaN | NaN | 788.7197 |
| Sf3b2 | GIEKPPFELPDFIK | -499 | -512 | 1 | 0 | 14 | NaN | NaN | 744.74805 | 908.5688 | 970.64435 | 1075.5349 |
| Sf3b2 | KIEEAMDGSETPQLF | -753 | -767 | 1 | 0 | 15 | NaN | NaN | 363.836 | NaN | 499.09113 | 534.5964 |
| Caprin1 | ELSLLDEFYK | -180 | -189 | 0 | 0.006677519 | 10 | NaN | NaN | 298.86493 | NaN | NaN | NaN |
| Kcnab2 | YDSGIPPYSR | -255 | -264 | 0 | 0.000471409 | 10 | NaN | NaN | 344.38953 | NaN | NaN | NaN |
| Csnk2b | FNLTGLNEQVPHYR | -34 | -47 | 0 | 0.004099215 | 14 | NaN | NaN | 1605.186 | NaN | NaN | NaN |
| Pitpnb | ELANTPDCPR | -180 | -189 | 0 | 0.003521747 | 10 | NaN | NaN | 295.19302 | NaN | NaN | 753.71246 |
| Otud6b | QADAPPIIVGEEYPR | -247 | -261 | 0 | 0.001610812 | 15 | NaN | NaN | 879.2675 | 922.3713 | 970.81757 | 1087.2355 |
| Xpo6 | ELDDETLDDDQQTEWQR | -472 | -488 | 0 | 0.002534621 | 17 | NaN | NaN | 322.42938 | NaN | NaN | NaN |
| Gmps | THHNDTELIR | -390 | -399 | 0 | 0 | 10 | NaN | NaN | 1405.2051 | 1264.1075 | NaN | NaN |
| Rps8 | LTPEEEEILNKK | -129 | -140 | 1 | 0 | 12 | NaN | NaN | 1002.10065 | NaN | 954.67914 | 938.1367 |
| Rps8 | SNNELVR | -86 | -92 | 0 | 0.00025015 | 7 | NaN | NaN | 678.2095 | 671.2747 | NaN | 1081.4586 |
| Ddx42 | KSEYTQPTPIQCQ | -270 | -282 | 1 | 0 | 13 | NaN | NaN | 779.6398 | NaN | NaN | NaN |
| Cyfip2 | NFDTNFEDR | -41 | -49 | 0 | 0 | 9 | NaN | NaN | 798.64594 | NaN | 543.3803 | 537.48944 |
| Cyfip2 | TDVFQSLR | -990 | -997 | 0 | 0.000298805 | 8 | NaN | NaN | 1086.2925 | NaN | NaN | NaN |
| Dhx8 | GIYYVVDPGFVK | -874 | -885 | 0 | 0.006624942 | 12 | NaN | NaN | 667.147 | NaN | NaN | NaN |
| Ipo9 | WTNIPLLVK | -908 | -916 | 0 | 0 | 9 | NaN | NaN | 488.22415 | 585.00824 | 812.02704 | 718.80786 |
| Armc10 | TDGSYDDILNAEQLK | -48 | -62 | 0 | 0.001868731 | 15 | NaN | NaN | 591.47406 | NaN | NaN | NaN |
| Nsa2 | ELGLVTQGGK | -226 | -235 | 0 | 0.009985598 | 10 | NaN | NaN | 3625.6912 | 922.4792 | NaN | NaN |
| Nsa2 | PQNEYIELHR | -2 | -11 | 0 | 0.006417541 | 10 | NaN | NaN | 217.98338 | NaN | NaN | NaN |
| Arfip1 | ELHEEFGYNADTQK | -209 | -222 | 0 | 0.000230707 | 14 | NaN | NaN | 328.9568 | NaN | 503.44284 | 443.90723 |
| Mapk1 | ELIFEETAR | -343 | -351 | 0 | 0.000627943 | 9 | NaN | NaN | 77.61724 | 2087.43 | NaN | NaN |
| Lnpep | QSPDGTCSLPSAR | -97 | -109 | 0 | 0 | 13 | NaN | NaN | 1696.1558 | 711.9416 | NaN | 1656.7593 |
| Arhgap30 | IIGEPGKDEER | -739 | -749 | 1 | 0 | 11 | NaN | NaN | 786.08185 | 706.37915 | 905.2494 | NaN |
| Map4k2 | ELPQVCVGAEGPEGPGCR | -661 | -678 | 0 | 0.001392869 | 18 | NaN | NaN | 140.79178 | NaN | NaN | NaN |
| Spr | ELPRPEGLQR | -86 | -95 | 1 | 0.001023236 | 10 | NaN | NaN | 2185.891 | 2192.0596 | NaN | NaN |
| Nup88 | WLPNHVVFLR | -17 | -26 | 0 | 0.003681885 | 10 | NaN | NaN | 332.6595 | NaN | NaN | NaN |
| Def6 | SSFTGFQPPPLAR | -565 | -577 | 0 | 0 | 13 | NaN | NaN | 644.0213 | 813.48047 | NaN | 1069.1815 |
| Def6 | IAELEEMQER | -428 | -437 | 0 | 0.00201047 | 10 | NaN | NaN | 924.23425 | NaN | NaN | NaN |
| Gsk3b | QTLPVIYVK | -151 | -159 | 0 | 0.004350979 | 9 | NaN | NaN | 5392.6606 | NaN | NaN | NaN |

| | | | | | | | | | | | | |
|---|---|---|---|---|---|---|---|---|---|---|---|---|
| Pdcd11 | SLRPIFLHPGR | -344 | -354 | 1 | 0.00204437 | 11 | NaN | NaN | 257.90884 | NaN | NaN | 181.72719 |
| Eif3l | IDESIHLQ | -382 | -389 | 0 | 0.001708008 | 8 | NaN | NaN | 1654.3381 | 1067.4963 | 1015.8286 | 977.8428 |
| Gar1 | ENKVPYFNAPVYLENK | -103 | -118 | 1 | 0.007374243 | 16 | NaN | NaN | 525.017 | NaN | NaN | NaN |
| Atad2 | GCLFYGPPGTGK | -117 | -128 | 0 | 0 | 12 | NaN | NaN | 511.049 | NaN | NaN | 636.52216 |
| Atad2 | ENLNFLHLNR | -422 | -431 | 0 | 0.000591124 | 10 | NaN | NaN | 1449.104 | NaN | NaN | NaN |
| Eif4g2 | KEDITQEFPGK | -869 | -879 | 1 | 0.000471409 | 11 | NaN | NaN | 2064.5151 | 1791.6624 | 1623.0747 | 1680.7191 |
| Ipo11 | FFTYPTLTEICR | -356 | -367 | 0 | 0 | 12 | NaN | NaN | 794.65845 | 1364.8237 | NaN | 1326.6316 |
| Mta2 | LADQGEIR | -141 | -148 | 0 | 0.002099344 | 8 | NaN | NaN | 403.73325 | 490.88724 | NaN | 1154.0094 |
| Mta2 | ENSSSNPYLVR | -15 | -25 | 0 | 0.000471409 | 11 | NaN | NaN | 2765.2192 | NaN | NaN | NaN |
| Smpd4 | ENVLPDSPLYHNK | -124 | -136 | 0 | 0.001708008 | 13 | NaN | NaN | 2204.5464 | NaN | 2697.5474 | 1093.3413 |
| Prpf40a | SGNWDTSGSELSEGELEK | -923 | -940 | 0 | 0.008075282 | 18 | NaN | NaN | 388.3667 | NaN | NaN | NaN |
| Ccdc88b | LEHLQEELEQAALER | -893 | -907 | 0 | 0.001092529 | 15 | NaN | NaN | 444.69193 | NaN | 507.5932 | 405.4458 |
| Ccdc88b | EPAHLLLPEAPANAS | -239 | -253 | 0 | 0.000431655 | 15 | NaN | NaN | 489.15582 | NaN | 833.48236 | NaN |
| Hspbp1 | TEHSPFHEHVLGAL | -281 | -294 | 0 | 0.00025015 | 14 | NaN | NaN | 514.3166 | 936.0921 | 5223.298 | 3435.0808 |
| Larp4 | VPDELGLENR | -489 | -498 | 0 | 0 | 10 | NaN | NaN | 475.3932 | 822.0465 | 428.496 | 364.53275 |
| Add3 | NTNPTEVLEK | -495 | -504 | 0 | 0 | 10 | NaN | NaN | 2460.1743 | 2492.918 | NaN | 942.2683 |
| Rpl17 | INPYMSSPCHIE | -136 | -147 | 0 | 0.00111006 | 12 | NaN | NaN | 215.91183 | NaN | NaN | 309.8686 |
| Tax1bp1 | VPSWEDNVVCSQPAR | -691 | -705 | 0 | 0.007987481 | 15 | NaN | NaN | 620.9155 | NaN | NaN | NaN |
| Golga5 | TLSNSSQSELESR | -576 | -588 | 0 | 0 | 13 | NaN | NaN | 2651.321 | 2979.5085 | NaN | NaN |
| Ints5 | EQPLLFELLK | -852 | -861 | 0 | 0.003102268 | 10 | NaN | NaN | 210.26666 | NaN | NaN | NaN |
| Nt5c | QYGALRPDLAEK | -51 | -62 | 1 | 0.000550307 | 12 | NaN | NaN | 1104.0198 | NaN | NaN | NaN |
| Cyba | ERPQVGGTIK | -140 | -149 | 1 | 0.001055699 | 10 | NaN | NaN | 7083.3354 | NaN | 6104.5405 | NaN |
| Psmc2 | PDYLGADQR | -2 | -10 | 0 | 0.00025015 | 9 | NaN | NaN | 738.70746 | 476.5907 | 534.4703 | NaN |
| Creb1 | SSPALPTQPAEEAAR | -256 | -270 | 0 | 0 | 15 | NaN | NaN | 1745.0908 | 1305.4313 | 1388.1555 | 317.0589 |
| Creb1 | ESEDSQESVDSVTDSQK | -93 | -109 | 0 | 0.000230707 | 17 | NaN | NaN | 250.02768 | 282.58533 | NaN | NaN |
| Tbca | MKAEDGENYAIK | -40 | -51 | 1 | 0.00111006 | 12 | NaN | NaN | 661.5124 | 547.72705 | NaN | NaN |
| Exoc4 | NKEIEGVTK | -546 | -554 | 1 | 0.00504706 | 9 | NaN | NaN | 1926.9242 | NaN | NaN | NaN |
| Exoc4 | ESEVLIGNLGDK | -683 | -694 | 0 | 0.000394361 | 12 | NaN | NaN | 616.1321 | NaN | NaN | NaN |
| Ptpn2 | ESGCLTPDHGPAVIH | -201 | -215 | 0 | 0.00078661 | 15 | NaN | NaN | 543.0945 | 594.8115 | NaN | NaN |
| Pak1ip1 | ESGDTVQEETSEPNSEK | -330 | -346 | 0 | 0 | 17 | NaN | NaN | 192.07379 | 432.61005 | 225.65796 | 245.87897 |
| Capg | ESHGIFFSGD | -37 | -46 | 0 | 0.00111006 | 10 | NaN | NaN | 480.12366 | NaN | 220.52072 | NaN |
| Ppa2 | ESNVEEEVWHF | -317 | -327 | 0 | 0 | 11 | NaN | NaN | 360.9972 | NaN | NaN | NaN |
| Hook3 | ESPVSAGHDAYVDLDR | -161 | -176 | 0 | 0.00025015 | 16 | NaN | NaN | 427.1076 | NaN | 347.2599 | NaN |
| Elob | LYKDDQLLDDGK | -44 | -55 | 1 | 0 | 12 | NaN | NaN | 2783.3975 | 642.9812 | 3279.6265 | 3447.9998 |
| Camk2d | FTDEYQLFEELGK | -10 | -22 | 0 | 0 | 13 | NaN | NaN | 974.53564 | 281.97247 | 1043.0465 | 1174.5938 |
| Dbnl | TYEGNSNDIR | -32 | -41 | 0 | 0 | 10 | NaN | NaN | 14862.24 | 11503.23 | NaN | 12918.342 |
| Dnpep | LVHIERPILR | -152 | -161 | 1 | 0.000869338 | 10 | NaN | NaN | 594.79083 | 551.4818 | 554.6958 | 782.11163 |
| Dnpep | LVHIERPILR | -152 | -161 | 1 | 0.000869338 | 10 | NaN | NaN | 594.79083 | 551.4818 | 554.6958 | 782.11163 |
| Dnpep | FELFPSVSR | -460 | -468 | 0 | 0.00111006 | 9 | NaN | NaN | 423.8587 | 662.9053 | 699.3746 | 641.4171 |
| Znf207 | LIHPDEDISLE | -380 | -390 | 0 | 0 | 11 | NaN | NaN | 1048.8567 | NaN | NaN | NaN |
| Gm12250 | SMSVEELESTIK | -333 | -344 | 0 | 0 | 12 | NaN | NaN | 1173.8164 | 920.327 | 584.6744 | 748.2496 |
| Pcbp1 | PATQCGSLIGK | -105 | -115 | 0 | 0.00450079 | 11 | NaN | NaN | 1027.9077 | NaN | 1319.9143 | 1275.2592 |
| Pcbp1 | TLSQSPQGR | -169 | -177 | 0 | 0.005176406 | 9 | NaN | NaN | 1009.4161 | NaN | NaN | NaN |
| Snrpd3 | VLHEAEGHIVTC | -9 | -20 | 0 | 0.000230707 | 12 | NaN | NaN | 4144.725 | 1890.6117 | 4429.7163 | 4190.9893 |
| Edc4 | LDHSDPITR | -1352 | -1360 | 0 | 0 | 9 | NaN | NaN | 995.63104 | 1040.5991 | 1531.9911 | 1264.5797 |
| Arpc1a | EVDEKPASTPWGSK | -175 | -188 | 1 | 0.000230707 | 14 | NaN | NaN | 10883.043 | NaN | NaN | NaN |
| Nrdc | EVEAVDSEYQLAR | -318 | -330 | 0 | 0 | 13 | NaN | NaN | 1084.0311 | 557.9427 | NaN | 814.1752 |
| Nrdc | LKECEDTHLGEEVDR | -1035 | -1049 | 1 | 0 | 15 | NaN | NaN | 921.6692 | 796.08936 | NaN | 343.96512 |
| Nrdc | LKECEDTHLGEEVDR | -1035 | -1049 | 1 | 0 | 15 | NaN | NaN | 921.6692 | 796.08936 | NaN | 343.96512 |
| Nrdc | SKETLDTLEK | -396 | -405 | 1 | 0.00025015 | 10 | NaN | NaN | 385.48013 | NaN | 588.2864 | 578.86224 |
| Nrdc | VVELPSGHHLCK | -907 | -918 | 0 | 0.009569685 | 12 | NaN | NaN | 655.0561 | NaN | NaN | NaN |
| Cul1 | LLETHIHNQ | -339 | -347 | 0 | 0.008919338 | 9 | NaN | NaN | 1824.3505 | 2252.8801 | NaN | NaN |
| Pon2 | FQEEENSLLHLK | -145 | -156 | 0 | 0 | 12 | NaN | NaN | 1461.3389 | NaN | 8709.579 | NaN |
| Ppp3ca | EVFDNDGKPR | -33 | -42 | 1 | 0 | 10 | NaN | NaN | 858.53674 | 1027.5409 | NaN | 886.3241 |
| NaN | SEEDKWPEGSPK | (100);(100) | (111);(111) | 1 | 0.001636916 | 12 | NaN | NaN | 523.5 | 702.4121 | NaN | NaN |
| Rpl15 | GATYGKPVHHGVNQLK | -78 | -93 | 1 | 0.00025015 | 16 | NaN | NaN | 508.50467 | NaN | NaN | NaN |
| Rpl15 | GATYGKPVHHGVNQLK | -78 | -93 | 1 | 0.00025015 | 16 | NaN | NaN | 508.50467 | NaN | NaN | NaN |
| Pck2 | RLEGEDSAQETPIGLVPK | -561 | -578 | 1 | 0 | 18 | NaN | NaN | 1099.6919 | 806.7722 | 1347.2399 | 739.3904 |
| Stk39 | SVHDSQAQPNANEDYR | -442 | -457 | 0 | 0.002880184 | 16 | NaN | NaN | 349.5752 | NaN | NaN | NaN |
| Gcsh | SCYEDGWLIK | -134 | -143 | 0 | 0 | 10 | NaN | NaN | 518.2408 | NaN | NaN | NaN |
| Usp39 | NFPITNVDLR | -471 | -480 | 0 | 0.00025015 | 10 | NaN | NaN | 5387.5664 | NaN | 3797.1472 | NaN |
| Flnb | EVTYDDVPIPNSPFK | -1305 | -1319 | 0 | 0.001463897 | 15 | NaN | NaN | 394.0522 | NaN | NaN | NaN |
| Ndufs5 | YTPPPHHSGR | -92 | -101 | 0 | 0.004224523 | 10 | NaN | NaN | 388.48007 | NaN | NaN | NaN |
| Stk4 | KLDEDSLTK | -16 | -24 | 1 | 0.000907206 | 9 | NaN | NaN | 267.7042 | NaN | NaN | NaN |
| Chchd3 | VAEELALEQAK | -66 | -76 | 0 | 0.00025015 | 11 | NaN | NaN | 244.43037 | 472.13232 | NaN | 302.00302 |
| Cycs | TGPNLHGL | -29 | -36 | 0 | 0.001392869 | 8 | NaN | NaN | 1540.0358 | 1204.8461 | 1010.9154 | NaN |
| Pdcd4 | EYLLSGDISEAEHCLK | -337 | -352 | 0 | 0 | 16 | NaN | NaN | 363.61993 | NaN | 2093.374 | 2110.7383 |
| Rasal3 | EYLQDTLGQVVR | -545 | -556 | 0 | 0 | 12 | NaN | NaN | 539.0295 | 567.4564 | 542.1598 | 844.70056 |
| Lat | SILIKPPQITVPR | -49 | -61 | 1 | 0 | 13 | NaN | NaN | 634.95825 | 565.8523 | 677.8531 | 742.3242 |
| Elavl1 | YLPQNMTQEELR | -26 | -37 | 0 | 0 | 12 | NaN | NaN | 634.20825 | 352.16827 | 648.8366 | 768.48517 |
| Uba6 | HCFPLETR | -720 | -727 | 0 | 0.00025015 | 8 | NaN | NaN | 831.49805 | NaN | 645.76184 | NaN |
| Rif1 | TSGEPSVNAEIDQNR | -1474 | -1488 | 0 | 0.00025015 | 15 | NaN | NaN | 296.4801 | 462.3888 | NaN | 422.16977 |
| Ndufs1 | KVAVTPPGLAR | -607 | -617 | 1 | 0.00272025 | 11 | NaN | NaN | 218.71451 | 362.10333 | NaN | NaN |
| Ndufs1 | KSWLHNDLK | -420 | -428 | 1 | 0.001776473 | 9 | NaN | NaN | 1850.7743 | NaN | NaN | NaN |
| Isyna1 | HVFVGGDDFK | -296 | -305 | 0 | 0.001023236 | 10 | NaN | NaN | 1278.8033 | 633.40845 | 1280.3649 | 876.01556 |
| Isyna1 | SPDVVYSPETIEAR | -11 | -24 | 0 | 0 | 14 | NaN | NaN | 621.98 | NaN | NaN | NaN |
| Mical1 | LLEVVNQR | -1024 | -1031 | 0 | 0.000347792 | 8 | NaN | NaN | 499.24127 | NaN | NaN | NaN |
| Themis | HFLIPISYK | -332 | -340 | 0 | 0 | 9 | NaN | NaN | 2663.6936 | 1706.6874 | 2455.734 | NaN |

| | | | | | | | | | | | | |
|---|---|---|---|---|---|---|---|---|---|---|---|---|
| Usp7 | IQSLLDIQEK | -1025 | -1034 | 0 | 0 | 10 | NaN | NaN | 1075.1138 | NaN | NaN | NaN |
| Fnbp1 | GVTPEDFSNFPPEQR | -395 | -409 | 0 | 0.00266297 | 15 | NaN | NaN | 586.5155 | NaN | NaN | NaN |
| Fnbp1 | RNEDEEGYVPTSYVEVY | -592 | -608 | 1 | 0.000712251 | 17 | NaN | NaN | 303.80905 | NaN | NaN | NaN |
| Atp5pf | FDDPKFEVIDKPQS | -95 | -108 | 2 | 0 | 14 | NaN | NaN | 1223.4885 | 1503.081 | 1547.6617 | 2319.2905 |
| Ddx46 | RVTYVVLDEADR | -522 | -533 | 1 | 0 | 12 | NaN | NaN | 493.22086 | NaN | 321.63333 | 413.867 |
| Ddx46 | SCPNHYEDYVHR | -695 | -706 | 0 | 0 | 12 | NaN | NaN | 504.12756 | NaN | NaN | NaN |
| Ddx46 | IEYEPFR | -326 | -332 | 0 | 0.00223472 | 7 | NaN | NaN | 1577.2245 | NaN | NaN | NaN |
| Aldh3a2 | FDHILYTGNTAVGK | -178 | -191 | 0 | 0 | 14 | NaN | NaN | 2499.8826 | NaN | 810.9783 | 310.93863 |
| Acbd3 | NANKPLLDEIVPVYR | -467 | -481 | 1 | 0.00450079 | 15 | NaN | NaN | 880.2699 | NaN | 935.96027 | NaN |
| Cc2d1b | NTNSPEFEQVFK | -738 | -749 | 0 | 0 | 12 | NaN | NaN | 8573.64 | 7387.471 | 8062.9126 | 7730.075 |
| Bola2 | LRQDLEAEHVEVE | -12 | -24 | 1 | 0.001941082 | 13 | NaN | NaN | 778.6838 | NaN | NaN | NaN |
| Setd7 | SGSSVYHFDK | -186 | -195 | 0 | 0 | 10 | NaN | NaN | 245.09898 | NaN | 202.07257 | 816.27234 |
| Setd7 | FEGNFVHGEK | -36 | -45 | 0 | 0.002752526 | 10 | NaN | NaN | 1469.1934 | NaN | 1397.0239 | NaN |
| Pdpk1 | FETITWENLHQQTPPK | -345 | -360 | 0 | 0.007343498 | 16 | NaN | NaN | 878.2565 | NaN | 1573.88 | NaN |
| Ctps1 | GVVDYFLR | -261 | -268 | 0 | 0.006164361 | 8 | NaN | NaN | 472.62836 | NaN | 409.59415 | NaN |
| Pum1 | FFEFGSHEQK | -906 | -915 | 0 | 0 | 10 | NaN | NaN | 2798.7268 | NaN | NaN | 1889.9363 |
| Atp6v0d1 | RADDYEQVK | -257 | -265 | 1 | 0.005698678 | 9 | NaN | NaN | 742.8482 | NaN | NaN | NaN |
| Ndufa11 | FFESYHEVPDGTQCHR | -5 | -20 | 0 | 0 | 16 | NaN | NaN | 477.4632 | 337.518 | 465.8123 | 434.9476 |
| Ndufa11 | FFESYHEVPDGTQCHR | -5 | -20 | 0 | 0 | 16 | NaN | NaN | 477.4632 | 337.518 | 465.8123 | 434.9476 |
| Glyr1 | TSFFLGEVGNAAK | -416 | -428 | 0 | 0.00025015 | 13 | NaN | NaN | 3313.217 | NaN | NaN | 623.8602 |
| Glyr1 | KWQPTATEPVK | -200 | -210 | 1 | 0.000230707 | 11 | NaN | NaN | 618.4059 | NaN | NaN | NaN |
| Pigx | FGESIEDLQTCR | -63 | -74 | 0 | 0 | 12 | NaN | NaN | 811.45654 | 243.88048 | 1223.8138 | NaN |
| Ddx1 | WQMNPYDR | -91 | -98 | 0 | 0.002199523 | 8 | NaN | NaN | 438.7532 | NaN | NaN | NaN |
| Tardbp | GISVHISNAEPK | -252 | -263 | 0 | 0 | 12 | NaN | NaN | 420.9803 | 333.1498 | 861.38513 | 462.52542 |
| Pals2 | FGTTVPFTSR | -379 | -388 | 0 | 0.005176406 | 10 | NaN | NaN | 1136.1604 | NaN | NaN | NaN |
| Nt5c3b | FHHYYPIEIDPHR | -81 | -93 | 0 | 0.000230707 | 13 | NaN | NaN | 1547.8365 | 1783.0005 | NaN | NaN |
| Cox6a1 | HEEHERPPFVAYPHLR | -65 | -80 | 1 | 0 | 16 | NaN | NaN | 7660.373 | 675.11865 | 5531.6743 | 6097.129 |
| Cox6a1 | HEEHERPPFVAYPHLR | -65 | -80 | 1 | 0 | 16 | NaN | NaN | 7660.373 | 675.11865 | 5531.6743 | 6097.129 |
| Cox6a1 | TKPFPWGDGNHT | -83 | -94 | 1 | 0.00025015 | 12 | NaN | NaN | 956.23926 | NaN | NaN | NaN |
| Acp5 | FVAVGDWGGVPNAPFHTAR | -30 | -48 | 0 | 0 | 19 | NaN | NaN | 456.95358 | 279.39996 | 2340.4468 | 1120.5522 |
| Acp5 | FVAVGDWGGVPNAPFHTAR | -30 | -48 | 0 | 0 | 19 | NaN | NaN | 456.95358 | 279.39996 | 2340.4468 | 1120.5522 |
| Acp5 | FHYGSEDSLGGFTHVEISPK | -283 | -302 | 0 | 0 | 20 | NaN | NaN | 430.6142 | NaN | 293.24045 | 382.61377 |
| Chchd4 | FHYSTEDIK | -75 | -83 | 0 | 0.001370415 | 9 | NaN | NaN | 625.4087 | NaN | NaN | NaN |
| Afg3l2 | FKDVAGCEEAK | -306 | -316 | 1 | 0 | 11 | NaN | NaN | 370.1959 | NaN | 1330.1818 | 629.67395 |
| Afg3l2 | WYLEHADPLLK | -582 | -592 | 0 | 0 | 11 | NaN | NaN | 1033.6868 | NaN | 1258.8116 | NaN |
| Afg3l2 | FIDEIDAVGR | -404 | -413 | 0 | 0.000907206 | 10 | NaN | NaN | 887.69666 | NaN | NaN | NaN |
| Timm8a1 | GAVDPQLQHFIEVETQK | -13 | -29 | 0 | 0 | 17 | NaN | NaN | 523.88385 | 510.20197 | 666.24915 | 692.8841 |
| Slfn1 | GIGEDLETSFK | -70 | -80 | 0 | 0 | 11 | NaN | NaN | 858.4643 | 745.63855 | 974.2686 | 891.2906 |
| Septin9 | THCEFAYLR | -527 | -535 | 0 | 0 | 9 | NaN | NaN | 503.0314 | 192.95937 | 831.4513 | 793.1364 |
| Tram1 | FINFQLR | -315 | -321 | 0 | 0.00025015 | 7 | NaN | NaN | 782.2392 | 834.2973 | 770.735 | 885.9199 |
| Tapbp | YPAGASAPHCEMSR | -85 | -98 | 0 | 0 | 14 | NaN | NaN | 964.8529 | 813.71216 | 759.4809 | 1059.6652 |
| Tapbp | QHGVHYVCR | -378 | -386 | 0 | 0 | 9 | NaN | NaN | 532.301 | 156.14479 | 698.97705 | 727.0958 |
| Tapbp | RFIPFPASAK | -98 | -107 | 1 | 0.007670622 | 10 | NaN | NaN | 250.30716 | NaN | NaN | NaN |
| Eif2b2 | FISNIGGNAPSYIYR | -324 | -338 | 0 | 0.00025015 | 15 | NaN | NaN | 1642.8214 | NaN | NaN | NaN |
| Vapa | HEQILVLDPPSDLK | -11 | -24 | 0 | 0 | 14 | NaN | NaN | 1747.9617 | 1882.3131 | 2639.2903 | 2295.9224 |
| Vapa | FKGPFTDVVTTN | -25 | -36 | 1 | 0.002810811 | 12 | NaN | NaN | 960.2202 | 5864.009 | 7071.879 | NaN |
| Sec62 | FKLEPHDDQVFLD | -168 | -180 | 1 | 0.00025015 | 13 | NaN | NaN | 1201.4977 | 1260.9359 | NaN | 784.8116 |
| Arhgap1 | WDDPYYDIAR | -60 | -69 | 0 | 0.000550307 | 10 | NaN | NaN | 445.92044 | NaN | NaN | NaN |
| Arhgap1 | TSDNKPSLSWLR | -130 | -141 | 1 | 0.002534621 | 12 | NaN | NaN | 322.9303 | NaN | NaN | NaN |
| Gclc | FKPPPPNSDIGWR | -411 | -423 | 1 | 0.004658491 | 13 | NaN | NaN | 687.4187 | NaN | NaN | NaN |
| Thop1 | NLNEDTTFLPFTR | -181 | -193 | 0 | 0 | 13 | NaN | NaN | 1443.1077 | 1177.907 | 2543.3123 | NaN |
| Rab27a | FLALGDSGVGK | -12 | -22 | 0 | 0.00025015 | 11 | NaN | NaN | 558.2935 | NaN | NaN | NaN |
| Ubash3a | GFFINDSPADAIR | -158 | -170 | 0 | 0.001055699 | 13 | NaN | NaN | 649.4552 | 468.60754 | NaN | 427.12946 |
| Ppp2r5a | GSQAELHPLPQLK | -48 | -60 | 0 | 0 | 13 | NaN | NaN | 4947.677 | NaN | 6992.422 | 7192.6 |
| Mettl1 | FLFPDPHFK | -153 | -161 | 0 | 0.006568856 | 9 | NaN | NaN | 529.25226 | NaN | 618.0722 | NaN |
| Pold2 | TDPFIFPECPHVY | -390 | -402 | 0 | 0 | 13 | NaN | NaN | 2720.3906 | 2298.598 | 3153.6836 | NaN |
| Oga | SFVQWLGCR | -656 | -664 | 0 | 0 | 9 | NaN | NaN | 744.7952 | 748.4338 | 836.1562 | 779.7951 |
| Oga | NDNQILTEIVEAK | -479 | -491 | 0 | 0 | 13 | NaN | NaN | 576.5252 | NaN | 1129.034 | NaN |
| Arhgef2 | FLNQLLER | -294 | -301 | 0 | 0 | 8 | NaN | NaN | 2290.7024 | NaN | 2137.014 | 2385.766 |
| Man2b1 | FLQDTFGSDGLPR | -178 | -190 | 0 | 0 | 13 | NaN | NaN | 907.4212 | NaN | 995.9033 | 603.34296 |
| Akap13 | FLSHSTDSLNK | -1888 | -1898 | 0 | 0.00078661 | 11 | NaN | NaN | 414.17575 | NaN | NaN | NaN |
| Rabggta | FLVENSVLK | -429 | -437 | 0 | 0.006677519 | 9 | NaN | NaN | 536.46387 | NaN | NaN | NaN |
| Ercc6l | FLVESESQNTVQR | -709 | -721 | 0 | 0.001055699 | 13 | NaN | NaN | 550.4089 | 271.7437 | NaN | 1158.7201 |
| Pgm1 | FNISNGGPAPEAITDK | -131 | -146 | 0 | 0.007905138 | 16 | NaN | NaN | 562.757 | NaN | NaN | NaN |
| Pigt | FNTLFPVSDGSSYFVR | -496 | -511 | 0 | 0.003490217 | 16 | NaN | NaN | 4377.565 | NaN | NaN | NaN |
| Cdc42 | FPSEYVPTVFDNY | -28 | -40 | 0 | 0.001636916 | 13 | NaN | NaN | 441.72296 | 481.06705 | 511.8074 | NaN |
| Larp1 | KFEPEYSQIK | -671 | -680 | 1 | 0.000431655 | 10 | NaN | NaN | 3514.3203 | 4612.2466 | 4273.576 | 2405.0781 |
| Larp1 | WHDQDETSSVK | -291 | -301 | 0 | 0 | 11 | NaN | NaN | 2869.7107 | 919.54065 | 1377.9944 | 1454.6189 |
| Larp1 | NWPTPGEIAHK | -148 | -158 | 0 | 0.000394361 | 11 | NaN | NaN | 697.3308 | NaN | NaN | 1159.4672 |
| Dpp4 | VEPHLPSHR | -170 | -178 | 0 | 0.000550307 | 9 | NaN | NaN | 763.9 | 818.815 | 635.4861 | 708.99365 |
| Dpp4 | GYHSVSPDR | -97 | -105 | 0 | 0.00078661 | 9 | NaN | NaN | 804.3696 | NaN | NaN | NaN |
| Dpp4 | FRPAEPHFT | -351 | -359 | 1 | 0.00272025 | 9 | NaN | NaN | 662.0354 | NaN | NaN | NaN |
| Dpp4 | SSAAPIQIPAPASVAR | -274 | -289 | 0 | 0.000826806 | 16 | NaN | NaN | 586.3678 | NaN | NaN | NaN |
| Kif5b | FRPLNESEVNR | -15 | -25 | 1 | 0.00383302 | 11 | NaN | NaN | 1710.618 | NaN | 1904.2548 | 1986.189 |
| Ints3 | YQDWFQR | -327 | -333 | 0 | 0.001868731 | 7 | NaN | NaN | 484.0981 | NaN | 353.76462 | 547.60486 |
| Ints3 | VLAHLAPLFDNPK | -467 | -479 | 0 | 0 | 13 | NaN | NaN | 1207.9922 | NaN | NaN | NaN |
| Ints3 | FSDDEEDLNSK | -534 | -544 | 0 | 0.00111006 | 11 | NaN | NaN | 617.47144 | NaN | NaN | NaN |
| Chp1 | FSEGEDQVNFR | -71 | -81 | 0 | 0 | 11 | NaN | NaN | 1509.3442 | NaN | NaN | NaN |

| | | | | | | | | | | | | |
|---|---|---|---|---|---|---|---|---|---|---|---|---|
| Oas3 | VDLSVLPAFDAVGPLK | -592 | -607 | 0 | 0.000550307 | 16 | NaN | NaN | 413.0621 | NaN | NaN | 486.75183 |
| Oas3 | LVPAFDVLGQPR | -145 | -156 | 0 | 0.00025015 | 12 | NaN | NaN | 1198.0166 | NaN | NaN | NaN |
| Jpt1 | FSLGFDEPAEQPVR | -30 | -43 | 0 | 0.00149795 | 14 | NaN | NaN | 11203.255 | 7148.2915 | 7740.082 | NaN |
| Il2rg | FSLPSVDELK | -209 | -218 | 0 | 0.00025015 | 10 | NaN | NaN | 297.8783 | NaN | 264.38678 | 236.55815 |
| Celf2 | NALHNIK | -101 | -107 | 0 | 0.00450079 | 7 | NaN | NaN | 1486.11 | 2011.0364 | NaN | NaN |
| Celf2 | MFVGQIPR | -42 | -49 | 0 | 0.005459848 | 8 | NaN | NaN | 718.31 | NaN | NaN | NaN |
| Snrpb2 | FSQFGHVVDIVALK | -31 | -44 | 0 | 0 | 14 | NaN | NaN | 1108.3035 | NaN | NaN | NaN |
| Trip12 | GNEETLGGFPVK | -474 | -485 | 0 | 0.001291103 | 12 | NaN | NaN | 1518.2714 | NaN | NaN | NaN |
| Zc3h15 | KEEKEDEISLEDLIER | -223 | -238 | 2 | 0.002068213 | 16 | NaN | NaN | 3179.097 | NaN | NaN | 3655.7632 |
| Gm2a | FSWDNCDEGKDPAVIK | -34 | -49 | 1 | 0.00025015 | 16 | NaN | NaN | 568.81067 | 477.28918 | 613.71747 | 606.26874 |
| Ncf1 | FTEIYEFHK | -44 | -52 | 0 | 0 | 9 | NaN | NaN | 514.80634 | NaN | 732.7405 | 891.00586 |
| Rbm3 | FTNPEHASDAMR | -54 | -65 | 0 | 0 | 12 | NaN | NaN | 3980.8853 | 3484.8918 | 4486.686 | 4761.0005 |
| Rbm3 | GPISEVVVVK | -30 | -39 | 0 | 0.000591124 | 10 | NaN | NaN | 883.2268 | NaN | 887.06757 | NaN |
| Suox | FVDLHPGGPSK | -115 | -125 | 0 | 0 | 11 | NaN | NaN | 1683.7904 | NaN | NaN | NaN |
| Gmppb | FVEKPQVFVS | -159 | -168 | 1 | 0.009688493 | 10 | NaN | NaN | 2133.192 | NaN | NaN | NaN |
| Gmppb | SIGESVPEPR | -348 | -357 | 0 | 0.000394361 | 10 | NaN | NaN | 2095.329 | NaN | NaN | NaN |
| Ptprcap | LSHASGGYYHPA | -57 | -68 | 0 | 0.00025015 | 12 | NaN | NaN | 10638.101 | 864.11786 | 1401.4895 | 1567.5248 |
| Ptprcap | LSHASGGYYHPAR | -57 | -69 | 0 | 0 | 13 | NaN | NaN | 485.23254 | 785.4454 | 1557.1746 | 1279.0372 |
| Ptprcap | FVIDGGPEEAAAK | -115 | -127 | 0 | 0.008222027 | 13 | NaN | NaN | 500.19104 | NaN | NaN | NaN |
| Ap2a2 | IIHHDSEPNLLVR | -316 | -328 | 0 | 0.00122419 | 13 | NaN | NaN | 478.185 | NaN | NaN | NaN |
| Las1l | GNEEVASHPEPSSR | -232 | -245 | 0 | 0 | 14 | NaN | NaN | 326.18643 | 521.558 | 520.87946 | NaN |
| Pdia6 | KDVVELTDDTFDK | -160 | -172 | 1 | 0 | 13 | NaN | NaN | 558.1442 | 1108.0754 | 775.7538 | 827.1552 |
| Csnk2a2 | LIDTVKDPVSK | -98 | -108 | 1 | 0 | 11 | NaN | NaN | 760.79535 | 801.84753 | 920.2257 | 820.0901 |
| Csnk2a2 | HLVSPEALDLLDK | -292 | -304 | 0 | 0 | 13 | NaN | NaN | 496.51685 | 427.19754 | 648.9564 | 672.3802 |
| Atp5mg | LATFWHYAK | -27 | -35 | 0 | 0 | 9 | NaN | NaN | 475.54044 | NaN | 1340.1984 | 1339.9907 |
| Thoc2 | YDNLITPVVDSLK | -590 | -602 | 0 | 0 | 13 | NaN | NaN | 227.88576 | 317.6556 | NaN | 301.69873 |
| Bcap31 | KYMEENDQLK | -149 | -158 | 1 | 0.00450079 | 10 | NaN | NaN | 897.61694 | NaN | NaN | NaN |
| Rps26 | GAAPRPPPKPM | -105 | -115 | 2 | 0 | 11 | NaN | NaN | 845.8307 | 560.556 | 677.5525 | 888.3085 |
| Mtpn | GADINAPDKHHITPL | -58 | -72 | 1 | 0.00025015 | 15 | NaN | NaN | 344.154 | 314.36108 | 529.60736 | 482.44122 |
| Nufip2 | SLDPSHIGDLQK | -589 | -600 | 0 | 0 | 12 | NaN | NaN | 235.72742 | NaN | NaN | NaN |
| Luc7l3 | GAFLIVGDAQSR | -196 | -207 | 0 | 0.00025015 | 12 | NaN | NaN | 1354.1467 | 581.04767 | NaN | 1570.2793 |
| Snrpa1 | GATLDQFDAIDFSDNEIR | -38 | -55 | 0 | 0.00078661 | 18 | NaN | NaN | 279.10065 | NaN | 562.083 | NaN |
| Tapbpl | KGATLEPEELLR | -256 | -267 | 1 | 0.000431655 | 12 | NaN | NaN | 2348.035 | NaN | NaN | NaN |
| Rcc1 | GCGEQGQLGR | -197 | -206 | 0 | 0 | 10 | NaN | NaN | 2296.545 | 3155.7058 | 571.37427 | 2862.2776 |
| Pus7 | HGLTELDVGICK | -106 | -117 | 0 | 0.002972566 | 12 | NaN | NaN | 857.22314 | NaN | NaN | NaN |
| Psap | PGVSDICK | -234 | -241 | 0 | 0 | 8 | NaN | NaN | 326.37152 | 438.00317 | 605.7541 | NaN |
| Vps29 | RGDFDENLNYPEQK | -60 | -73 | 1 | 0.00025015 | 14 | NaN | NaN | 404.77994 | 720.6513 | 738.07806 | 782.71655 |
| Il12rb1 | GDGTESPQAAPECALDTR | -649 | -666 | 0 | 0.00111006 | 18 | NaN | NaN | 1106.2241 | 1231.9084 | 1264.5555 | NaN |
| Sec11a | RVEDPIR | -72 | -78 | 1 | 0.002595029 | 7 | NaN | NaN | 2502.9172 | NaN | 77.983986 | 3277.0256 |
| Sec11a | GQHWLEK | -129 | -135 | 0 | 0.00644558 | 7 | NaN | NaN | 1528.0273 | NaN | NaN | 1328.0115 |
| Helb | GDSVLDAQDGDDHVR | -405 | -419 | 0 | 0 | 15 | NaN | NaN | 608.3336 | NaN | NaN | NaN |
| Atp5pb | GEFIDKLNEEK | -121 | -131 | 1 | 0.000431655 | 11 | NaN | NaN | 736.252 | 677.7978 | 926.2664 | 1044.4502 |
| Glipr2 | QFNNEVLK | -8 | -15 | 0 | 0.000907206 | 8 | NaN | NaN | 1337.8086 | 1762.8303 | 1245.9485 | NaN |
| Etf1 | TELSQSDMFDQR | -234 | -245 | 0 | 0.000230707 | 12 | NaN | NaN | 498.93988 | 588.7958 | 550.3287 | 580.4185 |
| Etf1 | YLTPEQEKDK | -345 | -354 | 1 | 0.000230707 | 10 | NaN | NaN | 780.81165 | NaN | NaN | 683.019 |
| Etf1 | SHFTDKETGQEHELIE | -355 | -370 | 1 | 0.001636916 | 16 | NaN | NaN | 639.724 | NaN | NaN | NaN |
| Rpl13 | TIGISVDPR | -93 | -101 | 0 | 0.00122419 | 9 | NaN | NaN | 859.2747 | NaN | NaN | NaN |
| Slc16a3 | KRPEVTEPEEVASEEK | -418 | -433 | 2 | 0 | 16 | NaN | NaN | 433.3069 | 489.35498 | 750.5287 | 682.5206 |
| Klhdc4 | TEQHWLK | -539 | -545 | 0 | 0.00437931 | 7 | NaN | NaN | 1324.845 | NaN | NaN | 1345.56 |
| Prep | GGGEYGETWHK | -506 | -516 | 0 | 0 | 11 | NaN | NaN | 387.3004 | NaN | 391.65732 | 573.8012 |
| Cnp | RLDEDLAGYCR | -102 | -112 | 1 | 0.00659934 | 11 | NaN | NaN | 350.53476 | NaN | NaN | NaN |
| Plpbp | GHDLSQGPNPDFQR | -187 | -200 | 0 | 0 | 14 | NaN | NaN | 1367.2197 | 1177.0961 | 1091.0017 | 1083.6005 |
| Lman1 | GHPDLQGQPADDIFESIGDR | -318 | -337 | 0 | 0 | 20 | NaN | NaN | 1570.2461 | 491.92926 | 444.09595 | 1667.7217 |
| Lman1 | TFQLTEPGKEPPTAEK | -272 | -287 | 1 | 0.001392869 | 16 | NaN | NaN | 397.53235 | NaN | NaN | 466.80768 |
| Aldh18a1 | GHVITDIVEGK | -337 | -347 | 0 | 0.00051306 | 11 | NaN | NaN | 455.54358 | NaN | 946.3865 | 1004.86584 |
| Smc4 | VHDLFQK | -574 | -580 | 0 | 0.00025015 | 7 | NaN | NaN | 945.681 | 899.9636 | 1097.9666 | 1525.0935 |
| Smc4 | LHPVEDNPVETVA | -1042 | -1054 | 0 | 0.002068213 | 13 | NaN | NaN | 1009.2118 | NaN | NaN | NaN |
| Smc4 | SHGIDLDHNR | -213 | -222 | 0 | 0.005946382 | 10 | NaN | NaN | 605.22736 | NaN | NaN | NaN |
| Rab21 | GIEELFLDLCK | -166 | -176 | 0 | 0 | 11 | NaN | NaN | 634.4491 | NaN | 767.55255 | 827.0155 |
| Rab21 | VNLAIWDTAGQER | -66 | -78 | 0 | 0.008014596 | 13 | NaN | NaN | 658.07184 | 963.1455 | NaN | NaN |
| Spag9 | NVSTESGENEEKSEVQ | -327 | -342 | 1 | 0.007224385 | 16 | NaN | NaN | 842.4144 | NaN | NaN | NaN |
| Golm1 | GIINVPVGSER | -367 | -377 | 0 | 0 | 11 | NaN | NaN | 430.423 | NaN | 509.6151 | NaN |
| Vps4b | SLSSTKPTVNEQDLLK | -417 | -432 | 1 | 0.00223472 | 16 | NaN | NaN | 412.7295 | NaN | NaN | NaN |
| Nsf | LEGPPHSGK | -541 | -549 | 0 | 0.003360691 | 9 | NaN | NaN | 368.59448 | NaN | 377.97186 | 486.9356 |
| Mad2l1 | WQFDIECDK | -100 | -108 | 0 | 0.00149795 | 9 | NaN | NaN | 633.12756 | 566.0736 | NaN | 705.6547 |
| Arcn1 | QHTFVETESVR | -45 | -55 | 0 | 0.009272078 | 11 | NaN | NaN | 780.939 | NaN | NaN | NaN |
| Add1 | GKNPTGLLALQ | -82 | -92 | 1 | 0.007987481 | 11 | NaN | NaN | 271.8609 | NaN | 254.00087 | NaN |
| Taf6 | GLAELGHDVIK | -379 | -389 | 0 | 0 | 11 | NaN | NaN | 1823.2771 | NaN | NaN | 511.6202 |
| Ly6c2 | QCLSFCPAGVPIKDPNIR | -73 | -90 | 1 | 0 | 18 | NaN | NaN | 1296.3422 | 501.45633 | 680.00446 | 652.1512 |
| Lpcat3 | GLWPGVEDLSLNK | -18 | -30 | 0 | 0.001811524 | 13 | NaN | NaN | 331.03912 | NaN | 1243.9337 | NaN |
| Gpn1 | GNASPVLDPSDLILTR | -311 | -326 | 0 | 0.00025015 | 16 | NaN | NaN | 907.55804 | NaN | NaN | 1196.0349 |
| Lgals3 | VAVNDAHLL | -225 | -233 | 0 | 0.000230707 | 9 | NaN | NaN | 3446.4438 | NaN | NaN | NaN |
| Srprb | GNSLTLIDLPGHESLR | -108 | -123 | 0 | 0 | 16 | NaN | NaN | 410.0104 | NaN | NaN | NaN |
| Fam98b | GPLLEEQALSK | -27 | -37 | 0 | 0 | 11 | NaN | NaN | 608.998 | 459.33966 | NaN | NaN |
| Rnaset2a;Rnas | GPLWSGSHEWK | (30);(30) | (40);(40) | 0 | 0 | 11 | NaN | NaN | 1251.264 | NaN | NaN | NaN |
| Nmt1 | LGEVVNTHGPVEPDKDNIR | -128 | -146 | 1 | 0 | 19 | NaN | NaN | 436.70428 | NaN | NaN | NaN |
| Ascc3 | GPVLESIVAR | -622 | -631 | 0 | 0 | 10 | NaN | NaN | 3696.4507 | NaN | 4085.0615 | NaN |
| Asah1 | GQFESHLR | -378 | -385 | 0 | 0.000712251 | 8 | NaN | NaN | 379.97345 | 1454.0773 | 1015.4015 | 389.84216 |

| | | | | | | | | | | | | |
|---|---|---|---|---|---|---|---|---|---|---|---|---|
| Smc6 | TLEDEAEENKIK | -746 | -757 | 1 | 0.00078661 | 12 | NaN | NaN | 486.63046 | 524.0974 | NaN | NaN |
| Smc6 | IEAENKYDTIK | -780 | -790 | 1 | 0.00051306 | 11 | NaN | NaN | 689.44385 | NaN | NaN | NaN |
| Gsto1 | NKPEWFFEK | -58 | -66 | 1 | 0.00078661 | 9 | NaN | NaN | 1712.0376 | 1734.1576 | 481.66248 | 2028.2655 |
| Lap3 | LHGSGDLEAWEK | -177 | -188 | 0 | 0 | 12 | NaN | NaN | 1106.6824 | 980.41235 | 1445.5658 | 896.6347 |
| Lap3 | HQDFPSVVVVGLGK | -90 | -103 | 0 | 0 | 14 | NaN | NaN | 635.13806 | 289.63504 | 624.72186 | 695.6204 |
| Lap3 | KLHGSGDLEAWEK | -176 | -188 | 1 | 0.000627943 | 13 | NaN | NaN | 938.80426 | NaN | NaN | 783.0827 |
| Lap3 | RFAEIIEK | -214 | -221 | 1 | 0.007523243 | 8 | NaN | NaN | 1164.9116 | NaN | NaN | NaN |
| Abce1 | LDLTHLK | -204 | -210 | 0 | 0.007546675 | 7 | NaN | NaN | 620.72675 | NaN | NaN | NaN |
| Emc4 | GSGQGDSLYPVGYLDK | -35 | -50 | 0 | 0.001092529 | 16 | NaN | NaN | 731.3847 | NaN | NaN | NaN |
| Sod2 | GSGWGWLGFNK | -144 | -154 | 0 | 0 | 11 | NaN | NaN | 1120.099 | 910.78845 | 192.81772 | 102.84675 |
| Vps36 | QGDVTEDETIR | -206 | -216 | 0 | 0 | 11 | NaN | NaN | 985.21564 | NaN | NaN | 1038.615 |
| Ybx3 | LAGSPGGDAAPGPAPASSAPAGGEDAEKK | -49 | -77 | 1 | 0.001023236 | 29 | NaN | NaN | 942.22906 | NaN | 1098.3325 | 1248.8792 |
| Ybx3 | SRPLNAVSQDGK | -328 | -339 | 1 | 0.001023236 | 12 | NaN | NaN | 572.9282 | NaN | 823.36755 | 1177.7657 |
| Ybx3 | TGGTTLPQAAADAPAAAPPDPAPK | -9 | -32 | 0 | 0.007791272 | 24 | NaN | NaN | 2024.6422 | NaN | NaN | NaN |
| Bri3bp | SVSSLFGEDNVR | -62 | -73 | 0 | 0.000826806 | 12 | NaN | NaN | 1416.9486 | 1127.597 | NaN | NaN |
| Nit1 | GSVTGPAHWEVLLR | -226 | -239 | 0 | 0 | 14 | NaN | NaN | 1048.9119 | NaN | NaN | NaN |
| Nup43 | HPDQQHVVAT | -226 | -235 | 0 | 0.002068213 | 10 | NaN | NaN | 777.9915 | 508.9092 | 1041.718 | 2311.696 |
| Nup43 | SSLFHQGGR | -299 | -307 | 0 | 0.00383302 | 9 | NaN | NaN | 769.8312 | 307.8251 | NaN | 761.3098 |
| Vdac1 | KLTFDSSFSPNTGK | -109 | -122 | 1 | 0.00025015 | 14 | NaN | NaN | 1438.8347 | 1185.4956 | 1292.176 | 1386.3927 |
| Cd3e | GTSVELTCPLDSDENLK | -35 | -51 | 0 | 0.002168954 | 17 | NaN | NaN | 473.95758 | NaN | NaN | NaN |
| Tigar | GVDAPLSETGFR | -26 | -37 | 0 | 0.001392869 | 12 | NaN | NaN | 1498.0054 | NaN | NaN | NaN |
| Sh3kbp1 | QLLSELDEEKK | -680 | -690 | 1 | 0.00025015 | 11 | NaN | NaN | 705.91125 | NaN | 739.1048 | NaN |
| Gbp7 | VITADEYLENALK | -193 | -205 | 0 | 0.000230707 | 13 | NaN | NaN | 2331.7104 | NaN | 5700.0874 | 1504.009 |
| Trp53i11 | GVGGEDDDGEVHR | -29 | -41 | 0 | 0 | 13 | NaN | NaN | 1389.1019 | NaN | NaN | 1089.7542 |
| Srpk1 | ITQLEESIR | -437 | -445 | 0 | 0.001023236 | 9 | NaN | NaN | 2115.9263 | NaN | 2084.3193 | 1799.0233 |
| Srpk1 | SAEQSLTR | -427 | -434 | 0 | 0.009569685 | 8 | NaN | NaN | 4126.5024 | NaN | NaN | NaN |
| Srpk1 | HKEDLHNANDCDVHTLK | -376 | -392 | 1 | 0.002972566 | 17 | NaN | NaN | 321.4585 | NaN | NaN | NaN |
| Coro7 | GVISLEGHEENKR | -56 | -68 | 1 | 0 | 13 | NaN | NaN | 479.38406 | 1371.313 | 1016.9253 | 800.54913 |
| FAM120A | VEGSSTASSGSQLAEGK | -501 | -517 | 0 | 0 | 17 | NaN | NaN | 496.1563 | NaN | 477.0515 | 482.12573 |
| FAM120A | GVQGFQDYIEK | -2 | -12 | 0 | 0.001463897 | 11 | NaN | NaN | 1635.455 | 2087.4092 | NaN | NaN |
| Aldoc | PHSYPALSAEQK | -2 | -13 | 0 | 0 | 12 | NaN | NaN | 3423.601 | 9019.573 | 2946.588 | NaN |
| Cbfb | RQQDPSPGSNLGGGDDLK | -168 | -185 | 1 | 0 | 18 | NaN | NaN | 808.4649 | 673.137 | 501.84027 | 464.3623 |
| Pus1 | GYAPESVLER | -303 | -312 | 0 | 0.00025015 | 10 | NaN | NaN | 4820.908 | NaN | NaN | NaN |
| Sart3 | KVPEKPEVR | -882 | -890 | 2 | 0.003102268 | 9 | NaN | NaN | 517.3521 | 340.75934 | 416.22357 | 583.5021 |
| Ap1g1 | IEFTFER | -717 | -723 | 0 | 0.002068213 | 7 | NaN | NaN | 1141.5588 | 1138.2747 | NaN | NaN |
| Trnt1 | IVDRPGDHDHETLEAIAE | -208 | -225 | 1 | 0.000230707 | 18 | NaN | NaN | 3820.746 | 397.25574 | 286.24524 | 252.98422 |
| Trnt1 | HAEVEFTTDWQK | -135 | -146 | 0 | 0.00025015 | 12 | NaN | NaN | 912.1385 | 1235.4136 | 528.54016 | 1741.3052 |
| Trnt1 | LHEENFEVT | -115 | -123 | 0 | 0.005605646 | 9 | NaN | NaN | 751.6135 | NaN | NaN | NaN |
| Hccs | HAHECPCGPSLVR | -176 | -188 | 0 | 0 | 13 | NaN | NaN | 537.94824 | 390.00314 | 730.09656 | 524.35706 |
| Cr1l | HCPAPSQLPSAK | -84 | -95 | 0 | 0 | 12 | NaN | NaN | 764.46747 | 1041.5681 | 1026.7238 | 1112.5757 |
| Cr1l | WEPELPSCFK | -329 | -338 | 0 | 0.00025015 | 10 | NaN | NaN | 892.9411 | NaN | 933.7229 | 786.8556 |
| Cr1l | SDGSWNPLLAK | -387 | -397 | 0 | 0.00383302 | 11 | NaN | NaN | 887.5954 | NaN | NaN | NaN |
| Dguok | HEDWFINK | -226 | -233 | 0 | 0.00111006 | 8 | NaN | NaN | 276.4693 | 987.2305 | 1268.7844 | NaN |
| Tor1aip1 | SAIHHPNEPSVK | -336 | -347 | 0 | 0 | 12 | NaN | NaN | 525.19855 | 1284.112 | 610.8641 | NaN |
| Sfxn2 | HFFNITDPR | -26 | -34 | 0 | 0.000230707 | 9 | NaN | NaN | 322.122 | NaN | NaN | NaN |
| Hvcn1 | HFTVVGDDYHTW | -27 | -38 | 0 | 0.002068213 | 12 | NaN | NaN | 673.328 | NaN | NaN | NaN |
| Ppp6r3 | WGILDPPVGNTR | -333 | -344 | 0 | 0 | 12 | NaN | NaN | 278.2981 | NaN | 539.4491 | 1455.8579 |
| Ppp6r3 | HIAFTPESQR | -598 | -607 | 0 | 0.000591124 | 10 | NaN | NaN | 3007.8547 | NaN | NaN | NaN |
| Ppp6r3 | NSTEPDPLLATLEK | -231 | -244 | 0 | 0.00122419 | 14 | NaN | NaN | 335.3785 | NaN | NaN | NaN |
| Eif5b | SEAPLSGSEDADDSNK | -132 | -147 | 0 | 0.001392869 | 16 | NaN | NaN | 259.84647 | NaN | NaN | NaN |
| Utp4 | HLHTFQPQSGTVE | -473 | -485 | 0 | 0.00713295 | 13 | NaN | NaN | 809.8242 | NaN | 2710.9626 | NaN |
| Utp4 | TVLYNPLPPK | -618 | -627 | 0 | 0.000826806 | 10 | NaN | NaN | 252.13037 | NaN | NaN | NaN |
| Sdhc | NTSSNRPLSPHLTIY | -45 | -59 | 1 | 0 | 15 | NaN | NaN | 1334.8248 | 953.4831 | 1449.289 | 1337.7417 |
| Foxp1 | HLNSEHALDDR | -354 | -364 | 0 | 0 | 11 | NaN | NaN | 592.78564 | 222.86554 | 944.37585 | 645.36365 |
| Nup85 | LTDLLDHCR | -377 | -385 | 0 | 0.002296977 | 9 | NaN | NaN | 2125.8506 | NaN | NaN | 475.1514 |
| Nup85 | HLTDLLDHCR | -376 | -385 | 0 | 0.000591124 | 10 | NaN | NaN | 341.74454 | NaN | NaN | 613.19183 |
| Ints14 | HLVLPIALNK | -271 | -280 | 0 | 0.003900536 | 10 | NaN | NaN | 190.35106 | NaN | NaN | NaN |
| H2-Ab1 | HNYEGPETHTSLR | -108 | -120 | 0 | 0.000230707 | 13 | NaN | NaN | 727.80566 | 778.69073 | NaN | 356.13016 |
| Cdk1 | SPEVLLGSAR | -171 | -180 | 0 | 0 | 10 | NaN | NaN | 354.26025 | NaN | 1007.4422 | NaN |
| Nip7 | VWVKPGAEQSFLYGN | -97 | -111 | 1 | 0.001868731 | 15 | NaN | NaN | 747.3303 | NaN | NaN | 682.48865 |
| Bzw2 | NEAGDDLEAVAK | -39 | -50 | 0 | 0 | 12 | NaN | NaN | 1543.145 | NaN | NaN | NaN |
| Sec63 | HSLLHFL | -421 | -427 | 0 | 0.00644558 | 7 | NaN | NaN | 426.16876 | NaN | NaN | 429.19525 |
| Hmgcs1 | YTIGLGQAR | -47 | -55 | 0 | 0.006788754 | 9 | NaN | NaN | 775.82544 | NaN | NaN | NaN |
| Nup188 | HTLSPEETSSVVR | -331 | -343 | 0 | 0.000394361 | 13 | NaN | NaN | 1486.4819 | NaN | 942.35657 | NaN |
| Nup188 | QLFLDVLDGTK | -1131 | -1141 | 0 | 0 | 11 | NaN | NaN | 899.59357 | NaN | NaN | NaN |
| Utp11 | HVFFFDTK | -136 | -143 | 0 | 0.00111006 | 8 | NaN | NaN | 1256.8275 | NaN | NaN | NaN |
| Gtf2h2 | HVILDETHYK | -220 | -229 | 0 | 0.000394361 | 10 | NaN | NaN | 1833.6707 | NaN | NaN | NaN |
| Smarcd2 | KLDQTIAR | -168 | -175 | 1 | 0.002595029 | 8 | NaN | NaN | 375.90976 | NaN | NaN | NaN |
| Dhx30 | QSLLELWR | -438 | -445 | 0 | 0.001868731 | 8 | NaN | NaN | 123.893776 | NaN | 194.14377 | NaN |
| Sh2d2a | VEGPSGTAPIGHPILR | -328 | -343 | 0 | 0.000712251 | 16 | NaN | NaN | 609.887 | NaN | NaN | NaN |
| Elp2 | HWQPSFYK | -289 | -296 | 0 | 0 | 8 | NaN | NaN | 744.057 | 1142.1769 | 991.84717 | 203.98093 |
| Ints7 | VFSVIHSNDPVAR | -107 | -119 | 0 | 0.00025015 | 13 | NaN | NaN | 455.1875 | 569.3475 | 467.01624 | 443.7397 |
| Wdhd1 | SEEEKEEDFR | -821 | -830 | 1 | 0.00078661 | 10 | NaN | NaN | 956.08563 | NaN | NaN | 343.51816 |
| Snx12 | IAGHPLAQNER | -129 | -139 | 0 | 0 | 11 | NaN | NaN | 961.58136 | NaN | 1007.7769 | 897.3074 |
| Gdi1 | ICDPSYIPDR | -281 | -290 | 0 | 0.000230707 | 10 | NaN | NaN | 3293.1375 | 3870.2473 | NaN | NaN |
| Atp2b4 | TSPIEGLSGNPADLEK | -60 | -75 | 0 | 0.008498216 | 16 | NaN | NaN | 1099.4388 | NaN | NaN | NaN |
| Ndufa10 | VVEDIEYLK | -269 | -277 | 0 | 0 | 9 | NaN | NaN | 807.7848 | 1566.1913 | 458.13623 | NaN |
| Ndufa10 | QDDWTFHYLR | -286 | -295 | 0 | 0 | 10 | NaN | NaN | 908.7422 | NaN | NaN | NaN |

| | | | | | | | | | | | | |
|---|---|---|---|---|---|---|---|---|---|---|---|---|
| Parn | IEDFLQSEEKR | -181 | -191 | 1 | 0 | 11 | NaN | NaN | 690.0373 | 418.03125 | 578.3933 | 795.67 |
| Parn | YCPLPADLNEFK | -294 | -305 | 0 | 0 | 12 | NaN | NaN | 542.5226 | NaN | NaN | NaN |
| Kif4 | IETELKEELVK | -913 | -923 | 1 | 0.008919338 | 11 | NaN | NaN | 1034.0876 | NaN | NaN | NaN |
| Dync1li1 | IGILHENFQTL | -333 | -343 | 0 | 0.00122419 | 11 | NaN | NaN | 64.89579 | NaN | 58.276123 | NaN |
| Get4 | IGQLFFGVPPK | -279 | -289 | 0 | 0.000550307 | 11 | NaN | NaN | 753.10364 | NaN | NaN | NaN |
| Eif4e3 | IHQLLPHIAFK | -177 | -187 | 0 | 0 | 11 | NaN | NaN | 640.07196 | 1041.8833 | NaN | NaN |
| Cyp51a1 | IHTPENPVIR | -488 | -497 | 0 | 0.000627943 | 10 | NaN | NaN | 217.28351 | 932.451 | NaN | NaN |
| Coasy | IIDSDHLGHR | -384 | -393 | 0 | 0.003745449 | 10 | NaN | NaN | 1887.5438 | NaN | NaN | NaN |
| Ints4 | IIEPAGESDNPLR | -825 | -837 | 0 | 0.00025015 | 13 | NaN | NaN | 941.09705 | NaN | NaN | NaN |
| Gnai3 | IQCQFEDLNR | -303 | -312 | 0 | 0 | 10 | NaN | NaN | 1596.2302 | NaN | 469.35223 | 350.0026 |
| LRWD1 | IILWDIGVPNQDYK | -419 | -432 | 0 | 0 | 14 | NaN | NaN | 408.35428 | NaN | 322.54306 | NaN |
| Synj2bp | IKEDGAAAQDGR | -47 | -58 | 1 | 0 | 12 | NaN | NaN | 319.5699 | NaN | NaN | NaN |
| Nup160 | KVDAAELLR | -1297 | -1305 | 1 | 0.004255025 | 9 | NaN | NaN | 1559.6621 | NaN | 1685.5522 | 1796.2466 |
| Dis3l2 | ILEEWFGR | -474 | -481 | 0 | 0.002068213 | 8 | NaN | NaN | 1185.0857 | NaN | NaN | NaN |
| Rpap3 | ILHDFYIER | -592 | -600 | 0 | 0 | 9 | NaN | NaN | 2321.6553 | 1933.5586 | 858.0468 | 534.3525 |
| Ciapin1 | NCTCGLAEELER | -246 | -257 | 0 | 0 | 12 | NaN | NaN | 2404.6733 | NaN | NaN | 813.471 |
| Gid8 | QQHLIELIR | -106 | -114 | 0 | 0 | 9 | NaN | NaN | 354.43228 | NaN | 2291.0198 | NaN |
| Gid8 | INSLHPELLDTNR | -86 | -98 | 0 | 0 | 13 | NaN | NaN | 1423.8936 | NaN | NaN | NaN |
| Lsm4 | TAQNHPMLVELK | -9 | -20 | 0 | 0 | 12 | NaN | NaN | 841.1843 | 1098.388 | 631.154 | NaN |
| Gns | IQEPYTFPAILK | -118 | -129 | 0 | 0.001392869 | 12 | NaN | NaN | 849.9145 | NaN | NaN | NaN |
| Fgb | IRPFFPQQ | -474 | -481 | 1 | 0.000672736 | 8 | NaN | NaN | 272.57245 | 892.082 | NaN | 1142.7014 |
| Fgb | QQTLLNQERPIK | -103 | -114 | 1 | 0.004439092 | 12 | NaN | NaN | 722.1407 | NaN | NaN | NaN |
| Cmtr1 | IRPVLEYR | -584 | -591 | 1 | 0 | 8 | NaN | NaN | 270.44943 | 246.88719 | 527.02454 | 601.9207 |
| Lrch3 | SVPSIHVPSPAVPK | -692 | -705 | 0 | 0.007546675 | 14 | NaN | NaN | 586.9904 | NaN | NaN | NaN |
| Rnf40 | ISLEYSELQDK | -254 | -264 | 0 | 0 | 11 | NaN | NaN | 482.65936 | NaN | 483.0176 | NaN |
| Rbm27 | ISQLKDELK | -874 | -882 | 1 | 0.00078661 | 9 | NaN | NaN | 105.474556 | NaN | NaN | 406.84863 |
| Acads | SVNNSLYLGPILK | -117 | -129 | 0 | 0 | 13 | NaN | NaN | 693.2834 | 372.23102 | 1589.1123 | 1709.625 |
| Utp15 | LWDIPNSK | -148 | -155 | 0 | 0.00437931 | 8 | NaN | NaN | 1040.1361 | 1232.5906 | NaN | NaN |
| Gpd1l | IVGSGNWGSAVAK | -10 | -22 | 0 | 0.004628201 | 13 | NaN | NaN | 2542.8093 | NaN | NaN | 3185.5173 |
| Eps15l1 | LSQLQESHLEAHR | -545 | -557 | 0 | 0 | 13 | NaN | NaN | 921.0893 | NaN | NaN | NaN |
| Ppp2r2a | SDYETYLSADDLR | -187 | -199 | 0 | 0 | 13 | NaN | NaN | 1145.0925 | 799.2327 | NaN | NaN |
| Mydgf | IWRPQGK | -87 | -93 | 1 | 0 | 7 | NaN | NaN | 1122.9884 | NaN | 1380.7794 | 1349.1333 |
| Ddx18 | VSDIQSQLEK | -548 | -557 | 0 | 0.00111006 | 10 | NaN | NaN | 1466.2887 | NaN | NaN | NaN |
| Dnttip2 | SIDPGLNIK | -565 | -573 | 0 | 0.003360691 | 9 | NaN | NaN | 3343.2517 | 2090.6948 | NaN | NaN |
| Nifk | SFPSLVLPKPK | -186 | -196 | 1 | 0 | 11 | NaN | NaN | 490.55215 | NaN | NaN | NaN |
| Grhpr | VFVTGPLPAEGR | -9 | -20 | 0 | 0.00383302 | 12 | NaN | NaN | 477.64767 | NaN | NaN | NaN |
| Fam210a | KEETDPLQDK | -110 | -119 | 1 | 0 | 10 | NaN | NaN | 678.7712 | NaN | 886.4365 | 694.39856 |
| Hmga1 | KLEKEEEEGISQ | -89 | -100 | 2 | 0.000627943 | 12 | NaN | NaN | 1015.5722 | 988.95374 | 1366.387 | NaN |
| Hmga1 | QKEPSEVPTPK | -45 | -55 | 1 | 0.005176406 | 11 | NaN | NaN | 1407.0237 | NaN | NaN | NaN |
| Hmga1 | KLEKEEEEGISQE | -89 | -101 | 2 | 0.00149795 | 13 | NaN | NaN | 686.26776 | NaN | NaN | NaN |
| Ifng | QAFNELIR | -129 | -136 | 0 | 0 | 8 | NaN | NaN | 1213.1473 | 937.83746 | NaN | 932.462 |
| Fkbp15 | KGEPAEAEVPSEIK | -1002 | -1015 | 1 | 0.000431655 | 14 | NaN | NaN | 968.6303 | NaN | NaN | 1012.65283 |
| Eif3j1;Eif3j2 | SRDDFTEFGK | (155);(157) | (164);(166) | 1 | 0.002629273 | 10 | NaN | NaN | 360.1354 | NaN | NaN | NaN |
| Kdm2a | KIEESDEEAVQAK | -688 | -700 | 1 | 0 | 13 | NaN | NaN | 804.8544 | 788.53284 | 710.1994 | NaN |
| Kdm2a | LTDQTLFFLR | -1106 | -1115 | 0 | 0.00025015 | 10 | NaN | NaN | 780.1745 | 596.7612 | NaN | NaN |
| Brox | KIPTEAPQLELK | -334 | -345 | 1 | 0.00025015 | 12 | NaN | NaN | 912.2473 | 546.5803 | 833.69324 | 927.53357 |
| H1-1 | KKPAGPSVSELIVQA | -35 | -49 | 2 | 0.00659934 | 15 | NaN | NaN | 100.22238 | NaN | 1009.012 | 377.43124 |
| Slamf1 | KLHDALTDQDPCTTIY | -300 | -315 | 1 | 0.001463897 | 16 | NaN | NaN | 228.11475 | NaN | NaN | NaN |
| Psme3 | LDECEEAFQGTK | -89 | -100 | 0 | 0.000627943 | 12 | NaN | NaN | 2736.9993 | NaN | NaN | NaN |
| Cmpk1 | KSVDEVFGEVVK | -179 | -190 | 1 | 0 | 12 | NaN | NaN | 565.21216 | 698.75146 | 1104.426 | 1020.3373 |
| Lztfl1 | VLQDLQLDQENQQDLLK | -188 | -204 | 0 | 0 | 17 | NaN | NaN | 314.23816 | 367.10712 | 275.256 | 414.21982 |
| Lztfl1 | KPIIDITKPK | -128 | -137 | 2 | 0.003966045 | 10 | NaN | NaN | 238.59352 | NaN | NaN | NaN |
| Clptm1l | SVAHLPWK | -461 | -468 | 0 | 0.00111006 | 8 | NaN | NaN | 1898.7793 | NaN | NaN | NaN |
| Tnfrsf8 | KQCAPDYYVNEDGK | -67 | -80 | 1 | 0.001291103 | 14 | NaN | NaN | 2715.4907 | 3162.6074 | 3508.1965 | 3010.0461 |
| Tnfrsf8 | LENLPLLDDSPAGNPFSPR | -383 | -401 | 0 | 0.000471409 | 19 | NaN | NaN | 354.9994 | NaN | 357.83563 | 326.23877 |
| Lsm6 | LEQTEEYVNGQLK | -45 | -57 | 0 | 0.00025015 | 13 | NaN | NaN | 857.89935 | 884.4885 | 586.144 | NaN |
| Pkn1 | SLAPVELLLR | -69 | -78 | 0 | 0 | 10 | NaN | NaN | 303.98163 | 300.68454 | 285.49805 | 291.8916 |
| Pkn1 | KSPLTLEDFK | -611 | -620 | 1 | 0.000230707 | 10 | NaN | NaN | 3778.9739 | NaN | NaN | NaN |
| Dtymk | KTELEDHSVHLL | -60 | -71 | 1 | 0.001868731 | 12 | NaN | NaN | 486.86224 | 436.3137 | 525.97296 | 535.0273 |
| Ndufc2 | KTYAEILEPFHPVR | -107 | -120 | 1 | 0 | 14 | NaN | NaN | 1027.3486 | NaN | NaN | NaN |
| Sec22b | NIEEVLQR | -152 | -159 | 0 | 0.00437931 | 8 | NaN | NaN | 1702.2256 | 970.52515 | NaN | NaN |
| Sf3b5 | SQLEHLQSK | -9 | -17 | 0 | 0 | 9 | NaN | NaN | 4539.034 | NaN | 2799.4104 | 1284.5189 |
| Nipsnap3b | KYDNFAHR | -89 | -96 | 1 | 0.007343498 | 8 | NaN | NaN | 1185.0002 | 749.71454 | 972.56537 | NaN |
| Nipsnap3b | TNEFLENFK | -49 | -57 | 0 | 0 | 9 | NaN | NaN | 511.41705 | NaN | 1250.8881 | 1348.7125 |
| Dynll1 | SVECATQALEK | -21 | -31 | 0 | 0.003620005 | 11 | NaN | NaN | 628.46466 | 599.78656 | NaN | NaN |
| Pitpna | WVDLTMDDIR | -236 | -245 | 0 | 0 | 10 | NaN | NaN | 1160.5314 | 644.8902 | NaN | 1029.8752 |
| Pitpna | WWGLQNK | -203 | -209 | 0 | 0.003490217 | 7 | NaN | NaN | 431.2657 | 610.4011 | 4439.9673 | NaN |
| Arfgap2 | LCIDCSGVHR | -45 | -54 | 0 | 0.00025015 | 10 | NaN | NaN | 1851.4264 | NaN | NaN | 358.26587 |
| Pmpcb | STLGEAEIER | -164 | -173 | 0 | 0.000471409 | 10 | NaN | NaN | 824.4531 | 860.7543 | 1037.8009 | 786.5331 |
| Samsn1 | LDDDSPYSGPFCGR | -154 | -167 | 0 | 0 | 14 | NaN | NaN | 151.13864 | 489.44244 | NaN | NaN |
| Isy1 | LDEDDGVIVPLEQEYEK | -163 | -179 | 0 | 0 | 17 | NaN | NaN | 485.6585 | NaN | 642.0953 | NaN |
| Arfgef2;Arfgef | LDGNAIVDFVR | (1089);(1132) | (1099);(1142) | 0 | 0 | 11 | NaN | NaN | 181.85048 | 1130.914 | 206.44638 | NaN |
| Nae1 | LWGDHGQEALESAHVC | -19 | -34 | 0 | 0.000471409 | 16 | NaN | NaN | 813.2659 | NaN | 947.03815 | 274.23245 |
| Prkch | LDNVLLDHEGHCK | -482 | -494 | 0 | 0 | 13 | NaN | NaN | 18059.482 | NaN | NaN | NaN |
| Kifc1 | LDPGLHLGPGER | -573 | -584 | 0 | 0.000230707 | 12 | NaN | NaN | 1579.3849 | NaN | 628.90454 | 569.4029 |
| Sympk | LDSDNDGIR | -172 | -180 | 0 | 0.005668207 | 9 | NaN | NaN | 828.76483 | NaN | NaN | 755.4039 |
| Pnpt1 | LDTEEHLK | -318 | -325 | 0 | 0.006164361 | 8 | NaN | NaN | 1549.6842 | NaN | 1527.7113 | NaN |
| Etnk1 | LDVTVQDQEEQR | -17 | -28 | 0 | 0 | 12 | NaN | NaN | 520.1898 | 550.1842 | NaN | NaN |

| | | | | | | | | | | | | |
|---|---|---|---|---|---|---|---|---|---|---|---|---|
| Noc2l | LKDRDPEFYK | -78 | -87 | 2 | 0.000230707 | 10 | NaN | NaN | 3950.2468 | NaN | 1209.7812 | NaN |
| Noc2l | LPEAHQGLK | -690 | -698 | 0 | 0.000230707 | 9 | NaN | NaN | 1615.4276 | NaN | NaN | NaN |
| Dgka | LEGGRPEDKLEFTFK | -107 | -121 | 2 | 0.00025015 | 15 | NaN | NaN | 527.2494 | NaN | NaN | NaN |
| Psd4 | TWHQSPQTLDSPLQK | -234 | -248 | 0 | 0.001023236 | 15 | NaN | NaN | 3054.1973 | NaN | NaN | 624.0559 |
| Psd4 | LEWAIDEEDADRPEK | -688 | -702 | 1 | 0 | 15 | NaN | NaN | 388.40213 | NaN | NaN | NaN |
| Prpf38b | LEWFSTLFPR | -212 | -221 | 0 | 0 | 10 | NaN | NaN | 688.02966 | 244.5864 | 1049.5917 | 746.7557 |
| Was | LFELLGR | -37 | -43 | 0 | 0.00897527 | 7 | NaN | NaN | 466.90625 | NaN | NaN | NaN |
| Rpf1 | LFPHNPQFIGR | -266 | -276 | 0 | 0.001846793 | 11 | NaN | NaN | 1666.7749 | 1841.3925 | NaN | NaN |
| Mrps9 | LFPSGLFEK | -111 | -119 | 0 | 0.001636916 | 9 | NaN | NaN | 479.28882 | NaN | NaN | NaN |
| Snx17 | LFSLTPAEVEQR | -64 | -75 | 0 | 0.00025015 | 12 | NaN | NaN | 279.10336 | 341.91266 | 593.35596 | 462.24524 |
| Gphn | TLIINLPGSK | -139 | -148 | 0 | 0.00025015 | 10 | NaN | NaN | 246.61124 | NaN | 269.67203 | 268.15015 |
| Capns1 | SHYSNIEANESEEVR | -85 | -99 | 0 | 0 | 15 | NaN | NaN | 842.4337 | 613.6285 | 951.74805 | 953.9391 |
| Riox1 | LGHFAPVDAVADQR | -450 | -463 | 0 | 0 | 14 | NaN | NaN | 713.36633 | 748.97046 | NaN | 701.7849 |
| Mrps16 | LGSYDPLPNSHGEK | -51 | -64 | 0 | 0.001636916 | 14 | NaN | NaN | 1630.881 | NaN | NaN | NaN |
| Ikbkb | QYLAPELLEQQK | -187 | -198 | 0 | 0.002199523 | 12 | NaN | NaN | 529.5307 | NaN | NaN | NaN |
| Acy1 | SHTDVVPVFK | -79 | -88 | 0 | 0.00025015 | 10 | NaN | NaN | 1683.6691 | NaN | NaN | NaN |
| Mtatp6 | LHSFQHWLVK | -42 | -51 | 0 | 0 | 10 | NaN | NaN | 1733.3104 | 3564.3384 | 2230.7588 | 1195.6958 |
| Trappc3 | LIEDFLAR | -55 | -62 | 0 | 0 | 8 | NaN | NaN | 612.6314 | 631.0004 | 525.7054 | NaN |
| Ptk2b | NLDHPHIVK | -480 | -488 | 0 | 0.00149795 | 9 | NaN | NaN | 1474.3737 | NaN | NaN | NaN |
| Ptk2b | LPKPELCPPVLYTL | -644 | -657 | 1 | 0.005605646 | 14 | NaN | NaN | 651.2306 | NaN | NaN | NaN |
| Ptk2b | QPFFWLENK | -624 | -632 | 0 | 0.000550307 | 9 | NaN | NaN | 538.427 | NaN | NaN | NaN |
| Iigp1 | SPAVFKPTDEETIQER | -347 | -362 | 1 | 0.00025015 | 16 | NaN | NaN | 635.01874 | 660.53143 | 686.7239 | 703.2421 |
| Iigp1 | SLPNITDSVIEK | -262 | -273 | 0 | 0 | 12 | NaN | NaN | 604.3875 | 763.0239 | 728.50555 | 798.89056 |
| Iigp1 | VDSDITNEADGKPQTFDK | -185 | -202 | 1 | 0.00142999 | 18 | NaN | NaN | 230.6855 | 406.50006 | 466.60562 | 285.07642 |
| Gsk3a | TSSFAEPGGGGGGGGGGPGGSASGPGGTGGGK | -19 | -50 | 0 | 0.002752526 | 32 | NaN | NaN | 158.48132 | NaN | NaN | NaN |
| Ldhb | LKDDEVAQLR | -309 | -318 | 1 | 0.00078661 | 10 | NaN | NaN | 983.4965 | NaN | NaN | 1293.2423 |
| Erbin | LKDEETNEESGR | -505 | -516 | 1 | 0 | 12 | NaN | NaN | 250.34656 | NaN | 334.82254 | NaN |
| Epc1 | LKEDDELIR | -164 | -172 | 1 | 0.00111006 | 9 | NaN | NaN | 947.0447 | NaN | NaN | NaN |
| Man2a1 | YSYDIPHTCGPDPK | -351 | -364 | 0 | 0 | 14 | NaN | NaN | 588.3383 | NaN | NaN | 247.77011 |
| Polr2l | LLNYAPLEK | -59 | -67 | 0 | 0.000394361 | 9 | NaN | NaN | 532.27765 | 555.2124 | 490.53137 | 523.946 |
| Carnmt1 | LLPQFPLHLDK | -92 | -102 | 0 | 0 | 11 | NaN | NaN | 676.2869 | 529.2372 | 839.32574 | 380.80066 |
| Mtx1 | TSNPWQSPSGTLPALR | -42 | -57 | 0 | 0 | 16 | NaN | NaN | 978.4899 | 852.3906 | 1049.082 | 985.3614 |
| Nudt21 | LMTEILGR | -124 | -131 | 0 | 0.002697088 | 8 | NaN | NaN | 737.0786 | NaN | NaN | NaN |
| Eps15 | LNDPFQPFPGNDSPK | -804 | -818 | 0 | 0.000627943 | 15 | NaN | NaN | 695.9135 | 588.0086 | NaN | NaN |
| Rwdd1 | SEAGENDETVQTTLK | -39 | -53 | 0 | 0 | 15 | NaN | NaN | 316.92838 | 453.5247 | 501.1388 | 508.4618 |
| Exoc1 | LNHFFEGVEAR | -778 | -788 | 0 | 0.000431655 | 11 | NaN | NaN | 323.39456 | NaN | NaN | NaN |
| Ppp1ca | SFQILKPADK | -292 | -301 | 1 | 0 | 10 | NaN | NaN | 735.92206 | NaN | NaN | NaN |
| Ints10 | LPANLLYK | -190 | -197 | 0 | 0 | 8 | NaN | NaN | 607.2052 | 513.0092 | 484.0341 | NaN |
| Uba3 | LPEHCIEYVR | -245 | -254 | 0 | 0 | 10 | NaN | NaN | 2644.108 | 3015.297 | 3082.3591 | 2879.0215 |
| Ssu72 | LPGPAPDKPNVYDFK | -44 | -58 | 1 | 0.00078661 | 15 | NaN | NaN | 399.61285 | 265.0852 | NaN | 251.48134 |
| Ngly1 | LPLEQSEDPAGLIR | -131 | -144 | 0 | 0.00025015 | 14 | NaN | NaN | 778.7427 | 635.0076 | NaN | NaN |
| Thumpd3 | QWDICNLPLR | -389 | -398 | 0 | 0.005301614 | 10 | NaN | NaN | 407.30783 | NaN | NaN | 161.65968 |
| Parp3 | LQLLDSGESEYK | -326 | -337 | 0 | 0.00025015 | 12 | NaN | NaN | 2459.4775 | NaN | NaN | 1099.6503 |
| Mlec | LYIEFVK | -190 | -196 | 0 | 0.005758933 | 7 | NaN | NaN | 611.1563 | NaN | NaN | NaN |
| Gga1 | LQPPSGTELPAFNPIVHPS | -568 | -586 | 0 | 0.00025015 | 19 | NaN | NaN | 387.08353 | NaN | 521.9341 | 679.5525 |
| Lrrc59 | VTGLHQQPLCT | -268 | -278 | 0 | 0.000230707 | 11 | NaN | NaN | 1055.7607 | 349.99203 | NaN | NaN |
| Maz | SRPDHLNSHVR | -347 | -357 | 1 | 0.004350979 | 11 | NaN | NaN | 452.0415 | NaN | NaN | NaN |
| Epb41 | LSTHSPFR | -702 | -709 | 0 | 0.000471409 | 8 | NaN | NaN | 1316.6113 | NaN | NaN | NaN |
| Mapk14 | SQERPTFYR | -2 | -10 | 1 | 0.001708008 | 9 | NaN | NaN | 1794.6345 | NaN | 1010.2398 | 1262.7428 |
| Eef1b | VFEAVSGPPPADLCHALR | -37 | -54 | 0 | 0 | 18 | NaN | NaN | 812.5052 | 272.44077 | 895.3797 | 992.47534 |
| Prf1 | SPHSGFHEVTCELNHGR | -499 | -515 | 0 | 0 | 17 | NaN | NaN | 281.63013 | NaN | 321.2711 | 533.1634 |
| Prf1 | LVQKPPLHLDFK | -182 | -193 | 1 | 0 | 12 | NaN | NaN | 298.6178 | NaN | 537.60095 | 590.31573 |
| Prkag1 | VVVDEHDVVK | -300 | -309 | 0 | 0 | 10 | NaN | NaN | 4286.995 | 6124.5933 | 4060.8516 | 3693.3682 |
| U2af1 | MHLKPISR | -172 | -179 | 1 | 0.005605646 | 8 | NaN | NaN | 4901.897 | NaN | NaN | NaN |
| Cdipt | MIDLSGNPVLR | -122 | -132 | 0 | 0 | 11 | NaN | NaN | 476.1717 | NaN | 543.2264 | NaN |
| Sfr1 | NDENNETLSEPGESSK | -155 | -170 | 0 | 0 | 16 | NaN | NaN | 729.0358 | NaN | NaN | NaN |
| Ufm1 | NDGIGINPAQTAGNVFLK | -52 | -69 | 0 | 0 | 18 | NaN | NaN | 2235.8413 | NaN | 926.303 | 879.2457 |
| Hnrnpll | VEFESVLCAQK | -259 | -269 | 0 | 0.008624882 | 11 | NaN | NaN | 826.25757 | NaN | NaN | NaN |
| Hnrnpll | TLSGLLEWK | -545 | -553 | 0 | 0.003133457 | 9 | NaN | NaN | 506.2118 | NaN | NaN | NaN |
| Atp2a3 | NEITAMTGDGVNDAPALK | -695 | -712 | 0 | 0.00897527 | 18 | NaN | NaN | 4905.6626 | 2481.0583 | 3347.905 | NaN |
| Vti1a | NELLGDAGNSSENQR | -101 | -115 | 0 | 0 | 15 | NaN | NaN | 694.1188 | 702.61597 | NaN | NaN |
| Ubr7 | NEPCSSSSSESDLQTVFK | -257 | -274 | 0 | 0.00051306 | 18 | NaN | NaN | 471.91803 | NaN | NaN | NaN |
| Cops4 | VGIPLETGQK | -128 | -137 | 0 | 0.003521747 | 10 | NaN | NaN | 839.64545 | NaN | 286.8021 | NaN |
| Pafah1b3 | NFGIGGDSTQHVLWR | -70 | -84 | 0 | 0.00025015 | 15 | NaN | NaN | 5337.6655 | 4952.0146 | 8818.635 | 11674.038 |
| Rbm22 | NHFYQFGEIR | -250 | -259 | 0 | 0 | 10 | NaN | NaN | 754.6654 | NaN | 923.23126 | 1322.9756 |
| Riok1 | NHIADPDIDKK | -521 | -531 | 1 | 0 | 11 | NaN | NaN | 861.4072 | NaN | 992.94104 | 1042.8706 |
| Vcpip1 | PGLIPAEK | -323 | -330 | 0 | 0.004065182 | 8 | NaN | NaN | 434.46185 | 302.28583 | 297.763 | 378.5186 |
| Pold3 | NIDEFVTDQNK | -9 | -19 | 0 | 0.006505855 | 11 | NaN | NaN | 500.0021 | NaN | NaN | NaN |
| Nop14 | NIYNLNEDEELTHYGQ | -117 | -132 | 0 | 0.008436628 | 16 | NaN | NaN | 309.88348 | NaN | 354.87546 | NaN |
| Copg2 | NLDLENLITDSNR | -326 | -338 | 0 | 0.001636916 | 13 | NaN | NaN | 493.1974 | NaN | NaN | 525.47156 |
| Mat2b | NLPSSHLRPITDSPVIGAQRPK | -270 | -291 | 2 | 0.00111006 | 22 | NaN | NaN | 563.7051 | NaN | NaN | NaN |
| Tm9sf3 | YLDPSFFQHR | -209 | -218 | 0 | 0 | 10 | NaN | NaN | 3325.9253 | 2485.4116 | 2639.1257 | 2380.6096 |
| Ufl1 | NNPVHLITEEDLK | -390 | -402 | 0 | 0 | 13 | NaN | NaN | 479.83148 | 314.04254 | NaN | 553.54333 |
| Gmppa | SPEALKPLR | -187 | -195 | 1 | 0.001190916 | 9 | NaN | NaN | 713.1678 | NaN | 1501.3228 | 1593.9991 |
| Cep55 | NQQWLVYDQQR | -181 | -191 | 0 | 0 | 11 | NaN | NaN | 947.27435 | 607.465 | 820.433 | 1024.1436 |
| Cep55 | SEELLSQVR | -324 | -332 | 0 | 0.00025015 | 9 | NaN | NaN | 889.82935 | NaN | NaN | NaN |
| Helz2 | NSLLSPGAPTSEFFTEPR | -1125 | -1142 | 0 | 0 | 18 | NaN | NaN | 1341.0597 | 1319.7455 | NaN | NaN |
| Helz2 | TVDPQGACNLDDALSVR | -1577 | -1593 | 0 | 0.00122419 | 17 | NaN | NaN | 361.11224 | NaN | NaN | 540.19385 |

| | | | | | | | | | | | | |
|---|---|---|---|---|---|---|---|---|---|---|---|---|
| Ctss | NSWGLNFGDQGYIR | -307 | -320 | 0 | 0 | 14 | NaN | NaN | 299.2818 | 439.44162 | 784.91626 | 586.6277 |
| Ctss | PDTVDWR | -124 | -130 | 0 | 0.00416551 | 7 | NaN | NaN | 798.7285 | NaN | 508.00714 | 585.1665 |
| Mrpl27 | VHVVPAKPEGTFK | -131 | -143 | 1 | 0.001291103 | 13 | NaN | NaN | 931.70514 | NaN | 487.89395 | NaN |
| Pcif1 | YSCVDDSAFER | -452 | -462 | 0 | 0 | 11 | NaN | NaN | 369.76538 | 123.44287 | NaN | NaN |
| Suv39h1 | NVFIDNLDER | -334 | -343 | 0 | 0.000230707 | 10 | NaN | NaN | 1022.6974 | NaN | NaN | 1776.607 |
| Dcaf13 | NWGFDSISSVK | -193 | -203 | 0 | 0 | 11 | NaN | NaN | 1447.1042 | NaN | NaN | NaN |
| Polr2c | SSVLHDEFIAHR | -56 | -67 | 0 | 0 | 12 | NaN | NaN | 612.48865 | NaN | 603.9582 | 190.5304 |
| Mrpl58 | QASSYIPLDR | -67 | -76 | 0 | 0.002296977 | 10 | NaN | NaN | 349.36853 | NaN | NaN | NaN |
| Slc2a1 | VTILELFR | -257 | -264 | 0 | 0.000627943 | 8 | NaN | NaN | 302.47644 | 293.16898 | 121.85753 | NaN |
| Arpp19 | TEVTGDHIPTPQDLPQR | -85 | -101 | 0 | 0.000230707 | 17 | NaN | NaN | 759.591 | 186.87227 | 1011.077 | 203.70284 |
| Thumpd1 | QQSPQPVAGK | -6 | -15 | 0 | 0.00644558 | 10 | NaN | NaN | 4116.2803 | NaN | NaN | NaN |
| Eif4ebp1 | QSQLPSSPEDKR | -94 | -105 | 1 | 0.00122419 | 12 | NaN | NaN | 1649.0272 | NaN | NaN | NaN |
| Eif4ebp1 | QSQLPSSPEDK | -94 | -104 | 0 | 0.005946382 | 11 | NaN | NaN | 1080.6144 | NaN | NaN | NaN |
| Cwc15 | QTTQDAPEEVR | -45 | -55 | 0 | 0 | 11 | NaN | NaN | 840.59326 | NaN | NaN | NaN |
| Pno1 | QVEDALALIR | -141 | -150 | 0 | 0.004412728 | 10 | NaN | NaN | 532.51526 | NaN | NaN | NaN |
| Ak3 | TQCSWLLDGFPR | -83 | -94 | 0 | 0 | 12 | NaN | NaN | 672.2768 | 1118.2852 | 1556.4028 | 1561.28 |
| Srsf2 | YGRPPDSHHS | -92 | -101 | 1 | 0 | 10 | NaN | NaN | 568.06195 | 658.707 | 746.4709 | 867.76013 |
| Casp6 | VDFCKDPDAIGK | -243 | -254 | 1 | 0.001776473 | 12 | NaN | NaN | 229.02502 | NaN | NaN | NaN |
| Eif2b4 | RVPEHTPADDPTLLR | -143 | -157 | 1 | 0.002068213 | 15 | NaN | NaN | 902.99536 | NaN | NaN | NaN |
| Yipf4 | SADAEDLSGSIAAPDVK | -22 | -38 | 0 | 0.002697088 | 17 | NaN | NaN | 2162.8853 | NaN | NaN | NaN |
| Sh3gl1 | SALVDAQLDYHR | -216 | -227 | 0 | 0 | 12 | NaN | NaN | 824.62335 | 2465.0667 | 3458.5725 | 3146.6707 |
| Polr1a | SAQEHLFAIWK | -267 | -277 | 0 | 0.001868731 | 11 | NaN | NaN | 912.01825 | NaN | NaN | NaN |
| Bltp3b | SDDGSFHIR | -1350 | -1358 | 0 | 0 | 9 | NaN | NaN | 1130.983 | NaN | NaN | NaN |
| Smn1 | SDDSDIWDDTALIK | -25 | -38 | 0 | 0.001463897 | 14 | NaN | NaN | 187.6359 | 397.2067 | NaN | NaN |
| Rbm15 | SDEAVEDGLFHEFK | -181 | -194 | 0 | 0 | 14 | NaN | NaN | 409.60544 | 102.752144 | 543.4919 | 777.9692 |
| Mapk1ip1l | SDEFSLADALPEQSSAKPPAVTNTK | -2 | -26 | 1 | 0.000550307 | 25 | NaN | NaN | 232.18958 | 306.73672 | 751.4559 | 553.2371 |
| Slc38a2 | YELPLVIK | -171 | -178 | 0 | 0 | 8 | NaN | NaN | 482.19363 | 469.37564 | 512.92676 | 540.5513 |
| Slc38a2 | SDFNYSYPTK | -24 | -33 | 0 | 0.009242859 | 10 | NaN | NaN | 2165.3572 | 2606.1821 | NaN | NaN |
| Emc8 | SDWTNPEINK | -192 | -201 | 0 | 0.001055699 | 10 | NaN | NaN | 1253.8263 | NaN | NaN | NaN |
| Scyl3 | SEDEKTEFFK | -263 | -272 | 1 | 0.003620005 | 10 | NaN | NaN | 6552.475 | NaN | NaN | NaN |
| Ndufa9 | SSEIEETKPAK | -363 | -373 | 1 | 0.001291103 | 11 | NaN | NaN | 579.4137 | NaN | 685.62463 | NaN |
| Pcnt | SELTEQLASSESR | -676 | -688 | 0 | 0.002534621 | 13 | NaN | NaN | 899.8687 | NaN | 1901.889 | NaN |
| Akap8 | SFEDEEIQK | -396 | -404 | 0 | 0.00111006 | 9 | NaN | NaN | 651.67114 | NaN | NaN | NaN |
| Brix1 | SFDPAFDDLPHYALLK | -167 | -182 | 0 | 0.00897527 | 16 | NaN | NaN | 815.1578 | NaN | NaN | NaN |
| Elovl5 | SFPSLENSVKPR | -283 | -294 | 1 | 0 | 12 | NaN | NaN | 9290.385 | 9824.378 | NaN | NaN |
| Yars2 | SFSELVLDPGTSVIDTCR | -392 | -409 | 0 | 0.000712251 | 18 | NaN | NaN | 319.1802 | NaN | NaN | NaN |
| Rtf2 | SGKPEEADPDPR | -226 | -237 | 1 | 0 | 12 | NaN | NaN | 262.4958 | NaN | 569.11456 | 541.6133 |
| Timmdc1 | SGVSDPGSLESGWDR | -43 | -57 | 0 | 0.000230707 | 15 | NaN | NaN | 1735.9995 | 993.53595 | 968.45013 | 790.3992 |
| Coro2a | SKENCYDSVPITR | -21 | -33 | 1 | 0 | 13 | NaN | NaN | 550.5364 | 729.2087 | 750.74445 | 776.27686 |
| Capn7 | SKIPSPYTLSK | -680 | -690 | 1 | 0.003003218 | 11 | NaN | NaN | 830.83954 | NaN | 904.8168 | NaN |
| Ercc2 | SLEQLQSEETLQR | -740 | -752 | 0 | 0.003745449 | 13 | NaN | NaN | 2791.7793 | NaN | NaN | NaN |
| Krt19 | TKFETEHALR | -170 | -179 | 1 | 0.00025015 | 10 | NaN | NaN | 3047.9937 | NaN | 240.14285 | 889.3252 |
| Larp4b | SLPLVQVDEK | -218 | -227 | 0 | 0.005522803 | 10 | NaN | NaN | 1188.7231 | NaN | NaN | NaN |
| Usp15 | SLPPVLVVHLK | -826 | -836 | 0 | 0.000591124 | 11 | NaN | NaN | 589.50146 | NaN | NaN | NaN |
| Ubl7 | SLSDWHLAVK | -2 | -11 | 0 | 0 | 10 | NaN | NaN | 587.7342 | NaN | 766.45575 | 431.65045 |
| Twf1 | SPLLEIVER | -277 | -285 | 0 | 0 | 9 | NaN | NaN | 713.57574 | 459.7903 | 603.44196 | 586.89044 |
| Comt | SQDLIPQLK | -162 | -170 | 0 | 0 | 9 | NaN | NaN | 919.3207 | 955.644 | NaN | NaN |
| Usp3 | SQGDSNVSLVEEFR | -210 | -223 | 0 | 0 | 14 | NaN | NaN | 591.0362 | 189.20111 | 290.63815 | 552.355 |
| Ablim1 | TLSPTPSAEGYQDVR | -473 | -487 | 0 | 0.000431655 | 15 | NaN | NaN | 563.4067 | NaN | NaN | NaN |
| Cnot8 | SSIDYQYQLLR | -56 | -66 | 0 | 0 | 11 | NaN | NaN | 869.19574 | NaN | NaN | NaN |
| Pigu | VEVVSPLSSWK | -32 | -42 | 0 | 0.000471409 | 11 | NaN | NaN | 661.4745 | NaN | 566.049 | NaN |
| Hgs | SSPVNSSAPLAEDIDPELAR | -309 | -328 | 0 | 0 | 20 | NaN | NaN | 1240.476 | NaN | 1529.1866 | 1561.5398 |
| Ap1g2 | STVVECLQETDASLSR | -349 | -364 | 0 | 0.001708008 | 16 | NaN | NaN | 554.27496 | NaN | 393.58545 | 1069.5481 |
| Polr1h | SVDEGPELQGPVIDR | -68 | -82 | 0 | 0 | 15 | NaN | NaN | 801.2412 | NaN | 1121.7616 | NaN |
| Sema7a | SVLQSINPAEPHR | -525 | -537 | 0 | 0.001092529 | 13 | NaN | NaN | 691.39246 | NaN | NaN | 360.05554 |
| Taf6l | SVSHDLEQLHR | -205 | -215 | 0 | 0 | 11 | NaN | NaN | 624.6233 | 433.94116 | NaN | NaN |
| Lasp1 | YKEEFEK | -86 | -92 | 1 | 0.001610812 | 7 | NaN | NaN | 3456.6292 | NaN | 3902.722 | NaN |
| Hspbap1 | SWIELEEDHLAR | -271 | -282 | 0 | 0.00078661 | 12 | NaN | NaN | 1766.5164 | NaN | NaN | 2116.5798 |
| Ube2o | SYGFYPGQVLIGPAK | -252 | -266 | 0 | 0.000298805 | 15 | NaN | NaN | 592.4802 | NaN | NaN | NaN |
| Zc3h18 | TTAPGPEPAKPGDLR | -648 | -662 | 1 | 0 | 15 | NaN | NaN | 404.94214 | 480.60953 | NaN | NaN |
| Gimap1 | TDPGCVETAR | -70 | -79 | 0 | 0 | 10 | NaN | NaN | 251.17447 | NaN | NaN | NaN |
| Sars2 | TELGLHFR | -380 | -387 | 0 | 0.00025015 | 8 | NaN | NaN | 3000.884 | NaN | NaN | NaN |
| Ddx24 | TESGVLPEEAR | -294 | -304 | 0 | 0 | 11 | NaN | NaN | 890.9928 | NaN | NaN | NaN |
| Nol12 | TESVQYDHPNHTVTVT | -94 | -109 | 0 | 0.000826806 | 16 | NaN | NaN | 737.3888 | 492.46695 | 388.88873 | NaN |
| Galnt2 | TFLDSHCECNER | -220 | -231 | 0 | 0 | 12 | NaN | NaN | 394.18845 | NaN | NaN | 330.88632 |
| Sh3glb1 | TGSLVITCPSNLNDLK | -287 | -302 | 0 | 0.008014596 | 16 | NaN | NaN | 616.4408 | NaN | NaN | NaN |
| Pnpo | THHGEEDWVYER | -247 | -258 | 0 | 0 | 12 | NaN | NaN | 2348.338 | NaN | NaN | 2852.1636 |
| Pnpo | THHGEEDWVYER | -247 | -258 | 0 | 0 | 12 | NaN | NaN | 2348.338 | NaN | NaN | 2852.1636 |
| Nedd4 | THLAEEFNTR | -352 | -361 | 0 | 0 | 10 | NaN | NaN | 1047.2831 | NaN | 2418.6458 | 1141.7463 |
| Cpsf2 | VCNNQVAVR | -743 | -751 | 0 | 0.004983786 | 9 | NaN | NaN | 2883.8743 | NaN | NaN | NaN |
| Nup214 | TTLLEGFAGVEEAR | -764 | -777 | 0 | 0.003360691 | 14 | NaN | NaN | 2429.1646 | NaN | NaN | NaN |
| Nisch | TLLTTFPDVAPR | -99 | -110 | 0 | 0 | 12 | NaN | NaN | 193.51822 | NaN | NaN | NaN |
| Wdr5 | TLPAHSDPVSAVHFNR | -166 | -181 | 0 | 0 | 16 | NaN | NaN | 867.36145 | 917.84424 | 1261.2633 | 1190.4325 |
| Gtf2f1 | TLTAEEAEEEWER | -154 | -166 | 0 | 0.000230707 | 13 | NaN | NaN | 86.73678 | NaN | NaN | 4080.7078 |
| Chid1 | TQFSDKPVQDR | -38 | -48 | 1 | 0.00111006 | 11 | NaN | NaN | 486.98425 | NaN | NaN | NaN |
| Yy1 | TREEVVGGDDSDGLR | -110 | -124 | 1 | 0.006850902 | 15 | NaN | NaN | 906.843 | NaN | NaN | NaN |
| Nsun5 | TRPEDAIDYFK | -145 | -155 | 1 | 0 | 11 | NaN | NaN | 337.77576 | NaN | 597.9567 | 579.8027 |
| Kif15 | YNASLADKEESK | -1019 | -1030 | 1 | 0.00025015 | 12 | NaN | NaN | 1643.9153 | NaN | NaN | NaN |

| | | | | | | | | | | | | |
|---|---|---|---|---|---|---|---|---|---|---|---|---|
| Tbrg4 | WEFPNFNSR | -565 | -573 | 0 | 0.004472887 | 9 | NaN | NaN | 395.08292 | NaN | NaN | NaN |
| Pogz | TSTDPLPVFLYPPVQR | -716 | -731 | 0 | 0 | 16 | NaN | NaN | 355.14426 | 364.2465 | 532.0001 | 399.29428 |
| Slc25a17 | TVFFPLDTAR | -25 | -34 | 0 | 0.00111006 | 10 | NaN | NaN | 758.41284 | NaN | NaN | NaN |
| Galm | TVFGELPSGGGTVEK | -7 | -21 | 0 | 0.00204437 | 15 | NaN | NaN | 624.1934 | NaN | NaN | 1824.2415 |
| Kcnh1 | TYCDLHVIK | -665 | -673 | 0 | 0.009153908 | 9 | NaN | NaN | 871.4471 | NaN | 1286.5028 | NaN |
| Abhd17b | TYCFDAFPNIDK | -208 | -219 | 0 | 0.000230707 | 12 | NaN | NaN | 737.71045 | NaN | NaN | NaN |
| Osgep | VTAPGTGFLPGDTAR | -33 | -47 | 0 | 0 | 15 | NaN | NaN | 5362.0547 | 2152.8438 | NaN | NaN |
| Mrfap1 | VEASEESALNHLQGAGGAEPR | -71 | -91 | 0 | 0.000712251 | 21 | NaN | NaN | 695.8988 | 1552.2842 | 245.55995 | 668.07587 |
| Mrpl30 | YGGDPQNPHK | -56 | -65 | 0 | 0.000712251 | 10 | NaN | NaN | 508.6886 | NaN | NaN | 360.67343 |
| Exog | VIGEDNVAVPSHLYK | -216 | -230 | 0 | 0.000230707 | 15 | NaN | NaN | 628.64514 | NaN | 696.08295 | 758.8461 |
| Tmed3 | VITGGHYDVDCYVEDPR | -61 | -77 | 0 | 0.00111006 | 17 | NaN | NaN | 275.09027 | NaN | NaN | NaN |
| Polr1d | VLHEEDHTLGNCLR | -43 | -56 | 0 | 0.000230707 | 14 | NaN | NaN | 529.4844 | NaN | 532.76587 | NaN |
| Gyg1 | VLPLLQHHGLVK | -257 | -268 | 0 | 0.00025015 | 12 | NaN | NaN | 60.378902 | NaN | NaN | NaN |
| Atp6v1d | VNAIEHVIIPR | -169 | -179 | 0 | 0.001291103 | 11 | NaN | NaN | 273.54034 | NaN | NaN | NaN |
| Wdr74 | WNHVWVGTET | -8 | -17 | 0 | 0.009242859 | 10 | NaN | NaN | 2582.9458 | NaN | NaN | NaN |
| Wdr74 | YFNAEEGTFLSQR | -69 | -81 | 0 | 0.001055699 | 13 | NaN | NaN | 837.9014 | NaN | NaN | NaN |
| Rpp38 | VPSLNVPWLPDR | -195 | -206 | 0 | 0.000591124 | 12 | NaN | NaN | 419.18832 | 780.5117 | 463.35956 | NaN |
| Vwa5a | VQSNCSLSPIQYLTDDK | -195 | -211 | 0 | 0.00078661 | 17 | NaN | NaN | 505.0969 | NaN | NaN | 1193.4775 |
| Cul5 | VSEDLNQAFK | -502 | -511 | 0 | 0.000431655 | 10 | NaN | NaN | 1583.1688 | NaN | NaN | NaN |
| Amdhd2 | VSLGHSVADLR | -207 | -217 | 0 | 0 | 11 | NaN | NaN | 1571.6447 | NaN | NaN | NaN |
| Psmb5 | VSSDNVADLHDK | -246 | -257 | 0 | 0.000591124 | 12 | NaN | NaN | 562.27844 | NaN | NaN | NaN |
| Mavs | VTTGGNQTGPNSSIR | -372 | -386 | 0 | 0.00025015 | 15 | NaN | NaN | 1132.5024 | 1999.0948 | NaN | NaN |
| Nom1 | VVEFSELDKPR | -761 | -771 | 1 | 0 | 11 | NaN | NaN | 650.8905 | NaN | NaN | NaN |
| Sra1 | WDAADDIHR | -159 | -167 | 0 | 0.000550307 | 9 | NaN | NaN | 1181.863 | 310.38467 | NaN | NaN |
| Bub1b | WDGLLWK | -963 | -969 | 0 | 0.00676572 | 7 | NaN | NaN | 357.6748 | NaN | 343.68176 | 174.0905 |
| Ubfd1 | WVPTQYVDAIK | -348 | -358 | 0 | 0 | 11 | NaN | NaN | 383.2463 | 453.37103 | NaN | 883.6653 |
| Dnajc21 | YDVLSDPQER | -53 | -62 | 0 | 0.000550307 | 10 | NaN | NaN | 1290.6665 | NaN | NaN | NaN |
| Mrps5 | YEGHTIFHDISLR | -287 | -299 | 0 | 0.000230707 | 13 | NaN | NaN | 11187.1455 | NaN | NaN | 383.38824 |
| Scyl2 | YFDTLFQR | -337 | -344 | 0 | 0.00025015 | 8 | NaN | NaN | 2093.4058 | 449.2736 | 289.86465 | 627.3072 |
| Prkcd | YFLEDGDCK | -120 | -128 | 0 | 0 | 9 | NaN | NaN | 439.592 | NaN | 544.13104 | NaN |
| Aars2 | YFSGDSQTGLDPDLETR | -166 | -182 | 0 | 0 | 17 | NaN | NaN | 756.59125 | NaN | NaN | NaN |
| Tra2a | YGPLSGVNVVYDQR | -140 | -153 | 0 | 0.001023236 | 14 | NaN | NaN | 7239.473 | NaN | NaN | NaN |
| Mrps18b | YGSRPVWAD | -66 | -74 | 1 | 0.000230707 | 9 | NaN | NaN | 543.3616 | NaN | 626.6352 | NaN |
| Tmem209 | YLDLTPNQEYLFER | -400 | -413 | 0 | 0.00025015 | 14 | NaN | NaN | 355.148 | 384.49496 | 597.0297 | 372.98346 |
| Ccnd3 | YLSCVPTR | -88 | -95 | 0 | 0.004439092 | 8 | NaN | NaN | 264.64145 | NaN | NaN | NaN |
| Tmem254 | YLVDHYHTFLR | -52 | -62 | 0 | 0 | 11 | NaN | NaN | 4252.1104 | NaN | 2694.1262 | 4070.0808 |
| Hmmr | YNDTAQSLR | (445)(466)(487)(508)(5 | (453)(474)(495)(516)( | 0 | 0 | 9 | NaN | NaN | 692.1298 | NaN | NaN | NaN |
| Tyms | YSLRDEFPLLTTK | -59 | -71 | 1 | 0.00051306 | 13 | NaN | NaN | 731.4708 | NaN | 1370.9862 | 518.25366 |
| Mef2a;Mef2d; | YTEYNEPHESR | (69);(69);(69) | (79);(79);(79) | 0 | 0.00025015 | 11 | NaN | NaN | 573.48096 | NaN | NaN | NaN |
| Dus3l | YVPVGLLERPPQR | -572 | -584 | 1 | 0.00078661 | 13 | NaN | NaN | 560.7274 | NaN | 1014.2944 | NaN |
| Trappc12 | YYPEQEPQLLSGIGR | -636 | -650 | 0 | 0 | 15 | NaN | NaN | 536.2169 | 390.6314 | 591.2035 | 464.35004 |
| Copb1 | LVEKPSPLTLAPHDFANIK | -768 | -786 | 1 | 0 | 19 | NaN | NaN | NaN | 2027.5566 | 1496.2208 | 1183.9844 |
| Copb1 | AAADVLEFVR | -416 | -425 | 0 | 0.000550307 | 10 | NaN | NaN | NaN | 712.144 | 931.54956 | 770.5739 |
| Naca | NILFVITKPDVY | (2073);(101) | (2084);(112) | 1 | 0.007546675 | 12 | NaN | NaN | NaN | 656.632 | 778.3014 | NaN |
| Mcm6 | SEDASQETVSKPSLR | -700 | -714 | 1 | 0 | 15 | NaN | NaN | NaN | 6413.233 | 6617.505 | 6768.195 |
| Mcm6 | TTGEGTSLR | -379 | -387 | 0 | 0.003745449 | 9 | NaN | NaN | NaN | 1518.8115 | 1140.1257 | NaN |
| Mcm6 | MEEEEDESALKR | -734 | -745 | 1 | 0.006624942 | 12 | NaN | NaN | NaN | 1936.9047 | NaN | NaN |
| Fh | SKEFAQVIK | -219 | -227 | 1 | 0.001392869 | 9 | NaN | NaN | NaN | 671.81006 | NaN | NaN |
| Smad4 | AAAGIGVDDLR | -485 | -495 | 0 | 0.000755556 | 11 | NaN | NaN | NaN | 673.46484 | NaN | NaN |
| Ddx20 | TQHLQELFSK | -298 | -307 | 0 | 0.00111006 | 10 | NaN | NaN | NaN | 1021.4948 | NaN | NaN |
| Lcp1 | HVVNIGAEDLK | -207 | -217 | 0 | 0 | 11 | NaN | NaN | NaN | 8496.898 | 8237.332 | 7888.794 |
| Lcp1 | EITENLMATGDLDQDGK | -52 | -68 | 0 | 0 | 17 | NaN | NaN | NaN | 1683.228 | 1852.5975 | 1832.8297 |
| Lcp1 | SAIGCHVVNIGAEDLK | -202 | -217 | 0 | 0 | 16 | NaN | NaN | NaN | 1402.6459 | 1109.1063 | 1209.6156 |
| Lcp1 | LREGESLEDLMK | -252 | -263 | 1 | 0.00142999 | 12 | NaN | NaN | NaN | 491.8505 | 698.96136 | 593.95264 |
| Lcp1 | IKVPVDWN | -433 | -440 | 1 | 0.00897527 | 8 | NaN | NaN | NaN | 1196.169 | 637.43555 | 1038.1173 |
| Lcp1 | SASAIGCHVVNIGAEDLK | -200 | -217 | 0 | 0 | 18 | NaN | NaN | NaN | 833.3063 | 1138.749 | 1011.84595 |
| Lcp1 | VDTDGNGYISCNELNDLFK | -21 | -39 | 0 | 0 | 19 | NaN | NaN | NaN | 531.34784 | 668.7203 | 516.10474 |
| Lcp1 | ANYHLENAGCTK | -274 | -285 | 0 | 0 | 12 | NaN | NaN | NaN | 314.56934 | 306.1671 | 410.09473 |
| Lcp1 | VNDDIIVNWVNTTLK | -516 | -530 | 0 | 0 | 15 | NaN | NaN | NaN | 271.12915 | 412.15207 | 277.2006 |
| Lcp1 | NYHLENAGCTK | -275 | -285 | 0 | 0.001708008 | 11 | NaN | NaN | NaN | 547.6437 | 696.0347 | NaN |
| Lcp1 | LLEQVAPK | -302 | -309 | 0 | 0.001904399 | 8 | NaN | NaN | NaN | 779.7542 | NaN | NaN |
| Lcp1 | VVIDMSGLR | -318 | -326 | 0 | 0.001610812 | 9 | NaN | NaN | NaN | 631.23676 | NaN | NaN |
| Lcp1 | NKYPALHKPENQDIDWGALEGETR | -372 | -395 | 2 | 0.00111006 | 24 | NaN | NaN | NaN | 414.93472 | NaN | NaN |
| Hmgb1 | FKDPNAPK | -89 | -96 | 1 | 0 | 8 | NaN | NaN | NaN | 5441.6733 | 5897.104 | 5745.537 |
| Hmgb1 | GKFEDMAK | -58 | -65 | 1 | 0.002265552 | 8 | NaN | NaN | NaN | 217.7631 | 488.84402 | 334.9013 |
| Hmgb1 | FLFCSEYRPK | -103 | -112 | 1 | 0 | 10 | NaN | NaN | NaN | 2319.1477 | 2362.712 | 2275.8691 |
| Nap1l1 | EFHFEPNDYFTNEVLTK | -215 | -231 | 0 | 0 | 17 | NaN | NaN | NaN | 332.72147 | 433.14954 | 434.7359 |
| Nap1l1 | KYAVLYQPLFDK | -105 | -116 | 1 | 0 | 12 | NaN | NaN | NaN | 510.61108 | 682.5383 | 615.845 |
| Nap1l1 | KNVDLLSDMVQEHDEPILK | -176 | -194 | 1 | 0 | 19 | NaN | NaN | NaN | 393.0255 | 218.1772 | 411.48523 |
| Nap1l1 | GIPEFWLT | -166 | -173 | 0 | 0.003299042 | 8 | NaN | NaN | NaN | 655.6313 | 1150.8496 | NaN |
| Hspa9 | GKDSETGENIR | -624 | -634 | 1 | 0.00025015 | 11 | NaN | NaN | NaN | 1573.9341 | 1786.7314 | 1956.511 |
| Hspa9 | AVNPDEAVAIGA | -410 | -421 | 0 | 0.005605646 | 12 | NaN | NaN | NaN | 1056.8402 | NaN | NaN |
| Hspa9 | TIAPCQK | -362 | -368 | 0 | 0.006164361 | 7 | NaN | NaN | NaN | 853.44464 | NaN | NaN |
| Hspa9 | SCVAVMEGK | -65 | -73 | 0 | 0.001868731 | 9 | NaN | NaN | NaN | 842.3854 | NaN | NaN |
| Hspa9 | EFKDQLPADECNK | -598 | -610 | 1 | 0.00025015 | 13 | NaN | NaN | NaN | 490.3169 | NaN | NaN |
| Hspa9 | MKETAENYLGHTA | -174 | -186 | 1 | 0.000627943 | 13 | NaN | NaN | NaN | 481.2422 | NaN | NaN |
| Hspa9 | GDTFLGGEDFDQALLR | -269 | -284 | 0 | 0.001055699 | 16 | NaN | NaN | NaN | 444.27243 | NaN | NaN |
| Ybx1 | DNQGAGEQGRPVR | -220 | -232 | 1 | 0.002168954 | 13 | NaN | NaN | NaN | 583.32465 | 618.3027 | NaN |

| | | | | | | | | | | | | |
|---|---|---|---|---|---|---|---|---|---|---|---|---|
| Ybx1 | SSEAETQQPPAAPAAALS | -2 | -19 | 0 | 0.005605646 | 18 | NaN | NaN | NaN | 108.179474 | NaN | NaN |
| Rfc1 | LHEDEEFAR | -218 | -226 | 0 | 0 | 9 | NaN | NaN | NaN | 2457.0134 | NaN | NaN |
| Pkm | LAPITSDPTEAAAV | -401 | -414 | 0 | 0 | 14 | NaN | NaN | NaN | 1375.4083 | 328.91733 | 1006.20325 |
| Pkm | PKPHSEAGTAF | -2 | -12 | 1 | 0 | 11 | NaN | NaN | NaN | 272.84384 | 5205.141 | 5137.188 |
| Pkm | NFSHGTHEYHAETIK | -75 | -89 | 0 | 0 | 15 | NaN | NaN | NaN | 2382.833 | 657.38666 | 672.62506 |
| Pkm | FSHGTHEYHAETIK | -76 | -89 | 0 | 0 | 14 | NaN | NaN | NaN | 1431.5251 | 1026.547 | 1194.3105 |
| Pkm | FSHGTHEYHAETIK | -76 | -89 | 0 | 0 | 14 | NaN | NaN | NaN | 1431.5251 | 1026.547 | 1194.3105 |
| Pkm | NTGIICTIGPASR | -44 | -56 | 0 | 0 | 13 | NaN | NaN | NaN | 1056.9397 | 990.3944 | 1438.9586 |
| Pkm | LQLFEELR | -392 | -399 | 0 | 0 | 8 | NaN | NaN | NaN | 765.5823 | 834.60254 | 856.6568 |
| Pkm | CRLDIDSAPITAR | -31 | -43 | 1 | 0.001092529 | 13 | NaN | NaN | NaN | 372.54257 | 2556.1572 | 2403.5417 |
| Pkm | LPGAAVDLPAVSEK | -211 | -224 | 0 | 0 | 14 | NaN | NaN | NaN | 1231.6237 | 1492.8096 | NaN |
| Pkm | LAPITSDPTEAAAVGA | -401 | -416 | 0 | 0.00149795 | 16 | NaN | NaN | NaN | 990.5978 | NaN | 169.99304 |
| Pkm | RFDEILEASDGIMVAR | -279 | -294 | 1 | 0.005522803 | 16 | NaN | NaN | NaN | 562.81335 | NaN | NaN |
| Pkm | RFDEILEASDGIMVA | -279 | -293 | 1 | 0.00204437 | 15 | NaN | NaN | NaN | 384.78326 | NaN | NaN |
| Pkm | ILWLDYK | -156 | -162 | 0 | 0.007670622 | 7 | NaN | NaN | NaN | 197.05267 | NaN | NaN |
| Pdcd6ip | KSLLSNLDEIKK | -553 | -564 | 2 | 0 | 12 | NaN | NaN | NaN | 2066.8936 | 483.30908 | 598.61053 |
| Pdcd6ip | LSNLDEIKK | -556 | -564 | 1 | 0 | 9 | NaN | NaN | NaN | 1280.4385 | 456.1487 | 1466.4736 |
| Pdcd6ip | TYPSGGEEQAQYCR | -28 | -41 | 0 | 0.000550307 | 14 | NaN | NaN | NaN | 363.77454 | NaN | 340.7654 |
| Pdcd6ip | KFGEEIAR | -269 | -276 | 1 | 0.001023236 | 8 | NaN | NaN | NaN | 1039.0267 | NaN | NaN |
| Pdcd6ip | YDEYVNVK | -291 | -298 | 0 | 0.003299042 | 8 | NaN | NaN | NaN | 1005.40344 | NaN | NaN |
| Pdcd6ip | QSNNEANLR | -641 | -649 | 0 | 0.000627943 | 9 | NaN | NaN | NaN | 739.9618 | NaN | NaN |
| Cct2 | GGSLADSYLDEGFLLDKK | -206 | -223 | 1 | 0 | 18 | NaN | NaN | NaN | 447.33365 | 1210.8438 | 1284.3938 |
| Cct2 | LIEEVMIGEDK | -348 | -358 | 0 | 0 | 11 | NaN | NaN | NaN | 1242.8439 | NaN | 729.57526 |
| Cct2 | KLIEEVMIGEDK | -347 | -358 | 1 | 0 | 12 | NaN | NaN | NaN | 3700.9583 | NaN | 4165.796 |
| Cct2 | QILDEAER | -381 | -388 | 0 | 0.006735683 | 8 | NaN | NaN | NaN | 666.9957 | NaN | NaN |
| Dhx9 | RDVVLAYPEVR | -530 | -540 | 1 | 0 | 11 | NaN | NaN | NaN | 6831.572 | 7020.2363 | 7020.2637 |
| Dhx9 | VIVDEIHER | -510 | -518 | 0 | 0.000230707 | 9 | NaN | NaN | NaN | 1357.2463 | 1581.8856 | 1132.6654 |
| Dhx9 | HLENNSHFGSH | -671 | -681 | 0 | 0.00111006 | 11 | NaN | NaN | NaN | 409.08044 | 473.27585 | 3239.559 |
| Dhx9 | HVIVDEIHER | -509 | -518 | 0 | 0.000712251 | 10 | NaN | NaN | NaN | 341.81564 | 154.53911 | 1472.2981 |
| Dhx9 | YQILPLHSQIPR | -683 | -694 | 0 | 0 | 12 | NaN | NaN | NaN | 400.9891 | 239.3459 | NaN |
| Dhx9 | AHFEPSQR | -318 | -325 | 0 | 0.007405213 | 8 | NaN | NaN | NaN | 1248.554 | NaN | NaN |
| Dhx9 | SGVESSGYGSPGPTWDR | -127 | -143 | 0 | 0.003069674 | 17 | NaN | NaN | NaN | 81.84595 | NaN | NaN |
| Pafah1b2 | AAEDIQGDDR | -14 | -23 | 0 | 0.00025015 | 10 | NaN | NaN | NaN | 3128.733 | 3544.031 | 3653.8167 |
| Pafah1b2 | NGELENIKPK | -87 | -96 | 1 | 0.000394361 | 10 | NaN | NaN | NaN | 1129.2147 | 2278.7668 | 1890.4818 |
| Tbcd | LPESSHTSLR | -285 | -294 | 0 | 0.000347792 | 10 | NaN | NaN | NaN | 1145.0167 | NaN | 1509.647 |
| Tbcd | AAEEEVEDDALVR | -11 | -23 | 0 | 0.001291103 | 13 | NaN | NaN | NaN | 2842.169 | NaN | NaN |
| Tbcd | AVSVGGLTESTVR | -990 | -1002 | 0 | 0 | 13 | NaN | NaN | NaN | 1431.2845 | NaN | NaN |
| Tbcd | DHETWETR | -143 | -150 | 0 | 0.00025015 | 8 | NaN | NaN | NaN | 747.6525 | NaN | NaN |
| Tbcd | QWLINDTLR | -706 | -714 | 0 | 0.002534621 | 9 | NaN | NaN | NaN | 365.8508 | NaN | NaN |
| Tbcd | GALPGFLLR | -782 | -790 | 0 | 0.001392869 | 9 | NaN | NaN | NaN | 289.66495 | NaN | NaN |
| Hspa4 | FLEMCDDLLAR | -306 | -316 | 0 | 0 | 11 | NaN | NaN | NaN | 790.2961 | 1422.3922 | 810.4569 |
| Hspa4 | RWNSPAEEGLSDCEVFPK | -405 | -422 | 1 | 0 | 18 | NaN | NaN | NaN | 677.5819 | 722.68677 | 702.80597 |
| Hspa4 | KNKEDQYEHLDAADVTK | -718 | -734 | 2 | 0.00025015 | 17 | NaN | NaN | NaN | 393.8726 | 648.07007 | 413.69617 |
| Hspa4 | HAEQNGPVDGQGDNPGSQA | -800 | -818 | 0 | 0.000230707 | 19 | NaN | NaN | NaN | 248.98459 | 368.15866 | 221.32639 |
| Hspa4 | PFVEAEK | -78 | -84 | 0 | 0.002168954 | 7 | NaN | NaN | NaN | 421.996 | NaN | NaN |
| Gvin1 | SLETCHVESIK | -1968 | -1978 | 0 | 0 | 11 | NaN | NaN | NaN | 3983.4602 | 1236.8956 | 1599.1279 |
| Gvin1 | ILSEHLCGCWK | -2283 | -2293 | 0 | 0 | 11 | NaN | NaN | NaN | 1843.617 | 2267.8518 | 2351.3594 |
| Gvin1 | SVLELLQDYSETHNK | -1337 | -1351 | 0 | 0.00078661 | 15 | NaN | NaN | NaN | 749.5504 | 915.802 | 914.90875 |
| Gvin1 | KDSLISDTR | -1835 | -1843 | 1 | 0 | 9 | NaN | NaN | NaN | 638.12854 | 786.2788 | NaN |
| Gvin1 | ELPFYFLQK | -905 | -913 | 0 | 0.005605646 | 9 | NaN | NaN | NaN | 493.39594 | 522.23346 | NaN |
| Gvin1 | HGEHHDWESLENDLR | -799 | -813 | 0 | 0.005605646 | 15 | NaN | NaN | NaN | 437.12918 | NaN | 874.60645 |
| Gvin1 | HGEHHDWESLENDLR | -799 | -813 | 0 | 0.005605646 | 15 | NaN | NaN | NaN | 437.12918 | NaN | 874.60645 |
| Gvin1 | LWHLWCK | -1284 | -1290 | 0 | 0.002629273 | 7 | NaN | NaN | NaN | 2726.6555 | NaN | NaN |
| Gvin1 | NDPVNYLER | -2060 | -2068 | 0 | 0.00111006 | 9 | NaN | NaN | NaN | 2030.8547 | NaN | NaN |
| Gvin1 | SEHLCGCWK | -2285 | -2293 | 0 | 0.009808007 | 9 | NaN | NaN | NaN | 927.2049 | NaN | NaN |
| Tln1 | VLGEAMTGISQNAK | -1402 | -1415 | 0 | 0.000230707 | 14 | NaN | NaN | NaN | 2323.9026 | 2229.7375 | 407.21964 |
| Tln1 | NLGTALAELR | -1026 | -1035 | 0 | 0 | 10 | NaN | NaN | NaN | 1966.4105 | 2071.9128 | 1099.99 |
| Tln1 | ATLDDFETLPPLGQDAASK | -511 | -529 | 0 | 0 | 19 | NaN | NaN | NaN | 560.80414 | 565.09717 | 538.10297 |
| Tln1 | KISIGNVVK | -7 | -15 | 1 | 0.001392869 | 9 | NaN | NaN | NaN | 540.08167 | 567.5843 | 496.8626 |
| Tln1 | AVAEQIPLLVQGVR | -958 | -971 | 0 | 0 | 14 | NaN | NaN | NaN | 413.69287 | 463.53497 | 430.84018 |
| Tln1 | LEAGVEMAGQAPSQEDR | -1282 | -1298 | 0 | 0.00025015 | 17 | NaN | NaN | NaN | 2038.192 | 1940.6556 | NaN |
| Tln1 | LLSDSLPPSTGTFQEAQSR | -1223 | -1241 | 0 | 0.001868731 | 19 | NaN | NaN | NaN | 884.38416 | NaN | 1019.79205 |
| Tln1 | KGIWLEAGK | -58 | -66 | 1 | 0.00111006 | 9 | NaN | NaN | NaN | 559.3422 | 490.58682 | NaN |
| Tln1 | ERIPEALAGPPNDFGLF | -34 | -50 | 1 | 0 | 17 | NaN | NaN | NaN | 271.98523 | 332.02625 | NaN |
| Tln1 | KEAAFHPEVAPDVR | -2220 | -2233 | 1 | 0.000230707 | 14 | NaN | NaN | NaN | 185.55962 | 989.7342 | NaN |
| Tln1 | EGAETFADHR | -2007 | -2016 | 0 | 0.001708008 | 10 | NaN | NaN | NaN | 1578.1016 | NaN | NaN |
| Tln1 | SNTSPEELGPLANQLTSDYGR | -1875 | -1895 | 0 | 0.000907206 | 21 | NaN | NaN | NaN | 431.45575 | NaN | NaN |
| Stat1 | FTYEPDPITK | -287 | -296 | 0 | 0 | 10 | NaN | NaN | NaN | 4204.379 | 5129.4775 | 4974.9224 |
| Stat1 | TELISVSEVHPSR | -704 | -716 | 0 | 0 | 13 | NaN | NaN | NaN | 1295.7703 | 580.1238 | 545.675 |
| Stat1 | NILGTHTK | -381 | -388 | 0 | 0.00025015 | 8 | NaN | NaN | NaN | 1716.5635 | 1784.4545 | NaN |
| Stat1 | FLEQVHQ | -14 | -20 | 0 | 0.000712251 | 7 | NaN | NaN | NaN | 359.36426 | NaN | 757.1798 |
| Stat1 | VHQLYDDSFPMEIR | -18 | -31 | 0 | 0.002434616 | 14 | NaN | NaN | NaN | 1042.8417 | NaN | NaN |
| Stat1 | ELLNSIELTQN | -219 | -229 | 0 | 0 | 11 | NaN | NaN | NaN | 578.8153 | NaN | NaN |
| Stat1 | EAPEPMELDDPK | -686 | -697 | 0 | 0.006100831 | 12 | NaN | NaN | NaN | 537.25134 | NaN | NaN |
| Plrg1 | SLAVALPSQTR | -125 | -135 | 0 | 0.000907206 | 11 | NaN | NaN | NaN | 520.83813 | NaN | 439.60303 |
| Nup93 | FDEYHSGHIDR | -699 | -709 | 0 | 0 | 11 | NaN | NaN | NaN | 785.898 | 455.89786 | 385.6423 |
| Nup93 | DKTEDYLWLK | -408 | -417 | 1 | 0.000230707 | 10 | NaN | NaN | NaN | 1429.5128 | 1592.3883 | 1383.5151 |
| Nup93 | PLNQESVEER | -721 | -730 | 0 | 0.00223472 | 10 | NaN | NaN | NaN | 160.82742 | 863.4636 | 945.499 |

| | | | | | | | | | | | | |
|---|---|---|---|---|---|---|---|---|---|---|---|---|
| Nup93 | HQLGEFK | -343 | -349 | 0 | 0.002068213 | 7 | NaN | NaN | NaN | 3411.2656 | 4194.275 | NaN |
| Nup93 | ESMLVEWEQVK | -131 | -141 | 0 | 0 | 11 | NaN | NaN | NaN | 1126.5035 | NaN | NaN |
| Nup93 | EALQYFYFLR | -546 | -555 | 0 | 0.000230707 | 10 | NaN | NaN | NaN | 272.03555 | NaN | NaN |
| Nup93 | TDVVLTPATDALK | -231 | -243 | 0 | 0 | 13 | NaN | NaN | NaN | 269.76865 | NaN | NaN |
| Ganab | MLDYLQGSGETPQTDIR | -338 | -354 | 0 | 0 | 17 | NaN | NaN | NaN | 1144.8501 | 928.9151 | 1307.2021 |
| Ganab | KNPEPELLVR | -653 | -662 | 1 | 0 | 10 | NaN | NaN | NaN | 492.7691 | 591.767 | 396.5867 |
| Ganab | HHGPQTLYLPVTL | -785 | -797 | 0 | 0 | 13 | NaN | NaN | NaN | 477.92886 | 749.3976 | 1668.8877 |
| Ganab | GRDDNSVELTVAEGPYK | -132 | -148 | 1 | 0.001846793 | 17 | NaN | NaN | NaN | 206.77296 | 518.74567 | 457.57162 |
| Ganab | LMLGPSVFDVFR | -366 | -377 | 0 | 0.001978766 | 12 | NaN | NaN | NaN | 1478.7544 | NaN | NaN |
| Ganab | NHGLYVK | -485 | -491 | 0 | 0.001092529 | 7 | NaN | NaN | NaN | 779.78125 | NaN | NaN |
| Snrnp200 | HHEDNLLR | -1862 | -1869 | 0 | 0.000947459 | 8 | NaN | NaN | NaN | 921.04675 | NaN | NaN |
| Snrnp200 | HDDLKGDPLLDQR | -940 | -952 | 1 | 0.000431655 | 13 | NaN | NaN | NaN | 2168.186 | 777.86176 | 1055.1262 |
| Snrnp200 | HINMDGTINVDDFK | -524 | -537 | 0 | 0.000591124 | 14 | NaN | NaN | NaN | 625.8965 | 548.34265 | 722.159 |
| Snrnp200 | LDFVAPATGGHNYTL | -2092 | -2106 | 0 | 0.000431655 | 15 | NaN | NaN | NaN | 201.6032 | 1264.9138 | 481.5985 |
| Snrnp200 | VHLQPITR | -1188 | -1195 | 0 | 0.003490217 | 8 | NaN | NaN | NaN | 138.15749 | 517.57996 | 516.37244 |
| Snrnp200 | HLSDHLSELVEQTL | -1780 | -1793 | 0 | 0.008436628 | 14 | NaN | NaN | NaN | 4112.574 | NaN | NaN |
| Snrnp200 | STFNFHPNVR | -1510 | -1519 | 0 | 0.00025015 | 10 | NaN | NaN | NaN | 2449.8538 | NaN | NaN |
| Dld | EANLAAAFGKPINF | -496 | -509 | 1 | 0.001392869 | 14 | NaN | NaN | NaN | 441.92493 | NaN | 506.71625 |
| Dld | FPFAANSR | -421 | -428 | 0 | 0.003998192 | 8 | NaN | NaN | NaN | 1504.158 | NaN | NaN |
| Dld | ALTGGIAHLF | -133 | -142 | 0 | 0.00272025 | 10 | NaN | NaN | NaN | 251.66608 | NaN | NaN |
| Tomm22 | PVVFETEK | -98 | -105 | 0 | 0.003745449 | 8 | NaN | NaN | NaN | 319.31396 | 625.06146 | 742.0991 |
| Dnajc2 | PVNTPER | -570 | -576 | 0 | 0.001811524 | 7 | NaN | NaN | NaN | 2587.4944 | NaN | NaN |
| Nolc1 | RDSQLSEVASK | -23 | -33 | 1 | 0.00111006 | 11 | NaN | NaN | NaN | 4473.4316 | 6456.197 | 5340.965 |
| Nolc1 | VREEEIEVDSR | -631 | -641 | 1 | 0 | 11 | NaN | NaN | NaN | 616.2973 | 651.0749 | 705.0817 |
| Nolc1 | RVVPSDLYPLVL | -8 | -19 | 1 | 0.000230707 | 12 | NaN | NaN | NaN | 373.0648 | 345.11493 | 483.04562 |
| Nolc1 | KESEEEEEEETEEK | -561 | -575 | 1 | 0.007343498 | 15 | NaN | NaN | NaN | 324.61646 | 638.78357 | 481.39996 |
| Nolc1 | PPPSVPLPK | -302 | -310 | 0 | 0.002972566 | 9 | NaN | NaN | NaN | 1172.1204 | NaN | 1695.0708 |
| Nolc1 | KQNETADEATTPQAK | -590 | -604 | 1 | 0.000230707 | 15 | NaN | NaN | NaN | 332.24805 | 471.11777 | NaN |
| Uqcrc1 | IPLAEWESR | -424 | -432 | 0 | 0 | 9 | NaN | NaN | NaN | 1440.5297 | 1560.414 | 550.32355 |
| Uqcrc1 | AFQGTPLAQAVEGPSENVR | -191 | -209 | 0 | 0.00111006 | 19 | NaN | NaN | NaN | 370.3625 | 739.53906 | 439.06647 |
| Uqcrc1 | FLEHLAFK | -92 | -99 | 0 | 0.002972566 | 8 | NaN | NaN | NaN | 1249.3132 | NaN | NaN |
| Uqcrc1 | LDGTTPVCEDIGR | -403 | -415 | 0 | 0.000230707 | 13 | NaN | NaN | NaN | 1247.6143 | NaN | NaN |
| Uqcrc1 | YFYDQCPAVA | -448 | -457 | 0 | 0.001392869 | 10 | NaN | NaN | NaN | 314.88718 | NaN | NaN |
| Parp14 | SGGGECEVVPEPGNPAR | -37 | -53 | 0 | 0 | 17 | NaN | NaN | NaN | 519.79443 | NaN | 538.11694 |
| Parp14 | AAGPELQTECDQIVK | -851 | -865 | 0 | 0.000471409 | 15 | NaN | NaN | NaN | 1187.2759 | NaN | NaN |
| Eef1g | NSSSISGVWVFR | -367 | -378 | 0 | 0 | 12 | NaN | NaN | NaN | 6568.522 | 6041.586 | 7521.223 |
| Eef1g | AAGTLYTYPENWR | -2 | -14 | 0 | 0 | 13 | NaN | NaN | NaN | 1393.5944 | 2066.8384 | 1973.1681 |
| Eef1g | WFLTCINQPQFR | -190 | -201 | 0 | 0 | 12 | NaN | NaN | NaN | 783.87897 | 1370.7975 | 1076.4318 |
| Eef1g | GTNNSSSISGVWVFR | -364 | -378 | 0 | 0 | 15 | NaN | NaN | NaN | 688.3419 | 1548.3267 | 1737.9845 |
| Eef1g | NNSSSISGVWVFR | -366 | -378 | 0 | 0 | 13 | NaN | NaN | NaN | 560.7477 | 692.265 | 280.34018 |
| Eef1g | VPAFEGDDGFCVFESN | -58 | -73 | 0 | 0 | 16 | NaN | NaN | NaN | 334.4859 | 362.1546 | 422.2752 |
| Eef1g | QYSGAQVR | -23 | -30 | 0 | 0.007609801 | 8 | NaN | NaN | NaN | 1283.4753 | NaN | NaN |
| Eef1g | VLDEFKR | -289 | -295 | 1 | 0.003133457 | 7 | NaN | NaN | NaN | 995.8031 | NaN | NaN |
| Gosr1 | AAGTSNYWEDLRK | -2 | -14 | 1 | 0 | 13 | NaN | NaN | NaN | 1304.7286 | 1338.5413 | 1185.2375 |
| Eef1a1 | GYAPVLDCHTAHIACK | -356 | -371 | 0 | 0 | 16 | NaN | NaN | NaN | 578.8759 | 1097.1748 | 798.30725 |
| Eef1a1 | GTTLLEALDCILPPTRPTDKPLR | -225 | -247 | 2 | 0.000591124 | 23 | NaN | NaN | NaN | 1561.2177 | 2139.6257 | 2004.0248 |
| Eef1a1 | DGSASGTTLLEAL | -220 | -232 | 0 | 0 | 13 | NaN | NaN | NaN | 1487.3849 | 2211.0376 | 1365.718 |
| Eef1a1 | DGSASGTTLLEAL | -220 | -232 | 0 | 0 | 13 | NaN | NaN | NaN | 1487.3849 | 2211.0376 | 1365.718 |
| Eef1a1 | NHPGQISAGYAPVLDCHTAHIACK | -348 | -371 | 0 | 0 | 24 | NaN | NaN | NaN | 1390.9061 | 2356.3425 | 2233.6704 |
| Eef1a1 | NHPGQISAGYAPVLDCHTAHIACK | -348 | -371 | 0 | 0 | 24 | NaN | NaN | NaN | 1390.9061 | 2356.3425 | 2233.6704 |
| Eef1a1 | AGYAPVLDCHTAHIACK | -355 | -371 | 0 | 0 | 17 | NaN | NaN | NaN | 1271.8594 | 1137.7415 | 1147.4331 |
| Eef1a1 | RYEEIVKEVSTY | -166 | -177 | 2 | 0.000550307 | 12 | NaN | NaN | NaN | 985.2389 | 1260.3373 | 1368.6324 |
| Eef1a1 | KYYVTIIDAPGHR | -84 | -96 | 1 | 0 | 13 | NaN | NaN | NaN | 752.9265 | 586.3183 | 966.54456 |
| Eef1a1 | KYYVTIIDAPGHR | -84 | -96 | 1 | 0 | 13 | NaN | NaN | NaN | 752.9265 | 586.3183 | 966.54456 |
| Eef1a1 | APVLDCHTAHIACK | -358 | -371 | 0 | 0 | 14 | NaN | NaN | NaN | 638.6124 | 1200.4414 | 1135.053 |
| Eef1a1 | APVLDCHTAHIACK | -358 | -371 | 0 | 0 | 14 | NaN | NaN | NaN | 638.6124 | 1200.4414 | 1135.053 |
| Eef1a1 | VNVTTEVK | -283 | -290 | 0 | 0.000550307 | 8 | NaN | NaN | NaN | 467.48166 | 608.69366 | 805.133 |
| Eef1a1 | IGYNPDTVAFVPISGW | -181 | -196 | 0 | 0.004224523 | 16 | NaN | NaN | NaN | 525.48096 | 461.99475 | NaN |
| Rplp1 | ANVNIGSLICNVGAGGPAPAAGAAPAGGAAPSTAA | -52 | -92 | 0 | 0 | 41 | NaN | NaN | NaN | 353.76617 | 472.13226 | 428.40442 |
| Rplp1 | ICNVGAGGPAPAAGAAPAGGAAPSTAAAPAEEK | -60 | -92 | 0 | 0.003490217 | 33 | NaN | NaN | NaN | 1138.7319 | NaN | 2053.2 |
| Pa2g4 | NCVCHFSPLK | -84 | -93 | 0 | 0 | 10 | NaN | NaN | NaN | 2619.0425 | 2347.486 | 627.55566 |
| Pa2g4 | SISVNNCVCHFSPLK | -79 | -93 | 0 | 0 | 15 | NaN | NaN | NaN | 2229.4163 | 2801.0234 | 2660.479 |
| Pa2g4 | AAHLCAEAALR | -145 | -155 | 0 | 0 | 11 | NaN | NaN | NaN | 1710.1683 | NaN | 1687.5677 |
| Fus | APKPDGPGGGPGGSHM | -442 | -457 | 1 | 0 | 16 | NaN | NaN | NaN | 299.23737 | 611.975 | 828.82104 |
| Fus | APKPDGPGGGPGGSHM | -442 | -457 | 1 | 0 | 16 | NaN | NaN | NaN | 299.23737 | 611.975 | 828.82104 |
| Fus | CPNPTCENMNFSWR | -421 | -434 | 0 | 0.00025015 | 14 | NaN | NaN | NaN | 546.02057 | 549.31433 | 537.50323 |
| Fus | LKGEATVSFDDPPSAK | -326 | -341 | 1 | 0.00025015 | 16 | NaN | NaN | NaN | 491.27292 | 597.55035 | 798.0348 |
| Phb1 | KLEAAEDIAYQLSR | -240 | -253 | 1 | 0 | 14 | NaN | NaN | NaN | 1921.6582 | 1786.7192 | 1904.507 |
| Phb1 | GLILDDVSLTHLTFGK | -162 | -177 | 0 | 0 | 16 | NaN | NaN | NaN | 808.1407 | 1083.8552 | 1007.97205 |
| Phb1 | NITYLPAGQSVLLQLPQ | -256 | -272 | 0 | 0 | 17 | NaN | NaN | NaN | 214.47662 | 285.24185 | 249.41304 |
| Phb1 | GTHFLIPWVQK | -53 | -63 | 0 | 0.00272025 | 11 | NaN | NaN | NaN | 1915.4476 | NaN | NaN |
| Phb1 | IGEDYDER | -110 | -117 | 0 | 0.001708008 | 8 | NaN | NaN | NaN | 518.0747 | NaN | NaN |
| Phb1 | AAIISAEGDSK | -209 | -219 | 0 | 0.000431655 | 11 | NaN | NaN | NaN | 438.3716 | NaN | NaN |
| Fbl | AAILGGVDQIHIKPGAK | -152 | -168 | 1 | 0.00111006 | 17 | NaN | NaN | NaN | 375.13647 | 439.82605 | 449.34033 |
| Fbl | VDQIHIKPGAK | -158 | -168 | 1 | 0.00025015 | 11 | NaN | NaN | NaN | 363.35492 | 315.87155 | 615.945 |
| Fbl | RVSISEGDDKIEYR | -128 | -141 | 2 | 0.003166358 | 14 | NaN | NaN | NaN | 1846.4341 | NaN | NaN |
| Fbl | GKEDALVTK | -107 | -115 | 1 | 0.00025015 | 9 | NaN | NaN | NaN | 1080.0469 | NaN | NaN |
| Ddx21 | LGSDGAEESMETLPKPSEK | -11 | -29 | 1 | 0 | 19 | NaN | NaN | NaN | 659.2088 | 639.49365 | 455.64737 |

| | | | | | | | | | | | | |
|---|---|---|---|---|---|---|---|---|---|---|---|---|
| Ddx21 | GLSQPSEEEVDIPKPK | -187 | -202 | 1 | 0 | 16 | NaN | NaN | NaN | 1697.8368 | 1702.1504 | 1736.3783 |
| Ddx21 | SKTEEATEGMEEAVSS | -38 | -53 | 1 | 0.002566359 | 16 | NaN | NaN | NaN | 602.7069 | 568.14966 | 478.29572 |
| Ddx21 | EIITEEPSEEEADMPKPK | -111 | -128 | 1 | 0.001708008 | 18 | NaN | NaN | NaN | 531.74365 | 521.657 | 581.37384 |
| Ddx21 | SVACFYGGTPYGGQIER | -360 | -376 | 0 | 0.00025015 | 17 | NaN | NaN | NaN | 329.20816 | NaN | NaN |
| Ddx21 | SSDAPGEESSSETEK | -235 | -249 | 0 | 0.001092529 | 15 | NaN | NaN | NaN | 200.9499 | NaN | NaN |
| Cap1 | DSLLANPVAEYLK | -47 | -59 | 0 | 0.000230707 | 13 | NaN | NaN | NaN | 1696.3594 | 2052.255 | 1992.6641 |
| Cap1 | KTDGCHAYLSK | -411 | -421 | 1 | 0.00078661 | 11 | NaN | NaN | NaN | 981.79346 | 1329.7434 | 1720.6824 |
| Cap1 | ALLATASQCQQPAGNK | -84 | -99 | 0 | 0.000550307 | 16 | NaN | NaN | NaN | 497.0102 | 620.1926 | NaN |
| Cap1 | AKPGPFVK | -147 | -154 | 1 | 0.007546675 | 8 | NaN | NaN | NaN | 1303.7415 | NaN | NaN |
| Cap1 | KEIGGDVQK | -62 | -70 | 1 | 0.00383302 | 9 | NaN | NaN | NaN | 507.38074 | NaN | NaN |
| Arpc4 | AALCLENFSSQVVER | -18 | -32 | 0 | 0.00025015 | 15 | NaN | NaN | NaN | 1374.3949 | 469.33975 | 1739.7053 |
| Arpc4 | TATLRPYLSAV | -2 | -12 | 1 | 0 | 11 | NaN | NaN | NaN | 1311.6417 | 1392.6136 | 443.3228 |
| Arpc4 | LCLENFSSQVVER | -20 | -32 | 0 | 0 | 13 | NaN | NaN | NaN | 688.29504 | 865.1824 | 952.86957 |
| Arpc4 | ELLLQPVTIS | -45 | -54 | 0 | 0.00025015 | 10 | NaN | NaN | NaN | 396.26028 | 655.97363 | 731.9355 |
| Arpc4 | NFHTEQMYK | -120 | -128 | 0 | 0 | 9 | NaN | NaN | NaN | 1197.7423 | NaN | NaN |
| Coro1c | AALEAEEWFEGK | -370 | -381 | 0 | 0 | 12 | NaN | NaN | NaN | 514.267 | 512.03064 | 415.08725 |
| Hars1 | IGDYVQQHGGVSLVEQLLQDPK | -267 | -288 | 0 | 0 | 22 | NaN | NaN | NaN | 495.42773 | 569.12006 | NaN |
| Hars1 | HGGVSLVEQLLQDPK | -274 | -288 | 0 | 0 | 15 | NaN | NaN | NaN | 169.43309 | NaN | 865.5434 |
| Hars1 | IGDYVQQH | -267 | -274 | 0 | 0.002780285 | 8 | NaN | NaN | NaN | 865.07837 | NaN | NaN |
| Rplp2 | KEESEESDDDMGFGL | -99 | -113 | 1 | 0.000431655 | 15 | NaN | NaN | NaN | 736.29675 | 721.352 | 663.78467 |
| Rplp2 | YLLAALGGNSSPSAK | -7 | -21 | 0 | 0 | 15 | NaN | NaN | NaN | 920.8967 | NaN | 858.5146 |
| Rplp2 | YVASYLLAALGGNSSPSAK | -3 | -21 | 0 | 0 | 19 | NaN | NaN | NaN | 255.019 | 587.32666 | NaN |
| Mcm3 | TAIHEVMEQGR | -420 | -430 | 0 | 0 | 11 | NaN | NaN | NaN | 4802.753 | 281.091 | 754.3271 |
| Mcm3 | TDDSQETQDSQKVELSEPR | -729 | -747 | 1 | 0.004923919 | 19 | NaN | NaN | NaN | 1447.5935 | 131.82112 | 1494.1783 |
| Mcm3 | VTTDQETGER | -382 | -391 | 0 | 0.000550307 | 10 | NaN | NaN | NaN | 776.8303 | 315.64166 | 520.0681 |
| Mcm3 | SKDIQPAFSADDIAK | -269 | -283 | 1 | 0 | 15 | NaN | NaN | NaN | 3240.7722 | 3634.7693 | NaN |
| Mcm3 | HLTESINR | -771 | -778 | 0 | 0.000431655 | 8 | NaN | NaN | NaN | 1404.0668 | NaN | 2017.6257 |
| Mcm3 | AALLEVFQEAHEQ | -753 | -765 | 0 | 0.003235556 | 13 | NaN | NaN | NaN | 1021.41583 | NaN | NaN |
| Mcm3 | LHLTESINR | -770 | -778 | 0 | 0 | 9 | NaN | NaN | NaN | 746.6954 | NaN | NaN |
| Mcm3 | ASEDESDLEDEEEKSQEDTEQK | -667 | -688 | 1 | 0.00025015 | 22 | NaN | NaN | NaN | 290.7327 | NaN | NaN |
| Mcm5 | ELPDAVPHGEMPR | -231 | -243 | 0 | 0 | 13 | NaN | NaN | NaN | 4273.051 | 4121.196 | 585.45416 |
| Mcm5 | ASFDEELADHLHK | -78 | -90 | 0 | 0 | 13 | NaN | NaN | NaN | 1172.1611 | 1478.362 | 75.677795 |
| Mcm5 | TTQEDQEMLSR | -664 | -674 | 0 | 0.002265552 | 11 | NaN | NaN | NaN | 123.994865 | 1062.8628 | NaN |
| Rps25 | GGDAPAAGEDA | -115 | -125 | 0 | 0.006100831 | 11 | NaN | NaN | NaN | 971.8743 | NaN | NaN |
| Rps25 | AALQELLSKG | -86 | -95 | 1 | 0.00383302 | 10 | NaN | NaN | NaN | 359.7484 | NaN | NaN |
| Rps25 | TKGGDAPAAGEDA | -113 | -125 | 1 | 0.001776473 | 13 | NaN | NaN | NaN | 190.90599 | NaN | NaN |
| Hadhb | AALSGLLHR | -83 | -91 | 0 | 0.004224523 | 9 | NaN | NaN | NaN | 780.1687 | NaN | NaN |
| Hadhb | KVGSPPLEK | -409 | -417 | 1 | 0 | 9 | NaN | NaN | NaN | 707.3225 | NaN | NaN |
| Cad | ASLGTADFYTEHGVK | -1344 | -1358 | 0 | 0 | 15 | NaN | NaN | NaN | 1747.208 | 570.9364 | 1045.8501 |
| Cad | GSGGLSIGQAGEFDYSGSQAIK | -400 | -421 | 0 | 0 | 22 | NaN | NaN | NaN | 279.55685 | 383.43555 | 336.4471 |
| Cad | RKEEILLIK | -1617 | -1625 | 2 | 0.004065182 | 9 | NaN | NaN | NaN | 371.53024 | NaN | 279.82724 |
| Cad | GIPGLQGVDTR | -112 | -122 | 0 | 0 | 11 | NaN | NaN | NaN | 1009.4967 | NaN | NaN |
| Lmnb1 | LSDYEQLLDVK | -358 | -368 | 0 | 0 | 11 | NaN | NaN | NaN | 4020.173 | 3341.631 | 2881.0378 |
| Lmnb1 | MYEEEINETR | -212 | -221 | 0 | 0 | 10 | NaN | NaN | NaN | 116.88798 | 839.03784 | 837.70526 |
| Lmnb1 | SQGEEVAQR | -535 | -543 | 0 | 0 | 9 | NaN | NaN | NaN | 680.4001 | 653.9255 | 501.19925 |
| Lmnb1 | SLETENSALQLQVTER | -53 | -68 | 0 | 0.000431655 | 16 | NaN | NaN | NaN | 1165.9874 | NaN | NaN |
| Lmnb1 | TEDLEFR | -203 | -209 | 0 | 0.008129024 | 7 | NaN | NaN | NaN | 740.7255 | NaN | NaN |
| Lmnb1 | RVDVEESEASSS | -419 | -430 | 1 | 0.001291103 | 12 | NaN | NaN | NaN | 431.8081 | NaN | NaN |
| Lmnb1 | RVDVEESEAS | -419 | -428 | 1 | 0 | 10 | NaN | NaN | NaN | 415.15775 | NaN | NaN |
| Ftsj3 | KADEDGFEVVPIQDPVK | -634 | -650 | 1 | 0.000230707 | 17 | NaN | NaN | NaN | 921.91016 | 639.7008 | 555.8533 |
| Ftsj3 | DEDGFEVVPIQDPVK | -636 | -650 | 0 | 0 | 15 | NaN | NaN | NaN | 588.75073 | 709.14716 | 605.849 |
| Ftsj3 | SEISIDDEELAQHPATTEDIR | -271 | -291 | 0 | 0 | 21 | NaN | NaN | NaN | 167.94946 | 219.03117 | 261.99826 |
| Ftsj3 | DTFLSDLPR | -442 | -450 | 0 | 0.000550307 | 9 | NaN | NaN | NaN | 520.1553 | NaN | 524.825 |
| Ftsj3 | AEGDLTLYHR | -242 | -251 | 0 | 0.009092375 | 10 | NaN | NaN | NaN | 1160.2682 | NaN | NaN |
| Tpi1 | LVGGASLKPEFVDIINAK | -231 | -248 | 1 | 0 | 18 | NaN | NaN | NaN | 657.16406 | 821.4048 | 858.2792 |
| Tpi1 | IIYGGSVTGATCK | -207 | -219 | 0 | 0 | 13 | NaN | NaN | NaN | 317.96542 | 2303.054 | 499.82153 |
| Tpi1 | DLGATWVVLGHSER | -86 | -99 | 0 | 0 | 14 | NaN | NaN | NaN | 831.6041 | 837.19666 | 130.92915 |
| Tpi1 | PVWAIGTGK | -167 | -175 | 0 | 0.001150701 | 9 | NaN | NaN | NaN | 769.6538 | 984.8047 | 1243.9291 |
| Tpi1 | PVWAIGTGK | -167 | -175 | 0 | 0.001150701 | 9 | NaN | NaN | NaN | 769.6538 | 984.8047 | 1243.9291 |
| Tpi1 | SNVNDGVAQST | -195 | -205 | 0 | 0.001392869 | 11 | NaN | NaN | NaN | 739.9672 | NaN | NaN |
| Ppp1r14b | ELLVDCYKPTEAFIS | -115 | -129 | 1 | 0.002845514 | 15 | NaN | NaN | NaN | 2289.279 | 3224.1619 | NaN |
| Nptn | AEDSGEYHCVYHFVSAPK | -209 | -226 | 0 | 0 | 18 | NaN | NaN | NaN | 1026.498 | 271.4651 | 282.4618 |
| Ctdp1 | AAPEHSGPQDTSGGR | -484 | -498 | 0 | 0 | 15 | NaN | NaN | NaN | 156.89003 | 248.3404 | 140.3447 |
| Me2 | SIVDNWPENHVK | -145 | -156 | 0 | 0 | 12 | NaN | NaN | NaN | 1235.5259 | 1238.4247 | 528.31177 |
| Me2 | FASGSPFEPVK | -443 | -453 | 0 | 0.00025015 | 11 | NaN | NaN | NaN | 1396.537 | NaN | NaN |
| Uqcrc2 | QIDSVADADVVK | -415 | -426 | 0 | 0 | 12 | NaN | NaN | NaN | 1302.3445 | NaN | NaN |
| Nfatc2 | AAPLGLADAHR | -774 | -784 | 0 | 0.00437931 | 11 | NaN | NaN | NaN | 30119.936 | 22166.582 | 24625.812 |
| Nfatc2 | VEFLGPCEQEER | -351 | -362 | 0 | 0.004255025 | 12 | NaN | NaN | NaN | 1270.9315 | NaN | NaN |
| H1-2 | AAPPAEKAPAK | -11 | -21 | 1 | 0.000591124 | 11 | NaN | NaN | NaN | 574.9448 | NaN | 571.84186 |
| H1-2 | SEAAPAAPAAAPPAEKAPAK | -2 | -21 | 1 | 0.001092529 | 20 | NaN | NaN | NaN | 316.3327 | NaN | 316.7682 |
| Wars1 | IGHPKPALL | -331 | -339 | 1 | 0.004224523 | 9 | NaN | NaN | NaN | 6363.7847 | NaN | NaN |
| Tcp1 | EQLAIAEFAR | -434 | -443 | 0 | 0.00078661 | 10 | NaN | NaN | NaN | 1486.8774 | 1170.7968 | 280.54565 |
| Tcp1 | AVNAAQDSTDLVAK | -453 | -466 | 0 | 0 | 14 | NaN | NaN | NaN | 791.2842 | 593.2892 | 501.5126 |
| Tcp1 | DLVHGKPR | -489 | -496 | 1 | 0.00383302 | 8 | NaN | NaN | NaN | 93.82101 | 708.2939 | 963.75366 |
| Tcp1 | ITDPEKLDQIR | -254 | -264 | 1 | 0.001868731 | 11 | NaN | NaN | NaN | 225.63806 | NaN | 1205.7273 |
| Tcp1 | ENLIINTDELGR | -134 | -145 | 0 | 0.00051306 | 12 | NaN | NaN | NaN | 919.87744 | NaN | NaN |
| Rps14 | FNDTFVHVTDLSGK | -37 | -50 | 0 | 0 | 14 | NaN | NaN | NaN | 1842.4156 | 3031.3616 | 2683.271 |
| Rps14 | ASFNDTFVHVTDLSGK | -35 | -50 | 0 | 0 | 16 | NaN | NaN | NaN | 1114.4985 | 1333.8828 | 1351.3823 |

| | | | | | | | | | | | | |
|---|---|---|---|---|---|---|---|---|---|---|---|---|
| Dpp3 | ETVINPETGEQIQSWYR | -474 | -490 | 0 | 0 | 17 | NaN | NaN | NaN | 310.39938 | 283.4715 | 311.90674 |
| Dpp3 | RDTVLLR | -659 | -665 | 1 | 0.001708008 | 7 | NaN | NaN | NaN | 515.6961 | 639.3219 | NaN |
| Slc16a1 | EGKEDEASTDVDEKPK | -454 | -469 | 2 | 0 | 16 | NaN | NaN | NaN | 924.96735 | 907.9996 | 702.7929 |
| Esd | VYLPPQAESGK | -34 | -44 | 0 | 0.004224523 | 11 | NaN | NaN | NaN | 698.32623 | 2316.4368 | NaN |
| Esd | YVNATEDPWK | -106 | -115 | 0 | 0 | 10 | NaN | NaN | NaN | 332.58493 | NaN | NaN |
| Esd | GEDDSWDFGTGAGF | -92 | -105 | 0 | 0.001291103 | 14 | NaN | NaN | NaN | 250.73117 | NaN | NaN |
| Pfdn1 | TLVDETNMYEGVGR | -53 | -66 | 0 | 0.002845514 | 14 | NaN | NaN | NaN | 1084.6642 | NaN | NaN |
| Hadh | AATGHTVVLVDQTEDILAK | -47 | -65 | 0 | 0.000431655 | 19 | NaN | NaN | NaN | 331.787 | 473.45007 | 666.5474 |
| Hadh | FAAEHTIFAS | -137 | -146 | 0 | 0.00078661 | 10 | NaN | NaN | NaN | 1114.8756 | 1211.2437 | NaN |
| Nosip | FWIPSLTPEAK | -162 | -172 | 0 | 0 | 11 | NaN | NaN | NaN | 100.12393 | 698.4612 | 103.96402 |
| Nosip | TLPNTEGEQPGPSVGPVGK | -135 | -153 | 0 | 0.00025015 | 19 | NaN | NaN | NaN | 1233.6738 | NaN | NaN |
| Got1 | WSNPPAQGAR | -296 | -305 | 0 | 0 | 10 | NaN | NaN | NaN | 1075.6501 | 5217.395 | NaN |
| Nop2 | AATPTEPDLKDQVTPK | -586 | -601 | 1 | 0.001636916 | 16 | NaN | NaN | NaN | 877.5997 | 1034.5023 | 820.50793 |
| Erap1 | VEDYFFGK | -385 | -392 | 0 | 0.00078661 | 8 | NaN | NaN | NaN | 4044.3704 | 3941.5122 | 5372.4604 |
| Erap1 | AVGAQNTEGWDFLYSK | -762 | -777 | 0 | 0 | 16 | NaN | NaN | NaN | 1246.4387 | 1420.2289 | 1235.0854 |
| Erap1 | EAHTTISSNDR | -615 | -625 | 0 | 0.002068213 | 11 | NaN | NaN | NaN | 607.0426 | 656.41064 | NaN |
| Dync1h1 | TKPVTGNLRPEEALQAL | -1262 | -1278 | 2 | 0.001463897 | 17 | NaN | NaN | NaN | 3041.5269 | 3803.6091 | 4823.8936 |
| Dync1h1 | SRQELEQHSVDTASTS | -1147 | -1162 | 1 | 0.00383302 | 16 | NaN | NaN | NaN | 355.76508 | 274.54166 | 99.28363 |
| Dync1h1 | ETVDQVEELRR | -3208 | -3218 | 1 | 0 | 11 | NaN | NaN | NaN | 76.46096 | 7620.6396 | NaN |
| Dync1h1 | GLTVEQAEAVVR | -3924 | -3935 | 0 | 0 | 12 | NaN | NaN | NaN | 1856.6077 | NaN | NaN |
| Dync1h1 | LHSLQPHACFR | -4111 | -4121 | 0 | 0 | 11 | NaN | NaN | NaN | 1827.5919 | NaN | NaN |
| Dync1h1 | LHSLQPHACFR | -4111 | -4121 | 0 | 0 | 11 | NaN | NaN | NaN | 1827.5919 | NaN | NaN |
| Dync1h1 | TFSEILNR | -871 | -878 | 0 | 0 | 8 | NaN | NaN | NaN | 1014.4361 | NaN | NaN |
| Dync1h1 | SIPLDEGEDEAQRR | -2382 | -2395 | 1 | 0.001610812 | 14 | NaN | NaN | NaN | 556.6479 | NaN | NaN |
| Etfa | APSSSSVGISEWLDQK | -188 | -203 | 0 | 0.001776473 | 16 | NaN | NaN | NaN | 1399.6333 | 1472.8785 | 1432.3315 |
| Etfa | LGGEVSCLVAGTK | -47 | -59 | 0 | 0 | 13 | NaN | NaN | NaN | 226.7544 | 610.4127 | 600.0008 |
| Etfa | AAVDAGFVPNDMQVGQTGK | -250 | -268 | 0 | 0.000471409 | 19 | NaN | NaN | NaN | 585.90393 | 219.386 | NaN |
| Etfa | VAPVSDIIEIK | -129 | -139 | 0 | 0 | 11 | NaN | NaN | NaN | 494.72638 | NaN | 644.65393 |
| Ets1 | QWTETHVR | -71 | -78 | 0 | 0.006735683 | 8 | NaN | NaN | NaN | 1605.2225 | NaN | NaN |
| Acat1 | EVYMGNVIQGGEGQAPTR | -85 | -102 | 0 | 0 | 18 | NaN | NaN | NaN | 609.97284 | 475.49478 | 573.0985 |
| Acat1 | AFADAAVDPIDFPLAPAYAVPK | -311 | -332 | 0 | 0.001868731 | 22 | NaN | NaN | NaN | 142.32288 | NaN | 269.34634 |
| Rrp12 | ALTHLLFEFK | -924 | -933 | 0 | 0.001392869 | 10 | NaN | NaN | NaN | 187.625 | 194.89693 | NaN |
| Rrp12 | VEEEEEEEEEEPVQSK | -1054 | -1070 | 0 | 0 | 17 | NaN | NaN | NaN | 179.12723 | NaN | 290.9118 |
| Rrp12 | SHLDDLKK | -813 | -820 | 1 | 0.005979575 | 8 | NaN | NaN | NaN | 1879.3202 | NaN | NaN |
| Kpnb1 | SVKPQILSVF | -708 | -717 | 1 | 0.007493591 | 10 | NaN | NaN | NaN | 678.26373 | 770.85114 | 894.9539 |
| Kpnb1 | TLVEVLGGEFLK | -634 | -645 | 0 | 0 | 12 | NaN | NaN | NaN | 526.30255 | 887.35126 | 526.4656 |
| Kpnb1 | TVSPDRLELEAA | -10 | -21 | 1 | 0 | 12 | NaN | NaN | NaN | 1316.3344 | 1953.2173 | NaN |
| Kpnb1 | QVVCEATQCPDTR | -220 | -232 | 0 | 0.000591124 | 13 | NaN | NaN | NaN | 702.17926 | NaN | 739.91486 |
| Kpnb1 | TKQDENDDDDDWNPCK | -331 | -346 | 1 | 0.003490217 | 16 | NaN | NaN | NaN | 64.887215 | 280.20996 | NaN |
| Kpnb1 | NVTNPNSTEHMK | -136 | -147 | 0 | 0.00025015 | 12 | NaN | NaN | NaN | 323.866 | NaN | NaN |
| Kpnb1 | GLKGDQENVHPDVMLVQPR | -775 | -793 | 1 | 0.003745449 | 19 | NaN | NaN | NaN | 308.92862 | NaN | NaN |
| Kpnb1 | SNEILTAIIQ | -170 | -179 | 0 | 0.000230707 | 10 | NaN | NaN | NaN | 298.5606 | NaN | NaN |
| Kpnb1 | ASEAAEQGRPPEHTSK | -297 | -312 | 1 | 0.00025015 | 16 | NaN | NaN | NaN | 244.1872 | NaN | NaN |
| Stip1 | YKDAIHF | -316 | -322 | 1 | 0.00025015 | 7 | NaN | NaN | NaN | 1497.3132 | 2447.774 | 2388.3342 |
| Stip1 | SAGNIDDALQCYSEAIK | -16 | -32 | 0 | 0 | 17 | NaN | NaN | NaN | 796.4323 | 967.41113 | 835.0893 |
| Stip1 | TVDLKPDWGKG | -64 | -74 | 2 | 0.00025015 | 11 | NaN | NaN | NaN | 1616.8558 | NaN | NaN |
| Stip1 | KFMNPFNLPNLYQK | -123 | -136 | 1 | 0.003037016 | 14 | NaN | NaN | NaN | 1113.0593 | NaN | NaN |
| Stip1 | INPDLALEEK | -355 | -364 | 0 | 0.002401448 | 10 | NaN | NaN | NaN | 936.5147 | NaN | NaN |
| Ndufv2 | NYPEGHQAAAVLPVLDLAQR | -68 | -87 | 0 | 0 | 20 | NaN | NaN | NaN | 330.3279 | 730.42377 | 1785.8938 |
| Ndufv2 | NYPEGHQAAAVLPVLDLAQR | -68 | -87 | 0 | 0 | 20 | NaN | NaN | NaN | 330.3279 | 730.42377 | 1785.8938 |
| Hsp90ab1 | KLGIHEDSTNR | -438 | -448 | 1 | 0 | 11 | NaN | NaN | NaN | 304.26926 | 699.57446 | 822.56433 |
| Hsp90ab1 | LELFSELAEDKENYKK | -413 | -428 | 2 | 0 | 16 | NaN | NaN | NaN | 164.66202 | 947.05945 | 889.4998 |
| Hsp90ab1 | FSELAEDKENYK | -416 | -427 | 1 | 0 | 12 | NaN | NaN | NaN | 1947.4989 | 1744.1532 | 1791.5027 |
| Hsp90ab1 | TSQSGDEMTSLSEYVSR | -459 | -475 | 0 | 0 | 17 | NaN | NaN | NaN | 1729.8217 | 2173.919 | 2073.7986 |
| Hsp90ab1 | RNPDDITQEEYGEFYK | -291 | -306 | 1 | 0 | 16 | NaN | NaN | NaN | 1221.3397 | 665.38556 | 192.00174 |
| Hsp90ab1 | RNPDDITQEEYGEFYK | -291 | -306 | 1 | 0 | 16 | NaN | NaN | NaN | 1221.3397 | 665.38556 | 192.00174 |
| Hsp90ab1 | GIDEDEVTAEEPSAAVPDEIPPLEGDEDASR | -689 | -719 | 0 | 0.00201047 | 31 | NaN | NaN | NaN | 640.6814 | 706.7994 | 634.1938 |
| Hsp90ab1 | LVSSPCCIVTSTY | -584 | -596 | 0 | 0.00078661 | 13 | NaN | NaN | NaN | 795.5564 | 1017.137 | NaN |
| Eno1 | YITPDQLADLY | -270 | -280 | 0 | 0.00025015 | 11 | NaN | NaN | NaN | 1017.65283 | 1045.187 | 984.61804 |
| Eno1 | EFMILPVGASSFR | -167 | -179 | 0 | 0 | 13 | NaN | NaN | NaN | 1112.447 | 1489.9031 | 1320.2045 |
| Eno1 | EFMILPVGASSFR | -167 | -179 | 0 | 0 | 13 | NaN | NaN | NaN | 1112.447 | 1489.9031 | 1320.2045 |
| Eno1 | SAGIQVVGDDLTVTNPK | -310 | -326 | 0 | 0 | 17 | NaN | NaN | NaN | 1044.5095 | 1268.5885 | 1200.1178 |
| Eno1 | GAEVYHNLK | -185 | -193 | 0 | 0.00111006 | 9 | NaN | NaN | NaN | 986.8501 | 483.00995 | 1269.81 |
| Eno1 | AREIFDSR | -8 | -15 | 1 | 0 | 8 | NaN | NaN | NaN | 558.511 | 858.5541 | 882.96643 |
| Eno1 | MIEMDGTENK | -94 | -103 | 0 | 0.000431655 | 10 | NaN | NaN | NaN | 446.51797 | 167.24426 | 518.86755 |
| Eno1 | SIEDPFDQDDWGAWQK | -291 | -306 | 0 | 0 | 16 | NaN | NaN | NaN | 365.07135 | 436.2383 | 348.58856 |
| Eno1 | VGDEGGFAPNILENK | -207 | -221 | 0 | 0 | 15 | NaN | NaN | NaN | 1860.2059 | 2149.2007 | NaN |
| Opa3 | AAWNALQDEVGR | -117 | -128 | 0 | 0.00111006 | 12 | NaN | NaN | NaN | 4187.759 | NaN | NaN |
| Eprs1 | NPDVGLKPVWYSPK | -536 | -549 | 1 | 0 | 14 | NaN | NaN | NaN | 2827.6135 | 3098.724 | 2707.7236 |
| Eprs1 | AINELNHCLSLR | -98 | -109 | 0 | 0 | 12 | NaN | NaN | NaN | 5587.84 | 5705.098 | 5588.232 |
| Eprs1 | EFLWQEGHSAFAT | -1166 | -1178 | 0 | 0 | 13 | NaN | NaN | NaN | 2354.8618 | 2338.1843 | 2693.7212 |
| Eprs1 | TELAEPIAIRPT | -1110 | -1121 | 1 | 0 | 12 | NaN | NaN | NaN | 1326.958 | 2385.1333 | 2227.8828 |
| Eprs1 | LTSAINELNHCLSLR | -95 | -109 | 0 | 0 | 15 | NaN | NaN | NaN | 869.03894 | 1064.0745 | 747.7927 |
| Eprs1 | KEENLAEWYSQ | -1020 | -1030 | 1 | 0.000471409 | 11 | NaN | NaN | NaN | 763.3839 | 298.9683 | 880.6703 |
| Eprs1 | EHLITKPVLGKDEDFK | -627 | -642 | 2 | 0.000230707 | 16 | NaN | NaN | NaN | 355.60748 | 447.25977 | 618.5827 |
| Eprs1 | QIPFCGEIDCEDWIK | -1444 | -1458 | 0 | 0.000550307 | 15 | NaN | NaN | NaN | 344.5611 | 306.5527 | 322.22815 |
| Eprs1 | SHSNPVSNAQPAGAEKPEAK | -883 | -902 | 1 | 0 | 20 | NaN | NaN | NaN | 299.7911 | 465.4215 | 435.31198 |
| Eprs1 | INELNHCLSLR | -99 | -109 | 0 | 0 | 11 | NaN | NaN | NaN | 1961.9618 | 1034.2417 | NaN |

| | | | | | | | | | | | | |
|---|---|---|---|---|---|---|---|---|---|---|---|---|
| Eprs1 | ESIKDFFDAEIK | -1058 | -1069 | 1 | 0.00111006 | 12 | NaN | NaN | NaN | 1666.1401 | NaN | 1336.7515 |
| Eprs1 | AIQGATSHHLGQNF | -1235 | -1248 | 0 | 0.00025015 | 14 | NaN | NaN | NaN | 1009.485 | 2653.254 | NaN |
| Eprs1 | GSGLSSGGAGEGQGPK | -994 | -1009 | 0 | 0 | 16 | NaN | NaN | NaN | 593.91077 | NaN | 633.64276 |
| Eprs1 | GDVSISVEEGKENLLR | -26 | -41 | 1 | 0.00149795 | 16 | NaN | NaN | NaN | 1932.2283 | NaN | NaN |
| Eprs1 | HEELMLGDPCLK | -651 | -662 | 0 | 0.001392869 | 12 | NaN | NaN | NaN | 767.91 | NaN | NaN |
| Rpl5 | YLMEEDEDAYKK | -210 | -221 | 1 | 0 | 12 | NaN | NaN | NaN | 89.97674 | NaN | NaN |
| Slc25a5 | YFGIYDTAK | -191 | -199 | 0 | 0 | 9 | NaN | NaN | NaN | 586.50543 | 763.01495 | 631.72327 |
| Slc25a5 | GTDIMYTGTLDCWR | -246 | -259 | 0 | 0 | 14 | NaN | NaN | NaN | 835.0567 | 749.44336 | NaN |
| Slc25a4 | AAYFGVYDTAK | -189 | -199 | 0 | 0 | 11 | NaN | NaN | NaN | 904.011 | 2068.8367 | NaN |
| Ccar2 | RDDGEDEFAGAK | -655 | -666 | 1 | 0.000230707 | 12 | NaN | NaN | NaN | 1936.4427 | 2647.7424 | 2586.2646 |
| Ccar2 | AAYNPGQAVPWNAVK | -98 | -112 | 0 | 0.002296977 | 15 | NaN | NaN | NaN | 592.0215 | NaN | NaN |
| Rars1 | EFAGYDVLR | -226 | -234 | 0 | 0.001846793 | 9 | NaN | NaN | NaN | 633.39935 | NaN | NaN |
| Vps35 | EDLPNLESSEETEQINK | -752 | -768 | 0 | 0 | 17 | NaN | NaN | NaN | 529.4243 | 674.61255 | 659.7907 |
| Vps35 | CFSEENHEPLR | -640 | -650 | 0 | 0 | 11 | NaN | NaN | NaN | 876.1721 | 527.9474 | NaN |
| Vps35 | FEYFDYESR | -425 | -433 | 0 | 0.000230707 | 9 | NaN | NaN | NaN | 491.457 | NaN | 543.16296 |
| Gars1 | GEFTIETEGK | -528 | -537 | 0 | 0.00025015 | 10 | NaN | NaN | NaN | 2020.3737 | 2069.6414 | 635.31995 |
| Gars1 | HVEEVVPSVIEPSFGLGR | -556 | -573 | 0 | 0 | 18 | NaN | NaN | NaN | 527.5417 | 754.6054 | 579.03186 |
| Gars1 | HVEEVVPSVIEPSFGLGR | -556 | -573 | 0 | 0 | 18 | NaN | NaN | NaN | 527.5417 | 754.6054 | 579.03186 |
| Gars1 | KVPLVAEKPLK | -467 | -477 | 2 | 0.00078661 | 11 | NaN | NaN | NaN | 350.22018 | NaN | NaN |
| Itgal | QIEDGVVVGAPGEGDNTGGLYHCR | -48 | -71 | 0 | 0 | 24 | NaN | NaN | NaN | 459.6882 | 472.856 | 355.31378 |
| Itgal | QVQHVYQVR | -944 | -952 | 0 | 0 | 9 | NaN | NaN | NaN | 575.8964 | NaN | 1426.4261 |
| Itgal | TQQVQHVYQVR | -942 | -952 | 0 | 0 | 11 | NaN | NaN | NaN | 302.47516 | 398.43246 | NaN |
| C3 | SSVAVPYVIVPLK | -892 | -904 | 0 | 0 | 13 | NaN | NaN | NaN | 583.28644 | 412.74542 | 371.00595 |
| C3 | VAVPYVIVPLK | -894 | -904 | 0 | 0.001636916 | 11 | NaN | NaN | NaN | 363.9886 | 404.3616 | 237.50029 |
| C3 | NVNFHLR | -472 | -478 | 0 | 0.000550307 | 7 | NaN | NaN | NaN | 4134.1763 | 3408.9941 | NaN |
| Lasp1 | ACFHCETCK | -28 | -36 | 0 | 0 | 9 | NaN | NaN | NaN | 320.75217 | 406.78015 | NaN |
| Gnb2 | TFVSGACDASIK | -198 | -209 | 0 | 0 | 12 | NaN | NaN | NaN | 2846.9607 | 1158.5206 | 1068.435 |
| Lgals1 | EPAFPFQPGSITEV | -75 | -88 | 0 | 0.000431655 | 14 | NaN | NaN | NaN | 509.56036 | 612.0346 | 583.08325 |
| Lgals1 | ACGLVASNLNLKPGECLK | -2 | -19 | 1 | 0.000230707 | 18 | NaN | NaN | NaN | 505.4393 | 664.43225 | 572.9019 |
| Lgals1 | EPAFPFQPGSITEVC | -75 | -89 | 0 | 0.000591124 | 15 | NaN | NaN | NaN | 407.63562 | 409.55423 | 371.80173 |
| Paics | IFQLLQEAGIK | -64 | -74 | 0 | 0 | 11 | NaN | NaN | NaN | 990.694 | 1175.7582 | 1082.4598 |
| Paics | AEYEGDGIPTVF | -314 | -325 | 0 | 0.000627943 | 12 | NaN | NaN | NaN | 177.29163 | 329.17126 | 374.72287 |
| Paics | LPSGIGCSTILSPEGSAQ | -368 | -385 | 0 | 0.000431655 | 18 | NaN | NaN | NaN | 1290.7418 | 1620.9824 | NaN |
| Paics | NNHLVWAK | -394 | -401 | 0 | 0.005333786 | 8 | NaN | NaN | NaN | 716.757 | 846.79944 | NaN |
| Paics | IFGLNNHLVWAK | -390 | -401 | 0 | 0.00025015 | 12 | NaN | NaN | NaN | 672.7611 | NaN | NaN |
| Aco2 | LQLLEPFDKWDGK | -565 | -577 | 1 | 0 | 13 | NaN | NaN | NaN | 624.0959 | 859.90393 | 919.2152 |
| Aco2 | MSHFEPSEYIR | -34 | -44 | 0 | 0 | 11 | NaN | NaN | NaN | 3077.8274 | 531.7356 | 411.2126 |
| Aco2 | HTFNETQIEWFR | -756 | -767 | 0 | 0 | 12 | NaN | NaN | NaN | 816.7152 | 556.8708 | 1646.0985 |
| Aco2 | VAVPSTIHCDHLIEAQVGGEK | -118 | -138 | 0 | 0 | 21 | NaN | NaN | NaN | 501.62622 | 622.53345 | 587.235 |
| Aco2 | TDHISAAGPWLK | -594 | -605 | 0 | 0 | 12 | NaN | NaN | NaN | 3482.194 | NaN | 616.05634 |
| Aco2 | EKEGWPLDIR | -369 | -378 | 1 | 0 | 10 | NaN | NaN | NaN | 2301.2827 | 4692.4106 | NaN |
| Aco2 | TFNETQIEWFR | -757 | -767 | 0 | 0 | 11 | NaN | NaN | NaN | 757.35034 | 88.777275 | NaN |
| Aco2 | QFISSGLPK | -109 | -117 | 0 | 0 | 9 | NaN | NaN | NaN | 541.2816 | 518.2668 | NaN |
| Aco2 | TDIANLAEEFK | -313 | -323 | 0 | 0.00025015 | 11 | NaN | NaN | NaN | 4521.5503 | NaN | NaN |
| Aco2 | HISAAGPWLK | -596 | -605 | 0 | 0 | 10 | NaN | NaN | NaN | 2351.8345 | NaN | NaN |
| Aco2 | NDANPETHAFVT | -480 | -491 | 0 | 0 | 12 | NaN | NaN | NaN | 1236.4182 | NaN | NaN |
| Aco2 | NHTFNETQIEWFR | -755 | -767 | 0 | 0.00025015 | 13 | NaN | NaN | NaN | 756.87964 | NaN | NaN |
| Eif4a3 | ATLPHEILEMTNK | -219 | -231 | 0 | 0 | 13 | NaN | NaN | NaN | 967.3353 | 2614.9653 | 2417.9316 |
| Eif4a3 | ACIGGTNVGEDIR | -139 | -151 | 0 | 0 | 13 | NaN | NaN | NaN | 369.50995 | 2021.7544 | 797.2394 |
| Cct3 | EIQVQHPAAK | -69 | -78 | 0 | 0 | 10 | NaN | NaN | NaN | 8849.79 | 11728.679 | 11065.041 |
| Cct3 | AMTGVEQWPYR | -428 | -438 | 0 | 0 | 11 | NaN | NaN | NaN | 1076.0625 | 1316.8274 | 186.44858 |
| Cct3 | RIDDIVSGHK | -518 | -527 | 1 | 0.001092529 | 10 | NaN | NaN | NaN | 792.3123 | 1572.4613 | 713.96735 |
| Cct3 | GISDLAQHYLMR | -295 | -306 | 0 | 0.000431655 | 12 | NaN | NaN | NaN | 882.74097 | 286.26672 | 700.37006 |
| Cct3 | NVLLDPQLVPGGGAS | -400 | -414 | 0 | 0 | 15 | NaN | NaN | NaN | 624.0247 | 766.27386 | 763.1007 |
| Cct3 | SSLACNIALDAVK | -169 | -181 | 0 | 0 | 13 | NaN | NaN | NaN | 538.3643 | 261.56964 | 343.98578 |
| Cct3 | TAVETAVLLLR | -508 | -518 | 0 | 0 | 11 | NaN | NaN | NaN | 511.85587 | 526.3013 | 562.5488 |
| Cct3 | WSSLACNIALDAVK | -168 | -181 | 0 | 0 | 14 | NaN | NaN | NaN | 214.96904 | 374.56482 | 320.42053 |
| Cct3 | KIGDEYFTFIT | -354 | -364 | 1 | 0.001291103 | 11 | NaN | NaN | NaN | 1495.5665 | NaN | 748.8142 |
| Cct3 | MMGHRPVLVL | -1 | -10 | 1 | 0.001868731 | 10 | NaN | NaN | NaN | 951.4898 | NaN | 322.3429 |
| Cct3 | KELGIWEPLAVK | -491 | -502 | 1 | 0 | 12 | NaN | NaN | NaN | 658.06335 | NaN | NaN |
| Cct3 | LLDSSLEYK | -240 | -248 | 0 | 0.003299042 | 9 | NaN | NaN | NaN | 441.2688 | NaN | NaN |
| Cct3 | IDDIVSGHKK | -519 | -528 | 1 | 0.007791272 | 10 | NaN | NaN | NaN | 432.64218 | NaN | NaN |
| Itgb2 | ASECIQEQSFVIR | -418 | -430 | 0 | 0 | 13 | NaN | NaN | NaN | 1260.75 | 1445.9373 | 1447.7047 |
| Itgb2 | EIFGQYCECDNVNCER | -529 | -544 | 0 | 0 | 16 | NaN | NaN | NaN | 360.72174 | 456.6765 | 459.21927 |
| Itgb2 | GDCDGVQINNPVTFQVK | -399 | -415 | 0 | 0 | 17 | NaN | NaN | NaN | 2502.43 | NaN | 3207.175 |
| Itgb2 | VTLYLRPGQAAAF | -104 | -116 | 1 | 0.000471409 | 13 | NaN | NaN | NaN | 1753.458 | NaN | 2157.41 |
| Itgb2 | YNSQVCGGSDR | -545 | -555 | 0 | 0 | 11 | NaN | NaN | NaN | 309.6364 | NaN | NaN |
| Prkcsh | TCLDGTATIPFDQVNDDYCDCK | -38 | -59 | 0 | 0.002595029 | 22 | NaN | NaN | NaN | 194.29514 | NaN | 214.8699 |
| Rpa2 | SQLQHMPVPSIK | -232 | -243 | 0 | 0.002099344 | 12 | NaN | NaN | NaN | 563.90485 | 484.0957 | NaN |
| Rpa2 | EKAPTNIVYK | -84 | -93 | 1 | 0.006293098 | 10 | NaN | NaN | NaN | 361.88013 | NaN | NaN |
| Hnrnpm | GDFFPPERPQQLPHGL | -285 | -300 | 1 | 0.00204437 | 16 | NaN | NaN | NaN | 2443.9038 | 2677.995 | 2133.1987 |
| Hnrnpm | GDFFPPERPQQLPH | -285 | -298 | 1 | 0.001023236 | 14 | NaN | NaN | NaN | 269.81778 | NaN | 399.8386 |
| Hnrnpm | GDFFPPERPQQLPH | -285 | -298 | 1 | 0.001023236 | 14 | NaN | NaN | NaN | 269.81778 | NaN | 399.8386 |
| Hnrnpm | MGPAMGPALGAGIER | -591 | -605 | 0 | 0.00025015 | 15 | NaN | NaN | NaN | 517.0038 | NaN | 968.6276 |
| Rps3a | NCLTNFHGMDLTR | -95 | -107 | 0 | 0 | 13 | NaN | NaN | NaN | 2138.6934 | 2456.9973 | 2380.9841 |
| Rps3a | KLITEDVQGK | -85 | -94 | 1 | 0 | 10 | NaN | NaN | NaN | 573.0035 | 754.93835 | 721.0995 |
| Plec | EGVVGPELHHK | -2900 | -2910 | 0 | 0.002168954 | 11 | NaN | NaN | NaN | 720.51056 | NaN | 738.65063 |
| Plec | QEEVYSELQAR | -3365 | -3375 | 0 | 0.00025015 | 11 | NaN | NaN | NaN | 2118.6204 | NaN | NaN |

| | | | | | | | | | | | | |
|---|---|---|---|---|---|---|---|---|---|---|---|---|
| Plec | VATGGIIDPVHSHR | -3613 | -3626 | 0 | 0.001055699 | 14 | NaN | NaN | NaN | 1543.638 | NaN | NaN |
| Plec | VATGGIIDPVHSHR | -3613 | -3626 | 0 | 0.001055699 | 14 | NaN | NaN | NaN | 1543.638 | NaN | NaN |
| Plec | NDESQLSPATR | -722 | -732 | 0 | 0.00025015 | 11 | NaN | NaN | NaN | 1117.0914 | NaN | NaN |
| Plec | ACTGGIIDPSTGER | -4460 | -4473 | 0 | 0.00051306 | 14 | NaN | NaN | NaN | 829.5288 | NaN | NaN |
| Plec | RHISDLYEDLR | -209 | -219 | 1 | 0.002595029 | 11 | NaN | NaN | NaN | 566.4205 | NaN | NaN |
| Plec | LAEDEAFQR | -2014 | -2022 | 0 | 0.00025015 | 9 | NaN | NaN | NaN | 330.44135 | NaN | NaN |
| Fasn | EHDLVLPMR | -2163 | -2171 | 0 | 0.00025015 | 9 | NaN | NaN | NaN | 6010.4976 | 415.26367 | NaN |
| Fasn | GTNDTVIGGTLPQR | -2067 | -2080 | 0 | 0 | 14 | NaN | NaN | NaN | 1078.5144 | 1044.1865 | 1113.1356 |
| Fasn | VAEVLAGEGHLYSR | -1237 | -1250 | 0 | 0 | 14 | NaN | NaN | NaN | 740.5192 | 529.67896 | 1939.7477 |
| Fasn | FLEIGKF | -1759 | -1765 | 1 | 0.007905138 | 7 | NaN | NaN | NaN | 672.03 | 583.27075 | 605.7212 |
| Fasn | FQGICEATLEGEQGK | -1006 | -1020 | 0 | 0 | 15 | NaN | NaN | NaN | 634.9094 | 674.42957 | 799.06555 |
| Fasn | TGGTYGEDLGADYNLSQVCDGK | -2443 | -2464 | 0 | 0 | 22 | NaN | NaN | NaN | 191.91411 | 263.48447 | 237.91125 |
| Fasn | EDTSFQWVDSLK | -1412 | -1423 | 0 | 0 | 12 | NaN | NaN | NaN | 155.77132 | 614.4243 | 691.28796 |
| Fasn | QVQPEGPYR | -2288 | -2296 | 0 | 0 | 9 | NaN | NaN | NaN | 2065.307 | NaN | NaN |
| Fasn | LKEDTQVADVT | -1070 | -1080 | 1 | 0.008045078 | 11 | NaN | NaN | NaN | 1512.3643 | NaN | NaN |
| Fasn | RHDGLPGLAVQ | -2042 | -2052 | 1 | 0.007905138 | 11 | NaN | NaN | NaN | 1498.3002 | NaN | NaN |
| Fasn | SIPEAQWQSSLAR | -716 | -728 | 0 | 0.00078661 | 13 | NaN | NaN | NaN | 1295.8999 | NaN | NaN |
| Fasn | YEAIVDGGINPASLR | -87 | -101 | 0 | 0 | 15 | NaN | NaN | NaN | 668.2355 | NaN | NaN |
| Fasn | LIGGVDMVTDDDRR | -26 | -39 | 1 | 0.000947459 | 14 | NaN | NaN | NaN | 564.5847 | NaN | NaN |
| Fasn | VIFPGTGYLC | -884 | -893 | 0 | 0.004658491 | 10 | NaN | NaN | NaN | 407.04895 | NaN | NaN |
| Fasn | GYDYGPQFQGICEATLEGEQGK | -999 | -1020 | 0 | 0.000230707 | 22 | NaN | NaN | NaN | 320.91672 | NaN | NaN |
| Fasn | VAEVLAGEGHLY | -1237 | -1248 | 0 | 0.00025015 | 12 | NaN | NaN | NaN | 246.94542 | NaN | NaN |
| Cct8 | AHEILPELVCCSAK | -139 | -152 | 0 | 0 | 14 | NaN | NaN | NaN | 1136.3269 | 1121.257 | 1054.3213 |
| Cct8 | FAEAFEAIPR | -441 | -450 | 0 | 0 | 10 | NaN | NaN | NaN | 9278.959 | 10852.726 | 10844.3955 |
| Cct8 | SKGEENLMDAQVK | -269 | -281 | 1 | 0 | 13 | NaN | NaN | NaN | 501.17365 | 915.30743 | 888.0922 |
| Cct8 | SKGEENLMDAQVK | -269 | -281 | 1 | 0 | 13 | NaN | NaN | NaN | 501.17365 | 915.30743 | 888.0922 |
| Cct8 | SCPFDGMITETK | -243 | -254 | 0 | 0.00078661 | 12 | NaN | NaN | NaN | 770.6523 | 486.45047 | 552.18756 |
| Cct8 | KAHEILPELVCCSAK | -138 | -152 | 1 | 0 | 15 | NaN | NaN | NaN | 181.49481 | 470.00128 | 556.6336 |
| Cct8 | VADIALHYANK | -297 | -307 | 0 | 0.000230707 | 11 | NaN | NaN | NaN | 2501.718 | 2701.0781 | NaN |
| Cct8 | SYGETCPGLEQYAIK | -425 | -439 | 0 | 0.00025015 | 15 | NaN | NaN | NaN | 448.97626 | 356.72812 | NaN |
| Cct8 | KFAEAFEAIPR | -440 | -450 | 1 | 0 | 11 | NaN | NaN | NaN | 687.8292 | NaN | NaN |
| Cct8 | KDWDDDQND | -540 | -548 | 1 | 0.001708008 | 9 | NaN | NaN | NaN | 601.03094 | NaN | NaN |
| Atp1a1 | PVNQVNPR | -651 | -658 | 0 | 0.001392869 | 8 | NaN | NaN | NaN | 778.38336 | 761.17456 | 825.0981 |
| Atp1a1 | AVFQANQENLPILK | -431 | -444 | 0 | 0 | 14 | NaN | NaN | NaN | 471.96332 | 498.8156 | 527.7663 |
| Atp1a1 | TSATWFALSR | -414 | -423 | 0 | 0 | 10 | NaN | NaN | NaN | 315.31424 | 521.1621 | 504.72568 |
| Atp1a1 | DKYEPAAVSEHGDKK | -8 | -22 | 2 | 0.000431655 | 15 | NaN | NaN | NaN | 240.5982 | 467.7208 | NaN |
| Atp1a1 | DVEDSYGQQWTYEQR | -897 | -911 | 0 | 0.004321969 | 15 | NaN | NaN | NaN | 546.0594 | NaN | NaN |
| Atp1a1 | EADTTENQSGVSFDK | -399 | -413 | 0 | 0.00266297 | 15 | NaN | NaN | NaN | 541.1328 | NaN | NaN |
| Prkar1a | HNIQALLK | -25 | -32 | 0 | 0.000471409 | 8 | NaN | NaN | NaN | 544.9663 | 1387.9508 | 1359.3026 |
| Prkar1a | ADALEPVQFEDGQK | -268 | -281 | 0 | 0.000471409 | 14 | NaN | NaN | NaN | 410.39856 | 680.6957 | 650.0186 |
| Cct6a | NSGFDLQETLVK | -454 | -465 | 0 | 0.000230707 | 12 | NaN | NaN | NaN | 4584.326 | 1442.8596 | 5328.5425 |
| Cct6a | LVYEYTLGEEK | -349 | -359 | 0 | 0 | 11 | NaN | NaN | NaN | 802.80273 | NaN | NaN |
| Api5 | FSPSDPDCVDR | -227 | -237 | 0 | 0 | 11 | NaN | NaN | NaN | 4642.448 | 4371.9194 | 4517.0884 |
| Api5 | GILADATEQVGQHK | -12 | -25 | 0 | 0 | 14 | NaN | NaN | NaN | 1781.8324 | 1675.588 | 1611.9342 |
| Ppa2 | ADCKEEHDIPR | -65 | -75 | 1 | 0.00025015 | 11 | NaN | NaN | NaN | 545.96124 | 917.7774 | 610.14667 |
| Mcm4 | FSNKVEAIDVEEAK | -748 | -761 | 1 | 0 | 14 | NaN | NaN | NaN | 2028.5409 | 587.81323 | 1170.411 |
| Mcm4 | APSIYEHEDIKK | -466 | -477 | 1 | 0 | 12 | NaN | NaN | NaN | 828.8519 | 994.39655 | 946.3515 |
| Mcm4 | LAPSIYEHEDIKK | -465 | -477 | 1 | 0 | 13 | NaN | NaN | NaN | 4940.8804 | 791.2554 | 814.3458 |
| Mcm4 | LAPSIYEHEDIKK | -465 | -477 | 1 | 0 | 13 | NaN | NaN | NaN | 4940.8804 | 791.2554 | 814.3458 |
| Mcm4 | TIENIQLPHTLLSR | -629 | -642 | 0 | 0 | 14 | NaN | NaN | NaN | 1359.4459 | 1388.7 | 1550.3452 |
| Mcm4 | VNEIFFDR | -240 | -247 | 0 | 0 | 8 | NaN | NaN | NaN | 812.22534 | 866.3889 | 746.75446 |
| Mcm4 | KTTIENIQLPHTL | -627 | -639 | 1 | 0 | 13 | NaN | NaN | NaN | 1718.6956 | NaN | 1112.0023 |
| Mcm4 | VEAIDVEEAK | -752 | -761 | 0 | 0 | 10 | NaN | NaN | NaN | 1802.817 | NaN | NaN |
| Mcm4 | FLQCFTDPLAK | -168 | -178 | 0 | 0.002972566 | 11 | NaN | NaN | NaN | 957.6111 | NaN | NaN |
| Mcm4 | HIDVIHYR | -414 | -421 | 0 | 0 | 8 | NaN | NaN | NaN | 281.16803 | NaN | NaN |
| Ppia | FDITADDEPLGR | -8 | -19 | 0 | 0 | 12 | NaN | NaN | NaN | 300.26266 | 16015.611 | 15655.762 |
| Ppia | TAKTEWLDGKHVVFGK | -116 | -131 | 2 | 0 | 16 | NaN | NaN | NaN | 633.0589 | 1145.1248 | 1128.7437 |
| Ppia | TAKTEWLDGKHVVFGK | -116 | -131 | 2 | 0 | 16 | NaN | NaN | NaN | 633.0589 | 1145.1248 | 1128.7437 |
| Ppia | TAKTEWLDGKHVVFGK | -116 | -131 | 2 | 0 | 16 | NaN | NaN | NaN | 633.0589 | 1145.1248 | 1128.7437 |
| Ppia | IIPGFMCQGGDFTR | -56 | -69 | 0 | 0.004199202 | 14 | NaN | NaN | NaN | 1304.2948 | 132.90965 | NaN |
| Ppia | VKEGMNIVEAM | -132 | -142 | 1 | 0.00025015 | 11 | NaN | NaN | NaN | 140.78825 | NaN | 319.45535 |
| Aldoa | VNPCIGGVIL | -70 | -79 | 0 | 0 | 10 | NaN | NaN | NaN | 840.88165 | 428.5912 | 993.39685 |
| Aldoa | KADDGRPFPQVIK | -87 | -99 | 2 | 0 | 13 | NaN | NaN | NaN | 1155.7217 | 502.9259 | 1375.2207 |
| Aldoa | ADESTGSIAK | -33 | -42 | 0 | 0 | 10 | NaN | NaN | NaN | 738.10944 | 345.17142 | 602.98334 |
| Aldoa | VNPCIGGVILFHETLYQK | -70 | -87 | 0 | 0 | 18 | NaN | NaN | NaN | 636.2488 | 994.0594 | 975.1803 |
| Aldoa | LSGGQSEEEASINLNAINK | -271 | -289 | 0 | 0 | 19 | NaN | NaN | NaN | 178.0058 | 1224.5197 | 1081.971 |
| Aldoa | VNPCIGGVILF | -70 | -80 | 0 | 0.002068213 | 11 | NaN | NaN | NaN | 174.6355 | 337.36655 | 219.3064 |
| Aldoa | AIMENANVLAR | -163 | -173 | 0 | 0.000471409 | 11 | NaN | NaN | NaN | 501.67618 | 1174.062 | NaN |
| Aldoa | YTPSGQSGAA | -343 | -352 | 0 | 0.00676572 | 10 | NaN | NaN | NaN | 984.1631 | NaN | NaN |
| Drg1 | ADDLIDVVEGNR | -227 | -238 | 0 | 0 | 12 | NaN | NaN | NaN | 3199.3652 | NaN | 3414.7402 |
| Sart1 | ADDLLPLGDQTQDGDFGSR | -420 | -438 | 0 | 0.002780285 | 19 | NaN | NaN | NaN | 963.2229 | 1334.4938 | 1288.255 |
| Sart1 | DGYKPDVK | -708 | -715 | 1 | 0.001392869 | 8 | NaN | NaN | NaN | 1143.9169 | 542.4766 | NaN |
| Snx27 | ADEDGEGIHPSAPHR | -2 | -16 | 0 | 0 | 15 | NaN | NaN | NaN | 528.23047 | 5523.095 | 5856.014 |
| Ipo7 | EYNEFAEVFLK | -284 | -294 | 0 | 0 | 11 | NaN | NaN | NaN | 1563.5437 | 643.8924 | 2840.5522 |
| Ipo7 | SDQNLQTALELTR | -490 | -502 | 0 | 0 | 13 | NaN | NaN | NaN | 1390.3302 | 1198.1884 | 1134.4691 |
| Ipo7 | VLTGVAGEDAECHAA | -725 | -739 | 0 | 0 | 15 | NaN | NaN | NaN | 309.8746 | 728.9789 | NaN |
| Ipo7 | ENIVEAIIH | -93 | -101 | 0 | 0.00383302 | 9 | NaN | NaN | NaN | 464.8538 | NaN | NaN |
| Prdx1 | HGEVCPAGWKPGSDTIKPDVNK | -169 | -190 | 2 | 0 | 22 | NaN | NaN | NaN | 5564.3174 | 5649.3496 | 4545.9272 |

| | | | | | | | | | | | | |
|---|---|---|---|---|---|---|---|---|---|---|---|---|
| Prdx1 | SVDSHFCHLAWINTPK | -77 | -92 | 0 | 0 | 16 | NaN | NaN | NaN | 788.87036 | 662.528 | 1508.7297 |
| Prdx1 | SVDSHFCHLAWINTPK | -77 | -92 | 0 | 0 | 16 | NaN | NaN | NaN | 788.87036 | 662.528 | 1508.7297 |
| Prdx1 | HGEVCPAGWKPGSDTIKPD | -169 | -187 | 2 | 0.00025015 | 19 | NaN | NaN | NaN | 317.0329 | 673.5385 | 511.54935 |
| Ldha | MMDLQHGSLFLK | -62 | -73 | 0 | 0 | 12 | NaN | NaN | NaN | 1214.1866 | 1308.7583 | 1340.6248 |
| Ldha | GEHGDSSVPVWSGVNVAGVSLK | -191 | -212 | 0 | 0 | 22 | NaN | NaN | NaN | 858.64856 | 816.7392 | 636.76636 |
| Ldha | GEHGDSSVPVWSGVNVAGVSLK | -191 | -212 | 0 | 0 | 22 | NaN | NaN | NaN | 858.64856 | 816.7392 | 636.76636 |
| Ldha | RNVNIFK | -112 | -118 | 1 | 0.002168954 | 7 | NaN | NaN | NaN | 690.3684 | 935.51013 | 858.2877 |
| Ldha | NLLKEEQAPQNK | -11 | -22 | 1 | 0.000471409 | 12 | NaN | NaN | NaN | 450.84216 | 453.88348 | 390.15088 |
| Ldha | ITVVGVGAVGMAC | -23 | -35 | 0 | 0.002168954 | 13 | NaN | NaN | NaN | 5896.86 | NaN | 165.15921 |
| Ldha | MKDLADELALVDVMEDK | -41 | -57 | 1 | 0.001092529 | 17 | NaN | NaN | NaN | 3027.9827 | NaN | 4366.836 |
| Myh9 | MEDGVGCLETAEEAK | -1373 | -1387 | 0 | 0 | 15 | NaN | NaN | NaN | 4018.7026 | 237.30373 | 3785.0884 |
| Myh9 | KMEDGVGCLETAEEAK | -1372 | -1387 | 1 | 0 | 16 | NaN | NaN | NaN | 3132.163 | 1830.2399 | 1305.8148 |
| Myh9 | TDLLLEPYNK | -290 | -299 | 0 | 0 | 10 | NaN | NaN | NaN | 454.39722 | 3212.747 | 137.9857 |
| Myh9 | RALEEAMEQK | -1483 | -1492 | 1 | 0.00111006 | 10 | NaN | NaN | NaN | 310.07297 | NaN | 502.28827 |
| Myh9 | SIRHEDELLAK | -840 | -850 | 1 | 0 | 11 | NaN | NaN | NaN | 554.53784 | 565.38904 | 571.9388 |
| Myh9 | QLEEAEEEAQR | -1878 | -1888 | 0 | 0 | 11 | NaN | NaN | NaN | 1233.9916 | 990.25726 | 1120.0469 |
| Myh9 | EAEMIQLQEELAAAER | -1679 | -1694 | 0 | 0 | 16 | NaN | NaN | NaN | 1137.6882 | 1203.8685 | 1061.9186 |
| Myh9 | AQQERDELADEIANSSGK | -1699 | -1716 | 1 | 0 | 18 | NaN | NaN | NaN | 548.41797 | 601.1809 | 2216.541 |
| Myh9 | AQQERDELADEIANSSGK | -1699 | -1716 | 1 | 0 | 18 | NaN | NaN | NaN | 548.41797 | 601.1809 | 2216.541 |
| Myh9 | KGTGDCSDEEVD | -1937 | -1948 | 1 | 0.00111006 | 12 | NaN | NaN | NaN | 330.19177 | 422.51544 | 404.4328 |
| Myh9 | EDGVGCLETAEEAKR | -1374 | -1388 | 1 | 0.001055699 | 15 | NaN | NaN | NaN | 1746.2139 | 423.5257 | NaN |
| Myh9 | KQELEEICHDLEA | -910 | -922 | 1 | 0 | 13 | NaN | NaN | NaN | 756.2553 | NaN | 691.7948 |
| Myh9 | EEDQIIMEDQNCK | -977 | -989 | 0 | 0.00078661 | 13 | NaN | NaN | NaN | 712.01953 | NaN | 310.05945 |
| Myh9 | ALEQQVEEMKTQ | -1529 | -1540 | 1 | 0.001392869 | 12 | NaN | NaN | NaN | 657.0735 | NaN | 1457.8336 |
| Myh9 | MIQLQEELAAAER | -1682 | -1694 | 0 | 0 | 13 | NaN | NaN | NaN | 585.9028 | 787.50775 | NaN |
| Myh9 | EDGVGCLETAEEAK | -1374 | -1387 | 0 | 0.001023236 | 14 | NaN | NaN | NaN | 499.57758 | 474.27258 | NaN |
| Myh9 | TEMEDLMSSK | -1504 | -1513 | 0 | 0.000431655 | 10 | NaN | NaN | NaN | 497.80646 | 453.84174 | NaN |
| Myh9 | VSHLLGINVT | -374 | -383 | 0 | 0.000471409 | 10 | NaN | NaN | NaN | 333.67404 | NaN | 391.01697 |
| Myh9 | LKDVLLQ | -1844 | -1850 | 1 | 0.005587349 | 7 | NaN | NaN | NaN | 802.7143 | NaN | NaN |
| Hnrnpul1 | ADFECGNDVELSFTK | -334 | -348 | 0 | 0 | 15 | NaN | NaN | NaN | 1848.7385 | 2031.7998 | 1903.9333 |
| Slc3a2 | NQKDEINETDLK | -160 | -171 | 1 | 0 | 12 | NaN | NaN | NaN | 1989.6708 | 2027.9027 | 2083.4849 |
| Slc3a2 | NAWFLPAQADIVATK | -209 | -223 | 0 | 0 | 15 | NaN | NaN | NaN | 447.44693 | 478.89832 | 459.6779 |
| Slc3a2 | LLLSTDSAR | -488 | -496 | 0 | 0.002466863 | 9 | NaN | NaN | NaN | 634.6826 | NaN | NaN |
| Icam1 | LFCSLEGLFPASEAR | -237 | -251 | 0 | 0.000230707 | 15 | NaN | NaN | NaN | 221.4176 | 290.15494 | 144.30692 |
| Icam1 | SQLEVSEGSQVTVK | -320 | -333 | 0 | 0.002629273 | 14 | NaN | NaN | NaN | 774.70074 | NaN | 323.69092 |
| Adss2 | DGETIPHFPANQEVLNK | -381 | -397 | 0 | 0 | 17 | NaN | NaN | NaN | 493.22818 | 543.5693 | 649.97705 |
| Adss2 | ADGIQEQQR | -143 | -151 | 0 | 0 | 9 | NaN | NaN | NaN | 416.58163 | 264.84003 | 658.05304 |
| Adss2 | PGWNTDISNAR | -406 | -416 | 0 | 0.001092529 | 11 | NaN | NaN | NaN | 310.72595 | 856.2323 | 751.7918 |
| Mrpl24 | KPTEIQWR | -129 | -136 | 1 | 0.000591124 | 8 | NaN | NaN | NaN | 1014.2357 | 319.81134 | NaN |
| Exosc4 | ADGSAYIEQGNTK | -35 | -47 | 0 | 0.001291103 | 13 | NaN | NaN | NaN | 3026.5596 | NaN | NaN |
| Pfn1 | EGVHGGLINK | -117 | -126 | 0 | 0 | 10 | NaN | NaN | NaN | 11075.377 | 9349.776 | 10280.6045 |
| Pfn1 | DSPSVWAAVPGK | -27 | -38 | 0 | 0 | 12 | NaN | NaN | NaN | 7634.5513 | 5952.1914 | 6442.0767 |
| Pfn1 | MADGTCQDAAIVGYK | -12 | -26 | 0 | 0 | 15 | NaN | NaN | NaN | 809.4943 | 833.16736 | 960.9433 |
| Pfn1 | ADGTCQDAAIVGYK | -13 | -26 | 0 | 0 | 14 | NaN | NaN | NaN | 878.54645 | 949.58704 | 943.50684 |
| Pfn1 | VSITPAEVGVLVGKDR | -41 | -56 | 1 | 0.00051306 | 16 | NaN | NaN | NaN | 459.89587 | 735.2446 | 583.36346 |
| Pfn1 | STGGAPTFNVTVT | -92 | -104 | 0 | 0.00025015 | 13 | NaN | NaN | NaN | 349.08636 | 320.23563 | 311.8553 |
| Pfn1 | IDSLMADGTCQDAAIVGYK | -8 | -26 | 0 | 0 | 19 | NaN | NaN | NaN | 1493.493 | NaN | 148.72195 |
| Pfn1 | GKEGVHGGLINKK | -115 | -127 | 2 | 0.00111006 | 13 | NaN | NaN | NaN | 376.84027 | NaN | NaN |
| Pfn1 | YIDSLMADGTCQDAAIVGYK | -7 | -26 | 0 | 0 | 20 | NaN | NaN | NaN | 348.21216 | NaN | NaN |
| Rpsa | ADHQPLTEAS | -129 | -138 | 0 | 0 | 10 | NaN | NaN | NaN | 693.4206 | 527.89557 | 680.83606 |
| Rpsa | EHPWEVMPDLY | -192 | -202 | 0 | 0 | 11 | NaN | NaN | NaN | 404.7829 | 349.60422 | 446.62494 |
| Rpsa | YVNLPTIALCNTDSPLR | -139 | -155 | 0 | 0 | 17 | NaN | NaN | NaN | 392.26675 | 552.48413 | 489.72095 |
| Rpsa | ADHQPLTEASYVNLPTIALC | -129 | -148 | 0 | 0 | 20 | NaN | NaN | NaN | 260.37366 | 282.96027 | 299.4193 |
| Rpsa | ADHQPLTEASYVNLPTIAL | -129 | -147 | 0 | 0.00025015 | 19 | NaN | NaN | NaN | 245.82837 | 297.75098 | 397.36536 |
| Rpsa | IENPADVSVISSR | -68 | -80 | 0 | 0 | 13 | NaN | NaN | NaN | 1184.1592 | NaN | 658.6824 |
| Rpsa | SYVNLPTIALCNTDSPLR | -138 | -155 | 0 | 0.000627943 | 18 | NaN | NaN | NaN | 170.62648 | 300.47574 | NaN |
| Hnrnpk | RPAEDMEEEQAFK | -22 | -34 | 1 | 0.00025015 | 13 | NaN | NaN | NaN | 6059.615 | 3550.6687 | 3899.897 |
| Hnrnpk | RTDYNASVSVPDSSGPER | -69 | -86 | 1 | 0 | 18 | NaN | NaN | NaN | 584.72485 | 1300.4294 | 1287.0304 |
| Hnrnpk | AVECLNYQHYK | -129 | -139 | 0 | 0 | 11 | NaN | NaN | NaN | 1607.9457 | 1498.7087 | 697.77924 |
| Hnrnpk | RNTDEMVELR | -37 | -46 | 1 | 0.001463897 | 10 | NaN | NaN | NaN | 891.8676 | 883.531 | 2064.7773 |
| Hnrnpk | PDSSGPER | -79 | -86 | 0 | 0.006788754 | 8 | NaN | NaN | NaN | 859.71436 | 1058.1895 | 804.4664 |
| Hnrnpk | IIPTLEEGLQLPSPTATS | -104 | -121 | 0 | 0 | 18 | NaN | NaN | NaN | 396.30145 | 373.33057 | 298.5784 |
| Hnrnpk | SQLPLESDAVECLNYQHYK | -121 | -139 | 0 | 0 | 19 | NaN | NaN | NaN | 318.51547 | 390.4801 | 371.80038 |
| Hnrnpk | AQPYDPNFYDETYDYGGFT | -222 | -240 | 0 | 0 | 19 | NaN | NaN | NaN | 309.4014 | 295.20587 | 294.22223 |
| Hnrnpk | AQPYDPNFYDETYDYGGF | -222 | -239 | 0 | 0.000712251 | 18 | NaN | NaN | NaN | 282.0022 | 318.0798 | 270.2868 |
| Hnrnpk | GTQDQIQNAQYLLQNSVK | -439 | -456 | 0 | 0 | 18 | NaN | NaN | NaN | 101.074554 | 398.11304 | 455.33795 |
| Hnrnpk | LFQECCPH | -180 | -187 | 0 | 0.001055699 | 8 | NaN | NaN | NaN | 779.19604 | 730.7615 | NaN |
| Hnrnpk | LLIHQSLAGGIIGVK | -149 | -163 | 0 | 0 | 15 | NaN | NaN | NaN | 556.212 | NaN | NaN |
| Hnrnpk | KIIPTLEEGLQLPSPT | -103 | -118 | 1 | 0.007670622 | 16 | NaN | NaN | NaN | 206.13666 | NaN | NaN |
| Csnk2a1 | QLYQTLTDYDIR | -123 | -134 | 0 | 0 | 12 | NaN | NaN | NaN | 1953.5813 | 2293.8726 | 2410.3013 |
| Csnk2a1 | EHVNNTDFK | -114 | -122 | 0 | 0.00025015 | 9 | NaN | NaN | NaN | 3220.29 | NaN | NaN |
| Npm1 | LDEDDEDDDEDDEDDEDDDDDDFDEEETEEKVPVK | -158 | -192 | 1 | 0.003620005 | 35 | NaN | NaN | NaN | 837.64526 | 677.5288 | 596.4528 |
| Npm1 | KADKDYHFK | -24 | -32 | 2 | 0.003299042 | 9 | NaN | NaN | NaN | 759.2032 | NaN | NaN |
| Npm1 | AESEDEDEEDVK | -123 | -134 | 0 | 0.002595029 | 12 | NaN | NaN | NaN | 213.94963 | NaN | NaN |
| Rbbp4 | SASDDHTICLWDISAVPK | -195 | -212 | 0 | 0.001291103 | 18 | NaN | NaN | NaN | 367.5236 | 431.49792 | 429.0633 |
| Rbbp4 | WDISAVPK | -205 | -212 | 0 | 0.00051306 | 8 | NaN | NaN | NaN | 761.89105 | 516.39374 | NaN |
| Tmsb10 | ADKPDMGEIA | -2 | -11 | 1 | 0.003003218 | 10 | NaN | NaN | NaN | 2151.2761 | 2370.5564 | NaN |
| Themis | LGHPCFYHLK | -113 | -122 | 0 | 0 | 10 | NaN | NaN | NaN | 1891.2102 | NaN | 560.72205 |

| | | | | | | | | | | | | |
|---|---|---|---|---|---|---|---|---|---|---|---|---|
| Themis | ADKTPYLSIEEITR | -91 | -104 | 1 | 0.00025015 | 14 | NaN | NaN | NaN | 71.580795 | NaN | 190.41785 |
| Capza2 | EATDPRPYEAENAIESW | -104 | -120 | 1 | 0.000230707 | 17 | NaN | NaN | NaN | 122.04462 | 425.16592 | 154.05856 |
| Capza2 | KVDGQQTIIAC | -147 | -157 | 1 | 0 | 11 | NaN | NaN | NaN | 1143.2749 | NaN | 1060.7098 |
| Sptbn1 | SHAYQQFLR | -1168 | -1176 | 0 | 0 | 9 | NaN | NaN | NaN | 1945.2764 | 1421.4607 | 444.235 |
| Sptbn1 | SQNIITDSSSLNAEAIR | -1548 | -1564 | 0 | 0 | 17 | NaN | NaN | NaN | 885.508 | 551.2531 | 617.33014 |
| Sptbn1 | DLDDFQSWLSR | -1070 | -1080 | 0 | 0 | 11 | NaN | NaN | NaN | 485.60553 | 412.4439 | 570.1982 |
| Sptbn1 | HLLGVEDLLQKH | -559 | -570 | 1 | 0.00025015 | 12 | NaN | NaN | NaN | 428.7993 | 418.61407 | 368.72906 |
| Ap1s1 | AIEQADLLQEEDESPR | -134 | -149 | 0 | 0.000755556 | 16 | NaN | NaN | NaN | 504.1831 | NaN | NaN |
| Efhd2 | SRFEEEIKAEQEER | -205 | -218 | 2 | 0.000591124 | 14 | NaN | NaN | NaN | 483.92465 | NaN | 3659.2288 |
| Efhd2 | LSEIDVSTEGVK | -177 | -188 | 0 | 0.00272025 | 12 | NaN | NaN | NaN | 6556.391 | NaN | NaN |
| Efhd2 | RVFNPYTEFK | -78 | -87 | 1 | 0.000947459 | 10 | NaN | NaN | NaN | 2247.3313 | NaN | NaN |
| Adsl | EMCFLFSDR | -25 | -33 | 0 | 0.001868731 | 9 | NaN | NaN | NaN | 714.0768 | 545.86255 | 758.12177 |
| Adsl | AFDLLLPK | -127 | -134 | 0 | 0 | 8 | NaN | NaN | NaN | 391.62653 | 423.80746 | 290.80173 |
| Adsl | QLFEGDHQK | -210 | -218 | 0 | 0 | 9 | NaN | NaN | NaN | 1031.561 | NaN | NaN |
| Adsl | FLQLFEGDHQK | -208 | -218 | 0 | 0.001055699 | 11 | NaN | NaN | NaN | 909.309 | NaN | NaN |
| Mical1 | RQDLSETDQLLGK | -310 | -322 | 1 | 0.00122419 | 13 | NaN | NaN | NaN | 759.12756 | 671.1394 | 716.37286 |
| Mical1 | TEADLQSENQVLR | -1010 | -1022 | 0 | 0 | 13 | NaN | NaN | NaN | 214.36386 | 504.06165 | 544.27704 |
| Mdh2 | HTPGVAADLSHIETR | -60 | -74 | 0 | 0 | 15 | NaN | NaN | NaN | 1281.4326 | 413.2156 | 1732.8282 |
| Mdh2 | DIAHTPGVAADLSHIETR | -57 | -74 | 0 | 0 | 18 | NaN | NaN | NaN | 1101.7996 | 1337.7604 | 1278.3722 |
| Mdh2 | PVIGGHAGK | -195 | -203 | 0 | 0.001392869 | 9 | NaN | NaN | NaN | 692.9847 | 2448.4421 | 2172.9365 |
| Mdh2 | KGEDFVKN | -329 | -336 | 2 | 0.001092529 | 8 | NaN | NaN | NaN | 1205.6207 | 434.9147 | 575.4675 |
| Mdh2 | LTLYDIAHTPGVAA | -53 | -66 | 0 | 0 | 14 | NaN | NaN | NaN | 632.41907 | 1477.3335 | 1486.8915 |
| Mdh2 | TLYDIAHTPGVAADLSHIETR | -54 | -74 | 0 | 0.000230707 | 21 | NaN | NaN | NaN | 479.76535 | 541.99677 | 165.6934 |
| Mdh2 | LYDIAHTPGVAADLSHIETR | -55 | -74 | 0 | 0.00025015 | 20 | NaN | NaN | NaN | 157.86421 | 999.67175 | 918.43445 |
| Ndufs3 | ILTDYGFEGHPFR | -186 | -198 | 0 | 0 | 13 | NaN | NaN | NaN | 1091.2802 | 233.02457 | 1338.5718 |
| Ndufs3 | IHPDGVIPTLTFLR | -87 | -100 | 0 | 0 | 14 | NaN | NaN | NaN | 382.72757 | 441.9747 | 459.16644 |
| Ndufs3 | TYADELTPIDSIVSVH | -144 | -159 | 0 | 0.00025015 | 16 | NaN | NaN | NaN | 656.1415 | 357.6181 | NaN |
| Bax | ADMFADGNFNWGR | -97 | -109 | 0 | 0 | 13 | NaN | NaN | NaN | 410.7836 | NaN | 381.4843 |
| Hdgfl2 | PHAFKPGDLVFAK | -2 | -14 | 1 | 0 | 13 | NaN | NaN | NaN | 507.62396 | 697.139 | 527.9981 |
| Hdgfl2 | PHAFKPGDLVFAK | -2 | -14 | 1 | 0 | 13 | NaN | NaN | NaN | 507.62396 | 697.139 | 527.9981 |
| Phb2 | QDTILAEGLHFR | -60 | -71 | 0 | 0 | 12 | NaN | NaN | NaN | 786.5057 | 1319.6678 | 1148.8674 |
| Phb2 | IGGVQQDTILAEGLHFR | -55 | -71 | 0 | 0 | 17 | NaN | NaN | NaN | 487.19598 | 895.0315 | 499.5202 |
| Phb2 | IGGVQQDTILAEGLHFR | -55 | -71 | 0 | 0 | 17 | NaN | NaN | NaN | 487.19598 | 895.0315 | 499.5202 |
| Phb2 | QVAQQEAQR | -201 | -209 | 0 | 0.00025015 | 9 | NaN | NaN | NaN | 429.844 | NaN | NaN |
| Gtf2e1 | ERPIWLR | -276 | -282 | 1 | 0.002780285 | 7 | NaN | NaN | NaN | 259.17035 | NaN | NaN |
| Gtf2e1 | ADPDVLTEVPAALK | -2 | -15 | 0 | 0.00111006 | 14 | NaN | NaN | NaN | 254.35295 | NaN | NaN |
| Stk24 | SQACGGNLGSIEELR | -379 | -393 | 0 | 0.001392869 | 15 | NaN | NaN | NaN | 782.7686 | NaN | NaN |
| Arhgdia | DYMVGSYGPR | -143 | -152 | 0 | 0.000591124 | 10 | NaN | NaN | NaN | 2730.3535 | 2659.8494 | 2422.159 |
| Arhgdia | EIQELDKDDESLRK | -37 | -50 | 2 | 0.000471409 | 14 | NaN | NaN | NaN | 614.60675 | 173.38806 | 529.26373 |
| Coro1a | EPVITLEGHTK | -122 | -132 | 0 | 0 | 11 | NaN | NaN | NaN | 9375.079 | 8606.875 | 8431.715 |
| Coro1a | KCEPIAMTVPR | -344 | -354 | 1 | 0 | 11 | NaN | NaN | NaN | 946.4816 | 2882.8188 | 2381.6072 |
| Coro1a | ICEASGGGAFLVLPLGK | -50 | -66 | 0 | 0 | 17 | NaN | NaN | NaN | 531.999 | 718.6617 | 741.4298 |
| Coro1a | EASGGGAFLVLPLGK | -52 | -66 | 0 | 0 | 15 | NaN | NaN | NaN | 447.28598 | 354.9814 | 547.44775 |
| Coro1a | GGGAFLVLPLGK | -55 | -66 | 0 | 0 | 12 | NaN | NaN | NaN | 221.90202 | 411.00006 | 378.87317 |
| Coro1a | GTPSSDTVSR | -423 | -432 | 0 | 0.000431655 | 10 | NaN | NaN | NaN | 3297.2703 | 498.51593 | NaN |
| Coro1a | DGALICTSCR | -187 | -196 | 0 | 0.000591124 | 10 | NaN | NaN | NaN | 1233.6091 | 1725.6805 | NaN |
| Coro1a | VDKNVPLVC | -70 | -78 | 1 | 0.001055699 | 9 | NaN | NaN | NaN | 284.94366 | NaN | NaN |
| Coro1a | SGGGAFLVLPLGK | -54 | -66 | 0 | 0.001190916 | 13 | NaN | NaN | NaN | 155.13992 | NaN | NaN |
| Nol11 | SEPFLLPHLK | -601 | -610 | 0 | 0 | 10 | NaN | NaN | NaN | 340.82352 | 815.4769 | NaN |
| Nol11 | ADQFLVTDSGR | -31 | -41 | 0 | 0 | 11 | NaN | NaN | NaN | 938.50836 | NaN | NaN |
| Mcm7 | EFYYENELGK | -19 | -28 | 0 | 0 | 10 | NaN | NaN | NaN | 1636.3129 | 1554.2865 | 1361.9548 |
| Mcm7 | WLIQDRPDR | -519 | -527 | 1 | 0.002199523 | 9 | NaN | NaN | NaN | 1540.0863 | 2329.6501 | 1906.7538 |
| Mcm7 | SVHFSEAEQR | -674 | -683 | 0 | 0 | 10 | NaN | NaN | NaN | 16465.727 | 19586.283 | 21132.72 |
| Mcm7 | QIAEEDFYEKL | -326 | -336 | 1 | 0 | 11 | NaN | NaN | NaN | 557.5851 | 291.8553 | 362.2829 |
| Mcm7 | ADQGVCCIDEFDK | -437 | -449 | 0 | 0 | 13 | NaN | NaN | NaN | 500.33936 | NaN | NaN |
| Calm1;Calm2;( | VFDKDGNGYISAAELR | (92);(92);(92) | (107);(107);(107) | 1 | 0.000712251 | 16 | NaN | NaN | NaN | 502.86337 | 665.2652 | 647.59424 |
| Calm1;Calm2;( | HVMTNLGEK | (108);(108);(108) | (116);(116);(116) | 0 | 0 | 9 | NaN | NaN | NaN | 460.08475 | 836.1322 | NaN |
| Nup133 | ISVDLIDDYPASDPR | -563 | -577 | 0 | 0.001636916 | 15 | NaN | NaN | NaN | 479.1402 | NaN | 727.97864 |
| Rab1a | EFADSLGIPFLETSAK | -141 | -156 | 0 | 0 | 16 | NaN | NaN | NaN | 234.49655 | 463.22247 | 347.92505 |
| Rab1a | SSMNPEYDYLFK | -2 | -13 | 0 | 0.000394361 | 12 | NaN | NaN | NaN | 375.99927 | NaN | 328.19135 |
| Rab1b | EFADSLGVPFLETSAK | -138 | -153 | 0 | 0 | 16 | NaN | NaN | NaN | 648.86237 | 502.42743 | 603.43085 |
| Pfkp | QQGGAPSPFDR | -672 | -682 | 0 | 0.00025015 | 11 | NaN | NaN | NaN | 1390.5333 | 1394.3948 | 1016.6719 |
| Pfkp | VSDDSICVLGICK | -711 | -723 | 0 | 0 | 13 | NaN | NaN | NaN | 1084.4498 | 93.62014 | 1141.7076 |
| Pfkp | NGDIDNDTVQK | -150 | -160 | 0 | 0.00025015 | 11 | NaN | NaN | NaN | 573.0588 | NaN | NaN |
| Nucb1 | YTEEELKR | -341 | -348 | 1 | 0 | 8 | NaN | NaN | NaN | 594.7427 | 1860.8414 | 1721.329 |
| Nucb1 | EHLDPQNQHTFEAR | -138 | -151 | 0 | 0.004658491 | 14 | NaN | NaN | NaN | 550.23926 | 943.4184 | NaN |
| Tars1 | DQELYFFHELSPGSC | -328 | -342 | 0 | 0 | 15 | NaN | NaN | NaN | 314.3672 | 359.6225 | NaN |
| Psmc4 | IQDEIPALSVSRPQTGL | -11 | -27 | 1 | 0 | 17 | NaN | NaN | NaN | 516.528 | 495.93436 | 494.09613 |
| Psmc4 | IQDEIPALSVSRPQTGLSF | -11 | -29 | 1 | 0.00437931 | 19 | NaN | NaN | NaN | 462.72147 | 439.41965 | 564.968 |
| Psmc4 | IEFPLPDRR | -331 | -339 | 1 | 0.001811524 | 9 | NaN | NaN | NaN | 1078.8434 | NaN | NaN |
| Aldh4a1 | STGSVVGQQPFGGAR | -509 | -523 | 0 | 0 | 15 | NaN | NaN | NaN | 1395.647 | 972.08374 | NaN |
| Aldh4a1 | ADVDSVVSGTLR | -326 | -337 | 0 | 0.008129024 | 12 | NaN | NaN | NaN | 2179.6445 | NaN | NaN |
| Sun2 | TNWGHPEYTCIYR | -710 | -722 | 0 | 0 | 13 | NaN | NaN | NaN | 338.48187 | 342.10242 | 299.71304 |
| Sun2 | HHQDSEDLFK | -380 | -389 | 0 | 0 | 10 | NaN | NaN | NaN | 427.89194 | 1109.3627 | NaN |
| Sun2 | SHEDALSLLEGLVSR | -338 | -352 | 0 | 0 | 15 | NaN | NaN | NaN | 426.65088 | NaN | 261.54065 |
| Sun2 | QKEGIVGVTEEQVHR | -530 | -544 | 1 | 0.006537474 | 15 | NaN | NaN | NaN | 1226.1278 | NaN | NaN |
| Sun2 | IRPTAVTLEHVPK | -633 | -645 | 1 | 0.005301614 | 13 | NaN | NaN | NaN | 275.39856 | NaN | NaN |
| Sptan1 | LQQSHPLSASQIQVK | -321 | -335 | 0 | 0 | 15 | NaN | NaN | NaN | 1003.0631 | 783.5781 | 1197.0598 |

| | | | | | | | | | | | | |
|---|---|---|---|---|---|---|---|---|---|---|---|---|
| Sptan1 | HQWIQETR | -2218 | -2225 | 0 | 0.004412728 | 8 | NaN | NaN | NaN | 4749.37 | NaN | NaN |
| Sptan1 | KLDPAQSASR | -1023 | -1032 | 1 | 0 | 10 | NaN | NaN | NaN | 735.75244 | NaN | NaN |
| Sptan1 | EKEPIAASTNR | -802 | -812 | 1 | 0.003133457 | 11 | NaN | NaN | NaN | 392.61597 | NaN | NaN |
| Gbp2 | LQTDQSLTEAAK | -474 | -485 | 0 | 0 | 12 | NaN | NaN | NaN | 2255.7578 | 2236.4287 | 2271.0193 |
| Gbp2 | QLLAEQENIIAA | -539 | -550 | 0 | 0 | 12 | NaN | NaN | NaN | 553.1314 | 194.16919 | 329.923 |
| Gbp2 | LGTFSKPGGYYLFLQ | -422 | -436 | 1 | 0.00111006 | 15 | NaN | NaN | NaN | 1336.474 | 1837.9045 | 1918.0782 |
| Gbp2 | FQTELGNLL | -378 | -386 | 0 | 0.002466863 | 9 | NaN | NaN | NaN | 677.45917 | 1108.6565 | 1032.1295 |
| Gbp2 | CFIFDRPAQR | -233 | -242 | 1 | 0.000431655 | 10 | NaN | NaN | NaN | 1134.6766 | 1385.479 | NaN |
| Gbp2 | LGTFSKPGGY | -422 | -431 | 1 | 0.003620005 | 10 | NaN | NaN | NaN | 1590.9108 | NaN | NaN |
| Iars2 | FLLGNLTGFNPETDSVPVK | -732 | -750 | 0 | 0.00025015 | 19 | NaN | NaN | NaN | 580.82336 | 790.9853 | 727.0351 |
| Iars2 | SGVDLEGGTCSHPLTPDK | -355 | -372 | 0 | 0 | 18 | NaN | NaN | NaN | 482.91245 | 3075.4556 | 261.0945 |
| Iars2 | ADWNNCYYTFDPK | -206 | -218 | 0 | 0 | 13 | NaN | NaN | NaN | 436.3558 | 390.15195 | 516.69147 |
| Iars2 | RDSFLGSIPGK | -886 | -896 | 1 | 0.006788754 | 11 | NaN | NaN | NaN | 1167.0952 | NaN | 1002.5925 |
| Erh | SHTILLVQPTK | -2 | -12 | 0 | 0 | 11 | NaN | NaN | NaN | 2434.3875 | 3106.1401 | 2548.6562 |
| Erh | MYEEHLK | -35 | -41 | 0 | 0 | 7 | NaN | NaN | NaN | 272.38345 | 1353.9691 | 1000.65826 |
| Erh | TQTYQPYNK | -76 | -84 | 0 | 0.009631128 | 9 | NaN | NaN | NaN | 723.02637 | NaN | NaN |
| Gcn1 | ASVQNPALCLVTR | -748 | -760 | 0 | 0 | 13 | NaN | NaN | NaN | 785.16034 | 753.26746 | NaN |
| Gcn1 | TLGTLVSHVTLR | -915 | -926 | 0 | 0 | 12 | NaN | NaN | NaN | 370.47708 | 174.21236 | NaN |
| Gcn1 | ALCDPLEEVR | -2013 | -2022 | 0 | 0.000591124 | 10 | NaN | NaN | NaN | 1771.4507 | NaN | NaN |
| Fubp1 | VPDGMVGFIIGR | -103 | -114 | 0 | 0 | 12 | NaN | NaN | NaN | 572.2126 | 632.5767 | 671.8205 |
| Fubp1 | EMVLELIR | -245 | -252 | 0 | 0.00025015 | 8 | NaN | NaN | NaN | 668.7362 | 452.1741 | 674.965 |
| Fubp1 | GTPESVQSAK | -148 | -157 | 0 | 0.002434616 | 10 | NaN | NaN | NaN | 322.91983 | NaN | NaN |
| Tufm | LDAVDTYIPVPTR | -240 | -252 | 0 | 0 | 13 | NaN | NaN | NaN | 1123.2488 | 801.06256 | 725.8068 |
| Tufm | DLDKPFLLPVESVYSIPGR | -253 | -271 | 1 | 0 | 19 | NaN | NaN | NaN | 436.37396 | 479.745 | 421.53827 |
| Tufm | AHTDCPGHADYVK | -123 | -135 | 0 | 0 | 13 | NaN | NaN | NaN | 296.5423 | 340.76154 | 306.2301 |
| Tufm | GDECELLGHNK | -287 | -297 | 0 | 0.00111006 | 11 | NaN | NaN | NaN | 754.1698 | NaN | NaN |
| Tufm | AEAGDNLGALVR | -316 | -327 | 0 | 0 | 12 | NaN | NaN | NaN | 567.6813 | NaN | NaN |
| Tkt | SSDLDGHPVPK | -104 | -114 | 0 | 0 | 11 | NaN | NaN | NaN | 3389.7253 | 11508.538 | 682.58014 |
| Tkt | TSRPENAIIYSNNEDFQVGQAK | -472 | -493 | 1 | 0 | 22 | NaN | NaN | NaN | 2684.5767 | 1944.4889 | 2454.8333 |
| Tkt | DPFTIKPLDR | -533 | -542 | 1 | 0.002752526 | 10 | NaN | NaN | NaN | 1441.4856 | 1547.8296 | 1702.574 |
| Tkt | QAFTDVATGSLGQGLGA | -115 | -131 | 0 | 0 | 17 | NaN | NaN | NaN | 678.2656 | 854.70526 | 2266.724 |
| Tkt | KLDNLVAIFDINR | -174 | -186 | 1 | 0 | 13 | NaN | NaN | NaN | 385.45505 | 571.9509 | 527.457 |
| Tkt | NMAEQIIQEIYSQ | -265 | -277 | 0 | 0.00122419 | 13 | NaN | NaN | NaN | 323.14178 | 394.63803 | 365.49316 |
| Tkt | KILATPPQEDAPSVDIA | -283 | -299 | 1 | 0.00149795 | 17 | NaN | NaN | NaN | 586.5438 | 299.65686 | NaN |
| Tkt | ILTVEDHYYEGGIGEAVSA | -556 | -574 | 0 | 0.009745776 | 19 | NaN | NaN | NaN | 2765.0989 | NaN | NaN |
| Hopx | AEAGLTEEQTQK | -38 | -49 | 0 | 0 | 12 | NaN | NaN | NaN | 867.2202 | NaN | 1258.3816 |
| Hopx | VNKHPDPTTL | -24 | -33 | 1 | 0.002168954 | 10 | NaN | NaN | NaN | 256.26816 | NaN | NaN |
| Taldo1 | DWHVANTDKK | -195 | -204 | 1 | 0.000230707 | 10 | NaN | NaN | NaN | 514.19446 | 524.9252 | 468.40558 |
| Taldo1 | LHNEDQMAVEK | -297 | -307 | 0 | 0 | 11 | NaN | NaN | NaN | 267.31152 | 369.2395 | 663.0368 |
| Taldo1 | LHNEDQMAVEK | -297 | -307 | 0 | 0 | 11 | NaN | NaN | NaN | 267.31152 | 369.2395 | 663.0368 |
| Taldo1 | PQEDPGVK | -208 | -215 | 0 | 0.002199523 | 8 | NaN | NaN | NaN | 1775.9084 | 1909.0753 | 649.6911 |
| Taldo1 | DWHVANTDK | -195 | -203 | 0 | 0.00078661 | 9 | NaN | NaN | NaN | 1551.9119 | 1014.0541 | 1009.4794 |
| Taldo1 | AEAGVTLISPFVGR | -179 | -192 | 0 | 0 | 14 | NaN | NaN | NaN | 1205.8972 | NaN | NaN |
| Taldo1 | GGPQEEQIK | -73 | -81 | 0 | 0 | 9 | NaN | NaN | NaN | 1099.9984 | NaN | NaN |
| Eif4a2 | AEAPHIVVGTPGR | -150 | -162 | 0 | 0.00025015 | 13 | NaN | NaN | NaN | 1185.3308 | NaN | NaN |
| Vdac2 | VNNSSLIGVGYTQTLRPGVK | -249 | -268 | 1 | 0 | 20 | NaN | NaN | NaN | 420.30994 | 383.31818 | 501.993 |
| Ptma | SDAAVDTSSEITTK | -2 | -15 | 0 | 0 | 14 | NaN | NaN | NaN | 5822.9185 | 5647.587 | 5892.552 |
| Ptma | EVVEEAENGR | -22 | -31 | 0 | 0 | 10 | NaN | NaN | NaN | 3488.0183 | 5682.136 | 7817.666 |
| Copa | TLIDKFDEHDGPVR | -42 | -55 | 1 | 0 | 14 | NaN | NaN | NaN | 625.5241 | 566.5202 | 625.7865 |
| Copa | IDKFDEHDGPVR | -44 | -55 | 1 | 0 | 12 | NaN | NaN | NaN | 1072.3406 | 1528.0226 | 1490.6903 |
| Copa | KETFDPEKETIPDIDPNAK | -775 | -793 | 2 | 0 | 19 | NaN | NaN | NaN | 541.1094 | 612.71625 | 488.81125 |
| Copa | YAVTTGDHGIIR | -551 | -562 | 0 | 0 | 12 | NaN | NaN | NaN | 2278.6572 | 2688.3494 | 1666.4604 |
| Copa | LDEDGFVEAPEGLGEDVLGK | -853 | -872 | 0 | 0 | 20 | NaN | NaN | NaN | 365.81815 | 450.70813 | 343.49118 |
| Copa | KLDALCNIHENIR | -517 | -529 | 1 | 0.00111006 | 13 | NaN | NaN | NaN | 682.7909 | 1108.3945 | NaN |
| Copa | TVGKFEEAVEK | -1018 | -1028 | 1 | 0.00149795 | 11 | NaN | NaN | NaN | 459.71094 | 775.6023 | NaN |
| Copa | TIDPTEFK | -591 | -598 | 0 | 0.006677519 | 8 | NaN | NaN | NaN | 1666.8251 | NaN | NaN |
| Anxa2 | GVDEVTIVNILTNR | -50 | -63 | 0 | 0.009897819 | 14 | NaN | NaN | NaN | 623.71674 | NaN | NaN |
| Rps3 | DEILPTTPISEQK | -215 | -227 | 0 | 0 | 13 | NaN | NaN | NaN | 12376.306 | 5132.9146 | 4763.3154 |
| Rps3 | RFIMESGAK | -124 | -132 | 1 | 0.00025015 | 9 | NaN | NaN | NaN | 1156.0962 | 519.6546 | 485.3625 |
| Rps3 | GDPVNYYVDTAVR | -161 | -173 | 0 | 0 | 13 | NaN | NaN | NaN | 1277.3453 | 307.78854 | 523.48444 |
| Rps3 | VTPTRTEIIIL | -41 | -51 | 1 | 0.001846793 | 11 | NaN | NaN | NaN | 540.44727 | 446.45065 | 486.5549 |
| Hdlbp | LGQALTEVYAK | -350 | -360 | 0 | 0.004824802 | 11 | NaN | NaN | NaN | 928.6052 | NaN | NaN |
| Rtcb | RTNLDESDVQPVK | -128 | -140 | 1 | 0.00078661 | 13 | NaN | NaN | NaN | 960.52155 | 2374.5 | NaN |
| Rtcb | NLDFQDVLDKL | -442 | -452 | 1 | 0.001978766 | 11 | NaN | NaN | NaN | 129.18314 | NaN | 50.53884 |
| Canx | IPDEEATKPEGW | -313 | -324 | 1 | 0.000230707 | 12 | NaN | NaN | NaN | 1066.701 | 1325.3508 | 1283.0557 |
| Canx | EVNFQNGIECGGAYVK | -152 | -167 | 0 | 0 | 16 | NaN | NaN | NaN | 735.9739 | 541.34247 | 410.83148 |
| Canx | APVPTGEVYFADSFDR | -63 | -78 | 0 | 0 | 16 | NaN | NaN | NaN | 702.56964 | 783.0997 | 811.12524 |
| Canx | LNKPFLFDTKPL | -136 | -147 | 2 | 0 | 12 | NaN | NaN | NaN | 2710.4077 | NaN | 4127.397 |
| Canx | RVVDDWANDGWGLK | -446 | -459 | 1 | 0.00051306 | 14 | NaN | NaN | NaN | 1692.0942 | NaN | NaN |
| Canx | YEVNFQNGIECGGAYVK | -151 | -167 | 0 | 0.001868731 | 17 | NaN | NaN | NaN | 723.7793 | NaN | NaN |
| Canx | TPYTIMFGPDK | -184 | -194 | 0 | 0.003360691 | 11 | NaN | NaN | NaN | 406.08676 | NaN | NaN |
| Canx | PDFFEDLEPFK | -406 | -416 | 0 | 0.009955823 | 11 | NaN | NaN | NaN | 383.45767 | NaN | NaN |
| P4ha1 | AEEDKLEQIK | -48 | -57 | 1 | 0.00111006 | 10 | NaN | NaN | NaN | 1720.8118 | NaN | NaN |
| Cdc37 | YLSDNVHLVC | -175 | -184 | 0 | 0 | 10 | NaN | NaN | NaN | 602.9021 | 391.1016 | NaN |
| Cdc37 | LGPGGLDPVEVY | -288 | -299 | 0 | 0.00025015 | 12 | NaN | NaN | NaN | 324.82587 | 411.32275 | NaN |
| Cdc37 | DVQMLQDAISK | -314 | -324 | 0 | 0.00051306 | 11 | NaN | NaN | NaN | 1424.4254 | NaN | NaN |
| Dbnl | LTQPETSYGR | -298 | -307 | 0 | 0 | 10 | NaN | NaN | NaN | 1110.141 | NaN | NaN |
| Numa1 | AEELGQELK | -1318 | -1326 | 0 | 0.006788754 | 9 | NaN | NaN | NaN | 550.2638 | NaN | NaN |

| | | | | | | | | | | | | |
|---|---|---|---|---|---|---|---|---|---|---|---|---|
| Clic1 | KFLDGNELTLADCNLLPK | -166 | -183 | 1 | 0 | 18 | NaN | NaN | NaN | 358.19608 | 368.8093 | 427.85266 |
| Clic1 | LTSPLPEEVDETSAEDEGISQR | -144 | -165 | 0 | 0 | 22 | NaN | NaN | NaN | 325.14432 | 441.01483 | 419.89023 |
| Clic1 | SNPALNDNLEK | -121 | -131 | 0 | 0.003490217 | 11 | NaN | NaN | NaN | 1506.5054 | NaN | NaN |
| Clic1 | KFLDGNELTLADC | -166 | -178 | 1 | 0.005459848 | 13 | NaN | NaN | NaN | 675.02814 | NaN | NaN |
| Ppp4r2 | NKHPDEDAVESEEHEVK | -230 | -246 | 1 | 0 | 17 | NaN | NaN | NaN | 361.14035 | 480.7268 | 131.85965 |
| Ipo5 | TDFAPGFQK | -428 | -436 | 0 | 0 | 9 | NaN | NaN | NaN | 1513.8839 | 1389.6207 | 3121.0103 |
| Ipo5 | SYLCDLIESNHPIVLGPNNTNLPK | -1014 | -1037 | 0 | 0.000471409 | 24 | NaN | NaN | NaN | 603.9319 | 214.81233 | NaN |
| Ipo5 | FILANEHNVALFK | -200 | -212 | 0 | 0.001291103 | 13 | NaN | NaN | NaN | 418.42978 | NaN | NaN |
| Ipo5 | LIPYLDNLVK | -478 | -487 | 0 | 0.002265552 | 10 | NaN | NaN | NaN | 398.7727 | NaN | NaN |
| Cd2bp2 | TWVSEGYFPDGVYCR | -307 | -321 | 0 | 0.00025015 | 15 | NaN | NaN | NaN | 638.2997 | 445.95316 | 937.4082 |
| Cd2bp2 | AEEVAEGELETPTPTQR | -253 | -269 | 0 | 0.002136192 | 17 | NaN | NaN | NaN | 187.10321 | NaN | NaN |
| Dhx15 | EEIDEACKR | -360 | -368 | 1 | 0 | 9 | NaN | NaN | NaN | 1104.7477 | 1214.7239 | NaN |
| Dhx15 | MAEFPLDPQLAK | -562 | -573 | 0 | 0.00025015 | 12 | NaN | NaN | NaN | 420.3313 | 429.6985 | NaN |
| Dhx15 | GQEEIDEACK | -358 | -367 | 0 | 0.001092529 | 10 | NaN | NaN | NaN | 1440.2633 | NaN | NaN |
| Dhx15 | RIFEPPPPK | -399 | -407 | 1 | 0.008284062 | 9 | NaN | NaN | NaN | 818.229 | NaN | NaN |
| Dhx15 | LQLPVWEY | -135 | -142 | 0 | 0.001708008 | 8 | NaN | NaN | NaN | 474.99652 | NaN | NaN |
| Dhx15 | LISAGLPPLK | -69 | -78 | 0 | 0.007819401 | 10 | NaN | NaN | NaN | 115.4309 | NaN | NaN |
| Rpl8 | RAVDFAER | -30 | -37 | 1 | 0.002910424 | 8 | NaN | NaN | NaN | 821.2837 | 1044.9828 | 828.0813 |
| Rpl8 | GIHTGQFVYCGK | -81 | -92 | 0 | 0 | 12 | NaN | NaN | NaN | 791.52185 | NaN | 1266.375 |
| Rpl8 | AEGIHTGQFVYCGK | -79 | -92 | 0 | 0.000230707 | 14 | NaN | NaN | NaN | 2373.5708 | NaN | NaN |
| Tnpo1 | ECLNNIGDSSPLIR | -102 | -115 | 0 | 0 | 14 | NaN | NaN | NaN | 15335.252 | 16246.719 | 16494.094 |
| Tnpo1 | NWPDLLPK | -133 | -140 | 0 | 0.001575548 | 8 | NaN | NaN | NaN | 735.41174 | 842.6432 | 882.80164 |
| Tnpo1 | SITCWTLSR | -464 | -472 | 0 | 0.00025015 | 9 | NaN | NaN | NaN | 241.18588 | 513.72375 | 503.82333 |
| Tnpo1 | VYRDELLPHILPLLK | -397 | -411 | 1 | 0.001291103 | 15 | NaN | NaN | NaN | 979.25055 | 1070.6906 | NaN |
| Tnpo1 | RDELLPHILPLLK | -399 | -411 | 1 | 0.000230707 | 13 | NaN | NaN | NaN | 94.36936 | 405.89777 | NaN |
| Flna | FNEEHIPDSPFVVPVASPSGDAR | -2311 | -2333 | 0 | 0 | 23 | NaN | NaN | NaN | 721.27325 | 806.321 | 887.11835 |
| Flna | TGVAVNKPAEFTVDAK | -685 | -700 | 1 | 0.00025015 | 16 | NaN | NaN | NaN | 697.7149 | 721.74884 | 714.6212 |
| Flna | TVPQHATSGPGPADVSK | -2543 | -2559 | 0 | 0 | 17 | NaN | NaN | NaN | 1261.0774 | 265.25818 | 175.4666 |
| Flna | VHGPGIQSGTTNKPNK | -1360 | -1375 | 1 | 0.00025015 | 16 | NaN | NaN | NaN | 718.523 | 427.08868 | 611.7482 |
| Flna | YIPVQQGPVGVNVTYGGDHIPK | -937 | -958 | 0 | 0 | 22 | NaN | NaN | NaN | 888.1365 | 958.78046 | 922.4823 |
| Flna | VSGQGLHEGHTFEPA | -2052 | -2066 | 0 | 0.00078661 | 15 | NaN | NaN | NaN | 431.84048 | NaN | NaN |
| Flna | VHSPSGALEECYVTEIDQDK | -2368 | -2387 | 0 | 0 | 20 | NaN | NaN | NaN | 360.1719 | 917.58563 | 1473.4473 |
| Flna | VLYGEEEVPR | -1523 | -1532 | 0 | 0 | 10 | NaN | NaN | NaN | 1324.2809 | 1167.8025 | 1185.5243 |
| Flna | AEVGVPAEFGIWTR | -2251 | -2264 | 0 | 0 | 14 | NaN | NaN | NaN | 1169.5164 | 1331.0273 | 1349.7784 |
| Flna | TVDCTEAGQGDVSIGIK | -793 | -809 | 0 | 0 | 17 | NaN | NaN | NaN | 735.26794 | 619.208 | 604.2777 |
| Flna | GVAVNKPAEFTVDAK | -686 | -700 | 1 | 0 | 15 | NaN | NaN | NaN | 678.7081 | 742.67126 | 838.42633 |
| Flna | SLHETSSVFVDSLTK | -2526 | -2540 | 0 | 0 | 15 | NaN | NaN | NaN | 518.82513 | 625.89996 | 589.2351 |
| Flna | VHSPSGALEECYVTEIDQDKYAVR | -2368 | -2391 | 1 | 0 | 24 | NaN | NaN | NaN | 415.63068 | 408.49393 | 545.1341 |
| Flna | YGGDEIPFSPY | -1622 | -1632 | 0 | 0 | 11 | NaN | NaN | NaN | 345.33994 | 381.36688 | 340.8057 |
| Flna | AGNNMLLVGVHGPR | -2585 | -2598 | 0 | 0.001055699 | 14 | NaN | NaN | NaN | 2109.9954 | NaN | 492.48914 |
| Flna | LPQLPITNF | -180 | -188 | 0 | 0.003998192 | 9 | NaN | NaN | NaN | 547.1309 | 732.1873 | NaN |
| Flna | YDGSPVPSSPFQVPVTEGCDPSR | -1335 | -1357 | 0 | 0.003003218 | 23 | NaN | NaN | NaN | 396.8385 | NaN | 430.4618 |
| Flna | KVEPGLGADNSVVR | -1019 | -1032 | 1 | 0.000230707 | 14 | NaN | NaN | NaN | 4204.429 | NaN | NaN |
| Flna | GPGLEPSGNIANK | -388 | -400 | 0 | 0 | 13 | NaN | NaN | NaN | 690.8651 | NaN | NaN |
| Rexo2 | WYPEDYEFAPK | -179 | -189 | 0 | 0 | 11 | NaN | NaN | NaN | 1201.7301 | 1179.0154 | 308.12952 |
| Rpl13a | AEGQVLVLDGR | -2 | -12 | 0 | 0 | 11 | NaN | NaN | NaN | 2009.5322 | 178.46693 | NaN |
| Kif5b | WLENELNR | -360 | -367 | 0 | 0.003102268 | 8 | NaN | NaN | NaN | 1159.5468 | NaN | NaN |
| Tgtp2;Tgtp1 | DYCSNHLQESLDSEPPVFLVS | (198);(198) | (218);(218) | 0 | 0.000591124 | 21 | NaN | NaN | NaN | 228.41876 | 265.60376 | 239.7893 |
| Eif2s3x | LDDPSCPRPECYR | -91 | -103 | 1 | 0 | 13 | NaN | NaN | NaN | 1774.5839 | 1858.3224 | 1813.99 |
| Eif2s3x | SHEVISR | -32 | -38 | 0 | 0.001636916 | 7 | NaN | NaN | NaN | 600.08984 | NaN | NaN |
| Lactb2 | ILEYISHR | -210 | -217 | 0 | 0 | 8 | NaN | NaN | NaN | 647.09106 | 687.4847 | 685.592 |
| Lactb2 | AEHNLLLHLR | -257 | -266 | 0 | 0.002168954 | 10 | NaN | NaN | NaN | 2912.972 | NaN | NaN |
| Lactb2 | AEHNLLLHLR | -257 | -266 | 0 | 0.002168954 | 10 | NaN | NaN | NaN | 2912.972 | NaN | NaN |
| Nop9 | GDTHGSFVVR | -199 | -208 | 0 | 0 | 10 | NaN | NaN | NaN | 942.7978 | 604.3808 | 1238.8027 |
| Nop9 | LQDLSACFLK | -257 | -266 | 0 | 0.002534621 | 10 | NaN | NaN | NaN | 936.569 | NaN | NaN |
| Nop9 | KVAPDAAEER | -73 | -82 | 1 | 0.001708008 | 10 | NaN | NaN | NaN | 647.358 | NaN | NaN |
| Psph | EHPPHLTPGIR | -82 | -92 | 0 | 0 | 11 | NaN | NaN | NaN | 1000.59454 | 658.3178 | 675.00824 |
| Psph | AEHPPHLTPGIR | -81 | -92 | 0 | 0 | 12 | NaN | NaN | NaN | 821.1369 | 814.82666 | 758.87244 |
| Mthfd1l | DAVPCYHWSAGGK | -823 | -835 | 0 | 0 | 13 | NaN | NaN | NaN | 6369.888 | 7192.8115 | 950.25146 |
| Mthfd1l | KTDPSTLTEEEVR | -546 | -558 | 1 | 0 | 13 | NaN | NaN | NaN | 1152.1346 | 1359.6311 | 1424.4501 |
| Mthfd1l | TDPSTLTEEEVRK | -547 | -559 | 1 | 0 | 13 | NaN | NaN | NaN | 1070.9138 | 675.97186 | 918.6582 |
| Mthfd1l | FLYDVQLPIVEK | -856 | -867 | 0 | 0 | 12 | NaN | NaN | NaN | 704.82007 | 773.3714 | 715.3084 |
| Mthfd1l | EYTEENIQLVADGCCNLQK | -764 | -782 | 0 | 0 | 19 | NaN | NaN | NaN | 433.043 | 482.53897 | 501.60077 |
| Vps25 | AEIITVSDGR | -162 | -171 | 0 | 0.00025015 | 10 | NaN | NaN | NaN | 1014.28345 | NaN | NaN |
| Anxa6 | GTVCAANDFNPDADAK | -355 | -370 | 0 | 0 | 16 | NaN | NaN | NaN | 3994.4902 | 4767.6274 | 4827.7124 |
| Anxa6 | DAISGVGTDEK | -103 | -113 | 0 | 0.000431655 | 11 | NaN | NaN | NaN | 1591.8967 | NaN | NaN |
| Anxa6 | DAFVAIVQSVK | -588 | -598 | 0 | 0 | 11 | NaN | NaN | NaN | 1041.2881 | 490.47314 | 1138.4601 |
| Anxa6 | RSEIDLLNIR | -629 | -638 | 1 | 0 | 10 | NaN | NaN | NaN | 967.3109 | 1621.3165 | 2139.778 |
| Anxa6 | TLIEILATR | -457 | -465 | 0 | 0.00025015 | 9 | NaN | NaN | NaN | 699.59753 | 785.54706 | 724.26764 |
| Anxa6 | YQMWELSAVSR | -340 | -350 | 0 | 0 | 11 | NaN | NaN | NaN | 589.5416 | 879.8409 | 875.65045 |
| Anxa6 | LCGGDDDAAGQFFPEAAQVA | -320 | -339 | 0 | 0.00025015 | 20 | NaN | NaN | NaN | 385.65295 | 505.32455 | 452.00644 |
| Anxa6 | GSVHDFPEFDANQDAEALYTA | -12 | -32 | 0 | 0.001868731 | 21 | NaN | NaN | NaN | 253.28812 | 280.9282 | 142.00053 |
| Anxa6 | MLVVLLQGTR | -157 | -166 | 0 | 0 | 10 | NaN | NaN | NaN | 1766.5074 | NaN | 1062.1508 |
| Anxa6 | TNEQMHQLVAAYK | -123 | -135 | 0 | 0.001291103 | 13 | NaN | NaN | NaN | 1682.8403 | NaN | 369.7829 |
| Anxa6 | ILISLATGNR | -500 | -509 | 0 | 0 | 10 | NaN | NaN | NaN | 1279.3156 | 1059.0194 | NaN |
| Anxa6 | VFQEFIK | -562 | -568 | 0 | 0.000826806 | 7 | NaN | NaN | NaN | 812.4485 | 655.6094 | NaN |
| Anxa6 | WGTDEAQFIYILGNR | -192 | -206 | 0 | 0 | 15 | NaN | NaN | NaN | 359.2659 | 362.27307 | NaN |
| Anxa6 | LCGGDDDAAGQFFPEAAQVAYQ | -320 | -341 | 0 | 0.004133093 | 22 | NaN | NaN | NaN | 175.73642 | NaN | NaN |

| | | | | | | | | | | | | |
|---|---|---|---|---|---|---|---|---|---|---|---|---|
| Gnl3 | AEISDVAPVEETR | -490 | -502 | 0 | 0 | 13 | NaN | NaN | NaN | 2927.2073 | 2559.5725 | 2677.5088 |
| Gnl3 | ELSPEQSTAGKPSDGSSA | -503 | -520 | 1 | 0.001055699 | 18 | NaN | NaN | NaN | 660.36285 | 582.4519 | NaN |
| Gnl3 | SPASIEELRPL | -319 | -329 | 1 | 0 | 11 | NaN | NaN | NaN | 3088.6387 | NaN | NaN |
| Eef2 | LKDLEEDHACIPIK | -558 | -571 | 1 | 0 | 14 | NaN | NaN | NaN | 1142.438 | 1324.0177 | 1331.4635 |
| Eef2 | EGALCEENMR | -689 | -698 | 0 | 0 | 10 | NaN | NaN | NaN | 319.1644 | 960.8088 | 879.3662 |
| Eef2 | YVEPIEDVPCGN | -457 | -468 | 0 | 0 | 12 | NaN | NaN | NaN | 176.47163 | 168.84511 | 206.98108 |
| Eef2 | GVQYLNEIK | -668 | -676 | 0 | 0.001291103 | 9 | NaN | NaN | NaN | 3892.4583 | 3439.6904 | 3422.3142 |
| Eef2 | WLPAGDALLQ | -343 | -352 | 0 | 0 | 10 | NaN | NaN | NaN | 2714.0015 | NaN | 4987.2324 |
| Eef2 | AYLPVNESFGFTA | -786 | -798 | 0 | 0.00025015 | 13 | NaN | NaN | NaN | 1098.6809 | 1139.579 | 1013.2195 |
| Eef2 | DSVVAGFQWATK | -677 | -688 | 0 | 0 | 12 | NaN | NaN | NaN | 996.11584 | 1198.2543 | 1222.0359 |
| Eef2 | ETVSEESNVLCLSK | -581 | -594 | 0 | 0 | 14 | NaN | NaN | NaN | 641.06995 | 570.90045 | 527.3225 |
| Eef2 | YFDPANGKF | -265 | -273 | 1 | 0.00122419 | 9 | NaN | NaN | NaN | 540.0173 | 547.41724 | 535.41675 |
| Eef2 | IWCFGPDGTGPN | -649 | -660 | 0 | 0.000230707 | 12 | NaN | NaN | NaN | 70.292206 | NaN | 445.94156 |
| Eef2 | GVQYLNEIKDSVVAGF | -668 | -683 | 1 | 0.00025015 | 16 | NaN | NaN | NaN | 344.1192 | NaN | NaN |
| Got2 | DDNGKPYVLPSVR | -69 | -81 | 1 | 0 | 13 | NaN | NaN | NaN | 960.3074 | 2038.0356 | 1911.9526 |
| Got2 | KASAELALGENNEVLK | -107 | -122 | 1 | 0 | 16 | NaN | NaN | NaN | 2345.8774 | 467.15668 | 963.2144 |
| Got2 | AYQGFASGDGDKDAWAVR | -245 | -262 | 1 | 0 | 18 | NaN | NaN | NaN | 595.18695 | 721.2597 | 652.8772 |
| Got2 | DFSGALEDISK | -190 | -200 | 0 | 0 | 11 | NaN | NaN | NaN | 161.14317 | NaN | 604.57495 |
| Got2 | KQWLQEVKG | -338 | -346 | 2 | 0.00223472 | 9 | NaN | NaN | NaN | 3353.7358 | NaN | NaN |
| Sf1 | AELGEAPVPASVGSTSGPATTPLASAPR | -318 | -345 | 0 | 0 | 28 | NaN | NaN | NaN | 251.61368 | 295.14487 | 274.21396 |
| Sf1 | ILRPWQSSETR | -261 | -271 | 1 | 0 | 11 | NaN | NaN | NaN | 3692.4565 | NaN | NaN |
| Sf1 | SVGSTSGPATTPLASAPR | -328 | -345 | 0 | 0 | 18 | NaN | NaN | NaN | 1503.7766 | NaN | NaN |
| Grap2 | QFHPHQQPSPQFTPGPQPPQQQR | -195 | -217 | 0 | 0.00272025 | 23 | NaN | NaN | NaN | 624.9367 | NaN | 549.0008 |
| Eif2s2 | DYTYEELLNR | -171 | -180 | 0 | 0 | 10 | NaN | NaN | NaN | 573.4056 | 2246.7852 | 1998.2065 |
| Eif2s2 | DASDDLDDLNFFNQ | -65 | -78 | 0 | 0.000230707 | 14 | NaN | NaN | NaN | 341.38785 | 124.97993 | 337.34467 |
| Eif2s2 | FVNFTDICK | -217 | -225 | 0 | 0.00078661 | 9 | NaN | NaN | NaN | 3783.9312 | NaN | NaN |
| Eif2s2 | EVEPEPTEEKDVDADEEDSRK | -43 | -63 | 2 | 0.005789881 | 21 | NaN | NaN | NaN | 864.01825 | NaN | NaN |
| Eif2s2 | FLQCETCHSR | -297 | -306 | 0 | 0.000230707 | 10 | NaN | NaN | NaN | 697.82385 | NaN | NaN |
| Umps | SWADIVNAHVVPGSGVVK | -335 | -352 | 0 | 0 | 18 | NaN | NaN | NaN | 1077.6873 | 1246.7726 | 1455.7488 |
| Umps | RAELPGTHPLASK | -230 | -242 | 1 | 0.000591124 | 13 | NaN | NaN | NaN | 1811.2568 | NaN | NaN |
| Psmd11 | YQEALHLGSQLLR | -143 | -155 | 0 | 0 | 13 | NaN | NaN | NaN | 1503.8677 | 1100.2405 | 748.8461 |
| Psmd11 | AELRDDPIISTH | -311 | -322 | 1 | 0.00025015 | 12 | NaN | NaN | NaN | 775.4608 | 91.10623 | NaN |
| Psmd11 | HLGSQLLR | -148 | -155 | 0 | 0 | 8 | NaN | NaN | NaN | 768.64246 | NaN | NaN |
| Psmd11 | DDPIISTHLAK | -315 | -325 | 0 | 0 | 11 | NaN | NaN | NaN | 368.68613 | NaN | NaN |
| Sin3a | LDGNIDSSQYEDSLR | -971 | -985 | 0 | 0.00025015 | 15 | NaN | NaN | NaN | 1197.6289 | 1299.048 | 1274.5795 |
| Sin3a | LHQILCLR | -899 | -906 | 0 | 0.003620005 | 8 | NaN | NaN | NaN | 269.0772 | 395.3441 | NaN |
| Nasp | AFQEAASLLGK | -65 | -75 | 0 | 0 | 11 | NaN | NaN | NaN | 659.2135 | 555.87305 | 766.3344 |
| Nasp | HLVMGDIPAAVNAF | -53 | -66 | 0 | 0.00223472 | 14 | NaN | NaN | NaN | 342.72574 | 342.34027 | NaN |
| Nasp | EIEELKELLPEIR | -624 | -636 | 1 | 0.000230707 | 13 | NaN | NaN | NaN | 521.5718 | NaN | NaN |
| Sra1 | AELYVKPGNK | -2 | -11 | 1 | 0.00078661 | 10 | NaN | NaN | NaN | 2791.4258 | 945.7036 | NaN |
| Cct4 | GLNPISTVTELR | -470 | -481 | 0 | 0 | 12 | NaN | NaN | NaN | 579.79517 | 815.8778 | 727.0401 |
| Cct4 | KAVADAIR | -42 | -49 | 1 | 0.003102268 | 8 | NaN | NaN | NaN | 673.60516 | NaN | 541.74176 |
| Cct4 | AYILNLVK | -281 | -288 | 0 | 0 | 8 | NaN | NaN | NaN | 486.87155 | 326.6814 | NaN |
| Cct4 | PVQLSDR | -154 | -160 | 0 | 0.005522803 | 7 | NaN | NaN | NaN | 690.19 | NaN | NaN |
| Lrrfip1 | GETSDTVNDVGYQAPTK | -207 | -223 | 0 | 0.002972566 | 17 | NaN | NaN | NaN | 396.76392 | NaN | NaN |
| Mybbp1a | LVQLADMLLNHNR | -543 | -555 | 0 | 0 | 13 | NaN | NaN | NaN | 1610.4944 | 1108.0819 | 1175.5657 |
| Mybbp1a | EHKLPNVALDLLR | -259 | -271 | 1 | 0 | 13 | NaN | NaN | NaN | 639.0474 | 652.501 | 739.0221 |
| Mybbp1a | GENMVISKPQNLFK | -332 | -345 | 1 | 0 | 14 | NaN | NaN | NaN | 205.35028 | 266.46545 | 186.08437 |
| Mybbp1a | GENMVISKPQNLFK | -332 | -345 | 1 | 0 | 14 | NaN | NaN | NaN | 205.35028 | 266.46545 | 186.08437 |
| Mybbp1a | DMFLQPDLNSL | -410 | -420 | 0 | 0 | 11 | NaN | NaN | NaN | 1064.956 | NaN | 1193.8685 |
| Mybbp1a | YCHEVGPC | -889 | -896 | 0 | 0.000431655 | 8 | NaN | NaN | NaN | 367.2503 | NaN | 425.90283 |
| Mybbp1a | FSFPLDDR | -499 | -506 | 0 | 0.002595029 | 8 | NaN | NaN | NaN | 243.84053 | NaN | 320.8298 |
| Mybbp1a | SLPLLSEEQLQLVMR | -310 | -324 | 0 | 0 | 15 | NaN | NaN | NaN | 1414.1561 | NaN | NaN |
| Mybbp1a | FCLFHAFFK | -476 | -484 | 0 | 0.00051306 | 9 | NaN | NaN | NaN | 655.5378 | NaN | NaN |
| Trap1 | TQEELVSNLGTIAR | -166 | -179 | 0 | 0 | 14 | NaN | NaN | NaN | 600.83167 | NaN | 1089.2053 |
| Trap1 | AEQDIKEDIAK | -458 | -468 | 1 | 0.00025015 | 11 | NaN | NaN | NaN | 235.7985 | 997.15204 | NaN |
| Trap1 | FEDTSPADER | -566 | -575 | 0 | 0 | 10 | NaN | NaN | NaN | 490.82526 | NaN | NaN |
| Csk | EYLEGNNFVHR | -303 | -313 | 0 | 0 | 11 | NaN | NaN | NaN | 2021.5316 | 1857.2228 | 2885.3164 |
| Csk | NFHGTAEQDLPFCK | -19 | -32 | 0 | 0 | 14 | NaN | NaN | NaN | 274.8127 | 372.0725 | 227.33264 |
| Pole3 | AERPEDLNLPNAVIT | -2 | -16 | 1 | 0.001392869 | 15 | NaN | NaN | NaN | 943.7742 | 1410.8275 | 158.19315 |
| Psmd12 | YVILAPFDNEQSDLVHR | -273 | -289 | 0 | 0 | 17 | NaN | NaN | NaN | 1675.8417 | 1442.5609 | 2002.3262 |
| Psmd12 | STLVEDYGVELR | -318 | -329 | 0 | 0 | 12 | NaN | NaN | NaN | 379.865 | 1512.2273 | 1216.2435 |
| Psmd12 | EVIETLLSLEK | -42 | -52 | 0 | 0.003490217 | 11 | NaN | NaN | NaN | 593.1923 | NaN | NaN |
| Sec23b | AESEEGPDVLR | -536 | -546 | 0 | 0.008680443 | 11 | NaN | NaN | NaN | 566.77515 | NaN | NaN |
| Ostf1 | TGLIPSNYVAEQAESIDNPLHEAAK | -58 | -82 | 0 | 0 | 25 | NaN | NaN | NaN | 332.79514 | 460.35098 | 369.32202 |
| Ostf1 | DMSDTSWWK | -43 | -51 | 0 | 0.00025015 | 9 | NaN | NaN | NaN | 331.59805 | NaN | NaN |
| Smc1a | ESLIGVYPEQGDCVISK | -1198 | -1214 | 0 | 0.002329279 | 17 | NaN | NaN | NaN | 1430.2372 | NaN | NaN |
| Smc1a | TALFEEISR | -152 | -160 | 0 | 0 | 9 | NaN | NaN | NaN | 702.84106 | NaN | NaN |
| Smc1a | TVALDGTLFQK | -638 | -648 | 0 | 0.001023236 | 11 | NaN | NaN | NaN | 548.7312 | NaN | NaN |
| Dis3 | YDDEIPSLR | -887 | -895 | 0 | 0 | 9 | NaN | NaN | NaN | 340.8047 | 444.94672 | 499.4589 |
| Dis3 | SVVLDDEGQNEDDVEKDEER | -307 | -326 | 1 | 0.000471409 | 20 | NaN | NaN | NaN | 314.34283 | 512.70557 | 534.82684 |
| Dis3 | IHEEFSEHALLR | -678 | -689 | 0 | 0.00025015 | 12 | NaN | NaN | NaN | 540.3718 | NaN | 1381.9526 |
| Dis3 | IHEEFSEHALLR | -678 | -689 | 0 | 0.00025015 | 12 | NaN | NaN | NaN | 540.3718 | NaN | 1381.9526 |
| Rpn1 | EGNHYFYSPYPTK | -158 | -170 | 0 | 0 | 13 | NaN | NaN | NaN | 1212.2781 | 825.699 | 812.1293 |
| Rpn1 | TILPAAAQDVYYR | -283 | -295 | 0 | 0 | 13 | NaN | NaN | NaN | 1009.7576 | 542.82587 | 465.40002 |
| Rpn1 | PAYSQDTFK | -208 | -216 | 0 | 0.002810811 | 9 | NaN | NaN | NaN | 780.26025 | 1169.335 | 952.4871 |
| Rpn1 | NLPSYEYLYNLGDQYALK | -336 | -353 | 0 | 0 | 18 | NaN | NaN | NaN | 349.30035 | 422.05576 | 357.53384 |
| Rpn1 | IDHILDAL | -601 | -608 | 0 | 0.009362332 | 8 | NaN | NaN | NaN | 764.0286 | NaN | NaN |

| | | | | | | | | | | | | |
|---|---|---|---|---|---|---|---|---|---|---|---|---|
| Eif3e | WIVNLIR | -370 | -376 | 0 | 0.001463897 | 7 | NaN | NaN | NaN | 1043.806 | 1477.4326 | 1574.1545 |
| Eif3e | LFIFETFCR | -338 | -346 | 0 | 0.000471409 | 9 | NaN | NaN | NaN | 863.66406 | 873.31946 | 805.18353 |
| Eif3e | MLFDYLADK | -112 | -120 | 0 | 0.00025015 | 9 | NaN | NaN | NaN | 1137.4153 | NaN | 133.76857 |
| Eif3e | LNMTPEEAER | -360 | -369 | 0 | 0 | 10 | NaN | NaN | NaN | 762.8333 | NaN | 324.83762 |
| Eif3e | LKETIDNNSVSSPL | -192 | -205 | 1 | 0.002265552 | 14 | NaN | NaN | NaN | 934.14856 | NaN | NaN |
| Ado | AEYTEASGPCVLTPHR | -160 | -175 | 0 | 0 | 16 | NaN | NaN | NaN | 1199.1091 | 1248.725 | 1269.4628 |
| Rangap1 | TAVLDAIDALMK | -490 | -501 | 0 | 0 | 12 | NaN | NaN | NaN | 1517.3976 | 1632.7451 | NaN |
| Rangap1 | AIPSLHGPLMVL | -533 | -544 | 0 | 0.001392869 | 12 | NaN | NaN | NaN | 1427.1211 | 1284.1271 | NaN |
| Rangap1 | VLDAIDALMK | -492 | -501 | 0 | 0.001055699 | 10 | NaN | NaN | NaN | 368.81546 | 322.78244 | NaN |
| Ddx27 | SADFNPDFVFTEK | -48 | -60 | 0 | 0 | 13 | NaN | NaN | NaN | 1616.2693 | 2108.6157 | NaN |
| Ddx27 | SVFDEELTNTSK | -711 | -722 | 0 | 0.000550307 | 12 | NaN | NaN | NaN | 517.87695 | NaN | 1414.1886 |
| Pin4 | AFALPVSGMDKPVFTDPPVK | -98 | -117 | 1 | 0.000471409 | 20 | NaN | NaN | NaN | 299.8059 | 371.2123 | 436.12866 |
| Ywhag | YYEIQNAPEQACHLAK | -183 | -198 | 0 | 0 | 16 | NaN | NaN | NaN | 673.3272 | 689.6983 | 693.1016 |
| Ywhaq | YLAEVACGDDR | -128 | -138 | 0 | 0 | 11 | NaN | NaN | NaN | 1072.977 | 1610.4215 | 231.54376 |
| Ywhaq | KQTIENSQGAYQEAFDISK | -139 | -157 | 1 | 0 | 19 | NaN | NaN | NaN | 242.159 | 295.5192 | 328.62897 |
| Ywhaq | YLIANATNPESK | -104 | -115 | 0 | 0 | 12 | NaN | NaN | NaN | 3513.4238 | NaN | NaN |
| Ywhaq | KQTIENSQGAYQ | -139 | -150 | 1 | 0.00025015 | 12 | NaN | NaN | NaN | 1112.3466 | NaN | NaN |
| Ywhaq | TVLELLDK | -96 | -103 | 0 | 0.004199202 | 8 | NaN | NaN | NaN | 938.8674 | NaN | NaN |
| Ywhab | NSQQAYQEAFEISK | -146 | -159 | 0 | 0.00025015 | 14 | NaN | NaN | NaN | 7010.139 | NaN | NaN |
| Ywhab | NDVLELLDK | -97 | -105 | 0 | 0.00111006 | 9 | NaN | NaN | NaN | 924.1361 | NaN | NaN |
| Ywhab | GHELSNEER | -35 | -43 | 0 | 0.00266297 | 9 | NaN | NaN | NaN | 240.24849 | NaN | NaN |
| Ywhaz | PNASQPESK | -107 | -115 | 0 | 0.002780285 | 9 | NaN | NaN | NaN | 749.2878 | 730.5733 | 1432.5575 |
| Ywhaz | IPNASQPESK | -106 | -115 | 0 | 0.00223472 | 10 | NaN | NaN | NaN | 2494.9094 | 160.58286 | 1287.3881 |
| Ywhaz | EKFLIPNASQPESK | -102 | -115 | 1 | 0 | 14 | NaN | NaN | NaN | 1803.4011 | 2187.3467 | 1886.4918 |
| Ywhaz | GIVDQSQQAYQEAF | -140 | -153 | 0 | 0.000230707 | 14 | NaN | NaN | NaN | 748.2576 | 412.3048 | 349.90585 |
| Ywhaz | DTQGDEAEAGEGGEN | -231 | -245 | 0 | 0.00025015 | 15 | NaN | NaN | NaN | 376.8078 | 294.17035 | NaN |
| Ywhaz | RYLAEVAAGDDKK | -127 | -139 | 2 | 0.001264841 | 13 | NaN | NaN | NaN | 1200.9844 | NaN | NaN |
| Pgm2 | AFDLAPPEAVPQQK | -277 | -290 | 0 | 0.00025015 | 14 | NaN | NaN | NaN | 3460.918 | NaN | NaN |
| Pgm2 | QKDPDPEFPTVK | -289 | -300 | 1 | 0.004350979 | 12 | NaN | NaN | NaN | 1029.9377 | NaN | NaN |
| Pgm2 | IAGGNKEELR | -47 | -56 | 1 | 0.001708008 | 10 | NaN | NaN | NaN | 471.05475 | NaN | NaN |
| Hsph1 | KPVTDCVISVPSFFTDAER | -135 | -153 | 1 | 0 | 19 | NaN | NaN | NaN | 615.9228 | 603.6274 | 941.6356 |
| Hsph1 | DVSTTLNADEAVAR | -361 | -374 | 0 | 0.005363933 | 14 | NaN | NaN | NaN | 1176.7723 | 572.26526 | 1153.2235 |
| Hsph1 | GIETIANEFSDR | -22 | -33 | 0 | 0.00078661 | 12 | NaN | NaN | NaN | 945.7031 | NaN | 1653.0894 |
| Hsph1 | NQQITHANNTVSSF | -54 | -67 | 0 | 0.002296977 | 14 | NaN | NaN | NaN | 1022.8396 | NaN | NaN |
| Eif3i | GHFGPINSVAFHPDGK | -283 | -298 | 0 | 0 | 16 | NaN | NaN | NaN | 1125.3441 | 3270.8877 | 3594.4702 |
| Eif3i | LFDSTTLEHQ | -214 | -223 | 0 | 0 | 10 | NaN | NaN | NaN | 353.44363 | 383.5695 | 868.37915 |
| Erp44 | KTPADCPVIAIDSFR | -313 | -327 | 1 | 0.00025015 | 15 | NaN | NaN | NaN | 882.949 | 2344.4219 | 1767.7366 |
| Erp44 | VFDLHSGK | -348 | -355 | 0 | 0.000550307 | 8 | NaN | NaN | NaN | 601.8047 | NaN | NaN |
| Ilf3 | EDITQSAQHALR | -312 | -323 | 0 | 0 | 12 | NaN | NaN | NaN | 794.0403 | 1184.7083 | 833.1445 |
| Cul4b | PEMVKLQEIFK | -756 | -766 | 1 | 0.009955823 | 11 | NaN | NaN | NaN | 837.96277 | NaN | NaN |
| Rpl29 | ASAPAQAPK | -137 | -145 | 0 | 0.00025015 | 9 | NaN | NaN | NaN | 538.3144 | 771.708 | 676.9772 |
| Actn1 | MHGTLEDQLSHLR | -660 | -672 | 0 | 0 | 13 | NaN | NaN | NaN | 852.3557 | 978.9176 | 1143.4158 |
| Actn1 | HTNYNMEHIR | -705 | -714 | 0 | 0 | 10 | NaN | NaN | NaN | 1482.6731 | 623.4128 | 634.4115 |
| Actn1 | GTQIENIEEDFRDGLK | -49 | -64 | 1 | 0 | 16 | NaN | NaN | NaN | 498.2419 | 746.3653 | 835.57965 |
| Actn1 | IDQLECDHQLIQEAL | -685 | -699 | 0 | 0.000907206 | 15 | NaN | NaN | NaN | 390.59122 | 642.42 | 753.81866 |
| Actn1 | KICDQWDNLGALTQK | -478 | -492 | 1 | 0.003490217 | 15 | NaN | NaN | NaN | 7021.8716 | NaN | NaN |
| Actn1 | IAQELNELDYYDSPSVNAR | -457 | -475 | 0 | 0 | 19 | NaN | NaN | NaN | 214.72296 | NaN | NaN |
| Lck | LHSYEPSHDGDLGFEK | -69 | -84 | 0 | 0.00025015 | 16 | NaN | NaN | NaN | 1886.8492 | 609.87634 | 1511.9553 |
| Lck | IEEQNYIHR | -355 | -363 | 0 | 0 | 9 | NaN | NaN | NaN | 3724.2832 | 4218.5083 | 4087.821 |
| Snrnp70 | HHNQPYCGIAPYIR | -33 | -46 | 0 | 0 | 14 | NaN | NaN | NaN | 2090.8672 | 1006.5403 | 1199.7051 |
| Snrnp70 | RVLVDVER | -173 | -180 | 1 | 0.00025015 | 8 | NaN | NaN | NaN | 951.0911 | 1878.922 | 886.5082 |
| Syncrip | WADPIEDPDPEVMAK | -322 | -336 | 0 | 0 | 15 | NaN | NaN | NaN | 619.9224 | 1312.1617 | 1086.5989 |
| Syncrip | GPIWDLR | -186 | -192 | 0 | 0.009242859 | 7 | NaN | NaN | NaN | 544.3697 | 520.0426 | 454.7666 |
| Syncrip | DLEGENIEIVFAKPPDQK | -395 | -412 | 1 | 0 | 18 | NaN | NaN | NaN | 509.5753 | 3229.013 | 3685.719 |
| Syncrip | MDPLTGLNR | -195 | -203 | 0 | 0.00111006 | 9 | NaN | NaN | NaN | 1447.1456 | NaN | NaN |
| Ogt | QYNPDLYCVR | -151 | -160 | 0 | 0.008919338 | 10 | NaN | NaN | NaN | 7127.9316 | NaN | NaN |
| Ogt | GHIYDNR | -727 | -733 | 0 | 0.009569685 | 7 | NaN | NaN | NaN | 1729.2521 | NaN | NaN |
| Ogt | SRQEYEDIAVK | -972 | -982 | 1 | 0 | 11 | NaN | NaN | NaN | 1318.3849 | NaN | NaN |
| Ogt | EYQAGDFEAAER | -31 | -42 | 0 | 0.00025015 | 12 | NaN | NaN | NaN | 632.2498 | NaN | NaN |
| Ogt | AIELQPHFPDAYCN | -285 | -298 | 0 | 0.006654467 | 14 | NaN | NaN | NaN | 623.639 | NaN | NaN |
| Gbp6 | GEDITSDDYLENALK | -189 | -203 | 0 | 0.000627943 | 15 | NaN | NaN | NaN | 1020.13196 | 266.2619 | 395.3803 |
| Vars1 | SKPEGHESLVDR | -973 | -984 | 1 | 0 | 12 | NaN | NaN | NaN | 837.2212 | 1755.7429 | 1478.5244 |
| Vars1 | VTEELFQR | -1066 | -1073 | 0 | 0 | 8 | NaN | NaN | NaN | 1558.7375 | 1142.1056 | 1067.6222 |
| Vars1 | FLEEVWK | -411 | -417 | 0 | 0.002068213 | 7 | NaN | NaN | NaN | 915.2013 | 882.50226 | 917.02954 |
| Vars1 | ATSKPEGHESLVDR | -971 | -984 | 1 | 0 | 14 | NaN | NaN | NaN | 361.73724 | 609.4822 | 542.6097 |
| Vars1 | AFLEEVWK | -410 | -417 | 0 | 0 | 8 | NaN | NaN | NaN | 691.5063 | 991.9452 | NaN |
| Vars1 | YVLDPSAR | -174 | -181 | 0 | 0.006078721 | 8 | NaN | NaN | NaN | 3952.3699 | NaN | NaN |
| Vars1 | KVPLEVQEADEAK | -1229 | -1241 | 1 | 0.00111006 | 13 | NaN | NaN | NaN | 1262.6561 | NaN | NaN |
| Vars1 | YDQLLNSNLDPSEVEK | -885 | -900 | 0 | 0 | 16 | NaN | NaN | NaN | 573.38824 | NaN | NaN |
| Vars1 | PPGEDPDGR | -743 | -751 | 0 | 0.005758933 | 9 | NaN | NaN | NaN | 475.68927 | NaN | NaN |
| Rpl34 | CGVCPGR | -46 | -52 | 0 | 0.001636916 | 7 | NaN | NaN | NaN | 502.2535 | 463.23666 | 443.58612 |
| Nfkb1 | RLEPVVSDAIYDS | -228 | -240 | 1 | 0.00122419 | 13 | NaN | NaN | NaN | 4900.925 | NaN | NaN |
| Nfkb1 | FDYAVTGDVK | -513 | -522 | 0 | 0.003069674 | 10 | NaN | NaN | NaN | 1307.6552 | NaN | NaN |
| Hnrnpu | FEYIEENKYSR | -234 | -244 | 1 | 0 | 11 | NaN | NaN | NaN | 1096.0835 | 1127.634 | 1074.614 |
| Hnrnpu | APQCLGK | -535 | -541 | 0 | 0.003745449 | 7 | NaN | NaN | NaN | 1068.1777 | 431.90668 | NaN |
| Hnrnpu | NFILDQTNVSAAAQR | -552 | -566 | 0 | 0 | 15 | NaN | NaN | NaN | 572.52386 | 702.48645 | NaN |
| Hnrnpu | SQHYHQGYY | -792 | -800 | 0 | 0.002810811 | 9 | NaN | NaN | NaN | 1173.0796 | NaN | NaN |
| Hnrnpu | TKDIDIHEVR | -327 | -336 | 1 | 0.000431655 | 10 | NaN | NaN | NaN | 377.88962 | NaN | NaN |

| | | | | | | | | | | | | |
|---|---|---|---|---|---|---|---|---|---|---|---|---|
| Hbb-b1 | VHLTDAEKA | -2 | -10 | 1 | 0.000230707 | 9 | NaN | NaN | NaN | 2261.166 | 757.0894 | 3082.6204 |
| Hbb-b1 | FNDGLNHLDSLK | -72 | -83 | 0 | 0 | 12 | NaN | NaN | NaN | 2481.1274 | NaN | NaN |
| Actr2 | HLWDYTFGPEK | -87 | -97 | 0 | 0 | 11 | NaN | NaN | NaN | 6041.5166 | 7710.3687 | 7487.0103 |
| Actr2 | VLVESYTLPDGR | -239 | -250 | 0 | 0.00025015 | 12 | NaN | NaN | NaN | 2564.1067 | 2856.9236 | 2276.737 |
| Actr2 | GYAFNHSADFETVR | -201 | -214 | 0 | 0 | 14 | NaN | NaN | NaN | 1683.6227 | 1063.8638 | 1075.3568 |
| Actr2 | HIVLSGGSTMYPGLPSR | -300 | -316 | 0 | 0.000907206 | 17 | NaN | NaN | NaN | 1065.4364 | 1283.8019 | 1358.9437 |
| Actr2 | KVVVCDNGTGFVK | -7 | -19 | 1 | 0 | 13 | NaN | NaN | NaN | 885.66876 | 984.35754 | 945.6656 |
| Actr2 | CGYAGSNFPEHIFPA | -20 | -34 | 0 | 0 | 15 | NaN | NaN | NaN | 794.71265 | 1130.913 | 1313.6176 |
| Actr2 | DLMVGDEASELR | -54 | -65 | 0 | 0 | 12 | NaN | NaN | NaN | 846.659 | NaN | NaN |
| Mettl26 | AFPNAEWQPSDVDQR | -48 | -62 | 0 | 0.00122419 | 15 | NaN | NaN | NaN | 297.7402 | NaN | NaN |
| Uggt1 | ILETTTFFQR | -659 | -668 | 0 | 0 | 10 | NaN | NaN | NaN | 1357.1807 | 1430.9673 | 1315.892 |
| Uggt1 | QQIAVDEPPPEGCK | -135 | -148 | 0 | 0.001868731 | 14 | NaN | NaN | NaN | 770.4663 | 810.9431 | NaN |
| Uggt1 | YSEIGHEEFSNIHHQLISK | -199 | -217 | 0 | 0.00025015 | 19 | NaN | NaN | NaN | 43.971344 | NaN | NaN |
| Prdx2 | AFQYTDEHGEVCPAGWKPGSDTIKPNVDDSK | -161 | -191 | 2 | 0.001463897 | 31 | NaN | NaN | NaN | 291.7603 | NaN | NaN |
| Pes1 | DHPTDVDYR | -210 | -218 | 0 | 0.00025015 | 9 | NaN | NaN | NaN | 1063.968 | NaN | 882.7422 |
| Pes1 | KGIYPHEPK | -41 | -49 | 1 | 0 | 9 | NaN | NaN | NaN | 1014.69055 | NaN | 670.21027 |
| Pes1 | ERYPTFIDAL | -122 | -131 | 1 | 0.002329279 | 10 | NaN | NaN | NaN | 632.0115 | NaN | NaN |
| Sfpq | GNLPADITEDEFKR | -294 | -307 | 1 | 0 | 14 | NaN | NaN | NaN | 524.914 | 304.48956 | 660.0129 |
| Sfpq | QHGTFEYEYSQR | -474 | -485 | 0 | 0 | 12 | NaN | NaN | NaN | 2936.83 | NaN | 3222.415 |
| Sfpq | EEYEGPNK | -688 | -695 | 0 | 0.001392869 | 8 | NaN | NaN | NaN | 490.56146 | 524.1274 | NaN |
| Sfpq | NMGDPYGSGGQK | -611 | -622 | 0 | 0.000907206 | 12 | NaN | NaN | NaN | 255.05284 | NaN | NaN |
| Rpl7a | YNDRYDEIR | -226 | -234 | 1 | 0.00025015 | 9 | NaN | NaN | NaN | 2454.9556 | 345.65308 | 2546.5195 |
| Rpl7a | QTATQLLK | -90 | -97 | 0 | 0.003490217 | 8 | NaN | NaN | NaN | 9047.178 | NaN | NaN |
| Acadl | PGAGSDLQGVR | -175 | -185 | 0 | 0 | 11 | NaN | NaN | NaN | 1500.4683 | 968.61774 | 1138.0068 |
| Acadl | FFQEEVIPH | -67 | -75 | 0 | 0 | 9 | NaN | NaN | NaN | 251.87607 | NaN | NaN |
| C1qbp | FLTDEIKEEK | -82 | -91 | 1 | 0 | 10 | NaN | NaN | NaN | 286.87387 | 549.50867 | 4868.6943 |
| Vav1 | AGAEGILTNR | -679 | -688 | 0 | 0.00025015 | 10 | NaN | NaN | NaN | 4720.7207 | 5846.384 | 6397.149 |
| Vav1 | SIENLDQSLANYGR | -389 | -402 | 0 | 0 | 14 | NaN | NaN | NaN | 497.60608 | 750.6494 | 639.6887 |
| Vav1 | EAEHNWWEGR | -631 | -640 | 0 | 0 | 10 | NaN | NaN | NaN | 1624.2958 | NaN | 1639.8727 |
| Atp5mk | YFNSYTLTGR | -18 | -27 | 0 | 0 | 10 | NaN | NaN | NaN | 1100.8955 | 912.1295 | 1155.2666 |
| Mri1 | AGAGGPGLAALVAFVR | -65 | -80 | 0 | 0 | 16 | NaN | NaN | NaN | 407.10333 | 260.64877 | 356.00293 |
| Mri1 | EGATEETVR | -114 | -122 | 0 | 0 | 9 | NaN | NaN | NaN | 1274.7128 | NaN | NaN |
| Rbm42 | ALRPPLEEPAAPR | -221 | -233 | 1 | 0 | 13 | NaN | NaN | NaN | 3974.7766 | 3477.3816 | NaN |
| Rbm42 | TYQQVQQTLEAR | -89 | -100 | 0 | 0 | 12 | NaN | NaN | NaN | 1782.8165 | 372.34888 | 808.875 |
| Rbm42 | GHLDSPEAR | -129 | -137 | 0 | 0 | 9 | NaN | NaN | NaN | 1178.6743 | NaN | 1231.1067 |
| Rplp0 | GHLENNPALEK | -67 | -77 | 0 | 0 | 11 | NaN | NaN | NaN | 5662.741 | 6628.9307 | 6464.8013 |
| Rplp0 | SVETEYTFPLTEKVK | -252 | -266 | 1 | 0 | 15 | NaN | NaN | NaN | 1606.5818 | 1579.4749 | 1716.7325 |
| Rplp0 | LENNPALEK | -69 | -77 | 0 | 0 | 9 | NaN | NaN | NaN | 839.3189 | 795.3622 | 1301.268 |
| Rplp0 | TSFFQALGITTK | -135 | -146 | 0 | 0 | 12 | NaN | NaN | NaN | 360.81335 | 330.01678 | 423.2597 |
| Cct5 | LGFAGVVQEISFGTTK | -353 | -368 | 0 | 0 | 16 | NaN | NaN | NaN | 446.6138 | 485.05627 | 491.61853 |
| Cct5 | DVDFELIK | -203 | -210 | 0 | 0 | 8 | NaN | NaN | NaN | 443.13358 | 168.41844 | 594.92834 |
| Cct5 | VLVDINNPEPLIQTAK | -155 | -170 | 0 | 0.002595029 | 16 | NaN | NaN | NaN | 400.28043 | 930.62256 | NaN |
| Cct5 | RDVDFELIK | -202 | -210 | 1 | 0.00383302 | 9 | NaN | NaN | NaN | 1677.0211 | NaN | NaN |
| Cct5 | FSELTSEK | -345 | -352 | 0 | 0.000431655 | 8 | NaN | NaN | NaN | 378.37735 | NaN | NaN |
| Hspa1l | VAMNPQNTVFDAK | -61 | -73 | 0 | 0.006472924 | 13 | NaN | NaN | NaN | 3957.685 | NaN | NaN |
| Fermt3 | KSQDEAPGDPTQQLNLK | -385 | -401 | 1 | 0.000230707 | 17 | NaN | NaN | NaN | 1061.1718 | 1258.1923 | 1063.6389 |
| Fermt3 | KSQDEAPGDPTQQLNLK | -385 | -401 | 1 | 0.000230707 | 17 | NaN | NaN | NaN | 1061.1718 | 1258.1923 | 1063.6389 |
| Fermt3 | LLVPSPEGMSEIYLR | -420 | -434 | 0 | 0 | 15 | NaN | NaN | NaN | 656.84216 | 841.08496 | 871.7163 |
| Fermt3 | VTGESHIGGVLLK | -36 | -48 | 0 | 0 | 13 | NaN | NaN | NaN | 643.3553 | 1059.1018 | 1134.8031 |
| Fermt3 | DWSDHAIWWEQK | -58 | -69 | 0 | 0 | 12 | NaN | NaN | NaN | 1152.0603 | 664.1202 | NaN |
| Ybx3 | PAPAIGEAEDKENQQAA | -290 | -306 | 1 | 0.00266297 | 17 | NaN | NaN | NaN | 819.46783 | 816.43335 | 958.93066 |
| Xpo1 | SAFPHLQDAQVK | -984 | -995 | 0 | 0 | 12 | NaN | NaN | NaN | 1318.7308 | 571.11957 | 1542.7561 |
| Xpo1 | VDTILEFSQNMNTK | -63 | -76 | 0 | 0 | 14 | NaN | NaN | NaN | 1322.2576 | NaN | 768.56104 |
| Xpo1 | KEHPDAWTR | -54 | -62 | 1 | 0 | 9 | NaN | NaN | NaN | 517.1248 | 1155.549 | NaN |
| Xpo1 | AVGHPFVIQLGR | -701 | -712 | 0 | 0 | 12 | NaN | NaN | NaN | 2134.0835 | NaN | NaN |
| Xpo1 | SLNQDIPAFK | -1003 | -1012 | 0 | 0 | 10 | NaN | NaN | NaN | 630.58813 | NaN | NaN |
| Xpo1 | LISTLIYK | -246 | -253 | 0 | 0 | 8 | NaN | NaN | NaN | 477.96884 | NaN | NaN |
| Sdha | SIAESCRPGDKVPSIK | -470 | -485 | 2 | 0.001055699 | 16 | NaN | NaN | NaN | 486.8829 | 1148.0044 | 1166.4548 |
| Sdha | KPFGEHW | -616 | -622 | 1 | 0.003966045 | 7 | NaN | NaN | NaN | 1219.6393 | 457.7345 | 736.2582 |
| Sdha | VRVDEYDYSKPIQGQ | -599 | -613 | 2 | 0.00149795 | 15 | NaN | NaN | NaN | 572.0707 | 568.0223 | 570.0792 |
| Rps21 | NDAGEFVDLYVPR | -3 | -15 | 0 | 0 | 13 | NaN | NaN | NaN | 1265.0405 | 1311.7612 | NaN |
| Tfrc | LDTYEALTQK | -573 | -582 | 0 | 0 | 10 | NaN | NaN | NaN | 2737.1833 | 3016.8105 | 3202.4878 |
| Tfrc | SAFSNLFGGEPLSYTR | -7 | -22 | 0 | 0 | 16 | NaN | NaN | NaN | 493.76514 | 599.56116 | 511.40063 |
| Tfrc | LAADEEENADNNMK | -40 | -53 | 0 | 0 | 14 | NaN | NaN | NaN | 315.27252 | NaN | 301.05215 |
| Rps18 | SLVIPEKFQHIL | -2 | -13 | 1 | 0.00025015 | 12 | NaN | NaN | NaN | 1061.7904 | 476.18378 | 569.4992 |
| Rps18 | GELTEDEVER | -57 | -66 | 0 | 0 | 10 | NaN | NaN | NaN | 587.9755 | 600.984 | 1200.0752 |
| Rps18 | GLDNKLREDLER | -102 | -113 | 2 | 0.00078661 | 12 | NaN | NaN | NaN | 326.94254 | 666.09717 | 709.81396 |
| Arpc5l | GEPGPDPCEVDGLLR | -35 | -49 | 0 | 0 | 15 | NaN | NaN | NaN | 665.2634 | 1038.034 | 1093.7953 |
| Arpc5l | AGEPGPDPCEVDGLLR | -34 | -49 | 0 | 0 | 16 | NaN | NaN | NaN | 722.6277 | NaN | NaN |
| Vcp | ATNRPNSIDPALR | -346 | -358 | 1 | 0.000230707 | 13 | NaN | NaN | NaN | 349.34128 | 764.0546 | 768.5029 |
| Vcp | WEDIGGLEDVK | -476 | -486 | 0 | 0 | 11 | NaN | NaN | NaN | 1143.7549 | 1293.5189 | 1198.0691 |
| Vcp | DHFEEAMR | -734 | -741 | 0 | 0 | 8 | NaN | NaN | NaN | 561.2636 | 504.92606 | 372.33716 |
| Vcp | DREVDIGIPDATGR | -364 | -377 | 1 | 0.004224523 | 14 | NaN | NaN | NaN | 1377.7184 | 1422.3505 | NaN |
| Vcp | RIHVLPIDDTVEGITGN | -113 | -129 | 1 | 0.004321969 | 17 | NaN | NaN | NaN | 232.16435 | NaN | 550.05133 |
| Vcp | VINQILTEMDGMSTK | -600 | -614 | 0 | 0.008105996 | 15 | NaN | NaN | NaN | 410.2034 | NaN | NaN |
| Vcp | VVEVPQVTWEDIGGLEDVKR | -468 | -487 | 1 | 0 | 20 | NaN | NaN | NaN | 379.5792 | NaN | NaN |
| Dyrk1a | AGESGHTVADYLK | -439 | -451 | 0 | 0.00025015 | 13 | NaN | NaN | NaN | 5365.0225 | NaN | NaN |
| Nono | KVELDNMPLR | -128 | -137 | 1 | 0.001392869 | 10 | NaN | NaN | NaN | 1895.0608 | 1845.1133 | 1880.0801 |

| | | | | | | | | | | | | |
|---|---|---|---|---|---|---|---|---|---|---|---|---|
| Nono | SQNEGLTIDLK | -52 | -62 | 0 | 0.00025015 | 11 | NaN | NaN | NaN | 1435.7211 | 1402.6973 | 1871.2043 |
| Nono | GIVEFSGKPAAR | -193 | -204 | 1 | 0 | 12 | NaN | NaN | NaN | 596.99365 | NaN | 1150.9037 |
| Dnmt1 | LHKEELSEEGYLAK | -68 | -81 | 1 | 0.000230707 | 14 | NaN | NaN | NaN | 952.2812 | 2230.604 | 2440.0347 |
| Dnmt1 | LHKEELSEEGYLA | -68 | -80 | 1 | 0 | 13 | NaN | NaN | NaN | 1031.3738 | 1455.4894 | 1232.387 |
| Dnmt1 | AGEVTNSLGQR | -1203 | -1213 | 0 | 0.001636916 | 11 | NaN | NaN | NaN | 1644.3699 | NaN | NaN |
| Dnmt1 | FLQNNPDAVYEDLINK | -521 | -536 | 0 | 0.002136192 | 16 | NaN | NaN | NaN | 456.23193 | NaN | NaN |
| Dnmt1 | QLGPEEPCEQEDDNR | -225 | -239 | 0 | 0.002265552 | 15 | NaN | NaN | NaN | 230.37497 | NaN | NaN |
| Matn4 | AGFALQQDQR | -286 | -295 | 0 | 0.000230707 | 10 | NaN | NaN | NaN | 1231.638 | NaN | NaN |
| Nat10 | ESLQDTQPVGVLVDCCK | -240 | -256 | 0 | 0.001868731 | 17 | NaN | NaN | NaN | 772.45264 | 697.9942 | 674.0005 |
| Nat10 | PGDAVEK | -473 | -479 | 0 | 0 | 7 | NaN | NaN | NaN | 112.42868 | 399.7334 | 464.9756 |
| Cyb5a | KFLEEHPGGEEVLR | -39 | -52 | 1 | 0 | 14 | NaN | NaN | NaN | 705.0364 | NaN | 1237.4824 |
| Cyb5a | KFLEEHPGGEEVLR | -39 | -52 | 1 | 0 | 14 | NaN | NaN | NaN | 705.0364 | NaN | 1237.4824 |
| Cyb5a | AGQSDKDVK | -2 | -10 | 1 | 0.00149795 | 9 | NaN | NaN | NaN | 678.33624 | NaN | 743.40436 |
| Ppp2r1a | EHSPSDLEAHFVPLVK | -118 | -133 | 0 | 0.00025015 | 16 | NaN | NaN | NaN | 577.16534 | 630.29504 | 658.44446 |
| Ppp2r1a | VKEFCENLSADCR | -306 | -318 | 1 | 0 | 13 | NaN | NaN | NaN | 360.95352 | 382.8965 | 497.09268 |
| Ppp2r1a | EWAHATIIPK | -476 | -485 | 0 | 0 | 10 | NaN | NaN | NaN | 1171.8748 | 776.80194 | NaN |
| Ppp2r1a | AISHEHSPSDLEAHFVPLVK | -114 | -133 | 0 | 0 | 20 | NaN | NaN | NaN | 693.575 | NaN | 795.86914 |
| Hnrnpl | QEFGPISYVVVMPK | -120 | -133 | 0 | 0 | 14 | NaN | NaN | NaN | 532.2214 | 2345.0972 | 2241.5112 |
| Hnrnpl | PASPVVHIR | -96 | -104 | 0 | 0 | 9 | NaN | NaN | NaN | 3826.9363 | NaN | NaN |
| Hnrnpl | AGGGGGENYDDPHK | -81 | -94 | 0 | 0 | 14 | NaN | NaN | NaN | 393.076 | 908.3066 | 1530.1794 |
| Hnrnpl | DVLYTICNPCGPVQR | -206 | -220 | 0 | 0.00025015 | 15 | NaN | NaN | NaN | 2779.7517 | NaN | 3274.743 |
| Hnrnpl | GHPAFVNYSTSQK | -163 | -175 | 0 | 0 | 13 | NaN | NaN | NaN | 441.04187 | NaN | 607.38605 |
| Hnrnpl | GGGGENYDDPHK | -83 | -94 | 0 | 0.001291103 | 12 | NaN | NaN | NaN | 410.6588 | 434.30197 | NaN |
| Hnrnpl | IQHPSNVLHF | -493 | -502 | 0 | 0.000947459 | 10 | NaN | NaN | NaN | 363.64935 | 4061.5015 | NaN |
| Hnrnpl | SVNSVLLF | -187 | -194 | 0 | 0.001708008 | 8 | NaN | NaN | NaN | 4529.995 | NaN | NaN |
| Hnrnpl | AITHLNNNFMFGQ | -432 | -444 | 0 | 0.002068213 | 13 | NaN | NaN | NaN | 1023.9878 | NaN | NaN |
| Hnrnpl | YKDFSESR | -471 | -478 | 1 | 0.006677519 | 8 | NaN | NaN | NaN | 872.72485 | NaN | NaN |
| Ttll12 | AYGEADPLIR | -238 | -247 | 0 | 0.000550307 | 10 | NaN | NaN | NaN | 1010.70544 | 794.39355 | NaN |
| Slc25a12 | QDLGLFGLYK | -473 | -482 | 0 | 0 | 10 | NaN | NaN | NaN | 504.2258 | 807.0343 | 507.5648 |
| Slc25a12 | DLPPANPDHIGGYR | -627 | -640 | 0 | 0.00830926 | 14 | NaN | NaN | NaN | 792.57623 | NaN | NaN |
| Rps2 | EDKEWIPVTK | -56 | -65 | 1 | 0.00025015 | 10 | NaN | NaN | NaN | 4358.1895 | 4497.0093 | 4569.683 |
| Rps2 | YQEFTDHLVK | -266 | -275 | 0 | 0.00025015 | 10 | NaN | NaN | NaN | 3920.3994 | 2781.598 | 2920.5303 |
| Rps2 | GKAEDKEWIPVTK | -53 | -65 | 2 | 0 | 13 | NaN | NaN | NaN | 3173.1426 | 448.9628 | 696.368 |
| Rps2 | GASLKDEVLK | -99 | -108 | 1 | 0.000230707 | 10 | NaN | NaN | NaN | 2104.4036 | 3328.5757 | 3548.694 |
| Rps2 | KSPYQEFTDHLVK | -263 | -275 | 1 | 0.00025015 | 13 | NaN | NaN | NaN | 1182.6122 | 1622.606 | 117.072266 |
| Rps2 | SLEEIYLF | -77 | -84 | 0 | 0.00051306 | 8 | NaN | NaN | NaN | 129.25719 | 601.198 | 723.1453 |
| Rps2 | SPYQEFTDH | -264 | -272 | 0 | 0.000431655 | 9 | NaN | NaN | NaN | 70.14204 | 917.5419 | 874.7861 |
| Rps2 | IGKPHTVPC | -174 | -182 | 1 | 0.000591124 | 9 | NaN | NaN | NaN | 3472.4954 | NaN | NaN |
| Rps2 | ASLKDEVLK | -100 | -108 | 1 | 0.006788754 | 9 | NaN | NaN | NaN | 809.67377 | NaN | NaN |
| Rps2 | TKSPYQEFTDHLVK | -262 | -275 | 1 | 0.005698678 | 14 | NaN | NaN | NaN | 346.1279 | NaN | NaN |
| Rps2 | GHVGLGVK | -135 | -142 | 0 | 0.003360691 | 8 | NaN | NaN | NaN | 333.33963 | NaN | NaN |
| Ahcy | HFEQMKDDAIVC | -286 | -297 | 1 | 0 | 12 | NaN | NaN | NaN | 700.58563 | 613.6056 | 686.7944 |
| Ahcy | KLDEAVAEAHLGK | -389 | -401 | 1 | 0 | 13 | NaN | NaN | NaN | 902.76306 | 879.4555 | 774.5853 |
| Ahcy | THPDKYPVGVHFLPK | -374 | -388 | 1 | 0 | 15 | NaN | NaN | NaN | 172.07529 | NaN | 604.9169 |
| Ahcy | VNIKPQVDRY | -319 | -328 | 2 | 0 | 10 | NaN | NaN | NaN | 1132.8855 | 1335.7141 | 1156.2352 |
| Ahcy | VIITEIDPINAL | -239 | -250 | 0 | 0.000712251 | 12 | NaN | NaN | NaN | 263.1109 | 259.86142 | NaN |
| Ahcy | GISEETTTGVHNL | -152 | -164 | 0 | 0.002099344 | 13 | NaN | NaN | NaN | 2472.1372 | NaN | NaN |
| Ahcy | LPYKVADIGL | -5 | -14 | 1 | 0.008528992 | 10 | NaN | NaN | NaN | 622.26996 | NaN | NaN |
| Ahcy | RIILLAEGR | -335 | -343 | 1 | 0.000471409 | 9 | NaN | NaN | NaN | 363.14987 | NaN | NaN |
| Rpl10a | AVAVGHVK | -166 | -173 | 0 | 0.003521747 | 8 | NaN | NaN | NaN | 546.5542 | NaN | 465.21307 |
| Rpl10a | VLCLAVAVGHVK | -162 | -173 | 0 | 0 | 12 | NaN | NaN | NaN | 339.85782 | 1069.7136 | NaN |
| Nifk | LALNPQEDSQFEK | -12 | -24 | 0 | 0.00951004 | 13 | NaN | NaN | NaN | 1327.349 | NaN | NaN |
| Timeless | AGLGPTEEEATGEEEWNSAPK | -1138 | -1158 | 0 | 0.003681885 | 21 | NaN | NaN | NaN | 191.33722 | 246.93663 | NaN |
| Fdxr | TEKPGVEEAAR | -288 | -298 | 1 | 0.00025015 | 11 | NaN | NaN | NaN | 3123.713 | 4484.0415 | 4325.4126 |
| Eif2s1 | RNECVVVIR | -67 | -75 | 1 | 0.00149795 | 9 | NaN | NaN | NaN | 3838.0972 | 3653.3062 | 3617.4197 |
| Eif2s1 | AWVFDDKYK | -135 | -143 | 1 | 0 | 9 | NaN | NaN | NaN | 1206.2936 | 1469.0719 | 1585.7534 |
| Eif2s1 | RPGYGAYDAF | -144 | -153 | 1 | 0.000550307 | 10 | NaN | NaN | NaN | 1131.0343 | 1495.846 | 258.64554 |
| Eif2s1 | AGLNCSTETMPIK | -214 | -226 | 0 | 0 | 13 | NaN | NaN | NaN | 908.5486 | 870.5461 | 1036.6593 |
| Eif2s1 | GRNECVVVIR | -66 | -75 | 1 | 0.001811524 | 10 | NaN | NaN | NaN | 1997.3438 | NaN | NaN |
| Ddx54 | SLDLQGLHR | -503 | -511 | 0 | 0 | 9 | NaN | NaN | NaN | 550.1398 | 766.37665 | 438.47424 |
| Ddx54 | GFAEQLQEIIGR | -255 | -266 | 0 | 0.000230707 | 12 | NaN | NaN | NaN | 2067.5454 | NaN | 1857.979 |
| Runx3 | VTPSTPSPR | -206 | -214 | 0 | 0.008014596 | 9 | NaN | NaN | NaN | 2127.4233 | NaN | NaN |
| Pcna | ALVFEAPNQEK | -100 | -110 | 0 | 0 | 11 | NaN | NaN | NaN | 3229.2117 | 550.3008 | 1951.5328 |
| Pcna | RSEGFDTYR | -53 | -61 | 1 | 0.000550307 | 9 | NaN | NaN | NaN | 5012.5864 | 5633.9834 | 713.0591 |
| Pcna | SMDSSHVSLVQLTLR | -39 | -53 | 0 | 0.00111006 | 15 | NaN | NaN | NaN | 607.66766 | 811.3534 | 780.77185 |
| Pcna | SMDSSHVSLVQLTLR | -39 | -53 | 0 | 0.00111006 | 15 | NaN | NaN | NaN | 607.66766 | 811.3534 | 780.77185 |
| Pcna | SMSADVPLVVEYK | -228 | -240 | 0 | 0 | 13 | NaN | NaN | NaN | 2438.5144 | 1695.0267 | NaN |
| Ndufb3 | KIEGTPLETVQK | -29 | -40 | 1 | 0.00025015 | 12 | NaN | NaN | NaN | 1045.709 | 1132.0798 | 322.46045 |
| Samm50 | GEEAEFVEVEPEAK | -23 | -36 | 0 | 0 | 14 | NaN | NaN | NaN | 2491.626 | 2728.092 | 2446.8516 |
| Samm50 | ICDGVQFGAGIR | -456 | -467 | 0 | 0 | 12 | NaN | NaN | NaN | 726.9647 | 491.30383 | 705.1009 |
| Samm50 | AGNLCNLNYGEGPK | -399 | -412 | 0 | 0 | 14 | NaN | NaN | NaN | 1109.8132 | NaN | NaN |
| Samm50 | SLEPLPSSGTDFGAL | -8 | -22 | 0 | 0.004439092 | 15 | NaN | NaN | NaN | 419.2555 | NaN | NaN |
| Ubqln4 | AGNSDNSSSQPLR | -305 | -317 | 0 | 0.001092529 | 13 | NaN | NaN | NaN | 868.40106 | NaN | NaN |
| Arpc5 | LKNPPINTK | -59 | -67 | 1 | 0.001636916 | 9 | NaN | NaN | NaN | 752.85956 | 697.7409 | NaN |
| Sart3 | VSNLPYSIEEPEVK | -708 | -721 | 0 | 0 | 14 | NaN | NaN | NaN | 1392.01 | 1396.8823 | 1044.2751 |
| Sart3 | GPGWDLQR | -52 | -59 | 0 | 0.001150701 | 8 | NaN | NaN | NaN | 169.9704 | NaN | NaN |
| Ddx6 | KGNEFEDYCLK | -94 | -104 | 1 | 0 | 11 | NaN | NaN | NaN | 923.8082 | 933.8996 | 281.64423 |
| Ddx6 | VCTDLFTR | -389 | -396 | 0 | 0 | 8 | NaN | NaN | NaN | 1785.6221 | NaN | 2039.8558 |

| | | | | | | | | | | | | |
|---|---|---|---|---|---|---|---|---|---|---|---|---|
| Sod1 | GDGPVQGTIHFEQK | -11 | -24 | 0 | 0 | 14 | NaN | NaN | NaN | 228.65495 | 532.8238 | 6705.033 |
| Sod1 | ASGEPVVLSGQITGLTEGQHGFHVH | -25 | -49 | 0 | 0.000550307 | 25 | NaN | NaN | NaN | 1027.9976 | 529.4041 | 550.1564 |
| Pnp | VFGFSLITNK | -235 | -244 | 0 | 0 | 10 | NaN | NaN | NaN | 1835.7578 | 2819.4624 | 2886.3909 |
| Pnp | GSGLGGLTAHLK | -32 | -43 | 0 | 0 | 12 | NaN | NaN | NaN | 744.9388 | 693.093 | 752.6124 |
| Nipsnap1 | AGPNIYELR | -180 | -188 | 0 | 0.00111006 | 9 | NaN | NaN | NaN | 685.11395 | NaN | 873.34283 |
| Bhmt | TFYASEDKLENR | -75 | -86 | 1 | 0 | 12 | NaN | NaN | NaN | 643.4801 | 751.1184 | 820.3996 |
| Pgd | GILFVGSGVSGGEEGAR | -120 | -136 | 0 | 0 | 17 | NaN | NaN | NaN | 320.02216 | 962.0888 | 653.4231 |
| Pgd | YGPSLMPGGNK | -137 | -147 | 0 | 0.000869338 | 11 | NaN | NaN | NaN | 1267.1406 | NaN | NaN |
| Pgd | QAVDDFIEK | -79 | -87 | 0 | 0 | 9 | NaN | NaN | NaN | 234.35678 | NaN | NaN |
| Acaca | RVDWQENDFSK | -265 | -275 | 1 | 0 | 11 | NaN | NaN | NaN | 1653.6987 | 641.61456 | 639.78064 |
| Tmpo | PLELSDFR | -373 | -380 | 0 | 0 | 8 | NaN | NaN | NaN | 953.0985 | 993.07855 | 1045.8761 |
| Clic4 | HPESNTAGMDIFAK | -111 | -124 | 0 | 0.001392869 | 14 | NaN | NaN | NaN | 1072.7717 | 2007.9333 | NaN |
| Arhgap45 | LSTPDGVPFIVK | -765 | -776 | 0 | 0 | 12 | NaN | NaN | NaN | 473.84283 | 130.39752 | 1011.54816 |
| Arhgap45 | RQIQEVIR | -488 | -495 | 1 | 0.00051306 | 8 | NaN | NaN | NaN | 1768.7788 | NaN | 1643.961 |
| Arhgap45 | KGSFNPGDASGPEAA | -575 | -589 | 1 | 0.001904399 | 15 | NaN | NaN | NaN | 384.9796 | NaN | NaN |
| Eef1d | GVVQDLQQAISK | -96 | -107 | 0 | 0.00025015 | 12 | NaN | NaN | NaN | 1937.2269 | 1395.9326 | 1560.7896 |
| Eef1d | SGPGGDHSELIVR | -71 | -83 | 0 | 0 | 13 | NaN | NaN | NaN | 1441.1145 | 1431.0796 | 1690.9086 |
| Eef1d | GSDEEEEDKEAAR | -161 | -173 | 1 | 0.00149795 | 13 | NaN | NaN | NaN | 169.27621 | 403.67505 | 291.65186 |
| Tbx21 | AGSSLGTPYSGGALVPAAPGR | -49 | -69 | 0 | 0 | 21 | NaN | NaN | NaN | 306.6966 | 255.17578 | 509.2555 |
| Lman2 | AGTGDLSDNHDIISIK | -259 | -274 | 0 | 0 | 16 | NaN | NaN | NaN | 3702.4304 | 4290.4863 | 4098.7847 |
| Lman2 | NCIDITGVR | -240 | -248 | 0 | 0 | 9 | NaN | NaN | NaN | 1625.5405 | 760.8667 | 1095.6685 |
| Baz1b | AVGEECDFEVGK | -115 | -126 | 0 | 0 | 12 | NaN | NaN | NaN | 4834.872 | 3323.6855 | 327.11118 |
| Phgdh | LDVFTEEPPR | -259 | -268 | 0 | 0 | 10 | NaN | NaN | NaN | 746.77563 | 138.64139 | 735.01575 |
| Phgdh | VTADVINAAEKL | -59 | -70 | 1 | 0 | 12 | NaN | NaN | NaN | 2580.5593 | 3360.9534 | NaN |
| Actn4 | TIPWLEDRVPQK | -313 | -324 | 1 | 0 | 12 | NaN | NaN | NaN | 12790.441 | 5147.02 | 944.0776 |
| Actn4 | HRPELIEYD | -206 | -214 | 1 | 0.002780285 | 9 | NaN | NaN | NaN | 1306.8209 | 1404.4517 | NaN |
| Actn4 | DHALLEEQSK | -635 | -644 | 0 | 0.000431655 | 10 | NaN | NaN | NaN | 1217.8405 | NaN | NaN |
| Actn4 | QGDAEFNR | -807 | -814 | 0 | 0.001743443 | 8 | NaN | NaN | NaN | 891.03217 | NaN | NaN |
| Actn4 | SDINNGWQHLEQAEK | -382 | -396 | 0 | 0.00383302 | 15 | NaN | NaN | NaN | 826.17474 | NaN | NaN |
| Pgk1 | VADKIQLINNMLDK | -217 | -230 | 1 | 0 | 14 | NaN | NaN | NaN | 5982.432 | 590.06946 | 8382.982 |
| Pgk1 | VLPGVDALSNV | -407 | -417 | 0 | 0 | 11 | NaN | NaN | NaN | 1499.073 | 3551.401 | NaN |
| Pgk1 | NNMEIGTSLYDEEGAK | -249 | -264 | 0 | 0 | 16 | NaN | NaN | NaN | 3955.962 | 2800.7188 | 2842.1292 |
| Pgk1 | ALESPERPFLAILGGAK | -200 | -216 | 1 | 0 | 17 | NaN | NaN | NaN | 610.1184 | 774.2929 | 820.1874 |
| Pgk1 | ALESPERPFLAILGGAK | -200 | -216 | 1 | 0 | 17 | NaN | NaN | NaN | 610.1184 | 774.2929 | 820.1874 |
| Pgk1 | DCVGPEVENACANPAAGTVILL | -98 | -119 | 0 | 0 | 22 | NaN | NaN | NaN | 243.28915 | 272.2797 | 389.8899 |
| Pgk1 | EPVAAELK | -79 | -86 | 0 | 0.007987481 | 8 | NaN | NaN | NaN | 352.7989 | 644.4059 | NaN |
| Pgk1 | GCITIIGGGDTATC | -366 | -379 | 0 | 0 | 14 | NaN | NaN | NaN | 1641.12 | NaN | NaN |
| Sfxn3 | YVYDSAFHPDTGEK | -72 | -85 | 0 | 0 | 14 | NaN | NaN | NaN | 439.5261 | 555.3655 | 442.99774 |
| Sfxn3 | AGVATPGLTEDQLWR | -55 | -69 | 0 | 0.00111006 | 15 | NaN | NaN | NaN | 1352.9388 | NaN | 1259.6525 |
| Sfxn3 | KTPTVVFW | -112 | -119 | 1 | 0.008075282 | 8 | NaN | NaN | NaN | 966.2348 | NaN | NaN |
| Prkcb | EHNVAYPK | -555 | -562 | 0 | 0.006260252 | 8 | NaN | NaN | NaN | 1562.7692 | NaN | NaN |
| Prkcb | ENIWDGVTTK | -490 | -499 | 0 | 0.006677519 | 10 | NaN | NaN | NaN | 582.6483 | NaN | NaN |
| Fto | GKDVLTPVSR | -87 | -96 | 1 | 0 | 10 | NaN | NaN | NaN | 1144.492 | 606.67993 | 634.9016 |
| Macroh2a1 | FVIHCNSPVWGADK | -269 | -282 | 0 | 0 | 14 | NaN | NaN | NaN | 845.8235 | 432.18893 | 1042.0494 |
| Metap1 | SCCTSVNEVICH | -192 | -203 | 0 | 0.000947459 | 12 | NaN | NaN | NaN | 450.82996 | NaN | 505.4516 |
| Metap1 | LQCPTCIK | -20 | -27 | 0 | 0.002972566 | 8 | NaN | NaN | NaN | 2816.9077 | NaN | NaN |
| Metap1 | AIDAVKPGVR | -259 | -268 | 1 | 0.001291103 | 10 | NaN | NaN | NaN | 770.1403 | NaN | NaN |
| Prmt5 | RVPLVAPEDLR | -154 | -164 | 1 | 0 | 11 | NaN | NaN | NaN | 1324.592 | 1434.4266 | 1240.0575 |
| Rack1 | DESHSEWVSCVR | -144 | -155 | 0 | 0 | 12 | NaN | NaN | NaN | 379.44934 | 3395.2522 | 3447.917 |
| Rack1 | LWDLNEGK | -218 | -225 | 0 | 0.003900536 | 8 | NaN | NaN | NaN | 7576.7266 | 7246.829 | 6380.0156 |
| Rack1 | SSNPIIVSCGWDK | -160 | -172 | 0 | 0 | 13 | NaN | NaN | NaN | 923.68896 | 983.8292 | 1078.2295 |
| Rack1 | FSPNSSNPIIVSCGWDK | -156 | -172 | 0 | 0 | 17 | NaN | NaN | NaN | 751.58295 | 1266.2821 | 828.3835 |
| Rack1 | AWSADGQTLFAGYTDNLVR | -290 | -308 | 0 | 0 | 19 | NaN | NaN | NaN | 427.3224 | 581.6663 | 547.7678 |
| Rack1 | DGQAMLWDLNEGK | -213 | -225 | 0 | 0 | 13 | NaN | NaN | NaN | 783.52026 | 587.8893 | NaN |
| Rack1 | YWLCAATGPSIK | -246 | -257 | 0 | 0 | 12 | NaN | NaN | NaN | 1277.978 | NaN | NaN |
| Rack1 | RDETNYGIPQR | -47 | -57 | 1 | 0.00142999 | 11 | NaN | NaN | NaN | 1153.179 | NaN | NaN |
| Tmed2 | HEQEYMEVR | -146 | -154 | 0 | 0 | 9 | NaN | NaN | NaN | 675.7539 | NaN | NaN |
| Pds5a | KFNQVLGDDEK | -557 | -567 | 1 | 0.004321969 | 11 | NaN | NaN | NaN | 1504.0521 | NaN | NaN |
| Pds5a | LIPILHQK | -719 | -726 | 0 | 0.009416377 | 8 | NaN | NaN | NaN | 400.84695 | NaN | NaN |
| Pds5a | KFFTQPEK | -1105 | -1112 | 1 | 0.00111006 | 8 | NaN | NaN | NaN | 261.80847 | NaN | NaN |
| Cebpz | SLIEEDEPEKK | -99 | -109 | 1 | 0 | 11 | NaN | NaN | NaN | 196.73032 | 2478.132 | 2134.043 |
| Cebpz | AIVSSGTLADR | -237 | -247 | 0 | 0.005698678 | 11 | NaN | NaN | NaN | 1057.3627 | NaN | NaN |
| Park7 | VTVAGLAGKDPVQCSR | -33 | -48 | 1 | 0 | 16 | NaN | NaN | NaN | 1556.2888 | 1575.629 | NaN |
| Park7 | AHEVGFGCK | -114 | -122 | 0 | 0.00201047 | 9 | NaN | NaN | NaN | 796.0404 | NaN | NaN |
| Prmt1 | TGFSTSPESPYTH | -299 | -311 | 0 | 0 | 13 | NaN | NaN | NaN | 1078.3774 | 298.31808 | NaN |
| Prmt1 | VEDLTFTSPF | -262 | -271 | 0 | 0.002099344 | 10 | NaN | NaN | NaN | 435.94562 | 647.56415 | NaN |
| Atp5f1b | LVLEVAQHLGESTVR | -95 | -109 | 0 | 0 | 15 | NaN | NaN | NaN | 5815.415 | 6934.3525 | 541.9008 |
| Atp5f1b | AHLDATTVLSR | -377 | -387 | 0 | 0 | 11 | NaN | NaN | NaN | 3628.616 | 4276.3623 | 4200.584 |
| Atp5f1b | IMDPNIVGNEHYDVARG | -407 | -423 | 1 | 0 | 17 | NaN | NaN | NaN | 1229.2223 | 382.48245 | 612.3175 |
| Atp5f1b | IMDPNIVGNEHYDVA | -407 | -421 | 0 | 0.00025015 | 15 | NaN | NaN | NaN | 787.50366 | 340.84476 | 402.9181 |
| Atp5f1b | EGNDLYHEMIESGVINLK | -242 | -259 | 0 | 0.00025015 | 18 | NaN | NaN | NaN | 566.123 | 478.70853 | 352.23187 |
| Atp5f1b | TVLIMELINNVAK | -213 | -225 | 0 | 0.00025015 | 13 | NaN | NaN | NaN | 422.48743 | NaN | 382.45895 |
| Atp5f1b | GGYSVFAGVGER | -228 | -239 | 0 | 0.000947459 | 12 | NaN | NaN | NaN | 3646.0327 | NaN | NaN |
| Atp5f1b | EGNDLYHE | -242 | -249 | 0 | 0.001392869 | 8 | NaN | NaN | NaN | 747.555 | NaN | NaN |
| Atp5f1b | TIAMDGTEGLVRGQ | -110 | -123 | 1 | 0.004791731 | 14 | NaN | NaN | NaN | 542.0389 | NaN | NaN |
| Atp5f1b | PLDSTSR | -400 | -406 | 0 | 0.002972566 | 7 | NaN | NaN | NaN | 376.9902 | NaN | NaN |
| Tgm2 | SVDSLTFGAVTGPDPSEEAGTK | -53 | -74 | 0 | 0.00025015 | 22 | NaN | NaN | NaN | 568.37085 | 782.7191 | 749.1635 |
| Tgm2 | LDPYSENSIPLR | -533 | -544 | 0 | 0 | 12 | NaN | NaN | NaN | 509.32315 | 1115.676 | 834.5641 |

| | | | | | | | | | | | | |
|---|---|---|---|---|---|---|---|---|---|---|---|---|
| Tgm2 | RNEFGELESNK | -317 | -327 | 1 | 0 | 11 | NaN | NaN | NaN | 447.81693 | 504.97186 | 543.7808 |
| Tgm2 | WCPADDVYLDSEEERR | -142 | -157 | 1 | 0 | 16 | NaN | NaN | NaN | 428.2343 | 389.86612 | 520.91547 |
| Gart | GFKDPLLASGTDGVGTK | -483 | -499 | 1 | 0 | 17 | NaN | NaN | NaN | 732.2044 | 912.5864 | 866.3174 |
| Gart | ISNAAVSVNDHSALAQ | -45 | -60 | 0 | 0 | 16 | NaN | NaN | NaN | 799.6148 | 958.0511 | 563.20605 |
| Gart | FGDPECQVILPL | -293 | -304 | 0 | 0 | 12 | NaN | NaN | NaN | 211.91455 | 308.2249 | 312.91058 |
| Gart | LLDGDEGPNTGGMGAYCPAPQVSK | -221 | -244 | 0 | 0.000471409 | 24 | NaN | NaN | NaN | 266.36334 | NaN | 231.71011 |
| Gart | YSSPAPGGCGDQTLGDLLLTPTR | -638 | -660 | 0 | 0 | 23 | NaN | NaN | NaN | 194.2193 | NaN | 262.22134 |
| Polr2a | HVIAEDVK | -301 | -308 | 0 | 0.00149795 | 8 | NaN | NaN | NaN | 8575.855 | NaN | NaN |
| Polr2a | EGGIKYPETTEGGRPK | -38 | -53 | 2 | 0.008680443 | 16 | NaN | NaN | NaN | 532.7827 | NaN | NaN |
| Ndufa2 | AHPNLPILIR | -47 | -56 | 0 | 0.00025015 | 10 | NaN | NaN | NaN | 2773.809 | NaN | NaN |
| Snrpg | TSGQQNNIGMVVIR | -50 | -63 | 0 | 0 | 14 | NaN | NaN | NaN | 1462.3932 | 1414.355 | 1450.3752 |
| Nmi | CHSVAVSPCIER | -231 | -242 | 0 | 0 | 12 | NaN | NaN | NaN | 1570.5066 | NaN | 1474.7621 |
| Nmi | NGGGEVESVDYDRK | -187 | -200 | 1 | 0 | 14 | NaN | NaN | NaN | 275.79788 | NaN | NaN |
| Qars1 | NFPAPKPLDIR | -581 | -591 | 1 | 0 | 11 | NaN | NaN | NaN | 641.7305 | 571.1476 | 607.8302 |
| Qars1 | AHQILGSTIDK | -40 | -50 | 0 | 0 | 11 | NaN | NaN | NaN | 1544.7235 | NaN | 1354.4294 |
| Qars1 | GFHQVPFAST | -602 | -611 | 0 | 0.001055699 | 10 | NaN | NaN | NaN | 179.46419 | 4108.5894 | NaN |
| Cpsf3 | RLYEALTPVH | -675 | -684 | 1 | 0.000907206 | 10 | NaN | NaN | NaN | 2127.0605 | 785.8787 | 1581.6543 |
| Cpsf3 | AHTDYQQTSEFIR | -395 | -407 | 0 | 0.001463897 | 13 | NaN | NaN | NaN | 1329.2021 | 1484.092 | 368.7501 |
| Dock8 | APDFNEEHLR | -1563 | -1572 | 0 | 0 | 10 | NaN | NaN | NaN | 734.84686 | 456.5629 | 1229.0977 |
| Dock8 | ISENFHCDLNSDQFK | -300 | -314 | 0 | 0.000989247 | 15 | NaN | NaN | NaN | 457.39294 | 478.72986 | NaN |
| Dock8 | TFESETLECSEPDTQTGPR | -162 | -180 | 0 | 0 | 19 | NaN | NaN | NaN | 278.2937 | 290.3261 | 266.33197 |
| Dock8 | LVIPILEAH | -1755 | -1763 | 0 | 0.001636916 | 9 | NaN | NaN | NaN | 120.55079 | 650.8702 | 617.73694 |
| Dock8 | LNILPTLISMR | -1084 | -1094 | 0 | 0.00051306 | 11 | NaN | NaN | NaN | 3937.471 | NaN | 4001.906 |
| Dock8 | IPELYKPIFR | -2066 | -2075 | 1 | 0 | 10 | NaN | NaN | NaN | 248.2857 | NaN | 314.9171 |
| Dock8 | GPLTSCDFDLR | -193 | -203 | 0 | 0 | 11 | NaN | NaN | NaN | 534.31287 | NaN | NaN |
| Mdh1 | NVIIWGNHSSTQYPDVNHAK | -180 | -199 | 0 | 0 | 20 | NaN | NaN | NaN | 1238.3251 | 1586.8531 | 1693.8046 |
| Mdh1 | ETAFEFLSSA | -325 | -334 | 0 | 0 | 10 | NaN | NaN | NaN | 470.78186 | 94.63476 | 756.7527 |
| Mdh1 | DIWFGTPEGEFVSM | -256 | -269 | 0 | 0.00272025 | 14 | NaN | NaN | NaN | 208.28958 | 292.20703 | 331.89667 |
| Mdh1 | DIWFGTPEGEFVSMG | -256 | -270 | 0 | 0.009955823 | 15 | NaN | NaN | NaN | 1606.6617 | NaN | NaN |
| Nvl | AIAGELDLPILK | -316 | -327 | 0 | 0 | 12 | NaN | NaN | NaN | 591.95844 | 533.2836 | 633.9034 |
| Mthfd1 | QVGDRDDSNLYINVK | -42 | -56 | 1 | 0.004439092 | 15 | NaN | NaN | NaN | 1562.5729 | 1479.5446 | 1389.651 |
| Mthfd1 | SQAPSSFQLLYDLK | -806 | -819 | 0 | 0 | 14 | NaN | NaN | NaN | 230.85095 | 451.41342 | NaN |
| Mthfd1 | TEEVELYGETK | -342 | -352 | 0 | 0 | 11 | NaN | NaN | NaN | 2970.0042 | NaN | NaN |
| Mthfd1 | DVDGLTSVSAGKL | -123 | -135 | 1 | 0.001811524 | 13 | NaN | NaN | NaN | 1208.9244 | NaN | NaN |
| Mthfd1 | ALGAHLR | -396 | -402 | 0 | 0.007224385 | 7 | NaN | NaN | NaN | 480.29797 | NaN | NaN |
| St13 | AIEINPDSAQPY | -173 | -184 | 0 | 0 | 12 | NaN | NaN | NaN | 576.21643 | 184.46472 | 216.47203 |
| St13 | LLGHWEEAAHDL | -193 | -204 | 0 | 0.000471409 | 12 | NaN | NaN | NaN | 2139.856 | 1782.4929 | 1904.2631 |
| St13 | KGAAIEALNDGELQK | -117 | -131 | 1 | 0.00204437 | 15 | NaN | NaN | NaN | 770.1694 | NaN | NaN |
| St13 | GAAIEALNDGELQK | -118 | -131 | 0 | 0 | 14 | NaN | NaN | NaN | 352.83795 | NaN | NaN |
| Rbm14 | AIEALHGHELRPGR | -51 | -64 | 1 | 0 | 14 | NaN | NaN | NaN | 1322.8551 | 349.24445 | 740.20123 |
| Rbm14 | IFVGNVDGADTTPEELAAL | -3 | -21 | 0 | 0.003166358 | 19 | NaN | NaN | NaN | 100.246475 | 241.66682 | NaN |
| Rbm14 | FDYQQAFGNSTGGFDGQAR | -185 | -203 | 0 | 0.001291103 | 19 | NaN | NaN | NaN | 362.20352 | NaN | NaN |
| Cyfip2 | DCPGTAEEYER | -426 | -436 | 0 | 0.002168954 | 11 | NaN | NaN | NaN | 711.782 | 635.54333 | 977.09076 |
| Cyfip2 | AIEDFHK | -590 | -596 | 0 | 0.00051306 | 7 | NaN | NaN | NaN | 210.91336 | 596.9544 | 596.8687 |
| Larp7 | GPQFVTGVIVK | -436 | -446 | 0 | 0 | 11 | NaN | NaN | NaN | 397.7494 | NaN | NaN |
| Fgfr1op2 | AIEIDEQQGCK | -189 | -199 | 0 | 0.001536401 | 11 | NaN | NaN | NaN | 191.87039 | NaN | NaN |
| Oxsr1 | SGVLDEPTIATILR | -115 | -128 | 0 | 0 | 14 | NaN | NaN | NaN | 448.57062 | 484.23825 | 558.7836 |
| Hnrnpab | ITFKEEDPVKK | -205 | -215 | 2 | 0 | 11 | NaN | NaN | NaN | 1594.4109 | 2939.524 | 3094.0305 |
| Pak2 | AVVTEEDDDDEDAAPPVIAPRPDHTK | -166 | -191 | 1 | 0 | 26 | NaN | NaN | NaN | 530.5642 | 533.0191 | 531.4568 |
| Ripk1 | AIFDNTTSLTDEHLNPIR | -559 | -576 | 0 | 0 | 18 | NaN | NaN | NaN | 644.5022 | 603.50464 | 705.5509 |
| Cblb | ALEIAQNNVEVAR | -952 | -964 | 0 | 0 | 13 | NaN | NaN | NaN | 1337.1971 | 1495.7083 | 703.16003 |
| Cblb | AIFPNGQFQGDNFR | -159 | -172 | 0 | 0.006735683 | 14 | NaN | NaN | NaN | 640.44604 | NaN | NaN |
| Cblb | GIIDAIQDAVGPPK | -24 | -37 | 0 | 0.00930008 | 14 | NaN | NaN | NaN | 189.79901 | NaN | NaN |
| Dlat | KILVPEGTR | (146)(273) | (154)(281) | 1 | 0.001291103 | 9 | NaN | NaN | NaN | 911.3611 | NaN | NaN |
| Ppm1g | SYDHKPEDEVELAR | -360 | -373 | 1 | 0.00025015 | 14 | NaN | NaN | NaN | 2062.9868 | NaN | 1952.9805 |
| Ppm1g | ATMTIEELLTR | -149 | -159 | 0 | 0.005668207 | 11 | NaN | NaN | NaN | 599.78015 | NaN | NaN |
| Sgta | AIGIDPGYSK | -152 | -161 | 0 | 0.006568856 | 10 | NaN | NaN | NaN | 430.37247 | NaN | 1283.6787 |
| Asns | TTAAHGLELR | -408 | -417 | 0 | 0 | 10 | NaN | NaN | NaN | 857.11316 | 1118.567 | 982.3278 |
| Kpna2 | NLTWTLSNLCR | -228 | -238 | 0 | 0 | 11 | NaN | NaN | NaN | 783.7315 | 632.4716 | 786.7003 |
| Kpna2 | IVTGTDEQTQK | -320 | -330 | 0 | 0 | 11 | NaN | NaN | NaN | 604.0446 | 550.9961 | 461.8339 |
| Kpna2 | GINSNNLESQLQAT | -84 | -97 | 0 | 0.007905138 | 14 | NaN | NaN | NaN | 659.75354 | NaN | NaN |
| Sf3b1 | LDDLVRPYVHK | -563 | -573 | 1 | 0 | 11 | NaN | NaN | NaN | 5856.229 | 6275.828 | 6062.5195 |
| Sf3b1 | LDDLVRPYVHK | -563 | -573 | 1 | 0 | 11 | NaN | NaN | NaN | 5856.229 | 6275.828 | 6062.5195 |
| Sf3b1 | LVDVDESTLSPEEQKER | -479 | -495 | 1 | 0 | 17 | NaN | NaN | NaN | 3370.929 | 3674.2888 | 3442.0527 |
| Sf3b1 | ELEDDDDDYSSSTSLLGQK | -62 | -80 | 0 | 0 | 19 | NaN | NaN | NaN | 471.66656 | 615.54364 | 592.2682 |
| Sf3b1 | GGDSIGETPTPGASK | -319 | -333 | 0 | 0 | 15 | NaN | NaN | NaN | 5171.3105 | 5282.0166 | NaN |
| Sf3b1 | IADREDEYK | -112 | -120 | 1 | 0.000712251 | 9 | NaN | NaN | NaN | 730.7931 | 966.8409 | NaN |
| Sf3b1 | TADQTPGATPK | -203 | -213 | 0 | 0.001055699 | 11 | NaN | NaN | NaN | 582.7388 | 610.2243 | NaN |
| Sf3b1 | VEIIEHGLVDEQQK | -687 | -700 | 0 | 0.000627943 | 14 | NaN | NaN | NaN | 1448.747 | NaN | NaN |
| Sf3b1 | LNDIPQSTEQYDPFAEHRPPK | -91 | -111 | 1 | 0.000230707 | 21 | NaN | NaN | NaN | 717.2736 | NaN | NaN |
| Psma5 | ITSPLMEPSSIEK | -54 | -66 | 0 | 0 | 13 | NaN | NaN | NaN | 590.785 | 1135.3002 | 1059.6829 |
| Psma5 | QVEYAIEAIK | -23 | -32 | 0 | 0 | 10 | NaN | NaN | NaN | 288.67395 | 513.7467 | 419.5122 |
| Psma5 | ALQFGEEDADPGAMSR | -120 | -135 | 0 | 0.00111006 | 16 | NaN | NaN | NaN | 555.7852 | NaN | NaN |
| Impdh2 | GAAIGTHEDDKYR | -247 | -259 | 1 | 0 | 13 | NaN | NaN | NaN | 377.71124 | 2056.6958 | 809.7072 |
| Impdh2 | RFGVPVIADGGIQNVGHIA | -356 | -374 | 1 | 0.00111006 | 19 | NaN | NaN | NaN | 414.3256 | NaN | 508.27478 |
| Impdh2 | TSSAQVEGGVH | -494 | -504 | 0 | 0 | 11 | NaN | NaN | NaN | 304.68057 | 446.39865 | NaN |
| Impdh2 | TADQVDLTSALTK | -45 | -57 | 0 | 0.001392869 | 13 | NaN | NaN | NaN | 1744.3997 | NaN | NaN |
| Sf3b3 | LETDEDMVTEIR | -315 | -326 | 0 | 0.00025015 | 12 | NaN | NaN | NaN | 1037.8489 | 1058.6846 | 1124.5248 |

| | | | | | | | | | | | | |
|---|---|---|---|---|---|---|---|---|---|---|---|---|
| Sf3b3 | ILELLRPDPNTGK | -37 | -49 | 1 | 0.00266297 | 13 | NaN | NaN | NaN | 602.834 | 642.3958 | 702.2339 |
| Sf3b3 | ASEQCPEGIVAISTNTLR | -746 | -763 | 0 | 0.00201047 | 18 | NaN | NaN | NaN | 672.7723 | 1269.1053 | NaN |
| Sf3b3 | FHLTPLSYETL | -729 | -739 | 0 | 0.00201047 | 11 | NaN | NaN | NaN | 572.4309 | NaN | 652.4622 |
| Sf3b3 | IHQETFGK | -102 | -109 | 0 | 0.00025015 | 8 | NaN | NaN | NaN | 6296.004 | NaN | NaN |
| Prrc2c | VAAPAVLNDISK | -1948 | -1959 | 0 | 0 | 12 | NaN | NaN | NaN | 1491.6521 | 895.7353 | NaN |
| Prrc2c | AIHNFPAVQHQELAK | -2441 | -2455 | 0 | 0.000431655 | 15 | NaN | NaN | NaN | 1836.0597 | NaN | NaN |
| Prrc2c | TEHVPSGPLR | -1201 | -1210 | 0 | 0.00111006 | 10 | NaN | NaN | NaN | 1261.351 | NaN | NaN |
| Prrc2c | VIEDPQSNLNDDGFTEVVSK | -1641 | -1660 | 0 | 0.001776473 | 20 | NaN | NaN | NaN | 402.6566 | NaN | NaN |
| Ncapd2 | GNAIYNLLPDIISR | -1150 | -1163 | 0 | 0 | 14 | NaN | NaN | NaN | 464.1006 | 426.7946 | 419.9463 |
| Ncapd2 | LGFDWEEER | -158 | -166 | 0 | 0 | 9 | NaN | NaN | NaN | 507.29385 | 653.1506 | NaN |
| Ncapd2 | EATSHFQESEPF | -536 | -547 | 0 | 0.000471409 | 12 | NaN | NaN | NaN | 504.99158 | NaN | 526.98444 |
| Rps7 | AQQNNVEHKVE | -161 | -171 | 1 | 0.00111006 | 11 | NaN | NaN | NaN | 262.48126 | NaN | NaN |
| Cstf3 | IDYLSHLNEDNNTR | -451 | -464 | 0 | 0.00025015 | 14 | NaN | NaN | NaN | 506.76706 | 1118.122 | 1150.4613 |
| Msto1 | EDEESPGELCPDVLYR | -37 | -52 | 0 | 0 | 16 | NaN | NaN | NaN | 2225.4788 | 2784.3154 | 2740.8696 |
| Rcc2 | DVACGANHTLVLDSQK | -332 | -347 | 0 | 0.001055699 | 16 | NaN | NaN | NaN | 1157.462 | 1206.0659 | 1327.0056 |
| Rcc2 | VFSWGFGGYGR | -349 | -359 | 0 | 0.00122419 | 11 | NaN | NaN | NaN | 885.7166 | 941.7279 | NaN |
| Rcc2 | IITEPEHTK | -82 | -90 | 0 | 0.000471409 | 9 | NaN | NaN | NaN | 887.4628 | NaN | NaN |
| Ppp4c | AILELDEHLQK | -271 | -281 | 0 | 0 | 11 | NaN | NaN | NaN | 1751.5956 | 433.74255 | 850.1531 |
| Lta4h | FSYQSVTTDDWK | -415 | -426 | 0 | 0 | 12 | NaN | NaN | NaN | 1132.1871 | 688.8909 | 1139.9891 |
| Lta4h | SANEFSETESMLK | -231 | -243 | 0 | 0 | 13 | NaN | NaN | NaN | 181.25175 | 261.28165 | 321.97717 |
| Lta4h | SANEFSETESMLK | -231 | -243 | 0 | 0 | 13 | NaN | NaN | NaN | 181.25175 | 261.28165 | 321.97717 |
| Lta4h | TVQSQEENLR | -42 | -51 | 0 | 0.00025015 | 10 | NaN | NaN | NaN | 344.50397 | 292.38623 | 576.82605 |
| Lta4h | LTYTAEVSVPK | -155 | -165 | 0 | 0.000550307 | 11 | NaN | NaN | NaN | 1413.708 | NaN | 1359.8573 |
| Lta4h | GSPMEISLPIALSK | -84 | -97 | 0 | 0 | 14 | NaN | NaN | NaN | 88.09084 | 756.4763 | NaN |
| Lta4h | GSPMEISLPIALSK | -84 | -97 | 0 | 0 | 14 | NaN | NaN | NaN | 88.09084 | 756.4763 | NaN |
| Lta4h | TWDHFWLNEGHTV | -311 | -323 | 0 | 0.00025015 | 13 | NaN | NaN | NaN | 158.59576 | 6307.643 | NaN |
| Lta4h | SKWEEAIPLALK | -546 | -557 | 1 | 0 | 12 | NaN | NaN | NaN | 467.8035 | NaN | NaN |
| Tubb5 | VTFIGNSTAIQELFK | -365 | -379 | 0 | 0 | 15 | NaN | NaN | NaN | 599.8774 | 611.6481 | 531.51355 |
| Tubb5 | AVTFIGNSTAIQELFK | -364 | -379 | 0 | 0 | 16 | NaN | NaN | NaN | 482.00278 | 611.44415 | 410.12277 |
| Tubb5 | TVPELTQQVFDAK | -285 | -297 | 0 | 0 | 13 | NaN | NaN | NaN | 562.0604 | 138.90297 | NaN |
| Pds5b | TTILHSDGDLTEQGK | -855 | -869 | 0 | 0 | 15 | NaN | NaN | NaN | 451.2406 | 240.2931 | 394.19968 |
| Iqgap1 | RDHINDIIK | -833 | -841 | 1 | 0.00025015 | 9 | NaN | NaN | NaN | 1456.0596 | 742.9092 | 787.5384 |
| Iqgap1 | ITLQDVVSH | -925 | -933 | 0 | 0.00025015 | 9 | NaN | NaN | NaN | 1066.1344 | 1190.37 | 1204.4725 |
| Iqgap1 | IIGNLLYYR | -1186 | -1194 | 0 | 0 | 9 | NaN | NaN | NaN | 824.1277 | 740.9946 | 451.21127 |
| Khsrp | MILIQDGSQNTNVDKPLR | -268 | -285 | 1 | 0.000230707 | 18 | NaN | NaN | NaN | 24536.943 | 27666.062 | 26896.59 |
| Khsrp | SVSLTGAPESVQK | -192 | -204 | 0 | 0 | 13 | NaN | NaN | NaN | 2990.9429 | 2696.1907 | 2090.2778 |
| Khsrp | IQDGSQNTNVDKPLR | -271 | -285 | 1 | 0.004224523 | 15 | NaN | NaN | NaN | 363.23624 | NaN | 439.3822 |
| Khsrp | QAQVATGGGPGAPPGSQPDYSAA | -656 | -678 | 0 | 0.001392869 | 23 | NaN | NaN | NaN | 169.59572 | NaN | 300.81027 |
| Khsrp | SGPPGPPGAPGMPPGGR | -396 | -412 | 0 | 0.00272025 | 17 | NaN | NaN | NaN | 618.263 | NaN | NaN |
| Nmt1 | GPVEPDKDNIR | -136 | -146 | 1 | 0 | 11 | NaN | NaN | NaN | 389.5888 | 903.0428 | 914.324 |
| Nmt1 | RLPETPK | -328 | -334 | 1 | 0.009745776 | 7 | NaN | NaN | NaN | 1036.579 | 1345.3975 | NaN |
| Rfk | AIQGDIEEAKK | -118 | -128 | 1 | 0.00025015 | 11 | NaN | NaN | NaN | 495.37277 | 1764.0405 | 1427.6278 |
| H2ac20;Hist2h | NDEELNKLLGK | (90);(90) | (100);(100) | 1 | 0.006820063 | 11 | NaN | NaN | NaN | 846.68713 | NaN | NaN |
| Eml4 | AISEDHVASVK | -53 | -63 | 0 | 0 | 11 | NaN | NaN | NaN | 2137.586 | 1292.5303 | 2378.8518 |
| Wdr1 | KVFASLPQVER | -7 | -17 | 1 | 0 | 11 | NaN | NaN | NaN | 401.8941 | 268.1393 | 2240.122 |
| Wdr1 | IKDIAWTEDSKR | -105 | -116 | 2 | 0 | 12 | NaN | NaN | NaN | 717.76715 | 852.66455 | 841.3138 |
| Wdr1 | GPVTDVAYSHDGAFLA | -490 | -505 | 0 | 0 | 16 | NaN | NaN | NaN | 902.59656 | 914.70776 | 1117.4924 |
| Wdr1 | WSPDSTHLLSASGDK | -242 | -256 | 0 | 0 | 15 | NaN | NaN | NaN | 887.0454 | 381.27405 | 1122.3787 |
| Wdr1 | WSPDSTHLLSASGDK | -242 | -256 | 0 | 0 | 15 | NaN | NaN | NaN | 887.0454 | 381.27405 | 1122.3787 |
| Wdr1 | WDTGSSVGEITGHNK | -133 | -147 | 0 | 0 | 15 | NaN | NaN | NaN | 879.43207 | 630.9068 | 701.27594 |
| Wdr1 | AISWSPDSTHLLSASGDK | -239 | -256 | 0 | 0 | 18 | NaN | NaN | NaN | 503.63943 | 203.70717 | 513.9476 |
| Wdr1 | AISWSPDSTHLLSASGDK | -239 | -256 | 0 | 0 | 18 | NaN | NaN | NaN | 503.63943 | 203.70717 | 513.9476 |
| Wdr1 | NIDNPAIADIYTEHAHQVVVA | -44 | -64 | 0 | 0 | 21 | NaN | NaN | NaN | 85.05125 | 185.582 | 305.91684 |
| Ppa1 | GQYISPFHDVPIYADK | -26 | -41 | 0 | 0 | 16 | NaN | NaN | NaN | 1959.0608 | 2313.7332 | 2327.951 |
| Ppa1 | NVDDPDAANYK | -160 | -170 | 0 | 0 | 11 | NaN | NaN | NaN | 802.47644 | NaN | NaN |
| Ddx39b | EDDEVETAAGADGTEAPAK | -14 | -32 | 0 | 0.001636916 | 19 | NaN | NaN | NaN | 8395.879 | NaN | 8726.367 |
| Rpl14 | VAIVDVIDQNR | -25 | -35 | 0 | 0 | 11 | NaN | NaN | NaN | 3711.9004 | 4725.6064 | 3922.9646 |
| Bin2 | EKEGAGSSGPEEPR | -457 | -470 | 1 | 0.00025015 | 14 | NaN | NaN | NaN | 511.1926 | 611.3703 | 625.55133 |
| Psmc1 | IETLDPALIRPG | -334 | -345 | 1 | 0 | 12 | NaN | NaN | NaN | 1620.1165 | 2253.3257 | 2041.0083 |
| Psmc1 | SFVDKDLLEPGCSVLLNHK | -137 | -155 | 1 | 0 | 19 | NaN | NaN | NaN | 521.5497 | 213.09752 | 593.7401 |
| Psmc1 | KYLGDGPK | -258 | -265 | 1 | 0 | 8 | NaN | NaN | NaN | 1617.4045 | NaN | 1105.2457 |
| Psmc1 | ESVELPLTHPE | -199 | -209 | 0 | 0.000550307 | 11 | NaN | NaN | NaN | 3267.7107 | NaN | NaN |
| Psmc1 | EEHAPSIVFIDEIDAIGTK | -275 | -293 | 0 | 0.00025015 | 19 | NaN | NaN | NaN | 259.10693 | NaN | NaN |
| Ezr | RKEDEVEEW | -437 | -445 | 2 | 0.001291103 | 9 | NaN | NaN | NaN | 314.82474 | 1188.9684 | 1166.3586 |
| Ezr | ALQLEEERR | -372 | -380 | 1 | 0.001868731 | 9 | NaN | NaN | NaN | 920.5958 | NaN | NaN |
| Ezr | SNELSQAR | -535 | -542 | 0 | 0.00830926 | 8 | NaN | NaN | NaN | 793.9187 | NaN | NaN |
| Ezr | THNDIIHNE | -548 | -556 | 0 | 0.004983786 | 9 | NaN | NaN | NaN | 383.96222 | NaN | NaN |
| Prkaca | AKEDFLK | -23 | -29 | 1 | 0.006568856 | 7 | NaN | NaN | NaN | 645.81445 | NaN | NaN |
| Prkacb | ILQAVEFPFLVR | -95 | -106 | 0 | 0 | 12 | NaN | NaN | NaN | 282.96646 | 295.90732 | 429.49362 |
| Gng2 | AKEDPLLTPVPASENPFR | -45 | -62 | 1 | 0 | 18 | NaN | NaN | NaN | 437.9915 | 446.8087 | 559.0702 |
| Tpr | AKEELEAEKR | -190 | -199 | 2 | 0.000431655 | 10 | NaN | NaN | NaN | 301.87994 | 429.48962 | NaN |
| Tpr | TDELLALGR | -279 | -287 | 0 | 0.00111006 | 9 | NaN | NaN | NaN | 204.8578 | 715.609 | NaN |
| Tpr | LQNEQLEK | -766 | -773 | 0 | 0.008222027 | 8 | NaN | NaN | NaN | 1195.577 | NaN | NaN |
| Tpr | APPQELGPPVQR | -2157 | -2168 | 0 | 0.001291103 | 12 | NaN | NaN | NaN | 746.64185 | NaN | NaN |
| Eif3k | VVGITYQHIDR | -145 | -155 | 0 | 0 | 11 | NaN | NaN | NaN | 2939.3257 | 3155.7837 | 2845.5981 |
| Eif3k | AKENAYDLEANLAVLK | -37 | -52 | 1 | 0.00078661 | 16 | NaN | NaN | NaN | 649.24274 | 726.9728 | 881.88965 |
| Eif3k | ALTNLPHTDFTL | -73 | -84 | 0 | 0.007224385 | 12 | NaN | NaN | NaN | 1361.9723 | NaN | NaN |
| Eif3k | KFICHVVGITY | -140 | -150 | 1 | 0.001190916 | 11 | NaN | NaN | NaN | 575.8 | NaN | NaN |

| | | | | | | | | | | | | |
|---|---|---|---|---|---|---|---|---|---|---|---|---|
| Hspa5 | ELEEIVQPIIS | -623 | -633 | 0 | 0 | 11 | NaN | NaN | NaN | 1108.9427 | 1467.3689 | 2389.192 |
| Hspa5 | ELEEIVQPIIS | -623 | -633 | 0 | 0 | 11 | NaN | NaN | NaN | 1108.9427 | 1467.3689 | 2389.192 |
| Hspa5 | QLTSNPENTVFDAK | -84 | -97 | 0 | 0 | 14 | NaN | NaN | NaN | 997.2314 | 1454.1027 | 1351.3599 |
| Hspa5 | KLYGSGGPPPTGEEDTSEKDEL | -634 | -655 | 2 | 0 | 22 | NaN | NaN | NaN | 863.29034 | 709.78284 | 673.53906 |
| Hspa5 | DNHLLGTFDLTGIPPAPR | -476 | -493 | 0 | 0 | 18 | NaN | NaN | NaN | 822.51685 | 925.38245 | 1038.532 |
| Hspa5 | DNHLLGTFDLTGIPPAPR | -476 | -493 | 0 | 0 | 18 | NaN | NaN | NaN | 822.51685 | 925.38245 | 1038.532 |
| Hspa5 | KFEELNMDLFR | -327 | -337 | 1 | 0 | 11 | NaN | NaN | NaN | 771.71954 | 808.355 | 780.22076 |
| Hspa5 | GSGGPPPTGEEDTSEKDEL | -637 | -655 | 1 | 0.00025015 | 19 | NaN | NaN | NaN | 423.21475 | 394.76392 | 263.2403 |
| Hspa5 | LTSNPENTVFDAK | -85 | -97 | 0 | 0.00025015 | 13 | NaN | NaN | NaN | 171.04654 | 3467.119 | 3425.7712 |
| Hspa5 | KKELEEIVQPIIS | -621 | -633 | 2 | 0.00111006 | 13 | NaN | NaN | NaN | 721.9037 | 1035.5001 | NaN |
| Hspa5 | TVPAYFNDAQR | -172 | -182 | 0 | 0.002068213 | 11 | NaN | NaN | NaN | 181.50163 | NaN | NaN |
| Top2a | AKGEEQDFPVDLEDTIAPR | -1487 | -1505 | 1 | 0 | 19 | NaN | NaN | NaN | 607.7381 | 623.7884 | 533.3408 |
| Top2a | AKGEEQDFPVDLEDTIAPR | -1487 | -1505 | 1 | 0 | 19 | NaN | NaN | NaN | 607.7381 | 623.7884 | 533.3408 |
| Top2a | DKVDETGNSLK | -276 | -286 | 1 | 0.00025015 | 11 | NaN | NaN | NaN | 818.17914 | NaN | NaN |
| Top2a | VNALIENPTFDSQTK | -363 | -377 | 0 | 0.00025015 | 15 | NaN | NaN | NaN | 620.7103 | NaN | NaN |
| Tmem43 | TATAPFVQIGR | -179 | -189 | 0 | 0.00713295 | 11 | NaN | NaN | NaN | 3745.714 | 1399.458 | NaN |
| Tmem43 | AKLEDPHVDIIR | -209 | -220 | 1 | 0 | 12 | NaN | NaN | NaN | 2114.586 | 2130.3945 | NaN |
| Tmem43 | LVHIIGALR | -88 | -96 | 0 | 0.002910424 | 9 | NaN | NaN | NaN | 573.00085 | NaN | NaN |
| Tmem43 | LVHIIGAL | -88 | -95 | 0 | 0.007987481 | 8 | NaN | NaN | NaN | 284.09558 | NaN | NaN |
| Grpel1 | TLRPALVGVVK | -205 | -215 | 1 | 0.002697088 | 11 | NaN | NaN | NaN | 2731.25 | NaN | NaN |
| Mapre1 | ENEGENDPVLQR | -230 | -241 | 0 | 0 | 12 | NaN | NaN | NaN | 955.417 | NaN | 1278.1576 |
| Ncl | SFNITEDELK | -403 | -412 | 0 | 0 | 10 | NaN | NaN | NaN | 3827.968 | 2549.4958 | 3313.0127 |
| Ncl | EVFEDAMEIR | -413 | -422 | 0 | 0 | 10 | NaN | NaN | NaN | 3042.1167 | 1080.2225 | 499.56125 |
| Ncl | IEFASFEDAK | -529 | -538 | 0 | 0 | 10 | NaN | NaN | NaN | 1814.0707 | 1965.127 | 2014.1958 |
| Ncl | KEAMEDGEIDGNK | -624 | -636 | 1 | 0 | 13 | NaN | NaN | NaN | 536.64264 | 633.5802 | 574.8558 |
| Ncl | KEAMEDGEIDGNK | -624 | -636 | 1 | 0 | 13 | NaN | NaN | NaN | 536.64264 | 633.5802 | 574.8558 |
| Ncl | KALVPTPGK | -116 | -124 | 1 | 0.002068213 | 9 | NaN | NaN | NaN | 1877.8993 | 1736.0527 | 1857.5249 |
| Ncl | PTTPFNLFIGNLNPNK | -305 | -320 | 0 | 0.00078661 | 16 | NaN | NaN | NaN | 282.5324 | 282.5433 | 184.76903 |
| Ncl | GGGGDFKPQ | -692 | -700 | 1 | 0.00025015 | 9 | NaN | NaN | NaN | 725.40326 | NaN | NaN |
| Cs | AKNPWPNVDAHSGVLL | -394 | -409 | 1 | 0 | 16 | NaN | NaN | NaN | 422.75644 | 616.74054 | 709.09375 |
| Cs | GYTDPQFTELMR | -245 | -256 | 0 | 0 | 12 | NaN | NaN | NaN | 302.86432 | 1364.1876 | 1325.8073 |
| Dlst | VEGGTPLFTL | -135 | -144 | 0 | 0.00025015 | 10 | NaN | NaN | NaN | 503.4367 | NaN | NaN |
| Rrp9 | EEEEEEELEETAQEK | -64 | -78 | 0 | 0 | 15 | NaN | NaN | NaN | 773.1979 | 688.4271 | 245.21626 |
| Hmgn2 | DANNPAENGDAK | -65 | -76 | 0 | 0.001392869 | 12 | NaN | NaN | NaN | 739.75446 | 589.968 | NaN |
| Fabp5 | QDGALVQHQQWDGK | -90 | -103 | 0 | 0 | 14 | NaN | NaN | NaN | 1135.2151 | NaN | NaN |
| Pygb | RVEDVEALDQK | -715 | -725 | 1 | 0.000230707 | 11 | NaN | NaN | NaN | 944.42334 | 1384.563 | 1126.3948 |
| Pygb | IGEGFLTDLSQLK | -508 | -520 | 0 | 0 | 13 | NaN | NaN | NaN | 790.3677 | 743.60913 | NaN |
| Pdia4 | DVQGSTEASAIKDYVVK | -368 | -384 | 1 | 0 | 17 | NaN | NaN | NaN | 4367.89 | 834.6358 | 5142.6597 |
| Pdia4 | AKVDATEQTDLAK | -226 | -238 | 1 | 0 | 13 | NaN | NaN | NaN | 1727.718 | 1874.9672 | 1457.3284 |
| Pdia4 | KFDVSGYPTIK | -124 | -134 | 1 | 0 | 11 | NaN | NaN | NaN | 905.3697 | 1257.6532 | 918.154 |
| Pdia3 | LKSEPIPESNEGPVK | -365 | -379 | 1 | 0.000230707 | 15 | NaN | NaN | NaN | 992.3589 | 803.0652 | 515.7283 |
| Pdia3 | FSDGHSEFLK | -164 | -173 | 0 | 0 | 10 | NaN | NaN | NaN | 972.4076 | 895.12775 | 656.358 |
| Pdia3 | DGIVSHLK | -122 | -129 | 0 | 0.000230707 | 8 | NaN | NaN | NaN | 775.3123 | 478.9669 | 466.12637 |
| Pdia3 | ELNDFISYLQR | -472 | -482 | 0 | 0 | 11 | NaN | NaN | NaN | 607.8783 | 565.2483 | 371.65555 |
| Pdia3 | FISDKDASVVGFF | -148 | -160 | 1 | 0 | 13 | NaN | NaN | NaN | 592.26306 | 451.94885 | 307.37598 |
| Pdia3 | AKVDCTANTNTCNK | -81 | -94 | 1 | 0.00025015 | 14 | NaN | NaN | NaN | 557.14844 | 613.6779 | 801.9112 |
| Pdia3 | SHELSDFGLESTTGEVPVVAIR | -308 | -329 | 0 | 0 | 22 | NaN | NaN | NaN | 469.54074 | 347.91473 | 340.43613 |
| Pdia3 | SHELSDFGLESTTGEVPVVAIR | -308 | -329 | 0 | 0 | 22 | NaN | NaN | NaN | 469.54074 | 347.91473 | 340.43613 |
| Pdia3 | QAGPASVPLR | -131 | -140 | 0 | 0 | 10 | NaN | NaN | NaN | 1350.6265 | 1116.6626 | NaN |
| Pdia3 | KFLDAGHKL | -289 | -297 | 2 | 0.001904399 | 9 | NaN | NaN | NaN | 629.0341 | NaN | 2433.3538 |
| Pdia3 | YYDVDYEK | -264 | -271 | 0 | 0.00078661 | 8 | NaN | NaN | NaN | 214.06592 | NaN | 199.19827 |
| Pdia3 | KFIQDSIFGL | -234 | -243 | 1 | 0.001610812 | 10 | NaN | NaN | NaN | 165.08954 | NaN | 253.35355 |
| Pdia3 | ISDKDASVVGFFR | -149 | -161 | 1 | 0.000431655 | 13 | NaN | NaN | NaN | 714.6735 | NaN | NaN |
| Rpl10 | NADEFEDMVAEK | -177 | -188 | 0 | 0 | 12 | NaN | NaN | NaN | 1318.223 | 1417.4048 | 1483.8276 |
| Serbp1 | KPANDITSQ | -345 | -353 | 1 | 0.002168954 | 9 | NaN | NaN | NaN | 1081.7588 | 975.0955 | 940.76855 |
| Serbp1 | SEEAHAEDSVMDHHFR | -329 | -344 | 0 | 0 | 16 | NaN | NaN | NaN | 504.71872 | 724.22003 | 710.7787 |
| Serbp1 | SEEAHAEDSVMDHHFR | -329 | -344 | 0 | 0 | 16 | NaN | NaN | NaN | 504.71872 | 724.22003 | 710.7787 |
| Serbp1 | SEEAHAEDSVMDHHFR | -329 | -344 | 0 | 0 | 16 | NaN | NaN | NaN | 504.71872 | 724.22003 | 710.7787 |
| Serbp1 | SEEAHAEDSVMDHHFR | -329 | -344 | 0 | 0 | 16 | NaN | NaN | NaN | 504.71872 | 724.22003 | 710.7787 |
| Serbp1 | VDRPIIER | -153 | -160 | 1 | 0.00223472 | 8 | NaN | NaN | NaN | 1069.9272 | 1001.78107 | 1386.4846 |
| Serbp1 | SAPDVDDPEAFPALA | -393 | -407 | 0 | 0.007224385 | 15 | NaN | NaN | NaN | 474.28113 | 190.7791 | 448.64822 |
| Serbp1 | HNWGTVK | -222 | -228 | 0 | 0 | 7 | NaN | NaN | NaN | 542.3908 | 816.25397 | NaN |
| Luc7l2 | LAETQEEISAEVAAK | -108 | -122 | 0 | 0 | 15 | NaN | NaN | NaN | 712.77856 | 605.7743 | 681.91144 |
| Naa15 | EALEHLCTYEK | -208 | -218 | 0 | 0 | 11 | NaN | NaN | NaN | 786.2329 | 739.5645 | 677.20276 |
| Naa15 | AKVETPLEEAIK | -648 | -659 | 1 | 0.00025015 | 12 | NaN | NaN | NaN | 1056.7318 | 671.0881 | NaN |
| Naa15 | WDKDNLQIL | -109 | -117 | 1 | 0.004224523 | 9 | NaN | NaN | NaN | 849.71704 | 822.6135 | NaN |
| H1-5 | SETAPAETAAPAPVEKSPAK | -2 | -21 | 1 | 0.000989247 | 20 | NaN | NaN | NaN | 424.00977 | 234.83934 | 305.8228 |
| H1-5 | TGPPVSELITK | -36 | -46 | 0 | 0 | 11 | NaN | NaN | NaN | 494.98972 | 562.18677 | NaN |
| H2-D1 | WEQSGAAEHY | -171 | -180 | 0 | 0.000230707 | 10 | NaN | NaN | NaN | 136.1976 | 616.96747 | 583.2433 |
| H2-D1 | KWEQSGAAEHY | -170 | -180 | 1 | 0 | 11 | NaN | NaN | NaN | 4627.6133 | NaN | 3355.9343 |
| H2-D1 | VSRPGLEEPR | -36 | -45 | 1 | 0.000431655 | 10 | NaN | NaN | NaN | 433.78305 | NaN | NaN |
| H2-D1 | KWEQSGAAEHYK | -170 | -181 | 1 | 0.000471409 | 12 | NaN | NaN | NaN | 419.0959 | NaN | NaN |
| H2-K1 | GGSHTIQVISGCEVGSDGR | -111 | -129 | 0 | 0 | 19 | NaN | NaN | NaN | 925.0415 | 1049.408 | 1044.567 |
| H2-K1 | KWEQAGEAER | -167 | -176 | 1 | 0 | 10 | NaN | NaN | NaN | 351.47345 | NaN | NaN |
| Nop56 | SSENSSTPEECEEVNEK | -461 | -477 | 0 | 0 | 17 | NaN | NaN | NaN | 272.7918 | 211.09589 | 235.40111 |
| Nop56 | STPEECEEVNEK | -466 | -477 | 0 | 0.000431655 | 12 | NaN | NaN | NaN | 207.18925 | NaN | NaN |
| Nudcd2 | ALAVGGREILK | -53 | -63 | 1 | 0.006100831 | 11 | NaN | NaN | NaN | 225.48764 | 836.8039 | NaN |
| Atp2a2 | TVEEVLGHFGVN | -8 | -19 | 0 | 0.000591124 | 12 | NaN | NaN | NaN | 764.93866 | NaN | 835.35223 |

| | | | | | | | | | | | | |
|---|---|---|---|---|---|---|---|---|---|---|---|---|
| Xrn2 | NSPGCQVASNPR | -447 | -458 | 0 | 0 | 12 | NaN | NaN | NaN | 356.67072 | 824.6942 | 619.29517 |
| Xrn2 | LEEVYPDLTPEENRR | -662 | -676 | 1 | 0.00142999 | 15 | NaN | NaN | NaN | 635.7285 | 1268.6859 | NaN |
| Xrn2 | TSFASDGSPSPLGGIK | -478 | -493 | 0 | 0 | 16 | NaN | NaN | NaN | 1534.5658 | NaN | NaN |
| Xrn2 | FKDPQFAEDYVFK | -755 | -767 | 1 | 0.000471409 | 13 | NaN | NaN | NaN | 683.18604 | NaN | NaN |
| Rnh1 | ELVLSNNDLHEPGVR | -169 | -183 | 0 | 0.00025015 | 15 | NaN | NaN | NaN | 772.2869 | 907.70435 | 964.859 |
| Rnh1 | SCPYFCSVLTK | -324 | -334 | 0 | 0 | 11 | NaN | NaN | NaN | 651.1105 | 317.1203 | 654.38257 |
| Rnh1 | SLQELDLSSNK | -223 | -233 | 0 | 0.00051306 | 11 | NaN | NaN | NaN | 828.66223 | 1195.4248 | NaN |
| Rnh1 | MSSNPLGDEGVQELCK | -343 | -358 | 0 | 0.00111006 | 16 | NaN | NaN | NaN | 543.1831 | NaN | 539.7584 |
| Mcm2 | VDSHGHNVFK | -207 | -216 | 0 | 0.00025015 | 10 | NaN | NaN | NaN | 1245.254 | 1150.3845 | 1007.6024 |
| Mcm2 | ALDDEDVEELTASQR | -96 | -110 | 0 | 0 | 15 | NaN | NaN | NaN | 2622.127 | 1996.5024 | 2188.998 |
| Mcm2 | ESMATGSIPITVR | -753 | -765 | 0 | 0 | 13 | NaN | NaN | NaN | 213.27936 | 803.83167 | NaN |
| Mcm2 | QDTIEIPEK | -855 | -863 | 0 | 0.001392869 | 9 | NaN | NaN | NaN | 721.15094 | NaN | NaN |
| Mcm2 | FVVGSHVR | -678 | -685 | 0 | 0.00025015 | 8 | NaN | NaN | NaN | 717.9698 | NaN | NaN |
| Prpf8 | TILQHLSEAWR | -708 | -718 | 0 | 0 | 11 | NaN | NaN | NaN | 1136.0593 | 1066.2219 | 2025.9199 |
| Prpf8 | RALDIPLVK | -420 | -428 | 1 | 0.00078661 | 9 | NaN | NaN | NaN | 1157.0472 | 634.8249 | 1380.995 |
| Prpf8 | GGVEGILEHTLFK | -1479 | -1491 | 0 | 0 | 13 | NaN | NaN | NaN | 1057.4282 | 1124.2433 | 911.6961 |
| Prpf8 | RFSPIPFPPLSY | -821 | -832 | 1 | 0 | 12 | NaN | NaN | NaN | 613.76654 | 421.5634 | 567.2081 |
| Prpf8 | TTVTEPHHIWPT | -1959 | -1970 | 0 | 0.001023236 | 12 | NaN | NaN | NaN | 848.4647 | 936.7881 | NaN |
| Prpf8 | ISLIQIFR | -1571 | -1578 | 0 | 0.001092529 | 8 | NaN | NaN | NaN | 573.8454 | 703.016 | NaN |
| Prpf8 | PVEEQPK | -1853 | -1859 | 0 | 0.003299042 | 7 | NaN | NaN | NaN | 18591.812 | NaN | NaN |
| Prpf8 | DIILGMEISAPSQQR | -2009 | -2023 | 0 | 0.001392869 | 15 | NaN | NaN | NaN | 1535.6896 | NaN | NaN |
| Prpf8 | EFYHEVHRPSHFLNF | -2303 | -2317 | 1 | 0.002199523 | 15 | NaN | NaN | NaN | 236.47847 | NaN | NaN |
| Lrpprc | ALDKDVASAK | -1327 | -1336 | 1 | 0 | 10 | NaN | NaN | NaN | 877.1425 | 385.9757 | 1827.1771 |
| Lrpprc | DESSDNFGSFFLR | -365 | -377 | 0 | 0 | 13 | NaN | NaN | NaN | 117.34491 | 198.60962 | 452.92145 |
| Lrpprc | PSSNISFPLVTVH | -819 | -831 | 0 | 0 | 13 | NaN | NaN | NaN | 775.0463 | 409.702 | NaN |
| Lrpprc | SKDESSDNFGSFFLR | -363 | -377 | 1 | 0.000627943 | 15 | NaN | NaN | NaN | 210.5552 | NaN | 249.21582 |
| Srsf6 | DADDAVYELNSK | -47 | -58 | 0 | 0 | 12 | NaN | NaN | NaN | 609.6706 | NaN | 603.92535 |
| Hk1 | TSDVAAIETDKEGVQNAK | -392 | -409 | 1 | 0.000431655 | 18 | NaN | NaN | NaN | 484.74704 | 623.60223 | 581.2936 |
| Hk1 | SDVAAIETDKEGVQNAK | -393 | -409 | 1 | 0.001023236 | 17 | NaN | NaN | NaN | 274.9703 | 448.68832 | 511.64413 |
| Hk1 | VGVDGSLYK | -466 | -474 | 0 | 0.000431655 | 9 | NaN | NaN | NaN | 411.49292 | 337.75098 | NaN |
| Hk1 | QLGLNSTCDDSILVK | -862 | -876 | 0 | 0.001636916 | 15 | NaN | NaN | NaN | 1017.8641 | NaN | NaN |
| Gusb | ALDLTRPVTF | -468 | -477 | 1 | 0.00051306 | 10 | NaN | NaN | NaN | 1059.4938 | NaN | 1309.5846 |
| Nup210 | ATGYQQSHLSEAR | -860 | -872 | 0 | 0 | 13 | NaN | NaN | NaN | 587.22815 | 541.1404 | 769.136 |
| Rps17 | QEIIEVDPDTK | -93 | -103 | 0 | 0 | 11 | NaN | NaN | NaN | 232.68674 | NaN | 195.51062 |
| Rps17 | EEIAIIPSKK | -36 | -45 | 1 | 0.00025015 | 10 | NaN | NaN | NaN | 567.28815 | 475.19354 | NaN |
| Rps17 | DNYVPEVSAL | -82 | -91 | 0 | 0.000431655 | 10 | NaN | NaN | NaN | 1985.169 | NaN | NaN |
| Rps17 | EEIAIIPSK | -36 | -44 | 0 | 0.00025015 | 9 | NaN | NaN | NaN | 1596.5542 | NaN | NaN |
| Uba6 | LETGQFLTFR | -239 | -248 | 0 | 0 | 10 | NaN | NaN | NaN | 777.8592 | 595.845 | 608.2363 |
| Uba6 | FWQAYPSAEDVLQK | -659 | -672 | 0 | 0 | 14 | NaN | NaN | NaN | 1316.2848 | 1437.7528 | NaN |
| Uba6 | ALDQFQENYNR | -331 | -341 | 0 | 0 | 11 | NaN | NaN | NaN | 520.13275 | NaN | NaN |
| Uba6 | FFLCEDDVVNER | -105 | -116 | 0 | 0.003333688 | 12 | NaN | NaN | NaN | 469.48468 | NaN | NaN |
| Myg1 | ALDQVSGIPGCIF | -329 | -341 | 0 | 0.00111006 | 13 | NaN | NaN | NaN | 874.66925 | 1090.6356 | NaN |
| Exosc5 | ALDSDGNLVLDPTTK | -158 | -172 | 0 | 0.000550307 | 15 | NaN | NaN | NaN | 1132.3997 | NaN | NaN |
| Rab11b | ALDSTNVEEAFK | -155 | -166 | 0 | 0.00635324 | 12 | NaN | NaN | NaN | 3518.94 | NaN | NaN |
| Pabpc1 | NLDDGIDDER | -300 | -309 | 0 | 0 | 10 | NaN | NaN | NaN | 2928.6792 | 2353.2124 | 2416.2222 |
| Pabpc1 | PAGPILSIR | -33 | -41 | 0 | 0 | 9 | NaN | NaN | NaN | 324.13913 | 751.6451 | 590.0147 |
| Pabpc1 | AVPNPVINPYQPAPPSGYF | -391 | -409 | 0 | 0.00025015 | 19 | NaN | NaN | NaN | 90.91279 | 304.34882 | 275.886 |
| Pabpc1 | ALDTMNFDVIK | -68 | -78 | 0 | 0 | 11 | NaN | NaN | NaN | 976.9206 | 996.2123 | NaN |
| Pabpc1 | SKVDEAVAVLQAHQAK | -605 | -620 | 1 | 0.001291103 | 16 | NaN | NaN | NaN | 466.1413 | 525.6258 | NaN |
| Pabpc1 | YVNFQQPADAER | -56 | -67 | 0 | 0.005176406 | 12 | NaN | NaN | NaN | 1898.1895 | NaN | NaN |
| Pabpc1 | EFSPFGTIT | -313 | -321 | 0 | 0.00122419 | 9 | NaN | NaN | NaN | 1076.9435 | NaN | NaN |
| Pabpc1 | ALYDTFSAF | -114 | -122 | 0 | 0.002168954 | 9 | NaN | NaN | NaN | 577.49023 | NaN | NaN |
| Pabpc1 | EREAELGAR | -178 | -186 | 1 | 0.002136192 | 9 | NaN | NaN | NaN | 422.6334 | NaN | NaN |
| Gspt1 | DAQIVIIEHK | -531 | -540 | 0 | 0 | 10 | NaN | NaN | NaN | 1070.6605 | 923.0094 | 935.3905 |
| Hprt1 | DLNHVCVISETGK | -201 | -213 | 0 | 0 | 13 | NaN | NaN | NaN | 2782.3708 | 2864.8809 | 2946.6963 |
| Hprt1 | SVGYRPDFVGFEIPDK | -171 | -186 | 1 | 0 | 16 | NaN | NaN | NaN | 705.2452 | 742.9423 | 884.63306 |
| Hprt1 | VFIPHGLIMDR | -35 | -45 | 0 | 0 | 11 | NaN | NaN | NaN | 789.79663 | 500.32437 | 928.3939 |
| Hprt1 | VFIPHGLIMDR | -35 | -45 | 0 | 0 | 11 | NaN | NaN | NaN | 789.79663 | 500.32437 | 928.3939 |
| Hprt1 | SIPMTVDFIR | -92 | -101 | 0 | 0 | 10 | NaN | NaN | NaN | 1523.5764 | 1004.88446 | 1247.3042 |
| Hprt1 | TMQTLLSLVK | -142 | -151 | 0 | 0.000230707 | 10 | NaN | NaN | NaN | 139.11252 | 915.75867 | 835.13055 |
| Hprt1 | PMTVDFIR | -94 | -101 | 0 | 0.000627943 | 8 | NaN | NaN | NaN | 118.62267 | 204.24413 | 1066.992 |
| Cndp2 | NVMLLPVGSADDGAHSQNEK | -431 | -450 | 0 | 0 | 20 | NaN | NaN | NaN | 490.53036 | 647.7627 | 651.79974 |
| Cndp2 | EGGSIPVTLTFQ | -414 | -425 | 0 | 0.00111006 | 12 | NaN | NaN | NaN | 7605.5347 | NaN | 8381.951 |
| Cndp2 | LYEVSQLKN | -467 | -475 | 1 | 0.00111006 | 9 | NaN | NaN | NaN | 988.13513 | 574.47534 | NaN |
| Cndp2 | VFQYIDENQDR | -7 | -17 | 0 | 0 | 11 | NaN | NaN | NaN | 1595.5044 | NaN | NaN |
| Cndp2 | LYEVSQLK | -467 | -474 | 0 | 0 | 8 | NaN | NaN | NaN | 897.4164 | NaN | NaN |
| Imp4 | ALEENQLIPTELR | -35 | -47 | 0 | 0 | 13 | NaN | NaN | NaN | 1942.1555 | 3292.769 | NaN |
| Imp4 | GTPVGLIVSHLPFGPT | -144 | -159 | 0 | 0 | 16 | NaN | NaN | NaN | 222.09308 | 339.1461 | NaN |
| Scrib | ALEGPYPVEEICLPR | -981 | -995 | 0 | 0.00025015 | 15 | NaN | NaN | NaN | 598.0993 | NaN | NaN |
| Scrib | VSLVGADDLR | -1388 | -1397 | 0 | 0.003966045 | 10 | NaN | NaN | NaN | 519.3948 | NaN | NaN |
| Cltc | ISGETIFVTAPHEATAGIIGVNR | -298 | -320 | 0 | 0 | 23 | NaN | NaN | NaN | 500.07257 | 610.49854 | 213.24973 |
| Cltc | GGLLDVDCSEDVIK | -817 | -830 | 0 | 0 | 14 | NaN | NaN | NaN | 392.74326 | 653.2904 | 697.329 |
| Cltc | GQCDLELINVCNENSLFK | -924 | -941 | 0 | 0 | 18 | NaN | NaN | NaN | 215.60275 | 258.57056 | 220.10706 |
| Cltc | VIQCFAETGQVQK | -488 | -500 | 0 | 0 | 13 | NaN | NaN | NaN | 757.3201 | 1242.9707 | NaN |
| Cltc | HSSLAGCQIINY | -145 | -156 | 0 | 0 | 12 | NaN | NaN | NaN | 734.19165 | 665.4321 | NaN |
| Cltc | LYDLETGTCIYMNR | -284 | -297 | 0 | 0 | 14 | NaN | NaN | NaN | 298.61957 | NaN | 302.41443 |
| Cltc | LSETQDPEEVSVTVK | -979 | -993 | 0 | 0.001392869 | 15 | NaN | NaN | NaN | 1864.4448 | NaN | NaN |
| Cltc | LAGAEELFAR | -357 | -366 | 0 | 0.00025015 | 10 | NaN | NaN | NaN | 741.58044 | NaN | NaN |

| | | | | | | | | | | | | |
|---|---|---|---|---|---|---|---|---|---|---|---|---|
| Cltc | ESDKFICIR | -33 | -41 | 1 | 0.002595029 | 9 | NaN | NaN | NaN | 567.7563 | NaN | NaN |
| Cltc | LEAALGLER | -1303 | -1311 | 0 | 0.005270854 | 9 | NaN | NaN | NaN | 478.47433 | NaN | NaN |
| Prpf6 | QELCEEALR | -695 | -703 | 0 | 0.002629273 | 9 | NaN | NaN | NaN | 541.203 | NaN | NaN |
| Fkbp4 | GEHSIVYLKPSYAFGSVGK | -214 | -232 | 1 | 0 | 19 | NaN | NaN | NaN | 236.32301 | 299.54456 | 309.7477 |
| Fkbp4 | DKFSFDLGKG | -75 | -84 | 2 | 0.002099344 | 10 | NaN | NaN | NaN | 949.6827 | 495.94955 | NaN |
| Fkbp4 | ALELDSNNEK | -345 | -354 | 0 | 0.002434616 | 10 | NaN | NaN | NaN | 2191.0183 | NaN | NaN |
| Msn | KTANDMIHAE | -538 | -547 | 1 | 0 | 10 | NaN | NaN | NaN | 467.83813 | NaN | 936.8437 |
| Msn | LNKDQWEER | -163 | -171 | 1 | 0 | 9 | NaN | NaN | NaN | 1167.7208 | NaN | 1563.5105 |
| Msn | LREDAVLEYLK | -183 | -193 | 1 | 0.00830926 | 11 | NaN | NaN | NaN | 943.4793 | NaN | NaN |
| Msn | ALELEQERK | -372 | -380 | 1 | 0.007493591 | 9 | NaN | NaN | NaN | 514.8311 | NaN | NaN |
| Itih3 | EVSFDVELPK | -76 | -85 | 0 | 0.00025015 | 10 | NaN | NaN | NaN | 949.6265 | 564.9915 | 732.7729 |
| Itih3 | FITNDLLGSALTK | -203 | -215 | 0 | 0 | 13 | NaN | NaN | NaN | 486.499 | 543.5226 | 544.93365 |
| Itih3 | FAPQGLPVVPK | -273 | -283 | 0 | 0.000298805 | 11 | NaN | NaN | NaN | 790.5163 | NaN | 754.17883 |
| Ubac2 | ASPPLEVSEEQVAR | -300 | -313 | 0 | 0.00719662 | 14 | NaN | NaN | NaN | 1064.7556 | NaN | NaN |
| Gbp7 | ALEQLHYVTELTER | -138 | -151 | 0 | 0 | 14 | NaN | NaN | NaN | 824.0493 | 821.4355 | 825.7981 |
| Gbp7 | ALEQLHYVTELTER | -138 | -151 | 0 | 0 | 14 | NaN | NaN | NaN | 824.0493 | 821.4355 | 825.7981 |
| Gbp7 | EAAEKEQELLR | -496 | -506 | 1 | 0.000431655 | 11 | NaN | NaN | NaN | 607.741 | NaN | NaN |
| Rbm39 | FVDVSSVPLAIGLTGQR | -201 | -217 | 0 | 0 | 17 | NaN | NaN | NaN | 371.79163 | 452.74042 | 334.0114 |
| Sugt1 | GICEYHEK | -86 | -93 | 0 | 0 | 8 | NaN | NaN | NaN | 756.7103 | 422.02252 | 992.75 |
| Sugt1 | EQNPDDAQYYCQR | -39 | -51 | 0 | 0.003966045 | 13 | NaN | NaN | NaN | 840.4448 | NaN | NaN |
| Nans | SASLEPGELAELVR | -251 | -264 | 0 | 0 | 14 | NaN | NaN | NaN | 7728.5435 | 6842.158 | 6444.971 |
| Nans | GYPPEDIFNLAGK | -321 | -333 | 0 | 0 | 13 | NaN | NaN | NaN | 426.2137 | NaN | 369.62778 |
| Kif11 | SVVECDHAR | -38 | -46 | 0 | 0.009569685 | 9 | NaN | NaN | NaN | 1033.3119 | 1382.8911 | NaN |
| Kif11 | TLHQIFEK | -138 | -145 | 0 | 0.00111006 | 8 | NaN | NaN | NaN | 12657.221 | NaN | NaN |
| Dkc1 | ALETLTGALFQRPPLIA | -172 | -188 | 1 | 0.000230707 | 17 | NaN | NaN | NaN | 205.35815 | 199.39072 | 227.47015 |
| Dkc1 | HGKPTDNTPATW | -401 | -412 | 1 | 0.00025015 | 12 | NaN | NaN | NaN | 1279.1785 | NaN | 1380.4619 |
| Dkc1 | DSAVNAICYGAK | -303 | -314 | 0 | 0.00025015 | 12 | NaN | NaN | NaN | 1553.1538 | NaN | NaN |
| Psmd2 | GTLTLCPYHSDR | -774 | -785 | 0 | 0 | 12 | NaN | NaN | NaN | 707.267 | 9205.413 | 9356.8955 |
| Psmd2 | EDVLTLLLPVMGDSK | -495 | -509 | 0 | 0 | 15 | NaN | NaN | NaN | 506.491 | 411.24396 | 399.09717 |
| Psmd2 | KEWQELDDAEK | -168 | -178 | 1 | 0 | 11 | NaN | NaN | NaN | 888.12823 | 1388.4788 | NaN |
| Psmd2 | AVPLALALISVSNPR | -689 | -703 | 0 | 0.00025015 | 15 | NaN | NaN | NaN | 197.14867 | 179.7023 | NaN |
| Abcf2 | EVPIPEHIDIYHLTR | -140 | -154 | 0 | 0 | 15 | NaN | NaN | NaN | 473.8694 | 497.44894 | 378.1134 |
| Abcf2 | LLTGELLPTDGMIR | -447 | -460 | 0 | 0 | 14 | NaN | NaN | NaN | 504.5456 | 672.6402 | NaN |
| Cdk2 | FQDVTKPVPHLR | -334 | -345 | 1 | 0 | 12 | NaN | NaN | NaN | 2173.9924 | NaN | NaN |
| Ftl1 | NLNQALLDLHALGSAR | -106 | -121 | 0 | 0 | 16 | NaN | NaN | NaN | 828.8794 | 710.95996 | 565.1555 |
| Raver1 | RPAEPPLPSPAVPGGGSGS | -543 | -561 | 1 | 0.000712251 | 19 | NaN | NaN | NaN | 744.1823 | NaN | NaN |
| Raver1 | YVHWTDAGQLTPALLHSR | -204 | -221 | 0 | 0.000431655 | 18 | NaN | NaN | NaN | 535.3347 | NaN | NaN |
| Dcps | RTDDPLLQLLQK | -321 | -332 | 1 | 0.001463897 | 12 | NaN | NaN | NaN | 501.00778 | 829.799 | 976.29126 |
| Dcps | TPFQVEHVAQLLTGSPELK | -86 | -104 | 0 | 0.00051306 | 19 | NaN | NaN | NaN | 318.59485 | 278.60178 | 295.11957 |
| Dnajc7 | TEPELLEDEDAKR | -14 | -26 | 1 | 0.00025015 | 13 | NaN | NaN | NaN | 7334.5513 | NaN | NaN |
| Dnajc7 | GDFDANNIFK | -461 | -470 | 0 | 0.005394768 | 10 | NaN | NaN | NaN | 296.85345 | NaN | NaN |
| Lrrc47 | LHDDLCEK | -361 | -368 | 0 | 0.000431655 | 8 | NaN | NaN | NaN | 1372.7156 | NaN | NaN |
| Cnot3 | DNDSILVDAIR | -191 | -201 | 0 | 0 | 11 | NaN | NaN | NaN | 452.9932 | 313.25394 | 2098.3848 |
| Cnot3 | GIEDPVPTLHLTDR | -553 | -566 | 0 | 0.006735683 | 14 | NaN | NaN | NaN | 1303.7233 | NaN | NaN |
| Apex1 | PLADSFR | -247 | -253 | 0 | 0.003966045 | 7 | NaN | NaN | NaN | 426.94144 | 343.21832 | 350.56232 |
| Ago2 | QLVVVILPGK | -517 | -526 | 0 | 0.008129024 | 10 | NaN | NaN | NaN | 268.56723 | 286.99866 | NaN |
| Ago2 | HYELDIKPEK | -57 | -66 | 1 | 0.001708008 | 10 | NaN | NaN | NaN | 163.92348 | NaN | 585.0196 |
| Cers2 | HDASDYLLESAK | -239 | -250 | 0 | 0.00025015 | 12 | NaN | NaN | NaN | 1139.3143 | 1144.3451 | 948.7473 |
| Dars1 | RETLINK | -207 | -213 | 1 | 0.001023236 | 7 | NaN | NaN | NaN | 622.10657 | NaN | NaN |
| Dars1 | KADDVVWVR | -55 | -63 | 1 | 0.00025015 | 9 | NaN | NaN | NaN | 362.70087 | NaN | NaN |
| Ptbp1 | GAGNSVLLVSNLNPER | -357 | -372 | 0 | 0 | 16 | NaN | NaN | NaN | 1560.6398 | 1573.5396 | 1578.8273 |
| Ptbp1 | VNYYTSVAPVLR | -110 | -121 | 0 | 0 | 12 | NaN | NaN | NaN | 1078.7428 | 900.14734 | 1232.9995 |
| Ptbp1 | LSLDGQNIYNACCTLR | -238 | -253 | 0 | 0 | 16 | NaN | NaN | NaN | 659.65234 | 806.06305 | 966.0029 |
| Ptbp1 | LSLDGQNIYNAC | -238 | -249 | 0 | 0.003102268 | 12 | NaN | NaN | NaN | 132.8336 | 1443.3639 | NaN |
| Bak1 | VGCDESPSPSEQQVAQDTEEVFR | -12 | -34 | 0 | 0.00111006 | 23 | NaN | NaN | NaN | 218.65933 | 269.03174 | NaN |
| Bak1 | LALYVYQR | -136 | -143 | 0 | 0.000431655 | 8 | NaN | NaN | NaN | 349.59503 | NaN | NaN |
| Rps12 | GGVMDVNTALQEVLK | -9 | -23 | 0 | 0 | 15 | NaN | NaN | NaN | 1196.5432 | 893.3326 | 1455.0112 |
| Rps12 | LVEALCAEHQINLIK | -64 | -78 | 0 | 0 | 15 | NaN | NaN | NaN | 554.3104 | 764.95154 | 738.793 |
| Rps12 | VVGCSCVVVK | -103 | -112 | 0 | 0 | 10 | NaN | NaN | NaN | 2440.5889 | NaN | NaN |
| Rrm1 | DFSYNYFGFK | -140 | -149 | 0 | 0 | 10 | NaN | NaN | NaN | 507.1093 | 490.38153 | 491.45157 |
| Pola1 | ALKPDLVNVIK | -524 | -534 | 1 | 0.00025015 | 11 | NaN | NaN | NaN | 947.2289 | 1102.0392 | 1024.1611 |
| Gltp | EHLLKPLPADR | -6 | -16 | 1 | 0.001708008 | 11 | NaN | NaN | NaN | 1385.8647 | NaN | NaN |
| Eif3d | FHEEDETSFQLVDTAR | -72 | -87 | 0 | 0.00111006 | 16 | NaN | NaN | NaN | 645.11536 | NaN | NaN |
| Psmb8 | DLSPEEAYDLGR | -218 | -229 | 0 | 0.004658491 | 12 | NaN | NaN | NaN | 1270.7737 | NaN | NaN |
| Psmb8 | KFQHGVIVAVDSR | -79 | -91 | 1 | 0.00025015 | 13 | NaN | NaN | NaN | 566.99445 | NaN | NaN |
| Lonp1 | ELGLEKDDKDAIEEK | -395 | -409 | 2 | 0.000712251 | 15 | NaN | NaN | NaN | 1798.9321 | 1370.2239 | 1615.7417 |
| Lonp1 | FLDHYLDVPVDLSK | -608 | -621 | 0 | 0 | 14 | NaN | NaN | NaN | 719.5824 | 948.73083 | 221.40929 |
| Lonp1 | ALLDNHSSEFNVTR | -435 | -448 | 0 | 0.00111006 | 14 | NaN | NaN | NaN | 102.16214 | NaN | NaN |
| Eif3h | SAVADKHELLSLA | -222 | -234 | 1 | 0.001023236 | 13 | NaN | NaN | NaN | 2109.6328 | NaN | 1826.0651 |
| Set | SKEFHLNESGDPSSK | -152 | -166 | 1 | 0 | 15 | NaN | NaN | NaN | 977.6622 | 1677.4642 | 1079.7859 |
| Glod4 | SLNYWSNLLGMK | -151 | -162 | 0 | 0 | 12 | NaN | NaN | NaN | 538.13666 | NaN | 1188.5703 |
| Glod4 | FVGDEAFR | -255 | -262 | 0 | 0 | 8 | NaN | NaN | NaN | 626.8462 | NaN | NaN |
| Glod4 | RHEEFEEGCK | -33 | -42 | 1 | 0.006100831 | 10 | NaN | NaN | NaN | 310.8263 | NaN | NaN |
| Gimap4 | FFQEVMHEFQNR | -174 | -185 | 0 | 0 | 12 | NaN | NaN | NaN | 2284.5212 | 1042.8132 | 1334.1499 |
| Gimap4 | TSPGPHALLLVVPLGR | -110 | -125 | 0 | 0 | 16 | NaN | NaN | NaN | 395.74213 | 518.2562 | 555.73364 |
| Gimap4 | TSPGPHALLLVVPLGR | -110 | -125 | 0 | 0 | 16 | NaN | NaN | NaN | 395.74213 | 518.2562 | 555.73364 |
| Gimap4 | RKDDLEDTDIHEYLEK | -155 | -170 | 2 | 0.00025015 | 16 | NaN | NaN | NaN | 210.64046 | 506.64197 | 556.9992 |
| Gimap4 | DTPGIFDTEVPDADTQR | -85 | -101 | 0 | 0 | 17 | NaN | NaN | NaN | 325.01718 | 586.3463 | NaN |

| | | | | | | | | | | | | |
|---|---|---|---|---|---|---|---|---|---|---|---|---|
| Gimap4 | REYEEQIK | -257 | -264 | 1 | 0.001868731 | 8 | NaN | NaN | NaN | 6606.054 | NaN | NaN |
| Gpi | FVSNIDGTHIAK | -183 | -194 | 0 | 0 | 12 | NaN | NaN | NaN | 4224.373 | 4811.157 | 4676.459 |
| Gpi | ETITNAETAK | -217 | -226 | 0 | 0 | 10 | NaN | NaN | NaN | 1626.0732 | 847.04846 | 1766.4471 |
| Gpi | KIEPELEGSSAVT | -524 | -536 | 1 | 0.000431655 | 13 | NaN | NaN | NaN | 3624.4155 | 2394.9792 | NaN |
| Gpi | SAVTSHDSSTNGLISFIK | -533 | -550 | 0 | 0.00025015 | 18 | NaN | NaN | NaN | 599.97565 | 525.7612 | 447.08725 |
| Gpi | EGNRPTNSIVFTK | -469 | -481 | 1 | 0.007905138 | 13 | NaN | NaN | NaN | 358.82135 | 534.8852 | NaN |
| Gpi | VSNIDGTHIAK | -184 | -194 | 0 | 0.00272025 | 11 | NaN | NaN | NaN | 1772.5067 | NaN | NaN |
| Shmt2 | KYSEGYPGK | -95 | -103 | 1 | 0.003461185 | 9 | NaN | NaN | NaN | 406.6539 | NaN | 602.54205 |
| Ube2n | IYHPNVDKL | -75 | -83 | 1 | 0 | 9 | NaN | NaN | NaN | 883.1831 | NaN | 1446.7734 |
| Ube2n | GPQDSPFEGGTFK | -41 | -53 | 0 | 0 | 13 | NaN | NaN | NaN | 336.24347 | NaN | NaN |
| Hsp90aa1 | KSLTNDWEEHLAVK | -315 | -328 | 1 | 0 | 14 | NaN | NaN | NaN | 102.13473 | 966.2064 | 912.2827 |
| Hsp90aa1 | TELAEDKENYKK | -426 | -437 | 2 | 0 | 12 | NaN | NaN | NaN | 739.1682 | 643.8104 | 468.07327 |
| Hsp90aa1 | PDDITNEEYGEFYK | -302 | -315 | 0 | 0 | 14 | NaN | NaN | NaN | 454.5746 | 578.8249 | NaN |
| Hsp90aa1 | LGLGIDEDDPTVDDTSAAVT | -695 | -714 | 0 | 0.00078661 | 20 | NaN | NaN | NaN | 351.2621 | NaN | 436.83344 |
| Hsp90aa1 | DQVANSAFVER | -501 | -511 | 0 | 0 | 11 | NaN | NaN | NaN | 1877.4596 | NaN | NaN |
| Hsp90aa1 | LELFTELAEDKENYKK | -422 | -437 | 2 | 0 | 16 | NaN | NaN | NaN | 688.86176 | NaN | NaN |
| Gsr | ALLTPVAIAAGR | -358 | -369 | 0 | 0 | 12 | NaN | NaN | NaN | 539.15784 | 408.24472 | 457.40405 |
| Sh3glb1 | ALNFLTPLR | -140 | -148 | 0 | 0 | 9 | NaN | NaN | NaN | 296.81918 | 354.72903 | 406.4856 |
| Gtpbp1 | TPEYLHIDQR | -547 | -556 | 0 | 0 | 10 | NaN | NaN | NaN | 2885.2273 | NaN | NaN |
| Wipf1 | PATPQLPSR | -386 | -394 | 0 | 0.001291103 | 9 | NaN | NaN | NaN | 921.6878 | 1106.4714 | 391.47296 |
| Wipf1 | PPLPPIPR | -486 | -493 | 0 | 0.002972566 | 8 | NaN | NaN | NaN | 718.3159 | NaN | 1157.1115 |
| Mphosph10 | SNFHFIPKPPVPEIK | -512 | -526 | 1 | 0.000627943 | 15 | NaN | NaN | NaN | 519.54944 | 585.2853 | 750.598 |
| Mphosph10 | KPWQLQGEVTAQ | -383 | -394 | 1 | 0.00078661 | 12 | NaN | NaN | NaN | 519.81995 | 918.04047 | NaN |
| Il16 | SFENFGSSQLPDR | -839 | -851 | 0 | 0 | 13 | NaN | NaN | NaN | 2665.284 | NaN | 687.2566 |
| Uba1 | NFPNAIEHTLQ | -636 | -646 | 0 | 0 | 11 | NaN | NaN | NaN | 2815.4307 | 2853.3362 | 2718.541 |
| Uba1 | NFPNAIEHTLQWAR | -636 | -649 | 0 | 0 | 14 | NaN | NaN | NaN | 1144.5372 | 1853.5391 | 1808.1299 |
| Uba1 | GCGEGGEVVVTDMDTIEK | -493 | -510 | 0 | 0 | 18 | NaN | NaN | NaN | 2218.3386 | 1163.857 | 1045.7944 |
| Uba1 | LFRPWDVTK | -518 | -526 | 1 | 0 | 9 | NaN | NaN | NaN | 1750.5327 | 1274.7385 | 1755.0984 |
| Uba1 | QTSSVLVSGLR | -71 | -81 | 0 | 0 | 11 | NaN | NaN | NaN | 861.9046 | 880.43195 | 1660.9911 |
| Uba1 | GSEADIDESLYSR | -45 | -57 | 0 | 0 | 13 | NaN | NaN | NaN | 624.4032 | 415.1403 | 563.56976 |
| Uba1 | VVIPFLTESYSSSQDPPEK | -609 | -627 | 0 | 0 | 19 | NaN | NaN | NaN | 430.24374 | 503.29135 | 508.61566 |
| Uba1 | VVQGHQQLDSYKN | -912 | -924 | 1 | 0.000431655 | 13 | NaN | NaN | NaN | 387.924 | 595.41 | 517.81506 |
| Uba1 | DESGEDVEVPYVR | -1042 | -1054 | 0 | 0.000230707 | 13 | NaN | NaN | NaN | 631.5372 | 566.2953 | NaN |
| Uba1 | NDESGEDVEVPYVR | -1041 | -1054 | 0 | 0 | 14 | NaN | NaN | NaN | 930.5717 | NaN | NaN |
| Uba1 | NEEDATELVGLAQ | -351 | -363 | 0 | 0.000627943 | 13 | NaN | NaN | NaN | 324.71155 | NaN | NaN |
| Gstp1 | YTIVYFPVR | -4 | -12 | 0 | 0 | 9 | NaN | NaN | NaN | 768.17365 | 619.6507 | 574.23676 |
| Tap1 | VWVVGLSR | -40 | -47 | 0 | 0.002199523 | 8 | NaN | NaN | NaN | 1020.8552 | 900.4 | 1134.9277 |
| Tap1 | QETGFFLK | -247 | -254 | 0 | 0.001291103 | 8 | NaN | NaN | NaN | 768.85504 | 771.777 | 796.4596 |
| Tap1 | WAILGLGVR | -48 | -56 | 0 | 0 | 9 | NaN | NaN | NaN | 279.4275 | 391.92896 | 391.95963 |
| Tap1 | GQEPLLFGR | -561 | -569 | 0 | 0.00025015 | 9 | NaN | NaN | NaN | 325.66064 | NaN | NaN |
| Bcas2 | TIVQLENEIYQIK | -198 | -210 | 0 | 0 | 13 | NaN | NaN | NaN | 527.8831 | 890.77234 | NaN |
| Cse1l | LLQPVLWER | -689 | -697 | 0 | 0.00025015 | 9 | NaN | NaN | NaN | 944.268 | 630.4721 | 1123.8062 |
| Cse1l | FEDQNTLTSICEK | -334 | -346 | 0 | 0 | 13 | NaN | NaN | NaN | 400.33154 | 550.2154 | 555.7807 |
| Cse1l | SLLLETHK | -668 | -675 | 0 | 0 | 8 | NaN | NaN | NaN | 151.19415 | 1127.959 | 1096.1718 |
| Cse1l | IWNLLVTTGR | -297 | -306 | 0 | 0 | 10 | NaN | NaN | NaN | 2513.1523 | 3267.3901 | NaN |
| Cse1l | SNELWTEIK | -166 | -174 | 0 | 0.00025015 | 9 | NaN | NaN | NaN | 1559.885 | NaN | NaN |
| Igtp | YRDPLEIISQVCDK | -266 | -279 | 1 | 0.000230707 | 14 | NaN | NaN | NaN | 632.1892 | 504.42923 | 1475.2704 |
| Igtp | KVIGIVKDEIQSK | -68 | -80 | 2 | 0.000627943 | 13 | NaN | NaN | NaN | 1233.6128 | NaN | 1102.7776 |
| Igtp | SSSPEVIEDIGK | -48 | -59 | 0 | 0.001023236 | 12 | NaN | NaN | NaN | 344.48645 | 1585.4347 | NaN |
| Selenoh | ALQLEAPELPVQVNPSKPR | -51 | -69 | 1 | 0 | 19 | NaN | NaN | NaN | 346.9016 | 302.09406 | 391.37238 |
| Gatad2b | QQEQEIEQR | -471 | -479 | 0 | 0 | 9 | NaN | NaN | NaN | 1778.4928 | NaN | 1095.7872 |
| Gatad2b | TDFTPHWK | -426 | -433 | 0 | 0.003102268 | 8 | NaN | NaN | NaN | 865.4308 | NaN | NaN |
| Copg1 | RVVLEHEEVR | -479 | -488 | 1 | 0 | 10 | NaN | NaN | NaN | 527.3893 | 1527.3121 | 1211.8508 |
| Copg1 | SVPLATTPMAEQRPESTATAA | -578 | -598 | 1 | 0.001536401 | 21 | NaN | NaN | NaN | 859.1833 | 1089.7954 | 639.70844 |
| Copg1 | ILYLINQGEHLGTT | -51 | -64 | 0 | 0.000471409 | 14 | NaN | NaN | NaN | 439.5602 | 978.44653 | 689.817 |
| Copg1 | ILHLLGQEGPK | -455 | -465 | 0 | 0 | 11 | NaN | NaN | NaN | 452.36612 | NaN | NaN |
| Copg1 | QLEEEDGSR | -239 | -247 | 0 | 0.00025015 | 9 | NaN | NaN | NaN | 343.5307 | NaN | NaN |
| Lypla1 | FSQGPINSANR | -152 | -162 | 0 | 0 | 11 | NaN | NaN | NaN | 1317.8531 | NaN | 1801.7856 |
| Shmt1 | ALVEPHGR | -124 | -131 | 0 | 0.001392869 | 8 | NaN | NaN | NaN | 1739.4514 | 922.2083 | NaN |
| Uhrf1 | LNDTIQLLVR | -65 | -74 | 0 | 0 | 10 | NaN | NaN | NaN | 3515.619 | 1891.1938 | 280.84985 |
| Uhrf1 | TLSEQQANLIK | -669 | -679 | 0 | 0 | 11 | NaN | NaN | NaN | 1465.8451 | 890.4759 | 716.48425 |
| Uhrf1 | RELYGNIR | -249 | -256 | 1 | 0 | 8 | NaN | NaN | NaN | 568.4242 | 780.98676 | 699.5653 |
| Uhrf1 | IEEVFHVEPQLQ | -32 | -43 | 0 | 0.000298805 | 12 | NaN | NaN | NaN | 1849.9802 | NaN | NaN |
| Psmb10 | PGTTPVLTR | -245 | -253 | 0 | 0.001463897 | 9 | NaN | NaN | NaN | 868.55554 | 1158.7942 | 974.8377 |
| Psmb10 | TTIAGLVFR | -40 | -48 | 0 | 0.000550307 | 9 | NaN | NaN | NaN | 516.3142 | 573.54553 | 702.3894 |
| Psmb10 | SDLGSGGNVDACVITAGGAK | -204 | -223 | 0 | 0 | 20 | NaN | NaN | NaN | 446.12592 | 534.76825 | 510.7392 |
| Psmb10 | ATNDSVVADK | -59 | -68 | 0 | 0 | 10 | NaN | NaN | NaN | 397.39807 | NaN | NaN |
| Psmb10 | VVGGVDLNGPQLYEVHPHGSYSR | -138 | -160 | 0 | 0.00201047 | 23 | NaN | NaN | NaN | 310.0313 | NaN | NaN |
| Elmo1 | TFNLLEDR | -301 | -308 | 0 | 0 | 8 | NaN | NaN | NaN | 1281.0258 | 176.61365 | 526.3065 |
| Elmo1 | YIILTHVIR | -273 | -281 | 0 | 0 | 9 | NaN | NaN | NaN | 492.29556 | 1238.5157 | 753.8217 |
| Elmo1 | HYGDLEESPQGEVPHDSLQDKLPVADIK | -587 | -614 | 1 | 0.001392869 | 28 | NaN | NaN | NaN | 604.4513 | NaN | 771.4258 |
| Elmo1 | IEWPGAYPK | -12 | -20 | 0 | 0.002265552 | 9 | NaN | NaN | NaN | 469.88538 | NaN | 300.0949 |
| Elmo1 | RLTTSPAQNAQQLHER | -76 | -91 | 1 | 0.002296977 | 16 | NaN | NaN | NaN | 10428.458 | NaN | NaN |
| Hnrnpll | ALVEFENIDSAK | -163 | -174 | 0 | 0 | 12 | NaN | NaN | NaN | 3707.7446 | NaN | 2486.5215 |
| Hnrnpll | HDGYGSHGPLLPLPSR | -337 | -352 | 0 | 0 | 16 | NaN | NaN | NaN | 914.51733 | NaN | NaN |
| Hnrnpll | ITRPGNTDDPSGGNK | -202 | -216 | 1 | 0.003102268 | 15 | NaN | NaN | NaN | 493.46655 | NaN | NaN |
| Ubr4 | NVLEINHEDLK | -2088 | -2098 | 0 | 0 | 11 | NaN | NaN | NaN | 932.4228 | 1066.6091 | 894.87067 |
| Ubr4 | FADKETLIQFLR | -3399 | -3410 | 1 | 0.00025015 | 12 | NaN | NaN | NaN | 3247.037 | 3928.4675 | NaN |

| | | | | | | | | | | | | |
|---|---|---|---|---|---|---|---|---|---|---|---|---|
| Ubr4 | LENPDEACAVSQK | -115 | -127 | 0 | 0 | 13 | NaN | NaN | NaN | 1227.1709 | 346.5669 | NaN |
| Ubr4 | VSLDEHFSK | -1132 | -1140 | 0 | 0.001636916 | 9 | NaN | NaN | NaN | 1213.0145 | NaN | 1360.4586 |
| Ubr4 | PDGGHIR | -2744 | -2750 | 0 | 0.002168954 | 7 | NaN | NaN | NaN | 482.80084 | 358.15686 | NaN |
| Ubr4 | VCLNLPYFLR | -259 | -268 | 0 | 0.001291103 | 10 | NaN | NaN | NaN | 463.73233 | 545.93427 | NaN |
| Ubr4 | HTLLSLVR | -1285 | -1292 | 0 | 0.00051306 | 8 | NaN | NaN | NaN | 259.87033 | 431.75073 | NaN |
| Ubr4 | YSGDQGQTIR | -1875 | -1884 | 0 | 0 | 10 | NaN | NaN | NaN | 250.27803 | NaN | NaN |
| Upf1 | GHEVEDVVIK | -453 | -462 | 0 | 0.000471409 | 10 | NaN | NaN | NaN | 2430.3486 | 2365.4512 | 2384.3096 |
| Upf1 | YKGDLAPLWK | -372 | -381 | 1 | 0 | 10 | NaN | NaN | NaN | 277.51614 | 568.80585 | 222.31415 |
| Upf1 | EAIDSPVSFL | -556 | -565 | 0 | 0.00025015 | 10 | NaN | NaN | NaN | 432.54977 | NaN | NaN |
| Upf1 | PLSLIQGPPGTGK | -492 | -504 | 0 | 0.00025015 | 13 | NaN | NaN | NaN | 339.05487 | NaN | NaN |
| Upf1 | VKLEADYDKK | -305 | -314 | 2 | 0.000431655 | 10 | NaN | NaN | NaN | 277.32986 | NaN | NaN |
| Psme2 | YAELYHIISSNLEK | -213 | -226 | 0 | 0 | 14 | NaN | NaN | NaN | 1839.2911 | 2248.1206 | 2205.8735 |
| Psme2 | TKVEAFQTTISK | -155 | -166 | 1 | 0 | 12 | NaN | NaN | NaN | 2292.2754 | 1951.6294 | 1876.5203 |
| Slc25a3 | TQPGYANTLR | -187 | -196 | 0 | 0 | 10 | NaN | NaN | NaN | 1532.7854 | NaN | NaN |
| Slc25a3 | GWAPTLIGYSMQ | -117 | -128 | 0 | 0.004350979 | 12 | NaN | NaN | NaN | 831.39 | NaN | NaN |
| Slc25a3 | LKEDGVR | -106 | -112 | 1 | 0.00025015 | 7 | NaN | NaN | NaN | 381.95563 | NaN | NaN |
| Rbm8a | MREDYDSVEQDGDEPGPQR | -50 | -68 | 1 | 0.00025015 | 19 | NaN | NaN | NaN | 301.3056 | 370.81522 | 372.4773 |
| Rbm8a | AMDEDGDESIHK | -16 | -27 | 0 | 0.001743443 | 12 | NaN | NaN | NaN | 5233 | NaN | NaN |
| Ptprc | DETVDDFWR | -719 | -727 | 0 | 0 | 9 | NaN | NaN | NaN | 2949.405 | 3773.6167 | 3809.1167 |
| Ptprc | DGSQQTGLFCAL | -1159 | -1170 | 0 | 0 | 12 | NaN | NaN | NaN | 932.68024 | 1143.9175 | 1221.2863 |
| Ptprc | LGIPETPKPSCGDPAAR | -374 | -390 | 1 | 0.000230707 | 17 | NaN | NaN | NaN | 395.46716 | 558.2046 | 570.8429 |
| Ptprc | ESAETEDVVDVFQVVK | -1175 | -1190 | 0 | 0.00025015 | 16 | NaN | NaN | NaN | 1326.8596 | 1603.8721 | 1073.9304 |
| Ptprc | VDVYGYVVK | -868 | -876 | 0 | 0.000230707 | 9 | NaN | NaN | NaN | 1227.9275 | 1689.2175 | 1717.55 |
| Ptprc | DEQQELVERDDEK | -602 | -614 | 1 | 0 | 13 | NaN | NaN | NaN | 1059.6638 | 841.01196 | 880.8951 |
| Ptprc | NLDEQQELVER | -600 | -610 | 0 | 0 | 11 | NaN | NaN | NaN | 149.74825 | 522.16754 | 118.05609 |
| Ptprc | LAEFQSIPR | -643 | -651 | 0 | 0.000230707 | 9 | NaN | NaN | NaN | 956.8519 | 869.62616 | NaN |
| Ptprc | NLDEQQELVERDDEK | -600 | -614 | 1 | 0.002910424 | 15 | NaN | NaN | NaN | 758.3776 | NaN | 776.0282 |
| Ptprc | NVINVQTDLGIPETPKPSCGDPAAR | -366 | -390 | 1 | 0.00051306 | 25 | NaN | NaN | NaN | 441.7657 | 360.9832 | NaN |
| Ptprc | VNAFSNFFSGPIVVH | -827 | -841 | 0 | 0.006850902 | 15 | NaN | NaN | NaN | 1245.4961 | NaN | NaN |
| Ptprc | DIVVTINDHK | -767 | -776 | 0 | 0.00025015 | 10 | NaN | NaN | NaN | 1178.0568 | NaN | NaN |
| Ptprc | AMLEGLEAEGK | -857 | -867 | 0 | 0.003745449 | 11 | NaN | NaN | NaN | 1119.1659 | NaN | NaN |
| Ptprc | ASYIDGFKEPR | -699 | -709 | 1 | 0 | 11 | NaN | NaN | NaN | 947.2226 | NaN | NaN |
| Ptprc | DLVSMIQDLK | -1125 | -1134 | 0 | 0.008436628 | 10 | NaN | NaN | NaN | 457.59277 | NaN | NaN |
| Yars1 | AMLESIGVPLEK | -103 | -114 | 0 | 0 | 12 | NaN | NaN | NaN | 2248.434 | NaN | NaN |
| Satb1 | LKDNSGLEVDVA | -707 | -718 | 1 | 0.004983786 | 12 | NaN | NaN | NaN | 1185.6195 | NaN | 1151.562 |
| Yme1l1 | QELQEVVEFLK | -291 | -301 | 0 | 0 | 11 | NaN | NaN | NaN | 160.42146 | 669.37115 | 577.0468 |
| Trir | FANDGSFLELFK | -55 | -66 | 0 | 0.000347792 | 12 | NaN | NaN | NaN | 318.04678 | 281.73843 | 405.0316 |
| Trir | KTEDEVLTSK | -134 | -143 | 1 | 0.00111006 | 10 | NaN | NaN | NaN | 513.8261 | 1152.5931 | NaN |
| Pdap1 | DSDESEDEDDDYQQK | -59 | -73 | 0 | 0 | 15 | NaN | NaN | NaN | 473.38416 | 466.52103 | 452.8692 |
| Prdx3 | DYGVLLESAGIALR | -172 | -185 | 0 | 0.000550307 | 14 | NaN | NaN | NaN | 363.5807 | NaN | NaN |
| Eif3a | IHEPIMLK | -46 | -53 | 0 | 0.001463897 | 8 | NaN | NaN | NaN | 921.35583 | 1371.8082 | NaN |
| Eif3a | LSIPITPER | -327 | -335 | 0 | 0.002629273 | 9 | NaN | NaN | NaN | 6059.116 | 4933.8154 | NaN |
| Eif3a | LYHDIAQQAFK | -173 | -183 | 0 | 0.00025015 | 11 | NaN | NaN | NaN | 3953.6448 | 4521.8384 | NaN |
| Eif3a | LQNNTILR | -431 | -438 | 0 | 0 | 8 | NaN | NaN | NaN | 632.13275 | 776.9382 | NaN |
| Eif3a | FLWESYR | -152 | -158 | 0 | 0.000298805 | 7 | NaN | NaN | NaN | 682.067 | NaN | NaN |
| Eif3a | TKNETDEDGWTT | -1330 | -1341 | 1 | 0.002265552 | 12 | NaN | NaN | NaN | 484.04102 | NaN | NaN |
| Eif3a | SVLQYVVPEVK | -378 | -388 | 0 | 0.000591124 | 11 | NaN | NaN | NaN | 381.385 | NaN | NaN |
| Ptbp3 | GVYGDVHR | -351 | -358 | 0 | 0.005758933 | 8 | NaN | NaN | NaN | 1623.5612 | 2333.2087 | NaN |
| Ptbp3 | KIPCDVTEAEVISL | -36 | -49 | 1 | 0.001291103 | 14 | NaN | NaN | NaN | 432.3401 | NaN | 1200.329 |
| Psat1 | GGGSGQFSAVPLNLIGLK | -77 | -94 | 0 | 0 | 18 | NaN | NaN | NaN | 421.28632 | 479.4403 | 482.4354 |
| Psat1 | ELLAVPNNYK | -62 | -71 | 0 | 0.00025015 | 10 | NaN | NaN | NaN | 1073.2496 | NaN | 1038.0166 |
| Hnrnpul2 | KANFSLPEK | -591 | -599 | 1 | 0 | 9 | NaN | NaN | NaN | 608.9541 | 1039.8848 | NaN |
| Pdcd11 | LQTGDTLHR | -683 | -691 | 0 | 0 | 9 | NaN | NaN | NaN | 1588.238 | NaN | 1574.8553 |
| Pdcd11 | LHLADIYTK | -1686 | -1694 | 0 | 0.001868731 | 9 | NaN | NaN | NaN | 1803.1019 | NaN | NaN |
| Sae1 | VEKEEAGGGGGGGISEEEAAQYDR | -2 | -25 | 1 | 0 | 24 | NaN | NaN | NaN | 595.80084 | 625.1504 | 554.13873 |
| Sae1 | AQNLNPMVDVK | -103 | -113 | 0 | 0 | 11 | NaN | NaN | NaN | 491.5282 | 1969.0173 | NaN |
| Septin7 | ANWEAQQR | -408 | -415 | 0 | 0 | 8 | NaN | NaN | NaN | 891.8151 | 555.35156 | 959.30396 |
| Septin7 | DVTNNVHYENY | -298 | -308 | 0 | 0 | 11 | NaN | NaN | NaN | 1547.2881 | 1526.2062 | NaN |
| Septin7 | ILEQQNSSR | -416 | -424 | 0 | 0 | 9 | NaN | NaN | NaN | 687.02844 | NaN | NaN |
| Septin7 | EFPETDDEEENK | -223 | -234 | 0 | 0.003133457 | 12 | NaN | NaN | NaN | 241.56418 | NaN | NaN |
| Cd247 | QDPNQLYNELNLGR | -66 | -79 | 0 | 0 | 14 | NaN | NaN | NaN | 1911.8694 | 1494.8558 | 1361.6819 |
| Ncbp1 | VIEENLHCIIK | -313 | -323 | 0 | 0 | 11 | NaN | NaN | NaN | 3744.9663 | 3601.3342 | 3619.8838 |
| Ncbp1 | TTLVGLLNAR | -87 | -96 | 0 | 0 | 10 | NaN | NaN | NaN | 380.79428 | 400.27063 | 445.09503 |
| Ncbp1 | LQQIFLQHHQTIQ | -747 | -759 | 0 | 0.000431655 | 13 | NaN | NaN | NaN | 598.67755 | NaN | 983.1561 |
| Tra2b | ANPDPNCCLGVF | -112 | -123 | 0 | 0 | 12 | NaN | NaN | NaN | 900.3752 | 120.08486 | 137.96101 |
| Smarca5 | RTEQEEDEELLTESSK | -144 | -159 | 1 | 0.00025015 | 16 | NaN | NaN | NaN | 671.1095 | 682.15015 | 721.01044 |
| Smarca5 | KESEITDEDIDGILER | -664 | -679 | 1 | 0.00111006 | 16 | NaN | NaN | NaN | 514.2874 | 653.85913 | 629.91565 |
| Smarca5 | LHMVLRPF | -383 | -390 | 1 | 0.002466863 | 8 | NaN | NaN | NaN | 639.6895 | 381.94894 | NaN |
| Smarca5 | LHMVLRPF | -383 | -390 | 1 | 0.002466863 | 8 | NaN | NaN | NaN | 639.6895 | 381.94894 | NaN |
| Smarca5 | SSAVEPPPPPPPESAPSKPSAAGAGGSSSGNK | -2 | -33 | 1 | 0.003681885 | 32 | NaN | NaN | NaN | 757.7391 | NaN | NaN |
| Serpinc1 | TSDQIHFFF | -148 | -156 | 0 | 0.000627943 | 9 | NaN | NaN | NaN | 376.69873 | 426.3841 | 487.955 |
| Atp5pd | LASLSEKPPAIDWAYYR | -42 | -58 | 1 | 0 | 17 | NaN | NaN | NaN | 4626.285 | 4225.3496 | 4997.753 |
| Atp5pd | NETFHAR | -35 | -41 | 0 | 0.003745449 | 7 | NaN | NaN | NaN | 2267.4248 | NaN | NaN |
| Acly | GFIGHYLDQK | -1058 | -1067 | 0 | 0.00025015 | 10 | NaN | NaN | NaN | 712.3931 | NaN | NaN |
| Acly | SFDELGEIIQS | -779 | -789 | 0 | 0.004031417 | 11 | NaN | NaN | NaN | 217.30707 | NaN | NaN |
| Vim | IDVDVSKPDLTAALR | -256 | -270 | 1 | 0 | 15 | NaN | NaN | NaN | 1187.7753 | 1370.2769 | 1416.3799 |
| Vim | QVDQLTNDK | -160 | -168 | 0 | 0 | 9 | NaN | NaN | NaN | 2619.6729 | 1950.0771 | NaN |
| Vim | ILLAELEQLKGQGK | -130 | -143 | 1 | 0 | 14 | NaN | NaN | NaN | 437.8915 | 487.32593 | 448.2879 |

| | | | | | | | | | | | | |
|---|---|---|---|---|---|---|---|---|---|---|---|---|
| Vim | KGTNESLER | -334 | -342 | 1 | 0.002265552 | 9 | NaN | NaN | NaN | 1635.9109 | NaN | NaN |
| Vim | QVQSLTCEVDALK | -322 | -334 | 0 | 0 | 13 | NaN | NaN | NaN | 1426.8367 | NaN | NaN |
| Vim | EEAESTLQSFR | -197 | -207 | 0 | 0 | 11 | NaN | NaN | NaN | 844.97455 | NaN | NaN |
| Mars1 | KQPQPQPPPPEGR | -204 | -216 | 1 | 0 | 13 | NaN | NaN | NaN | 2136.8743 | 2601.723 | NaN |
| Mars1 | KQPQPQPPPPEGR | -204 | -216 | 1 | 0 | 13 | NaN | NaN | NaN | 2136.8743 | 2601.723 | NaN |
| Mars1 | RDTVEQLR | -383 | -390 | 1 | 0.009871162 | 8 | NaN | NaN | NaN | 868.29785 | NaN | NaN |
| Mars1 | KWGTPVPLEGFEDK | -502 | -515 | 1 | 0.002595029 | 14 | NaN | NaN | NaN | 381.83194 | NaN | NaN |
| Mars1 | LTHIDHSLSR | -121 | -130 | 0 | 0.003003218 | 10 | NaN | NaN | NaN | 269.7602 | NaN | NaN |
| Prpf4 | APDDLVAPVVK | -15 | -25 | 0 | 0 | 11 | NaN | NaN | NaN | 777.72845 | 1089.0903 | NaN |
| Gtf2b | RLDALPR | -6 | -12 | 1 | 0.008624882 | 7 | NaN | NaN | NaN | 524.48065 | NaN | 651.97144 |
| Rps13 | APDLPEDLYHLIK | -81 | -93 | 0 | 0 | 13 | NaN | NaN | NaN | 517.4328 | 588.7474 | 510.74576 |
| Rpl23a | YDALDVANK | -144 | -152 | 0 | 0 | 9 | NaN | NaN | NaN | 1285.9088 | 460.26352 | NaN |
| Actr1a | SEEHPVLLTEAPLNPR | -103 | -118 | 0 | 0.000712251 | 16 | NaN | NaN | NaN | 1734.1295 | 3204.998 | 1515.6445 |
| Actr1a | SEEHPVLLTEAPLNPR | -103 | -118 | 0 | 0.000712251 | 16 | NaN | NaN | NaN | 1734.1295 | 3204.998 | 1515.6445 |
| Actr1a | APELLFRPD | -258 | -266 | 1 | 0.00201047 | 9 | NaN | NaN | NaN | 401.21335 | NaN | 542.5324 |
| Cfl1 | ASGVAVSDGVIKVF | -2 | -15 | 1 | 0.001904399 | 14 | NaN | NaN | NaN | 649.5647 | 493.2849 | 495.53418 |
| Cfl1 | HELQANCYEEVK | -133 | -144 | 0 | 0 | 12 | NaN | NaN | NaN | 399.75708 | 277.92087 | 382.64355 |
| Cfl1 | LPDKDCR | -75 | -81 | 1 | 0.008680443 | 7 | NaN | NaN | NaN | 156.81743 | NaN | 141.02095 |
| Cfl1 | STPEEVK | -24 | -30 | 0 | 0.004690496 | 7 | NaN | NaN | NaN | 187.93358 | NaN | NaN |
| G3bp2 | LPNFGFVVFDDSEPVQR | -371 | -387 | 0 | 0.000431655 | 17 | NaN | NaN | NaN | 298.29514 | 294.03287 | 330.10883 |
| Hspd1 | LGDFVNMVEK | -507 | -516 | 0 | 0 | 10 | NaN | NaN | NaN | 2206.383 | 2233.6675 | 3131.3223 |
| Hspd1 | VGGTSDVEVNEKK | -406 | -418 | 1 | 0 | 13 | NaN | NaN | NaN | 2152.318 | 3267.5981 | 3257.923 |
| Hspd1 | IQEITEQLDITTSEYEK | -371 | -387 | 0 | 0 | 17 | NaN | NaN | NaN | 842.6041 | 881.1899 | 945.43884 |
| Hspd1 | EDVDGEALSTLVLNR | -276 | -290 | 0 | 0 | 15 | NaN | NaN | NaN | 506.92606 | 680.56165 | 647.9602 |
| Hspd1 | GGGCALLR | -439 | -446 | 0 | 0.00149795 | 8 | NaN | NaN | NaN | 557.8764 | NaN | NaN |
| Hspd1 | PAMTIAK | -475 | -481 | 0 | 0.001904399 | 7 | NaN | NaN | NaN | 302.70978 | NaN | NaN |
| Mki67 | LESSQAEPVR | -2301 | -2310 | 0 | 0 | 10 | NaN | NaN | NaN | 1365.3345 | 715.01843 | 297.02713 |
| Mki67 | RYCSDPLIGNK | -1686 | -1696 | 1 | 0.00025015 | 11 | NaN | NaN | NaN | 1183.1432 | 367.2314 | 1163.5564 |
| Mki67 | NIVEPTSGDLFK | -187 | -198 | 0 | 0 | 12 | NaN | NaN | NaN | 1107.624 | NaN | 1706.7334 |
| Mki67 | QTPAGANDSVTVEESAK | -1334 | -1350 | 0 | 0.000591124 | 17 | NaN | NaN | NaN | 1027.2654 | NaN | NaN |
| Mki67 | QTPAGASDPVSVEESAK | -1923 | -1939 | 0 | 0.008653659 | 17 | NaN | NaN | NaN | 718.4903 | NaN | NaN |
| Mki67 | IALQSPQPGHIINPA | -1821 | -1835 | 0 | 0.002068213 | 15 | NaN | NaN | NaN | 435.55243 | NaN | NaN |
| Eif4a1 | GFKDQIYDIF | -191 | -200 | 1 | 0.000431655 | 10 | NaN | NaN | NaN | 1076.9181 | 1123.8956 | 1189.4297 |
| Eif4a1 | GFKDQIYDIFQK | -191 | -202 | 1 | 0.000230707 | 12 | NaN | NaN | NaN | 967.6855 | 2537.5537 | 2991.0627 |
| Eif4a1 | MEAPHIIVGTPGR | -149 | -161 | 0 | 0 | 13 | NaN | NaN | NaN | 792.0657 | 954.9984 | NaN |
| Eif4a1 | KDQIYDIFQK | -193 | -202 | 1 | 0.00025015 | 10 | NaN | NaN | NaN | 759.7147 | 912.02075 | NaN |
| Eif4a1 | TMPSDVLEVTK | -215 | -225 | 0 | 0 | 11 | NaN | NaN | NaN | 1266.4608 | NaN | NaN |
| Tbl2 | APIINIGIADTGK | -185 | -197 | 0 | 0.00223472 | 13 | NaN | NaN | NaN | 688.2726 | NaN | 874.18945 |
| Tbl2 | VELDHATLVR | -130 | -139 | 0 | 0 | 10 | NaN | NaN | NaN | 1658.5039 | NaN | NaN |
| Ddx17 | TFFTPGNLK | -441 | -449 | 0 | 0.001868731 | 9 | NaN | NaN | NaN | 310.6536 | 266.3548 | 831.20105 |
| Ddx17 | AYQYPPPPPPPPPSR | -635 | -649 | 0 | 0.000230707 | 15 | NaN | NaN | NaN | 2381.7146 | NaN | 2183.3606 |
| Rpl4 | ELAGHQTSAESWGTGR | -56 | -71 | 0 | 0.00025015 | 16 | NaN | NaN | NaN | 1202.4624 | 669.22235 | 387.32794 |
| Srrm1 | SAQENIAGIPSAFLELK | -111 | -127 | 0 | 0 | 17 | NaN | NaN | NaN | 1778.0481 | 2672.9548 | 2369.5898 |
| Sema4d | APLAFCEK | -498 | -505 | 0 | 0.000431655 | 8 | NaN | NaN | NaN | 1531.4757 | 1625.0378 | 1583.511 |
| Sema4d | DPYCAWSPAIK | -517 | -527 | 0 | 0 | 11 | NaN | NaN | NaN | 1424.299 | 1338.7723 | 1466.9137 |
| Sema4d | PGACIDSEAR | -367 | -376 | 0 | 0 | 10 | NaN | NaN | NaN | 995.525 | 1915.7693 | 898.66364 |
| Btf3 | APLATGEDDDDEVPDLVENFDEASKN | -176 | -201 | 1 | 0.005732205 | 26 | NaN | NaN | NaN | 255.3912 | 324.55307 | NaN |
| Psme1 | KAPLDIPVPDPVK | -58 | -70 | 1 | 0.000230707 | 13 | NaN | NaN | NaN | 1070.9427 | 1215.2319 | 1154.473 |
| Psme1 | LEGFHTQISK | -167 | -176 | 0 | 0.00025015 | 10 | NaN | NaN | NaN | 3195.8457 | 3480.4165 | 1178.6707 |
| Psme1 | VFELMTNLHTK | -156 | -166 | 0 | 0 | 11 | NaN | NaN | NaN | 1152.0664 | 369.54486 | 221.92944 |
| Psme1 | IVVLLQR | -110 | -116 | 0 | 0.007343498 | 7 | NaN | NaN | NaN | 321.41086 | 272.80463 | 277.60086 |
| Nckap1l | LTEEFGPHTK | -190 | -199 | 0 | 0 | 10 | NaN | NaN | NaN | 694.9909 | NaN | 698.2443 |
| Nckap1l | EKAEIIR | -80 | -86 | 1 | 0.004350979 | 7 | NaN | NaN | NaN | 174.69583 | NaN | NaN |
| Iars1 | NYPDPVSIIDKY | -607 | -618 | 1 | 0.002910424 | 12 | NaN | NaN | NaN | 756.9127 | 917.75507 | 1001.9996 |
| Iars1 | YNEHTVR | -680 | -686 | 0 | 0.005270854 | 7 | NaN | NaN | NaN | 492.06107 | 350.96814 | 482.12146 |
| Iars1 | VREELIDK | -809 | -816 | 1 | 0.000431655 | 8 | NaN | NaN | NaN | 883.27295 | 574.7227 | NaN |
| Iars1 | DKDTLSIH | -795 | -802 | 1 | 0.000627943 | 8 | NaN | NaN | NaN | 3791.4258 | NaN | NaN |
| Gm12250 | APLNIAVTGETGAGK | -67 | -81 | 0 | 0 | 15 | NaN | NaN | NaN | 1180.7573 | 1230.215 | 1308.0153 |
| Gm12250 | KIDSDLWNEK | -183 | -192 | 1 | 0 | 10 | NaN | NaN | NaN | 636.1659 | 1596.8851 | 1157.37 |
| Gm12250 | VISIINAALTDIEK | -53 | -66 | 0 | 0 | 14 | NaN | NaN | NaN | 243.3538 | 255.26988 | NaN |
| Gm12250 | LLEELPAHK | -246 | -254 | 0 | 0.000627943 | 9 | NaN | NaN | NaN | 870.17584 | NaN | NaN |
| Gm12250 | FLDTVTEDAK | -392 | -401 | 0 | 0.00025015 | 10 | NaN | NaN | NaN | 428.25412 | NaN | NaN |
| Gm12250 | EELPAHK | -248 | -254 | 0 | 0.000230707 | 7 | NaN | NaN | NaN | 155.18707 | NaN | NaN |
| Leo1 | TDLGNDLYFVK | -368 | -378 | 0 | 0.003003218 | 11 | NaN | NaN | NaN | 1084.6478 | 245.42206 | NaN |
| Chd4 | TEVEKVEEK | -1642 | -1650 | 1 | 0.00025015 | 9 | NaN | NaN | NaN | 567.96643 | 533.77527 | 591.7068 |
| Chd4 | SLKEEESTEGEKEVK | -1563 | -1577 | 2 | 0.000591124 | 15 | NaN | NaN | NaN | 387.6823 | 1160.3464 | 1068.3689 |
| Chd4 | TAPEDDKAPAEPPEGEEK | -1603 | -1620 | 1 | 0.000230707 | 18 | NaN | NaN | NaN | 338.0225 | 803.57513 | 738.16205 |
| Rpl18a | TKRPNTFF | -169 | -176 | 2 | 0.006654467 | 8 | NaN | NaN | NaN | 778.23553 | 1250.2582 | 1226.2954 |
| Rpl18a | IKFPLPHR | -150 | -157 | 1 | 0.001291103 | 8 | NaN | NaN | NaN | 725.92395 | 490.15356 | 467.06824 |
| Rpl18a | SSGEIVYCGQVFEK | -57 | -70 | 0 | 0 | 14 | NaN | NaN | NaN | 1383.8965 | NaN | NaN |
| Ctla2a | APPPDPSLDNEWK | -29 | -41 | 0 | 0 | 13 | NaN | NaN | NaN | 1019.2745 | 756.88965 | 824.1179 |
| Cdk4 | EHIDQDLR | -94 | -101 | 0 | 0.006677519 | 8 | NaN | NaN | NaN | 842.8469 | NaN | NaN |
| Cope | YGVVLDEIKPSSAPELQ | -66 | -82 | 1 | 0.00025015 | 17 | NaN | NaN | NaN | 344.2921 | 380.55496 | 343.65768 |
| Chchd2 | APSAVGSPAAAPR | -39 | -51 | 0 | 0.000550307 | 13 | NaN | NaN | NaN | 17328.607 | NaN | NaN |
| Capn1 | GPWSDSSYEWNK | -305 | -316 | 0 | 0 | 12 | NaN | NaN | NaN | 686.1548 | 609.17 | 659.2126 |
| Capn1 | SAQGNEFWSALLEK | -180 | -193 | 0 | 0.00025015 | 14 | NaN | NaN | NaN | 615.1241 | 674.1453 | 754.1822 |
| Eif2a | SWQPFLDGIFPAK | -412 | -424 | 0 | 0.00149795 | 13 | NaN | NaN | NaN | 120.144966 | 138.78906 | NaN |
| Eif2a | WQPFLDGIFPAK | -413 | -424 | 0 | 0.00051306 | 12 | NaN | NaN | NaN | 1132.2457 | NaN | NaN |

| | | | | | | | | | | | | |
|---|---|---|---|---|---|---|---|---|---|---|---|---|
| Dido1 | APTGSTPGPQGTLPAR | -1574 | -1589 | 0 | 0.001291103 | 16 | NaN | NaN | NaN | 744.4388 | 827.9178 | NaN |
| Hint1 | IIFEDDR | -31 | -37 | 0 | 0.006078721 | 7 | NaN | NaN | NaN | 837.136 | NaN | NaN |
| Hint1 | CLAFHDISPQAPTHF | -38 | -52 | 0 | 0.001636916 | 15 | NaN | NaN | NaN | 754.881 | NaN | NaN |
| Cars1 | AVHEALCDNIDTR | -583 | -595 | 0 | 0 | 13 | NaN | NaN | NaN | 4556.1523 | 7606.494 | 7838.2505 |
| Cars1 | SPNDFALWK | -399 | -407 | 0 | 0.002199523 | 9 | NaN | NaN | NaN | 1107.5548 | NaN | NaN |
| Cars1 | REQKPPATQLLK | -199 | -210 | 2 | 0.002972566 | 12 | NaN | NaN | NaN | 509.49008 | NaN | NaN |
| Necap2 | APVDQFPGTAVESVTDSSR | -71 | -89 | 0 | 0.001636916 | 19 | NaN | NaN | NaN | 342.89313 | NaN | 803.8677 |
| Pfas | FLRPEDSTRPS | -283 | -293 | 2 | 0.005112649 | 11 | NaN | NaN | NaN | 2348.5806 | 2629.0046 | 2485.575 |
| Pfas | FGEPVLAGFAR | -396 | -406 | 0 | 0 | 11 | NaN | NaN | NaN | 443.52063 | 610.9095 | 372.82306 |
| Pfas | GSIDILAEGRPALEK | -172 | -186 | 1 | 0 | 15 | NaN | NaN | NaN | 357.62518 | 384.97855 | 292.4375 |
| Pfas | WAAETLPWAEEMK | -46 | -58 | 0 | 0 | 13 | NaN | NaN | NaN | 272.91632 | 717.1519 | 651.10803 |
| Pfas | KAVVVEEPVGELR | -995 | -1007 | 1 | 0 | 13 | NaN | NaN | NaN | 176.45192 | 1324.1277 | 1336.9628 |
| Pfas | GSIDILAEGR | -172 | -181 | 0 | 0 | 10 | NaN | NaN | NaN | 1252.4613 | 1004.3881 | NaN |
| Pfas | HITQGLLK | -892 | -899 | 0 | 0.003360691 | 8 | NaN | NaN | NaN | 728.235 | NaN | NaN |
| Plac8 | LFCPVCSVCQLK | -89 | -100 | 0 | 0 | 12 | NaN | NaN | NaN | 1353.7614 | 1527.7206 | 1305.5739 |
| Plac8 | FCPVCSVCQLK | -90 | -100 | 0 | 0 | 11 | NaN | NaN | NaN | 746.07697 | 1466.9288 | 1666.8452 |
| Plac8 | VIVTQPGFVR | -7 | -16 | 0 | 0.00025015 | 10 | NaN | NaN | NaN | 353.307 | NaN | NaN |
| Ctsc | SWGSNWGESGYFR | -427 | -439 | 0 | 0 | 13 | NaN | NaN | NaN | 684.50195 | 475.71146 | 698.9488 |
| Ctsc | AQDFGVVEESCFPYTAK | -310 | -326 | 0 | 0 | 17 | NaN | NaN | NaN | 240.74295 | 379.49332 | 403.65796 |
| Ctsc | GCDGGFPYLIAGK | -296 | -308 | 0 | 0.001291103 | 13 | NaN | NaN | NaN | 257.5802 | NaN | NaN |
| Esyt1 | VFDKDLDKDDFLGR | -707 | -720 | 2 | 0 | 14 | NaN | NaN | NaN | 917.1738 | 213.017 | 199.94492 |
| Esyt1 | GGPTHPASPVLEVR | -930 | -943 | 0 | 0 | 14 | NaN | NaN | NaN | 736.1453 | 941.6237 | 1197.5764 |
| Esyt1 | KPHAESLELQVR | -842 | -853 | 1 | 0 | 12 | NaN | NaN | NaN | 3509.8499 | 722.1731 | 647.32666 |
| Esyt1 | KPHAESLELQVR | -842 | -853 | 1 | 0 | 12 | NaN | NaN | NaN | 3509.8499 | 722.1731 | 647.32666 |
| Esyt1 | VWPFLGQYMEK | -139 | -149 | 0 | 0.00025015 | 11 | NaN | NaN | NaN | 317.41382 | 421.5844 | 372.17764 |
| Esyt1 | VWPFLGQYMEK | -139 | -149 | 0 | 0.00025015 | 11 | NaN | NaN | NaN | 317.41382 | 421.5844 | 372.17764 |
| Esyt1 | TRVELGEKPLR | -172 | -182 | 2 | 0 | 11 | NaN | NaN | NaN | 1413.6146 | 410.55746 | NaN |
| Esyt1 | VLEAQDLIAK | -640 | -649 | 0 | 0.002972566 | 10 | NaN | NaN | NaN | 1322.3384 | NaN | NaN |
| Esyt1 | LDQVLQWNR | -448 | -456 | 0 | 0 | 9 | NaN | NaN | NaN | 825.14856 | NaN | NaN |
| Esyt1 | SAPVWEESASFLIR | -828 | -841 | 0 | 0.00025015 | 14 | NaN | NaN | NaN | 685.2892 | NaN | NaN |
| Nsun2 | LFEHYYQEL | -47 | -55 | 0 | 0.00025015 | 9 | NaN | NaN | NaN | 12900.527 | 14326.505 | 14360.866 |
| Nsun2 | AQEGIYTLYPFINSR | -603 | -617 | 0 | 0 | 15 | NaN | NaN | NaN | 1113.6238 | 1505.6943 | 1497.2977 |
| Nsun2 | QEGIYTLYPFINSR | -604 | -617 | 0 | 0 | 14 | NaN | NaN | NaN | 586.01227 | 704.67957 | 567.1314 |
| Nsun2 | YEPDSANPDTLQCPIVLC | -660 | -677 | 0 | 0.000431655 | 18 | NaN | NaN | NaN | 510.4493 | 543.0635 | 616.7764 |
| Nsun2 | WEGGYPEIVK | -34 | -43 | 0 | 0 | 10 | NaN | NaN | NaN | 1043.5663 | 516.7262 | NaN |
| Nsun2 | ILCDVPCSGDGTMR | -265 | -278 | 0 | 0 | 14 | NaN | NaN | NaN | 419.5498 | NaN | NaN |
| Cat | GYFEVTHDITR | -83 | -93 | 0 | 0 | 11 | NaN | NaN | NaN | 3113.339 | 1098.5852 | NaN |
| Cat | AQEDPDYGLR | -254 | -263 | 0 | 0.001904399 | 10 | NaN | NaN | NaN | 596.61426 | NaN | NaN |
| Cat | DVVFTDEMAHFDR | -54 | -66 | 0 | 0.00111006 | 13 | NaN | NaN | NaN | 341.36423 | NaN | NaN |
| Ppid | HVVFGQVIK | -146 | -154 | 0 | 0.000431655 | 9 | NaN | NaN | NaN | 1175.9283 | 780.11414 | 712.7401 |
| Echs1 | AQFGQPEILLGTIPGAGGTQ | -158 | -177 | 0 | 0 | 20 | NaN | NaN | NaN | 120.680725 | 285.51202 | 282.2362 |
| Atp5f1a | DTSVDLEETGR | -63 | -73 | 0 | 0 | 11 | NaN | NaN | NaN | 1215.4131 | 930.4061 | 1116.1078 |
| Atp5f1a | HALIIYDDLSKQ | -306 | -317 | 1 | 0 | 12 | NaN | NaN | NaN | 1127.3751 | 2315.61 | 3187.5864 |
| Atp5f1a | KAPGIIPR | -175 | -182 | 1 | 0.000471409 | 8 | NaN | NaN | NaN | 514.97955 | 668.68726 | 619.9083 |
| Atp5f1a | QYLAPYSGCSMGEYFR | -286 | -301 | 0 | 0 | 16 | NaN | NaN | NaN | 465.72363 | 320.55286 | 472.09207 |
| Atp5f1a | FGSDLDAATQQLLSR | -449 | -463 | 0 | 0 | 15 | NaN | NaN | NaN | 427.7064 | 422.60748 | 321.93274 |
| Atp5f1a | LKEIVTNFLA | -540 | -549 | 1 | 0.001055699 | 10 | NaN | NaN | NaN | 179.19958 | 386.0206 | 158.04776 |
| Atp5f1a | TGTAEMSSILEER | -46 | -58 | 0 | 0 | 13 | NaN | NaN | NaN | 1000.7545 | 835.0309 | NaN |
| Atp5f1a | FNDGTDEK | -232 | -239 | 0 | 0 | 8 | NaN | NaN | NaN | 351.8377 | NaN | 317.33575 |
| Atp5f1a | FNDGTDEK | -232 | -239 | 0 | 0 | 8 | NaN | NaN | NaN | 351.8377 | NaN | 317.33575 |
| Atp5f1a | SGCSMGEYFR | -292 | -301 | 0 | 0 | 10 | NaN | NaN | NaN | 315.39438 | 252.71425 | NaN |
| Atp5f1a | FENAFLSHVIS | -507 | -517 | 0 | 0.003998192 | 11 | NaN | NaN | NaN | 994.792 | NaN | NaN |
| Bin1 | HEPEPASGASPGATIPK | -315 | -331 | 0 | 0.00025015 | 17 | NaN | NaN | NaN | 1706.6831 | 609.6582 | 602.1506 |
| Bin1 | HDYTATDTDELQLK | -524 | -537 | 0 | 0.00204437 | 14 | NaN | NaN | NaN | 510.84143 | NaN | NaN |
| Gsn | YLWVGAGASEAEK | -601 | -613 | 0 | 0 | 13 | NaN | NaN | NaN | 1499.1116 | 1332.3387 | NaN |
| Gsn | VPFDAATLHTS | -418 | -428 | 0 | 0.001868731 | 11 | NaN | NaN | NaN | 10982.434 | NaN | NaN |
| Pola2 | DVHHEPVYPQPPFTF | -428 | -442 | 0 | 0.00111006 | 15 | NaN | NaN | NaN | 824.14795 | 901.6567 | NaN |
| Pola2 | EIFSSSGTSDR | -484 | -494 | 0 | 0.002810811 | 11 | NaN | NaN | NaN | 1559.2646 | NaN | NaN |
| Gapdh | DAGAGIALNDNFVK | -294 | -307 | 0 | 0 | 14 | NaN | NaN | NaN | 4105.2183 | 5082.8164 | 4620.827 |
| Gapdh | RVIISAPSADAPMFVM | -116 | -131 | 1 | 0 | 16 | NaN | NaN | NaN | 1385.5016 | 1664.6438 | 1673.3304 |
| Gapdh | PASTGAAK | -206 | -213 | 0 | 0.002780285 | 8 | NaN | NaN | NaN | 4116.9707 | 5226.398 | 3610.2148 |
| Gapdh | GVNHEKYDNSLK | -132 | -143 | 1 | 0 | 12 | NaN | NaN | NaN | 1139.0176 | 1623.847 | 1766.3132 |
| Gapdh | MTTVHAITATQK | -173 | -184 | 0 | 0 | 12 | NaN | NaN | NaN | 862.3478 | 573.4956 | 614.34125 |
| Gapdh | GKPITIFQER | -69 | -78 | 1 | 0 | 10 | NaN | NaN | NaN | 830.1922 | 603.47864 | 693.0924 |
| Gapdh | GAGIALNDNFVK | -296 | -307 | 0 | 0 | 12 | NaN | NaN | NaN | 546.80975 | 783.38354 | 917.0921 |
| Gapdh | GILGYTEDQVVSCDFNSNSHSS | -270 | -291 | 0 | 0.001055699 | 22 | NaN | NaN | NaN | 440.91904 | 573.24817 | 546.02875 |
| Mpst | HWLNQNLPISSGK | -134 | -146 | 0 | 0 | 13 | NaN | NaN | NaN | 2591.6482 | 2207.3574 | NaN |
| Mpst | THVVIYDDSDQGLYSAPR | -95 | -112 | 0 | 0.00025015 | 18 | NaN | NaN | NaN | 238.55496 | 276.59363 | NaN |
| Sp110 | TKNDAVDFSPTLPVTCGK | -351 | -368 | 1 | 0.00025015 | 18 | NaN | NaN | NaN | 413.78174 | 225.76024 | 371.09412 |
| Sp110 | SLEACQNLVPLSK | -51 | -63 | 0 | 0 | 13 | NaN | NaN | NaN | 2463.48 | NaN | NaN |
| Sp110 | DEQPSPSPR | -171 | -179 | 0 | 0 | 9 | NaN | NaN | NaN | 859.09656 | NaN | NaN |
| Fkbp8 | SQVLEHQPDNIK | -286 | -297 | 0 | 0 | 12 | NaN | NaN | NaN | 715.2253 | 1032.4927 | 156.03366 |
| Stat5a | VFPDRPKDEVFAK | -669 | -681 | 2 | 0 | 13 | NaN | NaN | NaN | 627.31366 | 581.89905 | NaN |
| Stat5a | HHKPHWNDGAIL | -584 | -595 | 1 | 0.002595029 | 12 | NaN | NaN | NaN | 125.06704 | 705.09863 | NaN |
| Eif3b | SPSQEPSAPGK | -88 | -98 | 0 | 0 | 11 | NaN | NaN | NaN | 405.2449 | 479.84912 | 345.0138 |
| Eif3b | IFLEYASPAHAVDAVK | -222 | -237 | 0 | 0.001092529 | 16 | NaN | NaN | NaN | 1087.888 | NaN | NaN |
| Eif3b | SAHWPIFK | -412 | -419 | 0 | 0.009897819 | 8 | NaN | NaN | NaN | 930.58276 | NaN | NaN |
| Eif3b | IDFSPCER | -368 | -375 | 0 | 0.006472924 | 8 | NaN | NaN | NaN | 849.62115 | NaN | NaN |

| | | | | | | | | | | | | |
|---|---|---|---|---|---|---|---|---|---|---|---|---|
| Eif3b | VSFYHVK | -578 | -584 | 0 | 0 | 7 | NaN | NaN | NaN | 770.2982 | NaN | NaN |
| Eif3b | LEEAECR | -286 | -292 | 0 | 0.00201047 | 7 | NaN | NaN | NaN | 684.1531 | NaN | NaN |
| Atp5f1d | ASHVPTLQVLRPGLVVVHTEDGTTTK | -71 | -96 | 1 | 0 | 26 | NaN | NaN | NaN | 595.8945 | 5124.8926 | 3661.246 |
| Atp5f1d | ASHVPTLQVLRPGLVVVHTEDGTTTK | -71 | -96 | 1 | 0 | 26 | NaN | NaN | NaN | 595.8945 | 5124.8926 | 3661.246 |
| Atp5f1d | SFTFASPTQVFFDSANVK | -39 | -56 | 0 | 0.00025015 | 18 | NaN | NaN | NaN | 316.05478 | 317.46835 | 285.24142 |
| Atp5f1d | FASPTQVFFDSANVK | -42 | -56 | 0 | 0 | 15 | NaN | NaN | NaN | 279.92273 | 845.18274 | 863.29755 |
| Rap1b | VKDTDDVPMILVGN | -103 | -116 | 1 | 0 | 14 | NaN | NaN | NaN | 1080.1869 | 907.7678 | 830.1448 |
| Rpl12 | QAQIEVVPSASAL | -68 | -80 | 0 | 0 | 13 | NaN | NaN | NaN | 170.84016 | 167.94196 | 156.94556 |
| Rpl12 | QAQIEVVPSASAL | -68 | -80 | 0 | 0 | 13 | NaN | NaN | NaN | 170.84016 | 167.94196 | 156.94556 |
| Rpl12 | QAQIEVVPSASALIIK | -68 | -83 | 0 | 0 | 16 | NaN | NaN | NaN | 543.06055 | 644.3138 | 1151.4226 |
| Ruvbl1 | EIIQDVTLHDLDVANARPQ | -233 | -251 | 1 | 0 | 19 | NaN | NaN | NaN | 355.1308 | 390.54956 | 393.40167 |
| Ruvbl1 | TEGINISEEALNHLGEIGTK | -381 | -400 | 0 | 0.001092529 | 20 | NaN | NaN | NaN | 306.1765 | 479.3112 | 372.38208 |
| Ruvbl1 | AQTEGINISEEALNHLGEIGTK | -379 | -400 | 0 | 0.000471409 | 22 | NaN | NaN | NaN | 291.07462 | 327.09494 | 327.36078 |
| Ruvbl1 | DIECFTYLHR | -308 | -317 | 0 | 0 | 10 | NaN | NaN | NaN | 126.41788 | 2385.6917 | 1800.8766 |
| Grwd1 | ELHWHPQCPGVLIST | -419 | -433 | 0 | 0.000394361 | 15 | NaN | NaN | NaN | 1166.691 | 1183.8857 | 1083.3048 |
| Stat3 | LLGPGVNYSGCQITWAK | -532 | -548 | 0 | 0.00025015 | 17 | NaN | NaN | NaN | 614.2211 | 466.16772 | 772.25775 |
| Stat3 | LGEIDQQYSR | -61 | -70 | 0 | 0.00025015 | 10 | NaN | NaN | NaN | 2501.4216 | 5081.9565 | NaN |
| Rpl35a | RDETEFYLGK | -36 | -45 | 1 | 0.001392869 | 10 | NaN | NaN | NaN | 1688.416 | 1581.7946 | 197.9727 |
| Hspa8 | LDKNQTAEKEEFEHQQK | -581 | -597 | 2 | 0 | 17 | NaN | NaN | NaN | 3377.4934 | 5305.1816 | 5070.2607 |
| Hspa8 | SFYPEEVSSMVLTK | -113 | -126 | 0 | 0 | 14 | NaN | NaN | NaN | 1331.1309 | 1241.6823 | 820.3951 |
| Hspa8 | MVVNDAGRPK | -93 | -102 | 1 | 0 | 10 | NaN | NaN | NaN | 271.70996 | 320.71973 | 984.496 |
| Hspa8 | DAGTIAGLNVLR | -160 | -171 | 0 | 0 | 12 | NaN | NaN | NaN | 865.70935 | 4806.3364 | 1733.717 |
| Hspa8 | LYQSAGGMPGGMPGGFPGGGAPPSGGAS | -610 | -637 | 0 | 0.00025015 | 28 | NaN | NaN | NaN | 482.2672 | 529.11285 | 523.8554 |
| Hspa8 | VTNAVVTVPAYFNDSQR | -139 | -155 | 0 | 0.002199523 | 17 | NaN | NaN | NaN | 1137.1506 | 1260.6299 | NaN |
| Hspa8 | GPAVGIDLGTTY | -4 | -15 | 0 | 0.002972566 | 12 | NaN | NaN | NaN | 652.50653 | NaN | NaN |
| Fkbp3 | QDHGSDSFLAEHK | -30 | -42 | 0 | 0 | 13 | NaN | NaN | NaN | 528.63477 | NaN | 797.4823 |
| Mrpl48 | ARPELEELLAK | -199 | -209 | 1 | 0.00025015 | 11 | NaN | NaN | NaN | 1167.0538 | 474.9012 | 1253.5913 |
| Ndufa8 | ARPEPNPVIEGDLKPA | -146 | -161 | 2 | 0.001055699 | 16 | NaN | NaN | NaN | 629.6565 | 702.0586 | 724.87195 |
| Hba | TYFPHFDVSHGSAQVK | -42 | -57 | 0 | 0 | 16 | NaN | NaN | NaN | 399.42358 | 1542.5969 | 1356.0836 |
| Hba | ASHHPADFTPAVHASLDK | -111 | -128 | 0 | 0 | 18 | NaN | NaN | NaN | 1031.2622 | 750.8705 | 1038.7913 |
| Hba | ASHHPADFTPAVHASLDK | -111 | -128 | 0 | 0 | 18 | NaN | NaN | NaN | 1031.2622 | 750.8705 | 1038.7913 |
| Hba | MFASFPTTK | -33 | -41 | 0 | 0 | 9 | NaN | NaN | NaN | 979.8674 | 1619.8905 | 1553.9277 |
| Hba | ASAAGHLDDLPGALSALSDLHAHK | -68 | -91 | 0 | 0.007670622 | 24 | NaN | NaN | NaN | 589.7706 | 615.9505 | NaN |
| Hba | LSDLHAHK | -84 | -91 | 0 | 0.00078661 | 8 | NaN | NaN | NaN | 404.03943 | 1003.60266 | NaN |
| Hba | SALSDLHAHK | -82 | -91 | 0 | 0 | 10 | NaN | NaN | NaN | 1534.7574 | NaN | NaN |
| Add1 | ASAGGPDNLVLLDPGK | -333 | -348 | 0 | 0 | 16 | NaN | NaN | NaN | 363.5954 | 1737.2339 | 1673.1073 |
| Sdf2l1 | EGIFIKPGADLSTGHDEL | -204 | -221 | 1 | 0.00025015 | 18 | NaN | NaN | NaN | 323.25217 | 375.44077 | 396.75317 |
| Sdf2l1 | ASAGLVTCGSVLK | -31 | -43 | 0 | 0.002329279 | 13 | NaN | NaN | NaN | 904.18225 | NaN | NaN |
| Sdf2l1 | QSVTGVEESDDANSYWR | -67 | -83 | 0 | 0.002972566 | 17 | NaN | NaN | NaN | 643.3618 | NaN | NaN |
| Glipr2 | EAQQYSEALASTR | -38 | -50 | 0 | 0 | 13 | NaN | NaN | NaN | 652.89685 | 125.22173 | 508.7669 |
| Glipr2 | ASDGSSFVVAR | -122 | -132 | 0 | 0 | 11 | NaN | NaN | NaN | 1434.4884 | 2492.9043 | NaN |
| Rpl22l1 | ASDKETYELR | -98 | -107 | 1 | 0 | 10 | NaN | NaN | NaN | 1459.0648 | 3144.8032 | NaN |
| Hsp90b1 | FQSSHHSTDITSLDQYVER | -512 | -530 | 0 | 0 | 19 | NaN | NaN | NaN | 588.9941 | 575.74634 | 531.6589 |
| Hsp90b1 | FQSSHHSTDITSLDQYVER | -512 | -530 | 0 | 0 | 19 | NaN | NaN | NaN | 588.9941 | 575.74634 | 531.6589 |
| Hsp90b1 | GVIEDHSNR | -495 | -503 | 0 | 0 | 9 | NaN | NaN | NaN | 1140.2457 | 469.72855 | 445.5145 |
| Hsp90b1 | RSGYLLPDTK | -724 | -733 | 1 | 0.001392869 | 10 | NaN | NaN | NaN | 570.7587 | 774.82434 | 1995.9744 |
| Hsp90b1 | EKEESDDEAAVEEEEEK | -302 | -319 | 1 | 0.00025015 | 18 | NaN | NaN | NaN | 527.5931 | 701.9993 | 614.5226 |
| Hsp90b1 | LIINSLYK | -88 | -95 | 0 | 0 | 8 | NaN | NaN | NaN | 964.52545 | NaN | 673.7407 |
| Hsp90b1 | KGYEVIYLTEPVDEYCIQ | -561 | -578 | 1 | 0.00051306 | 18 | NaN | NaN | NaN | 481.48627 | 684.8273 | NaN |
| Hsp90b1 | EEEAIQLDGLNASQIR | -52 | -67 | 0 | 0.000230707 | 16 | NaN | NaN | NaN | 1373.2568 | NaN | NaN |
| Hsp90b1 | KGYEVIYL | -561 | -568 | 1 | 0.006677519 | 8 | NaN | NaN | NaN | 788.58405 | NaN | NaN |
| Hsp90b1 | GVVDSDDLPLNVS | -435 | -447 | 0 | 0 | 13 | NaN | NaN | NaN | 192.57217 | NaN | NaN |
| Dck | KQASEDWEVVPEPVAR | -42 | -57 | 1 | 0.000471409 | 16 | NaN | NaN | NaN | 940.65985 | 986.98663 | 888.10315 |
| Dck | KQASEDWEVVPEPVAR | -42 | -57 | 1 | 0.000471409 | 16 | NaN | NaN | NaN | 940.65985 | 986.98663 | 888.10315 |
| Chaf1b | ASEEPSPELVNLPYR | -309 | -323 | 0 | 0 | 15 | NaN | NaN | NaN | 1487.9552 | NaN | 1300.9186 |
| Gapvd1 | DQVLHEHIQR | -1321 | -1330 | 0 | 0 | 10 | NaN | NaN | NaN | 202.46167 | 808.48206 | 884.87335 |
| Sf3a3 | ELNAISGPNEFAEFYNR | -71 | -87 | 0 | 0.000826806 | 17 | NaN | NaN | NaN | 238.18553 | 776.262 | NaN |
| Sf3a3 | KWDNGTFPGWPK | -220 | -231 | 1 | 0.003490217 | 12 | NaN | NaN | NaN | 1174.612 | NaN | NaN |
| Sf3a3 | KEELNAISGPNEFAEFYNR | -69 | -87 | 1 | 0.002502116 | 19 | NaN | NaN | NaN | 449.3956 | NaN | NaN |
| Ranbp1 | EEDEEELFK | -42 | -50 | 0 | 0 | 9 | NaN | NaN | NaN | 1505.6757 | 1055.8059 | 1291.3547 |
| Ranbp1 | NADESNHDPQFEPIVSLPEQEIK | -17 | -39 | 0 | 0 | 23 | NaN | NaN | NaN | 1417.7006 | 2912.27 | 1731.5071 |
| Ranbp1 | SHEDHDTSTENADESNHDPQFEPIVSLPEQEIK | -7 | -39 | 0 | 0.000471409 | 33 | NaN | NaN | NaN | 408.10498 | 841.83716 | 869.1349 |
| Ranbp1 | HADFADECPKPELLAIR | -125 | -141 | 1 | 0 | 17 | NaN | NaN | NaN | 530.0537 | 790.32434 | NaN |
| Eif6 | FENNCEVGCFAK | -7 | -18 | 0 | 0 | 12 | NaN | NaN | NaN | 377.24243 | 589.9802 | 559.81396 |
| Eif6 | ASFENNCEVGCFAK | -5 | -18 | 0 | 0.000550307 | 14 | NaN | NaN | NaN | 380.39737 | NaN | NaN |
| Rad23b | TTTTSGGHPLEFLR | -267 | -280 | 0 | 0.000431655 | 14 | NaN | NaN | NaN | 1249.7635 | NaN | NaN |
| Tnpo3 | FYELPTDSHASLR | -75 | -87 | 0 | 0.003133457 | 13 | NaN | NaN | NaN | 1552.2516 | NaN | NaN |
| Tnpo3 | NDEVIHSIFK | -357 | -366 | 0 | 0.00025015 | 10 | NaN | NaN | NaN | 371.54022 | NaN | NaN |
| Cox5b | KEDPNLVPSISNK | -73 | -85 | 1 | 0.002466863 | 13 | NaN | NaN | NaN | 1504.7543 | NaN | 1208.092 |
| Exosc10 | HSLDHLLR | -401 | -408 | 0 | 0 | 8 | NaN | NaN | NaN | 726.24384 | 1059.0906 | NaN |
| Exosc10 | STDGTLPLQK | -624 | -633 | 0 | 0.00266297 | 10 | NaN | NaN | NaN | 4581.822 | NaN | NaN |
| Atp5pf | ASGGPVDIGPEYQQDLDR | -56 | -73 | 0 | 0 | 18 | NaN | NaN | NaN | 1697.8761 | 551.83966 | 2525.3748 |
| Atp5pf | GGPVDIGPEYQQDLDR | -58 | -73 | 0 | 0.001868731 | 16 | NaN | NaN | NaN | 219.32448 | NaN | NaN |
| Akr1b1 | AYSPLGSPDRPWAK | -209 | -222 | 1 | 0 | 14 | NaN | NaN | NaN | 1179.1376 | 971.93243 | 1398.1414 |
| Akr1b1 | EQLVDEGLVK | -146 | -155 | 0 | 0.00025015 | 10 | NaN | NaN | NaN | 822.35986 | 1682.6611 | 1830.6105 |
| Arpc2 | ELQAHGADELLK | -66 | -77 | 0 | 0 | 12 | NaN | NaN | NaN | 3153.4167 | 3585.3853 | 3770.6318 |
| Arpc2 | DNTINLIHTF | -238 | -247 | 0 | 0 | 10 | NaN | NaN | NaN | 2661.95 | 375.79077 | 544.92224 |
| Arpc2 | ELQAHGADELLKR | -66 | -78 | 1 | 0 | 13 | NaN | NaN | NaN | 634.9162 | 859.28326 | 692.3195 |

| | | | | | | | | | | | | |
|---|---|---|---|---|---|---|---|---|---|---|---|---|
| Sf3a1 | ASKPLPPAPAPDEYLVSPITGEK | -395 | -417 | 1 | 0 | 23 | NaN | NaN | NaN | 856.70703 | 975.10254 | 978.0956 |
| Sf3a1 | QNEINNPK | -73 | -80 | 0 | 0.00025015 | 8 | NaN | NaN | NaN | 673.17 | 1068.2777 | 981.809 |
| Sf3a1 | VTWDGHSGSMAR | -498 | -509 | 0 | 0.00025015 | 12 | NaN | NaN | NaN | 444.94556 | 451.05347 | NaN |
| Rpn2 | SNLDPSNVDSLFYAA | -77 | -91 | 0 | 0.004321969 | 15 | NaN | NaN | NaN | 450.07285 | 463.966 | 317.7287 |
| Rpn2 | SLGVQVPDVK | -60 | -69 | 0 | 0.001392869 | 10 | NaN | NaN | NaN | 428.41217 | NaN | 448.66837 |
| Rpn2 | KIEFDSASGTYTLY | -469 | -482 | 1 | 0.006654467 | 14 | NaN | NaN | NaN | 3267.1099 | NaN | NaN |
| Psmd1 | DTSPGSAYQEGGGLYAL | -430 | -446 | 0 | 0.00025015 | 17 | NaN | NaN | NaN | 190.66408 | 544.09906 | 573.31995 |
| Psmd1 | KKEPEPNFQLL | -868 | -878 | 2 | 0.004439092 | 11 | NaN | NaN | NaN | 686.9208 | 1177.8743 | NaN |
| Lmnb2 | VKDLESLFHR | -127 | -136 | 1 | 0.002199523 | 10 | NaN | NaN | NaN | 973.00635 | 328.3961 | NaN |
| Lmnb2 | ENHIHELEEALAGER | -310 | -324 | 0 | 0.007670622 | 15 | NaN | NaN | NaN | 978.176 | 1762.1292 | NaN |
| Lmnb2 | ENHIHELEEALAGER | -310 | -324 | 0 | 0.007670622 | 15 | NaN | NaN | NaN | 978.176 | 1762.1292 | NaN |
| Lmnb2 | TNWGPGESFR | -528 | -537 | 0 | 0.000947459 | 10 | NaN | NaN | NaN | 223.51515 | 781.33704 | NaN |
| Lmnb2 | VLEGEDIAYK | -483 | -492 | 0 | 0.000627943 | 10 | NaN | NaN | NaN | 1069.4933 | NaN | NaN |
| Rad21 | FFLPAQPLWNNR | -373 | -384 | 0 | 0 | 12 | NaN | NaN | NaN | 316.56027 | 577.4217 | 487.29126 |
| Rad21 | LADCNEAFIK | -75 | -84 | 0 | 0.001846793 | 10 | NaN | NaN | NaN | 175.41983 | NaN | NaN |
| Cryl1 | ASPAAGGVVIVGSGLIGR | -2 | -19 | 0 | 0 | 18 | NaN | NaN | NaN | 561.7879 | 503.94873 | 671.0765 |
| Hnrnpa0 | LFIGGLNVQTSESGLR | -9 | -24 | 0 | 0 | 16 | NaN | NaN | NaN | 1018.739 | 1277.9681 | 1485.2825 |
| Hnrnpa0 | GLNVQTSESGLR | -13 | -24 | 0 | 0.00025015 | 12 | NaN | NaN | NaN | 1017.9956 | NaN | NaN |
| Nceh1 | VYFPEQIHDVIR | -153 | -164 | 0 | 0.00383302 | 12 | NaN | NaN | NaN | 313.38483 | NaN | NaN |
| Pcbp2 | PQSIIECVK | -152 | -160 | 0 | 0.003102268 | 9 | NaN | NaN | NaN | 1677.2776 | 1001.49817 | 922.5439 |
| Pcbp2 | LAQYLINVR | -343 | -351 | 0 | 0 | 9 | NaN | NaN | NaN | 558.2215 | 786.80884 | NaN |
| Ddost | SFFPDKPITQYPHAVGR | -208 | -224 | 1 | 0 | 17 | NaN | NaN | NaN | 426.57938 | 556.07007 | 629.0951 |
| Arpc1b | WVSHDSTVCLVDADKK | -219 | -234 | 1 | 0.00025015 | 16 | NaN | NaN | NaN | 415.03732 | 920.49695 | 1098.6764 |
| Arpc1b | VAWVSHDSTVCLVDADKK | -217 | -234 | 1 | 0 | 18 | NaN | NaN | NaN | 967.5053 | 916.0793 | 916.38745 |
| Arpc1b | VAWVSHDSTVCLVDADKK | -217 | -234 | 1 | 0 | 18 | NaN | NaN | NaN | 967.5053 | 916.0793 | 916.38745 |
| Arpc1b | WVSHDSTVCLVDADK | -219 | -233 | 0 | 0.001708008 | 15 | NaN | NaN | NaN | 1063.5555 | 915.6105 | NaN |
| Arpc1b | VAWVSHDSTVC | -217 | -227 | 0 | 0.001370415 | 11 | NaN | NaN | NaN | 137.63972 | 988.3083 | NaN |
| Arpc1b | SSEGGAATGAGLDSLHK | -310 | -326 | 0 | 0.00078661 | 17 | NaN | NaN | NaN | 3626.3298 | NaN | NaN |
| Arpc1b | TGAGLDSLHK | -317 | -326 | 0 | 0 | 10 | NaN | NaN | NaN | 1167.8861 | NaN | NaN |
| Ssrp1 | KAGEIWK | -580 | -586 | 1 | 0.004350979 | 7 | NaN | NaN | NaN | 846.88336 | NaN | 469.67264 |
| Ssrp1 | KITVPGNFQGH | -325 | -335 | 1 | 0.001868731 | 11 | NaN | NaN | NaN | 351.46506 | NaN | 1388.9042 |
| Ssrp1 | ITVPGNFQGH | -326 | -335 | 0 | 0.000627943 | 10 | NaN | NaN | NaN | 600.9261 | NaN | NaN |
| Ssrp1 | PTFLHLHGK | -220 | -228 | 0 | 0.003133457 | 9 | NaN | NaN | NaN | 445.14838 | NaN | NaN |
| Aimp1 | KHPDADSLYVEEVDVGEAAPR | -165 | -185 | 1 | 0.00201047 | 21 | NaN | NaN | NaN | 133.56644 | NaN | 345.9246 |
| Aimp1 | ASTDSKPIDASR | -141 | -152 | 1 | 0.00078661 | 12 | NaN | NaN | NaN | 403.37592 | NaN | NaN |
| Rpl3 | ASTDYDLSDK | -303 | -312 | 0 | 0 | 10 | NaN | NaN | NaN | 769.761 | 979.38135 | 961.1327 |
| Rpl3 | DDASKPVHLTA | -35 | -45 | 1 | 0.001291103 | 11 | NaN | NaN | NaN | 4883.313 | 4014.3 | 4380.0625 |
| Rpl3 | YGEVTNDFIMLK | -323 | -334 | 0 | 0 | 12 | NaN | NaN | NaN | 916.5008 | 948.58905 | 407.35324 |
| Rpl3 | VFGQDEMIDVIGVTK | -210 | -224 | 0 | 0 | 15 | NaN | NaN | NaN | 209.75569 | 341.41507 | 631.0553 |
| Rpl3 | GGFVHYGEVTNDFIMLK | -318 | -334 | 0 | 0.00025015 | 17 | NaN | NaN | NaN | 5786.9067 | 6275.967 | NaN |
| Rpl3 | ERLEQQVPVN | -199 | -208 | 1 | 0.001776473 | 10 | NaN | NaN | NaN | 17411.795 | NaN | NaN |
| Cacybp | EKPSYDTEADPSEGLMN | -178 | -194 | 1 | 0.00122419 | 17 | NaN | NaN | NaN | 104.209076 | 437.92838 | 3959.3215 |
| Mif | SGTNDPCALCSLHSIGK | -51 | -67 | 0 | 0 | 17 | NaN | NaN | NaN | 5742.4736 | 4554.91 | 4401.6743 |
| Mif | PMFIVNTNVPR | -2 | -12 | 0 | 0 | 11 | NaN | NaN | NaN | 1985.6246 | 2642.8374 | 2584.9524 |
| Mif | NDPCALCSLHSIGK | -54 | -67 | 0 | 0 | 14 | NaN | NaN | NaN | 882.1501 | 1252.0006 | 920.8127 |
| Mif | GTNDPCALCSLHSIGK | -52 | -67 | 0 | 0 | 16 | NaN | NaN | NaN | 364.52393 | 753.327 | 855.7784 |
| Mif | ASVPEGFLSELTQQLAQ | -13 | -29 | 0 | 0.001392869 | 17 | NaN | NaN | NaN | 375.3831 | 453.0733 | NaN |
| Mif | LHISPDRVYIN | -88 | -98 | 1 | 0.00111006 | 11 | NaN | NaN | NaN | 1580.8647 | NaN | NaN |
| Incenp | VLSPILLNNILPTTAK | -257 | -272 | 0 | 0.00078661 | 16 | NaN | NaN | NaN | 324.60358 | NaN | NaN |
| Rab14 | ATAPYNYSYIFK | -2 | -13 | 0 | 0.00025015 | 12 | NaN | NaN | NaN | 756.21564 | 373.13828 | 1193.8069 |
| Rab14 | IYQNIQDGSLDLNAAESGVQHKPSAPQGGR | -172 | -201 | 1 | 0.000712251 | 30 | NaN | NaN | NaN | 415.8305 | 358.1458 | 361.2203 |
| Rab14 | YIIIGDMGVGK | -14 | -24 | 0 | 0.00025015 | 11 | NaN | NaN | NaN | 984.373 | NaN | 2158.9912 |
| Rab14 | SCLLHQFTEK | -25 | -34 | 0 | 0 | 10 | NaN | NaN | NaN | 2455.4556 | NaN | NaN |
| Rab14 | LLHQFTEK | -27 | -34 | 0 | 0.001636916 | 8 | NaN | NaN | NaN | 579.6144 | NaN | NaN |
| Aars1 | HNDLDDVGKDVYHHT | -82 | -96 | 1 | 0 | 15 | NaN | NaN | NaN | 697.4961 | 731.50146 | 722.35266 |
| Aars1 | DDSSEDKTEFTVK | -552 | -564 | 1 | 0 | 13 | NaN | NaN | NaN | 888.7491 | 20446.338 | 20038.682 |
| Aars1 | QLIDSNPNQPLVIL | -847 | -860 | 0 | 0.00025015 | 14 | NaN | NaN | NaN | 225.78984 | 1274.3049 | 1055.9852 |
| Aars1 | VFDETYPDPVR | -685 | -695 | 0 | 0 | 11 | NaN | NaN | NaN | 3089.015 | 1924.7673 | NaN |
| Aars1 | SEIHYDR | -188 | -194 | 0 | 0.005145856 | 7 | NaN | NaN | NaN | 820.09875 | NaN | NaN |
| Aars1 | ATAVIPQWQK | -803 | -812 | 0 | 0.003620005 | 10 | NaN | NaN | NaN | 433.52716 | NaN | NaN |
| Aars1 | FDETYPDPVR | -686 | -695 | 0 | 0.007987481 | 10 | NaN | NaN | NaN | 426.31766 | NaN | NaN |
| Hp1bp3 | LSFPYYPSPGVLFPK | -411 | -425 | 0 | 0 | 15 | NaN | NaN | NaN | 459.94736 | 347.6008 | 314.3948 |
| Trim28 | KDHQYQFLEDAVR | -239 | -251 | 1 | 0 | 13 | NaN | NaN | NaN | 617.50616 | 792.6675 | 59.337772 |
| Trim28 | KDHQYQFLEDAVR | -239 | -251 | 1 | 0 | 13 | NaN | NaN | NaN | 617.50616 | 792.6675 | 59.337772 |
| Trim28 | TEGPGAEGPR | -582 | -591 | 0 | 0 | 10 | NaN | NaN | NaN | 556.68994 | 561.5444 | 602.5861 |
| Trim28 | TAASAASGSPGSGEGSAGGEK | -15 | -35 | 0 | 0.000550307 | 21 | NaN | NaN | NaN | 479.054 | 322.4553 | 568.49664 |
| Trim28 | WDLNAWTK | -394 | -401 | 0 | 0 | 8 | NaN | NaN | NaN | 434.56808 | 525.5155 | 482.54694 |
| Trim28 | ATDSTFSMEQPGGTLDLTLIR | -724 | -744 | 0 | 0 | 21 | NaN | NaN | NaN | 306.84763 | 383.93207 | 451.4628 |
| Cstf2 | GALPTNVPTPR | -309 | -319 | 0 | 0.001743443 | 11 | NaN | NaN | NaN | 1504.7728 | NaN | NaN |
| Hypk | KHDSGAADLER | -27 | -37 | 1 | 0.00025015 | 11 | NaN | NaN | NaN | 375.2143 | 261.1847 | NaN |
| Hnrnph1 | STAGASGGAYEHR | -363 | -375 | 0 | 0 | 13 | NaN | NaN | NaN | 485.7524 | 541.3171 | 457.45416 |
| Hnrnph1 | EIVPNGITLPVDFQGR | -135 | -150 | 0 | 0.001392869 | 16 | NaN | NaN | NaN | 690.1848 | NaN | NaN |
| Apoh | EHSSLAFWK | -328 | -336 | 0 | 0.001463897 | 9 | NaN | NaN | NaN | 298.02444 | NaN | NaN |
| Rbm12 | ATGEGFVEFR | -470 | -479 | 0 | 0.009060424 | 10 | NaN | NaN | NaN | 1642.5781 | NaN | NaN |
| Prpsap2 | ATHGLLSSDAPR | -297 | -308 | 0 | 0 | 12 | NaN | NaN | NaN | 1058.5278 | NaN | NaN |
| Rad50 | NFHELVK | -395 | -401 | 0 | 0.005270854 | 7 | NaN | NaN | NaN | 2458.2869 | NaN | NaN |
| Rab7a | SKNNIPYFETSAK | -145 | -157 | 1 | 0.00201047 | 13 | NaN | NaN | NaN | 1095.9716 | NaN | NaN |
| Gnpnat1 | GQLTETGVVSPEQFMK | -58 | -73 | 0 | 0 | 16 | NaN | NaN | NaN | 343.87503 | 1011.72174 | 291.52432 |

| | | | | | | | | | | | | |
|---|---|---|---|---|---|---|---|---|---|---|---|---|
| Mtch2 | KEYSSSFDR | -111 | -119 | 1 | 0 | 9 | NaN | NaN | NaN | 759.2852 | 799.1288 | 786.8315 |
| Ppp2r5a | KIEEPLFK | -347 | -354 | 1 | 0.002199523 | 8 | NaN | NaN | NaN | 503.11798 | 347.42365 | NaN |
| Gnb1 | LFVSGACDASAK | -198 | -209 | 0 | 0 | 12 | NaN | NaN | NaN | 2010.0767 | NaN | NaN |
| Psmc6 | HGEIDYEAIVK | -323 | -333 | 0 | 0 | 11 | NaN | NaN | NaN | 1604.5186 | 11271.796 | 3014.8396 |
| Psmc6 | REVIELPLTNPELFQR | -146 | -161 | 1 | 0 | 16 | NaN | NaN | NaN | 467.47787 | 560.43475 | 552.17773 |
| Supt6h | ATPANINLADLTR | -1538 | -1550 | 0 | 0 | 13 | NaN | NaN | NaN | 2198.1377 | 2921.7622 | 2874.342 |
| Supt6h | IVHELDQGQQL | -867 | -877 | 0 | 0 | 11 | NaN | NaN | NaN | 7105.378 | 7986.0693 | NaN |
| Supt6h | HITLYDIR | -1120 | -1127 | 0 | 0.000591124 | 8 | NaN | NaN | NaN | 780.0098 | NaN | NaN |
| Supt6h | FHNINFK | -1335 | -1341 | 0 | 0.005605646 | 7 | NaN | NaN | NaN | 381.6844 | NaN | NaN |
| Tcof1 | TSSSPAPTQTLPNSITQR | -1125 | -1142 | 0 | 0 | 18 | NaN | NaN | NaN | 751.0482 | 1797.5299 | 1534.925 |
| Tcof1 | KAEDDETLQAK | -64 | -74 | 1 | 0 | 11 | NaN | NaN | NaN | 400.43698 | 685.7946 | 562.0471 |
| Fmnl1 | RELETSLR | -122 | -129 | 1 | 0 | 8 | NaN | NaN | NaN | 1983.9194 | 4731.2104 | 4302.8564 |
| Fmnl1 | DGAIEDIITDLR | -1052 | -1063 | 0 | 0 | 12 | NaN | NaN | NaN | 2822.0803 | 3130.739 | NaN |
| Fmnl1 | ATPSGDDAQAPR | -508 | -519 | 0 | 0 | 12 | NaN | NaN | NaN | 567.5554 | 781.4597 | NaN |
| Fmnl1 | VPVPTVVRPS | -464 | -473 | 1 | 0.002199523 | 10 | NaN | NaN | NaN | 956.8147 | NaN | NaN |
| Cmas | WVSTDHDEIENVAK | -87 | -100 | 0 | 0 | 14 | NaN | NaN | NaN | 1023.793 | 1260.9816 | 725.26855 |
| Cmas | EVAYLGNEVSDEECLK | -365 | -380 | 0 | 0.00025015 | 16 | NaN | NaN | NaN | 1885.5417 | NaN | NaN |
| Cmas | GNEVSDEECLK | -370 | -380 | 0 | 0.000907206 | 11 | NaN | NaN | NaN | 500.7762 | NaN | NaN |
| Xpo5 | ATTQLCWPLLK | -1047 | -1057 | 0 | 0 | 11 | NaN | NaN | NaN | 645.4917 | 687.62 | NaN |
| Psma2 | QEPIPTAQLVQR | -102 | -113 | 0 | 0 | 12 | NaN | NaN | NaN | 2189.3262 | 1786.8062 | 209.98503 |
| Psma2 | YLVYQEPIPTAQLVQR | -98 | -113 | 0 | 0 | 16 | NaN | NaN | NaN | 368.76266 | 526.8323 | 449.4478 |
| Septin11 | SYELQESNVR | -84 | -93 | 0 | 0 | 10 | NaN | NaN | NaN | 968.87225 | NaN | 1754.969 |
| Septin11 | KKELEEEVSNF | -385 | -395 | 2 | 0.00122419 | 11 | NaN | NaN | NaN | 1428.4309 | NaN | NaN |
| Psmd14 | AVEEEDKMTPEQLAIK | -258 | -273 | 1 | 0.00051306 | 16 | NaN | NaN | NaN | 809.68085 | 800.66626 | 771.0588 |
| Psmd14 | TLQDYSEHCK | -230 | -239 | 0 | 0.00025015 | 10 | NaN | NaN | NaN | 1539.9237 | 1249.057 | NaN |
| Psmd14 | LGEFVDDYTVR | -58 | -68 | 0 | 0 | 11 | NaN | NaN | NaN | 805.8799 | NaN | NaN |
| Cyp51a1 | AVCGEDLPPLTYDQLK | -343 | -358 | 0 | 0.001636916 | 16 | NaN | NaN | NaN | 882.8615 | NaN | 1106.6041 |
| Ddx19a | LVDNTNQVEVLQR | -69 | -81 | 0 | 0 | 13 | NaN | NaN | NaN | 741.9077 | 801.7676 | 773.87915 |
| Ddx19a | REEETLDTIK | -298 | -307 | 1 | 0.002595029 | 10 | NaN | NaN | NaN | 2192.5408 | NaN | NaN |
| Ankrd44 | AVDIVGCTALHR | -698 | -709 | 0 | 0.00930008 | 12 | NaN | NaN | NaN | 3561.9944 | NaN | NaN |
| Ankrd44 | LINHGAEVTCK | -192 | -202 | 0 | 0 | 11 | NaN | NaN | NaN | 2205.9277 | NaN | NaN |
| Rpl6 | PQLQGYLR | -273 | -280 | 0 | 0.001636916 | 8 | NaN | NaN | NaN | 960.20715 | 645.86755 | 778.57306 |
| Lgals3bp | GGYSVVDVTSFEGSK | -519 | -533 | 0 | 0 | 15 | NaN | NaN | NaN | 126.74043 | 1224.7649 | NaN |
| Poglut2 | AVDTSGEQFTSSPGEK | -54 | -69 | 0 | 0.000591124 | 16 | NaN | NaN | NaN | 161.22554 | NaN | NaN |
| Psmb9 | YREDLLAHLIVA | -110 | -121 | 1 | 0.001055699 | 12 | NaN | NaN | NaN | 485.2687 | 409.93906 | NaN |
| Grk2 | FIESDKFTR | -165 | -173 | 1 | 0.00025015 | 9 | NaN | NaN | NaN | 347.18365 | NaN | NaN |
| Psma3 | CKDGVVFGVEK | -42 | -52 | 1 | 0 | 11 | NaN | NaN | NaN | 1150.362 | NaN | NaN |
| Sh3bgrl3 | LVDISQDNALRDEMR | -37 | -51 | 1 | 0.006100831 | 15 | NaN | NaN | NaN | 798.77264 | NaN | NaN |
| Sh3bgrl3 | VEAVEQDTLQEFLK | -78 | -91 | 0 | 0 | 14 | NaN | NaN | NaN | 164.21735 | 316.88297 | 447.479 |
| Ssb | HQIEYYFGDFNLPR | -19 | -32 | 0 | 0 | 14 | NaN | NaN | NaN | 1047.1478 | 871.69714 | 715.3781 |
| Ssb | ITDDQQESLNKW | -317 | -328 | 1 | 0 | 12 | NaN | NaN | NaN | 574.0253 | 655.60065 | 491.46008 |
| Ssb | AVFDSIQSAK | -156 | -165 | 0 | 0.001708008 | 10 | NaN | NaN | NaN | 100.029526 | 618.9988 | NaN |
| Ssb | GDLDDQTCR | -237 | -245 | 0 | 0.00830926 | 9 | NaN | NaN | NaN | 551.3269 | NaN | NaN |
| Ssb | KEDYFAK | -185 | -191 | 1 | 0.003490217 | 7 | NaN | NaN | NaN | 422.78067 | NaN | NaN |
| Cstf1 | AVFNHTEDYILLPDER | -359 | -374 | 0 | 0 | 16 | NaN | NaN | NaN | 408.94263 | 530.4209 | 547.2515 |
| Msh2 | IILEFLSK | -883 | -890 | 0 | 0.004595494 | 8 | NaN | NaN | NaN | 249.9618 | NaN | 467.57245 |
| Hcls1 | EVQQPSMPVEEPAAPAQLPK | -277 | -296 | 0 | 0.000431655 | 20 | NaN | NaN | NaN | 88.312744 | 537.9946 | 480.4075 |
| Hcls1 | GDDWDTDPDFVNDISEK | -18 | -34 | 0 | 0 | 17 | NaN | NaN | NaN | 395.8112 | 274.5752 | 145.30151 |
| Wrnip1 | YSEPVDQDYLPEELR | -636 | -650 | 0 | 0 | 15 | NaN | NaN | NaN | 623.8309 | 1882.5071 | 2271.7566 |
| Pex11b | GHPPLLLDVLR | -203 | -213 | 0 | 0.00025015 | 11 | NaN | NaN | NaN | 555.9914 | 321.7835 | NaN |
| Cops4 | VISFEEQVASIR | -96 | -107 | 0 | 0.000826806 | 12 | NaN | NaN | NaN | 1778.4843 | NaN | NaN |
| Cert1 | KIPALTENDPETWIVC | -514 | -529 | 1 | 0.000431655 | 16 | NaN | NaN | NaN | 412.02505 | 342.58325 | NaN |
| Rac2 | LAPITYPQGLAL | -134 | -145 | 0 | 0 | 12 | NaN | NaN | NaN | 588.9419 | 702.7062 | 659.3579 |
| Rac2 | FSLVSPASYENVR | -82 | -94 | 0 | 0 | 13 | NaN | NaN | NaN | 1792.036 | 2144.779 | 2032.4661 |
| Rac2 | LCPQPTR | -177 | -183 | 0 | 0.003681885 | 7 | NaN | NaN | NaN | 2231.129 | 6798.643 | NaN |
| Hyou1 | EEAEPPAEETSQPPPSEPK | -624 | -642 | 0 | 0.001868731 | 19 | NaN | NaN | NaN | 499.0858 | NaN | 785.19403 |
| Hyou1 | YFQHLLGK | -99 | -106 | 0 | 0.00025015 | 8 | NaN | NaN | NaN | 1566.1199 | NaN | NaN |
| Hyou1 | KYPDYESK | -530 | -537 | 1 | 0.00644558 | 8 | NaN | NaN | NaN | 1170.6069 | NaN | NaN |
| Hyou1 | QKLEELTLR | -721 | -729 | 1 | 0.005522803 | 9 | NaN | NaN | NaN | 1083.0482 | NaN | NaN |
| Hyou1 | LCQGLFFR | -804 | -811 | 0 | 0.001055699 | 8 | NaN | NaN | NaN | 801.3539 | NaN | NaN |
| Hyou1 | YSHDFNFHIN | -487 | -496 | 0 | 0.00025015 | 10 | NaN | NaN | NaN | 150.50256 | NaN | NaN |
| Gemin5 | TVVTEHTLHQHTISAL | -100 | -115 | 0 | 0.000431655 | 16 | NaN | NaN | NaN | 1311.8517 | 1125.97 | 315.71616 |
| Gemin5 | GHLESGHPELFHQL | -911 | -924 | 0 | 0.001610812 | 14 | NaN | NaN | NaN | 4048.5845 | NaN | NaN |
| Gemin5 | VVGELVGHTER | -56 | -66 | 0 | 0 | 11 | NaN | NaN | NaN | 1325.5869 | NaN | NaN |
| Gemin5 | HWSPTVK | -116 | -122 | 0 | 0 | 7 | NaN | NaN | NaN | 601.4136 | NaN | NaN |
| Gemin5 | AWSPHHDGR | -646 | -654 | 0 | 0.005587349 | 9 | NaN | NaN | NaN | 489.37143 | NaN | NaN |
| Calr | FGPDICGPGTK | -132 | -142 | 0 | 0 | 11 | NaN | NaN | NaN | 388.87738 | 9378.794 | 9341.722 |
| Calr | NSQVESGSLEDDWDFLPPKK | -188 | -207 | 1 | 0 | 20 | NaN | NaN | NaN | 1013.7633 | 1246.4562 | 1203.629 |
| Calr | VKHEQNIDCGGGYVK | -97 | -111 | 1 | 0 | 15 | NaN | NaN | NaN | 470.8991 | 1097.2623 | 1118.3475 |
| Calr | GKFYGDLEK | -54 | -62 | 1 | 0 | 9 | NaN | NaN | NaN | 1057.4672 | 693.20123 | 592.728 |
| Calr | RPDNTYEVK | -177 | -185 | 1 | 0.008014596 | 9 | NaN | NaN | NaN | 1046.958 | 1825.5494 | 1955.9427 |
| Calr | IDNSQVESGSLEDDWDFLPPKK | -186 | -207 | 1 | 0 | 22 | NaN | NaN | NaN | 838.7956 | 818.4004 | 821.6526 |
| Calr | IDNSQVESGSLEDDWDFLPPKK | -186 | -207 | 1 | 0 | 22 | NaN | NaN | NaN | 838.7956 | 818.4004 | 821.6526 |
| Calr | PDAAKPEDWDER | -211 | -222 | 1 | 0.003966045 | 12 | NaN | NaN | NaN | 465.1353 | 904.88165 | NaN |
| Calr | DWDKPEHIPDPDAK | -235 | -248 | 1 | 0.002168954 | 14 | NaN | NaN | NaN | 645.6913 | NaN | NaN |
| Cyb5r3 | AVLKDPNDHTVC | -193 | -204 | 1 | 0.00025015 | 12 | NaN | NaN | NaN | 661.94995 | 867.239 | 689.2666 |
| Rhog | LKEQGQAPITPQQGQ | -129 | -143 | 1 | 0.00659934 | 15 | NaN | NaN | NaN | 205.8689 | 300.01263 | NaN |
| Rhog | YLECSALQQDGVK | -154 | -166 | 0 | 0 | 13 | NaN | NaN | NaN | 1318.1456 | NaN | NaN |

| | | | | | | | | | | | | |
|---|---|---|---|---|---|---|---|---|---|---|---|---|
| Trmt5 | AVLPEGQDVTSGFSR | -181 | -195 | 0 | 0 | 15 | NaN | NaN | NaN | 1108.9166 | 564.89233 | 1199.5187 |
| Trmt5 | ENNYTYEFDFSK | -260 | -271 | 0 | 0.00149795 | 12 | NaN | NaN | NaN | 763.3127 | 523.09534 | NaN |
| Mlec | KEEEEEEEEYDEGSNLK | -229 | -245 | 1 | 0.003490217 | 17 | NaN | NaN | NaN | 381.93497 | 579.0887 | NaN |
| Mlec | AVNAGGEAHVDVHGIHFR | -50 | -67 | 0 | 0.000471409 | 18 | NaN | NaN | NaN | 332.16837 | NaN | NaN |
| Zc3h4 | LADDAEAGAEDEKEVEELKK | -481 | -500 | 2 | 0.001610812 | 20 | NaN | NaN | NaN | 1563.4181 | NaN | NaN |
| Zc3h4 | GGSGPGDSGPSDPR | -887 | -900 | 0 | 0 | 14 | NaN | NaN | NaN | 386.12134 | NaN | NaN |
| Cpne1 | LLLTDGAVTDVEATCK | -417 | -432 | 0 | 0 | 16 | NaN | NaN | NaN | 2398.4136 | 2938.523 | 3051.352 |
| Cpne1 | LLTDGAVTDVEATCK | -418 | -432 | 0 | 0.00149795 | 15 | NaN | NaN | NaN | 648.26013 | 497.65063 | NaN |
| Aip | PVIHQEGNR | -180 | -188 | 0 | 0 | 9 | NaN | NaN | NaN | 737.77136 | 458.96136 | NaN |
| Cnot1 | NKPILHTDLDVK | -1206 | -1217 | 1 | 0 | 12 | NaN | NaN | NaN | 489.6631 | 466.72495 | NaN |
| Eif4b | KPEENPASKF | -578 | -587 | 2 | 0.000471409 | 10 | NaN | NaN | NaN | 1024.5295 | NaN | 413.59518 |
| Vtn | DVWGIEGPIDAAFTR | -197 | -211 | 0 | 0 | 15 | NaN | NaN | NaN | 549.5139 | 802.7592 | 470.41208 |
| Tcerg1 | ETEEEDPKEEPVK | -490 | -502 | 1 | 0 | 13 | NaN | NaN | NaN | 224.91505 | 452.56912 | NaN |
| Tcerg1 | AVSEWTEYK | -433 | -441 | 0 | 0.00450079 | 9 | NaN | NaN | NaN | 229.10645 | NaN | NaN |
| Itgb1 | FCECDNFNCDR | -552 | -562 | 0 | 0 | 11 | NaN | NaN | NaN | 516.37604 | 418.475 | 89.473495 |
| Itgb1 | EKLPQPVQVDPVTHCK | -677 | -692 | 1 | 0.002780285 | 16 | NaN | NaN | NaN | 459.73553 | 617.12146 | 601.5455 |
| Ywhah | NSVVEASEAAYK | -144 | -155 | 0 | 0 | 12 | NaN | NaN | NaN | 466.59058 | 899.86017 | NaN |
| H4c1 | KTVTAMDVVYAL | -80 | -91 | 1 | 0.001023236 | 12 | NaN | NaN | NaN | 1912.1946 | 2305.2166 | 2269.5576 |
| H4c1 | KTVTAMDVVYAL | -80 | -91 | 1 | 0.001023236 | 12 | NaN | NaN | NaN | 1912.1946 | 2305.2166 | 2269.5576 |
| H4c1 | VTYTEHAK | -71 | -78 | 0 | 0.000471409 | 8 | NaN | NaN | NaN | 1578.7233 | 192.0843 | 1564.9224 |
| H4c1 | TVTAMDVVYAL | -81 | -91 | 0 | 0.001811524 | 11 | NaN | NaN | NaN | 1510.1892 | 2144.5012 | 1696.4395 |
| H4c1 | TYTEHAK | -72 | -78 | 0 | 0.007546675 | 7 | NaN | NaN | NaN | 120.371605 | 3896.234 | 3418.8809 |
| H4c1 | LENVIRDAVTYTEHAKR | -63 | -79 | 2 | 0 | 17 | NaN | NaN | NaN | 437.63284 | 1170.4967 | 2175.652 |
| H4c1 | GLIYEETRGVLK | -49 | -60 | 1 | 0 | 12 | NaN | NaN | NaN | 2222.6345 | NaN | NaN |
| H4c1 | DVVYALKR | -86 | -93 | 1 | 0.007987481 | 8 | NaN | NaN | NaN | 1012.14154 | NaN | NaN |
| Adk | GCTFPEKPDFH | -351 | -361 | 1 | 0 | 11 | NaN | NaN | NaN | 1826.5343 | 2098.7407 | 1768.7565 |
| Dnajc9 | KISLEDIQAF | -107 | -116 | 1 | 0.007105559 | 10 | NaN | NaN | NaN | 728.40216 | 791.88794 | 904.0001 |
| Lbr | EIESEEQNLVTK | -189 | -200 | 0 | 0 | 12 | NaN | NaN | NaN | 2980.6228 | 3523.2349 | 3521.6953 |
| Lbr | LHELWEPR | -261 | -268 | 0 | 0 | 8 | NaN | NaN | NaN | 970.29224 | 3496.496 | 3610.5588 |
| Lbr | SGNAVYDFFIGR | -372 | -383 | 0 | 0 | 12 | NaN | NaN | NaN | 941.13416 | 992.28815 | 949.5589 |
| Lbr | HFPPPLPALHELWEPR | -253 | -268 | 0 | 0 | 16 | NaN | NaN | NaN | 1928.1506 | 509.70474 | NaN |
| Arl6ip5 | RAWDDFFPGSDR | -9 | -20 | 1 | 0.001055699 | 12 | NaN | NaN | NaN | 854.1274 | NaN | NaN |
| Htatsf1 | EAFLNAPEASR | -371 | -381 | 0 | 0.00078661 | 11 | NaN | NaN | NaN | 1143.1637 | NaN | NaN |
| Sars1 | GIPYHIVNIVSGSLNHAASK | -356 | -375 | 0 | 0 | 20 | NaN | NaN | NaN | 1100.0878 | 869.6082 | 878.4172 |
| Sars1 | REIGNLLHPSVPISNDEDADNKVER | -133 | -157 | 2 | 0.005605646 | 25 | NaN | NaN | NaN | 1265.0022 | NaN | NaN |
| Sars1 | ELVSCSNCTDYQAR | -391 | -404 | 0 | 0 | 14 | NaN | NaN | NaN | 602.0581 | NaN | NaN |
| Cyrib | GLLGALTSTPYSPT | -106 | -119 | 0 | 0.00025015 | 14 | NaN | NaN | NaN | 387.99054 | 426.25363 | 484.712 |
| Cyrib | FYEFSQR | -93 | -99 | 0 | 0.00204437 | 7 | NaN | NaN | NaN | 1199.2156 | NaN | NaN |
| Sting1 | NLCCLAEELCHVQSR | -62 | -76 | 0 | 0.00025015 | 15 | NaN | NaN | NaN | 512.2395 | 588.86145 | 619.2068 |
| Sting1 | AWSYYIGYLR | -159 | -168 | 0 | 0 | 10 | NaN | NaN | NaN | 276.13583 | 2671.198 | NaN |
| Sting1 | HLASHELGLLLK | -50 | -61 | 0 | 0.00025015 | 12 | NaN | NaN | NaN | 525.8148 | NaN | NaN |
| Sting1 | HLASHELGLLLK | -50 | -61 | 0 | 0.00025015 | 12 | NaN | NaN | NaN | 525.8148 | NaN | NaN |
| Sting1 | KLNVAHGL | -151 | -158 | 1 | 0 | 8 | NaN | NaN | NaN | 503.99 | NaN | NaN |
| Dock11 | NLINDYISGFSPK | -1045 | -1057 | 0 | 0.000431655 | 13 | NaN | NaN | NaN | 1128.1246 | 1411.8228 | 1349.3184 |
| Gdi2 | FVSISDLFVPK | -380 | -390 | 0 | 0 | 11 | NaN | NaN | NaN | 1789.64 | 2485.427 | 2601.2192 |
| Gdi2 | VLHMDQNPYYGGESA | -30 | -44 | 0 | 0.00383302 | 15 | NaN | NaN | NaN | 924.315 | 736.4752 | 734.7197 |
| Gdi2 | AYDATTHFETTCDDIK | -403 | -418 | 0 | 0.00025015 | 16 | NaN | NaN | NaN | 335.72678 | 280.65262 | 288.1909 |
| Gdi2 | QLICDPSYVK | -279 | -288 | 0 | 0 | 10 | NaN | NaN | NaN | 552.55316 | NaN | NaN |
| Camk2g | FYFENLLSK | -448 | -456 | 0 | 0.009388464 | 9 | NaN | NaN | NaN | 2506.6614 | NaN | NaN |
| Stxbp2 | REPIPSLEAIYL | -65 | -76 | 1 | 0 | 12 | NaN | NaN | NaN | 346.54584 | 157.93732 | 328.8768 |
| Tmed10 | ITDSAGHILYAK | -76 | -87 | 0 | 0 | 12 | NaN | NaN | NaN | 1085.7925 | 1287.6804 | 1251.9005 |
| Tmed10 | IPDQLVILDMK | -117 | -127 | 0 | 0 | 11 | NaN | NaN | NaN | 950.2183 | 1319.592 | 1402.4314 |
| Kars1 | TELNDPVR | -498 | -505 | 0 | 0.000550307 | 8 | NaN | NaN | NaN | 1659.11 | NaN | 2040.0793 |
| Kars1 | GEDPYPHK | -89 | -96 | 0 | 0.000550307 | 8 | NaN | NaN | NaN | 386.36765 | NaN | NaN |
| Atp1b1 | AYGENIGYSEK | -279 | -289 | 0 | 0.000907206 | 11 | NaN | NaN | NaN | 566.00507 | NaN | NaN |
| Nup98 | KAEVTLDGVWPTDK | -818 | -831 | 1 | 0.001636916 | 14 | NaN | NaN | NaN | 1975.5331 | NaN | NaN |
| Nup98 | SFTAAPTGTTIK | -152 | -163 | 0 | 0.002910424 | 12 | NaN | NaN | NaN | 1135.902 | NaN | NaN |
| Nup98 | AYHIHPAGIVLTK | -726 | -738 | 0 | 0.001023236 | 13 | NaN | NaN | NaN | 297.0716 | NaN | NaN |
| Vma21 | AYIFEGALGMSNR | -50 | -62 | 0 | 0.000826806 | 13 | NaN | NaN | NaN | 2297.5054 | NaN | 1392.5994 |
| Bclaf1 | FYPEGDQETAK | -320 | -330 | 0 | 0.00078661 | 11 | NaN | NaN | NaN | 342.95462 | NaN | NaN |
| Ddb1 | KLPSFELLHK | -570 | -579 | 1 | 0 | 10 | NaN | NaN | NaN | 613.0737 | 353.53162 | 354.18655 |
| Ddb1 | PLYESPR | -716 | -722 | 0 | 0.002265552 | 7 | NaN | NaN | NaN | 972.98126 | 1157.5895 | 1371.2473 |
| Ddb1 | NSTLTIGTIDEIQK | -696 | -709 | 0 | 0 | 14 | NaN | NaN | NaN | 501.26007 | 381.99173 | 876.6329 |
| Ddb1 | RLYDGLFK | -134 | -141 | 1 | 0 | 8 | NaN | NaN | NaN | 495.36517 | 1578.555 | 2708.5583 |
| Ddb1 | VVTAEGLRPVK | -43 | -53 | 1 | 0 | 11 | NaN | NaN | NaN | 485.06122 | 417.66922 | 465.0797 |
| Ddb1 | AYKPMEGNFEEIAR | -934 | -947 | 1 | 0 | 14 | NaN | NaN | NaN | 210.24039 | 502.90125 | 625.78406 |
| Ddb1 | IHPQELR | -521 | -527 | 0 | 0.001291103 | 7 | NaN | NaN | NaN | 1482.2463 | NaN | NaN |
| Acss1 | AYPGYYFTGDGAHR | -517 | -530 | 0 | 0 | 14 | NaN | NaN | NaN | 669.1459 | NaN | 453.5007 |
| Snu13 | LLDLVQQSCNYK | -22 | -33 | 0 | 0 | 12 | NaN | NaN | NaN | 3891.936 | 3168.0369 | 1468.2642 |
| Snu13 | QQIQSIQQSIER | -114 | -125 | 0 | 0 | 12 | NaN | NaN | NaN | 1304.8308 | 1811.4135 | 1301.391 |
| Psmd6 | AYSQLLESYR | -303 | -312 | 0 | 0 | 10 | NaN | NaN | NaN | 714.1043 | 482.19614 | 468.73608 |
| Psmd6 | IHAYSQLLESYR | -301 | -312 | 0 | 0 | 12 | NaN | NaN | NaN | 836.65936 | 731.1348 | NaN |
| Psmd6 | LFSLYECR | -260 | -267 | 0 | 0.001868731 | 8 | NaN | NaN | NaN | 504.16794 | NaN | 555.0266 |
| Gc | EFPTYVEPTNDEICEAFR | -132 | -149 | 0 | 0 | 18 | NaN | NaN | NaN | 422.7405 | 528.4106 | 504.80878 |
| Gc | CADYSENTFTEYK | -407 | -419 | 0 | 0 | 13 | NaN | NaN | NaN | 572.4151 | NaN | 555.74005 |
| Gc | VEPTNDEICEAFR | -137 | -149 | 0 | 0.009657961 | 13 | NaN | NaN | NaN | 713.3998 | NaN | NaN |
| Alb | LGEYGFQNAILV | -422 | -433 | 0 | 0.003003218 | 12 | NaN | NaN | NaN | 399.37326 | 609.6657 | 615.46484 |
| Alb | RPCFSALTVDETYVPK | -509 | -524 | 1 | 0.00025015 | 16 | NaN | NaN | NaN | 7564.718 | NaN | NaN |

| | | | | | | | | | | | | |
|---|---|---|---|---|---|---|---|---|---|---|---|---|
| Alb | YNDLGEQHFK | -35 | -44 | 0 | 0.001636916 | 10 | NaN | NaN | NaN | 1000.93274 | NaN | NaN |
| Alb | LQTCCDKPLLK | -299 | -309 | 1 | 0.000826806 | 11 | NaN | NaN | NaN | 737.3758 | NaN | NaN |
| Rps27a | TYCFNKPEDK | -147 | -156 | 1 | 0.00025015 | 10 | NaN | NaN | NaN | 2443.1157 | NaN | NaN |
| Rps27a | CCLTYCFNKPED | -144 | -155 | 1 | 0.009657961 | 12 | NaN | NaN | NaN | 94.44804 | NaN | NaN |
| Crip1 | TLTSGGHAEHEGKPY | -37 | -51 | 1 | 0 | 15 | NaN | NaN | NaN | 1195.1078 | 479.39233 | 642.96326 |
| Crip1 | TLTSGGHAEHEGKPYCNHPCYSA | -37 | -59 | 1 | 0.000230707 | 23 | NaN | NaN | NaN | 1128.1162 | 1454.13 | 1180.7555 |
| Cast | DKDGKPLLPEPEETSK | -426 | -441 | 2 | 0.00111006 | 16 | NaN | NaN | NaN | 543.213 | 636.6553 | 442.06992 |
| Cast | SPVQSTPSKPSDK | -219 | -231 | 1 | 0 | 13 | NaN | NaN | NaN | 843.9211 | 1890.3588 | 1856.3569 |
| Cast | CGEDEDTVPAEYR | -408 | -420 | 0 | 0.009031675 | 13 | NaN | NaN | NaN | 416.66504 | NaN | NaN |
| Cnbp | CGESGHLAR | -162 | -170 | 0 | 0 | 9 | NaN | NaN | NaN | 414.33105 | 452.94354 | 740.07837 |
| Cnbp | CGETGHVAINCSK | -141 | -153 | 0 | 0 | 13 | NaN | NaN | NaN | 373.3892 | 631.0179 | 705.99176 |
| Dnaja2 | SVCSDCNGEGEVINEK | -184 | -199 | 0 | 0 | 16 | NaN | NaN | NaN | 482.63867 | 564.51587 | 625.4143 |
| Dnaja2 | CGFQFTFK | -280 | -287 | 0 | 0.006677519 | 8 | NaN | NaN | NaN | 237.78334 | NaN | NaN |
| Dpp4 | WEYYDSVYTER | -653 | -663 | 0 | 0 | 11 | NaN | NaN | NaN | 971.24884 | 1025.4431 | 864.76636 |
| Dpp4 | IISDKDGYK | -368 | -376 | 1 | 0.000550307 | 9 | NaN | NaN | NaN | 809.31104 | NaN | NaN |
| Dpp4 | FRPAEPHFTSDGSSF | -351 | -365 | 1 | 0.001092529 | 15 | NaN | NaN | NaN | 411.62375 | NaN | NaN |
| Ampd2 | PGNAPCLK | -47 | -54 | 0 | 0.004983786 | 8 | NaN | NaN | NaN | 162.46101 | NaN | NaN |
| Tnfrsf18 | VESQGDIVFGFR | -88 | -99 | 0 | 0.00111006 | 12 | NaN | NaN | NaN | 1905.1663 | NaN | NaN |
| Ppp1r7 | CIENLEELQSLR | -113 | -124 | 0 | 0.006915721 | 12 | NaN | NaN | NaN | 360.7636 | NaN | NaN |
| Gzmb | DQQPEAICGGF | -42 | -52 | 0 | 0.00051306 | 11 | NaN | NaN | NaN | 1899.1058 | 2183.4983 | 2524.9429 |
| Gzmb | NQICAGDPK | -185 | -193 | 0 | 0.00025015 | 9 | NaN | NaN | NaN | 1675.6534 | NaN | 1316.7174 |
| Gzmb | RNVNVKPGDVCYV | -132 | -144 | 2 | 0 | 13 | NaN | NaN | NaN | 689.03125 | 1008.5353 | 1273.6293 |
| Gzmb | NVNVKPGDVCYV | -133 | -144 | 1 | 0.001055699 | 12 | NaN | NaN | NaN | 1422.6802 | NaN | 1434.7788 |
| Gzmb | TFSNDIMLL | -104 | -112 | 0 | 0 | 9 | NaN | NaN | NaN | 738.00903 | 156.96246 | NaN |
| Gzmb | NVNVKPGDVCY | -133 | -143 | 1 | 0.00025015 | 11 | NaN | NaN | NaN | 1102.589 | NaN | NaN |
| Gzmb | ECESYFKN | -172 | -179 | 1 | 0.006788754 | 8 | NaN | NaN | NaN | 81.602776 | NaN | NaN |
| Rps8 | LTPEEEEILNK | -129 | -139 | 0 | 0 | 11 | NaN | NaN | NaN | 2478.378 | 4394.951 | 3921.5193 |
| Rps8 | TPEEEEILNK | -130 | -139 | 0 | 0 | 10 | NaN | NaN | NaN | 991.87976 | 944.2168 | 955.05164 |
| Ube2i | FKDDYPSSPPK | -64 | -74 | 1 | 0 | 11 | NaN | NaN | NaN | 2459.7366 | 2913.4353 | 2799.7664 |
| Ube2i | SILEEDKDWRPAITIK | -95 | -110 | 2 | 0 | 16 | NaN | NaN | NaN | 838.99384 | 579.8765 | 651.52563 |
| Ube2i | SILEEDKDWRPAITIK | -95 | -110 | 2 | 0 | 16 | NaN | NaN | NaN | 838.99384 | 579.8765 | 651.52563 |
| Ube2i | FEPPLFHPNVYPSGTVC | -77 | -93 | 0 | 0 | 17 | NaN | NaN | NaN | 587.52264 | 420.6777 | 654.7739 |
| Ube2i | PWEGGLFK | -52 | -59 | 0 | 0.001708008 | 8 | NaN | NaN | NaN | 576.8331 | 952.80725 | 141.44788 |
| Rps5 | VNQAIWLLCTGAR | -147 | -159 | 0 | 0 | 13 | NaN | NaN | NaN | 1221.9657 | 1567.8585 | 1151.4268 |
| Rps5 | DISLQDYIAVK | -32 | -42 | 0 | 0 | 11 | NaN | NaN | NaN | 248.39787 | 992.69104 | 989.063 |
| Nhp2 | CLEEVQALPTPL | -142 | -153 | 0 | 0.001055699 | 12 | NaN | NaN | NaN | 372.77237 | 545.5433 | NaN |
| Tpt1 | TEGAIDDSLIGGNA | -39 | -52 | 0 | 0.00078661 | 14 | NaN | NaN | NaN | 8800.111 | NaN | 15429.414 |
| Rbm22 | RGEECPYR | -175 | -182 | 1 | 0 | 8 | NaN | NaN | NaN | 1178.3518 | 793.72797 | 1310.2582 |
| Rbm22 | CLGENPYIR | -27 | -35 | 0 | 0.001868731 | 9 | NaN | NaN | NaN | 1670.5933 | NaN | NaN |
| Rpl11 | KVLEQLTGQTPVFSK | -38 | -52 | 1 | 0 | 15 | NaN | NaN | NaN | 1685.0585 | 2219.025 | 2387.662 |
| Rpl11 | QKYDGIILPGK | -168 | -178 | 1 | 0 | 11 | NaN | NaN | NaN | 537.2818 | 275.18948 | 880.369 |
| Pebp1 | GNDISSGTVLSDYVGSGPPSGTGLHR | -94 | -119 | 0 | 0 | 26 | NaN | NaN | NaN | 603.3944 | 832.0263 | 1342.7495 |
| Npm3 | SFFFGCELSGHTR | -36 | -48 | 0 | 0 | 13 | NaN | NaN | NaN | 997.1703 | 785.0015 | 475.2644 |
| Npm3 | LCGILPAK | -162 | -169 | 0 | 0.00051306 | 8 | NaN | NaN | NaN | 431.89407 | NaN | 400.2984 |
| Nutf2 | IDASCLTWEGQQFQGK | -34 | -49 | 0 | 0 | 16 | NaN | NaN | NaN | 768.5 | 1188.6776 | 981.53894 |
| Nutf2 | IQHSITAQDHQPTPDSCIIS | -64 | -83 | 0 | 0 | 20 | NaN | NaN | NaN | 354.64935 | 476.71033 | 419.68387 |
| Naga | CMGYPGTTLDK | -127 | -137 | 0 | 0.002434616 | 11 | NaN | NaN | NaN | 499.4946 | 363.64496 | 507.98206 |
| Txn | EAFQEALAAAGDK | -9 | -21 | 0 | 0 | 13 | NaN | NaN | NaN | 7224.171 | 7372.059 | 7248.132 |
| Txn | DCQDVAADCEVK | -61 | -72 | 0 | 0.000230707 | 12 | NaN | NaN | NaN | 419.30756 | NaN | 357.45508 |
| Itga2b | NRPPLEEDEEEE | -1022 | -1033 | 1 | 0 | 12 | NaN | NaN | NaN | 784.78107 | 824.6436 | 371.47427 |
| Itga2b | CNPLLFDLR | -96 | -104 | 0 | 0 | 9 | NaN | NaN | NaN | 827.49927 | 815.98694 | NaN |
| Itga2b | HGSVSIVVGAPR | -61 | -72 | 0 | 0 | 12 | NaN | NaN | NaN | 534.77454 | 501.68747 | NaN |
| Itga2b | DCGEDDLCVPQLR | -631 | -643 | 0 | 0.000591124 | 13 | NaN | NaN | NaN | 1332.3147 | NaN | NaN |
| Itga2b | DKHGSVSIVVGAPR | -59 | -72 | 1 | 0.001868731 | 14 | NaN | NaN | NaN | 218.23932 | NaN | NaN |
| Zyx | CNTCGQPITDR | -436 | -446 | 0 | 0.000431655 | 11 | NaN | NaN | NaN | 394.12787 | NaN | NaN |
| Vdac3 | SCSGVEFSTSGHAYTDTGK | -35 | -53 | 0 | 0 | 19 | NaN | NaN | NaN | 1384.7399 | 1569.64 | 1425.6638 |
| Vdac3 | VCNYGLTFTQK | -64 | -74 | 0 | 0 | 11 | NaN | NaN | NaN | 1146.3198 | 1208.7323 | 1237.6608 |
| Inhca | CQLCAGK | -185 | -191 | 0 | 0.005758933 | 7 | NaN | NaN | NaN | 1066.8851 | 906.70917 | NaN |
| Inhca | CQLCAGK | -185 | -191 | 0 | 0.005758933 | 7 | NaN | NaN | NaN | 1066.8851 | 906.70917 | NaN |
| Tagln2 | PEGQAPVK | -71 | -78 | 0 | 0.003998192 | 8 | NaN | NaN | NaN | 689.23413 | 950.0295 | 830.23444 |
| Tagln2 | TLMNLGGLAVAR | -128 | -139 | 0 | 0 | 12 | NaN | NaN | NaN | 864.3494 | 1337.4014 | 1075.0287 |
| Tagln2 | EQISQFLQAAER | -91 | -102 | 0 | 0 | 12 | NaN | NaN | NaN | 1529.7313 | 1399.2535 | 1541.0593 |
| Tagln2 | QMEQISQFLQAAER | -89 | -102 | 0 | 0 | 14 | NaN | NaN | NaN | 1564.3777 | 1649.424 | NaN |
| Ptges3 | LTFSCLGGSDNFK | -36 | -48 | 0 | 0 | 13 | NaN | NaN | NaN | 612.3393 | 831.4753 | 723.55853 |
| Ptges3 | NEIDLFHCIDPNDSK | -51 | -65 | 0 | 0.00122419 | 15 | NaN | NaN | NaN | 428.57324 | 720.03455 | 757.25464 |
| Ptges3 | DVNVNFEK | -26 | -33 | 0 | 0.001904399 | 8 | NaN | NaN | NaN | 971.94714 | NaN | NaN |
| Psmb2 | CVGEAGDTVQFAEYIQK | -46 | -62 | 0 | 0 | 17 | NaN | NaN | NaN | 614.9211 | 634.46747 | 660.06256 |
| Glud1 | IIAEGANGPTTPEADKIFLER | -400 | -420 | 1 | 0.00025015 | 21 | NaN | NaN | NaN | 511.33752 | 342.59467 | 496.08713 |
| Glud1 | PTAEFQDR | -489 | -496 | 0 | 0.00676572 | 8 | NaN | NaN | NaN | 449.91498 | 547.1672 | 588.6898 |
| Phf5a | HHPDLIFCR | -4 | -12 | 0 | 0.000907206 | 9 | NaN | NaN | NaN | 3492.9207 | 641.9463 | 1386.4932 |
| Phf5a | CVICDSYVRPC | -30 | -40 | 1 | 0 | 11 | NaN | NaN | NaN | 1043.887 | 828.3296 | 214.33417 |
| Phf5a | CVICGGPGVSDAYYCK | -58 | -73 | 0 | 0 | 16 | NaN | NaN | NaN | 741.22314 | 812.4041 | 763.2019 |
| Phf5a | GGPGVSDAYYCK | -62 | -73 | 0 | 0 | 12 | NaN | NaN | NaN | 669.1392 | NaN | 876.81 |
| Phf5a | HHPDLIFC | -4 | -11 | 0 | 0.006624942 | 8 | NaN | NaN | NaN | 949.21326 | NaN | NaN |
| Phf5a | KTDLFYER | -95 | -102 | 1 | 0.007105559 | 8 | NaN | NaN | NaN | 268.22684 | NaN | NaN |
| Strap | GHFGPIHCVR | -263 | -272 | 0 | 0 | 10 | NaN | NaN | NaN | 853.93823 | 936.25 | 804.97754 |
| Sdhd | CYFNYHDVGICR | -140 | -151 | 0 | 0 | 12 | NaN | NaN | NaN | 317.5149 | 364.09695 | 195.95497 |
| Sdhd | SLHWTSER | -63 | -70 | 0 | 0.00149795 | 8 | NaN | NaN | NaN | 816.0194 | NaN | NaN |

| | | | | | | | | | | | | |
|---|---|---|---|---|---|---|---|---|---|---|---|---|
| Alad | DEQGSAADSEDSPTIEAVR | -88 | -106 | 0 | 0 | 19 | NaN | NaN | NaN | 2320.0972 | 2779.3413 | 2761.3762 |
| Alad | SVLHSGYFHPLLR | -5 | -17 | 0 | 0 | 13 | NaN | NaN | NaN | 682.3331 | 798.3407 | 554.3474 |
| Alad | FASCFYGPFR | -200 | -209 | 0 | 0 | 10 | NaN | NaN | NaN | 365.64932 | 436.97266 | NaN |
| Alad | DAAQSSPAFGDR | -210 | -221 | 0 | 0 | 12 | NaN | NaN | NaN | 1034.129 | NaN | NaN |
| Tm9sf2 | VHGDIFRPPR | -350 | -359 | 1 | 0 | 10 | NaN | NaN | NaN | 625.65906 | 779.66144 | NaN |
| Tm9sf2 | IYYHVVETGSMGAR | -215 | -228 | 0 | 0.000471409 | 14 | NaN | NaN | NaN | 1971.9153 | NaN | NaN |
| Tm9sf2 | EDAQEEFGWK | -339 | -348 | 0 | 0 | 10 | NaN | NaN | NaN | 93.904686 | NaN | NaN |
| Srsf5 | EKDVERF | -17 | -23 | 2 | 0.005176406 | 7 | NaN | NaN | NaN | 292.17145 | 326.5879 | 2294.3464 |
| Srsf5 | VTFADAHRPK | -134 | -143 | 1 | 0 | 10 | NaN | NaN | NaN | 2923.2573 | NaN | NaN |
| Safb | EGQHYPER | -798 | -805 | 0 | 0 | 8 | NaN | NaN | NaN | 227.81769 | NaN | 331.65793 |
| Glyr1 | VDAVEEFLR | -83 | -91 | 0 | 0.000431655 | 9 | NaN | NaN | NaN | 2325.65 | NaN | NaN |
| Glyr1 | FKPDFYLK | -487 | -494 | 1 | 0.002168954 | 8 | NaN | NaN | NaN | 837.07916 | NaN | NaN |
| Glyr1 | DLVLGPSGVLQGIRPGK | -332 | -348 | 1 | 0.000907206 | 17 | NaN | NaN | NaN | 425.0539 | NaN | NaN |
| Npepps | LGLQNDLFSLAR | -617 | -628 | 0 | 0 | 12 | NaN | NaN | NaN | 955.52606 | 1223.8208 | 1193.2446 |
| Npepps | VTLSFPSTLQTGTGTLK | -130 | -146 | 0 | 0.00025015 | 17 | NaN | NaN | NaN | 336.50436 | 380.6757 | 468.39694 |
| Npepps | DADSIHQYLLQ | -902 | -912 | 0 | 0.00122419 | 11 | NaN | NaN | NaN | 770.2139 | NaN | NaN |
| Npepps | VLGATLSPELIQK | -783 | -795 | 0 | 0 | 13 | NaN | NaN | NaN | 446.35388 | NaN | NaN |
| Srsf2 | SYGRPPPDVEGMTSL | -2 | -16 | 1 | 0.003620005 | 15 | NaN | NaN | NaN | 393.36743 | 387.66318 | 664.36505 |
| Srsf2 | YGRPPDSHHS | -92 | -101 | 1 | 0 | 10 | NaN | NaN | NaN | 658.707 | 746.4709 | 867.76013 |
| Srsf1 | TKDIEDVFYK | -29 | -38 | 1 | 0.00025015 | 10 | NaN | NaN | NaN | 4416.1216 | 5130.146 | 4495.506 |
| Srsf1 | DGYDYDGYR | -75 | -83 | 0 | 0 | 9 | NaN | NaN | NaN | 1208.1986 | 1252.1061 | 1293.4766 |
| Srsf1 | RDGTGVVEFVR | -154 | -164 | 1 | 0.00142999 | 11 | NaN | NaN | NaN | 319.69897 | 1204.1813 | 1170.407 |
| Srsf1 | TKDIEDVFYKY | -29 | -39 | 2 | 0.001776473 | 11 | NaN | NaN | NaN | 36987.5 | NaN | NaN |
| Nudc | LVTSDPEINTK | -258 | -268 | 0 | 0 | 11 | NaN | NaN | NaN | 3222.3542 | 3701.176 | 3668.248 |
| Akr1a1 | KHHPEDVEPALR | -85 | -96 | 1 | 0.00025015 | 12 | NaN | NaN | NaN | 910.84076 | NaN | NaN |
| Mvp | SFFLQPGER | -307 | -315 | 0 | 0.007374243 | 9 | NaN | NaN | NaN | 289.22827 | NaN | NaN |
| Mvp | DAGPDGALLPR | -611 | -621 | 0 | 0.009031675 | 11 | NaN | NaN | NaN | 196.85333 | NaN | NaN |
| Eif3c | TCHSFIINEK | -749 | -758 | 0 | 0 | 10 | NaN | NaN | NaN | 1280.815 | 636.5083 | 1455.0952 |
| Eif3c | SEQDQAENEGEDSAVLMER | -530 | -548 | 0 | 0.000230707 | 19 | NaN | NaN | NaN | 276.7396 | 321.9513 | 277.0357 |
| Prpf38a | RVCDIILPR | -156 | -164 | 1 | 0.001636916 | 9 | NaN | NaN | NaN | 644.5757 | NaN | NaN |
| Glg1 | HLYFACR | -307 | -313 | 0 | 0.001463897 | 7 | NaN | NaN | NaN | 1532.0076 | NaN | NaN |
| Glg1 | DAHSQGEVVSCLEK | -255 | -268 | 0 | 0.00078661 | 14 | NaN | NaN | NaN | 530.13794 | NaN | NaN |
| Nudt9 | WVDISDQLK | -311 | -319 | 0 | 0 | 9 | NaN | NaN | NaN | 1228.108 | 1797.8278 | NaN |
| Pgam1 | FSGWYDADLSPAGHEEAK | -22 | -39 | 0 | 0 | 18 | NaN | NaN | NaN | 2908.511 | 3834.0503 | 3751.9478 |
| Pgam1 | FSGWYDADLSPAGHEEAKR | -22 | -40 | 1 | 0 | 19 | NaN | NaN | NaN | 846.0142 | 144.6947 | 925.82574 |
| Pgam1 | SYDVPPPPMEPDHPFYSNISK | -118 | -138 | 0 | 0 | 21 | NaN | NaN | NaN | 333.1914 | 646.33044 | 560.0402 |
| Pgam1 | GESAWNLENR | -12 | -21 | 0 | 0 | 10 | NaN | NaN | NaN | 1680.6134 | 1005.46356 | NaN |
| P4hb | LEPDMEEDDDQK | -492 | -503 | 0 | 0 | 12 | NaN | NaN | NaN | 2245.4385 | 2799.467 | 2747.3455 |
| P4hb | LITLEEEMTK | -319 | -328 | 0 | 0 | 10 | NaN | NaN | NaN | 823.7952 | NaN | NaN |
| P4hb | KVDATEESDLAQQYGVR | -83 | -99 | 1 | 0.000298805 | 17 | NaN | NaN | NaN | 669.2527 | 668.09564 | 690.4916 |
| P4hb | FLESGGQDGAGDDEDLDLE | -471 | -489 | 0 | 0.00078661 | 19 | NaN | NaN | NaN | 341.5026 | 303.20325 | 376.64886 |
| P4hb | FEEVAFDEKK | -379 | -388 | 1 | 0 | 10 | NaN | NaN | NaN | 1463.9839 | NaN | NaN |
| Galk2 | DALHPEPYSR | -288 | -297 | 0 | 0.003069674 | 10 | NaN | NaN | NaN | 1223.5322 | 1261.366 | NaN |
| Adprh | LAQMGDLEAIDVAR | -42 | -55 | 0 | 0 | 14 | NaN | NaN | NaN | 915.51434 | 1014.3673 | NaN |
| Adprh | DALLAAGDSWK | -285 | -295 | 0 | 0.005917757 | 11 | NaN | NaN | NaN | 562.4414 | NaN | NaN |
| Thoc2 | DALLDHDLALPL | -731 | -742 | 0 | 0.001055699 | 12 | NaN | NaN | NaN | 760.31146 | NaN | NaN |
| Rrbp1 | DALNQATSQVESK | -1010 | -1022 | 0 | 0 | 13 | NaN | NaN | NaN | 1628.0496 | NaN | NaN |
| Ikzf3 | SHTGERPFQCNQC | -138 | -150 | 1 | 0.000471409 | 13 | NaN | NaN | NaN | 413.1934 | 464.5949 | 455.32727 |
| Arhgdib | GPRPEEYEFLTPVEEAPK | -146 | -163 | 1 | 0 | 18 | NaN | NaN | NaN | 568.92456 | 702.1496 | 897.8764 |
| Arhgdib | TLLGDVPVVADPTVPNVTVT | -50 | -69 | 0 | 0.000230707 | 20 | NaN | NaN | NaN | 413.42197 | 364.46103 | 505.86176 |
| Arhgdib | MVGSYGPRPEEYEFLTPVEEAPK | -141 | -163 | 1 | 0 | 23 | NaN | NaN | NaN | 3569.106 | 479.6971 | NaN |
| Gfpt1 | VDSTTCLFPVEEK | -257 | -269 | 0 | 0 | 13 | NaN | NaN | NaN | 934.50543 | 1449.7902 | NaN |
| Gfpt1 | LNYHVPR | -8 | -14 | 0 | 0.00272025 | 7 | NaN | NaN | NaN | 579.19006 | NaN | NaN |
| Sp100 | RDHEFITGK | -51 | -59 | 1 | 0.001636916 | 9 | NaN | NaN | NaN | 942.616 | 331.90045 | NaN |
| Sp100 | KGVLQDPPR | -455 | -463 | 1 | 0.00201047 | 9 | NaN | NaN | NaN | 832.1689 | NaN | NaN |
| Eftud2 | SIVIRPLEPQPAPHLAR | -914 | -930 | 1 | 0 | 17 | NaN | NaN | NaN | 611.20605 | 811.0247 | 780.7538 |
| Eftud2 | DAVVAQEPLHR | -792 | -802 | 0 | 0 | 11 | NaN | NaN | NaN | 866.5377 | 1106.9591 | NaN |
| Eftud2 | SGTIHAGQPVK | -497 | -507 | 0 | 0.001636916 | 11 | NaN | NaN | NaN | 670.58344 | 942.45544 | NaN |
| Eftud2 | PVNPSELPK | -593 | -601 | 0 | 0.005270854 | 9 | NaN | NaN | NaN | 515.71246 | NaN | 1839.2157 |
| Eftud2 | VEESGEHVILGT | -620 | -631 | 0 | 0.001055699 | 12 | NaN | NaN | NaN | 1251.5356 | NaN | NaN |
| Atl3 | EQLQALIPYVLNPSK | -291 | -305 | 0 | 0.000431655 | 15 | NaN | NaN | NaN | 500.29507 | 535.1332 | 298.95325 |
| Atl3 | LKDIASEFK | -282 | -290 | 1 | 0 | 9 | NaN | NaN | NaN | 396.5176 | 531.64215 | 511.95248 |
| Abce1 | DAYTHPQFVTDVMKPL | -434 | -449 | 1 | 0.001392869 | 16 | NaN | NaN | NaN | 273.4519 | NaN | 645.2695 |
| Ero1a | SGYLDDCTCDVETIDK | -40 | -55 | 0 | 0.000431655 | 16 | NaN | NaN | NaN | 400.61655 | 303.9232 | 376.3863 |
| Ero1a | NMPESGPSYEFQLTR | -419 | -433 | 0 | 0.001092529 | 15 | NaN | NaN | NaN | 726.6543 | NaN | NaN |
| Ero1a | LLEILHEIK | -340 | -348 | 0 | 0.001708008 | 9 | NaN | NaN | NaN | 672.5542 | NaN | NaN |
| Ero1a | RDCAVKPCHSDEVPDGIK | -97 | -114 | 2 | 0.00437931 | 18 | NaN | NaN | NaN | 307.32114 | NaN | NaN |
| Banf1 | GEKPVGSLAGIGDVLSK | -16 | -32 | 1 | 0 | 17 | NaN | NaN | NaN | 1695.5361 | 2151.6746 | 2073.671 |
| Samhd1 | SFLYEIVSNKR | -327 | -337 | 1 | 0 | 11 | NaN | NaN | NaN | 900.9097 | 2252.569 | NaN |
| Smc3 | LPIDKEEVSLR | -89 | -99 | 1 | 0 | 11 | NaN | NaN | NaN | 304.08124 | 314.07288 | 326.3248 |
| Smc3 | RVDALNDEIR | -796 | -805 | 1 | 0.005333786 | 10 | NaN | NaN | NaN | 1794.257 | NaN | NaN |
| Cul1 | LHESTQDELAR | -279 | -289 | 0 | 0 | 11 | NaN | NaN | NaN | 4665.798 | NaN | NaN |
| Cul1 | DCLFRPL | -169 | -175 | 1 | 0.003681885 | 7 | NaN | NaN | NaN | 695.85645 | NaN | NaN |
| Hpcal1 | LKDCPTGHLTVDEFK | -35 | -49 | 1 | 0 | 15 | NaN | NaN | NaN | 629.55664 | 567.00696 | 412.47437 |
| Thbs1 | TIVTTLQDSIR | -289 | -299 | 0 | 0.00025015 | 11 | NaN | NaN | NaN | 895.9836 | 324.29538 | 987.83374 |
| Thbs1 | FQDLLDAVWADK | -75 | -86 | 0 | 0 | 12 | NaN | NaN | NaN | 713.4568 | 1007.5132 | 980.51074 |
| Hnrnpc | RVPPPPPIAR | -142 | -151 | 1 | 0.00025015 | 10 | NaN | NaN | NaN | 1125.9126 | 1549.5232 | 1318.9448 |
| Hnrnpc | QVLDINLAAEPK | -78 | -89 | 0 | 0 | 12 | NaN | NaN | NaN | 775.70245 | 1006.8315 | 816.6265 |

| | | | | | | | | | | | | |
|---|---|---|---|---|---|---|---|---|---|---|---|---|
| Hnrnpc | KVDSLLESLEK | -207 | -217 | 1 | 0.00025015 | 11 | NaN | NaN | NaN | 833.168 | NaN | 1337.6694 |
| Hnrnpc | DSAEEGDLLDDDDNEDRGDDQLELK | -267 | -291 | 1 | 0.00025015 | 25 | NaN | NaN | NaN | 413.4853 | NaN | 368.67926 |
| Hnrnpc | VPPPPPIAR | -143 | -151 | 0 | 0.002595029 | 9 | NaN | NaN | NaN | 8676.385 | NaN | NaN |
| Hnrnpc | LDYDFQR | -124 | -130 | 0 | 0.00025015 | 7 | NaN | NaN | NaN | 627.2407 | NaN | NaN |
| Psma1 | QTCPSANYFDCR | -146 | -157 | 0 | 0 | 12 | NaN | NaN | NaN | 753.58673 | 705.2748 | 716.49445 |
| Psma1 | QAAEEPAEK | -248 | -256 | 0 | 0 | 9 | NaN | NaN | NaN | 589.0418 | 303.57504 | NaN |
| Psma1 | TQIPTQR | -116 | -122 | 0 | 0.008253452 | 7 | NaN | NaN | NaN | 616.41003 | NaN | NaN |
| Hnrnpa2b1 | FDDHDPVDK | -160 | -168 | 0 | 0 | 9 | NaN | NaN | NaN | 8912.32 | 10106.232 | 10222.71 |
| Hnrnpa2b1 | TIEIITDRQSGK | -140 | -151 | 1 | 0.000550307 | 12 | NaN | NaN | NaN | 2625.9233 | 643.7926 | 743.1416 |
| Rbbp7 | YMPQNPHIIATK | -131 | -142 | 0 | 0 | 12 | NaN | NaN | NaN | 755.91425 | 681.33466 | 735.35266 |
| Rbbp7 | GKIECEIK | -112 | -119 | 1 | 0.00025015 | 8 | NaN | NaN | NaN | 1353.3954 | 583.4618 | NaN |
| Rbbp7 | PSHLVDAHTA | -264 | -273 | 0 | 0.004983786 | 10 | NaN | NaN | NaN | 842.82666 | NaN | NaN |
| Suclg2 | SHNGPVIVGSPQGGVDIEEVAAS | -162 | -184 | 0 | 0.000431655 | 23 | NaN | NaN | NaN | 598.2823 | 966.84644 | 607.07587 |
| Sub1 | SRDDNMFQIGK | -58 | -68 | 1 | 0 | 11 | NaN | NaN | NaN | 859.55426 | 195.29117 | 2407.4158 |
| Sub1 | KGISLNMEQWSQLK | -101 | -114 | 1 | 0 | 14 | NaN | NaN | NaN | 792.0354 | 3249.0823 | 532.69684 |
| Cdc42 | FDTAGQEDYDR | -56 | -66 | 0 | 0 | 11 | NaN | NaN | NaN | 179.29608 | 355.7772 | 412.54932 |
| Cdc42 | WVPEITHH | -97 | -104 | 0 | 0.000712251 | 8 | NaN | NaN | NaN | 460.87564 | 337.86218 | NaN |
| Cdc42 | DDPSTIEKL | -121 | -129 | 1 | 0.004321969 | 9 | NaN | NaN | NaN | 3524.3499 | NaN | NaN |
| Cdc42 | LFDTAGQEDYDR | -55 | -66 | 0 | 0 | 12 | NaN | NaN | NaN | 1438.8344 | NaN | NaN |
| Cdc42 | NVFDEAILAA | -167 | -176 | 0 | 0.000431655 | 10 | NaN | NaN | NaN | 439.4681 | NaN | NaN |
| Ptprcap | LSHASGGYYHPA | -57 | -68 | 0 | 0.00025015 | 12 | NaN | NaN | NaN | 864.11786 | 1401.4895 | 1567.5248 |
| Ptprcap | LSHASGGYYHPAR | -57 | -69 | 0 | 0 | 13 | NaN | NaN | NaN | 785.4454 | 1557.1746 | 1279.0372 |
| Ptprcap | LGAALWGHTCR | -70 | -80 | 0 | 0 | 11 | NaN | NaN | NaN | 1987.3342 | 1380.0746 | 1458.2588 |
| Srrt | FKGPEFVR | -721 | -728 | 1 | 0 | 8 | NaN | NaN | NaN | 1290.6251 | 1256.9653 | 1180.889 |
| Srrt | EICWNLQNIR | -476 | -485 | 0 | 0 | 10 | NaN | NaN | NaN | 963.3233 | 1294.1931 | 1333.9528 |
| Srrt | TQLWASEPGTPPVPTSLPSQNPILK | -534 | -558 | 0 | 0 | 25 | NaN | NaN | NaN | 333.27292 | 443.1941 | NaN |
| Srrt | LRECELSPGVN | -486 | -496 | 1 | 0 | 11 | NaN | NaN | NaN | 1112.0043 | NaN | NaN |
| Srrt | LDDSVDETEAVKR | -159 | -171 | 1 | 0.002068213 | 13 | NaN | NaN | NaN | 316.3866 | NaN | NaN |
| Hsf1 | DDTEFQHPCFLR | -95 | -106 | 0 | 0.000230707 | 12 | NaN | NaN | NaN | 949.0273 | 742.874 | 700.7084 |
| Bub3 | DLNTDQENLVGTHDAPIR | -83 | -100 | 0 | 0 | 18 | NaN | NaN | NaN | 730.1561 | 519.09717 | NaN |
| Bub3 | ENNIEQIYPVNAISF | -228 | -242 | 0 | 0.00437931 | 15 | NaN | NaN | NaN | 233.22156 | 417.39084 | NaN |
| Bub3 | TPCNAGTFSQPEK | -127 | -139 | 0 | 0.00025015 | 13 | NaN | NaN | NaN | 5414.11 | NaN | NaN |
| Uqcrb | DDTLHETEDVK | -35 | -45 | 0 | 0.00078661 | 11 | NaN | NaN | NaN | 197.53171 | 219.6784 | 1208.6711 |
| Fdps | QNFIQHFSQIVK | -15 | -26 | 0 | 0.001023236 | 12 | NaN | NaN | NaN | 446.3961 | 1167.9567 | 1301.5347 |
| Fdps | DLMTAPQGHVDLGR | -174 | -187 | 0 | 0 | 14 | NaN | NaN | NaN | 568.0712 | 483.01044 | NaN |
| Psmc3 | VSPDVNYEELAR | -378 | -389 | 0 | 0.000627943 | 12 | NaN | NaN | NaN | 3258.8142 | NaN | NaN |
| Nsfl1c | SSILINEAEPTTNIQIR | -285 | -301 | 0 | 0.000230707 | 17 | NaN | NaN | NaN | 519.7516 | 436.21606 | 378.67844 |
| Nsfl1c | HSGQDVHVV | -175 | -183 | 0 | 0.005758933 | 9 | NaN | NaN | NaN | 743.0846 | NaN | NaN |
| Nsfl1c | QDPSNAQFLESIR | -202 | -214 | 0 | 0.00025015 | 13 | NaN | NaN | NaN | 315.68433 | NaN | NaN |
| Dock2 | KEDGTTLHDGYHELVVLK | -536 | -553 | 1 | 0 | 18 | NaN | NaN | NaN | 1245.1909 | 272.91003 | 474.64645 |
| Dock2 | KEDGTTLHDGYHELVVLK | -536 | -553 | 1 | 0 | 18 | NaN | NaN | NaN | 1245.1909 | 272.91003 | 474.64645 |
| Dock2 | SVENFVSLVK | -1169 | -1178 | 0 | 0 | 10 | NaN | NaN | NaN | 360.1168 | 731.42114 | 511.32294 |
| Dock2 | LLHTWLLK | -1244 | -1251 | 0 | 0.00453359 | 8 | NaN | NaN | NaN | 990.6396 | NaN | NaN |
| Dock2 | KSVENFVSLVK | -1168 | -1178 | 1 | 0.002780285 | 11 | NaN | NaN | NaN | 967.71094 | NaN | NaN |
| Mtrex | EVALPADEDYIPLKPR | -111 | -126 | 1 | 0 | 16 | NaN | NaN | NaN | 784.9725 | 745.9713 | 739.9317 |
| Mtrex | LPQVEHVLPLLK | -436 | -447 | 0 | 0.00025015 | 12 | NaN | NaN | NaN | 322.0067 | 339.82266 | 345.84418 |
| Mtrex | THEVALPADEDYIPLKPR | -109 | -126 | 1 | 0 | 18 | NaN | NaN | NaN | 434.58786 | 599.00885 | NaN |
| Ranbp2 | FKTPEEAALFK | -1279 | -1289 | 1 | 0.000471409 | 11 | NaN | NaN | NaN | 806.0641 | NaN | 817.49146 |
| Ranbp2 | QNQPTSCVSAPASSETSR | -1587 | -1604 | 0 | 0.00025015 | 18 | NaN | NaN | NaN | 405.77655 | 327.82544 | NaN |
| Ranbp2 | KAEDIENDALSPEEQEECK | -690 | -708 | 1 | 0.001868731 | 19 | NaN | NaN | NaN | 222.9067 | NaN | 230.32277 |
| Ranbp2 | NSIPEPIDPLFK | -620 | -631 | 0 | 0.00122419 | 12 | NaN | NaN | NaN | 1268.7048 | NaN | NaN |
| Galnt6 | NLGTNQCLDVGENNR | -503 | -517 | 0 | 0.000394361 | 15 | NaN | NaN | NaN | 753.24603 | 592.89716 | 793.75684 |
| Galnt6 | DEEWELTQDQLIR | -578 | -590 | 0 | 0.000627943 | 13 | NaN | NaN | NaN | 1103.302 | NaN | NaN |
| Rsl1d1 | VKVPLPHSIL | -80 | -89 | 1 | 0.00025015 | 10 | NaN | NaN | NaN | 559.13196 | 537.8417 | NaN |
| Rsl1d1 | DEFDSPEQTEGFYK | -101 | -114 | 0 | 0 | 14 | NaN | NaN | NaN | 458.05884 | 454.32184 | NaN |
| Psmc2 | ESDTGLAPPALWDLAADKQ | -67 | -85 | 1 | 0.009002904 | 19 | NaN | NaN | NaN | 433.95804 | 458.47995 | NaN |
| Psmc2 | REVVETPLLHPER | -188 | -200 | 1 | 0.00025015 | 13 | NaN | NaN | NaN | 415.26645 | NaN | NaN |
| Psmc2 | DEGDIALLK | -26 | -34 | 0 | 0.003620005 | 9 | NaN | NaN | NaN | 399.4303 | NaN | NaN |
| Thy1 | VTSLTACLVNQNLR | -22 | -35 | 0 | 0 | 14 | NaN | NaN | NaN | 614.6372 | 830.5067 | 755.2991 |
| Usp4 | DEGLSHITPR | -550 | -559 | 0 | 0.000550307 | 10 | NaN | NaN | NaN | 2044.3344 | NaN | NaN |
| Cwc22 | FFGLLAGR | -505 | -512 | 0 | 0.003490217 | 8 | NaN | NaN | NaN | 541.75024 | 502.7263 | NaN |
| Parp1 | NELIWNIKDELKK | -242 | -254 | 2 | 0 | 13 | NaN | NaN | NaN | 464.66364 | 558.30255 | 578.05096 |
| Eif5b | SIDALKDWFR | -1186 | -1195 | 1 | 0.00025015 | 10 | NaN | NaN | NaN | 2386.436 | 2598.4321 | 711.33356 |
| Eif5b | HFEATDILVSK | -1171 | -1181 | 0 | 0.00025015 | 11 | NaN | NaN | NaN | 10796.442 | 7666.5664 | NaN |
| Eif5b | HTHVQDGEAGGIT | -649 | -661 | 0 | 0.000230707 | 13 | NaN | NaN | NaN | 2222.5332 | NaN | NaN |
| Cdv3 | DENEWKEFEQR | -106 | -116 | 1 | 0.00025015 | 11 | NaN | NaN | NaN | 819.1775 | 719.11584 | NaN |
| Rps19 | LKVPEWVDTV | -28 | -37 | 1 | 0.000712251 | 10 | NaN | NaN | NaN | 939.5778 | 1091.8354 | 1106.1803 |
| Rps19 | VLQALEGLK | -103 | -111 | 0 | 0.001055699 | 9 | NaN | NaN | NaN | 635.47833 | 389.6187 | 407.57333 |
| Rps19 | VPEWVDTVKL | -30 | -39 | 1 | 0 | 10 | NaN | NaN | NaN | 623.6516 | NaN | 1064.6667 |
| Rfc2 | DEPHPLLVK | -251 | -259 | 0 | 0.004983786 | 9 | NaN | NaN | NaN | 1305.9702 | NaN | NaN |
| Rfc2 | GHYELPWVEK | -27 | -36 | 0 | 0.002810811 | 10 | NaN | NaN | NaN | 610.6785 | NaN | NaN |
| Pml | LQDFISCITQR | -387 | -397 | 0 | 0.00111006 | 11 | NaN | NaN | NaN | 761.5285 | NaN | NaN |
| Pml | SNPESTVPEKK | -536 | -546 | 1 | 0.001392869 | 11 | NaN | NaN | NaN | 226.36316 | NaN | NaN |
| Anxa11 | LLISLSQGNR | -335 | -344 | 0 | 0 | 10 | NaN | NaN | NaN | 612.5791 | 604.8334 | 503.40692 |
| Anxa11 | GFGTDEQAIIDCLGSR | -213 | -228 | 0 | 0.00025015 | 16 | NaN | NaN | NaN | 234.34326 | NaN | NaN |
| Capzb | SGSGTMNLGGSLTR | -182 | -195 | 0 | 0 | 14 | NaN | NaN | NaN | 751.9473 | 1756.5458 | 312.54373 |
| Capzb | LCEDLLSSVDQPLK | -35 | -48 | 0 | 0 | 14 | NaN | NaN | NaN | 884.30817 | 442.4162 | 805.7925 |
| Supt16h | SINLGFSDLTNK | -378 | -389 | 0 | 0 | 12 | NaN | NaN | NaN | 1176.0284 | 2089.3635 | 2095.1482 |

| | | | | | | | | | | | | |
|---|---|---|---|---|---|---|---|---|---|---|---|---|
| Supt16h | NEGNIFPNPEAT | -582 | -593 | 0 | 0.002566359 | 12 | NaN | NaN | NaN | 257.44907 | NaN | NaN |
| Pkn2 | DFDLEPEPPPAPPR | -566 | -579 | 0 | 0.00025015 | 14 | NaN | NaN | NaN | 340.0652 | NaN | NaN |
| Pdlim1 | HPECYVCTDCGINLK | -278 | -292 | 0 | 0.00025015 | 15 | NaN | NaN | NaN | 449.7569 | 470.83823 | 584.3362 |
| Pdlim1 | TTQQIVLQGPGPWGFR | -2 | -17 | 0 | 0.000471409 | 16 | NaN | NaN | NaN | 255.05365 | 439.6699 | 331.5449 |
| Pdlim1 | SEPQEVLHIGSAHNR | -108 | -122 | 0 | 0.001636916 | 15 | NaN | NaN | NaN | 183.67412 | NaN | NaN |
| G3bp1 | HPDSHQLFIGNLPHEVDK | -334 | -351 | 0 | 0 | 18 | NaN | NaN | NaN | 4925.4727 | 1611.928 | 1462.762 |
| G3bp1 | NLPPSGAVPVTGTPPHVVK | -256 | -274 | 0 | 0.00025015 | 19 | NaN | NaN | NaN | 1513.2683 | 2064.7375 | 1867.9034 |
| G3bp1 | FGFVVFDDSEPVQK | -378 | -391 | 0 | 0.00111006 | 14 | NaN | NaN | NaN | 1197.2429 | NaN | NaN |
| G3bp1 | VPASQPRPE | -275 | -283 | 1 | 0.004690496 | 9 | NaN | NaN | NaN | 1049.3389 | NaN | NaN |
| G3bp1 | YQDEVFGGF | -133 | -141 | 0 | 0.005668207 | 9 | NaN | NaN | NaN | 956.75 | NaN | NaN |
| Was | DFIEDQGGLEAVR | -294 | -306 | 0 | 0 | 13 | NaN | NaN | NaN | 568.1568 | 343.56503 | 615.995 |
| Was | GGPPPPPPPATGR | -382 | -394 | 0 | 0.003521747 | 13 | NaN | NaN | NaN | 331.707 | NaN | NaN |
| Fkbp5 | VHLEGCCGGR | -170 | -179 | 0 | 0.000471409 | 10 | NaN | NaN | NaN | 559.71063 | NaN | 788.02344 |
| Psip1 | SNASSDVEVEEK | -102 | -113 | 0 | 0 | 12 | NaN | NaN | NaN | 510.31848 | NaN | NaN |
| H3c1 | QEACEAYLVGLFEDTNLCAIHAK | -94 | -116 | 0 | 0 | 23 | NaN | NaN | NaN | 363.76508 | 401.60492 | 482.9289 |
| Ebna1bp2 | SDQELQDAFSR | -19 | -29 | 0 | 0 | 11 | NaN | NaN | NaN | 1666.8496 | NaN | NaN |
| Ap2a1 | LLQCYPPPEDAAVK | -264 | -277 | 0 | 0.007987481 | 14 | NaN | NaN | NaN | 454.1866 | NaN | NaN |
| Aqr | HLHIIPTEPFPTSR | -1333 | -1346 | 0 | 0 | 14 | NaN | NaN | NaN | 907.2799 | 1500.8792 | NaN |
| Aqr | TVVEVAKPNIGENWPTR | -518 | -534 | 1 | 0 | 17 | NaN | NaN | NaN | 894.2071 | 463.51846 | 430.94702 |
| Aqr | HTQIEAIR | -804 | -811 | 0 | 0.00111006 | 8 | NaN | NaN | NaN | 198.21036 | NaN | NaN |
| Rpl9 | RDFNHINVELS | -36 | -46 | 1 | 0 | 11 | NaN | NaN | NaN | 990.7283 | 1489.2091 | 1699.6633 |
| Rpl9 | TICSHVQN | -72 | -79 | 0 | 0.006654467 | 8 | NaN | NaN | NaN | 3644.6123 | NaN | NaN |
| Son | SHHDGNLESDSFLK | -142 | -155 | 0 | 0 | 14 | NaN | NaN | NaN | 291.98462 | 391.3647 | NaN |
| Son | DFPESSTTDVQR | -322 | -333 | 0 | 0.000712251 | 12 | NaN | NaN | NaN | 1161.8789 | NaN | NaN |
| Son | SDLYENEYDR | -1550 | -1559 | 0 | 0.00204437 | 10 | NaN | NaN | NaN | 119.53258 | NaN | NaN |
| Mfap1a;Mfap1 | ESSDEEDEEFQFIK | (51);(51) | (64);(64) | 0 | 0 | 14 | NaN | NaN | NaN | 368.05518 | NaN | NaN |
| Acaa2 | LCGSGFQSIVSGCQEICSK | -91 | -109 | 0 | 0 | 19 | NaN | NaN | NaN | 389.4543 | 393.8571 | 414.5318 |
| Larp1 | DFSQLLNCPEFVPR | -472 | -485 | 0 | 0.001392869 | 14 | NaN | NaN | NaN | 1270.2227 | NaN | NaN |
| Larp1 | HFDYQFGYR | -333 | -341 | 0 | 0.002434616 | 9 | NaN | NaN | NaN | 388.22562 | NaN | NaN |
| Larp1 | FEPEYSQIK | -672 | -680 | 0 | 0.00450079 | 9 | NaN | NaN | NaN | 146.53922 | NaN | NaN |
| Atp5mf | DFTPSGIAGAF | -28 | -38 | 0 | 0.000230707 | 11 | NaN | NaN | NaN | 590.5337 | 1007.9915 | 876.8886 |
| Prdx6 | PGGLLLGDEAPNFEANTTIGR | -2 | -22 | 0 | 0 | 21 | NaN | NaN | NaN | 425.9318 | 546.3602 | 525.7147 |
| Rnf213 | TYSPGLQGQEPVR | -4156 | -4168 | 0 | 0.00025015 | 13 | NaN | NaN | NaN | 1883.5957 | 757.76355 | NaN |
| Rnf213 | LVEIGFPEK | -915 | -923 | 0 | 0.001636916 | 9 | NaN | NaN | NaN | 1333.9603 | NaN | NaN |
| Rnf213 | DGGLHDSVSR | -957 | -966 | 0 | 0.000230707 | 10 | NaN | NaN | NaN | 719.33154 | NaN | NaN |
| Rnf213 | DFVDKYDELK | -1272 | -1281 | 1 | 0 | 10 | NaN | NaN | NaN | 635.58685 | NaN | NaN |
| Cdc123 | DYTQYYDHISK | -194 | -204 | 0 | 0 | 11 | NaN | NaN | NaN | 310.3653 | NaN | 677.77435 |
| Irf3 | NFLNPAPQENPLK | -176 | -188 | 0 | 0.00025015 | 13 | NaN | NaN | NaN | 1614.771 | NaN | 1198.886 |
| Irf3 | DGAVFDLRPF | -309 | -318 | 1 | 0 | 10 | NaN | NaN | NaN | 671.26654 | 306.80194 | NaN |
| Zranb2 | DGDWICPDKK | -10 | -19 | 1 | 0 | 10 | NaN | NaN | NaN | 432.35434 | NaN | 374.81323 |
| Zranb2 | ESEGEEEDEDEDLSKY | -152 | -167 | 1 | 0.003393785 | 16 | NaN | NaN | NaN | 368.9323 | NaN | NaN |
| Blmh | ICFVNDPRPQHK | -268 | -279 | 1 | 0 | 12 | NaN | NaN | NaN | 761.34265 | 821.00574 | 719.91724 |
| Blmh | YNNQPIDFLK | -300 | -309 | 0 | 0.00025015 | 10 | NaN | NaN | NaN | 844.5249 | 1331.104 | 1548.075 |
| Blmh | TFTAVSEKDNQEGTFVK | -375 | -391 | 1 | 0.001092529 | 17 | NaN | NaN | NaN | 573.69116 | 469.94858 | 533.44666 |
| Myl12b | FTDEEVDELYR | -134 | -144 | 0 | 0 | 11 | NaN | NaN | NaN | 3117.0442 | 3117.5579 | 3048.4868 |
| Myl12b | GNFNYIEFT | -152 | -160 | 0 | 0.001610812 | 9 | NaN | NaN | NaN | 692.39856 | NaN | NaN |
| Macf1 | LLLEECEQR | -1058 | -1066 | 0 | 0.002199523 | 9 | NaN | NaN | NaN | 2116.1853 | NaN | NaN |
| Hdac1 | QCGSDSLSGDR | -260 | -270 | 0 | 0 | 11 | NaN | NaN | NaN | 400.05542 | 702.8386 | 461.49185 |
| Hdac1 | DGIDDESYEAIFKPVMS | -230 | -246 | 1 | 0.002780285 | 17 | NaN | NaN | NaN | 508.22852 | NaN | NaN |
| Psmb7 | IHFISPNIYCC | -77 | -87 | 0 | 0 | 11 | NaN | NaN | NaN | 771.2423 | 999.39386 | 1077.8492 |
| Usp5 | EEDTSAGTGDPPR | -81 | -93 | 0 | 0.000230707 | 13 | NaN | NaN | NaN | 726.93634 | NaN | NaN |
| Cmss1 | DGLHLNPLK | -220 | -228 | 0 | 0.00111006 | 9 | NaN | NaN | NaN | 692.3148 | 356.92596 | NaN |
| Hadha | MLSSCTTPQEATR | -106 | -118 | 0 | 0.007819401 | 13 | NaN | NaN | NaN | 517.79193 | NaN | NaN |
| Tcea1 | DGPSTDKDPEEK | -78 | -89 | 1 | 0.00713295 | 12 | NaN | NaN | NaN | 593.9955 | 625.3788 | 584.3154 |
| Xpot | SIDQEEDLDFVAR | -279 | -291 | 0 | 0.000907206 | 13 | NaN | NaN | NaN | 738.5698 | NaN | NaN |
| Xpot | EEACDCLFEIVNK | -242 | -254 | 0 | 0.000230707 | 13 | NaN | NaN | NaN | 626.9405 | NaN | NaN |
| Atic | RLDFNLVR | -90 | -97 | 1 | 0.007851348 | 8 | NaN | NaN | NaN | 1466.4275 | 1766.4142 | 1659.6006 |
| Atic | THTAQYDEAISDYFR | -180 | -194 | 0 | 0.001610812 | 15 | NaN | NaN | NaN | 823.12646 | 559.9352 | 716.65497 |
| Atic | NAWQLVTELR | -245 | -254 | 0 | 0 | 10 | NaN | NaN | NaN | 528.9374 | 340.71814 | 1366.8805 |
| Atic | LSGVSVSSDAFFPFR | -531 | -545 | 0 | 0 | 15 | NaN | NaN | NaN | 364.88788 | 956.2729 | 887.77167 |
| Atic | YKPDENEVR | -371 | -379 | 1 | 0.008014596 | 9 | NaN | NaN | NaN | 2765.2546 | NaN | NaN |
| Gmps | HNDTELIR | -392 | -399 | 0 | 0 | 8 | NaN | NaN | NaN | 2121.007 | 2385.4902 | 2399.3464 |
| Gmps | VIEPLKDFHKDEVR | -407 | -420 | 2 | 0.000230707 | 14 | NaN | NaN | NaN | 320.2125 | 754.71594 | NaN |
| Gmps | QADFEAHNILR | -590 | -600 | 0 | 0 | 11 | NaN | NaN | NaN | 8222.621 | NaN | NaN |
| Ptpn6 | WYHGHISGGQAESLLQAK | -110 | -127 | 0 | 0.000471409 | 18 | NaN | NaN | NaN | 618.845 | 401.31018 | 357.8465 |
| Ptpn6 | GEPWTFLVR | -128 | -136 | 0 | 0.005522803 | 9 | NaN | NaN | NaN | 3665.7798 | 1522.7821 | NaN |
| Ptpn6 | VRVDDQVTHIR | -43 | -53 | 1 | 0.000907206 | 11 | NaN | NaN | NaN | 343.9053 | NaN | NaN |
| Psmd3 | QLDSPEDAEFIVAK | -423 | -436 | 0 | 0 | 14 | NaN | NaN | NaN | 1585.6053 | 1990.024 | 2018.6353 |
| Psmd3 | LHYSLYDQAEK | -255 | -265 | 0 | 0 | 11 | NaN | NaN | NaN | 919.08813 | 845.0636 | 962.2549 |
| Psmd3 | DGVIEASINHEK | -440 | -451 | 0 | 0 | 12 | NaN | NaN | NaN | 873.6602 | NaN | 624.20575 |
| Psmd3 | EPQLAFHQ | -467 | -474 | 0 | 0.009060424 | 8 | NaN | NaN | NaN | 1066.5068 | NaN | NaN |
| Ddx5 | IVHINHQPFLER | -153 | -164 | 0 | 0 | 12 | NaN | NaN | NaN | 1584.8206 | 1847.4882 | 1554.142 |
| Ddx5 | SYLLPAIVHINHQPFLER | -147 | -164 | 0 | 0 | 18 | NaN | NaN | NaN | 1244.2734 | 786.38007 | 680.6964 |
| Ddx5 | SYLLPAIVHINHQPFLER | -147 | -164 | 0 | 0 | 18 | NaN | NaN | NaN | 1244.2734 | 786.38007 | 680.6964 |
| Ddx5 | RTTYLVLDEADR | -241 | -252 | 1 | 0.000627943 | 12 | NaN | NaN | NaN | 846.6981 | 961.8593 | 2243.0913 |
| Ddx5 | HNILQIVDVCHDVEK | -311 | -325 | 0 | 0.000230707 | 15 | NaN | NaN | NaN | 407.32193 | NaN | 203.57278 |
| Ddx5 | RCDELTR | -353 | -359 | 1 | 0.001392869 | 7 | NaN | NaN | NaN | 3804.9932 | NaN | NaN |
| Ddx5 | QPFLERGDGPICLVLAPTR | -159 | -177 | 1 | 0.006100831 | 19 | NaN | NaN | NaN | 90.139725 | NaN | NaN |

| | | | | | | | | | | | | |
|---|---|---|---|---|---|---|---|---|---|---|---|---|
| Pgls | EGTLPAALVQPR | -220 | -231 | 0 | 0.005946382 | 12 | NaN | NaN | NaN | 626.9005 | 701.5804 | 316.9165 |
| Arhgap25 | DHEVLFPK | -349 | -356 | 0 | 0.001868731 | 8 | NaN | NaN | NaN | 2957.4238 | 3084.282 | 3179.1145 |
| Arhgap25 | RDAFDAGERPSFDR | -212 | -225 | 2 | 0.001610812 | 14 | NaN | NaN | NaN | 789.65247 | 627.2902 | 627.2638 |
| Ube3a | ESEDYSPLIR | -137 | -146 | 0 | 0.00025015 | 10 | NaN | NaN | NaN | 737.63257 | NaN | NaN |
| Nup155 | TLTLVHVR | -384 | -391 | 0 | 0 | 8 | NaN | NaN | NaN | 542.7828 | 364.31174 | 259.7214 |
| Nup155 | KFQEAQLSEK | -766 | -775 | 1 | 0.002199523 | 10 | NaN | NaN | NaN | 950.5388 | 303.2635 | NaN |
| Nup155 | LVSNVGGDGEEIER | -524 | -537 | 0 | 0.000550307 | 14 | NaN | NaN | NaN | 2544.0793 | NaN | NaN |
| Eif3m | FIDISEEDQAAELR | -6 | -19 | 0 | 0 | 14 | NaN | NaN | NaN | 951.1605 | 657.9 | 1031.4729 |
| Eif3m | YTVYCSLIK | -121 | -129 | 0 | 0 | 9 | NaN | NaN | NaN | 1040.8523 | 488.19174 | NaN |
| Eif3m | ISEEDQAAELR | -9 | -19 | 0 | 0 | 11 | NaN | NaN | NaN | 635.62244 | NaN | 1094.7671 |
| Eif3m | KDPNAFLFDHLLTLKPVK | -213 | -230 | 2 | 0.00025015 | 18 | NaN | NaN | NaN | 100.79426 | NaN | NaN |
| Celf1 | DHPDQPDLDAIK | -6 | -17 | 0 | 0.000471409 | 12 | NaN | NaN | NaN | 1624.0325 | 646.9044 | 1798.5173 |
| Celf1 | FSSFGQIEECR | -128 | -138 | 0 | 0 | 11 | NaN | NaN | NaN | 1017.2335 | 1024.082 | 713.5954 |
| Celf1 | LDHPDQPDLDAIK | -5 | -17 | 0 | 0 | 13 | NaN | NaN | NaN | 291.37115 | 425.70654 | 606.7265 |
| Nol10 | LIEEEEEKQK | -457 | -466 | 1 | 0.001055699 | 10 | NaN | NaN | NaN | 517.1184 | 620.23895 | 354.08228 |
| Aldh18a1 | IEVVDSVQEAIDHIHK | -675 | -690 | 0 | 0 | 16 | NaN | NaN | NaN | 827.9073 | 897.14844 | 1054.4132 |
| Aldh18a1 | GSSHTDVIVTENEK | -692 | -705 | 0 | 0 | 14 | NaN | NaN | NaN | 2321.1157 | 2785.4158 | 501.05084 |
| Aldh18a1 | DHVVSDFSEHGSLK | -768 | -781 | 0 | 0 | 14 | NaN | NaN | NaN | 420.5014 | 174.68172 | 755.5338 |
| Aldh18a1 | DHVVSDFSEHGSLK | -768 | -781 | 0 | 0 | 14 | NaN | NaN | NaN | 420.5014 | 174.68172 | 755.5338 |
| Aldh18a1 | HTDVIVTENEK | -695 | -705 | 0 | 0.00025015 | 11 | NaN | NaN | NaN | 798.94507 | 1262.353 | 951.9153 |
| Aldh18a1 | LIDIFYPGDQQ | -283 | -293 | 0 | 0 | 11 | NaN | NaN | NaN | 685.0954 | 729.8684 | NaN |
| Aldh18a1 | EAIQLVNTR | -531 | -539 | 0 | 0.00025015 | 9 | NaN | NaN | NaN | 667.2408 | NaN | NaN |
| Ctsd | ILDIACWVHHK | -112 | -122 | 0 | 0 | 11 | NaN | NaN | NaN | 3835.9678 | 2281.2285 | 2076.286 |
| Ctsd | ILDIACWVHHK | -112 | -122 | 0 | 0 | 11 | NaN | NaN | NaN | 3835.9678 | 2281.2285 | 2076.286 |
| Ctsd | DTGSSNLWVPSIHCK | -97 | -111 | 0 | 0 | 15 | NaN | NaN | NaN | 3489.549 | 4369.1934 | 4314.561 |
| Puf60 | GLPPLTPEQQEALQK | -60 | -74 | 0 | 0 | 15 | NaN | NaN | NaN | 665.05853 | 840.7588 | NaN |
| Puf60 | DIDDDLEGEVTEECGKF | -479 | -495 | 1 | 0.000550307 | 17 | NaN | NaN | NaN | 115.20836 | NaN | 275.4787 |
| Sms | QVLIEDCIPVLK | -250 | -261 | 0 | 0.004439092 | 12 | NaN | NaN | NaN | 335.75443 | 376.50574 | NaN |
| Sms | ILSGDVNLAESDLAYTR | -163 | -179 | 0 | 0.001055699 | 17 | NaN | NaN | NaN | 200.50429 | 54.980545 | NaN |
| Zap70 | KGPEVLDFIK | -543 | -552 | 1 | 0.001636916 | 10 | NaN | NaN | NaN | 860.0024 | 925.0524 | NaN |
| Zap70 | KEEIPVSNVAELL | -429 | -441 | 1 | 0.003166358 | 13 | NaN | NaN | NaN | 1079.258 | NaN | NaN |
| Pus10 | FLEPEKPELILEAPNPPLKK | -51 | -70 | 2 | 0.001463897 | 20 | NaN | NaN | NaN | 529.09235 | 670.72577 | 880.09827 |
| Edf1 | PNNQVLGK | -116 | -123 | 0 | 0.001636916 | 8 | NaN | NaN | NaN | 1260.7638 | 1097.3175 | 1062.9614 |
| Usp47 | LEDSNNVDDSVHDR | -139 | -152 | 0 | 0 | 14 | NaN | NaN | NaN | 258.27563 | NaN | NaN |
| Stk38 | IGAPGVEEIK | -368 | -377 | 0 | 0 | 10 | NaN | NaN | NaN | 5707.4863 | NaN | 1193.5658 |
| Pitrm1 | GFEDDRIEAL | -419 | -428 | 1 | 0.000230707 | 10 | NaN | NaN | NaN | 995.79285 | 1053.368 | 1021.0813 |
| Rps6ka1 | YVDESGNPECLR | -532 | -543 | 0 | 0 | 12 | NaN | NaN | NaN | 1646.6476 | 1610.6296 | 1657.4832 |
| Pwp1 | NAHNDEISGLDLSSQIK | -381 | -397 | 0 | 0.00025015 | 17 | NaN | NaN | NaN | 490.2014 | 980.7212 | 274.50934 |
| Rbm25 | IPTAKPELFAYPL | -740 | -752 | 1 | 0.00025015 | 13 | NaN | NaN | NaN | 1926.8413 | 2354.7932 | 2466.215 |
| Rps6 | KLIEVDDER | -14 | -22 | 1 | 0 | 9 | NaN | NaN | NaN | 2207.6606 | 1664.5577 | 1804.1263 |
| Rps6 | NKEEAAEY | -202 | -209 | 1 | 0.000989247 | 8 | NaN | NaN | NaN | 521.0294 | NaN | NaN |
| Rps6 | NKEEAAEY | -202 | -209 | 1 | 0.000989247 | 8 | NaN | NaN | NaN | 521.0294 | NaN | NaN |
| Akap13 | TGDEVDLLSR | -1317 | -1326 | 0 | 0.000550307 | 10 | NaN | NaN | NaN | 139.21983 | NaN | NaN |
| Cybc1 | KDIQDVNVEEEK | -105 | -116 | 1 | 0.003133457 | 12 | NaN | NaN | NaN | 2020.3888 | NaN | NaN |
| Snd1 | RADDADEFGYSR | -899 | -910 | 1 | 0 | 12 | NaN | NaN | NaN | 2948.7402 | 5294.7046 | 3348.985 |
| Snd1 | DVETGTQLEK | -700 | -709 | 0 | 0 | 10 | NaN | NaN | NaN | 781.20105 | NaN | NaN |
| Snd1 | YEERPVEEVMPVLEEK | -661 | -676 | 1 | 0 | 16 | NaN | NaN | NaN | 401.45044 | 475.60556 | 496.01385 |
| Snd1 | DTPDEPWAFPA | -72 | -82 | 0 | 0.00051306 | 11 | NaN | NaN | NaN | 369.66156 | 385.32892 | 411.45233 |
| Snd1 | NLPGLVQEGEPF | -566 | -577 | 0 | 0.00025015 | 12 | NaN | NaN | NaN | 238.03104 | 242.53102 | 101.44093 |
| Snd1 | QVLNADAIVVK | -349 | -359 | 0 | 0 | 11 | NaN | NaN | NaN | 1262.384 | NaN | NaN |
| Snd1 | SEAVVEYVFSGSR | -527 | -539 | 0 | 0.00025015 | 13 | NaN | NaN | NaN | 641.95374 | NaN | NaN |
| Snd1 | LAGIECPR | -555 | -562 | 0 | 0.003299042 | 8 | NaN | NaN | NaN | 358.1075 | NaN | NaN |
| Itk | DISIPCHYK | -69 | -77 | 0 | 0.002099344 | 9 | NaN | NaN | NaN | 221.02924 | 235.17032 | NaN |
| Aprt | KDALEPGQR | -114 | -122 | 1 | 0.001291103 | 9 | NaN | NaN | NaN | 1366.2286 | NaN | NaN |
| Arih1 | DISQDSLQDIK | -510 | -520 | 0 | 0 | 11 | NaN | NaN | NaN | 382.84976 | NaN | NaN |
| Rab5a | IVVYDITNEESFAR | -97 | -110 | 0 | 0.00025015 | 14 | NaN | NaN | NaN | 878.88116 | NaN | 552.12274 |
| Eif4g1 | LSVDEVEKK | -1237 | -1245 | 1 | 0.003299042 | 9 | NaN | NaN | NaN | 605.1226 | 283.7965 | NaN |
| Ash2l | EHPDPGSKDPEEDYPK | -233 | -248 | 1 | 0.002068213 | 16 | NaN | NaN | NaN | 350.68088 | NaN | 479.61765 |
| Csde1 | YTSEDVEGNVQLETGDK | -141 | -157 | 0 | 0.000550307 | 17 | NaN | NaN | NaN | 316.372 | 632.9746 | NaN |
| Wdr75 | DVPESFTSETHQWLNR | -637 | -652 | 0 | 0 | 16 | NaN | NaN | NaN | 651.5458 | NaN | 574.73706 |
| Nipsnap3b | DKEWQER | -106 | -112 | 1 | 0.007009655 | 7 | NaN | NaN | NaN | 474.03943 | NaN | NaN |
| Itih2 | MLADSPPQDHSCCSGAL | -639 | -655 | 0 | 0.00025015 | 17 | NaN | NaN | NaN | 881.84106 | 967.08295 | 967.9097 |
| Itih2 | MLADSPPQDHSCCSGAL | -639 | -655 | 0 | 0.00025015 | 17 | NaN | NaN | NaN | 881.84106 | 967.08295 | 967.9097 |
| Itih2 | DKHADPDFTK | -572 | -581 | 1 | 0.00111006 | 10 | NaN | NaN | NaN | 845.6464 | NaN | NaN |
| Ogdh | DKICEEAFTR | -563 | -572 | 1 | 0 | 10 | NaN | NaN | NaN | 1359.2122 | 1471.8365 | 1489.4675 |
| Ogdh | TCPSTGLEEDVLFHIGK | -603 | -619 | 0 | 0.00025015 | 17 | NaN | NaN | NaN | 725.54333 | NaN | 966.573 |
| Ogdh | VVVNNQIGFTTDPR | -440 | -453 | 0 | 0 | 14 | NaN | NaN | NaN | 501.0905 | 369.92087 | NaN |
| Ogdh | VFHLPTTTFIGGQEPALPLR | -185 | -204 | 0 | 0 | 20 | NaN | NaN | NaN | 979.3589 | NaN | NaN |
| Ogdh | PYPTDVAR | -459 | -466 | 0 | 0.000394361 | 8 | NaN | NaN | NaN | 583.24634 | NaN | NaN |
| Tmed9 | VEQVEQIQK | -172 | -180 | 0 | 0.002068213 | 9 | NaN | NaN | NaN | 1224.2332 | NaN | NaN |
| Cbl | GIFPSGLFQGDTFR | -165 | -178 | 0 | 0 | 14 | NaN | NaN | NaN | 631.0755 | 523.3046 | 554.4489 |
| Cbl | SSPVAGPESEHPK | -649 | -661 | 0 | 0.00111006 | 13 | NaN | NaN | NaN | 595.49164 | 847.2521 | NaN |
| Cbl | GTEPIVVDPFDPR | -423 | -435 | 0 | 0 | 13 | NaN | NaN | NaN | 3197.182 | NaN | NaN |
| Cbl | DKPLPIPPTLR | -528 | -538 | 1 | 0.00078661 | 11 | NaN | NaN | NaN | 91.27195 | NaN | NaN |
| Elp1 | LWESPSLAVEPWK | -583 | -595 | 0 | 0.004983786 | 13 | NaN | NaN | NaN | 636.4826 | 896.17316 | NaN |
| Hmgb2 | KHPDSSVNFA | -30 | -39 | 1 | 0 | 10 | NaN | NaN | NaN | 3443.924 | 4341.5933 | 4657.3535 |
| Hmgb2 | FCSENRPK | -105 | -112 | 1 | 0 | 8 | NaN | NaN | NaN | 782.7407 | 843.00464 | 687.07007 |
| Hmgb2 | IKIEHPGLSIGDTA | -113 | -126 | 1 | 0 | 14 | NaN | NaN | NaN | 762.0699 | 1060.9285 | 1025.5201 |

| | | | | | | | | | | | | |
|---|---|---|---|---|---|---|---|---|---|---|---|---|
| Hmgb2 | IEHPGLSIGDTA | -115 | -126 | 0 | 0.00111006 | 12 | NaN | NaN | NaN | 3967.2869 | NaN | 3456.1018 |
| Hmgb2 | IKIEHPGLSIGD | -113 | -124 | 1 | 0.000394361 | 12 | NaN | NaN | NaN | 1655.4062 | 1938.4441 | NaN |
| Ube2f | EHSIDGTGWAPTR | -122 | -134 | 0 | 0 | 13 | NaN | NaN | NaN | 152.28043 | NaN | 167.87488 |
| Ruvbl2 | SLHEIDVINSR | -243 | -253 | 0 | 0 | 11 | NaN | NaN | NaN | 479.99597 | 1466.6494 | 1294.0438 |
| Ruvbl2 | IKEETEIIEGEVVEIQ | -131 | -146 | 1 | 0.00111006 | 16 | NaN | NaN | NaN | 5654.545 | 6056.196 | 6360.776 |
| Ruvbl2 | KFVQCPDGELQK | -223 | -234 | 1 | 0.006195994 | 12 | NaN | NaN | NaN | 484.01688 | NaN | NaN |
| Npc2 | DKVYSYLNK | -104 | -112 | 1 | 0.00051306 | 9 | NaN | NaN | NaN | 3456.5046 | 3236.28 | 3008.5571 |
| Npc2 | LNKLPVK | -110 | -116 | 1 | 0.007819401 | 7 | NaN | NaN | NaN | 328.95807 | NaN | NaN |
| Ccdc25 | FPDLAAEK | -141 | -148 | 0 | 0.002534621 | 8 | NaN | NaN | NaN | 356.91837 | 313.8509 | NaN |
| Noc2l | DLCGCLQK | -215 | -222 | 0 | 0.002296977 | 8 | NaN | NaN | NaN | 571.8852 | 565.911 | NaN |
| Tmod3 | LVEVNLNNIK | -200 | -209 | 0 | 0 | 10 | NaN | NaN | NaN | 798.75995 | 789.2814 | NaN |
| Flnb | DLDIIDNYDYSHTVK | -895 | -909 | 0 | 0 | 15 | NaN | NaN | NaN | 127.41368 | 860.6189 | 447.29382 |
| Baz1a | IQVPDYYDIIK | -1460 | -1470 | 0 | 0.007546675 | 11 | NaN | NaN | NaN | 852.94745 | NaN | NaN |
| Bag6 | HYLGGQEPTPSNIR | -814 | -827 | 0 | 0.00201047 | 14 | NaN | NaN | NaN | 1091.9902 | 215.98643 | NaN |
| Bag6 | MEQAPIQSEPR | -153 | -163 | 0 | 0.005668207 | 11 | NaN | NaN | NaN | 1298.1235 | NaN | NaN |
| Bag6 | THTLITGLEEYVR | -830 | -842 | 0 | 0.000471409 | 13 | NaN | NaN | NaN | 1187.7491 | NaN | NaN |
| Anp32a | DLFNCEVTNLNAYR | -119 | -132 | 0 | 0.00051306 | 14 | NaN | NaN | NaN | 1086.603 | 1242.8795 | 1221.5627 |
| Anp32a | SLDLFNCEVTNLNAYR | -117 | -132 | 0 | 0 | 16 | NaN | NaN | NaN | 107.90777 | 298.89914 | 436.11017 |
| Anp32a | SLDLFNCEVTNL | -117 | -128 | 0 | 0.000230707 | 12 | NaN | NaN | NaN | 265.11517 | 306.0167 | NaN |
| Hdgf | KGFSEGLWEIENNPTVK | -80 | -96 | 1 | 0 | 17 | NaN | NaN | NaN | 969.5343 | 1355.2404 | 1236.192 |
| Bag5 | DLGQNHSIIK | -269 | -278 | 0 | 0 | 10 | NaN | NaN | NaN | 994.9972 | NaN | NaN |
| Rps20 | LIDLHSPSEIVK | -88 | -99 | 0 | 0 | 12 | NaN | NaN | NaN | 7010.558 | 8489.925 | 8739.08 |
| Rps20 | DLHSPSEIVK | -90 | -99 | 0 | 0.005363933 | 10 | NaN | NaN | NaN | 916.63055 | 1899.1368 | 1678.7903 |
| Rps20 | VCADLIR | -35 | -41 | 0 | 0 | 7 | NaN | NaN | NaN | 601.4825 | NaN | NaN |
| Seh1l | DLIHDVSFDFHGR | -13 | -25 | 0 | 0 | 13 | NaN | NaN | NaN | 317.87402 | 470.51212 | 87.57645 |
| Seh1l | SLQHEVSCK | -154 | -162 | 0 | 0.000550307 | 9 | NaN | NaN | NaN | 661.656 | NaN | 1052.0742 |
| Eif4g2 | WKEDITQEFPGK | -868 | -879 | 1 | 0.001055699 | 12 | NaN | NaN | NaN | 2755.5474 | 2983.5276 | 2852.8132 |
| Eif4g2 | KLQDEFENR | -170 | -178 | 1 | 0.004439092 | 9 | NaN | NaN | NaN | 1205.1168 | 1210.7202 | 351.51746 |
| Eif4g2 | DLIHESILHK | -221 | -230 | 0 | 0 | 10 | NaN | NaN | NaN | 1092.453 | 514.152 | 1631.8612 |
| Eif4g2 | EAVVTDYLNSGNANDAVSGVR | -548 | -568 | 0 | 0.001291103 | 21 | NaN | NaN | NaN | 303.12207 | NaN | 326.2793 |
| Eif4g2 | LQDTVELR | -298 | -305 | 0 | 0.001941082 | 8 | NaN | NaN | NaN | 1877.3872 | NaN | NaN |
| Tpp2 | STLIDALCR | -1131 | -1139 | 0 | 0 | 9 | NaN | NaN | NaN | 1012.07056 | 1500.6783 | 1489.8152 |
| Tpp2 | ETYPAYLPLYVAR | -1060 | -1072 | 0 | 0 | 13 | NaN | NaN | NaN | 618.0487 | 598.23486 | 510.36172 |
| Tpp2 | SYGEATHWPNSGR | -328 | -340 | 0 | 0 | 13 | NaN | NaN | NaN | 380.7328 | 542.00415 | 799.85693 |
| Tpp2 | LKDLPFIVSH | -903 | -912 | 1 | 0.001055699 | 10 | NaN | NaN | NaN | 1962.6456 | 1975.2043 | NaN |
| Tpp2 | KWTDLFDTK | -1185 | -1193 | 1 | 0.00025015 | 9 | NaN | NaN | NaN | 1004.7239 | 367.25793 | NaN |
| Tpp2 | YEDLAPCITLK | -781 | -791 | 0 | 0 | 11 | NaN | NaN | NaN | 702.6833 | NaN | 627.7355 |
| Tpp2 | KYEDLAPCITLK | -780 | -791 | 1 | 0.00025015 | 12 | NaN | NaN | NaN | 3844.9336 | NaN | NaN |
| Tpp2 | LVEEKPTK | -1218 | -1225 | 1 | 0.003998192 | 8 | NaN | NaN | NaN | 302.04117 | NaN | NaN |
| Tpp2 | LKDLPFIVSHR | -903 | -913 | 1 | 0.006164361 | 11 | NaN | NaN | NaN | 217.56746 | NaN | NaN |
| Prp4k | DVEEEDEEALIEQR | -528 | -541 | 0 | 0.00025015 | 14 | NaN | NaN | NaN | 984.8501 | 1043.9138 | 654.9035 |
| Prp4k | DLLADLIGCQR | -954 | -964 | 0 | 0 | 11 | NaN | NaN | NaN | 350.09872 | NaN | NaN |
| Prp4k | TKLDDLALLEDLEK | -99 | -112 | 1 | 0.002810811 | 14 | NaN | NaN | NaN | 194.8926 | NaN | NaN |
| Prp4k | DNWTDAEGYYR | -666 | -676 | 0 | 0.001743443 | 11 | NaN | NaN | NaN | 100.35387 | NaN | NaN |
| Esyt2 | TQWYDLTEDGTRPQVIT | -829 | -845 | 1 | 0.001743443 | 17 | NaN | NaN | NaN | 912.1008 | 1180.0378 | 945.39294 |
| Esyt2 | RQDLEVEVK | -521 | -529 | 1 | 0.001636916 | 9 | NaN | NaN | NaN | 2043.645 | 1853.4436 | NaN |
| Pdcd4 | ELEVPHFHHE | -353 | -362 | 0 | 0.003681885 | 10 | NaN | NaN | NaN | 773.7575 | NaN | NaN |
| Ddx3x | GGAEIGQQIR | -302 | -311 | 0 | 0 | 10 | NaN | NaN | NaN | 491.0079 | NaN | NaN |
| Il2 | DLQCLEDELGPLR | -89 | -101 | 0 | 0 | 13 | NaN | NaN | NaN | 502.906 | 482.2827 | NaN |
| Bop1 | LALPGHAESYNPPPPEYLPTEEER | -291 | -313 | 0 | 0.00025015 | 23 | NaN | NaN | NaN | 571.1651 | 1063.2245 | NaN |
| Mrpl11 | DLSAEELEAFQK | -158 | -169 | 0 | 0 | 12 | NaN | NaN | NaN | 831.0849 | 377.2722 | 737.62866 |
| Mrpl11 | QDVPLSSVVR | -133 | -142 | 0 | 0 | 10 | NaN | NaN | NaN | 347.18878 | NaN | NaN |
| Gtpbp4 | VVDTPGILDHPLEDR | -219 | -233 | 0 | 0 | 15 | NaN | NaN | NaN | 185.06656 | NaN | 645.4194 |
| Flii | SNKLDFDGLPSGIGK | -301 | -315 | 1 | 0.000230707 | 15 | NaN | NaN | NaN | 991.72546 | 1013.76624 | 759.01965 |
| Lig1 | FCEVPDLDR | -505 | -513 | 0 | 0.002810811 | 9 | NaN | NaN | NaN | 2785.212 | 2538.163 | NaN |
| Map4 | SLDNVGHLPAGGAVK | -1046 | -1060 | 0 | 0 | 15 | NaN | NaN | NaN | 595.86536 | 634.8963 | 649.716 |
| Map4 | PPEDVLLTK | -361 | -369 | 0 | 0.00201047 | 9 | NaN | NaN | NaN | 504.5551 | NaN | 752.5301 |
| Map4 | SGSVPAAPHK | -716 | -725 | 0 | 0.000431655 | 10 | NaN | NaN | NaN | 473.43283 | 711.5979 | NaN |
| Map4 | MDLAPPEDVLLTK | -357 | -369 | 0 | 0.001846793 | 13 | NaN | NaN | NaN | 897.94446 | NaN | NaN |
| Dpysl2 | SITIANQTNCPLYVTK | -239 | -254 | 0 | 0 | 16 | NaN | NaN | NaN | 1087.1525 | NaN | NaN |
| Dpysl2 | SDADLVIWDPDSVK | -405 | -418 | 0 | 0.000989247 | 14 | NaN | NaN | NaN | 362.42313 | NaN | NaN |
| Clk2 | DNNYLPYPIHQVR | -255 | -267 | 0 | 0.000627943 | 13 | NaN | NaN | NaN | 557.9297 | NaN | NaN |
| Emg1 | SVEDISDIR | -179 | -187 | 0 | 0.000627943 | 9 | NaN | NaN | NaN | 1792.0175 | 1621.502 | 1967.007 |
| Emg1 | EQDWETTPPK | -19 | -28 | 0 | 0.001868731 | 10 | NaN | NaN | NaN | 430.55862 | NaN | NaN |
| Khdrbs1 | KDDEENYLDLF | -139 | -149 | 1 | 0.00025015 | 11 | NaN | NaN | NaN | 2645.2852 | 3047.163 | 777.97876 |
| Ctsz | DQDCDKFNQCGTCTEFK | -163 | -179 | 1 | 0.001370415 | 17 | NaN | NaN | NaN | 275.82587 | NaN | 340.88284 |
| Ppp6r3 | LGEDESLLMK | -96 | -105 | 0 | 0.009185548 | 10 | NaN | NaN | NaN | 1857.7173 | NaN | NaN |
| Ppp6r3 | DQDSTGDNLLLK | -414 | -425 | 0 | 0.00025015 | 12 | NaN | NaN | NaN | 313.8438 | NaN | NaN |
| Cd27 | HCNSGFLIR | -86 | -94 | 0 | 0.00111006 | 9 | NaN | NaN | NaN | 1840.7412 | 1863.8169 | NaN |
| Cd27 | HYWTGGGLCCR | -31 | -41 | 0 | 0.000394361 | 11 | NaN | NaN | NaN | 444.5421 | NaN | NaN |
| Farsa | KCWELTTEGEEIAR | -59 | -72 | 1 | 0 | 14 | NaN | NaN | NaN | 477.19473 | 521.77637 | 522.35565 |
| Farsa | SELMHLPSGK | -92 | -101 | 0 | 0.00272025 | 10 | NaN | NaN | NaN | 1773.2522 | NaN | 2104.9036 |
| Farsa | GVLPDSGHLHPL | -216 | -227 | 0 | 0 | 12 | NaN | NaN | NaN | 1590.139 | 1769.0466 | NaN |
| Cd48 | LEVFDPVPK | -122 | -130 | 0 | 0.000431655 | 9 | NaN | NaN | NaN | 1476.3328 | 611.79944 | 839.7998 |
| Cd48 | STFYTCQVSNPVSSK | -188 | -202 | 0 | 0 | 15 | NaN | NaN | NaN | 1410.6536 | 2254.5337 | 1734.5515 |
| Cd48 | EASTDSCHLR | -139 | -148 | 0 | 0.002910424 | 10 | NaN | NaN | NaN | 815.537 | NaN | 1316.5907 |
| Cd48 | HVDYTWYESSGPFPK | -157 | -171 | 0 | 0.00025015 | 15 | NaN | NaN | NaN | 728.57007 | 264.34528 | NaN |
| Col1a1 | DRDLEVDTTLK | -1215 | -1225 | 1 | 0 | 11 | NaN | NaN | NaN | 283.90848 | 677.3 | 109.12897 |

| | | | | | | | | | | | | |
|---|---|---|---|---|---|---|---|---|---|---|---|---|
| Actr3 | GYAGNTEPQFIIPSCIAIK | -20 | -38 | 0 | 0 | 19 | NaN | NaN | NaN | 278.44373 | 298.87677 | 328.91663 |
| Senp3 | DRLDFHQGWK | -501 | -510 | 1 | 0.002595029 | 10 | NaN | NaN | NaN | 412.5133 | NaN | NaN |
| Apobec3 | NKDNIHAEIC | -77 | -86 | 1 | 0.00025015 | 10 | NaN | NaN | NaN | 1898.6964 | NaN | NaN |
| Nme2 | PFFPGLVK | -59 | -66 | 0 | 0.005605646 | 8 | NaN | NaN | NaN | 292.12982 | NaN | NaN |
| Nme1 | FLQASEDLLK | -40 | -49 | 0 | 0 | 10 | NaN | NaN | NaN | 30803.479 | 30858.367 | 30774.59 |
| Nme1 | QASEDLLK | -42 | -49 | 0 | 0.001776473 | 8 | NaN | NaN | NaN | 270.28265 | 613.8981 | NaN |
| Cd5 | DRPLGLGNLICK | -191 | -202 | 1 | 0.00051306 | 12 | NaN | NaN | NaN | 1210.5355 | 1020.3712 | NaN |
| Cd5 | SQWEALCDSSAAR | -295 | -307 | 0 | 0 | 13 | NaN | NaN | NaN | 809.3388 | NaN | NaN |
| Cd5 | EQQCGDLISFH | -318 | -328 | 0 | 0.001392869 | 11 | NaN | NaN | NaN | 481.4538 | NaN | NaN |
| Mbp | DRPSESDELQTIQ | -109 | -121 | 1 | 0.00111006 | 13 | NaN | NaN | NaN | 589.3762 | NaN | 721.358 |
| Wnk1 | VESGYVCEGDHK | -541 | -552 | 0 | 0.000230707 | 12 | NaN | NaN | NaN | 1322.4756 | 1795.746 | NaN |
| Wnk1 | EGPVTSPPFR | -1968 | -1977 | 0 | 0 | 10 | NaN | NaN | NaN | 1122.4127 | 2016.6831 | NaN |
| Wnk1 | VPPAVIIPPAAPL | -2103 | -2115 | 0 | 0.001291103 | 13 | NaN | NaN | NaN | 253.13994 | 509.83276 | NaN |
| Brcc3;C6.1al | DRVEISPEQL | (88);(88) | (97);(97) | 1 | 0.006100831 | 10 | NaN | NaN | NaN | 4945.8877 | NaN | NaN |
| Mrpl4 | IGLAELHPDVFATAPR | -77 | -92 | 0 | 0 | 16 | NaN | NaN | NaN | 660.71967 | 337.03616 | 616.10767 |
| Mrpl4 | IGLAELHPDVFATAPR | -77 | -92 | 0 | 0 | 16 | NaN | NaN | NaN | 660.71967 | 337.03616 | 616.10767 |
| Plg | DSCQGDSGGPLVCFEK | -756 | -771 | 0 | 0.000989247 | 16 | NaN | NaN | NaN | 322.1029 | 348.23517 | 323.62927 |
| Prss2 | DSCQGDSGGPVVCNGELQGIV | -194 | -214 | 0 | 0.000431655 | 21 | NaN | NaN | NaN | 523.51953 | 561.6183 | 469.0871 |
| Try5;Try4 | DSCQGDSGGPVVCNGQLQGIVSW | (194);(194) | (216);(216) | 0 | 0.00201047 | 23 | NaN | NaN | NaN | 149.31036 | 305.8498 | 227.51093 |
| Cbx3 | KVEEAEPEEFVVE | -21 | -33 | 1 | 0.00025015 | 13 | NaN | NaN | NaN | 6047.6113 | 5216.5815 | 5220.9062 |
| Cbx3 | WHSCPEDEAQ | -174 | -183 | 0 | 0.000298805 | 10 | NaN | NaN | NaN | 777.1724 | 933.7053 | 1024.8293 |
| Supt5h | DSDQNNIHVK | -586 | -595 | 0 | 0.00025015 | 10 | NaN | NaN | NaN | 204.18108 | NaN | NaN |
| Nars1 | NYSDAIEWLK | -403 | -412 | 0 | 0 | 10 | NaN | NaN | NaN | 4317.2637 | NaN | 3672.6377 |
| Nars1 | NYSDAIEWLK | -403 | -412 | 0 | 0 | 10 | NaN | NaN | NaN | 4317.2637 | NaN | 3672.6377 |
| Nars1 | LMTDTINEPILLC | -437 | -449 | 0 | 0 | 13 | NaN | NaN | NaN | 1040.2048 | 1365.6998 | NaN |
| Nars1 | INEESDVDVQLNNR | -231 | -244 | 0 | 0.001463897 | 14 | NaN | NaN | NaN | 513.8507 | NaN | 1328.7446 |
| Nars1 | HLAEFTHVE | -342 | -350 | 0 | 0.00025015 | 9 | NaN | NaN | NaN | 266.62274 | NaN | NaN |
| Mat2a | KYLDEDTIYHLQPSGR | -234 | -249 | 1 | 0.00051306 | 16 | NaN | NaN | NaN | 516.6278 | 615.1043 | 490.696 |
| Mat2a | HDEEVCLDEMR | -209 | -219 | 0 | 0 | 11 | NaN | NaN | NaN | 546.55884 | 366.53613 | 453.0745 |
| Mat2a | HIGYDDSSK | -89 | -97 | 0 | 0.001092529 | 9 | NaN | NaN | NaN | 597.6409 | NaN | NaN |
| Rmdn3 | DSLDLDVEAASSPAAAALE | -211 | -229 | 0 | 0.007905138 | 19 | NaN | NaN | NaN | 1161.6849 | NaN | NaN |
| Purb | GGGGGGGGGPGGFQPAPRG | -11 | -29 | 1 | 0.00025015 | 19 | NaN | NaN | NaN | 978.6593 | NaN | 957.009 |
| Gls | VADYIPQLAK | -251 | -260 | 0 | 0 | 10 | NaN | NaN | NaN | 541.5912 | 561.7412 | 624.16656 |
| Gls | NDLGTEYVHR | -303 | -312 | 0 | 0 | 10 | NaN | NaN | NaN | 1497.5259 | 739.9092 | NaN |
| Sdcbp | LVQANSPASLVGLR | -141 | -154 | 0 | 0 | 14 | NaN | NaN | NaN | 290.9956 | 330.19397 | NaN |
| Wdr76 | DSSSISSQLPLLSTIR | -521 | -536 | 0 | 0 | 16 | NaN | NaN | NaN | 142.45456 | 818.03625 | 3153.5247 |
| Smchd1 | VEGDGVVEGVIR | -417 | -428 | 0 | 0 | 12 | NaN | NaN | NaN | 6423.324 | 703.68524 | 762.32007 |
| Smchd1 | GNDKEDGCFYFR | -1439 | -1450 | 1 | 0 | 12 | NaN | NaN | NaN | 1151.9979 | 1699.3008 | NaN |
| Smchd1 | LVPDSQPATPAVSNVR | -1492 | -1507 | 0 | 0.000550307 | 16 | NaN | NaN | NaN | 429.54364 | NaN | NaN |
| Arhgef6 | GPLEPPQIIKPW | -577 | -588 | 1 | 0 | 12 | NaN | NaN | NaN | 2842.3762 | 3091.9883 | 3086.0256 |
| Arhgef6 | DSVPQVLLPEEEK | -678 | -690 | 0 | 0 | 13 | NaN | NaN | NaN | 1396.4791 | 2901.1416 | NaN |
| Arhgef6 | QCLEEELK | -732 | -739 | 0 | 0.006677519 | 8 | NaN | NaN | NaN | 951.2209 | NaN | NaN |
| Nhlrc2 | LQHPLGVAWDEER | -474 | -486 | 0 | 0.001092529 | 13 | NaN | NaN | NaN | 788.5087 | NaN | 644.9873 |
| Hmgn5 | LHEEDGEEGQPK | -293 | -304 | 0 | 0 | 12 | NaN | NaN | NaN | 209.35199 | 355.16663 | NaN |
| Wdr43 | PGIPGHHAPIK | -405 | -415 | 0 | 0.001868731 | 11 | NaN | NaN | NaN | 895.41156 | 674.3987 | NaN |
| Wdr43 | VNCIQWHQDNDCLYSC | -129 | -144 | 0 | 0.000431655 | 16 | NaN | NaN | NaN | 233.68953 | 263.9106 | NaN |
| Wdr43 | HFTGHATPVSSL | -203 | -214 | 0 | 0.00025015 | 12 | NaN | NaN | NaN | 1206.0991 | NaN | NaN |
| Wdr43 | LHQEYVPSAHLSGTC | -48 | -62 | 0 | 0.002810811 | 15 | NaN | NaN | NaN | 926.3906 | NaN | NaN |
| Wdr43 | VTPKPIPIL | -327 | -335 | 1 | 0.006100831 | 9 | NaN | NaN | NaN | 335.05234 | NaN | NaN |
| Gna13 | GIHEYDFEIK | -205 | -214 | 0 | 0 | 10 | NaN | NaN | NaN | 1090.3234 | 1959.9243 | 1381.9478 |
| Wdr48 | DTNNNVAYWDVLK | -379 | -391 | 0 | 0 | 13 | NaN | NaN | NaN | 335.5977 | 332.85837 | NaN |
| Xpnpep1 | GGVTEISAADKAEEFR | -355 | -370 | 1 | 0.00025015 | 16 | NaN | NaN | NaN | 1600.2189 | 1620.2556 | 1756.6309 |
| Xpnpep1 | SLDEVYLIDSGAQYK | -407 | -421 | 0 | 0.00025015 | 15 | NaN | NaN | NaN | 800.4774 | 973.5341 | 1228.5741 |
| Xpnpep1 | GSLTFEPLTLVPIQTK | -555 | -570 | 0 | 0 | 16 | NaN | NaN | NaN | 307.7847 | NaN | 311.19574 |
| Stim1 | YVELPQYEETFR | -144 | -155 | 0 | 0 | 12 | NaN | NaN | NaN | 956.8851 | 1738.89 | NaN |
| Smarcc1 | SQKEEDEQEDLTK | -356 | -368 | 1 | 0 | 13 | NaN | NaN | NaN | 371.77167 | 561.60626 | 532.3124 |
| Smarcc1 | VREEVPLELVEAH | -724 | -736 | 1 | 0.00025015 | 13 | NaN | NaN | NaN | 597.0383 | 467.5316 | NaN |
| Smarcc1 | NTEKEQDSDVSEDVKPEEK | -814 | -832 | 2 | 0.000230707 | 19 | NaN | NaN | NaN | 361.47617 | NaN | 525.7604 |
| Notch1 | QDVNECSQNPGLCR | -177 | -190 | 0 | 0.00272025 | 14 | NaN | NaN | NaN | 437.44592 | NaN | NaN |
| Iah1 | QHVPLDEYSANLR | -100 | -112 | 0 | 0.00644558 | 13 | NaN | NaN | NaN | 486.59665 | NaN | NaN |
| Smu1 | YIHLENLLAR | -114 | -123 | 0 | 0 | 10 | NaN | NaN | NaN | 339.1177 | 6084.6255 | 6125.1055 |
| Uba2 | ESVLQFHPQ | -78 | -86 | 0 | 0.00051306 | 9 | NaN | NaN | NaN | 347.29294 | 536.501 | NaN |
| Uba2 | GEANADQQNEPQLGLK | -291 | -306 | 0 | 0.000230707 | 16 | NaN | NaN | NaN | 4163.0293 | NaN | NaN |
| Uba2 | SIMNPDYNVEFFR | -95 | -107 | 0 | 0.005698678 | 13 | NaN | NaN | NaN | 670.937 | NaN | NaN |
| Uba2 | VAPDVQIEDGK | -474 | -484 | 0 | 0 | 11 | NaN | NaN | NaN | 160.21469 | NaN | NaN |
| Pdia6 | DVVELTDDTFDKN | -161 | -173 | 1 | 0 | 13 | NaN | NaN | NaN | 1900.831 | 570.1949 | 2557.319 |
| Pdia6 | FSEQGINEFLR | -376 | -386 | 0 | 0 | 11 | NaN | NaN | NaN | 2655.7346 | 1607.8352 | NaN |
| Pdia6 | KDVVELTDDTFDKN | -160 | -173 | 2 | 0.002752526 | 14 | NaN | NaN | NaN | 1121.3035 | NaN | NaN |
| Pdia6 | TGEAIVDAALSALR | -119 | -132 | 0 | 0.00025015 | 14 | NaN | NaN | NaN | 895.0926 | NaN | NaN |
| Rpl27 | NIDDGTSDRPYSHALVA | -28 | -44 | 1 | 0.00025015 | 17 | NaN | NaN | NaN | 812.552 | 1215.9302 | 350.27164 |
| Evl | QEILEEVVR | -383 | -391 | 0 | 0.003003218 | 9 | NaN | NaN | NaN | 811.007 | NaN | 907.52637 |
| Nucks1 | SQEDSEDSEEKDVK | -54 | -67 | 1 | 0.000550307 | 14 | NaN | NaN | NaN | 264.4118 | 315.54984 | 299.5376 |
| Nucks1 | EEDEEAESPPEKK | -207 | -219 | 1 | 0 | 13 | NaN | NaN | NaN | 519.0592 | NaN | 687.1287 |
| Cpsf1 | RVEPAVGWTGGK | -423 | -434 | 1 | 0.000712251 | 12 | NaN | NaN | NaN | 299.65768 | 425.0594 | 771.4702 |
| Cpsf1 | DVIEVVPEPGQPLTK | -1141 | -1155 | 0 | 0.001868731 | 15 | NaN | NaN | NaN | 1111.0171 | NaN | 1477.1731 |
| Asl | SSVEWQIR | -446 | -453 | 0 | 0 | 8 | NaN | NaN | NaN | 724.5868 | 443.3789 | 489.00354 |
| Asl | INVLPLGSGAIAGNPLGVDR | -194 | -213 | 0 | 0.00450079 | 20 | NaN | NaN | NaN | 354.83035 | NaN | NaN |
| Ppp6r1 | LLTCVERPQLR | -169 | -179 | 1 | 0.001055699 | 11 | NaN | NaN | NaN | 1138.8691 | NaN | 1865.9191 |

| | | | | | | | | | | | | |
|---|---|---|---|---|---|---|---|---|---|---|---|---|
| Ppp6r1 | SDGEYGAWQGSQPVR | -636 | -650 | 0 | 0.001023236 | 15 | NaN | NaN | NaN | 3932.9307 | NaN | NaN |
| Ppp6r1 | NTVDLVNTH | -517 | -525 | 0 | 0.001291103 | 9 | NaN | NaN | NaN | 1221.5869 | NaN | NaN |
| Letm1 | KLEEGGPVYSPPAQVV | -132 | -147 | 1 | 0 | 16 | NaN | NaN | NaN | 671.0282 | 1779.7454 | 2434.244 |
| Diaph1 | LIEECVSQIVLHK | -424 | -436 | 0 | 0 | 13 | NaN | NaN | NaN | 809.0681 | 1688.2521 | 1479.9595 |
| Diaph1 | GFNISFLCK | -981 | -989 | 0 | 0.000394361 | 9 | NaN | NaN | NaN | 269.40442 | NaN | 331.44412 |
| Hk2 | LSPELLTTGSFETK | -324 | -337 | 0 | 0 | 14 | NaN | NaN | NaN | 1005.80914 | NaN | 674.60815 |
| Hk2 | SHEQLLEVK | -480 | -488 | 0 | 0.00025015 | 9 | NaN | NaN | NaN | 1980.2185 | NaN | NaN |
| Manf | LCYYIGATDDAATK | -71 | -84 | 0 | 0 | 14 | NaN | NaN | NaN | 1190.6893 | 191.79915 | 737.91516 |
| Manf | DVTFSPATIEEEIK | -45 | -59 | 0 | 0 | 15 | NaN | NaN | NaN | 616.0625 | 754.72144 | 611.4327 |
| Manf | KDSQICELK | -109 | -117 | 1 | 0 | 9 | NaN | NaN | NaN | 418.4194 | 557.8048 | 156.7128 |
| Calu | VHHEPQLSDK | -28 | -37 | 0 | 0.000627943 | 10 | NaN | NaN | NaN | 797.3208 | 781.1488 | NaN |
| Dazap1 | GGIPHNCGETELR | -118 | -130 | 0 | 0 | 13 | NaN | NaN | NaN | 1659.8694 | NaN | 1732.7465 |
| Dazap1 | NIDPKPCTPR | -79 | -88 | 1 | 0.002466863 | 10 | NaN | NaN | NaN | 1128.5372 | NaN | NaN |
| Nelfe | DWSYEEHGSAR | -167 | -177 | 0 | 0 | 11 | NaN | NaN | NaN | 361.0889 | NaN | 139.80397 |
| Irf4 | YPGLVWENEEK | -37 | -47 | 0 | 0 | 11 | NaN | NaN | NaN | 433.509 | 1083.0226 | 742.10535 |
| Irf4 | NLDQVLFPYPDDNGQR | -286 | -301 | 0 | 0 | 16 | NaN | NaN | NaN | 374.98 | 522.2603 | 467.87018 |
| Irf4 | DYAPDQSHPEIPYQCPVTF | -180 | -198 | 0 | 0.002296977 | 19 | NaN | NaN | NaN | 422.90945 | NaN | NaN |
| Lss | DYASIDWPAQR | -268 | -278 | 0 | 0 | 11 | NaN | NaN | NaN | 910.4316 | 1434.0005 | 1444.4076 |
| Rab35 | WLHEINQNCDDVCR | -102 | -115 | 0 | 0.00025015 | 14 | NaN | NaN | NaN | 1829.176 | 242.42249 | 45.044273 |
| Rras2 | QVTQEEGQQLAR | -136 | -147 | 0 | 0.00025015 | 12 | NaN | NaN | NaN | 1562.6187 | NaN | NaN |
| Dpp9 | DYDWTEPLSPTEDEFK | -464 | -479 | 0 | 0.00025015 | 16 | NaN | NaN | NaN | 630.0205 | 736.4641 | 595.28125 |
| Pgm1 | SIPYFQQTGVR | -319 | -329 | 0 | 0 | 11 | NaN | NaN | NaN | 1444.1743 | 456.70947 | 1531.3455 |
| Pgm1 | KADNFEYSDPVDGSISK | -470 | -486 | 1 | 0 | 17 | NaN | NaN | NaN | 825.26953 | 770.50574 | 775.34753 |
| Pgm1 | HGVVGPYVK | -226 | -234 | 0 | 0.002329279 | 9 | NaN | NaN | NaN | 773.9827 | NaN | NaN |
| Pgm1 | DYEEVEAEGANK | -429 | -440 | 0 | 0 | 12 | NaN | NaN | NaN | 658.72144 | NaN | NaN |
| Pgm1 | EAIQLIVR | -69 | -76 | 0 | 0.003133457 | 8 | NaN | NaN | NaN | 616.1417 | NaN | NaN |
| Pgm1 | YDYEEVEAEGANK | -428 | -440 | 0 | 0 | 13 | NaN | NaN | NaN | 561.3729 | NaN | NaN |
| Morf4l1 | HGPLLYEAK | -21 | -29 | 0 | 0.003521747 | 9 | NaN | NaN | NaN | 5510.8184 | 6801.276 | 6676.708 |
| Hnrnpa1 | GLSFETTDESLR | -20 | -31 | 0 | 0.000230707 | 12 | NaN | NaN | NaN | 1808.6504 | 1692.932 | 1797.4498 |
| Hnrnpa1 | NDFGNYNNQSSNFGPMK | -261 | -277 | 0 | 0.000627943 | 17 | NaN | NaN | NaN | 690.4837 | 836.1261 | 817.5886 |
| Pitpna | DYKAEEDPAK | -154 | -163 | 1 | 0 | 10 | NaN | NaN | NaN | 380.52396 | 487.3427 | 8252.705 |
| Pitpna | WKQELVNQK | -178 | -186 | 1 | 0.002466863 | 9 | NaN | NaN | NaN | 489.07892 | NaN | NaN |
| Rps10 | HPELADK | -32 | -38 | 0 | 0.000431655 | 7 | NaN | NaN | NaN | 893.83563 | 699.9099 | 859.39 |
| Rps10 | IYELLFK | -11 | -17 | 0 | 0.004224523 | 7 | NaN | NaN | NaN | 529.77875 | 525.26794 | NaN |
| Rps10 | GPEGERPA | -108 | -115 | 1 | 0.005668207 | 8 | NaN | NaN | NaN | 483.37463 | NaN | NaN |
| Cox4i1 | KWDYDKNEWK | -159 | -168 | 2 | 0.00025015 | 10 | NaN | NaN | NaN | 985.04346 | 1517.447 | 828.55493 |
| Erbin | DYSHCSLEQVPK | -28 | -39 | 0 | 0 | 12 | NaN | NaN | NaN | 1462.5067 | 2189.353 | 2109.1025 |
| Erbin | WLSDNQSKPLIPLQK | -397 | -411 | 1 | 0.000471409 | 15 | NaN | NaN | NaN | 2434.743 | NaN | 3230.0903 |
| Erbin | HIVNHDDVFEE | -584 | -594 | 0 | 0.002296977 | 11 | NaN | NaN | NaN | 3107.491 | NaN | NaN |
| Pfkl | SPVTELKK | -720 | -727 | 1 | 0.00272025 | 8 | NaN | NaN | NaN | 546.36914 | 450.68124 | NaN |
| Rpl13 | EAAEQDVEK | -201 | -209 | 0 | 0.005668207 | 9 | NaN | NaN | NaN | 110.03512 | NaN | 424.59863 |
| Cpox | REEDGDELAR | -105 | -114 | 1 | 0.005605646 | 10 | NaN | NaN | NaN | 431.5962 | 841.9313 | 424.21454 |
| Cpox | EACDQHGPDIYPK | -291 | -303 | 0 | 0.00111006 | 13 | NaN | NaN | NaN | 342.79108 | 793.00287 | 498.08942 |
| Ccar1 | STVMDDIHTVLK | -1115 | -1126 | 0 | 0.00272025 | 12 | NaN | NaN | NaN | 1272.1531 | 1520.0612 | 1443.1567 |
| Ccar1 | EDDIDEPKPK | -816 | -825 | 1 | 0 | 10 | NaN | NaN | NaN | 1259.3009 | NaN | 2371.6714 |
| Psmd7 | SVVALHNLINNK | -268 | -279 | 0 | 0 | 12 | NaN | NaN | NaN | 698.35376 | 641.1089 | 935.31476 |
| Psmd7 | SEIGAEEAEEVGVEHLLR | -160 | -177 | 0 | 0.00025015 | 18 | NaN | NaN | NaN | 231.01837 | 368.1636 | 366.61554 |
| Edc4 | SALHLLSPR | -779 | -787 | 0 | 0 | 9 | NaN | NaN | NaN | 879.6818 | NaN | NaN |
| Edc4 | HNTWPVDVSQIK | -276 | -287 | 0 | 0.001291103 | 12 | NaN | NaN | NaN | 680.3666 | NaN | NaN |
| Vapa | LQNPSDR | -39 | -45 | 0 | 0 | 7 | NaN | NaN | NaN | 785.7998 | NaN | 608.50616 |
| Vapa | EAKPDELMDSK | -115 | -125 | 1 | 0.000550307 | 11 | NaN | NaN | NaN | 397.60513 | NaN | NaN |
| Prpf19 | SVPNTSCVQVVR | -292 | -303 | 0 | 0 | 12 | NaN | NaN | NaN | 2213.6184 | 1141.5873 | 1048.9475 |
| Prpf19 | EALATLKPQAGLIVPQ | -129 | -144 | 1 | 0.001636916 | 16 | NaN | NaN | NaN | 602.1327 | 669.0048 | 649.6759 |
| Prpf19 | IAENGTDPINNQPLSEEQLIDIK | -34 | -56 | 0 | 0.001636916 | 23 | NaN | NaN | NaN | 241.28258 | 220.0564 | 244.05438 |
| Prpf19 | KVTSVVFHPSQELVFSA | -266 | -282 | 1 | 0.00025015 | 17 | NaN | NaN | NaN | 348.51007 | NaN | 395.5391 |
| Prpf19 | ENGTDPINNQPLSEEQLIDIK | -36 | -56 | 0 | 0.007670622 | 21 | NaN | NaN | NaN | 181.46689 | 136.53435 | NaN |
| Chmp4b | REALENANTNTEVLK | -93 | -107 | 1 | 0.007609801 | 15 | NaN | NaN | NaN | 1896.3926 | 2385.7886 | 2242.1335 |
| Chmp4b | GGPTPQEAIQR | -18 | -28 | 0 | 0.00025015 | 11 | NaN | NaN | NaN | 3213.953 | NaN | NaN |
| Chmp4b | LEELEQEELDK | -168 | -178 | 0 | 0 | 11 | NaN | NaN | NaN | 639.6998 | NaN | NaN |
| Chmp4b | NLLEISGPETVPLPNVPSVALPSKPAK | -179 | -205 | 1 | 0 | 27 | NaN | NaN | NaN | 279.57687 | NaN | NaN |
| Osgep | EALIVGGVGCNLR | -256 | -268 | 0 | 0.001023236 | 13 | NaN | NaN | NaN | 1522.2407 | NaN | NaN |
| Nherf1 | EALVEPASESPRPALA | -266 | -281 | 1 | 0.003069674 | 16 | NaN | NaN | NaN | 294.42404 | 322.03967 | NaN |
| Nherf1 | SEQAEPPAAADTH | -117 | -129 | 0 | 0.001743443 | 13 | NaN | NaN | NaN | 1293.5021 | NaN | NaN |
| Nherf1 | VVDPETDER | -90 | -98 | 0 | 0 | 9 | NaN | NaN | NaN | 1066.4222 | NaN | NaN |
| Zc3h11a | TLEEIKLEK | -449 | -457 | 1 | 0 | 9 | NaN | NaN | NaN | 1010.3014 | 327.3396 | NaN |
| Zc3h11a | GIGDSILNVK | -608 | -617 | 0 | 0.00051306 | 10 | NaN | NaN | NaN | 1370.9451 | NaN | NaN |
| Zc3h11a | SVLQQPQPNPGPEK | -241 | -254 | 0 | 0.002629273 | 14 | NaN | NaN | NaN | 498.40152 | NaN | NaN |
| Tdp2 | EAPDSTTVIFAGDTNLR | -260 | -276 | 0 | 0.000230707 | 17 | NaN | NaN | NaN | 3758.3157 | 6189.197 | NaN |
| Eif3f | VIGTLLGTVDKH | -120 | -131 | 1 | 0.000230707 | 12 | NaN | NaN | NaN | 782.66754 | 796.1461 | 974.33276 |
| Eif3f | YLANLTQSQIALNEK | -343 | -357 | 0 | 0 | 15 | NaN | NaN | NaN | 0 | 3555.6614 | 4093.38 |
| Eif3f | QTPAPSQPGPALPGPFPGGR | -75 | -94 | 0 | 0.001811524 | 20 | NaN | NaN | NaN | 406.22934 | NaN | 1531.1719 |
| Eif3f | SVLIHEYYSR | -187 | -196 | 0 | 0 | 10 | NaN | NaN | NaN | 690.6621 | NaN | NaN |
| Slc12a7 | EAQLIHDR | -949 | -956 | 0 | 0 | 8 | NaN | NaN | NaN | 1996.2056 | NaN | 1897.9612 |
| Slc12a7 | GVVEHEEDEDSR | -98 | -109 | 0 | 0.009985598 | 12 | NaN | NaN | NaN | 215.13684 | NaN | 220.52885 |
| Sh3kbp1 | VSSQAAIEELK | -647 | -657 | 0 | 0 | 11 | NaN | NaN | NaN | 457.67328 | 566.331 | NaN |
| Rnmt | LQPQDDEVPQK | -116 | -126 | 0 | 0.000394361 | 11 | NaN | NaN | NaN | 964.7103 | NaN | NaN |
| Rnmt | GNYPLFGCK | -338 | -346 | 0 | 0.002595029 | 9 | NaN | NaN | NaN | 822.75305 | NaN | NaN |
| Jpt1 | GFDEPAEQPVRK | -33 | -44 | 1 | 0 | 12 | NaN | NaN | NaN | 3221.1865 | 2608.8123 | 1833.9689 |

| | | | | | | | | | | | | |
|---|---|---|---|---|---|---|---|---|---|---|---|---|
| Jpt1 | SSEASSGDFLDLK | -87 | -99 | 0 | 0 | 13 | NaN | NaN | NaN | 366.36234 | 368.64014 | 502.3933 |
| Jpt1 | EASSGDFLDLK | -89 | -99 | 0 | 0 | 11 | NaN | NaN | NaN | 902.4822 | 1144.4274 | NaN |
| Jpt1 | GFDEPAEQPVR | -33 | -43 | 0 | 0.00025015 | 11 | NaN | NaN | NaN | 604.09546 | NaN | 2158.209 |
| Ahctf1 | SGPTLNEVIPDSYNR | -506 | -520 | 0 | 0 | 15 | NaN | NaN | NaN | 679.532 | NaN | 3881.5586 |
| Ahctf1 | TSVLDGIVPIESR | -1430 | -1442 | 0 | 0.00025015 | 13 | NaN | NaN | NaN | 676.2846 | NaN | NaN |
| Ahctf1 | EATPVAAGEAGDEDGAVIVSK | -1679 | -1699 | 0 | 0.002629273 | 21 | NaN | NaN | NaN | 326.87457 | NaN | NaN |
| Acadvl | EATQAVLDKPETL | -44 | -56 | 1 | 0.002099344 | 13 | NaN | NaN | NaN | 4326.086 | 4347.3374 | NaN |
| Inpp4b | TLEFVPVNLHLQ | -319 | -330 | 0 | 0 | 12 | NaN | NaN | NaN | 1598.6266 | 538.9031 | NaN |
| Inpp4b | EATSNDVLPVLTGR | -684 | -697 | 0 | 0.000471409 | 14 | NaN | NaN | NaN | 913.7078 | NaN | NaN |
| Eef1b | SILLDVKPWDDETDMTK | -141 | -157 | 1 | 0.008528992 | 17 | NaN | NaN | NaN | 1547.5905 | 2024.9324 | NaN |
| Eef1b | SIQADGLVWGSSK | -164 | -176 | 0 | 0 | 13 | NaN | NaN | NaN | 757.0129 | NaN | NaN |
| Tsn | EILTLLQGVHQGTGFQDIPK | -37 | -56 | 0 | 0.000550307 | 20 | NaN | NaN | NaN | 228.17574 | 223.68065 | 224.69562 |
| Tsn | INELDSGFR | -174 | -182 | 0 | 0.00051306 | 9 | NaN | NaN | NaN | 88.46566 | 649.3846 | NaN |
| Tsn | EILTLLQGVHQ | -37 | -47 | 0 | 0.00025015 | 11 | NaN | NaN | NaN | 385.1459 | NaN | NaN |
| Stt3b | FIAIVPGYISR | -201 | -211 | 0 | 0 | 11 | NaN | NaN | NaN | 452.8579 | 523.97766 | 539.18054 |
| Stt3a | SKLNPQQFEVLFR | -288 | -300 | 1 | 0.00025015 | 13 | NaN | NaN | NaN | 400.88245 | 776.04407 | 1521.3025 |
| Stt3a | SKLNPQQFEVLFR | -288 | -300 | 1 | 0.00025015 | 13 | NaN | NaN | NaN | 400.88245 | 776.04407 | 1521.3025 |
| Stt3a | VIFGGLTGYSSDDINK | -580 | -595 | 0 | 0 | 16 | NaN | NaN | NaN | 1206.537 | 1175.1421 | NaN |
| Stt3a | RPPGFDR | -656 | -662 | 1 | 0.001291103 | 7 | NaN | NaN | NaN | 807.22107 | NaN | 1526.4868 |
| Stt3a | RFLAEEGFYK | -58 | -67 | 1 | 0.001055699 | 10 | NaN | NaN | NaN | 2085.3206 | NaN | NaN |
| Adh5 | PFQLVTGR | -305 | -312 | 0 | 0.003102268 | 8 | NaN | NaN | NaN | 1282.4044 | 968.0166 | 981.2854 |
| Adh5 | PQCGECK | -95 | -101 | 0 | 0.00149795 | 7 | NaN | NaN | NaN | 378.94016 | 645.1939 | 560.9407 |
| Cdk2ap1;Cdk2 | ECLAETER | (103);(116) | (110);(123) | 0 | 0.000230707 | 8 | NaN | NaN | NaN | 662.0733 | NaN | NaN |
| Elac2 | NVSSTLVNLSPDK | -494 | -506 | 0 | 0.001636916 | 13 | NaN | NaN | NaN | 4066.1921 | NaN | NaN |
| Huwe1 | VGEGGSGIDFK | -3561 | -3571 | 0 | 0 | 11 | NaN | NaN | NaN | 1259.9241 | 1429.832 | 945.1662 |
| Huwe1 | LLECYFTR | -4126 | -4133 | 0 | 0.002199523 | 8 | NaN | NaN | NaN | 814.03015 | 638.90094 | 693.8989 |
| Rpl36a | ECVEPNCR | -71 | -78 | 0 | 0 | 8 | NaN | NaN | NaN | 537.58124 | NaN | NaN |
| Suz12 | HGGGGGGGSGPSAGSGGGGFGGSA | -6 | -29 | 0 | 0.002566359 | 24 | NaN | NaN | NaN | 394.46527 | NaN | NaN |
| Ddx47 | IEHLIGK | -378 | -384 | 0 | 0.005394768 | 7 | NaN | NaN | NaN | 249.84293 | NaN | 638.37854 |
| Rala | EDENVPFLLVGNK | -116 | -128 | 0 | 0.005605646 | 13 | NaN | NaN | NaN | 2629.0144 | NaN | NaN |
| Ola1 | KYEDFKDEGSENAVK | -349 | -363 | 2 | 0.00025015 | 15 | NaN | NaN | NaN | 527.4838 | 376.6578 | 1385.7279 |
| Arhgef2 | EDFPLIETEDK | -578 | -588 | 0 | 0 | 11 | NaN | NaN | NaN | 501.03256 | 713.83044 | NaN |
| Ubash3a | EDGKWDLVNPPVK | -593 | -605 | 1 | 0.00025015 | 13 | NaN | NaN | NaN | 3299.6147 | NaN | NaN |
| Ubash3a | EDQKVECLYEALR | -120 | -132 | 1 | 0.001392869 | 13 | NaN | NaN | NaN | 876.0433 | NaN | NaN |
| Casp3 | SVDSGIYLDSSYK | -26 | -38 | 0 | 0 | 13 | NaN | NaN | NaN | 662.6343 | 783.00543 | 1085.6136 |
| Casp3 | EDILELMDSVSK | -94 | -105 | 0 | 0 | 12 | NaN | NaN | NaN | 120.13889 | 267.21408 | 187.9826 |
| Stk17b | EDKENIPEDGSLVSK | -338 | -352 | 1 | 0 | 15 | NaN | NaN | NaN | 906.67487 | 478.04556 | 389.90332 |
| Rpl7 | HFVEGGDAGNREDQINR | -248 | -264 | 1 | 0 | 17 | NaN | NaN | NaN | 2845.5642 | 3625.0957 | 3535.6401 |
| Rpl7 | MEDLIHEIYTVGK | -209 | -221 | 0 | 0 | 13 | NaN | NaN | NaN | 4267.0225 | 5122.2295 | 4568.708 |
| Rpl7 | RNFAELK | -45 | -51 | 1 | 0.00025015 | 7 | NaN | NaN | NaN | 801.5304 | NaN | 835.9801 |
| Eif5a | NDFQLIGIQDGYLS | -87 | -100 | 0 | 0.000627943 | 14 | NaN | NaN | NaN | 485.85873 | 524.5901 | 531.6344 |
| Prex1 | SHYEFHNICDTK | -905 | -916 | 0 | 0.001610812 | 12 | NaN | NaN | NaN | 214.30196 | NaN | NaN |
| Dock10 | KYEQYSGDIR | -126 | -135 | 1 | 0 | 10 | NaN | NaN | NaN | 1221.2529 | NaN | NaN |
| Ly9 | EDTHLNTYDGSHTLR | -194 | -208 | 0 | 0.000230707 | 15 | NaN | NaN | NaN | 702.6241 | 706.14545 | 399.73975 |
| Hnrnpa3 | EDSVKPGAHLT | -114 | -124 | 1 | 0.000230707 | 11 | NaN | NaN | NaN | 4458.704 | 2154.934 | 5385.421 |
| Hnrnpa3 | KYHTINGHNCEVK | -187 | -199 | 1 | 0 | 13 | NaN | NaN | NaN | 1552.4225 | 1953.5278 | 1996.9381 |
| Hnrnpa3 | YGGGYGSGGGSGGYGSR | -361 | -377 | 0 | 0 | 17 | NaN | NaN | NaN | 404.82642 | 774.03296 | 890.1416 |
| Hnrnpa3 | KIFVGGIKEDTEEYNLR | -127 | -143 | 2 | 0.00025015 | 17 | NaN | NaN | NaN | 260.8042 | NaN | 2838.1292 |
| Hnrnpa3 | KIFVGGIKEDTEEYNLR | -127 | -143 | 2 | 0.00025015 | 17 | NaN | NaN | NaN | 260.8042 | NaN | 2838.1292 |
| Anxa1 | EDTLIEILTTR | -134 | -144 | 0 | 0.007609801 | 11 | NaN | NaN | NaN | 1389.497 | NaN | NaN |
| Psap | TCEWIHDSSLSASCK | -93 | -107 | 0 | 0 | 15 | NaN | NaN | NaN | 1247.7032 | 1437.6335 | 1423.275 |
| Psap | KLVLYLEHNLEK | -447 | -458 | 1 | 0 | 12 | NaN | NaN | NaN | 921.231 | 1719.8196 | 810.48035 |
| Psap | EEILAALEK | -463 | -471 | 0 | 0.000471409 | 9 | NaN | NaN | NaN | 792.8482 | 879.86176 | 615.5389 |
| Psap | GLSNACALLPDPAR | -343 | -356 | 0 | 0 | 14 | NaN | NaN | NaN | 496.5687 | 1625.9927 | 1437.4603 |
| Psap | EVVDSYLPVILDMIK | -108 | -122 | 0 | 0.00025015 | 15 | NaN | NaN | NaN | 220.59967 | NaN | 275.69797 |
| Psap | GGFCEVCK | -439 | -446 | 0 | 0 | 8 | NaN | NaN | NaN | 344.58167 | NaN | NaN |
| Psap | EDVCQDCMK | -194 | -202 | 0 | 0.00383302 | 9 | NaN | NaN | NaN | 291.2409 | NaN | NaN |
| Ppme1 | GIIEEEEEDEEGSESVNKR | -253 | -271 | 1 | 0.00025015 | 19 | NaN | NaN | NaN | 344.2329 | 375.97586 | 401.4968 |
| Ppme1 | LPSRPPLPGSGGSQ | -13 | -26 | 1 | 0.000431655 | 14 | NaN | NaN | NaN | 15660.135 | NaN | 15830.967 |
| Fxyd5 | EEATGSQTAAQTETQQLTK | -64 | -82 | 0 | 0.000431655 | 19 | NaN | NaN | NaN | 644.16095 | 617.70746 | NaN |
| Lars1 | EECCPGKPLNVFR | -1052 | -1064 | 1 | 0 | 13 | NaN | NaN | NaN | 724.0423 | 1255.4633 | 1153.8387 |
| Lars1 | GCPPDFPEEEEEEEESSAKPGDIVVR | -114 | -139 | 1 | 0.001846793 | 26 | NaN | NaN | NaN | 683.6392 | 659.9445 | 715.62006 |
| Lars1 | TEPGVPVSLVNPQPSSGHFSTK | -1065 | -1086 | 0 | 0.000471409 | 22 | NaN | NaN | NaN | 303.85007 | 377.18768 | 333.83734 |
| Lars1 | NFEASLDWLQEHACSR | -562 | -577 | 0 | 0 | 16 | NaN | NaN | NaN | 252.4811 | 229.7003 | 355.58072 |
| Lars1 | LHLGHTF | -61 | -67 | 0 | 0.00051306 | 7 | NaN | NaN | NaN | 580.15955 | NaN | NaN |
| Lars1 | TNCWVRPDMK | -305 | -314 | 1 | 0.00713295 | 10 | NaN | NaN | NaN | 272.66855 | NaN | NaN |
| Atp1b3 | EVTVECHIAGTR | -244 | -255 | 0 | 0 | 12 | NaN | NaN | NaN | 2853.3145 | NaN | NaN |
| Sec13 | LTYTGEGQWEVK | -132 | -143 | 0 | 0 | 12 | NaN | NaN | NaN | 989.1907 | 1237.2838 | NaN |
| Sec13 | SVDGQWVCISDVNK | -292 | -305 | 0 | 0 | 14 | NaN | NaN | NaN | 385.4807 | NaN | 622.17456 |
| Sec13 | REEEDGQWK | -195 | -203 | 1 | 0.001023236 | 9 | NaN | NaN | NaN | 325.4961 | NaN | NaN |
| Xrcc5 | EEEDIVEDLFPTSK | -468 | -481 | 0 | 0 | 14 | NaN | NaN | NaN | 605.02594 | 436.1114 | 561.36346 |
| Eif5 | VLCPECENPETDLHVNPK | -97 | -114 | 0 | 0.000230707 | 18 | NaN | NaN | NaN | 566.5147 | 650.1865 | 609.0574 |
| Eif5 | EEEEDDDWGEDTTEEAQR | -194 | -211 | 0 | 0 | 18 | NaN | NaN | NaN | 239.94424 | 380.16687 | 363.0001 |
| Mbd3 | SAFDIAEELVR | -147 | -157 | 0 | 0 | 11 | NaN | NaN | NaN | 697.18604 | 820.8971 | 924.7614 |
| Snrpd2 | HCNMVLENVK | -62 | -71 | 0 | 0 | 10 | NaN | NaN | NaN | 1605.6138 | 683.07623 | 1464.9222 |
| Degs1 | EEFEWVYTDQPHAA | -8 | -21 | 0 | 0 | 14 | NaN | NaN | NaN | 437.35724 | 531.8936 | 580.7661 |
| Sash3 | ETHLNELNIMDPQHR | -289 | -303 | 0 | 0 | 15 | NaN | NaN | NaN | 571.0648 | 501.12592 | 581.6423 |
| Sash3 | EFNLDDNIPEDDSGVLTPEDSGK | -45 | -67 | 0 | 0.00025015 | 23 | NaN | NaN | NaN | 263.42996 | 208.6814 | 230.34085 |

| | | | | | | | | | | | | |
|---|---|---|---|---|---|---|---|---|---|---|---|---|
| Sash3 | FIYVDVLPEEAVGPVRPS | -226 | -243 | 1 | 0.001023236 | 18 | NaN | NaN | NaN | 263.51233 | NaN | NaN |
| U2af2 | KYCDPDSYHR | -462 | -471 | 1 | 0 | 10 | NaN | NaN | NaN | 175.60966 | 784.9843 | 557.2437 |
| Acadm | GGYGFNTEYPVEK | -377 | -389 | 0 | 0.002880184 | 13 | NaN | NaN | NaN | 785.22406 | NaN | NaN |
| Casp8 | EIHEILEGYQSADHK | -293 | -307 | 0 | 0 | 15 | NaN | NaN | NaN | 1816.7026 | 534.71924 | 1539.1774 |
| Lpxn | EELGSSPFFER | -183 | -193 | 0 | 0 | 11 | NaN | NaN | NaN | 129.71599 | NaN | NaN |
| Utp6 | YLDWAYR | -481 | -487 | 0 | 0.005758933 | 7 | NaN | NaN | NaN | 1072.8368 | NaN | 1008.4108 |
| Acad9 | GVFPFPEVSQHELSEIN | -56 | -72 | 0 | 0.00025015 | 17 | NaN | NaN | NaN | 370.02206 | 339.79355 | 427.76544 |
| Acad9 | EENVHYFGR | -506 | -514 | 0 | 0 | 9 | NaN | NaN | NaN | 415.77698 | 445.43573 | NaN |
| Slc29a1 | WLPVLVASR | -360 | -368 | 0 | 0 | 9 | NaN | NaN | NaN | 560.1702 | 718.5272 | 703.3876 |
| Slc29a1 | EESGVPGPNSPPTNR | -264 | -278 | 0 | 0.001743443 | 15 | NaN | NaN | NaN | 1097.9214 | NaN | NaN |
| Spag7 | HGEEWDPQKAEEK | -123 | -135 | 1 | 0.000627943 | 13 | NaN | NaN | NaN | 850.2938 | 1062.3093 | 1104.1469 |
| Spag7 | SFSFGEDDDCR | -92 | -102 | 0 | 0.000431655 | 11 | NaN | NaN | NaN | 396.6525 | NaN | NaN |
| Spag7 | SFGEDDDCR | -94 | -102 | 0 | 0 | 9 | NaN | NaN | NaN | 198.8792 | NaN | NaN |
| Usp24 | NQLETAPPHELK | -2436 | -2447 | 0 | 0.001868731 | 12 | NaN | NaN | NaN | 620.56036 | 465.77325 | NaN |
| Usp24 | EFDYLPPVDSR | -1670 | -1680 | 0 | 0.004983786 | 11 | NaN | NaN | NaN | 1612.361 | NaN | NaN |
| Mta2 | LNPADAPNPVVF | -596 | -607 | 0 | 0 | 12 | NaN | NaN | NaN | 1013.48486 | 1561.1097 | 524.3736 |
| Mta2 | EFEEESKQPGVSEQQ | -65 | -79 | 1 | 0 | 15 | NaN | NaN | NaN | 450.25375 | 521.01086 | 604.261 |
| Mta2 | LAEGESDNR | -161 | -169 | 0 | 0 | 9 | NaN | NaN | NaN | 586.3248 | NaN | 610.31635 |
| Mta2 | EFEEESKQPGVSEQQR | -65 | -80 | 1 | 0 | 16 | NaN | NaN | NaN | 291.18964 | NaN | 358.8766 |
| Rps15a | TTSAGIMDHEEAR | -105 | -117 | 0 | 0 | 13 | NaN | NaN | NaN | 1282.1195 | 1135.4932 | 1408.1261 |
| Rps15a | SAGIMDHEEAR | -107 | -117 | 0 | 0 | 11 | NaN | NaN | NaN | 410.43164 | 592.9405 | 715.5195 |
| Rps15a | KCGVISPR | -71 | -78 | 1 | 0.005605646 | 8 | NaN | NaN | NaN | 3835.4282 | NaN | NaN |
| Ndufb5 | GDGPWYQFPTPEK | -164 | -176 | 0 | 0 | 13 | NaN | NaN | NaN | 877.62744 | NaN | 1189.2294 |
| Atxn2l | STSTPTSPGPR | -681 | -691 | 0 | 0 | 11 | NaN | NaN | NaN | 644.14667 | NaN | NaN |
| Cd69 | EFNSWFNLTGSGR | -160 | -172 | 0 | 0 | 13 | NaN | NaN | NaN | 477.65436 | 483.94037 | 598.78174 |
| Map2k1 | KKPTPIQLNPAPDGSAVN | -4 | -21 | 2 | 0.00149795 | 18 | NaN | NaN | NaN | 211.55898 | 618.4081 | 638.646 |
| Gtf2f1 | EFRPEDQPWLL | -90 | -100 | 1 | 0.000230707 | 11 | NaN | NaN | NaN | 643.9873 | 654.1134 | 340.2463 |
| Gfm1 | RYDEIPAGLR | -235 | -244 | 1 | 0.00025015 | 10 | NaN | NaN | NaN | 2293.8528 | 1256.7307 | 1501.4669 |
| Fbln1 | EFTRPEEIIFL | -610 | -620 | 1 | 0.005821388 | 11 | NaN | NaN | NaN | 1778.367 | 1813.9362 | NaN |
| Fbln1 | EFTRPEEIIFLR | -610 | -621 | 1 | 0 | 12 | NaN | NaN | NaN | 2874.827 | NaN | NaN |
| Fbln1 | EFTRPEEIIFLR | -610 | -621 | 1 | 0 | 12 | NaN | NaN | NaN | 2874.827 | NaN | NaN |
| Ecpas | GHAETDEQLQNIISK | -18 | -32 | 0 | 0 | 15 | NaN | NaN | NaN | 947.3316 | 521.68884 | 739.183 |
| Ecpas | TIAVLLPCLLDK | -1245 | -1256 | 0 | 0 | 12 | NaN | NaN | NaN | 658.43713 | 751.1186 | NaN |
| Ehmt2 | EGDTAWDLTPER | -936 | -947 | 0 | 0.000755556 | 12 | NaN | NaN | NaN | 274.7235 | 4414.121 | NaN |
| Rps28 | FMDDTSR | -34 | -40 | 0 | 0.002972566 | 7 | NaN | NaN | NaN | 144.22957 | 437.62708 | NaN |
| Cdk11b | SVEEFQCLNR | -423 | -432 | 0 | 0 | 10 | NaN | NaN | NaN | 380.32114 | NaN | NaN |
| Smarcc2 | IEESGTEEARPEGQAA | -751 | -766 | 1 | 0.00111006 | 16 | NaN | NaN | NaN | 344.8634 | NaN | 150.51306 |
| Ctcf | FHCPHCDTVIAR | -437 | -448 | 0 | 0 | 12 | NaN | NaN | NaN | 520.7913 | 691.1587 | NaN |
| Ctcf | TLPLPEGFQVVK | -157 | -168 | 0 | 0.001055699 | 12 | NaN | NaN | NaN | 442.80444 | 857.25397 | NaN |
| Mogs | HLWSPFGLR | -718 | -726 | 0 | 0.001055699 | 9 | NaN | NaN | NaN | 614.34955 | 1014.4404 | 864.1035 |
| Mogs | SGHTSELGDFR | -230 | -240 | 0 | 0.000550307 | 11 | NaN | NaN | NaN | 998.58325 | 659.33246 | NaN |
| Prkcq | SPYDCSNFDKEFLSEKPR | -657 | -674 | 2 | 0.00025015 | 18 | NaN | NaN | NaN | 438.67728 | 450.43732 | NaN |
| Nup37 | EGQELASVSDDHTCR | -136 | -150 | 0 | 0.002629273 | 15 | NaN | NaN | NaN | 190.5562 | NaN | NaN |
| Srrm2 | SVVQTTPVAGSQSLSSGTVAK | -2416 | -2436 | 0 | 0.000591124 | 21 | NaN | NaN | NaN | 419.94678 | 843.7034 | 762.78577 |
| Srrm2 | PASVNLADSR | -2203 | -2212 | 0 | 0.00201047 | 10 | NaN | NaN | NaN | 501.63406 | 994.22253 | NaN |
| Srrm2 | LGLIQEDVASSCIPR | -1133 | -1147 | 0 | 0 | 15 | NaN | NaN | NaN | 2998.6052 | NaN | NaN |
| Cnot2 | FFDCLNWR | -501 | -508 | 0 | 0.004350979 | 8 | NaN | NaN | NaN | 581.7043 | 489.549 | NaN |
| Cct7 | QVKPYVEEGLHPQIIIR | -107 | -123 | 1 | 0 | 17 | NaN | NaN | NaN | 1521.3435 | 168.563 | 1588.4193 |
| Cct7 | VWEPAMVR | -492 | -499 | 0 | 0 | 8 | NaN | NaN | NaN | 259.0871 | 1146.1938 | 880.76685 |
| Cct7 | VWEPAMVR | -492 | -499 | 0 | 0 | 8 | NaN | NaN | NaN | 259.0871 | 1146.1938 | 880.76685 |
| Cct7 | GGALEESQLVAGVAFK | -202 | -217 | 0 | 0 | 16 | NaN | NaN | NaN | 1093.9966 | NaN | NaN |
| Cct7 | VKPYVEEGLHPQIIIR | -108 | -123 | 1 | 0 | 16 | NaN | NaN | NaN | 496.2143 | NaN | NaN |
| Enpp4 | EGVLVEHVK | -58 | -66 | 0 | 0 | 9 | NaN | NaN | NaN | 1295.1886 | 854.0371 | NaN |
| Traf1 | EHAIDAFRPDLSSA | -349 | -362 | 1 | 0.000907206 | 14 | NaN | NaN | NaN | 592.40265 | NaN | NaN |
| Smc2 | SDYDWINAEK | -936 | -945 | 0 | 0.00025015 | 10 | NaN | NaN | NaN | 1372.3169 | NaN | 1384.2045 |
| Smc2 | ELDHSISK | -912 | -919 | 0 | 0 | 8 | NaN | NaN | NaN | 1080.7792 | NaN | NaN |
| Smc2 | QQEELDALKK | -741 | -750 | 1 | 0.00078661 | 10 | NaN | NaN | NaN | 907.8402 | NaN | NaN |
| Parp3 | QAAPGEEEEKVEEVPHPLDR | -298 | -317 | 1 | 0.00025015 | 20 | NaN | NaN | NaN | 121.720375 | NaN | 511.2981 |
| Cops2 | IDQVNQLLELDHQK | -402 | -415 | 0 | 0 | 14 | NaN | NaN | NaN | 690.9186 | 572.52997 | 557.78735 |
| Chordc1 | TTDFSDFLSIVGCTK | -47 | -61 | 0 | 0 | 15 | NaN | NaN | NaN | 454.4137 | 465.41202 | 525.465 |
| Pnkp | EHLDPALQR | -506 | -514 | 0 | 0 | 9 | NaN | NaN | NaN | 828.2801 | 381.02933 | 598.784 |
| Pnkp | STFIQEHLVSA | -378 | -388 | 0 | 0.000907206 | 11 | NaN | NaN | NaN | 831.81134 | NaN | NaN |
| Eef1e1 | SLGLKPGNK | -13 | -21 | 1 | 0.000591124 | 9 | NaN | NaN | NaN | 741.064 | NaN | NaN |
| Ndufa10 | SAYLQDIENAYK | -231 | -242 | 0 | 0 | 12 | NaN | NaN | NaN | 1271.5652 | NaN | 1077.4971 |
| Ndufa10 | EHLLSTGQGVVLER | -148 | -161 | 0 | 0 | 14 | NaN | NaN | NaN | 1558.0256 | NaN | NaN |
| Orc4 | EHLQQLELIKPVER | -374 | -387 | 1 | 0 | 14 | NaN | NaN | NaN | 279.34415 | 665.4705 | NaN |
| Paox | EHQDTFFEPPLPAK | -287 | -300 | 0 | 0 | 14 | NaN | NaN | NaN | 2307.3584 | NaN | NaN |
| Por | EHTQAVDYVK | -655 | -664 | 0 | 0.00025015 | 10 | NaN | NaN | NaN | 1399.4863 | 1880.426 | NaN |
| Por | KKEEIPEFSK | -47 | -56 | 2 | 0.000394361 | 10 | NaN | NaN | NaN | 673.63947 | NaN | 617.16626 |
| Por | QYASEPSEQEHLHK | -391 | -404 | 0 | 0.001846793 | 14 | NaN | NaN | NaN | 273.7828 | NaN | NaN |
| Capn2 | SAEGSEFWSALLEK | -170 | -183 | 0 | 0 | 14 | NaN | NaN | NaN | 414.6208 | 603.3349 | 453.6417 |
| Capn2 | LEEEDEDEEDGER | -391 | -403 | 0 | 0 | 13 | NaN | NaN | NaN | 346.81638 | NaN | NaN |
| Mrps34 | EIEQVMYHDWR | -130 | -140 | 0 | 0 | 11 | NaN | NaN | NaN | 313.03433 | NaN | 156.25102 |
| Mrps34 | LNSVPYPPLLR | -161 | -171 | 0 | 0.002168954 | 11 | NaN | NaN | NaN | 1301.075 | NaN | NaN |
| Tuba4a | KEIIDPVLDR | -112 | -121 | 1 | 0.00025015 | 10 | NaN | NaN | NaN | 543.8287 | 525.9168 | 411.58484 |
| Arhgef1 | SPNLAAPER | -905 | -913 | 0 | 0 | 9 | NaN | NaN | NaN | 16639.36 | 21221.553 | NaN |
| Arhgef1 | LVTEAAHVR | -424 | -432 | 0 | 0.000431655 | 9 | NaN | NaN | NaN | 784.9656 | 934.56903 | NaN |
| Arhgef1 | KLVHEGPLTW | -646 | -655 | 1 | 0.00951004 | 10 | NaN | NaN | NaN | 673.95044 | NaN | 186.61873 |

| | | | | | | | | | | | | |
|---|---|---|---|---|---|---|---|---|---|---|---|---|
| Fga | EINLQDYEGHQK | -192 | -203 | 0 | 0 | 12 | NaN | NaN | NaN | 5874.59 | 6309.605 | NaN |
| Pdk1 | TIYEFTDTVIR | -134 | -144 | 0 | 0 | 11 | NaN | NaN | NaN | 876.13 | NaN | 548.4935 |
| Vps26a | EITGIGPSTTTETETIAK | -215 | -232 | 0 | 0.00078661 | 18 | NaN | NaN | NaN | 584.57935 | NaN | NaN |
| Vps26a | HYLFYDGESVSGK | -39 | -51 | 0 | 0 | 13 | NaN | NaN | NaN | 92.53366 | NaN | NaN |
| Hnrnpf | RTEMDWVLK | -90 | -98 | 1 | 0.00383302 | 9 | NaN | NaN | NaN | 1322.8035 | 1540.4617 | 684.89465 |
| Hnrnpf | MLGPEGGEGYVVK | -2 | -14 | 0 | 0 | 13 | NaN | NaN | NaN | 687.15875 | NaN | NaN |
| Tm9sf4 | VHWEESDIK | -255 | -263 | 0 | 0 | 9 | NaN | NaN | NaN | 2165.9993 | NaN | NaN |
| Txlna | FEEFQNTLSK | -394 | -403 | 0 | 0 | 10 | NaN | NaN | NaN | 465.17078 | 533.9636 | 523.28503 |
| Txlna | KFEEFQNTLSK | -393 | -403 | 1 | 0.00383302 | 11 | NaN | NaN | NaN | 1760.6185 | NaN | NaN |
| Rpl17 | EKEQIVPKPEEEVAQK | -152 | -167 | 2 | 0.000431655 | 16 | NaN | NaN | NaN | 1089.6958 | 571.13184 | 884.51373 |
| Rpl17 | TEKEQIVPKPEEEVAQK | -151 | -167 | 2 | 0 | 17 | NaN | NaN | NaN | 512.8284 | 581.68005 | 514.4757 |
| Rpl17 | KEQIVPKPEEEVAQK | -153 | -167 | 2 | 0.000471409 | 15 | NaN | NaN | NaN | 613.5291 | 437.06207 | NaN |
| Rpl17 | NAESNAELK | -97 | -105 | 0 | 0.001868731 | 9 | NaN | NaN | NaN | 435.86642 | NaN | NaN |
| Rpl17 | EQIVPKPEEEVAQKK | -154 | -168 | 2 | 0 | 15 | NaN | NaN | NaN | 169.55699 | NaN | NaN |
| Thrap3 | GLTHEELK | -687 | -694 | 0 | 0 | 8 | NaN | NaN | NaN | 1196.318 | NaN | 2070.6482 |
| Thrap3 | YKDDPVDLRL | -707 | -716 | 2 | 0.00201047 | 10 | NaN | NaN | NaN | 1089.7893 | NaN | NaN |
| Sf3b2 | TKTEEEEIDR | -679 | -688 | 1 | 0 | 10 | NaN | NaN | NaN | 565.9789 | 1700.8726 | NaN |
| Eif3g | PTGDFDSKPS | -2 | -11 | 1 | 0.000230707 | 10 | NaN | NaN | NaN | 820.23975 | 658.48584 | 789.75464 |
| Eif3g | GIPLPTGDTSPEPELLPGDPLPPPK | -33 | -57 | 0 | 0.00051306 | 25 | NaN | NaN | NaN | 425.0578 | 366.5029 | 400.94244 |
| Eif3g | SKEDLNCQEEEDPMNK | -133 | -148 | 1 | 0.000947459 | 16 | NaN | NaN | NaN | 366.078 | 279.5725 | 267.3649 |
| Eif3g | SKEDLNCQEEEDPMNK | -133 | -148 | 1 | 0.000947459 | 16 | NaN | NaN | NaN | 366.078 | 279.5725 | 267.3649 |
| Ddx56 | PAEQPFLGK | -391 | -399 | 0 | 0.009745776 | 9 | NaN | NaN | NaN | 475.06024 | 562.89294 | 329.5606 |
| Ddx56 | GIDFHHVSAVL | -348 | -358 | 0 | 0.00383302 | 11 | NaN | NaN | NaN | 1057.2275 | NaN | NaN |
| Caprin1 | SEGYTASQPLYQPSHATEQRPQK | -437 | -459 | 1 | 0 | 23 | NaN | NaN | NaN | 572.4065 | 909.03406 | 530.1763 |
| Caprin1 | STSEGYTASQPLYQPSHATEQRPQK | -435 | -459 | 1 | 0.00025015 | 25 | NaN | NaN | NaN | 473.0382 | 528.7627 | 479.59714 |
| Caprin1 | LNQDQLDAVSKY | -86 | -97 | 1 | 0.00025015 | 12 | NaN | NaN | NaN | 673.9234 | NaN | 419.48053 |
| Tes | HFCCFDCDHILAGK | -327 | -340 | 0 | 0 | 14 | NaN | NaN | NaN | 496.3529 | 924.7563 | 755.5974 |
| Tes | EKQPVAGSEGAQY | -138 | -150 | 1 | 0.000826806 | 13 | NaN | NaN | NaN | 2351.1606 | 870.1723 | 581.05145 |
| Tes | VTDKPVCKPCYVK | -345 | -357 | 2 | 0 | 13 | NaN | NaN | NaN | 714.2909 | 621.0791 | 623.3955 |
| Hacd3 | EKRPLFLAPDFDR | -97 | -109 | 2 | 0 | 13 | NaN | NaN | NaN | 4186.499 | 98.67189 | 2867.853 |
| Hacd3 | PLFLAPDFDR | -100 | -109 | 0 | 0.001811524 | 10 | NaN | NaN | NaN | 618.05634 | NaN | 626.3392 |
| Tpm4 | SEKEDKYEEEIK | -179 | -190 | 2 | 0.000230707 | 12 | NaN | NaN | NaN | 2019.4888 | NaN | 691.7259 |
| Tpm4 | QNLEEAEK | -75 | -82 | 0 | 0 | 8 | NaN | NaN | NaN | 1063.7241 | NaN | NaN |
| Top2b | KVIHELANER | -296 | -305 | 1 | 0 | 10 | NaN | NaN | NaN | 560.87616 | 995.3139 | NaN |
| Csnk2b | VDEDYIQDK | -25 | -33 | 0 | 0.00122419 | 9 | NaN | NaN | NaN | 852.1688 | NaN | NaN |
| Atp5me | ELAEAQDDSILK | -60 | -71 | 0 | 0 | 12 | NaN | NaN | NaN | 826.5736 | 1556.2295 | 1371.8596 |
| S100a13 | KGSLNINEFK | -30 | -39 | 1 | 0.001055699 | 10 | NaN | NaN | NaN | 973.61304 | 1096.4337 | 1828.6053 |
| S100a13 | ELATQQLPHLLK | -40 | -51 | 0 | 0 | 12 | NaN | NaN | NaN | 1596.4777 | NaN | NaN |
| Eif2b4 | SHLPQYSR | -182 | -189 | 0 | 0 | 8 | NaN | NaN | NaN | 2004.1986 | NaN | NaN |
| Eif2b4 | NWQSHPSLR | -476 | -484 | 0 | 0.000907206 | 9 | NaN | NaN | NaN | 1017.1721 | NaN | NaN |
| Eif2b4 | ELDDPDDLQCK | -457 | -467 | 0 | 0.00111006 | 11 | NaN | NaN | NaN | 606.17236 | NaN | NaN |
| Tbl3 | VAFHPDPTR | -157 | -165 | 0 | 0.003360691 | 9 | NaN | NaN | NaN | 3263.3032 | NaN | NaN |
| Tbl3 | STLLATGGCDGAVR | -121 | -134 | 0 | 0.006677519 | 14 | NaN | NaN | NaN | 1895.5535 | NaN | NaN |
| Immt | ERPPEEVAAR | -195 | -204 | 1 | 0.00025015 | 10 | NaN | NaN | NaN | 1712.4421 | 2779.9915 | 3841.5876 |
| Immt | SAEMPTIPLGSAVEAIR | -583 | -599 | 0 | 0.000947459 | 17 | NaN | NaN | NaN | 338.53357 | 1357.1447 | 587.30145 |
| Rgcc | ELEDFIADLDR | -122 | -132 | 0 | 0.000230707 | 11 | NaN | NaN | NaN | 296.57358 | NaN | NaN |
| Prpf40a | ELEDLEGYQNTIVAGGLITK | -214 | -233 | 0 | 0.001055699 | 20 | NaN | NaN | NaN | 485.25793 | NaN | 637.7619 |
| Prpf40a | RTLLEQLDDDQ | -943 | -953 | 1 | 0.009122264 | 11 | NaN | NaN | NaN | 616.5244 | NaN | NaN |
| Cotl1 | IQQCTDDVR | -49 | -57 | 0 | 0 | 9 | NaN | NaN | NaN | 592.987 | NaN | NaN |
| Pdcl3 | GKDYVQEVTK | -100 | -109 | 1 | 0.003360691 | 10 | NaN | NaN | NaN | 743.84924 | NaN | NaN |
| Rfc4 | LHDAIIENENLSDK | -310 | -323 | 0 | 0.002466863 | 14 | NaN | NaN | NaN | 1656.2819 | 1748.573 | NaN |
| Dhx8 | IFDPAPPGSR | -848 | -857 | 0 | 0.000550307 | 10 | NaN | NaN | NaN | 1715.447 | NaN | 417.62146 |
| Fth1 | YFLHQSHEER | -55 | -64 | 0 | 0 | 10 | NaN | NaN | NaN | 136.48502 | 1157.9675 | NaN |
| Ipo9 | FRPPETTER | -96 | -104 | 1 | 0.003360691 | 9 | NaN | NaN | NaN | 1331.1271 | 943.83624 | NaN |
| Acap1 | YFQQGHEELNR | -202 | -212 | 0 | 0.00025015 | 11 | NaN | NaN | NaN | 671.04913 | 1353.6252 | 891.72974 |
| Acap1 | LGSCLLESGQHYLAA | -41 | -55 | 0 | 0.00025015 | 15 | NaN | NaN | NaN | 433.94522 | 541.9287 | 1007.7431 |
| Armc10 | NNLSVNVENQTK | -126 | -137 | 0 | 0 | 12 | NaN | NaN | NaN | 696.5691 | 666.7914 | 468.36414 |
| Npepl1 | VHLDIAAPVHAGER | -461 | -474 | 0 | 0 | 14 | NaN | NaN | NaN | 838.02625 | NaN | NaN |
| Blvra | YICTESSSHEDYIR | -71 | -84 | 0 | 0 | 14 | NaN | NaN | NaN | 964.6118 | 94.51366 | 853.0289 |
| Zc3hc1 | SFFDPTSQHR | -416 | -425 | 0 | 0 | 10 | NaN | NaN | NaN | 2238.9392 | 668.7051 | NaN |
| Mapk1 | FRHENIIGIN | -76 | -85 | 1 | 0.002595029 | 10 | NaN | NaN | NaN | 1914.7211 | NaN | NaN |
| Acd | ELILGSETLSSPR | -15 | -27 | 0 | 0 | 13 | NaN | NaN | NaN | 600.65607 | NaN | NaN |
| Spc25 | ELKEEYAR | -107 | -114 | 1 | 0.002068213 | 8 | NaN | NaN | NaN | 495.97534 | NaN | NaN |
| Dnpep | LTAFEEAIPK | -328 | -337 | 0 | 0 | 10 | NaN | NaN | NaN | 581.914 | 639.5154 | 587.0655 |
| Dnpep | ELKETEGWDIVPENK | -47 | -61 | 1 | 0.000550307 | 15 | NaN | NaN | NaN | 1856.0507 | NaN | NaN |
| Dnpep | GTPEPGPLGATDER | -200 | -213 | 0 | 0.00078661 | 14 | NaN | NaN | NaN | 1017.7667 | NaN | NaN |
| Cdc16 | ELLDSLPLNK | -183 | -192 | 0 | 0.00025015 | 10 | NaN | NaN | NaN | 6382.262 | 7073.403 | 6311.3804 |
| Isg15 | ELLGEYGLKPQCTVIK | -133 | -148 | 1 | 0 | 16 | NaN | NaN | NaN | 10557.914 | 11263.353 | NaN |
| Isg15 | GHSNIYEVFLTQ | -89 | -100 | 0 | 0.00122419 | 12 | NaN | NaN | NaN | 332.9838 | NaN | 257.33905 |
| Rps9 | VNIPSFIVR | -142 | -150 | 0 | 0 | 9 | NaN | NaN | NaN | 1276.62 | 1491.1075 | 1388.2487 |
| Rps9 | KGQGGAGAGDDEEED | -180 | -194 | 1 | 0.000550307 | 15 | NaN | NaN | NaN | 143.344 | NaN | NaN |
| Snx9 | ELLTVGQEHWK | -415 | -425 | 0 | 0 | 11 | NaN | NaN | NaN | 10888.432 | NaN | NaN |
| Lnpep | SIVAGSTHLFSTK | -965 | -977 | 0 | 0.000230707 | 13 | NaN | NaN | NaN | 792.3344 | 712.5119 | 738.67346 |
| Lnpep | SVPAEEGLIQDEFSESVK | -336 | -353 | 0 | 0 | 18 | NaN | NaN | NaN | 637.35144 | 1006.038 | 660.2946 |
| Lnpep | NSYYGFYGITYTDK | -269 | -282 | 0 | 0.000550307 | 14 | NaN | NaN | NaN | 845.9708 | NaN | NaN |
| Mavs | ELPGLADQVTR | -80 | -90 | 0 | 0 | 11 | NaN | NaN | NaN | 402.09943 | 724.1477 | 1256.1895 |
| Mtfr1l | ELQDELSHLR | -127 | -136 | 0 | 0 | 10 | NaN | NaN | NaN | 431.8263 | NaN | 5574.0728 |
| Zc3h15 | FGQQNPR | -52 | -58 | 0 | 0.007546675 | 7 | NaN | NaN | NaN | 75.50719 | NaN | NaN |

| | | | | | | | | | | | | |
|---|---|---|---|---|---|---|---|---|---|---|---|---|
| Eea1 | ELQEQEEVVSCTK | -773 | -785 | 0 | 0 | 13 | NaN | NaN | NaN | 1240.8179 | NaN | NaN |
| Def6 | KLQELELLQEAQR | -360 | -372 | 1 | 0 | 13 | NaN | NaN | NaN | 599.29694 | 412.68628 | 447.49457 |
| Cox5a | NKPDIDAWELR | -57 | -67 | 1 | 0 | 11 | NaN | NaN | NaN | 1943.4387 | 1899.9304 | 1835.0851 |
| Cox5a | FNKPDIDAWELR | -56 | -67 | 1 | 0 | 12 | NaN | NaN | NaN | 258.02026 | 2208.844 | 1388.6941 |
| Cox5a | ELRPTLNELGISTPEELGLDKV | -125 | -146 | 2 | 0.000627943 | 22 | NaN | NaN | NaN | 606.0573 | NaN | NaN |
| Rbm10 | ELSQGLLPLPQPY | -264 | -276 | 0 | 0 | 13 | NaN | NaN | NaN | 703.0943 | NaN | 382.96313 |
| Rbm10 | VEFSHLQDATR | -176 | -186 | 0 | 0.000550307 | 11 | NaN | NaN | NaN | 8732.885 | NaN | NaN |
| Rnf40 | NAELEENQELANSR | -336 | -349 | 0 | 0.000627943 | 14 | NaN | NaN | NaN | 116.10325 | 661.42926 | 705.66016 |
| Txnl1 | GCECLNESDEHGFDNCLR | -134 | -151 | 0 | 0.001463897 | 18 | NaN | NaN | NaN | 299.32602 | 124.82408 | 333.14642 |
| Txnl1 | TNNISATPTFLFFR | -70 | -83 | 0 | 0.000230707 | 14 | NaN | NaN | NaN | 790.82 | 886.43115 | NaN |
| Txnl1 | FVQSNQGEEETTR | -247 | -259 | 0 | 0.00437931 | 13 | NaN | NaN | NaN | 172.3131 | NaN | NaN |
| Pak1ip1 | ENAHIVEWSPSGGK | -165 | -178 | 0 | 0 | 14 | NaN | NaN | NaN | 602.96497 | 1060.301 | 1066.0183 |
| Pak1ip1 | ESGDTVQEETSEPNSEKS | -330 | -347 | 1 | 0.001868731 | 18 | NaN | NaN | NaN | 224.47797 | 296.0737 | 305.24582 |
| Hnrnph2 | ENDIYNFFSPLNPMR | -302 | -316 | 0 | 0 | 15 | NaN | NaN | NaN | 408.55472 | 719.33014 | 599.3795 |
| Mrpl18 | VYQPTPWEASSDSIK | -145 | -159 | 0 | 0 | 15 | NaN | NaN | NaN | 835.41644 | 202.37482 | 747.03784 |
| Mrpl18 | ENEAVAPEFTNR | -40 | -51 | 0 | 0 | 12 | NaN | NaN | NaN | 7995.3135 | NaN | NaN |
| Msh6 | LCAPLCSPSAISDR | -775 | -788 | 0 | 0.008129024 | 14 | NaN | NaN | NaN | 737.3435 | NaN | NaN |
| Msh6 | GLLEEVAGGFTSK | -870 | -882 | 0 | 0.00201047 | 13 | NaN | NaN | NaN | 242.02652 | NaN | NaN |
| Dtymk | ENFSLDWCK | -110 | -118 | 0 | 0 | 9 | NaN | NaN | NaN | 478.84387 | 674.267 | NaN |
| Dtymk | RWEQVPLIK | -76 | -84 | 1 | 0.00383302 | 9 | NaN | NaN | NaN | 1339.8295 | NaN | NaN |
| Skic8 | ENIETVVTGSLDDLVK | -30 | -45 | 0 | 0.00201047 | 16 | NaN | NaN | NaN | 677.3157 | NaN | NaN |
| Trmt1 | ENLAPGDWPR | -132 | -141 | 0 | 0 | 10 | NaN | NaN | NaN | 310.98862 | 1329.951 | NaN |
| Sumo2 | TENNDHINLK | -12 | -21 | 0 | 0 | 10 | NaN | NaN | NaN | 5612.4194 | 5635.9478 | 6184.6416 |
| Sumo2 | ENNDHINLK | -13 | -21 | 0 | 0.000431655 | 9 | NaN | NaN | NaN | 1219.5929 | NaN | NaN |
| Nelfa | ENPCPEQGDVIQIK | -470 | -483 | 0 | 0 | 14 | NaN | NaN | NaN | 475.80695 | 514.7205 | 549.2706 |
| Bola1 | ENPQLDISPPCLGGSK | -116 | -131 | 0 | 0.00111006 | 16 | NaN | NaN | NaN | 895.7463 | NaN | NaN |
| Fmr1 | QIPFHDVR | -41 | -48 | 0 | 0.007523243 | 8 | NaN | NaN | NaN | 801.63617 | NaN | NaN |
| Ndufs8 | RYPSGEER | -105 | -112 | 1 | 0.001392869 | 8 | NaN | NaN | NaN | 281.12292 | 550.5199 | 805.25006 |
| Ndufs8 | STETHEELLYNK | -176 | -187 | 0 | 0.000394361 | 12 | NaN | NaN | NaN | 971.8854 | NaN | 791.5197 |
| Ndufs8 | FRGEHALR | -97 | -104 | 1 | 0.002595029 | 8 | NaN | NaN | NaN | 461.75806 | NaN | NaN |
| Lsp1 | LEESNLSYR | -133 | -141 | 0 | 0.00025015 | 9 | NaN | NaN | NaN | 513.8878 | 1269.5574 | 443.87405 |
| Lsp1 | FWGNEGTAEGTEPSQSERPEEK | -99 | -120 | 1 | 0 | 22 | NaN | NaN | NaN | 465.29306 | 560.1797 | 501.55243 |
| Lsp1 | TLWETGEVQSQ | -256 | -266 | 0 | 0 | 11 | NaN | NaN | NaN | 57.47217 | 208.93765 | 247.95029 |
| Lsp1 | KSSELDEDEGFGDWSQKPEPR | -76 | -96 | 2 | 0.002780285 | 21 | NaN | NaN | NaN | 487.25812 | NaN | NaN |
| Hspbp1 | EQVLGLGALR | -179 | -188 | 0 | 0 | 10 | NaN | NaN | NaN | 763.8681 | 761.30096 | 694.301 |
| Ift25 | EPLDFEQWVEK | -81 | -91 | 0 | 0.00078661 | 11 | NaN | NaN | NaN | 476.3685 | 567.35974 | 350.31433 |
| Larp4 | HNPTVTGQQEQTYLPK | -408 | -423 | 0 | 0 | 16 | NaN | NaN | NaN | 1214.2966 | 1208.918 | 1176.254 |
| Trmt112 | VLEGTLQCPESGR | -93 | -105 | 0 | 0 | 13 | NaN | NaN | NaN | 1378.6301 | NaN | NaN |
| Add3 | VGEIEFEGLMR | -366 | -376 | 0 | 0.001291103 | 11 | NaN | NaN | NaN | 558.1076 | 495.8709 | NaN |
| Uso1 | SVPVEGESEHVSAA | -828 | -841 | 0 | 0.000826806 | 14 | NaN | NaN | NaN | 24267.803 | NaN | NaN |
| Tax1bp1 | ETDLDSLKDK | -249 | -258 | 1 | 0.00025015 | 10 | NaN | NaN | NaN | 532.13293 | 789.17957 | 868.7644 |
| Ifit1 | HIEQDIHLR | -364 | -372 | 0 | 0 | 9 | NaN | NaN | NaN | 467.59363 | 77.692955 | NaN |
| Ifit1 | EQVQFLDIK | -41 | -49 | 0 | 0 | 9 | NaN | NaN | NaN | 1128.5258 | NaN | NaN |
| Ctsb | WSNCPTIGQIR | -90 | -100 | 0 | 0 | 11 | NaN | NaN | NaN | 1795.4286 | 1232.1108 | NaN |
| Rps4x | NKPWISLPR | -232 | -240 | 1 | 0 | 9 | NaN | NaN | NaN | 925.95953 | 1342.4409 | 73.8261 |
| Rps4x | RITPEEAKYK | -113 | -122 | 2 | 0.000826806 | 10 | NaN | NaN | NaN | 423.57608 | 103.45383 | NaN |
| Rps4x | KPWISLPR | -233 | -240 | 1 | 0 | 8 | NaN | NaN | NaN | 332.99704 | NaN | 388.32892 |
| Rps4x | LTGDEVKK | -56 | -63 | 1 | 0.003620005 | 8 | NaN | NaN | NaN | 397.49786 | NaN | NaN |
| Akt2 | TFHVDSPDER | -87 | -96 | 0 | 0.000550307 | 10 | NaN | NaN | NaN | 2258.8296 | 2293.0989 | NaN |
| Emc2 | ERYDDAIQLYDR | -101 | -112 | 1 | 0.00078661 | 12 | NaN | NaN | NaN | 1480.7494 | 1889.8132 | NaN |
| Pf4 | HCAVPQLIATLK | -70 | -81 | 0 | 0.001868731 | 12 | NaN | NaN | NaN | 552.7816 | NaN | 439.58972 |
| Pf4 | HITSLEVIK | -58 | -66 | 0 | 0.00201047 | 9 | NaN | NaN | NaN | 1518.9165 | NaN | NaN |
| Grb2 | KVLNEECDQNWYK | -26 | -38 | 1 | 0.00051306 | 13 | NaN | NaN | NaN | 940.063 | 1055.6492 | 1108.7451 |
| Ptpn2 | TISHFHYT | -171 | -178 | 0 | 0.002296977 | 8 | NaN | NaN | NaN | 4099.9976 | NaN | NaN |
| Ptpn2 | LLQLENINTGETR | -158 | -170 | 0 | 0 | 13 | NaN | NaN | NaN | 815.47687 | NaN | NaN |
| Ptpn2 | LVDIEEAQR | -73 | -81 | 0 | 0.000431655 | 9 | NaN | NaN | NaN | 390.01 | NaN | NaN |
| Ptpn1 | KDPSSVDIK | -239 | -247 | 1 | 0 | 9 | NaN | NaN | NaN | 2517.4697 | NaN | NaN |
| U2surp | NLLALIHR | -426 | -433 | 0 | 0.000230707 | 8 | NaN | NaN | NaN | 6802.1567 | 532.7319 | 544.3559 |
| Ubxn1 | ESLIEMGFPR | -8 | -17 | 0 | 0.002199523 | 10 | NaN | NaN | NaN | 399.12463 | 769.3645 | NaN |
| Rad23a | SDDVPIKEYHIDEK | -56 | -69 | 1 | 0.001811524 | 14 | NaN | NaN | NaN | 479.90637 | 592.01373 | NaN |
| Ptpn11 | ESQSHPGDFVLSVR | -139 | -152 | 0 | 0 | 14 | NaN | NaN | NaN | 987.3049 | 1754.9194 | 457.76273 |
| Ptpn11 | VVLHDGDPNEPVSDYINAN | -290 | -308 | 0 | 0.000907206 | 19 | NaN | NaN | NaN | 398.06 | 244.12912 | NaN |
| Nol9 | ESSPLPVFTGHVL | -501 | -513 | 0 | 0 | 13 | NaN | NaN | NaN | 471.19284 | 633.7165 | 679.95 |
| Nploc4 | ESSSEQYVPDVFYK | -414 | -427 | 0 | 0 | 14 | NaN | NaN | NaN | 586.3687 | 137.87517 | NaN |
| Nploc4 | GAPNVVEDEIDQYLSK | -101 | -116 | 0 | 0.00025015 | 16 | NaN | NaN | NaN | 476.71075 | NaN | 484.757 |
| Elob | KESSTVFELK | -19 | -28 | 1 | 0.00025015 | 10 | NaN | NaN | NaN | 3548.7915 | 3350.889 | 3508.4548 |
| Tbc1d10c | ESTKPEIPR | -383 | -391 | 1 | 0 | 9 | NaN | NaN | NaN | 2302.0974 | 2077.3896 | 2353.6152 |
| Tbc1d10c | SLPGPAPLPQAR | -354 | -365 | 0 | 0 | 12 | NaN | NaN | NaN | 757.2336 | 679.8435 | 710.0781 |
| Ociad1 | FDSNVSGQSSFGTSPAADNIEK | -134 | -155 | 0 | 0.001023236 | 22 | NaN | NaN | NaN | 88.84762 | 194.61346 | 150.84106 |
| Usp14 | KLEAIEDDSGR | -214 | -224 | 1 | 0 | 11 | NaN | NaN | NaN | 1437.8253 | 1436.4683 | 2105.0896 |
| Usp14 | SSSSGHYVSW | -429 | -438 | 0 | 0 | 10 | NaN | NaN | NaN | 526.6564 | NaN | 664.95337 |
| Lrrc8c | IPHAVFSLL | -604 | -612 | 0 | 0.006384543 | 9 | NaN | NaN | NaN | 327.40906 | 690.9722 | 271.3896 |
| Lrrc8c | ETGIDDIPDVK | -357 | -367 | 0 | 0.00204437 | 11 | NaN | NaN | NaN | 905.1647 | NaN | NaN |
| Ppt1 | YWHDPIK | -185 | -191 | 0 | 0 | 7 | NaN | NaN | NaN | 1310.543 | NaN | 1367.5022 |
| Pcbp1 | IITLTGPTNAIFK | -58 | -70 | 0 | 0 | 13 | NaN | NaN | NaN | 4156.5566 | 4821.737 | 4937.526 |
| Pcbp1 | TTHELTIPNNLIGCIIGR | -280 | -297 | 0 | 0 | 18 | NaN | NaN | NaN | 422.89215 | 540.52704 | 486.32016 |
| Snrpd3 | MSNITVTYR | -43 | -51 | 0 | 0.007374243 | 9 | NaN | NaN | NaN | 352.1234 | NaN | NaN |
| Basp1 | ETPAASEAPSSAAK | -182 | -195 | 0 | 0 | 14 | NaN | NaN | NaN | 875.8595 | 894.0937 | NaN |

| | | | | | | | | | | | | |
|---|---|---|---|---|---|---|---|---|---|---|---|---|
| Ltn1 | ETPDTLSDPQTVPEEER | -201 | -217 | 0 | 0.001055699 | 17 | NaN | NaN | NaN | 1265.4519 | 1246.2169 | 1141.6609 |
| Ltn1 | HALVTPWLK | -729 | -737 | 0 | 0 | 9 | NaN | NaN | NaN | 381.5472 | NaN | NaN |
| Tsfm | VICETPEQIANLEEVGR | -237 | -253 | 0 | 0 | 17 | NaN | NaN | NaN | 1307.6849 | 632.13824 | 654.6046 |
| Tsfm | SSELSELAAGPDR | -165 | -177 | 0 | 0 | 13 | NaN | NaN | NaN | 2031.768 | NaN | NaN |
| Elof1 | ETQFTCPFCNHEK | -21 | -33 | 0 | 0 | 13 | NaN | NaN | NaN | 48.2304 | 960.43243 | 928.7911 |
| Vps26b | LVLIDEEERR | -274 | -283 | 1 | 0.000591124 | 10 | NaN | NaN | NaN | 1360.0447 | 1250.0667 | 1257.6237 |
| Vps26b | ETTGTGPNVYHENDTIAK | -213 | -230 | 0 | 0.000907206 | 18 | NaN | NaN | NaN | 1630.3761 | NaN | NaN |
| Atg3 | FKETGVITPEEFVAA | -28 | -42 | 1 | 0.002265552 | 15 | NaN | NaN | NaN | 2343.9646 | 817.58435 | NaN |
| Atg3 | EVAEYLTPVLK | -14 | -24 | 0 | 0.00025015 | 11 | NaN | NaN | NaN | 811.11505 | NaN | NaN |
| Parvg | WSVETIFNK | -126 | -134 | 0 | 0 | 9 | NaN | NaN | NaN | 956.75635 | 1064.3315 | 1089.3734 |
| Parvg | EVAQPTEEELPR | -15 | -26 | 0 | 0 | 12 | NaN | NaN | NaN | 901.3738 | 580.10425 | NaN |
| Tor1aip1 | EVDQELTDGFK | -472 | -482 | 0 | 0 | 11 | NaN | NaN | NaN | 1745.7212 | 1918.4839 | NaN |
| Smarca4 | GPTPFNQNQLHQL | -168 | -180 | 0 | 0 | 13 | NaN | NaN | NaN | 3479.1528 | NaN | NaN |
| Hectd1 | KEDLPGFVFESNR | -783 | -795 | 1 | 0.000550307 | 13 | NaN | NaN | NaN | 1076.7081 | NaN | NaN |
| Hectd1 | HGLTEELLSR | -230 | -239 | 0 | 0.006100831 | 10 | NaN | NaN | NaN | 791.1774 | NaN | NaN |
| Cdca8 | EVENLYNIEILR | -49 | -60 | 0 | 0 | 12 | NaN | NaN | NaN | 1196.5828 | NaN | NaN |
| Tbc1d10a | PALQPSPSIR | -402 | -411 | 0 | 0.005605646 | 10 | NaN | NaN | NaN | 584.7462 | NaN | NaN |
| Pck2 | LTEQVNQDLPK | -614 | -624 | 0 | 0 | 11 | NaN | NaN | NaN | 2189.0854 | 2149.6655 | 1711.9835 |
| Pck2 | LWPGFGENAR | -544 | -553 | 0 | 0 | 10 | NaN | NaN | NaN | 1051.4788 | NaN | NaN |
| Ppp1r18 | LSPGETPEESLR | -193 | -204 | 0 | 0.001743443 | 12 | NaN | NaN | NaN | 3971.3264 | NaN | NaN |
| Pfdn2 | IIETLSQQLQAK | -100 | -111 | 0 | 0 | 12 | NaN | NaN | NaN | 423.7242 | 338.05917 | 435.10077 |
| Atad2 | IIFFDEIDGLAPVR | -182 | -195 | 0 | 0.006788754 | 14 | NaN | NaN | NaN | 130.0906 | NaN | NaN |
| Csrp1 | TVYFAEEVQCEGNSFHK | -16 | -32 | 0 | 0 | 17 | NaN | NaN | NaN | 610.8605 | 646.0128 | 759.9348 |
| Urb1 | STVENFSQLGR | -936 | -946 | 0 | 0 | 11 | NaN | NaN | NaN | 2768.0205 | NaN | NaN |
| Septin1 | EVTHDLLYEGY | -278 | -288 | 0 | 0.001190916 | 11 | NaN | NaN | NaN | 1006.3513 | 830.0657 | 570.3618 |
| Septin1 | VHCCLYFISPFGR | -134 | -146 | 0 | 0 | 13 | NaN | NaN | NaN | 2773.3162 | 4361.7695 | 5354.7573 |
| Septin1 | VDTPGFGDSVDCSDCWLPVVR | -88 | -108 | 0 | 0.001846793 | 21 | NaN | NaN | NaN | 258.1744 | NaN | 340.6829 |
| Atp13a1 | RDLEEESTPIVK | -935 | -946 | 1 | 0.00897527 | 12 | NaN | NaN | NaN | 3544.8706 | 4216.4194 | NaN |
| Cul3 | EVVTEHLINK | -79 | -88 | 0 | 0 | 10 | NaN | NaN | NaN | 2530.9666 | 1721.688 | 1848.303 |
| Lactb | EVWSEGLGYADVENR | -135 | -149 | 0 | 0 | 15 | NaN | NaN | NaN | 1859.762 | 2275.8042 | 2206.5151 |
| Dek | LADANLEEVTMK | -338 | -349 | 0 | 0.00025015 | 12 | NaN | NaN | NaN | 761.72003 | NaN | NaN |
| Colgalt1 | WSPESPLQAPR | -38 | -48 | 0 | 0.001636916 | 11 | NaN | NaN | NaN | 1301.2119 | NaN | NaN |
| Ckap5 | FGQYAGHVVPTILEK | -355 | -369 | 0 | 0.000431655 | 15 | NaN | NaN | NaN | 450.98303 | NaN | NaN |
| Hnrnpd | FKESESVDK | -145 | -153 | 1 | 0 | 9 | NaN | NaN | NaN | 2183.623 | 2283.4158 | 2480.4592 |
| Hnrnpd | SPDTPEEK | -190 | -197 | 0 | 0.005176406 | 8 | NaN | NaN | NaN | 4130.8164 | NaN | NaN |
| Hnrnpd | GGLSPDTPEEK | -187 | -197 | 0 | 0.000627943 | 11 | NaN | NaN | NaN | 1052.8406 | NaN | NaN |
| Vrk1 | YCPDGVHK | -204 | -211 | 0 | 0.000431655 | 8 | NaN | NaN | NaN | 264.27042 | 803.08075 | 819.6699 |
| Rasal3 | VVLELDEVSIPR | -409 | -420 | 0 | 0 | 12 | NaN | NaN | NaN | 490.3805 | NaN | NaN |
| Rasal3 | TQDSLEPLPTILR | -748 | -760 | 0 | 0.00142999 | 13 | NaN | NaN | NaN | 249.27164 | NaN | NaN |
| Ikbkg | SLLGELQESQSR | -148 | -159 | 0 | 0.00025015 | 12 | NaN | NaN | NaN | 1356.9354 | 1759.0874 | NaN |
| Ikbkg | LFQDYDSHIK | -237 | -246 | 0 | 0 | 10 | NaN | NaN | NaN | 1281.9501 | NaN | 578.24994 |
| Mvb12a | EYNYGFVVEK | -251 | -260 | 0 | 0 | 10 | NaN | NaN | NaN | 640.7627 | NaN | NaN |
| Ap3d1 | ILIEDSDQNLK | -310 | -320 | 0 | 0.001868731 | 11 | NaN | NaN | NaN | 2105.708 | NaN | NaN |
| Lmf2 | FAHHLPVWLHK | -215 | -225 | 0 | 0.000591124 | 11 | NaN | NaN | NaN | 304.08722 | NaN | NaN |
| Hsd17b4 | NAVFEWHITK | -646 | -655 | 0 | 0 | 10 | NaN | NaN | NaN | 1377.7373 | NaN | NaN |
| Hsd17b4 | FAKPVYPGQTLQTE | -562 | -575 | 1 | 0.00111006 | 14 | NaN | NaN | NaN | 773.0235 | NaN | NaN |
| Isyna1 | VIVLWTANTER | -223 | -233 | 0 | 0.000230707 | 11 | NaN | NaN | NaN | 557.1656 | NaN | NaN |
| Rfc5 | FCLICNYLSK | -150 | -159 | 0 | 0 | 10 | NaN | NaN | NaN | 723.8367 | NaN | 687.172 |
| Rfc5 | FGPLTPELMVPR | -173 | -184 | 0 | 0 | 12 | NaN | NaN | NaN | 521.5355 | NaN | NaN |
| Pmpca | FCPVENIAK | -223 | -231 | 0 | 0.000471409 | 9 | NaN | NaN | NaN | 1250.3667 | NaN | 733.8943 |
| Pmpca | NVKPEDIK | -470 | -477 | 1 | 0.002168954 | 8 | NaN | NaN | NaN | 1282.2427 | NaN | NaN |
| Pmpca | PVIFEDVGR | -442 | -450 | 0 | 0.001055699 | 9 | NaN | NaN | NaN | 538.5991 | NaN | NaN |
| Pmpca | TVDLVELER | -417 | -425 | 0 | 0.001610812 | 9 | NaN | NaN | NaN | 374.23737 | NaN | NaN |
| Nudt16l1 | FDGLLGFPGGFVDR | -56 | -69 | 0 | 0 | 14 | NaN | NaN | NaN | 743.8886 | 82.27893 | 808.3813 |
| Nudt16l1 | LYAANPGQLFGR | -32 | -43 | 0 | 0.000431655 | 12 | NaN | NaN | NaN | 613.1925 | NaN | NaN |
| Ppig | FDIAINNQPAGR | -12 | -23 | 0 | 0.002401448 | 12 | NaN | NaN | NaN | 1572.7773 | NaN | NaN |
| Hm13 | GSGENKEEIINYEFDTK | -145 | -161 | 1 | 0 | 17 | NaN | NaN | NaN | 587.13293 | 638.3081 | 458.81506 |
| Rcsd1 | FDPAALLPGASPK | -95 | -107 | 0 | 0.00635324 | 13 | NaN | NaN | NaN | 1053.357 | NaN | NaN |
| Gbe1 | FDRDMNRLEER | -583 | -593 | 2 | 0.008014596 | 11 | NaN | NaN | NaN | 3170.9895 | NaN | NaN |
| Srsf7 | GHYAYDCHRY | -142 | -151 | 1 | 0.001636916 | 10 | NaN | NaN | NaN | 66.73153 | NaN | 362.44754 |
| Gnai2 | SCAAEEQGMLPEDLSGVIR | -111 | -129 | 0 | 0 | 19 | NaN | NaN | NaN | 282.3983 | 946.80365 | 457.3753 |
| Gnai2 | LWADHGVQACFGR | -131 | -143 | 0 | 0 | 13 | NaN | NaN | NaN | 1025.0499 | 1070.4067 | 545.6787 |
| Cc2d1b | NLPAPPGVTPDDLDAFVR | -700 | -717 | 0 | 0.00025015 | 18 | NaN | NaN | NaN | 445.3589 | 455.0462 | 485.03912 |
| Cox7a2 | LFQEDNGMPVHLK | -34 | -46 | 0 | 0 | 13 | NaN | NaN | NaN | 530.7084 | 570.69183 | 841.22363 |
| Fxr2 | FENNWQSER | -42 | -50 | 0 | 0 | 9 | NaN | NaN | NaN | 548.4648 | 647.59045 | 548.5016 |
| Srp68 | FESFCLDPSLVTK | -556 | -568 | 0 | 0 | 13 | NaN | NaN | NaN | 621.99493 | NaN | 457.6774 |
| Srp68 | WSEALVLYDR | -478 | -487 | 0 | 0.000627943 | 10 | NaN | NaN | NaN | 533.33527 | NaN | NaN |
| Psmd5 | LLQAVEPIHLAR | -71 | -82 | 0 | 0 | 12 | NaN | NaN | NaN | 411.8703 | 493.52853 | 625.6943 |
| Ctps1 | NVLGWQDANSTEFDPK | -412 | -427 | 0 | 0.00025015 | 16 | NaN | NaN | NaN | 403.0134 | 431.30826 | 427.56436 |
| Ctps1 | KLYGDTDYLEER | -466 | -477 | 1 | 0 | 12 | NaN | NaN | NaN | 144.03918 | 900.296 | 942.5187 |
| Nanp | YHYKEEAEIICDK | -39 | -51 | 1 | 0.000947459 | 13 | NaN | NaN | NaN | 419.37003 | NaN | NaN |
| Nanp | FFDLDNTLIDTAGASR | -10 | -25 | 0 | 0 | 16 | NaN | NaN | NaN | 331.73773 | NaN | NaN |
| Otub1 | FFEHFIEGGR | -189 | -198 | 0 | 0 | 10 | NaN | NaN | NaN | 1325.849 | 313.63745 | NaN |
| Otub1 | GFSHLEALLDDSK | -97 | -109 | 0 | 0.000431655 | 13 | NaN | NaN | NaN | 1070.4614 | 319.33243 | 997.2715 |
| Ndufa6 | FFHETETPRPK | -109 | -119 | 1 | 0 | 11 | NaN | NaN | NaN | 437.31296 | NaN | NaN |
| Txndc5 | VDCTQHYAVCSEHQVR | -231 | -246 | 0 | 0 | 16 | NaN | NaN | NaN | 374.8641 | 423.3313 | 396.7796 |
| Ddt | LCAATATILDKPEDR | -23 | -37 | 1 | 0.000230707 | 15 | NaN | NaN | NaN | 1160.7953 | 551.71674 | 611.0947 |
| Ddt | STEPCAHLLVSSIGVVGTAEQNR | -53 | -75 | 0 | 0 | 23 | NaN | NaN | NaN | 396.34192 | NaN | NaN |

| | | | | | | | | | | | | |
|---|---|---|---|---|---|---|---|---|---|---|---|---|
| Oxct1 | FFSSDESFAMIR | -379 | -390 | 0 | 0 | 12 | NaN | NaN | NaN | 1379.0216 | 654.75696 | NaN |
| Chd8 | FFVENPSEEDAAIVDK | -633 | -648 | 0 | 0.001092529 | 16 | NaN | NaN | NaN | 308.74466 | NaN | NaN |
| Nop14 | YIGDLATDSTPDLK | -483 | -496 | 0 | 0 | 14 | NaN | NaN | NaN | 844.04486 | 467.166 | 663.30225 |
| Ddx1 | SKAPDNYIVK | -259 | -268 | 1 | 0 | 10 | NaN | NaN | NaN | 8585.619 | NaN | NaN |
| Ddx1 | KAPDNYIVK | -260 | -268 | 1 | 0.003620005 | 9 | NaN | NaN | NaN | 2704.1682 | NaN | NaN |
| As3mt | SLQNVHEDVSSR | -47 | -58 | 0 | 0.006537474 | 12 | NaN | NaN | NaN | 885.15576 | NaN | NaN |
| As3mt | FGFQAPNVTFLHGR | -123 | -136 | 0 | 0.006568856 | 14 | NaN | NaN | NaN | 756.5697 | NaN | NaN |
| Tardbp | TVTAQFPGACGLR | -30 | -42 | 0 | 0.002466863 | 13 | NaN | NaN | NaN | 114.16373 | NaN | NaN |
| Prkch | TNKPTYNEEFCAN | -63 | -75 | 1 | 0.001868731 | 13 | NaN | NaN | NaN | 793.8793 | NaN | NaN |
| Cd47 | FGILTLK | -158 | -164 | 0 | 0.001636916 | 7 | NaN | NaN | NaN | 233.13841 | 213.8434 | 173.85612 |
| Scfd1 | VVGGGNYIEYQNLVDYIK | -592 | -609 | 0 | 0.00025015 | 18 | NaN | NaN | NaN | 787.3517 | NaN | NaN |
| Pmpcb | FHFGDSLCSHK | -258 | -268 | 0 | 0 | 11 | NaN | NaN | NaN | 635.4532 | 654.91736 | 374.43298 |
| Nipsnap2 | FHNVKPECLDAYNK | -73 | -86 | 1 | 0 | 14 | NaN | NaN | NaN | 354.11346 | NaN | 670.8532 |
| Cbr1 | KGVHAEEGWPNSAYGVT | -181 | -197 | 1 | 0.002972566 | 17 | NaN | NaN | NaN | 896.8867 | 912.4146 | NaN |
| Igf2r | HDLNPLIK | -179 | -186 | 0 | 0.001291103 | 8 | NaN | NaN | NaN | 789.624 | 991.11206 | 186.95135 |
| Timm8a1 | SKPVFSESLSD | -87 | -97 | 1 | 0.003461185 | 11 | NaN | NaN | NaN | 798.69183 | 677.6096 | 521.1933 |
| Slfn1 | IKNPDYILSK | -59 | -68 | 1 | 0.000230707 | 10 | NaN | NaN | NaN | 617.22485 | 511.01566 | 578.40076 |
| Sbds | FILPVNEGK | -176 | -184 | 0 | 0.000230707 | 9 | NaN | NaN | NaN | 936.5884 | 1007.5296 | 1003.44684 |
| Sbds | GTDDQTEICK | -76 | -85 | 0 | 0.000230707 | 10 | NaN | NaN | NaN | 128.81299 | 749.4373 | NaN |
| Pacs1 | VSIDGVEWSDIK | -924 | -935 | 0 | 0.006788754 | 12 | NaN | NaN | NaN | 1213.3904 | NaN | NaN |
| Tapbp | QHGVHYVCR | -378 | -386 | 0 | 0 | 9 | NaN | NaN | NaN | 156.14479 | 698.97705 | 727.0958 |
| Tapbp | HHSDGSVSQSGHLQLPPVTAK | -357 | -377 | 0 | 0 | 21 | NaN | NaN | NaN | 1216.0425 | 1770.331 | NaN |
| Nfu1 | IRPTVQEDGGDVIYR | -181 | -195 | 1 | 0.008828666 | 15 | NaN | NaN | NaN | 619.5416 | NaN | 936.29224 |
| Taf10 | SVAGPAPLVSAPAALPTSTAAESK | -19 | -42 | 0 | 0 | 24 | NaN | NaN | NaN | 205.95613 | 238.58575 | 221.84755 |
| Mad2l1 | FITNCEEVR | -174 | -182 | 0 | 0 | 9 | NaN | NaN | NaN | 1101.1603 | 1247.5851 | NaN |
| Hnrnpdl | GEIENIELPMDTK | -138 | -150 | 0 | 0 | 13 | NaN | NaN | NaN | 788.7248 | 881.50165 | 1029.191 |
| Ubr5 | GLIYIDPSNLR | -1737 | -1747 | 0 | 0.00025015 | 11 | NaN | NaN | NaN | 357.8647 | NaN | NaN |
| Idh2 | FKDIFQEIFDK | -262 | -272 | 1 | 0 | 11 | NaN | NaN | NaN | 520.12933 | 539.1665 | 231.10143 |
| Nob1 | TPLPNIDR | -163 | -170 | 0 | 0.00025015 | 8 | NaN | NaN | NaN | 1872.0692 | 1876.1948 | NaN |
| Nob1 | FKEPLPEYVR | -55 | -64 | 1 | 0.002566359 | 10 | NaN | NaN | NaN | 1562.8712 | 990.7086 | NaN |
| Sec62 | TDGDCDEEDDDKDGEVPK | -375 | -392 | 1 | 0.009657961 | 18 | NaN | NaN | NaN | 216.38054 | NaN | NaN |
| Nudt21 | QTKPLTLER | -27 | -35 | 1 | 0.001636916 | 9 | NaN | NaN | NaN | 676.86444 | 664.36646 | 727.98505 |
| Nudt21 | FKLPGGELNPGEDEVEGLKR | -104 | -123 | 2 | 0 | 20 | NaN | NaN | NaN | 442.5077 | 459.36774 | 100.88989 |
| Cfb | KIVLDPSGSMNIY | -257 | -269 | 1 | 0.001708008 | 13 | NaN | NaN | NaN | 1193.2966 | NaN | NaN |
| Thop1 | RNILDFPQHVSPCK | -79 | -92 | 1 | 0.00051306 | 14 | NaN | NaN | NaN | 929.57 | 614.8268 | 494.75854 |
| Thop1 | TSQTVATFLDELAQK | -287 | -301 | 0 | 0 | 15 | NaN | NaN | NaN | 778.24634 | 456.94464 | 828.89655 |
| Thop1 | QTVATFLDELAQK | -289 | -301 | 0 | 0.002910424 | 13 | NaN | NaN | NaN | 505.01343 | NaN | 505.49304 |
| Thop1 | TGSEAPQDLLEK | -527 | -538 | 0 | 0 | 12 | NaN | NaN | NaN | 1702.263 | NaN | NaN |
| Thop1 | NILDFPQHVSPCK | -80 | -92 | 0 | 0 | 13 | NaN | NaN | NaN | 1368.0951 | NaN | NaN |
| Thop1 | TDADPAEEYAR | -568 | -578 | 0 | 0.00025015 | 11 | NaN | NaN | NaN | 375.21118 | NaN | NaN |
| Dcun1d1 | KLEQLYTR | -64 | -71 | 1 | 0.002136192 | 8 | NaN | NaN | NaN | 614.33405 | 458.49768 | NaN |
| Nup107 | VMTQDDSEELPR | -470 | -481 | 0 | 0.007523243 | 12 | NaN | NaN | NaN | 214.52284 | NaN | NaN |
| Ubtf | HPELNISEEGITK | -267 | -279 | 0 | 0.00025015 | 13 | NaN | NaN | NaN | 1727.7885 | 1639.5006 | NaN |
| Smarce1 | TVEEPPTDPVPEDEKKE | -395 | -411 | 2 | 0.000627943 | 17 | NaN | NaN | NaN | 143.22144 | 594.40454 | 560.82404 |
| Kdm3a | FLGDQQSVFVSK | -133 | -144 | 0 | 0.00025015 | 12 | NaN | NaN | NaN | 354.78754 | 828.46234 | 708.0611 |
| Kdm3a | VAEWPWLSGTIR | -38 | -49 | 0 | 0.00025015 | 12 | NaN | NaN | NaN | 469.23422 | 446.57068 | NaN |
| Kdm3a | GQCEQEEEVLR | -1140 | -1150 | 0 | 0.009745776 | 11 | NaN | NaN | NaN | 2288.8447 | NaN | NaN |
| Apip | FLIPELCK | -25 | -32 | 0 | 0.000712251 | 8 | NaN | NaN | NaN | 811.80896 | 690.3236 | 620.6908 |
| Alg3 | VHSIFVLR | -164 | -171 | 0 | 0.006472924 | 8 | NaN | NaN | NaN | 910.66516 | NaN | NaN |
| Hspe1 | PEYGGTK | -74 | -80 | 0 | 0.00383302 | 7 | NaN | NaN | NaN | 785.1342 | 1347.0901 | 1253.3435 |
| Hspe1 | VLQATVVAVGSGGK | -41 | -54 | 0 | 0 | 14 | NaN | NaN | NaN | 553.43896 | NaN | NaN |
| Gar1 | QDQGPPER | -73 | -80 | 0 | 0.00025015 | 8 | NaN | NaN | NaN | 313.39844 | 409.52518 | NaN |
| Gar1 | GEFMHPCEDDIVCK | -85 | -98 | 0 | 0.001092529 | 14 | NaN | NaN | NaN | 312.81992 | NaN | NaN |
| Snx3 | FLQDEIIDK | -144 | -152 | 0 | 0 | 9 | NaN | NaN | NaN | 11144.862 | 2358.533 | 8433.135 |
| Snx3 | KQGLEQFINK | -119 | -128 | 1 | 0.000298805 | 10 | NaN | NaN | NaN | 2579.446 | 976.653 | 2848.6494 |
| Ctu2 | FLQQQQQQQR | -181 | -191 | 0 | 0 | 11 | NaN | NaN | NaN | 416.42084 | NaN | NaN |
| Eif3l | QYEQQTYQVIPEVIK | -39 | -53 | 0 | 0 | 15 | NaN | NaN | NaN | 964.9045 | 1124.5392 | 1111.5023 |
| Eif3l | YDPYAYPGDYDMHTGDPK | -14 | -31 | 0 | 0.00111006 | 18 | NaN | NaN | NaN | 358.97415 | 375.7889 | 424.0471 |
| Rtf1 | KFNDQDIEEIVK | -491 | -502 | 1 | 0 | 12 | NaN | NaN | NaN | 1060.0868 | 869.00256 | 669.9973 |
| Emb | NDPDAGKEFEQIEQL | -293 | -307 | 1 | 0.00025015 | 15 | NaN | NaN | NaN | 734.993 | 854.8499 | 749.029 |
| Emb | LEEDGGSYWCR | -231 | -241 | 0 | 0 | 11 | NaN | NaN | NaN | 407.98795 | NaN | NaN |
| Emb | FNIHVPK | -156 | -162 | 0 | 0.003102268 | 7 | NaN | NaN | NaN | 294.79926 | NaN | NaN |
| Lsm14a | TIDTENSTVALAK | -28 | -40 | 0 | 0 | 13 | NaN | NaN | NaN | 745.887 | 387.89856 | NaN |
| Selenof | GCCQEEAQFETK | -69 | -80 | 0 | 0 | 12 | NaN | NaN | NaN | 416.5036 | 451.01 | 487.6585 |
| Selenof | FNLLPLDPVCR | -58 | -68 | 0 | 0.001708008 | 11 | NaN | NaN | NaN | 377.14328 | NaN | NaN |
| Ppif | VYLDVGADGQPLGR | -47 | -60 | 0 | 0.000550307 | 14 | NaN | NaN | NaN | 671.49615 | NaN | NaN |
| Vps4b | WSDVAGLEGAK | -133 | -143 | 0 | 0 | 11 | NaN | NaN | NaN | 1029.8076 | 712.937 | NaN |
| Vps4b | NLFQLAR | -218 | -224 | 0 | 0.00713295 | 7 | NaN | NaN | NaN | 392.22733 | NaN | NaN |
| Vps4b | KFTEDFGQEG | -435 | -444 | 1 | 0.000989247 | 10 | NaN | NaN | NaN | 274.55615 | NaN | NaN |
| Ufl1 | FQDEIEDCLR | -478 | -487 | 0 | 0.00111006 | 10 | NaN | NaN | NaN | 146.7448 | NaN | NaN |
| Pigt | FQDTVTDVDK | -120 | -129 | 0 | 0 | 10 | NaN | NaN | NaN | 710.5006 | NaN | NaN |
| Tap2 | GGQLLLDGEPLTEYDHHYLHR | -523 | -543 | 0 | 0 | 21 | NaN | NaN | NaN | 440.95465 | 66.4563 | NaN |
| Tap2 | VVLVGQEPVLFSGSVK | -545 | -560 | 0 | 0 | 16 | NaN | NaN | NaN | 391.4151 | NaN | NaN |
| Ncstn | FQGETFDYIGSSR | -329 | -341 | 0 | 0 | 13 | NaN | NaN | NaN | 526.9736 | 331.3523 | 507.96866 |
| Ncstn | SFFWNVAPGAESAVAS | -285 | -300 | 0 | 0.00025015 | 16 | NaN | NaN | NaN | 135.77269 | NaN | NaN |
| Cab39 | SAEFLEQHYDR | -199 | -209 | 0 | 0 | 11 | NaN | NaN | NaN | 966.0471 | 760.9723 | 472.19064 |
| Ppp1r8 | FSEDDEIINPEDVDPSVGR | -203 | -221 | 0 | 0 | 19 | NaN | NaN | NaN | 1055.2499 | 1627.8732 | 1496.6141 |
| Ilk | WQGNDIVVK | -212 | -220 | 0 | 0.00025015 | 9 | NaN | NaN | NaN | 1623.3363 | NaN | 4892.1987 |

| | | | | | | | | | | | | |
|---|---|---|---|---|---|---|---|---|---|---|---|---|
| Ilk | LNENHSGELWK | -199 | -209 | 0 | 0.00025015 | 11 | NaN | NaN | NaN | 2994.6843 | NaN | NaN |
| NaN | FSGSGSGTDFTLK | -67 | -79 | 0 | 0.00025015 | 13 | NaN | NaN | NaN | 305.88477 | NaN | 1877.3545 |
| Celf2 | LDHSDQPDPDAIK | -29 | -41 | 0 | 0.00025015 | 13 | NaN | NaN | NaN | 638.59985 | 1154.2286 | 1068.3748 |
| Celf2 | HSDQPDPDAIK | -31 | -41 | 0 | 0 | 11 | NaN | NaN | NaN | 1721.025 | 326.75473 | NaN |
| Trip12 | SVYHLEDIVR | -1781 | -1790 | 0 | 0.000230707 | 10 | NaN | NaN | NaN | 11131.067 | 7168.28 | 8904.835 |
| Trip12 | SHDLFDIDPVVAR | -1768 | -1780 | 0 | 0 | 13 | NaN | NaN | NaN | 1763.7123 | 2099.415 | NaN |
| Trip12 | GLPPHLFGPLGPR | -416 | -428 | 0 | 0 | 13 | NaN | NaN | NaN | 1035.2404 | NaN | NaN |
| Trip12 | NTAGAQPQDDSIGGR | -19 | -33 | 0 | 0.001291103 | 15 | NaN | NaN | NaN | 696.0551 | NaN | NaN |
| Ddx52 | GFKPPVLVF | -411 | -419 | 1 | 0.00025015 | 9 | NaN | NaN | NaN | 435.89127 | 778.1011 | NaN |
| Rela | GSFSQADVHR | -237 | -246 | 0 | 0 | 10 | NaN | NaN | NaN | 389.92352 | 743.16486 | 605.7363 |
| Aldh9a1 | YINNYNVSPVELPFGGYK | -444 | -461 | 0 | 0.000431655 | 18 | NaN | NaN | NaN | 276.20953 | 378.46143 | 265.70447 |
| Ctnnbl1 | FVDILGLR | -373 | -380 | 0 | 0.00025015 | 8 | NaN | NaN | NaN | 122.26888 | 163.15952 | 117.625404 |
| Nae1 | FVEESPENLLDNDPSFFCR | -104 | -122 | 0 | 0.00025015 | 19 | NaN | NaN | NaN | 171.17644 | NaN | 232.40323 |
| Nae1 | QLLESTLLR | -130 | -138 | 0 | 0 | 9 | NaN | NaN | NaN | 2144.3315 | NaN | NaN |
| Snrpa | FVEFDNEVQAGAAR | -253 | -266 | 0 | 0.000550307 | 14 | NaN | NaN | NaN | 3453.3535 | NaN | NaN |
| Fau | RFVNVVPTF | -115 | -123 | 1 | 0.001023236 | 9 | NaN | NaN | NaN | 259.3462 | 346.93677 | 133.87361 |
| Fau | RFVNVVPTFG | -115 | -124 | 1 | 0.00122419 | 10 | NaN | NaN | NaN | 317.01212 | NaN | 1044.8737 |
| Stoml2 | VLIPVLDR | -65 | -72 | 0 | 0 | 8 | NaN | NaN | NaN | 372.7383 | NaN | NaN |
| Dynlt1 | FWDSSTDGSCTVR | -84 | -96 | 0 | 0.000550307 | 13 | NaN | NaN | NaN | 339.79688 | NaN | NaN |
| Rer1 | SEGDSVGDSVHGK | -2 | -14 | 0 | 0 | 13 | NaN | NaN | NaN | 697.59973 | 785.8661 | 874.82294 |
| Rer1 | VDPSLMEDSDDGPSLPTK | -87 | -104 | 0 | 0 | 18 | NaN | NaN | NaN | 805.2226 | 1376.0503 | 695.4617 |
| Snrpc | FYCDYCDTYLTH | -4 | -15 | 0 | 0 | 12 | NaN | NaN | NaN | 357.62793 | 383.11963 | 344.9389 |
| Atp5mg | VELVPPTPAEIPTAIQ | -36 | -51 | 0 | 0.00025015 | 16 | NaN | NaN | NaN | 764.45 | 822.84534 | 849.7215 |
| Bcap31 | KLKDELASTK | -189 | -198 | 2 | 0.00025015 | 10 | NaN | NaN | NaN | 520.5048 | 532.436 | 828.18274 |
| Rps26 | LHYCVSCAIHSK | -71 | -82 | 0 | 0 | 12 | NaN | NaN | NaN | 612.5376 | 1110.8875 | 1077.292 |
| Rps26 | LHYCVSCAIHSK | -71 | -82 | 0 | 0 | 12 | NaN | NaN | NaN | 612.5376 | 1110.8875 | 1077.292 |
| Rps26 | YCVSCAIHSK | -73 | -82 | 0 | 0 | 10 | NaN | NaN | NaN | 383.72772 | 172.93845 | 700.23865 |
| Rps26 | LHYCVSCAIHS | -71 | -81 | 0 | 0.000230707 | 11 | NaN | NaN | NaN | 494.66455 | NaN | 693.2537 |
| Mtpn | GPDGLTALEATDNQAIK | -98 | -114 | 0 | 0.000230707 | 17 | NaN | NaN | NaN | 1539.4504 | 1271.3553 | 1495.4156 |
| Mtpn | GADINAPDK | -58 | -66 | 0 | 0 | 9 | NaN | NaN | NaN | 443.17056 | NaN | NaN |
| Mtpn | KNGDLDEVKDYVAK | -11 | -24 | 2 | 0.005605646 | 14 | NaN | NaN | NaN | 293.61206 | NaN | NaN |
| Svil | GAFEEEEEPSYPILGR | -926 | -941 | 0 | 0 | 16 | NaN | NaN | NaN | 459.80182 | 434.66748 | 352.77582 |
| Svil | TIYPLADLLAR | -2111 | -2121 | 0 | 0.003299042 | 11 | NaN | NaN | NaN | 258.53018 | NaN | NaN |
| Luc7l3 | IHDENLR | -61 | -67 | 0 | 0 | 7 | NaN | NaN | NaN | 1514.867 | NaN | NaN |
| Luc7l3 | LSQNQQSSGAAGPTGK | -109 | -124 | 0 | 0.001023236 | 16 | NaN | NaN | NaN | 296.75693 | NaN | NaN |
| Anxa5 | SEIDLFNIR | -275 | -283 | 0 | 0 | 9 | NaN | NaN | NaN | 676.4785 | 827.4165 | 863.28406 |
| Anxa5 | GTDEDSILNLLTSR | -30 | -43 | 0 | 0 | 14 | NaN | NaN | NaN | 390.33923 | 322.37686 | 234.19824 |
| Anxa5 | GLGTDEDSILNLLTSR | -28 | -43 | 0 | 0 | 16 | NaN | NaN | NaN | 374.02362 | 708.88226 | 685.364 |
| Anxa5 | GTDDHTLIR | -261 | -269 | 0 | 0 | 9 | NaN | NaN | NaN | 1113.7484 | NaN | 976.7313 |
| Matr3 | NLPHSGYSDSAVLK | -502 | -515 | 0 | 0 | 14 | NaN | NaN | NaN | 1456.425 | 2597.6523 | 2523.822 |
| Matr3 | ITPENLPQILLQ | -133 | -144 | 0 | 0.001868731 | 12 | NaN | NaN | NaN | 754.8884 | 834.80115 | 742.1503 |
| Matr3 | SQESGYYDR | -208 | -216 | 0 | 0.00111006 | 9 | NaN | NaN | NaN | 803.1479 | NaN | NaN |
| Wdr89 | LFVIGGTNTGK | -288 | -298 | 0 | 0.00025015 | 11 | NaN | NaN | NaN | 2152.517 | 322.89536 | NaN |
| Wdr89 | GASEKPVQLFK | -101 | -111 | 1 | 0 | 11 | NaN | NaN | NaN | 490.0924 | 1398.4019 | NaN |
| Psmb4 | TQNPMVTGTSVLGVK | -46 | -60 | 0 | 0 | 15 | NaN | NaN | NaN | 518.5092 | 899.3001 | 376.36273 |
| Psmb4 | VNDSTMLGASGDYADFQYLK | -90 | -109 | 0 | 0 | 20 | NaN | NaN | NaN | 292.98294 | 290.3314 | 277.62408 |
| Itgb7 | SVDSCVSPEGGLCSGHGYCK | -601 | -620 | 0 | 0.001904399 | 20 | NaN | NaN | NaN | 405.58447 | 371.55008 | 452.85825 |
| Itgb7 | GATLPVYQELR | -350 | -360 | 0 | 0 | 11 | NaN | NaN | NaN | 382.08896 | NaN | NaN |
| Itgb7 | LCECDDASCER | -558 | -568 | 0 | 0 | 11 | NaN | NaN | NaN | 202.74901 | NaN | NaN |
| Abcf1 | SIDALGEAINDYK | -769 | -781 | 0 | 0 | 13 | NaN | NaN | NaN | 617.42975 | NaN | 462.77942 |
| Abcf1 | VLKPAKPEK | -149 | -157 | 2 | 0.000550307 | 9 | NaN | NaN | NaN | 265.09265 | NaN | 190.6428 |
| Rcc1 | TLGQGDVGQLGLGESVLER | -39 | -57 | 0 | 0 | 19 | NaN | NaN | NaN | 2059.5254 | 3020.4814 | 2963.5364 |
| Rcc1 | VSSGGQHTVLLVK | -404 | -416 | 0 | 0.000471409 | 13 | NaN | NaN | NaN | 702.2387 | 602.9071 | 606.30786 |
| Rcc1 | SAGDSHTAALTEDGR | -125 | -139 | 0 | 0.001392869 | 15 | NaN | NaN | NaN | 1627.9858 | NaN | NaN |
| Anxa4 | GCLIEILASR | -107 | -116 | 0 | 0.001023236 | 10 | NaN | NaN | NaN | 326.66095 | 395.75168 | 545.25745 |
| Snx12 | GDEGIFEESFIEER | -105 | -118 | 0 | 0.00025015 | 14 | NaN | NaN | NaN | 773.72485 | NaN | NaN |
| Triap1 | GDGSGDPCTDLFK | -30 | -42 | 0 | 0 | 13 | NaN | NaN | NaN | 451.80408 | NaN | NaN |
| Lsm3 | NTVEEPLDLIR | -12 | -22 | 0 | 0 | 11 | NaN | NaN | NaN | 1543.5878 | 286.91763 | 2195.8765 |
| Sec11a | NRVEDPIR | -71 | -78 | 1 | 0.001291103 | 8 | NaN | NaN | NaN | 459.70654 | 354.5571 | 584.86084 |
| Pgrmc1 | GDNDDDEPPPLPR | -55 | -67 | 0 | 0 | 13 | NaN | NaN | NaN | 1010.30743 | 492.15762 | 430.2033 |
| Thumpd3 | GDNNPLAVNR | -351 | -360 | 0 | 0.004224523 | 10 | NaN | NaN | NaN | 2571.9731 | NaN | NaN |
| Gmpr2 | GDVEHTIR | -299 | -306 | 0 | 0.00025015 | 8 | NaN | NaN | NaN | 842.73773 | 276.6525 | NaN |
| Gmpr2 | VGIGPGSVCTTR | -178 | -189 | 0 | 0 | 12 | NaN | NaN | NaN | 756.0883 | NaN | NaN |
| Ttc27 | QSLPWWTLR | -183 | -191 | 0 | 0.00416551 | 9 | NaN | NaN | NaN | 79.20892 | 452.69617 | 537.2145 |
| Spn | GSEVLETEGSGQRPTLTT | -319 | -336 | 1 | 0.001868731 | 18 | NaN | NaN | NaN | 687.51843 | NaN | NaN |
| Atp5pb | RKEEEHMIDWVEK | -209 | -221 | 2 | 0.001092529 | 13 | NaN | NaN | NaN | 614.29407 | 569.20605 | 546.0277 |
| Atp5pb | PPLPEYGGK | -45 | -53 | 0 | 0 | 9 | NaN | NaN | NaN | 1020.52234 | 568.63 | 1299.3741 |
| Sod2 | HHAAYVNNLNATEEK | -54 | -68 | 0 | 0 | 15 | NaN | NaN | NaN | 1742.0455 | 492.29813 | 510.57764 |
| Sod2 | GELLEAIKR | -115 | -123 | 1 | 0 | 9 | NaN | NaN | NaN | 820.5187 | 621.5356 | 515.6262 |
| Sod2 | YVNNLNATEEK | -58 | -68 | 0 | 0 | 11 | NaN | NaN | NaN | 3884.6228 | 606.76044 | NaN |
| Lims1 | NNSWHPECFR | -87 | -96 | 0 | 0 | 10 | NaN | NaN | NaN | 767.77026 | NaN | NaN |
| Vamp8 | NKTEDLEATSEHFK | -46 | -59 | 1 | 0 | 14 | NaN | NaN | NaN | 638.2483 | 1096.5282 | 788.37695 |
| Timm44 | GEVYDGDPDKVQR | -406 | -418 | 1 | 0.000550307 | 13 | NaN | NaN | NaN | 1245.4976 | NaN | NaN |
| Timm44 | VTDLLGGLFSK | -283 | -293 | 0 | 0.000431655 | 11 | NaN | NaN | NaN | 348.6696 | NaN | NaN |
| Fn1 | GEWTCKPIAEK | -176 | -186 | 1 | 0.000431655 | 11 | NaN | NaN | NaN | 423.80713 | 795.2693 | NaN |
| Nup160 | SLFTPGHFIN | -426 | -435 | 0 | 0.003490217 | 10 | NaN | NaN | NaN | 438.77853 | NaN | NaN |
| Polr2b | GFDQEEVFEKPTR | -822 | -834 | 1 | 0 | 13 | NaN | NaN | NaN | 1233.44 | 1362.9819 | 1300.3882 |
| Polr2b | IPQIGDKF | -928 | -935 | 1 | 0.006788754 | 8 | NaN | NaN | NaN | 730.30994 | NaN | 1184.178 |

| | | | | | | | | | | | | |
|---|---|---|---|---|---|---|---|---|---|---|---|---|
| Rabl6 | GFQDDVDIEDQTHGK | -452 | -466 | 0 | 0.00025015 | 15 | NaN | NaN | NaN | 941.6032 | NaN | NaN |
| Slc16a3 | PTAGFITGLK | -283 | -292 | 0 | 0 | 10 | NaN | NaN | NaN | 909.4331 | 1154.0321 | 984.4146 |
| Slc16a3 | LHKPPVDVGVD | -434 | -444 | 1 | 0.000471409 | 11 | NaN | NaN | NaN | 746.18994 | NaN | 608.854 |
| Mre11 | GGDLFHENKPSR | -58 | -69 | 1 | 0.00025015 | 12 | NaN | NaN | NaN | 370.53986 | 591.8787 | 680.97046 |
| Klhdc4 | QETGGSSAPEPLEVIK | -371 | -386 | 0 | 0 | 16 | NaN | NaN | NaN | 1158.0504 | 967.8572 | 642.29364 |
| Junb | GGGSGGGTGGGVTEEQEGFADGFVK | -114 | -138 | 0 | 0.000627943 | 25 | NaN | NaN | NaN | 45.130653 | 281.58542 | 328.23956 |
| Wapl | GGLDHIVDK | -828 | -836 | 0 | 0 | 9 | NaN | NaN | NaN | 998.7864 | 690.4197 | 774.1509 |
| Pum3 | KFQPDGESDESGAK | -92 | -105 | 1 | 0 | 14 | NaN | NaN | NaN | 555.688 | 414.54785 | 424.15213 |
| Suox | GGLTWQEAELEGEEQCPR | -467 | -484 | 0 | 0.007343498 | 18 | NaN | NaN | NaN | 405.71667 | NaN | NaN |
| Haus3 | GGNQLLLCDQELK | -517 | -529 | 0 | 0.000230707 | 13 | NaN | NaN | NaN | 3839.0967 | NaN | NaN |
| Mmtag2 | GGQDQFNWEDVK | -11 | -22 | 0 | 0.00078661 | 12 | NaN | NaN | NaN | 957.1116 | NaN | NaN |
| Cnp | GQVFLEELGNHK | -205 | -216 | 0 | 0 | 12 | NaN | NaN | NaN | 285.63394 | NaN | 1111.9724 |
| Gmip | GGVDAALEYAK | -96 | -106 | 0 | 0.003360691 | 11 | NaN | NaN | NaN | 1034.8855 | NaN | NaN |
| Keap1 | LNSAECYYPER | -484 | -494 | 0 | 0.00025015 | 11 | NaN | NaN | NaN | 605.03894 | 475.62634 | 625.8228 |
| Keap1 | LADLQVPR | -355 | -362 | 0 | 0 | 8 | NaN | NaN | NaN | 822.4637 | NaN | NaN |
| Plpbp | VQINTSGEDSK | -143 | -153 | 0 | 0.00111006 | 11 | NaN | NaN | NaN | 812.1229 | NaN | NaN |
| Rngtt | GHGECPTTENTETFIR | -93 | -108 | 0 | 0.007987481 | 16 | NaN | NaN | NaN | 1029.7975 | NaN | NaN |
| Hdac3 | GHGECVEYVK | -275 | -284 | 0 | 0 | 10 | NaN | NaN | NaN | 450.7339 | 959.50806 | 586.967 |
| Lman1 | YVSSLTEEISR | -370 | -380 | 0 | 0 | 11 | NaN | NaN | NaN | 385.2265 | NaN | 1469.6832 |
| Smc4 | SIDTAQECVNFLK | -645 | -657 | 0 | 0 | 13 | NaN | NaN | NaN | 639.97363 | NaN | NaN |
| Smc4 | YLEDIIGCGR | -251 | -260 | 0 | 0.007905138 | 10 | NaN | NaN | NaN | 408.63995 | NaN | NaN |
| Jpt2 | GIFDESTPVQTR | -70 | -81 | 0 | 0 | 12 | NaN | NaN | NaN | 20135.926 | 21017.482 | 16105.002 |
| Sh3bp1 | THLQDLGR | -280 | -287 | 0 | 0 | 8 | NaN | NaN | NaN | 988.4226 | NaN | 668.00165 |
| Iigp1 | GIGNEEEGAAK | -91 | -101 | 0 | 0 | 11 | NaN | NaN | NaN | 520.3897 | NaN | NaN |
| Psma6 | HITIFSPEGR | -12 | -21 | 0 | 0 | 10 | NaN | NaN | NaN | 1462.9937 | 1720.8242 | 1076.173 |
| Psma6 | SIDFKPSEIEVGVVTVENPK | -207 | -226 | 1 | 0 | 20 | NaN | NaN | NaN | 631.0417 | 240.87595 | 512.634 |
| Psma6 | QTESTSFLEK | -172 | -181 | 0 | 0.00025015 | 10 | NaN | NaN | NaN | 774.6618 | NaN | NaN |
| Opa1 | WNDFAEDSLR | -714 | -723 | 0 | 0 | 10 | NaN | NaN | NaN | 1108.2546 | 1231.1208 | 1107.8282 |
| Opa1 | GKEHDDIFDK | -690 | -699 | 1 | 0.001636916 | 10 | NaN | NaN | NaN | 732.65265 | NaN | 1095.7819 |
| Prmt3 | GKIEEVSLPVEK | -308 | -319 | 1 | 0.00204437 | 12 | NaN | NaN | NaN | 2203.758 | 2924.6191 | 2977.181 |
| Prmt3 | KLEDTIVLIK | -298 | -307 | 1 | 0.00025015 | 10 | NaN | NaN | NaN | 334.82446 | 285.5055 | NaN |
| Dnajc3 | VDGDPDNYIAYYR | -64 | -76 | 0 | 0.001463897 | 13 | NaN | NaN | NaN | 410.91617 | NaN | NaN |
| Impdh1 | GKLPIVNDQDELVAIIA | -207 | -223 | 1 | 0.00272025 | 17 | NaN | NaN | NaN | 520.0981 | NaN | NaN |
| Mbd2 | SASDVTEQIIK | -294 | -304 | 0 | 0 | 11 | NaN | NaN | NaN | 4385.7095 | NaN | 2200.5376 |
| Psme3ip1 | GLDEDETNFLDEVSR | -81 | -95 | 0 | 0.00111006 | 15 | NaN | NaN | NaN | 476.25922 | NaN | NaN |
| Farsb | NIFIECTGTDFTK | -250 | -262 | 0 | 0 | 13 | NaN | NaN | NaN | 1671.0507 | NaN | NaN |
| Mtor | GLLGALDPYK | -889 | -898 | 0 | 0.001092529 | 10 | NaN | NaN | NaN | 684.9362 | NaN | NaN |
| Snrpa1 | KTFNPGAGLPTDK | -179 | -191 | 1 | 0 | 13 | NaN | NaN | NaN | 1051.6921 | 1064.4034 | 1059.4489 |
| Cyc1 | GLLSSLDHTSIR | -100 | -111 | 0 | 0 | 12 | NaN | NaN | NaN | 1470.5852 | 909.2362 | 577.27356 |
| Cyc1 | SLDHTSIR | -104 | -111 | 0 | 0.001868731 | 8 | NaN | NaN | NaN | 892.3556 | 4852.7456 | 5397.0547 |
| Rhbdd1 | GLSEEEQLER | -267 | -276 | 0 | 0 | 10 | NaN | NaN | NaN | 7881.214 | NaN | NaN |
| Eefsec | SVDHCFSIK | -213 | -221 | 0 | 0 | 9 | NaN | NaN | NaN | 931.2143 | 288.5684 | NaN |
| Eefsec | IHIPGGLSPESK | -517 | -528 | 0 | 0.00025015 | 12 | NaN | NaN | NaN | 1683.9945 | NaN | NaN |
| Eefsec | VEFEKPVTCPR | -401 | -411 | 1 | 0.00111006 | 11 | NaN | NaN | NaN | 1180.6345 | NaN | NaN |
| Lgals3 | TVPYDLPLPGGVMPR | -129 | -143 | 0 | 0 | 15 | NaN | NaN | NaN | 1373.3146 | 1788.5924 | 1828.1511 |
| Fam118b | GNDYADLPEYFK | -304 | -315 | 0 | 0 | 12 | NaN | NaN | NaN | 798.96924 | 739.08734 | NaN |
| Acot7 | MIEEAGAIISTR | -80 | -91 | 0 | 0.00025015 | 12 | NaN | NaN | NaN | 515.45575 | 1318.791 | 1369.8783 |
| Ppih | GPFADENFK | -93 | -101 | 0 | 0.00025015 | 9 | NaN | NaN | NaN | 592.0474 | 523.37695 | 524.2 |
| Oxsm | GPHEGQFNEENFVSK | -95 | -109 | 0 | 0.000394361 | 15 | NaN | NaN | NaN | 1679.5094 | NaN | 643.2657 |
| Fam98b | INDALSCEYECR | -210 | -221 | 0 | 0 | 12 | NaN | NaN | NaN | 559.92706 | 450.1268 | 541.09393 |
| Rtn4 | LPEDDEPPARPPAPAGASPL | -128 | -147 | 1 | 0.00111006 | 20 | NaN | NaN | NaN | 3170.517 | NaN | 3267.655 |
| Rtn4 | HQAQIDHYLGLANK | -1128 | -1141 | 0 | 0 | 14 | NaN | NaN | NaN | 715.87823 | NaN | NaN |
| Ctss | QYIIDNGGIEADASYPYK | -199 | -216 | 0 | 0.00025015 | 18 | NaN | NaN | NaN | 561.2018 | 553.2531 | 631.182 |
| Ctss | GPVSVGIDASHS | -255 | -266 | 0 | 0.000431655 | 12 | NaN | NaN | NaN | 21653.555 | NaN | NaN |
| Mdp1 | GQNIQLYPEVPEVLGR | -42 | -57 | 0 | 0 | 16 | NaN | NaN | NaN | 610.52094 | 490.35892 | 518.6289 |
| Fen1 | RLDPSKYPVPENWLHK | -260 | -275 | 2 | 0.001023236 | 16 | NaN | NaN | NaN | 559.6042 | 1882.2139 | 1785.855 |
| Fen1 | GSDYCESIR | -229 | -237 | 0 | 0 | 9 | NaN | NaN | NaN | 310.51965 | NaN | NaN |
| Sacm1l | GSEKPLEQTFAN | -312 | -323 | 1 | 0.00142999 | 12 | NaN | NaN | NaN | 566.26184 | NaN | 937.6841 |
| Zfr | HATIYPTEEELQAVQK | -731 | -746 | 0 | 0.000230707 | 16 | NaN | NaN | NaN | 503.1365 | NaN | NaN |
| Zfr | SDVQPVGHDYVEEVR | -559 | -573 | 0 | 0.002265552 | 15 | NaN | NaN | NaN | 444.30392 | NaN | NaN |
| Mycbp2 | HGTDFLEYK | -4661 | -4669 | 0 | 0.000826806 | 9 | NaN | NaN | NaN | 308.77863 | NaN | NaN |
| Mrps16 | GSYDPLPNSHGEK | -52 | -64 | 0 | 0 | 13 | NaN | NaN | NaN | 3815.9702 | 4328.849 | 4309.0674 |
| Mapk3 | IADPEHDHTGFLT | -191 | -203 | 0 | 0.000471409 | 13 | NaN | NaN | NaN | 243.32771 | NaN | NaN |
| Slc7a5 | RGDGADPEGEGVTLQR | -34 | -49 | 1 | 0 | 16 | NaN | NaN | NaN | 756.8646 | 647.6197 | 620.0479 |
| Slc7a5 | KPELERPIK | -427 | -435 | 2 | 0.00051306 | 9 | NaN | NaN | NaN | 691.97534 | 394.87686 | 353.15198 |
| Slc7a5 | GTIIGSGIFVTPTGVLK | -62 | -78 | 0 | 0.001291103 | 17 | NaN | NaN | NaN | 247.23477 | NaN | NaN |
| Fxn | GTLDNPSSLDETAYER | -79 | -94 | 0 | 0.001636916 | 16 | NaN | NaN | NaN | 1032.9543 | 938.03754 | 899.5426 |
| Lsm2 | VQLPADEVDTQLLQDAAR | -70 | -87 | 0 | 0 | 18 | NaN | NaN | NaN | 410.40494 | 489.48132 | 571.282 |
| Tex10 | GTVDEDLSSPENLK | -304 | -317 | 0 | 0 | 14 | NaN | NaN | NaN | 895.38043 | NaN | 689.8168 |
| Tex10 | LLLQHVPLR | -865 | -873 | 0 | 0 | 9 | NaN | NaN | NaN | 57.323036 | 464.58395 | NaN |
| Ptpn7 | GTVEPICSVNTPR | -56 | -68 | 0 | 0.000550307 | 13 | NaN | NaN | NaN | 331.43176 | NaN | NaN |
| Mov10 | VGSVEEFQGQER | -863 | -874 | 0 | 0 | 12 | NaN | NaN | NaN | 936.3757 | 276.79993 | 439.46143 |
| Mov10 | SSETQQDDPVTYK | -403 | -415 | 0 | 0 | 13 | NaN | NaN | NaN | 4415.3975 | NaN | 2897.4177 |
| Mov10 | LSSETQQDDPVTYK | -402 | -415 | 0 | 0 | 14 | NaN | NaN | NaN | 498.26923 | NaN | NaN |
| Fkbp1b | GVEIETISPGDGR | -2 | -14 | 0 | 0.006735683 | 13 | NaN | NaN | NaN | 762.21954 | NaN | NaN |
| Gatd3 | GVTEAHVDQK | -222 | -231 | 0 | 0.000347792 | 10 | NaN | NaN | NaN | 686.0691 | 429.8616 | 499.5755 |
| Exosc3 | HKEPSGGGGGVY | -88 | -99 | 1 | 0.000471409 | 12 | NaN | NaN | NaN | 1459.9705 | NaN | NaN |
| Coro7 | RPHPCFTSS | -390 | -398 | 1 | 0.005758933 | 9 | NaN | NaN | NaN | 138.12878 | NaN | NaN |

| | | | | | | | | | | | | |
|---|---|---|---|---|---|---|---|---|---|---|---|---|
| Nbeal2 | SEESISPQLCQGLYK | -1211 | -1225 | 0 | 0.000550307 | 15 | NaN | NaN | NaN | 531.44244 | NaN | NaN |
| Cpsf6 | ISSLQDCLHGIESK | -470 | -483 | 0 | 0 | 14 | NaN | NaN | NaN | 1334.25 | 782.9027 | 337.23682 |
| Orc3 | GVNVTDHDVILR | -115 | -126 | 0 | 0 | 12 | NaN | NaN | NaN | 3190.6394 | 2136.314 | 1895.0494 |
| Nubp2 | SSGSGEELAR | -211 | -220 | 0 | 0.006677519 | 10 | NaN | NaN | NaN | 516.8828 | NaN | NaN |
| Prdx5 | PGAFTPGCSK | -89 | -98 | 0 | 0.000347792 | 10 | NaN | NaN | NaN | 225.24779 | 326.96353 | 212.51497 |
| Prdx5 | SVNDVFVIEEWGR | -123 | -135 | 0 | 0.007819401 | 13 | NaN | NaN | NaN | 761.8118 | NaN | NaN |
| Ly6c2 | PIKDPNIR | -83 | -90 | 1 | 0.00266297 | 8 | NaN | NaN | NaN | 1558.7314 | 1895.0876 | 1988.2952 |
| Ly6c2 | QCLSFCPAGVPIKDPNIR | -73 | -90 | 1 | 0 | 18 | NaN | NaN | NaN | 501.45633 | 680.00446 | 652.1512 |
| FAM120A | VDADNCLHR | -48 | -56 | 0 | 0.000627943 | 9 | NaN | NaN | NaN | 503.16135 | NaN | NaN |
| Rabgap1 | GVSDEDTDEEKETLK | -985 | -999 | 1 | 0.005854446 | 15 | NaN | NaN | NaN | 2888.93 | NaN | NaN |
| Man2b1 | WHIDPFGHSR | -193 | -202 | 0 | 0.00025015 | 10 | NaN | NaN | NaN | 497.90247 | 632.2775 | 596.86957 |
| Man2b1 | LPVYEGNFIVK | -528 | -538 | 0 | 0 | 11 | NaN | NaN | NaN | 473.589 | 550.0462 | NaN |
| Septin2 | QDLQEVTQDLHYENFR | -285 | -300 | 0 | 0.00201047 | 16 | NaN | NaN | NaN | 496.30286 | 498.98444 | 540.08997 |
| Septin2 | NLPNQVHR | -22 | -29 | 0 | 0.00111006 | 8 | NaN | NaN | NaN | 831.049 | 1098.3033 | NaN |
| Septin2 | QQPTQFINPETPGYVGF | -4 | -20 | 0 | 0.001708008 | 17 | NaN | NaN | NaN | 466.7727 | 577.4774 | NaN |
| Pik3cd | RHEVQEHFPEALAR | -534 | -547 | 1 | 0.00078661 | 14 | NaN | NaN | NaN | 787.9071 | NaN | NaN |
| Pik3cd | RHEVQEHFPEALAR | -534 | -547 | 1 | 0.00078661 | 14 | NaN | NaN | NaN | 787.9071 | NaN | NaN |
| Cbfb | KGWIDLHR | -111 | -118 | 1 | 0.000394361 | 8 | NaN | NaN | NaN | 694.6766 | 938.2484 | NaN |
| Prdx4 | IPLLSDLNHQISK | -177 | -189 | 0 | 0 | 13 | NaN | NaN | NaN | 1054.885 | NaN | 1192.805 |
| Prdx4 | LQGLESDER | -38 | -46 | 0 | 0.00111006 | 9 | NaN | NaN | NaN | 510.1311 | NaN | NaN |
| Gmds | GYEVHGIVR | -47 | -55 | 0 | 0.00025015 | 9 | NaN | NaN | NaN | 2757.251 | NaN | NaN |
| Gmds | LHYGDLTDSTCLVK | -82 | -95 | 0 | 0.00201047 | 14 | NaN | NaN | NaN | 1926.7036 | NaN | NaN |
| Vdac1 | SENGLEFTSSGSANTETTK | -48 | -66 | 0 | 0.000431655 | 19 | NaN | NaN | NaN | 449.43112 | 460.0668 | 352.1618 |
| Vdac1 | WTEYGLTF | -77 | -84 | 0 | 0.001636916 | 8 | NaN | NaN | NaN | 519.4597 | NaN | NaN |
| Vdac1 | VNNSSLIGLGYTQTLKPGIK | -250 | -269 | 1 | 0.00025015 | 20 | NaN | NaN | NaN | 478.6604 | NaN | NaN |
| Morf4l2 | GYLHDFLK | -254 | -261 | 0 | 0.003966045 | 8 | NaN | NaN | NaN | 234.45848 | NaN | NaN |
| Ap1g1 | VLAINILGR | -307 | -315 | 0 | 0 | 9 | NaN | NaN | NaN | 444.0935 | 459.46716 | 485.00177 |
| Trnt1 | LQSPEFQSLFTEGLK | -32 | -46 | 0 | 0 | 15 | NaN | NaN | NaN | 418.48996 | 718.48444 | 838.1112 |
| Cr1l | RDIVEFR | -302 | -308 | 1 | 0.000230707 | 7 | NaN | NaN | NaN | 946.7493 | 545.4077 | 508.56686 |
| Cr1l | SGPPPQCIELNK | -268 | -279 | 0 | 0.00025015 | 12 | NaN | NaN | NaN | 696.5702 | NaN | NaN |
| Rcn2 | VIDFDENTALDDTEEGSFR | -133 | -151 | 0 | 0 | 19 | NaN | NaN | NaN | 369.89258 | 418.9111 | 418.0948 |
| Rcn2 | HDDYFYHDEL | -311 | -320 | 0 | 0.000826806 | 10 | NaN | NaN | NaN | 396.69598 | NaN | NaN |
| Kifc1 | HDFSFDR | -371 | -377 | 0 | 0.004224523 | 7 | NaN | NaN | NaN | 564.5052 | NaN | NaN |
| Git2 | SVPSPTLPSTEDVIR | -580 | -594 | 0 | 0 | 15 | NaN | NaN | NaN | 942.92285 | NaN | 921.438 |
| Fam162a | HESLTSLNLER | -130 | -140 | 0 | 0 | 11 | NaN | NaN | NaN | 2529.1277 | 780.85065 | 2218.646 |
| Fam162a | RHESLTSLNLER | -129 | -140 | 1 | 0 | 12 | NaN | NaN | NaN | 391.93152 | 603.8105 | NaN |
| Fam162a | RHGVPLHKPTDFEK | -56 | -69 | 2 | 0.002810811 | 14 | NaN | NaN | NaN | 363.96375 | NaN | 590.86816 |
| Gcsh | SGEVTEVNEALAENPGLVNK | -114 | -133 | 0 | 0 | 20 | NaN | NaN | NaN | 295.95935 | 352.24265 | 312.49762 |
| Afg3l2 | IFIGPPDIK | -472 | -480 | 0 | 0.002752526 | 9 | NaN | NaN | NaN | 992.29297 | NaN | NaN |
| Pwp2 | QQGHFNSMVALAYSPDGQYI | -369 | -388 | 0 | 0.009928578 | 20 | NaN | NaN | NaN | 414.5599 | 655.13586 | NaN |
| Rpl26 | TVHVGIHPSK | -94 | -103 | 0 | 0 | 10 | NaN | NaN | NaN | 5196.5615 | 5398.5864 | 6014.564 |
| Ndufv1 | VVNADEGEPGTCK | -114 | -126 | 0 | 0 | 13 | NaN | NaN | NaN | 467.2514 | 455.58115 | 503.09622 |
| Ndufv1 | TSFGSLKDEDR | -30 | -40 | 1 | 0.001055699 | 11 | NaN | NaN | NaN | 766.5784 | NaN | NaN |
| Srpk1 | NYPENSNNETLR | -364 | -375 | 0 | 0.006100831 | 12 | NaN | NaN | NaN | 2213.316 | NaN | NaN |
| Srpk1 | IIQQVLQGLDYLHTK | -192 | -206 | 0 | 0.001868731 | 15 | NaN | NaN | NaN | 324.00607 | NaN | NaN |
| Toe1 | NTQPAEEFVVR | -23 | -33 | 0 | 0 | 11 | NaN | NaN | NaN | 1577.5646 | NaN | NaN |
| Toe1 | HGFNFNR | -144 | -150 | 0 | 0.007374243 | 7 | NaN | NaN | NaN | 870.9157 | NaN | NaN |
| Sec24a | HGNFFVR | -760 | -766 | 0 | 0.005242018 | 7 | NaN | NaN | NaN | 1000.95654 | NaN | 776.7259 |
| Sec24a | SLLDNLDLLPGNTR | -525 | -538 | 0 | 0.001291103 | 14 | NaN | NaN | NaN | 490.99707 | NaN | NaN |
| Agap2 | HGPLDTSVEDPQLR | -1073 | -1086 | 0 | 0.007343498 | 14 | NaN | NaN | NaN | 918.7493 | NaN | NaN |
| Apeh | VTSLVVDIVPR | -340 | -350 | 0 | 0 | 11 | NaN | NaN | NaN | 384.98502 | NaN | 490.3705 |
| Apeh | SRQDLFAVDTQTGSVT | -384 | -399 | 1 | 0.00025015 | 16 | NaN | NaN | NaN | 267.73163 | 863.9482 | NaN |
| Ddx46 | VIVIEEEKK | -597 | -605 | 1 | 0.000431655 | 9 | NaN | NaN | NaN | 1042.9166 | 905.79926 | 890.30756 |
| Ddx46 | LWSDFKDQQK | -752 | -761 | 1 | 0.00149795 | 10 | NaN | NaN | NaN | 796.7518 | NaN | NaN |
| Ddx46 | KVVTVVTTK | -263 | -271 | 1 | 0.004791731 | 9 | NaN | NaN | NaN | 328.4535 | NaN | NaN |
| Mycbp | YEEPEKPTSALDFLK | -35 | -49 | 1 | 0.000431655 | 15 | NaN | NaN | NaN | 2131.3757 | 605.7623 | 474.05664 |
| Mycbp | LYEEPEKPTSALDFLK | -34 | -49 | 1 | 0.002099344 | 16 | NaN | NaN | NaN | 398.7821 | NaN | 478.25754 |
| Far1 | HIDEVVYPPPVDPK | -162 | -175 | 0 | 0 | 14 | NaN | NaN | NaN | 853.4924 | 675.311 | 713.19727 |
| Gabpa | TISDETSEQVTR | -154 | -165 | 0 | 0.003620005 | 12 | NaN | NaN | NaN | 607.6997 | NaN | NaN |
| Dhx36 | HIVLDEIHER | -323 | -332 | 0 | 0 | 10 | NaN | NaN | NaN | 616.07996 | 441.08707 | 364.8516 |
| Dhx36 | IESPHPVDWDDTK | -954 | -966 | 0 | 0 | 13 | NaN | NaN | NaN | 1477.4749 | 935.61865 | NaN |
| Nup188 | SDGPEGQGQGQLLIK | -798 | -812 | 0 | 0.00025015 | 15 | NaN | NaN | NaN | 470.63696 | 207.03125 | 437.82755 |
| Nup188 | VKPIIDLVHK | -565 | -574 | 1 | 0.000431655 | 10 | NaN | NaN | NaN | 218.40797 | NaN | NaN |
| Nup188 | HKPHDVLSHEDGTLWR | -480 | -495 | 1 | 0.004658491 | 16 | NaN | NaN | NaN | 262.61334 | NaN | NaN |
| Dcaf7 | HLEHSTIIYEDPQHHPLLR | -208 | -226 | 0 | 0 | 19 | NaN | NaN | NaN | 139.90776 | NaN | NaN |
| Sdhc | SLCLGPTLIYSAK | -105 | -117 | 0 | 0 | 13 | NaN | NaN | NaN | 738.49744 | 935.5487 | 984.9154 |
| Cisd2 | QLPVPDSITGFAR | -20 | -32 | 0 | 0 | 13 | NaN | NaN | NaN | 661.7516 | 774.18787 | 769.7796 |
| Msra | YYAEDYHQQYLSK | -200 | -212 | 0 | 0 | 13 | NaN | NaN | NaN | 503.98456 | 606.2542 | 487.7597 |
| H2-Ab1 | HNYEGPETHTSLR | -108 | -120 | 0 | 0.000230707 | 13 | NaN | NaN | NaN | 778.69073 | NaN | 356.13016 |
| Nup43 | TIDNADSSTLHAV | -166 | -178 | 0 | 0 | 13 | NaN | NaN | NaN | 1275.2489 | 1271.1204 | NaN |
| Plaa | SSHLINLLNPK | -639 | -649 | 0 | 0.003966045 | 11 | NaN | NaN | NaN | 431.2453 | NaN | 568.66656 |
| Cdk1 | NLLIDDKGTIK | -133 | -143 | 1 | 0 | 11 | NaN | NaN | NaN | 2354.6958 | NaN | NaN |
| Twf1 | HQTLQGVAFPISR | -172 | -184 | 0 | 0 | 13 | NaN | NaN | NaN | 1494.7776 | NaN | 2980.9795 |
| Mrps22 | KPINDVLAEDK | -135 | -145 | 1 | 0 | 11 | NaN | NaN | NaN | 1376.8888 | 1588.7935 | 1670.1323 |
| Mrps22 | HQTYEDIDR | -246 | -254 | 0 | 0.00930008 | 9 | NaN | NaN | NaN | 947.6801 | NaN | NaN |
| Usp7 | VTDHDIPQQLVER | -532 | -544 | 0 | 0.000230707 | 13 | NaN | NaN | NaN | 514.4969 | NaN | 599.1792 |
| Vac14 | HSELQVPTSGR | -492 | -502 | 0 | 0 | 11 | NaN | NaN | NaN | 3932.9434 | 3900.4128 | 3831.3872 |
| Vac14 | LVTPEDDEPDEPK | -331 | -343 | 0 | 0.00025015 | 13 | NaN | NaN | NaN | 1660.1337 | 3002.6423 | NaN |

| | | | | | | | | | | | | |
|---|---|---|---|---|---|---|---|---|---|---|---|---|
| Hmgcs1 | PGSLPLNAEACWPK | -2 | -15 | 0 | 0.000550307 | 14 | NaN | NaN | NaN | 12626.958 | NaN | NaN |
| Hscb | HSTLVNDAYK | -119 | -128 | 0 | 0.000431655 | 10 | NaN | NaN | NaN | 220.9953 | NaN | NaN |
| Ccnt1 | HTCLHPQESLPDTR | -109 | -122 | 0 | 0.00025015 | 14 | NaN | NaN | NaN | 825.1394 | NaN | NaN |
| Nmd3 | HTLGEVWIQ | -340 | -348 | 0 | 0.008045078 | 9 | NaN | NaN | NaN | 461.94284 | NaN | NaN |
| Abhd12 | HVFDWIK | -226 | -232 | 0 | 0.001868731 | 7 | NaN | NaN | NaN | 5673.549 | NaN | NaN |
| Galnt2 | HVGSNLCLDSR | -532 | -542 | 0 | 0.002780285 | 11 | NaN | NaN | NaN | 1767.3702 | 2111.1353 | 2138.8547 |
| Galnt2 | LYFEELGK | -315 | -322 | 0 | 0.000591124 | 8 | NaN | NaN | NaN | 537.543 | NaN | NaN |
| Pnn | HVIAEQEVMETN | -397 | -408 | 0 | 0.000550307 | 12 | NaN | NaN | NaN | 3892.1833 | NaN | NaN |
| Mthfd2 | NAVSPDKDVDGFHVINVGR | -146 | -164 | 1 | 0.001463897 | 19 | NaN | NaN | NaN | 1516.1829 | NaN | NaN |
| Smarcd2 | RPGDLNVK | -290 | -297 | 1 | 0.009601182 | 8 | NaN | NaN | NaN | 679.9443 | 612.5729 | NaN |
| Smarcd2 | QELEQVLGIR | -520 | -529 | 0 | 0.00025015 | 10 | NaN | NaN | NaN | 2279.6538 | NaN | NaN |
| A2m | VCGIYTYGK | -256 | -264 | 0 | 0.001392869 | 9 | NaN | NaN | NaN | 1069.007 | NaN | NaN |
| Nup50 | TSEDADELHK | -449 | -458 | 0 | 0 | 10 | NaN | NaN | NaN | 462.627 | 531.4536 | 581.01434 |
| Nup50 | SFSAGSSSLFGK | -292 | -303 | 0 | 0.000230707 | 12 | NaN | NaN | NaN | 647.86566 | NaN | NaN |
| Wdr77 | HVVPLEPLPNPGPDSVVE | -325 | -342 | 0 | 0.00025015 | 18 | NaN | NaN | NaN | 340.45807 | 337.24304 | NaN |
| Apaf1 | HVVPVESGLGR | -278 | -288 | 0 | 0 | 11 | NaN | NaN | NaN | 3477.3774 | 3269.8374 | 3247.6497 |
| Apaf1 | RLPLNIEEAK | -215 | -224 | 1 | 0.005884925 | 10 | NaN | NaN | NaN | 488.31873 | 440.79303 | 456.1053 |
| Apaf1 | LNGEPGWVTIYGMAGCGK | -143 | -160 | 0 | 0.007343498 | 18 | NaN | NaN | NaN | 534.2596 | NaN | NaN |
| Apaf1 | SQLQDLHR | -451 | -458 | 0 | 0.002296977 | 8 | NaN | NaN | NaN | 485.31396 | NaN | NaN |
| Enoph1 | HWEEEECQQDVSLLR | -46 | -60 | 0 | 0.00025015 | 15 | NaN | NaN | NaN | 342.9232 | 426.525 | 399.2037 |
| H6pd | HWVPEQSR | -382 | -389 | 0 | 0.000989247 | 8 | NaN | NaN | NaN | 5734.124 | NaN | NaN |
| Ints7 | STPLNPLSFQCEFVK | -608 | -622 | 0 | 0.000230707 | 15 | NaN | NaN | NaN | 1933.705 | 2437.4673 | NaN |
| Ints7 | ILNVDEFVK | -97 | -105 | 0 | 0 | 9 | NaN | NaN | NaN | 801.3722 | 462.12387 | NaN |
| Zmpste24 | WHYSHPPLLER | -455 | -465 | 0 | 0 | 11 | NaN | NaN | NaN | 123.02316 | 893.699 | 112.90439 |
| Map2k2 | LPSGVFSSDFQEFVNK | -334 | -349 | 0 | 0.00025015 | 16 | NaN | NaN | NaN | 172.6616 | 549.90753 | 305.925 |
| Elmo2 | IAFDAESDPSNVPGSGTEK | -334 | -352 | 0 | 0 | 19 | NaN | NaN | NaN | 5506.3213 | NaN | NaN |
| Ube2l3 | LVNDPQPEHPLR | -111 | -122 | 0 | 0.000471409 | 12 | NaN | NaN | NaN | 681.8693 | 2482.5073 | 753.2858 |
| Bag1 | IANHLQELNK | -253 | -262 | 0 | 0.000230707 | 10 | NaN | NaN | NaN | 1755.0359 | NaN | 1886.5282 |
| Bag1 | LKDLEVSAEK | -243 | -252 | 1 | 0.009657961 | 10 | NaN | NaN | NaN | 910.5051 | NaN | NaN |
| Gpd1l | ICDEITGR | -103 | -110 | 0 | 0.00025015 | 8 | NaN | NaN | NaN | 666.9832 | 422.75464 | 694.0351 |
| Gpd1l | KFCETTIGSK | -162 | -171 | 1 | 0.00025015 | 10 | NaN | NaN | NaN | 575.27576 | NaN | NaN |
| Gdi1 | NPYYGGESSSITPLEELYK | -36 | -54 | 0 | 0 | 19 | NaN | NaN | NaN | 553.58545 | 682.53186 | NaN |
| Gdi1 | STTVETAEPEKEVEPALELLEPIDQK | -354 | -379 | 1 | 0 | 26 | NaN | NaN | NaN | 634.6622 | NaN | NaN |
| Prep | WMGGAELSDDGR | -234 | -245 | 0 | 0 | 12 | NaN | NaN | NaN | 509.10526 | NaN | 355.7324 |
| Prep | VLYVQDSLEGEAR | -99 | -111 | 0 | 0.000947459 | 13 | NaN | NaN | NaN | 1502.5826 | NaN | NaN |
| Wdr5 | IWDTASGQCLKT | -197 | -208 | 1 | 0.000947459 | 12 | NaN | NaN | NaN | 972.66425 | 1099.2968 | 894.0576 |
| Wdr5 | IDDDNPPVSFVK | -210 | -221 | 0 | 0.001636916 | 12 | NaN | NaN | NaN | 572.349 | NaN | NaN |
| Napa | LLEAHEEQNVDSYTE | -247 | -261 | 0 | 0.00025015 | 15 | NaN | NaN | NaN | 1195.9972 | 923.4519 | NaN |
| Rpl38 | IEEIKDFLLTA | -5 | -15 | 1 | 0.002296977 | 11 | NaN | NaN | NaN | 1440.5574 | 1656.2911 | 1457.5602 |
| Nudcd1 | YQDSVYYIDNLGR | -70 | -82 | 0 | 0.00025015 | 13 | NaN | NaN | NaN | 921.84546 | NaN | NaN |
| Mtx2 | TTQLTSDELSEK | -224 | -235 | 0 | 0.007880195 | 12 | NaN | NaN | NaN | 161.19485 | NaN | NaN |
| Pdcd10 | IERPEPEFQDLNEK | -93 | -106 | 1 | 0 | 14 | NaN | NaN | NaN | 651.0522 | NaN | 409.21048 |
| Slc38a1 | VCHPSVLPIYSELK | -285 | -298 | 0 | 0 | 14 | NaN | NaN | NaN | 1615.7103 | 1838.0928 | 644.45166 |
| Slc38a1 | VQSDLLHKY | -335 | -343 | 1 | 0.000230707 | 9 | NaN | NaN | NaN | 1111.4062 | 1009.8563 | 1022.0813 |
| Slc38a1 | IFILPSSLYLK | -428 | -438 | 0 | 0 | 11 | NaN | NaN | NaN | 640.3356 | 714.98566 | 664.1553 |
| Slc38a1 | LGEQVFGTTGK | -137 | -147 | 0 | 0 | 11 | NaN | NaN | NaN | 1681.71 | NaN | 1999.4647 |
| Slc38a1 | VQSDLLHK | -335 | -342 | 0 | 0.004824802 | 8 | NaN | NaN | NaN | 1086.1189 | NaN | NaN |
| Cul2 | SVLEVHGK | -342 | -349 | 0 | 0 | 8 | NaN | NaN | NaN | 664.5872 | NaN | 1341.7393 |
| Chd1 | IGDHITECLK | -1454 | -1463 | 0 | 0.00025015 | 10 | NaN | NaN | NaN | 1361.8711 | NaN | 1073.0223 |
| Pmm2 | LQEQLGNDVVEK | -48 | -59 | 0 | 0.008222027 | 12 | NaN | NaN | NaN | 617.87744 | NaN | NaN |
| Eif4e3 | RPLWEEESNAK | -95 | -105 | 1 | 0.002810811 | 11 | NaN | NaN | NaN | 1811.0518 | 1572.9583 | NaN |
| Eif4e3 | IHQLLPHIA | -177 | -185 | 0 | 0.002068213 | 9 | NaN | NaN | NaN | 254.507 | NaN | NaN |
| Rae1 | IHYINPPNPAK | -242 | -252 | 0 | 0 | 11 | NaN | NaN | NaN | 1460.3638 | 1293.5322 | 1237.5334 |
| Idi1 | IIADTFLFK | -199 | -207 | 0 | 0.001776473 | 9 | NaN | NaN | NaN | 699.5904 | 808.542 | 868.5588 |
| Idi1 | WWDNLNHLSPFVDHEK | -208 | -223 | 0 | 0.00149795 | 16 | NaN | NaN | NaN | 64.56853 | NaN | NaN |
| Magoh;Magol | IIDDSEITKEDDALWPPPDR | (63);(63) | (82);(82) | 1 | 0.00025015 | 20 | NaN | NaN | NaN | 321.20172 | 327.69424 | 825.59186 |
| Magoh;Magol | KIGSLIDVNQSK | (103);(103) | (114);(114) | 1 | 0.00025015 | 12 | NaN | NaN | NaN | 662.28406 | 613.1803 | 591.84534 |
| Magoh;Magol | VFYYLVQDLK | (121);(121) | (130);(130) | 0 | 0.000230707 | 10 | NaN | NaN | NaN | 635.31415 | 73.16814 | 833.2627 |
| Bud31 | IIECTHCGCR | -131 | -140 | 0 | 0 | 10 | NaN | NaN | NaN | 756.38074 | 986.6084 | 959.89307 |
| Ephx1 | NQYPHFK | -111 | -117 | 0 | 0.004595494 | 7 | NaN | NaN | NaN | 1670.8335 | NaN | NaN |
| Rpl19 | KDGLIIR | -46 | -52 | 1 | 0.009928578 | 7 | NaN | NaN | NaN | 339.6449 | 540.9602 | NaN |
| Rpl19 | VWLDPNETNEIANA | -22 | -35 | 0 | 0.001636916 | 14 | NaN | NaN | NaN | 892.1487 | NaN | NaN |
| Vps28 | IKEDRPITIK | -110 | -119 | 2 | 0.001610812 | 10 | NaN | NaN | NaN | 229.58894 | NaN | NaN |
| Micos13 | LVYDQELLGPSDK | -24 | -36 | 0 | 0 | 13 | NaN | NaN | NaN | 4025.8093 | NaN | NaN |
| Pycr1 | ILDEIGANIEDR | -78 | -89 | 0 | 0.00133086 | 12 | NaN | NaN | NaN | 1176.14 | NaN | NaN |
| Efl1 | ILEENSDLIR | -765 | -774 | 0 | 0.005242018 | 10 | NaN | NaN | NaN | 1347.4318 | NaN | 2227.5593 |
| Rpap3 | VLELEPDNFEATNELR | -228 | -243 | 0 | 0 | 16 | NaN | NaN | NaN | 810.6388 | 823.07184 | 915.3534 |
| Rpap3 | SDLKEDSVEELK | -644 | -655 | 1 | 0.000394361 | 12 | NaN | NaN | NaN | 4662.572 | NaN | NaN |
| Dpm1 | NYTIGEVPISFVDR | -221 | -234 | 0 | 0 | 14 | NaN | NaN | NaN | 888.81177 | 330.45544 | 807.8135 |
| Slk | SSEVGTDEALDDTQK | -542 | -556 | 0 | 0.000627943 | 15 | NaN | NaN | NaN | 999.7206 | 957.6634 | NaN |
| Trpv2 | YLTDSAYTEGSTGK | -100 | -113 | 0 | 0.001392869 | 14 | NaN | NaN | NaN | 821.26166 | NaN | 368.89874 |
| Padi2 | ILSNESLTQENQYFQR | -539 | -554 | 0 | 0 | 16 | NaN | NaN | NaN | 453.52548 | 780.2038 | 1035.01 |
| Etf1 | TEEEKILYLTPEQEK | -338 | -352 | 1 | 0.001023236 | 15 | NaN | NaN | NaN | 476.50256 | NaN | NaN |
| RTRAF | YKDLVPDNR | -99 | -107 | 1 | 0.000550307 | 9 | NaN | NaN | NaN | 1116.7957 | 1629.6646 | 511.27576 |
| RTRAF | NIHSSDWPK | -50 | -58 | 0 | 0.00025015 | 9 | NaN | NaN | NaN | 631.6094 | 321.89508 | 583.307 |
| RTRAF | INLDVNNPDFK | -121 | -131 | 0 | 0 | 11 | NaN | NaN | NaN | 376.50125 | 507.01785 | 440.69147 |
| RTRAF | RLLHIEELR | -205 | -213 | 1 | 0 | 9 | NaN | NaN | NaN | 387.02054 | NaN | NaN |
| Ikbkb | INYETQITPR | -395 | -404 | 0 | 0.003998192 | 10 | NaN | NaN | NaN | 957.634 | NaN | NaN |

| | | | | | | | | | | | | |
|---|---|---|---|---|---|---|---|---|---|---|---|---|
| Morc3 | KLPDGIDQLPEK | -421 | -432 | 1 | 0 | 12 | NaN | NaN | NaN | 3501.3623 | 3521.0735 | 3025.361 |
| Morc3 | IPEDLDETAGR | -221 | -231 | 0 | 0 | 11 | NaN | NaN | NaN | 2990.8699 | NaN | 1425.6132 |
| Septin9 | KADTLTLEER | -443 | -452 | 1 | 0.000907206 | 10 | NaN | NaN | NaN | 2036.2614 | NaN | 2781.2302 |
| Gatad2a | TLEPDLTEDDVENK | -14 | -27 | 0 | 0.00111006 | 14 | NaN | NaN | NaN | 1057.6355 | NaN | NaN |
| Stk4 | IPQDGDYEFLK | -427 | -437 | 0 | 0 | 11 | NaN | NaN | NaN | 2513.5876 | 2982.8577 | 2714.2043 |
| Stk4 | QLLQHPFVK | -274 | -282 | 0 | 0.00025015 | 9 | NaN | NaN | NaN | 470.58154 | NaN | NaN |
| Slc1a5 | IPQSCIQR | -218 | -225 | 0 | 0.002265552 | 8 | NaN | NaN | NaN | 518.32684 | 2443.6836 | 630.66 |
| Slc1a5 | LGPEGELLIR | -260 | -269 | 0 | 0 | 10 | NaN | NaN | NaN | 2493.7068 | NaN | NaN |
| Psme3 | KYPHVEDYR | -195 | -203 | 1 | 0 | 9 | NaN | NaN | NaN | 1575.6885 | 1648.3635 | 1996.7957 |
| Psme3 | KYPHVEDYR | -195 | -203 | 1 | 0 | 9 | NaN | NaN | NaN | 1575.6885 | 1648.3635 | 1996.7957 |
| Psme3 | IQEETVAELR | -157 | -166 | 0 | 0.001636916 | 10 | NaN | NaN | NaN | 823.02856 | 1509.4854 | NaN |
| Psme3 | NQQLVDIIEK | -112 | -121 | 0 | 0.004595494 | 10 | NaN | NaN | NaN | 362.04263 | 1545.158 | NaN |
| Psme3 | RLDECEEAF | -88 | -96 | 1 | 0.004658491 | 9 | NaN | NaN | NaN | 133.03334 | NaN | NaN |
| Znf638 | IQENDPEANIDK | -991 | -1002 | 0 | 0.00025015 | 12 | NaN | NaN | NaN | 6287.08 | NaN | NaN |
| Znf638 | SENLEDDTLPEGK | -641 | -653 | 0 | 0.000627943 | 13 | NaN | NaN | NaN | 346.73157 | NaN | NaN |
| Fgb | TPCTVSCNIPVVSGK | -215 | -229 | 0 | 0 | 15 | NaN | NaN | NaN | 2055.9233 | 1252.6241 | 1818.4246 |
| Fgb | QDGSVDFGR | -276 | -284 | 0 | 0.000347792 | 9 | NaN | NaN | NaN | 283.4019 | NaN | NaN |
| Frmd8 | ISLEGVHVIDNR | -297 | -308 | 0 | 0 | 12 | NaN | NaN | NaN | 1237.3766 | NaN | NaN |
| Frmd8 | LDSFLPAHLCK | -197 | -207 | 0 | 0.005698678 | 11 | NaN | NaN | NaN | 749.8783 | NaN | NaN |
| Nr2c2ap | IVDFYPEDTNALQTF | -89 | -103 | 0 | 0.00078661 | 15 | NaN | NaN | NaN | 348.46558 | 549.5332 | 362.10678 |
| Ppat | YCHEVKPGEIVEIS | -246 | -259 | 1 | 0.00025015 | 14 | NaN | NaN | NaN | 985.9603 | NaN | NaN |
| Ehd3 | IVNTPEVIR | -243 | -251 | 0 | 0.008736733 | 9 | NaN | NaN | NaN | 889.71014 | NaN | NaN |
| Ndufs7 | IVPVDIYVPGCPPTAEAL | -184 | -201 | 0 | 0 | 18 | NaN | NaN | NaN | 443.92886 | 489.99164 | 513.1189 |
| Uck2 | LFVDTDADTR | -153 | -162 | 0 | 0.001092529 | 10 | NaN | NaN | NaN | 1189.755 | NaN | NaN |
| Rab5b | IVVYDITNQETFAR | -97 | -110 | 0 | 0 | 14 | NaN | NaN | NaN | 397.8126 | 410.35828 | 556.0871 |
| Ppp2r2a | TTNNLYIFQDK | -435 | -445 | 0 | 0 | 11 | NaN | NaN | NaN | 2585.2825 | NaN | 576.8958 |
| Rps24 | VIDVLHPGK | -24 | -32 | 0 | 0.001055699 | 9 | NaN | NaN | NaN | 329.6988 | 1295.5986 | NaN |
| Memo1 | TPLYDLR | -98 | -104 | 0 | 0.00223472 | 7 | NaN | NaN | NaN | 294.61322 | NaN | NaN |
| Uqcrq | LIYTWGNQEFEQSK | -58 | -71 | 0 | 0 | 14 | NaN | NaN | NaN | 1246.6279 | 1435.7654 | 1415.1305 |
| Papss1 | IYWNDGLDQYR | -387 | -397 | 0 | 0 | 11 | NaN | NaN | NaN | 1394.7788 | NaN | NaN |
| Dnm1l | KAEELLAEEK | -597 | -606 | 1 | 0.003490217 | 10 | NaN | NaN | NaN | 341.0036 | 1556.2013 | NaN |
| Dnm1l | SVTDSIRDEYAFLQ | -257 | -270 | 1 | 0.002810811 | 14 | NaN | NaN | NaN | 255.4881 | NaN | NaN |
| Zc3h18 | KAVEDAIAR | -931 | -939 | 1 | 0.000826806 | 9 | NaN | NaN | NaN | 317.1725 | 746.05066 | NaN |
| Lamp1 | KCNTEEHIFVSK | -326 | -337 | 1 | 0.00025015 | 12 | NaN | NaN | NaN | 1277.6885 | NaN | 698.68994 |
| Thumpd1 | YAETFLEPWFK | -179 | -189 | 0 | 0.009388464 | 11 | NaN | NaN | NaN | 310.6639 | 560.4029 | 657.5169 |
| Thumpd1 | KCVEEAYSL | -56 | -64 | 1 | 0.003360691 | 9 | NaN | NaN | NaN | 653.5602 | 770.30585 | NaN |
| Polr2e | KEEVTELLAR | -153 | -162 | 1 | 0.000431655 | 10 | NaN | NaN | NaN | 4833.424 | NaN | NaN |
| Hmga1 | KQPPVSPGTA | -31 | -40 | 1 | 0.00272025 | 10 | NaN | NaN | NaN | 895.1684 | NaN | 1625.3698 |
| Myh10 | KFDQLLAEEKG | -1452 | -1462 | 2 | 0.000591124 | 11 | NaN | NaN | NaN | 6485.353 | 7267.8745 | 7287.1553 |
| Strn4 | NAALDVEPIHAF | -483 | -494 | 0 | 0.001392869 | 12 | NaN | NaN | NaN | 857.0015 | NaN | 368.83838 |
| Strn4 | KFGTDLNQGEK | -136 | -146 | 1 | 0.000431655 | 11 | NaN | NaN | NaN | 679.36664 | NaN | NaN |
| Strn4 | KNAALDVEPIHAF | -482 | -494 | 1 | 0.007987481 | 13 | NaN | NaN | NaN | 308.79443 | NaN | NaN |
| Ldhb | KFIIPQIVK | -119 | -127 | 1 | 0.000591124 | 9 | NaN | NaN | NaN | 548.31494 | NaN | 525.75726 |
| Supv3l1 | KFNDPNDPCK | -410 | -419 | 1 | 0 | 10 | NaN | NaN | NaN | 805.025 | NaN | NaN |
| Fis1 | LQTEPQNNQAK | -98 | -108 | 0 | 0.003360691 | 11 | NaN | NaN | NaN | 339.3573 | NaN | NaN |
| Prkca | KGTEELYAIK | -359 | -368 | 1 | 0.001392869 | 10 | NaN | NaN | NaN | 4626.5825 | NaN | NaN |
| Ppwd1 | KIEEGIEFVK | -118 | -127 | 1 | 0 | 10 | NaN | NaN | NaN | 2602.7468 | 1536.3292 | NaN |
| Ppwd1 | YWTGPPHEYK | -249 | -258 | 0 | 0 | 10 | NaN | NaN | NaN | 1043.3384 | NaN | NaN |
| Cd2 | NIPNFQMTDDIDEVR | -39 | -53 | 0 | 0.001392869 | 15 | NaN | NaN | NaN | 619.213 | NaN | NaN |
| Ints1 | TDQGLDEEGEDER | -2019 | -2031 | 0 | 0 | 13 | NaN | NaN | NaN | 700.065 | NaN | NaN |
| C7orf50 | KLDEAEPEDSGGSPGK | -168 | -183 | 1 | 0.001392869 | 16 | NaN | NaN | NaN | 561.5084 | 721.10785 | 824.66315 |
| Hook3 | KLEDLGDLR | -326 | -334 | 1 | 0.00450079 | 9 | NaN | NaN | NaN | 4021.2646 | NaN | NaN |
| Tpx2 | STAEQEAEELEK | -385 | -396 | 0 | 0 | 12 | NaN | NaN | NaN | 880.8434 | 495.9678 | NaN |
| Tpx2 | SSELPLTVPVSPK | -727 | -739 | 0 | 0.008253452 | 13 | NaN | NaN | NaN | 641.2916 | NaN | NaN |
| Mrpl54 | KLEELEPESR | -103 | -112 | 1 | 0.00025015 | 10 | NaN | NaN | NaN | 939.1927 | NaN | 1220.7649 |
| Pyhin1 | RPSSSILDPPVSSGT | -182 | -196 | 1 | 0.001636916 | 15 | NaN | NaN | NaN | 3698.886 | 1790.7402 | NaN |
| Gpaa1 | KLNHLLER | -345 | -352 | 1 | 0.009153908 | 8 | NaN | NaN | NaN | 550.3047 | NaN | NaN |
| Krr1 | KNIHPIYNIK | -210 | -219 | 1 | 0 | 10 | NaN | NaN | NaN | 1092.7351 | 386.51495 | 346.5354 |
| Krr1 | TFDPYIIIR | -106 | -114 | 0 | 0 | 9 | NaN | NaN | NaN | 699.89874 | 684.2677 | NaN |
| Bsg | SEHSSEGELAK | -229 | -239 | 0 | 0.009031675 | 11 | NaN | NaN | NaN | 428.08514 | NaN | NaN |
| Iqgap2 | KPEAQLPAVYPF | -328 | -339 | 1 | 0.00383302 | 12 | NaN | NaN | NaN | 2401.5579 | 375.12744 | 2821.5815 |
| Iqgap2 | VDQVQDIVTGNPTVIK | -951 | -966 | 0 | 0.00025015 | 16 | NaN | NaN | NaN | 1004.0235 | NaN | NaN |
| Mrpl43 | KPFHTDNPSIQ | -117 | -127 | 1 | 0.003521747 | 11 | NaN | NaN | NaN | 890.10944 | NaN | NaN |
| Cd8a | LPQPTFVVY | -74 | -82 | 0 | 0.00122419 | 9 | NaN | NaN | NaN | 723.8608 | 1204.3372 | NaN |
| Dnajb11 | QEVVCDECPNVK | -189 | -200 | 0 | 0.00025015 | 12 | NaN | NaN | NaN | 444.29108 | 429.15256 | 367.19183 |
| Dnajb11 | QYDTYGEEGLKDGHQS | -85 | -100 | 1 | 0.001055699 | 16 | NaN | NaN | NaN | 355.3391 | 403.24622 | 343.63635 |
| Dnajb11 | TLEVEIEPGVR | -207 | -217 | 0 | 0 | 11 | NaN | NaN | NaN | 1220.4457 | 1444.7073 | NaN |
| Dnajb11 | LHPDRNPDDPQAQEK | -52 | -66 | 1 | 0 | 15 | NaN | NaN | NaN | 577.3468 | NaN | NaN |
| Rbm17 | SDSNPLTEILK | -295 | -305 | 0 | 0.00204437 | 11 | NaN | NaN | NaN | 216.17746 | NaN | NaN |
| RtcA | KSEEEEDATKDTYVIE | -340 | -355 | 2 | 0.002265552 | 16 | NaN | NaN | NaN | 850.483 | 930.4214 | NaN |
| Ptk2b | RYIEDEDYYK | -572 | -581 | 1 | 0.00025015 | 10 | NaN | NaN | NaN | 606.3186 | 941.17505 | 881.2478 |
| Clybl | KTLEDFDLGTTEK | -90 | -102 | 1 | 0 | 13 | NaN | NaN | NaN | 515.1997 | 558.42694 | 625.8586 |
| Trim25 | VACDHCLTEIAVK | -104 | -116 | 0 | 0 | 13 | NaN | NaN | NaN | 660.50653 | NaN | NaN |
| Trim25 | KTPVAPGPPSHFSPN | -392 | -406 | 1 | 0.000907206 | 15 | NaN | NaN | NaN | 313.66452 | NaN | NaN |
| H1-4 | SETAPAAPAAPAPAEK | -2 | -17 | 0 | 0 | 16 | NaN | NaN | NaN | 2625.3977 | 1939.2856 | 1863.2377 |
| Utp14a | TLEELEELGKEDSLPNK | -509 | -525 | 1 | 0.000712251 | 17 | NaN | NaN | NaN | 3272.6453 | 3399.3977 | 3339.3142 |
| Utp14a | KVDPDAALEELEK | -273 | -285 | 1 | 0.002296977 | 13 | NaN | NaN | NaN | 476.0936 | NaN | NaN |
| Tmx1 | TSHSYFVHDL | -164 | -173 | 0 | 0.00111006 | 10 | NaN | NaN | NaN | 752.58966 | NaN | NaN |

| | | | | | | | | | | | | |
|---|---|---|---|---|---|---|---|---|---|---|---|---|
| Tpd52l2 | TQETLSQAGQK | -129 | -139 | 0 | 0 | 11 | NaN | NaN | NaN | 495.99786 | 770.11646 | 739.0446 |
| Eci1 | KVGVVDEVVPEDQVHSK | -206 | -222 | 1 | 0.00111006 | 17 | NaN | NaN | NaN | 426.5022 | 583.43536 | 616.9167 |
| Xrcc6 | LFTNEDDPHGR | -166 | -176 | 0 | 0 | 11 | NaN | NaN | NaN | 1154.1512 | 1318.4674 | 1408.992 |
| Sec22b | VANIEEVLQR | -150 | -159 | 0 | 0 | 10 | NaN | NaN | NaN | 986.73236 | 1035.2635 | 973.86694 |
| Arrb2 | KVQFAPETPGPQPSAETT | -171 | -188 | 1 | 0.000627943 | 18 | NaN | NaN | NaN | 654.32135 | 808.33325 | NaN |
| Polr1e | KVTSEDILK | -300 | -308 | 1 | 0.003745449 | 9 | NaN | NaN | NaN | 2327.2085 | NaN | NaN |
| Rab21 | VVLLGEGCVGK | -20 | -30 | 0 | 0.00201047 | 11 | NaN | NaN | NaN | 2445.4216 | 4068.5764 | NaN |
| Rab21 | KVVLLGEGCVGK | -19 | -30 | 1 | 0.007343498 | 12 | NaN | NaN | NaN | 356.28513 | NaN | NaN |
| Phpt1 | KWAEYHADIYDK | -47 | -58 | 1 | 0 | 12 | NaN | NaN | NaN | 997.7714 | 2606.8994 | 4306.395 |
| Slc38a2 | SSVTHLLCPTK | -393 | -403 | 0 | 0 | 11 | NaN | NaN | NaN | 1527.8928 | 1710.5911 | 1605.0583 |
| Slc38a2 | TANEGGSLLYEQLGHK | -125 | -140 | 0 | 0 | 16 | NaN | NaN | NaN | 238.82156 | 1132.8164 | 1048.0482 |
| Slc38a2 | KYELPLVIK | -170 | -178 | 1 | 0 | 9 | NaN | NaN | NaN | 1009.57416 | NaN | 1458.2545 |
| Slc38a2 | LYEQLGHK | -133 | -140 | 0 | 0.002880184 | 8 | NaN | NaN | NaN | 566.0704 | NaN | NaN |
| Cks2 | KYFDEHYEYR | -11 | -20 | 1 | 0 | 10 | NaN | NaN | NaN | 400.12177 | 1650.3833 | 1685.5648 |
| Tnfrsf9 | NSCDNCQPGTFCR | -26 | -38 | 0 | 0.000394361 | 13 | NaN | NaN | NaN | 363.67816 | NaN | NaN |
| QNG1 | KYSDELLK | -251 | -258 | 1 | 0 | 8 | NaN | NaN | NaN | 1946.2017 | 3693.466 | 4800.901 |
| Rprd1b | LLNIWQER | -107 | -114 | 0 | 0.000230707 | 8 | NaN | NaN | NaN | 819.4584 | 1052.1119 | 1095.3674 |
| Rprd1b | SLPQEVQDVSLLEK | -203 | -216 | 0 | 0 | 14 | NaN | NaN | NaN | 514.9385 | NaN | NaN |
| Luc7l | SHLLDCCPHDILAGTR | -38 | -53 | 0 | 0 | 16 | NaN | NaN | NaN | 956.43677 | 1127.9159 | 1543.424 |
| Vbp1 | LADNLYCK | -106 | -113 | 0 | 0 | 8 | NaN | NaN | NaN | 739.69 | 604.44635 | 548.4979 |
| Dennd10 | LADSEITIAPLAK | -255 | -267 | 0 | 0.001291103 | 13 | NaN | NaN | NaN | 454.86975 | NaN | NaN |
| Samsn1 | SDDSIEVHDR | -38 | -47 | 0 | 0.000471409 | 10 | NaN | NaN | NaN | 858.7702 | 1550.5586 | NaN |
| Arfgef2;Arfgef | SWHEILK | (995);(1040) | (1001);(1046) | 0 | 0.004224523 | 7 | NaN | NaN | NaN | 3399.0425 | NaN | NaN |
| Nup54 | YYIDADLLR | -456 | -464 | 0 | 0 | 9 | NaN | NaN | NaN | 419.48026 | 363.16086 | NaN |
| Ints3 | LDLLSELYQK | -672 | -681 | 0 | 0 | 10 | NaN | NaN | NaN | 882.01965 | NaN | NaN |
| Tor1a | LDYYDVVDEVSYQK | -188 | -201 | 0 | 0.001291103 | 14 | NaN | NaN | NaN | 528.14343 | NaN | NaN |
| Cggbp1 | LEANIPLEK | -112 | -120 | 0 | 0.000230707 | 9 | NaN | NaN | NaN | 815.0754 | 1054.1533 | 1078.2601 |
| Nub1 | LECCENEVENIIEEIR | -49 | -64 | 0 | 0 | 16 | NaN | NaN | NaN | 773.17444 | 1250.7969 | NaN |
| Nub1 | LWKPPYTK | -22 | -29 | 1 | 0.008653659 | 8 | NaN | NaN | NaN | 865.58923 | NaN | NaN |
| Ppp1r12a | WIGSETDLEPPVVK | -17 | -30 | 0 | 0 | 14 | NaN | NaN | NaN | 876.5128 | 8097.5664 | 8553.321 |
| Ppp1r12a | QGVDIEAAR | -162 | -170 | 0 | 0.002068213 | 9 | NaN | NaN | NaN | 1162.9045 | NaN | NaN |
| Ppp1r12a | NKETLIIEPEK | -319 | -329 | 1 | 0.006472924 | 11 | NaN | NaN | NaN | 1012.3285 | NaN | NaN |
| Ppp1r12a | LTPVRDEESESQR | -668 | -680 | 1 | 0.005301614 | 13 | NaN | NaN | NaN | 719.5473 | NaN | NaN |
| Ppp1r12a | LEKDDSTDFK | -923 | -932 | 1 | 0.000907206 | 10 | NaN | NaN | NaN | 539.93536 | NaN | NaN |
| Syf2 | LEWELQEEEK | -74 | -83 | 0 | 0.00122419 | 10 | NaN | NaN | NaN | 1159.7072 | NaN | NaN |
| Prpf38b | TNILSSPYFK | -65 | -74 | 0 | 0.00025015 | 10 | NaN | NaN | NaN | 654.961 | 879.16315 | NaN |
| Rab27a | SWIPEGVVR | -192 | -200 | 0 | 0.006654467 | 9 | NaN | NaN | NaN | 876.98975 | NaN | NaN |
| Nrbp1 | LTSLLEETLNK | -507 | -517 | 0 | 0.000230707 | 11 | NaN | NaN | NaN | 1920.1167 | 560.33234 | NaN |
| Nrbp1 | LFEVPSLK | -327 | -334 | 0 | 0 | 8 | NaN | NaN | NaN | 349.08325 | NaN | 302.23938 |
| Cnot7 | LFFPVIYDVK | -187 | -196 | 0 | 0 | 10 | NaN | NaN | NaN | 153.03607 | NaN | NaN |
| Ndufa9 | SVFPEAIIIRPS | -193 | -204 | 1 | 0.002099344 | 12 | NaN | NaN | NaN | 797.08655 | NaN | 184.78253 |
| Gclm | LFIVGSNSSSSTR | -95 | -107 | 0 | 0 | 13 | NaN | NaN | NaN | 2324.0217 | NaN | NaN |
| Ric8a | LFQELHGVR | -182 | -190 | 0 | 0 | 9 | NaN | NaN | NaN | 1063.7822 | NaN | NaN |
| Cdipt | SVISVIHLITAA | -187 | -198 | 0 | 0 | 12 | NaN | NaN | NaN | 399.6646 | 456.14618 | NaN |
| Cdipt | LFVPNLIGYAR | -8 | -18 | 0 | 0.009569685 | 11 | NaN | NaN | NaN | 185.60197 | NaN | NaN |
| Cdc42se1 | LGCCVVEKPQPK | -8 | -19 | 1 | 0.00025015 | 12 | NaN | NaN | NaN | 542.9183 | 535.9502 | 399.40448 |
| Hsd17b10 | TIAPGLFATPLLTTLPEK | -195 | -212 | 0 | 0.00025015 | 18 | NaN | NaN | NaN | 176.11601 | 189.52342 | 229.39926 |
| Vps16 | LGDTPGVSYSDIAAR | -511 | -525 | 0 | 0 | 15 | NaN | NaN | NaN | 957.43066 | NaN | 1539.9755 |
| Capns1 | TDGFGIDTCR | -136 | -145 | 0 | 0.000431655 | 10 | NaN | NaN | NaN | 150.74048 | NaN | NaN |
| Gfer | LGKPDFDCSR | -174 | -183 | 1 | 0.000431655 | 10 | NaN | NaN | NaN | 2220.7354 | NaN | NaN |
| Kdm3b | LGLGSVVPVEYLVDR | -130 | -144 | 0 | 0 | 15 | NaN | NaN | NaN | 377.42776 | 232.54407 | 300.03702 |
| Polr1c | SQCFSPGVIEVEEVQGK | -254 | -270 | 0 | 0.003681885 | 17 | NaN | NaN | NaN | 2897.7954 | NaN | NaN |
| Ifrd1 | LGPGFESEEILK | -154 | -165 | 0 | 0.00025015 | 12 | NaN | NaN | NaN | 1302.9042 | NaN | NaN |
| Sar1a | LVDCADHSR | -99 | -107 | 0 | 0.00025015 | 9 | NaN | NaN | NaN | 1033.3119 | NaN | NaN |
| Pde12 | SQHDISFQEALK | -394 | -405 | 0 | 0.00025015 | 12 | NaN | NaN | NaN | 777.09296 | NaN | NaN |
| Sh2d2a | LHQAPPINPIYQEPDEPIAF | -290 | -309 | 0 | 0.00025015 | 20 | NaN | NaN | NaN | 316.4796 | NaN | 412.94525 |
| Synj2bp | NLLHQDAVDLFR | -75 | -86 | 0 | 0 | 12 | NaN | NaN | NaN | 444.14453 | 576.7709 | 541.9048 |
| Synj2bp | LHQDAVDLFR | -77 | -86 | 0 | 0 | 10 | NaN | NaN | NaN | 1627.4338 | NaN | NaN |
| Ppp1r2 | LHYNEGLNIK | -147 | -156 | 0 | 0 | 10 | NaN | NaN | NaN | 1874.8364 | NaN | 2263.7163 |
| Srpra | VGNEAVDQLVK | -553 | -563 | 0 | 0.00383302 | 11 | NaN | NaN | NaN | 466.6543 | NaN | NaN |
| Ppil4 | LILDEQGEDSK | -365 | -375 | 0 | 0.005242018 | 11 | NaN | NaN | NaN | 772.6903 | NaN | NaN |
| Smurf1 | LIPQHLLK | -588 | -595 | 0 | 0.001392869 | 8 | NaN | NaN | NaN | 400.84695 | NaN | 476.63297 |
| Ubxn4 | LISNFLFSNPPPAQTS | -426 | -441 | 0 | 0.000471409 | 16 | NaN | NaN | NaN | 459.1457 | 611.48694 | 560.5392 |
| Elavl1 | LIVNYLPQNMTQEELR | -22 | -37 | 0 | 0 | 16 | NaN | NaN | NaN | 482.5756 | 721.1754 | 595.8492 |
| Elavl1 | SEAEEAITSFNGHKPPGSSEPITVK | -158 | -182 | 1 | 0.000298805 | 25 | NaN | NaN | NaN | 568.9068 | 744.0548 | 712.9177 |
| Ndufaf1 | LKDEIVAHLR | -98 | -107 | 1 | 0.001023236 | 10 | NaN | NaN | NaN | 34.54983 | NaN | NaN |
| Itsn2 | LKEQLDALEK | -596 | -605 | 1 | 0.00078661 | 10 | NaN | NaN | NaN | 281.39468 | NaN | NaN |
| Snrpe | VLDDAEEIHSK | -57 | -67 | 0 | 0 | 11 | NaN | NaN | NaN | 4993.556 | 4908.5513 | 5081.9263 |
| Snrpe | LKGDNITLLQSVSN | -79 | -92 | 1 | 0.004412728 | 14 | NaN | NaN | NaN | 1956.1819 | NaN | NaN |
| Cd3d | VVSSVQVHYR | -82 | -91 | 0 | 0.00111006 | 10 | NaN | NaN | NaN | 793.5634 | NaN | NaN |
| Gimap1 | LLAGLWGWR | -233 | -241 | 0 | 0.00122419 | 9 | NaN | NaN | NaN | 252.62735 | NaN | NaN |
| Psma4 | QLYQSDPSGNYGGWK | -146 | -160 | 0 | 0 | 15 | NaN | NaN | NaN | 544.8485 | 487.03778 | NaN |
| Alg8 | WSSFSVVR | -230 | -237 | 0 | 0.001868731 | 8 | NaN | NaN | NaN | 284.02203 | 335.07217 | NaN |
| Ap3b1 | WLIGENCER | -502 | -510 | 0 | 0.000471409 | 9 | NaN | NaN | NaN | 438.5395 | 517.58875 | NaN |
| Ktn1 | LLEEQLQHEVASK | -687 | -699 | 0 | 0.00111006 | 13 | NaN | NaN | NaN | 928.94946 | NaN | NaN |
| Ppp1ca | LLEVQGSRPGK | -16 | -26 | 1 | 0 | 11 | NaN | NaN | NaN | 471.438 | 788.9957 | 964.3469 |
| Ndufa13 | VGESVFHTTR | -106 | -115 | 0 | 0 | 10 | NaN | NaN | NaN | 515.1846 | NaN | 732.9336 |
| Carnmt1 | VNILVPGAGLGR | -193 | -204 | 0 | 0.000431655 | 12 | NaN | NaN | NaN | 478.96213 | NaN | NaN |

| | | | | | | | | | | | | |
|---|---|---|---|---|---|---|---|---|---|---|---|---|
| Focad | TVAGIPNFILK | -806 | -816 | 0 | 0 | 11 | NaN | NaN | NaN | 568.22437 | 520.93726 | 427.4261 |
| Focad | LLSLTEGQK | -335 | -343 | 0 | 0.000550307 | 9 | NaN | NaN | NaN | 659.05164 | NaN | NaN |
| Cdk9 | VDKYELFEK | -256 | -264 | 1 | 0 | 9 | NaN | NaN | NaN | 2574.3699 | NaN | NaN |
| Osbpl8 | VKFEEQDDFESEK | -673 | -685 | 1 | 0 | 13 | NaN | NaN | NaN | 1254.6532 | 1209.237 | 1193.319 |
| Osbpl8 | LNACEIIERPSK | -193 | -204 | 1 | 0.000230707 | 12 | NaN | NaN | NaN | 1031.0159 | 541.5495 | 582.72906 |
| Osbpl8 | SLSAILEGEAR | -546 | -556 | 0 | 0.001610812 | 11 | NaN | NaN | NaN | 2624.5806 | NaN | NaN |
| Sco2 | LNPDGLFTDYYGR | -220 | -232 | 0 | 0 | 13 | NaN | NaN | NaN | 579.05804 | NaN | 685.12067 |
| Ablim1 | LNSGLGQLILK | -694 | -704 | 0 | 0 | 11 | NaN | NaN | NaN | 688.71857 | 422.19662 | NaN |
| Gga1 | LPEDAIFPLPPPRPK | -151 | -165 | 1 | 0.00025015 | 15 | NaN | NaN | NaN | 502.56854 | NaN | NaN |
| Cnot8 | LPEEEHEFFHIL | -174 | -185 | 0 | 0 | 12 | NaN | NaN | NaN | 443.6915 | 334.53656 | 494.96292 |
| Uba3 | VIGAGGLGCELLK | -74 | -86 | 0 | 0 | 13 | NaN | NaN | NaN | 685.40283 | 899.4012 | 899.7659 |
| Scly | LPLEHLVENQ | -138 | -147 | 0 | 0.00201047 | 10 | NaN | NaN | NaN | 727.7302 | NaN | 755.7928 |
| Cpm | LPLFWNDNK | -293 | -301 | 0 | 0.001055699 | 9 | NaN | NaN | NaN | 1065.6908 | NaN | 486.71423 |
| Blvrb | LQDVTDDHIR | -125 | -134 | 0 | 0 | 10 | NaN | NaN | NaN | 1951.907 | 2657.427 | NaN |
| Ccdc124 | LQEEGSVEAR | -131 | -140 | 0 | 0.000230707 | 10 | NaN | NaN | NaN | 1713.2141 | NaN | NaN |
| Eps15l1 | LSQLQESHLEAH | -545 | -556 | 0 | 0.001392869 | 12 | NaN | NaN | NaN | 482.30264 | NaN | NaN |
| Clns1a | LQQPDTEAVLNGK | -18 | -30 | 0 | 0 | 13 | NaN | NaN | NaN | 1268.2097 | NaN | 673.22015 |
| Tyms | LRDEFPLLTTK | -61 | -71 | 1 | 0.007224385 | 11 | NaN | NaN | NaN | 309.80505 | NaN | NaN |
| Pus1 | LSAETLQQVNR | -220 | -230 | 0 | 0 | 11 | NaN | NaN | NaN | 9782.16 | NaN | NaN |
| Skic3 | LSDGASSEEAVR | -363 | -374 | 0 | 0.003998192 | 12 | NaN | NaN | NaN | 908.49475 | NaN | NaN |
| Hexb | LSHVYTPNDVR | -241 | -251 | 0 | 0.00111006 | 11 | NaN | NaN | NaN | 1116.3375 | NaN | NaN |
| Ints9 | LSNLPGWSLK | -49 | -58 | 0 | 0 | 10 | NaN | NaN | NaN | 1514.0809 | 1187.7028 | 1825.1051 |
| Taf6 | LSNTVLPSESMKVV | -9 | -22 | 1 | 0.009416377 | 14 | NaN | NaN | NaN | 404.5708 | NaN | NaN |
| Nqo1 | LSPDIVAEHK | -81 | -90 | 0 | 0.000431655 | 10 | NaN | NaN | NaN | 802.22363 | NaN | NaN |
| Mia2 | LSPPTLLEGPLR | -1130 | -1141 | 0 | 0.000550307 | 12 | NaN | NaN | NaN | 193.96063 | 365.91046 | 322.9132 |
| Eif2b2 | SELYHPDDHVL | -341 | -351 | 0 | 0.00051306 | 11 | NaN | NaN | NaN | 1359.632 | NaN | NaN |
| Mad1l1 | LSQALEELGVQK | -456 | -467 | 0 | 0.00111006 | 12 | NaN | NaN | NaN | 436.61542 | NaN | NaN |
| Sgpp1 | LSSIPCHDIR | -383 | -392 | 0 | 0 | 10 | NaN | NaN | NaN | 2431.2 | NaN | NaN |
| Mrpl21 | LSSPPWQEVVLPDPVEETR | -60 | -78 | 0 | 0 | 19 | NaN | NaN | NaN | 231.97995 | 211.72404 | 264.9847 |
| Lrrc59 | STLPSDFCGLTHLVK | -52 | -66 | 0 | 0 | 15 | NaN | NaN | NaN | 1004.28906 | 1162.6996 | 1050.0974 |
| Lrrc59 | LVTLPVSFAQLK | -97 | -108 | 0 | 0 | 12 | NaN | NaN | NaN | 420.70602 | 418.7573 | 358.31943 |
| Lrrc59 | LSTLPSDFCGLTHLVK | -51 | -66 | 0 | 0 | 16 | NaN | NaN | NaN | 330.38068 | 452.50964 | 524.5227 |
| Lrrc59 | QHLDLLNNR | -88 | -96 | 0 | 0.00025015 | 9 | NaN | NaN | NaN | 644.1196 | NaN | NaN |
| Tmem33 | SGQGSLLQPFIYYR | -177 | -190 | 0 | 0 | 14 | NaN | NaN | NaN | 401.40128 | 437.18875 | NaN |
| Tbrg4 | LSYLLSEKPK | -105 | -114 | 1 | 0.00111006 | 10 | NaN | NaN | NaN | 1265.6268 | NaN | NaN |
| Rif1 | SEPYSESTENSQLNVK | -976 | -991 | 0 | 0.005884925 | 16 | NaN | NaN | NaN | 950.4341 | 663.181 | NaN |
| Rif1 | VYPEELKPILR | -943 | -953 | 1 | 0.000394361 | 11 | NaN | NaN | NaN | 233.08487 | 639.83417 | NaN |
| Rif1 | SLEDLTTEEK | -1351 | -1360 | 0 | 0 | 10 | NaN | NaN | NaN | 1976.5548 | NaN | NaN |
| Ergic1 | VVNELYVDDPDKDSGGK | -52 | -68 | 1 | 0.003490217 | 17 | NaN | NaN | NaN | 477.36975 | 468.67914 | NaN |
| Slc25a17 | LTTPLWVVNTR | -118 | -128 | 0 | 0.000712251 | 11 | NaN | NaN | NaN | 6081.7646 | NaN | NaN |
| Ctdspl2 | LVELNEDVRPHIR | -441 | -453 | 1 | 0.002566359 | 13 | NaN | NaN | NaN | 484.57132 | 210.32356 | 143.2851 |
| Ighm | SILEGSDEYLVCK | -77 | -89 | 0 | 0 | 13 | NaN | NaN | NaN | 686.5572 | 1018.9069 | NaN |
| Chaf1a | LVGGQGPIDSFLR | -86 | -98 | 0 | 0 | 13 | NaN | NaN | NaN | 746.757 | 1282.2517 | 822.9049 |
| Kif23 | LVQAPLDADGDNVLQEK | -309 | -325 | 0 | 0 | 17 | NaN | NaN | NaN | 676.30054 | 650.1947 | 859.1971 |
| Kif23 | VAATQLEMQNKL | -634 | -645 | 1 | 0.007880195 | 12 | NaN | NaN | NaN | 762.85455 | NaN | NaN |
| Comp | LWHTGDTASQVR | -652 | -663 | 0 | 0 | 12 | NaN | NaN | NaN | 849.8428 | 926.9293 | NaN |
| Rab4b | MGSGIQYGDISLR | -183 | -195 | 0 | 0.00078661 | 13 | NaN | NaN | NaN | 760.21326 | NaN | NaN |
| Lyar | NCECLSCIDCGK | -27 | -38 | 0 | 0 | 12 | NaN | NaN | NaN | 767.1518 | 651.57117 | 602.6548 |
| Rbp4 | MVGTFTDTEDPAK | -91 | -103 | 0 | 0.001392869 | 13 | NaN | NaN | NaN | 487.24872 | NaN | NaN |
| Sf3b6 | NACDHLSGFNVCNR | -72 | -85 | 0 | 0.00111006 | 14 | NaN | NaN | NaN | 3675.2773 | 2983.5369 | 1064.3035 |
| Hmgcl | NANCSIEESFQR | -138 | -149 | 0 | 0.00025015 | 12 | NaN | NaN | NaN | 2401.4531 | NaN | NaN |
| Timm13 | NAQELLQR | -34 | -41 | 0 | 0.001055699 | 8 | NaN | NaN | NaN | 535.0269 | NaN | NaN |
| Itgb3 | VTFDCDCACQAFAQPSSPR | -454 | -472 | 0 | 0 | 19 | NaN | NaN | NaN | 234.59135 | 337.92355 | 308.95334 |
| Itgb3 | NDASHLLVF | -265 | -273 | 0 | 0.001811524 | 9 | NaN | NaN | NaN | 463.09473 | NaN | NaN |
| Rel | NDPYKPHPHDLVGK | -69 | -82 | 1 | 0.00111006 | 14 | NaN | NaN | NaN | 419.5304 | 864.8737 | 514.08136 |
| Psmd9 | NEPLVDCEGYPR | -53 | -64 | 0 | 0.000230707 | 12 | NaN | NaN | NaN | 3106.1548 | 760.0382 | NaN |
| Psmd9 | YDVLESQK | -41 | -48 | 0 | 0.001055699 | 8 | NaN | NaN | NaN | 635.55164 | NaN | 867.0312 |
| Wdr33 | NFETILQAHDSPVR | -152 | -165 | 0 | 0 | 14 | NaN | NaN | NaN | 1195.3522 | NaN | 1175.2101 |
| Atp6v1b2 | YAEIVHL | -68 | -74 | 0 | 0.002502116 | 7 | NaN | NaN | NaN | 603.4084 | NaN | NaN |
| Ssr3 | NFNPTVNYIL | -159 | -168 | 0 | 0.000712251 | 10 | NaN | NaN | NaN | 456.32858 | 468.536 | 487.2427 |
| Nup214 | NHDSGYLHLL | -783 | -792 | 0 | 0.001610812 | 10 | NaN | NaN | NaN | 609.13855 | NaN | NaN |
| Rab8b | TIELDGKK | -49 | -56 | 1 | 0.001023236 | 8 | NaN | NaN | NaN | 879.61316 | NaN | NaN |
| Wdr37 | NINIPALLQEK | -486 | -496 | 0 | 0 | 11 | NaN | NaN | NaN | 430.82114 | 553.55756 | 527.9508 |
| Vps36 | SLTEVYCLVNR | -265 | -275 | 0 | 0.002099344 | 11 | NaN | NaN | NaN | 290.59653 | NaN | NaN |
| Recql | NKEDFPWFGK | -69 | -78 | 1 | 0 | 10 | NaN | NaN | NaN | 410.81198 | 310.79022 | NaN |
| Tmprss13 | NKPGVYTK | -515 | -522 | 1 | 0 | 8 | NaN | NaN | NaN | 5770.3286 | 5185.2764 | 5649.4673 |
| Psmf1 | NLDDYIDAEDLSDFHR | -107 | -122 | 0 | 0 | 16 | NaN | NaN | NaN | 554.4973 | 717.10834 | 649.1839 |
| Ctsl | NLDHGVLLVGY | -273 | -283 | 0 | 0.001291103 | 11 | NaN | NaN | NaN | 799.88727 | NaN | NaN |
| Exosc2 | NLEPVALSDR | -222 | -231 | 0 | 0 | 10 | NaN | NaN | NaN | 2592.959 | NaN | NaN |
| Trim30a | NLHVANIVERLK | -70 | -81 | 1 | 0.001636916 | 12 | NaN | NaN | NaN | 959.0716 | NaN | NaN |
| Etfdh | VDHTVGWPL | -290 | -298 | 0 | 0.002972566 | 9 | NaN | NaN | NaN | 438.98776 | NaN | NaN |
| Tk1 | NLVPLAESVVK | -138 | -148 | 0 | 0.001392869 | 11 | NaN | NaN | NaN | 841.23346 | NaN | NaN |
| Ankfy1 | NNLQIIPDFSLK | -574 | -585 | 0 | 0.00025015 | 12 | NaN | NaN | NaN | 408.88025 | 464.8905 | NaN |
| Itpr2 | NNPALFWFSR | -2215 | -2224 | 0 | 0.009745776 | 10 | NaN | NaN | NaN | 727.1497 | NaN | NaN |
| Rpia | NNQVLGIGSGSTIVHAVQR | -89 | -107 | 0 | 0 | 19 | NaN | NaN | NaN | 635.3362 | 1598.7925 | 1082.0399 |
| Uqcc1 | NPQSILKPHAPTYNDEGL | -278 | -295 | 1 | 0.00025015 | 18 | NaN | NaN | NaN | 279.601 | 581.8129 | 517.58875 |
| Arcn1 | TTFLVDKYEIL | -501 | -511 | 1 | 0 | 11 | NaN | NaN | NaN | 888.3141 | 255.11533 | 1252.8914 |
| Arcn1 | RNTLEWCLPVIDAK | -435 | -448 | 1 | 0.00670918 | 14 | NaN | NaN | NaN | 535.86755 | 587.9539 | NaN |

| | | | | | | | | | | | | |
|---|---|---|---|---|---|---|---|---|---|---|---|---|
| Lmna | NSNLVGAAHEELQQSR | -281 | -296 | 0 | 0 | 16 | NaN | NaN | NaN | 643.8556 | 622.3627 | NaN |
| Ssbp3 | NSPNNISGISNPPGTPR | -346 | -362 | 0 | 0.002099344 | 17 | NaN | NaN | NaN | 647.3442 | NaN | 717.46466 |
| Hmg20a | NTVLQQHLETLR | -277 | -288 | 0 | 0 | 12 | NaN | NaN | NaN | 1270.1348 | 1459.3655 | 1498.3839 |
| Golga2 | NVDDSASEESNQALHVA | -785 | -801 | 0 | 0.005605646 | 17 | NaN | NaN | NaN | 683.9811 | NaN | NaN |
| Dnajc8 | QREEEIEAQ | -191 | -199 | 1 | 0.002434616 | 9 | NaN | NaN | NaN | 8699.11 | NaN | NaN |
| Pgam5 | NVESGEDELTSR | -76 | -87 | 0 | 0 | 12 | NaN | NaN | NaN | 3885.2617 | NaN | NaN |
| Anapc1 | NVESHLLNK | -718 | -726 | 0 | 0.00111006 | 9 | NaN | NaN | NaN | 682.4924 | NaN | NaN |
| Anks1a | NVIAEHEIR | -1015 | -1023 | 0 | 0 | 9 | NaN | NaN | NaN | 391.41208 | NaN | NaN |
| Pigk | NVLITDFFGSVR | -297 | -308 | 0 | 0 | 12 | NaN | NaN | NaN | 762.3685 | 1049.9092 | 1158.831 |
| Unc45a | NWFEGHGLAGK | -401 | -411 | 0 | 0 | 11 | NaN | NaN | NaN | 929.43335 | NaN | 1907.2205 |
| Nit1 | NYTKPGGTLEPPVK | -175 | -188 | 1 | 0.006472924 | 14 | NaN | NaN | NaN | 2142.3267 | 1180.2953 | NaN |
| Srp9 | PQFQTWEEFSR | -2 | -12 | 0 | 0 | 11 | NaN | NaN | NaN | 1577.3354 | NaN | NaN |
| Kif2a | PQRDDLK | -586 | -592 | 1 | 0.008653659 | 7 | NaN | NaN | NaN | 3209.0742 | NaN | NaN |
| Apoa1 | PYLDEFQK | -122 | -129 | 0 | 0 | 8 | NaN | NaN | NaN | 618.9172 | 785.2906 | 774.87476 |
| Tmco1 | QAGGFLGPPPPSGK | -173 | -186 | 0 | 0.00167576 | 14 | NaN | NaN | NaN | 4432.8257 | NaN | 3586.6917 |
| Iap | QCADSFIPR | -200 | -208 | 0 | 0.004595494 | 9 | NaN | NaN | NaN | 610.84894 | NaN | NaN |
| Traf3ip3 | SSETQDLQDQLK | -390 | -401 | 0 | 0 | 12 | NaN | NaN | NaN | 670.3892 | NaN | NaN |
| Ccnb1 | QLEEEQSVRPK | -177 | -187 | 1 | 0 | 11 | NaN | NaN | NaN | 456.40546 | 230.74957 | 970.31726 |
| Itsn1 | QLELEKQL | -405 | -412 | 1 | 0.001055699 | 8 | NaN | NaN | NaN | 825.51556 | 823.8952 | 905.0013 |
| Aup1 | QLGEESEEFALR | -232 | -243 | 0 | 0.001055699 | 12 | NaN | NaN | NaN | 781.484 | NaN | NaN |
| Dffb | QLLSDEQAPLR | -108 | -118 | 0 | 0.000550307 | 11 | NaN | NaN | NaN | 7652.5264 | NaN | NaN |
| Arpp19 | YPHLGQK | -36 | -42 | 0 | 0.007546675 | 7 | NaN | NaN | NaN | 227.14255 | NaN | 1339.528 |
| Apool | QLVRPDQLPIYTAPPLHSK | -34 | -52 | 1 | 0 | 19 | NaN | NaN | NaN | 1206.4861 | NaN | 527.83905 |
| Ctc1 | QQFQCGELPLLTR | -1172 | -1184 | 0 | 0.005363933 | 13 | NaN | NaN | NaN | 17090.795 | 15059.257 | 13993.656 |
| Rassf2 | QRLEEIAETPETI | -314 | -326 | 1 | 0.004133093 | 13 | NaN | NaN | NaN | 266.88 | 469.0799 | NaN |
| Aspscr1 | QSTSEPASAPFVPFSGGGQR | -247 | -266 | 0 | 0.000230707 | 20 | NaN | NaN | NaN | 510.17804 | NaN | NaN |
| Ubap2 | TTEDWTEDLSETK | -259 | -271 | 0 | 0 | 13 | NaN | NaN | NaN | 1113.5682 | NaN | NaN |
| Ubap2 | SQPEPSPVLSQLSQR | -428 | -442 | 0 | 0.000230707 | 15 | NaN | NaN | NaN | 651.0117 | NaN | NaN |
| Imp3 | RDLPERDPFR | -63 | -72 | 2 | 0.000627943 | 10 | NaN | NaN | NaN | 1600.9456 | 1009.42596 | NaN |
| Tma16 | REWDFDLK | -144 | -151 | 1 | 0.00025015 | 8 | NaN | NaN | NaN | 1758.5071 | 1914.9768 | 1957.1243 |
| Hells | TVVEGNIPIESEVNLK | -527 | -542 | 0 | 0 | 16 | NaN | NaN | NaN | 622.0688 | NaN | 400.519 |
| Hells | RFTPEIPTLL | -275 | -284 | 1 | 0.002068213 | 10 | NaN | NaN | NaN | 409.24817 | NaN | 480.05237 |
| Hells | SLCVEDIQK | -150 | -158 | 0 | 0.002566359 | 9 | NaN | NaN | NaN | 753.9407 | NaN | NaN |
| Tm9sf1 | WSETSVEHR | -208 | -216 | 0 | 0.00644558 | 9 | NaN | NaN | NaN | 1752.9824 | NaN | NaN |
| Epb41 | RGEEPAEPAEPEPTEAWK | -592 | -609 | 1 | 0.001392869 | 18 | NaN | NaN | NaN | 789.77094 | NaN | NaN |
| Mat2b | RIPVLYGEVEK | -181 | -191 | 1 | 0.000431655 | 11 | NaN | NaN | NaN | 4418.3774 | 5875.376 | 3726.08 |
| Mrpl19 | RLDDNLLYLR | -182 | -191 | 1 | 0.001636916 | 10 | NaN | NaN | NaN | 1059.3123 | NaN | NaN |
| Ythdf1 | RQEEEEVVR | -542 | -550 | 1 | 0.000627943 | 9 | NaN | NaN | NaN | 817.17773 | NaN | NaN |
| Ythdf1 | RLDGAFR | -420 | -426 | 1 | 0.000230707 | 7 | NaN | NaN | NaN | 735.96216 | NaN | NaN |
| Mtif2 | RPGTDPALLKPR | -168 | -179 | 2 | 0.00204437 | 12 | NaN | NaN | NaN | 182.77583 | NaN | NaN |
| Prrc2a | RPPTAPENTPSVPSGVK | -124 | -140 | 1 | 0.000627943 | 17 | NaN | NaN | NaN | 591.6738 | 580.7407 | 568.98315 |
| Coro2a | RSESYQEDIYPPTAA | -360 | -374 | 1 | 0.004224523 | 15 | NaN | NaN | NaN | 447.11896 | 357.73138 | NaN |
| Nkiras2 | RVDPDVAQHW | -129 | -138 | 1 | 0.001023236 | 10 | NaN | NaN | NaN | 2697.3843 | NaN | NaN |
| Arglu1 | RVEEELEKR | -139 | -147 | 2 | 0.000471409 | 9 | NaN | NaN | NaN | 896.89026 | NaN | NaN |
| Gpd2 | RVPIPVDR | -715 | -722 | 1 | 0.001708008 | 8 | NaN | NaN | NaN | 215.05307 | 336.7886 | 398.6476 |
| Mtco1 | SATMIIAIPTGVK | -307 | -319 | 0 | 0.001636916 | 13 | NaN | NaN | NaN | 569.6526 | 390.2154 | NaN |
| Impa2 | SAADLVTETDHR | -51 | -62 | 0 | 0 | 12 | NaN | NaN | NaN | 314.17703 | 664.9275 | NaN |
| Rrp1b | SDTGSFSQEELLK | -41 | -53 | 0 | 0 | 13 | NaN | NaN | NaN | 504.6536 | NaN | NaN |
| Mrpl58 | SADWIEEPVR | -107 | -116 | 0 | 0.001023236 | 10 | NaN | NaN | NaN | 586.3821 | NaN | NaN |
| Nelfb | SALPFLDLHGTPR | -46 | -58 | 0 | 0 | 13 | NaN | NaN | NaN | 2639.784 | NaN | NaN |
| M6pr | SNDKETVVGR | -97 | -106 | 1 | 0 | 10 | NaN | NaN | NaN | 646.59314 | NaN | 727.79114 |
| M6pr | SCDLVGEKDK | -32 | -41 | 1 | 0.000627943 | 10 | NaN | NaN | NaN | 366.80246 | 491.58536 | NaN |
| Niban1 | SCPGPIEEPFEAQEPAEK | -801 | -818 | 0 | 0 | 18 | NaN | NaN | NaN | 514.4275 | 527.1721 | NaN |
| Ascc3 | SDAEEYTDLPVR | -1851 | -1862 | 0 | 0 | 12 | NaN | NaN | NaN | 2921.761 | 2676.0571 | 2588.8352 |
| Mapk1ip1l | SDEFSLADALPEQSSAKPPAVTNTK | -2 | -26 | 1 | 0.000550307 | 25 | NaN | NaN | NaN | 306.73672 | 751.4559 | 553.2371 |
| Ddx31 | SDGPYALVLVPTR | -176 | -188 | 0 | 0 | 13 | NaN | NaN | NaN | 2926.1108 | 3017.5237 | 3180.4417 |
| Rbm28 | VVLHPDTEHSK | -356 | -366 | 0 | 0 | 11 | NaN | NaN | NaN | 319.46478 | NaN | 665.4937 |
| Gins4 | SEGEPSSLSPEEFVFAK | -115 | -131 | 0 | 0 | 17 | NaN | NaN | NaN | 2245.631 | 2493.1848 | NaN |
| Pik3r1 | VLSEIFSPVLFR | -263 | -274 | 0 | 0.007224385 | 12 | NaN | NaN | NaN | 376.67322 | NaN | NaN |
| Copg2 | SEITLVTPKPEK | -591 | -602 | 1 | 0 | 12 | NaN | NaN | NaN | 825.90204 | 453.6181 | NaN |
| Cdca7 | SELESFPGLFSGR | -140 | -152 | 0 | 0 | 13 | NaN | NaN | NaN | 1582.9987 | NaN | NaN |
| Ikzf1 | YEFSSHITR | -501 | -509 | 0 | 0.000431655 | 9 | NaN | NaN | NaN | 1715.9838 | NaN | 1754.5166 |
| Ggps1 | SFCEDLTEGK | -203 | -212 | 0 | 0 | 10 | NaN | NaN | NaN | 457.13596 | 540.57245 | 647.2765 |
| Kras | SFEDIHHYR | -89 | -97 | 0 | 0 | 9 | NaN | NaN | NaN | 1186.3793 | 1701.3 | NaN |
| Ugp2 | SFENSLGINVPR | -378 | -389 | 0 | 0 | 12 | NaN | NaN | NaN | 1260.6743 | 1321.5475 | NaN |
| Ugp2 | SFNTDEDTKK | -164 | -173 | 1 | 0.000230707 | 10 | NaN | NaN | NaN | 1284.2931 | NaN | NaN |
| Tor1aip2 | SFPLEWAPQNLTR | -67 | -79 | 0 | 0 | 13 | NaN | NaN | NaN | 147.84988 | 523.1422 | NaN |
| Gid8 | SYAEKPDEITK | -2 | -12 | 1 | 0.00025015 | 11 | NaN | NaN | NaN | 2973.5808 | 1499.0077 | 1511.9587 |
| Gid8 | SLHPELLDTNR | -88 | -98 | 0 | 0 | 11 | NaN | NaN | NaN | 2930.6155 | NaN | NaN |
| Strn | SVIDTSTIVR | -259 | -268 | 0 | 0.003102268 | 10 | NaN | NaN | NaN | 2787.6133 | 4433.486 | NaN |
| Strn | SHFDGIR | -461 | -467 | 0 | 0.000550307 | 7 | NaN | NaN | NaN | 2855.6975 | NaN | NaN |
| Atrip | SHFLLEQEK | -162 | -170 | 0 | 0.00111006 | 9 | NaN | NaN | NaN | 1211.7642 | NaN | NaN |
| Ube2z | SHWDPTLSSDWDGER | -83 | -97 | 0 | 0.000755556 | 15 | NaN | NaN | NaN | 167.6874 | NaN | NaN |
| Bloc1s6 | SHYLPDLQR | -58 | -66 | 0 | 0.000431655 | 9 | NaN | NaN | NaN | 3895.2676 | NaN | NaN |
| Ciao1 | SIAFDPSGQR | -199 | -208 | 0 | 0.000431655 | 10 | NaN | NaN | NaN | 307.7636 | NaN | NaN |
| Naxe | SIDIPSGWDVEK | -210 | -221 | 0 | 0 | 12 | NaN | NaN | NaN | 1027.3701 | NaN | NaN |
| Nisch | SIGSLPCLER | -399 | -408 | 0 | 0.003133457 | 10 | NaN | NaN | NaN | 829.6093 | NaN | NaN |
| Scrn2 | SPQVLSPTFGAQDPVR | -328 | -343 | 0 | 0.00025015 | 16 | NaN | NaN | NaN | 708.9844 | 992.23175 | 841.23456 |

| | | | | | | | | | | | | |
|---|---|---|---|---|---|---|---|---|---|---|---|---|
| Edc3 | SINCGDSLGVYQGR | -11 | -24 | 0 | 0 | 14 | NaN | NaN | NaN | 2518.4104 | NaN | NaN |
| Ciao2b | SINDPEHPLTLEEL | -52 | -65 | 0 | 0.000394361 | 14 | NaN | NaN | NaN | 1231.7677 | NaN | 1467.8484 |
| Nkap | SKAEEPSDLIGPEAPK | -282 | -297 | 1 | 0 | 16 | NaN | NaN | NaN | 3498.7026 | NaN | NaN |
| Grhpr | STVGIVGLGR | -154 | -163 | 0 | 0.000712251 | 10 | NaN | NaN | NaN | 523.839 | NaN | NaN |
| Ankrd17 | SLAASGGYVNIIK | -1184 | -1196 | 0 | 0.001904399 | 13 | NaN | NaN | NaN | 473.41815 | NaN | NaN |
| Cpsf2 | SLCHGLSDLAR | -309 | -319 | 0 | 0 | 11 | NaN | NaN | NaN | 473.545 | 254.15155 | 1849.7639 |
| Ifnar2 | SLEEDTILLDEGPQR | -448 | -462 | 0 | 0.000550307 | 15 | NaN | NaN | NaN | 541.73047 | NaN | NaN |
| Eif2b5 | SLFQGSGDGVEIR | -225 | -237 | 0 | 0 | 13 | NaN | NaN | NaN | 543.66785 | NaN | 738.0644 |
| Eif2b5 | TDNFDYQTR | -257 | -265 | 0 | 0.000471409 | 9 | NaN | NaN | NaN | 479.85754 | NaN | NaN |
| Cytip | SLGDFSWSQR | -66 | -75 | 0 | 0.00025015 | 10 | NaN | NaN | NaN | 901.92737 | NaN | 1224.2439 |
| Atrx | WAEEFNDETNVR | -2081 | -2092 | 0 | 0 | 12 | NaN | NaN | NaN | 303.70905 | NaN | 625.59937 |
| Rbms1 | SNSGWDQLSK | -52 | -61 | 0 | 0 | 10 | NaN | NaN | NaN | 891.4949 | 1088.938 | NaN |
| Atf1 | SNTTETASQPGSTVAGPHVS | -7 | -26 | 0 | 0.00078661 | 20 | NaN | NaN | NaN | 636.8255 | NaN | 480.90726 |
| Etv6 | WAENEFSLRPIESNK | -73 | -87 | 1 | 0 | 15 | NaN | NaN | NaN | 1289.9761 | 1643.787 | 1507.9805 |
| Etv6 | SPHSGDVLYELL | -107 | -118 | 0 | 0.00149795 | 12 | NaN | NaN | NaN | 1042.8855 | NaN | NaN |
| Tmem214 | SQSVFTGNPSVWLK | -130 | -143 | 0 | 0 | 14 | NaN | NaN | NaN | 1456.1692 | NaN | NaN |
| Htra2 | SQYNFIADVVEK | -145 | -156 | 0 | 0 | 12 | NaN | NaN | NaN | 5466.7534 | NaN | NaN |
| Brd4 | SREDEDALEQAR | -1298 | -1309 | 1 | 0 | 12 | NaN | NaN | NaN | 355.27585 | NaN | 649.31055 |
| Pcbd2 | SSDAQWLTAEER | -35 | -46 | 0 | 0.001636916 | 12 | NaN | NaN | NaN | 720.463 | NaN | 607.30334 |
| Cwc27 | SSHDLLKDDPHLS | -231 | -243 | 1 | 0 | 13 | NaN | NaN | NaN | 997.23303 | NaN | 4620.9297 |
| Cwc27 | VEEGREEEEAAPDGAVA | -339 | -355 | 1 | 0.003745449 | 17 | NaN | NaN | NaN | 703.8112 | NaN | NaN |
| Dus3l | SSPVPTCGPLTDEDVIR | -263 | -279 | 0 | 0.007343498 | 17 | NaN | NaN | NaN | 862.2939 | NaN | NaN |
| Tgfb1 | SSVEQHVELYQK | -162 | -173 | 0 | 0.00149795 | 12 | NaN | NaN | NaN | 498.27396 | NaN | 273.9819 |
| Pogz | STDPLPVFLYPPVQR | -717 | -731 | 0 | 0.00383302 | 15 | NaN | NaN | NaN | 204.66039 | 251.379 | NaN |
| Pogz | VPEPNENAGDAVQTK | -447 | -461 | 0 | 0.008222027 | 15 | NaN | NaN | NaN | 655.6906 | NaN | NaN |
| Gtse1 | STNEDDEVFFGPVGHK | -28 | -43 | 0 | 0.003133457 | 16 | NaN | NaN | NaN | 983.95215 | NaN | NaN |
| Pom121 | STTPTFKPIFDK | -738 | -749 | 1 | 0.000712251 | 12 | NaN | NaN | NaN | 3794.6484 | NaN | NaN |
| Actr10 | SVLCPSHFR | -94 | -102 | 0 | 0.000471409 | 9 | NaN | NaN | NaN | 705.70483 | NaN | NaN |
| Ufm1 | SVPESTPFTAVLK | -22 | -34 | 0 | 0.002168954 | 13 | NaN | NaN | NaN | 1211.9163 | NaN | NaN |
| Ighv3-6; | SVTTEDTATYYCAR | (103);(103) | (116);(116) | 0 | 0.000627943 | 14 | NaN | NaN | NaN | 253.90222 | NaN | NaN |
| Abhd16a | SWDTYYQPR | -43 | -51 | 0 | 0.001392869 | 9 | NaN | NaN | NaN | 3319.9954 | NaN | NaN |
| Sumf2 | VFEDFVSPELR | -103 | -113 | 0 | 0 | 11 | NaN | NaN | NaN | 2231.8076 | 2476.0386 | 2464.9587 |
| Sumf2 | SWIDTADGSANHR | -263 | -275 | 0 | 0 | 13 | NaN | NaN | NaN | 656.046 | 468.3238 | 428.59256 |
| Vps33b | SWQGLDEVVR | -533 | -542 | 0 | 0 | 10 | NaN | NaN | NaN | 3907.7368 | 1362.8715 | 1606.8799 |
| Golm1 | SYSSLQQDIFQFQK | -130 | -143 | 0 | 0.00272025 | 14 | NaN | NaN | NaN | 922.85785 | NaN | 1094.3448 |
| Ptms | TAEEEDEADPK | -81 | -91 | 0 | 0.00142999 | 11 | NaN | NaN | NaN | 392.45932 | NaN | NaN |
| Magt1 | TALQLHR | -79 | -85 | 0 | 0.001392869 | 7 | NaN | NaN | NaN | 200.07422 | NaN | NaN |
| Dap3 | TAYLPHELLGK | -314 | -324 | 0 | 0 | 11 | NaN | NaN | NaN | 1600.3177 | NaN | NaN |
| Ighg1 | TCNVAHPASSTK | (81);(81) | (92);(92) | 0 | 0 | 12 | NaN | NaN | NaN | 662.8006 | NaN | 518.3411 |
| Telo2 | TCVAEFEQLR | -568 | -577 | 0 | 0 | 10 | NaN | NaN | NaN | 178.65897 | NaN | NaN |
| Arfip1 | TDLEELNLGPR | -272 | -282 | 0 | 0 | 11 | NaN | NaN | NaN | 743.59375 | 776.7114 | 772.1653 |
| Ccdc134 | TDQTEFIPSTDPFQK | -184 | -198 | 0 | 0 | 15 | NaN | NaN | NaN | 422.6286 | 535.4904 | 679.19556 |
| Gpx4 | TDVNYTQLVDLHAR | -76 | -89 | 0 | 0 | 14 | NaN | NaN | NaN | 1047.1296 | NaN | NaN |
| Timm8b | TENCLSSCVDR | -52 | -62 | 0 | 0 | 11 | NaN | NaN | NaN | 663.00305 | 659.37146 | 729.60876 |
| Ermp1 | TFHNLEGSVVK | -688 | -698 | 0 | 0 | 11 | NaN | NaN | NaN | 779.2 | NaN | NaN |
| Tmx2 | TFSEENVIR | -245 | -253 | 0 | 0.000471409 | 9 | NaN | NaN | NaN | 1805.4344 | 1937.1456 | 1846.2162 |
| Pold3 | TFVDEEGCIVTEK | -387 | -399 | 0 | 0 | 13 | NaN | NaN | NaN | 580.77277 | NaN | NaN |
| Ptgr2 | TGNVAEQLR | -210 | -218 | 0 | 0.000230707 | 9 | NaN | NaN | NaN | 1474.4016 | NaN | NaN |
| Capn7 | TGPLLIELR | -724 | -732 | 0 | 0.001291103 | 9 | NaN | NaN | NaN | 5021.756 | NaN | NaN |
| Prf1 | TGVVWNNNNPR | -449 | -459 | 0 | 0.000230707 | 11 | NaN | NaN | NaN | 916.11615 | NaN | 1072.119 |
| Prf1 | VFFGGQEFR | -440 | -448 | 0 | 0 | 9 | NaN | NaN | NaN | 637.26276 | 941.20905 | NaN |
| Ghitm | VHSISYEQSPGPK | -178 | -190 | 0 | 0 | 13 | NaN | NaN | NaN | 1356.3821 | NaN | NaN |
| Atg12 | TIQGLIDFIK | -81 | -90 | 0 | 0.001370415 | 10 | NaN | NaN | NaN | 336.42624 | 428.8808 | 430.74646 |
| Mapk14 | TLFPGTDHIDQLK | -221 | -233 | 0 | 0.00025015 | 13 | NaN | NaN | NaN | 952.15466 | 1912.8038 | NaN |
| Mapk14 | TIWEVPER | -16 | -23 | 0 | 0.00201047 | 8 | NaN | NaN | NaN | 434.2004 | NaN | 320.6303 |
| Smarcad1 | TKPADEQSIYEK | -689 | -700 | 1 | 0.001291103 | 12 | NaN | NaN | NaN | 1071.1105 | NaN | NaN |
| Dhrs7b | YFGPVALTK | -165 | -173 | 0 | 0.005605646 | 9 | NaN | NaN | NaN | 274.27832 | NaN | 349.29892 |
| Fgg | VAQLEAQCQEPCK | -153 | -165 | 0 | 0.00122419 | 13 | NaN | NaN | NaN | 496.0831 | 352.13477 | 575.92896 |
| Fgg | YLQEIYNSNNQK | -134 | -145 | 0 | 0 | 12 | NaN | NaN | NaN | 362.58185 | NaN | NaN |
| Rbm26 | VIQPLVQQPILPVVK | -624 | -638 | 0 | 0.001708008 | 15 | NaN | NaN | NaN | 210.56429 | 247.63019 | 209.02072 |
| Kpna1 | TNIASGNSLQTR | -152 | -163 | 0 | 0.002910424 | 12 | NaN | NaN | NaN | 1955.0299 | NaN | NaN |
| Exoc5 | TNLPLGPSIDTHGETF | -395 | -410 | 0 | 0.003299042 | 16 | NaN | NaN | NaN | 942.46277 | 1784.2908 | 1166.8677 |
| Igfbp2 | TPCQQELDQVLER | -205 | -217 | 0 | 0 | 13 | NaN | NaN | NaN | 1224.8096 | NaN | NaN |
| Eef1akmt4-Ec | TPVSPPELPEK | (6);(6) | (16);(16) | 0 | 0.00111006 | 11 | NaN | NaN | NaN | 1143.1947 | NaN | NaN |
| Eef1akmt1 | TSETIQFLTR | -153 | -162 | 0 | 0.00078661 | 10 | NaN | NaN | NaN | 4455.6553 | NaN | NaN |
| Riok1 | TSQDHVDEEVFKQ | -414 | -426 | 1 | 0.000431655 | 13 | NaN | NaN | NaN | 717.81055 | 327.34 | 536.388 |
| Papola | TSSTDLSDIPALPANPIPVIK | -710 | -730 | 0 | 0.001868731 | 21 | NaN | NaN | NaN | 574.6217 | 409.0872 | NaN |
| Bcl7b | TTEEEEDSGAPPLK | -173 | -186 | 0 | 0.005270854 | 14 | NaN | NaN | NaN | 2932.2815 | NaN | NaN |
| Cfap418 | TVEELTDLQTDHQLR | -189 | -203 | 0 | 0.00122419 | 15 | NaN | NaN | NaN | 1048.4277 | NaN | 1673.1472 |
| Rabepk | TWSQPETHGSPPSPR | -187 | -201 | 0 | 0.009388464 | 15 | NaN | NaN | NaN | 342.25018 | NaN | NaN |
| Asf1b | TYHGQEFIR | -100 | -108 | 0 | 0.002296977 | 9 | NaN | NaN | NaN | 17761.133 | NaN | NaN |
| Tmem14c | VATGGIIGYAK | -22 | -32 | 0 | 0.00122419 | 11 | NaN | NaN | NaN | 240.84282 | NaN | NaN |
| Cd6 | VCDSEWYPSEAK | -288 | -299 | 0 | 0 | 12 | NaN | NaN | NaN | 1009.6218 | 531.67847 | NaN |
| Emc7 | VDGEEHVGFLK | -68 | -78 | 0 | 0.000471409 | 11 | NaN | NaN | NaN | 470.93286 | NaN | NaN |
| Zcchc17 | VDKPSEIVDVGDK | -54 | -66 | 1 | 0.00111006 | 13 | NaN | NaN | NaN | 916.9078 | NaN | NaN |
| Rap1gds1 | VDQAGGAQIVIDHLR | -127 | -141 | 0 | 0 | 15 | NaN | NaN | NaN | 3206.0537 | 2135.667 | 2484.839 |
| Ppp1r9b | VFQPPPPPPAPSGDGATEK | -250 | -268 | 0 | 0.009745776 | 19 | NaN | NaN | NaN | 490.06433 | NaN | NaN |
| Ssbp1 | VGQDPVMR | -39 | -46 | 0 | 0 | 8 | NaN | NaN | NaN | 465.9947 | NaN | 818.1899 |

| | | | | | | | | | | | | |
|---|---|---|---|---|---|---|---|---|---|---|---|---|
| Syvn1 | VHTFPLFAIR | -237 | -246 | 0 | 0.002972566 | 10 | NaN | NaN | NaN | 482.08673 | 436.08725 | 361.88345 |
| Actg1 | VIDNGSGMCK | -9 | -18 | 0 | 0 | 10 | NaN | NaN | NaN | 229.39172 | 258.79672 | 300.68112 |
| Paip2 | VISDGSSLEDLVVK | -95 | -108 | 0 | 0.00025015 | 14 | NaN | NaN | NaN | 410.18292 | 733.01636 | NaN |
| Rnasek | VLIEDVPFTEK | -39 | -49 | 0 | 0 | 11 | NaN | NaN | NaN | 1068.3591 | NaN | NaN |
| Utp25 | WPVLGHL | -234 | -240 | 0 | 0.002972566 | 7 | NaN | NaN | NaN | 410.93655 | 410.6045 | NaN |
| Tsr3 | VLSPVGTEYVSPADR | -97 | -111 | 0 | 0.001636916 | 15 | NaN | NaN | NaN | 566.2462 | NaN | NaN |
| Rabl3 | VLVLGDSGVGK | -9 | -19 | 0 | 0.002595029 | 11 | NaN | NaN | NaN | 2263.0625 | NaN | NaN |
| Lamp2 | YWGIHLQ | -164 | -170 | 0 | 0.005789881 | 7 | NaN | NaN | NaN | 892.50134 | 825.86426 | NaN |
| Sp3 | VVDEEGDQQHQEGK | -582 | -595 | 0 | 0.007343498 | 14 | NaN | NaN | NaN | 431.5745 | NaN | NaN |
| Helb | VSIPDEYTK | -60 | -68 | 0 | 0.00078661 | 9 | NaN | NaN | NaN | 553.5117 | NaN | NaN |
| LRWD1 | VSPEEETDKVR | -156 | -166 | 1 | 0.00025015 | 11 | NaN | NaN | NaN | 2748.9785 | NaN | NaN |
| Itga4 | VTLTIPLR | -771 | -778 | 0 | 0.00437931 | 8 | NaN | NaN | NaN | 336.3209 | NaN | NaN |
| Ubr7 | VVCEDWFHGR | -156 | -165 | 0 | 0 | 10 | NaN | NaN | NaN | 1786.642 | 2493.919 | NaN |
| Prrc1 | VVEDPVHGIR | -348 | -357 | 0 | 0 | 10 | NaN | NaN | NaN | 421.90427 | 1123.8042 | 3363.5989 |
| Cmtm7 | VYPLTHGALFK | -31 | -41 | 0 | 0 | 11 | NaN | NaN | NaN | 957.1621 | NaN | NaN |
| FRRS1 | VYWDAPSPAPDHIR | -128 | -141 | 0 | 0 | 14 | NaN | NaN | NaN | 428.1512 | 782.9973 | NaN |
| Prpf39 | WENEQGNLR | -213 | -221 | 0 | 0.00025015 | 9 | NaN | NaN | NaN | 393.18744 | NaN | NaN |
| Clgn | YEVNFQDGIDCGGAYIK | -141 | -157 | 0 | 0.00025015 | 17 | NaN | NaN | NaN | 313.80164 | 727.1434 | 486.16275 |
| Ncbp3 | YHEELPGLK | -368 | -376 | 0 | 0.00025015 | 9 | NaN | NaN | NaN | 712.9743 | 1132.6887 | 371.12082 |
| Arl2bp | YIEEQLLER | -76 | -84 | 0 | 0 | 9 | NaN | NaN | NaN | 565.19196 | 306.9675 | NaN |
| Arfgap2 | YKDNPFSLGETF | -357 | -368 | 1 | 0 | 12 | NaN | NaN | NaN | 598.47284 | 397.7076 | 414.46964 |
| Stat4 | YLEDLQDEFDYR | -163 | -174 | 0 | 0 | 12 | NaN | NaN | NaN | 444.3496 | 570.57697 | 520.1275 |
| Virma | YLITFNPVGR | -799 | -808 | 0 | 0.008736733 | 10 | NaN | NaN | NaN | 596.4961 | NaN | NaN |
| Cwf19l1 | YVGSFGNSSGEVDTK | -160 | -174 | 0 | 0.000591124 | 15 | NaN | NaN | NaN | 707.4181 | NaN | NaN |
| Copb1 | FVLPLQDHTIK | -65 | -75 | 0 | 0 | 11 | NaN | NaN | NaN | NaN | 1781.5349 | 1783.0995 |
| Copb1 | EEDRPPLR | -536 | -543 | 1 | 0.00149795 | 8 | NaN | NaN | NaN | NaN | 2794.711 | 3136.0903 |
| Copb1 | LPGLLMTIIR | -55 | -64 | 0 | 0.00025015 | 10 | NaN | NaN | NaN | NaN | 235.26883 | 1879.2733 |
| Mcm6 | LKEIESEIDSEEELINKK | -753 | -770 | 2 | 0 | 18 | NaN | NaN | NaN | NaN | 3024.8027 | NaN |
| Mcm6 | ACHVAPTNPR | -300 | -309 | 0 | 0.00111006 | 10 | NaN | NaN | NaN | NaN | 840.7931 | NaN |
| Mcm6 | PGAGSQHPEVR | -9 | -19 | 0 | 0.002068213 | 11 | NaN | NaN | NaN | NaN | 698.621 | NaN |
| Fh | TAEQFDEWVKPK | -490 | -501 | 1 | 0.00204437 | 12 | NaN | NaN | NaN | NaN | 2132.2449 | NaN |
| Fh | LNPHIGYDK | -456 | -464 | 0 | 0.009569685 | 9 | NaN | NaN | NaN | NaN | 1028.0162 | NaN |
| Uqcrc1 | QSVPETQVSILDNGLR | -43 | -58 | 0 | 0.004983786 | 16 | NaN | NaN | NaN | NaN | 1343.1803 | 921.1242 |
| Uqcrc1 | AAAGGVEHQQLLDLAQK | -232 | -248 | 0 | 0.000230707 | 17 | NaN | NaN | NaN | NaN | 1320.6095 | 1738.8811 |
| Uqcrc1 | ALVSHLDGTTPVCEDIGR | -398 | -415 | 0 | 0.000550307 | 18 | NaN | NaN | NaN | NaN | 1065.4773 | 1166.5562 |
| Snd1 | VDSHHQKPVNAIIEHVR | -187 | -203 | 1 | 0 | 17 | NaN | NaN | NaN | NaN | 4088.2888 | 3839.6423 |
| Snd1 | VDSHHQKPVNAIIEHVR | -187 | -203 | 1 | 0 | 17 | NaN | NaN | NaN | NaN | 4088.2888 | 3839.6423 |
| Snd1 | HFVDSHHQKPVNAIIEHVR | -185 | -203 | 1 | 0.00111006 | 19 | NaN | NaN | NaN | NaN | 706.9139 | NaN |
| Snd1 | AGNFIGWLH | -604 | -612 | 0 | 0 | 9 | NaN | NaN | NaN | NaN | 677.5274 | NaN |
| Snd1 | PVNAIIEHVR | -194 | -203 | 0 | 0.00025015 | 10 | NaN | NaN | NaN | NaN | 510.5752 | NaN |
| Snd1 | AAATQPDGKDTPDEPWAFPA | -63 | -82 | 1 | 0.009002904 | 20 | NaN | NaN | NaN | NaN | 362.55093 | NaN |
| Stt3b | AACFIAIVPGYISR | -198 | -211 | 0 | 0 | 14 | NaN | NaN | NaN | NaN | 246.75919 | 391.0387 |
| Lcp1 | YPALHKPENQDIDWGALEGETR | -374 | -395 | 1 | 0 | 22 | NaN | NaN | NaN | NaN | 1883.0695 | 1831.4072 |
| Lcp1 | YPALHKPENQDIDWGAL | -374 | -390 | 1 | 0.000431655 | 17 | NaN | NaN | NaN | NaN | 827.18567 | 684.8598 |
| Lcp1 | SASAIGCHVVNIGAEDLK | -200 | -217 | 0 | 0 | 18 | NaN | NaN | NaN | NaN | 1138.749 | 1011.84595 |
| Lcp1 | VNHLYSDLSDALVIFQ | -413 | -428 | 0 | 0.00201047 | 16 | NaN | NaN | NaN | NaN | 198.70161 | 158.55952 |
| Lcp1 | GCHVVNIGAEDLK | -205 | -217 | 0 | 0.001463897 | 13 | NaN | NaN | NaN | NaN | 1505.2988 | NaN |
| Lcp1 | EQSSVGTQHSYSEEEK | -108 | -123 | 0 | 0.008014596 | 16 | NaN | NaN | NaN | NaN | 389.45956 | NaN |
| Apex1 | EGYSGVGLLSR | -125 | -135 | 0 | 0.00025015 | 11 | NaN | NaN | NaN | NaN | 1237.5557 | NaN |
| Nap1l1 | NVDLLSDMVQEHDEPILK | -177 | -194 | 0 | 0 | 18 | NaN | NaN | NaN | NaN | 545.8987 | 470.61182 |
| Nap1l1 | LEDVEEVEEEETGEETK | -16 | -32 | 0 | 0.00025015 | 17 | NaN | NaN | NaN | NaN | 315.4563 | 130.99118 |
| Hspa9 | MEEFKDQLPADECNK | -596 | -610 | 1 | 0 | 15 | NaN | NaN | NaN | NaN | 735.1141 | 722.23456 |
| Hspa9 | KVLENAEGAR | -76 | -85 | 1 | 0.00111006 | 10 | NaN | NaN | NaN | NaN | 2054.567 | 1880.4851 |
| Prpf3 | DQTKPTPLILDEQGR | -249 | -263 | 1 | 0 | 15 | NaN | NaN | NaN | NaN | 495.43958 | 603.9684 |
| Stxbp2 | KLWPFVSDPAPVPSSQ | -494 | -509 | 1 | 0.003620005 | 16 | NaN | NaN | NaN | NaN | 433.1464 | 169.31096 |
| Rfc1 | GVLESIERDEAK | -413 | -424 | 1 | 0.003681885 | 12 | NaN | NaN | NaN | NaN | 1438.349 | NaN |
| Trmt112 | AADTLNLAEVPKEPTEGYEHDETFLR | -57 | -82 | 1 | 0.001291103 | 26 | NaN | NaN | NaN | NaN | 355.94348 | 308.7757 |
| Pkm | KITLDNAYMEK | -141 | -151 | 1 | 0 | 11 | NaN | NaN | NaN | NaN | 1704.4441 | 1721.866 |
| Pkm | HLQLFEELRR | -391 | -400 | 1 | 0 | 10 | NaN | NaN | NaN | NaN | 1927.5309 | 2709.5474 |
| Pkm | NFSHGTHEYHAETIK | -75 | -89 | 0 | 0 | 15 | NaN | NaN | NaN | NaN | 657.38666 | 672.62506 |
| Pkm | NFSHGTHEYHAETIK | -75 | -89 | 0 | 0 | 15 | NaN | NaN | NaN | NaN | 657.38666 | 672.62506 |
| Pkm | KIENHEGVR | -270 | -278 | 1 | 0.004412728 | 9 | NaN | NaN | NaN | NaN | 355.867 | 350.15912 |
| Pkm | IISKIENHEGVR | -267 | -278 | 1 | 0.00111006 | 12 | NaN | NaN | NaN | NaN | 809.9997 | 667.57874 |
| Pkm | AEGSDVANAVLDGADCIML | -343 | -361 | 0 | 0 | 19 | NaN | NaN | NaN | NaN | 279.28003 | 275.46317 |
| Pkm | RLNFSHGTHEYHAETIK | -73 | -89 | 1 | 0.005605646 | 17 | NaN | NaN | NaN | NaN | 716.6824 | NaN |
| Pdcd6ip | LCKPEPELNAAIPSANPAK | -523 | -541 | 1 | 0.001636916 | 19 | NaN | NaN | NaN | NaN | 422.7237 | 465.84183 |
| Pdcd6ip | ALGRPLDK | -53 | -60 | 1 | 0 | 8 | NaN | NaN | NaN | NaN | 420.89294 | 560.5823 |
| Cct2 | IIADNAGYDSADLVAQLR | -449 | -466 | 0 | 0.00272025 | 18 | NaN | NaN | NaN | NaN | 402.6178 | NaN |
| Dhx9 | KVFDPVPDGVTK | -699 | -710 | 1 | 0 | 12 | NaN | NaN | NaN | NaN | 3186.1013 | 3282.5322 |
| Dhx9 | LDPVNEHMLNTIR | -1125 | -1137 | 0 | 0 | 13 | NaN | NaN | NaN | NaN | 966.7742 | 806.7631 |
| Dhx9 | SEEVPAVGIVPPPPILSD | -77 | -94 | 0 | 0.000298805 | 18 | NaN | NaN | NaN | NaN | 248.80492 | 148.31815 |
| Dhx9 | TFPVQEYFLEDCIQ | -569 | -582 | 0 | 0.000907206 | 14 | NaN | NaN | NaN | NaN | 544.02423 | NaN |
| Dhx9 | SHVIVDEIHER | -508 | -518 | 0 | 0.00294445 | 11 | NaN | NaN | NaN | NaN | 536.5831 | NaN |
| Dhx9 | RTPLHEIALSIK | -797 | -808 | 1 | 0.001776473 | 12 | NaN | NaN | NaN | NaN | 417.2356 | NaN |
| Dhx9 | RYQILPLHSQIPR | -682 | -694 | 1 | 0.001708008 | 13 | NaN | NaN | NaN | NaN | 411.16684 | NaN |
| Pafah1b2 | QQYEIWR | -54 | -60 | 0 | 0.000627943 | 7 | NaN | NaN | NaN | NaN | 1010.81006 | NaN |
| Hspa4 | RWNSPAEEGLSDCEVFPK | -405 | -422 | 1 | 0 | 18 | NaN | NaN | NaN | NaN | 722.68677 | 702.80597 |
| Hspa4 | KNKEDQYEHLDAADVTK | -718 | -734 | 2 | 0.00025015 | 17 | NaN | NaN | NaN | NaN | 648.07007 | 413.69617 |

| | | | | | | | | | | | | | |
|---|---|---|---|---|---|---|---|---|---|---|---|---|---|
| Stat1 | DQKQEQLLLHK | -191 | -201 | 1 | | 0.00025015 | 11 | NaN | NaN | NaN | NaN | 4241.323 | 4353.463 |
| Stat1 | LINDELVEWK | -231 | -240 | 0 | | 0.003620005 | 10 | NaN | NaN | NaN | NaN | 512.46484 | NaN |
| Nup93 | DNALLSAIEESR | -107 | -118 | 0 | | 0.001055699 | 12 | NaN | NaN | NaN | NaN | 477.8956 | 569.06604 |
| Sarnp | SGIPQTER | -95 | -102 | 0 | | 0.001868731 | 8 | NaN | NaN | NaN | NaN | 631.5442 | NaN |
| Wdr12 | GAEEWILSGSYDK | -109 | -121 | 0 | | 0 | 13 | NaN | NaN | NaN | NaN | 1115.6747 | 2353.0898 |
| Wdr12 | AAEVADLSNIINK | -27 | -39 | 0 | | 0.000550307 | 13 | NaN | NaN | NaN | NaN | 1461.561 | NaN |
| Snrnp200 | HDDLKGDPLLDQR | -940 | -952 | 1 | | 0.000431655 | 13 | NaN | NaN | NaN | NaN | 777.86176 | 1055.1262 |
| Snrnp200 | ALETDENLLLCAPTGAGK | -492 | -509 | 0 | | 0.000431655 | 18 | NaN | NaN | NaN | NaN | 585.86597 | 490.06012 |
| Snrnp200 | SVHLQPITR | -1187 | -1195 | 0 | | 0 | 9 | NaN | NaN | NaN | NaN | 582.4409 | 614.73975 |
| Snrnp200 | LEREEEVTGPVIAPLFPQK | -2041 | -2059 | 1 | | 0.00453359 | 19 | NaN | NaN | NaN | NaN | 458.8565 | 410.64734 |
| Snrnp200 | QLPHFTSEHIK | -1962 | -1972 | 0 | | 0.00111006 | 11 | NaN | NaN | NaN | NaN | 1312.9224 | NaN |
| Snrnp200 | NHNEIGELIR | -1157 | -1166 | 0 | | 0 | 10 | NaN | NaN | NaN | NaN | 888.53546 | NaN |
| Snrnp200 | GYEEVHVPALKPKPF | -441 | -455 | 2 | | 0.00025015 | 15 | NaN | NaN | NaN | NaN | 539.34045 | NaN |
| Snrnp200 | RNSAFESLYQDK | -1313 | -1324 | 1 | | 0.00025015 | 12 | NaN | NaN | NaN | NaN | 367.4401 | NaN |
| Tln1 | AKPAAVAAENEEIGAHIK | -1900 | -1917 | 1 | | 0 | 18 | NaN | NaN | NaN | NaN | 3507.4956 | 3199.0828 |
| Tln1 | TLDDFETLPPLGQDAASK | -512 | -529 | 0 | | 0 | 18 | NaN | NaN | NaN | NaN | 2384.912 | 462.3451 |
| Tln1 | DAEGESDLENSRK | -843 | -855 | 1 | | 0 | 13 | NaN | NaN | NaN | NaN | 786.8664 | 1011.83453 |
| Tln1 | REQGVEEHETLLLR | -181 | -194 | 1 | | 0.00025015 | 14 | NaN | NaN | NaN | NaN | 739.66675 | 117.71411 |
| Tln1 | GALQCSPSDVYTK | -1935 | -1947 | 0 | | 0 | 13 | NaN | NaN | NaN | NaN | 734.246 | 627.5574 |
| Tln1 | AVTQALNELLQHVK | -773 | -786 | 0 | | 0 | 14 | NaN | NaN | NaN | NaN | 278.20544 | 492.7291 |
| Tln1 | SPVCQEQLVEAGR | -729 | -741 | 0 | | 0 | 13 | NaN | NaN | NaN | NaN | 1133.2782 | NaN |
| Tln1 | DDILNGSHPVSFDK | -221 | -234 | 0 | | 0.00078661 | 14 | NaN | NaN | NaN | NaN | 800.3592 | NaN |
| Tln1 | VGAIPANALDDGQWSQGLIS | -2376 | -2395 | 0 | | 0.004224523 | 20 | NaN | NaN | NaN | NaN | 662.3812 | NaN |
| Dld | SEEQLKEEGIEFK | -405 | -417 | 1 | | 0 | 13 | NaN | NaN | NaN | NaN | 2586.8254 | 2578.8354 |
| Srm | SLFPVVDYAYC | -226 | -236 | 0 | | 0 | 11 | NaN | NaN | NaN | NaN | 152.48485 | NaN |
| Tomm22 | QILLGPNTGLSGGMPGALPPLPGK | -118 | -141 | 0 | | 0.000947459 | 24 | NaN | NaN | NaN | NaN | 372.29782 | 407.95035 |
| Tomm22 | ILVLPVVFETEK | -94 | -105 | 0 | | 0.00450079 | 12 | NaN | NaN | NaN | NaN | 312.73993 | NaN |
| Dnajc2 | SYLDEEEKEK | -228 | -237 | 1 | | 0 | 10 | NaN | NaN | NaN | NaN | 245.46265 | 1010.86487 |
| Ilf2 | RWFEENASQSTVK | -209 | -221 | 1 | | 0.009657961 | 13 | NaN | NaN | NaN | NaN | 2362.7988 | NaN |
| Cnot3 | SGIEDPVPTLHLTDR | -552 | -566 | 0 | | 0 | 15 | NaN | NaN | NaN | NaN | 991.93256 | 988.72205 |
| Cnot3 | GTLLNGPPQFSTTPEIK | -517 | -533 | 0 | | 0.002296977 | 17 | NaN | NaN | NaN | NaN | 608.55176 | NaN |
| Cnot3 | AAGTLLNGPPQFSTTPEIK | -515 | -533 | 0 | | 0.001291103 | 19 | NaN | NaN | NaN | NaN | 595.3162 | NaN |
| Eef1g | VPAFEGDDGFCVF | -58 | -70 | 0 | | 0 | 13 | NaN | NaN | NaN | NaN | 1780.5616 | 1571.0828 |
| Eef1g | KLDPGSEETQTL | -401 | -412 | 1 | | 0.000627943 | 12 | NaN | NaN | NaN | NaN | 221.33328 | 3487.8066 |
| Eef1g | PPHFHFGQTNR | -35 | -45 | 0 | | 0 | 11 | NaN | NaN | NaN | NaN | 1554.2467 | 233.77191 |
| Eef1g | RILGLLDTHLK | -137 | -147 | 1 | | 0.00025015 | 11 | NaN | NaN | NaN | NaN | 242.13206 | NaN |
| Gosr1 | DILQDYTHEFHK | -113 | -124 | 0 | | 0.000907206 | 12 | NaN | NaN | NaN | NaN | 419.97745 | 467.0596 |
| Eef1a1 | VETGVLKPGMVVT | -267 | -279 | 1 | | 0 | 13 | NaN | NaN | NaN | NaN | 4824.0063 | 4364.203 |
| Eef1a1 | IGYNPDTVAFVPIS | -181 | -194 | 0 | | 0 | 14 | NaN | NaN | NaN | NaN | 1557.6075 | 1542.3475 |
| Eef1a1 | AGYAPVLDCHTAHIACK | -355 | -371 | 0 | | 0 | 17 | NaN | NaN | NaN | NaN | 1137.7415 | 1147.4331 |
| Eef1a1 | VNVTTEVK | -283 | -290 | 0 | | 0.000550307 | 8 | NaN | NaN | NaN | NaN | 608.69366 | 805.133 |
| Eef1a1 | KIGYNPDTVAFVPIS | -180 | -194 | 1 | | 0.001392869 | 15 | NaN | NaN | NaN | NaN | 906.9341 | 772.8197 |
| Eef1a1 | VIILNHPGQISAGYAPVLDCHTAHIACK | -344 | -371 | 0 | | 0 | 28 | NaN | NaN | NaN | NaN | 658.7677 | 761.6793 |
| Eef1a1 | TLLEALDCILPPTRPTDK | -227 | -244 | 1 | | 0 | 18 | NaN | NaN | NaN | NaN | 373.31387 | 347.81085 |
| Eef1a1 | DSTEPPYSQKR | -156 | -166 | 1 | | 0.008680443 | 11 | NaN | NaN | NaN | NaN | 699.1737 | NaN |
| Rplp1 | AAPAGGAAPSTAAAPAEEK | -74 | -92 | 0 | | 0.00025015 | 19 | NaN | NaN | NaN | NaN | 877.24316 | 1056.8677 |
| Rplp1 | VNIGSLICNVGAGGPAPAAGAAPAGGAAPSTAAAP[illegible] | -54 | -92 | 0 | | 0 | 39 | NaN | NaN | NaN | NaN | 319.16153 | 323.93335 |
| Rplp1 | EPFWPGLFAK | -40 | -49 | 0 | | 0 | 10 | NaN | NaN | NaN | NaN | 615.0764 | NaN |
| Mogs | SWFQHRPPGASPDR | -282 | -295 | 1 | | 0.00051306 | 14 | NaN | NaN | NaN | NaN | 584.90063 | 922.8196 |
| Mogs | LVGSQLTQALESHAA | -349 | -363 | 0 | | 0.00149795 | 15 | NaN | NaN | NaN | NaN | 823.4704 | NaN |
| Fus | LKGEATVSFDDPPSAK | -326 | -341 | 1 | | 0.00025015 | 16 | NaN | NaN | NaN | NaN | 597.55035 | 798.0348 |
| Fbl | GGVDQIHIKPGAK | -156 | -168 | 1 | | 0 | 13 | NaN | NaN | NaN | NaN | 2801.884 | 2846.3254 |
| Fbl | AAILGGVDQIHIKPGAK | -152 | -168 | 1 | | 0.00111006 | 17 | NaN | NaN | NaN | NaN | 439.82605 | 449.34033 |
| Fbl | VDQIHIKPGAK | -158 | -168 | 1 | | 0.00025015 | 11 | NaN | NaN | NaN | NaN | 315.87155 | 615.945 |
| Ddx21 | LKDGLSQPSEPK | -221 | -232 | 1 | | 0 | 12 | NaN | NaN | NaN | NaN | 267.3604 | 3105.1587 |
| Ddx21 | LGSDGAEESMETLPKPSEK | -11 | -29 | 1 | | 0 | 19 | NaN | NaN | NaN | NaN | 639.49365 | 455.64737 |
| Ddx21 | AHISGATSVDQR | -688 | -699 | 0 | | 0 | 12 | NaN | NaN | NaN | NaN | 564.90576 | 1212.6373 |
| Ddx21 | GLSQPSEEEADIPKPK | -150 | -165 | 1 | | 0.000431655 | 16 | NaN | NaN | NaN | NaN | 529.211 | 617.26154 |
| Ddx21 | GSDGAEESMETLPKPSEK | -12 | -29 | 1 | | 0 | 18 | NaN | NaN | NaN | NaN | 496.97763 | 600.5256 |
| Ddx21 | DSEDNPQTLLFSA | -436 | -448 | 0 | | 0 | 13 | NaN | NaN | NaN | NaN | 294.3785 | 363.7929 |
| Ddx21 | DSEDNPQTLLF | -436 | -446 | 0 | | 0.000627943 | 11 | NaN | NaN | NaN | NaN | 464.97366 | NaN |
| Cap1 | HAEMVHTGLK | -71 | -80 | 0 | | 0.000550307 | 10 | NaN | NaN | NaN | NaN | 841.79645 | 1171.9385 |
| Arpc4 | KELLLQPVTISR | -44 | -55 | 1 | | 0.00383302 | 12 | NaN | NaN | NaN | NaN | 436.6911 | 307.42487 |
| Arpc4 | RKPVEGYDIS | -106 | -115 | 2 | | 0.002780285 | 10 | NaN | NaN | NaN | NaN | 5555.057 | NaN |
| Shmt2 | SLVSGGTDTHLVLVDLRPK | -371 | -389 | 1 | | 0.00025015 | 19 | NaN | NaN | NaN | NaN | 270.25558 | 267.74323 |
| Shmt2 | AALEALGSCLNNK | -83 | -95 | 0 | | 0.00025015 | 13 | NaN | NaN | NaN | NaN | 2155.3638 | NaN |
| Hars1 | DKVSWEEVK | -242 | -250 | 1 | | 0.002296977 | 9 | NaN | NaN | NaN | NaN | 1315.292 | NaN |
| Rplp2 | KEESEESDDDMGFGL | -99 | -113 | 1 | | 0.000431655 | 15 | NaN | NaN | NaN | NaN | 721.352 | 663.78467 |
| Mcm5 | CPLDPYFIMPDK | -207 | -218 | 0 | | 0 | 12 | NaN | NaN | NaN | NaN | 456.2217 | NaN |
| Mcm5 | AIACLLFGGSR | -352 | -362 | 0 | | 0.001708008 | 11 | NaN | NaN | NaN | NaN | 1332.9512 | 1368.908 |
| Mcm5 | PQEEEEFR | -316 | -323 | 0 | | 0.00111006 | 8 | NaN | NaN | NaN | NaN | 502.59323 | 549.4492 |
| Septin1 | SATEIPLPMLPLADTEK | -314 | -330 | 0 | | 0.000230707 | 17 | NaN | NaN | NaN | NaN | 2777.7483 | 1389.5939 |
| Septin1 | IEEQFEQYLR | -110 | -119 | 0 | | 0.00025015 | 10 | NaN | NaN | NaN | NaN | 664.9928 | 359.40497 |
| Eif4g1 | LGEESEAPGQR | -1426 | -1436 | 0 | | 0 | 11 | NaN | NaN | NaN | NaN | 809.14197 | 660.7045 |
| Eif4g1 | ITKPGSIDSNNQ | -1076 | -1087 | 1 | | 0.005884925 | 12 | NaN | NaN | NaN | NaN | 1360.4194 | NaN |
| Eif4g1 | DKDDDEVFEKK | -858 | -868 | 2 | | 0 | 11 | NaN | NaN | NaN | NaN | 434.13614 | NaN |
| Cad | GTPVETIELTEDR | -503 | -515 | 0 | | 0.00635324 | 13 | NaN | NaN | NaN | NaN | 3656.5518 | 2825.4014 |
| Cad | FALGGLGSGFASTK | -558 | -571 | 0 | | 0 | 14 | NaN | NaN | NaN | NaN | 981.0912 | 1800.3187 |

| | | | | | | | | | | | | |
|---|---|---|---|---|---|---|---|---|---|---|---|---|
| Cad | RLCPPELPIPGSGLPPPR | -377 | -394 | 1 | 0.000298805 | 18 | NaN | NaN | NaN | NaN | 778.5178 | 1016.57837 |
| Cad | GLPVTCEVAPHHLFLNR | -1631 | -1647 | 0 | 0.00025015 | 17 | NaN | NaN | NaN | NaN | 509.12476 | 321.4903 |
| Cad | RIDCWFLHR | -831 | -839 | 1 | 0.001636916 | 9 | NaN | NaN | NaN | NaN | 381.9465 | 531.7938 |
| Cad | WFESSEIHVA | -75 | -84 | 0 | 0 | 10 | NaN | NaN | NaN | NaN | 243.63768 | 841.8495 |
| Cad | YASLGTADFYTEHGVK | -1343 | -1358 | 0 | 0 | 16 | NaN | NaN | NaN | NaN | 1823.7786 | NaN |
| Lmnb1 | FKAEHDQLLLNYA | -111 | -123 | 1 | 0.00111006 | 13 | NaN | NaN | NaN | NaN | 345.3334 | 1272.1227 |
| Grhpr | VTGPLPAEGR | -11 | -20 | 0 | 0 | 10 | NaN | NaN | NaN | NaN | 748.14197 | 1140.7742 |
| Grhpr | AANNLLAGLR | -309 | -318 | 0 | 0 | 10 | NaN | NaN | NaN | NaN | 206.86996 | 721.92163 |
| Tpi1 | YEPVWAIGTGK | -165 | -175 | 0 | 0.001150701 | 11 | NaN | NaN | NaN | NaN | 1864.5393 | 6229.581 |
| Tpi1 | SLKPEFVDIINAKQ | -236 | -249 | 2 | 0 | 14 | NaN | NaN | NaN | NaN | 540.8886 | 1040.1613 |
| Vim | VESLQEEIAFLKK | -224 | -236 | 1 | 0.00025015 | 13 | NaN | NaN | NaN | NaN | 1547.8945 | 1289.6038 |
| Vim | NLRETNLESLPLVDTHSKR | -422 | -440 | 2 | 0.000712251 | 19 | NaN | NaN | NaN | NaN | 558.6647 | NaN |
| Nptn | SVGYPHPEWIWR | -260 | -271 | 0 | 0.001023236 | 12 | NaN | NaN | NaN | NaN | 4186.597 | NaN |
| Wars1 | AIDQDPYFR | -314 | -322 | 0 | 0.000431655 | 9 | NaN | NaN | NaN | NaN | 1633.6425 | 1650.4869 |
| Wars1 | ISFPAVQAAPSFSNSFPK | -282 | -299 | 0 | 0 | 18 | NaN | NaN | NaN | NaN | 524.6226 | 502.1828 |
| Wars1 | PALQGAQTK | -345 | -353 | 0 | 0.002099344 | 9 | NaN | NaN | NaN | NaN | 487.2149 | 727.924 |
| Wars1 | TLIDVLQPLIAEHQAR | -437 | -452 | 0 | 0.006788754 | 16 | NaN | NaN | NaN | NaN | 877.2734 | NaN |
| N6amt1 | VFNPPYVVTPPEEVGSR | -120 | -136 | 0 | 0.000907206 | 17 | NaN | NaN | NaN | NaN | 1042.5691 | NaN |
| Tcp1 | WIGLDLVHGKPR | -485 | -496 | 1 | 0 | 12 | NaN | NaN | NaN | NaN | 1222.8784 | 1546.4143 |
| Tcp1 | YPVNSVNILK | -190 | -199 | 0 | 0.000591124 | 10 | NaN | NaN | NaN | NaN | 15728.018 | 16089.226 |
| Tcp1 | FGPVGLDK | -36 | -43 | 0 | 0.006624942 | 8 | NaN | NaN | NaN | NaN | 619.8991 | 810.88306 |
| Tcp1 | CELADLQDK | -76 | -84 | 0 | 0.000431655 | 9 | NaN | NaN | NaN | NaN | 732.5742 | NaN |
| Nup58 | AAQLQSIHENVK | -388 | -399 | 0 | 0.001636916 | 12 | NaN | NaN | NaN | NaN | 605.01465 | NaN |
| Ywhae | YLAEFATGNDRK | -131 | -142 | 1 | 0 | 12 | NaN | NaN | NaN | NaN | 1527.6675 | 1673.4219 |
| Esd | KVFEHSSVELK | -17 | -27 | 1 | 0 | 11 | NaN | NaN | NaN | NaN | 1049.2264 | 376.8913 |
| Esd | GYQQAASEHGLVVIAPDTSPR | -66 | -86 | 0 | 0.000755556 | 21 | NaN | NaN | NaN | NaN | 334.1557 | 418.54938 |
| Anxa4 | AASGFNATEDAQTLR | -10 | -24 | 0 | 0.000591124 | 15 | NaN | NaN | NaN | NaN | 3709.7107 | NaN |
| Anxa4 | SGFNATEDAQTLR | -12 | -24 | 0 | 0 | 13 | NaN | NaN | NaN | NaN | 1385.4998 | NaN |
| Plbd2 | AASGPTWDQCPPFQWSK | -553 | -569 | 0 | 0.00025015 | 17 | NaN | NaN | NaN | NaN | 618.1911 | NaN |
| Thop1 | QANAGLFNLR | -544 | -553 | 0 | 0 | 10 | NaN | NaN | NaN | NaN | 4608.782 | NaN |
| Trim28 | SCDTLTCR | -224 | -231 | 0 | 0.000471409 | 8 | NaN | NaN | NaN | NaN | 558.923 | 498.9962 |
| Trim28 | ASSPAGGGGEAQELLEHCGVCR | -49 | -70 | 0 | 0 | 22 | NaN | NaN | NaN | NaN | 428.45737 | 699.7935 |
| Trim28 | LTEDKADVQ | -775 | -783 | 1 | 0.001392869 | 9 | NaN | NaN | NaN | NaN | 3412.4834 | NaN |
| Trim28 | AATAASAASGSPGSGEGSAGGEK | -13 | -35 | 0 | 0 | 23 | NaN | NaN | NaN | NaN | 500.74088 | NaN |
| Trim28 | NSGDGGSAGDGAMVDCPVCK | -103 | -122 | 0 | 0.002329279 | 20 | NaN | NaN | NaN | NaN | 437.02524 | NaN |
| Trim28 | NNSGDGGSAGDGAMVDCPVCK | -102 | -122 | 0 | 0.000347792 | 21 | NaN | NaN | NaN | NaN | 132.82283 | NaN |
| Dock2 | VPAESFDLESAPPK | -1660 | -1673 | 0 | 0.000712251 | 14 | NaN | NaN | NaN | NaN | 2419.2595 | 1623.9216 |
| Dock2 | LFESINNLMK | -773 | -782 | 0 | 0.00051306 | 10 | NaN | NaN | NaN | NaN | 133.03223 | 138.49464 |
| Dock2 | LFESINNLMK | -773 | -782 | 0 | 0.00051306 | 10 | NaN | NaN | NaN | NaN | 133.03223 | 138.49464 |
| Dock2 | SSPTHSLYVFVR | -217 | -228 | 0 | 0 | 12 | NaN | NaN | NaN | NaN | 873.53143 | 611.6154 |
| Dock2 | EHPEDQDKLSHLK | -1557 | -1569 | 1 | 0 | 13 | NaN | NaN | NaN | NaN | 429.31342 | 377.64172 |
| Dock2 | LFHAHEEATYK | -183 | -193 | 0 | 0 | 11 | NaN | NaN | NaN | NaN | 1853.7238 | NaN |
| Dock2 | AATESDLKR | -1712 | -1720 | 1 | 0.000947459 | 9 | NaN | NaN | NaN | NaN | 333.08188 | NaN |
| Hadh | KTFESLVDFCK | -192 | -202 | 1 | 0.000230707 | 11 | NaN | NaN | NaN | NaN | 690.9138 | 671.8283 |
| Aagab | AATLPHEQR | -277 | -285 | 0 | 0.00025015 | 9 | NaN | NaN | NaN | NaN | 1410.7367 | NaN |
| Got1 | NLDYVATSIHEAVTK | -397 | -411 | 0 | 0 | 15 | NaN | NaN | NaN | NaN | 699.4828 | 806.5595 |
| Got1 | VNLGVGAYR | -34 | -42 | 0 | 0 | 9 | NaN | NaN | NaN | NaN | 1364.5243 | NaN |
| Txnl1 | AATNNISATPTFLFFR | -68 | -83 | 0 | 0.003620005 | 16 | NaN | NaN | NaN | NaN | 346.1556 | 224.52734 |
| Vcp | ETVVEVPQVTWEDIGGLEDVKR | -466 | -487 | 1 | 0.000627943 | 22 | NaN | NaN | NaN | NaN | 454.02747 | NaN |
| Nop2 | VEDEEAFVLPPAGETDQDGQAPDLQR | -198 | -223 | 0 | 0.00025015 | 26 | NaN | NaN | NaN | NaN | 368.68732 | 371.01355 |
| Nop2 | GPSAPFVSPH | -760 | -769 | 0 | 0 | 10 | NaN | NaN | NaN | NaN | 1297.4618 | NaN |
| Nop2 | RGVNLDPLGK | -316 | -325 | 1 | 0.003490217 | 10 | NaN | NaN | NaN | NaN | 354.927 | NaN |
| Erap1 | RDYLSADTFK | -438 | -447 | 1 | 0.000230707 | 10 | NaN | NaN | NaN | NaN | 520.1876 | 746.90564 |
| Erap1 | GLTTYRESSLLYDK | -312 | -325 | 1 | 0.007880195 | 14 | NaN | NaN | NaN | NaN | 801.1299 | NaN |
| Dync1h1 | FNEVLDHVLR | -2910 | -2919 | 0 | 0 | 10 | NaN | NaN | NaN | NaN | 4021.1494 | 1458.3376 |
| Dync1h1 | RAPVIDADKPVSSQ | -119 | -132 | 2 | 0.001291103 | 14 | NaN | NaN | NaN | NaN | 602.2941 | 560.3209 |
| Dync1h1 | INEWLTLVEK | -1696 | -1705 | 0 | 0 | 10 | NaN | NaN | NaN | NaN | 424.81387 | 619.35333 |
| Dync1h1 | IKETVDQVEELR | -3206 | -3217 | 1 | 0.000230707 | 12 | NaN | NaN | NaN | NaN | 2171.3206 | NaN |
| Dync1h1 | RAPVIDADKPVSSQL | -119 | -133 | 2 | 0.002780285 | 15 | NaN | NaN | NaN | NaN | 1400.2388 | NaN |
| Dync1h1 | VPQIEVETH | -2550 | -2558 | 0 | 0.002099344 | 9 | NaN | NaN | NaN | NaN | 802.9949 | NaN |
| Dync1h1 | VIYLNPPIEECR | -966 | -977 | 0 | 0.00025015 | 12 | NaN | NaN | NaN | NaN | 768.63184 | NaN |
| Dync1h1 | TQEIFDDWAR | -693 | -702 | 0 | 0.000627943 | 10 | NaN | NaN | NaN | NaN | 738.4126 | NaN |
| Dync1h1 | QNLDGLLNQLK | -1359 | -1369 | 0 | 0.000230707 | 11 | NaN | NaN | NaN | NaN | 550.79047 | NaN |
| Dync1h1 | RAPVIDADKPVSSQLR | -119 | -134 | 2 | 0.002199523 | 16 | NaN | NaN | NaN | NaN | 528.23773 | NaN |
| Dync1h1 | DHYCEYR | -2634 | -2640 | 0 | 0.005145856 | 7 | NaN | NaN | NaN | NaN | 383.27042 | NaN |
| Dync1h1 | VTDFGDKVEDPTFLNQL | -227 | -243 | 1 | 0.00416551 | 17 | NaN | NaN | NaN | NaN | 237.31534 | NaN |
| Sri | ELWAVLNGWR | -97 | -106 | 0 | 0 | 10 | NaN | NaN | NaN | NaN | 712.0589 | 652.45496 |
| Sri | RQHFISFDSDR | -106 | -116 | 1 | 0.00149795 | 11 | NaN | NaN | NaN | NaN | 700.3067 | NaN |
| Etfa | TIYAGNALCTVK | -147 | -158 | 0 | 0.00897527 | 12 | NaN | NaN | NaN | NaN | 2280.673 | NaN |
| Etfa | TIVAINKDPEAPIFQ | -295 | -309 | 1 | 0.00025015 | 15 | NaN | NaN | NaN | NaN | 1178.0845 | NaN |
| Rrp12 | AVPLEIDGSEETLDFPR | -551 | -567 | 0 | 0.00025015 | 17 | NaN | NaN | NaN | NaN | 6622.3423 | 6826.2695 |
| Rrp12 | PEVILCTK | -861 | -868 | 0 | 0.002910424 | 8 | NaN | NaN | NaN | NaN | 793.34784 | 988.5592 |
| Rrp12 | VLEEVCASSQGPAAR | -795 | -809 | 0 | 0 | 15 | NaN | NaN | NaN | NaN | 1447.6017 | NaN |
| Hspd1 | KNAGVEGSLIVEK | -481 | -493 | 1 | 0.00025015 | 13 | NaN | NaN | NaN | NaN | 5261.5938 | 4989.434 |
| Hspd1 | KVGGTSDVEVNEK | -405 | -417 | 1 | 0 | 13 | NaN | NaN | NaN | NaN | 3276.4158 | 3253.5688 |
| Hspd1 | CEFQDAYVLL | -237 | -246 | 0 | 0.001868731 | 10 | NaN | NaN | NaN | NaN | 201.84311 | 178.87866 |
| Hspd1 | AAVEEGIVLGGGCALLR | -430 | -446 | 0 | 0.007343498 | 17 | NaN | NaN | NaN | NaN | 195.84016 | NaN |
| Kpnb1 | SEAAEQGRPPEHTSK | -298 | -312 | 1 | 0 | 15 | NaN | NaN | NaN | NaN | 435.56952 | 411.71515 |

| | | | | | | | | | | | | |
|---|---|---|---|---|---|---|---|---|---|---|---|---|
| Ewsr1 | QDHPSSMGVYGQESGGFSGPGENR | -269 | -292 | 0 | 0.00149795 | 24 | NaN | NaN | NaN | NaN | 286.28122 | NaN |
| Stip1 | KEPKPEPMEEDLPENK | -207 | -222 | 2 | 0.001868731 | 16 | NaN | NaN | NaN | NaN | 584.61206 | 474.29477 |
| Stip1 | PQALSEHLK | -515 | -523 | 0 | 0.004758481 | 9 | NaN | NaN | NaN | NaN | 1052.6309 | 931.77814 |
| Ndufv2 | VGETTPDKLFTL | -159 | -170 | 1 | 0.003490217 | 12 | NaN | NaN | NaN | NaN | 356.24722 | NaN |
| Hsp90ab1 | SELAEDKENYK | -417 | -427 | 1 | 0 | 11 | NaN | NaN | NaN | NaN | 4139.412 | 4001.9385 |
| Hsp90ab1 | AEEPSAAVPDEIPPLEGDEDASR | -697 | -719 | 0 | 0 | 23 | NaN | NaN | NaN | NaN | 1635.5262 | 1515.3972 |
| Hsp90ab1 | IEDVGSDEEDDSGKDK | -250 | -265 | 1 | 0.00025015 | 16 | NaN | NaN | NaN | NaN | 1368.534 | 1253.136 |
| Hsp90ab1 | ELFSELAEDKENYK | -414 | -427 | 1 | 0.000230707 | 14 | NaN | NaN | NaN | NaN | 519.3484 | 1143.0049 |
| Hsp90ab1 | ELKIDIIPNPQER | -70 | -82 | 1 | 0 | 13 | NaN | NaN | NaN | NaN | 1813.0951 | NaN |
| Hsp90ab1 | FSLEDPQTHSNR | -668 | -679 | 0 | 0.00025015 | 12 | NaN | NaN | NaN | NaN | 1126.0406 | NaN |
| Hsp90ab1 | KCLELFSELAEDKENYK | -411 | -427 | 2 | 0.001092529 | 17 | NaN | NaN | NaN | NaN | 387.13934 | NaN |
| Hsp90ab1 | DVGSDEEDDSGK | -252 | -263 | 0 | 0.00111006 | 12 | NaN | NaN | NaN | NaN | 273.86377 | NaN |
| Snx5 | EVDDFFEQEK | -204 | -213 | 0 | 0 | 10 | NaN | NaN | NaN | NaN | 646.4071 | 396.48047 |
| Eno1 | AEVYHNLK | -186 | -193 | 0 | 0.000550307 | 8 | NaN | NaN | NaN | NaN | 1943.8223 | 3618.1414 |
| Eno1 | IGAEVYHNLKN | -184 | -194 | 1 | 0.000230707 | 11 | NaN | NaN | NaN | NaN | 1529.2181 | 1565.3906 |
| Eno1 | VNVVEQEKIDKL | -82 | -93 | 2 | 0 | 12 | NaN | NaN | NaN | NaN | 1098.998 | 1090.8604 |
| Eno1 | TPDQLADLYK | -272 | -281 | 0 | 0.002466863 | 10 | NaN | NaN | NaN | NaN | 408.44034 | 643.6975 |
| Eno1 | AGIQVVGDDLTVTNPKR | -311 | -327 | 1 | 0.00025015 | 17 | NaN | NaN | NaN | NaN | 321.25116 | 323.39972 |
| Eno1 | QEFMILPVGASSFR | -166 | -179 | 0 | 0.003299042 | 14 | NaN | NaN | NaN | NaN | 1584.2633 | NaN |
| Eprs1 | GFFICDQPYEPVSPY | -676 | -690 | 0 | 0 | 15 | NaN | NaN | NaN | NaN | 92.76477 | 356.47772 |
| Eprs1 | EAPCILIYIPDGHTK | -694 | -708 | 0 | 0 | 15 | NaN | NaN | NaN | NaN | 2543.1467 | NaN |
| Eprs1 | TELAEPIAIRPTSETVMYPA | -1110 | -1129 | 1 | 0.005605646 | 20 | NaN | NaN | NaN | NaN | 702.7462 | NaN |
| Rpl5 | RYLMEEDEDAYKK | -209 | -221 | 2 | 0 | 13 | NaN | NaN | NaN | NaN | 982.11884 | 228.4907 |
| Rpl5 | YLMEEDEDAYK | -210 | -220 | 0 | 0.00122419 | 11 | NaN | NaN | NaN | NaN | 530.28815 | NaN |
| Slc25a5 | VLYDEIKKYT | -289 | -298 | 2 | 0.000230707 | 10 | NaN | NaN | NaN | NaN | 1389.577 | 1189.039 |
| Slc25a4 | REFNGLGDCLTK | -152 | -163 | 1 | 0.005605646 | 12 | NaN | NaN | NaN | NaN | 1073.7231 | 1012.8998 |
| Slc25a4 | VAGLVSYPFDTVR | -223 | -235 | 0 | 0.009272078 | 13 | NaN | NaN | NaN | NaN | 2349.4153 | NaN |
| Rars1 | KRFDADEEFK | -278 | -287 | 2 | 0.004439092 | 10 | NaN | NaN | NaN | NaN | 663.66406 | 565.5255 |
| Rars1 | LFEFAGYDVLR | -224 | -234 | 0 | 0 | 11 | NaN | NaN | NaN | NaN | 435.25797 | 507.94543 |
| Vps18 | AAYTDHPPPFR | -249 | -259 | 0 | 0.001092529 | 11 | NaN | NaN | NaN | NaN | 450.14667 | NaN |
| Vps35 | IKAELAELPLR | -572 | -582 | 1 | 0.001291103 | 11 | NaN | NaN | NaN | NaN | 2507.474 | 1188.1553 |
| Vps35 | SEENHEPLR | -642 | -650 | 0 | 0 | 9 | NaN | NaN | NaN | NaN | 687.2868 | 437.6736 |
| Sacm1l | RELSAQPEVHR | -174 | -184 | 1 | 0 | 11 | NaN | NaN | NaN | NaN | 1462.1146 | NaN |
| Sacm1l | DVPPSAVTRPIFGIL | -47 | -61 | 1 | 0.008498216 | 15 | NaN | NaN | NaN | NaN | 570.4817 | NaN |
| Sucla2 | AQGGVNIEDVAAENPEAIVK | -186 | -205 | 0 | 0.005301614 | 20 | NaN | NaN | NaN | NaN | 344.96307 | 318.61398 |
| Sucla2 | ACDDLDEAAK | -429 | -438 | 0 | 0.00025015 | 10 | NaN | NaN | NaN | NaN | 632.5768 | NaN |
| Esyt2 | SSSSLLASPSHIAAK | -660 | -674 | 0 | 0.008045078 | 15 | NaN | NaN | NaN | NaN | 573.9838 | NaN |
| Esyt2 | ACDLPAWVHFPDTER | -104 | -118 | 0 | 0 | 15 | NaN | NaN | NaN | NaN | 432.992 | NaN |
| Ttc38 | DALKPNSPLTER | -442 | -453 | 1 | 0.00025015 | 12 | NaN | NaN | NaN | NaN | 1499.717 | 1586.9453 |
| C3 | FYHPEKDDGMLSK | -1492 | -1504 | 1 | 0 | 13 | NaN | NaN | NaN | NaN | 324.58313 | 687.76624 |
| C3 | QSGYLFIQTDK | -127 | -137 | 0 | 0 | 11 | NaN | NaN | NaN | NaN | 719.9162 | NaN |
| Lasp1 | KTQDQISNIK | -112 | -121 | 1 | 0.007905138 | 10 | NaN | NaN | NaN | NaN | 5386.5376 | NaN |
| Lasp1 | MGPSGGEGVEPER | -131 | -143 | 0 | 0 | 13 | NaN | NaN | NaN | NaN | 449.13254 | NaN |
| Paics | SHATQAIFEILEK | -166 | -178 | 0 | 0.002199523 | 13 | NaN | NaN | NaN | NaN | 1426.0966 | NaN |
| Paics | AEYEGDGIPTVFVS | -314 | -327 | 0 | 0.007987481 | 14 | NaN | NaN | NaN | NaN | 1077.802 | NaN |
| Uba1 | VVQGHQQLDSY | -912 | -922 | 0 | 0 | 11 | NaN | NaN | NaN | NaN | 1243.8263 | 1140.1176 |
| Uba1 | FKQPAENVNQYLTDSK | -656 | -671 | 1 | 0.000550307 | 16 | NaN | NaN | NaN | NaN | 491.5955 | 533.0042 |
| Uba1 | YYNQEWTLWDR | -947 | -957 | 0 | 0 | 11 | NaN | NaN | NaN | NaN | 4730.1636 | 4631.583 |
| Uba1 | LAELNSYVPVTAY | -135 | -147 | 0 | 0 | 13 | NaN | NaN | NaN | NaN | 2458.0432 | 2412.5571 |
| Uba1 | AVTLHDQGTTQWADLSSQ | -98 | -115 | 0 | 0.001868731 | 18 | NaN | NaN | NaN | NaN | 2263.3914 | 2235.085 |
| Uba1 | ACHHWHTQYCNNIR | -710 | -723 | 0 | 0 | 14 | NaN | NaN | NaN | NaN | 497.25433 | 614.6093 |
| Uba1 | ACHHWHTQYCNNIR | -710 | -723 | 0 | 0 | 14 | NaN | NaN | NaN | NaN | 497.25433 | 614.6093 |
| Uba1 | VLELCCNDESGEDVEVPYVR | -1035 | -1054 | 0 | 0 | 20 | NaN | NaN | NaN | NaN | 309.0417 | 391.99655 |
| Uba1 | IGGLAAQEVMK | -401 | -411 | 0 | 0.003235556 | 11 | NaN | NaN | NaN | NaN | 855.31354 | NaN |
| Uba1 | SLLQSVQVPEFTPK | -789 | -802 | 0 | 0 | 14 | NaN | NaN | NaN | NaN | 660.96643 | NaN |
| Uba1 | IHVSDQELQSANAS | -807 | -820 | 0 | 0.003620005 | 14 | NaN | NaN | NaN | NaN | 428.17572 | NaN |
| Uba1 | SLPASLVEPDFVMTDFAK | -305 | -322 | 0 | 0.004658491 | 18 | NaN | NaN | NaN | NaN | 399.61966 | NaN |
| Eif4a3 | YLPPATQVVLIS | -207 | -218 | 0 | 0.000431655 | 12 | NaN | NaN | NaN | NaN | 179.21597 | 1463.7891 |
| Eif4a3 | ATLPHEILEMTNK | -219 | -231 | 0 | 0 | 13 | NaN | NaN | NaN | NaN | 2614.9653 | 2417.9316 |
| Eif4a3 | QALILAPTR | -108 | -116 | 0 | 0.001846793 | 9 | NaN | NaN | NaN | NaN | 1211.9542 | NaN |
| Eif3e | ACLEDFIENAR | -327 | -337 | 0 | 0 | 11 | NaN | NaN | NaN | NaN | 497.78192 | 569.07074 |
| Cnot1 | ACLFGGIIEK | -913 | -922 | 0 | 0 | 10 | NaN | NaN | NaN | NaN | 830.29034 | 738.92993 |
| Cnot1 | FWNHEFVH | -2336 | -2343 | 0 | 0.000230707 | 8 | NaN | NaN | NaN | NaN | 1492.8014 | NaN |
| Cct3 | EDDVGTGAGLLEIK | -340 | -353 | 0 | 0.000230707 | 14 | NaN | NaN | NaN | NaN | 1362.7205 | 1383.1284 |
| Cct3 | IVSRPEELREDDVGTGAGLLEIK | -331 | -353 | 2 | 0.001868731 | 23 | NaN | NaN | NaN | NaN | 467.45184 | 543.9756 |
| Cct3 | PGGIIEDSCVLR | -205 | -216 | 0 | 0.006624942 | 12 | NaN | NaN | NaN | NaN | 777.16675 | NaN |
| Cct3 | NVLLDPQLVPGGGASEM | -400 | -416 | 0 | 0.000755556 | 17 | NaN | NaN | NaN | NaN | 532.8205 | NaN |
| Cct3 | HTQESCETWGVN | -470 | -481 | 0 | 0.001291103 | 12 | NaN | NaN | NaN | NaN | 501.7744 | NaN |
| Itgb2 | SNEFDYPSVGQLAH | -297 | -310 | 0 | 0.00025015 | 14 | NaN | NaN | NaN | NaN | 810.9574 | 803.5413 |
| Itgb2 | LDHSTLPDTLK | -370 | -380 | 0 | 0.003620005 | 11 | NaN | NaN | NaN | NaN | 1086.1614 | NaN |
| Itgb2 | NIHVEDSLECVK | -688 | -699 | 0 | 0.003360691 | 12 | NaN | NaN | NaN | NaN | 545.3097 | NaN |
| Itgb2 | PEEIGWR | -248 | -254 | 0 | 0.00111006 | 7 | NaN | NaN | NaN | NaN | 191.0767 | NaN |
| Prkcsh | SQCYELTTNEYVYR | -406 | -419 | 0 | 0 | 14 | NaN | NaN | NaN | NaN | 1757.4216 | 1708.3802 |
| Znf330 | ACPLTDAECVECER | -121 | -134 | 0 | 0.00025015 | 14 | NaN | NaN | NaN | NaN | 333.31696 | 296.70383 |
| Rpa2 | ACPRPEGLNFQDLR | -218 | -231 | 1 | 0 | 14 | NaN | NaN | NaN | NaN | 1779.9359 | 842.9836 |
| Rpa2 | PPETYVK | -121 | -127 | 0 | 0.00504706 | 7 | NaN | NaN | NaN | NaN | 967.7815 | 953.14954 |
| Rps3a | ACQSIYPLHDVF | -200 | -211 | 0 | 0 | 12 | NaN | NaN | NaN | NaN | 2229.9646 | 299.691 |
| Rps3a | PLHDVFVR | -206 | -213 | 0 | 0.003003218 | 8 | NaN | NaN | NaN | NaN | 1361.121 | 1232.0911 |

| | | | | | | | | | | | | |
|---|---|---|---|---|---|---|---|---|---|---|---|---|
| Rps3a | PPVQESV | -258 | -264 | 0 | 0.004567621 | 7 | NaN | NaN | NaN | NaN | 1263.9656 | 1351.0347 |
| Bcat2 | ACQVCPVHQILYEGK | -342 | -356 | 0 | 0.001092529 | 15 | NaN | NaN | NaN | NaN | 875.5963 | NaN |
| Fasn | STSIPEAQWQSSLAR | -714 | -728 | 0 | 0.00111006 | 15 | NaN | NaN | NaN | NaN | 13475.81 | 10571.617 |
| Fasn | LDALFEEANDSWR | -1789 | -1801 | 0 | 0.00201047 | 13 | NaN | NaN | NaN | NaN | 1086.2004 | 537.1351 |
| Fasn | GLAPESLEYIEAHGTGTK | -281 | -298 | 0 | 0 | 18 | NaN | NaN | NaN | NaN | 574.723 | 485.58096 |
| Fasn | SFGFGGSNVHVILQPNTR | -392 | -409 | 0 | 0 | 18 | NaN | NaN | NaN | NaN | 411.4971 | 477.4948 |
| Fasn | QQEQLVPTLEK | -1107 | -1117 | 0 | 0 | 11 | NaN | NaN | NaN | NaN | 331.1181 | 501.09027 |
| Fasn | RHFQLEQDKPK | -1508 | -1518 | 2 | 0.000230707 | 11 | NaN | NaN | NaN | NaN | 309.3462 | 645.7764 |
| Fasn | MLENQTPELFQDVNKPK | -1972 | -1988 | 1 | 0.006100831 | 17 | NaN | NaN | NaN | NaN | 1382.1239 | NaN |
| Fasn | EAMGTNDTVIGGTLPQR | -2064 | -2080 | 0 | 0.001023236 | 17 | NaN | NaN | NaN | NaN | 1312.9431 | NaN |
| Fasn | EFVEQLKQEGVFAK | -660 | -673 | 1 | 0.000394361 | 14 | NaN | NaN | NaN | NaN | 988.02844 | NaN |
| Fasn | WDHSQTW | -836 | -842 | 0 | 0.004957095 | 7 | NaN | NaN | NaN | NaN | 676.4031 | NaN |
| Fasn | VHVILQPNTR | -400 | -409 | 0 | 0.001776473 | 10 | NaN | NaN | NaN | NaN | 441.8157 | NaN |
| Fasn | VAASVDLITK | -2390 | -2399 | 0 | 0 | 10 | NaN | NaN | NaN | NaN | 347.52728 | NaN |
| Fasn | HAFEVSDTGNLIVSGK | -942 | -957 | 0 | 0.001776473 | 16 | NaN | NaN | NaN | NaN | 279.22476 | NaN |
| Fasn | LKEDTQVADVTTSR | -1070 | -1083 | 1 | 0.001023236 | 14 | NaN | NaN | NaN | NaN | 236.10822 | NaN |
| Cct8 | LTPPVQEEMGHCD | -336 | -348 | 0 | 0.001190916 | 13 | NaN | NaN | NaN | NaN | 936.0719 | 876.5695 |
| Cct8 | SGLEEAVYR | -23 | -31 | 0 | 0 | 9 | NaN | NaN | NaN | NaN | 801.48364 | 1153.3833 |
| Cct8 | PGGGATEIELAK | -410 | -421 | 0 | 0.001392869 | 12 | NaN | NaN | NaN | NaN | 801.2002 | 1356.6553 |
| Cct8 | AKPAGGPKPPSGK | -527 | -539 | 2 | 0.003037016 | 13 | NaN | NaN | NaN | NaN | 566.39545 | 644.4906 |
| Cct8 | AKPAGGPKPPSGK | -527 | -539 | 2 | 0.003037016 | 13 | NaN | NaN | NaN | NaN | 566.39545 | 644.4906 |
| Cct8 | DMLEASILDTYLGK | -491 | -504 | 0 | 0 | 14 | NaN | NaN | NaN | NaN | 394.72205 | 293.17596 |
| Cct8 | KHEKEDGAISTIVLR | -364 | -378 | 2 | 0 | 15 | NaN | NaN | NaN | NaN | 329.10443 | NaN |
| Sema4d | ACVTLHQEEASSR | -528 | -540 | 0 | 0 | 13 | NaN | NaN | NaN | NaN | 316.97018 | 884.5473 |
| Sema4d | ACVTLHQEEASSR | -528 | -540 | 0 | 0 | 13 | NaN | NaN | NaN | NaN | 316.97018 | 884.5473 |
| Sema4d | CPFDPAHSY | -156 | -164 | 0 | 0.007493591 | 9 | NaN | NaN | NaN | NaN | 241.73929 | NaN |
| Atp1a1 | DKYEPAAVSEHGDKK | -8 | -22 | 2 | 0.000431655 | 15 | NaN | NaN | NaN | NaN | 467.7208 | NaN |
| Yipf4 | ADAEDLSGSIAAPDVK | -23 | -38 | 0 | 0 | 16 | NaN | NaN | NaN | NaN | 1304.6436 | NaN |
| Atg3 | GLLETDEATLDTR | -164 | -176 | 0 | 0 | 13 | NaN | NaN | NaN | NaN | 919.78235 | NaN |
| Prkar1a | AEVYTEEDAASYVR | -102 | -115 | 0 | 0 | 14 | NaN | NaN | NaN | NaN | 1405.458 | NaN |
| Rps6 | MATEVAADALGEEWK | -32 | -46 | 0 | 0 | 15 | NaN | NaN | NaN | NaN | 1349.9423 | 1461.1194 |
| Rps6 | GEKDIPGLTDTTVPR | -117 | -131 | 1 | 0.003069674 | 15 | NaN | NaN | NaN | NaN | 853.1309 | NaN |
| Api5 | SFHQLGR | -349 | -355 | 0 | 0.002068213 | 7 | NaN | NaN | NaN | NaN | 1744.7709 | 1697.8885 |
| Api5 | PQFATGENLPR | -87 | -97 | 0 | 0.00111006 | 11 | NaN | NaN | NaN | NaN | 1121.8507 | 1194.7917 |
| Ppa2 | ADCKEEHDIPR | -65 | -75 | 1 | 0.00025015 | 11 | NaN | NaN | NaN | NaN | 917.7774 | 610.14667 |
| Ppa2 | GTLALIDQSETDWK | -185 | -198 | 0 | 0 | 14 | NaN | NaN | NaN | NaN | 344.38733 | 300.50812 |
| Mcm4 | YPDSILEHQIQVRPF | -248 | -262 | 1 | 0.00025015 | 15 | NaN | NaN | NaN | NaN | 1245.7313 | 1715.3988 |
| Mcm4 | TTIENIQLPHTL | -628 | -639 | 0 | 0.00025015 | 12 | NaN | NaN | NaN | NaN | 2468.822 | 2010.05 |
| Mcm4 | PQDEAYDR | -652 | -659 | 0 | 0.00272025 | 8 | NaN | NaN | NaN | NaN | 558.74426 | 562.29736 |
| Mcm4 | SVLHEVMEQQ | -584 | -593 | 0 | 0.007343498 | 10 | NaN | NaN | NaN | NaN | 844.9287 | NaN |
| Mcm4 | IAEPCSCVHCH | -321 | -331 | 0 | 0.000471409 | 11 | NaN | NaN | NaN | NaN | 323.5238 | NaN |
| Samhd1 | NGIDVDKWDYFAR | -338 | -350 | 1 | 0 | 13 | NaN | NaN | NaN | NaN | 1303.7207 | 1241.8301 |
| Samhd1 | NDPIHGHIEFHPLLIR | -151 | -166 | 0 | 0 | 16 | NaN | NaN | NaN | NaN | 762.2 | 748.31134 |
| Samhd1 | NDPIHGHIEFHPLLIR | -151 | -166 | 0 | 0 | 16 | NaN | NaN | NaN | NaN | 762.2 | 748.31134 |
| Samhd1 | VINVDYGMEDKNPIDR | -544 | -559 | 1 | 0 | 16 | NaN | NaN | NaN | NaN | 4107.844 | 4131.723 |
| Samhd1 | WEPEDVCSFLENR | -77 | -89 | 0 | 0 | 13 | NaN | NaN | NaN | NaN | 2283.8167 | 2593.8792 |
| Samhd1 | FEHSLGVGYLAGC | -197 | -209 | 0 | 0 | 13 | NaN | NaN | NaN | NaN | 417.93393 | 406.04504 |
| Fgb | ADDDYDEPTDSLDAR | -20 | -34 | 0 | 0.006788754 | 15 | NaN | NaN | NaN | NaN | 5223.241 | 5347.441 |
| Fgb | EDGGGWWYNR | -417 | -426 | 0 | 0 | 10 | NaN | NaN | NaN | NaN | 456.34955 | 476.95233 |
| Pds5a | VDESGPPAPSKPR | -1255 | -1267 | 1 | 0 | 13 | NaN | NaN | NaN | NaN | 868.4102 | 814.4229 |
| Pds5a | FNDIHVPVRLE | -331 | -341 | 1 | 0 | 11 | NaN | NaN | NaN | NaN | 1009.1333 | 2071.8657 |
| Pds5a | DLALVNDQLLGFVR | -390 | -403 | 0 | 0.00025015 | 14 | NaN | NaN | NaN | NaN | 387.8894 | 171.2053 |
| Pds5a | IAPVHIDSEAISAL | -617 | -630 | 0 | 0.00635324 | 14 | NaN | NaN | NaN | NaN | 1900.2272 | NaN |
| Pds5a | LVSEGDLTEQKR | -867 | -878 | 1 | 0.00111006 | 12 | NaN | NaN | NaN | NaN | 560.4415 | NaN |
| Ppia | YGEKFEDENFILK | -79 | -91 | 1 | 0 | 13 | NaN | NaN | NaN | NaN | 2671.3667 | 3215.6157 |
| Ppia | TEWLDGKHVVFGK | -119 | -131 | 1 | 0 | 13 | NaN | NaN | NaN | NaN | 1063.9225 | NaN |
| Ppia | TEWLDGKHVVFGK | -119 | -131 | 1 | 0 | 13 | NaN | NaN | NaN | NaN | 1063.9225 | NaN |
| Psme2 | ELYHIISSNLEK | -215 | -226 | 0 | 0.00111006 | 12 | NaN | NaN | NaN | NaN | 1048.5438 | 1146.0576 |
| Psme2 | DTHVMDYR | -181 | -188 | 0 | 0.001291103 | 8 | NaN | NaN | NaN | NaN | 186.35268 | NaN |
| Psme2 | QEADDFLCTFLPR | -26 | -38 | 0 | 0.000550307 | 13 | NaN | NaN | NaN | NaN | 496.07928 | NaN |
| Aldoa | VNPCIGGVILF | -70 | -80 | 0 | 0.002068213 | 11 | NaN | NaN | NaN | NaN | 337.36655 | 219.3064 |
| Uhrf1 | ALNCHSPINEK | -514 | -524 | 0 | 0 | 11 | NaN | NaN | NaN | NaN | 1288.5609 | NaN |
| Arsb | ADDLGWNDLGFHGSVIR | -53 | -69 | 0 | 0.005605646 | 17 | NaN | NaN | NaN | NaN | 970.7208 | NaN |
| Ncapd2 | HSQELSSILDDAALSGSDR | -92 | -110 | 0 | 0 | 19 | NaN | NaN | NaN | NaN | 834.31305 | 989.17535 |
| Ncapd2 | LSDIDLAGPLQK | -441 | -452 | 0 | 0 | 12 | NaN | NaN | NaN | NaN | 913.2502 | NaN |
| Ncapd2 | ADDTEAELIR | -978 | -987 | 0 | 0 | 10 | NaN | NaN | NaN | NaN | 725.4186 | NaN |
| Ncapd2 | NFFNELSHK | -1141 | -1149 | 0 | 0.001092529 | 9 | NaN | NaN | NaN | NaN | 392.04895 | NaN |
| Pitpnb | ADEDPALFHSVK | -156 | -167 | 0 | 0.00025015 | 12 | NaN | NaN | NaN | NaN | 1505.6404 | NaN |
| Ipo7 | AFAVGVQQVLLK | -295 | -306 | 0 | 0 | 12 | NaN | NaN | NaN | NaN | 318.582 | 299.32318 |
| Ipo7 | QILTEPNADPR | -417 | -427 | 0 | 0.001092529 | 11 | NaN | NaN | NaN | NaN | 2296.8699 | NaN |
| Slc25a20 | FFGFGLGK | -90 | -97 | 0 | 0.003745449 | 8 | NaN | NaN | NaN | NaN | 268.90536 | 318.88083 |
| Myh9 | LQQELDDLLVDLDHQR | -1418 | -1433 | 0 | 0 | 16 | NaN | NaN | NaN | NaN | 1431.9995 | 1413.8995 |
| Myh9 | HQSSDKFVSELWK | -601 | -613 | 1 | 0.000431655 | 13 | NaN | NaN | NaN | NaN | 1292.0541 | 1196.6722 |
| Myh9 | KTDLLLEPYNK | -289 | -299 | 1 | 0 | 11 | NaN | NaN | NaN | NaN | 1281.9401 | 1388.1621 |
| Myh9 | KMEDGVGCLETAEEAKR | -1372 | -1388 | 2 | 0.00025015 | 17 | NaN | NaN | NaN | NaN | 531.54193 | 140.91393 |
| Myh9 | SLKEEVGEEAIVELVENGKK | -45 | -64 | 2 | 0 | 20 | NaN | NaN | NaN | NaN | 231.46829 | 696.19525 |
| Myh9 | LEEELEEEQGNTELINDR | -1734 | -1751 | 0 | 0.000230707 | 18 | NaN | NaN | NaN | NaN | 1760.4891 | NaN |
| Myh9 | RQLEEAEEEAQRA | -1877 | -1889 | 2 | 0.004983786 | 13 | NaN | NaN | NaN | NaN | 1497.3302 | NaN |

| | | | | | | | | | | | | |
|---|---|---|---|---|---|---|---|---|---|---|---|---|
| Myh9 | NGHVTIPGQQDK | -305 | -316 | 0 | 0.00111006 | 12 | NaN | NaN | NaN | NaN | 1006.3409 | NaN |
| Gsr | ADFDNTVAIHPTSSEELVTLR | -480 | -500 | 0 | 0 | 21 | NaN | NaN | NaN | NaN | 291.26138 | 686.412 |
| Gsr | SNCTEELENAGVEVLK | -254 | -269 | 0 | 0.000712251 | 16 | NaN | NaN | NaN | NaN | 3910.183 | 3623.651 |
| Hnrnpul1 | VYHPELK | -137 | -143 | 0 | 0.003360691 | 7 | NaN | NaN | NaN | NaN | 660.7732 | NaN |
| Slc3a2 | ASNLPAGISLPASAK | -473 | -487 | 0 | 0 | 15 | NaN | NaN | NaN | NaN | 4224.504 | 3761.39 |
| Slc3a2 | VLGPIHK | -153 | -159 | 0 | 0.002752526 | 7 | NaN | NaN | NaN | NaN | 320.8111 | NaN |
| Sptan1 | TALLELWELR | -450 | -459 | 0 | 0 | 10 | NaN | NaN | NaN | NaN | 746.9702 | 1088.061 |
| Sptan1 | ADFNQLAELDR | -2145 | -2155 | 0 | 0 | 11 | NaN | NaN | NaN | NaN | 597.8733 | 625.1594 |
| Sptan1 | QWELLLEK | -130 | -137 | 0 | 0.005789881 | 8 | NaN | NaN | NaN | NaN | 970.8296 | NaN |
| Sptan1 | SDDNTIGQEEIQQR | -1831 | -1844 | 0 | 0 | 14 | NaN | NaN | NaN | NaN | 877.1719 | NaN |
| Sptan1 | QKLEDSYR | -38 | -45 | 1 | 0.00504706 | 8 | NaN | NaN | NaN | NaN | 831.615 | NaN |
| Sptan1 | EAFETDFTVHK | -1913 | -1923 | 0 | 0 | 11 | NaN | NaN | NaN | NaN | 548.2434 | NaN |
| Sptan1 | QFQDAGHFDAENIKK | -743 | -757 | 1 | 0.00670918 | 15 | NaN | NaN | NaN | NaN | 496.20203 | NaN |
| Sptan1 | SAQADFNQLAELDR | -2142 | -2155 | 0 | 0.00025015 | 14 | NaN | NaN | NaN | NaN | 452.80685 | NaN |
| Sptan1 | LQTASDESYKDPTNIQSK | -1571 | -1588 | 1 | 0.00437931 | 18 | NaN | NaN | NaN | NaN | 310.764 | NaN |
| Sptan1 | DLSSVQTLLTK | -2007 | -2017 | 0 | 0.00272025 | 11 | NaN | NaN | NaN | NaN | 268.43948 | NaN |
| Vars1 | ATSKPEGHESLVDR | -971 | -984 | 1 | 0 | 14 | NaN | NaN | NaN | NaN | 609.4822 | 542.6097 |
| Vars1 | VQGSDSDEEVVVATTR | -521 | -536 | 0 | 0 | 16 | NaN | NaN | NaN | NaN | 2140.658 | 2615.05 |
| Vars1 | KDVSGAMPDSYSPQ | -289 | -302 | 1 | 0.006100831 | 14 | NaN | NaN | NaN | NaN | 718.8588 | 597.32825 |
| Vars1 | SPHPDAFPSLR | -7 | -17 | 0 | 0.00025015 | 11 | NaN | NaN | NaN | NaN | 1821.2648 | NaN |
| Vars1 | TLNSAISDIEVDKK | -479 | -492 | 1 | 0 | 14 | NaN | NaN | NaN | NaN | 790.784 | NaN |
| Vars1 | LWPAGLGGPGGSR | -79 | -91 | 0 | 0.00437931 | 13 | NaN | NaN | NaN | NaN | 409.55457 | NaN |
| Vars1 | ADTELIPAACGATLPALGLR | -103 | -122 | 0 | 0.005605646 | 20 | NaN | NaN | NaN | NaN | 361.83298 | NaN |
| Nup155 | ADGEFLHELEEK | -1136 | -1147 | 0 | 0 | 12 | NaN | NaN | NaN | NaN | 1407.2512 | 1572.2786 |
| Nup155 | YVENPSLVLNCER | -1334 | -1346 | 0 | 0 | 13 | NaN | NaN | NaN | NaN | 1180.8909 | 365.8645 |
| Nup155 | LQIQETLQR | -1155 | -1163 | 0 | 0.00025015 | 9 | NaN | NaN | NaN | NaN | 846.60364 | NaN |
| Ppp1cb | GDIHGQYTDLLR | -62 | -73 | 0 | 0 | 12 | NaN | NaN | NaN | NaN | 1452.9628 | NaN |
| Adss2 | VGIGAFPTEQDNEIGELL | -304 | -321 | 0 | 0.00204437 | 18 | NaN | NaN | NaN | NaN | 349.82178 | NaN |
| Pfn1 | PSVWAAVPGK | -29 | -38 | 0 | 0.00142999 | 10 | NaN | NaN | NaN | NaN | 1176.86 | 1158.2904 |
| Pfn1 | PAEVGVLVGK | -45 | -54 | 0 | 0.001463897 | 10 | NaN | NaN | NaN | NaN | 714.0607 | 753.4495 |
| Pfn1 | CYEMASHLR | -128 | -136 | 0 | 0.000471409 | 9 | NaN | NaN | NaN | NaN | 1613.759 | NaN |
| Hnrnpk | EPLEGSEDR | -425 | -433 | 0 | 0.003299042 | 9 | NaN | NaN | NaN | NaN | 630.85864 | NaN |
| Hnrnpm | GLDHMASSIER | -460 | -470 | 0 | 0.00025015 | 11 | NaN | NaN | NaN | NaN | 322.68906 | 650.576 |
| Csnk2a1 | GGPNIITLADIVK | -90 | -102 | 0 | 0 | 13 | NaN | NaN | NaN | NaN | 338.63193 | 348.7048 |
| Npm1 | MTDQEAIQDLWQWR | -276 | -289 | 0 | 0 | 14 | NaN | NaN | NaN | NaN | 758.7341 | 668.50507 |
| Npm1 | SPLRPQNYLFGCELK | -10 | -24 | 1 | 0.001868731 | 15 | NaN | NaN | NaN | NaN | 927.83057 | 895.8695 |
| Npm1 | CGSGPVHISG | -104 | -113 | 0 | 0.000550307 | 10 | NaN | NaN | NaN | NaN | 14375.386 | NaN |
| Rbbp4 | SNNTSKPSHSVDAHTAEVN | -259 | -277 | 1 | 0 | 19 | NaN | NaN | NaN | NaN | 353.90967 | 393.0263 |
| Rbbp4 | SNNTSKPSHSVDAHTAEVN | -259 | -277 | 1 | 0 | 19 | NaN | NaN | NaN | NaN | 353.90967 | 393.0263 |
| Rbbp4 | GKIEIEIK | -113 | -120 | 1 | 0 | 8 | NaN | NaN | NaN | NaN | 259.64154 | 239.43791 |
| Tmsb10 | EKNTLPTK | -25 | -32 | 1 | 0.001776473 | 8 | NaN | NaN | NaN | NaN | 1411.7535 | 1374.1416 |
| Ranbp3 | LNDMASTDDGTLQSR | -352 | -366 | 0 | 0.00025015 | 15 | NaN | NaN | NaN | NaN | 401.25195 | NaN |
| Sptbn1 | KLLDPEDISVDHPDEK | -249 | -264 | 1 | 0.003745449 | 16 | NaN | NaN | NaN | NaN | 3104.44 | NaN |
| Sptbn1 | EGMQLISEKPETEAVVK | -1336 | -1352 | 1 | 0.002068213 | 17 | NaN | NaN | NaN | NaN | 1120.8604 | 253.96376 |
| Sptbn1 | LIEETEKR | -1577 | -1584 | 1 | 0.00111006 | 8 | NaN | NaN | NaN | NaN | 576.0768 | 517.6155 |
| Sptbn1 | HENIKNEIDNYEEDYQK | -1104 | -1120 | 1 | 0.00635324 | 17 | NaN | NaN | NaN | NaN | 1393.4948 | NaN |
| Sptbn1 | VADSHPESER | -1659 | -1668 | 0 | 0.000550307 | 10 | NaN | NaN | NaN | NaN | 1153.3474 | NaN |
| Efhd2 | RADLNQGIGEPQ | -62 | -73 | 1 | 0.004350979 | 12 | NaN | NaN | NaN | NaN | 2611.37 | NaN |
| Efhd2 | SRFEEEIK | -205 | -212 | 1 | 0.005917757 | 8 | NaN | NaN | NaN | NaN | 2606.8066 | NaN |
| Efhd2 | VFNPYTEFKEF | -79 | -89 | 1 | 0 | 11 | NaN | NaN | NaN | NaN | 346.28552 | NaN |
| Adsl | VKQEGGDNDLIER | -414 | -426 | 1 | 0.000394361 | 13 | NaN | NaN | NaN | NaN | 1031.9249 | 1023.2151 |
| Adsl | FLEEEVRPLLKPY | -460 | -472 | 2 | 0.00025015 | 13 | NaN | NaN | NaN | NaN | 429.53937 | 450.4319 |
| Adsl | ADLPTLGFTH | -150 | -159 | 0 | 0.002168954 | 10 | NaN | NaN | NaN | NaN | 421.22852 | 1105.0369 |
| Mdh2 | GYLGPEQLPDCLK | -79 | -91 | 0 | 0 | 13 | NaN | NaN | NaN | NaN | 5766.647 | 5452.714 |
| Mdh2 | VDFPQDQLATLT | -216 | -227 | 0 | 0.000431655 | 12 | NaN | NaN | NaN | NaN | 1742.7814 | 2784.5823 |
| Mdh2 | VDFPQDQLATL | -216 | -226 | 0 | 0.000627943 | 11 | NaN | NaN | NaN | NaN | 5348.519 | 5121.901 |
| Mdh2 | YDIAHTPGVAADLSHIETR | -56 | -74 | 0 | 0.000298805 | 19 | NaN | NaN | NaN | NaN | 1183.0321 | NaN |
| Cct6a | AEMGVWDNYCVK | -490 | -501 | 0 | 0.008947895 | 12 | NaN | NaN | NaN | NaN | 695.4645 | NaN |
| Phb2 | IGGVQQDTILAEGL | -55 | -68 | 0 | 0.00122419 | 14 | NaN | NaN | NaN | NaN | 377.21613 | 397.88116 |
| Prpf8 | LKDLILADYGK | -1983 | -1993 | 1 | 0.00025015 | 11 | NaN | NaN | NaN | NaN | 192.0569 | 772.9702 |
| Prpf8 | KFICISDLR | -2113 | -2121 | 1 | 0.008919338 | 9 | NaN | NaN | NaN | NaN | 1109.4333 | NaN |
| Prpf8 | AQIAGYLYGVSPPDNPQVK | -2122 | -2140 | 0 | 0.00025015 | 19 | NaN | NaN | NaN | NaN | 570.0421 | NaN |
| Glod4 | ELPDLEDLMK | -210 | -219 | 0 | 0.001743443 | 10 | NaN | NaN | NaN | NaN | 1153.5416 | NaN |
| Stk24 | GGSDSGDWIFTIR | -318 | -330 | 0 | 0 | 13 | NaN | NaN | NaN | NaN | 883.61884 | NaN |
| Arhgdia | STAPGPLELDLTGDLESFKK | -80 | -99 | 1 | 0 | 20 | NaN | NaN | NaN | NaN | 502.83234 | 475.9908 |
| Copb2 | GSSSPNFTLEGHEK | -173 | -186 | 0 | 0 | 14 | NaN | NaN | NaN | NaN | 2622.9321 | 2713.0435 |
| Copb2 | LWDWDKK | -125 | -131 | 1 | 0.000627943 | 7 | NaN | NaN | NaN | NaN | 1881.6497 | 1611.2534 |
| Copb2 | CIAVHPTQPFIL | -104 | -115 | 0 | 0 | 12 | NaN | NaN | NaN | NaN | 1553.608 | NaN |
| Coro1a | DRPHEGTRPVHAVF | -215 | -228 | 2 | 0.002629273 | 14 | NaN | NaN | NaN | NaN | 1324.23 | 419.70978 |
| Coro1a | DRPHEGTRPVHAVFVSEGK | -215 | -233 | 2 | 0.00025015 | 19 | NaN | NaN | NaN | NaN | 402.13913 | 1388.9989 |
| Coro1a | KSDLFQEDLYPPTAGPDPA | -355 | -373 | 1 | 0.00272025 | 19 | NaN | NaN | NaN | NaN | 425.71277 | 589.1051 |
| Coro1a | DRPHEGTRPVHA | -215 | -226 | 2 | 0.000755556 | 12 | NaN | NaN | NaN | NaN | 858.4818 | NaN |
| Coro1a | SDLFQEDLYPPTAGPDPALTAE | -356 | -377 | 0 | 0.00504706 | 22 | NaN | NaN | NaN | NaN | 184.0106 | NaN |
| Calm1;Calm2; | HVMTNLGEK | (108);(108);(108) | (116);(116);(116) | 0 | 0 | 9 | NaN | NaN | NaN | NaN | 836.1322 | NaN |
| Calm1;Calm2; | KEAFSLFDKDGDGTITTK | (14);(14);(14) | (31);(31);(31) | 2 | 0.002068213 | 18 | NaN | NaN | NaN | NaN | 468.9032 | NaN |
| Plcg1 | YQQPFEDFR | -1253 | -1261 | 0 | 0 | 9 | NaN | NaN | NaN | NaN | 632.26105 | 1606.4899 |
| Plcg1 | LSEPVPQTNAHESK | -653 | -666 | 0 | 0.001023236 | 14 | NaN | NaN | NaN | NaN | 1504.7751 | NaN |
| Pfkp | DLLFQPVAELK | -725 | -735 | 0 | 0 | 11 | NaN | NaN | NaN | NaN | 4925.829 | 4954.9673 |

| | | | | | | | | | | | | |
|---|---|---|---|---|---|---|---|---|---|---|---|---|
| Pfkp | SEGAIDMQNKPITSEK | -271 | -286 | 1 | 0.006915721 | 16 | NaN | NaN | NaN | NaN | 711.886 | 609.0503 |
| Pfkp | FVSDDSICVLGICK | -710 | -723 | 0 | 0.00078661 | 14 | NaN | NaN | NaN | NaN | 1342.1912 | NaN |
| Tars1 | ADTDLDPGCTLNKK | -647 | -660 | 1 | 0 | 14 | NaN | NaN | NaN | NaN | 1781.5688 | NaN |
| Tars1 | DQELYFFHEL | -328 | -337 | 0 | 0.004350979 | 10 | NaN | NaN | NaN | NaN | 440.7605 | NaN |
| Psmc4 | EFLHAQEEVKR | -71 | -81 | 1 | 0.001055699 | 11 | NaN | NaN | NaN | NaN | 2590.6692 | 2151.5242 |
| Psmc4 | FLEAVDQNTAIVGSTTGSNYYVR | -92 | -114 | 0 | 0.001092529 | 23 | NaN | NaN | NaN | NaN | 133.03308 | 156.07245 |
| Ldha | AIGLSVADLAESIMK | -251 | -265 | 0 | 0 | 15 | NaN | NaN | NaN | NaN | 1526.3893 | 1317.5831 |
| Ldha | RVHPISTMIK | -269 | -278 | 1 | 0 | 10 | NaN | NaN | NaN | NaN | 432.1757 | 469.5753 |
| Ldha | GEMMDLQHGSLFLK | -60 | -73 | 0 | 0 | 14 | NaN | NaN | NaN | NaN | 1364.4857 | 1253.7855 |
| Ldha | VADLAESIMK | -256 | -265 | 0 | 0.001463897 | 10 | NaN | NaN | NaN | NaN | 1267.193 | 2646.4832 |
| Smarcc1 | FWESPDTVSQLDSVR | -41 | -55 | 0 | 0.009416377 | 15 | NaN | NaN | NaN | NaN | 3677.968 | NaN |
| Smarcc1 | DKQVLVHW | -225 | -232 | 1 | 0.005789881 | 8 | NaN | NaN | NaN | NaN | 2635.206 | NaN |
| Smarcc1 | ADTPSGLVPLHLR | -559 | -571 | 0 | 0.008736733 | 13 | NaN | NaN | NaN | NaN | 2391.2454 | NaN |
| Smarcc1 | WESPDTVSQLDSVR | -42 | -55 | 0 | 0.00111006 | 14 | NaN | NaN | NaN | NaN | 520.8254 | NaN |
| Sun2 | KQPEVWESR | -270 | -278 | 1 | 0.001708008 | 9 | NaN | NaN | NaN | NaN | 1150.5883 | NaN |
| Sun2 | DEMHAQLQELENK | -493 | -505 | 0 | 0.001392869 | 13 | NaN | NaN | NaN | NaN | 361.56964 | NaN |
| Pcna | GNEDIITLR | -83 | -91 | 0 | 0 | 9 | NaN | NaN | NaN | NaN | 1237.6105 | 1331.6561 |
| Pcna | PLVVEYK | -234 | -240 | 0 | 0.004350979 | 7 | NaN | NaN | NaN | NaN | 412.28146 | 309.14365 |
| Pcna | PLVVEYK | -234 | -240 | 0 | 0.004350979 | 7 | NaN | NaN | NaN | NaN | 412.28146 | 309.14365 |
| Pcna | DLSHIGDAVVISCAK | -150 | -164 | 0 | 0.00025015 | 15 | NaN | NaN | NaN | NaN | 1086.1616 | NaN |
| Gbp2 | KAITHYEEQMNQK | -328 | -340 | 1 | 0 | 13 | NaN | NaN | NaN | NaN | 687.85425 | 838.9446 |
| Iars2 | FSGVDLEGGTCSHPLTPDK | -354 | -372 | 0 | 0 | 19 | NaN | NaN | NaN | NaN | 498.0327 | 480.0457 |
| Iars2 | YRDTVLLPQ | -60 | -68 | 1 | 0.00450079 | 9 | NaN | NaN | NaN | NaN | 872.357 | NaN |
| Gcn1 | LFLDHPHR | -549 | -556 | 0 | 0 | 8 | NaN | NaN | NaN | NaN | 1100.219 | 1901.7512 |
| Gcn1 | LTEVLTDSHVK | -1543 | -1553 | 0 | 0.000471409 | 11 | NaN | NaN | NaN | NaN | 914.8712 | NaN |
| Fubp1 | CQHAAEIITDLLR | -328 | -340 | 0 | 0 | 13 | NaN | NaN | NaN | NaN | 689.3362 | NaN |
| Fubp1 | LLDQIVEK | -159 | -166 | 0 | 0 | 8 | NaN | NaN | NaN | NaN | 623.45905 | NaN |
| Fubp1 | AEIITDLLR | -332 | -340 | 0 | 0 | 9 | NaN | NaN | NaN | NaN | 241.54459 | NaN |
| Tkt | SKDDQVTVIGAGVTLH | -498 | -513 | 1 | 0 | 16 | NaN | NaN | NaN | NaN | 940.80756 | 734.5092 |
| Tkt | KLDNLVAIFDINR | -174 | -186 | 1 | 0 | 13 | NaN | NaN | NaN | NaN | 571.9509 | 527.457 |
| Tkt | TIIVDGHSVEELCK | -213 | -226 | 0 | 0 | 14 | NaN | NaN | NaN | NaN | 949.43964 | NaN |
| Tkt | PFTIKPLDR | -534 | -542 | 1 | 0.003428289 | 9 | NaN | NaN | NaN | NaN | 841.49524 | NaN |
| Tkt | QHQVDIYQK | -196 | -204 | 0 | 0.00272025 | 9 | NaN | NaN | NaN | NaN | 390.70645 | NaN |
| Surf4 | LCLISTFLEDGIR | -31 | -43 | 0 | 0 | 13 | NaN | NaN | NaN | NaN | 433.8922 | NaN |
| Copa | IQVPNCDEIFYA | -448 | -459 | 0 | 0.005758933 | 12 | NaN | NaN | NaN | NaN | 400.5434 | 550.35645 |
| Copa | KGYPEVALH | -632 | -640 | 1 | 0.007040624 | 9 | NaN | NaN | NaN | NaN | 8019.259 | NaN |
| Copa | ASNLENSTYDLYTIPK | -383 | -398 | 0 | 0.006820063 | 16 | NaN | NaN | NaN | NaN | 3170.5857 | NaN |
| Copa | GYPEVALHFVK | -633 | -643 | 0 | 0.00383302 | 11 | NaN | NaN | NaN | NaN | 1214.6183 | NaN |
| Copa | KLNDLIQR | -1002 | -1009 | 1 | 0 | 8 | NaN | NaN | NaN | NaN | 1061.5789 | NaN |
| Copa | ETIPDIDPNAK | -783 | -793 | 0 | 0 | 11 | NaN | NaN | NaN | NaN | 1035.3705 | NaN |
| Copa | KIQVPNCDEIFYA | -447 | -459 | 1 | 0.002752526 | 13 | NaN | NaN | NaN | NaN | 507.22314 | NaN |
| Anxa2 | CIQNKPLYFADR | -262 | -273 | 1 | 0 | 12 | NaN | NaN | NaN | NaN | 862.54456 | 767.0268 |
| Anxa2 | DLEKDIISDTSGDFR | -154 | -168 | 1 | 0 | 15 | NaN | NaN | NaN | NaN | 1379.9238 | NaN |
| Anxa2 | QDIAFAYQR | -69 | -77 | 0 | 0 | 9 | NaN | NaN | NaN | NaN | 1266.7169 | NaN |
| Rps3 | LPTTPISEQK | -218 | -227 | 0 | 0.004350979 | 10 | NaN | NaN | NaN | NaN | 432.827 | 618.6175 |
| Ranbp2 | VELVTGEEDEKVLYSQ | -1860 | -1875 | 1 | 0.007546675 | 16 | NaN | NaN | NaN | NaN | 996.6422 | 1294.0481 |
| Ranbp2 | FESPATGILSPR | -945 | -956 | 0 | 0 | 12 | NaN | NaN | NaN | NaN | 704.37335 | 202.52698 |
| Ranbp2 | EGHWDCSVCLVR | -1411 | -1422 | 0 | 0 | 12 | NaN | NaN | NaN | NaN | 427.10593 | 435.26868 |
| Ranbp2 | SYQPLCLPLPVCR | -509 | -521 | 0 | 0.00272025 | 13 | NaN | NaN | NaN | NaN | 685.19745 | NaN |
| Ranbp2 | AEDIENDALSPEEQEECK | -691 | -708 | 0 | 0.000471409 | 18 | NaN | NaN | NaN | NaN | 340.03687 | NaN |
| Hdlbp | IQIPRPDDPSNQIK | -184 | -197 | 1 | 0.00025015 | 14 | NaN | NaN | NaN | NaN | 950.3624 | 839.9885 |
| Hdlbp | SSVAVLTQESFAEHR | -2 | -16 | 0 | 0.000230707 | 15 | NaN | NaN | NaN | NaN | 1995.2145 | 1835.2561 |
| Hdlbp | RCDIIIISGR | -947 | -956 | 1 | 0.006472924 | 10 | NaN | NaN | NaN | NaN | 885.8622 | 728.0614 |
| Rtcb | NVTDVVNTCHDAGISK | -477 | -492 | 0 | 0 | 16 | NaN | NaN | NaN | NaN | 352.96866 | NaN |
| Trmt2a | WTVPYTEQLEQK | -166 | -177 | 0 | 0.000471409 | 12 | NaN | NaN | NaN | NaN | 3677.3196 | NaN |
| Trmt2a | EHQEFQTPR | -590 | -598 | 0 | 0 | 9 | NaN | NaN | NaN | NaN | 1948.7781 | NaN |
| Arfgap1 | AEDSSDRPLEGH | -298 | -309 | 1 | 0.004595494 | 12 | NaN | NaN | NaN | NaN | 550.7941 | NaN |
| Canx | TDAPQPDVKDEEGKEEEK | -518 | -535 | 2 | 0 | 18 | NaN | NaN | NaN | NaN | 575.212 | 621.3272 |
| Canx | PGCGVWQR | -365 | -372 | 0 | 0.000712251 | 8 | NaN | NaN | NaN | NaN | 665.62146 | 1187.3619 |
| Canx | SDIFFDNFIISGDR | -432 | -445 | 0 | 0.002329279 | 14 | NaN | NaN | NaN | NaN | 554.39136 | NaN |
| P4ha1 | NYGVGGQYEPHFDFAR | -419 | -434 | 0 | 0 | 16 | NaN | NaN | NaN | NaN | 1324.886 | 1305.8318 |
| P4ha1 | LTQLEEGELSTVDK | -191 | -204 | 0 | 0.004412728 | 14 | NaN | NaN | NaN | NaN | 513.61755 | NaN |
| Cdc37 | LGPGGLDPVEVY | -288 | -299 | 0 | 0.00025015 | 12 | NaN | NaN | NaN | NaN | 411.32275 | NaN |
| Acin1 | AEEEEDQETPSR | -245 | -256 | 0 | 0 | 12 | NaN | NaN | NaN | NaN | 2235.4695 | 2292.462 |
| Acin1 | LVDRPSETK | -1098 | -1106 | 1 | 0.00025015 | 9 | NaN | NaN | NaN | NaN | 365.5804 | 486.52646 |
| Supt6h | FVEEEDDDEEEEEENLDDQDER | -30 | -51 | 0 | 0.003102268 | 22 | NaN | NaN | NaN | NaN | 164.2343 | 158.47566 |
| Supt6h | AEESEEEYNHEGEVVPR | -9 | -25 | 0 | 0.001811524 | 17 | NaN | NaN | NaN | NaN | 558.47504 | NaN |
| Ipo5 | SIIAEGEMHEAIK | -1040 | -1052 | 0 | 0 | 13 | NaN | NaN | NaN | NaN | 543.40045 | 484.53574 |
| Ipo5 | FLFDSVSSQNMGLR | -135 | -148 | 0 | 0 | 14 | NaN | NaN | NaN | NaN | 456.71014 | 375.9684 |
| Ipo5 | EFQQYLPVVMGPL | -618 | -630 | 0 | 0.000591124 | 13 | NaN | NaN | NaN | NaN | 307.89056 | 432.0201 |
| Ipo5 | LLQTMEDQGNQR | -446 | -457 | 0 | 0.003037016 | 12 | NaN | NaN | NaN | NaN | 1275.8173 | NaN |
| Pa2g4 | PFEPDLYK | -337 | -344 | 0 | 0 | 8 | NaN | NaN | NaN | NaN | 872.68195 | 565.3496 |
| Phf6 | RAEFGDFDIK | -257 | -266 | 1 | 0.001743443 | 10 | NaN | NaN | NaN | NaN | 692.9914 | NaN |
| Twf2 | GIHATEELK | -6 | -14 | 0 | 0.00025015 | 9 | NaN | NaN | NaN | NaN | 914.7869 | 973.5441 |
| Twf2 | GIHATEELKEFFAK | -6 | -19 | 1 | 0.00025015 | 14 | NaN | NaN | NaN | NaN | 930.9055 | NaN |
| Dhx15 | YTPEPERDYLEAAIR | -323 | -337 | 1 | 0.00078661 | 15 | NaN | NaN | NaN | NaN | 1748.609 | 1094.9502 |
| Mars1 | YHAIHADIYR | -338 | -347 | 0 | 0 | 10 | NaN | NaN | NaN | NaN | 566.4454 | 608.0989 |
| Mars1 | WELQHYHQLLEK | -689 | -700 | 0 | 0 | 12 | NaN | NaN | NaN | NaN | 960.94885 | 1078.7867 |

| | | | | | | | | | | | | |
|---|---|---|---|---|---|---|---|---|---|---|---|---|
| Mars1 | PGSDWTPNAR | -472 | -481 | 0 | 0.000230707 | 10 | NaN | NaN | NaN | NaN | 1081.2655 | 1320.5558 |
| Mars1 | AEGKPIETPK | -888 | -897 | 1 | 0.000431655 | 10 | NaN | NaN | NaN | NaN | 2089.654 | NaN |
| Mars1 | YHAIHADIY | -338 | -346 | 0 | 0.003360691 | 9 | NaN | NaN | NaN | NaN | 128.6453 | NaN |
| Tnpo1 | DSDVLDRPLNIMIPK | -172 | -186 | 1 | 0.00025015 | 15 | NaN | NaN | NaN | NaN | 227.75331 | 426.32898 |
| Ddb1 | KICYQEVSQCF | -723 | -733 | 1 | 0.000627943 | 11 | NaN | NaN | NaN | NaN | 513.0552 | 954.557 |
| Ddb1 | ANNSTLTIGTIDEIQK | -694 | -709 | 0 | 0 | 16 | NaN | NaN | NaN | NaN | 349.92896 | 329.08496 |
| Ddb1 | GLNIETGLLSDR | -615 | -626 | 0 | 0.00051306 | 12 | NaN | NaN | NaN | NaN | 1011.2417 | NaN |
| Flna | VAQPSITDNKDGTVTVR | -1815 | -1831 | 1 | 0 | 17 | NaN | NaN | NaN | NaN | 1981.4298 | 1911.8617 |
| Flna | YSPSEAGLHEMDIR | -1832 | -1845 | 0 | 0 | 14 | NaN | NaN | NaN | NaN | 1708.7625 | 1678.4205 |
| Flna | GLVEPVDVVDNADGTQTVN | -1492 | -1510 | 0 | 0.005605646 | 19 | NaN | NaN | NaN | NaN | 10200.453 | 9160.648 |
| Flna | EAEIVEGENHTYCIR | -2187 | -2201 | 0 | 0 | 15 | NaN | NaN | NaN | NaN | 733.0886 | 482.47894 |
| Flna | CSGPGLER | -1165 | -1172 | 0 | 0.001023236 | 8 | NaN | NaN | NaN | NaN | 537.838 | 853.34326 |
| Flna | WDEEEDEEAK | -155 | -164 | 0 | 0.00149795 | 10 | NaN | NaN | NaN | NaN | 313.01733 | 372.47192 |
| Flna | YIPVQQGPVGVN | -937 | -948 | 0 | 0.003490217 | 12 | NaN | NaN | NaN | NaN | 117.18193 | NaN |
| Flna | VAQPSITDNK | -1815 | -1824 | 0 | 0.001055699 | 10 | NaN | NaN | NaN | NaN | 6184.043 | NaN |
| Flna | RAEFTVETR | -301 | -309 | 1 | 0.001776473 | 9 | NaN | NaN | NaN | NaN | 1406.0018 | NaN |
| Flna | LPQLPITNFSR | -180 | -190 | 0 | 0 | 11 | NaN | NaN | NaN | NaN | 1175.5754 | NaN |
| Flna | VSGLGDKVDVGKDQEFTVK | -976 | -994 | 2 | 0.000431655 | 19 | NaN | NaN | NaN | NaN | 612.76996 | NaN |
| Flna | PQHATSGPGPADVSK | -2545 | -2559 | 0 | 0.00025015 | 15 | NaN | NaN | NaN | NaN | 346.2829 | NaN |
| Rpl13a | RLAHEVGWK | -140 | -148 | 1 | 0 | 9 | NaN | NaN | NaN | NaN | 683.726 | 716.0845 |
| Rpl13a | RLAHEVGWK | -140 | -148 | 1 | 0 | 9 | NaN | NaN | NaN | NaN | 683.726 | 716.0845 |
| Ppp5c | HGGLFSEDGVTLDDIR | -352 | -367 | 0 | 0.00111006 | 16 | NaN | NaN | NaN | NaN | 875.8388 | NaN |
| Ppp5c | HGGLFSEDGVTLDDIR | -352 | -367 | 0 | 0.00111006 | 16 | NaN | NaN | NaN | NaN | 875.8388 | NaN |
| Ipo4 | ICSLILPDNR | -1027 | -1036 | 0 | 0 | 10 | NaN | NaN | NaN | NaN | 1054.5211 | 1021.37463 |
| Ipo4 | AEHGGCPAQDHFPK | -928 | -941 | 0 | 0 | 14 | NaN | NaN | NaN | NaN | 575.40967 | NaN |
| Eif2s3x | KLDDPSCPRPECYR | -90 | -103 | 2 | 0.00025015 | 14 | NaN | NaN | NaN | NaN | 806.885 | 184.66144 |
| Eif2s3x | STPDEFPTDIPGTK | -108 | -121 | 0 | 0.000907206 | 14 | NaN | NaN | NaN | NaN | 868.0629 | NaN |
| Uchl5 | MLIEEEVQK | -269 | -277 | 0 | 0 | 9 | NaN | NaN | NaN | NaN | 2538.3489 | 652.6822 |
| Mthfd1l | THLSLSHEPDKK | -909 | -920 | 1 | 0 | 12 | NaN | NaN | NaN | NaN | 1088.5099 | 794.31903 |
| Mthfd1l | YDVQLPIVEK | -858 | -867 | 0 | 0 | 10 | NaN | NaN | NaN | NaN | 1681.2983 | 384.85614 |
| Mthfd1l | KLQPLSPVPSDIEISR | -351 | -366 | 1 | 0.002595029 | 16 | NaN | NaN | NaN | NaN | 1437.7548 | 296.12302 |
| Mthfd1l | LQDGQTFSSHGPGNPEAPGMDSIVR | -51 | -75 | 0 | 0.002434616 | 25 | NaN | NaN | NaN | NaN | 500.08435 | 421.02402 |
| Ap2m1 | AEIELLPTNDK | -368 | -378 | 0 | 0.000230707 | 11 | NaN | NaN | NaN | NaN | 3731.1 | NaN |
| Ganab | YRVPDVLVADPPTAR | -113 | -127 | 1 | 0 | 15 | NaN | NaN | NaN | NaN | 902.1506 | 823.66547 |
| Ganab | DAVHYGGWEHR | -559 | -569 | 0 | 0 | 11 | NaN | NaN | NaN | NaN | 575.3749 | 885.63403 |
| Ganab | DWSIHLR | -938 | -944 | 0 | 0.008222027 | 7 | NaN | NaN | NaN | NaN | 776.67285 | NaN |
| Ganab | GELFLDDGHTFNYQTR | -842 | -857 | 0 | 0.00142999 | 16 | NaN | NaN | NaN | NaN | 392.69037 | NaN |
| Eef2 | DIKLDSEDKDK | -320 | -330 | 2 | 0 | 11 | NaN | NaN | NaN | NaN | 849.63855 | 853.5166 |
| Eef2 | AYLPVNESFGFT | -786 | -797 | 0 | 0.000394361 | 12 | NaN | NaN | NaN | NaN | 953.4485 | 776.6435 |
| Eef2 | GEGQLSAAER | -240 | -249 | 0 | 0 | 10 | NaN | NaN | NaN | NaN | 3817.8223 | 4368.8086 |
| Eef2 | PFPDGLAEDIDKGEVSAR | -608 | -625 | 1 | 0 | 18 | NaN | NaN | NaN | NaN | 3611.6985 | 6074.1963 |
| Eef2 | RIMGPNYTPGK | -428 | -438 | 1 | 0.007791272 | 11 | NaN | NaN | NaN | NaN | 1890.032 | NaN |
| Eef2 | EGIPALDNFLDKL | -846 | -858 | 1 | 0 | 13 | NaN | NaN | NaN | NaN | 407.72766 | NaN |
| Got2 | EGSSHNWQHITDQIGMF | -365 | -381 | 0 | 0.00025015 | 17 | NaN | NaN | NaN | NaN | 205.1734 | 361.8684 |
| Sf1 | DGQMLPGEDEPLHAL | -192 | -206 | 0 | 0 | 15 | NaN | NaN | NaN | NaN | 517.8417 | 481.32837 |
| Grap2 | GGSLDINDGHCGLGSEVNAT | -228 | -247 | 0 | 0.00149795 | 20 | NaN | NaN | NaN | NaN | 407.45575 | 338.9198 |
| Eif2s2 | EYVTCHTCR | -275 | -283 | 0 | 0 | 9 | NaN | NaN | NaN | NaN | 504.295 | 436.14075 |
| Umps | SWADIVNAHVVPGSGVVK | -335 | -352 | 0 | 0 | 18 | NaN | NaN | NaN | NaN | 1246.7726 | 1455.7488 |
| Psmd11 | SILELGSLLAK | -49 | -59 | 0 | 0.00025015 | 11 | NaN | NaN | NaN | NaN | 238.9033 | 250.0997 |
| Psmd11 | TGQAAELGGLLK | -60 | -71 | 0 | 0.007224385 | 12 | NaN | NaN | NaN | NaN | 2173.5823 | NaN |
| Sin3a | TQYEEHIYR | -604 | -612 | 0 | 0.00025015 | 9 | NaN | NaN | NaN | NaN | 5775.55 | 6121.628 |
| Nasp | ESLDVDSEAKK | -35 | -45 | 1 | 0.00025015 | 11 | NaN | NaN | NaN | NaN | 2767.568 | 2060.0295 |
| Nasp | SNCVTDISHLVR | -692 | -703 | 0 | 0.002365814 | 12 | NaN | NaN | NaN | NaN | 1366.4146 | NaN |
| Sra1 | FTVEPENQTIPGFQQPS | -204 | -220 | 0 | 0.000230707 | 17 | NaN | NaN | NaN | NaN | 1389.4456 | NaN |
| Gvin1 | ILSEHLCGCWK | -2283 | -2293 | 0 | 0 | 11 | NaN | NaN | NaN | NaN | 2267.8518 | 2351.3594 |
| Gvin1 | LLDFSQPNSVAELQ | -78 | -91 | 0 | 0.001055699 | 14 | NaN | NaN | NaN | NaN | 580.588 | 550.4291 |
| Gvin1 | RTTVEITDISTEEKR | -760 | -774 | 2 | 0.00111006 | 15 | NaN | NaN | NaN | NaN | 556.75635 | 413.20435 |
| Gvin1 | TTVEITDISTEEK | -761 | -773 | 0 | 0.00078661 | 13 | NaN | NaN | NaN | NaN | 527.5542 | 512.25946 |
| Gvin1 | GKELSDEELHEK | -1883 | -1894 | 1 | 0 | 12 | NaN | NaN | NaN | NaN | 448.04245 | 663.04333 |
| Gvin1 | VSTPIVSFIR | -1053 | -1062 | 0 | 0.003521747 | 10 | NaN | NaN | NaN | NaN | 423.67343 | 221.26172 |
| Gvin1 | NEAELTEELTNAIK | -1205 | -1218 | 0 | 0 | 14 | NaN | NaN | NaN | NaN | 2714.2473 | NaN |
| Gvin1 | LLIEGDYK | -814 | -821 | 0 | 0.004439092 | 8 | NaN | NaN | NaN | NaN | 856.8311 | NaN |
| Kpna3 | NCDVLSHFPNLLSHPK | -324 | -339 | 0 | 0.001868731 | 16 | NaN | NaN | NaN | NaN | 600.0611 | 504.63254 |
| Fga | IDEANQDFTNR | -75 | -85 | 0 | 0.00111006 | 11 | NaN | NaN | NaN | NaN | 435.41345 | NaN |
| Mybbp1a | LVQLADMLLNHNR | -543 | -555 | 0 | 0 | 13 | NaN | NaN | NaN | NaN | 1108.0819 | 1175.5657 |
| Mybbp1a | GLPVMPTFWR | -382 | -391 | 0 | 0 | 10 | NaN | NaN | NaN | NaN | 653.4808 | 667.5708 |
| Mybbp1a | PSDTQSPVSTK | -1159 | -1169 | 0 | 0.001708008 | 11 | NaN | NaN | NaN | NaN | 1276.9142 | 839.6124 |
| Mybbp1a | VDILSEVPESMFQEILPK | -180 | -197 | 0 | 0.001023236 | 18 | NaN | NaN | NaN | NaN | 370.4535 | 248.89674 |
| Mybbp1a | RYCHEVGPCAEAL | -888 | -900 | 1 | 0.001636916 | 13 | NaN | NaN | NaN | NaN | 5726.207 | NaN |
| Mybbp1a | LEGCQDDPKR | -358 | -367 | 1 | 0.00111006 | 10 | NaN | NaN | NaN | NaN | 2185.0535 | NaN |
| Mybbp1a | VYSASLESLLTK | -972 | -983 | 0 | 0 | 12 | NaN | NaN | NaN | NaN | 590.02997 | NaN |
| Mybbp1a | CLDLDFVTR | -963 | -971 | 0 | 0.001904399 | 9 | NaN | NaN | NaN | NaN | 511.95007 | NaN |
| Mybbp1a | DTEDSEDQAANCLDLDFVTR | -952 | -971 | 0 | 0.000471409 | 20 | NaN | NaN | NaN | NaN | 318.84357 | NaN |
| Trap1 | EQDIKEDIAK | -459 | -468 | 1 | 0 | 10 | NaN | NaN | NaN | NaN | 761.26697 | NaN |
| Ogdh | PENGPAAQDPHK | -886 | -897 | 0 | 0.00111006 | 12 | NaN | NaN | NaN | NaN | 3453.6335 | 13634.933 |
| Ogdh | TKAEQFYCGDTEGK | -388 | -401 | 1 | 0 | 14 | NaN | NaN | NaN | NaN | 514.88306 | 480.01343 |
| Ogdh | GLHESDLDK | -176 | -184 | 0 | 0.002296977 | 9 | NaN | NaN | NaN | NaN | 7920.55 | NaN |
| Ogdh | NAGAPPGTAYQSPLSLSR | -84 | -101 | 0 | 0.008222027 | 18 | NaN | NaN | NaN | NaN | 5033.1133 | NaN |

| | | | | | | | | | | | | |
|---|---|---|---|---|---|---|---|---|---|---|---|---|
| Ogdh | AEQFYCGDTEGKK | -390 | -402 | 1 | 0.001636916 | 13 | NaN | NaN | NaN | NaN | 1025.856 | NaN |
| Ogdh | AFDLDAFKK | -1013 | -1021 | 1 | 0.001291103 | 9 | NaN | NaN | NaN | NaN | 798.8032 | NaN |
| Luc7l | SHLLDCCPHDILAGTR | -38 | -53 | 0 | 0 | 16 | NaN | NaN | NaN | NaN | 1127.9159 | 1543.424 |
| Luc7l | LAETQEEISAEVSAK | -108 | -122 | 0 | 0.001868731 | 15 | NaN | NaN | NaN | NaN | 763.92334 | 1105.6279 |
| Ubqln1 | DQDTLSQHGIHDGLTVH | -78 | -94 | 0 | 0 | 17 | NaN | NaN | NaN | NaN | 605.15326 | 501.52148 |
| Ubqln1 | EKEEFAVPENSSVQQFK | -37 | -53 | 1 | 0.002296977 | 17 | NaN | NaN | NaN | NaN | 456.64105 | 505.1048 |
| Ubqln1 | ALSNLESIPGGYNAL | -249 | -263 | 0 | 0.001610812 | 15 | NaN | NaN | NaN | NaN | 995.57263 | NaN |
| Ubqln1 | LNNPLFAGNPQLQEQMR | -413 | -429 | 0 | 0.00025015 | 17 | NaN | NaN | NaN | NaN | 766.83075 | NaN |
| Psmd12 | LDQHEGSYLSICK | -233 | -245 | 0 | 0 | 13 | NaN | NaN | NaN | NaN | 679.3812 | 865.18176 |
| Asf1a | AESEEYDQVLDSVLVGPVPAGR | -48 | -69 | 0 | 0.00025015 | 22 | NaN | NaN | NaN | NaN | 374.67468 | 341.98502 |
| Smc1a | FESVATNIDEIYK | -1074 | -1086 | 0 | 0.00025015 | 13 | NaN | NaN | NaN | NaN | 1387.1287 | NaN |
| Smc1a | HEYSEELEK | -115 | -123 | 0 | 0.000627943 | 9 | NaN | NaN | NaN | NaN | 942.4136 | NaN |
| Smc1a | QKEVTAIETK | -909 | -918 | 1 | 0 | 10 | NaN | NaN | NaN | NaN | 271.55115 | NaN |
| Dis3 | SVVLDDEGQNEDDVEKDEER | -307 | -326 | 1 | 0.000471409 | 20 | NaN | NaN | NaN | NaN | 512.70557 | 534.82684 |
| Dis3 | AVQEGIPAFTCEEYVK | -184 | -199 | 0 | 0.000431655 | 16 | NaN | NaN | NaN | NaN | 1584.658 | 2714.0842 |
| Dis3 | QIDVLEHPAIR | -76 | -86 | 0 | 0.000230707 | 11 | NaN | NaN | NaN | NaN | 1580.7102 | 1177.8362 |
| Dis3 | DVSHFIR | -508 | -514 | 0 | 0.003461185 | 7 | NaN | NaN | NaN | NaN | 600.6688 | 197.60187 |
| Bzw1 | QSVEHFTK | -224 | -231 | 0 | 0.00078661 | 8 | NaN | NaN | NaN | NaN | 2151.109 | NaN |
| Gars1 | YTILEHTFHVR | -576 | -586 | 0 | 0 | 11 | NaN | NaN | NaN | NaN | 4952.139 | 4608.134 |
| Gars1 | SEKGEFTIETEGK | -525 | -537 | 1 | 0.004595494 | 13 | NaN | NaN | NaN | NaN | 274.39493 | 1130.7737 |
| Gars1 | SPTTGNDLSPPVPF | -255 | -268 | 0 | 0.001868731 | 14 | NaN | NaN | NaN | NaN | 1001.5282 | NaN |
| Gars1 | PQVDVDR | -77 | -83 | 0 | 0.008253452 | 7 | NaN | NaN | NaN | NaN | 967.2653 | NaN |
| Gars1 | ITWADVEAR | -704 | -712 | 0 | 0.00025015 | 9 | NaN | NaN | NaN | NaN | 637.5251 | NaN |
| Rpn1 | ASSFVLALEPELESR | -67 | -81 | 0 | 0 | 15 | NaN | NaN | NaN | NaN | 643.7155 | 736.82214 |
| Rpn1 | KIDHILDAL | -600 | -608 | 1 | 0.001291103 | 9 | NaN | NaN | NaN | NaN | 542.3975 | 474.55194 |
| Rpn1 | IILPEGAKNI | -373 | -382 | 1 | 0.009897819 | 10 | NaN | NaN | NaN | NaN | 6578.074 | NaN |
| Rpn1 | HFDETVNR | -496 | -503 | 0 | 0.006100831 | 8 | NaN | NaN | NaN | NaN | 2273.6187 | NaN |
| Rpn1 | LHPYPTQITQSEK | -140 | -152 | 0 | 0.00204437 | 13 | NaN | NaN | NaN | NaN | 796.9614 | NaN |
| Cct7 | QVKPYVEEGLHPQIIIR | -107 | -123 | 1 | 0 | 17 | NaN | NaN | NaN | NaN | 168.563 | 1588.4193 |
| Cct7 | QVKPYVEEGLHPQIIIR | -107 | -123 | 1 | 0 | 17 | NaN | NaN | NaN | NaN | 168.563 | 1588.4193 |
| Cct7 | IVSVDETIKNPR | -513 | -524 | 1 | 0.00025015 | 12 | NaN | NaN | NaN | NaN | 779.73926 | 833.74664 |
| Cct7 | EGTDSSQGIPQLVSNISAC | -11 | -29 | 0 | 0.000550307 | 19 | NaN | NaN | NaN | NaN | 152.20654 | 333.60342 |
| Cct7 | AEWNILYDKLEK | -269 | -280 | 1 | 0 | 12 | NaN | NaN | NaN | NaN | 556.79425 | NaN |
| Psmd6 | VYQGLYCVAIR | -182 | -192 | 0 | 0 | 11 | NaN | NaN | NaN | NaN | 2009.627 | 1464.2461 |
| Utp3 | AFAKPVPQVDEAETR | -184 | -198 | 1 | 0.000471409 | 15 | NaN | NaN | NaN | NaN | 436.25464 | 463.5447 |
| Utp3 | AKPVPQVDEAETR | -186 | -198 | 1 | 0.00272025 | 13 | NaN | NaN | NaN | NaN | 3061.1138 | 2555.9346 |
| Utp3 | KKEENSAEEQALEEQNAK | -381 | -398 | 2 | 0 | 18 | NaN | NaN | NaN | NaN | 395.63602 | 379.8907 |
| Ddx27 | SLVGEEERK | -531 | -539 | 1 | 0.007766418 | 9 | NaN | NaN | NaN | NaN | 265.5687 | NaN |
| Pin4 | QGGDLGWMTR | -78 | -87 | 0 | 0 | 10 | NaN | NaN | NaN | NaN | 611.9566 | NaN |
| Ywhag | WTSDQQDDDGGEGNN | -233 | -247 | 0 | 0.001023236 | 15 | NaN | NaN | NaN | NaN | 240.3527 | NaN |
| Ywhaq | YYEILNNPELACTLAK | -178 | -193 | 0 | 0.000907206 | 16 | NaN | NaN | NaN | NaN | 879.1307 | 1108.8158 |
| Ywhab | RYLSEVASGENK | -129 | -140 | 1 | 0.000550307 | 12 | NaN | NaN | NaN | NaN | 999.5533 | 868.97003 |
| Ywhab | QTTVSNSQQAYQEAFEISK | -141 | -159 | 0 | 0 | 19 | NaN | NaN | NaN | NaN | 568.5562 | 604.9977 |
| Ywhaz | GIVDQSQQAYQE | -140 | -151 | 0 | 0 | 12 | NaN | NaN | NaN | NaN | 255.79578 | 248.34985 |
| Cct5 | GVIVDKDFSHPQMPK | -227 | -241 | 1 | 0.00025015 | 15 | NaN | NaN | NaN | NaN | 1167.07 | 961.67755 |
| Cct5 | GVIVDKDFSHPQMPK | -227 | -241 | 1 | 0.00025015 | 15 | NaN | NaN | NaN | NaN | 1167.07 | 961.67755 |
| Cct5 | GSNDMQYQHVIETLIGK | -497 | -513 | 0 | 0.000712251 | 17 | NaN | NaN | NaN | NaN | 871.57587 | 213.29874 |
| Cct5 | LIIKDQDR | -17 | -24 | 1 | 0 | 8 | NaN | NaN | NaN | NaN | 396.3212 | 466.30212 |
| Pccb | AFDNDVDALCNLR | -262 | -274 | 0 | 0.00676572 | 13 | NaN | NaN | NaN | NaN | 1240.3643 | 1186.958 |
| Hsph1 | NHDSEETEGVHEVFSR | -407 | -422 | 0 | 0 | 16 | NaN | NaN | NaN | NaN | 223.84023 | 154.85033 |
| Hsph1 | NAVEECVYEFR | -637 | -647 | 0 | 0 | 11 | NaN | NaN | NaN | NaN | 1037.2097 | 528.85834 |
| Hsph1 | ETEDWLYEEGEDQAK | -672 | -686 | 0 | 0 | 15 | NaN | NaN | NaN | NaN | 297.46066 | 278.46985 |
| Rnpep | AFFPCFDTPAVK | -177 | -188 | 0 | 0 | 12 | NaN | NaN | NaN | NaN | 1207.717 | NaN |
| Rnpep | LAHLVGDQDQFDK | -424 | -436 | 0 | 0.00051306 | 13 | NaN | NaN | NaN | NaN | 1027.9131 | NaN |
| Rnpep | VKIEPGVDPDDTYNETPYEK | -397 | -416 | 1 | 0.001743443 | 20 | NaN | NaN | NaN | NaN | 203.39052 | NaN |
| Usf2 | AFGDHNIQYQFR | -58 | -69 | 0 | 0.00078661 | 12 | NaN | NaN | NaN | NaN | 686.69147 | NaN |
| Actn1 | MHGTLEDQLSHLR | -660 | -672 | 0 | 0 | 13 | NaN | NaN | NaN | NaN | 978.9176 | 1143.4158 |
| Actn1 | GTLEDQLSHLR | -662 | -672 | 0 | 0 | 11 | NaN | NaN | NaN | NaN | 1547.2 | NaN |
| Actn1 | IEMHGTLEDQLSHLR | -658 | -672 | 0 | 0.00294445 | 15 | NaN | NaN | NaN | NaN | 838.40247 | NaN |
| Actn1 | DDPLTNLNTAFDVAER | -199 | -214 | 0 | 0.001636916 | 16 | NaN | NaN | NaN | NaN | 664.32556 | NaN |
| Actn1 | HGTLEDQLSHLR | -661 | -672 | 0 | 0.000591124 | 12 | NaN | NaN | NaN | NaN | 442.9068 | NaN |
| Snrnp70 | DPIPYLPPLEK | -17 | -27 | 0 | 0 | 11 | NaN | NaN | NaN | NaN | 3333.6792 | 3505.617 |
| Snrnp70 | EFEVYGPIKR | -122 | -131 | 1 | 0.002566359 | 10 | NaN | NaN | NaN | NaN | 825.8812 | NaN |
| Syncrip | YGGPPPDSVYSGQQPSVGTEIFVGK | -144 | -168 | 0 | 0.002296977 | 25 | NaN | NaN | NaN | NaN | 1411.7817 | 1385.9546 |
| Eftud2 | FVDCLIEQTHPEIR | -144 | -157 | 0 | 0 | 14 | NaN | NaN | NaN | NaN | 1132.8519 | 1286.8372 |
| Eftud2 | PVTVVLPDTK | -184 | -193 | 0 | 0 | 10 | NaN | NaN | NaN | NaN | 1035.134 | 329.86008 |
| Eftud2 | SVFHHWQIVPGDPLDK | -898 | -913 | 0 | 0 | 16 | NaN | NaN | NaN | NaN | 669.0945 | 459.50974 |
| Eftud2 | AFIPAIDSF | -872 | -880 | 0 | 0.001868731 | 9 | NaN | NaN | NaN | NaN | 874.77716 | NaN |
| Eftud2 | RYHIEVNR | -535 | -542 | 1 | 0.00266297 | 8 | NaN | NaN | NaN | NaN | 318.32126 | NaN |
| Gbp6 | LPTETLQELLDVHAA | -342 | -356 | 0 | 0.004255025 | 15 | NaN | NaN | NaN | NaN | 292.9317 | NaN |
| Ilk | AFLHTLEPLIPR | -306 | -317 | 0 | 0 | 12 | NaN | NaN | NaN | NaN | 356.1139 | 749.54517 |
| Nfkb1 | KQEDVVEDLLR | -589 | -599 | 1 | 0.00051306 | 11 | NaN | NaN | NaN | NaN | 2510.371 | 2468.485 |
| Nfkb1 | PDSTGSFTR | -219 | -227 | 0 | 0.00383302 | 9 | NaN | NaN | NaN | NaN | 369.1092 | 513.0463 |
| Nfkb1 | GYNPGLLVHSDLAYLQ | -162 | -177 | 0 | 0.002910424 | 16 | NaN | NaN | NaN | NaN | 343.7333 | 483.953 |
| Nfkb1 | SDLETSEPKPF | -335 | -345 | 1 | 0.002629273 | 11 | NaN | NaN | NaN | NaN | 1166.961 | NaN |
| Actr2 | IKDLMVGDEASELR | -52 | -65 | 1 | 0.001392869 | 14 | NaN | NaN | NaN | NaN | 2476.9277 | 2292.0608 |
| Actr2 | LEVNYPMENGIVR | -68 | -80 | 0 | 0.000907206 | 13 | NaN | NaN | NaN | NaN | 1296.5804 | NaN |
| Actr2 | GYAGSNFPEHIFPALVGR | -21 | -38 | 0 | 0.001023236 | 18 | NaN | NaN | NaN | NaN | 932.3246 | NaN |

| | | | | | | | | | | | | |
|---|---|---|---|---|---|---|---|---|---|---|---|---|
| Krtcap2 | TAFNNLENLVFGK | -53 | -65 | 0 | 0 | 13 | NaN | NaN | NaN | NaN | 319.1334 | NaN |
| Aimp1 | LCNLKPAK | -208 | -215 | 1 | 0 | 8 | NaN | NaN | NaN | NaN | 1169.7583 | NaN |
| Lancl1 | FLADLLVPTK | -382 | -391 | 0 | 0.00025015 | 10 | NaN | NaN | NaN | NaN | 476.40744 | 366.84308 |
| Gzma | SINKEPEQQILTVK | -84 | -97 | 1 | 0 | 14 | NaN | NaN | NaN | NaN | 1425.4061 | 1489.0469 |
| Uggt1 | LRAEVEENQK | -369 | -378 | 1 | 0.002534621 | 10 | NaN | NaN | NaN | NaN | 2218.6716 | 2194.2153 |
| Uggt1 | SILGIDSAYDQNRK | -604 | -617 | 1 | 0.001291103 | 14 | NaN | NaN | NaN | NaN | 296.46243 | 649.7846 |
| Pes1 | AFSHDHPTDVDYR | -206 | -218 | 0 | 0 | 13 | NaN | NaN | NaN | NaN | 718.10834 | 684.4846 |
| Pes1 | IVDRPGQQTPIIGR | -376 | -389 | 1 | 0.000471409 | 14 | NaN | NaN | NaN | NaN | 928.32965 | 949.74164 |
| Ndufs1 | AFTARPWETR | -237 | -246 | 1 | 0.005394768 | 10 | NaN | NaN | NaN | NaN | 1137.1575 | 147.57158 |
| Prpf31 | TVEIENELNIIHK | -100 | -112 | 0 | 0 | 13 | NaN | NaN | NaN | NaN | 3564.6611 | 3591.9214 |
| Rps2 | KAEDKEWIPVTK | -54 | -65 | 2 | 0 | 12 | NaN | NaN | NaN | NaN | 1116.5475 | 1115.1272 |
| Rps2 | AIGDYNGHVGLGVK | -129 | -142 | 0 | 0.006100831 | 14 | NaN | NaN | NaN | NaN | 1672.0239 | NaN |
| Rps2 | LGASLKDEVLK | -98 | -108 | 1 | 0 | 11 | NaN | NaN | NaN | NaN | 904.5401 | NaN |
| Rps2 | SPYQEFTDHL | -264 | -273 | 0 | 0.004439092 | 10 | NaN | NaN | NaN | NaN | 769.06836 | NaN |
| Slc38a1 | VCHPSVLPIYSELK | -285 | -298 | 0 | 0 | 14 | NaN | NaN | NaN | NaN | 1838.0928 | 644.45166 |
| Slc38a1 | AFVCHPSVLPIYSELK | -283 | -298 | 0 | 0.000230707 | 16 | NaN | NaN | NaN | NaN | 1772.8196 | NaN |
| Acadl | KFFQEEVIPHHTEWEK | -66 | -81 | 1 | 0 | 16 | NaN | NaN | NaN | NaN | 921.11523 | 664.4708 |
| Acadl | QGLLGINIAEK | -96 | -106 | 0 | 0.009242859 | 11 | NaN | NaN | NaN | NaN | 1116.0275 | NaN |
| Upf2 | HCGDDIAGLVPR | -313 | -324 | 0 | 0 | 12 | NaN | NaN | NaN | NaN | 1795.4846 | NaN |
| Upf2 | EGLSLIYEQLK | -281 | -291 | 0 | 0.000230707 | 11 | NaN | NaN | NaN | NaN | 604.75073 | NaN |
| Oat | AFYNNVLGEYEEYITK | -114 | -129 | 0 | 0.001150701 | 16 | NaN | NaN | NaN | NaN | 305.70355 | NaN |
| Pcnp | GGPEEEAEKPVK | -21 | -32 | 1 | 0.000591124 | 12 | NaN | NaN | NaN | NaN | 1442.3269 | 1285.7498 |
| Cyb5b | GEKVEGSEPSVTYYR | -10 | -24 | 1 | 0.007288197 | 15 | NaN | NaN | NaN | NaN | 332.7047 | NaN |
| Ddx39b | AGADGTEAPAK | -22 | -32 | 0 | 0 | 11 | NaN | NaN | NaN | NaN | 680.12317 | NaN |
| Mri1 | APSSSCDLHLETGK | -280 | -293 | 0 | 0 | 14 | NaN | NaN | NaN | NaN | 598.52783 | 380.7177 |
| Mri1 | YSPGSLQILDQL | -8 | -19 | 0 | 0 | 12 | NaN | NaN | NaN | NaN | 486.39792 | 429.97864 |
| Rplp0 | PAAAAAPAK | -289 | -297 | 0 | 0 | 9 | NaN | NaN | NaN | NaN | 531.3822 | 610.5195 |
| Aldh9a1 | AGAPPGLFNVVQGGAATGQ | -199 | -217 | 0 | 0.000394361 | 19 | NaN | NaN | NaN | NaN | 1239.5043 | 458.08453 |
| Aldh9a1 | AGEHIQLPGGSFGYTR | -128 | -143 | 0 | 0.00025015 | 16 | NaN | NaN | NaN | NaN | 1187.7124 | 1574.6365 |
| Aldh9a1 | MGPLINAPHLER | -327 | -338 | 0 | 0.00025015 | 12 | NaN | NaN | NaN | NaN | 1080.3557 | 990.38025 |
| Aldh9a1 | AGAPPGLFNVVQGGAAT | -199 | -215 | 0 | 0.003166358 | 17 | NaN | NaN | NaN | NaN | 771.29614 | NaN |
| Taldo1 | MESALDQLK | -11 | -19 | 0 | 0 | 9 | NaN | NaN | NaN | NaN | 776.778 | 1694.3523 |
| Taldo1 | SSTWEGIQAGK | -144 | -154 | 0 | 0.00025015 | 11 | NaN | NaN | NaN | NaN | 1195.4377 | 903.4816 |
| Qtrt2 | EALKNDTLAQLK | -396 | -407 | 1 | 0 | 12 | NaN | NaN | NaN | NaN | 2307.862 | NaN |
| Qtrt2 | AGDCTVDIPGCLLYTR | -23 | -38 | 0 | 0.00111006 | 16 | NaN | NaN | NaN | NaN | 761.10364 | NaN |
| Acap1 | YFQQGHEELNR | -202 | -212 | 0 | 0.00025015 | 11 | NaN | NaN | NaN | NaN | 1353.6252 | 891.72974 |
| Acap1 | LTHNAEVPR | -138 | -146 | 0 | 0.002265552 | 9 | NaN | NaN | NaN | NaN | 781.42426 | NaN |
| Xpo1 | LVSEVEETEIFK | -356 | -367 | 0 | 0.001776473 | 12 | NaN | NaN | NaN | NaN | 2156.0452 | 1879.0425 |
| Xpo1 | TSSDPTCVEK | -113 | -122 | 0 | 0.00025015 | 10 | NaN | NaN | NaN | NaN | 560.4923 | NaN |
| Xpo1 | AVGHPFVIQL | -701 | -710 | 0 | 0.007987481 | 10 | NaN | NaN | NaN | NaN | 399.15436 | NaN |
| Rps18 | LREDLERL | -107 | -114 | 2 | 0.006260252 | 8 | NaN | NaN | NaN | NaN | 861.97906 | NaN |
| Ssrp1 | EFNDIFQEVK | -6 | -15 | 0 | 0 | 10 | NaN | NaN | NaN | NaN | 2032.4543 | 2076.1472 |
| Numa1 | EAEQMGGELER | -960 | -970 | 0 | 0.002780285 | 11 | NaN | NaN | NaN | NaN | 1208.5533 | NaN |
| Numa1 | KVIEGSEMELAK | -86 | -97 | 1 | 0.000431655 | 12 | NaN | NaN | NaN | NaN | 1207.434 | NaN |
| Numa1 | KHPSSTQCLVSVQ | -73 | -85 | 1 | 0 | 13 | NaN | NaN | NaN | NaN | 908.2506 | NaN |
| Nono | TFPRPVTVEPMDQLDDEEGLPEK | -219 | -241 | 1 | 0.00025015 | 23 | NaN | NaN | NaN | NaN | 3907.405 | 31.894033 |
| Nono | EGIGAIGGTPPAFNR | -444 | -458 | 0 | 0.007546675 | 15 | NaN | NaN | NaN | NaN | 813.63495 | 874.2987 |
| Nono | FACHSASLTVR | -145 | -155 | 0 | 0 | 11 | NaN | NaN | NaN | NaN | 39020.62 | NaN |
| Actr1b | DIIANQPVVIDNGSGVIK | -5 | -22 | 0 | 0.002329279 | 18 | NaN | NaN | NaN | NaN | 1098.3107 | NaN |
| Actr1b | SINPQKDEALETEK | -225 | -238 | 1 | 0.001055699 | 14 | NaN | NaN | NaN | NaN | 938.32434 | NaN |
| Dnajc11 | AGFFGTIVEYGAER | -328 | -341 | 0 | 0 | 14 | NaN | NaN | NaN | NaN | 568.71106 | NaN |
| Ly6e | SAAAGFGNVNLGYTLNK | -58 | -74 | 0 | 0.002910424 | 17 | NaN | NaN | NaN | NaN | 553.94324 | NaN |
| Pdk1 | SPELELEELNAK | -245 | -256 | 0 | 0 | 12 | NaN | NaN | NaN | NaN | 2909.5662 | 2923.3086 |
| Pdk1 | KEISLLPDNLLR | -92 | -103 | 1 | 0.000627943 | 12 | NaN | NaN | NaN | NaN | 658.55927 | 615.1385 |
| Pdk1 | HEADDWCVPSR | -413 | -423 | 0 | 0.000431655 | 11 | NaN | NaN | NaN | NaN | 464.26227 | NaN |
| Eif3l | SLLGDYYQAIK | -282 | -292 | 0 | 0.00025015 | 11 | NaN | NaN | NaN | NaN | 2109.7585 | 2253.0042 |
| Eif3l | FSLVGLLR | -272 | -279 | 0 | 0.003620005 | 8 | NaN | NaN | NaN | NaN | 136.79707 | 272.1018 |
| Eif3l | AGFLDLTEQEFR | -476 | -487 | 0 | 0.007987481 | 12 | NaN | NaN | NaN | NaN | 1138.6344 | NaN |
| Eif3l | YEQQTYQVIPEVIK | -40 | -53 | 0 | 0.001868731 | 14 | NaN | NaN | NaN | NaN | 649.50024 | NaN |
| Syne3 | AGFLINPQDPIPR | -775 | -787 | 0 | 0.001463897 | 13 | NaN | NaN | NaN | NaN | 875.1973 | 1116.6509 |
| Slc25a12 | DIEVTKEEFAQSAIR | -245 | -259 | 1 | 0.000431655 | 15 | NaN | NaN | NaN | NaN | 1212.2048 | 476.9857 |
| Slc25a12 | TAGALAGVPAASLVTPADVIK | -523 | -543 | 0 | 0.00025015 | 21 | NaN | NaN | NaN | NaN | 291.39606 | 324.67856 |
| Slc25a12 | NSLLNNMELVR | -223 | -233 | 0 | 0.006677519 | 11 | NaN | NaN | NaN | NaN | 869.926 | NaN |
| Ppp2r1a | EHSPSDLEAHFVPLVK | -118 | -133 | 0 | 0.00025015 | 16 | NaN | NaN | NaN | NaN | 630.29504 | 658.44446 |
| Hnrnpl | TPASPVVHIRG | -95 | -105 | 1 | 0 | 11 | NaN | NaN | NaN | NaN | 1218.2683 | 1205.5331 |
| Hnrnpl | QPAIMPGQSYGLEDGSCSYK | -453 | -472 | 0 | 0.001023236 | 20 | NaN | NaN | NaN | NaN | 751.41516 | 671.602 |
| Hnrnpl | RFSTPEQAAK | -481 | -490 | 1 | 0.001392869 | 10 | NaN | NaN | NaN | NaN | 747.931 | 824.0038 |
| Hnrnpl | IMPGQSYGLEDGSCSYK | -456 | -472 | 0 | 0.000826806 | 17 | NaN | NaN | NaN | NaN | 731.4825 | NaN |
| Hnrnpl | DQDTWDYTNPNLSGQGDPGSNPNK | -276 | -299 | 0 | 0 | 24 | NaN | NaN | NaN | NaN | 151.26564 | NaN |
| Ttll12 | LLDEFVSLHGPTLR | -22 | -35 | 0 | 0.000431655 | 14 | NaN | NaN | NaN | NaN | 1059.2162 | 998.70306 |
| Ttll12 | IESPVLFLR | -448 | -456 | 0 | 0.002296977 | 9 | NaN | NaN | NaN | NaN | 4501.735 | NaN |
| Ttll12 | LPLAISPVARPQ | -285 | -296 | 1 | 0 | 12 | NaN | NaN | NaN | NaN | 298.1574 | NaN |
| Ahcy | ALDIAENEMPGLMR | -21 | -34 | 0 | 0.00051306 | 14 | NaN | NaN | NaN | NaN | 882.64764 | NaN |
| Gsn | GWDDNYWSVDPLDR | -760 | -773 | 0 | 0.00025015 | 14 | NaN | NaN | NaN | NaN | 463.72937 | NaN |
| Rpl10a | AVDIPHMDIEALK | -79 | -91 | 0 | 0 | 13 | NaN | NaN | NaN | NaN | 2792.062 | 2670.7668 |
| Timeless | NILHVPANLEQEK | -165 | -177 | 0 | 0 | 13 | NaN | NaN | NaN | NaN | 381.29214 | 233.91205 |
| Timeless | SIDDDASIHDR | -178 | -188 | 0 | 0.002296977 | 11 | NaN | NaN | NaN | NaN | 8616.303 | NaN |
| Cd6 | EDAGVVCSEHQSWR | -252 | -265 | 0 | 0.00122419 | 14 | NaN | NaN | NaN | NaN | 357.3446 | NaN |

| | | | | | | | | | | | | |
|---|---|---|---|---|---|---|---|---|---|---|---|---|
| Eif2s1 | RVSPEEAIK | -89 | -97 | 1 | 0.003360691 | 9 | NaN | NaN | NaN | NaN | 871.8863 | NaN |
| Sdha | VDEYDYSKPIQGQ | -601 | -613 | 1 | 0.00025015 | 13 | NaN | NaN | NaN | NaN | 2977.828 | 3182.0293 |
| Sdha | QYPVVDHEFDAVVVGAGGAGLR | -54 | -75 | 0 | 0.00122419 | 22 | NaN | NaN | NaN | NaN | 622.6166 | 523.603 |
| Sdha | EPIPVLPTVH | -398 | -407 | 0 | 0.009745776 | 10 | NaN | NaN | NaN | NaN | 669.04755 | NaN |
| Sdha | LGNMEEDNWR | -111 | -120 | 0 | 0.00383302 | 10 | NaN | NaN | NaN | NaN | 538.75604 | NaN |
| H2ac4;H2ac6;I | GNYSERVGAGAPVYLA | (38);(38);(38);(38);(38); | (53);(53);(53);(53);(53 | 1 | 0.007670622 | 16 | NaN | NaN | NaN | NaN | 891.5936 | 768.3449 |
| Pdhb | IMEGPAFNFLDAPAVR | -309 | -324 | 0 | 0.000230707 | 16 | NaN | NaN | NaN | NaN | 397.04276 | 449.86368 |
| Ddx54 | SVEYVVFDEADR | -239 | -250 | 0 | 0 | 12 | NaN | NaN | NaN | NaN | 916.48145 | NaN |
| Plec | AVTGYKDPYTGEQISLF | -2917 | -2933 | 1 | 0.001055699 | 17 | NaN | NaN | NaN | NaN | 3944.325 | 3135.866 |
| Plec | EDHPARPTVESF | -830 | -841 | 1 | 0.00025015 | 12 | NaN | NaN | NaN | NaN | 1034.2334 | 892.2293 |
| Plec | LKAEAELLQQQK | -2470 | -2481 | 1 | 0 | 12 | NaN | NaN | NaN | NaN | 879.9795 | 162.11613 |
| Plec | QYDIDDAITK | -4346 | -4355 | 0 | 0.006788754 | 10 | NaN | NaN | NaN | NaN | 1424.1141 | NaN |
| Plec | CITDPQTGLCLLPLK | -4252 | -4266 | 0 | 0 | 15 | NaN | NaN | NaN | NaN | 559.3848 | NaN |
| Plec | FLVPPPNQEAQEAVAR | -1001 | -1016 | 0 | 0.005176406 | 16 | NaN | NaN | NaN | NaN | 157.4737 | NaN |
| Top1 | SGDHLHNDSQIEADFR | -2 | -17 | 0 | 0.000394361 | 16 | NaN | NaN | NaN | NaN | 452.88684 | 327.42123 |
| Top1 | RDEDDADYKPK | -142 | -152 | 2 | 0.001636916 | 11 | NaN | NaN | NaN | NaN | 188.98557 | NaN |
| Sart3 | AGPEAEGEEDEAKPAGVQR | -18 | -36 | 1 | 0.00025015 | 19 | NaN | NaN | NaN | NaN | 780.6262 | 533.5331 |
| Sart3 | RLEGELSR | -124 | -131 | 1 | 0.001610812 | 8 | NaN | NaN | NaN | NaN | 1222.4254 | 595.85425 |
| Ddx6 | ITYDDRFNLK | -438 | -447 | 1 | 0.001708008 | 10 | NaN | NaN | NaN | NaN | 1047.8833 | 931.8956 |
| Ddx6 | VHCLNTLFSR | -322 | -331 | 0 | 0 | 10 | NaN | NaN | NaN | NaN | 1549.2758 | NaN |
| Ddx6 | GAYLIPLLER | -148 | -157 | 0 | 0.007288197 | 10 | NaN | NaN | NaN | NaN | 416.2541 | NaN |
| Sod1 | ASGEPVVLSGQITGLTEGQHGFHVH | -25 | -49 | 0 | 0.000550307 | 25 | NaN | NaN | NaN | NaN | 529.4041 | 550.1564 |
| Sod1 | ISLSGEHSIIGR | -105 | -116 | 0 | 0.00025015 | 12 | NaN | NaN | NaN | NaN | 1766.4473 | 1768.1577 |
| Sod1 | SLSGEHSIIGR | -106 | -116 | 0 | 0.004690496 | 11 | NaN | NaN | NaN | NaN | 1532.1536 | 1635.0812 |
| Sod1 | ASGEPVVLSGQITGL | -25 | -39 | 0 | 0.002566359 | 15 | NaN | NaN | NaN | NaN | 1269.0947 | 259.20105 |
| Sod1 | GLTEGQHGFHVHQYGDNTQGCTSAGPHFNPHSK | -38 | -70 | 0 | 0.00025015 | 33 | NaN | NaN | NaN | NaN | 817.091 | NaN |
| Sod1 | GLTEGQHGFHVHQYGDNTQGCTSAGPHFNPHSK | -38 | -70 | 0 | 0.00025015 | 33 | NaN | NaN | NaN | NaN | 817.091 | NaN |
| Pnp | LVFGLLNGR | -68 | -76 | 0 | 0.004857192 | 9 | NaN | NaN | NaN | NaN | 251.74675 | 220.07309 |
| Pnp | PNFETVAESR | -198 | -207 | 0 | 0.003299042 | 10 | NaN | NaN | NaN | NaN | 6169.849 | NaN |
| Pnp | WLLQHTEYRPQ | -16 | -26 | 1 | 0 | 11 | NaN | NaN | NaN | NaN | 1383.564 | NaN |
| Nudt9 | AGPQWADPQISESNFSPK | -106 | -123 | 0 | 0 | 18 | NaN | NaN | NaN | NaN | 613.5309 | NaN |
| Pgd | CDWVGDEGAGHFVK | -171 | -184 | 0 | 0.006164361 | 14 | NaN | NaN | NaN | NaN | 620.30383 | NaN |
| Acaca | FADLHDTPGR | -2185 | -2194 | 0 | 0.009002904 | 10 | NaN | NaN | NaN | NaN | 737.7984 | NaN |
| Acaca | ATLVEHGIR | -1316 | -1324 | 0 | 0.00437931 | 9 | NaN | NaN | NaN | NaN | 455.43805 | NaN |
| Clic4 | FLDGDEMTLADCNLLPK | -178 | -194 | 0 | 0 | 17 | NaN | NaN | NaN | NaN | 401.12085 | 420.27615 |
| Clic4 | RKPADLQNLAPGTHPPFITF | -61 | -80 | 2 | 0.00450079 | 20 | NaN | NaN | NaN | NaN | 586.40356 | NaN |
| Arhgap45 | LKECVLQDDLLEAR | -132 | -145 | 1 | 0 | 14 | NaN | NaN | NaN | NaN | 1322.1886 | 1213.532 |
| Arhgap45 | SSLVDYPHQAR | -948 | -958 | 0 | 0.00078661 | 11 | NaN | NaN | NaN | NaN | 350.75336 | NaN |
| Eef1d | PGGDHSELIVR | -73 | -83 | 0 | 0.001392869 | 11 | NaN | NaN | NaN | NaN | 386.12543 | 486.94778 |
| Tbx21 | VDVVLVDQHHWR | -186 | -197 | 0 | 0 | 12 | NaN | NaN | NaN | NaN | 597.8804 | 652.5067 |
| Psmg1 | AGTEEEEEEEEQSR | -16 | -29 | 0 | 0.00025015 | 14 | NaN | NaN | NaN | NaN | 1428.436 | 317.3834 |
| Psmg1 | SGVWEEVGCAK | -91 | -101 | 0 | 0.002265552 | 11 | NaN | NaN | NaN | NaN | 4217.8643 | NaN |
| Lman2 | DITDGNSEHLKR | -47 | -58 | 1 | 0 | 12 | NaN | NaN | NaN | NaN | 1374.6842 | 1644.2686 |
| Phgdh | GTGVDNVDLEAATR | -77 | -90 | 0 | 0 | 14 | NaN | NaN | NaN | NaN | 448.75473 | 453.4301 |
| Phgdh | GAALDVFTEEPPR | -256 | -268 | 0 | 0.007791272 | 13 | NaN | NaN | NaN | NaN | 998.2773 | NaN |
| Actn4 | GTQIENIDEDFRDGLK | -69 | -84 | 1 | 0 | 16 | NaN | NaN | NaN | NaN | 937.2199 | 1509.1139 |
| Actn4 | GTLEDQLSHLK | -682 | -692 | 0 | 0 | 11 | NaN | NaN | NaN | NaN | 710.80707 | 2134.0522 |
| Actn4 | KAGTQIENIDEDFRDGLK | -67 | -84 | 2 | 0.000431655 | 18 | NaN | NaN | NaN | NaN | 370.20883 | 598.72186 |
| Actn4 | PGALDYK | -894 | -900 | 0 | 0 | 7 | NaN | NaN | NaN | NaN | 187.40182 | NaN |
| Pgk1 | VADKIQLINNMLDK | -217 | -230 | 1 | 0 | 14 | NaN | NaN | NaN | NaN | 590.06946 | 8382.982 |
| Pgk1 | SGIPAGWMGLDCGTESSKK | -305 | -323 | 1 | 0 | 19 | NaN | NaN | NaN | NaN | 214.90656 | 503.2648 |
| Pgk1 | DKIQLINNMLDK | -219 | -230 | 1 | 0.00272025 | 12 | NaN | NaN | NaN | NaN | 2992.9856 | 772.09674 |
| Pgk1 | WNTEDKVSHVSTGGGASLELLEGK | -383 | -406 | 1 | 0.003299042 | 24 | NaN | NaN | NaN | NaN | 1325.9384 | 748.40027 |
| Pgk1 | HSSMVGVNLPQK | -173 | -184 | 0 | 0.000230707 | 12 | NaN | NaN | NaN | NaN | 547.66 | 452.5416 |
| Pgk1 | DCVGPEVENACANPAAGTVIL | -98 | -118 | 0 | 0.000431655 | 21 | NaN | NaN | NaN | NaN | 243.27206 | 337.5392 |
| Pgk1 | AKQIVWNGPVGVF | -331 | -343 | 1 | 0.00383302 | 13 | NaN | NaN | NaN | NaN | 808.1142 | NaN |
| Mcm3 | TDDSQEKTDDSQETQDSQK | -722 | -740 | 1 | 0.001092529 | 19 | NaN | NaN | NaN | NaN | 179.83786 | 280.9922 |
| Mcm3 | DFSEAETQMPQVHTPK | -706 | -721 | 0 | 0.000394361 | 16 | NaN | NaN | NaN | NaN | 1301.104 | NaN |
| Fto | VSWHHDENLVDR | -225 | -236 | 0 | 0 | 12 | NaN | NaN | NaN | NaN | 346.67462 | NaN |
| Pdcd6 | AGVNFSEFTGVWK | -78 | -90 | 0 | 0.000550307 | 13 | NaN | NaN | NaN | NaN | 1494.3667 | 1438.125 |
| Pdcd6 | RLTDIFR | -160 | -166 | 1 | 0.002068213 | 7 | NaN | NaN | NaN | NaN | 914.95795 | 610.5985 |
| Pdcd6 | ALPDQSFLWNVFQR | -21 | -34 | 0 | 0.00025015 | 14 | NaN | NaN | NaN | NaN | 156.68744 | NaN |
| Slc2a1 | AGVQQPVYATIGSGIVN | -301 | -317 | 0 | 0.007343498 | 17 | NaN | NaN | NaN | NaN | 607.7394 | NaN |
| Slc2a1 | SFEMLILGR | -118 | -126 | 0 | 0.00272025 | 9 | NaN | NaN | NaN | NaN | 215.3884 | NaN |
| Metap1 | HTAPNVPHYAK | -301 | -311 | 0 | 0.001055699 | 11 | NaN | NaN | NaN | NaN | 856.47626 | NaN |
| Metap1 | NCYPSPLNYYNFPK | -178 | -191 | 0 | 0.000907206 | 14 | NaN | NaN | NaN | NaN | 474.8244 | NaN |
| Prmt5 | DWNTLIVGK | -77 | -85 | 0 | 0.00930008 | 9 | NaN | NaN | NaN | NaN | 1564.4048 | NaN |
| Ankrd44 | DKDLNTPLHLAISK | -903 | -916 | 1 | 0 | 14 | NaN | NaN | NaN | NaN | 386.25345 | 366.15076 |
| Ankrd44 | NSALQTPLHIAAR | -939 | -951 | 0 | 0 | 13 | NaN | NaN | NaN | NaN | 5699.362 | 5299.623 |
| Ankrd44 | NSALQTPLHIA | -939 | -949 | 0 | 0.00025015 | 11 | NaN | NaN | NaN | NaN | 1985.798 | NaN |
| Ankrd44 | GAIDPSIVSCR | -825 | -835 | 0 | 0.00025015 | 11 | NaN | NaN | NaN | NaN | 1107.5204 | NaN |
| Ankrd44 | AHDNSEELER | -493 | -502 | 0 | 0.00025015 | 10 | NaN | NaN | NaN | NaN | 435.34885 | NaN |
| Preb | KAHEGEIGDLTLGPDGK | -193 | -209 | 1 | 0.001743443 | 17 | NaN | NaN | NaN | NaN | 332.69815 | NaN |
| Prmt1 | FYESMLNTVLHAR | -169 | -181 | 0 | 0 | 13 | NaN | NaN | NaN | NaN | 1637.4784 | 1795.5406 |
| Prmt1 | FYESMLNTVLHAR | -169 | -181 | 0 | 0 | 13 | NaN | NaN | NaN | NaN | 1637.4784 | 1795.5406 |
| Prmt1 | VEDLTFTSPFCLQ | -262 | -274 | 0 | 0.00025015 | 13 | NaN | NaN | NaN | NaN | 253.72253 | 78.79932 |
| Ap1m1 | ILQEYITQEGHK | -130 | -141 | 0 | 0 | 12 | NaN | NaN | NaN | NaN | 483.5289 | 538.7887 |
| Atp5f1b | PNIVGNEHYDVAR | -410 | -422 | 0 | 0.00025015 | 13 | NaN | NaN | NaN | NaN | 237.24176 | 369.06137 |

| | | | | | | | | | | | | |
|---|---|---|---|---|---|---|---|---|---|---|---|---|
| Atp5f1b | IMDPNIVGNEHYDVARG | -407 | -423 | 1 | 0 | 17 | NaN | NaN | NaN | NaN | 382.48245 | 612.3175 |
| Atp5f1b | NVIGEPIDERGPIKTK | -146 | -161 | 2 | 0 | 16 | NaN | NaN | NaN | NaN | 499.13596 | 80.84526 |
| Atp5f1b | VALTGLTVAEYFR | -282 | -294 | 0 | 0 | 13 | NaN | NaN | NaN | NaN | 1126.1642 | NaN |
| Atp5f1b | DQEGQDVLLF | -295 | -304 | 0 | 0.002972566 | 10 | NaN | NaN | NaN | NaN | 358.61118 | NaN |
| Psmc5 | VPDSTYEMIGGLDK | -143 | -156 | 0 | 0 | 14 | NaN | NaN | NaN | NaN | 788.02924 | 2144.9353 |
| Psmc5 | EELQLLQEQGSYVGEVVR | -61 | -78 | 0 | 0.000550307 | 18 | NaN | NaN | NaN | NaN | 281.0443 | 263.3553 |
| Psmc5 | ILPNKVDPLVSL | -126 | -137 | 1 | 0.00111006 | 12 | NaN | NaN | NaN | NaN | 463.33282 | NaN |
| Tgm2 | WCPADDVYLDSEEERR | -142 | -157 | 1 | 0 | 16 | NaN | NaN | NaN | NaN | 389.86612 | 520.91547 |
| Tgm2 | AWCPADDVYLDSEEERR | -141 | -157 | 1 | 0 | 17 | NaN | NaN | NaN | NaN | 484.0669 | 217.90396 |
| Tgm2 | HDQNSNLLIEYFR | -305 | -317 | 0 | 0 | 13 | NaN | NaN | NaN | NaN | 3005.0073 | NaN |
| Tgm2 | AWIGSVDILR | -253 | -262 | 0 | 0 | 10 | NaN | NaN | NaN | NaN | 1061.0908 | NaN |
| Gart | SWLQQEGELSEEEMAR | -714 | -729 | 0 | 0.00051306 | 16 | NaN | NaN | NaN | NaN | 679.0341 | NaN |
| Cnpy2 | SQDLHCGACR | -23 | -32 | 0 | 0.000627943 | 10 | NaN | NaN | NaN | NaN | 192.24783 | 813.18286 |
| Nampt | KYDGHLPIEVK | -107 | -117 | 1 | 0.009242859 | 11 | NaN | NaN | NaN | NaN | 587.6427 | NaN |
| Snrpg | GNSIIMLEALER | -64 | -75 | 0 | 0 | 12 | NaN | NaN | NaN | NaN | 5362.886 | NaN |
| Rrbp1 | SKCEELSSLHGQLK | -1134 | -1147 | 1 | 0.002697088 | 14 | NaN | NaN | NaN | NaN | 2346.2087 | NaN |
| Rrbp1 | KLQEQLGK | -1585 | -1592 | 1 | 0.006624942 | 8 | NaN | NaN | NaN | NaN | 598.0184 | NaN |
| Iars1 | LYLINSPVVR | -625 | -634 | 0 | 0 | 10 | NaN | NaN | NaN | NaN | 566.18976 | 512.08966 |
| Iars1 | AHQSGFHVDR | -72 | -81 | 0 | 0.00025015 | 10 | NaN | NaN | NaN | NaN | 2493.9438 | NaN |
| Iars1 | AHQSGFHVDR | -72 | -81 | 0 | 0.00025015 | 10 | NaN | NaN | NaN | NaN | 2493.9438 | NaN |
| Iars1 | YKQESDYEILER | -267 | -278 | 1 | 0.00025015 | 12 | NaN | NaN | NaN | NaN | 1206.574 | NaN |
| Iars1 | FGWDCHGLPVEYEIDK | -83 | -98 | 0 | 0 | 16 | NaN | NaN | NaN | NaN | 701.2767 | NaN |
| Iars1 | LFLNETQTQEIT | -1227 | -1238 | 0 | 0.008712551 | 12 | NaN | NaN | NaN | NaN | 534.5438 | NaN |
| Cpsf3 | KQININNPFVF | -295 | -305 | 1 | 0.00450079 | 11 | NaN | NaN | NaN | NaN | 527.9459 | NaN |
| Cpsf3 | GLIPVFALGR | -236 | -245 | 0 | 0.003521747 | 10 | NaN | NaN | NaN | NaN | 200.25674 | NaN |
| B2m | TEFTPTETDTYACR | -88 | -101 | 0 | 0.00025015 | 14 | NaN | NaN | NaN | NaN | 93.381966 | 2021.774 |
| Dock8 | SPDFYEEVK | -625 | -633 | 0 | 0.00025015 | 9 | NaN | NaN | NaN | NaN | 664.96783 | 673.2179 |
| Dock8 | EVYVPHTVYR | -552 | -561 | 0 | 0 | 10 | NaN | NaN | NaN | NaN | 724.25586 | NaN |
| Dock8 | EILEFPIR | -544 | -551 | 0 | 0.001610812 | 8 | NaN | NaN | NaN | NaN | 717.16785 | NaN |
| Dock8 | HPLNVLCDVSGK | -181 | -192 | 0 | 0.008680443 | 12 | NaN | NaN | NaN | NaN | 494.85184 | NaN |
| Dock8 | ALYPPVDEEDAVEIRPVPECPK | -241 | -262 | 1 | 0.00025015 | 22 | NaN | NaN | NaN | NaN | 204.19139 | NaN |
| Smchd1 | GDFESDHSGYVRPLPVPR | -209 | -226 | 1 | 0.000230707 | 18 | NaN | NaN | NaN | NaN | 1590.6394 | 1568.0237 |
| Smchd1 | AHVEGDGVVEGVIR | -415 | -428 | 0 | 0.00078661 | 14 | NaN | NaN | NaN | NaN | 1639.1263 | NaN |
| Smchd1 | SLPFMFYNDVK | -1609 | -1619 | 0 | 0.000627943 | 11 | NaN | NaN | NaN | NaN | 1138.0669 | NaN |
| Smchd1 | RIEIPSCK | -1013 | -1020 | 1 | 0.001392869 | 8 | NaN | NaN | NaN | NaN | 983.90656 | NaN |
| Smchd1 | FTVRPLPDDPK | -1140 | -1150 | 1 | 0 | 11 | NaN | NaN | NaN | NaN | 94.497116 | NaN |
| Htt | DASGWFDVLQK | -1366 | -1376 | 0 | 0.003900536 | 11 | NaN | NaN | NaN | NaN | 799.7591 | NaN |
| Mdh1 | NVIIWGNHSSTQYPDVNHA | -180 | -198 | 0 | 0.00025015 | 19 | NaN | NaN | NaN | NaN | 1218.065 | 1143.7264 |
| Mdh1 | KLGVTADDVK | -170 | -179 | 1 | 0.000907206 | 10 | NaN | NaN | NaN | NaN | 847.2372 | NaN |
| Psmb1 | NMQNVEHVPLTLDR | -184 | -197 | 0 | 0 | 14 | NaN | NaN | NaN | NaN | 3600.257 | 3434.6848 |
| Psmb1 | SAMLQPLLDNQVGFK | -169 | -183 | 0 | 0.001846793 | 15 | NaN | NaN | NaN | NaN | 3795.1206 | 3909.1235 |
| Nvl | APEIVSGVSGESEQK | -330 | -344 | 0 | 0 | 15 | NaN | NaN | NaN | NaN | 785.9274 | 1187.7961 |
| Nvl | KELEDGHLAK | -76 | -85 | 1 | 0.000394361 | 10 | NaN | NaN | NaN | NaN | 1473.3604 | NaN |
| Mthfd1 | ASQAPSSFQLLYDLK | -805 | -819 | 0 | 0.00025015 | 15 | NaN | NaN | NaN | NaN | 495.7813 | 160.11948 |
| Mtrex | AIDCLSDEDKK | -425 | -435 | 1 | 0 | 11 | NaN | NaN | NaN | NaN | 1067.2504 | NaN |
| Gatd1 | LIPSCPGALTDLASSGSLAR | -90 | -109 | 0 | 0 | 20 | NaN | NaN | NaN | NaN | 685.391 | 701.612 |
| Gatd1 | AIDFVDVTESNAR | -50 | -62 | 0 | 0.001291103 | 13 | NaN | NaN | NaN | NaN | 896.9759 | NaN |
| Ndufs6 | ITHTGQVYDEK | -31 | -41 | 0 | 0 | 11 | NaN | NaN | NaN | NaN | 947.6482 | 669.14056 |
| Ndufs6 | IIACDGGGGALGHPK | -76 | -90 | 0 | 0 | 15 | NaN | NaN | NaN | NaN | 490.9945 | 521.4419 |
| Ndufs6 | AIDLIAQQPVNEVEHR | -60 | -75 | 0 | 0.001636916 | 16 | NaN | NaN | NaN | NaN | 1717.0609 | NaN |
| Ndufs6 | TGTCGYCGLQFK | -101 | -112 | 0 | 0.001023236 | 12 | NaN | NaN | NaN | NaN | 339.95917 | NaN |
| Larp7 | LLEGDTECHAR | -474 | -484 | 0 | 0 | 11 | NaN | NaN | NaN | NaN | 2176.5483 | NaN |
| Set | QHEEPESFFTW | -194 | -204 | 0 | 0.001291103 | 11 | NaN | NaN | NaN | NaN | 42.016853 | 4237.0986 |
| Fxr2 | GGLASGGDVEPGLPVEVR | -2 | -19 | 0 | 0.001092529 | 18 | NaN | NaN | NaN | NaN | 2700.8193 | NaN |
| Hnrnpab | FKEEDPVKK | -207 | -215 | 2 | 0 | 9 | NaN | NaN | NaN | NaN | 5105.749 | 5316.9214 |
| Hnrnpab | AIELPIDPKLNK | -187 | -198 | 1 | 0.006735683 | 12 | NaN | NaN | NaN | NaN | 436.15567 | NaN |
| Etf1 | AIELSTEVLSNVK | -267 | -279 | 0 | 0 | 13 | NaN | NaN | NaN | NaN | 2498.2095 | 595.6135 |
| Etf1 | YFDEISQDTGKY | -290 | -301 | 1 | 0.000869338 | 12 | NaN | NaN | NaN | NaN | 12116.753 | NaN |
| Etf1 | EYQGGDDEFFDLDDY | -423 | -437 | 0 | 0.000230707 | 15 | NaN | NaN | NaN | NaN | 332.59897 | NaN |
| Otulin | PQPGAWPGASCAETPAR | -7 | -23 | 0 | 0.004255025 | 17 | NaN | NaN | NaN | NaN | 1110.3827 | NaN |
| Jak1 | LSDPGIPVSVLTR | -736 | -748 | 0 | 0.00025015 | 13 | NaN | NaN | NaN | NaN | 424.1581 | NaN |
| Ripk1 | KDLKPENILVDR | -137 | -148 | 2 | 0.001291103 | 12 | NaN | NaN | NaN | NaN | 251.1297 | 412.46774 |
| Ripk1 | EYPDQSPVLQR | -308 | -318 | 0 | 0.004690496 | 11 | NaN | NaN | NaN | NaN | 1347.0336 | NaN |
| Ap3d1 | SYLITTPCYSDAFAK | -1092 | -1106 | 0 | 0 | 15 | NaN | NaN | NaN | NaN | 743.92676 | 685.9251 |
| Ap3d1 | ALDIDLDKPLADSEK | -797 | -811 | 1 | 0.006788754 | 15 | NaN | NaN | NaN | NaN | 714.8389 | NaN |
| Ppm1g | KALQDAFLAIDAK | -92 | -104 | 1 | 0.000471409 | 13 | NaN | NaN | NaN | NaN | 2432.8442 | 2546.8188 |
| Ppm1g | KYLPDIIK | -73 | -80 | 1 | 0 | 8 | NaN | NaN | NaN | NaN | 1368.8003 | 1211.7838 |
| Asns | LFPGFDLETVK | -222 | -232 | 0 | 0 | 11 | NaN | NaN | NaN | NaN | 1878.5284 | 1825.274 |
| Kpna2 | SFPDDATSPLQENR | -55 | -68 | 0 | 0.000869338 | 14 | NaN | NaN | NaN | NaN | 14068.408 | NaN |
| Impdh2 | GVPVIADGGIQNVGHIAK | -358 | -375 | 0 | 0 | 18 | NaN | NaN | NaN | NaN | 2408.9565 | NaN |
| Sf3b3 | PGQFLAVDPK | -117 | -126 | 0 | 0.003745449 | 10 | NaN | NaN | NaN | NaN | 877.52496 | NaN |
| Sf3b3 | FSNTGEDWYVLVGVAK | -896 | -911 | 0 | 0.00025015 | 16 | NaN | NaN | NaN | NaN | 596.7951 | NaN |
| Sf3b3 | TPVEEVPAAIAPF | -943 | -955 | 0 | 0.005015524 | 13 | NaN | NaN | NaN | NaN | 485.4848 | NaN |
| Tex10 | TEHISPFFPL | -131 | -140 | 0 | 0.001392869 | 10 | NaN | NaN | NaN | NaN | 164.88347 | 288.83627 |
| Tex10 | AIHLPEQLKEDR | -37 | -48 | 1 | 0.006472924 | 12 | NaN | NaN | NaN | NaN | 447.4586 | NaN |
| Psmb4 | MVIDEELLGDGHSYSPR | -115 | -131 | 0 | 0 | 17 | NaN | NaN | NaN | NaN | 725.2472 | 909.53204 |
| Rpl30 | IDPGDSDIIR | -97 | -106 | 0 | 0.001904399 | 10 | NaN | NaN | NaN | NaN | 1533.0457 | 1823.4005 |
| Rps7 | IVKPNGEKPDEFESGIS | -8 | -24 | 2 | 0.00025015 | 17 | NaN | NaN | NaN | NaN | 884.1402 | 363.49173 |

| | | | | | | | | | | | | |
|---|---|---|---|---|---|---|---|---|---|---|---|---|
| Rps7 | TAVHDAILEDLVFPSEIVGK | -123 | -142 | 0 | 0.004255025 | 20 | NaN | NaN | NaN | NaN | 388.50842 | 314.33722 |
| Cstf3 | IDYLSHLNEDNNTR | -451 | -464 | 0 | 0.00025015 | 14 | NaN | NaN | NaN | NaN | 1118.122 | 1150.4613 |
| Msto1 | WWNQQDAALGR | -24 | -34 | 0 | 0.000298805 | 11 | NaN | NaN | NaN | NaN | 762.86053 | 779.7757 |
| Msto1 | EAIIPFPLVR | -357 | -366 | 0 | 0.003069674 | 10 | NaN | NaN | NaN | NaN | 463.4288 | 606.09894 |
| Rcc2 | NLGQNLWGPH | -129 | -138 | 0 | 0.005015524 | 10 | NaN | NaN | NaN | NaN | 185.60597 | 864.017 |
| Rcc2 | HTLVLDSQK | -339 | -347 | 0 | 0.003360691 | 9 | NaN | NaN | NaN | NaN | 3076.5889 | NaN |
| Sars1 | AILENYQAEK | -439 | -448 | 0 | 0.00149795 | 10 | NaN | NaN | NaN | NaN | 888.9444 | 990.0691 |
| Sars1 | FKDPGLVDQL | -27 | -36 | 1 | 0.002972566 | 10 | NaN | NaN | NaN | NaN | 3681.6824 | NaN |
| Sars1 | LHPSVPISNDEDADNK | -139 | -154 | 0 | 0.001868731 | 16 | NaN | NaN | NaN | NaN | 441.00748 | NaN |
| Sec23b | LGETIAQWR | -662 | -670 | 0 | 0.00122419 | 9 | NaN | NaN | NaN | NaN | 645.99243 | NaN |
| Lta4h | QWLTPEQTSGK | -117 | -127 | 0 | 0 | 11 | NaN | NaN | NaN | NaN | 196.63585 | NaN |
| Lta4h | QHPYLFSQC | -128 | -136 | 0 | 0.00025015 | 9 | NaN | NaN | NaN | NaN | 1460.4818 | 1191.0813 |
| Lta4h | PTLLAGDK | -281 | -288 | 0 | 0.002168954 | 8 | NaN | NaN | NaN | NaN | 1214.8115 | 604.1806 |
| Lta4h | EISHSWTGNLVTNK | -297 | -310 | 0 | 0 | 14 | NaN | NaN | NaN | NaN | 995.66327 | 1099.5278 |
| Lta4h | PCQDTPSVK | -146 | -154 | 0 | 0.00383302 | 9 | NaN | NaN | NaN | NaN | 815.3263 | 818.32104 |
| Lta4h | ENPCLTFVTPTLLAGDK | -272 | -288 | 0 | 0.00142999 | 17 | NaN | NaN | NaN | NaN | 298.3936 | 267.80206 |
| Tubb5 | YHGDSDLQLDR | -36 | -46 | 0 | 0 | 11 | NaN | NaN | NaN | NaN | 3263.363 | 3087.1228 |
| Tubb5 | EEEDFGEEAEEEA | -432 | -444 | 0 | 0 | 13 | NaN | NaN | NaN | NaN | 110.21688 | NaN |
| Iqgap1 | HYQDTLIR | -578 | -585 | 0 | 0.00450079 | 8 | NaN | NaN | NaN | NaN | 4988.2334 | 4848.8696 |
| Iqgap1 | FYGEQVDYYK | -1519 | -1528 | 0 | 0 | 10 | NaN | NaN | NaN | NaN | 843.75165 | 1091.9358 |
| Iqgap1 | VIAINEAIDR | -221 | -230 | 0 | 0 | 10 | NaN | NaN | NaN | NaN | 1179.4875 | NaN |
| Aim2 | VATETDFFFVK | -187 | -197 | 0 | 0 | 11 | NaN | NaN | NaN | NaN | 998.66064 | NaN |
| Aim2 | AINPFECETQEGR | -167 | -179 | 0 | 0.004224523 | 13 | NaN | NaN | NaN | NaN | 169.75621 | NaN |
| Nmt1 | GFTWDALDLGDR | -155 | -166 | 0 | 0 | 12 | NaN | NaN | NaN | NaN | 996.0724 | 614.4699 |
| Rangap1 | SFNSNTFLTR | -508 | -517 | 0 | 0.003461185 | 10 | NaN | NaN | NaN | NaN | 1028.2356 | NaN |
| Rangap1 | EIEEFDGLEAL | -44 | -54 | 0 | 0.002265552 | 11 | NaN | NaN | NaN | NaN | 81.62883 | NaN |
| Huwe1 | DHVFEDSYR | -4027 | -4035 | 0 | 0 | 9 | NaN | NaN | NaN | NaN | 2584.9773 | 3190.4583 |
| Huwe1 | LEHEVDLCR | -460 | -468 | 0 | 0.001846793 | 9 | NaN | NaN | NaN | NaN | 163.31525 | 413.30463 |
| Huwe1 | AIQDPAFSDGIR | -518 | -529 | 0 | 0.001092529 | 12 | NaN | NaN | NaN | NaN | 996.9707 | NaN |
| Psmd3 | VYEFLDKLDVVR | -213 | -224 | 1 | 0 | 12 | NaN | NaN | NaN | NaN | 1838.9081 | 766.9115 |
| Psmd3 | ELDTVTLEDIKEHVR | -62 | -76 | 1 | 0.001023236 | 15 | NaN | NaN | NaN | NaN | 281.8567 | 249.14548 |
| Psmd3 | FQTDGTYTLIIR | -382 | -393 | 0 | 0 | 12 | NaN | NaN | NaN | NaN | 325.5947 | NaN |
| H2ac20;Hist2h | QLAIRNDEELNKLLGK | (85);(85) | (100);(100) | 2 | 0.00670918 | 16 | NaN | NaN | NaN | NaN | 312.74182 | 337.97394 |
| Macroh2a1 | EFVEAVLELR | -237 | -246 | 0 | 0 | 10 | NaN | NaN | NaN | NaN | 500.4178 | 394.54767 |
| Macroh2a1 | TIASGGVLPNIHPELLAK | -99 | -116 | 0 | 0 | 18 | NaN | NaN | NaN | NaN | 414.22595 | 535.1576 |
| Macroh2a1 | FVIHCNSPVWGADKCEELLEK | -269 | -289 | 1 | 0.003521747 | 21 | NaN | NaN | NaN | NaN | 4654.71 | NaN |
| Macroh2a1 | FVIHCNSPVWGADKC | -269 | -283 | 1 | 0.005758933 | 15 | NaN | NaN | NaN | NaN | 1007.81586 | NaN |
| Eml4 | HLDWSPDNK | -737 | -745 | 0 | 0.001846793 | 9 | NaN | NaN | NaN | NaN | 765.28516 | NaN |
| Wdr1 | NIDNPAIADIYTEHAHQVVVAK | -44 | -65 | 0 | 0 | 22 | NaN | NaN | NaN | NaN | 840.02563 | 800.01025 |
| Cdv3 | ETPEPAMPSGVYRPPGAR | -181 | -198 | 1 | 0.000627943 | 18 | NaN | NaN | NaN | NaN | 605.99506 | NaN |
| Rpl14 | SFGPHAGK | -16 | -23 | 0 | 0.005758933 | 8 | NaN | NaN | NaN | NaN | 1743.649 | NaN |
| Ablim1 | SSGREEDEEELLR | -670 | -682 | 1 | 0 | 13 | NaN | NaN | NaN | NaN | 727.8162 | 406.817 |
| Ablim1 | AIYDIERPDLITYEPF | -438 | -453 | 1 | 0.005605646 | 16 | NaN | NaN | NaN | NaN | 372.2647 | NaN |
| Rpa1 | KSAEDVGLK | -103 | -111 | 1 | 0 | 9 | NaN | NaN | NaN | NaN | 529.18286 | NaN |
| Atp2a2 | AKDIVPGDIVEIA | -142 | -154 | 1 | 0.002752526 | 13 | NaN | NaN | NaN | NaN | 1024.7949 | 979.51825 |
| Atp2a2 | EEMHLEDSANFIK | -573 | -585 | 0 | 0.00111006 | 13 | NaN | NaN | NaN | NaN | 346.12186 | 411.77014 |
| Atp2a2 | ALATHDNPLK | -562 | -571 | 0 | 0.007320451 | 10 | NaN | NaN | NaN | NaN | 848.99756 | NaN |
| Mia2 | AKDLEEELER | -1030 | -1039 | 1 | 0 | 10 | NaN | NaN | NaN | NaN | 1133.5917 | 1333.0768 |
| Mia2 | DPYALDVPNTAFGR | -1096 | -1109 | 0 | 0.000230707 | 14 | NaN | NaN | NaN | NaN | 267.33527 | NaN |
| Prkacb | EFPFLVR | -100 | -106 | 0 | 0.001392869 | 7 | NaN | NaN | NaN | NaN | 7494.799 | NaN |
| Prkacb | DLKPENLLIDHQGYIQ | -167 | -182 | 1 | 0.002099344 | 16 | NaN | NaN | NaN | NaN | 539.3011 | NaN |
| Prkacb | KWENPPPSNAGLEDFER | -30 | -46 | 1 | 0.001904399 | 17 | NaN | NaN | NaN | NaN | 322.86615 | NaN |
| Nxn | AKEEEAPLLFF | -361 | -371 | 1 | 0.002068213 | 11 | NaN | NaN | NaN | NaN | 352.85898 | NaN |
| Tpr | QLQEIFENYKK | -748 | -758 | 1 | 0.001392869 | 11 | NaN | NaN | NaN | NaN | 5243.3325 | 6265.252 |
| Tpr | QLQEIFENYK | -748 | -757 | 0 | 0 | 10 | NaN | NaN | NaN | NaN | 1021.5894 | 1131.6882 |
| Tpr | SSTSQTVSTPAPEPVIDSTEAIEAK | -719 | -743 | 0 | 0.002099344 | 25 | NaN | NaN | NaN | NaN | 285.6836 | 369.02402 |
| Tpr | QVTEEVHK | -1055 | -1062 | 0 | 0 | 8 | NaN | NaN | NaN | NaN | 687.2197 | NaN |
| Tpr | FLAEQQSEIDCLK | -102 | -114 | 0 | 0 | 13 | NaN | NaN | NaN | NaN | 619.8437 | NaN |
| Tpr | DPDTEEYRK | -1422 | -1430 | 1 | 0.00383302 | 9 | NaN | NaN | NaN | NaN | 581.25354 | NaN |
| Tpr | HFFDDEDR | -2193 | -2200 | 0 | 0.000394361 | 8 | NaN | NaN | NaN | NaN | 353.93 | NaN |
| Pabpc1 | AKEFTNVYIK | -187 | -196 | 1 | 0 | 10 | NaN | NaN | NaN | NaN | 463.37976 | 231.17212 |
| Pabpc1 | GFGFVSFER | -232 | -240 | 0 | 0.000230707 | 9 | NaN | NaN | NaN | NaN | 174.30672 | 603.90735 |
| Eif3k | ICSQEESIKPK | -189 | -199 | 1 | 0 | 11 | NaN | NaN | NaN | NaN | 1049.0352 | 1005.72064 |
| Eif3k | LYQFNPAFFQT | -53 | -63 | 0 | 0.002910424 | 11 | NaN | NaN | NaN | NaN | 875.5919 | 242.44305 |
| Eif3k | HVVGITYQHIDR | -144 | -155 | 0 | 0 | 12 | NaN | NaN | NaN | NaN | 519.47394 | NaN |
| Hspa5 | AKFEELNMDLFR | -326 | -337 | 1 | 0 | 12 | NaN | NaN | NaN | NaN | 2151.6777 | 702.7682 |
| Hspa5 | KNQLTSNPENTVFDAK | -82 | -97 | 1 | 0.000431655 | 16 | NaN | NaN | NaN | NaN | 736.08167 | 141.1639 |
| Hspa5 | KLYGSGGPPPTGEEDTSEKDEL | -634 | -655 | 2 | 0 | 22 | NaN | NaN | NaN | NaN | 709.78284 | 673.53906 |
| Hspa5 | WNDPSVQQDIK | -104 | -114 | 0 | 0.00111006 | 11 | NaN | NaN | NaN | NaN | 1070.0354 | 965.5388 |
| Hspa5 | FFEGEDFSETLTR | -313 | -325 | 0 | 0 | 13 | NaN | NaN | NaN | NaN | 955.55237 | 1024.466 |
| Top2a | PEFEEWK | -592 | -598 | 0 | 0.001636916 | 7 | NaN | NaN | NaN | NaN | 544.66437 | 459.66455 |
| Top2a | WVYDEDVGINYR | -61 | -72 | 0 | 0.008653659 | 12 | NaN | NaN | NaN | NaN | 410.0606 | 649.8201 |
| Calr | YTLIVRPDNTYEVK | -172 | -185 | 1 | 0.00025015 | 14 | NaN | NaN | NaN | NaN | 2184.154 | 2288.7837 |
| Calr | AKIDDPTDSKPE | -223 | -234 | 2 | 0.002296977 | 12 | NaN | NaN | NaN | NaN | 1418.7653 | NaN |
| Tmem43 | SLVVSPDSIHSVAPENEGR | -69 | -87 | 0 | 0 | 19 | NaN | NaN | NaN | NaN | 692.3413 | 584.00903 |
| Tmem43 | SSDDPDLGPAHVVTVIAR | -249 | -266 | 0 | 0.000550307 | 18 | NaN | NaN | NaN | NaN | 3094.9243 | NaN |
| Tmem43 | HHGDFSAEEVFR | -287 | -298 | 0 | 0.001636916 | 12 | NaN | NaN | NaN | NaN | 403.96246 | NaN |
| Ncl | FEDAMEIR | -415 | -422 | 0 | 0.001868731 | 8 | NaN | NaN | NaN | NaN | 585.843 | 573.22 |

| | | | | | | | | | | | | |
|---|---|---|---|---|---|---|---|---|---|---|---|---|
| Ncl | KVTLDWAK | -636 | -643 | 1 | 0.00025015 | 8 | NaN | NaN | NaN | NaN | 343.65964 | 1863.1367 |
| Ncl | TTPFNLFIGNLNPNK | -306 | -320 | 0 | 0.001743443 | 15 | NaN | NaN | NaN | NaN | 615.74554 | NaN |
| Ncl | VEGSEPTTPFNLFIGN | -300 | -315 | 0 | 0.001708008 | 16 | NaN | NaN | NaN | NaN | 414.59885 | NaN |
| Ncl | SVAEEEDDEEEDEDDEDEDDEEEDDEDDDEEEEEEEP | -238 | -276 | 0 | 0.003745449 | 39 | NaN | NaN | NaN | NaN | 162.28561 | NaN |
| Cs | STNLKDVLSNLIPK | -30 | -43 | 1 | 0.001868731 | 14 | NaN | NaN | NaN | NaN | 419.1019 | 550.405 |
| Cs | DYIWNTLNSGR | -330 | -340 | 0 | 0 | 11 | NaN | NaN | NaN | NaN | 85.788574 | NaN |
| Rrp9 | EAHGLHGEPGLEQPFWVS | -363 | -380 | 0 | 0.00383302 | 18 | NaN | NaN | NaN | NaN | 667.10114 | 2084.6448 |
| Pygb | HLDHVAALFPGDVDR | -411 | -425 | 0 | 0 | 15 | NaN | NaN | NaN | NaN | 696.41504 | 801.36694 |
| Pygb | HLDHVAALFPGDVDR | -411 | -425 | 0 | 0 | 15 | NaN | NaN | NaN | NaN | 696.41504 | 801.36694 |
| Pygb | SVIEEGDCKR | -430 | -439 | 1 | 0 | 10 | NaN | NaN | NaN | NaN | 514.5111 | 1260.3507 |
| Galnt7 | AKPLVLGPEYK | -153 | -163 | 1 | 0 | 11 | NaN | NaN | NaN | NaN | 1692.6528 | 1594.9968 |
| Galnt7 | EKLDEYIK | -256 | -263 | 1 | 0.004983786 | 8 | NaN | NaN | NaN | NaN | 813.7814 | 787.5685 |
| Pdia4 | RFDVSGYPTLK | -239 | -249 | 1 | 0.004595494 | 11 | NaN | NaN | NaN | NaN | 2323.0466 | NaN |
| Pdia4 | NDGNFDNFVADK | -61 | -72 | 0 | 0.008129024 | 12 | NaN | NaN | NaN | NaN | 439.62308 | NaN |
| Pdia3 | HLANKFEDK | -210 | -218 | 1 | 0.001092529 | 9 | NaN | NaN | NaN | NaN | 4354.0894 | 742.25616 |
| Pdia3 | LSKDPNIVIA | -423 | -432 | 1 | 0.00830926 | 10 | NaN | NaN | NaN | NaN | 2991.3657 | 2683.1104 |
| Pdia3 | PEYEAAATR | -65 | -73 | 0 | 0.00025015 | 9 | NaN | NaN | NaN | NaN | 852.02075 | 680.26855 |
| Pdia3 | ALEQFLQEYFDGNLKR | -348 | -363 | 1 | 0.001055699 | 16 | NaN | NaN | NaN | NaN | 449.1127 | 380.13092 |
| Pdia3 | ALEQFLQEYFDGNLK | -348 | -362 | 0 | 0 | 15 | NaN | NaN | NaN | NaN | 358.3577 | 350.7499 |
| Pdia3 | TFSHELSDFGLESTTGEVPVVAIR | -306 | -329 | 0 | 0 | 24 | NaN | NaN | NaN | NaN | 467.51053 | NaN |
| Rpl10 | NADEFEDMVAEKR | -177 | -189 | 1 | 0.002534621 | 13 | NaN | NaN | NaN | NaN | 776.2118 | NaN |
| Serbp1 | NVTEETPEGEEHPVADTENKENEVEEVKEEGPK | -253 | -285 | 2 | 0.00025015 | 33 | NaN | NaN | NaN | NaN | 2274.462 | 2098.9744 |
| Serbp1 | ENKENEVEEVKEEGPK | -270 | -285 | 2 | 0.000298805 | 16 | NaN | NaN | NaN | NaN | 1039.068 | 1102.1859 |
| Serbp1 | SEEAHAEDSVMDHHFR | -329 | -344 | 0 | 0 | 16 | NaN | NaN | NaN | NaN | 724.22003 | 710.7787 |
| Serbp1 | GGSGSHNWGTVKDELTESPK | -217 | -236 | 1 | 0.00111006 | 20 | NaN | NaN | NaN | NaN | 244.36797 | NaN |
| Tspan3 | AKVENEVDR | -109 | -117 | 1 | 0.00025015 | 9 | NaN | NaN | NaN | NaN | 413.8159 | NaN |
| Luc7l2 | SHLLNCCPHDVLSGT | -38 | -52 | 0 | 0.001291103 | 15 | NaN | NaN | NaN | NaN | 2085.2087 | NaN |
| Dbi | TDRPGLLDLK | -42 | -51 | 1 | 0.00111006 | 10 | NaN | NaN | NaN | NaN | 3292.535 | 2800.8271 |
| Dbi | SQAEFDKAAEEVKR | -2 | -15 | 2 | 0 | 14 | NaN | NaN | NaN | NaN | 737.2077 | NaN |
| Gabarapl2 | VSGSQIVDIDKR | -36 | -47 | 1 | 0.000230707 | 12 | NaN | NaN | NaN | NaN | 747.7041 | 643.29834 |
| Gabarapl2 | AKYPDRVPVIVEK | -23 | -35 | 2 | 0 | 13 | NaN | NaN | NaN | NaN | 490.6929 | 279.00427 |
| Gabarapl2 | AKYPDRVPVIVE | -23 | -34 | 2 | 0.00149795 | 12 | NaN | NaN | NaN | NaN | 322.2612 | NaN |
| Nop56 | VALENANAVSEGVVHEDLR | -59 | -77 | 0 | 0.001392869 | 19 | NaN | NaN | NaN | NaN | 249.48492 | 345.91144 |
| Arpc5l | GFEKPTENSSAVLLQ | -115 | -129 | 1 | 0.007493591 | 15 | NaN | NaN | NaN | NaN | 1178.957 | NaN |
| Tomm34 | LPPIPVVPVSAQ | -144 | -155 | 0 | 0.001610812 | 12 | NaN | NaN | NaN | NaN | 711.72675 | 811.2124 |
| Tomm34 | VPSAGDVER | -184 | -192 | 0 | 0.001978766 | 9 | NaN | NaN | NaN | NaN | 3958.1216 | NaN |
| Tomm34 | VSDSVEQLR | -5 | -13 | 0 | 0.002910424 | 9 | NaN | NaN | NaN | NaN | 763.3209 | NaN |
| Xrn2 | LEEVYPDLTPEENRR | -662 | -676 | 1 | 0.00142999 | 15 | NaN | NaN | NaN | NaN | 1268.6859 | NaN |
| Xrn2 | YAWQGVALLPFVDER | -642 | -656 | 0 | 0.00025015 | 15 | NaN | NaN | NaN | NaN | 734.1382 | 709.1303 |
| Xrn2 | SSPSISPNTSFASDGSPSPLGGIK | -470 | -493 | 0 | 0.00051306 | 24 | NaN | NaN | NaN | NaN | 896.54675 | NaN |
| Xrn2 | VREEILAK | -135 | -142 | 1 | 0.003360691 | 8 | NaN | NaN | NaN | NaN | 721.58026 | NaN |
| Xrn2 | SVGAEPLLPWNR | -884 | -895 | 0 | 0.005270854 | 12 | NaN | NaN | NaN | NaN | 647.5658 | NaN |
| Rnh1 | WLWECDITAEGCK | -257 | -269 | 0 | 0.00025015 | 13 | NaN | NaN | NaN | NaN | 2402.0474 | 2368.674 |
| Rnh1 | LCPGLLLPSCK | -242 | -252 | 0 | 0.00122419 | 11 | NaN | NaN | NaN | NaN | 1206.5242 | NaN |
| Rnh1 | NCGLTEAGCGILPGMLR | -89 | -105 | 0 | 0.002068213 | 17 | NaN | NaN | NaN | NaN | 539.76495 | NaN |
| Mcm2 | PSIYGHEDIKR | -483 | -493 | 1 | 0.00025015 | 11 | NaN | NaN | NaN | NaN | 703.61456 | 925.3898 |
| Mcm2 | IAPSIYGHEDIK | -481 | -492 | 0 | 0.000431655 | 12 | NaN | NaN | NaN | NaN | 1168.1394 | NaN |
| Lrpprc | CSSGSPGSNQALLLLR | -112 | -127 | 0 | 0.003299042 | 16 | NaN | NaN | NaN | NaN | 3509.9646 | NaN |
| Lrpprc | NTYHVPELIK | -630 | -639 | 0 | 0.00025015 | 10 | NaN | NaN | NaN | NaN | 744.1603 | NaN |
| Prpf19 | KVTSVVFHPSQELVF | -266 | -280 | 1 | 0.00025015 | 15 | NaN | NaN | NaN | NaN | 3569.3936 | 2915.8687 |
| Prpf19 | TNVANFPGHSGPITSIA | -383 | -399 | 0 | 0.002972566 | 17 | NaN | NaN | NaN | NaN | 2669.8027 | NaN |
| Prpf19 | PNTSCVQVVR | -294 | -303 | 0 | 0.00930008 | 10 | NaN | NaN | NaN | NaN | 755.9146 | NaN |
| Gusb | AVLENYHSVLDQK | -560 | -572 | 0 | 0 | 13 | NaN | NaN | NaN | NaN | 881.146 | 907.4662 |
| Gusb | ALDLTRPVTFVSN | -468 | -480 | 1 | 0.005605646 | 13 | NaN | NaN | NaN | NaN | 1144.1168 | NaN |
| Gusb | DTSFDFFNYAGLHR | -202 | -215 | 0 | 0.00142999 | 14 | NaN | NaN | NaN | NaN | 747.30133 | NaN |
| Rnf213 | HSDLSPVLR | -632 | -640 | 0 | 0 | 9 | NaN | NaN | NaN | NaN | 659.302 | 303.65845 |
| Rnf213 | LDLLNEDNTR | -3510 | -3519 | 0 | 0.001291103 | 10 | NaN | NaN | NaN | NaN | 1452.9341 | NaN |
| Rnf213 | SYVGFHGGLWR | -3387 | -3397 | 0 | 0.00122419 | 11 | NaN | NaN | NaN | NaN | 616.9061 | NaN |
| Rnf213 | LEDGCIEDDPAPYKK | -2825 | -2839 | 1 | 0.003393785 | 15 | NaN | NaN | NaN | NaN | 377.90207 | NaN |
| Rnf213 | NIDFDNLPR | -2323 | -2331 | 0 | 0.00078661 | 9 | NaN | NaN | NaN | NaN | 175.31192 | NaN |
| Cmc1 | YNDPAFYEECKLEYLK | -68 | -83 | 1 | 0 | 16 | NaN | NaN | NaN | NaN | 270.80127 | 638.55615 |
| Nup210 | LSNHYTSSEVK | -1700 | -1710 | 0 | 0 | 11 | NaN | NaN | NaN | NaN | 591.5096 | 610.6574 |
| Nup210 | SVGNRPSLSNPFPAVEPTVVK | -723 | -743 | 1 | 0.000230707 | 21 | NaN | NaN | NaN | NaN | 408.30865 | 458.5471 |
| Nup210 | LPPGPEHCSGVK | -598 | -609 | 0 | 0.001392869 | 12 | NaN | NaN | NaN | NaN | 4572.758 | NaN |
| Nup210 | NSVLNFATNR | -1420 | -1429 | 0 | 0.000591124 | 10 | NaN | NaN | NaN | NaN | 1604.3574 | NaN |
| Nup210 | ALDPTAGQLHGLGK | -1259 | -1272 | 0 | 0.00122419 | 14 | NaN | NaN | NaN | NaN | 1366.0516 | NaN |
| Nup210 | LDLSCPLLQQNK | -763 | -774 | 0 | 0 | 12 | NaN | NaN | NaN | NaN | 786.7113 | NaN |
| Nup210 | DVQNPLHFGEMK | -518 | -529 | 0 | 0.000230707 | 12 | NaN | NaN | NaN | NaN | 443.87143 | NaN |
| Hprt1 | SYCNDQSTGDIK | -104 | -115 | 0 | 0 | 12 | NaN | NaN | NaN | NaN | 1727.6353 | 1823.4443 |
| Hprt1 | VFIPHGLIMDR | -35 | -45 | 0 | 0 | 11 | NaN | NaN | NaN | NaN | 500.32437 | 928.3939 |
| Hprt1 | SIPMTVDFIR | -92 | -101 | 0 | 0 | 10 | NaN | NaN | NaN | NaN | 1004.88446 | 1247.3042 |
| Hprt1 | NHVCVISETGK | -203 | -213 | 0 | 0 | 11 | NaN | NaN | NaN | NaN | 867.71387 | NaN |
| Cndp2 | NVMLLPVGSADDGAHSQNEK | -431 | -450 | 0 | 0 | 20 | NaN | NaN | NaN | NaN | 647.7627 | 651.79974 |
| Cndp2 | RLGGSVELVDIGK | -54 | -66 | 1 | 0.006735683 | 13 | NaN | NaN | NaN | NaN | 802.6312 | NaN |
| Smc2 | TVTLGGDVFDPHGTLSGGAR | -649 | -668 | 0 | 0 | 20 | NaN | NaN | NaN | NaN | 782.2946 | 756.24066 |
| Smc2 | ALEEELAGLK | -697 | -706 | 0 | 0 | 10 | NaN | NaN | NaN | NaN | 22162.176 | NaN |
| Imp4 | NVELTEVGPR | -241 | -250 | 0 | 0.001055699 | 10 | NaN | NaN | NaN | NaN | 12843.353 | 12900.601 |
| Cltc | GSIVNFSQDPDVHFK | -716 | -730 | 0 | 0 | 15 | NaN | NaN | NaN | NaN | 1418.992 | 2263.4016 |

| | | | | | | | | | | | | |
|---|---|---|---|---|---|---|---|---|---|---|---|---|
| Cltc | KAVDVFFPPEAQ | -246 | -257 | 1 | 0.003299042 | 12 | NaN | NaN | NaN | NaN | 1109.1885 | 609.0603 |
| Cltc | FSTDELVAEVEKR | -840 | -852 | 1 | 0.000394361 | 13 | NaN | NaN | NaN | NaN | 655.00946 | 1325.5786 |
| Cltc | KVSQPIEGH | -189 | -197 | 1 | 0.001291103 | 9 | NaN | NaN | NaN | NaN | 423.6905 | 462.4948 |
| Cltc | LKEDKLECSEELGDLVK | -452 | -468 | 2 | 0 | 17 | NaN | NaN | NaN | NaN | 374.92007 | 627.0685 |
| Cltc | LKEDKLECSEELGDLVK | -452 | -468 | 2 | 0 | 17 | NaN | NaN | NaN | NaN | 374.92007 | 627.0685 |
| Cltc | EHLELFWSR | -1334 | -1342 | 0 | 0.001392869 | 9 | NaN | NaN | NaN | NaN | 378.47867 | 152.74673 |
| Cltc | TMTDDVTFWK | -103 | -112 | 0 | 0 | 10 | NaN | NaN | NaN | NaN | 70.64943 | 350.03098 |
| Sgta | LGNYVGAVQDCER | -139 | -151 | 0 | 0.001291103 | 13 | NaN | NaN | NaN | NaN | 2244.8123 | 2324.332 |
| Pcmt1 | LILPVGPAGGNQMLEQYDK | -179 | -197 | 0 | 0 | 19 | NaN | NaN | NaN | NaN | 2541.8882 | NaN |
| Itih3 | HNVVTTR | -60 | -66 | 0 | 0.008680443 | 7 | NaN | NaN | NaN | NaN | 549.1111 | 680.0131 |
| Itih3 | FFAPQGLPVVPK | -272 | -283 | 0 | 0 | 12 | NaN | NaN | NaN | NaN | 7099.3765 | NaN |
| Sugt1 | DSTDTNFDTWIK | -109 | -120 | 0 | 0 | 12 | NaN | NaN | NaN | NaN | 1413.8201 | 1287.4843 |
| Sugt1 | KLDSTDTNFDTWIK | -107 | -120 | 1 | 0.00078661 | 14 | NaN | NaN | NaN | NaN | 989.8728 | NaN |
| Emc1 | RPEIPTEQSR | -886 | -895 | 1 | 0.003620005 | 10 | NaN | NaN | NaN | NaN | 430.64413 | NaN |
| Dkc1 | EEFLIKPESK | -34 | -43 | 1 | 0.001811524 | 10 | NaN | NaN | NaN | NaN | 6415.372 | 5770.0938 |
| Dkc1 | TGFINLDKPSNPSSHEVVAW | -89 | -108 | 1 | 0.00025015 | 20 | NaN | NaN | NaN | NaN | 1217.9972 | 1195.1923 |
| Psmd2 | EWQELDDAEK | -169 | -178 | 0 | 0 | 10 | NaN | NaN | NaN | NaN | 651.3389 | NaN |
| Psmd2 | DTSLYRPALEELR | -67 | -79 | 1 | 0 | 13 | NaN | NaN | NaN | NaN | 856.93805 | 2029.5918 |
| Psmd2 | GLGLAYAGSNR | -484 | -494 | 0 | 0.001463897 | 11 | NaN | NaN | NaN | NaN | 2751.4465 | 2664.9749 |
| Psmd2 | SRFPEALR | -260 | -267 | 1 | 0.007040624 | 8 | NaN | NaN | NaN | NaN | 1453.9514 | 1361.8982 |
| Atic | MDQSYKPDENEVR | -367 | -379 | 1 | 0 | 13 | NaN | NaN | NaN | NaN | 431.04364 | 517.82043 |
| Atic | VGVPLSEDEAR | -275 | -285 | 0 | 0.000947459 | 11 | NaN | NaN | NaN | NaN | 2467.6768 | NaN |
| Abcf2 | AVTGVLASHPNSTDVHIINL | -77 | -96 | 0 | 0.001291103 | 20 | NaN | NaN | NaN | NaN | 992.0499 | NaN |
| Bysl | GVDHQAEVIVDPEDER | -115 | -130 | 0 | 0.001055699 | 16 | NaN | NaN | NaN | NaN | 2751.8877 | 2964.1611 |
| Ndufa4 | FYSVNVDYSK | -65 | -74 | 0 | 0.00025015 | 10 | NaN | NaN | NaN | NaN | 2794.3853 | NaN |
| Ndufv1 | ALGDGAAWPVQGLIR | -426 | -440 | 0 | 0 | 15 | NaN | NaN | NaN | NaN | 1208.5149 | 778.5658 |
| Dcps | ENPDPSDGFVLIPDLK | -191 | -206 | 0 | 0.001023236 | 16 | NaN | NaN | NaN | NaN | 882.9119 | 910.4503 |
| Myl6 | DQGTYEDYVEGL | -82 | -93 | 0 | 0 | 12 | NaN | NaN | NaN | NaN | 671.2631 | 640.2729 |
| Dars1 | HHGIDLEK | -444 | -451 | 0 | 0.000826806 | 8 | NaN | NaN | NaN | NaN | 2757.2178 | 2578.7493 |
| Dars1 | FGAPPHAGGGIGL | -461 | -473 | 0 | 0.000550307 | 13 | NaN | NaN | NaN | NaN | 391.7631 | NaN |
| Nup188 | ALHTVQYGLLK | -1556 | -1566 | 0 | 0.006537474 | 11 | NaN | NaN | NaN | NaN | 387.22852 | NaN |
| Ptbp1 | LSLDGQNIYNAC | -238 | -249 | 0 | 0.003102268 | 12 | NaN | NaN | NaN | NaN | 1443.3639 | NaN |
| Ptbp1 | SLDGQNIYNACCTLR | -239 | -253 | 0 | 0 | 15 | NaN | NaN | NaN | NaN | 313.00635 | 254.45792 |
| Rps12 | LVEALCAEHQIN | -64 | -75 | 0 | 0.002972566 | 12 | NaN | NaN | NaN | NaN | 872.73334 | NaN |
| Diaph1 | LQDLQGEK | -504 | -511 | 0 | 0.006100831 | 8 | NaN | NaN | NaN | NaN | 912.4403 | 787.6411 |
| Diaph1 | ALITPAEELDFR | -311 | -322 | 0 | 0 | 12 | NaN | NaN | NaN | NaN | 744.96967 | 822.596 |
| Diaph1 | DVQNFPAATDEKDKFVEK | -1055 | -1072 | 2 | 0.00078661 | 18 | NaN | NaN | NaN | NaN | 854.6807 | NaN |
| Rrm1 | PEIPDDLK | -670 | -677 | 0 | 0.002595029 | 8 | NaN | NaN | NaN | NaN | 1572.6108 | NaN |
| Rrm1 | GNNESIEPYTSNIYTR | -612 | -627 | 0 | 0.003814256 | 16 | NaN | NaN | NaN | NaN | 902.617 | NaN |
| Rrm1 | DDSIEGIYDTLK | -225 | -236 | 0 | 0 | 12 | NaN | NaN | NaN | NaN | 507.08054 | NaN |
| Pola1 | IVIPTENIR | -725 | -733 | 0 | 0.000471409 | 9 | NaN | NaN | NaN | NaN | 709.72455 | 813.5814 |
| Pola1 | EIFDDDLEDDALDTCGK | -99 | -115 | 0 | 0 | 17 | NaN | NaN | NaN | NaN | 266.85202 | 431.13568 |
| Pola1 | WLGLDSTQFR | -1242 | -1251 | 0 | 0 | 10 | NaN | NaN | NaN | NaN | 1042.1774 | NaN |
| Acsl5 | RADIPVVICDTPQK | -177 | -190 | 1 | 0.000591124 | 14 | NaN | NaN | NaN | NaN | 4187.703 | 2992.756 |
| Acsl5 | SSQDQFVGIF | -129 | -138 | 0 | 0.007791272 | 10 | NaN | NaN | NaN | NaN | 191.9208 | NaN |
| Eif3h | HVNIDHLHVGWYQ | -109 | -121 | 0 | 0.002099344 | 13 | NaN | NaN | NaN | NaN | 1301.5914 | NaN |
| Eif3h | HYQEEGQGTEVVQGVLLGL | -52 | -70 | 0 | 0 | 19 | NaN | NaN | NaN | NaN | 345.3379 | NaN |
| Ide | QDDKFFLPK | -563 | -571 | 1 | 0.00078661 | 9 | NaN | NaN | NaN | NaN | 656.5359 | 1063.6891 |
| Ide | GLPLFPLVKPH | -1001 | -1011 | 1 | 0.00025015 | 11 | NaN | NaN | NaN | NaN | 422.8121 | 412.10092 |
| Ide | TEQWYGTQYK | -492 | -501 | 0 | 0.001743443 | 10 | NaN | NaN | NaN | NaN | 5409.834 | NaN |
| Ero1a | KPCPFWNDINQCGR | -83 | -96 | 1 | 0.000431655 | 14 | NaN | NaN | NaN | NaN | 1472.9244 | 1390.9132 |
| Ero1a | LDDCTCDVETIDK | -43 | -55 | 0 | 0 | 13 | NaN | NaN | NaN | NaN | 1162.9377 | 997.43567 |
| Ero1a | LIANMPESGPSYEFQLTR | -416 | -433 | 0 | 0 | 18 | NaN | NaN | NaN | NaN | 566.4559 | 611.07465 |
| Ero1a | SFPLHFDENSFFAGDKNEAHK | -349 | -369 | 1 | 0.003681885 | 21 | NaN | NaN | NaN | NaN | 404.51138 | 453.33038 |
| Ero1a | KGPDAWR | -191 | -197 | 1 | 0.002329279 | 7 | NaN | NaN | NaN | NaN | 434.26605 | NaN |
| Ero1a | ALLLEILHEIK | -338 | -348 | 0 | 0.001392869 | 11 | NaN | NaN | NaN | NaN | 327.26953 | NaN |
| Gimap4 | HELGNQDQGIPQLR | -19 | -32 | 0 | 0.000298805 | 14 | NaN | NaN | NaN | NaN | 1766.6398 | 1716.5747 |
| Gpi | SLNLNTNHGHILVDYSK | -41 | -57 | 0 | 0.00025015 | 17 | NaN | NaN | NaN | NaN | 637.9484 | NaN |
| Gpi | SLNLNTNHGHILVDYSK | -41 | -57 | 0 | 0.00025015 | 17 | NaN | NaN | NaN | NaN | 637.9484 | NaN |
| Gpi | HDSSTNGLISFIK | -538 | -550 | 0 | 0.004439092 | 13 | NaN | NaN | NaN | NaN | 508.7106 | NaN |
| Gpi | TTQETITNAETAK | -214 | -226 | 0 | 0.000230707 | 13 | NaN | NaN | NaN | NaN | 469.24103 | NaN |
| Gpi | FEADPER | -30 | -36 | 0 | 0.003299042 | 7 | NaN | NaN | NaN | NaN | 317.67505 | NaN |
| Ube2n | KLELFLPEEYPMAAPK | -53 | -68 | 1 | 0 | 16 | NaN | NaN | NaN | NaN | 1708.7329 | NaN |
| Hsp90aa1 | LGIHEDSQNRK | -448 | -458 | 1 | 0 | 11 | NaN | NaN | NaN | NaN | 708.80414 | 740.9856 |
| Hsp90aa1 | VTEEMPPLEGDDDTSR | -713 | -728 | 0 | 0.00025015 | 16 | NaN | NaN | NaN | NaN | 313.92285 | NaN |
| Lrpap1 | ERDELNWK | -90 | -97 | 1 | 0.00437931 | 8 | NaN | NaN | NaN | NaN | 1465.2764 | NaN |
| Gtpbp1 | TPEYLHID | -547 | -554 | 0 | 0.00111006 | 8 | NaN | NaN | NaN | NaN | 982.7413 | NaN |
| Tap1 | ALPAAALWHK | -123 | -132 | 0 | 0 | 10 | NaN | NaN | NaN | NaN | 1088.5155 | NaN |
| Tap1 | KIFEYLDR | -453 | -460 | 1 | 0.002068213 | 8 | NaN | NaN | NaN | NaN | 914.07324 | NaN |
| Gtf2i | KQVEELFER | -363 | -371 | 1 | 0 | 9 | NaN | NaN | NaN | NaN | 6226.981 | 5247.474 |
| Gtf2i | ILDSAEFIKF | -929 | -938 | 1 | 0 | 10 | NaN | NaN | NaN | NaN | 521.08435 | 469.22928 |
| Mphosph10 | SNFHFIPKPPVPEIK | -512 | -526 | 1 | 0.000627943 | 15 | NaN | NaN | NaN | NaN | 585.2853 | 750.598 |
| Il16 | KGPPVAPKPA | -778 | -787 | 2 | 0.00111006 | 10 | NaN | NaN | NaN | NaN | 1003.5677 | 992.1635 |
| Pgam1 | SYDVPPPPMEPDHPFYSNISK | -118 | -138 | 0 | 0 | 21 | NaN | NaN | NaN | NaN | 646.33044 | 560.0402 |
| Pgam1 | RYADLTEDQLPSCESLK | -141 | -157 | 1 | 0.000591124 | 17 | NaN | NaN | NaN | NaN | 1351.3646 | NaN |
| Pgam1 | FSGWYDADLSPA | -22 | -33 | 0 | 0.00025015 | 12 | NaN | NaN | NaN | NaN | 259.81265 | NaN |
| Cse1l | ALTLPGSSENEYIMK | -560 | -574 | 0 | 0 | 15 | NaN | NaN | NaN | NaN | 2456.6328 | 2395.732 |
| Cse1l | ALPLTNLFK | -181 | -189 | 0 | 0.006654467 | 9 | NaN | NaN | NaN | NaN | 820.93915 | 707.703 |

| | | | | | | | | | | | | |
|---|---|---|---|---|---|---|---|---|---|---|---|---|
| Cse1l | SNNVNEFPVLK | -471 | -481 | 0 | 0 | 11 | NaN | NaN | NaN | NaN | 2244.7795 | NaN |
| Cse1l | EHDPVGQMVNNPK | -913 | -925 | 0 | 0.002752526 | 13 | NaN | NaN | NaN | NaN | 441.07434 | NaN |
| Cse1l | WERTGNIPALVR | -695 | -706 | 1 | 0.00025015 | 12 | NaN | NaN | NaN | NaN | 391.10052 | NaN |
| Igtp | CFSPSFHDFPELR | -242 | -254 | 0 | 0 | 13 | NaN | NaN | NaN | NaN | 834.66125 | 859.31384 |
| Igtp | GVDDGSLQQVAR | -323 | -334 | 0 | 0 | 12 | NaN | NaN | NaN | NaN | 1830.8877 | 1581.6157 |
| Igtp | SPSFHDFPELR | -244 | -254 | 0 | 0 | 11 | NaN | NaN | NaN | NaN | 1088.7705 | 1472.3931 |
| Igtp | YRDPLEIISQV | -266 | -276 | 1 | 0.002168954 | 11 | NaN | NaN | NaN | NaN | 257.56943 | NaN |
| Copg1 | CNLDLENLVTDSNR | -325 | -338 | 0 | 0 | 14 | NaN | NaN | NaN | NaN | 818.8778 | 759.56805 |
| Copg1 | VVVVQAISALCQK | -377 | -389 | 0 | 0.00111006 | 13 | NaN | NaN | NaN | NaN | 4355.6143 | NaN |
| Shmt1 | MLSQPLKDSDAEVY | -15 | -28 | 1 | 0.007987481 | 14 | NaN | NaN | NaN | NaN | 1345.6405 | NaN |
| Shmt1 | DATLWASHEK | -5 | -14 | 0 | 0.005270854 | 10 | NaN | NaN | NaN | NaN | 897.41095 | NaN |
| Rack1 | TVQDESHSEWVSCVR | -141 | -155 | 0 | 0 | 15 | NaN | NaN | NaN | NaN | 1547.1772 | 1438.3712 |
| Rack1 | DGQTLFAGYTDNLVR | -294 | -308 | 0 | 0 | 15 | NaN | NaN | NaN | NaN | 2127.2961 | 1980.5244 |
| Rack1 | ALSGSWDGTLR | -78 | -88 | 0 | 0.001846793 | 11 | NaN | NaN | NaN | NaN | 23082.064 | NaN |
| Rack1 | GYLNTVTVSPDGSLCASGGK | -193 | -212 | 0 | 0 | 20 | NaN | NaN | NaN | NaN | 2669.2644 | NaN |
| Rab4b | SCLLHQFIENK | -22 | -32 | 0 | 0 | 11 | NaN | NaN | NaN | NaN | 458.9219 | 1389.6539 |
| Cand1 | PQLTSPR | -182 | -188 | 0 | 0.007437881 | 7 | NaN | NaN | NaN | NaN | 499.69586 | NaN |
| Elmo1 | TSPAQNAQQLHER | -79 | -91 | 0 | 0 | 13 | NaN | NaN | NaN | NaN | 711.687 | 696.2621 |
| Elmo1 | SAIDISILQR | -191 | -200 | 0 | 0 | 10 | NaN | NaN | NaN | NaN | 1845.1836 | 2152.0781 |
| Elmo1 | GDLEESPQGEVPHDSLQDK | -589 | -607 | 0 | 0.001708008 | 19 | NaN | NaN | NaN | NaN | 1059.0996 | 1152.5946 |
| Elmo1 | HYGDLEESPQGEVPHDSLQDK | -587 | -607 | 0 | 0.00051306 | 21 | NaN | NaN | NaN | NaN | 633.9765 | 146.19653 |
| Elmo1 | DLENIQIPDAPPPIPK | -700 | -715 | 0 | 0 | 16 | NaN | NaN | NaN | NaN | 288.9726 | 1114.2638 |
| Map4k1 | GLPVDIEDEEPEPPPAIPR | -297 | -315 | 0 | 0.000431655 | 19 | NaN | NaN | NaN | NaN | 404.7547 | 1226.9167 |
| Map4k1 | SDPSLWNPAAPEPGQPPLVPPR | -452 | -473 | 0 | 0.00122419 | 22 | NaN | NaN | NaN | NaN | 311.30637 | 249.45026 |
| Upf1 | GLPDLNHSQVYAVK | -473 | -486 | 0 | 0 | 14 | NaN | NaN | NaN | NaN | 834.98987 | 916.6333 |
| Upf1 | DETGELSSADEKR | -588 | -600 | 1 | 0 | 13 | NaN | NaN | NaN | NaN | 215.61404 | 1311.8147 |
| Upf1 | LKESQTQDNITVR | -315 | -327 | 1 | 0.002534621 | 13 | NaN | NaN | NaN | NaN | 2475.3352 | NaN |
| Rnf20 | EHLLQSNIGTGEK | -807 | -819 | 0 | 0 | 13 | NaN | NaN | NaN | NaN | 1005.05554 | 1023.53406 |
| Rnf20 | ALVVPEPEPDSDSNQER | -126 | -142 | 0 | 0.008680443 | 17 | NaN | NaN | NaN | NaN | 792.4534 | NaN |
| Slc25a3 | LPRPPPPEMPESL | -337 | -349 | 1 | 0.002466863 | 13 | NaN | NaN | NaN | NaN | 1532.5123 | NaN |
| Slc25a3 | TALQWFIYDSVK | -321 | -332 | 0 | 0.00025015 | 12 | NaN | NaN | NaN | NaN | 791.962 | NaN |
| Satb1 | DCKEEHAEFVLVR | -90 | -102 | 1 | 0 | 13 | NaN | NaN | NaN | NaN | 736.3118 | 594.39 |
| Satb1 | SMNKPLEQQVSTN | -356 | -368 | 1 | 0.00450079 | 13 | NaN | NaN | NaN | NaN | 971.0802 | NaN |
| Satb1 | ASIYDEIQQEMK | -496 | -507 | 0 | 0.002972566 | 12 | NaN | NaN | NaN | NaN | 415.54108 | NaN |
| Myo1g | ELYERPPHLYAVA | -76 | -88 | 1 | 0.00025015 | 13 | NaN | NaN | NaN | NaN | 602.16187 | 1546.982 |
| Pfkl | TGISEGHTVYIVHDGFEGLAK | -424 | -444 | 0 | 0 | 21 | NaN | NaN | NaN | NaN | 1426.8188 | 1277.9235 |
| Pfkl | VQEVGWHDVAGWLGR | -447 | -461 | 0 | 0.001092529 | 15 | NaN | NaN | NaN | NaN | 1389.3536 | 1300.0594 |
| Pfkl | MADYVSGELEHVTR | -758 | -771 | 0 | 0.00051306 | 14 | NaN | NaN | NaN | NaN | 598.8246 | 683.4689 |
| Pfkl | CHEHYTTEFLYN | -630 | -641 | 0 | 0.003681885 | 12 | NaN | NaN | NaN | NaN | 714.9967 | NaN |
| Pfkl | YIVHDGFEGLAK | -433 | -444 | 0 | 0.005758933 | 12 | NaN | NaN | NaN | NaN | 533.92 | NaN |
| Psmd4 | ITFCTGIR | -84 | -91 | 0 | 0.002502116 | 8 | NaN | NaN | NaN | NaN | 1474.8839 | NaN |
| Tbc1d15 | VTNYIFDSLR | -236 | -245 | 0 | 0 | 10 | NaN | NaN | NaN | NaN | 446.201 | NaN |
| Prdx3 | TPAVTQHAPYFK | -63 | -74 | 0 | 0 | 12 | NaN | NaN | NaN | NaN | 1523.9917 | 857.88794 |
| Prdx3 | FIIDPNGVVK | -188 | -197 | 0 | 0 | 10 | NaN | NaN | NaN | NaN | 837.53326 | 635.2608 |
| Ptbp3 | LQYADPVNAQYAK | -195 | -207 | 0 | 0 | 13 | NaN | NaN | NaN | NaN | 794.81696 | 852.2304 |
| Psat1 | NETVHGVEFDFVPDVK | -154 | -169 | 0 | 0 | 16 | NaN | NaN | NaN | NaN | 520.68585 | 1425.8334 |
| Psat1 | GMVLEWIK | -252 | -259 | 0 | 0.001023236 | 8 | NaN | NaN | NaN | NaN | 512.0171 | 609.18304 |
| Psat1 | SNFLSRPVDVSK | -179 | -190 | 1 | 0.001868731 | 12 | NaN | NaN | NaN | NaN | 871.5062 | NaN |
| Psat1 | DDLLGFSLR | -214 | -222 | 0 | 0.001055699 | 9 | NaN | NaN | NaN | NaN | 496.3419 | NaN |
| Psat1 | VCDMSSNFLSR | -174 | -184 | 0 | 0.001291103 | 11 | NaN | NaN | NaN | NaN | 443.67157 | NaN |
| Psat1 | PSVLDYK | -225 | -231 | 0 | 0.00025015 | 7 | NaN | NaN | NaN | NaN | 284.018 | NaN |
| Hnrnpu | AVVVCPKDEDYK | -579 | -590 | 1 | 0.002972566 | 12 | NaN | NaN | NaN | NaN | 417.70084 | NaN |
| Hnrnpu | NFETDEVELSYAK | -387 | -399 | 0 | 0 | 13 | NaN | NaN | NaN | NaN | 415.6293 | NaN |
| Lap3 | SAGVDDQENWHEGK | -105 | -118 | 0 | 0 | 14 | NaN | NaN | NaN | NaN | 455.21008 | 464.53937 |
| Acbd3 | IRQDADSVITVR | -384 | -395 | 1 | 0.00025015 | 12 | NaN | NaN | NaN | NaN | 705.1037 | 717.05347 |
| Acbd3 | ANKPLLDEIVPVYR | -468 | -481 | 1 | 0.009928578 | 14 | NaN | NaN | NaN | NaN | 506.67276 | NaN |
| Agfg1 | IWLGLFDDR | -92 | -100 | 0 | 0 | 9 | NaN | NaN | NaN | NaN | 430.27325 | 250.83403 |
| Sae1 | GRDPTSESYKEDAELLL | -258 | -274 | 2 | 0.00272025 | 17 | NaN | NaN | NaN | NaN | 695.2981 | 491.00568 |
| Sae1 | FFTGDVFGYHGY | -157 | -168 | 0 | 0.000947459 | 12 | NaN | NaN | NaN | NaN | 277.34702 | 265.7463 |
| Vwa5a | ANLGGTEILTPLCNIYK | -362 | -378 | 0 | 0 | 17 | NaN | NaN | NaN | NaN | 227.95206 | 446.51062 |
| Birc6 | DLEELGANPSLTNSK | -537 | -551 | 0 | 0 | 15 | NaN | NaN | NaN | NaN | 17653.266 | NaN |
| Birc6 | VLCELFQTAPQR | -3807 | -3818 | 0 | 0.000230707 | 12 | NaN | NaN | NaN | NaN | 2674.172 | NaN |
| Birc6 | ANLIRPGDAK | -2921 | -2930 | 1 | 0.001463897 | 10 | NaN | NaN | NaN | NaN | 866.27637 | NaN |
| Ttn | GVGEPTETTEPVR | -22737 | -22749 | 0 | 0.001023236 | 13 | NaN | NaN | NaN | NaN | 1167.4409 | 1110.002 |
| Septin7 | DNSNCWQPVIDYIDSK | -121 | -136 | 0 | 0.00078661 | 16 | NaN | NaN | NaN | NaN | 359.12628 | NaN |
| Ncbp1 | FLSDLVNCH | -130 | -138 | 0 | 0.003900536 | 9 | NaN | NaN | NaN | NaN | 2837.3503 | NaN |
| Smarca5 | RLDGQTPHDER | -527 | -537 | 1 | 0.005363933 | 11 | NaN | NaN | NaN | NaN | 847.065 | NaN |
| Serpinc1 | YQHLADSK | -96 | -103 | 0 | 0 | 8 | NaN | NaN | NaN | NaN | 681.31104 | 855.8039 |
| Serpinc1 | NDNDNIFLSPL | -104 | -114 | 0 | 0.006624942 | 11 | NaN | NaN | NaN | NaN | 210.9612 | NaN |
| Nudt5 | ANSKPFEVPFLK | -206 | -217 | 1 | 0.005459848 | 12 | NaN | NaN | NaN | NaN | 3800.2478 | NaN |
| Atp5pd | WNETFHAR | -34 | -41 | 0 | 0.005145856 | 8 | NaN | NaN | NaN | NaN | 1126.932 | NaN |
| Atp5pd | ANVAKPGLVDDFEKK | -59 | -73 | 2 | 0 | 15 | NaN | NaN | NaN | NaN | 559.5272 | NaN |
| Zcchc9 | QPGHGIADCPAVLESQD | -135 | -151 | 0 | 0.005522803 | 17 | NaN | NaN | NaN | NaN | 814.2718 | NaN |
| Acly | LNASGSTSTPAPSR | -439 | -452 | 0 | 0 | 14 | NaN | NaN | NaN | NaN | 295.0871 | 2433.8318 |
| Ada | GLQEGEQAFGIK | -136 | -147 | 0 | 0.000550307 | 12 | NaN | NaN | NaN | NaN | 2946.0894 | NaN |
| Ada | VGHGYHTIEDEALYNR | -236 | -251 | 0 | 0.00025015 | 16 | NaN | NaN | NaN | NaN | 1133.678 | NaN |
| Ada | DMGFTEEEFKR | -314 | -324 | 1 | 0.000431655 | 11 | NaN | NaN | NaN | NaN | 723.9487 | NaN |
| H1-2 | PAAAPPAEK | -9 | -17 | 0 | 0.006100831 | 9 | NaN | NaN | NaN | NaN | 970.48596 | NaN |

| | | | | | | | | | | | | |
|---|---|---|---|---|---|---|---|---|---|---|---|---|
| H1-3 | PAAPAAPAPVEK | -6 | -17 | 0 | 0.00078661 | 12 | NaN | NaN | NaN | NaN | 606.22656 | 593.2275 |
| H1-3 | APAAPAAPAPVEK | -5 | -17 | 0 | 0.002068213 | 13 | NaN | NaN | NaN | NaN | 845.7762 | 664.5209 |
| Prpf4 | FHQDGSLAGTGGLDAFGR | -372 | -389 | 0 | 0.006735683 | 18 | NaN | NaN | NaN | NaN | 1368.0764 | NaN |
| Sdhaf2 | APDLEYLFEKPH | -153 | -164 | 1 | 0.00078661 | 12 | NaN | NaN | NaN | NaN | 1176.8052 | 2268.8608 |
| Ddx1 | APDNYIVK | -261 | -268 | 0 | 0.004983786 | 8 | NaN | NaN | NaN | NaN | 496.64914 | NaN |
| Nif3l1 | APDYPTEGAHR | -157 | -167 | 0 | 0 | 11 | NaN | NaN | NaN | NaN | 367.54947 | NaN |
| Anp32e | IKDLSTVEALQNLK | -100 | -113 | 1 | 0.00025015 | 14 | NaN | NaN | NaN | NaN | 583.9499 | 835.9008 |
| Anp32e | SLDLFNCEIT | -117 | -126 | 0 | 0.002466863 | 10 | NaN | NaN | NaN | NaN | 216.04431 | 361.50073 |
| Gins1 | APEGQLPAFNEDGLR | -16 | -30 | 0 | 0 | 15 | NaN | NaN | NaN | NaN | 627.58826 | 388.28003 |
| Cfl1 | NIILEEGKEILVGDVGQTV | -46 | -64 | 1 | 0.00025015 | 19 | NaN | NaN | NaN | NaN | 693.2167 | 659.56934 |
| Cfl1 | HELQANCYEEVKDR | -133 | -146 | 1 | 0 | 14 | NaN | NaN | NaN | NaN | 260.01477 | 645.2266 |
| Cfl1 | SGVAVSDGVIK | -3 | -13 | 0 | 0.007343498 | 11 | NaN | NaN | NaN | NaN | 2309.6575 | NaN |
| Supt16h | APGEQTVPALNLQNAF | -607 | -622 | 0 | 0.000627943 | 16 | NaN | NaN | NaN | NaN | 938.44885 | NaN |
| Supt16h | ICDVYNSVMDVVK | -322 | -334 | 0 | 0.001370415 | 13 | NaN | NaN | NaN | NaN | 823.6699 | NaN |
| Mki67 | QTPAGASDSVTVEESAK | (1214)(2161) | (1230)(2177) | 0 | 0.000230707 | 17 | NaN | NaN | NaN | NaN | 1276.9747 | 1040.2635 |
| Mki67 | LESSQAEPVK | (1233)(1353)(2063) | (1242)(1362)(2072) | 0 | 0.00025015 | 10 | NaN | NaN | NaN | NaN | 977.2076 | 3510.3494 |
| Mki67 | KIDPVASVPVSK | -2713 | -2724 | 1 | 0 | 12 | NaN | NaN | NaN | NaN | 960.7011 | 2681.4668 |
| Mki67 | RSGDDGAHFPL | -12 | -22 | 1 | 0.003521747 | 11 | NaN | NaN | NaN | NaN | 1312.7509 | NaN |
| Mki67 | KLDTEVNVPR | -2491 | -2500 | 1 | 0.000394361 | 10 | NaN | NaN | NaN | NaN | 1240.1068 | NaN |
| Mki67 | QEATDEKPVPEK | -2958 | -2969 | 1 | 0.00111006 | 12 | NaN | NaN | NaN | NaN | 696.4936 | NaN |
| Mki67 | RENPVTPDQNSR | -3016 | -3027 | 1 | 0.001023236 | 12 | NaN | NaN | NaN | NaN | 378.009 | NaN |
| Rpl18 | TFDQLALESPKGR | -122 | -134 | 1 | 0 | 13 | NaN | NaN | NaN | NaN | 1235.0361 | NaN |
| Eif4a1 | DDMNLSESLLR | -35 | -45 | 0 | 0.003299042 | 11 | NaN | NaN | NaN | NaN | 1491.4838 | NaN |
| Rpl27a | APIIDVVR | -98 | -105 | 0 | 0.004224523 | 8 | NaN | NaN | NaN | NaN | 2466.3958 | 2630.9333 |
| Rpl27a | PIIDVVR | -99 | -105 | 0 | 0.006164361 | 7 | NaN | NaN | NaN | NaN | 323.43967 | 259.24258 |
| Idh3a | KTPYTDVNIVTIR | -134 | -146 | 1 | 0.00025015 | 13 | NaN | NaN | NaN | NaN | 1125.0605 | 1103.3894 |
| Rpl4 | LAGHQTSAESWGTGR | -57 | -71 | 0 | 0.001463897 | 15 | NaN | NaN | NaN | NaN | 449.19095 | 418.78885 |
| Srrm1 | KFAECLEK | -29 | -36 | 1 | 0.002566359 | 8 | NaN | NaN | NaN | NaN | 628.6248 | 920.6343 |
| Srrm1 | KAPKPEPVPEPK | -246 | -257 | 2 | 0.002466863 | 12 | NaN | NaN | NaN | NaN | 607.9544 | 1349.2318 |
| Psme1 | KGDEDDKGPPCGPVNCNEK | -91 | -109 | 2 | 0.00025015 | 19 | NaN | NaN | NaN | NaN | 296.2772 | 821.5481 |
| Psme1 | NAYAVLYDIILK | -221 | -232 | 0 | 0 | 12 | NaN | NaN | NaN | NaN | 215.07495 | NaN |
| Ppp6c | APLDLDKYVEIAR | -2 | -14 | 1 | 0 | 13 | NaN | NaN | NaN | NaN | 1066.3975 | 839.5242 |
| Nckap1l | EILRPSELLAGVK | -733 | -745 | 1 | 0 | 13 | NaN | NaN | NaN | NaN | 818.2184 | 814.4944 |
| Nckap1l | IFPSEYLSSHLEAR | -703 | -716 | 0 | 0 | 14 | NaN | NaN | NaN | NaN | 1958.9348 | NaN |
| Nckap1l | SVSLLQLGQETDKLK | -1064 | -1078 | 1 | 0.000230707 | 15 | NaN | NaN | NaN | NaN | 1062.6161 | NaN |
| Gm12250 | CLPTITEPAIDR | -261 | -272 | 0 | 0 | 12 | NaN | NaN | NaN | NaN | 4508.3 | 4045.5564 |
| Gm12250 | IDSDLWNEKK | -184 | -193 | 1 | 0 | 10 | NaN | NaN | NaN | NaN | 494.67914 | 394.71164 |
| Gm12250 | MSFFNDDIEEFEK | -297 | -309 | 0 | 0.003745449 | 13 | NaN | NaN | NaN | NaN | 194.37396 | 703.7474 |
| Gm12250 | IQEGDIQK | -45 | -52 | 0 | 0.001708008 | 8 | NaN | NaN | NaN | NaN | 1948.0691 | NaN |
| Leo1 | VNTDLGNDLYFVK | -366 | -378 | 0 | 0 | 13 | NaN | NaN | NaN | NaN | 919.27313 | NaN |
| Leo1 | LKVENTIR | -415 | -422 | 1 | 0.00272025 | 8 | NaN | NaN | NaN | NaN | 251.44604 | NaN |
| Chd4 | SLKEEESTEGEKEVK | -1563 | -1577 | 2 | 0.000591124 | 15 | NaN | NaN | NaN | NaN | 1160.3464 | 1068.3689 |
| Chd4 | DGGELLCCDTCPSSYHIH | -450 | -467 | 0 | 0.00025015 | 18 | NaN | NaN | NaN | NaN | 4523.5415 | 5718.754 |
| Chd4 | SEEGDEDFDER | -1363 | -1373 | 0 | 0.003360691 | 11 | NaN | NaN | NaN | NaN | 221.84152 | NaN |
| Crkl | KGLFPFTHVK | -283 | -292 | 1 | 0.000712251 | 10 | NaN | NaN | NaN | NaN | 198.7242 | 401.94495 |
| Crkl | LYDFPGNDAEDLPFK | -131 | -145 | 0 | 0.000431655 | 15 | NaN | NaN | NaN | NaN | 427.97714 | NaN |
| Cope | SQHLGKPPEVTNR | -260 | -272 | 1 | 0 | 13 | NaN | NaN | NaN | NaN | 962.08514 | 848.6882 |
| Gnl2 | FIGYPNVGK | -315 | -323 | 0 | 0.000230707 | 9 | NaN | NaN | NaN | NaN | 835.6914 | 750.9224 |
| Adpgk | APQEFTTSHLESGSR | -429 | -443 | 0 | 0 | 15 | NaN | NaN | NaN | NaN | 730.65295 | 561.47345 |
| Pdcd4 | APQLVGQFIAR | -257 | -267 | 0 | 0 | 11 | NaN | NaN | NaN | NaN | 1323.7836 | 1255.1549 |
| Pdcd4 | PELALDTPR | -248 | -256 | 0 | 0.005821388 | 9 | NaN | NaN | NaN | NaN | 635.9413 | 540.76605 |
| Pdcd4 | AVGDGILCNTYIDSYK | -268 | -283 | 0 | 0 | 16 | NaN | NaN | NaN | NaN | 1139.4764 | NaN |
| Pdcd4 | FVEECFQAGIISK | -429 | -441 | 0 | 0.001291103 | 13 | NaN | NaN | NaN | NaN | 728.3288 | NaN |
| Capn1 | SEHFINLR | -473 | -480 | 0 | 0 | 8 | NaN | NaN | NaN | NaN | 1556.3208 | NaN |
| Dnmt1 | LPLFPEPLHVFAPR | -1327 | -1340 | 0 | 0 | 14 | NaN | NaN | NaN | NaN | 701.54395 | 697.2791 |
| Dnmt1 | VSEPKEPEAAIK | -1127 | -1138 | 1 | 0.000431655 | 12 | NaN | NaN | NaN | NaN | 1117.8566 | 1009.6094 |
| Dnmt1 | LEEVTAGTQLGPEEPCEQEDDNR | -217 | -239 | 0 | 0.002566359 | 23 | NaN | NaN | NaN | NaN | 333.77264 | NaN |
| Dnmt1 | APSENWAMEGGTDPETTLPGAEDGK | -841 | -865 | 0 | 0.002199523 | 25 | NaN | NaN | NaN | NaN | 181.70833 | NaN |
| Psmc3 | TNRVDILDPALLR | -335 | -347 | 1 | 0 | 13 | NaN | NaN | NaN | NaN | 595.142 | 577.3086 |
| Psmc3 | APSIIFIDELDAIGTK | -282 | -297 | 0 | 0.000230707 | 16 | NaN | NaN | NaN | NaN | 231.55937 | 272.74127 |
| Psmc3 | ATVWDEAEQDGIGEEVLK | -21 | -38 | 0 | 0.000431655 | 18 | NaN | NaN | NaN | NaN | 1048.6061 | NaN |
| Apbb1ip | YKAPTDHCFVL | -377 | -387 | 1 | 0.005854446 | 11 | NaN | NaN | NaN | NaN | 1297.8944 | NaN |
| Rrm2 | PSVEDEPLLR | -66 | -75 | 0 | 0.006164361 | 10 | NaN | NaN | NaN | NaN | 1610.1365 | NaN |
| Cars1 | SKPGEPSWPCPWGK | -409 | -422 | 1 | 0.002265552 | 14 | NaN | NaN | NaN | NaN | 684.2258 | 144.25378 |
| Cars1 | SKFDENGLPTHDTEGK | -782 | -797 | 1 | 0.000431655 | 16 | NaN | NaN | NaN | NaN | 1498.5021 | 1510.2667 |
| Cars1 | SKFDENGLPTHDTEGK | -782 | -797 | 1 | 0.000431655 | 16 | NaN | NaN | NaN | NaN | 1498.5021 | 1510.2667 |
| Cars1 | NKDVFIPQDGK | -121 | -131 | 1 | 0 | 11 | NaN | NaN | NaN | NaN | 678.0837 | 747.5346 |
| Cars1 | VSEYVPEIVNFVQK | -324 | -337 | 0 | 0 | 14 | NaN | NaN | NaN | NaN | 431.83246 | 610.79376 |
| Nob1 | APVEHVVADAGAF | -2 | -14 | 0 | 0.009897819 | 13 | NaN | NaN | NaN | NaN | 1746.5781 | NaN |
| Pfas | RFQELQR | -206 | -212 | 1 | 0.00111006 | 7 | NaN | NaN | NaN | NaN | 1791.9529 | NaN |
| Pfas | NPSTVEVFDLAQSNSEHSR | -213 | -231 | 0 | 0.001868731 | 19 | NaN | NaN | NaN | NaN | 376.34604 | NaN |
| Dhx33 | EHKDDQLTLTPIGR | -473 | -486 | 1 | 0 | 14 | NaN | NaN | NaN | NaN | 475.25583 | 479.79486 |
| Dhx33 | APVLYLEGR | -237 | -245 | 0 | 0.00111006 | 9 | NaN | NaN | NaN | NaN | 953.46106 | NaN |
| Yars1 | SSSEEESKIDLLDR | -224 | -237 | 1 | 0.001392869 | 14 | NaN | NaN | NaN | NaN | 828.9184 | 779.6926 |
| Yars1 | RNLQEVLGEEK | -16 | -26 | 1 | 0.00025015 | 11 | NaN | NaN | NaN | NaN | 739.95123 | 721.57733 |
| H2-D1 | YFETAVSRPGLEEPR | -31 | -45 | 1 | 0.000298805 | 15 | NaN | NaN | NaN | NaN | 172.20752 | 833.9363 |
| Hadhb | LRDFIYVSQDPK | -338 | -349 | 1 | 0.00111006 | 12 | NaN | NaN | NaN | NaN | 467.61633 | 195.38216 |
| Ctsc | NLPESWDWR | -229 | -237 | 0 | 0.007343498 | 9 | NaN | NaN | NaN | NaN | 513.9692 | NaN |

| | | | | | | | | | | | | |
|---|---|---|---|---|---|---|---|---|---|---|---|---|
| Esyt1 | ELPAWVSFPDVEKA | -116 | -129 | 1 | 0 | 14 | NaN | NaN | NaN | NaN | 390.10583 | 348.7411 |
| Nsun2 | DGQWFADWHEVPQGR | -370 | -384 | 0 | 0 | 15 | NaN | NaN | NaN | NaN | 1706.5758 | 1617.479 |
| Nsun2 | QQPPQPEGEEDASDGGR | -11 | -27 | 0 | 0 | 17 | NaN | NaN | NaN | NaN | 579.62646 | 752.44434 |
| Nsun2 | DGVCGPPPSK | -498 | -507 | 0 | 0.00122419 | 10 | NaN | NaN | NaN | NaN | 1303.0867 | NaN |
| Ppid | ECGELKEGDDWGIFPK | -180 | -195 | 1 | 0.00078661 | 16 | NaN | NaN | NaN | NaN | 876.7917 | 894.2462 |
| Atp5f1a | GADTSVDLEETGR | -61 | -73 | 0 | 0.000471409 | 13 | NaN | NaN | NaN | NaN | 5256.1006 | 3087.236 |
| Atp5f1a | SQHQSLLGNIR | -517 | -527 | 0 | 0.00025015 | 11 | NaN | NaN | NaN | NaN | 1834.9414 | 692.9518 |
| Atp5f1a | ENAFLSHVISQHQSLLGNIR | -508 | -527 | 0 | 0 | 20 | NaN | NaN | NaN | NaN | 424.33466 | 456.90363 |
| Pcnt | LLDLFGDTLK | -939 | -948 | 0 | 0.00025015 | 10 | NaN | NaN | NaN | NaN | 543.6573 | 618.1832 |
| Pcnt | AQHESELEHLR | -390 | -400 | 0 | 0 | 11 | NaN | NaN | NaN | NaN | 330.5765 | NaN |
| Rpl8 | AQLNIGNVLPVGTMPEGTIV | -94 | -113 | 0 | 0.005112649 | 20 | NaN | NaN | NaN | NaN | 288.0611 | 212.70071 |
| Rpl8 | AQLNIGNVLPVGTMPEGTIVC | -94 | -114 | 0 | 0.00383302 | 21 | NaN | NaN | NaN | NaN | 245.93135 | NaN |
| Pola2 | SSGSHLVFVPSLR | -415 | -427 | 0 | 0 | 13 | NaN | NaN | NaN | NaN | 1318.9564 | 1296.1252 |
| Pola2 | KIEELGSELK | -223 | -232 | 1 | 0.000627943 | 10 | NaN | NaN | NaN | NaN | 110.187996 | 746.92725 |
| Gapdh | VYMFQYDSTHGK | -42 | -53 | 0 | 0 | 12 | NaN | NaN | NaN | NaN | 1406.2751 | 1586.0817 |
| Gapdh | MTTVHAITATQK | -173 | -184 | 0 | 0 | 12 | NaN | NaN | NaN | NaN | 573.4956 | 614.34125 |
| Gapdh | TVHAITATQK | -175 | -184 | 0 | 0 | 10 | NaN | NaN | NaN | NaN | 4985.234 | 5317.824 |
| Gapdh | LEKPAKYDDIK | -247 | -257 | 2 | 0 | 11 | NaN | NaN | NaN | NaN | 591.83875 | 1179.9268 |
| Gapdh | LVINGKPITIFQER | -65 | -78 | 1 | 0 | 14 | NaN | NaN | NaN | NaN | 418.14075 | 399.57346 |
| Gapdh | GILGYTEDQVVSCDFNSNSHSSTF | -270 | -293 | 0 | 0.00078661 | 24 | NaN | NaN | NaN | NaN | 156.92725 | 197.74866 |
| Gapdh | SSTFDAGAGIALNDNFVK | -290 | -307 | 0 | 0.000712251 | 18 | NaN | NaN | NaN | NaN | 1444.8302 | NaN |
| Sfxn3 | VATPGLTEDQLWR | -57 | -69 | 0 | 0.000230707 | 13 | NaN | NaN | NaN | NaN | 1397.5432 | 1387.238 |
| Sfxn3 | NLLLSGEQLEAS | -35 | -46 | 0 | 0.005270854 | 12 | NaN | NaN | NaN | NaN | 198.81345 | NaN |
| Sp110 | TKNDAVDFSPTLPVTCGK | -351 | -368 | 1 | 0.00025015 | 18 | NaN | NaN | NaN | NaN | 225.76024 | 371.09412 |
| Msn | QVWHEEHR | -173 | -180 | 0 | 0.00659934 | 8 | NaN | NaN | NaN | NaN | 10361.003 | NaN |
| Msn | KESEAVEWQQK | -438 | -448 | 1 | 0 | 11 | NaN | NaN | NaN | NaN | 2022.5775 | NaN |
| Eif3b | GFHCESSAHWPIFK | -406 | -419 | 0 | 0.002780285 | 14 | NaN | NaN | NaN | NaN | 622.62177 | 575.31604 |
| Eif3b | SIYETPSMGLLDK | -436 | -448 | 0 | 0 | 13 | NaN | NaN | NaN | NaN | 4716.879 | 3094.3127 |
| Atp5f1d | AQSELSGAADEAAR | -137 | -150 | 0 | 0 | 14 | NaN | NaN | NaN | NaN | 1710.5688 | 7940.0786 |
| Atp5f1d | TFASPTQVFFDSANVK | -41 | -56 | 0 | 0.00025015 | 16 | NaN | NaN | NaN | NaN | 519.13336 | 414.6354 |
| Ruvbl1 | EIIQDVTLHDLDVANARPQ | -233 | -251 | 1 | 0 | 19 | NaN | NaN | NaN | NaN | 390.54956 | 393.40167 |
| Ruvbl1 | EHVEEISELFYDAK | -428 | -441 | 0 | 0 | 14 | NaN | NaN | NaN | NaN | 932.7069 | 780.31 |
| Capza1 | AQYNMDQFTPVK | -55 | -66 | 0 | 0.001392869 | 12 | NaN | NaN | NaN | NaN | 570.7907 | NaN |
| Eif3a | AREESWGPPR | -1145 | -1154 | 1 | 0.00025015 | 10 | NaN | NaN | NaN | NaN | 371.25095 | NaN |
| Hspa8 | QVAMNPTNTVFDAK | -58 | -71 | 0 | 0 | 14 | NaN | NaN | NaN | NaN | 1370.3928 | 1204.3457 |
| Hspa8 | MKEIAEAYLGK | -127 | -137 | 1 | 0 | 11 | NaN | NaN | NaN | NaN | 785.3431 | 636.17786 |
| Hspa8 | SFYPEEVSSM | -113 | -122 | 0 | 0.007009655 | 10 | NaN | NaN | NaN | NaN | 865.1385 | NaN |
| Hspa8 | HWPFMVVNDAGR | -89 | -100 | 0 | 0 | 12 | NaN | NaN | NaN | NaN | 811.7182 | NaN |
| Baz1b | KDHSNLPDDDYCPR | -943 | -956 | 1 | 0 | 14 | NaN | NaN | NaN | NaN | 465.46332 | 446.66687 |
| Baz1b | RFEDQELGGR | -580 | -589 | 1 | 0.001055699 | 10 | NaN | NaN | NaN | NaN | 7799.9067 | 7525.7593 |
| Baz1b | ELDELLSCLHPQ | -999 | -1010 | 0 | 0.000431655 | 12 | NaN | NaN | NaN | NaN | 1162.1073 | NaN |
| Baz1b | SNEVPGLFIEK | -917 | -927 | 0 | 0 | 11 | NaN | NaN | NaN | NaN | 1060.4287 | NaN |
| Baz1b | QGQNLWFLCDSQK | -986 | -998 | 0 | 0.002780285 | 13 | NaN | NaN | NaN | NaN | 966.69037 | NaN |
| Ndufa8 | LGWVRPDLGQLSK | -115 | -127 | 1 | 0 | 13 | NaN | NaN | NaN | NaN | 677.6639 | 1231.9135 |
| Ptprc | SNFFSGPIVVHCSAGVGR | -831 | -848 | 0 | 0.00025015 | 18 | NaN | NaN | NaN | NaN | 373.51483 | 347.19095 |
| Ptprc | ARPGVVCSYEQY | -1195 | -1206 | 1 | 0.00025015 | 12 | NaN | NaN | NaN | NaN | 11731.203 | NaN |
| Ptprc | KRDPPSDPSPLEAEYQR | -923 | -939 | 2 | 0.004199202 | 17 | NaN | NaN | NaN | NaN | 564.4878 | NaN |
| Ptprc | WKGEELPAEPK | -1114 | -1124 | 1 | 0.00051306 | 11 | NaN | NaN | NaN | NaN | 558.71655 | NaN |
| Kbtbd11 | TPGEDSLPLPAEEEGAASTA | -18 | -37 | 0 | 0.003745449 | 20 | NaN | NaN | NaN | NaN | 1025.1771 | NaN |
| Hba | LSALSDLHAHK | -81 | -91 | 0 | 0.00713295 | 11 | NaN | NaN | NaN | NaN | 478.68604 | NaN |
| Hba | HHPADFTPAVHASLDK | -113 | -128 | 0 | 0 | 16 | NaN | NaN | NaN | NaN | 208.87738 | NaN |
| Cluh | RVVEEPYTVR | -117 | -126 | 1 | 0 | 10 | NaN | NaN | NaN | NaN | 1305.1997 | 1882.1809 |
| Cluh | ASDAFHFF | -977 | -984 | 0 | 0.00078661 | 8 | NaN | NaN | NaN | NaN | 103.8617 | NaN |
| Glipr2 | SYNFQQPGFTSGTGHFTA | -89 | -106 | 0 | 0.00111006 | 18 | NaN | NaN | NaN | NaN | 644.26776 | NaN |
| Hsp90b1 | NLLHVTDTGVGMTR | -143 | -156 | 0 | 0 | 14 | NaN | NaN | NaN | NaN | 976.6044 | 473.18567 |
| Hsp90b1 | IKEDEDDKTVMDLA | -701 | -714 | 2 | 0.000627943 | 14 | NaN | NaN | NaN | NaN | 742.7611 | 670.7149 |
| Hsp90b1 | FAFQAEVNR | -76 | -84 | 0 | 0 | 9 | NaN | NaN | NaN | NaN | 1372.6494 | NaN |
| Hsp90b1 | EATEKEFEPLLN | -609 | -620 | 1 | 0.001708008 | 12 | NaN | NaN | NaN | NaN | 887.09467 | NaN |
| Ist1 | ASEDIDFDDLSR | -342 | -353 | 0 | 0 | 12 | NaN | NaN | NaN | NaN | 655.65906 | 578.5785 |
| Arl3 | ASEDISHITPTQGFNIK | -38 | -54 | 0 | 0.00025015 | 17 | NaN | NaN | NaN | NaN | 431.03644 | NaN |
| Dck | DVNEDFKDKHESLVEK | -237 | -252 | 2 | 0.000431655 | 16 | NaN | NaN | NaN | NaN | 953.23535 | 796.1673 |
| Gapvd1 | DQVLHEHIQR | -1321 | -1330 | 0 | 0 | 10 | NaN | NaN | NaN | NaN | 808.48206 | 884.87335 |
| Gapvd1 | GFQETAYGEFLSR | -90 | -102 | 0 | 0.002068213 | 13 | NaN | NaN | NaN | NaN | 1274.572 | 738.4099 |
| Ranbp1 | MELKPNAGSDR | -108 | -118 | 1 | 0 | 11 | NaN | NaN | NaN | NaN | 1156.0953 | 1487.6505 |
| Ranbp1 | NHDPQFEPIVSLPEQEIK | -22 | -39 | 0 | 0.00025015 | 18 | NaN | NaN | NaN | NaN | 559.6742 | 622.985 |
| Ranbp1 | NHDPQFEPIVSLPEQEIK | -22 | -39 | 0 | 0.00025015 | 18 | NaN | NaN | NaN | NaN | 559.6742 | 622.985 |
| Srrt | PSQNPILK | -551 | -558 | 0 | 0.005522803 | 8 | NaN | NaN | NaN | NaN | 656.1285 | 385.01633 |
| Srrt | GDSDDEYDRR | -2 | -11 | 1 | 0.000431655 | 10 | NaN | NaN | NaN | NaN | 584.4323 | 498.00385 |
| Srrt | ASEPGTPPVPTSLPSQNPILK | -538 | -558 | 0 | 0.000230707 | 21 | NaN | NaN | NaN | NaN | 383.01285 | 377.25403 |
| Itgal | IQGAQVFPGIR | -567 | -577 | 0 | 0 | 11 | NaN | NaN | NaN | NaN | 1570.5602 | 1365.5514 |
| Itgal | TCDQNTYLSGL | -115 | -125 | 0 | 0.001023236 | 11 | NaN | NaN | NaN | NaN | 207.06772 | 177.43137 |
| Itgal | HFHLYGSK | -1061 | -1068 | 0 | 0.00025015 | 8 | NaN | NaN | NaN | NaN | 1013.91187 | NaN |
| Itgal | TDPLVTCHSEDLK | -988 | -1000 | 0 | 0 | 13 | NaN | NaN | NaN | NaN | 884.9681 | NaN |
| Hddc2 | LQDFYDSTAGKF | -159 | -170 | 1 | 0.004923919 | 12 | NaN | NaN | NaN | NaN | 1237.5242 | 1324.6259 |
| Rad23b | QIIQQNPSLLPALLQQIGR | -290 | -308 | 0 | 0.000550307 | 19 | NaN | NaN | NaN | NaN | 205.50433 | 159.27864 |
| Adh5 | ASGEEISTRPFQLVTGR | -296 | -312 | 1 | 0.007040624 | 17 | NaN | NaN | NaN | NaN | 2158.2354 | NaN |
| Adh5 | EAGKPLSIEEIEVAPPK | -15 | -31 | 1 | 0.002068213 | 17 | NaN | NaN | NaN | NaN | 724.03107 | NaN |
| Adh5 | EAGKPLSIEEIEVAPPK | -15 | -31 | 1 | 0.002068213 | 17 | NaN | NaN | NaN | NaN | 724.03107 | NaN |

| | | | | | | | | | | | | |
|---|---|---|---|---|---|---|---|---|---|---|---|---|
| Dtx3l | FGGPANFGYPDPDYLK | -721 | -736 | 0 | 0.001868731 | 16 | NaN | NaN | NaN | NaN | 2108.8774 | 2761.1997 |
| Dtx3l | LLEPELLQEISEIEQK | -391 | -406 | 0 | 0.000550307 | 16 | NaN | NaN | NaN | NaN | 425.00092 | NaN |
| Cox5b | CPNCGTHYK | -112 | -120 | 0 | 0.006195994 | 9 | NaN | NaN | NaN | NaN | 329.70737 | NaN |
| Exosc10 | TEDFIVDTLELR | -335 | -346 | 0 | 0 | 12 | NaN | NaN | NaN | NaN | 528.3095 | 681.9033 |
| Exosc10 | SGGLPQFGDEYDFYR | -50 | -64 | 0 | 0.004224523 | 15 | NaN | NaN | NaN | NaN | 507.77094 | NaN |
| Akr1b1 | SPLGSPDRPWAKPEDPSLLEDPR | -211 | -233 | 2 | 0 | 23 | NaN | NaN | NaN | NaN | 400.98892 | 412.23425 |
| Arpc2 | DTDAAVGDNIGYIT | -211 | -224 | 0 | 0 | 14 | NaN | NaN | NaN | NaN | 888.398 | NaN |
| Arpc2 | YRDDETMYVESK | -146 | -157 | 1 | 0.00025015 | 12 | NaN | NaN | NaN | NaN | 852.19446 | 830.08105 |
| Arpc2 | SHTAPQVLFSHR | -192 | -203 | 0 | 0.00025015 | 12 | NaN | NaN | NaN | NaN | 767.0471 | 3171.421 |
| Arpc2 | MILLEVNNR | -1 | -9 | 0 | 0.00142999 | 9 | NaN | NaN | NaN | NaN | 360.37967 | 753.72845 |
| Uba2 | LFGEEDADQEVSPDR | -196 | -210 | 0 | 0.000627943 | 15 | NaN | NaN | NaN | NaN | 1301.5491 | 1330.0952 |
| Uba2 | HSEDLGKDVEFEVVGDSPEK | -532 | -551 | 1 | 0.00078661 | 20 | NaN | NaN | NaN | NaN | 293.45932 | 1190.9259 |
| Sf3a1 | ITLQEQIEAIHK | -518 | -529 | 0 | 0 | 12 | NaN | NaN | NaN | NaN | 294.97427 | 347.26266 |
| Sf3a1 | PLTDQVSVIK | -730 | -739 | 0 | 0.005668207 | 10 | NaN | NaN | NaN | NaN | 697.494 | 673.8608 |
| Rpn2 | GLPLASNEALGALTAR | -136 | -151 | 0 | 0 | 16 | NaN | NaN | NaN | NaN | 1447.3866 | 1231.5055 |
| Rpn2 | NPILWNVADVVIK | -492 | -504 | 0 | 0 | 13 | NaN | NaN | NaN | NaN | 318.69614 | 299.9436 |
| Rpn2 | SNLDPSNVDSLFYAAQ | -77 | -92 | 0 | 0.001536401 | 16 | NaN | NaN | NaN | NaN | 268.8849 | 1474.1957 |
| Rpn2 | GFGLPLASNEALGALTAR | -134 | -151 | 0 | 0.000471409 | 18 | NaN | NaN | NaN | NaN | 216.05237 | NaN |
| Psmd1 | IYMNLEKPDFINVC | -204 | -217 | 1 | 0.005758933 | 14 | NaN | NaN | NaN | NaN | 2587.944 | NaN |
| Psmd1 | EPEPNFQLLDNPAR | -870 | -883 | 0 | 0.00025015 | 14 | NaN | NaN | NaN | NaN | 1397.44 | NaN |
| Psmd1 | QCVENADLPEGEK | -111 | -123 | 0 | 0.000591124 | 13 | NaN | NaN | NaN | NaN | 808.998 | NaN |
| Hexim1 | ASLQSGSRPGQEGEGGLK | -51 | -68 | 1 | 0.00122419 | 18 | NaN | NaN | NaN | NaN | 8039.3706 | 7693.3394 |
| Tram1 | ASLYHYGIK | -70 | -78 | 0 | 0 | 9 | NaN | NaN | NaN | NaN | 2341.3298 | 2385.7686 |
| Rad21 | KGGEADNLDEFLK | -406 | -418 | 1 | 0.000627943 | 13 | NaN | NaN | NaN | NaN | 1969.391 | 1790.4714 |
| Acadvl | SLSEGYPTAQHEK | -589 | -601 | 0 | 0.000394361 | 13 | NaN | NaN | NaN | NaN | 473.20007 | 540.37744 |
| Acadvl | EATQAVLDKPETLSSDASTR | -44 | -63 | 1 | 0.000826806 | 20 | NaN | NaN | NaN | NaN | 379.1783 | NaN |
| Clint1 | ENYHFVDEHGK | -120 | -130 | 0 | 0 | 11 | NaN | NaN | NaN | NaN | 4978.108 | 5159.1587 |
| Clint1 | IGSTIDDTISK | -216 | -226 | 0 | 0.000298805 | 11 | NaN | NaN | NaN | NaN | 990.6543 | 946.197 |
| Clint1 | IGSTIDDTISKF | -216 | -227 | 1 | 0 | 12 | NaN | NaN | NaN | NaN | 908.2305 | NaN |
| Szrd1 | ASPEEEQEKPILDRPTR | -106 | -122 | 2 | 0.00122419 | 17 | NaN | NaN | NaN | NaN | 440.46954 | 499.54825 |
| Szrd1 | EEAADSGEIDRR | -11 | -22 | 1 | 0 | 12 | NaN | NaN | NaN | NaN | 661.9832 | NaN |
| Strn4 | ASPGPGGLSGGESLLVK | -222 | -238 | 0 | 0 | 17 | NaN | NaN | NaN | NaN | 1402.4502 | 1655.9612 |
| Hnrnpa0 | KLFVGGLKG | -99 | -107 | 2 | 0.006624942 | 9 | NaN | NaN | NaN | NaN | 2995.2515 | NaN |
| Nceh1 | EGSPKPEEPLRR | -96 | -107 | 2 | 0.001055699 | 12 | NaN | NaN | NaN | NaN | 80.48942 | NaN |
| Wdr46 | SGLDHQLK | -376 | -383 | 0 | 0.002329279 | 8 | NaN | NaN | NaN | NaN | 457.0312 | NaN |
| Zc3hav1 | ICGQPQPCER | -161 | -170 | 0 | 0.003360691 | 10 | NaN | NaN | NaN | NaN | 8294.0205 | NaN |
| Cat | NSANEDNVTQVR | -433 | -444 | 0 | 0 | 12 | NaN | NaN | NaN | NaN | 1002.63763 | 1088.8754 |
| Cat | KEAETFPFNPFDLTK | -287 | -301 | 1 | 0.00383302 | 15 | NaN | NaN | NaN | NaN | 569.78503 | NaN |
| Cat | STVTGESGSADTVR | -114 | -127 | 0 | 0.00078661 | 14 | NaN | NaN | NaN | NaN | 290.5475 | NaN |
| Ppp6r1 | WEAFVSGPLAETNKK | -502 | -516 | 1 | 0.001023236 | 15 | NaN | NaN | NaN | NaN | 356.98727 | 394.55 |
| Ppp6r1 | ASQSLCDIIR | -211 | -220 | 0 | 0.005176406 | 10 | NaN | NaN | NaN | NaN | 1017.6688 | NaN |
| Ppp6r1 | WEAFVSGPLAETNK | -502 | -515 | 0 | 0.001610812 | 14 | NaN | NaN | NaN | NaN | 792.87476 | NaN |
| Ppp6r1 | SGDSLNPLLASFFSK | -112 | -126 | 0 | 0 | 15 | NaN | NaN | NaN | NaN | 417.45065 | NaN |
| Arpc1b | LVEPISCHAWNK | -7 | -18 | 0 | 0 | 12 | NaN | NaN | NaN | NaN | 1512.2289 | 1066.3967 |
| Arpc1b | GGAATGAGLDSLHK | -313 | -326 | 0 | 0 | 14 | NaN | NaN | NaN | NaN | 132.62636 | 1066.6571 |
| Arpc1b | VAWVSHDSTVCLVDADK | -217 | -233 | 0 | 0.00025015 | 17 | NaN | NaN | NaN | NaN | 257.84015 | NaN |
| Ogt | LKPDFIDGYIN | -118 | -128 | 1 | 0.00078661 | 11 | NaN | NaN | NaN | NaN | 739.7924 | NaN |
| Rpl3 | VFGQDEMIDVIGVTK | -210 | -224 | 0 | 0 | 15 | NaN | NaN | NaN | NaN | 341.41507 | 631.0553 |
| Rpl3 | KTVFAEHISDECK | -103 | -115 | 1 | 0.001846793 | 13 | NaN | NaN | NaN | NaN | 1176.7308 | 1265.6191 |
| Rpl3 | KAFMGPLK | -386 | -393 | 1 | 0 | 8 | NaN | NaN | NaN | NaN | 476.40524 | NaN |
| Ndufa6 | RFFHETETPRPK | -108 | -119 | 2 | 0.002752526 | 12 | NaN | NaN | NaN | NaN | 632.5643 | NaN |
| Ndufa6 | RFFHETETPRPK | -108 | -119 | 2 | 0.002752526 | 12 | NaN | NaN | NaN | NaN | 632.5643 | NaN |
| Ndufa6 | ASTSVKPIFSR | -16 | -26 | 1 | 0 | 11 | NaN | NaN | NaN | NaN | 575.8655 | NaN |
| Pepd | NPSVLLTLR | -140 | -148 | 0 | 0.00025015 | 9 | NaN | NaN | NaN | NaN | 1412.2142 | 1662.9381 |
| Pepd | YCTDTSIIFR | -57 | -66 | 0 | 0 | 10 | NaN | NaN | NaN | NaN | 709.3774 | 717.6183 |
| Pepd | NVNNTILHPEIVECR | -170 | -184 | 0 | 0.00133086 | 15 | NaN | NaN | NaN | NaN | 3987.414 | NaN |
| Pepd | NTILHPEIVECR | -173 | -184 | 0 | 0.001846793 | 12 | NaN | NaN | NaN | NaN | 613.9466 | NaN |
| Cacybp | NYGWDQSDKFVK | -78 | -89 | 1 | 0 | 12 | NaN | NaN | NaN | NaN | 318.0996 | 542.4024 |
| Mif | PAQYIAVH | -34 | -41 | 0 | 0.000712251 | 8 | NaN | NaN | NaN | NaN | 3016.0908 | 2368.043 |
| Mif | FIVNTNVPR | -4 | -12 | 0 | 0 | 9 | NaN | NaN | NaN | NaN | 1498.1129 | NaN |
| Mif | GLLSDRLHISPDR | -82 | -94 | 1 | 0 | 13 | NaN | NaN | NaN | NaN | 470.5128 | NaN |
| Incenp | LAEQQLQEEQAK | -735 | -746 | 0 | 0.00025015 | 12 | NaN | NaN | NaN | NaN | 2781.8718 | NaN |
| Phb1 | LIPWVQKPIIFDCR | -57 | -70 | 1 | 0 | 14 | NaN | NaN | NaN | NaN | 343.62955 | 341.89557 |
| Phb1 | AVAGGVVNSALYNVDAGHR | -17 | -35 | 0 | 0 | 19 | NaN | NaN | NaN | NaN | 822.9276 | 1236.1921 |
| Rab14 | STYNHLSSWLTDAR | -97 | -110 | 0 | 0 | 14 | NaN | NaN | NaN | NaN | 494.34702 | 462.8638 |
| Rab14 | KIYQNIQDGSLDLNAAESGVQHKPSAPQGGR | -171 | -201 | 2 | 0.00201047 | 31 | NaN | NaN | NaN | NaN | 540.4299 | NaN |
| Drg1 | DVLKPLGHK | -150 | -158 | 1 | 0.005145856 | 9 | NaN | NaN | NaN | NaN | 1275.8848 | 1192.4509 |
| Drg1 | TCPQSELDAETVK | -194 | -206 | 0 | 0.004595494 | 13 | NaN | NaN | NaN | NaN | 662.83984 | 679.5216 |
| Denr | KWPEVDDDSIEDLGEVKK | -181 | -198 | 2 | 0.009897819 | 18 | NaN | NaN | NaN | NaN | 4685.059 | NaN |
| Ubap2l | DWNEDLSETK | -268 | -277 | 0 | 0 | 10 | NaN | NaN | NaN | NaN | 1844.3455 | NaN |
| Ubap2l | ISDHNDADFEEK | -42 | -53 | 0 | 0 | 12 | NaN | NaN | NaN | NaN | 672.119 | NaN |
| Ubap2l | ATQLQTTQSVEGATGSAVK | -646 | -664 | 0 | 0.000431655 | 19 | NaN | NaN | NaN | NaN | 256.22052 | NaN |
| Sf3b1 | EFQSPDEEMKK | -776 | -786 | 1 | 0 | 11 | NaN | NaN | NaN | NaN | 967.38965 | 824.23016 |
| Sf3b1 | DSTGYYDQEIYGGSDSR | -34 | -50 | 0 | 0 | 17 | NaN | NaN | NaN | NaN | 482.7695 | 539.2932 |
| Dnajc7 | QLEDAIEDCTNAVK | -309 | -322 | 0 | 0 | 14 | NaN | NaN | NaN | NaN | 847.17334 | 960.0093 |
| Qars1 | QHLEITGGQVR | -255 | -265 | 0 | 0.00555405 | 11 | NaN | NaN | NaN | NaN | 3244.1033 | 3233.9062 |
| Qars1 | IHWVSQPLVCEIR | -678 | -690 | 0 | 0.000826806 | 13 | NaN | NaN | NaN | NaN | 347.65686 | 394.61 |
| Qars1 | LAKPFDK | -734 | -740 | 1 | 0.006850902 | 7 | NaN | NaN | NaN | NaN | 1846.6736 | NaN |

| | | | | | | | | | | | | |
|---|---|---|---|---|---|---|---|---|---|---|---|---|
| Qars1 | RFDDTNPEKEEAK | -301 | -313 | 2 | 0.00025015 | 13 | NaN | NaN | NaN | NaN | 507.20612 | NaN |
| Rad50 | LKEIEHNLSK | -255 | -264 | 1 | 0.00025015 | 10 | NaN | NaN | NaN | NaN | 610.4237 | 237.54303 |
| Rab7a | NEFPEPIKLDK | -184 | -194 | 1 | 0.00111006 | 11 | NaN | NaN | NaN | NaN | 11496.933 | NaN |
| Gnpnat1 | ITLECLPQNVGFYK | -153 | -166 | 0 | 0.00051306 | 14 | NaN | NaN | NaN | NaN | 1359.8586 | 1452.5302 |
| Ndufb4 | YKPAPLATLPSTLDPAEYDVSPETR | -6 | -30 | 1 | 0.00025015 | 25 | NaN | NaN | NaN | NaN | 393.91602 | NaN |
| Mrps18a | ATPDPPNPSGQCPICR | -59 | -74 | 0 | 0.001392869 | 16 | NaN | NaN | NaN | NaN | 505.24753 | 1411.6731 |
| Ap1b1 | SEIAESHPSSNLLDLNPQSINK | -185 | -206 | 0 | 0 | 22 | NaN | NaN | NaN | NaN | 542.3848 | NaN |
| Larp1 | ATQVEPLLPAGAPLLQAEEHGLAR | -2 | -25 | 0 | 0.008014596 | 24 | NaN | NaN | NaN | NaN | 300.544 | 288.44748 |
| Larp1 | FNKKMYEEFK | -904 | -913 | 2 | 0.008014596 | 10 | NaN | NaN | NaN | NaN | 2557.6497 | NaN |
| Larp1 | KFDGTEGPR | -342 | -350 | 1 | 0.000907206 | 9 | NaN | NaN | NaN | NaN | 1381.1993 | NaN |
| Cmas | GNEVSDEECLKR | -370 | -381 | 1 | 0 | 12 | NaN | NaN | NaN | NaN | 1163.2671 | NaN |
| Acp1 | ATSTYEVGNPPDYR | -46 | -59 | 0 | 0 | 14 | NaN | NaN | NaN | NaN | 809.5893 | NaN |
| Xpo5 | LLDNLLALVR | -772 | -781 | 0 | 0.002265552 | 10 | NaN | NaN | NaN | NaN | 544.50323 | NaN |
| Psma2 | VYQEPIPTAQLVQR | -100 | -113 | 0 | 0.00025015 | 14 | NaN | NaN | NaN | NaN | 609.78705 | NaN |
| Suox | DHGFPVR | -361 | -367 | 0 | 0.002068213 | 7 | NaN | NaN | NaN | NaN | 762.21716 | NaN |
| Suox | AVDDSYNVQPDTVAPIWNLR | -511 | -530 | 0 | 0 | 20 | NaN | NaN | NaN | NaN | 257.9091 | NaN |
| Usp15 | AVDENLSSIMER | -535 | -546 | 0 | 0.002910424 | 12 | NaN | NaN | NaN | NaN | 921.1697 | NaN |
| Stat5a | NLWNLKPFTTR | -639 | -649 | 1 | 0.008075282 | 11 | NaN | NaN | NaN | NaN | 1712.8989 | NaN |
| Rpl6 | TSTKVDISDVKIPK | -205 | -218 | 2 | 0.002810811 | 14 | NaN | NaN | NaN | NaN | 465.45508 | NaN |
| Lgals3bp | STETIASHEDIER | -329 | -341 | 0 | 0 | 13 | NaN | NaN | NaN | NaN | 1461.4816 | 1460.9573 |
| Lgals3bp | TPSLFPCASGAFSSFR | -547 | -562 | 0 | 0 | 16 | NaN | NaN | NaN | NaN | 579.59155 | 468.65118 |
| Lgals3bp | TSSYSNPTIGYENR | -500 | -513 | 0 | 0.001463897 | 14 | NaN | NaN | NaN | NaN | 542.39703 | NaN |
| Slc1a5 | MPSSEPELIQVK | -503 | -514 | 0 | 0.000907206 | 12 | NaN | NaN | NaN | NaN | 1521.7772 | NaN |
| Grk2 | IEEICQNLR | -150 | -158 | 0 | 0.00078661 | 9 | NaN | NaN | NaN | NaN | 4367.2983 | NaN |
| Psmd5 | ENSEAVTEILNNAELLK | -112 | -128 | 0 | 0.00025015 | 17 | NaN | NaN | NaN | NaN | 787.881 | NaN |
| Aars1 | HNDLDDVGKDVYHHT | -82 | -96 | 1 | 0 | 15 | NaN | NaN | NaN | NaN | 731.50146 | 722.35266 |
| Aars1 | CLCQVPQNAANR | -901 | -912 | 0 | 0.00025015 | 12 | NaN | NaN | NaN | NaN | 1117.7612 | 1078.6359 |
| Aars1 | KVDDSSEDKTEFTVK | -550 | -564 | 2 | 0.000550307 | 15 | NaN | NaN | NaN | NaN | 975.4093 | 878.941 |
| Aars1 | YAEQGGQIYDEGYLVK | -535 | -550 | 0 | 0.00383302 | 16 | NaN | NaN | NaN | NaN | 1612.2318 | NaN |
| Tuba4a | KLSDQCTGLQGFLVFH | -124 | -139 | 1 | 0.002880184 | 16 | NaN | NaN | NaN | NaN | 4096.929 | NaN |
| Msh2 | FDPNLSELR | -474 | -482 | 0 | 0.00450079 | 9 | NaN | NaN | NaN | NaN | 4078.6177 | NaN |
| Gale | AVGESVQKPLDYYR | -92 | -105 | 1 | 0.001023236 | 14 | NaN | NaN | NaN | NaN | 458.2839 | 379.6728 |
| Gale | DYIHVVDLAK | -239 | -248 | 0 | 0.000230707 | 10 | NaN | NaN | NaN | NaN | 580.8161 | NaN |
| Hcls1 | TEHINIHQLR | -49 | -58 | 0 | 0 | 10 | NaN | NaN | NaN | NaN | 518.591 | 615.7607 |
| Hcls1 | SAVGHEYVADVEKH | -97 | -110 | 1 | 0 | 14 | NaN | NaN | NaN | NaN | 505.76846 | 2153.404 |
| Acsl4 | LKDWQEGGYTVHDKPNPR | -499 | -516 | 2 | 0.00111006 | 18 | NaN | NaN | NaN | NaN | 547.8432 | 522.35266 |
| Ube2b;Ube2a | AVIFGPEGTPFEDGTFK | (38);(38) | (54);(54) | 0 | 0.009897819 | 17 | NaN | NaN | NaN | NaN | 699.5426 | NaN |
| Prpf6 | ELQHGTEEQQEEVR | -885 | -898 | 0 | 0.000907206 | 14 | NaN | NaN | NaN | NaN | 502.41028 | 525.1711 |
| Prpf6 | HGELWCAVSK | -908 | -917 | 0 | 0.008129024 | 10 | NaN | NaN | NaN | NaN | 502.545 | NaN |
| Ghitm | AVIWPQYVK | -114 | -122 | 0 | 0.008129024 | 9 | NaN | NaN | NaN | NaN | 1441.7482 | 768.88855 |
| Rac1 | HHCPNTPIIL | -103 | -112 | 0 | 0.005605646 | 10 | NaN | NaN | NaN | NaN | 2160.838 | 1912.0245 |
| Rac2 | PASYENVR | -87 | -94 | 0 | 0.001776473 | 8 | NaN | NaN | NaN | NaN | 861.9095 | 606.3906 |
| Hyou1 | SRFPEHELIVDPQR | -118 | -131 | 1 | 0 | 14 | NaN | NaN | NaN | NaN | 3009.7036 | 3122.2668 |
| Hyou1 | GDLGFLGPEDLR | -498 | -509 | 0 | 0.000627943 | 12 | NaN | NaN | NaN | NaN | 560.2786 | 294.9865 |
| Hyou1 | GLMDDVDFK | -332 | -340 | 0 | 0.000431655 | 9 | NaN | NaN | NaN | NaN | 841.53455 | NaN |
| Cmpk2 | HFVLGDATDHR | -48 | -58 | 0 | 0 | 11 | NaN | NaN | NaN | NaN | 849.5807 | NaN |
| Cmpk2 | KIFDDEPTIIR | -293 | -303 | 1 | 0 | 11 | NaN | NaN | NaN | NaN | 541.8365 | 511.3166 |
| Cmpk2 | ENPSCHLVDASPSR | -418 | -431 | 0 | 0 | 14 | NaN | NaN | NaN | NaN | 788.5726 | NaN |
| Iba57 | AVLPCVPQTSETAPLEER | -162 | -179 | 0 | 0 | 18 | NaN | NaN | NaN | NaN | 464.8661 | NaN |
| Pdia6 | AVLPHILDTGAAGR | -300 | -313 | 0 | 0 | 14 | NaN | NaN | NaN | NaN | 413.56546 | NaN |
| Cpne1 | LQHFCGGDLSTPIQVR | -195 | -210 | 0 | 0.000431655 | 16 | NaN | NaN | NaN | NaN | 268.7672 | NaN |
| Aip | EYYEVLDHCSSILNK | -246 | -260 | 0 | 0.00025015 | 15 | NaN | NaN | NaN | NaN | 3251.7954 | NaN |
| Igf2r | AVQDCEYTFSWPTPSACPVK | -1483 | -1502 | 0 | 0.003003218 | 20 | NaN | NaN | NaN | NaN | 1478.315 | 1121.0454 |
| Igf2r | DKEVPCYAFDDK | -164 | -175 | 1 | 0.000298805 | 12 | NaN | NaN | NaN | NaN | 1303.8706 | NaN |
| Nol6 | SREELYKEPTNEELNR | -65 | -80 | 2 | 0.004690496 | 16 | NaN | NaN | NaN | NaN | 161.80025 | NaN |
| Eif4b | GDDSFGDKYR | -216 | -225 | 1 | 0.001636916 | 10 | NaN | NaN | NaN | NaN | 773.14655 | NaN |
| Ago2 | QVLHHELLAIR | -679 | -689 | 0 | 0.005605646 | 11 | NaN | NaN | NaN | NaN | 274.49518 | NaN |
| Serpinf1 | AVSNFGYDLYR | -56 | -66 | 0 | 0.000471409 | 11 | NaN | NaN | NaN | NaN | 2245.8684 | 1763.0488 |
| Itgb1 | CNCQSHGIPASPK | -464 | -476 | 0 | 0.001392869 | 13 | NaN | NaN | NaN | NaN | 324.66708 | NaN |
| Tsn | AVTEILGIEPDREK | -118 | -131 | 1 | 0.002329279 | 14 | NaN | NaN | NaN | NaN | 1628.7074 | 911.35364 |
| Tsn | EILTLLQGVHQGTGF | -37 | -51 | 0 | 0.001291103 | 15 | NaN | NaN | NaN | NaN | 837.4868 | 152.71472 |
| Ywhah | KNSVVEASEAAYK | -143 | -155 | 1 | 0 | 13 | NaN | NaN | NaN | NaN | 267.05737 | 566.00256 |
| Snrnp40 | QGNVHNFEK | -279 | -287 | 0 | 0.005605646 | 9 | NaN | NaN | NaN | NaN | 661.9452 | NaN |
| H4c1 | LENVIRDAVTYTEHAK | -63 | -78 | 1 | 0 | 16 | NaN | NaN | NaN | NaN | 3395.4473 | 3238.9565 |
| H4c1 | KTVTAMDVVYALK | -80 | -92 | 1 | 0 | 13 | NaN | NaN | NaN | NaN | 2475.2764 | 2274.9265 |
| Glud1 | IIAEGANGPTTPEADKIFLER | -400 | -420 | 1 | 0.00025015 | 21 | NaN | NaN | NaN | NaN | 342.59467 | 496.08713 |
| Emc7 | AVVPGVKPQDWISAA | -50 | -64 | 1 | 0.00111006 | 15 | NaN | NaN | NaN | NaN | 1265.6161 | 800.0347 |
| Lbr | VSGWWGFVR | -547 | -555 | 0 | 0 | 9 | NaN | NaN | NaN | NaN | 607.55695 | 681.5709 |
| Lbr | LTPLVLKPFGNSVS | -127 | -140 | 1 | 0.002199523 | 14 | NaN | NaN | NaN | NaN | 237.59062 | NaN |
| Mtpn | AVYEGHVSCVK | -75 | -85 | 0 | 0.000431655 | 11 | NaN | NaN | NaN | NaN | 1347.2521 | NaN |
| Mtpn | HHITPLLSAVYEGHVSCVK | -67 | -85 | 0 | 0.00111006 | 19 | NaN | NaN | NaN | NaN | 1139.7219 | NaN |
| Cyrib | GLLGALTSTPYSPTQHLER | -106 | -124 | 0 | 0.00025015 | 19 | NaN | NaN | NaN | NaN | 493.2455 | 396.69153 |
| Cyrib | LDFENAQPTESEK | -21 | -33 | 0 | 0 | 13 | NaN | NaN | NaN | NaN | 1446.107 | NaN |
| Cyrib | MTNPAIQNDFSYYR | -147 | -160 | 0 | 0.000471409 | 14 | NaN | NaN | NaN | NaN | 156.27722 | NaN |
| Otud6b | AWGGQLELR | -225 | -233 | 0 | 0 | 9 | NaN | NaN | NaN | NaN | 1296.9247 | NaN |
| Kin | AWLQPGIVVK | -277 | -286 | 0 | 0 | 10 | NaN | NaN | NaN | NaN | 402.59045 | NaN |
| Kin | LDQTHLETVIPAPGK | -325 | -339 | 0 | 0.000471409 | 15 | NaN | NaN | NaN | NaN | 359.8129 | NaN |
| Tspo | LQKPSWHPPR | -37 | -46 | 1 | 0 | 10 | NaN | NaN | NaN | NaN | 216.52946 | 364.35666 |

| | | | | | | | | | | | | |
|---|---|---|---|---|---|---|---|---|---|---|---|---|
| Tspo | SLQKPSWHPPR | -36 | -46 | 1 | 0.002466863 | 11 | NaN | NaN | NaN | NaN | 451.81427 | NaN |
| Dock11 | RYEVISEISK | -1711 | -1720 | 1 | 0.00025015 | 10 | NaN | NaN | NaN | NaN | 8209.308 | 395.37585 |
| Dock11 | DFSGIEPDVKPFEEK | -348 | -362 | 1 | 0.005821388 | 15 | NaN | NaN | NaN | NaN | 675.8066 | 1157.186 |
| Dock11 | GSLIPEGATGFPDPGSTSENTR | -1239 | -1260 | 0 | 0.00149795 | 22 | NaN | NaN | NaN | NaN | 386.21173 | 548.99774 |
| Dock11 | DTVEAIQEEETSSQGK | -284 | -299 | 0 | 0.00025015 | 16 | NaN | NaN | NaN | NaN | 1566.3201 | NaN |
| Coasy | AYAPGGPAYQPVVEAF | -394 | -409 | 0 | 0 | 16 | NaN | NaN | NaN | NaN | 451.0696 | 331.01962 |
| Gdi2 | KFVSISDLFVPK | -379 | -390 | 1 | 0.00078661 | 12 | NaN | NaN | NaN | NaN | 180.55714 | 649.5154 |
| Gdi2 | AYDATTHFETTCDDIKDIYK | -403 | -422 | 1 | 0 | 20 | NaN | NaN | NaN | NaN | 415.93158 | NaN |
| Usp9x | SHTNDTIGSVR | -906 | -916 | 0 | 0.00051306 | 11 | NaN | NaN | NaN | NaN | 1760.2933 | 1429.7808 |
| Dnaja1 | CGFQKPISTLDNR | -271 | -283 | 1 | 0.000471409 | 13 | NaN | NaN | NaN | NaN | 382.42993 | NaN |
| Tmed10 | GISFHLPVNSR | -31 | -41 | 0 | 0 | 11 | NaN | NaN | NaN | NaN | 3075.3315 | 3355.4673 |
| Tmed10 | SFHLPVNSR | -33 | -41 | 0 | 0.003360691 | 9 | NaN | NaN | NaN | NaN | 473.12796 | 337.62564 |
| Tmed10 | DLLVTGAYEITDQSGGAGGLR | -51 | -71 | 0 | 0.001055699 | 21 | NaN | NaN | NaN | NaN | 754.2552 | NaN |
| Kars1 | KLPETSLFETEETRK | -405 | -419 | 2 | 0 | 15 | NaN | NaN | NaN | NaN | 2754.3418 | NaN |
| Kars1 | LTMFLTDSNNIK | -552 | -563 | 0 | 0.001392869 | 12 | NaN | NaN | NaN | NaN | 1229.2615 | NaN |
| Kars1 | YHPDGPEGQAYEVDFTPPFR | -371 | -390 | 0 | 0.002780285 | 20 | NaN | NaN | NaN | NaN | 73.75301 | NaN |
| Hspa14 | QVVLQDLDKK | -460 | -469 | 1 | 0 | 10 | NaN | NaN | NaN | NaN | 590.6031 | 900.49725 |
| Bclaf1 | FHDSEGDDTEETEDYR | -392 | -407 | 0 | 0 | 16 | NaN | NaN | NaN | NaN | 520.7349 | 498.59375 |
| Bclaf1 | DTFEHDPSESIDEFNKS | -189 | -205 | 1 | 0.009060424 | 17 | NaN | NaN | NaN | NaN | 729.4022 | NaN |
| Bclaf1 | LKELFDYSPPLHK | -503 | -515 | 1 | 0 | 13 | NaN | NaN | NaN | NaN | 417.27118 | NaN |
| Hsdl1 | AYLDHFSR | -223 | -230 | 0 | 0.00051306 | 8 | NaN | NaN | NaN | NaN | 565.3866 | 354.1232 |
| Sec22b | AYLEDLHSEFDEQHGK | -86 | -101 | 0 | 0 | 16 | NaN | NaN | NaN | NaN | 292.0406 | 427.1119 |
| Sec22b | AYLEDLHSEFDEQHGK | -86 | -101 | 0 | 0 | 16 | NaN | NaN | NaN | NaN | 292.0406 | 427.1119 |
| Sec22b | NLGSINTELQDVQR | -134 | -147 | 0 | 0 | 14 | NaN | NaN | NaN | NaN | 463.12427 | NaN |
| Snu13 | KLLDLVQQSC | -21 | -30 | 1 | 0.002068213 | 10 | NaN | NaN | NaN | NaN | 463.1103 | 2672.4468 |
| Snu13 | PLADAHLTK | -12 | -20 | 0 | 0.001941082 | 9 | NaN | NaN | NaN | NaN | 341.27295 | 414.46207 |
| Micos10 | AYSNCQHDFQAPYLLHGK | -54 | -71 | 0 | 0.00025015 | 18 | NaN | NaN | NaN | NaN | 914.2275 | NaN |
| Alb | HPYFYAPELLY | -170 | -180 | 0 | 0 | 11 | NaN | NaN | NaN | NaN | 1612.8263 | 1581.2191 |
| Alb | RHPYFYAPELLYY | -169 | -181 | 1 | 0 | 13 | NaN | NaN | NaN | NaN | 946.87823 | 540.48804 |
| Alb | RHPYFYAPELLYYA | -169 | -182 | 1 | 0 | 14 | NaN | NaN | NaN | NaN | 833.9126 | 890.25995 |
| Alb | LGEYGFQN | -422 | -429 | 0 | 0 | 8 | NaN | NaN | NaN | NaN | 30675.244 | 30357.547 |
| Alb | LECADDR | -275 | -281 | 0 | 0.00383302 | 7 | NaN | NaN | NaN | NaN | 504.66138 | NaN |
| Pml | NHSHLHCDIGEEIQQW | -225 | -240 | 0 | 0.001150701 | 16 | NaN | NaN | NaN | NaN | 1072.9343 | NaN |
| Pml | DNVFFESLQR | -110 | -119 | 0 | 0.007493591 | 10 | NaN | NaN | NaN | NaN | 669.118 | NaN |
| Pml | CCTCALLDR | -216 | -224 | 0 | 0.000627943 | 9 | NaN | NaN | NaN | NaN | 190.47638 | NaN |
| Gspt1 | CDLNNLCHSGR | -518 | -528 | 0 | 0 | 11 | NaN | NaN | NaN | NaN | 1803.2335 | 1079.0941 |
| Ergic3 | CESCYGAESEDIK | -142 | -154 | 0 | 0.000712251 | 13 | NaN | NaN | NaN | NaN | 162.882 | 245.99391 |
| Ergic3 | NPDTIEQCR | -175 | -183 | 0 | 0.00149795 | 9 | NaN | NaN | NaN | NaN | 3627.7566 | NaN |
| Tfrc | HANFGTK | -236 | -242 | 0 | 0.002199523 | 7 | NaN | NaN | NaN | NaN | 557.1742 | 487.03177 |
| Tfrc | SGDIWNIDNEF | -753 | -763 | 0 | 0.00051306 | 11 | NaN | NaN | NaN | NaN | 140.49141 | 232.97354 |
| Igkc | RADAAPTVSIFPPSSEQLTSGGASVV | -1 | -26 | 1 | 0.000627943 | 26 | NaN | NaN | NaN | NaN | 604.0976 | 521.8103 |
| Cnbp | CYSCGEFGHIQK | -120 | -131 | 0 | 0 | 12 | NaN | NaN | NaN | NaN | 1725.9783 | 1679.0273 |
| Arhgap15 | SSSSELPSHCHIDR | -216 | -229 | 0 | 0.002365814 | 14 | NaN | NaN | NaN | NaN | 267.4882 | NaN |
| Timm13 | LDPGAIMEQVK | -17 | -27 | 0 | 0 | 11 | NaN | NaN | NaN | NaN | 442.26816 | 63.637493 |
| Pebp1 | QVMNRPSSISWDGLDPGK | -45 | -62 | 1 | 0.002434616 | 18 | NaN | NaN | NaN | NaN | 1368.6548 | 1594.2463 |
| Pebp1 | VLTPTQVMNRPSSISWDGLDPGK | -40 | -62 | 1 | 0.002780285 | 23 | NaN | NaN | NaN | NaN | 773.5923 | 721.28 |
| Pebp1 | CYQAEWDDYVPK | -168 | -179 | 0 | 0.00204437 | 12 | NaN | NaN | NaN | NaN | 226.76219 | 361.28397 |
| Pebp1 | YAGVTVDELGK | -29 | -39 | 0 | 0.007343498 | 11 | NaN | NaN | NaN | NaN | 2451.5288 | NaN |
| Nutf2 | IQHSITAQDHQPTPDSCIIS | -64 | -83 | 0 | 0 | 20 | NaN | NaN | NaN | NaN | 476.71033 | 419.68387 |
| Naga | LNIDDCWIGGR | -75 | -85 | 0 | 0 | 11 | NaN | NaN | NaN | NaN | 908.8644 | 381.0326 |
| Txn | MIKPFFHSLCDK | -37 | -48 | 1 | 0.00025015 | 12 | NaN | NaN | NaN | NaN | 757.2295 | 910.553 |
| Itga2b | DVQPQGGLLCSTQPSPK | -847 | -863 | 0 | 0.000431655 | 17 | NaN | NaN | NaN | NaN | 7152.574 | 853.7103 |
| Itga2b | IFLGQSEGLSPR | -421 | -432 | 0 | 0.008222027 | 12 | NaN | NaN | NaN | NaN | 718.65985 | NaN |
| Itga2b | EEGDREQEDLDR | -794 | -805 | 1 | 0.008680443 | 12 | NaN | NaN | NaN | NaN | 165.34898 | NaN |
| Ubl5 | CNTDDTIGDLKK | -18 | -29 | 1 | 0 | 12 | NaN | NaN | NaN | NaN | 3969.5396 | NaN |
| Lamp1 | CNTEEHIFVSK | -327 | -337 | 0 | 0.00078661 | 11 | NaN | NaN | NaN | NaN | 865.6949 | NaN |
| Hectd1 | SCGLFTAPFPQDSDELER | -2228 | -2245 | 0 | 0.000550307 | 18 | NaN | NaN | NaN | NaN | 203.80045 | 382.16714 |
| Hectd1 | GSPLVTHDLLR | -289 | -299 | 0 | 0 | 11 | NaN | NaN | NaN | NaN | 1155.4839 | NaN |
| Hectd1 | TWPLDPAKDEK | -1202 | -1212 | 1 | 0.004224523 | 11 | NaN | NaN | NaN | NaN | 902.4254 | NaN |
| Hectd1 | DLYDDHFK | -840 | -847 | 0 | 0.00078661 | 8 | NaN | NaN | NaN | NaN | 838.4426 | NaN |
| Slc44a2 | CQFAFYGGESTYHR | -427 | -440 | 0 | 0 | 14 | NaN | NaN | NaN | NaN | 3403.6406 | 3180.2258 |
| Inhca | LFNQQELFGR | -623 | -632 | 0 | 0.000471409 | 10 | NaN | NaN | NaN | NaN | 1321.6818 | 790.7599 |
| Tagln2 | WITTQCR | -33 | -39 | 0 | 0.002697088 | 7 | NaN | NaN | NaN | NaN | 1029.2803 | NaN |
| Tagln2 | ENPRNFSDNQLQEGK | -157 | -171 | 1 | 0.00713295 | 15 | NaN | NaN | NaN | NaN | 742.04376 | NaN |
| Tagln2 | QMEQISQFLQAA | -89 | -100 | 0 | 0.006788754 | 12 | NaN | NaN | NaN | NaN | 488.00922 | NaN |
| Mier1 | FSSEIEDLAR | -51 | -60 | 0 | 0.001868731 | 10 | NaN | NaN | NaN | NaN | 4661.9805 | NaN |
| Mier1 | CSFDTEEALR | -262 | -271 | 0 | 0.00713295 | 10 | NaN | NaN | NaN | NaN | 336.61166 | NaN |
| Eif3i | LWDCETGK | -78 | -85 | 0 | 0.002466863 | 8 | NaN | NaN | NaN | NaN | 842.8021 | 933.9578 |
| Ptges3 | FCVEDSKDVNVNFEK | -19 | -33 | 1 | 0 | 15 | NaN | NaN | NaN | NaN | 2040.2972 | 905.64417 |
| Exosc7 | CVEVETDVVSNTSGSAR | -34 | -50 | 0 | 0.001190916 | 17 | NaN | NaN | NaN | NaN | 1180.5789 | NaN |
| Psmb2 | LTLSILDR | -138 | -145 | 0 | 0.006568856 | 8 | NaN | NaN | NaN | NaN | 490.09732 | 427.11813 |
| Rbm17 | PGAPDDEAVR | -349 | -358 | 0 | 0.00051306 | 10 | NaN | NaN | NaN | NaN | 4036.4583 | NaN |
| Rbm17 | CVIFEIPGAPDDEAVR | -343 | -358 | 0 | 0 | 16 | NaN | NaN | NaN | NaN | 1070.5782 | NaN |
| Rbm17 | PPTSLVEK | -177 | -184 | 0 | 0.006195994 | 8 | NaN | NaN | NaN | NaN | 610.715 | NaN |
| Strap | KCVLPEEDSGELAKPK | -304 | -319 | 2 | 0.00025015 | 16 | NaN | NaN | NaN | NaN | 530.3235 | NaN |
| Elac2 | CVLSGPPQLEK | -128 | -138 | 0 | 0 | 11 | NaN | NaN | NaN | NaN | 12602.455 | 10374.872 |
| Elac2 | ELPSFQESVEEYRK | -448 | -461 | 1 | 0 | 14 | NaN | NaN | NaN | NaN | 618.7285 | NaN |
| Xab2 | RSHPLPETAVR | -157 | -167 | 1 | 0.000394361 | 11 | NaN | NaN | NaN | NaN | 805.85693 | NaN |

| | | | | | | | | | | | | |
|---|---|---|---|---|---|---|---|---|---|---|---|---|
| Xab2 | IINTYTEAVQTVDPFK | -374 | -389 | 0 | 0.007224385 | 16 | NaN | NaN | NaN | NaN | 416.46008 | NaN |
| Emc4 | CWDIALGPLK | -69 | -78 | 0 | 0 | 10 | NaN | NaN | NaN | NaN | 955.5744 | 484.01538 |
| Glyr1 | TVDADTVTELAQVIVSR | -355 | -371 | 0 | 0 | 17 | NaN | NaN | NaN | NaN | 145.84721 | 295.38156 |
| Srsf2 | DAEDAMDAMDGAVLDGR | -67 | -83 | 0 | 0 | 17 | NaN | NaN | NaN | NaN | 152.70291 | 121.216515 |
| Srsf1 | KEDMTYAVR | -165 | -173 | 1 | 0.00025015 | 9 | NaN | NaN | NaN | NaN | 546.6592 | 1101.0211 |
| Nudc | KTDFFIGGEEGMAEK | -39 | -53 | 1 | 0 | 15 | NaN | NaN | NaN | NaN | 902.9648 | NaN |
| Ppp1r7 | RNIEGIDK | -156 | -163 | 1 | 0.007343498 | 8 | NaN | NaN | NaN | NaN | 872.3578 | 1186.9277 |
| Ppp1r7 | IENISNLHQLQ | -180 | -190 | 0 | 0.002629273 | 11 | NaN | NaN | NaN | NaN | 1610.2267 | NaN |
| Ppp1r7 | GIEVIEGLENNNK | -264 | -276 | 0 | 0.000230707 | 13 | NaN | NaN | NaN | NaN | 1592.637 | NaN |
| Akr1a1 | TASSVLLHTGQK | -2 | -13 | 0 | 0 | 12 | NaN | NaN | NaN | NaN | 2136.3872 | NaN |
| Eif3c | NILEESENLHNFDQPLR | -421 | -437 | 0 | 0 | 17 | NaN | NaN | NaN | NaN | 825.3343 | 801.23895 |
| Eif3c | QNPEQSADEDAEKN | -161 | -174 | 1 | 0.000471409 | 14 | NaN | NaN | NaN | NaN | 497.60724 | 239.56007 |
| Eif3c | EESENLHNFDQPLR | -424 | -437 | 0 | 0.007040624 | 14 | NaN | NaN | NaN | NaN | 935.5512 | NaN |
| Snrpa1 | VELGDLDPLASLK | -101 | -113 | 0 | 0.001291103 | 13 | NaN | NaN | NaN | NaN | 722.037 | NaN |
| Snrpa1 | DAIDFSDNEIR | -45 | -55 | 0 | 0.00933166 | 11 | NaN | NaN | NaN | NaN | 682.85547 | NaN |
| Snrpa1 | LTAELIEQAAQYTNAVR | -4 | -20 | 0 | 0.00025015 | 17 | NaN | NaN | NaN | NaN | 299.83487 | NaN |
| Pycard | DAILDALENLSGDELKK | -6 | -22 | 1 | 0 | 17 | NaN | NaN | NaN | NaN | 556.37354 | 549.65045 |
| P4hb | YKPESDELTAEK | -329 | -340 | 1 | 0 | 12 | NaN | NaN | NaN | NaN | 3869.166 | 3888.381 |
| P4hb | VDATEESDLAQQYGVR | -84 | -99 | 0 | 0 | 16 | NaN | NaN | NaN | NaN | 998.93494 | 340.42322 |
| Gbf1 | LLEVFTEHWR | -801 | -810 | 0 | 0.001291103 | 10 | NaN | NaN | NaN | NaN | 3154.294 | NaN |
| Gbf1 | DALEWESCFNK | -1583 | -1593 | 0 | 0.008653659 | 11 | NaN | NaN | NaN | NaN | 315.20694 | NaN |
| Galk2 | ILTPNTQDELTFK | -315 | -327 | 0 | 0 | 13 | NaN | NaN | NaN | NaN | 4220.2573 | 4080.537 |
| Galk2 | GIQEHFGLSK | -117 | -126 | 0 | 0.00025015 | 10 | NaN | NaN | NaN | NaN | 718.394 | 933.0394 |
| Galk2 | GLQWDNVLR | -259 | -267 | 0 | 0.004439092 | 9 | NaN | NaN | NaN | NaN | 1544.6482 | NaN |
| Galk2 | VNIIGEHIDYCGY | -44 | -56 | 0 | 0.003428289 | 13 | NaN | NaN | NaN | NaN | 1060.8821 | NaN |
| Kpna4 | LDKGDFGTQK | -377 | -386 | 1 | 0.000907206 | 10 | NaN | NaN | NaN | NaN | 1768.3423 | 1715.8359 |
| Kpna4 | GFNSSTNVPTEGFQF | -507 | -521 | 0 | 0 | 15 | NaN | NaN | NaN | NaN | 417.78568 | 449.79623 |
| Ikzf3 | SLPKPHEIENVDSR | -30 | -43 | 1 | 0.000394361 | 14 | NaN | NaN | NaN | NaN | 1292.3237 | 1405.8435 |
| Ctbp1 | IGSGFDNIDIK | -98 | -108 | 0 | 0.00025015 | 11 | NaN | NaN | NaN | NaN | 3373.226 | 557.4845 |
| Brd4 | SDSFEHFR | -1257 | -1264 | 0 | 0.000230707 | 8 | NaN | NaN | NaN | NaN | 385.09357 | NaN |
| Arhgdib | VGSYGPRPEEYEFLTPVEEAPK | -142 | -163 | 1 | 0 | 22 | NaN | NaN | NaN | NaN | 1861.1548 | 2019.3641 |
| Arhgdib | MVGSYGPRPEEYEFLTPVEEAPK | -141 | -163 | 1 | 0 | 23 | NaN | NaN | NaN | NaN | 479.6971 | NaN |
| Arhgdib | DAQPQLEEADDDLDSKLNYKPPPQK | -5 | -29 | 2 | 0.005668207 | 25 | NaN | NaN | NaN | NaN | 1215.3474 | NaN |
| Arhgdib | LEEADDDLDSKLNYKPPPQK | -10 | -29 | 2 | 0 | 20 | NaN | NaN | NaN | NaN | 704.9973 | NaN |
| Nt5c2 | LVDFFTNCPR | -168 | -177 | 0 | 0 | 10 | NaN | NaN | NaN | NaN | 747.59607 | NaN |
| Eif4e | SFDDYSDDVCGAVVNVR | -141 | -157 | 0 | 0 | 17 | NaN | NaN | NaN | NaN | 294.5355 | NaN |
| Atl3 | EHQHEEIQNVR | -240 | -250 | 0 | 0 | 11 | NaN | NaN | NaN | NaN | 585.40173 | 437.28198 |
| Pak1ip1 | SGHQETWTPVADFTHHSHTA | -24 | -43 | 0 | 0 | 20 | NaN | NaN | NaN | NaN | 518.7708 | 824.7406 |
| Pak1ip1 | DCDSLECLCEFR | -231 | -242 | 0 | 0 | 12 | NaN | NaN | NaN | NaN | 389.4525 | 376.94293 |
| Apobec3 | DCDSPVSLHHGVFK | -63 | -76 | 0 | 0 | 14 | NaN | NaN | NaN | NaN | 500.59076 | 1026.3293 |
| Banf1 | DFVAEPMGEKPVGSLAGIGDVL | -9 | -30 | 1 | 0.00437931 | 22 | NaN | NaN | NaN | NaN | 245.40916 | 244.99533 |
| Mad2l1bp | DCGEDWFQPK | -222 | -231 | 0 | 0 | 10 | NaN | NaN | NaN | NaN | 233.07237 | 354.90063 |
| Gabpa | ELCSLNQEDFFQR | -221 | -233 | 0 | 0 | 13 | NaN | NaN | NaN | NaN | 495.97375 | 485.0105 |
| Smc3 | NLPGEVTFLPLNK | -580 | -592 | 0 | 0 | 13 | NaN | NaN | NaN | NaN | 475.53534 | 488.38788 |
| Smc3 | IYNQELNETR | -227 | -236 | 0 | 0.003299042 | 10 | NaN | NaN | NaN | NaN | 888.0658 | NaN |
| Smc3 | EKINELLK | -187 | -194 | 1 | 0.002265552 | 8 | NaN | NaN | NaN | NaN | 499.20325 | NaN |
| Smc3 | TMDCITLEGDQVSHR | -647 | -661 | 0 | 0.00025015 | 15 | NaN | NaN | NaN | NaN | 385.9979 | NaN |
| Hpcal1 | LKDCPTGHLTVDEFK | -35 | -49 | 1 | 0 | 15 | NaN | NaN | NaN | NaN | 567.00696 | 412.47437 |
| Bnip1 | DCSGPLSELTELNTK | -32 | -46 | 0 | 0.000230707 | 15 | NaN | NaN | NaN | NaN | 514.40436 | 539.9087 |
| Thbs1 | DIDGDGILNER | -800 | -810 | 0 | 0.000431655 | 11 | NaN | NaN | NaN | NaN | 1034.9353 | 1043.6178 |
| Hnrnpc | VFIGNLNTLVVK | -18 | -29 | 0 | 0 | 12 | NaN | NaN | NaN | NaN | 11462.38 | 11633.53 |
| Hnrnpc | GDDLQAIK | -191 | -198 | 0 | 0.00111006 | 8 | NaN | NaN | NaN | NaN | 1633.982 | NaN |
| Hnrnpc | SEAGADDSAEEGDLLDDDDNEDRGDDQLELK | -261 | -291 | 1 | 0.004031417 | 31 | NaN | NaN | NaN | NaN | 308.76205 | NaN |
| Psma1 | PAEQDLTTK | -200 | -208 | 0 | 0.00272025 | 9 | NaN | NaN | NaN | NaN | 1235.4106 | 941.25037 |
| Mesd | EQWEKDDDIEEGDLPEHK | -59 | -76 | 1 | 0 | 18 | NaN | NaN | NaN | NaN | 216.4685 | 694.1593 |
| Vrk1 | DDGKLDFSAVENGSVK | -335 | -350 | 1 | 0.003299042 | 16 | NaN | NaN | NaN | NaN | 465.1828 | 541.1998 |
| Hnrnpa2b1 | SFETTEESLR | -29 | -38 | 0 | 0 | 10 | NaN | NaN | NaN | NaN | 8988.932 | 8529.249 |
| Hnrnpa2b1 | TFDDHDPVDK | -159 | -168 | 0 | 0.000431655 | 10 | NaN | NaN | NaN | NaN | 654.4822 | NaN |
| Emd | VRDDIFSSLEEEGKDR | -135 | -150 | 2 | 0.008129024 | 16 | NaN | NaN | NaN | NaN | 1085.1727 | NaN |
| Emd | DDIFSSLEEEGKDR | -137 | -150 | 1 | 0.000591124 | 14 | NaN | NaN | NaN | NaN | 241.7391 | NaN |
| Tsr1 | VADFVPSEESDLVGTLK | -257 | -273 | 0 | 0.007987481 | 17 | NaN | NaN | NaN | NaN | 1191.8146 | NaN |
| Sub1 | EQISDIDDAVR | -115 | -125 | 0 | 0 | 11 | NaN | NaN | NaN | NaN | 6624.486 | 6791.52 |
| Sub1 | QAVPEKPVKK | -30 | -39 | 2 | 0 | 10 | NaN | NaN | NaN | NaN | 357.20786 | 331.90356 |
| Sub1 | WMDSEGEMKPGR | -89 | -100 | 1 | 0.001636916 | 12 | NaN | NaN | NaN | NaN | 774.19104 | 495.67484 |
| Ptpn6 | KYPLNCSDPTSER | -97 | -109 | 1 | 0 | 13 | NaN | NaN | NaN | NaN | 1122.0276 | 1261.7115 |
| Ptpn6 | KLEIIQSQK | -522 | -530 | 1 | 0.000471409 | 9 | NaN | NaN | NaN | NaN | 506.57562 | NaN |
| Bub3 | FSPNTSQFLLVS | -22 | -33 | 0 | 0.001392869 | 12 | NaN | NaN | NaN | NaN | 174.3631 | NaN |
| Bub3 | LKENNIEQIYPVNAISF | -226 | -242 | 1 | 0.001708008 | 17 | NaN | NaN | NaN | NaN | 688.08673 | 69.596695 |
| Uqcrb | LPEDLYNDR | -51 | -59 | 0 | 0.006100831 | 9 | NaN | NaN | NaN | NaN | 668.3 | NaN |
| Nsfl1c | DLIHDQDEEEEEEEGQR | -77 | -93 | 0 | 0.00025015 | 17 | NaN | NaN | NaN | NaN | 782.04175 | 719.208 |
| Nsfl1c | KSPNELVDDLFK | -113 | -124 | 1 | 0.00078661 | 12 | NaN | NaN | NaN | NaN | 1537.1493 | 1462.1993 |
| Nsfl1c | SSPAQQAENEAK | -271 | -282 | 0 | 0.002296977 | 12 | NaN | NaN | NaN | NaN | 1215.2113 | 1621.3431 |
| Ftsj3 | QVDDNKEEEGENPLLVPLEEK | -494 | -514 | 1 | 0.000550307 | 21 | NaN | NaN | NaN | NaN | 106.61064 | NaN |
| Mycbp2 | DEDLEEELDEKDEK | -737 | -750 | 1 | 0.002566359 | 14 | NaN | NaN | NaN | NaN | 1106.6333 | NaN |
| Gmfb | DEELEGVSPDELKDELPER | -46 | -64 | 1 | 0.001055699 | 19 | NaN | NaN | NaN | NaN | 601.97455 | 668.53674 |
| Thy1 | DEGDYFCELQVS | -99 | -110 | 0 | 0.001776473 | 12 | NaN | NaN | NaN | NaN | 834.29895 | 30.893145 |
| Thy1 | KHVLSGTLGIPEHTYR | -61 | -76 | 1 | 0 | 16 | NaN | NaN | NaN | NaN | 807.2788 | 787.9168 |
| Dctn1 | KFDLSENCSER | -629 | -639 | 1 | 0.002466863 | 11 | NaN | NaN | NaN | NaN | 2291.7258 | 1657.34 |

| | | | | | | | | | | | | |
|---|---|---|---|---|---|---|---|---|---|---|---|---|
| Dctn1 | QADIDQLEAEK | -1020 | -1030 | 0 | 0.001904399 | 11 | NaN | NaN | NaN | NaN | 590.5531 | NaN |
| Dctn1 | QLQHENSILR | -1106 | -1115 | 0 | 0.009242859 | 10 | NaN | NaN | NaN | NaN | 225.75212 | NaN |
| Dnajc3 | GKLDEAEDDFKK | -119 | -130 | 2 | 0 | 12 | NaN | NaN | NaN | NaN | 511.32062 | 533.94763 |
| Dnajc3 | SKDEKPVEAIR | -317 | -327 | 2 | 0.008075282 | 11 | NaN | NaN | NaN | NaN | 348.75867 | NaN |
| Parp1 | FYTLIPHDFGMK | -735 | -746 | 0 | 0 | 12 | NaN | NaN | NaN | NaN | 676.457 | 611.9678 |
| Parp1 | VPHWYHF | -48 | -54 | 0 | 0.004595494 | 7 | NaN | NaN | NaN | NaN | 489.46524 | 1524.1929 |
| Eif5b | PIPGESPK | -1159 | -1166 | 0 | 0.000230707 | 8 | NaN | NaN | NaN | NaN | 544.417 | 173.06088 |
| Chp1 | LYDLDKDDK | -121 | -129 | 1 | 0 | 9 | NaN | NaN | NaN | NaN | 1568.5034 | 795.4464 |
| Cxcr4 | DENVHFNR | -33 | -40 | 0 | 0.00025015 | 8 | NaN | NaN | NaN | NaN | 486.23593 | 368.5258 |
| Xpnpep1 | DEPGYYEDGAFGIR | -522 | -535 | 0 | 0.00383302 | 14 | NaN | NaN | NaN | NaN | 792.14703 | NaN |
| Xpnpep1 | IENVVLVVPAK | -536 | -546 | 0 | 0 | 11 | NaN | NaN | NaN | NaN | 237.98022 | NaN |
| Scrn2 | DEVQEVVFIPAGTHVPGSR | -38 | -56 | 0 | 0 | 19 | NaN | NaN | NaN | NaN | 625.0135 | 500.7488 |
| Aco2 | TIHCDHLIEAQVGGEK | -123 | -138 | 0 | 0.000550307 | 16 | NaN | NaN | NaN | NaN | 3774.796 | 973.7295 |
| Aco2 | NDANPETHAFVTSPEIVTA | -480 | -498 | 0 | 0.000431655 | 19 | NaN | NaN | NaN | NaN | 426.32443 | 334.21927 |
| Unc119b | DFDFGFCIPSSR | -187 | -198 | 0 | 0 | 12 | NaN | NaN | NaN | NaN | 679.372 | NaN |
| Unc119b | DTIRPEHVLR | -66 | -75 | 1 | 0.000394361 | 10 | NaN | NaN | NaN | NaN | 252.901 | NaN |
| G3bp1 | HPDSHQLFIGNLPHEVDK | -334 | -351 | 0 | 0 | 18 | NaN | NaN | NaN | NaN | 1611.928 | 1462.762 |
| G3bp1 | YTLLNQAPDMLHR | -20 | -32 | 0 | 0.00111006 | 13 | NaN | NaN | NaN | NaN | 2069.7405 | 2127.6958 |
| G3bp1 | SELKDFFQNF | -352 | -361 | 1 | 0.00111006 | 10 | NaN | NaN | NaN | NaN | 1421.7289 | 1185.8264 |
| G3bp1 | HPDSHQLFIGNLPHEVDKSELK | -334 | -355 | 1 | 0 | 22 | NaN | NaN | NaN | NaN | 841.1479 | 631.24695 |
| G3bp1 | SYAHGGLDSNGKPADAVYGQK | -39 | -59 | 1 | 0.000230707 | 21 | NaN | NaN | NaN | NaN | 754.3432 | NaN |
| Matr3 | NDENKEDYTIPDEYR | -765 | -779 | 1 | 0 | 15 | NaN | NaN | NaN | NaN | 758.694 | 754.03235 |
| Nubp2 | DFIQEFPK | -245 | -252 | 0 | 0.001846793 | 8 | NaN | NaN | NaN | NaN | 914.8702 | 932.6527 |
| Tyms | DFLDSLGFSAR | -110 | -120 | 0 | 0 | 11 | NaN | NaN | NaN | NaN | 168.67076 | 192.42366 |
| Nsun5 | DFLTVSPSDQR | -285 | -295 | 0 | 0.007905138 | 11 | NaN | NaN | NaN | NaN | 2300.1802 | NaN |
| Pi4ka | LDPGAVSDVPEAIK | -1581 | -1594 | 0 | 0 | 14 | NaN | NaN | NaN | NaN | 712.59753 | NaN |
| Aqr | TVVEVAKPNIGENWPTR | -518 | -534 | 1 | 0 | 17 | NaN | NaN | NaN | NaN | 463.51846 | 430.94702 |
| Aqr | SNQALNQLFEK | -855 | -865 | 0 | 0.009955823 | 11 | NaN | NaN | NaN | NaN | 859.98083 | 967.5375 |
| Aqr | FLTLHDYLLR | -459 | -468 | 0 | 0 | 10 | NaN | NaN | NaN | NaN | 814.1369 | 626.9263 |
| Ndufs3 | VVAEPVELAQEFR | -218 | -230 | 0 | 0 | 13 | NaN | NaN | NaN | NaN | 1425.6089 | NaN |
| Dync1li1 | HFDFIQSHIR | -263 | -272 | 0 | 0.00025015 | 10 | NaN | NaN | NaN | NaN | 571.3508 | 69.90393 |
| Dync1li1 | KTGSPGGPGVGGSPGGGAA | -455 | -473 | 1 | 0.003966045 | 19 | NaN | NaN | NaN | NaN | 544.68146 | NaN |
| Acaa2 | NEAGYFNEEMAPIEVK | -194 | -209 | 0 | 0.006195994 | 16 | NaN | NaN | NaN | NaN | 381.5521 | NaN |
| Top2b | LHGLPEQFLY | -686 | -695 | 0 | 0.003521747 | 10 | NaN | NaN | NaN | NaN | 1607.38 | 1170.9482 |
| Top2b | KVIHELANER | -296 | -305 | 1 | 0 | 10 | NaN | NaN | NaN | NaN | 995.3139 | NaN |
| Top2b | DFTSEPPALPR | -1576 | -1586 | 0 | 0.001868731 | 11 | NaN | NaN | NaN | NaN | 391.46573 | NaN |
| Psmd13 | DRLELWCTDVK | -351 | -361 | 1 | 0.00025015 | 11 | NaN | NaN | NaN | NaN | 878.746 | 1065.1653 |
| Txnrd1 | IGEHMEEHGIK | -350 | -360 | 0 | 0.001392869 | 11 | NaN | NaN | NaN | NaN | 279.76422 | NaN |
| Blmh | HVPEEVLAVL | -428 | -437 | 0 | 0.000755556 | 10 | NaN | NaN | NaN | NaN | 935.95605 | NaN |
| Macf1 | KHINDLYEDLR | -97 | -107 | 1 | 0.000230707 | 11 | NaN | NaN | NaN | NaN | 210.89346 | NaN |
| Hdac1 | KVCYYYDGDVGNY | -10 | -22 | 1 | 0.001868731 | 13 | NaN | NaN | NaN | NaN | 345.9781 | 374.86826 |
| Cnn3 | DGIILCELINK | -54 | -64 | 0 | 0 | 11 | NaN | NaN | NaN | NaN | 749.1874 | NaN |
| Usp5 | VLFSIPDFQR | -343 | -352 | 0 | 0.00078661 | 10 | NaN | NaN | NaN | NaN | 926.87646 | NaN |
| Usp5 | EDKFEFDEDVK | -111 | -121 | 1 | 0.005605646 | 11 | NaN | NaN | NaN | NaN | 323.51935 | NaN |
| Rbm10 | ELSQGLLPLPQPYQAQ | -264 | -279 | 0 | 0 | 16 | NaN | NaN | NaN | NaN | 2951.914 | 2726.1768 |
| Rbm10 | DGLGSDNIGSR | -854 | -864 | 0 | 0.001868731 | 11 | NaN | NaN | NaN | NaN | 263.22757 | 1046.3369 |
| Rbm10 | QALHPPLTIDGK | -364 | -375 | 0 | 0.000550307 | 12 | NaN | NaN | NaN | NaN | 714.4497 | NaN |
| Ubap2 | SVEEWTTEDWTEDLSETK | -254 | -271 | 0 | 0.009122264 | 18 | NaN | NaN | NaN | NaN | 255.80426 | NaN |
| Oas3 | DGNPIKPWPVK | -1125 | -1135 | 1 | 0.00025015 | 11 | NaN | NaN | NaN | NaN | 1297.9878 | 1427.2998 |
| Nfkb2 | VDLVTHSDPPR | -93 | -103 | 0 | 0.001023236 | 11 | NaN | NaN | NaN | NaN | 983.35626 | 943.36945 |
| Nfkb2 | DGSFSLPLKPVISQPIHDSK | -202 | -221 | 1 | 0.001092529 | 20 | NaN | NaN | NaN | NaN | 198.52312 | NaN |
| Afg2a | DGSGLGLEESTGLK | -276 | -289 | 0 | 0.007880195 | 14 | NaN | NaN | NaN | NaN | 5068.174 | NaN |
| Ddx5 | HNILQIVDVCHDVEKDEK | -311 | -328 | 1 | 0 | 18 | NaN | NaN | NaN | NaN | 1681.5652 | 1417.6111 |
| Ddx5 | FYQEHPDLAR | -58 | -67 | 0 | 0 | 10 | NaN | NaN | NaN | NaN | 558.0184 | 7455.875 |
| Ddx5 | FYQEHPDLAR | -58 | -67 | 0 | 0 | 10 | NaN | NaN | NaN | NaN | 558.0184 | 7455.875 |
| Ddx5 | IVHINHQPFLER | -153 | -164 | 0 | 0 | 12 | NaN | NaN | NaN | NaN | 1847.4882 | 1554.142 |
| Ddx5 | GHNCPKPVLNFY | -86 | -97 | 1 | 0 | 12 | NaN | NaN | NaN | NaN | 695.24493 | NaN |
| Rela | LLTPVLSHPIFDNR | -174 | -187 | 0 | 0.000550307 | 14 | NaN | NaN | NaN | NaN | 404.05927 | NaN |
| Ints11 | DHCVQHLPDGSVTVE | -540 | -554 | 0 | 0.000712251 | 15 | NaN | NaN | NaN | NaN | 549.3584 | NaN |
| Ube3a | EFCASCPTFLR | -50 | -60 | 0 | 0 | 11 | NaN | NaN | NaN | NaN | 602.11005 | NaN |
| Eif3m | VAASCGAIQYIPTELDQVR | -130 | -148 | 0 | 0 | 19 | NaN | NaN | NaN | NaN | 358.21606 | NaN |
| Gnpda1 | HIDIHPENTHILDGN | -96 | -110 | 0 | 0.001370415 | 15 | NaN | NaN | NaN | NaN | 1397.5859 | 714.0674 |
| Gnpda1 | DHPESYHSF | -80 | -88 | 0 | 0.002972566 | 9 | NaN | NaN | NaN | NaN | 2527.3582 | 2446.4436 |
| Ncoa5 | DHRDPVYDR | -101 | -109 | 1 | 0.007609801 | 9 | NaN | NaN | NaN | NaN | 630.8667 | 489.97168 |
| Aldh18a1 | GQDHVVSDFSEHGSLK | -766 | -781 | 0 | 0 | 16 | NaN | NaN | NaN | NaN | 560.0849 | 487.15817 |
| Aldh18a1 | TQEALSIHGVK | -520 | -530 | 0 | 0 | 11 | NaN | NaN | NaN | NaN | 1225.2504 | 1311.2653 |
| Aldh18a1 | WSNIPFITVPLSR | -43 | -55 | 0 | 0.00111006 | 13 | NaN | NaN | NaN | NaN | 471.51755 | NaN |
| Ctsd | FDTGSSNLWVPSIHCK | -96 | -111 | 0 | 0.000627943 | 16 | NaN | NaN | NaN | NaN | 1275.4512 | NaN |
| Ube2o | DIADHPDFR | -639 | -647 | 0 | 0.005946382 | 9 | NaN | NaN | NaN | NaN | 1584.7391 | NaN |
| Acot9 | VCFTQDNYIQVR | -361 | -372 | 0 | 0.00025015 | 12 | NaN | NaN | NaN | NaN | 809.99493 | NaN |
| Npc1 | DICVAPLSPYNK | -466 | -477 | 0 | 0 | 12 | NaN | NaN | NaN | NaN | 852.0169 | 249.95334 |
| Puf60 | NIGQAQPIIDQLAEEAR | -212 | -228 | 0 | 0.000230707 | 17 | NaN | NaN | NaN | NaN | 625.4088 | 828.1319 |
| Puf60 | SIYYELGEDTIR | -140 | -151 | 0 | 0 | 12 | NaN | NaN | NaN | NaN | 376.96008 | NaN |
| Atl2 | GCFLLPHPGLK | -292 | -302 | 0 | 0 | 11 | NaN | NaN | NaN | NaN | 371.1097 | NaN |
| Atl2 | DIDEDFKR | -315 | -322 | 1 | 0.00272025 | 8 | NaN | NaN | NaN | NaN | 263.21402 | NaN |
| Ublcp1 | DIEDEVVEVENR | -100 | -111 | 0 | 0.000826806 | 12 | NaN | NaN | NaN | NaN | 2182.43 | NaN |
| Zap70 | ESPYSDPEELKDKK | -315 | -328 | 2 | 0.00383302 | 14 | NaN | NaN | NaN | NaN | 1856.647 | NaN |
| Tgtp2;Tgtp1 | DIESAPLHIAVTGETGAGK | (51);(51) | (69);(69) | 0 | 0.00122419 | 19 | NaN | NaN | NaN | NaN | 2660.1987 | 2899.9568 |

| | | | | | | | | | | | | |
|---|---|---|---|---|---|---|---|---|---|---|---|---|
| Lonp1 | TKTENPLVLIDEVDK | -569 | -583 | 1 | 0.002752526 | 15 | NaN | NaN | NaN | NaN | 2798.1208 | 2608.724 |
| Lonp1 | GLEVHFVEHYR | -919 | -929 | 0 | 0.00025015 | 11 | NaN | NaN | NaN | NaN | 430.08618 | 569.7169 |
| Usp47 | GIPLEDIEFAK | -1265 | -1275 | 0 | 0.00025015 | 11 | NaN | NaN | NaN | NaN | 717.8316 | 644.8179 |
| Parn | DIINNTSLAELEK | -329 | -341 | 0 | 0 | 13 | NaN | NaN | NaN | NaN | 1138.1097 | NaN |
| Pitrm1 | LEQSTAVPAQPHWDKPR | -291 | -307 | 1 | 0 | 17 | NaN | NaN | NaN | NaN | 656.1809 | 844.13696 |
| Tnfaip8l2 | DILLELVEHHLTPK | -125 | -138 | 0 | 0.000712251 | 14 | NaN | NaN | NaN | NaN | 386.53827 | NaN |
| Rbm25 | LKEFLEDYDDDRDDPK | -491 | -506 | 2 | 0.000471409 | 16 | NaN | NaN | NaN | NaN | 395.9576 | 445.2575 |
| Rbm25 | EGHPDPDAELQR | -548 | -559 | 0 | 0.00025015 | 12 | NaN | NaN | NaN | NaN | 1081.2524 | 377.0074 |
| Arhgef1 | RQLEDFR | -150 | -156 | 1 | 0.006195994 | 7 | NaN | NaN | NaN | NaN | 1958.7625 | 1699.2052 |
| Cast | DKDGKPLLPK | -566 | -575 | 2 | 0.00201047 | 10 | NaN | NaN | NaN | NaN | 556.5136 | NaN |
| Csde1 | SKVTLLEGDHVR | -306 | -317 | 1 | 0.001776473 | 12 | NaN | NaN | NaN | NaN | 5579.573 | NaN |
| Fkbp4 | LEYESSFSGEEMQK | -300 | -313 | 0 | 0.00051306 | 14 | NaN | NaN | NaN | NaN | 1320.151 | 1368.1261 |
| Srrm2 | VSSPVLETVQQR | -1358 | -1369 | 0 | 0.00025015 | 12 | NaN | NaN | NaN | NaN | 2302.7134 | 1491.5541 |
| Srrm2 | GPVSEVNTGFSPEVK | -1295 | -1309 | 0 | 0.001092529 | 15 | NaN | NaN | NaN | NaN | 707.7136 | NaN |
| Usp4 | ETLGEHDPWYCPTCK | -788 | -802 | 0 | 0.000591124 | 15 | NaN | NaN | NaN | NaN | 2109.8806 | NaN |
| Tmed9 | VEVKDPEDK | -87 | -95 | 1 | 0.001291103 | 9 | NaN | NaN | NaN | NaN | 666.6788 | NaN |
| Msmo1 | DKPETFEGQWK | -86 | -96 | 1 | 0.00167576 | 11 | NaN | NaN | NaN | NaN | 1074.9546 | NaN |
| Elp1 | LWDPVTPCR | -350 | -358 | 0 | 0.00078661 | 9 | NaN | NaN | NaN | NaN | 655.9958 | NaN |
| Rasa3 | HVDADSEVQGK | -116 | -126 | 0 | 0 | 11 | NaN | NaN | NaN | NaN | 1159.9453 | NaN |
| Hmgb2 | KKNEPEDEEEEEEEEEEDDEEEEEDE | -183 | -209 | 2 | 0.002780285 | 27 | NaN | NaN | NaN | NaN | 702.79205 | 605.86505 |
| Ube2c | FLTPCYHPNVDTQ | -98 | -110 | 0 | 0 | 13 | NaN | NaN | NaN | NaN | 4003.1748 | NaN |
| Ube2c | DKWSALYDVR | -120 | -129 | 1 | 0.000471409 | 10 | NaN | NaN | NaN | NaN | 1272.6296 | NaN |
| Atad3a | FVDEADAFLR | -408 | -417 | 0 | 0 | 10 | NaN | NaN | NaN | NaN | 484.05682 | 626.19226 |
| Noc2l | KEAFLGPILK | -326 | -335 | 1 | 0.002910424 | 10 | NaN | NaN | NaN | NaN | 1513.8549 | 1435.5026 |
| Noc2l | LKDRDPEFYK | -78 | -87 | 2 | 0.000230707 | 10 | NaN | NaN | NaN | NaN | 1209.7812 | NaN |
| Tmod3 | ETVLDDLDPENALLPAGFR | -33 | -51 | 0 | 0.000550307 | 19 | NaN | NaN | NaN | NaN | 187.06729 | NaN |
| Bag6 | FHQHYLGGQEPTPSNIR | -811 | -827 | 0 | 0.001055699 | 17 | NaN | NaN | NaN | NaN | 1482.0385 | 269.1249 |
| Bag6 | ENASPAPGTTAEEAMSR | -992 | -1008 | 0 | 0.002566359 | 17 | NaN | NaN | NaN | NaN | 1677.1617 | NaN |
| Bag6 | HGGAPLPGTR | -109 | -118 | 0 | 0.001636916 | 10 | NaN | NaN | NaN | NaN | 1244.3909 | NaN |
| Bag6 | PLTSPESLSR | -1100 | -1109 | 0 | 0 | 10 | NaN | NaN | NaN | NaN | 801.33215 | NaN |
| Ephx1 | KVETSDEEIKDLHQR | -52 | -66 | 2 | 0.000550307 | 15 | NaN | NaN | NaN | NaN | 496.16046 | 609.9215 |
| Ephx1 | DLELLYPFK | -278 | -286 | 0 | 0.00025015 | 9 | NaN | NaN | NaN | NaN | 630.8226 | NaN |
| Lmnb2 | VATGVVNIDEVDPEGR | -445 | -460 | 0 | 0.000230707 | 16 | NaN | NaN | NaN | NaN | 1539.6656 | 1324.8477 |
| Lmnb2 | LAEYQELLDIK | -349 | -359 | 0 | 0 | 11 | NaN | NaN | NaN | NaN | 1172.9945 | 926.9345 |
| Lmnb2 | RLVEVDSSR | -218 | -226 | 1 | 0.00272025 | 9 | NaN | NaN | NaN | NaN | 6188.7866 | NaN |
| Anp32a | IKDLSTIEPLKK | -100 | -111 | 2 | 0 | 12 | NaN | NaN | NaN | NaN | 825.6073 | 938.19525 |
| Slc12a7 | DLFSLKPDQSNVR | -1003 | -1015 | 1 | 0.000627943 | 13 | NaN | NaN | NaN | NaN | 328.88162 | 459.19916 |
| Slc12a7 | EADNPFSWK | -801 | -809 | 0 | 0.00078661 | 9 | NaN | NaN | NaN | NaN | 390.103 | NaN |
| Gls | GGTPPQQQQQQQQQPGASPPAAPGPK | -86 | -111 | 0 | 0.005605646 | 26 | NaN | NaN | NaN | NaN | 266.81668 | NaN |
| Gls | DLGTEYVHR | -304 | -312 | 0 | 0.000471409 | 9 | NaN | NaN | NaN | NaN | 1437.4832 | NaN |
| Rps20 | DTGKTPVEPEVAIH | -5 | -18 | 1 | 0.00149795 | 14 | NaN | NaN | NaN | NaN | 1364.1971 | 1551.6946 |
| Seh1l | DLIHDVSFDFHGR | -13 | -25 | 0 | 0 | 13 | NaN | NaN | NaN | NaN | 470.51212 | 87.57645 |
| Eif4g2 | YISSEVSPPSDETDSSSAPSK | -778 | -798 | 0 | 0.002595029 | 21 | NaN | NaN | NaN | NaN | 421.8249 | NaN |
| Lars1 | RFDDPLLGPR | -1119 | -1128 | 1 | 0.001023236 | 10 | NaN | NaN | NaN | NaN | 4174.0786 | 1942.0707 |
| Lars1 | TEPGVPVSLVNPQPSSGHFS | -1065 | -1084 | 0 | 0.004224523 | 20 | NaN | NaN | NaN | NaN | 449.87997 | NaN |
| Lars1 | LSAPLTCYK | -364 | -372 | 0 | 0.005145856 | 9 | NaN | NaN | NaN | NaN | 354.59363 | NaN |
| Tpp2 | IPANWTNPLGK | -99 | -109 | 0 | 0.00025015 | 11 | NaN | NaN | NaN | NaN | 505.7689 | NaN |
| Tpp2 | VHFKPGQIR | -652 | -660 | 1 | 0.003592055 | 9 | NaN | NaN | NaN | NaN | 393.83188 | NaN |
| A2m | DLKPAIVK | -1441 | -1448 | 1 | 0 | 8 | NaN | NaN | NaN | NaN | 4152.5674 | NaN |
| Prkaca | QAVNFPFLVK | -97 | -106 | 0 | 0 | 10 | NaN | NaN | NaN | NaN | 344.66357 | 313.9949 |
| Prkaca | VNFPFLVK | -99 | -106 | 0 | 0 | 8 | NaN | NaN | NaN | NaN | 305.47968 | 174.60239 |
| Mapk14 | DLLIDEWK | -331 | -338 | 0 | 0.00383302 | 8 | NaN | NaN | NaN | NaN | 413.721 | NaN |
| Imp3 | DLPERDPFR | -64 | -72 | 1 | 0.001055699 | 9 | NaN | NaN | NaN | NaN | 1249.6241 | NaN |
| Bop1 | IHPVTNRPADK | -219 | -229 | 1 | 0.00025015 | 11 | NaN | NaN | NaN | NaN | 254.4802 | 261.40167 |
| Bop1 | LALPGHAESYNPPPEYLPTEEER | -291 | -313 | 0 | 0.00025015 | 23 | NaN | NaN | NaN | NaN | 1063.2245 | NaN |
| Bop1 | STDQLLEAFTPPEEPALQPAR | -480 | -500 | 0 | 0.000230707 | 21 | NaN | NaN | NaN | NaN | 325.85452 | NaN |
| G6pdx | SYVAGQYDDAASYK | -106 | -119 | 0 | 0.00025015 | 14 | NaN | NaN | NaN | NaN | 1150.1633 | 1476.9095 |
| Mrpl11 | QAIPGPPLGPILGQR | -28 | -42 | 0 | 0.00111006 | 15 | NaN | NaN | NaN | NaN | 715.24207 | NaN |
| Ngdn | SLQGHPAVLR | -79 | -88 | 0 | 0.00122419 | 10 | NaN | NaN | NaN | NaN | 496.74643 | NaN |
| Flii | NQLTSLPSAICK | -279 | -290 | 0 | 0.000431655 | 12 | NaN | NaN | NaN | NaN | 766.7136 | NaN |
| Lig1 | NYHPIEDACWK | -270 | -280 | 0 | 0 | 11 | NaN | NaN | NaN | NaN | 3257.1157 | 3567.9556 |
| Lig1 | TKTENIPVCK | -195 | -204 | 1 | 0.000712251 | 10 | NaN | NaN | NaN | NaN | 6931.38 | NaN |
| Xpo6 | DLSSLLQAVGR | -556 | -566 | 0 | 0.000826806 | 11 | NaN | NaN | NaN | NaN | 1128.5225 | 1401.6271 |
| Irgm1 | SLDPLLYDFPK | -233 | -243 | 0 | 0 | 11 | NaN | NaN | NaN | NaN | 1509.9172 | 1468.9817 |
| Hnrnpf | DLSYCLSGMYDHR | -263 | -275 | 0 | 0.000230707 | 13 | NaN | NaN | NaN | NaN | 249.7048 | 280.41058 |
| Anxa5 | VYEEEYGSNLEDDVVGDTSGYYQR | -126 | -149 | 0 | 0 | 24 | NaN | NaN | NaN | NaN | 178.96562 | NaN |
| Gnai2 | LWADHGVQACFGR | -131 | -143 | 0 | 0 | 13 | NaN | NaN | NaN | NaN | 1070.4067 | 545.6787 |
| Ogfod1 | DLYDTDEHLQPK | -183 | -194 | 0 | 0.00078661 | 12 | NaN | NaN | NaN | NaN | 6303.713 | NaN |
| RtcA | LDPINLTDR | -178 | -186 | 0 | 0.00111006 | 9 | NaN | NaN | NaN | NaN | 1671.4528 | NaN |
| RtcA | DLYVSIQPVQEAR | -224 | -236 | 0 | 0.00897527 | 13 | NaN | NaN | NaN | NaN | 1540.7095 | NaN |
| Map4 | DVAPPMEEEIVPGNDTTSPK | -328 | -347 | 0 | 0.001868731 | 20 | NaN | NaN | NaN | NaN | 500.6156 | 515.75244 |
| Polr2c | VYPKPEEWPK | -196 | -205 | 1 | 0.00025015 | 10 | NaN | NaN | NaN | NaN | 805.5785 | 613.3815 |
| Polr2c | DNDPNDYVEQDDILIVK | -136 | -152 | 0 | 0 | 17 | NaN | NaN | NaN | NaN | 1206.4369 | NaN |
| Cdc5l | QVVQTPNTVLSTPFR | -400 | -414 | 0 | 0.00450079 | 15 | NaN | NaN | NaN | NaN | 763.2317 | NaN |
| Lrrc59 | LVNLQHLDLLNNR | -84 | -96 | 0 | 0 | 13 | NaN | NaN | NaN | NaN | 440.5125 | 281.4684 |
| Npepps | DNWEELHNR | -834 | -842 | 0 | 0 | 9 | NaN | NaN | NaN | NaN | 601.65125 | 432.97964 |
| Npepps | IKDNWEELHNR | -832 | -842 | 1 | 0.000431655 | 11 | NaN | NaN | NaN | NaN | 1431.0868 | NaN |
| Npepps | IKDNWEELHNR | -832 | -842 | 1 | 0.000431655 | 11 | NaN | NaN | NaN | NaN | 1431.0868 | NaN |

| | | | | | | | | | | | | |
|---|---|---|---|---|---|---|---|---|---|---|---|---|
| Npepps | LGWDPKPGEGHLDALLR | -691 | -707 | 1 | 0 | 17 | NaN | NaN | NaN | NaN | 1168.1207 | NaN |
| Uqcrfs1 | PDFSDYR | -86 | -92 | 0 | 0 | 7 | NaN | NaN | NaN | NaN | 480.49112 | 497.62527 |
| Khdrbs1 | SKDPSGAHPSVR | -20 | -31 | 1 | 0.001868731 | 12 | NaN | NaN | NaN | NaN | 367.4026 | 1013.4053 |
| Ppat | HVFTEDNLK | -72 | -80 | 0 | 0.000230707 | 9 | NaN | NaN | NaN | NaN | 1783.5212 | NaN |
| Ppat | DPYGNRPLCIGR | -199 | -210 | 1 | 0.006260252 | 12 | NaN | NaN | NaN | NaN | 506.9163 | NaN |
| Dda1 | DQEQVEAEGESSAPPR | -73 | -88 | 0 | 0 | 16 | NaN | NaN | NaN | NaN | 559.6838 | NaN |
| Rap1gds1 | KVDSNKEDDVAELK | -277 | -290 | 2 | 0 | 14 | NaN | NaN | NaN | NaN | 484.70444 | 494.03238 |
| Farsa | QALGEVIEAELR | -45 | -56 | 0 | 0 | 12 | NaN | NaN | NaN | NaN | 1605.9694 | 1580.1786 |
| Farsa | GEVIEAELR | -48 | -56 | 0 | 0.000826806 | 9 | NaN | NaN | NaN | NaN | 1070.4635 | NaN |
| Steep1 | DREEYDDGEKPLHVY | -16 | -30 | 2 | 0.001392869 | 15 | NaN | NaN | NaN | NaN | 235.94974 | 803.6249 |
| Steep1 | TIDEEEEEIEAR | -161 | -172 | 0 | 0.00025015 | 12 | NaN | NaN | NaN | NaN | 841.82227 | NaN |
| Actr3 | LKPKPIDVQVITH | -358 | -370 | 2 | 0 | 13 | NaN | NaN | NaN | NaN | 2467.189 | 524.21326 |
| Actr3 | LKPKPIDVQVITH | -358 | -370 | 2 | 0 | 13 | NaN | NaN | NaN | NaN | 2467.189 | 524.21326 |
| Actr3 | FIIPSCIAIK | -29 | -38 | 0 | 0.002099344 | 10 | NaN | NaN | NaN | NaN | 590.90814 | NaN |
| Actr3 | LKPKPIDVQVITHH | -358 | -371 | 2 | 0.002972566 | 14 | NaN | NaN | NaN | NaN | 395.74603 | NaN |
| Aarsd1 | HVVLEDTLLFPEGGGQPDDR | -42 | -61 | 0 | 0.002265552 | 20 | NaN | NaN | NaN | NaN | 473.4926 | 499.04706 |
| Nme2 | PEELIDYK | -136 | -143 | 0 | 0.00504706 | 8 | NaN | NaN | NaN | NaN | 728.6873 | 790.68604 |
| Nme1 | PEELVEYK | -136 | -143 | 0 | 0.007009655 | 8 | NaN | NaN | NaN | NaN | 832.2716 | 2382.293 |
| Cd5 | SLQCGSFLTHL | -203 | -213 | 0 | 0.007791272 | 11 | NaN | NaN | NaN | NaN | 459.42993 | 358.88412 |
| Mark2 | DRVETLRPH | -651 | -659 | 2 | 0.005176406 | 9 | NaN | NaN | NaN | NaN | 280.39993 | 327.90836 |
| Farsb | SAGSAFFPGR | -538 | -547 | 0 | 0.00272025 | 10 | NaN | NaN | NaN | NaN | 551.5592 | 639.4483 |
| Farsb | KLVITEETAK | -107 | -116 | 1 | 0.002296977 | 10 | NaN | NaN | NaN | NaN | 444.73328 | 534.5749 |
| Prss2 | DSCQGDSGGPVVCNGELQGIVSW | -194 | -216 | 0 | 0 | 23 | NaN | NaN | NaN | NaN | 244.92586 | 231.48734 |
| Try5;Try4 | DSCQGDSGGPVVCNGQLQGIV | (194);(194) | (214);(214) | 0 | 0.00025015 | 21 | NaN | NaN | NaN | NaN | 502.28854 | 494.97186 |
| Cbx3 | WHSCPEDEAQ | -174 | -183 | 0 | 0.000298805 | 10 | NaN | NaN | NaN | NaN | 933.7053 | 1024.8293 |
| Mat2a | ICDQISDAVLDAHLQQDPDAK | -33 | -53 | 0 | 0 | 21 | NaN | NaN | NaN | NaN | 509.87122 | 440.00452 |
| Mat2a | GTLPWLRPDSK | -171 | -181 | 1 | 0.000230707 | 11 | NaN | NaN | NaN | NaN | 1061.5702 | 1232.0472 |
| Sash3 | ETHLNELNIMDPQHR | -289 | -303 | 0 | 0 | 15 | NaN | NaN | NaN | NaN | 501.12592 | 581.6423 |
| Sash3 | FLEESPAPQYTGPFCGR | -161 | -177 | 0 | 0 | 17 | NaN | NaN | NaN | NaN | 370.5467 | 287.51636 |
| Eif4g3 | DSHADDPESIK | -735 | -745 | 0 | 0.00111006 | 11 | NaN | NaN | NaN | NaN | 901.5791 | NaN |
| Siae | DSPFGSIHPR | -396 | -405 | 0 | 0 | 10 | NaN | NaN | NaN | NaN | 1911.8191 | NaN |
| Trnt1 | DSSEPLKPYQDFVIDSR | -338 | -354 | 1 | 0.000591124 | 17 | NaN | NaN | NaN | NaN | 396.5072 | 1037.7372 |
| Trnt1 | RIDVTTDGR | -126 | -134 | 1 | 0.001846793 | 9 | NaN | NaN | NaN | NaN | 1659.4828 | 1420.8094 |
| Prdx6 | SKDINAYNGETPTEK | -83 | -97 | 1 | 0.002168954 | 15 | NaN | NaN | NaN | NaN | 1909.5431 | 852.4937 |
| Arpc3 | EKVFDPQSDKPSK | -146 | -158 | 2 | 0 | 13 | NaN | NaN | NaN | NaN | 4918.735 | 3093.0508 |
| Arpc3 | GITNFPIPGEPGFPLNAIYAK | -99 | -119 | 0 | 0.006788754 | 21 | NaN | NaN | NaN | NaN | 222.00638 | NaN |
| Wdr43 | LHQEYVPSAHLSGTCTCL | -48 | -65 | 0 | 0.000471409 | 18 | NaN | NaN | NaN | NaN | 830.92523 | 1010.4871 |
| Wdr43 | TGHATPVSSLR | -205 | -215 | 0 | 0 | 11 | NaN | NaN | NaN | NaN | 705.334 | NaN |
| Ddost | SSDIGDPLR | -116 | -124 | 0 | 0.000627943 | 9 | NaN | NaN | NaN | NaN | 1284.385 | 1275.5562 |
| Gna13 | WEDSGIQNAYDR | -153 | -164 | 0 | 0 | 12 | NaN | NaN | NaN | NaN | 2582.3765 | NaN |
| Atp6v1f | PAVLEIPSK | -86 | -94 | 0 | 0.002099344 | 9 | NaN | NaN | NaN | NaN | 1187.5612 | 1330.0605 |
| Xrcc6 | SFENPVLQQHFR | -475 | -486 | 0 | 0 | 12 | NaN | NaN | NaN | NaN | 1424.6144 | 1270.2994 |
| Xrcc6 | RVLELDQFK | -113 | -121 | 1 | 0.00025015 | 9 | NaN | NaN | NaN | NaN | 661.73944 | 1259.1062 |
| Xrcc6 | VFLPYADDKR | -433 | -442 | 1 | 0.001055699 | 10 | NaN | NaN | NaN | NaN | 980.50885 | NaN |
| Xrcc6 | ELVYPPGYNPEGK | -525 | -537 | 0 | 0.001868731 | 13 | NaN | NaN | NaN | NaN | 955.59973 | NaN |
| Xrcc6 | QIVLEKEETEELKR | -324 | -337 | 2 | 0.000550307 | 14 | NaN | NaN | NaN | NaN | 533.4586 | NaN |
| Xrcc6 | DTVGHGSDYSLSEVL | -131 | -145 | 0 | 0.002697088 | 15 | NaN | NaN | NaN | NaN | 442.92004 | NaN |
| Supt5h | LHSTCQTISVDR | -730 | -741 | 0 | 0 | 12 | NaN | NaN | NaN | NaN | 4845.079 | 7208.9336 |
| Supt5h | QFTDTPLQIK | -202 | -211 | 0 | 0.002595029 | 10 | NaN | NaN | NaN | NaN | 5751.146 | NaN |
| Supt5h | DTYLDTQIVGQTGVIR | -984 | -999 | 0 | 0 | 16 | NaN | NaN | NaN | NaN | 1361.6641 | NaN |
| Mcmbp | LSAVLPSVLNK | -540 | -550 | 0 | 0.000431655 | 11 | NaN | NaN | NaN | NaN | 715.00824 | 739.7726 |
| Mcmbp | SLLPADCQIHLQPQ | -513 | -526 | 0 | 0.007987481 | 14 | NaN | NaN | NaN | NaN | 941.30396 | NaN |
| Sec13 | DVAWAPSIGLPTSTIA | -217 | -232 | 0 | 0.00142999 | 16 | NaN | NaN | NaN | NaN | 144.99765 | NaN |
| Mthfd2 | RVQDPVTAK | -278 | -286 | 1 | 0.00025015 | 9 | NaN | NaN | NaN | NaN | 839.98474 | 541.5006 |
| Mthfd2 | DVDGFHVINVGR | -153 | -164 | 0 | 0.000431655 | 12 | NaN | NaN | NaN | NaN | 2236.4316 | NaN |
| Prp4k | HADIKPDNILVNESK | -813 | -827 | 1 | 0.00078661 | 15 | NaN | NaN | NaN | NaN | 1216.4672 | NaN |
| Smu1 | YIHLENLLAR | -114 | -123 | 0 | 0 | 10 | NaN | NaN | NaN | NaN | 6084.6255 | 6125.1055 |
| Smu1 | VKDVEEEKFPTQLSR | -193 | -207 | 2 | 0.00111006 | 15 | NaN | NaN | NaN | NaN | 2278.1658 | 183.8027 |
| Smu1 | YSEDGLLK | -502 | -509 | 0 | 0.007987481 | 8 | NaN | NaN | NaN | NaN | 737.8979 | NaN |
| Nucks1 | EDDDDSDYGSSK | -139 | -150 | 0 | 0.00676572 | 12 | NaN | NaN | NaN | NaN | 145.42882 | NaN |
| Uba6 | LHCFPLETR | -719 | -727 | 0 | 0.00025015 | 9 | NaN | NaN | NaN | NaN | 1785.434 | NaN |
| Uba6 | DVHGIWSR | -187 | -194 | 0 | 0.001743443 | 8 | NaN | NaN | NaN | NaN | 281.93878 | NaN |
| Nck1 | DVIEKPENDPEWWK | -218 | -231 | 1 | 0.004857192 | 14 | NaN | NaN | NaN | NaN | 1190.0045 | NaN |
| Pold1 | GIFPEPERDPVIQIC | -322 | -336 | 1 | 0.00383302 | 15 | NaN | NaN | NaN | NaN | 2664.6453 | NaN |
| Ppwd1 | TDKPYEDVSIINIT | -631 | -644 | 1 | 0.001291103 | 14 | NaN | NaN | NaN | NaN | 1333.5659 | 1423.3717 |
| Ppwd1 | DVITHVVCTK | -91 | -100 | 0 | 0.002780285 | 10 | NaN | NaN | NaN | NaN | 1251.19 | NaN |
| Mcm7 | LLVGGVDQSPQGMK | -357 | -370 | 0 | 0.00025015 | 14 | NaN | NaN | NaN | NaN | 684.573 | NaN |
| Park7 | GAEEMETVIPVDVMR | -13 | -27 | 0 | 0.00025015 | 15 | NaN | NaN | NaN | NaN | 1358.2516 | NaN |
| Park7 | TQGPYDVVVLPGGNLGAQNL | -63 | -82 | 0 | 0.002068213 | 20 | NaN | NaN | NaN | NaN | 708.02313 | 621.6814 |
| Rps19 | LKVPEWVDTVK | -28 | -38 | 1 | 0 | 11 | NaN | NaN | NaN | NaN | 6382.23 | 6488.93 |
| Epb41 | SQVTEGPGAPIKK | -568 | -580 | 1 | 0.000431655 | 13 | NaN | NaN | NaN | NaN | 591.3137 | NaN |
| Hmbs | DVPTILPPGFTIGAICK | -99 | -115 | 0 | 0.00078661 | 17 | NaN | NaN | NaN | NaN | 94.90642 | NaN |
| Dnaja2 | LYDILGVPPGASENELKK | -9 | -26 | 1 | 0.003133457 | 18 | NaN | NaN | NaN | NaN | 609.7638 | NaN |
| Hk2 | YVCATPDGTEK | -515 | -525 | 0 | 0 | 11 | NaN | NaN | NaN | NaN | 154.17886 | NaN |
| Manf | LRPGDCEVCISYLGR | -22 | -36 | 1 | 0 | 15 | NaN | NaN | NaN | NaN | 658.88226 | 577.8082 |
| Tfip11 | DWDLQNEFNPNR | -29 | -40 | 0 | 0.00025015 | 12 | NaN | NaN | NaN | NaN | 629.42554 | NaN |
| Calu | LHPEEYDYMK | -180 | -189 | 0 | 0 | 10 | NaN | NaN | NaN | NaN | 767.4139 | 776.2689 |
| Lss | FSNLYPDNTLAGHI | -720 | -733 | 0 | 0 | 14 | NaN | NaN | NaN | NaN | 2102.841 | 2181.434 |

| | | | | | | | | | | | | |
|---|---|---|---|---|---|---|---|---|---|---|---|---|
| Rab35 | LHEINQNCDDVCR | -103 | -115 | 0 | 0.000627943 | 13 | NaN | NaN | NaN | NaN | 561.89636 | 590.9692 |
| Rras2 | MNVDQAFHELVR | -162 | -173 | 0 | 0 | 12 | NaN | NaN | NaN | NaN | 804.17523 | 1689.5172 |
| Rras2 | SYFVTDYDPTIEDSYTK | -37 | -53 | 0 | 0.00051306 | 17 | NaN | NaN | NaN | NaN | 1313.1907 | 1225.5961 |
| Morf4l1 | NVDSILEDYANYK | -228 | -240 | 0 | 0.00025015 | 13 | NaN | NaN | NaN | NaN | 469.72958 | NaN |
| Prdx4 | DYGVYLEDSGHTLR | -190 | -203 | 0 | 0 | 14 | NaN | NaN | NaN | NaN | 12986.327 | 15156.688 |
| Prdx4 | HGEVCPAGWKPGSETIIPDPAGK | -244 | -266 | 1 | 0 | 23 | NaN | NaN | NaN | NaN | 456.0397 | 467.83752 |
| Prdx4 | ISKPAPYWEGTAVIN | -84 | -98 | 1 | 0 | 15 | NaN | NaN | NaN | NaN | 1845.9141 | NaN |
| F2 | YGFYTHVFR | -597 | -605 | 0 | 0 | 9 | NaN | NaN | NaN | NaN | 1112.282 | 1014.8156 |
| F2 | VLQVVNLPIVERPVCK | -519 | -534 | 1 | 0.00025015 | 16 | NaN | NaN | NaN | NaN | 431.53052 | 429.42087 |
| Pitpna | NETGGGEGVEVLVNEPYEK | -32 | -50 | 0 | 0.002595029 | 19 | NaN | NaN | NaN | NaN | 418.22632 | NaN |
| Rbm3 | ITFTNPEHASDAMR | -52 | -65 | 0 | 0 | 14 | NaN | NaN | NaN | NaN | 884.01276 | 556.96906 |
| Rbm3 | ITFTNPEHASDAMR | -52 | -65 | 0 | 0 | 14 | NaN | NaN | NaN | NaN | 884.01276 | 556.96906 |
| Rbm3 | GFITFTNPEHASDAMR | -50 | -65 | 0 | 0 | 16 | NaN | NaN | NaN | NaN | 116.767105 | 159.71889 |
| Rbm3 | DYSGSQGGYDRYSGGNYR | -131 | -148 | 1 | 0.003133457 | 18 | NaN | NaN | NaN | NaN | 198.66222 | NaN |
| Rpl13 | LILFPRKPSAPK | -124 | -135 | 2 | 0.000230707 | 12 | NaN | NaN | NaN | NaN | 385.85968 | 406.82074 |
| Rpl13 | EAAEQDVEKK | -201 | -210 | 1 | 0 | 10 | NaN | NaN | NaN | NaN | 395.47467 | NaN |
| Ccar1 | FTWVDAFPLSR | -396 | -406 | 0 | 0.00025015 | 11 | NaN | NaN | NaN | NaN | 442.26282 | 514.8918 |
| Ccar1 | SHCGYLLEK | -931 | -939 | 0 | 0.001610812 | 9 | NaN | NaN | NaN | NaN | 1049.4564 | NaN |
| Pafah1b1 | EAELDMNEELDKK | -34 | -46 | 1 | 0.00025015 | 13 | NaN | NaN | NaN | NaN | 599.559 | 721.8609 |
| Ttc27 | EAELQDLSNQLR | -833 | -844 | 0 | 0 | 12 | NaN | NaN | NaN | NaN | 498.30533 | NaN |
| Ncdn | EAEPDLLAVLR | -213 | -223 | 0 | 0.000550307 | 11 | NaN | NaN | NaN | NaN | 552.96027 | 527.76294 |
| Inpp5b | QIILTLPSFHK | -918 | -928 | 0 | 0.004439092 | 11 | NaN | NaN | NaN | NaN | 561.917 | NaN |
| Edc4 | SLDLQEVEPLGLPQASPSR | -712 | -730 | 0 | 0.003620005 | 19 | NaN | NaN | NaN | NaN | 503.50693 | NaN |
| Edc4 | ELAELWHNQEELLQR | -965 | -979 | 0 | 0.009657961 | 15 | NaN | NaN | NaN | NaN | 254.00833 | NaN |
| Edc4 | LVPDPLSSGDGNSTNKPGIR | -48 | -67 | 1 | 0 | 20 | NaN | NaN | NaN | NaN | 216.42975 | NaN |
| Fdx2 | EAHLDLLPPPEER | -111 | -123 | 0 | 0.000431655 | 13 | NaN | NaN | NaN | NaN | 909.5954 | NaN |
| Vasp | QVGEKPPKDESASEESEAR | -275 | -293 | 2 | 0 | 19 | NaN | NaN | NaN | NaN | 1068.4575 | 1143.2507 |
| Vasp | MKEEIIEVF | -357 | -365 | 1 | 0.008680443 | 9 | NaN | NaN | NaN | NaN | 729.8573 | NaN |
| Nemf | EAKPSLDADKPAEDILTY | -253 | -270 | 2 | 0.007343498 | 18 | NaN | NaN | NaN | NaN | 661.08966 | NaN |
| Nemf | KLPCESGDLEVIEEK | -817 | -831 | 1 | 0.00025015 | 15 | NaN | NaN | NaN | NaN | 510.9252 | NaN |
| Zc3h11a | LKAEEPSGADDSPSGTK | -356 | -372 | 1 | 0.00025015 | 17 | NaN | NaN | NaN | NaN | 532.32697 | 720.5236 |
| Zc3h11a | SSKPEEPLVR | -267 | -276 | 1 | 0.001392869 | 10 | NaN | NaN | NaN | NaN | 3417.3147 | NaN |
| Zc3h11a | SEIPCYWENQPVGCQK | -62 | -77 | 0 | 0 | 16 | NaN | NaN | NaN | NaN | 282.99933 | NaN |
| Sh3kbp1 | EAQNDDELTIK | -323 | -333 | 0 | 0 | 11 | NaN | NaN | NaN | NaN | 2628.7627 | 2677.4292 |
| Ak3 | REDDKPETVIK | -161 | -171 | 2 | 0.00078661 | 11 | NaN | NaN | NaN | NaN | 570.1721 | 991.03485 |
| Ak3 | EAQTEPVLQYYQK | -177 | -189 | 0 | 0.00830926 | 13 | NaN | NaN | NaN | NaN | 726.3007 | NaN |
| Ak3 | PQAEALDK | -97 | -104 | 0 | 0.000471409 | 8 | NaN | NaN | NaN | NaN | 332.54416 | NaN |
| Rnmt | QALEQYPAHENSK | -405 | -417 | 0 | 0 | 13 | NaN | NaN | NaN | NaN | 711.49493 | NaN |
| Tomm6 | SEASEVPDNVGDWLR | -15 | -29 | 0 | 0 | 15 | NaN | NaN | NaN | NaN | 838.11884 | NaN |
| Eef1b | TPAGLQVLNDYLADKS | -8 | -23 | 1 | 0.00111006 | 16 | NaN | NaN | NaN | NaN | 915.01636 | 304.456 |
| Rpl23 | FEDNAGVIVNNK | -98 | -109 | 0 | 0.00025015 | 12 | NaN | NaN | NaN | NaN | 1918.1287 | 2290.7126 |
| Rpl23 | GLPVGAVINCADNTGAK | -19 | -35 | 0 | 0.00644558 | 17 | NaN | NaN | NaN | NaN | 1087.9989 | NaN |
| Ifrd1 | GPEVLDEEGTQEDLEYK | -65 | -81 | 0 | 0.005522803 | 17 | NaN | NaN | NaN | NaN | 675.7704 | 352.33755 |
| Ifrd1 | ECFENIFTK | -211 | -219 | 0 | 0.009122264 | 9 | NaN | NaN | NaN | NaN | 1127.1743 | NaN |
| Mipep | ECFLQDPFNR | -646 | -655 | 0 | 0.000826806 | 10 | NaN | NaN | NaN | NaN | 383.84634 | NaN |
| Sema7a | ECPNPKPDEAPLQK | -538 | -551 | 1 | 0.005821388 | 14 | NaN | NaN | NaN | NaN | 1036.7886 | NaN |
| Ddx47 | QQYYLFIPSK | -239 | -248 | 0 | 0.001636916 | 10 | NaN | NaN | NaN | NaN | 1160.0685 | 1222.9636 |
| Ddx47 | REDAGDDDDKEGAIGVR | -425 | -441 | 2 | 0 | 17 | NaN | NaN | NaN | NaN | 247.68668 | NaN |
| Ndufab1 | EIVDYIADK | -143 | -151 | 0 | 0.00383302 | 9 | NaN | NaN | NaN | NaN | 2113.2544 | NaN |
| Ndufab1 | KLYDKIDPEK | -88 | -97 | 2 | 0.003069674 | 10 | NaN | NaN | NaN | NaN | 1632.8632 | NaN |
| Ndufab1 | PPLTLDGIK | -72 | -80 | 0 | 0.001846793 | 9 | NaN | NaN | NaN | NaN | 594.7839 | NaN |
| Ola1 | FYHDWNDKEIEVL | -202 | -214 | 1 | 0.00025015 | 13 | NaN | NaN | NaN | NaN | 349.3154 | 511.6855 |
| Ola1 | KIGIVGLPNVGK | -24 | -35 | 1 | 0.00133086 | 12 | NaN | NaN | NaN | NaN | 420.30084 | 422.02252 |
| Arhgef2 | LIHEGCLLWK | -474 | -483 | 0 | 0.00122419 | 10 | NaN | NaN | NaN | NaN | 386.62793 | NaN |
| Cdc40 | EWDIPVDFK | -446 | -454 | 0 | 0 | 9 | NaN | NaN | NaN | NaN | 622.34814 | NaN |
| Dock10 | WVDGGKPLFK | -835 | -844 | 1 | 0 | 10 | NaN | NaN | NaN | NaN | 621.5007 | 463.97003 |
| Dock10 | QSEEPHIK | -474 | -481 | 0 | 0.004031417 | 8 | NaN | NaN | NaN | NaN | 1371.619 | NaN |
| Dock10 | YELIADVNKPIIA | -1826 | -1838 | 1 | 0.001392869 | 13 | NaN | NaN | NaN | NaN | 391.4431 | NaN |
| Mkln1 | EDSCNAGPEDIQSR | -443 | -456 | 0 | 0 | 14 | NaN | NaN | NaN | NaN | 295.62985 | NaN |
| Hnrnpa3 | KYHTINGHNCEVK | -187 | -199 | 1 | 0 | 13 | NaN | NaN | NaN | NaN | 1953.5278 | 1996.9381 |
| Hnrnpa3 | SREDSVKPGAHLTVK | -112 | -126 | 2 | 0.001575548 | 15 | NaN | NaN | NaN | NaN | 1355.807 | NaN |
| Hnrnpa3 | GSYGGGDGGYNGFG | -258 | -271 | 0 | 0.009031675 | 14 | NaN | NaN | NaN | NaN | 297.45905 | NaN |
| Dis3l2 | EDVSEGLKR | -57 | -65 | 1 | 0.006624942 | 9 | NaN | NaN | NaN | NaN | 350.20322 | NaN |
| Rbm8a | MREDYDSVEQDGDEPGPQR | -50 | -68 | 1 | 0.00025015 | 19 | NaN | NaN | NaN | NaN | 370.81522 | 372.4773 |
| Rbm8a | HEAGGEDFAMDEDGDESIHK | -8 | -27 | 0 | 0.006100831 | 20 | NaN | NaN | NaN | NaN | 155.94025 | NaN |
| Oxct1 | EEAITGDFALVK | -200 | -211 | 0 | 0.00450079 | 12 | NaN | NaN | NaN | NaN | 670.4273 | NaN |
| Atp1b3 | KSFHQSLAEWK | -7 | -17 | 1 | 0.00078661 | 11 | NaN | NaN | NaN | NaN | 1250.9688 | NaN |
| Atp1b3 | KSFHQSLAEWK | -7 | -17 | 1 | 0.00078661 | 11 | NaN | NaN | NaN | NaN | 1250.9688 | NaN |
| Xrcc5 | GDLDSGLDHLKPSFPQK | -179 | -195 | 1 | 0 | 17 | NaN | NaN | NaN | NaN | 503.20328 | 531.0707 |
| Psmc2 | LREVVETPLLHPER | -187 | -200 | 1 | 0.00025015 | 14 | NaN | NaN | NaN | NaN | 2350.128 | NaN |
| Fam50a | EEELEREEITTK | -141 | -152 | 1 | 0.001708008 | 12 | NaN | NaN | NaN | NaN | 1018.554 | NaN |
| Fam50a | YEEELEREEITTK | -140 | -152 | 1 | 0.002434616 | 13 | NaN | NaN | NaN | NaN | 147.70302 | NaN |
| Tpx2 | SLAENPSGSLVQEPFQLATER | -645 | -665 | 0 | 0.00111006 | 21 | NaN | NaN | NaN | NaN | 261.94907 | 374.24805 |
| Degs1 | YDFVTDDTISPYSR | -298 | -311 | 0 | 0.00025015 | 14 | NaN | NaN | NaN | NaN | 725.7017 | 653.976 |
| Rcn2 | EELHYPQGEHR | -29 | -39 | 0 | 0.001291103 | 11 | NaN | NaN | NaN | NaN | 598.2103 | NaN |
| Rcn2 | RDPTANEDPEWIL | -220 | -232 | 1 | 0.000947459 | 13 | NaN | NaN | NaN | NaN | 263.97424 | NaN |
| Utp6 | QWIDLLLR | -375 | -382 | 0 | 0.001868731 | 8 | NaN | NaN | NaN | NaN | 307.90057 | 270.99005 |
| Caprin1 | EELSLLDEFYK | -179 | -189 | 0 | 0.005605646 | 11 | NaN | NaN | NaN | NaN | 638.3845 | NaN |

| | | | | | | | | | | | | |
|---|---|---|---|---|---|---|---|---|---|---|---|---|
| Caprin1 | QQQPQAASPSVPEPHSLTPVAQSDPLVR | -326 | -353 | 0 | 0 | 28 | NaN | NaN | NaN | NaN | 404.85458 | NaN |
| Pdcl3 | EENEDEFSEEDER | -58 | -70 | 0 | 0.008284062 | 13 | NaN | NaN | NaN | NaN | 252.25531 | NaN |
| Acad9 | LFIALTGLQHAGR | -438 | -450 | 0 | 0.00025015 | 13 | NaN | NaN | NaN | NaN | 3213.9539 | 3305.4478 |
| Acad9 | LSSGEHIAAF | -173 | -182 | 0 | 0.002972566 | 10 | NaN | NaN | NaN | NaN | 1266.8309 | 1250.6652 |
| Gnl1 | LCDCPGLIFPSLLPR | -409 | -423 | 0 | 0.00111006 | 15 | NaN | NaN | NaN | NaN | 255.08073 | NaN |
| Atp6v1a | LAEMPADSGYPAYLGAR | -365 | -381 | 0 | 0.004224523 | 17 | NaN | NaN | NaN | NaN | 1339.4554 | NaN |
| Ppil4 | IGADEEIDDFK | -191 | -201 | 0 | 0 | 11 | NaN | NaN | NaN | NaN | 761.94135 | 2413.8853 |
| Ppil4 | IHVDFSQSVAK | -311 | -321 | 0 | 0 | 11 | NaN | NaN | NaN | NaN | 3834.192 | NaN |
| Ppil4 | EEVEEIKAEK | -206 | -215 | 1 | 0.00111006 | 10 | NaN | NaN | NaN | NaN | 324.09732 | NaN |
| Sfpq | EEYEGPNK | -688 | -695 | 0 | 0.001392869 | 8 | NaN | NaN | NaN | NaN | 524.1274 | NaN |
| Sfpq | GPGTPAGYGR | -676 | -685 | 0 | 0.000230707 | 10 | NaN | NaN | NaN | NaN | 865.2794 | NaN |
| Mien1 | SAVKEEYPGIEIESR | -44 | -58 | 1 | 0.00051306 | 15 | NaN | NaN | NaN | NaN | 529.0401 | 457.9973 |
| Atp5f1c | KPPTFGDASVIAL | -159 | -171 | 1 | 0.000471409 | 13 | NaN | NaN | NaN | NaN | 2482.2195 | 2422.544 |
| Cherp | VKLEDHEYKPLDPK | -668 | -681 | 2 | 0.00122419 | 14 | NaN | NaN | NaN | NaN | 839.96484 | NaN |
| Cherp | VKLEDHEYKPLDPK | -668 | -681 | 2 | 0.00122419 | 14 | NaN | NaN | NaN | NaN | 839.96484 | NaN |
| Cherp | YSPPSHDRPR | -703 | -712 | 1 | 0 | 10 | NaN | NaN | NaN | NaN | 626.56696 | NaN |
| Mta2 | GHLSRPEAQ | -424 | -432 | 1 | 0.00111006 | 9 | NaN | NaN | NaN | NaN | 692.0152 | 623.5371 |
| Rps15a | TTSAGIMDHEEAR | -105 | -117 | 0 | 0 | 13 | NaN | NaN | NaN | NaN | 1135.4932 | 1408.1261 |
| Rps15a | SAGIMDHEEAR | -107 | -117 | 0 | 0 | 11 | NaN | NaN | NaN | NaN | 592.9405 | 715.5195 |
| Rps15a | KHGYIGEFEIIDDHR | -43 | -57 | 1 | 0.001392869 | 15 | NaN | NaN | NaN | NaN | 1098.5563 | 712.59235 |
| Csnk2b | EFFCEVDEDYIQDK | -20 | -33 | 0 | 0 | 14 | NaN | NaN | NaN | NaN | 217.57996 | NaN |
| Kdm3b | EFGDQDVDLVNCR | -1438 | -1450 | 0 | 0.000471409 | 13 | NaN | NaN | NaN | NaN | 504.6383 | NaN |
| Ndufb5 | LFVVKPSLYYDAR | -54 | -66 | 1 | 0 | 13 | NaN | NaN | NaN | NaN | 403.78668 | NaN |
| Tmx2 | EFNLNELYQR | -254 | -263 | 0 | 0 | 10 | NaN | NaN | NaN | NaN | 1041.3983 | NaN |
| Gc | HQPQEFPTYVEPTNDEICEAFR | -128 | -149 | 0 | 0 | 22 | NaN | NaN | NaN | NaN | 899.28656 | 932.20624 |
| Map2k1 | TPGRPLSSY | -292 | -300 | 1 | 0.005301614 | 9 | NaN | NaN | NaN | NaN | 1155.262 | 955.8488 |
| Gfm1 | ETIVAPVPFDFTHK | -527 | -540 | 0 | 0.004983786 | 14 | NaN | NaN | NaN | NaN | 1658.8662 | NaN |
| Anxa7 | GTDEQAIVDVVSNR | -177 | -190 | 0 | 0 | 14 | NaN | NaN | NaN | NaN | 310.6709 | NaN |
| Maip1 | EFTQGVKPDWTIAR | -270 | -283 | 1 | 0.001463897 | 14 | NaN | NaN | NaN | NaN | 529.9551 | 652.5902 |
| Fbln1 | TCINTEGSYTCQK | -326 | -338 | 0 | 0.000591124 | 13 | NaN | NaN | NaN | NaN | 313.5382 | 336.35767 |
| Sec16a | EFVQNQEVLPR | -478 | -488 | 0 | 0.000627943 | 11 | NaN | NaN | NaN | NaN | 412.85272 | NaN |
| Ecpas | YQFDPNLGIR | -1125 | -1134 | 0 | 0.00383302 | 10 | NaN | NaN | NaN | NaN | 1718.7454 | NaN |
| Ehmt2 | VHSSLGDTPQSEETLPK | -94 | -110 | 0 | 0.002972566 | 17 | NaN | NaN | NaN | NaN | 461.12546 | 497.20728 |
| Pck2 | EGEDSAQETPIGLVPK | -563 | -578 | 0 | 0 | 16 | NaN | NaN | NaN | NaN | 2565.811 | 2298.1533 |
| Pck2 | HSVGQPLTGHGDPVGQWPCNPEK | -212 | -234 | 0 | 0.00051306 | 23 | NaN | NaN | NaN | NaN | 531.5816 | 635.2573 |
| Pck2 | IGHVPDQR | -237 | -244 | 0 | 0.008890469 | 8 | NaN | NaN | NaN | NaN | 1505.747 | NaN |
| Pck2 | LEGEDSAQETPIGLVPK | -562 | -578 | 0 | 0 | 17 | NaN | NaN | NaN | NaN | 529.50616 | NaN |
| Ndufb10 | TPAPSPQTSLPNPITYL | -17 | -33 | 0 | 0.00111006 | 17 | NaN | NaN | NaN | NaN | 376.3489 | 369.51767 |
| Prdx2 | EGGLGPLNIPLLADVTK | -93 | -109 | 0 | 0.009688493 | 17 | NaN | NaN | NaN | NaN | 239.34735 | NaN |
| Ctcf | VEDQNTGAIENIIVEVK | -682 | -698 | 0 | 0.004595494 | 17 | NaN | NaN | NaN | NaN | 6737.6167 | 6230.075 |
| Ctcf | IVEESETFIK | -9 | -18 | 0 | 0.001291103 | 10 | NaN | NaN | NaN | NaN | 984.69196 | NaN |
| Sdf2l1 | NLHTHHFPSPLSNNQEVSAF | -114 | -133 | 0 | 0.002099344 | 20 | NaN | NaN | NaN | NaN | 287.05798 | 579.76227 |
| Ensa | NLVTGDHIPTPQDLPQR | -90 | -106 | 0 | 0.009031675 | 17 | NaN | NaN | NaN | NaN | 625.9211 | NaN |
| Prkcq | NETRPPCVPTPGK | -326 | -338 | 1 | 0.006100831 | 13 | NaN | NaN | NaN | NaN | 608.0207 | NaN |
| Nudt21 | EGVLIVHEHR | -81 | -90 | 0 | 0.001291103 | 10 | NaN | NaN | NaN | NaN | 871.8656 | 768.0118 |
| Nudt21 | FKLPGGELNPGEDEVEGLK | -104 | -122 | 1 | 0.00111006 | 19 | NaN | NaN | NaN | NaN | 308.0398 | NaN |
| Pym1 | VSLGDTAQIPSALQGPQATPL | -115 | -135 | 0 | 0 | 21 | NaN | NaN | NaN | NaN | 375.7668 | 197.74005 |
| Atp11b | IHVDEFALK | -614 | -622 | 0 | 0 | 9 | NaN | NaN | NaN | NaN | 1587.1056 | NaN |
| Parp9 | LPCHLIIHAVGPR | -190 | -202 | 0 | 0.005522803 | 13 | NaN | NaN | NaN | NaN | 348.95282 | NaN |
| Rb1 | EHECNIDEVK | -274 | -283 | 0 | 0.001868731 | 10 | NaN | NaN | NaN | NaN | 861.79376 | NaN |
| Rb1 | TLQTDPIDSFETER | -336 | -349 | 0 | 0 | 14 | NaN | NaN | NaN | NaN | 825.6912 | NaN |
| Parp3 | EHHITIDDPSLK | -449 | -460 | 0 | 0 | 12 | NaN | NaN | NaN | NaN | 729.9232 | 838.1844 |
| Cops2 | SGINPFDSQEAKPY | -289 | -302 | 1 | 0 | 14 | NaN | NaN | NaN | NaN | 1528.1278 | NaN |
| Chordc1 | RPSPDEPMTNLELK | -108 | -121 | 1 | 0.008014596 | 14 | NaN | NaN | NaN | NaN | 2208.1648 | NaN |
| Acads | ENLLGEPGMGFK | -251 | -262 | 0 | 0.008105996 | 12 | NaN | NaN | NaN | NaN | 895.91174 | NaN |
| Pum3 | GSKPQVAEIIR | -225 | -235 | 1 | 0 | 11 | NaN | NaN | NaN | NaN | 2294.7651 | 2163.9502 |
| Pum3 | WLLEQDKK | -559 | -566 | 1 | 0 | 8 | NaN | NaN | NaN | NaN | 980.5069 | 966.0902 |
| Pum3 | FIQYGNEEQR | -185 | -194 | 0 | 0 | 10 | NaN | NaN | NaN | NaN | 923.5255 | NaN |
| Pecam1 | EHTESYVIPQAR | -79 | -90 | 0 | 0.00078661 | 12 | NaN | NaN | NaN | NaN | 1024.2168 | NaN |
| Malt1 | EHTEVLPLLNPPGLK | -120 | -134 | 0 | 0.000471409 | 15 | NaN | NaN | NaN | NaN | 513.8023 | NaN |
| Por | HPFPCPTTYR | -359 | -368 | 0 | 0.006735683 | 10 | NaN | NaN | NaN | NaN | 1082.7507 | NaN |
| Pdcd11 | LPELFSPGMLVR | -139 | -150 | 0 | 0 | 12 | NaN | NaN | NaN | NaN | 1269.3174 | 858.9306 |
| Capn2 | VVPPDQSFQENYAGIF | -124 | -139 | 0 | 0.000230707 | 16 | NaN | NaN | NaN | NaN | 645.5718 | 419.05582 |
| Capn2 | RPTEICADPQFIIGGATR | -77 | -94 | 1 | 0.00025015 | 18 | NaN | NaN | NaN | NaN | 3834.1558 | NaN |
| Capn2 | FQDPSFPALPSSLGYK | -46 | -61 | 0 | 0.001868731 | 16 | NaN | NaN | NaN | NaN | 759.35944 | NaN |
| Capn2 | NPWGQVEWTGK | -286 | -296 | 0 | 0 | 11 | NaN | NaN | NaN | NaN | 655.56366 | NaN |
| Capn2 | EIDANEEDIDDGFR | -522 | -535 | 0 | 0 | 14 | NaN | NaN | NaN | NaN | 570.9819 | NaN |
| Lsm8 | EIDEETDSALDLGNIR | -72 | -87 | 0 | 0.000230707 | 16 | NaN | NaN | NaN | NaN | 3218.714 | 3322.4263 |
| Etfb | VISVEEPPQR | -224 | -233 | 0 | 0.005605646 | 10 | NaN | NaN | NaN | NaN | 3827.8274 | 4171.8325 |
| Nedd8 | EIEIDIEPTDK | -12 | -22 | 0 | 0.001291103 | 11 | NaN | NaN | NaN | NaN | 850.9258 | NaN |
| Impa1 | EKYPCHSFIGEESVAAGEK | -60 | -78 | 1 | 0.00111006 | 19 | NaN | NaN | NaN | NaN | 479.358 | 1136.3683 |
| Impa1 | SSPADLVTVTDQKVEK | -37 | -52 | 1 | 0.000907206 | 16 | NaN | NaN | NaN | NaN | 4553.823 | NaN |
| Ciao2b | EIFDLIR | -45 | -51 | 0 | 0.00025015 | 7 | NaN | NaN | NaN | NaN | 6689.459 | 5927.9463 |
| Casp8 | GMLEEGNLSFLK | -52 | -63 | 0 | 0 | 12 | NaN | NaN | NaN | NaN | 1749.5063 | 440.2389 |
| Casp8 | TMLAENNLETLK | -150 | -161 | 0 | 0.002972566 | 12 | NaN | NaN | NaN | NaN | 1552.36 | 1636.8805 |
| Pdlim1 | MNLASEPQEVLHIG | -104 | -117 | 0 | 0 | 14 | NaN | NaN | NaN | NaN | 1050.2352 | 1065.1711 |
| Pdlim1 | YNSPTGLYSSENISNFNNAVESK | -142 | -164 | 0 | 0.003620005 | 23 | NaN | NaN | NaN | NaN | 2168.775 | NaN |
| Sh3bp1 | EIQADYHR | -232 | -239 | 0 | 0.002752526 | 8 | NaN | NaN | NaN | NaN | 1653.6361 | NaN |

| | | | | | | | | | | | | |
|---|---|---|---|---|---|---|---|---|---|---|---|---|
| Sh3bp1 | GPNLLWPPEK | -428 | -437 | 0 | 0.007880195 | 10 | NaN | NaN | NaN | NaN | 405.99976 | NaN |
| Naa25 | FGDKPCCFTDLK | -359 | -370 | 1 | 0.001636916 | 12 | NaN | NaN | NaN | NaN | 566.2871 | NaN |
| Tm9sf4 | EKDVQFEHGYR | -159 | -169 | 1 | 0 | 11 | NaN | NaN | NaN | NaN | 1659.2946 | NaN |
| Tm9sf4 | TQLPYEYYSLPFCQPIK | -52 | -68 | 0 | 0.000431655 | 17 | NaN | NaN | NaN | NaN | 303.39633 | 330.06845 |
| Tm9sf4 | PFCQPIK | -62 | -68 | 0 | 0.008222027 | 7 | NaN | NaN | NaN | NaN | 535.1652 | NaN |
| Czib | EKDWTDYDEK | -134 | -143 | 1 | 0.00025015 | 10 | NaN | NaN | NaN | NaN | 818.56934 | NaN |
| Snx17 | EKEDGAFSFVR | -167 | -177 | 1 | 0.001392869 | 11 | NaN | NaN | NaN | NaN | 1319.3877 | 1304.9714 |
| Ik | EKEEEELMEKPQK | -182 | -194 | 2 | 0.00025015 | 13 | NaN | NaN | NaN | NaN | 718.55896 | NaN |
| Ik | EYNEDEDPAAR | -63 | -73 | 0 | 0 | 11 | NaN | NaN | NaN | NaN | 515.4802 | 557.83374 |
| Kif22 | ISPELLAHGR | -575 | -584 | 0 | 0.00025015 | 10 | NaN | NaN | NaN | NaN | 846.66156 | 737.91907 |
| Kif22 | EKENTPTILQPPASYSGT | -501 | -518 | 1 | 0.000550307 | 18 | NaN | NaN | NaN | NaN | 267.9654 | NaN |
| Rpl17 | TEKEQIVPKPEEEVAQK | -151 | -167 | 2 | 0 | 17 | NaN | NaN | NaN | NaN | 581.68005 | 514.4757 |
| Rpl17 | EKEQIVPKPEEEVAQK | -152 | -167 | 2 | 0.000431655 | 16 | NaN | NaN | NaN | NaN | 571.13184 | 884.51373 |
| Rpl17 | PKPEEEVAQK | -158 | -167 | 1 | 0.00437931 | 10 | NaN | NaN | NaN | NaN | 1570.5251 | 2109.038 |
| Rpl17 | KEQIVPKPEEEVAQK | -153 | -167 | 2 | 0.000471409 | 15 | NaN | NaN | NaN | NaN | 437.06207 | NaN |
| Hsd17b12 | SQDKLNQVSNNIK | -83 | -95 | 1 | 0.001811524 | 13 | NaN | NaN | NaN | NaN | 756.80945 | NaN |
| Thrap3 | MADFHKEELDEHDKDK | -419 | -434 | 2 | 0.00025015 | 16 | NaN | NaN | NaN | NaN | 850.55695 | 445.58548 |
| Thrap3 | KYYLHDDR | -875 | -882 | 1 | 0.001776473 | 8 | NaN | NaN | NaN | NaN | 283.44104 | 843.20557 |
| Eif3g | QVEEEGEDDKCVTSELLK | -15 | -32 | 1 | 0.003745449 | 18 | NaN | NaN | NaN | NaN | 394.3859 | 480.08673 |
| Ddx56 | ELVLHNPVTLK | -209 | -219 | 0 | 0 | 11 | NaN | NaN | NaN | NaN | 474.09332 | 343.4511 |
| Tes | QLPAHDQDPSK | -158 | -168 | 0 | 0 | 11 | NaN | NaN | NaN | NaN | 746.2501 | 901.2493 |
| Tes | VTDKPVCKPCYVK | -345 | -357 | 2 | 0 | 13 | NaN | NaN | NaN | NaN | 621.0791 | 623.3955 |
| Tes | NHAVVCQGCHNAIDPEVQ | -358 | -375 | 0 | 0.002434616 | 18 | NaN | NaN | NaN | NaN | 2373.381 | 3220.4893 |
| Hacd3 | LSDVQNPAISITDNVLHFK | -25 | -43 | 0 | 0 | 19 | NaN | NaN | NaN | NaN | 538.3231 | NaN |
| Tpm4 | LLEEELDR | -58 | -65 | 0 | 0.002296977 | 8 | NaN | NaN | NaN | NaN | 688.88025 | NaN |
| Tpm4 | TLDQTLNELNCI | -237 | -248 | 0 | 0.001055699 | 12 | NaN | NaN | NaN | NaN | 642.021 | NaN |
| Kcnab2 | EKVEVQLPELFHK | -223 | -235 | 1 | 0 | 13 | NaN | NaN | NaN | NaN | 261.76862 | 1022.4798 |
| Lamtor3 | KLPSVEGLHAIVVSDR | -12 | -27 | 1 | 0 | 16 | NaN | NaN | NaN | NaN | 1640.8198 | 2251.0469 |
| Ddx52 | ELASQIHR | -246 | -253 | 0 | 0 | 8 | NaN | NaN | NaN | NaN | 762.8816 | NaN |
| Polr2b | ELDDRDHYGNK | -381 | -391 | 1 | 0.000431655 | 11 | NaN | NaN | NaN | NaN | 477.45798 | 418.1264 |
| Polr2b | SYTGYNQEDSVIMNR | -784 | -798 | 0 | 0.001776473 | 15 | NaN | NaN | NaN | NaN | 841.81555 | NaN |
| Tbl3 | QFAGYSEEVLDVR | -329 | -341 | 0 | 0.001370415 | 13 | NaN | NaN | NaN | NaN | 1043.3156 | 643.021 |
| Tbl3 | ESFLVTGSQDCTVK | -434 | -447 | 0 | 0 | 14 | NaN | NaN | NaN | NaN | 443.06027 | NaN |
| Osbpl8 | ELDPLTGEWHYK | -733 | -744 | 0 | 0 | 12 | NaN | NaN | NaN | NaN | 2591.6743 | 1742.0721 |
| Osbpl8 | NLHSGDNFQLNDSEIER | -302 | -318 | 0 | 0.00383302 | 17 | NaN | NaN | NaN | NaN | 289.87634 | NaN |
| Immt | VVSQYHELVVQAR | -353 | -365 | 0 | 0 | 13 | NaN | NaN | NaN | NaN | 4029.7878 | NaN |
| Immt | KELDSITPDITPGWK | -370 | -384 | 1 | 0.000230707 | 15 | NaN | NaN | NaN | NaN | 2221.4465 | NaN |
| Eps15 | SGPSVAELPWAVK | -113 | -125 | 0 | 0 | 13 | NaN | NaN | NaN | NaN | 465.2833 | NaN |
| Nup62 | GGPADTSDPLQQICK | -466 | -480 | 0 | 0.00272025 | 15 | NaN | NaN | NaN | NaN | 7520.474 | NaN |
| Ddx42 | ERPGLGSENSDR | -662 | -673 | 1 | 0 | 12 | NaN | NaN | NaN | NaN | 120.48846 | 633.9754 |
| Ddx42 | SAPPQLPSFYK | -51 | -61 | 0 | 0.001868731 | 11 | NaN | NaN | NaN | NaN | 1165.8495 | NaN |
| Tomm40 | GSVNSNWIVGATLEK | -316 | -330 | 0 | 0 | 15 | NaN | NaN | NaN | NaN | 421.47806 | 604.0135 |
| Tomm40 | ELFPVQMEGVK | -92 | -102 | 0 | 0.001291103 | 11 | NaN | NaN | NaN | NaN | 146.46834 | 803.55634 |
| Npepl1 | ELGITPTIIRDEQLK | -207 | -221 | 1 | 0.003360691 | 15 | NaN | NaN | NaN | NaN | 983.65 | NaN |
| Arfip1 | IEQSQHLFQIH | -290 | -300 | 0 | 0.001811524 | 11 | NaN | NaN | NaN | NaN | 1121.3451 | NaN |
| Zc3hc1 | SWESSSPVDRPELEAA | -353 | -368 | 1 | 0.000471409 | 16 | NaN | NaN | NaN | NaN | 568.4748 | NaN |
| Nop14 | ELIEELIAK | -189 | -197 | 0 | 0.000230707 | 9 | NaN | NaN | NaN | NaN | 370.959 | 377.13538 |
| Nop14 | QYIGDLATDSTPDLK | -482 | -496 | 0 | 0.000431655 | 15 | NaN | NaN | NaN | NaN | 3618.5457 | NaN |
| Crlf3 | FFYPGWK | -432 | -438 | 0 | 0.003037016 | 7 | NaN | NaN | NaN | NaN | 433.72635 | NaN |
| Cebpz | LITDLLPDSR | -293 | -302 | 0 | 0.002329279 | 10 | NaN | NaN | NaN | NaN | 1784.7881 | NaN |
| Cebpz | TLDENEEVVDGGK | -60 | -72 | 0 | 0.004595494 | 13 | NaN | NaN | NaN | NaN | 406.6052 | NaN |
| Gpd1l | ELLQTPNFR | -181 | -189 | 0 | 0.002534621 | 9 | NaN | NaN | NaN | NaN | 843.3576 | NaN |
| Psmd8 | ILFAEATR | -285 | -292 | 0 | 0.005946382 | 8 | NaN | NaN | NaN | NaN | 624.5695 | NaN |
| Utp20 | ELNLTEHFGK | -57 | -66 | 0 | 0 | 10 | NaN | NaN | NaN | NaN | 3436.1606 | 3578.2368 |
| Utp20 | SFKEEIVHF | -984 | -992 | 1 | 0.007546675 | 9 | NaN | NaN | NaN | NaN | 690.8046 | 843.1202 |
| Utp20 | YLLELESGNK | -2565 | -2574 | 0 | 0.003681885 | 10 | NaN | NaN | NaN | NaN | 1664.389 | NaN |
| Utp20 | LGNVNIDIIHR | -23 | -33 | 0 | 0.001776473 | 11 | NaN | NaN | NaN | NaN | 693.1933 | NaN |
| Utp20 | LGNVNIDIIHR | -23 | -33 | 0 | 0.001776473 | 11 | NaN | NaN | NaN | NaN | 693.1933 | NaN |
| Utp20 | EVMEANLPSILLK | -1837 | -1849 | 0 | 0.002566359 | 13 | NaN | NaN | NaN | NaN | 537.9125 | NaN |
| Lnpep | GTELLLQQER | -625 | -634 | 0 | 0.000298805 | 10 | NaN | NaN | NaN | NaN | 652.8871 | NaN |
| Arhgap30 | EVLQELPIQNYR | -132 | -143 | 0 | 0 | 12 | NaN | NaN | NaN | NaN | 5213.3745 | NaN |
| Abr | ELPEPLLTDR | -727 | -736 | 0 | 0.007493591 | 10 | NaN | NaN | NaN | NaN | 5300.539 | NaN |
| Zc3h15 | KLEEVVNK | -157 | -164 | 1 | 0.001291103 | 8 | NaN | NaN | NaN | NaN | 15628.882 | 14442.119 |
| Paox | ELSEENQLVDTGGHVALPS | -86 | -104 | 0 | 0.000627943 | 19 | NaN | NaN | NaN | NaN | 259.405 | 5173.6846 |
| Anxa11 | EMSGDLEQGMLAVVK | -411 | -425 | 0 | 0.00025015 | 15 | NaN | NaN | NaN | NaN | 1522.4531 | NaN |
| Smarcb1 | ENASQPEVLVPIR | -178 | -190 | 0 | 0 | 13 | NaN | NaN | NaN | NaN | 1805.9095 | NaN |
| Msh6 | RWDTAFDHEK | -908 | -917 | 1 | 0 | 10 | NaN | NaN | NaN | NaN | 569.22723 | 668.5264 |
| Msh6 | YHETLEWLKPEK | -365 | -376 | 1 | 0 | 12 | NaN | NaN | NaN | NaN | 2570.671 | 2631.1013 |
| Msh6 | FFDDSPTR | -133 | -140 | 0 | 0.003681885 | 8 | NaN | NaN | NaN | NaN | 450.7665 | 376.2165 |
| Rpl9 | SVYAHFPINVV | -94 | -104 | 0 | 0.005459848 | 11 | NaN | NaN | NaN | NaN | 342.20355 | NaN |
| Trmt1 | WAEPIHDLDFVGR | -397 | -409 | 0 | 0 | 13 | NaN | NaN | NaN | NaN | 341.19873 | 318.95972 |
| Rwdd1 | ENLEDNDVSDILK | -73 | -85 | 0 | 0 | 13 | NaN | NaN | NaN | NaN | 2963.6855 | 3821.957 |
| Rwdd1 | TDYGEEQRNELEAL | -2 | -15 | 1 | 0.005015524 | 14 | NaN | NaN | NaN | NaN | 1112.4622 | NaN |
| Cstb | ENQKFDVF | -31 | -38 | 1 | 0.001463897 | 8 | NaN | NaN | NaN | NaN | 458.4309 | NaN |
| Prpf40a | QSTWEKPDDLK | -165 | -175 | 1 | 0 | 11 | NaN | NaN | NaN | NaN | 286.6889 | 3323.3853 |
| Prpf40a | QATPPIELDAVWEDIR | -752 | -767 | 0 | 0.00365438 | 16 | NaN | NaN | NaN | NaN | 592.87445 | NaN |
| Ccdc88b | LGEAHADLDSLR | -403 | -414 | 0 | 0.00025015 | 12 | NaN | NaN | NaN | NaN | 920.78796 | NaN |
| Ccdc88b | RVLEPGPLPR | -1305 | -1314 | 1 | 0.003003218 | 10 | NaN | NaN | NaN | NaN | 794.16565 | NaN |

| | | | | | | | | | | | | |
|---|---|---|---|---|---|---|---|---|---|---|---|---|
| Lsp1 | KVHLEESNLSYR | -130 | -141 | 1 | 0 | 12 | NaN | NaN | NaN | NaN | 851.03235 | 668.88544 |
| Lsp1 | SSELDEDEGFGDW | -77 | -89 | 0 | 0.00025015 | 13 | NaN | NaN | NaN | NaN | 331.5446 | NaN |
| Kpna1 | EPNPPIDEVINTPGVVAR | -113 | -130 | 0 | 0.001463897 | 18 | NaN | NaN | NaN | NaN | 1291.528 | NaN |
| Larp4 | QLEFCFSR | -122 | -129 | 0 | 0.000755556 | 8 | NaN | NaN | NaN | NaN | 420.21545 | NaN |
| Larp4 | EYEVMYSPSCETTR | -63 | -76 | 0 | 0.008161009 | 14 | NaN | NaN | NaN | NaN | 117.91874 | NaN |
| Add3 | KEQDHIIIIPR | -167 | -177 | 1 | 0 | 11 | NaN | NaN | NaN | NaN | 357.2078 | 336.16205 |
| Tmx3 | LLYELGDTGK | -241 | -250 | 0 | 0 | 10 | NaN | NaN | NaN | NaN | 673.20496 | 168.00162 |
| Tmx3 | TKDDIIEFAHR | -121 | -131 | 1 | 0.000712251 | 11 | NaN | NaN | NaN | NaN | 712.90204 | NaN |
| Txlna | LIEQYELR | -312 | -319 | 0 | 0.00051306 | 8 | NaN | NaN | NaN | NaN | 1092.8046 | 1291.8708 |
| Txlna | HKDLQQQLVDAK | -329 | -340 | 1 | 0.000627943 | 12 | NaN | NaN | NaN | NaN | 791.47437 | 365.42703 |
| Gripap1 | EQLQQAQEER | -535 | -544 | 0 | 0.00111006 | 10 | NaN | NaN | NaN | NaN | 2115.4893 | NaN |
| Itgb3 | SKVELEVR | -378 | -385 | 1 | 0.006384543 | 8 | NaN | NaN | NaN | NaN | 790.317 | NaN |
| Pcm1 | EQWQEQINQLK | -994 | -1004 | 0 | 0 | 11 | NaN | NaN | NaN | NaN | 2958.336 | NaN |
| Nt5c | HLALPPTR | -172 | -179 | 0 | 0.003003218 | 8 | NaN | NaN | NaN | NaN | 1383.1708 | NaN |
| Rps4x | TDITYPAGFMDVISIDK | -78 | -94 | 0 | 0 | 17 | NaN | NaN | NaN | NaN | 462.4114 | NaN |
| Rps4x | HWMLDKL | -17 | -23 | 1 | 0.003461185 | 7 | NaN | NaN | NaN | NaN | 955.04767 | 861.76636 |
| Pam16 | ERLDEELR | -103 | -110 | 1 | 0.002566359 | 8 | NaN | NaN | NaN | NaN | 799.1997 | 913.1708 |
| Akt2 | YDSLDPLELDQR | -456 | -467 | 0 | 0.001941082 | 12 | NaN | NaN | NaN | NaN | 695.60284 | NaN |
| Dpy30 | TQVAENPHSEYGLTDSVER | -11 | -29 | 0 | 0.001708008 | 19 | NaN | NaN | NaN | NaN | 365.46646 | 388.66443 |
| Akt1 | ERPQDVDQR | -40 | -48 | 1 | 0.003490217 | 9 | NaN | NaN | NaN | NaN | 747.8112 | 393.93634 |
| Otub1 | LLYRPGHYDILYK | -259 | -271 | 1 | 0.00078661 | 13 | NaN | NaN | NaN | NaN | 883.2052 | 928.8293 |
| Hbs1l | ESDVAFIPTSGLSGENLTAR | -434 | -453 | 0 | 0 | 20 | NaN | NaN | NaN | NaN | 1093.4412 | 971.0794 |
| Tbca | KQAEILQESR | -52 | -61 | 1 | 0.002068213 | 10 | NaN | NaN | NaN | NaN | 698.1799 | 670.78784 |
| Ptpn1 | FSYLAVIEGAK | -269 | -279 | 0 | 0 | 11 | NaN | NaN | NaN | NaN | 532.1727 | NaN |
| Cdca8 | ESIKEEEEEEEGGGGGGR | -121 | -139 | 1 | 0.003133457 | 19 | NaN | NaN | NaN | NaN | 513.2558 | NaN |
| U2surp | SEEQETEAFVEEPSK | -516 | -530 | 0 | 0.001392869 | 15 | NaN | NaN | NaN | NaN | 1257.078 | NaN |
| U2surp | TIQGHLQSENFK | -636 | -647 | 0 | 0.000471409 | 12 | NaN | NaN | NaN | NaN | 542.7659 | NaN |
| Ifit3 | ESLEAILPQLK | -7 | -17 | 0 | 0 | 11 | NaN | NaN | NaN | NaN | 808.6892 | 646.71783 |
| Ubxn1 | SSPPATDPGPVPSSPSQEPPTK | -187 | -208 | 0 | 0.004595494 | 22 | NaN | NaN | NaN | NaN | 453.68173 | NaN |
| Samsn1 | KGDIIDIICK | -188 | -197 | 1 | 0.000591124 | 10 | NaN | NaN | NaN | NaN | 436.32904 | 659.5687 |
| Samsn1 | ESLLDEETTVEHEK | -300 | -313 | 0 | 0.00078661 | 14 | NaN | NaN | NaN | NaN | 643.8493 | NaN |
| Son | SHHDGNLESDSFLK | -142 | -155 | 0 | 0 | 14 | NaN | NaN | NaN | NaN | 391.3647 | NaN |
| Rasa2 | PDANAIFR | -389 | -396 | 0 | 0.008919338 | 8 | NaN | NaN | NaN | NaN | 394.6025 | 514.56494 |
| Rasa2 | ESSGTSEPVHLK | -595 | -606 | 0 | 0.00025015 | 12 | NaN | NaN | NaN | NaN | 1804.4749 | NaN |
| Nploc4 | PDGHFGSK | -359 | -366 | 0 | 0.00437931 | 8 | NaN | NaN | NaN | NaN | 479.7002 | NaN |
| Dbnl | PEQEAVSR | -243 | -250 | 0 | 0.00383302 | 8 | NaN | NaN | NaN | NaN | 4950.737 | NaN |
| Dbnl | GYGPDGHFGMFPA | -417 | -429 | 0 | 0.004439092 | 13 | NaN | NaN | NaN | NaN | 693.7301 | NaN |
| Ints3 | ESVLDILTEQR | -184 | -194 | 0 | 0.001092529 | 11 | NaN | NaN | NaN | NaN | 571.0318 | 658.3766 |
| Ints3 | SVLDILTEQR | -185 | -194 | 0 | 0.002099344 | 10 | NaN | NaN | NaN | NaN | 717.52246 | NaN |
| Ociad1 | GILSSHPK | -65 | -72 | 0 | 0.00272025 | 8 | NaN | NaN | NaN | NaN | 981.76465 | 835.55457 |
| Ociad1 | VSRPIPDIGGGYIPTEEEWR | -14 | -33 | 1 | 0.001392869 | 20 | NaN | NaN | NaN | NaN | 737.5437 | NaN |
| Eif6 | ETEEILADVLKVE | -124 | -136 | 1 | 0.00204437 | 13 | NaN | NaN | NaN | NaN | 140.33038 | 944.38525 |
| Ahsa1 | LKETFLTSPEELYR | -211 | -224 | 1 | 0 | 14 | NaN | NaN | NaN | NaN | 1485.335 | 1516.3748 |
| Gimap8 | KSDDDVFSIIK | -376 | -386 | 1 | 0.001023236 | 11 | NaN | NaN | NaN | NaN | 1451.7726 | 1314.743 |
| Gimap8 | ETGEDQETQVK | -642 | -652 | 0 | 0.001392869 | 11 | NaN | NaN | NaN | NaN | 818.1304 | NaN |
| Arcn1 | MITETIIETDKPK | -199 | -211 | 1 | 0 | 13 | NaN | NaN | NaN | NaN | 573.89996 | NaN |
| Arcn1 | GVQLQTHPNVDKK | -324 | -336 | 1 | 0 | 13 | NaN | NaN | NaN | NaN | 476.94293 | NaN |
| Hmox2 | KYFFGENWEEQVK | -113 | -125 | 1 | 0.00142999 | 13 | NaN | NaN | NaN | NaN | 772.65045 | 420.15176 |
| Hmox2 | IHYVGQNEPELLVAH | -137 | -151 | 0 | 0 | 15 | NaN | NaN | NaN | NaN | 569.6717 | 572.1936 |
| Hmox2 | IHYVGQNEPELLVAH | -137 | -151 | 0 | 0 | 15 | NaN | NaN | NaN | NaN | 569.6717 | 572.1936 |
| Pcbp1 | MHGGTGFAGIDSSSPEVK | -251 | -268 | 0 | 0.00025015 | 18 | NaN | NaN | NaN | NaN | 437.77582 | 584.3791 |
| Pcbp1 | IITLTGPTNAIF | -58 | -69 | 0 | 0.005363933 | 12 | NaN | NaN | NaN | NaN | 79.61569 | NaN |
| Eif4a2 | ETQALVLAPTR | -101 | -111 | 0 | 0 | 11 | NaN | NaN | NaN | NaN | 503.04785 | 468.0921 |
| Mcm10 | EVCLSIDHPQK | -383 | -393 | 0 | 0 | 11 | NaN | NaN | NaN | NaN | 626.56195 | NaN |
| Frg1 | KKDDIPEED | -190 | -198 | 2 | 0.00272025 | 9 | NaN | NaN | NaN | NaN | 4872.8857 | NaN |
| Tor1aip1 | NEDPISNLCRPPLR | -187 | -200 | 1 | 0.000627943 | 14 | NaN | NaN | NaN | NaN | 1304.7502 | 1212.281 |
| Tor1aip1 | HLVLPVQPENTLK | -578 | -590 | 0 | 0.004350979 | 13 | NaN | NaN | NaN | NaN | 991.3135 | 1063.7087 |
| Gigyf2 | EVESPYEVHDYTR | -1169 | -1181 | 0 | 0.005522803 | 13 | NaN | NaN | NaN | NaN | 774.3117 | NaN |
| Pon2 | VVNHPQFK | -130 | -137 | 0 | 0.00025015 | 8 | NaN | NaN | NaN | NaN | 5206.591 | NaN |
| Hadha | MQLLEIITTDK | -506 | -516 | 0 | 0.00025015 | 11 | NaN | NaN | NaN | NaN | 5077.5254 | NaN |
| Casp6 | SVAEGYYSHR | -193 | -202 | 0 | 0.006384543 | 10 | NaN | NaN | NaN | NaN | 999.0629 | NaN |
| Casp6 | EVFDPAEQYK | -12 | -21 | 0 | 0.002068213 | 10 | NaN | NaN | NaN | NaN | 991.39374 | NaN |
| Ppp1r18 | PGETPEESLR | -195 | -204 | 0 | 0.00025015 | 10 | NaN | NaN | NaN | NaN | 544.6615 | 556.87396 |
| Cwc27 | RLTEVDIDDEERPR | -143 | -156 | 2 | 0.002534621 | 14 | NaN | NaN | NaN | NaN | 485.674 | 586.0653 |
| Aup1 | EVLPHVPLNVIQR | -306 | -318 | 0 | 0 | 13 | NaN | NaN | NaN | NaN | 434.35327 | 588.46747 |
| Aup1 | EVLPHVPLNVIQ | -306 | -317 | 0 | 0.00025015 | 12 | NaN | NaN | NaN | NaN | 602.57654 | NaN |
| Rnaset2a;Rnas | EVNSCQDSLDYWTIH | (55);(55) | (69);(69) | 0 | 0 | 15 | NaN | NaN | NaN | NaN | 605.51843 | 583.58466 |
| Usp39 | TLDLPTAPLYK | -391 | -401 | 0 | 0 | 11 | NaN | NaN | NaN | NaN | 3014.172 | NaN |
| Atp13a1 | SPQENLVPCDVLLLR | -325 | -339 | 0 | 0.000627943 | 15 | NaN | NaN | NaN | NaN | 710.9718 | NaN |
| Cul3 | SKEIESGHIF | -650 | -659 | 1 | 0.002296977 | 10 | NaN | NaN | NaN | NaN | 1529.1349 | 655.6831 |
| Dnm2 | GMEELIPLVNK | -5 | -15 | 0 | 0 | 11 | NaN | NaN | NaN | NaN | 1751.2996 | NaN |
| Nup160 | EYEFSQDEFR | -478 | -487 | 0 | 0 | 10 | NaN | NaN | NaN | NaN | 627.9572 | 671.03723 |
| Nup160 | YSLDHISSL | -347 | -355 | 0 | 0.002566359 | 9 | NaN | NaN | NaN | NaN | 370.26855 | 426.99292 |
| Nup160 | LHNEVVGIIESR | -1024 | -1035 | 0 | 0.00025015 | 12 | NaN | NaN | NaN | NaN | 245.97945 | 228.7541 |
| Nup160 | NLDLSWNELKK | -452 | -462 | 1 | 0.000431655 | 11 | NaN | NaN | NaN | NaN | 4098.688 | NaN |
| Nup160 | LILPHPSR | -130 | -137 | 0 | 0.001846793 | 8 | NaN | NaN | NaN | NaN | 478.39832 | NaN |
| Chchd3 | YEYHPVCADLQTK | -177 | -189 | 0 | 0.001636916 | 13 | NaN | NaN | NaN | NaN | 468.12247 | NaN |
| Rasal3 | FFSGEKPGFLAPR | -787 | -799 | 1 | 0 | 13 | NaN | NaN | NaN | NaN | 5020.7417 | 4963.923 |

| | | | | | | | | | | | | |
|---|---|---|---|---|---|---|---|---|---|---|---|---|
| Ikbkg | VTSLLGELQESQSR | -146 | -159 | 0 | 0 | 14 | NaN | NaN | NaN | NaN | 546.0075 | NaN |
| Prkaa1 | YLNPSVISLLK | -247 | -257 | 0 | 0.000431655 | 11 | NaN | NaN | NaN | NaN | 305.65845 | 264.1033 |
| Ubr4 | TPVFIFER | -4324 | -4331 | 0 | 0.002265552 | 8 | NaN | NaN | NaN | NaN | 878.78107 | 848.70984 |
| Ubr4 | LEKDPQQEDFLQGR | -4350 | -4363 | 1 | 0.000230707 | 14 | NaN | NaN | NaN | NaN | 2474.3171 | NaN |
| Hdac3 | FAPDFTLHPDVSTR | -332 | -345 | 0 | 0.00025015 | 14 | NaN | NaN | NaN | NaN | 1183.9113 | 847.839 |
| Tmem160 | FAPGHGPR | -37 | -44 | 0 | 0.000591124 | 8 | NaN | NaN | NaN | NaN | 632.45306 | NaN |
| Pspc1 | FAQPGTFEFEYASR | -264 | -277 | 0 | 0.00025015 | 14 | NaN | NaN | NaN | NaN | 1967.2048 | 595.0353 |
| Ldah | FATPFLCQFR | -179 | -188 | 0 | 0 | 10 | NaN | NaN | NaN | NaN | 748.4962 | NaN |
| Isyna1 | NHLGNNDGQNLSAPLQFR | -333 | -350 | 0 | 0 | 18 | NaN | NaN | NaN | NaN | 412.77475 | 473.948 |
| Raver1 | SFSHLLPSPEPSPEGSYVGQ | -709 | -728 | 0 | 0.000431655 | 20 | NaN | NaN | NaN | NaN | 945.24805 | NaN |
| Raver1 | RAPEQELPPLDPEEIR | -30 | -45 | 1 | 0 | 16 | NaN | NaN | NaN | NaN | 749.69696 | NaN |
| Mical1 | GLVSQPSGTPSAILFLGK | -618 | -635 | 0 | 0.001392869 | 18 | NaN | NaN | NaN | NaN | 379.20245 | 467.32666 |
| Mical1 | NIIPDSGAEPPPKPPR | -804 | -819 | 1 | 0.005605646 | 16 | NaN | NaN | NaN | NaN | 406.26038 | NaN |
| Themis | TPYLSIEEITR | -94 | -104 | 0 | 0 | 11 | NaN | NaN | NaN | NaN | 5500.953 | 5369.409 |
| Usp7 | VDVIFCDK | -795 | -802 | 0 | 0.009416377 | 8 | NaN | NaN | NaN | NaN | 702.9348 | NaN |
| Spcs2 | RLHDSLATER | -214 | -223 | 1 | 0.00383302 | 10 | NaN | NaN | NaN | NaN | 771.6805 | NaN |
| Spcs2 | SGGGGGGSSGAGGGPSCGTSSSR | -11 | -32 | 0 | 0.002168954 | 22 | NaN | NaN | NaN | NaN | 200.39352 | NaN |
| Eps15l1 | FDEIFLK | -277 | -283 | 0 | 0.008653659 | 7 | NaN | NaN | NaN | NaN | 430.16873 | NaN |
| Ubtf | FDGRPTKPPPNSY | -293 | -305 | 2 | 0.00111006 | 13 | NaN | NaN | NaN | NaN | 765.0338 | 760.55237 |
| Ubtf | SQELLSNGELNHLPLK | -579 | -594 | 0 | 0.003299042 | 16 | NaN | NaN | NaN | NaN | 299.09235 | 271.44043 |
| Ubtf | KHPDFPK | -107 | -113 | 1 | 0.000230707 | 7 | NaN | NaN | NaN | NaN | 813.6487 | NaN |
| Aldh3a2 | YLAPTILTDVDPNSK | -312 | -326 | 0 | 0.001868731 | 15 | NaN | NaN | NaN | NaN | 605.39343 | 558.63873 |
| Hm13 | SSSDMPETITSR | -62 | -73 | 0 | 0.004595494 | 12 | NaN | NaN | NaN | NaN | 403.34848 | NaN |
| Abcb7 | FDNVHFEYIEGQK | -474 | -486 | 0 | 0 | 13 | NaN | NaN | NaN | NaN | 16471.037 | 16109.564 |
| Abcb7 | LSYVWPEDRPDLR | -125 | -137 | 1 | 0.00025015 | 13 | NaN | NaN | NaN | NaN | 100.44284 | 307.164 |
| Abcb7 | ISLGFLGGAK | -142 | -151 | 0 | 0.009388464 | 10 | NaN | NaN | NaN | NaN | 361.67163 | 341.8747 |
| Abcb7 | RYDGFLK | -355 | -361 | 1 | 0.003745449 | 7 | NaN | NaN | NaN | NaN | 2144.396 | NaN |
| Abcb7 | GQNLQDVSLESLR | -533 | -545 | 0 | 0.000230707 | 13 | NaN | NaN | NaN | NaN | 1199.8503 | NaN |
| Abcb7 | GLHDAILR | -586 | -593 | 0 | 0.004350979 | 8 | NaN | NaN | NaN | NaN | 696.99506 | NaN |
| Cdc42 | IGGEPYTLGLFDTAGQEDYDR | -46 | -66 | 0 | 0.009745776 | 21 | NaN | NaN | NaN | NaN | 176.69527 | 211.80817 |
| Cc2d1b | GQPVDLSGMPPAPADLK | -323 | -339 | 0 | 0.009242859 | 17 | NaN | NaN | NaN | NaN | 317.41052 | NaN |
| Fnbp4 | KLQDAAEQLK | -589 | -598 | 1 | 0.006788754 | 10 | NaN | NaN | NaN | NaN | 1581.6261 | NaN |
| Dnpep | NINENFGPNTEIHLVPILAT | -172 | -191 | 0 | 0.002329279 | 20 | NaN | NaN | NaN | NaN | 224.53143 | 225.03783 |
| Dnpep | NDSPCGTTIGPILASR | -407 | -422 | 0 | 0 | 16 | NaN | NaN | NaN | NaN | 23607.44 | NaN |
| Poldip2 | LENLDSDVVQLR | -269 | -280 | 0 | 0 | 12 | NaN | NaN | NaN | NaN | 712.4889 | 585.02014 |
| Pdpk1 | LEYHFPEK | -300 | -307 | 0 | 0.002595029 | 8 | NaN | NaN | NaN | NaN | 899.02344 | 966.7972 |
| Pgm1 | FFGNLMDASK | -361 | -370 | 0 | 0.00025015 | 10 | NaN | NaN | NaN | NaN | 407.68805 | NaN |
| Nomo1 | SSIDSEPALVLGPLK | -678 | -692 | 0 | 0 | 15 | NaN | NaN | NaN | NaN | 8796.294 | 879.9824 |
| Nomo1 | VQVVVPEAETR | -456 | -466 | 0 | 0.002595029 | 11 | NaN | NaN | NaN | NaN | 10926.916 | NaN |
| Nomo1 | FFHFPPLLR | -1089 | -1097 | 0 | 0.000394361 | 9 | NaN | NaN | NaN | NaN | 1052.5099 | NaN |
| Bacc1 | FGDDLNHISCVIK | -57 | -69 | 0 | 0 | 13 | NaN | NaN | NaN | NaN | 3745.61 | 3561.251 |
| Naf1 | FGGSGQGSAAPQPPEDR | -15 | -31 | 0 | 0.001023236 | 17 | NaN | NaN | NaN | NaN | 774.4296 | NaN |
| Nudt19 | FGLGPEPPR | -74 | -82 | 0 | 0.000431655 | 9 | NaN | NaN | NaN | NaN | 505.3195 | NaN |
| Pofut1 | FGNQADHFLGSL | -49 | -60 | 0 | 0 | 12 | NaN | NaN | NaN | NaN | 401.8917 | 360.29358 |
| Scfd1 | QAPSEVDLEQYK | -463 | -474 | 0 | 0.00025015 | 12 | NaN | NaN | NaN | NaN | 2027.1101 | 1919.9635 |
| Scfd1 | FVMPTEENIDR | -88 | -98 | 0 | 0.00383302 | 11 | NaN | NaN | NaN | NaN | 708.9214 | NaN |
| Mrps30 | SVDFYWLR | -199 | -206 | 0 | 0.005176406 | 8 | NaN | NaN | NaN | NaN | 116.56357 | NaN |
| Nrdc | TDPFDTPAFNK | -428 | -438 | 0 | 0.003620005 | 11 | NaN | NaN | NaN | NaN | 461.91617 | NaN |
| Cox6a1 | TKPFPWGDGNHTLF | -83 | -96 | 1 | 0.005605646 | 14 | NaN | NaN | NaN | NaN | 1314.67 | 827.2817 |
| Lpxn | TWHPEHFFCS | -229 | -238 | 0 | 0.004983786 | 10 | NaN | NaN | NaN | NaN | 364.53723 | NaN |
| Exosc8 | SQFIADVVDNSQVIK | -106 | -120 | 0 | 0 | 15 | NaN | NaN | NaN | NaN | 1002.7936 | NaN |
| Slfn1 | SQSPDVSVGKPATIATN | -104 | -120 | 1 | 0.003745449 | 17 | NaN | NaN | NaN | NaN | 724.3363 | NaN |
| Prdx1 | PGSDTIKPDVNK | -179 | -190 | 1 | 0 | 12 | NaN | NaN | NaN | NaN | 2364.2942 | 2113.5605 |
| Prdx1 | GGLGPMNIPLISDPK | -95 | -109 | 0 | 0 | 15 | NaN | NaN | NaN | NaN | 1220.0558 | 1251.7689 |
| Prdx1 | KLNCQVIGASVDSHFCHL | -68 | -85 | 1 | 0.000550307 | 18 | NaN | NaN | NaN | NaN | 875.51105 | NaN |
| Atad1 | SEESHDEDEIRPVQQQDLHR | -319 | -338 | 1 | 0.002566359 | 20 | NaN | NaN | NaN | NaN | 482.13617 | 292.98462 |
| Nfu1 | LQGSCTSCPSSIITLK | -206 | -221 | 0 | 0.002780285 | 16 | NaN | NaN | NaN | NaN | 1434.7478 | NaN |
| Nfu1 | PGKPVLETR | -74 | -82 | 1 | 0.001776473 | 9 | NaN | NaN | NaN | NaN | 606.68286 | NaN |
| Cops8 | SEPAPVPPIPNEQQLAR | -183 | -199 | 0 | 0.00025015 | 17 | NaN | NaN | NaN | NaN | 575.3199 | NaN |
| Sdr39u1 | FIPVPSTVVR | -242 | -251 | 0 | 0 | 10 | NaN | NaN | NaN | NaN | 695.889 | 665.6887 |
| Paf1 | FITYPFDQNR | -48 | -57 | 0 | 0.00122419 | 10 | NaN | NaN | NaN | NaN | 416.7064 | NaN |
| Ubr5 | LLCDSAVLQPYLR | -1085 | -1097 | 0 | 0.003620005 | 13 | NaN | NaN | NaN | NaN | 967.30316 | NaN |
| Ubr5 | IGHLLPEEQVYLN | -1354 | -1366 | 0 | 0.001776473 | 13 | NaN | NaN | NaN | NaN | 727.68976 | NaN |
| Idh2 | LNEHFLNTTDFLDTIK | -427 | -442 | 0 | 0.00266297 | 16 | NaN | NaN | NaN | NaN | 757.99506 | 960.89825 |
| Dazap1 | NHNVQGFHPYR | -395 | -405 | 0 | 0 | 11 | NaN | NaN | NaN | NaN | 656.9492 | NaN |
| Afg3l2 | GSLDEDTSLPEGLQDWNK | -763 | -780 | 0 | 0 | 18 | NaN | NaN | NaN | NaN | 848.46497 | 852.7625 |
| Afg3l2 | TVAYHEAGHAVAG | -569 | -581 | 0 | 0.00122419 | 13 | NaN | NaN | NaN | NaN | 911.15985 | NaN |
| Hnrnpd | KIFVGGLSPDTPEEK | -183 | -197 | 1 | 0.005394768 | 15 | NaN | NaN | NaN | NaN | 708.8781 | NaN |
| Polr1a | FKEEVNHY | -872 | -879 | 1 | 0.001392869 | 8 | NaN | NaN | NaN | NaN | 2909.9937 | NaN |
| Polr1a | SLPCFEPYEFTPR | -941 | -953 | 0 | 0.00025015 | 13 | NaN | NaN | NaN | NaN | 785.3579 | NaN |
| Mon2 | FLDSDKPQWLR | -341 | -351 | 1 | 0 | 11 | NaN | NaN | NaN | NaN | 902.23157 | 152.4095 |
| Rars2 | FLEDVLNEVQSR | -392 | -403 | 0 | 0 | 12 | NaN | NaN | NaN | NaN | 924.8044 | NaN |
| Rars2 | SDYQFSWDR | -443 | -451 | 0 | 0.002502116 | 9 | NaN | NaN | NaN | NaN | 260.1059 | NaN |
| Med23 | FLTEALLPVVK | -1230 | -1240 | 0 | 0 | 11 | NaN | NaN | NaN | NaN | 378.88315 | 327.77664 |
| Uchl3 | MHFESGSTLK | -111 | -120 | 0 | 0 | 10 | NaN | NaN | NaN | NaN | 785.8983 | 1375.4038 |
| Smarce1 | FLESTDSFNNELKR | -259 | -272 | 1 | 0 | 14 | NaN | NaN | NaN | NaN | 399.50348 | 2124.9822 |
| Mettl1 | HFEEHPLFER | -204 | -213 | 0 | 0.00025015 | 10 | NaN | NaN | NaN | NaN | 807.229 | 409.7225 |
| Mettl1 | HSGAQVEFADIGCGY | -67 | -81 | 0 | 0 | 15 | NaN | NaN | NaN | NaN | 504.26712 | 504.33734 |

| | | | | | | | | | | | | |
|---|---|---|---|---|---|---|---|---|---|---|---|---|
| Pold2 | YIHPDDELVLEDELQR | -126 | -141 | 0 | 0.005605646 | 16 | NaN | NaN | NaN | NaN | 763.5968 | 598.5424 |
| Pold2 | HISPTAPDTLGCYPF | -373 | -387 | 0 | 0.00149795 | 15 | NaN | NaN | NaN | NaN | 736.0258 | 461.3603 |
| Snrpd3 | VLHEAEGHIVTCETNTGEVYR | -9 | -29 | 0 | 0.00383302 | 21 | NaN | NaN | NaN | NaN | 332.57715 | NaN |
| Sympk | FLIPVLNGLEK | -890 | -900 | 0 | 0.000627943 | 11 | NaN | NaN | NaN | NaN | 454.10223 | NaN |
| Nup35 | FLPGFLMGDLPAPVTPQPR | -34 | -52 | 0 | 0 | 19 | NaN | NaN | NaN | NaN | 550.0606 | NaN |
| Gar1 | KFYIDPYK | -154 | -161 | 1 | 0.00025015 | 8 | NaN | NaN | NaN | NaN | 122.759125 | 674.04425 |
| Gar1 | RFLPRPPGEK | -167 | -176 | 2 | 0.005605646 | 10 | NaN | NaN | NaN | NaN | 440.65747 | NaN |
| Man2b1 | HVLVIENK | -604 | -611 | 0 | 0.007880195 | 8 | NaN | NaN | NaN | NaN | 978.64703 | NaN |
| Man2b1 | GVSEPLLETDTGDKVR | -825 | -840 | 1 | 0.00025015 | 16 | NaN | NaN | NaN | NaN | 915.96027 | NaN |
| Stat3 | KLEELQQK | -283 | -290 | 1 | 0.001463897 | 8 | NaN | NaN | NaN | NaN | 846.4699 | 855.79736 |
| Stat3 | TLTDEELADWKR | -234 | -245 | 1 | 0.00051306 | 12 | NaN | NaN | NaN | NaN | 2668.483 | NaN |
| Sf3b4 | VEFLSEEDADYAIK | -59 | -72 | 0 | 0 | 14 | NaN | NaN | NaN | NaN | 630.9367 | 483.2674 |
| Gcc1 | FLTLPDSLGR | -733 | -742 | 0 | 0.000947459 | 10 | NaN | NaN | NaN | NaN | 344.29343 | NaN |
| Cdk11b | FLTYYPGR | -692 | -699 | 0 | 0.00149795 | 8 | NaN | NaN | NaN | NaN | 356.3146 | NaN |
| Ssr1 | YPQDYQFYIQ | -126 | -135 | 0 | 0.00111006 | 10 | NaN | NaN | NaN | NaN | 218.13327 | 176.00717 |
| Rpl26 | FNAPSHIR | -19 | -26 | 0 | 0.008890469 | 8 | NaN | NaN | NaN | NaN | 531.00116 | 2377.2014 |
| Gmeb1 | FNDQLISPK | -113 | -121 | 0 | 0.00111006 | 9 | NaN | NaN | NaN | NaN | 595.4214 | NaN |
| Clptm1 | VAGIFPCPTFK | -448 | -458 | 0 | 0 | 11 | NaN | NaN | NaN | NaN | 2528.832 | 2368.5986 |
| Clptm1 | GPSPQDQSGPGGAPR | -76 | -90 | 0 | 0 | 15 | NaN | NaN | NaN | NaN | 1776.1593 | 1178.8721 |
| Nhlrc2 | FSDKDGLLIVGVH | -107 | -119 | 1 | 0.00025015 | 13 | NaN | NaN | NaN | NaN | 1299.727 | 1512.4443 |
| Nhlrc2 | FNEPGGLCIGESGR | -530 | -543 | 0 | 0.005112649 | 14 | NaN | NaN | NaN | NaN | 491.20398 | NaN |
| Cisd1 | FPFCDGAHIK | -80 | -89 | 0 | 0 | 10 | NaN | NaN | NaN | NaN | 974.7523 | NaN |
| Ints14 | FPLPFPFPSK | -142 | -151 | 0 | 0.00025015 | 10 | NaN | NaN | NaN | NaN | 181.6888 | 138.58272 |
| Ints14 | IDIADISSPPVLSR | -257 | -270 | 0 | 0.00078661 | 14 | NaN | NaN | NaN | NaN | 308.02173 | NaN |
| Ncstn | LENIDSFVELGQVALR | -355 | -370 | 0 | 0.00111006 | 16 | NaN | NaN | NaN | NaN | 303.48166 | NaN |
| Derl1 | FQIWRPF | -50 | -56 | 1 | 0.003681885 | 7 | NaN | NaN | NaN | NaN | 417.07373 | NaN |
| Cab39 | LLSAEFLEQHYDR | -197 | -209 | 0 | 0 | 13 | NaN | NaN | NaN | NaN | 3964.9983 | NaN |
| Cab39 | LLHSENYVTK | -217 | -226 | 0 | 0.00025015 | 10 | NaN | NaN | NaN | NaN | 1458.3228 | NaN |
| Ufd1 | TTGDVIAINYNEK | -153 | -165 | 0 | 0.007523243 | 13 | NaN | NaN | NaN | NaN | 1194.01 | NaN |
| Chchd4 | FSCFHYSTEDIK | -72 | -83 | 0 | 0 | 12 | NaN | NaN | NaN | NaN | 386.7982 | 418.97427 |
| Prps2 | KFSNQETSVEIGESVR | -34 | -49 | 1 | 0.008129024 | 16 | NaN | NaN | NaN | NaN | 546.05725 | 6870.5205 |
| Bri3bp | FSQSVSSLFGEDNVR | -59 | -73 | 0 | 0 | 15 | NaN | NaN | NaN | NaN | 647.6169 | 572.5205 |
| Tomm70 | WKQDLDK | -526 | -532 | 1 | 0.001291103 | 7 | NaN | NaN | NaN | NaN | 733.1895 | 649.04956 |
| Tomm70 | FSSFTDDIISQPMLK | -264 | -278 | 0 | 0 | 15 | NaN | NaN | NaN | NaN | 983.8091 | NaN |
| Trip12 | SSAVVVDAIPVFLEK | -521 | -535 | 0 | 0.005242018 | 15 | NaN | NaN | NaN | NaN | 2184.4578 | NaN |
| Trip12 | WALNEGVCR | -1861 | -1869 | 0 | 0.005459848 | 9 | NaN | NaN | NaN | NaN | 841.04364 | NaN |
| Nmd3 | FSVEIVPICK | -242 | -251 | 0 | 0 | 10 | NaN | NaN | NaN | NaN | 484.29196 | NaN |
| Nmd3 | VPDVVLIK | -398 | -405 | 0 | 0.006788754 | 8 | NaN | NaN | NaN | NaN | 332.3221 | NaN |
| Ncaph | FTNTQITEHYSTCIK | -97 | -111 | 0 | 0.001463897 | 15 | NaN | NaN | NaN | NaN | 850.13947 | NaN |
| Acot7 | FTYVSLNQEGK | -331 | -341 | 0 | 0.00294445 | 11 | NaN | NaN | NaN | NaN | 1426.5968 | 1263.062 |
| Rheb | RVISYEEGK | -127 | -135 | 1 | 0.000907206 | 9 | NaN | NaN | NaN | NaN | 5463.2437 | 5213.75 |
| Rheb | FVEGQFVDSYDPTIENTFTK | -26 | -45 | 0 | 0.00204437 | 20 | NaN | NaN | NaN | NaN | 379.3517 | 358.9575 |
| Rbm39 | TGIDLGTTGR | -347 | -356 | 0 | 0 | 10 | NaN | NaN | NaN | NaN | 1700.299 | NaN |
| Tapbp | YPAGASAPHCEMSR | -85 | -98 | 0 | 0 | 14 | NaN | NaN | NaN | NaN | 759.4809 | 1059.6652 |
| Tapbp | HFFPAEGLEVK | -322 | -332 | 0 | 0 | 11 | NaN | NaN | NaN | NaN | 203.76392 | NaN |
| H2bc3 | PEPSKSAPAPK | -2 | -12 | 1 | 0.00025015 | 11 | NaN | NaN | NaN | NaN | 869.24805 | NaN |
| Spart | FVNPAGEVSAPSYPGYLR | -232 | -249 | 0 | 0.000394361 | 18 | NaN | NaN | NaN | NaN | 959.0941 | NaN |
| Ndufb6 | IFPGDTILETGEVIPPMR | -104 | -121 | 0 | 0 | 18 | NaN | NaN | NaN | NaN | 261.56537 | 617.63574 |
| Txndc5 | NEEPATPEPEAEPPR | -155 | -169 | 0 | 0.00025015 | 15 | NaN | NaN | NaN | NaN | 396.81323 | 435.24243 |
| Snrpc | FYCDYCDTYLTHDSPSVR | -4 | -21 | 0 | 0 | 18 | NaN | NaN | NaN | NaN | 231.36292 | NaN |
| Csnk2a2 | WENFIHSENR | -282 | -291 | 0 | 0 | 10 | NaN | NaN | NaN | NaN | 190.10422 | NaN |
| Glt8d1 | FYLPILVPSAK | -155 | -165 | 0 | 0.000230707 | 11 | NaN | NaN | NaN | NaN | 600.12384 | NaN |
| Bcap31 | RLVTLISQQ | -118 | -126 | 1 | 0.001868731 | 9 | NaN | NaN | NaN | NaN | 697.6149 | 560.29706 |
| Bcap31 | GAAEDGDKLDIGNTEMK | -160 | -176 | 1 | 0.004255025 | 17 | NaN | NaN | NaN | NaN | 1231.5159 | NaN |
| Rps26 | LHYCVSCAIH | -71 | -80 | 0 | 0.00025015 | 10 | NaN | NaN | NaN | NaN | 721.6605 | 409.65747 |
| Svil | SKEEEDTPGSSLEK | -56 | -69 | 1 | 0.000550307 | 14 | NaN | NaN | NaN | NaN | 971.6987 | 952.2042 |
| Aacs | LNYAENLLR | -99 | -107 | 0 | 0.004857192 | 9 | NaN | NaN | NaN | NaN | 472.7701 | 369.90695 |
| Lman1 | GAGTPGQPGQVSQQELDTVVK | -382 | -402 | 0 | 0.001291103 | 21 | NaN | NaN | NaN | NaN | 546.98047 | 702.8358 |
| Lman1 | GPHLVQSDGTVPF | -62 | -74 | 0 | 0.005605646 | 13 | NaN | NaN | NaN | NaN | 500.6822 | 1021.08 |
| Nmt2 | GAIEPDKDNIR | -169 | -179 | 1 | 0.000431655 | 11 | NaN | NaN | NaN | NaN | 1458.4365 | NaN |
| Adrm1 | SAAVTPSSSTSSAR | -213 | -226 | 0 | 0 | 14 | NaN | NaN | NaN | NaN | 689.4536 | NaN |
| Erp29 | GALPLDTVTFY | -39 | -49 | 0 | 0.00450079 | 11 | NaN | NaN | NaN | NaN | 161.44902 | NaN |
| Rcc1 | SWVGFSGGQHH | -295 | -305 | 0 | 0.00111006 | 11 | NaN | NaN | NaN | NaN | 6993.9614 | NaN |
| Polr2h | RIEGDETSTEAATR | -98 | -111 | 1 | 0.001055699 | 14 | NaN | NaN | NaN | NaN | 798.843 | 2541.9888 |
| Eif1 | HSFDPFADASK | -8 | -18 | 0 | 0.000712251 | 11 | NaN | NaN | NaN | NaN | 5223.8247 | 4744.891 |
| Il12rb1 | KWHLPAQELTETR | -353 | -365 | 1 | 0.000550307 | 13 | NaN | NaN | NaN | NaN | 667.3294 | NaN |
| Txndc12 | GFGDHIHWR | -31 | -39 | 0 | 0.000230707 | 9 | NaN | NaN | NaN | NaN | 7312.7427 | NaN |
| Adi1 | GDMITLPAGIYHR | -122 | -134 | 0 | 0.005394768 | 13 | NaN | NaN | NaN | NaN | 4969.6943 | NaN |
| Naa40 | GDPLEAFPVFK | -36 | -46 | 0 | 0 | 11 | NaN | NaN | NaN | NaN | 791.3317 | 851.3055 |
| Plaa | GDQFSSHLINLLNPK | -635 | -649 | 0 | 0.00078661 | 15 | NaN | NaN | NaN | NaN | 1366.0592 | 1414.1217 |
| Plaa | LSCSLPGHELDVR | -10 | -22 | 0 | 0 | 13 | NaN | NaN | NaN | NaN | 1228.7689 | 868.1369 |
| Mrps9 | GEDPETFAEEDIDR | -93 | -106 | 0 | 0.003966045 | 14 | NaN | NaN | NaN | NaN | 357.81335 | NaN |
| Mrps9 | GVDYLIYFPITQDR | -291 | -304 | 0 | 0.00025015 | 14 | NaN | NaN | NaN | NaN | 341.74036 | NaN |
| Chd1 | GEDVPQPEDEEFETIER | -258 | -274 | 0 | 0.000907206 | 17 | NaN | NaN | NaN | NaN | 417.09677 | 159.3656 |
| Sod2 | HHAAYVNNLNATEEK | -54 | -68 | 0 | 0 | 15 | NaN | NaN | NaN | NaN | 492.29813 | 510.57764 |
| Lims1 | NDPYHPDHFNCANCGK | -151 | -166 | 0 | 0.000712251 | 16 | NaN | NaN | NaN | NaN | 245.97415 | NaN |
| Vamp8 | NKTEDLEATSEHFK | -46 | -59 | 1 | 0 | 14 | NaN | NaN | NaN | NaN | 1096.5282 | 788.37695 |
| Fn1 | SVNIPDLLPGR | -773 | -783 | 0 | 0.00025015 | 11 | NaN | NaN | NaN | NaN | 744.7224 | NaN |

| | | | | | | | | | | | | |
|---|---|---|---|---|---|---|---|---|---|---|---|---|
| Man2a1 | GFDCQFSSR | -1078 | -1086 | 0 | 0.009242859 | 9 | NaN | NaN | NaN | NaN | 400.58768 | NaN |
| Abcf1 | LEQGDDTAAEKLEK | -393 | -406 | 1 | 0 | 14 | NaN | NaN | NaN | NaN | 678.66095 | 448.2257 |
| Cbx5 | HAYPEDAENK | -175 | -184 | 0 | 0.00025015 | 10 | NaN | NaN | NaN | NaN | 722.458 | NaN |
| Pde12 | GGAACEATGPEPVATCEPVVK | -103 | -123 | 0 | 0.00025015 | 21 | NaN | NaN | NaN | NaN | 419.2057 | 641.6988 |
| Klhdc4 | SRKEEEDLEAL | -29 | -39 | 2 | 0.001776473 | 11 | NaN | NaN | NaN | NaN | 1446.8737 | NaN |
| Prep | SNPLLIHVDTK | -667 | -677 | 0 | 0.00025015 | 11 | NaN | NaN | NaN | NaN | 921.6931 | NaN |
| Prep | VLVPEHEKDVLEWVAC | -328 | -343 | 1 | 0.003102268 | 16 | NaN | NaN | NaN | NaN | 771.2324 | NaN |
| Prep | GLSASGSDWVTIK | -142 | -154 | 0 | 0.000431655 | 13 | NaN | NaN | NaN | NaN | 469.61728 | NaN |
| Prep | VLVPEHEKDVLEWVA | -328 | -342 | 1 | 0.00644558 | 15 | NaN | NaN | NaN | NaN | 296.8696 | NaN |
| Prep | IDFTDPDESKWK | -316 | -327 | 1 | 0 | 12 | NaN | NaN | NaN | NaN | 195.6086 | NaN |
| Ptprcap | LGAALWGHTCR | -70 | -80 | 0 | 0 | 11 | NaN | NaN | NaN | NaN | 1380.0746 | 1458.2588 |
| B4galnt1 | SCSDVVVDHASK | -475 | -486 | 0 | 0 | 12 | NaN | NaN | NaN | NaN | 1168.5962 | 221.71832 |
| B4galnt1 | VGSCSDVVVDHASK | -473 | -486 | 0 | 0.00149795 | 14 | NaN | NaN | NaN | NaN | 1001.34393 | NaN |
| Plpbp | SFGHDLSQGPNPDFQR | -185 | -200 | 0 | 0.00266297 | 16 | NaN | NaN | NaN | NaN | 1521.6462 | NaN |
| Zmpste24 | HYSHPPLLER | -456 | -465 | 0 | 0.00142999 | 10 | NaN | NaN | NaN | NaN | 1000.66437 | 54.14399 |
| Zmpste24 | WHYSHPPLLER | -455 | -465 | 0 | 0 | 11 | NaN | NaN | NaN | NaN | 893.699 | 112.90439 |
| Zmpste24 | GHELGHWK | -334 | -341 | 0 | 0 | 8 | NaN | NaN | NaN | NaN | 1321.7489 | NaN |
| Pex11b | NACDLFIPLDK | -214 | -224 | 0 | 0 | 11 | NaN | NaN | NaN | NaN | 600.58 | NaN |
| Rpap3 | GHWDDVFLDSTQR | -403 | -415 | 0 | 0.002972566 | 13 | NaN | NaN | NaN | NaN | 1080.0546 | NaN |
| Actbl2 | GIHETTFNSIM | -274 | -284 | 0 | 0 | 11 | NaN | NaN | NaN | NaN | 90.93809 | 1388.1487 |
| Fyn | TEDDLSFHKGEK | -97 | -108 | 1 | 0 | 12 | NaN | NaN | NaN | NaN | 1738.914 | NaN |
| Yme1l1 | NFPEALDNALIRPGR | -425 | -439 | 1 | 0.001743443 | 15 | NaN | NaN | NaN | NaN | 821.88043 | 837.874 |
| Nsf | KLFADAEEEQR | -304 | -314 | 1 | 0 | 11 | NaN | NaN | NaN | NaN | 1084.3593 | 1539.6382 |
| Nsf | GVHDTVVNQLLSK | -345 | -357 | 0 | 0.001291103 | 13 | NaN | NaN | NaN | NaN | 1129.841 | NaN |
| Pcbp2 | GVTIPYRPKPS | -173 | -183 | 2 | 0.001092529 | 11 | NaN | NaN | NaN | NaN | 909.0425 | 1007.4852 |
| Isoc1 | LFAEEWGQYVDLPK | -71 | -84 | 0 | 0 | 14 | NaN | NaN | NaN | NaN | 181.71239 | NaN |
| Psma4 | SDANVLTNELR | -81 | -91 | 0 | 0 | 11 | NaN | NaN | NaN | NaN | 873.94214 | 937.1395 |
| Qtrt1 | GITTEQLDSLGCR | -56 | -68 | 0 | 0.002199523 | 13 | NaN | NaN | NaN | NaN | 439.32593 | NaN |
| Psma6 | SIDFKPSEIEVGVVTVENPK | -207 | -226 | 1 | 0 | 20 | NaN | NaN | NaN | NaN | 240.87595 | 512.634 |
| Rpl21 | VYNVTQHAVGIIVNK | -64 | -78 | 0 | 0 | 15 | NaN | NaN | NaN | NaN | 1081.9952 | 675.7601 |
| Spc25 | GKEQESEELTAK | -93 | -104 | 1 | 0.001291103 | 12 | NaN | NaN | NaN | NaN | 1199.9481 | NaN |
| Smarca2 | ISPIQKPQGLDPVEIL | -333 | -348 | 1 | 0 | 16 | NaN | NaN | NaN | NaN | 158.02554 | 239.3167 |
| Abcf3 | GKLEEIEADKAPAR | -293 | -306 | 2 | 0.000907206 | 14 | NaN | NaN | NaN | NaN | 363.9698 | 217.65213 |
| Abcf3 | GKLEEIEADKAPAR | -293 | -306 | 2 | 0.000907206 | 14 | NaN | NaN | NaN | NaN | 363.9698 | 217.65213 |
| Ppil1 | GKQFEDELHPDLK | -79 | -91 | 1 | 0 | 13 | NaN | NaN | NaN | NaN | 963.8638 | 972.0036 |
| Ppil1 | GKQFEDELHPDLK | -79 | -91 | 1 | 0 | 13 | NaN | NaN | NaN | NaN | 963.8638 | 972.0036 |
| Acot2 | LVELEVLDGHEPDGGQR | -144 | -160 | 0 | 0.00025015 | 17 | NaN | NaN | NaN | NaN | 720.262 | 606.6144 |
| Malsu1 | GLEDDTSSLTPVEFK | -212 | -226 | 0 | 0.00111006 | 15 | NaN | NaN | NaN | NaN | 580.7445 | NaN |
| Dpm3 | TFHDCEDAAR | -63 | -72 | 0 | 0.000591124 | 10 | NaN | NaN | NaN | NaN | 585.7085 | 510.9111 |
| Ddx18 | GLEQGYVVCPSEK | -389 | -401 | 0 | 0 | 13 | NaN | NaN | NaN | NaN | 5509.3984 | 2489.5403 |
| Rbm12b2;Rbn | GLPFIAGPVDIR | (9);(9) | (20);(20) | 0 | 0 | 12 | NaN | NaN | NaN | NaN | 379.781 | NaN |
| Rad51 | GLSGSDVLDNVAYAR | -179 | -193 | 0 | 0.000627943 | 15 | NaN | NaN | NaN | NaN | 1026.0657 | NaN |
| Lgals3 | RGNDVAFHFNPR | -165 | -176 | 1 | 0.003069674 | 12 | NaN | NaN | NaN | NaN | 615.33655 | NaN |
| Cmtm7 | GPGQPSEGLLDR | -19 | -30 | 0 | 0 | 12 | NaN | NaN | NaN | NaN | 758.2904 | 940.95105 |
| Cmtm7 | SHGSGLVR | -2 | -9 | 0 | 0.002780285 | 8 | NaN | NaN | NaN | NaN | 672.5614 | NaN |
| Rtn4 | SAAPVPPAAAPLLDFSSDSVPPAPR | -65 | -89 | 0 | 0.00111006 | 25 | NaN | NaN | NaN | NaN | 297.29337 | 434.86908 |
| Rtn4 | SADSPPRPPPAFK | -13 | -25 | 1 | 0 | 13 | NaN | NaN | NaN | NaN | 796.70636 | NaN |
| Ube2r2 | GPPNTLYEGGYFK | -47 | -59 | 0 | 0.000431655 | 13 | NaN | NaN | NaN | NaN | 1175.9386 | 883.09015 |
| Cpsf6 | TPLSEAEFEEIMNR | -407 | -420 | 0 | 0 | 14 | NaN | NaN | NaN | NaN | 378.12106 | 262.6117 |
| Cpsf6 | GPPPTDPYGRPPPYDR | -377 | -392 | 1 | 0.002265552 | 16 | NaN | NaN | NaN | NaN | 1138.3854 | 327.00723 |
| Memo1 | GPSHHVPLSR | -78 | -87 | 0 | 0 | 10 | NaN | NaN | NaN | NaN | 108.736664 | 653.66254 |
| Ppm1b | GPTEQLVSPEPEVYEIV | -214 | -230 | 0 | 0.00078661 | 17 | NaN | NaN | NaN | NaN | 418.72968 | NaN |
| Cmtr1 | GQENLQEILSK | -370 | -380 | 0 | 0 | 11 | NaN | NaN | NaN | NaN | 1575.0969 | NaN |
| Asah1 | IYDVLSTKPVLNK | -353 | -365 | 1 | 0.001291103 | 13 | NaN | NaN | NaN | NaN | 70.21393 | 548.92725 |
| Fen1 | LDPSKYPVPENWLHK | -261 | -275 | 1 | 0 | 15 | NaN | NaN | NaN | NaN | 919.1631 | 982.598 |
| Fen1 | LIADVAPSAIR | -9 | -19 | 0 | 0 | 11 | NaN | NaN | NaN | NaN | 922.0449 | NaN |
| Prrc2a | LKAEPAAPPVTPAAPALPPVVPK | -502 | -524 | 1 | 0.000431655 | 23 | NaN | NaN | NaN | NaN | 224.67976 | 619.0987 |
| Prrc2a | SLEIQKPAVAPR | -30 | -41 | 1 | 0.006100831 | 12 | NaN | NaN | NaN | NaN | 296.06427 | NaN |
| Ccnd3 | GSSSQGPSQTSTPTDVTAIHL | -272 | -292 | 0 | 0.001636916 | 21 | NaN | NaN | NaN | NaN | 539.8624 | 544.8566 |
| Nup107 | VLEENQEHYHIVQK | -505 | -518 | 0 | 0.000989247 | 14 | NaN | NaN | NaN | NaN | 774.514 | 785.6859 |
| Nup107 | GSTRPLVTELDPDAPIR | -315 | -331 | 1 | 0.000431655 | 17 | NaN | NaN | NaN | NaN | 498.24634 | NaN |
| Med18 | GSVLDHSLESLIHR | -25 | -38 | 0 | 0.00025015 | 14 | NaN | NaN | NaN | NaN | 549.55115 | 575.61694 |
| Mix23 | GSVNCEEFAEFQELLK | -6 | -21 | 0 | 0 | 16 | NaN | NaN | NaN | NaN | 645.6642 | 661.7015 |
| Mapk3 | HENVIGIR | -98 | -105 | 0 | 0.00078661 | 8 | NaN | NaN | NaN | NaN | 1479.5259 | 1380.1056 |
| Cwf19l1 | GTESLDEPVPAHSFSPK | -115 | -131 | 0 | 0.002068213 | 17 | NaN | NaN | NaN | NaN | 1576.5886 | 2131.0938 |
| Padi2 | GTQLWTDVYSAAPAGAK | -30 | -46 | 0 | 0.00025015 | 17 | NaN | NaN | NaN | NaN | 1317.4694 | NaN |
| Snrpd1 | GVDVSMNTHLK | -31 | -41 | 0 | 0 | 11 | NaN | NaN | NaN | NaN | 831.17883 | 628.37115 |
| Gatd3 | IFAPDVPQMHVIDHTK | -77 | -92 | 0 | 0 | 16 | NaN | NaN | NaN | NaN | 661.7896 | 681.5856 |
| Pmpca | LVGAEPAWGAPGTVDVDR | -270 | -287 | 0 | 0.000591124 | 18 | NaN | NaN | NaN | NaN | 403.33014 | NaN |
| Amdhd2 | GVHLEGPFISR | -139 | -149 | 0 | 0 | 11 | NaN | NaN | NaN | NaN | 1908.2317 | NaN |
| Nbeal2 | LLFSGGHWDGSLR | -2461 | -2473 | 0 | 0.00078661 | 13 | NaN | NaN | NaN | NaN | 397.1545 | 408.6367 |
| Septin2 | ILDEIEEH | -199 | -206 | 0 | 0.00383302 | 8 | NaN | NaN | NaN | NaN | 1484.1995 | NaN |
| Septin2 | QQPTQFINPETPGYVGFA | -4 | -21 | 0 | 0.003037016 | 18 | NaN | NaN | NaN | NaN | 459.68817 | NaN |
| Pik3cd | HEYLYGNYPLCHF | -247 | -259 | 0 | 0.00025015 | 13 | NaN | NaN | NaN | NaN | 289.87616 | NaN |
| Rel | GYNPYVEIIEQPR | -5 | -17 | 0 | 0.001743443 | 13 | NaN | NaN | NaN | NaN | 860.77704 | 1109.6105 |
| Ap1g1 | KLVPQLVR | -214 | -221 | 1 | 0.00025015 | 8 | NaN | NaN | NaN | NaN | 395.8267 | 436.5011 |
| Vdac1 | NTDNTLGTEITVEDQLAR | -89 | -106 | 0 | 0.002845514 | 18 | NaN | NaN | NaN | NaN | 972.95953 | 1062.7618 |
| Vdac1 | TFDSSFSPNTGK | -111 | -122 | 0 | 0.00025015 | 12 | NaN | NaN | NaN | NaN | 528.8984 | 556.8093 |

| | | | | | | | | | | | |
|---|---|---|---|---|---|---|---|---|---|---|---|
| Vdac1 | NVNAGGHKL | -280 | -288 | 1 | 0.00078661 | 9 | NaN | NaN | NaN | NaN | 1029.7218 | NaN |
| FAM120A | VAQSIEDHHQEVIGF | -159 | -173 | 0 | 0.001291103 | 15 | NaN | NaN | NaN | NaN | 437.12314 | NaN |
| Ca2 | HDPALQPLLISYDK | -40 | -53 | 0 | 0.000431655 | 14 | NaN | NaN | NaN | NaN | 385.60974 | 618.32776 |
| S100a11 | HDSFIQTSQK | -87 | -96 | 0 | 0.00025015 | 10 | NaN | NaN | NaN | NaN | 4037.3533 | 1546.5771 |
| Rpusd2 | IDGIAEAAPQHLDTPEK | -477 | -493 | 0 | 0.001868731 | 17 | NaN | NaN | NaN | NaN | 824.5132 | NaN |
| Apeh | VGWWHEPFR | -261 | -269 | 0 | 0 | 9 | NaN | NaN | NaN | NaN | 1026.7452 | NaN |
| Mycbp | HHLGAATPENPEIELL | -50 | -65 | 0 | 0.00111006 | 16 | NaN | NaN | NaN | NaN | 632.13293 | NaN |
| Dap3 | HIPDAHLWVK | -147 | -156 | 0 | 0.007880195 | 10 | NaN | NaN | NaN | NaN | 387.2486 | NaN |
| Swi5 | HISLLHEYNDIK | -47 | -58 | 0 | 0.001023236 | 12 | NaN | NaN | NaN | NaN | 947.3717 | NaN |
| Stim2 | HITVEDLWK | -118 | -126 | 0 | 0 | 9 | NaN | NaN | NaN | NaN | 658.9039 | NaN |
| Dhx36 | SVSHIVLDEIHER | -320 | -332 | 0 | 0 | 13 | NaN | NaN | NaN | NaN | 715.40625 | NaN |
| Cdc73 | NVHLSHPVYVR | -66 | -76 | 0 | 0 | 11 | NaN | NaN | NaN | NaN | 758.5432 | 697.8556 |
| Suclg1 | ITEGIPQQDMVR | -143 | -154 | 0 | 0.000230707 | 12 | NaN | NaN | NaN | NaN | 1125.5934 | NaN |
| Cbr4 | HLGPVNFLVN | -73 | -82 | 0 | 0.008129024 | 10 | NaN | NaN | NaN | NaN | 1344.1803 | NaN |
| Utp4 | SNQGSLHIVHLSEGSFK | -456 | -472 | 0 | 0.00078661 | 17 | NaN | NaN | NaN | NaN | 1072.2208 | 293.22083 |
| Thoc6 | HWQLSGELK | -282 | -290 | 0 | 0.00025015 | 9 | NaN | NaN | NaN | NaN | 1263.5728 | NaN |
| Nup85 | RIPLNTEQK | -441 | -449 | 1 | 0.000550307 | 9 | NaN | NaN | NaN | NaN | 879.5344 | NaN |
| Nipbl | SEEDSSYKPVK | -1952 | -1962 | 1 | 0.00122419 | 11 | NaN | NaN | NaN | NaN | 2242.8965 | NaN |
| Nipbl | KYEESLADSDNK | -1978 | -1989 | 1 | 0.009388464 | 12 | NaN | NaN | NaN | NaN | 744.2502 | NaN |
| Rassf2 | MDYTHQPALIPCGQDK | -1 | -16 | 0 | 0.005522803 | 16 | NaN | NaN | NaN | NaN | 314.8487 | NaN |
| Supt7l | HSDPESDFYR | -129 | -138 | 0 | 0.000394361 | 10 | NaN | NaN | NaN | NaN | 487.49564 | 467.4215 |
| M6pr | HTLAANFNPVSEER | -149 | -162 | 0 | 0 | 14 | NaN | NaN | NaN | NaN | 369.03915 | 1346.4022 |
| M6pr | SFESTVGQGSDTYSYIF | -60 | -76 | 0 | 0.000947459 | 17 | NaN | NaN | NaN | NaN | 271.3817 | NaN |
| Ak2 | LVSDEMVVELIEK | -73 | -85 | 0 | 0 | 13 | NaN | NaN | NaN | NaN | 972.9667 | 377.65274 |
| Eif2d | HTYQDHLWR | -170 | -178 | 0 | 0 | 9 | NaN | NaN | NaN | NaN | 423.0975 | 472.49246 |
| Eif2d | SSPPAIAPLDPTDSCEEK | -183 | -200 | 0 | 0.00025015 | 18 | NaN | NaN | NaN | NaN | 1634.1323 | NaN |
| Tbc1d1;Tbc1d | HVDHFGFICR | (320);(401) | (329);(410) | 0 | 0.00025015 | 10 | NaN | NaN | NaN | NaN | 449.5752 | NaN |
| Smarcd2 | IITDVIGNPEEER | -481 | -493 | 0 | 0 | 13 | NaN | NaN | NaN | NaN | 877.2362 | 708.1925 |
| Nup50 | LQQESPFSFHGN | -230 | -241 | 0 | 0 | 12 | NaN | NaN | NaN | NaN | 1574.9423 | 326.7044 |
| Nup50 | KAEDTSEKVEFTAEK | -242 | -256 | 2 | 0.001776473 | 15 | NaN | NaN | NaN | NaN | 2799.1904 | NaN |
| Hexb | TVTDLENAYK | -497 | -506 | 0 | 0 | 10 | NaN | NaN | NaN | NaN | 1322.6985 | 853.0377 |
| Hexb | ILEIISSLK | -370 | -378 | 0 | 0 | 9 | NaN | NaN | NaN | NaN | 252.61913 | NaN |
| Nanp | HWEEAIQETK | -75 | -84 | 0 | 0.000431655 | 10 | NaN | NaN | NaN | NaN | 301.966 | NaN |
| Enoph1 | STEGDILELIDGHFDTK | -164 | -180 | 0 | 0.00122419 | 17 | NaN | NaN | NaN | NaN | 581.5893 | NaN |
| Tecr | RLLETLFVHR | -141 | -150 | 1 | 0 | 10 | NaN | NaN | NaN | NaN | 981.1499 | 204.71013 |
| Il2rb | HYIEPYLEFEAR | -160 | -171 | 0 | 0.001190916 | 12 | NaN | NaN | NaN | NaN | 253.20946 | NaN |
| Wdhd1 | NQVLHGDPLPL | -604 | -614 | 0 | 0.00122419 | 11 | NaN | NaN | NaN | NaN | 716.4617 | NaN |
| Cox6b1 | SLCPVSWVSAWDDR | -63 | -76 | 0 | 0 | 14 | NaN | NaN | NaN | NaN | 340.51013 | NaN |
| Vdac2 | IAIEDQICQGLK | -97 | -108 | 0 | 0.00025015 | 12 | NaN | NaN | NaN | NaN | 5313.639 | NaN |
| Tex30 | ICISYPLHHPK | -128 | -138 | 0 | 0.000627943 | 11 | NaN | NaN | NaN | NaN | 423.7132 | NaN |
| Dnm1l | ICYIFHETFGR | -366 | -376 | 0 | 0 | 11 | NaN | NaN | NaN | NaN | 787.93414 | NaN |
| Atp2b4 | LDIQEGNPELVSK | -1147 | -1159 | 0 | 0.007819401 | 13 | NaN | NaN | NaN | NaN | 6007.405 | NaN |
| Bpnt2 | IDPLDATQEYTEDLRK | -171 | -186 | 1 | 0.00025015 | 16 | NaN | NaN | NaN | NaN | 804.4521 | NaN |
| Opa1 | SDYDASYNTQDHLPR | -276 | -290 | 0 | 0.00450079 | 15 | NaN | NaN | NaN | NaN | 608.4535 | NaN |
| Ndufa10 | QDDWTFHYL | -286 | -294 | 0 | 0 | 9 | NaN | NaN | NaN | NaN | 1254.6487 | 819.9207 |
| Napa | SKIEEACEIYAR | -36 | -47 | 1 | 0 | 12 | NaN | NaN | NaN | NaN | 351.16327 | 2053.7893 |
| Rpl38 | KIEEIKDFL | -4 | -12 | 2 | 0.000394361 | 9 | NaN | NaN | NaN | NaN | 603.49 | NaN |
| Rpl38 | IEEIKDFLL | -5 | -13 | 1 | 0.007880195 | 9 | NaN | NaN | NaN | NaN | 556.9607 | NaN |
| Nudcd1 | PDNINIK | -307 | -313 | 0 | 0.003360691 | 7 | NaN | NaN | NaN | NaN | 353.65643 | NaN |
| Pno1 | IFTPIVEHL | -91 | -99 | 0 | 0.00078661 | 9 | NaN | NaN | NaN | NaN | 10438.093 | NaN |
| Eed | KIHFPDFSTR | -293 | -302 | 1 | 0.000947459 | 10 | NaN | NaN | NaN | NaN | 855.03375 | 825.83954 |
| Eed | SGTNTERPDTPTNTPNAPGR | -46 | -65 | 1 | 0 | 20 | NaN | NaN | NaN | NaN | 338.10352 | NaN |
| Sdk1 | IHLEGNR | -97 | -103 | 0 | 0.000712251 | 7 | NaN | NaN | NaN | NaN | 1511.9595 | 1353.4034 |
| Hsd17b10 | LVAGEMGQNEPDQGGQR | -131 | -147 | 0 | 0 | 17 | NaN | NaN | NaN | NaN | 731.89166 | 813.91693 |
| Idi1 | SVFLFNTENK | -55 | -64 | 0 | 0 | 10 | NaN | NaN | NaN | NaN | 656.9603 | NaN |
| Ints4 | LYPESIVPIPSSNEEIR | -263 | -279 | 0 | 0.003490217 | 17 | NaN | NaN | NaN | NaN | 509.02045 | NaN |
| Ttf2 | IILDEAHNVK | -710 | -719 | 0 | 0.00111006 | 10 | NaN | NaN | NaN | NaN | 1242.1133 | NaN |
| LRWD1 | LCPVATCPDDFLLA | -450 | -463 | 0 | 0.006677519 | 14 | NaN | NaN | NaN | NaN | 308.9304 | NaN |
| Rab18 | KILIIGESGVGK | -10 | -21 | 1 | 0.00122419 | 12 | NaN | NaN | NaN | NaN | 1183.5417 | NaN |
| Rpl19 | ILMEHIHK | -137 | -144 | 0 | 0.008284062 | 8 | NaN | NaN | NaN | NaN | 605.2616 | NaN |
| Cd82 | YIHSEDYSK | -254 | -262 | 0 | 0 | 9 | NaN | NaN | NaN | NaN | 375.4782 | 437.99353 |
| Gpx4 | ILAFPCNQFGR | -97 | -107 | 0 | 0 | 11 | NaN | NaN | NaN | NaN | 513.95404 | 439.64606 |
| Elp3 | SPSDLIELVAR | -341 | -351 | 0 | 0.00025015 | 11 | NaN | NaN | NaN | NaN | 2786.217 | 2574.8809 |
| Dennd2d | TSEQINEHVSGPFVQ | -367 | -381 | 0 | 0.001708008 | 15 | NaN | NaN | NaN | NaN | 642.62244 | NaN |
| Mmut | ILFDGIPLEK | -170 | -179 | 0 | 0.00204437 | 10 | NaN | NaN | NaN | NaN | 311.63684 | NaN |
| Rcn1 | ILPQDYDHAQAEAR | -267 | -280 | 0 | 0 | 14 | NaN | NaN | NaN | NaN | 690.3505 | 877.8535 |
| Ppp3cb | ILYPSTLFLLR | -147 | -157 | 0 | 0 | 11 | NaN | NaN | NaN | NaN | 429.28918 | 600.9505 |
| Edf1 | INEKPQVIADYESGR | -99 | -113 | 1 | 0 | 15 | NaN | NaN | NaN | NaN | 658.4696 | 751.0433 |
| Ptcd3 | WNDILDLLK | -312 | -320 | 0 | 0.001636916 | 9 | NaN | NaN | NaN | NaN | 208.06215 | NaN |
| Mpo | IPCFLAGDMR | -370 | -379 | 0 | 0 | 10 | NaN | NaN | NaN | NaN | 667.3922 | 717.6789 |
| Mpo | TSFVDASGVYGSEDPLAR | -308 | -325 | 0 | 0 | 18 | NaN | NaN | NaN | NaN | 618.24054 | 612.19885 |
| Lrrc8c | YLDLSYNDIR | -687 | -696 | 0 | 0 | 10 | NaN | NaN | NaN | NaN | 617.6798 | 346.95972 |
| Stk4 | RQPILDAIEAK | -470 | -480 | 1 | 0.000431655 | 11 | NaN | NaN | NaN | NaN | 1952.9376 | 1644.7003 |
| Cactin | IQEDNRLELQ | -258 | -267 | 1 | 0.001708008 | 10 | NaN | NaN | NaN | NaN | 798.5867 | NaN |
| Psme3 | NSHDGLDGPTYK | -74 | -85 | 0 | 0.003102268 | 12 | NaN | NaN | NaN | NaN | 1260.8872 | NaN |
| Rab2a | KEEGEAFAR | -131 | -139 | 1 | 0.005363933 | 9 | NaN | NaN | NaN | NaN | 1207.2311 | 1311.0498 |
| Gga1 | IQSPQEWEAIQAL | -55 | -67 | 0 | 0.004224523 | 13 | NaN | NaN | NaN | NaN | 1134.9001 | NaN |
| Hsd17b4 | ISDEDWDIIHR | -111 | -121 | 0 | 0 | 11 | NaN | NaN | NaN | NaN | 1355.6041 | NaN |

| | | | | | | | | | | | | |
|---|---|---|---|---|---|---|---|---|---|---|---|---|
| Cnpy4 | ISDGQEEADDEEEEEEEEITK | -213 | -233 | 0 | 0.007987481 | 21 | NaN | NaN | NaN | NaN | 132.01523 | NaN |
| Znf638 | IYDPEIPTDEVK | -228 | -239 | 0 | 0.000550307 | 12 | NaN | NaN | NaN | NaN | 1408.7255 | NaN |
| Znf638 | SKEEPLFTF | -1525 | -1533 | 1 | 0.002910424 | 9 | NaN | NaN | NaN | NaN | 304.8954 | NaN |
| Rbm15 | ISHLSGSGSGDER | -203 | -215 | 0 | 0.00111006 | 13 | NaN | NaN | NaN | NaN | 1519.3236 | NaN |
| Ylpm1 | ITDESGHLAER | -1368 | -1378 | 0 | 0 | 11 | NaN | NaN | NaN | NaN | 1463.7291 | 1906.2816 |
| Ylpm1 | STFDADHAGQR | -915 | -925 | 0 | 0 | 11 | NaN | NaN | NaN | NaN | 1332.6116 | 1314.5083 |
| Utp15 | RNDSDPVPEHVPAELPEEK | -496 | -514 | 1 | 0.006624942 | 19 | NaN | NaN | NaN | NaN | 652.80273 | NaN |
| Rps24 | MVIDVLHPGK | -23 | -32 | 0 | 0 | 10 | NaN | NaN | NaN | NaN | 1642.1045 | 1148.5676 |
| Uqcrq | TWGNQEFEQSK | -61 | -71 | 0 | 0 | 11 | NaN | NaN | NaN | NaN | 1110.9635 | NaN |
| Tnfrsf8 | KADTVIVGSVK | -424 | -434 | 1 | 0.009897819 | 11 | NaN | NaN | NaN | NaN | 1600.2083 | NaN |
| Nedd4 | VDVPLYPLPTENPR | -165 | -178 | 0 | 0.00025015 | 14 | NaN | NaN | NaN | NaN | 809.5434 | 958.00055 |
| Nedd4 | WILENDPTELDLR | -674 | -686 | 0 | 0.002068213 | 13 | NaN | NaN | NaN | NaN | 626.12476 | NaN |
| Nedd4 | KDILGASDPYVR | -93 | -104 | 1 | 0 | 12 | NaN | NaN | NaN | NaN | 515.9148 | NaN |
| Hsdl2 | KDSLSDEVVR | -390 | -399 | 1 | 0.003133457 | 10 | NaN | NaN | NaN | NaN | 1257.1055 | NaN |
| Mlec | KEEEEEEEEYDEGSNLKR | -229 | -246 | 2 | 0.00025015 | 18 | NaN | NaN | NaN | NaN | 296.29138 | 340.45425 |
| Mlec | KEEEEEEEEYDEGS | -229 | -242 | 1 | 0.002780285 | 14 | NaN | NaN | NaN | NaN | 242.24532 | 445.5721 |
| Ppil3 | VIDGLETLDELEKLPVNEK | -121 | -139 | 1 | 0 | 19 | NaN | NaN | NaN | NaN | 259.53415 | 272.7411 |
| Dnajb11 | KQYDTYGEEGLKDGHQS | -84 | -100 | 2 | 0.008680443 | 17 | NaN | NaN | NaN | NaN | 378.98224 | 435.4383 |
| Smyd5 | KGETIFIERPLVAAQ | -44 | -58 | 2 | 0.00111006 | 15 | NaN | NaN | NaN | NaN | 340.64255 | 432.7095 |
| Trip4 | KGQFIEDLITK | -60 | -70 | 1 | 0.00025015 | 11 | NaN | NaN | NaN | NaN | 328.98203 | NaN |
| Prkcb | LDNVMLDSEGHIK | -469 | -481 | 0 | 0.001291103 | 13 | NaN | NaN | NaN | NaN | 261.1194 | NaN |
| Ppip5k2 | KIPDIYDCIK | -728 | -737 | 1 | 0 | 10 | NaN | NaN | NaN | NaN | 501.57596 | NaN |
| Sgpp1 | KITIPLACK | -374 | -382 | 1 | 0.005979575 | 9 | NaN | NaN | NaN | NaN | 2622.5461 | NaN |
| Rbm34 | KLDDDDDDEEESVSQTK | -81 | -97 | 1 | 0.001291103 | 17 | NaN | NaN | NaN | NaN | 284.62683 | 233.8461 |
| C7orf50 | VQEASPELSPEER | -44 | -56 | 0 | 0 | 13 | NaN | NaN | NaN | NaN | 976.16125 | 904.8353 |
| Nxf1 | NELKPEQIEQLK | -201 | -212 | 1 | 0.00025015 | 12 | NaN | NaN | NaN | NaN | 2061.1738 | NaN |
| Morc3 | SAVAELIDNAYDPDVNAK | -31 | -48 | 0 | 0.00025015 | 18 | NaN | NaN | NaN | NaN | 1998.0807 | NaN |
| Def6 | YILDKVEEGAFVK | -75 | -87 | 1 | 0.00025015 | 13 | NaN | NaN | NaN | NaN | 633.0708 | 956.9006 |
| Def6 | LLEEEEEKLK | -462 | -471 | 1 | 0.00025015 | 10 | NaN | NaN | NaN | NaN | 665.8253 | NaN |
| Krr1 | KNIHPIYNIK | -210 | -219 | 1 | 0 | 10 | NaN | NaN | NaN | NaN | 386.51495 | 346.5354 |
| Bsg | RKPDQTLDEDDPGAAPL | -353 | -369 | 2 | 0.000471409 | 17 | NaN | NaN | NaN | NaN | 946.7392 | 1135.5265 |
| Bsg | RKPDQTLDEDDPGAAP | -353 | -368 | 2 | 0.001291103 | 16 | NaN | NaN | NaN | NaN | 613.59845 | 2974.865 |
| Iqgap2 | LGIAPQIQDLLGK | -159 | -171 | 0 | 0 | 13 | NaN | NaN | NaN | NaN | 172.4076 | NaN |
| Raly | VFIGNLNTAVVK | -23 | -34 | 0 | 0 | 12 | NaN | NaN | NaN | NaN | 963.42303 | NaN |
| Tpd52l2 | SQDINLNSPNK | -5 | -15 | 0 | 0.00025015 | 11 | NaN | NaN | NaN | NaN | 7681.475 | NaN |
| Eci1 | KVGVVDEVVPEDQVHSK | -206 | -222 | 1 | 0.00111006 | 17 | NaN | NaN | NaN | NaN | 583.43536 | 616.9167 |
| Eci1 | NVQELWLR | -114 | -121 | 0 | 0.00025015 | 8 | NaN | NaN | NaN | NaN | 2085.2534 | 2166.609 |
| Mtmr1 | KVYDPVSEYK | -232 | -241 | 1 | 0.00111006 | 10 | NaN | NaN | NaN | NaN | 3082.6467 | NaN |
| Phpt1 | KWAEYHADIYDK | -47 | -58 | 1 | 0 | 12 | NaN | NaN | NaN | NaN | 2606.8994 | 4306.395 |
| Zmat2 | KWDKDEYEK | -15 | -23 | 2 | 0.00025015 | 9 | NaN | NaN | NaN | NaN | 833.3603 | NaN |
| Sf3b5 | YTIHSQLEHLQSK | -5 | -17 | 0 | 0.001610812 | 13 | NaN | NaN | NaN | NaN | 3910.3518 | NaN |
| Sf3b5 | YIGTGHADTTK | -18 | -28 | 0 | 0.002780285 | 11 | NaN | NaN | NaN | NaN | 587.9661 | NaN |
| Nipsnap3b | KYDNFAHR | -89 | -96 | 1 | 0.007343498 | 8 | NaN | NaN | NaN | NaN | 972.56537 | NaN |
| Cks2 | THLMSEEEWR | -35 | -44 | 0 | 0 | 10 | NaN | NaN | NaN | NaN | 3068.9028 | NaN |
| Pxn | YAHQQPPSPLPVYSSS | -76 | -91 | 0 | 0.008129024 | 16 | NaN | NaN | NaN | NaN | 1267.1919 | NaN |
| Usp24 | TDHWSIPYK | -134 | -142 | 0 | 0.002199523 | 9 | NaN | NaN | NaN | NaN | 2501.642 | 2928.7114 |
| Usp24 | LADSSPSNLQIITK | -1641 | -1654 | 0 | 0 | 14 | NaN | NaN | NaN | NaN | 445.907 | 538.3293 |
| Ube2g2 | LAEPNDESGANVDASK | -127 | -142 | 0 | 0.003490217 | 16 | NaN | NaN | NaN | NaN | 409.60474 | NaN |
| Scly | LANNETGVIMPVSEISR | -183 | -199 | 0 | 0.004658491 | 17 | NaN | NaN | NaN | NaN | 460.17795 | NaN |
| Rpia | WSEVNTAIK | -261 | -269 | 0 | 0.00025015 | 9 | NaN | NaN | NaN | NaN | 1207.3778 | 647.1228 |
| Rpia | LASHTAVENHVK | -77 | -88 | 0 | 0.00635324 | 12 | NaN | NaN | NaN | NaN | 419.7356 | NaN |
| Soat1 | LCAPLFR | -334 | -340 | 0 | 0.00111006 | 7 | NaN | NaN | NaN | NaN | 638.4529 | NaN |
| Kifc1 | LCDLNEELKR | -144 | -153 | 1 | 0 | 10 | NaN | NaN | NaN | NaN | 990.88666 | NaN |
| Ndufs8 | LCEAICPAQAITIEAEPR | -118 | -135 | 0 | 0.001023236 | 18 | NaN | NaN | NaN | NaN | 909.79395 | NaN |
| Pcbd1 | LDHHPEWFNVYNK | -60 | -72 | 0 | 0.000627943 | 13 | NaN | NaN | NaN | NaN | 289.75006 | NaN |
| Rasgrp2 | LDQALVAEHIEK | -418 | -429 | 0 | 0 | 12 | NaN | NaN | NaN | NaN | 662.81854 | NaN |
| Tsg101 | SENNDIDEVIIPTAPLYK | -310 | -327 | 0 | 0.00025015 | 18 | NaN | NaN | NaN | NaN | 248.96094 | NaN |
| Atad2 | TKPVDPDEVPDYVTVIK | -659 | -675 | 1 | 0.000550307 | 17 | NaN | NaN | NaN | NaN | 691.6249 | NaN |
| Vti1b | LEDLQGVPER | -22 | -31 | 0 | 0.001636916 | 10 | NaN | NaN | NaN | NaN | 4842.1714 | NaN |
| Vti1b | TLENEHLNR | -116 | -124 | 0 | 0 | 9 | NaN | NaN | NaN | NaN | 1166.7307 | NaN |
| Rab8a | NIEEHASADVEK | -105 | -116 | 0 | 0 | 12 | NaN | NaN | NaN | NaN | 2085.9045 | 2175.0735 |
| Wdr4 | PVGGEWQAAPDGAVSPR | -321 | -337 | 0 | 0 | 17 | NaN | NaN | NaN | NaN | 1185.8799 | 1279.1249 |
| Ints12 | LELDPIFLK | -7 | -15 | 0 | 0.000230707 | 9 | NaN | NaN | NaN | NaN | 433.78046 | NaN |
| Ube2s | LENYEEYAAR | -138 | -147 | 0 | 0 | 10 | NaN | NaN | NaN | NaN | 81.3711 | NaN |
| Mnat1 | LFEDPTVDKEVEIR | -62 | -75 | 1 | 0 | 14 | NaN | NaN | NaN | NaN | 632.6834 | NaN |
| Pld3 | LTHGVLHTK | -193 | -201 | 0 | 0.000907206 | 9 | NaN | NaN | NaN | NaN | 316.51904 | 276.1912 |
| Atp6v1h | LGESVQDLSSFDEYSSELK | -342 | -360 | 0 | 0.00122419 | 19 | NaN | NaN | NaN | NaN | 222.54382 | NaN |
| Polr1c | NNTSIVQDEILAHR | -96 | -109 | 0 | 0.000471409 | 14 | NaN | NaN | NaN | NaN | 731.3237 | NaN |
| Ctps2 | RVPLLLEEQGVVK | -252 | -264 | 1 | 0.000627943 | 13 | NaN | NaN | NaN | NaN | 900.51196 | 1023.1094 |
| Tmem209 | LGSPDSTSPSTSPTFWNY | -258 | -275 | 0 | 0 | 18 | NaN | NaN | NaN | NaN | 337.26068 | 185.11884 |
| Tmem209 | SQAPCANKDEADLISK | -297 | -312 | 1 | 0.000712251 | 16 | NaN | NaN | NaN | NaN | 1879.6176 | NaN |
| Mtx2 | RIEQHYFEDWGK | -248 | -259 | 1 | 0 | 12 | NaN | NaN | NaN | NaN | 460.89053 | 341.29398 |
| Ppp1r12a | RLASTSDIEEK | -504 | -514 | 1 | 0.003166358 | 11 | NaN | NaN | NaN | NaN | 2184.473 | 2379.0247 |
| Exosc4 | LHEDHLEQVLEAAAQ | -205 | -219 | 0 | 0.007493591 | 15 | NaN | NaN | NaN | NaN | 3976.1987 | NaN |
| Ifrd2 | LHFVEGGECEEETVR | -336 | -350 | 0 | 0.003133457 | 15 | NaN | NaN | NaN | NaN | 827.3115 | NaN |
| Pts | LHSPSLSDEENLR | -25 | -37 | 0 | 0 | 13 | NaN | NaN | NaN | NaN | 1206.9347 | 1222.698 |
| Adar | LIDQSGPPHEPK | -695 | -706 | 0 | 0 | 12 | NaN | NaN | NaN | NaN | 281.47916 | NaN |
| Dhfr | QEFESDTFFPEIDLGK | -141 | -156 | 0 | 0.00025015 | 16 | NaN | NaN | NaN | NaN | 351.12753 | NaN |

| | | | | | | | | | | | | |
|---|---|---|---|---|---|---|---|---|---|---|---|---|
| Atxn3 | LIGEELAHLK | -191 | -200 | 0 | 0 | 10 | NaN | NaN | NaN | NaN | 426.7443 | 409.5014 |
| Iigp1 | TKVDSDITNEADGKPQ | -183 | -198 | 2 | 0.003133457 | 16 | NaN | NaN | NaN | NaN | 823.29987 | 825.84906 |
| Prim1 | LKDDIYIR | -49 | -56 | 1 | 0.00383302 | 8 | NaN | NaN | NaN | NaN | 607.82837 | 891.0492 |
| Rrp1b | LKDSGSPLQFDYK | -282 | -294 | 1 | 0.00025015 | 13 | NaN | NaN | NaN | NaN | 2004.4146 | 2085.7065 |
| Phf23 | LKDSLFDLDGPK | -133 | -144 | 1 | 0.00025015 | 12 | NaN | NaN | NaN | NaN | 856.06995 | NaN |
| Lzic | REELDADEYEETKK | -34 | -47 | 2 | 0.00025015 | 14 | NaN | NaN | NaN | NaN | 2245.1443 | NaN |
| Rabgap1 | LLDPQTNTEIANYPIYK | -193 | -209 | 0 | 0 | 17 | NaN | NaN | NaN | NaN | 188.56812 | NaN |
| Cfap54 | LLIDMLIK | -1690 | -1697 | 0 | 0.002595029 | 8 | NaN | NaN | NaN | NaN | 2034.9727 | NaN |
| Rcl1 | VETKPCGEELK | -341 | -351 | 1 | 0.002595029 | 11 | NaN | NaN | NaN | NaN | 503.1446 | NaN |
| Chd1l | LLLTGTPIQNSLR | -193 | -205 | 0 | 0 | 13 | NaN | NaN | NaN | NaN | 580.7881 | 536.4973 |
| Stat6 | SLPLVVIVHGNQDNNAK | -407 | -423 | 0 | 0.001463897 | 17 | NaN | NaN | NaN | NaN | 1623.1736 | NaN |
| Ndufs2 | TQDIVFGEIDR | -453 | -463 | 0 | 0.000755556 | 11 | NaN | NaN | NaN | NaN | 1614.7534 | 1537.8081 |
| Ehd3 | LLPLEEYYR | -33 | -41 | 0 | 0.000712251 | 9 | NaN | NaN | NaN | NaN | 416.93805 | NaN |
| Cdk9 | VRDPYALDLIDK | -283 | -294 | 1 | 0.00951004 | 12 | NaN | NaN | NaN | NaN | 769.5529 | NaN |
| Cdk9 | QYDSVECPFCDEVTK | -4 | -18 | 0 | 0.00659934 | 15 | NaN | NaN | NaN | NaN | 757.50586 | NaN |
| Cdk9 | WYRPPELLLGER | -193 | -204 | 1 | 0.000907206 | 12 | NaN | NaN | NaN | NaN | 148.28812 | NaN |
| Alg10b | LNIPLPPISR | -428 | -437 | 0 | 0 | 10 | NaN | NaN | NaN | NaN | 514.3658 | 642.32385 |
| Bst2 | VSQALEQQAR | -111 | -120 | 0 | 0.003900536 | 10 | NaN | NaN | NaN | NaN | 511.70248 | NaN |
| Virma | LNSILTQPWR | -978 | -987 | 0 | 0.00025015 | 10 | NaN | NaN | NaN | NaN | 764.171 | NaN |
| Ints8 | LPDIPLR | -650 | -656 | 0 | 0.005605646 | 7 | NaN | NaN | NaN | NaN | 315.17557 | NaN |
| Cul4a | LPDNYTQDTWR | -54 | -64 | 0 | 0.00111006 | 11 | NaN | NaN | NaN | NaN | 1154.9878 | 1160.616 |
| Rnaseh2b | LPEPPASLTNPPSK | -231 | -244 | 0 | 0.001636916 | 14 | NaN | NaN | NaN | NaN | 1400.5752 | NaN |
| Uba7 | LPKPWDPDDAETVVEL | -298 | -313 | 1 | 0.001392869 | 16 | NaN | NaN | NaN | NaN | 2417.2253 | NaN |
| Atxn7l3b | LPLCSLPGDPGDGPQTELQR | -72 | -91 | 0 | 0 | 20 | NaN | NaN | NaN | NaN | 338.60327 | 350.09326 |
| Blvrb | LPSEGPQPAHVVVGDVR | -40 | -56 | 0 | 0.001055699 | 17 | NaN | NaN | NaN | NaN | 1223.0771 | 1224.124 |
| Ergic2 | YEYEVDKDFSSK | -62 | -73 | 1 | 0.008436628 | 12 | NaN | NaN | NaN | NaN | 414.59488 | NaN |
| Plin3 | TPQDAEKDPAKPEQVEAR | -312 | -329 | 2 | 0.002068213 | 18 | NaN | NaN | NaN | NaN | 646.5009 | 655.195 |
| Plin3 | LQESLPILQQPTEK | -103 | -116 | 0 | 0.000431655 | 14 | NaN | NaN | NaN | NaN | 661.3995 | NaN |
| Dcaf8 | VVVWDWVR | -211 | -218 | 0 | 0.000989247 | 8 | NaN | NaN | NaN | NaN | 333.2625 | 420.36118 |
| Dcaf8 | LQHGLEGHTGCVN | -179 | -191 | 0 | 0.002265552 | 13 | NaN | NaN | NaN | NaN | 489.2807 | NaN |
| Nfkbia | LQPQEAPLAAEPWK | -54 | -67 | 0 | 0.00078661 | 14 | NaN | NaN | NaN | NaN | 727.2565 | 873.1128 |
| Ccna2 | LSLHQEDQENVNPEK | -17 | -31 | 0 | 0 | 15 | NaN | NaN | NaN | NaN | 882.8967 | NaN |
| Tsc1 | LSQAPLLPSLLK | -107 | -118 | 0 | 0.001392869 | 12 | NaN | NaN | NaN | NaN | 372.31598 | NaN |
| Mrpl21 | TSLSSPPWQEVVLPDPVEETR | -58 | -78 | 0 | 0.000431655 | 21 | NaN | NaN | NaN | NaN | 201.97801 | 163.26402 |
| Mms19 | LTDPSTQLQLVGIR | -453 | -466 | 0 | 0 | 14 | NaN | NaN | NaN | NaN | 542.1898 | 456.4462 |
| Ints5 | LTGVDPILGHQLSAR | -39 | -53 | 0 | 0.000907206 | 15 | NaN | NaN | NaN | NaN | 560.4626 | 305.1133 |
| Ufsp2 | LVDAGDKPATFVGSR | -326 | -340 | 1 | 0.000947459 | 15 | NaN | NaN | NaN | NaN | 2124.577 | NaN |
| Morf4l2 | LVEDWDLVTR | -133 | -142 | 0 | 0.00201047 | 10 | NaN | NaN | NaN | NaN | 2795.8499 | NaN |
| Cdk4 | LVFEHIDQDLR | -91 | -101 | 0 | 0 | 11 | NaN | NaN | NaN | NaN | 2105.5742 | 3379.3477 |
| Pdlim5 | SVGPPQPSDQDTLVQR | -387 | -402 | 0 | 0.002972566 | 16 | NaN | NaN | NaN | NaN | 937.7616 | NaN |
| Pih1d1 | LVIGGPQQLYHLD | -244 | -256 | 0 | 0 | 13 | NaN | NaN | NaN | NaN | 994.87445 | 1074.217 |
| Snrpe | LYEQVNMR | -35 | -42 | 0 | 0.00051306 | 8 | NaN | NaN | NaN | NaN | 519.99097 | 1605.2272 |
| Snrpe | VWLYEQVNMR | -33 | -42 | 0 | 0.002199523 | 10 | NaN | NaN | NaN | NaN | 579.04425 | NaN |
| Bcap29 | LVTLITQLAK | -120 | -129 | 0 | 0 | 10 | NaN | NaN | NaN | NaN | 488.08618 | 514.58136 |
| Uprt | LVVEEGLNQLPYK | -147 | -159 | 0 | 0.00051306 | 13 | NaN | NaN | NaN | NaN | 420.78516 | NaN |
| Elp5 | LVYHDLFR | -64 | -71 | 0 | 0.001291103 | 8 | NaN | NaN | NaN | NaN | 1918.1494 | 67.58254 |
| H6pd | VVFEKPFGHDHLSAQ | -167 | -181 | 1 | 0 | 15 | NaN | NaN | NaN | NaN | 570.308 | 411.5122 |
| Thoc2 | LYDQCHDTLVQF | -769 | -780 | 0 | 0.003745449 | 12 | NaN | NaN | NaN | NaN | 868.902 | 653.0011 |
| Slamf1 | SWSDEAGTHLLSR | -178 | -190 | 0 | 0 | 13 | NaN | NaN | NaN | NaN | 433.2824 | 1478.5281 |
| Mtco2 | LYMMDEINNPVLTVK | -84 | -98 | 0 | 0.0074703 | 15 | NaN | NaN | NaN | NaN | 760.0033 | NaN |
| Gmppb | LYSGPGIVGNVLVDPSAR | -244 | -261 | 0 | 0.004224523 | 18 | NaN | NaN | NaN | NaN | 2206.6453 | 1203.005 |
| Hlcs | LYVGSGSEEALGR | -165 | -177 | 0 | 0 | 13 | NaN | NaN | NaN | NaN | 255.1249 | NaN |
| Oga | RAPVIWDNIHAN | -273 | -284 | 1 | 0.001055699 | 12 | NaN | NaN | NaN | NaN | 2978.0022 | NaN |
| Rpl31 | NEDEDSPNKLY | -93 | -103 | 1 | 0 | 11 | NaN | NaN | NaN | NaN | 872.6824 | 979.6366 |
| Cyp51a1 | SPIEFLENAYEK | -80 | -91 | 0 | 0 | 12 | NaN | NaN | NaN | NaN | 397.1657 | NaN |
| Atp6v1b2 | TVSGVNGPLVILDHVK | -49 | -64 | 0 | 0.001941082 | 16 | NaN | NaN | NaN | NaN | 411.79953 | NaN |
| Riok1 | TSQDHVDEEVFK | -414 | -425 | 0 | 0 | 12 | NaN | NaN | NaN | NaN | 591.8384 | 416.28677 |
| Exoc4 | NITVIFHPLLR | -496 | -506 | 0 | 0 | 11 | NaN | NaN | NaN | NaN | 503.5852 | NaN |
| Myo5a | NKDTVFEEQIK | -568 | -578 | 1 | 0.00111006 | 11 | NaN | NaN | NaN | NaN | 882.2722 | NaN |
| Krt7 | NKYEEEINR | -172 | -180 | 1 | 0.000712251 | 9 | NaN | NaN | NaN | NaN | 3866.88 | 4063.3618 |
| Mecr | NLELTAVEGSDVHVR | -62 | -76 | 0 | 0 | 15 | NaN | NaN | NaN | NaN | 1461.6995 | 1455.7119 |
| Ggh | NLGGISHAPNAVK | -253 | -265 | 0 | 0 | 13 | NaN | NaN | NaN | NaN | 593.2669 | NaN |
| Slc7a1 | SATFDELIGKPIGEFSR | -133 | -149 | 1 | 0 | 17 | NaN | NaN | NaN | NaN | 937.36334 | 729.9598 |
| Slc7a1 | NLLGLGQQMLR | -5 | -15 | 0 | 0 | 11 | NaN | NaN | NaN | NaN | 827.74744 | NaN |
| Cox7c | PFSVENK | -28 | -34 | 0 | 0.009897819 | 7 | NaN | NaN | NaN | NaN | 319.74094 | NaN |
| Nit2 | TLSPGDSFSTFDTPYCK | -131 | -147 | 0 | 0 | 17 | NaN | NaN | NaN | NaN | 368.3242 | 223.03914 |
| Rfc3 | YHLEVNPSDAGNSDR | -91 | -105 | 0 | 0.005015524 | 15 | NaN | NaN | NaN | NaN | 749.70776 | NaN |
| Tf | NQQEGVCPEGSIDNSPVK | -344 | -361 | 0 | 0.001055699 | 18 | NaN | NaN | NaN | NaN | 817.2477 | NaN |
| Otulinl | NSLFSDAILECK | -224 | -235 | 0 | 0.004199202 | 12 | NaN | NaN | NaN | NaN | 966.2069 | 376.05753 |
| Niban1 | NSQPFVLLPK | -150 | -159 | 0 | 0.00078661 | 10 | NaN | NaN | NaN | NaN | 789.124 | NaN |
| Man1a1 | NTEAHPFPILR | -634 | -644 | 0 | 0.000471409 | 11 | NaN | NaN | NaN | NaN | 733.5441 | NaN |
| Gtf2f2 | RLQIEESSKPVR | -129 | -140 | 2 | 0 | 12 | NaN | NaN | NaN | NaN | 188.82814 | NaN |
| Gfus | NVHINDNVLHS | -90 | -100 | 0 | 0.001023236 | 11 | NaN | NaN | NaN | NaN | 3735.3713 | NaN |
| Htatip2 | NVNQEVVDFEKL | -64 | -75 | 1 | 0.005242018 | 12 | NaN | NaN | NaN | NaN | 364.1034 | 2388.6653 |
| Htatip2 | VDRDYVLK | -107 | -114 | 1 | 0.00025015 | 8 | NaN | NaN | NaN | NaN | 883.8151 | NaN |
| Htatip2 | PGEWLAR | -179 | -185 | 0 | 0.008436628 | 7 | NaN | NaN | NaN | NaN | 337.009 | NaN |
| Pcca | PGFDGVVK | -189 | -196 | 0 | 0.00437931 | 8 | NaN | NaN | NaN | NaN | 644.8321 | 525.64246 |
| Dnajb6 | PGPAPAPVR | -255 | -263 | 0 | 0.006624942 | 9 | NaN | NaN | NaN | NaN | 161.60414 | 250.81111 |

| | | | | | | | | | | | | |
|---|---|---|---|---|---|---|---|---|---|---|---|---|
| Vwf | TIDVPWNVVQEK | -1740 | -1751 | 0 | 0.002068213 | 12 | NaN | NaN | NaN | NaN | 470.3414 | NaN |
| Clpp | VLVHPPQDGEDEPELVQK | -240 | -257 | 0 | 0.000550307 | 18 | NaN | NaN | NaN | NaN | 833.66187 | 1228.1533 |
| Ppp1r10 | STGLELDTPSLVPVK | -210 | -224 | 0 | 0 | 15 | NaN | NaN | NaN | NaN | 4784.395 | 4540.387 |
| Meaf6 | PPQIPDTR | -9 | -16 | 0 | 0.001291103 | 8 | NaN | NaN | NaN | NaN | 441.97702 | NaN |
| Apoa1 | PYLDEFQK | -122 | -129 | 0 | 0 | 8 | NaN | NaN | NaN | NaN | 785.2906 | 774.87476 |
| Apoa1 | QKVQPYLDEFQK | -118 | -129 | 1 | 0.004255025 | 12 | NaN | NaN | NaN | NaN | 476.85913 | NaN |
| Scamp2 | QAQEEFSQGIFSSR | -299 | -312 | 0 | 0.000230707 | 14 | NaN | NaN | NaN | NaN | 2380.8293 | NaN |
| Rmdn3 | QELPDVTNEDSAFQK | -446 | -460 | 0 | 0.008129024 | 15 | NaN | NaN | NaN | NaN | 222.96077 | NaN |
| Snx2 | QFLESSELPR | -260 | -269 | 0 | 0.00025015 | 10 | NaN | NaN | NaN | NaN | 4334.8555 | NaN |
| Gnb1 | QITNNIDPVGR | -32 | -42 | 0 | 0.003620005 | 11 | NaN | NaN | NaN | NaN | 320.20346 | NaN |
| Ptpn2 | QLENINTGETR | -160 | -170 | 0 | 0.002329279 | 11 | NaN | NaN | NaN | NaN | 1523.6307 | NaN |
| Ctc1 | TVSPGPLIPIPVLYPEK | -192 | -208 | 0 | 0.00025015 | 17 | NaN | NaN | NaN | NaN | 175.09747 | 218.92838 |
| Golga2 | VQDNESLSHLNR | -487 | -498 | 0 | 0 | 12 | NaN | NaN | NaN | NaN | 1065.7783 | 1172.3523 |
| Golga2 | QQNQDTLDQLEK | -185 | -196 | 0 | 0.00383302 | 12 | NaN | NaN | NaN | NaN | 1789.8756 | NaN |
| Eif2b4 | QQVIQDYTTPPSEELSR | -235 | -251 | 0 | 0.00025015 | 17 | NaN | NaN | NaN | NaN | 288.08722 | NaN |
| Tiprl | QSSHQDFSFGPWK | -7 | -19 | 0 | 0.002434616 | 13 | NaN | NaN | NaN | NaN | 336.58652 | 297.22998 |
| Vps28 | QVQGSEISSIDEFCR | -83 | -97 | 0 | 0.000550307 | 15 | NaN | NaN | NaN | NaN | 1214.4797 | NaN |
| Vps33b | QVSPIESLR | -390 | -398 | 0 | 0 | 9 | NaN | NaN | NaN | NaN | 907.81354 | NaN |
| Nsd2 | RADDPAEDVDVEDAPR | -531 | -546 | 1 | 0.00201047 | 16 | NaN | NaN | NaN | NaN | 910.16016 | NaN |
| Nsd2 | YAHENDITHFY | -1108 | -1118 | 0 | 0.001636916 | 11 | NaN | NaN | NaN | NaN | 239.33786 | NaN |
| Eif4ebp1 | RAGGEESQFEMDI | -105 | -117 | 1 | 0.00025015 | 13 | NaN | NaN | NaN | NaN | 537.0332 | NaN |
| Cox7b | RAPSFHDK | -29 | -36 | 1 | 0.004983786 | 8 | NaN | NaN | NaN | NaN | 584.5439 | NaN |
| Dglucy | RDLTGPSHAGAY | -137 | -148 | 1 | 0.000712251 | 12 | NaN | NaN | NaN | NaN | 903.8361 | 566.26263 |
| Dglucy | SLAPAFESFCQGNR | -45 | -58 | 0 | 0.00078661 | 14 | NaN | NaN | NaN | NaN | 418.00037 | NaN |
| Plp2 | RHTAAPTDPTDGP | -140 | -152 | 1 | 0 | 13 | NaN | NaN | NaN | NaN | 592.08887 | 932.7008 |
| Rcor1 | VGPQYQAAVPDFDPAK | -101 | -116 | 0 | 0.00122419 | 16 | NaN | NaN | NaN | NaN | 563.5724 | NaN |
| Orai1 | RLQDQLDHR | -284 | -292 | 1 | 0.001392869 | 9 | NaN | NaN | NaN | NaN | 696.44836 | 674.5544 |
| Mrps17 | RLVLDPYLLK | -32 | -41 | 1 | 0.000591124 | 10 | NaN | NaN | NaN | NaN | 214.05829 | 498.658 |
| Prkag1 | RLVVVDEHDVVK | -298 | -309 | 1 | 0.003620005 | 12 | NaN | NaN | NaN | NaN | 328.86072 | NaN |
| Lxn | SVAENCINYQQGTPHK | -15 | -30 | 0 | 0 | 16 | NaN | NaN | NaN | NaN | 718.5244 | 721.0881 |
| Lxn | RPLEAQDIPDNFGNVSPQ | -115 | -132 | 1 | 0.003900536 | 18 | NaN | NaN | NaN | NaN | 543.4196 | NaN |
| Coro2a | SLIEPVSMIVPR | -348 | -359 | 0 | 0 | 12 | NaN | NaN | NaN | NaN | 701.9379 | NaN |
| Mtco1 | WLFSTNHK | -6 | -13 | 0 | 0.003299042 | 8 | NaN | NaN | NaN | NaN | 2085.6929 | NaN |
| Sco2 | SAEQIVESIR | -235 | -244 | 0 | 0 | 10 | NaN | NaN | NaN | NaN | 4648.043 | NaN |
| Rnf114 | YTFPCPYCPEK | -140 | -150 | 0 | 0 | 11 | NaN | NaN | NaN | NaN | 896.9091 | NaN |
| Dnajc15 | SDADIDHTAGR | -26 | -36 | 0 | 0.00111006 | 11 | NaN | NaN | NaN | NaN | 565.6654 | NaN |
| Mettl3 | SDLALTLPTDAVSIR | -87 | -101 | 0 | 0 | 15 | NaN | NaN | NaN | NaN | 2299.1567 | NaN |
| Psd4 | SDLTQPILEDLQR | -357 | -369 | 0 | 0.000298805 | 13 | NaN | NaN | NaN | NaN | 5923.461 | NaN |
| Ykt6 | YTGLDDHLSK | -120 | -129 | 0 | 0 | 10 | NaN | NaN | NaN | NaN | 420.87057 | NaN |
| Atp5if1 | SDSSDSMDTGAGSIR | -27 | -41 | 0 | 0 | 15 | NaN | NaN | NaN | NaN | 324.48022 | NaN |
| Eef1akmt2 | SEGFELFEELPTPK | -204 | -217 | 0 | 0.00025015 | 14 | NaN | NaN | NaN | NaN | 3408.9795 | NaN |
| Copg2 | SSEPVQLTEAETEYFVR | -630 | -646 | 0 | 0.00111006 | 17 | NaN | NaN | NaN | NaN | 533.6056 | NaN |
| Copg2 | SIPLAMAPVFEQK | -578 | -590 | 0 | 0.000627943 | 13 | NaN | NaN | NaN | NaN | 503.67273 | NaN |
| Klc1 | SEQSVAQLEEEKK | -134 | -146 | 1 | 0.00025015 | 13 | NaN | NaN | NaN | NaN | 647.71136 | NaN |
| Rbl1 | TPNCSSEPVKEER | -910 | -922 | 1 | 0.003360691 | 13 | NaN | NaN | NaN | NaN | 1134.1324 | NaN |
| Wdr82 | SFDKGPFATFK | -178 | -188 | 1 | 0 | 11 | NaN | NaN | NaN | NaN | 1183.6073 | NaN |
| Cirbp | SFDTNEQALEQVFSK | -14 | -28 | 0 | 0 | 15 | NaN | NaN | NaN | NaN | 1592.7079 | 1423.198 |
| Tsnax | VYDGFSFIGNTGPYEVSK | -224 | -241 | 0 | 0 | 18 | NaN | NaN | NaN | NaN | 383.9979 | NaN |
| Phf14 | SFVPEEEKHEER | -828 | -839 | 1 | 0 | 12 | NaN | NaN | NaN | NaN | 1372.561 | 1345.4861 |
| Pars2 | SHEFQLPVDIGEDR | -256 | -269 | 0 | 0.000627943 | 14 | NaN | NaN | NaN | NaN | 460.09058 | 594.58765 |
| Riok2 | SHPNAEWYFDR | -254 | -264 | 0 | 0.000712251 | 11 | NaN | NaN | NaN | NaN | 562.4171 | NaN |
| Pbk | SHSPWAVK | -56 | -63 | 0 | 0 | 8 | NaN | NaN | NaN | NaN | 1082.5157 | NaN |
| Lmf2 | SHVANYPFHERPPTYLR | -541 | -557 | 1 | 0.00078661 | 17 | NaN | NaN | NaN | NaN | 1811.1954 | NaN |
| Lsm1 | SIDQFANLVLHQ | -34 | -45 | 0 | 0.00025015 | 12 | NaN | NaN | NaN | NaN | 384.73343 | NaN |
| Tmem254 | YLVDHYHTFLR | -52 | -62 | 0 | 0 | 11 | NaN | NaN | NaN | NaN | 2694.1262 | 4070.0808 |
| Tmem254 | SIPYQSLGPLGPFTK | -37 | -51 | 0 | 0.003003218 | 15 | NaN | NaN | NaN | NaN | 625.77966 | NaN |
| Lrrc58 | SISYTPYDLPGNLLR | -269 | -283 | 0 | 0 | 15 | NaN | NaN | NaN | NaN | 2653.3403 | 166.57079 |
| S100a9 | SITTIIDTFHQYSR | -12 | -25 | 0 | 0.00025015 | 14 | NaN | NaN | NaN | NaN | 1206.4648 | NaN |
| Uxs1 | SKIEEIVEPLR | -53 | -63 | 1 | 0.000431655 | 11 | NaN | NaN | NaN | NaN | 500.96704 | 525.8715 |
| Tmem167a | TGLLGIFWK | -37 | -45 | 0 | 0.00266297 | 9 | NaN | NaN | NaN | NaN | 340.60028 | NaN |
| Tssc4 | SLEDVSEASEQSNR | -163 | -176 | 0 | 0.002068213 | 14 | NaN | NaN | NaN | NaN | 810.6608 | NaN |
| Gpatch4 | SLGDELLGHTDR | -224 | -235 | 0 | 0 | 12 | NaN | NaN | NaN | NaN | 937.0205 | 851.28766 |
| Ptpn7 | SLGTVEPICSVNTPR | -54 | -68 | 0 | 0.001392869 | 15 | NaN | NaN | NaN | NaN | 1177.4908 | 849.8192 |
| Scamp4 | SLPEYPTVPSYSSSGGHWP | -212 | -230 | 0 | 0.00111006 | 19 | NaN | NaN | NaN | NaN | 170.80423 | NaN |
| Hsd11b1 | SLSLFHDDIHSVR | -125 | -137 | 0 | 0 | 13 | NaN | NaN | NaN | NaN | 586.76996 | NaN |
| Vapb | TSPLDDTEVKK | -158 | -168 | 1 | 0.000431655 | 11 | NaN | NaN | NaN | NaN | 1602.9292 | 932.34076 |
| Rbm26 | SLVDDSLLQDDDEEEEDNESR | -988 | -1008 | 0 | 0.007224385 | 21 | NaN | NaN | NaN | NaN | 222.28201 | 201.19362 |
| Gyg1 | SVNCDSQDPTVSHPEFLNL | -230 | -248 | 0 | 0.00025015 | 19 | NaN | NaN | NaN | NaN | 348.19992 | 366.5566 |
| Gyg1 | SNIDDLFER | -109 | -117 | 0 | 0 | 9 | NaN | NaN | NaN | NaN | 738.7746 | NaN |
| Magt1 | SNMEFLFNK | -208 | -216 | 0 | 0.000230707 | 9 | NaN | NaN | NaN | NaN | 218.75104 | NaN |
| Twf1 | WSPDHSHVR | -88 | -96 | 0 | 0.000712251 | 9 | NaN | NaN | NaN | NaN | 1378.1505 | NaN |
| Syvn1 | SPNISWLFHCR | -130 | -140 | 0 | 0 | 11 | NaN | NaN | NaN | NaN | 1822.3302 | 162.8601 |
| Tsfm | YVQPQGVTVVDFVR | -298 | -311 | 0 | 0 | 14 | NaN | NaN | NaN | NaN | 583.8216 | 536.332 |
| Tsfm | SPSLQNLVLGK | -222 | -232 | 0 | 0.004983786 | 11 | NaN | NaN | NaN | NaN | 454.96533 | NaN |
| Mrpl40 | SQELIPIEDFITPVR | -79 | -93 | 0 | 0 | 15 | NaN | NaN | NaN | NaN | 473.93396 | 700.50323 |
| Tmem165 | SQLTTIVLAAR | -252 | -262 | 0 | 0.00025015 | 11 | NaN | NaN | NaN | NaN | 250.1689 | NaN |
| Cuta | SSLVPALTEFVR | -125 | -136 | 0 | 0.00201047 | 12 | NaN | NaN | NaN | NaN | 846.9674 | NaN |
| Znf148 | STADFLDQVTSQK | -701 | -713 | 0 | 0.000431655 | 13 | NaN | NaN | NaN | NaN | 348.4593 | NaN |

| | | | | | | | | | | | | |
|---|---|---|---|---|---|---|---|---|---|---|---|---|
| Lpp | STGEPLGHVPAR | -11 | -22 | 0 | 0 | 12 | NaN | NaN | NaN | NaN | 127.483765 | 1264.0552 |
| Ktn1 | TAEHEAAQQDLQSK | -511 | -524 | 0 | 0.000989247 | 14 | NaN | NaN | NaN | NaN | 769.64166 | NaN |
| Ktn1 | STPQFEELESVLK | -904 | -916 | 0 | 0 | 13 | NaN | NaN | NaN | NaN | 473.4571 | NaN |
| Acot8 | SVLNLEPLDEDLYR | -30 | -43 | 0 | 0.001846793 | 14 | NaN | NaN | NaN | NaN | 935.61975 | NaN |
| Ano6 | SVSDDKLVFVK | -117 | -127 | 1 | 0 | 11 | NaN | NaN | NaN | NaN | 575.79596 | NaN |
| Lgals9 | SVWIICEGHCFK | -305 | -316 | 0 | 0 | 12 | NaN | NaN | NaN | NaN | 790.9172 | 882.38965 |
| Rpap1 | SWSPIGVGPTPSLYK | -617 | -631 | 0 | 0.006260252 | 15 | NaN | NaN | NaN | NaN | 1182.2969 | 402.52853 |
| Rpap1 | TLLVAPGDEELLDR | -484 | -497 | 0 | 0 | 14 | NaN | NaN | NaN | NaN | 1171.5425 | 778.062 |
| Fcho1 | SYFGEHFWGDK | -2 | -12 | 0 | 0.001743443 | 11 | NaN | NaN | NaN | NaN | 278.53888 | NaN |
| Nfrkb | TADKIELGDSDLK | -246 | -258 | 1 | 0.000471409 | 13 | NaN | NaN | NaN | NaN | 822.37244 | NaN |
| Grsf1 | TAFESDKEIELPK | -369 | -381 | 1 | 0 | 13 | NaN | NaN | NaN | NaN | 750.9924 | 242.02426 |
| Rbpj | YLHVEGGNFHAS | -228 | -239 | 0 | 0.002068213 | 12 | NaN | NaN | NaN | NaN | 1794.8156 | 1446.8105 |
| Efemp1 | TCQDINECETTNECR | -331 | -345 | 0 | 0.000591124 | 15 | NaN | NaN | NaN | NaN | 666.4241 | 623.8647 |
| Srbd1 | TDFDKPDFK | -886 | -894 | 1 | 0.00078661 | 9 | NaN | NaN | NaN | NaN | 965.36584 | 456.8617 |
| Adam10 | YDLEECTCASSDGKDDK | -576 | -592 | 1 | 0.000627943 | 17 | NaN | NaN | NaN | NaN | 387.54913 | 337.935 |
| Cd1d1 | TDSVVWLGDLQTHR | -44 | -57 | 0 | 0 | 14 | NaN | NaN | NaN | NaN | 1426.3599 | NaN |
| Atp6v1c1 | TEFWLISAPGEK | -2 | -13 | 0 | 0 | 12 | NaN | NaN | NaN | NaN | 597.0677 | 620.5874 |
| Pcyt1a;Pcyt1b | TEGISTSDIITR | (197);(197) | (208);(208) | 0 | 0.00078661 | 12 | NaN | NaN | NaN | NaN | 1140.7917 | NaN |
| Vps26c | TEVPFEFPLLVK | -89 | -100 | 0 | 0.000627943 | 12 | NaN | NaN | NaN | NaN | 933.7036 | 315.0985 |
| Yy1 | TGDRPYVCPFDGCNK | -378 | -392 | 1 | 0 | 15 | NaN | NaN | NaN | NaN | 412.79883 | 521.96576 |
| Exosc2 | THFHDLPCGA | -184 | -193 | 0 | 0.002168954 | 10 | NaN | NaN | NaN | NaN | 533.6747 | 171.91452 |
| Rock2 | THGSEIINDLQGR | -635 | -647 | 0 | 0.006100831 | 13 | NaN | NaN | NaN | NaN | 1074.097 | NaN |
| Polr2j | TINKEDHTLGNIIK | -34 | -47 | 1 | 0.00025015 | 14 | NaN | NaN | NaN | NaN | 89.86151 | NaN |
| Arpc1a | TQIALSPNNHEVHIY | -21 | -35 | 0 | 0.000869338 | 15 | NaN | NaN | NaN | NaN | 1158.9972 | 1519.6432 |
| Ahctf1 | VTDLPSAVTADQESHK | -1634 | -1649 | 0 | 0.00025015 | 16 | NaN | NaN | NaN | NaN | 660.4173 | NaN |
| Ahctf1 | YTLDLAGGTFPLR | -331 | -343 | 0 | 0 | 13 | NaN | NaN | NaN | NaN | 476.2758 | NaN |
| Ebp | TTNTVPLHPYWPR | -2 | -14 | 0 | 0 | 13 | NaN | NaN | NaN | NaN | 1022.26355 | 957.68506 |
| Macrod1 | TVGPIAVGGQPTASQAAELR | -224 | -242 | 0 | 0.000907206 | 19 | NaN | NaN | NaN | NaN | 287.9344 | 618.28107 |
| Adnp | TWEDVGLWDPSLTK | -52 | -65 | 0 | 0 | 14 | NaN | NaN | NaN | NaN | 281.41736 | 209.9435 |
| Etv6 | TYQEDEPTIASPVGWPR | -442 | -458 | 0 | 0 | 17 | NaN | NaN | NaN | NaN | 926.8387 | NaN |
| Gmds | YYRPTEVDFLQGDCSK | -323 | -338 | 1 | 0 | 16 | NaN | NaN | NaN | NaN | 359.84488 | 832.7385 |
| Lancl2 | VCPLPTTEDPGLPFHPN | -50 | -66 | 0 | 0.000826806 | 17 | NaN | NaN | NaN | NaN | 299.66806 | NaN |
| Qrich1 | VDDIFSDLYYVR | -566 | -577 | 0 | 0 | 12 | NaN | NaN | NaN | NaN | 631.2237 | NaN |
| Acox3 | VDVIAPSDFVLNSPIAK | -641 | -657 | 0 | 0 | 17 | NaN | NaN | NaN | NaN | 359.04752 | 396.78674 |
| Mrfap1 | VEASEESALNHLQGAGGAEPR | -71 | -91 | 0 | 0.000712251 | 21 | NaN | NaN | NaN | NaN | 245.55995 | 668.07587 |
| Ei24 | VFIPVLQSVTAR | -92 | -103 | 0 | 0 | 12 | NaN | NaN | NaN | NaN | 1480.8539 | 45.72984 |
| Mrpl10 | VFPSQVLKPF | -125 | -134 | 1 | 0.001291103 | 10 | NaN | NaN | NaN | NaN | 389.079 | NaN |
| Mrpl30 | VFQPKPEDHEK | -45 | -55 | 1 | 0.000230707 | 11 | NaN | NaN | NaN | NaN | 1077.1066 | 1153.211 |
| Mrpl30 | WHLKPVEQEAK | -149 | -159 | 1 | 0 | 11 | NaN | NaN | NaN | NaN | 421.04208 | 431.6423 |
| Tmem14c | VGISLLSSPHP | -104 | -114 | 0 | 0 | 11 | NaN | NaN | NaN | NaN | 932.8959 | 886.5642 |
| Jak3 | VGLPGAQEEPGLLR | -256 | -269 | 0 | 0 | 14 | NaN | NaN | NaN | NaN | 6951.359 | 6642.653 |
| Gstt2 | VHEYLGWHADNIR | -95 | -107 | 0 | 0.000471409 | 13 | NaN | NaN | NaN | NaN | 752.3972 | NaN |
| Herc4 | WEVFHELPLEK | -973 | -983 | 0 | 0 | 11 | NaN | NaN | NaN | NaN | 645.8693 | 330.62936 |
| Iah1 | VILITPPPLCEAA | -128 | -140 | 0 | 0.000550307 | 13 | NaN | NaN | NaN | NaN | 286.27972 | NaN |
| Rala | VKEDENVPFLLVGNK | -114 | -128 | 1 | 0 | 15 | NaN | NaN | NaN | NaN | 407.83734 | 417.5043 |
| Asf1b | VLENPSPFHSPFR | -11 | -23 | 0 | 0.000230707 | 13 | NaN | NaN | NaN | NaN | 484.60904 | 566.19086 |
| Slc35c1 | YVGVPFYNVGR | -136 | -146 | 0 | 0 | 11 | NaN | NaN | NaN | NaN | 623.2234 | 681.12225 |
| Mcm3ap | VLVLPDVEEQTPESPGR | -1355 | -1371 | 0 | 0.003133457 | 17 | NaN | NaN | NaN | NaN | 916.43604 | NaN |
| Acsf2 | VNPAYQSSELEYVLR | -154 | -168 | 0 | 0 | 15 | NaN | NaN | NaN | NaN | 1512.993 | NaN |
| Golt1b | VPVLGSLLNLPGIR | -112 | -125 | 0 | 0 | 14 | NaN | NaN | NaN | NaN | 189.044 | 151.19049 |
| Sp3 | WHSGERPFIC | -646 | -655 | 1 | 0.004628201 | 10 | NaN | NaN | NaN | NaN | 1577.8591 | NaN |
| Ubfd1 | VVGSTINDVLAVNTPK | -211 | -226 | 0 | 0.002810811 | 16 | NaN | NaN | NaN | NaN | 1266.059 | NaN |
| Mrpl37 | VYEIPGLEPITYEGK | -49 | -63 | 0 | 0.000431655 | 15 | NaN | NaN | NaN | NaN | 1024.7717 | 1075.1619 |
| Glb1 | WLAVLLPK | -162 | -169 | 0 | 0.000550307 | 8 | NaN | NaN | NaN | NaN | 482.69037 | NaN |
| Bles03 | WLIHLSPGFK | -176 | -185 | 0 | 0 | 10 | NaN | NaN | NaN | NaN | 456.91507 | 1055.9742 |
| Strip1 | WTELDTNQHR | -114 | -123 | 0 | 0 | 10 | NaN | NaN | NaN | NaN | 768.146 | NaN |
| Mad1l1 | YAEHQTDCLIFK | -654 | -665 | 0 | 0.000627943 | 12 | NaN | NaN | NaN | NaN | 687.1357 | NaN |
| Kat8 | YFSPFPEDYGK | -193 | -203 | 0 | 0 | 11 | NaN | NaN | NaN | NaN | 4842.714 | NaN |
| Alcam | YLDLSLNPSGEVTK | -331 | -344 | 0 | 0 | 14 | NaN | NaN | NaN | NaN | 761.24084 | 725.482 |
| Ubxn6 | YLDNIHLHPEEEK | -181 | -193 | 0 | 0.00025015 | 13 | NaN | NaN | NaN | NaN | 1045.7822 | NaN |
| Aco1 | YTINIPEDLKPR | -835 | -846 | 1 | 0 | 12 | NaN | NaN | NaN | NaN | 626.4179 | NaN |
| Pdxdc1 | YVDDPNWPGIGVVR | -511 | -524 | 0 | 0.001291103 | 14 | NaN | NaN | NaN | NaN | 1699.6062 | NaN |
| Heatr5b | YVGGIGSGQHLK | -922 | -933 | 0 | 0.00111006 | 12 | NaN | NaN | NaN | NaN | 4838.6143 | NaN |
| Copb1 | LVEKPSPLTLAPHDFANIK | -768 | -786 | 1 | 0 | 19 | NaN | NaN | NaN | NaN | NaN | 1183.9844 |
| Copb1 | LPGLLMTIIR | -55 | -64 | 0 | 0.00025015 | 10 | NaN | NaN | NaN | NaN | NaN | 1879.2733 |
| Copb1 | ILVCDAYR | -99 | -106 | 0 | 0.008222027 | 8 | NaN | NaN | NaN | NaN | NaN | 852.26605 |
| Mcm6 | CTSLFPTIHGNDEVK | -351 | -365 | 0 | 0.00025015 | 15 | NaN | NaN | NaN | NaN | NaN | 785.7179 |
| Mcm6 | FLDFLEEFQGSDGEIK | -30 | -45 | 0 | 0.002566359 | 16 | NaN | NaN | NaN | NaN | NaN | 404.90103 |
| Uqcrc1 | NNGAGYFLEHL | -86 | -96 | 0 | 0.003428289 | 11 | NaN | NaN | NaN | NaN | NaN | 825.5258 |
| Uqcrc1 | ALQSVPETQVSILDNGLR | -41 | -58 | 0 | 0.005459848 | 18 | NaN | NaN | NaN | NaN | NaN | 284.84192 |
| Ddx20 | QEAPETQVK | -713 | -721 | 0 | 0.009631128 | 9 | NaN | NaN | NaN | NaN | NaN | 1550.1205 |
| Blvra | AAAQELWELAAQK | -104 | -116 | 0 | 0.002752526 | 13 | NaN | NaN | NaN | NaN | NaN | 712.0122 |
| Lcp1 | ENLDDEEKLNNAK | -572 | -584 | 1 | 0.00025015 | 13 | NaN | NaN | NaN | NaN | NaN | 2233.5425 |
| Lcp1 | NSASAIGCHVVNIGAEDLK | -199 | -217 | 0 | 0.009031675 | 19 | NaN | NaN | NaN | NaN | NaN | 1853.5543 |
| Hmgb1 | TAADDKQPYEK | -136 | -146 | 1 | 0 | 11 | NaN | NaN | NaN | NaN | NaN | 1246.0656 |
| Nap1l1 | KNVDLLSDMVQEHDEPILK | -176 | -194 | 1 | 0 | 19 | NaN | NaN | NaN | NaN | NaN | 411.48523 |
| Ddrgk1 | AADGEPLHNEEER | -30 | -42 | 0 | 0 | 13 | NaN | NaN | NaN | NaN | NaN | 482.61118 |
| Prpf3 | VLGTEAVQDPTK | -484 | -495 | 0 | 0 | 12 | NaN | NaN | NaN | NaN | NaN | 1015.749 |

| | | | | | | | | | | | | |
|---|---|---|---|---|---|---|---|---|---|---|---|---|
| Entpd5 | AADTHLIDYEK | -335 | -345 | 0 | 0.00025015 | 11 | NaN | NaN | NaN | NaN | NaN | 1871.871 |
| Trmt112 | EVDVLEGTLQCPESGR | -90 | -105 | 0 | 0 | 16 | NaN | NaN | NaN | NaN | NaN | 773.9257 |
| Pkm | CRLDIDSAPITAR | -31 | -43 | 1 | 0.001092529 | 13 | NaN | NaN | NaN | NaN | NaN | 2403.5417 |
| Pkm | EKGADFLVTEVENGGSLGSK | -187 | -206 | 1 | 0.006384543 | 20 | NaN | NaN | NaN | NaN | NaN | 1242.3527 |
| Pkm | LAPITSDPTEAAAVGAV | -401 | -417 | 0 | 0.006624942 | 17 | NaN | NaN | NaN | NaN | NaN | 1077.4664 |
| Pkm | SKIENHEGVR | -269 | -278 | 1 | 0.00676572 | 10 | NaN | NaN | NaN | NaN | NaN | 424.61258 |
| Pkm | GIFPVLCKDAVLN | -468 | -480 | 1 | 0.00111006 | 13 | NaN | NaN | NaN | NaN | NaN | 377.48322 |
| Pdcd6ip | KDNDFIYHDR | -313 | -322 | 1 | 0 | 10 | NaN | NaN | NaN | NaN | NaN | 460.28296 |
| Pdcd6ip | PLDKHEGALETLLR | -57 | -70 | 1 | 0 | 14 | NaN | NaN | NaN | NaN | NaN | 591.5917 |
| Dhx9 | HLENNSHFGSH | -671 | -681 | 0 | 0.00111006 | 11 | NaN | NaN | NaN | NaN | NaN | 3239.559 |
| Dhx9 | ELDALDANDELTPLGR | -840 | -855 | 0 | 0.00111006 | 16 | NaN | NaN | NaN | NaN | NaN | 2749.9497 |
| Gspt1 | AAEFVPSFLR | -82 | -91 | 0 | 0 | 10 | NaN | NaN | NaN | NaN | NaN | 666.8622 |
| Hspa4 | LSGEYEKF | -633 | -640 | 1 | 0.00713295 | 8 | NaN | NaN | NaN | NaN | NaN | 648.6526 |
| Hspa4 | MQVDQEEPHTEEQQQQPQTPAENK | -522 | -545 | 0 | 0.004321969 | 24 | NaN | NaN | NaN | NaN | NaN | 409.5276 |
| Tln1 | ADAEGESDLENSRK | -842 | -855 | 1 | 0 | 14 | NaN | NaN | NaN | NaN | NaN | 744.2823 |
| Tln1 | AFHPEVAPDVR | -2223 | -2233 | 0 | 0 | 11 | NaN | NaN | NaN | NaN | NaN | 711.7796 |
| Tln1 | ISSPVCQEQLVEAGR | -727 | -741 | 0 | 0.00272025 | 15 | NaN | NaN | NaN | NaN | NaN | 676.6392 |
| Stat1 | LGPNAGPDGLIPWTR | -527 | -541 | 0 | 0.000230707 | 15 | NaN | NaN | NaN | NaN | NaN | 370.21353 |
| Smyd5 | AAEQYHQILCPGPSHDPR | -127 | -144 | 0 | 0.000431655 | 18 | NaN | NaN | NaN | NaN | NaN | 516.2465 |
| Nup93 | QMTDVVLTPATDALK | -229 | -243 | 0 | 0.00294445 | 15 | NaN | NaN | NaN | NaN | NaN | 265.4805 |
| Snrnp200 | YEVVDKDSIR | -2021 | -2030 | 1 | 0.001392869 | 10 | NaN | NaN | NaN | NaN | NaN | 1599.8754 |
| Snrnp200 | SDTEEILSK | -1910 | -1918 | 0 | 0.004983786 | 9 | NaN | NaN | NaN | NaN | NaN | 918.14264 |
| Snrnp200 | NSDDNVFVGAPTGSGK | -1341 | -1356 | 0 | 0.00025015 | 16 | NaN | NaN | NaN | NaN | NaN | 760.4721 |
| Tomm22 | EEELEEDDDDELDETLSER | -29 | -47 | 0 | 0.00078661 | 19 | NaN | NaN | NaN | NaN | NaN | 191.64368 |
| Gart | SSSHIVLVISNK | -833 | -844 | 0 | 0.000431655 | 12 | NaN | NaN | NaN | NaN | NaN | 685.44775 |
| Gart | QEEAWVIGSVVACPEDSPR | -757 | -775 | 0 | 0.001868731 | 19 | NaN | NaN | NaN | NaN | NaN | 407.0681 |
| Nolc1 | PAVTPKPSAA | -397 | -406 | 1 | 0.005394768 | 10 | NaN | NaN | NaN | NaN | NaN | 2417.3691 |
| Kpnb1 | GDQENVHPDVMLVQPR | -778 | -793 | 0 | 0 | 16 | NaN | NaN | NaN | NaN | NaN | 724.7852 |
| Kpnb1 | YMEAFKPFLGIGLK | -646 | -659 | 1 | 0 | 14 | NaN | NaN | NaN | NaN | NaN | 642.7343 |
| Kpnb1 | AKDCYPAVQK | -540 | -549 | 1 | 0.003102268 | 10 | NaN | NaN | NaN | NaN | NaN | 2683.8406 |
| Eef1g | YFSWEGTFQHVGK | -416 | -428 | 0 | 0.00025015 | 13 | NaN | NaN | NaN | NaN | NaN | 1153.3679 |
| Eef1a1 | PVNVTTEVK | -282 | -290 | 0 | 0 | 9 | NaN | NaN | NaN | NaN | NaN | 3402.758 |
| Rplp1 | ANVNIGSLICNVGAGGPAPAAGAAPAGGAAPSTAA | -52 | -92 | 0 | 0 | 41 | NaN | NaN | NaN | NaN | NaN | 428.40442 |
| Mogs | VPPEFLVQR | -495 | -503 | 0 | 0.002068213 | 9 | NaN | NaN | NaN | NaN | NaN | 126.934814 |
| Pa2g4 | DHEKAEFEVHEVY | -212 | -224 | 1 | 0.00025015 | 13 | NaN | NaN | NaN | NaN | NaN | 584.10333 |
| Impdh2 | RFGVPVIADGGIQNVGHIA | -356 | -374 | 1 | 0.00111006 | 19 | NaN | NaN | NaN | NaN | NaN | 508.27478 |
| Impdh2 | AAIGTHEDDKYR | -248 | -259 | 1 | 0.002780285 | 12 | NaN | NaN | NaN | NaN | NaN | 1201.457 |
| Impdh2 | REDLVVAPAGVTL | -182 | -194 | 1 | 0.001868731 | 13 | NaN | NaN | NaN | NaN | NaN | 714.6658 |
| Ddx21 | SKTEEATEGMEEAVSS | -38 | -53 | 1 | 0.002566359 | 16 | NaN | NaN | NaN | NaN | NaN | 478.29572 |
| Ddx21 | LLDSVPPTAISHF | -654 | -666 | 0 | 0.001023236 | 13 | NaN | NaN | NaN | NaN | NaN | 1493.7723 |
| Vta1 | QYEDVGTAVQNLQK | -286 | -299 | 0 | 0 | 14 | NaN | NaN | NaN | NaN | NaN | 1074.5245 |
| Arpc4 | VLIEGSINSVR | -61 | -71 | 0 | 0.00111006 | 11 | NaN | NaN | NaN | NaN | NaN | 543.81976 |
| Mcm5 | VSEHSIVQDFTK | -689 | -700 | 0 | 0.002810811 | 12 | NaN | NaN | NaN | NaN | NaN | 1233.5032 |
| Cad | VLGTSPEAIDSAENR | -1034 | -1048 | 0 | 0.00025015 | 15 | NaN | NaN | NaN | NaN | NaN | 1913.1052 |
| Cad | ALWAGYSVER | -816 | -825 | 0 | 0.00719662 | 10 | NaN | NaN | NaN | NaN | NaN | 1249.4757 |
| Cad | AFEPSLDYCVVK | -728 | -739 | 0 | 0.00025015 | 12 | NaN | NaN | NaN | NaN | NaN | 911.8793 |
| Lmnb1 | APATPLSPTR | -18 | -27 | 0 | 0.001392869 | 10 | NaN | NaN | NaN | NaN | NaN | 528.6118 |
| Ftsj3 | KSPQGQNEVPK | -575 | -585 | 1 | 0.00025015 | 11 | NaN | NaN | NaN | NaN | NaN | 812.1383 |
| Tpi1 | NVPAGTEVVCAPPTAYIDFAR | -33 | -53 | 0 | 0 | 21 | NaN | NaN | NaN | NaN | NaN | 1916.9105 |
| Tpi1 | RIIYGGSVTGATCK | -206 | -219 | 1 | 0.009060424 | 14 | NaN | NaN | NaN | NaN | NaN | 438.14706 |
| Vim | FADLSEAANRNNDALR | -295 | -310 | 1 | 0.006677519 | 16 | NaN | NaN | NaN | NaN | NaN | 860.05646 |
| Dek | LKKPPTDEELK | -321 | -331 | 2 | 0 | 11 | NaN | NaN | NaN | NaN | NaN | 1005.57623 |
| Dek | AAPAAEGEDAPVPPSSEKEPEMPGPR | -5 | -30 | 1 | 0.00025015 | 26 | NaN | NaN | NaN | NaN | NaN | 844.859 |
| Mrpl12 | VQGINLVQAK | -156 | -165 | 0 | 0.000471409 | 10 | NaN | NaN | NaN | NaN | NaN | 835.38074 |
| Me2 | HISDTVFLEAAK | -485 | -496 | 0 | 0.001463897 | 12 | NaN | NaN | NaN | NaN | NaN | 11467.962 |
| Me2 | NTLIQFEDFGNHN | -249 | -261 | 0 | 0.000471409 | 13 | NaN | NaN | NaN | NaN | NaN | 429.88922 |
| Nfatc2 | DAGLSPEQPALAL | -132 | -144 | 0 | 0.001291103 | 13 | NaN | NaN | NaN | NaN | NaN | 1122.979 |
| H1-2 | APAAAPPAEKAPAK | -8 | -21 | 1 | 0.000471409 | 14 | NaN | NaN | NaN | NaN | NaN | 1259.2267 |
| Lsm12 | AAPPGEYFSVGSQVSCR | -2 | -18 | 0 | 0.002068213 | 17 | NaN | NaN | NaN | NaN | NaN | 828.2206 |
| Wars1 | SDLEYMGQSPGFYR | -240 | -253 | 0 | 0.002401448 | 14 | NaN | NaN | NaN | NaN | NaN | 557.1654 |
| Wars1 | HLGHLVPFIFTK | -174 | -185 | 0 | 0.000907206 | 12 | NaN | NaN | NaN | NaN | NaN | 535.96564 |
| Wars1 | TFFLEDDDRLEQIR | -408 | -421 | 1 | 0.002068213 | 14 | NaN | NaN | NaN | NaN | NaN | 227.49036 |
| Tcp1 | GLDLVHGKPR | -487 | -496 | 1 | 0.005758933 | 10 | NaN | NaN | NaN | NaN | NaN | 399.89026 |
| Txlna | AAQPGALCDVSEELSR | -61 | -76 | 0 | 0 | 16 | NaN | NaN | NaN | NaN | NaN | 3096.8064 |
| Esd | SGSQIDILIDQGK | -212 | -224 | 0 | 0.007987481 | 13 | NaN | NaN | NaN | NaN | NaN | 4014.0037 |
| Erap1 | NTEGWDFLYSK | -767 | -777 | 0 | 0 | 11 | NaN | NaN | NaN | NaN | NaN | 1304.403 |
| Erap1 | TDVLILPEAVQWIK | -575 | -588 | 0 | 0.00383302 | 14 | NaN | NaN | NaN | NaN | NaN | 303.8438 |
| Dync1h1 | VFEPPPGVK | -4144 | -4152 | 0 | 0.002265552 | 9 | NaN | NaN | NaN | NaN | NaN | 792.97144 |
| Sri | FDDYIACCVK | -156 | -165 | 0 | 0.002566359 | 10 | NaN | NaN | NaN | NaN | NaN | 527.4079 |
| Acat1 | ASEITPITISVK | -229 | -240 | 0 | 0 | 12 | NaN | NaN | NaN | NaN | NaN | 818.0028 |
| Rrp12 | EIDGSEETLDFPR | -555 | -567 | 0 | 0.00025015 | 13 | NaN | NaN | NaN | NaN | NaN | 793.3561 |
| Rrp12 | LYSTIRPYLESK | -773 | -784 | 1 | 0.002780285 | 12 | NaN | NaN | NaN | NaN | NaN | 714.3083 |
| Hsp90ab1 | GIDEDEVTAEEPSAAVPDEIPPLEGDEDASR | -689 | -719 | 0 | 0.00201047 | 31 | NaN | NaN | NaN | NaN | NaN | 634.1938 |
| Hsp90ab1 | DHGEPIGR | -170 | -177 | 0 | 0.002099344 | 8 | NaN | NaN | NaN | NaN | NaN | 633.5802 |
| Hsp90ab1 | LEINPDHPIVETLR | -626 | -639 | 0 | 0.001846793 | 14 | NaN | NaN | NaN | NaN | NaN | 563.97076 |
| Mcm3 | AAVTTDQETGER | -380 | -391 | 0 | 0 | 12 | NaN | NaN | NaN | NaN | NaN | 843.1129 |
| Eprs1 | KEENLAEWYSQVITK | -1020 | -1034 | 1 | 0.00025015 | 15 | NaN | NaN | NaN | NaN | NaN | 775.46423 |
| Eprs1 | SHSNPVSNAQPAGAEKPEAK | -883 | -902 | 1 | 0 | 20 | NaN | NaN | NaN | NaN | NaN | 435.31198 |

| | | | | | | | | | | | | |
|---|---|---|---|---|---|---|---|---|---|---|---|---|
| Eprs1 | EENLAEWYSQVITK | -1021 | -1034 | 0 | 0 | 14 | NaN | NaN | NaN | NaN | NaN | 559.3244 |
| Eprs1 | EELLAIPVVR | -1198 | -1207 | 0 | 0.002265552 | 10 | NaN | NaN | NaN | NaN | NaN | 161.21255 |
| Pdhb | AAWYGHCPGLK | -163 | -173 | 0 | 0.004791731 | 11 | NaN | NaN | NaN | NaN | NaN | 1370.1798 |
| Rpl5 | KQFSQYIK | -221 | -228 | 1 | 0.001868731 | 8 | NaN | NaN | NaN | NaN | NaN | 2832.4268 |
| Rars1 | GFDILGIKPVQR | -648 | -659 | 1 | 0.000550307 | 12 | NaN | NaN | NaN | NaN | NaN | 415.00058 |
| Vps35 | IREDLPNLESSEETEQINK | -750 | -768 | 1 | 0.000298805 | 19 | NaN | NaN | NaN | NaN | NaN | 111.90813 |
| Sacm1l | VVTNQDGVFR | -377 | -386 | 0 | 0.006677519 | 10 | NaN | NaN | NaN | NaN | NaN | 1539.2545 |
| Sacm1l | TYDLTHTLQR | -136 | -145 | 0 | 0.001392869 | 10 | NaN | NaN | NaN | NaN | NaN | 791.6545 |
| Itgal | QIEDGVVVGAPGEGDNTGGLYHCR | -48 | -71 | 0 | 0 | 24 | NaN | NaN | NaN | NaN | NaN | 355.31378 |
| C3 | IWDVVEKA | -615 | -622 | 1 | 0.004439092 | 8 | NaN | NaN | NaN | NaN | NaN | 1687.5205 |
| Vars1 | SILYVSPHPDAFPSLR | -2 | -17 | 0 | 0 | 16 | NaN | NaN | NaN | NaN | NaN | 422.58133 |
| Vars1 | VRQPEFR | -195 | -201 | 1 | 0.001023236 | 7 | NaN | NaN | NaN | NaN | NaN | 2226.7336 |
| Vars1 | ACGATLPALGLR | -111 | -122 | 0 | 0.00025015 | 12 | NaN | NaN | NaN | NaN | NaN | 498.02313 |
| Gnb2 | GYDDFNCNIWDAMK | -288 | -301 | 0 | 0.00111006 | 14 | NaN | NaN | NaN | NaN | NaN | 162.50078 |
| Paics | FKDDANNDPQWSEEQLIAAK | -130 | -149 | 1 | 0 | 20 | NaN | NaN | NaN | NaN | NaN | 441.07602 |
| Paics | SAHKGPDETLR | -301 | -311 | 1 | 0.004439092 | 11 | NaN | NaN | NaN | NaN | NaN | 713.1285 |
| Uba1 | GLQPNGEEMTLK | -962 | -973 | 0 | 0.007987481 | 12 | NaN | NaN | NaN | NaN | NaN | 423.50192 |
| Eif4a3 | FMTDPIR | -232 | -238 | 0 | 0.00025015 | 7 | NaN | NaN | NaN | NaN | NaN | 2535.584 |
| Cnot1 | HYAWHGQGQSPSIR | -522 | -535 | 0 | 0 | 14 | NaN | NaN | NaN | NaN | NaN | 559.6607 |
| Cnot1 | YSNFLDTLK | -1148 | -1156 | 0 | 0.001708008 | 9 | NaN | NaN | NaN | NaN | NaN | 370.0982 |
| Nt5c2 | ACLVDFFTNCPR | -166 | -177 | 0 | 0.003521747 | 12 | NaN | NaN | NaN | NaN | NaN | 251.9476 |
| Cct3 | IVSRPEELREDDVGTGAGLLEIK | -331 | -353 | 2 | 0.001868731 | 23 | NaN | NaN | NaN | NaN | NaN | 543.9756 |
| Cct3 | AQALEVIPR | -441 | -449 | 0 | 0.000431655 | 9 | NaN | NaN | NaN | NaN | NaN | 1662.3505 |
| Cct3 | LCEDIIQLKPDVVITEK | -278 | -294 | 1 | 0.000947459 | 17 | NaN | NaN | NaN | NaN | NaN | 412.33075 |
| Cct3 | KIGDEYFTFITDCK | -354 | -367 | 1 | 0.003745449 | 14 | NaN | NaN | NaN | NaN | NaN | 341.34277 |
| Itgb2 | DSEGCWITYTLQQK | -668 | -681 | 0 | 0 | 14 | NaN | NaN | NaN | NaN | NaN | 637.9862 |
| Itgb2 | CHLEDNMYK | -287 | -295 | 0 | 0.00025015 | 9 | NaN | NaN | NaN | NaN | NaN | 357.41428 |
| Itgb2 | SNEFDYPSVGQL | -297 | -308 | 0 | 0.00025015 | 12 | NaN | NaN | NaN | NaN | NaN | 127.978264 |
| Prkcsh | KILIEEWK | -142 | -149 | 1 | 0.005270854 | 8 | NaN | NaN | NaN | NaN | NaN | 547.4332 |
| Rps3a | IEAHVDVK | -121 | -128 | 0 | 0.00025015 | 8 | NaN | NaN | NaN | NaN | NaN | 755.54626 |
| Fasn | EDPPQHGLPR | -1166 | -1175 | 0 | 0.007343498 | 10 | NaN | NaN | NaN | NaN | NaN | 3222.8015 |
| Cct8 | VYSCPFDGMITETK | -241 | -254 | 0 | 0 | 14 | NaN | NaN | NaN | NaN | NaN | 1048.0195 |
| Cct8 | AHEILPELVCC | -139 | -149 | 0 | 0 | 11 | NaN | NaN | NaN | NaN | NaN | 739.91534 |
| Cct8 | HEKEDGAISTIVLR | -365 | -378 | 1 | 0 | 14 | NaN | NaN | NaN | NaN | NaN | 347.9542 |
| Sema4d | SNSGVVQAPLAFCEK | -491 | -505 | 0 | 0 | 15 | NaN | NaN | NaN | NaN | NaN | 752.62115 |
| Sema4d | SPDGPEGEDDKVYFF | -220 | -234 | 1 | 0.001392869 | 15 | NaN | NaN | NaN | NaN | NaN | 465.78467 |
| Cct6a | VHAELADVLTE | -160 | -170 | 0 | 0.00078661 | 11 | NaN | NaN | NaN | NaN | NaN | 651.492 |
| Cct6a | EMGVWDNYCVK | -491 | -501 | 0 | 0.00025015 | 11 | NaN | NaN | NaN | NaN | NaN | 495.52194 |
| Api5 | IKDLFHIPPSYK | -425 | -436 | 1 | 0.001904399 | 12 | NaN | NaN | NaN | NaN | NaN | 1986.2488 |
| Api5 | DLFHIPPSYK | -427 | -436 | 0 | 0.004350979 | 10 | NaN | NaN | NaN | NaN | NaN | 769.6814 |
| Ppa2 | NYGALPQTWEDPHLR | -136 | -150 | 0 | 0 | 15 | NaN | NaN | NaN | NaN | NaN | 1579.3966 |
| Ppa2 | STDCCGDNDPIDVCEIGSK | -153 | -171 | 0 | 0.00025015 | 19 | NaN | NaN | NaN | NaN | NaN | 366.7359 |
| Fgb | YDRDNDGWVTTDPR | -398 | -411 | 1 | 0.002780285 | 14 | NaN | NaN | NaN | NaN | NaN | 431.8578 |
| Tubg1;Tubg2 | ADDEHYIPR | (55);(55) | (63);(63) | 0 | 0 | 9 | NaN | NaN | NaN | NaN | NaN | 878.25244 |
| Ppia | IYGEKFEDENFILK | -78 | -91 | 1 | 0.00644558 | 14 | NaN | NaN | NaN | NaN | NaN | 1006.4198 |
| Psme2 | VEAFQTTISK | -157 | -166 | 0 | 0.00025015 | 10 | NaN | NaN | NaN | NaN | NaN | 2405.2961 |
| Aldoa | AGTNGETTTQGLDGLSER | -117 | -134 | 0 | 0.000550307 | 18 | NaN | NaN | NaN | NaN | NaN | 2366.647 |
| Aldoa | IVAPGKGILAADESTGSIA | -23 | -41 | 1 | 0.005884925 | 19 | NaN | NaN | NaN | NaN | NaN | 334.66086 |
| Aldoa | PYPALTPEQK | -4 | -13 | 0 | 0.006677519 | 10 | NaN | NaN | NaN | NaN | NaN | 153.50993 |
| Sart1 | TILDEEPIVNR | -632 | -642 | 0 | 0.001055699 | 11 | NaN | NaN | NaN | NaN | NaN | 407.5126 |
| Mrrf | TAPGSLDHITVVTADGK | -111 | -127 | 0 | 0.00078661 | 17 | NaN | NaN | NaN | NaN | NaN | 951.6787 |
| Commd3 | SLPHITDVSWR | -121 | -131 | 0 | 0.003360691 | 11 | NaN | NaN | NaN | NaN | NaN | 2615.3188 |
| Pin1 | NHITNASQWER | -26 | -36 | 0 | 0.001463897 | 11 | NaN | NaN | NaN | NaN | NaN | 1677.4308 |
| Ipo7 | KVLTGVAGEDAECHAAK | -724 | -740 | 1 | 0.002595029 | 17 | NaN | NaN | NaN | NaN | NaN | 15675.727 |
| Ipo7 | MSEQLDLPVR | -44 | -53 | 0 | 0.002068213 | 10 | NaN | NaN | NaN | NaN | NaN | 1376.1964 |
| Ipo7 | WALHILAR | -263 | -270 | 0 | 0.009242859 | 8 | NaN | NaN | NaN | NaN | NaN | 288.92505 |
| Prdx1 | HGEVCPAGWKPGSDTIKPDVNK | -169 | -190 | 2 | 0 | 22 | NaN | NaN | NaN | NaN | NaN | 4545.9272 |
| Prdx1 | HGEVCPAGWKPGSDTIKPD | -169 | -187 | 2 | 0.00025015 | 19 | NaN | NaN | NaN | NaN | NaN | 511.54935 |
| Prdx1 | AFQFTDK | -162 | -168 | 0 | 0.008828666 | 7 | NaN | NaN | NaN | NaN | NaN | 538.4198 |
| Ldha | LLKEEQAPQNK | -12 | -22 | 1 | 0 | 11 | NaN | NaN | NaN | NaN | NaN | 3815.5176 |
| Ldha | RVHPISTMIK | -269 | -278 | 1 | 0 | 10 | NaN | NaN | NaN | NaN | NaN | 469.5753 |
| Ldha | VADLAESIMK | -256 | -265 | 0 | 0.001463897 | 10 | NaN | NaN | NaN | NaN | NaN | 2646.4832 |
| Ldha | GEMMDLQHGSLFLK | -60 | -73 | 0 | 0 | 14 | NaN | NaN | NaN | NaN | NaN | 1253.7855 |
| Rps5 | WSTDDVQINDISLQ | -23 | -36 | 0 | 0.00416551 | 14 | NaN | NaN | NaN | NaN | NaN | 1064.4702 |
| Ranbp1 | ADESNHDPQFEPIVSLPEQEIK | -18 | -39 | 0 | 0.00025015 | 22 | NaN | NaN | NaN | NaN | NaN | 638.3419 |
| Rcc2 | TKDGQILPVPN | -317 | -327 | 1 | 0.003069674 | 11 | NaN | NaN | NaN | NaN | NaN | 4732.3115 |
| Rcc2 | RLPEYTPR | -511 | -518 | 1 | 0.002752526 | 8 | NaN | NaN | NaN | NaN | NaN | 281.845 |
| Slc3a2 | QINPTLGSQEDFKDLLQSAK | -172 | -191 | 1 | 0 | 20 | NaN | NaN | NaN | NaN | NaN | 743.6344 |
| Sptan1 | QFQDAGHFDAENIK | -743 | -756 | 0 | 0.00122419 | 14 | NaN | NaN | NaN | NaN | NaN | 796.1616 |
| Sptan1 | EANQQQQFNR | -676 | -685 | 0 | 0.002434616 | 10 | NaN | NaN | NaN | NaN | NaN | 287.32288 |
| Nup155 | SIAADGEFLHELEEK | -1133 | -1147 | 0 | 0.000591124 | 15 | NaN | NaN | NaN | NaN | NaN | 3572.1868 |
| Pfn1 | IDSLMADGTCQDAAIVGYK | -8 | -26 | 0 | 0 | 19 | NaN | NaN | NaN | NaN | NaN | 148.72195 |
| Pfn1 | YKDSPSVWAAVPGK | -25 | -38 | 1 | 0 | 14 | NaN | NaN | NaN | NaN | NaN | 1168.3346 |
| Rpsa | RDPEEIEK | -205 | -212 | 1 | 0.00111006 | 8 | NaN | NaN | NaN | NaN | NaN | 2090.7722 |
| Ppp1cc | WSDPDKDVLGWGENDR | -206 | -221 | 1 | 0.00111006 | 16 | NaN | NaN | NaN | NaN | NaN | 267.4682 |
| Ppp1cc | WSDPDKDVLGWGENDR | -206 | -221 | 1 | 0.00111006 | 16 | NaN | NaN | NaN | NaN | NaN | 267.4682 |
| Hnrnpk | IDEPLEGSEDR | -423 | -433 | 0 | 0 | 11 | NaN | NaN | NaN | NaN | NaN | 6637.4214 |
| Hnrnpk | ADIETIGEILKK | -92 | -103 | 1 | 0 | 12 | NaN | NaN | NaN | NaN | NaN | 2294.1858 |

| | | | | | | | | | | | | |
|---|---|---|---|---|---|---|---|---|---|---|---|---|
| Hnrnpk | RNTDEMVELR | -37 | -46 | 1 | 0.001463897 | 10 | NaN | NaN | NaN | NaN | NaN | 2064.7773 |
| Hnrnpm | GPGGQPIDANHLSK | -308 | -321 | 0 | 0.000431655 | 14 | NaN | NaN | NaN | NaN | NaN | 376.42593 |
| Hnrnpm | GLDHMASSIER | -460 | -470 | 0 | 0.00025015 | 11 | NaN | NaN | NaN | NaN | NaN | 650.576 |
| Hnrnpm | MGPLGLDHMASSIER | -456 | -470 | 0 | 0.00051306 | 15 | NaN | NaN | NaN | NaN | NaN | 510.83627 |
| Csnk2a1 | EWGNQDDYQLVR | -32 | -43 | 0 | 0.003166358 | 12 | NaN | NaN | NaN | NaN | NaN | 1131.7854 |
| Npm1 | DELHIVEAEAM | -55 | -65 | 0 | 0 | 11 | NaN | NaN | NaN | NaN | NaN | 1927.8208 |
| Npm1 | VKLDEDDEDDDEDDEDDEDDDDDDFDEEETEEKVP[illegible] | -156 | -192 | 2 | 0.006654467 | 37 | NaN | NaN | NaN | NaN | NaN | 1152.6998 |
| Npm1 | SPLRPQNYLFGCELK | -10 | -24 | 1 | 0.001868731 | 15 | NaN | NaN | NaN | NaN | NaN | 895.8695 |
| Npm1 | IQDLWQWR | -282 | -289 | 0 | 0.005758933 | 8 | NaN | NaN | NaN | NaN | NaN | 768.3251 |
| Npm1 | DQEAIQDLWQWRK | -278 | -290 | 1 | 0.000230707 | 13 | NaN | NaN | NaN | NaN | NaN | 372.71756 |
| Themis | LGHPCFYHLK | -113 | -122 | 0 | 0 | 10 | NaN | NaN | NaN | NaN | NaN | 560.72205 |
| Themis | ELSPVSVGDQFLVHH | -380 | -394 | 0 | 0.001636916 | 15 | NaN | NaN | NaN | NaN | NaN | 864.18494 |
| Capza2 | EAENAIESWR | -112 | -121 | 0 | 0.009955823 | 10 | NaN | NaN | NaN | NaN | NaN | 611.4048 |
| Hprt1 | ADLLDYIK | -76 | -83 | 0 | 0.008947895 | 8 | NaN | NaN | NaN | NaN | NaN | 299.79538 |
| Sptbn1 | STDHGHNLQTVQLLIK | -1510 | -1525 | 0 | 0.000826806 | 16 | NaN | NaN | NaN | NaN | NaN | 3035.548 |
| Sptbn1 | ITDLYTDLR | -75 | -83 | 0 | 0 | 9 | NaN | NaN | NaN | NaN | NaN | 2858.8071 |
| Sptbn1 | EAEKLESEHPDQAQAILSR | -1018 | -1036 | 1 | 0 | 19 | NaN | NaN | NaN | NaN | NaN | 425.33063 |
| Adsl | DRADLPTLGFTHFQPA | -148 | -163 | 1 | 0.001092529 | 16 | NaN | NaN | NaN | NaN | NaN | 1221.5748 |
| Glod4 | DRSPSQSDPVLK | -129 | -140 | 1 | 0.001636916 | 12 | NaN | NaN | NaN | NaN | NaN | 1549.6177 |
| Uba2 | HSEDLGKDVEFEVVGDSPEK | -532 | -551 | 1 | 0.00078661 | 20 | NaN | NaN | NaN | NaN | NaN | 1190.9259 |
| Arhgdia | TAPGPLELDLTGDLESFK | -81 | -98 | 0 | 0.001636916 | 18 | NaN | NaN | NaN | NaN | NaN | 367.70654 |
| Rps6ka3 | SFFSTIDWNK | -325 | -334 | 0 | 0 | 10 | NaN | NaN | NaN | NaN | NaN | 651.0601 |
| Coro1a | LDRLEETVQAK | -451 | -461 | 1 | 0 | 11 | NaN | NaN | NaN | NaN | NaN | 10670.336 |
| Coro1a | VDKNVPLVCGH | -70 | -80 | 1 | 0.002199523 | 11 | NaN | NaN | NaN | NaN | NaN | 373.79773 |
| Coro1a | PHEGTRPVHAV | -217 | -227 | 1 | 0.006100831 | 11 | NaN | NaN | NaN | NaN | NaN | 138.37953 |
| Smc1a | KLEGELTEEVEMAK | -454 | -467 | 1 | 0.000755556 | 14 | NaN | NaN | NaN | NaN | NaN | 484.51596 |
| Smc1a | ADQDRLDLEER | -402 | -412 | 1 | 0 | 11 | NaN | NaN | NaN | NaN | NaN | 151.02365 |
| Nup133 | AEDLEESLTSSVGGR | -482 | -496 | 0 | 0 | 15 | NaN | NaN | NaN | NaN | NaN | 840.14966 |
| Nup133 | DQNFSDFLFR | -893 | -902 | 0 | 0.009217481 | 10 | NaN | NaN | NaN | NaN | NaN | 724.90643 |
| Cdk6 | DQQYECVAEIGEGAYGK | -10 | -26 | 0 | 0 | 17 | NaN | NaN | NaN | NaN | NaN | 489.40366 |
| Tars1 | LTYVSHDGDDK | -572 | -582 | 0 | 0.001392869 | 11 | NaN | NaN | NaN | NaN | NaN | 1065.7549 |
| Tars1 | DFSSLETLCK | -197 | -206 | 0 | 0.002972566 | 10 | NaN | NaN | NaN | NaN | NaN | 235.30586 |
| Psmc4 | SEEVDLEDYVARPDK | -355 | -369 | 1 | 0.003003218 | 15 | NaN | NaN | NaN | NaN | NaN | 847.4256 |
| Psmc4 | IEFPLPDR | -331 | -338 | 0 | 0.007546675 | 8 | NaN | NaN | NaN | NaN | NaN | 530.9903 |
| Sun2 | EAAASLGQILQK | -520 | -531 | 0 | 0.003620005 | 12 | NaN | NaN | NaN | NaN | NaN | 484.82538 |
| Gbp2 | LGTFSKPGGYY | -422 | -432 | 1 | 0.008828666 | 11 | NaN | NaN | NaN | NaN | NaN | 1467.378 |
| Gbp2 | TKAEAAEAANR | -492 | -502 | 1 | 0.002099344 | 11 | NaN | NaN | NaN | NaN | NaN | 189.53397 |
| Kars1 | ADYHDLMEITEK | -343 | -354 | 0 | 0.003299042 | 12 | NaN | NaN | NaN | NaN | NaN | 868.64185 |
| Timm8a1 | AEACFVNCVER | -59 | -69 | 0 | 0 | 11 | NaN | NaN | NaN | NaN | NaN | 176.65659 |
| Tkt | WAEAGFLPEAELLNLR | -86 | -101 | 0 | 0.001868731 | 16 | NaN | NaN | NaN | NaN | NaN | 406.09747 |
| Copa | VTTGDHGIIR | -553 | -562 | 0 | 0.007343498 | 10 | NaN | NaN | NaN | NaN | NaN | 1050.0824 |
| Anxa2 | TDLEKDIISDTSGDFR | -153 | -168 | 1 | 0 | 16 | NaN | NaN | NaN | NaN | NaN | 1327.0529 |
| Hdlbp | DKFPEVIINFPDPAQK | -535 | -550 | 1 | 0.005301614 | 16 | NaN | NaN | NaN | NaN | NaN | 563.5252 |
| Rps2 | AEDKEWIPVTK | -55 | -65 | 1 | 0.00025015 | 11 | NaN | NaN | NaN | NaN | NaN | 186.24525 |
| Canx | KIPNPDFFEDLEPF | -402 | -415 | 1 | 0.00266297 | 14 | NaN | NaN | NaN | NaN | NaN | 333.01172 |
| P4ha1 | DTYNLDTNTISK | -139 | -150 | 0 | 0.009031675 | 12 | NaN | NaN | NaN | NaN | NaN | 4479.6206 |
| Cdc37 | IDSGLWVPNSK | -338 | -348 | 0 | 0.007987481 | 11 | NaN | NaN | NaN | NaN | NaN | 1315.1776 |
| Cdc37 | RLGPGGLDPVEVY | -287 | -299 | 1 | 0.000826806 | 13 | NaN | NaN | NaN | NaN | NaN | 723.3317 |
| Dbnl | QEWESAGQQAPHPR | -253 | -266 | 0 | 0 | 14 | NaN | NaN | NaN | NaN | NaN | 1932.6855 |
| Dbnl | ARAEEDVEPECIMEK | -117 | -131 | 1 | 0 | 15 | NaN | NaN | NaN | NaN | NaN | 564.5062 |
| Acin1 | VVDLHADDSR | -887 | -896 | 0 | 0.009242859 | 10 | NaN | NaN | NaN | NaN | NaN | 1152.6117 |
| Acin1 | DYAEQDELDYHR | -1084 | -1095 | 0 | 0 | 12 | NaN | NaN | NaN | NaN | NaN | 663.36115 |
| Clic1 | PALNDNLEK | -123 | -131 | 0 | 0.000230707 | 9 | NaN | NaN | NaN | NaN | NaN | 3276.1594 |
| Clic1 | TCPDDEEIELAYEQVAR | -222 | -238 | 0 | 0 | 17 | NaN | NaN | NaN | NaN | NaN | 774.87573 |
| Exosc8 | GYVVPNVDLPPLCSSR | -77 | -92 | 0 | 0.000394361 | 16 | NaN | NaN | NaN | NaN | NaN | 2264.4744 |
| Twf2 | TAEFLYDEVHPK | -304 | -315 | 0 | 0.001776473 | 12 | NaN | NaN | NaN | NaN | NaN | 2093.445 |
| Twf2 | VHTEPTNVAQLPSR | -215 | -228 | 0 | 0.00272025 | 14 | NaN | NaN | NaN | NaN | NaN | 705.38635 |
| Twf2 | HLSSCAAPAPLTSAER | -137 | -152 | 0 | 0.000471409 | 16 | NaN | NaN | NaN | NaN | NaN | 592.356 |
| Dhx15 | MNDPLLER | -246 | -253 | 0 | 0 | 8 | NaN | NaN | NaN | NaN | NaN | 1262.5955 |
| Arhgap30 | AEFVQEYGVVDGIYR | -41 | -55 | 0 | 0.00025015 | 15 | NaN | NaN | NaN | NaN | NaN | 397.08582 |
| Tnpo1 | ESQSPDTTIQR | -29 | -39 | 0 | 0 | 11 | NaN | NaN | NaN | NaN | NaN | 12017.063 |
| Tnpo1 | DVEEDETIPDSEQDIRPR | -329 | -346 | 1 | 0.003393785 | 18 | NaN | NaN | NaN | NaN | NaN | 365.19666 |
| Ddb1 | EMLGGEIIPR | -580 | -589 | 0 | 0 | 10 | NaN | NaN | NaN | NaN | NaN | 1252.9869 |
| Ddb1 | EEQMDGTVTLK | -288 | -298 | 0 | 0.008129024 | 11 | NaN | NaN | NaN | NaN | NaN | 560.2034 |
| Ddb1 | GHFTSAEDLNLLIAK | -21 | -35 | 0 | 0.00051306 | 15 | NaN | NaN | NaN | NaN | NaN | 469.09314 |
| Rexo2 | QAEYEFLSFVR | -119 | -129 | 0 | 0.00025015 | 11 | NaN | NaN | NaN | NaN | NaN | 1447.9152 |
| Ppp5c | LLYPDHFHLLR | -291 | -301 | 0 | 0 | 11 | NaN | NaN | NaN | NaN | NaN | 724.3511 |
| Tgtp2;Tgtp1 | TKIDSDLDNEQK | (170);(170) | (181);(181) | 1 | 0 | 12 | NaN | NaN | NaN | NaN | NaN | 2459.4668 |
| Psph | LNIPTTNVFAN | -123 | -133 | 0 | 0.005605646 | 11 | NaN | NaN | NaN | NaN | NaN | 560.8798 |
| Psph | EACPPADAFIGFGGNVIR | -185 | -202 | 0 | 0.001463897 | 18 | NaN | NaN | NaN | NaN | NaN | 398.89584 |
| Uchl5 | KWQPGEEPAGSVVQDSR | -57 | -73 | 1 | 0.00111006 | 17 | NaN | NaN | NaN | NaN | NaN | 800.90875 |
| Stat5b | HLKPHWNDGAIL | -584 | -595 | 1 | 0.00272025 | 12 | NaN | NaN | NaN | NaN | NaN | 343.4449 |
| Anxa6 | TNEQMHQLVAAYK | -123 | -135 | 0 | 0.001291103 | 13 | NaN | NaN | NaN | NaN | NaN | 369.7829 |
| Anxa6 | MWELSAVSR | -342 | -350 | 0 | 0.00149795 | 9 | NaN | NaN | NaN | NaN | NaN | 1698.417 |
| Anxa6 | ISGVGTDEK | -105 | -113 | 0 | 0.006100831 | 9 | NaN | NaN | NaN | NaN | NaN | 704.3478 |
| Anxa6 | AEILEIADTPSGDKTSLETR | -527 | -546 | 1 | 0.00365438 | 20 | NaN | NaN | NaN | NaN | NaN | 549.4934 |
| Anxa6 | RSYPHLR | -554 | -560 | 1 | 0.00830926 | 7 | NaN | NaN | NaN | NaN | NaN | 310.72357 |
| Flna | AGNNMLLVGVHGPR | -2585 | -2598 | 0 | 0.001055699 | 14 | NaN | NaN | NaN | NaN | NaN | 492.48914 |

| | | | | | | | | | | | | |
|---|---|---|---|---|---|---|---|---|---|---|---|---|
| Flna | KSNFTVDCSK | -2575 | -2584 | 1 | 0.003490217 | 10 | NaN | NaN | NaN | NaN | NaN | 4169.4575 |
| Flna | ENGIYLIDVK | -2396 | -2405 | 0 | 0.002810811 | 10 | NaN | NaN | NaN | NaN | NaN | 2324.4155 |
| Flna | EAGAGGLAIAVEGPSK | -2265 | -2280 | 0 | 0.00025015 | 16 | NaN | NaN | NaN | NaN | NaN | 1179.6606 |
| Flna | CYGPGIEGQGVFR | -1260 | -1272 | 0 | 0.002068213 | 13 | NaN | NaN | NaN | NaN | NaN | 850.8895 |
| Flna | GGHQVPGSPFK | -1429 | -1439 | 0 | 0.00111006 | 11 | NaN | NaN | NaN | NaN | NaN | 579.46326 |
| Eef2 | SVIAHVDHGK | -23 | -32 | 0 | 0 | 10 | NaN | NaN | NaN | NaN | NaN | 2484.622 |
| Eef2 | PFPDGLAEDIDKGEVSAR | -608 | -625 | 1 | 0 | 18 | NaN | NaN | NaN | NaN | NaN | 6074.1963 |
| Eef2 | IWCFGPDGTGPN | -649 | -660 | 0 | 0.000230707 | 12 | NaN | NaN | NaN | NaN | NaN | 445.94156 |
| Eef2 | VEAKNPADLPK | -509 | -519 | 1 | 0.005301614 | 11 | NaN | NaN | NaN | NaN | NaN | 1698.141 |
| Got2 | IPEQSVLLLHA | -201 | -211 | 0 | 0.00025015 | 11 | NaN | NaN | NaN | NaN | NaN | 510.7364 |
| Sin3a | RWSDYVER | -1098 | -1105 | 1 | 0.008828666 | 8 | NaN | NaN | NaN | NaN | NaN | 581.9994 |
| Eloc | SSDGHEFIVK | -23 | -32 | 0 | 0.006624942 | 10 | NaN | NaN | NaN | NaN | NaN | 1927.9137 |
| Hexim1 | ELELELDR | -321 | -328 | 0 | 0.006472924 | 8 | NaN | NaN | NaN | NaN | NaN | 1149.888 |
| Fga | NFDNTYGQVSEDLR | -123 | -136 | 0 | 0.00025015 | 14 | NaN | NaN | NaN | NaN | NaN | 465.18665 |
| Gatad2a | AEPAAPHPTLK | -486 | -496 | 0 | 0.003780189 | 11 | NaN | NaN | NaN | NaN | NaN | 659.9298 |
| Sp2 | AEQQQVVQIPQQALR | -307 | -321 | 0 | 0 | 15 | NaN | NaN | NaN | NaN | NaN | 310.8573 |
| Psmd12 | LQEVIETLLSLEK | -40 | -52 | 0 | 0 | 13 | NaN | NaN | NaN | NaN | NaN | 418.03766 |
| Ostf1 | ESIDNPLHEAAK | -71 | -82 | 0 | 0.00025015 | 12 | NaN | NaN | NaN | NaN | NaN | 3037.7778 |
| Ostf1 | NYVAEQAESIDNPLHEAAK | -64 | -82 | 0 | 0.00078661 | 19 | NaN | NaN | NaN | NaN | NaN | 1371.9347 |
| Dis3 | LHQIDVLEHPAIR | -74 | -86 | 0 | 0 | 13 | NaN | NaN | NaN | NaN | NaN | 319.6048 |
| Dis3 | ETYIEQEQGENANDR | -130 | -144 | 0 | 0 | 15 | NaN | NaN | NaN | NaN | NaN | 1232.1178 |
| Dis3 | VAIGADCTYPELTDK | -793 | -807 | 0 | 0.000550307 | 15 | NaN | NaN | NaN | NaN | NaN | 1103.1255 |
| Dmap1 | AETDHLFDLSR | -155 | -165 | 0 | 0 | 11 | NaN | NaN | NaN | NaN | NaN | 2601.3477 |
| Morc3 | AEYPVNLPVEDIQK | -391 | -404 | 0 | 0.00383302 | 14 | NaN | NaN | NaN | NaN | NaN | 1877.168 |
| Ado | RPPPEQQFEPPLQPLER | -132 | -148 | 1 | 0.00025015 | 17 | NaN | NaN | NaN | NaN | NaN | 603.09406 |
| Utp3 | KKEENSAEEQALEEQNAK | -381 | -398 | 2 | 0 | 18 | NaN | NaN | NaN | NaN | NaN | 379.8907 |
| Ddx27 | GLQVGELHGNLSQTQR | -453 | -468 | 0 | 0.00111006 | 16 | NaN | NaN | NaN | NaN | NaN | 263.56262 |
| Ywhaq | TTVLELLDK | -95 | -103 | 0 | 0.002566359 | 9 | NaN | NaN | NaN | NaN | NaN | 327.05933 |
| Nol6 | LLPCPPLDFFRPC | -262 | -274 | 1 | 0.00201047 | 13 | NaN | NaN | NaN | NaN | NaN | 553.35455 |
| Nol6 | AFDHVVHLHPLSR | -475 | -487 | 0 | 0.006735683 | 13 | NaN | NaN | NaN | NaN | NaN | 368.69662 |
| Pccb | SVTNEDVTQEQLGGAK | -235 | -250 | 0 | 0.000230707 | 16 | NaN | NaN | NaN | NaN | NaN | 1099.159 |
| Hsph1 | KQDLPNAEEKPR | -185 | -196 | 2 | 0 | 12 | NaN | NaN | NaN | NaN | NaN | 572.153 |
| Hsph1 | TDGQQTSQSPPSPELTSEESK | -547 | -567 | 0 | 0.001610812 | 21 | NaN | NaN | NaN | NaN | NaN | 209.76968 |
| Nrm | SLRPLLGGIPESGGPDAR | -30 | -47 | 1 | 0.000471409 | 18 | NaN | NaN | NaN | NaN | NaN | 180.64249 |
| Actn1 | RDQALTEEHAR | -614 | -624 | 1 | 0 | 11 | NaN | NaN | NaN | NaN | NaN | 904.9793 |
| Actn1 | DYETATLSEIK | -421 | -431 | 0 | 0 | 11 | NaN | NaN | NaN | NaN | NaN | 850.3533 |
| Snrnp70 | QQEVETELK | -79 | -87 | 0 | 0.002780285 | 9 | NaN | NaN | NaN | NaN | NaN | 577.4676 |
| Ilk | AFLHTLEPLIPR | -306 | -317 | 0 | 0 | 12 | NaN | NaN | NaN | NaN | NaN | 749.54517 |
| Nfkb1 | EGFGDFSPTDVHR | -293 | -305 | 0 | 0.008736733 | 13 | NaN | NaN | NaN | NaN | NaN | 1112.5128 |
| Nfkb1 | NDLYQTPLHL | -575 | -584 | 0 | 0.00201047 | 10 | NaN | NaN | NaN | NaN | NaN | 909.8052 |
| Hnrnpu | CHNCAVEFNFGQK | -426 | -438 | 0 | 0.00078661 | 13 | NaN | NaN | NaN | NaN | NaN | 2248.47 |
| Hnrnpu | CAVEFNFGQK | -429 | -438 | 0 | 0.003681885 | 10 | NaN | NaN | NaN | NaN | NaN | 778.1626 |
| Galk1 | AFMEEFGAEPELAVSAPGR | -19 | -37 | 0 | 0 | 19 | NaN | NaN | NaN | NaN | NaN | 707.7442 |
| Rpa1 | VMDVKPVDFR | -600 | -609 | 1 | 0 | 10 | NaN | NaN | NaN | NaN | NaN | 594.9724 |
| Actr2 | LTEPPMNPTK | -109 | -118 | 0 | 0.003998192 | 10 | NaN | NaN | NaN | NaN | NaN | 524.47723 |
| Gzma | IIGGDTVVPHSRPY | -29 | -42 | 1 | 0 | 14 | NaN | NaN | NaN | NaN | NaN | 731.4859 |
| Paxbp1 | AFQNSEYGDDSIR | -813 | -825 | 0 | 0.002329279 | 13 | NaN | NaN | NaN | NaN | NaN | 715.93146 |
| Uggt1 | SEIGHEEFSNIHHQLISK | -200 | -217 | 0 | 0.000471409 | 18 | NaN | NaN | NaN | NaN | NaN | 239.00928 |
| Mars1 | TVGPEECVVPFLTR | -32 | -45 | 0 | 0.001392869 | 14 | NaN | NaN | NaN | NaN | NaN | 541.9582 |
| C1qbp | EHQEYITFLEDLK | -260 | -272 | 0 | 0 | 13 | NaN | NaN | NaN | NaN | NaN | 782.89496 |
| C1qbp | GVDNTFADELVEL | -243 | -255 | 0 | 0.008947895 | 13 | NaN | NaN | NaN | NaN | NaN | 100.3976 |
| Bub3 | AFYDPTHAWSGGLDHQLK | -63 | -80 | 0 | 0 | 18 | NaN | NaN | NaN | NaN | NaN | 523.2238 |
| Ppp2r2a | AGAGGGNDIQWCFSQVK | -2 | -18 | 0 | 0.000230707 | 17 | NaN | NaN | NaN | NaN | NaN | 355.97794 |
| Rplp0 | CFIVGADNVGSK | -27 | -38 | 0 | 0 | 12 | NaN | NaN | NaN | NaN | NaN | 2798.9324 |
| Taldo1 | PQEDPGVK | -208 | -215 | 0 | 0.002199523 | 8 | NaN | NaN | NaN | NaN | NaN | 649.6911 |
| Rpf2 | NITRPFEDQTSL | -67 | -78 | 1 | 0.006677519 | 12 | NaN | NaN | NaN | NaN | NaN | 744.4712 |
| Uqcrfs1 | RDPQHDLDR | -196 | -204 | 1 | 0.001636916 | 9 | NaN | NaN | NaN | NaN | NaN | 784.3685 |
| Uqcrfs1 | AGDFGGYYCPCHGSHYDASGR | -228 | -248 | 0 | 0.00025015 | 21 | NaN | NaN | NaN | NaN | NaN | 140.70453 |
| Top2a | AGDMELKPF | -200 | -208 | 1 | 0.002697088 | 9 | NaN | NaN | NaN | NaN | NaN | 325.84677 |
| Fermt3 | QDWSDHAIWWEQK | -57 | -69 | 0 | 0.00025015 | 13 | NaN | NaN | NaN | NaN | NaN | 221.79472 |
| Xpo1 | KDFEEYPEHR | -836 | -845 | 1 | 0.000712251 | 10 | NaN | NaN | NaN | NaN | NaN | 389.22247 |
| Numa1 | QTMQEQEQAAQGLR | -531 | -544 | 0 | 0 | 14 | NaN | NaN | NaN | NaN | NaN | 715.41565 |
| Nono | TTFPRPVTVEPMDQLDDEEGLPEK | -218 | -241 | 1 | 0.001392869 | 24 | NaN | NaN | NaN | NaN | NaN | 596.14935 |
| Actbl2 | GIHETTFNSIMKC | -274 | -286 | 1 | 0.00111006 | 13 | NaN | NaN | NaN | NaN | NaN | 1975.855 |
| Nat10 | AVSLLEEVITPR | -677 | -688 | 0 | 0 | 12 | NaN | NaN | NaN | NaN | NaN | 608.48737 |
| Ppp2r1a | AMSGDPNYLHR | -488 | -498 | 0 | 0 | 11 | NaN | NaN | NaN | NaN | NaN | 3602.0098 |
| Ppp2r1a | TQDQDVDVK | -568 | -576 | 0 | 0.003166358 | 9 | NaN | NaN | NaN | NaN | NaN | 393.94937 |
| Hnrnpl | AGGGGGENYDDPHKTPASPVVHIR | -81 | -104 | 1 | 0.000550307 | 24 | NaN | NaN | NaN | NaN | NaN | 424.78113 |
| Hnrnpl | GGGGGENYDDPHKTPASPVVHIR | -82 | -104 | 1 | 0.001392869 | 23 | NaN | NaN | NaN | NaN | NaN | 389.79437 |
| H2az1;H2az2 | TIAGGGVIPHIHK | (104);(104) | (116);(116) | 0 | 0 | 13 | NaN | NaN | NaN | NaN | NaN | 1683.3745 |
| Rpl4 | AGHQTSAESWGTGR | -58 | -71 | 0 | 0.005333786 | 14 | NaN | NaN | NaN | NaN | NaN | 1902.148 |
| Rpl10a | KAVDIPHMDIE | -78 | -88 | 1 | 0.000230707 | 11 | NaN | NaN | NaN | NaN | NaN | 930.63354 |
| Sdha | LCIEDGSIHR | -237 | -246 | 0 | 0.002329279 | 10 | NaN | NaN | NaN | NaN | NaN | 734.8634 |
| Htra2 | AGLRPGDVILAIGEK | -405 | -419 | 1 | 0.007040624 | 15 | NaN | NaN | NaN | NaN | NaN | 331.25693 |
| Top1 | WWEEERYPEGIK | -207 | -218 | 1 | 0.001463897 | 12 | NaN | NaN | NaN | NaN | NaN | 8394.009 |
| Top1 | SGDHLHNDSQIEADFR | -2 | -17 | 0 | 0.000394361 | 16 | NaN | NaN | NaN | NaN | NaN | 327.42123 |
| Vps4b | AGNYEEALQLYQH | -24 | -36 | 0 | 0.002099344 | 13 | NaN | NaN | NaN | NaN | NaN | 771.34937 |
| Aarsd1 | LEDTLLFPEGGGQPDDR | -45 | -61 | 0 | 0.004439092 | 17 | NaN | NaN | NaN | NaN | NaN | 289.98145 |

| | | | | | | | | | | | | |
|---|---|---|---|---|---|---|---|---|---|---|---|---|
| Pnp | DYNEIPNFPQSTVQGHAGR | -49 | -67 | 0 | 0.00025015 | 19 | NaN | NaN | NaN | NaN | NaN | 1910.6267 |
| Nipsnap1 | EFSFWNEPQPR | -169 | -179 | 0 | 0.00025015 | 11 | NaN | NaN | NaN | NaN | NaN | 1756.6553 |
| Pgd | VDDFLANEAK | -39 | -48 | 0 | 0.00111006 | 10 | NaN | NaN | NaN | NaN | NaN | 4354.6465 |
| Arid1a | NDPFVVDCSDK | -1818 | -1828 | 0 | 0.00149795 | 11 | NaN | NaN | NaN | NaN | NaN | 428.78 |
| Acaca | LYDEDKVPR | -1273 | -1281 | 1 | 0.000431655 | 9 | NaN | NaN | NaN | NaN | NaN | 1030.2446 |
| Arhgap45 | GAECEECCLACHK | -722 | -734 | 0 | 0 | 13 | NaN | NaN | NaN | NaN | NaN | 291.00943 |
| Baz1b | LSEIPLTLHSV | -679 | -689 | 0 | 0.002265552 | 11 | NaN | NaN | NaN | NaN | NaN | 3489.6658 |
| Phgdh | EELIAELQDCEGLIVR | -39 | -54 | 0 | 0.00025015 | 16 | NaN | NaN | NaN | NaN | NaN | 261.4021 |
| Actn4 | FNHFDKDHGGALGPEEFK | -775 | -792 | 1 | 0 | 18 | NaN | NaN | NaN | NaN | NaN | 109.28101 |
| Pgk1 | ITLPVDFVTADK | -280 | -291 | 0 | 0 | 12 | NaN | NaN | NaN | NaN | NaN | 2064.2979 |
| Pgk1 | NNMEIGTSLYDEEGAK | -249 | -264 | 0 | 0 | 16 | NaN | NaN | NaN | NaN | NaN | 2842.1292 |
| Pgk1 | DKIQLINNMLDK | -219 | -230 | 1 | 0.00272025 | 12 | NaN | NaN | NaN | NaN | NaN | 772.09674 |
| Pgk1 | VLNNMEIGTSLYDEEGAK | -247 | -264 | 0 | 0.004791731 | 18 | NaN | NaN | NaN | NaN | NaN | 2240.6506 |
| Pgk1 | VNEMIIGGGMAF | -231 | -242 | 0 | 0.00025015 | 12 | NaN | NaN | NaN | NaN | NaN | 79.9572 |
| Son | AGVPLPPNLKPAPPPTIEEK | -2091 | -2110 | 1 | 0.000550307 | 20 | NaN | NaN | NaN | NaN | NaN | 341.37726 |
| Preb | EAHGIVVTDVTFLPEK | -340 | -355 | 0 | 0.001708008 | 16 | NaN | NaN | NaN | NaN | NaN | 984.19696 |
| Preb | LLGPHETALF | -361 | -370 | 0 | 0.004255025 | 10 | NaN | NaN | NaN | NaN | NaN | 200.52527 |
| Prmt1 | DHVVTIIK | -136 | -143 | 0 | 0.003490217 | 8 | NaN | NaN | NaN | NaN | NaN | 296.60678 |
| Ap1m1 | ILQEYITQEGHKL | -130 | -142 | 1 | 0.001708008 | 13 | NaN | NaN | NaN | NaN | NaN | 972.2591 |
| Atp5f1b | KVLDSGAPIKIPVGPETLGR | -124 | -143 | 2 | 0.00025015 | 20 | NaN | NaN | NaN | NaN | NaN | 760.2097 |
| Atp5f1b | FAHLDATTVLSR | -376 | -387 | 0 | 0.001868731 | 12 | NaN | NaN | NaN | NaN | NaN | 11645.631 |
| Atp5f1b | PVGPETLGR | -135 | -143 | 0 | 0.00266297 | 9 | NaN | NaN | NaN | NaN | NaN | 1296.1847 |
| Atp5f1b | QFAPIHAEAPEFIEMS | -162 | -177 | 0 | 0.006735683 | 16 | NaN | NaN | NaN | NaN | NaN | 480.8591 |
| Psmc5 | KIEELQLIVNDK | -27 | -38 | 1 | 0.001392869 | 12 | NaN | NaN | NaN | NaN | NaN | 557.92505 |
| Tgm2 | AWCPADDVYLDSEEERR | -141 | -157 | 1 | 0 | 17 | NaN | NaN | NaN | NaN | NaN | 217.90396 |
| Nampt | EKYDGHLPIEVK | -106 | -117 | 1 | 0.001941082 | 12 | NaN | NaN | NaN | NaN | NaN | 1505.4679 |
| Cand1 | ALVPPVVACVGDPFYK | -519 | -534 | 0 | 0.002168954 | 16 | NaN | NaN | NaN | NaN | NaN | 1027.2042 |
| Cand1 | DLLDSVLPHLYNETK | -1043 | -1057 | 0 | 0.000431655 | 15 | NaN | NaN | NaN | NaN | NaN | 286.96008 |
| Polr2a | PVFHVGFL | -93 | -100 | 0 | 0.004690496 | 8 | NaN | NaN | NaN | NaN | NaN | 261.17004 |
| Qars1 | GHNPLPSPWR | -367 | -376 | 0 | 0.007609801 | 10 | NaN | NaN | NaN | NaN | NaN | 1876.1886 |
| Qars1 | WVSQPLVCEIR | -680 | -690 | 0 | 0.009122264 | 11 | NaN | NaN | NaN | NaN | NaN | 863.24927 |
| Tufm | DLDKPFLLPVESVYSIPGR | -253 | -271 | 1 | 0 | 19 | NaN | NaN | NaN | NaN | NaN | 421.53827 |
| Htt | LGEEEALEDDSESR | -424 | -437 | 0 | 0.005301614 | 14 | NaN | NaN | NaN | NaN | NaN | 669.15936 |
| Htt | VGDWLGNIR | -767 | -775 | 0 | 0 | 9 | NaN | NaN | NaN | NaN | NaN | 109.47722 |
| Ptpn1 | AHWPSHWKPFLVN | -400 | -412 | 1 | 0.002068213 | 13 | NaN | NaN | NaN | NaN | NaN | 417.67307 |
| Psmb1 | GFHGDCLTLTK | -83 | -93 | 0 | 0.001868731 | 11 | NaN | NaN | NaN | NaN | NaN | 263.422 |
| Nvl | KKPEIEGLPSEGDQEER | -498 | -514 | 2 | 0.00272025 | 17 | NaN | NaN | NaN | NaN | NaN | 247.24887 |
| Bpnt1 | AIAGIINQPYYNYQAGPDAALGR | -145 | -167 | 0 | 0.002199523 | 23 | NaN | NaN | NaN | NaN | NaN | 352.30783 |
| Mthfd1 | GEWIKPGAVVIDCGIN | -224 | -239 | 1 | 0.00051306 | 16 | NaN | NaN | NaN | NaN | NaN | 2594.4592 |
| Mthfd1 | ANIAHGNSSIIADR | -640 | -653 | 0 | 0 | 14 | NaN | NaN | NaN | NaN | NaN | 509.66675 |
| Mthfd1 | RGDLNDCFIPCTPK | -137 | -150 | 1 | 0.001291103 | 14 | NaN | NaN | NaN | NaN | NaN | 435.37808 |
| Gcn1 | PNDASYDAVR | -1307 | -1316 | 0 | 0.000907206 | 10 | NaN | NaN | NaN | NaN | NaN | 486.28354 |
| Ier3ip1 | NIGWGTDQGIGGFGEEPGIK | -30 | -49 | 0 | 0.000230707 | 20 | NaN | NaN | NaN | NaN | NaN | 761.53424 |
| St13 | DYDEDASAMLR | -211 | -221 | 0 | 0.006788754 | 11 | NaN | NaN | NaN | NaN | NaN | 240.99197 |
| Nop56 | AIDLINIESFSSR | -258 | -270 | 0 | 0.00025015 | 13 | NaN | NaN | NaN | NaN | NaN | 3830.5088 |
| Nop56 | VREWYGYHFPELVK | -189 | -202 | 1 | 0.001868731 | 14 | NaN | NaN | NaN | NaN | NaN | 341.02988 |
| Etf1 | GTIVTEEGKEK | -98 | -108 | 1 | 0.00111006 | 11 | NaN | NaN | NaN | NaN | NaN | 829.91724 |
| Txnrd2 | AIGDVAEGRPELTPTAIK | -356 | -373 | 1 | 0.001291103 | 18 | NaN | NaN | NaN | NaN | NaN | 889.5749 |
| Sgta | TPSASHEEQQE | -305 | -315 | 0 | 0.001868731 | 11 | NaN | NaN | NaN | NaN | NaN | 409.05887 |
| Kpna2 | AVPDLSTLACGYLR | -214 | -227 | 0 | 0.002265552 | 14 | NaN | NaN | NaN | NaN | NaN | 524.49084 |
| Sf3b1 | AIVNVIGMHK | -999 | -1008 | 0 | 0 | 10 | NaN | NaN | NaN | NaN | NaN | 1002.3998 |
| Sf3b1 | VDVDESTLSPEEQK | -480 | -493 | 0 | 0 | 14 | NaN | NaN | NaN | NaN | NaN | 556.6603 |
| Sf3b3 | VIVSDVQESFIWVR | -1000 | -1013 | 0 | 0.00025015 | 14 | NaN | NaN | NaN | NaN | NaN | 2187.787 |
| Sf3b3 | TSHEDHDFFQHVEMHLR | -1133 | -1149 | 0 | 0.003069674 | 17 | NaN | NaN | NaN | NaN | NaN | 377.33414 |
| Rpl30 | SGNNIELGTACGK | -75 | -87 | 0 | 0 | 13 | NaN | NaN | NaN | NaN | NaN | 508.3066 |
| Rps7 | AQQNNVEHKVETF | -161 | -173 | 1 | 0 | 13 | NaN | NaN | NaN | NaN | NaN | 778.2068 |
| Itprip | AIINHPLLFPR | -15 | -25 | 0 | 0.00025015 | 11 | NaN | NaN | NaN | NaN | NaN | 248.78781 |
| Sars1 | HRDEWLRPEDLPIK | -280 | -293 | 2 | 0.001055699 | 14 | NaN | NaN | NaN | NaN | NaN | 2530.1328 |
| Sars1 | VKLEAERF | -122 | -129 | 2 | 0.009955823 | 8 | NaN | NaN | NaN | NaN | NaN | 950.8768 |
| Sars1 | LHRDEWLRPEDLPIK | -279 | -293 | 2 | 0.00111006 | 15 | NaN | NaN | NaN | NaN | NaN | 119.58976 |
| Copz1 | ILEPSLYTVK | -5 | -14 | 0 | 0.000627943 | 10 | NaN | NaN | NaN | NaN | NaN | 1645.4222 |
| Tubb5 | HGDSDLQLDR | -37 | -46 | 0 | 0 | 10 | NaN | NaN | NaN | NaN | NaN | 11305.686 |
| Pds5b | HLASDFFLK | -62 | -70 | 0 | 0.001291103 | 9 | NaN | NaN | NaN | NaN | NaN | 311.56897 |
| Iqgap1 | WLDEIQGGIWQSNK | -602 | -615 | 0 | 0.003620005 | 14 | NaN | NaN | NaN | NaN | NaN | 1731.8802 |
| Iqgap1 | YLDELMK | -497 | -503 | 0 | 0.003966045 | 7 | NaN | NaN | NaN | NaN | NaN | 814.5066 |
| Khsrp | SQGDSIGSQLGPIHPPPR | -126 | -143 | 0 | 0.005270854 | 18 | NaN | NaN | NaN | NaN | NaN | 267.93942 |
| H2ac20;Hist2h | AIRNDEELNKLLGK | (87);(87) | (100);(100) | 2 | 0.000230707 | 14 | NaN | NaN | NaN | NaN | NaN | 1032.4524 |
| Zpr1 | LVQIPHFK | -272 | -279 | 0 | 0.003966045 | 8 | NaN | NaN | NaN | NaN | NaN | 909.9426 |
| Nop58 | IEEVDKEDEITEK | -445 | -457 | 1 | 0 | 13 | NaN | NaN | NaN | NaN | NaN | 5587.5503 |
| Wdr1 | VHPGGDTVAVGGTDGNVR | -453 | -470 | 0 | 0.00025015 | 18 | NaN | NaN | NaN | NaN | NaN | 357.75113 |
| Psma7 | YVAEIEKEKEENEK | -228 | -241 | 2 | 0.000550307 | 14 | NaN | NaN | NaN | NaN | NaN | 454.8956 |
| Rbm19 | AIVEFLEPLEAR | -622 | -633 | 0 | 0.001392869 | 12 | NaN | NaN | NaN | NaN | NaN | 331.3835 |
| Rdh11 | AKDFLAEEK | -108 | -116 | 1 | 0.00025015 | 9 | NaN | NaN | NaN | NaN | NaN | 799.28827 |
| Rps29 | AKDIGFIK | -47 | -54 | 1 | 0.004031417 | 8 | NaN | NaN | NaN | NaN | NaN | 772.0107 |
| Ezr | THNDIIHNENMR | -548 | -559 | 0 | 0 | 12 | NaN | NaN | NaN | NaN | NaN | 764.18414 |
| Pabpc1 | AVPNPVINPYQPAPPSGYFMAA | -391 | -412 | 0 | 0.00133086 | 22 | NaN | NaN | NaN | NaN | NaN | 334.62402 |
| Pabpc1 | AVPNPVINPYQPAPPSGYFM | -391 | -410 | 0 | 0.001291103 | 20 | NaN | NaN | NaN | NaN | NaN | 234.38995 |
| Eif3k | ENAYDLEANLAVLK | -39 | -52 | 0 | 0.00111006 | 14 | NaN | NaN | NaN | NaN | NaN | 391.00644 |

| | | | | | | | | | | | | |
|---|---|---|---|---|---|---|---|---|---|---|---|---|
| Hspa5 | HAVVTVPAYFNDAQR | -168 | -182 | 0 | 0.007493591 | 15 | NaN | NaN | NaN | NaN | NaN | 1050.7399 |
| Mapre1 | AKLEHEYIQNFK | -65 | -76 | 1 | 0 | 12 | NaN | NaN | NaN | NaN | NaN | 1042.9558 |
| Plec | QKVPAQQLQEAGIL | -2795 | -2808 | 1 | 0.001291103 | 14 | NaN | NaN | NaN | NaN | NaN | 584.1963 |
| Ncl | IKVPQNPHGKPK | -513 | -524 | 2 | 0 | 12 | NaN | NaN | NaN | NaN | NaN | 372.06903 |
| Ncl | PFNLFIGNLNPNK | -308 | -320 | 0 | 0.001291103 | 13 | NaN | NaN | NaN | NaN | NaN | 853.6357 |
| Cs | EVGKDVSDEK | -318 | -327 | 1 | 0.000907206 | 10 | NaN | NaN | NaN | NaN | NaN | 558.4329 |
| Hmgn2 | SAKPAPPKPEPK | -29 | -40 | 2 | 0 | 12 | NaN | NaN | NaN | NaN | NaN | 193.97308 |
| Fabp5 | KTETVCTFQDGALVQHQ | -82 | -98 | 1 | 0.004983786 | 17 | NaN | NaN | NaN | NaN | NaN | 2994.9895 |
| Pdia4 | NPIKFEGGNR | -606 | -615 | 1 | 0.002168954 | 10 | NaN | NaN | NaN | NaN | NaN | 822.86755 |
| Pdia3 | TADGIVSHLKK | -120 | -130 | 1 | 0.000431655 | 11 | NaN | NaN | NaN | NaN | NaN | 724.294 |
| Pdia3 | TGEVPVVAIR | -320 | -329 | 0 | 0 | 10 | NaN | NaN | NaN | NaN | NaN | 305.49588 |
| Pdia3 | ELSDFGLESTTGEVPVVAIR | -310 | -329 | 0 | 0.001868731 | 20 | NaN | NaN | NaN | NaN | NaN | 290.0229 |
| Rpl10 | FNADEFEDMVAEK | -176 | -188 | 0 | 0.000394361 | 13 | NaN | NaN | NaN | NaN | NaN | 308.44855 |
| Serbp1 | EETPEGEEHPVADTENK | -256 | -272 | 0 | 0.009002904 | 17 | NaN | NaN | NaN | NaN | NaN | 647.16626 |
| Esyt2 | ALALLEDEEQAVR | -87 | -99 | 0 | 0.002168954 | 13 | NaN | NaN | NaN | NaN | NaN | 417.53033 |
| Xrn2 | RNSSPSISPNTSFASDGSPSPLGGIK | -468 | -493 | 1 | 0.001610812 | 26 | NaN | NaN | NaN | NaN | NaN | 787.15955 |
| Rnh1 | NDLHEPGVR | -175 | -183 | 0 | 0.003998192 | 9 | NaN | NaN | NaN | NaN | NaN | 3099.9004 |
| Rnh1 | DISSAVQANPALTELSLR | -42 | -59 | 0 | 0.002068213 | 18 | NaN | NaN | NaN | NaN | NaN | 1414.9893 |
| Lrpprc | VFPCFDSAQSVR | -480 | -491 | 0 | 0.00025015 | 12 | NaN | NaN | NaN | NaN | NaN | 1243.1506 |
| Dctn2 | LLGPDAAINLADPDGALAK | -157 | -175 | 0 | 0.008768524 | 19 | NaN | NaN | NaN | NaN | NaN | 1188.3944 |
| Dctn2 | ALDLAVLDQVEAR | -271 | -283 | 0 | 0.001610812 | 13 | NaN | NaN | NaN | NaN | NaN | 514.76843 |
| Nup210 | VKQPHDPLVPVSA | -873 | -885 | 1 | 0.00272025 | 13 | NaN | NaN | NaN | NaN | NaN | 530.7704 |
| Acadvl | IFEGANDILR | -461 | -470 | 0 | 0.00025015 | 10 | NaN | NaN | NaN | NaN | NaN | 1605.8214 |
| Acadvl | ALDQFATVVEAK | -540 | -551 | 0 | 0 | 12 | NaN | NaN | NaN | NaN | NaN | 682.05347 |
| Myg1 | SSLCPGKPWQTK | -116 | -127 | 1 | 0 | 12 | NaN | NaN | NaN | NaN | NaN | 1023.2319 |
| Myg1 | NFYQHSWLPAR | -232 | -242 | 0 | 0.000627943 | 11 | NaN | NaN | NaN | NaN | NaN | 672.405 |
| Stim1 | ALDTVLFGPPLLTR | -194 | -207 | 0 | 0 | 14 | NaN | NaN | NaN | NaN | NaN | 2789.1504 |
| Cndp2 | DLHSGVYGGSVHEA | -226 | -239 | 0 | 0.005522803 | 14 | NaN | NaN | NaN | NaN | NaN | 2771.4739 |
| Cndp2 | DLHSGVYGGSVH | -226 | -237 | 0 | 0.004350979 | 12 | NaN | NaN | NaN | NaN | NaN | 796.1806 |
| Krt10 | QSLEASLAETEGR | -385 | -397 | 0 | 0.007288197 | 13 | NaN | NaN | NaN | NaN | NaN | 1009.09576 |
| Cltc | RALEHFTDLYDIK | -625 | -637 | 1 | 0 | 13 | NaN | NaN | NaN | NaN | NaN | 555.27216 |
| Cltc | RDPHLACVA | -912 | -920 | 1 | 0.003133457 | 9 | NaN | NaN | NaN | NaN | NaN | 3456.8699 |
| Cltc | IIEVGTPPTGNQPFPK | -230 | -245 | 0 | 0.003360691 | 16 | NaN | NaN | NaN | NaN | NaN | 725.2888 |
| Aars1 | ALELLTQEFGIPVER | -118 | -132 | 0 | 0.007670622 | 15 | NaN | NaN | NaN | NaN | NaN | 353.58286 |
| Pycard | ALENLSGDELKK | -11 | -22 | 1 | 0.00122419 | 12 | NaN | NaN | NaN | NaN | NaN | 705.451 |
| Gbp7 | ISIEDSIMQSDK | -468 | -479 | 0 | 0.004350979 | 12 | NaN | NaN | NaN | NaN | NaN | 364.18347 |
| Sugt1 | NMYPSSSHYTR | -246 | -256 | 0 | 0 | 11 | NaN | NaN | NaN | NaN | NaN | 723.0066 |
| Sugt1 | GDPQAALEELTK | -25 | -36 | 0 | 0.00025015 | 12 | NaN | NaN | NaN | NaN | NaN | 397.3336 |
| Dkc1 | TGFINLDKPSNPSSHEVVAW | -89 | -108 | 1 | 0.00025015 | 20 | NaN | NaN | NaN | NaN | NaN | 1195.1923 |
| Rpl7l1 | ALEVPDKHPLAFVIR | -78 | -92 | 1 | 0.004350979 | 15 | NaN | NaN | NaN | NaN | NaN | 175.77727 |
| Psmd2 | DAGEKDKEQELSEEDK | -35 | -50 | 2 | 0.009362332 | 16 | NaN | NaN | NaN | NaN | NaN | 936.05194 |
| Ftl1 | QNYSTEVEAAVNR | -7 | -19 | 0 | 0 | 13 | NaN | NaN | NaN | NaN | NaN | 509.34955 |
| Baz1a | ALFQIEQGIER | -1084 | -1094 | 0 | 0 | 11 | NaN | NaN | NaN | NaN | NaN | 728.1482 |
| Ndufv1 | GDGAAWPVQGLIR | -428 | -440 | 0 | 0 | 13 | NaN | NaN | NaN | NaN | NaN | 801.21405 |
| Raver1 | WTDAGQLTPALLHSR | -207 | -221 | 0 | 0 | 15 | NaN | NaN | NaN | NaN | NaN | 951.5696 |
| Dcps | TPFQVEHVAQLLTGSPELK | -86 | -104 | 0 | 0.00051306 | 19 | NaN | NaN | NaN | NaN | NaN | 295.11957 |
| Lrrc47 | TDLITLPPHVTVVR | -568 | -581 | 0 | 0.006735683 | 14 | NaN | NaN | NaN | NaN | NaN | 1121.6025 |
| Apex1 | GDLNVAHEEIDLR | -208 | -220 | 0 | 0 | 13 | NaN | NaN | NaN | NaN | NaN | 921.48584 |
| Apex1 | LPAELQELPGLTH | -103 | -115 | 0 | 0.00272025 | 13 | NaN | NaN | NaN | NaN | NaN | 1413.8811 |
| Dars1 | LQSGICHLFR | -198 | -207 | 0 | 0 | 10 | NaN | NaN | NaN | NaN | NaN | 927.0966 |
| Ptbp1 | MDGIVPDIAVGTK | -1 | -13 | 0 | 0.008498216 | 13 | NaN | NaN | NaN | NaN | NaN | 1272.7737 |
| Plcl2 | GAAGGALPTSPSPALGAK | -7 | -24 | 0 | 0.001023236 | 18 | NaN | NaN | NaN | NaN | NaN | 539.41187 |
| Bak1 | LEQLQPTAGNAYELFTK | -95 | -111 | 0 | 0.006100831 | 17 | NaN | NaN | NaN | NaN | NaN | 704.0383 |
| Rps12 | VLASNCDEPMYVK | -51 | -63 | 0 | 0 | 13 | NaN | NaN | NaN | NaN | NaN | 2401.9316 |
| Rps12 | KTALIHDGLAR | -23 | -33 | 1 | 0.00051306 | 11 | NaN | NaN | NaN | NaN | NaN | 443.12207 |
| Rrm1 | IIDINYYPIPEAHLS | -480 | -494 | 0 | 0.004224523 | 15 | NaN | NaN | NaN | NaN | NaN | 543.29266 |
| Eif3h | DFSPEALKK | -181 | -189 | 1 | 0.000431655 | 9 | NaN | NaN | NaN | NaN | NaN | 578.01666 |
| Set | EQQEAIEHIDEVQ | -39 | -51 | 0 | 0.001776473 | 13 | NaN | NaN | NaN | NaN | NaN | 665.16925 |
| Xpot | LFVTEYLTK | -112 | -120 | 0 | 0.00142999 | 9 | NaN | NaN | NaN | NaN | NaN | 357.96484 |
| Sp110 | ALLQHFIY | -11 | -18 | 0 | 0.006384543 | 8 | NaN | NaN | NaN | NaN | NaN | 6872.0117 |
| Shmt2 | INFAVFPSLQGGPHNHAIAA | -315 | -334 | 0 | 0 | 20 | NaN | NaN | NaN | NaN | NaN | 663.1204 |
| Shmt2 | GYSLVSGGTDTHLVLVDLRPK | -369 | -389 | 1 | 0 | 21 | NaN | NaN | NaN | NaN | NaN | 621.12366 |
| Shmt2 | ESLSDSDPEMWELLQR | -47 | -62 | 0 | 0 | 16 | NaN | NaN | NaN | NaN | NaN | 724.7488 |
| Shmt2 | ESLSDSDPEMWELLQR | -47 | -62 | 0 | 0 | 16 | NaN | NaN | NaN | NaN | NaN | 724.7488 |
| Hsp90aa1 | KHSQFIGYPITLF | -209 | -221 | 1 | 0.00078661 | 13 | NaN | NaN | NaN | NaN | NaN | 308.22928 |
| Hsp90aa1 | TELAEDKENYKK | -426 | -437 | 2 | 0 | 12 | NaN | NaN | NaN | NaN | NaN | 468.07327 |
| Hsp90aa1 | EEMPPLEGDDDTSRMEEVD | -715 | -733 | 1 | 0.005668207 | 19 | NaN | NaN | NaN | NaN | NaN | 368.305 |
| Hsp90aa1 | FYEQFSK | -438 | -444 | 0 | 0.000431655 | 7 | NaN | NaN | NaN | NaN | NaN | 361.38638 |
| Gtf2i | QKVENLFNEK | -735 | -744 | 1 | 0.00201047 | 10 | NaN | NaN | NaN | NaN | NaN | 587.7711 |
| Chd1l | KDLDAFENETAK | -290 | -301 | 1 | 0.001868731 | 12 | NaN | NaN | NaN | NaN | NaN | 940.1509 |
| Mphosph10 | DAALLAPEEIKEK | -546 | -558 | 1 | 0.00051306 | 13 | NaN | NaN | NaN | NaN | NaN | 784.4962 |
| Mphosph10 | KRPENSLLEETLHFDHAVR | -395 | -413 | 2 | 0.00111006 | 19 | NaN | NaN | NaN | NaN | NaN | 344.5345 |
| Utp14a | EAFAGDDVIK | -594 | -603 | 0 | 0.002910424 | 10 | NaN | NaN | NaN | NaN | NaN | 6643.2505 |
| Igtp | YRDPLEIISQVCDK | -266 | -279 | 1 | 0.000230707 | 14 | NaN | NaN | NaN | NaN | NaN | 1475.2704 |
| Igtp | YSSQNSSSPEVIEDIGK | -43 | -59 | 0 | 0 | 17 | NaN | NaN | NaN | NaN | NaN | 1528.786 |
| Uba6 | ALQLLHCFPLETR | -715 | -727 | 0 | 0.00383302 | 13 | NaN | NaN | NaN | NaN | NaN | 572.5491 |
| Rpn2 | ALQTASHLSQQADLR | -164 | -178 | 0 | 0.000471409 | 15 | NaN | NaN | NaN | NaN | NaN | 482.67563 |
| Rpn2 | SGFGLPLASNEALGALTAR | -133 | -151 | 0 | 0.00025015 | 19 | NaN | NaN | NaN | NaN | NaN | 255.11874 |

| | | | | | | | | | | | | |
|---|---|---|---|---|---|---|---|---|---|---|---|---|
| Uhrf1 | TVWEDTDLGLYK | -121 | -132 | 0 | 0.005821388 | 12 | NaN | NaN | NaN | NaN | NaN | 1176.5679 |
| Ubr4 | LDSEACEVLFSKPVK | -948 | -962 | 1 | 0 | 15 | NaN | NaN | NaN | NaN | NaN | 3158.524 |
| Scfd1 | LFTGDPLGTGQFSFQR | -237 | -252 | 0 | 0.006624942 | 16 | NaN | NaN | NaN | NaN | NaN | 2090.2073 |
| Map4k1 | PVVVETRPTDDPTAPSNL | -806 | -823 | 1 | 0.000907206 | 18 | NaN | NaN | NaN | NaN | NaN | 517.8612 |
| Prim1 | ALVPETIHDELQR | -245 | -257 | 0 | 0 | 13 | NaN | NaN | NaN | NaN | NaN | 969.1245 |
| Prim1 | VFEQFLENLDK | -391 | -401 | 0 | 0.000550307 | 11 | NaN | NaN | NaN | NaN | NaN | 481.4039 |
| Psmd14 | QTGRPEMVVGW | -101 | -111 | 1 | 0.008828666 | 11 | NaN | NaN | NaN | NaN | NaN | 662.2282 |
| Psmd14 | AMPQSGTGVSVEAVDPVFQAK | -74 | -94 | 0 | 0.002466863 | 21 | NaN | NaN | NaN | NaN | NaN | 433.5521 |
| Ppp2r1b | FTNLASDEQDSVR | -221 | -233 | 0 | 0.001392869 | 13 | NaN | NaN | NaN | NaN | NaN | 973.90857 |
| Psap | ANEDVCQDCMK | -192 | -202 | 0 | 0 | 11 | NaN | NaN | NaN | NaN | NaN | 382.982 |
| Eif3a | NNPESQSMHLETR | -221 | -233 | 0 | 0.002780285 | 13 | NaN | NaN | NaN | NaN | NaN | 1397.9777 |
| Eif3a | RHCDLQVR | -476 | -483 | 1 | 0.001392869 | 8 | NaN | NaN | NaN | NaN | NaN | 422.69028 |
| Lactb2 | ANIIYPGHGPVIHNAEAK | -192 | -209 | 0 | 0.000627943 | 18 | NaN | NaN | NaN | NaN | NaN | 286.14783 |
| Lap3 | YMGSPNATEAPLVFVGK | -266 | -282 | 0 | 0.000826806 | 17 | NaN | NaN | NaN | NaN | NaN | 493.56543 |
| Pdcd11 | ELYNGCLPEGK | -1376 | -1386 | 0 | 0.008890469 | 11 | NaN | NaN | NaN | NaN | NaN | 1395.1664 |
| Idh3a | ENTEGEYSGIEHVIVD | -147 | -162 | 0 | 0.002099344 | 16 | NaN | NaN | NaN | NaN | NaN | 2039.1014 |
| Idh3a | ANVRPCVSIEGYK | -122 | -134 | 1 | 0.008284062 | 13 | NaN | NaN | NaN | NaN | NaN | 655.6717 |
| Acly | NFLLNASGSTSTPAPSR | -436 | -452 | 0 | 0.001636916 | 17 | NaN | NaN | NaN | NaN | NaN | 1421.5056 |
| Gtf2b | KAVELDLVPGR | -238 | -248 | 1 | 0 | 11 | NaN | NaN | NaN | NaN | NaN | 13832.319 |
| Anp32e | ELSDNIISGGLEVLAEK | -70 | -86 | 0 | 0.00025015 | 17 | NaN | NaN | NaN | NaN | NaN | 178.98737 |
| Gins1 | GGDEGLDITQDVKPPK | -129 | -144 | 1 | 0.000550307 | 16 | NaN | NaN | NaN | NaN | NaN | 398.4412 |
| Hspd1 | AVEEGIVLGGGCALLR | -431 | -446 | 0 | 0 | 16 | NaN | NaN | NaN | NaN | NaN | 1102.6725 |
| Hspd1 | VTEIPKEEK | -546 | -554 | 1 | 0.009002904 | 9 | NaN | NaN | NaN | NaN | NaN | 374.81824 |
| Rpl27a | QSFCPTVNLDK | -67 | -77 | 0 | 0.000431655 | 11 | NaN | NaN | NaN | NaN | NaN | 946.0805 |
| Rpl27a | GVAPIIDVVR | -96 | -105 | 0 | 0.005301614 | 10 | NaN | NaN | NaN | NaN | NaN | 282.33405 |
| Ddx17 | ELAQQVQQVADDYGK | -176 | -190 | 0 | 0.00111006 | 15 | NaN | NaN | NaN | NaN | NaN | 1207.0222 |
| Psme1 | APLDIPVPDPVKEK | -59 | -72 | 1 | 0 | 14 | NaN | NaN | NaN | NaN | NaN | 1891.8877 |
| Psme1 | EPALNEANLSNL | -46 | -57 | 0 | 0.005758933 | 12 | NaN | NaN | NaN | NaN | NaN | 288.32 |
| Iars1 | VQQVPENISFPAEEEK | -2 | -17 | 0 | 0 | 16 | NaN | NaN | NaN | NaN | NaN | 966.0042 |
| Gm12250 | FGLDDESLENMAK | -318 | -330 | 0 | 0 | 13 | NaN | NaN | NaN | NaN | NaN | 1041.4047 |
| Isyna1 | APLVPPGSPVVNALFR | -466 | -481 | 0 | 0.002972566 | 16 | NaN | NaN | NaN | NaN | NaN | 293.18616 |
| Ubap2 | APPNLPQGVPPLLHNQ | -807 | -822 | 0 | 0.005605646 | 16 | NaN | NaN | NaN | NaN | NaN | 741.5186 |
| Gars1 | YTILEHTFHVR | -576 | -586 | 0 | 0 | 11 | NaN | NaN | NaN | NaN | NaN | 4608.134 |
| Ighg2b | APQVYILPPPAEQLSR | -233 | -248 | 0 | 0.001636916 | 16 | NaN | NaN | NaN | NaN | NaN | 2013.3817 |
| Etfa | GISGAIQHLAGMK | -279 | -291 | 0 | 0.003745449 | 13 | NaN | NaN | NaN | NaN | NaN | 2806.1736 |
| Etfa | YDLADQLHAAVGASR | -235 | -249 | 0 | 0.009272078 | 15 | NaN | NaN | NaN | NaN | NaN | 508.92584 |
| Eif2a | GPPHFTESTVLPR | -21 | -33 | 0 | 0 | 13 | NaN | NaN | NaN | NaN | NaN | 454.47522 |
| Rrm2 | EYLFNAIETMPCVK | -192 | -205 | 0 | 0.000627943 | 14 | NaN | NaN | NaN | NaN | NaN | 654.8346 |
| Oga | APVIWDNIHAN | -274 | -284 | 0 | 0.000591124 | 11 | NaN | NaN | NaN | NaN | NaN | 1815.145 |
| Oga | TIRPYFPK | -716 | -723 | 1 | 0.006978206 | 8 | NaN | NaN | NaN | NaN | NaN | 737.44165 |
| Yars1 | PEEVIPSR | -360 | -367 | 0 | 0.002099344 | 8 | NaN | NaN | NaN | NaN | NaN | 1774.0323 |
| H2-D1 | YFETAVSRPGLEEPR | -31 | -45 | 1 | 0.000298805 | 15 | NaN | NaN | NaN | NaN | NaN | 833.9363 |
| Exosc4 | AQADGSAYIEQGNTK | -33 | -47 | 0 | 0.007343498 | 15 | NaN | NaN | NaN | NaN | NaN | 481.44537 |
| Esyt1 | GDSPSEAPVGPLGQVK | -949 | -964 | 0 | 0.000230707 | 16 | NaN | NaN | NaN | NaN | NaN | 557.0285 |
| Ppid | SHPDFPEDADIDLK | -201 | -214 | 0 | 0.001904399 | 14 | NaN | NaN | NaN | NaN | NaN | 966.3124 |
| Atp5f1a | QFGSDLDAATQQLLSR | -448 | -463 | 0 | 0.00078661 | 16 | NaN | NaN | NaN | NaN | NaN | 837.3988 |
| Atp5f1a | EAYPGDVFYL | -335 | -344 | 0 | 0.002697088 | 10 | NaN | NaN | NaN | NaN | NaN | 327.44952 |
| Atp5f1a | LSHVISQHQSLLGNIR | -512 | -527 | 0 | 0.004224523 | 16 | NaN | NaN | NaN | NaN | NaN | 286.81732 |
| Bin1 | LNQNLNDVLVSLEK | -259 | -272 | 0 | 0.000627943 | 14 | NaN | NaN | NaN | NaN | NaN | 616.271 |
| Gsn | VADENPFAQGALR | -313 | -325 | 0 | 0 | 13 | NaN | NaN | NaN | NaN | NaN | 2163.9636 |
| Rpl8 | CCLEEKPGDR | -114 | -123 | 1 | 0.001811524 | 10 | NaN | NaN | NaN | NaN | NaN | 3457.9004 |
| Msn | SKYGDFNK | -132 | -139 | 1 | 0.000869338 | 8 | NaN | NaN | NaN | NaN | NaN | 19298.404 |
| Msn | KTQEQLASEMAELTAR | -412 | -427 | 1 | 0.007343498 | 16 | NaN | NaN | NaN | NaN | NaN | 1254.9652 |
| Eif3b | VDNAYWLWTFQGR | -666 | -678 | 0 | 0.000627943 | 13 | NaN | NaN | NaN | NaN | NaN | 1087.1124 |
| Eif3b | SDVEWDPTGR | -644 | -653 | 0 | 0.000550307 | 10 | NaN | NaN | NaN | NaN | NaN | 482.32803 |
| Ruvbl1 | EVYEGEVTELTPCETENPMGGYGK | -129 | -152 | 0 | 0.001392869 | 24 | NaN | NaN | NaN | NaN | NaN | 256.91293 |
| Eftud2 | AQVVGDVDTSLPR | -381 | -393 | 0 | 0.00025015 | 13 | NaN | NaN | NaN | NaN | NaN | 819.6091 |
| Add1 | YHGILVDEEEK | -273 | -283 | 0 | 0.00025015 | 11 | NaN | NaN | NaN | NaN | NaN | 1758.3838 |
| Nup98 | ASDAIINENYDYLK | -1654 | -1667 | 0 | 0.00450079 | 14 | NaN | NaN | NaN | NaN | NaN | 1216.097 |
| Capza1 | FTITPPSAQVVGVLK | -179 | -193 | 0 | 0.009122264 | 15 | NaN | NaN | NaN | NaN | NaN | 1469.2665 |
| Hsp90b1 | SLTDENALAGNEELTVK | -119 | -135 | 0 | 0.002466863 | 17 | NaN | NaN | NaN | NaN | NaN | 931.2973 |
| Hsp90b1 | NDIKPIWQRPSK | -337 | -348 | 2 | 0.000230707 | 12 | NaN | NaN | NaN | NaN | NaN | 787.3721 |
| Dck | GRNEEQGIPLEYLEK | -193 | -207 | 1 | 0.004439092 | 15 | NaN | NaN | NaN | NaN | NaN | 1014.39716 |
| Tnpo3 | DEVIHSIFK | -358 | -366 | 0 | 0.002780285 | 9 | NaN | NaN | NaN | NaN | NaN | 1455.1205 |
| Grap2 | ELGSQEGYVPK | -38 | -48 | 0 | 0.002566359 | 11 | NaN | NaN | NaN | NaN | NaN | 521.24994 |
| Grap2 | SQGGPHPSGTVGEEIRPSVN | -163 | -182 | 1 | 0.000907206 | 20 | NaN | NaN | NaN | NaN | NaN | 108.962105 |
| Dtx3l | KPVEDLRPR | -90 | -98 | 2 | 0.007791272 | 9 | NaN | NaN | NaN | NaN | NaN | 328.76456 |
| Exosc10 | KFVKPIFTDE | -480 | -489 | 2 | 0.00111006 | 10 | NaN | NaN | NaN | NaN | NaN | 838.16187 |
| Exosc10 | QYELDHFTPPQSVLQR | -259 | -274 | 0 | 0.001868731 | 16 | NaN | NaN | NaN | NaN | NaN | 356.19055 |
| Hba | DFTPAVHASLDK | -117 | -128 | 0 | 0 | 12 | NaN | NaN | NaN | NaN | NaN | 1335.4261 |
| Akr1b1 | MEQLVDEGLVK | -145 | -155 | 0 | 0.00078661 | 11 | NaN | NaN | NaN | NaN | NaN | 4136.882 |
| Arpc2 | SHTAPQVLFSHR | -192 | -203 | 0 | 0.00025015 | 12 | NaN | NaN | NaN | NaN | NaN | 3171.421 |
| Arpc2 | YRDDETMYVESK | -146 | -157 | 1 | 0.00025015 | 12 | NaN | NaN | NaN | NaN | NaN | 830.08105 |
| Supt16h | ASITSEVFNK | -187 | -196 | 0 | 0 | 10 | NaN | NaN | NaN | NaN | NaN | 6754.6753 |
| Supt16h | NISMSVEGDYTYLR | -556 | -569 | 0 | 0 | 14 | NaN | NaN | NaN | NaN | NaN | 967.7321 |
| Sf3a1 | TEDSLMPEEEFLR | -685 | -697 | 0 | 0.000550307 | 13 | NaN | NaN | NaN | NaN | NaN | 793.14856 |
| Psmd1 | SQQFLSSVIQNLR | -257 | -269 | 0 | 0.000712251 | 13 | NaN | NaN | NaN | NaN | NaN | 785.40283 |
| Pak1ip1 | ASLPPAAEPCPDQPK | -311 | -325 | 0 | 0.00025015 | 15 | NaN | NaN | NaN | NaN | NaN | 1338.7217 |

| | | | | | | | | | | | | |
|---|---|---|---|---|---|---|---|---|---|---|---|---|
| Cryl1 | VPDDPEHLAAR | -288 | -298 | 0 | 0 | 11 | NaN | NaN | NaN | NaN | NaN | 427.24448 |
| Hnrnpa0 | MAASPHAVDGNTVELK | -65 | -80 | 0 | 0.00025015 | 16 | NaN | NaN | NaN | NaN | NaN | 773.955 |
| Gba1 | VDSPIIVDIPK | -416 | -426 | 0 | 0.000907206 | 11 | NaN | NaN | NaN | NaN | NaN | 851.57794 |
| Zc3hav1 | YSHDVLSEQNFQVL | -108 | -121 | 0 | 0.001776473 | 14 | NaN | NaN | NaN | NaN | NaN | 848.25244 |
| Cat | ASQRPDVLTTGGGNPIGDK | -20 | -38 | 1 | 0 | 19 | NaN | NaN | NaN | NaN | NaN | 484.0715 |
| Cat | SALEHSVQCAVDVK | -417 | -430 | 0 | 0.00025015 | 14 | NaN | NaN | NaN | NaN | NaN | 1359.3881 |
| Dazap1 | FVGGLDWSTTQETLR | -13 | -27 | 0 | 0 | 15 | NaN | NaN | NaN | NaN | NaN | 717.64923 |
| Dazap1 | ASRPHTLDGR | -69 | -78 | 1 | 0.002752526 | 10 | NaN | NaN | NaN | NaN | NaN | 441.0599 |
| Pcbp2 | NPVEGSTDR | -322 | -330 | 0 | 0.004255025 | 9 | NaN | NaN | NaN | NaN | NaN | 9910.525 |
| Pcbp2 | THGNTGFSGIESSSPEVK | -255 | -272 | 0 | 0.000471409 | 18 | NaN | NaN | NaN | NaN | NaN | 939.6317 |
| Mki67 | RHSVNLDEGESA | -335 | -346 | 1 | 0.001055699 | 12 | NaN | NaN | NaN | NaN | NaN | 1010.7375 |
| Mki67 | ASSAHLVTIK | -2 | -11 | 0 | 0 | 10 | NaN | NaN | NaN | NaN | NaN | 596.8686 |
| Mki67 | ELFQTPASGKDPVTVDETTK | -1801 | -1820 | 1 | 0.002566359 | 20 | NaN | NaN | NaN | NaN | NaN | 332.29413 |
| Serpina3f | ASSNTDFAFSLYK | -38 | -50 | 0 | 0.00025015 | 13 | NaN | NaN | NaN | NaN | NaN | 1526.2756 |
| Trim28 | SMEQPGGTLDLTLIR | -730 | -744 | 0 | 0.00450079 | 15 | NaN | NaN | NaN | NaN | NaN | 390.8451 |
| Sh3kbp1 | ASSPSLFSTEGKPK | -629 | -642 | 1 | 0.00078661 | 14 | NaN | NaN | NaN | NaN | NaN | 9862.928 |
| Sh3kbp1 | LSELDEEKK | -682 | -690 | 1 | 0.008736733 | 9 | NaN | NaN | NaN | NaN | NaN | 813.8844 |
| Rgs10 | ASSQVNVEGQSR | -102 | -113 | 0 | 0 | 12 | NaN | NaN | NaN | NaN | NaN | 1712.2711 |
| Ogt | GQLQEAIEHYR | -103 | -113 | 0 | 0 | 11 | NaN | NaN | NaN | NaN | NaN | 1971.7103 |
| Tpr | QQVLERPELNK | -80 | -90 | 1 | 0.004031417 | 11 | NaN | NaN | NaN | NaN | NaN | 2942.5664 |
| Tpr | KKLENEVEQR | -927 | -936 | 2 | 0.00078661 | 10 | NaN | NaN | NaN | NaN | NaN | 2471.2544 |
| Tpr | ELLHNQNSWLNTELK | -262 | -276 | 0 | 0.00025015 | 15 | NaN | NaN | NaN | NaN | NaN | 915.5009 |
| Pepd | GLDVHDVGGYPEGVER | -373 | -388 | 0 | 0.00025015 | 16 | NaN | NaN | NaN | NaN | NaN | 635.959 |
| Mif | ELTQQLAQATGK | -22 | -33 | 0 | 0.004224523 | 12 | NaN | NaN | NaN | NaN | NaN | 490.49512 |
| Phb1 | GGVVNSALYNVDAGHR | -20 | -35 | 0 | 0 | 16 | NaN | NaN | NaN | NaN | NaN | 760.29034 |
| Stk10 | AVPPLVNHTQDSANVTQPSLDSNK | -333 | -356 | 0 | 0 | 24 | NaN | NaN | NaN | NaN | NaN | 459.2958 |
| Eif5a | EDLRLPEGDLGK | -110 | -121 | 1 | 0.00025015 | 12 | NaN | NaN | NaN | NaN | NaN | 12207.456 |
| Bag6 | SFFHQHYLGGQEPT | -809 | -822 | 0 | 0.00830926 | 14 | NaN | NaN | NaN | NaN | NaN | 5597.0923 |
| Bag6 | ATHGGAPLPGTR | -107 | -118 | 0 | 0.00025015 | 12 | NaN | NaN | NaN | NaN | NaN | 1677.6896 |
| Rad50 | ILELDQELTK | -474 | -483 | 0 | 0.000627943 | 10 | NaN | NaN | NaN | NaN | NaN | 1494.622 |
| Rad50 | VFQGTDEQLNDLYHNH | -296 | -311 | 0 | 0.001392869 | 16 | NaN | NaN | NaN | NaN | NaN | 1099.3849 |
| Rad50 | VFQGTDEQLNDLYHNHQR | -296 | -313 | 0 | 0 | 18 | NaN | NaN | NaN | NaN | NaN | 318.7846 |
| Nomo1 | FLLFSSLVNK | -227 | -236 | 0 | 0.002845514 | 10 | NaN | NaN | NaN | NaN | NaN | 254.80144 |
| Cotl1 | ATKIDKEACR | -2 | -11 | 2 | 0.004857192 | 10 | NaN | NaN | NaN | NaN | NaN | 981.5435 |
| Ndufb4 | SANIYPNFRPTPK | -73 | -85 | 1 | 0.005176406 | 13 | NaN | NaN | NaN | NaN | NaN | 489.42972 |
| Psmc6 | IHIDLPNEQAR | -299 | -309 | 0 | 0 | 11 | NaN | NaN | NaN | NaN | NaN | 34341.215 |
| Supt6h | IKEDGDEEGEGEEAEDEEQRGPELK | -492 | -516 | 2 | 0.007791272 | 25 | NaN | NaN | NaN | NaN | NaN | 593.3537 |
| Supt6h | SLGDSTDSYIEVLDGSR | -1041 | -1057 | 0 | 0.003333688 | 17 | NaN | NaN | NaN | NaN | NaN | 492.72754 |
| Ap1b1 | ESHPSSNLLDLNPQSINK | -189 | -206 | 0 | 0.00659934 | 18 | NaN | NaN | NaN | NaN | NaN | 439.12292 |
| Larp1 | WHDQDETSSVK | -291 | -301 | 0 | 0 | 11 | NaN | NaN | NaN | NaN | NaN | 1454.6189 |
| Larp1 | KMDADGFLPITLIA | -404 | -417 | 1 | 0.005605646 | 14 | NaN | NaN | NaN | NaN | NaN | 265.51877 |
| Septin11 | LTIVDTVGFGDQINK | -96 | -110 | 0 | 0.00078661 | 15 | NaN | NaN | NaN | NaN | NaN | 491.91632 |
| Cyp51a1 | LKDSWAER | -411 | -418 | 1 | 0.00644558 | 8 | NaN | NaN | NaN | NaN | NaN | 423.33832 |
| Grk2 | QLGHEEDYALGK | -546 | -557 | 0 | 0 | 12 | NaN | NaN | NaN | NaN | NaN | 1431.2816 |
| Sh3bgrl3 | AVEQDTLQEFLK | -80 | -91 | 0 | 0 | 12 | NaN | NaN | NaN | NaN | NaN | 5145.393 |
| Tuba4a | TIGGGDDSFTTFFCETGAGK | -41 | -60 | 0 | 0.000230707 | 20 | NaN | NaN | NaN | NaN | NaN | 431.61075 |
| Msh2 | VFVTPLIDLR | -435 | -444 | 0 | 0.009657961 | 10 | NaN | NaN | NaN | NaN | NaN | 294.07703 |
| Cars1 | FENDCWVR | -468 | -475 | 0 | 0.002296977 | 8 | NaN | NaN | NaN | NaN | NaN | 510.3735 |
| Capg | AVHLNTLLGER | -81 | -91 | 0 | 0.001023236 | 11 | NaN | NaN | NaN | NaN | NaN | 2245.0833 |
| Bop1 | LTDEQVALVH | -182 | -191 | 0 | 0.00149795 | 10 | NaN | NaN | NaN | NaN | NaN | 1595.5372 |
| Bop1 | DELDQFLDK | -153 | -161 | 0 | 0.006677519 | 9 | NaN | NaN | NaN | NaN | NaN | 1431.9628 |
| Prpf6 | DANDPVDDR | -39 | -47 | 0 | 0.001392869 | 9 | NaN | NaN | NaN | NaN | NaN | 313.6833 |
| Cmpk2 | SQGLLWQR | -168 | -175 | 0 | 0.00025015 | 8 | NaN | NaN | NaN | NaN | NaN | 853.77106 |
| Gemin5 | TVVTEHTLHQHTISAL | -100 | -115 | 0 | 0.000431655 | 16 | NaN | NaN | NaN | NaN | NaN | 315.71616 |
| Calr | FTVKHEQNIDCGGGYVK | -95 | -111 | 1 | 0.002068213 | 17 | NaN | NaN | NaN | NaN | NaN | 693.2085 |
| Rhog | FSIASPPSYENVR | -82 | -94 | 0 | 0 | 13 | NaN | NaN | NaN | NaN | NaN | 1511.0875 |
| Noc4l | VHRPQGPELDADPYDPTEKDPAR | -405 | -427 | 2 | 0 | 23 | NaN | NaN | NaN | NaN | NaN | 674.89813 |
| Cpne1 | QIEYHFETVQK | -62 | -72 | 0 | 0 | 11 | NaN | NaN | NaN | NaN | NaN | 679.0405 |
| Aip | RNIAEGKDPLEGQR | -106 | -119 | 2 | 0.000550307 | 14 | NaN | NaN | NaN | NaN | NaN | 658.51495 |
| Metap2 | AVQQELDKESGALVDEVAK | -51 | -69 | 1 | 0 | 19 | NaN | NaN | NaN | NaN | NaN | 456.29367 |
| Metap2 | AVQQELDKESGALVDEVAK | -51 | -69 | 1 | 0 | 19 | NaN | NaN | NaN | NaN | NaN | 456.29367 |
| Ppp1r12a | QWLNSGHIS | -183 | -191 | 0 | 0.00555405 | 9 | NaN | NaN | NaN | NaN | NaN | 603.94666 |
| Ppp1r12a | ILVDNLCDMETVNK | -250 | -263 | 0 | 0.000298805 | 14 | NaN | NaN | NaN | NaN | NaN | 329.219 |
| Ywhah | ELETVCNDVLAL | -92 | -103 | 0 | 0.002199523 | 12 | NaN | NaN | NaN | NaN | NaN | 640.33405 |
| H4c1 | NVIRDAVTYTEHAK | -65 | -78 | 1 | 0.00078661 | 14 | NaN | NaN | NaN | NaN | NaN | 760.1445 |
| Trip12 | GNIPTWLTELGK | -1558 | -1569 | 0 | 0.001708008 | 12 | NaN | NaN | NaN | NaN | NaN | 1199.5376 |
| Trip12 | SADLGSPSLQHSR | -1005 | -1017 | 0 | 0.001868731 | 13 | NaN | NaN | NaN | NaN | NaN | 1109.0841 |
| Trip12 | AVVVDAIPVFLEK | -523 | -535 | 0 | 0 | 13 | NaN | NaN | NaN | NaN | NaN | 592.9049 |
| Pfas | GWVGSDPSEEQAEPGQDSQPTQPGLLLR | -1167 | -1194 | 0 | 0 | 28 | NaN | NaN | NaN | NaN | NaN | 789.443 |
| Pfas | DSWDLDFYTK | -196 | -205 | 0 | 0 | 10 | NaN | NaN | NaN | NaN | NaN | 432.87268 |
| Dnajc9 | SLQVHPDRVEEDQKEDATR | -39 | -57 | 2 | 0.005605646 | 19 | NaN | NaN | NaN | NaN | NaN | 641.139 |
| Rack1 | HIGHTGYLNTVTVSPDGSLCASGGK | -188 | -212 | 0 | 0.000471409 | 25 | NaN | NaN | NaN | NaN | NaN | 374.9891 |
| Dock11 | YHVSCEINTK | -733 | -742 | 0 | 0.004099215 | 10 | NaN | NaN | NaN | NaN | NaN | 742.55 |
| Glo1 | FLAYEDKNDIPK | -72 | -83 | 1 | 0.00025015 | 12 | NaN | NaN | NaN | NaN | NaN | 1166.2069 |
| Prrc2c | NRPLEKPYIR | -1288 | -1297 | 2 | 0.002329279 | 10 | NaN | NaN | NaN | NaN | NaN | 437.93018 |
| Psmd6 | KVNEIVETNRPDSK | -349 | -362 | 2 | 0.000431655 | 14 | NaN | NaN | NaN | NaN | NaN | 559.647 |
| Psmd6 | QYLFSLYECR | -258 | -267 | 0 | 0.002199523 | 10 | NaN | NaN | NaN | NaN | NaN | 1071.7155 |
| Psmd6 | KDWLFAPH | -284 | -291 | 1 | 0.003166358 | 8 | NaN | NaN | NaN | NaN | NaN | 646.8043 |

| | | | | | | | | | | | | |
|---|---|---|---|---|---|---|---|---|---|---|---|---|
| Ptprc | TLVSWPEPVSKPESASKPHGY | -392 | -412 | 2 | 0 | 21 | NaN | NaN | NaN | NaN | NaN | 3838.1038 |
| Ptprc | TLVSWPEPVSKPESASKPHGYVLCYK | -392 | -417 | 2 | 0.009985598 | 26 | NaN | NaN | NaN | NaN | NaN | 668.48804 |
| Alb | PCFSALTVDETYVPK | -510 | -524 | 0 | 0.000907206 | 15 | NaN | NaN | NaN | NaN | NaN | 781.735 |
| Rps27a | CLTYCFNKPEDK | -145 | -156 | 1 | 0.00111006 | 12 | NaN | NaN | NaN | NaN | NaN | 409.5706 |
| Lsp1 | CEEQEELHAE | -10 | -19 | 0 | 0.00111006 | 10 | NaN | NaN | NaN | NaN | NaN | 100.06917 |
| Tfrc | VPQLNQMVR | -583 | -591 | 0 | 0.008014596 | 9 | NaN | NaN | NaN | NaN | NaN | 11631.896 |
| Tfrc | FCFCEDADYPYLGTR | -558 | -572 | 0 | 0 | 15 | NaN | NaN | NaN | NaN | NaN | 260.48828 |
| Igkc | HNSYTCEATHK | -82 | -92 | 0 | 0.000712251 | 11 | NaN | NaN | NaN | NaN | NaN | 265.57977 |
| Rhoa | KQEPVKPEEGR | -135 | -145 | 2 | 0.000431655 | 11 | NaN | NaN | NaN | NaN | NaN | 351.35004 |
| S100a11 | CIESLIAVFQK | -8 | -18 | 0 | 0 | 11 | NaN | NaN | NaN | NaN | NaN | 2836.785 |
| Gzmb | SNTLQEVELTVQK | -157 | -169 | 0 | 0.00204437 | 13 | NaN | NaN | NaN | NaN | NaN | 1906.9921 |
| Slfn1 | FIEVHDSGVVC | -283 | -293 | 0 | 0.000431655 | 11 | NaN | NaN | NaN | NaN | NaN | 1745.0269 |
| Slfn1 | CIQQLPVTHFCEER | -261 | -274 | 0 | 0.002068213 | 14 | NaN | NaN | NaN | NaN | NaN | 215.86082 |
| Notch2 | CPEGFLGEYCQHR | -53 | -65 | 0 | 0.00025015 | 13 | NaN | NaN | NaN | NaN | NaN | 646.4406 |
| Txndc17 | CQVGDKPYWK | -69 | -78 | 1 | 0.00025015 | 10 | NaN | NaN | NaN | NaN | NaN | 321.1628 |
| Tagln2 | YPEGQAPVK | -70 | -78 | 0 | 0.006472924 | 9 | NaN | NaN | NaN | NaN | NaN | 1651.0914 |
| Tagln2 | NFSDNQLQEGKNVIGL | -161 | -176 | 1 | 0.005946382 | 16 | NaN | NaN | NaN | NaN | NaN | 493.55045 |
| Tagln2 | MEQISQFLQAAER | -90 | -102 | 0 | 0.007546675 | 13 | NaN | NaN | NaN | NaN | NaN | 169.02245 |
| Itgb3 | GTLHEENTCSR | -648 | -658 | 0 | 0 | 11 | NaN | NaN | NaN | NaN | NaN | 417.11478 |
| Eif6 | CVFSNQGGLVHPK | -152 | -164 | 0 | 0.001023236 | 13 | NaN | NaN | NaN | NaN | NaN | 953.05334 |
| Psmb2 | DGIHNLENIAFPK | -186 | -198 | 0 | 0 | 13 | NaN | NaN | NaN | NaN | NaN | 1239.7571 |
| Try5;Try4 | CVGFLEGGK | (185);(185) | (193);(193) | 0 | 0.002534621 | 9 | NaN | NaN | NaN | NaN | NaN | 587.56323 |
| Glud1 | VSYFEWLK | -437 | -444 | 0 | 0.008680443 | 8 | NaN | NaN | NaN | NaN | NaN | 227.30453 |
| Elac2 | SVLLDCGEGTFGQLCR | -507 | -522 | 0 | 0 | 16 | NaN | NaN | NaN | NaN | NaN | 912.5232 |
| Elac2 | GIELAVRPH | -152 | -160 | 1 | 0.00201047 | 9 | NaN | NaN | NaN | NaN | NaN | 423.2539 |
| Xab2 | YAQLEEEWGLAR | -599 | -610 | 0 | 0 | 12 | NaN | NaN | NaN | NaN | NaN | 398.74567 |
| Emc4 | GDSLYPVGYLDK | -39 | -50 | 0 | 0.001092529 | 12 | NaN | NaN | NaN | NaN | NaN | 619.4971 |
| Usp5 | DAALNKEELLEYEEK | -494 | -508 | 1 | 0.008653659 | 15 | NaN | NaN | NaN | NaN | NaN | 327.40244 |
| Tm9sf2 | FYLPGLAPVNFCAEEK | -36 | -51 | 0 | 0.001291103 | 16 | NaN | NaN | NaN | NaN | NaN | 364.00143 |
| Srsf10 | YLRPPNTSLF | -4 | -13 | 1 | 0.005270854 | 10 | NaN | NaN | NaN | NaN | NaN | 1915.0493 |
| Srsf10 | DAEDALHNLDRK | -63 | -74 | 1 | 0 | 12 | NaN | NaN | NaN | NaN | NaN | 829.6059 |
| Srsf1 | EAGDVCYADVYR | -143 | -154 | 0 | 0.001291103 | 12 | NaN | NaN | NaN | NaN | NaN | 2095.247 |
| Srsf1 | RGGPPFAFVEFEDPR | -51 | -65 | 1 | 0.00111006 | 15 | NaN | NaN | NaN | NaN | NaN | 472.33398 |
| Ppp1r7 | IENISNLHQL | -180 | -189 | 0 | 0.001868731 | 10 | NaN | NaN | NaN | NaN | NaN | 1891.7434 |
| Eif3c | RLDEEEEDNEGGEWER | -286 | -301 | 1 | 0.000230707 | 16 | NaN | NaN | NaN | NaN | NaN | 384.35208 |
| P4hb | KMDSTANEVEAVK | -426 | -438 | 1 | 0.002697088 | 13 | NaN | NaN | NaN | NaN | NaN | 822.49036 |
| Nudt5 | HANSKPFEVPFLK | -205 | -217 | 1 | 0.000989247 | 13 | NaN | NaN | NaN | NaN | NaN | 748.1666 |
| Ikzf3 | SLPKPHEIENVDSR | -30 | -43 | 1 | 0.000394361 | 14 | NaN | NaN | NaN | NaN | NaN | 1405.8435 |
| Ikzf3 | SHTGERPFQCNQCGASF | -138 | -154 | 1 | 0.00078661 | 17 | NaN | NaN | NaN | NaN | NaN | 2411.0354 |
| Ikzf3 | YEFSSHIAR | -491 | -499 | 0 | 0.001636916 | 9 | NaN | NaN | NaN | NaN | NaN | 1295.9785 |
| Arhgdib | PEEYEFLTPVEEAPK | -149 | -163 | 0 | 0.003360691 | 15 | NaN | NaN | NaN | NaN | NaN | 2010.6296 |
| Isoc2a | ILNTALDLLHR | -115 | -125 | 0 | 0 | 11 | NaN | NaN | NaN | NaN | NaN | 888.0665 |
| Wdr77 | DATWSPLNHSLLTTV | -302 | -316 | 0 | 0.000907206 | 15 | NaN | NaN | NaN | NaN | NaN | 738.0073 |
| Atl3 | GGSDPETTGIQIW | -110 | -122 | 0 | 0.00142999 | 13 | NaN | NaN | NaN | NaN | NaN | 244.24193 |
| Ero1a | KLLESDYFR | -67 | -75 | 1 | 0.002136192 | 9 | NaN | NaN | NaN | NaN | NaN | 31223.738 |
| Ero1a | GYLDDCTCDVETIDK | -41 | -55 | 0 | 0.001868731 | 15 | NaN | NaN | NaN | NaN | NaN | 683.3527 |
| Ngp | DCDFLEDGEER | -86 | -96 | 0 | 0.002752526 | 11 | NaN | NaN | NaN | NaN | NaN | 275.17114 |
| Serinc3 | WLAVWFK | -439 | -445 | 0 | 0.009808007 | 7 | NaN | NaN | NaN | NaN | NaN | 340.5012 |
| Creld2 | KNFGGGNTAWEEK | -49 | -61 | 1 | 0.002595029 | 13 | NaN | NaN | NaN | NaN | NaN | 5653.7876 |
| Bnip1 | TLLDANEEFK | -169 | -178 | 0 | 0.000826806 | 10 | NaN | NaN | NaN | NaN | NaN | 544.21857 |
| Thbs1 | LWHTGNTPGQVR | -1080 | -1091 | 0 | 0.002168954 | 12 | NaN | NaN | NaN | NaN | NaN | 1162.3124 |
| Magoh;Magol | YYLVQDLK | (123);(123) | (130);(130) | 0 | 0.005605646 | 8 | NaN | NaN | NaN | NaN | NaN | 301.627 |
| Hnrnpc | SFDLDYDFQR | -121 | -130 | 0 | 0.00025015 | 10 | NaN | NaN | NaN | NaN | NaN | 1679.6409 |
| Hnrnpc | QKVDSLLESL | -206 | -215 | 1 | 0.007905138 | 10 | NaN | NaN | NaN | NaN | NaN | 1717.8342 |
| Usp14 | FDDDKVSIVTPEDILR | -449 | -464 | 1 | 0 | 16 | NaN | NaN | NaN | NaN | NaN | 780.1827 |
| Usp14 | SSGHYVSWVR | -431 | -440 | 0 | 0.000907206 | 10 | NaN | NaN | NaN | NaN | NaN | 1190.3643 |
| Sub1 | GISLNMEQWSQLK | -102 | -114 | 0 | 0 | 13 | NaN | NaN | NaN | NaN | NaN | 1289.6702 |
| Fdps | KQDAESLQR | -76 | -84 | 1 | 0.003069674 | 9 | NaN | NaN | NaN | NaN | NaN | 368.1505 |
| Dock2 | DEDGNILDPDKTSVIS | -167 | -182 | 1 | 0.003299042 | 16 | NaN | NaN | NaN | NaN | NaN | 1101.501 |
| Ranbp2 | QLFHHLPQETSR | -459 | -470 | 0 | 0 | 12 | NaN | NaN | NaN | NaN | NaN | 5111.8403 |
| Ranbp2 | EGQWDCSLCFVR | -1559 | -1570 | 0 | 0.00111006 | 12 | NaN | NaN | NaN | NaN | NaN | 198.78604 |
| Alad | SVLHSGYFHPLLR | -5 | -17 | 0 | 0 | 13 | NaN | NaN | NaN | NaN | NaN | 554.3474 |
| Pml | ERELLEAVNDR | -298 | -308 | 1 | 0.00111006 | 11 | NaN | NaN | NaN | NaN | NaN | 896.694 |
| Scrn2 | SIGTDISAEHPELR | -196 | -209 | 0 | 0 | 14 | NaN | NaN | NaN | NaN | NaN | 403.89795 |
| Aco2 | GHLDDPANQEIER | -72 | -84 | 0 | 0.00201047 | 13 | NaN | NaN | NaN | NaN | NaN | 3418.3647 |
| Aco2 | VAVPSTIHCDHLIEA | -118 | -132 | 0 | 0.00025015 | 15 | NaN | NaN | NaN | NaN | NaN | 208.29814 |
| Eif5b | RLEELEAK | -367 | -374 | 1 | 0.001291103 | 8 | NaN | NaN | NaN | NaN | NaN | 3326.066 |
| Eif5b | KDDEIPVLKDELIHELK | -948 | -964 | 2 | 0.001636916 | 17 | NaN | NaN | NaN | NaN | NaN | 521.2173 |
| Eif5b | RDAAAVKPTENNEEESASK | -76 | -94 | 2 | 0.002068213 | 19 | NaN | NaN | NaN | NaN | NaN | 261.27252 |
| Matr3 | GYPHLCSICDLPVHSNK | -288 | -304 | 0 | 0.000550307 | 17 | NaN | NaN | NaN | NaN | NaN | 174.62608 |
| Gvin1 | ISPSCLFVHQ | -1610 | -1619 | 0 | 0 | 10 | NaN | NaN | NaN | NaN | NaN | 1454.2086 |
| Gvin1 | LLEEPLSR | -1791 | -1798 | 0 | 0.006735683 | 8 | NaN | NaN | NaN | NaN | NaN | 417.21454 |
| Fkbp5 | REEQCILYLGPR | -211 | -222 | 1 | 0.00078661 | 12 | NaN | NaN | NaN | NaN | NaN | 1801.2351 |
| Ebna1bp2 | DVTLGPVPEVSETQPTPQNQDQK | -71 | -93 | 0 | 0.00450079 | 23 | NaN | NaN | NaN | NaN | NaN | 173.98633 |
| Aqr | LGHGEEELETEKDFSR | -879 | -894 | 1 | 0.000826806 | 16 | NaN | NaN | NaN | NaN | NaN | 689.28033 |
| Aqr | QDIEDSVSR | -483 | -491 | 0 | 0.004983786 | 9 | NaN | NaN | NaN | NaN | NaN | 631.28107 |
| Aqr | LLQNPQDGFSR | -1081 | -1091 | 0 | 0.002595029 | 11 | NaN | NaN | NaN | NaN | NaN | 197.99063 |
| Eif4h | VGNLPFNTVQGDIDAIFK | -46 | -63 | 0 | 0.00272025 | 18 | NaN | NaN | NaN | NaN | NaN | 157.48207 |

| | | | | | | | | | | | | |
|---|---|---|---|---|---|---|---|---|---|---|---|---|
| Mfap1a;Mfap1 | MDEDEEVYKR | (349);(349) | (358);(358) | 1 | 0.005605646 | 10 | NaN | NaN | NaN | NaN | NaN | 464.3142 |
| Mfap1a;Mfap1 | EQEAEPEEQEEDSSSDPR | (68);(68) | (85);(85) | 0 | 0.001291103 | 18 | NaN | NaN | NaN | NaN | NaN | 209.72858 |
| Samhd1 | FEHLVNSNELK | -273 | -283 | 0 | 0.00025015 | 11 | NaN | NaN | NaN | NaN | NaN | 1102.1499 |
| Samhd1 | GNLYDMFHTR | -400 | -409 | 0 | 0.009657961 | 10 | NaN | NaN | NaN | NaN | NaN | 1028.9264 |
| Samhd1 | HFVQWCALR | -610 | -618 | 0 | 0 | 9 | NaN | NaN | NaN | NaN | NaN | 975.2446 |
| Prdx6 | VVDSLQLTGTKPVATPVDWK | -163 | -182 | 1 | 0 | 20 | NaN | NaN | NaN | NaN | NaN | 478.38806 |
| Psmd13 | IGNHASYYK | -166 | -174 | 0 | 0.00204437 | 9 | NaN | NaN | NaN | NaN | NaN | 1170.1128 |
| Zranb2 | NLDASEEEDSNK | -184 | -195 | 0 | 0 | 12 | NaN | NaN | NaN | NaN | NaN | 771.08203 |
| Wdr89 | DPLGAYSETHSDDITQVR | -159 | -176 | 0 | 0.002068213 | 18 | NaN | NaN | NaN | NaN | NaN | 1203.4478 |
| Wdr89 | DGHLDYLIGGLYHEK | -270 | -284 | 0 | 0.00025015 | 15 | NaN | NaN | NaN | NaN | NaN | 237.65265 |
| Wdr89 | IYDKETLHLLR | -54 | -64 | 1 | 0.002534621 | 11 | NaN | NaN | NaN | NaN | NaN | 159.46138 |
| Hdac1 | LHISPSNMTNQNTNEYLEK | -343 | -361 | 0 | 0 | 19 | NaN | NaN | NaN | NaN | NaN | 473.91177 |
| Tcea1 | KLLDGPSTDKDPEEK | -75 | -89 | 2 | 0.000826806 | 15 | NaN | NaN | NaN | NaN | NaN | 1111.1506 |
| Naa50 | DGTFDNIYLHVQ | -103 | -114 | 0 | 0.003393785 | 12 | NaN | NaN | NaN | NaN | NaN | 800.1936 |
| Ptpn6 | RVNAADIENR | -217 | -226 | 1 | 0.002329279 | 10 | NaN | NaN | NaN | NaN | NaN | 1330.7847 |
| Ptpn6 | TQQQGILQDR | -80 | -89 | 0 | 0.002068213 | 10 | NaN | NaN | NaN | NaN | NaN | 533.1197 |
| Celf2 | DHSDQPDPDAIK | -30 | -41 | 0 | 0.000471409 | 12 | NaN | NaN | NaN | NaN | NaN | 395.38577 |
| Huwe1 | NAEDKDTPALAR | -2122 | -2133 | 1 | 0.00051306 | 12 | NaN | NaN | NaN | NaN | NaN | 1854.0289 |
| Huwe1 | HFEFYADPGAEVK | -199 | -211 | 0 | 0 | 13 | NaN | NaN | NaN | NaN | NaN | 804.86786 |
| Huwe1 | TTPLKPSSLPVIPDTIK | -1927 | -1943 | 1 | 0.001392869 | 17 | NaN | NaN | NaN | NaN | NaN | 356.78186 |
| Aldh18a1 | VGTFFSEVKPAGPTVEQQ | -349 | -366 | 1 | 0.000947459 | 18 | NaN | NaN | NaN | NaN | NaN | 280.29602 |
| Aldh18a1 | KVGTFFSEVKPAGPTVEQQ | -348 | -366 | 2 | 0.001708008 | 19 | NaN | NaN | NaN | NaN | NaN | 278.06186 |
| Nploc4 | DIDKFGNEITQL | -428 | -439 | 1 | 0.00450079 | 12 | NaN | NaN | NaN | NaN | NaN | 886.899 |
| Zap70 | PDPAAHLPFFYGSIS | -2 | -16 | 0 | 0.00713295 | 15 | NaN | NaN | NaN | NaN | NaN | 731.4759 |
| Zap70 | KWEDRPDFLTVEQR | -578 | -591 | 2 | 0.001392869 | 14 | NaN | NaN | NaN | NaN | NaN | 291.81943 |
| Lonp1 | FLDHYLDVPVDLSK | -608 | -621 | 0 | 0 | 14 | NaN | NaN | NaN | NaN | NaN | 221.40929 |
| Stk38 | LFFEGVDWEHIR | -380 | -391 | 0 | 0 | 12 | NaN | NaN | NaN | NaN | NaN | 516.14514 |
| Rps6ka1 | DKLPQSQLSHQDLQLVK | -668 | -684 | 1 | 0.001291103 | 17 | NaN | NaN | NaN | NaN | NaN | 496.02826 |
| Dhrs7 | DILVLPLDLTDTSSHEAATK | -104 | -123 | 0 | 0.00051306 | 20 | NaN | NaN | NaN | NaN | NaN | 275.22446 |
| Dhrs7 | EKDILVLPLDLTDTSSHEAATK | -102 | -123 | 1 | 0.005605646 | 22 | NaN | NaN | NaN | NaN | NaN | 129.74785 |
| Rbm25 | KLVPLDYGEDDKNATK | -706 | -721 | 2 | 0 | 16 | NaN | NaN | NaN | NaN | NaN | 1359.1324 |
| Rbm25 | KFEDEDSDDVPR | -692 | -703 | 1 | 0.000471409 | 12 | NaN | NaN | NaN | NaN | NaN | 1360.3617 |
| Akap13 | SLSEEHLEPDQR | -1611 | -1622 | 0 | 0.005668207 | 12 | NaN | NaN | NaN | NaN | NaN | 288.837 |
| Psmd11 | LRDDPIISTHLAK | -313 | -325 | 1 | 0 | 13 | NaN | NaN | NaN | NaN | NaN | 734.84393 |
| Snd1 | YEERPVEEVMPVLEEK | -661 | -676 | 1 | 0 | 16 | NaN | NaN | NaN | NaN | NaN | 496.01385 |
| Ehd1 | LLDTVDDMLANDIAR | -378 | -392 | 0 | 0.002972566 | 15 | NaN | NaN | NaN | NaN | NaN | 1207.9023 |
| Ankrd44 | TNTGFEESDGGALK | -555 | -568 | 0 | 0.001463897 | 14 | NaN | NaN | NaN | NaN | NaN | 713.5872 |
| Epb41l2 | VRPAELEQF | -452 | -460 | 1 | 0.009242859 | 9 | NaN | NaN | NaN | NaN | NaN | 1668.0303 |
| Wdr75 | TSETHQWLNR | -643 | -652 | 0 | 0.005176406 | 10 | NaN | NaN | NaN | NaN | NaN | 11702.14 |
| Fkbp4 | RGEAHLAVNDFDLAR | -359 | -373 | 1 | 0.001463897 | 15 | NaN | NaN | NaN | NaN | NaN | 459.83618 |
| Fkbp4 | DLTEEEDGGIIR | -141 | -152 | 0 | 0 | 12 | NaN | NaN | NaN | NaN | NaN | 109.10704 |
| Srrm2 | SLLPNSSQDELMEVEK | -1188 | -1203 | 0 | 0.000431655 | 16 | NaN | NaN | NaN | NaN | NaN | 653.3121 |
| Srrm2 | DKFSPTQDRPESSTVL | -1148 | -1163 | 2 | 0.002752526 | 16 | NaN | NaN | NaN | NaN | NaN | 527.8748 |
| Mrpl38 | VAEGQETCPYLPPFPAR | -245 | -261 | 0 | 0.000471409 | 17 | NaN | NaN | NaN | NaN | NaN | 872.4448 |
| Usp19 | LYLPVPLPQK | -712 | -721 | 0 | 0.00644558 | 10 | NaN | NaN | NaN | NaN | NaN | 73.89508 |
| Sfpq | GQGGAGPVGGQGPR | -660 | -673 | 0 | 0.00025015 | 14 | NaN | NaN | NaN | NaN | NaN | 2359.7031 |
| Sfpq | DKLESEMEDAYHEHQANLLR | -509 | -528 | 1 | 0.00051306 | 20 | NaN | NaN | NaN | NaN | NaN | 921.9131 |
| Hmgb2 | IKIEHPGLSIGDTAKK | -113 | -128 | 2 | 0.001743443 | 16 | NaN | NaN | NaN | NaN | NaN | 243.94998 |
| Ube2n | LAEPVPGIK | -16 | -24 | 0 | 0.002810811 | 9 | NaN | NaN | NaN | NaN | NaN | 237.18755 |
| Ccnk | DLAHTPSQLEGLDPATEAR | -30 | -48 | 0 | 0.001092529 | 19 | NaN | NaN | NaN | NaN | NaN | 544.33813 |
| Ik | VVDLDDEYADTDIPTTLIR | -240 | -258 | 0 | 0.00149795 | 19 | NaN | NaN | NaN | NaN | NaN | 458.6718 |
| Flnb | SPFTVGVAAPLDLSK | -932 | -946 | 0 | 0.00111006 | 15 | NaN | NaN | NaN | NaN | NaN | 1009.40967 |
| Anp32a | LLPQVMYLDGYDR | -138 | -150 | 0 | 0.000755556 | 13 | NaN | NaN | NaN | NaN | NaN | 2415.0918 |
| Sf3a3 | DLYDDKDGLRK | -59 | -69 | 2 | 0 | 11 | NaN | NaN | NaN | NaN | NaN | 733.74976 |
| Sf3a3 | WQPDTEEEYEDSSGNVV | -471 | -487 | 0 | 0.000627943 | 17 | NaN | NaN | NaN | NaN | NaN | 682.4097 |
| Rps20 | LIDLHSPSEIVKQ | -88 | -100 | 1 | 0.00025015 | 13 | NaN | NaN | NaN | NaN | NaN | 5989.53 |
| Rps20 | IDLHSPSEIVK | -89 | -99 | 0 | 0.00025015 | 11 | NaN | NaN | NaN | NaN | NaN | 1314.3052 |
| Seh1l | IFTLKPLR | -247 | -254 | 1 | 0.00051306 | 8 | NaN | NaN | NaN | NaN | NaN | 201.6245 |
| Ripk3 | TWNHDVAVK | -43 | -51 | 0 | 0 | 9 | NaN | NaN | NaN | NaN | NaN | 1198.7023 |
| Pcif1 | DLPEELVHAGWEK | -39 | -51 | 0 | 0 | 13 | NaN | NaN | NaN | NaN | NaN | 746.29865 |
| Ndufa9 | DLPGLEDLGVQPTPLELK | -331 | -348 | 0 | 0.001023236 | 18 | NaN | NaN | NaN | NaN | NaN | 257.7258 |
| Cdc5l | DLPRPSEVNETILRPL | -544 | -559 | 2 | 0.00111006 | 16 | NaN | NaN | NaN | NaN | NaN | 542.26886 |
| Gnai2 | SDYIPTQQDVLR | -166 | -177 | 0 | 0.001904399 | 12 | NaN | NaN | NaN | NaN | NaN | 1700.9113 |
| Gnai2 | IIHEDGYSEEECRQ | -55 | -68 | 1 | 0.002845514 | 14 | NaN | NaN | NaN | NaN | NaN | 611.8056 |
| Ddi2 | NPPLAEALLSGDLEK | -156 | -170 | 0 | 0.002099344 | 15 | NaN | NaN | NaN | NaN | NaN | 274.86102 |
| Tfdp1 | VFIDQNLSPGK | -16 | -26 | 0 | 0.008947895 | 11 | NaN | NaN | NaN | NaN | NaN | 3419.8274 |
| Tfdp1 | NSAQECQNLEVER | -197 | -209 | 0 | 0.004690496 | 13 | NaN | NaN | NaN | NaN | NaN | 864.19604 |
| Lrrc59 | LSTLPSDFCGLTHL | -51 | -64 | 0 | 0.002534621 | 14 | NaN | NaN | NaN | NaN | NaN | 1133.2439 |
| Wbp4 | DNRPSVEFHER | -22 | -32 | 1 | 0.001392869 | 11 | NaN | NaN | NaN | NaN | NaN | 589.432 |
| Eif3i | LFDSTTLEHQK | -214 | -224 | 0 | 0 | 11 | NaN | NaN | NaN | NaN | NaN | 2541.3928 |
| Emb | DPTDPTFTSLPVR | -34 | -46 | 0 | 0.000550307 | 13 | NaN | NaN | NaN | NaN | NaN | 655.6931 |
| Elf1 | DPVEVGQPSEVLR | -357 | -369 | 0 | 0.003521747 | 13 | NaN | NaN | NaN | NaN | NaN | 411.57874 |
| Tpp2 | ICEVINEAVWK | -341 | -351 | 0 | 0.000712251 | 11 | NaN | NaN | NaN | NaN | NaN | 2578.1008 |
| Tpp2 | HFVEVPEGATWAEVT | -662 | -676 | 0 | 0.005394768 | 15 | NaN | NaN | NaN | NaN | NaN | 709.53424 |
| Tpp2 | KTDPRPDAATIK | -1112 | -1123 | 2 | 0.007987481 | 12 | NaN | NaN | NaN | NaN | NaN | 312.46338 |
| Hars1 | LVSELWDAGIK | -427 | -437 | 0 | 0 | 11 | NaN | NaN | NaN | NaN | NaN | 1817.4039 |
| Hars1 | IGDYVQQHGGVSL | -267 | -279 | 0 | 0.004857192 | 13 | NaN | NaN | NaN | NaN | NaN | 628.71967 |
| Farsa | LPDSGHLHPLLK | -218 | -229 | 0 | 0.00078661 | 12 | NaN | NaN | NaN | NaN | NaN | 152.19304 |

| | | | | | | | | | | | |
|---|---|---|---|---|---|---|---|---|---|---|---|
| Ddx3x | VRPCVVYGGAEIGQQ | -295 | -309 | 1 | 0.002780285 | 15 | NaN | NaN | NaN | NaN | NaN | 1797.6188 |
| Steep1 | NAPVTFIVDGAVVK | -108 | -121 | 0 | 0.001055699 | 14 | NaN | NaN | NaN | NaN | NaN | 725.36334 |
| Actr3 | KDYEEIGPSIC | -398 | -408 | 1 | 0 | 11 | NaN | NaN | NaN | NaN | NaN | 462.45636 |
| Actr3 | LASTPEFYQVCHTK | -384 | -397 | 0 | 0.000230707 | 14 | NaN | NaN | NaN | NaN | NaN | 1758.2347 |
| Actr3 | STPEFYQVCHTK | -386 | -397 | 0 | 0 | 12 | NaN | NaN | NaN | NaN | NaN | 603.2722 |
| Rasal3 | DSCEEVFENIIHSY | -586 | -599 | 0 | 0.00122419 | 14 | NaN | NaN | NaN | NaN | NaN | 381.94708 |
| Nars1 | RSWDSEEILEGYK | -489 | -501 | 1 | 0.003360691 | 13 | NaN | NaN | NaN | NaN | NaN | 833.2859 |
| Ndufb7 | HEQHDWDYCEHLDYVK | -82 | -97 | 0 | 0 | 16 | NaN | NaN | NaN | NaN | NaN | 470.5545 |
| Atl2 | DSQSWIGGNNEPLTGF | -122 | -137 | 0 | 0.002199523 | 16 | NaN | NaN | NaN | NaN | NaN | 227.59573 |
| Smchd1 | QGDFESDHSGYVRPLPVPR | -208 | -226 | 1 | 0.002265552 | 19 | NaN | NaN | NaN | NaN | NaN | 710.686 |
| Ipo11 | GFLHGIFER | -237 | -245 | 0 | 0.001291103 | 9 | NaN | NaN | NaN | NaN | NaN | 1718.5017 |
| Ipo11 | DTAVLKPAEEQLK | -21 | -33 | 1 | 0.002168954 | 13 | NaN | NaN | NaN | NaN | NaN | 1046.5691 |
| Mcmbp | GGLHWCGEPK | -195 | -204 | 0 | 0.001636916 | 10 | NaN | NaN | NaN | NaN | NaN | 2054.181 |
| Mcmbp | KLQHINPLLPTCLNK | -314 | -328 | 1 | 0.001023236 | 15 | NaN | NaN | NaN | NaN | NaN | 889.15497 |
| Smarcc1 | VREEVPLELVEAHVK | -724 | -738 | 1 | 0 | 15 | NaN | NaN | NaN | NaN | NaN | 4308.2603 |
| Snx6 | KDLNFHVF | -159 | -166 | 1 | 0.007493591 | 8 | NaN | NaN | NaN | NaN | NaN | 194.35606 |
| Evl | VKPAGSVNDVGLDALDLDR | -362 | -380 | 1 | 0 | 19 | NaN | NaN | NaN | NaN | NaN | 416.34436 |
| Mcm7 | IAPEIYGHEDVKK | -340 | -352 | 1 | 0 | 13 | NaN | NaN | NaN | NaN | NaN | 6825.2554 |
| Rps19 | HKELAPYDENWFYTR | -42 | -56 | 1 | 0 | 15 | NaN | NaN | NaN | NaN | NaN | 710.66644 |
| Rps19 | NQQEFVR | -10 | -16 | 0 | 0.003299042 | 7 | NaN | NaN | NaN | NaN | NaN | 880.3256 |
| Dpp9 | TKDYDWTEPLSPTEDEFK | -462 | -479 | 1 | 0.002265552 | 18 | NaN | NaN | NaN | NaN | NaN | 248.92441 |
| F2 | VVNLPIVERPVCK | -522 | -534 | 1 | 0 | 13 | NaN | NaN | NaN | NaN | NaN | 563.3992 |
| Hcls1 | REVQQPSMPVEEPAAPAQLPK | -276 | -296 | 1 | 0.000431655 | 21 | NaN | NaN | NaN | NaN | NaN | 500.42242 |
| Hcls1 | YEPEPETEPDYEDVGELDR | -378 | -396 | 0 | 0.002296977 | 19 | NaN | NaN | NaN | NaN | NaN | 292.86633 |
| Capzb | SPWSNKYDPPLEDGAMPSAR | -73 | -92 | 1 | 0 | 20 | NaN | NaN | NaN | NaN | NaN | 1119.2369 |
| Capzb | SPWSNKYDPPLEDGAMPSAR | -73 | -92 | 1 | 0 | 20 | NaN | NaN | NaN | NaN | NaN | 1119.2369 |
| Anxa3 | DYPGFSPSVDAEAIR | -14 | -28 | 0 | 0.00025015 | 15 | NaN | NaN | NaN | NaN | NaN | 677.88416 |
| Ccar1 | TVPAHVETVVL | -558 | -568 | 0 | 0.003814256 | 11 | NaN | NaN | NaN | NaN | NaN | 2271.3467 |
| Ubxn7 | EAGLCPQETVFVQER | -452 | -466 | 0 | 0.00025015 | 15 | NaN | NaN | NaN | NaN | NaN | 631.7643 |
| Vasp | NGPSPEELEQQKR | -131 | -143 | 1 | 0.002401448 | 13 | NaN | NaN | NaN | NaN | NaN | 721.19226 |
| Sp1 | KFACPECPK | -686 | -694 | 1 | 0.001392869 | 9 | NaN | NaN | NaN | NaN | NaN | 645.21655 |
| Naa15 | NSDSLPHR | -757 | -764 | 0 | 0.002780285 | 8 | NaN | NaN | NaN | NaN | NaN | 894.1483 |
| Nherf1 | LVVDPETDER | -89 | -98 | 0 | 0.00383302 | 10 | NaN | NaN | NaN | NaN | NaN | 1585.4475 |
| Eif3f | GHDITEHSVLIHEYYSR | -180 | -196 | 0 | 0.001463897 | 17 | NaN | NaN | NaN | NaN | NaN | 610.0084 |
| Eef1b | SYIEGYVPSQADVA | -23 | -36 | 0 | 0.00025015 | 14 | NaN | NaN | NaN | NaN | NaN | 154.56615 |
| Eef1b | SSILLDVKPWDDETDMTK | -140 | -157 | 1 | 0.002595029 | 18 | NaN | NaN | NaN | NaN | NaN | 1007.5402 |
| Uba7 | ECLPEDETLLPSPEDCQPR | -380 | -398 | 0 | 0.003069674 | 19 | NaN | NaN | NaN | NaN | NaN | 299.07416 |
| Mob4 | QIFDEYENETFLCH | -176 | -189 | 0 | 0.00025015 | 14 | NaN | NaN | NaN | NaN | NaN | 411.67032 |
| Lig3 | EDEQQQALQDIASR | -395 | -408 | 0 | 0.00025015 | 14 | NaN | NaN | NaN | NaN | NaN | 1691.5432 |
| Ola1 | FDFLCQYHKPASK | -71 | -83 | 1 | 0.000591124 | 13 | NaN | NaN | NaN | NaN | NaN | 711.3496 |
| Prex1 | FSNIEDILEVHK | -95 | -106 | 0 | 0.002068213 | 12 | NaN | NaN | NaN | NaN | NaN | 414.7123 |
| Dock10 | GFFEEEEGKEYIYK | -1886 | -1899 | 1 | 0.001776473 | 14 | NaN | NaN | NaN | NaN | NaN | 1144.7593 |
| Wdr3 | TREPLILEEER | -704 | -714 | 1 | 0.009060424 | 11 | NaN | NaN | NaN | NaN | NaN | 691.39844 |
| Hnrnpa3 | YSCVEEVDAAMCAR | -83 | -96 | 0 | 0.00025015 | 14 | NaN | NaN | NaN | NaN | NaN | 330.20032 |
| Ppme1 | VLPQCGHAVHEDAPDK | -343 | -358 | 0 | 0.002099344 | 16 | NaN | NaN | NaN | NaN | NaN | 414.49435 |
| Rbm8a | GVHEEATEEDIHDKFAEYGEIK | -79 | -100 | 1 | 0.00025015 | 22 | NaN | NaN | NaN | NaN | NaN | 365.57077 |
| Sec13 | IDQPSGQKPNYIK | -168 | -180 | 1 | 0 | 13 | NaN | NaN | NaN | NaN | NaN | 614.7709 |
| Eif5 | KFVLCPECENPETDLHVNPK | -95 | -114 | 1 | 0.002099344 | 20 | NaN | NaN | NaN | NaN | NaN | 508.86353 |
| Tpx2 | SLAENPSGSLVQEPFQLA | -645 | -662 | 0 | 0.009897819 | 18 | NaN | NaN | NaN | NaN | NaN | 545.02747 |
| Degs1 | NVGYHNEHHDFPNVPGK | -255 | -271 | 0 | 0 | 17 | NaN | NaN | NaN | NaN | NaN | 700.66254 |
| Slc25a11 | IFLEQMNK | -298 | -305 | 0 | 0.005270854 | 8 | NaN | NaN | NaN | NaN | NaN | 3596.0295 |
| U2af2 | ENPSVYVPGVVSTVVPDSAHK | -240 | -260 | 0 | 0.00025015 | 21 | NaN | NaN | NaN | NaN | NaN | 643.7941 |
| Acadm | YCVTEPSAGSDVAAIK | -158 | -173 | 0 | 0.009631128 | 16 | NaN | NaN | NaN | NaN | NaN | 581.6348 |
| Gmps | FHPEVGLTENGK | -189 | -200 | 0 | 0.002466863 | 12 | NaN | NaN | NaN | NaN | NaN | 1192.9171 |
| Gmps | EEIVLLTHGDSVDK | -146 | -159 | 0 | 0.002534621 | 14 | NaN | NaN | NaN | NaN | NaN | 1031.7574 |
| Gtpbp4 | LHLAVPNKR | -360 | -368 | 1 | 0.00025015 | 9 | NaN | NaN | NaN | NaN | NaN | 405.61154 |
| Atp2a3 | EFDDLSPEQQR | -657 | -667 | 0 | 0 | 11 | NaN | NaN | NaN | NaN | NaN | 419.97482 |
| Atp5f1c | LKESTTSEQSAR | -242 | -253 | 1 | 0 | 12 | NaN | NaN | NaN | NaN | NaN | 240.18446 |
| Ablim1 | EFDKLPLWR | -842 | -850 | 1 | 0.00025015 | 9 | NaN | NaN | NaN | NaN | NaN | 626.2271 |
| Ablim1 | SGLGQLILK | -696 | -704 | 0 | 0.003966045 | 9 | NaN | NaN | NaN | NaN | NaN | 218.12671 |
| Mta2 | EFEEESKQPGVSEQQR | -65 | -80 | 1 | 0 | 16 | NaN | NaN | NaN | NaN | NaN | 358.8766 |
| Stk26 | EFIDACLNKDPSFRPT | -249 | -264 | 2 | 0.000591124 | 16 | NaN | NaN | NaN | NaN | NaN | 557.0899 |
| Ubl3 | EFLFSPNDSASDIAK | -23 | -37 | 0 | 0 | 15 | NaN | NaN | NaN | NaN | NaN | 1525.732 |
| Atxn2l | SSKFELAVDAVHR | -151 | -163 | 1 | 0 | 13 | NaN | NaN | NaN | NaN | NaN | 426.62454 |
| Map2k1 | HVEGDAAETPPRPR | -278 | -291 | 1 | 0 | 14 | NaN | NaN | NaN | NaN | NaN | 385.55774 |
| Ptgr2 | EFSLPDALNEGQVQVR | -27 | -42 | 0 | 0.006100831 | 16 | NaN | NaN | NaN | NaN | NaN | 4369.819 |
| Casp6 | RVDFCKDPDAIGK | -242 | -254 | 2 | 0.000431655 | 13 | NaN | NaN | NaN | NaN | NaN | 1026.7026 |
| Casp6 | EFTELLTLVNR | -226 | -236 | 0 | 0 | 11 | NaN | NaN | NaN | NaN | NaN | 345.62875 |
| Prkd2 | EFVLLPAASELAH | -56 | -68 | 0 | 0.00111006 | 13 | NaN | NaN | NaN | NaN | NaN | 907.80414 |
| Zyx | SPGAPSGPGPQPNQK | -272 | -286 | 0 | 0.001610812 | 15 | NaN | NaN | NaN | NaN | NaN | 561.56104 |
| Eif3m | GAELSEENSEGGLHVDLA | -26 | -43 | 0 | 0.001636916 | 18 | NaN | NaN | NaN | NaN | NaN | 366.16724 |
| Nup37 | EGHSDFINDLVFHPK | -121 | -135 | 0 | 0.00025015 | 15 | NaN | NaN | NaN | NaN | NaN | 2001.8475 |
| Nup37 | EGHSDFINDLVFHPK | -121 | -135 | 0 | 0.00025015 | 15 | NaN | NaN | NaN | NaN | NaN | 2001.8475 |
| Sdf2l1 | NLHTHHFPSPLSNNQEVSAF | -114 | -133 | 0 | 0.002099344 | 20 | NaN | NaN | NaN | NaN | NaN | 579.76227 |
| Smc2 | EHASNEQQLDAVNEAIK | -832 | -848 | 0 | 0.00025015 | 17 | NaN | NaN | NaN | NaN | NaN | 759.71484 |
| Mtap | TSLRPQTFYDGSHCSAR | -117 | -133 | 1 | 0.00122419 | 17 | NaN | NaN | NaN | NaN | NaN | 357.46826 |
| Lag3 | KWTPPGGGPELPVAGK | -291 | -306 | 1 | 0 | 16 | NaN | NaN | NaN | NaN | NaN | 494.04944 |
| Nmt2 | EHIIDTFVVENPSGK | -412 | -426 | 0 | 0.001291103 | 15 | NaN | NaN | NaN | NaN | NaN | 2190.9907 |

| | | | | | | | | | | | | |
|---|---|---|---|---|---|---|---|---|---|---|---|---|
| Chordc1 | HSGVPIFHEGMK | -179 | -190 | 0 | 0.003102268 | 12 | NaN | NaN | NaN | NaN | NaN | 368.33914 |
| Clint1 | SKWDEEWDK | -190 | -198 | 1 | 0.004857192 | 9 | NaN | NaN | NaN | NaN | NaN | 1489.5917 |
| Arsb | TISEGHPSPR | -380 | -389 | 0 | 0.001708008 | 10 | NaN | NaN | NaN | NaN | NaN | 1519.4763 |
| Arsb | EHPHIVQNLLSR | -491 | -502 | 0 | 0.001370415 | 12 | NaN | NaN | NaN | NaN | NaN | 1413.0463 |
| Pofut1 | EHPVLALPGAPAQFPVLEEHR | -191 | -211 | 0 | 0 | 21 | NaN | NaN | NaN | NaN | NaN | 492.7935 |
| Wdr18 | EHSFQPEQNTGK | -252 | -263 | 0 | 0.00142999 | 12 | NaN | NaN | NaN | NaN | NaN | 498.01035 |
| Arhgap15 | HNVESVDLCGAHIE | -140 | -153 | 0 | 0.00025015 | 14 | NaN | NaN | NaN | NaN | NaN | 754.5422 |
| Impa1 | KLQVSQQEDITK | -145 | -156 | 1 | 0.001092529 | 12 | NaN | NaN | NaN | NaN | NaN | 297.185 |
| Mrps34 | LPLFGLGR | -71 | -78 | 0 | 0.002434616 | 8 | NaN | NaN | NaN | NaN | NaN | 383.79184 |
| Nmd3 | EIHDGLDFYYSSK | -190 | -202 | 0 | 0.001868731 | 13 | NaN | NaN | NaN | NaN | NaN | 1340.5333 |
| Casp8 | KFLCLDYIPHK | -23 | -33 | 1 | 0.00025015 | 11 | NaN | NaN | NaN | NaN | NaN | 589.26697 |
| Sec24a | EIHLTPSTDFYK | -660 | -671 | 0 | 0.003360691 | 12 | NaN | NaN | NaN | NaN | NaN | 964.3337 |
| Galnt6 | EIILVDDASTDEHLK | -211 | -225 | 0 | 0.001023236 | 15 | NaN | NaN | NaN | NaN | NaN | 1675.3025 |
| Arhgef1 | EVISELLVTEAAHVR | -418 | -432 | 0 | 0 | 15 | NaN | NaN | NaN | NaN | NaN | 417.13724 |
| Mlx | EIVIGVLHQLK | -284 | -294 | 0 | 0 | 11 | NaN | NaN | NaN | NaN | NaN | 511.97607 |
| Pwp2 | HFKPEIR | -698 | -704 | 1 | 0.00111006 | 7 | NaN | NaN | NaN | NaN | NaN | 1086.843 |
| Hmmr | EKEVELEK | -422 | -429 | 1 | 0.008498216 | 8 | NaN | NaN | NaN | NaN | NaN | 1269.2651 |
| Thrap3 | KAEEMEDEPFTER | -481 | -493 | 1 | 0 | 13 | NaN | NaN | NaN | NaN | NaN | 2041.7982 |
| Thrap3 | KYYLHDDR | -875 | -882 | 1 | 0.001776473 | 8 | NaN | NaN | NaN | NaN | NaN | 843.20557 |
| Sf3b2 | SSVPAGMETPELIELR | -735 | -750 | 0 | 0 | 16 | NaN | NaN | NaN | NaN | NaN | 801.675 |
| Tecr | THPQWYPAR | -39 | -47 | 0 | 0.00294445 | 9 | NaN | NaN | NaN | NaN | NaN | 1604.2593 |
| Tecr | EKLCFLDK | -15 | -22 | 1 | 0.000230707 | 8 | NaN | NaN | NaN | NaN | NaN | 822.48737 |
| Eif3g | RADDNATIR | -234 | -242 | 1 | 0.000627943 | 9 | NaN | NaN | NaN | NaN | NaN | 2674.0579 |
| Bcat1 | HYAVELFEGLK | -89 | -99 | 0 | 0.007493591 | 11 | NaN | NaN | NaN | NaN | NaN | 1075.3628 |
| Bcat1 | GTACVVCPVSDILYK | -332 | -346 | 0 | 0.002199523 | 15 | NaN | NaN | NaN | NaN | NaN | 929.5145 |
| Tes | EYTQAENQNWHLK | -314 | -326 | 0 | 0 | 13 | NaN | NaN | NaN | NaN | NaN | 1284.8608 |
| Top2b | VEELDKVEAQER | -1181 | -1192 | 1 | 0.000471409 | 12 | NaN | NaN | NaN | NaN | NaN | 764.967 |
| Kcnab2 | VEVQLPELFHK | -225 | -235 | 0 | 0 | 11 | NaN | NaN | NaN | NaN | NaN | 766.58466 |
| Capn1 | NPWGEVEWK | -296 | -304 | 0 | 0.007405213 | 9 | NaN | NaN | NaN | NaN | NaN | 1671.4727 |
| Lamtor3 | VANDSAPEHALRPGFLSTF | -35 | -53 | 1 | 0.00025015 | 19 | NaN | NaN | NaN | NaN | NaN | 453.42526 |
| Rcl1 | ELASNPQGQGAAVLPEDLGR | -258 | -277 | 0 | 0.000298805 | 20 | NaN | NaN | NaN | NaN | NaN | 362.65317 |
| Unc13d | IFHNVLK | -511 | -517 | 0 | 0.006788754 | 7 | NaN | NaN | NaN | NaN | NaN | 379.604 |
| Polr2b | LSDYGYHLR | -1015 | -1023 | 0 | 0 | 9 | NaN | NaN | NaN | NaN | NaN | 988.9343 |
| Polr2b | HASGEVEEPPR | -73 | -83 | 0 | 0.000230707 | 11 | NaN | NaN | NaN | NaN | NaN | 485.47095 |
| Polr2b | TYSAPLYVDITK | -119 | -130 | 0 | 0.001055699 | 12 | NaN | NaN | NaN | NaN | NaN | 470.69897 |
| Pitrm1 | LGCGILNYR | -625 | -633 | 0 | 0.000230707 | 9 | NaN | NaN | NaN | NaN | NaN | 1029.2292 |
| Tbl3 | SSGLHFLTAGDQGILR | -257 | -272 | 0 | 0 | 16 | NaN | NaN | NaN | NaN | NaN | 665.69434 |
| Tbl3 | SVLLPEQPAPALGVK | -242 | -256 | 0 | 0.00111006 | 15 | NaN | NaN | NaN | NaN | NaN | 319.724 |
| Immt | KAVDEAADALLK | -285 | -296 | 1 | 0.000550307 | 12 | NaN | NaN | NaN | NaN | NaN | 1822.3965 |
| Immt | LSTDDLNSLIAHAHR | -395 | -409 | 0 | 0.00111006 | 15 | NaN | NaN | NaN | NaN | NaN | 385.21683 |
| Prpf40a | FTNMLGQPGSTALDLFK | -642 | -658 | 0 | 0 | 17 | NaN | NaN | NaN | NaN | NaN | 760.3231 |
| Alyref | WQHDLFDSGFGGGA | -86 | -99 | 0 | 0.00383302 | 14 | NaN | NaN | NaN | NaN | NaN | 240.71605 |
| Rfc4 | LVNQLHDAIIENENLSDK | -306 | -323 | 0 | 0.006164361 | 18 | NaN | NaN | NaN | NaN | NaN | 257.9033 |
| Tomm40 | WIVGATLEK | -322 | -330 | 0 | 0.000755556 | 9 | NaN | NaN | NaN | NaN | NaN | 1042.9501 |
| Arfip1 | IEQSQHLFQIHK | -290 | -301 | 0 | 0.00383302 | 12 | NaN | NaN | NaN | NaN | NaN | 455.68484 |
| Zc3hc1 | LQPTFDFGR | -128 | -136 | 0 | 0 | 9 | NaN | NaN | NaN | NaN | NaN | 1090.3463 |
| Dhcr24 | ELILADGSFVR | -191 | -201 | 0 | 0.00025015 | 11 | NaN | NaN | NaN | NaN | NaN | 844.349 |
| Cdc16 | NTEEAGLAPLQNSTK | -574 | -588 | 0 | 0 | 15 | NaN | NaN | NaN | NaN | NaN | 522.1985 |
| Arhgef2 | ELLSNVDQDVHELEK | -425 | -439 | 0 | 0.00025015 | 15 | NaN | NaN | NaN | NaN | NaN | 455.24945 |
| Def6 | SLQHAQQQLEEVR | -504 | -516 | 0 | 0 | 13 | NaN | NaN | NaN | NaN | NaN | 410.76266 |
| Msh6 | SCLQEGLIPGSQFWDATK | -614 | -631 | 0 | 0.002099344 | 18 | NaN | NaN | NaN | NaN | NaN | 300.36197 |
| Pdcd10 | IERPEPEFQDLNEK | -93 | -106 | 1 | 0 | 14 | NaN | NaN | NaN | NaN | NaN | 409.21048 |
| Asf1b | NVAVLENPSPFHSPFR | -8 | -23 | 0 | 0.007791272 | 16 | NaN | NaN | NaN | NaN | NaN | 739.8446 |
| Hspbp1 | TEHSPFHEHVLGALC | -281 | -295 | 0 | 0.00051306 | 15 | NaN | NaN | NaN | NaN | NaN | 4724.235 |
| Hspbp1 | REPELGLEELLR | -309 | -320 | 1 | 0.002199523 | 12 | NaN | NaN | NaN | NaN | NaN | 1469.9492 |
| Add3 | VNIIGEVVDQGSTDLK | -192 | -207 | 0 | 0.000298805 | 16 | NaN | NaN | NaN | NaN | NaN | 427.8429 |
| Cd3e | ERPPPVPNPDYEPIRK | -160 | -175 | 2 | 0.009185548 | 16 | NaN | NaN | NaN | NaN | NaN | 327.05292 |
| Psmc2 | EVVETPLLHPE | -189 | -199 | 0 | 0.001776473 | 11 | NaN | NaN | NaN | NaN | NaN | 3043.2534 |
| Psmc2 | GVLLFGPPGTGK | -211 | -222 | 0 | 0.009541249 | 12 | NaN | NaN | NaN | NaN | NaN | 1765.7941 |
| Sephs1 | LGIGMDTCVIPLR | -64 | -76 | 0 | 0.000826806 | 13 | NaN | NaN | NaN | NaN | NaN | 1433.6372 |
| Ifit3 | YCNLQEYHR | -365 | -373 | 0 | 0 | 9 | NaN | NaN | NaN | NaN | NaN | 19378.385 |
| As3mt | FAPDFLFTPVDASLPAPQGR | -316 | -335 | 0 | 0.00025015 | 20 | NaN | NaN | NaN | NaN | NaN | 280.3416 |
| Elob | TLGECGFTSQTAR | -56 | -68 | 0 | 0.002534621 | 13 | NaN | NaN | NaN | NaN | NaN | 560.26996 |
| Gfus | ETCTWFTDNYEQAR | -307 | -320 | 0 | 0.001392869 | 14 | NaN | NaN | NaN | NaN | NaN | 819.32434 |
| Dnpep | EVAGQVGVPLQDLMVR | -391 | -406 | 0 | 0.00149795 | 16 | NaN | NaN | NaN | NaN | NaN | 267.97278 |
| Ahsa1 | VDGNVTGEFTDLVPEK | -253 | -268 | 0 | 0.001636916 | 16 | NaN | NaN | NaN | NaN | NaN | 930.6554 |
| Ahsa1 | FYEWTIK | -80 | -86 | 0 | 0.008222027 | 7 | NaN | NaN | NaN | NaN | NaN | 428.50085 |
| Nol11 | GVLGAGPEGFLGVEQTDK | -13 | -30 | 0 | 0.002972566 | 18 | NaN | NaN | NaN | NaN | NaN | 1887.2339 |
| Pcbp1 | QICLVMLETLSQSPQGR | -161 | -177 | 0 | 0.001392869 | 17 | NaN | NaN | NaN | NaN | NaN | 917.8804 |
| Sec16a | WSQDGTEPQQCR | -1909 | -1920 | 0 | 0.001291103 | 12 | NaN | NaN | NaN | NaN | NaN | 272.91177 |
| Ltn1 | THLPDFLICK | -860 | -869 | 0 | 0 | 10 | NaN | NaN | NaN | NaN | NaN | 883.2196 |
| Smarca4 | EVEAQLPEKVEYVIK | -983 | -997 | 1 | 0 | 15 | NaN | NaN | NaN | NaN | NaN | 696.10114 |
| Nrdc | EVEAVDSEYQLARPS | -318 | -332 | 1 | 0.007546675 | 15 | NaN | NaN | NaN | NaN | NaN | 1254.43 |
| Nrdc | TYLPPSPVLAESTTPITDIR | -1126 | -1145 | 0 | 0.001868731 | 20 | NaN | NaN | NaN | NaN | NaN | 456.63745 |
| Cul1 | KCEQVLIEK | -290 | -298 | 1 | 0.00149795 | 9 | NaN | NaN | NaN | NaN | NaN | 415.3723 |
| NaN | ILYEILLGK | (143);(143) | (151);(151) | 0 | 0.007791272 | 9 | NaN | NaN | NaN | NaN | NaN | 376.29367 |
| Ppp4r3b | EVIPITDSELR | -250 | -260 | 0 | 0.001055699 | 11 | NaN | NaN | NaN | NaN | NaN | 2355.2815 |
| Oat | FADGNFWGR | -172 | -180 | 0 | 0.004255025 | 9 | NaN | NaN | NaN | NaN | NaN | 1009.7535 |

| | | | | | | | | | | | | |
|---|---|---|---|---|---|---|---|---|---|---|---|---|
| Aurka | EVLEHPWIK | -367 | -375 | 0 | 0.00025015 | 9 | NaN | NaN | NaN | NaN | NaN | 1469.9032 |
| Septin1 | LVDTPGFGDSVDCSDCWLPVVR | -87 | -108 | 0 | 0.001868731 | 22 | NaN | NaN | NaN | NaN | NaN | 165.26237 |
| Gnai3 | KIIHEDGYSEDECK | -54 | -67 | 1 | 0.00025015 | 14 | NaN | NaN | NaN | NaN | NaN | 496.43735 |
| Gnai3 | EVYTHFTCATDTK | -318 | -330 | 0 | 0.000550307 | 13 | NaN | NaN | NaN | NaN | NaN | 271.8212 |
| Stk4 | FLEYFEQK | -395 | -402 | 0 | 0.00223472 | 8 | NaN | NaN | NaN | NaN | NaN | 793.8071 |
| Cd3g | EYDQYSHLQGNQLR | -167 | -180 | 0 | 0.000989247 | 14 | NaN | NaN | NaN | NaN | NaN | 310.57285 |
| Gsto1 | EYLDEAYPEKK | -91 | -101 | 1 | 0 | 11 | NaN | NaN | NaN | NaN | NaN | 882.9358 |
| Pmpca | YLSGIAHFLEK | -103 | -113 | 0 | 0 | 11 | NaN | NaN | NaN | NaN | NaN | 624.2995 |
| Fnbp1 | SGFEPPGDIEFEDYTQPMK | -274 | -292 | 0 | 0.00294445 | 19 | NaN | NaN | NaN | NaN | NaN | 177.13974 |
| Ppp2r5c | QINNIFYR | -203 | -210 | 0 | 0.001023236 | 8 | NaN | NaN | NaN | NaN | NaN | 814.3613 |
| Ppp2r5c | LFDDCTQQFK | -404 | -413 | 0 | 0.001023236 | 10 | NaN | NaN | NaN | NaN | NaN | 506.23816 |
| Ubtf | WVEISNEVR | -72 | -80 | 0 | 0.00025015 | 9 | NaN | NaN | NaN | NaN | NaN | 4131.4355 |
| Ubtf | TPQQLWYTHE | -201 | -210 | 0 | 0.000712251 | 10 | NaN | NaN | NaN | NaN | NaN | 536.2456 |
| Ubtf | KLHPEMSNLDLTK | -132 | -144 | 1 | 0.007819401 | 13 | NaN | NaN | NaN | NaN | NaN | 523.4859 |
| Rpa3 | YIDRPVCFVGK | -20 | -30 | 1 | 0.000230707 | 11 | NaN | NaN | NaN | NaN | NaN | 1328.4081 |
| Cdc42 | QKPITPETAEKL | -134 | -145 | 2 | 0.001610812 | 12 | NaN | NaN | NaN | NaN | NaN | 2614.037 |
| Cdc42 | SVVSPSSFENVK | -83 | -94 | 0 | 0 | 12 | NaN | NaN | NaN | NaN | NaN | 2122.7607 |
| Cdc42 | TCLLISYTTNK | -17 | -27 | 0 | 0.001776473 | 11 | NaN | NaN | NaN | NaN | NaN | 712.1821 |
| Tor1aip1 | VLPVQPENTLK | -580 | -590 | 0 | 0.006078721 | 11 | NaN | NaN | NaN | NaN | NaN | 924.8353 |
| Ctps1 | LYGDTDYLEER | -467 | -477 | 0 | 0 | 11 | NaN | NaN | NaN | NaN | NaN | 1154.0336 |
| Ctps1 | LCSAHGVLVPGGFGVR | -361 | -376 | 0 | 0.003620005 | 16 | NaN | NaN | NaN | NaN | NaN | 282.23785 |
| Txndc5 | VDCTADSDVCSAQGVR | -105 | -120 | 0 | 0.001868731 | 16 | NaN | NaN | NaN | NaN | NaN | 509.78845 |
| Scamp3 | SGELDNPFQDPAVIQHR | -14 | -30 | 0 | 0 | 17 | NaN | NaN | NaN | NaN | NaN | 1143.6805 |
| Rasa3 | LDIDGDRETER | -722 | -732 | 1 | 0.001055699 | 11 | NaN | NaN | NaN | NaN | NaN | 2369.3557 |
| Pus1 | FGGDGLHEPLDWTQEEGK | -343 | -360 | 0 | 0.001708008 | 18 | NaN | NaN | NaN | NaN | NaN | 228.47546 |
| Tardbp | LVYVVNYPK | -71 | -79 | 0 | 0.00111006 | 9 | NaN | NaN | NaN | NaN | NaN | 1330.7578 |
| Cd47 | FVASNQR | -289 | -295 | 0 | 0.003620005 | 7 | NaN | NaN | NaN | NaN | NaN | 840.91565 |
| Sacs | FGQTTPPLVDFLK | -89 | -101 | 0 | 0 | 13 | NaN | NaN | NaN | NaN | NaN | 452.5134 |
| Trim33 | FHCDPTFWAK | -475 | -484 | 0 | 0 | 10 | NaN | NaN | NaN | NaN | NaN | 546.53955 |
| Clpb | LAAGADPNLGDEFSSVYK | -187 | -204 | 0 | 0.004957095 | 18 | NaN | NaN | NaN | NaN | NaN | 6290.456 |
| Clpb | VDGYNVHYGAR | -580 | -590 | 0 | 0.00025015 | 11 | NaN | NaN | NaN | NaN | NaN | 1009.4993 |
| Clpb | FIGSPPGYIGHEEGGQLTK | -393 | -411 | 0 | 0.002566359 | 19 | NaN | NaN | NaN | NaN | NaN | 280.79758 |
| Ldhb | SADTLWDIQK | -320 | -329 | 0 | 0.001610812 | 10 | NaN | NaN | NaN | NaN | NaN | 1059.4331 |
| Tapbp | PFPASAK | -101 | -107 | 0 | 0.003299042 | 7 | NaN | NaN | NaN | NaN | NaN | 788.57 |
| Tapbp | SPEQNCPR | -113 | -120 | 0 | 0.001023236 | 8 | NaN | NaN | NaN | NaN | NaN | 682.94305 |
| Sdr39u1 | RLDTTHLLAK | -82 | -91 | 1 | 0.005270854 | 10 | NaN | NaN | NaN | NaN | NaN | 247.48415 |
| Shcbp1 | FIQHDSVEGILIIHHGK | -438 | -454 | 0 | 0.007546675 | 17 | NaN | NaN | NaN | NaN | NaN | 972.48004 |
| Hnrnpd | FKEEEPVKK | -230 | -238 | 2 | 0 | 9 | NaN | NaN | NaN | NaN | NaN | 2645.6807 |
| Yars2 | FKEFFPESGTK | -42 | -52 | 1 | 0.00930008 | 11 | NaN | NaN | NaN | NaN | NaN | 1598.7996 |
| Vapa | HLRDEGLR | -195 | -202 | 1 | 0.00635324 | 8 | NaN | NaN | NaN | NaN | NaN | 357.3597 |
| Adprm | FLEDQIAQHPETTPSENYYAY | -130 | -150 | 0 | 0.000431655 | 21 | NaN | NaN | NaN | NaN | NaN | 267.48715 |
| Hspe1 | PEYGGTK | -74 | -80 | 0 | 0.00383302 | 7 | NaN | NaN | NaN | NaN | NaN | 1253.3435 |
| Ccdc88b | QLEEELIQLR | -952 | -961 | 0 | 0.005176406 | 10 | NaN | NaN | NaN | NaN | NaN | 5955.6846 |
| Ccz1 | FLTGPLNLNDPEAK | -280 | -293 | 0 | 0.00122419 | 14 | NaN | NaN | NaN | NaN | NaN | 2917.9517 |
| Ssr1 | QATFEYSFIPAEPMGGRPF | -151 | -169 | 1 | 0.00272025 | 19 | NaN | NaN | NaN | NaN | NaN | 2741.806 |
| Cox5a | YFNKPDIDAWELR | -55 | -67 | 1 | 0.000591124 | 13 | NaN | NaN | NaN | NaN | NaN | 950.92084 |
| Cluh | FPESCQDEVR | -730 | -739 | 0 | 0.006260252 | 10 | NaN | NaN | NaN | NaN | NaN | 505.33014 |
| Cdc34 | FPIDYPYSPPAFR | -64 | -76 | 0 | 0.001868731 | 13 | NaN | NaN | NaN | NaN | NaN | 1604.1925 |
| Tap2 | KPNLPQPGILAPPWLEGR | -449 | -466 | 1 | 0 | 18 | NaN | NaN | NaN | NaN | NaN | 2692.7795 |
| Usp24 | FQPALVTTVDALR | -372 | -384 | 0 | 0.003133457 | 13 | NaN | NaN | NaN | NaN | NaN | 432.36658 |
| Ufd1 | IIMPPSALDQLSR | -46 | -58 | 0 | 0 | 13 | NaN | NaN | NaN | NaN | NaN | 2449.4421 |
| Dpp4 | YNIYDVNKR | -126 | -134 | 1 | 0.004255025 | 9 | NaN | NaN | NaN | NaN | NaN | 2327.11 |
| Kif5b | TSQEQVYNDCAK | -56 | -67 | 0 | 0.00025015 | 12 | NaN | NaN | NaN | NaN | NaN | 1040.4843 |
| Nhlrc2 | LQHPLGVAWDEER | -474 | -486 | 0 | 0.001092529 | 13 | NaN | NaN | NaN | NaN | NaN | 644.9873 |
| Nhlrc2 | GNLLFSLIGEGHR | -172 | -184 | 0 | 0 | 13 | NaN | NaN | NaN | NaN | NaN | 856.47363 |
| Acot7 | HVSAEITYTSK | -124 | -134 | 0 | 0.001023236 | 11 | NaN | NaN | NaN | NaN | NaN | 1751.2048 |
| Atad3a | WSNFDPTGLER | -46 | -56 | 0 | 0 | 11 | NaN | NaN | NaN | NaN | NaN | 1415.1926 |
| Ctnnbl1 | LDESVREEADGVHNTL | -204 | -219 | 1 | 0.000591124 | 16 | NaN | NaN | NaN | NaN | NaN | 961.5783 |
| Nae1 | QSVGQAPESISEK | -366 | -378 | 0 | 0.000471409 | 13 | NaN | NaN | NaN | NaN | NaN | 2380.9504 |
| Emc1 | WVEHLPESDSILYQ | -164 | -177 | 0 | 0.009002904 | 14 | NaN | NaN | NaN | NaN | NaN | 926.4699 |
| Emc1 | TEVQKPVSAGDGSVA | -341 | -355 | 1 | 0.003490217 | 15 | NaN | NaN | NaN | NaN | NaN | 539.88983 |
| Txn2 | FVGIKDEDQLEAFLK | -148 | -162 | 1 | 0.002752526 | 15 | NaN | NaN | NaN | NaN | NaN | 648.9419 |
| Ube2h | FYGPQGTPYEGGVWK | -38 | -52 | 0 | 0.00025015 | 15 | NaN | NaN | NaN | NaN | NaN | 2202.479 |
| Samm50 | FYLGGPTSVR | -337 | -346 | 0 | 0.004595494 | 10 | NaN | NaN | NaN | NaN | NaN | 813.8464 |
| Mtpn | HHITPLLSAVYEGHVS | -67 | -82 | 0 | 0.003360691 | 16 | NaN | NaN | NaN | NaN | NaN | 894.2282 |
| Mtpn | GADINAPDKHHITPLLSAVYEGHVSCVK | -58 | -85 | 1 | 0.000550307 | 28 | NaN | NaN | NaN | NaN | NaN | 755.1584 |
| Ankfy1 | GANPNLQTEEALPVPK | -514 | -529 | 0 | 0.00078661 | 16 | NaN | NaN | NaN | NaN | NaN | 82.71665 |
| Rcc1 | TVSSGGQHTVLLVK | -403 | -416 | 0 | 0 | 14 | NaN | NaN | NaN | NaN | NaN | 678.9265 |
| Marcksl1 | GDAIEPAPPSQEAEAK | -62 | -77 | 0 | 0.00122419 | 16 | NaN | NaN | NaN | NaN | NaN | 4420.4165 |
| Ogg1 | GDDSQVSRPTLEELETLHK | -80 | -98 | 1 | 0.001291103 | 19 | NaN | NaN | NaN | NaN | NaN | 1015.214 |
| Ppwd1 | GDNQPLHIFDK | -210 | -220 | 0 | 0.00051306 | 11 | NaN | NaN | NaN | NaN | NaN | 259.4459 |
| Fbxo3 | GDVEEVQGPGVVGEFPIISPGR | -338 | -359 | 0 | 0 | 22 | NaN | NaN | NaN | NaN | NaN | 430.82526 |
| Sod2 | HTIFWTNLSPK | -98 | -108 | 0 | 0.001868731 | 11 | NaN | NaN | NaN | NaN | NaN | 2562.8008 |
| Sod2 | VWEHAYYLQYK | -184 | -194 | 0 | 0.000907206 | 11 | NaN | NaN | NaN | NaN | NaN | 1460.0549 |
| Sod2 | GDVTTQVALQPALK | -76 | -89 | 0 | 0 | 14 | NaN | NaN | NaN | NaN | NaN | 1263.9894 |
| Sod2 | GVQGSGWGWLGFNK | -141 | -154 | 0 | 0.00383302 | 14 | NaN | NaN | NaN | NaN | NaN | 625.58325 |
| Otub1 | YRPGHYDILYK | -261 | -271 | 1 | 0 | 11 | NaN | NaN | NaN | NaN | NaN | 1789.7393 |
| Ccna2 | LHQEDQENVNPEK | -19 | -31 | 0 | 0.00272025 | 13 | NaN | NaN | NaN | NaN | NaN | 387.2249 |

| | | | | | | | | | | | | |
|---|---|---|---|---|---|---|---|---|---|---|---|---|
| Ccna2 | GELSLIDADPYLK | -327 | -339 | 0 | 0 | 13 | NaN | NaN | NaN | NaN | NaN | 303.11475 |
| Oxsm | GPHEGQFNEENFVSK | -95 | -109 | 0 | 0.000394361 | 15 | NaN | NaN | NaN | NaN | NaN | 643.2657 |
| Oxsm | GESGIVSVVGDEYK | -69 | -82 | 0 | 0.002910424 | 14 | NaN | NaN | NaN | NaN | NaN | 1178.6838 |
| Golt1b | GFFPVVVGFIR | -100 | -110 | 0 | 0 | 11 | NaN | NaN | NaN | NaN | NaN | 409.91666 |
| Dhodh | SETGGLSGKPLR | -298 | -309 | 1 | 0.003102268 | 12 | NaN | NaN | NaN | NaN | NaN | 709.5188 |
| Hdhd5 | QMLVSGQGPLVENAR | -133 | -147 | 0 | 0.001811524 | 15 | NaN | NaN | NaN | NaN | NaN | 177.50764 |
| Xpo5 | GFVVGYTPSGNPIFR | -748 | -762 | 0 | 0.009955823 | 15 | NaN | NaN | NaN | NaN | NaN | 1811.2516 |
| Exosc7 | GGDDLGTEIANTLYR | -97 | -111 | 0 | 0.00111006 | 15 | NaN | NaN | NaN | NaN | NaN | 1589.6676 |
| Mre11 | NHPNLFNPDNPK | -316 | -327 | 0 | 0.00051306 | 12 | NaN | NaN | NaN | NaN | NaN | 491.2146 |
| Fam3c | GGDVAPFIEFLK | -126 | -137 | 0 | 0.005668207 | 12 | NaN | NaN | NaN | NaN | NaN | 380.1952 |
| Bub1b | GIAADPEGLLQEEDLDGK | -623 | -640 | 0 | 0.004690496 | 18 | NaN | NaN | NaN | NaN | NaN | 976.505 |
| G3bp2 | GIEEPLEESSHEPEPEPESETK | -184 | -205 | 0 | 0.008129024 | 22 | NaN | NaN | NaN | NaN | NaN | 879.38135 |
| Sh3bp1 | KVEQCKDEYLADLY | -201 | -214 | 2 | 0.003745449 | 14 | NaN | NaN | NaN | NaN | NaN | 639.1889 |
| Sh3bp1 | SSPPAPSLPPGSVSPGTPQALPR | -576 | -598 | 0 | 0.007791272 | 23 | NaN | NaN | NaN | NaN | NaN | 303.7503 |
| Sar1a | GIVFLVDCADHSR | -95 | -107 | 0 | 0 | 13 | NaN | NaN | NaN | NaN | NaN | 324.4011 |
| Rpl21 | HAVGIIVNK | -70 | -78 | 0 | 0.00437931 | 9 | NaN | NaN | NaN | NaN | NaN | 310.70624 |
| Smg1 | GLEDLHLDER | -2154 | -2163 | 0 | 0.005668207 | 10 | NaN | NaN | NaN | NaN | NaN | 737.7375 |
| Crkl | IHYLDTTTLIEPAPR | -90 | -104 | 0 | 0.001092529 | 15 | NaN | NaN | NaN | NaN | NaN | 738.08563 |
| Dohh | GLGGPDAISWISR | -33 | -45 | 0 | 0.00078661 | 13 | NaN | NaN | NaN | NaN | NaN | 1156.3849 |
| Fis1 | GLLQTEPQNNQAK | -96 | -108 | 0 | 0.00111006 | 13 | NaN | NaN | NaN | NaN | NaN | 578.64514 |
| Khnyn | GLLSPQFQGVR | -211 | -221 | 0 | 0.000826806 | 11 | NaN | NaN | NaN | NaN | NaN | 1233.7338 |
| Cyc1 | HLVGVCYTEEEAK | -134 | -146 | 0 | 0.000471409 | 13 | NaN | NaN | NaN | NaN | NaN | 288.78513 |
| Ecsit | KDEPWLVPRPPEPQR | -49 | -63 | 2 | 0.00111006 | 15 | NaN | NaN | NaN | NaN | NaN | 2861.873 |
| Ecsit | GLQETNPTLAQIPVVFR | -376 | -392 | 0 | 0.009242859 | 17 | NaN | NaN | NaN | NaN | NaN | 290.747 |
| Kif15 | GPSDSDNFSHNLR | -121 | -133 | 0 | 0.00111006 | 13 | NaN | NaN | NaN | NaN | NaN | 11662.834 |
| Asah1 | TVNLDFQR | -185 | -192 | 0 | 0.001092529 | 8 | NaN | NaN | NaN | NaN | NaN | 1360.7146 |
| Ndufs2 | NFGPQHPAAHGVLR | -83 | -96 | 0 | 0.000989247 | 14 | NaN | NaN | NaN | NaN | NaN | 2724.5498 |
| Ndufs2 | NFGPQHPAAHGVLR | -83 | -96 | 0 | 0.000989247 | 14 | NaN | NaN | NaN | NaN | NaN | 2724.5498 |
| Mix23 | IVHELNTTVPTASFAGK | -30 | -46 | 0 | 0.002068213 | 17 | NaN | NaN | NaN | NaN | NaN | 919.45325 |
| Git2 | LQPFPTHIGR | -490 | -499 | 0 | 0.003681885 | 10 | NaN | NaN | NaN | NaN | NaN | 4031.4395 |
| Gbe1 | GTHDLWDSR | -317 | -325 | 0 | 0.007987481 | 9 | NaN | NaN | NaN | NaN | NaN | 519.82635 |
| Lsm2 | LHSVDQYLNIK | -29 | -39 | 0 | 0.006326148 | 11 | NaN | NaN | NaN | NaN | NaN | 1681.8531 |
| Agpat3 | KWEEDRDTVIEGLR | -147 | -160 | 2 | 0.000230707 | 14 | NaN | NaN | NaN | NaN | NaN | 1338.4849 |
| Nbeal2 | NLENVEAGWGQVLVPR | -91 | -106 | 0 | 0.002068213 | 16 | NaN | NaN | NaN | NaN | NaN | 404.45663 |
| Nbeal2 | QWLPALPTAELR | -456 | -467 | 0 | 0.001708008 | 12 | NaN | NaN | NaN | NaN | NaN | 339.77313 |
| Atad1 | GVLLYGPPGCGK | -128 | -139 | 0 | 0.006677519 | 12 | NaN | NaN | NaN | NaN | NaN | 1873.4856 |
| Atad1 | KDTVILPIK | -104 | -112 | 1 | 0.00078661 | 9 | NaN | NaN | NaN | NaN | NaN | 420.3108 |
| Abcf1 | VLKPAKPEK | -149 | -157 | 2 | 0.000550307 | 9 | NaN | NaN | NaN | NaN | NaN | 190.6428 |
| Abcf1 | LEQGDDTAAEK | -393 | -403 | 0 | 0.001291103 | 11 | NaN | NaN | NaN | NaN | NaN | 830.9764 |
| Vdac1 | TQTLKPGIK | -261 | -269 | 1 | 0.002068213 | 9 | NaN | NaN | NaN | NaN | NaN | 763.03625 |
| Rnaseh2c | HDADGLQASFR | -54 | -64 | 0 | 0.00111006 | 11 | NaN | NaN | NaN | NaN | NaN | 1330.9956 |
| Cdk9 | HENVVNLIEICR | -75 | -86 | 0 | 0.008105996 | 12 | NaN | NaN | NaN | NaN | NaN | 569.6759 |
| Mettl1 | HFEEHPLFER | -204 | -213 | 0 | 0.00025015 | 10 | NaN | NaN | NaN | NaN | NaN | 409.7225 |
| Pdcd5 | VSEQGLIEILEK | -87 | -98 | 0 | 0.004031417 | 12 | NaN | NaN | NaN | NaN | NaN | 273.01627 |
| Hsd17b4 | HGEQYLELYKPLPR | -405 | -418 | 1 | 0 | 14 | NaN | NaN | NaN | NaN | NaN | 956.6174 |
| Pelo | HINFEVVK | -196 | -203 | 0 | 0.000550307 | 8 | NaN | NaN | NaN | NaN | NaN | 1235.4437 |
| Psma6 | ILIGIDEEQGPQVYK | -139 | -153 | 0 | 0.00025015 | 15 | NaN | NaN | NaN | NaN | NaN | 291.6878 |
| Pygb | LVDDEAFIR | -525 | -533 | 0 | 0.000989247 | 9 | NaN | NaN | NaN | NaN | NaN | 12671.342 |
| Ezh2 | NTETALDNKPCGPQ | -310 | -323 | 1 | 0.00204437 | 14 | NaN | NaN | NaN | NaN | NaN | 150.1602 |
| Cisd2 | KVQLPAYLK | -11 | -19 | 1 | 0.00383302 | 9 | NaN | NaN | NaN | NaN | NaN | 12073.661 |
| Msra | VISAEEALPGR | -28 | -38 | 0 | 0 | 11 | NaN | NaN | NaN | NaN | NaN | 1105.8079 |
| Nup43 | LVCGTDAEAIYVTR | -363 | -376 | 0 | 0 | 14 | NaN | NaN | NaN | NaN | NaN | 1982.0137 |
| Atp6v1f | HPNFLVVEK | -33 | -41 | 0 | 0.002068213 | 9 | NaN | NaN | NaN | NaN | NaN | 971.548 |
| Hgh1 | HSWEPEPDVR | -326 | -335 | 0 | 0.000431655 | 10 | NaN | NaN | NaN | NaN | NaN | 599.3523 |
| Apaf1 | TVLCEGGVPQRPVIF | -112 | -126 | 1 | 0.00204437 | 15 | NaN | NaN | NaN | NaN | NaN | 923.9724 |
| Wdhd1 | NISDQTCAVSWPVLQK | -164 | -179 | 0 | 0 | 16 | NaN | NaN | NaN | NaN | NaN | 1628.026 |
| Cox6b1 | KTAPFDSR | -13 | -20 | 1 | 0.003037016 | 8 | NaN | NaN | NaN | NaN | NaN | 1440.3335 |
| U2af1 | IALLNIYR | -46 | -53 | 0 | 0 | 8 | NaN | NaN | NaN | NaN | NaN | 891.74994 |
| U2af1 | WFNGQPIHAELSPVTDFR | -134 | -151 | 0 | 0 | 18 | NaN | NaN | NaN | NaN | NaN | 96.32353 |
| Thumpd3 | IALTEESLHR | -285 | -294 | 0 | 0.000431655 | 10 | NaN | NaN | NaN | NaN | NaN | 42445.133 |
| Gbf1 | IDCFLPHLR | -1707 | -1715 | 0 | 0 | 9 | NaN | NaN | NaN | NaN | NaN | 3758.5688 |
| Rtf1 | SQPVSLPEELNR | -355 | -366 | 0 | 0.006788754 | 12 | NaN | NaN | NaN | NaN | NaN | 732.28796 |
| Sar1b | LVDCADHER | -99 | -107 | 0 | 0.000550307 | 9 | NaN | NaN | NaN | NaN | NaN | 737.00507 |
| Rpl38 | YTLVITDK | -43 | -50 | 0 | 0.002329279 | 8 | NaN | NaN | NaN | NaN | NaN | 1051.2211 |
| Pmm2 | IEFYELDKK | -138 | -146 | 1 | 0.004224523 | 9 | NaN | NaN | NaN | NaN | NaN | 2675.3096 |
| Ccdc47 | IESVHFSDQFSGPK | -328 | -341 | 0 | 0.00635324 | 14 | NaN | NaN | NaN | NaN | NaN | 231.1721 |
| Nfkb2 | IEVDLVTHSDPPR | -91 | -103 | 0 | 0.00223472 | 13 | NaN | NaN | NaN | NaN | NaN | 635.7823 |
| Cog4 | IFPLLGLHEDGLSK | -221 | -234 | 0 | 0 | 14 | NaN | NaN | NaN | NaN | NaN | 963.6758 |
| Zc3h18 | KEQEPDFEEK | -315 | -324 | 1 | 0.000627943 | 10 | NaN | NaN | NaN | NaN | NaN | 1572.4849 |
| Zc3h18 | KPAPPPAPPQAT | -635 | -646 | 1 | 0.002810811 | 12 | NaN | NaN | NaN | NaN | NaN | 489.64478 |
| Nt5c3b | NSSVCENSSYFQQLQNK | -206 | -222 | 0 | 0 | 17 | NaN | NaN | NaN | NaN | NaN | 229.31544 |
| Trrap | IHTIPTVPIVDCFQK | -1752 | -1766 | 0 | 0.000627943 | 15 | NaN | NaN | NaN | NaN | NaN | 2509.826 |
| Idi1 | KNCHLNENIDK | -37 | -47 | 1 | 0.000591124 | 11 | NaN | NaN | NaN | NaN | NaN | 445.84378 |
| Rpl19 | VWLDPNETNEIA | -22 | -33 | 0 | 0 | 12 | NaN | NaN | NaN | NaN | NaN | 1519.9053 |
| Ckap2 | IKDPDLTTPDSK | -554 | -565 | 1 | 0.000947459 | 12 | NaN | NaN | NaN | NaN | NaN | 2562.4016 |
| Mmrn1 | IKEDDAVAPDFSK | -1062 | -1074 | 1 | 0.00051306 | 13 | NaN | NaN | NaN | NaN | NaN | 1198.2216 |
| Eed | IKESYDYNPNK | -274 | -284 | 1 | 0.000627943 | 11 | NaN | NaN | NaN | NaN | NaN | 1297.2935 |
| Dlg3;Dlg4;Dlg | ILAGGPADLSGELR | (432);(341);(494) | (445);(354);(507) | 0 | 0.00111006 | 14 | NaN | NaN | NaN | NaN | NaN | 708.7665 |

| | | | | | | | | | | | | |
|---|---|---|---|---|---|---|---|---|---|---|---|---|
| Mrps2 | ILDTPLQHSDFFNVK | -59 | -73 | 0 | 0 | 15 | NaN | NaN | NaN | NaN | NaN | 859.4115 |
| Carnmt1 | ILKPGGIWIN | -322 | -331 | 1 | 0.003102268 | 10 | NaN | NaN | NaN | NaN | NaN | 530.5536 |
| Mrpl15 | YGFNEGHSF | -80 | -88 | 0 | 0.006677519 | 9 | NaN | NaN | NaN | NaN | NaN | 355.80005 |
| Nif3l1 | INIILSETDRDPLR | -361 | -374 | 1 | 0.00142999 | 14 | NaN | NaN | NaN | NaN | NaN | 632.6588 |
| Myo18a | ISELTSELTDER | -1287 | -1298 | 0 | 0 | 12 | NaN | NaN | NaN | NaN | NaN | 1015.64124 |
| Kif22 | LLDLLNEGSAR | -587 | -597 | 0 | 0.000627943 | 11 | NaN | NaN | NaN | NaN | NaN | 586.0339 |
| Ylpm1 | WDTNDDQGLNSEFK | -825 | -838 | 0 | 0.000471409 | 14 | NaN | NaN | NaN | NaN | NaN | 369.42236 |
| Stag1 | TLDKEYDVAVEAIR | -382 | -395 | 1 | 0.000230707 | 14 | NaN | NaN | NaN | NaN | NaN | 1273.109 |
| Stag1 | ITDGSPSKEDLLVLR | -752 | -766 | 1 | 0.00025015 | 15 | NaN | NaN | NaN | NaN | NaN | 524.27655 |
| Utp15 | NVLCVEHGQPVESVLLFPS | -199 | -217 | 0 | 0.001868731 | 19 | NaN | NaN | NaN | NaN | NaN | 652.36725 |
| Uck2 | TVQIPVYDFVSHS | -106 | -118 | 0 | 0.000627943 | 13 | NaN | NaN | NaN | NaN | NaN | 726.9795 |
| Eps15 | IWDLADTDGK | -56 | -65 | 0 | 0 | 10 | NaN | NaN | NaN | NaN | NaN | 1726.1029 |
| Mtatp8 | IYLPHSLPQQ | -58 | -67 | 0 | 0.007343498 | 10 | NaN | NaN | NaN | NaN | NaN | 6046.8022 |
| Msmo1 | KDKPETFEGQWK | -85 | -96 | 2 | 0.000627943 | 12 | NaN | NaN | NaN | NaN | NaN | 745.4334 |
| Ptrhd1 | KVVLEAADETTLK | -80 | -92 | 1 | 0.00025015 | 13 | NaN | NaN | NaN | NaN | NaN | 632.96936 |
| Ptrhd1 | KDLSQAPFSWPTGALVAQ | -33 | -50 | 1 | 0.003360691 | 18 | NaN | NaN | NaN | NaN | NaN | 474.54126 |
| Gtf2e2 | KDYSDITPGK | -283 | -292 | 1 | 0.002168954 | 10 | NaN | NaN | NaN | NaN | NaN | 799.7135 |
| Gimap8 | YMIVLFTR | -596 | -603 | 0 | 0.00635324 | 8 | NaN | NaN | NaN | NaN | NaN | 1058.8041 |
| Hmga1 | SQKEPSEVPTPK | -44 | -55 | 1 | 0 | 12 | NaN | NaN | NaN | NaN | NaN | 1102.2113 |
| Nit2 | SIYLIGGSIPEEDAGK | -77 | -92 | 0 | 0.005758933 | 16 | NaN | NaN | NaN | NaN | NaN | 328.00726 |
| Nit2 | KIHLFDIDVPGK | -112 | -123 | 1 | 0.002265552 | 12 | NaN | NaN | NaN | NaN | NaN | 257.77844 |
| H1-1 | SETAPVAQAASTATEKPAAAK | -2 | -22 | 1 | 0.005605646 | 21 | NaN | NaN | NaN | NaN | NaN | 1497.1888 |
| Krr1 | QDDVACDIIK | -136 | -145 | 0 | 0.001610812 | 10 | NaN | NaN | NaN | NaN | NaN | 2676.7715 |
| Ncln | KRDPEFVFYDQLK | -499 | -511 | 2 | 0.001708008 | 13 | NaN | NaN | NaN | NaN | NaN | 586.88965 |
| Mrps31 | LWEIEFAK | -266 | -273 | 0 | 0.00025015 | 8 | NaN | NaN | NaN | NaN | NaN | 673.5723 |
| Sumo1 | KVIGQDSSEIHFK | -25 | -37 | 1 | 0.00051306 | 13 | NaN | NaN | NaN | NaN | NaN | 927.4739 |
| Mtarc2 | RVGDPVYR | -328 | -335 | 1 | 0.001536401 | 8 | NaN | NaN | NaN | NaN | NaN | 308.08923 |
| Phpt1 | WAEYHADIYDK | -48 | -58 | 0 | 0 | 11 | NaN | NaN | NaN | NaN | NaN | 512.5825 |
| Slc38a2 | SFVCHPAVLPIYEELK | -299 | -314 | 0 | 0 | 16 | NaN | NaN | NaN | NaN | NaN | 3576.984 |
| Pik3c3 | KYENFDDIK | -404 | -412 | 1 | 0.00111006 | 9 | NaN | NaN | NaN | NaN | NaN | 4210.371 |
| Rigi | SECLINQECEEIR | -128 | -140 | 0 | 0.001291103 | 13 | NaN | NaN | NaN | NaN | NaN | 1450.215 |
| Emc3 | KYYFNNPEDGFFK | -77 | -89 | 1 | 0.003490217 | 13 | NaN | NaN | NaN | NaN | NaN | 349.66733 |
| Pxn | VLDPLDQWQPSGSR | -62 | -75 | 0 | 0.003620005 | 14 | NaN | NaN | NaN | NaN | NaN | 4487.5176 |
| Pxn | TWHPEHFFCAQ | -435 | -445 | 0 | 0.005522803 | 11 | NaN | NaN | NaN | NaN | NaN | 1063.7456 |
| Fcho1 | LALCHLELTR | -81 | -90 | 0 | 0.002265552 | 10 | NaN | NaN | NaN | NaN | NaN | 678.9472 |
| Nup54 | SFSAPTNTGSTGLLGGTQNK | -40 | -59 | 0 | 0.004224523 | 20 | NaN | NaN | NaN | NaN | NaN | 303.13226 |
| Pla2g12a | LDLLGGEDGLCQYK | -56 | -69 | 0 | 0 | 14 | NaN | NaN | NaN | NaN | NaN | 6803.197 |
| Cggbp1 | SLQCNSPAQTEK | -89 | -100 | 0 | 0.003461185 | 12 | NaN | NaN | NaN | NaN | NaN | 657.06384 |
| Niban1 | LEAVQFFR | -213 | -220 | 0 | 0.00025015 | 8 | NaN | NaN | NaN | NaN | NaN | 687.29315 |
| Tmed5 | SGHIQTLLR | -170 | -178 | 0 | 0.005698678 | 9 | NaN | NaN | NaN | NaN | NaN | 234.01588 |
| Tmem43 | YPEVGDVR | -233 | -240 | 0 | 0.00111006 | 8 | NaN | NaN | NaN | NaN | NaN | 1766.7938 |
| Paip2 | LEEEEEHEWFIPAR | -65 | -78 | 0 | 0.000471409 | 14 | NaN | NaN | NaN | NaN | NaN | 236.3288 |
| Racgap1 | LEQQIQLIR | -97 | -105 | 0 | 0.006260252 | 9 | NaN | NaN | NaN | NaN | NaN | 417.718 |
| Kif4 | LFEEQNHFAK | -903 | -912 | 0 | 0.000394361 | 10 | NaN | NaN | NaN | NaN | NaN | 864.0062 |
| Scyl3 | LFEVHEEHVR | -345 | -354 | 0 | 0.000230707 | 10 | NaN | NaN | NaN | NaN | NaN | 381.5322 |
| Washc5 | LFLEDYVR | -1127 | -1134 | 0 | 0.000627943 | 8 | NaN | NaN | NaN | NaN | NaN | 996.0751 |
| Gmppa | LFSPEALKPLR | -185 | -195 | 1 | 0.000230707 | 11 | NaN | NaN | NaN | NaN | NaN | 532.09766 |
| Smc4 | LPQTQQELK | -546 | -554 | 0 | 0.002534621 | 9 | NaN | NaN | NaN | NaN | NaN | 2340.1252 |
| Ikbkb | YLNQFENCCGLR | -107 | -118 | 0 | 0.009871162 | 12 | NaN | NaN | NaN | NaN | NaN | 655.5294 |
| Acy1 | TVQPNPDYGGAITFLEER | -24 | -41 | 0 | 0.008436628 | 18 | NaN | NaN | NaN | NaN | NaN | 550.52136 |
| Prkag1 | LHETLETIINR | -280 | -290 | 0 | 0 | 11 | NaN | NaN | NaN | NaN | NaN | 987.26404 |
| Ppp1r2 | TSAASPPVVPSAEQPRPIVEEELSK | -19 | -43 | 1 | 0.001708008 | 25 | NaN | NaN | NaN | NaN | NaN | 634.4409 |
| Mrpl44 | LIAEGPGETVLVAEEEAAR | -280 | -298 | 0 | 0.00025015 | 19 | NaN | NaN | NaN | NaN | NaN | 457.5524 |
| Syf2 | LKLPANWEAK | -61 | -70 | 1 | 0.001055699 | 10 | NaN | NaN | NaN | NaN | NaN | 7119.8276 |
| Ube2v2 | WQNSYSIK | -109 | -116 | 0 | 0.000431655 | 8 | NaN | NaN | NaN | NaN | NaN | 1152.7241 |
| Orc5 | LLLEQILNK | -71 | -79 | 0 | 0.00025015 | 9 | NaN | NaN | NaN | NaN | NaN | 939.03894 |
| Fam169b | LLPVLDTVFVR | -183 | -193 | 0 | 0.001743443 | 11 | NaN | NaN | NaN | NaN | NaN | 180.65616 |
| Rabep2 | LNEENQGLR | -365 | -373 | 0 | 0.009688493 | 9 | NaN | NaN | NaN | NaN | NaN | 2001.6882 |
| Birc6 | LPCPEGLDPDIEDASPVCR | -4836 | -4854 | 0 | 0.003966045 | 19 | NaN | NaN | NaN | NaN | NaN | 388.65457 |
| Ctss | LPDTVDWR | -123 | -130 | 0 | 0.008045078 | 8 | NaN | NaN | NaN | NaN | NaN | 4778.9014 |
| Flii | LVTLPEAIHFL | -352 | -362 | 0 | 0.006568856 | 11 | NaN | NaN | NaN | NaN | NaN | 173.06767 |
| Sec23a | LQAPVDDAQEILHSR | -685 | -699 | 0 | 0.001463897 | 15 | NaN | NaN | NaN | NaN | NaN | 819.6358 |
| Sema7a | LQDVFLLPDPSGQWR | -301 | -315 | 0 | 0.002099344 | 15 | NaN | NaN | NaN | NaN | NaN | 334.45584 |
| Mff | LQLLEEENKER | -256 | -266 | 1 | 0.000471409 | 11 | NaN | NaN | NaN | NaN | NaN | 497.40668 |
| Etv6 | LQQENNHQETYPL | -225 | -237 | 0 | 0.00111006 | 13 | NaN | NaN | NaN | NaN | NaN | 449.61972 |
| Vdac2 | SFNAGGHKL | -279 | -287 | 1 | 0.002068213 | 9 | NaN | NaN | NaN | NaN | NaN | 2717.5227 |
| Ddx24 | LTLVHQAPAR | -510 | -519 | 0 | 0.001868731 | 10 | NaN | NaN | NaN | NaN | NaN | 321.92853 |
| Gtf3c4 | LVDLTEIYGDR | -201 | -211 | 0 | 0.001023236 | 11 | NaN | NaN | NaN | NaN | NaN | 859.76276 |
| Cd5 | LVGGSSVCEGIAEVR | -278 | -292 | 0 | 0.00025015 | 15 | NaN | NaN | NaN | NaN | NaN | 4167.8096 |
| Cd5 | PGHEGLR | -163 | -169 | 0 | 0.002534621 | 7 | NaN | NaN | NaN | NaN | NaN | 918.7146 |
| Pdlim5 | SWHPEEFNCAHCK | -433 | -445 | 0 | 0.008129024 | 13 | NaN | NaN | NaN | NaN | NaN | 230.79523 |
| Vps26b | YFLFYDGETVSGK | -37 | -49 | 0 | 0 | 13 | NaN | NaN | NaN | NaN | NaN | 589.00604 |
| Gpd2 | LVQDYGLESEVAQHLAK | -483 | -499 | 0 | 0 | 17 | NaN | NaN | NaN | NaN | NaN | 505.47983 |
| Cd2 | VQPKPPCGSGDGVSLPPPN | -326 | -344 | 1 | 0.002099344 | 19 | NaN | NaN | NaN | NaN | NaN | 811.103 |
| Vps28 | MFHGIPATPGVGAPGNKPELYEEVK | -1 | -25 | 1 | 0.003681885 | 25 | NaN | NaN | NaN | NaN | NaN | 351.81177 |
| Mtco1 | SWLATLHGGNIK | -322 | -333 | 0 | 0 | 12 | NaN | NaN | NaN | NaN | NaN | 1613.6924 |
| Sf3b6 | YGINTDPPK | -117 | -125 | 0 | 0.00201047 | 9 | NaN | NaN | NaN | NaN | NaN | 833.407 |
| Rpl31 | RNEDEDSPNKLY | -92 | -103 | 2 | 0.00025015 | 12 | NaN | NaN | NaN | NaN | NaN | 6425.892 |

| | | | | | | | | | | | | |
|---|---|---|---|---|---|---|---|---|---|---|---|---|
| Armt1 | NELQTDKPITPLVDK | -67 | -81 | 1 | 0.001636916 | 15 | NaN | NaN | NaN | NaN | NaN | 1408.7311 |
| Plaat3 | NKHDEEYTPLPLSK | -78 | -91 | 1 | 0.00025015 | 14 | NaN | NaN | NaN | NaN | NaN | 387.80295 |
| Usp3 | NKVDTYVQFPLR | -422 | -433 | 1 | 0.000431655 | 12 | NaN | NaN | NaN | NaN | NaN | 443.18402 |
| Dimt1 | NLPYQISSPFVFK | -128 | -140 | 0 | 0.00383302 | 13 | NaN | NaN | NaN | NaN | NaN | 371.09085 |
| Atxn10 | SEPHTEDKEALVTIR | -291 | -305 | 1 | 0.00078661 | 15 | NaN | NaN | NaN | NaN | NaN | 1044.7076 |
| Rfc3 | SNYHLEVNPSDAGNSDR | -89 | -105 | 0 | 0.00078661 | 17 | NaN | NaN | NaN | NaN | NaN | 268.00278 |
| Hgfac | NPDKDERPWCYVVK | -337 | -350 | 2 | 0.00111006 | 14 | NaN | NaN | NaN | NaN | NaN | 238.86948 |
| Ntaq1 | NSCYCEENIWK | -26 | -36 | 0 | 0.001636916 | 11 | NaN | NaN | NaN | NaN | NaN | 69.401634 |
| Lsm14a | SSFQSVGSYGPFGR | -103 | -116 | 0 | 0.004224523 | 14 | NaN | NaN | NaN | NaN | NaN | 1042.1119 |
| Znf593 | PKPEPDAEPDPDLPGGGLHR | -43 | -62 | 1 | 0.001291103 | 20 | NaN | NaN | NaN | NaN | NaN | 861.80634 |
| Gins4 | QENILVEPEADEQR | -179 | -192 | 0 | 0.00025015 | 14 | NaN | NaN | NaN | NaN | NaN | 1304.2363 |
| Faf1 | QILENELQIPVPK | -126 | -138 | 0 | 0.00025015 | 13 | NaN | NaN | NaN | NaN | NaN | 275.9407 |
| Arpp19 | TEVTGDHIPTPQDLPQR | -85 | -101 | 0 | 0.000230707 | 17 | NaN | NaN | NaN | NaN | NaN | 203.70284 |
| Pycr3 | TDVLTPAGTTIHGLHALER | -232 | -250 | 0 | 0.001092529 | 19 | NaN | NaN | NaN | NaN | NaN | 6090.443 |
| Capn2 | RPTEICADPQFIIGGAT | -77 | -93 | 1 | 0.00025015 | 17 | NaN | NaN | NaN | NaN | NaN | 1533.5144 |
| Srprb | SAAPSTLDSSSTAPAQLGK | -207 | -225 | 0 | 0.001636916 | 19 | NaN | NaN | NaN | NaN | NaN | 1037.9694 |
| Mtmr3 | SEAVLYPVCHVR | -556 | -567 | 0 | 0.000431655 | 12 | NaN | NaN | NaN | NaN | NaN | 563.3243 |
| Igf2 | SCDLALLETYCATPAK | -74 | -89 | 0 | 0 | 16 | NaN | NaN | NaN | NaN | NaN | 1199.6464 |
| Ddx31 | WLIVDEADR | -257 | -265 | 0 | 0.00078661 | 9 | NaN | NaN | NaN | NaN | NaN | 970.74585 |
| Plekho2 | SDISEDQPQEPPR | -190 | -202 | 0 | 0.004350979 | 13 | NaN | NaN | NaN | NaN | NaN | 1968.163 |
| Rbm28 | VVLHPDTEHSK | -356 | -366 | 0 | 0 | 11 | NaN | NaN | NaN | NaN | NaN | 665.4937 |
| Maz | SFSRPDHLNSHVR | -345 | -357 | 1 | 0.001023236 | 13 | NaN | NaN | NaN | NaN | NaN | 610.1431 |
| Atad5 | SGLCDEFSLENR | -1668 | -1679 | 0 | 0.000431655 | 12 | NaN | NaN | NaN | NaN | NaN | 409.72034 |
| Tsc2 | SHLEELAAGGIPIER | -1348 | -1362 | 0 | 0.00025015 | 15 | NaN | NaN | NaN | NaN | NaN | 707.1673 |
| Ccdc124 | SHLELPLEENLNR | -117 | -129 | 0 | 0 | 13 | NaN | NaN | NaN | NaN | NaN | 1191.8146 |
| Spr | SLCALQPYK | -158 | -166 | 0 | 0.007987481 | 9 | NaN | NaN | NaN | NaN | NaN | 4225.7227 |
| Wdr73 | SLGSDGQLCLLDSR | -243 | -256 | 0 | 0.00111006 | 14 | NaN | NaN | NaN | NaN | NaN | 1492.7106 |
| Arpc1a | SLHQFLLEPITCH | -2 | -14 | 0 | 0.000431655 | 13 | NaN | NaN | NaN | NaN | NaN | 208.96852 |
| Slamf1 | SNHCQPPVEEK | -273 | -283 | 0 | 0.001055699 | 11 | NaN | NaN | NaN | NaN | NaN | 1778.4485 |
| Vwf | SQEEVDIHYCQGK | -2741 | -2753 | 0 | 0 | 13 | NaN | NaN | NaN | NaN | NaN | 294.6867 |
| Fli1 | SQYWTSPTAGIYPNPSVPR | -420 | -438 | 0 | 0.00025015 | 19 | NaN | NaN | NaN | NaN | NaN | 319.12878 |
| Znf280c | SSIFSDVETHFR | -429 | -440 | 0 | 0.000907206 | 12 | NaN | NaN | NaN | NaN | NaN | 543.838 |
| Fn1 | TFYQIGDSWEK | -568 | -578 | 0 | 0.00025015 | 11 | NaN | NaN | NaN | NaN | NaN | 721.95746 |
| Enpp4 | SYDLPHLQNFIK | -46 | -57 | 0 | 0.000230707 | 12 | NaN | NaN | NaN | NaN | NaN | 820.5349 |
| Ndufaf5 | TAVNDLGHLLGR | -220 | -231 | 0 | 0.00025015 | 12 | NaN | NaN | NaN | NaN | NaN | 2196.005 |
| Lipa | TGQEQIYYVGHSQ | -161 | -173 | 0 | 0.001776473 | 13 | NaN | NaN | NaN | NaN | NaN | 555.3846 |
| Timm50 | TPEQVTEIANR | -33 | -43 | 0 | 0.000230707 | 11 | NaN | NaN | NaN | NaN | NaN | 849.5782 |
| Ublcp1 | TLSEDDTVLDLK | -18 | -29 | 0 | 0.00223472 | 12 | NaN | NaN | NaN | NaN | NaN | 781.4633 |
| Rbms1 | TPPGVSAPTEPLLCK | -208 | -222 | 0 | 0.002780285 | 15 | NaN | NaN | NaN | NaN | NaN | 4116.2524 |
| Ercc2 | TPTIANPVLHF | -425 | -435 | 0 | 0.00025015 | 11 | NaN | NaN | NaN | NaN | NaN | 692.9867 |
| Rad51 | TSVEEESFGPQPISR | -13 | -27 | 0 | 0.001392869 | 15 | NaN | NaN | NaN | NaN | NaN | 653.4279 |
| Focad | TSVPTDYSYLPEGSFIR | -1346 | -1362 | 0 | 0.002502116 | 17 | NaN | NaN | NaN | NaN | NaN | 1196.5793 |
| Trappc1 | TVHNLYLFDR | -2 | -11 | 0 | 0 | 10 | NaN | NaN | NaN | NaN | NaN | 950.6431 |
| Get4 | TVLCEQYQPSLR | -256 | -267 | 0 | 0.002780285 | 12 | NaN | NaN | NaN | NaN | NaN | 464.656 |
| Nudcd3 | VDEEEQAVLDR | -299 | -309 | 0 | 0 | 11 | NaN | NaN | NaN | NaN | NaN | 6478.068 |
| Gpatch8 | VLEVEKEDTEELR | -100 | -112 | 1 | 0.00025015 | 13 | NaN | NaN | NaN | NaN | NaN | 3059.702 |
| Nemf | VLNPLLPYGPALIEHC | -186 | -201 | 0 | 0.001392869 | 16 | NaN | NaN | NaN | NaN | NaN | 243.8555 |
| Wdr74 | VYDPVSPQR | -209 | -217 | 0 | 0.004595494 | 9 | NaN | NaN | NaN | NaN | NaN | 822.49036 |
| Itch | VTEENKEEYIR | -690 | -700 | 1 | 0 | 11 | NaN | NaN | NaN | NaN | NaN | 1026.5509 |
| Bles03 | YGDPGSPNSEPVGWIA | -112 | -127 | 0 | 0.00383302 | 16 | NaN | NaN | NaN | NaN | NaN | 294.68173 |
| Nkg7 | WSQTPFSQVQ | -117 | -126 | 0 | 0.000712251 | 10 | NaN | NaN | NaN | NaN | NaN | 472.14966 |
| Dis3l2 | YEADIPEEGCGHHPLQQSR | -137 | -155 | 0 | 0.003900536 | 19 | NaN | NaN | NaN | NaN | NaN | 819.43 |
| Urb1 | YHDLTFLK | -1439 | -1446 | 0 | 0.001291103 | 8 | NaN | NaN | NaN | NaN | NaN | 778.9863 |
| Crk | YLDTTTLIEPVAR | -108 | -120 | 0 | 0.00111006 | 13 | NaN | NaN | NaN | NaN | NaN | 11240.958 |
| Ogfod1 | YLVPSWDR | -169 | -176 | 0 | 0.001811524 | 8 | NaN | NaN | NaN | NaN | NaN | 577.5016 |
