## Supplementary material for "Maleic acid adjuvates BCG to induce CD4^+^CD8^+^ T cells and sterilize tuberculosis": Table S3

| REAGENT or RESOURCE | SOURCE | IDENTIFIER |
|---|---|---|
| **Antibodies for flow cytometry** | | |
| Hamster monoclonal PE Anti-Mouse CD3e (clone 145-2C11) | BD Biosciences | Cat# 553063; RRID: AB_394597 |
| Rat monoclonal Alexa Fluor® 700 anti-mouse CD3 Antibody (clone 17A2) | Biolegend | Cat# 100216; RRID: AB_493697 |
| Rat monoclonal BV605 Anti-Mouse CD4 (clone RM4-5) | BD Biosciences | Cat# 563151; RRID: |
| Rat monoclonal APC/Cyanine7 anti-mouse CD4 Antibody (clone RM4-5) | Biolegend | Cat# 553046; RRID: AB_394582 |
| Rat monoclonal BV421 Anti-Mouse CD4 (clone GK1.5) | BD Biosciences | Cat# 562891; RRID: |
| Rat monoclonal FITC Anti-Mouse CD4 (clone GK1.5) | BD Biosciences | Cat# 100414; RRID: AB_312699 |
| Rat monoclonal PerCP-Cy™5.5 Anti-Mouse CD8a (clone 53-6.7) | BD Biosciences | Cat# 551162; RRID: AB_394081 |
| Rat monoclonal APC-Cy™7 Anti-Mouse CD8a (clone 53-6.7) | BD Biosciences | Cat# 557654; RRID: AB_396769 |
| Rat monoclonal PE-Cy™7 Anti-Mouse CD8a (clone 53-6.7) | BD Biosciences | Cat# 552877; RRID: AB_394506 |
| Rat monoclonal FITC Anti-Mouse CD8a (clone 53-6.7) | BD Biosciences | Cat# 561966; RRID: AB_10896291 |
| Rat monoclonal PE Anti-Mouse CD44 (clone IM7) | BD Biosciences | Cat# 553134; RRID: AB_394649 |
| Rat monoclonal FITC Anti-Mouse CD44 (clone IM7) | BD Biosciences | Cat# 553133; RRID: |
| Rat monoclonal APC anti-mouse CD62L (clone MEL-14) | Biolegend | Cat# 104412; RRID: AB_313099 |
| Rat monoclonal Brilliant Violet 650™ anti-mouse CD62L (clone MEL-14) | Biolegend | Cat# 104453; RRID: |
| Rat monoclonal APC/Cyanine7 anti-mouse CD62L (clone MEL-14) | Biolegend | Cat# 104428; RRID: AB_830799 |
| Rat monoclonal PE-Cy™7 Anti-Mouse CD62L (clone MEL-14) | BD Biosciences | Cat# 560516; RRID: |
| Armenian Hamster monoclonal anti-mouse CD69 (H1.2F3), FITC (clone H1.2F3) | Thermofisher | Cat# 11-0691-85; RRID: AB_465120 |
| Armenian Hamster monoclonal PerCP/Cyanine5.5 anti-mouse CD103 (clone 2E7 ) | Biolegend | Cat# 121416; RRID: |
| Rat monoclonal BV421 Anti-Mouse CD127 (clone A7R34) | BD Biosciences | Cat# 566377; RRID: |
| Syrian hamster monoclonal anti-KLRG1 (2F1), APC (clone 2F1) | Thermofisher | Cat# 17-5893-82; RRID: AB_469469 |
| Rat monoclonal APC-Cy™7 Anti-Mouse CD45 (clone 30-F11) | BD Biosciences | Cat# 557659; RRID: AB_396774 |
| Rat monoclonal Alexa Fluor® 700 anti-mouse CD11b (clone M1/70) | BD Biosciences | Cat# 557960; RRID: AB_396960 |
| Rat monoclonal PE-Cy™7 Anti-Mouse Ly-6C (clone AL-21) | BD Biosciences | Cat# 560593; RRID: |
| Rat monoclonal APC Anti-Mouse F4/80 (clone T45-2342) | BD Biosciences | Cat# 566787; RRID: |
| Hamster monoclonal BV605 Anti-Mouse CD11c (clone HL3) | BD Biosciences | Cat# 563057; RRID: |
| Rat monoclonal PerCP-Cy™5.5 Anti-Mouse CD103 (clone M290 ) | BD Biosciences | Cat# 563637; RRID: |
| Mouse monoclonal APC Anti-Mouse NK-1.1 (clone PK136) | BD Biosciences | Cat# 550627; RRID: AB_398463 |
| Rat monoclonal FITC anti-mouse CD86 (clone GL-1) | Biolegend | Cat# 105006; RRID: AB_313149 |
| Rat monoclonal Brilliant Violet 421 anti-mouse CD206 MMR (clone C068C2) | Biolegend | Cat# 141717; RRID: |
| Rat monoclonal PE anti-mouse I-A/I-E (clone M5/114.15.2) | Biolegend | Cat# 107608; RRID: AB_313323 |
| Armenian Hamster monoclonal APC anti-mouse CD152 (clone UC10-4B9) | Biolegend | Cat# 106310; RRID: |
| Rat monoclonal PerCP/Cyanine5.5 anti-mouse/human CD45R/B220 (clone RA3-6B2) | Biolegend | Cat# 103236; RRID: AB_893354 |
| Mouse monoclonal BV421 Anti-Human CD4 (clone L200) | BD Biosciences | Cat# 562842; RRID: |
| Mouse monoclonal BV605 Anti-Human CD45RA (clone HI100) | BD Biosciences | Cat# 562886; RRID: |
| Mouse monoclonal APC-Cy™7 Anti-Human CD8 (clone RPA-T8) | BD Biosciences | Cat# 557760; RRID: AB_396865 |
| Rat monoclonal PE Anti-Human CCR7 (CD197) (clone 3D12) | BD Biosciences | Cat# 552176; RRID: AB_394354 |
| Rat monoclonal PerCP/Cyanine5.5 anti-mouse CD150 (SLAM) Antibody | Biolegend | Cat# 115921; RRID: |
| Mouse monoclonal BV650 Mouse Anti-Human CD4 | BD Biosciences | Cat# 563737; RRID: |
| Mouse monoclonal RB705 Mouse Anti-Human CD8 | BD Biosciences | Cat# 757674; RRID: |
| PE Donkey anti-rabbit IgG (minimal x-reactivity) (clone Poly4064) | BioLegend | Cat# 406421; RRID |
| Phospho-STA5B-Ser127 Rabbit Polyclonal Antibody | ABclonal | N/A |
| Mouse monoclonal anti-Ets-1 (clone C-4) | Santa Cruz | Cat# sc-55581PE |
| Mouse monoclonal Alexa Fluor™ 488 Anti-RUNX3 (clone R3-5G4) | BD Biosciences | Cat# 565740; RRID: AB_2916367 |
| Rat monoclonal Alexa Fluor® 647 Anti-Mouse Zbtb7b (ThPok) (clone T43-94) | BD Biosciences | Cat# 565500; RRID: AB_2739268 |

| | | |
|---|---|---|
| Rat monoclonal R718 Anti-Mouse Zbtb7b (ThPok) (clone T43-94) | BD Biosciences | Cat# 566942; RRID: AB_2869959 |
| **Antibodies for Immunofluorescence** | | |
| Rat monoclonal Purified anti-mouse/human CD45R/B220 (clone RA3-6B2) | Biolegend | Cat# 103201; RRID: AB_312986 |
| Rabbit recombinant multiclonal Anti-human/mouse CD4 (clone RM1013) | Abcam | Cat# ab288724 |
| Rabbit recombinant monoclonal Anti-mouse CD8 alpha (clone EPR21769) | Abcam | Cat# ab217344 |
| Rabbit Recombinant Monoclonal Anti-human/mouse CD44 (clone EPR18668) | Abcam | Cat# ab232556 |
| **Antibodies for CUT&Tag** | | |
| Rabbit Monoclonal anti-TCF1/TCF7 (clone C63D9) | Cell Signaling | Cat# 2203 |
| Rabbit Monoclonal anti-ETS-1 (clone D8O8A) | Cell Signaling | Cat# 14069 |
| Rabbit monoclonal Anti-c-Myb (phospho S11) (clone EP769Y) | Abcam | Cat# ab45150 |
| Rabbit polyclonal anti-SOX4 | Diagenode | Cat# C15310129 |
| Rabbit Polyclonal anti-TCF12 | Bethyl | Cat# A300-754A |
| Mouse monoclonal PE anti-Ets-2 (clone E-5) | Santa Cruz | Cat# sc-365666PE |
| **Bacterial** | | |
| *M. tuberculosis* H37Rv | Laboratory stock | N/A |
| Bacillus Calmette-Guerin, BCG | Laboratory stock | N/A |
| **Chemicals** | | |
| Middlebrook 7H9 broth | BD Biosciences | Cat# 271310 |
| OADC | BD Biosciences | Cat# 212351 |
| Tween-80 | Sigma | Cat# 9005-65-6 |
| Middlebrook 7H10 agar | Becton Dickson | Cat# 262710 |
| Penicillin | Sigma | Cat# P4333 |
| Heat-inactivated fetal bovine serum | ThermoFisher | Cat# 26010066 |
| Penicillin and streptomycin | ThermoFisher | Cat# 15140148 |
| RPMI-1640 | ThermoFisher | Cat# C11875500 |
| Red Blood Cell Lysing Buffer | Beyotime | Cat# C3702 |
| Macrophage colony-stimulating factor (M-CSF) | ABclonal | Cat# RP01216 |
| 4% paraformaldehyde fixative | Beyotime | Cat# P0099 |
| BD Pharmingen™ Stain Buffer (FBS) | BD Pharmingen™ | Cat# 554656; RRID: |
| Cell Trace Violet | Thermofisher | Cat# C34557 |
| Collagenase II | Diamond | Cat# A004174 |
| DNase I | Sigma | Cat# 10104159001 |
| Incomplete Freund's adjuvant (IFA) | MedChemExpress | Cat# HY-153808A |
| Complete Freund's adjuvant(CFA) | MedChemExpress | Cat# HY-153808 |
| Purified Protein Derivative of Tuberculin (TB-PPD) | Simcere | |
| Pyruvic acid | MedChemExpress | Cat# HY-Y0781 |
| Fumaric acid | MedChemExpress | Cat# HY-W015883 |
| Malic acid | MedChemExpress | Cat# HY-Y1311 |
| Butyric acid | MedChemExpress | Cat# HY-B0350 |
| β-Muricholic acid | MedChemExpress | Cat# HY-133707 |
| Orotic acid | MedChemExpress | Cat# HY-N0157 |
| Cafestol | MedChemExpress | Cat# HY-N6257 |
| Maleic Acid | MedChemExpress | Cat# HY-Y0367 |
| 16:1 (DELTA 9-Cis) PC | MedChemExpress | Cat# HY-157423 |
| α-Linolenic acid | MedChemExpress | Cat# HY-N0728 |
| Gamma-linolenic acid | MedChemExpress | Cat# HY-N7140 |
| Vancomycin | MedChemExpress | Cat# HY-B0671 |
| Neomycin(sulfate) | MedChemExpress | Cat# HY-B0470 |
| Metronidazole | MedChemExpress | Cat# HY-B0318 |

| | | |
|---|---|---|
| Ampicillin | MedChemExpress | Cat# HY-B0522 |
| Parasutterella excrementihominis | HZBio,Microbial Conservation | Cat# HZB904599 |
| Bacteroides fragilis | HZBio,Microbial Conservation | Cat# HZB358365 |
| Rikenella microfusus | HZBio,Microbial Conservation | Cat# HZB721060 |
| Phocaeicola vulgatus | HZBio,Microbial Conservation | Cat# HZB358368 |
| Clostridium butyricum | HZBio,Microbial Conservation | Cat# HZB133180 |
| **Critical commercial assays** | | |
| fixation/permeabilization kit | BD Biosciences | Cat# 554714, RRID: |
| CD4 (L3T4) MicroBeads, mouse | Miltenyi Biotec | Cat# 130-117-043 |
| CD8 (TIL) MicroBeads, mouse | Miltenyi Biotec | Cat# 130-116-478 |
| MojoSort™ Mouse CD8 T Cell Isolation Kit | BioLegend | Cat# 480035 |
| ProteanFect Max Mouse Immunocyte Transfection Kit | Nanoportal Biotech | Cat# PTO3-0010A |
| Easy editing$^{TM}$ kit | Tongxin Technology, Beijing, China | Cat# TX-EE12 |
| InVivoMAb anti-mouse CD3ε (clone 145-2C11) | BioXcell | Cat# BE0001-1 |
| InVivoMAb anti-mouse CD28 (clone 37.51) | BioXcell | Cat# BE0015-1 |
| InVivoMAb anti-mouse Ly6G (clone 1A8) | BioXcell | Cat# BE0075-1 |
| InVivoMAb rat IgG2a isotype control (clone 2A3) | BioXcell | Cat# BE0089 |
| Recombinant Murine IL-2 | Peprotech | Cat# 212-12 |
| Recombinant Human PAK2 | Finetest | Cat# P5717 |
| **Experimental models: Organisms/strains** | | |
| SCID mice | Shanghai Model Organisms Center | Cat# SM-015 |
| C57BL/6 mice | Cyagen Biosciences | N/A |
| CD45.1 mice | Cyagen Biosciences | N/A |
| *Rag1-/-* mice | Cyagen Biosciences | N/A |
| SLAMF1-deficient (*Slamf1* $^{-/-}$) mice | Anhui Medical University, Hefei, China | N/A |
| *Stat5b* $^{S127A}$ | Cyagen Biosciences | N/A |
| **Software and algorithms** | | |
| GraphPad Prism 10.1.2 | Tongji University | https://nic.tongji.edu.cn/zbrjpt/list.htm |
| LAS X | Leica microsystems | https://www.leicamicrosystems.com.cn/cn/ |
| FlowJo 10 | TreeStar | N/A |
| BioRender | BioRender | www.biorender.com |
